## Supplementary material for "A handle on mass coincidence errors in *de novo* sequencing of antibodies by bottom-up proteomics": Combined_027.html

Details Combined\_027 | Stitch OverviewUndefined

### Read Combined\_027

#### Sequence (length=13)

TVLHQDWLDGKEY

#### Spectrum 7243? Spectrum 7243 The raw spectrum of this peptide as annotated by Hecklib. The fragments are coloured according to ion type (see legend). Any peaks with a star '\*' as text can be hovered over to see the full details, first the ion type second the mass shift type. By hovering over the amino acids in the peptide or ions in the legend the corresponding peaks are highlighted. By toggling the 'Unassigned' label you can turn the background (unassigned) peaks on or off in the plot. By updating the slider in the Ion legend you can update the spectrum to only show the top X% of the peaks with labels. The top X% means any peak that is within X% of the highest intensity. By dragging in the spectrum you can zoom in to a specific part of the spectrum and use 'Zoom Out' to get back to the original zoom level. The annotation of the spectrum is based on the given sequence in the peptides file and is done with different software so inconsistencies are likely. The peaks are annotated based on the given sequence, with 20 ppm tolerance.

Copy Data

##### Spectrum 7243 (TSV)

###### Preview

```
Loading example...
```

*Click on the button to copy the data to your clipboard.*

Mz MinMz MaxIntensity Max

WidthHeightPeptide font sizePeptide stroke widthSpectrum font sizeSpectrum stroke widthCompact peptide

Ion legend

wxyz

abcd

OtherUnassignedIon^Charge^_Position_Show for top:%

TVLHQDWLDGKEY

04.57e+59.15e+51.37e+61.83e+6

Zoom Out

y^+1^_2_y^+1^_2_y^+1^_3_c^+2^_7_c^+1^_4_y^+1^_4_c^+1^_5_c^+1^_5_c^+1^_5_y^+1^_5_y^+2^_10_c^+2^_11_y^+2^_11_c^+1^_6_y^+2^_11_c^+1^_6_c^+2^_12_y^+1^_6_y^+2^_12_c^+1^_7_y^+1^_7_z^+1^_7_c^+1^_7_y^+1^_7_c^+1^_8_c^+1^_8_z^+1^_8_c^+1^_8_y^+1^_8_w^+1^_9_c^+1^_9_c^+1^_9_c^+1^_9_y^+1^_9_z^+1^_9_y^+1^_9_c^+1^_10_c^+1^_10_z^+1^_10_y^+1^_10_c^+1^_11_c^+1^_11_w^+1^_11_z^+1^_11_y^+1^_11_c^+1^_12_z^+1^_12_

0778155623343112

Fragment Matches Table

Show background peaks

| Position | Ion type | Intensity | mz Theoretical | mz Error (Th) | mz Error (ppm) | Charge | Series Number |
| --- | --- | --- | --- | --- | --- | --- | --- |
| - | - | 1618 | 122.9 | - | - | 0 | - |
| - | - | 1681 | 126.5 | - | - | 0 | - |
| - | - | 1537 | 127.7 | - | - | 0 | - |
| - | - | 2.987E+04 | 129.1 | - | - | 0 | - |
| - | - | 2464 | 130.1 | - | - | 0 | - |
| - | - | 1800 | 139.3 | - | - | 0 | - |
| - | - | 2134 | 145.6 | - | - | 0 | - |
| - | - | 2206 | 148.9 | - | - | 0 | - |
| - | - | 2565 | 148.9 | - | - | 0 | - |
| - | - | 1796 | 148.9 | - | - | 0 | - |
| - | - | 2697 | 148.9 | - | - | 0 | - |
| - | - | 3279 | 148.9 | - | - | 0 | - |
| - | - | 3579 | 148.9 | - | - | 0 | - |
| - | - | 5176 | 148.9 | - | - | 0 | - |
| - | - | 5307 | 148.9 | - | - | 0 | - |
| - | - | 1.392E+04 | 148.9 | - | - | 0 | - |
| - | - | 2.303E+04 | 148.9 | - | - | 0 | - |
| - | - | 1.712E+04 | 149 | - | - | 0 | - |
| - | - | 8608 | 149 | - | - | 0 | - |
| - | - | 5345 | 149 | - | - | 0 | - |
| - | - | 5102 | 149 | - | - | 0 | - |
| - | - | 4507 | 149 | - | - | 0 | - |
| - | - | 2217 | 149 | - | - | 0 | - |
| - | - | 2383 | 149 | - | - | 0 | - |
| - | - | 1969 | 149.1 | - | - | 0 | - |
| - | - | 1911 | 149.2 | - | - | 0 | - |
| - | - | 1848 | 150.9 | - | - | 0 | - |
| - | - | 2047 | 163.1 | - | - | 0 | - |
| - | - | 1.788E+04 | 173.1 | - | - | 0 | - |
| - | - | 2010 | 186.1 | - | - | 0 | - |
| - | - | 2.732E+04 | 186.1 | - | - | 0 | - |
| - | - | 2130 | 191.4 | - | - | 0 | - |
| - | - | 2497 | 195.1 | - | - | 0 | - |
| - | - | 3.301E+04 | 201.1 | - | - | 0 | - |
| - | - | 2712 | 202.1 | - | - | 0 | - |
| - | - | 2319 | 247.3 | - | - | 0 | - |
| - | - | 4256 | 251.1 | - | - | 0 | - |
| - | - | 2044 | 263.3 | - | - | 0 | - |
| - | - | 2419 | 267.3 | - | - | 0 | - |
| - | - | 1.06E+04 | 283.1 | - | - | 0 | - |
| 12 | y | 3.146E+04 | 293.1 | 0.000283 | 0.9657 | +1 | 2 |
| - | - | 4965 | 294.1 | - | - | 0 | - |
| - | - | 2851 | 297.2 | - | - | 0 | - |
| - | - | 1.843E+04 | 301.2 | - | - | 0 | - |
| 12 | y | 1.047E+04 | 311.1 | 0.0003192 | 1.026 | +1 | 2 |
| - | - | 8473 | 314.2 | - | - | 0 | - |
| - | - | 6823 | 315.2 | - | - | 0 | - |
| - | - | 2760 | 376.9 | - | - | 0 | - |
| - | - | 2715 | 383.9 | - | - | 0 | - |
| - | - | 4220 | 396.2 | - | - | 0 | - |
| - | - | 2415 | 414.2 | - | - | 0 | - |
| - | - | 8067 | 423.3 | - | - | 0 | - |
| - | - | 2984 | 430.2 | - | - | 0 | - |
| 11 | y | 9236 | 439.2 | 0.0009218 | 2.099 | +1 | 3 |
| 7 | c | 2749 | 440.2 | 0.004014 | 9.118 | +2 | 7 |
| - | - | 3.326E+04 | 451.3 | - | - | 0 | - |
| - | - | 8422 | 452.3 | - | - | 0 | - |
| - | - | 2945 | 458.1 | - | - | 0 | - |
| 4 | c | 3275 | 468.3 | 0.0005964 | 1.274 | +1 | 4 |
| 10 | y | 6.557E+04 | 496.2 | 0.000535 | 1.078 | +1 | 4 |
| - | - | 1.439E+04 | 497.2 | - | - | 0 | - |
| - | - | 3641 | 534.3 | - | - | 0 | - |
| - | - | 3156 | 543.3 | - | - | 0 | - |
| - | - | 1.064E+04 | 552.3 | - | - | 0 | - |
| - | - | 6321 | 553.3 | - | - | 0 | - |
| - | - | 3068 | 555.3 | - | - | 0 | - |
| - | - | 8042 | 577.3 | - | - | 0 | - |
| 5 | c | 3018 | 578.3 | 0.007472 | 12.92 | +1 | 5 |
| 5 | c | 2.065E+04 | 579.3 | 0.0002731 | 0.4714 | +1 | 5 |
| - | - | 7619 | 580.3 | - | - | 0 | - |
| - | - | 1.46E+04 | 595.3 | - | - | 0 | - |
| 5 | c | 3.072E+04 | 596.4 | 0.0008243 | 1.382 | +1 | 5 |
| - | - | 8912 | 597.4 | - | - | 0 | - |
| - | - | 2932 | 600.3 | - | - | 0 | - |
| 9 | y | 3.59E+04 | 611.3 | 0.00047 | 0.7688 | +1 | 5 |
| - | - | 1.273E+04 | 611.8 | - | - | 0 | - |
| - | - | 9321 | 612.3 | - | - | 0 | - |
| - | - | 1.163E+04 | 638.3 | - | - | 0 | - |
| - | - | 9659 | 638.8 | - | - | 0 | - |
| 4 | y | 4.55E+04 | 645.8 | 0.0005208 | 0.8064 | +2 | 10 |
| - | - | 2.949E+04 | 646.3 | - | - | 0 | - |
| - | - | 1.233E+04 | 646.8 | - | - | 0 | - |
| 11 | c | 2.48E+04 | 647.3 | 0.0005467 | 0.8446 | +2 | 11 |
| - | - | 1.853E+04 | 647.8 | - | - | 0 | - |
| - | - | 8028 | 648.3 | - | - | 0 | - |
| - | - | 2.345E+04 | 667.4 | - | - | 0 | - |
| - | - | 2.325E+04 | 668.4 | - | - | 0 | - |
| - | - | 7833 | 669.4 | - | - | 0 | - |
| - | - | 2422 | 688.8 | - | - | 0 | - |
| 3 | y | 5513 | 693.8 | 0.01326 | 19.12 | +2 | 11 |
| 6 | c | 3.924E+04 | 694.4 | 0.001401 | 2.018 | +1 | 6 |
| - | - | 1.174E+04 | 695.4 | - | - | 0 | - |
| - | - | 1.082E+04 | 697.9 | - | - | 0 | - |
| - | - | 3571 | 698.9 | - | - | 0 | - |
| - | - | 3263 | 702.3 | - | - | 0 | - |
| 3 | y | 6.709E+04 | 702.3 | 0.0007437 | 1.059 | +2 | 11 |
| - | - | 6.482E+04 | 702.8 | - | - | 0 | - |
| - | - | 3.622E+04 | 703.3 | - | - | 0 | - |
| - | - | 7786 | 703.8 | - | - | 0 | - |
| - | - | 3131 | 704.3 | - | - | 0 | - |
| - | - | 7141 | 710.4 | - | - | 0 | - |
| 6 | c | 5.699E+04 | 711.4 | 0.001644 | 2.311 | +1 | 6 |
| 12 | c | 1.468E+05 | 711.9 | 0.0001148 | 0.1612 | +2 | 12 |
| - | - | 1.104E+05 | 712.4 | - | - | 0 | - |
| - | - | 4.759E+04 | 712.9 | - | - | 0 | - |
| - | - | 3934 | 713.4 | - | - | 0 | - |
| - | - | 5369 | 720.9 | - | - | 0 | - |
| - | - | 3250 | 721.4 | - | - | 0 | - |
| - | - | 2636 | 721.9 | - | - | 0 | - |
| 8 | y | 1.558E+04 | 724.4 | 0.0003665 | 0.5059 | +1 | 6 |
| - | - | 6240 | 725.4 | - | - | 0 | - |
| 2 | y | 4002 | 751.9 | 0.0002666 | 0.3546 | +2 | 12 |
| - | - | 5123 | 752.4 | - | - | 0 | - |
| - | - | 5173 | 779.4 | - | - | 0 | - |
| - | - | 5061 | 779.9 | - | - | 0 | - |
| - | - | 7121 | 780.4 | - | - | 0 | - |
| - | - | 4008 | 780.9 | - | - | 0 | - |
| - | - | 4142 | 781.4 | - | - | 0 | - |
| - | - | 3729 | 784.9 | - | - | 0 | - |
| - | - | 5.025E+04 | 793.4 | - | - | 0 | - |
| - | - | 5.666E+04 | 793.9 | - | - | 0 | - |
| - | - | 3.306E+04 | 794.4 | - | - | 0 | - |
| - | - | 5611 | 794.9 | - | - | 0 | - |
| - | - | 2.204E+04 | 801.9 | - | - | 0 | - |
| - | - | 1.076E+06 | 802.4 | - | - | 0 | - |
| - | - | 9.753E+05 | 802.9 | - | - | 0 | - |
| - | - | 4.227E+05 | 803.4 | - | - | 0 | - |
| - | - | 5.523E+04 | 803.9 | - | - | 0 | - |
| - | - | 2680 | 852.4 | - | - | 0 | - |
| - | - | 2.275E+04 | 853.4 | - | - | 0 | - |
| - | - | 2.714E+04 | 854.4 | - | - | 0 | - |
| - | - | 9311 | 855.5 | - | - | 0 | - |
| - | - | 3279 | 878.4 | - | - | 0 | - |
| 7 | c | 3.268E+04 | 880.4 | 0.001002 | 1.139 | +1 | 7 |
| - | - | 1.441E+04 | 881.4 | - | - | 0 | - |
| - | - | 2995 | 882.4 | - | - | 0 | - |
| 7 | y | 2915 | 893.4 | 0.01206 | 13.49 | +1 | 7 |
| 7 | z | 1.96E+04 | 894.4 | 0.0004085 | 0.4567 | +1 | 7 |
| - | - | 1.28E+04 | 895.4 | - | - | 0 | - |
| - | - | 2.311E+04 | 896.4 | - | - | 0 | - |
| 7 | c | 9.562E+04 | 897.5 | 0.0008181 | 0.9116 | +1 | 7 |
| - | - | 4.789E+04 | 898.5 | - | - | 0 | - |
| - | - | 7918 | 899.5 | - | - | 0 | - |
| - | - | 5192 | 908.4 | - | - | 0 | - |
| - | - | 5449 | 909.4 | - | - | 0 | - |
| 7 | y | 4.586E+04 | 910.4 | 0.0006999 | 0.7688 | +1 | 7 |
| - | - | 1.821E+04 | 911.4 | - | - | 0 | - |
| - | - | 3993 | 912.4 | - | - | 0 | - |
| - | - | 1.12E+04 | 965.4 | - | - | 0 | - |
| - | - | 5334 | 966.4 | - | - | 0 | - |
| - | - | 1.312E+04 | 966.5 | - | - | 0 | - |
| - | - | 1.634E+04 | 967.5 | - | - | 0 | - |
| - | - | 5916 | 968.5 | - | - | 0 | - |
| - | - | 4337 | 975.5 | - | - | 0 | - |
| - | - | 3097 | 980.5 | - | - | 0 | - |
| - | - | 3.606E+04 | 991.5 | - | - | 0 | - |
| 8 | c | 1.607E+04 | 992.5 | 0.005592 | 5.634 | +1 | 8 |
| 8 | c | 6.239E+04 | 993.5 | 0.0009322 | 0.9382 | +1 | 8 |
| - | - | 3.177E+04 | 994.5 | - | - | 0 | - |
| - | - | 8220 | 995.5 | - | - | 0 | - |
| 6 | z | 3.054E+04 | 1009 | 0.002754 | 2.729 | +1 | 8 |
| - | - | 7.786E+04 | 1010 | - | - | 0 | - |
| - | - | 3.337E+04 | 1010 | - | - | 0 | - |
| 8 | c | 1.889E+05 | 1011 | 0.001081 | 1.07 | +1 | 8 |
| - | - | 1.268E+04 | 1011 | - | - | 0 | - |
| - | - | 9.705E+04 | 1012 | - | - | 0 | - |
| - | - | 2.362E+04 | 1013 | - | - | 0 | - |
| - | - | 6363 | 1024 | - | - | 0 | - |
| 6 | y | 4.009E+04 | 1025 | 0.0001891 | 0.1844 | +1 | 8 |
| - | - | 1.975E+04 | 1026 | - | - | 0 | - |
| - | - | 6071 | 1027 | - | - | 0 | - |
| - | - | 6896 | 1064 | - | - | 0 | - |
| - | - | 4056 | 1065 | - | - | 0 | - |
| - | - | 4357 | 1068 | - | - | 0 | - |
| - | - | 3926 | 1069 | - | - | 0 | - |
| 5 | w | 3516 | 1079 | 0.005371 | 4.975 | +1 | 9 |
| - | - | 4.553E+04 | 1082 | - | - | 0 | - |
| - | - | 4.798E+04 | 1083 | - | - | 0 | - |
| - | - | 2.009E+04 | 1084 | - | - | 0 | - |
| - | - | 4894 | 1085 | - | - | 0 | - |
| - | - | 6602 | 1094 | - | - | 0 | - |
| 9 | c | 7007 | 1108 | 0.003848 | 3.475 | +1 | 9 |
| 9 | c | 8.555E+04 | 1109 | 0.0004173 | 0.3765 | +1 | 9 |
| - | - | 5.374E+04 | 1110 | - | - | 0 | - |
| - | - | 1.371E+04 | 1111 | - | - | 0 | - |
| - | - | 3.417E+04 | 1125 | - | - | 0 | - |
| 9 | c | 4.007E+04 | 1126 | 0.003305 | 2.936 | +1 | 9 |
| - | - | 2.117E+04 | 1127 | - | - | 0 | - |
| - | - | 5999 | 1128 | - | - | 0 | - |
| 5 | y | 9394 | 1136 | 0.008932 | 7.859 | +1 | 9 |
| 5 | z | 2.399E+05 | 1137 | 0.000846 | 0.7437 | +1 | 9 |
| - | - | 1.789E+05 | 1139 | - | - | 0 | - |
| - | - | 6.104E+04 | 1140 | - | - | 0 | - |
| - | - | 8898 | 1141 | - | - | 0 | - |
| - | - | 5379 | 1152 | - | - | 0 | - |
| - | - | 3.527E+04 | 1153 | - | - | 0 | - |
| 5 | y | 1.19E+05 | 1154 | 0.001382 | 1.198 | +1 | 9 |
| - | - | 6.064E+04 | 1155 | - | - | 0 | - |
| - | - | 1.466E+04 | 1156 | - | - | 0 | - |
| - | - | 3151 | 1164 | - | - | 0 | - |
| 10 | c | 3.369E+04 | 1166 | 0.001659 | 1.423 | +1 | 10 |
| - | - | 2.123E+04 | 1167 | - | - | 0 | - |
| - | - | 7581 | 1168 | - | - | 0 | - |
| - | - | 4.694E+04 | 1182 | - | - | 0 | - |
| 10 | c | 3.559E+05 | 1183 | 0.0004763 | 0.4028 | +1 | 10 |
| - | - | 2.271E+05 | 1184 | - | - | 0 | - |
| - | - | 6.695E+04 | 1185 | - | - | 0 | - |
| - | - | 6953 | 1186 | - | - | 0 | - |
| - | - | 3814 | 1204 | - | - | 0 | - |
| - | - | 3720 | 1224 | - | - | 0 | - |
| - | - | 3484 | 1232 | - | - | 0 | - |
| - | - | 3304 | 1267 | - | - | 0 | - |
| - | - | 6166 | 1268 | - | - | 0 | - |
| 4 | z | 2.053E+05 | 1275 | 0.001774 | 1.392 | +1 | 10 |
| - | - | 1.432E+05 | 1276 | - | - | 0 | - |
| - | - | 4.923E+04 | 1277 | - | - | 0 | - |
| - | - | 6312 | 1278 | - | - | 0 | - |
| - | - | 9558 | 1290 | - | - | 0 | - |
| 4 | y | 6.255E+04 | 1291 | 0.002066 | 1.601 | +1 | 10 |
| - | - | 4.529E+04 | 1292 | - | - | 0 | - |
| - | - | 1.363E+04 | 1293 | - | - | 0 | - |
| 11 | c | 2.554E+04 | 1294 | 0.002484 | 1.92 | +1 | 11 |
| - | - | 1.616E+04 | 1295 | - | - | 0 | - |
| - | - | 4163 | 1296 | - | - | 0 | - |
| - | - | 1.946E+04 | 1310 | - | - | 0 | - |
| 11 | c | 1.505E+05 | 1311 | 0.001567 | 1.196 | +1 | 11 |
| - | - | 1.184E+05 | 1312 | - | - | 0 | - |
| - | - | 3.579E+04 | 1313 | - | - | 0 | - |
| - | - | 5744 | 1314 | - | - | 0 | - |
| - | - | 1.628E+04 | 1337 | - | - | 0 | - |
| - | - | 9329 | 1338 | - | - | 0 | - |
| - | - | 4826 | 1344 | - | - | 0 | - |
| 3 | w | 1.688E+04 | 1345 | 0.008854 | 6.585 | +1 | 11 |
| - | - | 1.507E+04 | 1346 | - | - | 0 | - |
| - | - | 4086 | 1347 | - | - | 0 | - |
| - | - | 3239 | 1352 | - | - | 0 | - |
| 3 | z | 1.081E+05 | 1388 | 0.001122 | 0.8083 | +1 | 11 |
| - | - | 9.977E+04 | 1389 | - | - | 0 | - |
| - | - | 3.624E+04 | 1390 | - | - | 0 | - |
| - | - | 4986 | 1391 | - | - | 0 | - |
| - | - | 3.214E+04 | 1396 | - | - | 0 | - |
| - | - | 2.823E+04 | 1397 | - | - | 0 | - |
| - | - | 7946 | 1398 | - | - | 0 | - |
| 3 | y | 4.901E+04 | 1404 | 0.002268 | 1.615 | +1 | 11 |
| - | - | 4.676E+04 | 1405 | - | - | 0 | - |
| - | - | 1.839E+04 | 1406 | - | - | 0 | - |
| - | - | 4355 | 1425 | - | - | 0 | - |
| - | - | 4736 | 1426 | - | - | 0 | - |
| - | - | 4650 | 1431 | - | - | 0 | - |
| - | - | 1.465E+04 | 1439 | - | - | 0 | - |
| 12 | c | 5.181E+05 | 1440 | 0.00168 | 1.167 | +1 | 12 |
| - | - | 3.929E+05 | 1441 | - | - | 0 | - |
| - | - | 1.505E+05 | 1442 | - | - | 0 | - |
| - | - | 1.482E+04 | 1443 | - | - | 0 | - |
| - | - | 4210 | 1475 | - | - | 0 | - |
| 2 | z | 2.226E+04 | 1487 | 0.001909 | 1.284 | +1 | 12 |
| - | - | 2.152E+04 | 1488 | - | - | 0 | - |
| - | - | 9579 | 1489 | - | - | 0 | - |
| - | - | 3443 | 1506 | - | - | 0 | - |
| - | - | 6467 | 1516 | - | - | 0 | - |
| - | - | 1.645E+04 | 1517 | - | - | 0 | - |
| - | - | 1.444E+04 | 1518 | - | - | 0 | - |
| - | - | 7972 | 1519 | - | - | 0 | - |
| - | - | 4110 | 1527 | - | - | 0 | - |
| - | - | 6495 | 1528 | - | - | 0 | - |
| - | - | 5255 | 1529 | - | - | 0 | - |
| - | - | 5459 | 1532 | - | - | 0 | - |
| - | - | 9770 | 1533 | - | - | 0 | - |
| - | - | 1.288E+04 | 1534 | - | - | 0 | - |
| - | - | 8161 | 1535 | - | - | 0 | - |
| - | - | 6316 | 1541 | - | - | 0 | - |
| - | - | 1.261E+04 | 1542 | - | - | 0 | - |
| - | - | 4.301E+04 | 1543 | - | - | 0 | - |
| - | - | 5.089E+04 | 1544 | - | - | 0 | - |
| - | - | 1.159E+05 | 1545 | - | - | 0 | - |
| - | - | 9.233E+04 | 1546 | - | - | 0 | - |
| - | - | 3.674E+04 | 1547 | - | - | 0 | - |
| - | - | 4733 | 1548 | - | - | 0 | - |
| - | - | 1.073E+04 | 1549 | - | - | 0 | - |
| - | - | 1.178E+04 | 1550 | - | - | 0 | - |
| - | - | 8308 | 1551 | - | - | 0 | - |
| - | - | 5.658E+04 | 1559 | - | - | 0 | - |
| - | - | 1.267E+05 | 1560 | - | - | 0 | - |
| - | - | 1.147E+05 | 1561 | - | - | 0 | - |
| - | - | 6.387E+04 | 1562 | - | - | 0 | - |
| - | - | 2.173E+04 | 1563 | - | - | 0 | - |
| - | - | 1.221E+04 | 1569 | - | - | 0 | - |
| - | - | 1.694E+04 | 1570 | - | - | 0 | - |
| - | - | 9192 | 1571 | - | - | 0 | - |
| - | - | 3585 | 1572 | - | - | 0 | - |
| - | - | 3176 | 1576 | - | - | 0 | - |
| - | - | 1.76E+05 | 1577 | - | - | 0 | - |
| - | - | 1.544E+05 | 1578 | - | - | 0 | - |
| - | - | 5308 | 1578 | - | - | 0 | - |
| - | - | 7.211E+04 | 1579 | - | - | 0 | - |
| - | - | 9178 | 1580 | - | - | 0 | - |
| - | - | 1.653E+05 | 1587 | - | - | 0 | - |
| - | - | 4.813E+05 | 1588 | - | - | 0 | - |
| - | - | 3.906E+05 | 1589 | - | - | 0 | - |
| - | - | 1.505E+05 | 1590 | - | - | 0 | - |
| - | - | 1.958E+04 | 1591 | - | - | 0 | - |
| - | - | 2.126E+04 | 1603 | - | - | 0 | - |
| - | - | 7.578E+05 | 1604 | - | - | 0 | - |
| - | - | 1.812E+06 | 1605 | - | - | 0 | - |
| - | - | 1.349E+06 | 1606 | - | - | 0 | - |
| - | - | 5.245E+05 | 1607 | - | - | 0 | - |
| - | - | 6.429E+04 | 1608 | - | - | 0 | - |
| - | - | 3907 | 3081 | - | - | 0 | - |

m/z Charge Intensity FragmentType MassShift Position
122.93534851074219 0 1617.7184
126.45089721679688 0 1681.311
127.67321014404297 0 1536.764
129.10211181640625 0 29872.287
130.10585021972656 0 2464.1118
139.26576232910156 0 1799.7001
145.59568786621094 0 2133.5798
148.8660430908203 0 2206.0464
148.87306213378906 0 2564.812
148.88034057617188 0 1796.4637
148.90185546875 0 2696.5718
148.9088897705078 0 3279.0964
148.91624450683594 0 3579.427
148.92347717285156 0 5175.8213
148.93081665039062 0 5307.4507
148.9381866455078 0 13920.432
148.9458770751953 0 23030.695
148.9624786376953 0 17115.715
148.9701385498047 0 8608.251
148.97732543945312 0 5344.6816
148.98414611816406 0 5102.4897
148.99156188964844 0 4506.977
149.01332092285156 0 2217.163
149.0352325439453 0 2383.324
149.1077117919922 0 1969.4014
149.2371368408203 0 1911.3608
150.8721466064453 0 1848.2457
163.0591583251953 0 2046.7974
173.12828063964844 0 17880.564
186.11618041992188 0 2010.2964
186.12339782714844 0 27315.773
191.39801025390625 0 2129.6353
195.11293029785156 0 2497.4941
201.12301635742188 0 33008.285
202.1261444091797 0 2712.0269
247.31971740722656 0 2319.1045
251.14883422851562 0 4256.4927
263.28558349609375 0 2043.8104
267.26800537109375 0 2418.668
283.1395263671875 0 10603.049
293.1129150390625 0 31457.666 y Water loss 11
294.1164245605469 0 4965.1187
297.15631103515625 0 2850.9744
301.15008544921875 0 18434.91
311.1234436035156 0 10467.84 y 11
314.2078857421875 0 8472.839
315.16656494140625 0 6823.3677
376.89044189453125 0 2760.256
383.9182434082031 0 2715.4736
396.2227783203125 0 4220.0063
414.23284912109375 0 2415.3352
423.27130126953125 0 8067.324
430.1943054199219 0 2984.1055
439.2178039550781 0 9236.299 y 10
440.22320556640625 0 2748.8801 c Water loss 6
451.26593017578125 0 33262.715
452.2705993652344 0 8421.94
458.1185302734375 0 2944.615
468.29229736328125 0 3274.5566 c 3
496.2396545410156 0 65572.055 y 9
497.24334716796875 0 14393.621
534.3260498046875 0 3641.2852
543.2762451171875 0 3156.3645
552.3367309570312 0 10637.914
553.340576171875 0 6320.7656
555.252685546875 0 3067.933
577.3338012695312 0 8042.0503
578.3334350585938 0 3018.3625 c Water loss 4
579.3251953125 0 20650.053 c Ammonia loss 4
580.3265380859375 0 7619.489
595.34375 0 14603.237
596.3506469726562 0 30722.908 c 4
597.35400390625 0 8912.423
600.3136596679688 0 2931.7627
611.2666625976562 0 35904.105 y 8
611.7962036132812 0 12727.103
612.2709350585938 0 9320.837
638.3272705078125 0 11625.174
638.8283081054688 0 9659.245
645.7905883789062 0 45501.465 y 3
646.2916870117188 0 29493.467
646.793701171875 0 12334.028
647.3323974609375 0 24802.043 c Ammonia loss 10
647.8341064453125 0 18532.195
648.3353271484375 0 8027.6406
667.36376953125 0 23448.965
668.3707275390625 0 23245.977
669.3734741210938 0 7833.042
688.8466796875 0 2421.844
693.8331298828125 0 5513.3364 y Ammonia loss 2
694.3504638671875 0 39235.55 c Ammonia loss 5
695.3549194335938 0 11744.003
697.8555297851562 0 10817.7295
698.8596801757812 0 3570.5752
702.2626953125 0 3263.3386
702.3323974609375 0 67094.98 y 2
702.8374633789062 0 64815.465
703.3408813476562 0 36224.168
703.8478393554688 0 7785.534
704.3443603515625 0 3130.9092
710.3707275390625 0 7141.384
711.3767700195312 0 56993.637 c 5
711.8541259765625 0 146756.86 c Ammonia loss 11
712.3565063476562 0 110375.11
712.8565673828125 0 47593.117
713.35595703125 0 3934.2324
720.8593139648438 0 5369.2476
721.3650512695312 0 3249.5415
721.86474609375 0 2636.2166
724.350830078125 0 15575.596 y 7
725.353271484375 0 6240.376
751.8676147460938 0 4002.2542 y 1
752.3698120117188 0 5122.772
779.3860473632812 0 5173.303
779.8873901367188 0 5060.5713
780.3797607421875 0 7121.4595
780.8836059570312 0 4008.387
781.378662109375 0 4142.239
784.8828735351562 0 3729.268
793.3858642578125 0 50254.797
793.884521484375 0 56664.332
794.3853149414062 0 33058.023
794.8837280273438 0 5610.837
801.8981323242188 0 22038.965
802.3909301757812 0 1076178.8
802.892333984375 0 975332.9
803.3939819335938 0 422729.9
803.8943481445312 0 55230.555
852.4362182617188 0 2679.8735
853.4425048828125 0 22746.39
854.4491577148438 0 27137.617
855.4530029296875 0 9311.351
878.4386596679688 0 3278.9758
880.43017578125 0 32682.36 c Ammonia loss 6
881.4346313476562 0 14406.033
882.4376831054688 0 2994.9248
893.416015625 0 2915.2031 y Ammonia loss 6
894.411376953125 0 19599.123 z 6
895.4169311523438 0 12804.64
896.4454956054688 0 23106.645
897.4569091796875 0 95616.55 c 6
898.4607543945312 0 47890.914
899.4607543945312 0 7918.2563
908.4241333007812 0 5191.9893
909.4258422851562 0 5448.9053
910.4298095703125 0 45859.957 y 6
911.4327392578125 0 18213.209
912.4351196289062 0 3992.5046
965.447021484375 0 11195.561
966.4396362304688 0 5333.7754
966.5316772460938 0 13119.694
967.53271484375 0 16344.382
968.5369262695312 0 5915.7695
975.501220703125 0 4336.739
980.4652099609375 0 3096.783
991.5232543945312 0 36064.742
992.525634765625 0 16072.156 c Water loss 7
993.5161743164062 0 62386.188 c Ammonia loss 7
994.5180053710938 0 31770.102
995.5182495117188 0 8219.935
1009.4359741210938 0 30542.133 z 5
1009.5350341796875 0 77856.516
1010.43994140625 0 33368.54
1010.5407104492188 0 188921.27 c 7
1011.44580078125 0 12678.417
1011.5441284179688 0 97053.414
1012.5480346679688 0 23618.637
1024.4552001953125 0 6362.5547
1025.4576416015625 0 40088.81 y 5
1026.4610595703125 0 19747.227
1027.463623046875 0 6071.3364
1063.54541015625 0 6895.944
1064.539306640625 0 4056.4875
1067.5343017578125 0 4356.665
1068.52978515625 0 3925.7395
1079.462646484375 0 3516.3955 w 4
1081.55615234375 0 45531.527
1082.5599365234375 0 47978.316
1083.5645751953125 0 20089.027
1084.5740966796875 0 4893.938
1093.5279541015625 0 6602.371
1107.5543212890625 0 7006.6733 c Water loss 8
1108.5426025390625 0 85554.914 c Ammonia loss 8
1109.5435791015625 0 53737.72
1110.5455322265625 0 13706.178
1124.560791015625 0 34172.086
1125.5654296875 0 40065.547 c 8
1126.5692138671875 0 21167.295
1127.5706787109375 0 5999.3135
1136.4984130859375 0 9393.876 y Ammonia loss 4
1137.4964599609375 0 239934.8 z 4
1138.50146484375 0 178903.47
1139.50537109375 0 61041.613
1140.51171875 0 8897.685
1151.5166015625 0 5379.1934
1152.508544921875 0 35274.195
1153.5146484375 0 118954.59 y 4
1154.5185546875 0 60638.004
1155.52001953125 0 14664.137
1163.5704345703125 0 3151.0059
1165.5653076171875 0 33692.37 c Ammonia loss 9
1166.565673828125 0 21226.742
1167.5740966796875 0 7581.078
1181.5845947265625 0 46937.586
1182.5897216796875 0 355920.9 c 9
1183.5926513671875 0 227094.67
1184.594482421875 0 66953.13
1185.602294921875 0 6953.211
1203.5157470703125 0 3814.3433
1223.5782470703125 0 3719.977
1231.5589599609375 0 3484.0742
1266.673583984375 0 3303.6978
1267.6688232421875 0 6165.5522
1274.554443359375 0 205269.75 z 3
1275.557861328125 0 143157.61
1276.560546875 0 49229.883
1277.575927734375 0 6311.801
1289.56591796875 0 9557.687
1290.5728759765625 0 62551.81 y 3
1291.5771484375 0 45292.918
1292.5797119140625 0 13631.681
1293.6561279296875 0 25535.084 c Ammonia loss 10
1294.660400390625 0 16157.729
1295.6507568359375 0 4163.225
1309.6798095703125 0 19457.236
1310.68359375 0 150479.56 c 10
1311.6864013671875 0 118418.59
1312.6888427734375 0 35789.195
1313.6875 0 5743.537
1336.699462890625 0 16284.648
1337.7022705078125 0 9328.998
1343.6358642578125 0 4825.747
1344.5943603515625 0 16880.031 w 2
1345.59326171875 0 15073.348
1346.5889892578125 0 4086.3394
1351.727294921875 0 3239.0422
1387.63916015625 0 108061.664 z 2
1388.642333984375 0 99773.49
1389.6439208984375 0 36238.31
1390.6468505859375 0 4986.1987
1395.7130126953125 0 32141.014
1396.7137451171875 0 28232.5
1397.7203369140625 0 7946.0396
1403.65673828125 0 49005.945 y 2
1404.659912109375 0 46760.516
1405.664306640625 0 18385.178
1424.7122802734375 0 4354.511
1425.709716796875 0 4735.904
1430.6502685546875 0 4650.2646
1438.7381591796875 0 14653.824
1439.72607421875 0 518083.9 c 11
1440.7294921875 0 392887.53
1441.731689453125 0 150502.75
1442.73046875 0 14821.219
1474.7313232421875 0 4209.805
1486.706787109375 0 22255.936 z 1
1487.7119140625 0 21516.031
1488.7161865234375 0 9579.238
1505.7449951171875 0 3442.9268
1515.763916015625 0 6466.788
1516.7261962890625 0 16453.46
1517.7249755859375 0 14442.368
1518.72265625 0 7971.601
1526.74609375 0 4110.3193
1527.7633056640625 0 6494.5454
1528.7579345703125 0 5255.318
1531.7259521484375 0 5459.131
1532.7611083984375 0 9769.686
1533.7491455078125 0 12881.745
1534.742919921875 0 8161.3804
1540.7750244140625 0 6316.281
1541.76123046875 0 12612.843
1542.768310546875 0 43012.08
1543.7706298828125 0 50887.08
1544.7626953125 0 115868.67
1545.7633056640625 0 92334.02
1546.7645263671875 0 36744.906
1547.7603759765625 0 4732.6323
1548.718505859375 0 10728.954
1549.726806640625 0 11777.881
1550.7264404296875 0 8307.716
1558.7757568359375 0 56584.652
1559.767578125 0 126688.73
1560.7744140625 0 114732.73
1561.7764892578125 0 63868.742
1562.78271484375 0 21732.127
1568.759765625 0 12214.863
1569.7587890625 0 16938.764
1570.7669677734375 0 9191.965
1571.7593994140625 0 3585.096
1575.78759765625 0 3176.4517
1576.7862548828125 0 176016.98
1577.7882080078125 0 154411.08
1578.0335693359375 0 5307.5186
1578.7911376953125 0 72111.81
1579.7952880859375 0 9178.407
1586.769775390625 0 165330.97
1587.759033203125 0 481295.75
1588.7607421875 0 390550.88
1589.76123046875 0 150485.88
1590.7633056640625 0 19578.736
1602.7921142578125 0 21260.82
1603.77392578125 0 757821.3
1604.7796630859375 0 1811625.4
1605.78369140625 0 1349095
1606.787109375 0 524460.94
1607.7890625 0 64287.51
3081.027099609375 0 3906.5366

Spectrum Details

|  |  |
| --- | --- |
| Matched peaks? Matched peaks The total absolute number of peaks matched. Additionally in brackets the total fraction of peaks matched and the total number of peaks is shown. | 47 (15.26% of 308) |
| FDR? FDR The false discovery rate estimated for this peptide. It is calculated by matching all theoretical fragments with a non-integer shift with the raw peaks for this spectrum. This is done with 40 different shifts. The resulting percentage is the average number of annotated peaks over the number of annotated peaks with the correct spectrum. | 3.19% |
| Satellite FDR? Satellite FDR See the FDR for details on its calculation. This satellite ion specific FDR only contains the satellite ions (d/w) for I/L/J positions. | 0.00% |
| PSM Score? PSM Score The PSM Score as given by Hecklib to this annotated spectrum. It is shown with three significant figures. | 493 |

#### Spectrum 7331? Spectrum 7331 The raw spectrum of this peptide as annotated by Hecklib. The fragments are coloured according to ion type (see legend). Any peaks with a star '\*' as text can be hovered over to see the full details, first the ion type second the mass shift type. By hovering over the amino acids in the peptide or ions in the legend the corresponding peaks are highlighted. By toggling the 'Unassigned' label you can turn the background (unassigned) peaks on or off in the plot. By updating the slider in the Ion legend you can update the spectrum to only show the top X% of the peaks with labels. The top X% means any peak that is within X% of the highest intensity. By dragging in the spectrum you can zoom in to a specific part of the spectrum and use 'Zoom Out' to get back to the original zoom level. The annotation of the spectrum is based on the given sequence in the peptides file and is done with different software so inconsistencies are likely. The peaks are annotated based on the given sequence, with 20 ppm tolerance.

Copy Data

##### Spectrum 7331 (TSV)

###### Preview

```
Loading example...
```

*Click on the button to copy the data to your clipboard.*

Mz MinMz MaxIntensity Max

WidthHeightPeptide font sizePeptide stroke widthSpectrum font sizeSpectrum stroke widthCompact peptide

Ion legend

wxyz

abcd

OtherUnassignedIon^Charge^_Position_Show for top:%

TVLHQDWLDGKEY

02.15e+54.30e+56.45e+58.60e+5

Zoom Out

a^+1^_2_y^+3^_4_a^+1^_2_y^+1^_1_b^+1^_2_b^+1^_2_y^+2^_3_y^+2^_4_b^+2^_5_y^+1^_2_b^+1^_3_y^+2^_5_y^+3^_7_y^+2^_5_y^+1^_2_b^+1^_3_b^+2^_6_b^+2^_6_b^+2^_6_y^+2^_6_y^+2^_6_y^+3^_9_b^+3^_10_y^+1^_3_y^+1^_3_b^+2^_7_b^+2^_7_b^+1^_4_y^+1^_3_b^+2^_7_y^+2^_7_b^+1^_4_y^+2^_7_y^+1^_4_b^+2^_8_b^+2^_8_y^+1^_4_\*\*\*b^+2^_9_b^+2^_9_b^+1^_5_y^+2^_9_y^+2^_9_b^+1^_5_y^+1^_5_y^+1^_5_y^+2^_10_y^+2^_10_y^+2^_10_b^+1^_6_b^+1^_6_y^+2^_11_y^+2^_11_b^+1^_6_y^+2^_11_b^+2^_12_b^+2^_12_y^+1^_6_b^+2^_12_y^+1^_6_y^+2^_12_y^+2^_12_y^+2^_12_\*b^+1^_7_b^+1^_7_b^+1^_7_y^+1^_7_b^+1^_8_b^+1^_8_b^+1^_8_y^+1^_8_y^+1^_8_b^+1^_9_b^+1^_9_y^+1^_9_y^+1^_9_y^+1^_9_b^+1^_10_

0778155723353114

Fragment Matches Table

Show background peaks

| Position | Ion type | Intensity | mz Theoretical | mz Error (Th) | mz Error (ppm) | Charge | Series Number |
| --- | --- | --- | --- | --- | --- | --- | --- |
| - | - | 8650 | 120.1 | - | - | 0 | - |
| - | - | 1005 | 121 | - | - | 0 | - |
| - | - | 1069 | 121.1 | - | - | 0 | - |
| - | - | 4219 | 122.1 | - | - | 0 | - |
| - | - | 4883 | 123 | - | - | 0 | - |
| - | - | 1507 | 123.1 | - | - | 0 | - |
| - | - | 948.9 | 125.1 | - | - | 0 | - |
| - | - | 761.9 | 125.4 | - | - | 0 | - |
| - | - | 1827 | 126.1 | - | - | 0 | - |
| - | - | 7478 | 127.1 | - | - | 0 | - |
| - | - | 1423 | 127.1 | - | - | 0 | - |
| - | - | 2911 | 127.1 | - | - | 0 | - |
| - | - | 1037 | 128 | - | - | 0 | - |
| - | - | 922.6 | 128.1 | - | - | 0 | - |
| - | - | 5.511E+04 | 128.1 | - | - | 0 | - |
| - | - | 3311 | 129.1 | - | - | 0 | - |
| - | - | 5.652E+05 | 129.1 | - | - | 0 | - |
| - | - | 2.382E+05 | 130.1 | - | - | 0 | - |
| - | - | 3574 | 130.1 | - | - | 0 | - |
| - | - | 3.574E+04 | 130.1 | - | - | 0 | - |
| - | - | 2.291E+04 | 131.1 | - | - | 0 | - |
| - | - | 5.45E+04 | 132.1 | - | - | 0 | - |
| - | - | 891.2 | 133.1 | - | - | 0 | - |
| - | - | 3894 | 133.1 | - | - | 0 | - |
| - | - | 1006 | 135.2 | - | - | 0 | - |
| - | - | 6.728E+04 | 136.1 | - | - | 0 | - |
| - | - | 5881 | 137.1 | - | - | 0 | - |
| - | - | 1088 | 137.2 | - | - | 0 | - |
| - | - | 6234 | 138.1 | - | - | 0 | - |
| - | - | 4064 | 138.1 | - | - | 0 | - |
| - | - | 1151 | 139.1 | - | - | 0 | - |
| - | - | 5655 | 139.1 | - | - | 0 | - |
| - | - | 1.004E+04 | 141.1 | - | - | 0 | - |
| - | - | 5138 | 142.1 | - | - | 0 | - |
| - | - | 3458 | 143 | - | - | 0 | - |
| - | - | 2311 | 144.1 | - | - | 0 | - |
| - | - | 4474 | 145.1 | - | - | 0 | - |
| - | - | 1386 | 146.1 | - | - | 0 | - |
| - | - | 3055 | 147 | - | - | 0 | - |
| - | - | 1.375E+04 | 149 | - | - | 0 | - |
| - | - | 3414 | 150.1 | - | - | 0 | - |
| - | - | 1.11E+04 | 151.1 | - | - | 0 | - |
| - | - | 1110 | 152.1 | - | - | 0 | - |
| - | - | 1037 | 154.1 | - | - | 0 | - |
| - | - | 2585 | 155 | - | - | 0 | - |
| - | - | 2058 | 155.1 | - | - | 0 | - |
| - | - | 3.481E+04 | 155.1 | - | - | 0 | - |
| 2 | a | 2.403E+04 | 155.1 | 0.0003966 | 2.557 | +1 | 2 |
| - | - | 1149 | 156 | - | - | 0 | - |
| - | - | 2875 | 156.1 | - | - | 0 | - |
| - | - | 1764 | 156.1 | - | - | 0 | - |
| - | - | 1451 | 156.1 | - | - | 0 | - |
| - | - | 2184 | 157.1 | - | - | 0 | - |
| - | - | 1113 | 157.1 | - | - | 0 | - |
| - | - | 3018 | 158.1 | - | - | 0 | - |
| - | - | 4.239E+05 | 159.1 | - | - | 0 | - |
| - | - | 4522 | 160.1 | - | - | 0 | - |
| - | - | 2756 | 160.1 | - | - | 0 | - |
| - | - | 4.24E+04 | 160.1 | - | - | 0 | - |
| - | - | 1818 | 161.1 | - | - | 0 | - |
| - | - | 1153 | 162.7 | - | - | 0 | - |
| - | - | 3.203E+04 | 165.1 | - | - | 0 | - |
| - | - | 1333 | 165.1 | - | - | 0 | - |
| - | - | 7.089E+04 | 166.1 | - | - | 0 | - |
| 10 | y | 3243 | 166.1 | 0.001969 | 11.86 | +3 | 4 |
| - | - | 5192 | 166.1 | - | - | 0 | - |
| - | - | 1.368E+04 | 167.1 | - | - | 0 | - |
| - | - | 4712 | 167.1 | - | - | 0 | - |
| - | - | 1101 | 167.1 | - | - | 0 | - |
| - | - | 6946 | 168.1 | - | - | 0 | - |
| - | - | 5292 | 168.1 | - | - | 0 | - |
| - | - | 5467 | 168.1 | - | - | 0 | - |
| - | - | 7294 | 169.1 | - | - | 0 | - |
| - | - | 1106 | 169.1 | - | - | 0 | - |
| - | - | 1.594E+04 | 169.1 | - | - | 0 | - |
| - | - | 1.825E+04 | 170.1 | - | - | 0 | - |
| - | - | 2023 | 171.1 | - | - | 0 | - |
| - | - | 2578 | 171.1 | - | - | 0 | - |
| - | - | 2174 | 171.1 | - | - | 0 | - |
| - | - | 6226 | 172.1 | - | - | 0 | - |
| - | - | 1678 | 172.1 | - | - | 0 | - |
| - | - | 8185 | 173.1 | - | - | 0 | - |
| 2 | a | 8.519E+05 | 173.1 | 0.0004978 | 2.875 | +1 | 2 |
| - | - | 2427 | 173.5 | - | - | 0 | - |
| - | - | 7.427E+04 | 174.1 | - | - | 0 | - |
| - | - | 4728 | 175.1 | - | - | 0 | - |
| - | - | 2135 | 177.1 | - | - | 0 | - |
| - | - | 2.656E+04 | 178.1 | - | - | 0 | - |
| - | - | 1488 | 179.1 | - | - | 0 | - |
| - | - | 5853 | 181.1 | - | - | 0 | - |
| 13 | y | 7.922E+04 | 182.1 | 0.0004954 | 2.721 | +1 | 1 |
| - | - | 6802 | 183.1 | - | - | 0 | - |
| 2 | b | 2.202E+04 | 183.1 | 0.0004008 | 2.189 | +1 | 2 |
| - | - | 2599 | 184.1 | - | - | 0 | - |
| - | - | 1201 | 185.1 | - | - | 0 | - |
| - | - | 3392 | 185.2 | - | - | 0 | - |
| - | - | 1.837E+05 | 186.1 | - | - | 0 | - |
| - | - | 3.825E+04 | 187.1 | - | - | 0 | - |
| - | - | 1.599E+04 | 187.1 | - | - | 0 | - |
| - | - | 1456 | 187.1 | - | - | 0 | - |
| - | - | 4542 | 188.1 | - | - | 0 | - |
| - | - | 4317 | 188.1 | - | - | 0 | - |
| - | - | 6685 | 190.1 | - | - | 0 | - |
| - | - | 4997 | 190.1 | - | - | 0 | - |
| - | - | 3914 | 194.1 | - | - | 0 | - |
| - | - | 4.161E+04 | 195.1 | - | - | 0 | - |
| - | - | 2.959E+04 | 196.1 | - | - | 0 | - |
| - | - | 2706 | 196.1 | - | - | 0 | - |
| - | - | 1667 | 197.1 | - | - | 0 | - |
| - | - | 3215 | 197.1 | - | - | 0 | - |
| - | - | 1455 | 197.2 | - | - | 0 | - |
| - | - | 5290 | 198.1 | - | - | 0 | - |
| - | - | 1.525E+04 | 198.1 | - | - | 0 | - |
| - | - | 3526 | 199.1 | - | - | 0 | - |
| - | - | 3673 | 199.1 | - | - | 0 | - |
| - | - | 1548 | 199.1 | - | - | 0 | - |
| - | - | 2508 | 200.1 | - | - | 0 | - |
| 2 | b | 2.27E+05 | 201.1 | 0.0004257 | 2.117 | +1 | 2 |
| - | - | 2121 | 202.1 | - | - | 0 | - |
| - | - | 2.161E+04 | 202.1 | - | - | 0 | - |
| - | - | 2696 | 204.1 | - | - | 0 | - |
| - | - | 1.708E+04 | 205.1 | - | - | 0 | - |
| - | - | 1651 | 205.1 | - | - | 0 | - |
| - | - | 5441 | 205.1 | - | - | 0 | - |
| - | - | 2179 | 206.1 | - | - | 0 | - |
| - | - | 4475 | 206.1 | - | - | 0 | - |
| - | - | 7481 | 207.2 | - | - | 0 | - |
| - | - | 1782 | 209.1 | - | - | 0 | - |
| - | - | 6253 | 210.1 | - | - | 0 | - |
| 11 | y | 6215 | 211.1 | 0.0005118 | 2.424 | +2 | 3 |
| - | - | 5798 | 212.1 | - | - | 0 | - |
| - | - | 1.956E+04 | 213.1 | - | - | 0 | - |
| - | - | 2111 | 213.2 | - | - | 0 | - |
| - | - | 1587 | 214.1 | - | - | 0 | - |
| - | - | 1183 | 215.1 | - | - | 0 | - |
| - | - | 1707 | 215.1 | - | - | 0 | - |
| - | - | 2254 | 215.1 | - | - | 0 | - |
| - | - | 4183 | 216.1 | - | - | 0 | - |
| - | - | 1251 | 221.1 | - | - | 0 | - |
| - | - | 2352 | 221.1 | - | - | 0 | - |
| - | - | 7312 | 221.1 | - | - | 0 | - |
| - | - | 4076 | 221.1 | - | - | 0 | - |
| - | - | 2459 | 221.1 | - | - | 0 | - |
| - | - | 2678 | 221.1 | - | - | 0 | - |
| - | - | 2052 | 222.1 | - | - | 0 | - |
| - | - | 6066 | 222.1 | - | - | 0 | - |
| - | - | 2951 | 223.1 | - | - | 0 | - |
| - | - | 1.169E+04 | 223.1 | - | - | 0 | - |
| - | - | 1994 | 223.1 | - | - | 0 | - |
| - | - | 7.524E+04 | 223.2 | - | - | 0 | - |
| - | - | 3399 | 224.1 | - | - | 0 | - |
| - | - | 7707 | 224.2 | - | - | 0 | - |
| - | - | 8065 | 225 | - | - | 0 | - |
| - | - | 1441 | 226 | - | - | 0 | - |
| - | - | 1.511E+04 | 226.1 | - | - | 0 | - |
| - | - | 1549 | 226.2 | - | - | 0 | - |
| - | - | 1.158E+04 | 227.1 | - | - | 0 | - |
| - | - | 3133 | 227.2 | - | - | 0 | - |
| - | - | 2776 | 228.1 | - | - | 0 | - |
| - | - | 2.048E+04 | 229.1 | - | - | 0 | - |
| - | - | 7156 | 229.2 | - | - | 0 | - |
| - | - | 1550 | 230.1 | - | - | 0 | - |
| - | - | 2885 | 230.1 | - | - | 0 | - |
| - | - | 2467 | 230.2 | - | - | 0 | - |
| - | - | 1.101E+04 | 233.1 | - | - | 0 | - |
| - | - | 8.248E+04 | 234.1 | - | - | 0 | - |
| - | - | 1.154E+04 | 235.1 | - | - | 0 | - |
| - | - | 2957 | 235.2 | - | - | 0 | - |
| - | - | 2417 | 237.1 | - | - | 0 | - |
| - | - | 8837 | 238.1 | - | - | 0 | - |
| - | - | 2876 | 238.1 | - | - | 0 | - |
| - | - | 2065 | 239.1 | - | - | 0 | - |
| - | - | 1.529E+04 | 239.1 | - | - | 0 | - |
| - | - | 1978 | 239.1 | - | - | 0 | - |
| - | - | 4276 | 240.1 | - | - | 0 | - |
| - | - | 2206 | 240.1 | - | - | 0 | - |
| - | - | 3.201E+04 | 240.1 | - | - | 0 | - |
| - | - | 2514 | 241.1 | - | - | 0 | - |
| - | - | 4121 | 241.1 | - | - | 0 | - |
| - | - | 2021 | 241.1 | - | - | 0 | - |
| - | - | 4307 | 241.2 | - | - | 0 | - |
| - | - | 1582 | 242.1 | - | - | 0 | - |
| - | - | 6282 | 243.1 | - | - | 0 | - |
| - | - | 3.015E+04 | 244.1 | - | - | 0 | - |
| - | - | 3647 | 245.1 | - | - | 0 | - |
| - | - | 4.643E+04 | 247.1 | - | - | 0 | - |
| - | - | 2708 | 247.1 | - | - | 0 | - |
| - | - | 1816 | 247.2 | - | - | 0 | - |
| - | - | 3017 | 248.1 | - | - | 0 | - |
| - | - | 2.011E+04 | 248.1 | - | - | 0 | - |
| 10 | y | 1207 | 248.6 | 3.105E-05 | 0.1249 | +2 | 4 |
| - | - | 2.404E+04 | 249.1 | - | - | 0 | - |
| - | - | 1949 | 249.1 | - | - | 0 | - |
| - | - | 6909 | 249.1 | - | - | 0 | - |
| - | - | 3170 | 250.1 | - | - | 0 | - |
| - | - | 1948 | 250.1 | - | - | 0 | - |
| - | - | 1299 | 250.2 | - | - | 0 | - |
| - | - | 2.79E+05 | 251.2 | - | - | 0 | - |
| - | - | 1901 | 252.1 | - | - | 0 | - |
| - | - | 3.599E+04 | 252.2 | - | - | 0 | - |
| - | - | 1622 | 252.6 | - | - | 0 | - |
| - | - | 1830 | 253.2 | - | - | 0 | - |
| - | - | 1408 | 254.2 | - | - | 0 | - |
| - | - | 1.548E+04 | 255.1 | - | - | 0 | - |
| - | - | 3678 | 256.1 | - | - | 0 | - |
| - | - | 1874 | 256.2 | - | - | 0 | - |
| - | - | 8665 | 257.1 | - | - | 0 | - |
| - | - | 1.215E+04 | 258.1 | - | - | 0 | - |
| - | - | 1628 | 259.1 | - | - | 0 | - |
| - | - | 1963 | 259.1 | - | - | 0 | - |
| - | - | 5544 | 261.1 | - | - | 0 | - |
| - | - | 2135 | 261.1 | - | - | 0 | - |
| - | - | 3724 | 262.1 | - | - | 0 | - |
| - | - | 3.06E+04 | 265.1 | - | - | 0 | - |
| - | - | 2313 | 265.1 | - | - | 0 | - |
| - | - | 7.648E+04 | 266.1 | - | - | 0 | - |
| - | - | 8367 | 267.1 | - | - | 0 | - |
| - | - | 1383 | 268.1 | - | - | 0 | - |
| - | - | 4046 | 268.1 | - | - | 0 | - |
| - | - | 6056 | 268.2 | - | - | 0 | - |
| - | - | 2468 | 269.2 | - | - | 0 | - |
| - | - | 3677 | 269.2 | - | - | 0 | - |
| - | - | 1.041E+04 | 270.1 | - | - | 0 | - |
| - | - | 2059 | 270.1 | - | - | 0 | - |
| - | - | 4283 | 270.6 | - | - | 0 | - |
| - | - | 1.87E+04 | 272.2 | - | - | 0 | - |
| - | - | 3510 | 273.2 | - | - | 0 | - |
| - | - | 1.555E+04 | 274.1 | - | - | 0 | - |
| - | - | 5420 | 274.1 | - | - | 0 | - |
| - | - | 4950 | 275.1 | - | - | 0 | - |
| - | - | 3021 | 275.1 | - | - | 0 | - |
| - | - | 2256 | 275.2 | - | - | 0 | - |
| - | - | 4061 | 276.1 | - | - | 0 | - |
| - | - | 2577 | 276.1 | - | - | 0 | - |
| - | - | 3182 | 276.2 | - | - | 0 | - |
| - | - | 2130 | 276.2 | - | - | 0 | - |
| - | - | 6042 | 279.1 | - | - | 0 | - |
| - | - | 7980 | 280.1 | - | - | 0 | - |
| - | - | 1842 | 280.1 | - | - | 0 | - |
| - | - | 1.123E+04 | 281.1 | - | - | 0 | - |
| - | - | 9141 | 282.2 | - | - | 0 | - |
| - | - | 1.995E+05 | 283.1 | - | - | 0 | - |
| - | - | 1358 | 283.4 | - | - | 0 | - |
| - | - | 1787 | 284.1 | - | - | 0 | - |
| - | - | 8713 | 284.1 | - | - | 0 | - |
| - | - | 2541 | 284.1 | - | - | 0 | - |
| - | - | 2.737E+04 | 284.1 | - | - | 0 | - |
| - | - | 2340 | 285.1 | - | - | 0 | - |
| - | - | 1670 | 285.2 | - | - | 0 | - |
| - | - | 1.491E+04 | 286.1 | - | - | 0 | - |
| - | - | 1764 | 286.2 | - | - | 0 | - |
| - | - | 1554 | 287.1 | - | - | 0 | - |
| - | - | 3686 | 287.2 | - | - | 0 | - |
| - | - | 5657 | 288.6 | - | - | 0 | - |
| 5 | b | 2956 | 290.2 | 0.0001753 | 0.604 | +2 | 5 |
| 12 | y | 1.132E+05 | 293.1 | 0.000663 | 2.262 | +1 | 2 |
| - | - | 2.038E+04 | 294.1 | - | - | 0 | - |
| - | - | 1404 | 295.1 | - | - | 0 | - |
| - | - | 2142 | 295.1 | - | - | 0 | - |
| - | - | 1306 | 296.1 | - | - | 0 | - |
| 3 | b | 3650 | 296.2 | 0.001038 | 3.506 | +1 | 3 |
| - | - | 2283 | 297.1 | - | - | 0 | - |
| 9 | y | 4489 | 297.1 | 0.001226 | 4.126 | +2 | 5 |
| - | - | 3.809E+04 | 297.2 | - | - | 0 | - |
| - | - | 1703 | 297.2 | - | - | 0 | - |
| - | - | 1728 | 297.6 | - | - | 0 | - |
| 7 | y | 4574 | 298.1 | 0.005032 | 16.88 | +3 | 7 |
| - | - | 4439 | 298.2 | - | - | 0 | - |
| - | - | 8.948E+04 | 299.1 | - | - | 0 | - |
| - | - | 2486 | 300.1 | - | - | 0 | - |
| - | - | 1.8E+04 | 300.2 | - | - | 0 | - |
| - | - | 1562 | 301.1 | - | - | 0 | - |
| - | - | 1.035E+05 | 301.2 | - | - | 0 | - |
| - | - | 1.993E+04 | 302.1 | - | - | 0 | - |
| - | - | 1.502E+04 | 302.2 | - | - | 0 | - |
| - | - | 2764 | 303.1 | - | - | 0 | - |
| - | - | 2866 | 305.2 | - | - | 0 | - |
| - | - | 2452 | 305.6 | - | - | 0 | - |
| 9 | y | 1.802E+04 | 306.1 | 0.0008875 | 2.899 | +2 | 5 |
| - | - | 1983 | 306.2 | - | - | 0 | - |
| - | - | 6637 | 306.6 | - | - | 0 | - |
| - | - | 3678 | 307.1 | - | - | 0 | - |
| - | - | 1798 | 309.2 | - | - | 0 | - |
| 12 | y | 1.521E+04 | 311.1 | 0.0005353 | 1.721 | +1 | 2 |
| - | - | 2120 | 312.1 | - | - | 0 | - |
| - | - | 3415 | 314.2 | - | - | 0 | - |
| 3 | b | 9154 | 314.2 | 0.0006971 | 2.218 | +1 | 3 |
| - | - | 6.307E+04 | 315.2 | - | - | 0 | - |
| - | - | 1.19E+04 | 316.2 | - | - | 0 | - |
| - | - | 4019 | 319.2 | - | - | 0 | - |
| - | - | 3923 | 325.2 | - | - | 0 | - |
| - | - | 1.396E+04 | 326.7 | - | - | 0 | - |
| - | - | 3753 | 327.2 | - | - | 0 | - |
| - | - | 1461 | 330.2 | - | - | 0 | - |
| - | - | 1637 | 331.7 | - | - | 0 | - |
| - | - | 6730 | 332.2 | - | - | 0 | - |
| - | - | 1.113E+04 | 333.2 | - | - | 0 | - |
| - | - | 2061 | 334.2 | - | - | 0 | - |
| - | - | 1711 | 338.2 | - | - | 0 | - |
| 6 | b | 2100 | 338.7 | 0.0006688 | 1.975 | +2 | 6 |
| 6 | b | 1342 | 339.2 | 0.004144 | 12.22 | +2 | 6 |
| - | - | 2405 | 339.7 | - | - | 0 | - |
| - | - | 3086 | 340.7 | - | - | 0 | - |
| - | - | 1977 | 341.2 | - | - | 0 | - |
| - | - | 5219 | 342.2 | - | - | 0 | - |
| - | - | 4721 | 343.2 | - | - | 0 | - |
| - | - | 1422 | 343.2 | - | - | 0 | - |
| - | - | 5018 | 344.2 | - | - | 0 | - |
| - | - | 1.138E+04 | 345 | - | - | 0 | - |
| - | - | 8565 | 345.1 | - | - | 0 | - |
| - | - | 2.975E+04 | 346.1 | - | - | 0 | - |
| - | - | 1763 | 346.1 | - | - | 0 | - |
| - | - | 5039 | 347.1 | - | - | 0 | - |
| 6 | b | 1.44E+04 | 347.7 | 0.000727 | 2.091 | +2 | 6 |
| - | - | 3745 | 348.2 | - | - | 0 | - |
| - | - | 1588 | 348.7 | - | - | 0 | - |
| - | - | 1739 | 349.2 | - | - | 0 | - |
| - | - | 2.496E+04 | 350.2 | - | - | 0 | - |
| - | - | 4269 | 351.2 | - | - | 0 | - |
| - | - | 2952 | 352.7 | - | - | 0 | - |
| 8 | y | 5242 | 353.7 | 0.0005148 | 1.456 | +2 | 6 |
| - | - | 1760 | 354.2 | - | - | 0 | - |
| - | - | 2821 | 355.1 | - | - | 0 | - |
| - | - | 1.177E+04 | 359 | - | - | 0 | - |
| - | - | 6442 | 360.7 | - | - | 0 | - |
| - | - | 1.072E+04 | 361.2 | - | - | 0 | - |
| - | - | 1686 | 361.7 | - | - | 0 | - |
| - | - | 9645 | 362.2 | - | - | 0 | - |
| 8 | y | 3.992E+04 | 362.7 | 0.0008783 | 2.422 | +2 | 6 |
| - | - | 4.678E+04 | 363.1 | - | - | 0 | - |
| - | - | 1.528E+04 | 363.2 | - | - | 0 | - |
| - | - | 1500 | 363.7 | - | - | 0 | - |
| - | - | 8181 | 364.1 | - | - | 0 | - |
| - | - | 7058 | 364.1 | - | - | 0 | - |
| - | - | 3348 | 366.2 | - | - | 0 | - |
| - | - | 4234 | 367.1 | - | - | 0 | - |
| - | - | 2241 | 367.2 | - | - | 0 | - |
| - | - | 5842 | 367.2 | - | - | 0 | - |
| - | - | 1468 | 367.7 | - | - | 0 | - |
| - | - | 3719 | 373.2 | - | - | 0 | - |
| - | - | 1634 | 373.7 | - | - | 0 | - |
| - | - | 2308 | 376.2 | - | - | 0 | - |
| - | - | 7160 | 376.2 | - | - | 0 | - |
| - | - | 2236 | 376.7 | - | - | 0 | - |
| - | - | 3199 | 377.1 | - | - | 0 | - |
| - | - | 2158 | 377.3 | - | - | 0 | - |
| - | - | 1122 | 378.2 | - | - | 0 | - |
| - | - | 1570 | 378.2 | - | - | 0 | - |
| - | - | 1.659E+04 | 378.2 | - | - | 0 | - |
| - | - | 7.731E+04 | 379.2 | - | - | 0 | - |
| - | - | 3508 | 380.2 | - | - | 0 | - |
| - | - | 1.622E+04 | 380.2 | - | - | 0 | - |
| - | - | 1.315E+05 | 381.2 | - | - | 0 | - |
| - | - | 4751 | 381.7 | - | - | 0 | - |
| - | - | 2.527E+04 | 382.2 | - | - | 0 | - |
| - | - | 4594 | 382.2 | - | - | 0 | - |
| - | - | 3157 | 383.2 | - | - | 0 | - |
| - | - | 6328 | 384.2 | - | - | 0 | - |
| - | - | 4213 | 384.2 | - | - | 0 | - |
| - | - | 9788 | 385.2 | - | - | 0 | - |
| 5 | y | 3743 | 385.2 | 0.00407 | 10.57 | +3 | 9 |
| - | - | 2154 | 386.2 | - | - | 0 | - |
| 10 | b | 1960 | 389.2 | 0.004715 | 12.11 | +3 | 10 |
| - | - | 8780 | 390.2 | - | - | 0 | - |
| - | - | 2611 | 390.7 | - | - | 0 | - |
| - | - | 6477 | 391.1 | - | - | 0 | - |
| - | - | 2622 | 391.2 | - | - | 0 | - |
| - | - | 3197 | 392.2 | - | - | 0 | - |
| - | - | 2012 | 394.1 | - | - | 0 | - |
| - | - | 1.871E+04 | 394.2 | - | - | 0 | - |
| - | - | 7163 | 395.1 | - | - | 0 | - |
| - | - | 5259 | 395.2 | - | - | 0 | - |
| - | - | 2269 | 395.2 | - | - | 0 | - |
| - | - | 3279 | 395.2 | - | - | 0 | - |
| - | - | 1948 | 396.2 | - | - | 0 | - |
| - | - | 2288 | 396.2 | - | - | 0 | - |
| - | - | 2.681E+04 | 396.2 | - | - | 0 | - |
| - | - | 2407 | 397.2 | - | - | 0 | - |
| - | - | 5531 | 397.2 | - | - | 0 | - |
| - | - | 1414 | 397.7 | - | - | 0 | - |
| - | - | 1.22E+04 | 398.2 | - | - | 0 | - |
| - | - | 1921 | 399.2 | - | - | 0 | - |
| - | - | 5400 | 402.2 | - | - | 0 | - |
| - | - | 1.189E+04 | 402.2 | - | - | 0 | - |
| - | - | 4571 | 403.2 | - | - | 0 | - |
| - | - | 3644 | 404.7 | - | - | 0 | - |
| - | - | 4231 | 405.3 | - | - | 0 | - |
| - | - | 2761 | 406.2 | - | - | 0 | - |
| - | - | 2034 | 407.2 | - | - | 0 | - |
| - | - | 1.062E+04 | 409.1 | - | - | 0 | - |
| - | - | 1723 | 409.2 | - | - | 0 | - |
| - | - | 2472 | 409.2 | - | - | 0 | - |
| - | - | 1820 | 409.7 | - | - | 0 | - |
| - | - | 6418 | 411.2 | - | - | 0 | - |
| - | - | 8.469E+04 | 412.2 | - | - | 0 | - |
| - | - | 7933 | 413.1 | - | - | 0 | - |
| - | - | 4099 | 413.2 | - | - | 0 | - |
| - | - | 1.225E+04 | 413.2 | - | - | 0 | - |
| - | - | 2032 | 413.3 | - | - | 0 | - |
| - | - | 1291 | 414.2 | - | - | 0 | - |
| - | - | 1498 | 414.2 | - | - | 0 | - |
| - | - | 2.526E+04 | 414.2 | - | - | 0 | - |
| - | - | 2116 | 414.3 | - | - | 0 | - |
| - | - | 2.243E+04 | 415 | - | - | 0 | - |
| - | - | 1.016E+04 | 415.2 | - | - | 0 | - |
| - | - | 6712 | 415.2 | - | - | 0 | - |
| - | - | 2310 | 416.2 | - | - | 0 | - |
| - | - | 1.702E+04 | 417.7 | - | - | 0 | - |
| - | - | 2.656E+04 | 418.2 | - | - | 0 | - |
| - | - | 1.297E+04 | 418.7 | - | - | 0 | - |
| - | - | 3968 | 419.2 | - | - | 0 | - |
| 11 | y | 2875 | 421.2 | 0.001891 | 4.49 | +1 | 3 |
| 11 | y | 2399 | 422.2 | 0.0003127 | 0.7406 | +1 | 3 |
| - | - | 2339 | 422.7 | - | - | 0 | - |
| - | - | 5735 | 423.2 | - | - | 0 | - |
| - | - | 3.297E+04 | 423.3 | - | - | 0 | - |
| - | - | 2305 | 423.7 | - | - | 0 | - |
| - | - | 1741 | 424.2 | - | - | 0 | - |
| - | - | 8756 | 424.3 | - | - | 0 | - |
| - | - | 2815 | 424.7 | - | - | 0 | - |
| - | - | 2.171E+05 | 426.7 | - | - | 0 | - |
| - | - | 1.086E+05 | 427.2 | - | - | 0 | - |
| - | - | 3.01E+04 | 427.7 | - | - | 0 | - |
| - | - | 3805 | 428.2 | - | - | 0 | - |
| - | - | 2.926E+04 | 429.2 | - | - | 0 | - |
| - | - | 2.143E+05 | 430.2 | - | - | 0 | - |
| - | - | 4.639E+04 | 431.2 | - | - | 0 | - |
| 7 | b | 2.559E+04 | 431.7 | 0.001112 | 2.577 | +2 | 7 |
| 7 | b | 1.963E+04 | 432.2 | 0.006663 | 15.42 | +2 | 7 |
| - | - | 4814 | 432.7 | - | - | 0 | - |
| - | - | 4025 | 433.2 | - | - | 0 | - |
| 4 | b | 6126 | 433.3 | 0.0003235 | 0.7467 | +1 | 4 |
| - | - | 2011 | 434.3 | - | - | 0 | - |
| - | - | 1368 | 437.2 | - | - | 0 | - |
| - | - | 4420 | 438.2 | - | - | 0 | - |
| - | - | 3669 | 438.7 | - | - | 0 | - |
| 11 | y | 5.65E+04 | 439.2 | 0.001001 | 2.279 | +1 | 3 |
| - | - | 2324 | 440.2 | - | - | 0 | - |
| - | - | 1.289E+04 | 440.2 | - | - | 0 | - |
| 7 | b | 1.083E+05 | 440.7 | 0.001079 | 2.448 | +2 | 7 |
| - | - | 6.018E+04 | 441.2 | - | - | 0 | - |
| - | - | 1561 | 441.3 | - | - | 0 | - |
| - | - | 1.744E+04 | 441.7 | - | - | 0 | - |
| - | - | 5005 | 442.2 | - | - | 0 | - |
| - | - | 2948 | 443.2 | - | - | 0 | - |
| 7 | y | 1.008E+04 | 447.2 | 0.001382 | 3.091 | +2 | 7 |
| - | - | 1699 | 447.2 | - | - | 0 | - |
| - | - | 4050 | 447.7 | - | - | 0 | - |
| - | - | 1.1E+04 | 448.2 | - | - | 0 | - |
| - | - | 1561 | 448.8 | - | - | 0 | - |
| - | - | 3518 | 449.2 | - | - | 0 | - |
| - | - | 1807 | 450.2 | - | - | 0 | - |
| - | - | 1538 | 451.2 | - | - | 0 | - |
| - | - | 2080 | 451.2 | - | - | 0 | - |
| 4 | b | 6.004E+04 | 451.3 | 0.001142 | 2.53 | +1 | 4 |
| - | - | 1.629E+04 | 452.3 | - | - | 0 | - |
| - | - | 2743 | 453.3 | - | - | 0 | - |
| - | - | 3443 | 455.3 | - | - | 0 | - |
| 7 | y | 6840 | 455.7 | 0.0004063 | 0.8917 | +2 | 7 |
| - | - | 6091 | 459.2 | - | - | 0 | - |
| - | - | 6750 | 460.8 | - | - | 0 | - |
| - | - | 6201 | 461.2 | - | - | 0 | - |
| - | - | 1.042E+04 | 461.3 | - | - | 0 | - |
| - | - | 2072 | 462.2 | - | - | 0 | - |
| - | - | 1689 | 462.2 | - | - | 0 | - |
| - | - | 1480 | 464.2 | - | - | 0 | - |
| - | - | 1669 | 465.7 | - | - | 0 | - |
| - | - | 2888 | 466.2 | - | - | 0 | - |
| - | - | 1.505E+04 | 468.3 | - | - | 0 | - |
| - | - | 2793 | 469.3 | - | - | 0 | - |
| - | - | 1361 | 470.2 | - | - | 0 | - |
| - | - | 1.402E+04 | 474.3 | - | - | 0 | - |
| - | - | 1.811E+04 | 474.8 | - | - | 0 | - |
| - | - | 9133 | 475.3 | - | - | 0 | - |
| - | - | 3579 | 475.8 | - | - | 0 | - |
| - | - | 4908 | 476.2 | - | - | 0 | - |
| - | - | 1.158E+04 | 477.2 | - | - | 0 | - |
| 10 | y | 1.275E+04 | 478.2 | 0.0004683 | 0.9793 | +1 | 4 |
| - | - | 6480 | 478.3 | - | - | 0 | - |
| - | - | 1547 | 479.2 | - | - | 0 | - |
| - | - | 1688 | 479.3 | - | - | 0 | - |
| - | - | 2078 | 480.2 | - | - | 0 | - |
| - | - | 2634 | 481.7 | - | - | 0 | - |
| - | - | 4.308E+04 | 483.3 | - | - | 0 | - |
| - | - | 1.838E+04 | 483.8 | - | - | 0 | - |
| - | - | 2183 | 484.2 | - | - | 0 | - |
| - | - | 7974 | 484.3 | - | - | 0 | - |
| - | - | 2306 | 484.8 | - | - | 0 | - |
| 8 | b | 9451 | 488.3 | 0.001561 | 3.197 | +2 | 8 |
| 8 | b | 9542 | 488.7 | 0.007905 | 16.17 | +2 | 8 |
| - | - | 3278 | 489.3 | - | - | 0 | - |
| - | - | 1499 | 489.8 | - | - | 0 | - |
| - | - | 1.431E+05 | 494.2 | - | - | 0 | - |
| - | - | 3.568E+04 | 495.2 | - | - | 0 | - |
| 10 | y | 1.761E+05 | 496.2 | 0.001113 | 2.243 | +1 | 4 |
| - | - | 4.415E+04 | 497.2 | - | - | 0 | - |
| - | - | 1.821E+04 | 497.8 | - | - | 0 | - |
| - | - | 5975 | 498.2 | - | - | 0 | - |
| - | - | 1558 | 502.2 | - | - | 0 | - |
| - | - | 2.052E+04 | 503.1 | - | - | 0 | - |
| - | - | 3922 | 504.2 | - | - | 0 | - |
| - | - | 5166 | 506.2 | - | - | 0 | - |
| - | - | 2898 | 506.3 | - | - | 0 | - |
| - | - | 2269 | 507.3 | - | - | 0 | - |
| - | - | 1.366E+04 | 511.3 | - | - | 0 | - |
| - | - | 2039 | 512.3 | - | - | 0 | - |
| - | - | 2398 | 513.2 | - | - | 0 | - |
| - | - | 2497 | 514.2 | - | - | 0 | - |
| - | - | 2125 | 515.3 | - | - | 0 | - |
| - | - | 6572 | 519.1 | - | - | 0 | - |
| - | - | 5058 | 523.3 | - | - | 0 | - |
| - | - | 2.301E+04 | 525.3 | - | - | 0 | - |
| - | - | 3583 | 526.2 | - | - | 0 | - |
| - | - | 6507 | 526.3 | - | - | 0 | - |
| - | - | 1899 | 527.3 | - | - | 0 | - |
| - | - | 2267 | 528.2 | - | - | 0 | - |
| 0 | Precursor | 4731 | 529.3 | 0.001474 | 2.785 | +3 | -1 |
| 0 | Precursor | 1.001E+04 | 529.6 | 0.007453 | 14.07 | +3 | -1 |
| - | - | 3344 | 529.9 | - | - | 0 | - |
| - | - | 2266 | 530.2 | - | - | 0 | - |
| - | - | 2006 | 531.2 | - | - | 0 | - |
| - | - | 5447 | 532.2 | - | - | 0 | - |
| - | - | 1649 | 534.9 | - | - | 0 | - |
| 0 | Precursor | 3522 | 535.3 | 0.00137 | 2.56 | +3 | -1 |
| - | - | 8788 | 535.6 | - | - | 0 | - |
| - | - | 3795 | 536.2 | - | - | 0 | - |
| - | - | 1951 | 537.7 | - | - | 0 | - |
| - | - | 1698 | 538.8 | - | - | 0 | - |
| - | - | 2.712E+04 | 539.2 | - | - | 0 | - |
| - | - | 6487 | 540.2 | - | - | 0 | - |
| - | - | 5746 | 542.3 | - | - | 0 | - |
| - | - | 5.335E+04 | 543.3 | - | - | 0 | - |
| - | - | 1.428E+04 | 544.3 | - | - | 0 | - |
| - | - | 2766 | 545.3 | - | - | 0 | - |
| 9 | b | 1953 | 545.8 | 0.0006931 | 1.27 | +2 | 9 |
| - | - | 6515 | 546.3 | - | - | 0 | - |
| - | - | 2739 | 546.7 | - | - | 0 | - |
| - | - | 1844 | 547.3 | - | - | 0 | - |
| - | - | 1873 | 548.2 | - | - | 0 | - |
| - | - | 1736 | 548.3 | - | - | 0 | - |
| - | - | 7358 | 549.2 | - | - | 0 | - |
| - | - | 2062 | 550.2 | - | - | 0 | - |
| - | - | 6199 | 551.3 | - | - | 0 | - |
| - | - | 3248 | 551.3 | - | - | 0 | - |
| - | - | 2322 | 552.3 | - | - | 0 | - |
| - | - | 1845 | 553.3 | - | - | 0 | - |
| - | - | 1593 | 554.3 | - | - | 0 | - |
| 9 | b | 5723 | 554.8 | 0.003246 | 5.852 | +2 | 9 |
| - | - | 1.584E+04 | 555.3 | - | - | 0 | - |
| - | - | 8004 | 555.8 | - | - | 0 | - |
| - | - | 2606 | 556.3 | - | - | 0 | - |
| 5 | b | 8027 | 561.3 | 0.000706 | 1.258 | +1 | 5 |
| - | - | 2115 | 562.3 | - | - | 0 | - |
| - | - | 5515 | 563.3 | - | - | 0 | - |
| - | - | 1739 | 564.3 | - | - | 0 | - |
| - | - | 3527 | 564.3 | - | - | 0 | - |
| - | - | 1.014E+04 | 565.3 | - | - | 0 | - |
| - | - | 2.887E+04 | 567.2 | - | - | 0 | - |
| - | - | 6324 | 568.2 | - | - | 0 | - |
| 5 | y | 3018 | 568.7 | 0.009922 | 17.45 | +2 | 9 |
| - | - | 1644 | 569.2 | - | - | 0 | - |
| - | - | 6623 | 575.2 | - | - | 0 | - |
| - | - | 1746 | 575.3 | - | - | 0 | - |
| - | - | 4274 | 576.2 | - | - | 0 | - |
| 5 | y | 1778 | 577.3 | 0.007207 | 12.48 | +2 | 9 |
| - | - | 1769 | 577.8 | - | - | 0 | - |
| - | - | 1478 | 578.3 | - | - | 0 | - |
| - | - | 2166 | 579.3 | - | - | 0 | - |
| 5 | b | 4.086E+04 | 579.3 | 0.001067 | 1.841 | +1 | 5 |
| - | - | 1.199E+04 | 580.3 | - | - | 0 | - |
| - | - | 2660 | 581.3 | - | - | 0 | - |
| - | - | 4432 | 592.3 | - | - | 0 | - |
| - | - | 4262 | 592.3 | - | - | 0 | - |
| 9 | y | 4.564E+04 | 593.3 | 0.0008174 | 1.378 | +1 | 5 |
| - | - | 1.838E+04 | 593.3 | - | - | 0 | - |
| - | - | 2710 | 593.8 | - | - | 0 | - |
| - | - | 1.47E+04 | 594.3 | - | - | 0 | - |
| - | - | 3104 | 594.3 | - | - | 0 | - |
| - | - | 6751 | 595.3 | - | - | 0 | - |
| - | - | 1608 | 596.3 | - | - | 0 | - |
| - | - | 1866 | 597.8 | - | - | 0 | - |
| - | - | 2018 | 598.3 | - | - | 0 | - |
| - | - | 4988 | 600.3 | - | - | 0 | - |
| - | - | 7570 | 602.8 | - | - | 0 | - |
| - | - | 1.056E+04 | 603.3 | - | - | 0 | - |
| - | - | 2420 | 603.8 | - | - | 0 | - |
| - | - | 2103 | 604.3 | - | - | 0 | - |
| - | - | 7.167E+04 | 610.3 | - | - | 0 | - |
| - | - | 4506 | 610.3 | - | - | 0 | - |
| 9 | y | 5.474E+05 | 611.3 | 0.001544 | 2.526 | +1 | 5 |
| - | - | 3.522E+04 | 611.8 | - | - | 0 | - |
| - | - | 1.758E+05 | 612.3 | - | - | 0 | - |
| - | - | 1.072E+04 | 612.8 | - | - | 0 | - |
| - | - | 4.362E+04 | 613.3 | - | - | 0 | - |
| - | - | 6275 | 614.3 | - | - | 0 | - |
| - | - | 3202 | 617.3 | - | - | 0 | - |
| - | - | 3776 | 620.8 | - | - | 0 | - |
| - | - | 6132 | 621.3 | - | - | 0 | - |
| - | - | 3799 | 621.3 | - | - | 0 | - |
| - | - | 2144 | 622.3 | - | - | 0 | - |
| - | - | 2858 | 622.4 | - | - | 0 | - |
| - | - | 3068 | 628.3 | - | - | 0 | - |
| - | - | 1725 | 630.3 | - | - | 0 | - |
| - | - | 5533 | 634.3 | - | - | 0 | - |
| - | - | 9762 | 635.3 | - | - | 0 | - |
| - | - | 4842 | 636.3 | - | - | 0 | - |
| 4 | y | 5522 | 636.8 | 0.001344 | 2.11 | +2 | 10 |
| 4 | y | 3254 | 637.3 | 0.009946 | 15.61 | +2 | 10 |
| - | - | 1772 | 637.8 | - | - | 0 | - |
| - | - | 5078 | 640.3 | - | - | 0 | - |
| - | - | 2521 | 641.3 | - | - | 0 | - |
| - | - | 6141 | 645.3 | - | - | 0 | - |
| 4 | y | 3.198E+04 | 645.8 | 0.001982 | 3.069 | +2 | 10 |
| - | - | 2.691E+04 | 646.3 | - | - | 0 | - |
| - | - | 9354 | 646.8 | - | - | 0 | - |
| - | - | 2695 | 648.3 | - | - | 0 | - |
| - | - | 3128 | 650.3 | - | - | 0 | - |
| - | - | 1709 | 651.3 | - | - | 0 | - |
| - | - | 5.6E+04 | 652.3 | - | - | 0 | - |
| - | - | 2.331E+04 | 653.3 | - | - | 0 | - |
| - | - | 3186 | 654.3 | - | - | 0 | - |
| - | - | 4086 | 658.3 | - | - | 0 | - |
| - | - | 1863 | 659.3 | - | - | 0 | - |
| - | - | 2303 | 660.9 | - | - | 0 | - |
| - | - | 2699 | 661.3 | - | - | 0 | - |
| - | - | 2.053E+04 | 662.3 | - | - | 0 | - |
| - | - | 1.191E+04 | 663.3 | - | - | 0 | - |
| - | - | 4139 | 664.3 | - | - | 0 | - |
| - | - | 1690 | 666.4 | - | - | 0 | - |
| - | - | 2105 | 668.3 | - | - | 0 | - |
| - | - | 1696 | 675.4 | - | - | 0 | - |
| 6 | b | 2.637E+04 | 676.3 | 0.001046 | 1.546 | +1 | 6 |
| 6 | b | 1.679E+04 | 677.3 | 0.008912 | 13.16 | +1 | 6 |
| - | - | 9466 | 678.3 | - | - | 0 | - |
| - | - | 5403 | 679.3 | - | - | 0 | - |
| - | - | 1532 | 679.8 | - | - | 0 | - |
| - | - | 9.325E+04 | 680.3 | - | - | 0 | - |
| - | - | 4.103E+04 | 681.3 | - | - | 0 | - |
| - | - | 8329 | 682.3 | - | - | 0 | - |
| - | - | 2548 | 684.3 | - | - | 0 | - |
| - | - | 8893 | 684.8 | - | - | 0 | - |
| - | - | 6568 | 685.3 | - | - | 0 | - |
| - | - | 2675 | 685.8 | - | - | 0 | - |
| - | - | 2815 | 688.3 | - | - | 0 | - |
| - | - | 3311 | 692.8 | - | - | 0 | - |
| 3 | y | 2.729E+04 | 693.3 | 0.001731 | 2.497 | +2 | 11 |
| 3 | y | 2.853E+04 | 693.8 | 0.008014 | 11.55 | +2 | 11 |
| 6 | b | 2.779E+05 | 694.4 | 0.0003687 | 0.5309 | +1 | 6 |
| - | - | 4801 | 694.8 | - | - | 0 | - |
| - | - | 1.048E+05 | 695.4 | - | - | 0 | - |
| - | - | 2.307E+04 | 696.4 | - | - | 0 | - |
| - | - | 1705 | 697.3 | - | - | 0 | - |
| - | - | 5017 | 697.4 | - | - | 0 | - |
| - | - | 1874 | 697.9 | - | - | 0 | - |
| - | - | 2512 | 698.3 | - | - | 0 | - |
| - | - | 2645 | 700.9 | - | - | 0 | - |
| - | - | 2092 | 701.3 | - | - | 0 | - |
| - | - | 2.902E+04 | 701.8 | - | - | 0 | - |
| - | - | 2563 | 701.9 | - | - | 0 | - |
| 3 | y | 2.168E+05 | 702.3 | 0.002186 | 3.112 | +2 | 11 |
| 12 | b | 1.818E+05 | 702.8 | 0.01253 | 17.83 | +2 | 12 |
| 12 | b | 8.905E+04 | 703.3 | 0.00277 | 3.939 | +2 | 12 |
| - | - | 2.177E+04 | 703.8 | - | - | 0 | - |
| - | - | 7010 | 704.3 | - | - | 0 | - |
| 8 | y | 1.928E+04 | 706.3 | 0.002508 | 3.55 | +1 | 6 |
| - | - | 9166 | 707.3 | - | - | 0 | - |
| - | - | 1898 | 708.3 | - | - | 0 | - |
| - | - | 1.095E+04 | 711.4 | - | - | 0 | - |
| 12 | b | 1.82E+04 | 711.9 | 0.001838 | 2.582 | +2 | 12 |
| - | - | 1.44E+04 | 712.4 | - | - | 0 | - |
| - | - | 5423 | 712.9 | - | - | 0 | - |
| - | - | 2530 | 713.4 | - | - | 0 | - |
| - | - | 2106 | 720.4 | - | - | 0 | - |
| - | - | 1.184E+04 | 721.4 | - | - | 0 | - |
| - | - | 4158 | 722.4 | - | - | 0 | - |
| - | - | 2.115E+04 | 723.4 | - | - | 0 | - |
| 8 | y | 1.818E+05 | 724.4 | 0.00177 | 2.443 | +1 | 6 |
| - | - | 6.997E+04 | 725.4 | - | - | 0 | - |
| - | - | 1.492E+04 | 726.4 | - | - | 0 | - |
| - | - | 2106 | 727.4 | - | - | 0 | - |
| - | - | 9043 | 729.4 | - | - | 0 | - |
| - | - | 2128 | 729.9 | - | - | 0 | - |
| - | - | 3913 | 730.4 | - | - | 0 | - |
| - | - | 6889 | 733.4 | - | - | 0 | - |
| - | - | 6259 | 734.4 | - | - | 0 | - |
| - | - | 2610 | 735.4 | - | - | 0 | - |
| - | - | 1933 | 737.9 | - | - | 0 | - |
| - | - | 2256 | 738.4 | - | - | 0 | - |
| 2 | y | 3716 | 742.9 | 0.004023 | 5.416 | +2 | 12 |
| 2 | y | 4902 | 743.4 | 0.01415 | 19.04 | +2 | 12 |
| - | - | 2139 | 743.9 | - | - | 0 | - |
| - | - | 1762 | 745.4 | - | - | 0 | - |
| - | - | 3951 | 747.4 | - | - | 0 | - |
| - | - | 1841 | 748.4 | - | - | 0 | - |
| - | - | 2401 | 749.3 | - | - | 0 | - |
| - | - | 3.184E+04 | 751.4 | - | - | 0 | - |
| 2 | y | 1.643E+04 | 751.9 | 0.001304 | 1.735 | +2 | 12 |
| - | - | 2.82E+04 | 752.4 | - | - | 0 | - |
| - | - | 1.022E+04 | 752.9 | - | - | 0 | - |
| - | - | 6427 | 753.4 | - | - | 0 | - |
| - | - | 2794 | 760.3 | - | - | 0 | - |
| - | - | 2810 | 761.4 | - | - | 0 | - |
| - | - | 4582 | 762.4 | - | - | 0 | - |
| - | - | 1.671E+04 | 763.4 | - | - | 0 | - |
| - | - | 6690 | 764.4 | - | - | 0 | - |
| - | - | 5894 | 765.4 | - | - | 0 | - |
| - | - | 2441 | 766.4 | - | - | 0 | - |
| - | - | 6990 | 767.3 | - | - | 0 | - |
| - | - | 2688 | 768.3 | - | - | 0 | - |
| - | - | 2084 | 769.4 | - | - | 0 | - |
| - | - | 3277 | 775.4 | - | - | 0 | - |
| - | - | 2689 | 776.4 | - | - | 0 | - |
| - | - | 1.742E+04 | 779.4 | - | - | 0 | - |
| - | - | 9306 | 780.4 | - | - | 0 | - |
| - | - | 7570 | 781.4 | - | - | 0 | - |
| - | - | 4247 | 782.4 | - | - | 0 | - |
| 0 | Precursor | 2.87E+04 | 793.4 | 0.01412 | 17.8 | +2 | -1 |
| - | - | 1.113E+04 | 794.4 | - | - | 0 | - |
| - | - | 9594 | 795.3 | - | - | 0 | - |
| - | - | 3802 | 795.4 | - | - | 0 | - |
| - | - | 4022 | 796.3 | - | - | 0 | - |
| - | - | 6342 | 807.4 | - | - | 0 | - |
| - | - | 5187 | 807.4 | - | - | 0 | - |
| - | - | 2141 | 808.4 | - | - | 0 | - |
| - | - | 3574 | 808.4 | - | - | 0 | - |
| - | - | 4972 | 818.4 | - | - | 0 | - |
| - | - | 3608 | 819.4 | - | - | 0 | - |
| - | - | 1929 | 820.4 | - | - | 0 | - |
| - | - | 2212 | 825.4 | - | - | 0 | - |
| - | - | 3103 | 826.4 | - | - | 0 | - |
| - | - | 3432 | 827.4 | - | - | 0 | - |
| - | - | 6313 | 834.4 | - | - | 0 | - |
| - | - | 8224 | 835.4 | - | - | 0 | - |
| - | - | 1.678E+04 | 836.4 | - | - | 0 | - |
| - | - | 1.07E+04 | 837.5 | - | - | 0 | - |
| - | - | 2445 | 838.5 | - | - | 0 | - |
| - | - | 3942 | 843.4 | - | - | 0 | - |
| - | - | 1.232E+04 | 844.4 | - | - | 0 | - |
| - | - | 4614 | 845.4 | - | - | 0 | - |
| - | - | 8912 | 846.4 | - | - | 0 | - |
| - | - | 6247 | 847.4 | - | - | 0 | - |
| - | - | 3293 | 848.4 | - | - | 0 | - |
| - | - | 6.196E+04 | 852.4 | - | - | 0 | - |
| - | - | 2093 | 853.4 | - | - | 0 | - |
| - | - | 3.299E+04 | 853.4 | - | - | 0 | - |
| - | - | 1.07E+04 | 854.4 | - | - | 0 | - |
| 7 | b | 4.201E+04 | 862.4 | 0.001139 | 1.321 | +1 | 7 |
| 7 | b | 3.155E+04 | 863.4 | 0.01175 | 13.61 | +1 | 7 |
| - | - | 5.354E+04 | 864.5 | - | - | 0 | - |
| - | - | 2.27E+04 | 865.5 | - | - | 0 | - |
| - | - | 6345 | 866.5 | - | - | 0 | - |
| - | - | 1829 | 867.5 | - | - | 0 | - |
| - | - | 1966 | 869.4 | - | - | 0 | - |
| - | - | 6088 | 876.4 | - | - | 0 | - |
| - | - | 2427 | 877.4 | - | - | 0 | - |
| 7 | b | 3.256E+05 | 880.4 | 0.0008897 | 1.011 | +1 | 7 |
| - | - | 1.676E+05 | 881.4 | - | - | 0 | - |
| - | - | 4.917E+04 | 882.4 | - | - | 0 | - |
| - | - | 9036 | 883.4 | - | - | 0 | - |
| - | - | 2423 | 890.4 | - | - | 0 | - |
| - | - | 3954 | 891.4 | - | - | 0 | - |
| - | - | 4710 | 892.5 | - | - | 0 | - |
| - | - | 1913 | 893.5 | - | - | 0 | - |
| - | - | 3485 | 894.5 | - | - | 0 | - |
| - | - | 2079 | 895.4 | - | - | 0 | - |
| - | - | 2081 | 907.4 | - | - | 0 | - |
| - | - | 2.799E+04 | 908.4 | - | - | 0 | - |
| - | - | 1.65E+04 | 909.4 | - | - | 0 | - |
| 7 | y | 2.756E+04 | 910.4 | 0.001558 | 1.712 | +1 | 7 |
| - | - | 1.366E+04 | 911.4 | - | - | 0 | - |
| - | - | 3734 | 912.4 | - | - | 0 | - |
| - | - | 3695 | 936.4 | - | - | 0 | - |
| - | - | 4689 | 937.4 | - | - | 0 | - |
| - | - | 2794 | 947.4 | - | - | 0 | - |
| - | - | 4567 | 954.4 | - | - | 0 | - |
| - | - | 1.019E+04 | 955.4 | - | - | 0 | - |
| - | - | 5964 | 956.4 | - | - | 0 | - |
| - | - | 2372 | 962.4 | - | - | 0 | - |
| - | - | 1686 | 964.5 | - | - | 0 | - |
| - | - | 1.088E+04 | 965.5 | - | - | 0 | - |
| - | - | 5189 | 966.5 | - | - | 0 | - |
| - | - | 2403 | 967.5 | - | - | 0 | - |
| - | - | 7750 | 972.4 | - | - | 0 | - |
| - | - | 4237 | 973.4 | - | - | 0 | - |
| 8 | b | 5062 | 975.5 | 0.001504 | 1.541 | +1 | 8 |
| 8 | b | 4363 | 976.5 | 0.0118 | 12.08 | +1 | 8 |
| - | - | 4897 | 980.5 | - | - | 0 | - |
| - | - | 4531 | 982.5 | - | - | 0 | - |
| - | - | 2080 | 983.5 | - | - | 0 | - |
| 8 | b | 1.556E+04 | 993.5 | 0.0005049 | 0.5082 | +1 | 8 |
| - | - | 1.132E+04 | 994.5 | - | - | 0 | - |
| - | - | 3200 | 995.5 | - | - | 0 | - |
| - | - | 2473 | 1007 | - | - | 0 | - |
| 6 | y | 1706 | 1008 | 0.01539 | 15.26 | +1 | 8 |
| - | - | 1615 | 1010 | - | - | 0 | - |
| - | - | 1890 | 1024 | - | - | 0 | - |
| 6 | y | 1.528E+04 | 1025 | 0.001288 | 1.256 | +1 | 8 |
| - | - | 1.046E+04 | 1026 | - | - | 0 | - |
| - | - | 3257 | 1027 | - | - | 0 | - |
| - | - | 2418 | 1076 | - | - | 0 | - |
| - | - | 2734 | 1077 | - | - | 0 | - |
| 9 | b | 1571 | 1092 | 0.02089 | 19.13 | +1 | 9 |
| - | - | 8192 | 1094 | - | - | 0 | - |
| - | - | 6978 | 1095 | - | - | 0 | - |
| - | - | 2103 | 1096 | - | - | 0 | - |
| 9 | b | 4365 | 1109 | 0.002879 | 2.597 | +1 | 9 |
| - | - | 2089 | 1110 | - | - | 0 | - |
| - | - | 2506 | 1118 | - | - | 0 | - |
| 5 | y | 1.034E+04 | 1136 | 0.001249 | 1.1 | +1 | 9 |
| 5 | y | 1.758E+04 | 1136 | 0.00881 | 7.752 | +1 | 9 |
| - | - | 1.017E+04 | 1137 | - | - | 0 | - |
| - | - | 3162 | 1139 | - | - | 0 | - |
| - | - | 4879 | 1153 | - | - | 0 | - |
| 5 | y | 3.103E+04 | 1154 | 0.001304 | 1.13 | +1 | 9 |
| - | - | 2.049E+04 | 1155 | - | - | 0 | - |
| - | - | 5828 | 1156 | - | - | 0 | - |
| - | - | 1594 | 1165 | - | - | 0 | - |
| 10 | b | 3954 | 1166 | 0.003734 | 3.204 | +1 | 10 |
| - | - | 1810 | 1167 | - | - | 0 | - |
| - | - | 1535 | 1495 | - | - | 0 | - |
| - | - | 1735 | 1711 | - | - | 0 | - |
| - | - | 1371 | 1720 | - | - | 0 | - |
| - | - | 1348 | 1952 | - | - | 0 | - |
| - | - | 1585 | 2769 | - | - | 0 | - |
| - | - | 1386 | 3083 | - | - | 0 | - |

m/z Charge Intensity FragmentType MassShift Position
120.0811767578125 0 8649.986
121.04016876220703 0 1004.66547
121.0845718383789 0 1069.1127
122.07170104980469 0 4218.539
123.0444564819336 0 4882.5874
123.05550384521484 0 1507.0856
125.10821533203125 0 948.9379
125.37583923339844 0 761.8791
126.09159851074219 0 1826.6792
127.05060577392578 0 7478.4526
127.08692932128906 0 1422.9551
127.12323760986328 0 2910.801
128.04901123046875 0 1037.3254
128.0541534423828 0 922.6204
128.10740661621094 0 55114.438
129.06625366210938 0 3310.992
129.10269165039062 0 565230.94
130.06553649902344 0 238166.84
130.09991455078125 0 3574.3108
130.1060028076172 0 35743.445
131.06886291503906 0 22908.508
132.08116149902344 0 54500.523
133.06085205078125 0 891.21844
133.0846405029297 0 3893.977
135.1682586669922 0 1006.20526
136.0760955810547 0 67277.75
137.0793914794922 0 5881.4478
137.16400146484375 0 1087.6134
138.0666046142578 0 6233.968
138.091796875 0 4064.2314
139.05067443847656 0 1151.0259
139.0869598388672 0 5654.8076
141.1026611328125 0 10035.268
142.0654754638672 0 5138.098
143.0457763671875 0 3458.1775
144.08151245117188 0 2310.6348
145.06130981445312 0 4474.455
146.06044006347656 0 1385.8187
147.04440307617188 0 3055.1697
149.04527282714844 0 13746.018
150.0667266845703 0 3413.6245
151.08706665039062 0 11096.127
152.09136962890625 0 1110.4309
154.09823608398438 0 1037.4235
155.0452880859375 0 2585.2375
155.08216857910156 0 2058.2966
155.09315490722656 0 34811.6
155.1182861328125 0 24034.566 a Water loss 1
156.02987670898438 0 1148.7776
156.07681274414062 0 2875.1064
156.0963134765625 0 1763.5522
156.12156677246094 0 1450.6041
157.09765625 0 2184.4023
157.13427734375 0 1113.0303
158.0841064453125 0 3018.3223
159.09214782714844 0 423888.97
160.07603454589844 0 4522.1953
160.08941650390625 0 2755.5054
160.09547424316406 0 42395.215
161.09881591796875 0 1817.5886
162.66802978515625 0 1152.6813
165.05503845214844 0 32029.95
165.0778045654297 0 1332.5521
166.06150817871094 0 70886.375
166.08688354492188 0 3243.0261 y 9
166.09788513183594 0 5191.5933
167.055908203125 0 13676.412
167.06472778320312 0 4712.211
167.11862182617188 0 1100.6453
168.08119201660156 0 6945.633
168.1023712158203 0 5291.71
168.11363220214844 0 5466.9023
169.07643127441406 0 7293.8228
169.08340454101562 0 1106.2295
169.0976104736328 0 15942.399
170.06045532226562 0 18247.807
171.0636444091797 0 2022.851
171.092041015625 0 2578.1753
171.1492919921875 0 2174.4216
172.07215881347656 0 6226.1704
172.10867309570312 0 1677.5334
173.05616760253906 0 8184.721
173.1289520263672 0 851863.94 a 1
173.4528350830078 0 2426.961
174.1322784423828 0 74271.516
175.13426208496094 0 4727.738
177.10272216796875 0 2134.5732
178.1343231201172 0 26559.402
179.137939453125 0 1487.5388
181.06117248535156 0 5853.0967
182.0816650390625 0 79219.055 y 12
183.0856475830078 0 6801.941
183.1132049560547 0 22020.197 b Water loss 1
184.11695861816406 0 2599.1616
185.1288604736328 0 1200.8776
185.1651611328125 0 3391.5015
186.12416076660156 0 183741.56
187.0870361328125 0 38245.766
187.12759399414062 0 15985.395
187.14474487304688 0 1456.1244
188.07081604003906 0 4541.703
188.090576171875 0 4316.8716
190.08287048339844 0 6684.8887
190.13430786132812 0 4997.4956
194.0928497314453 0 3913.8901
195.11326599121094 0 41609.277
196.1084747314453 0 29591.084
196.1165313720703 0 2705.6453
197.11181640625 0 1666.761
197.12855529785156 0 3215.187
197.16529846191406 0 1454.5344
198.0878448486328 0 5290.221
198.128173828125 0 15249.846
199.07167053222656 0 3525.7158
199.08712768554688 0 3673.3594
199.1315460205078 0 1547.8448
200.1435089111328 0 2507.6626
201.12379455566406 0 226962.03 b 1
202.08670043945312 0 2120.6191
202.12701416015625 0 21611.203
204.11407470703125 0 2696.1213
205.09767150878906 0 17080.656
205.10748291015625 0 1650.909
205.1451873779297 0 5441.0635
206.10107421875 0 2178.7527
206.1293487548828 0 4474.8203
207.1608123779297 0 7481.2734
209.05662536621094 0 1782.2583
210.12753295898438 0 6253.4873
211.1082305908203 0 6215.3994 y Water loss 10
212.1398468017578 0 5798.4443
213.123779296875 0 19562.82
213.1601104736328 0 2111.4792
214.11834716796875 0 1586.8372
215.10414123535156 0 1183.4669
215.11741638183594 0 1706.5869
215.13931274414062 0 2253.795
216.0981903076172 0 4183.23
221.0594482421875 0 1251.2124
221.07260131835938 0 2352.3157
221.08482360839844 0 7312.3154
221.1042022705078 0 4075.5671
221.12913513183594 0 2458.6863
221.1400604248047 0 2677.8174
222.08535766601562 0 2052.4993
222.12396240234375 0 6066.0693
223.0641326904297 0 2951.1707
223.10809326171875 0 11693.582
223.11891174316406 0 1993.9552
223.15582275390625 0 75239.73
224.1032257080078 0 3398.5684
224.15911865234375 0 7707.326
225.04342651367188 0 8065.1147
226.04339599609375 0 1440.955
226.082763671875 0 15113.314
226.15538024902344 0 1549.0907
227.06666564941406 0 11579.124
227.1546173095703 0 3133.229
228.13442993164062 0 2775.969
229.11883544921875 0 20483.316
229.1553192138672 0 7155.6333
230.1041717529297 0 1550.1643
230.1221923828125 0 2885.342
230.1503143310547 0 2466.5532
233.14028930664062 0 11008.17
234.124267578125 0 82477.46
235.12750244140625 0 11538.525
235.1560821533203 0 2957.132
237.13555908203125 0 2417.435
238.11907958984375 0 8837.379
238.1307830810547 0 2876.1523
239.08241271972656 0 2065.4917
239.0955352783203 0 15294.385
239.11480712890625 0 1978.463
240.09603881835938 0 4275.852
240.1210174560547 0 2205.924
240.13482666015625 0 32013.293
241.09336853027344 0 2514.1797
241.1188507080078 0 4120.8433
241.13812255859375 0 2021.4436
241.1915283203125 0 4306.7925
242.09280395507812 0 1581.9354
243.10975646972656 0 6281.717
244.09341430664062 0 30150.691
245.09686279296875 0 3646.9453
247.1083526611328 0 46431.645
247.14385986328125 0 2708.319
247.1563720703125 0 1815.801
248.10348510742188 0 3017.218
248.11436462402344 0 20112.25
248.62376403808594 0 1207.121 y 9
249.09873962402344 0 24038.305
249.11260986328125 0 1949.1505
249.1352996826172 0 6909.0986
250.1022186279297 0 3170.4368
250.11842346191406 0 1947.713
250.16680908203125 0 1299.0415
251.15087890625 0 279046.75
252.13409423828125 0 1901.0453
252.15399169921875 0 35988.406
252.60317993164062 0 1621.9037
253.1563262939453 0 1829.9108
254.15155029296875 0 1407.8213
255.14840698242188 0 15481.656
256.1082458496094 0 3677.991
256.1520080566406 0 1873.8695
257.1135559082031 0 8664.929
258.14520263671875 0 12146.988
259.12908935546875 0 1627.9259
259.1498107910156 0 1962.7513
261.1195983886719 0 5544.141
261.13421630859375 0 2134.5793
262.1195983886719 0 3724.315
265.1297912597656 0 30596.39
265.14373779296875 0 2312.7688
266.1253662109375 0 76481.12
267.12823486328125 0 8367.188
268.1142578125 0 1383.0647
268.1300964355469 0 4046.3118
268.1774597167969 0 6056.2275
269.1610412597656 0 2468.2922
269.1856994628906 0 3677.2961
270.1221008300781 0 10410.24
270.1453857421875 0 2059.066
270.6234130859375 0 4283.3438
272.1761779785156 0 18702.582
273.1793518066406 0 3510.1172
274.118896484375 0 15549.645
274.1316223144531 0 5419.9956
275.1029357910156 0 4950.032
275.1212158203125 0 3020.952
275.1500244140625 0 2255.501
276.10906982421875 0 4061.492
276.1260986328125 0 2576.5789
276.1554870605469 0 3182.1873
276.1705017089844 0 2130.1523
279.14544677734375 0 6041.971
280.1298828125 0 7980.1646
280.1466369628906 0 1842.2621
281.0517883300781 0 11227.749
282.1565246582031 0 9140.938
283.14068603515625 0 199512.16
283.35296630859375 0 1357.5121
284.0877380371094 0 1787.0176
284.1039733886719 0 8713.042
284.12408447265625 0 2541.2292
284.14385986328125 0 27365.418
285.14422607421875 0 2339.692
285.15924072265625 0 1670.0244
286.1402587890625 0 14908.315
286.15643310546875 0 1763.9647
287.1428527832031 0 1553.6853
287.172119140625 0 3685.7412
288.619384765625 0 5656.748
290.1659240722656 0 2956.2358 b 4
293.1138610839844 0 113156.05 y Water loss 11
294.1179504394531 0 20382.018
295.1033020019531 0 1403.7208
295.12017822265625 0 2142.4712
296.10614013671875 0 1305.502
296.1979064941406 0 3649.6238 b Water loss 2
297.0835876464844 0 2282.9473
297.1331481933594 0 4489.3374 y Water loss 8
297.15643310546875 0 38087.766
297.1993103027344 0 1702.8221
297.63372802734375 0 1728.1488
298.1398010253906 0 4573.9897 y Water loss 6
298.1595153808594 0 4438.879
299.0624084472656 0 89479.42
300.0634765625 0 2486.2405
300.1688537597656 0 17997.21
301.0589904785156 0 1562.3153
301.1513366699219 0 103467.91
302.1142578125 0 19932.973
302.1545715332031 0 15017.131
303.1180419921875 0 2763.718
305.178466796875 0 2865.513
305.6473388671875 0 2451.8813
306.1380920410156 0 18021.04 y 8
306.2295227050781 0 1982.7198
306.6397399902344 0 6636.575
307.14044189453125 0 3677.948
309.1680603027344 0 1798.0121
311.1242980957031 0 15206.751 y 11
312.1273498535156 0 2119.8662
314.18646240234375 0 3415.4834
314.2081298828125 0 9153.521 b 2
315.16705322265625 0 63073.477
316.17022705078125 0 11901.33
319.1524963378906 0 4018.9739
325.1514892578125 0 3922.6035
326.6644287109375 0 13958.244
327.1661071777344 0 3752.976
330.1664733886719 0 1460.5361
331.6578369140625 0 1637.235
332.2088623046875 0 6729.5435
333.1776123046875 0 11126.287
334.184326171875 0 2060.9346
338.1827697753906 0 1711.1094
338.6749572753906 0 2100.459 b Water loss 5
339.1704406738281 0 1341.5834 b Ammonia loss 5
339.6762390136719 0 2405.4028
340.6619873046875 0 3086.1497
341.1626892089844 0 1977.3928
342.1822204589844 0 5219.4043
343.1619873046875 0 4720.6445
343.1838073730469 0 1422.131
344.1715393066406 0 5018.4297
344.9771423339844 0 11378.182
345.13140869140625 0 8564.724
346.11541748046875 0 29748.762
346.1372375488281 0 1763.0566
347.1178894042969 0 5039.068
347.6802978515625 0 14403.483 b 5
348.1817932128906 0 3745.0684
348.6849060058594 0 1587.8402
349.15130615234375 0 1738.9587
350.219482421875 0 24956.393
351.2216491699219 0 4268.987
352.67437744140625 0 2952.4622
353.6744689941406 0 5242.2505 y Water loss 7
354.1766662597656 0 1760.1376
355.0698547363281 0 2820.6858
359.02899169921875 0 11769.362
360.69561767578125 0 6442.378
361.19781494140625 0 10718.569
361.68914794921875 0 1686.2834
362.1844177246094 0 9645.143
362.68011474609375 0 39921.2 y 7
363.1418151855469 0 46782.58
363.1820068359375 0 15276.644
363.6815185546875 0 1499.6288
364.1256103515625 0 8181.0444
364.1457824707031 0 7058.483
366.17852783203125 0 3348.169
367.1404724121094 0 4233.8315
367.1635437011719 0 2240.9568
367.24615478515625 0 5842.261
367.68505859375 0 1467.6685
373.1856994628906 0 3718.9006
373.6866760253906 0 1633.8551
376.1633605957031 0 2308.12
376.1979675292969 0 7160.2573
376.7003479003906 0 2235.7852
377.146240234375 0 3199.419
377.266357421875 0 2158.149
378.1795654296875 0 1121.8375
378.1847839355469 0 1570.3175
378.2142639160156 0 16587.22
379.20977783203125 0 77314.91
380.1641845703125 0 3507.7815
380.2126159667969 0 16216.327
381.1524658203125 0 131502.98
381.71209716796875 0 4751.2236
382.15557861328125 0 25269.773
382.18878173828125 0 4593.7837
383.1564025878906 0 3157.386
384.167236328125 0 6327.8804
384.1883544921875 0 4212.6963
385.15087890625 0 9787.503
385.17279052734375 0 3742.5852 y 4
386.1794738769531 0 2153.924
389.19744873046875 0 1960.0223 b 9
390.1960144042969 0 8780.034
390.6959533691406 0 2611.0918
391.1368408203125 0 6477.4585
391.1849670410156 0 2622.1384
392.2093811035156 0 3197.059
394.1490783691406 0 2011.8701
394.1728820800781 0 18713.766
395.1344909667969 0 7162.824
395.1575012207031 0 5258.711
395.1790771484375 0 2268.8328
395.2406311035156 0 3279.004
396.1616516113281 0 1948.4733
396.1947326660156 0 2287.5498
396.225341796875 0 26812.95
397.2023620605469 0 2406.5483
397.2283630371094 0 5530.597
397.704833984375 0 1414.4988
398.1771545410156 0 12203.812
399.18353271484375 0 1921.2289
402.1763000488281 0 5399.509
402.1997375488281 0 11887.847
403.20159912109375 0 4571.2705
404.7105712890625 0 3644.384
405.26275634765625 0 4230.9814
406.2453308105469 0 2760.8198
407.2033996582031 0 2034.1193
409.1473083496094 0 10621.621
409.17315673828125 0 1722.9512
409.205322265625 0 2471.7822
409.7015686035156 0 1820.1904
411.1998596191406 0 6417.8677
412.18365478515625 0 84689.67
413.145263671875 0 7933.0005
413.1622009277344 0 4099.319
413.18701171875 0 12253.522
413.25177001953125 0 2032.489
414.1934814453125 0 1290.6696
414.2010498046875 0 1498.1451
414.2355651855469 0 25264.059
414.2646484375 0 2115.6106
415.0377197265625 0 22428.598
415.1980285644531 0 10158.18
415.2389831542969 0 6712.1616
416.2018737792969 0 2310.4897
417.71746826171875 0 17020.293
418.21087646484375 0 26556.04
418.71142578125 0 12970.441
419.2143249511719 0 3968.2769
421.2100524902344 0 2874.7705 y Water loss 10
422.1918640136719 0 2399.2249 y Ammonia loss 10
422.70965576171875 0 2338.667
423.20306396484375 0 5734.7783
423.2723693847656 0 32973.973
423.70513916015625 0 2305.0574
424.2252502441406 0 1740.943
424.2748107910156 0 8756.298
424.73284912109375 0 2814.6138
426.72271728515625 0 217115.17
427.2239990234375 0 108594.195
427.7255859375 0 30098.105
428.2259521484375 0 3805.074
429.2101745605469 0 29257.686
430.1944274902344 0 214305.64
431.1972961425781 0 46385.57
431.7150573730469 0 25587.629 b Water loss 6
432.2126159667969 0 19628.977 b Ammonia loss 6
432.71160888671875 0 4813.6313
433.2102966308594 0 4024.502
433.256103515625 0 6125.7495 b Water loss 3
434.25946044921875 0 2010.6709
437.18719482421875 0 1367.5396
438.1732482910156 0 4420.1777
438.7261962890625 0 3668.9233
439.2197265625 0 56500.68 y 10
440.1855163574219 0 2323.5686
440.2230529785156 0 12892.369
440.7203063964844 0 108304.375 b 6
441.22161865234375 0 60179.426
441.2574462890625 0 1560.6243
441.7230224609375 0 17442.6
442.2249450683594 0 5004.7
443.23968505859375 0 2948.145
447.2070007324219 0 10080.205 y Ammonia loss 6
447.23907470703125 0 1698.9529
447.70977783203125 0 4050.404
448.2054748535156 0 10999.819
448.7676086425781 0 1561.1622
449.21148681640625 0 3518.3774
450.2359313964844 0 1807.1024
451.1749572753906 0 1537.7747
451.23681640625 0 2080.1042
451.2674865722656 0 60044.875 b 3
452.2705993652344 0 16288.472
453.27215576171875 0 2742.6873
455.266845703125 0 3442.6829
455.71929931640625 0 6840.4834 y 6
459.19976806640625 0 6091.238
460.75408935546875 0 6749.801
461.2032470703125 0 6200.9224
461.2539367675781 0 10420.655
462.20794677734375 0 2072.2852
462.2468566894531 0 1689.468
464.2168273925781 0 1480.139
465.7471618652344 0 1669.0934
466.244140625 0 2887.9688
468.2938232421875 0 15053.043
469.2971496582031 0 2793.4373
470.2467956542969 0 1360.7737
474.260009765625 0 14020.09
474.75341796875 0 18108.193
475.2549133300781 0 9133.27
475.7550964355469 0 3578.7976
476.22637939453125 0 4908.3804
477.2107238769531 0 11576.515
478.2291564941406 0 12750.043 y Water loss 9
478.2775573730469 0 6479.942
479.24176025390625 0 1546.7664
479.2817077636719 0 1687.97
480.2215881347656 0 2078.0977
481.7289733886719 0 2633.6492
483.2649841308594 0 43083.66
483.7662048339844 0 18378.623
484.2228088378906 0 2182.9941
484.26751708984375 0 7973.8525
484.7705078125 0 2305.6968
488.2575378417969 0 9451.316 b Water loss 7
488.7558898925781 0 9542.388 b Ammonia loss 7
489.25927734375 0 3277.798
489.7533264160156 0 1498.9905
494.23681640625 0 143069.69
495.23968505859375 0 35682.203
496.2413024902344 0 176143.77 y 9
497.2466125488281 0 44146.414
497.76361083984375 0 18206.107
498.2449645996094 0 5975.4697
502.2296447753906 0 1558.3413
503.10821533203125 0 20518.766
504.2016906738281 0 3921.7468
506.2268371582031 0 5166.152
506.2732238769531 0 2898.0493
507.26068115234375 0 2269.4268
511.2633056640625 0 13659.001
512.2660522460938 0 2038.73
513.207763671875 0 2398.098
514.18994140625 0 2497.409
515.2811279296875 0 2124.668
519.1400756835938 0 6572.224
523.259521484375 0 5057.9634
525.2677001953125 0 23011.053
526.230712890625 0 3583.0117
526.271240234375 0 6506.6025
527.27197265625 0 1898.7697
528.2383422851562 0 2267.385
529.2611694335938 0 4730.62 Precursor Water loss
529.5951538085938 0 10014.822 Precursor Ammonia loss
529.9292602539062 0 3344.4995
530.22216796875 0 2265.8655
531.2203979492188 0 2006.1747
532.194580078125 0 5447.453
534.888671875 0 1649.1097
535.2645874023438 0 3522.0254 Precursor
535.5987548828125 0 8787.811
536.16650390625 0 3795.1206
537.7337036132812 0 1950.8851
538.773193359375 0 1698.2753
539.237060546875 0 27119.035
540.2390747070312 0 6487.0093
542.2933349609375 0 5746.4053
543.278076171875 0 53345.82
544.2816162109375 0 14284.284
545.2830810546875 0 2766.277
545.7701416015625 0 1952.8391 b Water loss 8
546.2503051757812 0 6515.443
546.7495727539062 0 2738.611
547.2811889648438 0 1844.4703
548.2396240234375 0 1873.1737
548.2950439453125 0 1736.4001
549.2210083007812 0 7358.329
550.2114868164062 0 2061.967
551.2584838867188 0 6199.086
551.33056640625 0 3247.6309
552.2586669921875 0 2322.1843
553.2628173828125 0 1844.8689
554.2821044921875 0 1592.5065
554.771484375 0 5723.129 b 8
555.2571411132812 0 15843.161
555.7577514648438 0 8004.3203
556.2559204101562 0 2606.4607
561.3150634765625 0 8027.305 b Water loss 4
562.3214111328125 0 2114.7783
563.2582397460938 0 5515.3853
564.2608642578125 0 1739.046
564.33154296875 0 3526.653
565.2639770507812 0 10141.219
567.2319946289062 0 28874.146
568.235595703125 0 6324.458
568.75830078125 0 3018.2346 y Ammonia loss 4
569.2337036132812 0 1643.9319
575.2465209960938 0 6623.1777
575.2946166992188 0 1746.0106
576.2339477539062 0 4274.159
577.2688598632812 0 1777.5295 y 4
577.7615356445312 0 1768.6211
578.2713623046875 0 1478.217
579.274169921875 0 2165.727
579.3259887695312 0 40856.1 b 4
580.328857421875 0 11989.5
581.3303833007812 0 2659.717
592.2732543945312 0 4431.734
592.3251342773438 0 4261.935
593.2573852539062 0 45639.766 y Water loss 8
593.306396484375 0 18382.926
593.7868041992188 0 2709.674
594.2591552734375 0 14698.167
594.3081665039062 0 3104.4133
595.256591796875 0 6751.47
596.2574462890625 0 1608.0154
597.7977294921875 0 1865.5125
598.300537109375 0 2018.1671
600.3157348632812 0 4988.3037
602.7937622070312 0 7569.8804
603.2904663085938 0 10559.814
603.7963256835938 0 2419.9124
604.2900390625 0 2102.99
610.2840576171875 0 71672.586
610.3302001953125 0 4505.896
611.2686767578125 0 547396.4 y 8
611.798095703125 0 35215.242
612.2718505859375 0 175818.47
612.7998046875 0 10719.093
613.2739868164062 0 43619.633
614.2765502929688 0 6274.9634
617.2845458984375 0 3201.6138
620.8034057617188 0 3775.9407
621.251220703125 0 6131.8325
621.301513671875 0 3799.0994
622.2577514648438 0 2143.9795
622.368896484375 0 2857.7158
628.2748413085938 0 3068.0098
630.2864379882812 0 1724.59
634.3055419921875 0 5533.147
635.2973022460938 0 9762.058
636.3291625976562 0 4842.266
636.7871704101562 0 5521.894 y Water loss 3
637.2877807617188 0 3254.3608 y Ammonia loss 3
637.7892456054688 0 1771.6068
640.2745971679688 0 5077.6523
641.2681884765625 0 2520.891
645.296142578125 0 6141.371
645.7930908203125 0 31983.67 y 3
646.29345703125 0 26910.686
646.795166015625 0 9354.105
648.342529296875 0 2695.4946
650.2694091796875 0 3127.6003
651.2765502929688 0 1709.2915
652.3212890625 0 55998.7
653.3236083984375 0 23312.209
654.3256225585938 0 3185.5173
658.2848510742188 0 4086.3586
659.3157958984375 0 1863.4033
660.8705444335938 0 2302.9216
661.3280639648438 0 2698.957
662.3062133789062 0 20530.627
663.29931640625 0 11905.117
664.3419189453125 0 4138.5537
666.3629760742188 0 1690.0663
668.2807006835938 0 2104.9656
675.3798217773438 0 1696.0261
676.3423461914062 0 26367.28 b Water loss 5
677.334228515625 0 16793.549 b Ammonia loss 5
678.3362426757812 0 9465.974
679.3341064453125 0 5402.622
679.83154296875 0 1531.7565
680.3164672851562 0 93251.64
681.3190307617188 0 41032.285
682.3223876953125 0 8328.831
684.322265625 0 2547.7007
684.8179931640625 0 8893.42
685.318603515625 0 6567.509
685.8224487304688 0 2674.5647
688.3336181640625 0 2815.0442
692.8370971679688 0 3310.8303
693.32958984375 0 27293.14 y Water loss 2
693.827880859375 0 28532.209 y Ammonia loss 2
694.3522338867188 0 277917.8 b 5
694.8287353515625 0 4800.8013
695.3555297851562 0 104772.836
696.3583374023438 0 23072.5
697.290283203125 0 1704.6381
697.3563232421875 0 5017.481
697.8602294921875 0 1873.7523
698.345703125 0 2512.36
700.8926391601562 0 2645.2979
701.337890625 0 2091.625
701.8419799804688 0 29015.934
701.9065551757812 0 2563.3547
702.3353271484375 0 216794.2 y 2
702.83642578125 0 181764.25 b Water loss 11
703.3381958007812 0 89051.76 b Ammonia loss 11
703.8388671875 0 21773.62
704.3384399414062 0 7010.0093
706.3431396484375 0 19284.852 y Water loss 7
707.3446655273438 0 9166.4795
708.3386840820312 0 1897.9121
711.3700561523438 0 10949.913
711.8560791015625 0 18199.287 b 11
712.3591918945312 0 14395.347
712.8599853515625 0 5422.782
713.3562622070312 0 2530.008
720.3742065429688 0 2106.4763
721.36376953125 0 11836.194
722.3670654296875 0 4157.708
723.3685913085938 0 21151.77
724.3529663085938 0 181803.19 y 7
725.3557739257812 0 69974.52
726.3580932617188 0 14922.069
727.3623657226562 0 2106.4082
729.3584594726562 0 9043.174
729.8616333007812 0 2128.1914
730.361572265625 0 3912.634
733.3648681640625 0 6889.1206
734.3563232421875 0 6259.3735
735.3562622070312 0 2610.0176
737.8616943359375 0 1932.892
738.3515625 0 2256.451
742.8660888671875 0 3715.6218 y Water loss 1
743.3682250976562 0 4901.969 y Ammonia loss 1
743.8583374023438 0 2139.32
745.35986328125 0 1762.1047
747.3889770507812 0 3951.0981
748.3795776367188 0 1841.3551
749.33154296875 0 2400.9573
751.3770141601562 0 31843.773
751.86865234375 0 16428.988 y 1
752.3748779296875 0 28202.846
752.87158203125 0 10216.986
753.376220703125 0 6426.9263
760.3053588867188 0 2794.0015
761.3783569335938 0 2810.424
762.4161987304688 0 4582.103
763.3646240234375 0 16709.984
764.3662109375 0 6690.2876
765.4020385742188 0 5894.168
766.4075927734375 0 2440.6992
767.347412109375 0 6989.7173
768.3475341796875 0 2687.755
769.405029296875 0 2084.4397
775.3907470703125 0 3276.9587
776.3707275390625 0 2689.2356
779.3841552734375 0 17423.277
780.38671875 0 9305.569
781.3672485351562 0 7570.317
782.3641967773438 0 4247.024
793.4000244140625 0 28699.365 Precursor Water loss
794.4024658203125 0 11128.257
795.3418579101562 0 9594.383
795.4095458984375 0 3802.3755
796.345703125 0 4021.5242
807.37744140625 0 6342.0723
807.4411010742188 0 5187.069
808.3701782226562 0 2141.4255
808.4314575195312 0 3574.1067
818.4392700195312 0 4971.8145
819.4442138671875 0 3607.8813
820.4362182617188 0 1928.887
825.3858642578125 0 2212.4065
826.377197265625 0 3102.7566
827.3712768554688 0 3431.9387
834.4260864257812 0 6313.227
835.4161987304688 0 8223.879
836.4461059570312 0 16777.463
837.4544677734375 0 10704.716
838.4544067382812 0 2445.0408
843.3966064453125 0 3942.0276
844.388671875 0 12317.377
845.3946533203125 0 4613.7637
846.447509765625 0 8912.421
847.443603515625 0 6246.7197
848.4468994140625 0 3293.4072
852.437255859375 0 61956.465
853.3524169921875 0 2093.4395
853.4403076171875 0 32991.41
854.4420776367188 0 10700.521
862.4217529296875 0 42005.684 b Water loss 6
863.4163818359375 0 31545.443 b Ammonia loss 6
864.4559326171875 0 53540.484
865.4605712890625 0 22700.668
866.4632568359375 0 6345.4004
867.4620971679688 0 1828.7144
869.38525390625 0 1966.4895
876.4365844726562 0 6088.231
877.4367065429688 0 2426.9907
880.4320678710938 0 325623.03 b 6
881.4344482421875 0 167566.64
882.43701171875 0 49170.566
883.4407958984375 0 9036.021
890.4137573242188 0 2422.9172
891.4091796875 0 3954.0725
892.4699096679688 0 4709.5234
893.4727783203125 0 1912.9083
894.453125 0 3484.628
895.447265625 0 2079.1685
907.4397583007812 0 2081.148
908.4276733398438 0 27991.398
909.4326171875 0 16497.074
910.4320678710938 0 27564.443 y 6
911.434814453125 0 13664.713
912.436279296875 0 3733.9658
936.4237670898438 0 3695.4983
937.4144897460938 0 4688.876
947.4387817382812 0 2793.8994
954.4354858398438 0 4566.6655
955.4212646484375 0 10191.582
956.421875 0 5963.827
962.4472045898438 0 2372.3694
964.4705810546875 0 1685.7017
965.4500732421875 0 10882.73
966.4517211914062 0 5189.293
967.4525146484375 0 2402.6418
972.4434204101562 0 7749.5557
973.4470825195312 0 4237.0815
975.503173828125 0 5062.142 b Water loss 7
976.50048828125 0 4362.7065 b Ammonia loss 7
980.4603881835938 0 4897.3623
982.472900390625 0 4531.485
983.4744262695312 0 2079.9275
993.5157470703125 0 15555.412 b 7
994.5202026367188 0 11323.308
995.5261840820312 0 3199.995
1007.4842529296875 0 2473.2666
1008.4462890625 0 1706.1749 y Ammonia loss 5
1009.5018310546875 0 1615.0729
1024.472900390625 0 1889.5896
1025.458740234375 0 15281.217 y 5
1026.4620361328125 0 10463.129
1027.4659423828125 0 3256.6985
1075.541748046875 0 2418.23
1076.5286865234375 0 2733.7312
1091.4947509765625 0 1571.1759 b Ammonia loss 8
1093.544189453125 0 8191.6084
1094.5430908203125 0 6978.2144
1095.5458984375 0 2102.9048
1108.539306640625 0 4365.052 b 8
1109.5416259765625 0 2089.4868
1118.48974609375 0 2506.0698
1135.5067138671875 0 10338.336 y Water loss 4
1136.498291015625 0 17579.895 y Ammonia loss 4
1137.4986572265625 0 10174.933
1138.50048828125 0 3162.262
1152.5313720703125 0 4878.8525
1153.517333984375 0 31025.19 y 4
1154.5198974609375 0 20490.166
1155.522216796875 0 5828.452
1164.5938720703125 0 1594.4017
1165.5673828125 0 3953.8123 b 9
1166.559326171875 0 1809.5311
1494.9261474609375 0 1535.155
1710.91162109375 0 1735.2775
1720.029052734375 0 1371.2191
1951.8946533203125 0 1348.0876
2768.503173828125 0 1584.9862
3082.859619140625 0 1385.8217

Spectrum Details

|  |  |
| --- | --- |
| Matched peaks? Matched peaks The total absolute number of peaks matched. Additionally in brackets the total fraction of peaks matched and the total number of peaks is shown. | 81 (9.82% of 825) |
| FDR? FDR The false discovery rate estimated for this peptide. It is calculated by matching all theoretical fragments with a non-integer shift with the raw peaks for this spectrum. This is done with 40 different shifts. The resulting percentage is the average number of annotated peaks over the number of annotated peaks with the correct spectrum. | 0.62% |
| Satellite FDR? Satellite FDR See the FDR for details on its calculation. This satellite ion specific FDR only contains the satellite ions (d/w) for I/L/J positions. | ∞ |
| PSM Score? PSM Score The PSM Score as given by Hecklib to this annotated spectrum. It is shown with three significant figures. | 604 |

#### Spectrum 7274? Spectrum 7274 The raw spectrum of this peptide as annotated by Hecklib. The fragments are coloured according to ion type (see legend). Any peaks with a star '\*' as text can be hovered over to see the full details, first the ion type second the mass shift type. By hovering over the amino acids in the peptide or ions in the legend the corresponding peaks are highlighted. By toggling the 'Unassigned' label you can turn the background (unassigned) peaks on or off in the plot. By updating the slider in the Ion legend you can update the spectrum to only show the top X% of the peaks with labels. The top X% means any peak that is within X% of the highest intensity. By dragging in the spectrum you can zoom in to a specific part of the spectrum and use 'Zoom Out' to get back to the original zoom level. The annotation of the spectrum is based on the given sequence in the peptides file and is done with different software so inconsistencies are likely. The peaks are annotated based on the given sequence, with 20 ppm tolerance.

Copy Data

##### Spectrum 7274 (TSV)

###### Preview

```
Loading example...
```

*Click on the button to copy the data to your clipboard.*

Mz MinMz MaxIntensity Max

WidthHeightPeptide font sizePeptide stroke widthSpectrum font sizeSpectrum stroke widthCompact peptide

Ion legend

wxyz

abcd

OtherUnassignedIon^Charge^_Position_Show for top:%

TVLHQDWLDGKEY

04.87e+59.75e+51.46e+61.95e+6

Zoom Out

a^+1^_2_a^+1^_2_y^+1^_1_b^+1^_2_b^+1^_2_y^+2^_3_y^+2^_4_b^+2^_5_y^+1^_2_b^+1^_3_y^+2^_5_y^+3^_7_y^+2^_5_y^+1^_2_b^+1^_3_b^+2^_6_b^+2^_6_y^+2^_6_y^+2^_6_y^+3^_9_b^+3^_10_y^+1^_3_y^+1^_3_b^+2^_7_b^+2^_7_b^+1^_4_y^+1^_3_b^+2^_7_y^+2^_7_b^+1^_4_y^+2^_7_y^+1^_4_b^+2^_8_b^+2^_8_y^+1^_4_\*\*\*b^+2^_9_b^+1^_5_b^+1^_5_y^+2^_9_b^+1^_5_b^+2^_10_y^+1^_5_y^+1^_5_y^+2^_10_y^+2^_10_b^+2^_11_y^+2^_10_b^+1^_6_b^+1^_6_y^+2^_11_y^+2^_11_b^+1^_6_y^+2^_11_b^+2^_12_b^+2^_12_y^+1^_6_b^+2^_12_y^+1^_6_y^+2^_12_y^+2^_12_y^+2^_12_\*b^+1^_7_b^+1^_7_b^+1^_7_y^+1^_7_b^+1^_8_b^+1^_8_b^+1^_8_y^+1^_8_b^+1^_9_b^+1^_9_y^+1^_9_y^+1^_9_y^+1^_9_b^+1^_10_

0799159823963195

Fragment Matches Table

Show background peaks

| Position | Ion type | Intensity | mz Theoretical | mz Error (Th) | mz Error (ppm) | Charge | Series Number |
| --- | --- | --- | --- | --- | --- | --- | --- |
| - | - | 5636 | 120.1 | - | - | 0 | - |
| - | - | 1.026E+04 | 122.1 | - | - | 0 | - |
| - | - | 1.179E+04 | 123 | - | - | 0 | - |
| - | - | 2414 | 123.1 | - | - | 0 | - |
| - | - | 3293 | 125.1 | - | - | 0 | - |
| - | - | 1535 | 125.6 | - | - | 0 | - |
| - | - | 1601 | 126 | - | - | 0 | - |
| - | - | 3884 | 126.1 | - | - | 0 | - |
| - | - | 1.455E+04 | 127.1 | - | - | 0 | - |
| - | - | 1538 | 127.1 | - | - | 0 | - |
| - | - | 5685 | 127.1 | - | - | 0 | - |
| - | - | 1642 | 128.1 | - | - | 0 | - |
| - | - | 3597 | 128.1 | - | - | 0 | - |
| - | - | 1.205E+05 | 128.1 | - | - | 0 | - |
| - | - | 1.29E+06 | 129.1 | - | - | 0 | - |
| - | - | 5.292E+05 | 130.1 | - | - | 0 | - |
| - | - | 9603 | 130.1 | - | - | 0 | - |
| - | - | 8.435E+04 | 130.1 | - | - | 0 | - |
| - | - | 5.087E+04 | 131.1 | - | - | 0 | - |
| - | - | 1948 | 131.1 | - | - | 0 | - |
| - | - | 1566 | 131.2 | - | - | 0 | - |
| - | - | 1.257E+05 | 132.1 | - | - | 0 | - |
| - | - | 2446 | 133.1 | - | - | 0 | - |
| - | - | 1.384E+04 | 133.1 | - | - | 0 | - |
| - | - | 2131 | 133.1 | - | - | 0 | - |
| - | - | 1.579E+05 | 136.1 | - | - | 0 | - |
| - | - | 1623 | 136.2 | - | - | 0 | - |
| - | - | 1.377E+04 | 137.1 | - | - | 0 | - |
| - | - | 1.542E+04 | 138.1 | - | - | 0 | - |
| - | - | 5696 | 138.1 | - | - | 0 | - |
| - | - | 1768 | 139.1 | - | - | 0 | - |
| - | - | 1.541E+04 | 139.1 | - | - | 0 | - |
| - | - | 1932 | 139.2 | - | - | 0 | - |
| - | - | 1.983E+04 | 141.1 | - | - | 0 | - |
| - | - | 7678 | 142.1 | - | - | 0 | - |
| - | - | 2321 | 142.1 | - | - | 0 | - |
| - | - | 1814 | 142.6 | - | - | 0 | - |
| - | - | 8487 | 143 | - | - | 0 | - |
| - | - | 2102 | 143.1 | - | - | 0 | - |
| - | - | 7782 | 144.1 | - | - | 0 | - |
| - | - | 1483 | 144.1 | - | - | 0 | - |
| - | - | 9071 | 145.1 | - | - | 0 | - |
| - | - | 2211 | 146.1 | - | - | 0 | - |
| - | - | 6829 | 147 | - | - | 0 | - |
| - | - | 2463 | 148.1 | - | - | 0 | - |
| - | - | 2381 | 149 | - | - | 0 | - |
| - | - | 1.757E+04 | 149 | - | - | 0 | - |
| - | - | 6623 | 150.1 | - | - | 0 | - |
| - | - | 2.476E+04 | 151.1 | - | - | 0 | - |
| - | - | 2716 | 153.1 | - | - | 0 | - |
| - | - | 2752 | 154.1 | - | - | 0 | - |
| - | - | 2700 | 154.1 | - | - | 0 | - |
| - | - | 4053 | 155 | - | - | 0 | - |
| - | - | 3701 | 155.1 | - | - | 0 | - |
| - | - | 7.863E+04 | 155.1 | - | - | 0 | - |
| 2 | a | 5.665E+04 | 155.1 | 0.0003966 | 2.557 | +1 | 2 |
| - | - | 4035 | 156.1 | - | - | 0 | - |
| - | - | 5106 | 156.1 | - | - | 0 | - |
| - | - | 2808 | 156.1 | - | - | 0 | - |
| - | - | 3274 | 156.1 | - | - | 0 | - |
| - | - | 1947 | 157.1 | - | - | 0 | - |
| - | - | 7224 | 157.1 | - | - | 0 | - |
| - | - | 4496 | 157.1 | - | - | 0 | - |
| - | - | 6481 | 158.1 | - | - | 0 | - |
| - | - | 9.717E+05 | 159.1 | - | - | 0 | - |
| - | - | 9117 | 160.1 | - | - | 0 | - |
| - | - | 7912 | 160.1 | - | - | 0 | - |
| - | - | 1.054E+05 | 160.1 | - | - | 0 | - |
| - | - | 5466 | 161.1 | - | - | 0 | - |
| - | - | 8.799E+04 | 165.1 | - | - | 0 | - |
| - | - | 4322 | 165.1 | - | - | 0 | - |
| - | - | 1.617E+05 | 166.1 | - | - | 0 | - |
| - | - | 1.445E+04 | 166.1 | - | - | 0 | - |
| - | - | 1.099E+04 | 167.1 | - | - | 0 | - |
| - | - | 1.09E+04 | 167.1 | - | - | 0 | - |
| - | - | 3045 | 167.1 | - | - | 0 | - |
| - | - | 6454 | 167.1 | - | - | 0 | - |
| - | - | 1.635E+04 | 168.1 | - | - | 0 | - |
| - | - | 1.046E+04 | 168.1 | - | - | 0 | - |
| - | - | 1.261E+04 | 168.1 | - | - | 0 | - |
| - | - | 1.406E+04 | 169.1 | - | - | 0 | - |
| - | - | 2866 | 169.1 | - | - | 0 | - |
| - | - | 3.83E+04 | 169.1 | - | - | 0 | - |
| - | - | 4.701E+04 | 170.1 | - | - | 0 | - |
| - | - | 5061 | 171.1 | - | - | 0 | - |
| - | - | 5164 | 171.1 | - | - | 0 | - |
| - | - | 1.532E+04 | 172.1 | - | - | 0 | - |
| - | - | 1.711E+04 | 173.1 | - | - | 0 | - |
| 2 | a | 1.93E+06 | 173.1 | 0.0004825 | 2.787 | +1 | 2 |
| - | - | 3533 | 173.5 | - | - | 0 | - |
| - | - | 1.696E+05 | 174.1 | - | - | 0 | - |
| - | - | 1.106E+04 | 175.1 | - | - | 0 | - |
| - | - | 2726 | 176.1 | - | - | 0 | - |
| - | - | 3156 | 177.1 | - | - | 0 | - |
| - | - | 5.843E+04 | 178.1 | - | - | 0 | - |
| - | - | 4126 | 179.1 | - | - | 0 | - |
| - | - | 5476 | 179.1 | - | - | 0 | - |
| - | - | 8064 | 181.1 | - | - | 0 | - |
| 13 | y | 1.805E+05 | 182.1 | 0.0004954 | 2.721 | +1 | 1 |
| - | - | 1.64E+04 | 183.1 | - | - | 0 | - |
| 2 | b | 5.006E+04 | 183.1 | 0.0003855 | 2.105 | +1 | 2 |
| - | - | 5178 | 184.1 | - | - | 0 | - |
| - | - | 1.517E+04 | 185.2 | - | - | 0 | - |
| - | - | 4.215E+05 | 186.1 | - | - | 0 | - |
| - | - | 8.119E+04 | 187.1 | - | - | 0 | - |
| - | - | 3.519E+04 | 187.1 | - | - | 0 | - |
| - | - | 7923 | 188.1 | - | - | 0 | - |
| - | - | 2154 | 188.1 | - | - | 0 | - |
| - | - | 7398 | 188.1 | - | - | 0 | - |
| - | - | 2152 | 188.1 | - | - | 0 | - |
| - | - | 1.673E+04 | 190.1 | - | - | 0 | - |
| - | - | 1.412E+04 | 190.1 | - | - | 0 | - |
| - | - | 1.024E+04 | 194.1 | - | - | 0 | - |
| - | - | 9.932E+04 | 195.1 | - | - | 0 | - |
| - | - | 6.703E+04 | 196.1 | - | - | 0 | - |
| - | - | 8471 | 196.1 | - | - | 0 | - |
| - | - | 4574 | 197.1 | - | - | 0 | - |
| - | - | 5261 | 197.1 | - | - | 0 | - |
| - | - | 1.016E+04 | 198.1 | - | - | 0 | - |
| - | - | 3.366E+04 | 198.1 | - | - | 0 | - |
| - | - | 8966 | 199.1 | - | - | 0 | - |
| - | - | 7084 | 199.1 | - | - | 0 | - |
| - | - | 5360 | 199.1 | - | - | 0 | - |
| - | - | 5695 | 200.1 | - | - | 0 | - |
| 2 | b | 5.181E+05 | 201.1 | 0.0003952 | 1.965 | +1 | 2 |
| - | - | 4231 | 202.1 | - | - | 0 | - |
| - | - | 5.208E+04 | 202.1 | - | - | 0 | - |
| - | - | 4338 | 203.1 | - | - | 0 | - |
| - | - | 1957 | 204.1 | - | - | 0 | - |
| - | - | 4458 | 204.1 | - | - | 0 | - |
| - | - | 3313 | 204.1 | - | - | 0 | - |
| - | - | 4.105E+04 | 205.1 | - | - | 0 | - |
| - | - | 6818 | 205.1 | - | - | 0 | - |
| - | - | 3837 | 206.1 | - | - | 0 | - |
| - | - | 9102 | 206.1 | - | - | 0 | - |
| - | - | 1.329E+04 | 207.2 | - | - | 0 | - |
| - | - | 1.645E+04 | 210.1 | - | - | 0 | - |
| 11 | y | 8375 | 211.1 | 0.0003897 | 1.846 | +2 | 3 |
| - | - | 1.258E+04 | 212.1 | - | - | 0 | - |
| - | - | 4.528E+04 | 213.1 | - | - | 0 | - |
| - | - | 6471 | 213.2 | - | - | 0 | - |
| - | - | 3882 | 215.1 | - | - | 0 | - |
| - | - | 2803 | 215.1 | - | - | 0 | - |
| - | - | 1.253E+04 | 216.1 | - | - | 0 | - |
| - | - | 2016 | 218 | - | - | 0 | - |
| - | - | 2332 | 219.1 | - | - | 0 | - |
| - | - | 5395 | 221.1 | - | - | 0 | - |
| - | - | 4247 | 221.1 | - | - | 0 | - |
| - | - | 4034 | 221.1 | - | - | 0 | - |
| - | - | 1.033E+04 | 221.1 | - | - | 0 | - |
| - | - | 2526 | 221.1 | - | - | 0 | - |
| - | - | 5488 | 221.1 | - | - | 0 | - |
| - | - | 1.713E+04 | 222.1 | - | - | 0 | - |
| - | - | 3292 | 222.1 | - | - | 0 | - |
| - | - | 3049 | 223.1 | - | - | 0 | - |
| - | - | 2.449E+04 | 223.1 | - | - | 0 | - |
| - | - | 2552 | 223.1 | - | - | 0 | - |
| - | - | 1.736E+05 | 223.2 | - | - | 0 | - |
| - | - | 7502 | 224.1 | - | - | 0 | - |
| - | - | 1.734E+04 | 224.2 | - | - | 0 | - |
| - | - | 5749 | 225 | - | - | 0 | - |
| - | - | 3491 | 225.1 | - | - | 0 | - |
| - | - | 3798 | 225.2 | - | - | 0 | - |
| - | - | 3.87E+04 | 226.1 | - | - | 0 | - |
| - | - | 3.348E+04 | 227.1 | - | - | 0 | - |
| - | - | 5089 | 227.1 | - | - | 0 | - |
| - | - | 9511 | 227.2 | - | - | 0 | - |
| - | - | 4489 | 228.1 | - | - | 0 | - |
| - | - | 3016 | 228.1 | - | - | 0 | - |
| - | - | 5.383E+04 | 229.1 | - | - | 0 | - |
| - | - | 5221 | 230.1 | - | - | 0 | - |
| - | - | 4105 | 230.2 | - | - | 0 | - |
| - | - | 2.977E+04 | 233.1 | - | - | 0 | - |
| - | - | 1.891E+05 | 234.1 | - | - | 0 | - |
| - | - | 3518 | 234.7 | - | - | 0 | - |
| - | - | 2269 | 235.1 | - | - | 0 | - |
| - | - | 2.874E+04 | 235.1 | - | - | 0 | - |
| - | - | 4875 | 235.2 | - | - | 0 | - |
| - | - | 2317 | 235.6 | - | - | 0 | - |
| - | - | 3679 | 237.1 | - | - | 0 | - |
| - | - | 2.329E+04 | 238.1 | - | - | 0 | - |
| - | - | 4019 | 238.1 | - | - | 0 | - |
| - | - | 3394 | 239.1 | - | - | 0 | - |
| - | - | 1.314E+04 | 239.1 | - | - | 0 | - |
| - | - | 3627 | 239.1 | - | - | 0 | - |
| - | - | 5106 | 240.1 | - | - | 0 | - |
| - | - | 7.171E+04 | 240.1 | - | - | 0 | - |
| - | - | 2615 | 241.1 | - | - | 0 | - |
| - | - | 1.385E+04 | 241.1 | - | - | 0 | - |
| - | - | 1.035E+04 | 241.1 | - | - | 0 | - |
| - | - | 9202 | 241.2 | - | - | 0 | - |
| - | - | 2318 | 242.2 | - | - | 0 | - |
| - | - | 1.156E+04 | 243.1 | - | - | 0 | - |
| - | - | 7.729E+04 | 244.1 | - | - | 0 | - |
| - | - | 6824 | 245.1 | - | - | 0 | - |
| - | - | 1.103E+05 | 247.1 | - | - | 0 | - |
| - | - | 4057 | 247.1 | - | - | 0 | - |
| - | - | 4571 | 248.1 | - | - | 0 | - |
| - | - | 4.13E+04 | 248.1 | - | - | 0 | - |
| 10 | y | 2430 | 248.6 | 0.0001531 | 0.6159 | +2 | 4 |
| - | - | 6.302E+04 | 249.1 | - | - | 0 | - |
| - | - | 4165 | 249.1 | - | - | 0 | - |
| - | - | 4141 | 249.1 | - | - | 0 | - |
| - | - | 1.518E+04 | 249.1 | - | - | 0 | - |
| - | - | 9602 | 250.1 | - | - | 0 | - |
| - | - | 6.869E+05 | 251.2 | - | - | 0 | - |
| - | - | 4442 | 252.1 | - | - | 0 | - |
| - | - | 8.252E+04 | 252.2 | - | - | 0 | - |
| - | - | 6010 | 253.2 | - | - | 0 | - |
| - | - | 3006 | 254.1 | - | - | 0 | - |
| - | - | 2.963E+04 | 255.1 | - | - | 0 | - |
| - | - | 8697 | 256.1 | - | - | 0 | - |
| - | - | 5068 | 256.2 | - | - | 0 | - |
| - | - | 2.032E+04 | 257.1 | - | - | 0 | - |
| - | - | 2.651E+04 | 258.1 | - | - | 0 | - |
| - | - | 3534 | 259.1 | - | - | 0 | - |
| - | - | 4281 | 259.1 | - | - | 0 | - |
| - | - | 1.236E+04 | 261.1 | - | - | 0 | - |
| - | - | 7914 | 261.1 | - | - | 0 | - |
| - | - | 8424 | 262.1 | - | - | 0 | - |
| - | - | 2468 | 263.1 | - | - | 0 | - |
| - | - | 2139 | 263.1 | - | - | 0 | - |
| - | - | 6.084E+04 | 265.1 | - | - | 0 | - |
| - | - | 1.741E+05 | 266.1 | - | - | 0 | - |
| - | - | 1.968E+04 | 267.1 | - | - | 0 | - |
| - | - | 3919 | 267.1 | - | - | 0 | - |
| - | - | 3931 | 267.6 | - | - | 0 | - |
| - | - | 8793 | 268.1 | - | - | 0 | - |
| - | - | 1.479E+04 | 268.2 | - | - | 0 | - |
| - | - | 6328 | 269.2 | - | - | 0 | - |
| - | - | 8709 | 269.2 | - | - | 0 | - |
| - | - | 2.793E+04 | 270.1 | - | - | 0 | - |
| - | - | 4513 | 270.1 | - | - | 0 | - |
| - | - | 2434 | 270.2 | - | - | 0 | - |
| - | - | 4278 | 270.6 | - | - | 0 | - |
| - | - | 3.61E+04 | 272.2 | - | - | 0 | - |
| - | - | 7503 | 273.2 | - | - | 0 | - |
| - | - | 3.375E+04 | 274.1 | - | - | 0 | - |
| - | - | 1.338E+04 | 274.1 | - | - | 0 | - |
| - | - | 5815 | 275.1 | - | - | 0 | - |
| - | - | 5663 | 275.1 | - | - | 0 | - |
| - | - | 1.016E+04 | 276.1 | - | - | 0 | - |
| - | - | 7098 | 276.1 | - | - | 0 | - |
| - | - | 7743 | 276.2 | - | - | 0 | - |
| - | - | 2369 | 276.7 | - | - | 0 | - |
| - | - | 1.517E+04 | 279.1 | - | - | 0 | - |
| - | - | 2702 | 279.6 | - | - | 0 | - |
| - | - | 1.817E+04 | 280.1 | - | - | 0 | - |
| - | - | 3048 | 280.1 | - | - | 0 | - |
| - | - | 1.399E+04 | 281.1 | - | - | 0 | - |
| - | - | 3677 | 281.1 | - | - | 0 | - |
| - | - | 1.509E+04 | 282.2 | - | - | 0 | - |
| - | - | 4.291E+05 | 283.1 | - | - | 0 | - |
| - | - | 3052 | 284.1 | - | - | 0 | - |
| - | - | 1.531E+04 | 284.1 | - | - | 0 | - |
| - | - | 3191 | 284.1 | - | - | 0 | - |
| - | - | 6.142E+04 | 284.1 | - | - | 0 | - |
| - | - | 2707 | 285.1 | - | - | 0 | - |
| - | - | 3793 | 285.1 | - | - | 0 | - |
| - | - | 4645 | 285.2 | - | - | 0 | - |
| - | - | 3.106E+04 | 286.1 | - | - | 0 | - |
| - | - | 3900 | 287.1 | - | - | 0 | - |
| - | - | 9098 | 287.2 | - | - | 0 | - |
| - | - | 4592 | 287.2 | - | - | 0 | - |
| - | - | 6310 | 288.6 | - | - | 0 | - |
| 5 | b | 4218 | 290.2 | 0.0004961 | 1.71 | +2 | 5 |
| - | - | 2487 | 290.7 | - | - | 0 | - |
| 12 | y | 2.564E+05 | 293.1 | 0.0006325 | 2.158 | +1 | 2 |
| - | - | 6782 | 293.2 | - | - | 0 | - |
| - | - | 5.494E+04 | 294.1 | - | - | 0 | - |
| - | - | 3451 | 295.1 | - | - | 0 | - |
| 3 | b | 8294 | 296.2 | 0.0007942 | 2.681 | +1 | 3 |
| - | - | 3350 | 296.7 | - | - | 0 | - |
| - | - | 2737 | 297.1 | - | - | 0 | - |
| 9 | y | 9000 | 297.1 | 0.001348 | 4.537 | +2 | 5 |
| - | - | 7.842E+04 | 297.2 | - | - | 0 | - |
| - | - | 3315 | 297.6 | - | - | 0 | - |
| 7 | y | 8725 | 298.1 | 0.004818 | 16.16 | +3 | 7 |
| - | - | 1.336E+04 | 298.2 | - | - | 0 | - |
| - | - | 8.491E+04 | 299.1 | - | - | 0 | - |
| - | - | 2.519E+04 | 300.2 | - | - | 0 | - |
| - | - | 2.583E+05 | 301.2 | - | - | 0 | - |
| - | - | 4.48E+04 | 302.1 | - | - | 0 | - |
| - | - | 3.596E+04 | 302.2 | - | - | 0 | - |
| - | - | 1.116E+04 | 303.1 | - | - | 0 | - |
| - | - | 4091 | 303.2 | - | - | 0 | - |
| - | - | 6414 | 305.2 | - | - | 0 | - |
| - | - | 3335 | 305.7 | - | - | 0 | - |
| 9 | y | 4.319E+04 | 306.1 | 0.0006434 | 2.102 | +2 | 5 |
| - | - | 4091 | 306.2 | - | - | 0 | - |
| - | - | 1.807E+04 | 306.6 | - | - | 0 | - |
| - | - | 8462 | 307.1 | - | - | 0 | - |
| - | - | 4116 | 309.2 | - | - | 0 | - |
| - | - | 3237 | 310.2 | - | - | 0 | - |
| 12 | y | 2.986E+04 | 311.1 | 0.0004743 | 1.524 | +1 | 2 |
| - | - | 1.481E+04 | 311.1 | - | - | 0 | - |
| - | - | 2631 | 311.7 | - | - | 0 | - |
| - | - | 7266 | 312.1 | - | - | 0 | - |
| - | - | 7958 | 314.2 | - | - | 0 | - |
| 3 | b | 2.309E+04 | 314.2 | 0.0007886 | 2.51 | +1 | 3 |
| - | - | 1.388E+05 | 315.2 | - | - | 0 | - |
| - | - | 4641 | 315.2 | - | - | 0 | - |
| - | - | 2.304E+04 | 316.2 | - | - | 0 | - |
| - | - | 3538 | 317.7 | - | - | 0 | - |
| - | - | 9810 | 319.2 | - | - | 0 | - |
| - | - | 2489 | 320.2 | - | - | 0 | - |
| - | - | 3803 | 322.2 | - | - | 0 | - |
| - | - | 1.038E+04 | 325.2 | - | - | 0 | - |
| - | - | 2330 | 325.2 | - | - | 0 | - |
| - | - | 3.168E+04 | 326.7 | - | - | 0 | - |
| - | - | 1.486E+04 | 327.2 | - | - | 0 | - |
| - | - | 4926 | 327.7 | - | - | 0 | - |
| - | - | 3282 | 331.2 | - | - | 0 | - |
| - | - | 4678 | 331.7 | - | - | 0 | - |
| - | - | 1.099E+04 | 332.2 | - | - | 0 | - |
| - | - | 1.976E+04 | 333.2 | - | - | 0 | - |
| - | - | 3847 | 334.2 | - | - | 0 | - |
| - | - | 2917 | 335.2 | - | - | 0 | - |
| - | - | 3775 | 338.2 | - | - | 0 | - |
| 6 | b | 4311 | 338.7 | 0.0008519 | 2.515 | +2 | 6 |
| - | - | 7622 | 339.2 | - | - | 0 | - |
| - | - | 4224 | 339.7 | - | - | 0 | - |
| - | - | 9754 | 340.7 | - | - | 0 | - |
| - | - | 3092 | 341.2 | - | - | 0 | - |
| - | - | 1.118E+04 | 342.2 | - | - | 0 | - |
| - | - | 1.116E+04 | 343.2 | - | - | 0 | - |
| - | - | 2507 | 343.2 | - | - | 0 | - |
| - | - | 1.422E+04 | 344.2 | - | - | 0 | - |
| - | - | 1.158E+04 | 345 | - | - | 0 | - |
| - | - | 1.789E+04 | 345.1 | - | - | 0 | - |
| - | - | 2834 | 345.2 | - | - | 0 | - |
| - | - | 7.541E+04 | 346.1 | - | - | 0 | - |
| - | - | 4612 | 346.1 | - | - | 0 | - |
| - | - | 3322 | 346.7 | - | - | 0 | - |
| - | - | 1.335E+04 | 347.1 | - | - | 0 | - |
| 6 | b | 4.05E+04 | 347.7 | 0.0006965 | 2.003 | +2 | 6 |
| - | - | 1.477E+04 | 348.2 | - | - | 0 | - |
| - | - | 2777 | 348.7 | - | - | 0 | - |
| - | - | 3119 | 349.1 | - | - | 0 | - |
| - | - | 4007 | 349.2 | - | - | 0 | - |
| - | - | 5.894E+04 | 350.2 | - | - | 0 | - |
| - | - | 3676 | 351.1 | - | - | 0 | - |
| - | - | 4270 | 351.2 | - | - | 0 | - |
| - | - | 7500 | 351.2 | - | - | 0 | - |
| - | - | 4809 | 352.2 | - | - | 0 | - |
| - | - | 4731 | 352.7 | - | - | 0 | - |
| 8 | y | 9196 | 353.7 | 0.0005148 | 1.456 | +2 | 6 |
| - | - | 4243 | 354.2 | - | - | 0 | - |
| - | - | 3164 | 357.2 | - | - | 0 | - |
| - | - | 1.082E+04 | 359 | - | - | 0 | - |
| - | - | 3946 | 360.2 | - | - | 0 | - |
| - | - | 1.832E+04 | 360.7 | - | - | 0 | - |
| - | - | 2.324E+04 | 361.2 | - | - | 0 | - |
| - | - | 2.084E+04 | 362.2 | - | - | 0 | - |
| 8 | y | 1.06E+05 | 362.7 | 0.0007257 | 2.001 | +2 | 6 |
| - | - | 9.942E+04 | 363.1 | - | - | 0 | - |
| - | - | 4.002E+04 | 363.2 | - | - | 0 | - |
| - | - | 7706 | 363.7 | - | - | 0 | - |
| - | - | 1.986E+04 | 364.1 | - | - | 0 | - |
| - | - | 1.883E+04 | 364.1 | - | - | 0 | - |
| - | - | 3500 | 365.1 | - | - | 0 | - |
| - | - | 2932 | 366.2 | - | - | 0 | - |
| - | - | 6505 | 366.2 | - | - | 0 | - |
| - | - | 9709 | 367.1 | - | - | 0 | - |
| - | - | 6685 | 367.2 | - | - | 0 | - |
| - | - | 9698 | 367.2 | - | - | 0 | - |
| - | - | 2887 | 368.1 | - | - | 0 | - |
| - | - | 3891 | 368.2 | - | - | 0 | - |
| - | - | 3726 | 369.2 | - | - | 0 | - |
| - | - | 3342 | 370.2 | - | - | 0 | - |
| - | - | 6082 | 373.2 | - | - | 0 | - |
| - | - | 6273 | 376.2 | - | - | 0 | - |
| - | - | 1.438E+04 | 376.2 | - | - | 0 | - |
| - | - | 4765 | 376.7 | - | - | 0 | - |
| - | - | 5831 | 377.1 | - | - | 0 | - |
| - | - | 4.399E+04 | 378.2 | - | - | 0 | - |
| - | - | 1.695E+05 | 379.2 | - | - | 0 | - |
| - | - | 6681 | 380.2 | - | - | 0 | - |
| - | - | 3.238E+04 | 380.2 | - | - | 0 | - |
| - | - | 2.949E+05 | 381.2 | - | - | 0 | - |
| - | - | 9830 | 381.7 | - | - | 0 | - |
| - | - | 4.717E+04 | 382.2 | - | - | 0 | - |
| - | - | 1.146E+04 | 382.2 | - | - | 0 | - |
| - | - | 4313 | 382.2 | - | - | 0 | - |
| - | - | 2794 | 382.7 | - | - | 0 | - |
| - | - | 8787 | 383.2 | - | - | 0 | - |
| - | - | 3680 | 383.2 | - | - | 0 | - |
| - | - | 1.259E+04 | 384.2 | - | - | 0 | - |
| - | - | 7825 | 384.2 | - | - | 0 | - |
| - | - | 2.698E+04 | 385.2 | - | - | 0 | - |
| 5 | y | 4278 | 385.2 | 0.006512 | 16.91 | +3 | 9 |
| - | - | 4076 | 386.2 | - | - | 0 | - |
| - | - | 4369 | 386.2 | - | - | 0 | - |
| 10 | b | 4310 | 389.2 | 0.000778 | 1.999 | +3 | 10 |
| - | - | 6754 | 390.2 | - | - | 0 | - |
| - | - | 5562 | 390.7 | - | - | 0 | - |
| - | - | 1.124E+04 | 391.1 | - | - | 0 | - |
| - | - | 4361 | 392.2 | - | - | 0 | - |
| - | - | 5017 | 393.2 | - | - | 0 | - |
| - | - | 4.029E+04 | 394.2 | - | - | 0 | - |
| - | - | 2.13E+04 | 395.1 | - | - | 0 | - |
| - | - | 8077 | 395.2 | - | - | 0 | - |
| - | - | 4443 | 395.2 | - | - | 0 | - |
| - | - | 3586 | 395.2 | - | - | 0 | - |
| - | - | 5313 | 396.1 | - | - | 0 | - |
| - | - | 4713 | 396.2 | - | - | 0 | - |
| - | - | 6.534E+04 | 396.2 | - | - | 0 | - |
| - | - | 5273 | 397.2 | - | - | 0 | - |
| - | - | 1.282E+04 | 397.2 | - | - | 0 | - |
| - | - | 3028 | 397.7 | - | - | 0 | - |
| - | - | 2.909E+04 | 398.2 | - | - | 0 | - |
| - | - | 3240 | 398.2 | - | - | 0 | - |
| - | - | 6758 | 399.2 | - | - | 0 | - |
| - | - | 1.612E+04 | 402.2 | - | - | 0 | - |
| - | - | 2.609E+04 | 402.2 | - | - | 0 | - |
| - | - | 2964 | 403.1 | - | - | 0 | - |
| - | - | 1.236E+04 | 403.2 | - | - | 0 | - |
| - | - | 3801 | 404.7 | - | - | 0 | - |
| - | - | 1.181E+04 | 405.3 | - | - | 0 | - |
| - | - | 6326 | 406.2 | - | - | 0 | - |
| - | - | 4062 | 407.2 | - | - | 0 | - |
| - | - | 2.557E+04 | 409.1 | - | - | 0 | - |
| - | - | 4845 | 409.2 | - | - | 0 | - |
| - | - | 3408 | 410.1 | - | - | 0 | - |
| - | - | 1.002E+04 | 411.2 | - | - | 0 | - |
| - | - | 1.921E+05 | 412.2 | - | - | 0 | - |
| - | - | 2.818E+04 | 413.1 | - | - | 0 | - |
| - | - | 3.101E+04 | 413.2 | - | - | 0 | - |
| - | - | 4795 | 414.1 | - | - | 0 | - |
| - | - | 6035 | 414.2 | - | - | 0 | - |
| - | - | 6.01E+04 | 414.2 | - | - | 0 | - |
| - | - | 2.108E+04 | 415 | - | - | 0 | - |
| - | - | 2.455E+04 | 415.2 | - | - | 0 | - |
| - | - | 1.176E+04 | 415.2 | - | - | 0 | - |
| - | - | 6564 | 416.2 | - | - | 0 | - |
| - | - | 3.618E+04 | 417.7 | - | - | 0 | - |
| - | - | 5.146E+04 | 418.2 | - | - | 0 | - |
| - | - | 2.536E+04 | 418.7 | - | - | 0 | - |
| - | - | 3797 | 419.2 | - | - | 0 | - |
| 11 | y | 9610 | 421.2 | 0.0018 | 4.273 | +1 | 3 |
| 11 | y | 4548 | 422.2 | 0.0003892 | 0.9219 | +1 | 3 |
| - | - | 2797 | 422.3 | - | - | 0 | - |
| - | - | 3089 | 422.7 | - | - | 0 | - |
| - | - | 1.381E+04 | 423.2 | - | - | 0 | - |
| - | - | 6.982E+04 | 423.3 | - | - | 0 | - |
| - | - | 4828 | 423.7 | - | - | 0 | - |
| - | - | 3320 | 424.2 | - | - | 0 | - |
| - | - | 2.119E+04 | 424.3 | - | - | 0 | - |
| - | - | 8772 | 424.7 | - | - | 0 | - |
| - | - | 3138 | 425.2 | - | - | 0 | - |
| - | - | 5.155E+05 | 426.7 | - | - | 0 | - |
| - | - | 2.639E+05 | 427.2 | - | - | 0 | - |
| - | - | 7.862E+04 | 427.7 | - | - | 0 | - |
| - | - | 1.253E+04 | 428.2 | - | - | 0 | - |
| - | - | 3.217E+04 | 429.2 | - | - | 0 | - |
| - | - | 3443 | 429.7 | - | - | 0 | - |
| - | - | 5.309E+05 | 430.2 | - | - | 0 | - |
| - | - | 1.097E+05 | 431.2 | - | - | 0 | - |
| 7 | b | 6.066E+04 | 431.7 | 0.0007157 | 1.658 | +2 | 7 |
| 7 | b | 3.063E+04 | 432.2 | 0.00706 | 16.33 | +2 | 7 |
| - | - | 1.134E+04 | 432.7 | - | - | 0 | - |
| - | - | 9996 | 433.2 | - | - | 0 | - |
| 4 | b | 2.06E+04 | 433.3 | 0.0005066 | 1.169 | +1 | 4 |
| - | - | 3924 | 434.3 | - | - | 0 | - |
| - | - | 4013 | 437.2 | - | - | 0 | - |
| - | - | 9008 | 438.2 | - | - | 0 | - |
| - | - | 3749 | 438.7 | - | - | 0 | - |
| 11 | y | 1.235E+05 | 439.2 | 0.001092 | 2.487 | +1 | 3 |
| - | - | 6288 | 440.2 | - | - | 0 | - |
| - | - | 2.938E+04 | 440.2 | - | - | 0 | - |
| 7 | b | 2.452E+05 | 440.7 | 0.001079 | 2.448 | +2 | 7 |
| - | - | 1.403E+05 | 441.2 | - | - | 0 | - |
| - | - | 3.924E+04 | 441.7 | - | - | 0 | - |
| - | - | 7294 | 442.2 | - | - | 0 | - |
| - | - | 2088 | 442.3 | - | - | 0 | - |
| - | - | 2652 | 444.2 | - | - | 0 | - |
| 7 | y | 2.622E+04 | 447.2 | 0.0006194 | 1.385 | +2 | 7 |
| - | - | 5147 | 447.2 | - | - | 0 | - |
| - | - | 9110 | 447.7 | - | - | 0 | - |
| - | - | 2.325E+04 | 448.2 | - | - | 0 | - |
| - | - | 1.025E+04 | 449.2 | - | - | 0 | - |
| - | - | 3876 | 450.2 | - | - | 0 | - |
| - | - | 7001 | 450.2 | - | - | 0 | - |
| - | - | 6252 | 450.3 | - | - | 0 | - |
| 4 | b | 1.421E+05 | 451.3 | 0.001142 | 2.53 | +1 | 4 |
| - | - | 3364 | 452.2 | - | - | 0 | - |
| - | - | 3.421E+04 | 452.3 | - | - | 0 | - |
| - | - | 4676 | 453.3 | - | - | 0 | - |
| - | - | 5224 | 455.2 | - | - | 0 | - |
| - | - | 7060 | 455.3 | - | - | 0 | - |
| 7 | y | 1.61E+04 | 455.7 | 0.001505 | 3.302 | +2 | 7 |
| - | - | 3503 | 456.2 | - | - | 0 | - |
| - | - | 1.181E+04 | 459.2 | - | - | 0 | - |
| - | - | 1.94E+04 | 460.8 | - | - | 0 | - |
| - | - | 1.084E+04 | 461.2 | - | - | 0 | - |
| - | - | 1.274E+04 | 461.3 | - | - | 0 | - |
| - | - | 5866 | 461.8 | - | - | 0 | - |
| - | - | 2895 | 462.2 | - | - | 0 | - |
| - | - | 5146 | 464.2 | - | - | 0 | - |
| - | - | 2559 | 464.2 | - | - | 0 | - |
| - | - | 3290 | 465.8 | - | - | 0 | - |
| - | - | 4684 | 466.2 | - | - | 0 | - |
| - | - | 4.144E+04 | 468.3 | - | - | 0 | - |
| - | - | 8606 | 469.3 | - | - | 0 | - |
| - | - | 2.592E+04 | 474.3 | - | - | 0 | - |
| - | - | 4.039E+04 | 474.8 | - | - | 0 | - |
| - | - | 2.083E+04 | 475.3 | - | - | 0 | - |
| - | - | 4349 | 475.8 | - | - | 0 | - |
| - | - | 1.339E+04 | 476.2 | - | - | 0 | - |
| - | - | 2.686E+04 | 477.2 | - | - | 0 | - |
| 10 | y | 2.563E+04 | 478.2 | 0.0002031 | 0.4246 | +1 | 4 |
| - | - | 1.735E+04 | 478.3 | - | - | 0 | - |
| - | - | 4245 | 479.2 | - | - | 0 | - |
| - | - | 3617 | 479.3 | - | - | 0 | - |
| - | - | 4049 | 479.7 | - | - | 0 | - |
| - | - | 3088 | 480.2 | - | - | 0 | - |
| - | - | 4651 | 481.7 | - | - | 0 | - |
| - | - | 8.649E+04 | 483.3 | - | - | 0 | - |
| - | - | 5.931E+04 | 483.8 | - | - | 0 | - |
| - | - | 6228 | 484.2 | - | - | 0 | - |
| - | - | 1.763E+04 | 484.3 | - | - | 0 | - |
| - | - | 2416 | 484.8 | - | - | 0 | - |
| 8 | b | 2.704E+04 | 488.3 | 0.001439 | 2.947 | +2 | 8 |
| 8 | b | 1.711E+04 | 488.7 | 0.009126 | 18.67 | +2 | 8 |
| - | - | 1.128E+04 | 489.3 | - | - | 0 | - |
| - | - | 3.138E+05 | 494.2 | - | - | 0 | - |
| - | - | 8.139E+04 | 495.2 | - | - | 0 | - |
| 10 | y | 4.226E+05 | 496.2 | 0.001052 | 2.12 | +1 | 4 |
| - | - | 1.121E+05 | 497.2 | - | - | 0 | - |
| - | - | 3.427E+04 | 497.8 | - | - | 0 | - |
| - | - | 1.879E+04 | 498.2 | - | - | 0 | - |
| - | - | 3246 | 502.2 | - | - | 0 | - |
| - | - | 2.037E+04 | 503.1 | - | - | 0 | - |
| - | - | 1.439E+04 | 504.2 | - | - | 0 | - |
| - | - | 4209 | 505.2 | - | - | 0 | - |
| - | - | 2549 | 505.2 | - | - | 0 | - |
| - | - | 8247 | 506.2 | - | - | 0 | - |
| - | - | 4287 | 506.3 | - | - | 0 | - |
| - | - | 7221 | 507.3 | - | - | 0 | - |
| - | - | 2.156E+04 | 511.3 | - | - | 0 | - |
| - | - | 4166 | 512.3 | - | - | 0 | - |
| - | - | 4851 | 513.2 | - | - | 0 | - |
| - | - | 6199 | 514.2 | - | - | 0 | - |
| - | - | 9976 | 515.3 | - | - | 0 | - |
| - | - | 6616 | 519.1 | - | - | 0 | - |
| - | - | 4080 | 519.3 | - | - | 0 | - |
| - | - | 5975 | 522.2 | - | - | 0 | - |
| - | - | 1.099E+04 | 523.3 | - | - | 0 | - |
| - | - | 4384 | 523.6 | - | - | 0 | - |
| - | - | 3098 | 524.3 | - | - | 0 | - |
| - | - | 5.616E+04 | 525.3 | - | - | 0 | - |
| - | - | 1.124E+04 | 526.2 | - | - | 0 | - |
| - | - | 1.249E+04 | 526.3 | - | - | 0 | - |
| - | - | 3380 | 527.2 | - | - | 0 | - |
| 0 | Precursor | 1.64E+04 | 529.3 | 0.001901 | 3.592 | +3 | -1 |
| 0 | Precursor | 2.269E+04 | 529.6 | 0.006782 | 12.81 | +3 | -1 |
| - | - | 1.11E+04 | 529.9 | - | - | 0 | - |
| - | - | 4273 | 530.2 | - | - | 0 | - |
| - | - | 3510 | 530.3 | - | - | 0 | - |
| - | - | 4593 | 531.2 | - | - | 0 | - |
| - | - | 1.438E+04 | 532.2 | - | - | 0 | - |
| - | - | 2988 | 533.2 | - | - | 0 | - |
| 0 | Precursor | 1.334E+04 | 535.3 | 0.0003938 | 0.7357 | +3 | -1 |
| - | - | 1.559E+04 | 535.6 | - | - | 0 | - |
| - | - | 8951 | 535.9 | - | - | 0 | - |
| - | - | 3511 | 536.2 | - | - | 0 | - |
| - | - | 6629 | 536.3 | - | - | 0 | - |
| - | - | 3485 | 537.2 | - | - | 0 | - |
| - | - | 1.157E+04 | 537.7 | - | - | 0 | - |
| - | - | 9512 | 538.3 | - | - | 0 | - |
| - | - | 6391 | 538.8 | - | - | 0 | - |
| - | - | 5.873E+04 | 539.2 | - | - | 0 | - |
| - | - | 4749 | 539.3 | - | - | 0 | - |
| - | - | 1.843E+04 | 540.2 | - | - | 0 | - |
| - | - | 7036 | 541.3 | - | - | 0 | - |
| - | - | 4281 | 542.3 | - | - | 0 | - |
| - | - | 1.288E+05 | 543.3 | - | - | 0 | - |
| - | - | 3.753E+04 | 544.3 | - | - | 0 | - |
| - | - | 7692 | 545.3 | - | - | 0 | - |
| - | - | 1.639E+04 | 546.2 | - | - | 0 | - |
| - | - | 1.043E+04 | 546.8 | - | - | 0 | - |
| - | - | 4509 | 547.2 | - | - | 0 | - |
| - | - | 5655 | 547.8 | - | - | 0 | - |
| - | - | 3659 | 548.2 | - | - | 0 | - |
| - | - | 3257 | 548.3 | - | - | 0 | - |
| - | - | 1.298E+04 | 549.2 | - | - | 0 | - |
| - | - | 8457 | 550.2 | - | - | 0 | - |
| - | - | 1.352E+04 | 551.3 | - | - | 0 | - |
| - | - | 7131 | 551.3 | - | - | 0 | - |
| - | - | 2948 | 552.3 | - | - | 0 | - |
| - | - | 3042 | 552.3 | - | - | 0 | - |
| - | - | 3317 | 553.3 | - | - | 0 | - |
| 9 | b | 1.45E+04 | 554.8 | 0.0002557 | 0.461 | +2 | 9 |
| - | - | 2.99E+04 | 555.3 | - | - | 0 | - |
| - | - | 1.687E+04 | 555.8 | - | - | 0 | - |
| - | - | 5581 | 556.3 | - | - | 0 | - |
| 5 | b | 1.456E+04 | 561.3 | 0.000767 | 1.366 | +1 | 5 |
| 5 | b | 7375 | 562.3 | 0.003507 | 6.237 | +1 | 5 |
| - | - | 1.28E+04 | 563.3 | - | - | 0 | - |
| - | - | 5617 | 564.3 | - | - | 0 | - |
| - | - | 1.98E+04 | 565.3 | - | - | 0 | - |
| - | - | 3817 | 566.3 | - | - | 0 | - |
| - | - | 6.678E+04 | 567.2 | - | - | 0 | - |
| - | - | 1.81E+04 | 568.2 | - | - | 0 | - |
| - | - | 8310 | 568.8 | - | - | 0 | - |
| - | - | 7337 | 569.2 | - | - | 0 | - |
| - | - | 1.566E+04 | 575.2 | - | - | 0 | - |
| - | - | 3584 | 576.2 | - | - | 0 | - |
| - | - | 3284 | 576.3 | - | - | 0 | - |
| - | - | 4511 | 577.2 | - | - | 0 | - |
| 5 | y | 3494 | 577.3 | 0.003606 | 6.246 | +2 | 9 |
| - | - | 4351 | 577.8 | - | - | 0 | - |
| - | - | 4410 | 579.3 | - | - | 0 | - |
| 5 | b | 8.837E+04 | 579.3 | 0.0009445 | 1.63 | +1 | 5 |
| - | - | 2.625E+04 | 580.3 | - | - | 0 | - |
| - | - | 3260 | 581.3 | - | - | 0 | - |
| - | - | 4010 | 581.3 | - | - | 0 | - |
| - | - | 4446 | 582.3 | - | - | 0 | - |
| 10 | b | 4457 | 583.3 | 0.001647 | 2.823 | +2 | 10 |
| - | - | 2930 | 587.3 | - | - | 0 | - |
| - | - | 3660 | 588.8 | - | - | 0 | - |
| - | - | 2812 | 589.3 | - | - | 0 | - |
| - | - | 4100 | 592.3 | - | - | 0 | - |
| - | - | 1.147E+04 | 592.3 | - | - | 0 | - |
| 9 | y | 1.005E+05 | 593.3 | 0.0005733 | 0.9663 | +1 | 5 |
| - | - | 4.794E+04 | 593.3 | - | - | 0 | - |
| - | - | 6232 | 593.8 | - | - | 0 | - |
| - | - | 3.029E+04 | 594.3 | - | - | 0 | - |
| - | - | 1.016E+04 | 594.3 | - | - | 0 | - |
| - | - | 3970 | 594.8 | - | - | 0 | - |
| - | - | 6311 | 595.2 | - | - | 0 | - |
| - | - | 2801 | 596.4 | - | - | 0 | - |
| - | - | 3432 | 597.3 | - | - | 0 | - |
| - | - | 1.055E+04 | 597.8 | - | - | 0 | - |
| - | - | 3668 | 598.3 | - | - | 0 | - |
| - | - | 1.006E+04 | 600.3 | - | - | 0 | - |
| - | - | 2821 | 602.3 | - | - | 0 | - |
| - | - | 2.057E+04 | 602.8 | - | - | 0 | - |
| - | - | 1.956E+04 | 603.3 | - | - | 0 | - |
| - | - | 1.001E+04 | 603.8 | - | - | 0 | - |
| - | - | 4709 | 604.3 | - | - | 0 | - |
| - | - | 8.561E+04 | 610.3 | - | - | 0 | - |
| - | - | 7488 | 610.3 | - | - | 0 | - |
| 9 | y | 1.307E+06 | 611.3 | 0.001483 | 2.426 | +1 | 5 |
| - | - | 8.374E+04 | 611.8 | - | - | 0 | - |
| - | - | 4.078E+05 | 612.3 | - | - | 0 | - |
| - | - | 2.465E+04 | 612.8 | - | - | 0 | - |
| - | - | 8.348E+04 | 613.3 | - | - | 0 | - |
| - | - | 1.613E+04 | 614.3 | - | - | 0 | - |
| - | - | 5718 | 616.3 | - | - | 0 | - |
| - | - | 7229 | 617.3 | - | - | 0 | - |
| - | - | 3016 | 619.3 | - | - | 0 | - |
| - | - | 6376 | 620.8 | - | - | 0 | - |
| - | - | 1.414E+04 | 621.3 | - | - | 0 | - |
| - | - | 7976 | 621.3 | - | - | 0 | - |
| - | - | 3180 | 622.3 | - | - | 0 | - |
| - | - | 5587 | 622.4 | - | - | 0 | - |
| - | - | 2583 | 623.2 | - | - | 0 | - |
| - | - | 3566 | 628.3 | - | - | 0 | - |
| - | - | 3665 | 628.8 | - | - | 0 | - |
| - | - | 1.718E+04 | 634.3 | - | - | 0 | - |
| - | - | 2.155E+04 | 635.3 | - | - | 0 | - |
| - | - | 1.097E+04 | 636.3 | - | - | 0 | - |
| 4 | y | 7394 | 636.8 | 0.0001839 | 0.2888 | +2 | 10 |
| 4 | y | 4253 | 637.3 | 0.005002 | 7.849 | +2 | 10 |
| - | - | 4149 | 637.3 | - | - | 0 | - |
| - | - | 7062 | 637.8 | - | - | 0 | - |
| 11 | b | 6576 | 638.3 | 0.001684 | 2.638 | +2 | 11 |
| - | - | 1.644E+04 | 640.3 | - | - | 0 | - |
| - | - | 3773 | 641.3 | - | - | 0 | - |
| - | - | 1.216E+04 | 645.3 | - | - | 0 | - |
| 4 | y | 8.444E+04 | 645.8 | 0.001432 | 2.218 | +2 | 10 |
| - | - | 7.61E+04 | 646.3 | - | - | 0 | - |
| - | - | 2.432E+04 | 646.8 | - | - | 0 | - |
| - | - | 3501 | 647.3 | - | - | 0 | - |
| - | - | 5030 | 648.3 | - | - | 0 | - |
| - | - | 4264 | 649.3 | - | - | 0 | - |
| - | - | 6332 | 650.3 | - | - | 0 | - |
| - | - | 4144 | 650.3 | - | - | 0 | - |
| - | - | 4126 | 651.3 | - | - | 0 | - |
| - | - | 1.309E+05 | 652.3 | - | - | 0 | - |
| - | - | 4.819E+04 | 653.3 | - | - | 0 | - |
| - | - | 8889 | 654.3 | - | - | 0 | - |
| - | - | 1.079E+04 | 658.3 | - | - | 0 | - |
| - | - | 6748 | 659.3 | - | - | 0 | - |
| - | - | 2734 | 660.3 | - | - | 0 | - |
| - | - | 3382 | 660.9 | - | - | 0 | - |
| - | - | 9412 | 661.3 | - | - | 0 | - |
| - | - | 6311 | 661.8 | - | - | 0 | - |
| - | - | 3.784E+04 | 662.3 | - | - | 0 | - |
| - | - | 3.558E+04 | 663.3 | - | - | 0 | - |
| - | - | 3857 | 664.3 | - | - | 0 | - |
| - | - | 3658 | 664.3 | - | - | 0 | - |
| - | - | 2937 | 666.3 | - | - | 0 | - |
| - | - | 3352 | 666.4 | - | - | 0 | - |
| - | - | 5624 | 668.3 | - | - | 0 | - |
| - | - | 2717 | 670.8 | - | - | 0 | - |
| 6 | b | 5.67E+04 | 676.3 | 0.001046 | 1.546 | +1 | 6 |
| 6 | b | 3.798E+04 | 677.3 | 0.008607 | 12.71 | +1 | 6 |
| - | - | 1.567E+04 | 678.3 | - | - | 0 | - |
| - | - | 1.415E+04 | 679.3 | - | - | 0 | - |
| - | - | 5735 | 679.8 | - | - | 0 | - |
| - | - | 2.289E+05 | 680.3 | - | - | 0 | - |
| - | - | 9.301E+04 | 681.3 | - | - | 0 | - |
| - | - | 1.911E+04 | 682.3 | - | - | 0 | - |
| - | - | 3683 | 683.3 | - | - | 0 | - |
| - | - | 6793 | 684.3 | - | - | 0 | - |
| - | - | 2.107E+04 | 684.8 | - | - | 0 | - |
| - | - | 1.202E+04 | 685.3 | - | - | 0 | - |
| - | - | 8515 | 685.8 | - | - | 0 | - |
| - | - | 2944 | 686.3 | - | - | 0 | - |
| - | - | 4925 | 692.8 | - | - | 0 | - |
| 3 | y | 6.773E+04 | 693.3 | 0.00167 | 2.409 | +2 | 11 |
| 3 | y | 6.121E+04 | 693.8 | 0.008503 | 12.25 | +2 | 11 |
| 6 | b | 6.146E+05 | 694.4 | 0.0004297 | 0.6188 | +1 | 6 |
| - | - | 1.433E+04 | 694.8 | - | - | 0 | - |
| - | - | 2.343E+05 | 695.4 | - | - | 0 | - |
| - | - | 5.721E+04 | 696.4 | - | - | 0 | - |
| - | - | 3240 | 697.3 | - | - | 0 | - |
| - | - | 1.02E+04 | 697.4 | - | - | 0 | - |
| - | - | 4213 | 697.9 | - | - | 0 | - |
| - | - | 7129 | 698.4 | - | - | 0 | - |
| - | - | 3.112E+04 | 701.8 | - | - | 0 | - |
| 3 | y | 5.278E+05 | 702.3 | 0.001637 | 2.33 | +2 | 11 |
| 12 | b | 4.413E+05 | 702.8 | 0.01253 | 17.83 | +2 | 12 |
| 12 | b | 2.019E+05 | 703.3 | 0.003198 | 4.546 | +2 | 12 |
| - | - | 6.474E+04 | 703.8 | - | - | 0 | - |
| - | - | 2.12E+04 | 704.3 | - | - | 0 | - |
| - | - | 5078 | 705.4 | - | - | 0 | - |
| 8 | y | 4.332E+04 | 706.3 | 0.002325 | 3.291 | +1 | 6 |
| - | - | 1.557E+04 | 707.3 | - | - | 0 | - |
| - | - | 4217 | 708.3 | - | - | 0 | - |
| - | - | 1.472E+04 | 711.4 | - | - | 0 | - |
| 12 | b | 3.42E+04 | 711.9 | 0.001533 | 2.154 | +2 | 12 |
| - | - | 4.377E+04 | 712.4 | - | - | 0 | - |
| - | - | 1.624E+04 | 712.9 | - | - | 0 | - |
| - | - | 7745 | 713.4 | - | - | 0 | - |
| - | - | 3222 | 715.3 | - | - | 0 | - |
| - | - | 5148 | 720.4 | - | - | 0 | - |
| - | - | 3114 | 720.9 | - | - | 0 | - |
| - | - | 2.862E+04 | 721.4 | - | - | 0 | - |
| - | - | 9457 | 722.4 | - | - | 0 | - |
| - | - | 3.198E+04 | 723.4 | - | - | 0 | - |
| 8 | y | 4.081E+05 | 724.4 | 0.001648 | 2.275 | +1 | 6 |
| - | - | 1.737E+05 | 725.4 | - | - | 0 | - |
| - | - | 4.146E+04 | 726.4 | - | - | 0 | - |
| - | - | 4979 | 727.4 | - | - | 0 | - |
| - | - | 4929 | 728.9 | - | - | 0 | - |
| - | - | 1.898E+04 | 729.4 | - | - | 0 | - |
| - | - | 9587 | 730.4 | - | - | 0 | - |
| - | - | 1.603E+04 | 733.4 | - | - | 0 | - |
| - | - | 1.616E+04 | 734.4 | - | - | 0 | - |
| - | - | 4404 | 735.4 | - | - | 0 | - |
| - | - | 4897 | 737.9 | - | - | 0 | - |
| - | - | 4473 | 738.4 | - | - | 0 | - |
| 2 | y | 3695 | 742.9 | 0.005366 | 7.223 | +2 | 12 |
| 2 | y | 9679 | 743.4 | 0.01177 | 15.84 | +2 | 12 |
| - | - | 4146 | 743.9 | - | - | 0 | - |
| - | - | 3344 | 744.4 | - | - | 0 | - |
| - | - | 2873 | 746.4 | - | - | 0 | - |
| - | - | 9205 | 747.4 | - | - | 0 | - |
| - | - | 1.037E+04 | 748.4 | - | - | 0 | - |
| - | - | 5931 | 749.3 | - | - | 0 | - |
| - | - | 3427 | 750.4 | - | - | 0 | - |
| - | - | 6.021E+04 | 751.4 | - | - | 0 | - |
| 2 | y | 4.737E+04 | 751.9 | 0.001487 | 1.978 | +2 | 12 |
| - | - | 7.281E+04 | 752.4 | - | - | 0 | - |
| - | - | 1.614E+04 | 752.9 | - | - | 0 | - |
| - | - | 1.561E+04 | 753.4 | - | - | 0 | - |
| - | - | 4086 | 760.3 | - | - | 0 | - |
| - | - | 4698 | 761.4 | - | - | 0 | - |
| - | - | 8278 | 762.4 | - | - | 0 | - |
| - | - | 3.815E+04 | 763.4 | - | - | 0 | - |
| - | - | 1.097E+04 | 764.4 | - | - | 0 | - |
| - | - | 1.594E+04 | 765.4 | - | - | 0 | - |
| - | - | 7682 | 766.4 | - | - | 0 | - |
| - | - | 1.365E+04 | 767.3 | - | - | 0 | - |
| - | - | 4085 | 768.3 | - | - | 0 | - |
| - | - | 4253 | 769.4 | - | - | 0 | - |
| - | - | 7724 | 775.4 | - | - | 0 | - |
| - | - | 8410 | 776.4 | - | - | 0 | - |
| - | - | 4.167E+04 | 779.4 | - | - | 0 | - |
| - | - | 1.651E+04 | 780.4 | - | - | 0 | - |
| - | - | 1.425E+04 | 781.4 | - | - | 0 | - |
| - | - | 8373 | 782.4 | - | - | 0 | - |
| - | - | 2794 | 791.4 | - | - | 0 | - |
| - | - | 3242 | 792.4 | - | - | 0 | - |
| 0 | Precursor | 6.793E+04 | 793.4 | 0.0143 | 18.03 | +2 | -1 |
| - | - | 2.713E+04 | 794.4 | - | - | 0 | - |
| - | - | 2.809E+04 | 795.3 | - | - | 0 | - |
| - | - | 4431 | 795.4 | - | - | 0 | - |
| - | - | 1.031E+04 | 796.3 | - | - | 0 | - |
| - | - | 3608 | 797.3 | - | - | 0 | - |
| - | - | 1.043E+04 | 807.4 | - | - | 0 | - |
| - | - | 7560 | 807.4 | - | - | 0 | - |
| - | - | 9907 | 808.4 | - | - | 0 | - |
| - | - | 3678 | 817.4 | - | - | 0 | - |
| - | - | 7909 | 818.4 | - | - | 0 | - |
| - | - | 7650 | 819.4 | - | - | 0 | - |
| - | - | 3712 | 820.4 | - | - | 0 | - |
| - | - | 6137 | 825.4 | - | - | 0 | - |
| - | - | 1.235E+04 | 826.4 | - | - | 0 | - |
| - | - | 3135 | 827.4 | - | - | 0 | - |
| - | - | 1.211E+04 | 834.4 | - | - | 0 | - |
| - | - | 2.526E+04 | 835.4 | - | - | 0 | - |
| - | - | 5.264E+04 | 836.4 | - | - | 0 | - |
| - | - | 2.089E+04 | 837.5 | - | - | 0 | - |
| - | - | 6326 | 838.5 | - | - | 0 | - |
| - | - | 9180 | 843.4 | - | - | 0 | - |
| - | - | 2.044E+04 | 844.4 | - | - | 0 | - |
| - | - | 1.345E+04 | 845.4 | - | - | 0 | - |
| - | - | 2.038E+04 | 846.4 | - | - | 0 | - |
| - | - | 1.345E+04 | 847.4 | - | - | 0 | - |
| - | - | 1.005E+04 | 852.4 | - | - | 0 | - |
| - | - | 1.48E+05 | 852.4 | - | - | 0 | - |
| - | - | 5105 | 853.4 | - | - | 0 | - |
| - | - | 8.043E+04 | 853.4 | - | - | 0 | - |
| - | - | 1.76E+04 | 854.4 | - | - | 0 | - |
| 7 | b | 8.893E+04 | 862.4 | 0.0008343 | 0.9673 | +1 | 7 |
| 7 | b | 7.164E+04 | 863.4 | 0.01224 | 14.18 | +1 | 7 |
| - | - | 1.105E+05 | 864.5 | - | - | 0 | - |
| - | - | 5.575E+04 | 865.5 | - | - | 0 | - |
| - | - | 2.044E+04 | 866.5 | - | - | 0 | - |
| - | - | 2970 | 867.5 | - | - | 0 | - |
| - | - | 5143 | 869.4 | - | - | 0 | - |
| - | - | 9589 | 876.4 | - | - | 0 | - |
| - | - | 4519 | 877.4 | - | - | 0 | - |
| 7 | b | 7.167E+05 | 880.4 | 0.0007676 | 0.8719 | +1 | 7 |
| - | - | 3.809E+05 | 881.4 | - | - | 0 | - |
| - | - | 1.203E+05 | 882.4 | - | - | 0 | - |
| - | - | 2.156E+04 | 883.4 | - | - | 0 | - |
| - | - | 5099 | 890.4 | - | - | 0 | - |
| - | - | 9915 | 891.4 | - | - | 0 | - |
| - | - | 3760 | 892.4 | - | - | 0 | - |
| - | - | 9595 | 892.5 | - | - | 0 | - |
| - | - | 8167 | 893.5 | - | - | 0 | - |
| - | - | 6106 | 894.4 | - | - | 0 | - |
| - | - | 3284 | 907.4 | - | - | 0 | - |
| - | - | 8.022E+04 | 908.4 | - | - | 0 | - |
| - | - | 4.365E+04 | 909.4 | - | - | 0 | - |
| 7 | y | 7.049E+04 | 910.4 | 0.001436 | 1.578 | +1 | 7 |
| - | - | 3.031E+04 | 911.4 | - | - | 0 | - |
| - | - | 1.009E+04 | 912.4 | - | - | 0 | - |
| - | - | 7365 | 936.4 | - | - | 0 | - |
| - | - | 1.093E+04 | 937.4 | - | - | 0 | - |
| - | - | 4667 | 938.4 | - | - | 0 | - |
| - | - | 8082 | 947.4 | - | - | 0 | - |
| - | - | 5153 | 948.4 | - | - | 0 | - |
| - | - | 1.131E+04 | 954.4 | - | - | 0 | - |
| - | - | 2.485E+04 | 955.4 | - | - | 0 | - |
| - | - | 1.066E+04 | 956.4 | - | - | 0 | - |
| - | - | 4174 | 957.4 | - | - | 0 | - |
| - | - | 3496 | 962.4 | - | - | 0 | - |
| - | - | 4068 | 964.5 | - | - | 0 | - |
| - | - | 2.286E+04 | 965.4 | - | - | 0 | - |
| - | - | 1.226E+04 | 966.4 | - | - | 0 | - |
| - | - | 3326 | 967.5 | - | - | 0 | - |
| - | - | 2.027E+04 | 972.4 | - | - | 0 | - |
| - | - | 1.438E+04 | 973.4 | - | - | 0 | - |
| - | - | 6115 | 974.4 | - | - | 0 | - |
| 8 | b | 5650 | 975.5 | 0.003013 | 3.089 | +1 | 8 |
| 8 | b | 6362 | 976.5 | 0.01534 | 15.7 | +1 | 8 |
| - | - | 7625 | 980.5 | - | - | 0 | - |
| - | - | 3524 | 981.5 | - | - | 0 | - |
| - | - | 7102 | 982.5 | - | - | 0 | - |
| - | - | 4027 | 983.5 | - | - | 0 | - |
| 8 | b | 3.803E+04 | 993.5 | 0.0009322 | 0.9382 | +1 | 8 |
| - | - | 2.229E+04 | 994.5 | - | - | 0 | - |
| - | - | 6279 | 995.5 | - | - | 0 | - |
| - | - | 8483 | 1007 | - | - | 0 | - |
| - | - | 4663 | 1008 | - | - | 0 | - |
| 6 | y | 3.478E+04 | 1025 | 0.002386 | 2.327 | +1 | 8 |
| - | - | 2.241E+04 | 1026 | - | - | 0 | - |
| - | - | 5972 | 1027 | - | - | 0 | - |
| - | - | 3866 | 1076 | - | - | 0 | - |
| - | - | 5523 | 1077 | - | - | 0 | - |
| - | - | 3902 | 1078 | - | - | 0 | - |
| 9 | b | 3475 | 1092 | 0.0193 | 17.68 | +1 | 9 |
| - | - | 2.082E+04 | 1094 | - | - | 0 | - |
| - | - | 1.636E+04 | 1095 | - | - | 0 | - |
| - | - | 4285 | 1096 | - | - | 0 | - |
| - | - | 3250 | 1097 | - | - | 0 | - |
| 9 | b | 1.073E+04 | 1109 | 0.001292 | 1.165 | +1 | 9 |
| - | - | 6271 | 1110 | - | - | 0 | - |
| - | - | 4139 | 1111 | - | - | 0 | - |
| - | - | 4198 | 1118 | - | - | 0 | - |
| - | - | 4769 | 1119 | - | - | 0 | - |
| 5 | y | 2.533E+04 | 1136 | 0.002225 | 1.96 | +1 | 9 |
| 5 | y | 4.094E+04 | 1136 | 0.006613 | 5.819 | +1 | 9 |
| - | - | 2.331E+04 | 1137 | - | - | 0 | - |
| - | - | 1.063E+04 | 1138 | - | - | 0 | - |
| 5 | y | 6.758E+04 | 1154 | 0.00106 | 0.9188 | +1 | 9 |
| - | - | 5.009E+04 | 1155 | - | - | 0 | - |
| - | - | 1.817E+04 | 1156 | - | - | 0 | - |
| - | - | 6260 | 1157 | - | - | 0 | - |
| 10 | b | 6440 | 1166 | 0.0008042 | 0.69 | +1 | 10 |
| - | - | 4575 | 1167 | - | - | 0 | - |
| - | - | 3727 | 1223 | - | - | 0 | - |
| - | - | 3064 | 1287 | - | - | 0 | - |
| - | - | 2744 | 2013 | - | - | 0 | - |
| - | - | 2852 | 3163 | - | - | 0 | - |

m/z Charge Intensity FragmentType MassShift Position
120.08126068115234 0 5635.7485
122.07168579101562 0 10261.165
123.04442596435547 0 11790.01
123.05549621582031 0 2414.1987
125.10782623291016 0 3293.2954
125.5638656616211 0 1535.4692
125.96668243408203 0 1600.5402
126.09197235107422 0 3883.5344
127.05060577392578 0 14552.156
127.08658599853516 0 1537.7372
127.12328338623047 0 5684.993
128.05419921875 0 1641.6833
128.0819854736328 0 3596.9895
128.10739135742188 0 120450.555
129.10267639160156 0 1290455.2
130.06552124023438 0 529168.6
130.09983825683594 0 9603.356
130.10598754882812 0 84349.9
131.06884765625 0 50866.246
131.10824584960938 0 1947.5447
131.22203063964844 0 1566.4266
132.08114624023438 0 125666.04
133.06076049804688 0 2445.5981
133.08448791503906 0 13843.133
133.08934020996094 0 2130.6343
136.0760955810547 0 157948.22
136.20199584960938 0 1623.0112
137.0794219970703 0 13769.69
138.06661987304688 0 15416.894
138.09182739257812 0 5695.9185
139.05026245117188 0 1767.9458
139.0869140625 0 15414.989
139.18386840820312 0 1931.665
141.10269165039062 0 19825.426
142.0654754638672 0 7678.003
142.12351989746094 0 2320.5173
142.63534545898438 0 1814.4125
143.04547119140625 0 8487.125
143.11875915527344 0 2101.9663
144.08114624023438 0 7782.0205
144.08689880371094 0 1483.1349
145.06130981445312 0 9071.229
146.05996704101562 0 2210.7305
147.0446319580078 0 6829.418
148.0609130859375 0 2462.9167
148.95428466796875 0 2381.1191
149.04525756835938 0 17574.928
150.06654357910156 0 6622.8164
151.0869903564453 0 24760.453
153.10250854492188 0 2716.3274
154.06149291992188 0 2752.1409
154.09774780273438 0 2699.5234
155.0458221435547 0 4053.4917
155.0822296142578 0 3701.3684
155.09312438964844 0 78626.49
155.1182861328125 0 56645.918 a Water loss 1
156.07681274414062 0 4034.542
156.09634399414062 0 5106.2856
156.10263061523438 0 2807.5193
156.12156677246094 0 3274.0046
157.0599365234375 0 1947.248
157.09759521484375 0 7224.417
157.13357543945312 0 4496.201
158.08438110351562 0 6481.278
159.09213256835938 0 971727.1
160.07608032226562 0 9116.724
160.08917236328125 0 7912.2363
160.09544372558594 0 105384.086
161.09889221191406 0 5465.555
165.05502319335938 0 87992.23
165.0773162841797 0 4322.0225
166.06149291992188 0 161668.1
166.0979461669922 0 14449.401
167.05615234375 0 10987.563
167.0651092529297 0 10900.412
167.0814971923828 0 3045.445
167.1182403564453 0 6454.2397
168.0812530517578 0 16347.59
168.1022491455078 0 10457.823
168.1135711669922 0 12611.257
169.07640075683594 0 14061.676
169.08387756347656 0 2865.6921
169.09759521484375 0 38299.1
170.0604705810547 0 47005.168
171.0636444091797 0 5060.775
171.09205627441406 0 5164.354
172.0721435546875 0 15322.3125
173.0562286376953 0 17105.734
173.12893676757812 0 1929929.4 a 1
173.45223999023438 0 3533.0676
174.1322784423828 0 169557.77
175.13389587402344 0 11064.365
176.08187866210938 0 2726.2346
177.10279846191406 0 3155.6758
178.13426208496094 0 58432.945
179.11831665039062 0 4126.3164
179.1376190185547 0 5476.48
181.06114196777344 0 8063.5103
182.0816650390625 0 180457.19 y 12
183.08526611328125 0 16398.605
183.11318969726562 0 50058.492 b Water loss 1
184.11642456054688 0 5178.111
185.1652374267578 0 15165.821
186.12417602539062 0 421505.4
187.0870361328125 0 81185.28
187.12759399414062 0 35191.32
188.0709991455078 0 7922.571
188.0804443359375 0 2153.617
188.0906219482422 0 7397.6587
188.12921142578125 0 2152.4941
190.08273315429688 0 16729.09
190.1343536376953 0 14121.066
194.0928192138672 0 10237.709
195.1132049560547 0 99320.695
196.10845947265625 0 67025.45
196.11705017089844 0 8470.511
197.1114501953125 0 4573.7573
197.12881469726562 0 5261.1494
198.0879669189453 0 10164.778
198.12820434570312 0 33657.508
199.07159423828125 0 8966.42
199.08692932128906 0 7084.48
199.13150024414062 0 5360.0977
200.14361572265625 0 5695.326
201.12376403808594 0 518060.78 b 1
202.08677673339844 0 4230.729
202.12709045410156 0 52075.04
203.12918090820312 0 4337.8965
204.0772705078125 0 1956.855
204.11375427246094 0 4458.175
204.13442993164062 0 3312.855
205.09764099121094 0 41051.156
205.14520263671875 0 6817.7407
206.1011505126953 0 3836.58
206.12908935546875 0 9102.301
207.16098022460938 0 13288.499
210.1278076171875 0 16447.652
211.1081085205078 0 8374.921 y Water loss 10
212.13992309570312 0 12580.939
213.123779296875 0 45281.63
213.1602020263672 0 6470.721
215.1185760498047 0 3882.4548
215.1393585205078 0 2802.533
216.09829711914062 0 12533.913
218.02247619628906 0 2015.5652
219.1138458251953 0 2332.1785
221.07156372070312 0 5395.4766
221.0833282470703 0 4246.8477
221.09378051757812 0 4034.0933
221.10382080078125 0 10332.378
221.12921142578125 0 2525.7026
221.14051818847656 0 5488.0327
222.1240692138672 0 17133.121
222.13580322265625 0 3292.43
223.06370544433594 0 3049.2546
223.10812377929688 0 24487.049
223.11920166015625 0 2551.6448
223.15582275390625 0 173646.95
224.10386657714844 0 7501.543
224.15907287597656 0 17337.73
225.04391479492188 0 5748.9565
225.12344360351562 0 3491.0605
225.16090393066406 0 3797.9958
226.08273315429688 0 38699.24
227.0667266845703 0 33477.56
227.08592224121094 0 5088.643
227.1548309326172 0 9511.469
228.0696258544922 0 4488.782
228.13381958007812 0 3015.907
229.1188201904297 0 53830.633
230.12205505371094 0 5221.177
230.15098571777344 0 4104.954
233.14012145996094 0 29770.457
234.12425231933594 0 189114.89
234.65045166015625 0 3518.1655
235.1072235107422 0 2268.572
235.12767028808594 0 28738.246
235.15538024902344 0 4875.4834
235.60598754882812 0 2317.1555
237.1356658935547 0 3678.88
238.119140625 0 23290.645
238.1298828125 0 4018.617
239.08267211914062 0 3394.0652
239.0956573486328 0 13139.138
239.11392211914062 0 3626.8047
240.09609985351562 0 5105.962
240.13478088378906 0 71713.7
241.0967254638672 0 2614.785
241.11851501464844 0 13851.9
241.137939453125 0 10352.756
241.19131469726562 0 9202.253
242.19552612304688 0 2318.2256
243.1097412109375 0 11555.414
244.09335327148438 0 77287.2
245.096435546875 0 6824.371
247.10830688476562 0 110333.695
247.1441650390625 0 4057.05
248.10134887695312 0 4571.223
248.114013671875 0 41298.87
248.62388610839844 0 2429.8008 y 9
249.09877014160156 0 63018.91
249.11502075195312 0 4165.265
249.11892700195312 0 4141.416
249.13531494140625 0 15177.06
250.10208129882812 0 9602.359
251.15087890625 0 686858.94
252.13433837890625 0 4442.4727
252.15406799316406 0 82521.9
253.15606689453125 0 6009.633
254.0766143798828 0 3006.3213
255.14840698242188 0 29630.617
256.1083679199219 0 8697.295
256.1529235839844 0 5068.417
257.1135559082031 0 20322.227
258.1452941894531 0 26514.73
259.12884521484375 0 3534.2195
259.1484375 0 4280.7856
261.1195068359375 0 12364.267
261.1354675292969 0 7913.659
262.1192626953125 0 8423.698
263.1048889160156 0 2467.7522
263.1221923828125 0 2139.026
265.1297302246094 0 60842.684
266.1253662109375 0 174147.84
267.12847900390625 0 19679.088
267.1448974609375 0 3919.0825
267.61212158203125 0 3930.6187
268.1299133300781 0 8792.783
268.1771240234375 0 14790.106
269.1611328125 0 6328.4478
269.18560791015625 0 8708.975
270.1222229003906 0 27927.684
270.1455078125 0 4512.725
270.16015625 0 2433.7214
270.6245422363281 0 4277.939
272.1761474609375 0 36098.016
273.17919921875 0 7502.7666
274.1188049316406 0 33748.36
274.1316223144531 0 13381.351
275.10333251953125 0 5815.4126
275.1230163574219 0 5663.355
276.1092834472656 0 10159.993
276.1274719238281 0 7098.041
276.1558837890625 0 7743.24
276.6714172363281 0 2369.0647
279.145751953125 0 15168.433
279.6144104003906 0 2702.1396
280.12969970703125 0 18172.111
280.14593505859375 0 3048.1882
281.05181884765625 0 13986.056
281.1328430175781 0 3677.046
282.1570129394531 0 15092.552
283.140625 0 429112.25
284.0881652832031 0 3052.3381
284.1036682128906 0 15305.342
284.1239013671875 0 3191.1428
284.1438903808594 0 61422.395
285.0899658203125 0 2706.7603
285.1051940917969 0 3792.6094
285.1575622558594 0 4645.435
286.1403503417969 0 31057.785
287.14208984375 0 3899.7354
287.17181396484375 0 9097.788
287.1883239746094 0 4591.5415
288.61907958984375 0 6310.431
290.1665954589844 0 4218.004 b 4
290.6689758300781 0 2486.9949
293.11383056640625 0 256374.94 y Water loss 11
293.161865234375 0 6781.954
294.11773681640625 0 54942.387
295.1201477050781 0 3451.4082
296.1976623535156 0 8293.904 b Water loss 2
296.66583251953125 0 3349.523
297.0832824707031 0 2737.2043
297.1332702636719 0 8999.883 y Water loss 8
297.1564636230469 0 78424.375
297.6349182128906 0 3314.929
298.1400146484375 0 8725.087 y Water loss 6
298.15948486328125 0 13359.997
299.06231689453125 0 84912.61
300.1689147949219 0 25194.729
301.15130615234375 0 258331.05
302.1142883300781 0 44797.54
302.1546630859375 0 35959.746
303.1172180175781 0 11164.901
303.1682434082031 0 4090.868
305.17889404296875 0 6414.1533
305.6824951171875 0 3335.026
306.1378479003906 0 43194.887 y 8
306.22967529296875 0 4090.9243
306.6395568847656 0 18070.684
307.14093017578125 0 8462.19
309.16693115234375 0 4116.219
310.1536560058594 0 3237.0083
311.1242370605469 0 29864.066 y 11
311.1377868652344 0 14810.062
311.6888427734375 0 2630.9495
312.1280517578125 0 7266.1323
314.18731689453125 0 7957.995
314.2082214355469 0 23093.535 b 2
315.1670227050781 0 138819.75
315.2105407714844 0 4640.5845
316.1707763671875 0 23044.826
317.6594543457031 0 3538.065
319.1521301269531 0 9810.484
320.17169189453125 0 2489.0918
322.2238464355469 0 3802.805
325.1510925292969 0 10384.505
325.1717224121094 0 2330.016
326.66448974609375 0 31681.957
327.16558837890625 0 14864.687
327.6676025390625 0 4926.401
331.1623840332031 0 3282.011
331.6566467285156 0 4678.0513
332.20880126953125 0 10985.618
333.1774597167969 0 19761.562
334.1839599609375 0 3846.8342
335.208984375 0 2917.2148
338.1828918457031 0 3775.4783
338.6751403808594 0 4310.9624 b Water loss 5
339.1731262207031 0 7621.762
339.6765441894531 0 4224.0537
340.662109375 0 9753.731
341.1649169921875 0 3092.478
342.1822814941406 0 11180.487
343.1625061035156 0 11163.202
343.185791015625 0 2507.2375
344.1723937988281 0 14217.404
344.97686767578125 0 11578.654
345.13128662109375 0 17890.219
345.1744689941406 0 2833.9897
346.1153259277344 0 75413.98
346.1374206542969 0 4611.877
346.699462890625 0 3322.1191
347.1182861328125 0 13346.6455
347.6802673339844 0 40498.44 b 5
348.18212890625 0 14765.311
348.68231201171875 0 2776.5298
349.1291809082031 0 3119.0386
349.15130615234375 0 4007.3052
350.21942138671875 0 58943.46
351.1452331542969 0 3676.1487
351.1668395996094 0 4269.613
351.2224426269531 0 7500.1953
352.1630554199219 0 4809.4214
352.6722106933594 0 4730.6543
353.6744689941406 0 9195.754 y Water loss 7
354.1766662597656 0 4242.998
357.1751403808594 0 3163.6995
359.0292663574219 0 10815.1875
360.2027893066406 0 3945.862
360.6955871582031 0 18317.617
361.19805908203125 0 23236.613
362.18402099609375 0 20839.16
362.6799621582031 0 106038.85 y 7
363.1417236328125 0 99423.31
363.1819763183594 0 40018.395
363.6824035644531 0 7706.0747
364.12530517578125 0 19864.975
364.1456604003906 0 18834.537
365.12841796875 0 3499.814
366.1556396484375 0 2932.1843
366.1779479980469 0 6504.512
367.1400451660156 0 9709.202
367.1624450683594 0 6684.666
367.24566650390625 0 9697.977
368.1454772949219 0 2887.0962
368.1922607421875 0 3891.4885
369.1966552734375 0 3726.2588
370.175048828125 0 3342.2048
373.1846618652344 0 6082.232
376.16204833984375 0 6273.097
376.1984558105469 0 14378.553
376.70013427734375 0 4765.456
377.1452941894531 0 5831.146
378.2144775390625 0 43994.09
379.2097473144531 0 169471.6
380.1620788574219 0 6680.6724
380.2126770019531 0 32383.613
381.1524658203125 0 294878.28
381.71160888671875 0 9830.046
382.15521240234375 0 47168.453
382.1888427734375 0 11462.275
382.215087890625 0 4313.059
382.6896057128906 0 2793.775
383.15679931640625 0 8787.491
383.2057800292969 0 3679.9387
384.1663818359375 0 12590.22
384.1888427734375 0 7824.8003
385.15093994140625 0 26981.51
385.17034912109375 0 4278.369 y 4
386.154541015625 0 4075.8213
386.1794128417969 0 4368.7666
389.1935119628906 0 4310.46 b 9
390.197021484375 0 6753.8066
390.6968688964844 0 5561.777
391.135986328125 0 11235.425
392.20794677734375 0 4361.247
393.213134765625 0 5017.0996
394.1730651855469 0 40294.145
395.13543701171875 0 21301.438
395.1567687988281 0 8076.653
395.17889404296875 0 4443.3555
395.240966796875 0 3585.8643
396.1390075683594 0 5312.8906
396.1941223144531 0 4712.9043
396.22515869140625 0 65343.926
397.20306396484375 0 5272.861
397.2283020019531 0 12817.49
397.7044677734375 0 3027.711
398.17730712890625 0 29091.137
398.227294921875 0 3240.457
399.18133544921875 0 6757.9688
402.1766662597656 0 16116.7295
402.1998596191406 0 26094.809
403.136474609375 0 2963.67
403.19976806640625 0 12358.374
404.710205078125 0 3800.9768
405.26123046875 0 11810.484
406.2464904785156 0 6325.7534
407.20379638671875 0 4061.763
409.1475524902344 0 25567.438
409.2019348144531 0 4845.1577
410.1497497558594 0 3407.813
411.20037841796875 0 10024.981
412.1837463378906 0 192065.62
413.1454162597656 0 28182.56
413.1869201660156 0 31012.596
414.14764404296875 0 4794.8027
414.19561767578125 0 6034.902
414.2355651855469 0 60095.28
415.0375061035156 0 21076.773
415.198486328125 0 24548.605
415.23883056640625 0 11762.346
416.2010498046875 0 6564.146
417.717529296875 0 36181.824
418.21087646484375 0 51459.816
418.7117004394531 0 25358.217
419.2136535644531 0 3796.585
421.2099609375 0 9609.87 y Water loss 10
422.19256591796875 0 4547.568 y Ammonia loss 10
422.2516784667969 0 2797.3755
422.7088928222656 0 3088.627
423.2015686035156 0 13805.937
423.27252197265625 0 69818.61
423.70745849609375 0 4827.5273
424.2247009277344 0 3319.6208
424.27532958984375 0 21186.068
424.7353820800781 0 8771.639
425.2350769042969 0 3138.1816
426.7227478027344 0 515548.8
427.22406005859375 0 263857.22
427.7254943847656 0 78619.3
428.2261657714844 0 12531.824
429.210205078125 0 32173.291
429.7296447753906 0 3442.5598
430.19439697265625 0 530864
431.1973571777344 0 109708.484
431.71466064453125 0 60661.848 b Water loss 6
432.2130126953125 0 30629.432 b Ammonia loss 6
432.71307373046875 0 11339.6455
433.2092590332031 0 9996.013
433.25628662109375 0 20596.148 b Water loss 3
434.2600402832031 0 3924.3342
437.1905822753906 0 4013.042
438.1742858886719 0 9007.988
438.7291564941406 0 3749.092
439.2198181152344 0 123505.08 y 10
440.1785888671875 0 6287.754
440.22320556640625 0 29376.623
440.7203063964844 0 245160.11 b 6
441.2216796875 0 140318.12
441.7230224609375 0 39240.367
442.22332763671875 0 7293.611
442.2672119140625 0 2088.41
444.1806335449219 0 2652.4485
447.20623779296875 0 26222.412 y Ammonia loss 6
447.23919677734375 0 5146.5493
447.7077941894531 0 9109.514
448.20526123046875 0 23251.32
449.20880126953125 0 10250.422
450.17413330078125 0 3875.6382
450.2373962402344 0 7000.875
450.2831115722656 0 6251.6895
451.2674865722656 0 142051.72 b 3
452.2396545410156 0 3364.2725
452.2706298828125 0 34210.633
453.2709655761719 0 4675.7666
455.1961364746094 0 5223.9966
455.2673034667969 0 7060.1753
455.72039794921875 0 16095.871 y 6
456.2204284667969 0 3503.051
459.1997985839844 0 11810.153
460.754150390625 0 19403.29
461.20428466796875 0 10838.783
461.2535400390625 0 12742.561
461.7567138671875 0 5865.7427
462.2052001953125 0 2894.8384
464.21649169921875 0 5145.529
464.2497863769531 0 2558.8242
465.75042724609375 0 3289.7754
466.24481201171875 0 4684.1587
468.2939147949219 0 41439.047
469.2960205078125 0 8606.003
474.2594299316406 0 25915.234
474.7545166015625 0 40385.156
475.2540283203125 0 20826.824
475.75372314453125 0 4349.4316
476.2259826660156 0 13386.288
477.21051025390625 0 26857.447
478.2298278808594 0 25632.328 y Water loss 9
478.2777099609375 0 17348.74
479.2372741699219 0 4245.247
479.2841796875 0 3616.9712
479.7479553222656 0 4048.8384
480.2205505371094 0 3087.7405
481.7294616699219 0 4650.843
483.2649230957031 0 86487.37
483.766357421875 0 59314.477
484.22088623046875 0 6228.0923
484.26690673828125 0 17625.324
484.76947021484375 0 2416.3933
488.2574157714844 0 27040.18 b Water loss 7
488.7571105957031 0 17111.84 b Ammonia loss 7
489.2532653808594 0 11284.675
494.23675537109375 0 313827.75
495.2396545410156 0 81394.375
496.2412414550781 0 422584.56 y 9
497.246826171875 0 112124
497.763671875 0 34265.164
498.2433166503906 0 18793.229
502.2298583984375 0 3246.0518
503.1089172363281 0 20370.842
504.20111083984375 0 14389.786
505.19970703125 0 4208.9883
505.2481384277344 0 2549.1138
506.2270202636719 0 8246.837
506.2727966308594 0 4287.36
507.25927734375 0 7220.9395
511.2630920410156 0 21564.389
512.2653198242188 0 4166.4175
513.2123413085938 0 4851.15
514.1885986328125 0 6199.2144
515.282470703125 0 9975.669
519.1399536132812 0 6616.168
519.330810546875 0 4079.7878
522.2335205078125 0 5975.112
523.258544921875 0 10985.967
523.5888061523438 0 4383.5063
524.2518310546875 0 3097.711
525.2669677734375 0 56164.645
526.2314453125 0 11239.3
526.2709350585938 0 12489.303
527.2326049804688 0 3379.6194
529.2615966796875 0 16400.682 Precursor Water loss
529.594482421875 0 22692.879 Precursor Ammonia loss
529.9284057617188 0 11095.883
530.2237548828125 0 4273.4556
530.263916015625 0 3510.0522
531.2205810546875 0 4592.5337
532.1942749023438 0 14383.438
533.1966552734375 0 2988.2783
535.2636108398438 0 13341.861 Precursor
535.598876953125 0 15587.111
535.9334716796875 0 8950.504
536.1658325195312 0 3510.7942
536.2661743164062 0 6628.8203
537.2484130859375 0 3485.107
537.7374877929688 0 11571.005
538.2711791992188 0 9512.348
538.7691650390625 0 6391.2725
539.2368774414062 0 58729.617
539.2821044921875 0 4749.2827
540.2395629882812 0 18430.992
541.2525024414062 0 7035.8633
542.295166015625 0 4281.32
543.278076171875 0 128805.92
544.2807006835938 0 37530.05
545.2828979492188 0 7692.119
546.2496948242188 0 16389.424
546.7513427734375 0 10434.099
547.2452392578125 0 4509.1313
547.7770385742188 0 5654.6943
548.2393798828125 0 3659.304
548.2965087890625 0 3257.167
549.2212524414062 0 12977.8545
550.21044921875 0 8457.227
551.2578125 0 13516.654
551.3319702148438 0 7130.9736
552.2584228515625 0 2948.0266
552.3296508789062 0 3041.7454
553.263916015625 0 3316.629
554.7744750976562 0 14501.373 b 8
555.2576293945312 0 29903.322
555.757080078125 0 16866.686
556.2586669921875 0 5581.4272
561.3151245117188 0 14555.29 b Water loss 4
562.3018798828125 0 7374.7046 b Ammonia loss 4
563.258056640625 0 12803.434
564.2596435546875 0 5617.442
565.26318359375 0 19799.424
566.2679443359375 0 3816.7686
567.2316284179688 0 66776.41
568.23583984375 0 18096.318
568.7598876953125 0 8310.392
569.2401733398438 0 7336.838
575.2465209960938 0 15658.9
576.232177734375 0 3583.785
576.2823486328125 0 3283.5496
577.2139892578125 0 4510.639
577.2652587890625 0 3494.3264 y 4
577.7645874023438 0 4350.7246
579.27392578125 0 4409.6655
579.3258666992188 0 88371.56 b 4
580.3284912109375 0 26251.33
581.2703857421875 0 3259.869
581.332275390625 0 4009.6558
582.3084716796875 0 4445.583
583.287109375 0 4456.564 b 9
587.2547607421875 0 2930.266
588.7960205078125 0 3659.9912
589.306884765625 0 2812.2808
592.2769165039062 0 4099.8755
592.32568359375 0 11470.691
593.2571411132812 0 100535.51 y Water loss 8
593.3062133789062 0 47940.727
593.7844848632812 0 6232.2925
594.2567138671875 0 30290.979
594.3099975585938 0 10159.864
594.7808837890625 0 3970.229
595.2486572265625 0 6311.376
596.3529663085938 0 2800.5142
597.3008422851562 0 3431.782
597.8009643554688 0 10552.307
598.3026733398438 0 3668.174
600.3141479492188 0 10055.535
602.2992553710938 0 2820.7854
602.79150390625 0 20571.523
603.2926025390625 0 19561.312
603.7914428710938 0 10006.808
604.2867431640625 0 4709.241
610.283935546875 0 85613.234
610.3313598632812 0 7487.961
611.2686157226562 0 1306835.6 y 8
611.7976684570312 0 83740.805
612.2718505859375 0 407759.47
612.7996215820312 0 24650.21
613.2740478515625 0 83475.13
614.2767333984375 0 16126.709
616.298095703125 0 5718.4834
617.28515625 0 7228.879
619.2752075195312 0 3016.172
620.803466796875 0 6375.7417
621.252685546875 0 14135.559
621.3018798828125 0 7976.106
622.2552490234375 0 3180.356
622.370849609375 0 5586.721
623.2496948242188 0 2582.8657
628.2760009765625 0 3566.0784
628.777099609375 0 3665.2932
634.3076171875 0 17177.867
635.2969970703125 0 21554.17
636.328125 0 10968.377
636.7860107421875 0 7393.8857 y Water loss 3
637.2828369140625 0 4252.8447 y Ammonia loss 3
637.3394165039062 0 4148.892
637.7880249023438 0 7061.742
638.329345703125 0 6576.3813 b Water loss 10
640.2745361328125 0 16438.896
641.26318359375 0 3772.8604
645.2855834960938 0 12163.268
645.7925415039062 0 84437.37 y 3
646.2933349609375 0 76104.43
646.79541015625 0 24324.941
647.2891845703125 0 3500.7017
648.3444213867188 0 5029.559
649.3428955078125 0 4263.5405
650.2684936523438 0 6331.6953
650.3269653320312 0 4143.721
651.274658203125 0 4125.7
652.3212280273438 0 130906.84
653.32421875 0 48192.49
654.3258666992188 0 8888.792
658.2839965820312 0 10791.659
659.3182983398438 0 6748.3203
660.3198852539062 0 2733.906
660.871826171875 0 3382.1353
661.3300170898438 0 9412.403
661.8320922851562 0 6310.615
662.3060913085938 0 37843.754
663.2953491210938 0 35578.387
664.294189453125 0 3857.3938
664.349609375 0 3657.9363
666.2875366210938 0 2937.2402
666.3525390625 0 3352.083
668.2764282226562 0 5623.8643
670.8245849609375 0 2717.0332
676.3423461914062 0 56702.883 b Water loss 5
677.3339233398438 0 37979.24 b Ammonia loss 5
678.3336181640625 0 15670.672
679.3345336914062 0 14151.325
679.829833984375 0 5734.872
680.31640625 0 228895.25
681.3187255859375 0 93009.13
682.3214721679688 0 19112.572
683.3211669921875 0 3683.0325
684.3216552734375 0 6792.7607
684.8175659179688 0 21068.74
685.3175048828125 0 12015.21
685.8193359375 0 8515.44
686.3201904296875 0 2944.1099
692.8341674804688 0 4925.466
693.3295288085938 0 67730.914 y Water loss 2
693.828369140625 0 61213.77 y Ammonia loss 2
694.352294921875 0 614619.75 b 5
694.8289794921875 0 14331.693
695.355224609375 0 234307.67
696.3577880859375 0 57207.89
697.2879638671875 0 3240.1975
697.3519897460938 0 10198.61
697.8592529296875 0 4212.5786
698.355224609375 0 7129.128
701.842041015625 0 31122.012
702.3347778320312 0 527801.7 y 2
702.83642578125 0 441281.25 b Water loss 11
703.3377685546875 0 201892.69 b Ammonia loss 11
703.8394165039062 0 64742.824
704.3394165039062 0 21198.31
705.3553466796875 0 5077.97
706.3429565429688 0 43315.836 y Water loss 7
707.3452758789062 0 15568.916
708.306640625 0 4217.08
711.3749389648438 0 14723.596
711.8557739257812 0 34202.14 b 11
712.35791015625 0 43772.71
712.857666015625 0 16243.754
713.3598022460938 0 7745.3154
715.3053588867188 0 3221.8064
720.3854370117188 0 5147.53
720.86328125 0 3113.6106
721.3642578125 0 28616.525
722.365478515625 0 9457.404
723.3695068359375 0 31984.15
724.3528442382812 0 408124.16 y 7
725.3554077148438 0 173714.66
726.3582763671875 0 41460.58
727.3577270507812 0 4979.2817
728.8580322265625 0 4928.662
729.3577880859375 0 18983.658
730.3630981445312 0 9586.697
733.3641967773438 0 16027.551
734.3573608398438 0 16164.745
735.3563842773438 0 4403.745
737.8675537109375 0 4896.938
738.3605346679688 0 4473.41
742.867431640625 0 3695.4968 y Water loss 1
743.3658447265625 0 9679.032 y Ammonia loss 1
743.8636474609375 0 4145.725
744.3695068359375 0 3344.1436
746.3502807617188 0 2873.1704
747.3907470703125 0 9204.814
748.384033203125 0 10369.035
749.3372192382812 0 5930.8613
750.3944702148438 0 3427.4265
751.3770751953125 0 60211.027
751.8688354492188 0 47369.535 y 1
752.373291015625 0 72810.88
752.872314453125 0 16136.248
753.3775024414062 0 15607.787
760.3052978515625 0 4086.2163
761.3750610351562 0 4698.3647
762.41552734375 0 8277.626
763.3684692382812 0 38147.33
764.3651733398438 0 10966.632
765.4011840820312 0 15943.116
766.408203125 0 7682.3535
767.3482666015625 0 13650.566
768.34521484375 0 4084.8027
769.409912109375 0 4252.879
775.390380859375 0 7723.531
776.3793334960938 0 8410.489
779.3842163085938 0 41668.31
780.3850708007812 0 16514.338
781.369384765625 0 14248.011
782.3671264648438 0 8373.25
791.38623046875 0 2794.2058
792.437744140625 0 3241.8704
793.4002075195312 0 67930.164 Precursor Water loss
794.4036865234375 0 27130.191
795.342529296875 0 28092.502
795.40771484375 0 4430.781
796.3455200195312 0 10306.909
797.3447265625 0 3607.558
807.3767700195312 0 10426.152
807.4410400390625 0 7560.062
808.3824462890625 0 9906.703
817.40380859375 0 3677.615
818.4415893554688 0 7908.789
819.4359741210938 0 7649.7583
820.4398803710938 0 3711.787
825.3868408203125 0 6137.0684
826.3782348632812 0 12348.624
827.374267578125 0 3134.8672
834.4276733398438 0 12108.862
835.4185180664062 0 25260.057
836.4492797851562 0 52639.79
837.4528198242188 0 20889.791
838.4544677734375 0 6326.0386
843.4031372070312 0 9179.664
844.3892822265625 0 20439.16
845.3927612304688 0 13447.59
846.4466552734375 0 20381.885
847.444091796875 0 13448.598
852.363525390625 0 10047.463
852.4373168945312 0 148018.12
853.3524780273438 0 5105.4546
853.4398193359375 0 80431.836
854.4421997070312 0 17598.326
862.4214477539062 0 88932.73 b Water loss 6
863.4168701171875 0 71635.72 b Ammonia loss 6
864.4558715820312 0 110456.49
865.4601440429688 0 55745.574
866.4629516601562 0 20438.451
867.4647827148438 0 2969.681
869.3908081054688 0 5143.2476
876.436279296875 0 9588.575
877.4334106445312 0 4518.525
880.4319458007812 0 716659.94 b 6
881.4342651367188 0 380891.25
882.4370727539062 0 120344.63
883.43798828125 0 21559.885
890.4139404296875 0 5098.5884
891.4078979492188 0 9915.17
892.39794921875 0 3760.1282
892.4683227539062 0 9594.52
893.4723510742188 0 8166.8926
894.436279296875 0 6106.015
907.4415283203125 0 3283.8809
908.4271240234375 0 80218.79
909.4310913085938 0 43653.617
910.4319458007812 0 70490.01 y 6
911.4344482421875 0 30313.715
912.4384765625 0 10087.626
936.421630859375 0 7364.837
937.4119262695312 0 10927.078
938.416259765625 0 4667.098
947.4393310546875 0 8081.665
948.424072265625 0 5152.6895
954.4321899414062 0 11305.188
955.4207153320312 0 24846.883
956.4215698242188 0 10662.717
957.4220581054688 0 4173.8066
962.4488525390625 0 3496.1643
964.4551391601562 0 4068.311
965.448974609375 0 22861.277
966.4498291015625 0 12262.424
967.4500732421875 0 3325.671
972.4429931640625 0 20272.465
973.4448852539062 0 14377.27
974.4473876953125 0 6115.028
975.5076904296875 0 5649.596 b Water loss 7
976.5040283203125 0 6361.9663 b Ammonia loss 7
980.4602661132812 0 7624.5464
981.4566040039062 0 3524.0786
982.4725341796875 0 7102.12
983.4810791015625 0 4026.9482
993.5161743164062 0 38033.676 b 7
994.5180053710938 0 22286.793
995.5189208984375 0 6279.106
1007.4812622070312 0 8482.783
1008.4948120117188 0 4662.7563
1025.4598388671875 0 34778.51 y 5
1026.4610595703125 0 22409.604
1027.4710693359375 0 5972.21
1075.5318603515625 0 3866.3413
1076.5245361328125 0 5522.954
1077.5257568359375 0 3902.0984
1091.496337890625 0 3475.0796 b Ammonia loss 8
1093.54150390625 0 20819.71
1094.545654296875 0 16357.722
1095.5458984375 0 4284.5947
1096.87548828125 0 3250.4465
1108.5408935546875 0 10726.851 b 8
1109.527099609375 0 6270.9814
1110.544189453125 0 4138.584
1118.4857177734375 0 4198.354
1119.484375 0 4769.0093
1135.5076904296875 0 25331.803 y Water loss 4
1136.49609375 0 40938.836 y Ammonia loss 4
1137.4952392578125 0 23308.922
1138.49951171875 0 10627.299
1153.51708984375 0 67581.93 y 4
1154.5201416015625 0 50087.156
1155.520751953125 0 18168.5
1156.527587890625 0 6260.496
1165.564453125 0 6439.9116 b 9
1166.5711669921875 0 4574.7676
1222.57666015625 0 3726.537
1286.6905517578125 0 3063.8826
2012.9296875 0 2743.7783
3163.44189453125 0 2851.9692

Spectrum Details

|  |  |
| --- | --- |
| Matched peaks? Matched peaks The total absolute number of peaks matched. Additionally in brackets the total fraction of peaks matched and the total number of peaks is shown. | 79 (8.77% of 901) |
| FDR? FDR The false discovery rate estimated for this peptide. It is calculated by matching all theoretical fragments with a non-integer shift with the raw peaks for this spectrum. This is done with 40 different shifts. The resulting percentage is the average number of annotated peaks over the number of annotated peaks with the correct spectrum. | 0.72% |
| Satellite FDR? Satellite FDR See the FDR for details on its calculation. This satellite ion specific FDR only contains the satellite ions (d/w) for I/L/J positions. | ∞ |
| PSM Score? PSM Score The PSM Score as given by Hecklib to this annotated spectrum. It is shown with three significant figures. | 604 |

#### Spectrum 7394? Spectrum 7394 The raw spectrum of this peptide as annotated by Hecklib. The fragments are coloured according to ion type (see legend). Any peaks with a star '\*' as text can be hovered over to see the full details, first the ion type second the mass shift type. By hovering over the amino acids in the peptide or ions in the legend the corresponding peaks are highlighted. By toggling the 'Unassigned' label you can turn the background (unassigned) peaks on or off in the plot. By updating the slider in the Ion legend you can update the spectrum to only show the top X% of the peaks with labels. The top X% means any peak that is within X% of the highest intensity. By dragging in the spectrum you can zoom in to a specific part of the spectrum and use 'Zoom Out' to get back to the original zoom level. The annotation of the spectrum is based on the given sequence in the peptides file and is done with different software so inconsistencies are likely. The peaks are annotated based on the given sequence, with 20 ppm tolerance.

Copy Data

##### Spectrum 7394 (TSV)

###### Preview

```
Loading example...
```

*Click on the button to copy the data to your clipboard.*

Mz MinMz MaxIntensity Max

WidthHeightPeptide font sizePeptide stroke widthSpectrum font sizeSpectrum stroke widthCompact peptide

Ion legend

wxyz

abcd

OtherUnassignedIon^Charge^_Position_Show for top:%

TVLHQDWLDGKEY

01.24e+52.47e+53.71e+54.94e+5

Zoom Out

a^+1^_2_y^+3^_4_a^+1^_2_y^+1^_1_b^+1^_2_b^+1^_2_y^+2^_3_b^+2^_5_y^+1^_2_b^+1^_3_y^+2^_5_y^+3^_7_y^+2^_5_y^+1^_2_b^+1^_3_b^+2^_6_b^+2^_6_y^+2^_6_y^+2^_6_y^+3^_9_b^+3^_10_y^+1^_3_y^+1^_3_b^+2^_7_b^+2^_7_b^+1^_4_y^+1^_3_b^+2^_7_y^+2^_7_b^+1^_4_y^+2^_7_y^+1^_4_b^+2^_8_b^+2^_8_y^+1^_4_\*\*\*b^+2^_9_b^+2^_9_b^+1^_5_b^+1^_5_b^+1^_5_y^+1^_5_y^+1^_5_y^+2^_10_y^+2^_10_b^+2^_11_y^+2^_10_b^+1^_6_b^+1^_6_y^+2^_11_y^+2^_11_b^+1^_6_y^+2^_11_b^+2^_12_b^+2^_12_y^+1^_6_b^+2^_12_y^+1^_6_y^+2^_12_y^+2^_12_y^+2^_12_\*b^+1^_7_b^+1^_7_b^+1^_7_y^+1^_7_b^+1^_8_b^+1^_8_b^+1^_8_y^+1^_8_b^+1^_9_y^+1^_9_y^+1^_9_y^+1^_9_b^+1^_10_y^+1^_10_

0872174526173489

Fragment Matches Table

Show background peaks

| Position | Ion type | Intensity | mz Theoretical | mz Error (Th) | mz Error (ppm) | Charge | Series Number |
| --- | --- | --- | --- | --- | --- | --- | --- |
| - | - | 8446 | 120.1 | - | - | 0 | - |
| - | - | 569.9 | 121.1 | - | - | 0 | - |
| - | - | 469.6 | 121.9 | - | - | 0 | - |
| - | - | 2263 | 122.1 | - | - | 0 | - |
| - | - | 2485 | 123 | - | - | 0 | - |
| - | - | 769.5 | 123.1 | - | - | 0 | - |
| - | - | 460.7 | 125 | - | - | 0 | - |
| - | - | 902.9 | 125.1 | - | - | 0 | - |
| - | - | 1224 | 126.1 | - | - | 0 | - |
| - | - | 5867 | 127.1 | - | - | 0 | - |
| - | - | 953 | 127.1 | - | - | 0 | - |
| - | - | 1657 | 127.1 | - | - | 0 | - |
| - | - | 2.764E+04 | 128.1 | - | - | 0 | - |
| - | - | 1904 | 129.1 | - | - | 0 | - |
| - | - | 3.227E+05 | 129.1 | - | - | 0 | - |
| - | - | 1.403E+05 | 130.1 | - | - | 0 | - |
| - | - | 1983 | 130.1 | - | - | 0 | - |
| - | - | 1.861E+04 | 130.1 | - | - | 0 | - |
| - | - | 498.9 | 130.8 | - | - | 0 | - |
| - | - | 1.308E+04 | 131.1 | - | - | 0 | - |
| - | - | 704 | 131.1 | - | - | 0 | - |
| - | - | 3.061E+04 | 132.1 | - | - | 0 | - |
| - | - | 834.3 | 133.1 | - | - | 0 | - |
| - | - | 3231 | 133.1 | - | - | 0 | - |
| - | - | 578.8 | 134 | - | - | 0 | - |
| - | - | 4.319E+04 | 136.1 | - | - | 0 | - |
| - | - | 3302 | 137.1 | - | - | 0 | - |
| - | - | 3975 | 138.1 | - | - | 0 | - |
| - | - | 1406 | 138.1 | - | - | 0 | - |
| - | - | 539.8 | 139.1 | - | - | 0 | - |
| - | - | 3874 | 139.1 | - | - | 0 | - |
| - | - | 5368 | 141.1 | - | - | 0 | - |
| - | - | 1878 | 142.1 | - | - | 0 | - |
| - | - | 1971 | 143 | - | - | 0 | - |
| - | - | 636.7 | 143.1 | - | - | 0 | - |
| - | - | 1013 | 144.1 | - | - | 0 | - |
| - | - | 3393 | 145.1 | - | - | 0 | - |
| - | - | 1189 | 146.1 | - | - | 0 | - |
| - | - | 1789 | 147 | - | - | 0 | - |
| - | - | 1.13E+04 | 149 | - | - | 0 | - |
| - | - | 624.1 | 150 | - | - | 0 | - |
| - | - | 1788 | 150.1 | - | - | 0 | - |
| - | - | 7253 | 151.1 | - | - | 0 | - |
| - | - | 683.5 | 153.1 | - | - | 0 | - |
| - | - | 1070 | 154.1 | - | - | 0 | - |
| - | - | 1495 | 155.1 | - | - | 0 | - |
| - | - | 2.102E+04 | 155.1 | - | - | 0 | - |
| 2 | a | 1.529E+04 | 155.1 | 0.0003356 | 2.163 | +1 | 2 |
| - | - | 1288 | 156.1 | - | - | 0 | - |
| - | - | 760 | 156.1 | - | - | 0 | - |
| - | - | 1122 | 156.1 | - | - | 0 | - |
| - | - | 875.4 | 157.1 | - | - | 0 | - |
| - | - | 1900 | 157.1 | - | - | 0 | - |
| - | - | 3165 | 158.1 | - | - | 0 | - |
| - | - | 2.435E+05 | 159.1 | - | - | 0 | - |
| - | - | 2313 | 160.1 | - | - | 0 | - |
| - | - | 1672 | 160.1 | - | - | 0 | - |
| - | - | 2.772E+04 | 160.1 | - | - | 0 | - |
| - | - | 918.5 | 161.1 | - | - | 0 | - |
| - | - | 1.998E+04 | 165.1 | - | - | 0 | - |
| - | - | 1164 | 165.1 | - | - | 0 | - |
| - | - | 3.752E+04 | 166.1 | - | - | 0 | - |
| 10 | y | 3830 | 166.1 | 0.001786 | 10.76 | +3 | 4 |
| - | - | 2409 | 166.1 | - | - | 0 | - |
| - | - | 1.369E+04 | 167.1 | - | - | 0 | - |
| - | - | 2441 | 167.1 | - | - | 0 | - |
| - | - | 608.9 | 167.1 | - | - | 0 | - |
| - | - | 1369 | 167.1 | - | - | 0 | - |
| - | - | 917.7 | 168.1 | - | - | 0 | - |
| - | - | 3933 | 168.1 | - | - | 0 | - |
| - | - | 3618 | 168.1 | - | - | 0 | - |
| - | - | 3512 | 168.1 | - | - | 0 | - |
| - | - | 4239 | 169.1 | - | - | 0 | - |
| - | - | 1339 | 169.1 | - | - | 0 | - |
| - | - | 9594 | 169.1 | - | - | 0 | - |
| - | - | 1.322E+04 | 170.1 | - | - | 0 | - |
| - | - | 1479 | 171.1 | - | - | 0 | - |
| - | - | 2283 | 171.1 | - | - | 0 | - |
| - | - | 1597 | 171.1 | - | - | 0 | - |
| - | - | 4491 | 172.1 | - | - | 0 | - |
| - | - | 3987 | 173.1 | - | - | 0 | - |
| 2 | a | 4.894E+05 | 173.1 | 0.0004673 | 2.699 | +1 | 2 |
| - | - | 2575 | 173.4 | - | - | 0 | - |
| - | - | 4.308E+04 | 174.1 | - | - | 0 | - |
| - | - | 2110 | 175.1 | - | - | 0 | - |
| - | - | 847.9 | 176.1 | - | - | 0 | - |
| - | - | 1563 | 177.1 | - | - | 0 | - |
| - | - | 1.289E+04 | 178.1 | - | - | 0 | - |
| - | - | 1147 | 179.1 | - | - | 0 | - |
| - | - | 984.3 | 179.1 | - | - | 0 | - |
| - | - | 2833 | 181.1 | - | - | 0 | - |
| 13 | y | 4.778E+04 | 182.1 | 0.0004496 | 2.469 | +1 | 1 |
| - | - | 3756 | 183.1 | - | - | 0 | - |
| 2 | b | 1.283E+04 | 183.1 | 0.0004008 | 2.189 | +1 | 2 |
| - | - | 1265 | 183.1 | - | - | 0 | - |
| - | - | 1498 | 184.1 | - | - | 0 | - |
| - | - | 837.2 | 185.1 | - | - | 0 | - |
| - | - | 3145 | 185.2 | - | - | 0 | - |
| - | - | 1.09E+05 | 186.1 | - | - | 0 | - |
| - | - | 1.791E+04 | 187.1 | - | - | 0 | - |
| - | - | 764.4 | 187.1 | - | - | 0 | - |
| - | - | 8458 | 187.1 | - | - | 0 | - |
| - | - | 2327 | 188.1 | - | - | 0 | - |
| - | - | 2725 | 188.1 | - | - | 0 | - |
| - | - | 861 | 188.1 | - | - | 0 | - |
| - | - | 3984 | 190.1 | - | - | 0 | - |
| - | - | 3032 | 190.1 | - | - | 0 | - |
| - | - | 656.8 | 191.1 | - | - | 0 | - |
| - | - | 688.2 | 193.1 | - | - | 0 | - |
| - | - | 596.8 | 193.1 | - | - | 0 | - |
| - | - | 3969 | 194.1 | - | - | 0 | - |
| - | - | 799.5 | 195.1 | - | - | 0 | - |
| - | - | 2.596E+04 | 195.1 | - | - | 0 | - |
| - | - | 1.554E+04 | 196.1 | - | - | 0 | - |
| - | - | 2218 | 196.1 | - | - | 0 | - |
| - | - | 1394 | 197.1 | - | - | 0 | - |
| - | - | 1931 | 197.1 | - | - | 0 | - |
| - | - | 813.4 | 197.2 | - | - | 0 | - |
| - | - | 2814 | 198.1 | - | - | 0 | - |
| - | - | 8202 | 198.1 | - | - | 0 | - |
| - | - | 1574 | 199.1 | - | - | 0 | - |
| - | - | 1973 | 199.1 | - | - | 0 | - |
| - | - | 1838 | 199.1 | - | - | 0 | - |
| - | - | 726.9 | 200.1 | - | - | 0 | - |
| - | - | 1684 | 200.1 | - | - | 0 | - |
| 2 | b | 1.345E+05 | 201.1 | 0.0003647 | 1.813 | +1 | 2 |
| - | - | 1684 | 202.1 | - | - | 0 | - |
| - | - | 1.213E+04 | 202.1 | - | - | 0 | - |
| - | - | 1110 | 203.1 | - | - | 0 | - |
| - | - | 1604 | 204.1 | - | - | 0 | - |
| - | - | 1.122E+04 | 205.1 | - | - | 0 | - |
| - | - | 662.4 | 205.1 | - | - | 0 | - |
| - | - | 1985 | 205.1 | - | - | 0 | - |
| - | - | 839.9 | 206.1 | - | - | 0 | - |
| - | - | 2025 | 206.1 | - | - | 0 | - |
| - | - | 3520 | 207.2 | - | - | 0 | - |
| - | - | 4193 | 210.1 | - | - | 0 | - |
| 11 | y | 3145 | 211.1 | 0.0003745 | 1.774 | +2 | 3 |
| - | - | 836.8 | 211.1 | - | - | 0 | - |
| - | - | 637.1 | 212.1 | - | - | 0 | - |
| - | - | 2962 | 212.1 | - | - | 0 | - |
| - | - | 1.077E+04 | 213.1 | - | - | 0 | - |
| - | - | 1471 | 213.2 | - | - | 0 | - |
| - | - | 3756 | 215.1 | - | - | 0 | - |
| - | - | 2645 | 216.1 | - | - | 0 | - |
| - | - | 2428 | 221.1 | - | - | 0 | - |
| - | - | 1213 | 221.1 | - | - | 0 | - |
| - | - | 4770 | 221.1 | - | - | 0 | - |
| - | - | 2759 | 221.1 | - | - | 0 | - |
| - | - | 735.4 | 221.1 | - | - | 0 | - |
| - | - | 1664 | 221.1 | - | - | 0 | - |
| - | - | 2208 | 222.1 | - | - | 0 | - |
| - | - | 4160 | 222.1 | - | - | 0 | - |
| - | - | 3425 | 223.1 | - | - | 0 | - |
| - | - | 1337 | 223.1 | - | - | 0 | - |
| - | - | 9063 | 223.1 | - | - | 0 | - |
| - | - | 4.402E+04 | 223.2 | - | - | 0 | - |
| - | - | 3080 | 224.1 | - | - | 0 | - |
| - | - | 4885 | 224.2 | - | - | 0 | - |
| - | - | 7328 | 225 | - | - | 0 | - |
| - | - | 1359 | 225.1 | - | - | 0 | - |
| - | - | 668 | 225.2 | - | - | 0 | - |
| - | - | 2008 | 226 | - | - | 0 | - |
| - | - | 8920 | 226.1 | - | - | 0 | - |
| - | - | 785 | 226.2 | - | - | 0 | - |
| - | - | 879.4 | 227 | - | - | 0 | - |
| - | - | 8433 | 227.1 | - | - | 0 | - |
| - | - | 963.9 | 227.1 | - | - | 0 | - |
| - | - | 1788 | 227.2 | - | - | 0 | - |
| - | - | 3911 | 228.1 | - | - | 0 | - |
| - | - | 9730 | 229.1 | - | - | 0 | - |
| - | - | 864.4 | 230.1 | - | - | 0 | - |
| - | - | 1547 | 230.2 | - | - | 0 | - |
| - | - | 1023 | 231.1 | - | - | 0 | - |
| - | - | 6735 | 233.1 | - | - | 0 | - |
| - | - | 4.764E+04 | 234.1 | - | - | 0 | - |
| - | - | 1145 | 234.7 | - | - | 0 | - |
| - | - | 6337 | 235.1 | - | - | 0 | - |
| - | - | 1360 | 235.6 | - | - | 0 | - |
| - | - | 2055 | 237.1 | - | - | 0 | - |
| - | - | 5559 | 238.1 | - | - | 0 | - |
| - | - | 1001 | 238.1 | - | - | 0 | - |
| - | - | 1.427E+04 | 239.1 | - | - | 0 | - |
| - | - | 3787 | 240.1 | - | - | 0 | - |
| - | - | 1.806E+04 | 240.1 | - | - | 0 | - |
| - | - | 2690 | 241.1 | - | - | 0 | - |
| - | - | 2404 | 241.1 | - | - | 0 | - |
| - | - | 2381 | 241.1 | - | - | 0 | - |
| - | - | 2121 | 241.2 | - | - | 0 | - |
| - | - | 4268 | 243.1 | - | - | 0 | - |
| - | - | 1.943E+04 | 244.1 | - | - | 0 | - |
| - | - | 2143 | 245.1 | - | - | 0 | - |
| - | - | 2.611E+04 | 247.1 | - | - | 0 | - |
| - | - | 1660 | 247.1 | - | - | 0 | - |
| - | - | 1263 | 247.2 | - | - | 0 | - |
| - | - | 2670 | 248.1 | - | - | 0 | - |
| - | - | 9236 | 248.1 | - | - | 0 | - |
| - | - | 1.586E+04 | 249.1 | - | - | 0 | - |
| - | - | 4344 | 249.1 | - | - | 0 | - |
| - | - | 1844 | 250.1 | - | - | 0 | - |
| - | - | 1344 | 250.1 | - | - | 0 | - |
| - | - | 1.644E+05 | 251.2 | - | - | 0 | - |
| - | - | 1515 | 252.1 | - | - | 0 | - |
| - | - | 2.13E+04 | 252.2 | - | - | 0 | - |
| - | - | 708.9 | 253.1 | - | - | 0 | - |
| - | - | 1533 | 253.2 | - | - | 0 | - |
| - | - | 9198 | 255.1 | - | - | 0 | - |
| - | - | 1817 | 256.1 | - | - | 0 | - |
| - | - | 775.4 | 256.2 | - | - | 0 | - |
| - | - | 906.1 | 256.6 | - | - | 0 | - |
| - | - | 3975 | 257.1 | - | - | 0 | - |
| - | - | 8156 | 258.1 | - | - | 0 | - |
| - | - | 3983 | 261.1 | - | - | 0 | - |
| - | - | 1322 | 261.1 | - | - | 0 | - |
| - | - | 2218 | 262.1 | - | - | 0 | - |
| - | - | 1.893E+04 | 265.1 | - | - | 0 | - |
| - | - | 1010 | 265.1 | - | - | 0 | - |
| - | - | 888.8 | 265.2 | - | - | 0 | - |
| - | - | 4.535E+04 | 266.1 | - | - | 0 | - |
| - | - | 4571 | 267.1 | - | - | 0 | - |
| - | - | 1880 | 268.1 | - | - | 0 | - |
| - | - | 4038 | 268.2 | - | - | 0 | - |
| - | - | 2123 | 269.2 | - | - | 0 | - |
| - | - | 3331 | 269.2 | - | - | 0 | - |
| - | - | 6709 | 270.1 | - | - | 0 | - |
| - | - | 1044 | 270.1 | - | - | 0 | - |
| - | - | 2151 | 270.6 | - | - | 0 | - |
| - | - | 8455 | 272.2 | - | - | 0 | - |
| - | - | 1124 | 273.2 | - | - | 0 | - |
| - | - | 8395 | 274.1 | - | - | 0 | - |
| - | - | 3343 | 274.1 | - | - | 0 | - |
| - | - | 2075 | 275.1 | - | - | 0 | - |
| - | - | 999 | 275.1 | - | - | 0 | - |
| - | - | 1009 | 275.2 | - | - | 0 | - |
| - | - | 2488 | 276.1 | - | - | 0 | - |
| - | - | 1499 | 276.1 | - | - | 0 | - |
| - | - | 2490 | 276.2 | - | - | 0 | - |
| - | - | 3477 | 279.1 | - | - | 0 | - |
| - | - | 4034 | 280.1 | - | - | 0 | - |
| - | - | 1.201E+04 | 281.1 | - | - | 0 | - |
| - | - | 906.2 | 281.1 | - | - | 0 | - |
| - | - | 7639 | 282.2 | - | - | 0 | - |
| - | - | 1396 | 283 | - | - | 0 | - |
| - | - | 1.167E+05 | 283.1 | - | - | 0 | - |
| - | - | 5510 | 284.1 | - | - | 0 | - |
| - | - | 968.4 | 284.1 | - | - | 0 | - |
| - | - | 1.606E+04 | 284.1 | - | - | 0 | - |
| - | - | 969.3 | 285 | - | - | 0 | - |
| - | - | 944.1 | 285.1 | - | - | 0 | - |
| - | - | 1359 | 285.1 | - | - | 0 | - |
| - | - | 1185 | 285.2 | - | - | 0 | - |
| - | - | 6694 | 286.1 | - | - | 0 | - |
| - | - | 825.6 | 287.1 | - | - | 0 | - |
| - | - | 1743 | 287.2 | - | - | 0 | - |
| - | - | 1577 | 288.6 | - | - | 0 | - |
| - | - | 822 | 289.1 | - | - | 0 | - |
| 5 | b | 1130 | 290.2 | 0.0009844 | 3.393 | +2 | 5 |
| 12 | y | 6.687E+04 | 293.1 | 0.0005409 | 1.845 | +1 | 2 |
| - | - | 1.34E+04 | 294.1 | - | - | 0 | - |
| - | - | 1414 | 295.1 | - | - | 0 | - |
| - | - | 1069 | 295.1 | - | - | 0 | - |
| - | - | 1360 | 296.1 | - | - | 0 | - |
| 3 | b | 2163 | 296.2 | 0.0006416 | 2.166 | +1 | 3 |
| - | - | 1004 | 296.7 | - | - | 0 | - |
| - | - | 2032 | 297.1 | - | - | 0 | - |
| 9 | y | 1408 | 297.1 | 0.003759 | 12.65 | +2 | 5 |
| - | - | 2.108E+04 | 297.2 | - | - | 0 | - |
| 7 | y | 2010 | 298.1 | 0.005337 | 17.9 | +3 | 7 |
| - | - | 3614 | 298.2 | - | - | 0 | - |
| - | - | 8.637E+04 | 299.1 | - | - | 0 | - |
| - | - | 1869 | 300.1 | - | - | 0 | - |
| - | - | 1.225E+04 | 300.2 | - | - | 0 | - |
| - | - | 1284 | 301.1 | - | - | 0 | - |
| - | - | 6.11E+04 | 301.2 | - | - | 0 | - |
| - | - | 1.093E+04 | 302.1 | - | - | 0 | - |
| - | - | 7615 | 302.2 | - | - | 0 | - |
| - | - | 1276 | 303.1 | - | - | 0 | - |
| - | - | 1989 | 305.2 | - | - | 0 | - |
| - | - | 3388 | 305.6 | - | - | 0 | - |
| 9 | y | 1.126E+04 | 306.1 | 0.0006739 | 2.201 | +2 | 5 |
| - | - | 5445 | 306.6 | - | - | 0 | - |
| - | - | 1400 | 307.1 | - | - | 0 | - |
| - | - | 1024 | 309.2 | - | - | 0 | - |
| - | - | 803.1 | 310.2 | - | - | 0 | - |
| 12 | y | 1.089E+04 | 311.1 | 0.0009931 | 3.192 | +1 | 2 |
| - | - | 2230 | 311.1 | - | - | 0 | - |
| - | - | 1759 | 312.1 | - | - | 0 | - |
| - | - | 1982 | 314.2 | - | - | 0 | - |
| 3 | b | 4935 | 314.2 | 0.0008496 | 2.704 | +1 | 3 |
| - | - | 3.599E+04 | 315.2 | - | - | 0 | - |
| - | - | 1076 | 315.2 | - | - | 0 | - |
| - | - | 5772 | 316.2 | - | - | 0 | - |
| - | - | 1237 | 317.7 | - | - | 0 | - |
| - | - | 1551 | 319.2 | - | - | 0 | - |
| - | - | 2747 | 325.2 | - | - | 0 | - |
| - | - | 7661 | 326.7 | - | - | 0 | - |
| - | - | 3389 | 327.2 | - | - | 0 | - |
| - | - | 1314 | 327.7 | - | - | 0 | - |
| - | - | 3638 | 332.2 | - | - | 0 | - |
| - | - | 6514 | 333.2 | - | - | 0 | - |
| - | - | 1134 | 334.2 | - | - | 0 | - |
| - | - | 1343 | 335.2 | - | - | 0 | - |
| - | - | 1133 | 338.2 | - | - | 0 | - |
| 6 | b | 2261 | 338.7 | 8.894E-05 | 0.2626 | +2 | 6 |
| - | - | 2218 | 340.7 | - | - | 0 | - |
| - | - | 1231 | 341 | - | - | 0 | - |
| - | - | 1745 | 341.2 | - | - | 0 | - |
| - | - | 2707 | 342.2 | - | - | 0 | - |
| - | - | 3537 | 343.2 | - | - | 0 | - |
| - | - | 963 | 343.2 | - | - | 0 | - |
| - | - | 2374 | 344.2 | - | - | 0 | - |
| - | - | 1.178E+04 | 345 | - | - | 0 | - |
| - | - | 4775 | 345.1 | - | - | 0 | - |
| - | - | 1.729E+04 | 346.1 | - | - | 0 | - |
| - | - | 1462 | 346.7 | - | - | 0 | - |
| - | - | 2568 | 347.1 | - | - | 0 | - |
| 6 | b | 9952 | 347.7 | 0.0004829 | 1.389 | +2 | 6 |
| - | - | 2247 | 348.2 | - | - | 0 | - |
| - | - | 1.284E+04 | 350.2 | - | - | 0 | - |
| - | - | 945.5 | 351.2 | - | - | 0 | - |
| - | - | 2626 | 351.2 | - | - | 0 | - |
| - | - | 984.5 | 352.7 | - | - | 0 | - |
| 8 | y | 2631 | 353.7 | 0.001247 | 3.527 | +2 | 6 |
| - | - | 1086 | 354.2 | - | - | 0 | - |
| - | - | 1407 | 355.1 | - | - | 0 | - |
| - | - | 1222 | 356.2 | - | - | 0 | - |
| - | - | 917.3 | 357.2 | - | - | 0 | - |
| - | - | 9929 | 359 | - | - | 0 | - |
| - | - | 1070 | 360.2 | - | - | 0 | - |
| - | - | 4415 | 360.7 | - | - | 0 | - |
| - | - | 735.9 | 361.2 | - | - | 0 | - |
| - | - | 4450 | 361.2 | - | - | 0 | - |
| - | - | 5578 | 362.2 | - | - | 0 | - |
| 8 | y | 2.187E+04 | 362.7 | 0.00039 | 1.075 | +2 | 6 |
| - | - | 2.691E+04 | 363.1 | - | - | 0 | - |
| - | - | 8921 | 363.2 | - | - | 0 | - |
| - | - | 3075 | 363.7 | - | - | 0 | - |
| - | - | 5215 | 364.1 | - | - | 0 | - |
| - | - | 5517 | 364.1 | - | - | 0 | - |
| - | - | 2686 | 366.2 | - | - | 0 | - |
| - | - | 1684 | 367.1 | - | - | 0 | - |
| - | - | 1856 | 367.2 | - | - | 0 | - |
| - | - | 2529 | 367.2 | - | - | 0 | - |
| - | - | 1238 | 367.7 | - | - | 0 | - |
| - | - | 1128 | 368.2 | - | - | 0 | - |
| - | - | 1178 | 368.2 | - | - | 0 | - |
| - | - | 1070 | 369.1 | - | - | 0 | - |
| - | - | 1103 | 369.2 | - | - | 0 | - |
| - | - | 965.1 | 370.1 | - | - | 0 | - |
| - | - | 934.1 | 373.2 | - | - | 0 | - |
| - | - | 904.8 | 375.2 | - | - | 0 | - |
| - | - | 1708 | 376.2 | - | - | 0 | - |
| - | - | 4052 | 376.2 | - | - | 0 | - |
| - | - | 1827 | 377.1 | - | - | 0 | - |
| - | - | 1079 | 377.3 | - | - | 0 | - |
| - | - | 9326 | 378.2 | - | - | 0 | - |
| - | - | 4.532E+04 | 379.2 | - | - | 0 | - |
| - | - | 1169 | 380.2 | - | - | 0 | - |
| - | - | 9664 | 380.2 | - | - | 0 | - |
| - | - | 8.108E+04 | 381.2 | - | - | 0 | - |
| - | - | 2166 | 381.7 | - | - | 0 | - |
| - | - | 1.462E+04 | 382.2 | - | - | 0 | - |
| - | - | 2790 | 382.2 | - | - | 0 | - |
| - | - | 1193 | 382.2 | - | - | 0 | - |
| - | - | 1619 | 383.2 | - | - | 0 | - |
| - | - | 2831 | 384.2 | - | - | 0 | - |
| - | - | 3085 | 384.2 | - | - | 0 | - |
| - | - | 5670 | 385.2 | - | - | 0 | - |
| 5 | y | 1932 | 385.2 | 0.003948 | 10.25 | +3 | 9 |
| - | - | 910.4 | 386.2 | - | - | 0 | - |
| 10 | b | 1582 | 389.2 | 0.0005644 | 1.45 | +3 | 10 |
| - | - | 3664 | 390.2 | - | - | 0 | - |
| - | - | 2876 | 391.1 | - | - | 0 | - |
| - | - | 1570 | 391.2 | - | - | 0 | - |
| - | - | 1591 | 392.2 | - | - | 0 | - |
| - | - | 1322 | 394.1 | - | - | 0 | - |
| - | - | 1.032E+04 | 394.2 | - | - | 0 | - |
| - | - | 4185 | 395.1 | - | - | 0 | - |
| - | - | 2507 | 395.2 | - | - | 0 | - |
| - | - | 1308 | 395.2 | - | - | 0 | - |
| - | - | 1739 | 395.2 | - | - | 0 | - |
| - | - | 1.536E+04 | 396.2 | - | - | 0 | - |
| - | - | 2861 | 397.2 | - | - | 0 | - |
| - | - | 5493 | 398.2 | - | - | 0 | - |
| - | - | 1394 | 398.7 | - | - | 0 | - |
| - | - | 914.6 | 399.2 | - | - | 0 | - |
| - | - | 995.5 | 401.2 | - | - | 0 | - |
| - | - | 3713 | 402.2 | - | - | 0 | - |
| - | - | 6443 | 402.2 | - | - | 0 | - |
| - | - | 2341 | 403.2 | - | - | 0 | - |
| - | - | 1033 | 404.2 | - | - | 0 | - |
| - | - | 1273 | 404.7 | - | - | 0 | - |
| - | - | 841 | 405.2 | - | - | 0 | - |
| - | - | 2762 | 405.3 | - | - | 0 | - |
| - | - | 2063 | 406.2 | - | - | 0 | - |
| - | - | 1051 | 407.2 | - | - | 0 | - |
| - | - | 1114 | 407.2 | - | - | 0 | - |
| - | - | 6374 | 409.1 | - | - | 0 | - |
| - | - | 1898 | 409.2 | - | - | 0 | - |
| - | - | 1040 | 409.7 | - | - | 0 | - |
| - | - | 5107 | 411.2 | - | - | 0 | - |
| - | - | 4.745E+04 | 412.2 | - | - | 0 | - |
| - | - | 5193 | 413.1 | - | - | 0 | - |
| - | - | 2922 | 413.2 | - | - | 0 | - |
| - | - | 7484 | 413.2 | - | - | 0 | - |
| - | - | 1568 | 413.2 | - | - | 0 | - |
| - | - | 1218 | 414.1 | - | - | 0 | - |
| - | - | 1.319E+04 | 414.2 | - | - | 0 | - |
| - | - | 2.157E+04 | 415 | - | - | 0 | - |
| - | - | 3616 | 415.2 | - | - | 0 | - |
| - | - | 2877 | 415.2 | - | - | 0 | - |
| - | - | 1204 | 416.2 | - | - | 0 | - |
| - | - | 1.021E+04 | 417.7 | - | - | 0 | - |
| - | - | 1.323E+04 | 418.2 | - | - | 0 | - |
| - | - | 6109 | 418.7 | - | - | 0 | - |
| - | - | 1645 | 419.2 | - | - | 0 | - |
| 11 | y | 2361 | 421.2 | 0.0005791 | 1.375 | +1 | 3 |
| 11 | y | 1262 | 422.2 | 0.0005113 | 1.211 | +1 | 3 |
| - | - | 880.1 | 422.3 | - | - | 0 | - |
| - | - | 1760 | 422.7 | - | - | 0 | - |
| - | - | 2875 | 423.2 | - | - | 0 | - |
| - | - | 1.834E+04 | 423.3 | - | - | 0 | - |
| - | - | 1724 | 423.7 | - | - | 0 | - |
| - | - | 926.5 | 423.7 | - | - | 0 | - |
| - | - | 5227 | 424.3 | - | - | 0 | - |
| - | - | 1470 | 424.7 | - | - | 0 | - |
| - | - | 907.7 | 425.3 | - | - | 0 | - |
| - | - | 1.307E+05 | 426.7 | - | - | 0 | - |
| - | - | 6.524E+04 | 427.2 | - | - | 0 | - |
| - | - | 1.759E+04 | 427.7 | - | - | 0 | - |
| - | - | 3321 | 428.2 | - | - | 0 | - |
| - | - | 1290 | 429.1 | - | - | 0 | - |
| - | - | 2.667E+04 | 429.2 | - | - | 0 | - |
| - | - | 1.213E+05 | 430.2 | - | - | 0 | - |
| - | - | 2.591E+04 | 431.2 | - | - | 0 | - |
| 7 | b | 1.423E+04 | 431.7 | 0.0006241 | 1.446 | +2 | 7 |
| 7 | b | 6780 | 432.2 | 0.007274 | 16.83 | +2 | 7 |
| - | - | 3993 | 432.7 | - | - | 0 | - |
| - | - | 1052 | 433 | - | - | 0 | - |
| - | - | 1510 | 433.2 | - | - | 0 | - |
| 4 | b | 4886 | 433.3 | 0.0006592 | 1.522 | +1 | 4 |
| - | - | 1145 | 434.3 | - | - | 0 | - |
| - | - | 2762 | 438.2 | - | - | 0 | - |
| - | - | 1229 | 438.7 | - | - | 0 | - |
| 11 | y | 3.447E+04 | 439.2 | 0.0008787 | 2.001 | +1 | 3 |
| - | - | 1563 | 440.2 | - | - | 0 | - |
| - | - | 8359 | 440.2 | - | - | 0 | - |
| 7 | b | 6.405E+04 | 440.7 | 0.0006823 | 1.548 | +2 | 7 |
| - | - | 3.228E+04 | 441.2 | - | - | 0 | - |
| - | - | 9595 | 441.7 | - | - | 0 | - |
| - | - | 1640 | 442.2 | - | - | 0 | - |
| - | - | 1246 | 443.2 | - | - | 0 | - |
| - | - | 863.2 | 444.2 | - | - | 0 | - |
| - | - | 1233 | 446.7 | - | - | 0 | - |
| 7 | y | 4452 | 447.2 | 0.00126 | 2.818 | +2 | 7 |
| - | - | 1350 | 447.7 | - | - | 0 | - |
| - | - | 6524 | 448.2 | - | - | 0 | - |
| - | - | 1425 | 449.2 | - | - | 0 | - |
| - | - | 1602 | 450.2 | - | - | 0 | - |
| - | - | 1853 | 450.3 | - | - | 0 | - |
| 4 | b | 3.774E+04 | 451.3 | 0.0006841 | 1.516 | +1 | 4 |
| - | - | 1.011E+04 | 452.3 | - | - | 0 | - |
| - | - | 1430 | 453.3 | - | - | 0 | - |
| - | - | 1463 | 455.3 | - | - | 0 | - |
| 7 | y | 2103 | 455.7 | 0.0002538 | 0.5568 | +2 | 7 |
| - | - | 2102 | 456.2 | - | - | 0 | - |
| - | - | 3321 | 459.2 | - | - | 0 | - |
| - | - | 3149 | 460.8 | - | - | 0 | - |
| - | - | 3319 | 461.2 | - | - | 0 | - |
| - | - | 4018 | 461.3 | - | - | 0 | - |
| - | - | 984.8 | 462.2 | - | - | 0 | - |
| - | - | 1124 | 462.3 | - | - | 0 | - |
| - | - | 1096 | 464.2 | - | - | 0 | - |
| - | - | 949.7 | 465.7 | - | - | 0 | - |
| - | - | 1168 | 466.2 | - | - | 0 | - |
| - | - | 874 | 466.2 | - | - | 0 | - |
| - | - | 1129 | 468.3 | - | - | 0 | - |
| - | - | 9060 | 468.3 | - | - | 0 | - |
| - | - | 2170 | 469.3 | - | - | 0 | - |
| - | - | 1001 | 472.2 | - | - | 0 | - |
| - | - | 8221 | 474.3 | - | - | 0 | - |
| - | - | 9329 | 474.8 | - | - | 0 | - |
| - | - | 4615 | 475.3 | - | - | 0 | - |
| - | - | 1188 | 475.8 | - | - | 0 | - |
| - | - | 3076 | 476.2 | - | - | 0 | - |
| - | - | 7228 | 477.2 | - | - | 0 | - |
| 10 | y | 6179 | 478.2 | 0.0003768 | 0.7879 | +1 | 4 |
| - | - | 4591 | 478.3 | - | - | 0 | - |
| - | - | 1765 | 479.2 | - | - | 0 | - |
| - | - | 1437 | 479.7 | - | - | 0 | - |
| - | - | 1053 | 480.2 | - | - | 0 | - |
| - | - | 2.64E+04 | 483.3 | - | - | 0 | - |
| - | - | 1.671E+04 | 483.8 | - | - | 0 | - |
| - | - | 1375 | 484.2 | - | - | 0 | - |
| - | - | 3977 | 484.3 | - | - | 0 | - |
| - | - | 1507 | 484.8 | - | - | 0 | - |
| 8 | b | 7700 | 488.3 | 0.001378 | 2.822 | +2 | 8 |
| 8 | b | 3988 | 488.7 | 0.007447 | 15.24 | +2 | 8 |
| - | - | 1892 | 489.3 | - | - | 0 | - |
| - | - | 8.271E+04 | 494.2 | - | - | 0 | - |
| - | - | 840.3 | 495.2 | - | - | 0 | - |
| - | - | 2.229E+04 | 495.2 | - | - | 0 | - |
| 10 | y | 9.704E+04 | 496.2 | 0.0007468 | 1.505 | +1 | 4 |
| - | - | 2.52E+04 | 497.2 | - | - | 0 | - |
| - | - | 8318 | 497.8 | - | - | 0 | - |
| - | - | 4965 | 498.2 | - | - | 0 | - |
| - | - | 2.307E+04 | 503.1 | - | - | 0 | - |
| - | - | 3397 | 504.2 | - | - | 0 | - |
| - | - | 1958 | 506.2 | - | - | 0 | - |
| - | - | 1358 | 507.3 | - | - | 0 | - |
| - | - | 799.4 | 510.3 | - | - | 0 | - |
| - | - | 5106 | 511.3 | - | - | 0 | - |
| - | - | 1345 | 512.3 | - | - | 0 | - |
| - | - | 1317 | 515.3 | - | - | 0 | - |
| - | - | 7093 | 519.1 | - | - | 0 | - |
| - | - | 1119 | 522.2 | - | - | 0 | - |
| - | - | 2621 | 523.3 | - | - | 0 | - |
| - | - | 1935 | 523.6 | - | - | 0 | - |
| - | - | 1196 | 524.3 | - | - | 0 | - |
| - | - | 1.348E+04 | 525.3 | - | - | 0 | - |
| - | - | 2990 | 526.2 | - | - | 0 | - |
| - | - | 3089 | 526.3 | - | - | 0 | - |
| 0 | Precursor | 4634 | 529.3 | 0.001291 | 2.439 | +3 | -1 |
| 0 | Precursor | 5376 | 529.6 | 0.007575 | 14.3 | +3 | -1 |
| - | - | 2757 | 529.9 | - | - | 0 | - |
| - | - | 1087 | 531.2 | - | - | 0 | - |
| - | - | 3701 | 532.2 | - | - | 0 | - |
| - | - | 1354 | 533.2 | - | - | 0 | - |
| - | - | 1992 | 534.9 | - | - | 0 | - |
| 0 | Precursor | 5123 | 535.3 | 0.00076 | 1.42 | +3 | -1 |
| - | - | 2912 | 535.6 | - | - | 0 | - |
| - | - | 1830 | 535.9 | - | - | 0 | - |
| - | - | 4730 | 536.2 | - | - | 0 | - |
| - | - | 1805 | 537.2 | - | - | 0 | - |
| - | - | 2018 | 537.7 | - | - | 0 | - |
| - | - | 1091 | 538.3 | - | - | 0 | - |
| - | - | 1337 | 538.8 | - | - | 0 | - |
| - | - | 1.408E+04 | 539.2 | - | - | 0 | - |
| - | - | 4427 | 540.2 | - | - | 0 | - |
| - | - | 1412 | 541.3 | - | - | 0 | - |
| - | - | 4726 | 542.3 | - | - | 0 | - |
| - | - | 2.63E+04 | 543.3 | - | - | 0 | - |
| - | - | 9098 | 544.3 | - | - | 0 | - |
| - | - | 2021 | 545.3 | - | - | 0 | - |
| 9 | b | 947.2 | 545.8 | 0.004983 | 9.131 | +2 | 9 |
| - | - | 4722 | 546.2 | - | - | 0 | - |
| - | - | 1851 | 546.8 | - | - | 0 | - |
| - | - | 1016 | 547.3 | - | - | 0 | - |
| - | - | 956.4 | 547.8 | - | - | 0 | - |
| - | - | 1324 | 548.2 | - | - | 0 | - |
| - | - | 3125 | 549.2 | - | - | 0 | - |
| - | - | 1526 | 550.2 | - | - | 0 | - |
| - | - | 3830 | 551.3 | - | - | 0 | - |
| - | - | 2112 | 551.3 | - | - | 0 | - |
| - | - | 1918 | 552.3 | - | - | 0 | - |
| - | - | 1245 | 553.3 | - | - | 0 | - |
| 9 | b | 3478 | 554.8 | 0.000744 | 1.341 | +2 | 9 |
| - | - | 6402 | 555.3 | - | - | 0 | - |
| - | - | 5645 | 555.8 | - | - | 0 | - |
| - | - | 797.8 | 556.3 | - | - | 0 | - |
| - | - | 1270 | 560.2 | - | - | 0 | - |
| 5 | b | 3542 | 561.3 | 0.0008891 | 1.584 | +1 | 5 |
| 5 | b | 1218 | 562.3 | 0.003324 | 5.911 | +1 | 5 |
| - | - | 3484 | 563.3 | - | - | 0 | - |
| - | - | 2208 | 564.3 | - | - | 0 | - |
| - | - | 780.7 | 564.3 | - | - | 0 | - |
| - | - | 5852 | 565.3 | - | - | 0 | - |
| - | - | 1562 | 566.3 | - | - | 0 | - |
| - | - | 1.61E+04 | 567.2 | - | - | 0 | - |
| - | - | 938.6 | 567.3 | - | - | 0 | - |
| - | - | 5199 | 568.2 | - | - | 0 | - |
| - | - | 852.7 | 569.2 | - | - | 0 | - |
| - | - | 784.9 | 569.8 | - | - | 0 | - |
| - | - | 4020 | 575.2 | - | - | 0 | - |
| - | - | 2935 | 576.2 | - | - | 0 | - |
| 5 | b | 2.551E+04 | 579.3 | 0.0002731 | 0.4714 | +1 | 5 |
| - | - | 7366 | 580.3 | - | - | 0 | - |
| - | - | 1831 | 581.3 | - | - | 0 | - |
| - | - | 813.9 | 587.3 | - | - | 0 | - |
| - | - | 3424 | 592.3 | - | - | 0 | - |
| - | - | 3493 | 592.3 | - | - | 0 | - |
| 9 | y | 3.154E+04 | 593.3 | 0.0002681 | 0.4519 | +1 | 5 |
| - | - | 1.162E+04 | 593.3 | - | - | 0 | - |
| - | - | 8577 | 594.3 | - | - | 0 | - |
| - | - | 3152 | 594.3 | - | - | 0 | - |
| - | - | 1206 | 594.8 | - | - | 0 | - |
| - | - | 2640 | 595.2 | - | - | 0 | - |
| - | - | 1417 | 596.4 | - | - | 0 | - |
| - | - | 1337 | 597.8 | - | - | 0 | - |
| - | - | 1424 | 600.3 | - | - | 0 | - |
| - | - | 4763 | 602.8 | - | - | 0 | - |
| - | - | 5696 | 603.3 | - | - | 0 | - |
| - | - | 813.9 | 603.4 | - | - | 0 | - |
| - | - | 1903 | 603.8 | - | - | 0 | - |
| - | - | 6.16E+04 | 610.3 | - | - | 0 | - |
| - | - | 2222 | 610.3 | - | - | 0 | - |
| 9 | y | 3.189E+05 | 611.3 | 0.001117 | 1.827 | +1 | 5 |
| - | - | 1.607E+04 | 611.8 | - | - | 0 | - |
| - | - | 9.869E+04 | 612.3 | - | - | 0 | - |
| - | - | 4353 | 612.8 | - | - | 0 | - |
| - | - | 2.238E+04 | 613.3 | - | - | 0 | - |
| - | - | 2939 | 614.3 | - | - | 0 | - |
| - | - | 1106 | 615.3 | - | - | 0 | - |
| - | - | 1142 | 616.3 | - | - | 0 | - |
| - | - | 1896 | 617.3 | - | - | 0 | - |
| - | - | 865.5 | 619.3 | - | - | 0 | - |
| - | - | 4079 | 621.3 | - | - | 0 | - |
| - | - | 1365 | 621.3 | - | - | 0 | - |
| - | - | 1109 | 622.3 | - | - | 0 | - |
| - | - | 3181 | 634.3 | - | - | 0 | - |
| - | - | 4829 | 635.3 | - | - | 0 | - |
| - | - | 3169 | 636.3 | - | - | 0 | - |
| 4 | y | 3476 | 636.8 | 0.000306 | 0.4805 | +2 | 10 |
| 4 | y | 4358 | 637.3 | 0.01104 | 17.33 | +2 | 10 |
| - | - | 1573 | 637.8 | - | - | 0 | - |
| 11 | b | 1150 | 638.3 | 0.005773 | 9.044 | +2 | 11 |
| - | - | 2678 | 640.3 | - | - | 0 | - |
| - | - | 1250 | 641.3 | - | - | 0 | - |
| - | - | 3820 | 645.3 | - | - | 0 | - |
| 4 | y | 2.086E+04 | 645.8 | 0.00186 | 2.88 | +2 | 10 |
| - | - | 1.379E+04 | 646.3 | - | - | 0 | - |
| - | - | 7297 | 646.8 | - | - | 0 | - |
| - | - | 1363 | 647.3 | - | - | 0 | - |
| - | - | 1575 | 648.3 | - | - | 0 | - |
| - | - | 1842 | 650.3 | - | - | 0 | - |
| - | - | 1091 | 650.3 | - | - | 0 | - |
| - | - | 1108 | 651.3 | - | - | 0 | - |
| - | - | 3.316E+04 | 652.3 | - | - | 0 | - |
| - | - | 1160 | 652.8 | - | - | 0 | - |
| - | - | 1.283E+04 | 653.3 | - | - | 0 | - |
| - | - | 2188 | 654.3 | - | - | 0 | - |
| - | - | 2022 | 657.4 | - | - | 0 | - |
| - | - | 2050 | 658.3 | - | - | 0 | - |
| - | - | 1009 | 658.3 | - | - | 0 | - |
| - | - | 1241 | 659.3 | - | - | 0 | - |
| - | - | 1839 | 660.9 | - | - | 0 | - |
| - | - | 2211 | 661.3 | - | - | 0 | - |
| - | - | 1320 | 661.8 | - | - | 0 | - |
| - | - | 1.279E+04 | 662.3 | - | - | 0 | - |
| - | - | 929.8 | 662.8 | - | - | 0 | - |
| - | - | 8104 | 663.3 | - | - | 0 | - |
| - | - | 2774 | 664.3 | - | - | 0 | - |
| - | - | 1086 | 665.3 | - | - | 0 | - |
| - | - | 1570 | 666.4 | - | - | 0 | - |
| - | - | 1668 | 668.3 | - | - | 0 | - |
| 6 | b | 1.797E+04 | 676.3 | 0.0004353 | 0.6436 | +1 | 6 |
| 6 | b | 8643 | 677.3 | 0.006593 | 9.734 | +1 | 6 |
| - | - | 3794 | 678.3 | - | - | 0 | - |
| - | - | 3358 | 679.3 | - | - | 0 | - |
| - | - | 1091 | 679.8 | - | - | 0 | - |
| - | - | 5.895E+04 | 680.3 | - | - | 0 | - |
| - | - | 2.434E+04 | 681.3 | - | - | 0 | - |
| - | - | 4592 | 682.3 | - | - | 0 | - |
| - | - | 2155 | 684.3 | - | - | 0 | - |
| - | - | 3242 | 684.8 | - | - | 0 | - |
| - | - | 2685 | 685.3 | - | - | 0 | - |
| - | - | 1386 | 685.8 | - | - | 0 | - |
| - | - | 1269 | 688.3 | - | - | 0 | - |
| - | - | 959.3 | 692.3 | - | - | 0 | - |
| - | - | 2716 | 692.8 | - | - | 0 | - |
| 3 | y | 1.654E+04 | 693.3 | 0.001609 | 2.321 | +2 | 11 |
| 3 | y | 1.89E+04 | 693.8 | 0.007648 | 11.02 | +2 | 11 |
| 6 | b | 1.664E+05 | 694.4 | 0.0001196 | 0.1723 | +1 | 6 |
| - | - | 2845 | 694.8 | - | - | 0 | - |
| - | - | 6.322E+04 | 695.4 | - | - | 0 | - |
| - | - | 1.424E+04 | 696.4 | - | - | 0 | - |
| - | - | 2678 | 697.4 | - | - | 0 | - |
| - | - | 1852 | 698.4 | - | - | 0 | - |
| - | - | 1240 | 700.9 | - | - | 0 | - |
| - | - | 2.262E+04 | 701.8 | - | - | 0 | - |
| 3 | y | 1.151E+05 | 702.3 | 0.001698 | 2.417 | +2 | 11 |
| 12 | b | 9.805E+04 | 702.8 | 0.01302 | 18.53 | +2 | 12 |
| 12 | b | 4.415E+04 | 703.3 | 0.00332 | 4.72 | +2 | 12 |
| - | - | 1.27E+04 | 703.8 | - | - | 0 | - |
| - | - | 5388 | 704.3 | - | - | 0 | - |
| - | - | 2235 | 705.4 | - | - | 0 | - |
| 8 | y | 9844 | 706.3 | 0.001775 | 2.513 | +1 | 6 |
| - | - | 4260 | 707.3 | - | - | 0 | - |
| - | - | 921.7 | 709.3 | - | - | 0 | - |
| - | - | 6664 | 711.4 | - | - | 0 | - |
| 12 | b | 8696 | 711.9 | 0.003059 | 4.297 | +2 | 12 |
| - | - | 7959 | 712.4 | - | - | 0 | - |
| - | - | 3125 | 712.9 | - | - | 0 | - |
| - | - | 1070 | 719.4 | - | - | 0 | - |
| - | - | 1129 | 720.4 | - | - | 0 | - |
| - | - | 1124 | 720.9 | - | - | 0 | - |
| - | - | 7277 | 721.4 | - | - | 0 | - |
| - | - | 2674 | 722.4 | - | - | 0 | - |
| - | - | 1.606E+04 | 723.4 | - | - | 0 | - |
| 8 | y | 9.565E+04 | 724.4 | 0.001159 | 1.601 | +1 | 6 |
| - | - | 3.873E+04 | 725.4 | - | - | 0 | - |
| - | - | 1.116E+04 | 726.4 | - | - | 0 | - |
| - | - | 1614 | 727.4 | - | - | 0 | - |
| - | - | 888.8 | 728.4 | - | - | 0 | - |
| - | - | 4206 | 729.4 | - | - | 0 | - |
| - | - | 2316 | 730.4 | - | - | 0 | - |
| - | - | 4025 | 733.4 | - | - | 0 | - |
| - | - | 3441 | 734.4 | - | - | 0 | - |
| - | - | 1193 | 737.9 | - | - | 0 | - |
| - | - | 1302 | 738.4 | - | - | 0 | - |
| - | - | 1106 | 739.4 | - | - | 0 | - |
| 2 | y | 1084 | 742.9 | 0.003545 | 4.772 | +2 | 12 |
| 2 | y | 2844 | 743.4 | 0.01336 | 17.97 | +2 | 12 |
| - | - | 1108 | 744.4 | - | - | 0 | - |
| - | - | 1104 | 745.4 | - | - | 0 | - |
| - | - | 2106 | 747.4 | - | - | 0 | - |
| - | - | 1982 | 748.4 | - | - | 0 | - |
| - | - | 1392 | 749.3 | - | - | 0 | - |
| - | - | 2.03E+04 | 751.4 | - | - | 0 | - |
| 2 | y | 1.046E+04 | 751.9 | 0.001976 | 2.628 | +2 | 12 |
| - | - | 1.759E+04 | 752.4 | - | - | 0 | - |
| - | - | 3830 | 752.9 | - | - | 0 | - |
| - | - | 3622 | 753.4 | - | - | 0 | - |
| - | - | 1046 | 754.4 | - | - | 0 | - |
| - | - | 1854 | 760.3 | - | - | 0 | - |
| - | - | 1968 | 761.4 | - | - | 0 | - |
| - | - | 2334 | 762.4 | - | - | 0 | - |
| - | - | 9422 | 763.4 | - | - | 0 | - |
| - | - | 5318 | 764.4 | - | - | 0 | - |
| - | - | 4030 | 765.4 | - | - | 0 | - |
| - | - | 1786 | 766.4 | - | - | 0 | - |
| - | - | 4489 | 767.3 | - | - | 0 | - |
| - | - | 1169 | 768.3 | - | - | 0 | - |
| - | - | 1923 | 769.4 | - | - | 0 | - |
| - | - | 2603 | 776.4 | - | - | 0 | - |
| - | - | 9376 | 779.4 | - | - | 0 | - |
| - | - | 4467 | 780.4 | - | - | 0 | - |
| - | - | 4533 | 781.4 | - | - | 0 | - |
| - | - | 1480 | 782.4 | - | - | 0 | - |
| 0 | Precursor | 1.749E+04 | 793.4 | 0.01351 | 17.03 | +2 | -1 |
| - | - | 8704 | 794.4 | - | - | 0 | - |
| - | - | 6227 | 795.3 | - | - | 0 | - |
| - | - | 1408 | 795.4 | - | - | 0 | - |
| - | - | 1518 | 796.3 | - | - | 0 | - |
| - | - | 4651 | 796.5 | - | - | 0 | - |
| - | - | 1501 | 797.5 | - | - | 0 | - |
| - | - | 3880 | 807.4 | - | - | 0 | - |
| - | - | 2194 | 807.4 | - | - | 0 | - |
| - | - | 2493 | 808.4 | - | - | 0 | - |
| - | - | 2661 | 818.4 | - | - | 0 | - |
| - | - | 2051 | 819.4 | - | - | 0 | - |
| - | - | 1624 | 825.4 | - | - | 0 | - |
| - | - | 1909 | 826.4 | - | - | 0 | - |
| - | - | 1121 | 827.4 | - | - | 0 | - |
| - | - | 1340 | 828.4 | - | - | 0 | - |
| - | - | 4722 | 834.4 | - | - | 0 | - |
| - | - | 6119 | 835.4 | - | - | 0 | - |
| - | - | 1.168E+04 | 836.4 | - | - | 0 | - |
| - | - | 4699 | 837.5 | - | - | 0 | - |
| - | - | 2487 | 838.5 | - | - | 0 | - |
| - | - | 2132 | 843.4 | - | - | 0 | - |
| - | - | 4327 | 844.4 | - | - | 0 | - |
| - | - | 3955 | 845.4 | - | - | 0 | - |
| - | - | 6845 | 846.4 | - | - | 0 | - |
| - | - | 5098 | 847.4 | - | - | 0 | - |
| - | - | 1247 | 848.4 | - | - | 0 | - |
| - | - | 4.2E+04 | 852.4 | - | - | 0 | - |
| - | - | 1.993E+04 | 853.4 | - | - | 0 | - |
| - | - | 6018 | 854.4 | - | - | 0 | - |
| - | - | 963.9 | 857.4 | - | - | 0 | - |
| 7 | b | 2.516E+04 | 862.4 | 0.0001018 | 0.1181 | +1 | 7 |
| 7 | b | 1.859E+04 | 863.4 | 0.012 | 13.89 | +1 | 7 |
| - | - | 2.757E+04 | 864.5 | - | - | 0 | - |
| - | - | 1.456E+04 | 865.5 | - | - | 0 | - |
| - | - | 5112 | 866.5 | - | - | 0 | - |
| - | - | 4322 | 876.4 | - | - | 0 | - |
| - | - | 1343 | 877.4 | - | - | 0 | - |
| - | - | 1275 | 878.4 | - | - | 0 | - |
| 7 | b | 1.975E+05 | 880.4 | 0.0003921 | 0.4453 | +1 | 7 |
| - | - | 1.028E+05 | 881.4 | - | - | 0 | - |
| - | - | 2.999E+04 | 882.4 | - | - | 0 | - |
| - | - | 6238 | 883.4 | - | - | 0 | - |
| - | - | 1303 | 884.4 | - | - | 0 | - |
| - | - | 1734 | 890.4 | - | - | 0 | - |
| - | - | 2257 | 891.4 | - | - | 0 | - |
| - | - | 2663 | 892.5 | - | - | 0 | - |
| - | - | 1287 | 893.5 | - | - | 0 | - |
| - | - | 1198 | 894.4 | - | - | 0 | - |
| - | - | 2276 | 907.4 | - | - | 0 | - |
| - | - | 1.441E+04 | 908.4 | - | - | 0 | - |
| - | - | 8984 | 909.4 | - | - | 0 | - |
| 7 | y | 1.721E+04 | 910.4 | 0.001436 | 1.578 | +1 | 7 |
| - | - | 7636 | 911.4 | - | - | 0 | - |
| - | - | 2366 | 912.4 | - | - | 0 | - |
| - | - | 1054 | 936.4 | - | - | 0 | - |
| - | - | 2910 | 937.4 | - | - | 0 | - |
| - | - | 1378 | 947.4 | - | - | 0 | - |
| - | - | 4273 | 954.4 | - | - | 0 | - |
| - | - | 4942 | 955.4 | - | - | 0 | - |
| - | - | 2852 | 956.4 | - | - | 0 | - |
| - | - | 1279 | 964.5 | - | - | 0 | - |
| - | - | 6231 | 965.4 | - | - | 0 | - |
| - | - | 2483 | 966.4 | - | - | 0 | - |
| - | - | 3035 | 972.4 | - | - | 0 | - |
| - | - | 2674 | 973.4 | - | - | 0 | - |
| 8 | b | 3304 | 975.5 | 2.223E-05 | 0.02279 | +1 | 8 |
| 8 | b | 1903 | 976.5 | 0.01418 | 14.52 | +1 | 8 |
| - | - | 1654 | 980.5 | - | - | 0 | - |
| - | - | 1649 | 982.5 | - | - | 0 | - |
| - | - | 1106 | 990.5 | - | - | 0 | - |
| 8 | b | 1.001E+04 | 993.5 | 0.0004106 | 0.4133 | +1 | 8 |
| - | - | 7558 | 994.5 | - | - | 0 | - |
| - | - | 1165 | 995.5 | - | - | 0 | - |
| - | - | 1570 | 1007 | - | - | 0 | - |
| - | - | 1097 | 1009 | - | - | 0 | - |
| - | - | 1813 | 1024 | - | - | 0 | - |
| 6 | y | 8183 | 1025 | 0.0007994 | 0.7796 | +1 | 8 |
| - | - | 4824 | 1026 | - | - | 0 | - |
| - | - | 1428 | 1027 | - | - | 0 | - |
| - | - | 1282 | 1065 | - | - | 0 | - |
| - | - | 1324 | 1076 | - | - | 0 | - |
| - | - | 1232 | 1078 | - | - | 0 | - |
| - | - | 4226 | 1094 | - | - | 0 | - |
| - | - | 2760 | 1095 | - | - | 0 | - |
| - | - | 1123 | 1096 | - | - | 0 | - |
| 9 | b | 2124 | 1109 | 0.001048 | 0.9449 | +1 | 9 |
| - | - | 1485 | 1110 | - | - | 0 | - |
| - | - | 1491 | 1117 | - | - | 0 | - |
| - | - | 2147 | 1118 | - | - | 0 | - |
| 5 | y | 6228 | 1136 | 0.001249 | 1.1 | +1 | 9 |
| 5 | y | 1.054E+04 | 1136 | 0.006491 | 5.711 | +1 | 9 |
| - | - | 5000 | 1137 | - | - | 0 | - |
| - | - | 1586 | 1138 | - | - | 0 | - |
| - | - | 3930 | 1153 | - | - | 0 | - |
| 5 | y | 1.585E+04 | 1154 | 0.00106 | 0.9188 | +1 | 9 |
| - | - | 1.008E+04 | 1155 | - | - | 0 | - |
| - | - | 3795 | 1156 | - | - | 0 | - |
| - | - | 925 | 1157 | - | - | 0 | - |
| 10 | b | 2266 | 1166 | 0.005565 | 4.774 | +1 | 10 |
| - | - | 882.4 | 1205 | - | - | 0 | - |
| 4 | y | 973.7 | 1291 | 0.005118 | 3.965 | +1 | 10 |
| - | - | 963.5 | 3081 | - | - | 0 | - |
| - | - | 1132 | 3455 | - | - | 0 | - |

m/z Charge Intensity FragmentType MassShift Position
120.0811538696289 0 8445.756
121.08434295654297 0 569.93335
121.92815399169922 0 469.64844
122.07157135009766 0 2263.3984
123.04450988769531 0 2485.313
123.05561065673828 0 769.5132
124.99102783203125 0 460.74606
125.10752868652344 0 902.8716
126.09156799316406 0 1223.9608
127.05061340332031 0 5866.7773
127.08692932128906 0 952.99084
127.12342071533203 0 1656.7916
128.1073760986328 0 27643.783
129.0663604736328 0 1903.9429
129.10267639160156 0 322747.84
130.06552124023438 0 140265.5
130.09983825683594 0 1982.5854
130.10598754882812 0 18614.938
130.78900146484375 0 498.8818
131.06886291503906 0 13081.435
131.1181182861328 0 703.99445
132.08114624023438 0 30608.861
133.06103515625 0 834.34314
133.0845489501953 0 3230.5242
134.04495239257812 0 578.76447
136.07608032226562 0 43186.582
137.07949829101562 0 3302.4238
138.06666564941406 0 3975.2585
138.09158325195312 0 1406.3265
139.0819854736328 0 539.8432
139.0869903564453 0 3873.5156
141.1027069091797 0 5367.975
142.06546020507812 0 1878.4285
143.04568481445312 0 1971.0459
143.08164978027344 0 636.7323
144.08131408691406 0 1013.1926
145.06130981445312 0 3393.2761
146.0606231689453 0 1188.7456
147.04461669921875 0 1789.1641
149.04530334472656 0 11298.224
150.04534912109375 0 624.0867
150.066650390625 0 1787.8827
151.08705139160156 0 7253.494
153.1027069091797 0 683.5373
154.09771728515625 0 1070.3788
155.08180236816406 0 1495.269
155.0931396484375 0 21019.492
155.11822509765625 0 15292.282 a Water loss 1
156.07704162597656 0 1287.6495
156.09632873535156 0 759.9911
156.12149047851562 0 1121.7467
157.0609130859375 0 875.35986
157.09750366210938 0 1900.2062
158.08401489257812 0 3165.3108
159.0921173095703 0 243524.56
160.0758819580078 0 2313.475
160.08926391601562 0 1672.236
160.095458984375 0 27718.436
161.098876953125 0 918.5101
165.0550079345703 0 19975.197
165.0778045654297 0 1164.3998
166.06143188476562 0 37516.96
166.08670043945312 0 3829.7402 y 9
166.09786987304688 0 2409.378
167.05587768554688 0 13693.762
167.06471252441406 0 2440.5598
167.0819091796875 0 608.9237
167.11837768554688 0 1369.195
168.05552673339844 0 917.7028
168.081298828125 0 3933.2925
168.10220336914062 0 3618.0635
168.1135711669922 0 3512.3894
169.07640075683594 0 4239.1997
169.08444213867188 0 1339.4023
169.0976104736328 0 9594.41
170.0604705810547 0 13222.829
171.06370544433594 0 1479.4734
171.09225463867188 0 2282.9504
171.149658203125 0 1596.7542
172.07211303710938 0 4490.713
173.05616760253906 0 3986.6094
173.12892150878906 0 489409.5 a 1
173.4400177001953 0 2574.8074
174.13226318359375 0 43076.664
175.13412475585938 0 2110.0862
176.08290100097656 0 847.86835
177.1028594970703 0 1563.474
178.13426208496094 0 12888.948
179.11819458007812 0 1146.6942
179.13783264160156 0 984.31726
181.06124877929688 0 2832.6033
182.0816192626953 0 47776.492 y 12
183.0852508544922 0 3755.7383
183.1132049560547 0 12825.797 b Water loss 1
183.14930725097656 0 1264.5991
184.11643981933594 0 1497.881
185.09243774414062 0 837.2407
185.1652374267578 0 3145.3376
186.12413024902344 0 109009.36
187.08700561523438 0 17909.996
187.1082305908203 0 764.38635
187.12759399414062 0 8458.302
188.07093811035156 0 2327.0388
188.09071350097656 0 2724.6665
188.12892150878906 0 861.0169
190.08267211914062 0 3983.5593
190.13449096679688 0 3032.2502
191.11875915527344 0 656.841
193.0986785888672 0 688.2083
193.1083526611328 0 596.8116
194.0928192138672 0 3969.0408
195.0913543701172 0 799.4898
195.1132049560547 0 25959.953
196.10842895507812 0 15541.586
196.1168975830078 0 2217.5747
197.1081085205078 0 1394.3418
197.12876892089844 0 1930.7744
197.1649932861328 0 813.43866
198.08767700195312 0 2814.158
198.1280975341797 0 8202.475
199.07162475585938 0 1574.1251
199.0872039794922 0 1973.183
199.13131713867188 0 1837.7579
200.10313415527344 0 726.8924
200.1425323486328 0 1684.4058
201.1237335205078 0 134461.06 b 1
202.08682250976562 0 1683.722
202.12696838378906 0 12125.675
203.12832641601562 0 1110.2494
204.11322021484375 0 1603.8445
205.09756469726562 0 11223.787
205.10806274414062 0 662.4132
205.1450653076172 0 1984.778
206.10101318359375 0 839.8539
206.12896728515625 0 2024.712
207.160888671875 0 3520.2493
210.12753295898438 0 4193.018
211.10809326171875 0 3144.9885 y Water loss 10
211.13156127929688 0 836.84186
212.09046936035156 0 637.0537
212.13966369628906 0 2961.5713
213.12374877929688 0 10772.373
213.1595001220703 0 1471.4066
215.13943481445312 0 3756.1868
216.09815979003906 0 2644.5308
221.05938720703125 0 2427.9683
221.07235717773438 0 1212.8611
221.08474731445312 0 4769.8887
221.10389709472656 0 2758.9255
221.12831115722656 0 735.4187
221.13970947265625 0 1663.7867
222.08535766601562 0 2208.1104
222.12393188476562 0 4159.5605
223.0640106201172 0 3425.1833
223.08265686035156 0 1337.0625
223.1081085205078 0 9062.64
223.15579223632812 0 44021.06
224.10316467285156 0 3079.942
224.1592254638672 0 4885.4785
225.04330444335938 0 7328.4575
225.12368774414062 0 1359.2712
225.16123962402344 0 668.0013
226.0437469482422 0 2007.8872
226.082763671875 0 8919.999
226.1561279296875 0 785.03235
227.02304077148438 0 879.42456
227.0666961669922 0 8433.065
227.08575439453125 0 963.8738
227.1546630859375 0 1788.4202
228.13475036621094 0 3911.098
229.1187286376953 0 9729.828
230.12240600585938 0 864.35156
230.15037536621094 0 1547.4296
231.0878448486328 0 1023.01935
233.14010620117188 0 6735.1914
234.12420654296875 0 47640.88
234.65052795410156 0 1144.5433
235.12754821777344 0 6336.848
235.60604858398438 0 1360.4401
237.1348419189453 0 2054.8594
238.11898803710938 0 5559.115
238.13027954101562 0 1000.5619
239.09544372558594 0 14266.147
240.09609985351562 0 3787.2407
240.13473510742188 0 18060.773
241.09292602539062 0 2689.5803
241.1185760498047 0 2403.5874
241.13784790039062 0 2380.5437
241.19126892089844 0 2120.836
243.1094970703125 0 4267.7256
244.09326171875 0 19430.63
245.09669494628906 0 2142.915
247.1082000732422 0 26111.896
247.14422607421875 0 1659.7825
247.1566162109375 0 1263.0387
248.1031494140625 0 2669.9587
248.11415100097656 0 9235.666
249.0987091064453 0 15860.13
249.1351776123047 0 4343.726
250.10191345214844 0 1844.4573
250.11866760253906 0 1344.1747
251.1508026123047 0 164438.03
252.13377380371094 0 1515.141
252.15394592285156 0 21300.004
253.11851501464844 0 708.9018
253.15621948242188 0 1533.0533
255.1480255126953 0 9197.871
256.10870361328125 0 1816.6045
256.1512145996094 0 775.4055
256.6111755371094 0 906.1081
257.1134033203125 0 3975.2473
258.145263671875 0 8156.229
261.1196594238281 0 3983.247
261.1346740722656 0 1322.0095
262.1193542480469 0 2217.6702
265.12969970703125 0 18926.168
265.1436462402344 0 1010.44055
265.1539611816406 0 888.81104
266.1252746582031 0 45354.17
267.1286926269531 0 4570.989
268.1302185058594 0 1880.4202
268.17706298828125 0 4037.9158
269.16156005859375 0 2123.0986
269.1862487792969 0 3331.1082
270.1218566894531 0 6709.377
270.1457214355469 0 1044.4563
270.623779296875 0 2150.7954
272.1760559082031 0 8455.499
273.17919921875 0 1123.8682
274.11883544921875 0 8395.394
274.1316833496094 0 3343.0823
275.1036682128906 0 2075.0007
275.1206970214844 0 999.0261
275.15032958984375 0 1008.74133
276.1089782714844 0 2487.627
276.1263732910156 0 1498.8685
276.1557312011719 0 2489.805
279.1458740234375 0 3477.0867
280.129638671875 0 4034.4302
281.05181884765625 0 12013.122
281.1331787109375 0 906.22516
282.1567687988281 0 7639.4976
283.0310974121094 0 1395.9752
283.1405334472656 0 116698.81
284.1037292480469 0 5510.3975
284.1235046386719 0 968.4081
284.1434631347656 0 16060.44
285.01043701171875 0 969.3222
285.1062927246094 0 944.0579
285.1436462402344 0 1358.8219
285.1584777832031 0 1185.489
286.140380859375 0 6693.6953
287.14306640625 0 825.56793
287.1718444824219 0 1742.5576
288.6177673339844 0 1576.6993
289.1216735839844 0 821.98376
290.1670837402344 0 1129.5226 b 4
293.1137390136719 0 66870.99 y Water loss 11
294.11773681640625 0 13396.2705
295.1031799316406 0 1413.7975
295.1195373535156 0 1068.7191
296.10498046875 0 1360.0527
296.197509765625 0 2162.994 b Water loss 2
296.6667175292969 0 1004.4825
297.08319091796875 0 2031.9806
297.13568115234375 0 1407.5125 y Water loss 8
297.1562805175781 0 21079.316
298.1394958496094 0 2010.4225 y Water loss 6
298.15924072265625 0 3614.2346
299.062255859375 0 86371.48
300.0625915527344 0 1868.514
300.1680908203125 0 12251.259
301.0634460449219 0 1284.4789
301.1512451171875 0 61104.69
302.1141357421875 0 10927.018
302.1544189453125 0 7615.4917
303.1170349121094 0 1275.6858
305.1793212890625 0 1989.2097
305.64569091796875 0 3388.1597
306.13787841796875 0 11259.35 y 8
306.63916015625 0 5445.316
307.1401672363281 0 1400.4714
309.16748046875 0 1024.3795
310.1545715332031 0 803.085
311.124755859375 0 10894.18 y 11
311.137939453125 0 2229.6328
312.1265869140625 0 1758.9198
314.18701171875 0 1982.0403
314.2082824707031 0 4935.0483 b 2
315.16693115234375 0 35991.98
315.2107849121094 0 1075.5618
316.1704406738281 0 5772.2695
317.6587829589844 0 1237.0149
319.1518859863281 0 1551.4375
325.1508483886719 0 2746.7524
326.6643981933594 0 7661.363
327.1650390625 0 3388.7104
327.666748046875 0 1314.2164
332.2088623046875 0 3638.2734
333.17730712890625 0 6513.7266
334.1850280761719 0 1134.2422
335.207275390625 0 1343.4669
338.18267822265625 0 1133.0931
338.67437744140625 0 2261.182 b Water loss 5
340.6615295410156 0 2217.8657
341.0168762207031 0 1231.17
341.1641845703125 0 1745.2687
342.1820373535156 0 2707.425
343.16192626953125 0 3537.4468
343.1835632324219 0 963.03235
344.1714782714844 0 2374.0122
344.9769592285156 0 11782.902
345.1312561035156 0 4774.7314
346.11505126953125 0 17288.062
346.6983947753906 0 1461.5779
347.11822509765625 0 2567.7317
347.6800537109375 0 9952.201 b 5
348.181640625 0 2246.8599
350.21923828125 0 12838.136
351.1657409667969 0 945.5396
351.2209777832031 0 2626.442
352.671142578125 0 984.53723
353.6752014160156 0 2630.8513 y Water loss 7
354.1757507324219 0 1086.0375
355.0705261230469 0 1406.5674
356.1947326660156 0 1221.9929
357.1731262207031 0 917.2555
359.0286865234375 0 9928.546
360.2027282714844 0 1069.5807
360.69586181640625 0 4415.2104
361.1815490722656 0 735.8945
361.1980895996094 0 4450.495
362.1847229003906 0 5578.32
362.67962646484375 0 21874.56 y 7
363.1414489746094 0 26905.936
363.1816101074219 0 8920.823
363.68280029296875 0 3074.9243
364.124755859375 0 5214.5195
364.1455383300781 0 5516.9927
366.1781311035156 0 2685.9248
367.1401062011719 0 1684.144
367.1626281738281 0 1855.5101
367.24578857421875 0 2529.0715
367.6856384277344 0 1238.4989
368.1663513183594 0 1128.1968
368.1915588378906 0 1178.4526
369.12310791015625 0 1070.4965
369.17529296875 0 1103.3706
370.123291015625 0 965.1128
373.18365478515625 0 934.09265
375.2005920410156 0 904.78644
376.162109375 0 1707.9114
376.1983337402344 0 4052.0044
377.14532470703125 0 1826.5375
377.26708984375 0 1079.2366
378.2139587402344 0 9325.7705
379.20953369140625 0 45318.137
380.1651611328125 0 1169.1443
380.21185302734375 0 9663.879
381.1522521972656 0 81084.414
381.71240234375 0 2165.5576
382.15496826171875 0 14624.233
382.18768310546875 0 2790.3833
382.2137145996094 0 1193.1495
383.15704345703125 0 1618.9525
384.1664123535156 0 2831.2922
384.1886291503906 0 3085.418
385.1509704589844 0 5669.5664
385.17291259765625 0 1931.6393 y 4
386.178955078125 0 910.4105
389.19329833984375 0 1582.0292 b 9
390.1965637207031 0 3664.3745
391.13677978515625 0 2876.3362
391.18389892578125 0 1570.0464
392.2082824707031 0 1591.4364
394.1492614746094 0 1322.1206
394.1729431152344 0 10316.981
395.13494873046875 0 4185.4434
395.1572265625 0 2506.6997
395.18035888671875 0 1308.0968
395.24114990234375 0 1739.2191
396.2247314453125 0 15359.546
397.2275390625 0 2860.6497
398.17730712890625 0 5493.336
398.737548828125 0 1393.63
399.2393493652344 0 914.6188
401.2142639160156 0 995.4579
402.17694091796875 0 3712.621
402.1993713378906 0 6442.6714
403.19964599609375 0 2340.5684
404.2058410644531 0 1033.2229
404.7109680175781 0 1272.7458
405.2101745605469 0 841.0456
405.2613830566406 0 2761.5774
406.2452392578125 0 2062.6284
407.2037353515625 0 1051.4425
407.24365234375 0 1114.4965
409.1473083496094 0 6374.093
409.20513916015625 0 1898.2244
409.7008972167969 0 1040.0475
411.1993408203125 0 5106.958
412.1834716796875 0 47446.664
413.1451110839844 0 5192.964
413.16253662109375 0 2921.5027
413.1868896484375 0 7484.2935
413.2496032714844 0 1567.5194
414.14678955078125 0 1218.0779
414.2354736328125 0 13186.924
415.0372314453125 0 21571.25
415.1982421875 0 3616.3218
415.2391662597656 0 2877.4722
416.2008361816406 0 1204.2274
417.7171630859375 0 10209.638
418.21051025390625 0 13229.32
418.71099853515625 0 6109.433
419.21185302734375 0 1645.2509
421.208740234375 0 2360.721 y Water loss 10
422.19268798828125 0 1261.666 y Ammonia loss 10
422.2541198730469 0 880.06683
422.7098083496094 0 1760.3027
423.2012939453125 0 2874.805
423.2720642089844 0 18338.758
423.7037353515625 0 1723.6923
423.7326965332031 0 926.4551
424.27459716796875 0 5226.641
424.7338562011719 0 1470.3387
425.2746887207031 0 907.7457
426.72247314453125 0 130729.01
427.22381591796875 0 65235.03
427.7252197265625 0 17588.488
428.2256774902344 0 3320.667
429.0882568359375 0 1290.3757
429.2099304199219 0 26671.871
430.19415283203125 0 121285.95
431.19720458984375 0 25907.252
431.7145690917969 0 14228.095 b Water loss 6
432.2132263183594 0 6780.317 b Ammonia loss 6
432.7112731933594 0 3993.2583
433.0466613769531 0 1051.6028
433.20697021484375 0 1509.6208
433.2564392089844 0 4886.2446 b Water loss 3
434.2604064941406 0 1144.8733
438.17425537109375 0 2761.733
438.7320556640625 0 1228.6401
439.2196044921875 0 34472.65 y 10
440.18121337890625 0 1563.4625
440.2223815917969 0 8359.271
440.71990966796875 0 64045.047 b 6
441.22149658203125 0 32283.688
441.7225341796875 0 9594.716
442.22406005859375 0 1639.8827
443.23907470703125 0 1246.0997
444.1815490722656 0 863.2126
446.7414855957031 0 1233.3821
447.2068786621094 0 4451.597 y Ammonia loss 6
447.70489501953125 0 1349.7245
448.2049560546875 0 6523.6274
449.2064514160156 0 1424.5099
450.23583984375 0 1602.1708
450.2825927734375 0 1853.475
451.26702880859375 0 37740.516 b 3
452.27044677734375 0 10106.334
453.2734680175781 0 1430.3463
455.2646179199219 0 1463.2045
455.7191467285156 0 2103.2312 y 6
456.2225036621094 0 2102.1582
459.1994934082031 0 3321.2253
460.753662109375 0 3148.861
461.2046203613281 0 3318.8438
461.2525634765625 0 4017.7297
462.2069396972656 0 984.7635
462.2557678222656 0 1123.6077
464.2138977050781 0 1096.1826
465.74560546875 0 949.67377
466.16778564453125 0 1168.3604
466.2436828613281 0 873.9905
468.2539978027344 0 1129.3855
468.2935485839844 0 9060.382
469.2964172363281 0 2169.5173
472.2196350097656 0 1001.39435
474.25897216796875 0 8220.935
474.75433349609375 0 9329.255
475.2548522949219 0 4614.526
475.7545471191406 0 1187.7013
476.2257385253906 0 3075.5103
477.2108459472656 0 7228.1294
478.229248046875 0 6178.8228 y Water loss 9
478.27752685546875 0 4591.065
479.2356872558594 0 1764.7113
479.744140625 0 1437.3752
480.2251281738281 0 1053.3228
483.2644958496094 0 26400.982
483.76605224609375 0 16712.807
484.22161865234375 0 1374.9557
484.26715087890625 0 3977.0781
484.7692565917969 0 1507.4052
488.2573547363281 0 7700.015 b Water loss 7
488.75543212890625 0 3988.0015 b Ammonia loss 7
489.2544250488281 0 1892.1659
494.2364196777344 0 82707.81
495.2015686035156 0 840.32715
495.23944091796875 0 22286.078
496.2409362792969 0 97043.55 y 9
497.2464294433594 0 25201.246
497.763427734375 0 8317.625
498.24395751953125 0 4965.0073
503.1081848144531 0 23065.834
504.20208740234375 0 3396.7366
506.2259826660156 0 1958.0529
507.2611389160156 0 1357.6106
510.2628479003906 0 799.44696
511.2628479003906 0 5106.151
512.2650146484375 0 1344.8857
515.2842407226562 0 1316.8552
519.1394653320312 0 7093.146
522.2293090820312 0 1118.7424
523.2579956054688 0 2620.7979
523.5887451171875 0 1934.5999
524.2821044921875 0 1195.926
525.2667846679688 0 13484.517
526.2301025390625 0 2990.206
526.2686767578125 0 3089.0098
529.260986328125 0 4633.822 Precursor Water loss
529.5952758789062 0 5376.1147 Precursor Ammonia loss
529.9255981445312 0 2756.7024
531.2216186523438 0 1087.2583
532.1944580078125 0 3700.687
533.2447509765625 0 1353.6124
534.8868408203125 0 1992.0455
535.2639770507812 0 5122.7246 Precursor
535.5971069335938 0 2912.1382
535.9346313476562 0 1830.2985
536.166748046875 0 4729.506
537.2438354492188 0 1805.1626
537.7359008789062 0 2017.9132
538.2730712890625 0 1091.0101
538.77880859375 0 1336.6794
539.2366943359375 0 14078.367
540.2396240234375 0 4426.718
541.254638671875 0 1412.3044
542.2936401367188 0 4725.5376
543.277587890625 0 26303.865
544.2808227539062 0 9097.866
545.2809448242188 0 2021.1154
545.7644653320312 0 947.20044 b Water loss 8
546.25 0 4721.6953
546.7520141601562 0 1850.819
547.2516479492188 0 1015.9165
547.7769775390625 0 956.36957
548.2379760742188 0 1323.6646
549.2207641601562 0 3124.749
550.2072143554688 0 1526.2018
551.2581176757812 0 3830.2283
551.3299560546875 0 2111.9424
552.2581787109375 0 1917.5328
553.2633666992188 0 1244.8809
554.7739868164062 0 3477.6118 b 8
555.2564086914062 0 6402.0557
555.7569580078125 0 5645.3374
556.265869140625 0 797.75146
560.2483520507812 0 1270.318
561.3152465820312 0 3542.3923 b Water loss 4
562.3016967773438 0 1218.2703 b Ammonia loss 4
563.2578125 0 3483.8142
564.2601928710938 0 2208.1052
564.33544921875 0 780.72876
565.2634887695312 0 5852.0215
566.2647094726562 0 1562.0532
567.2313842773438 0 16099.674
567.2802124023438 0 938.60046
568.2347412109375 0 5199.4185
569.2415161132812 0 852.69037
569.8236083984375 0 784.9426
575.2471313476562 0 4020.308
576.2332153320312 0 2934.6484
579.3251953125 0 25514.217 b 4
580.3287353515625 0 7366.01
581.3314819335938 0 1830.9584
587.3340454101562 0 813.88873
592.2730712890625 0 3424.3796
592.3252563476562 0 3493.1714
593.2568359375 0 31537.45 y Water loss 8
593.30615234375 0 11618.547
594.2547607421875 0 8577.438
594.3073120117188 0 3151.767
594.7855224609375 0 1206.1965
595.25 0 2639.7988
596.3516845703125 0 1416.9683
597.7979736328125 0 1337.1938
600.3145751953125 0 1424.4491
602.793212890625 0 4763.014
603.2907104492188 0 5696.3457
603.3605346679688 0 813.8512
603.7915649414062 0 1903.4326
610.2835693359375 0 61602.3
610.3287353515625 0 2222.333
611.2682495117188 0 318883.78 y 8
611.7976684570312 0 16069.522
612.271484375 0 98694.65
612.8013916015625 0 4353.409
613.2733764648438 0 22384.09
614.275146484375 0 2938.7925
615.3176879882812 0 1106.2161
616.3006591796875 0 1141.659
617.2811889648438 0 1895.9087
619.2676391601562 0 865.4591
621.251953125 0 4078.5833
621.299560546875 0 1364.7186
622.2566528320312 0 1108.9961
634.3041381835938 0 3181.4485
635.2962036132812 0 4828.7725
636.3238525390625 0 3168.7659
636.7861328125 0 3476.163 y Water loss 3
637.2888793945312 0 4357.5767 y Ammonia loss 3
637.78662109375 0 1572.9272
638.3334350585938 0 1149.8179 b Water loss 10
640.27490234375 0 2677.5186
641.2759399414062 0 1249.7673
645.2973022460938 0 3820.1504
645.79296875 0 20860.723 y 3
646.2940673828125 0 13788.938
646.7947387695312 0 7296.596
647.2901000976562 0 1363.1769
648.3404541015625 0 1574.6715
650.2684326171875 0 1842.4385
650.3253173828125 0 1090.5322
651.2740478515625 0 1107.5215
652.3206787109375 0 33159.58
652.8245849609375 0 1159.6952
653.32373046875 0 12830.847
654.3252563476562 0 2188.0532
657.3602294921875 0 2021.6633
658.2830810546875 0 2050.228
658.33642578125 0 1008.83527
659.3182983398438 0 1241.2316
660.8706665039062 0 1839.0955
661.3261108398438 0 2211.4219
661.8289184570312 0 1320.3104
662.3056030273438 0 12790.13
662.8306274414062 0 929.84106
663.2957763671875 0 8104.335
664.295166015625 0 2774.205
665.3456420898438 0 1086.1282
666.3561401367188 0 1569.8293
668.2811889648438 0 1668.1078
676.3417358398438 0 17966.758 b Water loss 5
677.3319091796875 0 8642.967 b Ammonia loss 5
678.3360595703125 0 3794.3918
679.3301391601562 0 3358.0508
679.8316040039062 0 1091.0181
680.3157348632812 0 58952.23
681.318359375 0 24340.47
682.3204345703125 0 4592.4463
684.3223876953125 0 2155.2354
684.8176879882812 0 3241.8489
685.3153076171875 0 2685.4897
685.8195190429688 0 1386.1031
688.3336791992188 0 1269.2827
692.2831420898438 0 959.31494
692.8359985351562 0 2715.7212
693.3294677734375 0 16537.697 y Water loss 2
693.8275146484375 0 18904.584 y Ammonia loss 2
694.3517456054688 0 166421.4 b 5
694.8271484375 0 2845.1042
695.354736328125 0 63218.1
696.3568725585938 0 14240.719
697.3515014648438 0 2678.1885
698.3575439453125 0 1852.2484
700.89404296875 0 1240.3636
701.8414306640625 0 22617.705
702.3348388671875 0 115138.66 y 2
702.8359375 0 98048.32 b Water loss 11
703.337646484375 0 44145.887 b Ammonia loss 11
703.8394165039062 0 12704.667
704.3385009765625 0 5387.782
705.358154296875 0 2234.7043
706.3424072265625 0 9844.453 y Water loss 7
707.34326171875 0 4259.7266
709.3150024414062 0 921.659
711.3712768554688 0 6664.449
711.8572998046875 0 8695.627 b 11
712.3583984375 0 7959.3164
712.8592529296875 0 3124.7134
719.391357421875 0 1069.9177
720.3846435546875 0 1129.325
720.8611450195312 0 1124.2526
721.3634033203125 0 7277.0835
722.3648681640625 0 2674.2336
723.3679809570312 0 16063.704
724.3523559570312 0 95645.664 y 7
725.3553466796875 0 38731.03
726.3580322265625 0 11159.033
727.3578491210938 0 1614.3843
728.3726196289062 0 888.7997
729.3594360351562 0 4205.9756
730.3587646484375 0 2315.6929
733.3636474609375 0 4024.9958
734.3530883789062 0 3440.9937
737.8638305664062 0 1193.283
738.3634643554688 0 1302.2968
739.355224609375 0 1105.5901
742.8585205078125 0 1083.8541 y Water loss 1
743.367431640625 0 2844.483 y Ammonia loss 1
744.3560180664062 0 1107.7196
745.364013671875 0 1104.2567
747.3890380859375 0 2105.8162
748.3796997070312 0 1981.649
749.3306884765625 0 1391.6677
751.3768310546875 0 20299.156
751.8693237304688 0 10457.73 y 1
752.3735961914062 0 17594.992
752.8712158203125 0 3830.4717
753.3765258789062 0 3621.6233
754.3790893554688 0 1046.049
760.3064575195312 0 1853.8425
761.3726806640625 0 1967.8796
762.4152221679688 0 2333.7732
763.3658447265625 0 9421.984
764.3655395507812 0 5317.7134
765.4000854492188 0 4029.815
766.4072875976562 0 1786.411
767.3479614257812 0 4488.6143
768.3414306640625 0 1168.8715
769.3995361328125 0 1922.5521
776.3787231445312 0 2603.3481
779.3836669921875 0 9375.878
780.385009765625 0 4467.137
781.3643188476562 0 4533.1143
782.3717651367188 0 1479.5645
793.3994140625 0 17487.64 Precursor Water loss
794.4022827148438 0 8703.791
795.3412475585938 0 6226.7075
795.4085083007812 0 1408.1593
796.3405151367188 0 1517.861
796.468505859375 0 4651.0117
797.4722290039062 0 1501.4993
807.3785400390625 0 3880.4717
807.4453735351562 0 2194.2104
808.3826904296875 0 2493.1365
818.44189453125 0 2660.5557
819.4409790039062 0 2051.3367
825.3867797851562 0 1624.0992
826.3762817382812 0 1908.8374
827.3750610351562 0 1120.7592
828.4361572265625 0 1340.2349
834.4263305664062 0 4722.374
835.4141235351562 0 6118.542
836.4487915039062 0 11680.796
837.4532470703125 0 4698.948
838.4530639648438 0 2486.6875
843.4004516601562 0 2132.0056
844.3889770507812 0 4327.1743
845.3938598632812 0 3954.7764
846.4464111328125 0 6845.0312
847.4450073242188 0 5097.8975
848.4458618164062 0 1247.1047
852.436279296875 0 42002.027
853.4389038085938 0 19932.438
854.441650390625 0 6018.204
857.4461669921875 0 963.93274
862.4207153320312 0 25161.646 b Water loss 6
863.4166259765625 0 18585.193 b Ammonia loss 6
864.453857421875 0 27566.133
865.4584350585938 0 14559.622
866.4617919921875 0 5111.808
876.435302734375 0 4322.149
877.4346313476562 0 1343.2958
878.4368896484375 0 1275.2078
880.4307861328125 0 197536.7 b 6
881.433349609375 0 102810.54
882.4365844726562 0 29986.793
883.4387817382812 0 6238.448
884.4345092773438 0 1302.8961
890.4207153320312 0 1734.131
891.4127197265625 0 2257.266
892.46826171875 0 2663.0227
893.4719848632812 0 1287.3727
894.4432373046875 0 1197.8921
907.442138671875 0 2275.8655
908.4265747070312 0 14413.575
909.4324951171875 0 8984.065
910.4319458007812 0 17211.502 y 6
911.4334106445312 0 7636.333
912.4358520507812 0 2365.735
936.4158935546875 0 1054.3966
937.4164428710938 0 2910.0125
947.4361572265625 0 1378.0979
954.431884765625 0 4273.015
955.42138671875 0 4942.1943
956.4210815429688 0 2851.5276
964.4603271484375 0 1278.7819
965.4484252929688 0 6231.0596
966.4486083984375 0 2482.6846
972.4446411132812 0 3035.4854
973.4425048828125 0 2673.7349
975.5046997070312 0 3304.355 b Water loss 7
976.5028686523438 0 1902.8713 b Ammonia loss 7
980.458984375 0 1654.2672
982.4724731445312 0 1648.762
990.4514770507812 0 1106.4436
993.5148315429688 0 10005.013 b 7
994.5172119140625 0 7558.4365
995.5236206054688 0 1164.7736
1007.4766845703125 0 1570.3733
1008.5011596679688 0 1096.7313
1024.4796142578125 0 1813.26
1025.458251953125 0 8183.039 y 5
1026.4598388671875 0 4823.971
1027.4625244140625 0 1428.0602
1064.5174560546875 0 1282.486
1075.531982421875 0 1323.8494
1077.531494140625 0 1231.729
1093.5400390625 0 4226.4985
1094.55029296875 0 2759.5508
1095.5518798828125 0 1122.641
1108.5411376953125 0 2124.0352 b 8
1109.5313720703125 0 1485.422
1117.4962158203125 0 1491.1777
1118.48681640625 0 2147.119
1135.5067138671875 0 6228.0283 y Water loss 4
1136.4959716796875 0 10536.762 y Ammonia loss 4
1137.496337890625 0 4999.9473
1138.49462890625 0 1585.589
1152.5323486328125 0 3930.1401
1153.51708984375 0 15850.836 y 4
1154.5198974609375 0 10084.531
1155.523681640625 0 3795.4812
1156.530029296875 0 925.0428
1165.5692138671875 0 2265.6182 b 9
1204.5731201171875 0 882.375
1290.56982421875 0 973.7044 y 3
3081.43359375 0 963.4725
3454.69873046875 0 1132.4221

Spectrum Details

|  |  |
| --- | --- |
| Matched peaks? Matched peaks The total absolute number of peaks matched. Additionally in brackets the total fraction of peaks matched and the total number of peaks is shown. | 78 (9.36% of 833) |
| FDR? FDR The false discovery rate estimated for this peptide. It is calculated by matching all theoretical fragments with a non-integer shift with the raw peaks for this spectrum. This is done with 40 different shifts. The resulting percentage is the average number of annotated peaks over the number of annotated peaks with the correct spectrum. | 0.73% |
| Satellite FDR? Satellite FDR See the FDR for details on its calculation. This satellite ion specific FDR only contains the satellite ions (d/w) for I/L/J positions. | ∞ |
| PSM Score? PSM Score The PSM Score as given by Hecklib to this annotated spectrum. It is shown with three significant figures. | 604 |

#### Spectrum 6887? Spectrum 6887 The raw spectrum of this peptide as annotated by Hecklib. The fragments are coloured according to ion type (see legend). Any peaks with a star '\*' as text can be hovered over to see the full details, first the ion type second the mass shift type. By hovering over the amino acids in the peptide or ions in the legend the corresponding peaks are highlighted. By toggling the 'Unassigned' label you can turn the background (unassigned) peaks on or off in the plot. By updating the slider in the Ion legend you can update the spectrum to only show the top X% of the peaks with labels. The top X% means any peak that is within X% of the highest intensity. By dragging in the spectrum you can zoom in to a specific part of the spectrum and use 'Zoom Out' to get back to the original zoom level. The annotation of the spectrum is based on the given sequence in the peptides file and is done with different software so inconsistencies are likely. The peaks are annotated based on the given sequence, with 20 ppm tolerance.

Copy Data

##### Spectrum 6887 (TSV)

###### Preview

```
Loading example...
```

*Click on the button to copy the data to your clipboard.*

Mz MinMz MaxIntensity Max

WidthHeightPeptide font sizePeptide stroke widthSpectrum font sizeSpectrum stroke widthCompact peptide

Ion legend

wxyz

abcd

OtherUnassignedIon^Charge^_Position_Show for top:%

TVLHQDWLDGKEY

08.08e+61.62e+72.42e+73.23e+7

Zoom Out

y^+1^_1_c^+1^_2_c^+1^_2_y^+1^_2_y^+2^_5_y^+2^_5_y^+1^_2_c^+1^_3_c^+2^_6_y^+2^_6_y^+2^_6_c^+3^_10_y^+1^_3_z^+1^_3_y^+1^_3_c^+2^_7_c^+2^_7_c^+1^_4_c^+1^_4_z^+1^_4_y^+1^_4_w^+1^_5_z^+1^_5_c^+1^_5_y^+1^_5_z^+1^_5_c^+1^_5_y^+1^_5_y^+2^_10_z^+2^_10_y^+2^_10_c^+2^_11_c^+2^_11_c^+2^_11_w^+1^_6_w^+2^_11_y^+2^_11_y^+2^_11_c^+1^_6_y^+2^_11_y^+1^_6_z^+1^_6_c^+1^_6_c^+2^_12_c^+2^_12_y^+1^_6_w^+2^_12_y^+2^_12_z^+2^_12_y^+2^_12_c^+1^_7_y^+1^_7_z^+1^_7_c^+1^_7_y^+1^_7_c^+1^_8_c^+1^_8_z^+1^_8_c^+1^_8_y^+1^_8_w^+1^_9_c^+1^_9_c^+1^_9_z^+1^_9_y^+1^_9_c^+1^_10_y^+1^_10_z^+1^_10_y^+1^_10_c^+1^_11_c^+1^_11_w^+1^_11_y^+1^_11_z^+1^_11_y^+1^_11_c^+1^_12_c^+1^_12_z^+1^_12_

0881176126423523

Fragment Matches Table

Show background peaks

| Position | Ion type | Intensity | mz Theoretical | mz Error (Th) | mz Error (ppm) | Charge | Series Number |
| --- | --- | --- | --- | --- | --- | --- | --- |
| - | - | 3.91E+04 | 121.1 | - | - | 0 | - |
| - | - | 4.104E+04 | 127.2 | - | - | 0 | - |
| - | - | 9.743E+04 | 128.1 | - | - | 0 | - |
| - | - | 5.584E+05 | 129.1 | - | - | 0 | - |
| - | - | 1.056E+06 | 130.1 | - | - | 0 | - |
| - | - | 9.327E+04 | 131.1 | - | - | 0 | - |
| - | - | 1.54E+05 | 131.1 | - | - | 0 | - |
| - | - | 3.896E+04 | 134 | - | - | 0 | - |
| - | - | 1.295E+05 | 136.1 | - | - | 0 | - |
| - | - | 3.752E+04 | 138.1 | - | - | 0 | - |
| - | - | 4.628E+04 | 138.1 | - | - | 0 | - |
| - | - | 4.575E+04 | 140.6 | - | - | 0 | - |
| - | - | 3.849E+04 | 141.3 | - | - | 0 | - |
| - | - | 3.903E+04 | 148.8 | - | - | 0 | - |
| - | - | 4.968E+04 | 150.1 | - | - | 0 | - |
| - | - | 3.578E+04 | 152 | - | - | 0 | - |
| - | - | 2.303E+06 | 155.1 | - | - | 0 | - |
| - | - | 1.433E+05 | 156.1 | - | - | 0 | - |
| - | - | 3.954E+04 | 156.6 | - | - | 0 | - |
| - | - | 2.377E+05 | 157.1 | - | - | 0 | - |
| - | - | 9.913E+05 | 159.1 | - | - | 0 | - |
| - | - | 3.998E+04 | 161.3 | - | - | 0 | - |
| - | - | 8.433E+04 | 165.1 | - | - | 0 | - |
| - | - | 8.272E+04 | 171.1 | - | - | 0 | - |
| - | - | 3.864E+06 | 173.1 | - | - | 0 | - |
| - | - | 3.661E+05 | 174.1 | - | - | 0 | - |
| 13 | y | 7.621E+05 | 182.1 | 0.0004343 | 2.385 | +1 | 1 |
| - | - | 6.496E+04 | 183.1 | - | - | 0 | - |
| - | - | 6.473E+04 | 183.1 | - | - | 0 | - |
| - | - | 7.273E+04 | 184.1 | - | - | 0 | - |
| - | - | 2.627E+05 | 186.1 | - | - | 0 | - |
| - | - | 4.338E+04 | 186.6 | - | - | 0 | - |
| - | - | 7.685E+04 | 187.1 | - | - | 0 | - |
| - | - | 1.458E+05 | 188.1 | - | - | 0 | - |
| - | - | 4.534E+04 | 194.1 | - | - | 0 | - |
| - | - | 4.863E+04 | 196 | - | - | 0 | - |
| - | - | 9.743E+04 | 198.1 | - | - | 0 | - |
| 2 | c | 6.688E+04 | 200.1 | 0.0002036 | 1.017 | +1 | 2 |
| - | - | 2.723E+06 | 201.1 | - | - | 0 | - |
| - | - | 2.939E+05 | 202.1 | - | - | 0 | - |
| - | - | 4.873E+04 | 205.7 | - | - | 0 | - |
| - | - | 9.101E+04 | 207.1 | - | - | 0 | - |
| - | - | 9.82E+04 | 213.1 | - | - | 0 | - |
| - | - | 1.22E+06 | 215.1 | - | - | 0 | - |
| - | - | 5.699E+04 | 216.1 | - | - | 0 | - |
| 2 | c | 1.025E+06 | 218.1 | 0.000549 | 2.516 | +1 | 2 |
| - | - | 6.61E+04 | 219.2 | - | - | 0 | - |
| - | - | 1.533E+05 | 223.2 | - | - | 0 | - |
| - | - | 6.577E+04 | 225.1 | - | - | 0 | - |
| - | - | 2.345E+05 | 226.1 | - | - | 0 | - |
| - | - | 1.205E+05 | 228.1 | - | - | 0 | - |
| - | - | 5.742E+04 | 234.1 | - | - | 0 | - |
| - | - | 6.633E+04 | 240.1 | - | - | 0 | - |
| - | - | 2.253E+05 | 243.1 | - | - | 0 | - |
| - | - | 4.084E+05 | 245.1 | - | - | 0 | - |
| - | - | 6.473E+04 | 247.1 | - | - | 0 | - |
| - | - | 7.942E+04 | 251.1 | - | - | 0 | - |
| - | - | 6.338E+05 | 251.2 | - | - | 0 | - |
| - | - | 1.577E+06 | 258.1 | - | - | 0 | - |
| - | - | 1.054E+05 | 258.2 | - | - | 0 | - |
| - | - | 1.764E+05 | 259.1 | - | - | 0 | - |
| - | - | 3.233E+05 | 260.2 | - | - | 0 | - |
| - | - | 7.226E+04 | 266.1 | - | - | 0 | - |
| - | - | 6.845E+04 | 267.7 | - | - | 0 | - |
| - | - | 1.503E+06 | 268.2 | - | - | 0 | - |
| - | - | 1.182E+05 | 269.2 | - | - | 0 | - |
| - | - | 7.732E+04 | 272.2 | - | - | 0 | - |
| - | - | 3.688E+05 | 283.1 | - | - | 0 | - |
| - | - | 8.538E+04 | 285.1 | - | - | 0 | - |
| - | - | 9.925E+04 | 287.2 | - | - | 0 | - |
| - | - | 1.106E+05 | 288.2 | - | - | 0 | - |
| - | - | 9.485E+04 | 288.6 | - | - | 0 | - |
| 12 | y | 6.172E+05 | 293.1 | 0.0005104 | 1.741 | +1 | 2 |
| - | - | 1.29E+05 | 294.1 | - | - | 0 | - |
| 9 | y | 5.35E+04 | 297.1 | 0.0005852 | 1.969 | +2 | 5 |
| - | - | 1.013E+05 | 299.1 | - | - | 0 | - |
| - | - | 2.09E+05 | 300.2 | - | - | 0 | - |
| - | - | 3.315E+05 | 301.2 | - | - | 0 | - |
| - | - | 6.717E+04 | 302.2 | - | - | 0 | - |
| - | - | 4.98E+04 | 305.5 | - | - | 0 | - |
| - | - | 2.066E+05 | 305.6 | - | - | 0 | - |
| 9 | y | 1.972E+05 | 306.1 | 0.001162 | 3.796 | +2 | 5 |
| - | - | 5.865E+04 | 306.2 | - | - | 0 | - |
| - | - | 1.128E+05 | 306.6 | - | - | 0 | - |
| - | - | 1.706E+05 | 307.2 | - | - | 0 | - |
| - | - | 6.018E+04 | 308.2 | - | - | 0 | - |
| 12 | y | 3.452E+05 | 311.1 | 0.0006574 | 2.113 | +1 | 2 |
| - | - | 3.825E+05 | 314.2 | - | - | 0 | - |
| - | - | 2.116E+05 | 315.2 | - | - | 0 | - |
| - | - | 8.879E+04 | 315.2 | - | - | 0 | - |
| - | - | 2.76E+06 | 317.2 | - | - | 0 | - |
| - | - | 3.909E+05 | 318.2 | - | - | 0 | - |
| - | - | 5.266E+05 | 320.2 | - | - | 0 | - |
| - | - | 1.012E+05 | 321.2 | - | - | 0 | - |
| 3 | c | 1.498E+06 | 331.2 | 0.0007593 | 2.292 | +1 | 3 |
| - | - | 2.289E+05 | 332.2 | - | - | 0 | - |
| - | - | 7.331E+04 | 342.1 | - | - | 0 | - |
| - | - | 1.399E+05 | 343.1 | - | - | 0 | - |
| 6 | c | 2.468E+05 | 347.7 | 0.0001777 | 0.5111 | +2 | 6 |
| - | - | 5.873E+04 | 348.2 | - | - | 0 | - |
| - | - | 6.06E+04 | 350.2 | - | - | 0 | - |
| 8 | y | 6.826E+04 | 353.7 | 0.001308 | 3.699 | +2 | 6 |
| - | - | 1.172E+05 | 357.2 | - | - | 0 | - |
| - | - | 1.489E+05 | 362.2 | - | - | 0 | - |
| 8 | y | 6.29E+05 | 362.7 | 0.0007257 | 2.001 | +2 | 6 |
| - | - | 2.035E+05 | 363.2 | - | - | 0 | - |
| - | - | 3.824E+05 | 364.2 | - | - | 0 | - |
| - | - | 9.928E+04 | 365.2 | - | - | 0 | - |
| - | - | 3.263E+05 | 374.2 | - | - | 0 | - |
| - | - | 8.546E+04 | 375.2 | - | - | 0 | - |
| - | - | 1.953E+05 | 378.2 | - | - | 0 | - |
| - | - | 1.436E+05 | 379.2 | - | - | 0 | - |
| - | - | 1.013E+05 | 381.2 | - | - | 0 | - |
| 10 | c | 1.174E+05 | 389.2 | 0.002487 | 6.391 | +3 | 10 |
| - | - | 5.301E+04 | 396.2 | - | - | 0 | - |
| - | - | 8.876E+04 | 406.2 | - | - | 0 | - |
| - | - | 7.232E+04 | 408.3 | - | - | 0 | - |
| - | - | 1.19E+05 | 412.2 | - | - | 0 | - |
| - | - | 7.441E+04 | 417.2 | - | - | 0 | - |
| - | - | 1.094E+05 | 417.7 | - | - | 0 | - |
| - | - | 1.516E+05 | 418.2 | - | - | 0 | - |
| - | - | 6.897E+04 | 420.2 | - | - | 0 | - |
| 11 | y | 7.063E+04 | 421.2 | 0.003051 | 7.244 | +1 | 3 |
| 11 | z | 3.629E+05 | 423.2 | 0.00114 | 2.693 | +1 | 3 |
| - | - | 1.343E+06 | 423.3 | - | - | 0 | - |
| - | - | 8.194E+04 | 424.2 | - | - | 0 | - |
| - | - | 3.604E+05 | 424.3 | - | - | 0 | - |
| - | - | 2.283E+06 | 426.7 | - | - | 0 | - |
| - | - | 1.292E+06 | 427.2 | - | - | 0 | - |
| - | - | 4.065E+05 | 427.7 | - | - | 0 | - |
| - | - | 6.747E+04 | 428.2 | - | - | 0 | - |
| - | - | 3.616E+05 | 429.2 | - | - | 0 | - |
| - | - | 5.434E+05 | 430.2 | - | - | 0 | - |
| - | - | 1.212E+05 | 431.2 | - | - | 0 | - |
| - | - | 3.258E+05 | 431.7 | - | - | 0 | - |
| - | - | 2.513E+05 | 432.2 | - | - | 0 | - |
| - | - | 6.764E+04 | 432.7 | - | - | 0 | - |
| - | - | 8.737E+04 | 438.7 | - | - | 0 | - |
| 11 | y | 5.001E+05 | 439.2 | 0.001275 | 2.904 | +1 | 3 |
| 7 | c | 1.044E+05 | 440.2 | 0.004838 | 10.99 | +2 | 7 |
| 7 | c | 1.629E+06 | 440.7 | 0.000957 | 2.171 | +2 | 7 |
| - | - | 9.459E+05 | 441.2 | - | - | 0 | - |
| - | - | 1.912E+05 | 441.7 | - | - | 0 | - |
| - | - | 6.663E+04 | 448.2 | - | - | 0 | - |
| 4 | c | 2.263E+05 | 450.3 | 0.001393 | 3.093 | +1 | 4 |
| - | - | 3.186E+06 | 451.3 | - | - | 0 | - |
| - | - | 6.773E+05 | 452.3 | - | - | 0 | - |
| - | - | 1.011E+05 | 453.3 | - | - | 0 | - |
| 4 | c | 1.182E+07 | 468.3 | 0.0009905 | 2.115 | +1 | 4 |
| - | - | 3.012E+06 | 469.3 | - | - | 0 | - |
| - | - | 4.567E+05 | 470.3 | - | - | 0 | - |
| - | - | 1.967E+05 | 474.3 | - | - | 0 | - |
| - | - | 1.618E+05 | 474.8 | - | - | 0 | - |
| - | - | 7.743E+04 | 475.3 | - | - | 0 | - |
| 10 | z | 1.173E+06 | 480.2 | 0.001099 | 2.289 | +1 | 4 |
| - | - | 3.858E+05 | 481.2 | - | - | 0 | - |
| - | - | 1.12E+05 | 482.2 | - | - | 0 | - |
| - | - | 9.01E+05 | 483.3 | - | - | 0 | - |
| - | - | 4.212E+05 | 483.8 | - | - | 0 | - |
| - | - | 9.045E+04 | 484.3 | - | - | 0 | - |
| - | - | 1.935E+05 | 488.2 | - | - | 0 | - |
| - | - | 1.702E+05 | 488.3 | - | - | 0 | - |
| - | - | 1.081E+05 | 488.8 | - | - | 0 | - |
| - | - | 4.135E+05 | 489.2 | - | - | 0 | - |
| - | - | 1.333E+05 | 490.2 | - | - | 0 | - |
| - | - | 5.737E+04 | 492.2 | - | - | 0 | - |
| - | - | 1.899E+05 | 494.2 | - | - | 0 | - |
| - | - | 8.783E+05 | 494.3 | - | - | 0 | - |
| - | - | 3.223E+05 | 495.2 | - | - | 0 | - |
| - | - | 2.387E+05 | 495.3 | - | - | 0 | - |
| 10 | y | 2.88E+06 | 496.2 | 0.0008994 | 1.812 | +1 | 4 |
| - | - | 6.603E+05 | 497.2 | - | - | 0 | - |
| - | - | 2.697E+05 | 497.8 | - | - | 0 | - |
| - | - | 1.428E+05 | 498.2 | - | - | 0 | - |
| - | - | 5.774E+04 | 511.3 | - | - | 0 | - |
| - | - | 6.758E+04 | 522.2 | - | - | 0 | - |
| - | - | 2.491E+05 | 523.2 | - | - | 0 | - |
| - | - | 1.83E+05 | 524.2 | - | - | 0 | - |
| - | - | 8.114E+04 | 524.2 | - | - | 0 | - |
| - | - | 7.485E+04 | 525.3 | - | - | 0 | - |
| - | - | 9.578E+04 | 528.9 | - | - | 0 | - |
| - | - | 7.054E+04 | 529.3 | - | - | 0 | - |
| - | - | 1.552E+05 | 529.6 | - | - | 0 | - |
| - | - | 8.653E+04 | 529.9 | - | - | 0 | - |
| - | - | 7.218E+04 | 534.9 | - | - | 0 | - |
| - | - | 3.671E+05 | 535.3 | - | - | 0 | - |
| - | - | 1.74E+05 | 535.6 | - | - | 0 | - |
| - | - | 1.704E+05 | 535.9 | - | - | 0 | - |
| - | - | 6.809E+04 | 536.2 | - | - | 0 | - |
| - | - | 1.555E+06 | 538.3 | - | - | 0 | - |
| - | - | 4.393E+05 | 539.3 | - | - | 0 | - |
| - | - | 1.459E+05 | 540.3 | - | - | 0 | - |
| - | - | 8.959E+04 | 543.3 | - | - | 0 | - |
| 9 | w | 1.724E+06 | 550.3 | 0.0008937 | 1.624 | +1 | 5 |
| - | - | 9.424E+04 | 550.3 | - | - | 0 | - |
| - | - | 5.254E+05 | 551.3 | - | - | 0 | - |
| - | - | 1.474E+05 | 551.3 | - | - | 0 | - |
| - | - | 7.709E+04 | 552.3 | - | - | 0 | - |
| - | - | 9.631E+04 | 552.3 | - | - | 0 | - |
| - | - | 1.544E+05 | 558.3 | - | - | 0 | - |
| - | - | 9.966E+04 | 565.3 | - | - | 0 | - |
| - | - | 1.076E+05 | 569.3 | - | - | 0 | - |
| - | - | 8.513E+04 | 569.8 | - | - | 0 | - |
| - | - | 5.978E+04 | 575.2 | - | - | 0 | - |
| 9 | z | 6.199E+04 | 577.2 | 0.002696 | 4.67 | +1 | 5 |
| - | - | 6.837E+04 | 579.1 | - | - | 0 | - |
| 5 | c | 6.492E+05 | 579.3 | 0.0007004 | 1.209 | +1 | 5 |
| - | - | 1.599E+05 | 580.3 | - | - | 0 | - |
| - | - | 7.591E+04 | 590.8 | - | - | 0 | - |
| - | - | 9.982E+04 | 592.3 | - | - | 0 | - |
| 9 | y | 3.919E+05 | 593.3 | 0.003747 | 6.316 | +1 | 5 |
| - | - | 6.642E+05 | 594.3 | - | - | 0 | - |
| 9 | z | 1.666E+06 | 595.2 | 0.00208 | 3.494 | +1 | 5 |
| - | - | 8.206E+05 | 596.3 | - | - | 0 | - |
| 5 | c | 7.433E+06 | 596.4 | 0.0008847 | 1.483 | +1 | 5 |
| - | - | 1.445E+05 | 597.3 | - | - | 0 | - |
| - | - | 2.463E+06 | 597.4 | - | - | 0 | - |
| - | - | 6.843E+04 | 598.3 | - | - | 0 | - |
| - | - | 5.229E+05 | 598.4 | - | - | 0 | - |
| - | - | 1.037E+05 | 599.4 | - | - | 0 | - |
| - | - | 2.188E+05 | 601.3 | - | - | 0 | - |
| - | - | 1.676E+05 | 602.3 | - | - | 0 | - |
| - | - | 2.859E+05 | 608.3 | - | - | 0 | - |
| - | - | 8.948E+04 | 609.3 | - | - | 0 | - |
| - | - | 2.347E+05 | 609.3 | - | - | 0 | - |
| - | - | 2.484E+06 | 610.3 | - | - | 0 | - |
| - | - | 9.054E+04 | 610.3 | - | - | 0 | - |
| 9 | y | 5.388E+06 | 611.3 | 0.002094 | 3.425 | +1 | 5 |
| - | - | 2.518E+05 | 611.8 | - | - | 0 | - |
| - | - | 1.757E+06 | 612.3 | - | - | 0 | - |
| - | - | 3.581E+05 | 613.3 | - | - | 0 | - |
| - | - | 4.363E+05 | 621.3 | - | - | 0 | - |
| - | - | 1.321E+05 | 622.3 | - | - | 0 | - |
| - | - | 1.03E+05 | 629.8 | - | - | 0 | - |
| - | - | 1.302E+05 | 633.3 | - | - | 0 | - |
| - | - | 2.424E+05 | 633.8 | - | - | 0 | - |
| - | - | 1.181E+05 | 634.3 | - | - | 0 | - |
| - | - | 5.875E+04 | 634.8 | - | - | 0 | - |
| - | - | 6.245E+04 | 635.3 | - | - | 0 | - |
| 4 | y | 8.525E+04 | 637.3 | 0.009153 | 14.36 | +2 | 10 |
| 4 | z | 2.805E+05 | 637.8 | 0.004752 | 7.451 | +2 | 10 |
| - | - | 6.202E+04 | 638.3 | - | - | 0 | - |
| - | - | 2.486E+05 | 645.3 | - | - | 0 | - |
| 4 | y | 4.335E+05 | 645.8 | 0.002958 | 4.581 | +2 | 10 |
| - | - | 7.649E+04 | 645.8 | - | - | 0 | - |
| - | - | 2.201E+05 | 646.3 | - | - | 0 | - |
| - | - | 6.466E+04 | 646.4 | - | - | 0 | - |
| - | - | 1.086E+05 | 646.8 | - | - | 0 | - |
| 11 | c | 1.084E+05 | 646.8 | 0.007452 | 11.52 | +2 | 11 |
| 11 | c | 1.813E+05 | 647.3 | 0.00751 | 11.6 | +2 | 11 |
| - | - | 9.128E+04 | 647.8 | - | - | 0 | - |
| - | - | 1.548E+05 | 648.3 | - | - | 0 | - |
| - | - | 5.228E+04 | 649.3 | - | - | 0 | - |
| - | - | 9.806E+04 | 651.3 | - | - | 0 | - |
| - | - | 1.629E+05 | 652.3 | - | - | 0 | - |
| - | - | 5.85E+05 | 654.9 | - | - | 0 | - |
| - | - | 3.82E+06 | 655.4 | - | - | 0 | - |
| 11 | c | 1.299E+07 | 655.8 | 0.002231 | 3.402 | +2 | 11 |
| - | - | 1.011E+07 | 656.3 | - | - | 0 | - |
| - | - | 4.239E+06 | 656.9 | - | - | 0 | - |
| - | - | 8.338E+05 | 657.4 | - | - | 0 | - |
| - | - | 2.415E+05 | 657.9 | - | - | 0 | - |
| - | - | 8.63E+04 | 663.4 | - | - | 0 | - |
| - | - | 2.512E+06 | 664.3 | - | - | 0 | - |
| - | - | 3.471E+05 | 664.3 | - | - | 0 | - |
| 8 | w | 1.375E+06 | 665.3 | 0.008863 | 13.32 | +1 | 6 |
| - | - | 1.003E+05 | 665.4 | - | - | 0 | - |
| - | - | 3.75E+05 | 666.3 | - | - | 0 | - |
| - | - | 1.835E+05 | 666.4 | - | - | 0 | - |
| - | - | 1.458E+05 | 667.3 | - | - | 0 | - |
| - | - | 5.247E+05 | 667.4 | - | - | 0 | - |
| - | - | 1.033E+06 | 668.4 | - | - | 0 | - |
| - | - | 8.999E+05 | 668.9 | - | - | 0 | - |
| - | - | 4.625E+05 | 669.4 | - | - | 0 | - |
| - | - | 1.861E+05 | 669.9 | - | - | 0 | - |
| - | - | 7.521E+04 | 671.8 | - | - | 0 | - |
| - | - | 8.486E+05 | 672.3 | - | - | 0 | - |
| 3 | w | 1.496E+06 | 672.8 | 0.004512 | 6.706 | +2 | 11 |
| - | - | 1.158E+06 | 673.3 | - | - | 0 | - |
| - | - | 4.498E+05 | 673.8 | - | - | 0 | - |
| - | - | 1.622E+05 | 676.3 | - | - | 0 | - |
| - | - | 6.242E+04 | 677.3 | - | - | 0 | - |
| - | - | 1.479E+05 | 679.3 | - | - | 0 | - |
| - | - | 8.432E+04 | 679.8 | - | - | 0 | - |
| - | - | 4.991E+05 | 680.3 | - | - | 0 | - |
| - | - | 1.652E+05 | 681.3 | - | - | 0 | - |
| - | - | 9.925E+04 | 684.3 | - | - | 0 | - |
| - | - | 6.625E+04 | 684.8 | - | - | 0 | - |
| - | - | 9.22E+04 | 689.9 | - | - | 0 | - |
| - | - | 1.028E+05 | 692.8 | - | - | 0 | - |
| 3 | y | 3.611E+05 | 693.3 | 0.00167 | 2.409 | +2 | 11 |
| 3 | y | 8.508E+05 | 693.8 | 0.0118 | 17.01 | +2 | 11 |
| 6 | c | 1.869E+06 | 694.4 | 0.009275 | 13.36 | +1 | 6 |
| - | - | 7.892E+05 | 694.8 | - | - | 0 | - |
| - | - | 5.272E+05 | 695.4 | - | - | 0 | - |
| - | - | 2.991E+05 | 696.4 | - | - | 0 | - |
| - | - | 1.572E+05 | 697.4 | - | - | 0 | - |
| - | - | 3.584E+05 | 697.9 | - | - | 0 | - |
| - | - | 3.723E+05 | 698.4 | - | - | 0 | - |
| - | - | 2.019E+05 | 698.9 | - | - | 0 | - |
| - | - | 6.702E+04 | 699.4 | - | - | 0 | - |
| - | - | 1.485E+06 | 701.8 | - | - | 0 | - |
| 3 | y | 3.037E+06 | 702.3 | 0.004627 | 6.589 | +2 | 11 |
| - | - | 1.904E+06 | 702.8 | - | - | 0 | - |
| - | - | 8.001E+05 | 703.3 | - | - | 0 | - |
| - | - | 1.62E+05 | 703.8 | - | - | 0 | - |
| 8 | y | 1.383E+05 | 706.3 | 0.003301 | 4.674 | +1 | 6 |
| - | - | 1.619E+06 | 707.3 | - | - | 0 | - |
| 8 | z | 1.78E+06 | 708.3 | 0.004991 | 7.046 | +1 | 6 |
| - | - | 1.774E+05 | 708.9 | - | - | 0 | - |
| - | - | 7.485E+05 | 709.3 | - | - | 0 | - |
| - | - | 1.855E+05 | 710.3 | - | - | 0 | - |
| 6 | c | 6.596E+06 | 711.4 | 0.000675 | 0.9489 | +1 | 6 |
| 12 | c | 2.819E+05 | 711.9 | 0.005317 | 7.47 | +2 | 12 |
| - | - | 2.542E+06 | 712.4 | - | - | 0 | - |
| - | - | 1.383E+05 | 712.9 | - | - | 0 | - |
| - | - | 6.494E+05 | 713.4 | - | - | 0 | - |
| - | - | 7.948E+04 | 713.9 | - | - | 0 | - |
| - | - | 1.167E+05 | 714.4 | - | - | 0 | - |
| - | - | 1.155E+05 | 714.9 | - | - | 0 | - |
| - | - | 1.962E+05 | 715.4 | - | - | 0 | - |
| - | - | 2.213E+05 | 715.8 | - | - | 0 | - |
| - | - | 1.368E+05 | 716.3 | - | - | 0 | - |
| - | - | 8.878E+04 | 716.8 | - | - | 0 | - |
| - | - | 5.697E+06 | 719.9 | - | - | 0 | - |
| 12 | c | 8.615E+06 | 720.4 | 0.005593 | 7.764 | +2 | 12 |
| - | - | 5.956E+06 | 720.9 | - | - | 0 | - |
| - | - | 2.259E+06 | 721.4 | - | - | 0 | - |
| - | - | 5.377E+05 | 721.9 | - | - | 0 | - |
| - | - | 1.062E+05 | 722.4 | - | - | 0 | - |
| - | - | 8.486E+04 | 722.8 | - | - | 0 | - |
| - | - | 7.272E+05 | 723.4 | - | - | 0 | - |
| 8 | y | 1.881E+06 | 724.4 | 0.002868 | 3.96 | +1 | 6 |
| - | - | 7.971E+05 | 725.4 | - | - | 0 | - |
| - | - | 1.658E+05 | 726.4 | - | - | 0 | - |
| - | - | 8.06E+04 | 728.4 | - | - | 0 | - |
| - | - | 6.408E+04 | 728.9 | - | - | 0 | - |
| - | - | 4.293E+05 | 729.4 | - | - | 0 | - |
| - | - | 6.041E+05 | 729.9 | - | - | 0 | - |
| - | - | 3.587E+05 | 730.4 | - | - | 0 | - |
| - | - | 1.375E+05 | 730.9 | - | - | 0 | - |
| 2 | w | 2.354E+05 | 736.3 | 0.0033 | 4.481 | +2 | 12 |
| - | - | 5.251E+05 | 736.9 | - | - | 0 | - |
| - | - | 6.276E+05 | 737.4 | - | - | 0 | - |
| - | - | 3.756E+05 | 737.9 | - | - | 0 | - |
| - | - | 1.05E+05 | 738.4 | - | - | 0 | - |
| - | - | 8.337E+04 | 739.4 | - | - | 0 | - |
| - | - | 8.195E+04 | 740.4 | - | - | 0 | - |
| - | - | 1.291E+05 | 740.9 | - | - | 0 | - |
| - | - | 1.692E+05 | 741.4 | - | - | 0 | - |
| - | - | 8.203E+04 | 742.9 | - | - | 0 | - |
| 2 | y | 2.942E+05 | 743.4 | 0.01464 | 19.69 | +2 | 12 |
| 2 | z | 6.605E+05 | 743.9 | 0.007676 | 10.32 | +2 | 12 |
| - | - | 5.883E+05 | 744.4 | - | - | 0 | - |
| - | - | 2.876E+05 | 744.9 | - | - | 0 | - |
| - | - | 8.134E+04 | 745.4 | - | - | 0 | - |
| - | - | 1.987E+05 | 748.4 | - | - | 0 | - |
| - | - | 2.903E+06 | 748.9 | - | - | 0 | - |
| - | - | 2.336E+06 | 749.4 | - | - | 0 | - |
| - | - | 1.064E+06 | 749.9 | - | - | 0 | - |
| - | - | 5.135E+05 | 750.4 | - | - | 0 | - |
| - | - | 1.888E+05 | 750.9 | - | - | 0 | - |
| - | - | 4.423E+05 | 751.4 | - | - | 0 | - |
| 2 | y | 3.943E+05 | 751.9 | 0.002647 | 3.521 | +2 | 12 |
| - | - | 2.996E+05 | 752.4 | - | - | 0 | - |
| - | - | 8.353E+04 | 752.9 | - | - | 0 | - |
| - | - | 3.088E+05 | 757.9 | - | - | 0 | - |
| - | - | 3.979E+05 | 758.4 | - | - | 0 | - |
| - | - | 3.206E+05 | 758.9 | - | - | 0 | - |
| - | - | 1.432E+05 | 759.4 | - | - | 0 | - |
| - | - | 8.091E+04 | 759.9 | - | - | 0 | - |
| - | - | 1.058E+05 | 762.4 | - | - | 0 | - |
| - | - | 1.102E+05 | 762.9 | - | - | 0 | - |
| - | - | 1.027E+05 | 763.4 | - | - | 0 | - |
| - | - | 1.138E+05 | 764.4 | - | - | 0 | - |
| - | - | 2.649E+05 | 764.9 | - | - | 0 | - |
| - | - | 2.954E+05 | 765.4 | - | - | 0 | - |
| - | - | 1.794E+05 | 765.9 | - | - | 0 | - |
| - | - | 1.584E+05 | 766.4 | - | - | 0 | - |
| - | - | 8.016E+04 | 767.4 | - | - | 0 | - |
| - | - | 1.565E+05 | 770.9 | - | - | 0 | - |
| - | - | 2.13E+05 | 771.4 | - | - | 0 | - |
| - | - | 5.473E+05 | 771.9 | - | - | 0 | - |
| - | - | 1.209E+06 | 772.4 | - | - | 0 | - |
| - | - | 1.196E+06 | 772.9 | - | - | 0 | - |
| - | - | 1.004E+06 | 773.4 | - | - | 0 | - |
| - | - | 3.871E+05 | 773.9 | - | - | 0 | - |
| - | - | 6.506E+04 | 774.4 | - | - | 0 | - |
| - | - | 2.789E+05 | 779.4 | - | - | 0 | - |
| - | - | 1.577E+06 | 779.9 | - | - | 0 | - |
| - | - | 1.369E+06 | 780.4 | - | - | 0 | - |
| - | - | 8.321E+05 | 780.9 | - | - | 0 | - |
| - | - | 3.298E+05 | 781.4 | - | - | 0 | - |
| - | - | 7.43E+04 | 781.9 | - | - | 0 | - |
| - | - | 3.367E+05 | 785.4 | - | - | 0 | - |
| - | - | 5.461E+05 | 785.9 | - | - | 0 | - |
| - | - | 7.761E+05 | 786.4 | - | - | 0 | - |
| - | - | 7.533E+05 | 786.9 | - | - | 0 | - |
| - | - | 5.658E+05 | 787.4 | - | - | 0 | - |
| - | - | 1.819E+05 | 787.9 | - | - | 0 | - |
| - | - | 1.282E+05 | 788.4 | - | - | 0 | - |
| - | - | 4.165E+05 | 793.4 | - | - | 0 | - |
| - | - | 3.224E+06 | 793.9 | - | - | 0 | - |
| - | - | 4.584E+06 | 794.4 | - | - | 0 | - |
| - | - | 3.542E+06 | 794.9 | - | - | 0 | - |
| - | - | 1.614E+06 | 795.4 | - | - | 0 | - |
| - | - | 4.063E+05 | 795.9 | - | - | 0 | - |
| - | - | 9.157E+04 | 796.4 | - | - | 0 | - |
| - | - | 4.217E+06 | 801.9 | - | - | 0 | - |
| - | - | 2.522E+07 | 802.4 | - | - | 0 | - |
| - | - | 3.198E+07 | 802.9 | - | - | 0 | - |
| - | - | 2.084E+07 | 803.4 | - | - | 0 | - |
| - | - | 9.184E+06 | 803.9 | - | - | 0 | - |
| - | - | 2.688E+06 | 804.4 | - | - | 0 | - |
| - | - | 5.172E+05 | 804.9 | - | - | 0 | - |
| - | - | 6.236E+04 | 835.4 | - | - | 0 | - |
| - | - | 7.533E+04 | 836.5 | - | - | 0 | - |
| - | - | 4.149E+05 | 837.4 | - | - | 0 | - |
| - | - | 6.191E+04 | 837.4 | - | - | 0 | - |
| - | - | 9.261E+04 | 837.5 | - | - | 0 | - |
| - | - | 9.461E+05 | 838.4 | - | - | 0 | - |
| - | - | 3.385E+05 | 839.4 | - | - | 0 | - |
| - | - | 1.852E+05 | 840.4 | - | - | 0 | - |
| - | - | 1.089E+05 | 844.4 | - | - | 0 | - |
| - | - | 1.486E+05 | 849.4 | - | - | 0 | - |
| - | - | 1.603E+05 | 850.4 | - | - | 0 | - |
| - | - | 8.546E+04 | 851.4 | - | - | 0 | - |
| - | - | 4.226E+05 | 852.4 | - | - | 0 | - |
| - | - | 4.711E+05 | 853.4 | - | - | 0 | - |
| - | - | 1.769E+05 | 854.4 | - | - | 0 | - |
| - | - | 2.813E+05 | 862.4 | - | - | 0 | - |
| - | - | 1.84E+05 | 863.4 | - | - | 0 | - |
| - | - | 6.974E+05 | 864.5 | - | - | 0 | - |
| - | - | 2.6E+05 | 865.5 | - | - | 0 | - |
| - | - | 1.022E+05 | 876.4 | - | - | 0 | - |
| 7 | c | 4.012E+06 | 880.4 | 0.0008287 | 0.9412 | +1 | 7 |
| - | - | 1.991E+06 | 881.4 | - | - | 0 | - |
| - | - | 6.211E+05 | 882.4 | - | - | 0 | - |
| - | - | 1.12E+05 | 883.4 | - | - | 0 | - |
| 7 | y | 1.512E+05 | 892.4 | 0.01058 | 11.86 | +1 | 7 |
| - | - | 1.359E+06 | 893.4 | - | - | 0 | - |
| 7 | z | 2.49E+06 | 894.4 | 0.005939 | 6.64 | +1 | 7 |
| - | - | 1.008E+06 | 895.4 | - | - | 0 | - |
| - | - | 3.507E+05 | 896.4 | - | - | 0 | - |
| 7 | c | 9.577E+06 | 897.5 | 0.0008909 | 0.9927 | +1 | 7 |
| - | - | 4.862E+06 | 898.5 | - | - | 0 | - |
| - | - | 1.478E+06 | 899.5 | - | - | 0 | - |
| - | - | 3.045E+05 | 900.5 | - | - | 0 | - |
| - | - | 1.171E+05 | 902.4 | - | - | 0 | - |
| - | - | 6.366E+04 | 907.4 | - | - | 0 | - |
| - | - | 7.385E+05 | 909.4 | - | - | 0 | - |
| 7 | y | 9.312E+05 | 910.4 | 0.008577 | 9.421 | +1 | 7 |
| - | - | 4.446E+05 | 911.4 | - | - | 0 | - |
| - | - | 1.61E+05 | 912.4 | - | - | 0 | - |
| - | - | 1.251E+05 | 936.4 | - | - | 0 | - |
| - | - | 2.623E+05 | 937.4 | - | - | 0 | - |
| - | - | 2.059E+05 | 938.4 | - | - | 0 | - |
| - | - | 6.437E+05 | 964.5 | - | - | 0 | - |
| - | - | 1.031E+06 | 965.5 | - | - | 0 | - |
| - | - | 4.095E+05 | 966.5 | - | - | 0 | - |
| - | - | 2.283E+05 | 966.5 | - | - | 0 | - |
| - | - | 1.055E+05 | 967.4 | - | - | 0 | - |
| - | - | 1.894E+05 | 967.5 | - | - | 0 | - |
| - | - | 1.351E+05 | 991.5 | - | - | 0 | - |
| 8 | c | 8.394E+04 | 992.5 | 0.009437 | 9.508 | +1 | 8 |
| 8 | c | 5.943E+05 | 993.5 | 0.001543 | 1.553 | +1 | 8 |
| - | - | 2.891E+05 | 994.5 | - | - | 0 | - |
| - | - | 1.148E+05 | 995.5 | - | - | 0 | - |
| - | - | 2.794E+06 | 1008 | - | - | 0 | - |
| 6 | z | 3.41E+06 | 1009 | 0.00811 | 8.034 | +1 | 8 |
| - | - | 1.069E+06 | 1010 | - | - | 0 | - |
| 8 | c | 7.059E+06 | 1011 | 0.001727 | 1.709 | +1 | 8 |
| - | - | 3.833E+05 | 1011 | - | - | 0 | - |
| - | - | 3.968E+06 | 1012 | - | - | 0 | - |
| - | - | 9.913E+04 | 1012 | - | - | 0 | - |
| - | - | 1.204E+06 | 1013 | - | - | 0 | - |
| - | - | 3.14E+05 | 1014 | - | - | 0 | - |
| - | - | 1.063E+06 | 1024 | - | - | 0 | - |
| 6 | y | 8.367E+05 | 1025 | 0.01472 | 14.35 | +1 | 8 |
| - | - | 4.098E+05 | 1026 | - | - | 0 | - |
| - | - | 1.376E+05 | 1027 | - | - | 0 | - |
| - | - | 8.853E+04 | 1038 | - | - | 0 | - |
| - | - | 2.893E+05 | 1063 | - | - | 0 | - |
| - | - | 2.48E+05 | 1064 | - | - | 0 | - |
| - | - | 9.402E+04 | 1065 | - | - | 0 | - |
| - | - | 1.824E+06 | 1068 | - | - | 0 | - |
| - | - | 1.124E+06 | 1069 | - | - | 0 | - |
| - | - | 3.851E+05 | 1070 | - | - | 0 | - |
| - | - | 1.483E+05 | 1078 | - | - | 0 | - |
| 5 | w | 7.957E+05 | 1079 | 0.00354 | 3.28 | +1 | 9 |
| - | - | 4.167E+05 | 1080 | - | - | 0 | - |
| - | - | 6.293E+05 | 1081 | - | - | 0 | - |
| - | - | 1.379E+05 | 1081 | - | - | 0 | - |
| - | - | 7.809E+05 | 1082 | - | - | 0 | - |
| - | - | 3.212E+05 | 1083 | - | - | 0 | - |
| - | - | 1.456E+05 | 1084 | - | - | 0 | - |
| - | - | 6.686E+04 | 1093 | - | - | 0 | - |
| - | - | 3.701E+05 | 1094 | - | - | 0 | - |
| - | - | 2.672E+05 | 1095 | - | - | 0 | - |
| - | - | 1.018E+05 | 1096 | - | - | 0 | - |
| 9 | c | 1.163E+05 | 1109 | 0.003591 | 3.24 | +1 | 9 |
| - | - | 8.627E+04 | 1110 | - | - | 0 | - |
| - | - | 1.711E+05 | 1111 | - | - | 0 | - |
| - | - | 7.833E+04 | 1120 | - | - | 0 | - |
| - | - | 1.413E+05 | 1124 | - | - | 0 | - |
| - | - | 1.164E+06 | 1125 | - | - | 0 | - |
| 9 | c | 1.662E+06 | 1126 | 0.008658 | 7.692 | +1 | 9 |
| - | - | 8.002E+05 | 1127 | - | - | 0 | - |
| - | - | 2.37E+05 | 1128 | - | - | 0 | - |
| - | - | 2.623E+06 | 1137 | - | - | 0 | - |
| 5 | z | 1.776E+07 | 1137 | 0.002572 | 2.261 | +1 | 9 |
| - | - | 1.09E+07 | 1139 | - | - | 0 | - |
| - | - | 4.119E+06 | 1140 | - | - | 0 | - |
| - | - | 9.663E+05 | 1141 | - | - | 0 | - |
| - | - | 1.14E+05 | 1142 | - | - | 0 | - |
| - | - | 3.666E+05 | 1153 | - | - | 0 | - |
| 5 | y | 8.305E+05 | 1154 | 0.005821 | 5.046 | +1 | 9 |
| - | - | 5.054E+05 | 1155 | - | - | 0 | - |
| - | - | 1.691E+05 | 1156 | - | - | 0 | - |
| - | - | 9.114E+04 | 1169 | - | - | 0 | - |
| - | - | 9.382E+04 | 1170 | - | - | 0 | - |
| - | - | 4.556E+06 | 1182 | - | - | 0 | - |
| 10 | c | 1.047E+07 | 1183 | 0.00636 | 5.378 | +1 | 10 |
| - | - | 6.296E+06 | 1184 | - | - | 0 | - |
| - | - | 2.212E+06 | 1185 | - | - | 0 | - |
| - | - | 4.435E+05 | 1186 | - | - | 0 | - |
| - | - | 1.315E+05 | 1187 | - | - | 0 | - |
| - | - | 1.712E+05 | 1231 | - | - | 0 | - |
| - | - | 1.26E+05 | 1232 | - | - | 0 | - |
| - | - | 3.2E+05 | 1267 | - | - | 0 | - |
| - | - | 5.218E+05 | 1268 | - | - | 0 | - |
| - | - | 2.361E+05 | 1269 | - | - | 0 | - |
| 4 | y | 9.196E+05 | 1274 | 0.02338 | 18.36 | +1 | 10 |
| 4 | z | 3.276E+06 | 1275 | 0.005672 | 4.45 | +1 | 10 |
| - | - | 3.137E+06 | 1276 | - | - | 0 | - |
| - | - | 1.53E+06 | 1277 | - | - | 0 | - |
| - | - | 5E+05 | 1278 | - | - | 0 | - |
| - | - | 8.581E+04 | 1279 | - | - | 0 | - |
| - | - | 1.655E+05 | 1290 | - | - | 0 | - |
| 4 | y | 5.033E+05 | 1291 | 0.002695 | 2.088 | +1 | 10 |
| - | - | 2.954E+05 | 1292 | - | - | 0 | - |
| 11 | c | 1.32E+05 | 1294 | 0.008746 | 6.761 | +1 | 11 |
| - | - | 3.016E+05 | 1295 | - | - | 0 | - |
| - | - | 2.488E+05 | 1296 | - | - | 0 | - |
| - | - | 8.983E+04 | 1297 | - | - | 0 | - |
| - | - | 1.671E+06 | 1310 | - | - | 0 | - |
| 11 | c | 4.877E+06 | 1311 | 0.004048 | 3.088 | +1 | 11 |
| - | - | 3.943E+06 | 1312 | - | - | 0 | - |
| - | - | 1.733E+06 | 1313 | - | - | 0 | - |
| - | - | 4.788E+05 | 1314 | - | - | 0 | - |
| - | - | 9.451E+04 | 1315 | - | - | 0 | - |
| - | - | 1.236E+05 | 1344 | - | - | 0 | - |
| 3 | w | 3.262E+05 | 1345 | 0.005192 | 3.861 | +1 | 11 |
| - | - | 9.22E+04 | 1346 | - | - | 0 | - |
| - | - | 8.747E+04 | 1380 | - | - | 0 | - |
| - | - | 1.086E+05 | 1381 | - | - | 0 | - |
| 3 | y | 6.098E+05 | 1387 | 0.02245 | 16.19 | +1 | 11 |
| 3 | z | 2.851E+06 | 1388 | 0.007057 | 5.086 | +1 | 11 |
| - | - | 3.476E+06 | 1389 | - | - | 0 | - |
| - | - | 2.263E+06 | 1390 | - | - | 0 | - |
| - | - | 8.979E+05 | 1391 | - | - | 0 | - |
| - | - | 1.89E+05 | 1392 | - | - | 0 | - |
| - | - | 3.132E+05 | 1395 | - | - | 0 | - |
| - | - | 8.173E+05 | 1396 | - | - | 0 | - |
| - | - | 8.129E+05 | 1397 | - | - | 0 | - |
| - | - | 3.58E+05 | 1398 | - | - | 0 | - |
| - | - | 1.498E+05 | 1399 | - | - | 0 | - |
| 3 | y | 2.969E+05 | 1404 | 0.006888 | 4.907 | +1 | 11 |
| - | - | 2.811E+05 | 1405 | - | - | 0 | - |
| - | - | 2.217E+05 | 1413 | - | - | 0 | - |
| - | - | 9.504E+04 | 1414 | - | - | 0 | - |
| 12 | c | 3.399E+05 | 1423 | 0.0245 | 17.22 | +1 | 12 |
| - | - | 4.395E+05 | 1424 | - | - | 0 | - |
| - | - | 3.848E+05 | 1425 | - | - | 0 | - |
| - | - | 2.011E+05 | 1426 | - | - | 0 | - |
| - | - | 6.077E+05 | 1439 | - | - | 0 | - |
| 12 | c | 2.156E+06 | 1440 | 0.01382 | 9.601 | +1 | 12 |
| - | - | 2.318E+06 | 1441 | - | - | 0 | - |
| - | - | 1.095E+06 | 1442 | - | - | 0 | - |
| - | - | 4.789E+05 | 1443 | - | - | 0 | - |
| - | - | 1.152E+05 | 1444 | - | - | 0 | - |
| 2 | z | 3.048E+05 | 1487 | 0.01726 | 11.61 | +1 | 12 |
| - | - | 7.509E+05 | 1488 | - | - | 0 | - |
| - | - | 6.131E+05 | 1489 | - | - | 0 | - |
| - | - | 1.854E+05 | 1490 | - | - | 0 | - |
| - | - | 8.647E+04 | 1491 | - | - | 0 | - |
| - | - | 7.698E+04 | 1504 | - | - | 0 | - |
| - | - | 1.084E+05 | 1542 | - | - | 0 | - |
| - | - | 2.739E+05 | 1543 | - | - | 0 | - |
| - | - | 3.244E+05 | 1544 | - | - | 0 | - |
| - | - | 6.647E+05 | 1545 | - | - | 0 | - |
| - | - | 7.623E+05 | 1546 | - | - | 0 | - |
| - | - | 3.963E+05 | 1547 | - | - | 0 | - |
| - | - | 1.584E+05 | 1548 | - | - | 0 | - |
| - | - | 4.642E+05 | 1559 | - | - | 0 | - |
| - | - | 1.905E+06 | 1560 | - | - | 0 | - |
| - | - | 1.653E+06 | 1561 | - | - | 0 | - |
| - | - | 1.077E+06 | 1562 | - | - | 0 | - |
| - | - | 3.789E+05 | 1563 | - | - | 0 | - |
| - | - | 8.09E+04 | 1564 | - | - | 0 | - |
| - | - | 2.118E+05 | 1570 | - | - | 0 | - |
| - | - | 3.663E+05 | 1571 | - | - | 0 | - |
| - | - | 3.699E+05 | 1572 | - | - | 0 | - |
| - | - | 1.259E+05 | 1573 | - | - | 0 | - |
| - | - | 1.875E+05 | 1576 | - | - | 0 | - |
| - | - | 1.033E+06 | 1577 | - | - | 0 | - |
| - | - | 1.156E+06 | 1578 | - | - | 0 | - |
| - | - | 6.736E+05 | 1579 | - | - | 0 | - |
| - | - | 2.922E+05 | 1580 | - | - | 0 | - |
| - | - | 7.379E+05 | 1587 | - | - | 0 | - |
| - | - | 4.35E+06 | 1588 | - | - | 0 | - |
| - | - | 7.105E+06 | 1589 | - | - | 0 | - |
| - | - | 4.886E+06 | 1590 | - | - | 0 | - |
| - | - | 2.5E+06 | 1591 | - | - | 0 | - |
| - | - | 7.238E+05 | 1592 | - | - | 0 | - |
| - | - | 9.879E+04 | 1593 | - | - | 0 | - |
| - | - | 2.9E+05 | 1603 | - | - | 0 | - |
| - | - | 2.238E+06 | 1604 | - | - | 0 | - |
| - | - | 8.123E+06 | 1605 | - | - | 0 | - |
| - | - | 1.32E+07 | 1606 | - | - | 0 | - |
| - | - | 9.209E+06 | 1607 | - | - | 0 | - |
| - | - | 4.224E+06 | 1608 | - | - | 0 | - |
| - | - | 1.259E+06 | 1609 | - | - | 0 | - |
| - | - | 2.306E+05 | 1610 | - | - | 0 | - |
| - | - | 1.063E+05 | 2139 | - | - | 0 | - |
| - | - | 6.622E+04 | 3488 | - | - | 0 | - |

m/z Charge Intensity FragmentType MassShift Position
121.06500244140625 0 39100.88
127.2240982055664 0 41036.598
128.10740661621094 0 97431.91
129.1026153564453 0 558405.3
130.06549072265625 0 1055545.8
131.0689697265625 0 93270.414
131.11822509765625 0 154023.95
133.9840850830078 0 38958.605
136.07611083984375 0 129450.81
138.0660858154297 0 37523.742
138.0911865234375 0 46277.605
140.55209350585938 0 45751.605
141.32044982910156 0 38489.855
148.8430633544922 0 39034.465
150.06680297851562 0 49680.348
152.01950073242188 0 35783.805
155.0930938720703 0 2302751.8
156.09649658203125 0 143265.25
156.55613708496094 0 39538.414
157.13397216796875 0 237650.55
159.09205627441406 0 991254.75
161.2538299560547 0 39978.848
165.05491638183594 0 84326.24
171.11312866210938 0 82715.4
173.12884521484375 0 3863877
174.13230895996094 0 366139.44
182.08160400390625 0 762144.2 y 12
183.08462524414062 0 64958.234
183.11349487304688 0 64734.598
184.06138610839844 0 72729.914
186.12432861328125 0 262670.47
186.5674285888672 0 43384.066
187.08738708496094 0 76854.734
188.1396942138672 0 145754.66
194.09320068359375 0 45341.25
195.95751953125 0 48626.562
198.11317443847656 0 97433.52
200.13955688476562 0 66875.56 c Water loss 1
201.1237335205078 0 2723082.5
202.1270294189453 0 293899.62
205.69387817382812 0 48728.523
207.1130828857422 0 91013.04
213.09971618652344 0 98197.664
215.139404296875 0 1220202.6
216.1433563232422 0 56993.16
218.1504669189453 0 1024993.06 c 1
219.15379333496094 0 66100.305
223.15594482421875 0 153261.98
225.12374877929688 0 65773.414
226.10781860351562 0 234497.08
228.1343231201172 0 120530.84
234.12350463867188 0 57420.484
240.13465881347656 0 66326.42
243.13458251953125 0 225299.56
245.1375274658203 0 408446.53
247.10841369628906 0 64728.434
251.10313415527344 0 79420.02
251.15089416503906 0 633823.7
258.145263671875 0 1576712
258.1600036621094 0 105379.7
259.148193359375 0 176404.95
260.16094970703125 0 323312.62
266.127197265625 0 72259.62
267.6654968261719 0 68448.32
268.1771545410156 0 1502597
269.1808776855469 0 118218.22
272.16021728515625 0 77319.09
283.14068603515625 0 368762.6
285.13104248046875 0 85380.91
287.22064208984375 0 99250.53
288.228515625 0 110631.53
288.6190185546875 0 94854.28
293.11370849609375 0 617245.75 y Water loss 11
294.1177673339844 0 129002.625
297.13250732421875 0 53501.582 y Water loss 8
299.1481628417969 0 101338.266
300.1675109863281 0 209041.86
301.1513977050781 0 331494.72
302.1573486328125 0 67166.5
305.5427551269531 0 49797.1
305.64630126953125 0 206579.78
306.13836669921875 0 197201.61 y 8
306.1818542480469 0 58651.418
306.6392517089844 0 112814.69
307.1895751953125 0 170602.97
308.1947326660156 0 60184.117
311.1244201660156 0 345218.38 y 11
314.20819091796875 0 382530.7
315.1668701171875 0 211615.02
315.21124267578125 0 88787.03
317.1589050292969 0 2760183.5
318.1619873046875 0 390855.16
320.1973876953125 0 526635.5
321.2005920410156 0 101206.84
331.2347412109375 0 1498199 c 2
332.23773193359375 0 228853.03
342.1169128417969 0 73305.914
343.1019592285156 0 139895.88
347.67974853515625 0 246777.7 c Ammonia loss 5
348.18170166015625 0 58726.14
350.22076416015625 0 60597.5
353.6752624511719 0 68256.79 y Water loss 7
357.17742919921875 0 117205.8
362.1873474121094 0 148899.42
362.6799621582031 0 629040.9 y 7
363.1811218261719 0 203516.8
364.1866760253906 0 382439.25
365.1907043457031 0 99278.305
374.179931640625 0 326269.47
375.1824645996094 0 85463.69
378.2078857421875 0 195257.4
379.20855712890625 0 143628.56
381.2251281738281 0 101336.68
389.19024658203125 0 117401.48 c Ammonia loss 9
396.23248291015625 0 53008.74
406.2446594238281 0 88761.13
408.2615051269531 0 72321.59
412.1825256347656 0 118952.51
417.18701171875 0 74413.984
417.7169189453125 0 109390.8
418.2123107910156 0 151603.69
420.1774597167969 0 68968.375
421.2112121582031 0 70634.2 y Water loss 10
423.2011413574219 0 362857.84 z 10
423.272216796875 0 1342585.8
424.20361328125 0 81938.94
424.2747802734375 0 360449.97
426.7225646972656 0 2282589.5
427.22412109375 0 1291809.5
427.7250061035156 0 406518.6
428.22442626953125 0 67470.7
429.2100830078125 0 361575.9
430.1944274902344 0 543374.8
431.1968994140625 0 121239.93
431.7144470214844 0 325846.2
432.2134094238281 0 251335
432.7079772949219 0 67638.71
438.7301025390625 0 87374.82
439.2200012207031 0 500109.47 y 10
440.2223815917969 0 104356.45 c Water loss 6
440.7201843261719 0 1628676.5 c Ammonia loss 6
441.22174072265625 0 945865.2
441.7231140136719 0 191175.72
448.2038879394531 0 66633.32
450.2837219238281 0 226257.9 c Water loss 3
451.2674255371094 0 3185540.5
452.2702331542969 0 677310.56
453.272216796875 0 101096.945
468.29388427734375 0 11820204 c 3
469.29669189453125 0 3012379.2
470.29901123046875 0 456659.3
474.2597961425781 0 196743.9
474.7560729980469 0 161762.36
475.2568664550781 0 77426.69
480.2225646972656 0 1172644.5 z 9
481.2281494140625 0 385763.8
482.2311706542969 0 112022.55
483.2648620605469 0 901045.56
483.7662658691406 0 421201.22
484.2664794921875 0 90446.81
488.22357177734375 0 193505.19
488.25665283203125 0 170180.9
488.754150390625 0 108050.64
489.2080078125 0 413465.97
490.2105712890625 0 133322.08
492.2462463378906 0 57366.38
494.2371520996094 0 189873.97
494.30950927734375 0 878264.44
495.2344970703125 0 322328.56
495.3129577636719 0 238708.17
496.2410888671875 0 2880370.8 y 9
497.2458190917969 0 660339.1
497.7632141113281 0 269714.16
498.2447204589844 0 142769.12
511.2617492675781 0 57743.188
522.24267578125 0 67582
523.2271118164062 0 249107.62
524.1744995117188 0 183027.75
524.2264404296875 0 81138.58
525.2677001953125 0 74853.086
528.9324340820312 0 95782.85
529.2597045898438 0 70538.37
529.595947265625 0 155191.98
529.9267578125 0 86526.27
534.9368896484375 0 72182.51
535.265380859375 0 367071.38
535.60107421875 0 174000.75
535.9334106445312 0 170428.86
536.2330322265625 0 68089.56
538.2517700195312 0 1555226.2
539.25439453125 0 439348.7
540.255615234375 0 145899.3
543.2807006835938 0 89591.71
550.2516479492188 0 1724121.4 w 8
550.2994384765625 0 94237.96
551.2555541992188 0 525388.94
551.3306884765625 0 147358.77
552.2592163085938 0 77085.94
552.3374633789062 0 96307.984
558.3007202148438 0 154356.61
565.262939453125 0 99659.305
569.3112182617188 0 107626.195
569.8114013671875 0 85127.44
575.2454833984375 0 59778.39
577.2405395507812 0 61992.793 z Water loss 8
579.0914306640625 0 68370.67
579.3256225585938 0 649213.56 c Ammonia loss 4
580.3283081054688 0 159936.7
590.8158569335938 0 75905.65
592.2714233398438 0 99822.18
593.2603149414062 0 391856.88 y Water loss 8
594.2637329101562 0 664150.44
595.25048828125 0 1665870.8 z 8
596.2557373046875 0 820585.94
596.3523559570312 0 7433140.5 c 4
597.2633056640625 0 144528.66
597.3550415039062 0 2462519.8
598.262451171875 0 68426.67
598.3572998046875 0 522879.5
599.3606567382812 0 103677.3
601.308349609375 0 218795.02
602.2954711914062 0 167568.03
608.280517578125 0 285867.8
609.2843017578125 0 89480.266
609.337890625 0 234722.98
610.2838745117188 0 2484171.5
610.3299560546875 0 90540.055
611.2692260742188 0 5388244.5 y 8
611.80126953125 0 251788.52
612.272216796875 0 1756905
613.2742919921875 0 358080.8
621.2893676757812 0 436269.94
622.2916259765625 0 132122.94
629.8010864257812 0 102950.37
633.3474731445312 0 130221.91
633.842041015625 0 242377.72
634.3428955078125 0 118140.28
634.841796875 0 58748.9
635.3432006835938 0 62447.508
637.2869873046875 0 85252.36 y Ammonia loss 3
637.7864990234375 0 280529.25 z 3
638.283203125 0 62024.64
645.2984619140625 0 248599.38
645.7940673828125 0 433519.1 y 3
645.8453369140625 0 76493.914
646.2957153320312 0 220087.19
646.35009765625 0 64658.508
646.7920532226562 0 108591.41
646.848388671875 0 108420.664 c Water loss 10
647.3404541015625 0 181309.73 c Ammonia loss 10
647.8401489257812 0 91284.61
648.3406372070312 0 154761.8
649.3431396484375 0 52278.684
651.2904663085938 0 98056.83
652.3214111328125 0 162933.45
654.85107421875 0 584971.75
655.3541259765625 0 3819723.8
655.8484497070312 0 12989158 c 10
656.3494262695312 0 10106434
656.8502197265625 0 4238540.5
657.351806640625 0 833757.56
657.8529052734375 0 241526.47
663.3518676757812 0 86301.32
664.2945556640625 0 2512476.5
664.3448486328125 0 347146.6
665.2865600585938 0 1375051.6 w 7
665.3541259765625 0 100301.54
666.2879638671875 0 375033.5
666.3572998046875 0 183458.5
667.2894287109375 0 145840.6
667.365234375 0 524677.7
668.36328125 0 1032756.9
668.8617553710938 0 899892.06
669.3610229492188 0 462520.66
669.8612670898438 0 186098.27
671.8356323242188 0 75205.7
672.3057861328125 0 848572.4
672.8009033203125 0 1495644.1 w 2
673.3016967773438 0 1157840.2
673.8017578125 0 449834
676.3447875976562 0 162162.22
677.3438720703125 0 62421.164
679.3341674804688 0 147946.05
679.83056640625 0 84316.93
680.3168334960938 0 499124.97
681.3221435546875 0 165156.05
684.3314208984375 0 99246.31
684.8219604492188 0 66250.24
689.8621215820312 0 92201.04
692.8367309570312 0 102782.8
693.3295288085938 0 361050.12 y Water loss 2
693.8316650390625 0 850774.5 y Ammonia loss 2
694.3425903320312 0 1868738 c Ammonia loss 5
694.8284912109375 0 789222.7
695.3507690429688 0 527152.4
696.36328125 0 299086.72
697.3702392578125 0 157158.14
697.8692626953125 0 358372.44
698.3673706054688 0 372284.16
698.8682250976562 0 201914.2
699.3666381835938 0 67021.38
701.8416137695312 0 1485007.6
702.3377685546875 0 3036762.2 y 2
702.8381958007812 0 1903842.8
703.3388061523438 0 800099.56
703.8431396484375 0 162035.83
706.3439331054688 0 138269.77 y Water loss 7
707.3495483398438 0 1619427.1
708.3374633789062 0 1779968.2 z 7
708.8579711914062 0 177405.83
709.3391723632812 0 748521
710.3446044921875 0 185454.19
711.3790893554688 0 6596272 c 5
711.8595581054688 0 281878.03 c Ammonia loss 11
712.3811645507812 0 2542248
712.8550415039062 0 138286.45
713.3823852539062 0 649394.1
713.86669921875 0 79478.61
714.3917236328125 0 116662.625
714.8731689453125 0 115503.18
715.3599853515625 0 196234.42
715.8333129882812 0 221290.77
716.3338623046875 0 136825.56
716.8302612304688 0 88776.89
719.876220703125 0 5697334.5
720.3731079101562 0 8615027 c 11
720.8728637695312 0 5955931.5
721.374267578125 0 2259295.5
721.873291015625 0 537654
722.3814697265625 0 106206.76
722.8406372070312 0 84859.945
723.3668823242188 0 727211.5
724.3540649414062 0 1880994.8 y 7
725.355712890625 0 797132.06
726.3575439453125 0 165767.75
728.376220703125 0 80601.59
728.8746948242188 0 64079.65
729.3613891601562 0 429312.25
729.85888671875 0 604106.3
730.3585205078125 0 358686.6
730.86181640625 0 137528.53
736.3495483398438 0 235385.23 w 1
736.8604736328125 0 525083.1
737.3629760742188 0 627625.1
737.864990234375 0 375569.25
738.3619995117188 0 104960.95
739.352783203125 0 83367.086
740.3630981445312 0 81953.21
740.8657836914062 0 129074.12
741.3682250976562 0 169178.94
742.8788452148438 0 82027.63
743.3687133789062 0 294180.34 y Ammonia loss 1
743.8656616210938 0 660506.5 z 1
744.3646850585938 0 588337.9
744.8663330078125 0 287570.44
745.3628540039062 0 81340.04
748.3748168945312 0 198714.1
748.8676147460938 0 2902629
749.369384765625 0 2335566
749.8712768554688 0 1063633.6
750.3746948242188 0 513531.94
750.8779907226562 0 188838.31
751.3779296875 0 442319.56
751.8699951171875 0 394332.25 y 1
752.3675537109375 0 299553.06
752.8729858398438 0 83529.6
757.8914794921875 0 308813.28
758.38330078125 0 397902.7
758.8724975585938 0 320597
759.372802734375 0 143226.45
759.8680419921875 0 80905.58
762.37353515625 0 105765.49
762.8770141601562 0 110161.19
763.3680419921875 0 102744.516
764.3713989257812 0 113763.56
764.8775634765625 0 264931.3
765.37744140625 0 295445.88
765.8649291992188 0 179370.56
766.3622436523438 0 158374.05
767.3646240234375 0 80160.59
770.8861083984375 0 156492.98
771.3865966796875 0 213033.61
771.8912963867188 0 547264.3
772.3912963867188 0 1209031
772.8834838867188 0 1196308.1
773.3870849609375 0 1003902.3
773.8858642578125 0 387104.47
774.39501953125 0 65056.81
779.3955688476562 0 278895.6
779.894287109375 0 1576915.1
780.3937377929688 0 1368977
780.8947143554688 0 832051.75
781.3925170898438 0 329750.8
781.8953857421875 0 74301.97
785.38623046875 0 336709
785.8847045898438 0 546129.6
786.3822631835938 0 776087.9
786.8770751953125 0 753309.8
787.377197265625 0 565771.3
787.8779907226562 0 181867.22
788.3745727539062 0 128165.17
793.399169921875 0 416473.72
793.891357421875 0 3223657.8
794.3883666992188 0 4583616.5
794.8887939453125 0 3541619
795.3889770507812 0 1614260.2
795.889404296875 0 406311.25
796.385498046875 0 91565.22
801.900146484375 0 4216528.5
802.3972778320312 0 25220720
802.8970947265625 0 31984112
803.3982543945312 0 20836000
803.89990234375 0 9184434
804.4006958007812 0 2687734.2
804.901123046875 0 517199.06
835.4188232421875 0 62358.61
836.4511108398438 0 75325.61
837.3663330078125 0 414935.34
837.4392700195312 0 61911.285
837.4541625976562 0 92614.805
838.3543090820312 0 946071.1
839.3544921875 0 338539.4
840.354248046875 0 185178.53
844.4071655273438 0 108875.17
849.43359375 0 148621.4
850.425048828125 0 160251.47
851.43310546875 0 85458.86
852.4368286132812 0 422595.3
853.443115234375 0 471060.4
854.444580078125 0 176879.56
862.421630859375 0 281342.16
863.4168090820312 0 183953.39
864.45751953125 0 697421.25
865.4595336914062 0 259980.56
876.4360961914062 0 102177.13
880.4320068359375 0 4012432.5 c Ammonia loss 6
881.434814453125 0 1991236.5
882.4381103515625 0 621116.44
883.4400024414062 0 112023.39
892.4093627929688 0 151185.64 y Water loss 6
893.4281005859375 0 1359002.1
894.417724609375 0 2489807 z 6
895.4190063476562 0 1007634.2
896.4215698242188 0 350652.9
897.4586181640625 0 9576859 c 6
898.4614868164062 0 4862264.5
899.464111328125 0 1478089.5
900.468017578125 0 304526.3
902.4146728515625 0 117075.8
907.4469604492188 0 63657.305
909.4462280273438 0 738467.56
910.4390869140625 0 931235.44 y 6
911.4390869140625 0 444644.94
912.440185546875 0 161009.06
936.431396484375 0 125085.6
937.420166015625 0 262314.9
938.4219360351562 0 205916.45
964.4645385742188 0 643709.75
965.4561157226562 0 1030674.56
966.4557495117188 0 409546.75
966.5368041992188 0 228299.19
967.4439086914062 0 105478.68
967.5359497070312 0 189436.97
991.5188598632812 0 135071.27
992.5217895507812 0 83944.984 c Water loss 7
993.5167846679688 0 594300.4 c Ammonia loss 7
994.5184936523438 0 289064.4
995.5217895507812 0 114776.914
1008.455810546875 0 2794352.8
1009.4468383789062 0 3410002.5 z 5
1010.443603515625 0 1069031
1010.5435180664062 0 7059225 c 7
1011.4474487304688 0 383282.62
1011.5459594726562 0 3968137.5
1012.4470825195312 0 99126.15
1012.5486450195312 0 1204411.9
1013.5506591796875 0 313955.5
1024.4769287109375 0 1062516.2
1025.47216796875 0 836652.6 y 5
1026.4735107421875 0 409789.44
1027.4703369140625 0 137641.44
1037.557373046875 0 88528.195
1062.5623779296875 0 289322.5
1063.56103515625 0 247965.44
1064.553955078125 0 94018.875
1067.5396728515625 0 1824075.2
1068.543212890625 0 1123956.4
1069.546142578125 0 385101.12
1078.4884033203125 0 148276.39
1079.4715576171875 0 795738.8 w 4
1080.4678955078125 0 416733.97
1080.5733642578125 0 629283
1081.4595947265625 0 137923.58
1081.5714111328125 0 780929.4
1082.5706787109375 0 321169.16
1083.5718994140625 0 145602.77
1092.5228271484375 0 66859.75
1093.5152587890625 0 370101.1
1094.51953125 0 267194.53
1095.5308837890625 0 101828.76
1108.5457763671875 0 116330.2 c Ammonia loss 8
1109.5433349609375 0 86268.96
1110.5394287109375 0 171064.6
1119.5816650390625 0 78331.51
1123.5758056640625 0 141282.73
1124.58447265625 0 1164344.4
1125.577392578125 0 1662195.5 c 8
1126.57666015625 0 800249.3
1127.5772705078125 0 237032.17
1136.513916015625 0 2623059.8
1137.4998779296875 0 17755968 z 4
1138.502197265625 0 10897770
1139.505126953125 0 4118751.8
1140.5076904296875 0 966319.44
1141.50634765625 0 113992.016
1152.53515625 0 366576.3
1153.5218505859375 0 830541.2 y 4
1154.5233154296875 0 505391.06
1155.5235595703125 0 169130.94
1169.4913330078125 0 91141.04
1170.4921875 0 93819.234
1181.607177734375 0 4555861
1182.5965576171875 0 10472164 c 9
1183.59716796875 0 6296342.5
1184.6002197265625 0 2212332.8
1185.603515625 0 443530.94
1186.6024169921875 0 131520.69
1230.567138671875 0 171174.53
1231.5728759765625 0 126047.67
1266.6796875 0 320036.47
1267.6795654296875 0 521756.94
1268.684326171875 0 236147.14
1273.57177734375 0 919554.2 y Ammonia loss 3
1274.5618896484375 0 3276356.5 z 3
1275.563720703125 0 3136555.2
1276.5665283203125 0 1530059.8
1277.5709228515625 0 499994.1
1278.5675048828125 0 85807.42
1289.581298828125 0 165469.53
1290.57763671875 0 503250.34 y 3
1291.577880859375 0 295389.94
1293.6673583984375 0 131989.69 c Ammonia loss 10
1294.67431640625 0 301574.2
1295.6741943359375 0 248838.17
1296.6678466796875 0 89827.25
1309.6988525390625 0 1671023.4
1310.689208984375 0 4877362.5 c 10
1311.6907958984375 0 3943418
1312.69287109375 0 1733434.9
1313.695068359375 0 478765.25
1314.697021484375 0 94506.1
1343.656494140625 0 123569.76
1344.5906982421875 0 326239.34 w 2
1345.6002197265625 0 92195.8
1379.7115478515625 0 87470.57
1380.7203369140625 0 108583.06
1386.6549072265625 0 609816.2 y Ammonia loss 2
1387.6473388671875 0 2851337.8 z 2
1388.6475830078125 0 3475557.8
1389.65185546875 0 2263432
1390.6531982421875 0 897877.6
1391.6573486328125 0 189036.98
1394.727294921875 0 313193.8
1395.732421875 0 817324.25
1396.7305908203125 0 812892.8
1397.732666015625 0 358028.88
1398.731689453125 0 149774.19
1403.6658935546875 0 296931.28 y 2
1404.666015625 0 281114.7
1412.7489013671875 0 221676.02
1413.7589111328125 0 95039.99
1422.7257080078125 0 339885.25 c Ammonia loss 11
1423.7230224609375 0 439504.5
1424.7197265625 0 384792.9
1425.733642578125 0 201103.3
1438.7418212890625 0 607657.5
1439.7415771484375 0 2155713.5 c 11
1440.7388916015625 0 2317566.5
1441.7420654296875 0 1094883.8
1442.740966796875 0 478900.5
1443.7371826171875 0 115167.39
1486.7259521484375 0 304770.62 z 1
1487.722412109375 0 750941.94
1488.7200927734375 0 613101.7
1489.7236328125 0 185407.95
1490.7122802734375 0 86473.44
1503.7183837890625 0 76977.695
1541.771240234375 0 108423.664
1542.7659912109375 0 273874.8
1543.775634765625 0 324434.2
1544.7786865234375 0 664699.1
1545.774658203125 0 762341.1
1546.773681640625 0 396272.44
1547.780029296875 0 158381.75
1558.7821044921875 0 464205.97
1559.7821044921875 0 1904811.8
1560.7808837890625 0 1653270.8
1561.7838134765625 0 1077242.4
1562.7847900390625 0 378890.78
1563.8011474609375 0 80897.55
1569.7593994140625 0 211752.36
1570.763427734375 0 366283.16
1571.76220703125 0 369893.88
1572.7659912109375 0 125863.414
1575.8031005859375 0 187499.75
1576.8018798828125 0 1032573.75
1577.802978515625 0 1155898.4
1578.804443359375 0 673616.25
1579.8087158203125 0 292181.1
1586.777587890625 0 737915.2
1587.774658203125 0 4350252.5
1588.7706298828125 0 7104683.5
1589.77099609375 0 4886042
1590.771728515625 0 2500417.8
1591.775390625 0 723796.2
1592.771484375 0 98792.445
1602.7894287109375 0 290001.28
1603.7874755859375 0 2238439.5
1604.7928466796875 0 8122818.5
1605.792236328125 0 13202066
1606.7943115234375 0 9209164
1607.7969970703125 0 4223514
1608.8009033203125 0 1259183
1609.7999267578125 0 230590.55
2139.271240234375 0 106290.1
3487.907958984375 0 66217.664

Spectrum Details

|  |  |
| --- | --- |
| Matched peaks? Matched peaks The total absolute number of peaks matched. Additionally in brackets the total fraction of peaks matched and the total number of peaks is shown. | 78 (12.46% of 626) |
| FDR? FDR The false discovery rate estimated for this peptide. It is calculated by matching all theoretical fragments with a non-integer shift with the raw peaks for this spectrum. This is done with 40 different shifts. The resulting percentage is the average number of annotated peaks over the number of annotated peaks with the correct spectrum. | 3.21% |
| Satellite FDR? Satellite FDR See the FDR for details on its calculation. This satellite ion specific FDR only contains the satellite ions (d/w) for I/L/J positions. | 0.00% |
| PSM Score? PSM Score The PSM Score as given by Hecklib to this annotated spectrum. It is shown with three significant figures. | 724 |

#### Spectrum 7458? Spectrum 7458 The raw spectrum of this peptide as annotated by Hecklib. The fragments are coloured according to ion type (see legend). Any peaks with a star '\*' as text can be hovered over to see the full details, first the ion type second the mass shift type. By hovering over the amino acids in the peptide or ions in the legend the corresponding peaks are highlighted. By toggling the 'Unassigned' label you can turn the background (unassigned) peaks on or off in the plot. By updating the slider in the Ion legend you can update the spectrum to only show the top X% of the peaks with labels. The top X% means any peak that is within X% of the highest intensity. By dragging in the spectrum you can zoom in to a specific part of the spectrum and use 'Zoom Out' to get back to the original zoom level. The annotation of the spectrum is based on the given sequence in the peptides file and is done with different software so inconsistencies are likely. The peaks are annotated based on the given sequence, with 20 ppm tolerance.

Copy Data

##### Spectrum 7458 (TSV)

###### Preview

```
Loading example...
```

*Click on the button to copy the data to your clipboard.*

Mz MinMz MaxIntensity Max

WidthHeightPeptide font sizePeptide stroke widthSpectrum font sizeSpectrum stroke widthCompact peptide

Ion legend

wxyz

abcd

OtherUnassignedIon^Charge^_Position_Show for top:%

TVLHQDWLDGKEY

09.07e+41.81e+52.72e+53.63e+5

Zoom Out

y^+3^_3_a^+1^_2_y^+3^_4_a^+1^_2_y^+1^_1_b^+1^_2_b^+1^_2_y^+2^_3_y^+1^_2_b^+1^_3_y^+2^_5_y^+3^_7_y^+2^_5_y^+1^_2_b^+1^_3_b^+2^_6_b^+2^_6_b^+2^_6_y^+2^_6_y^+2^_6_y^+3^_9_y^+1^_3_y^+1^_3_b^+2^_7_b^+2^_7_b^+1^_4_y^+1^_3_b^+2^_7_y^+2^_7_b^+1^_4_y^+2^_7_y^+1^_4_b^+2^_8_b^+2^_8_y^+1^_4_\*\*\*b^+2^_9_b^+2^_9_b^+2^_9_b^+1^_5_b^+1^_5_b^+1^_5_y^+1^_5_y^+1^_5_y^+2^_10_y^+2^_10_y^+2^_10_b^+2^_11_b^+1^_6_b^+1^_6_y^+2^_11_y^+2^_11_b^+1^_6_y^+2^_11_b^+2^_12_b^+2^_12_y^+1^_6_b^+2^_12_y^+1^_6_y^+2^_12_y^+2^_12_y^+2^_12_\*b^+1^_7_b^+1^_7_b^+1^_7_y^+1^_7_b^+1^_8_b^+1^_8_b^+1^_8_y^+1^_8_b^+1^_9_y^+1^_9_y^+1^_9_y^+1^_9_b^+1^_10_

0778155623343112

Fragment Matches Table

Show background peaks

| Position | Ion type | Intensity | mz Theoretical | mz Error (Th) | mz Error (ppm) | Charge | Series Number |
| --- | --- | --- | --- | --- | --- | --- | --- |
| - | - | 4691 | 120.1 | - | - | 0 | - |
| - | - | 663.5 | 121.1 | - | - | 0 | - |
| - | - | 351 | 121.2 | - | - | 0 | - |
| - | - | 1573 | 122.1 | - | - | 0 | - |
| - | - | 454.6 | 122.1 | - | - | 0 | - |
| - | - | 2166 | 123 | - | - | 0 | - |
| - | - | 634.4 | 123.1 | - | - | 0 | - |
| - | - | 418.8 | 123.4 | - | - | 0 | - |
| - | - | 358.2 | 123.7 | - | - | 0 | - |
| - | - | 891 | 125.1 | - | - | 0 | - |
| - | - | 811.2 | 126.1 | - | - | 0 | - |
| - | - | 362 | 126.1 | - | - | 0 | - |
| - | - | 4845 | 127.1 | - | - | 0 | - |
| - | - | 444.7 | 127.1 | - | - | 0 | - |
| - | - | 905 | 127.1 | - | - | 0 | - |
| - | - | 2.243E+04 | 128.1 | - | - | 0 | - |
| - | - | 2.368E+05 | 129.1 | - | - | 0 | - |
| - | - | 1.054E+05 | 130.1 | - | - | 0 | - |
| - | - | 1434 | 130.1 | - | - | 0 | - |
| - | - | 1.564E+04 | 130.1 | - | - | 0 | - |
| - | - | 1.104E+04 | 131.1 | - | - | 0 | - |
| - | - | 635.9 | 131.1 | - | - | 0 | - |
| - | - | 2.301E+04 | 132.1 | - | - | 0 | - |
| - | - | 518.4 | 132.5 | - | - | 0 | - |
| - | - | 948.1 | 133.1 | - | - | 0 | - |
| - | - | 2257 | 133.1 | - | - | 0 | - |
| - | - | 3.074E+04 | 136.1 | - | - | 0 | - |
| - | - | 2618 | 137.1 | - | - | 0 | - |
| - | - | 3254 | 138.1 | - | - | 0 | - |
| - | - | 1602 | 138.1 | - | - | 0 | - |
| - | - | 501 | 139.1 | - | - | 0 | - |
| - | - | 3123 | 139.1 | - | - | 0 | - |
| - | - | 4555 | 141.1 | - | - | 0 | - |
| - | - | 1390 | 142.1 | - | - | 0 | - |
| - | - | 1758 | 143 | - | - | 0 | - |
| - | - | 1085 | 144.1 | - | - | 0 | - |
| - | - | 2304 | 145.1 | - | - | 0 | - |
| - | - | 501.3 | 145.1 | - | - | 0 | - |
| - | - | 1253 | 146.1 | - | - | 0 | - |
| - | - | 1494 | 147 | - | - | 0 | - |
| 11 | y | 425.5 | 147.1 | 0.0007637 | 5.193 | +3 | 3 |
| - | - | 814.7 | 148.9 | - | - | 0 | - |
| - | - | 1.25E+04 | 149 | - | - | 0 | - |
| - | - | 1614 | 150.1 | - | - | 0 | - |
| - | - | 5012 | 151.1 | - | - | 0 | - |
| - | - | 551.6 | 152.1 | - | - | 0 | - |
| - | - | 688.4 | 153.1 | - | - | 0 | - |
| - | - | 560.2 | 154.1 | - | - | 0 | - |
| - | - | 539.4 | 155.1 | - | - | 0 | - |
| - | - | 1.275E+04 | 155.1 | - | - | 0 | - |
| 2 | a | 1.122E+04 | 155.1 | 0.0003813 | 2.458 | +1 | 2 |
| - | - | 1150 | 156.1 | - | - | 0 | - |
| - | - | 988.5 | 156.1 | - | - | 0 | - |
| - | - | 644.7 | 156.1 | - | - | 0 | - |
| - | - | 788.3 | 157.1 | - | - | 0 | - |
| - | - | 1272 | 157.1 | - | - | 0 | - |
| - | - | 697.4 | 157.1 | - | - | 0 | - |
| - | - | 1282 | 158.1 | - | - | 0 | - |
| - | - | 1.815E+05 | 159.1 | - | - | 0 | - |
| - | - | 1453 | 160.1 | - | - | 0 | - |
| - | - | 1053 | 160.1 | - | - | 0 | - |
| - | - | 1.925E+04 | 160.1 | - | - | 0 | - |
| - | - | 438.1 | 161.1 | - | - | 0 | - |
| - | - | 1.633E+04 | 165.1 | - | - | 0 | - |
| - | - | 2.749E+04 | 166.1 | - | - | 0 | - |
| 10 | y | 2819 | 166.1 | 0.001802 | 10.85 | +3 | 4 |
| - | - | 1744 | 166.1 | - | - | 0 | - |
| - | - | 1.637E+04 | 167.1 | - | - | 0 | - |
| - | - | 2370 | 167.1 | - | - | 0 | - |
| - | - | 742.2 | 167.1 | - | - | 0 | - |
| - | - | 1234 | 167.1 | - | - | 0 | - |
| - | - | 566.6 | 167.3 | - | - | 0 | - |
| - | - | 429.2 | 167.7 | - | - | 0 | - |
| - | - | 971.2 | 168.1 | - | - | 0 | - |
| - | - | 3221 | 168.1 | - | - | 0 | - |
| - | - | 1575 | 168.1 | - | - | 0 | - |
| - | - | 2270 | 168.1 | - | - | 0 | - |
| - | - | 2220 | 169.1 | - | - | 0 | - |
| - | - | 6731 | 169.1 | - | - | 0 | - |
| - | - | 8240 | 170.1 | - | - | 0 | - |
| - | - | 677.3 | 171.1 | - | - | 0 | - |
| - | - | 733.7 | 171.1 | - | - | 0 | - |
| - | - | 1671 | 171.1 | - | - | 0 | - |
| - | - | 1966 | 171.1 | - | - | 0 | - |
| - | - | 3502 | 172.1 | - | - | 0 | - |
| - | - | 3993 | 173.1 | - | - | 0 | - |
| 2 | a | 3.592E+05 | 173.1 | 0.0004825 | 2.787 | +1 | 2 |
| - | - | 1112 | 173.4 | - | - | 0 | - |
| - | - | 627.8 | 174.1 | - | - | 0 | - |
| - | - | 3.082E+04 | 174.1 | - | - | 0 | - |
| - | - | 2470 | 175.1 | - | - | 0 | - |
| - | - | 466.9 | 176 | - | - | 0 | - |
| - | - | 1247 | 177.1 | - | - | 0 | - |
| - | - | 484.7 | 177.1 | - | - | 0 | - |
| - | - | 1.12E+04 | 178.1 | - | - | 0 | - |
| - | - | 886.7 | 179.1 | - | - | 0 | - |
| - | - | 1896 | 181.1 | - | - | 0 | - |
| 13 | y | 3.512E+04 | 182.1 | 0.0004496 | 2.469 | +1 | 1 |
| - | - | 2975 | 183.1 | - | - | 0 | - |
| 2 | b | 1.015E+04 | 183.1 | 0.0004466 | 2.439 | +1 | 2 |
| - | - | 1076 | 184.1 | - | - | 0 | - |
| - | - | 653 | 185.1 | - | - | 0 | - |
| - | - | 2506 | 185.2 | - | - | 0 | - |
| - | - | 7.231E+04 | 186.1 | - | - | 0 | - |
| - | - | 1.56E+04 | 187.1 | - | - | 0 | - |
| - | - | 666.5 | 187.1 | - | - | 0 | - |
| - | - | 6024 | 187.1 | - | - | 0 | - |
| - | - | 524.1 | 187.1 | - | - | 0 | - |
| - | - | 1606 | 188.1 | - | - | 0 | - |
| - | - | 1579 | 188.1 | - | - | 0 | - |
| - | - | 714.9 | 189.1 | - | - | 0 | - |
| - | - | 3424 | 190.1 | - | - | 0 | - |
| - | - | 2314 | 190.1 | - | - | 0 | - |
| - | - | 676.7 | 193.1 | - | - | 0 | - |
| - | - | 2322 | 194.1 | - | - | 0 | - |
| - | - | 1.839E+04 | 195.1 | - | - | 0 | - |
| - | - | 746.3 | 196.1 | - | - | 0 | - |
| - | - | 1.261E+04 | 196.1 | - | - | 0 | - |
| - | - | 1221 | 196.1 | - | - | 0 | - |
| - | - | 661.1 | 197.1 | - | - | 0 | - |
| - | - | 1006 | 197.1 | - | - | 0 | - |
| - | - | 834.6 | 197.1 | - | - | 0 | - |
| - | - | 2459 | 198.1 | - | - | 0 | - |
| - | - | 6403 | 198.1 | - | - | 0 | - |
| - | - | 1241 | 199.1 | - | - | 0 | - |
| - | - | 1438 | 199.1 | - | - | 0 | - |
| - | - | 1593 | 199.1 | - | - | 0 | - |
| - | - | 923.1 | 199.1 | - | - | 0 | - |
| - | - | 1194 | 200.1 | - | - | 0 | - |
| 2 | b | 9.768E+04 | 201.1 | 0.0003952 | 1.965 | +1 | 2 |
| - | - | 1240 | 202.1 | - | - | 0 | - |
| - | - | 9606 | 202.1 | - | - | 0 | - |
| - | - | 1271 | 203.1 | - | - | 0 | - |
| - | - | 1165 | 203.1 | - | - | 0 | - |
| - | - | 973.5 | 204.1 | - | - | 0 | - |
| - | - | 8332 | 205.1 | - | - | 0 | - |
| - | - | 885.1 | 205.1 | - | - | 0 | - |
| - | - | 1357 | 205.1 | - | - | 0 | - |
| - | - | 769.8 | 206.1 | - | - | 0 | - |
| - | - | 2204 | 206.1 | - | - | 0 | - |
| - | - | 3143 | 207.2 | - | - | 0 | - |
| - | - | 609.6 | 208.1 | - | - | 0 | - |
| - | - | 1045 | 209.1 | - | - | 0 | - |
| - | - | 665.1 | 209.1 | - | - | 0 | - |
| - | - | 3519 | 210.1 | - | - | 0 | - |
| 11 | y | 2318 | 211.1 | 0.0006186 | 2.93 | +2 | 3 |
| - | - | 2581 | 212.1 | - | - | 0 | - |
| - | - | 8387 | 213.1 | - | - | 0 | - |
| - | - | 1417 | 213.2 | - | - | 0 | - |
| - | - | 1024 | 214.1 | - | - | 0 | - |
| - | - | 633.9 | 215.1 | - | - | 0 | - |
| - | - | 1703 | 215.1 | - | - | 0 | - |
| - | - | 1773 | 216.1 | - | - | 0 | - |
| - | - | 1946 | 221.1 | - | - | 0 | - |
| - | - | 953.9 | 221.1 | - | - | 0 | - |
| - | - | 5825 | 221.1 | - | - | 0 | - |
| - | - | 1446 | 221.1 | - | - | 0 | - |
| - | - | 1026 | 221.1 | - | - | 0 | - |
| - | - | 2031 | 222.1 | - | - | 0 | - |
| - | - | 3002 | 222.1 | - | - | 0 | - |
| - | - | 3359 | 223.1 | - | - | 0 | - |
| - | - | 973.6 | 223.1 | - | - | 0 | - |
| - | - | 5381 | 223.1 | - | - | 0 | - |
| - | - | 1314 | 223.1 | - | - | 0 | - |
| - | - | 3.202E+04 | 223.2 | - | - | 0 | - |
| - | - | 1935 | 224.1 | - | - | 0 | - |
| - | - | 3743 | 224.2 | - | - | 0 | - |
| - | - | 7653 | 225 | - | - | 0 | - |
| - | - | 685.7 | 225.1 | - | - | 0 | - |
| - | - | 1852 | 226 | - | - | 0 | - |
| - | - | 8040 | 226.1 | - | - | 0 | - |
| - | - | 931.1 | 226.2 | - | - | 0 | - |
| - | - | 799.6 | 227 | - | - | 0 | - |
| - | - | 1447 | 227 | - | - | 0 | - |
| - | - | 6396 | 227.1 | - | - | 0 | - |
| - | - | 1034 | 227.2 | - | - | 0 | - |
| - | - | 704.3 | 228.1 | - | - | 0 | - |
| - | - | 682.5 | 228.1 | - | - | 0 | - |
| - | - | 2169 | 228.1 | - | - | 0 | - |
| - | - | 7332 | 229.1 | - | - | 0 | - |
| - | - | 1043 | 230.2 | - | - | 0 | - |
| - | - | 5059 | 233.1 | - | - | 0 | - |
| - | - | 3.8E+04 | 234.1 | - | - | 0 | - |
| - | - | 748.6 | 234.7 | - | - | 0 | - |
| - | - | 4662 | 235.1 | - | - | 0 | - |
| - | - | 977.1 | 235.6 | - | - | 0 | - |
| - | - | 1863 | 237.1 | - | - | 0 | - |
| - | - | 4252 | 238.1 | - | - | 0 | - |
| - | - | 880.5 | 238.1 | - | - | 0 | - |
| - | - | 563.5 | 239.1 | - | - | 0 | - |
| - | - | 1.415E+04 | 239.1 | - | - | 0 | - |
| - | - | 4865 | 240.1 | - | - | 0 | - |
| - | - | 1.375E+04 | 240.1 | - | - | 0 | - |
| - | - | 2851 | 241.1 | - | - | 0 | - |
| - | - | 1969 | 241.1 | - | - | 0 | - |
| - | - | 2131 | 241.1 | - | - | 0 | - |
| - | - | 1561 | 241.2 | - | - | 0 | - |
| - | - | 2994 | 243.1 | - | - | 0 | - |
| - | - | 1.551E+04 | 244.1 | - | - | 0 | - |
| - | - | 589.3 | 244.6 | - | - | 0 | - |
| - | - | 1066 | 245.1 | - | - | 0 | - |
| - | - | 1.81E+04 | 247.1 | - | - | 0 | - |
| - | - | 1413 | 247.1 | - | - | 0 | - |
| - | - | 2355 | 248.1 | - | - | 0 | - |
| - | - | 8123 | 248.1 | - | - | 0 | - |
| - | - | 1.18E+04 | 249.1 | - | - | 0 | - |
| - | - | 3041 | 249.1 | - | - | 0 | - |
| - | - | 1402 | 250.1 | - | - | 0 | - |
| - | - | 628.5 | 250.1 | - | - | 0 | - |
| - | - | 677.9 | 250.2 | - | - | 0 | - |
| - | - | 553.1 | 250.3 | - | - | 0 | - |
| - | - | 635.9 | 251 | - | - | 0 | - |
| - | - | 1.21E+05 | 251.2 | - | - | 0 | - |
| - | - | 632.5 | 252.1 | - | - | 0 | - |
| - | - | 1.364E+04 | 252.2 | - | - | 0 | - |
| - | - | 712.5 | 253.1 | - | - | 0 | - |
| - | - | 1571 | 253.2 | - | - | 0 | - |
| - | - | 6790 | 255.1 | - | - | 0 | - |
| - | - | 1617 | 256.1 | - | - | 0 | - |
| - | - | 907.4 | 256.2 | - | - | 0 | - |
| - | - | 1329 | 256.6 | - | - | 0 | - |
| - | - | 3205 | 257.1 | - | - | 0 | - |
| - | - | 5190 | 258.1 | - | - | 0 | - |
| - | - | 975.1 | 259.1 | - | - | 0 | - |
| - | - | 2862 | 261.1 | - | - | 0 | - |
| - | - | 919.8 | 261.1 | - | - | 0 | - |
| - | - | 1357 | 262.1 | - | - | 0 | - |
| - | - | 1.45E+04 | 265.1 | - | - | 0 | - |
| - | - | 3613 | 266.1 | - | - | 0 | - |
| - | - | 2.884E+04 | 266.1 | - | - | 0 | - |
| - | - | 759.4 | 267.1 | - | - | 0 | - |
| - | - | 3286 | 267.1 | - | - | 0 | - |
| - | - | 1015 | 267.1 | - | - | 0 | - |
| - | - | 1407 | 268.1 | - | - | 0 | - |
| - | - | 3414 | 268.2 | - | - | 0 | - |
| - | - | 1398 | 269.2 | - | - | 0 | - |
| - | - | 1733 | 269.2 | - | - | 0 | - |
| - | - | 4669 | 270.1 | - | - | 0 | - |
| - | - | 7326 | 272.2 | - | - | 0 | - |
| - | - | 1501 | 273.2 | - | - | 0 | - |
| - | - | 6925 | 274.1 | - | - | 0 | - |
| - | - | 1667 | 274.1 | - | - | 0 | - |
| - | - | 1577 | 275.1 | - | - | 0 | - |
| - | - | 1442 | 275.1 | - | - | 0 | - |
| - | - | 717.5 | 275.1 | - | - | 0 | - |
| - | - | 1424 | 276.1 | - | - | 0 | - |
| - | - | 2120 | 276.1 | - | - | 0 | - |
| - | - | 2139 | 276.2 | - | - | 0 | - |
| - | - | 2883 | 279.1 | - | - | 0 | - |
| - | - | 3105 | 280.1 | - | - | 0 | - |
| - | - | 9481 | 281.1 | - | - | 0 | - |
| - | - | 6666 | 282.2 | - | - | 0 | - |
| - | - | 829.4 | 282.7 | - | - | 0 | - |
| - | - | 1798 | 283 | - | - | 0 | - |
| - | - | 8.79E+04 | 283.1 | - | - | 0 | - |
| - | - | 3215 | 284.1 | - | - | 0 | - |
| - | - | 799.7 | 284.1 | - | - | 0 | - |
| - | - | 1.15E+04 | 284.1 | - | - | 0 | - |
| - | - | 694.2 | 285.1 | - | - | 0 | - |
| - | - | 1018 | 285.1 | - | - | 0 | - |
| - | - | 1153 | 285.1 | - | - | 0 | - |
| - | - | 728.7 | 285.2 | - | - | 0 | - |
| - | - | 5584 | 286.1 | - | - | 0 | - |
| - | - | 825.4 | 287.1 | - | - | 0 | - |
| - | - | 1761 | 287.2 | - | - | 0 | - |
| - | - | 571.4 | 288.1 | - | - | 0 | - |
| - | - | 1713 | 288.6 | - | - | 0 | - |
| 12 | y | 4.661E+04 | 293.1 | 0.000602 | 2.054 | +1 | 2 |
| - | - | 1477 | 293.2 | - | - | 0 | - |
| - | - | 9563 | 294.1 | - | - | 0 | - |
| - | - | 1245 | 295.1 | - | - | 0 | - |
| - | - | 1130 | 295.1 | - | - | 0 | - |
| - | - | 1186 | 296.1 | - | - | 0 | - |
| 3 | b | 1123 | 296.2 | 0.0008553 | 2.887 | +1 | 3 |
| - | - | 1636 | 297.1 | - | - | 0 | - |
| - | - | 834.9 | 297.1 | - | - | 0 | - |
| 9 | y | 1525 | 297.1 | 0.001989 | 6.694 | +2 | 5 |
| - | - | 1.732E+04 | 297.2 | - | - | 0 | - |
| - | - | 747.8 | 297.2 | - | - | 0 | - |
| 7 | y | 1499 | 298.1 | 0.005154 | 17.29 | +3 | 7 |
| - | - | 1603 | 298.2 | - | - | 0 | - |
| - | - | 9.137E+04 | 299.1 | - | - | 0 | - |
| - | - | 2548 | 300.1 | - | - | 0 | - |
| - | - | 1.173E+04 | 300.2 | - | - | 0 | - |
| - | - | 1624 | 301.1 | - | - | 0 | - |
| - | - | 999.4 | 301.1 | - | - | 0 | - |
| - | - | 4.339E+04 | 301.2 | - | - | 0 | - |
| - | - | 1466 | 301.2 | - | - | 0 | - |
| - | - | 8016 | 302.1 | - | - | 0 | - |
| - | - | 5699 | 302.2 | - | - | 0 | - |
| - | - | 1278 | 303.1 | - | - | 0 | - |
| - | - | 604.7 | 303.2 | - | - | 0 | - |
| - | - | 1021 | 305.2 | - | - | 0 | - |
| - | - | 1991 | 305.6 | - | - | 0 | - |
| 9 | y | 6905 | 306.1 | 0.0008265 | 2.7 | +2 | 5 |
| - | - | 642.8 | 306.2 | - | - | 0 | - |
| - | - | 3437 | 306.6 | - | - | 0 | - |
| - | - | 1857 | 307.1 | - | - | 0 | - |
| - | - | 534.8 | 307.2 | - | - | 0 | - |
| - | - | 600 | 309.2 | - | - | 0 | - |
| 12 | y | 6064 | 311.1 | 0.0003217 | 1.034 | +1 | 2 |
| - | - | 2905 | 311.1 | - | - | 0 | - |
| - | - | 608.4 | 311.7 | - | - | 0 | - |
| - | - | 1556 | 312.1 | - | - | 0 | - |
| - | - | 732.5 | 314.1 | - | - | 0 | - |
| - | - | 1880 | 314.2 | - | - | 0 | - |
| 3 | b | 3944 | 314.2 | 0.0009412 | 2.995 | +1 | 3 |
| - | - | 2.454E+04 | 315.2 | - | - | 0 | - |
| - | - | 4352 | 316.2 | - | - | 0 | - |
| - | - | 890.1 | 317.7 | - | - | 0 | - |
| - | - | 605.5 | 318.1 | - | - | 0 | - |
| - | - | 1797 | 319.2 | - | - | 0 | - |
| - | - | 595 | 324.2 | - | - | 0 | - |
| - | - | 1508 | 325.2 | - | - | 0 | - |
| - | - | 579.5 | 326 | - | - | 0 | - |
| - | - | 6704 | 326.7 | - | - | 0 | - |
| - | - | 628 | 327 | - | - | 0 | - |
| - | - | 2557 | 327.2 | - | - | 0 | - |
| - | - | 2564 | 332.2 | - | - | 0 | - |
| - | - | 4962 | 333.2 | - | - | 0 | - |
| - | - | 982.4 | 334.2 | - | - | 0 | - |
| - | - | 575.3 | 336 | - | - | 0 | - |
| 6 | b | 1092 | 338.7 | 0.0006383 | 1.885 | +2 | 6 |
| 6 | b | 1029 | 339.2 | 0.005335 | 15.73 | +2 | 6 |
| - | - | 1158 | 339.7 | - | - | 0 | - |
| - | - | 2796 | 340.7 | - | - | 0 | - |
| - | - | 704.3 | 341 | - | - | 0 | - |
| - | - | 1242 | 341.2 | - | - | 0 | - |
| - | - | 2445 | 342.2 | - | - | 0 | - |
| - | - | 1627 | 343.2 | - | - | 0 | - |
| - | - | 835.4 | 343.2 | - | - | 0 | - |
| - | - | 1965 | 344.2 | - | - | 0 | - |
| - | - | 1.098E+04 | 345 | - | - | 0 | - |
| - | - | 726.6 | 345 | - | - | 0 | - |
| - | - | 3800 | 345.1 | - | - | 0 | - |
| - | - | 1.34E+04 | 346.1 | - | - | 0 | - |
| - | - | 981 | 346.1 | - | - | 0 | - |
| - | - | 2581 | 347.1 | - | - | 0 | - |
| 6 | b | 7019 | 347.7 | 0.000727 | 2.091 | +2 | 6 |
| - | - | 2454 | 348.2 | - | - | 0 | - |
| - | - | 691 | 348.7 | - | - | 0 | - |
| - | - | 887.8 | 349.2 | - | - | 0 | - |
| - | - | 1.187E+04 | 350.2 | - | - | 0 | - |
| - | - | 974.2 | 351.2 | - | - | 0 | - |
| - | - | 2520 | 351.2 | - | - | 0 | - |
| - | - | 786.1 | 352.2 | - | - | 0 | - |
| - | - | 1817 | 352.7 | - | - | 0 | - |
| 8 | y | 2094 | 353.7 | 0.0004233 | 1.197 | +2 | 6 |
| - | - | 1680 | 354.2 | - | - | 0 | - |
| - | - | 1676 | 355.1 | - | - | 0 | - |
| - | - | 9757 | 359 | - | - | 0 | - |
| - | - | 871.6 | 359.1 | - | - | 0 | - |
| - | - | 709.8 | 360.2 | - | - | 0 | - |
| - | - | 2764 | 360.7 | - | - | 0 | - |
| - | - | 4300 | 361.2 | - | - | 0 | - |
| - | - | 915.4 | 361.7 | - | - | 0 | - |
| - | - | 5333 | 362.2 | - | - | 0 | - |
| 8 | y | 1.725E+04 | 362.7 | 0.0009088 | 2.506 | +2 | 6 |
| - | - | 2.059E+04 | 363.1 | - | - | 0 | - |
| - | - | 6536 | 363.2 | - | - | 0 | - |
| - | - | 1398 | 363.7 | - | - | 0 | - |
| - | - | 3516 | 364.1 | - | - | 0 | - |
| - | - | 3275 | 364.1 | - | - | 0 | - |
| - | - | 1885 | 366.2 | - | - | 0 | - |
| - | - | 1425 | 367.1 | - | - | 0 | - |
| - | - | 1516 | 367.2 | - | - | 0 | - |
| - | - | 1456 | 367.2 | - | - | 0 | - |
| - | - | 1085 | 368.2 | - | - | 0 | - |
| - | - | 1218 | 369.1 | - | - | 0 | - |
| - | - | 1141 | 370.1 | - | - | 0 | - |
| - | - | 802.1 | 370.2 | - | - | 0 | - |
| - | - | 1519 | 373.2 | - | - | 0 | - |
| - | - | 1845 | 376.2 | - | - | 0 | - |
| - | - | 4923 | 376.2 | - | - | 0 | - |
| - | - | 1334 | 376.7 | - | - | 0 | - |
| - | - | 1352 | 377.1 | - | - | 0 | - |
| - | - | 7963 | 378.2 | - | - | 0 | - |
| - | - | 3.071E+04 | 379.2 | - | - | 0 | - |
| - | - | 1098 | 380.2 | - | - | 0 | - |
| - | - | 7215 | 380.2 | - | - | 0 | - |
| - | - | 5.88E+04 | 381.2 | - | - | 0 | - |
| - | - | 1989 | 381.7 | - | - | 0 | - |
| - | - | 1.014E+04 | 382.2 | - | - | 0 | - |
| - | - | 2168 | 382.2 | - | - | 0 | - |
| - | - | 1008 | 382.2 | - | - | 0 | - |
| - | - | 1918 | 383.2 | - | - | 0 | - |
| - | - | 945 | 383.2 | - | - | 0 | - |
| - | - | 2008 | 384.2 | - | - | 0 | - |
| - | - | 3193 | 384.2 | - | - | 0 | - |
| - | - | 3608 | 385.2 | - | - | 0 | - |
| 5 | y | 1345 | 385.2 | 0.003735 | 9.696 | +3 | 9 |
| - | - | 1229 | 386.2 | - | - | 0 | - |
| - | - | 2190 | 390.2 | - | - | 0 | - |
| - | - | 807.6 | 390.7 | - | - | 0 | - |
| - | - | 3067 | 391.1 | - | - | 0 | - |
| - | - | 698.8 | 391.2 | - | - | 0 | - |
| - | - | 730.1 | 392.1 | - | - | 0 | - |
| - | - | 8377 | 394.2 | - | - | 0 | - |
| - | - | 2627 | 395.1 | - | - | 0 | - |
| - | - | 3226 | 395.2 | - | - | 0 | - |
| - | - | 621.9 | 395.2 | - | - | 0 | - |
| - | - | 1396 | 395.2 | - | - | 0 | - |
| - | - | 788.4 | 396.1 | - | - | 0 | - |
| - | - | 1.104E+04 | 396.2 | - | - | 0 | - |
| - | - | 742.1 | 397.2 | - | - | 0 | - |
| - | - | 1757 | 397.2 | - | - | 0 | - |
| - | - | 3560 | 398.2 | - | - | 0 | - |
| - | - | 717.8 | 399.2 | - | - | 0 | - |
| - | - | 702.2 | 401.2 | - | - | 0 | - |
| - | - | 690.6 | 401.2 | - | - | 0 | - |
| - | - | 3309 | 402.2 | - | - | 0 | - |
| - | - | 5203 | 402.2 | - | - | 0 | - |
| - | - | 2357 | 403.2 | - | - | 0 | - |
| - | - | 1747 | 404.7 | - | - | 0 | - |
| - | - | 2390 | 405.3 | - | - | 0 | - |
| - | - | 907.3 | 406.2 | - | - | 0 | - |
| - | - | 1159 | 406.2 | - | - | 0 | - |
| - | - | 1153 | 407.2 | - | - | 0 | - |
| - | - | 582.6 | 407.2 | - | - | 0 | - |
| - | - | 715.5 | 408.7 | - | - | 0 | - |
| - | - | 4271 | 409.1 | - | - | 0 | - |
| - | - | 987.7 | 409.2 | - | - | 0 | - |
| - | - | 1333 | 410.1 | - | - | 0 | - |
| - | - | 4728 | 411.2 | - | - | 0 | - |
| - | - | 3.417E+04 | 412.2 | - | - | 0 | - |
| - | - | 4530 | 413.1 | - | - | 0 | - |
| - | - | 1874 | 413.2 | - | - | 0 | - |
| - | - | 4057 | 413.2 | - | - | 0 | - |
| - | - | 921.9 | 413.3 | - | - | 0 | - |
| - | - | 1165 | 414.1 | - | - | 0 | - |
| - | - | 1212 | 414.2 | - | - | 0 | - |
| - | - | 859.1 | 414.2 | - | - | 0 | - |
| - | - | 8655 | 414.2 | - | - | 0 | - |
| - | - | 2.031E+04 | 415 | - | - | 0 | - |
| - | - | 3434 | 415.2 | - | - | 0 | - |
| - | - | 2120 | 415.2 | - | - | 0 | - |
| - | - | 750.9 | 416.2 | - | - | 0 | - |
| - | - | 6457 | 417.7 | - | - | 0 | - |
| - | - | 1.016E+04 | 418.2 | - | - | 0 | - |
| - | - | 4908 | 418.7 | - | - | 0 | - |
| - | - | 1916 | 419.2 | - | - | 0 | - |
| 11 | y | 1398 | 421.2 | 0.001769 | 4.201 | +1 | 3 |
| 11 | y | 938.4 | 422.2 | 0.001457 | 3.452 | +1 | 3 |
| - | - | 759 | 422.3 | - | - | 0 | - |
| - | - | 1044 | 422.7 | - | - | 0 | - |
| - | - | 1694 | 423.2 | - | - | 0 | - |
| - | - | 1.595E+04 | 423.3 | - | - | 0 | - |
| - | - | 747.3 | 423.7 | - | - | 0 | - |
| - | - | 4005 | 424.3 | - | - | 0 | - |
| - | - | 1450 | 424.7 | - | - | 0 | - |
| - | - | 9.478E+04 | 426.7 | - | - | 0 | - |
| - | - | 5.056E+04 | 427.2 | - | - | 0 | - |
| - | - | 707.6 | 427.3 | - | - | 0 | - |
| - | - | 1.299E+04 | 427.7 | - | - | 0 | - |
| - | - | 2996 | 428.2 | - | - | 0 | - |
| - | - | 1.947E+04 | 429.2 | - | - | 0 | - |
| - | - | 8.576E+04 | 430.2 | - | - | 0 | - |
| - | - | 1.915E+04 | 431.2 | - | - | 0 | - |
| 7 | b | 1.079E+04 | 431.7 | 0.0005326 | 1.234 | +2 | 7 |
| 7 | b | 6875 | 432.2 | 0.006968 | 16.12 | +2 | 7 |
| - | - | 2100 | 432.7 | - | - | 0 | - |
| - | - | 1056 | 433 | - | - | 0 | - |
| - | - | 2501 | 433.2 | - | - | 0 | - |
| 4 | b | 3596 | 433.3 | 0.0005677 | 1.31 | +1 | 4 |
| - | - | 1091 | 434.3 | - | - | 0 | - |
| - | - | 1574 | 438.2 | - | - | 0 | - |
| - | - | 1133 | 438.7 | - | - | 0 | - |
| 11 | y | 2.445E+04 | 439.2 | 0.001092 | 2.487 | +1 | 3 |
| - | - | 1003 | 440.2 | - | - | 0 | - |
| - | - | 5145 | 440.2 | - | - | 0 | - |
| 7 | b | 4.604E+04 | 440.7 | 0.001049 | 2.379 | +2 | 7 |
| - | - | 1143 | 440.9 | - | - | 0 | - |
| - | - | 2.612E+04 | 441.2 | - | - | 0 | - |
| - | - | 5873 | 441.7 | - | - | 0 | - |
| - | - | 1476 | 442.2 | - | - | 0 | - |
| - | - | 861.4 | 442.7 | - | - | 0 | - |
| 7 | y | 2672 | 447.2 | 0.002084 | 4.661 | +2 | 7 |
| - | - | 2098 | 447.7 | - | - | 0 | - |
| - | - | 5031 | 448.2 | - | - | 0 | - |
| - | - | 879.1 | 448.3 | - | - | 0 | - |
| - | - | 1775 | 449.2 | - | - | 0 | - |
| - | - | 1464 | 450.2 | - | - | 0 | - |
| - | - | 1159 | 450.3 | - | - | 0 | - |
| 4 | b | 2.392E+04 | 451.3 | 0.001081 | 2.395 | +1 | 4 |
| - | - | 6736 | 452.3 | - | - | 0 | - |
| - | - | 771.2 | 455.2 | - | - | 0 | - |
| - | - | 1865 | 455.3 | - | - | 0 | - |
| 7 | y | 1337 | 455.7 | 0.001719 | 3.771 | +2 | 7 |
| - | - | 912 | 456.2 | - | - | 0 | - |
| - | - | 2957 | 459.2 | - | - | 0 | - |
| - | - | 2053 | 460.8 | - | - | 0 | - |
| - | - | 3553 | 461.2 | - | - | 0 | - |
| - | - | 3268 | 461.3 | - | - | 0 | - |
| - | - | 937.7 | 462.2 | - | - | 0 | - |
| - | - | 841.8 | 464.2 | - | - | 0 | - |
| - | - | 968.3 | 466.2 | - | - | 0 | - |
| - | - | 7032 | 468.3 | - | - | 0 | - |
| - | - | 1655 | 469.3 | - | - | 0 | - |
| - | - | 6641 | 474.3 | - | - | 0 | - |
| - | - | 7534 | 474.8 | - | - | 0 | - |
| - | - | 2981 | 475.3 | - | - | 0 | - |
| - | - | 1682 | 475.8 | - | - | 0 | - |
| - | - | 2179 | 476.2 | - | - | 0 | - |
| - | - | 717.8 | 476.3 | - | - | 0 | - |
| - | - | 4897 | 477.2 | - | - | 0 | - |
| 10 | y | 4793 | 478.2 | 0.0003862 | 0.8075 | +1 | 4 |
| - | - | 3470 | 478.3 | - | - | 0 | - |
| - | - | 1147 | 479.2 | - | - | 0 | - |
| - | - | 1326 | 479.3 | - | - | 0 | - |
| - | - | 1.767E+04 | 483.3 | - | - | 0 | - |
| - | - | 1.282E+04 | 483.8 | - | - | 0 | - |
| - | - | 862.5 | 484.2 | - | - | 0 | - |
| - | - | 3726 | 484.3 | - | - | 0 | - |
| 8 | b | 4598 | 488.3 | 0.001652 | 3.384 | +2 | 8 |
| 8 | b | 2473 | 488.7 | 0.006684 | 13.68 | +2 | 8 |
| - | - | 1302 | 489.3 | - | - | 0 | - |
| - | - | 5.931E+04 | 494.2 | - | - | 0 | - |
| - | - | 1.413E+04 | 495.2 | - | - | 0 | - |
| 10 | y | 7.57E+04 | 496.2 | 0.001021 | 2.058 | +1 | 4 |
| - | - | 1.673E+04 | 497.2 | - | - | 0 | - |
| - | - | 7117 | 497.8 | - | - | 0 | - |
| - | - | 3426 | 498.2 | - | - | 0 | - |
| - | - | 699.5 | 499.9 | - | - | 0 | - |
| - | - | 2.174E+04 | 503.1 | - | - | 0 | - |
| - | - | 2371 | 504.2 | - | - | 0 | - |
| - | - | 2050 | 506.2 | - | - | 0 | - |
| - | - | 2064 | 507.3 | - | - | 0 | - |
| - | - | 3954 | 511.3 | - | - | 0 | - |
| - | - | 1255 | 512.3 | - | - | 0 | - |
| - | - | 759.2 | 513.2 | - | - | 0 | - |
| - | - | 1114 | 515.3 | - | - | 0 | - |
| - | - | 7656 | 519.1 | - | - | 0 | - |
| - | - | 713.1 | 519.3 | - | - | 0 | - |
| - | - | 866.9 | 520.3 | - | - | 0 | - |
| - | - | 786 | 521.2 | - | - | 0 | - |
| - | - | 1482 | 523.3 | - | - | 0 | - |
| - | - | 1451 | 524.3 | - | - | 0 | - |
| - | - | 1.182E+04 | 525.3 | - | - | 0 | - |
| - | - | 1110 | 526.2 | - | - | 0 | - |
| - | - | 2696 | 526.3 | - | - | 0 | - |
| 0 | Precursor | 2959 | 529.3 | 0.002512 | 4.745 | +3 | -1 |
| 0 | Precursor | 2387 | 529.6 | 0.007636 | 14.42 | +3 | -1 |
| - | - | 2141 | 529.9 | - | - | 0 | - |
| - | - | 676.5 | 531.2 | - | - | 0 | - |
| - | - | 2251 | 532.2 | - | - | 0 | - |
| - | - | 807.5 | 533.2 | - | - | 0 | - |
| - | - | 1055 | 534.3 | - | - | 0 | - |
| - | - | 769.5 | 534.6 | - | - | 0 | - |
| - | - | 881.9 | 534.6 | - | - | 0 | - |
| - | - | 1280 | 534.9 | - | - | 0 | - |
| - | - | 889.7 | 535.2 | - | - | 0 | - |
| 0 | Precursor | 3219 | 535.3 | 0.00076 | 1.42 | +3 | -1 |
| - | - | 1161 | 535.6 | - | - | 0 | - |
| - | - | 2575 | 535.6 | - | - | 0 | - |
| - | - | 2151 | 535.9 | - | - | 0 | - |
| - | - | 5160 | 536.2 | - | - | 0 | - |
| - | - | 739.7 | 536.3 | - | - | 0 | - |
| - | - | 993.1 | 537.2 | - | - | 0 | - |
| - | - | 1419 | 537.7 | - | - | 0 | - |
| - | - | 1037 | 538.3 | - | - | 0 | - |
| - | - | 1.143E+04 | 539.2 | - | - | 0 | - |
| - | - | 853 | 539.3 | - | - | 0 | - |
| - | - | 3005 | 540.2 | - | - | 0 | - |
| - | - | 795.9 | 541.8 | - | - | 0 | - |
| - | - | 3757 | 542.3 | - | - | 0 | - |
| - | - | 1.712E+04 | 543.3 | - | - | 0 | - |
| - | - | 5082 | 544.3 | - | - | 0 | - |
| - | - | 1218 | 545.3 | - | - | 0 | - |
| 9 | b | 1119 | 545.8 | 0.002237 | 4.098 | +2 | 9 |
| 9 | b | 2084 | 546.3 | 0.009625 | 17.62 | +2 | 9 |
| - | - | 977.8 | 547.2 | - | - | 0 | - |
| - | - | 2320 | 549.2 | - | - | 0 | - |
| - | - | 1060 | 550.2 | - | - | 0 | - |
| - | - | 2310 | 551.3 | - | - | 0 | - |
| - | - | 1866 | 551.3 | - | - | 0 | - |
| - | - | 1236 | 552.3 | - | - | 0 | - |
| - | - | 1016 | 553.3 | - | - | 0 | - |
| 9 | b | 2114 | 554.8 | 0.004101 | 7.392 | +2 | 9 |
| - | - | 4799 | 555.3 | - | - | 0 | - |
| - | - | 3492 | 555.8 | - | - | 0 | - |
| 5 | b | 2319 | 561.3 | 0.001499 | 2.671 | +1 | 5 |
| 5 | b | 815.8 | 562.3 | 0.006375 | 11.34 | +1 | 5 |
| - | - | 1889 | 563.3 | - | - | 0 | - |
| - | - | 1425 | 564.3 | - | - | 0 | - |
| - | - | 743.7 | 564.3 | - | - | 0 | - |
| - | - | 3841 | 565.3 | - | - | 0 | - |
| - | - | 1157 | 566.3 | - | - | 0 | - |
| - | - | 1.257E+04 | 567.2 | - | - | 0 | - |
| - | - | 3951 | 568.2 | - | - | 0 | - |
| - | - | 3956 | 575.2 | - | - | 0 | - |
| - | - | 760.7 | 575.3 | - | - | 0 | - |
| - | - | 1953 | 576.2 | - | - | 0 | - |
| 5 | b | 1.666E+04 | 579.3 | 0.0008224 | 1.42 | +1 | 5 |
| - | - | 5994 | 580.3 | - | - | 0 | - |
| - | - | 1189 | 581.3 | - | - | 0 | - |
| - | - | 637.5 | 589.3 | - | - | 0 | - |
| - | - | 2396 | 592.3 | - | - | 0 | - |
| - | - | 3403 | 592.3 | - | - | 0 | - |
| 9 | y | 2.122E+04 | 593.3 | 0.0006343 | 1.069 | +1 | 5 |
| - | - | 7055 | 593.3 | - | - | 0 | - |
| - | - | 6648 | 594.3 | - | - | 0 | - |
| - | - | 1601 | 594.3 | - | - | 0 | - |
| - | - | 855.5 | 595.2 | - | - | 0 | - |
| - | - | 802.1 | 595.3 | - | - | 0 | - |
| - | - | 947.1 | 596.4 | - | - | 0 | - |
| - | - | 900.7 | 597.3 | - | - | 0 | - |
| - | - | 1748 | 597.8 | - | - | 0 | - |
| - | - | 1146 | 600.3 | - | - | 0 | - |
| - | - | 1932 | 602.3 | - | - | 0 | - |
| - | - | 3297 | 602.8 | - | - | 0 | - |
| - | - | 4279 | 603.3 | - | - | 0 | - |
| - | - | 1216 | 603.8 | - | - | 0 | - |
| - | - | 4.745E+04 | 610.3 | - | - | 0 | - |
| - | - | 2090 | 610.3 | - | - | 0 | - |
| 9 | y | 2.295E+05 | 611.3 | 0.001727 | 2.826 | +1 | 5 |
| - | - | 1.182E+04 | 611.8 | - | - | 0 | - |
| - | - | 7.038E+04 | 612.3 | - | - | 0 | - |
| - | - | 2978 | 612.8 | - | - | 0 | - |
| - | - | 1.734E+04 | 613.3 | - | - | 0 | - |
| - | - | 2513 | 614.3 | - | - | 0 | - |
| - | - | 846.4 | 617.3 | - | - | 0 | - |
| - | - | 1632 | 620.8 | - | - | 0 | - |
| - | - | 3994 | 621.3 | - | - | 0 | - |
| - | - | 1820 | 621.3 | - | - | 0 | - |
| - | - | 1461 | 622.3 | - | - | 0 | - |
| - | - | 1336 | 622.4 | - | - | 0 | - |
| - | - | 622.8 | 629.3 | - | - | 0 | - |
| - | - | 777.4 | 630.3 | - | - | 0 | - |
| - | - | 2780 | 634.3 | - | - | 0 | - |
| - | - | 2757 | 635.3 | - | - | 0 | - |
| - | - | 1544 | 636.3 | - | - | 0 | - |
| 4 | y | 2660 | 636.8 | 0.001405 | 2.206 | +2 | 10 |
| 4 | y | 2173 | 637.3 | 0.009885 | 15.51 | +2 | 10 |
| - | - | 924.4 | 637.3 | - | - | 0 | - |
| - | - | 2389 | 640.3 | - | - | 0 | - |
| - | - | 746.4 | 644.3 | - | - | 0 | - |
| - | - | 4053 | 645.3 | - | - | 0 | - |
| 4 | y | 1.511E+04 | 645.8 | 0.002226 | 3.447 | +2 | 10 |
| - | - | 9223 | 646.3 | - | - | 0 | - |
| - | - | 2554 | 646.8 | - | - | 0 | - |
| - | - | 765.8 | 647.3 | - | - | 0 | - |
| 11 | b | 817.6 | 647.3 | 0.004275 | 6.604 | +2 | 11 |
| - | - | 913.9 | 648.3 | - | - | 0 | - |
| - | - | 1824 | 650.3 | - | - | 0 | - |
| - | - | 1460 | 650.3 | - | - | 0 | - |
| - | - | 2.435E+04 | 652.3 | - | - | 0 | - |
| - | - | 8705 | 653.3 | - | - | 0 | - |
| - | - | 1534 | 654.3 | - | - | 0 | - |
| - | - | 1023 | 658.3 | - | - | 0 | - |
| - | - | 2221 | 660.9 | - | - | 0 | - |
| - | - | 1166 | 661.3 | - | - | 0 | - |
| - | - | 1145 | 661.4 | - | - | 0 | - |
| - | - | 7492 | 662.3 | - | - | 0 | - |
| - | - | 743 | 662.8 | - | - | 0 | - |
| - | - | 6051 | 663.3 | - | - | 0 | - |
| - | - | 942.5 | 664.3 | - | - | 0 | - |
| - | - | 1136 | 664.3 | - | - | 0 | - |
| - | - | 661.9 | 666.4 | - | - | 0 | - |
| - | - | 834.8 | 668.3 | - | - | 0 | - |
| - | - | 836.8 | 675.8 | - | - | 0 | - |
| 6 | b | 1.202E+04 | 676.3 | 0.001107 | 1.636 | +1 | 6 |
| 6 | b | 7946 | 677.3 | 0.0108 | 15.95 | +1 | 6 |
| - | - | 1010 | 677.8 | - | - | 0 | - |
| - | - | 2240 | 678.3 | - | - | 0 | - |
| - | - | 1821 | 679.3 | - | - | 0 | - |
| - | - | 4.469E+04 | 680.3 | - | - | 0 | - |
| - | - | 1.842E+04 | 681.3 | - | - | 0 | - |
| - | - | 3980 | 682.3 | - | - | 0 | - |
| - | - | 1034 | 684.3 | - | - | 0 | - |
| - | - | 3436 | 684.8 | - | - | 0 | - |
| - | - | 1963 | 685.3 | - | - | 0 | - |
| - | - | 1060 | 685.8 | - | - | 0 | - |
| - | - | 1903 | 688.3 | - | - | 0 | - |
| - | - | 2709 | 692.8 | - | - | 0 | - |
| 3 | y | 1.352E+04 | 693.3 | 0.00228 | 3.289 | +2 | 11 |
| 3 | y | 1.141E+04 | 693.8 | 0.008319 | 11.99 | +2 | 11 |
| 6 | b | 1.194E+05 | 694.4 | 0.0003687 | 0.5309 | +1 | 6 |
| - | - | 2993 | 694.8 | - | - | 0 | - |
| - | - | 4.718E+04 | 695.4 | - | - | 0 | - |
| - | - | 1.076E+04 | 696.4 | - | - | 0 | - |
| - | - | 1612 | 697.4 | - | - | 0 | - |
| - | - | 901.7 | 697.9 | - | - | 0 | - |
| - | - | 1390 | 698.4 | - | - | 0 | - |
| - | - | 800 | 700.9 | - | - | 0 | - |
| - | - | 1489 | 701.3 | - | - | 0 | - |
| - | - | 2.187E+04 | 701.8 | - | - | 0 | - |
| 3 | y | 8.272E+04 | 702.3 | 0.002613 | 3.721 | +2 | 11 |
| 12 | b | 6.662E+04 | 702.8 | 0.01253 | 17.83 | +2 | 12 |
| 12 | b | 3.119E+04 | 703.3 | 0.003137 | 4.46 | +2 | 12 |
| - | - | 8579 | 703.8 | - | - | 0 | - |
| - | - | 4029 | 704.3 | - | - | 0 | - |
| - | - | 1424 | 705.4 | - | - | 0 | - |
| 8 | y | 7753 | 706.3 | 0.002508 | 3.55 | +1 | 6 |
| - | - | 4132 | 707.3 | - | - | 0 | - |
| - | - | 1023 | 708.3 | - | - | 0 | - |
| - | - | 1702 | 710.4 | - | - | 0 | - |
| - | - | 967 | 710.9 | - | - | 0 | - |
| - | - | 4982 | 711.4 | - | - | 0 | - |
| 12 | b | 6996 | 711.9 | 0.00312 | 4.383 | +2 | 12 |
| - | - | 7378 | 712.4 | - | - | 0 | - |
| - | - | 2789 | 712.9 | - | - | 0 | - |
| - | - | 1188 | 713.4 | - | - | 0 | - |
| - | - | 766.8 | 716.3 | - | - | 0 | - |
| - | - | 1009 | 718.4 | - | - | 0 | - |
| - | - | 1029 | 720.4 | - | - | 0 | - |
| - | - | 5552 | 721.4 | - | - | 0 | - |
| - | - | 2411 | 722.4 | - | - | 0 | - |
| - | - | 1.314E+04 | 723.4 | - | - | 0 | - |
| 8 | y | 7.082E+04 | 724.4 | 0.002075 | 2.865 | +1 | 6 |
| - | - | 2.943E+04 | 725.4 | - | - | 0 | - |
| - | - | 7366 | 726.4 | - | - | 0 | - |
| - | - | 808.4 | 727.4 | - | - | 0 | - |
| - | - | 1102 | 728.4 | - | - | 0 | - |
| - | - | 2159 | 729.4 | - | - | 0 | - |
| - | - | 745.2 | 729.5 | - | - | 0 | - |
| - | - | 1576 | 730.4 | - | - | 0 | - |
| - | - | 3761 | 733.4 | - | - | 0 | - |
| - | - | 2821 | 734.4 | - | - | 0 | - |
| 2 | y | 1115 | 742.9 | 0.002619 | 3.526 | +2 | 12 |
| 2 | y | 2380 | 743.4 | 0.01427 | 19.2 | +2 | 12 |
| - | - | 776.5 | 743.9 | - | - | 0 | - |
| - | - | 1003 | 744.4 | - | - | 0 | - |
| - | - | 1858 | 747.4 | - | - | 0 | - |
| - | - | 2462 | 748.4 | - | - | 0 | - |
| - | - | 1764 | 749.3 | - | - | 0 | - |
| - | - | 900.9 | 750.4 | - | - | 0 | - |
| - | - | 1.327E+04 | 751.4 | - | - | 0 | - |
| 2 | y | 6776 | 751.9 | 0.002647 | 3.521 | +2 | 12 |
| - | - | 1.36E+04 | 752.4 | - | - | 0 | - |
| - | - | 2934 | 752.9 | - | - | 0 | - |
| - | - | 2496 | 753.4 | - | - | 0 | - |
| - | - | 902.7 | 755.4 | - | - | 0 | - |
| - | - | 1350 | 761.4 | - | - | 0 | - |
| - | - | 1652 | 762.4 | - | - | 0 | - |
| - | - | 7072 | 763.4 | - | - | 0 | - |
| - | - | 3755 | 764.4 | - | - | 0 | - |
| - | - | 2708 | 765.4 | - | - | 0 | - |
| - | - | 1683 | 766.4 | - | - | 0 | - |
| - | - | 2959 | 767.3 | - | - | 0 | - |
| - | - | 929.2 | 767.4 | - | - | 0 | - |
| - | - | 1015 | 768.3 | - | - | 0 | - |
| - | - | 1466 | 769.4 | - | - | 0 | - |
| - | - | 1150 | 775.4 | - | - | 0 | - |
| - | - | 1097 | 776.4 | - | - | 0 | - |
| - | - | 8026 | 779.4 | - | - | 0 | - |
| - | - | 3507 | 780.4 | - | - | 0 | - |
| - | - | 3501 | 781.4 | - | - | 0 | - |
| - | - | 1898 | 782.4 | - | - | 0 | - |
| 0 | Precursor | 1.326E+04 | 793.4 | 0.01449 | 18.26 | +2 | -1 |
| - | - | 4548 | 794.4 | - | - | 0 | - |
| - | - | 3176 | 795.3 | - | - | 0 | - |
| - | - | 916.3 | 795.4 | - | - | 0 | - |
| - | - | 1247 | 796.3 | - | - | 0 | - |
| - | - | 3852 | 807.4 | - | - | 0 | - |
| - | - | 1715 | 807.4 | - | - | 0 | - |
| - | - | 1090 | 808.4 | - | - | 0 | - |
| - | - | 1765 | 818.4 | - | - | 0 | - |
| - | - | 1612 | 819.4 | - | - | 0 | - |
| - | - | 860.3 | 820.4 | - | - | 0 | - |
| - | - | 1252 | 825.4 | - | - | 0 | - |
| - | - | 1822 | 826.4 | - | - | 0 | - |
| - | - | 740.4 | 827.4 | - | - | 0 | - |
| - | - | 1826 | 834.4 | - | - | 0 | - |
| - | - | 4122 | 835.4 | - | - | 0 | - |
| - | - | 7887 | 836.5 | - | - | 0 | - |
| - | - | 4736 | 837.5 | - | - | 0 | - |
| - | - | 1799 | 838.5 | - | - | 0 | - |
| - | - | 1663 | 843.4 | - | - | 0 | - |
| - | - | 2972 | 844.4 | - | - | 0 | - |
| - | - | 2419 | 845.4 | - | - | 0 | - |
| - | - | 3481 | 846.4 | - | - | 0 | - |
| - | - | 2757 | 847.4 | - | - | 0 | - |
| - | - | 1575 | 848.4 | - | - | 0 | - |
| - | - | 2.837E+04 | 852.4 | - | - | 0 | - |
| - | - | 1.473E+04 | 853.4 | - | - | 0 | - |
| - | - | 4611 | 854.4 | - | - | 0 | - |
| 7 | b | 1.726E+04 | 862.4 | 0.000346 | 0.4012 | +1 | 7 |
| 7 | b | 1.34E+04 | 863.4 | 0.01297 | 15.03 | +1 | 7 |
| - | - | 2.055E+04 | 864.5 | - | - | 0 | - |
| - | - | 1.308E+04 | 865.5 | - | - | 0 | - |
| - | - | 3619 | 866.5 | - | - | 0 | - |
| - | - | 1116 | 869.4 | - | - | 0 | - |
| - | - | 2781 | 876.4 | - | - | 0 | - |
| - | - | 901.7 | 877.4 | - | - | 0 | - |
| 7 | b | 1.428E+05 | 880.4 | 0.0003404 | 0.3866 | +1 | 7 |
| - | - | 7.637E+04 | 881.4 | - | - | 0 | - |
| - | - | 2.448E+04 | 882.4 | - | - | 0 | - |
| - | - | 3727 | 883.4 | - | - | 0 | - |
| - | - | 1029 | 884.4 | - | - | 0 | - |
| - | - | 1129 | 890.4 | - | - | 0 | - |
| - | - | 1692 | 891.4 | - | - | 0 | - |
| - | - | 2775 | 892.5 | - | - | 0 | - |
| - | - | 1186 | 893.5 | - | - | 0 | - |
| - | - | 925.8 | 894.5 | - | - | 0 | - |
| - | - | 1427 | 895.5 | - | - | 0 | - |
| - | - | 1802 | 907.4 | - | - | 0 | - |
| - | - | 9881 | 908.4 | - | - | 0 | - |
| - | - | 7822 | 909.4 | - | - | 0 | - |
| 7 | y | 1.161E+04 | 910.4 | 0.002108 | 2.315 | +1 | 7 |
| - | - | 3272 | 911.4 | - | - | 0 | - |
| - | - | 1857 | 912.4 | - | - | 0 | - |
| - | - | 1104 | 936.4 | - | - | 0 | - |
| - | - | 1130 | 937.4 | - | - | 0 | - |
| - | - | 1162 | 938.4 | - | - | 0 | - |
| - | - | 987.6 | 947.4 | - | - | 0 | - |
| - | - | 2987 | 954.4 | - | - | 0 | - |
| - | - | 3493 | 955.4 | - | - | 0 | - |
| - | - | 1882 | 956.4 | - | - | 0 | - |
| - | - | 891.5 | 962.4 | - | - | 0 | - |
| - | - | 1095 | 964.5 | - | - | 0 | - |
| - | - | 3506 | 965.4 | - | - | 0 | - |
| - | - | 1813 | 966.5 | - | - | 0 | - |
| - | - | 928.3 | 971.5 | - | - | 0 | - |
| - | - | 2465 | 972.4 | - | - | 0 | - |
| - | - | 1356 | 973.4 | - | - | 0 | - |
| - | - | 916.1 | 974.4 | - | - | 0 | - |
| 8 | b | 1773 | 975.5 | 0.001365 | 1.399 | +1 | 8 |
| 8 | b | 1185 | 976.5 | 0.0179 | 18.33 | +1 | 8 |
| - | - | 814.3 | 981.5 | - | - | 0 | - |
| - | - | 1373 | 982.5 | - | - | 0 | - |
| 8 | b | 8568 | 993.5 | 0.0005659 | 0.5696 | +1 | 8 |
| - | - | 4537 | 994.5 | - | - | 0 | - |
| - | - | 2806 | 995.5 | - | - | 0 | - |
| - | - | 778.6 | 1007 | - | - | 0 | - |
| - | - | 1107 | 1024 | - | - | 0 | - |
| 6 | y | 4594 | 1025 | 0.003363 | 3.279 | +1 | 8 |
| - | - | 2996 | 1026 | - | - | 0 | - |
| - | - | 809 | 1027 | - | - | 0 | - |
| - | - | 935.4 | 1065 | - | - | 0 | - |
| - | - | 755.1 | 1076 | - | - | 0 | - |
| - | - | 1117 | 1077 | - | - | 0 | - |
| - | - | 3343 | 1094 | - | - | 0 | - |
| - | - | 2219 | 1095 | - | - | 0 | - |
| - | - | 946.4 | 1096 | - | - | 0 | - |
| 9 | b | 736.5 | 1109 | 0.002737 | 2.469 | +1 | 9 |
| - | - | 1109 | 1117 | - | - | 0 | - |
| - | - | 1611 | 1118 | - | - | 0 | - |
| - | - | 809.4 | 1119 | - | - | 0 | - |
| 5 | y | 5505 | 1136 | 0.001981 | 1.745 | +1 | 9 |
| 5 | y | 7009 | 1136 | 0.008322 | 7.322 | +1 | 9 |
| - | - | 3565 | 1137 | - | - | 0 | - |
| - | - | 1516 | 1139 | - | - | 0 | - |
| - | - | 2642 | 1153 | - | - | 0 | - |
| 5 | y | 1.091E+04 | 1154 | 0.001914 | 1.66 | +1 | 9 |
| - | - | 6003 | 1155 | - | - | 0 | - |
| - | - | 2527 | 1156 | - | - | 0 | - |
| - | - | 1017 | 1157 | - | - | 0 | - |
| 10 | b | 1961 | 1166 | 0.006053 | 5.193 | +1 | 10 |
| - | - | 1064 | 1183 | - | - | 0 | - |
| - | - | 659.2 | 3081 | - | - | 0 | - |

m/z Charge Intensity FragmentType MassShift Position
120.0811767578125 0 4691.317
121.0843505859375 0 663.5456
121.21499633789062 0 350.9592
122.0716323852539 0 1573.3623
122.14251708984375 0 454.56314
123.04443359375 0 2165.8308
123.0555648803711 0 634.40845
123.39301300048828 0 418.7689
123.74199676513672 0 358.2154
125.10765838623047 0 891.017
126.09171295166016 0 811.1962
126.1018295288086 0 362.02988
127.05062866210938 0 4844.75
127.08773040771484 0 444.74646
127.1233901977539 0 905.0133
128.1073760986328 0 22426.666
129.1026611328125 0 236773.33
130.06552124023438 0 105439.76
130.09982299804688 0 1433.76
130.10598754882812 0 15643.422
131.06886291503906 0 11040.29
131.11846923828125 0 635.88495
132.08116149902344 0 23011.816
132.53701782226562 0 518.3584
133.06112670898438 0 948.06683
133.08460998535156 0 2257.2227
136.0760955810547 0 30735.977
137.07948303222656 0 2617.5789
138.06658935546875 0 3254.1182
138.09176635742188 0 1601.6487
139.0508575439453 0 500.9774
139.08700561523438 0 3123.0786
141.10267639160156 0 4554.7734
142.0655059814453 0 1389.7577
143.04547119140625 0 1757.6964
144.0815887451172 0 1085.4393
145.06101989746094 0 2304.1836
145.08518981933594 0 501.33444
146.060302734375 0 1253.2208
147.04449462890625 0 1493.587
147.07699584960938 0 425.4708 y 10
148.94796752929688 0 814.6537
149.04525756835938 0 12498.76
150.066650390625 0 1614.0977
151.0869140625 0 5012.447
152.09080505371094 0 551.58276
153.10250854492188 0 688.4255
154.09750366210938 0 560.19366
155.08203125 0 539.42786
155.09310913085938 0 12753.498
155.11827087402344 0 11221.993 a Water loss 1
156.0769805908203 0 1150.0275
156.09637451171875 0 988.5332
156.12158203125 0 644.7496
157.06100463867188 0 788.26263
157.0975799560547 0 1271.6581
157.13356018066406 0 697.39734
158.08424377441406 0 1281.5643
159.09213256835938 0 181487.88
160.07598876953125 0 1452.7112
160.0890655517578 0 1053.0791
160.095458984375 0 19250.166
161.09881591796875 0 438.09744
165.0550079345703 0 16325.157
166.0614471435547 0 27493.16
166.0867156982422 0 2818.864 y 9
166.09783935546875 0 1744.3608
167.05589294433594 0 16373.978
167.06504821777344 0 2369.7212
167.0816192626953 0 742.1531
167.11842346191406 0 1233.5715
167.28375244140625 0 566.6411
167.6746063232422 0 429.2287
168.0552978515625 0 971.1789
168.08123779296875 0 3221.0488
168.10220336914062 0 1574.7073
168.11361694335938 0 2270.3835
169.07638549804688 0 2220.2173
169.0975799560547 0 6730.7764
170.0605010986328 0 8240.221
171.0640106201172 0 677.3386
171.0767822265625 0 733.72314
171.09205627441406 0 1671.3356
171.1497039794922 0 1966.1989
172.07211303710938 0 3501.8503
173.0560760498047 0 3992.9539
173.12893676757812 0 359225.72 a 1
173.43882751464844 0 1111.7494
174.05477905273438 0 627.7878
174.13226318359375 0 30821.621
175.13381958007812 0 2470.0542
176.0373077392578 0 466.93774
177.1029510498047 0 1247.0677
177.11183166503906 0 484.7291
178.13436889648438 0 11198.525
179.13751220703125 0 886.6557
181.06134033203125 0 1895.6725
182.0816192626953 0 35122.51 y 12
183.08517456054688 0 2975.005
183.11325073242188 0 10152.891 b Water loss 1
184.11679077148438 0 1075.8584
185.0924530029297 0 653.0006
185.1654510498047 0 2506.2925
186.1241455078125 0 72305.27
187.0869903564453 0 15599.751
187.1077117919922 0 666.4602
187.1276092529297 0 6023.566
187.14443969726562 0 524.0586
188.0710906982422 0 1605.9396
188.09046936035156 0 1579.208
189.0990753173828 0 714.8705
190.08267211914062 0 3424.2327
190.13441467285156 0 2313.6067
193.09754943847656 0 676.73895
194.09278869628906 0 2322.398
195.1132354736328 0 18389.865
196.09805297851562 0 746.2537
196.10845947265625 0 12606.412
196.11666870117188 0 1220.7552
197.07086181640625 0 661.1352
197.11158752441406 0 1005.62946
197.12876892089844 0 834.63544
198.08779907226562 0 2459.0017
198.12803649902344 0 6402.5083
199.07154846191406 0 1241.4612
199.0872039794922 0 1437.8384
199.13186645507812 0 1593.247
199.14437866210938 0 923.06024
200.1435546875 0 1193.7812
201.12376403808594 0 97681.805 b 1
202.0870819091797 0 1240.4355
202.12705993652344 0 9605.764
203.10304260253906 0 1271.3114
203.1294403076172 0 1164.9175
204.1129913330078 0 973.5487
205.09759521484375 0 8332.297
205.107666015625 0 885.0795
205.14547729492188 0 1357.0664
206.1015167236328 0 769.77844
206.12892150878906 0 2204.3062
207.160888671875 0 3142.5686
208.1087646484375 0 609.64026
209.05648803710938 0 1045.2213
209.0929718017578 0 665.0885
210.1275177001953 0 3518.5085
211.10833740234375 0 2318.2988 y Water loss 10
212.13980102539062 0 2581.098
213.1238250732422 0 8387.068
213.160400390625 0 1416.8756
214.1277618408203 0 1024.3645
215.11956787109375 0 633.8684
215.1393280029297 0 1703.0394
216.09820556640625 0 1773.4623
221.059326171875 0 1945.638
221.07220458984375 0 953.8733
221.08489990234375 0 5824.856
221.1039581298828 0 1445.8615
221.14002990722656 0 1025.6644
222.08541870117188 0 2031.082
222.12435913085938 0 3001.6365
223.0643310546875 0 3359.3044
223.08148193359375 0 973.5643
223.10806274414062 0 5381.135
223.1194305419922 0 1314.1958
223.1558074951172 0 32021.412
224.1034393310547 0 1935.0234
224.15919494628906 0 3742.5503
225.04344177246094 0 7653.168
225.1237335205078 0 685.74554
226.04345703125 0 1852.1412
226.08282470703125 0 8040.226
226.15602111816406 0 931.13245
227.02264404296875 0 799.58484
227.04010009765625 0 1447.0896
227.06668090820312 0 6396.3174
227.1537628173828 0 1033.8383
228.07080078125 0 704.3121
228.09808349609375 0 682.4842
228.13487243652344 0 2169.1335
229.11878967285156 0 7332.1064
230.15109252929688 0 1043.4242
233.1400909423828 0 5059.4053
234.12425231933594 0 38001.633
234.65093994140625 0 748.6178
235.12765502929688 0 4661.89
235.6050567626953 0 977.08545
237.1348114013672 0 1862.7429
238.11915588378906 0 4251.795
238.1300811767578 0 880.5254
239.08233642578125 0 563.5052
239.09547424316406 0 14149.678
240.09640502929688 0 4865.4995
240.13479614257812 0 13752.279
241.09329223632812 0 2850.6204
241.11862182617188 0 1968.6555
241.137939453125 0 2131.498
241.19125366210938 0 1561.0593
243.1098175048828 0 2993.7915
244.0933074951172 0 15507.599
244.61099243164062 0 589.3363
245.09642028808594 0 1066.0197
247.10841369628906 0 18098
247.14437866210938 0 1413.4443
248.10313415527344 0 2355.2695
248.11407470703125 0 8123.097
249.0987548828125 0 11800.523
249.13504028320312 0 3040.8203
250.1023712158203 0 1402.1151
250.11793518066406 0 628.4947
250.16795349121094 0 677.92346
250.3199005126953 0 553.0735
251.00486755371094 0 635.9418
251.15084838867188 0 121043.49
252.1363067626953 0 632.45715
252.15406799316406 0 13639.22
253.1192626953125 0 712.47284
253.1555938720703 0 1571.1523
255.1480255126953 0 6789.842
256.10833740234375 0 1617.4545
256.15203857421875 0 907.36017
256.6114196777344 0 1329.1244
257.1135559082031 0 3205.1196
258.1453857421875 0 5189.687
259.1480407714844 0 975.0636
261.1197204589844 0 2861.697
261.134521484375 0 919.8282
262.1195068359375 0 1357.1339
265.1297302246094 0 14502.981
266.1139831542969 0 3613.0432
266.1253662109375 0 28842.99
267.0770263671875 0 759.4328
267.12847900390625 0 3286.1665
267.1448059082031 0 1014.5262
268.1295471191406 0 1407.1243
268.1772766113281 0 3413.6892
269.16131591796875 0 1397.9792
269.18560791015625 0 1733.3582
270.1220397949219 0 4668.516
272.17620849609375 0 7325.5835
273.1800231933594 0 1500.5778
274.1188659667969 0 6925.006
274.1312561035156 0 1667.1814
275.10308837890625 0 1577.221
275.121337890625 0 1441.5986
275.1499328613281 0 717.536
276.10919189453125 0 1423.6339
276.1269226074219 0 2119.8604
276.155517578125 0 2138.6382
279.14581298828125 0 2882.5864
280.1298828125 0 3105.274
281.0517883300781 0 9480.836
282.1568908691406 0 6666.2886
282.6683044433594 0 829.44507
283.0312194824219 0 1798.4498
283.140625 0 87899.74
284.1036682128906 0 3215.4917
284.1233215332031 0 799.7218
284.1435852050781 0 11500.552
285.0887145996094 0 694.2304
285.1060485839844 0 1018.408
285.1431579589844 0 1152.7922
285.1585693359375 0 728.65405
286.14031982421875 0 5584.127
287.1418151855469 0 825.42944
287.1716613769531 0 1760.8793
288.1278991699219 0 571.38367
288.6188659667969 0 1712.9069
293.1138000488281 0 46607.74 y Water loss 11
293.1618957519531 0 1476.7555
294.1178283691406 0 9562.57
295.1036682128906 0 1245.078
295.1204833984375 0 1129.6914
296.10394287109375 0 1185.6123
296.1977233886719 0 1122.5758 b Water loss 2
297.0833740234375 0 1635.8042
297.1013488769531 0 834.9014
297.1339111328125 0 1525.2378 y Water loss 8
297.1564025878906 0 17315.883
297.20001220703125 0 747.7951
298.1396789550781 0 1499.0641 y Water loss 6
298.15863037109375 0 1602.9937
299.0623779296875 0 91366.23
300.06292724609375 0 2548.024
300.16815185546875 0 11734.954
301.06072998046875 0 1624.201
301.1170959472656 0 999.35144
301.1512756347656 0 43390.08
301.1683349609375 0 1465.5953
302.1143798828125 0 8016.492
302.1545715332031 0 5699.0234
303.11822509765625 0 1278.4045
303.1685791015625 0 604.6916
305.1799621582031 0 1021.44806
305.64617919921875 0 1991.286
306.1380310058594 0 6905.373 y 8
306.2287902832031 0 642.8047
306.6398010253906 0 3436.5747
307.14031982421875 0 1857.0289
307.2318420410156 0 534.8017
309.1667175292969 0 600.018
311.12408447265625 0 6064.489 y 11
311.1376647949219 0 2904.6445
311.6877746582031 0 608.39
312.1279296875 0 1555.9181
314.1143493652344 0 732.47925
314.18701171875 0 1880.0978
314.2083740234375 0 3944.3877 b 2
315.1669921875 0 24537.814
316.1706237792969 0 4352.4004
317.65948486328125 0 890.06604
318.1197509765625 0 605.4985
319.15191650390625 0 1797.1908
324.1706848144531 0 595.04755
325.15093994140625 0 1508.0294
326.0198974609375 0 579.53156
326.6645812988281 0 6703.885
326.9665222167969 0 628.00934
327.1658630371094 0 2557.387
332.208740234375 0 2564.0469
333.17742919921875 0 4962.449
334.18511962890625 0 982.4087
336.0015563964844 0 575.31586
338.6749267578125 0 1091.885 b Water loss 5
339.171630859375 0 1028.6539 b Ammonia loss 5
339.6761474609375 0 1157.5765
340.6617431640625 0 2795.66
341.0185852050781 0 704.27386
341.16387939453125 0 1242.2761
342.1820068359375 0 2445.038
343.1617736816406 0 1626.6083
343.1847839355469 0 835.3513
344.17218017578125 0 1965.3647
344.9772033691406 0 10977.75
344.9999084472656 0 726.5555
345.1313781738281 0 3800.396
346.1152648925781 0 13403.497
346.1374206542969 0 980.9954
347.1182861328125 0 2581.1921
347.6802978515625 0 7019.259 b 5
348.1818542480469 0 2453.9028
348.68218994140625 0 691.0358
349.1513366699219 0 887.80005
350.2193603515625 0 11865.747
351.1671447753906 0 974.172
351.2214050292969 0 2519.9802
352.1653747558594 0 786.06433
352.6713562011719 0 1816.6654
353.67437744140625 0 2093.7288 y Water loss 7
354.1762390136719 0 1679.5315
355.070068359375 0 1676.48
359.029052734375 0 9757.049
359.134765625 0 871.6104
360.203369140625 0 709.75464
360.6954040527344 0 2763.6042
361.1976013183594 0 4299.9336
361.6866455078125 0 915.4111
362.1856689453125 0 5333.4595
362.6801452636719 0 17248.738 y 7
363.1416931152344 0 20590.715
363.1819763183594 0 6536.4907
363.6824951171875 0 1397.94
364.12530517578125 0 3516.3267
364.1454772949219 0 3274.8118
366.17816162109375 0 1885.3181
367.1398620605469 0 1424.6761
367.1619567871094 0 1515.5074
367.24530029296875 0 1456.4658
368.1932067871094 0 1085.1375
369.12261962890625 0 1217.8967
370.1224365234375 0 1141.3043
370.17620849609375 0 802.12036
373.1854248046875 0 1519.0228
376.1619873046875 0 1845.1149
376.1985778808594 0 4922.664
376.69866943359375 0 1333.5302
377.14556884765625 0 1351.6747
378.2143859863281 0 7963.266
379.2097473144531 0 30706.723
380.1657409667969 0 1098.1349
380.2126159667969 0 7214.5874
381.15240478515625 0 58796.414
381.7120361328125 0 1988.8477
382.1551208496094 0 10143.747
382.18914794921875 0 2168.1733
382.2137145996094 0 1007.77405
383.1581726074219 0 1918.4303
383.20733642578125 0 945.0019
384.16619873046875 0 2007.9121
384.1883850097656 0 3192.758
385.1509704589844 0 3608.1963
385.1731262207031 0 1344.6729 y 4
386.1550598144531 0 1229.1395
390.196044921875 0 2190.3608
390.698486328125 0 807.6151
391.1363220214844 0 3066.5422
391.18670654296875 0 698.768
392.1195373535156 0 730.07526
394.173095703125 0 8377.457
395.13409423828125 0 2627.2866
395.15777587890625 0 3226.1975
395.2136535644531 0 621.8763
395.2415771484375 0 1395.651
396.1380920410156 0 788.3943
396.2250061035156 0 11042.991
397.2034912109375 0 742.1494
397.2283935546875 0 1756.9711
398.1782531738281 0 3560.0366
399.1814270019531 0 717.83093
401.1826477050781 0 702.2011
401.2146301269531 0 690.5999
402.1767883300781 0 3308.7158
402.1996154785156 0 5203.4087
403.201171875 0 2356.7227
404.7115783691406 0 1746.6919
405.2616271972656 0 2390.2268
406.2110595703125 0 907.2981
406.2451477050781 0 1158.6077
407.2043762207031 0 1153.0826
407.2437744140625 0 582.5587
408.71221923828125 0 715.51135
409.1475524902344 0 4271.1953
409.2075500488281 0 987.72003
410.14849853515625 0 1332.5809
411.199462890625 0 4727.58
412.18359375 0 34170.51
413.1454772949219 0 4529.697
413.1638488769531 0 1873.5231
413.18804931640625 0 4057.2334
413.2516174316406 0 921.9242
414.1488342285156 0 1164.9792
414.1942443847656 0 1211.7921
414.199951171875 0 859.0846
414.2354431152344 0 8655.301
415.0375671386719 0 20314.562
415.1988525390625 0 3434.0205
415.23883056640625 0 2119.9578
416.20086669921875 0 750.9443
417.71734619140625 0 6456.7715
418.21112060546875 0 10156.376
418.71240234375 0 4907.6094
419.2132568359375 0 1916.3959
421.2099304199219 0 1398.4022 y Water loss 10
422.1936340332031 0 938.4496 y Ammonia loss 10
422.2537536621094 0 759.0268
422.7073059082031 0 1043.6455
423.2041015625 0 1694.1802
423.2723083496094 0 15953.364
423.7030334472656 0 747.326
424.27557373046875 0 4004.5671
424.7351989746094 0 1449.7885
426.7226867675781 0 94782.89
427.2240295410156 0 50559.74
427.2586364746094 0 707.6397
427.7254638671875 0 12985.723
428.2253723144531 0 2995.5369
429.210205078125 0 19474.377
430.1943664550781 0 85758.8
431.1973876953125 0 19151.477
431.7144775390625 0 10790.941 b Water loss 6
432.2129211425781 0 6875.4185 b Ammonia loss 6
432.7146911621094 0 2100.2686
433.049560546875 0 1056.373
433.2091369628906 0 2500.7761
433.25634765625 0 3595.603 b Water loss 3
434.2597351074219 0 1090.5114
438.1727600097656 0 1574.1504
438.7308044433594 0 1133.4563
439.2198181152344 0 24451.572 y 10
440.18463134765625 0 1003.3677
440.2227478027344 0 5144.994
440.72027587890625 0 46039.99 b 6
440.91778564453125 0 1143.1488
441.2216491699219 0 26122.406
441.7222900390625 0 5873.0024
442.2244567871094 0 1475.8746
442.7210693359375 0 861.374
447.20770263671875 0 2672.4 y Ammonia loss 6
447.7081604003906 0 2098.034
448.20489501953125 0 5030.8037
448.2648010253906 0 879.0675
449.2091064453125 0 1775.352
450.23583984375 0 1464.4849
450.2826232910156 0 1159.0381
451.2674255371094 0 23918.236 b 3
452.2703552246094 0 6736.2397
455.19647216796875 0 771.24835
455.2660217285156 0 1865.2369
455.7206115722656 0 1337.4535 y 6
456.2191467285156 0 911.9896
459.1993103027344 0 2957.083
460.7528991699219 0 2053.1528
461.2043762207031 0 3553.2603
461.2539367675781 0 3268.077
462.20819091796875 0 937.73145
464.2171325683594 0 841.7606
466.24310302734375 0 968.2761
468.2939758300781 0 7032.3467
469.2977600097656 0 1654.5299
474.2599182128906 0 6641.436
474.75537109375 0 7533.6636
475.2546081542969 0 2981.1846
475.756103515625 0 1681.6414
476.22564697265625 0 2179.3557
476.2608642578125 0 717.80164
477.2111511230469 0 4897.418
478.2300109863281 0 4793.1504 y Water loss 9
478.27813720703125 0 3469.6145
479.23822021484375 0 1146.855
479.2834167480469 0 1326.1304
483.2648010253906 0 17674.494
483.7660827636719 0 12816.167
484.21966552734375 0 862.54553
484.26666259765625 0 3726.4702
488.25762939453125 0 4597.6343 b Water loss 7
488.7546691894531 0 2473.006 b Ammonia loss 7
489.25933837890625 0 1302.0481
494.2366638183594 0 59307.77
495.23956298828125 0 14134.481
496.2412109375 0 75698.56 y 9
497.24652099609375 0 16734.9
497.7634582519531 0 7117.119
498.245361328125 0 3425.9822
499.928955078125 0 699.5477
503.1085205078125 0 21744.572
504.2008972167969 0 2371.443
506.22900390625 0 2049.731
507.2580871582031 0 2064.3108
511.2642822265625 0 3954.4175
512.26806640625 0 1255.3501
513.2081298828125 0 759.19025
515.2857666015625 0 1114.2445
519.1397094726562 0 7656.27
519.3277587890625 0 713.14386
520.2661743164062 0 866.8682
521.2232666015625 0 786.0059
523.2603759765625 0 1481.5657
524.2835693359375 0 1451.09
525.267578125 0 11820.495
526.2335815429688 0 1109.7256
526.2694091796875 0 2696.3352
529.26220703125 0 2959.2917 Precursor Water loss
529.5953369140625 0 2387.231 Precursor Ammonia loss
529.929443359375 0 2141.4924
531.219970703125 0 676.522
532.1959838867188 0 2251.2368
533.1971435546875 0 807.4803
534.3058471679688 0 1055.0299
534.55419921875 0 769.5139
534.634765625 0 881.90485
534.8897094726562 0 1280.0117
535.2232055664062 0 889.6899
535.2639770507812 0 3218.688 Precursor
535.5562133789062 0 1160.8201
535.5992431640625 0 2575.3367
535.9354248046875 0 2150.9678
536.1659545898438 0 5159.9727
536.2637329101562 0 739.72095
537.2440795898438 0 993.1087
537.7384643554688 0 1418.7906
538.2724609375 0 1036.7245
539.2366333007812 0 11428.111
539.2820434570312 0 852.967
540.239990234375 0 3004.795
541.7594604492188 0 795.8975
542.2942504882812 0 3757.421
543.2781982421875 0 17117.324
544.2816772460938 0 5081.9434
545.284423828125 0 1218.4238
545.7672119140625 0 1118.983 b Water loss 8
546.2518310546875 0 2083.877 b Ammonia loss 8
547.2470703125 0 977.8203
549.2202758789062 0 2319.8464
550.2101440429688 0 1060.4606
551.2576904296875 0 2310.3457
551.3306884765625 0 1865.9608
552.2604370117188 0 1235.5045
553.2655639648438 0 1015.90967
554.7706298828125 0 2114.007 b 8
555.2568359375 0 4798.9624
555.7571411132812 0 3492.2603
561.3158569335938 0 2318.943 b Water loss 4
562.3047485351562 0 815.7724 b Ammonia loss 4
563.2588500976562 0 1888.8473
564.2637939453125 0 1425.4393
564.32763671875 0 743.67523
565.2644653320312 0 3841.438
566.2665405273438 0 1157.0275
567.2319946289062 0 12566.308
568.2369995117188 0 3951.4775
575.2481079101562 0 3956.084
575.2931518554688 0 760.65466
576.2385864257812 0 1953.4817
579.3257446289062 0 16657.945 b 4
580.3279418945312 0 5994.279
581.3309936523438 0 1188.8849
589.3094482421875 0 637.48267
592.2723388671875 0 2396.142
592.3253173828125 0 3402.5913
593.2572021484375 0 21224.691 y Water loss 8
593.3062133789062 0 7055.1245
594.2551879882812 0 6647.718
594.309814453125 0 1601.4973
595.2272338867188 0 855.45746
595.26611328125 0 802.07715
596.3533325195312 0 947.1164
597.2986450195312 0 900.72485
597.8027954101562 0 1748.4344
600.3159790039062 0 1145.7177
602.2994384765625 0 1932.4064
602.792724609375 0 3296.596
603.2924194335938 0 4279.4707
603.7935791015625 0 1216.1348
610.2838745117188 0 47453.79
610.3276977539062 0 2089.9348
611.2688598632812 0 229492.45 y 8
611.798828125 0 11816.659
612.2720947265625 0 70376.15
612.8002319335938 0 2977.6426
613.2739868164062 0 17337.744
614.2769165039062 0 2513.0286
617.288818359375 0 846.41144
620.804931640625 0 1632.2692
621.2522583007812 0 3994.1323
621.3033447265625 0 1820.4471
622.2559814453125 0 1461.1357
622.3685302734375 0 1336.0852
629.29931640625 0 622.78107
630.2901611328125 0 777.36694
634.3070068359375 0 2780.184
635.2965698242188 0 2756.8171
636.327392578125 0 1543.5651
636.7872314453125 0 2659.9424 y Water loss 3
637.2877197265625 0 2173.3862 y Ammonia loss 3
637.33984375 0 924.43304
640.2750244140625 0 2389.1067
644.29736328125 0 746.41486
645.296630859375 0 4052.7976
645.7933349609375 0 15108.249 y 3
646.2945556640625 0 9223.454
646.7942504882812 0 2554.0754
647.28662109375 0 765.81793
647.3372192382812 0 817.6249 b 10
648.3421630859375 0 913.9256
650.2703857421875 0 1823.9645
650.3272705078125 0 1459.6696
652.3209838867188 0 24350.855
653.323486328125 0 8705.275
654.3219604492188 0 1534.0266
658.2822875976562 0 1022.80396
660.8733520507812 0 2220.8428
661.3259887695312 0 1165.5272
661.38232421875 0 1144.7954
662.3057250976562 0 7491.7417
662.8274536132812 0 742.97266
663.2994384765625 0 6050.9053
664.2913208007812 0 942.4932
664.3472900390625 0 1136.1743
666.36083984375 0 661.9468
668.283447265625 0 834.823
675.8099975585938 0 836.7538
676.3424072265625 0 12016.918 b Water loss 5
677.3361206054688 0 7945.8447 b Ammonia loss 5
677.8064575195312 0 1010
678.3261108398438 0 2240.0547
679.3328857421875 0 1821.0642
680.3162841796875 0 44694.164
681.3184204101562 0 18416.137
682.3201904296875 0 3980.3896
684.3184814453125 0 1034.4318
684.8189697265625 0 3435.8142
685.3173828125 0 1962.661
685.8209228515625 0 1060.1323
688.3345336914062 0 1903.4294
692.8361206054688 0 2708.5203
693.3301391601562 0 13521.668 y Water loss 2
693.8281860351562 0 11414.457 y Ammonia loss 2
694.3522338867188 0 119359.08 b 5
694.8282470703125 0 2992.5771
695.3551025390625 0 47182.996
696.3579711914062 0 10759.138
697.3507690429688 0 1611.699
697.8607177734375 0 901.70197
698.3500366210938 0 1390.4531
700.8980102539062 0 800.01154
701.3405151367188 0 1488.9387
701.841796875 0 21874.967
702.3357543945312 0 82721.26 y 2
702.83642578125 0 66615.55 b Water loss 11
703.3378295898438 0 31187.377 b Ammonia loss 11
703.8396606445312 0 8578.54
704.3378295898438 0 4029.2336
705.3580322265625 0 1424.296
706.3431396484375 0 7752.591 y Water loss 7
707.3451538085938 0 4131.8296
708.307861328125 0 1022.5425
710.4055786132812 0 1701.9766
710.9064331054688 0 966.95294
711.370361328125 0 4981.6694
711.8573608398438 0 6995.5547 b 11
712.3583374023438 0 7378.275
712.8592529296875 0 2788.8806
713.3572387695312 0 1188.0854
716.34228515625 0 766.8417
718.387451171875 0 1009.4202
720.375 0 1028.969
721.3635864257812 0 5551.6294
722.3660888671875 0 2410.637
723.3684692382812 0 13142.235
724.353271484375 0 70819.7 y 7
725.3557739257812 0 29430.49
726.358642578125 0 7365.515
727.3579711914062 0 808.3893
728.3736572265625 0 1102.1869
729.3605346679688 0 2158.5803
729.4525146484375 0 745.2138
730.3617553710938 0 1575.7147
733.3639526367188 0 3761.2778
734.3565063476562 0 2820.932
742.8646850585938 0 1115.1709 y Water loss 1
743.3683471679688 0 2380.2979 y Ammonia loss 1
743.859375 0 776.4891
744.37939453125 0 1003.14154
747.3930053710938 0 1857.9452
748.3771362304688 0 2461.566
749.33740234375 0 1763.9866
750.391357421875 0 900.9304
751.376953125 0 13268.917
751.8699951171875 0 6775.8965 y 1
752.3744506835938 0 13596.129
752.8716430664062 0 2933.6855
753.3758544921875 0 2496.0984
755.4257202148438 0 902.73376
761.3776245117188 0 1350.0875
762.4169921875 0 1651.7123
763.3668212890625 0 7071.6997
764.3679809570312 0 3755.0706
765.4043579101562 0 2708.4912
766.4057006835938 0 1683.3633
767.3486328125 0 2959.1821
767.4187622070312 0 929.1736
768.3474731445312 0 1015.0852
769.3992919921875 0 1465.5759
775.3956909179688 0 1149.6986
776.378662109375 0 1097.1614
779.3840942382812 0 8025.8125
780.3869018554688 0 3507.2378
781.3700561523438 0 3501.1936
782.3683471679688 0 1897.8635
793.400390625 0 13260.637 Precursor Water loss
794.4019165039062 0 4548.2705
795.3434448242188 0 3176.1528
795.4168090820312 0 916.3455
796.3442993164062 0 1246.9287
807.3776245117188 0 3851.5
807.445068359375 0 1714.6438
808.3696899414062 0 1089.8291
818.44189453125 0 1765.0276
819.4417724609375 0 1612.3031
820.438232421875 0 860.274
825.3941650390625 0 1252.3431
826.3789672851562 0 1821.8444
827.3777465820312 0 740.37537
834.4254760742188 0 1826.1935
835.4174194335938 0 4122.381
836.4503173828125 0 7887.2524
837.4575805664062 0 4736.1294
838.4508056640625 0 1798.937
843.4019165039062 0 1663.4962
844.3912353515625 0 2971.6155
845.3955688476562 0 2419.4546
846.4474487304688 0 3480.8442
847.44482421875 0 2756.8767
848.4466552734375 0 1574.5845
852.4368896484375 0 28371.027
853.4391479492188 0 14734.347
854.4423217773438 0 4610.8022
862.4209594726562 0 17261.068 b Water loss 6
863.4176025390625 0 13398.28 b Ammonia loss 6
864.4564819335938 0 20548.17
865.45947265625 0 13079.098
866.4618530273438 0 3618.7058
869.4093017578125 0 1115.741
876.439208984375 0 2780.8013
877.4351196289062 0 901.6569
880.4315185546875 0 142830.61 b 6
881.43408203125 0 76366.06
882.43701171875 0 24475.56
883.437255859375 0 3726.643
884.426513671875 0 1028.5685
890.4122314453125 0 1128.82
891.4107666015625 0 1691.6223
892.4644165039062 0 2774.579
893.4801635742188 0 1185.9662
894.4519653320312 0 925.781
895.524658203125 0 1427.1833
907.4423217773438 0 1802.2798
908.4282836914062 0 9880.947
909.432861328125 0 7822.4004
910.4326171875 0 11609.628 y 6
911.434814453125 0 3272.0603
912.4390258789062 0 1857.3713
936.4203491210938 0 1104.397
937.4345092773438 0 1129.9828
938.4176635742188 0 1161.8003
947.4298095703125 0 987.6484
954.42919921875 0 2986.81
955.4208374023438 0 3493.4436
956.4249877929688 0 1881.5377
962.4432373046875 0 891.5178
964.4635620117188 0 1095.4558
965.4483032226562 0 3505.9055
966.452392578125 0 1813.3383
971.4573974609375 0 928.3464
972.4493408203125 0 2464.9414
973.443359375 0 1356.0236
974.4489135742188 0 916.1074
975.5060424804688 0 1772.7163 b Water loss 7
976.506591796875 0 1185.2943 b Ammonia loss 7
981.46337890625 0 814.34467
982.4736328125 0 1372.8744
993.5158081054688 0 8567.56 b 7
994.517578125 0 4536.682
995.5185546875 0 2805.561
1007.4802856445312 0 778.6046
1024.475341796875 0 1106.7285
1025.4608154296875 0 4594.1597 y 5
1026.4615478515625 0 2995.692
1027.4576416015625 0 808.9994
1064.5115966796875 0 935.4115
1075.532958984375 0 755.09045
1076.5406494140625 0 1117.3785
1093.5450439453125 0 3342.6106
1094.54833984375 0 2219.3535
1095.550048828125 0 946.4336
1108.544921875 0 736.45404 b 8
1117.49755859375 0 1108.547
1118.485107421875 0 1610.9995
1119.4857177734375 0 809.36414
1135.5074462890625 0 5504.5513 y Water loss 4
1136.497802734375 0 7008.837 y Ammonia loss 4
1137.49609375 0 3565.2854
1138.5028076171875 0 1516.157
1152.5362548828125 0 2641.5117
1153.5179443359375 0 10909.179 y 4
1154.5205078125 0 6002.5913
1155.5230712890625 0 2527.2986
1156.51708984375 0 1017.22504
1165.5697021484375 0 1961.4661 b 9
1182.590576171875 0 1063.5142
3081.01513671875 0 659.1839

Spectrum Details

|  |  |
| --- | --- |
| Matched peaks? Matched peaks The total absolute number of peaks matched. Additionally in brackets the total fraction of peaks matched and the total number of peaks is shown. | 78 (9.18% of 850) |
| FDR? FDR The false discovery rate estimated for this peptide. It is calculated by matching all theoretical fragments with a non-integer shift with the raw peaks for this spectrum. This is done with 40 different shifts. The resulting percentage is the average number of annotated peaks over the number of annotated peaks with the correct spectrum. | 1.01% |
| Satellite FDR? Satellite FDR See the FDR for details on its calculation. This satellite ion specific FDR only contains the satellite ions (d/w) for I/L/J positions. | ∞ |
| PSM Score? PSM Score The PSM Score as given by Hecklib to this annotated spectrum. It is shown with three significant figures. | 583 |

#### Spectrum 8106? Spectrum 8106 The raw spectrum of this peptide as annotated by Hecklib. The fragments are coloured according to ion type (see legend). Any peaks with a star '\*' as text can be hovered over to see the full details, first the ion type second the mass shift type. By hovering over the amino acids in the peptide or ions in the legend the corresponding peaks are highlighted. By toggling the 'Unassigned' label you can turn the background (unassigned) peaks on or off in the plot. By updating the slider in the Ion legend you can update the spectrum to only show the top X% of the peaks with labels. The top X% means any peak that is within X% of the highest intensity. By dragging in the spectrum you can zoom in to a specific part of the spectrum and use 'Zoom Out' to get back to the original zoom level. The annotation of the spectrum is based on the given sequence in the peptides file and is done with different software so inconsistencies are likely. The peaks are annotated based on the given sequence, with 20 ppm tolerance.

Copy Data

##### Spectrum 8106 (TSV)

###### Preview

```
Loading example...
```

*Click on the button to copy the data to your clipboard.*

Mz MinMz MaxIntensity Max

WidthHeightPeptide font sizePeptide stroke widthSpectrum font sizeSpectrum stroke widthCompact peptide

Ion legend

wxyz

abcd

OtherUnassignedIon^Charge^_Position_Show for top:%

TVLHQDWLDGKEY

01.90e+43.80e+45.70e+47.60e+4

Zoom Out

y^+1^_1_c^+1^_2_y^+1^_2_z^+1^_2_y^+1^_2_c^+1^_3_y^+2^_6_z^+1^_3_y^+1^_3_c^+2^_7_c^+1^_4_z^+1^_4_y^+1^_4_w^+1^_5_c^+1^_5_y^+1^_5_z^+1^_5_c^+1^_5_y^+1^_5_y^+2^_10_c^+2^_11_w^+1^_6_w^+2^_11_y^+2^_11_y^+2^_11_c^+1^_6_y^+2^_11_z^+1^_6_c^+1^_6_c^+2^_12_c^+2^_12_y^+1^_6_y^+2^_12_z^+2^_12_y^+2^_12_c^+1^_7_z^+1^_7_c^+1^_7_y^+1^_7_z^+1^_8_c^+1^_8_y^+1^_8_w^+1^_9_c^+1^_9_y^+1^_9_z^+1^_9_y^+1^_9_c^+1^_10_y^+1^_10_z^+1^_10_y^+1^_10_c^+1^_11_y^+1^_11_z^+1^_11_c^+1^_12_c^+1^_12_z^+1^_12_

0750150122513001

Fragment Matches Table

Show background peaks

| Position | Ion type | Intensity | mz Theoretical | mz Error (Th) | mz Error (ppm) | Charge | Series Number |
| --- | --- | --- | --- | --- | --- | --- | --- |
| - | - | 621.1 | 120.1 | - | - | 0 | - |
| - | - | 1361 | 129.1 | - | - | 0 | - |
| - | - | 2466 | 130.1 | - | - | 0 | - |
| - | - | 779.8 | 149 | - | - | 0 | - |
| - | - | 557.9 | 149 | - | - | 0 | - |
| - | - | 5279 | 149 | - | - | 0 | - |
| - | - | 476.5 | 151 | - | - | 0 | - |
| - | - | 427.9 | 155 | - | - | 0 | - |
| - | - | 515.6 | 155.1 | - | - | 0 | - |
| - | - | 6502 | 155.1 | - | - | 0 | - |
| - | - | 720.1 | 156.1 | - | - | 0 | - |
| - | - | 501.2 | 157.1 | - | - | 0 | - |
| - | - | 423.1 | 158.3 | - | - | 0 | - |
| - | - | 2299 | 159.1 | - | - | 0 | - |
| - | - | 473.2 | 159.1 | - | - | 0 | - |
| - | - | 478.7 | 160.9 | - | - | 0 | - |
| - | - | 465.5 | 166.5 | - | - | 0 | - |
| - | - | 2668 | 167.1 | - | - | 0 | - |
| - | - | 1.094E+04 | 173.1 | - | - | 0 | - |
| - | - | 606.3 | 174.1 | - | - | 0 | - |
| 13 | y | 2496 | 182.1 | 2.344E-05 | 0.1287 | +1 | 1 |
| - | - | 970.5 | 186.1 | - | - | 0 | - |
| - | - | 8072 | 201.1 | - | - | 0 | - |
| - | - | 895 | 202.1 | - | - | 0 | - |
| - | - | 2333 | 215.1 | - | - | 0 | - |
| 2 | c | 2110 | 218.1 | 3.086E-05 | 0.1415 | +1 | 2 |
| - | - | 9047 | 221.1 | - | - | 0 | - |
| - | - | 2447 | 222.1 | - | - | 0 | - |
| - | - | 2804 | 223.1 | - | - | 0 | - |
| - | - | 1230 | 223.1 | - | - | 0 | - |
| - | - | 2897 | 225 | - | - | 0 | - |
| - | - | 684.8 | 226 | - | - | 0 | - |
| - | - | 944.9 | 229.2 | - | - | 0 | - |
| - | - | 1.108E+04 | 239.1 | - | - | 0 | - |
| - | - | 3580 | 240.1 | - | - | 0 | - |
| - | - | 2634 | 241.1 | - | - | 0 | - |
| - | - | 540.8 | 242.1 | - | - | 0 | - |
| - | - | 1123 | 243.1 | - | - | 0 | - |
| - | - | 955.7 | 245.1 | - | - | 0 | - |
| - | - | 1520 | 251.2 | - | - | 0 | - |
| - | - | 4372 | 258.1 | - | - | 0 | - |
| - | - | 647.5 | 259.1 | - | - | 0 | - |
| - | - | 1056 | 260.2 | - | - | 0 | - |
| - | - | 3203 | 268.2 | - | - | 0 | - |
| - | - | 587.8 | 280.6 | - | - | 0 | - |
| - | - | 1.776E+04 | 281.1 | - | - | 0 | - |
| 12 | y | 1395 | 293.1 | 0.0003273 | 1.117 | +1 | 2 |
| 12 | z | 2709 | 295.1 | 0.001737 | 5.885 | +1 | 2 |
| - | - | 2650 | 296.1 | - | - | 0 | - |
| - | - | 4151 | 297.1 | - | - | 0 | - |
| - | - | 1612 | 297.1 | - | - | 0 | - |
| - | - | 7.526E+04 | 299.1 | - | - | 0 | - |
| - | - | 1223 | 300.1 | - | - | 0 | - |
| - | - | 842.6 | 301.1 | - | - | 0 | - |
| 12 | y | 965.5 | 311.1 | 0.0002912 | 0.9359 | +1 | 2 |
| - | - | 904.1 | 314.2 | - | - | 0 | - |
| - | - | 5816 | 317.2 | - | - | 0 | - |
| - | - | 1176 | 318.2 | - | - | 0 | - |
| - | - | 786.9 | 320.2 | - | - | 0 | - |
| 3 | c | 3292 | 331.2 | 3.654E-06 | 0.01103 | +1 | 3 |
| - | - | 1239 | 341 | - | - | 0 | - |
| - | - | 553.9 | 348.3 | - | - | 0 | - |
| - | - | 2011 | 355.1 | - | - | 0 | - |
| - | - | 708.5 | 356.1 | - | - | 0 | - |
| - | - | 6189 | 359 | - | - | 0 | - |
| 8 | y | 1090 | 362.7 | 0.001136 | 3.132 | +2 | 6 |
| - | - | 519.5 | 367.8 | - | - | 0 | - |
| - | - | 1498 | 369.1 | - | - | 0 | - |
| - | - | 2918 | 370.1 | - | - | 0 | - |
| - | - | 2225 | 371.1 | - | - | 0 | - |
| - | - | 714 | 374.2 | - | - | 0 | - |
| - | - | 626.8 | 378.2 | - | - | 0 | - |
| - | - | 2.534E+04 | 415 | - | - | 0 | - |
| - | - | 566.2 | 417.7 | - | - | 0 | - |
| - | - | 565.3 | 418.2 | - | - | 0 | - |
| 11 | z | 1168 | 423.2 | 0.0002546 | 0.6017 | +1 | 3 |
| - | - | 3568 | 423.3 | - | - | 0 | - |
| - | - | 1143 | 424.3 | - | - | 0 | - |
| - | - | 4451 | 426.7 | - | - | 0 | - |
| - | - | 2232 | 427.2 | - | - | 0 | - |
| - | - | 747.6 | 427.7 | - | - | 0 | - |
| - | - | 1223 | 429.1 | - | - | 0 | - |
| - | - | 825.2 | 429.2 | - | - | 0 | - |
| - | - | 1089 | 430.2 | - | - | 0 | - |
| - | - | 671.9 | 431.7 | - | - | 0 | - |
| - | - | 601.4 | 432.2 | - | - | 0 | - |
| - | - | 656.2 | 433 | - | - | 0 | - |
| - | - | 855.1 | 438.7 | - | - | 0 | - |
| 11 | y | 940.2 | 439.2 | 0.0007387 | 1.682 | +1 | 3 |
| 7 | c | 3611 | 440.7 | 1.955E-05 | 0.04437 | +2 | 7 |
| - | - | 1798 | 441.2 | - | - | 0 | - |
| - | - | 7668 | 451.3 | - | - | 0 | - |
| - | - | 1382 | 452.3 | - | - | 0 | - |
| 4 | c | 2.873E+04 | 468.3 | 0.0001665 | 0.3556 | +1 | 4 |
| - | - | 7414 | 469.3 | - | - | 0 | - |
| - | - | 1316 | 470.3 | - | - | 0 | - |
| - | - | 613.3 | 471.4 | - | - | 0 | - |
| 10 | z | 3123 | 480.2 | 0.0008856 | 1.844 | +1 | 4 |
| - | - | 789.9 | 481.2 | - | - | 0 | - |
| - | - | 950.6 | 483.3 | - | - | 0 | - |
| - | - | 658.2 | 483.8 | - | - | 0 | - |
| - | - | 671.7 | 489.2 | - | - | 0 | - |
| - | - | 3089 | 494.3 | - | - | 0 | - |
| - | - | 705 | 495.2 | - | - | 0 | - |
| - | - | 630.6 | 495.3 | - | - | 0 | - |
| 10 | y | 7996 | 496.2 | 0.0001993 | 0.4016 | +1 | 4 |
| - | - | 2317 | 497.2 | - | - | 0 | - |
| - | - | 707.2 | 497.8 | - | - | 0 | - |
| - | - | 6.45E+04 | 503.1 | - | - | 0 | - |
| - | - | 2.394E+04 | 519.1 | - | - | 0 | - |
| - | - | 1054 | 529.3 | - | - | 0 | - |
| - | - | 738 | 534.3 | - | - | 0 | - |
| - | - | 1.416E+04 | 536.2 | - | - | 0 | - |
| - | - | 2536 | 538.3 | - | - | 0 | - |
| - | - | 1097 | 539.3 | - | - | 0 | - |
| 9 | w | 3646 | 550.3 | 0.0006496 | 1.181 | +1 | 5 |
| - | - | 2602 | 551.3 | - | - | 0 | - |
| - | - | 723.5 | 551.3 | - | - | 0 | - |
| 5 | c | 996 | 579.3 | 0.0002152 | 0.3714 | +1 | 5 |
| 9 | y | 969.1 | 593.3 | 0.002234 | 3.766 | +1 | 5 |
| - | - | 1361 | 594.3 | - | - | 0 | - |
| 9 | z | 3115 | 595.2 | 0.0007981 | 1.341 | +1 | 5 |
| - | - | 1409 | 596.3 | - | - | 0 | - |
| 5 | c | 2.1E+04 | 596.4 | 3.087E-05 | 0.05177 | +1 | 5 |
| - | - | 5465 | 597.4 | - | - | 0 | - |
| - | - | 1152 | 598.4 | - | - | 0 | - |
| - | - | 4627 | 610.3 | - | - | 0 | - |
| 9 | y | 9747 | 611.3 | 0.0009339 | 1.528 | +1 | 5 |
| - | - | 3263 | 612.3 | - | - | 0 | - |
| - | - | 843.2 | 621.3 | - | - | 0 | - |
| - | - | 1221 | 633.8 | - | - | 0 | - |
| - | - | 775.6 | 645.3 | - | - | 0 | - |
| 4 | y | 860.6 | 645.8 | 0.00168 | 2.602 | +2 | 10 |
| - | - | 867.1 | 646.3 | - | - | 0 | - |
| - | - | 775.7 | 648.3 | - | - | 0 | - |
| - | - | 1750 | 654.8 | - | - | 0 | - |
| - | - | 6987 | 655.4 | - | - | 0 | - |
| 11 | c | 3.154E+04 | 655.8 | 0.001132 | 1.727 | +2 | 11 |
| - | - | 2.477E+04 | 656.3 | - | - | 0 | - |
| - | - | 8638 | 656.8 | - | - | 0 | - |
| - | - | 3472 | 657.4 | - | - | 0 | - |
| - | - | 5593 | 664.3 | - | - | 0 | - |
| - | - | 1035 | 664.3 | - | - | 0 | - |
| 8 | w | 5021 | 665.3 | 0.003126 | 4.698 | +1 | 6 |
| - | - | 1132 | 666.3 | - | - | 0 | - |
| - | - | 752.5 | 666.4 | - | - | 0 | - |
| - | - | 1426 | 667.4 | - | - | 0 | - |
| - | - | 981.3 | 668.4 | - | - | 0 | - |
| - | - | 1933 | 668.9 | - | - | 0 | - |
| - | - | 1355 | 669.9 | - | - | 0 | - |
| - | - | 2009 | 672.3 | - | - | 0 | - |
| 3 | w | 2775 | 672.8 | 0.003779 | 5.617 | +2 | 11 |
| - | - | 3246 | 673.3 | - | - | 0 | - |
| - | - | 1023 | 673.8 | - | - | 0 | - |
| - | - | 1224 | 680.3 | - | - | 0 | - |
| - | - | 621.3 | 688.3 | - | - | 0 | - |
| - | - | 671.9 | 690.3 | - | - | 0 | - |
| 3 | y | 830.4 | 693.3 | 0.006675 | 9.627 | +2 | 11 |
| 3 | y | 1448 | 693.8 | 0.005695 | 8.208 | +2 | 11 |
| 6 | c | 2920 | 694.4 | 0.01269 | 18.28 | +1 | 6 |
| - | - | 1503 | 694.8 | - | - | 0 | - |
| - | - | 2106 | 695.4 | - | - | 0 | - |
| - | - | 809.7 | 698.4 | - | - | 0 | - |
| - | - | 3091 | 701.8 | - | - | 0 | - |
| 3 | y | 7023 | 702.3 | 0.000477 | 0.6792 | +2 | 11 |
| - | - | 2980 | 702.8 | - | - | 0 | - |
| - | - | 1996 | 703.3 | - | - | 0 | - |
| - | - | 2797 | 707.3 | - | - | 0 | - |
| 8 | z | 4477 | 708.3 | 0.004991 | 7.046 | +1 | 6 |
| - | - | 1138 | 709.3 | - | - | 0 | - |
| 6 | c | 1.72E+04 | 711.4 | 0.0005457 | 0.7671 | +1 | 6 |
| 12 | c | 672.6 | 711.9 | 0.01203 | 16.9 | +2 | 12 |
| - | - | 5669 | 712.4 | - | - | 0 | - |
| - | - | 1792 | 713.4 | - | - | 0 | - |
| - | - | 2524 | 718.9 | - | - | 0 | - |
| - | - | 1032 | 719.4 | - | - | 0 | - |
| - | - | 457.5 | 719.8 | - | - | 0 | - |
| - | - | 1.143E+04 | 719.9 | - | - | 0 | - |
| - | - | 706.1 | 719.9 | - | - | 0 | - |
| 12 | c | 2.301E+04 | 720.4 | 0.003395 | 4.713 | +2 | 12 |
| - | - | 1.461E+04 | 720.9 | - | - | 0 | - |
| - | - | 7854 | 721.4 | - | - | 0 | - |
| - | - | 1530 | 721.9 | - | - | 0 | - |
| - | - | 1378 | 723.4 | - | - | 0 | - |
| 8 | y | 4299 | 724.4 | 0.00177 | 2.443 | +1 | 6 |
| - | - | 1554 | 725.4 | - | - | 0 | - |
| - | - | 843.3 | 726.4 | - | - | 0 | - |
| - | - | 1317 | 729.9 | - | - | 0 | - |
| - | - | 677.7 | 730.4 | - | - | 0 | - |
| - | - | 1847 | 736.9 | - | - | 0 | - |
| - | - | 5443 | 737.4 | - | - | 0 | - |
| - | - | 983 | 737.9 | - | - | 0 | - |
| - | - | 912.1 | 741.4 | - | - | 0 | - |
| - | - | 796 | 742.9 | - | - | 0 | - |
| 2 | y | 728.2 | 743.4 | 0.006522 | 8.774 | +2 | 12 |
| 2 | z | 1316 | 743.9 | 0.007248 | 9.744 | +2 | 12 |
| - | - | 1481 | 744.4 | - | - | 0 | - |
| - | - | 628.4 | 748.1 | - | - | 0 | - |
| - | - | 5326 | 748.9 | - | - | 0 | - |
| - | - | 5619 | 749.4 | - | - | 0 | - |
| - | - | 3213 | 749.9 | - | - | 0 | - |
| - | - | 596.8 | 750.4 | - | - | 0 | - |
| - | - | 1471 | 751.4 | - | - | 0 | - |
| 2 | y | 1610 | 751.9 | 0.001731 | 2.303 | +2 | 12 |
| - | - | 856.2 | 758.9 | - | - | 0 | - |
| - | - | 773.6 | 764.9 | - | - | 0 | - |
| - | - | 1187 | 765.4 | - | - | 0 | - |
| - | - | 700.6 | 771.4 | - | - | 0 | - |
| - | - | 1608 | 771.9 | - | - | 0 | - |
| - | - | 2100 | 772.4 | - | - | 0 | - |
| - | - | 4317 | 772.9 | - | - | 0 | - |
| - | - | 3109 | 773.4 | - | - | 0 | - |
| - | - | 878 | 773.9 | - | - | 0 | - |
| - | - | 3690 | 779.9 | - | - | 0 | - |
| - | - | 3472 | 780.4 | - | - | 0 | - |
| - | - | 1126 | 780.9 | - | - | 0 | - |
| - | - | 617.1 | 781.4 | - | - | 0 | - |
| - | - | 1422 | 785.4 | - | - | 0 | - |
| - | - | 1328 | 785.9 | - | - | 0 | - |
| - | - | 2248 | 786.4 | - | - | 0 | - |
| - | - | 1918 | 786.9 | - | - | 0 | - |
| - | - | 2256 | 787.4 | - | - | 0 | - |
| - | - | 1172 | 793.4 | - | - | 0 | - |
| - | - | 8276 | 793.9 | - | - | 0 | - |
| - | - | 1.239E+04 | 794.4 | - | - | 0 | - |
| - | - | 8933 | 794.9 | - | - | 0 | - |
| - | - | 4659 | 795.4 | - | - | 0 | - |
| - | - | 1377 | 795.9 | - | - | 0 | - |
| - | - | 1156 | 800.9 | - | - | 0 | - |
| - | - | 845.9 | 801.3 | - | - | 0 | - |
| - | - | 904.7 | 801.4 | - | - | 0 | - |
| - | - | 943.5 | 801.8 | - | - | 0 | - |
| - | - | 8993 | 801.9 | - | - | 0 | - |
| - | - | 5.343E+04 | 802.4 | - | - | 0 | - |
| - | - | 7.314E+04 | 802.9 | - | - | 0 | - |
| - | - | 4.981E+04 | 803.4 | - | - | 0 | - |
| - | - | 2.183E+04 | 803.9 | - | - | 0 | - |
| - | - | 6316 | 804.4 | - | - | 0 | - |
| - | - | 2787 | 804.9 | - | - | 0 | - |
| - | - | 1457 | 837.4 | - | - | 0 | - |
| - | - | 1836 | 838.4 | - | - | 0 | - |
| - | - | 1329 | 839.4 | - | - | 0 | - |
| - | - | 681.6 | 850.4 | - | - | 0 | - |
| - | - | 797.1 | 852.4 | - | - | 0 | - |
| - | - | 836.8 | 853.4 | - | - | 0 | - |
| - | - | 733.2 | 864.5 | - | - | 0 | - |
| 7 | c | 7084 | 880.4 | 0.001063 | 1.208 | +1 | 7 |
| - | - | 3074 | 881.4 | - | - | 0 | - |
| - | - | 880.8 | 882.4 | - | - | 0 | - |
| - | - | 3607 | 893.4 | - | - | 0 | - |
| 7 | z | 6288 | 894.4 | 0.001972 | 2.205 | +1 | 7 |
| - | - | 1816 | 895.4 | - | - | 0 | - |
| 7 | c | 2.158E+04 | 897.5 | 0.0002688 | 0.2995 | +1 | 7 |
| - | - | 1.001E+04 | 898.5 | - | - | 0 | - |
| - | - | 4039 | 899.5 | - | - | 0 | - |
| - | - | 887.5 | 909.4 | - | - | 0 | - |
| 7 | y | 2773 | 910.4 | 0.003756 | 4.125 | +1 | 7 |
| - | - | 1441 | 911.4 | - | - | 0 | - |
| - | - | 1662 | 964.5 | - | - | 0 | - |
| - | - | 2418 | 965.5 | - | - | 0 | - |
| - | - | 776.8 | 966.5 | - | - | 0 | - |
| - | - | 6115 | 1008 | - | - | 0 | - |
| 6 | z | 1.033E+04 | 1009 | 0.004692 | 4.648 | +1 | 8 |
| - | - | 614 | 1010 | - | - | 0 | - |
| - | - | 4143 | 1010 | - | - | 0 | - |
| 8 | c | 1.729E+04 | 1011 | 0.000384 | 0.38 | +1 | 8 |
| - | - | 947.7 | 1011 | - | - | 0 | - |
| - | - | 8613 | 1012 | - | - | 0 | - |
| - | - | 2720 | 1013 | - | - | 0 | - |
| - | - | 1671 | 1024 | - | - | 0 | - |
| 6 | y | 2911 | 1025 | 0.0113 | 11.02 | +1 | 8 |
| - | - | 1169 | 1026 | - | - | 0 | - |
| - | - | 603.6 | 1063 | - | - | 0 | - |
| - | - | 1079 | 1064 | - | - | 0 | - |
| - | - | 2195 | 1068 | - | - | 0 | - |
| - | - | 2241 | 1068 | - | - | 0 | - |
| - | - | 1333 | 1069 | - | - | 0 | - |
| - | - | 967.5 | 1070 | - | - | 0 | - |
| - | - | 902.6 | 1071 | - | - | 0 | - |
| - | - | 916.6 | 1072 | - | - | 0 | - |
| - | - | 1.106E+04 | 1072 | - | - | 0 | - |
| - | - | 5168 | 1073 | - | - | 0 | - |
| 5 | w | 1079 | 1079 | 0.008057 | 7.464 | +1 | 9 |
| - | - | 953.2 | 1080 | - | - | 0 | - |
| - | - | 1820 | 1081 | - | - | 0 | - |
| - | - | 1819 | 1082 | - | - | 0 | - |
| - | - | 1763 | 1083 | - | - | 0 | - |
| - | - | 904.6 | 1095 | - | - | 0 | - |
| - | - | 2530 | 1125 | - | - | 0 | - |
| 9 | c | 4285 | 1126 | 0.00402 | 3.571 | +1 | 9 |
| - | - | 3196 | 1127 | - | - | 0 | - |
| 5 | y | 5940 | 1136 | 0.02273 | 20 | +1 | 9 |
| 5 | z | 4.188E+04 | 1137 | 0.0001306 | 0.1148 | +1 | 9 |
| - | - | 2.368E+04 | 1139 | - | - | 0 | - |
| - | - | 9569 | 1140 | - | - | 0 | - |
| - | - | 2304 | 1141 | - | - | 0 | - |
| - | - | 896.2 | 1153 | - | - | 0 | - |
| 5 | y | 2129 | 1154 | 0.0046 | 3.988 | +1 | 9 |
| - | - | 1387 | 1155 | - | - | 0 | - |
| - | - | 9571 | 1182 | - | - | 0 | - |
| 10 | c | 2.402E+04 | 1183 | 0.003186 | 2.694 | +1 | 10 |
| - | - | 1.486E+04 | 1184 | - | - | 0 | - |
| - | - | 5647 | 1185 | - | - | 0 | - |
| - | - | 1435 | 1186 | - | - | 0 | - |
| - | - | 1023 | 1187 | - | - | 0 | - |
| - | - | 721.4 | 1224 | - | - | 0 | - |
| - | - | 659.9 | 1266 | - | - | 0 | - |
| - | - | 904 | 1267 | - | - | 0 | - |
| - | - | 869.2 | 1268 | - | - | 0 | - |
| 4 | y | 2012 | 1274 | 0.01606 | 12.61 | +1 | 10 |
| 4 | z | 8545 | 1275 | 0.002986 | 2.343 | +1 | 10 |
| - | - | 7168 | 1276 | - | - | 0 | - |
| - | - | 4165 | 1277 | - | - | 0 | - |
| - | - | 1476 | 1278 | - | - | 0 | - |
| 4 | y | 1024 | 1291 | 0.00123 | 0.9531 | +1 | 10 |
| - | - | 859.9 | 1295 | - | - | 0 | - |
| - | - | 860.5 | 1296 | - | - | 0 | - |
| - | - | 3891 | 1310 | - | - | 0 | - |
| 11 | c | 1.376E+04 | 1311 | 0.001851 | 1.412 | +1 | 11 |
| - | - | 1.16E+04 | 1312 | - | - | 0 | - |
| - | - | 4352 | 1313 | - | - | 0 | - |
| - | - | 1184 | 1314 | - | - | 0 | - |
| 3 | y | 1348 | 1387 | 0.007436 | 5.362 | +1 | 11 |
| 3 | z | 8100 | 1388 | 0.005836 | 4.206 | +1 | 11 |
| - | - | 8303 | 1389 | - | - | 0 | - |
| - | - | 5374 | 1390 | - | - | 0 | - |
| - | - | 1407 | 1391 | - | - | 0 | - |
| - | - | 739.6 | 1395 | - | - | 0 | - |
| - | - | 2093 | 1396 | - | - | 0 | - |
| - | - | 2359 | 1397 | - | - | 0 | - |
| - | - | 833.9 | 1398 | - | - | 0 | - |
| 12 | c | 854.3 | 1423 | 0.01217 | 8.557 | +1 | 12 |
| - | - | 1176 | 1424 | - | - | 0 | - |
| 12 | c | 6486 | 1440 | 0.01041 | 7.227 | +1 | 12 |
| - | - | 7676 | 1441 | - | - | 0 | - |
| - | - | 4460 | 1442 | - | - | 0 | - |
| - | - | 1605 | 1443 | - | - | 0 | - |
| 2 | z | 1052 | 1487 | 0.01567 | 10.54 | +1 | 12 |
| - | - | 1775 | 1488 | - | - | 0 | - |
| - | - | 1540 | 1489 | - | - | 0 | - |
| - | - | 908.6 | 1490 | - | - | 0 | - |
| - | - | 1426 | 1543 | - | - | 0 | - |
| - | - | 928.6 | 1544 | - | - | 0 | - |
| - | - | 2261 | 1545 | - | - | 0 | - |
| - | - | 2420 | 1546 | - | - | 0 | - |
| - | - | 3061 | 1547 | - | - | 0 | - |
| - | - | 828 | 1548 | - | - | 0 | - |
| - | - | 934.1 | 1549 | - | - | 0 | - |
| - | - | 1157 | 1559 | - | - | 0 | - |
| - | - | 4194 | 1560 | - | - | 0 | - |
| - | - | 5867 | 1561 | - | - | 0 | - |
| - | - | 3698 | 1562 | - | - | 0 | - |
| - | - | 1394 | 1563 | - | - | 0 | - |
| - | - | 1600 | 1571 | - | - | 0 | - |
| - | - | 936.7 | 1572 | - | - | 0 | - |
| - | - | 3017 | 1577 | - | - | 0 | - |
| - | - | 2705 | 1578 | - | - | 0 | - |
| - | - | 2657 | 1579 | - | - | 0 | - |
| - | - | 691.5 | 1580 | - | - | 0 | - |
| - | - | 1808 | 1587 | - | - | 0 | - |
| - | - | 1.365E+04 | 1588 | - | - | 0 | - |
| - | - | 2.121E+04 | 1589 | - | - | 0 | - |
| - | - | 960.2 | 1589 | - | - | 0 | - |
| - | - | 1.428E+04 | 1590 | - | - | 0 | - |
| - | - | 6910 | 1591 | - | - | 0 | - |
| - | - | 2652 | 1592 | - | - | 0 | - |
| - | - | 666 | 1603 | - | - | 0 | - |
| - | - | 5943 | 1604 | - | - | 0 | - |
| - | - | 2.158E+04 | 1605 | - | - | 0 | - |
| - | - | 3.848E+04 | 1606 | - | - | 0 | - |
| - | - | 2.616E+04 | 1607 | - | - | 0 | - |
| - | - | 1.228E+04 | 1608 | - | - | 0 | - |
| - | - | 5167 | 1609 | - | - | 0 | - |
| - | - | 2640 | 1610 | - | - | 0 | - |
| - | - | 740.9 | 2195 | - | - | 0 | - |
| - | - | 975.2 | 2963 | - | - | 0 | - |
| - | - | 746.8 | 2972 | - | - | 0 | - |

m/z Charge Intensity FragmentType MassShift Position
120.08094024658203 0 621.14087
129.10238647460938 0 1361.422
130.0651092529297 0 2466.352
148.95416259765625 0 779.7532
149.03929138183594 0 557.92035
149.04495239257812 0 5279.3135
151.0419921875 0 476.47693
155.0412139892578 0 427.90918
155.0870819091797 0 515.6322
155.09275817871094 0 6501.899
156.0960693359375 0 720.10846
157.13369750976562 0 501.1744
158.3282012939453 0 423.05515
159.0916290283203 0 2298.798
159.09869384765625 0 473.15762
160.94061279296875 0 478.68274
166.53457641601562 0 465.48373
167.05538940429688 0 2667.787
173.12844848632812 0 10936.795
174.1326904296875 0 606.3256
182.08114624023438 0 2495.5566 y 12
186.12388610839844 0 970.4724
201.12335205078125 0 8071.7925
202.12623596191406 0 895.01117
215.1389617919922 0 2332.7358
218.14988708496094 0 2110.2258 c 1
221.084228515625 0 9047.383
222.08497619628906 0 2447.0696
223.06346130371094 0 2804.4978
223.0817413330078 0 1229.7097
225.04295349121094 0 2896.766
226.042724609375 0 684.7641
229.15501403808594 0 944.8997
239.09490966796875 0 11084.145
240.09596252441406 0 3579.604
241.091796875 0 2634.4438
242.090576171875 0 540.78595
243.13328552246094 0 1122.6516
245.13645935058594 0 955.68506
251.15011596679688 0 1519.8904
258.1449890136719 0 4372.263
259.1487731933594 0 647.47125
260.15966796875 0 1056.098
268.17669677734375 0 3203.413
280.5548400878906 0 587.82654
281.0511169433594 0 17764.264
293.113525390625 0 1395.001 y Water loss 11
295.1033020019531 0 2708.9167 z 11
296.1038818359375 0 2650.0059
297.0824890136719 0 4151.0728
297.1006164550781 0 1612.125
299.06170654296875 0 75255.08
300.0625 0 1223.2744
301.05877685546875 0 842.6148
311.1240539550781 0 965.5139 y 11
314.2081298828125 0 904.0696
317.1579895019531 0 5816.2876
318.161376953125 0 1176.1477
320.1977844238281 0 786.88226
331.2339782714844 0 3292.1797 c 2
341.017333984375 0 1238.6904
348.253173828125 0 553.853
355.0694580078125 0 2010.816
356.0690002441406 0 708.46954
359.028076171875 0 6188.74
362.6781005859375 0 1089.6324 y 7
367.8006286621094 0 519.51324
369.1227111816406 0 1497.9724
370.1221008300781 0 2917.8037
371.119384765625 0 2224.6528
374.1773681640625 0 713.9685
378.20233154296875 0 626.7545
415.0366516113281 0 25338.28
417.7179260253906 0 566.23224
418.2086181640625 0 565.2646
423.20025634765625 0 1167.5164 z 10
423.2713317871094 0 3568.2817
424.27435302734375 0 1143.4944
426.72161865234375 0 4450.8164
427.2237243652344 0 2232.2603
427.7243957519531 0 747.6217
429.0895080566406 0 1222.8638
429.2096252441406 0 825.21814
430.1942443847656 0 1088.7137
431.7164611816406 0 671.92114
432.2105407714844 0 601.4256
433.04736328125 0 656.2428
438.7288818359375 0 855.05115
439.2179870605469 0 940.20056 y 10
440.7192077636719 0 3610.5679 c Ammonia loss 6
441.22076416015625 0 1797.9608
451.2662658691406 0 7668.2173
452.26898193359375 0 1382.0034
468.2930603027344 0 28728.6 c 3
469.29541015625 0 7414.015
470.2975158691406 0 1315.7174
471.38250732421875 0 613.343
480.22235107421875 0 3122.7783 z 9
481.2242736816406 0 789.8677
483.2633361816406 0 950.5949
483.7640686035156 0 658.15155
489.2073669433594 0 671.6925
494.3083801269531 0 3089.4946
495.2303466796875 0 705.0375
495.3094787597656 0 630.60266
496.239990234375 0 7995.584 y 9
497.2450866699219 0 2316.894
497.7626037597656 0 707.18256
503.1074523925781 0 64495.445
519.1387939453125 0 23936.16
529.2584838867188 0 1054.3909
534.3255004882812 0 738.02966
536.1649780273438 0 14162.294
538.2518310546875 0 2536.4624
539.25537109375 0 1097.0978
550.2514038085938 0 3646.4966 w 8
551.2572021484375 0 2602.1475
551.3276977539062 0 723.4972
579.32470703125 0 995.99493 c Ammonia loss 4
593.2543334960938 0 969.0909 y Water loss 8
594.2651977539062 0 1361.2754
595.2492065429688 0 3114.8906 z 8
596.2571411132812 0 1409.4967
596.3514404296875 0 21003.664 c 4
597.3539428710938 0 5465.4307
598.3532104492188 0 1151.7981
610.2828369140625 0 4627.096
611.26806640625 0 9747.26 y 8
612.2720947265625 0 3263.1257
621.28759765625 0 843.2265
633.837646484375 0 1220.6418
645.2957153320312 0 775.57574
645.7894287109375 0 860.60504 y 3
646.2909545898438 0 867.1252
648.33349609375 0 775.6712
654.847900390625 0 1750.1158
655.3519287109375 0 6986.9585
655.8473510742188 0 31537.262 c 10
656.3480834960938 0 24766.227
656.8499755859375 0 8638.34
657.350341796875 0 3472.3906
664.2930908203125 0 5592.8643
664.3468017578125 0 1035.2087
665.2808227539062 0 5020.783 w 7
666.28271484375 0 1131.7782
666.3556518554688 0 752.5094
667.3649291992188 0 1425.7759
668.3639526367188 0 981.3346
668.8593139648438 0 1933.2488
669.8555908203125 0 1354.5898
672.303955078125 0 2008.6418
672.8001708984375 0 2775.3037 w 2
673.2991943359375 0 3246.27
673.802734375 0 1023.16077
680.3168334960938 0 1223.923
688.3136596679688 0 621.2764
690.34130859375 0 671.8738
693.3345336914062 0 830.43243 y Water loss 2
693.8255615234375 0 1448.4445 y Ammonia loss 2
694.3391723632812 0 2919.632 c Ammonia loss 5
694.8270263671875 0 1502.913
695.3506469726562 0 2106.1714
698.3709106445312 0 809.67554
701.83984375 0 3091.015
702.3336181640625 0 7022.8623 y 2
702.8326416015625 0 2980.409
703.3358154296875 0 1996.1936
707.3462524414062 0 2797.2732
708.3374633789062 0 4477.047 z 7
709.3404541015625 0 1138.2189
711.3778686523438 0 17200.914 c 5
711.8662719726562 0 672.56244 c Ammonia loss 11
712.3801879882812 0 5669.4146
713.3807983398438 0 1791.9186
718.9205932617188 0 2523.6082
719.4235229492188 0 1031.773
719.8048095703125 0 457.5097
719.8751220703125 0 11428.415
719.935302734375 0 706.09454
720.3709106445312 0 23012.934 c 11
720.8706665039062 0 14614.185
721.371826171875 0 7853.788
721.8717041015625 0 1530.4551
723.3692626953125 0 1377.5012
724.3529663085938 0 4298.736 y 7
725.3539428710938 0 1553.5317
726.356689453125 0 843.27167
729.854736328125 0 1316.6348
730.3605346679688 0 677.72925
736.8599853515625 0 1847.3295
737.3602905273438 0 5442.7583
737.8658447265625 0 983
741.3591918945312 0 912.12213
742.8859252929688 0 796.03784
743.360595703125 0 728.21655 y Ammonia loss 1
743.865234375 0 1315.603 z 1
744.3629760742188 0 1480.8588
748.053466796875 0 628.40314
748.8662109375 0 5325.807
749.3684692382812 0 5618.922
749.8695678710938 0 3212.643
750.3905029296875 0 596.8154
751.382080078125 0 1471.0975
751.8690795898438 0 1610.3708 y 1
758.866455078125 0 856.1875
764.8727416992188 0 773.63116
765.3770141601562 0 1187.3181
771.3900146484375 0 700.5604
771.893310546875 0 1608.1562
772.386962890625 0 2099.6545
772.88671875 0 4316.6055
773.3819580078125 0 3108.8306
773.8654174804688 0 877.99744
779.892333984375 0 3690.0542
780.39208984375 0 3472.1282
780.895751953125 0 1125.5488
781.38818359375 0 617.0943
785.3864135742188 0 1422.213
785.880615234375 0 1328.1187
786.3844604492188 0 2247.5273
786.8768920898438 0 1918.0778
787.3773193359375 0 2255.6458
793.3960571289062 0 1171.548
793.8897094726562 0 8275.812
794.38623046875 0 12394.693
794.8861694335938 0 8933.475
795.39013671875 0 4658.708
795.8850708007812 0 1376.9095
800.925537109375 0 1156.1454
801.3196411132812 0 845.89075
801.3884887695312 0 904.6553
801.8263549804688 0 943.5378
801.8991088867188 0 8992.685
802.3955078125 0 53430.93
802.8958740234375 0 73140.414
803.3971557617188 0 49807.246
803.8989868164062 0 21832.271
804.3990478515625 0 6315.822
804.8963012695312 0 2787.0447
837.3638305664062 0 1457.4429
838.3560180664062 0 1836.2561
839.3505249023438 0 1329.1887
850.4237670898438 0 681.64325
852.4408569335938 0 797.05347
853.44189453125 0 836.78754
864.4591674804688 0 733.16046
880.4301147460938 0 7083.6187 c Ammonia loss 6
881.4356079101562 0 3073.6733
882.4453735351562 0 880.8276
893.4255981445312 0 3607.1218
894.4137573242188 0 6288.3994 z 6
895.4180908203125 0 1816.0309
897.4574584960938 0 21579.428 c 6
898.4607543945312 0 10007.014
899.4631958007812 0 4038.9104
909.4424438476562 0 887.5296
910.4342651367188 0 2772.8052 y 6
911.4307250976562 0 1441.0653
964.4627685546875 0 1662.3225
965.4541015625 0 2417.6562
966.5380249023438 0 776.8116
1008.4546508789062 0 6115.2207
1009.4434204101562 0 10325.928 z 5
1009.55126953125 0 614.03894
1010.4403686523438 0 4142.941
1010.5421752929688 0 17285.186 c 7
1011.4446411132812 0 947.65186
1011.544677734375 0 8613.1875
1012.5466918945312 0 2719.9316
1024.476318359375 0 1670.634
1025.46875 0 2911.2024 y 5
1026.4698486328125 0 1168.6865
1062.5797119140625 0 603.6489
1063.5614013671875 0 1079.3606
1067.5394287109375 0 2194.8337
1068.4326171875 0 2241.1875
1068.5455322265625 0 1333.0913
1070.4412841796875 0 967.5146
1071.432861328125 0 902.6486
1071.58984375 0 916.596
1072.4287109375 0 11063.785
1073.4300537109375 0 5168.195
1079.47607421875 0 1079.2964 w 4
1080.4598388671875 0 953.1874
1080.5733642578125 0 1819.5829
1081.564208984375 0 1819.123
1082.5606689453125 0 1763.3981
1094.5302734375 0 904.58777
1124.580322265625 0 2529.8892
1125.57275390625 0 4284.5312 c 8
1126.569091796875 0 3196.0232
1136.51220703125 0 5939.649 y Ammonia loss 4
1137.4974365234375 0 41882.676 z 4
1138.5008544921875 0 23684.604
1139.50341796875 0 9568.635
1140.505615234375 0 2303.7747
1152.5233154296875 0 896.18884
1153.5206298828125 0 2128.698 y 4
1154.5203857421875 0 1387.1295
1181.6048583984375 0 9570.851
1182.5933837890625 0 24022.799 c 9
1183.594970703125 0 14856.345
1184.5977783203125 0 5646.571
1185.604248046875 0 1435.182
1186.6124267578125 0 1023.0155
1223.592041015625 0 721.41235
1265.6732177734375 0 659.8577
1266.6739501953125 0 904.00214
1267.676513671875 0 869.1846
1273.564453125 0 2012.4465 y Ammonia loss 3
1274.5592041015625 0 8545.361 z 3
1275.564208984375 0 7168.4443
1276.564697265625 0 4164.689
1277.5653076171875 0 1475.9451
1290.576171875 0 1023.75885 y 3
1294.6759033203125 0 859.8926
1295.66259765625 0 860.49286
1309.69677734375 0 3891.389
1310.68701171875 0 13755.191 c 10
1311.689208984375 0 11599.92
1312.68994140625 0 4351.822
1313.6868896484375 0 1184.3188
1386.639892578125 0 1347.555 y Ammonia loss 2
1387.6461181640625 0 8099.7803 z 2
1388.6468505859375 0 8303.417
1389.6492919921875 0 5373.981
1390.6427001953125 0 1406.6887
1394.7215576171875 0 739.58374
1395.7376708984375 0 2092.8264
1396.7271728515625 0 2358.781
1397.726318359375 0 833.8799
1422.71337890625 0 854.3473 c Ammonia loss 11
1423.7130126953125 0 1175.52
1439.7381591796875 0 6485.7803 c 11
1440.7374267578125 0 7675.7285
1441.7376708984375 0 4460.4478
1442.7376708984375 0 1604.8807
1486.724365234375 0 1051.6776 z 1
1487.7177734375 0 1775.2706
1488.7181396484375 0 1540.1656
1489.73974609375 0 908.5528
1542.7764892578125 0 1426.3422
1543.76708984375 0 928.5679
1544.78125 0 2260.6855
1545.7742919921875 0 2420.3206
1546.7728271484375 0 3061.0564
1547.775634765625 0 828.01654
1548.7767333984375 0 934.06085
1558.79296875 0 1157.2347
1559.7806396484375 0 4194.1196
1560.7796630859375 0 5867.0884
1561.781005859375 0 3697.5208
1562.7861328125 0 1393.8876
1570.758056640625 0 1600.3005
1571.7449951171875 0 936.74023
1576.7940673828125 0 3016.811
1577.807373046875 0 2705.365
1578.7958984375 0 2657.2534
1579.79931640625 0 691.5069
1586.771240234375 0 1808.3694
1587.772216796875 0 13645.452
1588.767333984375 0 21212.773
1589.1182861328125 0 960.20667
1589.76904296875 0 14278.089
1590.773193359375 0 6909.9707
1591.7691650390625 0 2651.9546
1602.8043212890625 0 666.0165
1603.7857666015625 0 5942.504
1604.7900390625 0 21580.559
1605.7901611328125 0 38478.613
1606.7923583984375 0 26157.963
1607.7960205078125 0 12284.25
1608.798095703125 0 5166.955
1609.788330078125 0 2640.3914
2195.46435546875 0 740.8992
2963.473388671875 0 975.1853
2971.61083984375 0 746.79083

Spectrum Details

|  |  |
| --- | --- |
| Matched peaks? Matched peaks The total absolute number of peaks matched. Additionally in brackets the total fraction of peaks matched and the total number of peaks is shown. | 57 (15.12% of 377) |
| FDR? FDR The false discovery rate estimated for this peptide. It is calculated by matching all theoretical fragments with a non-integer shift with the raw peaks for this spectrum. This is done with 40 different shifts. The resulting percentage is the average number of annotated peaks over the number of annotated peaks with the correct spectrum. | 3.26% |
| Satellite FDR? Satellite FDR See the FDR for details on its calculation. This satellite ion specific FDR only contains the satellite ions (d/w) for I/L/J positions. | 1.19% |
| PSM Score? PSM Score The PSM Score as given by Hecklib to this annotated spectrum. It is shown with three significant figures. | 549 |

#### Spectrum 7713? Spectrum 7713 The raw spectrum of this peptide as annotated by Hecklib. The fragments are coloured according to ion type (see legend). Any peaks with a star '\*' as text can be hovered over to see the full details, first the ion type second the mass shift type. By hovering over the amino acids in the peptide or ions in the legend the corresponding peaks are highlighted. By toggling the 'Unassigned' label you can turn the background (unassigned) peaks on or off in the plot. By updating the slider in the Ion legend you can update the spectrum to only show the top X% of the peaks with labels. The top X% means any peak that is within X% of the highest intensity. By dragging in the spectrum you can zoom in to a specific part of the spectrum and use 'Zoom Out' to get back to the original zoom level. The annotation of the spectrum is based on the given sequence in the peptides file and is done with different software so inconsistencies are likely. The peaks are annotated based on the given sequence, with 20 ppm tolerance.

Copy Data

##### Spectrum 7713 (TSV)

###### Preview

```
Loading example...
```

*Click on the button to copy the data to your clipboard.*

Mz MinMz MaxIntensity Max

WidthHeightPeptide font sizePeptide stroke widthSpectrum font sizeSpectrum stroke widthCompact peptide

Ion legend

wxyz

abcd

OtherUnassignedIon^Charge^_Position_Show for top:%

TVLHQDWLDGKEY

03.51e+47.03e+41.05e+51.41e+5

Zoom Out

y^+1^_1_c^+1^_2_y^+1^_2_z^+1^_2_y^+1^_2_c^+1^_3_c^+2^_6_y^+2^_6_y^+1^_3_c^+2^_7_c^+1^_4_c^+1^_4_z^+1^_4_y^+1^_4_w^+1^_5_c^+1^_5_y^+1^_5_z^+1^_5_c^+1^_5_y^+1^_5_z^+2^_10_y^+2^_10_c^+2^_11_w^+1^_6_w^+2^_11_y^+2^_11_y^+2^_11_c^+1^_6_y^+2^_11_z^+1^_6_c^+1^_6_c^+2^_12_c^+2^_12_y^+1^_6_w^+2^_12_y^+2^_12_z^+2^_12_y^+2^_12_c^+1^_7_z^+1^_7_c^+1^_7_y^+1^_7_c^+1^_8_z^+1^_8_c^+1^_8_y^+1^_8_w^+1^_9_c^+1^_9_z^+1^_9_y^+1^_9_c^+1^_10_y^+1^_10_z^+1^_10_y^+1^_10_c^+1^_11_y^+1^_11_z^+1^_11_y^+1^_11_c^+1^_12_c^+1^_12_z^+1^_12_

040681312191626

Fragment Matches Table

Show background peaks

| Position | Ion type | Intensity | mz Theoretical | mz Error (Th) | mz Error (ppm) | Charge | Series Number |
| --- | --- | --- | --- | --- | --- | --- | --- |
| - | - | 668.8 | 120.1 | - | - | 0 | - |
| - | - | 485.2 | 128.1 | - | - | 0 | - |
| - | - | 3227 | 129.1 | - | - | 0 | - |
| - | - | 3541 | 130.1 | - | - | 0 | - |
| - | - | 436.6 | 131.1 | - | - | 0 | - |
| - | - | 1031 | 131.1 | - | - | 0 | - |
| - | - | 1084 | 136.1 | - | - | 0 | - |
| - | - | 489.6 | 139.5 | - | - | 0 | - |
| - | - | 588.2 | 148.9 | - | - | 0 | - |
| - | - | 5728 | 149 | - | - | 0 | - |
| - | - | 429.6 | 150.9 | - | - | 0 | - |
| - | - | 1.222E+04 | 155.1 | - | - | 0 | - |
| - | - | 495 | 156.1 | - | - | 0 | - |
| - | - | 1038 | 157.1 | - | - | 0 | - |
| - | - | 3875 | 159.1 | - | - | 0 | - |
| - | - | 502.6 | 160 | - | - | 0 | - |
| - | - | 724.3 | 165.1 | - | - | 0 | - |
| - | - | 1577 | 167.1 | - | - | 0 | - |
| - | - | 586.2 | 171.1 | - | - | 0 | - |
| - | - | 1.673E+04 | 173.1 | - | - | 0 | - |
| - | - | 1499 | 174.1 | - | - | 0 | - |
| - | - | 552.7 | 181.1 | - | - | 0 | - |
| 13 | y | 3263 | 182.1 | 6.812E-05 | 0.3741 | +1 | 1 |
| - | - | 792.6 | 184.1 | - | - | 0 | - |
| - | - | 1267 | 186.1 | - | - | 0 | - |
| - | - | 993.2 | 188.1 | - | - | 0 | - |
| - | - | 466 | 198.2 | - | - | 0 | - |
| - | - | 1.213E+04 | 201.1 | - | - | 0 | - |
| - | - | 992.3 | 202.1 | - | - | 0 | - |
| - | - | 1412 | 213.2 | - | - | 0 | - |
| - | - | 6556 | 215.1 | - | - | 0 | - |
| - | - | 809 | 216.1 | - | - | 0 | - |
| - | - | 520.8 | 216.2 | - | - | 0 | - |
| 2 | c | 3331 | 218.1 | 7.595E-05 | 0.3482 | +1 | 2 |
| - | - | 7143 | 221.1 | - | - | 0 | - |
| - | - | 2597 | 222.1 | - | - | 0 | - |
| - | - | 2621 | 223.1 | - | - | 0 | - |
| - | - | 1461 | 223.1 | - | - | 0 | - |
| - | - | 1113 | 223.2 | - | - | 0 | - |
| - | - | 2176 | 225 | - | - | 0 | - |
| - | - | 614.1 | 226 | - | - | 0 | - |
| - | - | 1027 | 226.1 | - | - | 0 | - |
| - | - | 1676 | 229.2 | - | - | 0 | - |
| - | - | 1.018E+04 | 239.1 | - | - | 0 | - |
| - | - | 4339 | 240.1 | - | - | 0 | - |
| - | - | 1588 | 241.1 | - | - | 0 | - |
| - | - | 1262 | 243.1 | - | - | 0 | - |
| - | - | 1511 | 245.1 | - | - | 0 | - |
| - | - | 2877 | 251.2 | - | - | 0 | - |
| - | - | 570.5 | 252.2 | - | - | 0 | - |
| - | - | 5995 | 258.1 | - | - | 0 | - |
| - | - | 1331 | 260.2 | - | - | 0 | - |
| - | - | 6460 | 268.2 | - | - | 0 | - |
| - | - | 598.6 | 269.2 | - | - | 0 | - |
| - | - | 1.709E+04 | 281.1 | - | - | 0 | - |
| - | - | 568.3 | 283 | - | - | 0 | - |
| - | - | 1551 | 283.1 | - | - | 0 | - |
| - | - | 769.2 | 287.2 | - | - | 0 | - |
| - | - | 960.9 | 288.2 | - | - | 0 | - |
| 12 | y | 2251 | 293.1 | 0.0003883 | 1.325 | +1 | 2 |
| - | - | 932.2 | 294.1 | - | - | 0 | - |
| 12 | z | 3189 | 295.1 | 0.001523 | 5.161 | +1 | 2 |
| - | - | 2243 | 296.1 | - | - | 0 | - |
| - | - | 3177 | 297.1 | - | - | 0 | - |
| - | - | 2139 | 297.1 | - | - | 0 | - |
| - | - | 824.9 | 298.1 | - | - | 0 | - |
| - | - | 6.748E+04 | 299.1 | - | - | 0 | - |
| - | - | 1723 | 300.1 | - | - | 0 | - |
| - | - | 1149 | 301.2 | - | - | 0 | - |
| - | - | 657.7 | 302.2 | - | - | 0 | - |
| - | - | 541.3 | 304.1 | - | - | 0 | - |
| - | - | 1293 | 305.1 | - | - | 0 | - |
| - | - | 811 | 306.1 | - | - | 0 | - |
| - | - | 744.4 | 307.2 | - | - | 0 | - |
| - | - | 557.7 | 308.2 | - | - | 0 | - |
| - | - | 507.4 | 309.2 | - | - | 0 | - |
| 12 | y | 2162 | 311.1 | 0.0004133 | 1.328 | +1 | 2 |
| - | - | 716.8 | 314.1 | - | - | 0 | - |
| - | - | 2057 | 314.2 | - | - | 0 | - |
| - | - | 595.9 | 315.2 | - | - | 0 | - |
| - | - | 1.102E+04 | 317.2 | - | - | 0 | - |
| - | - | 1842 | 318.2 | - | - | 0 | - |
| - | - | 1874 | 320.2 | - | - | 0 | - |
| - | - | 8654 | 323.1 | - | - | 0 | - |
| - | - | 773.8 | 324.1 | - | - | 0 | - |
| - | - | 2993 | 327.2 | - | - | 0 | - |
| 3 | c | 5873 | 331.2 | 0.0001795 | 0.5418 | +1 | 3 |
| - | - | 780.1 | 332.2 | - | - | 0 | - |
| - | - | 1139 | 341 | - | - | 0 | - |
| - | - | 643.5 | 343.1 | - | - | 0 | - |
| - | - | 528.2 | 346.3 | - | - | 0 | - |
| 6 | c | 587.4 | 347.7 | 0.0003411 | 0.9811 | +2 | 6 |
| - | - | 1493 | 355.1 | - | - | 0 | - |
| - | - | 680.7 | 356.1 | - | - | 0 | - |
| - | - | 806.5 | 357.1 | - | - | 0 | - |
| - | - | 5828 | 359 | - | - | 0 | - |
| 8 | y | 1705 | 362.7 | 0.0007257 | 2.001 | +2 | 6 |
| - | - | 1498 | 364.2 | - | - | 0 | - |
| - | - | 1751 | 369.1 | - | - | 0 | - |
| - | - | 3148 | 370.1 | - | - | 0 | - |
| - | - | 2771 | 371.1 | - | - | 0 | - |
| - | - | 762.1 | 372.1 | - | - | 0 | - |
| - | - | 2074 | 374.2 | - | - | 0 | - |
| - | - | 965.5 | 378.2 | - | - | 0 | - |
| - | - | 786.4 | 408.3 | - | - | 0 | - |
| - | - | 1188 | 412.2 | - | - | 0 | - |
| - | - | 2.187E+04 | 415 | - | - | 0 | - |
| - | - | 6178 | 423.3 | - | - | 0 | - |
| - | - | 1547 | 424.3 | - | - | 0 | - |
| - | - | 9214 | 426.7 | - | - | 0 | - |
| - | - | 3323 | 427.2 | - | - | 0 | - |
| - | - | 1442 | 427.7 | - | - | 0 | - |
| - | - | 1591 | 429.1 | - | - | 0 | - |
| - | - | 640.3 | 429.2 | - | - | 0 | - |
| - | - | 2352 | 430.2 | - | - | 0 | - |
| - | - | 1238 | 431.7 | - | - | 0 | - |
| - | - | 1006 | 432.2 | - | - | 0 | - |
| 11 | y | 2589 | 439.2 | 0.0006651 | 1.514 | +1 | 3 |
| 7 | c | 5000 | 440.7 | 0.0001416 | 0.3214 | +2 | 7 |
| - | - | 2338 | 441.2 | - | - | 0 | - |
| - | - | 863.6 | 444.2 | - | - | 0 | - |
| - | - | 994 | 445.2 | - | - | 0 | - |
| 4 | c | 670.8 | 450.3 | 0.0004382 | 0.9732 | +1 | 4 |
| - | - | 1.336E+04 | 451.3 | - | - | 0 | - |
| - | - | 3223 | 452.3 | - | - | 0 | - |
| - | - | 675.5 | 453.3 | - | - | 0 | - |
| 4 | c | 5.233E+04 | 468.3 | 0.0004412 | 0.9421 | +1 | 4 |
| - | - | 1.478E+04 | 469.3 | - | - | 0 | - |
| - | - | 2493 | 470.3 | - | - | 0 | - |
| - | - | 640 | 474.3 | - | - | 0 | - |
| - | - | 1035 | 474.8 | - | - | 0 | - |
| 10 | z | 5198 | 480.2 | 0.0005805 | 1.209 | +1 | 4 |
| - | - | 1934 | 481.2 | - | - | 0 | - |
| - | - | 2324 | 483.3 | - | - | 0 | - |
| - | - | 930.7 | 483.8 | - | - | 0 | - |
| - | - | 2264 | 489.2 | - | - | 0 | - |
| - | - | 949.2 | 494.2 | - | - | 0 | - |
| - | - | 3802 | 494.3 | - | - | 0 | - |
| - | - | 1592 | 495.2 | - | - | 0 | - |
| - | - | 1424 | 495.3 | - | - | 0 | - |
| 10 | y | 1.565E+04 | 496.2 | 0.0004721 | 0.9514 | +1 | 4 |
| - | - | 3276 | 497.2 | - | - | 0 | - |
| - | - | 646.6 | 497.8 | - | - | 0 | - |
| - | - | 5.492E+04 | 503.1 | - | - | 0 | - |
| - | - | 978 | 517.3 | - | - | 0 | - |
| - | - | 2.209E+04 | 519.1 | - | - | 0 | - |
| - | - | 1040 | 523.2 | - | - | 0 | - |
| - | - | 686.9 | 528.9 | - | - | 0 | - |
| - | - | 635.2 | 529.3 | - | - | 0 | - |
| - | - | 1138 | 529.6 | - | - | 0 | - |
| - | - | 753.2 | 529.9 | - | - | 0 | - |
| - | - | 673 | 534.3 | - | - | 0 | - |
| - | - | 3572 | 535.3 | - | - | 0 | - |
| - | - | 1.23E+04 | 536.2 | - | - | 0 | - |
| - | - | 1.026E+04 | 536.3 | - | - | 0 | - |
| - | - | 5113 | 538.3 | - | - | 0 | - |
| - | - | 1452 | 539.3 | - | - | 0 | - |
| 9 | w | 4954 | 550.3 | 0.0007717 | 1.402 | +1 | 5 |
| - | - | 4494 | 551.3 | - | - | 0 | - |
| - | - | 884.6 | 552.3 | - | - | 0 | - |
| 5 | c | 1961 | 579.3 | 0.0007034 | 1.214 | +1 | 5 |
| - | - | 808.3 | 580.3 | - | - | 0 | - |
| 9 | y | 902.3 | 593.3 | 0.0006475 | 1.091 | +1 | 5 |
| - | - | 1602 | 594.3 | - | - | 0 | - |
| 9 | z | 5278 | 595.2 | 0.0009201 | 1.546 | +1 | 5 |
| - | - | 637.2 | 595.3 | - | - | 0 | - |
| - | - | 2032 | 596.3 | - | - | 0 | - |
| 5 | c | 3.081E+04 | 596.4 | 0.0001522 | 0.2553 | +1 | 5 |
| - | - | 9995 | 597.4 | - | - | 0 | - |
| - | - | 2131 | 598.4 | - | - | 0 | - |
| - | - | 790.4 | 602.3 | - | - | 0 | - |
| - | - | 844.6 | 608.3 | - | - | 0 | - |
| - | - | 1234 | 609.3 | - | - | 0 | - |
| - | - | 7901 | 610.3 | - | - | 0 | - |
| 9 | y | 1.94E+04 | 611.3 | 0.001178 | 1.927 | +1 | 5 |
| - | - | 766 | 611.8 | - | - | 0 | - |
| - | - | 6165 | 612.3 | - | - | 0 | - |
| - | - | 1012 | 613.3 | - | - | 0 | - |
| - | - | 1438 | 621.3 | - | - | 0 | - |
| - | - | 1599 | 633.8 | - | - | 0 | - |
| - | - | 1109 | 634.3 | - | - | 0 | - |
| - | - | 1187 | 637.3 | - | - | 0 | - |
| 4 | z | 684.4 | 637.8 | 0.003653 | 5.728 | +2 | 10 |
| 4 | y | 1897 | 645.8 | 0.001493 | 2.312 | +2 | 10 |
| - | - | 1005 | 646.3 | - | - | 0 | - |
| - | - | 604.6 | 651.3 | - | - | 0 | - |
| - | - | 2821 | 654.8 | - | - | 0 | - |
| - | - | 1.401E+04 | 655.4 | - | - | 0 | - |
| 11 | c | 5.846E+04 | 655.8 | 0.001254 | 1.913 | +2 | 11 |
| - | - | 4.473E+04 | 656.3 | - | - | 0 | - |
| - | - | 1.837E+04 | 656.8 | - | - | 0 | - |
| - | - | 4226 | 657.4 | - | - | 0 | - |
| - | - | 958.5 | 657.9 | - | - | 0 | - |
| - | - | 7353 | 664.3 | - | - | 0 | - |
| - | - | 1534 | 664.3 | - | - | 0 | - |
| 8 | w | 9190 | 665.3 | 0.003187 | 4.79 | +1 | 6 |
| - | - | 1184 | 665.3 | - | - | 0 | - |
| - | - | 3472 | 666.3 | - | - | 0 | - |
| - | - | 1730 | 666.4 | - | - | 0 | - |
| - | - | 999 | 667.3 | - | - | 0 | - |
| - | - | 2408 | 667.4 | - | - | 0 | - |
| - | - | 3963 | 668.4 | - | - | 0 | - |
| - | - | 4530 | 668.9 | - | - | 0 | - |
| - | - | 2279 | 669.4 | - | - | 0 | - |
| - | - | 1412 | 669.9 | - | - | 0 | - |
| - | - | 3384 | 672.3 | - | - | 0 | - |
| 3 | w | 5621 | 672.8 | 0.003047 | 4.529 | +2 | 11 |
| - | - | 4469 | 673.3 | - | - | 0 | - |
| - | - | 2011 | 673.8 | - | - | 0 | - |
| - | - | 1342 | 680.3 | - | - | 0 | - |
| - | - | 709.5 | 690.8 | - | - | 0 | - |
| 3 | y | 1707 | 693.3 | 0.0006324 | 0.9121 | +2 | 11 |
| 3 | y | 2339 | 693.8 | 0.008747 | 12.61 | +2 | 11 |
| 6 | c | 5594 | 694.4 | 0.01092 | 15.73 | +1 | 6 |
| - | - | 2604 | 694.8 | - | - | 0 | - |
| - | - | 2579 | 695.4 | - | - | 0 | - |
| - | - | 897.4 | 697.9 | - | - | 0 | - |
| - | - | 1758 | 698.4 | - | - | 0 | - |
| - | - | 812.6 | 698.9 | - | - | 0 | - |
| - | - | 3342 | 701.8 | - | - | 0 | - |
| 3 | y | 9799 | 702.3 | 0.002735 | 3.895 | +2 | 11 |
| - | - | 6758 | 702.8 | - | - | 0 | - |
| - | - | 3533 | 703.3 | - | - | 0 | - |
| - | - | 1497 | 703.8 | - | - | 0 | - |
| - | - | 810.4 | 704.3 | - | - | 0 | - |
| - | - | 5842 | 707.3 | - | - | 0 | - |
| 8 | z | 7528 | 708.3 | 0.003465 | 4.892 | +1 | 6 |
| - | - | 2376 | 709.3 | - | - | 0 | - |
| - | - | 768.9 | 710.3 | - | - | 0 | - |
| 6 | c | 2.632E+04 | 711.4 | 0.0003626 | 0.5097 | +1 | 6 |
| 12 | c | 1459 | 711.9 | 0.004402 | 6.184 | +2 | 12 |
| - | - | 1.117E+04 | 712.4 | - | - | 0 | - |
| - | - | 2279 | 713.4 | - | - | 0 | - |
| - | - | 660.4 | 714.3 | - | - | 0 | - |
| - | - | 948.8 | 718.9 | - | - | 0 | - |
| - | - | 932.2 | 718.9 | - | - | 0 | - |
| - | - | 1234 | 719.4 | - | - | 0 | - |
| - | - | 815.5 | 719.8 | - | - | 0 | - |
| - | - | 2.16E+04 | 719.9 | - | - | 0 | - |
| 12 | c | 4.427E+04 | 720.4 | 0.002724 | 3.781 | +2 | 12 |
| - | - | 2.79E+04 | 720.9 | - | - | 0 | - |
| - | - | 1.292E+04 | 721.4 | - | - | 0 | - |
| - | - | 3326 | 721.9 | - | - | 0 | - |
| - | - | 795.8 | 722.4 | - | - | 0 | - |
| - | - | 2886 | 723.4 | - | - | 0 | - |
| 8 | y | 8369 | 724.4 | 0.001343 | 1.853 | +1 | 6 |
| - | - | 3918 | 725.4 | - | - | 0 | - |
| - | - | 2316 | 729.4 | - | - | 0 | - |
| - | - | 1854 | 729.9 | - | - | 0 | - |
| - | - | 1787 | 730.4 | - | - | 0 | - |
| - | - | 799.1 | 735.9 | - | - | 0 | - |
| 2 | w | 1014 | 736.3 | 0.001766 | 2.398 | +2 | 12 |
| - | - | 2364 | 736.9 | - | - | 0 | - |
| - | - | 5637 | 737.4 | - | - | 0 | - |
| - | - | 2514 | 737.9 | - | - | 0 | - |
| - | - | 902 | 738.4 | - | - | 0 | - |
| - | - | 755 | 741.4 | - | - | 0 | - |
| 2 | y | 878.8 | 743.4 | 0.006461 | 8.692 | +2 | 12 |
| 2 | z | 2671 | 743.9 | 0.004868 | 6.544 | +2 | 12 |
| - | - | 2314 | 744.4 | - | - | 0 | - |
| - | - | 1208 | 744.9 | - | - | 0 | - |
| - | - | 1.111E+04 | 748.9 | - | - | 0 | - |
| - | - | 8290 | 749.4 | - | - | 0 | - |
| - | - | 5458 | 749.9 | - | - | 0 | - |
| - | - | 876.1 | 750.4 | - | - | 0 | - |
| - | - | 2068 | 750.9 | - | - | 0 | - |
| - | - | 1843 | 751.4 | - | - | 0 | - |
| 2 | y | 1659 | 751.9 | 0.0003437 | 0.4572 | +2 | 12 |
| - | - | 1107 | 752.4 | - | - | 0 | - |
| - | - | 1205 | 757.9 | - | - | 0 | - |
| - | - | 1136 | 758.4 | - | - | 0 | - |
| - | - | 896.1 | 758.9 | - | - | 0 | - |
| - | - | 844.9 | 759.4 | - | - | 0 | - |
| - | - | 887 | 764.4 | - | - | 0 | - |
| - | - | 1550 | 764.9 | - | - | 0 | - |
| - | - | 1465 | 765.4 | - | - | 0 | - |
| - | - | 1179 | 766.4 | - | - | 0 | - |
| - | - | 913.6 | 771.4 | - | - | 0 | - |
| - | - | 2616 | 771.9 | - | - | 0 | - |
| - | - | 4507 | 772.4 | - | - | 0 | - |
| - | - | 9283 | 772.9 | - | - | 0 | - |
| - | - | 6820 | 773.4 | - | - | 0 | - |
| - | - | 1855 | 773.9 | - | - | 0 | - |
| - | - | 973 | 774.4 | - | - | 0 | - |
| - | - | 5454 | 779.9 | - | - | 0 | - |
| - | - | 6248 | 780.4 | - | - | 0 | - |
| - | - | 3351 | 780.9 | - | - | 0 | - |
| - | - | 2232 | 781.4 | - | - | 0 | - |
| - | - | 913 | 784.9 | - | - | 0 | - |
| - | - | 1687 | 785.4 | - | - | 0 | - |
| - | - | 3428 | 785.9 | - | - | 0 | - |
| - | - | 2582 | 786.4 | - | - | 0 | - |
| - | - | 4693 | 786.9 | - | - | 0 | - |
| - | - | 2910 | 787.4 | - | - | 0 | - |
| - | - | 1303 | 787.9 | - | - | 0 | - |
| - | - | 1305 | 793.4 | - | - | 0 | - |
| - | - | 1.246E+04 | 793.9 | - | - | 0 | - |
| - | - | 2.239E+04 | 794.4 | - | - | 0 | - |
| - | - | 1.771E+04 | 794.9 | - | - | 0 | - |
| - | - | 7543 | 795.4 | - | - | 0 | - |
| - | - | 1995 | 795.9 | - | - | 0 | - |
| - | - | 966.8 | 797.4 | - | - | 0 | - |
| - | - | 1772 | 801.4 | - | - | 0 | - |
| - | - | 938.4 | 801.5 | - | - | 0 | - |
| - | - | 2209 | 801.8 | - | - | 0 | - |
| - | - | 1.82E+04 | 801.9 | - | - | 0 | - |
| - | - | 9.133E+04 | 802.4 | - | - | 0 | - |
| - | - | 1.391E+05 | 802.9 | - | - | 0 | - |
| - | - | 8.988E+04 | 803.4 | - | - | 0 | - |
| - | - | 4.146E+04 | 803.9 | - | - | 0 | - |
| - | - | 1.634E+04 | 804.4 | - | - | 0 | - |
| - | - | 4061 | 804.9 | - | - | 0 | - |
| - | - | 2160 | 837.4 | - | - | 0 | - |
| - | - | 4693 | 838.4 | - | - | 0 | - |
| - | - | 1920 | 839.4 | - | - | 0 | - |
| - | - | 1256 | 850.4 | - | - | 0 | - |
| - | - | 1219 | 851.4 | - | - | 0 | - |
| - | - | 655.8 | 852.2 | - | - | 0 | - |
| - | - | 1825 | 853.4 | - | - | 0 | - |
| - | - | 1213 | 862.4 | - | - | 0 | - |
| - | - | 1772 | 864.5 | - | - | 0 | - |
| - | - | 830.3 | 865.5 | - | - | 0 | - |
| 7 | c | 1.298E+04 | 880.4 | 0.0005752 | 0.6533 | +1 | 7 |
| - | - | 6990 | 881.4 | - | - | 0 | - |
| - | - | 1602 | 882.4 | - | - | 0 | - |
| - | - | 926.9 | 883.4 | - | - | 0 | - |
| - | - | 5193 | 893.4 | - | - | 0 | - |
| 7 | z | 9944 | 894.4 | 0.002887 | 3.228 | +1 | 7 |
| - | - | 4166 | 895.4 | - | - | 0 | - |
| - | - | 837.1 | 896.4 | - | - | 0 | - |
| 7 | c | 3.891E+04 | 897.5 | 0.0002078 | 0.2315 | +1 | 7 |
| - | - | 2.186E+04 | 898.5 | - | - | 0 | - |
| - | - | 5253 | 899.5 | - | - | 0 | - |
| - | - | 1780 | 900.5 | - | - | 0 | - |
| - | - | 3232 | 909.4 | - | - | 0 | - |
| 7 | y | 5403 | 910.4 | 0.003634 | 3.991 | +1 | 7 |
| - | - | 2572 | 911.4 | - | - | 0 | - |
| - | - | 1345 | 937.4 | - | - | 0 | - |
| - | - | 817.9 | 938.4 | - | - | 0 | - |
| - | - | 2308 | 964.5 | - | - | 0 | - |
| - | - | 5066 | 965.5 | - | - | 0 | - |
| - | - | 1849 | 966.5 | - | - | 0 | - |
| - | - | 919 | 967.5 | - | - | 0 | - |
| - | - | 1105 | 968.5 | - | - | 0 | - |
| - | - | 690.3 | 991.5 | - | - | 0 | - |
| 8 | c | 1907 | 993.5 | 0.0003218 | 0.3239 | +1 | 8 |
| - | - | 1447 | 994.5 | - | - | 0 | - |
| - | - | 8978 | 1008 | - | - | 0 | - |
| 6 | z | 1.547E+04 | 1009 | 0.004753 | 4.709 | +1 | 8 |
| - | - | 972.3 | 1010 | - | - | 0 | - |
| - | - | 6244 | 1010 | - | - | 0 | - |
| 8 | c | 2.939E+04 | 1011 | 0.000384 | 0.38 | +1 | 8 |
| - | - | 1672 | 1011 | - | - | 0 | - |
| - | - | 1.647E+04 | 1012 | - | - | 0 | - |
| - | - | 1129 | 1012 | - | - | 0 | - |
| - | - | 6149 | 1013 | - | - | 0 | - |
| - | - | 819.1 | 1014 | - | - | 0 | - |
| - | - | 3587 | 1024 | - | - | 0 | - |
| 6 | y | 4586 | 1025 | 0.008368 | 8.16 | +1 | 8 |
| - | - | 2244 | 1026 | - | - | 0 | - |
| - | - | 728.1 | 1027 | - | - | 0 | - |
| - | - | 1532 | 1064 | - | - | 0 | - |
| - | - | 764.4 | 1065 | - | - | 0 | - |
| - | - | 4186 | 1068 | - | - | 0 | - |
| - | - | 3419 | 1069 | - | - | 0 | - |
| - | - | 689.5 | 1069 | - | - | 0 | - |
| - | - | 1143 | 1070 | - | - | 0 | - |
| - | - | 1162 | 1071 | - | - | 0 | - |
| 5 | w | 2773 | 1079 | 0.002564 | 2.375 | +1 | 9 |
| - | - | 1850 | 1080 | - | - | 0 | - |
| - | - | 2725 | 1081 | - | - | 0 | - |
| - | - | 743.8 | 1081 | - | - | 0 | - |
| - | - | 5722 | 1082 | - | - | 0 | - |
| - | - | 2378 | 1083 | - | - | 0 | - |
| - | - | 1030 | 1084 | - | - | 0 | - |
| - | - | 758.5 | 1093 | - | - | 0 | - |
| - | - | 1642 | 1094 | - | - | 0 | - |
| - | - | 805.7 | 1095 | - | - | 0 | - |
| - | - | 596.9 | 1109 | - | - | 0 | - |
| - | - | 643.1 | 1110 | - | - | 0 | - |
| - | - | 4579 | 1125 | - | - | 0 | - |
| 9 | c | 7672 | 1126 | 0.004508 | 4.005 | +1 | 9 |
| - | - | 4991 | 1127 | - | - | 0 | - |
| - | - | 1485 | 1128 | - | - | 0 | - |
| - | - | 1.04E+04 | 1137 | - | - | 0 | - |
| 5 | z | 7.661E+04 | 1137 | 0.0002526 | 0.2221 | +1 | 9 |
| - | - | 4.827E+04 | 1139 | - | - | 0 | - |
| - | - | 1.83E+04 | 1140 | - | - | 0 | - |
| - | - | 3441 | 1141 | - | - | 0 | - |
| - | - | 702.3 | 1142 | - | - | 0 | - |
| - | - | 1133 | 1153 | - | - | 0 | - |
| 5 | y | 5166 | 1154 | 0.001792 | 1.554 | +1 | 9 |
| - | - | 2468 | 1155 | - | - | 0 | - |
| - | - | 1334 | 1156 | - | - | 0 | - |
| - | - | 1.614E+04 | 1182 | - | - | 0 | - |
| 10 | c | 4.135E+04 | 1183 | 0.003308 | 2.797 | +1 | 10 |
| - | - | 2.375E+04 | 1184 | - | - | 0 | - |
| - | - | 1.094E+04 | 1185 | - | - | 0 | - |
| - | - | 2939 | 1186 | - | - | 0 | - |
| - | - | 1459 | 1187 | - | - | 0 | - |
| - | - | 1098 | 1224 | - | - | 0 | - |
| - | - | 1755 | 1267 | - | - | 0 | - |
| - | - | 1673 | 1268 | - | - | 0 | - |
| - | - | 1084 | 1269 | - | - | 0 | - |
| 4 | y | 3389 | 1274 | 0.02009 | 15.77 | +1 | 10 |
| 4 | z | 1.462E+04 | 1275 | 0.002376 | 1.864 | +1 | 10 |
| - | - | 1.533E+04 | 1276 | - | - | 0 | - |
| - | - | 6048 | 1277 | - | - | 0 | - |
| - | - | 1779 | 1278 | - | - | 0 | - |
| - | - | 675.1 | 1279 | - | - | 0 | - |
| - | - | 695.8 | 1290 | - | - | 0 | - |
| 4 | y | 2480 | 1291 | 0.004892 | 3.791 | +1 | 10 |
| - | - | 1160 | 1292 | - | - | 0 | - |
| - | - | 1422 | 1295 | - | - | 0 | - |
| - | - | 718.4 | 1296 | - | - | 0 | - |
| - | - | 5608 | 1310 | - | - | 0 | - |
| 11 | c | 2.561E+04 | 1311 | 0.001362 | 1.039 | +1 | 11 |
| - | - | 1.984E+04 | 1312 | - | - | 0 | - |
| - | - | 8289 | 1313 | - | - | 0 | - |
| - | - | 2319 | 1314 | - | - | 0 | - |
| - | - | 854.6 | 1344 | - | - | 0 | - |
| - | - | 768.1 | 1345 | - | - | 0 | - |
| 3 | y | 1442 | 1387 | 0.01171 | 8.444 | +1 | 11 |
| 3 | z | 1.36E+04 | 1388 | 0.003761 | 2.71 | +1 | 11 |
| - | - | 1.477E+04 | 1389 | - | - | 0 | - |
| - | - | 9024 | 1390 | - | - | 0 | - |
| - | - | 3719 | 1391 | - | - | 0 | - |
| - | - | 1070 | 1392 | - | - | 0 | - |
| - | - | 1008 | 1395 | - | - | 0 | - |
| - | - | 4601 | 1396 | - | - | 0 | - |
| - | - | 4618 | 1397 | - | - | 0 | - |
| - | - | 3199 | 1398 | - | - | 0 | - |
| - | - | 939.1 | 1399 | - | - | 0 | - |
| 3 | y | 1490 | 1404 | 0.00469 | 3.342 | +1 | 11 |
| - | - | 1592 | 1405 | - | - | 0 | - |
| - | - | 1043 | 1413 | - | - | 0 | - |
| 12 | c | 1514 | 1423 | 0.02389 | 16.79 | +1 | 12 |
| - | - | 2439 | 1424 | - | - | 0 | - |
| - | - | 2436 | 1425 | - | - | 0 | - |
| - | - | 1081 | 1426 | - | - | 0 | - |
| - | - | 3127 | 1439 | - | - | 0 | - |
| 12 | c | 9250 | 1440 | 0.007597 | 5.277 | +1 | 12 |
| - | - | 1.2E+04 | 1441 | - | - | 0 | - |
| - | - | 8151 | 1442 | - | - | 0 | - |
| - | - | 3197 | 1443 | - | - | 0 | - |
| - | - | 907.8 | 1444 | - | - | 0 | - |
| 2 | z | 1610 | 1487 | 0.01286 | 8.651 | +1 | 12 |
| - | - | 3198 | 1488 | - | - | 0 | - |
| - | - | 2072 | 1489 | - | - | 0 | - |
| - | - | 1372 | 1490 | - | - | 0 | - |
| - | - | 677.6 | 1491 | - | - | 0 | - |
| - | - | 880.7 | 1529 | - | - | 0 | - |
| - | - | 1664 | 1543 | - | - | 0 | - |
| - | - | 1437 | 1544 | - | - | 0 | - |
| - | - | 2154 | 1545 | - | - | 0 | - |
| - | - | 6370 | 1546 | - | - | 0 | - |
| - | - | 4642 | 1547 | - | - | 0 | - |
| - | - | 2201 | 1548 | - | - | 0 | - |
| - | - | 958.3 | 1549 | - | - | 0 | - |
| - | - | 2153 | 1559 | - | - | 0 | - |
| - | - | 7428 | 1560 | - | - | 0 | - |
| - | - | 9745 | 1561 | - | - | 0 | - |
| - | - | 5275 | 1562 | - | - | 0 | - |
| - | - | 2397 | 1563 | - | - | 0 | - |
| - | - | 1207 | 1570 | - | - | 0 | - |
| - | - | 2474 | 1571 | - | - | 0 | - |
| - | - | 1239 | 1572 | - | - | 0 | - |
| - | - | 983.8 | 1573 | - | - | 0 | - |
| - | - | 744.5 | 1576 | - | - | 0 | - |
| - | - | 4570 | 1577 | - | - | 0 | - |
| - | - | 6813 | 1578 | - | - | 0 | - |
| - | - | 4526 | 1579 | - | - | 0 | - |
| - | - | 2287 | 1580 | - | - | 0 | - |
| - | - | 3053 | 1587 | - | - | 0 | - |
| - | - | 1.903E+04 | 1588 | - | - | 0 | - |
| - | - | 3.59E+04 | 1589 | - | - | 0 | - |
| - | - | 2.567E+04 | 1590 | - | - | 0 | - |
| - | - | 1.163E+04 | 1591 | - | - | 0 | - |
| - | - | 4642 | 1592 | - | - | 0 | - |
| - | - | 1363 | 1593 | - | - | 0 | - |
| - | - | 999.7 | 1603 | - | - | 0 | - |
| - | - | 9743 | 1604 | - | - | 0 | - |
| - | - | 3.668E+04 | 1605 | - | - | 0 | - |
| - | - | 6.324E+04 | 1606 | - | - | 0 | - |
| - | - | 4.677E+04 | 1607 | - | - | 0 | - |
| - | - | 2.206E+04 | 1608 | - | - | 0 | - |
| - | - | 7884 | 1609 | - | - | 0 | - |
| - | - | 3423 | 1610 | - | - | 0 | - |

m/z Charge Intensity FragmentType MassShift Position
120.06583404541016 0 668.7516
128.1071319580078 0 485.2024
129.10238647460938 0 3226.6194
130.0653076171875 0 3540.651
131.06863403320312 0 436.60083
131.11817932128906 0 1030.6396
136.0758056640625 0 1084.2795
139.5156707763672 0 489.5873
148.94757080078125 0 588.1633
149.04502868652344 0 5728.491
150.85986328125 0 429.63507
155.09288024902344 0 12215.934
156.09629821777344 0 494.98016
157.1334228515625 0 1037.9093
159.09181213378906 0 3875.2993
160.02146911621094 0 502.5916
165.05455017089844 0 724.3431
167.0554962158203 0 1576.5376
171.1129913330078 0 586.1625
173.12860107421875 0 16726.037
174.13186645507812 0 1499.4857
181.10906982421875 0 552.6512
182.08123779296875 0 3262.5613 y 12
184.06016540527344 0 792.55225
186.12368774414062 0 1267.1764
188.13941955566406 0 993.16187
198.21083068847656 0 465.9615
201.12347412109375 0 12132.795
202.12677001953125 0 992.26056
213.1601104736328 0 1411.9236
215.13916015625 0 6556.314
216.1424560546875 0 808.9974
216.2119598388672 0 520.7579
218.14999389648438 0 3331.361 c 1
221.0845947265625 0 7142.9155
222.084716796875 0 2596.5144
223.06419372558594 0 2620.8164
223.08160400390625 0 1461.1029
223.15567016601562 0 1112.8208
225.04324340820312 0 2176.172
226.04380798339844 0 614.06494
226.10801696777344 0 1027.2568
229.15484619140625 0 1675.7449
239.09506225585938 0 10176.396
240.0957794189453 0 4338.822
241.0925750732422 0 1587.7162
243.13375854492188 0 1262.4827
245.13714599609375 0 1510.8966
251.15029907226562 0 2876.757
252.15664672851562 0 570.53326
258.14508056640625 0 5994.511
260.1597900390625 0 1330.8799
268.1767578125 0 6459.696
269.1814880371094 0 598.5638
281.0513000488281 0 17088.297
283.03021240234375 0 568.25397
283.1399841308594 0 1550.5684
287.2201232910156 0 769.18787
288.22723388671875 0 960.91376
293.11358642578125 0 2250.9558 y Water loss 11
294.11639404296875 0 932.18115
295.103515625 0 3189.4233 z 11
296.10394287109375 0 2242.9512
297.0826110839844 0 3176.5144
297.1007385253906 0 2138.5422
298.1009521484375 0 824.9013
299.0618896484375 0 67477.02
300.0632019042969 0 1722.8198
301.1507873535156 0 1149.029
302.1507568359375 0 657.7052
304.10809326171875 0 541.26935
305.124755859375 0 1292.9049
306.1075439453125 0 810.99316
307.189453125 0 744.4339
308.1934814453125 0 557.6687
309.2047424316406 0 507.41296
311.1241760253906 0 2161.824 y 11
314.1149597167969 0 716.82794
314.2080993652344 0 2057.2908
315.1658630371094 0 595.94617
317.15826416015625 0 11023.204
318.1617126464844 0 1841.6107
320.1966857910156 0 1873.8508
323.1352233886719 0 8653.754
324.1386413574219 0 773.77704
327.203125 0 2992.7815
331.2341613769531 0 5873.4863 c 2
332.23663330078125 0 780.14264
341.0180358886719 0 1139.2722
343.1023864746094 0 643.50494
346.2668151855469 0 528.24
347.6792297363281 0 587.4168 c Ammonia loss 5
355.06964111328125 0 1492.6672
356.0703125 0 680.6775
357.06854248046875 0 806.5323
359.0285949707031 0 5827.895
362.6799621582031 0 1704.8057 y 7
364.18621826171875 0 1497.7001
369.12164306640625 0 1750.6353
370.1229248046875 0 3147.9485
371.12054443359375 0 2770.6865
372.12054443359375 0 762.1432
374.1793518066406 0 2073.5586
378.2130432128906 0 965.5494
408.2606201171875 0 786.35455
412.183349609375 0 1187.58
415.03704833984375 0 21873.887
423.271728515625 0 6177.649
424.27471923828125 0 1546.6936
426.72198486328125 0 9214.431
427.2233581542969 0 3323.1094
427.7247009277344 0 1441.8191
429.0885314941406 0 1590.8389
429.21002197265625 0 640.34265
430.1940002441406 0 2351.7966
431.71368408203125 0 1238.4576
432.2138366699219 0 1006.40094
439.2193908691406 0 2589.328 y 10
440.7190856933594 0 4999.625 c Ammonia loss 6
441.2214050292969 0 2337.5513
444.2486572265625 0 863.6425
445.21673583984375 0 993.9998
450.2818908691406 0 670.81226 c Water loss 3
451.26678466796875 0 13357.775
452.2703857421875 0 3223.2473
453.27606201171875 0 675.484
468.2933349609375 0 52326.043 c 3
469.2961120605469 0 14780.352
470.2998046875 0 2492.582
474.2569274902344 0 640.0301
474.75567626953125 0 1035.0421
480.2220458984375 0 5198.4805 z 9
481.22467041015625 0 1934.337
483.26446533203125 0 2324.0498
483.764892578125 0 930.74976
489.20733642578125 0 2264.2825
494.2373046875 0 949.2243
494.30865478515625 0 3801.5203
495.2322998046875 0 1591.61
495.3121643066406 0 1424.4884
496.24066162109375 0 15649.24 y 9
497.2467346191406 0 3275.6553
497.7647399902344 0 646.5523
503.107666015625 0 54921.695
517.2781372070312 0 977.9643
519.1388549804688 0 22089.537
523.2299194335938 0 1039.7328
528.9321899414062 0 686.9133
529.2620239257812 0 635.2435
529.5934448242188 0 1137.8229
529.9268188476562 0 753.19275
534.3301391601562 0 673.0048
535.286865234375 0 3571.803
536.165283203125 0 12304.893
536.2695922851562 0 10262.868
538.2509155273438 0 5112.966
539.25439453125 0 1451.7719
550.2515258789062 0 4954.098 w 8
551.2560424804688 0 4494.442
552.2587890625 0 884.6097
579.32421875 0 1960.9045 c Ammonia loss 4
580.325439453125 0 808.2749
593.2559204101562 0 902.2893 y Water loss 8
594.2628784179688 0 1601.8708
595.2493286132812 0 5277.82 z 8
595.3029174804688 0 637.196
596.2567138671875 0 2031.5055
596.3516235351562 0 30812.543 c 4
597.3546142578125 0 9994.601
598.3572998046875 0 2130.658
602.2972412109375 0 790.4399
608.2784423828125 0 844.62915
609.3342895507812 0 1234.4562
610.282958984375 0 7901.022
611.268310546875 0 19396.332 y 8
611.7994384765625 0 766.04443
612.2706909179688 0 6165.2666
613.2711181640625 0 1011.8758
621.289306640625 0 1437.8275
633.8432006835938 0 1598.5354
634.3384399414062 0 1108.7334
637.2922973632812 0 1186.7931
637.785400390625 0 684.381 z 3
645.7926025390625 0 1897.0757 y 3
646.2880249023438 0 1004.89856
651.345947265625 0 604.59955
654.8496704101562 0 2821.1396
655.352783203125 0 14014.143
655.8474731445312 0 58459.906 c 10
656.3482055664062 0 44727.242
656.8494262695312 0 18374.248
657.3504638671875 0 4225.5684
657.8526611328125 0 958.5421
664.2933349609375 0 7353.479
664.3452758789062 0 1533.639
665.2808837890625 0 9189.574 w 7
665.3467407226562 0 1183.9695
666.282958984375 0 3471.7622
666.3558349609375 0 1730.3635
667.292236328125 0 998.9795
667.3644409179688 0 2408.1594
668.3645629882812 0 3963.1392
668.859375 0 4530.1064
669.3594970703125 0 2278.5298
669.8616333007812 0 1411.645
672.3031005859375 0 3383.6235
672.7994384765625 0 5621.375 w 2
673.2984619140625 0 4469.3286
673.7992553710938 0 2010.5262
680.3168334960938 0 1342.1836
690.8471069335938 0 709.5448
693.3284912109375 0 1707.0178 y Water loss 2
693.82861328125 0 2338.9854 y Ammonia loss 2
694.3409423828125 0 5594.2773 c Ammonia loss 5
694.8267211914062 0 2603.6294
695.353271484375 0 2578.6978
697.8689575195312 0 897.41785
698.3662719726562 0 1757.8788
698.8629150390625 0 812.61725
701.8388061523438 0 3341.616
702.3358764648438 0 9799.463 y 2
702.836669921875 0 6757.7856
703.3377685546875 0 3533.3962
703.838623046875 0 1496.8771
704.341064453125 0 810.4257
707.3493041992188 0 5842.342
708.3359375 0 7528.0835 z 7
709.3384399414062 0 2375.5745
710.3397827148438 0 768.948
711.3780517578125 0 26317.62 c 5
711.858642578125 0 1458.728 c Ammonia loss 11
712.3804321289062 0 11169.315
713.3828125 0 2278.6365
714.3435668945312 0 660.3691
718.8636474609375 0 948.81824
718.92626953125 0 932.2432
719.4251098632812 0 1234.4236
719.7992553710938 0 815.4514
719.8750610351562 0 21598.479
720.3702392578125 0 44272.19 c 11
720.8707885742188 0 27902.713
721.3719482421875 0 12918.871
721.8731689453125 0 3325.6428
722.3760375976562 0 795.7944
723.3648071289062 0 2885.951
724.3525390625 0 8368.912 y 7
725.355224609375 0 3918.275
729.3583984375 0 2315.562
729.8579711914062 0 1854.4149
730.358154296875 0 1787.236
735.8630981445312 0 799.1078
736.344482421875 0 1013.7441 w 1
736.8658447265625 0 2364.479
737.35888671875 0 5637.457
737.8636474609375 0 2513.621
738.3641967773438 0 902.0017
741.3693237304688 0 754.9649
743.3605346679688 0 878.82605 y Ammonia loss 1
743.8628540039062 0 2670.8025 z 1
744.3635864257812 0 2313.919
744.8643798828125 0 1208.4338
748.8666381835938 0 11106.579
749.3677978515625 0 8290.435
749.8677978515625 0 5457.8086
750.376953125 0 876.0781
750.8759765625 0 2068.3308
751.3773803710938 0 1842.5237
751.8670043945312 0 1658.563 y 1
752.3795166015625 0 1106.952
757.8914794921875 0 1205.4178
758.378173828125 0 1135.8964
758.86962890625 0 896.14734
759.3670043945312 0 844.93835
764.3776245117188 0 886.96564
764.8755493164062 0 1550.3652
765.36865234375 0 1464.7415
766.3595581054688 0 1178.7773
771.3936157226562 0 913.62866
771.8916625976562 0 2615.891
772.3861694335938 0 4506.99
772.8845825195312 0 9282.633
773.3853149414062 0 6820.2417
773.88037109375 0 1854.6072
774.3823852539062 0 972.98865
779.8931884765625 0 5453.5713
780.39306640625 0 6248.3706
780.8956298828125 0 3351.379
781.3907470703125 0 2232.1726
784.8809204101562 0 912.9579
785.3857421875 0 1687.4258
785.884521484375 0 3428.0164
786.3809814453125 0 2582.4607
786.8781127929688 0 4693.467
787.3744506835938 0 2909.9902
787.8737182617188 0 1303.0286
793.3908081054688 0 1305.3484
793.8897094726562 0 12461.018
794.385498046875 0 22385.027
794.8870849609375 0 17711.85
795.386474609375 0 7543.036
795.8912353515625 0 1995.2668
797.3831787109375 0 966.8176
801.3931274414062 0 1771.5359
801.4611206054688 0 938.3742
801.826904296875 0 2208.53
801.8991088867188 0 18202.002
802.3951416015625 0 91332.336
802.8956909179688 0 139122.27
803.3966674804688 0 89876.86
803.898681640625 0 41462.78
804.3992309570312 0 16341.057
804.8977661132812 0 4061.4846
837.3658447265625 0 2160.198
838.3540649414062 0 4693.124
839.3523559570312 0 1920.4636
850.4282836914062 0 1255.9581
851.406494140625 0 1219.4026
852.1968383789062 0 655.77576
853.4384155273438 0 1825.4209
862.4222412109375 0 1213.1014
864.455810546875 0 1772.2651
865.4559326171875 0 830.2891
880.4306030273438 0 12980.593 c Ammonia loss 6
881.4340209960938 0 6990.4478
882.4358520507812 0 1601.9729
883.4314575195312 0 926.9217
893.4271240234375 0 5192.7466
894.4146728515625 0 9943.777 z 6
895.418701171875 0 4165.5347
896.4201049804688 0 837.0564
897.45751953125 0 38908.953 c 6
898.4603271484375 0 21857.695
899.4625854492188 0 5252.777
900.4653930664062 0 1780.0673
909.4464111328125 0 3232.388
910.4341430664062 0 5402.748 y 6
911.4381713867188 0 2571.739
937.4203491210938 0 1345.1667
938.4232177734375 0 817.89197
964.4657592773438 0 2307.682
965.4545288085938 0 5065.774
966.45263671875 0 1849.3798
967.53955078125 0 918.9528
968.4857788085938 0 1104.8661
991.5180053710938 0 690.2919
993.5155639648438 0 1907.2988 c Ammonia loss 7
994.5184326171875 0 1446.6561
1008.45361328125 0 8977.743
1009.4434814453125 0 15474.314 z 5
1009.5524291992188 0 972.341
1010.4406127929688 0 6243.7876
1010.5421752929688 0 29388.59 c 7
1011.4452514648438 0 1671.562
1011.5445556640625 0 16468.023
1012.4456176757812 0 1129.2655
1012.5482788085938 0 6149.3926
1013.5567016601562 0 819.1036
1024.4736328125 0 3586.6138
1025.4658203125 0 4585.6616 y 5
1026.4697265625 0 2244.4932
1027.46533203125 0 728.0814
1063.5504150390625 0 1532.2644
1064.54296875 0 764.4427
1067.5350341796875 0 4186.1475
1068.5369873046875 0 3419.349
1068.6551513671875 0 689.4805
1069.5380859375 0 1143.1792
1070.5264892578125 0 1162.4412
1079.4705810546875 0 2773.0576 w 4
1080.470703125 0 1849.8038
1080.5704345703125 0 2725.4126
1081.449462890625 0 743.7657
1081.5625 0 5721.713
1082.56689453125 0 2378.4482
1083.565673828125 0 1030.4714
1092.50732421875 0 758.4836
1093.5174560546875 0 1641.9269
1094.516845703125 0 805.66705
1108.5196533203125 0 596.9434
1109.5313720703125 0 643.14575
1124.5830078125 0 4579.3354
1125.5732421875 0 7672.394 c 8
1126.5753173828125 0 4991.198
1127.57470703125 0 1484.7749
1136.5123291015625 0 10401.216
1137.49755859375 0 76605.11 z 4
1138.50048828125 0 48271.863
1139.5028076171875 0 18298.21
1140.5079345703125 0 3440.7656
1141.5009765625 0 702.3176
1152.537109375 0 1133.2249
1153.517822265625 0 5165.966 y 4
1154.5181884765625 0 2467.5703
1155.5279541015625 0 1334.3015
1181.60546875 0 16143.399
1182.593505859375 0 41345.457 c 9
1183.5955810546875 0 23747.377
1184.5972900390625 0 10940.251
1185.602294921875 0 2938.9634
1186.610107421875 0 1458.5243
1223.595458984375 0 1098.4172
1266.6739501953125 0 1754.6954
1267.678955078125 0 1672.794
1268.6859130859375 0 1083.6975
1273.5684814453125 0 3388.7488 y Ammonia loss 3
1274.55859375 0 14619.583 z 3
1275.5615234375 0 15326.243
1276.5655517578125 0 6048.2637
1277.5654296875 0 1778.6163
1278.5672607421875 0 675.0736
1289.5792236328125 0 695.79846
1290.579833984375 0 2479.5464 y 3
1291.5784912109375 0 1159.6869
1294.6708984375 0 1422.4883
1295.67041015625 0 718.40936
1309.6983642578125 0 5607.8525
1310.6865234375 0 25611.549 c 10
1311.6885986328125 0 19835.281
1312.690185546875 0 8289.02
1313.6939697265625 0 2319.2007
1343.626220703125 0 854.55414
1344.619873046875 0 768.14734
1386.6441650390625 0 1441.6262 y Ammonia loss 2
1387.64404296875 0 13602.451 z 2
1388.6463623046875 0 14768.205
1389.65087890625 0 9024.3545
1390.6524658203125 0 3719.276
1391.649658203125 0 1069.7156
1394.7244873046875 0 1008.30225
1395.7249755859375 0 4601.0166
1396.7274169921875 0 4617.7725
1397.7269287109375 0 3199.2676
1398.736083984375 0 939.0572
1403.6636962890625 0 1489.7805 y 2
1404.6668701171875 0 1591.7617
1412.74853515625 0 1042.701
1422.72509765625 0 1514.375 c Ammonia loss 11
1423.7213134765625 0 2439.2625
1424.714599609375 0 2436.1177
1425.7176513671875 0 1080.5751
1438.7451171875 0 3126.6382
1439.7353515625 0 9250.297 c 11
1440.7357177734375 0 11999.548
1441.739990234375 0 8151.2056
1442.7421875 0 3197.0015
1443.7425537109375 0 907.82294
1486.7215576171875 0 1609.6061 z 1
1487.7135009765625 0 3198.3665
1488.7159423828125 0 2071.9019
1489.7232666015625 0 1371.724
1490.7122802734375 0 677.6077
1528.7542724609375 0 880.66833
1542.7630615234375 0 1664.0923
1543.779052734375 0 1436.7167
1544.7706298828125 0 2153.519
1545.7711181640625 0 6370.474
1546.769775390625 0 4641.991
1547.7808837890625 0 2200.6785
1548.7691650390625 0 958.3491
1558.783447265625 0 2152.6775
1559.7833251953125 0 7428.2744
1560.78271484375 0 9745.256
1561.785888671875 0 5274.8086
1562.782958984375 0 2396.9636
1569.7763671875 0 1206.6993
1570.7607421875 0 2474.289
1571.7584228515625 0 1239.4469
1572.7623291015625 0 983.78265
1575.7822265625 0 744.4541
1576.79736328125 0 4570.398
1577.7974853515625 0 6813.281
1578.7972412109375 0 4526.4375
1579.80419921875 0 2287.1497
1586.780517578125 0 3053.169
1587.7723388671875 0 19029.785
1588.7681884765625 0 35900.438
1589.769775390625 0 25666.574
1590.771484375 0 11630.793
1591.7747802734375 0 4641.7485
1592.767578125 0 1363.0038
1602.79150390625 0 999.6787
1603.7861328125 0 9743.297
1604.7890625 0 36675.508
1605.78955078125 0 63241.72
1606.7921142578125 0 46771.05
1607.79443359375 0 22061.812
1608.7984619140625 0 7883.5347
1609.793701171875 0 3423.1448

Spectrum Details

|  |  |
| --- | --- |
| Matched peaks? Matched peaks The total absolute number of peaks matched. Additionally in brackets the total fraction of peaks matched and the total number of peaks is shown. | 61 (12.50% of 488) |
| FDR? FDR The false discovery rate estimated for this peptide. It is calculated by matching all theoretical fragments with a non-integer shift with the raw peaks for this spectrum. This is done with 40 different shifts. The resulting percentage is the average number of annotated peaks over the number of annotated peaks with the correct spectrum. | 3.75% |
| Satellite FDR? Satellite FDR See the FDR for details on its calculation. This satellite ion specific FDR only contains the satellite ions (d/w) for I/L/J positions. | 0.00% |
| PSM Score? PSM Score The PSM Score as given by Hecklib to this annotated spectrum. It is shown with three significant figures. | 567 |

#### Spectrum 7024? Spectrum 7024 The raw spectrum of this peptide as annotated by Hecklib. The fragments are coloured according to ion type (see legend). Any peaks with a star '\*' as text can be hovered over to see the full details, first the ion type second the mass shift type. By hovering over the amino acids in the peptide or ions in the legend the corresponding peaks are highlighted. By toggling the 'Unassigned' label you can turn the background (unassigned) peaks on or off in the plot. By updating the slider in the Ion legend you can update the spectrum to only show the top X% of the peaks with labels. The top X% means any peak that is within X% of the highest intensity. By dragging in the spectrum you can zoom in to a specific part of the spectrum and use 'Zoom Out' to get back to the original zoom level. The annotation of the spectrum is based on the given sequence in the peptides file and is done with different software so inconsistencies are likely. The peaks are annotated based on the given sequence, with 20 ppm tolerance.

Copy Data

##### Spectrum 7024 (TSV)

###### Preview

```
Loading example...
```

*Click on the button to copy the data to your clipboard.*

Mz MinMz MaxIntensity Max

WidthHeightPeptide font sizePeptide stroke widthSpectrum font sizeSpectrum stroke widthCompact peptide

Ion legend

wxyz

abcd

OtherUnassignedIon^Charge^_Position_Show for top:%

TVLHQDWLDGKEY

05.54e+51.11e+61.66e+62.21e+6

Zoom Out

a^+1^_2_a^+1^_2_y^+1^_1_b^+1^_2_b^+1^_2_y^+2^_3_b^+2^_4_y^+2^_4_b^+2^_5_y^+1^_2_b^+1^_3_y^+2^_5_y^+3^_7_y^+2^_5_y^+1^_2_b^+1^_3_b^+2^_6_b^+2^_6_b^+2^_6_y^+2^_6_y^+2^_6_b^+3^_9_y^+3^_9_y^+1^_3_y^+1^_3_b^+2^_7_b^+2^_7_b^+1^_4_y^+1^_3_b^+2^_7_y^+2^_7_y^+2^_7_b^+1^_4_y^+2^_7_y^+1^_4_b^+2^_8_b^+2^_8_y^+1^_4_\*\*\*b^+2^_9_b^+2^_9_b^+2^_9_b^+1^_5_b^+1^_5_y^+2^_9_y^+2^_9_b^+1^_5_y^+1^_5_y^+1^_5_y^+2^_10_y^+2^_10_b^+2^_11_y^+2^_10_b^+1^_6_b^+1^_6_y^+2^_11_y^+2^_11_b^+1^_6_y^+2^_11_b^+2^_12_b^+2^_12_y^+1^_6_b^+2^_12_y^+1^_6_y^+2^_12_y^+2^_12_y^+2^_12_\*b^+1^_7_b^+1^_7_b^+1^_7_y^+1^_7_b^+1^_8_b^+1^_8_b^+1^_8_y^+1^_8_y^+1^_8_b^+1^_9_b^+1^_9_y^+1^_9_y^+1^_9_y^+1^_9_b^+1^_10_y^+1^_10_

0874174726213494

Fragment Matches Table

Show background peaks

| Position | Ion type | Intensity | mz Theoretical | mz Error (Th) | mz Error (ppm) | Charge | Series Number |
| --- | --- | --- | --- | --- | --- | --- | --- |
| - | - | 7131 | 120.1 | - | - | 0 | - |
| - | - | 8729 | 122.1 | - | - | 0 | - |
| - | - | 1.022E+04 | 123 | - | - | 0 | - |
| - | - | 2693 | 123.1 | - | - | 0 | - |
| - | - | 2086 | 124.9 | - | - | 0 | - |
| - | - | 3802 | 125.1 | - | - | 0 | - |
| - | - | 7024 | 126.1 | - | - | 0 | - |
| - | - | 3.431E+04 | 127.1 | - | - | 0 | - |
| - | - | 3663 | 127.1 | - | - | 0 | - |
| - | - | 3889 | 127.1 | - | - | 0 | - |
| - | - | 1.296E+05 | 128.1 | - | - | 0 | - |
| - | - | 9255 | 129.1 | - | - | 0 | - |
| - | - | 1.454E+06 | 129.1 | - | - | 0 | - |
| - | - | 6.795E+05 | 130.1 | - | - | 0 | - |
| - | - | 1.093E+04 | 130.1 | - | - | 0 | - |
| - | - | 8.897E+04 | 130.1 | - | - | 0 | - |
| - | - | 6.908E+04 | 131.1 | - | - | 0 | - |
| - | - | 3459 | 131.1 | - | - | 0 | - |
| - | - | 3202 | 131.1 | - | - | 0 | - |
| - | - | 1.495E+05 | 132.1 | - | - | 0 | - |
| - | - | 4335 | 133.1 | - | - | 0 | - |
| - | - | 1.602E+04 | 133.1 | - | - | 0 | - |
| - | - | 2094 | 133.9 | - | - | 0 | - |
| - | - | 1785 | 136 | - | - | 0 | - |
| - | - | 1.755E+05 | 136.1 | - | - | 0 | - |
| - | - | 1.394E+04 | 137.1 | - | - | 0 | - |
| - | - | 1.72E+04 | 138.1 | - | - | 0 | - |
| - | - | 9131 | 138.1 | - | - | 0 | - |
| - | - | 2037 | 138.4 | - | - | 0 | - |
| - | - | 1.678E+04 | 139.1 | - | - | 0 | - |
| - | - | 2.604E+04 | 141.1 | - | - | 0 | - |
| - | - | 1.157E+04 | 142.1 | - | - | 0 | - |
| - | - | 2133 | 142.1 | - | - | 0 | - |
| - | - | 7148 | 143 | - | - | 0 | - |
| - | - | 3876 | 144.1 | - | - | 0 | - |
| - | - | 7656 | 144.1 | - | - | 0 | - |
| - | - | 1.57E+04 | 145.1 | - | - | 0 | - |
| - | - | 2885 | 146.1 | - | - | 0 | - |
| - | - | 1.059E+04 | 147 | - | - | 0 | - |
| - | - | 1.476E+04 | 149 | - | - | 0 | - |
| - | - | 1.098E+04 | 150.1 | - | - | 0 | - |
| - | - | 3.771E+04 | 151.1 | - | - | 0 | - |
| - | - | 4187 | 152.1 | - | - | 0 | - |
| - | - | 3804 | 153.1 | - | - | 0 | - |
| - | - | 6181 | 154.1 | - | - | 0 | - |
| - | - | 2223 | 154.1 | - | - | 0 | - |
| - | - | 4419 | 155 | - | - | 0 | - |
| - | - | 5389 | 155.1 | - | - | 0 | - |
| - | - | 9.558E+04 | 155.1 | - | - | 0 | - |
| 2 | a | 7.052E+04 | 155.1 | 0.0003508 | 2.262 | +1 | 2 |
| - | - | 5017 | 156.1 | - | - | 0 | - |
| - | - | 5642 | 156.1 | - | - | 0 | - |
| - | - | 5919 | 156.1 | - | - | 0 | - |
| - | - | 8595 | 157.1 | - | - | 0 | - |
| - | - | 1.098E+04 | 158.1 | - | - | 0 | - |
| - | - | 1.156E+06 | 159.1 | - | - | 0 | - |
| - | - | 8257 | 160.1 | - | - | 0 | - |
| - | - | 9588 | 160.1 | - | - | 0 | - |
| - | - | 1.257E+05 | 160.1 | - | - | 0 | - |
| - | - | 4396 | 161.1 | - | - | 0 | - |
| - | - | 9.229E+04 | 165.1 | - | - | 0 | - |
| - | - | 5576 | 165.1 | - | - | 0 | - |
| - | - | 1.782E+05 | 166.1 | - | - | 0 | - |
| - | - | 1.584E+04 | 166.1 | - | - | 0 | - |
| - | - | 9711 | 167.1 | - | - | 0 | - |
| - | - | 1.134E+04 | 167.1 | - | - | 0 | - |
| - | - | 3421 | 167.1 | - | - | 0 | - |
| - | - | 1.803E+04 | 168.1 | - | - | 0 | - |
| - | - | 1.183E+04 | 168.1 | - | - | 0 | - |
| - | - | 1.717E+04 | 168.1 | - | - | 0 | - |
| - | - | 1.823E+04 | 169.1 | - | - | 0 | - |
| - | - | 4124 | 169.1 | - | - | 0 | - |
| - | - | 4.112E+04 | 169.1 | - | - | 0 | - |
| - | - | 5.318E+04 | 170.1 | - | - | 0 | - |
| - | - | 3527 | 170.1 | - | - | 0 | - |
| - | - | 6346 | 171.1 | - | - | 0 | - |
| - | - | 4885 | 171.1 | - | - | 0 | - |
| - | - | 2.488E+04 | 171.1 | - | - | 0 | - |
| - | - | 2.988E+04 | 172.1 | - | - | 0 | - |
| - | - | 2352 | 172.1 | - | - | 0 | - |
| - | - | 2634 | 172.2 | - | - | 0 | - |
| - | - | 1.921E+04 | 173.1 | - | - | 0 | - |
| 2 | a | 2.193E+06 | 173.1 | 0.0004368 | 2.523 | +1 | 2 |
| - | - | 1.906E+05 | 174.1 | - | - | 0 | - |
| - | - | 1.205E+04 | 175.1 | - | - | 0 | - |
| - | - | 4175 | 176.1 | - | - | 0 | - |
| - | - | 5051 | 177.1 | - | - | 0 | - |
| - | - | 6.786E+04 | 178.1 | - | - | 0 | - |
| - | - | 7648 | 179.1 | - | - | 0 | - |
| - | - | 1.478E+04 | 181.1 | - | - | 0 | - |
| 13 | y | 2.057E+05 | 182.1 | 0.0004191 | 2.302 | +1 | 1 |
| - | - | 1.981E+04 | 183.1 | - | - | 0 | - |
| 2 | b | 6.032E+04 | 183.1 | 0.0003703 | 2.022 | +1 | 2 |
| - | - | 4779 | 183.1 | - | - | 0 | - |
| - | - | 4958 | 184.1 | - | - | 0 | - |
| - | - | 1.523E+04 | 185.2 | - | - | 0 | - |
| - | - | 4.557E+05 | 186.1 | - | - | 0 | - |
| - | - | 9.648E+04 | 187.1 | - | - | 0 | - |
| - | - | 2544 | 187.1 | - | - | 0 | - |
| - | - | 3387 | 187.1 | - | - | 0 | - |
| - | - | 3.584E+04 | 187.1 | - | - | 0 | - |
| - | - | 6646 | 188.1 | - | - | 0 | - |
| - | - | 1.158E+04 | 188.1 | - | - | 0 | - |
| - | - | 5654 | 189.1 | - | - | 0 | - |
| - | - | 1.906E+04 | 190.1 | - | - | 0 | - |
| - | - | 1.332E+04 | 190.1 | - | - | 0 | - |
| - | - | 1.624E+04 | 194.1 | - | - | 0 | - |
| - | - | 1.106E+05 | 195.1 | - | - | 0 | - |
| - | - | 6293 | 195.1 | - | - | 0 | - |
| - | - | 8.655E+04 | 196.1 | - | - | 0 | - |
| - | - | 9783 | 196.1 | - | - | 0 | - |
| - | - | 4045 | 197.1 | - | - | 0 | - |
| - | - | 1.005E+04 | 197.1 | - | - | 0 | - |
| - | - | 3602 | 197.2 | - | - | 0 | - |
| - | - | 2646 | 197.4 | - | - | 0 | - |
| - | - | 1.5E+04 | 198.1 | - | - | 0 | - |
| - | - | 3046 | 198.1 | - | - | 0 | - |
| - | - | 4.72E+04 | 198.1 | - | - | 0 | - |
| - | - | 6464 | 199.1 | - | - | 0 | - |
| - | - | 1.193E+04 | 199.1 | - | - | 0 | - |
| - | - | 4338 | 199.1 | - | - | 0 | - |
| - | - | 1.31E+04 | 199.1 | - | - | 0 | - |
| - | - | 5389 | 200.1 | - | - | 0 | - |
| - | - | 3161 | 200.3 | - | - | 0 | - |
| 2 | b | 5.98E+05 | 201.1 | 0.0003494 | 1.737 | +1 | 2 |
| - | - | 4613 | 202.1 | - | - | 0 | - |
| - | - | 5.297E+04 | 202.1 | - | - | 0 | - |
| - | - | 4910 | 203.1 | - | - | 0 | - |
| - | - | 3662 | 204.1 | - | - | 0 | - |
| - | - | 5.518E+04 | 205.1 | - | - | 0 | - |
| - | - | 1891 | 205.1 | - | - | 0 | - |
| - | - | 7953 | 205.1 | - | - | 0 | - |
| - | - | 2914 | 206.1 | - | - | 0 | - |
| - | - | 1.083E+04 | 206.1 | - | - | 0 | - |
| - | - | 1.723E+04 | 207.2 | - | - | 0 | - |
| - | - | 3873 | 208.1 | - | - | 0 | - |
| - | - | 4315 | 209.1 | - | - | 0 | - |
| - | - | 1.823E+04 | 210.1 | - | - | 0 | - |
| - | - | 2765 | 211.1 | - | - | 0 | - |
| 11 | y | 1.744E+04 | 211.1 | 0.000344 | 1.629 | +2 | 3 |
| - | - | 6104 | 211.1 | - | - | 0 | - |
| - | - | 1.392E+04 | 212.1 | - | - | 0 | - |
| - | - | 5.842E+04 | 213.1 | - | - | 0 | - |
| - | - | 4778 | 213.1 | - | - | 0 | - |
| - | - | 7456 | 213.2 | - | - | 0 | - |
| - | - | 5114 | 214.1 | - | - | 0 | - |
| - | - | 5209 | 215.1 | - | - | 0 | - |
| - | - | 1.436E+04 | 216.1 | - | - | 0 | - |
| - | - | 5783 | 221.1 | - | - | 0 | - |
| - | - | 8086 | 221.1 | - | - | 0 | - |
| - | - | 1.004E+04 | 221.1 | - | - | 0 | - |
| - | - | 7080 | 221.1 | - | - | 0 | - |
| - | - | 1.961E+04 | 222.1 | - | - | 0 | - |
| - | - | 2.958E+04 | 223.1 | - | - | 0 | - |
| - | - | 2.063E+05 | 223.2 | - | - | 0 | - |
| - | - | 1.149E+04 | 224.1 | - | - | 0 | - |
| - | - | 1.751E+04 | 224.2 | - | - | 0 | - |
| - | - | 5791 | 225 | - | - | 0 | - |
| - | - | 2393 | 225.1 | - | - | 0 | - |
| - | - | 3438 | 225.1 | - | - | 0 | - |
| - | - | 4892 | 225.2 | - | - | 0 | - |
| - | - | 3.871E+04 | 226.1 | - | - | 0 | - |
| - | - | 3753 | 226.1 | - | - | 0 | - |
| 4 | b | 3107 | 226.1 | 0.0005643 | 2.495 | +2 | 4 |
| - | - | 3.684E+04 | 227.1 | - | - | 0 | - |
| - | - | 4965 | 227.1 | - | - | 0 | - |
| - | - | 1.251E+04 | 227.2 | - | - | 0 | - |
| - | - | 2919 | 228.1 | - | - | 0 | - |
| - | - | 2.583E+04 | 228.1 | - | - | 0 | - |
| - | - | 2.874E+04 | 229.1 | - | - | 0 | - |
| - | - | 3447 | 230.1 | - | - | 0 | - |
| - | - | 4853 | 230.2 | - | - | 0 | - |
| - | - | 3.405E+04 | 233.1 | - | - | 0 | - |
| - | - | 2.218E+05 | 234.1 | - | - | 0 | - |
| - | - | 2988 | 234.7 | - | - | 0 | - |
| - | - | 3.462E+04 | 235.1 | - | - | 0 | - |
| - | - | 5388 | 235.2 | - | - | 0 | - |
| - | - | 5067 | 235.6 | - | - | 0 | - |
| - | - | 1.02E+04 | 237.1 | - | - | 0 | - |
| - | - | 3.442E+04 | 238.1 | - | - | 0 | - |
| - | - | 4776 | 238.1 | - | - | 0 | - |
| - | - | 5063 | 239.1 | - | - | 0 | - |
| - | - | 1.127E+04 | 239.1 | - | - | 0 | - |
| - | - | 5311 | 240.1 | - | - | 0 | - |
| - | - | 9.607E+04 | 240.1 | - | - | 0 | - |
| - | - | 4882 | 241.1 | - | - | 0 | - |
| - | - | 1.185E+04 | 241.1 | - | - | 0 | - |
| - | - | 1.194E+04 | 241.1 | - | - | 0 | - |
| - | - | 1.333E+04 | 241.2 | - | - | 0 | - |
| - | - | 1.838E+04 | 243.1 | - | - | 0 | - |
| - | - | 8.991E+04 | 244.1 | - | - | 0 | - |
| - | - | 1.105E+04 | 245.1 | - | - | 0 | - |
| - | - | 2703 | 245.1 | - | - | 0 | - |
| - | - | 1.287E+05 | 247.1 | - | - | 0 | - |
| - | - | 6766 | 247.1 | - | - | 0 | - |
| - | - | 3850 | 247.2 | - | - | 0 | - |
| - | - | 2.298E+04 | 248.1 | - | - | 0 | - |
| - | - | 4.038E+04 | 248.1 | - | - | 0 | - |
| - | - | 2863 | 248.1 | - | - | 0 | - |
| 10 | y | 4319 | 248.6 | 0.0002589 | 1.041 | +2 | 4 |
| - | - | 6.085E+04 | 249.1 | - | - | 0 | - |
| - | - | 3871 | 249.1 | - | - | 0 | - |
| - | - | 5595 | 249.1 | - | - | 0 | - |
| - | - | 2.432E+04 | 249.1 | - | - | 0 | - |
| - | - | 3110 | 249.6 | - | - | 0 | - |
| - | - | 1.112E+04 | 250.1 | - | - | 0 | - |
| - | - | 8364 | 250.1 | - | - | 0 | - |
| - | - | 3030 | 250.2 | - | - | 0 | - |
| - | - | 7.618E+05 | 251.2 | - | - | 0 | - |
| - | - | 6421 | 252.1 | - | - | 0 | - |
| - | - | 9.329E+04 | 252.2 | - | - | 0 | - |
| - | - | 7626 | 253.2 | - | - | 0 | - |
| - | - | 2758 | 254.1 | - | - | 0 | - |
| - | - | 4.395E+04 | 255.1 | - | - | 0 | - |
| - | - | 1.223E+04 | 256.1 | - | - | 0 | - |
| - | - | 5884 | 256.2 | - | - | 0 | - |
| - | - | 6893 | 256.6 | - | - | 0 | - |
| - | - | 2.314E+04 | 257.1 | - | - | 0 | - |
| - | - | 2610 | 258.1 | - | - | 0 | - |
| - | - | 3079 | 258.1 | - | - | 0 | - |
| - | - | 3.36E+04 | 258.1 | - | - | 0 | - |
| - | - | 5267 | 259.1 | - | - | 0 | - |
| - | - | 3668 | 259.1 | - | - | 0 | - |
| - | - | 1.645E+04 | 261.1 | - | - | 0 | - |
| - | - | 7146 | 261.1 | - | - | 0 | - |
| - | - | 1.066E+04 | 262.1 | - | - | 0 | - |
| - | - | 1.107E+05 | 265.1 | - | - | 0 | - |
| - | - | 2.818E+04 | 266.1 | - | - | 0 | - |
| - | - | 2.079E+05 | 266.1 | - | - | 0 | - |
| - | - | 2.232E+04 | 267.1 | - | - | 0 | - |
| - | - | 4696 | 267.6 | - | - | 0 | - |
| - | - | 8554 | 268.1 | - | - | 0 | - |
| - | - | 2.458E+04 | 268.2 | - | - | 0 | - |
| - | - | 8523 | 269.2 | - | - | 0 | - |
| - | - | 1.154E+04 | 269.2 | - | - | 0 | - |
| - | - | 3.23E+04 | 270.1 | - | - | 0 | - |
| - | - | 5740 | 270.1 | - | - | 0 | - |
| - | - | 3150 | 270.2 | - | - | 0 | - |
| - | - | 7026 | 270.6 | - | - | 0 | - |
| - | - | 3536 | 271.1 | - | - | 0 | - |
| - | - | 4.776E+04 | 272.2 | - | - | 0 | - |
| - | - | 8527 | 273.2 | - | - | 0 | - |
| - | - | 4.363E+04 | 274.1 | - | - | 0 | - |
| - | - | 1.222E+04 | 274.1 | - | - | 0 | - |
| - | - | 8759 | 275.1 | - | - | 0 | - |
| - | - | 6839 | 275.1 | - | - | 0 | - |
| - | - | 3801 | 275.2 | - | - | 0 | - |
| - | - | 9893 | 276.1 | - | - | 0 | - |
| - | - | 1.264E+04 | 276.1 | - | - | 0 | - |
| - | - | 1.52E+04 | 276.2 | - | - | 0 | - |
| - | - | 3325 | 278.2 | - | - | 0 | - |
| - | - | 3200 | 279.1 | - | - | 0 | - |
| - | - | 1.677E+04 | 279.1 | - | - | 0 | - |
| - | - | 1.546E+04 | 280.1 | - | - | 0 | - |
| - | - | 4150 | 280.1 | - | - | 0 | - |
| - | - | 1.046E+04 | 281.1 | - | - | 0 | - |
| - | - | 6.575E+04 | 282.2 | - | - | 0 | - |
| - | - | 5.971E+05 | 283.1 | - | - | 0 | - |
| - | - | 2.592E+04 | 284.1 | - | - | 0 | - |
| - | - | 4314 | 284.1 | - | - | 0 | - |
| - | - | 8.122E+04 | 284.1 | - | - | 0 | - |
| - | - | 4134 | 284.2 | - | - | 0 | - |
| - | - | 1.072E+04 | 285.2 | - | - | 0 | - |
| - | - | 3.045E+04 | 286.1 | - | - | 0 | - |
| - | - | 3827 | 286.2 | - | - | 0 | - |
| - | - | 3535 | 287.1 | - | - | 0 | - |
| - | - | 8828 | 287.2 | - | - | 0 | - |
| - | - | 3794 | 287.2 | - | - | 0 | - |
| - | - | 1.583E+04 | 288.6 | - | - | 0 | - |
| 5 | b | 4944 | 290.2 | 0.0005572 | 1.92 | +2 | 5 |
| 12 | y | 3.129E+05 | 293.1 | 0.0005409 | 1.845 | +1 | 2 |
| - | - | 7579 | 293.2 | - | - | 0 | - |
| - | - | 6.175E+04 | 294.1 | - | - | 0 | - |
| - | - | 5575 | 294.2 | - | - | 0 | - |
| - | - | 3291 | 295.1 | - | - | 0 | - |
| - | - | 5874 | 295.1 | - | - | 0 | - |
| - | - | 5522 | 296.2 | - | - | 0 | - |
| 3 | b | 1.216E+04 | 296.2 | 0.0005806 | 1.96 | +1 | 3 |
| - | - | 4566 | 296.6 | - | - | 0 | - |
| - | - | 4097 | 296.7 | - | - | 0 | - |
| 9 | y | 1.326E+04 | 297.1 | 0.001684 | 5.667 | +2 | 5 |
| - | - | 1.058E+05 | 297.2 | - | - | 0 | - |
| 7 | y | 1.097E+04 | 298.1 | 0.005215 | 17.49 | +3 | 7 |
| - | - | 1.439E+04 | 298.2 | - | - | 0 | - |
| - | - | 8.423E+04 | 299.1 | - | - | 0 | - |
| - | - | 3838 | 300.1 | - | - | 0 | - |
| - | - | 1.109E+05 | 300.2 | - | - | 0 | - |
| - | - | 1.033E+04 | 301.1 | - | - | 0 | - |
| - | - | 2.537E+05 | 301.2 | - | - | 0 | - |
| - | - | 1.772E+04 | 301.2 | - | - | 0 | - |
| - | - | 4.747E+04 | 302.1 | - | - | 0 | - |
| - | - | 3.705E+04 | 302.2 | - | - | 0 | - |
| - | - | 9973 | 303.1 | - | - | 0 | - |
| - | - | 3978 | 303.2 | - | - | 0 | - |
| - | - | 9636 | 305.2 | - | - | 0 | - |
| - | - | 2.411E+04 | 305.6 | - | - | 0 | - |
| 9 | y | 4.221E+04 | 306.1 | 0.000857 | 2.799 | +2 | 5 |
| - | - | 4040 | 306.2 | - | - | 0 | - |
| - | - | 1.54E+04 | 306.6 | - | - | 0 | - |
| - | - | 9294 | 307.1 | - | - | 0 | - |
| - | - | 4786 | 309.2 | - | - | 0 | - |
| - | - | 3830 | 310.2 | - | - | 0 | - |
| 12 | y | 5.124E+04 | 311.1 | 0.0007489 | 2.407 | +1 | 2 |
| - | - | 8457 | 311.1 | - | - | 0 | - |
| - | - | 9714 | 312.1 | - | - | 0 | - |
| - | - | 1.028E+04 | 314.2 | - | - | 0 | - |
| 3 | b | 2.857E+04 | 314.2 | 0.0006971 | 2.218 | +1 | 3 |
| - | - | 1.744E+05 | 315.2 | - | - | 0 | - |
| - | - | 7571 | 315.2 | - | - | 0 | - |
| - | - | 2.711E+04 | 316.2 | - | - | 0 | - |
| - | - | 3032 | 317.7 | - | - | 0 | - |
| - | - | 5631 | 318.1 | - | - | 0 | - |
| - | - | 1.114E+04 | 319.2 | - | - | 0 | - |
| - | - | 1.065E+04 | 325.2 | - | - | 0 | - |
| - | - | 4.393E+04 | 326.7 | - | - | 0 | - |
| - | - | 1.406E+04 | 327.2 | - | - | 0 | - |
| - | - | 6195 | 327.7 | - | - | 0 | - |
| - | - | 1.823E+04 | 332.2 | - | - | 0 | - |
| - | - | 3.585E+04 | 333.2 | - | - | 0 | - |
| - | - | 7346 | 334.2 | - | - | 0 | - |
| - | - | 3209 | 338.2 | - | - | 0 | - |
| 6 | b | 6519 | 338.7 | 0.002012 | 5.939 | +2 | 6 |
| 6 | b | 6048 | 339.2 | 0.004633 | 13.66 | +2 | 6 |
| - | - | 4688 | 339.7 | - | - | 0 | - |
| - | - | 1.377E+04 | 340.7 | - | - | 0 | - |
| - | - | 4402 | 341.2 | - | - | 0 | - |
| - | - | 1.206E+04 | 342.2 | - | - | 0 | - |
| - | - | 1.324E+04 | 343.2 | - | - | 0 | - |
| - | - | 4197 | 343.2 | - | - | 0 | - |
| - | - | 1.417E+04 | 344.2 | - | - | 0 | - |
| - | - | 3023 | 344.7 | - | - | 0 | - |
| - | - | 1.067E+04 | 345 | - | - | 0 | - |
| - | - | 2.772E+04 | 345.1 | - | - | 0 | - |
| - | - | 9.634E+04 | 346.1 | - | - | 0 | - |
| - | - | 6277 | 346.1 | - | - | 0 | - |
| - | - | 5311 | 346.7 | - | - | 0 | - |
| - | - | 1.622E+04 | 347.1 | - | - | 0 | - |
| 6 | b | 4.511E+04 | 347.7 | 0.0006355 | 1.828 | +2 | 6 |
| - | - | 1.603E+04 | 348.2 | - | - | 0 | - |
| - | - | 4080 | 349.2 | - | - | 0 | - |
| - | - | 6.375E+04 | 350.2 | - | - | 0 | - |
| - | - | 1.428E+04 | 351.2 | - | - | 0 | - |
| - | - | 6116 | 352.2 | - | - | 0 | - |
| - | - | 8582 | 352.7 | - | - | 0 | - |
| 8 | y | 1.67E+04 | 353.7 | 0.000698 | 1.973 | +2 | 6 |
| - | - | 4787 | 354.2 | - | - | 0 | - |
| - | - | 8584 | 359 | - | - | 0 | - |
| - | - | 6354 | 359.1 | - | - | 0 | - |
| - | - | 1.974E+04 | 360.7 | - | - | 0 | - |
| - | - | 1.99E+04 | 361.2 | - | - | 0 | - |
| - | - | 7443 | 361.7 | - | - | 0 | - |
| - | - | 4.21E+04 | 362.2 | - | - | 0 | - |
| 8 | y | 9.518E+04 | 362.7 | 0.0009393 | 2.59 | +2 | 6 |
| - | - | 1.379E+05 | 363.1 | - | - | 0 | - |
| - | - | 4.802E+04 | 363.2 | - | - | 0 | - |
| - | - | 7764 | 363.7 | - | - | 0 | - |
| - | - | 2.268E+04 | 364.1 | - | - | 0 | - |
| - | - | 1.942E+04 | 364.1 | - | - | 0 | - |
| 9 | b | 3557 | 364.2 | 0.004313 | 11.84 | +3 | 9 |
| - | - | 6480 | 365.1 | - | - | 0 | - |
| - | - | 3819 | 365.2 | - | - | 0 | - |
| - | - | 1.271E+04 | 366.2 | - | - | 0 | - |
| - | - | 1.074E+04 | 367.1 | - | - | 0 | - |
| - | - | 5482 | 367.2 | - | - | 0 | - |
| - | - | 8581 | 367.2 | - | - | 0 | - |
| - | - | 4316 | 368.2 | - | - | 0 | - |
| - | - | 8777 | 368.2 | - | - | 0 | - |
| - | - | 4054 | 370.2 | - | - | 0 | - |
| - | - | 3918 | 373.2 | - | - | 0 | - |
| - | - | 1.295E+04 | 376.2 | - | - | 0 | - |
| - | - | 2.196E+04 | 376.2 | - | - | 0 | - |
| - | - | 6223 | 376.7 | - | - | 0 | - |
| - | - | 3660 | 377.1 | - | - | 0 | - |
| - | - | 1.367E+04 | 377.1 | - | - | 0 | - |
| - | - | 5.488E+04 | 378.2 | - | - | 0 | - |
| - | - | 2.151E+05 | 379.2 | - | - | 0 | - |
| - | - | 5568 | 380.2 | - | - | 0 | - |
| - | - | 4.596E+04 | 380.2 | - | - | 0 | - |
| - | - | 4.081E+05 | 381.2 | - | - | 0 | - |
| - | - | 1.728E+04 | 381.7 | - | - | 0 | - |
| - | - | 7.994E+04 | 382.2 | - | - | 0 | - |
| - | - | 1.171E+04 | 382.2 | - | - | 0 | - |
| - | - | 5688 | 382.2 | - | - | 0 | - |
| - | - | 6034 | 382.7 | - | - | 0 | - |
| - | - | 8505 | 383.2 | - | - | 0 | - |
| - | - | 4267 | 383.2 | - | - | 0 | - |
| - | - | 1.43E+04 | 384.2 | - | - | 0 | - |
| - | - | 1.969E+04 | 384.2 | - | - | 0 | - |
| - | - | 2.355E+04 | 385.2 | - | - | 0 | - |
| 5 | y | 1.33E+04 | 385.2 | 0.004315 | 11.2 | +3 | 9 |
| - | - | 5959 | 386.2 | - | - | 0 | - |
| - | - | 1.79E+04 | 390.2 | - | - | 0 | - |
| - | - | 7901 | 390.7 | - | - | 0 | - |
| - | - | 1.229E+04 | 391.1 | - | - | 0 | - |
| - | - | 5430 | 391.2 | - | - | 0 | - |
| - | - | 6036 | 392.2 | - | - | 0 | - |
| - | - | 5159 | 393.2 | - | - | 0 | - |
| - | - | 6.283E+04 | 394.2 | - | - | 0 | - |
| - | - | 2.014E+04 | 395.1 | - | - | 0 | - |
| - | - | 1.955E+04 | 395.2 | - | - | 0 | - |
| - | - | 1.829E+04 | 395.2 | - | - | 0 | - |
| - | - | 5137 | 396.2 | - | - | 0 | - |
| - | - | 6.112E+04 | 396.2 | - | - | 0 | - |
| - | - | 4066 | 397.2 | - | - | 0 | - |
| - | - | 1.32E+04 | 397.2 | - | - | 0 | - |
| - | - | 3.121E+04 | 398.2 | - | - | 0 | - |
| - | - | 6539 | 399.2 | - | - | 0 | - |
| - | - | 1.054E+04 | 401.2 | - | - | 0 | - |
| - | - | 2.788E+04 | 402.2 | - | - | 0 | - |
| - | - | 2.92E+04 | 402.2 | - | - | 0 | - |
| - | - | 1.273E+04 | 403.2 | - | - | 0 | - |
| - | - | 3696 | 404.7 | - | - | 0 | - |
| - | - | 1.508E+04 | 405.3 | - | - | 0 | - |
| - | - | 7008 | 406.2 | - | - | 0 | - |
| - | - | 4790 | 407.2 | - | - | 0 | - |
| - | - | 3447 | 408.7 | - | - | 0 | - |
| - | - | 3.787E+04 | 409.1 | - | - | 0 | - |
| - | - | 6935 | 409.2 | - | - | 0 | - |
| - | - | 7591 | 409.7 | - | - | 0 | - |
| - | - | 3992 | 410.1 | - | - | 0 | - |
| - | - | 3.825E+04 | 411.2 | - | - | 0 | - |
| - | - | 2.409E+05 | 412.2 | - | - | 0 | - |
| - | - | 1.948E+04 | 413.1 | - | - | 0 | - |
| - | - | 2.12E+04 | 413.2 | - | - | 0 | - |
| - | - | 3.298E+04 | 413.2 | - | - | 0 | - |
| - | - | 1.104E+04 | 413.3 | - | - | 0 | - |
| - | - | 5496 | 414.1 | - | - | 0 | - |
| - | - | 4090 | 414.2 | - | - | 0 | - |
| - | - | 6.48E+04 | 414.2 | - | - | 0 | - |
| - | - | 1.882E+04 | 415 | - | - | 0 | - |
| - | - | 1.723E+04 | 415.2 | - | - | 0 | - |
| - | - | 1.215E+04 | 415.2 | - | - | 0 | - |
| - | - | 5822 | 416.2 | - | - | 0 | - |
| - | - | 4.556E+04 | 417.7 | - | - | 0 | - |
| - | - | 6.779E+04 | 418.2 | - | - | 0 | - |
| - | - | 3.055E+04 | 418.7 | - | - | 0 | - |
| - | - | 1.203E+04 | 419.2 | - | - | 0 | - |
| 11 | y | 1.286E+04 | 421.2 | 0.0007622 | 1.81 | +1 | 3 |
| - | - | 3562 | 422.2 | - | - | 0 | - |
| 11 | y | 5818 | 422.2 | 0.00161 | 3.813 | +1 | 3 |
| - | - | 5973 | 422.7 | - | - | 0 | - |
| - | - | 1.179E+04 | 423.2 | - | - | 0 | - |
| - | - | 1.019E+05 | 423.3 | - | - | 0 | - |
| - | - | 5130 | 423.7 | - | - | 0 | - |
| - | - | 8304 | 424.2 | - | - | 0 | - |
| - | - | 2.411E+04 | 424.3 | - | - | 0 | - |
| - | - | 5088 | 424.7 | - | - | 0 | - |
| - | - | 6.492E+05 | 426.7 | - | - | 0 | - |
| - | - | 3.328E+05 | 427.2 | - | - | 0 | - |
| - | - | 4882 | 427.3 | - | - | 0 | - |
| - | - | 9.486E+04 | 427.7 | - | - | 0 | - |
| - | - | 2.132E+04 | 428.2 | - | - | 0 | - |
| - | - | 3694 | 428.7 | - | - | 0 | - |
| - | - | 1.952E+05 | 429.2 | - | - | 0 | - |
| - | - | 4.966E+05 | 430.2 | - | - | 0 | - |
| - | - | 1.065E+05 | 431.2 | - | - | 0 | - |
| 7 | b | 7.057E+04 | 431.7 | 0.0008072 | 1.87 | +2 | 7 |
| 7 | b | 4.105E+04 | 432.2 | 0.007213 | 16.69 | +2 | 7 |
| - | - | 1.548E+04 | 432.7 | - | - | 0 | - |
| - | - | 7245 | 432.7 | - | - | 0 | - |
| - | - | 1.101E+04 | 433.2 | - | - | 0 | - |
| 4 | b | 2.3E+04 | 433.3 | 4.27E-05 | 0.09855 | +1 | 4 |
| - | - | 1.066E+04 | 438.2 | - | - | 0 | - |
| - | - | 8228 | 438.7 | - | - | 0 | - |
| 11 | y | 1.593E+05 | 439.2 | 0.001123 | 2.556 | +1 | 3 |
| - | - | 1.218E+04 | 440.2 | - | - | 0 | - |
| - | - | 3.681E+04 | 440.2 | - | - | 0 | - |
| 7 | b | 3.331E+05 | 440.7 | 0.0008349 | 1.894 | +2 | 7 |
| - | - | 1.782E+05 | 441.2 | - | - | 0 | - |
| - | - | 5.488E+04 | 441.7 | - | - | 0 | - |
| - | - | 1.065E+04 | 442.2 | - | - | 0 | - |
| - | - | 3838 | 444.2 | - | - | 0 | - |
| 7 | y | 4588 | 446.7 | 0.002368 | 5.301 | +2 | 7 |
| - | - | 4035 | 446.7 | - | - | 0 | - |
| 7 | y | 2.536E+04 | 447.2 | 0.002511 | 5.616 | +2 | 7 |
| - | - | 4447 | 447.7 | - | - | 0 | - |
| - | - | 3.017E+04 | 448.2 | - | - | 0 | - |
| - | - | 3040 | 448.2 | - | - | 0 | - |
| - | - | 8990 | 449.2 | - | - | 0 | - |
| - | - | 4135 | 450.2 | - | - | 0 | - |
| - | - | 7062 | 450.2 | - | - | 0 | - |
| - | - | 7078 | 450.3 | - | - | 0 | - |
| 4 | b | 1.864E+05 | 451.3 | 0.00102 | 2.26 | +1 | 4 |
| - | - | 4.806E+04 | 452.3 | - | - | 0 | - |
| - | - | 6282 | 453.3 | - | - | 0 | - |
| - | - | 3721 | 455.2 | - | - | 0 | - |
| - | - | 1.129E+04 | 455.3 | - | - | 0 | - |
| 7 | y | 1.11E+04 | 455.7 | 0.00532 | 11.67 | +2 | 7 |
| - | - | 3558 | 456.2 | - | - | 0 | - |
| - | - | 3069 | 456.3 | - | - | 0 | - |
| - | - | 1.79E+04 | 459.2 | - | - | 0 | - |
| - | - | 2.478E+04 | 460.8 | - | - | 0 | - |
| - | - | 1.328E+04 | 461.2 | - | - | 0 | - |
| - | - | 2.643E+04 | 461.3 | - | - | 0 | - |
| - | - | 8341 | 461.8 | - | - | 0 | - |
| - | - | 7039 | 462.2 | - | - | 0 | - |
| - | - | 3757 | 462.3 | - | - | 0 | - |
| - | - | 4120 | 465.7 | - | - | 0 | - |
| - | - | 4494 | 466.2 | - | - | 0 | - |
| - | - | 3416 | 466.7 | - | - | 0 | - |
| - | - | 4.944E+04 | 468.3 | - | - | 0 | - |
| - | - | 1.023E+04 | 469.3 | - | - | 0 | - |
| - | - | 3299 | 470.3 | - | - | 0 | - |
| - | - | 3.939E+04 | 474.3 | - | - | 0 | - |
| - | - | 5.59E+04 | 474.8 | - | - | 0 | - |
| - | - | 2.713E+04 | 475.3 | - | - | 0 | - |
| - | - | 5765 | 475.8 | - | - | 0 | - |
| - | - | 1.367E+04 | 476.2 | - | - | 0 | - |
| - | - | 2.983E+04 | 477.2 | - | - | 0 | - |
| 10 | y | 3.124E+04 | 478.2 | 0.0003556 | 0.7437 | +1 | 4 |
| - | - | 2.028E+04 | 478.3 | - | - | 0 | - |
| - | - | 6964 | 479.2 | - | - | 0 | - |
| - | - | 5723 | 479.3 | - | - | 0 | - |
| - | - | 3628 | 479.7 | - | - | 0 | - |
| - | - | 5069 | 480.2 | - | - | 0 | - |
| - | - | 3295 | 481.2 | - | - | 0 | - |
| - | - | 1.388E+05 | 483.3 | - | - | 0 | - |
| - | - | 7.767E+04 | 483.8 | - | - | 0 | - |
| - | - | 7559 | 484.2 | - | - | 0 | - |
| - | - | 2.111E+04 | 484.3 | - | - | 0 | - |
| - | - | 8919 | 484.8 | - | - | 0 | - |
| - | - | 3472 | 486.2 | - | - | 0 | - |
| - | - | 5200 | 488.2 | - | - | 0 | - |
| 8 | b | 4.331E+04 | 488.3 | 0.001439 | 2.947 | +2 | 8 |
| 8 | b | 2.404E+04 | 488.7 | 0.007569 | 15.49 | +2 | 8 |
| - | - | 1.344E+04 | 489.3 | - | - | 0 | - |
| - | - | 3.816E+05 | 494.2 | - | - | 0 | - |
| - | - | 3552 | 495.2 | - | - | 0 | - |
| - | - | 9.986E+04 | 495.2 | - | - | 0 | - |
| 10 | y | 5.421E+05 | 496.2 | 0.0009909 | 1.997 | +1 | 4 |
| - | - | 1.16E+05 | 497.2 | - | - | 0 | - |
| - | - | 5.006E+04 | 497.8 | - | - | 0 | - |
| - | - | 1.826E+04 | 498.2 | - | - | 0 | - |
| - | - | 3229 | 499.2 | - | - | 0 | - |
| - | - | 3146 | 501.2 | - | - | 0 | - |
| - | - | 4643 | 502.2 | - | - | 0 | - |
| - | - | 1.946E+04 | 503.1 | - | - | 0 | - |
| - | - | 3554 | 503.2 | - | - | 0 | - |
| - | - | 1.327E+04 | 504.2 | - | - | 0 | - |
| - | - | 4188 | 505.2 | - | - | 0 | - |
| - | - | 1.4E+04 | 506.2 | - | - | 0 | - |
| - | - | 6972 | 506.3 | - | - | 0 | - |
| - | - | 1.276E+04 | 507.3 | - | - | 0 | - |
| - | - | 2.907E+04 | 511.3 | - | - | 0 | - |
| - | - | 5837 | 512.3 | - | - | 0 | - |
| - | - | 5690 | 513.2 | - | - | 0 | - |
| - | - | 6660 | 515.3 | - | - | 0 | - |
| - | - | 4913 | 519.1 | - | - | 0 | - |
| - | - | 4776 | 519.3 | - | - | 0 | - |
| - | - | 7350 | 522.2 | - | - | 0 | - |
| - | - | 1.24E+04 | 523.3 | - | - | 0 | - |
| - | - | 4691 | 523.6 | - | - | 0 | - |
| - | - | 1.053E+04 | 524.3 | - | - | 0 | - |
| - | - | 6.259E+04 | 525.3 | - | - | 0 | - |
| - | - | 1.147E+04 | 526.2 | - | - | 0 | - |
| - | - | 1.767E+04 | 526.3 | - | - | 0 | - |
| - | - | 3465 | 527.2 | - | - | 0 | - |
| - | - | 6167 | 528.9 | - | - | 0 | - |
| 0 | Precursor | 2.22E+04 | 529.3 | 0.002878 | 5.437 | +3 | -1 |
| 0 | Precursor | 1.746E+04 | 529.6 | 0.007087 | 13.38 | +3 | -1 |
| - | - | 1.366E+04 | 529.9 | - | - | 0 | - |
| - | - | 4999 | 530.3 | - | - | 0 | - |
| - | - | 3457 | 531.2 | - | - | 0 | - |
| - | - | 4981 | 531.8 | - | - | 0 | - |
| - | - | 1.292E+04 | 532.2 | - | - | 0 | - |
| - | - | 4403 | 533.2 | - | - | 0 | - |
| - | - | 8560 | 534.9 | - | - | 0 | - |
| 0 | Precursor | 1.412E+04 | 535.3 | 0.0005769 | 1.078 | +3 | -1 |
| - | - | 1.45E+04 | 535.6 | - | - | 0 | - |
| - | - | 5818 | 535.9 | - | - | 0 | - |
| - | - | 4654 | 536.2 | - | - | 0 | - |
| - | - | 5817 | 537.2 | - | - | 0 | - |
| - | - | 7940 | 537.7 | - | - | 0 | - |
| - | - | 3876 | 537.8 | - | - | 0 | - |
| - | - | 9566 | 538.3 | - | - | 0 | - |
| - | - | 3419 | 538.8 | - | - | 0 | - |
| - | - | 7.955E+04 | 539.2 | - | - | 0 | - |
| - | - | 2.223E+04 | 540.2 | - | - | 0 | - |
| - | - | 4843 | 541.3 | - | - | 0 | - |
| - | - | 3.829E+04 | 542.3 | - | - | 0 | - |
| - | - | 8.99E+04 | 543.3 | - | - | 0 | - |
| - | - | 3.256E+04 | 544.3 | - | - | 0 | - |
| - | - | 5952 | 545.3 | - | - | 0 | - |
| 9 | b | 5931 | 545.8 | 0.003213 | 5.887 | +2 | 9 |
| 9 | b | 1.11E+04 | 546.3 | 0.008649 | 15.83 | +2 | 9 |
| - | - | 7516 | 546.7 | - | - | 0 | - |
| - | - | 4010 | 547.3 | - | - | 0 | - |
| - | - | 3981 | 548.3 | - | - | 0 | - |
| - | - | 1.617E+04 | 549.2 | - | - | 0 | - |
| - | - | 9303 | 550.2 | - | - | 0 | - |
| - | - | 1.886E+04 | 551.3 | - | - | 0 | - |
| - | - | 8162 | 551.3 | - | - | 0 | - |
| - | - | 8685 | 552.3 | - | - | 0 | - |
| - | - | 3410 | 552.3 | - | - | 0 | - |
| - | - | 5044 | 553.3 | - | - | 0 | - |
| - | - | 5421 | 554.3 | - | - | 0 | - |
| 9 | b | 2.306E+04 | 554.8 | 0.001415 | 2.551 | +2 | 9 |
| - | - | 2.41E+04 | 555.3 | - | - | 0 | - |
| - | - | 1.817E+04 | 555.8 | - | - | 0 | - |
| - | - | 6822 | 558.2 | - | - | 0 | - |
| - | - | 3891 | 559.3 | - | - | 0 | - |
| 5 | b | 1.556E+04 | 561.3 | 0.0002787 | 0.4965 | +1 | 5 |
| 5 | b | 8305 | 562.3 | 0.009366 | 16.66 | +1 | 5 |
| - | - | 1.273E+04 | 563.3 | - | - | 0 | - |
| - | - | 3996 | 564.3 | - | - | 0 | - |
| - | - | 2.684E+04 | 565.3 | - | - | 0 | - |
| - | - | 9629 | 566.3 | - | - | 0 | - |
| - | - | 8.342E+04 | 567.2 | - | - | 0 | - |
| - | - | 6129 | 567.8 | - | - | 0 | - |
| - | - | 2.669E+04 | 568.2 | - | - | 0 | - |
| 5 | y | 4852 | 568.7 | 0.01059 | 18.63 | +2 | 9 |
| - | - | 6892 | 569.2 | - | - | 0 | - |
| - | - | 3.4E+04 | 575.2 | - | - | 0 | - |
| - | - | 5382 | 575.3 | - | - | 0 | - |
| - | - | 1.541E+04 | 576.2 | - | - | 0 | - |
| - | - | 3747 | 576.8 | - | - | 0 | - |
| 5 | y | 7983 | 577.3 | 0.004765 | 8.255 | +2 | 9 |
| 5 | b | 1.204E+05 | 579.3 | 0.0007614 | 1.314 | +1 | 5 |
| - | - | 4.109E+04 | 580.3 | - | - | 0 | - |
| - | - | 7387 | 581.3 | - | - | 0 | - |
| - | - | 7033 | 582.3 | - | - | 0 | - |
| - | - | 4816 | 586.3 | - | - | 0 | - |
| - | - | 4546 | 588.8 | - | - | 0 | - |
| - | - | 8064 | 589.3 | - | - | 0 | - |
| - | - | 2.982E+04 | 592.3 | - | - | 0 | - |
| - | - | 1.738E+04 | 592.3 | - | - | 0 | - |
| 9 | y | 1.53E+05 | 593.3 | 0.0006953 | 1.172 | +1 | 5 |
| - | - | 4.934E+04 | 593.3 | - | - | 0 | - |
| - | - | 6875 | 593.8 | - | - | 0 | - |
| - | - | 5.708E+04 | 594.3 | - | - | 0 | - |
| - | - | 1.382E+04 | 594.3 | - | - | 0 | - |
| - | - | 1.391E+04 | 595.3 | - | - | 0 | - |
| - | - | 6974 | 596.4 | - | - | 0 | - |
| - | - | 6089 | 597.8 | - | - | 0 | - |
| - | - | 5578 | 598.3 | - | - | 0 | - |
| - | - | 4389 | 599.3 | - | - | 0 | - |
| - | - | 4118 | 600.3 | - | - | 0 | - |
| - | - | 7833 | 602.3 | - | - | 0 | - |
| - | - | 2.29E+04 | 602.8 | - | - | 0 | - |
| - | - | 1.653E+04 | 603.3 | - | - | 0 | - |
| - | - | 8772 | 603.8 | - | - | 0 | - |
| - | - | 5.127E+05 | 610.3 | - | - | 0 | - |
| 9 | y | 1.391E+06 | 611.3 | 0.001971 | 3.225 | +1 | 5 |
| - | - | 7.137E+04 | 611.8 | - | - | 0 | - |
| - | - | 4.436E+05 | 612.3 | - | - | 0 | - |
| - | - | 1.464E+04 | 612.8 | - | - | 0 | - |
| - | - | 1.079E+05 | 613.3 | - | - | 0 | - |
| - | - | 1.681E+04 | 614.3 | - | - | 0 | - |
| - | - | 4860 | 615.3 | - | - | 0 | - |
| - | - | 4864 | 616.3 | - | - | 0 | - |
| - | - | 1.009E+04 | 617.3 | - | - | 0 | - |
| - | - | 3765 | 619.3 | - | - | 0 | - |
| - | - | 4715 | 619.3 | - | - | 0 | - |
| - | - | 9441 | 620.8 | - | - | 0 | - |
| - | - | 2.507E+04 | 621.3 | - | - | 0 | - |
| - | - | 1.291E+04 | 621.3 | - | - | 0 | - |
| - | - | 8050 | 622.3 | - | - | 0 | - |
| - | - | 4539 | 622.4 | - | - | 0 | - |
| - | - | 3777 | 623.4 | - | - | 0 | - |
| - | - | 4536 | 626.3 | - | - | 0 | - |
| - | - | 4007 | 627.3 | - | - | 0 | - |
| - | - | 4166 | 627.8 | - | - | 0 | - |
| - | - | 8664 | 628.3 | - | - | 0 | - |
| - | - | 5708 | 628.8 | - | - | 0 | - |
| - | - | 5370 | 630.3 | - | - | 0 | - |
| - | - | 2.147E+04 | 634.3 | - | - | 0 | - |
| - | - | 2.762E+04 | 635.3 | - | - | 0 | - |
| - | - | 1.796E+04 | 636.3 | - | - | 0 | - |
| 4 | y | 1.638E+04 | 636.8 | 0.003846 | 6.04 | +2 | 10 |
| 4 | y | 1.295E+04 | 637.3 | 0.009763 | 15.32 | +2 | 10 |
| - | - | 5423 | 637.3 | - | - | 0 | - |
| - | - | 5557 | 637.8 | - | - | 0 | - |
| 11 | b | 3987 | 638.3 | 0.009496 | 14.88 | +2 | 11 |
| - | - | 5889 | 638.8 | - | - | 0 | - |
| - | - | 1.206E+04 | 640.3 | - | - | 0 | - |
| - | - | 3708 | 641.3 | - | - | 0 | - |
| - | - | 4575 | 644.3 | - | - | 0 | - |
| - | - | 3.038E+04 | 645.3 | - | - | 0 | - |
| 4 | y | 8.522E+04 | 645.8 | 0.003569 | 5.526 | +2 | 10 |
| - | - | 6.483E+04 | 646.3 | - | - | 0 | - |
| - | - | 2.421E+04 | 646.8 | - | - | 0 | - |
| - | - | 7785 | 647.3 | - | - | 0 | - |
| - | - | 8923 | 648.3 | - | - | 0 | - |
| - | - | 7029 | 650.3 | - | - | 0 | - |
| - | - | 5064 | 650.3 | - | - | 0 | - |
| - | - | 4740 | 651.8 | - | - | 0 | - |
| - | - | 1.638E+05 | 652.3 | - | - | 0 | - |
| - | - | 5.895E+04 | 653.3 | - | - | 0 | - |
| - | - | 1.337E+04 | 654.3 | - | - | 0 | - |
| - | - | 9407 | 658.3 | - | - | 0 | - |
| - | - | 4669 | 659.3 | - | - | 0 | - |
| - | - | 3951 | 660.8 | - | - | 0 | - |
| - | - | 7101 | 661.3 | - | - | 0 | - |
| - | - | 5378 | 661.8 | - | - | 0 | - |
| - | - | 5.107E+04 | 662.3 | - | - | 0 | - |
| - | - | 3.35E+04 | 663.3 | - | - | 0 | - |
| - | - | 5809 | 664.3 | - | - | 0 | - |
| - | - | 8547 | 664.3 | - | - | 0 | - |
| - | - | 5258 | 665.3 | - | - | 0 | - |
| - | - | 3732 | 666.4 | - | - | 0 | - |
| - | - | 4013 | 668.3 | - | - | 0 | - |
| 6 | b | 7.147E+04 | 676.3 | 0.0005574 | 0.8241 | +1 | 6 |
| 6 | b | 5.216E+04 | 677.3 | 0.01111 | 16.4 | +1 | 6 |
| - | - | 1.877E+04 | 678.3 | - | - | 0 | - |
| - | - | 1.553E+04 | 679.3 | - | - | 0 | - |
| - | - | 3824 | 679.8 | - | - | 0 | - |
| - | - | 2.874E+05 | 680.3 | - | - | 0 | - |
| - | - | 1.157E+05 | 681.3 | - | - | 0 | - |
| - | - | 2.757E+04 | 682.3 | - | - | 0 | - |
| - | - | 1.568E+04 | 684.3 | - | - | 0 | - |
| - | - | 1.879E+04 | 684.8 | - | - | 0 | - |
| - | - | 1.22E+04 | 685.3 | - | - | 0 | - |
| - | - | 5670 | 685.8 | - | - | 0 | - |
| - | - | 7058 | 688.3 | - | - | 0 | - |
| - | - | 4195 | 689.3 | - | - | 0 | - |
| - | - | 4552 | 690.3 | - | - | 0 | - |
| - | - | 1.725E+04 | 692.8 | - | - | 0 | - |
| 3 | y | 1.036E+05 | 693.3 | 0.002402 | 3.465 | +2 | 11 |
| 3 | y | 8.876E+04 | 693.8 | 0.01021 | 14.72 | +2 | 11 |
| 6 | b | 8.047E+05 | 694.4 | 0.0003076 | 0.443 | +1 | 6 |
| - | - | 1.734E+04 | 694.8 | - | - | 0 | - |
| - | - | 3.15E+05 | 695.4 | - | - | 0 | - |
| - | - | 7.503E+04 | 696.4 | - | - | 0 | - |
| - | - | 4538 | 697.3 | - | - | 0 | - |
| - | - | 1.72E+04 | 697.4 | - | - | 0 | - |
| - | - | 4246 | 697.9 | - | - | 0 | - |
| - | - | 4817 | 698.4 | - | - | 0 | - |
| - | - | 8733 | 700.8 | - | - | 0 | - |
| - | - | 1.706E+04 | 701.3 | - | - | 0 | - |
| - | - | 2.166E+05 | 701.8 | - | - | 0 | - |
| 3 | y | 5.314E+05 | 702.3 | 0.003651 | 5.198 | +2 | 11 |
| 12 | b | 4.072E+05 | 702.8 | 0.01131 | 16.09 | +2 | 12 |
| 12 | b | 1.877E+05 | 703.3 | 0.00277 | 3.939 | +2 | 12 |
| - | - | 4.985E+04 | 703.8 | - | - | 0 | - |
| - | - | 2.059E+04 | 704.3 | - | - | 0 | - |
| - | - | 1.014E+04 | 705.4 | - | - | 0 | - |
| 8 | y | 4.941E+04 | 706.3 | 0.002447 | 3.464 | +1 | 6 |
| - | - | 2.038E+04 | 707.3 | - | - | 0 | - |
| - | - | 3021 | 708.3 | - | - | 0 | - |
| - | - | 3.875E+04 | 711.4 | - | - | 0 | - |
| 12 | b | 4.254E+04 | 711.9 | 0.004707 | 6.612 | +2 | 12 |
| - | - | 3.611E+04 | 712.4 | - | - | 0 | - |
| - | - | 6161 | 712.9 | - | - | 0 | - |
| - | - | 4753 | 718.4 | - | - | 0 | - |
| - | - | 6031 | 720.4 | - | - | 0 | - |
| - | - | 3.672E+04 | 721.4 | - | - | 0 | - |
| - | - | 1.378E+04 | 722.4 | - | - | 0 | - |
| - | - | 1.532E+05 | 723.4 | - | - | 0 | - |
| 8 | y | 4.167E+05 | 724.4 | 0.002685 | 3.707 | +1 | 6 |
| - | - | 1.584E+05 | 725.4 | - | - | 0 | - |
| - | - | 4.869E+04 | 726.4 | - | - | 0 | - |
| - | - | 6679 | 727.4 | - | - | 0 | - |
| - | - | 8225 | 728.4 | - | - | 0 | - |
| - | - | 4769 | 728.9 | - | - | 0 | - |
| - | - | 1.478E+04 | 729.4 | - | - | 0 | - |
| - | - | 7757 | 730.4 | - | - | 0 | - |
| - | - | 2.386E+04 | 733.4 | - | - | 0 | - |
| - | - | 2.214E+04 | 734.4 | - | - | 0 | - |
| - | - | 5811 | 735.3 | - | - | 0 | - |
| - | - | 7715 | 737.9 | - | - | 0 | - |
| - | - | 5268 | 738.4 | - | - | 0 | - |
| - | - | 4720 | 739.4 | - | - | 0 | - |
| 2 | y | 7443 | 742.9 | 0.002497 | 3.362 | +2 | 12 |
| 2 | y | 1.49E+04 | 743.4 | 0.0122 | 16.41 | +2 | 12 |
| - | - | 4750 | 744.4 | - | - | 0 | - |
| - | - | 1.127E+04 | 747.4 | - | - | 0 | - |
| - | - | 9483 | 748.4 | - | - | 0 | - |
| - | - | 9130 | 749.3 | - | - | 0 | - |
| - | - | 7005 | 750.4 | - | - | 0 | - |
| - | - | 1.012E+05 | 751.4 | - | - | 0 | - |
| 2 | y | 4.286E+04 | 751.9 | 0.003257 | 4.332 | +2 | 12 |
| - | - | 6.598E+04 | 752.4 | - | - | 0 | - |
| - | - | 1.498E+04 | 752.9 | - | - | 0 | - |
| - | - | 2.012E+04 | 753.4 | - | - | 0 | - |
| - | - | 4597 | 760.3 | - | - | 0 | - |
| - | - | 6003 | 761.4 | - | - | 0 | - |
| - | - | 5237 | 762.4 | - | - | 0 | - |
| - | - | 1.037E+04 | 762.4 | - | - | 0 | - |
| - | - | 5.418E+04 | 763.4 | - | - | 0 | - |
| - | - | 2.475E+04 | 764.4 | - | - | 0 | - |
| - | - | 2.01E+04 | 765.4 | - | - | 0 | - |
| - | - | 8020 | 766.4 | - | - | 0 | - |
| - | - | 1.462E+04 | 767.3 | - | - | 0 | - |
| - | - | 5768 | 768.3 | - | - | 0 | - |
| - | - | 1.291E+04 | 769.4 | - | - | 0 | - |
| - | - | 8463 | 775.4 | - | - | 0 | - |
| - | - | 8314 | 776.4 | - | - | 0 | - |
| - | - | 4.759E+04 | 779.4 | - | - | 0 | - |
| - | - | 2.54E+04 | 780.4 | - | - | 0 | - |
| - | - | 2.697E+04 | 781.4 | - | - | 0 | - |
| - | - | 1.127E+04 | 782.4 | - | - | 0 | - |
| - | - | 4346 | 791.4 | - | - | 0 | - |
| 0 | Precursor | 9.083E+04 | 793.4 | 0.014 | 17.64 | +2 | -1 |
| - | - | 3.642E+04 | 794.4 | - | - | 0 | - |
| - | - | 1.337E+04 | 795.3 | - | - | 0 | - |
| - | - | 7832 | 795.4 | - | - | 0 | - |
| - | - | 8776 | 796.3 | - | - | 0 | - |
| - | - | 1.883E+04 | 807.4 | - | - | 0 | - |
| - | - | 1.071E+04 | 807.4 | - | - | 0 | - |
| - | - | 1.269E+04 | 808.4 | - | - | 0 | - |
| - | - | 4619 | 809.4 | - | - | 0 | - |
| - | - | 3629 | 817.4 | - | - | 0 | - |
| - | - | 1.854E+04 | 818.4 | - | - | 0 | - |
| - | - | 8384 | 819.4 | - | - | 0 | - |
| - | - | 3889 | 820.4 | - | - | 0 | - |
| - | - | 1.003E+04 | 825.4 | - | - | 0 | - |
| - | - | 1.125E+04 | 826.4 | - | - | 0 | - |
| - | - | 4601 | 827.4 | - | - | 0 | - |
| - | - | 1.495E+04 | 834.4 | - | - | 0 | - |
| - | - | 3.002E+04 | 835.4 | - | - | 0 | - |
| - | - | 5.978E+04 | 836.4 | - | - | 0 | - |
| - | - | 3.538E+04 | 837.5 | - | - | 0 | - |
| - | - | 1.15E+04 | 838.5 | - | - | 0 | - |
| - | - | 1.285E+04 | 843.4 | - | - | 0 | - |
| - | - | 1.785E+04 | 844.4 | - | - | 0 | - |
| - | - | 1.251E+04 | 845.4 | - | - | 0 | - |
| - | - | 2.348E+04 | 846.4 | - | - | 0 | - |
| - | - | 1.973E+04 | 847.4 | - | - | 0 | - |
| - | - | 7308 | 848.4 | - | - | 0 | - |
| - | - | 5729 | 851.4 | - | - | 0 | - |
| - | - | 2.114E+05 | 852.4 | - | - | 0 | - |
| - | - | 5398 | 853.3 | - | - | 0 | - |
| - | - | 1.127E+05 | 853.4 | - | - | 0 | - |
| - | - | 3.157E+04 | 854.4 | - | - | 0 | - |
| 7 | b | 1.328E+05 | 862.4 | 0.0006511 | 0.755 | +1 | 7 |
| 7 | b | 9.411E+04 | 863.4 | 0.01267 | 14.67 | +1 | 7 |
| - | - | 1.706E+05 | 864.5 | - | - | 0 | - |
| - | - | 8.708E+04 | 865.5 | - | - | 0 | - |
| - | - | 2.049E+04 | 866.5 | - | - | 0 | - |
| - | - | 4794 | 868.4 | - | - | 0 | - |
| - | - | 1.715E+04 | 876.4 | - | - | 0 | - |
| - | - | 8013 | 877.4 | - | - | 0 | - |
| 7 | b | 9.897E+05 | 880.4 | 0.0004014 | 0.4559 | +1 | 7 |
| - | - | 5.477E+05 | 881.4 | - | - | 0 | - |
| - | - | 1.623E+05 | 882.4 | - | - | 0 | - |
| - | - | 3.188E+04 | 883.4 | - | - | 0 | - |
| - | - | 7151 | 890.4 | - | - | 0 | - |
| - | - | 1.074E+04 | 891.4 | - | - | 0 | - |
| - | - | 1.487E+04 | 892.5 | - | - | 0 | - |
| - | - | 3976 | 893.5 | - | - | 0 | - |
| - | - | 4802 | 894.5 | - | - | 0 | - |
| - | - | 2.036E+04 | 907.4 | - | - | 0 | - |
| - | - | 5.798E+04 | 908.4 | - | - | 0 | - |
| - | - | 4.866E+04 | 909.4 | - | - | 0 | - |
| 7 | y | 7.01E+04 | 910.4 | 0.002108 | 2.315 | +1 | 7 |
| - | - | 3.279E+04 | 911.4 | - | - | 0 | - |
| - | - | 9109 | 912.4 | - | - | 0 | - |
| - | - | 9911 | 936.4 | - | - | 0 | - |
| - | - | 8710 | 937.4 | - | - | 0 | - |
| - | - | 3772 | 938.4 | - | - | 0 | - |
| - | - | 3989 | 946.5 | - | - | 0 | - |
| - | - | 4404 | 947.4 | - | - | 0 | - |
| - | - | 3638 | 953.4 | - | - | 0 | - |
| - | - | 1.603E+04 | 954.4 | - | - | 0 | - |
| - | - | 2.08E+04 | 955.4 | - | - | 0 | - |
| - | - | 8205 | 956.4 | - | - | 0 | - |
| - | - | 4350 | 962.5 | - | - | 0 | - |
| - | - | 9516 | 964.5 | - | - | 0 | - |
| - | - | 2.077E+04 | 965.5 | - | - | 0 | - |
| - | - | 1.104E+04 | 966.5 | - | - | 0 | - |
| - | - | 7788 | 971.5 | - | - | 0 | - |
| - | - | 1.369E+04 | 972.4 | - | - | 0 | - |
| - | - | 4362 | 973.4 | - | - | 0 | - |
| 8 | b | 1.352E+04 | 975.5 | 0.0002219 | 0.2275 | +1 | 8 |
| 8 | b | 1.037E+04 | 976.5 | 0.01076 | 11.02 | +1 | 8 |
| - | - | 4717 | 979.5 | - | - | 0 | - |
| - | - | 7981 | 980.5 | - | - | 0 | - |
| - | - | 6200 | 981.5 | - | - | 0 | - |
| - | - | 5943 | 982.5 | - | - | 0 | - |
| - | - | 5777 | 983.5 | - | - | 0 | - |
| 8 | b | 7.231E+04 | 993.5 | 0.0001997 | 0.201 | +1 | 8 |
| - | - | 3.799E+04 | 994.5 | - | - | 0 | - |
| - | - | 1.262E+04 | 995.5 | - | - | 0 | - |
| 6 | y | 8143 | 1007 | 0.002636 | 2.617 | +1 | 8 |
| - | - | 3629 | 1009 | - | - | 0 | - |
| - | - | 8467 | 1009 | - | - | 0 | - |
| - | - | 1.605E+04 | 1024 | - | - | 0 | - |
| 6 | y | 3.483E+04 | 1025 | 0.004217 | 4.113 | +1 | 8 |
| - | - | 1.747E+04 | 1026 | - | - | 0 | - |
| - | - | 3930 | 1027 | - | - | 0 | - |
| - | - | 9272 | 1076 | - | - | 0 | - |
| - | - | 8704 | 1077 | - | - | 0 | - |
| 9 | b | 5697 | 1092 | 0.01588 | 14.55 | +1 | 9 |
| - | - | 8430 | 1093 | - | - | 0 | - |
| - | - | 2.204E+04 | 1094 | - | - | 0 | - |
| - | - | 1.302E+04 | 1095 | - | - | 0 | - |
| - | - | 5049 | 1096 | - | - | 0 | - |
| - | - | 3636 | 1108 | - | - | 0 | - |
| 9 | b | 8043 | 1109 | 0.006907 | 6.231 | +1 | 9 |
| - | - | 7723 | 1110 | - | - | 0 | - |
| - | - | 4329 | 1117 | - | - | 0 | - |
| - | - | 7454 | 1118 | - | - | 0 | - |
| - | - | 3793 | 1119 | - | - | 0 | - |
| - | - | 6377 | 1135 | - | - | 0 | - |
| 5 | y | 3.284E+04 | 1136 | 0.001859 | 1.637 | +1 | 9 |
| 5 | y | 3.688E+04 | 1136 | 0.01003 | 8.826 | +1 | 9 |
| - | - | 1.787E+04 | 1137 | - | - | 0 | - |
| - | - | 5362 | 1138 | - | - | 0 | - |
| - | - | 2.605E+04 | 1153 | - | - | 0 | - |
| 5 | y | 6.598E+04 | 1154 | 0.005943 | 5.152 | +1 | 9 |
| - | - | 3.967E+04 | 1155 | - | - | 0 | - |
| - | - | 1.22E+04 | 1156 | - | - | 0 | - |
| - | - | 3858 | 1165 | - | - | 0 | - |
| 10 | b | 6807 | 1166 | 0.006053 | 5.193 | +1 | 10 |
| - | - | 4057 | 1206 | - | - | 0 | - |
| - | - | 4236 | 1207 | - | - | 0 | - |
| 4 | y | 4397 | 1291 | 0.00939 | 7.276 | +1 | 10 |
| - | - | 3248 | 1434 | - | - | 0 | - |
| - | - | 4157 | 3083 | - | - | 0 | - |
| - | - | 4604 | 3084 | - | - | 0 | - |
| - | - | 3626 | 3460 | - | - | 0 | - |

m/z Charge Intensity FragmentType MassShift Position
120.08129119873047 0 7130.725
122.071533203125 0 8729.244
123.04448699951172 0 10224.336
123.05551147460938 0 2693.2634
124.8956527709961 0 2086.4507
125.10771942138672 0 3801.787
126.09173583984375 0 7023.65
127.05057525634766 0 34307.74
127.08688354492188 0 3663.2551
127.123779296875 0 3888.8252
128.1073760986328 0 129605.305
129.06640625 0 9255.279
129.10263061523438 0 1453667.8
130.06549072265625 0 679489.75
130.09971618652344 0 10926.62
130.10592651367188 0 88968.09
131.06881713867188 0 69078.086
131.10690307617188 0 3458.6174
131.1182861328125 0 3202.2317
132.0811004638672 0 149538.92
133.06097412109375 0 4335.2373
133.08444213867188 0 16015.144
133.92430114746094 0 2093.8484
135.95419311523438 0 1785.2856
136.07606506347656 0 175484.36
137.0795135498047 0 13944.563
138.0665283203125 0 17201.973
138.09173583984375 0 9131.066
138.4046173095703 0 2036.6486
139.08692932128906 0 16778.715
141.10263061523438 0 26037.63
142.06549072265625 0 11567.292
142.12301635742188 0 2133.2146
143.04544067382812 0 7147.9404
144.07615661621094 0 3876.491
144.0814208984375 0 7655.8213
145.06121826171875 0 15699.85
146.0926055908203 0 2884.6213
147.0444793701172 0 10585.975
149.04518127441406 0 14761.888
150.0664825439453 0 10978.413
151.0869903564453 0 37712.574
152.10733032226562 0 4187.1396
153.10287475585938 0 3804.2905
154.0615234375 0 6181.26
154.09828186035156 0 2222.7295
155.04559326171875 0 4419.08
155.08224487304688 0 5388.694
155.09307861328125 0 95580.7
155.1182403564453 0 70517.43 a Water loss 1
156.07701110839844 0 5017.3745
156.09640502929688 0 5641.993
156.12156677246094 0 5918.917
157.09754943847656 0 8595.283
158.0843048095703 0 10976.055
159.0920867919922 0 1156464.9
160.07598876953125 0 8256.515
160.08892822265625 0 9588.308
160.0954132080078 0 125747.766
161.09860229492188 0 4395.7095
165.05499267578125 0 92291.05
165.0771942138672 0 5575.651
166.06143188476562 0 178205.4
166.09786987304688 0 15836.473
167.05621337890625 0 9710.914
167.06495666503906 0 11336.43
167.1184844970703 0 3420.7986
168.08111572265625 0 18032.156
168.10215759277344 0 11826.029
168.11354064941406 0 17171.76
169.0764617919922 0 18225.906
169.08401489257812 0 4123.9497
169.0975341796875 0 41124.414
170.06040954589844 0 53179.35
170.10079956054688 0 3526.5928
171.06370544433594 0 6346.046
171.09202575683594 0 4884.884
171.14955139160156 0 24880.23
172.0720977783203 0 29883.52
172.1458282470703 0 2352.3674
172.15333557128906 0 2633.6602
173.05621337890625 0 19210.68
173.12889099121094 0 2192835.2 a 1
174.13221740722656 0 190598.9
175.1342315673828 0 12048.853
176.0823516845703 0 4174.891
177.10281372070312 0 5051.0117
178.13424682617188 0 67855.72
179.1377410888672 0 7648.2954
181.06106567382812 0 14783.033
182.0815887451172 0 205740.69 y 12
183.08522033691406 0 19814.887
183.11317443847656 0 60320.527 b Water loss 1
183.12184143066406 0 4778.9097
184.1164093017578 0 4958.467
185.165283203125 0 15233.128
186.1240997314453 0 455716.28
187.08694458007812 0 96480.08
187.1075439453125 0 2543.7822
187.1207733154297 0 3387.4727
187.1275634765625 0 35837.664
188.07070922851562 0 6646.181
188.09024047851562 0 11584.724
189.09835815429688 0 5654.0356
190.08265686035156 0 19062.074
190.13412475585938 0 13317.253
194.09286499023438 0 16235.896
195.1131591796875 0 110561.1
195.1233367919922 0 6292.8696
196.1083221435547 0 86550.414
196.1166229248047 0 9783.166
197.1108856201172 0 4045.1147
197.1287384033203 0 10047.375
197.16490173339844 0 3602.4734
197.39039611816406 0 2646.0144
198.0876922607422 0 14996.17
198.11244201660156 0 3046.2268
198.1281280517578 0 47197.8
199.07142639160156 0 6464.475
199.0868377685547 0 11926.271
199.1317138671875 0 4337.816
199.14463806152344 0 13104.026
200.14288330078125 0 5389.3145
200.2943878173828 0 3160.8142
201.12371826171875 0 597981.1 b 1
202.08668518066406 0 4612.6733
202.12709045410156 0 52966.785
203.12799072265625 0 4909.8774
204.1129913330078 0 3662.1245
205.09751892089844 0 55181.035
205.10736083984375 0 1891.3009
205.14501953125 0 7952.631
206.10145568847656 0 2913.6255
206.12901306152344 0 10825.792
207.16065979003906 0 17233.424
208.10829162597656 0 3872.6936
209.05572509765625 0 4315.418
210.12718200683594 0 18230.158
211.08724975585938 0 2764.7039
211.10806274414062 0 17438.32 y Water loss 10
211.1304473876953 0 6104.389
212.13986206054688 0 13919.865
213.12376403808594 0 58421.867
213.1359100341797 0 4777.8936
213.1602783203125 0 7455.836
214.12770080566406 0 5113.893
215.1395263671875 0 5209.4136
216.0983123779297 0 14359.04
221.07142639160156 0 5782.7886
221.0924072265625 0 8086.3086
221.1037139892578 0 10043.822
221.1400146484375 0 7080.305
222.1241912841797 0 19612.25
223.10804748535156 0 29583.719
223.15576171875 0 206312.67
224.1034393310547 0 11489.607
224.15933227539062 0 17506.342
225.0438232421875 0 5790.744
225.09983825683594 0 2393.4236
225.1230010986328 0 3438.3794
225.1603240966797 0 4892.456
226.08261108398438 0 38709.105
226.09437561035156 0 3753.4822
226.1373748779297 0 3107.1152 b 3
227.06663513183594 0 36839.32
227.08583068847656 0 4965.114
227.154541015625 0 12509.578
228.06996154785156 0 2919.4307
228.1346893310547 0 25825.352
229.11874389648438 0 28735.623
230.12315368652344 0 3446.7722
230.15054321289062 0 4853.199
233.14007568359375 0 34053.082
234.12420654296875 0 221769.92
234.6510467529297 0 2987.7573
235.1276397705078 0 34617.91
235.1553497314453 0 5387.8174
235.60568237304688 0 5066.6914
237.1353759765625 0 10202.92
238.11904907226562 0 34416.56
238.1300048828125 0 4776.0366
239.0824432373047 0 5062.9067
239.09593200683594 0 11266.438
240.0967559814453 0 5310.6333
240.13478088378906 0 96073.09
241.09613037109375 0 4881.674
241.11874389648438 0 11850.189
241.1379852294922 0 11940.088
241.19142150878906 0 13329.975
243.1095428466797 0 18376.81
244.09326171875 0 89906.516
245.096923828125 0 11053.905
245.11224365234375 0 2703.3638
247.1082305908203 0 128700.58
247.14418029785156 0 6765.8096
247.1562957763672 0 3850.4563
248.10308837890625 0 22982.344
248.11424255371094 0 40379.043
248.1468048095703 0 2862.51
248.62347412109375 0 4319.059 y 9
249.09864807128906 0 60848.117
249.11305236816406 0 3871.0474
249.119140625 0 5595.2217
249.13514709472656 0 24315.174
249.6030731201172 0 3109.604
250.1018524169922 0 11116.098
250.11900329589844 0 8363.909
250.16600036621094 0 3030.2178
251.1508026123047 0 761779.25
252.13372802734375 0 6421.4614
252.15396118164062 0 93294.94
253.1558074951172 0 7625.723
254.07704162597656 0 2757.9365
255.14805603027344 0 43945.53
256.10845947265625 0 12231.743
256.15203857421875 0 5884.2915
256.6109924316406 0 6892.947
257.1132507324219 0 23138.742
258.1246032714844 0 2609.9602
258.12811279296875 0 3078.843
258.1451110839844 0 33597.24
259.1293640136719 0 5267.4585
259.1486511230469 0 3667.971
261.11907958984375 0 16448.514
261.13507080078125 0 7146.377
262.1197204589844 0 10660.039
265.1297912597656 0 110701.35
266.11376953125 0 28183.707
266.12530517578125 0 207880.11
267.12860107421875 0 22316.354
267.61285400390625 0 4695.9424
268.13006591796875 0 8553.67
268.1769714355469 0 24584.863
269.1611328125 0 8523.395
269.18560791015625 0 11536.026
270.1221008300781 0 32304.678
270.1455383300781 0 5740.14
270.15966796875 0 3149.9092
270.6227722167969 0 7026.3906
271.1249084472656 0 3535.5745
272.1762390136719 0 47757.156
273.1789245605469 0 8526.514
274.1187438964844 0 43634.3
274.13153076171875 0 12220.627
275.1031188964844 0 8758.848
275.1215515136719 0 6839.1665
275.1506652832031 0 3800.875
276.1089782714844 0 9892.5205
276.12701416015625 0 12635.426
276.15582275390625 0 15195.02
278.1602783203125 0 3325.1328
279.1298522949219 0 3199.8167
279.14556884765625 0 16768.102
280.12933349609375 0 15463.397
280.1448974609375 0 4150.429
281.05169677734375 0 10457.754
282.15667724609375 0 65751.04
283.14056396484375 0 597148.75
284.1036071777344 0 25917.441
284.1236267089844 0 4314.1206
284.1437072753906 0 81216.95
284.16204833984375 0 4133.708
285.15618896484375 0 10719.618
286.1402587890625 0 30453.143
286.1589660644531 0 3826.6985
287.14178466796875 0 3534.7349
287.1717529296875 0 8828.477
287.1875 0 3793.656
288.61944580078125 0 15833.336
290.1666564941406 0 4943.9546 b 4
293.1137390136719 0 312947.9 y Water loss 11
293.16259765625 0 7579.2695
294.1180114746094 0 61750.77
294.1812438964844 0 5575.3286
295.1029052734375 0 3291.0842
295.1200866699219 0 5874.483
296.17645263671875 0 5521.787
296.19744873046875 0 12164.746 b Water loss 2
296.6424865722656 0 4565.9263
296.6664733886719 0 4096.949
297.13360595703125 0 13263.542 y Water loss 8
297.1563720703125 0 105821.78
298.1396179199219 0 10969.534 y Water loss 6
298.1589660644531 0 14394.81
299.062255859375 0 84226.555
300.13134765625 0 3837.751
300.1677551269531 0 110871.24
301.1144104003906 0 10333.614
301.1512145996094 0 253653.98
301.1697692871094 0 17719.762
302.11419677734375 0 47466.223
302.15472412109375 0 37054.15
303.11712646484375 0 9973.158
303.1671447753906 0 3978.4238
305.17999267578125 0 9635.668
305.6456298828125 0 24105.15
306.1380615234375 0 42207.17 y 8
306.2299499511719 0 4039.6138
306.6399230957031 0 15402.481
307.1407470703125 0 9293.692
309.1669006347656 0 4786.47
310.1522521972656 0 3830.3777
311.12451171875 0 51243.906 y 11
311.1398620605469 0 8456.76
312.1277770996094 0 9714.103
314.1866760253906 0 10282.407
314.2081298828125 0 28570.85 b 2
315.1669616699219 0 174438.33
315.2112121582031 0 7571.408
316.1702880859375 0 27114.188
317.6589050292969 0 3032.133
318.12152099609375 0 5631.088
319.1513671875 0 11136.223
325.1507873535156 0 10648.628
326.6645812988281 0 43929.082
327.1666564941406 0 14062.474
327.6671142578125 0 6195.1157
332.2087097167969 0 18234.578
333.1772766113281 0 35847.293
334.18377685546875 0 7346.146
338.1830139160156 0 3208.8892
338.6763000488281 0 6518.597 b Water loss 5
339.1709289550781 0 6048.293 b Ammonia loss 5
339.6762390136719 0 4687.7793
340.66192626953125 0 13768.102
341.1643371582031 0 4401.7866
342.1819763183594 0 12055.05
343.16143798828125 0 13242.869
343.18402099609375 0 4196.636
344.1717834472656 0 14172.156
344.6711730957031 0 3023.0767
344.9771423339844 0 10674.517
345.1312255859375 0 27717.637
346.1152038574219 0 96339.68
346.1375732421875 0 6276.766
346.6988830566406 0 5311.27
347.1178283691406 0 16223.18
347.6802062988281 0 45107.32 b 5
348.18145751953125 0 16028.472
349.15069580078125 0 4080.0715
350.2192077636719 0 63747.582
351.2206726074219 0 14279.914
352.16461181640625 0 6116.0767
352.6732177734375 0 8582.391
353.6746520996094 0 16697.53 y Water loss 7
354.1757507324219 0 4786.746
359.02838134765625 0 8583.916
359.134521484375 0 6354.407
360.6956787109375 0 19737.977
361.1982116699219 0 19903.896
361.6852722167969 0 7442.991
362.1860046386719 0 42100.492
362.68017578125 0 95176.24 y 7
363.1416320800781 0 137883.75
363.1820373535156 0 48024.426
363.6834411621094 0 7764.2344
364.1249694824219 0 22683.94
364.1454772949219 0 19422.045
364.1863708496094 0 3556.791 b Water loss 8
365.1276550292969 0 6479.805
365.1500244140625 0 3818.8152
366.177978515625 0 12708.16
367.1399841308594 0 10743.69
367.16131591796875 0 5481.8315
367.245361328125 0 8581.403
368.1670227050781 0 4316.0127
368.1934509277344 0 8776.735
370.17401123046875 0 4054.0325
373.1854248046875 0 3917.7244
376.1619873046875 0 12951.343
376.19775390625 0 21957.768
376.697509765625 0 6222.677
377.1221618652344 0 3659.7534
377.1456298828125 0 13671.369
378.2142028808594 0 54882.445
379.2096862792969 0 215069.2
380.1662292480469 0 5568.3447
380.21240234375 0 45960.676
381.15234375 0 408107.28
381.7120361328125 0 17280.166
382.1551208496094 0 79935.625
382.1885681152344 0 11707.199
382.21429443359375 0 5687.892
382.68865966796875 0 6033.877
383.1581726074219 0 8504.769
383.20758056640625 0 4267.412
384.1655578613281 0 14301.048
384.1893615722656 0 19692.213
385.1506042480469 0 23546.941
385.17254638671875 0 13302.648 y 4
386.1795654296875 0 5958.934
390.1959533691406 0 17900.824
390.6957702636719 0 7901.0684
391.1365966796875 0 12292.677
391.1859130859375 0 5430.098
392.2074279785156 0 6035.983
393.18988037109375 0 5158.53
394.17279052734375 0 62825.72
395.13494873046875 0 20141.814
395.157470703125 0 19554.826
395.2409973144531 0 18289.766
396.1919250488281 0 5136.8936
396.2249450683594 0 61118.84
397.2055358886719 0 4066.0056
397.228515625 0 13200.185
398.177978515625 0 31205.705
399.1823425292969 0 6539.373
401.2152404785156 0 10535.233
402.17694091796875 0 27881.902
402.1998291015625 0 29204.836
403.2010803222656 0 12734.761
404.7124938964844 0 3695.5193
405.26141357421875 0 15084.68
406.2448425292969 0 7008.1807
407.24365234375 0 4789.8657
408.71148681640625 0 3446.8313
409.1474914550781 0 37869.49
409.20416259765625 0 6934.5273
409.7007751464844 0 7590.9414
410.1477355957031 0 3991.9375
411.19940185546875 0 38245.547
412.1835021972656 0 240889.3
413.1450500488281 0 19481.955
413.1644287109375 0 21199.352
413.18743896484375 0 32976.473
413.25103759765625 0 11038.926
414.1475524902344 0 5496.3203
414.1934814453125 0 4090.3633
414.23565673828125 0 64804.594
415.037353515625 0 18821.297
415.1984558105469 0 17227.895
415.2394714355469 0 12146.878
416.2020263671875 0 5821.6123
417.7174987792969 0 45556.4
418.2109680175781 0 67793.664
418.71209716796875 0 30554.87
419.2134094238281 0 12033.296
421.20892333984375 0 12864.239 y Water loss 10
422.16571044921875 0 3562.0999
422.19378662109375 0 5817.7715 y Ammonia loss 10
422.7093200683594 0 5973.4277
423.20306396484375 0 11789.817
423.2723083496094 0 101910.44
423.7039794921875 0 5130.2344
424.22161865234375 0 8304.045
424.27569580078125 0 24113.871
424.7384948730469 0 5087.814
426.72259521484375 0 649159.2
427.2240905761719 0 332780.72
427.2597351074219 0 4881.6216
427.7255554199219 0 94859.37
428.2259521484375 0 21321.86
428.7250061035156 0 3694.458
429.2101135253906 0 195167.34
430.1943054199219 0 496606.53
431.1971740722656 0 106523.86
431.7147521972656 0 70573.85 b Water loss 6
432.2131652832031 0 41049.93 b Ammonia loss 6
432.71099853515625 0 15479.906
432.74224853515625 0 7244.8486
433.21026611328125 0 11010.518
433.2557373046875 0 22999.738 b Water loss 3
438.1741027832031 0 10656.678
438.7312927246094 0 8227.708
439.2198486328125 0 159329.89 y 10
440.17877197265625 0 12181.42
440.2225646972656 0 36806.117
440.7200622558594 0 333067.3 b 6
441.2214660644531 0 178218.5
441.72283935546875 0 54875.836
442.2225036621094 0 10651.727
444.17498779296875 0 3838.1785
446.71124267578125 0 4588.018 y Water loss 6
446.7420654296875 0 4034.662
447.2081298828125 0 25360.992 y Ammonia loss 6
447.7041015625 0 4446.642
448.205078125 0 30173.408
448.2356872558594 0 3040.235
449.20672607421875 0 8989.688
450.1760559082031 0 4135.0513
450.2371826171875 0 7062.261
450.2836608886719 0 7078.2
451.2673645019531 0 186413 b 3
452.2702941894531 0 48055.043
453.2721862792969 0 6281.6475
455.23162841796875 0 3721.2183
455.2659912109375 0 11294.256
455.7242126464844 0 11103.746 y 6
456.2227783203125 0 3558.1667
456.271240234375 0 3068.599
459.1994934082031 0 17899.264
460.7536315917969 0 24783.748
461.2040100097656 0 13279.309
461.2532653808594 0 26432.914
461.7561950683594 0 8341.381
462.2112731933594 0 7039.407
462.2509765625 0 3756.8145
465.74859619140625 0 4120.0117
466.2438049316406 0 4493.7417
466.7449035644531 0 3415.5752
468.29376220703125 0 49443.227
469.2968444824219 0 10234.272
470.2998962402344 0 3299.261
474.25933837890625 0 39392.56
474.7547912597656 0 55896.6
475.25579833984375 0 27133.404
475.75640869140625 0 5765.207
476.22589111328125 0 13665.148
477.2106628417969 0 29826.922
478.22998046875 0 31238.314 y Water loss 9
478.2777404785156 0 20279.48
479.23675537109375 0 6963.971
479.28021240234375 0 5723.204
479.743896484375 0 3628.3533
480.2170104980469 0 5068.529
481.2362976074219 0 3294.5056
483.2646789550781 0 138779.7
483.7662658691406 0 77674.016
484.221923828125 0 7558.6484
484.2677307128906 0 21114.615
484.7689208984375 0 8918.996
486.1873779296875 0 3471.683
488.2205810546875 0 5199.748
488.2574157714844 0 43305.938 b Water loss 7
488.75555419921875 0 24035.744 b Ammonia loss 7
489.2572326660156 0 13438.867
494.236572265625 0 381607.25
495.2005920410156 0 3552.0264
495.239501953125 0 99863.55
496.2411804199219 0 542127.4 y 9
497.2463684082031 0 115968.625
497.76373291015625 0 50062.926
498.24658203125 0 18257.482
499.2462463378906 0 3228.7146
501.2449951171875 0 3146.0764
502.2300109863281 0 4642.7344
503.10845947265625 0 19456.191
503.2201232910156 0 3554.09
504.2001647949219 0 13274.299
505.23431396484375 0 4187.732
506.2270202636719 0 14003.666
506.27435302734375 0 6971.902
507.2585754394531 0 12758.886
511.2638854980469 0 29067.977
512.2656860351562 0 5837.1206
513.21044921875 0 5689.774
515.2868041992188 0 6660.368
519.1371459960938 0 4912.8438
519.3296508789062 0 4776.3726
522.2122192382812 0 7349.582
523.2601318359375 0 12395.622
523.5868530273438 0 4691.278
524.2799682617188 0 10533.474
525.2675170898438 0 62589.703
526.2301025390625 0 11466.151
526.2709350585938 0 17666.02
527.2337036132812 0 3464.7756
528.9329833984375 0 6167.4507
529.2625732421875 0 22200.355 Precursor Water loss
529.5947875976562 0 17461.68 Precursor Ammonia loss
529.9286499023438 0 13661.303
530.2651977539062 0 4999.4443
531.2154541015625 0 3456.8464
531.7770385742188 0 4981.065
532.1953735351562 0 12923.543
533.1998901367188 0 4402.86
534.9363403320312 0 8560.252
535.2637939453125 0 14115.178 Precursor
535.6005249023438 0 14502.673
535.9330444335938 0 5817.761
536.1646728515625 0 4653.7197
537.24560546875 0 5817.4795
537.7398681640625 0 7940.0547
537.7827758789062 0 3875.8408
538.269775390625 0 9566.332
538.775390625 0 3418.7283
539.2367553710938 0 79554.234
540.2396240234375 0 22227.25
541.2508544921875 0 4842.5537
542.2939453125 0 38286.344
543.2782592773438 0 89895.414
544.2816772460938 0 32560.258
545.2835083007812 0 5951.705
545.7662353515625 0 5931.212 b Water loss 8
546.2528076171875 0 11101.435 b Ammonia loss 8
546.7487182617188 0 7516.053
547.2811279296875 0 4010.4224
548.2927856445312 0 3980.8193
549.2213134765625 0 16166.166
550.2103881835938 0 9303.194
551.2568359375 0 18862.379
551.3299560546875 0 8162.197
552.2617797851562 0 8684.898
552.333251953125 0 3409.5374
553.2574462890625 0 5043.553
554.2801513671875 0 5421.072
554.7733154296875 0 23060.76 b 8
555.2589111328125 0 24101.26
555.7572631835938 0 18171.436
558.2201538085938 0 6821.749
559.2533569335938 0 3890.7861
561.3146362304688 0 15558.904 b Water loss 4
562.3077392578125 0 8305.423 b Ammonia loss 4
563.2581787109375 0 12726.095
564.26904296875 0 3995.5542
565.2628173828125 0 26841.633
566.2615966796875 0 9628.959
567.23193359375 0 83419.46
567.7685546875 0 6128.5054
568.2371215820312 0 26686.258
568.7589721679688 0 4852.2515 y Ammonia loss 4
569.2382202148438 0 6891.7534
575.2466430664062 0 33999.098
575.2927856445312 0 5381.833
576.2359619140625 0 15413.272
576.7677001953125 0 3747.2637
577.2664184570312 0 7983.0327 y 4
579.32568359375 0 120429.19 b 4
580.3286743164062 0 41091.465
581.3308715820312 0 7386.546
582.3035888671875 0 7032.6187
586.2655029296875 0 4816.316
588.794189453125 0 4546.1104
589.3074340820312 0 8064.0166
592.2723999023438 0 29817.127
592.3250122070312 0 17384.242
593.2572631835938 0 153023.4 y Water loss 8
593.3057861328125 0 49335.28
593.7847290039062 0 6874.8667
594.2569580078125 0 57080.37
594.3093872070312 0 13819.384
595.2572021484375 0 13905.101
596.3538208007812 0 6973.939
597.799072265625 0 6089.0806
598.3009643554688 0 5577.8687
599.3290405273438 0 4388.8896
600.3158569335938 0 4118.3228
602.2996215820312 0 7832.5723
602.793701171875 0 22904.865
603.2898559570312 0 16534.262
603.7903442382812 0 8772.042
610.2838745117188 0 512726.12
611.2691040039062 0 1390715.9 y 8
611.79931640625 0 71374
612.2721557617188 0 443601.06
612.7987060546875 0 14637.554
613.273681640625 0 107942.484
614.2750854492188 0 16808.523
615.3172607421875 0 4860.0835
616.3074340820312 0 4864.462
617.2846069335938 0 10085.932
619.28515625 0 3765.295
619.3412475585938 0 4715.3975
620.80322265625 0 9440.608
621.2515869140625 0 25074.348
621.3014526367188 0 12909.827
622.2538452148438 0 8050.2812
622.3701171875 0 4538.8096
623.3656005859375 0 3776.6875
626.2986450195312 0 4536.274
627.274658203125 0 4007.2415
627.7781982421875 0 4166.169
628.2784423828125 0 8664.334
628.7762451171875 0 5708.2666
630.2896118164062 0 5370.2305
634.30810546875 0 21467.496
635.2970581054688 0 27623.445
636.3282470703125 0 17956.629
636.7896728515625 0 16375.424 y Water loss 3
637.28759765625 0 12948.289 y Ammonia loss 3
637.3392944335938 0 5423.2524
637.7899780273438 0 5556.721
638.337158203125 0 3987.4646 b Water loss 10
638.83251953125 0 5888.9316
640.2748413085938 0 12060.232
641.2567138671875 0 3707.7163
644.29736328125 0 4574.731
645.2955322265625 0 30380.568
645.794677734375 0 85215.984 y 3
646.2942504882812 0 64826.816
646.7955322265625 0 24205.64
647.2926025390625 0 7785.2104
648.3443603515625 0 8922.686
650.2681274414062 0 7028.8516
650.3226318359375 0 5063.772
651.8040161132812 0 4740.058
652.32080078125 0 163798.53
653.3236694335938 0 58946.754
654.3256225585938 0 13366.252
658.2832641601562 0 9407.446
659.3211669921875 0 4669.0605
660.8309326171875 0 3951.052
661.330810546875 0 7100.805
661.8322143554688 0 5377.838
662.30615234375 0 51065.293
663.2975463867188 0 33503.37
664.2891235351562 0 5809.387
664.3416748046875 0 8546.531
665.3451538085938 0 5258.325
666.35009765625 0 3731.6394
668.2781982421875 0 4012.5083
676.3418579101562 0 71468.164 b Water loss 5
677.33642578125 0 52160.53 b Ammonia loss 5
678.3344116210938 0 18768.293
679.334228515625 0 15525.813
679.8314208984375 0 3823.7317
680.3162231445312 0 287380.03
681.31884765625 0 115735.06
682.3209838867188 0 27568.297
684.3262939453125 0 15676.871
684.820556640625 0 18794.145
685.3176879882812 0 12196.238
685.8230590820312 0 5670.4956
688.3333740234375 0 7058.422
689.3272705078125 0 4194.7827
690.3004150390625 0 4551.6665
692.8359375 0 17245.557
693.3302612304688 0 103595.77 y Water loss 2
693.830078125 0 88756.67 y Ammonia loss 2
694.3521728515625 0 804705.25 b 5
694.8291625976562 0 17340.65
695.3554077148438 0 314956.1
696.3585205078125 0 75030.13
697.2907104492188 0 4538.374
697.3564453125 0 17204.893
697.8598022460938 0 4246.3145
698.3599243164062 0 4816.777
700.8411254882812 0 8733.007
701.3424072265625 0 17062.238
701.8418579101562 0 216614.4
702.3367919921875 0 531384.7 y 2
702.837646484375 0 407174.84 b Water loss 11
703.3381958007812 0 187689.86 b Ammonia loss 11
703.8400268554688 0 49849.652
704.3389282226562 0 20590.312
705.3587036132812 0 10135.721
706.3430786132812 0 49412.72 y Water loss 7
707.3439331054688 0 20379.621
708.3346557617188 0 3021.0122
711.3681640625 0 38750.254
711.8589477539062 0 42542.035 b 11
712.36328125 0 36111.87
712.85888671875 0 6160.7773
718.3927001953125 0 4752.9146
720.3846435546875 0 6030.9385
721.3621215820312 0 36723.383
722.3654174804688 0 13776.002
723.3681030273438 0 153188.27
724.3538818359375 0 416726.72 y 7
725.3566284179688 0 158355.48
726.3582153320312 0 48689.344
727.360595703125 0 6679.0913
728.3707275390625 0 8225.318
728.8601684570312 0 4769.1636
729.35986328125 0 14777.89
730.3642578125 0 7756.57
733.3644409179688 0 23862.879
734.3546752929688 0 22140.977
735.3478393554688 0 5811.17
737.8679809570312 0 7714.5938
738.36279296875 0 5267.9727
739.3524169921875 0 4720.311
742.8645629882812 0 7442.675 y Water loss 1
743.3662719726562 0 14904.997 y Ammonia loss 1
744.36083984375 0 4749.8374
747.3898315429688 0 11269.92
748.3832397460938 0 9482.817
749.3355102539062 0 9130.364
750.3892211914062 0 7004.792
751.3770141601562 0 101223.234
751.87060546875 0 42861.613 y 1
752.3760986328125 0 65982.55
752.873291015625 0 14979.296
753.3762817382812 0 20122.156
760.3120727539062 0 4597.4966
761.3743286132812 0 6002.5454
762.3516845703125 0 5236.831
762.4141845703125 0 10368.793
763.3659057617188 0 54181.457
764.3663330078125 0 24751.91
765.40087890625 0 20098.975
766.40771484375 0 8019.6284
767.346435546875 0 14624.586
768.3465576171875 0 5767.589
769.40185546875 0 12909.912
775.3912353515625 0 8463.158
776.3814086914062 0 8313.889
779.3842163085938 0 47590.285
780.3856201171875 0 25395.342
781.3668212890625 0 26973.352
782.3670043945312 0 11266.62
791.3988037109375 0 4346.1235
793.39990234375 0 90834.95 Precursor Water loss
794.4019165039062 0 36423.164
795.3424682617188 0 13371.955
795.407958984375 0 7832.1953
796.3416748046875 0 8775.87
807.3779296875 0 18829.387
807.4405517578125 0 10714.049
808.382568359375 0 12694.997
809.420166015625 0 4619.128
817.4110107421875 0 3629.4304
818.4396362304688 0 18535.6
819.4407958984375 0 8384.207
820.4380493164062 0 3889.0415
825.391357421875 0 10027.667
826.3822021484375 0 11252.558
827.3800048828125 0 4600.8013
834.4254760742188 0 14946.229
835.415771484375 0 30019.715
836.4490356445312 0 59784.734
837.4518432617188 0 35376.426
838.4533081054688 0 11497.211
843.4043579101562 0 12853.66
844.3893432617188 0 17854.81
845.3941040039062 0 12505.648
846.4464721679688 0 23480.719
847.444091796875 0 19728.973
848.4479370117188 0 7307.686
851.3809204101562 0 5728.908
852.4367065429688 0 211437.75
853.3499145507812 0 5397.847
853.439697265625 0 112733.59
854.4429931640625 0 31565.3
862.4212646484375 0 132822.4 b Water loss 6
863.4172973632812 0 94112.69 b Ammonia loss 6
864.4562377929688 0 170601.56
865.4605102539062 0 87076.99
866.4620971679688 0 20490.05
868.4085693359375 0 4794.195
876.43798828125 0 17150.67
877.4420776367188 0 8012.6826
880.4315795898438 0 989745.75 b 6
881.4342651367188 0 547685.06
882.4375 0 162343.72
883.4389038085938 0 31875.342
890.4190063476562 0 7151.2026
891.41943359375 0 10739.655
892.461181640625 0 14870.554
893.478759765625 0 3976.3948
894.4564819335938 0 4802.32
907.4425048828125 0 20361.688
908.428955078125 0 57983.258
909.4362182617188 0 48663.094
910.4326171875 0 70098.87 y 6
911.4360961914062 0 32788.234
912.4353637695312 0 9108.593
936.4249877929688 0 9911.371
937.4231567382812 0 8710.052
938.41845703125 0 3772.211
946.4503784179688 0 3989.241
947.4298095703125 0 4404.0146
953.4456176757812 0 3638.445
954.4331665039062 0 16027.622
955.4247436523438 0 20797.125
956.426025390625 0 8205.249
962.4513549804688 0 4350.2065
964.4596557617188 0 9516.457
965.4534912109375 0 20768.314
966.4523315429688 0 11036.13
971.45654296875 0 7788.1934
972.4476318359375 0 13691.098
973.4472045898438 0 4361.8213
975.5044555664062 0 13517.197 b Water loss 7
976.4994506835938 0 10373.186 b Ammonia loss 7
979.48095703125 0 4716.811
980.4656982421875 0 7980.6973
981.4779663085938 0 6199.7383
982.4732055664062 0 5943.377
983.4752197265625 0 5776.527
993.5154418945312 0 72311.98 b 7
994.518310546875 0 37990.09
995.5183715820312 0 12617.169
1007.4495239257812 0 8142.6597 y Water loss 5
1008.5059204101562 0 3628.907
1009.436767578125 0 8466.502
1024.476806640625 0 16053.152
1025.461669921875 0 34829.547 y 5
1026.46533203125 0 17472.457
1027.464599609375 0 3930.259
1075.5291748046875 0 9271.503
1076.5396728515625 0 8703.941
1091.499755859375 0 5696.934 b Ammonia loss 8
1092.553466796875 0 8429.83
1093.547607421875 0 22040.764
1094.548828125 0 13022.685
1095.5357666015625 0 5049.293
1107.568115234375 0 3636.2024
1108.5352783203125 0 8043.182 b 8
1109.5224609375 0 7722.5293
1117.494384765625 0 4329.3335
1118.481201171875 0 7454.4595
1119.4835205078125 0 3793.297
1134.527587890625 0 6376.6196
1135.50732421875 0 32840.742 y Water loss 4
1136.49951171875 0 36878.477 y Ammonia loss 4
1137.4989013671875 0 17865.092
1138.497802734375 0 5362.071
1152.5303955078125 0 26053.123
1153.52197265625 0 65978.96 y 4
1154.521728515625 0 39669.562
1155.526611328125 0 12198.332
1164.57958984375 0 3857.9968
1165.5697021484375 0 6807.4556 b 9
1205.5787353515625 0 4056.9087
1206.567138671875 0 4236.31
1290.5655517578125 0 4396.6846 y 3
1434.2408447265625 0 3247.996
3082.6318359375 0 4156.7925
3084.28369140625 0 4604.37
3459.765380859375 0 3626.3994

Spectrum Details

|  |  |
| --- | --- |
| Matched peaks? Matched peaks The total absolute number of peaks matched. Additionally in brackets the total fraction of peaks matched and the total number of peaks is shown. | 86 (9.44% of 911) |
| FDR? FDR The false discovery rate estimated for this peptide. It is calculated by matching all theoretical fragments with a non-integer shift with the raw peaks for this spectrum. This is done with 40 different shifts. The resulting percentage is the average number of annotated peaks over the number of annotated peaks with the correct spectrum. | 0.66% |
| Satellite FDR? Satellite FDR See the FDR for details on its calculation. This satellite ion specific FDR only contains the satellite ions (d/w) for I/L/J positions. | - |
| PSM Score? PSM Score The PSM Score as given by Hecklib to this annotated spectrum. It is shown with three significant figures. | 647 |

#### Spectrum 7645? Spectrum 7645 The raw spectrum of this peptide as annotated by Hecklib. The fragments are coloured according to ion type (see legend). Any peaks with a star '\*' as text can be hovered over to see the full details, first the ion type second the mass shift type. By hovering over the amino acids in the peptide or ions in the legend the corresponding peaks are highlighted. By toggling the 'Unassigned' label you can turn the background (unassigned) peaks on or off in the plot. By updating the slider in the Ion legend you can update the spectrum to only show the top X% of the peaks with labels. The top X% means any peak that is within X% of the highest intensity. By dragging in the spectrum you can zoom in to a specific part of the spectrum and use 'Zoom Out' to get back to the original zoom level. The annotation of the spectrum is based on the given sequence in the peptides file and is done with different software so inconsistencies are likely. The peaks are annotated based on the given sequence, with 20 ppm tolerance.

Copy Data

##### Spectrum 7645 (TSV)

###### Preview

```
Loading example...
```

*Click on the button to copy the data to your clipboard.*

Mz MinMz MaxIntensity Max

WidthHeightPeptide font sizePeptide stroke widthSpectrum font sizeSpectrum stroke widthCompact peptide

Ion legend

wxyz

abcd

OtherUnassignedIon^Charge^_Position_Show for top:%

TVLHQDWLDGKEY

05.19e+41.04e+51.56e+52.07e+5

Zoom Out

a^+1^_2_y^+3^_4_a^+1^_2_y^+1^_1_b^+1^_2_b^+1^_2_y^+2^_3_b^+3^_6_y^+1^_2_b^+1^_3_y^+3^_7_y^+2^_5_y^+1^_2_b^+1^_3_b^+2^_6_b^+2^_6_y^+2^_6_y^+2^_6_y^+3^_9_y^+1^_3_y^+1^_3_b^+2^_7_b^+2^_7_b^+1^_4_y^+1^_3_b^+2^_7_y^+2^_7_b^+1^_4_y^+2^_7_y^+1^_4_b^+2^_8_b^+2^_8_y^+1^_4_\*\*b^+2^_9_b^+2^_9_b^+1^_5_b^+1^_5_y^+1^_5_y^+1^_5_y^+2^_10_y^+2^_10_y^+2^_10_b^+1^_6_b^+1^_6_y^+2^_11_y^+2^_11_b^+1^_6_y^+2^_11_b^+2^_12_b^+2^_12_y^+1^_6_b^+2^_12_y^+1^_6_y^+2^_12_y^+2^_12_\*b^+1^_7_b^+1^_7_b^+1^_7_y^+1^_7_b^+1^_8_b^+1^_8_y^+1^_8_b^+1^_9_y^+1^_9_y^+1^_9_y^+1^_9_

0778155523333111

Fragment Matches Table

Show background peaks

| Position | Ion type | Intensity | mz Theoretical | mz Error (Th) | mz Error (ppm) | Charge | Series Number |
| --- | --- | --- | --- | --- | --- | --- | --- |
| - | - | 4269 | 120.1 | - | - | 0 | - |
| - | - | 6490 | 120.1 | - | - | 0 | - |
| - | - | 471.5 | 121.1 | - | - | 0 | - |
| - | - | 851.8 | 122.1 | - | - | 0 | - |
| - | - | 955.7 | 123 | - | - | 0 | - |
| - | - | 551.5 | 126.1 | - | - | 0 | - |
| - | - | 3131 | 127.1 | - | - | 0 | - |
| - | - | 1.125E+04 | 128.1 | - | - | 0 | - |
| - | - | 1166 | 129.1 | - | - | 0 | - |
| - | - | 1.419E+05 | 129.1 | - | - | 0 | - |
| - | - | 6.318E+04 | 130.1 | - | - | 0 | - |
| - | - | 687.4 | 130.1 | - | - | 0 | - |
| - | - | 706.4 | 130.1 | - | - | 0 | - |
| - | - | 9185 | 130.1 | - | - | 0 | - |
| - | - | 6957 | 131.1 | - | - | 0 | - |
| - | - | 1.25E+04 | 132.1 | - | - | 0 | - |
| - | - | 425.1 | 132.1 | - | - | 0 | - |
| - | - | 572.4 | 133.1 | - | - | 0 | - |
| - | - | 1286 | 133.1 | - | - | 0 | - |
| - | - | 371.5 | 133.1 | - | - | 0 | - |
| - | - | 400.2 | 134 | - | - | 0 | - |
| - | - | 2.019E+04 | 136.1 | - | - | 0 | - |
| - | - | 1784 | 137.1 | - | - | 0 | - |
| - | - | 464.8 | 137.1 | - | - | 0 | - |
| - | - | 2006 | 138.1 | - | - | 0 | - |
| - | - | 738 | 138.1 | - | - | 0 | - |
| - | - | 1636 | 139.1 | - | - | 0 | - |
| - | - | 3164 | 141.1 | - | - | 0 | - |
| - | - | 517.8 | 142.1 | - | - | 0 | - |
| - | - | 862.4 | 143 | - | - | 0 | - |
| - | - | 585.2 | 143.1 | - | - | 0 | - |
| - | - | 655.9 | 144.1 | - | - | 0 | - |
| - | - | 1540 | 145.1 | - | - | 0 | - |
| - | - | 1005 | 146.1 | - | - | 0 | - |
| - | - | 806.2 | 147 | - | - | 0 | - |
| - | - | 982.2 | 149 | - | - | 0 | - |
| - | - | 1.559E+04 | 149 | - | - | 0 | - |
| - | - | 975.6 | 150.1 | - | - | 0 | - |
| - | - | 419 | 151 | - | - | 0 | - |
| - | - | 3913 | 151.1 | - | - | 0 | - |
| - | - | 460.2 | 152.1 | - | - | 0 | - |
| - | - | 1934 | 155.1 | - | - | 0 | - |
| - | - | 8438 | 155.1 | - | - | 0 | - |
| 2 | a | 6033 | 155.1 | 0.0002288 | 1.475 | +1 | 2 |
| - | - | 695.8 | 156.1 | - | - | 0 | - |
| - | - | 618.7 | 156.1 | - | - | 0 | - |
| - | - | 613.2 | 157.1 | - | - | 0 | - |
| - | - | 662.2 | 158.1 | - | - | 0 | - |
| - | - | 1.078E+05 | 159.1 | - | - | 0 | - |
| - | - | 971.7 | 160.1 | - | - | 0 | - |
| - | - | 441 | 160.1 | - | - | 0 | - |
| - | - | 1.142E+04 | 160.1 | - | - | 0 | - |
| - | - | 635.1 | 161.1 | - | - | 0 | - |
| - | - | 9220 | 165.1 | - | - | 0 | - |
| - | - | 887.9 | 165.1 | - | - | 0 | - |
| - | - | 1.595E+04 | 166.1 | - | - | 0 | - |
| 10 | y | 3080 | 166.1 | 0.001588 | 9.561 | +3 | 4 |
| - | - | 1267 | 166.1 | - | - | 0 | - |
| - | - | 1.948E+04 | 167.1 | - | - | 0 | - |
| - | - | 840.5 | 167.1 | - | - | 0 | - |
| - | - | 1045 | 168.1 | - | - | 0 | - |
| - | - | 2180 | 168.1 | - | - | 0 | - |
| - | - | 1308 | 168.1 | - | - | 0 | - |
| - | - | 1631 | 168.1 | - | - | 0 | - |
| - | - | 639.3 | 169.1 | - | - | 0 | - |
| - | - | 1402 | 169.1 | - | - | 0 | - |
| - | - | 4734 | 169.1 | - | - | 0 | - |
| - | - | 4989 | 170.1 | - | - | 0 | - |
| - | - | 646.9 | 171.1 | - | - | 0 | - |
| - | - | 844.5 | 171.1 | - | - | 0 | - |
| - | - | 1348 | 171.1 | - | - | 0 | - |
| - | - | 1939 | 172.1 | - | - | 0 | - |
| - | - | 1904 | 173.1 | - | - | 0 | - |
| 2 | a | 2.054E+05 | 173.1 | 0.0002842 | 1.641 | +1 | 2 |
| - | - | 3189 | 173.5 | - | - | 0 | - |
| - | - | 1.802E+04 | 174.1 | - | - | 0 | - |
| - | - | 1492 | 175.1 | - | - | 0 | - |
| - | - | 6139 | 178.1 | - | - | 0 | - |
| - | - | 999.3 | 179.1 | - | - | 0 | - |
| - | - | 1367 | 181.1 | - | - | 0 | - |
| 13 | y | 2.096E+04 | 182.1 | 0.0001902 | 1.045 | +1 | 1 |
| - | - | 2171 | 183.1 | - | - | 0 | - |
| 2 | b | 5234 | 183.1 | 0.0002482 | 1.356 | +1 | 2 |
| - | - | 627.6 | 183.1 | - | - | 0 | - |
| - | - | 733.3 | 185.1 | - | - | 0 | - |
| - | - | 1653 | 185.2 | - | - | 0 | - |
| - | - | 4.371E+04 | 186.1 | - | - | 0 | - |
| - | - | 8767 | 187.1 | - | - | 0 | - |
| - | - | 3910 | 187.1 | - | - | 0 | - |
| - | - | 949.8 | 187.1 | - | - | 0 | - |
| - | - | 1494 | 188.1 | - | - | 0 | - |
| - | - | 731.7 | 188.1 | - | - | 0 | - |
| - | - | 781.9 | 189.1 | - | - | 0 | - |
| - | - | 1972 | 190.1 | - | - | 0 | - |
| - | - | 1364 | 190.1 | - | - | 0 | - |
| - | - | 422.8 | 194.1 | - | - | 0 | - |
| - | - | 1212 | 194.1 | - | - | 0 | - |
| - | - | 1.083E+04 | 195.1 | - | - | 0 | - |
| - | - | 7430 | 196.1 | - | - | 0 | - |
| - | - | 745.9 | 196.1 | - | - | 0 | - |
| - | - | 1846 | 197.1 | - | - | 0 | - |
| - | - | 503 | 197.8 | - | - | 0 | - |
| - | - | 520.3 | 197.8 | - | - | 0 | - |
| - | - | 1270 | 198.1 | - | - | 0 | - |
| - | - | 3628 | 198.1 | - | - | 0 | - |
| - | - | 708.7 | 199.1 | - | - | 0 | - |
| - | - | 546.8 | 199.1 | - | - | 0 | - |
| - | - | 478.9 | 199.1 | - | - | 0 | - |
| - | - | 707.7 | 200.1 | - | - | 0 | - |
| 2 | b | 5.957E+04 | 201.1 | 0.0001205 | 0.5993 | +1 | 2 |
| - | - | 743 | 202.1 | - | - | 0 | - |
| - | - | 6425 | 202.1 | - | - | 0 | - |
| - | - | 748.9 | 203.1 | - | - | 0 | - |
| - | - | 4578 | 205.1 | - | - | 0 | - |
| - | - | 802.8 | 205.1 | - | - | 0 | - |
| - | - | 885.8 | 206.1 | - | - | 0 | - |
| - | - | 760.9 | 206.1 | - | - | 0 | - |
| - | - | 943.2 | 207.2 | - | - | 0 | - |
| - | - | 706.1 | 209.1 | - | - | 0 | - |
| - | - | 1836 | 210.1 | - | - | 0 | - |
| 11 | y | 1492 | 211.1 | 0.0001914 | 0.9065 | +2 | 3 |
| - | - | 789 | 211.1 | - | - | 0 | - |
| - | - | 505.6 | 211.5 | - | - | 0 | - |
| - | - | 508.5 | 212.1 | - | - | 0 | - |
| - | - | 1554 | 212.1 | - | - | 0 | - |
| - | - | 5971 | 213.1 | - | - | 0 | - |
| - | - | 854.3 | 213.2 | - | - | 0 | - |
| - | - | 1502 | 215.1 | - | - | 0 | - |
| - | - | 1444 | 216.1 | - | - | 0 | - |
| - | - | 1999 | 219.1 | - | - | 0 | - |
| - | - | 2861 | 221.1 | - | - | 0 | - |
| - | - | 998.5 | 221.1 | - | - | 0 | - |
| - | - | 7675 | 221.1 | - | - | 0 | - |
| - | - | 1173 | 221.1 | - | - | 0 | - |
| - | - | 1016 | 221.1 | - | - | 0 | - |
| - | - | 2547 | 222.1 | - | - | 0 | - |
| - | - | 1500 | 222.1 | - | - | 0 | - |
| - | - | 5593 | 223.1 | - | - | 0 | - |
| - | - | 1013 | 223.1 | - | - | 0 | - |
| - | - | 3727 | 223.1 | - | - | 0 | - |
| - | - | 1.968E+04 | 223.2 | - | - | 0 | - |
| - | - | 1263 | 224.1 | - | - | 0 | - |
| - | - | 1613 | 224.2 | - | - | 0 | - |
| - | - | 8602 | 225 | - | - | 0 | - |
| - | - | 591.7 | 225.2 | - | - | 0 | - |
| - | - | 2474 | 226 | - | - | 0 | - |
| - | - | 4368 | 226.1 | - | - | 0 | - |
| 6 | b | 3961 | 226.1 | 0.0002329 | 1.03 | +3 | 6 |
| - | - | 669.4 | 226.2 | - | - | 0 | - |
| - | - | 931.7 | 227 | - | - | 0 | - |
| - | - | 1650 | 227 | - | - | 0 | - |
| - | - | 3374 | 227.1 | - | - | 0 | - |
| - | - | 1213 | 227.2 | - | - | 0 | - |
| - | - | 2296 | 228.1 | - | - | 0 | - |
| - | - | 2916 | 229.1 | - | - | 0 | - |
| - | - | 3944 | 229.2 | - | - | 0 | - |
| - | - | 838.9 | 230.1 | - | - | 0 | - |
| - | - | 3956 | 233.1 | - | - | 0 | - |
| - | - | 2.004E+04 | 234.1 | - | - | 0 | - |
| - | - | 945 | 234.1 | - | - | 0 | - |
| - | - | 900.1 | 235.1 | - | - | 0 | - |
| - | - | 2119 | 235.1 | - | - | 0 | - |
| - | - | 561.1 | 235.6 | - | - | 0 | - |
| - | - | 1030 | 237.1 | - | - | 0 | - |
| - | - | 2961 | 238.1 | - | - | 0 | - |
| - | - | 715.3 | 238.1 | - | - | 0 | - |
| - | - | 1.958E+04 | 239.1 | - | - | 0 | - |
| - | - | 6815 | 240.1 | - | - | 0 | - |
| - | - | 9097 | 240.1 | - | - | 0 | - |
| - | - | 4007 | 241.1 | - | - | 0 | - |
| - | - | 1324 | 241.1 | - | - | 0 | - |
| - | - | 497.3 | 241.1 | - | - | 0 | - |
| - | - | 1238 | 241.2 | - | - | 0 | - |
| - | - | 1127 | 243.1 | - | - | 0 | - |
| - | - | 7896 | 244.1 | - | - | 0 | - |
| - | - | 1.15E+04 | 247.1 | - | - | 0 | - |
| - | - | 2280 | 248.1 | - | - | 0 | - |
| - | - | 4269 | 248.1 | - | - | 0 | - |
| - | - | 1009 | 249.1 | - | - | 0 | - |
| - | - | 7370 | 249.1 | - | - | 0 | - |
| - | - | 2404 | 249.1 | - | - | 0 | - |
| - | - | 1000 | 250.1 | - | - | 0 | - |
| - | - | 6.666E+04 | 251.2 | - | - | 0 | - |
| - | - | 8558 | 252.2 | - | - | 0 | - |
| - | - | 1004 | 253.2 | - | - | 0 | - |
| - | - | 4691 | 255.1 | - | - | 0 | - |
| - | - | 853.9 | 256.1 | - | - | 0 | - |
| - | - | 854.8 | 256.2 | - | - | 0 | - |
| - | - | 630.4 | 256.6 | - | - | 0 | - |
| - | - | 2093 | 257.1 | - | - | 0 | - |
| - | - | 3685 | 258.1 | - | - | 0 | - |
| - | - | 585 | 259.1 | - | - | 0 | - |
| - | - | 1740 | 261.1 | - | - | 0 | - |
| - | - | 896 | 261.1 | - | - | 0 | - |
| - | - | 679.6 | 261.2 | - | - | 0 | - |
| - | - | 704.6 | 262.1 | - | - | 0 | - |
| - | - | 9129 | 265.1 | - | - | 0 | - |
| - | - | 951.8 | 265.1 | - | - | 0 | - |
| - | - | 1.692E+04 | 266.1 | - | - | 0 | - |
| - | - | 2412 | 267.1 | - | - | 0 | - |
| - | - | 646.4 | 267.1 | - | - | 0 | - |
| - | - | 978 | 268.1 | - | - | 0 | - |
| - | - | 1443 | 268.2 | - | - | 0 | - |
| - | - | 1441 | 269.2 | - | - | 0 | - |
| - | - | 2963 | 270.1 | - | - | 0 | - |
| - | - | 4259 | 272.2 | - | - | 0 | - |
| - | - | 3635 | 274.1 | - | - | 0 | - |
| - | - | 1188 | 274.1 | - | - | 0 | - |
| - | - | 975.5 | 275.1 | - | - | 0 | - |
| - | - | 827.5 | 276.1 | - | - | 0 | - |
| - | - | 1323 | 276.2 | - | - | 0 | - |
| - | - | 1532 | 279.1 | - | - | 0 | - |
| - | - | 861.4 | 280.1 | - | - | 0 | - |
| - | - | 1.434E+04 | 281.1 | - | - | 0 | - |
| - | - | 4479 | 282.2 | - | - | 0 | - |
| - | - | 2236 | 283 | - | - | 0 | - |
| - | - | 5.448E+04 | 283.1 | - | - | 0 | - |
| - | - | 1567 | 284.1 | - | - | 0 | - |
| - | - | 766.9 | 284.1 | - | - | 0 | - |
| - | - | 7928 | 284.1 | - | - | 0 | - |
| - | - | 796 | 285 | - | - | 0 | - |
| - | - | 858.9 | 285.2 | - | - | 0 | - |
| - | - | 4682 | 286.1 | - | - | 0 | - |
| - | - | 815 | 286.2 | - | - | 0 | - |
| - | - | 530.8 | 287.1 | - | - | 0 | - |
| - | - | 636 | 287.2 | - | - | 0 | - |
| - | - | 760.4 | 288.6 | - | - | 0 | - |
| 12 | y | 2.873E+04 | 293.1 | 0.0001137 | 0.3878 | +1 | 2 |
| - | - | 668.9 | 293.2 | - | - | 0 | - |
| - | - | 5203 | 294.1 | - | - | 0 | - |
| - | - | 1560 | 295.1 | - | - | 0 | - |
| - | - | 875.7 | 295.1 | - | - | 0 | - |
| - | - | 1895 | 296.1 | - | - | 0 | - |
| 3 | b | 1068 | 296.2 | 9.231E-05 | 0.3117 | +1 | 3 |
| - | - | 2697 | 297.1 | - | - | 0 | - |
| - | - | 1041 | 297.1 | - | - | 0 | - |
| - | - | 8463 | 297.2 | - | - | 0 | - |
| - | - | 1460 | 297.2 | - | - | 0 | - |
| 7 | y | 1252 | 298.1 | 0.004665 | 15.65 | +3 | 7 |
| - | - | 1395 | 298.2 | - | - | 0 | - |
| - | - | 1.117E+05 | 299.1 | - | - | 0 | - |
| - | - | 3714 | 300.1 | - | - | 0 | - |
| - | - | 8122 | 300.2 | - | - | 0 | - |
| - | - | 2429 | 301.1 | - | - | 0 | - |
| - | - | 2.324E+04 | 301.2 | - | - | 0 | - |
| - | - | 958.5 | 301.2 | - | - | 0 | - |
| - | - | 4191 | 302.1 | - | - | 0 | - |
| - | - | 3745 | 302.2 | - | - | 0 | - |
| - | - | 762.3 | 303.1 | - | - | 0 | - |
| - | - | 1125 | 305.6 | - | - | 0 | - |
| 9 | y | 3158 | 306.1 | 0.0003382 | 1.105 | +2 | 5 |
| - | - | 1397 | 306.6 | - | - | 0 | - |
| - | - | 577.9 | 309.2 | - | - | 0 | - |
| - | - | 908.7 | 310.2 | - | - | 0 | - |
| 12 | y | 4382 | 311.1 | 0.0003827 | 1.23 | +1 | 2 |
| - | - | 795.7 | 312.1 | - | - | 0 | - |
| - | - | 726.7 | 314.1 | - | - | 0 | - |
| - | - | 566.2 | 314.2 | - | - | 0 | - |
| 3 | b | 2600 | 314.2 | 0.0004016 | 1.278 | +1 | 3 |
| - | - | 1.439E+04 | 315.2 | - | - | 0 | - |
| - | - | 1088 | 315.2 | - | - | 0 | - |
| - | - | 1174 | 316.2 | - | - | 0 | - |
| - | - | 2222 | 316.2 | - | - | 0 | - |
| - | - | 553.1 | 317.1 | - | - | 0 | - |
| - | - | 853.6 | 319.2 | - | - | 0 | - |
| - | - | 635.8 | 324.2 | - | - | 0 | - |
| - | - | 832 | 325.2 | - | - | 0 | - |
| - | - | 2124 | 325.2 | - | - | 0 | - |
| - | - | 4537 | 326.7 | - | - | 0 | - |
| - | - | 657.3 | 327 | - | - | 0 | - |
| - | - | 1798 | 327.2 | - | - | 0 | - |
| - | - | 1853 | 332.2 | - | - | 0 | - |
| - | - | 2708 | 333.2 | - | - | 0 | - |
| - | - | 525.3 | 338.2 | - | - | 0 | - |
| 6 | b | 622 | 338.7 | 0.0007045 | 2.08 | +2 | 6 |
| - | - | 1157 | 340.7 | - | - | 0 | - |
| - | - | 1084 | 341 | - | - | 0 | - |
| - | - | 1275 | 342.2 | - | - | 0 | - |
| - | - | 959.9 | 343.2 | - | - | 0 | - |
| - | - | 610.2 | 343.2 | - | - | 0 | - |
| - | - | 1166 | 344.2 | - | - | 0 | - |
| - | - | 1.442E+04 | 345 | - | - | 0 | - |
| - | - | 2455 | 345.1 | - | - | 0 | - |
| - | - | 793.3 | 345.2 | - | - | 0 | - |
| - | - | 9904 | 346.1 | - | - | 0 | - |
| - | - | 742.8 | 346.7 | - | - | 0 | - |
| - | - | 1114 | 347.1 | - | - | 0 | - |
| 6 | b | 4884 | 347.7 | 0.000219 | 0.63 | +2 | 6 |
| - | - | 840.5 | 348.2 | - | - | 0 | - |
| - | - | 712.6 | 349.2 | - | - | 0 | - |
| - | - | 1422 | 350.1 | - | - | 0 | - |
| - | - | 5848 | 350.2 | - | - | 0 | - |
| - | - | 1071 | 351.2 | - | - | 0 | - |
| - | - | 900.8 | 352.2 | - | - | 0 | - |
| 8 | y | 845.1 | 353.7 | 0.0003396 | 0.9603 | +2 | 6 |
| - | - | 2223 | 355.1 | - | - | 0 | - |
| - | - | 662.1 | 356.1 | - | - | 0 | - |
| - | - | 838.9 | 356.2 | - | - | 0 | - |
| - | - | 1.145E+04 | 359 | - | - | 0 | - |
| - | - | 1114 | 360.7 | - | - | 0 | - |
| - | - | 1958 | 361.2 | - | - | 0 | - |
| - | - | 3609 | 362.2 | - | - | 0 | - |
| 8 | y | 9803 | 362.7 | 0.0002509 | 0.6918 | +2 | 6 |
| - | - | 1.259E+04 | 363.1 | - | - | 0 | - |
| - | - | 3869 | 363.2 | - | - | 0 | - |
| - | - | 689.8 | 363.7 | - | - | 0 | - |
| - | - | 1843 | 364.1 | - | - | 0 | - |
| - | - | 1766 | 364.1 | - | - | 0 | - |
| - | - | 1221 | 366.2 | - | - | 0 | - |
| - | - | 1358 | 367.1 | - | - | 0 | - |
| - | - | 818.5 | 367.2 | - | - | 0 | - |
| - | - | 1023 | 369.1 | - | - | 0 | - |
| - | - | 2011 | 370.1 | - | - | 0 | - |
| - | - | 1114 | 371.1 | - | - | 0 | - |
| - | - | 710.6 | 373.2 | - | - | 0 | - |
| - | - | 599 | 374.2 | - | - | 0 | - |
| - | - | 1039 | 376.2 | - | - | 0 | - |
| - | - | 1232 | 376.2 | - | - | 0 | - |
| - | - | 997.5 | 377.1 | - | - | 0 | - |
| - | - | 5371 | 378.2 | - | - | 0 | - |
| - | - | 1.803E+04 | 379.2 | - | - | 0 | - |
| - | - | 787.3 | 380.2 | - | - | 0 | - |
| - | - | 4160 | 380.2 | - | - | 0 | - |
| - | - | 3.319E+04 | 381.2 | - | - | 0 | - |
| - | - | 1195 | 381.7 | - | - | 0 | - |
| - | - | 6739 | 382.2 | - | - | 0 | - |
| - | - | 1613 | 382.2 | - | - | 0 | - |
| - | - | 682.3 | 382.2 | - | - | 0 | - |
| - | - | 1181 | 384.2 | - | - | 0 | - |
| - | - | 1140 | 384.2 | - | - | 0 | - |
| - | - | 2602 | 385.1 | - | - | 0 | - |
| 5 | y | 862.4 | 385.2 | 0.003826 | 9.934 | +3 | 9 |
| - | - | 798.5 | 386.2 | - | - | 0 | - |
| - | - | 1108 | 390.2 | - | - | 0 | - |
| - | - | 671.8 | 391.1 | - | - | 0 | - |
| - | - | 5612 | 394.2 | - | - | 0 | - |
| - | - | 2259 | 395.1 | - | - | 0 | - |
| - | - | 1039 | 395.2 | - | - | 0 | - |
| - | - | 1865 | 395.2 | - | - | 0 | - |
| - | - | 934.3 | 396.1 | - | - | 0 | - |
| - | - | 6333 | 396.2 | - | - | 0 | - |
| - | - | 1058 | 397.2 | - | - | 0 | - |
| - | - | 3352 | 398.2 | - | - | 0 | - |
| - | - | 999.2 | 401.2 | - | - | 0 | - |
| - | - | 2572 | 402.2 | - | - | 0 | - |
| - | - | 3406 | 402.2 | - | - | 0 | - |
| - | - | 1442 | 403.2 | - | - | 0 | - |
| - | - | 804.4 | 405.3 | - | - | 0 | - |
| - | - | 2619 | 409.1 | - | - | 0 | - |
| - | - | 3096 | 411.2 | - | - | 0 | - |
| - | - | 1.962E+04 | 412.2 | - | - | 0 | - |
| - | - | 1412 | 413.1 | - | - | 0 | - |
| - | - | 2334 | 413.2 | - | - | 0 | - |
| - | - | 1810 | 413.2 | - | - | 0 | - |
| - | - | 1217 | 413.3 | - | - | 0 | - |
| - | - | 6088 | 414.2 | - | - | 0 | - |
| - | - | 2.418E+04 | 415 | - | - | 0 | - |
| - | - | 1871 | 415.2 | - | - | 0 | - |
| - | - | 1542 | 415.2 | - | - | 0 | - |
| - | - | 4019 | 417.7 | - | - | 0 | - |
| - | - | 6828 | 418.2 | - | - | 0 | - |
| - | - | 2914 | 418.7 | - | - | 0 | - |
| 11 | y | 748.7 | 421.2 | 0.00183 | 4.345 | +1 | 3 |
| 11 | y | 1226 | 422.2 | 2.302E-05 | 0.05452 | +1 | 3 |
| - | - | 1408 | 423.2 | - | - | 0 | - |
| - | - | 7751 | 423.3 | - | - | 0 | - |
| - | - | 2274 | 424.3 | - | - | 0 | - |
| - | - | 632 | 426.2 | - | - | 0 | - |
| - | - | 5.53E+04 | 426.7 | - | - | 0 | - |
| - | - | 2.931E+04 | 427.2 | - | - | 0 | - |
| - | - | 8248 | 427.7 | - | - | 0 | - |
| - | - | 1532 | 428.2 | - | - | 0 | - |
| - | - | 1403 | 429.1 | - | - | 0 | - |
| - | - | 1.47E+04 | 429.2 | - | - | 0 | - |
| - | - | 4.728E+04 | 430.2 | - | - | 0 | - |
| - | - | 9451 | 431.2 | - | - | 0 | - |
| 7 | b | 6182 | 431.7 | 0.0005356 | 1.241 | +2 | 7 |
| 7 | b | 3612 | 432.2 | 0.00645 | 14.92 | +2 | 7 |
| - | - | 1224 | 432.7 | - | - | 0 | - |
| - | - | 1058 | 433 | - | - | 0 | - |
| 4 | b | 1860 | 433.3 | 0.0005982 | 1.381 | +1 | 4 |
| - | - | 944.7 | 438.2 | - | - | 0 | - |
| 11 | y | 1.416E+04 | 439.2 | 0.0001463 | 0.333 | +1 | 3 |
| - | - | 640.1 | 440.2 | - | - | 0 | - |
| - | - | 3705 | 440.2 | - | - | 0 | - |
| 7 | b | 2.698E+04 | 440.7 | 1.955E-05 | 0.04437 | +2 | 7 |
| - | - | 919.1 | 440.9 | - | - | 0 | - |
| - | - | 1.297E+04 | 441.2 | - | - | 0 | - |
| - | - | 3402 | 441.7 | - | - | 0 | - |
| - | - | 750.8 | 442.2 | - | - | 0 | - |
| - | - | 4018 | 444.2 | - | - | 0 | - |
| 7 | y | 2141 | 447.2 | 0.001687 | 3.773 | +2 | 7 |
| - | - | 2973 | 448.2 | - | - | 0 | - |
| - | - | 1130 | 449.2 | - | - | 0 | - |
| - | - | 1121 | 450.2 | - | - | 0 | - |
| - | - | 1254 | 450.3 | - | - | 0 | - |
| 4 | b | 1.602E+04 | 451.3 | 0.0001043 | 0.2311 | +1 | 4 |
| - | - | 3376 | 452.3 | - | - | 0 | - |
| - | - | 991.7 | 455.3 | - | - | 0 | - |
| 7 | y | 925.6 | 455.7 | 0.0007115 | 1.561 | +2 | 7 |
| - | - | 1259 | 459.2 | - | - | 0 | - |
| - | - | 2095 | 460.8 | - | - | 0 | - |
| - | - | 1492 | 461.2 | - | - | 0 | - |
| - | - | 1520 | 461.3 | - | - | 0 | - |
| - | - | 592.5 | 463.2 | - | - | 0 | - |
| - | - | 724.1 | 464.2 | - | - | 0 | - |
| - | - | 3389 | 468.3 | - | - | 0 | - |
| - | - | 923.2 | 469.3 | - | - | 0 | - |
| - | - | 2168 | 474.3 | - | - | 0 | - |
| - | - | 4190 | 474.8 | - | - | 0 | - |
| - | - | 2840 | 475.3 | - | - | 0 | - |
| - | - | 938.5 | 475.8 | - | - | 0 | - |
| - | - | 1833 | 476.2 | - | - | 0 | - |
| - | - | 3066 | 477.2 | - | - | 0 | - |
| 10 | y | 3102 | 478.2 | 0.0003768 | 0.7879 | +1 | 4 |
| - | - | 1816 | 478.3 | - | - | 0 | - |
| - | - | 630.8 | 479.3 | - | - | 0 | - |
| - | - | 9461 | 483.3 | - | - | 0 | - |
| - | - | 6941 | 483.8 | - | - | 0 | - |
| - | - | 606.7 | 484.2 | - | - | 0 | - |
| - | - | 1415 | 484.3 | - | - | 0 | - |
| 8 | b | 3605 | 488.3 | 0.0001786 | 0.3658 | +2 | 8 |
| 8 | b | 2554 | 488.7 | 0.006959 | 14.24 | +2 | 8 |
| - | - | 1004 | 489.3 | - | - | 0 | - |
| - | - | 1260 | 494.2 | - | - | 0 | - |
| - | - | 3.342E+04 | 494.2 | - | - | 0 | - |
| - | - | 8263 | 495.2 | - | - | 0 | - |
| 10 | y | 4.244E+04 | 496.2 | 0.0001059 | 0.2134 | +1 | 4 |
| - | - | 1.027E+04 | 497.2 | - | - | 0 | - |
| - | - | 3701 | 497.8 | - | - | 0 | - |
| - | - | 1364 | 498.2 | - | - | 0 | - |
| - | - | 2.726E+04 | 503.1 | - | - | 0 | - |
| - | - | 1050 | 504.2 | - | - | 0 | - |
| - | - | 3174 | 511.3 | - | - | 0 | - |
| - | - | 1082 | 513.2 | - | - | 0 | - |
| - | - | 647.8 | 514.2 | - | - | 0 | - |
| - | - | 8959 | 519.1 | - | - | 0 | - |
| - | - | 831.6 | 520.3 | - | - | 0 | - |
| - | - | 638.7 | 521.2 | - | - | 0 | - |
| - | - | 790.6 | 522.2 | - | - | 0 | - |
| - | - | 1593 | 523.3 | - | - | 0 | - |
| - | - | 5685 | 525.3 | - | - | 0 | - |
| - | - | 824.6 | 526.2 | - | - | 0 | - |
| - | - | 1974 | 526.3 | - | - | 0 | - |
| - | - | 889.5 | 528.9 | - | - | 0 | - |
| 0 | Precursor | 3225 | 529.3 | 0.0009064 | 1.713 | +3 | -1 |
| 0 | Precursor | 1867 | 529.6 | 0.004707 | 8.887 | +3 | -1 |
| - | - | 1355 | 529.9 | - | - | 0 | - |
| - | - | 1114 | 532.2 | - | - | 0 | - |
| - | - | 1121 | 534.6 | - | - | 0 | - |
| - | - | 923.9 | 535.3 | - | - | 0 | - |
| - | - | 1214 | 535.6 | - | - | 0 | - |
| - | - | 919.9 | 535.9 | - | - | 0 | - |
| - | - | 6376 | 536.2 | - | - | 0 | - |
| - | - | 5803 | 536.3 | - | - | 0 | - |
| - | - | 885.3 | 538.3 | - | - | 0 | - |
| - | - | 6094 | 539.2 | - | - | 0 | - |
| - | - | 1883 | 540.2 | - | - | 0 | - |
| - | - | 3282 | 542.3 | - | - | 0 | - |
| - | - | 9142 | 543.3 | - | - | 0 | - |
| - | - | 1928 | 544.3 | - | - | 0 | - |
| 9 | b | 1002 | 545.8 | 0.0009548 | 1.75 | +2 | 9 |
| - | - | 1226 | 546.8 | - | - | 0 | - |
| - | - | 1467 | 549.2 | - | - | 0 | - |
| - | - | 1946 | 551.3 | - | - | 0 | - |
| - | - | 876.1 | 551.3 | - | - | 0 | - |
| 9 | b | 1117 | 554.8 | 0.001782 | 3.211 | +2 | 9 |
| - | - | 2561 | 555.3 | - | - | 0 | - |
| - | - | 1616 | 555.8 | - | - | 0 | - |
| 5 | b | 1379 | 561.3 | 0.003269 | 5.825 | +1 | 5 |
| - | - | 1720 | 563.3 | - | - | 0 | - |
| - | - | 1128 | 564.3 | - | - | 0 | - |
| - | - | 2447 | 565.3 | - | - | 0 | - |
| - | - | 885.1 | 566.3 | - | - | 0 | - |
| - | - | 5521 | 567.2 | - | - | 0 | - |
| - | - | 2574 | 568.2 | - | - | 0 | - |
| - | - | 2426 | 575.2 | - | - | 0 | - |
| - | - | 827 | 576.2 | - | - | 0 | - |
| 5 | b | 1.168E+04 | 579.3 | 0.0002762 | 0.4768 | +1 | 5 |
| - | - | 2938 | 580.3 | - | - | 0 | - |
| - | - | 2233 | 592.3 | - | - | 0 | - |
| - | - | 1627 | 592.3 | - | - | 0 | - |
| 9 | y | 1.351E+04 | 593.3 | 0.0007085 | 1.194 | +1 | 5 |
| - | - | 3216 | 593.3 | - | - | 0 | - |
| - | - | 4860 | 594.3 | - | - | 0 | - |
| - | - | 1578 | 595.2 | - | - | 0 | - |
| - | - | 1035 | 597.8 | - | - | 0 | - |
| - | - | 999.6 | 598.3 | - | - | 0 | - |
| - | - | 2320 | 602.8 | - | - | 0 | - |
| - | - | 3180 | 603.3 | - | - | 0 | - |
| - | - | 761.1 | 603.8 | - | - | 0 | - |
| - | - | 3.89E+04 | 610.3 | - | - | 0 | - |
| 9 | y | 1.313E+05 | 611.3 | 0.0003235 | 0.5292 | +1 | 5 |
| - | - | 5919 | 611.8 | - | - | 0 | - |
| - | - | 4.117E+04 | 612.3 | - | - | 0 | - |
| - | - | 2021 | 612.8 | - | - | 0 | - |
| - | - | 9795 | 613.3 | - | - | 0 | - |
| - | - | 2290 | 621.3 | - | - | 0 | - |
| - | - | 960.3 | 621.3 | - | - | 0 | - |
| - | - | 635.2 | 622.4 | - | - | 0 | - |
| - | - | 1552 | 626.3 | - | - | 0 | - |
| - | - | 929.1 | 628.3 | - | - | 0 | - |
| - | - | 1470 | 634.3 | - | - | 0 | - |
| - | - | 3139 | 635.3 | - | - | 0 | - |
| - | - | 1070 | 636.3 | - | - | 0 | - |
| 4 | y | 1252 | 636.8 | 0.0006095 | 0.9572 | +2 | 10 |
| 4 | y | 1164 | 637.3 | 0.004514 | 7.083 | +2 | 10 |
| - | - | 900.5 | 637.3 | - | - | 0 | - |
| - | - | 1213 | 640.3 | - | - | 0 | - |
| - | - | 2146 | 645.3 | - | - | 0 | - |
| 4 | y | 8384 | 645.8 | 0.001371 | 2.123 | +2 | 10 |
| - | - | 5073 | 646.3 | - | - | 0 | - |
| - | - | 2702 | 646.8 | - | - | 0 | - |
| - | - | 727.6 | 648.3 | - | - | 0 | - |
| - | - | 1134 | 650.3 | - | - | 0 | - |
| - | - | 1.359E+04 | 652.3 | - | - | 0 | - |
| - | - | 4169 | 653.3 | - | - | 0 | - |
| - | - | 1182 | 654.3 | - | - | 0 | - |
| - | - | 776.9 | 658.3 | - | - | 0 | - |
| - | - | 1483 | 660.9 | - | - | 0 | - |
| - | - | 1537 | 661.4 | - | - | 0 | - |
| - | - | 4175 | 662.3 | - | - | 0 | - |
| - | - | 2716 | 663.3 | - | - | 0 | - |
| - | - | 937.2 | 664.3 | - | - | 0 | - |
| - | - | 838.9 | 664.3 | - | - | 0 | - |
| - | - | 868.1 | 670.8 | - | - | 0 | - |
| 6 | b | 6825 | 676.3 | 0.0005413 | 0.8003 | +1 | 6 |
| 6 | b | 4071 | 677.3 | 0.004396 | 6.49 | +1 | 6 |
| - | - | 1766 | 678.3 | - | - | 0 | - |
| - | - | 1021 | 679.3 | - | - | 0 | - |
| - | - | 2.7E+04 | 680.3 | - | - | 0 | - |
| - | - | 9486 | 681.3 | - | - | 0 | - |
| - | - | 2269 | 682.3 | - | - | 0 | - |
| - | - | 1056 | 684.3 | - | - | 0 | - |
| - | - | 2351 | 684.8 | - | - | 0 | - |
| - | - | 1098 | 685.3 | - | - | 0 | - |
| - | - | 1158 | 685.8 | - | - | 0 | - |
| - | - | 1170 | 688.3 | - | - | 0 | - |
| - | - | 791.8 | 692.8 | - | - | 0 | - |
| 3 | y | 7612 | 693.3 | 3.899E-05 | 0.05624 | +2 | 11 |
| 3 | y | 9547 | 693.8 | 0.007282 | 10.5 | +2 | 11 |
| 6 | b | 7.061E+04 | 694.4 | 0.001218 | 1.755 | +1 | 6 |
| - | - | 751.8 | 694.8 | - | - | 0 | - |
| - | - | 2.531E+04 | 695.4 | - | - | 0 | - |
| - | - | 5601 | 696.4 | - | - | 0 | - |
| - | - | 1715 | 697.4 | - | - | 0 | - |
| - | - | 623.1 | 699.3 | - | - | 0 | - |
| - | - | 1276 | 701.3 | - | - | 0 | - |
| - | - | 1.234E+04 | 701.8 | - | - | 0 | - |
| 3 | y | 5.046E+04 | 702.3 | 0.001148 | 1.635 | +2 | 11 |
| 12 | b | 3.826E+04 | 702.8 | 0.01351 | 19.22 | +2 | 12 |
| 12 | b | 2.066E+04 | 703.3 | 0.00454 | 6.455 | +2 | 12 |
| - | - | 5581 | 703.8 | - | - | 0 | - |
| - | - | 3624 | 704.3 | - | - | 0 | - |
| - | - | 1115 | 705.4 | - | - | 0 | - |
| 8 | y | 5639 | 706.3 | 0.0001274 | 0.1804 | +1 | 6 |
| - | - | 2662 | 707.3 | - | - | 0 | - |
| - | - | 1411 | 710.4 | - | - | 0 | - |
| - | - | 679.2 | 710.9 | - | - | 0 | - |
| - | - | 4348 | 711.4 | - | - | 0 | - |
| 12 | b | 4236 | 711.9 | 0.002632 | 3.697 | +2 | 12 |
| - | - | 2883 | 712.4 | - | - | 0 | - |
| - | - | 1575 | 712.9 | - | - | 0 | - |
| - | - | 3598 | 721.4 | - | - | 0 | - |
| - | - | 1038 | 722.4 | - | - | 0 | - |
| - | - | 1.034E+04 | 723.4 | - | - | 0 | - |
| 8 | y | 4.125E+04 | 724.4 | 0.0001218 | 0.1682 | +1 | 6 |
| - | - | 1.747E+04 | 725.4 | - | - | 0 | - |
| - | - | 4045 | 726.4 | - | - | 0 | - |
| - | - | 1623 | 729.4 | - | - | 0 | - |
| - | - | 2387 | 733.4 | - | - | 0 | - |
| - | - | 1983 | 734.4 | - | - | 0 | - |
| - | - | 746.9 | 735.3 | - | - | 0 | - |
| 2 | y | 1555 | 743.4 | 0.01336 | 17.97 | +2 | 12 |
| - | - | 667.4 | 747.4 | - | - | 0 | - |
| - | - | 1976 | 748.4 | - | - | 0 | - |
| - | - | 8798 | 751.4 | - | - | 0 | - |
| 2 | y | 3751 | 751.9 | 0.0007549 | 1.004 | +2 | 12 |
| - | - | 5970 | 752.4 | - | - | 0 | - |
| - | - | 1711 | 752.9 | - | - | 0 | - |
| - | - | 1723 | 753.4 | - | - | 0 | - |
| - | - | 811.8 | 760.3 | - | - | 0 | - |
| - | - | 990 | 761.4 | - | - | 0 | - |
| - | - | 994.2 | 762.3 | - | - | 0 | - |
| - | - | 959.2 | 762.4 | - | - | 0 | - |
| - | - | 4389 | 763.4 | - | - | 0 | - |
| - | - | 2200 | 764.4 | - | - | 0 | - |
| - | - | 2650 | 765.4 | - | - | 0 | - |
| - | - | 1121 | 767.3 | - | - | 0 | - |
| - | - | 972.7 | 768.3 | - | - | 0 | - |
| - | - | 854.7 | 769.4 | - | - | 0 | - |
| - | - | 928.3 | 776.4 | - | - | 0 | - |
| - | - | 3801 | 779.4 | - | - | 0 | - |
| - | - | 1622 | 780.4 | - | - | 0 | - |
| - | - | 2184 | 781.4 | - | - | 0 | - |
| - | - | 1143 | 782.4 | - | - | 0 | - |
| 0 | Precursor | 7894 | 793.4 | 0.01156 | 14.57 | +2 | -1 |
| - | - | 3633 | 794.4 | - | - | 0 | - |
| - | - | 1442 | 795.3 | - | - | 0 | - |
| - | - | 751.4 | 795.4 | - | - | 0 | - |
| - | - | 1654 | 807.4 | - | - | 0 | - |
| - | - | 886.5 | 818.4 | - | - | 0 | - |
| - | - | 1124 | 825.4 | - | - | 0 | - |
| - | - | 1566 | 834.4 | - | - | 0 | - |
| - | - | 2370 | 835.4 | - | - | 0 | - |
| - | - | 4944 | 836.4 | - | - | 0 | - |
| - | - | 2526 | 837.5 | - | - | 0 | - |
| - | - | 1109 | 843.4 | - | - | 0 | - |
| - | - | 1805 | 844.4 | - | - | 0 | - |
| - | - | 1425 | 845.4 | - | - | 0 | - |
| - | - | 2346 | 846.4 | - | - | 0 | - |
| - | - | 1432 | 847.4 | - | - | 0 | - |
| - | - | 3116 | 851.4 | - | - | 0 | - |
| - | - | 1.618E+04 | 852.4 | - | - | 0 | - |
| - | - | 9483 | 853.4 | - | - | 0 | - |
| - | - | 2781 | 854.4 | - | - | 0 | - |
| 7 | b | 1.093E+04 | 862.4 | 0.001912 | 2.217 | +1 | 7 |
| 7 | b | 9101 | 863.4 | 0.01041 | 12.06 | +1 | 7 |
| - | - | 1.294E+04 | 864.5 | - | - | 0 | - |
| - | - | 6384 | 865.5 | - | - | 0 | - |
| - | - | 1965 | 866.5 | - | - | 0 | - |
| - | - | 1254 | 876.4 | - | - | 0 | - |
| - | - | 966.4 | 877.4 | - | - | 0 | - |
| - | - | 2224 | 878.4 | - | - | 0 | - |
| - | - | 788.6 | 879.4 | - | - | 0 | - |
| 7 | b | 8.287E+04 | 880.4 | 0.001735 | 1.97 | +1 | 7 |
| - | - | 4.519E+04 | 881.4 | - | - | 0 | - |
| - | - | 636.6 | 881.5 | - | - | 0 | - |
| - | - | 732 | 881.5 | - | - | 0 | - |
| - | - | 1.419E+04 | 882.4 | - | - | 0 | - |
| - | - | 3219 | 883.4 | - | - | 0 | - |
| - | - | 1528 | 892.5 | - | - | 0 | - |
| - | - | 850.2 | 893.5 | - | - | 0 | - |
| - | - | 921.5 | 907.4 | - | - | 0 | - |
| - | - | 5936 | 908.4 | - | - | 0 | - |
| - | - | 3999 | 909.4 | - | - | 0 | - |
| 7 | y | 6165 | 910.4 | 0.0009441 | 1.037 | +1 | 7 |
| - | - | 2758 | 911.4 | - | - | 0 | - |
| - | - | 796.6 | 936.4 | - | - | 0 | - |
| - | - | 1560 | 954.4 | - | - | 0 | - |
| - | - | 2193 | 955.4 | - | - | 0 | - |
| - | - | 831.1 | 964.5 | - | - | 0 | - |
| - | - | 2227 | 965.5 | - | - | 0 | - |
| - | - | 794.8 | 966.4 | - | - | 0 | - |
| - | - | 1699 | 972.4 | - | - | 0 | - |
| 8 | b | 993 | 975.5 | 0.001792 | 1.837 | +1 | 8 |
| 8 | b | 5275 | 993.5 | 0.0002275 | 0.229 | +1 | 8 |
| - | - | 3526 | 994.5 | - | - | 0 | - |
| - | - | 782 | 995.5 | - | - | 0 | - |
| - | - | 1452 | 1024 | - | - | 0 | - |
| 6 | y | 3622 | 1025 | 0.003485 | 3.398 | +1 | 8 |
| - | - | 1730 | 1026 | - | - | 0 | - |
| - | - | 1093 | 1076 | - | - | 0 | - |
| - | - | 2018 | 1094 | - | - | 0 | - |
| - | - | 879.8 | 1095 | - | - | 0 | - |
| - | - | 998.7 | 1096 | - | - | 0 | - |
| 9 | b | 1039 | 1109 | 0.007375 | 6.653 | +1 | 9 |
| - | - | 982.7 | 1118 | - | - | 0 | - |
| - | - | 681.1 | 1119 | - | - | 0 | - |
| 5 | y | 3043 | 1136 | 0.001925 | 1.696 | +1 | 9 |
| 5 | y | 3988 | 1136 | 0.007467 | 6.571 | +1 | 9 |
| - | - | 1903 | 1137 | - | - | 0 | - |
| - | - | 2140 | 1153 | - | - | 0 | - |
| 5 | y | 6197 | 1154 | 0.002969 | 2.573 | +1 | 9 |
| - | - | 3401 | 1155 | - | - | 0 | - |
| - | - | 1508 | 1156 | - | - | 0 | - |
| - | - | 638.9 | 1429 | - | - | 0 | - |
| - | - | 872.6 | 3080 | - | - | 0 | - |

m/z Charge Intensity FragmentType MassShift Position
120.06575775146484 0 4268.7593
120.08104705810547 0 6489.8516
121.08455657958984 0 471.49005
122.07140350341797 0 851.7858
123.04428100585938 0 955.6674
126.09154510498047 0 551.45636
127.05049896240234 0 3130.9563
128.1072235107422 0 11248.9795
129.0659942626953 0 1165.6168
129.10252380371094 0 141937.17
130.0653839111328 0 63176.203
130.06971740722656 0 687.3955
130.10006713867188 0 706.43665
130.10580444335938 0 9185.159
131.06874084472656 0 6957.2183
132.08099365234375 0 12501.938
132.10194396972656 0 425.07388
133.0610809326172 0 572.4028
133.08456420898438 0 1285.8226
133.14089965820312 0 371.52353
134.0269012451172 0 400.21393
136.075927734375 0 20185.307
137.0792236328125 0 1783.6339
137.0968475341797 0 464.7549
138.0663299560547 0 2006.3236
138.09173583984375 0 738.0401
139.08668518066406 0 1636.2133
141.10252380371094 0 3163.9448
142.06591796875 0 517.8014
143.04515075683594 0 862.38684
143.1181182861328 0 585.201
144.08119201660156 0 655.8961
145.0609130859375 0 1540.4952
146.06027221679688 0 1005.1711
147.044189453125 0 806.16064
149.0384521484375 0 982.18964
149.04510498046875 0 15591.563
150.06655883789062 0 975.5662
150.98753356933594 0 419.03207
151.08682250976562 0 3912.9573
152.08999633789062 0 460.18573
155.08160400390625 0 1933.9741
155.0929412841797 0 8437.807
155.1181182861328 0 6033.402 a Water loss 1
156.09632873535156 0 695.8425
156.12135314941406 0 618.67865
157.09762573242188 0 613.1927
158.0836639404297 0 662.2378
159.09188842773438 0 107793.57
160.07574462890625 0 971.6641
160.08946228027344 0 440.97742
160.09523010253906 0 11415.052
161.09799194335938 0 635.1415
165.05482482910156 0 9220.039
165.0774383544922 0 887.93585
166.06126403808594 0 15948.901
166.0865020751953 0 3079.935 y 9
166.0978546142578 0 1267.1276
167.0556182861328 0 19479.615
167.0639190673828 0 840.4685
168.0548095703125 0 1044.5466
168.08106994628906 0 2179.9353
168.10218811035156 0 1307.967
168.11351013183594 0 1631.0663
169.05224609375 0 639.3026
169.0760955810547 0 1402.358
169.09739685058594 0 4733.608
170.06021118164062 0 4988.945
171.07638549804688 0 646.90076
171.09193420410156 0 844.5407
171.14930725097656 0 1347.5104
172.07186889648438 0 1939.3265
173.05601501464844 0 1904.4738
173.1287384033203 0 205406.05 a 1
173.45034790039062 0 3189.198
174.1320343017578 0 18019.729
175.13427734375 0 1492.308
178.13404846191406 0 6138.828
179.13723754882812 0 999.30774
181.06077575683594 0 1367.4706
182.08135986328125 0 20963.43 y 12
183.08497619628906 0 2171.236
183.11305236816406 0 5233.641 b Water loss 1
183.14895629882812 0 627.6496
185.07217407226562 0 733.3496
185.16531372070312 0 1652.705
186.12391662597656 0 43714.34
187.0867919921875 0 8766.974
187.1272735595703 0 3910.3003
187.14459228515625 0 949.7676
188.0706024169922 0 1493.6407
188.09078979492188 0 731.7292
189.0984649658203 0 781.88245
190.08262634277344 0 1971.7063
190.13424682617188 0 1363.7935
194.0817413330078 0 422.77414
194.0924530029297 0 1211.6173
195.11294555664062 0 10830.098
196.108154296875 0 7429.7173
196.1165008544922 0 745.917
197.12860107421875 0 1845.7767
197.76683044433594 0 503.0285
197.79295349121094 0 520.2676
198.08749389648438 0 1270.3054
198.12771606445312 0 3628.1086
199.07142639160156 0 708.6999
199.0867156982422 0 546.7653
199.13063049316406 0 478.92444
200.1421661376953 0 707.74274
201.1234893798828 0 59570.918 b 1
202.08665466308594 0 742.9835
202.12681579589844 0 6424.9355
203.12841796875 0 748.8548
205.097412109375 0 4577.533
205.1450958251953 0 802.79535
206.10055541992188 0 885.8126
206.1284637451172 0 760.8899
207.16123962402344 0 943.20703
209.0926971435547 0 706.0729
210.12767028808594 0 1836.3407
211.10791015625 0 1492.4849 y Water loss 10
211.14413452148438 0 788.9959
211.46197509765625 0 505.64978
212.06748962402344 0 508.48325
212.1394500732422 0 1554.2369
213.12350463867188 0 5970.779
213.16055297851562 0 854.28845
215.1394500732422 0 1502.2134
216.09837341308594 0 1444.4932
219.13404846191406 0 1999.1842
221.05882263183594 0 2861.0022
221.07179260253906 0 998.5232
221.0845184326172 0 7675.154
221.10365295410156 0 1173.4662
221.14002990722656 0 1015.8088
222.08493041992188 0 2546.6045
222.1239013671875 0 1499.7345
223.0637664794922 0 5592.563
223.08201599121094 0 1012.8459
223.10781860351562 0 3727.4875
223.15548706054688 0 19680.506
224.10313415527344 0 1263.4943
224.15878295898438 0 1613.1967
225.04310607910156 0 8601.6
225.1601104736328 0 591.7161
226.04322814941406 0 2473.7896
226.08229064941406 0 4368.238
226.1188507080078 0 3961.0562 b Water loss 5
226.15394592285156 0 669.3574
227.02236938476562 0 931.6912
227.0403594970703 0 1649.8013
227.0664520263672 0 3374.2961
227.15524291992188 0 1212.9034
228.1344451904297 0 2295.9917
229.1182861328125 0 2915.9746
229.15480041503906 0 3943.697
230.12088012695312 0 838.877
233.13983154296875 0 3955.7063
234.12387084960938 0 20038.965
234.14361572265625 0 945.00555
235.10789489746094 0 900.06915
235.1271209716797 0 2119.0508
235.6051483154297 0 561.10913
237.13510131835938 0 1030.0485
238.11859130859375 0 2960.5122
238.1302032470703 0 715.2665
239.0950927734375 0 19578.26
240.0955047607422 0 6815.392
240.1343536376953 0 9096.802
241.09255981445312 0 4006.585
241.1182861328125 0 1323.7428
241.13729858398438 0 497.3406
241.19168090820312 0 1238.1943
243.1095733642578 0 1127.4015
244.09291076660156 0 7896.205
247.10789489746094 0 11499.096
248.10275268554688 0 2280.1848
248.11387634277344 0 4269.4683
249.05422973632812 0 1009.4653
249.09844970703125 0 7369.76
249.13470458984375 0 2404.009
250.10107421875 0 1000.0009
251.15040588378906 0 66656.94
252.15359497070312 0 8558.324
253.1552734375 0 1003.97577
255.14797973632812 0 4690.5654
256.1081237792969 0 853.9275
256.1526794433594 0 854.75165
256.61114501953125 0 630.4474
257.1129150390625 0 2093.4998
258.1448974609375 0 3685.4497
259.1450500488281 0 584.98804
261.1195068359375 0 1739.9274
261.13446044921875 0 896.0392
261.16015625 0 679.62134
262.11907958984375 0 704.57263
265.1294250488281 0 9128.63
265.1438293457031 0 951.84955
266.1249084472656 0 16918.688
267.1279296875 0 2412.2385
267.1445617675781 0 646.3682
268.1286926269531 0 978.04
268.17645263671875 0 1442.939
269.1856384277344 0 1441.4745
270.12176513671875 0 2963.1582
272.17584228515625 0 4258.629
274.118408203125 0 3635.1052
274.132080078125 0 1188.4587
275.1017150878906 0 975.485
276.10888671875 0 827.5051
276.15545654296875 0 1322.711
279.1454772949219 0 1532.2615
280.1297912597656 0 861.37787
281.05133056640625 0 14336.277
282.1564025878906 0 4478.9756
283.0299987792969 0 2236.0662
283.1402282714844 0 54478.45
284.10284423828125 0 1566.656
284.1243591308594 0 766.88495
284.14337158203125 0 7927.78
285.0101318359375 0 795.9696
285.15716552734375 0 858.88574
286.1398620605469 0 4682.327
286.156494140625 0 815.0403
287.1431884765625 0 530.82935
287.187744140625 0 635.99786
288.6192321777344 0 760.391
293.1133117675781 0 28727.756 y Water loss 11
293.1614074707031 0 668.8592
294.11724853515625 0 5202.85
295.1031188964844 0 1560.1072
295.1199035644531 0 875.73883
296.103759765625 0 1894.9579
296.19696044921875 0 1068.0297 b Water loss 2
297.0826110839844 0 2697.2534
297.1006164550781 0 1041.0786
297.1557922363281 0 8462.773
297.1928405761719 0 1459.9336
298.1401672363281 0 1252.2856 y Water loss 6
298.1586608886719 0 1395.1349
299.0618591308594 0 111692.55
300.0624084472656 0 3713.7349
300.16748046875 0 8121.8467
301.059326171875 0 2429.2332
301.1508483886719 0 23239.854
301.16912841796875 0 958.54395
302.1138000488281 0 4190.5034
302.1539611816406 0 3745.3582
303.1163635253906 0 762.34564
305.6448669433594 0 1125.4294
306.1375427246094 0 3157.785 y 8
306.6383361816406 0 1396.8392
309.1682434082031 0 577.8596
310.15289306640625 0 908.69775
311.1241455078125 0 4381.5503 y 11
312.12677001953125 0 795.7179
314.11578369140625 0 726.68286
314.18865966796875 0 566.2033
314.20703125 0 2599.5303 b 2
315.1664123535156 0 14388.84
315.1856689453125 0 1088.0999
316.1501770019531 0 1174.4418
316.1693420410156 0 2221.7297
317.13238525390625 0 553.1231
319.1531066894531 0 853.5799
324.1718444824219 0 635.8071
325.1514587402344 0 831.9722
325.1868591308594 0 2124.349
326.66400146484375 0 4537.3433
326.9669189453125 0 657.3145
327.1654052734375 0 1797.6628
332.207763671875 0 1853.2039
333.1770324707031 0 2707.5027
338.181884765625 0 525.31744
338.673583984375 0 622.02045 b Water loss 5
340.6605529785156 0 1157.1394
341.01800537109375 0 1083.697
342.18109130859375 0 1275.2467
343.1619873046875 0 959.92883
343.1846618652344 0 610.2498
344.1717224121094 0 1165.7988
344.97650146484375 0 14424.306
345.1310729980469 0 2455.1553
345.173828125 0 793.30524
346.1145935058594 0 9904.236
346.6963806152344 0 742.8481
347.117431640625 0 1113.5002
347.6793518066406 0 4884.353 b 5
348.181396484375 0 840.47174
349.1510314941406 0 712.63605
350.13446044921875 0 1421.7283
350.21875 0 5848.2393
351.222412109375 0 1071.3385
352.1632995605469 0 900.77844
353.6736145019531 0 845.09607 y Water loss 7
355.070068359375 0 2223.148
356.070068359375 0 662.12744
356.1910705566406 0 838.86554
359.0282287597656 0 11453.284
360.6949462890625 0 1114.4443
361.1966552734375 0 1957.5098
362.1848449707031 0 3609.1118
362.6789855957031 0 9803.115 y 7
363.1408996582031 0 12588.727
363.1814880371094 0 3869.3967
363.6801452636719 0 689.83923
364.1244812011719 0 1842.6335
364.1441955566406 0 1766.4539
366.17694091796875 0 1221.1387
367.13916015625 0 1358.2781
367.2438049316406 0 818.496
369.1226501464844 0 1023.4413
370.1233825683594 0 2011.2849
371.1205139160156 0 1114.1614
373.18560791015625 0 710.583
374.2022399902344 0 599.03815
376.1611633300781 0 1038.6844
376.19677734375 0 1232.4751
377.1443176269531 0 997.48816
378.21343994140625 0 5371.487
379.20867919921875 0 18026.129
380.1626892089844 0 787.26575
380.211669921875 0 4159.986
381.1517028808594 0 33189.625
381.71087646484375 0 1194.9213
382.1545715332031 0 6738.993
382.18914794921875 0 1613.4547
382.2149353027344 0 682.3059
384.164794921875 0 1181.4412
384.1890869140625 0 1140.3171
385.14990234375 0 2601.8003
385.17303466796875 0 862.4449 y 4
386.153564453125 0 798.5407
390.19439697265625 0 1108.439
391.1360168457031 0 671.8489
394.172119140625 0 5611.9067
395.13409423828125 0 2258.5432
395.1570739746094 0 1038.8837
395.2394714355469 0 1865.4528
396.1398010253906 0 934.2603
396.2240295410156 0 6333.117
397.22723388671875 0 1058.408
398.1775207519531 0 3351.9333
401.21331787109375 0 999.16016
402.1761169433594 0 2572.3176
402.19927978515625 0 3405.5376
403.1999816894531 0 1441.9296
405.25897216796875 0 804.4157
409.1467590332031 0 2618.8682
411.1993713378906 0 3095.8362
412.182861328125 0 19617.69
413.1439208984375 0 1412.2744
413.16644287109375 0 2334.4504
413.188720703125 0 1810.2611
413.25030517578125 0 1216.8749
414.23480224609375 0 6088.302
415.0367736816406 0 24179.764
415.1979064941406 0 1870.6567
415.2381896972656 0 1542.4331
417.716796875 0 4019.4053
418.20989990234375 0 6828.4033
418.71112060546875 0 2914.3804
421.2099914550781 0 748.67413 y Water loss 10
422.19219970703125 0 1226.017 y Ammonia loss 10
423.2030944824219 0 1407.8147
423.2714538574219 0 7751.4785
424.27288818359375 0 2274.0818
426.199951171875 0 631.9525
426.7218017578125 0 55300.074
427.2232360839844 0 29312.52
427.7244873046875 0 8247.666
428.2279968261719 0 1531.7051
429.08905029296875 0 1403.1904
429.20941162109375 0 14696.86
430.1934814453125 0 47283.58
431.1970520019531 0 9451.332
431.7134094238281 0 6181.769 b Water loss 6
432.21240234375 0 3612.4414 b Ammonia loss 6
432.7081298828125 0 1224.1953
433.04736328125 0 1058.2538
433.2563781738281 0 1859.5271 b Water loss 3
438.173095703125 0 944.7422
439.2188720703125 0 14164.483 y 10
440.1826477050781 0 640.06366
440.2201232910156 0 3704.7266
440.7192077636719 0 26979.986 b 6
440.9195556640625 0 919.08954
441.2209167480469 0 12973.788
441.7218933105469 0 3401.5903
442.2205505371094 0 750.82745
444.24505615234375 0 4018.3481
447.2073059082031 0 2140.8596 y Ammonia loss 6
448.2037658691406 0 2973.3833
449.20635986328125 0 1130.0745
450.23577880859375 0 1120.5443
450.28131103515625 0 1253.6196
451.2664489746094 0 16018.999 b 3
452.2692565917969 0 3375.938
455.2662048339844 0 991.71533
455.7196044921875 0 925.59064 y 6
459.1994323730469 0 1258.943
460.75347900390625 0 2094.6301
461.2027282714844 0 1491.6919
461.2532043457031 0 1520.0447
463.2197570800781 0 592.4706
464.2140808105469 0 724.1251
468.2928466796875 0 3388.8274
469.2947082519531 0 923.21844
474.2582702636719 0 2167.768
474.7536315917969 0 4190.0415
475.25579833984375 0 2839.9014
475.7586975097656 0 938.5251
476.224609375 0 1833.1796
477.2088928222656 0 3065.777
478.229248046875 0 3102.2866 y Water loss 9
478.2756042480469 0 1815.8463
479.2804260253906 0 630.7807
483.2639465332031 0 9460.604
483.7651062011719 0 6940.927
484.21942138671875 0 606.71387
484.265625 0 1415.123
488.25579833984375 0 3605.0823 b Water loss 7
488.75494384765625 0 2554.062 b Ammonia loss 7
489.2502136230469 0 1004.0805
494.1944274902344 0 1259.6368
494.2356872558594 0 33417.395
495.2386169433594 0 8263.053
496.24029541015625 0 42435.812 y 9
497.2458190917969 0 10270.946
497.76226806640625 0 3700.818
498.2402648925781 0 1363.981
503.1072998046875 0 27263.37
504.198486328125 0 1049.7506
511.2620544433594 0 3173.697
513.1998291015625 0 1081.791
514.1918334960938 0 647.7529
519.1384887695312 0 8958.586
520.267822265625 0 831.5847
521.2269897460938 0 638.6758
522.2125854492188 0 790.5814
523.2572631835938 0 1592.9421
525.2662353515625 0 5685.3857
526.2272338867188 0 824.6081
526.2686157226562 0 1973.5272
528.9290161132812 0 889.48016
529.2587890625 0 3224.879 Precursor Water loss
529.5924072265625 0 1867.1715 Precursor Ammonia loss
529.9271850585938 0 1355.0968
532.1929931640625 0 1113.5618
534.634521484375 0 1120.5778
535.3020629882812 0 923.86926
535.5968017578125 0 1214.1475
535.9307250976562 0 919.8636
536.1650390625 0 6376.1333
536.2686157226562 0 5802.8213
538.2701416015625 0 885.345
539.2357788085938 0 6094.0923
540.2380981445312 0 1883.4807
542.2921142578125 0 3282.3628
543.276611328125 0 9142.203
544.2804565429688 0 1928.0952
545.7684936523438 0 1002.2697 b Water loss 8
546.7506103515625 0 1225.5635
549.220703125 0 1466.998
551.2566528320312 0 1945.8263
551.3301391601562 0 876.0646
554.77294921875 0 1117.3541 b 8
555.2548217773438 0 2561.2554
555.7568969726562 0 1615.9612
561.317626953125 0 1379.1417 b Water loss 4
563.256591796875 0 1719.6827
564.2658081054688 0 1127.8875
565.2635498046875 0 2447.3315
566.2648315429688 0 885.11774
567.230712890625 0 5521.3877
568.2337646484375 0 2573.5342
575.246826171875 0 2425.7798
576.2310791015625 0 826.99835
579.3246459960938 0 11684.5 b 4
580.3262329101562 0 2938.2822
592.2715454101562 0 2232.7961
592.3236083984375 0 1626.8748
593.255859375 0 13509.741 y Water loss 8
593.3045654296875 0 3216.0308
594.25341796875 0 4859.712
595.2489624023438 0 1578.0347
597.799560546875 0 1034.5758
598.295654296875 0 999.64795
602.7926025390625 0 2319.8784
603.2886962890625 0 3179.9026
603.7936401367188 0 761.0757
610.28271484375 0 38899.453
611.2674560546875 0 131261.69 y 8
611.7981567382812 0 5918.5317
612.2703857421875 0 41168.723
612.7979125976562 0 2021.4347
613.272705078125 0 9794.601
621.2507934570312 0 2289.5874
621.3021850585938 0 960.264
622.3652954101562 0 635.24286
626.2813720703125 0 1551.9044
628.2775268554688 0 929.08734
634.3031005859375 0 1469.6322
635.2946166992188 0 3139.2168
636.2877197265625 0 1069.6249
636.7852172851562 0 1251.7312 y Water loss 3
637.2823486328125 0 1163.9921 y Ammonia loss 3
637.3438720703125 0 900.4908
640.2706298828125 0 1213.3905
645.2919311523438 0 2146.2349
645.79248046875 0 8383.535 y 3
646.2919921875 0 5072.9053
646.7947998046875 0 2701.811
648.3424072265625 0 727.5724
650.3240966796875 0 1133.7714
652.3189086914062 0 13585.741
653.322021484375 0 4169.249
654.3199462890625 0 1182.2839
658.2848510742188 0 776.92395
660.8704223632812 0 1483.4656
661.3727416992188 0 1537.0493
662.304443359375 0 4175.221
663.2982788085938 0 2715.5186
664.2902221679688 0 937.1543
664.34716796875 0 838.89496
670.7987060546875 0 868.0565
676.3407592773438 0 6824.687 b Water loss 5
677.3297119140625 0 4070.7458 b Ammonia loss 5
678.3043212890625 0 1765.8265
679.3339233398438 0 1021.1654
680.314697265625 0 26997.568
681.316650390625 0 9486.328
682.3192749023438 0 2269.054
684.3217163085938 0 1056.3138
684.81787109375 0 2351.4492
685.3226318359375 0 1097.6637
685.817138671875 0 1158.357
688.330078125 0 1170.4607
692.8368530273438 0 791.7701
693.3278198242188 0 7612.113 y Water loss 2
693.8271484375 0 9546.967 y Ammonia loss 2
694.3506469726562 0 70606.81 b 5
694.824951171875 0 751.8214
695.3535766601562 0 25314.201
696.3548583984375 0 5600.655
697.3566284179688 0 1714.9943
699.3411254882812 0 623.08215
701.3363037109375 0 1276.0345
701.83984375 0 12343.937
702.3342895507812 0 50461.625 y 2
702.83544921875 0 38256.73 b Water loss 11
703.33642578125 0 20662.613 b Ammonia loss 11
703.8379516601562 0 5580.907
704.3355712890625 0 3623.8247
705.3611450195312 0 1114.6986
706.3407592773438 0 5638.979 y Water loss 7
707.3433837890625 0 2662.3232
710.4044799804688 0 1411.186
710.9112548828125 0 679.2126
711.3682250976562 0 4347.6636
711.8568725585938 0 4236.088 b 11
712.3573608398438 0 2882.5925
712.8566284179688 0 1575.4873
721.3609619140625 0 3598.2827
722.36767578125 0 1037.5645
723.36669921875 0 10342.755
724.351318359375 0 41249.832 y 7
725.3538818359375 0 17474.584
726.3567504882812 0 4044.619
729.3600463867188 0 1622.6699
733.3602294921875 0 2386.757
734.35400390625 0 1983.3622
735.340576171875 0 746.9328
743.367431640625 0 1554.7852 y Ammonia loss 1
747.382568359375 0 667.4373
748.3822021484375 0 1976.2125
751.3753051757812 0 8798.318
751.8681030273438 0 3750.682 y 1
752.3727416992188 0 5970.0083
752.8707885742188 0 1710.5933
753.3721923828125 0 1722.5021
760.3071899414062 0 811.80994
761.3698120117188 0 990.02814
762.3494262695312 0 994.1813
762.4193115234375 0 959.2201
763.3629760742188 0 4389.0933
764.3659057617188 0 2199.8755
765.4033203125 0 2649.5303
767.3444213867188 0 1120.5502
768.3456420898438 0 972.7342
769.3963012695312 0 854.67267
776.3793334960938 0 928.28375
779.3828735351562 0 3801.1157
780.3743896484375 0 1622.3397
781.3653564453125 0 2183.557
782.36279296875 0 1142.925
793.3974609375 0 7894.337 Precursor Water loss
794.39990234375 0 3632.6958
795.3348999023438 0 1442.3619
795.4102172851562 0 751.38293
807.3779907226562 0 1653.5198
818.4463500976562 0 886.5283
825.38671875 0 1124.3721
834.421630859375 0 1566.2311
835.4129638671875 0 2370.1072
836.4458618164062 0 4943.975
837.4556884765625 0 2525.9229
843.3997802734375 0 1108.9669
844.3947143554688 0 1804.9075
845.38525390625 0 1424.8088
846.4462890625 0 2346.156
847.4435424804688 0 1432.0509
851.3906860351562 0 3116.4492
852.4337158203125 0 16180.922
853.4368286132812 0 9482.939
854.4411010742188 0 2780.9343
862.418701171875 0 10928.632 b Water loss 6
863.4150390625 0 9100.984 b Ammonia loss 6
864.4542236328125 0 12938.266
865.458251953125 0 6384.128
866.4610595703125 0 1964.7776
876.4376831054688 0 1254.4381
877.4339599609375 0 966.36743
878.41552734375 0 2224.1963
879.4212646484375 0 788.55237
880.429443359375 0 82872.14 b 6
881.4318237304688 0 45186.266
881.5088500976562 0 636.5915
881.5352783203125 0 732.0116
882.4348754882812 0 14186.282
883.4385986328125 0 3219.228
892.466064453125 0 1527.6106
893.4736328125 0 850.2358
907.44189453125 0 921.4672
908.4267578125 0 5935.589
909.4336547851562 0 3999.4526
910.4295654296875 0 6165.242 y 6
911.4330444335938 0 2758.4585
936.4194946289062 0 796.59436
954.4293823242188 0 1560.1069
955.425537109375 0 2192.5781
964.45751953125 0 831.1234
965.4501342773438 0 2226.5674
966.4439086914062 0 794.783
972.4381103515625 0 1698.9521
975.5064697265625 0 993.0094 b Water loss 7
993.5150146484375 0 5274.7925 b 7
994.5147094726562 0 3525.5962
995.5057373046875 0 782.0401
1024.4678955078125 0 1452.3369
1025.4609375 0 3621.7644 y 5
1026.4588623046875 0 1730.133
1075.5294189453125 0 1092.5552
1093.5450439453125 0 2018.269
1094.536376953125 0 879.79877
1095.537841796875 0 998.67163
1108.549560546875 0 1039.4375 b 8
1118.4873046875 0 982.65186
1119.478271484375 0 681.05383
1135.5035400390625 0 3042.577 y Water loss 4
1136.4969482421875 0 3988.484 y Ammonia loss 4
1137.48876953125 0 1903.1298
1152.5308837890625 0 2139.8586
1153.5130615234375 0 6196.909 y 4
1154.517578125 0 3401.191
1155.5185546875 0 1508.149
1428.8629150390625 0 638.86786
3080.07080078125 0 872.6028

Spectrum Details

|  |  |
| --- | --- |
| Matched peaks? Matched peaks The total absolute number of peaks matched. Additionally in brackets the total fraction of peaks matched and the total number of peaks is shown. | 69 (10.33% of 668) |
| FDR? FDR The false discovery rate estimated for this peptide. It is calculated by matching all theoretical fragments with a non-integer shift with the raw peaks for this spectrum. This is done with 40 different shifts. The resulting percentage is the average number of annotated peaks over the number of annotated peaks with the correct spectrum. | 0.90% |
| Satellite FDR? Satellite FDR See the FDR for details on its calculation. This satellite ion specific FDR only contains the satellite ions (d/w) for I/L/J positions. | ∞ |
| PSM Score? PSM Score The PSM Score as given by Hecklib to this annotated spectrum. It is shown with three significant figures. | 521 |

#### Spectrum 7777? Spectrum 7777 The raw spectrum of this peptide as annotated by Hecklib. The fragments are coloured according to ion type (see legend). Any peaks with a star '\*' as text can be hovered over to see the full details, first the ion type second the mass shift type. By hovering over the amino acids in the peptide or ions in the legend the corresponding peaks are highlighted. By toggling the 'Unassigned' label you can turn the background (unassigned) peaks on or off in the plot. By updating the slider in the Ion legend you can update the spectrum to only show the top X% of the peaks with labels. The top X% means any peak that is within X% of the highest intensity. By dragging in the spectrum you can zoom in to a specific part of the spectrum and use 'Zoom Out' to get back to the original zoom level. The annotation of the spectrum is based on the given sequence in the peptides file and is done with different software so inconsistencies are likely. The peaks are annotated based on the given sequence, with 20 ppm tolerance.

Copy Data

##### Spectrum 7777 (TSV)

###### Preview

```
Loading example...
```

*Click on the button to copy the data to your clipboard.*

Mz MinMz MaxIntensity Max

WidthHeightPeptide font sizePeptide stroke widthSpectrum font sizeSpectrum stroke widthCompact peptide

Ion legend

wxyz

abcd

OtherUnassignedIon^Charge^_Position_Show for top:%

TVLHQDWLDGKEY

04.43e+48.85e+41.33e+51.77e+5

Zoom Out

a^+1^_2_y^+3^_4_a^+1^_2_y^+1^_1_b^+1^_2_b^+1^_2_y^+2^_3_b^+3^_6_y^+1^_2_b^+1^_3_y^+3^_7_y^+2^_5_y^+1^_2_b^+1^_3_b^+2^_6_b^+2^_6_y^+2^_6_y^+2^_6_y^+3^_9_y^+1^_3_b^+2^_7_b^+2^_7_b^+1^_4_y^+1^_3_b^+2^_7_y^+2^_7_b^+1^_4_y^+1^_4_b^+2^_8_b^+2^_8_y^+1^_4_\*\*\*b^+2^_9_b^+1^_5_b^+1^_5_y^+1^_5_y^+1^_5_y^+2^_10_y^+2^_10_b^+1^_6_b^+1^_6_y^+2^_11_y^+2^_11_b^+1^_6_y^+2^_11_b^+2^_12_b^+2^_12_y^+1^_6_b^+2^_12_y^+1^_6_y^+2^_12_y^+2^_12_y^+2^_12_\*b^+1^_7_b^+1^_7_b^+1^_7_y^+1^_7_b^+1^_8_b^+1^_8_b^+1^_8_y^+1^_8_y^+1^_9_y^+1^_9_y^+1^_9_

0778155623343112

Fragment Matches Table

Show background peaks

| Position | Ion type | Intensity | mz Theoretical | mz Error (Th) | mz Error (ppm) | Charge | Series Number |
| --- | --- | --- | --- | --- | --- | --- | --- |
| - | - | 700.4 | 120.1 | - | - | 0 | - |
| - | - | 6911 | 120.1 | - | - | 0 | - |
| - | - | 605.6 | 121.1 | - | - | 0 | - |
| - | - | 615.1 | 122.1 | - | - | 0 | - |
| - | - | 388 | 122.9 | - | - | 0 | - |
| - | - | 758 | 123 | - | - | 0 | - |
| - | - | 395.5 | 124.4 | - | - | 0 | - |
| - | - | 616.6 | 125.1 | - | - | 0 | - |
| - | - | 587.4 | 126.1 | - | - | 0 | - |
| - | - | 2424 | 127.1 | - | - | 0 | - |
| - | - | 348.6 | 127.5 | - | - | 0 | - |
| - | - | 1.168E+04 | 128.1 | - | - | 0 | - |
| - | - | 1240 | 129.1 | - | - | 0 | - |
| - | - | 1.188E+05 | 129.1 | - | - | 0 | - |
| - | - | 5.076E+04 | 130.1 | - | - | 0 | - |
| - | - | 646.4 | 130.1 | - | - | 0 | - |
| - | - | 518.3 | 130.1 | - | - | 0 | - |
| - | - | 1059 | 130.1 | - | - | 0 | - |
| - | - | 9144 | 130.1 | - | - | 0 | - |
| - | - | 4845 | 131.1 | - | - | 0 | - |
| - | - | 417.6 | 131.1 | - | - | 0 | - |
| - | - | 1.196E+04 | 132.1 | - | - | 0 | - |
| - | - | 1545 | 133.1 | - | - | 0 | - |
| - | - | 596.3 | 134 | - | - | 0 | - |
| - | - | 955.1 | 136.1 | - | - | 0 | - |
| - | - | 1.777E+04 | 136.1 | - | - | 0 | - |
| - | - | 1687 | 137.1 | - | - | 0 | - |
| - | - | 1742 | 138.1 | - | - | 0 | - |
| - | - | 747.8 | 138.1 | - | - | 0 | - |
| - | - | 1378 | 139.1 | - | - | 0 | - |
| - | - | 3874 | 141.1 | - | - | 0 | - |
| - | - | 600.3 | 142.1 | - | - | 0 | - |
| - | - | 527.1 | 143 | - | - | 0 | - |
| - | - | 524.7 | 144.1 | - | - | 0 | - |
| - | - | 1465 | 145.1 | - | - | 0 | - |
| - | - | 1153 | 146.1 | - | - | 0 | - |
| - | - | 1021 | 147 | - | - | 0 | - |
| - | - | 526.9 | 147.1 | - | - | 0 | - |
| - | - | 1.39E+04 | 149 | - | - | 0 | - |
| - | - | 600.5 | 150 | - | - | 0 | - |
| - | - | 1149 | 150.1 | - | - | 0 | - |
| - | - | 790 | 151 | - | - | 0 | - |
| - | - | 2709 | 151.1 | - | - | 0 | - |
| - | - | 922.4 | 153.1 | - | - | 0 | - |
| - | - | 548.4 | 154.1 | - | - | 0 | - |
| - | - | 462.9 | 155 | - | - | 0 | - |
| - | - | 6314 | 155.1 | - | - | 0 | - |
| 2 | a | 5911 | 155.1 | 0.0002288 | 1.475 | +1 | 2 |
| - | - | 509.4 | 156.1 | - | - | 0 | - |
| - | - | 763.2 | 156.1 | - | - | 0 | - |
| - | - | 457.5 | 157.1 | - | - | 0 | - |
| - | - | 971.1 | 157.1 | - | - | 0 | - |
| - | - | 984.3 | 157.1 | - | - | 0 | - |
| - | - | 919.8 | 158.1 | - | - | 0 | - |
| - | - | 444.2 | 158.5 | - | - | 0 | - |
| - | - | 9.244E+04 | 159.1 | - | - | 0 | - |
| - | - | 729.4 | 160.1 | - | - | 0 | - |
| - | - | 525.9 | 160.1 | - | - | 0 | - |
| - | - | 1.172E+04 | 160.1 | - | - | 0 | - |
| - | - | 7615 | 165.1 | - | - | 0 | - |
| - | - | 1.355E+04 | 166.1 | - | - | 0 | - |
| 10 | y | 5065 | 166.1 | 0.001649 | 9.928 | +3 | 4 |
| - | - | 1007 | 166.1 | - | - | 0 | - |
| - | - | 1.953E+04 | 167.1 | - | - | 0 | - |
| - | - | 1177 | 167.1 | - | - | 0 | - |
| - | - | 933.3 | 168.1 | - | - | 0 | - |
| - | - | 1480 | 168.1 | - | - | 0 | - |
| - | - | 750.8 | 168.1 | - | - | 0 | - |
| - | - | 1144 | 168.1 | - | - | 0 | - |
| - | - | 553.6 | 169.1 | - | - | 0 | - |
| - | - | 1212 | 169.1 | - | - | 0 | - |
| - | - | 549.3 | 169.1 | - | - | 0 | - |
| - | - | 4051 | 169.1 | - | - | 0 | - |
| - | - | 4267 | 170.1 | - | - | 0 | - |
| - | - | 741.8 | 171.1 | - | - | 0 | - |
| - | - | 1126 | 171.1 | - | - | 0 | - |
| - | - | 1555 | 172.1 | - | - | 0 | - |
| - | - | 1476 | 173.1 | - | - | 0 | - |
| 2 | a | 1.753E+05 | 173.1 | 0.0003147 | 1.818 | +1 | 2 |
| - | - | 546.3 | 174.1 | - | - | 0 | - |
| - | - | 1.53E+04 | 174.1 | - | - | 0 | - |
| - | - | 528.2 | 177.1 | - | - | 0 | - |
| - | - | 493.7 | 177.1 | - | - | 0 | - |
| - | - | 5030 | 178.1 | - | - | 0 | - |
| - | - | 614.3 | 179.1 | - | - | 0 | - |
| - | - | 1145 | 181.1 | - | - | 0 | - |
| - | - | 923.7 | 182.1 | - | - | 0 | - |
| 13 | y | 1.884E+04 | 182.1 | 0.0002512 | 1.38 | +1 | 1 |
| - | - | 1542 | 183.1 | - | - | 0 | - |
| 2 | b | 5396 | 183.1 | 0.0001872 | 1.022 | +1 | 2 |
| - | - | 807.7 | 183.1 | - | - | 0 | - |
| - | - | 560.2 | 185.1 | - | - | 0 | - |
| - | - | 1483 | 185.2 | - | - | 0 | - |
| - | - | 3.398E+04 | 186.1 | - | - | 0 | - |
| - | - | 7511 | 187.1 | - | - | 0 | - |
| - | - | 2409 | 187.1 | - | - | 0 | - |
| - | - | 3837 | 187.1 | - | - | 0 | - |
| - | - | 1311 | 188.1 | - | - | 0 | - |
| - | - | 1054 | 190.1 | - | - | 0 | - |
| - | - | 899.6 | 190.1 | - | - | 0 | - |
| - | - | 1686 | 194.1 | - | - | 0 | - |
| - | - | 8871 | 195.1 | - | - | 0 | - |
| - | - | 5646 | 196.1 | - | - | 0 | - |
| - | - | 837.6 | 196.1 | - | - | 0 | - |
| - | - | 1278 | 197.1 | - | - | 0 | - |
| - | - | 1634 | 198.1 | - | - | 0 | - |
| - | - | 2754 | 198.1 | - | - | 0 | - |
| - | - | 866.8 | 199.1 | - | - | 0 | - |
| - | - | 961.2 | 199.1 | - | - | 0 | - |
| - | - | 733.7 | 200.1 | - | - | 0 | - |
| 2 | b | 4.827E+04 | 201.1 | 0.0001968 | 0.9787 | +1 | 2 |
| - | - | 1197 | 202.1 | - | - | 0 | - |
| - | - | 4858 | 202.1 | - | - | 0 | - |
| - | - | 515.9 | 203.1 | - | - | 0 | - |
| - | - | 4378 | 205.1 | - | - | 0 | - |
| - | - | 857 | 205.1 | - | - | 0 | - |
| - | - | 664.6 | 206.1 | - | - | 0 | - |
| - | - | 677.4 | 206.1 | - | - | 0 | - |
| - | - | 1434 | 207.2 | - | - | 0 | - |
| - | - | 536.2 | 208.1 | - | - | 0 | - |
| - | - | 728 | 209.1 | - | - | 0 | - |
| - | - | 1510 | 210.1 | - | - | 0 | - |
| 11 | y | 1056 | 211.1 | 0.0002982 | 1.412 | +2 | 3 |
| - | - | 779.2 | 212.1 | - | - | 0 | - |
| - | - | 4539 | 213.1 | - | - | 0 | - |
| - | - | 840.1 | 213.2 | - | - | 0 | - |
| - | - | 2118 | 215.1 | - | - | 0 | - |
| - | - | 1177 | 216.1 | - | - | 0 | - |
| - | - | 1931 | 221.1 | - | - | 0 | - |
| - | - | 911.2 | 221.1 | - | - | 0 | - |
| - | - | 7593 | 221.1 | - | - | 0 | - |
| - | - | 1149 | 221.1 | - | - | 0 | - |
| - | - | 601.7 | 221.1 | - | - | 0 | - |
| - | - | 596 | 221.6 | - | - | 0 | - |
| - | - | 531.3 | 222 | - | - | 0 | - |
| - | - | 1509 | 222.1 | - | - | 0 | - |
| - | - | 1501 | 222.1 | - | - | 0 | - |
| - | - | 4253 | 223.1 | - | - | 0 | - |
| - | - | 1148 | 223.1 | - | - | 0 | - |
| - | - | 2822 | 223.1 | - | - | 0 | - |
| - | - | 1.551E+04 | 223.2 | - | - | 0 | - |
| - | - | 969.8 | 224.1 | - | - | 0 | - |
| - | - | 1985 | 224.2 | - | - | 0 | - |
| - | - | 9682 | 225 | - | - | 0 | - |
| - | - | 2979 | 226 | - | - | 0 | - |
| - | - | 3036 | 226.1 | - | - | 0 | - |
| 6 | b | 745 | 226.1 | 4.177E-05 | 0.1847 | +3 | 6 |
| - | - | 899.8 | 226.2 | - | - | 0 | - |
| - | - | 2510 | 227 | - | - | 0 | - |
| - | - | 2468 | 227.1 | - | - | 0 | - |
| - | - | 718 | 227.2 | - | - | 0 | - |
| - | - | 1551 | 228.1 | - | - | 0 | - |
| - | - | 2624 | 229.1 | - | - | 0 | - |
| - | - | 3824 | 229.2 | - | - | 0 | - |
| - | - | 1536 | 233.1 | - | - | 0 | - |
| - | - | 575.6 | 233.2 | - | - | 0 | - |
| - | - | 1.697E+04 | 234.1 | - | - | 0 | - |
| - | - | 1845 | 235.1 | - | - | 0 | - |
| - | - | 1253 | 237.1 | - | - | 0 | - |
| - | - | 3274 | 238.1 | - | - | 0 | - |
| - | - | 1.498E+04 | 239.1 | - | - | 0 | - |
| - | - | 5274 | 240.1 | - | - | 0 | - |
| - | - | 7741 | 240.1 | - | - | 0 | - |
| - | - | 2984 | 241.1 | - | - | 0 | - |
| - | - | 862.7 | 241.1 | - | - | 0 | - |
| - | - | 797.7 | 241.1 | - | - | 0 | - |
| - | - | 717.6 | 241.2 | - | - | 0 | - |
| - | - | 801.4 | 243.1 | - | - | 0 | - |
| - | - | 7966 | 244.1 | - | - | 0 | - |
| - | - | 8373 | 247.1 | - | - | 0 | - |
| - | - | 620.1 | 247.1 | - | - | 0 | - |
| - | - | 1064 | 248.1 | - | - | 0 | - |
| - | - | 3693 | 248.1 | - | - | 0 | - |
| - | - | 939.4 | 249.1 | - | - | 0 | - |
| - | - | 5907 | 249.1 | - | - | 0 | - |
| - | - | 1803 | 249.1 | - | - | 0 | - |
| - | - | 622.9 | 250.1 | - | - | 0 | - |
| - | - | 608.9 | 250.2 | - | - | 0 | - |
| - | - | 5.664E+04 | 251.2 | - | - | 0 | - |
| - | - | 6675 | 252.2 | - | - | 0 | - |
| - | - | 1003 | 253.2 | - | - | 0 | - |
| - | - | 2844 | 255.1 | - | - | 0 | - |
| - | - | 1078 | 256.1 | - | - | 0 | - |
| - | - | 489 | 257.1 | - | - | 0 | - |
| - | - | 2180 | 257.1 | - | - | 0 | - |
| - | - | 1847 | 258.1 | - | - | 0 | - |
| - | - | 492.7 | 260.2 | - | - | 0 | - |
| - | - | 1433 | 261.1 | - | - | 0 | - |
| - | - | 580.6 | 262.1 | - | - | 0 | - |
| - | - | 6676 | 265.1 | - | - | 0 | - |
| - | - | 624.9 | 265.1 | - | - | 0 | - |
| - | - | 1728 | 266.1 | - | - | 0 | - |
| - | - | 1.452E+04 | 266.1 | - | - | 0 | - |
| - | - | 541.2 | 267 | - | - | 0 | - |
| - | - | 549.5 | 267 | - | - | 0 | - |
| - | - | 2258 | 267.1 | - | - | 0 | - |
| - | - | 600.6 | 267.1 | - | - | 0 | - |
| - | - | 934.3 | 268.1 | - | - | 0 | - |
| - | - | 1396 | 268.2 | - | - | 0 | - |
| - | - | 825.3 | 269.2 | - | - | 0 | - |
| - | - | 850.1 | 269.2 | - | - | 0 | - |
| - | - | 2658 | 270.1 | - | - | 0 | - |
| - | - | 918.6 | 270.6 | - | - | 0 | - |
| - | - | 3237 | 272.2 | - | - | 0 | - |
| - | - | 911 | 273.2 | - | - | 0 | - |
| - | - | 3012 | 274.1 | - | - | 0 | - |
| - | - | 790.5 | 274.1 | - | - | 0 | - |
| - | - | 820.4 | 275.1 | - | - | 0 | - |
| - | - | 974.2 | 276.2 | - | - | 0 | - |
| - | - | 1244 | 279.1 | - | - | 0 | - |
| - | - | 1738 | 280.1 | - | - | 0 | - |
| - | - | 1.287E+04 | 281.1 | - | - | 0 | - |
| - | - | 4212 | 282.2 | - | - | 0 | - |
| - | - | 1748 | 283 | - | - | 0 | - |
| - | - | 4.598E+04 | 283.1 | - | - | 0 | - |
| - | - | 1721 | 284.1 | - | - | 0 | - |
| - | - | 5810 | 284.1 | - | - | 0 | - |
| - | - | 819 | 285 | - | - | 0 | - |
| - | - | 1000 | 285.2 | - | - | 0 | - |
| - | - | 2617 | 286.1 | - | - | 0 | - |
| - | - | 928.5 | 287.2 | - | - | 0 | - |
| - | - | 773.2 | 288.6 | - | - | 0 | - |
| 12 | y | 2.147E+04 | 293.1 | 0.0002358 | 0.8043 | +1 | 2 |
| - | - | 650.3 | 293.2 | - | - | 0 | - |
| - | - | 4569 | 294.1 | - | - | 0 | - |
| - | - | 1882 | 295.1 | - | - | 0 | - |
| - | - | 705.1 | 295.1 | - | - | 0 | - |
| - | - | 1206 | 296.1 | - | - | 0 | - |
| 3 | b | 1253 | 296.2 | 0.0001533 | 0.5177 | +1 | 3 |
| - | - | 2080 | 297.1 | - | - | 0 | - |
| - | - | 1010 | 297.1 | - | - | 0 | - |
| - | - | 6947 | 297.2 | - | - | 0 | - |
| 7 | y | 819.5 | 298.1 | 0.00494 | 16.57 | +3 | 7 |
| - | - | 1186 | 298.2 | - | - | 0 | - |
| - | - | 1.073E+05 | 299.1 | - | - | 0 | - |
| - | - | 3043 | 300.1 | - | - | 0 | - |
| - | - | 7279 | 300.2 | - | - | 0 | - |
| - | - | 2529 | 301.1 | - | - | 0 | - |
| - | - | 1.834E+04 | 301.2 | - | - | 0 | - |
| - | - | 715.1 | 301.2 | - | - | 0 | - |
| - | - | 4811 | 302.1 | - | - | 0 | - |
| - | - | 2379 | 302.2 | - | - | 0 | - |
| - | - | 841.1 | 305.2 | - | - | 0 | - |
| - | - | 1653 | 305.6 | - | - | 0 | - |
| 9 | y | 5241 | 306.1 | 0.0003077 | 1.005 | +2 | 5 |
| - | - | 1668 | 306.6 | - | - | 0 | - |
| 12 | y | 3445 | 311.1 | 0.0001691 | 0.5435 | +1 | 2 |
| - | - | 832.7 | 313.1 | - | - | 0 | - |
| - | - | 796.8 | 314.2 | - | - | 0 | - |
| 3 | b | 2338 | 314.2 | 0.0003308 | 1.053 | +1 | 3 |
| - | - | 1.237E+04 | 315.2 | - | - | 0 | - |
| - | - | 648.1 | 315.2 | - | - | 0 | - |
| - | - | 2042 | 316.2 | - | - | 0 | - |
| - | - | 741.2 | 317.7 | - | - | 0 | - |
| - | - | 761.1 | 323.2 | - | - | 0 | - |
| - | - | 719.2 | 325.2 | - | - | 0 | - |
| - | - | 2501 | 326.7 | - | - | 0 | - |
| - | - | 865.9 | 327.2 | - | - | 0 | - |
| - | - | 634.6 | 332.2 | - | - | 0 | - |
| - | - | 3301 | 333.2 | - | - | 0 | - |
| 6 | b | 674.7 | 338.7 | 8.894E-05 | 0.2626 | +2 | 6 |
| - | - | 682.1 | 339.7 | - | - | 0 | - |
| - | - | 960 | 340.7 | - | - | 0 | - |
| - | - | 1368 | 341 | - | - | 0 | - |
| - | - | 1118 | 342.2 | - | - | 0 | - |
| - | - | 1026 | 343.2 | - | - | 0 | - |
| - | - | 1.3E+04 | 345 | - | - | 0 | - |
| - | - | 2162 | 345.1 | - | - | 0 | - |
| - | - | 6492 | 346.1 | - | - | 0 | - |
| - | - | 544.6 | 346.1 | - | - | 0 | - |
| - | - | 871.3 | 347.1 | - | - | 0 | - |
| 6 | b | 2947 | 347.7 | 5.412E-06 | 0.01557 | +2 | 6 |
| - | - | 743 | 349.2 | - | - | 0 | - |
| - | - | 4796 | 350.2 | - | - | 0 | - |
| - | - | 882.8 | 351.2 | - | - | 0 | - |
| 8 | y | 1560 | 353.7 | 0.0007059 | 1.996 | +2 | 6 |
| - | - | 686.2 | 354.2 | - | - | 0 | - |
| - | - | 2163 | 355.1 | - | - | 0 | - |
| - | - | 815.8 | 356.1 | - | - | 0 | - |
| - | - | 782.6 | 356.2 | - | - | 0 | - |
| - | - | 1.423E+04 | 359 | - | - | 0 | - |
| - | - | 503.6 | 359 | - | - | 0 | - |
| - | - | 1801 | 360.7 | - | - | 0 | - |
| - | - | 1896 | 361.2 | - | - | 0 | - |
| - | - | 2388 | 362.2 | - | - | 0 | - |
| 8 | y | 8615 | 362.7 | 9.831E-05 | 0.2711 | +2 | 6 |
| - | - | 1.08E+04 | 363.1 | - | - | 0 | - |
| - | - | 2314 | 363.2 | - | - | 0 | - |
| - | - | 1680 | 364.1 | - | - | 0 | - |
| - | - | 1265 | 364.1 | - | - | 0 | - |
| - | - | 1408 | 366.2 | - | - | 0 | - |
| - | - | 1597 | 367.1 | - | - | 0 | - |
| - | - | 863.8 | 369.1 | - | - | 0 | - |
| - | - | 1062 | 370.1 | - | - | 0 | - |
| - | - | 689.8 | 371.1 | - | - | 0 | - |
| - | - | 593.6 | 373.2 | - | - | 0 | - |
| - | - | 898 | 376.2 | - | - | 0 | - |
| - | - | 1644 | 376.2 | - | - | 0 | - |
| - | - | 898 | 377.1 | - | - | 0 | - |
| - | - | 3967 | 378.2 | - | - | 0 | - |
| - | - | 1.672E+04 | 379.2 | - | - | 0 | - |
| - | - | 2762 | 380.2 | - | - | 0 | - |
| - | - | 2.951E+04 | 381.2 | - | - | 0 | - |
| - | - | 664.2 | 381.7 | - | - | 0 | - |
| - | - | 5075 | 382.2 | - | - | 0 | - |
| - | - | 1104 | 382.2 | - | - | 0 | - |
| - | - | 1580 | 384.2 | - | - | 0 | - |
| - | - | 1508 | 384.2 | - | - | 0 | - |
| - | - | 1047 | 385.1 | - | - | 0 | - |
| 5 | y | 979.7 | 385.2 | 0.002484 | 6.448 | +3 | 9 |
| - | - | 1449 | 390.2 | - | - | 0 | - |
| - | - | 1120 | 391.1 | - | - | 0 | - |
| - | - | 796.2 | 392 | - | - | 0 | - |
| - | - | 5035 | 394.2 | - | - | 0 | - |
| - | - | 1407 | 395.1 | - | - | 0 | - |
| - | - | 941.8 | 395.2 | - | - | 0 | - |
| - | - | 1362 | 395.2 | - | - | 0 | - |
| - | - | 6145 | 396.2 | - | - | 0 | - |
| - | - | 916.7 | 397.2 | - | - | 0 | - |
| - | - | 2170 | 398.2 | - | - | 0 | - |
| - | - | 692.1 | 399 | - | - | 0 | - |
| - | - | 941.6 | 401.2 | - | - | 0 | - |
| - | - | 1688 | 402.2 | - | - | 0 | - |
| - | - | 1590 | 402.2 | - | - | 0 | - |
| - | - | 831.7 | 403.2 | - | - | 0 | - |
| - | - | 1121 | 405.3 | - | - | 0 | - |
| - | - | 1580 | 409.1 | - | - | 0 | - |
| - | - | 1236 | 409.2 | - | - | 0 | - |
| - | - | 1959 | 411.2 | - | - | 0 | - |
| - | - | 1.901E+04 | 412.2 | - | - | 0 | - |
| - | - | 1385 | 413.1 | - | - | 0 | - |
| - | - | 2145 | 413.2 | - | - | 0 | - |
| - | - | 4934 | 414.2 | - | - | 0 | - |
| - | - | 2.7E+04 | 415 | - | - | 0 | - |
| - | - | 1781 | 415.2 | - | - | 0 | - |
| - | - | 1180 | 415.2 | - | - | 0 | - |
| - | - | 1868 | 417.7 | - | - | 0 | - |
| - | - | 5842 | 418.2 | - | - | 0 | - |
| - | - | 2378 | 418.7 | - | - | 0 | - |
| 11 | y | 1002 | 421.2 | 0.000274 | 0.6504 | +1 | 3 |
| - | - | 714.4 | 422.7 | - | - | 0 | - |
| - | - | 1182 | 423.2 | - | - | 0 | - |
| - | - | 7488 | 423.3 | - | - | 0 | - |
| - | - | 1692 | 424.3 | - | - | 0 | - |
| - | - | 897.1 | 424.7 | - | - | 0 | - |
| - | - | 4.731E+04 | 426.7 | - | - | 0 | - |
| - | - | 2.651E+04 | 427.2 | - | - | 0 | - |
| - | - | 5775 | 427.7 | - | - | 0 | - |
| - | - | 1473 | 428.2 | - | - | 0 | - |
| - | - | 920.4 | 429.1 | - | - | 0 | - |
| - | - | 1.283E+04 | 429.2 | - | - | 0 | - |
| - | - | 655.5 | 429.6 | - | - | 0 | - |
| - | - | 4.028E+04 | 430.2 | - | - | 0 | - |
| - | - | 9716 | 431.2 | - | - | 0 | - |
| 7 | b | 4801 | 431.7 | 0.0002884 | 0.6681 | +2 | 7 |
| 7 | b | 2984 | 432.2 | 0.002665 | 6.167 | +2 | 7 |
| - | - | 611 | 432.7 | - | - | 0 | - |
| - | - | 673.6 | 432.7 | - | - | 0 | - |
| - | - | 1547 | 433 | - | - | 0 | - |
| - | - | 1011 | 433.2 | - | - | 0 | - |
| 4 | b | 1188 | 433.3 | 0.002338 | 5.396 | +1 | 4 |
| 11 | y | 1.217E+04 | 439.2 | 0.0002073 | 0.472 | +1 | 3 |
| - | - | 822 | 439.7 | - | - | 0 | - |
| - | - | 2604 | 440.2 | - | - | 0 | - |
| 7 | b | 2.22E+04 | 440.7 | 1.096E-05 | 0.02487 | +2 | 7 |
| - | - | 1.265E+04 | 441.2 | - | - | 0 | - |
| - | - | 983.5 | 441.3 | - | - | 0 | - |
| - | - | 2904 | 441.7 | - | - | 0 | - |
| 7 | y | 744 | 447.2 | 0.005014 | 11.21 | +2 | 7 |
| - | - | 2007 | 448.2 | - | - | 0 | - |
| - | - | 728.9 | 448.3 | - | - | 0 | - |
| - | - | 597.6 | 449.2 | - | - | 0 | - |
| - | - | 690.4 | 450.3 | - | - | 0 | - |
| - | - | 635 | 451.2 | - | - | 0 | - |
| 4 | b | 1.482E+04 | 451.3 | 1.778E-05 | 0.03941 | +1 | 4 |
| - | - | 2930 | 452.3 | - | - | 0 | - |
| - | - | 948.5 | 455.2 | - | - | 0 | - |
| - | - | 1265 | 455.3 | - | - | 0 | - |
| - | - | 1036 | 459.2 | - | - | 0 | - |
| - | - | 2211 | 460.8 | - | - | 0 | - |
| - | - | 1254 | 461.2 | - | - | 0 | - |
| - | - | 1224 | 461.3 | - | - | 0 | - |
| - | - | 666.1 | 464.2 | - | - | 0 | - |
| - | - | 3329 | 468.3 | - | - | 0 | - |
| - | - | 1025 | 469.3 | - | - | 0 | - |
| - | - | 3423 | 474.3 | - | - | 0 | - |
| - | - | 3897 | 474.8 | - | - | 0 | - |
| - | - | 1136 | 475.3 | - | - | 0 | - |
| - | - | 1000 | 475.3 | - | - | 0 | - |
| - | - | 1504 | 476.2 | - | - | 0 | - |
| - | - | 2150 | 477.2 | - | - | 0 | - |
| 10 | y | 1672 | 478.2 | 0.001384 | 2.894 | +1 | 4 |
| - | - | 1664 | 478.3 | - | - | 0 | - |
| - | - | 693.6 | 479.2 | - | - | 0 | - |
| - | - | 9716 | 483.3 | - | - | 0 | - |
| - | - | 5135 | 483.8 | - | - | 0 | - |
| - | - | 955.2 | 484.2 | - | - | 0 | - |
| - | - | 1614 | 484.3 | - | - | 0 | - |
| 8 | b | 2133 | 488.3 | 0.0009505 | 1.947 | +2 | 8 |
| 8 | b | 1711 | 488.7 | 0.00406 | 8.307 | +2 | 8 |
| - | - | 2.696E+04 | 494.2 | - | - | 0 | - |
| - | - | 7092 | 495.2 | - | - | 0 | - |
| 10 | y | 3.534E+04 | 496.2 | 0.000228 | 0.4594 | +1 | 4 |
| - | - | 8731 | 497.2 | - | - | 0 | - |
| - | - | 4553 | 497.8 | - | - | 0 | - |
| - | - | 1685 | 498.2 | - | - | 0 | - |
| - | - | 2.71E+04 | 503.1 | - | - | 0 | - |
| - | - | 1569 | 504.2 | - | - | 0 | - |
| - | - | 1649 | 506.2 | - | - | 0 | - |
| - | - | 935.7 | 507.3 | - | - | 0 | - |
| - | - | 2054 | 511.3 | - | - | 0 | - |
| - | - | 8751 | 519.1 | - | - | 0 | - |
| - | - | 812.6 | 521.8 | - | - | 0 | - |
| - | - | 1460 | 523.3 | - | - | 0 | - |
| - | - | 4911 | 525.3 | - | - | 0 | - |
| - | - | 837 | 526.3 | - | - | 0 | - |
| - | - | 780.3 | 528.9 | - | - | 0 | - |
| 0 | Precursor | 1399 | 529.3 | 0.002878 | 5.437 | +3 | -1 |
| 0 | Precursor | 1414 | 529.6 | 0.004768 | 9.002 | +3 | -1 |
| - | - | 1355 | 532.2 | - | - | 0 | - |
| - | - | 711.8 | 534.3 | - | - | 0 | - |
| - | - | 856.5 | 534.6 | - | - | 0 | - |
| 0 | Precursor | 1261 | 535.3 | 0.001254 | 2.343 | +3 | -1 |
| - | - | 1379 | 535.6 | - | - | 0 | - |
| - | - | 5862 | 536.2 | - | - | 0 | - |
| - | - | 1340 | 536.3 | - | - | 0 | - |
| - | - | 5996 | 539.2 | - | - | 0 | - |
| - | - | 2140 | 540.2 | - | - | 0 | - |
| - | - | 2443 | 542.3 | - | - | 0 | - |
| - | - | 7523 | 543.3 | - | - | 0 | - |
| - | - | 2672 | 544.3 | - | - | 0 | - |
| - | - | 945.3 | 545.3 | - | - | 0 | - |
| - | - | 1252 | 546.2 | - | - | 0 | - |
| - | - | 1576 | 549.2 | - | - | 0 | - |
| - | - | 1245 | 551.3 | - | - | 0 | - |
| 9 | b | 1070 | 554.8 | 0.00874 | 15.75 | +2 | 9 |
| - | - | 1738 | 555.3 | - | - | 0 | - |
| - | - | 1015 | 555.8 | - | - | 0 | - |
| 5 | b | 1956 | 561.3 | 0.001369 | 2.439 | +1 | 5 |
| - | - | 2105 | 565.3 | - | - | 0 | - |
| - | - | 5265 | 567.2 | - | - | 0 | - |
| - | - | 1747 | 568.2 | - | - | 0 | - |
| - | - | 2141 | 575.2 | - | - | 0 | - |
| - | - | 874.1 | 576.2 | - | - | 0 | - |
| 5 | b | 9043 | 579.3 | 2.897E-05 | 0.05001 | +1 | 5 |
| - | - | 3465 | 580.3 | - | - | 0 | - |
| - | - | 1070 | 582.3 | - | - | 0 | - |
| - | - | 697.4 | 589.3 | - | - | 0 | - |
| - | - | 1640 | 592.3 | - | - | 0 | - |
| - | - | 1591 | 592.3 | - | - | 0 | - |
| 9 | y | 1.256E+04 | 593.3 | 0.0002812 | 0.4741 | +1 | 5 |
| - | - | 3687 | 593.3 | - | - | 0 | - |
| - | - | 3935 | 594.3 | - | - | 0 | - |
| - | - | 1905 | 595.2 | - | - | 0 | - |
| - | - | 901.8 | 596.3 | - | - | 0 | - |
| - | - | 1242 | 602.8 | - | - | 0 | - |
| - | - | 1689 | 603.3 | - | - | 0 | - |
| - | - | 813 | 603.8 | - | - | 0 | - |
| - | - | 785.4 | 604.3 | - | - | 0 | - |
| - | - | 3.118E+04 | 610.3 | - | - | 0 | - |
| - | - | 1080 | 610.3 | - | - | 0 | - |
| 9 | y | 1.068E+05 | 611.3 | 0.0004456 | 0.7289 | +1 | 5 |
| - | - | 5172 | 611.8 | - | - | 0 | - |
| - | - | 3.511E+04 | 612.3 | - | - | 0 | - |
| - | - | 1258 | 612.8 | - | - | 0 | - |
| - | - | 7150 | 613.3 | - | - | 0 | - |
| - | - | 1351 | 614.3 | - | - | 0 | - |
| - | - | 797.5 | 620.8 | - | - | 0 | - |
| - | - | 1223 | 621.2 | - | - | 0 | - |
| - | - | 1004 | 622.4 | - | - | 0 | - |
| - | - | 1031 | 628.4 | - | - | 0 | - |
| - | - | 1025 | 634.3 | - | - | 0 | - |
| - | - | 1806 | 635.3 | - | - | 0 | - |
| 4 | y | 1562 | 636.8 | 0.001405 | 2.206 | +2 | 10 |
| - | - | 815.6 | 640.3 | - | - | 0 | - |
| - | - | 2697 | 645.3 | - | - | 0 | - |
| 4 | y | 6228 | 645.8 | 0.001676 | 2.596 | +2 | 10 |
| - | - | 4010 | 646.3 | - | - | 0 | - |
| - | - | 2309 | 646.8 | - | - | 0 | - |
| - | - | 765.1 | 650.3 | - | - | 0 | - |
| - | - | 1.138E+04 | 652.3 | - | - | 0 | - |
| - | - | 5222 | 653.3 | - | - | 0 | - |
| - | - | 984.3 | 659.3 | - | - | 0 | - |
| - | - | 2008 | 660.9 | - | - | 0 | - |
| - | - | 1100 | 661.4 | - | - | 0 | - |
| - | - | 3547 | 662.3 | - | - | 0 | - |
| - | - | 3315 | 663.3 | - | - | 0 | - |
| - | - | 870.3 | 664.3 | - | - | 0 | - |
| 6 | b | 5384 | 676.3 | 0.0002361 | 0.3491 | +1 | 6 |
| 6 | b | 2609 | 677.3 | 0.00525 | 7.751 | +1 | 6 |
| - | - | 1180 | 677.8 | - | - | 0 | - |
| - | - | 690.2 | 678.3 | - | - | 0 | - |
| - | - | 1210 | 679.3 | - | - | 0 | - |
| - | - | 2.032E+04 | 680.3 | - | - | 0 | - |
| - | - | 7697 | 681.3 | - | - | 0 | - |
| - | - | 2384 | 682.3 | - | - | 0 | - |
| - | - | 1622 | 684.3 | - | - | 0 | - |
| - | - | 1578 | 684.8 | - | - | 0 | - |
| - | - | 695.5 | 685.3 | - | - | 0 | - |
| - | - | 718.5 | 690.3 | - | - | 0 | - |
| - | - | 1560 | 692.8 | - | - | 0 | - |
| 3 | y | 6206 | 693.3 | 3.899E-05 | 0.05624 | +2 | 11 |
| 3 | y | 7737 | 693.8 | 0.007831 | 11.29 | +2 | 11 |
| 6 | b | 6.039E+04 | 694.4 | 0.001096 | 1.579 | +1 | 6 |
| - | - | 2.261E+04 | 695.4 | - | - | 0 | - |
| - | - | 1231 | 695.9 | - | - | 0 | - |
| - | - | 4995 | 696.4 | - | - | 0 | - |
| - | - | 1706 | 697.4 | - | - | 0 | - |
| - | - | 926.5 | 700.9 | - | - | 0 | - |
| - | - | 769 | 701.3 | - | - | 0 | - |
| - | - | 1.364E+04 | 701.8 | - | - | 0 | - |
| 3 | y | 4.228E+04 | 702.3 | 0.001759 | 2.504 | +2 | 11 |
| 12 | b | 2.691E+04 | 702.8 | 0.01381 | 19.65 | +2 | 12 |
| 12 | b | 1.394E+04 | 703.3 | 0.003686 | 5.241 | +2 | 12 |
| - | - | 3628 | 703.8 | - | - | 0 | - |
| - | - | 2717 | 704.3 | - | - | 0 | - |
| 8 | y | 4239 | 706.3 | 0.00147 | 2.081 | +1 | 6 |
| - | - | 809.8 | 707.3 | - | - | 0 | - |
| - | - | 1340 | 710.4 | - | - | 0 | - |
| - | - | 2590 | 711.4 | - | - | 0 | - |
| 12 | b | 3197 | 711.9 | 0.005012 | 7.041 | +2 | 12 |
| - | - | 2005 | 712.4 | - | - | 0 | - |
| - | - | 1159 | 712.4 | - | - | 0 | - |
| - | - | 1043 | 712.9 | - | - | 0 | - |
| - | - | 3014 | 721.4 | - | - | 0 | - |
| - | - | 1034 | 722.4 | - | - | 0 | - |
| - | - | 9105 | 723.4 | - | - | 0 | - |
| 8 | y | 3.016E+04 | 724.4 | 0.000488 | 0.6738 | +1 | 6 |
| - | - | 1.328E+04 | 725.4 | - | - | 0 | - |
| - | - | 3707 | 726.4 | - | - | 0 | - |
| - | - | 2009 | 729.4 | - | - | 0 | - |
| - | - | 936.1 | 730.4 | - | - | 0 | - |
| - | - | 1040 | 733.4 | - | - | 0 | - |
| - | - | 835.2 | 734.4 | - | - | 0 | - |
| 2 | y | 672.3 | 742.9 | 0.001887 | 2.54 | +2 | 12 |
| 2 | y | 1189 | 743.4 | 0.01385 | 18.63 | +2 | 12 |
| - | - | 939.4 | 748.4 | - | - | 0 | - |
| - | - | 6873 | 751.4 | - | - | 0 | - |
| 2 | y | 3506 | 751.9 | 0.0004497 | 0.5982 | +2 | 12 |
| - | - | 5368 | 752.4 | - | - | 0 | - |
| - | - | 1030 | 752.9 | - | - | 0 | - |
| - | - | 3798 | 763.4 | - | - | 0 | - |
| - | - | 1291 | 764.4 | - | - | 0 | - |
| - | - | 1785 | 765.4 | - | - | 0 | - |
| - | - | 1679 | 767.3 | - | - | 0 | - |
| - | - | 877.3 | 775.4 | - | - | 0 | - |
| - | - | 886.9 | 776.4 | - | - | 0 | - |
| - | - | 3924 | 779.4 | - | - | 0 | - |
| - | - | 1618 | 780.4 | - | - | 0 | - |
| - | - | 2597 | 781.4 | - | - | 0 | - |
| - | - | 737.6 | 782.4 | - | - | 0 | - |
| 0 | Precursor | 6848 | 793.4 | 0.01253 | 15.8 | +2 | -1 |
| - | - | 2511 | 794.4 | - | - | 0 | - |
| - | - | 2067 | 807.4 | - | - | 0 | - |
| - | - | 821.7 | 808.4 | - | - | 0 | - |
| - | - | 798 | 818.4 | - | - | 0 | - |
| - | - | 791 | 825.4 | - | - | 0 | - |
| - | - | 757.1 | 826.4 | - | - | 0 | - |
| - | - | 1506 | 835.4 | - | - | 0 | - |
| - | - | 4156 | 836.5 | - | - | 0 | - |
| - | - | 1700 | 837.5 | - | - | 0 | - |
| - | - | 1116 | 838.4 | - | - | 0 | - |
| - | - | 1249 | 839.5 | - | - | 0 | - |
| - | - | 1166 | 840.5 | - | - | 0 | - |
| - | - | 765.1 | 843.4 | - | - | 0 | - |
| - | - | 1353 | 844.4 | - | - | 0 | - |
| - | - | 967.3 | 845.4 | - | - | 0 | - |
| - | - | 2166 | 846.4 | - | - | 0 | - |
| - | - | 1748 | 847.4 | - | - | 0 | - |
| - | - | 792.3 | 851.4 | - | - | 0 | - |
| - | - | 995.8 | 852.3 | - | - | 0 | - |
| - | - | 1.562E+04 | 852.4 | - | - | 0 | - |
| - | - | 7324 | 853.4 | - | - | 0 | - |
| - | - | 2317 | 854.4 | - | - | 0 | - |
| 7 | b | 8560 | 862.4 | 0.00179 | 2.076 | +1 | 7 |
| 7 | b | 6713 | 863.4 | 0.01206 | 13.97 | +1 | 7 |
| - | - | 9829 | 864.5 | - | - | 0 | - |
| - | - | 5905 | 865.5 | - | - | 0 | - |
| - | - | 1940 | 866.5 | - | - | 0 | - |
| - | - | 1069 | 876.4 | - | - | 0 | - |
| 7 | b | 7.023E+04 | 880.4 | 0.001369 | 1.554 | +1 | 7 |
| - | - | 3.836E+04 | 881.4 | - | - | 0 | - |
| - | - | 1.14E+04 | 882.4 | - | - | 0 | - |
| - | - | 1958 | 883.4 | - | - | 0 | - |
| - | - | 949.3 | 892.5 | - | - | 0 | - |
| - | - | 701.8 | 902.4 | - | - | 0 | - |
| - | - | 4238 | 908.4 | - | - | 0 | - |
| - | - | 2904 | 909.4 | - | - | 0 | - |
| 7 | y | 5263 | 910.4 | 0.0002766 | 0.3039 | +1 | 7 |
| - | - | 2435 | 911.4 | - | - | 0 | - |
| - | - | 911 | 954.4 | - | - | 0 | - |
| - | - | 1480 | 955.4 | - | - | 0 | - |
| - | - | 1108 | 965.4 | - | - | 0 | - |
| - | - | 1273 | 972.4 | - | - | 0 | - |
| 8 | b | 1245 | 975.5 | 0.003152 | 3.231 | +1 | 8 |
| 8 | b | 844 | 976.5 | 0.008377 | 8.579 | +1 | 8 |
| - | - | 685.9 | 979.5 | - | - | 0 | - |
| 8 | b | 4999 | 993.5 | 0.00157 | 1.581 | +1 | 8 |
| - | - | 2877 | 994.5 | - | - | 0 | - |
| 6 | y | 1853 | 1025 | 0.0006654 | 0.6489 | +1 | 8 |
| - | - | 1323 | 1026 | - | - | 0 | - |
| - | - | 892.3 | 1076 | - | - | 0 | - |
| - | - | 1548 | 1094 | - | - | 0 | - |
| - | - | 1336 | 1095 | - | - | 0 | - |
| - | - | 671 | 1110 | - | - | 0 | - |
| - | - | 669.8 | 1119 | - | - | 0 | - |
| 5 | y | 2436 | 1136 | 0.004123 | 3.631 | +1 | 9 |
| 5 | y | 2825 | 1136 | 0.005636 | 4.959 | +1 | 9 |
| - | - | 1473 | 1137 | - | - | 0 | - |
| - | - | 1055 | 1138 | - | - | 0 | - |
| - | - | 1737 | 1153 | - | - | 0 | - |
| 5 | y | 4738 | 1154 | 0.0005715 | 0.4955 | +1 | 9 |
| - | - | 2639 | 1155 | - | - | 0 | - |
| - | - | 1035 | 1156 | - | - | 0 | - |
| - | - | 729.9 | 1167 | - | - | 0 | - |
| - | - | 858.8 | 3081 | - | - | 0 | - |

m/z Charge Intensity FragmentType MassShift Position
120.06581115722656 0 700.42285
120.08106231689453 0 6911.4478
121.08484649658203 0 605.5514
122.07136535644531 0 615.05194
122.88048553466797 0 388.00342
123.0443115234375 0 758.02344
124.4383544921875 0 395.45227
125.10746765136719 0 616.62994
126.0915756225586 0 587.40857
127.05050659179688 0 2423.731
127.47537231445312 0 348.62418
128.10728454589844 0 11677.117
129.0662078857422 0 1240.4596
129.1025390625 0 118791.055
130.06541442871094 0 50756.54
130.06971740722656 0 646.40137
130.086669921875 0 518.3208
130.10031127929688 0 1058.5771
130.10586547851562 0 9143.519
131.06874084472656 0 4844.906
131.10824584960938 0 417.55606
132.0810089111328 0 11960.443
133.08518981933594 0 1544.9015
134.0274200439453 0 596.32965
136.0701141357422 0 955.1229
136.0760040283203 0 17767.734
137.0795440673828 0 1686.581
138.0666046142578 0 1741.7639
138.09161376953125 0 747.7676
139.0867462158203 0 1378.1453
141.10252380371094 0 3874.3655
142.0657958984375 0 600.339
143.04542541503906 0 527.091
144.07644653320312 0 524.7182
145.0609893798828 0 1465.0093
146.06016540527344 0 1152.8105
147.04454040527344 0 1020.63306
147.11331176757812 0 526.9125
149.04513549804688 0 13901.03
150.04432678222656 0 600.46936
150.06658935546875 0 1148.7281
151.0417938232422 0 790.0218
151.08689880371094 0 2708.6006
153.10289001464844 0 922.42017
154.06121826171875 0 548.3755
155.04527282714844 0 462.87912
155.0929718017578 0 6314.1177
155.1181182861328 0 5910.9873 a Water loss 1
156.0767822265625 0 509.4229
156.09603881835938 0 763.1903
157.0609130859375 0 457.4521
157.09754943847656 0 971.1143
157.13381958007812 0 984.29266
158.084228515625 0 919.76245
158.5285186767578 0 444.21283
159.09194946289062 0 92438.46
160.075927734375 0 729.3668
160.089599609375 0 525.9312
160.09527587890625 0 11717.284
165.05487060546875 0 7614.6445
166.06124877929688 0 13552.717
166.08656311035156 0 5065.26 y 9
166.0976104736328 0 1007.2587
167.05567932128906 0 19531.586
167.06394958496094 0 1176.925
168.0548553466797 0 933.25586
168.08087158203125 0 1480.1908
168.1025848388672 0 750.7836
168.11325073242188 0 1143.8475
169.0524444580078 0 553.62787
169.07643127441406 0 1212.2601
169.08444213867188 0 549.2732
169.0973358154297 0 4050.6853
170.06031799316406 0 4266.5176
171.11306762695312 0 741.77057
171.14956665039062 0 1126.0752
172.0717315673828 0 1555.3535
173.0560302734375 0 1475.7272
173.12876892089844 0 175296.64 a 1
174.05540466308594 0 546.3446
174.13206481933594 0 15297.115
177.10311889648438 0 528.1579
177.1121826171875 0 493.74695
178.13412475585938 0 5029.6772
179.13731384277344 0 614.2932
181.06109619140625 0 1144.9474
182.072021484375 0 923.66235
182.0814208984375 0 18837.344 y 12
183.08517456054688 0 1541.9849
183.1129913330078 0 5396.0513 b Water loss 1
183.14894104003906 0 807.6742
185.091796875 0 560.22723
185.165283203125 0 1483.1941
186.12396240234375 0 33976.3
187.0868377685547 0 7511.2363
187.1274871826172 0 2409.1348
187.1442413330078 0 3837.3472
188.07115173339844 0 1310.6378
190.08221435546875 0 1053.9674
190.13438415527344 0 899.5783
194.0927734375 0 1685.7415
195.113037109375 0 8871.135
196.10824584960938 0 5645.6943
196.11636352539062 0 837.5702
197.12872314453125 0 1278.3745
198.0878143310547 0 1633.8623
198.12803649902344 0 2754.0505
199.07110595703125 0 866.8473
199.08721923828125 0 961.1601
200.1430206298828 0 733.7211
201.12356567382812 0 48268.97 b 1
202.08595275878906 0 1197.2603
202.12684631347656 0 4858.128
203.10275268554688 0 515.8814
205.0972442626953 0 4378.0176
205.1448516845703 0 857.02106
206.10043334960938 0 664.61017
206.12905883789062 0 677.41736
207.16061401367188 0 1434.211
208.1082305908203 0 536.23334
209.09161376953125 0 728.04565
210.12744140625 0 1510.3481
211.10801696777344 0 1055.7747 y Water loss 10
212.1396942138672 0 779.1601
213.12359619140625 0 4538.592
213.16006469726562 0 840.0707
215.13902282714844 0 2117.7566
216.09840393066406 0 1176.5671
221.05889892578125 0 1930.7987
221.07284545898438 0 911.1881
221.0845184326172 0 7592.739
221.10340881347656 0 1148.9905
221.14013671875 0 601.6715
221.57655334472656 0 596.0178
221.96128845214844 0 531.2524
222.08534240722656 0 1508.6307
222.1238555908203 0 1500.7872
223.0638427734375 0 4252.7417
223.08169555664062 0 1147.6204
223.1078338623047 0 2822.1711
223.1555938720703 0 15508.047
224.10304260253906 0 969.8311
224.15907287597656 0 1984.8447
225.0431365966797 0 9681.827
226.04342651367188 0 2978.6553
226.08224487304688 0 3036.0972
226.1185760498047 0 744.98303 b Water loss 5
226.15513610839844 0 899.8145
227.04000854492188 0 2510.114
227.0666046142578 0 2467.781
227.15492248535156 0 717.98834
228.13441467285156 0 1550.5485
229.11834716796875 0 2624.2058
229.1549072265625 0 3823.662
233.13998413085938 0 1536.4418
233.1656951904297 0 575.6426
234.12393188476562 0 16971.236
235.1273193359375 0 1844.9414
237.13522338867188 0 1252.7112
238.11886596679688 0 3274.1975
239.0951690673828 0 14981.495
240.0963134765625 0 5274.269
240.1344757080078 0 7741.0303
241.0924072265625 0 2983.649
241.11903381347656 0 862.731
241.13746643066406 0 797.65356
241.19143676757812 0 717.56775
243.10931396484375 0 801.37994
244.09303283691406 0 7966.3057
247.10787963867188 0 8372.576
247.14285278320312 0 620.1301
248.10116577148438 0 1064.0826
248.11367797851562 0 3693.4895
249.05337524414062 0 939.4121
249.09841918945312 0 5907.375
249.1345672607422 0 1802.7122
250.1026611328125 0 622.92017
250.16714477539062 0 608.87054
251.15048217773438 0 56636.81
252.1536102294922 0 6675.186
253.1565704345703 0 1002.95526
255.14756774902344 0 2844.1648
256.1086120605469 0 1077.807
257.0921936035156 0 489.00943
257.1128845214844 0 2180.0068
258.1448059082031 0 1847.4609
260.2461242675781 0 492.7149
261.1191711425781 0 1432.5698
262.1181945800781 0 580.6283
265.12939453125 0 6675.6865
265.1433410644531 0 624.8598
266.11395263671875 0 1727.5334
266.1250305175781 0 14520.444
266.9874267578125 0 541.1583
267.0030822753906 0 549.511
267.1279296875 0 2258.211
267.1449890136719 0 600.63257
268.1294250488281 0 934.284
268.1766357421875 0 1395.5308
269.1606140136719 0 825.3021
269.18634033203125 0 850.0864
270.12158203125 0 2657.7595
270.62384033203125 0 918.62897
272.17572021484375 0 3236.8818
273.17913818359375 0 910.9601
274.1184387207031 0 3011.8357
274.1327209472656 0 790.48865
275.102294921875 0 820.43097
276.1553649902344 0 974.2106
279.1449279785156 0 1243.9891
280.1289367675781 0 1738.4315
281.05145263671875 0 12868.266
282.15631103515625 0 4211.9043
283.0302429199219 0 1748.1385
283.1402587890625 0 45977.2
284.1037902832031 0 1721.0707
284.1435852050781 0 5809.645
285.0096435546875 0 819.0112
285.15765380859375 0 1000.0717
286.1400451660156 0 2616.5317
287.1714782714844 0 928.5037
288.6187438964844 0 773.16437
293.1134338378906 0 21470.053 y Water loss 11
293.1600341796875 0 650.3006
294.1173095703125 0 4569.1694
295.1031799316406 0 1881.7163
295.1194763183594 0 705.0863
296.1043701171875 0 1205.801
296.197021484375 0 1253.3425 b Water loss 2
297.082763671875 0 2080.3992
297.1008605957031 0 1009.5459
297.1559753417969 0 6946.606
298.139892578125 0 819.4638 y Water loss 6
298.15838623046875 0 1185.8188
299.06195068359375 0 107269.67
300.062255859375 0 3043.2014
300.1672058105469 0 7278.9814
301.0599365234375 0 2529.0208
301.1509704589844 0 18340.979
301.1695251464844 0 715.0736
302.11376953125 0 4811.255
302.15478515625 0 2378.986
305.1788024902344 0 841.0839
305.6457214355469 0 1652.7415
306.13751220703125 0 5240.9673 y 8
306.6396179199219 0 1667.575
311.1239318847656 0 3444.9746 y 11
313.1144714355469 0 832.6931
314.18768310546875 0 796.8337
314.207763671875 0 2337.6099 b 2
315.1663818359375 0 12372.75
315.1849670410156 0 648.1277
316.169677734375 0 2041.7104
317.659912109375 0 741.1755
323.2075500488281 0 761.0932
325.1504211425781 0 719.16394
326.6637268066406 0 2500.871
327.1659240722656 0 865.89246
332.2083740234375 0 634.62476
333.17694091796875 0 3300.6047
338.67437744140625 0 674.74176 b Water loss 5
339.6741943359375 0 682.0754
340.66015625 0 959.9924
341.0185241699219 0 1367.9288
342.1812438964844 0 1118.2693
343.1612854003906 0 1025.5847
344.9765319824219 0 13004.174
345.130859375 0 2162.1946
346.1148681640625 0 6492.2646
346.13690185546875 0 544.6471
347.1155090332031 0 871.3419
347.6795654296875 0 2947.3843 b 5
349.15216064453125 0 742.9604
350.2189025878906 0 4795.88
351.2218933105469 0 882.8106
353.6732482910156 0 1560.0394 y Water loss 7
354.1787109375 0 686.22186
355.06964111328125 0 2163.4717
356.0724182128906 0 815.7822
356.1927185058594 0 782.64844
359.0284118652344 0 14230.611
359.04901123046875 0 503.55823
360.6947021484375 0 1800.5728
361.1966247558594 0 1896.2843
362.185546875 0 2388.165
362.67913818359375 0 8614.707 y 7
363.1410827636719 0 10795.334
363.181640625 0 2314.3176
364.12371826171875 0 1679.6923
364.1453857421875 0 1264.9939
366.176513671875 0 1407.7703
367.1394958496094 0 1597.1106
369.12066650390625 0 863.7729
370.12213134765625 0 1062.4485
371.1202392578125 0 689.81024
373.18255615234375 0 593.6069
376.16229248046875 0 897.95123
376.1980895996094 0 1644.313
377.1452941894531 0 898.0037
378.2137756347656 0 3966.5156
379.2089538574219 0 16715.465
380.211669921875 0 2762.006
381.1518249511719 0 29511.465
381.7097473144531 0 664.1711
382.15447998046875 0 5074.9224
382.1869201660156 0 1104.2605
384.1664733886719 0 1579.6403
384.188720703125 0 1508.1216
385.148681640625 0 1047.2158
385.17437744140625 0 979.6722 y 4
390.1952209472656 0 1449.1151
391.1355895996094 0 1120.321
391.959228515625 0 796.1924
394.1725769042969 0 5034.5337
395.13446044921875 0 1406.8915
395.1786804199219 0 941.795
395.2400207519531 0 1362.02
396.2247009277344 0 6145.0415
397.22613525390625 0 916.7157
398.1769714355469 0 2169.93
399.00469970703125 0 692.06036
401.2161865234375 0 941.6164
402.1769104003906 0 1688.0292
402.19927978515625 0 1589.5454
403.1998291015625 0 831.728
405.2611999511719 0 1121.3618
409.1461181640625 0 1579.7318
409.2039489746094 0 1236.3291
411.1990661621094 0 1959.3605
412.18304443359375 0 19014.129
413.1436767578125 0 1384.8213
413.18756103515625 0 2144.7188
414.2347106933594 0 4934.12
415.0368347167969 0 26999.025
415.1988830566406 0 1781.2806
415.23907470703125 0 1179.9888
417.7170715332031 0 1868.0427
418.2100524902344 0 5842.095
418.7110290527344 0 2378.4473
421.20843505859375 0 1002.0705 y Water loss 10
422.7104187011719 0 714.35376
423.2019348144531 0 1182.2979
423.2716979980469 0 7487.677
424.27392578125 0 1692.4382
424.7342529296875 0 897.11884
426.721923828125 0 47313.965
427.22344970703125 0 26506.9
427.7252502441406 0 5775.3467
428.2261047363281 0 1472.7831
429.08935546875 0 920.3784
429.2095947265625 0 12828.977
429.5750427246094 0 655.48615
430.1935729980469 0 40275.12
431.197021484375 0 9716.117
431.7142333984375 0 4800.848 b Water loss 6
432.2086181640625 0 2983.679 b Ammonia loss 6
432.70819091796875 0 611.0171
432.74176025390625 0 673.57764
433.047119140625 0 1547.0101
433.2095947265625 0 1011.13446
433.25811767578125 0 1188.4192 b Water loss 3
439.21893310546875 0 12169.872 y 10
439.7136535644531 0 821.9717
440.22161865234375 0 2603.6914
440.71923828125 0 22199.914 b 6
441.2210693359375 0 12652.571
441.25384521484375 0 983.51276
441.72149658203125 0 2903.6863
447.21063232421875 0 743.99426 y Ammonia loss 6
448.2042541503906 0 2007.3633
448.2652282714844 0 728.9239
449.21197509765625 0 597.5523
450.2808532714844 0 690.41943
451.2348327636719 0 635.036
451.2663269042969 0 14824.694 b 3
452.2694396972656 0 2930.1343
455.22637939453125 0 948.5187
455.26312255859375 0 1264.9899
459.1986389160156 0 1036.0336
460.75335693359375 0 2211.0776
461.2027893066406 0 1254.2864
461.2518310546875 0 1223.7554
464.2237854003906 0 666.0657
468.292724609375 0 3329.1375
469.2959289550781 0 1025.4053
474.25946044921875 0 3423.4377
474.75286865234375 0 3897.1252
475.2536315917969 0 1136.1902
475.2942199707031 0 1000.23474
476.2251281738281 0 1503.539
477.2108154296875 0 2149.8818
478.2282409667969 0 1671.9736 y Water loss 9
478.2782287597656 0 1663.5679
479.23760986328125 0 693.60284
483.2638244628906 0 9716.266
483.7655334472656 0 5135.418
484.21929931640625 0 955.1538
484.2662048339844 0 1614.4513
488.2569274902344 0 2132.7268 b Water loss 7
488.7520446777344 0 1710.7397 b Ammonia loss 7
494.2356872558594 0 26961.51
495.2389221191406 0 7092.143
496.24041748046875 0 35343.957 y 9
497.2460632324219 0 8730.692
497.7631530761719 0 4552.748
498.2424621582031 0 1684.5374
503.10736083984375 0 27098.66
504.1996765136719 0 1569.294
506.22564697265625 0 1648.9769
507.2584533691406 0 935.7484
511.26214599609375 0 2054.339
519.1388549804688 0 8751.1045
521.7820434570312 0 812.64197
523.2572021484375 0 1459.8217
525.2666015625 0 4911.48
526.2686157226562 0 837.0073
528.9331665039062 0 780.25525
529.2625732421875 0 1399.1062 Precursor Water loss
529.5924682617188 0 1413.6455 Precursor Ammonia loss
532.1936645507812 0 1354.5157
534.2991333007812 0 711.7953
534.5562133789062 0 856.45795
535.261962890625 0 1260.5934 Precursor
535.5974731445312 0 1379.3779
536.1649780273438 0 5861.5527
536.2713012695312 0 1340.2578
539.2354125976562 0 5995.85
540.2385864257812 0 2139.6252
542.29296875 0 2442.875
543.2774047851562 0 7522.808
544.2791748046875 0 2671.583
545.2794799804688 0 945.34125
546.2465209960938 0 1251.6633
549.2214965820312 0 1575.8228
551.256103515625 0 1244.6414
554.7659912109375 0 1070.4524 b 8
555.2564697265625 0 1738.15
555.755859375 0 1014.76825
561.31298828125 0 1955.9021 b Water loss 4
565.2633666992188 0 2104.8723
567.2307739257812 0 5265.1616
568.2353515625 0 1747.1149
575.2451782226562 0 2140.687
576.2344970703125 0 874.0603
579.324951171875 0 9042.959 b 4
580.32763671875 0 3464.5432
582.3028564453125 0 1070.3314
589.2947387695312 0 697.3827
592.2725219726562 0 1639.8596
592.3248291015625 0 1590.7109
593.2562866210938 0 12558.372 y Water loss 8
593.304931640625 0 3687.002
594.2552490234375 0 3935.015
595.2483520507812 0 1904.8307
596.3490600585938 0 901.7947
602.793212890625 0 1241.8652
603.2906494140625 0 1689.0535
603.7943115234375 0 813.00745
604.28662109375 0 785.4054
610.28271484375 0 31183.367
610.3287963867188 0 1079.7333
611.267578125 0 106798.15 y 8
611.7982788085938 0 5172.219
612.270751953125 0 35113.715
612.7984619140625 0 1257.8158
613.2722778320312 0 7150.4014
614.2744750976562 0 1351.4629
620.8001708984375 0 797.48517
621.2491455078125 0 1223.064
622.3635864257812 0 1003.72095
628.38916015625 0 1030.9607
634.3062744140625 0 1024.7319
635.296142578125 0 1806.2548
636.7872314453125 0 1561.6715 y Water loss 3
640.2718505859375 0 815.6489
645.2969970703125 0 2697.202
645.7927856445312 0 6227.8135 y 3
646.2936401367188 0 4009.577
646.7942504882812 0 2309.4011
650.321533203125 0 765.0801
652.3192138671875 0 11376.656
653.32275390625 0 5221.8696
659.316162109375 0 984.2758
660.8724975585938 0 2008.0896
661.3760986328125 0 1099.7782
662.3038940429688 0 3547.217
663.2947998046875 0 3315.3835
664.2910766601562 0 870.25714
676.341064453125 0 5383.8223 b Water loss 5
677.33056640625 0 2609.2246 b Ammonia loss 5
677.8054809570312 0 1180.2397
678.3101196289062 0 690.23785
679.3267822265625 0 1209.6414
680.3148803710938 0 20322.309
681.3171997070312 0 7697.2075
682.32080078125 0 2383.9285
684.3171997070312 0 1622.2338
684.821533203125 0 1578.413
685.3172607421875 0 695.46405
690.2993774414062 0 718.48254
692.8346557617188 0 1560.1764
693.3278198242188 0 6205.5796 y Water loss 2
693.8276977539062 0 7736.628 y Ammonia loss 2
694.3507690429688 0 60388.227 b 5
695.3549194335938 0 22606.723
695.8779296875 0 1230.9088
696.358154296875 0 4994.8623
697.359375 0 1705.5758
700.8853759765625 0 926.47815
701.3310546875 0 768.95715
701.8405151367188 0 13638.915
702.3348999023438 0 42277.11 y 2
702.8351440429688 0 26906.162 b Water loss 11
703.3372802734375 0 13942.023 b Ammonia loss 11
703.838623046875 0 3628.0708
704.3375244140625 0 2716.8035
706.3421020507812 0 4238.6353 y Water loss 7
707.3396606445312 0 809.84436
710.4120483398438 0 1339.8839
711.3677368164062 0 2590.2632
711.8592529296875 0 3197.0347 b 11
712.359375 0 2005.4102
712.4313354492188 0 1159.419
712.854736328125 0 1042.9269
721.3624267578125 0 3013.935
722.3643798828125 0 1034.4478
723.3663330078125 0 9104.51
724.3516845703125 0 30162.326 y 7
725.354736328125 0 13276.261
726.356689453125 0 3707.3237
729.3582153320312 0 2008.9131
730.36279296875 0 936.0792
733.3671264648438 0 1040.3064
734.3599853515625 0 835.1849
742.8639526367188 0 672.3415 y Water loss 1
743.367919921875 0 1189.3812 y Ammonia loss 1
748.3756713867188 0 939.38965
751.376220703125 0 6872.9976
751.8677978515625 0 3506.3215 y 1
752.3735961914062 0 5368.1943
752.8674926757812 0 1029.7135
763.3605346679688 0 3798.2024
764.3687133789062 0 1291.0632
765.4008178710938 0 1784.685
767.3468017578125 0 1679.4546
775.3953247070312 0 877.3184
776.38623046875 0 886.8948
779.3836669921875 0 3924.3564
780.3795776367188 0 1617.9725
781.3642578125 0 2596.9902
782.36376953125 0 737.5843
793.3984375 0 6847.605 Precursor Water loss
794.3995361328125 0 2510.569
807.3739624023438 0 2066.6714
808.3781127929688 0 821.7439
818.4397583007812 0 797.97925
825.3841552734375 0 791.03784
826.3921508789062 0 757.1363
835.4151611328125 0 1506.3519
836.4501953125 0 4156.117
837.4506225585938 0 1699.8735
838.4439086914062 0 1115.5571
839.4783935546875 0 1249.4038
840.4751586914062 0 1166.3417
843.3967895507812 0 765.0713
844.39013671875 0 1353.0613
845.3914184570312 0 967.2577
846.44189453125 0 2166.3882
847.4456787109375 0 1748.0012
851.3867797851562 0 792.32574
852.3488159179688 0 995.8155
852.435302734375 0 15624.95
853.4374389648438 0 7324.452
854.441650390625 0 2316.9788
862.4188232421875 0 8559.716 b Water loss 6
863.4166870117188 0 6713.147 b Ammonia loss 6
864.4548950195312 0 9829.0625
865.4588012695312 0 5904.551
866.4623413085938 0 1940.1438
876.4391479492188 0 1068.7177
880.4298095703125 0 70233.555 b 6
881.4324951171875 0 38357.754
882.4349975585938 0 11402.526
883.4301147460938 0 1957.6777
892.4716186523438 0 949.2951
902.4459228515625 0 701.82465
908.427001953125 0 4238.4976
909.4341430664062 0 2904.3276
910.4307861328125 0 5262.752 y 6
911.4356689453125 0 2435.2122
954.4312133789062 0 911.0116
955.4169921875 0 1480.4587
965.445556640625 0 1107.7596
972.445556640625 0 1273.2483
975.5015258789062 0 1244.8643 b Water loss 7
976.4970703125 0 844.03845 b Ammonia loss 7
979.4757080078125 0 685.881
993.513671875 0 4998.814 b 7
994.5183715820312 0 2877.4048
1025.456787109375 0 1853.3119 y 5
1026.4521484375 0 1323.4196
1075.53125 0 892.28735
1093.5411376953125 0 1547.7405
1094.540283203125 0 1335.9475
1109.5120849609375 0 670.957
1119.495849609375 0 669.76605
1135.5013427734375 0 2436.1047 y Water loss 4
1136.4951171875 0 2824.8093 y Ammonia loss 4
1137.493408203125 0 1472.5297
1138.4920654296875 0 1055.2576
1152.5303955078125 0 1736.5234
1153.5166015625 0 4737.556 y 4
1154.521484375 0 2639.012
1155.5174560546875 0 1034.6836
1166.5677490234375 0 729.9399
3081.1044921875 0 858.8179

Spectrum Details

|  |  |
| --- | --- |
| Matched peaks? Matched peaks The total absolute number of peaks matched. Additionally in brackets the total fraction of peaks matched and the total number of peaks is shown. | 67 (10.88% of 616) |
| FDR? FDR The false discovery rate estimated for this peptide. It is calculated by matching all theoretical fragments with a non-integer shift with the raw peaks for this spectrum. This is done with 40 different shifts. The resulting percentage is the average number of annotated peaks over the number of annotated peaks with the correct spectrum. | 1.00% |
| Satellite FDR? Satellite FDR See the FDR for details on its calculation. This satellite ion specific FDR only contains the satellite ions (d/w) for I/L/J positions. | ∞ |
| PSM Score? PSM Score The PSM Score as given by Hecklib to this annotated spectrum. It is shown with three significant figures. | 501 |

#### Spectrum 6234? Spectrum 6234 The raw spectrum of this peptide as annotated by Hecklib. The fragments are coloured according to ion type (see legend). Any peaks with a star '\*' as text can be hovered over to see the full details, first the ion type second the mass shift type. By hovering over the amino acids in the peptide or ions in the legend the corresponding peaks are highlighted. By toggling the 'Unassigned' label you can turn the background (unassigned) peaks on or off in the plot. By updating the slider in the Ion legend you can update the spectrum to only show the top X% of the peaks with labels. The top X% means any peak that is within X% of the highest intensity. By dragging in the spectrum you can zoom in to a specific part of the spectrum and use 'Zoom Out' to get back to the original zoom level. The annotation of the spectrum is based on the given sequence in the peptides file and is done with different software so inconsistencies are likely. The peaks are annotated based on the given sequence, with 20 ppm tolerance.

Copy Data

##### Spectrum 6234 (TSV)

###### Preview

```
Loading example...
```

*Click on the button to copy the data to your clipboard.*

Mz MinMz MaxIntensity Max

WidthHeightPeptide font sizePeptide stroke widthSpectrum font sizeSpectrum stroke widthCompact peptide

Ion legend

wxyz

abcd

OtherUnassignedIon^Charge^_Position_Show for top:%

TVLHQDWLDGKEY

08.79e+41.76e+52.64e+53.52e+5

Zoom Out

y^+3^_3_a^+1^_2_d^+1^_2_y^+3^_4_a^+1^_2_y^+1^_1_b^+1^_2_b^+1^_2_y^+2^_3_y^+2^_4_y^+1^_2_b^+1^_3_y^+2^_5_y^+1^_2_b^+1^_3_b^+3^_8_y^+3^_8_b^+2^_6_y^+2^_6_y^+2^_6_y^+1^_3_y^+1^_3_y^+3^_10_b^+1^_4_y^+1^_3_b^+2^_7_b^+2^_7_b^+1^_4_y^+2^_7_y^+2^_7_y^+1^_4_b^+2^_8_b^+2^_8_\*\*\*b^+1^_5_b^+1^_5_b^+2^_9_y^+2^_9_b^+1^_5_y^+1^_5_y^+1^_5_y^+2^_10_y^+2^_10_b^+2^_11_b^+1^_6_b^+1^_6_b^+1^_6_y^+2^_11_y^+2^_11_y^+1^_6_y^+2^_11_b^+2^_12_b^+2^_12_b^+2^_12_y^+1^_6_y^+2^_12_y^+2^_12_y^+2^_12_\*b^+1^_7_b^+1^_7_b^+1^_7_y^+1^_7_y^+1^_7_b^+1^_8_b^+1^_8_b^+1^_8_y^+1^_8_y^+1^_8_b^+1^_9_b^+1^_9_y^+1^_9_y^+1^_9_y^+1^_9_b^+1^_10_

0778155723353114

Fragment Matches Table

Show background peaks

| Position | Ion type | Intensity | mz Theoretical | mz Error (Th) | mz Error (ppm) | Charge | Series Number |
| --- | --- | --- | --- | --- | --- | --- | --- |
| - | - | 1.251E+04 | 120.1 | - | - | 0 | - |
| - | - | 529.7 | 121.1 | - | - | 0 | - |
| - | - | 685.1 | 121.1 | - | - | 0 | - |
| - | - | 419.3 | 121.5 | - | - | 0 | - |
| - | - | 1589 | 122.1 | - | - | 0 | - |
| - | - | 1784 | 123 | - | - | 0 | - |
| - | - | 942.5 | 123.1 | - | - | 0 | - |
| - | - | 1436 | 125.1 | - | - | 0 | - |
| - | - | 960.7 | 126.1 | - | - | 0 | - |
| - | - | 1042 | 126.1 | - | - | 0 | - |
| - | - | 5841 | 127.1 | - | - | 0 | - |
| - | - | 745.5 | 127.1 | - | - | 0 | - |
| - | - | 2.281E+04 | 128.1 | - | - | 0 | - |
| - | - | 1605 | 129.1 | - | - | 0 | - |
| - | - | 2.15E+05 | 129.1 | - | - | 0 | - |
| - | - | 4.424E+04 | 130.1 | - | - | 0 | - |
| - | - | 828.6 | 130.1 | - | - | 0 | - |
| - | - | 1.461E+04 | 130.1 | - | - | 0 | - |
| - | - | 4402 | 131.1 | - | - | 0 | - |
| - | - | 7975 | 132 | - | - | 0 | - |
| - | - | 505.3 | 132.1 | - | - | 0 | - |
| - | - | 757.8 | 133 | - | - | 0 | - |
| - | - | 510.4 | 133.1 | - | - | 0 | - |
| - | - | 2.973E+04 | 136.1 | - | - | 0 | - |
| - | - | 2455 | 137.1 | - | - | 0 | - |
| - | - | 3699 | 138.1 | - | - | 0 | - |
| - | - | 1444 | 138.1 | - | - | 0 | - |
| - | - | 2243 | 139.1 | - | - | 0 | - |
| - | - | 1323 | 140.1 | - | - | 0 | - |
| - | - | 3558 | 141.1 | - | - | 0 | - |
| - | - | 477.3 | 143.4 | - | - | 0 | - |
| - | - | 825.5 | 144.1 | - | - | 0 | - |
| - | - | 3144 | 145.1 | - | - | 0 | - |
| - | - | 3411 | 146.1 | - | - | 0 | - |
| - | - | 1412 | 147 | - | - | 0 | - |
| 11 | y | 716.9 | 147.1 | 0.0007485 | 5.089 | +3 | 3 |
| - | - | 867.3 | 148.9 | - | - | 0 | - |
| - | - | 547.2 | 149 | - | - | 0 | - |
| - | - | 1608 | 150.1 | - | - | 0 | - |
| - | - | 6560 | 151.1 | - | - | 0 | - |
| - | - | 539.5 | 153.1 | - | - | 0 | - |
| - | - | 469.1 | 153.6 | - | - | 0 | - |
| - | - | 1104 | 154.1 | - | - | 0 | - |
| - | - | 560.9 | 155 | - | - | 0 | - |
| - | - | 965.6 | 155.1 | - | - | 0 | - |
| - | - | 1.457E+04 | 155.1 | - | - | 0 | - |
| 2 | a | 1.008E+04 | 155.1 | 0.0004271 | 2.753 | +1 | 2 |
| - | - | 846.8 | 156.1 | - | - | 0 | - |
| - | - | 1020 | 156.1 | - | - | 0 | - |
| - | - | 1088 | 156.1 | - | - | 0 | - |
| - | - | 956.9 | 156.1 | - | - | 0 | - |
| - | - | 1084 | 157.1 | - | - | 0 | - |
| - | - | 1.285E+04 | 157.1 | - | - | 0 | - |
| - | - | 1787 | 157.1 | - | - | 0 | - |
| - | - | 3603 | 157.1 | - | - | 0 | - |
| - | - | 652.6 | 157.1 | - | - | 0 | - |
| - | - | 4.027E+04 | 158.1 | - | - | 0 | - |
| - | - | 1256 | 158.1 | - | - | 0 | - |
| - | - | 1320 | 158.1 | - | - | 0 | - |
| - | - | 4304 | 159.1 | - | - | 0 | - |
| - | - | 2511 | 159.1 | - | - | 0 | - |
| 2 | d | 1613 | 159.1 | 0.0004466 | 2.807 | +1 | 2 |
| - | - | 1.532E+04 | 165.1 | - | - | 0 | - |
| - | - | 666.7 | 165.1 | - | - | 0 | - |
| - | - | 2.547E+04 | 166.1 | - | - | 0 | - |
| 10 | y | 7446 | 166.1 | 0.001802 | 10.85 | +3 | 4 |
| - | - | 2620 | 166.1 | - | - | 0 | - |
| - | - | 983.6 | 167.1 | - | - | 0 | - |
| - | - | 1819 | 167.1 | - | - | 0 | - |
| - | - | 1171 | 167.1 | - | - | 0 | - |
| - | - | 1922 | 168.1 | - | - | 0 | - |
| - | - | 2784 | 168.1 | - | - | 0 | - |
| - | - | 5908 | 169.1 | - | - | 0 | - |
| - | - | 2662 | 169.1 | - | - | 0 | - |
| - | - | 1996 | 171.1 | - | - | 0 | - |
| - | - | 2715 | 172.1 | - | - | 0 | - |
| - | - | 531.4 | 172.1 | - | - | 0 | - |
| - | - | 3742 | 173.1 | - | - | 0 | - |
| 2 | a | 3.482E+05 | 173.1 | 0.0005283 | 3.052 | +1 | 2 |
| - | - | 3.048E+04 | 174.1 | - | - | 0 | - |
| - | - | 2.111E+05 | 175.1 | - | - | 0 | - |
| - | - | 2536 | 175.1 | - | - | 0 | - |
| - | - | 1.987E+04 | 176.1 | - | - | 0 | - |
| - | - | 1376 | 177.1 | - | - | 0 | - |
| - | - | 821.1 | 177.1 | - | - | 0 | - |
| - | - | 1.102E+04 | 178.1 | - | - | 0 | - |
| - | - | 1207 | 179.1 | - | - | 0 | - |
| - | - | 1326 | 181.1 | - | - | 0 | - |
| 13 | y | 3.401E+04 | 182.1 | 0.0005106 | 2.804 | +1 | 1 |
| - | - | 3550 | 183.1 | - | - | 0 | - |
| 2 | b | 1.771E+04 | 183.1 | 0.0004618 | 2.522 | +1 | 2 |
| - | - | 1109 | 183.1 | - | - | 0 | - |
| - | - | 1306 | 184.1 | - | - | 0 | - |
| - | - | 2643 | 185.1 | - | - | 0 | - |
| - | - | 1362 | 185.1 | - | - | 0 | - |
| - | - | 822.7 | 185.1 | - | - | 0 | - |
| - | - | 2382 | 185.2 | - | - | 0 | - |
| - | - | 516.6 | 185.4 | - | - | 0 | - |
| - | - | 5.294E+04 | 186.1 | - | - | 0 | - |
| - | - | 2524 | 187.1 | - | - | 0 | - |
| - | - | 1873 | 187.1 | - | - | 0 | - |
| - | - | 4213 | 187.1 | - | - | 0 | - |
| - | - | 1227 | 187.1 | - | - | 0 | - |
| - | - | 6138 | 189.1 | - | - | 0 | - |
| - | - | 952.5 | 189.1 | - | - | 0 | - |
| - | - | 5086 | 190.1 | - | - | 0 | - |
| - | - | 2246 | 190.1 | - | - | 0 | - |
| - | - | 733.8 | 193.1 | - | - | 0 | - |
| - | - | 643.5 | 194.1 | - | - | 0 | - |
| - | - | 2955 | 194.1 | - | - | 0 | - |
| - | - | 1.679E+04 | 195.1 | - | - | 0 | - |
| - | - | 5364 | 196.1 | - | - | 0 | - |
| - | - | 1142 | 196.1 | - | - | 0 | - |
| - | - | 782.5 | 196.1 | - | - | 0 | - |
| - | - | 3459 | 197.1 | - | - | 0 | - |
| - | - | 2069 | 198.1 | - | - | 0 | - |
| - | - | 3187 | 199.1 | - | - | 0 | - |
| 2 | b | 1.071E+05 | 201.1 | 0.000441 | 2.193 | +1 | 2 |
| - | - | 842.4 | 202.1 | - | - | 0 | - |
| - | - | 9186 | 202.1 | - | - | 0 | - |
| - | - | 1208 | 203.1 | - | - | 0 | - |
| - | - | 1075 | 203.1 | - | - | 0 | - |
| - | - | 849.2 | 203.1 | - | - | 0 | - |
| - | - | 612 | 203.2 | - | - | 0 | - |
| - | - | 7546 | 205.1 | - | - | 0 | - |
| - | - | 1010 | 205.1 | - | - | 0 | - |
| - | - | 595 | 206.1 | - | - | 0 | - |
| - | - | 1630 | 206.1 | - | - | 0 | - |
| - | - | 3128 | 207.2 | - | - | 0 | - |
| - | - | 1483 | 210.1 | - | - | 0 | - |
| 11 | y | 2250 | 211.1 | 0.0005271 | 2.497 | +2 | 3 |
| - | - | 656.4 | 211.1 | - | - | 0 | - |
| - | - | 9987 | 211.2 | - | - | 0 | - |
| - | - | 979.7 | 212.1 | - | - | 0 | - |
| - | - | 2800 | 212.1 | - | - | 0 | - |
| - | - | 6865 | 213.1 | - | - | 0 | - |
| - | - | 1187 | 213.2 | - | - | 0 | - |
| - | - | 1059 | 214.1 | - | - | 0 | - |
| - | - | 3279 | 215.1 | - | - | 0 | - |
| - | - | 1315 | 215.1 | - | - | 0 | - |
| - | - | 1734 | 216.1 | - | - | 0 | - |
| - | - | 656.8 | 216.1 | - | - | 0 | - |
| - | - | 1.067E+04 | 217.1 | - | - | 0 | - |
| - | - | 1342 | 220.1 | - | - | 0 | - |
| - | - | 6050 | 221.1 | - | - | 0 | - |
| - | - | 1905 | 221.1 | - | - | 0 | - |
| - | - | 1095 | 221.1 | - | - | 0 | - |
| - | - | 848.9 | 222.1 | - | - | 0 | - |
| - | - | 3095 | 222.1 | - | - | 0 | - |
| - | - | 946.2 | 223.1 | - | - | 0 | - |
| - | - | 4317 | 223.1 | - | - | 0 | - |
| - | - | 2.958E+04 | 223.2 | - | - | 0 | - |
| - | - | 2818 | 224.2 | - | - | 0 | - |
| - | - | 5165 | 226.1 | - | - | 0 | - |
| - | - | 706.7 | 226.2 | - | - | 0 | - |
| - | - | 4909 | 227.1 | - | - | 0 | - |
| - | - | 757.6 | 227.1 | - | - | 0 | - |
| - | - | 1568 | 227.1 | - | - | 0 | - |
| - | - | 2291 | 228.1 | - | - | 0 | - |
| - | - | 2073 | 228.2 | - | - | 0 | - |
| - | - | 4016 | 229.1 | - | - | 0 | - |
| - | - | 694.6 | 230.2 | - | - | 0 | - |
| - | - | 1236 | 231.1 | - | - | 0 | - |
| - | - | 5684 | 233.1 | - | - | 0 | - |
| - | - | 943.3 | 233.2 | - | - | 0 | - |
| - | - | 3.463E+04 | 234.1 | - | - | 0 | - |
| - | - | 3256 | 235.1 | - | - | 0 | - |
| - | - | 2278 | 237.1 | - | - | 0 | - |
| - | - | 3845 | 238.1 | - | - | 0 | - |
| - | - | 1241 | 238.1 | - | - | 0 | - |
| - | - | 6688 | 239.2 | - | - | 0 | - |
| - | - | 1.34E+04 | 240.1 | - | - | 0 | - |
| - | - | 586.3 | 241.1 | - | - | 0 | - |
| - | - | 1588 | 241.1 | - | - | 0 | - |
| - | - | 1879 | 241.1 | - | - | 0 | - |
| - | - | 2574 | 241.2 | - | - | 0 | - |
| - | - | 2038 | 242.1 | - | - | 0 | - |
| - | - | 2281 | 243.1 | - | - | 0 | - |
| - | - | 713.8 | 243.1 | - | - | 0 | - |
| - | - | 5171 | 243.1 | - | - | 0 | - |
| - | - | 7939 | 244.1 | - | - | 0 | - |
| - | - | 1105 | 244.1 | - | - | 0 | - |
| - | - | 940.9 | 244.2 | - | - | 0 | - |
| - | - | 1.633E+04 | 247.1 | - | - | 0 | - |
| - | - | 1316 | 247.1 | - | - | 0 | - |
| - | - | 1051 | 248.1 | - | - | 0 | - |
| - | - | 7002 | 248.1 | - | - | 0 | - |
| 10 | y | 675.8 | 248.6 | 0.0005193 | 2.089 | +2 | 4 |
| - | - | 9959 | 249.1 | - | - | 0 | - |
| - | - | 1071 | 249.1 | - | - | 0 | - |
| - | - | 2802 | 249.1 | - | - | 0 | - |
| - | - | 1081 | 250.1 | - | - | 0 | - |
| - | - | 737.2 | 250.1 | - | - | 0 | - |
| - | - | 1.146E+05 | 251.2 | - | - | 0 | - |
| - | - | 767.4 | 252.1 | - | - | 0 | - |
| - | - | 1.483E+04 | 252.2 | - | - | 0 | - |
| - | - | 1618 | 253.2 | - | - | 0 | - |
| - | - | 747 | 254.1 | - | - | 0 | - |
| - | - | 3396 | 255.1 | - | - | 0 | - |
| - | - | 3246 | 255.1 | - | - | 0 | - |
| - | - | 2102 | 256.2 | - | - | 0 | - |
| - | - | 2456 | 257.1 | - | - | 0 | - |
| - | - | 4896 | 258.1 | - | - | 0 | - |
| - | - | 992 | 259.1 | - | - | 0 | - |
| - | - | 3159 | 261.1 | - | - | 0 | - |
| - | - | 1622 | 261.1 | - | - | 0 | - |
| - | - | 1273 | 262.1 | - | - | 0 | - |
| - | - | 895.4 | 262.2 | - | - | 0 | - |
| - | - | 688.4 | 263.1 | - | - | 0 | - |
| - | - | 653.6 | 263.1 | - | - | 0 | - |
| - | - | 1.686E+04 | 265.1 | - | - | 0 | - |
| - | - | 3.106E+04 | 266.1 | - | - | 0 | - |
| - | - | 3531 | 267.1 | - | - | 0 | - |
| - | - | 658.9 | 267.1 | - | - | 0 | - |
| - | - | 2516 | 268.1 | - | - | 0 | - |
| - | - | 2211 | 268.2 | - | - | 0 | - |
| - | - | 1424 | 269.1 | - | - | 0 | - |
| - | - | 1674 | 269.2 | - | - | 0 | - |
| - | - | 991.6 | 270.1 | - | - | 0 | - |
| - | - | 6418 | 270.2 | - | - | 0 | - |
| - | - | 900.8 | 271.2 | - | - | 0 | - |
| - | - | 910.3 | 272.1 | - | - | 0 | - |
| - | - | 4103 | 274.1 | - | - | 0 | - |
| - | - | 1247 | 275.1 | - | - | 0 | - |
| - | - | 1637 | 276.1 | - | - | 0 | - |
| - | - | 2688 | 276.2 | - | - | 0 | - |
| - | - | 1773 | 279.1 | - | - | 0 | - |
| - | - | 3588 | 280.1 | - | - | 0 | - |
| - | - | 4557 | 282.2 | - | - | 0 | - |
| - | - | 7.985E+04 | 283.1 | - | - | 0 | - |
| - | - | 1180 | 284.1 | - | - | 0 | - |
| - | - | 1.256E+04 | 284.1 | - | - | 0 | - |
| - | - | 697.8 | 284.2 | - | - | 0 | - |
| - | - | 866.6 | 285.1 | - | - | 0 | - |
| - | - | 648.8 | 285.2 | - | - | 0 | - |
| - | - | 4067 | 286.1 | - | - | 0 | - |
| - | - | 4628 | 286.2 | - | - | 0 | - |
| - | - | 1416 | 287.2 | - | - | 0 | - |
| - | - | 962.1 | 287.2 | - | - | 0 | - |
| - | - | 1.23E+04 | 288.2 | - | - | 0 | - |
| - | - | 989.2 | 288.6 | - | - | 0 | - |
| - | - | 2320 | 289.2 | - | - | 0 | - |
| - | - | 5665 | 290.1 | - | - | 0 | - |
| 12 | y | 4.083E+04 | 293.1 | 0.0006325 | 2.158 | +1 | 2 |
| - | - | 1008 | 293.2 | - | - | 0 | - |
| - | - | 8997 | 294.1 | - | - | 0 | - |
| - | - | 693.2 | 295.1 | - | - | 0 | - |
| 3 | b | 1871 | 296.2 | 0.001008 | 3.403 | +1 | 3 |
| - | - | 1.339E+04 | 297.2 | - | - | 0 | - |
| - | - | 1044 | 297.2 | - | - | 0 | - |
| - | - | 1447 | 298.1 | - | - | 0 | - |
| - | - | 3704 | 298.2 | - | - | 0 | - |
| - | - | 1983 | 299.1 | - | - | 0 | - |
| - | - | 1854 | 300.1 | - | - | 0 | - |
| - | - | 1126 | 300.1 | - | - | 0 | - |
| - | - | 8864 | 300.2 | - | - | 0 | - |
| - | - | 4335 | 301.1 | - | - | 0 | - |
| - | - | 3581 | 301.1 | - | - | 0 | - |
| - | - | 3.96E+04 | 301.2 | - | - | 0 | - |
| - | - | 1613 | 302.1 | - | - | 0 | - |
| - | - | 755.6 | 302.1 | - | - | 0 | - |
| - | - | 1466 | 302.1 | - | - | 0 | - |
| - | - | 5434 | 302.2 | - | - | 0 | - |
| - | - | 1565 | 303.1 | - | - | 0 | - |
| - | - | 981.3 | 304.1 | - | - | 0 | - |
| - | - | 662 | 304.7 | - | - | 0 | - |
| - | - | 3900 | 305.2 | - | - | 0 | - |
| - | - | 1267 | 305.6 | - | - | 0 | - |
| 9 | y | 9452 | 306.1 | 0.0006129 | 2.002 | +2 | 5 |
| - | - | 2641 | 306.6 | - | - | 0 | - |
| - | - | 1086 | 307.1 | - | - | 0 | - |
| - | - | 1323 | 309.2 | - | - | 0 | - |
| - | - | 1285 | 310.2 | - | - | 0 | - |
| 12 | y | 6539 | 311.1 | 0.0009321 | 2.996 | +1 | 2 |
| - | - | 753.7 | 311.7 | - | - | 0 | - |
| - | - | 1275 | 312.1 | - | - | 0 | - |
| - | - | 1.194E+04 | 312.2 | - | - | 0 | - |
| - | - | 833.5 | 313.2 | - | - | 0 | - |
| - | - | 2233 | 313.2 | - | - | 0 | - |
| - | - | 7108 | 314.1 | - | - | 0 | - |
| 3 | b | 3103 | 314.2 | 0.0008802 | 2.801 | +1 | 3 |
| - | - | 2.17E+04 | 315.2 | - | - | 0 | - |
| - | - | 2139 | 316.2 | - | - | 0 | - |
| - | - | 6087 | 316.2 | - | - | 0 | - |
| - | - | 9065 | 318.1 | - | - | 0 | - |
| - | - | 1510 | 319.1 | - | - | 0 | - |
| - | - | 1300 | 319.2 | - | - | 0 | - |
| - | - | 795.4 | 322.2 | - | - | 0 | - |
| - | - | 1278 | 325.2 | - | - | 0 | - |
| - | - | 1112 | 326.1 | - | - | 0 | - |
| - | - | 3265 | 327.3 | - | - | 0 | - |
| - | - | 4410 | 330.2 | - | - | 0 | - |
| 8 | b | 617.4 | 331.2 | 0.004207 | 12.7 | +3 | 8 |
| - | - | 2114 | 332.2 | - | - | 0 | - |
| - | - | 6729 | 333.2 | - | - | 0 | - |
| - | - | 1352 | 334.2 | - | - | 0 | - |
| - | - | 1966 | 334.7 | - | - | 0 | - |
| - | - | 1781 | 338.2 | - | - | 0 | - |
| - | - | 1500 | 341.2 | - | - | 0 | - |
| 6 | y | 2200 | 342.1 | 0.003795 | 11.09 | +3 | 8 |
| - | - | 2333 | 344.2 | - | - | 0 | - |
| - | - | 3853 | 345.1 | - | - | 0 | - |
| - | - | 1.244E+04 | 346.1 | - | - | 0 | - |
| - | - | 2279 | 347.1 | - | - | 0 | - |
| 6 | b | 4648 | 347.7 | 0.0004829 | 1.389 | +2 | 6 |
| - | - | 2200 | 348.2 | - | - | 0 | - |
| - | - | 1500 | 348.7 | - | - | 0 | - |
| - | - | 675.3 | 350.2 | - | - | 0 | - |
| - | - | 9947 | 350.2 | - | - | 0 | - |
| - | - | 1672 | 351.2 | - | - | 0 | - |
| 8 | y | 1234 | 353.7 | 0.000698 | 1.973 | +2 | 6 |
| - | - | 868.3 | 354.2 | - | - | 0 | - |
| - | - | 794.9 | 355.2 | - | - | 0 | - |
| - | - | 1440 | 355.2 | - | - | 0 | - |
| - | - | 866.1 | 357.2 | - | - | 0 | - |
| - | - | 2618 | 358.2 | - | - | 0 | - |
| - | - | 687.1 | 359.2 | - | - | 0 | - |
| - | - | 888.3 | 360.2 | - | - | 0 | - |
| - | - | 3381 | 361.2 | - | - | 0 | - |
| - | - | 3948 | 362.2 | - | - | 0 | - |
| 8 | y | 1.339E+04 | 362.7 | 0.0008783 | 2.422 | +2 | 6 |
| - | - | 2.084E+04 | 363.1 | - | - | 0 | - |
| - | - | 5254 | 363.2 | - | - | 0 | - |
| - | - | 1329 | 363.7 | - | - | 0 | - |
| - | - | 4369 | 364.1 | - | - | 0 | - |
| - | - | 2060 | 364.1 | - | - | 0 | - |
| - | - | 793.1 | 364.2 | - | - | 0 | - |
| - | - | 806.9 | 364.7 | - | - | 0 | - |
| - | - | 958.5 | 365.1 | - | - | 0 | - |
| - | - | 2084 | 366.2 | - | - | 0 | - |
| - | - | 1878 | 367.2 | - | - | 0 | - |
| - | - | 1583 | 367.2 | - | - | 0 | - |
| - | - | 1312 | 368.7 | - | - | 0 | - |
| - | - | 647.3 | 369.2 | - | - | 0 | - |
| - | - | 2471 | 373.2 | - | - | 0 | - |
| - | - | 910.1 | 373.7 | - | - | 0 | - |
| - | - | 1089 | 375.7 | - | - | 0 | - |
| - | - | 1042 | 376.7 | - | - | 0 | - |
| - | - | 1347 | 377.1 | - | - | 0 | - |
| - | - | 814.9 | 378.2 | - | - | 0 | - |
| - | - | 916.2 | 378.2 | - | - | 0 | - |
| - | - | 7319 | 378.2 | - | - | 0 | - |
| - | - | 2.833E+04 | 379.2 | - | - | 0 | - |
| - | - | 6069 | 380.2 | - | - | 0 | - |
| - | - | 5.714E+04 | 381.2 | - | - | 0 | - |
| - | - | 716.6 | 381.7 | - | - | 0 | - |
| - | - | 1.149E+04 | 382.2 | - | - | 0 | - |
| - | - | 1130 | 382.2 | - | - | 0 | - |
| - | - | 1924 | 383.1 | - | - | 0 | - |
| - | - | 1240 | 383.2 | - | - | 0 | - |
| - | - | 2361 | 383.2 | - | - | 0 | - |
| - | - | 742.9 | 383.7 | - | - | 0 | - |
| - | - | 5001 | 384.2 | - | - | 0 | - |
| - | - | 1352 | 384.7 | - | - | 0 | - |
| - | - | 1907 | 385.2 | - | - | 0 | - |
| - | - | 998.5 | 385.2 | - | - | 0 | - |
| - | - | 1258 | 389.2 | - | - | 0 | - |
| - | - | 690 | 389.7 | - | - | 0 | - |
| - | - | 2755 | 390.2 | - | - | 0 | - |
| - | - | 870.1 | 390.7 | - | - | 0 | - |
| - | - | 2733 | 391.1 | - | - | 0 | - |
| - | - | 876.6 | 392.2 | - | - | 0 | - |
| - | - | 7845 | 394.2 | - | - | 0 | - |
| - | - | 3294 | 395.2 | - | - | 0 | - |
| - | - | 3043 | 395.2 | - | - | 0 | - |
| - | - | 1281 | 396.2 | - | - | 0 | - |
| - | - | 8660 | 396.2 | - | - | 0 | - |
| - | - | 1572 | 397.2 | - | - | 0 | - |
| - | - | 2637 | 397.3 | - | - | 0 | - |
| - | - | 3410 | 398.2 | - | - | 0 | - |
| - | - | 775.9 | 398.2 | - | - | 0 | - |
| - | - | 1162 | 398.3 | - | - | 0 | - |
| - | - | 2001 | 398.7 | - | - | 0 | - |
| - | - | 2719 | 399.2 | - | - | 0 | - |
| - | - | 2200 | 400.2 | - | - | 0 | - |
| - | - | 1886 | 401.1 | - | - | 0 | - |
| - | - | 2474 | 401.2 | - | - | 0 | - |
| - | - | 5368 | 402.2 | - | - | 0 | - |
| - | - | 2580 | 403.2 | - | - | 0 | - |
| - | - | 2378 | 405.3 | - | - | 0 | - |
| - | - | 1296 | 406.2 | - | - | 0 | - |
| - | - | 687.6 | 407.2 | - | - | 0 | - |
| - | - | 2500 | 407.2 | - | - | 0 | - |
| - | - | 934.7 | 408.2 | - | - | 0 | - |
| - | - | 4675 | 409.1 | - | - | 0 | - |
| - | - | 1315 | 409.2 | - | - | 0 | - |
| - | - | 1831 | 410.1 | - | - | 0 | - |
| - | - | 973.9 | 410.2 | - | - | 0 | - |
| - | - | 2432 | 411.1 | - | - | 0 | - |
| - | - | 3277 | 411.2 | - | - | 0 | - |
| - | - | 3.378E+04 | 412.2 | - | - | 0 | - |
| - | - | 3322 | 412.3 | - | - | 0 | - |
| - | - | 4784 | 413.2 | - | - | 0 | - |
| - | - | 5303 | 413.2 | - | - | 0 | - |
| - | - | 983.9 | 413.3 | - | - | 0 | - |
| - | - | 1633 | 414.2 | - | - | 0 | - |
| - | - | 1.002E+04 | 414.2 | - | - | 0 | - |
| - | - | 1.052E+04 | 414.3 | - | - | 0 | - |
| - | - | 1087 | 414.7 | - | - | 0 | - |
| - | - | 2296 | 415.2 | - | - | 0 | - |
| - | - | 1141 | 416.7 | - | - | 0 | - |
| - | - | 2705 | 417.2 | - | - | 0 | - |
| - | - | 778.3 | 417.7 | - | - | 0 | - |
| - | - | 2439 | 418.2 | - | - | 0 | - |
| - | - | 849.7 | 418.7 | - | - | 0 | - |
| 11 | y | 1625 | 421.2 | 0.0008233 | 1.955 | +1 | 3 |
| 11 | y | 927.7 | 422.2 | 8.405E-05 | 0.1991 | +1 | 3 |
| - | - | 4316 | 423.2 | - | - | 0 | - |
| - | - | 1.311E+04 | 423.3 | - | - | 0 | - |
| - | - | 862.5 | 424.2 | - | - | 0 | - |
| - | - | 8423 | 424.3 | - | - | 0 | - |
| - | - | 3153 | 425.3 | - | - | 0 | - |
| - | - | 6826 | 425.7 | - | - | 0 | - |
| - | - | 9998 | 426.2 | - | - | 0 | - |
| - | - | 4311 | 426.7 | - | - | 0 | - |
| - | - | 7218 | 428.2 | - | - | 0 | - |
| - | - | 1.517E+04 | 429.1 | - | - | 0 | - |
| - | - | 1.403E+04 | 429.2 | - | - | 0 | - |
| - | - | 3473 | 430.1 | - | - | 0 | - |
| 4 | y | 8.59E+04 | 430.2 | 0.00454 | 10.55 | +3 | 10 |
| - | - | 2291 | 430.7 | - | - | 0 | - |
| - | - | 2.783E+04 | 431.2 | - | - | 0 | - |
| - | - | 2758 | 431.7 | - | - | 0 | - |
| - | - | 4715 | 432.2 | - | - | 0 | - |
| - | - | 684.2 | 432.2 | - | - | 0 | - |
| 4 | b | 3501 | 433.3 | 0.0008728 | 2.015 | +1 | 4 |
| - | - | 936.3 | 434.3 | - | - | 0 | - |
| - | - | 6.051E+04 | 434.7 | - | - | 0 | - |
| - | - | 2.973E+04 | 435.2 | - | - | 0 | - |
| - | - | 8070 | 435.7 | - | - | 0 | - |
| - | - | 1985 | 436.2 | - | - | 0 | - |
| 11 | y | 2.446E+04 | 439.2 | 0.001001 | 2.279 | +1 | 3 |
| - | - | 1246 | 439.3 | - | - | 0 | - |
| 7 | b | 2.699E+04 | 439.7 | 0.003274 | 7.446 | +2 | 7 |
| - | - | 1.754E+04 | 440.2 | - | - | 0 | - |
| - | - | 5603 | 440.7 | - | - | 0 | - |
| - | - | 973.8 | 440.7 | - | - | 0 | - |
| - | - | 817.2 | 441.2 | - | - | 0 | - |
| - | - | 1254 | 441.2 | - | - | 0 | - |
| - | - | 1405 | 441.3 | - | - | 0 | - |
| - | - | 2.55E+04 | 442.3 | - | - | 0 | - |
| - | - | 1.112E+04 | 446.2 | - | - | 0 | - |
| - | - | 1882 | 446.7 | - | - | 0 | - |
| - | - | 3010 | 447.2 | - | - | 0 | - |
| - | - | 1495 | 447.2 | - | - | 0 | - |
| - | - | 3493 | 448.2 | - | - | 0 | - |
| 7 | b | 3.854E+04 | 448.7 | 0.003363 | 7.495 | +2 | 7 |
| - | - | 2.44E+04 | 449.2 | - | - | 0 | - |
| - | - | 783.8 | 449.5 | - | - | 0 | - |
| - | - | 5922 | 449.7 | - | - | 0 | - |
| - | - | 1241 | 450.3 | - | - | 0 | - |
| 4 | b | 2.608E+04 | 451.3 | 0.001111 | 2.463 | +1 | 4 |
| - | - | 6173 | 452.3 | - | - | 0 | - |
| - | - | 1152 | 453.3 | - | - | 0 | - |
| - | - | 2764 | 454.2 | - | - | 0 | - |
| 7 | y | 9321 | 454.7 | 0.004036 | 8.876 | +2 | 7 |
| - | - | 5418 | 455.2 | - | - | 0 | - |
| - | - | 960.5 | 455.3 | - | - | 0 | - |
| - | - | 2337 | 455.7 | - | - | 0 | - |
| - | - | 789.5 | 456.2 | - | - | 0 | - |
| - | - | 698 | 458.2 | - | - | 0 | - |
| - | - | 822.2 | 458.2 | - | - | 0 | - |
| - | - | 2228 | 459.2 | - | - | 0 | - |
| - | - | 1961 | 461.2 | - | - | 0 | - |
| - | - | 1057 | 461.2 | - | - | 0 | - |
| - | - | 1548 | 461.3 | - | - | 0 | - |
| - | - | 753.4 | 463.2 | - | - | 0 | - |
| 7 | y | 2767 | 463.7 | 0.003545 | 7.644 | +2 | 7 |
| - | - | 3486 | 464.2 | - | - | 0 | - |
| - | - | 1365 | 466.2 | - | - | 0 | - |
| - | - | 6926 | 468.3 | - | - | 0 | - |
| - | - | 1007 | 468.8 | - | - | 0 | - |
| - | - | 1046 | 469.3 | - | - | 0 | - |
| - | - | 1974 | 469.3 | - | - | 0 | - |
| - | - | 916.1 | 470.2 | - | - | 0 | - |
| - | - | 1288 | 471.3 | - | - | 0 | - |
| - | - | 1976 | 473.3 | - | - | 0 | - |
| - | - | 972.3 | 473.7 | - | - | 0 | - |
| - | - | 798.1 | 474.2 | - | - | 0 | - |
| - | - | 753.1 | 475.2 | - | - | 0 | - |
| - | - | 842.4 | 475.3 | - | - | 0 | - |
| - | - | 2625 | 476.2 | - | - | 0 | - |
| - | - | 3646 | 477.2 | - | - | 0 | - |
| 10 | y | 4203 | 478.2 | 0.0002336 | 0.4884 | +1 | 4 |
| - | - | 3506 | 478.3 | - | - | 0 | - |
| - | - | 1174 | 478.3 | - | - | 0 | - |
| - | - | 1958 | 479.2 | - | - | 0 | - |
| - | - | 1.23E+04 | 482.3 | - | - | 0 | - |
| - | - | 9895 | 482.8 | - | - | 0 | - |
| - | - | 3761 | 483.3 | - | - | 0 | - |
| - | - | 5828 | 483.3 | - | - | 0 | - |
| - | - | 922 | 487.7 | - | - | 0 | - |
| - | - | 843.8 | 488.3 | - | - | 0 | - |
| - | - | 891 | 488.8 | - | - | 0 | - |
| - | - | 1286 | 490.2 | - | - | 0 | - |
| - | - | 2.027E+04 | 491.3 | - | - | 0 | - |
| - | - | 9173 | 491.8 | - | - | 0 | - |
| - | - | 2489 | 492.3 | - | - | 0 | - |
| - | - | 5.186E+04 | 494.2 | - | - | 0 | - |
| - | - | 1.503E+04 | 495.2 | - | - | 0 | - |
| - | - | 4660 | 495.3 | - | - | 0 | - |
| 8 | b | 6.813E+04 | 496.3 | 0.008728 | 17.59 | +2 | 8 |
| - | - | 6534 | 496.3 | - | - | 0 | - |
| - | - | 1.404E+04 | 496.8 | - | - | 0 | - |
| - | - | 1.592E+04 | 497.2 | - | - | 0 | - |
| - | - | 1032 | 497.3 | - | - | 0 | - |
| - | - | 3948 | 498.2 | - | - | 0 | - |
| - | - | 1319 | 500.2 | - | - | 0 | - |
| 8 | b | 9327 | 505.3 | 0.003445 | 6.819 | +2 | 8 |
| - | - | 6742 | 505.8 | - | - | 0 | - |
| - | - | 996 | 506.1 | - | - | 0 | - |
| - | - | 1963 | 506.3 | - | - | 0 | - |
| - | - | 2820 | 507.3 | - | - | 0 | - |
| - | - | 8560 | 511.3 | - | - | 0 | - |
| - | - | 7686 | 511.3 | - | - | 0 | - |
| - | - | 1333 | 512.2 | - | - | 0 | - |
| - | - | 1637 | 512.3 | - | - | 0 | - |
| - | - | 1.114E+05 | 513.4 | - | - | 0 | - |
| - | - | 715.8 | 513.7 | - | - | 0 | - |
| - | - | 1166 | 514.2 | - | - | 0 | - |
| - | - | 2350 | 520.2 | - | - | 0 | - |
| - | - | 5321 | 523.3 | - | - | 0 | - |
| - | - | 4638 | 524.3 | - | - | 0 | - |
| - | - | 1460 | 524.8 | - | - | 0 | - |
| - | - | 9873 | 525.3 | - | - | 0 | - |
| - | - | 3953 | 525.8 | - | - | 0 | - |
| - | - | 2751 | 526.3 | - | - | 0 | - |
| - | - | 3692 | 526.3 | - | - | 0 | - |
| - | - | 1612 | 526.8 | - | - | 0 | - |
| - | - | 815.1 | 529.2 | - | - | 0 | - |
| - | - | 1101 | 530.2 | - | - | 0 | - |
| 0 | Precursor | 5295 | 534.6 | 0.003139 | 5.872 | +3 | -1 |
| 0 | Precursor | 6066 | 534.9 | 0.00979 | 18.3 | +3 | -1 |
| - | - | 1647 | 535.3 | - | - | 0 | - |
| - | - | 4261 | 535.6 | - | - | 0 | - |
| - | - | 2398 | 535.9 | - | - | 0 | - |
| - | - | 1164 | 536.3 | - | - | 0 | - |
| - | - | 931.1 | 537.2 | - | - | 0 | - |
| - | - | 744.8 | 538.2 | - | - | 0 | - |
| - | - | 2.468E+04 | 539.8 | - | - | 0 | - |
| - | - | 1308 | 540.3 | - | - | 0 | - |
| - | - | 1.8E+04 | 540.3 | - | - | 0 | - |
| 0 | Precursor | 6205 | 540.6 | 0.003707 | 6.857 | +3 | -1 |
| - | - | 5650 | 540.8 | - | - | 0 | - |
| - | - | 5307 | 540.9 | - | - | 0 | - |
| - | - | 1.789E+05 | 541.3 | - | - | 0 | - |
| - | - | 6780 | 541.6 | - | - | 0 | - |
| - | - | 6594 | 541.9 | - | - | 0 | - |
| - | - | 1334 | 542.2 | - | - | 0 | - |
| - | - | 3353 | 542.3 | - | - | 0 | - |
| - | - | 866.9 | 542.3 | - | - | 0 | - |
| - | - | 1.349E+04 | 543.3 | - | - | 0 | - |
| - | - | 4888 | 544.3 | - | - | 0 | - |
| - | - | 1890 | 545.2 | - | - | 0 | - |
| - | - | 839.5 | 545.3 | - | - | 0 | - |
| - | - | 1184 | 545.7 | - | - | 0 | - |
| - | - | 1507 | 546.3 | - | - | 0 | - |
| - | - | 947.1 | 546.8 | - | - | 0 | - |
| - | - | 768.6 | 547.3 | - | - | 0 | - |
| - | - | 2643 | 548.2 | - | - | 0 | - |
| - | - | 1226 | 549.2 | - | - | 0 | - |
| - | - | 622.7 | 549.2 | - | - | 0 | - |
| - | - | 714.3 | 551.3 | - | - | 0 | - |
| - | - | 1696 | 551.3 | - | - | 0 | - |
| - | - | 6531 | 554.3 | - | - | 0 | - |
| - | - | 1485 | 554.7 | - | - | 0 | - |
| - | - | 3121 | 554.8 | - | - | 0 | - |
| - | - | 4787 | 555.2 | - | - | 0 | - |
| - | - | 1125 | 555.3 | - | - | 0 | - |
| - | - | 1479 | 556.2 | - | - | 0 | - |
| - | - | 741.5 | 557.2 | - | - | 0 | - |
| - | - | 1705 | 559.3 | - | - | 0 | - |
| 5 | b | 1071 | 561.3 | 0.007969 | 14.2 | +1 | 5 |
| 5 | b | 2069 | 562.3 | 0.0008212 | 1.461 | +1 | 5 |
| 9 | b | 1738 | 562.8 | 0.0002887 | 0.513 | +2 | 9 |
| - | - | 5227 | 563.3 | - | - | 0 | - |
| - | - | 2819 | 563.8 | - | - | 0 | - |
| - | - | 1809 | 565.2 | - | - | 0 | - |
| - | - | 5268 | 565.3 | - | - | 0 | - |
| - | - | 1215 | 566.2 | - | - | 0 | - |
| - | - | 1493 | 566.3 | - | - | 0 | - |
| - | - | 962.7 | 572.3 | - | - | 0 | - |
| - | - | 5126 | 573.8 | - | - | 0 | - |
| - | - | 2821 | 574.3 | - | - | 0 | - |
| - | - | 4314 | 575.2 | - | - | 0 | - |
| - | - | 1192 | 575.3 | - | - | 0 | - |
| 5 | y | 899.1 | 576.3 | 0.008939 | 15.51 | +2 | 9 |
| - | - | 838.6 | 576.3 | - | - | 0 | - |
| 5 | b | 1.888E+04 | 579.3 | 0.0005783 | 0.9982 | +1 | 5 |
| - | - | 1130 | 580.3 | - | - | 0 | - |
| - | - | 5589 | 580.3 | - | - | 0 | - |
| - | - | 1083 | 581.3 | - | - | 0 | - |
| - | - | 1.639E+04 | 583.2 | - | - | 0 | - |
| - | - | 4963 | 584.2 | - | - | 0 | - |
| - | - | 927.3 | 584.3 | - | - | 0 | - |
| - | - | 955.9 | 587.3 | - | - | 0 | - |
| - | - | 827.1 | 588.8 | - | - | 0 | - |
| - | - | 1172 | 589.3 | - | - | 0 | - |
| - | - | 1988 | 592.3 | - | - | 0 | - |
| 9 | y | 2.346E+04 | 593.3 | 0.0005733 | 0.9663 | +1 | 5 |
| - | - | 1.112E+04 | 593.3 | - | - | 0 | - |
| - | - | 9722 | 594.3 | - | - | 0 | - |
| - | - | 2659 | 594.3 | - | - | 0 | - |
| - | - | 2168 | 595.2 | - | - | 0 | - |
| - | - | 1276 | 595.3 | - | - | 0 | - |
| - | - | 1129 | 596.4 | - | - | 0 | - |
| - | - | 874.6 | 596.8 | - | - | 0 | - |
| - | - | 4.91E+04 | 597.8 | - | - | 0 | - |
| - | - | 2.68E+04 | 598.3 | - | - | 0 | - |
| - | - | 979.3 | 599 | - | - | 0 | - |
| - | - | 1059 | 602.3 | - | - | 0 | - |
| - | - | 1200 | 605.8 | - | - | 0 | - |
| - | - | 1070 | 606.3 | - | - | 0 | - |
| - | - | 2273 | 608.3 | - | - | 0 | - |
| - | - | 3.394E+04 | 610.3 | - | - | 0 | - |
| - | - | 3714 | 610.3 | - | - | 0 | - |
| - | - | 3378 | 610.8 | - | - | 0 | - |
| 9 | y | 2.306E+05 | 611.3 | 0.0013 | 2.127 | +1 | 5 |
| - | - | 1884 | 611.8 | - | - | 0 | - |
| - | - | 7.307E+04 | 612.3 | - | - | 0 | - |
| - | - | 808.9 | 613.2 | - | - | 0 | - |
| - | - | 1.447E+04 | 613.3 | - | - | 0 | - |
| - | - | 1860 | 614.3 | - | - | 0 | - |
| - | - | 899.3 | 618.8 | - | - | 0 | - |
| - | - | 4537 | 619.3 | - | - | 0 | - |
| - | - | 1.39E+04 | 619.8 | - | - | 0 | - |
| - | - | 6690 | 620.3 | - | - | 0 | - |
| - | - | 3265 | 620.8 | - | - | 0 | - |
| - | - | 2897 | 621.3 | - | - | 0 | - |
| - | - | 1826 | 621.3 | - | - | 0 | - |
| - | - | 1070 | 622.3 | - | - | 0 | - |
| - | - | 1781 | 628.8 | - | - | 0 | - |
| - | - | 2253 | 629.3 | - | - | 0 | - |
| - | - | 1166 | 629.8 | - | - | 0 | - |
| - | - | 1185 | 631.8 | - | - | 0 | - |
| - | - | 1397 | 633.3 | - | - | 0 | - |
| - | - | 1213 | 636.8 | - | - | 0 | - |
| - | - | 6184 | 638.3 | - | - | 0 | - |
| - | - | 4289 | 638.8 | - | - | 0 | - |
| 4 | y | 3164 | 644.8 | 0.004085 | 6.336 | +2 | 10 |
| - | - | 2669 | 645.3 | - | - | 0 | - |
| - | - | 808.8 | 645.8 | - | - | 0 | - |
| - | - | 804.6 | 646.3 | - | - | 0 | - |
| - | - | 5.088E+04 | 647.3 | - | - | 0 | - |
| - | - | 2.726E+04 | 647.8 | - | - | 0 | - |
| - | - | 2233 | 648.3 | - | - | 0 | - |
| - | - | 1383 | 649.3 | - | - | 0 | - |
| - | - | 2772 | 650.3 | - | - | 0 | - |
| - | - | 4732 | 651.3 | - | - | 0 | - |
| - | - | 1110 | 652.3 | - | - | 0 | - |
| - | - | 2070 | 652.3 | - | - | 0 | - |
| - | - | 3489 | 653.3 | - | - | 0 | - |
| - | - | 949.6 | 653.4 | - | - | 0 | - |
| 4 | y | 1.467E+04 | 653.8 | 0.004235 | 6.478 | +2 | 10 |
| - | - | 1.095E+04 | 654.3 | - | - | 0 | - |
| - | - | 4560 | 654.8 | - | - | 0 | - |
| 11 | b | 934.1 | 655.3 | 0.004026 | 6.144 | +2 | 11 |
| - | - | 2177 | 656.3 | - | - | 0 | - |
| - | - | 1294 | 658.3 | - | - | 0 | - |
| - | - | 860.1 | 658.8 | - | - | 0 | - |
| - | - | 2823 | 659.3 | - | - | 0 | - |
| - | - | 2106 | 660.3 | - | - | 0 | - |
| - | - | 1522 | 661.3 | - | - | 0 | - |
| - | - | 868.5 | 664.3 | - | - | 0 | - |
| - | - | 1080 | 666.3 | - | - | 0 | - |
| - | - | 2.198E+04 | 668.3 | - | - | 0 | - |
| - | - | 800.6 | 668.8 | - | - | 0 | - |
| - | - | 9716 | 669.3 | - | - | 0 | - |
| - | - | 852.8 | 669.8 | - | - | 0 | - |
| - | - | 1408 | 670.3 | - | - | 0 | - |
| - | - | 2640 | 671.3 | - | - | 0 | - |
| - | - | 1107 | 672.3 | - | - | 0 | - |
| - | - | 3465 | 672.8 | - | - | 0 | - |
| - | - | 1940 | 673.3 | - | - | 0 | - |
| - | - | 2865 | 674.3 | - | - | 0 | - |
| - | - | 759.9 | 674.8 | - | - | 0 | - |
| 6 | b | 1.494E+04 | 676.3 | 0.0004963 | 0.7338 | +1 | 6 |
| 6 | b | 9619 | 677.3 | 0.006715 | 9.914 | +1 | 6 |
| - | - | 1295 | 677.8 | - | - | 0 | - |
| - | - | 2090 | 678.3 | - | - | 0 | - |
| - | - | 3460 | 678.3 | - | - | 0 | - |
| - | - | 1190 | 678.8 | - | - | 0 | - |
| - | - | 5269 | 679.3 | - | - | 0 | - |
| - | - | 1269 | 680.3 | - | - | 0 | - |
| - | - | 1.181E+04 | 681.8 | - | - | 0 | - |
| - | - | 6741 | 682.3 | - | - | 0 | - |
| - | - | 1209 | 684.3 | - | - | 0 | - |
| - | - | 1154 | 687.3 | - | - | 0 | - |
| - | - | 933.7 | 687.8 | - | - | 0 | - |
| - | - | 1669 | 688.3 | - | - | 0 | - |
| - | - | 5856 | 689.4 | - | - | 0 | - |
| - | - | 1618 | 690.4 | - | - | 0 | - |
| - | - | 3234 | 692.3 | - | - | 0 | - |
| - | - | 4324 | 692.8 | - | - | 0 | - |
| - | - | 1749 | 693.3 | - | - | 0 | - |
| - | - | 794.9 | 693.8 | - | - | 0 | - |
| 6 | b | 1.198E+05 | 694.4 | 0.0006128 | 0.8825 | +1 | 6 |
| - | - | 4.523E+04 | 695.4 | - | - | 0 | - |
| - | - | 3.388E+04 | 696.3 | - | - | 0 | - |
| - | - | 5606 | 696.4 | - | - | 0 | - |
| - | - | 1.467E+04 | 697.3 | - | - | 0 | - |
| - | - | 3564 | 698.3 | - | - | 0 | - |
| - | - | 781.4 | 699.3 | - | - | 0 | - |
| - | - | 1914 | 700.3 | - | - | 0 | - |
| - | - | 4142 | 700.8 | - | - | 0 | - |
| 3 | y | 1.543E+04 | 701.3 | 0.003679 | 5.246 | +2 | 11 |
| 3 | y | 1.353E+04 | 701.8 | 0.01137 | 16.2 | +2 | 11 |
| - | - | 9049 | 702.3 | - | - | 0 | - |
| - | - | 6574 | 702.8 | - | - | 0 | - |
| - | - | 5970 | 703.3 | - | - | 0 | - |
| - | - | 856 | 705.4 | - | - | 0 | - |
| - | - | 1384 | 705.9 | - | - | 0 | - |
| 8 | y | 8022 | 706.3 | 0.002386 | 3.378 | +1 | 6 |
| - | - | 2883 | 707.3 | - | - | 0 | - |
| - | - | 1032 | 708.3 | - | - | 0 | - |
| - | - | 2595 | 709.3 | - | - | 0 | - |
| - | - | 2.152E+04 | 709.8 | - | - | 0 | - |
| 3 | y | 9.325E+04 | 710.3 | 0.0045 | 6.336 | +2 | 11 |
| 12 | b | 6.918E+04 | 710.8 | 0.01022 | 14.37 | +2 | 12 |
| 12 | b | 3.164E+04 | 711.3 | 0.000578 | 0.8126 | +2 | 12 |
| - | - | 7.838E+04 | 711.9 | - | - | 0 | - |
| - | - | 4.828E+04 | 712.4 | - | - | 0 | - |
| - | - | 1476 | 712.9 | - | - | 0 | - |
| - | - | 1189 | 713.3 | - | - | 0 | - |
| - | - | 2829 | 719.4 | - | - | 0 | - |
| 12 | b | 7820 | 719.8 | 0.006533 | 9.076 | +2 | 12 |
| - | - | 5710 | 720.4 | - | - | 0 | - |
| - | - | 3873 | 720.9 | - | - | 0 | - |
| - | - | 1072 | 721.4 | - | - | 0 | - |
| - | - | 1.119E+04 | 723.4 | - | - | 0 | - |
| 8 | y | 6.967E+04 | 724.4 | 0.001404 | 1.938 | +1 | 6 |
| - | - | 2.773E+04 | 725.4 | - | - | 0 | - |
| - | - | 7109 | 726.4 | - | - | 0 | - |
| - | - | 3044 | 727.4 | - | - | 0 | - |
| - | - | 1227 | 727.9 | - | - | 0 | - |
| - | - | 1513 | 728.4 | - | - | 0 | - |
| - | - | 1039 | 728.9 | - | - | 0 | - |
| - | - | 1317 | 729.4 | - | - | 0 | - |
| - | - | 1980 | 734.3 | - | - | 0 | - |
| - | - | 1002 | 737.4 | - | - | 0 | - |
| - | - | 855 | 737.9 | - | - | 0 | - |
| - | - | 2659 | 739.4 | - | - | 0 | - |
| - | - | 1959 | 740.4 | - | - | 0 | - |
| - | - | 1595 | 741.4 | - | - | 0 | - |
| - | - | 2137 | 745.4 | - | - | 0 | - |
| - | - | 1.243E+04 | 746.4 | - | - | 0 | - |
| - | - | 7873 | 746.9 | - | - | 0 | - |
| 2 | y | 2246 | 750.9 | 0.002859 | 3.807 | +2 | 12 |
| 2 | y | 3163 | 751.3 | 0.01317 | 17.53 | +2 | 12 |
| - | - | 1418 | 751.9 | - | - | 0 | - |
| - | - | 900.6 | 752.4 | - | - | 0 | - |
| - | - | 1.323E+05 | 755.4 | - | - | 0 | - |
| - | - | 9.106E+04 | 755.9 | - | - | 0 | - |
| - | - | 1194 | 756.4 | - | - | 0 | - |
| - | - | 4247 | 759.4 | - | - | 0 | - |
| 2 | y | 7323 | 759.9 | 0.004839 | 6.369 | +2 | 12 |
| - | - | 1.248E+04 | 760.4 | - | - | 0 | - |
| - | - | 5438 | 760.9 | - | - | 0 | - |
| - | - | 1687 | 762.3 | - | - | 0 | - |
| - | - | 1146 | 762.4 | - | - | 0 | - |
| - | - | 5005 | 763.4 | - | - | 0 | - |
| - | - | 2768 | 764.4 | - | - | 0 | - |
| - | - | 1214 | 765.3 | - | - | 0 | - |
| - | - | 1105 | 766.3 | - | - | 0 | - |
| - | - | 2446 | 767.4 | - | - | 0 | - |
| - | - | 2534 | 768.4 | - | - | 0 | - |
| - | - | 1149 | 769.4 | - | - | 0 | - |
| - | - | 6672 | 779.3 | - | - | 0 | - |
| - | - | 2476 | 780.3 | - | - | 0 | - |
| - | - | 3874 | 781.4 | - | - | 0 | - |
| - | - | 2086 | 782.4 | - | - | 0 | - |
| - | - | 2196 | 783.3 | - | - | 0 | - |
| - | - | 1268 | 784.3 | - | - | 0 | - |
| - | - | 1527 | 791.4 | - | - | 0 | - |
| - | - | 1573 | 792.4 | - | - | 0 | - |
| - | - | 9888 | 795.4 | - | - | 0 | - |
| - | - | 5165 | 796.4 | - | - | 0 | - |
| - | - | 6970 | 797.4 | - | - | 0 | - |
| - | - | 2126 | 798.4 | - | - | 0 | - |
| - | - | 928.4 | 806.4 | - | - | 0 | - |
| - | - | 1425 | 807.4 | - | - | 0 | - |
| - | - | 1.473E+04 | 809.4 | - | - | 0 | - |
| 0 | Precursor | 7878 | 810.4 | 0.008649 | 10.67 | +2 | -1 |
| - | - | 2401 | 811.3 | - | - | 0 | - |
| - | - | 1757 | 811.4 | - | - | 0 | - |
| - | - | 1107 | 812.3 | - | - | 0 | - |
| - | - | 6906 | 823.4 | - | - | 0 | - |
| - | - | 2425 | 824.4 | - | - | 0 | - |
| - | - | 1023 | 825.4 | - | - | 0 | - |
| - | - | 1298 | 828.4 | - | - | 0 | - |
| - | - | 1.381E+04 | 836.4 | - | - | 0 | - |
| - | - | 4400 | 837.4 | - | - | 0 | - |
| - | - | 791.6 | 840.4 | - | - | 0 | - |
| - | - | 1354 | 841.4 | - | - | 0 | - |
| - | - | 1763 | 842.4 | - | - | 0 | - |
| - | - | 1610 | 850.4 | - | - | 0 | - |
| - | - | 2207 | 851.4 | - | - | 0 | - |
| - | - | 3490 | 852.4 | - | - | 0 | - |
| - | - | 1123 | 853.4 | - | - | 0 | - |
| - | - | 1518 | 859.4 | - | - | 0 | - |
| - | - | 3114 | 860.4 | - | - | 0 | - |
| - | - | 2978 | 861.4 | - | - | 0 | - |
| - | - | 1363 | 862.4 | - | - | 0 | - |
| - | - | 777.1 | 864.5 | - | - | 0 | - |
| - | - | 1043 | 867.4 | - | - | 0 | - |
| - | - | 1.705E+04 | 868.4 | - | - | 0 | - |
| - | - | 7488 | 869.4 | - | - | 0 | - |
| - | - | 2952 | 870.4 | - | - | 0 | - |
| 7 | b | 1.595E+04 | 878.4 | 0.004731 | 5.386 | +1 | 7 |
| 7 | b | 1.476E+04 | 879.4 | 0.0151 | 17.17 | +1 | 7 |
| - | - | 5552 | 880.4 | - | - | 0 | - |
| - | - | 1233 | 881.4 | - | - | 0 | - |
| - | - | 814.6 | 887.4 | - | - | 0 | - |
| - | - | 830.6 | 890.4 | - | - | 0 | - |
| - | - | 1028 | 891.4 | - | - | 0 | - |
| - | - | 2926 | 892.4 | - | - | 0 | - |
| 7 | b | 1.446E+05 | 896.4 | 0.004908 | 5.475 | +1 | 7 |
| - | - | 7.395E+04 | 897.4 | - | - | 0 | - |
| - | - | 2.135E+04 | 898.4 | - | - | 0 | - |
| - | - | 4409 | 899.4 | - | - | 0 | - |
| - | - | 932.3 | 900.4 | - | - | 0 | - |
| - | - | 1328 | 906.4 | - | - | 0 | - |
| - | - | 1292 | 907.4 | - | - | 0 | - |
| 7 | y | 2909 | 908.4 | 0.008329 | 9.169 | +1 | 7 |
| - | - | 2023 | 909.4 | - | - | 0 | - |
| - | - | 1999 | 910.4 | - | - | 0 | - |
| - | - | 1398 | 923.4 | - | - | 0 | - |
| - | - | 8034 | 924.4 | - | - | 0 | - |
| - | - | 4572 | 925.4 | - | - | 0 | - |
| 7 | y | 7437 | 926.4 | 0.006859 | 7.403 | +1 | 7 |
| - | - | 3784 | 927.4 | - | - | 0 | - |
| - | - | 1742 | 952.4 | - | - | 0 | - |
| - | - | 1239 | 953.4 | - | - | 0 | - |
| - | - | 1527 | 963.4 | - | - | 0 | - |
| - | - | 2359 | 970.4 | - | - | 0 | - |
| - | - | 3159 | 971.4 | - | - | 0 | - |
| - | - | 993.2 | 972.4 | - | - | 0 | - |
| - | - | 1516 | 975.5 | - | - | 0 | - |
| - | - | 900.3 | 976.5 | - | - | 0 | - |
| - | - | 1880 | 980.5 | - | - | 0 | - |
| - | - | 3833 | 981.4 | - | - | 0 | - |
| - | - | 1430 | 982.4 | - | - | 0 | - |
| - | - | 1126 | 986.5 | - | - | 0 | - |
| - | - | 1554 | 987.5 | - | - | 0 | - |
| - | - | 3028 | 988.4 | - | - | 0 | - |
| - | - | 1158 | 989.4 | - | - | 0 | - |
| 8 | b | 967.9 | 991.5 | 0.0099 | 9.985 | +1 | 8 |
| 8 | b | 1182 | 992.5 | 0.01874 | 18.89 | +1 | 8 |
| - | - | 884.6 | 993.5 | - | - | 0 | - |
| - | - | 1065 | 996.5 | - | - | 0 | - |
| - | - | 916.5 | 997.5 | - | - | 0 | - |
| - | - | 1202 | 998.5 | - | - | 0 | - |
| - | - | 999.2 | 999.5 | - | - | 0 | - |
| 8 | b | 6947 | 1010 | 0.004523 | 4.481 | +1 | 8 |
| - | - | 4958 | 1011 | - | - | 0 | - |
| - | - | 1809 | 1012 | - | - | 0 | - |
| - | - | 1018 | 1014 | - | - | 0 | - |
| 6 | y | 1827 | 1023 | 0.01929 | 18.85 | +1 | 8 |
| - | - | 1399 | 1030 | - | - | 0 | - |
| - | - | 846.5 | 1031 | - | - | 0 | - |
| - | - | 951.6 | 1040 | - | - | 0 | - |
| 6 | y | 6648 | 1041 | 0.007992 | 7.674 | +1 | 8 |
| - | - | 3620 | 1042 | - | - | 0 | - |
| - | - | 1125 | 1074 | - | - | 0 | - |
| - | - | 1439 | 1092 | - | - | 0 | - |
| 9 | b | 895.8 | 1108 | 0.008566 | 7.734 | +1 | 9 |
| - | - | 3041 | 1110 | - | - | 0 | - |
| - | - | 1771 | 1111 | - | - | 0 | - |
| - | - | 1045 | 1124 | - | - | 0 | - |
| 9 | b | 1267 | 1125 | 0.008708 | 7.744 | +1 | 9 |
| - | - | 716.5 | 1126 | - | - | 0 | - |
| - | - | 789 | 1133 | - | - | 0 | - |
| - | - | 1666 | 1134 | - | - | 0 | - |
| - | - | 902.7 | 1135 | - | - | 0 | - |
| - | - | 1142 | 1151 | - | - | 0 | - |
| 5 | y | 5074 | 1151 | 0.004046 | 3.514 | +1 | 9 |
| 5 | y | 7698 | 1152 | 0.01112 | 9.648 | +1 | 9 |
| - | - | 4125 | 1153 | - | - | 0 | - |
| - | - | 1827 | 1155 | - | - | 0 | - |
| - | - | 924.5 | 1161 | - | - | 0 | - |
| - | - | 2036 | 1169 | - | - | 0 | - |
| 5 | y | 1.139E+04 | 1170 | 0.006055 | 5.177 | +1 | 9 |
| - | - | 7441 | 1171 | - | - | 0 | - |
| - | - | 2443 | 1172 | - | - | 0 | - |
| - | - | 3498 | 1178 | - | - | 0 | - |
| - | - | 1262 | 1179 | - | - | 0 | - |
| 10 | b | 2188 | 1182 | 0.01019 | 8.627 | +1 | 10 |
| - | - | 836.9 | 1689 | - | - | 0 | - |
| - | - | 712.9 | 2375 | - | - | 0 | - |
| - | - | 852.9 | 3083 | - | - | 0 | - |

m/z Charge Intensity FragmentType MassShift Position
120.08121490478516 0 12507.683
121.0652847290039 0 529.6981
121.08450317382812 0 685.13556
121.54315948486328 0 419.26355
122.0716781616211 0 1589.1208
123.04450225830078 0 1783.643
123.05594635009766 0 942.5289
125.10777282714844 0 1436.1276
126.05562591552734 0 960.6778
126.0918197631836 0 1041.5883
127.05071258544922 0 5840.716
127.12338256835938 0 745.5115
128.107421875 0 22811.146
129.0662078857422 0 1604.9385
129.10272216796875 0 214958.95
130.0655517578125 0 44242.93
130.10018920898438 0 828.61725
130.1060333251953 0 14605.463
131.06895446777344 0 4402.2783
132.04481506347656 0 7975.175
132.1018524169922 0 505.27994
133.04808044433594 0 757.76935
133.0615692138672 0 510.4088
136.07614135742188 0 29734.662
137.07958984375 0 2455.4197
138.06661987304688 0 3698.9084
138.0918731689453 0 1443.9684
139.08717346191406 0 2242.6992
140.08200073242188 0 1322.9518
141.1026153564453 0 3557.5771
143.43099975585938 0 477.33185
144.07717895507812 0 825.5498
145.06124877929688 0 3143.8386
146.06051635742188 0 3411.4558
147.04441833496094 0 1411.551
147.07701110839844 0 716.9268 y 10
148.94786071777344 0 867.3451
149.04556274414062 0 547.2171
150.0664825439453 0 1608.3156
151.08705139160156 0 6559.8154
153.1029510498047 0 539.547
153.61981201171875 0 469.1184
154.0502471923828 0 1104.4791
155.04527282714844 0 560.92145
155.08218383789062 0 965.63824
155.09320068359375 0 14565.755
155.11831665039062 0 10081.651 a Water loss 1
156.0772247314453 0 846.75684
156.0963134765625 0 1020.4389
156.10267639160156 0 1087.7467
156.12147521972656 0 956.8769
157.0614471435547 0 1083.6504
157.07647705078125 0 12852.812
157.09732055664062 0 1786.5535
157.10879516601562 0 3603.2925
157.13381958007812 0 652.64374
158.06045532226562 0 40268.05
158.08013916015625 0 1255.5784
158.09275817871094 0 1319.9092
159.06394958496094 0 4304.3364
159.092041015625 0 2510.6792
159.11325073242188 0 1612.6387 d 1
165.05506896972656 0 15322.928
165.07679748535156 0 666.6885
166.06149291992188 0 25468.172
166.0867156982422 0 7445.771 y 9
166.09796142578125 0 2619.6868
167.05642700195312 0 983.6166
167.06507873535156 0 1819.0195
167.1183624267578 0 1170.6592
168.10235595703125 0 1921.6066
168.11361694335938 0 2784.313
169.0976104736328 0 5908.2373
169.13397216796875 0 2661.592
171.076904296875 0 1996.111
172.0721893310547 0 2714.5032
172.10830688476562 0 531.35394
173.05616760253906 0 3742.4229
173.1289825439453 0 348238.66 a 1
174.13233947753906 0 30477.633
175.0870819091797 0 211144.98
175.11944580078125 0 2535.9204
176.0904083251953 0 19874.018
177.0938720703125 0 1376.1097
177.1025390625 0 821.1304
178.13430786132812 0 11022.537
179.1374969482422 0 1206.5371
181.06130981445312 0 1325.7358
182.08168029785156 0 34013.188 y 12
183.085205078125 0 3549.756
183.11326599121094 0 17706.096 b Water loss 1
183.14971923828125 0 1108.506
184.1168975830078 0 1305.9565
185.07131958007812 0 2642.505
185.09286499023438 0 1362.4812
185.12918090820312 0 822.6562
185.16546630859375 0 2382.0515
185.37356567382812 0 516.56287
186.12417602539062 0 52938.047
187.08697509765625 0 2523.7449
187.1079864501953 0 1872.5887
187.12765502929688 0 4213.2305
187.1444091796875 0 1226.7767
189.08749389648438 0 6138.391
189.09689331054688 0 952.4512
190.08277893066406 0 5086.3857
190.13421630859375 0 2246.121
193.0973663330078 0 733.80176
194.083740234375 0 643.51733
194.0926971435547 0 2955.4724
195.11325073242188 0 16793.389
196.10842895507812 0 5364.3203
196.11683654785156 0 1141.7797
196.14422607421875 0 782.47363
197.12884521484375 0 3458.535
198.0880889892578 0 2069.3481
199.0719451904297 0 3186.9832
201.12380981445312 0 107143.84 b 1
202.09774780273438 0 842.388
202.12713623046875 0 9185.568
203.06637573242188 0 1207.5868
203.0814208984375 0 1074.7368
203.12904357910156 0 849.1566
203.15176391601562 0 612.04
205.09759521484375 0 7545.914
205.1446533203125 0 1010.0031
206.10195922851562 0 594.9911
206.12908935546875 0 1629.5143
207.160888671875 0 3127.5403
210.12437438964844 0 1482.7152
211.10824584960938 0 2249.7964 y Water loss 10
211.1450958251953 0 656.44916
211.1558837890625 0 9987.179
212.1035614013672 0 979.73016
212.13986206054688 0 2799.5376
213.12376403808594 0 6864.563
213.1599578857422 0 1186.6179
214.12741088867188 0 1058.504
215.0818328857422 0 3278.9407
215.13963317871094 0 1315.116
216.09854125976562 0 1733.7467
216.1348114013672 0 656.80414
217.0823516845703 0 10670.601
220.10781860351562 0 1342.4298
221.09243774414062 0 6049.64
221.10400390625 0 1904.8855
221.14053344726562 0 1095.1667
222.09632873535156 0 848.9451
222.12425231933594 0 3095.1165
223.06419372558594 0 946.2136
223.10824584960938 0 4316.622
223.15582275390625 0 29579.066
224.1593475341797 0 2817.6016
226.08274841308594 0 5164.9546
226.1553192138672 0 706.7329
227.06695556640625 0 4908.6763
227.08399963378906 0 757.6435
227.11477661132812 0 1567.9071
228.1348419189453 0 2291.264
228.18272399902344 0 2072.5388
229.11875915527344 0 4016.0698
230.15060424804688 0 694.6259
231.11358642578125 0 1236.3088
233.14012145996094 0 5684.406
233.1649169921875 0 943.2672
234.12425231933594 0 34629.207
235.12754821777344 0 3255.673
237.13502502441406 0 2278.3076
238.1191864013672 0 3844.727
238.13095092773438 0 1240.8541
239.15093994140625 0 6688.1895
240.13479614257812 0 13398.337
241.09786987304688 0 586.30566
241.11883544921875 0 1587.9097
241.1373291015625 0 1879.2308
241.1920166015625 0 2573.8665
242.1145477294922 0 2038.2338
243.1094207763672 0 2280.9724
243.13710021972656 0 713.8471
243.1496124267578 0 5170.797
244.09327697753906 0 7939.338
244.13050842285156 0 1105.3606
244.15301513671875 0 940.8889
247.10829162597656 0 16331.335
247.14459228515625 0 1315.947
248.1012725830078 0 1050.6975
248.11398315429688 0 7001.6797
248.62425231933594 0 675.7886 y 9
249.0988006591797 0 9959.239
249.11270141601562 0 1070.7596
249.13491821289062 0 2802.0754
250.10269165039062 0 1080.5867
250.11788940429688 0 737.21063
251.15086364746094 0 114640.52
252.13641357421875 0 767.44745
252.15408325195312 0 14827.287
253.15594482421875 0 1618.0642
254.11390686035156 0 746.96423
255.109619140625 0 3395.841
255.1457977294922 0 3245.51
256.1772766113281 0 2101.814
257.1132507324219 0 2456.055
258.1454162597656 0 4896.3843
259.1281433105469 0 992.0389
261.1199035644531 0 3159.3276
261.1353759765625 0 1621.6488
262.1188659667969 0 1272.516
262.1519470214844 0 895.36945
263.1023864746094 0 688.4238
263.1395263671875 0 653.6152
265.1299133300781 0 16857.43
266.1253662109375 0 31060.102
267.1287536621094 0 3531.174
267.14349365234375 0 658.94684
268.1292419433594 0 2515.9111
268.17767333984375 0 2210.7673
269.1289367675781 0 1424.2368
269.1858215332031 0 1674.4318
270.1458740234375 0 991.5609
270.16058349609375 0 6417.6924
271.16387939453125 0 900.8306
272.1026916503906 0 910.2555
274.1302185058594 0 4103.099
275.1024169921875 0 1246.5183
276.10931396484375 0 1636.9265
276.15594482421875 0 2687.6865
279.14599609375 0 1772.8087
280.1298828125 0 3588.2385
282.1567077636719 0 4557.1943
283.140625 0 79849.234
284.1244812011719 0 1180.1819
284.1435546875 0 12556.088
284.1614685058594 0 697.8324
285.1435852050781 0 866.60046
285.15838623046875 0 648.78827
286.14013671875 0 4066.9717
286.1557922363281 0 4627.501
287.1727600097656 0 1415.9451
287.1878356933594 0 962.1043
288.1712646484375 0 12296.111
288.6202697753906 0 989.18475
289.17431640625 0 2319.537
290.1141052246094 0 5665.187
293.11383056640625 0 40831.082 y Water loss 11
293.1625061035156 0 1007.78467
294.11785888671875 0 8997.303
295.1195983886719 0 693.16986
296.1978759765625 0 1870.6289 b Water loss 2
297.1564025878906 0 13393.071
297.1734313964844 0 1044.4205
298.1393737792969 0 1446.8033
298.1573181152344 0 3703.9053
299.06182861328125 0 1982.8218
300.0638732910156 0 1853.6833
300.09832763671875 0 1125.5326
300.167236328125 0 8863.959
301.0600891113281 0 4334.968
301.0824890136719 0 3580.787
301.1512756347656 0 39602.094
302.0616149902344 0 1613.4396
302.0827941894531 0 755.567
302.12554931640625 0 1466.1808
302.1546630859375 0 5434.4863
303.0573425292969 0 1565.3285
304.1307373046875 0 981.25745
304.6639404296875 0 661.9993
305.1616516113281 0 3900.027
305.646240234375 0 1266.6279
306.1378173828125 0 9452.454 y 8
306.6388854980469 0 2640.9155
307.1403503417969 0 1085.893
309.167724609375 0 1323.4089
310.224365234375 0 1285.201
311.12469482421875 0 6538.6133 y 11
311.688232421875 0 753.73364
312.1267395019531 0 1274.5605
312.1559753417969 0 11944.283
313.1599426269531 0 833.49304
313.1874694824219 0 2233.2673
314.1497802734375 0 7108.4927
314.20831298828125 0 3103.2563 b 2
315.1670227050781 0 21700.629
316.15032958984375 0 2138.578
316.1679382324219 0 6087.2793
318.109130859375 0 9064.577
319.1126708984375 0 1509.7422
319.15277099609375 0 1300.4604
322.1872863769531 0 795.3766
325.15087890625 0 1277.6454
326.14874267578125 0 1112.4977
327.2511901855469 0 3265.212
330.16656494140625 0 4409.812
331.1739501953125 0 617.35913 b Water loss 7
332.2091064453125 0 2113.6013
333.177490234375 0 6728.5566
334.1809997558594 0 1352.4585
334.6615905761719 0 1966.408
338.2192687988281 0 1780.961
341.18267822265625 0 1500.4873
342.141357421875 0 2200.0413 y Ammonia loss 5
344.1721496582031 0 2333.1804
345.1312561035156 0 3852.936
346.1150817871094 0 12442.041
347.1180419921875 0 2279.4177
347.6800537109375 0 4647.6416 b 5
348.18157958984375 0 2200.1475
348.6586608886719 0 1500.2849
350.18109130859375 0 675.3093
350.21923828125 0 9946.542
351.22186279296875 0 1672.346
353.6746520996094 0 1233.96 y Water loss 7
354.1753845214844 0 868.29816
355.16595458984375 0 794.89746
355.24609375 0 1440.4958
357.17767333984375 0 866.10065
358.17779541015625 0 2618.266
359.1670837402344 0 687.14496
360.20391845703125 0 888.3328
361.19879150390625 0 3381.0862
362.18450927734375 0 3948.0952
362.68011474609375 0 13389.755 y 7
363.1415710449219 0 20839.35
363.1820373535156 0 5254.2803
363.6846618652344 0 1329.0526
364.1251525878906 0 4368.745
364.14581298828125 0 2059.5637
364.1778564453125 0 793.13165
364.6766357421875 0 806.91284
365.1271667480469 0 958.5336
366.1782531738281 0 2083.8442
367.1623229980469 0 1877.8141
367.245361328125 0 1582.783
368.6942138671875 0 1311.8787
369.232177734375 0 647.3046
373.1858215332031 0 2471.4543
373.6864929199219 0 910.09314
375.68310546875 0 1088.911
376.68011474609375 0 1041.8163
377.1448669433594 0 1346.7692
378.17718505859375 0 814.8966
378.1846008300781 0 916.21515
378.2142639160156 0 7318.554
379.2096252441406 0 28325.463
380.2115478515625 0 6068.7114
381.1523132324219 0 57137.844
381.7126159667969 0 716.57074
382.1548767089844 0 11494.948
382.1908874511719 0 1130.0784
383.1344299316406 0 1923.801
383.1595153808594 0 1239.5216
383.2292785644531 0 2361.0408
383.72802734375 0 742.9472
384.1914367675781 0 5001.3
384.6969909667969 0 1352.3501
385.1707458496094 0 1906.9319
385.19329833984375 0 998.49634
389.19061279296875 0 1258.339
389.6832275390625 0 690.0172
390.1788330078125 0 2754.801
390.6790771484375 0 870.0787
391.13665771484375 0 2733.0825
392.17376708984375 0 876.5548
394.17291259765625 0 7844.6094
395.15667724609375 0 3294.0054
395.24029541015625 0 3042.8035
396.1576232910156 0 1280.9884
396.2249450683594 0 8660.436
397.2273254394531 0 1572.3856
397.2572021484375 0 2637.3923
398.1787414550781 0 3410.1133
398.1995544433594 0 775.9456
398.2870788574219 0 1162.0553
398.6939392089844 0 2000.9229
399.1845397949219 0 2718.6567
400.162841796875 0 2200.121
401.14697265625 0 1886.2947
401.23968505859375 0 2473.683
402.1993713378906 0 5368.4204
403.199462890625 0 2580.0168
405.260986328125 0 2377.9622
406.2438659667969 0 1295.822
407.20758056640625 0 687.5813
407.24090576171875 0 2499.8535
408.2001037597656 0 934.7062
409.14739990234375 0 4674.9307
409.2209777832031 0 1314.959
410.14910888671875 0 1830.732
410.20306396484375 0 973.85156
411.1307067871094 0 2431.9373
411.19891357421875 0 3276.7717
412.183349609375 0 33783.73
412.266845703125 0 3321.7207
413.16571044921875 0 4784.315
413.1872863769531 0 5303.086
413.2519226074219 0 983.93085
414.1697082519531 0 1632.7666
414.2350769042969 0 10020.101
414.283203125 0 10520.451
414.7204895019531 0 1086.5149
415.2391052246094 0 2295.5928
416.70819091796875 0 1141.0144
417.20220947265625 0 2705.3896
417.7012939453125 0 778.3396
418.1732482910156 0 2438.5288
418.71771240234375 0 849.7108
421.208984375 0 1625.0392 y Water loss 10
422.1922607421875 0 927.6597 y Ammonia loss 10
423.234130859375 0 4315.943
423.2720947265625 0 13109.183
424.219970703125 0 862.512
424.2696533203125 0 8422.703
425.2513122558594 0 3152.883
425.7148132324219 0 6825.984
426.2080078125 0 9997.936
426.70916748046875 0 4310.736
428.15716552734375 0 7217.822
429.1413269042969 0 15170.482
429.2101135253906 0 14026.135
430.1427917480469 0 3473.3403
430.1941833496094 0 85904.55 y Water loss 3
430.7075500488281 0 2291.0737
431.197265625 0 27825.357
431.70220947265625 0 2758.2395
432.1993103027344 0 4714.6074
432.22869873046875 0 684.20416
433.25665283203125 0 3500.825 b Water loss 3
434.25604248046875 0 936.2893
434.72015380859375 0 60508.53
435.22161865234375 0 29732.17
435.7228088378906 0 8070.272
436.2264404296875 0 1985.0422
439.2197265625 0 24457.564 y 10
439.2547607421875 0 1245.945
439.71221923828125 0 26987.844 b Water loss 6
440.21307373046875 0 17537.98
440.7098083496094 0 5603.0225
440.7386169433594 0 973.8387
441.17791748046875 0 817.2243
441.2105407714844 0 1253.8922
441.2925109863281 0 1404.7877
442.2779846191406 0 25499.65
446.1678771972656 0 11124.723
446.71893310546875 0 1881.6068
447.17095947265625 0 3010.4805
447.21966552734375 0 1494.6902
448.2044982910156 0 3492.7966
448.71759033203125 0 38537.74 b 6
449.2185974121094 0 24398.406
449.50482177734375 0 783.8429
449.720458984375 0 5921.5356
450.28369140625 0 1240.9878
451.2674560546875 0 26077.414 b 3
452.2699279785156 0 6172.743
453.27191162109375 0 1151.6621
454.2200927734375 0 2764.259
454.712646484375 0 9321.297 y Water loss 6
455.2146911621094 0 5418.0913
455.2708740234375 0 960.5271
455.7169189453125 0 2336.544
456.21234130859375 0 789.4982
458.1886291503906 0 698.042
458.2364807128906 0 822.2048
459.19976806640625 0 2228.4453
461.2049560546875 0 1961.1481
461.24749755859375 0 1056.9744
461.2912902832031 0 1547.9176
463.22637939453125 0 753.4323
463.7174377441406 0 2767.1606 y 6
464.2184753417969 0 3486.0862
466.2439270019531 0 1364.6033
468.2936096191406 0 6926.0684
468.7514343261719 0 1007.1471
469.2527160644531 0 1046.4777
469.2944030761719 0 1973.7018
470.2068786621094 0 916.0624
471.260498046875 0 1288.1672
473.2530517578125 0 1975.9824
473.7492370605469 0 972.2501
474.249755859375 0 798.108
475.23687744140625 0 753.10834
475.2911071777344 0 842.39075
476.2270202636719 0 2625.3938
477.2110290527344 0 3646.1348
478.2298583984375 0 4203.3345 y Water loss 9
478.2777404785156 0 3506.4414
478.3134460449219 0 1174.188
479.228271484375 0 1957.6448
482.2568359375 0 12300.886
482.7547302246094 0 9895.174
483.25421142578125 0 3761.323
483.341064453125 0 5828.461
487.7438049316406 0 921.9703
488.2560729980469 0 843.8059
488.75653076171875 0 891.0279
490.2310791015625 0 1285.9385
491.26202392578125 0 20265.82
491.76373291015625 0 9173.112
492.26409912109375 0 2488.692
494.23651123046875 0 51860.74
495.2394714355469 0 15030.551
495.3406982421875 0 4659.7524
496.24224853515625 0 68134.99 b Water loss 7
496.3253479003906 0 6533.857
496.7659606933594 0 14043.759
497.2448425292969 0 15919.403
497.2788391113281 0 1032.3596
498.2485046386719 0 3947.6143
500.2153015136719 0 1319.0553
505.25970458984375 0 9327.054 b 7
505.7609558105469 0 6741.598
506.1066589355469 0 996.0371
506.26776123046875 0 1962.5262
507.2577819824219 0 2820.2915
511.2627868652344 0 8559.535
511.33587646484375 0 7686.142
512.177978515625 0 1332.6859
512.2685546875 0 1637.2306
513.3515014648438 0 111409.56
513.7290649414062 0 715.7844
514.2317504882812 0 1165.7188
520.1937255859375 0 2350.3525
523.335693359375 0 5320.7734
524.2637329101562 0 4637.6055
524.7656860351562 0 1460.263
525.267578125 0 9872.877
525.8129272460938 0 3953.32
526.27099609375 0 2750.886
526.3145141601562 0 3691.99
526.8140869140625 0 1611.7239
529.2039794921875 0 815.14905
530.1846313476562 0 1100.9042
534.5928344726562 0 5294.7007 Precursor Water loss
534.927490234375 0 6066.086 Precursor Ammonia loss
535.2588500976562 0 1646.6713
535.605224609375 0 4261.492
535.94091796875 0 2398.4253
536.26123046875 0 1163.7734
537.224853515625 0 931.05096
538.205810546875 0 744.8131
539.81005859375 0 24678.023
540.2684326171875 0 1307.536
540.3110961914062 0 17997.469
540.596923828125 0 6204.6733 Precursor
540.8126220703125 0 5649.6973
540.93115234375 0 5306.9424
541.3463134765625 0 178886.94
541.6092529296875 0 6780.1455
541.9434204101562 0 6593.8438
542.22509765625 0 1334.2421
542.2935791015625 0 3352.808
542.3441162109375 0 866.9043
543.2786254882812 0 13486.985
544.281494140625 0 4887.6113
545.24267578125 0 1889.9629
545.2881469726562 0 839.4785
545.7345581054688 0 1183.5054
546.2706909179688 0 1507.0563
546.767578125 0 947.1221
547.2584838867188 0 768.59045
548.18896484375 0 2642.5972
549.1934814453125 0 1225.8722
549.2496337890625 0 622.68524
551.2599487304688 0 714.2998
551.330078125 0 1696.4996
554.2808227539062 0 6530.9717
554.7424926757812 0 1484.5178
554.7833862304688 0 3121.1667
555.2322998046875 0 4787.0435
555.2760620117188 0 1125.0566
556.2344970703125 0 1478.7687
557.233642578125 0 741.48193
559.2514038085938 0 1704.7313
561.3223266601562 0 1071.4125 b Water loss 4
562.2991943359375 0 2069.007 b Ammonia loss 4
562.77001953125 0 1737.5437 b 8
563.2548828125 0 5226.5444
563.7549438476562 0 2818.7578
565.21435546875 0 1808.8457
565.2626342773438 0 5268.055
566.2064208984375 0 1215.2129
566.263427734375 0 1492.5869
572.2579345703125 0 962.73
573.7978515625 0 5126.3604
574.2982788085938 0 2821.3655
575.2470703125 0 4313.749
575.2930297851562 0 1192.1951
576.242431640625 0 899.1198 y Water loss 4
576.278076171875 0 838.60944
579.3255004882812 0 18877.293 b 4
580.2811889648438 0 1129.5477
580.3285522460938 0 5589.2397
581.3329467773438 0 1082.7875
583.2269287109375 0 16387.27
584.2296752929688 0 4963.3853
584.3145141601562 0 927.3395
587.2781982421875 0 955.8535
588.7943115234375 0 827.0715
589.2960205078125 0 1172.4608
592.2723388671875 0 1987.955
593.2571411132812 0 23456.533 y Water loss 8
593.3060913085938 0 11121.328
594.2551879882812 0 9721.841
594.3088989257812 0 2658.8909
595.2484130859375 0 2168.0347
595.317138671875 0 1276.1545
596.3558349609375 0 1129.356
596.7921142578125 0 874.6159
597.797119140625 0 49095.21
598.2986450195312 0 26802.55
599.0022583007812 0 979.2698
602.2799072265625 0 1058.7448
605.7974853515625 0 1200.1779
606.3023681640625 0 1069.9092
608.3184814453125 0 2273.415
610.2836303710938 0 33936.535
610.3321533203125 0 3714.2854
610.7902221679688 0 3378.4849
611.2684326171875 0 230621.44 y 8
611.7896728515625 0 1884.4014
612.2711181640625 0 73071.45
613.2183227539062 0 808.89
613.273681640625 0 14466.814
614.2767333984375 0 1859.7517
618.7906494140625 0 899.26324
619.3011474609375 0 4537.367
619.79541015625 0 13902.509
620.2975463867188 0 6689.994
620.7984008789062 0 3264.659
621.2521362304688 0 2896.8823
621.3007202148438 0 1826.3468
622.2548217773438 0 1069.876
628.799560546875 0 1781.07
629.3115234375 0 2253.389
629.811279296875 0 1165.8604
631.825439453125 0 1185.4069
633.274658203125 0 1397.1943
636.7755126953125 0 1212.5278
638.3192138671875 0 6184.16
638.8217163085938 0 4288.7847
644.784912109375 0 3164.419 y Water loss 3
645.2857666015625 0 2668.5942
645.7852172851562 0 808.79913
646.2881469726562 0 804.6036
647.33154296875 0 50875.49
647.8328857421875 0 27255.146
648.34228515625 0 2232.6377
649.3353271484375 0 1382.5538
650.3076171875 0 2771.8992
651.2932739257812 0 4732.3047
652.2871704101562 0 1109.9143
652.3405151367188 0 2070.0986
653.29638671875 0 3488.745
653.3529052734375 0 949.6314
653.7903442382812 0 14670.893 y 3
654.2914428710938 0 10949.846
654.7926025390625 0 4559.64
655.3319702148438 0 934.0938 b 10
656.269775390625 0 2176.679
658.3311157226562 0 1293.5625
658.7808837890625 0 860.07764
659.3200073242188 0 2822.7827
660.3221435546875 0 2105.9265
661.2726440429688 0 1522.3508
664.3330688476562 0 868.5276
666.2637329101562 0 1080.0575
668.3157348632812 0 21981.99
668.8215942382812 0 800.56775
669.3197021484375 0 9716.022
669.8240966796875 0 852.77124
670.3214721679688 0 1407.704
671.3462524414062 0 2639.6597
672.3396606445312 0 1106.7701
672.8281860351562 0 3464.858
673.3309936523438 0 1939.9452
674.3081665039062 0 2864.96
674.8259887695312 0 759.8806
676.341796875 0 14941.226 b Water loss 5
677.33203125 0 9618.82 b Ammonia loss 5
677.8276977539062 0 1295.0458
678.2971801757812 0 2089.6147
678.3391723632812 0 3460.3945
678.8267211914062 0 1189.603
679.2864379882812 0 5269.005
680.2919311523438 0 1269.061
681.8342895507812 0 11811.973
682.3350830078125 0 6741.3047
684.2738647460938 0 1208.7936
687.3311767578125 0 1153.9335
687.8329467773438 0 933.7261
688.331787109375 0 1668.5201
689.357177734375 0 5856.2915
690.3594360351562 0 1617.5631
692.32080078125 0 3233.5046
692.8162231445312 0 4323.596
693.3160400390625 0 1748.6276
693.8214111328125 0 794.87915
694.3524780273438 0 119839.484 b 5
695.3551025390625 0 45233.19
696.3104248046875 0 33877.566
696.3583374023438 0 5606.2275
697.313232421875 0 14670.042
698.3163452148438 0 3563.8333
699.3421630859375 0 781.4172
700.3231811523438 0 1913.5845
700.8314819335938 0 4141.525
701.3265380859375 0 15427.117 y Water loss 2
701.8262329101562 0 13526.407 y Ammonia loss 2
702.3270874023438 0 9049.344
702.8421630859375 0 6574.201
703.3450927734375 0 5970.091
705.3538818359375 0 856.00757
705.8572998046875 0 1383.7006
706.343017578125 0 8022.4336 y Water loss 7
707.3447265625 0 2882.7808
708.3488159179688 0 1032.2366
709.32861328125 0 2594.722
709.8389892578125 0 21519.623
710.3326416015625 0 93254.68 y 2
710.833740234375 0 69176.41 b Water loss 11
711.3353881835938 0 31641.266 b Ammonia loss 11
711.8510131835938 0 78377.16
712.3541870117188 0 48284.004
712.856201171875 0 1476.0305
713.3369750976562 0 1189.3861
719.358154296875 0 2829.2952
719.8557739257812 0 7820.115 b 11
720.3540649414062 0 5709.923
720.85498046875 0 3872.7192
721.358154296875 0 1071.537
723.3676147460938 0 11194.408
724.3526000976562 0 69670.36 y 7
725.35498046875 0 27734.543
726.3577880859375 0 7109.169
727.3562622070312 0 3043.6711
727.8612060546875 0 1227.1752
728.35595703125 0 1513.4517
728.859375 0 1039.3578
729.3589477539062 0 1316.5697
734.3370971679688 0 1979.8717
737.3614501953125 0 1001.52905
737.8593139648438 0 854.96783
739.3746337890625 0 2658.9382
740.3707885742188 0 1959.1337
741.35791015625 0 1595.2407
745.358154296875 0 2137.3599
746.3627319335938 0 12432.498
746.864501953125 0 7872.7134
750.8599243164062 0 2246.4414 y Water loss 1
751.3622436523438 0 3163.3264 y Ammonia loss 1
751.8615112304688 0 1418.038
752.3657836914062 0 900.5679
755.3685302734375 0 132282.17
755.8699951171875 0 91060.51
756.3690185546875 0 1193.8777
759.3702392578125 0 4246.6016
759.8671875 0 7322.932 y 1
760.3641967773438 0 12481.198
760.8657836914062 0 5437.9995
762.3206787109375 0 1686.9503
762.40966796875 0 1145.5753
763.38134765625 0 5004.802
764.3828125 0 2768.1018
765.326416015625 0 1214.43
766.3355102539062 0 1104.6774
767.3839721679688 0 2446.4092
768.3804321289062 0 2533.696
769.3713989257812 0 1148.6049
779.3485717773438 0 6672.1514
780.3466186523438 0 2476.4521
781.3995361328125 0 3873.5823
782.4004516601562 0 2085.676
783.3421020507812 0 2195.5393
784.3397216796875 0 1268.4956
791.3844604492188 0 1527.0155
792.3751831054688 0 1573.1842
795.3790283203125 0 9887.801
796.38232421875 0 5164.9565
797.3605346679688 0 6969.869
798.3652954101562 0 2125.523
806.3574829101562 0 928.3807
807.436767578125 0 1425.1927
809.394287109375 0 14733.525
810.3948364257812 0 7878.4097 Precursor
811.3378295898438 0 2401.1729
811.4028930664062 0 1757.2639
812.3370361328125 0 1107.2106
823.3717651367188 0 6906.3022
824.3768310546875 0 2424.6965
825.384521484375 0 1023.45416
828.43310546875 0 1298.3334
836.426513671875 0 13808.672
837.4303588867188 0 4399.632
840.3690185546875 0 791.6283
841.3832397460938 0 1353.837
842.37548828125 0 1762.9556
850.423828125 0 1610.3424
851.4090576171875 0 2207.2285
852.4141845703125 0 3490.0344
853.4278564453125 0 1123.2086
859.40283203125 0 1517.996
860.3896484375 0 3114.2708
861.3903198242188 0 2977.9583
862.3995361328125 0 1362.5652
864.4586791992188 0 777.14435
867.374267578125 0 1042.8685
868.4313354492188 0 17052.97
869.4337768554688 0 7488.0005
870.4370727539062 0 2951.6382
878.4153442382812 0 15953.892 b Water loss 6
879.4097290039062 0 14764.523 b Ammonia loss 6
880.4109497070312 0 5551.5093
881.4202880859375 0 1232.9248
887.4310913085938 0 814.5772
890.4146728515625 0 830.60815
891.4285888671875 0 1028.1439
892.4293823242188 0 2925.7234
896.4260864257812 0 144555.75 b 6
897.4288940429688 0 73946.984
898.4312744140625 0 21351.893
899.4367065429688 0 4408.5283
900.4432983398438 0 932.2817
906.4066772460938 0 1328.2177
907.4114379882812 0 1292.2155
908.4182739257812 0 2909.401 y Water loss 6
909.443115234375 0 2023.3365
910.4383544921875 0 1998.6194
923.4325561523438 0 1398.4026
924.4229736328125 0 8033.997
925.426513671875 0 4572.117
926.4273681640625 0 7437.1035 y 6
927.4327392578125 0 3784.1323
952.41943359375 0 1741.5734
953.405029296875 0 1239.1469
963.4346923828125 0 1527.4747
970.4259643554688 0 2359.4614
971.415771484375 0 3159.052
972.4098510742188 0 993.2322
975.5031127929688 0 1516.081
976.5093994140625 0 900.2929
980.45361328125 0 1879.9824
981.4478149414062 0 3833.0176
982.4412841796875 0 1429.7317
986.5007934570312 0 1126.1752
987.4908447265625 0 1554.1868
988.4385986328125 0 3028.1592
989.4357299804688 0 1158.1162
991.5045776367188 0 967.9341 b Water loss 7
992.4974365234375 0 1182.305 b Ammonia loss 7
993.5105590820312 0 884.57806
996.4564208984375 0 1065.24
997.463134765625 0 916.5245
998.4688720703125 0 1201.5328
999.4666137695312 0 999.15704
1009.509765625 0 6947.451 b 7
1010.5127563476562 0 4957.588
1011.5107421875 0 1808.7021
1014.4832153320312 0 1018.00195
1023.4561767578125 0 1826.5145 y Water loss 5
1030.4842529296875 0 1399.0728
1031.4935302734375 0 846.48553
1040.471923828125 0 951.62866
1041.4554443359375 0 6648.1934 y 5
1042.456298828125 0 3619.6294
1073.5223388671875 0 1125.4794
1091.517333984375 0 1439.0188
1107.4970703125 0 895.84576 b Ammonia loss 8
1109.5386962890625 0 3040.6892
1110.5404052734375 0 1770.7252
1123.550048828125 0 1044.5007
1124.5408935546875 0 1266.5266 b 8
1125.518310546875 0 716.462
1133.4859619140625 0 788.9556
1134.4830322265625 0 1665.5432
1135.4896240234375 0 902.66187
1150.506103515625 0 1142.0333
1151.49951171875 0 5073.8867 y Water loss 4
1152.4906005859375 0 7697.5654 y Ammonia loss 4
1153.491455078125 0 4124.6626
1154.5001220703125 0 1826.6091
1160.5435791015625 0 924.507
1168.5240478515625 0 2035.9751
1169.5120849609375 0 11389.636 y 4
1170.51416015625 0 7440.6587
1171.516357421875 0 2443.225
1177.5587158203125 0 3498.3176
1178.5634765625 0 1262.447
1181.5638427734375 0 2188.343 b 9
1688.857421875 0 836.85864
2375.11328125 0 712.9243
3083.054931640625 0 852.9032

Spectrum Details

|  |  |
| --- | --- |
| Matched peaks? Matched peaks The total absolute number of peaks matched. Additionally in brackets the total fraction of peaks matched and the total number of peaks is shown. | 77 (8.65% of 890) |
| FDR? FDR The false discovery rate estimated for this peptide. It is calculated by matching all theoretical fragments with a non-integer shift with the raw peaks for this spectrum. This is done with 40 different shifts. The resulting percentage is the average number of annotated peaks over the number of annotated peaks with the correct spectrum. | 1.39% |
| Satellite FDR? Satellite FDR See the FDR for details on its calculation. This satellite ion specific FDR only contains the satellite ions (d/w) for I/L/J positions. | ∞ |
| PSM Score? PSM Score The PSM Score as given by Hecklib to this annotated spectrum. It is shown with three significant figures. | 647 |

#### Spectrum 6988? Spectrum 6988 The raw spectrum of this peptide as annotated by Hecklib. The fragments are coloured according to ion type (see legend). Any peaks with a star '\*' as text can be hovered over to see the full details, first the ion type second the mass shift type. By hovering over the amino acids in the peptide or ions in the legend the corresponding peaks are highlighted. By toggling the 'Unassigned' label you can turn the background (unassigned) peaks on or off in the plot. By updating the slider in the Ion legend you can update the spectrum to only show the top X% of the peaks with labels. The top X% means any peak that is within X% of the highest intensity. By dragging in the spectrum you can zoom in to a specific part of the spectrum and use 'Zoom Out' to get back to the original zoom level. The annotation of the spectrum is based on the given sequence in the peptides file and is done with different software so inconsistencies are likely. The peaks are annotated based on the given sequence, with 20 ppm tolerance.

Copy Data

##### Spectrum 6988 (TSV)

###### Preview

```
Loading example...
```

*Click on the button to copy the data to your clipboard.*

Mz MinMz MaxIntensity Max

WidthHeightPeptide font sizePeptide stroke widthSpectrum font sizeSpectrum stroke widthCompact peptide

Ion legend

wxyz

abcd

OtherUnassignedIon^Charge^_Position_Show for top:%

TVLHQDWLDGKEY

07.01e+51.40e+62.10e+62.81e+6

Zoom Out

a^+1^_2_y^+3^_4_a^+1^_2_y^+1^_1_b^+1^_2_b^+1^_2_y^+2^_3_y^+2^_4_y^+1^_2_b^+1^_3_y^+2^_5_y^+3^_7_y^+2^_5_y^+1^_2_b^+1^_3_b^+2^_6_b^+2^_6_b^+2^_6_y^+2^_6_y^+2^_6_y^+3^_9_b^+3^_10_y^+1^_3_y^+1^_3_b^+2^_7_b^+2^_7_b^+1^_4_y^+1^_3_b^+2^_7_y^+2^_7_y^+2^_7_b^+1^_4_y^+2^_7_y^+1^_4_b^+2^_8_b^+2^_8_y^+1^_4_\*\*\*b^+2^_9_b^+2^_9_b^+2^_9_b^+1^_5_y^+2^_9_b^+1^_5_y^+1^_5_y^+1^_5_y^+2^_10_y^+2^_10_b^+2^_11_b^+2^_11_y^+2^_10_b^+1^_6_b^+1^_6_y^+2^_11_y^+2^_11_b^+1^_6_y^+2^_11_b^+2^_12_b^+2^_12_y^+1^_6_b^+2^_12_y^+1^_6_y^+2^_12_y^+2^_12_y^+2^_12_\*b^+1^_7_b^+1^_7_b^+1^_7_y^+1^_7_b^+1^_8_b^+1^_8_b^+1^_8_y^+1^_8_y^+1^_8_b^+1^_9_b^+1^_9_y^+1^_9_y^+1^_9_y^+1^_9_b^+1^_10_

0716143321492866

Fragment Matches Table

Show background peaks

| Position | Ion type | Intensity | mz Theoretical | mz Error (Th) | mz Error (ppm) | Charge | Series Number |
| --- | --- | --- | --- | --- | --- | --- | --- |
| - | - | 7388 | 120.1 | - | - | 0 | - |
| - | - | 1.409E+04 | 122.1 | - | - | 0 | - |
| - | - | 1.517E+04 | 123 | - | - | 0 | - |
| - | - | 4411 | 123.1 | - | - | 0 | - |
| - | - | 8772 | 125.1 | - | - | 0 | - |
| - | - | 7028 | 126.1 | - | - | 0 | - |
| - | - | 5.095E+04 | 127.1 | - | - | 0 | - |
| - | - | 7734 | 127.1 | - | - | 0 | - |
| - | - | 5935 | 128.1 | - | - | 0 | - |
| - | - | 1.819E+05 | 128.1 | - | - | 0 | - |
| - | - | 1.129E+04 | 129.1 | - | - | 0 | - |
| - | - | 1.808E+06 | 129.1 | - | - | 0 | - |
| - | - | 9.154E+05 | 130.1 | - | - | 0 | - |
| - | - | 1.38E+04 | 130.1 | - | - | 0 | - |
| - | - | 1.212E+05 | 130.1 | - | - | 0 | - |
| - | - | 8738 | 131.1 | - | - | 0 | - |
| - | - | 9.963E+04 | 131.1 | - | - | 0 | - |
| - | - | 8403 | 131.1 | - | - | 0 | - |
| - | - | 2.019E+05 | 132.1 | - | - | 0 | - |
| - | - | 4294 | 133.1 | - | - | 0 | - |
| - | - | 2.158E+04 | 133.1 | - | - | 0 | - |
| - | - | 2.336E+05 | 136.1 | - | - | 0 | - |
| - | - | 2883 | 137.1 | - | - | 0 | - |
| - | - | 2.402E+04 | 137.1 | - | - | 0 | - |
| - | - | 2.742E+04 | 138.1 | - | - | 0 | - |
| - | - | 7847 | 138.1 | - | - | 0 | - |
| - | - | 7125 | 139.1 | - | - | 0 | - |
| - | - | 2.072E+04 | 139.1 | - | - | 0 | - |
| - | - | 2.921E+04 | 141.1 | - | - | 0 | - |
| - | - | 1.39E+04 | 142.1 | - | - | 0 | - |
| - | - | 3415 | 142.1 | - | - | 0 | - |
| - | - | 1.245E+04 | 143 | - | - | 0 | - |
| - | - | 7730 | 144.1 | - | - | 0 | - |
| - | - | 9477 | 144.1 | - | - | 0 | - |
| - | - | 1.835E+04 | 145.1 | - | - | 0 | - |
| - | - | 1.037E+04 | 147 | - | - | 0 | - |
| - | - | 1.595E+04 | 149 | - | - | 0 | - |
| - | - | 1.462E+04 | 150.1 | - | - | 0 | - |
| - | - | 4.994E+04 | 151.1 | - | - | 0 | - |
| - | - | 4271 | 153.1 | - | - | 0 | - |
| - | - | 6798 | 154.1 | - | - | 0 | - |
| - | - | 5914 | 155 | - | - | 0 | - |
| - | - | 9235 | 155.1 | - | - | 0 | - |
| - | - | 1.308E+05 | 155.1 | - | - | 0 | - |
| 2 | a | 8.621E+04 | 155.1 | 0.0003508 | 2.262 | +1 | 2 |
| - | - | 8775 | 156.1 | - | - | 0 | - |
| - | - | 1.078E+04 | 156.1 | - | - | 0 | - |
| - | - | 1.034E+04 | 156.1 | - | - | 0 | - |
| - | - | 1.269E+04 | 157.1 | - | - | 0 | - |
| - | - | 1.473E+04 | 158.1 | - | - | 0 | - |
| - | - | 1.544E+06 | 159.1 | - | - | 0 | - |
| - | - | 7250 | 160.1 | - | - | 0 | - |
| - | - | 1.581E+05 | 160.1 | - | - | 0 | - |
| - | - | 8470 | 161.1 | - | - | 0 | - |
| - | - | 1.329E+05 | 165.1 | - | - | 0 | - |
| - | - | 5200 | 165.1 | - | - | 0 | - |
| - | - | 2.156E+05 | 166.1 | - | - | 0 | - |
| 10 | y | 5187 | 166.1 | 0.002168 | 13.05 | +3 | 4 |
| - | - | 1.72E+04 | 166.1 | - | - | 0 | - |
| - | - | 8779 | 167.1 | - | - | 0 | - |
| - | - | 1.284E+04 | 167.1 | - | - | 0 | - |
| - | - | 4604 | 167.1 | - | - | 0 | - |
| - | - | 2.997E+04 | 168.1 | - | - | 0 | - |
| - | - | 2.078E+04 | 168.1 | - | - | 0 | - |
| - | - | 2.073E+04 | 168.1 | - | - | 0 | - |
| - | - | 2.561E+04 | 169.1 | - | - | 0 | - |
| - | - | 4.996E+04 | 169.1 | - | - | 0 | - |
| - | - | 7.639E+04 | 170.1 | - | - | 0 | - |
| - | - | 9858 | 171.1 | - | - | 0 | - |
| - | - | 1.367E+04 | 171.1 | - | - | 0 | - |
| - | - | 4.805E+04 | 172.1 | - | - | 0 | - |
| - | - | 3947 | 172.1 | - | - | 0 | - |
| - | - | 1.574E+04 | 173.1 | - | - | 0 | - |
| 2 | a | 2.778E+06 | 173.1 | 0.000452 | 2.611 | +1 | 2 |
| - | - | 2.559E+05 | 174.1 | - | - | 0 | - |
| - | - | 1.473E+04 | 175.1 | - | - | 0 | - |
| - | - | 7735 | 176.1 | - | - | 0 | - |
| - | - | 6147 | 177.1 | - | - | 0 | - |
| - | - | 8.523E+04 | 178.1 | - | - | 0 | - |
| - | - | 4712 | 179.1 | - | - | 0 | - |
| - | - | 1.559E+04 | 181.1 | - | - | 0 | - |
| 13 | y | 2.631E+05 | 182.1 | 0.0004343 | 2.385 | +1 | 1 |
| - | - | 2.992E+04 | 183.1 | - | - | 0 | - |
| 2 | b | 8.103E+04 | 183.1 | 0.0003703 | 2.022 | +1 | 2 |
| - | - | 7750 | 184.1 | - | - | 0 | - |
| - | - | 2.318E+04 | 185.2 | - | - | 0 | - |
| - | - | 5.351E+05 | 186.1 | - | - | 0 | - |
| - | - | 1.34E+05 | 187.1 | - | - | 0 | - |
| - | - | 4524 | 187.1 | - | - | 0 | - |
| - | - | 4.657E+04 | 187.1 | - | - | 0 | - |
| - | - | 5008 | 187.1 | - | - | 0 | - |
| - | - | 7184 | 188.1 | - | - | 0 | - |
| - | - | 1.143E+04 | 188.1 | - | - | 0 | - |
| - | - | 4855 | 188.1 | - | - | 0 | - |
| - | - | 1.371E+04 | 189.1 | - | - | 0 | - |
| - | - | 3.231E+04 | 190.1 | - | - | 0 | - |
| - | - | 2.289E+04 | 190.1 | - | - | 0 | - |
| - | - | 8479 | 193.1 | - | - | 0 | - |
| - | - | 1.734E+04 | 194.1 | - | - | 0 | - |
| - | - | 1.288E+05 | 195.1 | - | - | 0 | - |
| - | - | 7829 | 195.1 | - | - | 0 | - |
| - | - | 1.267E+05 | 196.1 | - | - | 0 | - |
| - | - | 1.073E+04 | 196.1 | - | - | 0 | - |
| - | - | 8965 | 197.1 | - | - | 0 | - |
| - | - | 1.352E+04 | 197.1 | - | - | 0 | - |
| - | - | 5031 | 197.2 | - | - | 0 | - |
| - | - | 2.272E+04 | 198.1 | - | - | 0 | - |
| - | - | 5.617E+04 | 198.1 | - | - | 0 | - |
| - | - | 9413 | 199.1 | - | - | 0 | - |
| - | - | 1.672E+04 | 199.1 | - | - | 0 | - |
| - | - | 8202 | 199.1 | - | - | 0 | - |
| - | - | 4806 | 200.1 | - | - | 0 | - |
| - | - | 1.086E+04 | 200.1 | - | - | 0 | - |
| 2 | b | 7.963E+05 | 201.1 | 0.0003494 | 1.737 | +1 | 2 |
| - | - | 1.045E+04 | 202.1 | - | - | 0 | - |
| - | - | 6.477E+04 | 202.1 | - | - | 0 | - |
| - | - | 6338 | 203.1 | - | - | 0 | - |
| - | - | 4064 | 203.7 | - | - | 0 | - |
| - | - | 8374 | 204.1 | - | - | 0 | - |
| - | - | 4645 | 204.1 | - | - | 0 | - |
| - | - | 7.193E+04 | 205.1 | - | - | 0 | - |
| - | - | 1.086E+04 | 205.1 | - | - | 0 | - |
| - | - | 8413 | 206.1 | - | - | 0 | - |
| - | - | 1.571E+04 | 206.1 | - | - | 0 | - |
| - | - | 4325 | 207.2 | - | - | 0 | - |
| - | - | 2.585E+04 | 207.2 | - | - | 0 | - |
| - | - | 5986 | 208.1 | - | - | 0 | - |
| - | - | 4801 | 208.7 | - | - | 0 | - |
| - | - | 5728 | 209.1 | - | - | 0 | - |
| - | - | 2.437E+04 | 210.1 | - | - | 0 | - |
| 11 | y | 2.389E+04 | 211.1 | 0.0003592 | 1.702 | +2 | 3 |
| - | - | 2.218E+04 | 212.1 | - | - | 0 | - |
| - | - | 7.572E+04 | 213.1 | - | - | 0 | - |
| - | - | 5403 | 213.1 | - | - | 0 | - |
| - | - | 7074 | 213.2 | - | - | 0 | - |
| - | - | 4999 | 214.1 | - | - | 0 | - |
| - | - | 7586 | 215.1 | - | - | 0 | - |
| - | - | 1.95E+04 | 216.1 | - | - | 0 | - |
| - | - | 4865 | 221.1 | - | - | 0 | - |
| - | - | 8455 | 221.1 | - | - | 0 | - |
| - | - | 1.743E+04 | 221.1 | - | - | 0 | - |
| - | - | 8971 | 221.1 | - | - | 0 | - |
| - | - | 5332 | 222.1 | - | - | 0 | - |
| - | - | 3.153E+04 | 222.1 | - | - | 0 | - |
| - | - | 5828 | 223.1 | - | - | 0 | - |
| - | - | 4.319E+04 | 223.1 | - | - | 0 | - |
| - | - | 2.581E+05 | 223.2 | - | - | 0 | - |
| - | - | 1.671E+04 | 224.1 | - | - | 0 | - |
| - | - | 2.723E+04 | 224.2 | - | - | 0 | - |
| - | - | 6614 | 225 | - | - | 0 | - |
| - | - | 5508 | 225.1 | - | - | 0 | - |
| - | - | 3919 | 225.2 | - | - | 0 | - |
| - | - | 4.064E+04 | 226.1 | - | - | 0 | - |
| - | - | 4.801E+04 | 227.1 | - | - | 0 | - |
| - | - | 1.365E+04 | 227.2 | - | - | 0 | - |
| - | - | 4962 | 228.1 | - | - | 0 | - |
| - | - | 4.1E+04 | 228.1 | - | - | 0 | - |
| - | - | 1.316E+04 | 229.1 | - | - | 0 | - |
| - | - | 4833 | 230.7 | - | - | 0 | - |
| - | - | 4726 | 231.1 | - | - | 0 | - |
| - | - | 4.753E+04 | 233.1 | - | - | 0 | - |
| - | - | 2.853E+05 | 234.1 | - | - | 0 | - |
| - | - | 3.82E+04 | 235.1 | - | - | 0 | - |
| - | - | 9882 | 235.2 | - | - | 0 | - |
| - | - | 7084 | 235.6 | - | - | 0 | - |
| - | - | 1.779E+04 | 237.1 | - | - | 0 | - |
| - | - | 4.987E+04 | 238.1 | - | - | 0 | - |
| - | - | 7392 | 238.1 | - | - | 0 | - |
| - | - | 9242 | 239.1 | - | - | 0 | - |
| - | - | 8421 | 239.1 | - | - | 0 | - |
| - | - | 5524 | 239.1 | - | - | 0 | - |
| - | - | 1.257E+05 | 240.1 | - | - | 0 | - |
| - | - | 1.608E+04 | 241.1 | - | - | 0 | - |
| - | - | 1.288E+04 | 241.1 | - | - | 0 | - |
| - | - | 2.24E+04 | 241.2 | - | - | 0 | - |
| - | - | 2.267E+04 | 243.1 | - | - | 0 | - |
| - | - | 1.202E+05 | 244.1 | - | - | 0 | - |
| - | - | 4356 | 244.6 | - | - | 0 | - |
| - | - | 7858 | 245.1 | - | - | 0 | - |
| - | - | 1.524E+05 | 247.1 | - | - | 0 | - |
| - | - | 8531 | 247.1 | - | - | 0 | - |
| - | - | 6800 | 247.2 | - | - | 0 | - |
| - | - | 4.458E+04 | 248.1 | - | - | 0 | - |
| - | - | 5.835E+04 | 248.1 | - | - | 0 | - |
| 10 | y | 5059 | 248.6 | 0.0002294 | 0.9228 | +2 | 4 |
| - | - | 9.994E+04 | 249.1 | - | - | 0 | - |
| - | - | 3.064E+04 | 249.1 | - | - | 0 | - |
| - | - | 9365 | 250.1 | - | - | 0 | - |
| - | - | 1.067E+04 | 250.1 | - | - | 0 | - |
| - | - | 9.766E+05 | 251.2 | - | - | 0 | - |
| - | - | 1.325E+05 | 252.2 | - | - | 0 | - |
| - | - | 5598 | 252.2 | - | - | 0 | - |
| - | - | 9138 | 253.1 | - | - | 0 | - |
| - | - | 1.184E+04 | 253.2 | - | - | 0 | - |
| - | - | 6.295E+04 | 255.1 | - | - | 0 | - |
| - | - | 8314 | 256.1 | - | - | 0 | - |
| - | - | 7695 | 256.2 | - | - | 0 | - |
| - | - | 1.799E+04 | 256.6 | - | - | 0 | - |
| - | - | 2.7E+04 | 257.1 | - | - | 0 | - |
| - | - | 4267 | 258.1 | - | - | 0 | - |
| - | - | 5337 | 258.1 | - | - | 0 | - |
| - | - | 4.378E+04 | 258.1 | - | - | 0 | - |
| - | - | 5578 | 259.1 | - | - | 0 | - |
| - | - | 2.562E+04 | 261.1 | - | - | 0 | - |
| - | - | 1.192E+04 | 261.1 | - | - | 0 | - |
| - | - | 1.719E+05 | 265.1 | - | - | 0 | - |
| - | - | 5033 | 265.6 | - | - | 0 | - |
| - | - | 5.573E+04 | 266.1 | - | - | 0 | - |
| - | - | 2.787E+05 | 266.1 | - | - | 0 | - |
| - | - | 3.146E+04 | 267.1 | - | - | 0 | - |
| - | - | 1.016E+04 | 267.1 | - | - | 0 | - |
| - | - | 1.492E+04 | 268.1 | - | - | 0 | - |
| - | - | 2.363E+04 | 268.2 | - | - | 0 | - |
| - | - | 6697 | 269.2 | - | - | 0 | - |
| - | - | 1.55E+04 | 269.2 | - | - | 0 | - |
| - | - | 3.691E+04 | 270.1 | - | - | 0 | - |
| - | - | 5804 | 270.2 | - | - | 0 | - |
| - | - | 7399 | 270.6 | - | - | 0 | - |
| - | - | 6.408E+04 | 272.2 | - | - | 0 | - |
| - | - | 5996 | 273.2 | - | - | 0 | - |
| - | - | 4.781E+04 | 274.1 | - | - | 0 | - |
| - | - | 1.654E+04 | 274.1 | - | - | 0 | - |
| - | - | 1.56E+04 | 275.1 | - | - | 0 | - |
| - | - | 6416 | 275.1 | - | - | 0 | - |
| - | - | 1.502E+04 | 276.1 | - | - | 0 | - |
| - | - | 1.02E+04 | 276.1 | - | - | 0 | - |
| - | - | 1.682E+04 | 276.2 | - | - | 0 | - |
| - | - | 5943 | 277.1 | - | - | 0 | - |
| - | - | 1.899E+04 | 279.1 | - | - | 0 | - |
| - | - | 2.646E+04 | 280.1 | - | - | 0 | - |
| - | - | 1.605E+04 | 281.1 | - | - | 0 | - |
| - | - | 1.374E+05 | 282.2 | - | - | 0 | - |
| - | - | 8.396E+05 | 283.1 | - | - | 0 | - |
| - | - | 5255 | 284.1 | - | - | 0 | - |
| - | - | 2.221E+04 | 284.1 | - | - | 0 | - |
| - | - | 8542 | 284.1 | - | - | 0 | - |
| - | - | 1.05E+05 | 284.1 | - | - | 0 | - |
| - | - | 5982 | 285.1 | - | - | 0 | - |
| - | - | 4597 | 285.1 | - | - | 0 | - |
| - | - | 1.813E+04 | 285.2 | - | - | 0 | - |
| - | - | 4.349E+04 | 286.1 | - | - | 0 | - |
| - | - | 7036 | 287.1 | - | - | 0 | - |
| - | - | 1.474E+04 | 287.2 | - | - | 0 | - |
| - | - | 5186 | 287.2 | - | - | 0 | - |
| - | - | 7743 | 288.1 | - | - | 0 | - |
| - | - | 2.72E+04 | 288.6 | - | - | 0 | - |
| - | - | 5773 | 289.1 | - | - | 0 | - |
| 12 | y | 3.74E+05 | 293.1 | 0.0004494 | 1.533 | +1 | 2 |
| - | - | 1.122E+04 | 293.2 | - | - | 0 | - |
| - | - | 7.713E+04 | 294.1 | - | - | 0 | - |
| - | - | 6237 | 295.1 | - | - | 0 | - |
| 3 | b | 1.588E+04 | 296.2 | 0.0004585 | 1.548 | +1 | 3 |
| - | - | 1.11E+04 | 296.6 | - | - | 0 | - |
| 9 | y | 2.155E+04 | 297.1 | 0.001348 | 4.537 | +2 | 5 |
| - | - | 1.295E+05 | 297.2 | - | - | 0 | - |
| 7 | y | 1.444E+04 | 298.1 | 0.005062 | 16.98 | +3 | 7 |
| - | - | 1.993E+04 | 298.2 | - | - | 0 | - |
| - | - | 8.994E+04 | 299.1 | - | - | 0 | - |
| - | - | 4548 | 299.1 | - | - | 0 | - |
| - | - | 1.961E+05 | 300.2 | - | - | 0 | - |
| - | - | 9224 | 301.1 | - | - | 0 | - |
| - | - | 2.703E+05 | 301.2 | - | - | 0 | - |
| - | - | 2.082E+04 | 301.2 | - | - | 0 | - |
| - | - | 7.286E+04 | 302.1 | - | - | 0 | - |
| - | - | 4.418E+04 | 302.2 | - | - | 0 | - |
| - | - | 1.364E+04 | 303.1 | - | - | 0 | - |
| - | - | 7160 | 305.2 | - | - | 0 | - |
| - | - | 5.292E+04 | 305.6 | - | - | 0 | - |
| 9 | y | 6.207E+04 | 306.1 | 0.001101 | 3.597 | +2 | 5 |
| - | - | 1.075E+04 | 306.2 | - | - | 0 | - |
| - | - | 2.004E+04 | 306.6 | - | - | 0 | - |
| - | - | 1.471E+04 | 307.1 | - | - | 0 | - |
| - | - | 1.167E+04 | 310.2 | - | - | 0 | - |
| 12 | y | 4.654E+04 | 311.1 | 0.0005353 | 1.721 | +1 | 2 |
| - | - | 1.683E+04 | 311.1 | - | - | 0 | - |
| - | - | 9716 | 312.1 | - | - | 0 | - |
| - | - | 1.465E+04 | 314.2 | - | - | 0 | - |
| 3 | b | 3.628E+04 | 314.2 | 0.0009412 | 2.995 | +1 | 3 |
| - | - | 2.077E+05 | 315.2 | - | - | 0 | - |
| - | - | 8450 | 315.2 | - | - | 0 | - |
| - | - | 3.478E+04 | 316.2 | - | - | 0 | - |
| - | - | 7552 | 317.7 | - | - | 0 | - |
| - | - | 5499 | 318.1 | - | - | 0 | - |
| - | - | 1.364E+04 | 319.2 | - | - | 0 | - |
| - | - | 7501 | 322.2 | - | - | 0 | - |
| - | - | 1.343E+04 | 325.2 | - | - | 0 | - |
| - | - | 5967 | 326.2 | - | - | 0 | - |
| - | - | 5.914E+04 | 326.7 | - | - | 0 | - |
| - | - | 1.592E+04 | 327.2 | - | - | 0 | - |
| - | - | 5790 | 330.7 | - | - | 0 | - |
| - | - | 2.417E+04 | 332.2 | - | - | 0 | - |
| - | - | 4.912E+04 | 333.2 | - | - | 0 | - |
| - | - | 1.289E+04 | 334.2 | - | - | 0 | - |
| 6 | b | 7668 | 338.7 | 0.001188 | 3.507 | +2 | 6 |
| 6 | b | 1.175E+04 | 339.2 | 0.002313 | 6.821 | +2 | 6 |
| - | - | 6768 | 339.7 | - | - | 0 | - |
| - | - | 1.538E+04 | 340.7 | - | - | 0 | - |
| - | - | 8645 | 341.2 | - | - | 0 | - |
| - | - | 1.87E+04 | 342.2 | - | - | 0 | - |
| - | - | 1.384E+04 | 343.2 | - | - | 0 | - |
| - | - | 5152 | 343.2 | - | - | 0 | - |
| - | - | 1.877E+04 | 344.2 | - | - | 0 | - |
| - | - | 1.201E+04 | 345 | - | - | 0 | - |
| - | - | 3.205E+04 | 345.1 | - | - | 0 | - |
| - | - | 1.279E+05 | 346.1 | - | - | 0 | - |
| - | - | 7102 | 346.1 | - | - | 0 | - |
| - | - | 6670 | 346.7 | - | - | 0 | - |
| - | - | 1.903E+04 | 347.1 | - | - | 0 | - |
| 6 | b | 6.494E+04 | 347.7 | 0.0005134 | 1.477 | +2 | 6 |
| - | - | 2.204E+04 | 348.2 | - | - | 0 | - |
| - | - | 8872 | 349.2 | - | - | 0 | - |
| - | - | 6203 | 350.1 | - | - | 0 | - |
| - | - | 8.474E+04 | 350.2 | - | - | 0 | - |
| - | - | 1.641E+04 | 351.2 | - | - | 0 | - |
| - | - | 7683 | 352.2 | - | - | 0 | - |
| - | - | 5439 | 352.7 | - | - | 0 | - |
| 8 | y | 3.191E+04 | 353.7 | 0.0009421 | 2.664 | +2 | 6 |
| - | - | 7415 | 354.2 | - | - | 0 | - |
| - | - | 1.16E+04 | 359 | - | - | 0 | - |
| - | - | 1.021E+04 | 360.2 | - | - | 0 | - |
| - | - | 2.995E+04 | 360.7 | - | - | 0 | - |
| - | - | 3.09E+04 | 361.2 | - | - | 0 | - |
| - | - | 6637 | 361.7 | - | - | 0 | - |
| - | - | 6.79E+04 | 362.2 | - | - | 0 | - |
| 8 | y | 1.21E+05 | 362.7 | 0.0008477 | 2.337 | +2 | 6 |
| - | - | 1.914E+05 | 363.1 | - | - | 0 | - |
| - | - | 4.569E+04 | 363.2 | - | - | 0 | - |
| - | - | 1.65E+04 | 363.7 | - | - | 0 | - |
| - | - | 2.945E+04 | 364.1 | - | - | 0 | - |
| - | - | 2.116E+04 | 364.1 | - | - | 0 | - |
| - | - | 2.295E+04 | 366.2 | - | - | 0 | - |
| - | - | 1.906E+04 | 367.1 | - | - | 0 | - |
| - | - | 1.141E+04 | 367.2 | - | - | 0 | - |
| - | - | 1.171E+04 | 367.2 | - | - | 0 | - |
| - | - | 5984 | 367.7 | - | - | 0 | - |
| - | - | 7954 | 368.2 | - | - | 0 | - |
| - | - | 6665 | 374.2 | - | - | 0 | - |
| - | - | 2.04E+04 | 376.2 | - | - | 0 | - |
| - | - | 2.209E+04 | 376.2 | - | - | 0 | - |
| - | - | 1.125E+04 | 376.7 | - | - | 0 | - |
| - | - | 2.375E+04 | 377.1 | - | - | 0 | - |
| - | - | 5103 | 378.2 | - | - | 0 | - |
| - | - | 7.205E+04 | 378.2 | - | - | 0 | - |
| - | - | 2.678E+05 | 379.2 | - | - | 0 | - |
| - | - | 7253 | 380.2 | - | - | 0 | - |
| - | - | 5.166E+04 | 380.2 | - | - | 0 | - |
| - | - | 5.43E+05 | 381.2 | - | - | 0 | - |
| - | - | 1.422E+04 | 381.7 | - | - | 0 | - |
| - | - | 8.959E+04 | 382.2 | - | - | 0 | - |
| - | - | 1.755E+04 | 382.2 | - | - | 0 | - |
| - | - | 6960 | 382.2 | - | - | 0 | - |
| - | - | 6605 | 382.7 | - | - | 0 | - |
| - | - | 1.344E+04 | 383.2 | - | - | 0 | - |
| - | - | 2.637E+04 | 384.2 | - | - | 0 | - |
| - | - | 2.757E+04 | 384.2 | - | - | 0 | - |
| - | - | 4.106E+04 | 385.2 | - | - | 0 | - |
| 5 | y | 1.667E+04 | 385.2 | 0.003979 | 10.33 | +3 | 9 |
| - | - | 8367 | 386.2 | - | - | 0 | - |
| - | - | 1.066E+04 | 386.2 | - | - | 0 | - |
| 10 | b | 6506 | 389.2 | 0.003128 | 8.037 | +3 | 10 |
| - | - | 1.401E+04 | 390.2 | - | - | 0 | - |
| - | - | 1.285E+04 | 390.7 | - | - | 0 | - |
| - | - | 1.853E+04 | 391.1 | - | - | 0 | - |
| - | - | 4910 | 391.2 | - | - | 0 | - |
| - | - | 6765 | 392.1 | - | - | 0 | - |
| - | - | 5511 | 392.2 | - | - | 0 | - |
| - | - | 9593 | 393.2 | - | - | 0 | - |
| - | - | 8.955E+04 | 394.2 | - | - | 0 | - |
| - | - | 2.478E+04 | 395.1 | - | - | 0 | - |
| - | - | 3.008E+04 | 395.2 | - | - | 0 | - |
| - | - | 3.626E+04 | 395.2 | - | - | 0 | - |
| - | - | 8612 | 396.2 | - | - | 0 | - |
| - | - | 7.879E+04 | 396.2 | - | - | 0 | - |
| - | - | 2.074E+04 | 397.2 | - | - | 0 | - |
| - | - | 3.427E+04 | 398.2 | - | - | 0 | - |
| - | - | 8204 | 399.2 | - | - | 0 | - |
| - | - | 1.499E+04 | 401.2 | - | - | 0 | - |
| - | - | 2.859E+04 | 402.2 | - | - | 0 | - |
| - | - | 3.515E+04 | 402.2 | - | - | 0 | - |
| - | - | 5974 | 403.1 | - | - | 0 | - |
| - | - | 2.131E+04 | 403.2 | - | - | 0 | - |
| - | - | 5758 | 404.2 | - | - | 0 | - |
| - | - | 1.021E+04 | 404.7 | - | - | 0 | - |
| - | - | 1.626E+04 | 405.3 | - | - | 0 | - |
| - | - | 1.256E+04 | 406.2 | - | - | 0 | - |
| - | - | 6259 | 407.2 | - | - | 0 | - |
| - | - | 6181 | 407.2 | - | - | 0 | - |
| - | - | 4.573E+04 | 409.1 | - | - | 0 | - |
| - | - | 8170 | 409.2 | - | - | 0 | - |
| - | - | 6629 | 409.7 | - | - | 0 | - |
| - | - | 8212 | 410.1 | - | - | 0 | - |
| - | - | 7.839E+04 | 411.2 | - | - | 0 | - |
| - | - | 2.819E+05 | 412.2 | - | - | 0 | - |
| - | - | 2.485E+04 | 413.1 | - | - | 0 | - |
| - | - | 3.15E+04 | 413.2 | - | - | 0 | - |
| - | - | 2.883E+04 | 413.2 | - | - | 0 | - |
| - | - | 1.799E+04 | 413.3 | - | - | 0 | - |
| - | - | 8144 | 414.1 | - | - | 0 | - |
| - | - | 1.016E+04 | 414.2 | - | - | 0 | - |
| - | - | 6.763E+04 | 414.2 | - | - | 0 | - |
| - | - | 1.691E+04 | 415 | - | - | 0 | - |
| - | - | 1.716E+04 | 415.2 | - | - | 0 | - |
| - | - | 9277 | 415.2 | - | - | 0 | - |
| - | - | 4.883E+04 | 417.7 | - | - | 0 | - |
| - | - | 8.506E+04 | 418.2 | - | - | 0 | - |
| - | - | 3.89E+04 | 418.7 | - | - | 0 | - |
| - | - | 1.251E+04 | 419.2 | - | - | 0 | - |
| 11 | y | 1.508E+04 | 421.2 | 0.0005791 | 1.375 | +1 | 3 |
| 11 | y | 1.18E+04 | 422.2 | 8.405E-05 | 0.1991 | +1 | 3 |
| - | - | 7225 | 422.7 | - | - | 0 | - |
| - | - | 2.263E+04 | 423.2 | - | - | 0 | - |
| - | - | 1.355E+05 | 423.3 | - | - | 0 | - |
| - | - | 8468 | 423.7 | - | - | 0 | - |
| - | - | 7527 | 424.2 | - | - | 0 | - |
| - | - | 3.747E+04 | 424.3 | - | - | 0 | - |
| - | - | 1.535E+04 | 424.7 | - | - | 0 | - |
| - | - | 6470 | 425.2 | - | - | 0 | - |
| - | - | 9.181E+05 | 426.7 | - | - | 0 | - |
| - | - | 4.211E+05 | 427.2 | - | - | 0 | - |
| - | - | 1.163E+05 | 427.7 | - | - | 0 | - |
| - | - | 1.888E+04 | 428.2 | - | - | 0 | - |
| - | - | 3.565E+05 | 429.2 | - | - | 0 | - |
| - | - | 5.61E+05 | 430.2 | - | - | 0 | - |
| - | - | 1.145E+05 | 431.2 | - | - | 0 | - |
| 7 | b | 1.019E+05 | 431.7 | 0.0006241 | 1.446 | +2 | 7 |
| 7 | b | 4.577E+04 | 432.2 | 0.006083 | 14.08 | +2 | 7 |
| - | - | 2.286E+04 | 432.7 | - | - | 0 | - |
| - | - | 1.568E+04 | 433.2 | - | - | 0 | - |
| 4 | b | 2.478E+04 | 433.3 | 1.834E-05 | 0.04233 | +1 | 4 |
| - | - | 7580 | 434.3 | - | - | 0 | - |
| - | - | 1.569E+04 | 438.2 | - | - | 0 | - |
| - | - | 8451 | 438.7 | - | - | 0 | - |
| 11 | y | 2.259E+05 | 439.2 | 0.0008177 | 1.862 | +1 | 3 |
| - | - | 1.529E+04 | 440.2 | - | - | 0 | - |
| - | - | 6.325E+04 | 440.2 | - | - | 0 | - |
| 7 | b | 4.34E+05 | 440.7 | 0.0006518 | 1.479 | +2 | 7 |
| - | - | 2.53E+05 | 441.2 | - | - | 0 | - |
| - | - | 5.923E+04 | 441.7 | - | - | 0 | - |
| - | - | 1.815E+04 | 442.2 | - | - | 0 | - |
| 7 | y | 7050 | 446.7 | 0.002826 | 6.326 | +2 | 7 |
| - | - | 6907 | 446.7 | - | - | 0 | - |
| 7 | y | 2.434E+04 | 447.2 | 0.007211 | 16.12 | +2 | 7 |
| - | - | 5730 | 447.7 | - | - | 0 | - |
| - | - | 2.389E+04 | 448.2 | - | - | 0 | - |
| - | - | 1.073E+04 | 449.2 | - | - | 0 | - |
| - | - | 8192 | 450.2 | - | - | 0 | - |
| - | - | 1.104E+04 | 450.2 | - | - | 0 | - |
| - | - | 1.123E+04 | 450.3 | - | - | 0 | - |
| 4 | b | 2.586E+05 | 451.3 | 0.0007757 | 1.719 | +1 | 4 |
| - | - | 5.116E+04 | 452.3 | - | - | 0 | - |
| - | - | 9789 | 453.3 | - | - | 0 | - |
| - | - | 5368 | 455.2 | - | - | 0 | - |
| - | - | 1.334E+04 | 455.3 | - | - | 0 | - |
| 7 | y | 6222 | 455.7 | 0.007059 | 15.49 | +2 | 7 |
| - | - | 1.006E+04 | 456.2 | - | - | 0 | - |
| - | - | 7164 | 457.2 | - | - | 0 | - |
| - | - | 2.649E+04 | 459.2 | - | - | 0 | - |
| - | - | 3.034E+04 | 460.8 | - | - | 0 | - |
| - | - | 1.585E+04 | 461.2 | - | - | 0 | - |
| - | - | 3.21E+04 | 461.3 | - | - | 0 | - |
| - | - | 5812 | 464.2 | - | - | 0 | - |
| - | - | 9264 | 465.7 | - | - | 0 | - |
| - | - | 9116 | 466.2 | - | - | 0 | - |
| - | - | 6.718E+04 | 468.3 | - | - | 0 | - |
| - | - | 1.491E+04 | 469.3 | - | - | 0 | - |
| - | - | 5529 | 471.2 | - | - | 0 | - |
| - | - | 7355 | 474.2 | - | - | 0 | - |
| - | - | 4.761E+04 | 474.3 | - | - | 0 | - |
| - | - | 7.843E+04 | 474.8 | - | - | 0 | - |
| - | - | 2.855E+04 | 475.3 | - | - | 0 | - |
| - | - | 8717 | 475.8 | - | - | 0 | - |
| - | - | 1.869E+04 | 476.2 | - | - | 0 | - |
| - | - | 4.164E+04 | 477.2 | - | - | 0 | - |
| 10 | y | 3.896E+04 | 478.2 | 0.0001937 | 0.405 | +1 | 4 |
| - | - | 3.242E+04 | 478.3 | - | - | 0 | - |
| - | - | 7514 | 479.3 | - | - | 0 | - |
| - | - | 6576 | 479.7 | - | - | 0 | - |
| - | - | 7610 | 480.2 | - | - | 0 | - |
| - | - | 6233 | 481.2 | - | - | 0 | - |
| - | - | 1.847E+05 | 483.3 | - | - | 0 | - |
| - | - | 1.258E+05 | 483.8 | - | - | 0 | - |
| - | - | 1.246E+04 | 484.2 | - | - | 0 | - |
| - | - | 3.25E+04 | 484.3 | - | - | 0 | - |
| - | - | 1.171E+04 | 484.8 | - | - | 0 | - |
| 8 | b | 5.13E+04 | 488.3 | 0.0008895 | 1.822 | +2 | 8 |
| 8 | b | 3.301E+04 | 488.7 | 0.007844 | 16.05 | +2 | 8 |
| - | - | 1.204E+04 | 489.3 | - | - | 0 | - |
| - | - | 8360 | 489.8 | - | - | 0 | - |
| - | - | 6161 | 492.3 | - | - | 0 | - |
| - | - | 5.135E+05 | 494.2 | - | - | 0 | - |
| - | - | 1.224E+05 | 495.2 | - | - | 0 | - |
| 10 | y | 5.933E+05 | 496.2 | 0.0007162 | 1.443 | +1 | 4 |
| - | - | 1.508E+05 | 497.2 | - | - | 0 | - |
| - | - | 5.881E+04 | 497.8 | - | - | 0 | - |
| - | - | 2.687E+04 | 498.2 | - | - | 0 | - |
| - | - | 5629 | 498.8 | - | - | 0 | - |
| - | - | 8032 | 499.2 | - | - | 0 | - |
| - | - | 2.384E+04 | 503.1 | - | - | 0 | - |
| - | - | 2.224E+04 | 504.2 | - | - | 0 | - |
| - | - | 1.328E+04 | 506.2 | - | - | 0 | - |
| - | - | 1.109E+04 | 506.3 | - | - | 0 | - |
| - | - | 9946 | 507.3 | - | - | 0 | - |
| - | - | 4.071E+04 | 511.3 | - | - | 0 | - |
| - | - | 1.324E+04 | 512.3 | - | - | 0 | - |
| - | - | 1.376E+04 | 513.2 | - | - | 0 | - |
| - | - | 6482 | 514.2 | - | - | 0 | - |
| - | - | 6023 | 515.3 | - | - | 0 | - |
| - | - | 7326 | 516.2 | - | - | 0 | - |
| - | - | 5910 | 519.1 | - | - | 0 | - |
| - | - | 1.55E+04 | 523.3 | - | - | 0 | - |
| - | - | 6127 | 523.6 | - | - | 0 | - |
| - | - | 2.195E+04 | 524.3 | - | - | 0 | - |
| - | - | 8.324E+04 | 525.3 | - | - | 0 | - |
| - | - | 1.005E+04 | 526.2 | - | - | 0 | - |
| - | - | 1.478E+04 | 526.3 | - | - | 0 | - |
| - | - | 4857 | 528.2 | - | - | 0 | - |
| - | - | 1.616E+04 | 528.9 | - | - | 0 | - |
| 0 | Precursor | 3.757E+04 | 529.3 | 0.002512 | 4.745 | +3 | -1 |
| 0 | Precursor | 3.227E+04 | 529.6 | 0.008247 | 15.57 | +3 | -1 |
| - | - | 2.157E+04 | 529.9 | - | - | 0 | - |
| - | - | 8052 | 530.3 | - | - | 0 | - |
| - | - | 7149 | 531.2 | - | - | 0 | - |
| - | - | 5407 | 531.8 | - | - | 0 | - |
| - | - | 1.275E+04 | 532.2 | - | - | 0 | - |
| 0 | Precursor | 2.535E+04 | 535.3 | 0.001737 | 3.244 | +3 | -1 |
| - | - | 1.369E+04 | 535.6 | - | - | 0 | - |
| - | - | 1.247E+04 | 535.9 | - | - | 0 | - |
| - | - | 5828 | 536.3 | - | - | 0 | - |
| - | - | 1.034E+04 | 537.2 | - | - | 0 | - |
| - | - | 8947 | 537.7 | - | - | 0 | - |
| - | - | 9821 | 537.8 | - | - | 0 | - |
| - | - | 6756 | 538.3 | - | - | 0 | - |
| - | - | 8036 | 538.8 | - | - | 0 | - |
| - | - | 8.809E+04 | 539.2 | - | - | 0 | - |
| - | - | 3667 | 539.3 | - | - | 0 | - |
| - | - | 3.049E+04 | 540.2 | - | - | 0 | - |
| - | - | 6625 | 541.2 | - | - | 0 | - |
| - | - | 8.367E+04 | 542.3 | - | - | 0 | - |
| - | - | 8.848E+04 | 543.3 | - | - | 0 | - |
| - | - | 2.317E+04 | 544.3 | - | - | 0 | - |
| - | - | 1.006E+04 | 545.3 | - | - | 0 | - |
| 9 | b | 1.05E+04 | 545.8 | 0.004983 | 9.131 | +2 | 9 |
| 9 | b | 1.331E+04 | 546.3 | 0.005475 | 10.02 | +2 | 9 |
| - | - | 7264 | 546.8 | - | - | 0 | - |
| - | - | 7755 | 547.2 | - | - | 0 | - |
| - | - | 5308 | 547.8 | - | - | 0 | - |
| - | - | 4955 | 548.2 | - | - | 0 | - |
| - | - | 2.826E+04 | 549.2 | - | - | 0 | - |
| - | - | 6613 | 550.2 | - | - | 0 | - |
| - | - | 3.234E+04 | 551.3 | - | - | 0 | - |
| - | - | 1.511E+04 | 551.3 | - | - | 0 | - |
| - | - | 9776 | 552.3 | - | - | 0 | - |
| - | - | 5156 | 553.3 | - | - | 0 | - |
| - | - | 8636 | 554.3 | - | - | 0 | - |
| 9 | b | 2.533E+04 | 554.8 | 0.01069 | 19.27 | +2 | 9 |
| - | - | 3.193E+04 | 555.3 | - | - | 0 | - |
| - | - | 1.996E+04 | 555.8 | - | - | 0 | - |
| - | - | 5457 | 558.2 | - | - | 0 | - |
| 5 | b | 2.252E+04 | 561.3 | 0.002041 | 3.635 | +1 | 5 |
| - | - | 7216 | 562.3 | - | - | 0 | - |
| - | - | 1.819E+04 | 563.3 | - | - | 0 | - |
| - | - | 1.073E+04 | 564.3 | - | - | 0 | - |
| - | - | 6421 | 564.3 | - | - | 0 | - |
| - | - | 3.834E+04 | 565.3 | - | - | 0 | - |
| - | - | 5646 | 566.3 | - | - | 0 | - |
| - | - | 1.268E+05 | 567.2 | - | - | 0 | - |
| - | - | 2.036E+04 | 568.2 | - | - | 0 | - |
| - | - | 1.2E+04 | 569.2 | - | - | 0 | - |
| - | - | 5.388E+04 | 575.2 | - | - | 0 | - |
| - | - | 1.978E+04 | 576.2 | - | - | 0 | - |
| - | - | 6566 | 577.2 | - | - | 0 | - |
| 5 | y | 7170 | 577.3 | 0.008244 | 14.28 | +2 | 9 |
| 5 | b | 1.602E+05 | 579.3 | 0.0004562 | 0.7875 | +1 | 5 |
| - | - | 5.727E+04 | 580.3 | - | - | 0 | - |
| - | - | 4982 | 581.3 | - | - | 0 | - |
| - | - | 1.28E+04 | 581.3 | - | - | 0 | - |
| - | - | 1.048E+04 | 582.3 | - | - | 0 | - |
| - | - | 6580 | 583.3 | - | - | 0 | - |
| - | - | 6359 | 588.8 | - | - | 0 | - |
| - | - | 6219 | 589.3 | - | - | 0 | - |
| - | - | 6.027E+04 | 592.3 | - | - | 0 | - |
| - | - | 1.863E+04 | 592.3 | - | - | 0 | - |
| 9 | y | 2.109E+05 | 593.3 | 0.0005733 | 0.9663 | +1 | 5 |
| - | - | 8.349E+04 | 593.3 | - | - | 0 | - |
| - | - | 9224 | 593.8 | - | - | 0 | - |
| - | - | 7.297E+04 | 594.3 | - | - | 0 | - |
| - | - | 1.653E+04 | 594.3 | - | - | 0 | - |
| - | - | 1.995E+04 | 595.3 | - | - | 0 | - |
| - | - | 5239 | 596.3 | - | - | 0 | - |
| - | - | 8664 | 596.3 | - | - | 0 | - |
| - | - | 5106 | 597.3 | - | - | 0 | - |
| - | - | 6751 | 597.8 | - | - | 0 | - |
| - | - | 1.042E+04 | 602.3 | - | - | 0 | - |
| - | - | 2.396E+04 | 602.8 | - | - | 0 | - |
| - | - | 6965 | 603.2 | - | - | 0 | - |
| - | - | 2.768E+04 | 603.3 | - | - | 0 | - |
| - | - | 1.486E+04 | 603.8 | - | - | 0 | - |
| - | - | 9.542E+05 | 610.3 | - | - | 0 | - |
| 9 | y | 1.733E+06 | 611.3 | 0.002032 | 3.325 | +1 | 5 |
| - | - | 9.078E+04 | 611.8 | - | - | 0 | - |
| - | - | 5.149E+05 | 612.3 | - | - | 0 | - |
| - | - | 1.395E+04 | 612.8 | - | - | 0 | - |
| - | - | 1.294E+05 | 613.3 | - | - | 0 | - |
| - | - | 1.498E+04 | 614.3 | - | - | 0 | - |
| - | - | 1.537E+04 | 617.3 | - | - | 0 | - |
| - | - | 8022 | 620.3 | - | - | 0 | - |
| - | - | 5891 | 620.8 | - | - | 0 | - |
| - | - | 4.213E+04 | 621.3 | - | - | 0 | - |
| - | - | 1.13E+04 | 621.3 | - | - | 0 | - |
| - | - | 1.641E+04 | 622.3 | - | - | 0 | - |
| - | - | 6407 | 622.4 | - | - | 0 | - |
| - | - | 8089 | 623.3 | - | - | 0 | - |
| - | - | 7830 | 628.3 | - | - | 0 | - |
| - | - | 6888 | 628.8 | - | - | 0 | - |
| - | - | 2.338E+04 | 634.3 | - | - | 0 | - |
| - | - | 3.448E+04 | 635.3 | - | - | 0 | - |
| - | - | 2.062E+04 | 636.3 | - | - | 0 | - |
| 4 | y | 2.051E+04 | 636.8 | 0.002015 | 3.164 | +2 | 10 |
| 4 | y | 1.806E+04 | 637.3 | 0.0108 | 16.95 | +2 | 10 |
| - | - | 1.182E+04 | 637.8 | - | - | 0 | - |
| 11 | b | 1.081E+04 | 638.3 | 0.007665 | 12.01 | +2 | 11 |
| 11 | b | 7828 | 638.8 | 0.01181 | 18.49 | +2 | 11 |
| - | - | 7783 | 640.3 | - | - | 0 | - |
| - | - | 5583 | 644.3 | - | - | 0 | - |
| - | - | 7.305E+04 | 645.3 | - | - | 0 | - |
| 4 | y | 1.013E+05 | 645.8 | 0.004484 | 6.944 | +2 | 10 |
| - | - | 7.124E+04 | 646.3 | - | - | 0 | - |
| - | - | 1.769E+04 | 646.8 | - | - | 0 | - |
| - | - | 7123 | 648.3 | - | - | 0 | - |
| - | - | 1.338E+04 | 650.3 | - | - | 0 | - |
| - | - | 2.024E+05 | 652.3 | - | - | 0 | - |
| - | - | 7.909E+04 | 653.3 | - | - | 0 | - |
| - | - | 1.781E+04 | 654.3 | - | - | 0 | - |
| - | - | 1.191E+04 | 658.3 | - | - | 0 | - |
| - | - | 7601 | 659.3 | - | - | 0 | - |
| - | - | 5957 | 661.3 | - | - | 0 | - |
| - | - | 7.639E+04 | 662.3 | - | - | 0 | - |
| - | - | 4.416E+04 | 663.3 | - | - | 0 | - |
| - | - | 1.706E+04 | 664.3 | - | - | 0 | - |
| - | - | 1.418E+04 | 664.3 | - | - | 0 | - |
| - | - | 6502 | 665.3 | - | - | 0 | - |
| - | - | 6530 | 668.3 | - | - | 0 | - |
| 6 | b | 1.086E+05 | 676.3 | 0.0003132 | 0.4631 | +1 | 6 |
| 6 | b | 5.789E+04 | 677.3 | 0.009523 | 14.06 | +1 | 6 |
| - | - | 2.337E+04 | 678.3 | - | - | 0 | - |
| - | - | 2.39E+04 | 679.3 | - | - | 0 | - |
| - | - | 3.876E+05 | 680.3 | - | - | 0 | - |
| - | - | 1.676E+05 | 681.3 | - | - | 0 | - |
| - | - | 3.428E+04 | 682.3 | - | - | 0 | - |
| - | - | 3.354E+04 | 684.3 | - | - | 0 | - |
| - | - | 2.898E+04 | 684.8 | - | - | 0 | - |
| - | - | 1.436E+04 | 685.3 | - | - | 0 | - |
| - | - | 1.438E+04 | 688.3 | - | - | 0 | - |
| - | - | 4.475E+04 | 692.8 | - | - | 0 | - |
| 3 | y | 1.514E+05 | 693.3 | 0.00283 | 4.081 | +2 | 11 |
| 3 | y | 1.084E+05 | 693.8 | 0.009418 | 13.57 | +2 | 11 |
| 6 | b | 1.066E+06 | 694.4 | 0.0001807 | 0.2602 | +1 | 6 |
| - | - | 2.113E+04 | 694.8 | - | - | 0 | - |
| - | - | 3.818E+05 | 695.4 | - | - | 0 | - |
| - | - | 6097 | 696.3 | - | - | 0 | - |
| - | - | 8.469E+04 | 696.4 | - | - | 0 | - |
| - | - | 2.176E+04 | 697.4 | - | - | 0 | - |
| - | - | 3.842E+05 | 701.8 | - | - | 0 | - |
| 3 | y | 6.647E+05 | 702.3 | 0.004566 | 6.502 | +2 | 11 |
| 12 | b | 4.524E+05 | 702.8 | 0.01082 | 15.4 | +2 | 12 |
| - | - | 9319 | 702.9 | - | - | 0 | - |
| - | - | 5924 | 702.9 | - | - | 0 | - |
| 12 | b | 1.735E+05 | 703.3 | 0.001245 | 1.769 | +2 | 12 |
| - | - | 5.614E+04 | 703.8 | - | - | 0 | - |
| - | - | 2.085E+04 | 704.3 | - | - | 0 | - |
| - | - | 2.825E+04 | 705.4 | - | - | 0 | - |
| 8 | y | 6.364E+04 | 706.3 | 0.002264 | 3.205 | +1 | 6 |
| - | - | 3.456E+04 | 707.3 | - | - | 0 | - |
| - | - | 6839 | 708.4 | - | - | 0 | - |
| - | - | 5645 | 709.3 | - | - | 0 | - |
| - | - | 6.384E+04 | 711.4 | - | - | 0 | - |
| 12 | b | 5.78E+04 | 711.9 | 0.006721 | 9.442 | +2 | 12 |
| - | - | 3.693E+04 | 712.4 | - | - | 0 | - |
| - | - | 1.933E+04 | 712.9 | - | - | 0 | - |
| - | - | 7065 | 720.4 | - | - | 0 | - |
| - | - | 4.328E+04 | 721.4 | - | - | 0 | - |
| - | - | 1.909E+04 | 722.4 | - | - | 0 | - |
| - | - | 2.702E+05 | 723.4 | - | - | 0 | - |
| 8 | y | 4.937E+05 | 724.4 | 0.003113 | 4.297 | +1 | 6 |
| - | - | 2.114E+05 | 725.4 | - | - | 0 | - |
| - | - | 5.79E+04 | 726.4 | - | - | 0 | - |
| - | - | 6278 | 727.4 | - | - | 0 | - |
| - | - | 1.748E+04 | 728.4 | - | - | 0 | - |
| - | - | 1.825E+04 | 729.4 | - | - | 0 | - |
| - | - | 6278 | 730.4 | - | - | 0 | - |
| - | - | 2.862E+04 | 733.4 | - | - | 0 | - |
| - | - | 3.028E+04 | 734.3 | - | - | 0 | - |
| - | - | 1.29E+04 | 735.3 | - | - | 0 | - |
| - | - | 7486 | 739.4 | - | - | 0 | - |
| 2 | y | 1.321E+04 | 742.9 | 0.005366 | 7.223 | +2 | 12 |
| 2 | y | 2.493E+04 | 743.4 | 0.01238 | 16.66 | +2 | 12 |
| - | - | 8840 | 745.4 | - | - | 0 | - |
| - | - | 1E+04 | 747.4 | - | - | 0 | - |
| - | - | 1.622E+04 | 748.4 | - | - | 0 | - |
| - | - | 1.032E+04 | 749.3 | - | - | 0 | - |
| - | - | 1.373E+05 | 751.4 | - | - | 0 | - |
| 2 | y | 5.023E+04 | 751.9 | 0.00399 | 5.306 | +2 | 12 |
| - | - | 9.423E+04 | 752.4 | - | - | 0 | - |
| - | - | 1.208E+04 | 752.9 | - | - | 0 | - |
| - | - | 1.748E+04 | 753.4 | - | - | 0 | - |
| - | - | 6326 | 754.4 | - | - | 0 | - |
| - | - | 1.196E+04 | 761.4 | - | - | 0 | - |
| - | - | 7948 | 762.4 | - | - | 0 | - |
| - | - | 1.474E+04 | 762.4 | - | - | 0 | - |
| - | - | 7.269E+04 | 763.4 | - | - | 0 | - |
| - | - | 3.265E+04 | 764.4 | - | - | 0 | - |
| - | - | 3.83E+04 | 765.4 | - | - | 0 | - |
| - | - | 1.959E+04 | 766.4 | - | - | 0 | - |
| - | - | 3.192E+04 | 767.3 | - | - | 0 | - |
| - | - | 9921 | 769.4 | - | - | 0 | - |
| - | - | 9306 | 775.4 | - | - | 0 | - |
| - | - | 8160 | 776.4 | - | - | 0 | - |
| - | - | 6.903E+04 | 779.4 | - | - | 0 | - |
| - | - | 3.426E+04 | 780.4 | - | - | 0 | - |
| - | - | 2.99E+04 | 781.4 | - | - | 0 | - |
| - | - | 1.636E+04 | 782.4 | - | - | 0 | - |
| - | - | 5763 | 783.4 | - | - | 0 | - |
| 0 | Precursor | 1.12E+05 | 793.4 | 0.01381 | 17.41 | +2 | -1 |
| - | - | 5.714E+04 | 794.4 | - | - | 0 | - |
| - | - | 7958 | 795.3 | - | - | 0 | - |
| - | - | 8613 | 795.4 | - | - | 0 | - |
| - | - | 2.391E+04 | 807.4 | - | - | 0 | - |
| - | - | 1.528E+04 | 807.4 | - | - | 0 | - |
| - | - | 1.474E+04 | 808.4 | - | - | 0 | - |
| - | - | 1.476E+04 | 818.4 | - | - | 0 | - |
| - | - | 1.344E+04 | 819.4 | - | - | 0 | - |
| - | - | 9183 | 820.4 | - | - | 0 | - |
| - | - | 1.235E+04 | 825.4 | - | - | 0 | - |
| - | - | 1.032E+04 | 826.4 | - | - | 0 | - |
| - | - | 5199 | 834.3 | - | - | 0 | - |
| - | - | 2.782E+04 | 834.4 | - | - | 0 | - |
| - | - | 3.888E+04 | 835.4 | - | - | 0 | - |
| - | - | 8.813E+04 | 836.4 | - | - | 0 | - |
| - | - | 3.741E+04 | 837.5 | - | - | 0 | - |
| - | - | 8096 | 838.5 | - | - | 0 | - |
| - | - | 6121 | 842.4 | - | - | 0 | - |
| - | - | 1.167E+04 | 843.4 | - | - | 0 | - |
| - | - | 1.805E+04 | 844.4 | - | - | 0 | - |
| - | - | 2.156E+04 | 845.4 | - | - | 0 | - |
| - | - | 3.692E+04 | 846.4 | - | - | 0 | - |
| - | - | 3.058E+04 | 847.4 | - | - | 0 | - |
| - | - | 8088 | 848.4 | - | - | 0 | - |
| - | - | 9397 | 851.4 | - | - | 0 | - |
| - | - | 3.103E+05 | 852.4 | - | - | 0 | - |
| - | - | 1.412E+05 | 853.4 | - | - | 0 | - |
| - | - | 4.317E+04 | 854.4 | - | - | 0 | - |
| - | - | 1.403E+04 | 855.4 | - | - | 0 | - |
| 7 | b | 1.759E+05 | 862.4 | 4.08E-05 | 0.04731 | +1 | 7 |
| 7 | b | 1.102E+05 | 863.4 | 0.01126 | 13.05 | +1 | 7 |
| - | - | 2.179E+05 | 864.5 | - | - | 0 | - |
| - | - | 1.198E+05 | 865.5 | - | - | 0 | - |
| - | - | 3.039E+04 | 866.5 | - | - | 0 | - |
| - | - | 2.268E+04 | 876.4 | - | - | 0 | - |
| - | - | 1.242E+04 | 877.4 | - | - | 0 | - |
| 7 | b | 1.421E+06 | 880.4 | 0.0003921 | 0.4453 | +1 | 7 |
| - | - | 7.304E+05 | 881.4 | - | - | 0 | - |
| - | - | 2.113E+05 | 882.4 | - | - | 0 | - |
| - | - | 4.882E+04 | 883.4 | - | - | 0 | - |
| - | - | 9398 | 884.4 | - | - | 0 | - |
| - | - | 7973 | 890.4 | - | - | 0 | - |
| - | - | 8208 | 891.4 | - | - | 0 | - |
| - | - | 3.077E+04 | 892.5 | - | - | 0 | - |
| - | - | 1.385E+04 | 893.5 | - | - | 0 | - |
| - | - | 8071 | 894.5 | - | - | 0 | - |
| - | - | 3.373E+04 | 907.4 | - | - | 0 | - |
| - | - | 3.992E+04 | 908.4 | - | - | 0 | - |
| - | - | 6.771E+04 | 909.4 | - | - | 0 | - |
| 7 | y | 6.934E+04 | 910.4 | 0.004305 | 4.728 | +1 | 7 |
| - | - | 3.178E+04 | 911.4 | - | - | 0 | - |
| - | - | 1.041E+04 | 936.4 | - | - | 0 | - |
| - | - | 8440 | 937.4 | - | - | 0 | - |
| - | - | 1.474E+04 | 947.4 | - | - | 0 | - |
| - | - | 7795 | 953.4 | - | - | 0 | - |
| - | - | 2.094E+04 | 954.4 | - | - | 0 | - |
| - | - | 2.753E+04 | 955.4 | - | - | 0 | - |
| - | - | 1.034E+04 | 956.4 | - | - | 0 | - |
| - | - | 2.15E+04 | 964.5 | - | - | 0 | - |
| - | - | 2.691E+04 | 965.5 | - | - | 0 | - |
| - | - | 1.459E+04 | 966.5 | - | - | 0 | - |
| - | - | 1.347E+04 | 971.5 | - | - | 0 | - |
| - | - | 1.452E+04 | 972.4 | - | - | 0 | - |
| - | - | 9599 | 973.4 | - | - | 0 | - |
| 8 | b | 1.975E+04 | 975.5 | 0.003091 | 3.168 | +1 | 8 |
| 8 | b | 1.138E+04 | 976.5 | 0.009415 | 9.642 | +1 | 8 |
| - | - | 9079 | 979.5 | - | - | 0 | - |
| - | - | 7010 | 980.5 | - | - | 0 | - |
| - | - | 8347 | 981.5 | - | - | 0 | - |
| 8 | b | 1.153E+05 | 993.5 | 1.662E-05 | 0.01673 | +1 | 8 |
| - | - | 5.914E+04 | 994.5 | - | - | 0 | - |
| - | - | 2.69E+04 | 995.5 | - | - | 0 | - |
| 6 | y | 9814 | 1007 | 0.001575 | 1.564 | +1 | 8 |
| - | - | 2.665E+04 | 1024 | - | - | 0 | - |
| 6 | y | 3.24E+04 | 1025 | 0.006171 | 6.017 | +1 | 8 |
| - | - | 1.906E+04 | 1026 | - | - | 0 | - |
| - | - | 7081 | 1027 | - | - | 0 | - |
| - | - | 9262 | 1076 | - | - | 0 | - |
| - | - | 1.021E+04 | 1077 | - | - | 0 | - |
| 9 | b | 6432 | 1092 | 0.01161 | 10.63 | +1 | 9 |
| - | - | 1.72E+04 | 1093 | - | - | 0 | - |
| - | - | 2.524E+04 | 1094 | - | - | 0 | - |
| - | - | 1.516E+04 | 1095 | - | - | 0 | - |
| - | - | 9191 | 1108 | - | - | 0 | - |
| 9 | b | 8978 | 1109 | 0.002146 | 1.936 | +1 | 9 |
| - | - | 7306 | 1117 | - | - | 0 | - |
| - | - | 2.062E+04 | 1118 | - | - | 0 | - |
| - | - | 1.335E+04 | 1135 | - | - | 0 | - |
| 5 | y | 4.21E+04 | 1136 | 0.00308 | 2.712 | +1 | 9 |
| 5 | y | 4.704E+04 | 1136 | 0.009298 | 8.182 | +1 | 9 |
| - | - | 2.258E+04 | 1138 | - | - | 0 | - |
| - | - | 5.318E+04 | 1153 | - | - | 0 | - |
| 5 | y | 8.394E+04 | 1154 | 0.006675 | 5.787 | +1 | 9 |
| - | - | 3.896E+04 | 1155 | - | - | 0 | - |
| - | - | 1.297E+04 | 1156 | - | - | 0 | - |
| 10 | b | 6783 | 1166 | 0.004344 | 3.727 | +1 | 10 |
| - | - | 5537 | 1342 | - | - | 0 | - |
| - | - | 6735 | 1792 | - | - | 0 | - |
| - | - | 6566 | 2838 | - | - | 0 | - |

m/z Charge Intensity FragmentType MassShift Position
120.08114624023438 0 7387.6865
122.07164764404297 0 14088.247
123.04425811767578 0 15166.934
123.05562591552734 0 4410.727
125.10804748535156 0 8772.066
126.09165954589844 0 7028.4946
127.05061340332031 0 50948.312
127.1233901977539 0 7734.487
128.08213806152344 0 5934.6934
128.1073760986328 0 181905.22
129.066162109375 0 11288.354
129.1026611328125 0 1807507.2
130.0655059814453 0 915389.5
130.09983825683594 0 13802.821
130.10597229003906 0 121230.06
131.06329345703125 0 8738.468
131.06886291503906 0 99630.12
131.11822509765625 0 8403.456
132.0811309814453 0 201890.02
133.06106567382812 0 4293.7046
133.0845489501953 0 21583.365
136.07606506347656 0 233637.58
137.07237243652344 0 2882.5833
137.07940673828125 0 24021.184
138.06658935546875 0 27424.62
138.09176635742188 0 7847.486
139.05059814453125 0 7125.1274
139.08697509765625 0 20722.498
141.1026611328125 0 29208.355
142.06565856933594 0 13903.449
142.12307739257812 0 3415.3726
143.04559326171875 0 12452.911
144.07647705078125 0 7730.283
144.08181762695312 0 9476.505
145.06117248535156 0 18353.715
147.0443572998047 0 10367.494
149.0452117919922 0 15945.941
150.06649780273438 0 14619.44
151.0869903564453 0 49938.188
153.10252380371094 0 4271.187
154.0612335205078 0 6798.284
155.045654296875 0 5914.0522
155.0819854736328 0 9235.059
155.0930938720703 0 130818.04
155.1182403564453 0 86209.06 a Water loss 1
156.07705688476562 0 8774.655
156.0963592529297 0 10775.182
156.12185668945312 0 10336.222
157.09762573242188 0 12687.961
158.08436584472656 0 14730.004
159.09210205078125 0 1544315.9
160.07579040527344 0 7249.967
160.09544372558594 0 158093.64
161.0985107421875 0 8469.898
165.05494689941406 0 132912.33
165.0770263671875 0 5200.425
166.0614471435547 0 215588.81
166.0870819091797 0 5186.7266 y 9
166.09768676757812 0 17200.324
167.05624389648438 0 8778.6875
167.0648956298828 0 12843.992
167.1179656982422 0 4604.1504
168.0811309814453 0 29974.436
168.1022491455078 0 20782.62
168.113525390625 0 20725.723
169.0764617919922 0 25612.725
169.0975799560547 0 49959.883
170.06040954589844 0 76392.21
171.0638427734375 0 9857.505
171.09173583984375 0 13667.963
172.07212829589844 0 48050.61
172.10842895507812 0 3946.8079
173.05604553222656 0 15740.574
173.12890625 0 2777850.2 a 1
174.13223266601562 0 255873.3
175.1345672607422 0 14730.993
176.08251953125 0 7735.203
177.1024627685547 0 6146.727
178.13424682617188 0 85226.91
179.1375732421875 0 4712.158
181.06109619140625 0 15590.503
182.08160400390625 0 263096.56 y 12
183.08517456054688 0 29922.812
183.11317443847656 0 81033.664 b Water loss 1
184.11669921875 0 7750.1655
185.16519165039062 0 23175.812
186.12413024902344 0 535137.3
187.0869903564453 0 133950.06
187.12039184570312 0 4524.0107
187.12757873535156 0 46574.76
187.14480590820312 0 5008.204
188.0709991455078 0 7183.7407
188.09027099609375 0 11426.838
188.12860107421875 0 4854.8145
189.0988006591797 0 13714.054
190.0826873779297 0 32306.473
190.13424682617188 0 22892.164
193.09756469726562 0 8478.612
194.09283447265625 0 17335.11
195.1131591796875 0 128782.93
195.1234130859375 0 7829.324
196.10833740234375 0 126739.44
196.1165008544922 0 10731.957
197.11166381835938 0 8964.613
197.12887573242188 0 13516.322
197.16543579101562 0 5030.847
198.08787536621094 0 22715.227
198.1280975341797 0 56166.332
199.07138061523438 0 9413.277
199.0869598388672 0 16715.27
199.13150024414062 0 8201.785
200.08966064453125 0 4806.1816
200.1427459716797 0 10859.123
201.12371826171875 0 796276.06 b 1
202.08644104003906 0 10452.66
202.12698364257812 0 64772.363
203.1282501220703 0 6338.166
203.67442321777344 0 4063.9744
204.11370849609375 0 8373.719
204.1345672607422 0 4644.772
205.0975341796875 0 71933.734
205.14498901367188 0 10859.795
206.10079956054688 0 8412.951
206.12913513183594 0 15710.903
207.15110778808594 0 4325.176
207.16067504882812 0 25850.023
208.1077423095703 0 5986.136
208.67100524902344 0 4801.3945
209.09251403808594 0 5728.48
210.12738037109375 0 24371.348
211.1080780029297 0 23891.164 y Water loss 10
212.13978576660156 0 22184.979
213.1237335205078 0 75719.625
213.135986328125 0 5402.5137
213.16014099121094 0 7073.8765
214.12783813476562 0 4999.4673
215.13926696777344 0 7586.1914
216.09825134277344 0 19499.105
221.07273864746094 0 4865.348
221.0933074951172 0 8455.416
221.10354614257812 0 17429.69
221.14039611816406 0 8970.637
222.10714721679688 0 5332.098
222.12393188476562 0 31527.062
223.08706665039062 0 5828.463
223.108154296875 0 43189.066
223.15579223632812 0 258072.23
224.10353088378906 0 16708.467
224.15914916992188 0 27230.994
225.0437469482422 0 6613.722
225.1242218017578 0 5507.981
225.1608428955078 0 3918.745
226.08274841308594 0 40642.926
227.06658935546875 0 48013.99
227.15464782714844 0 13653.686
228.0699462890625 0 4962.008
228.13462829589844 0 41002.855
229.11874389648438 0 13158.956
230.70733642578125 0 4833.4614
231.0889892578125 0 4726.366
233.14013671875 0 47534.82
234.12416076660156 0 285316.1
235.1277313232422 0 38196.293
235.15567016601562 0 9882.332
235.6057586669922 0 7084.2593
237.13497924804688 0 17792.516
238.11880493164062 0 49866.684
238.1304168701172 0 7392.351
239.08197021484375 0 9242.121
239.0946502685547 0 8420.991
239.12283325195312 0 5523.7764
240.13473510742188 0 125744.84
241.11825561523438 0 16082.049
241.13754272460938 0 12877.8545
241.19131469726562 0 22397.463
243.1099395751953 0 22672.607
244.0932159423828 0 120178.99
244.61141967773438 0 4356.1963
245.0974578857422 0 7857.6406
247.10821533203125 0 152397.16
247.14407348632812 0 8531.459
247.15713500976562 0 6799.5273
248.10317993164062 0 44581.812
248.114501953125 0 58348.246
248.62396240234375 0 5059.4395 y 9
249.0986328125 0 99942.375
249.13526916503906 0 30636.246
250.1027374267578 0 9364.698
250.1194305419922 0 10671.589
251.1507568359375 0 976619.44
252.1539764404297 0 132460.72
252.16940307617188 0 5597.7915
253.1186981201172 0 9138.34
253.1562957763672 0 11842.166
255.1476287841797 0 62954.55
256.10845947265625 0 8314.261
256.15142822265625 0 7694.894
256.61102294921875 0 17993.86
257.1131286621094 0 27001.518
258.12371826171875 0 4266.56
258.1305847167969 0 5337.13
258.1451721191406 0 43781.38
259.1293029785156 0 5578.2563
261.11907958984375 0 25619.537
261.1349792480469 0 11923.82
265.1297302246094 0 171907.84
265.616943359375 0 5033.001
266.1133728027344 0 55729.44
266.12530517578125 0 278677.5
267.12823486328125 0 31459.385
267.1458740234375 0 10155.809
268.1294250488281 0 14918.503
268.1770935058594 0 23632.117
269.160888671875 0 6697.045
269.1859436035156 0 15499.989
270.1221618652344 0 36908.555
270.1612854003906 0 5803.6934
270.62451171875 0 7399.0635
272.17608642578125 0 64081.523
273.1785888671875 0 5996.4565
274.118896484375 0 47814.785
274.1317138671875 0 16540.87
275.10284423828125 0 15597.882
275.1199035644531 0 6416.134
276.10888671875 0 15019.081
276.1258850097656 0 10198.608
276.1556091308594 0 16823.812
277.1295166015625 0 5943.2144
279.1453857421875 0 18987.324
280.1296691894531 0 26463.594
281.0521545410156 0 16052.051
282.15655517578125 0 137449.95
283.1405334472656 0 839644
284.08843994140625 0 5254.6123
284.1034851074219 0 22213.596
284.1219482421875 0 8542.373
284.14373779296875 0 104969.78
285.1065979003906 0 5982.465
285.1268310546875 0 4596.7793
285.1560974121094 0 18131.738
286.1401672363281 0 43485.777
287.14208984375 0 7036.218
287.1717529296875 0 14744.935
287.1876525878906 0 5185.564
288.1272277832031 0 7743.1367
288.6191101074219 0 27201.15
289.1208801269531 0 5772.79
293.1136474609375 0 373965.62 y Water loss 11
293.162353515625 0 11221.711
294.1178283691406 0 77131.33
295.12109375 0 6237.1396
296.19732666015625 0 15879.035 b Water loss 2
296.6396789550781 0 11104.3545
297.1332702636719 0 21550.412 y Water loss 8
297.1562805175781 0 129489.55
298.1397705078125 0 14439.481 y Water loss 6
298.15887451171875 0 19932.854
299.0621337890625 0 89936.6
299.12506103515625 0 4547.9463
300.16741943359375 0 196137.03
301.1174011230469 0 9224.279
301.1511535644531 0 270306.03
301.1694641113281 0 20820.703
302.114013671875 0 72862.15
302.15460205078125 0 44184.527
303.117919921875 0 13641.102
305.1795959472656 0 7159.956
305.64569091796875 0 52921.906
306.1383056640625 0 62072.145 y 8
306.22943115234375 0 10750.763
306.6390380859375 0 20041.809
307.14117431640625 0 14710.672
310.154052734375 0 11670.425
311.1242980957031 0 46538.414 y 11
311.13812255859375 0 16827.252
312.1268615722656 0 9715.955
314.1864318847656 0 14649.367
314.2083740234375 0 36283.56 b 2
315.1669006347656 0 207676
315.21063232421875 0 8450.487
316.170166015625 0 34784.57
317.65875244140625 0 7552.409
318.1204833984375 0 5498.528
319.15240478515625 0 13642.899
322.22479248046875 0 7501.1494
325.15069580078125 0 13430.042
326.1541748046875 0 5967.2827
326.6640625 0 59136.875
327.16583251953125 0 15920.648
330.65869140625 0 5789.8086
332.20867919921875 0 24166.762
333.1773376464844 0 49116.832
334.1837463378906 0 12891.315
338.67547607421875 0 7668.0415 b Water loss 5
339.1686096191406 0 11751.477 b Ammonia loss 5
339.67706298828125 0 6768.1226
340.66229248046875 0 15383.16
341.1625061035156 0 8644.967
342.1819763183594 0 18701.592
343.1615905761719 0 13840.927
343.1840515136719 0 5151.5
344.1720886230469 0 18770.002
344.9764709472656 0 12006.883
345.1312255859375 0 32048.58
346.1151123046875 0 127865.02
346.1377258300781 0 7102.031
346.6984558105469 0 6669.6196
347.1178894042969 0 19034.982
347.6800842285156 0 64935.023 b 5
348.1815185546875 0 22039.645
349.1512756347656 0 8872.1045
350.1343688964844 0 6203.3794
350.2190246582031 0 84740.76
351.22161865234375 0 16405.633
352.1673583984375 0 7683.378
352.6728820800781 0 5439.2085
353.6748962402344 0 31914.453 y Water loss 7
354.17510986328125 0 7414.9326
359.029052734375 0 11600.533
360.20404052734375 0 10205.642
360.695556640625 0 29947.908
361.1972961425781 0 30901.982
361.68701171875 0 6637.061
362.18597412109375 0 67902.04
362.6800842285156 0 121043.984 y 7
363.1413879394531 0 191423.66
363.1825256347656 0 45689.348
363.6815490722656 0 16503.898
364.1250305175781 0 29452.033
364.1455078125 0 21162.418
366.1776123046875 0 22946.44
367.139892578125 0 19058.107
367.16229248046875 0 11411.626
367.24554443359375 0 11707.585
367.68377685546875 0 5984.0557
368.1943664550781 0 7953.8374
374.20062255859375 0 6664.6533
376.162109375 0 20399.055
376.1983947753906 0 22089.76
376.6976318359375 0 11246.166
377.14642333984375 0 23746.492
378.1510925292969 0 5102.967
378.2139587402344 0 72054.484
379.2095642089844 0 267759.78
380.16693115234375 0 7253.252
380.2121276855469 0 51657.695
381.1521301269531 0 542951.7
381.7115173339844 0 14215.237
382.15484619140625 0 89591.78
382.1886901855469 0 17554.97
382.2137451171875 0 6960.1587
382.6932373046875 0 6605.0024
383.1565246582031 0 13441.238
384.1664123535156 0 26372.945
384.1888427734375 0 27571.4
385.1507263183594 0 41063.78
385.1728820800781 0 16669.309 y 4
386.15423583984375 0 8366.92
386.1803894042969 0 10655.255
389.19586181640625 0 6505.819 b 9
390.1961364746094 0 14010.749
390.6961669921875 0 12854.691
391.1374206542969 0 18533.312
391.1858215332031 0 4910.2393
392.1180114746094 0 6765.2275
392.2091369628906 0 5511.392
393.1888732910156 0 9593.095
394.1727600097656 0 89550.98
395.1340637207031 0 24778.805
395.1576232910156 0 30077.668
395.2406921386719 0 36260.285
396.191650390625 0 8611.984
396.2248840332031 0 78786.46
397.2274169921875 0 20743.994
398.1783752441406 0 34269.734
399.1838684082031 0 8203.807
401.2153015136719 0 14989.94
402.17669677734375 0 28590.922
402.1994323730469 0 35153.613
403.1368408203125 0 5973.662
403.20062255859375 0 21308.285
404.2046813964844 0 5757.7134
404.7103576660156 0 10214.03
405.26171875 0 16258.059
406.2452087402344 0 12563.051
407.203857421875 0 6259.341
407.24664306640625 0 6180.9497
409.1473693847656 0 45731.598
409.2066345214844 0 8169.7144
409.7033386230469 0 6628.906
410.1487121582031 0 8211.856
411.1992492675781 0 78387.74
412.183349609375 0 281875.72
413.14459228515625 0 24853.959
413.1656799316406 0 31504.258
413.1881103515625 0 28829.303
413.2517395019531 0 17991.209
414.1468811035156 0 8143.9155
414.1915588378906 0 10158.186
414.23565673828125 0 67630.266
415.0372009277344 0 16909.676
415.1981201171875 0 17155.549
415.24041748046875 0 9276.592
417.7171325683594 0 48831.734
418.2109680175781 0 85055.28
418.71173095703125 0 38901.44
419.2134094238281 0 12510.077
421.208740234375 0 15079.1045 y Water loss 10
422.1922607421875 0 11797.308 y Ammonia loss 10
422.7099914550781 0 7225.4956
423.20367431640625 0 22630.488
423.27197265625 0 135482.78
423.7020263671875 0 8468.188
424.2226257324219 0 7526.7427
424.2747802734375 0 37470.703
424.73431396484375 0 15353.955
425.2383117675781 0 6469.502
426.722412109375 0 918131.94
427.22381591796875 0 421061.03
427.7251281738281 0 116291.33
428.2256164550781 0 18879.16
429.20989990234375 0 356530.75
430.1941223144531 0 560984.7
431.1972351074219 0 114536.79
431.7145690917969 0 101906.8 b Water loss 6
432.2120361328125 0 45765.18 b Ammonia loss 6
432.71160888671875 0 22855.355
433.2098388671875 0 15678.732
433.25579833984375 0 24777.406 b Water loss 3
434.2591857910156 0 7579.642
438.1737365722656 0 15686.554
438.7301025390625 0 8451.401
439.21954345703125 0 225872.02 y 10
440.1783142089844 0 15288.835
440.222412109375 0 63249.04
440.7198791503906 0 434023.03 b 6
441.2215270996094 0 252963.6
441.7227783203125 0 59232.848
442.2240295410156 0 18148.229
446.7107849121094 0 7050.261 y Water loss 6
446.74078369140625 0 6906.823
447.21282958984375 0 24335.35 y Ammonia loss 6
447.7100524902344 0 5729.6104
448.20416259765625 0 23894.855
449.208984375 0 10731.6875
450.1734924316406 0 8192.117
450.2373046875 0 11040.538
450.2821960449219 0 11231.018
451.2671203613281 0 258622.55 b 3
452.2701721191406 0 51164.34
453.27203369140625 0 9789.056
455.23272705078125 0 5368.39
455.26666259765625 0 13335.183
455.7259521484375 0 6222.1367 y 6
456.2218933105469 0 10064.085
457.2069396972656 0 7164.329
459.1989440917969 0 26489.273
460.75396728515625 0 30336.113
461.2057189941406 0 15847.311
461.2533264160156 0 32103.953
464.2156066894531 0 5811.583
465.748046875 0 9263.632
466.2406311035156 0 9115.623
468.2936096191406 0 67184.81
469.2967529296875 0 14905.476
471.236328125 0 5528.695
474.22393798828125 0 7355.113
474.2594299316406 0 47610.836
474.7542724609375 0 78428.01
475.2554931640625 0 28553.188
475.7548828125 0 8716.869
476.2259216308594 0 18694.152
477.2106018066406 0 41640.14
478.22943115234375 0 38963.16 y Water loss 9
478.2771301269531 0 32422.924
479.2818603515625 0 7514.302
479.74688720703125 0 6575.8574
480.22296142578125 0 7610.195
481.2325744628906 0 6232.6934
483.264404296875 0 184733.64
483.76617431640625 0 125796.78
484.22137451171875 0 12460.738
484.266845703125 0 32503.836
484.76617431640625 0 11709.207
488.2568664550781 0 51302.637 b Water loss 7
488.7558288574219 0 33005.895 b Ammonia loss 7
489.2514953613281 0 12041.387
489.7578430175781 0 8359.565
492.3361511230469 0 6160.7104
494.236328125 0 513536.06
495.2393493652344 0 122366.79
496.24090576171875 0 593320.9 y 9
497.2467346191406 0 150797.86
497.7637023925781 0 58809.945
498.24176025390625 0 26874.338
498.7671813964844 0 5628.7354
499.24224853515625 0 8031.7783
503.10784912109375 0 23843.207
504.20172119140625 0 22239.8
506.22607421875 0 13283.726
506.27288818359375 0 11094.891
507.2615661621094 0 9946.364
511.2628479003906 0 40705.99
512.262451171875 0 13235.291
513.2095947265625 0 13756.669
514.189208984375 0 6481.824
515.2799682617188 0 6022.715
516.22119140625 0 7326.279
519.1399536132812 0 5909.9814
523.2581176757812 0 15502.01
523.5902099609375 0 6127.214
524.2809448242188 0 21947.91
525.2669677734375 0 83237.21
526.2283935546875 0 10046.338
526.2689208984375 0 14775.324
528.2427368164062 0 4856.936
528.9332885742188 0 16157.726
529.26220703125 0 37571.84 Precursor Water loss
529.595947265625 0 32274.184 Precursor Ammonia loss
529.9314575195312 0 21568.734
530.265625 0 8051.727
531.2144165039062 0 7149.2124
531.7681274414062 0 5407.088
532.1937866210938 0 12748.42
535.2649536132812 0 25347.174 Precursor
535.5997924804688 0 13687.792
535.9354248046875 0 12468.516
536.2616577148438 0 5828.005
537.2437744140625 0 10343.608
537.7390747070312 0 8947.203
537.7809448242188 0 9820.994
538.2734375 0 6755.711
538.7703247070312 0 8036.4404
539.2364501953125 0 88092.766
539.2799072265625 0 3667.0532
540.2390747070312 0 30490.64
541.24658203125 0 6625.028
542.2938232421875 0 83672.48
543.2781982421875 0 88479.05
544.283203125 0 23168.137
545.282470703125 0 10056.907
545.7644653320312 0 10501.946 b Water loss 8
546.2559814453125 0 13306.307 b Ammonia loss 8
546.75048828125 0 7263.823
547.2479248046875 0 7755.499
547.7745361328125 0 5307.6895
548.2432250976562 0 4954.931
549.22119140625 0 28263.41
550.2071533203125 0 6613.0327
551.2579956054688 0 32336.125
551.3302001953125 0 15109.6875
552.2601318359375 0 9776.216
553.2650146484375 0 5156.33
554.2826538085938 0 8635.898
554.7640380859375 0 25328.707 b 8
555.2598266601562 0 31933.295
555.759765625 0 19962.393
558.2228393554688 0 5456.9683
561.3123168945312 0 22517.926 b Water loss 4
562.3115844726562 0 7215.6313
563.25634765625 0 18194.736
564.267578125 0 10732.208
564.3308715820312 0 6420.5845
565.2640991210938 0 38341.344
566.2681884765625 0 5646.1753
567.231201171875 0 126812.414
568.2369384765625 0 20356.889
569.2369995117188 0 12003.846
575.2466430664062 0 53882.418
576.2384033203125 0 19778.875
577.2206420898438 0 6566.216
577.2698974609375 0 7170.1104 y 4
579.3253784179688 0 160177.7 b 4
580.328125 0 57273.016
581.2634887695312 0 4982.2837
581.3302001953125 0 12799.246
582.3041381835938 0 10480.714
583.305908203125 0 6579.6045
588.7978515625 0 6358.5386
589.30810546875 0 6219.293
592.2730712890625 0 60270.32
592.325439453125 0 18626.008
593.2571411132812 0 210912.89 y Water loss 8
593.3057250976562 0 83487.34
593.7859497070312 0 9224.287
594.256103515625 0 72974.9
594.30859375 0 16533.402
595.2551879882812 0 19946.986
596.2582397460938 0 5238.626
596.3499145507812 0 8664.262
597.3087768554688 0 5105.901
597.8037719726562 0 6750.5093
602.3001098632812 0 10424.837
602.7944946289062 0 23957.81
603.24072265625 0 6965.269
603.2924194335938 0 27684.268
603.795166015625 0 14862.862
610.2835083007812 0 954237.25
611.2691650390625 0 1732930 y 8
611.80078125 0 90775.61
612.2720336914062 0 514948.84
612.8021240234375 0 13949.436
613.2736206054688 0 129436.89
614.2747192382812 0 14978.273
617.2817993164062 0 15372.725
620.2734985351562 0 8021.682
620.8079223632812 0 5891.3154
621.2509765625 0 42130.633
621.3023071289062 0 11298.865
622.255859375 0 16405.465
622.3677368164062 0 6407.013
623.2552490234375 0 8088.721
628.2799072265625 0 7829.841
628.7769775390625 0 6888.074
634.3068237304688 0 23379.217
635.2974853515625 0 34477.03
636.3290405273438 0 20619.314
636.787841796875 0 20509.498 y Water loss 3
637.2886352539062 0 18062.928 y Ammonia loss 3
637.7926635742188 0 11818.386
638.3353271484375 0 10810.068 b Water loss 10
638.8314819335938 0 7828.4146 b Ammonia loss 10
640.2839965820312 0 7783.4473
644.2941284179688 0 5582.9795
645.2963256835938 0 73049.69
645.7955932617188 0 101274.766 y 3
646.2950439453125 0 71241.9
646.7953491210938 0 17692.35
648.3416748046875 0 7123.2544
650.3258056640625 0 13377.985
652.3206176757812 0 202418.52
653.3234252929688 0 79090.39
654.3245239257812 0 17805.572
658.3333129882812 0 11913.753
659.3148803710938 0 7600.681
661.3289184570312 0 5956.914
662.3057250976562 0 76392.35
663.2933349609375 0 44164.87
664.2927856445312 0 17059.715
664.3411254882812 0 14183.407
665.3429565429688 0 6502.3857
668.2782592773438 0 6529.815
676.3416137695312 0 108569.516 b Water loss 5
677.3348388671875 0 57891.766 b Ammonia loss 5
678.3314819335938 0 23372.121
679.33349609375 0 23903.586
680.315673828125 0 387561.75
681.3182983398438 0 167557.9
682.3218994140625 0 34281.39
684.3228149414062 0 33539.75
684.8223876953125 0 28980.732
685.316650390625 0 14362.172
688.3339233398438 0 14378.772
692.8352661132812 0 44753.88
693.3306884765625 0 151376.05 y Water loss 2
693.8292846679688 0 108421.06 y Ammonia loss 2
694.3516845703125 0 1065576.4 b 5
694.8299560546875 0 21132.658
695.3547973632812 0 381825.4
696.2960815429688 0 6096.5776
696.35791015625 0 84693.12
697.3555297851562 0 21761.81
701.84130859375 0 384178.47
702.3377075195312 0 664686.7 y 2
702.838134765625 0 452385.47 b Water loss 11
702.923095703125 0 9318.948
702.9351196289062 0 5923.55
703.3397216796875 0 173477.06 b Ammonia loss 11
703.8408813476562 0 56144.74
704.3355712890625 0 20852.205
705.3554077148438 0 28251.637
706.3428955078125 0 63637.227 y Water loss 7
707.3441162109375 0 34562.445
708.3502197265625 0 6838.6357
709.327880859375 0 5645.3794
711.3673706054688 0 63841.484
711.8609619140625 0 57798.234 b 11
712.3643188476562 0 36930.066
712.8628540039062 0 19328.066
720.379638671875 0 7064.6035
721.3635864257812 0 43278.516
722.3653564453125 0 19087.357
723.3675537109375 0 270219.38
724.3543090820312 0 493727.06 y 7
725.3555908203125 0 211376.88
726.3575439453125 0 57896.46
727.3604736328125 0 6278.447
728.372314453125 0 17476.443
729.3633422851562 0 18246.162
730.3629760742188 0 6277.703
733.3630981445312 0 28623.137
734.34765625 0 30283.516
735.3497314453125 0 12902.838
739.35693359375 0 7485.8193
742.867431640625 0 13208.464 y Water loss 1
743.366455078125 0 24925.498 y Ammonia loss 1
745.3570556640625 0 8839.787
747.3939819335938 0 10002.624
748.379638671875 0 16218.828
749.336181640625 0 10315.559
751.3751831054688 0 137256.88
751.871337890625 0 50226.535 y 1
752.3759155273438 0 94232.234
752.8715209960938 0 12084.249
753.3784790039062 0 17482.184
754.3787841796875 0 6326.3716
761.3746337890625 0 11961.575
762.35205078125 0 7947.583
762.4133911132812 0 14735.276
763.3639526367188 0 72685.984
764.3656616210938 0 32652.938
765.40087890625 0 38298.715
766.406494140625 0 19589.111
767.3468017578125 0 31916.926
769.4024047851562 0 9921.107
775.3883666992188 0 9306.262
776.375 0 8160.0107
779.3842163085938 0 69028.55
780.3864135742188 0 34261.27
781.371826171875 0 29900.125
782.3663330078125 0 16356.143
783.3757934570312 0 5762.5156
793.3997192382812 0 111962.97 Precursor Water loss
794.4010009765625 0 57139.895
795.339111328125 0 7957.5737
795.4115600585938 0 8613.43
807.3775634765625 0 23909.135
807.4396362304688 0 15284.606
808.3834838867188 0 14735.802
818.4310302734375 0 14759.175
819.4384155273438 0 13443.004
820.43798828125 0 9182.685
825.3933715820312 0 12353.5625
826.3787231445312 0 10316.48
834.3489379882812 0 5198.901
834.42626953125 0 27815.201
835.4148559570312 0 38877.633
836.448486328125 0 88134.94
837.4522094726562 0 37412.215
838.4536743164062 0 8095.871
842.4076538085938 0 6121.163
843.3978881835938 0 11670.111
844.3968505859375 0 18050.262
845.3954467773438 0 21563.516
846.4459228515625 0 36919.43
847.4423828125 0 30578.893
848.4434814453125 0 8087.5522
851.37939453125 0 9397.275
852.4359741210938 0 310341.3
853.4385986328125 0 141195.27
854.4414672851562 0 43165.188
855.44189453125 0 14030.807
862.420654296875 0 175899.81 b Water loss 6
863.4158935546875 0 110237.88 b Ammonia loss 6
864.4552612304688 0 217861.45
865.4596557617188 0 119809.14
866.4613037109375 0 30387.73
876.4380493164062 0 22682.883
877.4400634765625 0 12420.295
880.4307861328125 0 1420555.2 b 6
881.4334716796875 0 730443.7
882.4367065429688 0 211322.17
883.43896484375 0 48819.38
884.43505859375 0 9398.015
890.4170532226562 0 7973.321
891.4200439453125 0 8207.572
892.4603271484375 0 30771.771
893.4695434570312 0 13850.494
894.4503784179688 0 8070.766
907.4418334960938 0 33725.703
908.4353637695312 0 39916.535
909.4421997070312 0 67711.43
910.434814453125 0 69341.77 y 6
911.4376831054688 0 31782.633
936.4267578125 0 10413.812
937.4234008789062 0 8440.479
947.4392700195312 0 14744.568
953.4472045898438 0 7795.223
954.4306030273438 0 20943.457
955.4248046875 0 27527.564
956.4267578125 0 10335.012
964.463623046875 0 21500.65
965.4544067382812 0 26907.32
966.4546508789062 0 14594.461
971.45751953125 0 13471.483
972.4487915039062 0 14515.3955
973.4495849609375 0 9598.721
975.5015869140625 0 19752.34 b Water loss 7
976.4981079101562 0 11383.959 b Ammonia loss 7
979.4713134765625 0 9078.735
980.4583129882812 0 7009.8643
981.4708251953125 0 8346.788
993.5152587890625 0 115319 b 7
994.5174560546875 0 59136.348
995.518310546875 0 26899.69
1007.4453125 0 9814.415 y Water loss 5
1024.4735107421875 0 26649.646
1025.463623046875 0 32397.889 y 5
1026.46728515625 0 19056.955
1027.47021484375 0 7080.5156
1075.530029296875 0 9261.665
1076.53857421875 0 10213.465
1091.5040283203125 0 6432.4575 b Ammonia loss 8
1092.5537109375 0 17199.697
1093.55126953125 0 25239.834
1094.5516357421875 0 15163.31
1107.548828125 0 9190.582
1108.5400390625 0 8977.796 b 8
1117.4874267578125 0 7305.853
1118.481689453125 0 20624.258
1134.521728515625 0 13347.473
1135.508544921875 0 42098.914 y Water loss 4
1136.498779296875 0 47040.242 y Ammonia loss 4
1137.5013427734375 0 22579.395
1152.5296630859375 0 53177.49
1153.522705078125 0 83939.03 y 4
1154.5213623046875 0 38958.62
1155.52099609375 0 12965.403
1165.5679931640625 0 6782.636 b 9
1341.54443359375 0 5537.173
1791.8846435546875 0 6734.711
2837.552978515625 0 6566.023

Spectrum Details

|  |  |
| --- | --- |
| Matched peaks? Matched peaks The total absolute number of peaks matched. Additionally in brackets the total fraction of peaks matched and the total number of peaks is shown. | 83 (10.10% of 822) |
| FDR? FDR The false discovery rate estimated for this peptide. It is calculated by matching all theoretical fragments with a non-integer shift with the raw peaks for this spectrum. This is done with 40 different shifts. The resulting percentage is the average number of annotated peaks over the number of annotated peaks with the correct spectrum. | 0.60% |
| Satellite FDR? Satellite FDR See the FDR for details on its calculation. This satellite ion specific FDR only contains the satellite ions (d/w) for I/L/J positions. | - |
| PSM Score? PSM Score The PSM Score as given by Hecklib to this annotated spectrum. It is shown with three significant figures. | 604 |

#### Spectrum 7091? Spectrum 7091 The raw spectrum of this peptide as annotated by Hecklib. The fragments are coloured according to ion type (see legend). Any peaks with a star '\*' as text can be hovered over to see the full details, first the ion type second the mass shift type. By hovering over the amino acids in the peptide or ions in the legend the corresponding peaks are highlighted. By toggling the 'Unassigned' label you can turn the background (unassigned) peaks on or off in the plot. By updating the slider in the Ion legend you can update the spectrum to only show the top X% of the peaks with labels. The top X% means any peak that is within X% of the highest intensity. By dragging in the spectrum you can zoom in to a specific part of the spectrum and use 'Zoom Out' to get back to the original zoom level. The annotation of the spectrum is based on the given sequence in the peptides file and is done with different software so inconsistencies are likely. The peaks are annotated based on the given sequence, with 20 ppm tolerance.

Copy Data

##### Spectrum 7091 (TSV)

###### Preview

```
Loading example...
```

*Click on the button to copy the data to your clipboard.*

Mz MinMz MaxIntensity Max

WidthHeightPeptide font sizePeptide stroke widthSpectrum font sizeSpectrum stroke widthCompact peptide

Ion legend

wxyz

abcd

OtherUnassignedIon^Charge^_Position_Show for top:%

TVLHQDWLDGKEY

09.12e+51.82e+62.74e+63.65e+6

Zoom Out

a^+1^_2_a^+1^_2_y^+1^_1_b^+1^_2_b^+1^_2_y^+2^_3_y^+2^_4_y^+2^_4_b^+2^_5_b^+2^_5_y^+1^_2_b^+1^_3_y^+2^_5_y^+3^_7_y^+2^_5_y^+1^_2_b^+1^_3_b^+2^_6_b^+2^_6_b^+2^_6_y^+2^_6_y^+2^_6_b^+3^_9_y^+3^_9_b^+3^_10_y^+1^_3_y^+1^_3_b^+2^_7_b^+2^_7_b^+1^_4_y^+1^_3_b^+2^_7_y^+2^_7_b^+1^_4_y^+2^_7_y^+3^_11_y^+3^_11_y^+1^_4_b^+2^_8_b^+2^_8_y^+1^_4_\*\*\*b^+2^_9_b^+1^_5_b^+1^_5_y^+2^_9_y^+2^_9_b^+1^_5_y^+1^_5_y^+1^_5_y^+2^_10_y^+2^_10_b^+2^_11_b^+2^_11_y^+2^_10_b^+1^_6_b^+1^_6_y^+2^_11_y^+2^_11_b^+1^_6_y^+2^_11_b^+2^_12_b^+2^_12_y^+1^_6_b^+2^_12_y^+1^_6_y^+2^_12_y^+2^_12_\*b^+1^_7_b^+1^_7_b^+1^_7_y^+1^_7_b^+1^_8_b^+1^_8_b^+1^_8_y^+1^_8_y^+1^_8_y^+1^_8_b^+1^_9_b^+1^_9_y^+1^_9_y^+1^_9_y^+1^_9_b^+1^_10_y^+1^_10_

0778155623343112

Fragment Matches Table

Show background peaks

| Position | Ion type | Intensity | mz Theoretical | mz Error (Th) | mz Error (ppm) | Charge | Series Number |
| --- | --- | --- | --- | --- | --- | --- | --- |
| - | - | 3544 | 120.1 | - | - | 0 | - |
| - | - | 8590 | 120.1 | - | - | 0 | - |
| - | - | 3742 | 121 | - | - | 0 | - |
| - | - | 1.754E+04 | 122.1 | - | - | 0 | - |
| - | - | 1.261E+04 | 123 | - | - | 0 | - |
| - | - | 4332 | 123.1 | - | - | 0 | - |
| - | - | 2916 | 123.1 | - | - | 0 | - |
| - | - | 6525 | 125.1 | - | - | 0 | - |
| - | - | 1.357E+04 | 126.1 | - | - | 0 | - |
| - | - | 7.674E+04 | 127.1 | - | - | 0 | - |
| - | - | 5544 | 127.1 | - | - | 0 | - |
| - | - | 7250 | 127.1 | - | - | 0 | - |
| - | - | 4433 | 128.1 | - | - | 0 | - |
| - | - | 2.185E+05 | 128.1 | - | - | 0 | - |
| - | - | 1.567E+04 | 129.1 | - | - | 0 | - |
| - | - | 2.599E+06 | 129.1 | - | - | 0 | - |
| - | - | 1.111E+06 | 130.1 | - | - | 0 | - |
| - | - | 1.95E+04 | 130.1 | - | - | 0 | - |
| - | - | 1.769E+05 | 130.1 | - | - | 0 | - |
| - | - | 3476 | 130.1 | - | - | 0 | - |
| - | - | 1.066E+05 | 131.1 | - | - | 0 | - |
| - | - | 2312 | 131.1 | - | - | 0 | - |
| - | - | 4495 | 131.1 | - | - | 0 | - |
| - | - | 2.803E+05 | 132.1 | - | - | 0 | - |
| - | - | 6550 | 133.1 | - | - | 0 | - |
| - | - | 4809 | 133.1 | - | - | 0 | - |
| - | - | 2.921E+04 | 133.1 | - | - | 0 | - |
| - | - | 2.418E+05 | 136.1 | - | - | 0 | - |
| - | - | 1.611E+04 | 137.1 | - | - | 0 | - |
| - | - | 3642 | 138.1 | - | - | 0 | - |
| - | - | 3.179E+04 | 138.1 | - | - | 0 | - |
| - | - | 1.174E+04 | 138.1 | - | - | 0 | - |
| - | - | 4379 | 139.1 | - | - | 0 | - |
| - | - | 2.674E+04 | 139.1 | - | - | 0 | - |
| - | - | 3.526E+04 | 141.1 | - | - | 0 | - |
| - | - | 2135 | 142.1 | - | - | 0 | - |
| - | - | 2.378E+04 | 142.1 | - | - | 0 | - |
| - | - | 1.535E+04 | 143 | - | - | 0 | - |
| - | - | 2584 | 143.1 | - | - | 0 | - |
| - | - | 1.25E+04 | 144.1 | - | - | 0 | - |
| - | - | 3.945E+04 | 145.1 | - | - | 0 | - |
| - | - | 3276 | 146.1 | - | - | 0 | - |
| - | - | 2860 | 146.1 | - | - | 0 | - |
| - | - | 1.492E+04 | 147 | - | - | 0 | - |
| - | - | 3583 | 148.1 | - | - | 0 | - |
| - | - | 2770 | 148.3 | - | - | 0 | - |
| - | - | 1.083E+04 | 149 | - | - | 0 | - |
| - | - | 3633 | 149.1 | - | - | 0 | - |
| - | - | 1.763E+04 | 150.1 | - | - | 0 | - |
| - | - | 5.078E+04 | 151.1 | - | - | 0 | - |
| - | - | 7784 | 152.1 | - | - | 0 | - |
| - | - | 2843 | 154.1 | - | - | 0 | - |
| - | - | 2554 | 154.2 | - | - | 0 | - |
| - | - | 4985 | 155 | - | - | 0 | - |
| - | - | 7602 | 155.1 | - | - | 0 | - |
| - | - | 1.564E+05 | 155.1 | - | - | 0 | - |
| 2 | a | 1.057E+05 | 155.1 | 0.0003966 | 2.557 | +1 | 2 |
| - | - | 1745 | 155.1 | - | - | 0 | - |
| - | - | 7617 | 156.1 | - | - | 0 | - |
| - | - | 3565 | 156.1 | - | - | 0 | - |
| - | - | 9769 | 156.1 | - | - | 0 | - |
| - | - | 2867 | 156.1 | - | - | 0 | - |
| - | - | 7931 | 156.1 | - | - | 0 | - |
| - | - | 8254 | 157.1 | - | - | 0 | - |
| - | - | 1.258E+04 | 157.1 | - | - | 0 | - |
| - | - | 2.149E+04 | 158.1 | - | - | 0 | - |
| - | - | 2.134E+06 | 159.1 | - | - | 0 | - |
| - | - | 1.596E+04 | 160.1 | - | - | 0 | - |
| - | - | 1.237E+04 | 160.1 | - | - | 0 | - |
| - | - | 2.213E+05 | 160.1 | - | - | 0 | - |
| - | - | 1.079E+04 | 161.1 | - | - | 0 | - |
| - | - | 2927 | 161.5 | - | - | 0 | - |
| - | - | 1.296E+05 | 165.1 | - | - | 0 | - |
| - | - | 6446 | 165.1 | - | - | 0 | - |
| - | - | 2.904E+05 | 166.1 | - | - | 0 | - |
| - | - | 2.171E+04 | 166.1 | - | - | 0 | - |
| - | - | 7351 | 167.1 | - | - | 0 | - |
| - | - | 2.462E+04 | 167.1 | - | - | 0 | - |
| - | - | 8857 | 167.1 | - | - | 0 | - |
| - | - | 3522 | 167.1 | - | - | 0 | - |
| - | - | 8185 | 167.1 | - | - | 0 | - |
| - | - | 3.263E+04 | 168.1 | - | - | 0 | - |
| - | - | 1.979E+04 | 168.1 | - | - | 0 | - |
| - | - | 1.991E+04 | 168.1 | - | - | 0 | - |
| - | - | 3.81E+04 | 169.1 | - | - | 0 | - |
| - | - | 6028 | 169.1 | - | - | 0 | - |
| - | - | 4.849E+04 | 169.1 | - | - | 0 | - |
| - | - | 1.035E+05 | 170.1 | - | - | 0 | - |
| - | - | 1.246E+04 | 171.1 | - | - | 0 | - |
| - | - | 5550 | 171.1 | - | - | 0 | - |
| - | - | 9837 | 171.1 | - | - | 0 | - |
| - | - | 1.644E+04 | 171.1 | - | - | 0 | - |
| - | - | 3.687E+04 | 172.1 | - | - | 0 | - |
| - | - | 2975 | 172.2 | - | - | 0 | - |
| - | - | 4.532E+04 | 173.1 | - | - | 0 | - |
| 2 | a | 3.613E+06 | 173.1 | 0.0005131 | 2.963 | +1 | 2 |
| - | - | 9402 | 173.5 | - | - | 0 | - |
| - | - | 3394 | 174.1 | - | - | 0 | - |
| - | - | 3.226E+05 | 174.1 | - | - | 0 | - |
| - | - | 1.579E+04 | 175.1 | - | - | 0 | - |
| - | - | 4267 | 176.1 | - | - | 0 | - |
| - | - | 6753 | 177.1 | - | - | 0 | - |
| - | - | 1.199E+05 | 178.1 | - | - | 0 | - |
| - | - | 1.295E+04 | 179.1 | - | - | 0 | - |
| - | - | 1.88E+04 | 181.1 | - | - | 0 | - |
| 13 | y | 2.842E+05 | 182.1 | 0.0005106 | 2.804 | +1 | 1 |
| - | - | 2.804E+04 | 183.1 | - | - | 0 | - |
| 2 | b | 8.92E+04 | 183.1 | 0.0004618 | 2.522 | +1 | 2 |
| - | - | 9835 | 184.1 | - | - | 0 | - |
| - | - | 4270 | 185.1 | - | - | 0 | - |
| - | - | 3.088E+04 | 185.2 | - | - | 0 | - |
| - | - | 5.374E+05 | 186.1 | - | - | 0 | - |
| - | - | 1.583E+05 | 187.1 | - | - | 0 | - |
| - | - | 4040 | 187.1 | - | - | 0 | - |
| - | - | 4.986E+04 | 187.1 | - | - | 0 | - |
| - | - | 1.131E+04 | 188.1 | - | - | 0 | - |
| - | - | 1.931E+04 | 188.1 | - | - | 0 | - |
| - | - | 5160 | 189.1 | - | - | 0 | - |
| - | - | 6.425E+04 | 190.1 | - | - | 0 | - |
| - | - | 2.625E+04 | 190.1 | - | - | 0 | - |
| - | - | 5278 | 191.1 | - | - | 0 | - |
| - | - | 5589 | 193.1 | - | - | 0 | - |
| - | - | 4044 | 193.1 | - | - | 0 | - |
| - | - | 1.826E+04 | 194.1 | - | - | 0 | - |
| - | - | 1.723E+05 | 195.1 | - | - | 0 | - |
| - | - | 3.282E+04 | 196.1 | - | - | 0 | - |
| - | - | 1.777E+04 | 196.1 | - | - | 0 | - |
| - | - | 5092 | 197.1 | - | - | 0 | - |
| - | - | 1.922E+04 | 197.1 | - | - | 0 | - |
| - | - | 5339 | 197.2 | - | - | 0 | - |
| - | - | 3.087E+04 | 198.1 | - | - | 0 | - |
| - | - | 6.931E+04 | 198.1 | - | - | 0 | - |
| - | - | 1.702E+04 | 199.1 | - | - | 0 | - |
| - | - | 2.056E+04 | 199.1 | - | - | 0 | - |
| - | - | 3309 | 199.1 | - | - | 0 | - |
| - | - | 4057 | 199.1 | - | - | 0 | - |
| - | - | 1.024E+04 | 199.1 | - | - | 0 | - |
| - | - | 7923 | 199.1 | - | - | 0 | - |
| - | - | 5281 | 200.1 | - | - | 0 | - |
| - | - | 1.218E+04 | 200.1 | - | - | 0 | - |
| 2 | b | 9.753E+05 | 201.1 | 0.0004562 | 2.268 | +1 | 2 |
| - | - | 6636 | 202.1 | - | - | 0 | - |
| - | - | 9.774E+04 | 202.1 | - | - | 0 | - |
| - | - | 5471 | 203.1 | - | - | 0 | - |
| - | - | 3574 | 203.2 | - | - | 0 | - |
| - | - | 7362 | 204.1 | - | - | 0 | - |
| - | - | 7580 | 204.1 | - | - | 0 | - |
| - | - | 8.085E+04 | 205.1 | - | - | 0 | - |
| - | - | 7115 | 205.1 | - | - | 0 | - |
| - | - | 1.416E+04 | 205.1 | - | - | 0 | - |
| - | - | 9947 | 206.1 | - | - | 0 | - |
| - | - | 1.37E+04 | 206.1 | - | - | 0 | - |
| - | - | 3.454E+04 | 207.2 | - | - | 0 | - |
| - | - | 8095 | 208.1 | - | - | 0 | - |
| - | - | 6413 | 209.1 | - | - | 0 | - |
| - | - | 3.609E+04 | 210.1 | - | - | 0 | - |
| - | - | 3787 | 211.1 | - | - | 0 | - |
| 11 | y | 2.346E+04 | 211.1 | 0.0004966 | 2.352 | +2 | 3 |
| - | - | 5749 | 211.1 | - | - | 0 | - |
| - | - | 3.192E+04 | 212.1 | - | - | 0 | - |
| - | - | 8.435E+04 | 213.1 | - | - | 0 | - |
| - | - | 1.243E+04 | 213.2 | - | - | 0 | - |
| - | - | 9414 | 214.1 | - | - | 0 | - |
| - | - | 4670 | 215.1 | - | - | 0 | - |
| - | - | 9330 | 215.1 | - | - | 0 | - |
| - | - | 3871 | 215.5 | - | - | 0 | - |
| - | - | 2.42E+04 | 216.1 | - | - | 0 | - |
| - | - | 1.356E+04 | 217.1 | - | - | 0 | - |
| - | - | 6342 | 221.1 | - | - | 0 | - |
| - | - | 5268 | 221.1 | - | - | 0 | - |
| - | - | 5856 | 221.1 | - | - | 0 | - |
| - | - | 1.71E+04 | 221.1 | - | - | 0 | - |
| - | - | 7427 | 221.1 | - | - | 0 | - |
| - | - | 4090 | 222.1 | - | - | 0 | - |
| - | - | 2.623E+04 | 222.1 | - | - | 0 | - |
| - | - | 4537 | 223.1 | - | - | 0 | - |
| - | - | 4.954E+04 | 223.1 | - | - | 0 | - |
| - | - | 3.338E+05 | 223.2 | - | - | 0 | - |
| - | - | 6688 | 224.1 | - | - | 0 | - |
| - | - | 4878 | 224.1 | - | - | 0 | - |
| - | - | 3.959E+04 | 224.2 | - | - | 0 | - |
| - | - | 6253 | 225.2 | - | - | 0 | - |
| - | - | 6.854E+04 | 226.1 | - | - | 0 | - |
| - | - | 6.514E+04 | 227.1 | - | - | 0 | - |
| - | - | 9245 | 227.1 | - | - | 0 | - |
| - | - | 1.968E+04 | 227.2 | - | - | 0 | - |
| - | - | 4280 | 228.1 | - | - | 0 | - |
| - | - | 1.208E+04 | 228.1 | - | - | 0 | - |
| - | - | 3.953E+04 | 229.1 | - | - | 0 | - |
| - | - | 4661 | 229.2 | - | - | 0 | - |
| - | - | 9098 | 230.2 | - | - | 0 | - |
| - | - | 4168 | 231.1 | - | - | 0 | - |
| - | - | 5.935E+04 | 233.1 | - | - | 0 | - |
| - | - | 3.652E+05 | 234.1 | - | - | 0 | - |
| - | - | 3962 | 234.6 | - | - | 0 | - |
| - | - | 4.522E+04 | 235.1 | - | - | 0 | - |
| - | - | 9377 | 235.2 | - | - | 0 | - |
| - | - | 4202 | 236.1 | - | - | 0 | - |
| - | - | 1.902E+04 | 237.1 | - | - | 0 | - |
| - | - | 2.753E+04 | 238.1 | - | - | 0 | - |
| - | - | 6334 | 238.1 | - | - | 0 | - |
| - | - | 9960 | 239.1 | - | - | 0 | - |
| - | - | 7453 | 239.1 | - | - | 0 | - |
| - | - | 4776 | 239.1 | - | - | 0 | - |
| 10 | y | 3245 | 239.6 | 0.001758 | 7.337 | +2 | 4 |
| - | - | 3866 | 240.1 | - | - | 0 | - |
| - | - | 1.6E+05 | 240.1 | - | - | 0 | - |
| - | - | 3483 | 241.1 | - | - | 0 | - |
| - | - | 2.084E+04 | 241.1 | - | - | 0 | - |
| - | - | 1.859E+04 | 241.1 | - | - | 0 | - |
| - | - | 1.839E+04 | 241.2 | - | - | 0 | - |
| - | - | 3.341E+04 | 243.1 | - | - | 0 | - |
| - | - | 4608 | 243.1 | - | - | 0 | - |
| - | - | 1.665E+05 | 244.1 | - | - | 0 | - |
| - | - | 1.557E+04 | 245.1 | - | - | 0 | - |
| - | - | 3782 | 246.1 | - | - | 0 | - |
| - | - | 1.835E+05 | 247.1 | - | - | 0 | - |
| - | - | 1.559E+04 | 247.1 | - | - | 0 | - |
| - | - | 5549 | 247.2 | - | - | 0 | - |
| - | - | 9.389E+04 | 248.1 | - | - | 0 | - |
| 10 | y | 6452 | 248.6 | 0.0002905 | 1.168 | +2 | 4 |
| - | - | 1.266E+05 | 249.1 | - | - | 0 | - |
| - | - | 6282 | 249.1 | - | - | 0 | - |
| - | - | 3.478E+04 | 249.1 | - | - | 0 | - |
| - | - | 1.814E+04 | 250.1 | - | - | 0 | - |
| - | - | 7627 | 250.1 | - | - | 0 | - |
| - | - | 4235 | 250.1 | - | - | 0 | - |
| - | - | 6093 | 250.2 | - | - | 0 | - |
| - | - | 1.262E+06 | 251.2 | - | - | 0 | - |
| - | - | 6191 | 252.1 | - | - | 0 | - |
| - | - | 6887 | 252.1 | - | - | 0 | - |
| - | - | 1.627E+05 | 252.2 | - | - | 0 | - |
| - | - | 5789 | 252.2 | - | - | 0 | - |
| - | - | 6357 | 253.1 | - | - | 0 | - |
| - | - | 1.997E+04 | 253.2 | - | - | 0 | - |
| - | - | 7.138E+04 | 255.1 | - | - | 0 | - |
| - | - | 3818 | 256.1 | - | - | 0 | - |
| - | - | 1.358E+04 | 256.1 | - | - | 0 | - |
| - | - | 6500 | 256.1 | - | - | 0 | - |
| - | - | 7240 | 256.2 | - | - | 0 | - |
| - | - | 4759 | 257.1 | - | - | 0 | - |
| - | - | 2.597E+04 | 257.1 | - | - | 0 | - |
| - | - | 1.076E+04 | 258.1 | - | - | 0 | - |
| - | - | 6.573E+04 | 258.1 | - | - | 0 | - |
| - | - | 7154 | 259.1 | - | - | 0 | - |
| - | - | 6616 | 259.1 | - | - | 0 | - |
| - | - | 2.832E+04 | 261.1 | - | - | 0 | - |
| - | - | 1.465E+04 | 261.1 | - | - | 0 | - |
| - | - | 1.41E+04 | 262.1 | - | - | 0 | - |
| - | - | 1.679E+05 | 265.1 | - | - | 0 | - |
| - | - | 3.66E+05 | 266.1 | - | - | 0 | - |
| - | - | 6872 | 267.1 | - | - | 0 | - |
| - | - | 4.066E+04 | 267.1 | - | - | 0 | - |
| - | - | 8266 | 267.1 | - | - | 0 | - |
| - | - | 1.577E+04 | 268.1 | - | - | 0 | - |
| - | - | 3.091E+04 | 268.2 | - | - | 0 | - |
| - | - | 1.019E+04 | 269.2 | - | - | 0 | - |
| - | - | 2.32E+04 | 269.2 | - | - | 0 | - |
| - | - | 4.247E+04 | 270.1 | - | - | 0 | - |
| - | - | 5349 | 270.1 | - | - | 0 | - |
| - | - | 4240 | 270.2 | - | - | 0 | - |
| - | - | 1.212E+04 | 270.6 | - | - | 0 | - |
| - | - | 7.377E+04 | 272.2 | - | - | 0 | - |
| - | - | 1.205E+04 | 273.2 | - | - | 0 | - |
| - | - | 6.007E+04 | 274.1 | - | - | 0 | - |
| - | - | 2.833E+04 | 274.1 | - | - | 0 | - |
| - | - | 1.762E+04 | 275.1 | - | - | 0 | - |
| - | - | 8696 | 275.1 | - | - | 0 | - |
| - | - | 1.044E+04 | 275.2 | - | - | 0 | - |
| - | - | 1.346E+04 | 276.1 | - | - | 0 | - |
| - | - | 6707 | 276.1 | - | - | 0 | - |
| - | - | 2.799E+04 | 276.2 | - | - | 0 | - |
| - | - | 4753 | 276.2 | - | - | 0 | - |
| - | - | 5822 | 277.1 | - | - | 0 | - |
| - | - | 2.559E+04 | 279.1 | - | - | 0 | - |
| - | - | 2.701E+04 | 280.1 | - | - | 0 | - |
| - | - | 1.018E+04 | 281.1 | - | - | 0 | - |
| 5 | b | 4533 | 281.2 | 0.002422 | 8.613 | +2 | 5 |
| - | - | 2.804E+04 | 282.2 | - | - | 0 | - |
| - | - | 6607 | 282.7 | - | - | 0 | - |
| - | - | 1.047E+06 | 283.1 | - | - | 0 | - |
| - | - | 5497 | 284.1 | - | - | 0 | - |
| - | - | 3.492E+04 | 284.1 | - | - | 0 | - |
| - | - | 1.635E+04 | 284.1 | - | - | 0 | - |
| - | - | 1.459E+05 | 284.1 | - | - | 0 | - |
| - | - | 7269 | 285.1 | - | - | 0 | - |
| - | - | 4709 | 285.1 | - | - | 0 | - |
| - | - | 1.308E+04 | 285.1 | - | - | 0 | - |
| - | - | 8450 | 285.2 | - | - | 0 | - |
| - | - | 5.858E+04 | 286.1 | - | - | 0 | - |
| - | - | 8833 | 287.1 | - | - | 0 | - |
| - | - | 1.413E+04 | 287.2 | - | - | 0 | - |
| - | - | 6199 | 287.2 | - | - | 0 | - |
| - | - | 3507 | 288.1 | - | - | 0 | - |
| - | - | 4085 | 288.6 | - | - | 0 | - |
| 5 | b | 6075 | 290.2 | 0.001015 | 3.498 | +2 | 5 |
| - | - | 3528 | 290.7 | - | - | 0 | - |
| - | - | 4103 | 291.2 | - | - | 0 | - |
| 12 | y | 4.286E+05 | 293.1 | 0.0007545 | 2.574 | +1 | 2 |
| - | - | 1.78E+04 | 293.2 | - | - | 0 | - |
| - | - | 8.91E+04 | 294.1 | - | - | 0 | - |
| - | - | 4582 | 294.2 | - | - | 0 | - |
| - | - | 8032 | 295.1 | - | - | 0 | - |
| 3 | b | 1.809E+04 | 296.2 | 0.0008247 | 2.784 | +1 | 3 |
| - | - | 7046 | 296.7 | - | - | 0 | - |
| 9 | y | 1.703E+04 | 297.1 | 0.001379 | 4.64 | +2 | 5 |
| - | - | 1.438E+05 | 297.2 | - | - | 0 | - |
| - | - | 7062 | 297.2 | - | - | 0 | - |
| - | - | 3709 | 297.6 | - | - | 0 | - |
| - | - | 3868 | 297.7 | - | - | 0 | - |
| 7 | y | 1.296E+04 | 298.1 | 0.004665 | 15.65 | +3 | 7 |
| - | - | 2.286E+04 | 298.2 | - | - | 0 | - |
| - | - | 5.868E+04 | 299.1 | - | - | 0 | - |
| - | - | 3808 | 300.1 | - | - | 0 | - |
| - | - | 6.804E+04 | 300.2 | - | - | 0 | - |
| - | - | 5.968E+05 | 301.2 | - | - | 0 | - |
| - | - | 7.654E+04 | 302.1 | - | - | 0 | - |
| - | - | 8.437E+04 | 302.2 | - | - | 0 | - |
| - | - | 1.55E+04 | 303.1 | - | - | 0 | - |
| - | - | 5929 | 303.2 | - | - | 0 | - |
| - | - | 1.284E+04 | 305.2 | - | - | 0 | - |
| - | - | 1.075E+04 | 305.6 | - | - | 0 | - |
| 9 | y | 1.507E+05 | 306.1 | 0.0008875 | 2.899 | +2 | 5 |
| - | - | 9143 | 306.2 | - | - | 0 | - |
| - | - | 5.654E+04 | 306.6 | - | - | 0 | - |
| - | - | 9352 | 307.1 | - | - | 0 | - |
| - | - | 9657 | 309.2 | - | - | 0 | - |
| 12 | y | 6.927E+04 | 311.1 | 0.0009626 | 3.094 | +1 | 2 |
| - | - | 5846 | 311.2 | - | - | 0 | - |
| - | - | 1.218E+04 | 312.1 | - | - | 0 | - |
| - | - | 2.23E+04 | 314.2 | - | - | 0 | - |
| 3 | b | 4.755E+04 | 314.2 | 0.0008802 | 2.801 | +1 | 3 |
| - | - | 2.67E+05 | 315.2 | - | - | 0 | - |
| - | - | 6118 | 316.1 | - | - | 0 | - |
| - | - | 3.611E+04 | 316.2 | - | - | 0 | - |
| - | - | 5346 | 317.7 | - | - | 0 | - |
| - | - | 5421 | 318.1 | - | - | 0 | - |
| - | - | 1.386E+04 | 319.2 | - | - | 0 | - |
| - | - | 4570 | 320.2 | - | - | 0 | - |
| - | - | 4681 | 324.2 | - | - | 0 | - |
| - | - | 7569 | 324.7 | - | - | 0 | - |
| - | - | 4296 | 325.2 | - | - | 0 | - |
| - | - | 6.554E+04 | 326.7 | - | - | 0 | - |
| - | - | 3.284E+04 | 327.2 | - | - | 0 | - |
| - | - | 3476 | 327.7 | - | - | 0 | - |
| - | - | 5283 | 329.2 | - | - | 0 | - |
| - | - | 4459 | 331.7 | - | - | 0 | - |
| - | - | 3999 | 332.2 | - | - | 0 | - |
| - | - | 2.72E+04 | 332.2 | - | - | 0 | - |
| - | - | 1.121E+05 | 333.2 | - | - | 0 | - |
| - | - | 6056 | 333.2 | - | - | 0 | - |
| - | - | 2.239E+04 | 334.2 | - | - | 0 | - |
| - | - | 3902 | 335.2 | - | - | 0 | - |
| - | - | 3798 | 337.2 | - | - | 0 | - |
| - | - | 7753 | 338.2 | - | - | 0 | - |
| 6 | b | 1.104E+04 | 338.7 | 0.000974 | 2.876 | +2 | 6 |
| 6 | b | 1.186E+04 | 339.2 | 0.006555 | 19.33 | +2 | 6 |
| - | - | 9774 | 339.7 | - | - | 0 | - |
| - | - | 4158 | 340.2 | - | - | 0 | - |
| - | - | 1.45E+04 | 340.7 | - | - | 0 | - |
| - | - | 1.071E+04 | 341.2 | - | - | 0 | - |
| - | - | 3.257E+04 | 342.2 | - | - | 0 | - |
| - | - | 5213 | 343.2 | - | - | 0 | - |
| - | - | 2.857E+04 | 344.2 | - | - | 0 | - |
| - | - | 1.028E+04 | 345 | - | - | 0 | - |
| - | - | 4.199E+04 | 345.1 | - | - | 0 | - |
| - | - | 6058 | 345.2 | - | - | 0 | - |
| - | - | 1.754E+05 | 346.1 | - | - | 0 | - |
| - | - | 8822 | 346.7 | - | - | 0 | - |
| - | - | 2.854E+04 | 347.1 | - | - | 0 | - |
| 6 | b | 6.694E+04 | 347.7 | 0.0009101 | 2.618 | +2 | 6 |
| - | - | 3.089E+04 | 348.2 | - | - | 0 | - |
| - | - | 8051 | 349.2 | - | - | 0 | - |
| - | - | 1.189E+05 | 350.2 | - | - | 0 | - |
| - | - | 2.35E+04 | 351.2 | - | - | 0 | - |
| - | - | 5271 | 352.2 | - | - | 0 | - |
| - | - | 1.244E+04 | 352.7 | - | - | 0 | - |
| 8 | y | 2.213E+04 | 353.7 | 0.000698 | 1.973 | +2 | 6 |
| - | - | 8153 | 354.2 | - | - | 0 | - |
| - | - | 5792 | 356.2 | - | - | 0 | - |
| - | - | 9793 | 357.2 | - | - | 0 | - |
| - | - | 5797 | 358.2 | - | - | 0 | - |
| - | - | 7443 | 359 | - | - | 0 | - |
| - | - | 8742 | 360.2 | - | - | 0 | - |
| - | - | 3.53E+04 | 360.7 | - | - | 0 | - |
| - | - | 4.179E+04 | 361.2 | - | - | 0 | - |
| - | - | 5268 | 361.7 | - | - | 0 | - |
| - | - | 3.248E+04 | 362.2 | - | - | 0 | - |
| 8 | y | 3.168E+05 | 362.7 | 0.0008477 | 2.337 | +2 | 6 |
| - | - | 2.305E+05 | 363.1 | - | - | 0 | - |
| - | - | 1.273E+05 | 363.2 | - | - | 0 | - |
| - | - | 2.824E+04 | 363.7 | - | - | 0 | - |
| - | - | 3.79E+04 | 364.1 | - | - | 0 | - |
| - | - | 3.037E+04 | 364.1 | - | - | 0 | - |
| 9 | b | 4278 | 364.2 | 0.001383 | 3.799 | +3 | 9 |
| - | - | 7758 | 365.1 | - | - | 0 | - |
| - | - | 6609 | 366.2 | - | - | 0 | - |
| - | - | 2.086E+04 | 366.2 | - | - | 0 | - |
| - | - | 1.654E+04 | 367.1 | - | - | 0 | - |
| - | - | 1.749E+04 | 367.2 | - | - | 0 | - |
| - | - | 2.046E+04 | 367.2 | - | - | 0 | - |
| - | - | 7000 | 368.1 | - | - | 0 | - |
| - | - | 1.138E+04 | 368.2 | - | - | 0 | - |
| - | - | 1.451E+04 | 373.2 | - | - | 0 | - |
| - | - | 1.332E+04 | 376.2 | - | - | 0 | - |
| - | - | 2.733E+04 | 376.2 | - | - | 0 | - |
| - | - | 1.433E+04 | 376.7 | - | - | 0 | - |
| - | - | 1.037E+04 | 377.1 | - | - | 0 | - |
| - | - | 4715 | 377.2 | - | - | 0 | - |
| - | - | 7.325E+04 | 378.2 | - | - | 0 | - |
| - | - | 3.474E+05 | 379.2 | - | - | 0 | - |
| - | - | 8910 | 380.2 | - | - | 0 | - |
| - | - | 5.947E+04 | 380.2 | - | - | 0 | - |
| - | - | 6.463E+05 | 381.2 | - | - | 0 | - |
| - | - | 2.579E+04 | 381.7 | - | - | 0 | - |
| - | - | 1.198E+05 | 382.2 | - | - | 0 | - |
| - | - | 2.644E+04 | 382.2 | - | - | 0 | - |
| - | - | 4914 | 382.2 | - | - | 0 | - |
| - | - | 4399 | 382.7 | - | - | 0 | - |
| - | - | 1.368E+04 | 383.2 | - | - | 0 | - |
| - | - | 4638 | 383.2 | - | - | 0 | - |
| - | - | 2.651E+04 | 384.2 | - | - | 0 | - |
| - | - | 5.137E+04 | 384.2 | - | - | 0 | - |
| - | - | 4.084E+04 | 385.2 | - | - | 0 | - |
| 5 | y | 3.082E+04 | 385.2 | 0.003735 | 9.696 | +3 | 9 |
| - | - | 5558 | 385.2 | - | - | 0 | - |
| - | - | 9071 | 386.2 | - | - | 0 | - |
| - | - | 1.183E+04 | 386.2 | - | - | 0 | - |
| - | - | 4847 | 387.2 | - | - | 0 | - |
| 10 | b | 8859 | 389.2 | 0.0009611 | 2.47 | +3 | 10 |
| - | - | 3.153E+04 | 390.2 | - | - | 0 | - |
| - | - | 1.394E+04 | 390.7 | - | - | 0 | - |
| - | - | 2.479E+04 | 391.1 | - | - | 0 | - |
| - | - | 6640 | 391.2 | - | - | 0 | - |
| - | - | 5902 | 391.7 | - | - | 0 | - |
| - | - | 5014 | 392.1 | - | - | 0 | - |
| - | - | 5250 | 392.1 | - | - | 0 | - |
| - | - | 9869 | 392.2 | - | - | 0 | - |
| - | - | 5901 | 393.2 | - | - | 0 | - |
| - | - | 9.464E+04 | 394.2 | - | - | 0 | - |
| - | - | 3.198E+04 | 395.1 | - | - | 0 | - |
| - | - | 3.174E+04 | 395.2 | - | - | 0 | - |
| - | - | 7975 | 395.2 | - | - | 0 | - |
| - | - | 5941 | 395.2 | - | - | 0 | - |
| - | - | 7459 | 395.2 | - | - | 0 | - |
| - | - | 5558 | 396.1 | - | - | 0 | - |
| - | - | 8313 | 396.2 | - | - | 0 | - |
| - | - | 7.767E+04 | 396.2 | - | - | 0 | - |
| - | - | 1.321E+04 | 397.2 | - | - | 0 | - |
| - | - | 1.865E+04 | 397.2 | - | - | 0 | - |
| - | - | 5700 | 397.7 | - | - | 0 | - |
| - | - | 5.475E+04 | 398.2 | - | - | 0 | - |
| - | - | 5760 | 399.2 | - | - | 0 | - |
| - | - | 2.83E+04 | 402.2 | - | - | 0 | - |
| - | - | 7.84E+04 | 402.2 | - | - | 0 | - |
| - | - | 5093 | 403.1 | - | - | 0 | - |
| - | - | 3.682E+04 | 403.2 | - | - | 0 | - |
| - | - | 6723 | 404.2 | - | - | 0 | - |
| - | - | 1.39E+04 | 404.7 | - | - | 0 | - |
| - | - | 1.823E+04 | 405.3 | - | - | 0 | - |
| - | - | 1.346E+04 | 406.2 | - | - | 0 | - |
| - | - | 7173 | 407.2 | - | - | 0 | - |
| - | - | 5624 | 407.2 | - | - | 0 | - |
| - | - | 7147 | 408.7 | - | - | 0 | - |
| - | - | 5.043E+04 | 409.1 | - | - | 0 | - |
| - | - | 5574 | 409.2 | - | - | 0 | - |
| - | - | 1.112E+04 | 409.2 | - | - | 0 | - |
| - | - | 8621 | 410.1 | - | - | 0 | - |
| - | - | 4439 | 410.2 | - | - | 0 | - |
| - | - | 1.459E+04 | 411.2 | - | - | 0 | - |
| - | - | 4.471E+05 | 412.2 | - | - | 0 | - |
| - | - | 2.4E+04 | 413.1 | - | - | 0 | - |
| - | - | 8.179E+04 | 413.2 | - | - | 0 | - |
| - | - | 4.493E+04 | 413.2 | - | - | 0 | - |
| - | - | 6944 | 413.2 | - | - | 0 | - |
| - | - | 1.602E+04 | 414.2 | - | - | 0 | - |
| - | - | 7179 | 414.2 | - | - | 0 | - |
| - | - | 1.637E+05 | 414.2 | - | - | 0 | - |
| - | - | 1.914E+04 | 415 | - | - | 0 | - |
| - | - | 2.164E+04 | 415.2 | - | - | 0 | - |
| - | - | 3.188E+04 | 415.2 | - | - | 0 | - |
| - | - | 4983 | 416.2 | - | - | 0 | - |
| - | - | 5.636E+04 | 417.7 | - | - | 0 | - |
| - | - | 1.183E+05 | 418.2 | - | - | 0 | - |
| - | - | 6.808E+04 | 418.7 | - | - | 0 | - |
| - | - | 1.877E+04 | 419.2 | - | - | 0 | - |
| - | - | 4821 | 420.2 | - | - | 0 | - |
| 11 | y | 1.804E+04 | 421.2 | 0.00122 | 2.896 | +1 | 3 |
| 11 | y | 1.964E+04 | 422.2 | 0.001488 | 3.524 | +1 | 3 |
| - | - | 9319 | 422.7 | - | - | 0 | - |
| - | - | 1.825E+04 | 423.2 | - | - | 0 | - |
| - | - | 1.7E+05 | 423.3 | - | - | 0 | - |
| - | - | 7604 | 423.7 | - | - | 0 | - |
| - | - | 3813 | 423.7 | - | - | 0 | - |
| - | - | 5719 | 424.2 | - | - | 0 | - |
| - | - | 3.614E+04 | 424.3 | - | - | 0 | - |
| - | - | 1.456E+04 | 424.7 | - | - | 0 | - |
| - | - | 9218 | 425.2 | - | - | 0 | - |
| - | - | 5892 | 426.2 | - | - | 0 | - |
| - | - | 1.06E+06 | 426.7 | - | - | 0 | - |
| - | - | 5.4E+05 | 427.2 | - | - | 0 | - |
| - | - | 1.639E+05 | 427.7 | - | - | 0 | - |
| - | - | 3.443E+04 | 428.2 | - | - | 0 | - |
| - | - | 9.216E+04 | 429.2 | - | - | 0 | - |
| - | - | 1.212E+06 | 430.2 | - | - | 0 | - |
| - | - | 2.618E+05 | 431.2 | - | - | 0 | - |
| 7 | b | 1.293E+05 | 431.7 | 0.001082 | 2.506 | +2 | 7 |
| 7 | b | 5.966E+04 | 432.2 | 0.007304 | 16.9 | +2 | 7 |
| - | - | 2.726E+04 | 432.7 | - | - | 0 | - |
| - | - | 2.236E+04 | 433.2 | - | - | 0 | - |
| 4 | b | 4.553E+04 | 433.3 | 0.0003846 | 0.8876 | +1 | 4 |
| - | - | 6029 | 433.7 | - | - | 0 | - |
| - | - | 5547 | 434.3 | - | - | 0 | - |
| - | - | 1.554E+04 | 438.2 | - | - | 0 | - |
| - | - | 6921 | 438.7 | - | - | 0 | - |
| 11 | y | 2.886E+05 | 439.2 | 0.001459 | 3.321 | +1 | 3 |
| - | - | 9037 | 440.2 | - | - | 0 | - |
| - | - | 6.68E+04 | 440.2 | - | - | 0 | - |
| 7 | b | 5.104E+05 | 440.7 | 0.001262 | 2.864 | +2 | 7 |
| - | - | 2.809E+05 | 441.2 | - | - | 0 | - |
| - | - | 7.812E+04 | 441.7 | - | - | 0 | - |
| - | - | 2.522E+04 | 442.2 | - | - | 0 | - |
| - | - | 5033 | 443.2 | - | - | 0 | - |
| - | - | 5546 | 446.7 | - | - | 0 | - |
| 7 | y | 5.11E+04 | 447.2 | 0.001108 | 2.477 | +2 | 7 |
| - | - | 6759 | 447.2 | - | - | 0 | - |
| - | - | 2.808E+04 | 447.7 | - | - | 0 | - |
| - | - | 6.466E+04 | 448.2 | - | - | 0 | - |
| - | - | 1.691E+04 | 449.2 | - | - | 0 | - |
| - | - | 1.026E+04 | 450.2 | - | - | 0 | - |
| - | - | 1.046E+04 | 450.2 | - | - | 0 | - |
| - | - | 1.218E+04 | 450.3 | - | - | 0 | - |
| 4 | b | 3.107E+05 | 451.3 | 0.001386 | 3.071 | +1 | 4 |
| - | - | 7.656E+04 | 452.3 | - | - | 0 | - |
| - | - | 1.186E+04 | 453.3 | - | - | 0 | - |
| - | - | 2.494E+04 | 455.3 | - | - | 0 | - |
| 7 | y | 2.516E+04 | 455.7 | 0.001291 | 2.834 | +2 | 7 |
| - | - | 1.299E+04 | 456.2 | - | - | 0 | - |
| - | - | 4973 | 456.3 | - | - | 0 | - |
| - | - | 2.973E+04 | 459.2 | - | - | 0 | - |
| - | - | 2.313E+04 | 460.8 | - | - | 0 | - |
| - | - | 2.1E+04 | 461.2 | - | - | 0 | - |
| - | - | 4.145E+04 | 461.3 | - | - | 0 | - |
| - | - | 9031 | 461.8 | - | - | 0 | - |
| - | - | 6097 | 462.2 | - | - | 0 | - |
| - | - | 5711 | 462.3 | - | - | 0 | - |
| 3 | y | 5730 | 462.6 | 0.001516 | 3.277 | +3 | 11 |
| 3 | y | 6026 | 462.9 | 0.005359 | 11.58 | +3 | 11 |
| - | - | 6197 | 464.2 | - | - | 0 | - |
| - | - | 5278 | 465.3 | - | - | 0 | - |
| - | - | 1.132E+04 | 465.7 | - | - | 0 | - |
| - | - | 6302 | 466.2 | - | - | 0 | - |
| - | - | 1.053E+04 | 466.2 | - | - | 0 | - |
| - | - | 8.088E+04 | 468.3 | - | - | 0 | - |
| - | - | 2.618E+04 | 469.3 | - | - | 0 | - |
| - | - | 5214 | 472.2 | - | - | 0 | - |
| - | - | 6.163E+04 | 474.3 | - | - | 0 | - |
| - | - | 6.702E+04 | 474.8 | - | - | 0 | - |
| - | - | 4.247E+04 | 475.3 | - | - | 0 | - |
| - | - | 1.002E+04 | 475.8 | - | - | 0 | - |
| - | - | 2.455E+04 | 476.2 | - | - | 0 | - |
| - | - | 4.778E+04 | 477.2 | - | - | 0 | - |
| 10 | y | 4.355E+04 | 478.2 | 0.0005082 | 1.063 | +1 | 4 |
| - | - | 4.087E+04 | 478.3 | - | - | 0 | - |
| - | - | 1.296E+04 | 479.2 | - | - | 0 | - |
| - | - | 7073 | 479.3 | - | - | 0 | - |
| - | - | 6944 | 480.2 | - | - | 0 | - |
| - | - | 1.7E+05 | 483.3 | - | - | 0 | - |
| - | - | 1.197E+05 | 483.8 | - | - | 0 | - |
| - | - | 1.126E+04 | 484.2 | - | - | 0 | - |
| - | - | 4.401E+04 | 484.3 | - | - | 0 | - |
| - | - | 6125 | 484.8 | - | - | 0 | - |
| - | - | 5796 | 485.2 | - | - | 0 | - |
| - | - | 4665 | 488.2 | - | - | 0 | - |
| 8 | b | 5.849E+04 | 488.3 | 0.002141 | 4.384 | +2 | 8 |
| 8 | b | 3.881E+04 | 488.7 | 0.007692 | 15.74 | +2 | 8 |
| - | - | 1.799E+04 | 489.3 | - | - | 0 | - |
| - | - | 4651 | 493.3 | - | - | 0 | - |
| - | - | 6.182E+05 | 494.2 | - | - | 0 | - |
| - | - | 4384 | 495.2 | - | - | 0 | - |
| - | - | 1.714E+05 | 495.2 | - | - | 0 | - |
| 10 | y | 6.514E+05 | 496.2 | 0.001296 | 2.612 | +1 | 4 |
| - | - | 1.515E+05 | 497.2 | - | - | 0 | - |
| - | - | 7.247E+04 | 497.8 | - | - | 0 | - |
| - | - | 3.084E+04 | 498.2 | - | - | 0 | - |
| - | - | 6360 | 502.2 | - | - | 0 | - |
| - | - | 1.973E+04 | 503.1 | - | - | 0 | - |
| - | - | 2.9E+04 | 504.2 | - | - | 0 | - |
| - | - | 9274 | 505.2 | - | - | 0 | - |
| - | - | 8682 | 506.2 | - | - | 0 | - |
| - | - | 8559 | 506.3 | - | - | 0 | - |
| - | - | 2.545E+04 | 507.3 | - | - | 0 | - |
| - | - | 7893 | 508.3 | - | - | 0 | - |
| - | - | 6.212E+04 | 511.3 | - | - | 0 | - |
| - | - | 1.727E+04 | 512.3 | - | - | 0 | - |
| - | - | 1.333E+04 | 513.2 | - | - | 0 | - |
| - | - | 7248 | 514.2 | - | - | 0 | - |
| - | - | 1.494E+04 | 515.3 | - | - | 0 | - |
| - | - | 4606 | 515.8 | - | - | 0 | - |
| - | - | 1.016E+04 | 516.3 | - | - | 0 | - |
| - | - | 5479 | 519.1 | - | - | 0 | - |
| - | - | 6559 | 520.3 | - | - | 0 | - |
| - | - | 7959 | 521.2 | - | - | 0 | - |
| - | - | 4570 | 521.3 | - | - | 0 | - |
| - | - | 6537 | 522.2 | - | - | 0 | - |
| - | - | 2.009E+04 | 523.3 | - | - | 0 | - |
| - | - | 1.194E+05 | 525.3 | - | - | 0 | - |
| - | - | 9018 | 526.2 | - | - | 0 | - |
| - | - | 3.541E+04 | 526.3 | - | - | 0 | - |
| 0 | Precursor | 3.211E+04 | 529.3 | 0.00184 | 3.477 | +3 | -1 |
| 0 | Precursor | 3.417E+04 | 529.6 | 0.007575 | 14.3 | +3 | -1 |
| - | - | 2.108E+04 | 529.9 | - | - | 0 | - |
| - | - | 9754 | 531.2 | - | - | 0 | - |
| - | - | 3.164E+04 | 532.2 | - | - | 0 | - |
| - | - | 6841 | 533.2 | - | - | 0 | - |
| - | - | 4840 | 533.3 | - | - | 0 | - |
| - | - | 5911 | 534.3 | - | - | 0 | - |
| 0 | Precursor | 3.36E+04 | 535.3 | 0.00076 | 1.42 | +3 | -1 |
| - | - | 3.867E+04 | 535.6 | - | - | 0 | - |
| - | - | 2.587E+04 | 535.9 | - | - | 0 | - |
| - | - | 5126 | 536.2 | - | - | 0 | - |
| - | - | 1.05E+04 | 536.3 | - | - | 0 | - |
| - | - | 6224 | 537.2 | - | - | 0 | - |
| - | - | 7494 | 537.7 | - | - | 0 | - |
| - | - | 5202 | 538.3 | - | - | 0 | - |
| - | - | 1.231E+05 | 539.2 | - | - | 0 | - |
| - | - | 4563 | 539.3 | - | - | 0 | - |
| - | - | 3.108E+04 | 540.2 | - | - | 0 | - |
| - | - | 1.306E+04 | 541.2 | - | - | 0 | - |
| - | - | 1.553E+04 | 542.3 | - | - | 0 | - |
| - | - | 2.388E+05 | 543.3 | - | - | 0 | - |
| - | - | 7.318E+04 | 544.3 | - | - | 0 | - |
| - | - | 1.261E+04 | 545.3 | - | - | 0 | - |
| - | - | 2.808E+04 | 546.2 | - | - | 0 | - |
| - | - | 1.174E+04 | 546.8 | - | - | 0 | - |
| - | - | 9485 | 547.2 | - | - | 0 | - |
| - | - | 8798 | 548.2 | - | - | 0 | - |
| - | - | 6919 | 548.3 | - | - | 0 | - |
| - | - | 3.177E+04 | 549.2 | - | - | 0 | - |
| - | - | 1.101E+04 | 550.2 | - | - | 0 | - |
| - | - | 6370 | 551.2 | - | - | 0 | - |
| - | - | 2.981E+04 | 551.3 | - | - | 0 | - |
| - | - | 1.942E+04 | 551.3 | - | - | 0 | - |
| - | - | 1.228E+04 | 552.3 | - | - | 0 | - |
| - | - | 6654 | 552.3 | - | - | 0 | - |
| - | - | 5167 | 553.3 | - | - | 0 | - |
| 9 | b | 1.2E+04 | 554.8 | 0.009228 | 16.63 | +2 | 9 |
| - | - | 3.516E+04 | 555.3 | - | - | 0 | - |
| - | - | 3.469E+04 | 555.8 | - | - | 0 | - |
| - | - | 1.165E+04 | 556.3 | - | - | 0 | - |
| - | - | 4897 | 557.2 | - | - | 0 | - |
| - | - | 5607 | 558.3 | - | - | 0 | - |
| - | - | 4667 | 559.7 | - | - | 0 | - |
| - | - | 8801 | 561.3 | - | - | 0 | - |
| 5 | b | 2.292E+04 | 561.3 | 0.00211 | 3.759 | +1 | 5 |
| 5 | b | 1.116E+04 | 562.3 | 0.004178 | 7.431 | +1 | 5 |
| - | - | 2.067E+04 | 563.3 | - | - | 0 | - |
| - | - | 7850 | 564.3 | - | - | 0 | - |
| - | - | 7598 | 564.3 | - | - | 0 | - |
| - | - | 7.607E+04 | 565.3 | - | - | 0 | - |
| - | - | 2.91E+04 | 566.3 | - | - | 0 | - |
| - | - | 1.333E+05 | 567.2 | - | - | 0 | - |
| - | - | 4.76E+04 | 568.2 | - | - | 0 | - |
| 5 | y | 8892 | 568.7 | 0.01114 | 19.59 | +2 | 9 |
| - | - | 1.023E+04 | 569.2 | - | - | 0 | - |
| - | - | 3.489E+04 | 575.2 | - | - | 0 | - |
| - | - | 2.084E+04 | 576.2 | - | - | 0 | - |
| - | - | 7342 | 576.3 | - | - | 0 | - |
| - | - | 8864 | 577.2 | - | - | 0 | - |
| 5 | y | 7539 | 577.3 | 0.007695 | 13.33 | +2 | 9 |
| 5 | b | 1.978E+05 | 579.3 | 0.001128 | 1.946 | +1 | 5 |
| - | - | 6.616E+04 | 580.3 | - | - | 0 | - |
| - | - | 5139 | 581.3 | - | - | 0 | - |
| - | - | 9198 | 581.3 | - | - | 0 | - |
| - | - | 7358 | 582.3 | - | - | 0 | - |
| - | - | 5647 | 586.3 | - | - | 0 | - |
| - | - | 1.161E+04 | 588.8 | - | - | 0 | - |
| - | - | 1.612E+04 | 589.3 | - | - | 0 | - |
| - | - | 1.044E+04 | 592.3 | - | - | 0 | - |
| - | - | 2.886E+04 | 592.3 | - | - | 0 | - |
| 9 | y | 2.605E+05 | 593.3 | 0.001062 | 1.789 | +1 | 5 |
| - | - | 8.048E+04 | 593.3 | - | - | 0 | - |
| - | - | 9211 | 593.8 | - | - | 0 | - |
| - | - | 9.597E+04 | 594.3 | - | - | 0 | - |
| - | - | 2.201E+04 | 594.3 | - | - | 0 | - |
| - | - | 6593 | 594.8 | - | - | 0 | - |
| - | - | 3.35E+04 | 595.3 | - | - | 0 | - |
| - | - | 4601 | 596.3 | - | - | 0 | - |
| - | - | 8583 | 596.4 | - | - | 0 | - |
| - | - | 7204 | 597.3 | - | - | 0 | - |
| - | - | 8272 | 597.8 | - | - | 0 | - |
| - | - | 1.002E+04 | 598.3 | - | - | 0 | - |
| - | - | 1.113E+04 | 600.3 | - | - | 0 | - |
| - | - | 5597 | 602.3 | - | - | 0 | - |
| - | - | 2.782E+04 | 602.8 | - | - | 0 | - |
| - | - | 3.203E+04 | 603.3 | - | - | 0 | - |
| - | - | 1.108E+04 | 603.8 | - | - | 0 | - |
| - | - | 9263 | 604.3 | - | - | 0 | - |
| - | - | 2.136E+05 | 610.3 | - | - | 0 | - |
| - | - | 2.564E+04 | 610.3 | - | - | 0 | - |
| 9 | y | 3.255E+06 | 611.3 | 0.001788 | 2.926 | +1 | 5 |
| - | - | 1.195E+05 | 611.8 | - | - | 0 | - |
| - | - | 1.014E+06 | 612.3 | - | - | 0 | - |
| - | - | 3.771E+04 | 612.8 | - | - | 0 | - |
| - | - | 2.316E+05 | 613.3 | - | - | 0 | - |
| - | - | 2.869E+04 | 614.3 | - | - | 0 | - |
| - | - | 1.272E+04 | 617.3 | - | - | 0 | - |
| - | - | 5638 | 618.3 | - | - | 0 | - |
| - | - | 1.243E+04 | 620.8 | - | - | 0 | - |
| - | - | 1.8E+04 | 621.3 | - | - | 0 | - |
| - | - | 1.721E+04 | 621.3 | - | - | 0 | - |
| - | - | 9273 | 622.3 | - | - | 0 | - |
| - | - | 1.023E+04 | 622.4 | - | - | 0 | - |
| - | - | 1.334E+04 | 628.3 | - | - | 0 | - |
| - | - | 5625 | 631.8 | - | - | 0 | - |
| - | - | 3E+04 | 634.3 | - | - | 0 | - |
| - | - | 3.969E+04 | 635.3 | - | - | 0 | - |
| - | - | 2.815E+04 | 636.3 | - | - | 0 | - |
| 4 | y | 2.348E+04 | 636.8 | 0.001588 | 2.493 | +2 | 10 |
| 4 | y | 1.917E+04 | 637.3 | 0.01068 | 16.76 | +2 | 10 |
| - | - | 9903 | 637.3 | - | - | 0 | - |
| - | - | 1.081E+04 | 637.8 | - | - | 0 | - |
| 11 | b | 7271 | 638.3 | 0.006506 | 10.19 | +2 | 11 |
| 11 | b | 4516 | 638.8 | 0.008577 | 13.43 | +2 | 11 |
| - | - | 1.708E+04 | 640.3 | - | - | 0 | - |
| - | - | 4821 | 640.3 | - | - | 0 | - |
| - | - | 2.014E+04 | 645.3 | - | - | 0 | - |
| 4 | y | 1.597E+05 | 645.8 | 0.002043 | 3.163 | +2 | 10 |
| - | - | 1.162E+05 | 646.3 | - | - | 0 | - |
| - | - | 4.332E+04 | 646.8 | - | - | 0 | - |
| - | - | 1.319E+04 | 647.3 | - | - | 0 | - |
| - | - | 6265 | 648.3 | - | - | 0 | - |
| - | - | 6407 | 649.3 | - | - | 0 | - |
| - | - | 1.226E+04 | 650.3 | - | - | 0 | - |
| - | - | 1.385E+04 | 650.3 | - | - | 0 | - |
| - | - | 7988 | 650.8 | - | - | 0 | - |
| - | - | 5580 | 651.3 | - | - | 0 | - |
| - | - | 2.692E+05 | 652.3 | - | - | 0 | - |
| - | - | 1.072E+05 | 653.3 | - | - | 0 | - |
| - | - | 2.239E+04 | 654.3 | - | - | 0 | - |
| - | - | 8581 | 658.3 | - | - | 0 | - |
| - | - | 1.166E+04 | 658.3 | - | - | 0 | - |
| - | - | 1.29E+04 | 659.3 | - | - | 0 | - |
| - | - | 1.303E+04 | 661.3 | - | - | 0 | - |
| - | - | 4614 | 661.8 | - | - | 0 | - |
| - | - | 8.75E+04 | 662.3 | - | - | 0 | - |
| - | - | 5.7E+04 | 663.3 | - | - | 0 | - |
| - | - | 1.668E+04 | 664.3 | - | - | 0 | - |
| - | - | 1.215E+04 | 664.3 | - | - | 0 | - |
| - | - | 6023 | 665.3 | - | - | 0 | - |
| - | - | 7060 | 666.4 | - | - | 0 | - |
| - | - | 9153 | 668.3 | - | - | 0 | - |
| - | - | 4337 | 670.3 | - | - | 0 | - |
| - | - | 5397 | 671.3 | - | - | 0 | - |
| 6 | b | 1.369E+05 | 676.3 | 0.001412 | 2.087 | +1 | 6 |
| 6 | b | 8.814E+04 | 677.3 | 0.008546 | 12.62 | +1 | 6 |
| - | - | 3.773E+04 | 678.3 | - | - | 0 | - |
| - | - | 2.348E+04 | 679.3 | - | - | 0 | - |
| - | - | 7287 | 679.8 | - | - | 0 | - |
| - | - | 4.935E+05 | 680.3 | - | - | 0 | - |
| - | - | 6669 | 680.9 | - | - | 0 | - |
| - | - | 1.956E+05 | 681.3 | - | - | 0 | - |
| - | - | 5E+04 | 682.3 | - | - | 0 | - |
| - | - | 2.097E+04 | 684.3 | - | - | 0 | - |
| - | - | 3.245E+04 | 684.8 | - | - | 0 | - |
| - | - | 2.881E+04 | 685.3 | - | - | 0 | - |
| - | - | 1.211E+04 | 685.8 | - | - | 0 | - |
| - | - | 8162 | 686.3 | - | - | 0 | - |
| - | - | 4515 | 687.3 | - | - | 0 | - |
| - | - | 1.512E+04 | 688.3 | - | - | 0 | - |
| - | - | 7781 | 688.9 | - | - | 0 | - |
| - | - | 5830 | 689.3 | - | - | 0 | - |
| - | - | 1.102E+04 | 692.8 | - | - | 0 | - |
| 3 | y | 1.15E+05 | 693.3 | 0.002463 | 3.553 | +2 | 11 |
| 3 | y | 1.28E+05 | 693.8 | 0.008136 | 11.73 | +2 | 11 |
| 6 | b | 1.37E+06 | 694.4 | 0.000979 | 1.41 | +1 | 6 |
| - | - | 1.877E+04 | 694.8 | - | - | 0 | - |
| - | - | 6766 | 694.9 | - | - | 0 | - |
| - | - | 5.224E+05 | 695.4 | - | - | 0 | - |
| - | - | 1.162E+05 | 696.4 | - | - | 0 | - |
| - | - | 2.579E+04 | 697.4 | - | - | 0 | - |
| - | - | 4993 | 698.4 | - | - | 0 | - |
| - | - | 1.985E+04 | 701.3 | - | - | 0 | - |
| - | - | 8.991E+04 | 701.8 | - | - | 0 | - |
| 3 | y | 8.187E+05 | 702.3 | 0.002491 | 3.547 | +2 | 11 |
| 12 | b | 6.744E+05 | 702.8 | 0.01217 | 17.31 | +2 | 12 |
| 12 | b | 3.346E+05 | 703.3 | 0.002709 | 3.852 | +2 | 12 |
| - | - | 1.026E+05 | 703.8 | - | - | 0 | - |
| - | - | 3.934E+04 | 704.3 | - | - | 0 | - |
| - | - | 1.079E+04 | 705.3 | - | - | 0 | - |
| 8 | y | 1.017E+05 | 706.3 | 0.001714 | 2.427 | +1 | 6 |
| - | - | 4.081E+04 | 707.3 | - | - | 0 | - |
| - | - | 5435 | 707.8 | - | - | 0 | - |
| - | - | 1.016E+04 | 708.3 | - | - | 0 | - |
| - | - | 6844 | 709.3 | - | - | 0 | - |
| - | - | 4508 | 710.4 | - | - | 0 | - |
| - | - | 4.466E+04 | 711.4 | - | - | 0 | - |
| 12 | b | 5.89E+04 | 711.9 | 0.002754 | 3.869 | +2 | 12 |
| - | - | 4.95E+04 | 712.4 | - | - | 0 | - |
| - | - | 1.366E+04 | 712.9 | - | - | 0 | - |
| - | - | 6424 | 713.4 | - | - | 0 | - |
| - | - | 6231 | 716.4 | - | - | 0 | - |
| - | - | 9620 | 720.4 | - | - | 0 | - |
| - | - | 9936 | 720.9 | - | - | 0 | - |
| - | - | 7.335E+04 | 721.4 | - | - | 0 | - |
| - | - | 2.002E+04 | 722.4 | - | - | 0 | - |
| - | - | 6.953E+04 | 723.4 | - | - | 0 | - |
| 8 | y | 1.096E+06 | 724.4 | 0.002014 | 2.78 | +1 | 6 |
| - | - | 4.427E+05 | 725.4 | - | - | 0 | - |
| - | - | 1.1E+05 | 726.4 | - | - | 0 | - |
| - | - | 1.421E+04 | 727.4 | - | - | 0 | - |
| - | - | 2.431E+04 | 729.4 | - | - | 0 | - |
| - | - | 1.316E+04 | 730.4 | - | - | 0 | - |
| - | - | 3.502E+04 | 733.4 | - | - | 0 | - |
| - | - | 3.144E+04 | 734.4 | - | - | 0 | - |
| - | - | 1.44E+04 | 735.4 | - | - | 0 | - |
| - | - | 5678 | 737.3 | - | - | 0 | - |
| - | - | 1.25E+04 | 737.9 | - | - | 0 | - |
| - | - | 8732 | 738.4 | - | - | 0 | - |
| - | - | 4714 | 738.9 | - | - | 0 | - |
| - | - | 1.616E+04 | 739.4 | - | - | 0 | - |
| - | - | 6344 | 739.9 | - | - | 0 | - |
| - | - | 7182 | 740.4 | - | - | 0 | - |
| 2 | y | 6191 | 742.9 | 0.002009 | 2.704 | +2 | 12 |
| - | - | 2.596E+04 | 743.4 | - | - | 0 | - |
| - | - | 6899 | 743.9 | - | - | 0 | - |
| - | - | 1.229E+04 | 744.4 | - | - | 0 | - |
| - | - | 5707 | 746.4 | - | - | 0 | - |
| - | - | 1.19E+04 | 747.4 | - | - | 0 | - |
| - | - | 2.573E+04 | 748.4 | - | - | 0 | - |
| - | - | 1.38E+04 | 749.3 | - | - | 0 | - |
| - | - | 1.352E+04 | 750.4 | - | - | 0 | - |
| - | - | 1.391E+05 | 751.4 | - | - | 0 | - |
| 2 | y | 5.9E+04 | 751.9 | 0.00222 | 2.952 | +2 | 12 |
| - | - | 1.218E+05 | 752.4 | - | - | 0 | - |
| - | - | 2.96E+04 | 752.9 | - | - | 0 | - |
| - | - | 6.973E+04 | 753.4 | - | - | 0 | - |
| - | - | 2.433E+04 | 753.9 | - | - | 0 | - |
| - | - | 1.934E+04 | 754.4 | - | - | 0 | - |
| - | - | 5650 | 754.9 | - | - | 0 | - |
| - | - | 5551 | 758.4 | - | - | 0 | - |
| - | - | 7199 | 760.3 | - | - | 0 | - |
| - | - | 1.167E+04 | 761.4 | - | - | 0 | - |
| - | - | 1.547E+04 | 762.4 | - | - | 0 | - |
| - | - | 8.274E+04 | 763.4 | - | - | 0 | - |
| - | - | 3.96E+04 | 764.4 | - | - | 0 | - |
| - | - | 3.864E+04 | 765.4 | - | - | 0 | - |
| - | - | 1.893E+04 | 766.4 | - | - | 0 | - |
| - | - | 3.464E+04 | 767.3 | - | - | 0 | - |
| - | - | 1.337E+04 | 768.4 | - | - | 0 | - |
| - | - | 7369 | 768.4 | - | - | 0 | - |
| - | - | 1.448E+04 | 769.4 | - | - | 0 | - |
| - | - | 9324 | 770.4 | - | - | 0 | - |
| - | - | 1.486E+04 | 775.4 | - | - | 0 | - |
| - | - | 1.832E+04 | 776.4 | - | - | 0 | - |
| - | - | 8730 | 777.4 | - | - | 0 | - |
| - | - | 8.377E+04 | 779.4 | - | - | 0 | - |
| - | - | 3.767E+04 | 780.4 | - | - | 0 | - |
| - | - | 4.275E+04 | 781.4 | - | - | 0 | - |
| - | - | 1.774E+04 | 782.4 | - | - | 0 | - |
| 0 | Precursor | 1.518E+05 | 793.4 | 0.01491 | 18.8 | +2 | -1 |
| - | - | 7.327E+04 | 794.4 | - | - | 0 | - |
| - | - | 1.936E+04 | 795.3 | - | - | 0 | - |
| - | - | 1.296E+04 | 795.4 | - | - | 0 | - |
| - | - | 9603 | 796.3 | - | - | 0 | - |
| - | - | 2.679E+04 | 807.4 | - | - | 0 | - |
| - | - | 1.814E+04 | 807.4 | - | - | 0 | - |
| - | - | 1.343E+04 | 808.4 | - | - | 0 | - |
| - | - | 6248 | 809.4 | - | - | 0 | - |
| - | - | 7553 | 817.4 | - | - | 0 | - |
| - | - | 2.109E+04 | 818.4 | - | - | 0 | - |
| - | - | 2.129E+04 | 819.4 | - | - | 0 | - |
| - | - | 7016 | 820.4 | - | - | 0 | - |
| - | - | 9236 | 825.4 | - | - | 0 | - |
| - | - | 1.191E+04 | 826.4 | - | - | 0 | - |
| - | - | 5732 | 827.4 | - | - | 0 | - |
| - | - | 6241 | 829.4 | - | - | 0 | - |
| - | - | 2.779E+04 | 834.4 | - | - | 0 | - |
| - | - | 4.531E+04 | 835.4 | - | - | 0 | - |
| - | - | 1.07E+05 | 836.5 | - | - | 0 | - |
| - | - | 4.205E+04 | 837.5 | - | - | 0 | - |
| - | - | 1.205E+04 | 838.5 | - | - | 0 | - |
| - | - | 9346 | 843.4 | - | - | 0 | - |
| - | - | 2.501E+04 | 844.4 | - | - | 0 | - |
| - | - | 2.379E+04 | 845.4 | - | - | 0 | - |
| - | - | 4.224E+04 | 846.4 | - | - | 0 | - |
| - | - | 2.873E+04 | 847.4 | - | - | 0 | - |
| - | - | 9748 | 848.4 | - | - | 0 | - |
| - | - | 3.299E+05 | 852.4 | - | - | 0 | - |
| - | - | 1.727E+05 | 853.4 | - | - | 0 | - |
| - | - | 5680 | 854.4 | - | - | 0 | - |
| - | - | 3.894E+04 | 854.4 | - | - | 0 | - |
| - | - | 9010 | 855.4 | - | - | 0 | - |
| 7 | b | 1.935E+05 | 862.4 | 0.001628 | 1.887 | +1 | 7 |
| 7 | b | 1.355E+05 | 863.4 | 0.01236 | 14.32 | +1 | 7 |
| - | - | 2.396E+05 | 864.5 | - | - | 0 | - |
| - | - | 1.152E+05 | 865.5 | - | - | 0 | - |
| - | - | 3.52E+04 | 866.5 | - | - | 0 | - |
| - | - | 6347 | 869.4 | - | - | 0 | - |
| - | - | 2.636E+04 | 876.4 | - | - | 0 | - |
| - | - | 1.402E+04 | 877.4 | - | - | 0 | - |
| - | - | 4723 | 878.4 | - | - | 0 | - |
| 7 | b | 1.576E+06 | 880.4 | 0.001439 | 1.634 | +1 | 7 |
| - | - | 8.066E+05 | 881.4 | - | - | 0 | - |
| - | - | 2.24E+05 | 882.4 | - | - | 0 | - |
| - | - | 5.384E+04 | 883.4 | - | - | 0 | - |
| - | - | 7716 | 884.4 | - | - | 0 | - |
| - | - | 1.05E+04 | 890.4 | - | - | 0 | - |
| - | - | 8225 | 891.4 | - | - | 0 | - |
| - | - | 3.027E+04 | 892.5 | - | - | 0 | - |
| - | - | 9509 | 893.5 | - | - | 0 | - |
| - | - | 1.381E+04 | 894.4 | - | - | 0 | - |
| - | - | 6510 | 895.5 | - | - | 0 | - |
| - | - | 5.855E+04 | 908.4 | - | - | 0 | - |
| - | - | 3.503E+04 | 909.4 | - | - | 0 | - |
| 7 | y | 9.638E+04 | 910.4 | 0.00223 | 2.449 | +1 | 7 |
| - | - | 4.71E+04 | 911.4 | - | - | 0 | - |
| - | - | 1.369E+04 | 912.4 | - | - | 0 | - |
| - | - | 8142 | 919.4 | - | - | 0 | - |
| - | - | 4636 | 936.4 | - | - | 0 | - |
| - | - | 1.693E+04 | 937.4 | - | - | 0 | - |
| - | - | 9438 | 938.4 | - | - | 0 | - |
| - | - | 1.065E+04 | 948.4 | - | - | 0 | - |
| - | - | 1.506E+04 | 954.4 | - | - | 0 | - |
| - | - | 2.952E+04 | 955.4 | - | - | 0 | - |
| - | - | 1.965E+04 | 956.4 | - | - | 0 | - |
| - | - | 1.04E+04 | 964.5 | - | - | 0 | - |
| - | - | 3.264E+04 | 965.4 | - | - | 0 | - |
| - | - | 2.143E+04 | 966.5 | - | - | 0 | - |
| - | - | 6318 | 971.5 | - | - | 0 | - |
| - | - | 1.687E+04 | 972.4 | - | - | 0 | - |
| - | - | 1.366E+04 | 973.4 | - | - | 0 | - |
| 8 | b | 1.229E+04 | 975.5 | 0.0008933 | 0.9157 | +1 | 8 |
| 8 | b | 9429 | 976.5 | 0.01869 | 19.14 | +1 | 8 |
| - | - | 6265 | 977.5 | - | - | 0 | - |
| - | - | 1.559E+04 | 980.5 | - | - | 0 | - |
| - | - | 7446 | 981.5 | - | - | 0 | - |
| - | - | 4954 | 982.5 | - | - | 0 | - |
| 8 | b | 8.022E+04 | 993.5 | 0.001359 | 1.368 | +1 | 8 |
| - | - | 5.043E+04 | 994.5 | - | - | 0 | - |
| - | - | 1.625E+04 | 995.5 | - | - | 0 | - |
| 6 | y | 1.307E+04 | 1007 | 0.006542 | 6.494 | +1 | 8 |
| 6 | y | 8985 | 1008 | 0.00678 | 6.723 | +1 | 8 |
| - | - | 7338 | 1024 | - | - | 0 | - |
| 6 | y | 6.803E+04 | 1025 | 0.002264 | 2.208 | +1 | 8 |
| - | - | 3.442E+04 | 1026 | - | - | 0 | - |
| - | - | 8894 | 1027 | - | - | 0 | - |
| - | - | 5635 | 1065 | - | - | 0 | - |
| - | - | 8522 | 1076 | - | - | 0 | - |
| - | - | 6025 | 1077 | - | - | 0 | - |
| 9 | b | 6986 | 1092 | 0.01918 | 17.57 | +1 | 9 |
| - | - | 5048 | 1093 | - | - | 0 | - |
| - | - | 2.491E+04 | 1094 | - | - | 0 | - |
| - | - | 1.801E+04 | 1095 | - | - | 0 | - |
| - | - | 7512 | 1096 | - | - | 0 | - |
| 9 | b | 6720 | 1109 | 0.01594 | 14.38 | +1 | 9 |
| - | - | 5193 | 1110 | - | - | 0 | - |
| - | - | 7984 | 1117 | - | - | 0 | - |
| - | - | 1.235E+04 | 1118 | - | - | 0 | - |
| - | - | 6148 | 1119 | - | - | 0 | - |
| 5 | y | 4.382E+04 | 1136 | 0.0007602 | 0.6695 | +1 | 9 |
| 5 | y | 7.362E+04 | 1136 | 0.005758 | 5.067 | +1 | 9 |
| - | - | 4.605E+04 | 1137 | - | - | 0 | - |
| - | - | 1.291E+04 | 1138 | - | - | 0 | - |
| - | - | 9435 | 1153 | - | - | 0 | - |
| 5 | y | 9.294E+04 | 1154 | 0.002281 | 1.977 | +1 | 9 |
| - | - | 6.351E+04 | 1155 | - | - | 0 | - |
| - | - | 2.555E+04 | 1156 | - | - | 0 | - |
| - | - | 5407 | 1157 | - | - | 0 | - |
| 10 | b | 1.17E+04 | 1166 | 0.01045 | 8.964 | +1 | 10 |
| - | - | 7704 | 1167 | - | - | 0 | - |
| - | - | 5399 | 1168 | - | - | 0 | - |
| 4 | y | 6718 | 1291 | 0.004751 | 3.682 | +1 | 10 |
| - | - | 5352 | 1292 | - | - | 0 | - |
| - | - | 3849 | 2538 | - | - | 0 | - |
| - | - | 3872 | 3081 | - | - | 0 | - |

m/z Charge Intensity FragmentType MassShift Position
120.05648040771484 0 3543.6804
120.08122253417969 0 8590.483
121.04010772705078 0 3741.868
122.07183837890625 0 17538.438
123.04458618164062 0 12610.971
123.05569458007812 0 4332.3223
123.09236907958984 0 2916.429
125.10774230957031 0 6524.672
126.09180450439453 0 13573.74
127.05064392089844 0 76737.76
127.0872802734375 0 5543.7354
127.1233139038086 0 7250.334
128.0541229248047 0 4432.6553
128.107421875 0 218484.7
129.06634521484375 0 15672.241
129.1027069091797 0 2599481.5
130.06553649902344 0 1110643
130.0998077392578 0 19497.12
130.1060028076172 0 176912
130.114013671875 0 3476.2366
131.06886291503906 0 106602.42
131.10720825195312 0 2311.5867
131.11805725097656 0 4494.7676
132.08116149902344 0 280334.06
133.0609588623047 0 6549.5312
133.0795440673828 0 4808.963
133.0845947265625 0 29207.332
136.0761260986328 0 241846.22
137.0795135498047 0 16111.188
138.05496215820312 0 3641.6526
138.06663513183594 0 31788.908
138.09178161621094 0 11739.522
139.05068969726562 0 4379.3696
139.08702087402344 0 26735.082
141.10264587402344 0 35256.727
142.06134033203125 0 2135.4187
142.06556701660156 0 23777.502
143.0456085205078 0 15348.663
143.1185760498047 0 2583.5793
144.08119201660156 0 12500.818
145.06117248535156 0 39446.473
146.0603790283203 0 3276.0657
146.09262084960938 0 2860.3425
147.04458618164062 0 14915.05
148.06101989746094 0 3583.4895
148.27313232421875 0 2769.6921
149.0452117919922 0 10825.682
149.10757446289062 0 3632.7085
150.0666961669922 0 17631.377
151.08705139160156 0 50780.46
152.10763549804688 0 7783.784
154.09796142578125 0 2843.179
154.18948364257812 0 2554.4834
155.04566955566406 0 4984.9194
155.0819091796875 0 7602.4985
155.0931854248047 0 156371.75
155.1182861328125 0 105739.16 a Water loss 1
155.12515258789062 0 1745.2225
156.076904296875 0 7616.8486
156.08999633789062 0 3564.9246
156.09658813476562 0 9769.385
156.10293579101562 0 2866.617
156.12156677246094 0 7931.072
157.06130981445312 0 8253.932
157.0977325439453 0 12576.48
158.084228515625 0 21493.287
159.0921630859375 0 2134174.8
160.07601928710938 0 15955.523
160.08961486816406 0 12374.954
160.09548950195312 0 221325.67
161.09869384765625 0 10789.227
161.5143280029297 0 2926.8076
165.0550537109375 0 129552.86
165.07728576660156 0 6445.811
166.06150817871094 0 290371.72
166.09796142578125 0 21709.51
167.05723571777344 0 7351.0615
167.0649871826172 0 24619.072
167.0820770263672 0 8857.275
167.09262084960938 0 3521.9216
167.11839294433594 0 8185.14
168.081298828125 0 32633.61
168.10232543945312 0 19791.334
168.1135711669922 0 19909.455
169.0764617919922 0 38101.805
169.08441162109375 0 6027.775
169.0976104736328 0 48488.016
170.06048583984375 0 103453.46
171.06394958496094 0 12455.956
171.07620239257812 0 5549.6235
171.09202575683594 0 9837.419
171.1497344970703 0 16441.549
172.07223510742188 0 36870.22
172.19496154785156 0 2975.2615
173.05628967285156 0 45318.895
173.12896728515625 0 3612609.5 a 1
173.4515838623047 0 9402.059
174.05975341796875 0 3394.0261
174.13230895996094 0 322577.53
175.134521484375 0 15793.469
176.08245849609375 0 4266.843
177.10281372070312 0 6752.96
178.1343994140625 0 119947.49
179.1378173828125 0 12948.147
181.0611572265625 0 18798.773
182.08168029785156 0 284203.75 y 12
183.0854949951172 0 28041.93
183.11326599121094 0 89201.2 b Water loss 1
184.11656188964844 0 9834.58
185.12904357910156 0 4270.06
185.1653594970703 0 30875.244
186.1242218017578 0 537407.56
187.0871124267578 0 158303.7
187.10797119140625 0 4040.0803
187.12762451171875 0 49855.305
188.07081604003906 0 11308.854
188.09056091308594 0 19311.45
189.0872802734375 0 5160.4546
190.08282470703125 0 64250.43
190.13427734375 0 26249.77
191.085693359375 0 5278.369
193.0977020263672 0 5588.6025
193.10850524902344 0 4043.613
194.09292602539062 0 18262.188
195.11326599121094 0 172348.86
196.10842895507812 0 32815.77
196.11688232421875 0 17767.293
197.10853576660156 0 5092.3916
197.1290283203125 0 19221.969
197.16502380371094 0 5339.226
198.08787536621094 0 30870.154
198.12815856933594 0 69310.01
199.07162475585938 0 17018.39
199.08700561523438 0 20562.824
199.09689331054688 0 3309.3975
199.10858154296875 0 4057.0024
199.1316680908203 0 10244.308
199.14427185058594 0 7923.168
200.09080505371094 0 5281.3755
200.14369201660156 0 12175.796
201.1238250732422 0 975347.4 b 1
202.0869903564453 0 6636.167
202.1271209716797 0 97744.29
203.1285858154297 0 5470.689
203.15110778808594 0 3573.655
204.11354064941406 0 7361.6543
204.1346435546875 0 7580.0225
205.09764099121094 0 80850.53
205.1085968017578 0 7114.6016
205.14520263671875 0 14160.341
206.10107421875 0 9947.204
206.12913513183594 0 13703.094
207.16094970703125 0 34537.766
208.10797119140625 0 8095.279
209.05674743652344 0 6412.7583
210.1265106201172 0 36089.176
211.0872344970703 0 3786.7678
211.10821533203125 0 23458.426 y Water loss 10
211.1317596435547 0 5749.076
212.1399688720703 0 31920.01
213.1239013671875 0 84345.72
213.1604461669922 0 12428.884
214.12713623046875 0 9413.696
215.1191864013672 0 4670.0737
215.13961791992188 0 9330.399
215.5143585205078 0 3871.0076
216.09848022460938 0 24201.584
217.08242797851562 0 13559.333
221.07211303710938 0 6342.2437
221.083251953125 0 5267.7183
221.09384155273438 0 5855.9106
221.1041259765625 0 17100.2
221.14013671875 0 7426.6206
222.10623168945312 0 4090.201
222.12425231933594 0 26226.777
223.08721923828125 0 4536.9746
223.10829162597656 0 49542.902
223.15589904785156 0 333800.03
224.1029510498047 0 6687.5664
224.12013244628906 0 4877.854
224.15939331054688 0 39590.78
225.16064453125 0 6253.1943
226.0828399658203 0 68539.234
227.06678771972656 0 65140.04
227.08660888671875 0 9244.9
227.1547393798828 0 19675.47
228.0701904296875 0 4280.4385
228.13461303710938 0 12082.647
229.11880493164062 0 39531.62
229.15550231933594 0 4661.459
230.1507568359375 0 9097.551
231.08741760253906 0 4168.2607
233.14031982421875 0 59350.242
234.12432861328125 0 365166.78
234.6499786376953 0 3961.647
235.12759399414062 0 45215.008
235.15586853027344 0 9377.432
236.12892150878906 0 4202.4756
237.13552856445312 0 19021.094
238.119384765625 0 27533.787
238.1299591064453 0 6334.268
239.08201599121094 0 9959.711
239.0950469970703 0 7452.574
239.11434936523438 0 4775.7754
239.62020874023438 0 3245.159 y Water loss 9
240.09634399414062 0 3866.3936
240.13485717773438 0 159970.67
241.09828186035156 0 3483.163
241.11874389648438 0 20838.701
241.13804626464844 0 18586.252
241.19166564941406 0 18392.053
243.10983276367188 0 33407.93
243.13388061523438 0 4607.7803
244.0934600830078 0 166516.3
245.0970458984375 0 15574.139
246.13462829589844 0 3782.072
247.10838317871094 0 183548.89
247.14486694335938 0 15588.879
247.15676879882812 0 5549.23
248.11436462402344 0 93892.95
248.6240234375 0 6451.9185 y 9
249.09884643554688 0 126614.17
249.1210479736328 0 6282.1836
249.1352996826172 0 34777.492
250.10227966308594 0 18137.396
250.11883544921875 0 7626.759
250.1385955810547 0 4234.5293
250.16665649414062 0 6092.884
251.15097045898438 0 1261514.5
252.1127166748047 0 6190.9585
252.1350860595703 0 6886.9087
252.15414428710938 0 162743.94
252.17002868652344 0 5789.091
253.11961364746094 0 6356.802
253.1561737060547 0 19966.879
255.14796447753906 0 71384.8
256.0950927734375 0 3818.3035
256.10894775390625 0 13575.084
256.1298522949219 0 6500.379
256.1513366699219 0 7239.669
257.0926513671875 0 4758.8667
257.11346435546875 0 25972.662
258.1239318847656 0 10764.324
258.14532470703125 0 65733.96
259.1300964355469 0 7153.855
259.1478271484375 0 6616.0884
261.1195983886719 0 28319.326
261.1351623535156 0 14647.134
262.12005615234375 0 14102.162
265.12994384765625 0 167870.89
266.12542724609375 0 366026.84
267.0778503417969 0 6871.8
267.12884521484375 0 40658.836
267.1453552246094 0 8265.744
268.1299743652344 0 15772.89
268.1770935058594 0 30907.04
269.1611633300781 0 10187.033
269.186279296875 0 23201.15
270.12255859375 0 42469.57
270.1458435058594 0 5348.729
270.1604919433594 0 4239.6187
270.6235046386719 0 12116.879
272.1763000488281 0 73771.32
273.179443359375 0 12047.004
274.11907958984375 0 60069.25
274.1316833496094 0 28333.04
275.10308837890625 0 17619.973
275.1212463378906 0 8695.726
275.1505432128906 0 10444.241
276.10931396484375 0 13456.85
276.1268615722656 0 6707.3574
276.1558532714844 0 27988.113
276.17071533203125 0 4752.641
277.0930480957031 0 5822.3076
279.1459655761719 0 25586.816
280.1299133300781 0 27005.783
281.0522155761719 0 10180.295
281.1632385253906 0 4532.931 b Water loss 4
282.1572265625 0 28041.482
282.6690368652344 0 6607.0815
283.1407470703125 0 1046788.7
284.0871276855469 0 5496.7485
284.10382080078125 0 34921.254
284.1238098144531 0 16352.826
284.14398193359375 0 145868.3
285.1089172363281 0 7269.3335
285.1286315917969 0 4709.4917
285.144775390625 0 13081.221
285.1585388183594 0 8449.585
286.14044189453125 0 58578.234
287.1427001953125 0 8832.9375
287.1721496582031 0 14126.955
287.18798828125 0 6199.126
288.1275939941406 0 3506.761
288.6192321777344 0 4085.3013
290.1671142578125 0 6074.805 b 4
290.6687927246094 0 3528.4875
291.2189025878906 0 4103.426
293.11395263671875 0 428629.97 y Water loss 11
293.16143798828125 0 17799.826
294.11810302734375 0 89102.78
294.18255615234375 0 4582.0845
295.120361328125 0 8032.0405
296.19769287109375 0 18091.145 b Water loss 2
296.666259765625 0 7045.99
297.13330078125 0 17030.213 y Water loss 8
297.1565246582031 0 143800.28
297.1756286621094 0 7061.644
297.6359558105469 0 3709.244
297.6568298339844 0 3867.7515
298.1401672363281 0 12955.89 y Water loss 6
298.15924072265625 0 22864.258
299.06243896484375 0 58683.35
300.111328125 0 3807.798
300.16888427734375 0 68037.83
301.15142822265625 0 596849.5
302.11431884765625 0 76539.75
302.1547546386719 0 84367.54
303.11785888671875 0 15502.668
303.1698913574219 0 5928.9517
305.17962646484375 0 12838.478
305.646240234375 0 10753.591
306.1380920410156 0 150739.12 y 8
306.229736328125 0 9142.635
306.63946533203125 0 56539.36
307.1408386230469 0 9351.7
309.1677551269531 0 9656.945
311.1247253417969 0 69271.65 y 11
311.1710205078125 0 5845.7397
312.1278076171875 0 12179.172
314.1869812011719 0 22302.291
314.20831298828125 0 47553.203 b 2
315.1672058105469 0 266978.84
316.1498718261719 0 6117.8667
316.1707458496094 0 36109.13
317.6590576171875 0 5346.4604
318.1203308105469 0 5421.0645
319.1522521972656 0 13860.43
320.15496826171875 0 4569.8457
324.19378662109375 0 4680.6865
324.6770324707031 0 7569.0337
325.15191650390625 0 4296.139
326.6647033691406 0 65541.984
327.16619873046875 0 32841.312
327.666259765625 0 3475.9858
329.1826477050781 0 5283.1436
331.65869140625 0 4458.935
332.1889343261719 0 3998.7432
332.2091979980469 0 27202.521
333.17767333984375 0 112065.63
333.198486328125 0 6056.047
334.1829833984375 0 22389.697
335.1838073730469 0 3901.6055
337.1624755859375 0 3798.0974
338.18328857421875 0 7752.99
338.6752624511719 0 11037.995 b Water loss 5
339.1728515625 0 11860.486 b Ammonia loss 5
339.67626953125 0 9774.309
340.1776428222656 0 4158.4097
340.66192626953125 0 14495.591
341.16357421875 0 10714.91
342.1822204589844 0 32574.016
343.1631164550781 0 5212.9473
344.1726989746094 0 28569.117
344.9770202636719 0 10280.648
345.13153076171875 0 41992.75
345.1760559082031 0 6058.4243
346.11541748046875 0 175388.25
346.6983947753906 0 8822.408
347.1187438964844 0 28537.584
347.68048095703125 0 66937.305 b 5
348.18182373046875 0 30892.283
349.1514892578125 0 8050.9653
350.2195129394531 0 118865.28
351.2220153808594 0 23500.34
352.1683349609375 0 5271.2515
352.67303466796875 0 12437.331
353.6746520996094 0 22127.197 y Water loss 7
354.1778869628906 0 8152.84
356.194580078125 0 5792.3384
357.17718505859375 0 9792.949
358.17938232421875 0 5796.5386
359.02783203125 0 7442.8994
360.2041015625 0 8742.174
360.6958923339844 0 35301.56
361.1980895996094 0 41789.426
361.6867370605469 0 5267.6387
362.1841735839844 0 32479.309
362.6800842285156 0 316754.03 y 7
363.14190673828125 0 230521.06
363.1817626953125 0 127312.02
363.68280029296875 0 28238.479
364.1253967285156 0 37902.51
364.145751953125 0 30367.24
364.1834411621094 0 4277.5996 b Water loss 8
365.1288146972656 0 7757.506
366.1567687988281 0 6609.43
366.1785583496094 0 20860.273
367.1400146484375 0 16544.148
367.1632080078125 0 17490.475
367.2457580566406 0 20464.084
368.1451416015625 0 7000.189
368.1919860839844 0 11376.142
373.1861877441406 0 14514.307
376.1625061035156 0 13322.41
376.1983947753906 0 27330.834
376.6997985839844 0 14329.4375
377.14642333984375 0 10371.251
377.20123291015625 0 4714.628
378.21453857421875 0 73254.766
379.2099609375 0 347365.4
380.1659240722656 0 8909.632
380.2127685546875 0 59474.695
381.152587890625 0 646274.1
381.7116394042969 0 25785.111
382.15533447265625 0 119834.83
382.19024658203125 0 26442.686
382.2117004394531 0 4913.699
382.691650390625 0 4399.2603
383.15740966796875 0 13675.898
383.2065734863281 0 4637.713
384.1663513183594 0 26506.99
384.18878173828125 0 51374.777
385.1507873535156 0 40840.086
385.1731262207031 0 30821.824 y 4
385.1960144042969 0 5557.8647
386.1548156738281 0 9071.085
386.1785888671875 0 11828.877
387.15606689453125 0 4846.5884
389.1936950683594 0 8859.157 b 9
390.19610595703125 0 31532.453
390.69757080078125 0 13936.509
391.1368103027344 0 24792.172
391.1846008300781 0 6640.307
391.6875 0 5901.8945
392.1191711425781 0 5014.225
392.1431884765625 0 5249.891
392.208740234375 0 9868.626
393.2135009765625 0 5900.9507
394.1733093261719 0 94643.9
395.1348876953125 0 31976.895
395.1576232910156 0 31743.953
395.1795959472656 0 7974.535
395.2022399902344 0 5941.285
395.2403259277344 0 7459.1167
396.13677978515625 0 5557.515
396.16046142578125 0 8313.193
396.22552490234375 0 77674.766
397.2033996582031 0 13205.048
397.22833251953125 0 18654.057
397.7059631347656 0 5700.4663
398.1788635253906 0 54753.56
399.1826477050781 0 5760.0225
402.1763916015625 0 28300.734
402.1998596191406 0 78398.555
403.1368103027344 0 5092.7153
403.20086669921875 0 36820.414
404.2122497558594 0 6723.351
404.71002197265625 0 13897.94
405.26171875 0 18230.158
406.2460632324219 0 13455.678
407.2047119140625 0 7173.135
407.24420166015625 0 5624.0938
408.7106018066406 0 7146.944
409.1477355957031 0 50428.566
409.17388916015625 0 5573.9517
409.2065124511719 0 11124.912
410.14959716796875 0 8621.39
410.1993713378906 0 4438.5205
411.19970703125 0 14585.846
412.18389892578125 0 447144.53
413.1439514160156 0 24000.727
413.1677551269531 0 81785.1
413.1898498535156 0 44926.73
413.2497863769531 0 6943.7397
414.171142578125 0 16017.761
414.1966552734375 0 7179.044
414.23583984375 0 163650.42
415.0377197265625 0 19140.006
415.1997375488281 0 21640.834
415.2394714355469 0 31878.941
416.20343017578125 0 4983.375
417.7177734375 0 56358.94
418.2107849121094 0 118349.89
418.7120361328125 0 68075.234
419.2130126953125 0 18774.639
420.16326904296875 0 4820.6797
421.2093811035156 0 18044.936 y Water loss 10
422.19366455078125 0 19638.045 y Ammonia loss 10
422.7104187011719 0 9319.29
423.2037353515625 0 18252.982
423.27252197265625 0 169987.69
423.7046203613281 0 7603.753
423.73565673828125 0 3812.5928
424.22625732421875 0 5719.0996
424.2755432128906 0 36137.07
424.7362365722656 0 14559.236
425.2338562011719 0 9217.535
426.19866943359375 0 5891.749
426.7229919433594 0 1060224.1
427.224365234375 0 540023.4
427.725830078125 0 163873.19
428.22674560546875 0 34428.04
429.2104187011719 0 92162.39
430.1945495605469 0 1212127.8
431.19744873046875 0 261808.48
431.71502685546875 0 129258.76 b Water loss 6
432.2132568359375 0 59663.875 b Ammonia loss 6
432.71356201171875 0 27255.396
433.2111511230469 0 22355.713
433.25616455078125 0 45534.793 b Water loss 3
433.7114562988281 0 6029.3613
434.26202392578125 0 5547.291
438.1746520996094 0 15543.777
438.7248229980469 0 6920.714
439.2201843261719 0 288629.84 y 10
440.1806640625 0 9036.878
440.22308349609375 0 66803.87
440.7204895019531 0 510383.44 b 6
441.22186279296875 0 280921.78
441.7232360839844 0 78117.73
442.2247619628906 0 25223.127
443.2402038574219 0 5032.9346
446.7405700683594 0 5546.4326
447.20672607421875 0 51099.57 y Ammonia loss 6
447.2398986816406 0 6758.8076
447.7079162597656 0 28081.484
448.2051696777344 0 64656.19
449.2090759277344 0 16911.533
450.1747741699219 0 10259.412
450.2359924316406 0 10459.765
450.2841796875 0 12183.423
451.2677307128906 0 310707.03 b 3
452.2705078125 0 76555.46
453.2729797363281 0 11863.271
455.2660827636719 0 24936.043
455.7201843261719 0 25159.627 y 6
456.22216796875 0 12991.647
456.26837158203125 0 4973.148
459.19921875 0 29725.496
460.7547302246094 0 23128.893
461.204833984375 0 20998.893
461.2535705566406 0 41453.168
461.7557373046875 0 9031.347
462.2085876464844 0 6096.9995
462.2508544921875 0 5710.9966
462.55584716796875 0 5730.155 y Water loss 2
462.8876953125 0 6026.4575 y Ammonia loss 2
464.2154846191406 0 6196.5527
465.2552795410156 0 5278.108
465.745361328125 0 11317.81
466.1683044433594 0 6301.892
466.243408203125 0 10529.015
468.2940673828125 0 80881.91
469.2973327636719 0 26179.701
472.2201843261719 0 5214.3716
474.2597351074219 0 61633.29
474.7558288574219 0 67017.805
475.2556457519531 0 42474.4
475.7570495605469 0 10016.6875
476.22662353515625 0 24550.656
477.21099853515625 0 47777.633
478.2301330566406 0 43546.336 y Water loss 9
478.2787170410156 0 40865.113
479.2344055175781 0 12955.054
479.2811279296875 0 7073.241
480.22265625 0 6944.1836
483.2650451660156 0 169954.06
483.7665710449219 0 119675.516
484.2205505371094 0 11259.865
484.2677917480469 0 44012.145
484.7705383300781 0 6125.285
485.2240905761719 0 5795.7456
488.221435546875 0 4665.4727
488.25811767578125 0 58493.75 b Water loss 7
488.75567626953125 0 38805.246 b Ammonia loss 7
489.2552185058594 0 17989.262
493.25238037109375 0 4650.9194
494.2370910644531 0 618151.5
495.2054443359375 0 4383.62
495.2399597167969 0 171368.34
496.2414855957031 0 651442.9 y 9
497.2467956542969 0 151455.47
497.76409912109375 0 72465.125
498.2457275390625 0 30840.887
502.2309265136719 0 6360.012
503.1083984375 0 19730.463
504.2010803222656 0 29001.932
505.201171875 0 9273.632
506.2312316894531 0 8682.182
506.2732238769531 0 8558.81
507.2574157714844 0 25451.84
508.2598571777344 0 7892.928
511.2635192871094 0 62119.926
512.2674560546875 0 17270.6
513.2105712890625 0 13333.721
514.1849365234375 0 7248.4053
515.2843017578125 0 14942.79
515.760498046875 0 4606.251
516.2869262695312 0 10161.266
519.14013671875 0 5478.728
520.2669677734375 0 6558.7754
521.2220458984375 0 7958.856
521.271484375 0 4570.0186
522.2123413085938 0 6537.2603
523.2604370117188 0 20091.37
525.2675170898438 0 119446.68
526.2322998046875 0 9018.263
526.2706909179688 0 35406.7
529.2615356445312 0 32106.898 Precursor Water loss
529.5952758789062 0 34173.656 Precursor Ammonia loss
529.9293823242188 0 21081.406
531.218505859375 0 9754.257
532.1956176757812 0 31643.535
533.2032470703125 0 6840.657
533.3185424804688 0 4840.4834
534.3031005859375 0 5911.1533
535.2639770507812 0 33598.9 Precursor
535.6004638671875 0 38672.625
535.937744140625 0 25867.61
536.1669921875 0 5125.912
536.2725830078125 0 10498.005
537.2461547851562 0 6223.9253
537.7374877929688 0 7493.6216
538.2749633789062 0 5201.8594
539.2371826171875 0 123076.61
539.2756958007812 0 4563.4136
540.2395629882812 0 31076.402
541.2485961914062 0 13064.323
542.2955322265625 0 15531.243
543.2781372070312 0 238815.27
544.2814331054688 0 73176.266
545.282958984375 0 12612.604
546.25 0 28078.676
546.75048828125 0 11740.942
547.2499389648438 0 9484.893
548.23681640625 0 8798.22
548.3010864257812 0 6918.7363
549.2225341796875 0 31770.41
550.2101440429688 0 11012.429
551.2145385742188 0 6370.248
551.2588500976562 0 29811.945
551.3312377929688 0 19415.312
552.2603759765625 0 12284.386
552.335205078125 0 6653.6396
553.2682495117188 0 5167.465
554.7655029296875 0 11999.995 b 8
555.25732421875 0 35159.477
555.7572021484375 0 34685.45
556.2609252929688 0 11649.944
557.2344970703125 0 4897.1616
558.2685546875 0 5607.3696
559.7487182617188 0 4666.918
561.2815551757812 0 8800.564
561.3164672851562 0 22924.297 b Water loss 4
562.3025512695312 0 11162.125 b Ammonia loss 4
563.2586059570312 0 20671.455
564.2644653320312 0 7849.7993
564.3327026367188 0 7598.1133
565.262939453125 0 76073.2
566.2640991210938 0 29098.543
567.2322998046875 0 133281.88
568.23681640625 0 47601.816
568.759521484375 0 8891.825 y Ammonia loss 4
569.2388916015625 0 10233.684
575.2471923828125 0 34887.934
576.234375 0 20837.084
576.2815551757812 0 7341.7646
577.2175903320312 0 8864.067
577.2693481445312 0 7538.8853 y 4
579.3260498046875 0 197772.98 b 4
580.3292846679688 0 66159.97
581.2703247070312 0 5139.3447
581.3335571289062 0 9197.876
582.3046264648438 0 7358.179
586.258056640625 0 5646.5254
588.7950439453125 0 11608.6045
589.3043823242188 0 16124.535
592.2738647460938 0 10442.946
592.3257446289062 0 28858.828
593.2576293945312 0 260482.34 y Water loss 8
593.306640625 0 80480.38
593.7875366210938 0 9210.675
594.2538452148438 0 95968.234
594.310791015625 0 22010.582
594.781982421875 0 6593.2314
595.2510986328125 0 33504.664
596.2543334960938 0 4600.5454
596.35302734375 0 8582.586
597.3065185546875 0 7203.6333
597.7999877929688 0 8271.616
598.3015747070312 0 10019.503
600.3157958984375 0 11129.259
602.301513671875 0 5596.9956
602.7933349609375 0 27815.48
603.290771484375 0 32025.41
603.7906494140625 0 11075.543
604.2880249023438 0 9263.484
610.2842407226562 0 213624.89
610.332763671875 0 25638.275
611.2689208984375 0 3254771.8 y 8
611.7977294921875 0 119540.24
612.2719116210938 0 1013553.8
612.8002319335938 0 37707.418
613.2739868164062 0 231642.4
614.2755737304688 0 28692.084
617.2845458984375 0 12719.553
618.3276977539062 0 5638.014
620.8032836914062 0 12432.584
621.2528076171875 0 17998.143
621.3008422851562 0 17209.256
622.2588500976562 0 9272.947
622.3689575195312 0 10230.37
628.2742309570312 0 13338.14
631.8369140625 0 5624.6836
634.3067626953125 0 30002.87
635.2980346679688 0 39690.855
636.3279418945312 0 28148.201
636.7874145507812 0 23483.072 y Water loss 3
637.2885131835938 0 19171.773 y Ammonia loss 3
637.3385009765625 0 9902.855
637.78955078125 0 10806.927
638.3341674804688 0 7270.7954 b Water loss 10
638.8282470703125 0 4515.6724 b Ammonia loss 10
640.2755737304688 0 17081.06
640.335693359375 0 4821.323
645.2870483398438 0 20139.91
645.7931518554688 0 159713.86 y 3
646.2938232421875 0 116151.01
646.7949829101562 0 43315.72
647.2952270507812 0 13185.179
648.3445434570312 0 6265.383
649.3389892578125 0 6406.6196
650.2687377929688 0 12263.303
650.3303833007812 0 13854.978
650.7872924804688 0 7988.244
651.273193359375 0 5579.748
652.3214721679688 0 269244.47
653.3241577148438 0 107229.016
654.326171875 0 22386.762
658.2799682617188 0 8581.293
658.3327026367188 0 11656.573
659.3154907226562 0 12902.417
661.3320922851562 0 13027.679
661.8308715820312 0 4614.131
662.3069458007812 0 87503.44
663.2970581054688 0 57002.98
664.2971801757812 0 16675.848
664.34521484375 0 12147.493
665.3458251953125 0 6023.1567
666.3535766601562 0 7059.9316
668.28271484375 0 9152.806
670.33203125 0 4337.286
671.3325805664062 0 5396.935
676.3427124023438 0 136872.6 b Water loss 5
677.3338623046875 0 88144.58 b Ammonia loss 5
678.3336181640625 0 37734.49
679.3347778320312 0 23484.04
679.83349609375 0 7286.6084
680.3167724609375 0 493480.4
680.8681640625 0 6669.371
681.3193969726562 0 195572.45
682.3214721679688 0 50004.848
684.3252563476562 0 20970.686
684.818359375 0 32452.477
685.318603515625 0 28807.838
685.8216552734375 0 12112.047
686.3253173828125 0 8162.3164
687.3079833984375 0 4514.9395
688.3360595703125 0 15123.383
688.8532104492188 0 7780.7783
689.3316650390625 0 5830.4717
692.8349609375 0 11019.415
693.330322265625 0 114955.84 y Water loss 2
693.8280029296875 0 128030.28 y Ammonia loss 2
694.3528442382812 0 1370389.2 b 5
694.826904296875 0 18773.832
694.8782958984375 0 6765.633
695.3563232421875 0 522406.28
696.359130859375 0 116248.125
697.3563842773438 0 25785.754
698.352783203125 0 4993.066
701.3432006835938 0 19845.102
701.8419799804688 0 89912.586
702.3356323242188 0 818668 y 2
702.8367919921875 0 674366.44 b Water loss 11
703.3382568359375 0 334574.4 b Ammonia loss 11
703.8389892578125 0 102599.62
704.3397216796875 0 39337.426
705.3449096679688 0 10789.55
706.3423461914062 0 101676.125 y Water loss 7
707.3443603515625 0 40805.65
707.829833984375 0 5435.4473
708.3394165039062 0 10158.498
709.3185424804688 0 6843.758
710.3603515625 0 4508.0073
711.375732421875 0 44660.445
711.8569946289062 0 58904.902 b 11
712.3606567382812 0 49495.19
712.8602905273438 0 13656.091
713.3677368164062 0 6423.5566
716.350830078125 0 6230.9116
720.3817138671875 0 9620.193
720.8591918945312 0 9936.458
721.3641357421875 0 73351.66
722.3668212890625 0 20021.281
723.36962890625 0 69527.11
724.3532104492188 0 1095648.5 y 7
725.3558349609375 0 442657.1
726.35791015625 0 110033.64
727.3612060546875 0 14212.301
729.3624267578125 0 24305.207
730.3636474609375 0 13162.674
733.3644409179688 0 35017.668
734.3547973632812 0 31435.795
735.3585205078125 0 14397.459
737.3387451171875 0 5677.5396
737.8641967773438 0 12501.527
738.3712158203125 0 8732.226
738.8663330078125 0 4714.191
739.3707275390625 0 16157.59
739.8734741210938 0 6344.304
740.3756713867188 0 7181.875
742.8640747070312 0 6190.7656 y Water loss 1
743.3693237304688 0 25956.207
743.8627319335938 0 6899.058
744.371337890625 0 12289.128
746.361083984375 0 5707.1187
747.3910522460938 0 11898.487
748.38623046875 0 25732.836
749.3382568359375 0 13800.937
750.3890380859375 0 13515.772
751.3775634765625 0 139082.69
751.8695678710938 0 59001.03 y 1
752.3753051757812 0 121763.445
752.8731079101562 0 29603.205
753.374267578125 0 69730
753.8743286132812 0 24328.877
754.3770141601562 0 19339.582
754.8795776367188 0 5650.3643
758.3652954101562 0 5551.3125
760.3035888671875 0 7199.0083
761.3758544921875 0 11673.892
762.4136352539062 0 15467.688
763.3665161132812 0 82744.86
764.3679809570312 0 39599.668
765.4043579101562 0 38642.695
766.4074096679688 0 18925.115
767.3475952148438 0 34639.664
768.3515625 0 13370.044
768.4163208007812 0 7368.784
769.4002075195312 0 14478.147
770.401611328125 0 9324.119
775.3912353515625 0 14861.395
776.3807983398438 0 18324.615
777.381591796875 0 8729.75
779.3848266601562 0 83771.5
780.3887329101562 0 37672.13
781.3671264648438 0 42749.082
782.367431640625 0 17743.678
793.4008178710938 0 151828.66 Precursor Water loss
794.4036254882812 0 73268.555
795.3421020507812 0 19360.75
795.408447265625 0 12959.449
796.3475952148438 0 9602.8125
807.3796997070312 0 26791.627
807.441650390625 0 18143.52
808.4126586914062 0 13426.572
809.4133911132812 0 6248.081
817.4031372070312 0 7552.554
818.4371948242188 0 21087.53
819.43896484375 0 21294.299
820.4357299804688 0 7015.515
825.3841552734375 0 9236.376
826.3779907226562 0 11908.086
827.3795166015625 0 5732.0645
829.4281616210938 0 6241.2983
834.4274291992188 0 27790.453
835.41650390625 0 45309.16
836.45068359375 0 107006.81
837.4512939453125 0 42045.637
838.4533081054688 0 12054.672
843.4006958007812 0 9345.951
844.3910522460938 0 25010.432
845.394775390625 0 23789.555
846.4482421875 0 42243.875
847.4472045898438 0 28733.47
848.4471435546875 0 9747.858
852.4378662109375 0 329879.6
853.4406127929688 0 172730.66
854.3668823242188 0 5680.0483
854.443359375 0 38943.18
855.4484252929688 0 9010.1045
862.4222412109375 0 193466.98 b Water loss 6
863.4169921875 0 135464.88 b Ammonia loss 6
864.456298828125 0 239592.28
865.4603271484375 0 115229.95
866.4638061523438 0 35197.066
869.3899536132812 0 6346.937
876.4408569335938 0 26356.207
877.4391479492188 0 14020.043
878.4321899414062 0 4723.2656
880.4326171875 0 1576009.4 b 6
881.43505859375 0 806623.2
882.438232421875 0 224003.4
883.4398803710938 0 53841.953
884.4398193359375 0 7715.9233
890.4182739257812 0 10499.787
891.4086303710938 0 8225.078
892.463623046875 0 30274.104
893.4798583984375 0 9508.644
894.4496459960938 0 13811.849
895.4510498046875 0 6510.056
908.4286499023438 0 58553.605
909.434814453125 0 35033.477
910.4327392578125 0 96377.13 y 6
911.4353637695312 0 47099.24
912.4369506835938 0 13692.082
919.4027099609375 0 8142.205
936.427490234375 0 4635.5864
937.4259643554688 0 16931.223
938.4119873046875 0 9437.541
948.4287719726562 0 10647.041
954.43310546875 0 15061.258
955.4215698242188 0 29518.11
956.4232177734375 0 19649.043
964.4620361328125 0 10395.665
965.4483642578125 0 32637.209
966.4515991210938 0 21430.842
971.4629516601562 0 6317.6304
972.443115234375 0 16872.637
973.4462280273438 0 13658.873
975.5037841796875 0 12289.19 b Water loss 7
976.5073852539062 0 9429.239 b Ammonia loss 7
977.501708984375 0 6264.83
980.459716796875 0 15594.244
981.4639282226562 0 7445.828
982.4736938476562 0 4954.441
993.5166015625 0 80217.875 b 7
994.5201416015625 0 50432.996
995.5239868164062 0 16249.038
1007.4534301757812 0 13074.881 y Water loss 5
1008.4376831054688 0 8984.99 y Ammonia loss 5
1024.4757080078125 0 7337.592
1025.459716796875 0 68034.44 y 5
1026.46240234375 0 34423.293
1027.458984375 0 8894.266
1064.5174560546875 0 5635.333
1075.5323486328125 0 8521.924
1076.5302734375 0 6025.0625
1091.4964599609375 0 6985.955 b Ammonia loss 8
1092.505126953125 0 5047.803
1093.545166015625 0 24909.348
1094.5472412109375 0 18013.805
1095.5428466796875 0 7512.4663
1108.5262451171875 0 6720.458 b 8
1109.518310546875 0 5193.2544
1117.4964599609375 0 7983.9624
1118.4874267578125 0 12347.667
1119.4840087890625 0 6148.438
1135.5062255859375 0 43820.23 y Water loss 4
1136.4952392578125 0 73623.63 y Ammonia loss 4
1137.495361328125 0 46045.164
1138.49658203125 0 12913.924
1152.5364990234375 0 9435.245
1153.518310546875 0 92938.62 y 4
1154.52099609375 0 63513.598
1155.521240234375 0 25552.195
1156.523193359375 0 5406.6807
1165.5740966796875 0 11700.773 b 9
1166.567626953125 0 7704.292
1167.568115234375 0 5399.496
1290.5701904296875 0 6718.036 y 3
1291.5673828125 0 5352.442
2537.555908203125 0 3849.1067
3081.468017578125 0 3872.2534

Spectrum Details

|  |  |
| --- | --- |
| Matched peaks? Matched peaks The total absolute number of peaks matched. Additionally in brackets the total fraction of peaks matched and the total number of peaks is shown. | 88 (9.02% of 976) |
| FDR? FDR The false discovery rate estimated for this peptide. It is calculated by matching all theoretical fragments with a non-integer shift with the raw peaks for this spectrum. This is done with 40 different shifts. The resulting percentage is the average number of annotated peaks over the number of annotated peaks with the correct spectrum. | 0.54% |
| Satellite FDR? Satellite FDR See the FDR for details on its calculation. This satellite ion specific FDR only contains the satellite ions (d/w) for I/L/J positions. | - |
| PSM Score? PSM Score The PSM Score as given by Hecklib to this annotated spectrum. It is shown with three significant figures. | 669 |

#### Spectrum 7158? Spectrum 7158 The raw spectrum of this peptide as annotated by Hecklib. The fragments are coloured according to ion type (see legend). Any peaks with a star '\*' as text can be hovered over to see the full details, first the ion type second the mass shift type. By hovering over the amino acids in the peptide or ions in the legend the corresponding peaks are highlighted. By toggling the 'Unassigned' label you can turn the background (unassigned) peaks on or off in the plot. By updating the slider in the Ion legend you can update the spectrum to only show the top X% of the peaks with labels. The top X% means any peak that is within X% of the highest intensity. By dragging in the spectrum you can zoom in to a specific part of the spectrum and use 'Zoom Out' to get back to the original zoom level. The annotation of the spectrum is based on the given sequence in the peptides file and is done with different software so inconsistencies are likely. The peaks are annotated based on the given sequence, with 20 ppm tolerance.

Copy Data

##### Spectrum 7158 (TSV)

###### Preview

```
Loading example...
```

*Click on the button to copy the data to your clipboard.*

Mz MinMz MaxIntensity Max

WidthHeightPeptide font sizePeptide stroke widthSpectrum font sizeSpectrum stroke widthCompact peptide

Ion legend

wxyz

abcd

OtherUnassignedIon^Charge^_Position_Show for top:%

TVLHQDWLDGKEY

02.02e+54.05e+56.07e+58.09e+5

Zoom Out

a^+1^_2_y^+3^_4_a^+1^_2_y^+1^_1_b^+1^_2_b^+1^_2_y^+2^_3_y^+2^_4_b^+2^_5_y^+1^_2_b^+1^_3_y^+2^_5_y^+3^_7_y^+2^_5_y^+1^_2_b^+1^_3_b^+2^_6_b^+2^_6_b^+2^_6_y^+2^_6_y^+2^_6_y^+3^_9_b^+3^_10_y^+1^_3_y^+1^_3_b^+2^_7_b^+2^_7_b^+1^_4_y^+1^_3_b^+2^_7_y^+2^_7_y^+2^_7_b^+1^_4_y^+2^_7_y^+3^_11_y^+1^_4_b^+2^_8_b^+2^_8_y^+1^_4_\*\*\*b^+2^_9_b^+2^_9_b^+2^_9_b^+1^_5_b^+1^_5_b^+1^_5_y^+1^_5_y^+1^_5_y^+2^_10_y^+2^_10_b^+2^_11_y^+2^_10_b^+1^_6_b^+1^_6_y^+2^_11_y^+2^_11_b^+1^_6_y^+2^_11_b^+2^_12_b^+2^_12_y^+1^_6_b^+2^_12_y^+1^_6_y^+2^_12_y^+2^_12_\*b^+1^_7_b^+1^_7_b^+1^_7_y^+1^_7_b^+1^_8_b^+1^_8_b^+1^_8_y^+1^_8_y^+1^_8_b^+1^_9_b^+1^_9_y^+1^_9_y^+1^_9_y^+1^_9_b^+1^_10_y^+1^_10_

0778155623343112

Fragment Matches Table

Show background peaks

| Position | Ion type | Intensity | mz Theoretical | mz Error (Th) | mz Error (ppm) | Charge | Series Number |
| --- | --- | --- | --- | --- | --- | --- | --- |
| - | - | 6800 | 120.1 | - | - | 0 | - |
| - | - | 3326 | 122.1 | - | - | 0 | - |
| - | - | 3338 | 123 | - | - | 0 | - |
| - | - | 2130 | 125.1 | - | - | 0 | - |
| - | - | 892.3 | 126.1 | - | - | 0 | - |
| - | - | 2425 | 126.1 | - | - | 0 | - |
| - | - | 1.326E+04 | 127.1 | - | - | 0 | - |
| - | - | 1600 | 127.1 | - | - | 0 | - |
| - | - | 1055 | 128.1 | - | - | 0 | - |
| - | - | 4.983E+04 | 128.1 | - | - | 0 | - |
| - | - | 3107 | 129.1 | - | - | 0 | - |
| - | - | 5.486E+05 | 129.1 | - | - | 0 | - |
| - | - | 2.467E+05 | 130.1 | - | - | 0 | - |
| - | - | 2652 | 130.1 | - | - | 0 | - |
| - | - | 3.516E+04 | 130.1 | - | - | 0 | - |
| - | - | 1880 | 131.1 | - | - | 0 | - |
| - | - | 2.156E+04 | 131.1 | - | - | 0 | - |
| - | - | 1939 | 131.1 | - | - | 0 | - |
| - | - | 5.576E+04 | 132.1 | - | - | 0 | - |
| - | - | 1.07E+04 | 133.1 | - | - | 0 | - |
| - | - | 940.3 | 134 | - | - | 0 | - |
| - | - | 6.009E+04 | 136.1 | - | - | 0 | - |
| - | - | 5537 | 137.1 | - | - | 0 | - |
| - | - | 7168 | 138.1 | - | - | 0 | - |
| - | - | 2459 | 138.1 | - | - | 0 | - |
| - | - | 888.7 | 138.7 | - | - | 0 | - |
| - | - | 1251 | 139.1 | - | - | 0 | - |
| - | - | 5583 | 139.1 | - | - | 0 | - |
| - | - | 8802 | 141.1 | - | - | 0 | - |
| - | - | 4977 | 142.1 | - | - | 0 | - |
| - | - | 907.1 | 142.3 | - | - | 0 | - |
| - | - | 3821 | 143 | - | - | 0 | - |
| - | - | 1152 | 143.1 | - | - | 0 | - |
| - | - | 1446 | 144.1 | - | - | 0 | - |
| - | - | 2483 | 144.1 | - | - | 0 | - |
| - | - | 7639 | 145.1 | - | - | 0 | - |
| - | - | 2404 | 146.1 | - | - | 0 | - |
| - | - | 2666 | 147 | - | - | 0 | - |
| - | - | 5561 | 147.1 | - | - | 0 | - |
| - | - | 894.3 | 148.2 | - | - | 0 | - |
| - | - | 1.328E+04 | 149 | - | - | 0 | - |
| - | - | 3034 | 150.1 | - | - | 0 | - |
| - | - | 1.32E+04 | 151.1 | - | - | 0 | - |
| - | - | 1396 | 153.1 | - | - | 0 | - |
| - | - | 1084 | 154.1 | - | - | 0 | - |
| - | - | 1340 | 155 | - | - | 0 | - |
| - | - | 2005 | 155.1 | - | - | 0 | - |
| - | - | 3.611E+04 | 155.1 | - | - | 0 | - |
| 2 | a | 2.348E+04 | 155.1 | 0.0003661 | 2.36 | +1 | 2 |
| - | - | 1474 | 156.1 | - | - | 0 | - |
| - | - | 1226 | 156.1 | - | - | 0 | - |
| - | - | 1521 | 156.1 | - | - | 0 | - |
| - | - | 1398 | 156.1 | - | - | 0 | - |
| - | - | 1400 | 157.1 | - | - | 0 | - |
| - | - | 2080 | 157.1 | - | - | 0 | - |
| - | - | 4346 | 158.1 | - | - | 0 | - |
| - | - | 4.34E+05 | 159.1 | - | - | 0 | - |
| - | - | 4211 | 160.1 | - | - | 0 | - |
| - | - | 2718 | 160.1 | - | - | 0 | - |
| - | - | 4.594E+04 | 160.1 | - | - | 0 | - |
| - | - | 1843 | 161.1 | - | - | 0 | - |
| - | - | 3.168E+04 | 165.1 | - | - | 0 | - |
| - | - | 1443 | 165.1 | - | - | 0 | - |
| - | - | 6.002E+04 | 166.1 | - | - | 0 | - |
| 10 | y | 2756 | 166.1 | 0.001496 | 9.01 | +3 | 4 |
| - | - | 5859 | 166.1 | - | - | 0 | - |
| - | - | 1.372E+04 | 167.1 | - | - | 0 | - |
| - | - | 4670 | 167.1 | - | - | 0 | - |
| - | - | 1668 | 167.1 | - | - | 0 | - |
| - | - | 1473 | 167.1 | - | - | 0 | - |
| - | - | 7902 | 168.1 | - | - | 0 | - |
| - | - | 5099 | 168.1 | - | - | 0 | - |
| - | - | 7278 | 168.1 | - | - | 0 | - |
| - | - | 6263 | 169.1 | - | - | 0 | - |
| - | - | 1678 | 169.1 | - | - | 0 | - |
| - | - | 1.571E+04 | 169.1 | - | - | 0 | - |
| - | - | 2.137E+04 | 170.1 | - | - | 0 | - |
| - | - | 3070 | 171.1 | - | - | 0 | - |
| - | - | 1090 | 171.1 | - | - | 0 | - |
| - | - | 1260 | 171.1 | - | - | 0 | - |
| - | - | 5646 | 171.1 | - | - | 0 | - |
| - | - | 9094 | 172.1 | - | - | 0 | - |
| - | - | 919.9 | 172.8 | - | - | 0 | - |
| - | - | 7537 | 173.1 | - | - | 0 | - |
| 2 | a | 8.012E+05 | 173.1 | 0.0004978 | 2.875 | +1 | 2 |
| - | - | 6.749E+04 | 174.1 | - | - | 0 | - |
| - | - | 3463 | 175.1 | - | - | 0 | - |
| - | - | 1189 | 177.1 | - | - | 0 | - |
| - | - | 3533 | 177.1 | - | - | 0 | - |
| - | - | 2.401E+04 | 178.1 | - | - | 0 | - |
| - | - | 2843 | 179.1 | - | - | 0 | - |
| - | - | 3795 | 181.1 | - | - | 0 | - |
| 13 | y | 7.482E+04 | 182.1 | 0.0004801 | 2.637 | +1 | 1 |
| - | - | 8033 | 183.1 | - | - | 0 | - |
| 2 | b | 2.245E+04 | 183.1 | 0.0004771 | 2.605 | +1 | 2 |
| - | - | 1043 | 184.1 | - | - | 0 | - |
| - | - | 2496 | 184.1 | - | - | 0 | - |
| - | - | 1684 | 185.1 | - | - | 0 | - |
| - | - | 7481 | 185.2 | - | - | 0 | - |
| - | - | 1.441E+05 | 186.1 | - | - | 0 | - |
| - | - | 3.626E+04 | 187.1 | - | - | 0 | - |
| - | - | 1213 | 187.1 | - | - | 0 | - |
| - | - | 1.428E+04 | 187.1 | - | - | 0 | - |
| - | - | 2044 | 187.1 | - | - | 0 | - |
| - | - | 2539 | 188.1 | - | - | 0 | - |
| - | - | 3814 | 188.1 | - | - | 0 | - |
| - | - | 1108 | 188.1 | - | - | 0 | - |
| - | - | 1101 | 189.1 | - | - | 0 | - |
| - | - | 1598 | 189.1 | - | - | 0 | - |
| - | - | 1649 | 189.1 | - | - | 0 | - |
| - | - | 1.051E+04 | 190.1 | - | - | 0 | - |
| - | - | 4753 | 190.1 | - | - | 0 | - |
| - | - | 1054 | 191.3 | - | - | 0 | - |
| - | - | 1444 | 192.1 | - | - | 0 | - |
| - | - | 1109 | 194.1 | - | - | 0 | - |
| - | - | 5897 | 194.1 | - | - | 0 | - |
| - | - | 3.943E+04 | 195.1 | - | - | 0 | - |
| - | - | 1972 | 195.1 | - | - | 0 | - |
| - | - | 2.314E+04 | 196.1 | - | - | 0 | - |
| - | - | 2945 | 196.1 | - | - | 0 | - |
| - | - | 3933 | 197.1 | - | - | 0 | - |
| - | - | 2081 | 197.2 | - | - | 0 | - |
| - | - | 3569 | 198.1 | - | - | 0 | - |
| - | - | 1.51E+04 | 198.1 | - | - | 0 | - |
| - | - | 3572 | 199.1 | - | - | 0 | - |
| - | - | 4345 | 199.1 | - | - | 0 | - |
| - | - | 3071 | 199.1 | - | - | 0 | - |
| - | - | 2381 | 199.1 | - | - | 0 | - |
| - | - | 1593 | 200.1 | - | - | 0 | - |
| 2 | b | 2.15E+05 | 201.1 | 0.0004257 | 2.117 | +1 | 2 |
| - | - | 2542 | 202.1 | - | - | 0 | - |
| - | - | 1.911E+04 | 202.1 | - | - | 0 | - |
| - | - | 2445 | 203.1 | - | - | 0 | - |
| - | - | 1486 | 203.3 | - | - | 0 | - |
| - | - | 1464 | 204.1 | - | - | 0 | - |
| - | - | 2.096E+04 | 205.1 | - | - | 0 | - |
| - | - | 1049 | 205.1 | - | - | 0 | - |
| - | - | 2774 | 205.1 | - | - | 0 | - |
| - | - | 2594 | 206.1 | - | - | 0 | - |
| - | - | 3795 | 206.1 | - | - | 0 | - |
| - | - | 8106 | 207.2 | - | - | 0 | - |
| - | - | 1876 | 208.1 | - | - | 0 | - |
| - | - | 1440 | 209.1 | - | - | 0 | - |
| - | - | 7632 | 210.1 | - | - | 0 | - |
| 11 | y | 5660 | 211.1 | 0.0006644 | 3.147 | +2 | 3 |
| - | - | 1253 | 211.1 | - | - | 0 | - |
| - | - | 1405 | 211.1 | - | - | 0 | - |
| - | - | 1316 | 212.1 | - | - | 0 | - |
| - | - | 5098 | 212.1 | - | - | 0 | - |
| - | - | 1.643E+04 | 213.1 | - | - | 0 | - |
| - | - | 1889 | 213.2 | - | - | 0 | - |
| - | - | 2021 | 214.1 | - | - | 0 | - |
| - | - | 3251 | 215.1 | - | - | 0 | - |
| - | - | 6183 | 216.1 | - | - | 0 | - |
| - | - | 1359 | 217.1 | - | - | 0 | - |
| - | - | 1962 | 221.1 | - | - | 0 | - |
| - | - | 6887 | 221.1 | - | - | 0 | - |
| - | - | 3759 | 221.1 | - | - | 0 | - |
| - | - | 4023 | 221.1 | - | - | 0 | - |
| - | - | 1475 | 222.1 | - | - | 0 | - |
| - | - | 6094 | 222.1 | - | - | 0 | - |
| - | - | 3641 | 223.1 | - | - | 0 | - |
| - | - | 1254 | 223.1 | - | - | 0 | - |
| - | - | 1.291E+04 | 223.1 | - | - | 0 | - |
| - | - | 7.169E+04 | 223.2 | - | - | 0 | - |
| - | - | 3203 | 224.1 | - | - | 0 | - |
| - | - | 9377 | 224.2 | - | - | 0 | - |
| - | - | 6749 | 225 | - | - | 0 | - |
| - | - | 1297 | 225.1 | - | - | 0 | - |
| - | - | 1352 | 225.1 | - | - | 0 | - |
| - | - | 2120 | 225.2 | - | - | 0 | - |
| - | - | 1254 | 226 | - | - | 0 | - |
| - | - | 1.574E+04 | 226.1 | - | - | 0 | - |
| - | - | 2062 | 226.2 | - | - | 0 | - |
| - | - | 1652 | 227 | - | - | 0 | - |
| - | - | 1.234E+04 | 227.1 | - | - | 0 | - |
| - | - | 4582 | 227.2 | - | - | 0 | - |
| - | - | 2236 | 228.1 | - | - | 0 | - |
| - | - | 9564 | 228.1 | - | - | 0 | - |
| - | - | 6036 | 229.1 | - | - | 0 | - |
| - | - | 1547 | 230.1 | - | - | 0 | - |
| - | - | 1683 | 230.2 | - | - | 0 | - |
| - | - | 1.012E+04 | 233.1 | - | - | 0 | - |
| - | - | 1222 | 233.2 | - | - | 0 | - |
| - | - | 8.54E+04 | 234.1 | - | - | 0 | - |
| - | - | 8499 | 235.1 | - | - | 0 | - |
| - | - | 1530 | 235.2 | - | - | 0 | - |
| - | - | 2024 | 235.6 | - | - | 0 | - |
| - | - | 1362 | 236.1 | - | - | 0 | - |
| - | - | 4377 | 237.1 | - | - | 0 | - |
| - | - | 1.128E+04 | 238.1 | - | - | 0 | - |
| - | - | 1881 | 238.1 | - | - | 0 | - |
| - | - | 2064 | 239.1 | - | - | 0 | - |
| - | - | 1.268E+04 | 239.1 | - | - | 0 | - |
| - | - | 3494 | 240.1 | - | - | 0 | - |
| - | - | 3.673E+04 | 240.1 | - | - | 0 | - |
| - | - | 2437 | 241.1 | - | - | 0 | - |
| - | - | 4655 | 241.1 | - | - | 0 | - |
| - | - | 3366 | 241.1 | - | - | 0 | - |
| - | - | 5326 | 241.2 | - | - | 0 | - |
| - | - | 6648 | 243.1 | - | - | 0 | - |
| - | - | 3.64E+04 | 244.1 | - | - | 0 | - |
| - | - | 1130 | 244.6 | - | - | 0 | - |
| - | - | 2761 | 245.1 | - | - | 0 | - |
| - | - | 4.148E+04 | 247.1 | - | - | 0 | - |
| - | - | 2514 | 247.1 | - | - | 0 | - |
| - | - | 1394 | 247.2 | - | - | 0 | - |
| - | - | 6471 | 248.1 | - | - | 0 | - |
| - | - | 1.89E+04 | 248.1 | - | - | 0 | - |
| 10 | y | 1361 | 248.6 | 0.0001673 | 0.6729 | +2 | 4 |
| - | - | 2.735E+04 | 249.1 | - | - | 0 | - |
| - | - | 1832 | 249.1 | - | - | 0 | - |
| - | - | 7458 | 249.1 | - | - | 0 | - |
| - | - | 1770 | 249.6 | - | - | 0 | - |
| - | - | 3963 | 250.1 | - | - | 0 | - |
| - | - | 2485 | 250.1 | - | - | 0 | - |
| - | - | 1372 | 250.2 | - | - | 0 | - |
| - | - | 2.775E+05 | 251.2 | - | - | 0 | - |
| - | - | 2066 | 252.1 | - | - | 0 | - |
| - | - | 3.92E+04 | 252.2 | - | - | 0 | - |
| - | - | 1542 | 252.2 | - | - | 0 | - |
| - | - | 1230 | 253.1 | - | - | 0 | - |
| - | - | 3631 | 253.2 | - | - | 0 | - |
| - | - | 1.654E+04 | 255.1 | - | - | 0 | - |
| - | - | 1642 | 256.1 | - | - | 0 | - |
| - | - | 3669 | 256.1 | - | - | 0 | - |
| - | - | 1502 | 256.2 | - | - | 0 | - |
| - | - | 1820 | 256.6 | - | - | 0 | - |
| - | - | 5960 | 257.1 | - | - | 0 | - |
| - | - | 1.26E+04 | 258.1 | - | - | 0 | - |
| - | - | 1800 | 259.1 | - | - | 0 | - |
| - | - | 1280 | 260.1 | - | - | 0 | - |
| - | - | 6304 | 261.1 | - | - | 0 | - |
| - | - | 3173 | 261.1 | - | - | 0 | - |
| - | - | 3181 | 262.1 | - | - | 0 | - |
| - | - | 1458 | 264.1 | - | - | 0 | - |
| - | - | 4.235E+04 | 265.1 | - | - | 0 | - |
| - | - | 1425 | 265.2 | - | - | 0 | - |
| - | - | 9904 | 266.1 | - | - | 0 | - |
| - | - | 7.786E+04 | 266.1 | - | - | 0 | - |
| - | - | 1.067E+04 | 267.1 | - | - | 0 | - |
| - | - | 1694 | 267.1 | - | - | 0 | - |
| - | - | 1898 | 267.6 | - | - | 0 | - |
| - | - | 4799 | 268.1 | - | - | 0 | - |
| - | - | 5544 | 268.2 | - | - | 0 | - |
| - | - | 2203 | 269.2 | - | - | 0 | - |
| - | - | 7062 | 269.2 | - | - | 0 | - |
| - | - | 9527 | 270.1 | - | - | 0 | - |
| - | - | 2414 | 270.6 | - | - | 0 | - |
| - | - | 1.587E+04 | 272.2 | - | - | 0 | - |
| - | - | 3145 | 273.2 | - | - | 0 | - |
| - | - | 1.466E+04 | 274.1 | - | - | 0 | - |
| - | - | 6831 | 274.1 | - | - | 0 | - |
| - | - | 2323 | 275.1 | - | - | 0 | - |
| - | - | 3314 | 275.1 | - | - | 0 | - |
| - | - | 2702 | 275.2 | - | - | 0 | - |
| - | - | 2348 | 276.1 | - | - | 0 | - |
| - | - | 2018 | 276.1 | - | - | 0 | - |
| - | - | 6262 | 276.2 | - | - | 0 | - |
| - | - | 1742 | 277.1 | - | - | 0 | - |
| - | - | 4311 | 279.1 | - | - | 0 | - |
| - | - | 7051 | 280.1 | - | - | 0 | - |
| - | - | 1495 | 280.1 | - | - | 0 | - |
| - | - | 1.102E+04 | 281.1 | - | - | 0 | - |
| - | - | 2.241E+04 | 282.2 | - | - | 0 | - |
| - | - | 2.293E+05 | 283.1 | - | - | 0 | - |
| - | - | 2565 | 283.6 | - | - | 0 | - |
| - | - | 9987 | 284.1 | - | - | 0 | - |
| - | - | 2958 | 284.1 | - | - | 0 | - |
| - | - | 3.298E+04 | 284.1 | - | - | 0 | - |
| - | - | 1954 | 285.1 | - | - | 0 | - |
| - | - | 3977 | 285.2 | - | - | 0 | - |
| - | - | 1.213E+04 | 286.1 | - | - | 0 | - |
| - | - | 2107 | 287.1 | - | - | 0 | - |
| - | - | 2844 | 287.2 | - | - | 0 | - |
| - | - | 1346 | 287.2 | - | - | 0 | - |
| - | - | 1323 | 288.1 | - | - | 0 | - |
| - | - | 2256 | 288.6 | - | - | 0 | - |
| - | - | 1448 | 289.1 | - | - | 0 | - |
| 5 | b | 2339 | 290.2 | 0.0005572 | 1.92 | +2 | 5 |
| 12 | y | 1.008E+05 | 293.1 | 0.0006325 | 2.158 | +1 | 2 |
| - | - | 2582 | 293.2 | - | - | 0 | - |
| - | - | 2.01E+04 | 294.1 | - | - | 0 | - |
| - | - | 1979 | 294.2 | - | - | 0 | - |
| - | - | 1396 | 295.1 | - | - | 0 | - |
| - | - | 3007 | 295.1 | - | - | 0 | - |
| 3 | b | 5341 | 296.2 | 0.0005501 | 1.857 | +1 | 3 |
| - | - | 1553 | 296.6 | - | - | 0 | - |
| - | - | 2124 | 297.1 | - | - | 0 | - |
| 9 | y | 4120 | 297.1 | 0.00205 | 6.899 | +2 | 5 |
| - | - | 3.118E+04 | 297.2 | - | - | 0 | - |
| 7 | y | 3049 | 298.1 | 0.005184 | 17.39 | +3 | 7 |
| - | - | 5619 | 298.2 | - | - | 0 | - |
| - | - | 8.133E+04 | 299.1 | - | - | 0 | - |
| - | - | 1603 | 300.1 | - | - | 0 | - |
| - | - | 3.703E+04 | 300.2 | - | - | 0 | - |
| - | - | 3208 | 301.1 | - | - | 0 | - |
| - | - | 9.923E+04 | 301.2 | - | - | 0 | - |
| - | - | 5080 | 301.2 | - | - | 0 | - |
| - | - | 1.827E+04 | 302.1 | - | - | 0 | - |
| - | - | 1.393E+04 | 302.2 | - | - | 0 | - |
| - | - | 2694 | 303.1 | - | - | 0 | - |
| - | - | 7152 | 305.6 | - | - | 0 | - |
| 9 | y | 2.415E+04 | 306.1 | 0.0007655 | 2.5 | +2 | 5 |
| - | - | 7449 | 306.6 | - | - | 0 | - |
| - | - | 4648 | 307.1 | - | - | 0 | - |
| - | - | 2281 | 309.2 | - | - | 0 | - |
| 12 | y | 1.4E+04 | 311.1 | 0.0009015 | 2.898 | +1 | 2 |
| - | - | 3099 | 311.1 | - | - | 0 | - |
| - | - | 2312 | 312.1 | - | - | 0 | - |
| - | - | 4824 | 314.2 | - | - | 0 | - |
| 3 | b | 9534 | 314.2 | 0.0007276 | 2.316 | +1 | 3 |
| - | - | 5.485E+04 | 315.2 | - | - | 0 | - |
| - | - | 1768 | 315.2 | - | - | 0 | - |
| - | - | 8564 | 316.2 | - | - | 0 | - |
| - | - | 1422 | 317.7 | - | - | 0 | - |
| - | - | 3588 | 319.2 | - | - | 0 | - |
| - | - | 2351 | 322.2 | - | - | 0 | - |
| - | - | 3456 | 325.2 | - | - | 0 | - |
| - | - | 1.361E+04 | 326.7 | - | - | 0 | - |
| - | - | 6200 | 327.2 | - | - | 0 | - |
| - | - | 5464 | 332.2 | - | - | 0 | - |
| - | - | 1.679E+04 | 333.2 | - | - | 0 | - |
| - | - | 3962 | 334.2 | - | - | 0 | - |
| 6 | b | 3876 | 338.7 | 0.001127 | 3.326 | +2 | 6 |
| 6 | b | 1876 | 339.2 | 0.005029 | 14.83 | +2 | 6 |
| - | - | 3375 | 339.7 | - | - | 0 | - |
| - | - | 4951 | 340.7 | - | - | 0 | - |
| - | - | 1335 | 341 | - | - | 0 | - |
| - | - | 2194 | 341.2 | - | - | 0 | - |
| - | - | 4513 | 342.2 | - | - | 0 | - |
| - | - | 2769 | 343.2 | - | - | 0 | - |
| - | - | 1581 | 343.2 | - | - | 0 | - |
| - | - | 4752 | 344.2 | - | - | 0 | - |
| - | - | 1.154E+04 | 345 | - | - | 0 | - |
| - | - | 8390 | 345.1 | - | - | 0 | - |
| - | - | 3.611E+04 | 346.1 | - | - | 0 | - |
| - | - | 1609 | 346.1 | - | - | 0 | - |
| - | - | 5177 | 347.1 | - | - | 0 | - |
| 6 | b | 1.409E+04 | 347.7 | 0.0006355 | 1.828 | +2 | 6 |
| - | - | 4891 | 348.2 | - | - | 0 | - |
| - | - | 2192 | 349.2 | - | - | 0 | - |
| - | - | 2.511E+04 | 350.2 | - | - | 0 | - |
| - | - | 4552 | 351.2 | - | - | 0 | - |
| - | - | 1449 | 352.2 | - | - | 0 | - |
| - | - | 3175 | 352.7 | - | - | 0 | - |
| - | - | 2045 | 353.2 | - | - | 0 | - |
| 8 | y | 3238 | 353.7 | 0.000759 | 2.146 | +2 | 6 |
| - | - | 3122 | 354.2 | - | - | 0 | - |
| - | - | 1629 | 355.1 | - | - | 0 | - |
| - | - | 1.29E+04 | 359 | - | - | 0 | - |
| - | - | 2043 | 360.2 | - | - | 0 | - |
| - | - | 9196 | 360.7 | - | - | 0 | - |
| - | - | 1939 | 361.2 | - | - | 0 | - |
| - | - | 8936 | 361.2 | - | - | 0 | - |
| - | - | 1.254E+04 | 362.2 | - | - | 0 | - |
| 8 | y | 4.709E+04 | 362.7 | 0.0009698 | 2.674 | +2 | 6 |
| - | - | 4.896E+04 | 363.1 | - | - | 0 | - |
| - | - | 1.805E+04 | 363.2 | - | - | 0 | - |
| - | - | 3845 | 363.7 | - | - | 0 | - |
| - | - | 8667 | 364.1 | - | - | 0 | - |
| - | - | 8487 | 364.1 | - | - | 0 | - |
| - | - | 1659 | 365.1 | - | - | 0 | - |
| - | - | 2965 | 366.2 | - | - | 0 | - |
| - | - | 4548 | 366.2 | - | - | 0 | - |
| - | - | 3599 | 367.1 | - | - | 0 | - |
| - | - | 2554 | 367.2 | - | - | 0 | - |
| - | - | 4071 | 367.2 | - | - | 0 | - |
| - | - | 2403 | 368.2 | - | - | 0 | - |
| - | - | 1456 | 370.1 | - | - | 0 | - |
| - | - | 1547 | 370.2 | - | - | 0 | - |
| - | - | 3264 | 373.2 | - | - | 0 | - |
| - | - | 4608 | 376.2 | - | - | 0 | - |
| - | - | 5234 | 376.2 | - | - | 0 | - |
| - | - | 2692 | 376.7 | - | - | 0 | - |
| - | - | 1693 | 377.1 | - | - | 0 | - |
| - | - | 5380 | 377.1 | - | - | 0 | - |
| - | - | 1.862E+04 | 378.2 | - | - | 0 | - |
| - | - | 7.245E+04 | 379.2 | - | - | 0 | - |
| - | - | 1895 | 380.2 | - | - | 0 | - |
| - | - | 1.515E+04 | 380.2 | - | - | 0 | - |
| - | - | 1.367E+05 | 381.2 | - | - | 0 | - |
| - | - | 5585 | 381.7 | - | - | 0 | - |
| - | - | 2.533E+04 | 382.2 | - | - | 0 | - |
| - | - | 4493 | 382.2 | - | - | 0 | - |
| - | - | 1486 | 382.2 | - | - | 0 | - |
| - | - | 3353 | 383.2 | - | - | 0 | - |
| - | - | 6484 | 384.2 | - | - | 0 | - |
| - | - | 6647 | 384.2 | - | - | 0 | - |
| - | - | 8030 | 385.2 | - | - | 0 | - |
| 5 | y | 5642 | 385.2 | 0.004193 | 10.88 | +3 | 9 |
| - | - | 1830 | 385.2 | - | - | 0 | - |
| - | - | 1825 | 386.2 | - | - | 0 | - |
| - | - | 2472 | 386.2 | - | - | 0 | - |
| - | - | 1562 | 386.2 | - | - | 0 | - |
| 10 | b | 3369 | 389.2 | 0.0003813 | 0.9797 | +3 | 10 |
| - | - | 5636 | 390.2 | - | - | 0 | - |
| - | - | 3225 | 390.7 | - | - | 0 | - |
| - | - | 5251 | 391.1 | - | - | 0 | - |
| - | - | 2099 | 392.2 | - | - | 0 | - |
| - | - | 1900 | 393.2 | - | - | 0 | - |
| - | - | 2.46E+04 | 394.2 | - | - | 0 | - |
| - | - | 5505 | 395.1 | - | - | 0 | - |
| - | - | 9836 | 395.2 | - | - | 0 | - |
| - | - | 4433 | 395.2 | - | - | 0 | - |
| - | - | 3482 | 396.1 | - | - | 0 | - |
| - | - | 2.369E+04 | 396.2 | - | - | 0 | - |
| - | - | 4565 | 397.2 | - | - | 0 | - |
| - | - | 1538 | 397.7 | - | - | 0 | - |
| - | - | 1.009E+04 | 398.2 | - | - | 0 | - |
| - | - | 2040 | 399.2 | - | - | 0 | - |
| - | - | 1469 | 401.2 | - | - | 0 | - |
| - | - | 1922 | 401.2 | - | - | 0 | - |
| - | - | 8237 | 402.2 | - | - | 0 | - |
| - | - | 1.294E+04 | 402.2 | - | - | 0 | - |
| - | - | 6393 | 403.2 | - | - | 0 | - |
| - | - | 1564 | 403.7 | - | - | 0 | - |
| - | - | 5930 | 405.3 | - | - | 0 | - |
| - | - | 2899 | 406.2 | - | - | 0 | - |
| - | - | 3538 | 407.2 | - | - | 0 | - |
| - | - | 1688 | 408.7 | - | - | 0 | - |
| - | - | 1.063E+04 | 409.1 | - | - | 0 | - |
| - | - | 2015 | 409.2 | - | - | 0 | - |
| - | - | 1395 | 409.7 | - | - | 0 | - |
| - | - | 1.299E+04 | 411.2 | - | - | 0 | - |
| - | - | 8.17E+04 | 412.2 | - | - | 0 | - |
| - | - | 6442 | 413.1 | - | - | 0 | - |
| - | - | 9453 | 413.2 | - | - | 0 | - |
| - | - | 1.37E+04 | 413.2 | - | - | 0 | - |
| - | - | 3024 | 413.3 | - | - | 0 | - |
| - | - | 2608 | 414.2 | - | - | 0 | - |
| - | - | 2.697E+04 | 414.2 | - | - | 0 | - |
| - | - | 2.109E+04 | 415 | - | - | 0 | - |
| - | - | 3948 | 415.2 | - | - | 0 | - |
| - | - | 6214 | 415.2 | - | - | 0 | - |
| - | - | 1.354E+04 | 417.7 | - | - | 0 | - |
| - | - | 2.482E+04 | 418.2 | - | - | 0 | - |
| - | - | 1.363E+04 | 418.7 | - | - | 0 | - |
| - | - | 4831 | 419.2 | - | - | 0 | - |
| 11 | y | 3268 | 421.2 | 0.001128 | 2.679 | +1 | 3 |
| 11 | y | 2100 | 422.2 | 3.802E-05 | 0.09005 | +1 | 3 |
| - | - | 2752 | 422.7 | - | - | 0 | - |
| - | - | 5912 | 423.2 | - | - | 0 | - |
| - | - | 4.009E+04 | 423.3 | - | - | 0 | - |
| - | - | 7858 | 424.3 | - | - | 0 | - |
| - | - | 2717 | 424.7 | - | - | 0 | - |
| - | - | 2363 | 426.2 | - | - | 0 | - |
| - | - | 2.492E+05 | 426.7 | - | - | 0 | - |
| - | - | 1.226E+05 | 427.2 | - | - | 0 | - |
| - | - | 3.75E+04 | 427.7 | - | - | 0 | - |
| - | - | 7159 | 428.2 | - | - | 0 | - |
| - | - | 2032 | 428.7 | - | - | 0 | - |
| - | - | 6.566E+04 | 429.2 | - | - | 0 | - |
| - | - | 2100 | 429.7 | - | - | 0 | - |
| - | - | 2.009E+05 | 430.2 | - | - | 0 | - |
| - | - | 3.968E+04 | 431.2 | - | - | 0 | - |
| 7 | b | 3.268E+04 | 431.7 | 0.001082 | 2.506 | +2 | 7 |
| 7 | b | 1.625E+04 | 432.2 | 0.004954 | 11.46 | +2 | 7 |
| - | - | 5757 | 432.7 | - | - | 0 | - |
| - | - | 2021 | 433.2 | - | - | 0 | - |
| 4 | b | 7050 | 433.3 | 0.0005371 | 1.24 | +1 | 4 |
| - | - | 1809 | 434.3 | - | - | 0 | - |
| - | - | 5066 | 438.2 | - | - | 0 | - |
| - | - | 3592 | 438.7 | - | - | 0 | - |
| 11 | y | 6.518E+04 | 439.2 | 0.001367 | 3.112 | +1 | 3 |
| - | - | 4092 | 440.2 | - | - | 0 | - |
| - | - | 1.489E+04 | 440.2 | - | - | 0 | - |
| 7 | b | 1.203E+05 | 440.7 | 0.001049 | 2.379 | +2 | 7 |
| - | - | 6.37E+04 | 441.2 | - | - | 0 | - |
| - | - | 1.858E+04 | 441.7 | - | - | 0 | - |
| - | - | 3326 | 442.2 | - | - | 0 | - |
| - | - | 1958 | 444.2 | - | - | 0 | - |
| 7 | y | 3572 | 446.7 | 0.000226 | 0.506 | +2 | 7 |
| - | - | 2462 | 446.7 | - | - | 0 | - |
| 7 | y | 7218 | 447.2 | 0.002725 | 6.094 | +2 | 7 |
| - | - | 2128 | 447.7 | - | - | 0 | - |
| - | - | 1.038E+04 | 448.2 | - | - | 0 | - |
| - | - | 1760 | 448.3 | - | - | 0 | - |
| - | - | 4099 | 449.2 | - | - | 0 | - |
| - | - | 2042 | 450.2 | - | - | 0 | - |
| - | - | 2682 | 450.2 | - | - | 0 | - |
| - | - | 3283 | 450.3 | - | - | 0 | - |
| 4 | b | 7.033E+04 | 451.3 | 0.001264 | 2.801 | +1 | 4 |
| - | - | 1.601E+04 | 452.3 | - | - | 0 | - |
| - | - | 1583 | 452.7 | - | - | 0 | - |
| - | - | 5402 | 455.3 | - | - | 0 | - |
| 7 | y | 3801 | 455.7 | 0.001627 | 3.57 | +2 | 7 |
| - | - | 6717 | 459.2 | - | - | 0 | - |
| - | - | 7841 | 460.8 | - | - | 0 | - |
| - | - | 4673 | 461.2 | - | - | 0 | - |
| - | - | 7120 | 461.3 | - | - | 0 | - |
| 3 | y | 1376 | 462.9 | 0.00664 | 14.35 | +3 | 11 |
| - | - | 1683 | 465.7 | - | - | 0 | - |
| - | - | 2885 | 466.2 | - | - | 0 | - |
| - | - | 2023 | 466.2 | - | - | 0 | - |
| - | - | 1.652E+04 | 468.3 | - | - | 0 | - |
| - | - | 3319 | 469.3 | - | - | 0 | - |
| - | - | 1.293E+04 | 474.3 | - | - | 0 | - |
| - | - | 1.865E+04 | 474.8 | - | - | 0 | - |
| - | - | 1.005E+04 | 475.3 | - | - | 0 | - |
| - | - | 1661 | 475.8 | - | - | 0 | - |
| - | - | 5945 | 476.2 | - | - | 0 | - |
| - | - | 1.148E+04 | 477.2 | - | - | 0 | - |
| 10 | y | 8796 | 478.2 | 0.0004378 | 0.9155 | +1 | 4 |
| - | - | 8747 | 478.3 | - | - | 0 | - |
| - | - | 2112 | 479.2 | - | - | 0 | - |
| - | - | 1882 | 479.7 | - | - | 0 | - |
| - | - | 1997 | 482.3 | - | - | 0 | - |
| - | - | 4.228E+04 | 483.3 | - | - | 0 | - |
| - | - | 3.057E+04 | 483.8 | - | - | 0 | - |
| - | - | 3229 | 484.2 | - | - | 0 | - |
| - | - | 6797 | 484.3 | - | - | 0 | - |
| - | - | 1927 | 484.8 | - | - | 0 | - |
| - | - | 1458 | 485.2 | - | - | 0 | - |
| - | - | 2071 | 488.1 | - | - | 0 | - |
| 8 | b | 1.303E+04 | 488.3 | 0.001652 | 3.384 | +2 | 8 |
| 8 | b | 8794 | 488.7 | 0.006959 | 14.24 | +2 | 8 |
| - | - | 2696 | 489.3 | - | - | 0 | - |
| - | - | 1830 | 490.2 | - | - | 0 | - |
| - | - | 1.474E+05 | 494.2 | - | - | 0 | - |
| - | - | 1620 | 495.2 | - | - | 0 | - |
| - | - | 3.379E+04 | 495.2 | - | - | 0 | - |
| 10 | y | 1.598E+05 | 496.2 | 0.001082 | 2.181 | +1 | 4 |
| - | - | 4.064E+04 | 497.2 | - | - | 0 | - |
| - | - | 1.823E+04 | 497.8 | - | - | 0 | - |
| - | - | 7749 | 498.2 | - | - | 0 | - |
| - | - | 2062 | 502.2 | - | - | 0 | - |
| - | - | 2.17E+04 | 503.1 | - | - | 0 | - |
| - | - | 6065 | 504.2 | - | - | 0 | - |
| - | - | 3517 | 505.2 | - | - | 0 | - |
| - | - | 2506 | 505.2 | - | - | 0 | - |
| - | - | 3702 | 506.2 | - | - | 0 | - |
| - | - | 2170 | 506.3 | - | - | 0 | - |
| - | - | 5428 | 507.3 | - | - | 0 | - |
| - | - | 1669 | 508.3 | - | - | 0 | - |
| - | - | 1.217E+04 | 511.3 | - | - | 0 | - |
| - | - | 2445 | 512.3 | - | - | 0 | - |
| - | - | 3222 | 513.2 | - | - | 0 | - |
| - | - | 1960 | 515.3 | - | - | 0 | - |
| - | - | 1.024E+04 | 517.3 | - | - | 0 | - |
| - | - | 3818 | 518.3 | - | - | 0 | - |
| - | - | 7873 | 519.1 | - | - | 0 | - |
| - | - | 2524 | 520.3 | - | - | 0 | - |
| - | - | 1633 | 522.2 | - | - | 0 | - |
| - | - | 3434 | 523.3 | - | - | 0 | - |
| - | - | 1474 | 523.6 | - | - | 0 | - |
| - | - | 3495 | 524.3 | - | - | 0 | - |
| - | - | 2.415E+04 | 525.3 | - | - | 0 | - |
| - | - | 2360 | 526.2 | - | - | 0 | - |
| - | - | 7831 | 526.3 | - | - | 0 | - |
| - | - | 1579 | 527.3 | - | - | 0 | - |
| - | - | 4253 | 528.9 | - | - | 0 | - |
| 0 | Precursor | 9639 | 529.3 | 0.003061 | 5.783 | +3 | -1 |
| 0 | Precursor | 9641 | 529.6 | 0.008186 | 15.46 | +3 | -1 |
| - | - | 5433 | 529.9 | - | - | 0 | - |
| - | - | 1440 | 530.2 | - | - | 0 | - |
| - | - | 1402 | 531.2 | - | - | 0 | - |
| - | - | 5961 | 532.2 | - | - | 0 | - |
| - | - | 9732 | 534.3 | - | - | 0 | - |
| - | - | 2238 | 534.9 | - | - | 0 | - |
| 0 | Precursor | 6360 | 535.3 | 0.002652 | 4.955 | +3 | -1 |
| - | - | 3094 | 535.4 | - | - | 0 | - |
| - | - | 6148 | 535.6 | - | - | 0 | - |
| - | - | 3202 | 535.9 | - | - | 0 | - |
| - | - | 4766 | 536.2 | - | - | 0 | - |
| - | - | 1756 | 537.2 | - | - | 0 | - |
| - | - | 2111 | 537.7 | - | - | 0 | - |
| - | - | 2484 | 538.3 | - | - | 0 | - |
| - | - | 2309 | 538.8 | - | - | 0 | - |
| - | - | 2.822E+04 | 539.2 | - | - | 0 | - |
| - | - | 8893 | 540.2 | - | - | 0 | - |
| - | - | 1578 | 541.2 | - | - | 0 | - |
| - | - | 1.524E+04 | 542.3 | - | - | 0 | - |
| - | - | 3.394E+04 | 543.3 | - | - | 0 | - |
| - | - | 9790 | 544.3 | - | - | 0 | - |
| - | - | 2147 | 545.3 | - | - | 0 | - |
| 9 | b | 2342 | 545.8 | 0.008035 | 14.72 | +2 | 9 |
| 9 | b | 5151 | 546.3 | 0.01048 | 19.18 | +2 | 9 |
| - | - | 2009 | 547.2 | - | - | 0 | - |
| - | - | 7383 | 549.2 | - | - | 0 | - |
| - | - | 2063 | 550.2 | - | - | 0 | - |
| - | - | 8025 | 551.3 | - | - | 0 | - |
| - | - | 1833 | 551.3 | - | - | 0 | - |
| - | - | 3867 | 552.3 | - | - | 0 | - |
| 9 | b | 7459 | 554.8 | 0.009533 | 17.18 | +2 | 9 |
| - | - | 7854 | 555.3 | - | - | 0 | - |
| - | - | 4246 | 555.8 | - | - | 0 | - |
| - | - | 1761 | 558.2 | - | - | 0 | - |
| 5 | b | 6777 | 561.3 | 0.000828 | 1.475 | +1 | 5 |
| 5 | b | 2193 | 562.3 | 0.0004605 | 0.819 | +1 | 5 |
| - | - | 3544 | 563.3 | - | - | 0 | - |
| - | - | 3514 | 564.3 | - | - | 0 | - |
| - | - | 2679 | 564.3 | - | - | 0 | - |
| - | - | 1.295E+04 | 565.3 | - | - | 0 | - |
| - | - | 2443 | 566.3 | - | - | 0 | - |
| - | - | 3.473E+04 | 567.2 | - | - | 0 | - |
| - | - | 1.034E+04 | 568.2 | - | - | 0 | - |
| - | - | 1870 | 569.2 | - | - | 0 | - |
| - | - | 1.106E+04 | 575.2 | - | - | 0 | - |
| - | - | 5136 | 576.2 | - | - | 0 | - |
| - | - | 1478 | 577.2 | - | - | 0 | - |
| - | - | 1342 | 577.8 | - | - | 0 | - |
| 5 | b | 4.843E+04 | 579.3 | 0.0009445 | 1.63 | +1 | 5 |
| - | - | 1.237E+04 | 580.3 | - | - | 0 | - |
| - | - | 1691 | 581.3 | - | - | 0 | - |
| - | - | 3179 | 581.3 | - | - | 0 | - |
| - | - | 1910 | 582.3 | - | - | 0 | - |
| - | - | 1815 | 589.3 | - | - | 0 | - |
| - | - | 1.03E+04 | 592.3 | - | - | 0 | - |
| - | - | 4968 | 592.3 | - | - | 0 | - |
| 9 | y | 6.226E+04 | 593.3 | 0.0008784 | 1.481 | +1 | 5 |
| - | - | 1.962E+04 | 593.3 | - | - | 0 | - |
| - | - | 2.351E+04 | 594.3 | - | - | 0 | - |
| - | - | 3603 | 594.3 | - | - | 0 | - |
| - | - | 4527 | 595.3 | - | - | 0 | - |
| - | - | 2751 | 596.2 | - | - | 0 | - |
| - | - | 2222 | 596.4 | - | - | 0 | - |
| - | - | 3332 | 597.8 | - | - | 0 | - |
| - | - | 2150 | 598.3 | - | - | 0 | - |
| - | - | 4024 | 602.3 | - | - | 0 | - |
| - | - | 9263 | 602.8 | - | - | 0 | - |
| - | - | 9849 | 603.3 | - | - | 0 | - |
| - | - | 3157 | 603.8 | - | - | 0 | - |
| - | - | 1.647E+05 | 610.3 | - | - | 0 | - |
| 9 | y | 5.782E+05 | 611.3 | 0.002032 | 3.325 | +1 | 5 |
| - | - | 2.88E+04 | 611.8 | - | - | 0 | - |
| - | - | 1.858E+05 | 612.3 | - | - | 0 | - |
| - | - | 8146 | 612.8 | - | - | 0 | - |
| - | - | 4.1E+04 | 613.3 | - | - | 0 | - |
| - | - | 6302 | 614.3 | - | - | 0 | - |
| - | - | 2746 | 617.3 | - | - | 0 | - |
| - | - | 2136 | 620.8 | - | - | 0 | - |
| - | - | 1.062E+04 | 621.3 | - | - | 0 | - |
| - | - | 3281 | 621.3 | - | - | 0 | - |
| - | - | 2564 | 622.3 | - | - | 0 | - |
| - | - | 1952 | 622.4 | - | - | 0 | - |
| - | - | 2379 | 628.3 | - | - | 0 | - |
| - | - | 2212 | 630.3 | - | - | 0 | - |
| - | - | 7679 | 634.3 | - | - | 0 | - |
| - | - | 9472 | 635.3 | - | - | 0 | - |
| - | - | 4659 | 636.3 | - | - | 0 | - |
| 4 | y | 7689 | 636.8 | 0.002015 | 3.164 | +2 | 10 |
| 4 | y | 3384 | 637.3 | 0.009519 | 14.94 | +2 | 10 |
| - | - | 1862 | 637.3 | - | - | 0 | - |
| - | - | 2177 | 637.8 | - | - | 0 | - |
| 11 | b | 3161 | 638.3 | 0.004369 | 6.845 | +2 | 11 |
| - | - | 3224 | 640.3 | - | - | 0 | - |
| - | - | 9431 | 645.3 | - | - | 0 | - |
| 4 | y | 3.011E+04 | 645.8 | 0.003446 | 5.337 | +2 | 10 |
| - | - | 2.342E+04 | 646.3 | - | - | 0 | - |
| - | - | 7777 | 646.8 | - | - | 0 | - |
| - | - | 2345 | 648.3 | - | - | 0 | - |
| - | - | 2883 | 650.3 | - | - | 0 | - |
| - | - | 3178 | 650.3 | - | - | 0 | - |
| - | - | 6.023E+04 | 652.3 | - | - | 0 | - |
| - | - | 2.602E+04 | 653.3 | - | - | 0 | - |
| - | - | 4558 | 654.3 | - | - | 0 | - |
| - | - | 2137 | 658.3 | - | - | 0 | - |
| - | - | 2122 | 659.3 | - | - | 0 | - |
| - | - | 1399 | 660.3 | - | - | 0 | - |
| - | - | 2235 | 661.3 | - | - | 0 | - |
| - | - | 2.234E+04 | 662.3 | - | - | 0 | - |
| - | - | 1427 | 662.8 | - | - | 0 | - |
| - | - | 1.368E+04 | 663.3 | - | - | 0 | - |
| - | - | 3426 | 664.3 | - | - | 0 | - |
| - | - | 2094 | 664.4 | - | - | 0 | - |
| - | - | 1670 | 666.3 | - | - | 0 | - |
| - | - | 1996 | 668.3 | - | - | 0 | - |
| - | - | 2265 | 670.3 | - | - | 0 | - |
| - | - | 1492 | 670.8 | - | - | 0 | - |
| - | - | 1963 | 675.8 | - | - | 0 | - |
| 6 | b | 3.155E+04 | 676.3 | 0.00129 | 1.907 | +1 | 6 |
| 6 | b | 1.657E+04 | 677.3 | 0.009462 | 13.97 | +1 | 6 |
| - | - | 8001 | 678.3 | - | - | 0 | - |
| - | - | 3566 | 679.3 | - | - | 0 | - |
| - | - | 2499 | 679.8 | - | - | 0 | - |
| - | - | 1.096E+05 | 680.3 | - | - | 0 | - |
| - | - | 4.316E+04 | 681.3 | - | - | 0 | - |
| - | - | 1.196E+04 | 682.3 | - | - | 0 | - |
| - | - | 1673 | 683.3 | - | - | 0 | - |
| - | - | 4147 | 684.3 | - | - | 0 | - |
| - | - | 1.028E+04 | 684.8 | - | - | 0 | - |
| - | - | 5794 | 685.3 | - | - | 0 | - |
| - | - | 1865 | 685.8 | - | - | 0 | - |
| - | - | 3403 | 688.3 | - | - | 0 | - |
| - | - | 1747 | 689.3 | - | - | 0 | - |
| - | - | 5328 | 692.8 | - | - | 0 | - |
| 3 | y | 3.249E+04 | 693.3 | 0.002891 | 4.169 | +2 | 11 |
| 3 | y | 3.479E+04 | 693.8 | 0.009235 | 13.31 | +2 | 11 |
| 6 | b | 2.954E+05 | 694.4 | 0.0004907 | 0.7067 | +1 | 6 |
| - | - | 4659 | 694.8 | - | - | 0 | - |
| - | - | 1.171E+05 | 695.4 | - | - | 0 | - |
| - | - | 2.723E+04 | 696.4 | - | - | 0 | - |
| - | - | 6091 | 697.4 | - | - | 0 | - |
| - | - | 2158 | 698.4 | - | - | 0 | - |
| - | - | 5656 | 701.3 | - | - | 0 | - |
| - | - | 2729 | 701.8 | - | - | 0 | - |
| - | - | 7.243E+04 | 701.8 | - | - | 0 | - |
| 3 | y | 1.832E+05 | 702.3 | 0.003834 | 5.459 | +2 | 11 |
| 12 | b | 1.448E+05 | 702.8 | 0.01168 | 16.62 | +2 | 12 |
| 12 | b | 6.136E+04 | 703.3 | 0.001611 | 2.29 | +2 | 12 |
| - | - | 1.769E+04 | 703.8 | - | - | 0 | - |
| - | - | 9272 | 704.3 | - | - | 0 | - |
| - | - | 5250 | 705.4 | - | - | 0 | - |
| 8 | y | 2.028E+04 | 706.3 | 0.002508 | 3.55 | +1 | 6 |
| - | - | 8450 | 707.3 | - | - | 0 | - |
| - | - | 2066 | 708.3 | - | - | 0 | - |
| - | - | 1895 | 710.9 | - | - | 0 | - |
| - | - | 1.549E+04 | 711.4 | - | - | 0 | - |
| 12 | b | 1.347E+04 | 711.9 | 0.007332 | 10.3 | +2 | 12 |
| - | - | 7630 | 712.4 | - | - | 0 | - |
| - | - | 3199 | 712.9 | - | - | 0 | - |
| - | - | 1916 | 713.4 | - | - | 0 | - |
| - | - | 1779 | 715.4 | - | - | 0 | - |
| - | - | 3428 | 720.4 | - | - | 0 | - |
| - | - | 1.445E+04 | 721.4 | - | - | 0 | - |
| - | - | 4661 | 722.4 | - | - | 0 | - |
| - | - | 4.414E+04 | 723.4 | - | - | 0 | - |
| 8 | y | 1.783E+05 | 724.4 | 0.002441 | 3.37 | +1 | 6 |
| - | - | 7.575E+04 | 725.4 | - | - | 0 | - |
| - | - | 1.914E+04 | 726.4 | - | - | 0 | - |
| - | - | 2259 | 727.4 | - | - | 0 | - |
| - | - | 2319 | 728.4 | - | - | 0 | - |
| - | - | 1616 | 728.9 | - | - | 0 | - |
| - | - | 5667 | 729.4 | - | - | 0 | - |
| - | - | 3531 | 730.4 | - | - | 0 | - |
| - | - | 8223 | 733.4 | - | - | 0 | - |
| - | - | 7187 | 734.4 | - | - | 0 | - |
| - | - | 2516 | 735.4 | - | - | 0 | - |
| - | - | 1771 | 737.9 | - | - | 0 | - |
| - | - | 2558 | 739.4 | - | - | 0 | - |
| 2 | y | 2151 | 742.9 | 0.0007377 | 0.993 | +2 | 12 |
| - | - | 5034 | 743.4 | - | - | 0 | - |
| - | - | 3574 | 744.4 | - | - | 0 | - |
| - | - | 3528 | 747.4 | - | - | 0 | - |
| - | - | 3286 | 748.4 | - | - | 0 | - |
| - | - | 3563 | 749.3 | - | - | 0 | - |
| - | - | 3.558E+04 | 751.4 | - | - | 0 | - |
| 2 | y | 1.562E+04 | 751.9 | 0.003074 | 4.089 | +2 | 12 |
| - | - | 2.464E+04 | 752.4 | - | - | 0 | - |
| - | - | 6773 | 752.9 | - | - | 0 | - |
| - | - | 8979 | 753.4 | - | - | 0 | - |
| - | - | 3237 | 753.9 | - | - | 0 | - |
| - | - | 1576 | 760.3 | - | - | 0 | - |
| - | - | 1981 | 762.4 | - | - | 0 | - |
| - | - | 2.289E+04 | 763.4 | - | - | 0 | - |
| - | - | 5666 | 764.4 | - | - | 0 | - |
| - | - | 7792 | 765.4 | - | - | 0 | - |
| - | - | 2663 | 766.4 | - | - | 0 | - |
| - | - | 6954 | 767.3 | - | - | 0 | - |
| - | - | 2321 | 769.4 | - | - | 0 | - |
| - | - | 2287 | 770.4 | - | - | 0 | - |
| - | - | 4986 | 775.4 | - | - | 0 | - |
| - | - | 3165 | 776.4 | - | - | 0 | - |
| - | - | 2083 | 778.4 | - | - | 0 | - |
| - | - | 1.775E+04 | 779.4 | - | - | 0 | - |
| - | - | 9070 | 780.4 | - | - | 0 | - |
| - | - | 9251 | 781.4 | - | - | 0 | - |
| - | - | 4970 | 782.4 | - | - | 0 | - |
| - | - | 1790 | 791.4 | - | - | 0 | - |
| 0 | Precursor | 3.352E+04 | 793.4 | 0.01369 | 17.26 | +2 | -1 |
| - | - | 1.478E+04 | 794.4 | - | - | 0 | - |
| - | - | 3590 | 795.3 | - | - | 0 | - |
| - | - | 7703 | 807.4 | - | - | 0 | - |
| - | - | 3721 | 807.4 | - | - | 0 | - |
| - | - | 4395 | 808.4 | - | - | 0 | - |
| - | - | 4146 | 818.4 | - | - | 0 | - |
| - | - | 3801 | 819.4 | - | - | 0 | - |
| - | - | 3137 | 825.4 | - | - | 0 | - |
| - | - | 2643 | 826.4 | - | - | 0 | - |
| - | - | 7742 | 834.4 | - | - | 0 | - |
| - | - | 1.187E+04 | 835.4 | - | - | 0 | - |
| - | - | 2.419E+04 | 836.4 | - | - | 0 | - |
| - | - | 1.041E+04 | 837.5 | - | - | 0 | - |
| - | - | 1546 | 838.3 | - | - | 0 | - |
| - | - | 1893 | 838.5 | - | - | 0 | - |
| - | - | 3903 | 843.4 | - | - | 0 | - |
| - | - | 4908 | 844.4 | - | - | 0 | - |
| - | - | 5398 | 845.4 | - | - | 0 | - |
| - | - | 1.006E+04 | 846.4 | - | - | 0 | - |
| - | - | 6822 | 847.4 | - | - | 0 | - |
| - | - | 2694 | 848.4 | - | - | 0 | - |
| - | - | 1946 | 851.4 | - | - | 0 | - |
| - | - | 7.624E+04 | 852.4 | - | - | 0 | - |
| - | - | 3.689E+04 | 853.4 | - | - | 0 | - |
| - | - | 1.056E+04 | 854.4 | - | - | 0 | - |
| 7 | b | 4.154E+04 | 862.4 | 0.000468 | 0.5427 | +1 | 7 |
| 7 | b | 3.309E+04 | 863.4 | 0.01249 | 14.46 | +1 | 7 |
| - | - | 6.13E+04 | 864.5 | - | - | 0 | - |
| - | - | 3.052E+04 | 865.5 | - | - | 0 | - |
| - | - | 9746 | 866.5 | - | - | 0 | - |
| - | - | 2185 | 867.5 | - | - | 0 | - |
| - | - | 5716 | 876.4 | - | - | 0 | - |
| - | - | 3110 | 877.4 | - | - | 0 | - |
| 7 | b | 3.828E+05 | 880.4 | 0.0006455 | 0.7332 | +1 | 7 |
| - | - | 1.951E+05 | 881.4 | - | - | 0 | - |
| - | - | 5.646E+04 | 882.4 | - | - | 0 | - |
| - | - | 9687 | 883.4 | - | - | 0 | - |
| - | - | 1943 | 884.4 | - | - | 0 | - |
| - | - | 1731 | 890.4 | - | - | 0 | - |
| - | - | 6567 | 892.5 | - | - | 0 | - |
| - | - | 3629 | 893.5 | - | - | 0 | - |
| - | - | 1847 | 894.5 | - | - | 0 | - |
| - | - | 2039 | 895.5 | - | - | 0 | - |
| - | - | 2034 | 896.5 | - | - | 0 | - |
| - | - | 3621 | 907.4 | - | - | 0 | - |
| - | - | 1.066E+04 | 908.4 | - | - | 0 | - |
| - | - | 1.127E+04 | 909.4 | - | - | 0 | - |
| 7 | y | 1.791E+04 | 910.4 | 0.004915 | 5.399 | +1 | 7 |
| - | - | 8497 | 911.4 | - | - | 0 | - |
| - | - | 3067 | 912.4 | - | - | 0 | - |
| - | - | 3519 | 936.4 | - | - | 0 | - |
| - | - | 3668 | 937.4 | - | - | 0 | - |
| - | - | 1861 | 947.4 | - | - | 0 | - |
| - | - | 1814 | 949.4 | - | - | 0 | - |
| - | - | 4419 | 954.4 | - | - | 0 | - |
| - | - | 6984 | 955.4 | - | - | 0 | - |
| - | - | 2181 | 956.4 | - | - | 0 | - |
| - | - | 4087 | 964.5 | - | - | 0 | - |
| - | - | 5636 | 965.5 | - | - | 0 | - |
| - | - | 2355 | 966.4 | - | - | 0 | - |
| - | - | 4942 | 972.4 | - | - | 0 | - |
| - | - | 2168 | 973.4 | - | - | 0 | - |
| 8 | b | 4037 | 975.5 | 0.0002664 | 0.2731 | +1 | 8 |
| 8 | b | 3621 | 976.5 | 0.01369 | 14.02 | +1 | 8 |
| - | - | 1875 | 977.5 | - | - | 0 | - |
| - | - | 1602 | 980.5 | - | - | 0 | - |
| - | - | 1988 | 981.5 | - | - | 0 | - |
| - | - | 1672 | 982.5 | - | - | 0 | - |
| 8 | b | 2.593E+04 | 993.5 | 0.000627 | 0.6311 | +1 | 8 |
| - | - | 1.716E+04 | 994.5 | - | - | 0 | - |
| - | - | 5402 | 995.5 | - | - | 0 | - |
| 6 | y | 2649 | 1007 | 0.01228 | 12.19 | +1 | 8 |
| - | - | 2427 | 1009 | - | - | 0 | - |
| - | - | 5596 | 1024 | - | - | 0 | - |
| 6 | y | 1.196E+04 | 1025 | 0.005194 | 5.065 | +1 | 8 |
| - | - | 6806 | 1026 | - | - | 0 | - |
| - | - | 1792 | 1027 | - | - | 0 | - |
| - | - | 2988 | 1076 | - | - | 0 | - |
| 9 | b | 1600 | 1092 | 0.01307 | 11.98 | +1 | 9 |
| - | - | 2079 | 1093 | - | - | 0 | - |
| - | - | 5286 | 1094 | - | - | 0 | - |
| - | - | 1981 | 1095 | - | - | 0 | - |
| 9 | b | 1840 | 1109 | 0.02094 | 18.89 | +1 | 9 |
| - | - | 2360 | 1117 | - | - | 0 | - |
| - | - | 5104 | 1118 | - | - | 0 | - |
| - | - | 1702 | 1135 | - | - | 0 | - |
| 5 | y | 1.187E+04 | 1136 | 0.001615 | 1.422 | +1 | 9 |
| 5 | y | 1.771E+04 | 1136 | 0.01027 | 9.041 | +1 | 9 |
| - | - | 6996 | 1137 | - | - | 0 | - |
| - | - | 1852 | 1138 | - | - | 0 | - |
| - | - | 1.182E+04 | 1153 | - | - | 0 | - |
| 5 | y | 2.523E+04 | 1154 | 0.0046 | 3.988 | +1 | 9 |
| - | - | 1.389E+04 | 1155 | - | - | 0 | - |
| - | - | 3927 | 1156 | - | - | 0 | - |
| 10 | b | 1823 | 1166 | 0.01924 | 16.5 | +1 | 10 |
| - | - | 1640 | 1167 | - | - | 0 | - |
| 4 | y | 2232 | 1291 | 0.01 | 7.749 | +1 | 10 |
| - | - | 1435 | 2084 | - | - | 0 | - |
| - | - | 1651 | 3081 | - | - | 0 | - |

m/z Charge Intensity FragmentType MassShift Position
120.0811767578125 0 6799.854
122.07160186767578 0 3325.8733
123.04454803466797 0 3338.2722
125.10771942138672 0 2130.2393
126.0669937133789 0 892.3032
126.09175109863281 0 2424.7383
127.05065155029297 0 13261.32
127.12360382080078 0 1600.3972
128.08236694335938 0 1055.1049
128.10740661621094 0 49825.42
129.06622314453125 0 3107.3486
129.10269165039062 0 548607.2
130.06553649902344 0 246690.06
130.10032653808594 0 2651.7256
130.1060028076172 0 35156.137
131.06324768066406 0 1879.9963
131.0688934326172 0 21559.393
131.11801147460938 0 1939.2285
132.08114624023438 0 55758.83
133.08572387695312 0 10699.404
134.0276641845703 0 940.3098
136.0760955810547 0 60092.36
137.0794677734375 0 5536.7085
138.06666564941406 0 7167.652
138.0918731689453 0 2458.6458
138.69801330566406 0 888.7123
139.05072021484375 0 1251.3474
139.0870819091797 0 5582.699
141.1026611328125 0 8801.852
142.0655517578125 0 4976.506
142.28421020507812 0 907.1138
143.045654296875 0 3821.0505
143.06883239746094 0 1152.25
144.0762939453125 0 1445.8473
144.08163452148438 0 2483.1094
145.0611114501953 0 7638.889
146.06039428710938 0 2404.2397
147.04457092285156 0 2666.4119
147.1020050048828 0 5561.3804
148.1792755126953 0 894.33276
149.04527282714844 0 13281.205
150.06668090820312 0 3033.5098
151.08706665039062 0 13201.001
153.1035614013672 0 1396.3434
154.06149291992188 0 1083.5009
155.04559326171875 0 1339.7935
155.08230590820312 0 2004.9412
155.0931854248047 0 36109.418
155.11825561523438 0 23475.135 a Water loss 1
156.0765380859375 0 1473.5044
156.09658813476562 0 1226.0314
156.10292053222656 0 1520.7335
156.12203979492188 0 1397.8774
157.0611572265625 0 1400.2726
157.0975341796875 0 2079.6575
158.08413696289062 0 4345.6997
159.09213256835938 0 434016.94
160.07611083984375 0 4210.7886
160.08909606933594 0 2717.8857
160.09547424316406 0 45942.215
161.0987091064453 0 1842.9171
165.0550537109375 0 31682.615
165.07730102539062 0 1443.4073
166.0614776611328 0 60018.16
166.08641052246094 0 2755.8716 y 9
166.0979461669922 0 5859.023
167.05592346191406 0 13723.815
167.06488037109375 0 4669.991
167.08189392089844 0 1668.1367
167.11827087402344 0 1473.4236
168.0812530517578 0 7902.121
168.10232543945312 0 5099.113
168.11361694335938 0 7277.582
169.07635498046875 0 6262.861
169.08387756347656 0 1678.3799
169.09764099121094 0 15706.903
170.06048583984375 0 21370.893
171.0635986328125 0 3069.9685
171.07667541503906 0 1090.241
171.0921173095703 0 1259.8044
171.1496124267578 0 5645.962
172.07223510742188 0 9093.79
172.82586669921875 0 919.9337
173.05621337890625 0 7536.876
173.1289520263672 0 801236.5 a 1
174.13226318359375 0 67485.09
175.13404846191406 0 3462.628
177.1038055419922 0 1189.4781
177.1126708984375 0 3532.6965
178.1343994140625 0 24010.81
179.13766479492188 0 2843.39
181.0609893798828 0 3795.492
182.08164978027344 0 74820.7 y 12
183.08522033691406 0 8033.1064
183.11328125 0 22451.404 b Water loss 1
184.08636474609375 0 1043.0187
184.11643981933594 0 2495.9248
185.12945556640625 0 1683.7324
185.16522216796875 0 7481.152
186.12417602539062 0 144062.7
187.08706665039062 0 36258.4
187.1204376220703 0 1212.5266
187.1276092529297 0 14284.111
187.1441650390625 0 2044.2866
188.07061767578125 0 2538.666
188.0904083251953 0 3813.8733
188.1304473876953 0 1108.212
189.0762176513672 0 1101.46
189.08770751953125 0 1597.5585
189.09848022460938 0 1648.9637
190.082763671875 0 10509.246
190.1343994140625 0 4752.7495
191.3355255126953 0 1053.5386
192.11407470703125 0 1444.4797
194.08316040039062 0 1108.77
194.09274291992188 0 5896.697
195.1132354736328 0 39430.523
195.1234130859375 0 1972.1396
196.10842895507812 0 23136.342
196.11651611328125 0 2944.5635
197.12896728515625 0 3932.9358
197.16517639160156 0 2081.2117
198.0878143310547 0 3569.0696
198.12811279296875 0 15101.497
199.0716094970703 0 3572.2517
199.08712768554688 0 4345.253
199.13150024414062 0 3071.4468
199.14492797851562 0 2380.8923
200.14060974121094 0 1593.3612
201.12379455566406 0 214987.05 b 1
202.08676147460938 0 2541.7651
202.12705993652344 0 19109.607
203.12832641601562 0 2444.7532
203.25453186035156 0 1485.5039
204.1131591796875 0 1464.2734
205.09759521484375 0 20961.596
205.1083221435547 0 1048.975
205.14553833007812 0 2773.9417
206.1017608642578 0 2594.386
206.12916564941406 0 3794.936
207.1609344482422 0 8106.1963
208.1088104248047 0 1876.4171
209.0923614501953 0 1440.0173
210.12753295898438 0 7631.985
211.10838317871094 0 5660.3794 y Water loss 10
211.13143920898438 0 1252.5588
211.143310546875 0 1404.8611
212.1126251220703 0 1315.8647
212.1399383544922 0 5098.01
213.123779296875 0 16431.268
213.1602783203125 0 1888.8087
214.1276092529297 0 2021.337
215.1392822265625 0 3251.4084
216.0984344482422 0 6183.2744
217.0828094482422 0 1359.298
221.07254028320312 0 1961.7664
221.0847930908203 0 6887.082
221.1036376953125 0 3758.577
221.1398468017578 0 4023.4897
222.08541870117188 0 1474.9866
222.1239776611328 0 6093.6133
223.0640411376953 0 3641.1133
223.08216857910156 0 1254.3905
223.10829162597656 0 12906.009
223.1558380126953 0 71690.305
224.1033935546875 0 3202.8296
224.1591339111328 0 9377.031
225.04318237304688 0 6748.811
225.09869384765625 0 1297.1499
225.1230010986328 0 1351.6575
225.16018676757812 0 2120.455
226.0433349609375 0 1253.6677
226.08277893066406 0 15743.653
226.15576171875 0 2062.2732
227.04006958007812 0 1651.8114
227.06668090820312 0 12343.01
227.15518188476562 0 4581.831
228.0989532470703 0 2236.0127
228.13462829589844 0 9564.001
229.11880493164062 0 6035.553
230.10311889648438 0 1547.338
230.1507568359375 0 1682.5795
233.1400909423828 0 10119.074
233.164306640625 0 1221.8551
234.12428283691406 0 85399.195
235.1276092529297 0 8499.478
235.15554809570312 0 1530.1995
235.60569763183594 0 2024.2214
236.13043212890625 0 1361.7861
237.13510131835938 0 4377.0195
238.11900329589844 0 11275.165
238.12986755371094 0 1881.4847
239.08192443847656 0 2063.752
239.09555053710938 0 12680.243
240.0972442626953 0 3493.7092
240.1348419189453 0 36734.297
241.0941619873047 0 2437.1133
241.1187286376953 0 4654.913
241.13784790039062 0 3366.0483
241.19146728515625 0 5325.5664
243.11001586914062 0 6647.903
244.0933837890625 0 36403.652
244.61141967773438 0 1130.2274
245.0974884033203 0 2761.3926
247.1083221435547 0 41484.95
247.14454650878906 0 2514.046
247.15701293945312 0 1394.4789
248.1032257080078 0 6470.9644
248.1142120361328 0 18899.477
248.62356567382812 0 1360.9799 y 9
249.09873962402344 0 27347.781
249.12037658691406 0 1832.4767
249.1353302001953 0 7458.259
249.60342407226562 0 1770.4536
250.10174560546875 0 3963.439
250.11856079101562 0 2485.107
250.166748046875 0 1372.179
251.15089416503906 0 277501.28
252.13633728027344 0 2065.7517
252.154052734375 0 39197.79
252.16928100585938 0 1541.7631
253.1200408935547 0 1229.5168
253.1563720703125 0 3631.0227
255.1475067138672 0 16537.03
256.09442138671875 0 1642.2585
256.10870361328125 0 3669.2075
256.1504821777344 0 1501.9148
256.61102294921875 0 1820.1481
257.1135559082031 0 5959.5073
258.14532470703125 0 12604.918
259.1473388671875 0 1800.4724
260.1398620605469 0 1280.4343
261.11981201171875 0 6304.2134
261.1351013183594 0 3173.2034
262.1208190917969 0 3181.485
264.1458740234375 0 1457.5771
265.1298522949219 0 42348.93
265.16485595703125 0 1425.4236
266.1142272949219 0 9903.652
266.1253662109375 0 77859.516
267.1286926269531 0 10669.966
267.1448669433594 0 1694.3668
267.6131591796875 0 1897.869
268.1297912597656 0 4798.5366
268.1774597167969 0 5543.8267
269.1614685058594 0 2202.7422
269.18634033203125 0 7062.063
270.1222229003906 0 9527.064
270.62408447265625 0 2414.186
272.1761169433594 0 15866.597
273.17901611328125 0 3144.9805
274.11883544921875 0 14664.462
274.13153076171875 0 6830.82
275.1039733886719 0 2323.1873
275.12237548828125 0 3314.4485
275.15045166015625 0 2701.9988
276.1097412109375 0 2348.22
276.1263732910156 0 2017.5428
276.1556091308594 0 6261.9995
277.0942077636719 0 1741.761
279.1459655761719 0 4310.8755
280.12945556640625 0 7051.374
280.14520263671875 0 1495.4058
281.0521240234375 0 11021.443
282.1567077636719 0 22414.049
283.140625 0 229261.81
283.6370544433594 0 2564.5667
284.1037292480469 0 9987.233
284.1238098144531 0 2958.0024
284.143798828125 0 32979.215
285.1429443359375 0 1953.7506
285.15673828125 0 3976.5952
286.140380859375 0 12132.637
287.14276123046875 0 2106.726
287.17169189453125 0 2844.2183
287.1885986328125 0 1345.9746
288.1275634765625 0 1322.898
288.61920166015625 0 2255.875
289.12103271484375 0 1447.6877
290.1666564941406 0 2338.6677 b 4
293.11383056640625 0 100810.805 y Water loss 11
293.16082763671875 0 2582.1057
294.1181335449219 0 20100.564
294.1819763183594 0 1979.437
295.10272216796875 0 1396.1371
295.119873046875 0 3006.7463
296.1974182128906 0 5341.475 b Water loss 2
296.6399841308594 0 1553.0487
297.0828857421875 0 2123.669
297.13397216796875 0 4120.067 y Water loss 8
297.1563720703125 0 31184.518
298.1396484375 0 3049.4111 y Water loss 6
298.1590576171875 0 5619.1206
299.06243896484375 0 81332.78
300.0621337890625 0 1603.3258
300.1679992675781 0 37030.17
301.1160888671875 0 3207.597
301.15130615234375 0 99233.66
301.1693420410156 0 5079.7275
302.1142883300781 0 18273.701
302.15472412109375 0 13929.89
303.116943359375 0 2694.2026
305.6452941894531 0 7151.974
306.1379699707031 0 24152.652 y 8
306.6395263671875 0 7449.1987
307.140380859375 0 4647.652
309.1670227050781 0 2281.1548
311.1246643066406 0 13995.376 y 11
311.1396484375 0 3098.8218
312.1280212402344 0 2312.4146
314.18695068359375 0 4823.9927
314.2081604003906 0 9533.944 b 2
315.1670837402344 0 54850.98
315.21063232421875 0 1767.5118
316.17022705078125 0 8563.804
317.6602478027344 0 1422.4442
319.1519775390625 0 3588.2957
322.2242126464844 0 2351.1375
325.1519470214844 0 3455.9583
326.6645812988281 0 13606.953
327.166259765625 0 6200.011
332.209228515625 0 5464.1255
333.177490234375 0 16787.059
334.18316650390625 0 3962.0881
338.6754150390625 0 3876.3682 b Water loss 5
339.17132568359375 0 1875.5121 b Ammonia loss 5
339.6772155761719 0 3374.7017
340.66204833984375 0 4950.78
341.0179443359375 0 1335.3737
341.1619873046875 0 2194.4226
342.181884765625 0 4512.596
343.16278076171875 0 2768.9248
343.18475341796875 0 1580.7792
344.1722412109375 0 4752.26
344.9771423339844 0 11536.291
345.1313171386719 0 8389.6875
346.1152648925781 0 36112.97
346.1340637207031 0 1608.9121
347.1182556152344 0 5177.3857
347.6802062988281 0 14089.153 b 5
348.181396484375 0 4890.7686
349.1513366699219 0 2192.1533
350.21929931640625 0 25112.742
351.2212829589844 0 4551.939
352.1643371582031 0 1449.3113
352.6724548339844 0 3175.4626
353.1770935058594 0 2044.821
353.6747131347656 0 3238.3716 y Water loss 7
354.17608642578125 0 3121.622
355.0693054199219 0 1629.0205
359.0295104980469 0 12899.358
360.2041931152344 0 2042.9269
360.6954040527344 0 9195.908
361.1815490722656 0 1938.7832
361.1982421875 0 8935.994
362.1851806640625 0 12538.64
362.6802062988281 0 47094.707 y 7
363.14166259765625 0 48960.297
363.18206787109375 0 18053.635
363.6824035644531 0 3845.0696
364.125 0 8667.472
364.1456298828125 0 8487.279
365.1492004394531 0 1658.6466
366.1558837890625 0 2965.2747
366.178466796875 0 4548.4053
367.1396789550781 0 3598.785
367.161376953125 0 2553.9226
367.24603271484375 0 4071.083
368.1947021484375 0 2402.712
370.1219482421875 0 1456.0939
370.1749267578125 0 1546.5613
373.18719482421875 0 3263.803
376.16265869140625 0 4607.886
376.1982116699219 0 5234.01
376.7003173828125 0 2692.2659
377.1234436035156 0 1693.0513
377.1465148925781 0 5379.5415
378.2145080566406 0 18623.133
379.2098083496094 0 72446.61
380.1663513183594 0 1895.4674
380.2125244140625 0 15151.223
381.1524353027344 0 136703.9
381.71209716796875 0 5584.944
382.15496826171875 0 25327.191
382.1861572265625 0 4493.044
382.2132263183594 0 1486.4011
383.1571960449219 0 3353.1877
384.1669921875 0 6483.971
384.1886291503906 0 6646.6333
385.1506652832031 0 8029.7124
385.17266845703125 0 5642.2524 y 4
385.19610595703125 0 1829.785
386.1539001464844 0 1824.5524
386.17852783203125 0 2472.4548
386.2039794921875 0 1562.041
389.193115234375 0 3368.7424 b 9
390.1966552734375 0 5636.4673
390.6969299316406 0 3225.2537
391.1368713378906 0 5251.0806
392.2084655761719 0 2098.9177
393.18914794921875 0 1899.8015
394.17291259765625 0 24603.838
395.1340637207031 0 5504.9253
395.1571350097656 0 9835.508
395.24127197265625 0 4432.8867
396.13897705078125 0 3482.3936
396.2253723144531 0 23685.08
397.2274475097656 0 4564.815
397.70440673828125 0 1538.4502
398.17913818359375 0 10088.994
399.18499755859375 0 2039.7107
401.1832275390625 0 1469.4592
401.2150573730469 0 1921.7424
402.17657470703125 0 8236.529
402.199951171875 0 12945
403.2007141113281 0 6392.559
403.71685791015625 0 1564.0995
405.2616271972656 0 5930.4717
406.2445983886719 0 2899.118
407.2047119140625 0 3537.7268
408.7099304199219 0 1687.5942
409.1475830078125 0 10629.976
409.20867919921875 0 2015.332
409.6966552734375 0 1395.3533
411.1995544433594 0 12994.059
412.1836242675781 0 81695.15
413.14434814453125 0 6441.501
413.1652526855469 0 9453.422
413.1880798339844 0 13704.446
413.25164794921875 0 3023.755
414.170166015625 0 2607.8792
414.2355041503906 0 26971.092
415.0373229980469 0 21089.422
415.19873046875 0 3948.4016
415.23968505859375 0 6214.194
417.7171630859375 0 13544.837
418.2109375 0 24822.814
418.71136474609375 0 13630.718
419.21136474609375 0 4831.29
421.20928955078125 0 3268.3896 y Water loss 10
422.192138671875 0 2100.2178 y Ammonia loss 10
422.7084655761719 0 2751.6343
423.2008361816406 0 5912.095
423.27227783203125 0 40088.94
424.27484130859375 0 7858.314
424.73345947265625 0 2717.2495
426.19744873046875 0 2362.6284
426.7227783203125 0 249236.69
427.2240905761719 0 122561.24
427.7253112792969 0 37502.99
428.22625732421875 0 7159.08
428.72930908203125 0 2032.335
429.2102355957031 0 65657.03
429.7291259765625 0 2100.1938
430.1944274902344 0 200904.75
431.19757080078125 0 39681.562
431.71502685546875 0 32679.12 b Water loss 6
432.2109069824219 0 16254.555 b Ammonia loss 6
432.71295166015625 0 5757.428
433.20965576171875 0 2020.8733
433.2563171386719 0 7049.9087 b Water loss 3
434.26068115234375 0 1808.8962
438.1741638183594 0 5065.672
438.72796630859375 0 3592.3225
439.2200927734375 0 65182.953 y 10
440.1780700683594 0 4091.6426
440.2230224609375 0 14886.95
440.72027587890625 0 120272.36 b 6
441.22174072265625 0 63700.883
441.722900390625 0 18576.531
442.2247619628906 0 3326.4985
444.1782531738281 0 1957.5405
446.7138366699219 0 3571.7537 y Water loss 6
446.7436218261719 0 2461.7017
447.2083435058594 0 7217.6094 y Ammonia loss 6
447.7074890136719 0 2128.4666
448.20513916015625 0 10381.036
448.2677307128906 0 1760.103
449.2086181640625 0 4099.1714
450.17633056640625 0 2042.0233
450.23870849609375 0 2682.0447
450.2824401855469 0 3283.1667
451.2676086425781 0 70332.96 b 3
452.2703857421875 0 16009.645
452.73828125 0 1582.8346
455.2668151855469 0 5401.9346
455.72052001953125 0 3800.7898 y 6
459.19952392578125 0 6717.0684
460.754150390625 0 7841.378
461.2041015625 0 4672.834
461.2537841796875 0 7119.92
462.88897705078125 0 1376.3075 y Ammonia loss 2
465.74737548828125 0 1683.4221
466.1692199707031 0 2884.9775
466.2449645996094 0 2022.7227
468.29364013671875 0 16515.686
469.2976379394531 0 3319.3582
474.25958251953125 0 12929.183
474.75494384765625 0 18645.3
475.254150390625 0 10051.36
475.75823974609375 0 1661.3789
476.22698974609375 0 5944.508
477.2110595703125 0 11479.2
478.22918701171875 0 8796.3125 y Water loss 9
478.2784118652344 0 8746.615
479.2364196777344 0 2112.0024
479.744384765625 0 1882.1113
482.2768859863281 0 1997.3934
483.2650146484375 0 42278.2
483.7662353515625 0 30566.75
484.2201843261719 0 3229.3118
484.26715087890625 0 6796.637
484.7656555175781 0 1926.7004
485.2216796875 0 1458.1646
488.12432861328125 0 2070.7925
488.25762939453125 0 13030.972 b Water loss 7
488.75494384765625 0 8793.595 b Ammonia loss 7
489.25408935546875 0 2696.483
490.2382507324219 0 1830.2433
494.2367858886719 0 147441.23
495.1997985839844 0 1619.9443
495.2396545410156 0 33787.84
496.24127197265625 0 159829.77 y 9
497.24652099609375 0 40637.426
497.76373291015625 0 18225.828
498.2447509765625 0 7748.72
502.2299499511719 0 2062.0457
503.1082763671875 0 21702.021
504.200439453125 0 6065.1313
505.15057373046875 0 3517.0945
505.2005615234375 0 2506.0886
506.2265319824219 0 3701.9358
506.2687072753906 0 2169.717
507.2570495605469 0 5427.7217
508.2520446777344 0 1668.6641
511.2626953125 0 12171.992
512.2650756835938 0 2444.9863
513.2103271484375 0 3222.382
515.2859497070312 0 1959.9755
517.322998046875 0 10244.122
518.3248901367188 0 3818.0142
519.139892578125 0 7873.049
520.2645874023438 0 2524.0625
522.2122802734375 0 1633.2415
523.2617797851562 0 3434.3223
523.58642578125 0 1473.6772
524.2841186523438 0 3494.8455
525.2671508789062 0 24151.045
526.2286376953125 0 2360.3044
526.2699584960938 0 7830.7476
527.2749633789062 0 1579.0999
528.933349609375 0 4252.915
529.2627563476562 0 9639.421 Precursor Water loss
529.5958862304688 0 9641.141 Precursor Ammonia loss
529.9287719726562 0 5432.794
530.2243041992188 0 1439.6161
531.2214965820312 0 1401.5366
532.1947021484375 0 5961.44
534.3490600585938 0 9731.888
534.9343872070312 0 2237.5256
535.265869140625 0 6359.861 Precursor
535.3511962890625 0 3093.6897
535.601806640625 0 6147.6963
535.9383544921875 0 3202.3486
536.1675415039062 0 4765.562
537.2452392578125 0 1755.8329
537.7406005859375 0 2111.2239
538.2730102539062 0 2484.3452
538.7750244140625 0 2308.5552
539.23681640625 0 28217.123
540.2409057617188 0 8893.3955
541.2451171875 0 1578.1351
542.2939453125 0 15240.2295
543.278076171875 0 33935.176
544.2816162109375 0 9789.592
545.2859497070312 0 2147.0986
545.7614135742188 0 2342.1938 b Water loss 8
546.2509765625 0 5150.8384 b Ammonia loss 8
547.2484130859375 0 2009.2738
549.2217407226562 0 7383.3906
550.211181640625 0 2062.9592
551.2578735351562 0 8024.983
551.33203125 0 1832.9032
552.259521484375 0 3866.8853
554.7651977539062 0 7458.9116 b 8
555.2573852539062 0 7854.089
555.7593994140625 0 4245.8574
558.2210693359375 0 1760.845
561.315185546875 0 6776.61 b Water loss 4
562.2979125976562 0 2192.6287 b Ammonia loss 4
563.2589111328125 0 3544.3481
564.2667236328125 0 3513.563
564.3298950195312 0 2679.4263
565.2636108398438 0 12953.378
566.2689208984375 0 2442.627
567.23193359375 0 34730.33
568.2357177734375 0 10337.08
569.2373046875 0 1869.6006
575.2473754882812 0 11060.754
576.23681640625 0 5136.2437
577.2267456054688 0 1478.4327
577.7557373046875 0 1342.0258
579.3258666992188 0 48431.918 b 4
580.3286743164062 0 12373.872
581.2719116210938 0 1690.8092
581.3300170898438 0 3178.9734
582.3037109375 0 1909.6605
589.3109741210938 0 1815.2528
592.272705078125 0 10299.793
592.3263549804688 0 4967.98
593.2574462890625 0 62262.457 y Water loss 8
593.3060302734375 0 19624.008
594.2568969726562 0 23513.275
594.3096313476562 0 3602.7383
595.256591796875 0 4526.845
596.2465209960938 0 2750.9272
596.3511352539062 0 2222.307
597.80029296875 0 3331.7102
598.2981567382812 0 2150.2786
602.3008422851562 0 4023.598
602.7939453125 0 9262.542
603.2921142578125 0 9848.599
603.7947998046875 0 3157.0457
610.2839965820312 0 164748.27
611.2691650390625 0 578151.2 y 8
611.8003540039062 0 28802.662
612.2720336914062 0 185788.11
612.8011474609375 0 8146.467
613.2738037109375 0 41000.715
614.275390625 0 6302.369
617.2871704101562 0 2746.0574
620.8052978515625 0 2136.1396
621.2520141601562 0 10622.36
621.3021240234375 0 3281.2607
622.2540283203125 0 2564.2063
622.369140625 0 1951.5714
628.2808227539062 0 2379.3992
630.28955078125 0 2212.43
634.3087158203125 0 7679.374
635.2974243164062 0 9471.618
636.3252563476562 0 4658.796
636.787841796875 0 7689.16 y Water loss 3
637.287353515625 0 3384.255 y Ammonia loss 3
637.344482421875 0 1861.775
637.786376953125 0 2176.9607
638.33203125 0 3161.2664 b Water loss 10
640.2753295898438 0 3224.4297
645.2950439453125 0 9430.992
645.7945556640625 0 30108.348 y 3
646.294189453125 0 23420.4
646.7953491210938 0 7776.7793
648.3457641601562 0 2345.4407
650.271240234375 0 2882.6309
650.3291015625 0 3177.5842
652.3209838867188 0 60234.29
653.3240966796875 0 26017.21
654.3262329101562 0 4557.799
658.3348388671875 0 2136.839
659.3153076171875 0 2121.6929
660.3284912109375 0 1398.7284
661.333251953125 0 2234.5872
662.3062744140625 0 22343.111
662.83984375 0 1427.3446
663.2987670898438 0 13681.448
664.2950439453125 0 3426.2524
664.3502197265625 0 2093.7373
666.3499755859375 0 1670.1598
668.284423828125 0 1995.6317
670.3283081054688 0 2264.5706
670.8245239257812 0 1491.9592
675.8079223632812 0 1963.3544
676.3425903320312 0 31549.842 b Water loss 5
677.3347778320312 0 16572.582 b Ammonia loss 5
678.3346557617188 0 8001.162
679.3322143554688 0 3566.2344
679.83935546875 0 2499.2834
680.31640625 0 109570.14
681.31884765625 0 43164.812
682.3213500976562 0 11963.141
683.3259887695312 0 1672.6738
684.3220825195312 0 4147.264
684.82080078125 0 10282.316
685.3197021484375 0 5793.9517
685.820556640625 0 1865.2583
688.3323974609375 0 3403.4004
689.3320922851562 0 1746.5751
692.8363037109375 0 5327.567
693.3307495117188 0 32492.902 y Water loss 2
693.8291015625 0 34793.293 y Ammonia loss 2
694.3523559570312 0 295370.4 b 5
694.83251953125 0 4659.4424
695.355712890625 0 117104.28
696.3577880859375 0 27232.014
697.3574829101562 0 6091.252
698.3626708984375 0 2158.3306
701.3397216796875 0 5655.9565
701.7718505859375 0 2729.0046
701.8419799804688 0 72433.01
702.3369750976562 0 183220.31 y 2
702.8372802734375 0 144783.25 b Water loss 11
703.33935546875 0 61363.31 b Ammonia loss 11
703.8407592773438 0 17694.268
704.3385620117188 0 9272.267
705.3569946289062 0 5249.664
706.3431396484375 0 20278.262 y Water loss 7
707.3425903320312 0 8449.634
708.3063354492188 0 2065.728
710.911376953125 0 1895.2784
711.3696899414062 0 15493.792
711.861572265625 0 13473.02 b 11
712.3634033203125 0 7629.9097
712.8609619140625 0 3198.6687
713.352783203125 0 1916.0756
715.3707885742188 0 1778.7415
720.3815307617188 0 3427.592
721.364013671875 0 14453.914
722.3654174804688 0 4660.618
723.3683471679688 0 44143.84
724.3536376953125 0 178337.62 y 7
725.3563232421875 0 75747.84
726.3582153320312 0 19138.607
727.3621826171875 0 2258.991
728.3731689453125 0 2319.0022
728.8595581054688 0 1615.5038
729.3621215820312 0 5667.4844
730.3681030273438 0 3531.2434
733.3644409179688 0 8222.691
734.3565063476562 0 7187.259
735.3592529296875 0 2515.5493
737.8646240234375 0 1771.2295
739.3590698242188 0 2558.1853
742.861328125 0 2151.4592 y Water loss 1
743.3704223632812 0 5034.204
744.3704223632812 0 3574.4458
747.3887329101562 0 3527.8982
748.3829956054688 0 3285.7231
749.337646484375 0 3562.8113
751.3767700195312 0 35581.6
751.8704223632812 0 15624.55 y 1
752.37646484375 0 24644.154
752.8721313476562 0 6773.278
753.375 0 8979.154
753.8734741210938 0 3236.8237
760.3037719726562 0 1576.0381
762.3563232421875 0 1980.7714
763.3665161132812 0 22892.17
764.365966796875 0 5665.866
765.4027709960938 0 7791.8755
766.4080810546875 0 2663.0332
767.3485107421875 0 6953.914
769.4095458984375 0 2320.676
770.4019775390625 0 2287.1838
775.3907470703125 0 4986.1025
776.3739624023438 0 3164.6116
778.39599609375 0 2083.1653
779.3843994140625 0 17750.059
780.3880004882812 0 9069.7295
781.3695068359375 0 9250.962
782.3660278320312 0 4969.9517
791.3848266601562 0 1789.6593
793.3995971679688 0 33518.29 Precursor Water loss
794.4024047851562 0 14784.706
795.34521484375 0 3590.278
807.3780517578125 0 7703.4653
807.445556640625 0 3720.6458
808.384765625 0 4394.5127
818.4360961914062 0 4146.0576
819.4378662109375 0 3800.6472
825.3902587890625 0 3136.97
826.3832397460938 0 2642.5347
834.42724609375 0 7742.3384
835.416015625 0 11865.962
836.448974609375 0 24194.484
837.4524536132812 0 10410.556
838.3197021484375 0 1546.0612
838.4557495117188 0 1892.9408
843.4022216796875 0 3902.6482
844.3936767578125 0 4908.2876
845.3953857421875 0 5397.6675
846.446533203125 0 10061.743
847.444580078125 0 6822.19
848.4461669921875 0 2693.778
851.384033203125 0 1946.143
852.4369506835938 0 76235.805
853.4397583007812 0 36885.324
854.4423217773438 0 10557.53
862.4210815429688 0 41543.645 b Water loss 6
863.4171142578125 0 33090.508 b Ammonia loss 6
864.4564819335938 0 61300.312
865.4610595703125 0 30515.152
866.4642333984375 0 9745.517
867.4637451171875 0 2184.8296
876.436279296875 0 5715.7563
877.4412231445312 0 3110.3691
880.4318237304688 0 382827.5 b 6
881.4343872070312 0 195136.17
882.437255859375 0 56462.812
883.4381103515625 0 9686.622
884.4400634765625 0 1943.3157
890.408203125 0 1730.5375
892.4639282226562 0 6567.262
893.4679565429688 0 3628.5288
894.4603271484375 0 1846.6066
895.5327758789062 0 2038.566
896.5303344726562 0 2033.6997
907.4389038085938 0 3620.8928
908.4315795898438 0 10656.085
909.4405517578125 0 11273.844
910.4354248046875 0 17909.336 y 6
911.4351196289062 0 8496.513
912.4346923828125 0 3067.255
936.4293212890625 0 3519.1167
937.42138671875 0 3668.4167
947.43115234375 0 1860.7498
949.4375 0 1813.6104
954.4299926757812 0 4419.3696
955.4208374023438 0 6983.604
956.42578125 0 2180.611
964.4659423828125 0 4087.293
965.454345703125 0 5635.6665
966.4457397460938 0 2355.421
972.4490356445312 0 4941.8633
973.440673828125 0 2167.6675
975.5049438476562 0 4036.7327 b Water loss 7
976.5023803710938 0 3621.0557 b Ammonia loss 7
977.500732421875 0 1875.1807
980.46337890625 0 1601.9546
981.4765625 0 1987.5854
982.4723510742188 0 1671.8243
993.515869140625 0 25928.957 b 7
994.5176391601562 0 17163.342
995.5225830078125 0 5402.0415
1007.4591674804688 0 2648.9216 y Water loss 5
1009.4348754882812 0 2427.149
1024.472412109375 0 5595.9404
1025.462646484375 0 11957.309 y 5
1026.4617919921875 0 6805.8657
1027.4573974609375 0 1791.6814
1075.5284423828125 0 2987.6865
1091.5025634765625 0 1600.0526 b Ammonia loss 8
1092.5653076171875 0 2079.0388
1093.5457763671875 0 5286.476
1094.5482177734375 0 1980.5715
1108.521240234375 0 1839.9607 b 8
1117.492431640625 0 2359.9778
1118.4810791015625 0 5104.1777
1134.5164794921875 0 1701.6589
1135.507080078125 0 11868.916 y Water loss 4
1136.499755859375 0 17713.164 y Ammonia loss 4
1137.4993896484375 0 6995.645
1138.4940185546875 0 1852.0079
1152.530029296875 0 11819.318
1153.5206298828125 0 25229.166 y 4
1154.523193359375 0 13888.159
1155.5233154296875 0 3926.9114
1165.5828857421875 0 1822.7478 b 9
1166.5733642578125 0 1639.7744
1290.56494140625 0 2231.8362 y 3
2084.44140625 0 1435.3265
3081.24853515625 0 1651.2219

Spectrum Details

|  |  |
| --- | --- |
| Matched peaks? Matched peaks The total absolute number of peaks matched. Additionally in brackets the total fraction of peaks matched and the total number of peaks is shown. | 84 (9.77% of 860) |
| FDR? FDR The false discovery rate estimated for this peptide. It is calculated by matching all theoretical fragments with a non-integer shift with the raw peaks for this spectrum. This is done with 40 different shifts. The resulting percentage is the average number of annotated peaks over the number of annotated peaks with the correct spectrum. | 0.82% |
| Satellite FDR? Satellite FDR See the FDR for details on its calculation. This satellite ion specific FDR only contains the satellite ions (d/w) for I/L/J positions. | ∞ |
| PSM Score? PSM Score The PSM Score as given by Hecklib to this annotated spectrum. It is shown with three significant figures. | 647 |

#### Spectrum 7841? Spectrum 7841 The raw spectrum of this peptide as annotated by Hecklib. The fragments are coloured according to ion type (see legend). Any peaks with a star '\*' as text can be hovered over to see the full details, first the ion type second the mass shift type. By hovering over the amino acids in the peptide or ions in the legend the corresponding peaks are highlighted. By toggling the 'Unassigned' label you can turn the background (unassigned) peaks on or off in the plot. By updating the slider in the Ion legend you can update the spectrum to only show the top X% of the peaks with labels. The top X% means any peak that is within X% of the highest intensity. By dragging in the spectrum you can zoom in to a specific part of the spectrum and use 'Zoom Out' to get back to the original zoom level. The annotation of the spectrum is based on the given sequence in the peptides file and is done with different software so inconsistencies are likely. The peaks are annotated based on the given sequence, with 20 ppm tolerance.

Copy Data

##### Spectrum 7841 (TSV)

###### Preview

```
Loading example...
```

*Click on the button to copy the data to your clipboard.*

Mz MinMz MaxIntensity Max

WidthHeightPeptide font sizePeptide stroke widthSpectrum font sizeSpectrum stroke widthCompact peptide

Ion legend

wxyz

abcd

OtherUnassignedIon^Charge^_Position_Show for top:%

TVLHQDWLDGKEY

04.06e+48.13e+41.22e+51.63e+5

Zoom Out

a^+1^_2_y^+3^_4_a^+1^_2_y^+1^_1_b^+1^_2_b^+1^_2_y^+2^_3_b^+3^_6_y^+1^_2_b^+1^_3_y^+2^_5_y^+3^_7_y^+2^_5_y^+1^_2_b^+1^_3_b^+2^_6_y^+2^_6_y^+2^_6_y^+1^_3_b^+2^_7_b^+2^_7_b^+1^_4_y^+1^_3_b^+2^_7_y^+2^_7_b^+1^_4_y^+1^_4_b^+2^_8_b^+2^_8_y^+1^_4_\*\*b^+2^_9_b^+1^_5_b^+1^_5_b^+1^_5_y^+1^_5_y^+1^_5_y^+1^_5_y^+2^_10_y^+2^_10_b^+1^_6_b^+1^_6_y^+2^_11_y^+2^_11_b^+1^_6_y^+2^_11_b^+2^_12_b^+2^_12_y^+1^_6_b^+2^_12_y^+1^_6_y^+2^_12_y^+2^_12_\*b^+1^_7_b^+1^_7_b^+1^_7_y^+1^_7_b^+1^_8_b^+1^_8_b^+1^_8_y^+1^_8_y^+1^_9_y^+1^_9_y^+1^_9_

0778155623343112

Fragment Matches Table

Show background peaks

| Position | Ion type | Intensity | mz Theoretical | mz Error (Th) | mz Error (ppm) | Charge | Series Number |
| --- | --- | --- | --- | --- | --- | --- | --- |
| - | - | 6723 | 120.1 | - | - | 0 | - |
| - | - | 371.8 | 120.1 | - | - | 0 | - |
| - | - | 704.9 | 121.1 | - | - | 0 | - |
| - | - | 1120 | 122.1 | - | - | 0 | - |
| - | - | 397.8 | 122.9 | - | - | 0 | - |
| - | - | 534.7 | 123 | - | - | 0 | - |
| - | - | 453.9 | 126.1 | - | - | 0 | - |
| - | - | 2524 | 127.1 | - | - | 0 | - |
| - | - | 907.1 | 128.1 | - | - | 0 | - |
| - | - | 1.006E+04 | 128.1 | - | - | 0 | - |
| - | - | 599 | 129.1 | - | - | 0 | - |
| - | - | 1.069E+05 | 129.1 | - | - | 0 | - |
| - | - | 4.735E+04 | 130.1 | - | - | 0 | - |
| - | - | 689.3 | 130.1 | - | - | 0 | - |
| - | - | 469.7 | 130.1 | - | - | 0 | - |
| - | - | 1105 | 130.1 | - | - | 0 | - |
| - | - | 6655 | 130.1 | - | - | 0 | - |
| - | - | 4794 | 131.1 | - | - | 0 | - |
| - | - | 393.7 | 131.1 | - | - | 0 | - |
| - | - | 1.151E+04 | 132.1 | - | - | 0 | - |
| - | - | 596.8 | 133.1 | - | - | 0 | - |
| - | - | 962.8 | 133.1 | - | - | 0 | - |
| - | - | 533.4 | 134 | - | - | 0 | - |
| - | - | 1.63E+04 | 136.1 | - | - | 0 | - |
| - | - | 1447 | 137.1 | - | - | 0 | - |
| - | - | 1378 | 138.1 | - | - | 0 | - |
| - | - | 1775 | 139.1 | - | - | 0 | - |
| - | - | 2697 | 141.1 | - | - | 0 | - |
| - | - | 409.5 | 141.7 | - | - | 0 | - |
| - | - | 583.7 | 142.1 | - | - | 0 | - |
| - | - | 569.9 | 142.1 | - | - | 0 | - |
| - | - | 721.8 | 143 | - | - | 0 | - |
| - | - | 609.6 | 144.1 | - | - | 0 | - |
| - | - | 802.4 | 145.1 | - | - | 0 | - |
| - | - | 1368 | 146.1 | - | - | 0 | - |
| - | - | 641.7 | 147 | - | - | 0 | - |
| - | - | 1076 | 149 | - | - | 0 | - |
| - | - | 1.504E+04 | 149 | - | - | 0 | - |
| - | - | 861.4 | 150.1 | - | - | 0 | - |
| - | - | 482.6 | 150.1 | - | - | 0 | - |
| - | - | 2106 | 151.1 | - | - | 0 | - |
| - | - | 447.5 | 151.3 | - | - | 0 | - |
| - | - | 794.3 | 155.1 | - | - | 0 | - |
| - | - | 7067 | 155.1 | - | - | 0 | - |
| 2 | a | 5617 | 155.1 | 0.0002288 | 1.475 | +1 | 2 |
| - | - | 692.6 | 156.1 | - | - | 0 | - |
| - | - | 741.5 | 157.1 | - | - | 0 | - |
| - | - | 646.9 | 158.1 | - | - | 0 | - |
| - | - | 7.829E+04 | 159.1 | - | - | 0 | - |
| - | - | 876.5 | 160.1 | - | - | 0 | - |
| - | - | 966.4 | 160.1 | - | - | 0 | - |
| - | - | 9194 | 160.1 | - | - | 0 | - |
| - | - | 472.2 | 160.8 | - | - | 0 | - |
| - | - | 504.2 | 161.1 | - | - | 0 | - |
| - | - | 8502 | 165.1 | - | - | 0 | - |
| - | - | 850.9 | 166.1 | - | - | 0 | - |
| - | - | 1.302E+04 | 166.1 | - | - | 0 | - |
| 10 | y | 3389 | 166.1 | 0.001542 | 9.285 | +3 | 4 |
| - | - | 1374 | 166.1 | - | - | 0 | - |
| - | - | 2.357E+04 | 167.1 | - | - | 0 | - |
| - | - | 1211 | 167.1 | - | - | 0 | - |
| - | - | 551.4 | 168.1 | - | - | 0 | - |
| - | - | 1526 | 168.1 | - | - | 0 | - |
| - | - | 755.7 | 168.1 | - | - | 0 | - |
| - | - | 1238 | 168.1 | - | - | 0 | - |
| - | - | 1385 | 169.1 | - | - | 0 | - |
| - | - | 3950 | 169.1 | - | - | 0 | - |
| - | - | 3723 | 170.1 | - | - | 0 | - |
| - | - | 534.1 | 171.1 | - | - | 0 | - |
| - | - | 560.4 | 171.1 | - | - | 0 | - |
| - | - | 2215 | 172.1 | - | - | 0 | - |
| - | - | 1487 | 173.1 | - | - | 0 | - |
| 2 | a | 1.609E+05 | 173.1 | 0.00033 | 1.906 | +1 | 2 |
| - | - | 2900 | 173.5 | - | - | 0 | - |
| - | - | 444.2 | 174.1 | - | - | 0 | - |
| - | - | 848 | 174.1 | - | - | 0 | - |
| - | - | 1.352E+04 | 174.1 | - | - | 0 | - |
| - | - | 642.3 | 175.1 | - | - | 0 | - |
| - | - | 4501 | 178.1 | - | - | 0 | - |
| - | - | 965.4 | 179.1 | - | - | 0 | - |
| - | - | 930.8 | 181.1 | - | - | 0 | - |
| 13 | y | 1.636E+04 | 182.1 | 0.0002512 | 1.38 | +1 | 1 |
| - | - | 1434 | 183.1 | - | - | 0 | - |
| 2 | b | 5108 | 183.1 | 0.0002787 | 1.522 | +1 | 2 |
| - | - | 1425 | 185.2 | - | - | 0 | - |
| - | - | 3.127E+04 | 186.1 | - | - | 0 | - |
| - | - | 6766 | 187.1 | - | - | 0 | - |
| - | - | 739.9 | 187.1 | - | - | 0 | - |
| - | - | 3311 | 187.1 | - | - | 0 | - |
| - | - | 5041 | 187.1 | - | - | 0 | - |
| - | - | 1645 | 188.1 | - | - | 0 | - |
| - | - | 1028 | 188.1 | - | - | 0 | - |
| - | - | 713.1 | 188.1 | - | - | 0 | - |
| - | - | 771.3 | 190.1 | - | - | 0 | - |
| - | - | 1246 | 190.1 | - | - | 0 | - |
| - | - | 1043 | 194.1 | - | - | 0 | - |
| - | - | 8366 | 195.1 | - | - | 0 | - |
| - | - | 5546 | 196.1 | - | - | 0 | - |
| - | - | 762.9 | 196.1 | - | - | 0 | - |
| - | - | 1009 | 197.1 | - | - | 0 | - |
| - | - | 1441 | 198.1 | - | - | 0 | - |
| - | - | 2483 | 198.1 | - | - | 0 | - |
| - | - | 593.7 | 199.1 | - | - | 0 | - |
| - | - | 930.6 | 199.1 | - | - | 0 | - |
| - | - | 513.3 | 200.1 | - | - | 0 | - |
| 2 | b | 4.668E+04 | 201.1 | 0.0001968 | 0.9787 | +1 | 2 |
| - | - | 567 | 201.6 | - | - | 0 | - |
| - | - | 677.6 | 202.1 | - | - | 0 | - |
| - | - | 4900 | 202.1 | - | - | 0 | - |
| - | - | 544 | 203.1 | - | - | 0 | - |
| - | - | 4204 | 205.1 | - | - | 0 | - |
| - | - | 1628 | 207.2 | - | - | 0 | - |
| - | - | 672 | 209.1 | - | - | 0 | - |
| - | - | 1220 | 210.1 | - | - | 0 | - |
| 11 | y | 815.5 | 211.1 | 0.0001609 | 0.762 | +2 | 3 |
| - | - | 1231 | 212.1 | - | - | 0 | - |
| - | - | 3936 | 213.1 | - | - | 0 | - |
| - | - | 1820 | 215.1 | - | - | 0 | - |
| - | - | 1268 | 216.1 | - | - | 0 | - |
| - | - | 755.4 | 216.1 | - | - | 0 | - |
| - | - | 1976 | 221.1 | - | - | 0 | - |
| - | - | 7191 | 221.1 | - | - | 0 | - |
| - | - | 941.9 | 221.1 | - | - | 0 | - |
| - | - | 613.7 | 221.1 | - | - | 0 | - |
| - | - | 2400 | 222.1 | - | - | 0 | - |
| - | - | 2410 | 222.1 | - | - | 0 | - |
| - | - | 4513 | 223.1 | - | - | 0 | - |
| - | - | 1296 | 223.1 | - | - | 0 | - |
| - | - | 2358 | 223.1 | - | - | 0 | - |
| - | - | 1.342E+04 | 223.2 | - | - | 0 | - |
| - | - | 668 | 224.1 | - | - | 0 | - |
| - | - | 2012 | 224.2 | - | - | 0 | - |
| - | - | 9605 | 225 | - | - | 0 | - |
| - | - | 2346 | 226 | - | - | 0 | - |
| - | - | 2301 | 226.1 | - | - | 0 | - |
| 6 | b | 737.3 | 226.1 | 0.000355 | 1.57 | +3 | 6 |
| - | - | 1692 | 226.2 | - | - | 0 | - |
| - | - | 1593 | 227 | - | - | 0 | - |
| - | - | 2365 | 227.1 | - | - | 0 | - |
| - | - | 1097 | 227.2 | - | - | 0 | - |
| - | - | 1629 | 228.1 | - | - | 0 | - |
| - | - | 2686 | 229.1 | - | - | 0 | - |
| - | - | 2552 | 233.1 | - | - | 0 | - |
| - | - | 1.463E+04 | 234.1 | - | - | 0 | - |
| - | - | 1549 | 235.1 | - | - | 0 | - |
| - | - | 1267 | 237.1 | - | - | 0 | - |
| - | - | 2616 | 238.1 | - | - | 0 | - |
| - | - | 2.265E+04 | 239.1 | - | - | 0 | - |
| - | - | 5763 | 240.1 | - | - | 0 | - |
| - | - | 6387 | 240.1 | - | - | 0 | - |
| - | - | 3343 | 241.1 | - | - | 0 | - |
| - | - | 1145 | 241.1 | - | - | 0 | - |
| - | - | 824.6 | 241.1 | - | - | 0 | - |
| - | - | 997.5 | 241.2 | - | - | 0 | - |
| - | - | 1052 | 243.1 | - | - | 0 | - |
| - | - | 6337 | 244.1 | - | - | 0 | - |
| - | - | 656.8 | 245.1 | - | - | 0 | - |
| - | - | 7881 | 247.1 | - | - | 0 | - |
| - | - | 1096 | 248.1 | - | - | 0 | - |
| - | - | 3513 | 248.1 | - | - | 0 | - |
| - | - | 793.6 | 249.1 | - | - | 0 | - |
| - | - | 4773 | 249.1 | - | - | 0 | - |
| - | - | 1681 | 249.1 | - | - | 0 | - |
| - | - | 674 | 250.1 | - | - | 0 | - |
| - | - | 5.289E+04 | 251.2 | - | - | 0 | - |
| - | - | 7466 | 252.2 | - | - | 0 | - |
| - | - | 707.4 | 253.2 | - | - | 0 | - |
| - | - | 3093 | 255.1 | - | - | 0 | - |
| - | - | 1560 | 257.1 | - | - | 0 | - |
| - | - | 3040 | 258.1 | - | - | 0 | - |
| - | - | 1343 | 261.1 | - | - | 0 | - |
| - | - | 706.5 | 261.1 | - | - | 0 | - |
| - | - | 1093 | 262.1 | - | - | 0 | - |
| - | - | 8211 | 265.1 | - | - | 0 | - |
| - | - | 1802 | 266.1 | - | - | 0 | - |
| - | - | 1.311E+04 | 266.1 | - | - | 0 | - |
| - | - | 1552 | 267.1 | - | - | 0 | - |
| - | - | 1415 | 268.1 | - | - | 0 | - |
| - | - | 1091 | 268.2 | - | - | 0 | - |
| - | - | 787.9 | 269.2 | - | - | 0 | - |
| - | - | 673 | 269.2 | - | - | 0 | - |
| - | - | 2639 | 270.1 | - | - | 0 | - |
| - | - | 3326 | 272.2 | - | - | 0 | - |
| - | - | 2835 | 274.1 | - | - | 0 | - |
| - | - | 872.9 | 274.1 | - | - | 0 | - |
| - | - | 1187 | 275.1 | - | - | 0 | - |
| - | - | 777.9 | 276.1 | - | - | 0 | - |
| - | - | 707.2 | 276.1 | - | - | 0 | - |
| - | - | 1191 | 276.2 | - | - | 0 | - |
| - | - | 1257 | 279.1 | - | - | 0 | - |
| - | - | 1174 | 280.1 | - | - | 0 | - |
| - | - | 1.503E+04 | 281.1 | - | - | 0 | - |
| - | - | 3754 | 282.2 | - | - | 0 | - |
| - | - | 1913 | 283 | - | - | 0 | - |
| - | - | 4.213E+04 | 283.1 | - | - | 0 | - |
| - | - | 1270 | 284.1 | - | - | 0 | - |
| - | - | 5243 | 284.1 | - | - | 0 | - |
| - | - | 711 | 285 | - | - | 0 | - |
| - | - | 1537 | 286.1 | - | - | 0 | - |
| 12 | y | 2.122E+04 | 293.1 | 0.0002358 | 0.8043 | +1 | 2 |
| - | - | 1113 | 293.1 | - | - | 0 | - |
| - | - | 863.4 | 293.2 | - | - | 0 | - |
| - | - | 4167 | 294.1 | - | - | 0 | - |
| - | - | 2029 | 295.1 | - | - | 0 | - |
| - | - | 1023 | 296.1 | - | - | 0 | - |
| 3 | b | 982 | 296.2 | 0.000518 | 1.749 | +1 | 3 |
| - | - | 2367 | 297.1 | - | - | 0 | - |
| - | - | 914.6 | 297.1 | - | - | 0 | - |
| 9 | y | 765 | 297.1 | 0.002599 | 8.748 | +2 | 5 |
| - | - | 5943 | 297.2 | - | - | 0 | - |
| 7 | y | 733.8 | 298.1 | 0.00436 | 14.62 | +3 | 7 |
| - | - | 1155 | 298.2 | - | - | 0 | - |
| - | - | 1.216E+05 | 299.1 | - | - | 0 | - |
| - | - | 3391 | 300.1 | - | - | 0 | - |
| - | - | 6133 | 300.2 | - | - | 0 | - |
| - | - | 1940 | 301.1 | - | - | 0 | - |
| - | - | 789.9 | 301.1 | - | - | 0 | - |
| - | - | 1.647E+04 | 301.2 | - | - | 0 | - |
| - | - | 1008 | 301.2 | - | - | 0 | - |
| - | - | 3231 | 302.1 | - | - | 0 | - |
| - | - | 2811 | 302.2 | - | - | 0 | - |
| - | - | 664.6 | 303.2 | - | - | 0 | - |
| - | - | 1029 | 305.6 | - | - | 0 | - |
| 9 | y | 3990 | 306.1 | 0.0009486 | 3.099 | +2 | 5 |
| - | - | 1791 | 306.6 | - | - | 0 | - |
| - | - | 860.2 | 307.1 | - | - | 0 | - |
| 12 | y | 3434 | 311.1 | 4.704E-05 | 0.1512 | +1 | 2 |
| - | - | 734.5 | 311.2 | - | - | 0 | - |
| - | - | 1168 | 313.1 | - | - | 0 | - |
| - | - | 997.3 | 314.1 | - | - | 0 | - |
| - | - | 1059 | 314.2 | - | - | 0 | - |
| 3 | b | 1918 | 314.2 | 0.0001783 | 0.5673 | +1 | 3 |
| - | - | 1.239E+04 | 315.2 | - | - | 0 | - |
| - | - | 876.9 | 315.2 | - | - | 0 | - |
| - | - | 1392 | 316.2 | - | - | 0 | - |
| - | - | 565.8 | 317.1 | - | - | 0 | - |
| - | - | 801.5 | 323.2 | - | - | 0 | - |
| - | - | 676.9 | 325.2 | - | - | 0 | - |
| - | - | 1784 | 326.7 | - | - | 0 | - |
| - | - | 1029 | 327 | - | - | 0 | - |
| - | - | 897.9 | 327.2 | - | - | 0 | - |
| - | - | 905.8 | 332.2 | - | - | 0 | - |
| - | - | 2121 | 333.2 | - | - | 0 | - |
| - | - | 599.4 | 339.7 | - | - | 0 | - |
| - | - | 968 | 340.7 | - | - | 0 | - |
| - | - | 1367 | 341 | - | - | 0 | - |
| - | - | 1245 | 342.2 | - | - | 0 | - |
| - | - | 539.1 | 343.9 | - | - | 0 | - |
| - | - | 1176 | 344.2 | - | - | 0 | - |
| - | - | 1.525E+04 | 345 | - | - | 0 | - |
| - | - | 1838 | 345.1 | - | - | 0 | - |
| - | - | 6831 | 346.1 | - | - | 0 | - |
| - | - | 589 | 347 | - | - | 0 | - |
| - | - | 1246 | 347.1 | - | - | 0 | - |
| 6 | b | 2877 | 347.7 | 0.0001472 | 0.4233 | +2 | 6 |
| - | - | 4269 | 350.2 | - | - | 0 | - |
| - | - | 935.3 | 351.2 | - | - | 0 | - |
| - | - | 619.9 | 352.2 | - | - | 0 | - |
| - | - | 958.7 | 352.7 | - | - | 0 | - |
| 8 | y | 1460 | 353.7 | 0.0005759 | 1.628 | +2 | 6 |
| - | - | 3379 | 355.1 | - | - | 0 | - |
| - | - | 1.524E+04 | 359 | - | - | 0 | - |
| - | - | 1155 | 361.2 | - | - | 0 | - |
| - | - | 1903 | 362.2 | - | - | 0 | - |
| 8 | y | 8252 | 362.7 | 5.428E-05 | 0.1497 | +2 | 6 |
| - | - | 9461 | 363.1 | - | - | 0 | - |
| - | - | 4177 | 363.2 | - | - | 0 | - |
| - | - | 1371 | 364.1 | - | - | 0 | - |
| - | - | 1637 | 364.1 | - | - | 0 | - |
| - | - | 728.8 | 366.2 | - | - | 0 | - |
| - | - | 752.9 | 367.1 | - | - | 0 | - |
| - | - | 747.4 | 367.2 | - | - | 0 | - |
| - | - | 794.2 | 368.2 | - | - | 0 | - |
| - | - | 1220 | 369.1 | - | - | 0 | - |
| - | - | 2480 | 370.1 | - | - | 0 | - |
| - | - | 764.6 | 371.1 | - | - | 0 | - |
| - | - | 1041 | 376.2 | - | - | 0 | - |
| - | - | 727.8 | 377.1 | - | - | 0 | - |
| - | - | 3872 | 378.2 | - | - | 0 | - |
| - | - | 1.348E+04 | 379.2 | - | - | 0 | - |
| - | - | 564.5 | 380.2 | - | - | 0 | - |
| - | - | 2418 | 380.2 | - | - | 0 | - |
| - | - | 2.558E+04 | 381.2 | - | - | 0 | - |
| - | - | 656.8 | 381.7 | - | - | 0 | - |
| - | - | 4836 | 382.2 | - | - | 0 | - |
| - | - | 1232 | 382.2 | - | - | 0 | - |
| - | - | 1065 | 384.2 | - | - | 0 | - |
| - | - | 1037 | 384.2 | - | - | 0 | - |
| - | - | 1489 | 385.1 | - | - | 0 | - |
| - | - | 1366 | 390.2 | - | - | 0 | - |
| - | - | 811 | 390.7 | - | - | 0 | - |
| - | - | 868.5 | 391.1 | - | - | 0 | - |
| - | - | 4598 | 394.2 | - | - | 0 | - |
| - | - | 1351 | 395.1 | - | - | 0 | - |
| - | - | 792 | 395.2 | - | - | 0 | - |
| - | - | 4891 | 396.2 | - | - | 0 | - |
| - | - | 695.1 | 397.2 | - | - | 0 | - |
| - | - | 2802 | 398.2 | - | - | 0 | - |
| - | - | 1541 | 402.2 | - | - | 0 | - |
| - | - | 2114 | 402.2 | - | - | 0 | - |
| - | - | 784 | 405.3 | - | - | 0 | - |
| - | - | 1976 | 409.1 | - | - | 0 | - |
| - | - | 635.4 | 409.2 | - | - | 0 | - |
| - | - | 2562 | 411.2 | - | - | 0 | - |
| - | - | 1.586E+04 | 412.2 | - | - | 0 | - |
| - | - | 2107 | 413.2 | - | - | 0 | - |
| - | - | 2211 | 413.2 | - | - | 0 | - |
| - | - | 919.1 | 413.3 | - | - | 0 | - |
| - | - | 4728 | 414.2 | - | - | 0 | - |
| - | - | 2.5E+04 | 415 | - | - | 0 | - |
| - | - | 1315 | 415.2 | - | - | 0 | - |
| - | - | 1071 | 415.2 | - | - | 0 | - |
| - | - | 3325 | 417.7 | - | - | 0 | - |
| - | - | 6181 | 418.2 | - | - | 0 | - |
| - | - | 2264 | 418.7 | - | - | 0 | - |
| 11 | y | 741.7 | 421.2 | 0.002563 | 6.084 | +1 | 3 |
| - | - | 6288 | 423.3 | - | - | 0 | - |
| - | - | 1365 | 424.3 | - | - | 0 | - |
| - | - | 4.722E+04 | 426.7 | - | - | 0 | - |
| - | - | 2.207E+04 | 427.2 | - | - | 0 | - |
| - | - | 5676 | 427.7 | - | - | 0 | - |
| - | - | 687 | 428.2 | - | - | 0 | - |
| - | - | 1366 | 429.1 | - | - | 0 | - |
| - | - | 1.174E+04 | 429.2 | - | - | 0 | - |
| - | - | 3.597E+04 | 430.2 | - | - | 0 | - |
| - | - | 8346 | 431.2 | - | - | 0 | - |
| 7 | b | 5156 | 431.7 | 4.727E-05 | 0.1095 | +2 | 7 |
| 7 | b | 3084 | 432.2 | 0.005076 | 11.75 | +2 | 7 |
| - | - | 1578 | 432.7 | - | - | 0 | - |
| - | - | 706.3 | 432.7 | - | - | 0 | - |
| - | - | 1198 | 433 | - | - | 0 | - |
| - | - | 863.8 | 433.2 | - | - | 0 | - |
| 4 | b | 2077 | 433.3 | 0.0009887 | 2.282 | +1 | 4 |
| - | - | 813.5 | 438.2 | - | - | 0 | - |
| - | - | 1131 | 438.7 | - | - | 0 | - |
| 11 | y | 1.116E+04 | 439.2 | 0.0004209 | 0.9584 | +1 | 3 |
| - | - | 2635 | 440.2 | - | - | 0 | - |
| 7 | b | 2.167E+04 | 440.7 | 7.2E-05 | 0.1634 | +2 | 7 |
| - | - | 1.169E+04 | 441.2 | - | - | 0 | - |
| - | - | 719.7 | 441.3 | - | - | 0 | - |
| - | - | 4512 | 441.7 | - | - | 0 | - |
| - | - | 871.3 | 442.2 | - | - | 0 | - |
| - | - | 941.1 | 442.7 | - | - | 0 | - |
| 7 | y | 1211 | 447.2 | 0.0008635 | 1.931 | +2 | 7 |
| - | - | 1571 | 448.2 | - | - | 0 | - |
| - | - | 1128 | 449.2 | - | - | 0 | - |
| - | - | 756.6 | 450.3 | - | - | 0 | - |
| 4 | b | 1.286E+04 | 451.3 | 0.0002264 | 0.5016 | +1 | 4 |
| - | - | 3578 | 452.3 | - | - | 0 | - |
| - | - | 919.1 | 455.2 | - | - | 0 | - |
| - | - | 791.4 | 455.3 | - | - | 0 | - |
| - | - | 745.8 | 459.2 | - | - | 0 | - |
| - | - | 1131 | 461.2 | - | - | 0 | - |
| - | - | 1629 | 461.3 | - | - | 0 | - |
| - | - | 2323 | 468.3 | - | - | 0 | - |
| - | - | 845.3 | 469.3 | - | - | 0 | - |
| - | - | 2319 | 474.3 | - | - | 0 | - |
| - | - | 3969 | 474.8 | - | - | 0 | - |
| - | - | 1056 | 475.3 | - | - | 0 | - |
| - | - | 1020 | 475.8 | - | - | 0 | - |
| - | - | 1023 | 476.2 | - | - | 0 | - |
| - | - | 1765 | 477.2 | - | - | 0 | - |
| 10 | y | 830.8 | 478.2 | 0.002004 | 4.19 | +1 | 4 |
| - | - | 1507 | 478.3 | - | - | 0 | - |
| - | - | 7875 | 483.3 | - | - | 0 | - |
| - | - | 6604 | 483.8 | - | - | 0 | - |
| - | - | 1059 | 484.3 | - | - | 0 | - |
| 8 | b | 2749 | 488.3 | 0.000859 | 1.759 | +2 | 8 |
| 8 | b | 1442 | 488.7 | 0.008485 | 17.36 | +2 | 8 |
| - | - | 2.438E+04 | 494.2 | - | - | 0 | - |
| - | - | 6362 | 495.2 | - | - | 0 | - |
| 10 | y | 2.947E+04 | 496.2 | 0.0003806 | 0.7669 | +1 | 4 |
| - | - | 7882 | 497.2 | - | - | 0 | - |
| - | - | 2891 | 497.8 | - | - | 0 | - |
| - | - | 1666 | 498.2 | - | - | 0 | - |
| - | - | 3.002E+04 | 503.1 | - | - | 0 | - |
| - | - | 1478 | 504.2 | - | - | 0 | - |
| - | - | 1398 | 506.1 | - | - | 0 | - |
| - | - | 1190 | 508.1 | - | - | 0 | - |
| - | - | 2375 | 511.3 | - | - | 0 | - |
| - | - | 1.129E+04 | 519.1 | - | - | 0 | - |
| - | - | 1481 | 521.8 | - | - | 0 | - |
| - | - | 1292 | 522.3 | - | - | 0 | - |
| - | - | 906.2 | 523.3 | - | - | 0 | - |
| - | - | 4396 | 525.3 | - | - | 0 | - |
| - | - | 743.4 | 526.2 | - | - | 0 | - |
| - | - | 1465 | 526.3 | - | - | 0 | - |
| 0 | Precursor | 2174 | 529.3 | 0.001718 | 3.246 | +3 | -1 |
| 0 | Precursor | 2080 | 529.6 | 0.007758 | 14.65 | +3 | -1 |
| - | - | 1168 | 529.9 | - | - | 0 | - |
| - | - | 1110 | 532.2 | - | - | 0 | - |
| - | - | 800.6 | 534.2 | - | - | 0 | - |
| - | - | 1735 | 534.6 | - | - | 0 | - |
| - | - | 1099 | 534.8 | - | - | 0 | - |
| - | - | 1551 | 534.9 | - | - | 0 | - |
| - | - | 1084 | 535.6 | - | - | 0 | - |
| - | - | 1020 | 535.9 | - | - | 0 | - |
| - | - | 6938 | 536.2 | - | - | 0 | - |
| - | - | 1583 | 536.3 | - | - | 0 | - |
| - | - | 739.5 | 537.7 | - | - | 0 | - |
| - | - | 4742 | 539.2 | - | - | 0 | - |
| - | - | 1627 | 540.2 | - | - | 0 | - |
| - | - | 2825 | 542.3 | - | - | 0 | - |
| - | - | 7010 | 543.3 | - | - | 0 | - |
| - | - | 1101 | 544.3 | - | - | 0 | - |
| - | - | 826.8 | 547.3 | - | - | 0 | - |
| - | - | 979 | 549.2 | - | - | 0 | - |
| - | - | 821.5 | 550.2 | - | - | 0 | - |
| - | - | 2181 | 551.3 | - | - | 0 | - |
| - | - | 815.3 | 551.3 | - | - | 0 | - |
| 9 | b | 1263 | 554.8 | 0.009655 | 17.4 | +2 | 9 |
| - | - | 1859 | 555.3 | - | - | 0 | - |
| - | - | 833 | 555.8 | - | - | 0 | - |
| 5 | b | 1308 | 561.3 | 0.002415 | 4.302 | +1 | 5 |
| 5 | b | 758.2 | 562.3 | 0.0003995 | 0.7104 | +1 | 5 |
| - | - | 1163 | 563.3 | - | - | 0 | - |
| - | - | 1648 | 565.3 | - | - | 0 | - |
| - | - | 5480 | 567.2 | - | - | 0 | - |
| - | - | 2659 | 568.2 | - | - | 0 | - |
| - | - | 1457 | 575.2 | - | - | 0 | - |
| - | - | 1093 | 576.2 | - | - | 0 | - |
| 5 | b | 7908 | 579.3 | 0.0002762 | 0.4768 | +1 | 5 |
| - | - | 2828 | 580.3 | - | - | 0 | - |
| - | - | 738.2 | 589.3 | - | - | 0 | - |
| - | - | 1775 | 592.3 | - | - | 0 | - |
| - | - | 1179 | 592.3 | - | - | 0 | - |
| 9 | y | 9813 | 593.3 | 0.0001592 | 0.2683 | +1 | 5 |
| - | - | 3586 | 593.3 | - | - | 0 | - |
| 9 | y | 1790 | 594.2 | 0.01125 | 18.93 | +1 | 5 |
| - | - | 1130 | 602.3 | - | - | 0 | - |
| - | - | 1753 | 602.8 | - | - | 0 | - |
| - | - | 1445 | 603.3 | - | - | 0 | - |
| - | - | 3.044E+04 | 610.3 | - | - | 0 | - |
| 9 | y | 9.882E+04 | 611.3 | 0.0007508 | 1.228 | +1 | 5 |
| - | - | 3966 | 611.8 | - | - | 0 | - |
| - | - | 3.09E+04 | 612.3 | - | - | 0 | - |
| - | - | 1016 | 612.8 | - | - | 0 | - |
| - | - | 7559 | 613.3 | - | - | 0 | - |
| - | - | 964.5 | 614.3 | - | - | 0 | - |
| - | - | 754.1 | 617.3 | - | - | 0 | - |
| - | - | 2391 | 621.3 | - | - | 0 | - |
| - | - | 1382 | 634.3 | - | - | 0 | - |
| - | - | 1413 | 635.3 | - | - | 0 | - |
| - | - | 597.5 | 636.3 | - | - | 0 | - |
| 4 | y | 1424 | 637.3 | 0.01007 | 15.8 | +2 | 10 |
| - | - | 2638 | 645.3 | - | - | 0 | - |
| 4 | y | 5836 | 645.8 | 0.002043 | 3.163 | +2 | 10 |
| - | - | 3739 | 646.3 | - | - | 0 | - |
| - | - | 1555 | 646.8 | - | - | 0 | - |
| - | - | 690.4 | 650.8 | - | - | 0 | - |
| - | - | 1.085E+04 | 652.3 | - | - | 0 | - |
| - | - | 4211 | 653.3 | - | - | 0 | - |
| - | - | 1475 | 654.3 | - | - | 0 | - |
| - | - | 1259 | 659.3 | - | - | 0 | - |
| - | - | 1180 | 660.9 | - | - | 0 | - |
| - | - | 1577 | 661.4 | - | - | 0 | - |
| - | - | 4048 | 662.3 | - | - | 0 | - |
| - | - | 2838 | 663.3 | - | - | 0 | - |
| 6 | b | 5891 | 676.3 | 0.001091 | 1.612 | +1 | 6 |
| 6 | b | 1755 | 677.3 | 0.00879 | 12.98 | +1 | 6 |
| - | - | 1127 | 677.8 | - | - | 0 | - |
| - | - | 1.871E+04 | 680.3 | - | - | 0 | - |
| - | - | 7455 | 681.3 | - | - | 0 | - |
| - | - | 1640 | 682.3 | - | - | 0 | - |
| - | - | 1029 | 684.3 | - | - | 0 | - |
| - | - | 1463 | 684.8 | - | - | 0 | - |
| - | - | 831.3 | 685.3 | - | - | 0 | - |
| - | - | 1097 | 692.8 | - | - | 0 | - |
| 3 | y | 6681 | 693.3 | 0.0001 | 0.1443 | +2 | 11 |
| 3 | y | 6997 | 693.8 | 0.007221 | 10.41 | +2 | 11 |
| 6 | b | 5.081E+04 | 694.4 | 0.001096 | 1.579 | +1 | 6 |
| - | - | 1365 | 694.8 | - | - | 0 | - |
| - | - | 1.939E+04 | 695.4 | - | - | 0 | - |
| - | - | 2271 | 695.9 | - | - | 0 | - |
| - | - | 4815 | 696.4 | - | - | 0 | - |
| - | - | 657.7 | 697.4 | - | - | 0 | - |
| - | - | 884.4 | 698.4 | - | - | 0 | - |
| - | - | 1.122E+04 | 701.8 | - | - | 0 | - |
| - | - | 688 | 701.9 | - | - | 0 | - |
| 3 | y | 3.381E+04 | 702.3 | 0.002064 | 2.939 | +2 | 11 |
| 12 | b | 2.495E+04 | 702.8 | 0.01327 | 18.87 | +2 | 12 |
| 12 | b | 1.216E+04 | 703.3 | 0.003747 | 5.327 | +2 | 12 |
| - | - | 2971 | 703.8 | - | - | 0 | - |
| - | - | 2182 | 704.3 | - | - | 0 | - |
| - | - | 801.3 | 705.4 | - | - | 0 | - |
| 8 | y | 4232 | 706.3 | 5.567E-05 | 0.07882 | +1 | 6 |
| - | - | 1212 | 707.3 | - | - | 0 | - |
| - | - | 1915 | 710.4 | - | - | 0 | - |
| - | - | 1061 | 710.9 | - | - | 0 | - |
| - | - | 2937 | 711.4 | - | - | 0 | - |
| 12 | b | 3319 | 711.9 | 0.001594 | 2.24 | +2 | 12 |
| - | - | 2382 | 712.4 | - | - | 0 | - |
| - | - | 2758 | 721.4 | - | - | 0 | - |
| - | - | 1282 | 722.4 | - | - | 0 | - |
| - | - | 8746 | 723.4 | - | - | 0 | - |
| 8 | y | 3.15E+04 | 724.4 | 0.0007322 | 1.011 | +1 | 6 |
| - | - | 1.138E+04 | 725.4 | - | - | 0 | - |
| - | - | 3024 | 726.4 | - | - | 0 | - |
| - | - | 1054 | 729.4 | - | - | 0 | - |
| - | - | 946.8 | 730.4 | - | - | 0 | - |
| - | - | 1827 | 733.4 | - | - | 0 | - |
| - | - | 1592 | 734.3 | - | - | 0 | - |
| 2 | y | 919.5 | 742.9 | 0.002985 | 4.019 | +2 | 12 |
| - | - | 707.7 | 746.4 | - | - | 0 | - |
| - | - | 6119 | 751.4 | - | - | 0 | - |
| 2 | y | 2883 | 751.9 | 0.001609 | 2.141 | +2 | 12 |
| - | - | 4552 | 752.4 | - | - | 0 | - |
| - | - | 720.7 | 752.9 | - | - | 0 | - |
| - | - | 3307 | 763.4 | - | - | 0 | - |
| - | - | 1542 | 764.4 | - | - | 0 | - |
| - | - | 1574 | 765.4 | - | - | 0 | - |
| - | - | 956.8 | 766.4 | - | - | 0 | - |
| - | - | 1154 | 767.3 | - | - | 0 | - |
| - | - | 3925 | 779.4 | - | - | 0 | - |
| - | - | 2602 | 780.4 | - | - | 0 | - |
| - | - | 1790 | 781.4 | - | - | 0 | - |
| 0 | Precursor | 6152 | 793.4 | 0.01302 | 16.41 | +2 | -1 |
| - | - | 2195 | 794.4 | - | - | 0 | - |
| - | - | 819.6 | 795.3 | - | - | 0 | - |
| - | - | 811.3 | 795.4 | - | - | 0 | - |
| - | - | 1303 | 834.4 | - | - | 0 | - |
| - | - | 1818 | 835.4 | - | - | 0 | - |
| - | - | 2509 | 836.5 | - | - | 0 | - |
| - | - | 1784 | 837.4 | - | - | 0 | - |
| - | - | 806.5 | 844.4 | - | - | 0 | - |
| - | - | 1061 | 845.4 | - | - | 0 | - |
| - | - | 1776 | 846.4 | - | - | 0 | - |
| - | - | 1628 | 847.4 | - | - | 0 | - |
| - | - | 721.6 | 848.5 | - | - | 0 | - |
| - | - | 1.143E+04 | 852.4 | - | - | 0 | - |
| - | - | 6738 | 853.4 | - | - | 0 | - |
| - | - | 1901 | 854.4 | - | - | 0 | - |
| 7 | b | 7947 | 862.4 | 0.001119 | 1.297 | +1 | 7 |
| 7 | b | 5702 | 863.4 | 0.01102 | 12.76 | +1 | 7 |
| - | - | 1.115E+04 | 864.5 | - | - | 0 | - |
| - | - | 5730 | 865.5 | - | - | 0 | - |
| - | - | 771.5 | 866.5 | - | - | 0 | - |
| - | - | 1216 | 876.4 | - | - | 0 | - |
| 7 | b | 6.247E+04 | 880.4 | 0.001369 | 1.554 | +1 | 7 |
| - | - | 3.695E+04 | 881.4 | - | - | 0 | - |
| - | - | 499.3 | 881.5 | - | - | 0 | - |
| - | - | 1.196E+04 | 882.4 | - | - | 0 | - |
| - | - | 2073 | 883.4 | - | - | 0 | - |
| - | - | 723 | 890.4 | - | - | 0 | - |
| - | - | 901.6 | 892.5 | - | - | 0 | - |
| - | - | 1129 | 907.4 | - | - | 0 | - |
| - | - | 2076 | 908.4 | - | - | 0 | - |
| - | - | 2589 | 909.4 | - | - | 0 | - |
| 7 | y | 4443 | 910.4 | 0.000948 | 1.041 | +1 | 7 |
| - | - | 2059 | 911.4 | - | - | 0 | - |
| - | - | 1421 | 954.4 | - | - | 0 | - |
| - | - | 1453 | 955.4 | - | - | 0 | - |
| - | - | 814.1 | 964.5 | - | - | 0 | - |
| - | - | 1228 | 965.5 | - | - | 0 | - |
| - | - | 1217 | 966.4 | - | - | 0 | - |
| 8 | b | 983.8 | 975.5 | 0.0007102 | 0.728 | +1 | 8 |
| 8 | b | 773.5 | 976.5 | 0.00911 | 9.329 | +1 | 8 |
| - | - | 654.4 | 982.5 | - | - | 0 | - |
| 8 | b | 4045 | 993.5 | 0.0005937 | 0.5976 | +1 | 8 |
| - | - | 2501 | 994.5 | - | - | 0 | - |
| - | - | 931 | 995.5 | - | - | 0 | - |
| 6 | y | 2456 | 1025 | 0.00202 | 1.97 | +1 | 8 |
| - | - | 1089 | 1026 | - | - | 0 | - |
| - | - | 748.9 | 1043 | - | - | 0 | - |
| - | - | 1252 | 1094 | - | - | 0 | - |
| 5 | y | 979.7 | 1136 | 0.0007046 | 0.6205 | +1 | 9 |
| 5 | y | 2777 | 1136 | 0.004782 | 4.207 | +1 | 9 |
| - | - | 1406 | 1137 | - | - | 0 | - |
| - | - | 1638 | 1153 | - | - | 0 | - |
| 5 | y | 3959 | 1154 | 0.003379 | 2.929 | +1 | 9 |
| - | - | 3084 | 1155 | - | - | 0 | - |
| - | - | 1312 | 1156 | - | - | 0 | - |
| - | - | 750.9 | 1711 | - | - | 0 | - |
| - | - | 677.2 | 2518 | - | - | 0 | - |
| - | - | 723.3 | 3080 | - | - | 0 | - |
| - | - | 884.6 | 3081 | - | - | 0 | - |

m/z Charge Intensity FragmentType MassShift Position
120.0810546875 0 6723.069
120.100341796875 0 371.844
121.08445739746094 0 704.86597
122.07176208496094 0 1119.5869
122.86956024169922 0 397.80304
123.04459381103516 0 534.7357
126.09160614013672 0 453.8635
127.0505142211914 0 2524.4934
128.10255432128906 0 907.0716
128.10733032226562 0 10056.545
129.06585693359375 0 599.0333
129.10255432128906 0 106938.15
130.0654296875 0 47350.05
130.0697021484375 0 689.29315
130.07127380371094 0 469.6861
130.1002197265625 0 1104.8434
130.10586547851562 0 6655.3477
131.0688018798828 0 4794.04
131.1072540283203 0 393.73267
132.08106994628906 0 11512.79
133.0609588623047 0 596.82135
133.08489990234375 0 962.7794
134.02783203125 0 533.40466
136.07598876953125 0 16300.441
137.07937622070312 0 1446.9677
138.06639099121094 0 1378.3588
139.08689880371094 0 1775.4353
141.10238647460938 0 2697.4006
141.72003173828125 0 409.45486
142.06564331054688 0 583.70215
142.1232147216797 0 569.86395
143.04551696777344 0 721.7762
144.08139038085938 0 609.60425
145.0609893798828 0 802.3665
146.0601806640625 0 1368.4922
147.044189453125 0 641.7048
148.9547576904297 0 1076.2795
149.04513549804688 0 15043.204
150.06639099121094 0 861.4127
150.09210205078125 0 482.64087
151.0870819091797 0 2105.7168
151.32376098632812 0 447.5137
155.08187866210938 0 794.27106
155.09304809570312 0 7067.3145
155.1181182861328 0 5616.5063 a Water loss 1
156.1211700439453 0 692.59467
157.09765625 0 741.45575
158.08416748046875 0 646.8726
159.09194946289062 0 78286.36
160.07638549804688 0 876.4552
160.08836364746094 0 966.42615
160.09535217285156 0 9193.549
160.8106689453125 0 472.23367
161.0989227294922 0 504.1932
165.05487060546875 0 8502.087
166.05360412597656 0 850.89166
166.06129455566406 0 13020.153
166.08645629882812 0 3389.0383 y 9
166.09767150878906 0 1373.836
167.0557098388672 0 23573.188
167.06385803222656 0 1210.5476
168.05487060546875 0 551.3661
168.0812530517578 0 1525.8107
168.10238647460938 0 755.7465
168.1134033203125 0 1238.2546
169.0760040283203 0 1385.2837
169.09732055664062 0 3949.9036
170.06028747558594 0 3723.208
171.06385803222656 0 534.1461
171.14947509765625 0 560.4032
172.072021484375 0 2215.045
173.0560302734375 0 1486.6123
173.1287841796875 0 160905.06 a 1
173.4506072998047 0 2899.8762
174.0556182861328 0 444.15427
174.12408447265625 0 848.0008
174.13214111328125 0 13515.4375
175.13406372070312 0 642.2942
178.13412475585938 0 4500.8545
179.13768005371094 0 965.4342
181.06088256835938 0 930.8034
182.0814208984375 0 16362.489 y 12
183.08494567871094 0 1433.6825
183.1130828857422 0 5108.408 b Water loss 1
185.16531372070312 0 1425.3948
186.12399291992188 0 31273.363
187.08685302734375 0 6765.842
187.1081085205078 0 739.88806
187.12762451171875 0 3310.8289
187.1443634033203 0 5040.6626
188.07069396972656 0 1644.9927
188.09042358398438 0 1028.0043
188.14720153808594 0 713.12335
190.08291625976562 0 771.2862
190.13394165039062 0 1246.0083
194.0924835205078 0 1043.1552
195.11309814453125 0 8366.5
196.10829162597656 0 5545.8086
196.11651611328125 0 762.89636
197.12847900390625 0 1008.80023
198.08770751953125 0 1441.3024
198.12794494628906 0 2482.9082
199.07086181640625 0 593.7243
199.08758544921875 0 930.6026
200.06594848632812 0 513.28253
201.12356567382812 0 46680.035 b 1
201.60848999023438 0 566.96875
202.08633422851562 0 677.6433
202.12681579589844 0 4900.1577
203.12911987304688 0 544.0172
205.0972900390625 0 4204.2134
207.16058349609375 0 1627.715
209.0924530029297 0 671.97614
210.12779235839844 0 1220.1396
211.10787963867188 0 815.53064 y Water loss 10
212.1395263671875 0 1230.9373
213.1236114501953 0 3936.3347
215.1390380859375 0 1820.189
216.09828186035156 0 1268.4315
216.13388061523438 0 755.3743
221.05917358398438 0 1976.2559
221.0846710205078 0 7191.212
221.10345458984375 0 941.9233
221.14002990722656 0 613.6568
222.08590698242188 0 2400.1497
222.1239776611328 0 2409.9348
223.06394958496094 0 4512.876
223.081787109375 0 1296.3772
223.10801696777344 0 2358.12
223.15560913085938 0 13420.446
224.10281372070312 0 667.9581
224.15908813476562 0 2012.3674
225.0431671142578 0 9604.636
226.04405212402344 0 2346.438
226.08253479003906 0 2300.6492
226.1189727783203 0 737.25275 b Water loss 5
226.15560913085938 0 1691.543
227.04029846191406 0 1592.7666
227.0664825439453 0 2364.9958
227.15501403808594 0 1097.0485
228.13467407226562 0 1629.1877
229.11865234375 0 2686.1443
233.13986206054688 0 2552.4727
234.1240234375 0 14633.739
235.12742614746094 0 1548.7825
237.13539123535156 0 1267.1932
238.1189422607422 0 2616.3428
239.09523010253906 0 22649.61
240.09593200683594 0 5762.9805
240.1344451904297 0 6387.06
241.09226989746094 0 3343.1104
241.11880493164062 0 1144.9877
241.13595581054688 0 824.61755
241.19107055664062 0 997.49316
243.10833740234375 0 1051.843
244.09295654296875 0 6336.97
245.0972137451172 0 656.7611
247.10806274414062 0 7881.31
248.1016082763672 0 1095.5446
248.11398315429688 0 3512.7717
249.0538787841797 0 793.5879
249.0984344482422 0 4773.1567
249.13502502441406 0 1680.968
250.10284423828125 0 673.9529
251.1505126953125 0 52892.305
252.15371704101562 0 7466.2744
253.1553955078125 0 707.35565
255.14791870117188 0 3093.4243
257.1131896972656 0 1559.5382
258.14495849609375 0 3039.6018
261.1188659667969 0 1342.9147
261.1339416503906 0 706.5332
262.1190185546875 0 1093.4635
265.1296691894531 0 8210.996
266.11376953125 0 1801.5405
266.1249694824219 0 13111.054
267.12847900390625 0 1552.1748
268.1290283203125 0 1415.3269
268.1769714355469 0 1090.554
269.1608581542969 0 787.8912
269.1865539550781 0 673.01013
270.1220397949219 0 2638.9275
272.1760559082031 0 3326.056
274.11871337890625 0 2834.5361
274.13299560546875 0 872.92084
275.1036682128906 0 1186.9177
276.1088562011719 0 777.9448
276.12701416015625 0 707.1873
276.15594482421875 0 1191.1499
279.14569091796875 0 1256.9574
280.1303405761719 0 1173.8534
281.0513916015625 0 15031.563
282.15631103515625 0 3753.8152
283.03070068359375 0 1912.7162
283.14031982421875 0 42132.52
284.1025390625 0 1269.6708
284.143310546875 0 5242.8657
285.010009765625 0 711.0121
286.1395263671875 0 1537.1511
293.1134338378906 0 21217.547 y Water loss 11
293.13079833984375 0 1112.6887
293.162353515625 0 863.4439
294.1177062988281 0 4167.2197
295.1034851074219 0 2028.737
296.1042175292969 0 1023.22925
296.19635009765625 0 981.95465 b Water loss 2
297.0823974609375 0 2367.286
297.09979248046875 0 914.59076
297.134521484375 0 765.02435 y Water loss 8
297.15594482421875 0 5942.8584
298.1404724121094 0 733.8383 y Water loss 6
298.1584167480469 0 1154.6099
299.0619812011719 0 121604.05
300.0623474121094 0 3390.8057
300.16741943359375 0 6132.7036
301.0601501464844 0 1940.3672
301.1164855957031 0 789.8652
301.15093994140625 0 16471.674
301.1692810058594 0 1008.3309
302.1140441894531 0 3231.397
302.1540222167969 0 2811.3682
303.1678771972656 0 664.5894
305.64605712890625 0 1028.8387
306.1381530761719 0 3989.6995 y 8
306.63922119140625 0 1791.4941
307.14013671875 0 860.22687
311.1238098144531 0 3433.7693 y 11
311.1535339355469 0 734.45776
313.11309814453125 0 1168.2594
314.1146240234375 0 997.32416
314.18798828125 0 1058.8682
314.2076110839844 0 1917.57 b 2
315.1665954589844 0 12391.527
315.1855773925781 0 876.8803
316.1697998046875 0 1392.3424
317.0749206542969 0 565.8242
323.20916748046875 0 801.4661
325.15118408203125 0 676.92194
326.6638488769531 0 1783.5663
326.9664001464844 0 1029.1517
327.1642150878906 0 897.9052
332.20953369140625 0 905.76917
333.1770935058594 0 2121.4082
339.6768798828125 0 599.3546
340.66259765625 0 967.9907
341.01806640625 0 1366.7814
342.1820068359375 0 1244.6843
343.8543701171875 0 539.1355
344.1721496582031 0 1175.557
344.9767150878906 0 15245.663
345.1312255859375 0 1837.7721
346.11480712890625 0 6830.7163
347.02886962890625 0 589.02374
347.11761474609375 0 1245.5146
347.6797180175781 0 2876.5903 b 5
350.21881103515625 0 4269.3213
351.2195129394531 0 935.3364
352.16741943359375 0 619.9063
352.672119140625 0 958.6871
353.6745300292969 0 1460.2971 y Water loss 7
355.070068359375 0 3379.2349
359.02838134765625 0 15240.127
361.1983947753906 0 1155.3256
362.1835632324219 0 1902.7847
362.6792907714844 0 8252.473 y 7
363.14105224609375 0 9460.953
363.18170166015625 0 4177.315
364.1238098144531 0 1370.7269
364.1446228027344 0 1636.8015
366.1779479980469 0 728.80475
367.1394958496094 0 752.86523
367.16253662109375 0 747.3714
368.18975830078125 0 794.18286
369.1219787597656 0 1219.8452
370.1219787597656 0 2480.267
371.1201477050781 0 764.5753
376.1983642578125 0 1040.8279
377.14544677734375 0 727.81714
378.21343994140625 0 3871.9673
379.2091064453125 0 13479.273
380.1650695800781 0 564.4904
380.2120056152344 0 2417.5864
381.1518859863281 0 25578.896
381.71026611328125 0 656.8002
382.15472412109375 0 4836.271
382.1889343261719 0 1231.7808
384.16400146484375 0 1064.7296
384.18988037109375 0 1036.8829
385.14996337890625 0 1488.5525
390.1953125 0 1365.9904
390.69586181640625 0 811.0014
391.13623046875 0 868.51434
394.1726379394531 0 4597.6685
395.1338195800781 0 1351.2816
395.15704345703125 0 792.0201
396.22412109375 0 4891.1353
397.22625732421875 0 695.05676
398.1772155761719 0 2801.8335
402.1764221191406 0 1540.7405
402.1989440917969 0 2113.9446
405.2603454589844 0 784.03754
409.1467590332031 0 1975.9275
409.2017822265625 0 635.3828
411.1991882324219 0 2561.7454
412.1829528808594 0 15857.14
413.1648864746094 0 2106.7478
413.1875305175781 0 2210.8594
413.2503967285156 0 919.0672
414.2354736328125 0 4728.199
415.0368347167969 0 25002.377
415.1999206542969 0 1314.6102
415.23858642578125 0 1070.6986
417.7168884277344 0 3325.4531
418.2109069824219 0 6181.1323
418.7103271484375 0 2263.8289
421.2107238769531 0 741.6954 y Water loss 10
423.2717590332031 0 6288.4053
424.2739562988281 0 1364.5157
426.72198486328125 0 47223.844
427.2236328125 0 22065.305
427.7247314453125 0 5675.772
428.22393798828125 0 687.0099
429.089111328125 0 1365.6434
429.2095947265625 0 11736.841
430.1935729980469 0 35974.99
431.197021484375 0 8345.755
431.7138977050781 0 5155.653 b Water loss 6
432.2110290527344 0 3083.8618 b Ammonia loss 6
432.71026611328125 0 1578.4358
432.74200439453125 0 706.26324
433.0479431152344 0 1198.1553
433.2091369628906 0 863.82306
433.2547912597656 0 2076.6504 b Water loss 3
438.17193603515625 0 813.51416
438.72955322265625 0 1130.8325
439.2191467285156 0 11157.089 y 10
440.2225646972656 0 2635.0425
440.71929931640625 0 21672.844 b 6
441.220947265625 0 11694.211
441.2518615722656 0 719.68335
441.7223815917969 0 4512.371
442.22564697265625 0 871.3059
442.7170104980469 0 941.0591
447.20648193359375 0 1210.8676 y Ammonia loss 6
448.20477294921875 0 1570.5696
449.2073669433594 0 1128.4277
450.28326416015625 0 756.5804
451.2665710449219 0 12860.948 b 3
452.269775390625 0 3578.0442
455.2281799316406 0 919.1244
455.26519775390625 0 791.4096
459.1978759765625 0 745.84204
461.20465087890625 0 1131.0055
461.25238037109375 0 1629.0216
468.29302978515625 0 2322.6123
469.29486083984375 0 845.3053
474.2579345703125 0 2318.9233
474.75421142578125 0 3968.8257
475.2530822753906 0 1055.685
475.7508850097656 0 1020.4314
476.2253723144531 0 1022.9793
477.2100524902344 0 1764.7266
478.23162841796875 0 830.7668 y Water loss 9
478.2779235839844 0 1507.3987
483.2640380859375 0 7875.365
483.7653503417969 0 6603.616
484.26611328125 0 1059.3698
488.2568359375 0 2748.5166 b Water loss 7
488.7564697265625 0 1442.0254 b Ammonia loss 7
494.23577880859375 0 24381.764
495.2385559082031 0 6361.869
496.2405700683594 0 29467.82 y 9
497.24652099609375 0 7881.527
497.763427734375 0 2890.5918
498.2425537109375 0 1665.5859
503.1075744628906 0 30016.316
504.1966857910156 0 1478.2096
506.1327209472656 0 1398.1008
508.0943298339844 0 1189.9865
511.2630920410156 0 2374.8823
519.138916015625 0 11287.688
521.78271484375 0 1480.9612
522.2850952148438 0 1291.9546
523.2572631835938 0 906.24567
525.26611328125 0 4396.126
526.2286376953125 0 743.44165
526.2696533203125 0 1464.7744
529.2614135742188 0 2174.196 Precursor Water loss
529.595458984375 0 2080.0022 Precursor Ammonia loss
529.9284057617188 0 1167.9951
532.1929931640625 0 1109.9974
534.1781616210938 0 800.587
534.5526733398438 0 1734.9443
534.826904296875 0 1098.9967
534.8889770507812 0 1551.1367
535.59814453125 0 1084.2977
535.9312133789062 0 1019.7546
536.1658935546875 0 6937.8257
536.2686767578125 0 1583.1011
537.7322998046875 0 739.4987
539.2359008789062 0 4742.3735
540.2395629882812 0 1626.8625
542.2940673828125 0 2824.772
543.277587890625 0 7010.4136
544.2817993164062 0 1101.2108
547.279541015625 0 826.80786
549.220703125 0 979.016
550.2100219726562 0 821.5343
551.257080078125 0 2181.1611
551.3302612304688 0 815.32104
554.7650756835938 0 1262.8577 b 8
555.25732421875 0 1859.0522
555.7562255859375 0 833.0441
561.3167724609375 0 1308.0427 b Water loss 4
562.2979736328125 0 758.22675 b Ammonia loss 4
563.2548217773438 0 1162.8193
565.26220703125 0 1647.551
567.2313232421875 0 5480.309
568.23486328125 0 2659.001
575.2449951171875 0 1457.3257
576.2321166992188 0 1093.4836
579.3246459960938 0 7908.406 b 4
580.326904296875 0 2828.1318
589.3076171875 0 738.2102
592.2717895507812 0 1774.6571
592.3232421875 0 1178.6532
593.2564086914062 0 9813.075 y Water loss 8
593.3045654296875 0 3586.0264
594.2518310546875 0 1789.8978 y Ammonia loss 8
602.2992553710938 0 1129.8278
602.7918090820312 0 1752.5054
603.2891845703125 0 1445.3606
610.2828979492188 0 30439.924
611.2678833007812 0 98820.77 y 8
611.7980346679688 0 3966.381
612.2706909179688 0 30897.031
612.8018188476562 0 1015.9387
613.2723388671875 0 7559.0903
614.2747192382812 0 964.54663
617.2830810546875 0 754.1073
621.2507934570312 0 2390.7942
634.3058471679688 0 1381.8721
635.294921875 0 1412.597
636.322998046875 0 597.45593
637.2879028320312 0 1423.7955 y Ammonia loss 3
645.2965087890625 0 2638.3364
645.7931518554688 0 5836.1787 y 3
646.2911987304688 0 3739.4927
646.7928466796875 0 1555.4856
650.7913208007812 0 690.3595
652.3198852539062 0 10853.017
653.322265625 0 4211.397
654.3284301757812 0 1474.7426
659.31494140625 0 1258.5332
660.8687133789062 0 1179.7587
661.37451171875 0 1577.2456
662.3043823242188 0 4047.6582
663.2985229492188 0 2838.0586
676.3402099609375 0 5891.488 b Water loss 5
677.3341064453125 0 1754.8254 b Ammonia loss 5
677.8099365234375 0 1126.7947
680.3151245117188 0 18710.621
681.3175659179688 0 7454.8677
682.3224487304688 0 1639.8865
684.3171997070312 0 1029.4911
684.8204345703125 0 1463.0505
685.3143310546875 0 831.28656
692.8350830078125 0 1097.081
693.3277587890625 0 6680.869 y Water loss 2
693.8270874023438 0 6996.6465 y Ammonia loss 2
694.3507690429688 0 50809.074 b 5
694.8240966796875 0 1364.6648
695.3549194335938 0 19390.95
695.8760986328125 0 2270.5378
696.3612670898438 0 4814.798
697.359375 0 657.7259
698.3697509765625 0 884.4071
701.8405151367188 0 11220.877
701.9047241210938 0 687.9632
702.335205078125 0 33810.387 y 2
702.835693359375 0 24945.035 b Water loss 11
703.3372192382812 0 12158.389 b Ammonia loss 11
703.8403930664062 0 2971.0386
704.3339233398438 0 2182.401
705.3521728515625 0 801.3288
706.340576171875 0 4231.76 y Water loss 7
707.3392944335938 0 1212.4594
710.4053955078125 0 1914.8005
710.906982421875 0 1061.3767
711.3709106445312 0 2937.4907
711.8558349609375 0 3319.483 b 11
712.3613891601562 0 2381.9014
721.3615112304688 0 2757.6243
722.3643798828125 0 1281.8821
723.3662109375 0 8746.476
724.3519287109375 0 31495.703 y 7
725.3545532226562 0 11377.837
726.3585205078125 0 3024.1965
729.3624267578125 0 1053.7758
730.3674926757812 0 946.80725
733.3600463867188 0 1826.96
734.3494262695312 0 1592.3171
742.8650512695312 0 919.5175 y Water loss 1
746.35546875 0 707.6519
751.37548828125 0 6118.872
751.8689575195312 0 2882.5667 y 1
752.3746337890625 0 4552.1177
752.8739624023438 0 720.6788
763.3656005859375 0 3307.0396
764.3653564453125 0 1541.8182
765.4000854492188 0 1574.0544
766.4066162109375 0 956.8202
767.3471069335938 0 1154.2311
779.3829345703125 0 3924.8494
780.3838500976562 0 2601.5579
781.3657836914062 0 1789.7169
793.39892578125 0 6152.1743 Precursor Water loss
794.4041748046875 0 2194.8506
795.3344116210938 0 819.6467
795.40771484375 0 811.3206
834.42138671875 0 1303.2289
835.4165649414062 0 1818.4408
836.4554443359375 0 2509.4421
837.446533203125 0 1783.5763
844.3858642578125 0 806.4761
845.3945922851562 0 1060.8181
846.4450073242188 0 1776.3582
847.445556640625 0 1628.0034
848.4564819335938 0 721.6221
852.4349365234375 0 11433.787
853.437255859375 0 6738.416
854.4405517578125 0 1900.7003
862.4194946289062 0 7946.6914 b Water loss 6
863.4156494140625 0 5702.0693 b Ammonia loss 6
864.4545288085938 0 11148.583
865.4575805664062 0 5729.7046
866.4653930664062 0 771.4993
876.4393310546875 0 1216.439
880.4298095703125 0 62469.848 b 6
881.4323120117188 0 36952.953
881.508056640625 0 499.30145
882.4357299804688 0 11956.148
883.4397583007812 0 2072.8623
890.4169311523438 0 722.95294
892.46923828125 0 901.5509
907.4415893554688 0 1128.7919
908.4251098632812 0 2076.2402
909.4354858398438 0 2589.044
910.4314575195312 0 4443.4907 y 6
911.4348754882812 0 2058.5503
954.4359741210938 0 1420.7994
955.4190063476562 0 1452.8718
964.4642944335938 0 814.10803
965.450439453125 0 1228.202
966.4475708007812 0 1217.3651
975.5039672851562 0 983.8197 b Water loss 7
976.497802734375 0 773.504 b Ammonia loss 7
982.4761352539062 0 654.4394
993.5146484375 0 4045.4272 b 7
994.5167236328125 0 2501.1897
995.5157470703125 0 931.0022
1025.45947265625 0 2456.4653 y 5
1026.4598388671875 0 1088.5668
1042.55810546875 0 748.8546
1093.5489501953125 0 1252.0172
1135.5047607421875 0 979.6597 y Water loss 4
1136.4942626953125 0 2776.6404 y Ammonia loss 4
1137.4957275390625 0 1406.3658
1152.5260009765625 0 1638.3429
1153.5194091796875 0 3959.0852 y 4
1154.520751953125 0 3084.2307
1155.520263671875 0 1312.386
1710.938720703125 0 750.93475
2518.10791015625 0 677.1838
3080.45703125 0 723.2729
3081.083251953125 0 884.5923

Spectrum Details

|  |  |
| --- | --- |
| Matched peaks? Matched peaks The total absolute number of peaks matched. Additionally in brackets the total fraction of peaks matched and the total number of peaks is shown. | 66 (11.46% of 576) |
| FDR? FDR The false discovery rate estimated for this peptide. It is calculated by matching all theoretical fragments with a non-integer shift with the raw peaks for this spectrum. This is done with 40 different shifts. The resulting percentage is the average number of annotated peaks over the number of annotated peaks with the correct spectrum. | 1.01% |
| Satellite FDR? Satellite FDR See the FDR for details on its calculation. This satellite ion specific FDR only contains the satellite ions (d/w) for I/L/J positions. | ∞ |
| PSM Score? PSM Score The PSM Score as given by Hecklib to this annotated spectrum. It is shown with three significant figures. | 541 |

#### Spectrum 7971? Spectrum 7971 The raw spectrum of this peptide as annotated by Hecklib. The fragments are coloured according to ion type (see legend). Any peaks with a star '\*' as text can be hovered over to see the full details, first the ion type second the mass shift type. By hovering over the amino acids in the peptide or ions in the legend the corresponding peaks are highlighted. By toggling the 'Unassigned' label you can turn the background (unassigned) peaks on or off in the plot. By updating the slider in the Ion legend you can update the spectrum to only show the top X% of the peaks with labels. The top X% means any peak that is within X% of the highest intensity. By dragging in the spectrum you can zoom in to a specific part of the spectrum and use 'Zoom Out' to get back to the original zoom level. The annotation of the spectrum is based on the given sequence in the peptides file and is done with different software so inconsistencies are likely. The peaks are annotated based on the given sequence, with 20 ppm tolerance.

Copy Data

##### Spectrum 7971 (TSV)

###### Preview

```
Loading example...
```

*Click on the button to copy the data to your clipboard.*

Mz MinMz MaxIntensity Max

WidthHeightPeptide font sizePeptide stroke widthSpectrum font sizeSpectrum stroke widthCompact peptide

Ion legend

wxyz

abcd

OtherUnassignedIon^Charge^_Position_Show for top:%

TVLHQDWLDGKEY

02.27e+44.54e+46.81e+49.08e+4

Zoom Out

y^+3^_4_y^+1^_1_c^+1^_2_y^+1^_2_z^+1^_2_y^+2^_5_y^+2^_5_y^+1^_2_c^+1^_3_c^+2^_6_y^+2^_6_z^+1^_3_y^+1^_3_c^+2^_7_c^+1^_4_c^+1^_4_z^+1^_4_y^+1^_4_w^+1^_5_c^+1^_5_y^+1^_5_z^+1^_5_c^+1^_5_y^+1^_5_y^+2^_10_c^+2^_11_w^+1^_6_w^+2^_11_y^+2^_11_c^+1^_6_y^+2^_11_z^+1^_6_c^+1^_6_c^+2^_12_c^+2^_12_y^+1^_6_w^+2^_12_y^+2^_12_z^+2^_12_c^+1^_7_z^+1^_7_c^+1^_7_y^+1^_7_z^+1^_8_c^+1^_8_y^+1^_8_w^+1^_9_c^+1^_9_y^+1^_9_z^+1^_9_y^+1^_9_c^+1^_10_y^+1^_10_z^+1^_10_y^+1^_10_c^+1^_11_z^+1^_11_y^+1^_11_c^+1^_12_

0739147822182957

Fragment Matches Table

Show background peaks

| Position | Ion type | Intensity | mz Theoretical | mz Error (Th) | mz Error (ppm) | Charge | Series Number |
| --- | --- | --- | --- | --- | --- | --- | --- |
| - | - | 467 | 120.1 | - | - | 0 | - |
| - | - | 572.2 | 120.1 | - | - | 0 | - |
| - | - | 654 | 128.1 | - | - | 0 | - |
| - | - | 1584 | 129.1 | - | - | 0 | - |
| - | - | 436.3 | 129.7 | - | - | 0 | - |
| - | - | 2398 | 130.1 | - | - | 0 | - |
| - | - | 408.8 | 131.1 | - | - | 0 | - |
| - | - | 1050 | 131.1 | - | - | 0 | - |
| - | - | 578.1 | 136.1 | - | - | 0 | - |
| - | - | 406.3 | 144.6 | - | - | 0 | - |
| - | - | 8621 | 147 | - | - | 0 | - |
| - | - | 464.9 | 148.9 | - | - | 0 | - |
| - | - | 6868 | 149 | - | - | 0 | - |
| - | - | 623.1 | 155.1 | - | - | 0 | - |
| - | - | 7890 | 155.1 | - | - | 0 | - |
| - | - | 750.6 | 156.1 | - | - | 0 | - |
| - | - | 737.3 | 157.1 | - | - | 0 | - |
| - | - | 2826 | 159.1 | - | - | 0 | - |
| - | - | 627 | 159.1 | - | - | 0 | - |
| - | - | 1847 | 161.1 | - | - | 0 | - |
| 10 | y | 698.4 | 166.1 | 0.001542 | 9.285 | +3 | 4 |
| - | - | 1778 | 167.1 | - | - | 0 | - |
| - | - | 469.5 | 167.5 | - | - | 0 | - |
| - | - | 448.1 | 168.9 | - | - | 0 | - |
| - | - | 685.4 | 173.1 | - | - | 0 | - |
| - | - | 1.349E+04 | 173.1 | - | - | 0 | - |
| - | - | 3047 | 173.5 | - | - | 0 | - |
| - | - | 1337 | 174.1 | - | - | 0 | - |
| 13 | y | 2729 | 182.1 | 8.178E-06 | 0.04492 | +1 | 1 |
| - | - | 1131 | 186.1 | - | - | 0 | - |
| - | - | 807.5 | 188.1 | - | - | 0 | - |
| - | - | 591.2 | 191.1 | - | - | 0 | - |
| - | - | 7249 | 191.1 | - | - | 0 | - |
| - | - | 8660 | 201.1 | - | - | 0 | - |
| - | - | 617.8 | 202.1 | - | - | 0 | - |
| - | - | 917.9 | 205.1 | - | - | 0 | - |
| - | - | 1.152E+04 | 205.1 | - | - | 0 | - |
| - | - | 989.7 | 206.1 | - | - | 0 | - |
| - | - | 4309 | 215.1 | - | - | 0 | - |
| 2 | c | 3326 | 218.1 | 9.121E-05 | 0.4181 | +1 | 2 |
| - | - | 8610 | 221.1 | - | - | 0 | - |
| - | - | 2402 | 222.1 | - | - | 0 | - |
| - | - | 2910 | 223.1 | - | - | 0 | - |
| - | - | 1853 | 223.1 | - | - | 0 | - |
| - | - | 2130 | 225 | - | - | 0 | - |
| - | - | 943.3 | 226.1 | - | - | 0 | - |
| - | - | 508 | 229.1 | - | - | 0 | - |
| - | - | 8544 | 235.1 | - | - | 0 | - |
| - | - | 1232 | 236.1 | - | - | 0 | - |
| - | - | 1288 | 237.1 | - | - | 0 | - |
| - | - | 1.247E+04 | 239.1 | - | - | 0 | - |
| - | - | 4140 | 240.1 | - | - | 0 | - |
| - | - | 1243 | 241.1 | - | - | 0 | - |
| - | - | 899.2 | 243.1 | - | - | 0 | - |
| - | - | 1372 | 245.1 | - | - | 0 | - |
| - | - | 2978 | 249.1 | - | - | 0 | - |
| - | - | 2380 | 251.2 | - | - | 0 | - |
| - | - | 4756 | 258.1 | - | - | 0 | - |
| - | - | 892.2 | 260.2 | - | - | 0 | - |
| - | - | 4697 | 268.2 | - | - | 0 | - |
| - | - | 537.9 | 269.2 | - | - | 0 | - |
| - | - | 539.4 | 277.1 | - | - | 0 | - |
| - | - | 3325 | 279.1 | - | - | 0 | - |
| - | - | 661.9 | 280.1 | - | - | 0 | - |
| - | - | 1.941E+04 | 281.1 | - | - | 0 | - |
| - | - | 908.4 | 281.1 | - | - | 0 | - |
| - | - | 1085 | 283.1 | - | - | 0 | - |
| 12 | y | 1540 | 293.1 | 0.0005714 | 1.95 | +1 | 2 |
| - | - | 1619 | 293.2 | - | - | 0 | - |
| 12 | z | 3597 | 295.1 | 0.001828 | 6.195 | +1 | 2 |
| - | - | 3486 | 296.1 | - | - | 0 | - |
| - | - | 3769 | 297.1 | - | - | 0 | - |
| - | - | 1595 | 297.1 | - | - | 0 | - |
| 9 | y | 950.9 | 297.1 | 0.004339 | 14.6 | +2 | 5 |
| - | - | 551.4 | 298.1 | - | - | 0 | - |
| - | - | 7.725E+04 | 299.1 | - | - | 0 | - |
| - | - | 1174 | 300.1 | - | - | 0 | - |
| - | - | 1091 | 301.1 | - | - | 0 | - |
| - | - | 667.3 | 301.2 | - | - | 0 | - |
| - | - | 613.3 | 301.8 | - | - | 0 | - |
| 9 | y | 783.1 | 306.1 | 0.0005824 | 1.902 | +2 | 5 |
| 12 | y | 713.6 | 311.1 | 0.0002301 | 0.7397 | +1 | 2 |
| - | - | 884.1 | 314.1 | - | - | 0 | - |
| - | - | 873.5 | 314.2 | - | - | 0 | - |
| - | - | 520.8 | 315.2 | - | - | 0 | - |
| - | - | 8550 | 317.2 | - | - | 0 | - |
| - | - | 1024 | 318.2 | - | - | 0 | - |
| - | - | 1301 | 320.2 | - | - | 0 | - |
| - | - | 749.3 | 323.2 | - | - | 0 | - |
| 3 | c | 4251 | 331.2 | 0.0001489 | 0.4496 | +1 | 3 |
| - | - | 1861 | 341 | - | - | 0 | - |
| - | - | 635 | 343.1 | - | - | 0 | - |
| 6 | c | 1018 | 347.7 | 0.000666 | 1.915 | +2 | 6 |
| - | - | 1156 | 355.1 | - | - | 0 | - |
| - | - | 513.8 | 357.1 | - | - | 0 | - |
| - | - | 6762 | 359 | - | - | 0 | - |
| 8 | y | 846.9 | 362.7 | 0.0006036 | 1.664 | +2 | 6 |
| - | - | 733.3 | 364.2 | - | - | 0 | - |
| - | - | 669.9 | 367.2 | - | - | 0 | - |
| - | - | 2076 | 369.1 | - | - | 0 | - |
| - | - | 3514 | 370.1 | - | - | 0 | - |
| - | - | 2580 | 371.1 | - | - | 0 | - |
| - | - | 1281 | 374.2 | - | - | 0 | - |
| - | - | 1455 | 381.2 | - | - | 0 | - |
| - | - | 559.1 | 388.7 | - | - | 0 | - |
| - | - | 6190 | 411.2 | - | - | 0 | - |
| - | - | 1396 | 412.2 | - | - | 0 | - |
| - | - | 2.159E+04 | 415 | - | - | 0 | - |
| 11 | z | 801.9 | 423.2 | 0.0006208 | 1.467 | +1 | 3 |
| - | - | 3723 | 423.3 | - | - | 0 | - |
| - | - | 1075 | 424.3 | - | - | 0 | - |
| - | - | 5411 | 426.7 | - | - | 0 | - |
| - | - | 2601 | 427.2 | - | - | 0 | - |
| - | - | 1113 | 427.7 | - | - | 0 | - |
| - | - | 1149 | 429.1 | - | - | 0 | - |
| - | - | 944 | 429.2 | - | - | 0 | - |
| - | - | 983.7 | 430.2 | - | - | 0 | - |
| 11 | y | 1898 | 439.2 | 0.001428 | 3.251 | +1 | 3 |
| 7 | c | 4558 | 440.7 | 0.0002246 | 0.5096 | +2 | 7 |
| - | - | 993 | 441.2 | - | - | 0 | - |
| - | - | 1031 | 441.7 | - | - | 0 | - |
| - | - | 845.7 | 444.2 | - | - | 0 | - |
| - | - | 799.3 | 445.2 | - | - | 0 | - |
| 4 | c | 561.9 | 450.3 | 0.0007825 | 1.738 | +1 | 4 |
| - | - | 8750 | 451.3 | - | - | 0 | - |
| - | - | 3113 | 452.3 | - | - | 0 | - |
| - | - | 1102 | 453.3 | - | - | 0 | - |
| - | - | 841.6 | 454.3 | - | - | 0 | - |
| 4 | c | 3.36E+04 | 468.3 | 0.0003496 | 0.7466 | +1 | 4 |
| - | - | 9235 | 469.3 | - | - | 0 | - |
| - | - | 1228 | 470.3 | - | - | 0 | - |
| - | - | 918.7 | 471.3 | - | - | 0 | - |
| 10 | z | 3808 | 480.2 | 0.0001227 | 0.2555 | +1 | 4 |
| - | - | 1498 | 481.2 | - | - | 0 | - |
| - | - | 1527 | 483.3 | - | - | 0 | - |
| - | - | 852.2 | 489.2 | - | - | 0 | - |
| - | - | 2077 | 494.3 | - | - | 0 | - |
| - | - | 1153 | 495.2 | - | - | 0 | - |
| 10 | y | 8474 | 496.2 | 0.000289 | 0.5824 | +1 | 4 |
| - | - | 2524 | 497.2 | - | - | 0 | - |
| - | - | 2.318E+04 | 499.3 | - | - | 0 | - |
| - | - | 8051 | 500.3 | - | - | 0 | - |
| - | - | 2722 | 501.3 | - | - | 0 | - |
| - | - | 6.07E+04 | 503.1 | - | - | 0 | - |
| - | - | 2.311E+04 | 519.1 | - | - | 0 | - |
| - | - | 637.6 | 524.2 | - | - | 0 | - |
| - | - | 5317 | 534.3 | - | - | 0 | - |
| - | - | 2670 | 535.3 | - | - | 0 | - |
| - | - | 1.379E+04 | 536.2 | - | - | 0 | - |
| - | - | 646.7 | 536.3 | - | - | 0 | - |
| - | - | 3881 | 538.3 | - | - | 0 | - |
| - | - | 1005 | 539.3 | - | - | 0 | - |
| 9 | w | 3809 | 550.3 | 0.0006496 | 1.181 | +1 | 5 |
| - | - | 2808 | 551.3 | - | - | 0 | - |
| - | - | 837.5 | 552.3 | - | - | 0 | - |
| 5 | c | 1632 | 579.3 | 0.0006393 | 1.104 | +1 | 5 |
| - | - | 671 | 583.7 | - | - | 0 | - |
| 9 | y | 976.2 | 593.3 | 0.001306 | 2.201 | +1 | 5 |
| - | - | 1530 | 594.3 | - | - | 0 | - |
| 9 | z | 3592 | 595.2 | 0.001958 | 3.289 | +1 | 5 |
| - | - | 1759 | 596.3 | - | - | 0 | - |
| 5 | c | 1.994E+04 | 596.4 | 0.0002133 | 0.3576 | +1 | 5 |
| - | - | 7560 | 597.4 | - | - | 0 | - |
| - | - | 1888 | 598.4 | - | - | 0 | - |
| - | - | 607.1 | 602.3 | - | - | 0 | - |
| - | - | 837 | 609.3 | - | - | 0 | - |
| - | - | 5998 | 610.3 | - | - | 0 | - |
| 9 | y | 1.393E+04 | 611.3 | 0.001544 | 2.526 | +1 | 5 |
| - | - | 1524 | 611.8 | - | - | 0 | - |
| - | - | 4321 | 612.3 | - | - | 0 | - |
| - | - | 583.1 | 621.3 | - | - | 0 | - |
| - | - | 705.5 | 627.3 | - | - | 0 | - |
| 4 | y | 1402 | 645.8 | 0.001925 | 2.98 | +2 | 10 |
| - | - | 1951 | 654.8 | - | - | 0 | - |
| - | - | 1.157E+04 | 655.4 | - | - | 0 | - |
| 11 | c | 3.884E+04 | 655.8 | 0.001132 | 1.727 | +2 | 11 |
| - | - | 2.634E+04 | 656.3 | - | - | 0 | - |
| - | - | 1.102E+04 | 656.8 | - | - | 0 | - |
| - | - | 2857 | 657.4 | - | - | 0 | - |
| - | - | 5565 | 664.3 | - | - | 0 | - |
| - | - | 829.8 | 664.3 | - | - | 0 | - |
| 8 | w | 6176 | 665.3 | 0.006116 | 9.194 | +1 | 6 |
| - | - | 2141 | 666.3 | - | - | 0 | - |
| - | - | 1019 | 666.4 | - | - | 0 | - |
| - | - | 1806 | 667.4 | - | - | 0 | - |
| - | - | 1802 | 668.4 | - | - | 0 | - |
| - | - | 1611 | 668.9 | - | - | 0 | - |
| - | - | 2113 | 669.4 | - | - | 0 | - |
| - | - | 671.8 | 669.9 | - | - | 0 | - |
| - | - | 2942 | 672.3 | - | - | 0 | - |
| 3 | w | 4026 | 672.8 | 0.005122 | 7.613 | +2 | 11 |
| - | - | 2614 | 673.3 | - | - | 0 | - |
| - | - | 624 | 673.8 | - | - | 0 | - |
| - | - | 1806 | 680.3 | - | - | 0 | - |
| - | - | 574.5 | 688.2 | - | - | 0 | - |
| 3 | y | 2648 | 693.8 | 0.01174 | 16.92 | +2 | 11 |
| 6 | c | 3837 | 694.4 | 0.01092 | 15.73 | +1 | 6 |
| - | - | 2030 | 694.8 | - | - | 0 | - |
| - | - | 1867 | 695.4 | - | - | 0 | - |
| - | - | 726.9 | 697.9 | - | - | 0 | - |
| - | - | 799.4 | 698.4 | - | - | 0 | - |
| - | - | 708.1 | 698.9 | - | - | 0 | - |
| - | - | 3711 | 701.8 | - | - | 0 | - |
| 3 | y | 6715 | 702.3 | 0.003468 | 4.937 | +2 | 11 |
| - | - | 4581 | 702.8 | - | - | 0 | - |
| - | - | 1996 | 703.3 | - | - | 0 | - |
| - | - | 3443 | 707.3 | - | - | 0 | - |
| 8 | z | 5799 | 708.3 | 0.004625 | 6.529 | +1 | 6 |
| - | - | 2483 | 709.3 | - | - | 0 | - |
| - | - | 705.4 | 710.9 | - | - | 0 | - |
| 6 | c | 1.891E+04 | 711.4 | 0.0004846 | 0.6813 | +1 | 6 |
| 12 | c | 822.8 | 711.9 | 0.003486 | 4.898 | +2 | 12 |
| - | - | 7222 | 712.4 | - | - | 0 | - |
| - | - | 802.1 | 713.4 | - | - | 0 | - |
| - | - | 783.9 | 716.3 | - | - | 0 | - |
| - | - | 1363 | 718.9 | - | - | 0 | - |
| - | - | 768.4 | 719.4 | - | - | 0 | - |
| - | - | 1.226E+04 | 719.9 | - | - | 0 | - |
| 12 | c | 2.627E+04 | 720.4 | 0.003517 | 4.883 | +2 | 12 |
| - | - | 1.997E+04 | 720.9 | - | - | 0 | - |
| - | - | 9381 | 721.4 | - | - | 0 | - |
| - | - | 2537 | 721.9 | - | - | 0 | - |
| - | - | 1231 | 723.4 | - | - | 0 | - |
| 8 | y | 4490 | 724.4 | 0.001831 | 2.528 | +1 | 6 |
| - | - | 2176 | 725.4 | - | - | 0 | - |
| - | - | 762.6 | 726.4 | - | - | 0 | - |
| - | - | 618 | 727.9 | - | - | 0 | - |
| - | - | 717.9 | 729.4 | - | - | 0 | - |
| - | - | 825.1 | 730.4 | - | - | 0 | - |
| - | - | 698.7 | 733.4 | - | - | 0 | - |
| 2 | w | 783 | 736.3 | 0.00256 | 3.476 | +2 | 12 |
| - | - | 2272 | 736.9 | - | - | 0 | - |
| - | - | 5489 | 737.4 | - | - | 0 | - |
| - | - | 1363 | 737.9 | - | - | 0 | - |
| - | - | 1138 | 741.4 | - | - | 0 | - |
| 2 | y | 912.1 | 743.4 | 0.01433 | 19.28 | +2 | 12 |
| 2 | z | 1733 | 743.9 | 0.008713 | 11.71 | +2 | 12 |
| - | - | 1223 | 744.4 | - | - | 0 | - |
| - | - | 857.6 | 744.9 | - | - | 0 | - |
| - | - | 7535 | 748.9 | - | - | 0 | - |
| - | - | 5990 | 749.4 | - | - | 0 | - |
| - | - | 2811 | 749.9 | - | - | 0 | - |
| - | - | 776.6 | 750.9 | - | - | 0 | - |
| - | - | 1502 | 752.4 | - | - | 0 | - |
| - | - | 920.5 | 757.9 | - | - | 0 | - |
| - | - | 674.3 | 759.4 | - | - | 0 | - |
| - | - | 1063 | 765.4 | - | - | 0 | - |
| - | - | 1738 | 771.9 | - | - | 0 | - |
| - | - | 4235 | 772.4 | - | - | 0 | - |
| - | - | 5474 | 772.9 | - | - | 0 | - |
| - | - | 3141 | 773.4 | - | - | 0 | - |
| - | - | 1861 | 773.9 | - | - | 0 | - |
| - | - | 755.7 | 774.4 | - | - | 0 | - |
| - | - | 4904 | 779.9 | - | - | 0 | - |
| - | - | 3931 | 780.4 | - | - | 0 | - |
| - | - | 2833 | 780.9 | - | - | 0 | - |
| - | - | 1553 | 781.4 | - | - | 0 | - |
| - | - | 959.2 | 785.4 | - | - | 0 | - |
| - | - | 1495 | 785.9 | - | - | 0 | - |
| - | - | 2725 | 786.4 | - | - | 0 | - |
| - | - | 2326 | 786.9 | - | - | 0 | - |
| - | - | 2371 | 787.4 | - | - | 0 | - |
| - | - | 939.4 | 787.9 | - | - | 0 | - |
| - | - | 1425 | 793.4 | - | - | 0 | - |
| - | - | 7895 | 793.9 | - | - | 0 | - |
| - | - | 1.431E+04 | 794.4 | - | - | 0 | - |
| - | - | 1.222E+04 | 794.9 | - | - | 0 | - |
| - | - | 3521 | 795.4 | - | - | 0 | - |
| - | - | 1432 | 795.9 | - | - | 0 | - |
| - | - | 1094 | 796.4 | - | - | 0 | - |
| - | - | 1228 | 800.4 | - | - | 0 | - |
| - | - | 897.4 | 800.9 | - | - | 0 | - |
| - | - | 1637 | 801.4 | - | - | 0 | - |
| - | - | 1181 | 801.8 | - | - | 0 | - |
| - | - | 1.332E+04 | 801.9 | - | - | 0 | - |
| - | - | 6.683E+04 | 802.4 | - | - | 0 | - |
| - | - | 8.989E+04 | 802.9 | - | - | 0 | - |
| - | - | 5.881E+04 | 803.4 | - | - | 0 | - |
| - | - | 2.691E+04 | 803.9 | - | - | 0 | - |
| - | - | 8719 | 804.4 | - | - | 0 | - |
| - | - | 2563 | 804.9 | - | - | 0 | - |
| - | - | 1925 | 837.4 | - | - | 0 | - |
| - | - | 3039 | 838.4 | - | - | 0 | - |
| - | - | 931.9 | 839.4 | - | - | 0 | - |
| - | - | 807.3 | 852.4 | - | - | 0 | - |
| - | - | 1093 | 853.4 | - | - | 0 | - |
| - | - | 1111 | 854.4 | - | - | 0 | - |
| - | - | 693.2 | 863.4 | - | - | 0 | - |
| - | - | 1579 | 864.5 | - | - | 0 | - |
| - | - | 857.4 | 865.5 | - | - | 0 | - |
| 7 | c | 8675 | 880.4 | 0.0005141 | 0.5839 | +1 | 7 |
| - | - | 4430 | 881.4 | - | - | 0 | - |
| - | - | 1412 | 882.4 | - | - | 0 | - |
| - | - | 3354 | 893.4 | - | - | 0 | - |
| 7 | z | 6721 | 894.4 | 0.004718 | 5.276 | +1 | 7 |
| - | - | 3735 | 895.4 | - | - | 0 | - |
| 7 | c | 2.428E+04 | 897.5 | 2.465E-05 | 0.02747 | +1 | 7 |
| - | - | 1.586E+04 | 898.5 | - | - | 0 | - |
| - | - | 4047 | 899.5 | - | - | 0 | - |
| - | - | 2460 | 909.4 | - | - | 0 | - |
| 7 | y | 3650 | 910.4 | 0.004305 | 4.728 | +1 | 7 |
| - | - | 726.3 | 911.4 | - | - | 0 | - |
| - | - | 658.1 | 937.4 | - | - | 0 | - |
| - | - | 1848 | 964.5 | - | - | 0 | - |
| - | - | 2801 | 965.5 | - | - | 0 | - |
| - | - | 822.6 | 965.6 | - | - | 0 | - |
| - | - | 950.5 | 966.4 | - | - | 0 | - |
| - | - | 972.2 | 994.5 | - | - | 0 | - |
| - | - | 6910 | 1008 | - | - | 0 | - |
| 6 | z | 1.141E+04 | 1009 | 0.004814 | 4.769 | +1 | 8 |
| - | - | 4270 | 1010 | - | - | 0 | - |
| 8 | c | 2.016E+04 | 1011 | 0.000323 | 0.3196 | +1 | 8 |
| - | - | 1677 | 1011 | - | - | 0 | - |
| - | - | 1.287E+04 | 1012 | - | - | 0 | - |
| - | - | 4059 | 1013 | - | - | 0 | - |
| - | - | 752.7 | 1014 | - | - | 0 | - |
| - | - | 2114 | 1024 | - | - | 0 | - |
| 6 | y | 3652 | 1025 | 0.007391 | 7.208 | +1 | 8 |
| - | - | 1839 | 1026 | - | - | 0 | - |
| - | - | 736.9 | 1027 | - | - | 0 | - |
| - | - | 612.3 | 1063 | - | - | 0 | - |
| - | - | 908.5 | 1064 | - | - | 0 | - |
| - | - | 716.2 | 1064 | - | - | 0 | - |
| - | - | 3931 | 1068 | - | - | 0 | - |
| - | - | 3020 | 1069 | - | - | 0 | - |
| - | - | 1027 | 1071 | - | - | 0 | - |
| - | - | 966.6 | 1071 | - | - | 0 | - |
| - | - | 1052 | 1072 | - | - | 0 | - |
| - | - | 690.4 | 1072 | - | - | 0 | - |
| 5 | w | 1714 | 1079 | 0.002686 | 2.488 | +1 | 9 |
| - | - | 1686 | 1081 | - | - | 0 | - |
| - | - | 3329 | 1082 | - | - | 0 | - |
| - | - | 2187 | 1083 | - | - | 0 | - |
| - | - | 678.1 | 1095 | - | - | 0 | - |
| - | - | 2434 | 1125 | - | - | 0 | - |
| 9 | c | 6679 | 1126 | 0.00463 | 4.113 | +1 | 9 |
| - | - | 3744 | 1127 | - | - | 0 | - |
| - | - | 811.1 | 1128 | - | - | 0 | - |
| 5 | y | 7275 | 1136 | 0.0209 | 18.39 | +1 | 9 |
| 5 | z | 4.951E+04 | 1137 | 0.0002526 | 0.2221 | +1 | 9 |
| - | - | 2.969E+04 | 1139 | - | - | 0 | - |
| - | - | 1.283E+04 | 1140 | - | - | 0 | - |
| - | - | 3303 | 1141 | - | - | 0 | - |
| - | - | 1122 | 1153 | - | - | 0 | - |
| 5 | y | 2420 | 1154 | 0.003745 | 3.247 | +1 | 9 |
| - | - | 2235 | 1155 | - | - | 0 | - |
| - | - | 1.144E+04 | 1182 | - | - | 0 | - |
| 10 | c | 2.698E+04 | 1183 | 0.003186 | 2.694 | +1 | 10 |
| - | - | 1.695E+04 | 1184 | - | - | 0 | - |
| - | - | 5591 | 1185 | - | - | 0 | - |
| - | - | 2224 | 1186 | - | - | 0 | - |
| - | - | 1111 | 1187 | - | - | 0 | - |
| - | - | 1121 | 1267 | - | - | 0 | - |
| - | - | 777.2 | 1268 | - | - | 0 | - |
| - | - | 1404 | 1269 | - | - | 0 | - |
| 4 | y | 1837 | 1274 | 0.01862 | 14.62 | +1 | 10 |
| 4 | z | 1.062E+04 | 1275 | 0.005184 | 4.067 | +1 | 10 |
| - | - | 9638 | 1276 | - | - | 0 | - |
| - | - | 4361 | 1277 | - | - | 0 | - |
| - | - | 2165 | 1278 | - | - | 0 | - |
| 4 | y | 1526 | 1291 | 0.0009673 | 0.7495 | +1 | 10 |
| - | - | 747.8 | 1293 | - | - | 0 | - |
| - | - | 3658 | 1310 | - | - | 0 | - |
| 11 | c | 1.627E+04 | 1311 | 0.001729 | 1.319 | +1 | 11 |
| - | - | 1.393E+04 | 1312 | - | - | 0 | - |
| - | - | 5025 | 1313 | - | - | 0 | - |
| - | - | 1861 | 1314 | - | - | 0 | - |
| 3 | z | 8743 | 1388 | 0.005104 | 3.678 | +1 | 11 |
| - | - | 9766 | 1389 | - | - | 0 | - |
| - | - | 6207 | 1390 | - | - | 0 | - |
| - | - | 2596 | 1391 | - | - | 0 | - |
| - | - | 2441 | 1396 | - | - | 0 | - |
| - | - | 2253 | 1397 | - | - | 0 | - |
| - | - | 1825 | 1398 | - | - | 0 | - |
| 3 | y | 1099 | 1404 | 0.01092 | 7.777 | +1 | 11 |
| - | - | 955.1 | 1405 | - | - | 0 | - |
| - | - | 706.7 | 1412 | - | - | 0 | - |
| - | - | 993.9 | 1413 | - | - | 0 | - |
| - | - | 2220 | 1424 | - | - | 0 | - |
| - | - | 1349 | 1425 | - | - | 0 | - |
| - | - | 1054 | 1426 | - | - | 0 | - |
| - | - | 1703 | 1439 | - | - | 0 | - |
| 12 | c | 6009 | 1440 | 0.009184 | 6.379 | +1 | 12 |
| - | - | 8122 | 1441 | - | - | 0 | - |
| - | - | 4055 | 1442 | - | - | 0 | - |
| - | - | 2032 | 1443 | - | - | 0 | - |
| - | - | 2258 | 1488 | - | - | 0 | - |
| - | - | 1876 | 1489 | - | - | 0 | - |
| - | - | 1180 | 1490 | - | - | 0 | - |
| - | - | 1567 | 1544 | - | - | 0 | - |
| - | - | 1554 | 1545 | - | - | 0 | - |
| - | - | 3659 | 1546 | - | - | 0 | - |
| - | - | 3131 | 1547 | - | - | 0 | - |
| - | - | 1078 | 1548 | - | - | 0 | - |
| - | - | 848.9 | 1559 | - | - | 0 | - |
| - | - | 4810 | 1560 | - | - | 0 | - |
| - | - | 6016 | 1561 | - | - | 0 | - |
| - | - | 3555 | 1562 | - | - | 0 | - |
| - | - | 1458 | 1563 | - | - | 0 | - |
| - | - | 767.5 | 1570 | - | - | 0 | - |
| - | - | 1768 | 1571 | - | - | 0 | - |
| - | - | 1212 | 1572 | - | - | 0 | - |
| - | - | 932.8 | 1573 | - | - | 0 | - |
| - | - | 4247 | 1577 | - | - | 0 | - |
| - | - | 3968 | 1578 | - | - | 0 | - |
| - | - | 2316 | 1579 | - | - | 0 | - |
| - | - | 1404 | 1580 | - | - | 0 | - |
| - | - | 2482 | 1587 | - | - | 0 | - |
| - | - | 1.56E+04 | 1588 | - | - | 0 | - |
| - | - | 2.407E+04 | 1589 | - | - | 0 | - |
| - | - | 1.655E+04 | 1590 | - | - | 0 | - |
| - | - | 8933 | 1591 | - | - | 0 | - |
| - | - | 3055 | 1592 | - | - | 0 | - |
| - | - | 939.7 | 1593 | - | - | 0 | - |
| - | - | 1235 | 1603 | - | - | 0 | - |
| - | - | 8985 | 1604 | - | - | 0 | - |
| - | - | 2.794E+04 | 1605 | - | - | 0 | - |
| - | - | 4.625E+04 | 1606 | - | - | 0 | - |
| - | - | 3.221E+04 | 1607 | - | - | 0 | - |
| - | - | 1.55E+04 | 1608 | - | - | 0 | - |
| - | - | 5674 | 1609 | - | - | 0 | - |
| - | - | 3341 | 1610 | - | - | 0 | - |
| - | - | 704.8 | 1913 | - | - | 0 | - |
| - | - | 698 | 2928 | - | - | 0 | - |

m/z Charge Intensity FragmentType MassShift Position
120.06597137451172 0 466.9846
120.08085632324219 0 572.18066
128.10711669921875 0 654.04865
129.10238647460938 0 1583.7207
129.71932983398438 0 436.30325
130.06521606445312 0 2398.0225
131.06930541992188 0 408.78687
131.1177978515625 0 1049.7625
136.07574462890625 0 578.05664
144.5670928955078 0 406.25146
147.04727172851562 0 8621.32
148.94802856445312 0 464.9392
149.04490661621094 0 6867.824
155.0860137939453 0 623.0976
155.09283447265625 0 7890.099
156.0963134765625 0 750.56995
157.13352966308594 0 737.3133
159.09165954589844 0 2826.1106
159.1017608642578 0 627.0335
161.08102416992188 0 1846.6921
166.08645629882812 0 698.3799 y 9
167.0557403564453 0 1778.0746
167.4801483154297 0 469.47028
168.87721252441406 0 448.12677
173.12039184570312 0 685.3784
173.12852478027344 0 13489.0205
173.45083618164062 0 3046.9404
174.13229370117188 0 1336.5537
182.08116149902344 0 2729.1943 y 12
186.12399291992188 0 1130.527
188.14002990722656 0 807.48584
191.06515502929688 0 591.1812
191.07342529296875 0 7249.312
201.1233673095703 0 8660.5
202.1262969970703 0 617.78705
205.0970001220703 0 917.90967
205.10711669921875 0 11517.05
206.1104736328125 0 989.6536
215.13905334472656 0 4309.4365
218.15000915527344 0 3325.7542 c 1
221.0844268798828 0 8609.775
222.08486938476562 0 2402.2988
223.06356811523438 0 2910.2812
223.08157348632812 0 1852.6097
225.0431671142578 0 2130.3364
226.10743713378906 0 943.33417
229.1195068359375 0 507.97992
235.099609375 0 8544.358
236.1016845703125 0 1231.7279
237.07945251464844 0 1287.6837
239.094970703125 0 12470.209
240.09547424316406 0 4139.9546
241.0916290283203 0 1243.1549
243.13412475585938 0 899.2264
245.1372833251953 0 1371.9916
249.1336212158203 0 2977.6938
251.1502685546875 0 2379.5562
258.1448974609375 0 4756.0293
260.1600646972656 0 892.2432
268.1766357421875 0 4696.898
269.1835021972656 0 537.875
277.1440734863281 0 539.38245
279.1258544921875 0 3325.436
280.12823486328125 0 661.9199
281.05126953125 0 19409.963
281.1051025390625 0 908.39307
283.1387023925781 0 1084.8881
293.11376953125 0 1540.116 y Water loss 11
293.1600646972656 0 1619.0161
295.10321044921875 0 3597.1262 z 11
296.1036376953125 0 3486.2715
297.0823669433594 0 3769.4905
297.1012878417969 0 1595.009
297.1362609863281 0 950.8569 y Water loss 8
298.10198974609375 0 551.39026
299.06182861328125 0 77253.766
300.060791015625 0 1173.9534
301.05999755859375 0 1091.1222
301.1506652832031 0 667.27795
301.8310852050781 0 613.29865
306.1377868652344 0 783.1322 y 8
311.1239929199219 0 713.6065 y 11
314.1172790527344 0 884.107
314.2070007324219 0 873.5462
315.1678771972656 0 520.7859
317.1583251953125 0 8549.525
318.1612243652344 0 1024.0978
320.197265625 0 1300.7329
323.152587890625 0 749.32825
331.234130859375 0 4250.6694 c 2
341.017578125 0 1860.5756
343.099609375 0 634.9603
347.68023681640625 0 1017.80945 c Ammonia loss 5
355.0697326660156 0 1156.0077
357.0672302246094 0 513.8378
359.0284423828125 0 6761.923
362.6798400878906 0 846.907 y 7
364.18597412109375 0 733.303
367.1791687011719 0 669.9098
369.122314453125 0 2076.4382
370.12249755859375 0 3514.1494
371.1198425292969 0 2579.7087
374.1800842285156 0 1281.4276
381.2123718261719 0 1454.9943
388.6943359375 0 559.09467
411.2044677734375 0 6190.481
412.20782470703125 0 1396.3086
415.03692626953125 0 21586.842
423.20062255859375 0 801.8791 z 10
423.2718505859375 0 3722.569
424.27362060546875 0 1074.9065
426.7220458984375 0 5411.4746
427.2222900390625 0 2600.7996
427.7235107421875 0 1113.4491
429.0888366699219 0 1148.7128
429.20880126953125 0 944.044
430.1930236816406 0 983.7091
439.22015380859375 0 1898.4617 y 10
440.7194519042969 0 4558.2974 c Ammonia loss 6
441.2208251953125 0 993.0154
441.721923828125 0 1031.4606
444.24566650390625 0 845.747
445.2163391113281 0 799.25696
450.2831115722656 0 561.86273 c Water loss 3
451.2667541503906 0 8749.592
452.269775390625 0 3112.531
453.25079345703125 0 1102.0762
454.2550964355469 0 841.61786
468.2932434082031 0 33601.188 c 3
469.2961120605469 0 9234.806
470.2971496582031 0 1227.8665
471.2618103027344 0 918.6522
480.2215881347656 0 3808.499 z 9
481.2261657714844 0 1497.6107
483.264892578125 0 1527.121
489.2052001953125 0 852.19635
494.308837890625 0 2077.2236
495.232177734375 0 1153.1063
496.240478515625 0 8473.594 y 9
497.2436218261719 0 2523.542
499.25732421875 0 23181.947
500.260009765625 0 8051.445
501.2601318359375 0 2722.3408
503.107666015625 0 60702.254
519.138916015625 0 23109.385
524.1781616210938 0 637.6149
534.29638671875 0 5317.345
535.2982788085938 0 2669.7515
536.1653442382812 0 13787.713
536.303466796875 0 646.69116
538.2503051757812 0 3881.1619
539.25439453125 0 1005.44977
550.2514038085938 0 3808.569 w 8
551.256103515625 0 2808.4185
552.2581787109375 0 837.51184
579.3255615234375 0 1632.4713 c Ammonia loss 4
583.711181640625 0 670.9657
593.2578735351562 0 976.1553 y Water loss 8
594.2630004882812 0 1529.71
595.2503662109375 0 3592.4187 z 8
596.2562866210938 0 1758.6382
596.3516845703125 0 19939.496 c 4
597.3544311523438 0 7559.6577
598.357666015625 0 1888.0056
602.2945556640625 0 607.1179
609.3328857421875 0 836.9685
610.2835693359375 0 5998.2935
611.2686767578125 0 13933.911 y 8
611.8004150390625 0 1524.3676
612.2721557617188 0 4321.4766
621.2911987304688 0 583.07166
627.3134765625 0 705.45825
645.7891845703125 0 1401.9878 y 3
654.8474731445312 0 1950.5255
655.352294921875 0 11568.844
655.8473510742188 0 38842.695 c 10
656.3483276367188 0 26336.7
656.8499145507812 0 11015.923
657.3504028320312 0 2857.4412
664.2932739257812 0 5564.592
664.3438110351562 0 829.8324
665.2838134765625 0 6175.947 w 7
666.2838134765625 0 2141.1594
666.3547973632812 0 1018.80023
667.365234375 0 1805.6084
668.3633422851562 0 1802.4744
668.8599853515625 0 1610.789
669.3587646484375 0 2112.5552
669.8535766601562 0 671.7683
672.3032836914062 0 2941.689
672.801513671875 0 4026.2346 w 2
673.2987060546875 0 2613.5037
673.80078125 0 623.9905
680.315673828125 0 1806.3378
688.2075805664062 0 574.4899
693.8316040039062 0 2648.2026 y Ammonia loss 2
694.3409423828125 0 3836.8467 c Ammonia loss 5
694.82666015625 0 2030.4563
695.35302734375 0 1867.0474
697.8668823242188 0 726.9289
698.3660888671875 0 799.3884
698.861083984375 0 708.08795
701.8410034179688 0 3711.2542
702.3366088867188 0 6715.2466 y 2
702.8370361328125 0 4581.257
703.3380126953125 0 1996.1243
707.3490600585938 0 3443.3557
708.3370971679688 0 5798.955 z 7
709.3377075195312 0 2483.3489
710.9081420898438 0 705.4111
711.3779296875 0 18907.9 c 5
711.8577270507812 0 822.76196 c Ammonia loss 11
712.380126953125 0 7221.842
713.3842163085938 0 802.1011
716.329345703125 0 783.9024
718.919921875 0 1362.7814
719.4185180664062 0 768.4122
719.875244140625 0 12256.168
720.3710327148438 0 26266.096 c 11
720.870849609375 0 19968.344
721.3721313476562 0 9380.7
721.8739013671875 0 2537.4563
723.364501953125 0 1230.8083
724.35302734375 0 4490.4946 y 7
725.3555908203125 0 2176.376
726.35791015625 0 762.5703
727.8754272460938 0 618.03186
729.362060546875 0 717.9278
730.3551635742188 0 825.05975
733.3628540039062 0 698.66754
736.3436889648438 0 782.95557 w 1
736.8638916015625 0 2272.224
737.359375 0 5488.909
737.8612060546875 0 1363.4491
741.3621215820312 0 1137.8577
743.368408203125 0 912.0656 y Ammonia loss 1
743.86669921875 0 1732.5685 z 1
744.355712890625 0 1223.3854
744.8662109375 0 857.6299
748.865966796875 0 7535.2856
749.3680419921875 0 5990.01
749.8705444335938 0 2810.9077
750.8749389648438 0 776.6346
752.371337890625 0 1501.9977
757.8943481445312 0 920.4522
759.3636474609375 0 674.26276
765.373291015625 0 1063.4456
771.8902587890625 0 1738.1848
772.3890380859375 0 4235.0986
772.8854370117188 0 5473.5747
773.382568359375 0 3140.95
773.8820190429688 0 1860.6074
774.391357421875 0 755.69525
779.8936767578125 0 4904.0205
780.3922119140625 0 3930.9229
780.893798828125 0 2833.4727
781.395751953125 0 1553.1243
785.384765625 0 959.246
785.8807983398438 0 1495.0488
786.3809814453125 0 2725.328
786.879150390625 0 2326.335
787.3728637695312 0 2371.4153
787.8732299804688 0 939.4008
793.3953857421875 0 1424.6589
793.890625 0 7894.848
794.3862915039062 0 14312.831
794.8857421875 0 12215.25
795.387451171875 0 3520.8264
795.888916015625 0 1432.0829
796.388671875 0 1094.3928
800.4051513671875 0 1227.558
800.8989868164062 0 897.3894
801.4452514648438 0 1637.0613
801.8275756835938 0 1181.2163
801.898681640625 0 13318.149
802.395751953125 0 66829.39
802.89599609375 0 89886.59
803.39697265625 0 58811.965
803.8984375 0 26908.04
804.3990478515625 0 8718.685
804.8982543945312 0 2562.7212
837.3648071289062 0 1924.7119
838.353515625 0 3039.254
839.357421875 0 931.8605
852.4404296875 0 807.3231
853.4390869140625 0 1093.4797
854.4437255859375 0 1111.1473
863.4209594726562 0 693.1806
864.4545288085938 0 1579.0004
865.4595336914062 0 857.36444
880.4306640625 0 8674.502 c Ammonia loss 6
881.4340209960938 0 4429.9697
882.4371337890625 0 1411.874
893.4275512695312 0 3353.6704
894.41650390625 0 6720.6797 z 6
895.4168090820312 0 3735.0015
897.4577026367188 0 24276.62 c 6
898.4612426757812 0 15857.455
899.4634399414062 0 4046.5657
909.4454345703125 0 2459.5876
910.434814453125 0 3649.6077 y 6
911.4373779296875 0 726.26166
937.4165649414062 0 658.1159
964.464111328125 0 1848.1372
965.4544067382812 0 2801.0435
965.6128540039062 0 822.6377
966.4472045898438 0 950.46094
994.5191040039062 0 972.1511
1008.4552612304688 0 6909.53
1009.4435424804688 0 11406.696 z 5
1010.4395141601562 0 4269.6963
1010.5421142578125 0 20156.164 c 7
1011.4436645507812 0 1676.7668
1011.5450439453125 0 12873.8545
1012.548095703125 0 4059.2485
1013.5392456054688 0 752.6573
1024.471923828125 0 2113.5527
1025.46484375 0 3651.92 y 5
1026.4749755859375 0 1839.4666
1027.470703125 0 736.94464
1062.58740234375 0 612.2643
1063.548583984375 0 908.54175
1064.1175537109375 0 716.1518
1067.5367431640625 0 3931.2407
1068.5421142578125 0 3020.4902
1070.62548828125 0 1027.0037
1071.1220703125 0 966.58435
1071.6282958984375 0 1051.6741
1072.4227294921875 0 690.41046
1079.470703125 0 1713.7341 w 4
1080.56884765625 0 1686.3079
1081.5645751953125 0 3329.3088
1082.5599365234375 0 2186.6455
1094.5093994140625 0 678.1453
1124.5838623046875 0 2433.745
1125.5733642578125 0 6678.5015 c 8
1126.5728759765625 0 3743.9043
1127.57275390625 0 811.1369
1136.5103759765625 0 7274.598 y Ammonia loss 4
1137.49755859375 0 49507.26 z 4
1138.5006103515625 0 29688.273
1139.5035400390625 0 12832.901
1140.5050048828125 0 3302.637
1152.525146484375 0 1121.8646
1153.519775390625 0 2419.6243 y 4
1154.5201416015625 0 2234.8708
1181.6055908203125 0 11442.932
1182.5933837890625 0 26975.426 c 9
1183.5941162109375 0 16950.766
1184.5958251953125 0 5591.364
1185.6064453125 0 2224.384
1186.6094970703125 0 1110.7616
1266.6695556640625 0 1120.7639
1267.6785888671875 0 777.1903
1268.6751708984375 0 1404.3547
1273.5670166015625 0 1837.0815 y Ammonia loss 3
1274.5614013671875 0 10623.361 z 3
1275.5621337890625 0 9638.056
1276.5635986328125 0 4360.6357
1277.5750732421875 0 2164.7593
1290.573974609375 0 1526.4669 y 3
1292.584228515625 0 747.82404
1309.6964111328125 0 3658.3274
1310.6868896484375 0 16272.632 c 10
1311.6890869140625 0 13930.542
1312.69140625 0 5024.826
1313.691650390625 0 1860.7949
1387.6453857421875 0 8743.23 z 2
1388.647216796875 0 9766.259
1389.6507568359375 0 6207.0015
1390.6544189453125 0 2596.265
1395.7298583984375 0 2440.9307
1396.72509765625 0 2252.7043
1397.72509765625 0 1824.9574
1403.669921875 0 1098.5507 y 2
1404.6622314453125 0 955.1255
1411.7303466796875 0 706.658
1412.7493896484375 0 993.874
1423.720703125 0 2220.2715
1424.7137451171875 0 1348.9122
1425.7283935546875 0 1054.2694
1438.7403564453125 0 1703.2511
1439.7369384765625 0 6009.3267 c 11
1440.7364501953125 0 8122.133
1441.734375 0 4054.7378
1442.7371826171875 0 2032.4381
1487.72216796875 0 2258.0137
1488.720947265625 0 1875.7255
1489.7178955078125 0 1179.8276
1543.777587890625 0 1567.1799
1544.7762451171875 0 1554.3058
1545.771728515625 0 3659.4731
1546.773193359375 0 3130.506
1547.783447265625 0 1077.513
1558.786865234375 0 848.9058
1559.781005859375 0 4809.712
1560.7786865234375 0 6016.1294
1561.7835693359375 0 3555.2437
1562.7830810546875 0 1458.381
1569.7587890625 0 767.53625
1570.7607421875 0 1768.4265
1571.7554931640625 0 1211.8677
1572.7425537109375 0 932.80743
1576.7999267578125 0 4246.8403
1577.7987060546875 0 3967.8977
1578.80322265625 0 2315.8162
1579.8070068359375 0 1404.342
1586.779052734375 0 2482.3452
1587.7728271484375 0 15602.505
1588.769775390625 0 24069.062
1589.77197265625 0 16546.627
1590.7701416015625 0 8932.56
1591.7750244140625 0 3054.922
1592.7642822265625 0 939.70123
1602.766845703125 0 1234.602
1603.7852783203125 0 8985.314
1604.791015625 0 27936
1605.7911376953125 0 46247.508
1606.79296875 0 32206.791
1607.7962646484375 0 15497.415
1608.7974853515625 0 5673.9917
1609.7882080078125 0 3340.812
1912.8067626953125 0 704.78754
2927.59521484375 0 698.0186

Spectrum Details

|  |  |
| --- | --- |
| Matched peaks? Matched peaks The total absolute number of peaks matched. Additionally in brackets the total fraction of peaks matched and the total number of peaks is shown. | 59 (13.92% of 424) |
| FDR? FDR The false discovery rate estimated for this peptide. It is calculated by matching all theoretical fragments with a non-integer shift with the raw peaks for this spectrum. This is done with 40 different shifts. The resulting percentage is the average number of annotated peaks over the number of annotated peaks with the correct spectrum. | 2.95% |
| Satellite FDR? Satellite FDR See the FDR for details on its calculation. This satellite ion specific FDR only contains the satellite ions (d/w) for I/L/J positions. | 0.00% |
| PSM Score? PSM Score The PSM Score as given by Hecklib to this annotated spectrum. It is shown with three significant figures. | 530 |

#### Spectrum 6245? Spectrum 6245 The raw spectrum of this peptide as annotated by Hecklib. The fragments are coloured according to ion type (see legend). Any peaks with a star '\*' as text can be hovered over to see the full details, first the ion type second the mass shift type. By hovering over the amino acids in the peptide or ions in the legend the corresponding peaks are highlighted. By toggling the 'Unassigned' label you can turn the background (unassigned) peaks on or off in the plot. By updating the slider in the Ion legend you can update the spectrum to only show the top X% of the peaks with labels. The top X% means any peak that is within X% of the highest intensity. By dragging in the spectrum you can zoom in to a specific part of the spectrum and use 'Zoom Out' to get back to the original zoom level. The annotation of the spectrum is based on the given sequence in the peptides file and is done with different software so inconsistencies are likely. The peaks are annotated based on the given sequence, with 20 ppm tolerance.

Copy Data

##### Spectrum 6245 (TSV)

###### Preview

```
Loading example...
```

*Click on the button to copy the data to your clipboard.*

Mz MinMz MaxIntensity Max

WidthHeightPeptide font sizePeptide stroke widthSpectrum font sizeSpectrum stroke widthCompact peptide

Ion legend

wxyz

abcd

OtherUnassignedIon^Charge^_Position_Show for top:%

TVLHQDWLDGKEY

06.20e+41.24e+51.86e+52.48e+5

Zoom Out

y^+3^_3_a^+1^_2_d^+1^_2_y^+3^_4_a^+1^_2_y^+1^_1_b^+1^_2_b^+1^_2_y^+2^_3_b^+3^_6_y^+1^_2_b^+1^_3_y^+2^_5_y^+1^_2_b^+1^_3_b^+3^_8_b^+2^_6_y^+3^_8_b^+2^_6_y^+2^_6_y^+1^_3_y^+3^_10_b^+1^_4_y^+1^_3_b^+2^_7_b^+2^_7_b^+1^_4_y^+2^_7_y^+2^_7_y^+1^_4_b^+2^_8_b^+2^_8_\*\*\*b^+1^_5_b^+2^_9_b^+1^_5_y^+1^_5_y^+1^_5_y^+2^_10_y^+2^_10_y^+2^_10_b^+2^_11_b^+1^_6_b^+1^_6_b^+1^_6_y^+2^_11_y^+2^_11_y^+1^_6_y^+2^_11_b^+2^_12_b^+2^_12_b^+2^_12_y^+1^_6_y^+2^_12_y^+2^_12_\*b^+1^_7_b^+1^_7_b^+1^_7_y^+1^_7_y^+1^_7_b^+1^_8_b^+1^_8_y^+1^_8_b^+1^_9_b^+1^_9_y^+1^_9_y^+1^_9_y^+1^_9_b^+1^_10_

0810162124313241

Fragment Matches Table

Show background peaks

| Position | Ion type | Intensity | mz Theoretical | mz Error (Th) | mz Error (ppm) | Charge | Series Number |
| --- | --- | --- | --- | --- | --- | --- | --- |
| - | - | 1.433E+04 | 120.1 | - | - | 0 | - |
| - | - | 715.6 | 121.1 | - | - | 0 | - |
| - | - | 1140 | 122.1 | - | - | 0 | - |
| - | - | 440.1 | 122.1 | - | - | 0 | - |
| - | - | 1058 | 123 | - | - | 0 | - |
| - | - | 1730 | 125.1 | - | - | 0 | - |
| - | - | 1331 | 126.1 | - | - | 0 | - |
| - | - | 755 | 126.1 | - | - | 0 | - |
| - | - | 4305 | 127.1 | - | - | 0 | - |
| - | - | 539 | 127.1 | - | - | 0 | - |
| - | - | 1.357E+04 | 128.1 | - | - | 0 | - |
| - | - | 848.2 | 129.1 | - | - | 0 | - |
| - | - | 1.469E+05 | 129.1 | - | - | 0 | - |
| - | - | 724.4 | 130.1 | - | - | 0 | - |
| - | - | 2.794E+04 | 130.1 | - | - | 0 | - |
| - | - | 1248 | 130.1 | - | - | 0 | - |
| - | - | 1.006E+04 | 130.1 | - | - | 0 | - |
| - | - | 538.3 | 130.4 | - | - | 0 | - |
| - | - | 445.4 | 131.1 | - | - | 0 | - |
| - | - | 2942 | 131.1 | - | - | 0 | - |
| - | - | 5625 | 132 | - | - | 0 | - |
| - | - | 1086 | 132.1 | - | - | 0 | - |
| - | - | 474.3 | 133.1 | - | - | 0 | - |
| - | - | 2.043E+04 | 136.1 | - | - | 0 | - |
| - | - | 2251 | 137.1 | - | - | 0 | - |
| - | - | 2596 | 138.1 | - | - | 0 | - |
| - | - | 1050 | 138.1 | - | - | 0 | - |
| - | - | 1030 | 139.1 | - | - | 0 | - |
| - | - | 1848 | 139.1 | - | - | 0 | - |
| - | - | 1401 | 140.1 | - | - | 0 | - |
| - | - | 2609 | 141.1 | - | - | 0 | - |
| - | - | 1622 | 145.1 | - | - | 0 | - |
| - | - | 3118 | 146.1 | - | - | 0 | - |
| - | - | 1041 | 147 | - | - | 0 | - |
| - | - | 681.9 | 147.1 | - | - | 0 | - |
| 11 | y | 1211 | 147.1 | 0.001008 | 6.853 | +3 | 3 |
| - | - | 850.1 | 149 | - | - | 0 | - |
| - | - | 2904 | 151.1 | - | - | 0 | - |
| - | - | 800.8 | 153.1 | - | - | 0 | - |
| - | - | 1927 | 154.1 | - | - | 0 | - |
| - | - | 712.5 | 154.1 | - | - | 0 | - |
| - | - | 8285 | 155.1 | - | - | 0 | - |
| 2 | a | 7948 | 155.1 | 0.0003813 | 2.458 | +1 | 2 |
| - | - | 864.1 | 156.1 | - | - | 0 | - |
| - | - | 1239 | 156.1 | - | - | 0 | - |
| - | - | 543 | 156.1 | - | - | 0 | - |
| - | - | 1169 | 157.1 | - | - | 0 | - |
| - | - | 8583 | 157.1 | - | - | 0 | - |
| - | - | 1883 | 157.1 | - | - | 0 | - |
| - | - | 4166 | 157.1 | - | - | 0 | - |
| - | - | 2.628E+04 | 158.1 | - | - | 0 | - |
| - | - | 1474 | 158.1 | - | - | 0 | - |
| - | - | 590.4 | 158.2 | - | - | 0 | - |
| - | - | 2877 | 159.1 | - | - | 0 | - |
| - | - | 3544 | 159.1 | - | - | 0 | - |
| 2 | d | 2002 | 159.1 | 0.000355 | 2.231 | +1 | 2 |
| - | - | 1.156E+04 | 165.1 | - | - | 0 | - |
| - | - | 1.909E+04 | 166.1 | - | - | 0 | - |
| 10 | y | 4807 | 166.1 | 0.001741 | 10.48 | +3 | 4 |
| - | - | 1262 | 166.1 | - | - | 0 | - |
| - | - | 651.6 | 167.1 | - | - | 0 | - |
| - | - | 923.5 | 167.1 | - | - | 0 | - |
| - | - | 1294 | 168.1 | - | - | 0 | - |
| - | - | 972.9 | 168.1 | - | - | 0 | - |
| - | - | 805.6 | 169.1 | - | - | 0 | - |
| - | - | 4029 | 169.1 | - | - | 0 | - |
| - | - | 2033 | 169.1 | - | - | 0 | - |
| - | - | 1842 | 171.1 | - | - | 0 | - |
| - | - | 614.7 | 171.1 | - | - | 0 | - |
| - | - | 2125 | 172.1 | - | - | 0 | - |
| - | - | 649.3 | 172.1 | - | - | 0 | - |
| - | - | 1998 | 173.1 | - | - | 0 | - |
| 2 | a | 2.457E+05 | 173.1 | 0.000452 | 2.611 | +1 | 2 |
| - | - | 916.2 | 173.4 | - | - | 0 | - |
| - | - | 1.914E+04 | 174.1 | - | - | 0 | - |
| - | - | 1.272E+05 | 175.1 | - | - | 0 | - |
| - | - | 1987 | 175.1 | - | - | 0 | - |
| - | - | 940.2 | 176.1 | - | - | 0 | - |
| - | - | 1.329E+04 | 176.1 | - | - | 0 | - |
| - | - | 803.1 | 177.1 | - | - | 0 | - |
| - | - | 927.3 | 177.1 | - | - | 0 | - |
| - | - | 6053 | 178.1 | - | - | 0 | - |
| - | - | 1140 | 179.1 | - | - | 0 | - |
| - | - | 781.9 | 181.1 | - | - | 0 | - |
| 13 | y | 2.362E+04 | 182.1 | 0.0004191 | 2.302 | +1 | 1 |
| - | - | 1308 | 182.1 | - | - | 0 | - |
| - | - | 2360 | 183.1 | - | - | 0 | - |
| 2 | b | 1.73E+04 | 183.1 | 0.0003703 | 2.022 | +1 | 2 |
| - | - | 1001 | 183.1 | - | - | 0 | - |
| - | - | 1163 | 184.1 | - | - | 0 | - |
| - | - | 2859 | 185.1 | - | - | 0 | - |
| - | - | 977.6 | 185.1 | - | - | 0 | - |
| - | - | 1361 | 185.1 | - | - | 0 | - |
| - | - | 1745 | 185.2 | - | - | 0 | - |
| - | - | 507.3 | 186.1 | - | - | 0 | - |
| - | - | 3.506E+04 | 186.1 | - | - | 0 | - |
| - | - | 1002 | 187.1 | - | - | 0 | - |
| - | - | 1340 | 187.1 | - | - | 0 | - |
| - | - | 2422 | 187.1 | - | - | 0 | - |
| - | - | 2276 | 187.1 | - | - | 0 | - |
| - | - | 1211 | 187.1 | - | - | 0 | - |
| - | - | 828.5 | 188.1 | - | - | 0 | - |
| - | - | 7969 | 189.1 | - | - | 0 | - |
| - | - | 554.8 | 189.4 | - | - | 0 | - |
| - | - | 2169 | 190.1 | - | - | 0 | - |
| - | - | 1856 | 190.1 | - | - | 0 | - |
| - | - | 2205 | 194.1 | - | - | 0 | - |
| - | - | 1.055E+04 | 195.1 | - | - | 0 | - |
| - | - | 4712 | 196.1 | - | - | 0 | - |
| - | - | 738.1 | 196.1 | - | - | 0 | - |
| - | - | 888.1 | 196.1 | - | - | 0 | - |
| - | - | 692.9 | 197.1 | - | - | 0 | - |
| - | - | 789 | 197.1 | - | - | 0 | - |
| - | - | 2879 | 197.1 | - | - | 0 | - |
| - | - | 582.1 | 197.2 | - | - | 0 | - |
| - | - | 628 | 198.1 | - | - | 0 | - |
| - | - | 4206 | 199.1 | - | - | 0 | - |
| - | - | 1281 | 199.1 | - | - | 0 | - |
| - | - | 581.3 | 200.1 | - | - | 0 | - |
| 2 | b | 8.236E+04 | 201.1 | 0.0003494 | 1.737 | +1 | 2 |
| - | - | 723.5 | 202.1 | - | - | 0 | - |
| - | - | 6216 | 202.1 | - | - | 0 | - |
| - | - | 2016 | 203.1 | - | - | 0 | - |
| - | - | 1008 | 203.1 | - | - | 0 | - |
| - | - | 4499 | 205.1 | - | - | 0 | - |
| - | - | 785.9 | 206.1 | - | - | 0 | - |
| - | - | 1150 | 206.1 | - | - | 0 | - |
| - | - | 1104 | 207.2 | - | - | 0 | - |
| - | - | 665.2 | 208.1 | - | - | 0 | - |
| - | - | 834.3 | 210.1 | - | - | 0 | - |
| - | - | 526.6 | 210.9 | - | - | 0 | - |
| 11 | y | 1461 | 211.1 | 3.879E-05 | 0.1837 | +2 | 3 |
| - | - | 905.4 | 211.1 | - | - | 0 | - |
| - | - | 1.203E+04 | 211.2 | - | - | 0 | - |
| - | - | 672.8 | 212.1 | - | - | 0 | - |
| - | - | 1623 | 212.1 | - | - | 0 | - |
| - | - | 1010 | 213.1 | - | - | 0 | - |
| - | - | 1019 | 213.1 | - | - | 0 | - |
| - | - | 4965 | 213.1 | - | - | 0 | - |
| - | - | 986.1 | 213.2 | - | - | 0 | - |
| - | - | 1370 | 215.1 | - | - | 0 | - |
| - | - | 590.6 | 215.1 | - | - | 0 | - |
| - | - | 1356 | 216.1 | - | - | 0 | - |
| - | - | 1262 | 216.1 | - | - | 0 | - |
| - | - | 1.214E+04 | 217.1 | - | - | 0 | - |
| - | - | 692.6 | 218.1 | - | - | 0 | - |
| - | - | 1494 | 219.1 | - | - | 0 | - |
| - | - | 709.3 | 220.1 | - | - | 0 | - |
| - | - | 3850 | 221.1 | - | - | 0 | - |
| - | - | 1010 | 221.1 | - | - | 0 | - |
| - | - | 681.2 | 221.1 | - | - | 0 | - |
| - | - | 596.7 | 222.1 | - | - | 0 | - |
| - | - | 1058 | 222.1 | - | - | 0 | - |
| - | - | 3017 | 222.1 | - | - | 0 | - |
| - | - | 617.5 | 223.1 | - | - | 0 | - |
| - | - | 3192 | 223.1 | - | - | 0 | - |
| - | - | 1.943E+04 | 223.2 | - | - | 0 | - |
| - | - | 860.8 | 224.1 | - | - | 0 | - |
| - | - | 2781 | 224.2 | - | - | 0 | - |
| - | - | 3957 | 226.1 | - | - | 0 | - |
| 6 | b | 1111 | 226.1 | 0.000416 | 1.84 | +3 | 6 |
| - | - | 562 | 226.2 | - | - | 0 | - |
| - | - | 2990 | 227.1 | - | - | 0 | - |
| - | - | 803.5 | 227.1 | - | - | 0 | - |
| - | - | 3175 | 227.1 | - | - | 0 | - |
| - | - | 1236 | 228.1 | - | - | 0 | - |
| - | - | 2127 | 228.2 | - | - | 0 | - |
| - | - | 3376 | 229.1 | - | - | 0 | - |
| - | - | 578.4 | 230.1 | - | - | 0 | - |
| - | - | 3858 | 233.1 | - | - | 0 | - |
| - | - | 1770 | 233.2 | - | - | 0 | - |
| - | - | 2.122E+04 | 234.1 | - | - | 0 | - |
| - | - | 3006 | 235.1 | - | - | 0 | - |
| - | - | 806.1 | 237.1 | - | - | 0 | - |
| - | - | 1517 | 237.1 | - | - | 0 | - |
| - | - | 557.8 | 238.1 | - | - | 0 | - |
| - | - | 1912 | 238.1 | - | - | 0 | - |
| - | - | 7687 | 239.2 | - | - | 0 | - |
| - | - | 1.062E+04 | 240.1 | - | - | 0 | - |
| - | - | 1270 | 241.2 | - | - | 0 | - |
| - | - | 3163 | 242.1 | - | - | 0 | - |
| - | - | 1773 | 243.1 | - | - | 0 | - |
| - | - | 2835 | 243.1 | - | - | 0 | - |
| - | - | 5854 | 244.1 | - | - | 0 | - |
| - | - | 686.6 | 245.1 | - | - | 0 | - |
| - | - | 732.4 | 245.1 | - | - | 0 | - |
| - | - | 1.196E+04 | 247.1 | - | - | 0 | - |
| - | - | 1099 | 247.1 | - | - | 0 | - |
| - | - | 1003 | 247.1 | - | - | 0 | - |
| - | - | 851.1 | 247.2 | - | - | 0 | - |
| - | - | 708.2 | 248.1 | - | - | 0 | - |
| - | - | 5243 | 248.1 | - | - | 0 | - |
| - | - | 1234 | 248.6 | - | - | 0 | - |
| - | - | 6194 | 249.1 | - | - | 0 | - |
| - | - | 2505 | 249.1 | - | - | 0 | - |
| - | - | 629.3 | 250.1 | - | - | 0 | - |
| - | - | 7.225E+04 | 251.2 | - | - | 0 | - |
| - | - | 9816 | 252.2 | - | - | 0 | - |
| - | - | 595.7 | 253.2 | - | - | 0 | - |
| - | - | 2780 | 255.1 | - | - | 0 | - |
| - | - | 2053 | 255.1 | - | - | 0 | - |
| - | - | 588.8 | 256.1 | - | - | 0 | - |
| - | - | 2593 | 256.2 | - | - | 0 | - |
| - | - | 1151 | 257.1 | - | - | 0 | - |
| - | - | 3837 | 258.1 | - | - | 0 | - |
| - | - | 2001 | 261.1 | - | - | 0 | - |
| - | - | 797.1 | 261.1 | - | - | 0 | - |
| - | - | 1130 | 262.1 | - | - | 0 | - |
| - | - | 773.2 | 262.2 | - | - | 0 | - |
| - | - | 8775 | 265.1 | - | - | 0 | - |
| - | - | 2084 | 266.1 | - | - | 0 | - |
| - | - | 1.714E+04 | 266.1 | - | - | 0 | - |
| - | - | 1256 | 267.1 | - | - | 0 | - |
| - | - | 709.3 | 268.1 | - | - | 0 | - |
| - | - | 1083 | 268.2 | - | - | 0 | - |
| - | - | 968.4 | 269.1 | - | - | 0 | - |
| - | - | 1045 | 269.2 | - | - | 0 | - |
| - | - | 1649 | 269.2 | - | - | 0 | - |
| - | - | 1176 | 270.1 | - | - | 0 | - |
| - | - | 4918 | 270.2 | - | - | 0 | - |
| - | - | 782.1 | 271.2 | - | - | 0 | - |
| - | - | 791.7 | 272.1 | - | - | 0 | - |
| - | - | 635.4 | 272.1 | - | - | 0 | - |
| - | - | 1811 | 274.1 | - | - | 0 | - |
| - | - | 848.7 | 275.1 | - | - | 0 | - |
| - | - | 817.2 | 276.1 | - | - | 0 | - |
| - | - | 1092 | 276.2 | - | - | 0 | - |
| - | - | 775.9 | 279.1 | - | - | 0 | - |
| - | - | 1303 | 280.1 | - | - | 0 | - |
| - | - | 3191 | 282.2 | - | - | 0 | - |
| - | - | 5.542E+04 | 283.1 | - | - | 0 | - |
| - | - | 966.8 | 284.1 | - | - | 0 | - |
| - | - | 7140 | 284.1 | - | - | 0 | - |
| - | - | 1122 | 284.2 | - | - | 0 | - |
| - | - | 3975 | 286.1 | - | - | 0 | - |
| - | - | 2602 | 286.2 | - | - | 0 | - |
| - | - | 819 | 287.2 | - | - | 0 | - |
| - | - | 1213 | 288.2 | - | - | 0 | - |
| - | - | 7514 | 288.2 | - | - | 0 | - |
| - | - | 1460 | 289.2 | - | - | 0 | - |
| - | - | 3068 | 290.1 | - | - | 0 | - |
| - | - | 639.3 | 291.1 | - | - | 0 | - |
| 12 | y | 3.144E+04 | 293.1 | 0.0003883 | 1.325 | +1 | 2 |
| - | - | 5966 | 294.1 | - | - | 0 | - |
| - | - | 1226 | 294.2 | - | - | 0 | - |
| 3 | b | 1792 | 296.2 | 0.0001228 | 0.4147 | +1 | 3 |
| - | - | 8524 | 297.2 | - | - | 0 | - |
| - | - | 2082 | 298.1 | - | - | 0 | - |
| - | - | 3262 | 298.2 | - | - | 0 | - |
| - | - | 1808 | 299.1 | - | - | 0 | - |
| - | - | 916.8 | 299.1 | - | - | 0 | - |
| - | - | 2209 | 300.1 | - | - | 0 | - |
| - | - | 4602 | 300.2 | - | - | 0 | - |
| - | - | 4524 | 301.1 | - | - | 0 | - |
| - | - | 2050 | 301.1 | - | - | 0 | - |
| - | - | 2.45E+04 | 301.2 | - | - | 0 | - |
| - | - | 1395 | 301.2 | - | - | 0 | - |
| - | - | 1823 | 302.1 | - | - | 0 | - |
| - | - | 1306 | 302.1 | - | - | 0 | - |
| - | - | 3200 | 302.2 | - | - | 0 | - |
| - | - | 3668 | 305.2 | - | - | 0 | - |
| - | - | 1551 | 305.6 | - | - | 0 | - |
| 9 | y | 5003 | 306.1 | 0.0007044 | 2.301 | +2 | 5 |
| - | - | 766.5 | 307.1 | - | - | 0 | - |
| - | - | 1256 | 310.2 | - | - | 0 | - |
| 12 | y | 4810 | 311.1 | 0.0005048 | 1.623 | +1 | 2 |
| - | - | 1.251E+04 | 312.2 | - | - | 0 | - |
| - | - | 1350 | 313.2 | - | - | 0 | - |
| - | - | 2391 | 313.2 | - | - | 0 | - |
| - | - | 5149 | 314.1 | - | - | 0 | - |
| 3 | b | 1826 | 314.2 | 0.0001269 | 0.4039 | +1 | 3 |
| - | - | 1.346E+04 | 315.2 | - | - | 0 | - |
| - | - | 1201 | 315.2 | - | - | 0 | - |
| - | - | 2762 | 316.2 | - | - | 0 | - |
| - | - | 3859 | 316.2 | - | - | 0 | - |
| - | - | 5971 | 318.1 | - | - | 0 | - |
| - | - | 710.2 | 319.1 | - | - | 0 | - |
| - | - | 963.9 | 319.2 | - | - | 0 | - |
| - | - | 1137 | 322.2 | - | - | 0 | - |
| - | - | 764.8 | 325.1 | - | - | 0 | - |
| - | - | 3874 | 327.3 | - | - | 0 | - |
| - | - | 5285 | 330.2 | - | - | 0 | - |
| 8 | b | 823.3 | 331.2 | 0.00204 | 6.16 | +3 | 8 |
| - | - | 1817 | 332.2 | - | - | 0 | - |
| - | - | 4912 | 333.2 | - | - | 0 | - |
| - | - | 907.2 | 334.2 | - | - | 0 | - |
| - | - | 1208 | 334.7 | - | - | 0 | - |
| - | - | 3105 | 338.2 | - | - | 0 | - |
| 6 | b | 948.9 | 338.7 | 0.0007655 | 2.26 | +2 | 6 |
| - | - | 641.6 | 339.2 | - | - | 0 | - |
| - | - | 2283 | 341.2 | - | - | 0 | - |
| 6 | y | 2905 | 342.1 | 0.004924 | 14.39 | +3 | 8 |
| - | - | 974.4 | 344.2 | - | - | 0 | - |
| - | - | 2918 | 345.1 | - | - | 0 | - |
| - | - | 6756 | 346.1 | - | - | 0 | - |
| - | - | 1093 | 347.1 | - | - | 0 | - |
| - | - | 1252 | 347.2 | - | - | 0 | - |
| 6 | b | 4553 | 347.7 | 0.0002692 | 0.7744 | +2 | 6 |
| - | - | 1413 | 348.2 | - | - | 0 | - |
| - | - | 1258 | 348.7 | - | - | 0 | - |
| - | - | 1210 | 349.2 | - | - | 0 | - |
| - | - | 7573 | 350.2 | - | - | 0 | - |
| - | - | 1241 | 351.2 | - | - | 0 | - |
| - | - | 1865 | 355.2 | - | - | 0 | - |
| - | - | 1253 | 357.2 | - | - | 0 | - |
| - | - | 1402 | 358.2 | - | - | 0 | - |
| - | - | 1870 | 359.2 | - | - | 0 | - |
| - | - | 675.8 | 360.2 | - | - | 0 | - |
| - | - | 1929 | 361.2 | - | - | 0 | - |
| - | - | 2794 | 362.2 | - | - | 0 | - |
| 8 | y | 1.013E+04 | 362.7 | 0.0006036 | 1.664 | +2 | 6 |
| - | - | 1.265E+04 | 363.1 | - | - | 0 | - |
| - | - | 3653 | 363.2 | - | - | 0 | - |
| - | - | 1349 | 363.7 | - | - | 0 | - |
| - | - | 2726 | 364.1 | - | - | 0 | - |
| - | - | 1461 | 364.1 | - | - | 0 | - |
| - | - | 1225 | 364.2 | - | - | 0 | - |
| - | - | 1025 | 364.7 | - | - | 0 | - |
| - | - | 930.3 | 365.1 | - | - | 0 | - |
| - | - | 1643 | 366.2 | - | - | 0 | - |
| - | - | 1066 | 367.2 | - | - | 0 | - |
| - | - | 986.9 | 367.2 | - | - | 0 | - |
| - | - | 1056 | 368.2 | - | - | 0 | - |
| - | - | 1036 | 368.7 | - | - | 0 | - |
| - | - | 804.8 | 369.2 | - | - | 0 | - |
| - | - | 1576 | 373.2 | - | - | 0 | - |
| - | - | 674.5 | 373.7 | - | - | 0 | - |
| - | - | 816.2 | 376.2 | - | - | 0 | - |
| - | - | 4667 | 378.2 | - | - | 0 | - |
| - | - | 1.974E+04 | 379.2 | - | - | 0 | - |
| - | - | 3880 | 380.2 | - | - | 0 | - |
| - | - | 3.645E+04 | 381.2 | - | - | 0 | - |
| - | - | 7456 | 382.2 | - | - | 0 | - |
| - | - | 953 | 382.7 | - | - | 0 | - |
| - | - | 1061 | 383.1 | - | - | 0 | - |
| - | - | 2735 | 383.2 | - | - | 0 | - |
| - | - | 1274 | 383.7 | - | - | 0 | - |
| - | - | 3491 | 384.2 | - | - | 0 | - |
| - | - | 1249 | 385.2 | - | - | 0 | - |
| - | - | 1957 | 390.2 | - | - | 0 | - |
| - | - | 1074 | 391.1 | - | - | 0 | - |
| - | - | 4487 | 394.2 | - | - | 0 | - |
| - | - | 2316 | 395.2 | - | - | 0 | - |
| - | - | 1824 | 395.2 | - | - | 0 | - |
| - | - | 1032 | 396.2 | - | - | 0 | - |
| - | - | 5980 | 396.2 | - | - | 0 | - |
| - | - | 1444 | 397.2 | - | - | 0 | - |
| - | - | 4167 | 397.3 | - | - | 0 | - |
| - | - | 2231 | 398.2 | - | - | 0 | - |
| - | - | 1197 | 398.3 | - | - | 0 | - |
| - | - | 1772 | 399.2 | - | - | 0 | - |
| - | - | 1046 | 400.2 | - | - | 0 | - |
| - | - | 1258 | 400.2 | - | - | 0 | - |
| - | - | 913.1 | 401.2 | - | - | 0 | - |
| - | - | 4396 | 402.2 | - | - | 0 | - |
| - | - | 1671 | 403.2 | - | - | 0 | - |
| - | - | 834.6 | 405.2 | - | - | 0 | - |
| - | - | 988.4 | 405.3 | - | - | 0 | - |
| - | - | 941 | 405.7 | - | - | 0 | - |
| - | - | 758.2 | 406.2 | - | - | 0 | - |
| - | - | 2353 | 407.2 | - | - | 0 | - |
| - | - | 3030 | 409.1 | - | - | 0 | - |
| - | - | 1641 | 409.2 | - | - | 0 | - |
| - | - | 957.3 | 410.1 | - | - | 0 | - |
| - | - | 1433 | 411.1 | - | - | 0 | - |
| - | - | 2883 | 411.2 | - | - | 0 | - |
| - | - | 2.296E+04 | 412.2 | - | - | 0 | - |
| - | - | 3044 | 412.3 | - | - | 0 | - |
| - | - | 2737 | 413.2 | - | - | 0 | - |
| - | - | 4333 | 413.2 | - | - | 0 | - |
| - | - | 1092 | 414.2 | - | - | 0 | - |
| - | - | 7351 | 414.2 | - | - | 0 | - |
| - | - | 1.217E+04 | 414.3 | - | - | 0 | - |
| - | - | 2051 | 414.7 | - | - | 0 | - |
| - | - | 1338 | 415.2 | - | - | 0 | - |
| - | - | 1192 | 417.2 | - | - | 0 | - |
| - | - | 1250 | 418 | - | - | 0 | - |
| - | - | 1402 | 418.2 | - | - | 0 | - |
| - | - | 865.4 | 418.2 | - | - | 0 | - |
| - | - | 1091 | 418.7 | - | - | 0 | - |
| 11 | y | 867.5 | 421.2 | 0.004943 | 11.74 | +1 | 3 |
| - | - | 4856 | 423.2 | - | - | 0 | - |
| - | - | 9490 | 423.3 | - | - | 0 | - |
| - | - | 1874 | 423.7 | - | - | 0 | - |
| - | - | 9482 | 424.3 | - | - | 0 | - |
| - | - | 3156 | 425.3 | - | - | 0 | - |
| - | - | 3912 | 425.7 | - | - | 0 | - |
| - | - | 6008 | 426.2 | - | - | 0 | - |
| - | - | 3440 | 426.7 | - | - | 0 | - |
| - | - | 3893 | 428.2 | - | - | 0 | - |
| - | - | 1062 | 428.2 | - | - | 0 | - |
| - | - | 9667 | 429.1 | - | - | 0 | - |
| - | - | 1.048E+04 | 429.2 | - | - | 0 | - |
| - | - | 1177 | 429.2 | - | - | 0 | - |
| - | - | 1810 | 430.1 | - | - | 0 | - |
| 4 | y | 5.519E+04 | 430.2 | 0.004265 | 9.915 | +3 | 10 |
| - | - | 1134 | 430.7 | - | - | 0 | - |
| - | - | 1.481E+04 | 431.2 | - | - | 0 | - |
| - | - | 2701 | 431.7 | - | - | 0 | - |
| - | - | 2091 | 432.2 | - | - | 0 | - |
| - | - | 897.5 | 432.2 | - | - | 0 | - |
| 4 | b | 1629 | 433.3 | 0.001117 | 2.578 | +1 | 4 |
| - | - | 3.863E+04 | 434.7 | - | - | 0 | - |
| - | - | 1.881E+04 | 435.2 | - | - | 0 | - |
| - | - | 3933 | 435.7 | - | - | 0 | - |
| - | - | 779.8 | 436.2 | - | - | 0 | - |
| 11 | y | 1.69E+04 | 439.2 | 0.000604 | 1.375 | +1 | 3 |
| 7 | b | 1.801E+04 | 439.7 | 0.002969 | 6.752 | +2 | 7 |
| - | - | 1.029E+04 | 440.2 | - | - | 0 | - |
| - | - | 3887 | 440.7 | - | - | 0 | - |
| - | - | 2136 | 441.3 | - | - | 0 | - |
| - | - | 3.085E+04 | 442.3 | - | - | 0 | - |
| - | - | 8927 | 446.2 | - | - | 0 | - |
| - | - | 1191 | 447.2 | - | - | 0 | - |
| - | - | 1137 | 447.2 | - | - | 0 | - |
| - | - | 2848 | 448.2 | - | - | 0 | - |
| 7 | b | 2.413E+04 | 448.7 | 0.003119 | 6.951 | +2 | 7 |
| - | - | 1.511E+04 | 449.2 | - | - | 0 | - |
| - | - | 2849 | 449.7 | - | - | 0 | - |
| - | - | 837.9 | 450.3 | - | - | 0 | - |
| 4 | b | 1.685E+04 | 451.3 | 0.0008977 | 1.989 | +1 | 4 |
| - | - | 4036 | 452.3 | - | - | 0 | - |
| - | - | 2647 | 454.2 | - | - | 0 | - |
| 7 | y | 8686 | 454.7 | 0.003517 | 7.735 | +2 | 7 |
| - | - | 4596 | 455.2 | - | - | 0 | - |
| - | - | 1433 | 455.3 | - | - | 0 | - |
| - | - | 707.5 | 455.7 | - | - | 0 | - |
| - | - | 1546 | 458.2 | - | - | 0 | - |
| - | - | 1278 | 459.2 | - | - | 0 | - |
| - | - | 1773 | 461.2 | - | - | 0 | - |
| - | - | 1686 | 461.3 | - | - | 0 | - |
| 7 | y | 3684 | 463.7 | 0.003575 | 7.71 | +2 | 7 |
| - | - | 1747 | 464.2 | - | - | 0 | - |
| - | - | 2472 | 466.2 | - | - | 0 | - |
| - | - | 801.5 | 466.3 | - | - | 0 | - |
| - | - | 5755 | 468.3 | - | - | 0 | - |
| - | - | 885.8 | 469.3 | - | - | 0 | - |
| - | - | 1230 | 469.3 | - | - | 0 | - |
| - | - | 930.8 | 473.7 | - | - | 0 | - |
| - | - | 791.1 | 474.8 | - | - | 0 | - |
| - | - | 810.3 | 475.8 | - | - | 0 | - |
| - | - | 1658 | 476.2 | - | - | 0 | - |
| - | - | 2981 | 477.2 | - | - | 0 | - |
| 10 | y | 2097 | 478.2 | 0.0006303 | 1.318 | +1 | 4 |
| - | - | 2875 | 478.3 | - | - | 0 | - |
| - | - | 889.1 | 478.3 | - | - | 0 | - |
| - | - | 2268 | 479.2 | - | - | 0 | - |
| - | - | 7040 | 482.3 | - | - | 0 | - |
| - | - | 4980 | 482.8 | - | - | 0 | - |
| - | - | 2351 | 483.3 | - | - | 0 | - |
| - | - | 6336 | 483.3 | - | - | 0 | - |
| - | - | 1794 | 488.3 | - | - | 0 | - |
| - | - | 7954 | 491.3 | - | - | 0 | - |
| - | - | 6479 | 491.8 | - | - | 0 | - |
| - | - | 2071 | 492.3 | - | - | 0 | - |
| - | - | 984.2 | 493.3 | - | - | 0 | - |
| - | - | 3.241E+04 | 494.2 | - | - | 0 | - |
| - | - | 1.007E+04 | 495.2 | - | - | 0 | - |
| - | - | 6703 | 495.3 | - | - | 0 | - |
| 8 | b | 4.381E+04 | 496.3 | 0.008667 | 17.47 | +2 | 8 |
| - | - | 7399 | 496.3 | - | - | 0 | - |
| - | - | 1.708E+04 | 496.8 | - | - | 0 | - |
| - | - | 8673 | 497.2 | - | - | 0 | - |
| - | - | 3994 | 497.3 | - | - | 0 | - |
| - | - | 827.7 | 497.9 | - | - | 0 | - |
| - | - | 3305 | 498.2 | - | - | 0 | - |
| - | - | 1281 | 504.3 | - | - | 0 | - |
| 8 | b | 7468 | 505.3 | 0.00488 | 9.658 | +2 | 8 |
| - | - | 3666 | 505.8 | - | - | 0 | - |
| - | - | 1446 | 506.1 | - | - | 0 | - |
| - | - | 825.8 | 506.2 | - | - | 0 | - |
| - | - | 2241 | 506.3 | - | - | 0 | - |
| - | - | 1189 | 507.3 | - | - | 0 | - |
| - | - | 4757 | 511.3 | - | - | 0 | - |
| - | - | 9394 | 511.3 | - | - | 0 | - |
| - | - | 1057 | 512.2 | - | - | 0 | - |
| - | - | 926.1 | 512.3 | - | - | 0 | - |
| - | - | 1.434E+05 | 513.4 | - | - | 0 | - |
| - | - | 1161 | 520.2 | - | - | 0 | - |
| - | - | 6336 | 523.3 | - | - | 0 | - |
| - | - | 4076 | 524.3 | - | - | 0 | - |
| - | - | 874.3 | 524.3 | - | - | 0 | - |
| - | - | 1696 | 524.8 | - | - | 0 | - |
| - | - | 5872 | 525.3 | - | - | 0 | - |
| - | - | 9482 | 525.8 | - | - | 0 | - |
| - | - | 1507 | 526.3 | - | - | 0 | - |
| - | - | 4462 | 526.3 | - | - | 0 | - |
| - | - | 1683 | 526.8 | - | - | 0 | - |
| - | - | 815.4 | 527.3 | - | - | 0 | - |
| - | - | 839.2 | 528.6 | - | - | 0 | - |
| - | - | 959.4 | 530.2 | - | - | 0 | - |
| - | - | 747.4 | 530.3 | - | - | 0 | - |
| - | - | 1002 | 532.3 | - | - | 0 | - |
| - | - | 973 | 534.3 | - | - | 0 | - |
| 0 | Precursor | 2284 | 534.6 | 0.003505 | 6.557 | +3 | -1 |
| 0 | Precursor | 1395 | 534.9 | 0.009851 | 18.42 | +3 | -1 |
| - | - | 1288 | 535.3 | - | - | 0 | - |
| - | - | 3939 | 535.6 | - | - | 0 | - |
| - | - | 1997 | 535.9 | - | - | 0 | - |
| - | - | 803.2 | 538.3 | - | - | 0 | - |
| - | - | 3.201E+04 | 539.8 | - | - | 0 | - |
| - | - | 834.9 | 540 | - | - | 0 | - |
| - | - | 1356 | 540.3 | - | - | 0 | - |
| - | - | 2.128E+04 | 540.3 | - | - | 0 | - |
| 0 | Precursor | 3176 | 540.6 | 0.0007771 | 1.437 | +3 | -1 |
| - | - | 6175 | 540.8 | - | - | 0 | - |
| - | - | 3434 | 540.9 | - | - | 0 | - |
| - | - | 2.178E+05 | 541.3 | - | - | 0 | - |
| - | - | 7755 | 541.6 | - | - | 0 | - |
| - | - | 7984 | 541.9 | - | - | 0 | - |
| - | - | 2755 | 542.3 | - | - | 0 | - |
| - | - | 784.2 | 542.3 | - | - | 0 | - |
| - | - | 8628 | 543.3 | - | - | 0 | - |
| - | - | 2713 | 544.3 | - | - | 0 | - |
| - | - | 1029 | 545.2 | - | - | 0 | - |
| - | - | 1092 | 545.8 | - | - | 0 | - |
| - | - | 1246 | 548.2 | - | - | 0 | - |
| - | - | 1527 | 551.3 | - | - | 0 | - |
| - | - | 7721 | 554.3 | - | - | 0 | - |
| - | - | 2411 | 554.8 | - | - | 0 | - |
| - | - | 4002 | 555.2 | - | - | 0 | - |
| - | - | 1136 | 555.3 | - | - | 0 | - |
| - | - | 882 | 556.2 | - | - | 0 | - |
| 5 | b | 1893 | 561.3 | 0.0005147 | 0.917 | +1 | 5 |
| 9 | b | 1681 | 562.8 | 0.008806 | 15.65 | +2 | 9 |
| - | - | 3328 | 563.3 | - | - | 0 | - |
| - | - | 1880 | 563.8 | - | - | 0 | - |
| - | - | 1813 | 564.8 | - | - | 0 | - |
| - | - | 1413 | 565.2 | - | - | 0 | - |
| - | - | 4400 | 565.3 | - | - | 0 | - |
| - | - | 877 | 566.2 | - | - | 0 | - |
| - | - | 1352 | 566.3 | - | - | 0 | - |
| - | - | 6163 | 573.8 | - | - | 0 | - |
| - | - | 2314 | 574.3 | - | - | 0 | - |
| - | - | 1892 | 575.2 | - | - | 0 | - |
| - | - | 1032 | 575.3 | - | - | 0 | - |
| 5 | b | 1.024E+04 | 579.3 | 0.0004562 | 0.7875 | +1 | 5 |
| - | - | 3522 | 580.3 | - | - | 0 | - |
| - | - | 967.4 | 582.8 | - | - | 0 | - |
| - | - | 1.059E+04 | 583.2 | - | - | 0 | - |
| - | - | 3180 | 584.2 | - | - | 0 | - |
| - | - | 999.7 | 587.3 | - | - | 0 | - |
| - | - | 1716 | 588.8 | - | - | 0 | - |
| - | - | 1390 | 589.3 | - | - | 0 | - |
| - | - | 1414 | 592.3 | - | - | 0 | - |
| 9 | y | 1.564E+04 | 593.3 | 0.0003901 | 0.6576 | +1 | 5 |
| - | - | 4962 | 593.3 | - | - | 0 | - |
| - | - | 4449 | 594.3 | - | - | 0 | - |
| - | - | 1245 | 594.3 | - | - | 0 | - |
| - | - | 1053 | 595.3 | - | - | 0 | - |
| - | - | 1231 | 595.3 | - | - | 0 | - |
| - | - | 1565 | 596.4 | - | - | 0 | - |
| - | - | 5.731E+04 | 597.8 | - | - | 0 | - |
| - | - | 2.937E+04 | 598.3 | - | - | 0 | - |
| - | - | 1882 | 598.7 | - | - | 0 | - |
| - | - | 1192 | 600.3 | - | - | 0 | - |
| - | - | 789 | 602.3 | - | - | 0 | - |
| - | - | 950 | 608.3 | - | - | 0 | - |
| - | - | 955 | 609.8 | - | - | 0 | - |
| - | - | 2.496E+04 | 610.3 | - | - | 0 | - |
| - | - | 2613 | 610.3 | - | - | 0 | - |
| - | - | 2888 | 610.8 | - | - | 0 | - |
| 9 | y | 1.415E+05 | 611.3 | 0.001056 | 1.727 | +1 | 5 |
| - | - | 1768 | 611.8 | - | - | 0 | - |
| - | - | 4.207E+04 | 612.3 | - | - | 0 | - |
| - | - | 9659 | 613.3 | - | - | 0 | - |
| - | - | 1391 | 614.3 | - | - | 0 | - |
| - | - | 1204 | 614.3 | - | - | 0 | - |
| - | - | 963.6 | 616.3 | - | - | 0 | - |
| - | - | 3309 | 619.3 | - | - | 0 | - |
| - | - | 8192 | 619.8 | - | - | 0 | - |
| - | - | 4658 | 620.3 | - | - | 0 | - |
| - | - | 2648 | 620.8 | - | - | 0 | - |
| - | - | 1760 | 621.3 | - | - | 0 | - |
| - | - | 1341 | 621.3 | - | - | 0 | - |
| - | - | 1571 | 628.3 | - | - | 0 | - |
| - | - | 2853 | 629.3 | - | - | 0 | - |
| - | - | 1734 | 629.8 | - | - | 0 | - |
| - | - | 3099 | 631.8 | - | - | 0 | - |
| - | - | 1918 | 632.3 | - | - | 0 | - |
| - | - | 1201 | 632.8 | - | - | 0 | - |
| - | - | 8748 | 638.3 | - | - | 0 | - |
| - | - | 5714 | 638.8 | - | - | 0 | - |
| - | - | 927.1 | 644.3 | - | - | 0 | - |
| 4 | y | 1763 | 644.8 | 0.001522 | 2.36 | +2 | 10 |
| 4 | y | 1398 | 645.3 | 0.01226 | 19 | +2 | 10 |
| - | - | 1395 | 645.8 | - | - | 0 | - |
| - | - | 5.786E+04 | 647.3 | - | - | 0 | - |
| - | - | 3.117E+04 | 647.8 | - | - | 0 | - |
| - | - | 1193 | 650.3 | - | - | 0 | - |
| - | - | 1869 | 651.3 | - | - | 0 | - |
| - | - | 2775 | 652.3 | - | - | 0 | - |
| - | - | 2306 | 653.3 | - | - | 0 | - |
| - | - | 1917 | 653.3 | - | - | 0 | - |
| - | - | 996.9 | 653.7 | - | - | 0 | - |
| 4 | y | 1.064E+04 | 653.8 | 0.003442 | 5.264 | +2 | 10 |
| - | - | 909.7 | 654 | - | - | 0 | - |
| - | - | 7033 | 654.3 | - | - | 0 | - |
| - | - | 3790 | 654.8 | - | - | 0 | - |
| 11 | b | 866.4 | 655.3 | 0.007204 | 10.99 | +2 | 11 |
| - | - | 1262 | 656.3 | - | - | 0 | - |
| - | - | 930.2 | 666.3 | - | - | 0 | - |
| - | - | 2706 | 667.3 | - | - | 0 | - |
| - | - | 1028 | 667.8 | - | - | 0 | - |
| - | - | 1.418E+04 | 668.3 | - | - | 0 | - |
| - | - | 5950 | 669.3 | - | - | 0 | - |
| - | - | 1401 | 670.3 | - | - | 0 | - |
| - | - | 2935 | 671.3 | - | - | 0 | - |
| - | - | 1714 | 672.3 | - | - | 0 | - |
| - | - | 4630 | 672.8 | - | - | 0 | - |
| - | - | 2634 | 673.3 | - | - | 0 | - |
| - | - | 1904 | 674.3 | - | - | 0 | - |
| 6 | b | 1.02E+04 | 676.3 | 0.0001301 | 0.1924 | +1 | 6 |
| 6 | b | 7262 | 677.3 | 0.008546 | 12.62 | +1 | 6 |
| - | - | 1307 | 677.8 | - | - | 0 | - |
| - | - | 1410 | 678.3 | - | - | 0 | - |
| - | - | 1072 | 678.3 | - | - | 0 | - |
| - | - | 2814 | 679.3 | - | - | 0 | - |
| - | - | 1.624E+04 | 681.8 | - | - | 0 | - |
| - | - | 1.019E+04 | 682.3 | - | - | 0 | - |
| - | - | 930.2 | 686.8 | - | - | 0 | - |
| - | - | 969.4 | 687.3 | - | - | 0 | - |
| - | - | 7505 | 689.4 | - | - | 0 | - |
| - | - | 892.5 | 690.4 | - | - | 0 | - |
| - | - | 1714 | 692.3 | - | - | 0 | - |
| - | - | 2727 | 692.8 | - | - | 0 | - |
| 6 | b | 7.524E+04 | 694.4 | 0.0001245 | 0.1793 | +1 | 6 |
| - | - | 2.806E+04 | 695.4 | - | - | 0 | - |
| - | - | 2.355E+04 | 696.3 | - | - | 0 | - |
| - | - | 3309 | 696.4 | - | - | 0 | - |
| - | - | 1.127E+04 | 697.3 | - | - | 0 | - |
| - | - | 2762 | 698.3 | - | - | 0 | - |
| - | - | 2473 | 699.3 | - | - | 0 | - |
| - | - | 2806 | 700.3 | - | - | 0 | - |
| - | - | 3301 | 700.8 | - | - | 0 | - |
| 3 | y | 1.157E+04 | 701.3 | 0.002336 | 3.332 | +2 | 11 |
| 3 | y | 1.085E+04 | 701.8 | 0.009596 | 13.67 | +2 | 11 |
| - | - | 6166 | 702.3 | - | - | 0 | - |
| - | - | 8337 | 702.8 | - | - | 0 | - |
| - | - | 5915 | 703.3 | - | - | 0 | - |
| - | - | 1297 | 705.4 | - | - | 0 | - |
| - | - | 1050 | 705.9 | - | - | 0 | - |
| 8 | y | 5250 | 706.3 | 0.0008599 | 1.217 | +1 | 6 |
| - | - | 2316 | 707.3 | - | - | 0 | - |
| - | - | 902.6 | 708.3 | - | - | 0 | - |
| - | - | 3592 | 709.3 | - | - | 0 | - |
| - | - | 1.501E+04 | 709.8 | - | - | 0 | - |
| 3 | y | 5.962E+04 | 710.3 | 0.004317 | 6.078 | +2 | 11 |
| 12 | b | 4.786E+04 | 710.8 | 0.01034 | 14.55 | +2 | 12 |
| 12 | b | 1.726E+04 | 711.3 | 0.0009442 | 1.327 | +2 | 12 |
| - | - | 8.902E+04 | 711.9 | - | - | 0 | - |
| - | - | 5.322E+04 | 712.4 | - | - | 0 | - |
| - | - | 1699 | 712.9 | - | - | 0 | - |
| - | - | 976.5 | 715.8 | - | - | 0 | - |
| - | - | 2143 | 719.4 | - | - | 0 | - |
| 12 | b | 5595 | 719.8 | 0.004946 | 6.871 | +2 | 12 |
| - | - | 2786 | 720.4 | - | - | 0 | - |
| - | - | 1989 | 720.9 | - | - | 0 | - |
| - | - | 848.9 | 722.4 | - | - | 0 | - |
| - | - | 7657 | 723.4 | - | - | 0 | - |
| 8 | y | 4.452E+04 | 724.4 | 0.0008543 | 1.179 | +1 | 6 |
| - | - | 1.704E+04 | 725.4 | - | - | 0 | - |
| - | - | 4247 | 726.4 | - | - | 0 | - |
| - | - | 1727 | 727.4 | - | - | 0 | - |
| - | - | 1237 | 728.9 | - | - | 0 | - |
| - | - | 1191 | 729.4 | - | - | 0 | - |
| - | - | 1142 | 734.3 | - | - | 0 | - |
| - | - | 1139 | 737.4 | - | - | 0 | - |
| - | - | 1051 | 737.9 | - | - | 0 | - |
| - | - | 2174 | 739.4 | - | - | 0 | - |
| - | - | 1563 | 740.4 | - | - | 0 | - |
| - | - | 911 | 740.9 | - | - | 0 | - |
| - | - | 1638 | 741.4 | - | - | 0 | - |
| - | - | 2327 | 745.4 | - | - | 0 | - |
| - | - | 1.242E+04 | 746.4 | - | - | 0 | - |
| - | - | 8442 | 746.9 | - | - | 0 | - |
| - | - | 825 | 747.4 | - | - | 0 | - |
| - | - | 986.6 | 750.4 | - | - | 0 | - |
| 2 | y | 2310 | 751.3 | 0.01238 | 16.47 | +2 | 12 |
| - | - | 1073 | 751.9 | - | - | 0 | - |
| - | - | 1.463E+05 | 755.4 | - | - | 0 | - |
| - | - | 1.012E+05 | 755.9 | - | - | 0 | - |
| - | - | 1821 | 756.4 | - | - | 0 | - |
| - | - | 2341 | 759.4 | - | - | 0 | - |
| 2 | y | 5166 | 759.9 | 0.002642 | 3.477 | +2 | 12 |
| - | - | 1.054E+04 | 760.4 | - | - | 0 | - |
| - | - | 5881 | 760.9 | - | - | 0 | - |
| - | - | 1255 | 761.4 | - | - | 0 | - |
| - | - | 3378 | 763.4 | - | - | 0 | - |
| - | - | 2077 | 764.4 | - | - | 0 | - |
| - | - | 871.3 | 765.3 | - | - | 0 | - |
| - | - | 1265 | 767.4 | - | - | 0 | - |
| - | - | 1102 | 768.4 | - | - | 0 | - |
| - | - | 1121 | 770.4 | - | - | 0 | - |
| - | - | 911.6 | 778.4 | - | - | 0 | - |
| - | - | 4262 | 779.3 | - | - | 0 | - |
| - | - | 1566 | 780.3 | - | - | 0 | - |
| - | - | 2446 | 781.4 | - | - | 0 | - |
| - | - | 1787 | 782.4 | - | - | 0 | - |
| - | - | 2112 | 783.3 | - | - | 0 | - |
| - | - | 1096 | 784.3 | - | - | 0 | - |
| - | - | 1111 | 787.4 | - | - | 0 | - |
| - | - | 976.7 | 791.5 | - | - | 0 | - |
| - | - | 912 | 792.4 | - | - | 0 | - |
| - | - | 972 | 793.4 | - | - | 0 | - |
| - | - | 5844 | 795.4 | - | - | 0 | - |
| - | - | 2691 | 796.4 | - | - | 0 | - |
| - | - | 2921 | 797.4 | - | - | 0 | - |
| - | - | 1786 | 798.4 | - | - | 0 | - |
| - | - | 9162 | 809.4 | - | - | 0 | - |
| 0 | Precursor | 5089 | 810.4 | 0.002973 | 3.668 | +2 | -1 |
| - | - | 1736 | 811.3 | - | - | 0 | - |
| - | - | 845.4 | 818.4 | - | - | 0 | - |
| - | - | 3017 | 823.4 | - | - | 0 | - |
| - | - | 1839 | 824.4 | - | - | 0 | - |
| - | - | 1.451E+04 | 836.4 | - | - | 0 | - |
| - | - | 3995 | 837.4 | - | - | 0 | - |
| - | - | 944.2 | 842.4 | - | - | 0 | - |
| - | - | 1550 | 850.4 | - | - | 0 | - |
| - | - | 2282 | 851.4 | - | - | 0 | - |
| - | - | 1767 | 852.4 | - | - | 0 | - |
| - | - | 1045 | 853.4 | - | - | 0 | - |
| - | - | 1027 | 859.4 | - | - | 0 | - |
| - | - | 1850 | 860.4 | - | - | 0 | - |
| - | - | 1276 | 861.4 | - | - | 0 | - |
| - | - | 1078 | 868.4 | - | - | 0 | - |
| - | - | 1.156E+04 | 868.4 | - | - | 0 | - |
| - | - | 795.3 | 869.4 | - | - | 0 | - |
| - | - | 6310 | 869.4 | - | - | 0 | - |
| - | - | 1932 | 870.4 | - | - | 0 | - |
| 7 | b | 1.06E+04 | 878.4 | 0.004487 | 5.108 | +1 | 7 |
| 7 | b | 8235 | 879.4 | 0.01602 | 18.21 | +1 | 7 |
| - | - | 3353 | 880.4 | - | - | 0 | - |
| - | - | 1231 | 881.4 | - | - | 0 | - |
| - | - | 1716 | 892.4 | - | - | 0 | - |
| - | - | 924.7 | 893.4 | - | - | 0 | - |
| 7 | b | 9.327E+04 | 896.4 | 0.00442 | 4.931 | +1 | 7 |
| - | - | 4.593E+04 | 897.4 | - | - | 0 | - |
| - | - | 1.492E+04 | 898.4 | - | - | 0 | - |
| - | - | 2778 | 899.4 | - | - | 0 | - |
| - | - | 872.5 | 906.4 | - | - | 0 | - |
| - | - | 840.5 | 907.4 | - | - | 0 | - |
| 7 | y | 3312 | 908.4 | 0.01364 | 15.01 | +1 | 7 |
| - | - | 1979 | 909.4 | - | - | 0 | - |
| - | - | 1661 | 910.4 | - | - | 0 | - |
| - | - | 909.4 | 915.4 | - | - | 0 | - |
| - | - | 1512 | 923.4 | - | - | 0 | - |
| - | - | 4414 | 924.4 | - | - | 0 | - |
| - | - | 3682 | 925.4 | - | - | 0 | - |
| 7 | y | 4622 | 926.4 | 0.006004 | 6.481 | +1 | 7 |
| - | - | 2301 | 927.4 | - | - | 0 | - |
| - | - | 1245 | 953.4 | - | - | 0 | - |
| - | - | 1362 | 970.4 | - | - | 0 | - |
| - | - | 1723 | 971.4 | - | - | 0 | - |
| - | - | 1326 | 975.5 | - | - | 0 | - |
| - | - | 807.2 | 976.5 | - | - | 0 | - |
| - | - | 2652 | 981.4 | - | - | 0 | - |
| - | - | 897.7 | 982.4 | - | - | 0 | - |
| - | - | 1204 | 986.5 | - | - | 0 | - |
| - | - | 1631 | 987.5 | - | - | 0 | - |
| - | - | 2151 | 988.4 | - | - | 0 | - |
| 8 | b | 1532 | 991.5 | 0.003675 | 3.706 | +1 | 8 |
| - | - | 938.3 | 996.5 | - | - | 0 | - |
| 8 | b | 5934 | 1010 | 0.003913 | 3.876 | +1 | 8 |
| - | - | 3385 | 1011 | - | - | 0 | - |
| - | - | 1292 | 1012 | - | - | 0 | - |
| - | - | 1031 | 1012 | - | - | 0 | - |
| - | - | 1357 | 1014 | - | - | 0 | - |
| - | - | 2071 | 1030 | - | - | 0 | - |
| - | - | 1087 | 1040 | - | - | 0 | - |
| 6 | y | 3043 | 1041 | 0.003964 | 3.806 | +1 | 8 |
| - | - | 1631 | 1042 | - | - | 0 | - |
| - | - | 885.6 | 1074 | - | - | 0 | - |
| - | - | 1013 | 1091 | - | - | 0 | - |
| - | - | 1474 | 1092 | - | - | 0 | - |
| - | - | 1373 | 1093 | - | - | 0 | - |
| - | - | 884.4 | 1102 | - | - | 0 | - |
| 9 | b | 777.4 | 1108 | 0.003195 | 2.885 | +1 | 9 |
| - | - | 2036 | 1110 | - | - | 0 | - |
| - | - | 1045 | 1112 | - | - | 0 | - |
| 9 | b | 1116 | 1125 | 0.003703 | 3.293 | +1 | 9 |
| - | - | 817.5 | 1135 | - | - | 0 | - |
| 5 | y | 3591 | 1151 | 0.002093 | 1.818 | +1 | 9 |
| 5 | y | 4753 | 1152 | 0.01075 | 9.331 | +1 | 9 |
| - | - | 2895 | 1153 | - | - | 0 | - |
| - | - | 1030 | 1155 | - | - | 0 | - |
| - | - | 2134 | 1169 | - | - | 0 | - |
| 5 | y | 7732 | 1170 | 0.005567 | 4.76 | +1 | 9 |
| - | - | 4009 | 1171 | - | - | 0 | - |
| - | - | 1249 | 1172 | - | - | 0 | - |
| - | - | 2729 | 1178 | - | - | 0 | - |
| - | - | 2167 | 1179 | - | - | 0 | - |
| 10 | b | 1453 | 1182 | 0.003114 | 2.635 | +1 | 10 |
| - | - | 835.1 | 1277 | - | - | 0 | - |
| - | - | 782.5 | 1721 | - | - | 0 | - |
| - | - | 713.4 | 1994 | - | - | 0 | - |
| - | - | 798.5 | 3070 | - | - | 0 | - |
| - | - | 838.7 | 3209 | - | - | 0 | - |

m/z Charge Intensity FragmentType MassShift Position
120.08116912841797 0 14331.761
121.08454132080078 0 715.61224
122.07148742675781 0 1140.4005
122.116943359375 0 440.05716
123.04461669921875 0 1058.404
125.10767364501953 0 1730.3588
126.05524444580078 0 1330.5542
126.0915756225586 0 754.95685
127.05057525634766 0 4304.618
127.0871353149414 0 539.03625
128.1073760986328 0 13569.351
129.06610107421875 0 848.24316
129.10264587402344 0 146904.77
130.05043029785156 0 724.43225
130.06549072265625 0 27939.172
130.1002197265625 0 1247.8767
130.10595703125 0 10059.489
130.43145751953125 0 538.2697
131.0638427734375 0 445.38638
131.06884765625 0 2942.463
132.04476928710938 0 5625.0405
132.10218811035156 0 1085.644
133.06150817871094 0 474.28183
136.0760955810547 0 20430.752
137.0793914794922 0 2251.397
138.0666046142578 0 2595.614
138.0917205810547 0 1049.9973
139.05062866210938 0 1030.1036
139.0871124267578 0 1848.4282
140.08229064941406 0 1401.4501
141.10260009765625 0 2609.431
145.0609893798828 0 1621.8574
146.06044006347656 0 3117.6882
147.04452514648438 0 1040.7262
147.06390380859375 0 681.8681
147.07675170898438 0 1211.1755 y 10
149.0448760986328 0 850.1456
151.0869903564453 0 2903.6797
153.1027374267578 0 800.8275
154.0501251220703 0 1927.2432
154.0618438720703 0 712.4583
155.09312438964844 0 8285.03
155.11827087402344 0 7948.353 a Water loss 1
156.07745361328125 0 864.1452
156.1027069091797 0 1239.0417
156.1215362548828 0 542.98395
157.06117248535156 0 1169.222
157.0764923095703 0 8583.457
157.0975799560547 0 1882.9365
157.10874938964844 0 4166.427
158.06039428710938 0 26278.299
158.0927734375 0 1474.2892
158.19668579101562 0 590.4237
159.063720703125 0 2876.8657
159.0921173095703 0 3544.1572
159.1131591796875 0 2001.5531 d 1
165.0550079345703 0 11560.958
166.06141662597656 0 19091.178
166.08665466308594 0 4806.5444 y 9
166.09786987304688 0 1262.0879
167.05645751953125 0 651.6246
167.082275390625 0 923.5042
168.10247802734375 0 1293.7189
168.11375427246094 0 972.9352
169.07656860351562 0 805.6068
169.0974578857422 0 4029.454
169.13389587402344 0 2033.2096
171.0767822265625 0 1842.3032
171.11276245117188 0 614.6972
172.0720977783203 0 2124.7927
172.1087646484375 0 649.2785
173.0562744140625 0 1997.8066
173.12890625 0 245650.52 a 1
173.43820190429688 0 916.199
174.13223266601562 0 19139.814
175.0869903564453 0 127189.46
175.11935424804688 0 1987.2014
176.082763671875 0 940.20233
176.0903778076172 0 13292.205
177.09390258789062 0 803.101
177.1021728515625 0 927.3301
178.13421630859375 0 6052.646
179.13778686523438 0 1139.9043
181.06129455566406 0 781.8704
182.0815887451172 0 23615.537 y 12
182.09117126464844 0 1308.2568
183.08538818359375 0 2360.2092
183.11317443847656 0 17297.475 b Water loss 1
183.14971923828125 0 1001.4142
184.11648559570312 0 1162.9849
185.0712432861328 0 2858.7214
185.09230041503906 0 977.642
185.12901306152344 0 1361.477
185.16514587402344 0 1744.9447
186.07473754882812 0 507.26605
186.1240692138672 0 35064.46
187.0714111328125 0 1001.8862
187.08663940429688 0 1339.6416
187.10800170898438 0 2422.4631
187.12754821777344 0 2275.6382
187.1449737548828 0 1211.0007
188.07093811035156 0 828.4511
189.08731079101562 0 7968.826
189.3579864501953 0 554.8376
190.08285522460938 0 2168.7761
190.13467407226562 0 1855.5391
194.0925750732422 0 2205.2678
195.11314392089844 0 10551.88
196.10853576660156 0 4711.5312
196.11692810058594 0 738.145
196.1449737548828 0 888.05853
197.0702362060547 0 692.8714
197.11181640625 0 788.97644
197.12889099121094 0 2878.6653
197.1656951904297 0 582.07446
198.08766174316406 0 627.98926
199.07176208496094 0 4206.0435
199.1448516845703 0 1281.3716
200.13917541503906 0 581.25275
201.12371826171875 0 82363.164 b 1
202.08721923828125 0 723.467
202.1269073486328 0 6215.787
203.06607055664062 0 2015.8854
203.0819854736328 0 1007.73395
205.0976104736328 0 4498.6567
206.10093688964844 0 785.8953
206.1290740966797 0 1149.5826
207.16107177734375 0 1104.0157
208.10865783691406 0 665.2344
210.12466430664062 0 834.32635
210.9343719482422 0 526.5783
211.10775756835938 0 1460.6041 y Water loss 10
211.14561462402344 0 905.4041
211.1557159423828 0 12025.854
212.10304260253906 0 672.8424
212.1396484375 0 1623.4729
213.06597900390625 0 1010.1157
213.08734130859375 0 1019.2875
213.12387084960938 0 4964.9736
213.15963745117188 0 986.08813
215.08299255371094 0 1369.8754
215.1392059326172 0 590.60846
216.09788513183594 0 1355.5586
216.1352081298828 0 1261.7998
217.08229064941406 0 12138.751
218.0861053466797 0 692.5964
219.124267578125 0 1493.9205
220.10842895507812 0 709.3421
221.092529296875 0 3850.2625
221.1034698486328 0 1009.6968
221.13986206054688 0 681.2065
222.08448791503906 0 596.67126
222.09619140625 0 1058.284
222.12396240234375 0 3016.6511
223.06517028808594 0 617.5132
223.10797119140625 0 3191.8103
223.15573120117188 0 19430.287
224.0635223388672 0 860.79205
224.1590576171875 0 2780.511
226.0825653076172 0 3956.938
226.11903381347656 0 1111.1409 b Water loss 5
226.15463256835938 0 561.99786
227.0663604736328 0 2990.034
227.10289001464844 0 803.5192
227.11428833007812 0 3174.838
228.13482666015625 0 1235.7496
228.18247985839844 0 2127.2842
229.11875915527344 0 3376.279
230.11346435546875 0 578.3713
233.13978576660156 0 3857.917
233.1653594970703 0 1770.1569
234.1241455078125 0 21222.22
235.12757873535156 0 3005.8052
237.09877014160156 0 806.12085
237.13458251953125 0 1516.9167
238.0842742919922 0 557.8429
238.1186065673828 0 1912.2025
239.15065002441406 0 7687.0215
240.13475036621094 0 10620.402
241.19122314453125 0 1269.7374
242.1139678955078 0 3163.1782
243.1092529296875 0 1772.9453
243.14956665039062 0 2834.9795
244.09341430664062 0 5853.837
245.09619140625 0 686.64105
245.12611389160156 0 732.41846
247.10813903808594 0 11962.173
247.1209716796875 0 1099.1707
247.143798828125 0 1003.0896
247.15664672851562 0 851.1456
248.10145568847656 0 708.23553
248.11380004882812 0 5243.4546
248.62966918945312 0 1233.5177
249.09864807128906 0 6194.3354
249.1349639892578 0 2505.311
250.10105895996094 0 629.2751
251.15074157714844 0 72251.055
252.1538543701172 0 9815.904
253.15675354003906 0 595.68097
255.1092529296875 0 2779.6355
255.14540100097656 0 2052.5457
256.1305847167969 0 588.7844
256.17718505859375 0 2592.768
257.1128234863281 0 1151.4441
258.1451416015625 0 3836.79
261.11962890625 0 2001.3682
261.13433837890625 0 797.09326
262.11859130859375 0 1130.0916
262.1546325683594 0 773.1691
265.1296691894531 0 8775.194
266.1142578125 0 2084.087
266.1251525878906 0 17139.738
267.12896728515625 0 1256.3248
268.1287536621094 0 709.3003
268.1773681640625 0 1083.4733
269.1292419433594 0 968.4373
269.1615905761719 0 1045.0919
269.1862487792969 0 1649.2949
270.1456604003906 0 1175.6752
270.160400390625 0 4917.7676
271.16278076171875 0 782.11145
272.1040344238281 0 791.6805
272.1352844238281 0 635.43506
274.13018798828125 0 1811.009
275.1035461425781 0 848.7209
276.1106872558594 0 817.18445
276.1548156738281 0 1091.906
279.1463317871094 0 775.9198
280.12860107421875 0 1303.3875
282.1563720703125 0 3191.2168
283.1404724121094 0 55415.44
284.12652587890625 0 966.76447
284.1435852050781 0 7140.493
284.160400390625 0 1121.8071
286.14013671875 0 3975.0896
286.1556091308594 0 2601.8142
287.1726989746094 0 819.02515
288.1546936035156 0 1212.583
288.171142578125 0 7513.5317
289.174560546875 0 1460.4628
290.1139831542969 0 3067.645
291.1188049316406 0 639.30945
293.11358642578125 0 31441.865 y Water loss 11
294.1174621582031 0 5965.7993
294.15704345703125 0 1225.5642
296.1969909667969 0 1791.8661 b Water loss 2
297.15643310546875 0 8524.183
298.1395263671875 0 2082.3596
298.1572570800781 0 3261.944
299.06195068359375 0 1807.7498
299.1368713378906 0 916.8057
300.0628662109375 0 2209.2197
300.16705322265625 0 4602.0747
301.0595703125 0 4523.7207
301.0826110839844 0 2049.7537
301.15106201171875 0 24495.166
301.1705627441406 0 1395.0894
302.0606384277344 0 1823.4016
302.1251220703125 0 1306.1512
302.1544189453125 0 3200.0295
305.1614990234375 0 3667.5273
305.64556884765625 0 1550.6211
306.1379089355469 0 5003.297 y 8
307.14056396484375 0 766.4557
310.2237243652344 0 1256.3079
311.124267578125 0 4809.594 y 11
312.1557922363281 0 12505.614
313.1573791503906 0 1350.2816
313.18780517578125 0 2391.3953
314.148681640625 0 5149.3945
314.2073059082031 0 1825.5192 b 2
315.1668701171875 0 13458.863
315.1856689453125 0 1201.0061
316.1503601074219 0 2762.4014
316.1687316894531 0 3858.8525
318.1089172363281 0 5970.906
319.1117248535156 0 710.19604
319.1506042480469 0 963.92816
322.1874694824219 0 1136.6373
325.14990234375 0 764.7994
327.2506408691406 0 3873.528
330.1662292480469 0 5284.514
331.1717834472656 0 823.3344 b Water loss 7
332.20892333984375 0 1817.4509
333.1772766113281 0 4911.8975
334.1823425292969 0 907.16895
334.663330078125 0 1208.3081
338.21917724609375 0 3105.0044
338.67352294921875 0 948.9244 b Water loss 5
339.1855773925781 0 641.57245
341.182861328125 0 2282.5457
342.1402282714844 0 2904.9878 y Ammonia loss 5
344.1724548339844 0 974.3788
345.1315612792969 0 2918.107
346.1150817871094 0 6755.9624
347.1182861328125 0 1092.6702
347.17364501953125 0 1252.229
347.6798400878906 0 4552.537 b 5
348.1817626953125 0 1412.5596
348.6589050292969 0 1258.2306
349.184326171875 0 1210.2784
350.21923828125 0 7572.922
351.2220153808594 0 1240.9753
355.2453308105469 0 1865.2618
357.1774597167969 0 1253.2539
358.1766052246094 0 1402.466
359.1680908203125 0 1869.597
360.2021484375 0 675.8473
361.19989013671875 0 1928.5339
362.18408203125 0 2794.4707
362.6798400878906 0 10130.394 y 7
363.1414489746094 0 12650.17
363.1822814941406 0 3653.0142
363.6809387207031 0 1348.9204
364.1253662109375 0 2726.3276
364.14691162109375 0 1461.4836
364.17401123046875 0 1224.6213
364.6722717285156 0 1024.6902
365.1280517578125 0 930.2999
366.1783142089844 0 1643.2719
367.15985107421875 0 1065.6843
367.24609375 0 986.8972
368.1634826660156 0 1056.1406
368.691162109375 0 1035.9427
369.1553649902344 0 804.79346
373.1851501464844 0 1575.6332
373.68670654296875 0 674.5011
376.1601257324219 0 816.185
378.2141418457031 0 4667.044
379.2093505859375 0 19737.883
380.2110595703125 0 3879.8123
381.1521301269531 0 36448.906
382.15496826171875 0 7456.219
382.69378662109375 0 952.9714
383.1338806152344 0 1061.0656
383.2281188964844 0 2735.2427
383.73046875 0 1273.6694
384.1918029785156 0 3491.0493
385.1707458496094 0 1249.1279
390.17730712890625 0 1956.5647
391.13616943359375 0 1074.2777
394.1720275878906 0 4487.433
395.1557312011719 0 2315.7878
395.2404479980469 0 1823.9896
396.1603088378906 0 1032.401
396.2253112792969 0 5980.016
397.22882080078125 0 1443.5607
397.25665283203125 0 4167.172
398.1784362792969 0 2230.9065
398.2892761230469 0 1197.491
399.1838073730469 0 1771.7079
400.1625061035156 0 1045.881
400.22021484375 0 1257.7175
401.2409973144531 0 913.05664
402.19854736328125 0 4395.974
403.20068359375 0 1670.6552
405.2015686035156 0 834.5776
405.2593994140625 0 988.3576
405.7169189453125 0 940.9868
406.24407958984375 0 758.17456
407.24151611328125 0 2353.4387
409.1474609375 0 3030.1733
409.2205810546875 0 1640.8347
410.149169921875 0 957.2518
411.1304016113281 0 1433.4417
411.19879150390625 0 2883.422
412.1831359863281 0 22957.348
412.2667236328125 0 3044.0862
413.1644592285156 0 2736.849
413.18658447265625 0 4332.6226
414.19036865234375 0 1092.0819
414.23443603515625 0 7350.6855
414.283203125 0 12167.996
414.7212829589844 0 2051.121
415.2394714355469 0 1338.1117
417.2062072753906 0 1191.51
418.0371398925781 0 1249.584
418.17144775390625 0 1401.5554
418.20172119140625 0 865.44495
418.7172546386719 0 1091.0493
421.2131042480469 0 867.49524 y Water loss 10
423.23358154296875 0 4856.2495
423.27197265625 0 9490.497
423.7350158691406 0 1874.1912
424.2690124511719 0 9481.651
425.2514953613281 0 3155.535
425.7144470214844 0 3912.2432
426.2082214355469 0 6007.5293
426.7086486816406 0 3439.7148
428.157470703125 0 3892.9846
428.2163391113281 0 1061.9485
429.1408996582031 0 9667.389
429.20989990234375 0 10476.255
429.23492431640625 0 1177.4736
430.1424560546875 0 1810.0768
430.19390869140625 0 55193.86 y Water loss 3
430.70550537109375 0 1133.7852
431.1970520019531 0 14813.058
431.7029724121094 0 2700.9255
432.1974792480469 0 2091.06
432.22955322265625 0 897.48804
433.25689697265625 0 1629.1053 b Water loss 3
434.7198181152344 0 38629.215
435.22125244140625 0 18810.125
435.7228698730469 0 3933.2698
436.22113037109375 0 779.77545
439.2193298339844 0 16903.32 y 10
439.7119140625 0 18010.996 b Water loss 6
440.21246337890625 0 10287.937
440.7118835449219 0 3887.1675
441.29388427734375 0 2135.8323
442.27777099609375 0 30853.969
446.1675109863281 0 8927.481
447.16949462890625 0 1190.553
447.2193298339844 0 1136.584
448.2042236328125 0 2847.9133
448.71734619140625 0 24130.402 b 6
449.21832275390625 0 15110.052
449.7203674316406 0 2849.3538
450.2829284667969 0 837.8651
451.2672424316406 0 16849.088 b 3
452.2698059082031 0 4035.6646
454.21917724609375 0 2646.777
454.7121276855469 0 8686.057 y Water loss 6
455.212646484375 0 4595.6323
455.2730407714844 0 1432.613
455.71612548828125 0 707.54175
458.23541259765625 0 1545.7057
459.19842529296875 0 1278.2506
461.2040100097656 0 1772.8362
461.291015625 0 1686.318
463.71746826171875 0 3684.0964 y 6
464.21966552734375 0 1747.4467
466.2417907714844 0 2472.0051
466.3138427734375 0 801.4733
468.293212890625 0 5755.4995
469.2514343261719 0 885.8069
469.2884216308594 0 1230.475
473.7472839355469 0 930.75806
474.79669189453125 0 791.0949
475.794921875 0 810.30963
476.2242431640625 0 1658.3434
477.2102355957031 0 2981.1174
478.2302551269531 0 2096.5293 y Water loss 9
478.2783508300781 0 2875.0454
478.3128967285156 0 889.1485
479.2253112792969 0 2268.3176
482.25677490234375 0 7039.57
482.7551574707031 0 4979.5977
483.25201416015625 0 2351.0586
483.3409423828125 0 6336.1685
488.25384521484375 0 1794.4843
491.26165771484375 0 7954.175
491.7631530761719 0 6478.7246
492.26458740234375 0 2070.8938
493.2535095214844 0 984.16656
494.2362976074219 0 32412.055
495.239013671875 0 10072.942
495.3404846191406 0 6702.5146
496.2423095703125 0 43805.535 b Water loss 7
496.32415771484375 0 7399.446
496.76702880859375 0 17084.604
497.24432373046875 0 8672.81
497.2741394042969 0 3994.082
497.9103698730469 0 827.69714
498.2463073730469 0 3304.9272
504.2540588378906 0 1280.9211
505.2611389160156 0 7467.9297 b 7
505.76153564453125 0 3666.2642
506.1064758300781 0 1446.3337
506.22637939453125 0 825.75684
506.26580810546875 0 2240.5076
507.2568664550781 0 1188.551
511.2627258300781 0 4757.332
511.33587646484375 0 9393.91
512.1762084960938 0 1057.4674
512.2651977539062 0 926.13855
513.3511962890625 0 143358.17
520.1943969726562 0 1161.1163
523.3352661132812 0 6336.1455
524.2644653320312 0 4076.131
524.3206787109375 0 874.34186
524.7633056640625 0 1695.533
525.2672119140625 0 5872.034
525.8124389648438 0 9481.96
526.2728881835938 0 1507.0914
526.3150024414062 0 4462.445
526.8139038085938 0 1682.5015
527.3138427734375 0 815.44934
528.5891723632812 0 839.21716
530.1868896484375 0 959.3772
530.270751953125 0 747.39996
532.2706298828125 0 1001.9509
534.303466796875 0 972.9543
534.5932006835938 0 2283.5269 Precursor Water loss
534.9275512695312 0 1395.1814 Precursor Ammonia loss
535.2596435546875 0 1287.8073
535.6055908203125 0 3939.387
535.9403686523438 0 1996.8489
538.345947265625 0 803.2086
539.8096923828125 0 32011.723
540.0474853515625 0 834.9067
540.2696533203125 0 1356.4498
540.31103515625 0 21284.79
540.593994140625 0 3176.2512 Precursor
540.8123779296875 0 6174.588
540.9304809570312 0 3433.893
541.3461303710938 0 217795.23
541.609130859375 0 7754.622
541.9437255859375 0 7983.611
542.29248046875 0 2755.2578
542.3436279296875 0 784.2342
543.2780151367188 0 8627.634
544.2802734375 0 2712.6453
545.2369995117188 0 1029.4691
545.7753295898438 0 1091.8682
548.1885375976562 0 1245.7229
551.3294677734375 0 1526.9565
554.2808837890625 0 7720.7896
554.7813720703125 0 2410.7432
555.2302856445312 0 4002.3242
555.2705688476562 0 1135.6193
556.234130859375 0 881.9648
561.3138427734375 0 1893.2563 b Water loss 4
562.7609252929688 0 1681.4067 b 8
563.2548217773438 0 3328.4326
563.7543334960938 0 1879.9379
564.7916870117188 0 1812.7402
565.2152709960938 0 1412.7396
565.261962890625 0 4400.065
566.1976928710938 0 877.0304
566.2660522460938 0 1352.0206
573.79638671875 0 6163.3584
574.2984008789062 0 2314.0596
575.2459106445312 0 1891.6393
575.292236328125 0 1031.7651
579.3253784179688 0 10235.098 b 4
580.3284912109375 0 3521.7722
582.78759765625 0 967.4138
583.2263793945312 0 10585.116
584.2305297851562 0 3180.2903
587.2769775390625 0 999.6779
588.7908935546875 0 1715.6345
589.2932739257812 0 1389.6399
592.2692260742188 0 1413.6755
593.2569580078125 0 15642.533 y Water loss 8
593.3055419921875 0 4961.9365
594.25341796875 0 4449.457
594.3092041015625 0 1245.2533
595.2542114257812 0 1052.8491
595.3191528320312 0 1231.1841
596.352783203125 0 1565.2748
597.796875 0 57305.945
598.2982177734375 0 29372.049
598.6708984375 0 1882.3391
600.2717895507812 0 1192.1459
602.2766723632812 0 788.95667
608.3206787109375 0 949.9757
609.7797241210938 0 955.043
610.283447265625 0 24955.629
610.3323974609375 0 2612.9634
610.78857421875 0 2887.7075
611.2681884765625 0 141530.58 y 8
611.791015625 0 1767.6306
612.270751953125 0 42073.145
613.27294921875 0 9659.134
614.2743530273438 0 1391.387
614.3306884765625 0 1203.9727
616.3059692382812 0 963.569
619.2994995117188 0 3309.228
619.7958374023438 0 8192.417
620.2949829101562 0 4658.0312
620.7993774414062 0 2648.248
621.2515869140625 0 1759.7402
621.3012084960938 0 1340.5875
628.2711181640625 0 1570.9343
629.314208984375 0 2853.281
629.8143310546875 0 1734.049
631.8230590820312 0 3099.1077
632.3189697265625 0 1918.3501
632.8242797851562 0 1201.0618
638.3198852539062 0 8747.738
638.8201293945312 0 5713.9004
644.3468017578125 0 927.0916
644.7823486328125 0 1763.4514 y Water loss 3
645.2850952148438 0 1398.2991 y Ammonia loss 3
645.7837524414062 0 1394.6606
647.3311157226562 0 57856.19
647.8326416015625 0 31166.525
650.3077392578125 0 1193.2246
651.2920532226562 0 1868.5057
652.3379516601562 0 2774.9338
653.2943725585938 0 2306.3909
653.3466186523438 0 1916.7382
653.6953735351562 0 996.9035
653.78955078125 0 10639.079 y 3
654.0294799804688 0 909.6594
654.2920532226562 0 7033.1533
654.7932739257812 0 3789.658
655.3207397460938 0 866.35223 b 10
656.2678833007812 0 1262.1642
666.26318359375 0 930.2419
667.3416748046875 0 2706.4902
667.8428955078125 0 1027.6129
668.3153076171875 0 14181.092
669.319091796875 0 5949.7114
670.3245239257812 0 1400.6909
671.3453979492188 0 2935.3767
672.3391723632812 0 1714.486
672.828857421875 0 4630.014
673.3303833007812 0 2633.903
674.30712890625 0 1903.9023
676.3414306640625 0 10195.719 b Water loss 5
677.3338623046875 0 7261.6655 b Ammonia loss 5
677.8276977539062 0 1306.554
678.2936401367188 0 1410.1364
678.3375854492188 0 1071.937
679.2868041992188 0 2813.721
681.8339233398438 0 16235.506
682.3345947265625 0 10194.6875
686.8211669921875 0 930.1558
687.328369140625 0 969.44666
689.3579711914062 0 7505.4375
690.3614501953125 0 892.5165
692.3213500976562 0 1713.8169
692.8143920898438 0 2727.385
694.3519897460938 0 75241.766 b 5
695.3546142578125 0 28057.48
696.3097534179688 0 23545.178
696.357666015625 0 3309.442
697.3128662109375 0 11271.231
698.3147583007812 0 2761.6309
699.339599609375 0 2473.1758
700.3201293945312 0 2805.8665
700.82421875 0 3300.943
701.3251953125 0 11567.278 y Water loss 2
701.824462890625 0 10852.897 y Ammonia loss 2
702.326171875 0 6166.04
702.8448486328125 0 8337.448
703.3469848632812 0 5914.538
705.3546752929688 0 1297.0731
705.8564453125 0 1050.3644
706.3414916992188 0 5250.121 y Water loss 7
707.3436279296875 0 2315.929
708.3495483398438 0 902.6243
709.3265991210938 0 3591.636
709.8372802734375 0 15009.685
710.3324584960938 0 59620.22 y 2
710.8336181640625 0 47864.617 b Water loss 11
711.3350219726562 0 17261.432 b Ammonia loss 11
711.8515014648438 0 89017.08
712.3538818359375 0 53224.004
712.8555297851562 0 1698.6711
715.8189697265625 0 976.52234
719.3580322265625 0 2143.4143
719.8541870117188 0 5594.885 b 11
720.3546752929688 0 2786.1643
720.8541870117188 0 1988.8469
722.3522338867188 0 848.9351
723.3663940429688 0 7657.4697
724.35205078125 0 44524.1 y 7
725.35498046875 0 17041.59
726.3560180664062 0 4247.006
727.3528442382812 0 1726.8483
728.8543090820312 0 1237.1328
729.356689453125 0 1191.4191
734.3411254882812 0 1142.1373
737.3623046875 0 1138.882
737.8507690429688 0 1051.4698
739.373779296875 0 2174.2344
740.3687133789062 0 1562.7692
740.8675537109375 0 910.95026
741.3560180664062 0 1638.4775
745.3560791015625 0 2327.3877
746.3627319335938 0 12423.73
746.8628540039062 0 8441.656
747.3611450195312 0 825.00195
750.368896484375 0 986.5977
751.3614501953125 0 2309.684 y Ammonia loss 1
751.8594360351562 0 1072.5582
755.3681640625 0 146291.8
755.8695068359375 0 101160.68
756.368408203125 0 1821.002
759.370361328125 0 2341.0315
759.864990234375 0 5166.406 y 1
760.3641357421875 0 10535.497
760.8618774414062 0 5880.8657
761.3709106445312 0 1255.4929
763.380126953125 0 3378.2637
764.3756713867188 0 2077.3962
765.3278198242188 0 871.2781
767.3859252929688 0 1265.1428
768.3731079101562 0 1102.3026
770.41357421875 0 1120.8877
778.361572265625 0 911.5612
779.3466796875 0 4261.743
780.3493041992188 0 1565.9059
781.396728515625 0 2445.6206
782.4004516601562 0 1786.7196
783.3397216796875 0 2112.3586
784.3422241210938 0 1096.3739
787.3941650390625 0 1111.0352
791.4631958007812 0 976.7456
792.3660888671875 0 912.00653
793.4202270507812 0 972.00195
795.3773193359375 0 5844.3257
796.380126953125 0 2691.4082
797.3606567382812 0 2921.3477
798.3590087890625 0 1785.974
809.3936157226562 0 9162.245
810.38916015625 0 5089.28 Precursor
811.3345336914062 0 1735.735
818.4159545898438 0 845.4381
823.3720703125 0 3016.7185
824.3770751953125 0 1838.9315
836.42529296875 0 14509.548
837.4281616210938 0 3995.2996
842.3825073242188 0 944.1594
850.4253540039062 0 1550.1333
851.4072875976562 0 2282.196
852.42138671875 0 1767.2201
853.4161987304688 0 1045.2225
859.4014282226562 0 1027.2528
860.3896484375 0 1849.7516
861.3893432617188 0 1275.55
868.361572265625 0 1077.878
868.4314575195312 0 11563.185
869.360595703125 0 795.2972
869.434814453125 0 6310.013
870.4365844726562 0 1931.9648
878.4151000976562 0 10597.906 b Water loss 6
879.41064453125 0 8234.695 b Ammonia loss 6
880.4143676757812 0 3352.6729
881.4360961914062 0 1231.2756
892.4293212890625 0 1716.1642
893.4268188476562 0 924.7274
896.4255981445312 0 93267.15 b 6
897.428466796875 0 45931.543
898.431640625 0 14919.502
899.4347534179688 0 2777.7693
906.4080810546875 0 872.4831
907.404296875 0 840.4768
908.423583984375 0 3311.8457 y Water loss 6
909.4270629882812 0 1978.5806
910.43408203125 0 1661.3682
915.4194946289062 0 909.44824
923.4396362304688 0 1511.5012
924.4232177734375 0 4414.2905
925.4284057617188 0 3682.255
926.426513671875 0 4621.6733 y 6
927.4324340820312 0 2301.19
953.39892578125 0 1245.321
970.4284057617188 0 1362.2749
971.4076538085938 0 1722.9827
975.4990234375 0 1326.3496
976.4848022460938 0 807.22546
981.4442749023438 0 2652.3984
982.4468994140625 0 897.71643
986.490966796875 0 1204.2864
987.4851684570312 0 1630.7631
988.4423828125 0 2151.3743
991.4983520507812 0 1531.9768 b Water loss 7
996.4673461914062 0 938.2839
1009.5091552734375 0 5933.976 b 7
1010.5115356445312 0 3385.1067
1011.5137939453125 0 1292.4889
1012.4905395507812 0 1031.3243
1014.4850463867188 0 1356.987
1030.49072265625 0 2071.3977
1040.472412109375 0 1086.7252
1041.451416015625 0 3042.6255 y 5
1042.459228515625 0 1630.632
1073.525146484375 0 885.5547
1090.513671875 0 1012.70184
1091.521484375 0 1474.138
1092.5321044921875 0 1373.4177
1101.5191650390625 0 884.4375
1107.50244140625 0 777.4281 b Ammonia loss 8
1109.5338134765625 0 2036.4585
1111.5447998046875 0 1044.8187
1124.535888671875 0 1116.1019 b 8
1135.470703125 0 817.4683
1151.49755859375 0 3590.7263 y Water loss 4
1152.490234375 0 4752.535 y Ammonia loss 4
1153.499267578125 0 2894.7825
1154.501708984375 0 1030.4124
1168.525390625 0 2134.073
1169.5115966796875 0 7732.3486 y 4
1170.510986328125 0 4008.5312
1171.5205078125 0 1249.2305
1177.5521240234375 0 2728.63
1178.562255859375 0 2167.4207
1181.5567626953125 0 1452.565 b 9
1276.6239013671875 0 835.0581
1720.5623779296875 0 782.48004
1993.904296875 0 713.35333
3069.5537109375 0 798.50934
3208.9931640625 0 838.67444

Spectrum Details

|  |  |
| --- | --- |
| Matched peaks? Matched peaks The total absolute number of peaks matched. Additionally in brackets the total fraction of peaks matched and the total number of peaks is shown. | 72 (9.02% of 798) |
| FDR? FDR The false discovery rate estimated for this peptide. It is calculated by matching all theoretical fragments with a non-integer shift with the raw peaks for this spectrum. This is done with 40 different shifts. The resulting percentage is the average number of annotated peaks over the number of annotated peaks with the correct spectrum. | 1.69% |
| Satellite FDR? Satellite FDR See the FDR for details on its calculation. This satellite ion specific FDR only contains the satellite ions (d/w) for I/L/J positions. | ∞ |
| PSM Score? PSM Score The PSM Score as given by Hecklib to this annotated spectrum. It is shown with three significant figures. | 562 |

#### Spectrum 7305? Spectrum 7305 The raw spectrum of this peptide as annotated by Hecklib. The fragments are coloured according to ion type (see legend). Any peaks with a star '\*' as text can be hovered over to see the full details, first the ion type second the mass shift type. By hovering over the amino acids in the peptide or ions in the legend the corresponding peaks are highlighted. By toggling the 'Unassigned' label you can turn the background (unassigned) peaks on or off in the plot. By updating the slider in the Ion legend you can update the spectrum to only show the top X% of the peaks with labels. The top X% means any peak that is within X% of the highest intensity. By dragging in the spectrum you can zoom in to a specific part of the spectrum and use 'Zoom Out' to get back to the original zoom level. The annotation of the spectrum is based on the given sequence in the peptides file and is done with different software so inconsistencies are likely. The peaks are annotated based on the given sequence, with 20 ppm tolerance.

Copy Data

##### Spectrum 7305 (TSV)

###### Preview

```
Loading example...
```

*Click on the button to copy the data to your clipboard.*

Mz MinMz MaxIntensity Max

WidthHeightPeptide font sizePeptide stroke widthSpectrum font sizeSpectrum stroke widthCompact peptide

Ion legend

wxyz

abcd

OtherUnassignedIon^Charge^_Position_Show for top:%

TVLHQDWLDGKEY

01.18e+52.37e+53.55e+54.74e+5

Zoom Out

a^+1^_2_a^+1^_2_y^+1^_1_b^+1^_2_b^+1^_2_y^+2^_3_b^+2^_4_y^+1^_2_b^+1^_3_y^+1^_2_b^+1^_3_y^+1^_3_b^+1^_4_y^+1^_3_b^+1^_4_y^+1^_4_b^+2^_8_y^+1^_4_b^+1^_5_b^+1^_5_b^+1^_5_b^+2^_10_y^+1^_5_y^+1^_5_y^+2^_10_y^+2^_10_b^+2^_11_b^+2^_11_y^+2^_10_b^+2^_11_b^+1^_6_b^+1^_6_y^+2^_11_y^+2^_11_b^+1^_6_y^+2^_11_b^+2^_12_b^+2^_12_y^+1^_6_b^+2^_12_y^+1^_6_y^+2^_12_\*\*\*b^+1^_7_b^+1^_7_b^+1^_7_y^+1^_7_y^+1^_7_y^+1^_7_b^+1^_8_b^+1^_8_b^+1^_8_y^+1^_8_y^+1^_8_b^+1^_9_b^+1^_9_b^+1^_9_y^+1^_9_y^+1^_9_b^+1^_10_b^+1^_10_y^+1^_9_b^+1^_10_y^+1^_10_y^+1^_10_b^+1^_11_b^+1^_11_y^+1^_10_b^+1^_11_y^+1^_11_y^+1^_11_y^+1^_11_b^+1^_12_

036873611051473

Fragment Matches Table

Show background peaks

| Position | Ion type | Intensity | mz Theoretical | mz Error (Th) | mz Error (ppm) | Charge | Series Number |
| --- | --- | --- | --- | --- | --- | --- | --- |
| - | - | 5603 | 120.1 | - | - | 0 | - |
| - | - | 621.9 | 121.1 | - | - | 0 | - |
| - | - | 2065 | 122.1 | - | - | 0 | - |
| - | - | 458.6 | 123 | - | - | 0 | - |
| - | - | 639.3 | 123.1 | - | - | 0 | - |
| - | - | 807 | 126.1 | - | - | 0 | - |
| - | - | 2059 | 127.1 | - | - | 0 | - |
| - | - | 763.5 | 127.1 | - | - | 0 | - |
| - | - | 595.1 | 128.1 | - | - | 0 | - |
| - | - | 5443 | 128.1 | - | - | 0 | - |
| - | - | 1.797E+05 | 129.1 | - | - | 0 | - |
| - | - | 1.007E+04 | 130.1 | - | - | 0 | - |
| - | - | 1477 | 130.1 | - | - | 0 | - |
| - | - | 1.035E+04 | 130.1 | - | - | 0 | - |
| - | - | 1056 | 131.1 | - | - | 0 | - |
| - | - | 2702 | 132.1 | - | - | 0 | - |
| - | - | 1.546E+04 | 136.1 | - | - | 0 | - |
| - | - | 1445 | 137.1 | - | - | 0 | - |
| - | - | 791.6 | 138.1 | - | - | 0 | - |
| - | - | 3427 | 138.1 | - | - | 0 | - |
| - | - | 568.8 | 138.1 | - | - | 0 | - |
| - | - | 1988 | 139.1 | - | - | 0 | - |
| - | - | 3873 | 141.1 | - | - | 0 | - |
| - | - | 479.3 | 142.1 | - | - | 0 | - |
| - | - | 1381 | 143 | - | - | 0 | - |
| - | - | 1463 | 145.1 | - | - | 0 | - |
| - | - | 464.5 | 146.1 | - | - | 0 | - |
| - | - | 572.8 | 147 | - | - | 0 | - |
| - | - | 2752 | 150.1 | - | - | 0 | - |
| - | - | 2517 | 151.1 | - | - | 0 | - |
| - | - | 610.4 | 153.1 | - | - | 0 | - |
| - | - | 503.8 | 153.1 | - | - | 0 | - |
| - | - | 504.7 | 154.1 | - | - | 0 | - |
| - | - | 957.2 | 155.1 | - | - | 0 | - |
| - | - | 1.697E+04 | 155.1 | - | - | 0 | - |
| 2 | a | 2366 | 155.1 | 0.000244 | 1.573 | +1 | 2 |
| - | - | 655.8 | 156.1 | - | - | 0 | - |
| - | - | 1416 | 156.1 | - | - | 0 | - |
| - | - | 1395 | 157.1 | - | - | 0 | - |
| - | - | 2.629E+04 | 159.1 | - | - | 0 | - |
| - | - | 563.4 | 160.1 | - | - | 0 | - |
| - | - | 2541 | 160.1 | - | - | 0 | - |
| - | - | 6197 | 165.1 | - | - | 0 | - |
| - | - | 1.16E+04 | 166.1 | - | - | 0 | - |
| - | - | 2412 | 166.1 | - | - | 0 | - |
| - | - | 3655 | 166.1 | - | - | 0 | - |
| - | - | 641.4 | 167.1 | - | - | 0 | - |
| - | - | 1401 | 167.1 | - | - | 0 | - |
| - | - | 762.7 | 167.1 | - | - | 0 | - |
| - | - | 1745 | 168.1 | - | - | 0 | - |
| - | - | 1536 | 168.1 | - | - | 0 | - |
| - | - | 640.4 | 169.1 | - | - | 0 | - |
| - | - | 6502 | 169.1 | - | - | 0 | - |
| - | - | 540.2 | 169.1 | - | - | 0 | - |
| - | - | 1493 | 170.1 | - | - | 0 | - |
| - | - | 472.2 | 170.1 | - | - | 0 | - |
| - | - | 532.8 | 171.1 | - | - | 0 | - |
| - | - | 886.7 | 171.1 | - | - | 0 | - |
| - | - | 490.4 | 171.1 | - | - | 0 | - |
| - | - | 1390 | 172.1 | - | - | 0 | - |
| - | - | 1536 | 173.1 | - | - | 0 | - |
| 2 | a | 1.045E+05 | 173.1 | 0.00033 | 1.906 | +1 | 2 |
| - | - | 3788 | 173.5 | - | - | 0 | - |
| - | - | 454.1 | 173.5 | - | - | 0 | - |
| - | - | 812.6 | 174.1 | - | - | 0 | - |
| - | - | 8858 | 174.1 | - | - | 0 | - |
| - | - | 1120 | 176.1 | - | - | 0 | - |
| - | - | 1136 | 177.1 | - | - | 0 | - |
| - | - | 4912 | 178.1 | - | - | 0 | - |
| - | - | 976.1 | 179.1 | - | - | 0 | - |
| - | - | 544.1 | 179.1 | - | - | 0 | - |
| - | - | 1175 | 181.1 | - | - | 0 | - |
| 13 | y | 1.822E+04 | 182.1 | 0.0003123 | 1.715 | +1 | 1 |
| - | - | 1701 | 183.1 | - | - | 0 | - |
| 2 | b | 1381 | 183.1 | 0.0001109 | 0.6055 | +1 | 2 |
| - | - | 848 | 183.2 | - | - | 0 | - |
| - | - | 640.5 | 185.1 | - | - | 0 | - |
| - | - | 527.7 | 185.1 | - | - | 0 | - |
| - | - | 888.1 | 185.2 | - | - | 0 | - |
| - | - | 7.027E+04 | 186.1 | - | - | 0 | - |
| - | - | 3872 | 187.1 | - | - | 0 | - |
| - | - | 475.3 | 187.1 | - | - | 0 | - |
| - | - | 697.3 | 187.1 | - | - | 0 | - |
| - | - | 5069 | 187.1 | - | - | 0 | - |
| - | - | 1370 | 188.1 | - | - | 0 | - |
| - | - | 1107 | 190.1 | - | - | 0 | - |
| - | - | 1385 | 190.1 | - | - | 0 | - |
| - | - | 982.2 | 191.1 | - | - | 0 | - |
| - | - | 689.3 | 193.1 | - | - | 0 | - |
| - | - | 3952 | 194.1 | - | - | 0 | - |
| - | - | 565.6 | 194.1 | - | - | 0 | - |
| - | - | 469.3 | 195.1 | - | - | 0 | - |
| - | - | 1.243E+04 | 195.1 | - | - | 0 | - |
| - | - | 4270 | 196.1 | - | - | 0 | - |
| - | - | 1019 | 196.1 | - | - | 0 | - |
| - | - | 656.7 | 197.1 | - | - | 0 | - |
| - | - | 1187 | 197.1 | - | - | 0 | - |
| - | - | 1905 | 198.1 | - | - | 0 | - |
| - | - | 1015 | 198.1 | - | - | 0 | - |
| - | - | 754.6 | 199.1 | - | - | 0 | - |
| - | - | 863.5 | 199.1 | - | - | 0 | - |
| - | - | 583.7 | 200.1 | - | - | 0 | - |
| 2 | b | 6.589E+04 | 201.1 | 0.0002579 | 1.282 | +1 | 2 |
| - | - | 986 | 202.1 | - | - | 0 | - |
| - | - | 5168 | 202.1 | - | - | 0 | - |
| - | - | 763.1 | 203.1 | - | - | 0 | - |
| - | - | 4245 | 205.1 | - | - | 0 | - |
| - | - | 729.8 | 205.1 | - | - | 0 | - |
| - | - | 835.2 | 206.1 | - | - | 0 | - |
| - | - | 527.6 | 207.1 | - | - | 0 | - |
| - | - | 1437 | 207.2 | - | - | 0 | - |
| - | - | 541.6 | 209.1 | - | - | 0 | - |
| 11 | y | 704.8 | 211.1 | 0.0009391 | 4.448 | +2 | 3 |
| - | - | 1191 | 212.1 | - | - | 0 | - |
| - | - | 5340 | 213.1 | - | - | 0 | - |
| - | - | 827.7 | 213.2 | - | - | 0 | - |
| - | - | 551.4 | 215.1 | - | - | 0 | - |
| - | - | 2297 | 216.1 | - | - | 0 | - |
| 4 | b | 767.7 | 217.1 | 0.002261 | 10.41 | +2 | 4 |
| - | - | 603 | 219.1 | - | - | 0 | - |
| - | - | 848.4 | 221.1 | - | - | 0 | - |
| - | - | 2783 | 221.1 | - | - | 0 | - |
| - | - | 5176 | 221.1 | - | - | 0 | - |
| - | - | 2430 | 221.1 | - | - | 0 | - |
| - | - | 1062 | 222.1 | - | - | 0 | - |
| - | - | 2434 | 223.1 | - | - | 0 | - |
| - | - | 2.139E+04 | 223.2 | - | - | 0 | - |
| - | - | 1062 | 224.1 | - | - | 0 | - |
| - | - | 3101 | 224.2 | - | - | 0 | - |
| - | - | 5517 | 226.1 | - | - | 0 | - |
| - | - | 2965 | 227.1 | - | - | 0 | - |
| - | - | 997.4 | 227.1 | - | - | 0 | - |
| - | - | 1688 | 227.1 | - | - | 0 | - |
| - | - | 1386 | 227.2 | - | - | 0 | - |
| - | - | 661.2 | 228.1 | - | - | 0 | - |
| - | - | 2034 | 229.1 | - | - | 0 | - |
| - | - | 625.9 | 229.2 | - | - | 0 | - |
| - | - | 644.2 | 231.1 | - | - | 0 | - |
| - | - | 2674 | 233.1 | - | - | 0 | - |
| - | - | 571.8 | 233.2 | - | - | 0 | - |
| - | - | 1.545E+04 | 234.1 | - | - | 0 | - |
| - | - | 781.6 | 234.1 | - | - | 0 | - |
| - | - | 1713 | 235.1 | - | - | 0 | - |
| - | - | 2502 | 238.1 | - | - | 0 | - |
| - | - | 850.6 | 238.1 | - | - | 0 | - |
| - | - | 844.8 | 239.1 | - | - | 0 | - |
| - | - | 8508 | 240.1 | - | - | 0 | - |
| - | - | 1208 | 241.1 | - | - | 0 | - |
| - | - | 1295 | 241.1 | - | - | 0 | - |
| - | - | 850.1 | 243.1 | - | - | 0 | - |
| - | - | 1.332E+04 | 244.1 | - | - | 0 | - |
| - | - | 1063 | 244.1 | - | - | 0 | - |
| - | - | 1503 | 245.1 | - | - | 0 | - |
| - | - | 517.4 | 246.1 | - | - | 0 | - |
| - | - | 1.92E+04 | 247.1 | - | - | 0 | - |
| - | - | 574.8 | 247.1 | - | - | 0 | - |
| - | - | 924.4 | 247.2 | - | - | 0 | - |
| - | - | 8178 | 248.1 | - | - | 0 | - |
| - | - | 531.2 | 248.2 | - | - | 0 | - |
| - | - | 1.165E+04 | 249.1 | - | - | 0 | - |
| - | - | 1435 | 249.1 | - | - | 0 | - |
| - | - | 5547 | 249.1 | - | - | 0 | - |
| - | - | 1177 | 250.1 | - | - | 0 | - |
| - | - | 5.023E+04 | 251.2 | - | - | 0 | - |
| - | - | 820.4 | 252.1 | - | - | 0 | - |
| - | - | 950.5 | 252.1 | - | - | 0 | - |
| - | - | 6117 | 252.2 | - | - | 0 | - |
| - | - | 1141 | 252.2 | - | - | 0 | - |
| - | - | 3007 | 255.1 | - | - | 0 | - |
| - | - | 834.3 | 256.1 | - | - | 0 | - |
| - | - | 2287 | 257.1 | - | - | 0 | - |
| - | - | 3111 | 258.1 | - | - | 0 | - |
| - | - | 1121 | 261.1 | - | - | 0 | - |
| - | - | 682.4 | 265.1 | - | - | 0 | - |
| - | - | 5659 | 265.1 | - | - | 0 | - |
| - | - | 687.3 | 265.1 | - | - | 0 | - |
| - | - | 3.245E+04 | 266.1 | - | - | 0 | - |
| - | - | 496.9 | 267.1 | - | - | 0 | - |
| - | - | 2326 | 267.1 | - | - | 0 | - |
| - | - | 1561 | 268.1 | - | - | 0 | - |
| - | - | 2163 | 268.2 | - | - | 0 | - |
| - | - | 631.7 | 268.5 | - | - | 0 | - |
| - | - | 741.6 | 269.2 | - | - | 0 | - |
| - | - | 1716 | 269.2 | - | - | 0 | - |
| - | - | 1214 | 270.1 | - | - | 0 | - |
| - | - | 4195 | 272.2 | - | - | 0 | - |
| - | - | 5917 | 274.1 | - | - | 0 | - |
| - | - | 1086 | 274.1 | - | - | 0 | - |
| - | - | 1481 | 275.1 | - | - | 0 | - |
| - | - | 866.9 | 275.1 | - | - | 0 | - |
| - | - | 1168 | 275.2 | - | - | 0 | - |
| - | - | 1776 | 276.1 | - | - | 0 | - |
| - | - | 662.6 | 276.2 | - | - | 0 | - |
| - | - | 1311 | 279.1 | - | - | 0 | - |
| - | - | 1501 | 280.1 | - | - | 0 | - |
| - | - | 896.5 | 281.1 | - | - | 0 | - |
| - | - | 1090 | 282.2 | - | - | 0 | - |
| - | - | 4.077E+04 | 283.1 | - | - | 0 | - |
| - | - | 2622 | 284.1 | - | - | 0 | - |
| - | - | 4247 | 284.1 | - | - | 0 | - |
| - | - | 834.2 | 285.2 | - | - | 0 | - |
| - | - | 4058 | 286.1 | - | - | 0 | - |
| - | - | 1235 | 287.2 | - | - | 0 | - |
| - | - | 908.9 | 292.1 | - | - | 0 | - |
| 12 | y | 4.562E+04 | 293.1 | 0.0003883 | 1.325 | +1 | 2 |
| - | - | 1529 | 293.2 | - | - | 0 | - |
| - | - | 8080 | 294.1 | - | - | 0 | - |
| - | - | 675.9 | 295.1 | - | - | 0 | - |
| 3 | b | 2163 | 296.2 | 0.0001533 | 0.5177 | +1 | 3 |
| - | - | 1.134E+04 | 297.2 | - | - | 0 | - |
| - | - | 1216 | 298.1 | - | - | 0 | - |
| - | - | 1549 | 298.2 | - | - | 0 | - |
| - | - | 3390 | 300.2 | - | - | 0 | - |
| - | - | 809.6 | 301.1 | - | - | 0 | - |
| - | - | 1.694E+04 | 301.2 | - | - | 0 | - |
| - | - | 1065 | 301.2 | - | - | 0 | - |
| - | - | 9618 | 302.1 | - | - | 0 | - |
| - | - | 699.9 | 302.1 | - | - | 0 | - |
| - | - | 2165 | 302.2 | - | - | 0 | - |
| - | - | 996.4 | 303.1 | - | - | 0 | - |
| - | - | 583.2 | 306 | - | - | 0 | - |
| - | - | 711.9 | 310.2 | - | - | 0 | - |
| 12 | y | 6933 | 311.1 | 0.0006574 | 2.113 | +1 | 2 |
| - | - | 970.9 | 312.1 | - | - | 0 | - |
| - | - | 619.3 | 314.1 | - | - | 0 | - |
| 3 | b | 8088 | 314.2 | 0.0006971 | 2.218 | +1 | 3 |
| - | - | 1.687E+04 | 315.2 | - | - | 0 | - |
| - | - | 1794 | 315.2 | - | - | 0 | - |
| - | - | 2185 | 316.2 | - | - | 0 | - |
| - | - | 1049 | 319.2 | - | - | 0 | - |
| - | - | 585.1 | 320.2 | - | - | 0 | - |
| - | - | 604.4 | 329.2 | - | - | 0 | - |
| - | - | 822 | 332.2 | - | - | 0 | - |
| - | - | 2417 | 333.2 | - | - | 0 | - |
| - | - | 1159 | 334.1 | - | - | 0 | - |
| - | - | 935.9 | 334.2 | - | - | 0 | - |
| - | - | 491.7 | 336.2 | - | - | 0 | - |
| - | - | 1082 | 340.2 | - | - | 0 | - |
| - | - | 1075 | 342.2 | - | - | 0 | - |
| - | - | 566.7 | 343.2 | - | - | 0 | - |
| - | - | 1097 | 344.2 | - | - | 0 | - |
| - | - | 2984 | 345.1 | - | - | 0 | - |
| - | - | 1.155E+04 | 346.1 | - | - | 0 | - |
| - | - | 710.9 | 346.1 | - | - | 0 | - |
| - | - | 1474 | 347.1 | - | - | 0 | - |
| - | - | 3251 | 350.2 | - | - | 0 | - |
| - | - | 959.7 | 352.2 | - | - | 0 | - |
| - | - | 923.8 | 361.2 | - | - | 0 | - |
| - | - | 1632 | 362.2 | - | - | 0 | - |
| - | - | 573.8 | 362.2 | - | - | 0 | - |
| - | - | 1.464E+04 | 363.1 | - | - | 0 | - |
| - | - | 1705 | 364.1 | - | - | 0 | - |
| - | - | 2378 | 364.1 | - | - | 0 | - |
| - | - | 623.8 | 365.1 | - | - | 0 | - |
| - | - | 1466 | 367.1 | - | - | 0 | - |
| - | - | 8735 | 378.2 | - | - | 0 | - |
| - | - | 1.558E+04 | 379.2 | - | - | 0 | - |
| - | - | 918.8 | 379.2 | - | - | 0 | - |
| - | - | 634.2 | 380.2 | - | - | 0 | - |
| - | - | 2580 | 380.2 | - | - | 0 | - |
| - | - | 3.413E+04 | 381.2 | - | - | 0 | - |
| - | - | 5098 | 382.2 | - | - | 0 | - |
| - | - | 951.5 | 383.2 | - | - | 0 | - |
| - | - | 2135 | 384.2 | - | - | 0 | - |
| - | - | 3151 | 385.2 | - | - | 0 | - |
| - | - | 923.2 | 386.2 | - | - | 0 | - |
| - | - | 2022 | 391.1 | - | - | 0 | - |
| - | - | 1601 | 394.2 | - | - | 0 | - |
| - | - | 2303 | 395.1 | - | - | 0 | - |
| - | - | 912.9 | 396.1 | - | - | 0 | - |
| - | - | 791.3 | 396.2 | - | - | 0 | - |
| - | - | 8224 | 396.2 | - | - | 0 | - |
| - | - | 1550 | 397.2 | - | - | 0 | - |
| - | - | 3301 | 398.2 | - | - | 0 | - |
| - | - | 1892 | 402.2 | - | - | 0 | - |
| - | - | 1078 | 402.2 | - | - | 0 | - |
| - | - | 747 | 403.2 | - | - | 0 | - |
| - | - | 1152 | 405.3 | - | - | 0 | - |
| - | - | 1784 | 406.2 | - | - | 0 | - |
| - | - | 947.3 | 407.2 | - | - | 0 | - |
| - | - | 2256 | 409.1 | - | - | 0 | - |
| - | - | 1858 | 412.2 | - | - | 0 | - |
| - | - | 7005 | 412.2 | - | - | 0 | - |
| - | - | 6497 | 413.1 | - | - | 0 | - |
| - | - | 1216 | 414.1 | - | - | 0 | - |
| - | - | 4752 | 414.2 | - | - | 0 | - |
| - | - | 5445 | 415.2 | - | - | 0 | - |
| - | - | 1031 | 415.2 | - | - | 0 | - |
| - | - | 618 | 418.2 | - | - | 0 | - |
| - | - | 681.4 | 421.2 | - | - | 0 | - |
| 11 | y | 723 | 421.2 | 0.0002143 | 0.5088 | +1 | 3 |
| - | - | 2.34E+04 | 423.3 | - | - | 0 | - |
| - | - | 4995 | 424.3 | - | - | 0 | - |
| - | - | 722.8 | 425.3 | - | - | 0 | - |
| - | - | 2765 | 426.7 | - | - | 0 | - |
| - | - | 1218 | 427.2 | - | - | 0 | - |
| - | - | 1168 | 429.2 | - | - | 0 | - |
| - | - | 5762 | 430.2 | - | - | 0 | - |
| - | - | 1.291E+04 | 430.2 | - | - | 0 | - |
| - | - | 1218 | 431.2 | - | - | 0 | - |
| - | - | 2198 | 431.2 | - | - | 0 | - |
| 4 | b | 2511 | 433.3 | 0.0003235 | 0.7467 | +1 | 4 |
| - | - | 680.9 | 434.3 | - | - | 0 | - |
| - | - | 789.4 | 438.2 | - | - | 0 | - |
| 11 | y | 1.456E+04 | 439.2 | 0.0007871 | 1.792 | +1 | 3 |
| - | - | 2716 | 440.2 | - | - | 0 | - |
| - | - | 1456 | 441.2 | - | - | 0 | - |
| - | - | 1302 | 448.2 | - | - | 0 | - |
| - | - | 848.6 | 449.2 | - | - | 0 | - |
| - | - | 956.9 | 450.2 | - | - | 0 | - |
| - | - | 855.2 | 450.3 | - | - | 0 | - |
| 4 | b | 3.88E+04 | 451.3 | 0.0006231 | 1.381 | +1 | 4 |
| - | - | 9672 | 452.3 | - | - | 0 | - |
| - | - | 1552 | 453.3 | - | - | 0 | - |
| - | - | 805.8 | 455.2 | - | - | 0 | - |
| - | - | 1211 | 459.2 | - | - | 0 | - |
| - | - | 882.7 | 461.2 | - | - | 0 | - |
| - | - | 658.1 | 466.2 | - | - | 0 | - |
| - | - | 1.005E+04 | 468.3 | - | - | 0 | - |
| - | - | 1782 | 469.3 | - | - | 0 | - |
| - | - | 767.9 | 472.2 | - | - | 0 | - |
| - | - | 1314 | 476.2 | - | - | 0 | - |
| - | - | 2129 | 477.2 | - | - | 0 | - |
| 10 | y | 3207 | 478.2 | 0.001231 | 2.575 | +1 | 4 |
| - | - | 678.6 | 478.3 | - | - | 0 | - |
| - | - | 794.9 | 479.2 | - | - | 0 | - |
| - | - | 967.1 | 480.2 | - | - | 0 | - |
| - | - | 974.5 | 482.2 | - | - | 0 | - |
| - | - | 652 | 483.3 | - | - | 0 | - |
| - | - | 733.7 | 484.2 | - | - | 0 | - |
| 8 | b | 743.1 | 488.3 | 0.00623 | 12.76 | +2 | 8 |
| - | - | 1.758E+04 | 494.2 | - | - | 0 | - |
| - | - | 3292 | 495.2 | - | - | 0 | - |
| 10 | y | 3.988E+04 | 496.2 | 0.0006857 | 1.382 | +1 | 4 |
| - | - | 1.15E+04 | 497.2 | - | - | 0 | - |
| - | - | 1168 | 498.2 | - | - | 0 | - |
| - | - | 2846 | 504.2 | - | - | 0 | - |
| - | - | 754.6 | 505.2 | - | - | 0 | - |
| - | - | 744.9 | 506.3 | - | - | 0 | - |
| - | - | 753.6 | 525.2 | - | - | 0 | - |
| - | - | 1899 | 525.3 | - | - | 0 | - |
| - | - | 1762 | 526.2 | - | - | 0 | - |
| - | - | 908.5 | 526.3 | - | - | 0 | - |
| - | - | 634.6 | 527.2 | - | - | 0 | - |
| - | - | 709.2 | 528.7 | - | - | 0 | - |
| - | - | 1001 | 529.2 | - | - | 0 | - |
| - | - | 1348 | 530.2 | - | - | 0 | - |
| - | - | 2484 | 532.2 | - | - | 0 | - |
| - | - | 1166 | 537.2 | - | - | 0 | - |
| - | - | 1102 | 537.7 | - | - | 0 | - |
| - | - | 6960 | 539.2 | - | - | 0 | - |
| - | - | 1642 | 540.2 | - | - | 0 | - |
| - | - | 685.8 | 541.8 | - | - | 0 | - |
| - | - | 4512 | 543.3 | - | - | 0 | - |
| - | - | 829.7 | 544.3 | - | - | 0 | - |
| - | - | 3334 | 546.2 | - | - | 0 | - |
| - | - | 1992 | 546.7 | - | - | 0 | - |
| - | - | 819.9 | 547.2 | - | - | 0 | - |
| - | - | 1707 | 549.2 | - | - | 0 | - |
| - | - | 1342 | 551.3 | - | - | 0 | - |
| - | - | 2003 | 551.3 | - | - | 0 | - |
| - | - | 767.3 | 552.3 | - | - | 0 | - |
| - | - | 1551 | 554.8 | - | - | 0 | - |
| - | - | 1.056E+04 | 555.3 | - | - | 0 | - |
| - | - | 5420 | 555.8 | - | - | 0 | - |
| - | - | 1342 | 556.3 | - | - | 0 | - |
| 5 | b | 2559 | 561.3 | 0.0006368 | 1.134 | +1 | 5 |
| 5 | b | 1423 | 562.3 | 0.01077 | 19.15 | +1 | 5 |
| - | - | 767.9 | 564.3 | - | - | 0 | - |
| - | - | 765.3 | 564.8 | - | - | 0 | - |
| - | - | 745.6 | 565.3 | - | - | 0 | - |
| - | - | 9223 | 567.2 | - | - | 0 | - |
| - | - | 2402 | 568.2 | - | - | 0 | - |
| 5 | b | 2.565E+04 | 579.3 | 0.0002731 | 0.4714 | +1 | 5 |
| - | - | 6880 | 580.3 | - | - | 0 | - |
| - | - | 1169 | 582.3 | - | - | 0 | - |
| 10 | b | 886.7 | 583.3 | 0.005919 | 10.15 | +2 | 10 |
| - | - | 898 | 587.2 | - | - | 0 | - |
| 9 | y | 3957 | 593.3 | 0.000146 | 0.2461 | +1 | 5 |
| - | - | 2140 | 593.3 | - | - | 0 | - |
| - | - | 782.5 | 594.3 | - | - | 0 | - |
| - | - | 644.9 | 596.2 | - | - | 0 | - |
| - | - | 648.6 | 597.3 | - | - | 0 | - |
| - | - | 2972 | 600.3 | - | - | 0 | - |
| - | - | 1160 | 601.2 | - | - | 0 | - |
| - | - | 2305 | 602.8 | - | - | 0 | - |
| - | - | 2915 | 603.3 | - | - | 0 | - |
| - | - | 2905 | 610.3 | - | - | 0 | - |
| 9 | y | 2.884E+04 | 611.3 | 0.0006897 | 1.128 | +1 | 5 |
| - | - | 8486 | 611.8 | - | - | 0 | - |
| - | - | 7537 | 612.3 | - | - | 0 | - |
| - | - | 1454 | 612.8 | - | - | 0 | - |
| - | - | 1555 | 613.3 | - | - | 0 | - |
| - | - | 1815 | 617.3 | - | - | 0 | - |
| - | - | 1344 | 621.3 | - | - | 0 | - |
| - | - | 887.3 | 625.3 | - | - | 0 | - |
| - | - | 898.7 | 628.3 | - | - | 0 | - |
| - | - | 952.6 | 629.8 | - | - | 0 | - |
| - | - | 724.5 | 630.3 | - | - | 0 | - |
| - | - | 2001 | 635.3 | - | - | 0 | - |
| - | - | 854 | 636.3 | - | - | 0 | - |
| 4 | y | 4019 | 636.8 | 0.0009774 | 1.535 | +2 | 10 |
| 4 | y | 1677 | 637.3 | 0.01001 | 15.7 | +2 | 10 |
| 11 | b | 6627 | 638.3 | 0.001623 | 2.542 | +2 | 11 |
| 11 | b | 4836 | 638.8 | 0.009981 | 15.62 | +2 | 11 |
| - | - | 1627 | 639.3 | - | - | 0 | - |
| - | - | 801 | 639.8 | - | - | 0 | - |
| - | - | 3127 | 640.3 | - | - | 0 | - |
| - | - | 799.8 | 640.7 | - | - | 0 | - |
| - | - | 1794 | 641.3 | - | - | 0 | - |
| - | - | 612.8 | 644.3 | - | - | 0 | - |
| - | - | 3378 | 645.3 | - | - | 0 | - |
| 4 | y | 2.515E+04 | 645.8 | 0.001127 | 1.745 | +2 | 10 |
| - | - | 1.712E+04 | 646.3 | - | - | 0 | - |
| - | - | 6917 | 646.8 | - | - | 0 | - |
| - | - | 928.6 | 647.2 | - | - | 0 | - |
| 11 | b | 1.049E+04 | 647.3 | 0.000674 | 1.041 | +2 | 11 |
| - | - | 6187 | 647.8 | - | - | 0 | - |
| - | - | 2494 | 648.3 | - | - | 0 | - |
| - | - | 7816 | 652.3 | - | - | 0 | - |
| - | - | 3115 | 653.3 | - | - | 0 | - |
| - | - | 1732 | 658.3 | - | - | 0 | - |
| - | - | 1450 | 659.3 | - | - | 0 | - |
| - | - | 1392 | 662.3 | - | - | 0 | - |
| - | - | 1495 | 663.3 | - | - | 0 | - |
| - | - | 688.5 | 664.3 | - | - | 0 | - |
| 6 | b | 4115 | 676.3 | 0.0003743 | 0.5534 | +1 | 6 |
| 6 | b | 1902 | 677.3 | 0.008607 | 12.71 | +1 | 6 |
| - | - | 757.7 | 678.3 | - | - | 0 | - |
| - | - | 1.586E+04 | 680.3 | - | - | 0 | - |
| - | - | 748 | 680.8 | - | - | 0 | - |
| - | - | 6953 | 681.3 | - | - | 0 | - |
| - | - | 959 | 684.8 | - | - | 0 | - |
| - | - | 614.9 | 689.3 | - | - | 0 | - |
| - | - | 869.7 | 690.3 | - | - | 0 | - |
| - | - | 718.3 | 692.8 | - | - | 0 | - |
| 3 | y | 3960 | 693.3 | 0.002769 | 3.993 | +2 | 11 |
| 3 | y | 2540 | 693.8 | 0.009723 | 14.01 | +2 | 11 |
| 6 | b | 4.838E+04 | 694.4 | 2.445E-06 | 0.003521 | +1 | 6 |
| - | - | 1594 | 694.5 | - | - | 0 | - |
| - | - | 761.4 | 694.8 | - | - | 0 | - |
| - | - | 1.741E+04 | 695.4 | - | - | 0 | - |
| - | - | 2776 | 696.4 | - | - | 0 | - |
| - | - | 5793 | 697.9 | - | - | 0 | - |
| - | - | 3864 | 698.4 | - | - | 0 | - |
| - | - | 1505 | 698.9 | - | - | 0 | - |
| - | - | 3006 | 701.8 | - | - | 0 | - |
| 3 | y | 2.789E+04 | 702.3 | 0.001759 | 2.504 | +2 | 11 |
| 12 | b | 3.234E+04 | 702.8 | 0.008504 | 12.1 | +2 | 12 |
| 12 | b | 1.75E+04 | 703.3 | 0.003089 | 4.392 | +2 | 12 |
| - | - | 6072 | 703.8 | - | - | 0 | - |
| 8 | y | 2551 | 706.3 | 0.001653 | 2.341 | +1 | 6 |
| - | - | 1056 | 707.8 | - | - | 0 | - |
| - | - | 8262 | 711.4 | - | - | 0 | - |
| 12 | b | 6.495E+04 | 711.9 | 0.001106 | 1.554 | +2 | 12 |
| - | - | 5.591E+04 | 712.4 | - | - | 0 | - |
| - | - | 2.038E+04 | 712.9 | - | - | 0 | - |
| - | - | 2909 | 713.4 | - | - | 0 | - |
| - | - | 839.1 | 715.3 | - | - | 0 | - |
| - | - | 4587 | 720.9 | - | - | 0 | - |
| - | - | 3739 | 721.4 | - | - | 0 | - |
| - | - | 1412 | 721.9 | - | - | 0 | - |
| - | - | 2168 | 723.4 | - | - | 0 | - |
| 8 | y | 1.17E+04 | 724.4 | 0.000366 | 0.5052 | +1 | 6 |
| - | - | 4639 | 725.4 | - | - | 0 | - |
| - | - | 662.5 | 726.4 | - | - | 0 | - |
| - | - | 5167 | 729.4 | - | - | 0 | - |
| - | - | 2570 | 730.4 | - | - | 0 | - |
| - | - | 962.1 | 734.4 | - | - | 0 | - |
| - | - | 1961 | 751.4 | - | - | 0 | - |
| 2 | y | 1492 | 751.9 | 0.00283 | 3.764 | +2 | 12 |
| - | - | 2797 | 752.4 | - | - | 0 | - |
| - | - | 907.4 | 753.4 | - | - | 0 | - |
| - | - | 1270 | 760.3 | - | - | 0 | - |
| - | - | 680 | 762.9 | - | - | 0 | - |
| - | - | 698.4 | 763.4 | - | - | 0 | - |
| - | - | 1735 | 765.4 | - | - | 0 | - |
| - | - | 701.7 | 766.4 | - | - | 0 | - |
| - | - | 921.2 | 775.9 | - | - | 0 | - |
| - | - | 747.6 | 777.3 | - | - | 0 | - |
| - | - | 961.3 | 778.3 | - | - | 0 | - |
| - | - | 3536 | 779.4 | - | - | 0 | - |
| - | - | 2175 | 779.9 | - | - | 0 | - |
| - | - | 5058 | 780.4 | - | - | 0 | - |
| - | - | 3916 | 780.9 | - | - | 0 | - |
| - | - | 2293 | 781.4 | - | - | 0 | - |
| - | - | 818.9 | 784.4 | - | - | 0 | - |
| - | - | 2034 | 784.9 | - | - | 0 | - |
| - | - | 920.6 | 785.4 | - | - | 0 | - |
| - | - | 2747 | 792.9 | - | - | 0 | - |
| 0 | Precursor | 3.068E+04 | 793.4 | 0.004232 | 5.334 | +2 | -1 |
| 0 | Precursor | 2.696E+04 | 793.9 | 0.007951 | 10.02 | +2 | -1 |
| - | - | 1.612E+04 | 794.4 | - | - | 0 | - |
| - | - | 4089 | 794.9 | - | - | 0 | - |
| - | - | 1.411E+04 | 795.3 | - | - | 0 | - |
| - | - | 5372 | 796.3 | - | - | 0 | - |
| - | - | 1025 | 797.3 | - | - | 0 | - |
| - | - | 853.1 | 801.4 | - | - | 0 | - |
| - | - | 5.341E+04 | 801.9 | - | - | 0 | - |
| 0 | Precursor | 4.692E+05 | 802.4 | 0.001574 | 1.961 | +2 | -1 |
| - | - | 4.257E+05 | 802.9 | - | - | 0 | - |
| - | - | 2.034E+05 | 803.4 | - | - | 0 | - |
| - | - | 2.458E+04 | 803.9 | - | - | 0 | - |
| - | - | 1335 | 807.4 | - | - | 0 | - |
| - | - | 1191 | 808.4 | - | - | 0 | - |
| - | - | 2785 | 825.4 | - | - | 0 | - |
| - | - | 3903 | 826.4 | - | - | 0 | - |
| - | - | 1890 | 827.4 | - | - | 0 | - |
| - | - | 787.6 | 834.3 | - | - | 0 | - |
| - | - | 1516 | 834.4 | - | - | 0 | - |
| - | - | 1805 | 835.4 | - | - | 0 | - |
| - | - | 2177 | 836.4 | - | - | 0 | - |
| - | - | 1015 | 837.5 | - | - | 0 | - |
| - | - | 2303 | 843.4 | - | - | 0 | - |
| - | - | 6728 | 844.4 | - | - | 0 | - |
| - | - | 3725 | 845.4 | - | - | 0 | - |
| - | - | 934.3 | 851.4 | - | - | 0 | - |
| - | - | 3421 | 852.4 | - | - | 0 | - |
| - | - | 1.114E+04 | 852.4 | - | - | 0 | - |
| - | - | 1926 | 853.4 | - | - | 0 | - |
| - | - | 4912 | 853.4 | - | - | 0 | - |
| - | - | 1111 | 854.4 | - | - | 0 | - |
| 7 | b | 7252 | 862.4 | 4.08E-05 | 0.04731 | +1 | 7 |
| 7 | b | 5741 | 863.4 | 0.012 | 13.89 | +1 | 7 |
| - | - | 3084 | 864.5 | - | - | 0 | - |
| - | - | 1135 | 865.5 | - | - | 0 | - |
| - | - | 2022 | 869.4 | - | - | 0 | - |
| - | - | 1441 | 876.4 | - | - | 0 | - |
| 7 | b | 3.346E+04 | 880.4 | 9.623E-05 | 0.1093 | +1 | 7 |
| - | - | 1.491E+04 | 881.4 | - | - | 0 | - |
| - | - | 2807 | 882.4 | - | - | 0 | - |
| - | - | 1251 | 890.4 | - | - | 0 | - |
| - | - | 772.9 | 891.4 | - | - | 0 | - |
| 7 | y | 1957 | 892.4 | 0.002174 | 2.436 | +1 | 7 |
| 7 | y | 1164 | 893.4 | 0.01389 | 15.54 | +1 | 7 |
| - | - | 8907 | 908.4 | - | - | 0 | - |
| - | - | 7076 | 909.4 | - | - | 0 | - |
| 7 | y | 2.029E+04 | 910.4 | 0.001436 | 1.578 | +1 | 7 |
| - | - | 9285 | 911.4 | - | - | 0 | - |
| - | - | 2169 | 912.4 | - | - | 0 | - |
| - | - | 838.2 | 918.4 | - | - | 0 | - |
| - | - | 2743 | 936.4 | - | - | 0 | - |
| - | - | 2630 | 937.4 | - | - | 0 | - |
| - | - | 1613 | 938.4 | - | - | 0 | - |
| - | - | 736.5 | 947.7 | - | - | 0 | - |
| - | - | 829.3 | 948.2 | - | - | 0 | - |
| - | - | 1458 | 949.5 | - | - | 0 | - |
| - | - | 2019 | 953.3 | - | - | 0 | - |
| - | - | 957 | 954.3 | - | - | 0 | - |
| - | - | 4133 | 954.4 | - | - | 0 | - |
| - | - | 7653 | 955.4 | - | - | 0 | - |
| - | - | 2915 | 956.4 | - | - | 0 | - |
| - | - | 1150 | 962.5 | - | - | 0 | - |
| - | - | 1400 | 963.4 | - | - | 0 | - |
| - | - | 2345 | 965.4 | - | - | 0 | - |
| - | - | 3783 | 965.5 | - | - | 0 | - |
| - | - | 1221 | 966.4 | - | - | 0 | - |
| - | - | 1883 | 966.5 | - | - | 0 | - |
| - | - | 5114 | 972.4 | - | - | 0 | - |
| - | - | 2076 | 973.4 | - | - | 0 | - |
| 8 | b | 4516 | 975.5 | 0.001992 | 2.042 | +1 | 8 |
| 8 | b | 5218 | 976.5 | 0.01241 | 12.7 | +1 | 8 |
| - | - | 1563 | 977.5 | - | - | 0 | - |
| - | - | 1588 | 979.5 | - | - | 0 | - |
| - | - | 4721 | 980.5 | - | - | 0 | - |
| - | - | 2960 | 981.5 | - | - | 0 | - |
| - | - | 1918 | 982.5 | - | - | 0 | - |
| 8 | b | 3.953E+04 | 993.5 | 0.0001997 | 0.201 | +1 | 8 |
| - | - | 2.188E+04 | 994.5 | - | - | 0 | - |
| - | - | 6191 | 995.5 | - | - | 0 | - |
| 6 | y | 2060 | 1007 | 0.009533 | 9.463 | +1 | 8 |
| - | - | 1395 | 1008 | - | - | 0 | - |
| - | - | 738.2 | 1009 | - | - | 0 | - |
| - | - | 882 | 1012 | - | - | 0 | - |
| - | - | 1277 | 1024 | - | - | 0 | - |
| 6 | y | 1.176E+04 | 1025 | 0.0006774 | 0.6606 | +1 | 8 |
| - | - | 6494 | 1026 | - | - | 0 | - |
| - | - | 1636 | 1027 | - | - | 0 | - |
| - | - | 824.6 | 1063 | - | - | 0 | - |
| - | - | 881.4 | 1065 | - | - | 0 | - |
| - | - | 821.1 | 1073 | - | - | 0 | - |
| - | - | 1076 | 1074 | - | - | 0 | - |
| - | - | 1611 | 1076 | - | - | 0 | - |
| - | - | 1212 | 1077 | - | - | 0 | - |
| 9 | b | 2231 | 1091 | 0.0009723 | 0.8915 | +1 | 9 |
| 9 | b | 6291 | 1092 | 0.01246 | 11.42 | +1 | 9 |
| - | - | 3910 | 1093 | - | - | 0 | - |
| - | - | 5191 | 1094 | - | - | 0 | - |
| - | - | 3154 | 1095 | - | - | 0 | - |
| - | - | 2925 | 1108 | - | - | 0 | - |
| 9 | b | 4.773E+04 | 1109 | 0.0005394 | 0.4866 | +1 | 9 |
| - | - | 2.836E+04 | 1110 | - | - | 0 | - |
| - | - | 9926 | 1111 | - | - | 0 | - |
| - | - | 781.4 | 1112 | - | - | 0 | - |
| - | - | 1023 | 1117 | - | - | 0 | - |
| - | - | 1639 | 1118 | - | - | 0 | - |
| - | - | 969.9 | 1119 | - | - | 0 | - |
| 5 | y | 6426 | 1136 | 0.0002163 | 0.1905 | +1 | 9 |
| 5 | y | 9512 | 1136 | 0.007711 | 6.785 | +1 | 9 |
| - | - | 5316 | 1137 | - | - | 0 | - |
| 10 | b | 3774 | 1148 | 1.635E-05 | 0.01425 | +1 | 10 |
| 10 | b | 3038 | 1149 | 0.01124 | 9.786 | +1 | 10 |
| - | - | 1627 | 1150 | - | - | 0 | - |
| - | - | 1622 | 1153 | - | - | 0 | - |
| 5 | y | 1.524E+04 | 1154 | 0.0002053 | 0.178 | +1 | 9 |
| - | - | 9122 | 1155 | - | - | 0 | - |
| - | - | 2601 | 1156 | - | - | 0 | - |
| - | - | 2773 | 1165 | - | - | 0 | - |
| 10 | b | 2.434E+04 | 1166 | 0.000438 | 0.3758 | +1 | 10 |
| - | - | 1.583E+04 | 1167 | - | - | 0 | - |
| - | - | 5240 | 1168 | - | - | 0 | - |
| - | - | 5565 | 1183 | - | - | 0 | - |
| - | - | 4196 | 1184 | - | - | 0 | - |
| - | - | 1390 | 1185 | - | - | 0 | - |
| - | - | 736.5 | 1198 | - | - | 0 | - |
| - | - | 1717 | 1205 | - | - | 0 | - |
| - | - | 1457 | 1206 | - | - | 0 | - |
| - | - | 2333 | 1223 | - | - | 0 | - |
| - | - | 2280 | 1224 | - | - | 0 | - |
| - | - | 819.1 | 1257 | - | - | 0 | - |
| 4 | y | 2030 | 1273 | 0.001907 | 1.499 | +1 | 10 |
| 4 | y | 2108 | 1274 | 0.01875 | 14.72 | +1 | 10 |
| - | - | 875.5 | 1275 | - | - | 0 | - |
| 11 | b | 1666 | 1276 | 0.005225 | 4.096 | +1 | 11 |
| 11 | b | 1195 | 1277 | 0.02455 | 19.23 | +1 | 11 |
| - | - | 1058 | 1278 | - | - | 0 | - |
| - | - | 1842 | 1290 | - | - | 0 | - |
| 4 | y | 2.048E+04 | 1291 | 0.001211 | 0.9387 | +1 | 10 |
| - | - | 1.219E+04 | 1292 | - | - | 0 | - |
| - | - | 3600 | 1293 | - | - | 0 | - |
| 11 | b | 1.118E+04 | 1294 | 0.002118 | 1.637 | +1 | 11 |
| - | - | 7363 | 1295 | - | - | 0 | - |
| - | - | 4056 | 1296 | - | - | 0 | - |
| 3 | y | 767.8 | 1386 | 0.01037 | 7.486 | +1 | 11 |
| 3 | y | 735.2 | 1387 | 0.02623 | 18.92 | +1 | 11 |
| - | - | 677.8 | 1388 | - | - | 0 | - |
| - | - | 772.9 | 1403 | - | - | 0 | - |
| 3 | y | 1.067E+04 | 1404 | 0.0005586 | 0.398 | +1 | 11 |
| - | - | 8277 | 1405 | - | - | 0 | - |
| 12 | b | 2864 | 1406 | 0.01596 | 11.36 | +1 | 12 |
| - | - | 803.1 | 1458 | - | - | 0 | - |

m/z Charge Intensity FragmentType MassShift Position
120.08106231689453 0 5602.668
121.08441162109375 0 621.8575
122.07154083251953 0 2065.4443
123.04436492919922 0 458.56042
123.0556640625 0 639.31744
126.05522918701172 0 806.9647
127.05046081542969 0 2059.1128
127.08672332763672 0 763.49524
128.0825653076172 0 595.11005
128.10731506347656 0 5442.9585
129.10256958007812 0 179687.16
130.06541442871094 0 10068.444
130.09976196289062 0 1477.0554
130.10586547851562 0 10350.976
131.06887817382812 0 1056.0237
132.0811309814453 0 2702.3313
136.07598876953125 0 15460.579
137.0792999267578 0 1444.973
138.05545043945312 0 791.55707
138.06651306152344 0 3426.5168
138.07167053222656 0 568.8369
139.08688354492188 0 1988.1326
141.1024932861328 0 3872.8474
142.1055145263672 0 479.26233
143.04554748535156 0 1380.7152
145.06124877929688 0 1463.4121
146.06031799316406 0 464.5424
147.04432678222656 0 572.77856
150.06651306152344 0 2751.812
151.08689880371094 0 2516.777
153.06622314453125 0 610.4203
153.1023406982422 0 503.7863
154.0980682373047 0 504.73715
155.08200073242188 0 957.19525
155.093017578125 0 16970.848
155.11813354492188 0 2365.635 a Water loss 1
156.07081604003906 0 655.7586
156.07730102539062 0 1416.4843
157.09735107421875 0 1394.7327
159.09197998046875 0 26290.785
160.0886993408203 0 563.4408
160.0954132080078 0 2540.9714
165.05490112304688 0 6197.4585
166.06138610839844 0 11595.822
166.0864715576172 0 2411.6497
166.0977783203125 0 3654.725
167.06471252441406 0 641.3982
167.08172607421875 0 1401.3013
167.11819458007812 0 762.733
168.10240173339844 0 1744.8267
168.11355590820312 0 1535.7806
169.07676696777344 0 640.37964
169.09747314453125 0 6502.4106
169.11756896972656 0 540.20166
170.060302734375 0 1492.9816
170.1005859375 0 472.18158
171.06369018554688 0 532.7637
171.09182739257812 0 886.6941
171.11187744140625 0 490.40878
172.07179260253906 0 1389.9043
173.0561065673828 0 1536.418
173.1287841796875 0 104499.414 a 1
173.45152282714844 0 3788.3481
173.54879760742188 0 454.13278
174.12503051757812 0 812.56647
174.1321563720703 0 8858.487
176.08201599121094 0 1120.0042
177.10260009765625 0 1136.3453
178.13417053222656 0 4911.8223
179.11851501464844 0 976.14075
179.13815307617188 0 544.07526
181.06126403808594 0 1174.5477
182.08148193359375 0 18221.562 y 12
183.0855255126953 0 1700.9618
183.1129150390625 0 1381.3766 b Water loss 1
183.1501007080078 0 848.01117
185.09219360351562 0 640.4931
185.12864685058594 0 527.70325
185.16542053222656 0 888.06934
186.1240234375 0 70266.26
187.08692932128906 0 3872.0178
187.09495544433594 0 475.2641
187.107421875 0 697.336
187.1273956298828 0 5068.894
188.0708770751953 0 1370.1813
190.08241271972656 0 1106.8362
190.13467407226562 0 1384.8868
191.1179962158203 0 982.18463
193.09747314453125 0 689.3158
194.0926971435547 0 3952.4033
194.12823486328125 0 565.59344
195.08718872070312 0 469.32385
195.11309814453125 0 12430.739
196.108154296875 0 4269.6816
196.11703491210938 0 1018.71844
197.1117706298828 0 656.66504
197.12867736816406 0 1186.5889
198.0873565673828 0 1904.6077
198.1276397705078 0 1015.4886
199.07156372070312 0 754.64545
199.1076202392578 0 863.5433
200.143310546875 0 583.69666
201.12362670898438 0 65888.51 b 1
202.08633422851562 0 986.0412
202.126953125 0 5167.865
203.1301727294922 0 763.12463
205.09750366210938 0 4245.0483
205.14476013183594 0 729.8029
206.12881469726562 0 835.21545
207.11228942871094 0 527.5935
207.16079711914062 0 1436.6088
209.0929412841797 0 541.5739
211.10865783691406 0 704.8186 y Water loss 10
212.1395721435547 0 1191.2173
213.12367248535156 0 5340.021
213.1600341796875 0 827.656
215.1387939453125 0 551.3774
216.0982666015625 0 2296.683
217.1337890625 0 767.7154 b Water loss 3
219.11376953125 0 602.9808
221.07106018066406 0 848.35944
221.1033935546875 0 2782.8872
221.12872314453125 0 5176.1436
221.14022827148438 0 2430.0928
222.12364196777344 0 1062.136
223.10780334472656 0 2433.5767
223.1556396484375 0 21389.533
224.10304260253906 0 1061.9569
224.15916442871094 0 3101.2524
226.08262634277344 0 5516.8633
227.06651306152344 0 2965.415
227.08653259277344 0 997.42413
227.10301208496094 0 1688.1132
227.15463256835938 0 1386.2615
228.13458251953125 0 661.23584
229.11880493164062 0 2034.2521
229.1553497314453 0 625.8834
231.08872985839844 0 644.20447
233.13986206054688 0 2674.1194
233.16400146484375 0 571.7997
234.12405395507812 0 15446.482
234.14405822753906 0 781.61176
235.12765502929688 0 1712.5568
238.11880493164062 0 2502.4644
238.13079833984375 0 850.5514
239.11434936523438 0 844.8038
240.1346893310547 0 8508.003
241.11956787109375 0 1207.7803
241.1378173828125 0 1294.7507
243.10948181152344 0 850.135
244.0930938720703 0 13315.684
244.10635375976562 0 1062.542
245.09707641601562 0 1503.4818
246.10044860839844 0 517.44415
247.108154296875 0 19200.736
247.14344787597656 0 574.8039
247.15591430664062 0 924.4262
248.11367797851562 0 8178.1484
248.15786743164062 0 531.1899
249.09852600097656 0 11654.862
249.1211395263672 0 1435.4855
249.13504028320312 0 5546.964
250.10183715820312 0 1176.7826
251.15065002441406 0 50226.47
252.11019897460938 0 820.37274
252.13478088378906 0 950.5114
252.15394592285156 0 6116.7935
252.16920471191406 0 1140.6787
255.1481170654297 0 3006.8428
256.1073913574219 0 834.30536
257.1136474609375 0 2287.1833
258.14508056640625 0 3111.1624
261.1352233886719 0 1120.5682
265.1169738769531 0 682.3943
265.1296691894531 0 5659.3223
265.14166259765625 0 687.30365
266.12506103515625 0 32454.713
267.1109619140625 0 496.87897
267.1280517578125 0 2325.851
268.1295166015625 0 1561.214
268.1768798828125 0 2163.1113
268.53363037109375 0 631.7423
269.1617736816406 0 741.641
269.18603515625 0 1715.508
270.12237548828125 0 1214.3757
272.1759033203125 0 4194.902
274.11859130859375 0 5916.9023
274.1325988769531 0 1086.0784
275.10272216796875 0 1480.9218
275.1215515136719 0 866.89404
275.1509704589844 0 1168.2708
276.1085510253906 0 1775.9661
276.1564025878906 0 662.5893
279.14544677734375 0 1310.7853
280.1292724609375 0 1501.1351
281.1255187988281 0 896.5405
282.1578674316406 0 1090.3436
283.1404724121094 0 40772.465
284.103515625 0 2622.0017
284.14306640625 0 4246.8506
285.1561279296875 0 834.1781
286.1399230957031 0 4058.4292
287.17169189453125 0 1235.4095
292.1288146972656 0 908.8731
293.11358642578125 0 45620.234 y Water loss 11
293.1619567871094 0 1528.8184
294.11737060546875 0 8080.2764
295.11962890625 0 675.8977
296.197021484375 0 2163.4714 b Water loss 2
297.1560363769531 0 11341.772
298.140380859375 0 1216.2253
298.1599426269531 0 1549.3204
300.1691589355469 0 3390.4333
301.1149597167969 0 809.62256
301.1510314941406 0 16944.105
301.1707458496094 0 1065.1008
302.114013671875 0 9618.405
302.1297912597656 0 699.88135
302.1554870605469 0 2165.1208
303.1156921386719 0 996.4013
305.9922180175781 0 583.1906
310.15478515625 0 711.863
311.1244201660156 0 6932.6494 y 11
312.1268615722656 0 970.9224
314.1144714355469 0 619.27075
314.2081298828125 0 8088.0005 b 2
315.1667175292969 0 16871.684
315.2114562988281 0 1793.817
316.170166015625 0 2184.5051
319.1515808105469 0 1048.9484
320.17108154296875 0 585.0522
329.1814270019531 0 604.44446
332.2097473144531 0 822.0053
333.17657470703125 0 2417.4832
334.13897705078125 0 1159.3038
334.18670654296875 0 935.90485
336.1709899902344 0 491.66113
340.1869812011719 0 1082.3145
342.15753173828125 0 1074.8137
343.160400390625 0 566.7309
344.1719970703125 0 1096.9359
345.1305847167969 0 2983.8142
346.1151428222656 0 11552.3955
346.1376037597656 0 710.8579
347.1177978515625 0 1474.0891
350.21905517578125 0 3250.8518
352.1656188964844 0 959.67896
361.1983337402344 0 923.78674
362.1819763183594 0 1632.4235
362.21826171875 0 573.75793
363.1415710449219 0 14642.687
364.1249694824219 0 1705.3546
364.14544677734375 0 2378.4092
365.1295166015625 0 623.8396
367.1400146484375 0 1465.7362
378.214111328125 0 8735.339
379.2094421386719 0 15579.925
379.243896484375 0 918.8066
380.1618347167969 0 634.1633
380.2117004394531 0 2579.8281
381.152099609375 0 34127.242
382.15478515625 0 5098.2617
383.1591796875 0 951.52625
384.1671447753906 0 2134.8904
385.1510314941406 0 3150.7756
386.1529541015625 0 923.23114
391.1361083984375 0 2022.4358
394.1733703613281 0 1601.3901
395.13519287109375 0 2303.3872
396.1404113769531 0 912.9042
396.1921691894531 0 791.25006
396.2248229980469 0 8223.585
397.2285461425781 0 1550.4437
398.1746520996094 0 3301.4727
402.1765441894531 0 1891.5944
402.20147705078125 0 1078.3617
403.19732666015625 0 747.0496
405.2625427246094 0 1151.7595
406.2459716796875 0 1784.3142
407.2432861328125 0 947.2853
409.1480712890625 0 2255.559
412.1607360839844 0 1858.2793
412.18341064453125 0 7005.3013
413.1458740234375 0 6496.9316
414.14971923828125 0 1215.6163
414.2352294921875 0 4752.426
415.1986083984375 0 5445.406
415.2383117675781 0 1031.4218
418.2091064453125 0 618.0166
421.17547607421875 0 681.40906
421.20794677734375 0 722.95013 y Water loss 10
423.2720642089844 0 23400.62
424.2745056152344 0 4994.842
425.27880859375 0 722.7986
426.7220764160156 0 2764.6514
427.2252502441406 0 1218.3629
429.2096862792969 0 1168.0741
430.17010498046875 0 5762.035
430.194580078125 0 12906.004
431.1709289550781 0 1217.9575
431.1976623535156 0 2198.1958
433.256103515625 0 2510.5703 b Water loss 3
434.2613525390625 0 680.88824
438.1754455566406 0 789.4454
439.2195129394531 0 14564.132 y 10
440.22265625 0 2715.9934
441.2215270996094 0 1455.631
448.2027587890625 0 1301.5348
449.2089538574219 0 848.58234
450.2358703613281 0 956.9478
450.2849426269531 0 855.20044
451.2669677734375 0 38797.492 b 3
452.2698974609375 0 9672.17
453.27276611328125 0 1551.6064
455.1933288574219 0 805.7927
459.1993713378906 0 1211.4155
461.2026062011719 0 882.7399
466.2381896972656 0 658.13776
468.2935485839844 0 10053.88
469.2941589355469 0 1782.4753
472.2215576171875 0 767.8742
476.2248229980469 0 1314.0654
477.2093505859375 0 2129.481
478.2283935546875 0 3206.831 y Water loss 9
478.2766418457031 0 678.5557
479.2325439453125 0 794.884
480.2254943847656 0 967.1029
482.2309265136719 0 974.4631
483.2658386230469 0 652.0437
484.2208251953125 0 733.6908
488.26220703125 0 743.1116 b Water loss 7
494.236328125 0 17581.576
495.2391052246094 0 3292.4883
496.2408752441406 0 39879.65 y 9
497.2447204589844 0 11503.036
498.2396545410156 0 1168.0665
504.2001037597656 0 2846.4365
505.2048645019531 0 754.55316
506.27215576171875 0 744.8865
525.2408447265625 0 753.5906
525.2667846679688 0 1899.0474
526.2308349609375 0 1761.7646
526.2720336914062 0 908.4795
527.2338256835938 0 634.5513
528.7299194335938 0 709.2465
529.23193359375 0 1000.9703
530.2242431640625 0 1348.2693
532.1942749023438 0 2483.5264
537.2444458007812 0 1165.8164
537.7372436523438 0 1101.5138
539.2365112304688 0 6960.492
540.2381591796875 0 1641.8214
541.7584228515625 0 685.8471
543.2769775390625 0 4511.851
544.2778930664062 0 829.7083
546.249267578125 0 3334.1943
546.7493286132812 0 1992.3175
547.2496337890625 0 819.86664
549.2207641601562 0 1707.4695
551.2581176757812 0 1341.6726
551.3300170898438 0 2003.2719
552.2528076171875 0 767.26404
554.7620239257812 0 1551.2384
555.2554931640625 0 10561.701
555.7556762695312 0 5420.4873
556.2567749023438 0 1342.4296
561.313720703125 0 2558.744 b Water loss 4
562.3091430664062 0 1422.5204 b Ammonia loss 4
564.2647094726562 0 767.909
564.7628784179688 0 765.29535
565.270263671875 0 745.5948
567.231689453125 0 9222.598
568.2348022460938 0 2402.306
579.3251953125 0 25652.768 b 4
580.3282470703125 0 6879.9043
582.3006591796875 0 1168.9635
583.2913818359375 0 886.67566 b 9
587.2478637695312 0 898.0441
593.2567138671875 0 3957.2197 y Water loss 8
593.3037109375 0 2140.4773
594.2650146484375 0 782.5064
596.2344970703125 0 644.8996
597.3002319335938 0 648.6137
600.31494140625 0 2972.3438
601.2319946289062 0 1159.8887
602.7904052734375 0 2304.545
603.29052734375 0 2914.565
610.284423828125 0 2904.893
611.267822265625 0 28837.336 y 8
611.7979125976562 0 8485.79
612.2727661132812 0 7536.9565
612.8026733398438 0 1453.9077
613.2735595703125 0 1554.8809
617.2846069335938 0 1815.0044
621.304443359375 0 1344.4758
625.2583618164062 0 887.3356
628.2777709960938 0 898.7123
629.8204345703125 0 952.6165
630.3294677734375 0 724.45483
635.295166015625 0 2000.5935
636.299560546875 0 854.00305
636.7868041992188 0 4018.6455 y Water loss 3
637.287841796875 0 1676.7048 y Ammonia loss 3
638.3292846679688 0 6627.1396 b Water loss 10
638.8296508789062 0 4835.616 b Ammonia loss 10
639.3280639648438 0 1627.3221
639.8321533203125 0 801.0456
640.2723999023438 0 3126.9617
640.720947265625 0 799.79834
641.2705688476562 0 1793.8635
644.3070678710938 0 612.84814
645.2962036132812 0 3378.2166
645.792236328125 0 25152.158 y 3
646.2931518554688 0 17122.229
646.7940673828125 0 6917.3325
647.2420043945312 0 928.6304
647.3336181640625 0 10485.941 b 10
647.8353881835938 0 6186.634
648.338134765625 0 2494.1953
652.3209838867188 0 7816.383
653.3228149414062 0 3115.0833
658.2830200195312 0 1732.3868
659.2882080078125 0 1449.7716
662.3049926757812 0 1392.2
663.2932739257812 0 1495.3346
664.3433837890625 0 688.465
676.3416748046875 0 4114.631 b Water loss 5
677.3339233398438 0 1902.442 b Ammonia loss 5
678.337646484375 0 757.6536
680.3158569335938 0 15861.77
680.8472900390625 0 748.0449
681.3186645507812 0 6952.9883
684.8179931640625 0 959.02606
689.3430786132812 0 614.85077
690.3403930664062 0 869.66235
692.8439331054688 0 718.3034
693.3306274414062 0 3959.8691 y Water loss 2
693.82958984375 0 2540.265 y Ammonia loss 2
694.3518676757812 0 48375.305 b 5
694.5438842773438 0 1594.0737
694.8324584960938 0 761.39575
695.3544921875 0 17410.602
696.3573608398438 0 2776.4287
697.85791015625 0 5792.7915
698.3553466796875 0 3863.9585
698.8607788085938 0 1504.9823
701.8414306640625 0 3006.3071
702.3348999023438 0 27888.947 y 2
702.8404541015625 0 32337.957 b Water loss 11
703.3440551757812 0 17502.59 b Ammonia loss 11
703.846923828125 0 6071.5737
706.34228515625 0 2550.7544 y Water loss 7
707.8291625976562 0 1056.1567
711.359130859375 0 8262.009
711.8553466796875 0 64951.992 b 11
712.3564453125 0 55913.805
712.8575439453125 0 20379.586
713.3560791015625 0 2908.5637
715.3021850585938 0 839.0629
720.860595703125 0 4586.5967
721.3614501953125 0 3739.398
721.86376953125 0 1412.2003
723.3676147460938 0 2167.5884
724.3515625 0 11697.433 y 7
725.3546142578125 0 4639.44
726.3534545898438 0 662.45215
729.35791015625 0 5166.9067
730.361572265625 0 2569.9697
734.3516235351562 0 962.06836
751.3792724609375 0 1960.533
751.8701782226562 0 1491.5226 y 1
752.3731079101562 0 2797.3718
753.3739013671875 0 907.3639
760.3054809570312 0 1269.8776
762.8651733398438 0 680.0227
763.3546752929688 0 698.40857
765.402587890625 0 1734.9695
766.4072875976562 0 701.68396
775.87255859375 0 921.1521
777.3250732421875 0 747.5586
778.3193359375 0 961.31683
779.3869018554688 0 3535.6333
779.8885498046875 0 2174.5972
780.3840942382812 0 5058.075
780.8797607421875 0 3915.884
781.3775634765625 0 2292.8955
784.386474609375 0 818.8893
784.8767700195312 0 2033.6133
785.3820190429688 0 920.64557
792.8933715820312 0 2746.8037
793.39013671875 0 30675.41 Precursor Water loss
793.8858642578125 0 26955.533 Precursor Ammonia loss
794.3885498046875 0 16117.025
794.8888549804688 0 4089.2068
795.3430786132812 0 14108.341
796.34619140625 0 5371.737
797.34765625 0 1024.5482
801.393798828125 0 853.0532
801.899658203125 0 53406.594
802.3927612304688 0 469244.1 Precursor
802.8941040039062 0 425650.75
803.3955078125 0 203354.97
803.897216796875 0 24580.027
807.3765258789062 0 1335.0897
808.3696899414062 0 1190.5258
825.3878173828125 0 2785.056
826.3764038085938 0 3902.5972
827.3772583007812 0 1889.5793
834.3464965820312 0 787.58765
834.4266967773438 0 1515.7617
835.41455078125 0 1805.2437
836.438720703125 0 2177.2356
837.45166015625 0 1014.98254
843.40087890625 0 2303.0708
844.3881225585938 0 6727.823
845.3886108398438 0 3724.9856
851.378662109375 0 934.25757
852.360595703125 0 3420.7568
852.4370727539062 0 11142.084
853.3656616210938 0 1926.2869
853.4386596679688 0 4912.3496
854.4472045898438 0 1111.4718
862.420654296875 0 7252.328 b Water loss 6
863.4166259765625 0 5740.689 b Ammonia loss 6
864.4540405273438 0 3083.8462
865.4556274414062 0 1135.0173
869.38916015625 0 2021.7938
876.439697265625 0 1441.4545
880.4312744140625 0 33458.195 b 6
881.4343872070312 0 14913.173
882.435546875 0 2806.8167
890.4196166992188 0 1251.0905
891.417724609375 0 772.8653
892.422119140625 0 1957.007 y Water loss 6
893.4178466796875 0 1164.3677 y Ammonia loss 6
908.4275512695312 0 8907.332
909.4346923828125 0 7076.0396
910.4319458007812 0 20294.408 y 6
911.4342041015625 0 9285.15
912.4375610351562 0 2168.8418
918.4165649414062 0 838.20874
936.421142578125 0 2742.935
937.4173583984375 0 2630.203
938.4102783203125 0 1613.4579
947.7109985351562 0 736.5275
948.2026977539062 0 829.2897
949.494140625 0 1457.9827
953.3453979492188 0 2018.6583
954.3431396484375 0 956.96606
954.4335327148438 0 4132.8804
955.4210815429688 0 7652.775
956.4212646484375 0 2914.9739
962.4522094726562 0 1150.2893
963.4481811523438 0 1399.8701
965.4398193359375 0 2344.534
965.5206909179688 0 3783.2393
966.4353637695312 0 1220.9641
966.5272827148438 0 1882.9027
972.4429931640625 0 5113.5654
973.4459838867188 0 2076.0078
975.502685546875 0 4515.946 b Water loss 7
976.5010986328125 0 5218.4673 b Ammonia loss 7
977.50048828125 0 1562.963
979.4747924804688 0 1588.41
980.458984375 0 4721.449
981.4637451171875 0 2960.2239
982.4636840820312 0 1918.3989
993.5154418945312 0 39526.645 b 7
994.51806640625 0 21880.424
995.5201416015625 0 6190.9233
1007.4564208984375 0 2059.6033 y Water loss 5
1008.4517211914062 0 1395.0281
1009.4528198242188 0 738.1981
1011.5321044921875 0 882.0318
1024.4725341796875 0 1277.241
1025.4581298828125 0 11763.171 y 5
1026.460693359375 0 6494.237
1027.4598388671875 0 1635.5349
1062.5484619140625 0 824.6036
1064.5196533203125 0 881.40857
1073.490234375 0 821.1052
1074.4893798828125 0 1076.2054
1075.5301513671875 0 1610.7603
1076.5350341796875 0 1211.8381
1090.5325927734375 0 2231.4097 b Water loss 8
1091.503173828125 0 6290.542 b Ammonia loss 8
1092.5054931640625 0 3910.4607
1093.5369873046875 0 5191.1274
1094.5418701171875 0 3154.4202
1107.5576171875 0 2925.0635
1108.542724609375 0 47726.777 b 8
1109.54345703125 0 28360.992
1110.544921875 0 9926.1045
1111.5439453125 0 781.4099
1117.4976806640625 0 1022.68506
1118.4813232421875 0 1638.6747
1119.486083984375 0 969.9073
1135.5052490234375 0 6425.898 y Water loss 4
1136.4971923828125 0 9511.729 y Ammonia loss 4
1137.494140625 0 5315.5317
1147.5531005859375 0 3774.276 b Water loss 9
1148.54833984375 0 3038.0059 b Ammonia loss 9
1149.5455322265625 0 1627.2957
1152.5361328125 0 1621.8367
1153.5162353515625 0 15236.756 y 4
1154.5194091796875 0 9121.676
1155.51953125 0 2600.8052
1164.577392578125 0 2773.1592
1165.5640869140625 0 24336.332 b 9
1166.566162109375 0 15832.792
1167.569091796875 0 5239.9873
1182.5927734375 0 5564.707
1183.5916748046875 0 4195.9443
1184.5926513671875 0 1389.6724
1198.2474365234375 0 736.4747
1204.577880859375 0 1717.4634
1205.5833740234375 0 1456.663
1222.5894775390625 0 2333.026
1223.5882568359375 0 2280.0361
1256.5430908203125 0 819.1219
1272.5662841796875 0 2030.031 y Water loss 3
1273.567138671875 0 2107.6758 y Ammonia loss 3
1274.5655517578125 0 875.51154
1275.642822265625 0 1665.7876 b Water loss 10
1276.6566162109375 0 1194.8073 b Ammonia loss 10
1277.6334228515625 0 1057.7369
1289.5921630859375 0 1842.1687
1290.57373046875 0 20478.041 y 3
1291.5770263671875 0 12192.055
1292.576904296875 0 3599.853
1293.656494140625 0 11182.495 b 10
1294.660400390625 0 7362.6943
1295.6639404296875 0 4056.357
1385.6588134765625 0 767.76556 y Water loss 2
1386.65869140625 0 735.22424 y Ammonia loss 2
1387.6495361328125 0 677.8204
1402.6719970703125 0 772.91724
1403.658447265625 0 10670.6455 y 2
1404.6619873046875 0 8277.45
1405.65869140625 0 2863.8142 b Ammonia loss 11
1458.199951171875 0 803.0669

Spectrum Details

|  |  |
| --- | --- |
| Matched peaks? Matched peaks The total absolute number of peaks matched. Additionally in brackets the total fraction of peaks matched and the total number of peaks is shown. | 75 (11.70% of 641) |
| FDR? FDR The false discovery rate estimated for this peptide. It is calculated by matching all theoretical fragments with a non-integer shift with the raw peaks for this spectrum. This is done with 40 different shifts. The resulting percentage is the average number of annotated peaks over the number of annotated peaks with the correct spectrum. | 0.83% |
| Satellite FDR? Satellite FDR See the FDR for details on its calculation. This satellite ion specific FDR only contains the satellite ions (d/w) for I/L/J positions. | - |
| PSM Score? PSM Score The PSM Score as given by Hecklib to this annotated spectrum. It is shown with three significant figures. | 952 |

#### Spectrum 7222? Spectrum 7222 The raw spectrum of this peptide as annotated by Hecklib. The fragments are coloured according to ion type (see legend). Any peaks with a star '\*' as text can be hovered over to see the full details, first the ion type second the mass shift type. By hovering over the amino acids in the peptide or ions in the legend the corresponding peaks are highlighted. By toggling the 'Unassigned' label you can turn the background (unassigned) peaks on or off in the plot. By updating the slider in the Ion legend you can update the spectrum to only show the top X% of the peaks with labels. The top X% means any peak that is within X% of the highest intensity. By dragging in the spectrum you can zoom in to a specific part of the spectrum and use 'Zoom Out' to get back to the original zoom level. The annotation of the spectrum is based on the given sequence in the peptides file and is done with different software so inconsistencies are likely. The peaks are annotated based on the given sequence, with 20 ppm tolerance.

Copy Data

##### Spectrum 7222 (TSV)

###### Preview

```
Loading example...
```

*Click on the button to copy the data to your clipboard.*

Mz MinMz MaxIntensity Max

WidthHeightPeptide font sizePeptide stroke widthSpectrum font sizeSpectrum stroke widthCompact peptide

Ion legend

wxyz

abcd

OtherUnassignedIon^Charge^_Position_Show for top:%

TVLHQDWLDGKEY

01.58e+63.17e+64.75e+66.33e+6

Zoom Out

a^+1^_2_a^+1^_2_y^+1^_1_b^+1^_2_b^+1^_2_y^+2^_3_b^+2^_4_y^+2^_4_y^+2^_4_a^+1^_3_b^+2^_5_b^+2^_5_y^+1^_2_b^+1^_3_y^+2^_5_y^+3^_7_y^+2^_5_y^+1^_2_b^+1^_3_b^+2^_6_b^+2^_6_y^+2^_6_y^+2^_6_b^+3^_9_y^+3^_9_b^+3^_10_y^+1^_3_y^+1^_3_b^+2^_7_b^+2^_7_b^+1^_4_y^+1^_3_b^+2^_7_y^+2^_7_b^+1^_4_y^+2^_7_y^+3^_11_b^+3^_12_b^+3^_12_y^+1^_4_b^+2^_8_y^+1^_4_\*\*\*b^+2^_9_b^+2^_9_b^+2^_9_b^+1^_5_b^+1^_5_y^+2^_9_b^+1^_5_b^+2^_10_y^+1^_5_y^+1^_5_y^+2^_10_b^+2^_11_b^+2^_11_y^+2^_10_b^+2^_11_b^+1^_6_b^+1^_6_y^+2^_11_y^+2^_11_b^+1^_6_y^+2^_11_b^+2^_12_b^+2^_12_y^+1^_6_b^+2^_12_y^+1^_6_y^+2^_12_y^+2^_12_b^+1^_7_b^+1^_7_y^+1^_7_y^+1^_7_y^+1^_7_b^+1^_8_b^+1^_8_y^+1^_8_b^+1^_9_b^+1^_9_y^+1^_9_y^+1^_9_y^+1^_9_b^+1^_10_y^+1^_10_

0778155623343112

Fragment Matches Table

Show background peaks

| Position | Ion type | Intensity | mz Theoretical | mz Error (Th) | mz Error (ppm) | Charge | Series Number |
| --- | --- | --- | --- | --- | --- | --- | --- |
| - | - | 1887 | 120.1 | - | - | 0 | - |
| - | - | 2907 | 120.1 | - | - | 0 | - |
| - | - | 1591 | 120.1 | - | - | 0 | - |
| - | - | 5805 | 121 | - | - | 0 | - |
| - | - | 2.906E+04 | 122.1 | - | - | 0 | - |
| - | - | 2654 | 123 | - | - | 0 | - |
| - | - | 2.967E+04 | 123 | - | - | 0 | - |
| - | - | 1.264E+04 | 123.1 | - | - | 0 | - |
| - | - | 5594 | 123.1 | - | - | 0 | - |
| - | - | 1440 | 125.1 | - | - | 0 | - |
| - | - | 1.121E+04 | 125.1 | - | - | 0 | - |
| - | - | 1.15E+04 | 126.1 | - | - | 0 | - |
| - | - | 1635 | 126.1 | - | - | 0 | - |
| - | - | 3.661E+04 | 127.1 | - | - | 0 | - |
| - | - | 8227 | 127.1 | - | - | 0 | - |
| - | - | 1.372E+04 | 127.1 | - | - | 0 | - |
| - | - | 2309 | 128 | - | - | 0 | - |
| - | - | 1741 | 128.1 | - | - | 0 | - |
| - | - | 1452 | 128.1 | - | - | 0 | - |
| - | - | 6996 | 128.1 | - | - | 0 | - |
| - | - | 3.817E+05 | 128.1 | - | - | 0 | - |
| - | - | 1.573E+04 | 129.1 | - | - | 0 | - |
| - | - | 3.946E+06 | 129.1 | - | - | 0 | - |
| - | - | 1.614E+06 | 130.1 | - | - | 0 | - |
| - | - | 2.587E+04 | 130.1 | - | - | 0 | - |
| - | - | 2.543E+05 | 130.1 | - | - | 0 | - |
| - | - | 6890 | 131.1 | - | - | 0 | - |
| - | - | 1.579E+05 | 131.1 | - | - | 0 | - |
| - | - | 5801 | 131.1 | - | - | 0 | - |
| - | - | 3.766E+05 | 132.1 | - | - | 0 | - |
| - | - | 7780 | 133.1 | - | - | 0 | - |
| - | - | 3.277E+04 | 133.1 | - | - | 0 | - |
| - | - | 4.917E+05 | 136.1 | - | - | 0 | - |
| - | - | 2075 | 137.1 | - | - | 0 | - |
| - | - | 4.256E+04 | 137.1 | - | - | 0 | - |
| - | - | 3.883E+04 | 138.1 | - | - | 0 | - |
| - | - | 2423 | 138.1 | - | - | 0 | - |
| - | - | 2.197E+04 | 138.1 | - | - | 0 | - |
| - | - | 6286 | 139.1 | - | - | 0 | - |
| - | - | 2282 | 139.1 | - | - | 0 | - |
| - | - | 4.749E+04 | 139.1 | - | - | 0 | - |
| - | - | 4033 | 140.1 | - | - | 0 | - |
| - | - | 3154 | 140.1 | - | - | 0 | - |
| - | - | 6.846E+04 | 141.1 | - | - | 0 | - |
| - | - | 3.012E+04 | 142.1 | - | - | 0 | - |
| - | - | 6071 | 142.1 | - | - | 0 | - |
| - | - | 5415 | 142.1 | - | - | 0 | - |
| - | - | 2.567E+04 | 143 | - | - | 0 | - |
| - | - | 3848 | 143.1 | - | - | 0 | - |
| - | - | 1946 | 143.1 | - | - | 0 | - |
| - | - | 2908 | 143.1 | - | - | 0 | - |
| - | - | 1690 | 144 | - | - | 0 | - |
| - | - | 1677 | 144.1 | - | - | 0 | - |
| - | - | 2.445E+04 | 144.1 | - | - | 0 | - |
| - | - | 3.798E+04 | 145.1 | - | - | 0 | - |
| - | - | 4100 | 146.1 | - | - | 0 | - |
| - | - | 5028 | 146.1 | - | - | 0 | - |
| - | - | 2.704E+04 | 147 | - | - | 0 | - |
| - | - | 3135 | 147.1 | - | - | 0 | - |
| - | - | 2005 | 148.1 | - | - | 0 | - |
| - | - | 6198 | 148.1 | - | - | 0 | - |
| - | - | 2486 | 148.1 | - | - | 0 | - |
| - | - | 8895 | 149 | - | - | 0 | - |
| - | - | 2.459E+04 | 150.1 | - | - | 0 | - |
| - | - | 4323 | 150.1 | - | - | 0 | - |
| - | - | 2045 | 150.1 | - | - | 0 | - |
| - | - | 1799 | 151.1 | - | - | 0 | - |
| - | - | 6.917E+04 | 151.1 | - | - | 0 | - |
| - | - | 2394 | 152.1 | - | - | 0 | - |
| - | - | 5846 | 152.1 | - | - | 0 | - |
| - | - | 6025 | 152.1 | - | - | 0 | - |
| - | - | 9163 | 153.1 | - | - | 0 | - |
| - | - | 6533 | 154.1 | - | - | 0 | - |
| - | - | 8510 | 154.1 | - | - | 0 | - |
| - | - | 7725 | 155 | - | - | 0 | - |
| - | - | 9336 | 155.1 | - | - | 0 | - |
| - | - | 2.537E+05 | 155.1 | - | - | 0 | - |
| 2 | a | 1.729E+05 | 155.1 | 0.0009459 | 6.098 | +1 | 2 |
| - | - | 1969 | 155.1 | - | - | 0 | - |
| - | - | 2280 | 156 | - | - | 0 | - |
| - | - | 1.605E+04 | 156.1 | - | - | 0 | - |
| - | - | 2426 | 156.1 | - | - | 0 | - |
| - | - | 1.297E+04 | 156.1 | - | - | 0 | - |
| - | - | 8459 | 156.1 | - | - | 0 | - |
| - | - | 1.756E+04 | 156.1 | - | - | 0 | - |
| - | - | 5140 | 157.1 | - | - | 0 | - |
| - | - | 3130 | 157.1 | - | - | 0 | - |
| - | - | 1.665E+04 | 157.1 | - | - | 0 | - |
| - | - | 7324 | 157.1 | - | - | 0 | - |
| - | - | 2.424E+04 | 158.1 | - | - | 0 | - |
| - | - | 1685 | 158.1 | - | - | 0 | - |
| - | - | 1618 | 158.8 | - | - | 0 | - |
| - | - | 3.089E+06 | 159.1 | - | - | 0 | - |
| - | - | 2.31E+04 | 160.1 | - | - | 0 | - |
| - | - | 2.016E+04 | 160.1 | - | - | 0 | - |
| - | - | 3.324E+05 | 160.1 | - | - | 0 | - |
| - | - | 2307 | 161.1 | - | - | 0 | - |
| - | - | 2786 | 161.1 | - | - | 0 | - |
| - | - | 1.535E+04 | 161.1 | - | - | 0 | - |
| - | - | 4540 | 164.1 | - | - | 0 | - |
| - | - | 2.506E+05 | 165.1 | - | - | 0 | - |
| - | - | 1.36E+04 | 165.1 | - | - | 0 | - |
| - | - | 1900 | 165.1 | - | - | 0 | - |
| - | - | 4.908E+05 | 166.1 | - | - | 0 | - |
| - | - | 3.594E+04 | 166.1 | - | - | 0 | - |
| - | - | 6193 | 167.1 | - | - | 0 | - |
| - | - | 3.81E+04 | 167.1 | - | - | 0 | - |
| - | - | 6825 | 167.1 | - | - | 0 | - |
| - | - | 4179 | 167.1 | - | - | 0 | - |
| - | - | 2594 | 167.1 | - | - | 0 | - |
| - | - | 1.192E+04 | 167.1 | - | - | 0 | - |
| - | - | 3203 | 168.1 | - | - | 0 | - |
| - | - | 4.667E+04 | 168.1 | - | - | 0 | - |
| - | - | 3.226E+04 | 168.1 | - | - | 0 | - |
| - | - | 4.419E+04 | 168.1 | - | - | 0 | - |
| - | - | 1316 | 168.1 | - | - | 0 | - |
| - | - | 1625 | 169.1 | - | - | 0 | - |
| - | - | 4.276E+04 | 169.1 | - | - | 0 | - |
| - | - | 4863 | 169.1 | - | - | 0 | - |
| - | - | 1.178E+05 | 169.1 | - | - | 0 | - |
| - | - | 1.574E+05 | 170.1 | - | - | 0 | - |
| - | - | 7278 | 170.1 | - | - | 0 | - |
| - | - | 3012 | 170.1 | - | - | 0 | - |
| - | - | 2.026E+04 | 171.1 | - | - | 0 | - |
| - | - | 4526 | 171.1 | - | - | 0 | - |
| - | - | 2.018E+04 | 171.1 | - | - | 0 | - |
| - | - | 1905 | 171.2 | - | - | 0 | - |
| - | - | 3.591E+04 | 172.1 | - | - | 0 | - |
| - | - | 2601 | 172.1 | - | - | 0 | - |
| - | - | 4163 | 172.1 | - | - | 0 | - |
| - | - | 6.896E+04 | 173.1 | - | - | 0 | - |
| 2 | a | 6.268E+06 | 173.1 | 0.001123 | 6.489 | +1 | 2 |
| - | - | 2474 | 174.1 | - | - | 0 | - |
| - | - | 5.577E+05 | 174.1 | - | - | 0 | - |
| - | - | 2384 | 175.1 | - | - | 0 | - |
| - | - | 2.978E+04 | 175.1 | - | - | 0 | - |
| - | - | 1.024E+04 | 176.1 | - | - | 0 | - |
| - | - | 1.381E+04 | 177.1 | - | - | 0 | - |
| - | - | 2.034E+05 | 178.1 | - | - | 0 | - |
| - | - | 1.062E+04 | 179.1 | - | - | 0 | - |
| - | - | 1745 | 179.1 | - | - | 0 | - |
| - | - | 2.303E+04 | 179.1 | - | - | 0 | - |
| - | - | 1892 | 180.1 | - | - | 0 | - |
| - | - | 2194 | 180.1 | - | - | 0 | - |
| - | - | 3.487E+04 | 181.1 | - | - | 0 | - |
| 13 | y | 5.639E+05 | 182.1 | 0.001151 | 6.324 | +1 | 1 |
| - | - | 3398 | 183 | - | - | 0 | - |
| - | - | 5.45E+04 | 183.1 | - | - | 0 | - |
| 2 | b | 1.615E+05 | 183.1 | 0.001118 | 6.105 | +1 | 2 |
| - | - | 2228 | 184.1 | - | - | 0 | - |
| - | - | 1.637E+04 | 184.1 | - | - | 0 | - |
| - | - | 2911 | 185.1 | - | - | 0 | - |
| - | - | 2595 | 185.1 | - | - | 0 | - |
| - | - | 1682 | 185.1 | - | - | 0 | - |
| - | - | 6149 | 185.1 | - | - | 0 | - |
| - | - | 1162 | 185.2 | - | - | 0 | - |
| - | - | 5.137E+04 | 185.2 | - | - | 0 | - |
| - | - | 1.518E+06 | 186.1 | - | - | 0 | - |
| - | - | 2.539E+05 | 187.1 | - | - | 0 | - |
| - | - | 1.235E+05 | 187.1 | - | - | 0 | - |
| - | - | 6404 | 187.1 | - | - | 0 | - |
| - | - | 2.608E+04 | 188.1 | - | - | 0 | - |
| - | - | 2.834E+04 | 188.1 | - | - | 0 | - |
| - | - | 7823 | 188.1 | - | - | 0 | - |
| - | - | 3708 | 189.1 | - | - | 0 | - |
| - | - | 4.619E+04 | 190.1 | - | - | 0 | - |
| - | - | 4.161E+04 | 190.1 | - | - | 0 | - |
| - | - | 2422 | 191.1 | - | - | 0 | - |
| - | - | 5077 | 191.1 | - | - | 0 | - |
| - | - | 1607 | 192.1 | - | - | 0 | - |
| - | - | 8815 | 193.1 | - | - | 0 | - |
| - | - | 4168 | 193.1 | - | - | 0 | - |
| - | - | 3.285E+04 | 194.1 | - | - | 0 | - |
| - | - | 3944 | 194.1 | - | - | 0 | - |
| - | - | 8663 | 195.1 | - | - | 0 | - |
| - | - | 3.315E+05 | 195.1 | - | - | 0 | - |
| - | - | 2.039E+05 | 196.1 | - | - | 0 | - |
| - | - | 1.906E+04 | 196.1 | - | - | 0 | - |
| - | - | 1760 | 197.1 | - | - | 0 | - |
| - | - | 2729 | 197.1 | - | - | 0 | - |
| - | - | 1.771E+04 | 197.1 | - | - | 0 | - |
| - | - | 1.993E+04 | 197.1 | - | - | 0 | - |
| - | - | 7059 | 197.2 | - | - | 0 | - |
| - | - | 3.933E+04 | 198.1 | - | - | 0 | - |
| - | - | 1.109E+05 | 198.1 | - | - | 0 | - |
| - | - | 2.037E+04 | 199.1 | - | - | 0 | - |
| - | - | 2.553E+04 | 199.1 | - | - | 0 | - |
| - | - | 2661 | 199.1 | - | - | 0 | - |
| - | - | 1.613E+04 | 199.1 | - | - | 0 | - |
| - | - | 3016 | 200.1 | - | - | 0 | - |
| - | - | 5089 | 200.1 | - | - | 0 | - |
| - | - | 2.465E+04 | 200.1 | - | - | 0 | - |
| 2 | b | 1.673E+06 | 201.1 | 0.001158 | 5.758 | +1 | 2 |
| - | - | 2.015E+04 | 202.1 | - | - | 0 | - |
| - | - | 1.693E+05 | 202.1 | - | - | 0 | - |
| - | - | 5486 | 203.1 | - | - | 0 | - |
| - | - | 1.727E+04 | 203.1 | - | - | 0 | - |
| - | - | 3975 | 203.2 | - | - | 0 | - |
| - | - | 5085 | 204.1 | - | - | 0 | - |
| - | - | 1.309E+04 | 204.1 | - | - | 0 | - |
| - | - | 5459 | 204.1 | - | - | 0 | - |
| - | - | 1.368E+05 | 205.1 | - | - | 0 | - |
| - | - | 2.468E+04 | 205.1 | - | - | 0 | - |
| - | - | 1.522E+04 | 206.1 | - | - | 0 | - |
| - | - | 2.8E+04 | 206.1 | - | - | 0 | - |
| - | - | 2951 | 206.1 | - | - | 0 | - |
| - | - | 7860 | 207.1 | - | - | 0 | - |
| - | - | 2552 | 207.1 | - | - | 0 | - |
| - | - | 3950 | 207.2 | - | - | 0 | - |
| - | - | 4.691E+04 | 207.2 | - | - | 0 | - |
| - | - | 5476 | 208.1 | - | - | 0 | - |
| - | - | 6635 | 208.2 | - | - | 0 | - |
| - | - | 8179 | 209.1 | - | - | 0 | - |
| - | - | 5360 | 209.1 | - | - | 0 | - |
| - | - | 5.031E+04 | 210.1 | - | - | 0 | - |
| - | - | 2616 | 211.1 | - | - | 0 | - |
| - | - | 3720 | 211.1 | - | - | 0 | - |
| 11 | y | 2.65E+04 | 211.1 | 0.001366 | 6.472 | +2 | 3 |
| - | - | 6557 | 211.1 | - | - | 0 | - |
| - | - | 2313 | 211.1 | - | - | 0 | - |
| - | - | 2937 | 212.1 | - | - | 0 | - |
| - | - | 3.141E+04 | 212.1 | - | - | 0 | - |
| - | - | 2314 | 212.2 | - | - | 0 | - |
| - | - | 1.585E+05 | 213.1 | - | - | 0 | - |
| - | - | 1.001E+04 | 213.1 | - | - | 0 | - |
| - | - | 1.485E+04 | 213.2 | - | - | 0 | - |
| - | - | 1.654E+04 | 214.1 | - | - | 0 | - |
| - | - | 2196 | 214.2 | - | - | 0 | - |
| - | - | 8918 | 215.1 | - | - | 0 | - |
| - | - | 1.533E+04 | 215.1 | - | - | 0 | - |
| - | - | 3.161E+04 | 216.1 | - | - | 0 | - |
| - | - | 2868 | 217.1 | - | - | 0 | - |
| - | - | 1.016E+04 | 221.1 | - | - | 0 | - |
| - | - | 4120 | 221.1 | - | - | 0 | - |
| - | - | 1.105E+04 | 221.1 | - | - | 0 | - |
| - | - | 3.134E+04 | 221.1 | - | - | 0 | - |
| - | - | 1.873E+04 | 221.1 | - | - | 0 | - |
| - | - | 1684 | 221.6 | - | - | 0 | - |
| - | - | 2919 | 222.1 | - | - | 0 | - |
| - | - | 3886 | 222.1 | - | - | 0 | - |
| - | - | 4.759E+04 | 222.1 | - | - | 0 | - |
| - | - | 2043 | 223.1 | - | - | 0 | - |
| - | - | 8613 | 223.1 | - | - | 0 | - |
| - | - | 8.87E+04 | 223.1 | - | - | 0 | - |
| - | - | 8976 | 223.1 | - | - | 0 | - |
| - | - | 5.53E+05 | 223.2 | - | - | 0 | - |
| - | - | 2.235E+04 | 224.1 | - | - | 0 | - |
| - | - | 6729 | 224.1 | - | - | 0 | - |
| - | - | 6.876E+04 | 224.2 | - | - | 0 | - |
| - | - | 2984 | 225 | - | - | 0 | - |
| - | - | 3512 | 225.1 | - | - | 0 | - |
| - | - | 8332 | 225.1 | - | - | 0 | - |
| - | - | 1.094E+04 | 225.2 | - | - | 0 | - |
| - | - | 2065 | 225.6 | - | - | 0 | - |
| - | - | 1.126E+05 | 226.1 | - | - | 0 | - |
| 4 | b | 3890 | 226.1 | 0.001159 | 5.127 | +2 | 4 |
| - | - | 9.168E+04 | 227.1 | - | - | 0 | - |
| - | - | 1.164E+04 | 227.1 | - | - | 0 | - |
| - | - | 2694 | 227.1 | - | - | 0 | - |
| - | - | 2.913E+04 | 227.2 | - | - | 0 | - |
| - | - | 7729 | 228.1 | - | - | 0 | - |
| - | - | 4493 | 228.1 | - | - | 0 | - |
| - | - | 3726 | 228.2 | - | - | 0 | - |
| - | - | 2.022E+05 | 229.1 | - | - | 0 | - |
| - | - | 2528 | 230.1 | - | - | 0 | - |
| - | - | 2.126E+04 | 230.1 | - | - | 0 | - |
| - | - | 1.138E+04 | 230.2 | - | - | 0 | - |
| - | - | 4180 | 230.2 | - | - | 0 | - |
| - | - | 9543 | 231.1 | - | - | 0 | - |
| - | - | 2270 | 232.1 | - | - | 0 | - |
| - | - | 2765 | 233.1 | - | - | 0 | - |
| - | - | 8.704E+04 | 233.1 | - | - | 0 | - |
| - | - | 6.421E+05 | 234.1 | - | - | 0 | - |
| - | - | 7508 | 234.7 | - | - | 0 | - |
| - | - | 3690 | 235.1 | - | - | 0 | - |
| - | - | 8.304E+04 | 235.1 | - | - | 0 | - |
| - | - | 1.525E+04 | 235.2 | - | - | 0 | - |
| - | - | 8535 | 235.6 | - | - | 0 | - |
| - | - | 3494 | 236.1 | - | - | 0 | - |
| - | - | 6124 | 236.1 | - | - | 0 | - |
| - | - | 1.533E+04 | 237.1 | - | - | 0 | - |
| - | - | 3947 | 238.1 | - | - | 0 | - |
| - | - | 6.554E+04 | 238.1 | - | - | 0 | - |
| - | - | 1.357E+04 | 238.1 | - | - | 0 | - |
| - | - | 2680 | 239.1 | - | - | 0 | - |
| - | - | 1.471E+04 | 239.1 | - | - | 0 | - |
| - | - | 8932 | 239.1 | - | - | 0 | - |
| - | - | 6511 | 239.1 | - | - | 0 | - |
| 10 | y | 2451 | 239.6 | 0.002948 | 12.3 | +2 | 4 |
| - | - | 2.106E+05 | 240.1 | - | - | 0 | - |
| - | - | 9407 | 241.1 | - | - | 0 | - |
| - | - | 2.972E+04 | 241.1 | - | - | 0 | - |
| - | - | 2.028E+04 | 241.1 | - | - | 0 | - |
| - | - | 2.871E+04 | 241.2 | - | - | 0 | - |
| - | - | 3091 | 242.2 | - | - | 0 | - |
| - | - | 4815 | 242.2 | - | - | 0 | - |
| - | - | 3.9E+04 | 243.1 | - | - | 0 | - |
| - | - | 3361 | 243.1 | - | - | 0 | - |
| - | - | 2.422E+05 | 244.1 | - | - | 0 | - |
| - | - | 3802 | 244.6 | - | - | 0 | - |
| - | - | 2.284E+04 | 245.1 | - | - | 0 | - |
| - | - | 3065 | 245.1 | - | - | 0 | - |
| - | - | 2878 | 246.1 | - | - | 0 | - |
| - | - | 3.464E+05 | 247.1 | - | - | 0 | - |
| - | - | 1.819E+04 | 247.1 | - | - | 0 | - |
| - | - | 7905 | 247.2 | - | - | 0 | - |
| - | - | 1.378E+05 | 248.1 | - | - | 0 | - |
| 10 | y | 4721 | 248.6 | 0.001435 | 5.771 | +2 | 4 |
| - | - | 2.031E+05 | 249.1 | - | - | 0 | - |
| - | - | 1.357E+04 | 249.1 | - | - | 0 | - |
| - | - | 5.092E+04 | 249.1 | - | - | 0 | - |
| - | - | 4829 | 249.6 | - | - | 0 | - |
| - | - | 2.256E+04 | 250.1 | - | - | 0 | - |
| - | - | 7982 | 250.1 | - | - | 0 | - |
| - | - | 3006 | 250.1 | - | - | 0 | - |
| - | - | 7122 | 250.2 | - | - | 0 | - |
| - | - | 2.101E+06 | 251.2 | - | - | 0 | - |
| - | - | 9074 | 252.1 | - | - | 0 | - |
| - | - | 2.79E+05 | 252.2 | - | - | 0 | - |
| - | - | 6423 | 252.6 | - | - | 0 | - |
| - | - | 8712 | 253.1 | - | - | 0 | - |
| - | - | 2657 | 253.1 | - | - | 0 | - |
| - | - | 2.202E+04 | 253.2 | - | - | 0 | - |
| - | - | 5359 | 254.1 | - | - | 0 | - |
| - | - | 1.026E+05 | 255.1 | - | - | 0 | - |
| - | - | 3893 | 256.1 | - | - | 0 | - |
| - | - | 2.291E+04 | 256.1 | - | - | 0 | - |
| - | - | 2478 | 256.1 | - | - | 0 | - |
| - | - | 1.986E+04 | 256.2 | - | - | 0 | - |
| - | - | 1.041E+04 | 256.6 | - | - | 0 | - |
| - | - | 5450 | 257.1 | - | - | 0 | - |
| - | - | 6.326E+04 | 257.1 | - | - | 0 | - |
| - | - | 1.014E+04 | 258.1 | - | - | 0 | - |
| - | - | 7.822E+04 | 258.1 | - | - | 0 | - |
| - | - | 1.086E+04 | 259.1 | - | - | 0 | - |
| - | - | 1.045E+04 | 259.1 | - | - | 0 | - |
| - | - | 4.037E+04 | 261.1 | - | - | 0 | - |
| - | - | 2.043E+04 | 261.1 | - | - | 0 | - |
| - | - | 2758 | 261.2 | - | - | 0 | - |
| - | - | 3.129E+04 | 262.1 | - | - | 0 | - |
| - | - | 4004 | 262.1 | - | - | 0 | - |
| - | - | 3671 | 263.1 | - | - | 0 | - |
| - | - | 2833 | 263.1 | - | - | 0 | - |
| - | - | 2300 | 263.2 | - | - | 0 | - |
| - | - | 1.731E+05 | 265.1 | - | - | 0 | - |
| - | - | 2262 | 265.6 | - | - | 0 | - |
| - | - | 5.315E+05 | 266.1 | - | - | 0 | - |
| - | - | 8194 | 267.1 | - | - | 0 | - |
| - | - | 6.628E+04 | 267.1 | - | - | 0 | - |
| - | - | 4676 | 267.1 | - | - | 0 | - |
| - | - | 6552 | 267.6 | - | - | 0 | - |
| - | - | 3751 | 268.1 | - | - | 0 | - |
| - | - | 2.51E+04 | 268.1 | - | - | 0 | - |
| - | - | 4.28E+04 | 268.2 | - | - | 0 | - |
| 3 | a | 4364 | 268.2 | 0.001751 | 6.53 | +1 | 3 |
| - | - | 3313 | 269.1 | - | - | 0 | - |
| - | - | 5061 | 269.1 | - | - | 0 | - |
| - | - | 2.009E+04 | 269.2 | - | - | 0 | - |
| - | - | 3.138E+04 | 269.2 | - | - | 0 | - |
| - | - | 8.598E+04 | 270.1 | - | - | 0 | - |
| - | - | 1.105E+04 | 270.1 | - | - | 0 | - |
| - | - | 7031 | 270.2 | - | - | 0 | - |
| - | - | 3268 | 270.2 | - | - | 0 | - |
| - | - | 2.857E+04 | 270.6 | - | - | 0 | - |
| - | - | 4140 | 271.1 | - | - | 0 | - |
| - | - | 2008 | 271.2 | - | - | 0 | - |
| - | - | 1.321E+05 | 272.2 | - | - | 0 | - |
| - | - | 2.583E+04 | 273.2 | - | - | 0 | - |
| - | - | 1.287E+05 | 274.1 | - | - | 0 | - |
| - | - | 2.884E+04 | 275.1 | - | - | 0 | - |
| - | - | 1.985E+04 | 275.1 | - | - | 0 | - |
| - | - | 1.049E+04 | 275.2 | - | - | 0 | - |
| - | - | 2.413E+04 | 276.1 | - | - | 0 | - |
| - | - | 1.341E+04 | 276.1 | - | - | 0 | - |
| - | - | 2.295E+04 | 276.2 | - | - | 0 | - |
| - | - | 6437 | 276.2 | - | - | 0 | - |
| - | - | 5075 | 276.6 | - | - | 0 | - |
| - | - | 2378 | 276.7 | - | - | 0 | - |
| - | - | 4174 | 277.1 | - | - | 0 | - |
| - | - | 4531 | 277.2 | - | - | 0 | - |
| - | - | 2164 | 277.2 | - | - | 0 | - |
| - | - | 4627 | 278.2 | - | - | 0 | - |
| - | - | 2130 | 278.2 | - | - | 0 | - |
| - | - | 4.602E+04 | 279.1 | - | - | 0 | - |
| - | - | 2654 | 279.2 | - | - | 0 | - |
| - | - | 3472 | 279.6 | - | - | 0 | - |
| - | - | 4.819E+04 | 280.1 | - | - | 0 | - |
| - | - | 7061 | 280.1 | - | - | 0 | - |
| - | - | 2.535E+04 | 281.1 | - | - | 0 | - |
| - | - | 3790 | 281.1 | - | - | 0 | - |
| - | - | 7342 | 281.1 | - | - | 0 | - |
| 5 | b | 4158 | 281.2 | 0.003429 | 12.19 | +2 | 5 |
| - | - | 2662 | 282.1 | - | - | 0 | - |
| - | - | 1.161E+04 | 282.2 | - | - | 0 | - |
| - | - | 6454 | 282.7 | - | - | 0 | - |
| - | - | 1.28E+06 | 283.1 | - | - | 0 | - |
| - | - | 6443 | 283.6 | - | - | 0 | - |
| - | - | 8435 | 284.1 | - | - | 0 | - |
| - | - | 6.554E+04 | 284.1 | - | - | 0 | - |
| - | - | 1.043E+04 | 284.1 | - | - | 0 | - |
| - | - | 1.81E+05 | 284.1 | - | - | 0 | - |
| - | - | 3007 | 284.6 | - | - | 0 | - |
| - | - | 6501 | 285.1 | - | - | 0 | - |
| - | - | 9643 | 285.1 | - | - | 0 | - |
| - | - | 3386 | 285.1 | - | - | 0 | - |
| - | - | 1.235E+04 | 285.1 | - | - | 0 | - |
| - | - | 1.444E+04 | 285.2 | - | - | 0 | - |
| - | - | 1.152E+05 | 286.1 | - | - | 0 | - |
| - | - | 1.836E+04 | 287.1 | - | - | 0 | - |
| - | - | 2.767E+04 | 287.2 | - | - | 0 | - |
| - | - | 1.058E+04 | 287.2 | - | - | 0 | - |
| - | - | 7304 | 288.1 | - | - | 0 | - |
| - | - | 3666 | 288.2 | - | - | 0 | - |
| - | - | 2.133E+04 | 288.6 | - | - | 0 | - |
| - | - | 6401 | 289.1 | - | - | 0 | - |
| - | - | 2898 | 289.2 | - | - | 0 | - |
| 5 | b | 1.876E+04 | 290.2 | 0.002236 | 7.705 | +2 | 5 |
| - | - | 5710 | 290.7 | - | - | 0 | - |
| - | - | 6830 | 291.2 | - | - | 0 | - |
| - | - | 2172 | 292.1 | - | - | 0 | - |
| - | - | 2671 | 292.2 | - | - | 0 | - |
| - | - | 2184 | 292.2 | - | - | 0 | - |
| 12 | y | 8.521E+05 | 293.1 | 0.002006 | 6.843 | +1 | 2 |
| - | - | 1.781E+04 | 293.2 | - | - | 0 | - |
| - | - | 1.61E+05 | 294.1 | - | - | 0 | - |
| - | - | 3079 | 294.2 | - | - | 0 | - |
| - | - | 3358 | 295.1 | - | - | 0 | - |
| - | - | 1.989E+04 | 295.1 | - | - | 0 | - |
| - | - | 2919 | 296.1 | - | - | 0 | - |
| - | - | 2777 | 296.1 | - | - | 0 | - |
| - | - | 3693 | 296.2 | - | - | 0 | - |
| 3 | b | 2.926E+04 | 296.2 | 0.002076 | 7.009 | +1 | 3 |
| - | - | 7260 | 296.7 | - | - | 0 | - |
| 9 | y | 2.104E+04 | 297.1 | 0.002935 | 9.878 | +2 | 5 |
| - | - | 2.702E+05 | 297.2 | - | - | 0 | - |
| - | - | 3772 | 297.6 | - | - | 0 | - |
| - | - | 3233 | 297.7 | - | - | 0 | - |
| 7 | y | 2.702E+04 | 298.1 | 0.003567 | 11.96 | +3 | 7 |
| - | - | 3.962E+04 | 298.2 | - | - | 0 | - |
| - | - | 1.917E+04 | 299.1 | - | - | 0 | - |
| - | - | 2822 | 299.1 | - | - | 0 | - |
| - | - | 5806 | 299.2 | - | - | 0 | - |
| - | - | 3301 | 300.1 | - | - | 0 | - |
| - | - | 3110 | 300.1 | - | - | 0 | - |
| - | - | 4.758E+04 | 300.2 | - | - | 0 | - |
| - | - | 1.901E+04 | 301.1 | - | - | 0 | - |
| - | - | 7.651E+05 | 301.2 | - | - | 0 | - |
| - | - | 1.411E+05 | 302.1 | - | - | 0 | - |
| - | - | 1.06E+05 | 302.2 | - | - | 0 | - |
| - | - | 2.011E+04 | 303.1 | - | - | 0 | - |
| - | - | 6471 | 303.2 | - | - | 0 | - |
| - | - | 1.027E+04 | 303.2 | - | - | 0 | - |
| - | - | 4281 | 304.2 | - | - | 0 | - |
| - | - | 2308 | 304.2 | - | - | 0 | - |
| - | - | 1.925E+04 | 305.2 | - | - | 0 | - |
| - | - | 3372 | 305.6 | - | - | 0 | - |
| - | - | 6977 | 305.7 | - | - | 0 | - |
| 9 | y | 1.424E+05 | 306.1 | 0.002139 | 6.986 | +2 | 5 |
| - | - | 1.327E+04 | 306.2 | - | - | 0 | - |
| - | - | 5.467E+04 | 306.6 | - | - | 0 | - |
| - | - | 2.203E+04 | 307.1 | - | - | 0 | - |
| - | - | 2735 | 307.2 | - | - | 0 | - |
| - | - | 3756 | 307.2 | - | - | 0 | - |
| - | - | 2660 | 308.1 | - | - | 0 | - |
| - | - | 2715 | 308.2 | - | - | 0 | - |
| - | - | 5090 | 309.1 | - | - | 0 | - |
| - | - | 8846 | 309.2 | - | - | 0 | - |
| - | - | 8862 | 310.2 | - | - | 0 | - |
| - | - | 2861 | 310.7 | - | - | 0 | - |
| 12 | y | 9.466E+04 | 311.1 | 0.001848 | 5.938 | +1 | 2 |
| - | - | 4.868E+04 | 311.1 | - | - | 0 | - |
| - | - | 2885 | 311.2 | - | - | 0 | - |
| - | - | 5563 | 311.7 | - | - | 0 | - |
| - | - | 1.573E+04 | 312.1 | - | - | 0 | - |
| - | - | 5902 | 312.1 | - | - | 0 | - |
| - | - | 2858 | 312.2 | - | - | 0 | - |
| - | - | 5300 | 313.1 | - | - | 0 | - |
| - | - | 4969 | 314.1 | - | - | 0 | - |
| - | - | 2.253E+04 | 314.2 | - | - | 0 | - |
| 3 | b | 6.334E+04 | 314.2 | 0.002223 | 7.075 | +1 | 3 |
| - | - | 4.34E+05 | 315.2 | - | - | 0 | - |
| - | - | 1.335E+04 | 315.2 | - | - | 0 | - |
| - | - | 6.874E+04 | 316.2 | - | - | 0 | - |
| - | - | 2758 | 317.1 | - | - | 0 | - |
| - | - | 5504 | 317.2 | - | - | 0 | - |
| - | - | 5494 | 317.7 | - | - | 0 | - |
| - | - | 5020 | 318.1 | - | - | 0 | - |
| - | - | 1.069E+04 | 318.2 | - | - | 0 | - |
| - | - | 3948 | 318.7 | - | - | 0 | - |
| - | - | 3562 | 319.1 | - | - | 0 | - |
| - | - | 2.662E+04 | 319.2 | - | - | 0 | - |
| - | - | 5826 | 319.2 | - | - | 0 | - |
| - | - | 4831 | 320.2 | - | - | 0 | - |
| - | - | 4323 | 320.2 | - | - | 0 | - |
| - | - | 4742 | 321.1 | - | - | 0 | - |
| - | - | 3919 | 321.2 | - | - | 0 | - |
| - | - | 5684 | 322.2 | - | - | 0 | - |
| - | - | 2084 | 323.1 | - | - | 0 | - |
| - | - | 2579 | 324.2 | - | - | 0 | - |
| - | - | 6016 | 324.2 | - | - | 0 | - |
| - | - | 4303 | 324.7 | - | - | 0 | - |
| - | - | 2.122E+04 | 325.2 | - | - | 0 | - |
| - | - | 3262 | 325.2 | - | - | 0 | - |
| - | - | 2470 | 326.2 | - | - | 0 | - |
| - | - | 4331 | 326.2 | - | - | 0 | - |
| - | - | 1.071E+05 | 326.7 | - | - | 0 | - |
| - | - | 3.897E+04 | 327.2 | - | - | 0 | - |
| - | - | 6594 | 327.7 | - | - | 0 | - |
| - | - | 2192 | 328.1 | - | - | 0 | - |
| - | - | 3983 | 330.2 | - | - | 0 | - |
| - | - | 3692 | 330.7 | - | - | 0 | - |
| - | - | 2507 | 331.1 | - | - | 0 | - |
| - | - | 6708 | 331.7 | - | - | 0 | - |
| - | - | 4.466E+04 | 332.2 | - | - | 0 | - |
| - | - | 5.179E+04 | 333.2 | - | - | 0 | - |
| - | - | 8277 | 333.2 | - | - | 0 | - |
| - | - | 4410 | 334.1 | - | - | 0 | - |
| - | - | 2434 | 334.1 | - | - | 0 | - |
| - | - | 1.189E+04 | 334.2 | - | - | 0 | - |
| - | - | 2534 | 335.1 | - | - | 0 | - |
| - | - | 1.806E+04 | 335.2 | - | - | 0 | - |
| - | - | 3860 | 336.2 | - | - | 0 | - |
| - | - | 3568 | 337.2 | - | - | 0 | - |
| - | - | 1.034E+04 | 338.2 | - | - | 0 | - |
| 6 | b | 2.021E+04 | 338.7 | 0.002469 | 7.291 | +2 | 6 |
| - | - | 1.711E+04 | 339.2 | - | - | 0 | - |
| - | - | 1.246E+04 | 339.7 | - | - | 0 | - |
| - | - | 2703 | 340.2 | - | - | 0 | - |
| - | - | 2.463E+04 | 340.7 | - | - | 0 | - |
| - | - | 3483 | 341 | - | - | 0 | - |
| - | - | 1.121E+04 | 341.2 | - | - | 0 | - |
| - | - | 3.588E+04 | 342.2 | - | - | 0 | - |
| - | - | 2601 | 342.7 | - | - | 0 | - |
| - | - | 4.294E+04 | 343.2 | - | - | 0 | - |
| - | - | 7179 | 343.2 | - | - | 0 | - |
| - | - | 3.757E+04 | 344.2 | - | - | 0 | - |
| - | - | 3114 | 345 | - | - | 0 | - |
| - | - | 5.822E+04 | 345.1 | - | - | 0 | - |
| - | - | 5251 | 345.2 | - | - | 0 | - |
| - | - | 2.14E+05 | 346.1 | - | - | 0 | - |
| - | - | 8820 | 346.7 | - | - | 0 | - |
| - | - | 3.868E+04 | 347.1 | - | - | 0 | - |
| 6 | b | 1.146E+05 | 347.7 | 0.002436 | 7.006 | +2 | 6 |
| - | - | 3667 | 348.1 | - | - | 0 | - |
| - | - | 4.284E+04 | 348.2 | - | - | 0 | - |
| - | - | 3469 | 348.2 | - | - | 0 | - |
| - | - | 1.318E+04 | 348.7 | - | - | 0 | - |
| - | - | 5512 | 349.1 | - | - | 0 | - |
| - | - | 9307 | 349.2 | - | - | 0 | - |
| - | - | 5425 | 349.2 | - | - | 0 | - |
| - | - | 2444 | 350.1 | - | - | 0 | - |
| - | - | 1.733E+05 | 350.2 | - | - | 0 | - |
| - | - | 5823 | 351.1 | - | - | 0 | - |
| - | - | 8564 | 351.2 | - | - | 0 | - |
| - | - | 3.529E+04 | 351.2 | - | - | 0 | - |
| - | - | 1.238E+04 | 352.2 | - | - | 0 | - |
| - | - | 3528 | 352.2 | - | - | 0 | - |
| - | - | 2624 | 352.2 | - | - | 0 | - |
| - | - | 1.485E+04 | 352.7 | - | - | 0 | - |
| - | - | 7506 | 353.2 | - | - | 0 | - |
| 8 | y | 2.604E+04 | 353.7 | 0.002193 | 6.202 | +2 | 6 |
| - | - | 1.46E+04 | 354.2 | - | - | 0 | - |
| - | - | 2724 | 354.7 | - | - | 0 | - |
| - | - | 2685 | 355.2 | - | - | 0 | - |
| - | - | 6516 | 356.2 | - | - | 0 | - |
| - | - | 2827 | 356.7 | - | - | 0 | - |
| - | - | 4737 | 357.2 | - | - | 0 | - |
| - | - | 3679 | 358.2 | - | - | 0 | - |
| - | - | 6380 | 359.1 | - | - | 0 | - |
| - | - | 3572 | 359.2 | - | - | 0 | - |
| - | - | 2323 | 360.2 | - | - | 0 | - |
| - | - | 1.063E+04 | 360.2 | - | - | 0 | - |
| - | - | 4.818E+04 | 360.7 | - | - | 0 | - |
| - | - | 5.732E+04 | 361.2 | - | - | 0 | - |
| - | - | 1.36E+04 | 361.7 | - | - | 0 | - |
| - | - | 5.365E+04 | 362.2 | - | - | 0 | - |
| 8 | y | 2.918E+05 | 362.7 | 0.002587 | 7.134 | +2 | 6 |
| - | - | 2.968E+05 | 363.1 | - | - | 0 | - |
| - | - | 1.247E+05 | 363.2 | - | - | 0 | - |
| - | - | 2.466E+04 | 363.7 | - | - | 0 | - |
| - | - | 5.146E+04 | 364.1 | - | - | 0 | - |
| - | - | 4.409E+04 | 364.1 | - | - | 0 | - |
| 9 | b | 5768 | 364.2 | 0.00416 | 11.42 | +3 | 9 |
| - | - | 4608 | 364.2 | - | - | 0 | - |
| - | - | 3080 | 364.7 | - | - | 0 | - |
| - | - | 1.079E+04 | 365.1 | - | - | 0 | - |
| - | - | 5668 | 365.2 | - | - | 0 | - |
| - | - | 8264 | 366.2 | - | - | 0 | - |
| - | - | 1.54E+04 | 366.2 | - | - | 0 | - |
| - | - | 3E+04 | 367.1 | - | - | 0 | - |
| - | - | 8526 | 367.2 | - | - | 0 | - |
| - | - | 2.644E+04 | 367.2 | - | - | 0 | - |
| - | - | 7749 | 367.7 | - | - | 0 | - |
| - | - | 5593 | 368.1 | - | - | 0 | - |
| - | - | 4937 | 368.1 | - | - | 0 | - |
| - | - | 3159 | 368.2 | - | - | 0 | - |
| - | - | 1.416E+04 | 368.2 | - | - | 0 | - |
| - | - | 2470 | 368.2 | - | - | 0 | - |
| - | - | 4879 | 368.3 | - | - | 0 | - |
| - | - | 3223 | 368.7 | - | - | 0 | - |
| - | - | 3095 | 369.2 | - | - | 0 | - |
| - | - | 7194 | 369.2 | - | - | 0 | - |
| - | - | 6796 | 369.2 | - | - | 0 | - |
| - | - | 7947 | 370.2 | - | - | 0 | - |
| - | - | 2863 | 370.2 | - | - | 0 | - |
| - | - | 1.495E+04 | 373.2 | - | - | 0 | - |
| - | - | 4028 | 373.7 | - | - | 0 | - |
| - | - | 4369 | 374.2 | - | - | 0 | - |
| - | - | 3938 | 374.7 | - | - | 0 | - |
| - | - | 2183 | 375.1 | - | - | 0 | - |
| - | - | 3855 | 375.2 | - | - | 0 | - |
| - | - | 4001 | 375.2 | - | - | 0 | - |
| - | - | 1.704E+04 | 376.2 | - | - | 0 | - |
| - | - | 4.098E+04 | 376.2 | - | - | 0 | - |
| - | - | 1.806E+04 | 376.7 | - | - | 0 | - |
| - | - | 7135 | 377.1 | - | - | 0 | - |
| - | - | 1.161E+04 | 377.1 | - | - | 0 | - |
| - | - | 2529 | 377.2 | - | - | 0 | - |
| - | - | 7027 | 377.3 | - | - | 0 | - |
| - | - | 1.056E+05 | 378.2 | - | - | 0 | - |
| - | - | 5.011E+05 | 379.2 | - | - | 0 | - |
| - | - | 1.793E+04 | 380.2 | - | - | 0 | - |
| - | - | 1.057E+05 | 380.2 | - | - | 0 | - |
| - | - | 8.539E+05 | 381.2 | - | - | 0 | - |
| - | - | 3.136E+04 | 381.7 | - | - | 0 | - |
| - | - | 1.528E+05 | 382.2 | - | - | 0 | - |
| - | - | 2.085E+04 | 382.2 | - | - | 0 | - |
| - | - | 1.553E+04 | 382.2 | - | - | 0 | - |
| - | - | 4660 | 382.3 | - | - | 0 | - |
| - | - | 1.551E+04 | 382.7 | - | - | 0 | - |
| - | - | 2.113E+04 | 383.2 | - | - | 0 | - |
| - | - | 8567 | 383.2 | - | - | 0 | - |
| - | - | 3743 | 383.7 | - | - | 0 | - |
| - | - | 3.871E+04 | 384.2 | - | - | 0 | - |
| - | - | 2.205E+04 | 384.2 | - | - | 0 | - |
| - | - | 6.208E+04 | 385.2 | - | - | 0 | - |
| 5 | y | 1.57E+04 | 385.2 | 0.002178 | 5.655 | +3 | 9 |
| - | - | 1.023E+04 | 386.2 | - | - | 0 | - |
| - | - | 1.256E+04 | 386.2 | - | - | 0 | - |
| - | - | 3193 | 386.2 | - | - | 0 | - |
| - | - | 3120 | 387.2 | - | - | 0 | - |
| 10 | b | 1.051E+04 | 389.2 | 0.002579 | 6.625 | +3 | 10 |
| - | - | 3.666E+04 | 390.2 | - | - | 0 | - |
| - | - | 1.517E+04 | 390.7 | - | - | 0 | - |
| - | - | 3.436E+04 | 391.1 | - | - | 0 | - |
| - | - | 1.107E+04 | 391.2 | - | - | 0 | - |
| - | - | 6184 | 391.7 | - | - | 0 | - |
| - | - | 9456 | 392.1 | - | - | 0 | - |
| - | - | 3772 | 392.1 | - | - | 0 | - |
| - | - | 1.393E+04 | 392.2 | - | - | 0 | - |
| - | - | 7169 | 393.2 | - | - | 0 | - |
| - | - | 9.647E+04 | 394.2 | - | - | 0 | - |
| - | - | 6.246E+04 | 395.1 | - | - | 0 | - |
| - | - | 2.206E+04 | 395.2 | - | - | 0 | - |
| - | - | 5136 | 395.2 | - | - | 0 | - |
| - | - | 1.337E+04 | 396.1 | - | - | 0 | - |
| - | - | 4578 | 396.2 | - | - | 0 | - |
| - | - | 6872 | 396.2 | - | - | 0 | - |
| - | - | 1.864E+05 | 396.2 | - | - | 0 | - |
| - | - | 7516 | 397.2 | - | - | 0 | - |
| - | - | 3.449E+04 | 397.2 | - | - | 0 | - |
| - | - | 7516 | 397.7 | - | - | 0 | - |
| - | - | 7.828E+04 | 398.2 | - | - | 0 | - |
| - | - | 5499 | 398.2 | - | - | 0 | - |
| - | - | 1.792E+04 | 399.2 | - | - | 0 | - |
| - | - | 4010 | 400.2 | - | - | 0 | - |
| - | - | 2779 | 400.7 | - | - | 0 | - |
| - | - | 3670 | 401.2 | - | - | 0 | - |
| - | - | 4.01E+04 | 402.2 | - | - | 0 | - |
| - | - | 7.385E+04 | 402.2 | - | - | 0 | - |
| - | - | 5649 | 403.1 | - | - | 0 | - |
| - | - | 2.732E+04 | 403.2 | - | - | 0 | - |
| - | - | 5943 | 404.2 | - | - | 0 | - |
| - | - | 2642 | 404.2 | - | - | 0 | - |
| - | - | 1.113E+04 | 404.7 | - | - | 0 | - |
| - | - | 5219 | 405.2 | - | - | 0 | - |
| - | - | 5576 | 405.2 | - | - | 0 | - |
| - | - | 3.202E+04 | 405.3 | - | - | 0 | - |
| - | - | 1.818E+04 | 406.2 | - | - | 0 | - |
| - | - | 1.777E+04 | 407.2 | - | - | 0 | - |
| - | - | 7824 | 407.2 | - | - | 0 | - |
| - | - | 2801 | 407.3 | - | - | 0 | - |
| - | - | 3224 | 408.2 | - | - | 0 | - |
| - | - | 3638 | 408.2 | - | - | 0 | - |
| - | - | 8296 | 408.7 | - | - | 0 | - |
| - | - | 6.147E+04 | 409.1 | - | - | 0 | - |
| - | - | 4611 | 409.2 | - | - | 0 | - |
| - | - | 1.593E+04 | 409.2 | - | - | 0 | - |
| - | - | 1.568E+04 | 409.7 | - | - | 0 | - |
| - | - | 1.232E+04 | 410.2 | - | - | 0 | - |
| - | - | 4712 | 410.2 | - | - | 0 | - |
| - | - | 3126 | 411.2 | - | - | 0 | - |
| - | - | 2892 | 411.2 | - | - | 0 | - |
| - | - | 5.509E+05 | 412.2 | - | - | 0 | - |
| - | - | 6.014E+04 | 413.1 | - | - | 0 | - |
| - | - | 8.483E+04 | 413.2 | - | - | 0 | - |
| - | - | 1.795E+04 | 414.2 | - | - | 0 | - |
| - | - | 1.706E+04 | 414.2 | - | - | 0 | - |
| - | - | 1.709E+05 | 414.2 | - | - | 0 | - |
| - | - | 8604 | 415 | - | - | 0 | - |
| - | - | 6.303E+04 | 415.2 | - | - | 0 | - |
| - | - | 3.42E+04 | 415.2 | - | - | 0 | - |
| - | - | 3104 | 416.1 | - | - | 0 | - |
| - | - | 1.357E+04 | 416.2 | - | - | 0 | - |
| - | - | 4051 | 416.2 | - | - | 0 | - |
| - | - | 2681 | 417.2 | - | - | 0 | - |
| - | - | 8.941E+04 | 417.7 | - | - | 0 | - |
| - | - | 1.517E+05 | 418.2 | - | - | 0 | - |
| - | - | 8.082E+04 | 418.7 | - | - | 0 | - |
| - | - | 1.913E+04 | 419.2 | - | - | 0 | - |
| - | - | 3188 | 419.7 | - | - | 0 | - |
| - | - | 3593 | 420.2 | - | - | 0 | - |
| - | - | 3687 | 421.1 | - | - | 0 | - |
| 11 | y | 1.905E+04 | 421.2 | 0.003814 | 9.055 | +1 | 3 |
| - | - | 3539 | 422.2 | - | - | 0 | - |
| 11 | y | 1.493E+04 | 422.2 | 0.00338 | 8.006 | +1 | 3 |
| - | - | 4789 | 422.3 | - | - | 0 | - |
| - | - | 1.211E+04 | 422.7 | - | - | 0 | - |
| - | - | 3.367E+04 | 423.2 | - | - | 0 | - |
| - | - | 2.049E+05 | 423.3 | - | - | 0 | - |
| - | - | 1.204E+04 | 423.7 | - | - | 0 | - |
| - | - | 5264 | 423.7 | - | - | 0 | - |
| - | - | 1.038E+04 | 424.2 | - | - | 0 | - |
| - | - | 5.53E+04 | 424.3 | - | - | 0 | - |
| - | - | 1.268E+04 | 424.7 | - | - | 0 | - |
| - | - | 5746 | 425.2 | - | - | 0 | - |
| - | - | 5120 | 425.3 | - | - | 0 | - |
| - | - | 3866 | 426.2 | - | - | 0 | - |
| - | - | 1.448E+06 | 426.7 | - | - | 0 | - |
| - | - | 7.528E+05 | 427.2 | - | - | 0 | - |
| - | - | 2.135E+05 | 427.7 | - | - | 0 | - |
| - | - | 3.583E+04 | 428.2 | - | - | 0 | - |
| - | - | 2471 | 428.7 | - | - | 0 | - |
| - | - | 1.49E+04 | 429.2 | - | - | 0 | - |
| - | - | 6346 | 429.7 | - | - | 0 | - |
| - | - | 1.44E+06 | 430.2 | - | - | 0 | - |
| - | - | 3.034E+05 | 431.2 | - | - | 0 | - |
| 7 | b | 1.588E+05 | 431.7 | 0.003218 | 7.454 | +2 | 7 |
| 7 | b | 8.954E+04 | 432.2 | 0.00764 | 17.68 | +2 | 7 |
| - | - | 3.466E+04 | 432.7 | - | - | 0 | - |
| - | - | 9702 | 432.7 | - | - | 0 | - |
| - | - | 2.473E+04 | 433.2 | - | - | 0 | - |
| 4 | b | 5.18E+04 | 433.3 | 0.002399 | 5.536 | +1 | 4 |
| - | - | 4469 | 433.7 | - | - | 0 | - |
| - | - | 3659 | 433.7 | - | - | 0 | - |
| - | - | 4678 | 434.2 | - | - | 0 | - |
| - | - | 1.338E+04 | 434.3 | - | - | 0 | - |
| - | - | 3469 | 437.2 | - | - | 0 | - |
| - | - | 3966 | 437.2 | - | - | 0 | - |
| - | - | 2.145E+04 | 438.2 | - | - | 0 | - |
| - | - | 2971 | 438.2 | - | - | 0 | - |
| - | - | 1.358E+04 | 438.7 | - | - | 0 | - |
| 11 | y | 3.045E+05 | 439.2 | 0.003351 | 7.629 | +1 | 3 |
| - | - | 1.429E+04 | 440.2 | - | - | 0 | - |
| - | - | 8.202E+04 | 440.2 | - | - | 0 | - |
| 7 | b | 7.011E+05 | 440.7 | 0.003185 | 7.226 | +2 | 7 |
| - | - | 3.56E+05 | 441.2 | - | - | 0 | - |
| - | - | 1.018E+05 | 441.7 | - | - | 0 | - |
| - | - | 2.385E+04 | 442.2 | - | - | 0 | - |
| - | - | 7725 | 443.2 | - | - | 0 | - |
| - | - | 4978 | 444.2 | - | - | 0 | - |
| - | - | 2800 | 444.2 | - | - | 0 | - |
| - | - | 2511 | 446.2 | - | - | 0 | - |
| - | - | 6135 | 446.2 | - | - | 0 | - |
| - | - | 7142 | 446.2 | - | - | 0 | - |
| - | - | 9001 | 446.7 | - | - | 0 | - |
| 7 | y | 6.407E+04 | 447.2 | 0.003183 | 7.117 | +2 | 7 |
| - | - | 6690 | 447.2 | - | - | 0 | - |
| - | - | 3809 | 447.3 | - | - | 0 | - |
| - | - | 3.016E+04 | 447.7 | - | - | 0 | - |
| - | - | 7.389E+04 | 448.2 | - | - | 0 | - |
| - | - | 2.437E+04 | 449.2 | - | - | 0 | - |
| - | - | 1.077E+04 | 450.2 | - | - | 0 | - |
| - | - | 1.773E+04 | 450.2 | - | - | 0 | - |
| - | - | 2.014E+04 | 450.3 | - | - | 0 | - |
| 4 | b | 3.789E+05 | 451.3 | 0.003339 | 7.4 | +1 | 4 |
| - | - | 2697 | 451.8 | - | - | 0 | - |
| - | - | 9.445E+04 | 452.3 | - | - | 0 | - |
| - | - | 6770 | 452.7 | - | - | 0 | - |
| - | - | 1536 | 453.3 | - | - | 0 | - |
| - | - | 1.241E+04 | 453.3 | - | - | 0 | - |
| - | - | 4935 | 454.7 | - | - | 0 | - |
| - | - | 9194 | 455.2 | - | - | 0 | - |
| - | - | 2.313E+04 | 455.3 | - | - | 0 | - |
| 7 | y | 3.74E+04 | 455.7 | 0.003519 | 7.722 | +2 | 7 |
| - | - | 2.278E+04 | 456.2 | - | - | 0 | - |
| - | - | 7372 | 456.3 | - | - | 0 | - |
| - | - | 5755 | 456.7 | - | - | 0 | - |
| - | - | 5046 | 458.2 | - | - | 0 | - |
| - | - | 3.22E+04 | 459.2 | - | - | 0 | - |
| - | - | 3682 | 460.2 | - | - | 0 | - |
| - | - | 2864 | 460.2 | - | - | 0 | - |
| - | - | 2938 | 460.3 | - | - | 0 | - |
| - | - | 3.423E+04 | 460.8 | - | - | 0 | - |
| - | - | 3.994E+04 | 461.2 | - | - | 0 | - |
| - | - | 4.071E+04 | 461.3 | - | - | 0 | - |
| - | - | 9041 | 461.8 | - | - | 0 | - |
| - | - | 1.192E+04 | 462.2 | - | - | 0 | - |
| - | - | 4074 | 462.2 | - | - | 0 | - |
| 3 | y | 7067 | 462.6 | 0.002462 | 5.322 | +3 | 11 |
| - | - | 1.559E+04 | 464.2 | - | - | 0 | - |
| - | - | 9549 | 464.3 | - | - | 0 | - |
| - | - | 9139 | 465.3 | - | - | 0 | - |
| - | - | 9836 | 465.8 | - | - | 0 | - |
| - | - | 5307 | 466.2 | - | - | 0 | - |
| - | - | 1.291E+04 | 466.2 | - | - | 0 | - |
| - | - | 3708 | 466.7 | - | - | 0 | - |
| - | - | 3208 | 468.2 | - | - | 0 | - |
| - | - | 9.22E+04 | 468.3 | - | - | 0 | - |
| 12 | b | 3011 | 468.9 | 0.0002723 | 0.5808 | +3 | 12 |
| - | - | 3302 | 469.2 | - | - | 0 | - |
| - | - | 2.093E+04 | 469.3 | - | - | 0 | - |
| - | - | 7648 | 472.2 | - | - | 0 | - |
| - | - | 5628 | 473.2 | - | - | 0 | - |
| - | - | 2855 | 474.2 | - | - | 0 | - |
| - | - | 2102 | 474.2 | - | - | 0 | - |
| - | - | 7.104E+04 | 474.3 | - | - | 0 | - |
| - | - | 8.72E+04 | 474.8 | - | - | 0 | - |
| 12 | b | 5091 | 474.9 | 0.003805 | 8.012 | +3 | 12 |
| - | - | 4.781E+04 | 475.3 | - | - | 0 | - |
| - | - | 1.33E+04 | 475.8 | - | - | 0 | - |
| - | - | 3.074E+04 | 476.2 | - | - | 0 | - |
| - | - | 6.384E+04 | 477.2 | - | - | 0 | - |
| - | - | 7620 | 477.7 | - | - | 0 | - |
| 10 | y | 6.889E+04 | 478.2 | 0.002126 | 4.445 | +1 | 4 |
| - | - | 4.842E+04 | 478.3 | - | - | 0 | - |
| - | - | 1.659E+04 | 479.2 | - | - | 0 | - |
| - | - | 8147 | 479.3 | - | - | 0 | - |
| - | - | 8842 | 479.7 | - | - | 0 | - |
| - | - | 1.314E+04 | 480.2 | - | - | 0 | - |
| - | - | 3014 | 481.2 | - | - | 0 | - |
| - | - | 3909 | 481.7 | - | - | 0 | - |
| - | - | 3268 | 482.3 | - | - | 0 | - |
| - | - | 2823 | 482.7 | - | - | 0 | - |
| - | - | 2.37E+05 | 483.3 | - | - | 0 | - |
| - | - | 1.525E+05 | 483.8 | - | - | 0 | - |
| - | - | 1.493E+04 | 484.2 | - | - | 0 | - |
| - | - | 4.441E+04 | 484.3 | - | - | 0 | - |
| - | - | 1.439E+04 | 484.8 | - | - | 0 | - |
| - | - | 3252 | 485.2 | - | - | 0 | - |
| - | - | 4023 | 486.2 | - | - | 0 | - |
| - | - | 3762 | 487.2 | - | - | 0 | - |
| - | - | 4526 | 488.2 | - | - | 0 | - |
| 8 | b | 7.804E+04 | 488.3 | 0.004033 | 8.26 | +2 | 8 |
| - | - | 5.021E+04 | 488.8 | - | - | 0 | - |
| - | - | 2.077E+04 | 489.3 | - | - | 0 | - |
| - | - | 7054 | 489.8 | - | - | 0 | - |
| - | - | 2819 | 491.3 | - | - | 0 | - |
| - | - | 5084 | 493.3 | - | - | 0 | - |
| - | - | 7.574E+05 | 494.2 | - | - | 0 | - |
| - | - | 4949 | 495.2 | - | - | 0 | - |
| - | - | 2.031E+05 | 495.2 | - | - | 0 | - |
| 10 | y | 1.092E+06 | 496.2 | 0.003707 | 7.47 | +1 | 4 |
| - | - | 2.712E+05 | 497.2 | - | - | 0 | - |
| - | - | 8.074E+04 | 497.8 | - | - | 0 | - |
| - | - | 4.288E+04 | 498.2 | - | - | 0 | - |
| - | - | 4537 | 498.8 | - | - | 0 | - |
| - | - | 1.176E+04 | 499.2 | - | - | 0 | - |
| - | - | 2540 | 499.7 | - | - | 0 | - |
| - | - | 8004 | 502.2 | - | - | 0 | - |
| - | - | 8903 | 503.1 | - | - | 0 | - |
| - | - | 4222 | 503.2 | - | - | 0 | - |
| - | - | 3.655E+04 | 504.2 | - | - | 0 | - |
| - | - | 9194 | 505.2 | - | - | 0 | - |
| - | - | 3371 | 505.2 | - | - | 0 | - |
| - | - | 2.152E+04 | 506.2 | - | - | 0 | - |
| - | - | 9141 | 506.3 | - | - | 0 | - |
| - | - | 1.578E+04 | 507.3 | - | - | 0 | - |
| - | - | 5414 | 508.2 | - | - | 0 | - |
| - | - | 5.621E+04 | 511.3 | - | - | 0 | - |
| - | - | 6025 | 512.2 | - | - | 0 | - |
| - | - | 1.403E+04 | 512.3 | - | - | 0 | - |
| - | - | 8368 | 513.2 | - | - | 0 | - |
| - | - | 1E+04 | 514.2 | - | - | 0 | - |
| - | - | 2707 | 514.3 | - | - | 0 | - |
| - | - | 2671 | 515.2 | - | - | 0 | - |
| - | - | 1.696E+04 | 515.3 | - | - | 0 | - |
| - | - | 3615 | 516.2 | - | - | 0 | - |
| - | - | 5522 | 516.3 | - | - | 0 | - |
| - | - | 5471 | 519.3 | - | - | 0 | - |
| - | - | 3424 | 520.3 | - | - | 0 | - |
| - | - | 6760 | 521.2 | - | - | 0 | - |
| - | - | 3020 | 521.3 | - | - | 0 | - |
| - | - | 1.333E+04 | 522.2 | - | - | 0 | - |
| - | - | 1.997E+04 | 523.3 | - | - | 0 | - |
| - | - | 9411 | 523.6 | - | - | 0 | - |
| - | - | 6839 | 523.9 | - | - | 0 | - |
| - | - | 6631 | 524.3 | - | - | 0 | - |
| - | - | 1.276E+05 | 525.3 | - | - | 0 | - |
| - | - | 2.193E+04 | 526.2 | - | - | 0 | - |
| - | - | 3.534E+04 | 526.3 | - | - | 0 | - |
| - | - | 7646 | 527.2 | - | - | 0 | - |
| - | - | 6885 | 527.3 | - | - | 0 | - |
| - | - | 8587 | 528.7 | - | - | 0 | - |
| 0 | Precursor | 3.381E+04 | 529.3 | 0.004221 | 7.974 | +3 | -1 |
| 0 | Precursor | 5.119E+04 | 529.6 | 0.008979 | 16.95 | +3 | -1 |
| - | - | 3789 | 529.8 | - | - | 0 | - |
| - | - | 2.863E+04 | 529.9 | - | - | 0 | - |
| - | - | 9645 | 530.2 | - | - | 0 | - |
| - | - | 8613 | 530.3 | - | - | 0 | - |
| - | - | 8397 | 531.2 | - | - | 0 | - |
| - | - | 3.211E+04 | 532.2 | - | - | 0 | - |
| - | - | 2519 | 532.2 | - | - | 0 | - |
| - | - | 1.079E+04 | 533.2 | - | - | 0 | - |
| - | - | 7488 | 533.3 | - | - | 0 | - |
| - | - | 6266 | 533.3 | - | - | 0 | - |
| - | - | 5269 | 534.2 | - | - | 0 | - |
| - | - | 4371 | 534.3 | - | - | 0 | - |
| 0 | Precursor | 4.196E+04 | 535.3 | 0.003385 | 6.323 | +3 | -1 |
| - | - | 4177 | 535.3 | - | - | 0 | - |
| - | - | 3.984E+04 | 535.6 | - | - | 0 | - |
| - | - | 2.173E+04 | 535.9 | - | - | 0 | - |
| - | - | 6668 | 536.3 | - | - | 0 | - |
| - | - | 6916 | 537.2 | - | - | 0 | - |
| - | - | 2.102E+04 | 537.7 | - | - | 0 | - |
| - | - | 9843 | 538.2 | - | - | 0 | - |
| - | - | 1.574E+04 | 538.3 | - | - | 0 | - |
| - | - | 1.285E+04 | 538.8 | - | - | 0 | - |
| - | - | 1.449E+05 | 539.2 | - | - | 0 | - |
| - | - | 7110 | 539.3 | - | - | 0 | - |
| - | - | 4.263E+04 | 540.2 | - | - | 0 | - |
| - | - | 4922 | 540.8 | - | - | 0 | - |
| - | - | 1.265E+04 | 541.3 | - | - | 0 | - |
| - | - | 5878 | 541.8 | - | - | 0 | - |
| - | - | 2876 | 542.3 | - | - | 0 | - |
| - | - | 3.3E+05 | 543.3 | - | - | 0 | - |
| - | - | 9.667E+04 | 544.3 | - | - | 0 | - |
| - | - | 2.349E+04 | 545.3 | - | - | 0 | - |
| 9 | b | 1.06E+04 | 545.8 | 0.002158 | 3.954 | +2 | 9 |
| 9 | b | 3.405E+04 | 546.3 | 0.00755 | 13.82 | +2 | 9 |
| - | - | 2.266E+04 | 546.8 | - | - | 0 | - |
| - | - | 9513 | 547.3 | - | - | 0 | - |
| - | - | 1.044E+04 | 547.3 | - | - | 0 | - |
| - | - | 1.175E+04 | 547.8 | - | - | 0 | - |
| - | - | 5847 | 548.2 | - | - | 0 | - |
| - | - | 3414 | 548.3 | - | - | 0 | - |
| - | - | 3.269E+04 | 549.2 | - | - | 0 | - |
| - | - | 1.69E+04 | 550.2 | - | - | 0 | - |
| - | - | 4877 | 551.2 | - | - | 0 | - |
| - | - | 2.839E+04 | 551.3 | - | - | 0 | - |
| - | - | 1.925E+04 | 551.3 | - | - | 0 | - |
| - | - | 1.555E+04 | 552.3 | - | - | 0 | - |
| - | - | 4438 | 552.3 | - | - | 0 | - |
| - | - | 6065 | 553.3 | - | - | 0 | - |
| 9 | b | 4.883E+04 | 554.8 | 0.003651 | 6.58 | +2 | 9 |
| - | - | 7.295E+04 | 555.3 | - | - | 0 | - |
| - | - | 5.529E+04 | 555.8 | - | - | 0 | - |
| - | - | 1.461E+04 | 556.3 | - | - | 0 | - |
| - | - | 3334 | 556.8 | - | - | 0 | - |
| - | - | 3197 | 557.2 | - | - | 0 | - |
| - | - | 4364 | 558.3 | - | - | 0 | - |
| - | - | 2614 | 559.8 | - | - | 0 | - |
| - | - | 7121 | 560.3 | - | - | 0 | - |
| 5 | b | 2.784E+04 | 561.3 | 0.003208 | 5.716 | +1 | 5 |
| 5 | b | 1.182E+04 | 562.3 | 0.007047 | 12.53 | +1 | 5 |
| - | - | 2.39E+04 | 563.3 | - | - | 0 | - |
| - | - | 3236 | 563.3 | - | - | 0 | - |
| - | - | 1.2E+04 | 564.3 | - | - | 0 | - |
| - | - | 9851 | 564.3 | - | - | 0 | - |
| - | - | 5453 | 564.8 | - | - | 0 | - |
| - | - | 4.105E+04 | 565.3 | - | - | 0 | - |
| - | - | 1.221E+04 | 566.3 | - | - | 0 | - |
| - | - | 1.426E+05 | 567.2 | - | - | 0 | - |
| - | - | 4.032E+04 | 568.2 | - | - | 0 | - |
| - | - | 9964 | 568.8 | - | - | 0 | - |
| - | - | 7940 | 569.2 | - | - | 0 | - |
| - | - | 4028 | 569.3 | - | - | 0 | - |
| - | - | 3263 | 569.8 | - | - | 0 | - |
| - | - | 2445 | 570.2 | - | - | 0 | - |
| - | - | 2.079E+04 | 575.3 | - | - | 0 | - |
| - | - | 6721 | 575.3 | - | - | 0 | - |
| - | - | 1.077E+04 | 576.2 | - | - | 0 | - |
| - | - | 8326 | 576.3 | - | - | 0 | - |
| - | - | 6627 | 577.2 | - | - | 0 | - |
| 5 | y | 1.145E+04 | 577.3 | 0.007146 | 12.38 | +2 | 9 |
| - | - | 9449 | 577.8 | - | - | 0 | - |
| - | - | 5230 | 578.3 | - | - | 0 | - |
| - | - | 4807 | 578.3 | - | - | 0 | - |
| 5 | b | 2.05E+05 | 579.3 | 0.004118 | 7.109 | +1 | 5 |
| - | - | 6.454E+04 | 580.3 | - | - | 0 | - |
| - | - | 5381 | 581.3 | - | - | 0 | - |
| - | - | 1.387E+04 | 581.3 | - | - | 0 | - |
| - | - | 6155 | 582.3 | - | - | 0 | - |
| 10 | b | 8297 | 583.3 | 0.005675 | 9.729 | +2 | 10 |
| - | - | 3151 | 585.3 | - | - | 0 | - |
| - | - | 4233 | 586.3 | - | - | 0 | - |
| - | - | 8625 | 587.3 | - | - | 0 | - |
| - | - | 4279 | 588.8 | - | - | 0 | - |
| - | - | 8961 | 589.3 | - | - | 0 | - |
| - | - | 2512 | 589.8 | - | - | 0 | - |
| - | - | 3864 | 592.3 | - | - | 0 | - |
| - | - | 2.547E+04 | 592.3 | - | - | 0 | - |
| 9 | y | 2.097E+05 | 593.3 | 0.003747 | 6.316 | +1 | 5 |
| - | - | 9.715E+04 | 593.3 | - | - | 0 | - |
| - | - | 8717 | 593.8 | - | - | 0 | - |
| - | - | 6.131E+04 | 594.3 | - | - | 0 | - |
| - | - | 2.517E+04 | 594.3 | - | - | 0 | - |
| - | - | 1.228E+04 | 594.8 | - | - | 0 | - |
| - | - | 1.893E+04 | 595.3 | - | - | 0 | - |
| - | - | 4823 | 596.3 | - | - | 0 | - |
| - | - | 9620 | 596.4 | - | - | 0 | - |
| - | - | 1.622E+04 | 597.8 | - | - | 0 | - |
| - | - | 1.4E+04 | 598.3 | - | - | 0 | - |
| - | - | 4486 | 598.8 | - | - | 0 | - |
| - | - | 2563 | 600.3 | - | - | 0 | - |
| - | - | 2.476E+04 | 600.3 | - | - | 0 | - |
| - | - | 9465 | 601.3 | - | - | 0 | - |
| - | - | 5.136E+04 | 602.8 | - | - | 0 | - |
| - | - | 6.202E+04 | 603.3 | - | - | 0 | - |
| - | - | 2.185E+04 | 603.8 | - | - | 0 | - |
| - | - | 1.228E+04 | 604.3 | - | - | 0 | - |
| - | - | 2995 | 605.3 | - | - | 0 | - |
| - | - | 3181 | 607.3 | - | - | 0 | - |
| - | - | 3054 | 609.4 | - | - | 0 | - |
| - | - | 3.561E+04 | 610.3 | - | - | 0 | - |
| - | - | 1.979E+04 | 610.3 | - | - | 0 | - |
| 9 | y | 2.833E+06 | 611.3 | 0.00423 | 6.92 | +1 | 5 |
| - | - | 1.843E+05 | 611.8 | - | - | 0 | - |
| - | - | 8.901E+05 | 612.3 | - | - | 0 | - |
| - | - | 5.199E+04 | 612.8 | - | - | 0 | - |
| - | - | 2.043E+05 | 613.3 | - | - | 0 | - |
| - | - | 2815 | 613.8 | - | - | 0 | - |
| - | - | 2.964E+04 | 614.3 | - | - | 0 | - |
| - | - | 2755 | 615.3 | - | - | 0 | - |
| - | - | 3509 | 616.3 | - | - | 0 | - |
| - | - | 1.421E+04 | 617.3 | - | - | 0 | - |
| - | - | 2389 | 618.3 | - | - | 0 | - |
| - | - | 3095 | 619.3 | - | - | 0 | - |
| - | - | 7037 | 619.3 | - | - | 0 | - |
| - | - | 3819 | 619.8 | - | - | 0 | - |
| - | - | 2492 | 620.3 | - | - | 0 | - |
| - | - | 1.466E+04 | 620.8 | - | - | 0 | - |
| - | - | 1.859E+04 | 621.3 | - | - | 0 | - |
| - | - | 2.399E+04 | 621.3 | - | - | 0 | - |
| - | - | 3392 | 621.8 | - | - | 0 | - |
| - | - | 1.009E+04 | 622.3 | - | - | 0 | - |
| - | - | 4150 | 622.3 | - | - | 0 | - |
| - | - | 8460 | 622.4 | - | - | 0 | - |
| - | - | 2875 | 622.8 | - | - | 0 | - |
| - | - | 5098 | 623.3 | - | - | 0 | - |
| - | - | 3103 | 626.3 | - | - | 0 | - |
| - | - | 3310 | 627.3 | - | - | 0 | - |
| - | - | 2944 | 627.8 | - | - | 0 | - |
| - | - | 1.299E+04 | 628.3 | - | - | 0 | - |
| - | - | 9476 | 628.8 | - | - | 0 | - |
| - | - | 5980 | 630.3 | - | - | 0 | - |
| - | - | 2.961E+04 | 634.3 | - | - | 0 | - |
| - | - | 4.244E+04 | 635.3 | - | - | 0 | - |
| - | - | 2.58E+04 | 636.3 | - | - | 0 | - |
| 4 | y | 3.224E+04 | 636.8 | 0.004762 | 7.477 | +2 | 10 |
| - | - | 2.577E+04 | 637.3 | - | - | 0 | - |
| - | - | 8581 | 637.3 | - | - | 0 | - |
| - | - | 1.166E+04 | 637.8 | - | - | 0 | - |
| 11 | b | 7674 | 638.3 | 0.006872 | 10.77 | +2 | 11 |
| 11 | b | 5999 | 638.8 | 0.009615 | 15.05 | +2 | 11 |
| - | - | 3.525E+04 | 640.3 | - | - | 0 | - |
| - | - | 1.055E+04 | 641.3 | - | - | 0 | - |
| - | - | 3875 | 642.3 | - | - | 0 | - |
| - | - | 5036 | 644.3 | - | - | 0 | - |
| - | - | 2.025E+04 | 645.3 | - | - | 0 | - |
| 4 | y | 1.815E+05 | 645.8 | 0.004423 | 6.849 | +2 | 10 |
| - | - | 1.401E+05 | 646.3 | - | - | 0 | - |
| - | - | 5.945E+04 | 646.8 | - | - | 0 | - |
| - | - | 1.17E+04 | 647.3 | - | - | 0 | - |
| 11 | b | 5840 | 647.3 | 0.007815 | 12.07 | +2 | 11 |
| - | - | 2957 | 647.4 | - | - | 0 | - |
| - | - | 9726 | 647.8 | - | - | 0 | - |
| - | - | 1.057E+04 | 648.3 | - | - | 0 | - |
| - | - | 6829 | 649.3 | - | - | 0 | - |
| - | - | 1.137E+04 | 650.3 | - | - | 0 | - |
| - | - | 1.059E+04 | 650.3 | - | - | 0 | - |
| - | - | 1.088E+04 | 650.8 | - | - | 0 | - |
| - | - | 9071 | 651.3 | - | - | 0 | - |
| - | - | 3085 | 651.3 | - | - | 0 | - |
| - | - | 2.512E+05 | 652.3 | - | - | 0 | - |
| - | - | 6199 | 652.8 | - | - | 0 | - |
| - | - | 9.079E+04 | 653.3 | - | - | 0 | - |
| - | - | 1.95E+04 | 654.3 | - | - | 0 | - |
| - | - | 2840 | 655.3 | - | - | 0 | - |
| - | - | 2683 | 657.9 | - | - | 0 | - |
| - | - | 2.731E+04 | 658.3 | - | - | 0 | - |
| - | - | 5643 | 658.3 | - | - | 0 | - |
| - | - | 9850 | 659.3 | - | - | 0 | - |
| - | - | 6154 | 660.3 | - | - | 0 | - |
| - | - | 2.421E+04 | 661.3 | - | - | 0 | - |
| - | - | 1.382E+04 | 661.8 | - | - | 0 | - |
| - | - | 9.142E+04 | 662.3 | - | - | 0 | - |
| - | - | 2799 | 662.8 | - | - | 0 | - |
| - | - | 6.088E+04 | 663.3 | - | - | 0 | - |
| - | - | 1.723E+04 | 664.3 | - | - | 0 | - |
| - | - | 1.175E+04 | 664.3 | - | - | 0 | - |
| - | - | 6661 | 665.3 | - | - | 0 | - |
| - | - | 5990 | 666.3 | - | - | 0 | - |
| - | - | 5658 | 666.4 | - | - | 0 | - |
| - | - | 8743 | 668.3 | - | - | 0 | - |
| - | - | 3904 | 669.3 | - | - | 0 | - |
| - | - | 3540 | 670.8 | - | - | 0 | - |
| - | - | 3279 | 671.3 | - | - | 0 | - |
| 6 | b | 1.236E+05 | 676.3 | 0.004342 | 6.419 | +1 | 6 |
| 6 | b | 7.249E+04 | 677.3 | 0.0127 | 18.75 | +1 | 6 |
| - | - | 3.482E+04 | 678.3 | - | - | 0 | - |
| - | - | 2.722E+04 | 679.3 | - | - | 0 | - |
| - | - | 1.203E+04 | 679.8 | - | - | 0 | - |
| - | - | 4.594E+05 | 680.3 | - | - | 0 | - |
| - | - | 1.822E+05 | 681.3 | - | - | 0 | - |
| - | - | 3.923E+04 | 682.3 | - | - | 0 | - |
| - | - | 6034 | 683.3 | - | - | 0 | - |
| - | - | 1.05E+04 | 684.3 | - | - | 0 | - |
| - | - | 3.615E+04 | 684.8 | - | - | 0 | - |
| - | - | 3.219E+04 | 685.3 | - | - | 0 | - |
| - | - | 1.073E+04 | 685.8 | - | - | 0 | - |
| - | - | 7098 | 686.3 | - | - | 0 | - |
| - | - | 7823 | 688.3 | - | - | 0 | - |
| - | - | 4080 | 689.3 | - | - | 0 | - |
| - | - | 6383 | 690.3 | - | - | 0 | - |
| - | - | 3497 | 692.3 | - | - | 0 | - |
| - | - | 2811 | 692.4 | - | - | 0 | - |
| 3 | y | 1.096E+05 | 693.3 | 0.004844 | 6.986 | +2 | 11 |
| 3 | y | 1.423E+05 | 693.8 | 0.01155 | 16.65 | +2 | 11 |
| 6 | b | 1.208E+06 | 694.4 | 0.003848 | 5.541 | +1 | 6 |
| - | - | 2.799E+04 | 694.8 | - | - | 0 | - |
| - | - | 4.761E+05 | 695.4 | - | - | 0 | - |
| - | - | 1.049E+05 | 696.4 | - | - | 0 | - |
| - | - | 1.662E+04 | 697.4 | - | - | 0 | - |
| - | - | 7669 | 697.9 | - | - | 0 | - |
| - | - | 2918 | 698.3 | - | - | 0 | - |
| - | - | 7771 | 698.4 | - | - | 0 | - |
| - | - | 3189 | 698.9 | - | - | 0 | - |
| - | - | 3499 | 701.4 | - | - | 0 | - |
| - | - | 1.94E+04 | 701.8 | - | - | 0 | - |
| 3 | y | 1.102E+06 | 702.3 | 0.004749 | 6.762 | +2 | 11 |
| 12 | b | 9.362E+05 | 702.8 | 0.009237 | 13.14 | +2 | 12 |
| 12 | b | 4.454E+05 | 703.3 | 9.826E-05 | 0.1397 | +2 | 12 |
| - | - | 1.228E+05 | 703.8 | - | - | 0 | - |
| - | - | 4.578E+04 | 704.3 | - | - | 0 | - |
| - | - | 5984 | 705.4 | - | - | 0 | - |
| 8 | y | 6.916E+04 | 706.3 | 0.004827 | 6.834 | +1 | 6 |
| - | - | 3.127E+04 | 707.3 | - | - | 0 | - |
| - | - | 1.143E+04 | 708.3 | - | - | 0 | - |
| - | - | 2884 | 709.3 | - | - | 0 | - |
| - | - | 2.555E+04 | 711.4 | - | - | 0 | - |
| 12 | b | 8.846E+04 | 711.9 | 0.004707 | 6.612 | +2 | 12 |
| - | - | 7.701E+04 | 712.4 | - | - | 0 | - |
| - | - | 2.747E+04 | 712.9 | - | - | 0 | - |
| - | - | 1.087E+04 | 713.4 | - | - | 0 | - |
| - | - | 3805 | 714.3 | - | - | 0 | - |
| - | - | 4992 | 715.3 | - | - | 0 | - |
| - | - | 3450 | 715.4 | - | - | 0 | - |
| - | - | 5598 | 716.3 | - | - | 0 | - |
| - | - | 4284 | 717.3 | - | - | 0 | - |
| - | - | 5327 | 718.4 | - | - | 0 | - |
| - | - | 9060 | 720.4 | - | - | 0 | - |
| - | - | 4763 | 720.9 | - | - | 0 | - |
| - | - | 5.59E+04 | 721.4 | - | - | 0 | - |
| - | - | 1.509E+04 | 722.4 | - | - | 0 | - |
| - | - | 1.882E+04 | 723.4 | - | - | 0 | - |
| 8 | y | 8.3E+05 | 724.4 | 0.004699 | 6.488 | +1 | 6 |
| - | - | 3.356E+05 | 725.4 | - | - | 0 | - |
| - | - | 9.719E+04 | 726.4 | - | - | 0 | - |
| - | - | 1.727E+04 | 727.4 | - | - | 0 | - |
| - | - | 4580 | 728.9 | - | - | 0 | - |
| - | - | 3.995E+04 | 729.4 | - | - | 0 | - |
| - | - | 2.02E+04 | 730.4 | - | - | 0 | - |
| - | - | 6465 | 731.4 | - | - | 0 | - |
| - | - | 3786 | 732.3 | - | - | 0 | - |
| - | - | 2.741E+04 | 733.4 | - | - | 0 | - |
| - | - | 2.651E+04 | 734.4 | - | - | 0 | - |
| - | - | 1.049E+04 | 735.4 | - | - | 0 | - |
| - | - | 3595 | 736.4 | - | - | 0 | - |
| - | - | 3728 | 737.3 | - | - | 0 | - |
| - | - | 1.409E+04 | 737.9 | - | - | 0 | - |
| - | - | 8025 | 738.4 | - | - | 0 | - |
| - | - | 6463 | 739.4 | - | - | 0 | - |
| 2 | y | 1.156E+04 | 742.9 | 0.003779 | 5.087 | +2 | 12 |
| - | - | 2.211E+04 | 743.4 | - | - | 0 | - |
| - | - | 7069 | 743.9 | - | - | 0 | - |
| - | - | 7972 | 744.4 | - | - | 0 | - |
| - | - | 6119 | 745.4 | - | - | 0 | - |
| - | - | 2540 | 746.3 | - | - | 0 | - |
| - | - | 1.472E+04 | 747.4 | - | - | 0 | - |
| - | - | 1.531E+04 | 748.4 | - | - | 0 | - |
| - | - | 1.724E+04 | 749.3 | - | - | 0 | - |
| - | - | 6371 | 750.3 | - | - | 0 | - |
| - | - | 5325 | 750.4 | - | - | 0 | - |
| - | - | 1.187E+05 | 751.4 | - | - | 0 | - |
| 2 | y | 7.837E+04 | 751.9 | 0.004356 | 5.794 | +2 | 12 |
| - | - | 1.354E+05 | 752.4 | - | - | 0 | - |
| - | - | 3.959E+04 | 752.9 | - | - | 0 | - |
| - | - | 2.811E+04 | 753.4 | - | - | 0 | - |
| - | - | 3827 | 756.9 | - | - | 0 | - |
| - | - | 4125 | 757.4 | - | - | 0 | - |
| - | - | 5766 | 758.4 | - | - | 0 | - |
| - | - | 1.172E+04 | 760.3 | - | - | 0 | - |
| - | - | 5430 | 761.3 | - | - | 0 | - |
| - | - | 8803 | 761.4 | - | - | 0 | - |
| - | - | 1.152E+04 | 762.4 | - | - | 0 | - |
| - | - | 7831 | 762.4 | - | - | 0 | - |
| - | - | 6.653E+04 | 763.4 | - | - | 0 | - |
| - | - | 2.536E+04 | 764.4 | - | - | 0 | - |
| - | - | 2.941E+04 | 765.4 | - | - | 0 | - |
| - | - | 1.522E+04 | 766.4 | - | - | 0 | - |
| - | - | 2.548E+04 | 767.4 | - | - | 0 | - |
| - | - | 3412 | 767.4 | - | - | 0 | - |
| - | - | 8783 | 768.4 | - | - | 0 | - |
| - | - | 3710 | 768.4 | - | - | 0 | - |
| - | - | 1.299E+04 | 769.4 | - | - | 0 | - |
| - | - | 5135 | 770.4 | - | - | 0 | - |
| - | - | 1.342E+04 | 775.4 | - | - | 0 | - |
| - | - | 1.048E+04 | 776.4 | - | - | 0 | - |
| - | - | 5064 | 777.3 | - | - | 0 | - |
| - | - | 3185 | 777.4 | - | - | 0 | - |
| - | - | 5007 | 778.3 | - | - | 0 | - |
| - | - | 3514 | 778.4 | - | - | 0 | - |
| - | - | 6.418E+04 | 779.4 | - | - | 0 | - |
| - | - | 2.945E+04 | 780.4 | - | - | 0 | - |
| - | - | 3.318E+04 | 781.4 | - | - | 0 | - |
| - | - | 1.61E+04 | 782.4 | - | - | 0 | - |
| - | - | 3840 | 783.4 | - | - | 0 | - |
| - | - | 3817 | 790.4 | - | - | 0 | - |
| - | - | 3456 | 791.4 | - | - | 0 | - |
| - | - | 3575 | 792.4 | - | - | 0 | - |
| - | - | 1.141E+05 | 793.4 | - | - | 0 | - |
| - | - | 5.129E+04 | 794.4 | - | - | 0 | - |
| - | - | 5.428E+04 | 795.3 | - | - | 0 | - |
| - | - | 8202 | 795.4 | - | - | 0 | - |
| - | - | 2.823E+04 | 796.3 | - | - | 0 | - |
| - | - | 3347 | 796.4 | - | - | 0 | - |
| - | - | 9492 | 797.4 | - | - | 0 | - |
| - | - | 2.073E+04 | 807.4 | - | - | 0 | - |
| - | - | 1.569E+04 | 807.4 | - | - | 0 | - |
| - | - | 1.481E+04 | 808.4 | - | - | 0 | - |
| - | - | 5164 | 809.4 | - | - | 0 | - |
| - | - | 3834 | 817.3 | - | - | 0 | - |
| - | - | 5692 | 817.4 | - | - | 0 | - |
| - | - | 1.203E+04 | 818.4 | - | - | 0 | - |
| - | - | 1.459E+04 | 819.4 | - | - | 0 | - |
| - | - | 8283 | 820.4 | - | - | 0 | - |
| - | - | 3386 | 824.4 | - | - | 0 | - |
| - | - | 9182 | 825.4 | - | - | 0 | - |
| - | - | 2.149E+04 | 826.4 | - | - | 0 | - |
| - | - | 1.04E+04 | 827.4 | - | - | 0 | - |
| - | - | 3877 | 829.4 | - | - | 0 | - |
| - | - | 3816 | 834.4 | - | - | 0 | - |
| - | - | 2.905E+04 | 834.4 | - | - | 0 | - |
| - | - | 3.412E+04 | 835.4 | - | - | 0 | - |
| - | - | 7.158E+04 | 836.5 | - | - | 0 | - |
| - | - | 3.457E+04 | 837.5 | - | - | 0 | - |
| - | - | 1.082E+04 | 838.5 | - | - | 0 | - |
| - | - | 1.416E+04 | 843.4 | - | - | 0 | - |
| - | - | 3.816E+04 | 844.4 | - | - | 0 | - |
| - | - | 2.788E+04 | 845.4 | - | - | 0 | - |
| - | - | 3.915E+04 | 846.5 | - | - | 0 | - |
| - | - | 2.346E+04 | 847.4 | - | - | 0 | - |
| - | - | 1.061E+04 | 848.5 | - | - | 0 | - |
| - | - | 2.576E+05 | 852.4 | - | - | 0 | - |
| - | - | 1.253E+05 | 853.4 | - | - | 0 | - |
| - | - | 3.751E+04 | 854.4 | - | - | 0 | - |
| - | - | 6490 | 855.5 | - | - | 0 | - |
| 7 | b | 1.511E+05 | 862.4 | 0.004679 | 5.426 | +1 | 7 |
| - | - | 1.171E+05 | 863.4 | - | - | 0 | - |
| - | - | 1.976E+05 | 864.5 | - | - | 0 | - |
| - | - | 1.025E+05 | 865.5 | - | - | 0 | - |
| - | - | 2.562E+04 | 866.5 | - | - | 0 | - |
| - | - | 5033 | 867.5 | - | - | 0 | - |
| - | - | 9335 | 869.4 | - | - | 0 | - |
| - | - | 4501 | 870.4 | - | - | 0 | - |
| - | - | 3583 | 873.4 | - | - | 0 | - |
| - | - | 2897 | 875.4 | - | - | 0 | - |
| - | - | 2.053E+04 | 876.4 | - | - | 0 | - |
| - | - | 9618 | 877.4 | - | - | 0 | - |
| 7 | b | 1.232E+06 | 880.4 | 0.004796 | 5.447 | +1 | 7 |
| - | - | 6.599E+05 | 881.4 | - | - | 0 | - |
| - | - | 1.937E+05 | 882.4 | - | - | 0 | - |
| - | - | 3.661E+04 | 883.4 | - | - | 0 | - |
| - | - | 3584 | 884.5 | - | - | 0 | - |
| - | - | 1.349E+04 | 890.4 | - | - | 0 | - |
| - | - | 2.236E+04 | 891.4 | - | - | 0 | - |
| 7 | y | 8128 | 892.4 | 0.009666 | 10.83 | +1 | 7 |
| - | - | 2.018E+04 | 892.5 | - | - | 0 | - |
| 7 | y | 5475 | 893.4 | 0.002107 | 2.358 | +1 | 7 |
| - | - | 7506 | 893.5 | - | - | 0 | - |
| - | - | 8485 | 894.5 | - | - | 0 | - |
| - | - | 4126 | 895.5 | - | - | 0 | - |
| - | - | 1.487E+05 | 908.4 | - | - | 0 | - |
| - | - | 8.685E+04 | 909.4 | - | - | 0 | - |
| 7 | y | 1.316E+05 | 910.4 | 0.005709 | 6.27 | +1 | 7 |
| - | - | 6.347E+04 | 911.4 | - | - | 0 | - |
| - | - | 1.708E+04 | 912.4 | - | - | 0 | - |
| - | - | 3565 | 913.4 | - | - | 0 | - |
| - | - | 5556 | 927.4 | - | - | 0 | - |
| - | - | 9713 | 936.4 | - | - | 0 | - |
| - | - | 1.772E+04 | 937.4 | - | - | 0 | - |
| - | - | 7650 | 938.4 | - | - | 0 | - |
| - | - | 4038 | 944.5 | - | - | 0 | - |
| - | - | 5444 | 947.4 | - | - | 0 | - |
| - | - | 8812 | 948.4 | - | - | 0 | - |
| - | - | 3684 | 949.4 | - | - | 0 | - |
| - | - | 2.269E+04 | 954.4 | - | - | 0 | - |
| - | - | 4.045E+04 | 955.4 | - | - | 0 | - |
| - | - | 2.244E+04 | 956.4 | - | - | 0 | - |
| - | - | 6840 | 957.4 | - | - | 0 | - |
| - | - | 4294 | 962.5 | - | - | 0 | - |
| - | - | 3019 | 963.4 | - | - | 0 | - |
| - | - | 5839 | 964.5 | - | - | 0 | - |
| - | - | 4.173E+04 | 965.5 | - | - | 0 | - |
| - | - | 2.605E+04 | 966.5 | - | - | 0 | - |
| - | - | 7918 | 967.5 | - | - | 0 | - |
| - | - | 3.135E+04 | 972.4 | - | - | 0 | - |
| - | - | 1.902E+04 | 973.4 | - | - | 0 | - |
| - | - | 6175 | 974.4 | - | - | 0 | - |
| 8 | b | 1.149E+04 | 975.5 | 0.001609 | 1.65 | +1 | 8 |
| - | - | 9220 | 976.5 | - | - | 0 | - |
| - | - | 3652 | 979.5 | - | - | 0 | - |
| - | - | 1.708E+04 | 980.5 | - | - | 0 | - |
| - | - | 5692 | 981.5 | - | - | 0 | - |
| - | - | 1.627E+04 | 982.5 | - | - | 0 | - |
| - | - | 6919 | 983.5 | - | - | 0 | - |
| - | - | 3215 | 984.5 | - | - | 0 | - |
| - | - | 4390 | 990.5 | - | - | 0 | - |
| - | - | 4283 | 991.5 | - | - | 0 | - |
| 8 | b | 5.886E+04 | 993.5 | 0.005449 | 5.484 | +1 | 8 |
| - | - | 3.293E+04 | 994.5 | - | - | 0 | - |
| - | - | 9472 | 995.5 | - | - | 0 | - |
| - | - | 1.19E+04 | 1007 | - | - | 0 | - |
| - | - | 7773 | 1008 | - | - | 0 | - |
| - | - | 2955 | 1009 | - | - | 0 | - |
| 6 | y | 6.264E+04 | 1025 | 0.00556 | 5.422 | +1 | 8 |
| - | - | 3.806E+04 | 1026 | - | - | 0 | - |
| - | - | 1.525E+04 | 1027 | - | - | 0 | - |
| - | - | 7139 | 1065 | - | - | 0 | - |
| - | - | 5566 | 1066 | - | - | 0 | - |
| - | - | 1.019E+04 | 1076 | - | - | 0 | - |
| - | - | 7264 | 1077 | - | - | 0 | - |
| - | - | 3337 | 1078 | - | - | 0 | - |
| 9 | b | 8540 | 1092 | 0.01368 | 12.54 | +1 | 9 |
| - | - | 6128 | 1093 | - | - | 0 | - |
| - | - | 4.256E+04 | 1094 | - | - | 0 | - |
| - | - | 2.764E+04 | 1095 | - | - | 0 | - |
| - | - | 1.022E+04 | 1096 | - | - | 0 | - |
| 9 | b | 1.737E+04 | 1109 | 0.004079 | 3.68 | +1 | 9 |
| - | - | 1.178E+04 | 1110 | - | - | 0 | - |
| - | - | 4110 | 1118 | - | - | 0 | - |
| - | - | 9657 | 1118 | - | - | 0 | - |
| - | - | 6071 | 1119 | - | - | 0 | - |
| 5 | y | 3.609E+04 | 1136 | 0.005399 | 4.755 | +1 | 9 |
| 5 | y | 7.24E+04 | 1136 | 0.01064 | 9.363 | +1 | 9 |
| - | - | 3.96E+04 | 1138 | - | - | 0 | - |
| - | - | 1.37E+04 | 1139 | - | - | 0 | - |
| - | - | 4419 | 1140 | - | - | 0 | - |
| 5 | y | 1.141E+05 | 1154 | 0.004966 | 4.305 | +1 | 9 |
| - | - | 7.646E+04 | 1155 | - | - | 0 | - |
| - | - | 2.787E+04 | 1156 | - | - | 0 | - |
| - | - | 7147 | 1157 | - | - | 0 | - |
| 10 | b | 1.464E+04 | 1166 | 0.006542 | 5.612 | +1 | 10 |
| - | - | 8615 | 1167 | - | - | 0 | - |
| - | - | 3343 | 1168 | - | - | 0 | - |
| - | - | 6565 | 1183 | - | - | 0 | - |
| - | - | 3573 | 1184 | - | - | 0 | - |
| - | - | 3369 | 1193 | - | - | 0 | - |
| - | - | 2958 | 1195 | - | - | 0 | - |
| - | - | 4148 | 1205 | - | - | 0 | - |
| - | - | 3206 | 1206 | - | - | 0 | - |
| - | - | 4541 | 1223 | - | - | 0 | - |
| - | - | 3324 | 1224 | - | - | 0 | - |
| 4 | y | 7347 | 1291 | 0.007578 | 5.872 | +1 | 10 |
| - | - | 6431 | 1292 | - | - | 0 | - |
| - | - | 2548 | 3081 | - | - | 0 | - |

m/z Charge Intensity FragmentType MassShift Position
120.05281066894531 0 1886.5111
120.0568618774414 0 2906.859
120.08171081542969 0 1590.5947
121.04045867919922 0 5805.1987
122.0721206665039 0 29060.215
123.04033660888672 0 2654.1323
123.04496002197266 0 29669.021
123.0560531616211 0 12642.961
123.09271240234375 0 5594.157
125.0716552734375 0 1439.9409
125.10819244384766 0 11214.299
126.09217071533203 0 11498.481
126.12843322753906 0 1635.2732
127.0511703491211 0 36614.71
127.08753967285156 0 8227.141
127.12393188476562 0 13717.899
128.0494842529297 0 2309.23
128.0543670654297 0 1740.8606
128.07215881347656 0 1452.415
128.082763671875 0 6995.533
128.1079559326172 0 381712.03
129.06727600097656 0 15731.774
129.1031951904297 0 3945991
130.06591796875 0 1614383.6
130.10018920898438 0 25869.85
130.10621643066406 0 254346.92
131.06356811523438 0 6889.739
131.06924438476562 0 157936.81
131.10862731933594 0 5800.5444
132.08157348632812 0 376594.75
133.0616455078125 0 7779.7554
133.0849609375 0 32766.531
136.07659912109375 0 491667.03
137.07423400878906 0 2075.3315
137.07997131347656 0 42555.42
138.06704711914062 0 38832.566
138.08319091796875 0 2422.9084
138.09225463867188 0 21971.146
139.05099487304688 0 6286.2275
139.0709991455078 0 2281.6423
139.08749389648438 0 47486.48
140.09095764160156 0 4033.4832
140.1444854736328 0 3154.332
141.1031951904297 0 68457.85
142.06610107421875 0 30118.904
142.1065673828125 0 6070.847
142.12347412109375 0 5414.6895
143.04615783691406 0 25671.309
143.06924438476562 0 3848.028
143.08290100097656 0 1945.9021
143.1192626953125 0 2908.0798
144.04885864257812 0 1689.5656
144.0771484375 0 1677.3717
144.0817108154297 0 24454.334
145.06173706054688 0 37976.125
146.06068420410156 0 4099.6343
146.0933380126953 0 5027.5283
147.0450439453125 0 27039.438
147.11376953125 0 3135.385
148.0508270263672 0 2004.8904
148.0614471435547 0 6197.78
148.08831787109375 0 2486.0864
149.0457763671875 0 8894.77
150.067138671875 0 24586.559
150.0924072265625 0 4323.292
150.10313415527344 0 2045.2258
151.07066345214844 0 1798.8855
151.0875701904297 0 69172.66
152.0717010498047 0 2393.676
152.09103393554688 0 5846.299
152.10797119140625 0 6024.5957
153.103271484375 0 9162.817
154.06248474121094 0 6532.7026
154.0984649658203 0 8510.118
155.0462188720703 0 7725.1055
155.08241271972656 0 9336.015
155.09371948242188 0 253667.47
155.11883544921875 0 172866.14 a Water loss 1
155.124755859375 0 1969.1671
156.03009033203125 0 2280.0996
156.0778045654297 0 16047.693
156.09060668945312 0 2426.0146
156.09698486328125 0 12968.714
156.10299682617188 0 8458.533
156.122314453125 0 17558.87
157.06166076660156 0 5140.0347
157.07699584960938 0 3130.484
157.09815979003906 0 16645.15
157.13473510742188 0 7324.048
158.08482360839844 0 24244.066
158.13815307617188 0 1685.3566
158.7574462890625 0 1617.6406
159.0926513671875 0 3089004.2
160.07655334472656 0 23099.98
160.08984375 0 20157.922
160.09597778320312 0 332381.6
161.08016967773438 0 2307.2454
161.09307861328125 0 2786.477
161.09939575195312 0 15348.912
164.1192169189453 0 4540.101
165.05563354492188 0 250576.58
165.07797241210938 0 13597.055
165.10250854492188 0 1900.2736
166.06204223632812 0 490842.62
166.0984344482422 0 35942.758
167.05860900878906 0 6193.013
167.0655975341797 0 38101.516
167.0825653076172 0 6825.37
167.0940704345703 0 4179.101
167.10166931152344 0 2594.382
167.11888122558594 0 11918.422
168.0669403076172 0 3202.7395
168.0818328857422 0 46669.81
168.10299682617188 0 32259.754
168.11416625976562 0 44189.984
168.12179565429688 0 1316.3925
169.0617218017578 0 1625.2003
169.07716369628906 0 42761.53
169.08497619628906 0 4862.9956
169.09825134277344 0 117770.18
170.06114196777344 0 157362.16
170.10157775878906 0 7278.4614
170.12940979003906 0 3012.0503
171.06463623046875 0 20258.63
171.07754516601562 0 4526.408
171.0927734375 0 20175.295
171.15005493164062 0 1905.4727
172.0729522705078 0 35914.22
172.0957489013672 0 2601.3071
172.10911560058594 0 4163.31
173.057373046875 0 68957.58
173.12957763671875 0 6268207.5 a 1
174.06028747558594 0 2473.8877
174.1327667236328 0 557666.7
175.0985107421875 0 2383.542
175.1349334716797 0 29775.455
176.0828857421875 0 10235.606
177.1033172607422 0 13805.006
178.13491821289062 0 203384.47
179.11900329589844 0 10620.659
179.13133239746094 0 1745.0498
179.13839721679688 0 23025.008
180.10243225097656 0 1891.7437
180.11375427246094 0 2194.1855
181.06185913085938 0 34870.67
182.0823211669922 0 563856.4 y 12
183.0417022705078 0 3398.215
183.08596801757812 0 54496.004
183.11392211914062 0 161486.58 b Water loss 1
184.0860595703125 0 2227.9072
184.1173553466797 0 16374.696
185.09291076660156 0 2911.2183
185.10873413085938 0 2595.0735
185.11874389648438 0 1681.9222
185.12973022460938 0 6149.2847
185.1588592529297 0 1162.1058
185.166015625 0 51367.496
186.12490844726562 0 1518361.9
187.08775329589844 0 253924.34
187.12823486328125 0 123507.07
187.14515686035156 0 6403.5225
188.0717010498047 0 26079.414
188.091064453125 0 28344.953
188.1302947998047 0 7822.7705
189.07554626464844 0 3708.4026
190.08340454101562 0 46191.88
190.13510131835938 0 41609.32
191.08607482910156 0 2421.7786
191.13865661621094 0 5076.9067
192.11373901367188 0 1606.6025
193.0983123779297 0 8814.729
193.1089324951172 0 4167.5156
194.093505859375 0 32847.04
194.1300048828125 0 3944.2527
195.09280395507812 0 8662.627
195.11399841308594 0 331466.6
196.10923767089844 0 203935.97
196.11740112304688 0 19055.604
197.05751037597656 0 1760.167
197.0723114013672 0 2729.145
197.1117706298828 0 17712.049
197.12973022460938 0 19929.379
197.16600036621094 0 7059.4663
198.08860778808594 0 39333.355
198.12890625 0 110883.81
199.0724639892578 0 20374.338
199.08802795410156 0 25525.535
199.1094970703125 0 2660.54
199.1322479248047 0 16129.305
200.09156799316406 0 3015.5388
200.10414123535156 0 5088.5566
200.1447296142578 0 24648.25
201.12452697753906 0 1672561 b 1
202.0873565673828 0 20150.43
202.12782287597656 0 169290.62
203.09283447265625 0 5485.618
203.1294403076172 0 17265.684
203.15138244628906 0 3974.7222
204.078125 0 5084.6655
204.11451721191406 0 13090.592
204.1358642578125 0 5459.371
205.09835815429688 0 136801.62
205.14599609375 0 24684.893
206.1019287109375 0 15220.094
206.12998962402344 0 27995.426
206.1487579345703 0 2950.7485
207.11402893066406 0 7859.8506
207.1338348388672 0 2551.6892
207.15109252929688 0 3950.0835
207.16171264648438 0 46909.145
208.10914611816406 0 5475.543
208.1651611328125 0 6635.357
209.05703735351562 0 8178.5405
209.09268188476562 0 5360.313
210.12896728515625 0 50305.426
211.07167053222656 0 2615.6604
211.08786010742188 0 3719.8271
211.1090850830078 0 26498.057 y Water loss 10
211.13270568847656 0 6556.7344
211.14427185058594 0 2312.8438
212.09336853027344 0 2937.0903
212.14080810546875 0 31410.256
212.176513671875 0 2313.6592
213.1246795654297 0 158539.2
213.14178466796875 0 10008.6
213.1612091064453 0 14851.019
214.12791442871094 0 16541.244
214.1654510498047 0 2196.4426
215.11907958984375 0 8918.442
215.14039611816406 0 15330.251
216.09922790527344 0 31608.578
217.10325622558594 0 2868.0974
221.07217407226562 0 10163.653
221.08404541015625 0 4119.71
221.09356689453125 0 11049.975
221.10479736328125 0 31338.863
221.1410369873047 0 18729.451
221.60870361328125 0 1684.1904
222.08721923828125 0 2918.692
222.10800170898438 0 3886.279
222.12510681152344 0 47586.207
223.06593322753906 0 2042.7208
223.08792114257812 0 8613.141
223.1092071533203 0 88695.33
223.1208953857422 0 8975.983
223.15676879882812 0 553032.1
224.1044158935547 0 22353.701
224.12022399902344 0 6729.039
224.16004943847656 0 68759.41
225.0443115234375 0 2983.6162
225.1027374267578 0 3511.726
225.1248016357422 0 8332.109
225.16172790527344 0 10938.538
225.64759826660156 0 2064.6797
226.083740234375 0 112598.96
226.13796997070312 0 3889.678 b 3
227.067626953125 0 91682.4
227.08743286132812 0 11636.785
227.14231872558594 0 2694.026
227.1557159423828 0 29130.646
228.071044921875 0 7729.0215
228.13629150390625 0 4493.4653
228.15902709960938 0 3726.0774
229.11976623535156 0 202230.33
230.1037139892578 0 2528.4224
230.1232147216797 0 21260.053
230.1514129638672 0 11376.173
230.1881103515625 0 4180.1304
231.08876037597656 0 9542.674
232.1466064453125 0 2269.9019
233.10601806640625 0 2764.8193
233.14129638671875 0 87036.04
234.12521362304688 0 642111.8
234.6513214111328 0 7507.989
235.10836791992188 0 3690.4197
235.12852478027344 0 83042.05
235.15655517578125 0 15245.605
235.60662841796875 0 8534.872
236.10906982421875 0 3494.4937
236.13157653808594 0 6124.439
237.13619995117188 0 15325.253
238.09945678710938 0 3946.8477
238.12020874023438 0 65538.164
238.13192749023438 0 13570.25
239.07041931152344 0 2680.4937
239.08290100097656 0 14711.219
239.10389709472656 0 8932.321
239.1156768798828 0 6511.185
239.62139892578125 0 2451.47 y Water loss 9
240.13589477539062 0 210622.31
241.09840393066406 0 9407.092
241.11990356445312 0 29719.012
241.1389617919922 0 20277.596
241.19271850585938 0 28709.693
242.15249633789062 0 3091.2563
242.1963348388672 0 4814.5615
243.11077880859375 0 38996.363
243.13430786132812 0 3360.8928
244.0944061279297 0 242223.28
244.61224365234375 0 3801.9636
245.0978240966797 0 22839.387
245.11346435546875 0 3065.4177
246.09925842285156 0 2878.3516
247.10939025878906 0 346392.22
247.14593505859375 0 18191.846
247.15731811523438 0 7904.6035
248.11505126953125 0 137815.08
248.6251678466797 0 4721.3003 y 9
249.09988403320312 0 203109.66
249.1180419921875 0 13573.108
249.13636779785156 0 50923.11
249.60435485839844 0 4829.0225
250.1032257080078 0 22555.615
250.1201934814453 0 7981.7466
250.13998413085938 0 3006.138
250.16775512695312 0 7122.461
251.15187072753906 0 2100508.5
252.13648986816406 0 9073.52
252.15504455566406 0 279019.72
252.6048583984375 0 6423.1104
253.1215057373047 0 8711.688
253.13916015625 0 2657.4673
253.1573486328125 0 22015.201
254.07875061035156 0 5359.1846
255.1497039794922 0 102644.42
256.0953674316406 0 3893.4653
256.10980224609375 0 22907.156
256.1319885253906 0 2478.11
256.1533508300781 0 19864.498
256.6125183105469 0 10410.278
257.09326171875 0 5450.232
257.1146545410156 0 63256.746
258.12353515625 0 10143.21
258.1465148925781 0 78224.25
259.1304626464844 0 10864.473
259.1495361328125 0 10445.67
261.1205749511719 0 40372.4
261.1363525390625 0 20433.701
261.16168212890625 0 2757.669
262.1205749511719 0 31289.832
262.138671875 0 4004.3457
263.1049499511719 0 3671.3237
263.12298583984375 0 2832.8574
263.1521301269531 0 2299.7068
265.1309814453125 0 173093.06
265.6158752441406 0 2262.1086
266.1265563964844 0 531523.44
267.0781555175781 0 8194.05
267.1296081542969 0 66275.03
267.1474304199219 0 4675.735
267.6153259277344 0 6552.2246
268.0945129394531 0 3751.2073
268.130859375 0 25095.871
268.1783447265625 0 42800.316
268.2037048339844 0 4363.8887 a Water loss 2
269.11419677734375 0 3312.8127
269.1337890625 0 5061.448
269.1624450683594 0 20088.352
269.18731689453125 0 31377.078
270.1236572265625 0 85975.86
270.1467590332031 0 11049.121
270.162353515625 0 7031.484
270.1900939941406 0 3268.144
270.62481689453125 0 28566.58
271.1258850097656 0 4139.5366
271.1513671875 0 2008.1415
272.1774597167969 0 132092.9
273.1806335449219 0 25828.168
274.120361328125 0 128719.586
275.1043395996094 0 28839.55
275.1234436035156 0 19848.006
275.1518249511719 0 10488.794
276.1101989746094 0 24134.398
276.1282043457031 0 13414.709
276.1571350097656 0 22954.66
276.1712951660156 0 6437.17
276.6305236816406 0 5075.105
276.6712646484375 0 2377.65
277.095458984375 0 4173.822
277.1590576171875 0 4530.624
277.2056884765625 0 2163.8914
278.1630554199219 0 4627.288
278.1900939941406 0 2130.1353
279.1470947265625 0 46020.867
279.1717834472656 0 2654.237
279.614501953125 0 3472.0664
280.13104248046875 0 48191.598
280.14971923828125 0 7060.614
281.0530700683594 0 25353.453
281.1156921386719 0 3790.3445
281.1344299316406 0 7341.9683
281.16424560546875 0 4158.2983 b Water loss 4
282.12060546875 0 2662.056
282.15960693359375 0 11607.968
282.6697082519531 0 6454.476
283.1419677734375 0 1280445
283.6381530761719 0 6443.0757
284.0890197753906 0 8435.316
284.10498046875 0 65540.88
284.124267578125 0 10433.767
284.1450500488281 0 181037.17
284.6203918457031 0 3007.1106
285.0891418457031 0 6501.254
285.1077575683594 0 9643.323
285.1287841796875 0 3386.466
285.1453857421875 0 12348.403
285.1581115722656 0 14439.645
286.1416320800781 0 115199.89
287.14404296875 0 18356.043
287.1731262207031 0 27669.89
287.1891784667969 0 10584.526
288.1297302246094 0 7303.657
288.17626953125 0 3666.4834
288.62066650390625 0 21332.66
289.1217346191406 0 6400.549
289.20367431640625 0 2897.7268
290.1683349609375 0 18763.354 b 4
290.66943359375 0 5709.5693
291.2192687988281 0 6829.711
292.1447448730469 0 2171.5461
292.179443359375 0 2670.9302
292.2229919433594 0 2184.0593
293.1152038574219 0 852131.1 y Water loss 11
293.1632080078125 0 17811.178
294.11920166015625 0 161036.95
294.16485595703125 0 3078.8809
295.1053771972656 0 3358.0054
295.1219177246094 0 19893.467
296.090087890625 0 2918.9663
296.125244140625 0 2777.3652
296.17822265625 0 3693.1023
296.1989440917969 0 29258.416 b Water loss 2
296.6679382324219 0 7260.2715
297.1348571777344 0 21043.543 y Water loss 8
297.1578063964844 0 270223.72
297.6356201171875 0 3772.0974
297.6603698730469 0 3233.4365
298.1412658691406 0 27018.666 y Water loss 6
298.1606750488281 0 39624.19
299.0637512207031 0 19173.062
299.1449279785156 0 2822.4148
299.1636962890625 0 5805.6333
300.11285400390625 0 3301.473
300.1315612792969 0 3109.5825
300.1721496582031 0 47579.156
301.1166076660156 0 19005.031
301.15264892578125 0 765123
302.11572265625 0 141128.23
302.1559753417969 0 106016.94
303.1190185546875 0 20105.049
303.15576171875 0 6470.641
303.1690368652344 0 10271.265
304.1687927246094 0 4281.4224
304.215087890625 0 2308.1084
305.18096923828125 0 19254.666
305.64862060546875 0 3372.2344
305.6825256347656 0 6976.693
306.13934326171875 0 142355.73 y 8
306.2308349609375 0 13266.819
306.6409912109375 0 54671.883
307.1421813964844 0 22025.766
307.1593933105469 0 2734.5618
307.2353515625 0 3755.8042
308.1441955566406 0 2659.5095
308.1777648925781 0 2714.5784
309.147216796875 0 5090.191
309.16925048828125 0 8845.759
310.1571960449219 0 8861.797
310.6659240722656 0 2861.3362
311.1256103515625 0 94663.85 y 11
311.1393737792969 0 48684.23
311.17022705078125 0 2884.9954
311.68963623046875 0 5563.001
312.1292724609375 0 15728.872
312.14215087890625 0 5901.7056
312.19110107421875 0 2857.6934
313.1157531738281 0 5299.8013
314.1170959472656 0 4968.529
314.18853759765625 0 22530.703
314.20965576171875 0 63335.52 b 2
315.16851806640625 0 434043.4
315.2126159667969 0 13346.516
316.1720886230469 0 68736.73
317.138427734375 0 2757.6362
317.1766052246094 0 5504.496
317.6613464355469 0 5494.4272
318.1219787597656 0 5019.665
318.15277099609375 0 10686.807
318.6518859863281 0 3948.125
319.1041259765625 0 3562.27
319.15386962890625 0 26617.19
319.2146911621094 0 5826.241
320.1571350097656 0 4831.2734
320.1739501953125 0 4323.2627
321.11004638671875 0 4742.043
321.1573181152344 0 3918.5117
322.2259216308594 0 5683.582
323.1475524902344 0 2084.273
324.172119140625 0 2579.1091
324.1969909667969 0 6016.1494
324.6790466308594 0 4302.57
325.1528015136719 0 21224.854
325.17254638671875 0 3261.772
326.1568908691406 0 2470.466
326.18719482421875 0 4330.939
326.666015625 0 107148.56
327.1675720214844 0 38973.387
327.6688232421875 0 6594.175
328.1158447265625 0 2192.1472
330.16473388671875 0 3983.016
330.660888671875 0 3691.532
331.14141845703125 0 2506.7434
331.6586608886719 0 6707.8667
332.2105407714844 0 44661.77
333.1791076660156 0 51787.45
333.2139587402344 0 8276.577
334.12060546875 0 4409.621
334.140869140625 0 2434.366
334.1866760253906 0 11885.007
335.123291015625 0 2533.7786
335.2106018066406 0 18061.021
336.21295166015625 0 3860.089
337.16619873046875 0 3567.739
338.1851501464844 0 10341.344
338.6767578125 0 20213.846 b Water loss 5
339.17523193359375 0 17113.008
339.67767333984375 0 12464.165
340.1802673339844 0 2703.3064
340.66363525390625 0 24634.555
341.0199279785156 0 3483.1384
341.165283203125 0 11213.168
342.18359375 0 35879.465
342.6733093261719 0 2600.8748
343.16363525390625 0 42942.707
343.1874084472656 0 7178.757
344.173583984375 0 37567.848
344.9779357910156 0 3113.9758
345.1331481933594 0 58220.3
345.1758728027344 0 5250.9395
346.1170654296875 0 213959.23
346.70074462890625 0 8820.304
347.1201477050781 0 38677.47
347.6820068359375 0 114643.54 b 5
348.120849609375 0 3666.6938
348.1834411621094 0 42843.953
348.2071838378906 0 3468.56
348.684326171875 0 13180.024
349.130615234375 0 5511.7
349.1536865234375 0 9306.884
349.1873779296875 0 5425.057
350.1142883300781 0 2444.488
350.2210388183594 0 173344.16
351.1463928222656 0 5823.286
351.1687927246094 0 8564.394
351.2232360839844 0 35288.285
352.1678466796875 0 12380.03
352.2019958496094 0 3528.3462
352.22528076171875 0 2623.99
352.6743469238281 0 14850.734
353.17486572265625 0 7505.508
353.6761474609375 0 26041.895 y Water loss 7
354.1785583496094 0 14597.556
354.6789855957031 0 2724.2087
355.1611633300781 0 2684.75
356.19573974609375 0 6515.5347
356.67120361328125 0 2826.877
357.1783752441406 0 4736.9663
358.1806640625 0 3678.8167
359.13739013671875 0 6379.685
359.187255859375 0 3571.5469
360.16900634765625 0 2323.3086
360.2052001953125 0 10631.846
360.697509765625 0 48176.133
361.1993103027344 0 57324.31
361.69000244140625 0 13595.986
362.185546875 0 53647.97
362.68182373046875 0 291789.53 y 7
363.14361572265625 0 296795.7
363.18359375 0 124696.04
363.6845703125 0 24661.13
364.1273498535156 0 51455.28
364.14727783203125 0 44085.918
364.18621826171875 0 5768.186 b Water loss 8
364.2137145996094 0 4608.482
364.6744079589844 0 3080.368
365.13037109375 0 10791.466
365.1507568359375 0 5668.1055
366.1578674316406 0 8264.341
366.18017578125 0 15398.775
367.1422119140625 0 30004.002
367.16522216796875 0 8526.245
367.2474365234375 0 26443.521
367.6866760253906 0 7749.4927
368.1253967285156 0 5593.2935
368.147705078125 0 4937.0728
368.1693115234375 0 3159.2659
368.194091796875 0 14158.155
368.2278137207031 0 2470.28
368.25213623046875 0 4879.004
368.6977233886719 0 3222.9187
369.1552429199219 0 3094.6814
369.17755126953125 0 7194.4834
369.1966552734375 0 6795.9604
370.1775817871094 0 7946.6694
370.22540283203125 0 2862.568
373.1876220703125 0 14950.628
373.68927001953125 0 4027.9626
374.20367431640625 0 4369.423
374.6964416503906 0 3937.936
375.1418151855469 0 2182.9502
375.1732177734375 0 3855.1672
375.2005310058594 0 4000.9368
376.1651306152344 0 17044.83
376.20068359375 0 40979.684
376.7020568847656 0 18059.15
377.1262512207031 0 7134.842
377.1484680175781 0 11605.928
377.1893005371094 0 2529.1462
377.2688293457031 0 7027.3447
378.21636962890625 0 105583.766
379.211669921875 0 501133.97
380.1648864746094 0 17927.861
380.2146911621094 0 105709.29
381.154296875 0 853867.9
381.7135314941406 0 31362.035
382.1572265625 0 152804.67
382.1904296875 0 20852.256
382.21600341796875 0 15532.199
382.2733459472656 0 4659.9023
382.6937255859375 0 15511.701
383.1588134765625 0 21134.295
383.2079772949219 0 8566.827
383.7093200683594 0 3742.5745
384.1684875488281 0 38705.67
384.1914978027344 0 22046.408
385.15283203125 0 62077.26
385.1746826171875 0 15700.087 y 4
386.1563415527344 0 10225.249
386.1820068359375 0 12564.139
386.2074890136719 0 3193.1853
387.21380615234375 0 3119.8699
389.1953125 0 10511.286 b 9
390.1985168457031 0 36658.03
390.6999206542969 0 15174.593
391.13885498046875 0 34363.875
391.18927001953125 0 11067.885
391.69036865234375 0 6184.036
392.1231384277344 0 9455.752
392.1468200683594 0 3772.0195
392.2105407714844 0 13933.36
393.21466064453125 0 7169.1943
394.175048828125 0 96473.04
395.1372985839844 0 62455.54
395.15997314453125 0 22059.002
395.18121337890625 0 5136.0435
396.1405334472656 0 13372.8545
396.164306640625 0 4578.334
396.1944580078125 0 6871.6245
396.2270812988281 0 186418.77
397.2052307128906 0 7515.838
397.2298583984375 0 34486.26
397.7074890136719 0 7515.5557
398.1787109375 0 78277.766
398.23187255859375 0 5499.134
399.18157958984375 0 17915.824
400.20220947265625 0 4010.4656
400.7025146484375 0 2779.3315
401.18499755859375 0 3670.048
402.1787414550781 0 40100.348
402.201416015625 0 73852.73
403.1374816894531 0 5648.982
403.2021484375 0 27321.225
404.18511962890625 0 5942.945
404.2163391113281 0 2641.8472
404.7124328613281 0 11134.433
405.2197570800781 0 5218.6987
405.2259521484375 0 5576.325
405.26416015625 0 32022.514
406.2480773925781 0 18182.299
407.2068176269531 0 17770.225
407.2446594238281 0 7824.0244
407.27508544921875 0 2800.7231
408.1647644042969 0 3224.0828
408.2124328613281 0 3638.1372
408.7137756347656 0 8295.923
409.1495361328125 0 61465.133
409.1777648925781 0 4610.582
409.20623779296875 0 15929.055
409.7054443359375 0 15684.747
410.15191650390625 0 12318.1455
410.20513916015625 0 4711.565
411.2047119140625 0 3126.2488
411.24005126953125 0 2891.8347
412.18572998046875 0 550936.44
413.1477966308594 0 60142.07
413.1890563964844 0 84830.86
414.1515808105469 0 17954.488
414.19580078125 0 17061.688
414.2377624511719 0 170911.39
415.0406188964844 0 8604.004
415.2006530761719 0 63034.45
415.24102783203125 0 34198.93
416.1410217285156 0 3104.029
416.2038269042969 0 13571.253
416.2325439453125 0 4051.088
417.2081604003906 0 2681.3257
417.71942138671875 0 89406.59
418.2133483886719 0 151718.66
418.71405029296875 0 80821.29
419.2147216796875 0 19125.514
419.7166748046875 0 3188.2864
420.172607421875 0 3593.1785
421.1499328613281 0 3686.716
421.21197509765625 0 19053.217 y Water loss 10
422.1668395996094 0 3538.807
422.195556640625 0 14933.818 y Ammonia loss 10
422.25439453125 0 4789.0635
422.71142578125 0 12107.64
423.205078125 0 33665.477
423.2745361328125 0 204948.2
423.706298828125 0 12040.447
423.7373046875 0 5263.524
424.2239074707031 0 10384.2
424.27740478515625 0 55303.168
424.7367248535156 0 12680.128
425.23583984375 0 5746.062
425.27960205078125 0 5119.847
426.1998596191406 0 3866.1768
426.7249450683594 0 1447729.5
427.22637939453125 0 752845.6
427.72772216796875 0 213513.5
428.22882080078125 0 35828.773
428.7296142578125 0 2471.0928
429.2132263183594 0 14898.463
429.72918701171875 0 6345.9927
430.1962890625 0 1439734.4
431.1993713378906 0 303358.5
431.7171630859375 0 158805.83 b Water loss 6
432.2135925292969 0 89544.46 b Ammonia loss 6
432.7151184082031 0 34663.875
432.7439880371094 0 9702.447
433.21209716796875 0 24728.467
433.2581787109375 0 51801.363 b Water loss 3
433.7154235839844 0 4468.6685
433.7440185546875 0 3658.638
434.2134704589844 0 4677.9116
434.2624816894531 0 13383.789
437.19305419921875 0 3468.566
437.2325134277344 0 3966.3728
438.17645263671875 0 21452.057
438.19989013671875 0 2971.424
438.7302551269531 0 13578.24
439.2220764160156 0 304471.44 y 10
440.1812438964844 0 14294.785
440.22528076171875 0 82017.14
440.722412109375 0 701125.2 b 6
441.2240295410156 0 356021.12
441.7252197265625 0 101840.83
442.2265930175781 0 23845.11
443.2422790527344 0 7725.3823
444.1814270019531 0 4977.612
444.2490539550781 0 2799.5938
446.1710510253906 0 2511.089
446.2064208984375 0 6134.7754
446.24334716796875 0 7142.402
446.7403259277344 0 9001.155
447.20880126953125 0 64065.746 y Ammonia loss 6
447.2423400878906 0 6690.161
447.2757568359375 0 3808.974
447.7109680175781 0 30161.19
448.20758056640625 0 73890.55
449.21124267578125 0 24373.23
450.1766357421875 0 10773.198
450.2388000488281 0 17729.494
450.28570556640625 0 20138.803
451.2696838378906 0 378893.12 b 3
451.75201416015625 0 2697.4087
452.2728271484375 0 94447.77
452.74273681640625 0 6770.1523
453.2505187988281 0 1535.7275
453.27471923828125 0 12408.675
454.71990966796875 0 4935.358
455.2016296386719 0 9194.412
455.2685852050781 0 23127.396
455.722412109375 0 37404.29 y 6
456.223876953125 0 22775.537
456.2721252441406 0 7372.3906
456.72674560546875 0 5754.512
458.1911315917969 0 5046.0093
459.20123291015625 0 32201.326
460.203125 0 3681.88
460.2272644042969 0 2864.276
460.2708435058594 0 2938.457
460.75653076171875 0 34231.746
461.20648193359375 0 39943.31
461.2557373046875 0 40714.785
461.75799560546875 0 9041.259
462.2092590332031 0 11918.242
462.2481384277344 0 4073.522
462.5567932128906 0 7067.451 y Water loss 2
464.2169189453125 0 15589.65
464.2553405761719 0 9548.955
465.2550354003906 0 9138.735
465.7506408691406 0 9836.366
466.1715087890625 0 5306.9805
466.24810791015625 0 12912.547
466.74420166015625 0 3708.2312
468.1853942871094 0 3207.7725
468.29638671875 0 92198.34
468.9014587402344 0 3011.0464 b Water loss 11
469.24395751953125 0 3302.459
469.29901123046875 0 20933.908
472.22216796875 0 7647.7295
473.2251892089844 0 5627.772
474.1669006347656 0 2855.0432
474.2299499511719 0 2101.5437
474.26202392578125 0 71043.99
474.7581481933594 0 87200.85
474.9090576171875 0 5091.21 b 11
475.2573547363281 0 47810.266
475.7585144042969 0 13297.791
476.2283630371094 0 30741.63
477.21319580078125 0 63836.53
477.72222900390625 0 7620.0117
478.23175048828125 0 68889.32 y Water loss 9
478.2806701660156 0 48421.562
479.23931884765625 0 16587.354
479.28387451171875 0 8147.4
479.7482604980469 0 8842.35
480.2284240722656 0 13142.226
481.23760986328125 0 3014.3235
481.7341003417969 0 3909.2495
482.27685546875 0 3268.2844
482.7361145019531 0 2823.0017
483.26751708984375 0 237041.31
483.76885986328125 0 152455.19
484.22283935546875 0 14932.011
484.27020263671875 0 44406.785
484.7712707519531 0 14392.82
485.2253112792969 0 3252.374
486.1702880859375 0 4023.01
487.1958923339844 0 3762.279
488.2245178222656 0 4526.068
488.260009765625 0 78038.6 b Water loss 7
488.75860595703125 0 50211.375
489.2573547363281 0 20774.936
489.7589111328125 0 7054.121
491.2787170410156 0 2818.9092
493.25537109375 0 5084.165
494.2394104003906 0 757432.06
495.20458984375 0 4948.982
495.2422790527344 0 203073.5
496.243896484375 0 1091719.5 y 9
497.2499084472656 0 271167.06
497.7666320800781 0 80735.6
498.2474670410156 0 42880.43
498.7723388671875 0 4537.484
499.2470703125 0 11758.0625
499.7411193847656 0 2540.3945
502.2348327636719 0 8003.5566
503.1103820800781 0 8902.924
503.2174987792969 0 4222.082
504.20355224609375 0 36547.56
505.2050476074219 0 9193.691
505.23834228515625 0 3371.0571
506.23052978515625 0 21521.086
506.2751770019531 0 9141.185
507.2604675292969 0 15782.766
508.2239990234375 0 5414.3223
511.2660827636719 0 56208.95
512.2189331054688 0 6024.758
512.2686767578125 0 14033.655
513.2144165039062 0 8368.154
514.1939086914062 0 10002.652
514.2647094726562 0 2707.037
515.1939697265625 0 2670.7307
515.2844848632812 0 16961.635
516.2255249023438 0 3615.408
516.2896118164062 0 5521.8423
519.3341674804688 0 5471.062
520.26953125 0 3423.638
521.225341796875 0 6759.9507
521.258544921875 0 3019.6824
522.228515625 0 13326.313
523.2632446289062 0 19968.916
523.5934448242188 0 9410.671
523.9241943359375 0 6838.8906
524.2518920898438 0 6630.519
525.2696533203125 0 127561.34
526.2337036132812 0 21925.447
526.2738037109375 0 35335.21
527.2352294921875 0 7645.7227
527.279052734375 0 6884.6055
528.7362060546875 0 8586.767
529.263916015625 0 33805.594 Precursor Water loss
529.5966796875 0 51187.164 Precursor Ammonia loss
529.7659912109375 0 3789.204
529.93017578125 0 28631.623
530.2290649414062 0 9645.081
530.263916015625 0 8612.963
531.2222290039062 0 8396.802
532.197509765625 0 32108.281
532.2406616210938 0 2519.2446
533.201171875 0 10789.097
533.2510986328125 0 7488.147
533.3223876953125 0 6265.9014
534.2391357421875 0 5269.149
534.3038330078125 0 4371.265
535.2666015625 0 41957.21 Precursor
535.310302734375 0 4177.225
535.6011962890625 0 39839.97
535.9359130859375 0 21729.994
536.2694702148438 0 6667.957
537.249267578125 0 6915.5337
537.7415161132812 0 21020.14
538.2391357421875 0 9842.856
538.2731323242188 0 15742.721
538.7741088867188 0 12852.552
539.2395629882812 0 144916.78
539.2772216796875 0 7109.613
540.2425537109375 0 42627.25
540.7809448242188 0 4922.242
541.254150390625 0 12646.448
541.7634887695312 0 5878.471
542.299560546875 0 2876.3528
543.28076171875 0 329987.4
544.2841186523438 0 96672.36
545.2864990234375 0 23493.322
545.7716064453125 0 10595.118 b Water loss 8
546.25390625 0 34053.77 b Ammonia loss 8
546.7547607421875 0 22663.252
547.2564697265625 0 9512.608
547.2796020507812 0 10438.789
547.7794799804688 0 11747.023
548.2413330078125 0 5847.2236
548.2933349609375 0 3414.1406
549.224609375 0 32692.486
550.2150268554688 0 16901.338
551.2135009765625 0 4877.293
551.26025390625 0 28391.453
551.334716796875 0 19254.469
552.2637329101562 0 15551.454
552.3319702148438 0 4438.095
553.2650146484375 0 6065.487
554.7783813476562 0 48828.65 b 8
555.2598876953125 0 72949.945
555.7604370117188 0 55285.93
556.26171875 0 14612.047
556.762451171875 0 3333.74
557.23974609375 0 3197.332
558.2714233398438 0 4364.1265
559.75146484375 0 2614.4426
560.251220703125 0 7121.476
561.3175659179688 0 27836.496 b Water loss 4
562.305419921875 0 11817.455 b Ammonia loss 4
563.2606201171875 0 23896.758
563.3045043945312 0 3236.4626
564.2625122070312 0 12002.238
564.336669921875 0 9851.381
564.7664184570312 0 5452.674
565.2662963867188 0 41052.29
566.267578125 0 12214.255
567.23486328125 0 142557.73
568.2406616210938 0 40320.098
568.7617797851562 0 9963.81
569.2421875 0 7940.4385
569.2931518554688 0 4028.1704
569.7578125 0 3263.3064
570.232421875 0 2444.7737
575.2501220703125 0 20790.506
575.2971801757812 0 6721.142
576.23779296875 0 10772.734
576.2835083007812 0 8325.518
577.2208251953125 0 6627.161
577.268798828125 0 11452.735 y 4
577.7660522460938 0 9448.864
578.2672729492188 0 5229.566
578.3468627929688 0 4806.866
579.3290405273438 0 204950.53 b 4
580.3316040039062 0 64535.562
581.2701416015625 0 5380.87
581.3341674804688 0 13871.188
582.3078002929688 0 6154.675
583.2911376953125 0 8297.015 b 9
585.2806396484375 0 3150.7856
586.2680053710938 0 4233.107
587.2504272460938 0 8624.8125
588.7990112304688 0 4279.1357
589.3125 0 8961.305
589.7947387695312 0 2511.513
592.2806396484375 0 3863.961
592.3289184570312 0 25472.445
593.2603149414062 0 209687.61 y Water loss 8
593.3099975585938 0 97147.43
593.7913818359375 0 8716.8125
594.2608642578125 0 61314.633
594.3123168945312 0 25168.633
594.7853393554688 0 12280.708
595.2573852539062 0 18925.691
596.2606201171875 0 4822.678
596.3566284179688 0 9619.899
597.8024291992188 0 16224.084
598.3040771484375 0 14002.789
598.806640625 0 4485.9023
600.2755737304688 0 2562.8845
600.3186645507812 0 24761.998
601.3199462890625 0 9465.434
602.795166015625 0 51355.883
603.2946166992188 0 62023.027
603.795654296875 0 21846.39
604.2908325195312 0 12275.111
605.3474731445312 0 2994.8206
607.3013916015625 0 3180.7405
609.3548583984375 0 3053.7314
610.2868041992188 0 35606.926
610.336669921875 0 19785.207
611.2713623046875 0 2832989.8 y 8
611.8004150390625 0 184280.05
612.2750854492188 0 890107.9
612.8026733398438 0 51988.63
613.27685546875 0 204303.73
613.8036499023438 0 2815.2312
614.2783203125 0 29644.863
615.322265625 0 2754.7998
616.31298828125 0 3508.7114
617.2884521484375 0 14214.124
618.3280639648438 0 2388.748
619.2755126953125 0 3095.3506
619.3279418945312 0 7036.644
619.7689819335938 0 3818.8027
620.2863159179688 0 2492.344
620.8062133789062 0 14663.589
621.254638671875 0 18591.92
621.3055419921875 0 23993.795
621.8098754882812 0 3391.6594
622.2616577148438 0 10089.892
622.3135375976562 0 4150.0522
622.3713989257812 0 8460.003
622.7884521484375 0 2874.7156
623.255859375 0 5097.9575
626.3025512695312 0 3103.2668
627.2793579101562 0 3310.263
627.787353515625 0 2943.7178
628.2791748046875 0 12985.261
628.780517578125 0 9475.889
630.2933349609375 0 5980.2715
634.3112182617188 0 29609.846
635.3007202148438 0 42441.605
636.3291625976562 0 25796.402
636.7905883789062 0 32239.375 y Water loss 3
637.2906494140625 0 25774.89
637.3408813476562 0 8581.006
637.792724609375 0 11656.923
638.3345336914062 0 7673.871 b Water loss 10
638.8292846679688 0 5999.3765 b Ammonia loss 10
640.2769775390625 0 35250.64
641.2684936523438 0 10548.634
642.26953125 0 3874.9512
644.3001708984375 0 5036.161
645.2843627929688 0 20247.078
645.7955322265625 0 181490.6 y 3
646.2965087890625 0 140068.86
646.7985229492188 0 59445.574
647.2968139648438 0 11703.835
647.3407592773438 0 5840.3 b 10
647.3952026367188 0 2957.1184
647.8375244140625 0 9726.299
648.3466186523438 0 10566.377
649.3438720703125 0 6829.3535
650.2716064453125 0 11365.467
650.3324584960938 0 10588.417
650.7869873046875 0 10876.889
651.2821655273438 0 9070.697
651.3358764648438 0 3084.8208
652.3246459960938 0 251241.52
652.828125 0 6199.386
653.3272705078125 0 90790.484
654.3292236328125 0 19504.963
655.3250732421875 0 2839.7612
657.859375 0 2683.257
658.2872924804688 0 27311.27
658.336669921875 0 5643.1436
659.3226318359375 0 9850.041
660.3204345703125 0 6153.5747
661.3349609375 0 24207.316
661.8350219726562 0 13822.316
662.3095092773438 0 91415.94
662.8379516601562 0 2798.8887
663.300048828125 0 60876.05
664.2987670898438 0 17230.8
664.3460083007812 0 11748.93
665.3458251953125 0 6660.581
666.2929077148438 0 5989.628
666.3591918945312 0 5658.341
668.283447265625 0 8742.569
669.287841796875 0 3904.06
670.8302612304688 0 3539.504
671.3348388671875 0 3278.5916
676.3456420898438 0 123570.53 b Water loss 5
677.3380126953125 0 72494.51 b Ammonia loss 5
678.3375854492188 0 34815.008
679.3358764648438 0 27218.547
679.8370361328125 0 12026.954
680.31982421875 0 459355.78
681.322509765625 0 182187.25
682.3245849609375 0 39226.445
683.3284301757812 0 6033.726
684.3287963867188 0 10497.031
684.820068359375 0 36146.01
685.3204956054688 0 32190.705
685.823974609375 0 10733.856
686.322509765625 0 7097.5903
688.3350830078125 0 7822.894
689.3328857421875 0 4079.9895
690.3067626953125 0 6383.3545
692.3317260742188 0 3496.9326
692.4110107421875 0 2811.4268
693.3327026367188 0 109553.66 y Water loss 2
693.8314208984375 0 142274.06 y Ammonia loss 2
694.355712890625 0 1208324.8 b 5
694.8311767578125 0 27991.932
695.3590698242188 0 476116.38
696.362060546875 0 104937.32
697.35791015625 0 16622.607
697.8591918945312 0 7668.8975
698.279296875 0 2917.5134
698.3544921875 0 7771.0913
698.8651123046875 0 3188.775
701.3740844726562 0 3498.853
701.8450927734375 0 19401.355
702.337890625 0 1101589.1 y 2
702.8397216796875 0 936166.4 b Water loss 11
703.341064453125 0 445372.56 b Ammonia loss 11
703.8427734375 0 122822.49
704.3424072265625 0 45781.723
705.3516845703125 0 5983.516
706.345458984375 0 69161.74 y Water loss 7
707.3472900390625 0 31270.201
708.3389892578125 0 11430.815
709.337646484375 0 2883.7188
711.3782958984375 0 25549.129
711.8589477539062 0 88459.36 b 11
712.3609008789062 0 77005.53
712.861572265625 0 27473.012
713.3607177734375 0 10874.148
714.3255004882812 0 3804.8767
715.3052978515625 0 4991.586
715.36083984375 0 3450.3108
716.341552734375 0 5598.3853
717.3444213867188 0 4284.4365
718.3930053710938 0 5326.774
720.3883056640625 0 9060.349
720.8635864257812 0 4763.466
721.3671264648438 0 55901.31
722.3694458007812 0 15091.695
723.3760375976562 0 18819.758
724.3558959960938 0 830026.1 y 7
725.3591918945312 0 335589.2
726.3614501953125 0 97189.06
727.36474609375 0 17266.84
728.8627319335938 0 4579.761
729.361572265625 0 39947.887
730.3643798828125 0 20203.656
731.3639526367188 0 6464.5444
732.3224487304688 0 3786.165
733.3681030273438 0 27408.65
734.35986328125 0 26510.727
735.3600463867188 0 10489.896
736.3959350585938 0 3595.1453
737.3411254882812 0 3727.536
737.8671875 0 14088.735
738.3657836914062 0 8025.2246
739.3558959960938 0 6462.7485
742.8658447265625 0 11564.6455 y Water loss 1
743.3717041015625 0 22109.307
743.8658447265625 0 7069.3467
744.37109375 0 7971.941
745.3607177734375 0 6119.2017
746.345947265625 0 2539.6157
747.3914184570312 0 14717.727
748.3876342773438 0 15309.18
749.3428344726562 0 17241.215
750.3381958007812 0 6371.2427
750.4000244140625 0 5324.8706
751.3810424804688 0 118702.9
751.8717041015625 0 78369.4 y 1
752.3770751953125 0 135441.36
752.8749389648438 0 39587.01
753.3810424804688 0 28114.195
756.8629760742188 0 3827.2412
757.3746948242188 0 4124.6494
758.3695678710938 0 5766.2183
760.3090209960938 0 11717.997
761.3099365234375 0 5430.2114
761.3779907226562 0 8803
762.3636474609375 0 11516.358
762.4222412109375 0 7831.366
763.3701171875 0 66528.51
764.3709106445312 0 25359.334
765.406494140625 0 29408.486
766.4099731445312 0 15220.605
767.3521728515625 0 25479.719
767.4193115234375 0 3411.7256
768.3529663085938 0 8782.617
768.4172973632812 0 3710.3047
769.4038696289062 0 12991.674
770.4038696289062 0 5134.825
775.3966064453125 0 13417.244
776.3851318359375 0 10475.002
777.3318481445312 0 5064.1997
777.3945922851562 0 3185.3547
778.3256225585938 0 5007.2534
778.4055786132812 0 3513.802
779.387939453125 0 64177.03
780.3916015625 0 29453.137
781.3712158203125 0 33176.957
782.3726196289062 0 16101.607
783.370849609375 0 3840.1055
790.3623046875 0 3817.2031
791.3897705078125 0 3456.2263
792.442626953125 0 3575.122
793.4041137695312 0 114139.72
794.4069213867188 0 51288.355
795.3466186523438 0 54281.594
795.4132690429688 0 8201.668
796.3494262695312 0 28230.236
796.418212890625 0 3346.6565
797.3532104492188 0 9491.763
807.3809204101562 0 20727.26
807.4440307617188 0 15686.324
808.384765625 0 14813.968
809.3659057617188 0 5164.497
817.3290405273438 0 3834.2678
817.4053344726562 0 5692.2744
818.4408569335938 0 12026.637
819.44287109375 0 14590.974
820.4419555664062 0 8282.73
824.3767700195312 0 3386.1594
825.3923950195312 0 9181.879
826.3807983398438 0 21486.908
827.3843994140625 0 10401.732
829.4341430664062 0 3877.4414
834.3593139648438 0 3815.7097
834.4303588867188 0 29046.908
835.4209594726562 0 34123.316
836.4539794921875 0 71579.125
837.455322265625 0 34565.688
838.4600219726562 0 10816.215
843.404052734375 0 14155.056
844.3936767578125 0 38157.887
845.3973999023438 0 27877.967
846.451416015625 0 39146.32
847.449462890625 0 23456.086
848.4505004882812 0 10610.516
852.4412841796875 0 257576
853.4437866210938 0 125345.89
854.44677734375 0 37513.805
855.4517822265625 0 6490.4604
862.42529296875 0 151106 b Water loss 6
863.4220581054688 0 117138.96
864.4603881835938 0 197644.81
865.4642333984375 0 102535.7
866.4674682617188 0 25622.068
867.4700317382812 0 5032.8774
869.3931884765625 0 9335.112
870.3935546875 0 4500.7954
873.3986206054688 0 3582.8586
875.4269409179688 0 2897.1133
876.4420166015625 0 20534.295
877.4439697265625 0 9617.818
880.4359741210938 0 1232254.8 b 6
881.4390869140625 0 659867.2
882.44189453125 0 193659.17
883.4429321289062 0 36613.535
884.4501342773438 0 3584.3938
890.4207153320312 0 13494.611
891.4107666015625 0 22362.105
892.4102783203125 0 8127.9453 y Water loss 6
892.4722290039062 0 20178.143
893.4060668945312 0 5474.7495 y Ammonia loss 6
893.4779663085938 0 7505.8857
894.4583740234375 0 8485.276
895.455810546875 0 4125.503
908.4312744140625 0 148739.78
909.43408203125 0 86853.58
910.4362182617188 0 131611.66 y 6
911.4382934570312 0 63471.918
912.4404907226562 0 17076.74
913.4422607421875 0 3564.5613
927.425537109375 0 5556.091
936.4287109375 0 9713.389
937.4202270507812 0 17722.957
938.4235229492188 0 7650.052
944.4583129882812 0 4037.8582
947.4435424804688 0 5443.847
948.4371337890625 0 8811.96
949.4340209960938 0 3683.8313
954.4369506835938 0 22694.379
955.42529296875 0 40445.38
956.426025390625 0 22436.227
957.4301147460938 0 6840.495
962.458740234375 0 4293.8125
963.4444580078125 0 3019.4
964.4602661132812 0 5839.3184
965.4534912109375 0 41731.117
966.455810546875 0 26054.38
967.4599609375 0 7917.615
972.447509765625 0 31345.426
973.4482421875 0 19018.002
974.448486328125 0 6175.0386
975.5062866210938 0 11486.098 b Water loss 7
976.5100708007812 0 9219.628
979.4872436523438 0 3652.4612
980.4663696289062 0 17083.152
981.4688110351562 0 5692.3564
982.4756469726562 0 16267.967
983.4810180664062 0 6919.039
984.4832153320312 0 3215.4998
990.4610595703125 0 4389.8555
991.4505004882812 0 4282.709
993.5206909179688 0 58858.137 b 7
994.52392578125 0 32933.332
995.5292358398438 0 9472.071
1007.4887084960938 0 11898.969
1008.486328125 0 7772.6265
1009.497802734375 0 2954.5972
1025.4630126953125 0 62644.016 y 5
1026.4659423828125 0 38060.742
1027.467529296875 0 15245.626
1064.527587890625 0 7139.1274
1065.526611328125 0 5566.4043
1075.53466796875 0 10188.7705
1076.53564453125 0 7263.694
1077.5318603515625 0 3336.5002
1091.501953125 0 8539.631 b Ammonia loss 8
1092.5068359375 0 6127.7197
1093.546630859375 0 42556.918
1094.550048828125 0 27644.908
1095.5506591796875 0 10219.459
1108.5462646484375 0 17374.168 b 8
1109.5419921875 0 11775.092
1117.5086669921875 0 4109.943
1118.4892578125 0 9656.772
1119.4891357421875 0 6070.932
1135.5108642578125 0 36085.664 y Water loss 4
1136.5001220703125 0 72395.59 y Ammonia loss 4
1137.5003662109375 0 39598.45
1138.5030517578125 0 13699.527
1139.5047607421875 0 4418.693
1153.52099609375 0 114085.05 y 4
1154.5235595703125 0 76459.625
1155.526611328125 0 27872.79
1156.5306396484375 0 7147.2446
1165.5701904296875 0 14643.668 b 9
1166.574951171875 0 8615.153
1167.580322265625 0 3342.7878
1182.594970703125 0 6565.2036
1183.589111328125 0 3573.3882
1192.612548828125 0 3369.052
1194.61865234375 0 2957.8303
1204.5811767578125 0 4148.2744
1205.5799560546875 0 3206.2117
1222.587646484375 0 4541.2573
1223.5887451171875 0 3323.6492
1290.58251953125 0 7346.6553 y 3
1291.580810546875 0 6430.787
3081.479248046875 0 2547.538

Spectrum Details

|  |  |
| --- | --- |
| Matched peaks? Matched peaks The total absolute number of peaks matched. Additionally in brackets the total fraction of peaks matched and the total number of peaks is shown. | 88 (6.44% of 1367) |
| FDR? FDR The false discovery rate estimated for this peptide. It is calculated by matching all theoretical fragments with a non-integer shift with the raw peaks for this spectrum. This is done with 40 different shifts. The resulting percentage is the average number of annotated peaks over the number of annotated peaks with the correct spectrum. | 1.03% |
| Satellite FDR? Satellite FDR See the FDR for details on its calculation. This satellite ion specific FDR only contains the satellite ions (d/w) for I/L/J positions. | - |
| PSM Score? PSM Score The PSM Score as given by Hecklib to this annotated spectrum. It is shown with three significant figures. | 647 |

#### Spectrum 8039? Spectrum 8039 The raw spectrum of this peptide as annotated by Hecklib. The fragments are coloured according to ion type (see legend). Any peaks with a star '\*' as text can be hovered over to see the full details, first the ion type second the mass shift type. By hovering over the amino acids in the peptide or ions in the legend the corresponding peaks are highlighted. By toggling the 'Unassigned' label you can turn the background (unassigned) peaks on or off in the plot. By updating the slider in the Ion legend you can update the spectrum to only show the top X% of the peaks with labels. The top X% means any peak that is within X% of the highest intensity. By dragging in the spectrum you can zoom in to a specific part of the spectrum and use 'Zoom Out' to get back to the original zoom level. The annotation of the spectrum is based on the given sequence in the peptides file and is done with different software so inconsistencies are likely. The peaks are annotated based on the given sequence, with 20 ppm tolerance.

Copy Data

##### Spectrum 8039 (TSV)

###### Preview

```
Loading example...
```

*Click on the button to copy the data to your clipboard.*

Mz MinMz MaxIntensity Max

WidthHeightPeptide font sizePeptide stroke widthSpectrum font sizeSpectrum stroke widthCompact peptide

Ion legend

wxyz

abcd

OtherUnassignedIon^Charge^_Position_Show for top:%

TVLHQDWLDGKEY

03.26e+46.53e+49.79e+41.31e+5

Zoom Out

a^+1^_2_y^+3^_4_a^+1^_2_y^+1^_1_b^+1^_2_b^+1^_2_y^+2^_3_b^+3^_6_y^+1^_2_b^+1^_3_y^+2^_5_y^+1^_2_b^+1^_3_b^+2^_6_y^+2^_6_y^+2^_6_y^+3^_9_b^+2^_7_b^+2^_7_b^+1^_4_y^+1^_3_b^+2^_7_y^+2^_7_b^+1^_4_y^+2^_7_y^+1^_4_b^+2^_8_b^+2^_8_y^+1^_4_\*\*\*b^+1^_5_b^+1^_5_y^+1^_5_y^+1^_5_y^+2^_10_y^+2^_10_y^+2^_10_b^+1^_6_b^+1^_6_y^+2^_11_y^+2^_11_b^+1^_6_y^+2^_11_b^+2^_12_b^+2^_12_y^+1^_6_b^+2^_12_y^+1^_6_y^+2^_12_y^+2^_12_\*b^+1^_7_b^+1^_7_b^+1^_7_y^+1^_7_b^+1^_8_y^+1^_8_y^+1^_9_y^+1^_9_y^+1^_9_

0850170125513401

Fragment Matches Table

Show background peaks

| Position | Ion type | Intensity | mz Theoretical | mz Error (Th) | mz Error (ppm) | Charge | Series Number |
| --- | --- | --- | --- | --- | --- | --- | --- |
| - | - | 5848 | 120.1 | - | - | 0 | - |
| - | - | 639.3 | 121.1 | - | - | 0 | - |
| - | - | 1073 | 123 | - | - | 0 | - |
| - | - | 481.6 | 125.1 | - | - | 0 | - |
| - | - | 420.6 | 126 | - | - | 0 | - |
| - | - | 1845 | 127.1 | - | - | 0 | - |
| - | - | 377 | 128.1 | - | - | 0 | - |
| - | - | 8348 | 128.1 | - | - | 0 | - |
| - | - | 1100 | 129.1 | - | - | 0 | - |
| - | - | 8.462E+04 | 129.1 | - | - | 0 | - |
| - | - | 3.609E+04 | 130.1 | - | - | 0 | - |
| - | - | 450 | 130.1 | - | - | 0 | - |
| - | - | 847.2 | 130.1 | - | - | 0 | - |
| - | - | 5629 | 130.1 | - | - | 0 | - |
| - | - | 584.6 | 131.1 | - | - | 0 | - |
| - | - | 3658 | 131.1 | - | - | 0 | - |
| - | - | 402.2 | 131.1 | - | - | 0 | - |
| - | - | 8495 | 132.1 | - | - | 0 | - |
| - | - | 1136 | 132.1 | - | - | 0 | - |
| - | - | 546.8 | 133.1 | - | - | 0 | - |
| - | - | 476.2 | 134 | - | - | 0 | - |
| - | - | 1.221E+04 | 136.1 | - | - | 0 | - |
| - | - | 1561 | 137.1 | - | - | 0 | - |
| - | - | 1019 | 138.1 | - | - | 0 | - |
| - | - | 987.3 | 139.1 | - | - | 0 | - |
| - | - | 551.3 | 140.7 | - | - | 0 | - |
| - | - | 2598 | 141.1 | - | - | 0 | - |
| - | - | 630.3 | 143.1 | - | - | 0 | - |
| - | - | 984.7 | 145.1 | - | - | 0 | - |
| - | - | 1095 | 146.1 | - | - | 0 | - |
| - | - | 1054 | 149 | - | - | 0 | - |
| - | - | 1.414E+04 | 149 | - | - | 0 | - |
| - | - | 964 | 149.1 | - | - | 0 | - |
| - | - | 796.1 | 150 | - | - | 0 | - |
| - | - | 539.3 | 150.1 | - | - | 0 | - |
| - | - | 676.7 | 151 | - | - | 0 | - |
| - | - | 1540 | 151.1 | - | - | 0 | - |
| - | - | 441.3 | 155 | - | - | 0 | - |
| - | - | 4729 | 155.1 | - | - | 0 | - |
| 2 | a | 3489 | 155.1 | 0.0003203 | 2.065 | +1 | 2 |
| - | - | 439.9 | 157.1 | - | - | 0 | - |
| - | - | 575.4 | 158.1 | - | - | 0 | - |
| - | - | 6.21E+04 | 159.1 | - | - | 0 | - |
| - | - | 407.2 | 159.3 | - | - | 0 | - |
| - | - | 891.2 | 160.1 | - | - | 0 | - |
| - | - | 6394 | 160.1 | - | - | 0 | - |
| - | - | 5351 | 165.1 | - | - | 0 | - |
| - | - | 9400 | 166.1 | - | - | 0 | - |
| 10 | y | 3089 | 166.1 | 0.001618 | 9.745 | +3 | 4 |
| - | - | 2.205E+04 | 167.1 | - | - | 0 | - |
| - | - | 1282 | 167.1 | - | - | 0 | - |
| - | - | 635.5 | 168.1 | - | - | 0 | - |
| - | - | 756 | 168.1 | - | - | 0 | - |
| - | - | 539 | 168.1 | - | - | 0 | - |
| - | - | 985.1 | 168.1 | - | - | 0 | - |
| - | - | 797.1 | 169.1 | - | - | 0 | - |
| - | - | 3027 | 169.1 | - | - | 0 | - |
| - | - | 2877 | 170.1 | - | - | 0 | - |
| - | - | 619.8 | 171.1 | - | - | 0 | - |
| - | - | 1635 | 172.1 | - | - | 0 | - |
| - | - | 897.3 | 173.1 | - | - | 0 | - |
| 2 | a | 1.235E+05 | 173.1 | 0.0003147 | 1.818 | +1 | 2 |
| - | - | 1235 | 173.5 | - | - | 0 | - |
| - | - | 1.202E+04 | 174.1 | - | - | 0 | - |
| - | - | 560.6 | 175.1 | - | - | 0 | - |
| - | - | 793 | 177.1 | - | - | 0 | - |
| - | - | 3473 | 178.1 | - | - | 0 | - |
| - | - | 544.3 | 179.1 | - | - | 0 | - |
| 13 | y | 1.131E+04 | 182.1 | 0.0002665 | 1.464 | +1 | 1 |
| - | - | 1507 | 183.1 | - | - | 0 | - |
| 2 | b | 3817 | 183.1 | 0.0001872 | 1.022 | +1 | 2 |
| - | - | 1181 | 183.1 | - | - | 0 | - |
| - | - | 943.9 | 185.2 | - | - | 0 | - |
| - | - | 2.446E+04 | 186.1 | - | - | 0 | - |
| - | - | 4703 | 187.1 | - | - | 0 | - |
| - | - | 490.9 | 187.1 | - | - | 0 | - |
| - | - | 1998 | 187.1 | - | - | 0 | - |
| - | - | 1086 | 187.1 | - | - | 0 | - |
| - | - | 650 | 188.1 | - | - | 0 | - |
| - | - | 827 | 190.1 | - | - | 0 | - |
| - | - | 1188 | 190.1 | - | - | 0 | - |
| - | - | 5946 | 195.1 | - | - | 0 | - |
| - | - | 4023 | 196.1 | - | - | 0 | - |
| - | - | 578.4 | 197.1 | - | - | 0 | - |
| - | - | 818 | 197.1 | - | - | 0 | - |
| - | - | 575.6 | 198.1 | - | - | 0 | - |
| - | - | 1855 | 198.1 | - | - | 0 | - |
| - | - | 717.3 | 199.1 | - | - | 0 | - |
| 2 | b | 3.309E+04 | 201.1 | 0.0001816 | 0.9028 | +1 | 2 |
| - | - | 3326 | 202.1 | - | - | 0 | - |
| - | - | 2206 | 205.1 | - | - | 0 | - |
| - | - | 508.6 | 206.1 | - | - | 0 | - |
| - | - | 1205 | 207.2 | - | - | 0 | - |
| - | - | 543.8 | 209.1 | - | - | 0 | - |
| - | - | 1183 | 210.1 | - | - | 0 | - |
| 11 | y | 563.7 | 211.1 | 0.0001914 | 0.9065 | +2 | 3 |
| - | - | 494.3 | 211.1 | - | - | 0 | - |
| - | - | 729.4 | 211.1 | - | - | 0 | - |
| - | - | 931.7 | 212.1 | - | - | 0 | - |
| - | - | 3618 | 213.1 | - | - | 0 | - |
| - | - | 692.5 | 213.2 | - | - | 0 | - |
| - | - | 572.1 | 215.1 | - | - | 0 | - |
| - | - | 671.2 | 216.1 | - | - | 0 | - |
| - | - | 2037 | 221.1 | - | - | 0 | - |
| - | - | 6474 | 221.1 | - | - | 0 | - |
| - | - | 854.1 | 221.1 | - | - | 0 | - |
| - | - | 847.2 | 221.1 | - | - | 0 | - |
| - | - | 1622 | 222.1 | - | - | 0 | - |
| - | - | 663.4 | 222.1 | - | - | 0 | - |
| - | - | 4657 | 223.1 | - | - | 0 | - |
| - | - | 1247 | 223.1 | - | - | 0 | - |
| - | - | 2615 | 223.1 | - | - | 0 | - |
| - | - | 1.071E+04 | 223.2 | - | - | 0 | - |
| - | - | 931.3 | 224.1 | - | - | 0 | - |
| - | - | 1336 | 224.2 | - | - | 0 | - |
| - | - | 9572 | 225 | - | - | 0 | - |
| - | - | 2807 | 226 | - | - | 0 | - |
| - | - | 2206 | 226.1 | - | - | 0 | - |
| 6 | b | 603.2 | 226.1 | 2.651E-05 | 0.1172 | +3 | 6 |
| - | - | 1040 | 227 | - | - | 0 | - |
| - | - | 1000 | 227 | - | - | 0 | - |
| - | - | 1840 | 227.1 | - | - | 0 | - |
| - | - | 1156 | 228.1 | - | - | 0 | - |
| - | - | 1824 | 229.1 | - | - | 0 | - |
| - | - | 805.7 | 229.2 | - | - | 0 | - |
| - | - | 1213 | 233.1 | - | - | 0 | - |
| - | - | 1.325E+04 | 234.1 | - | - | 0 | - |
| - | - | 1895 | 235.1 | - | - | 0 | - |
| - | - | 935.9 | 237.1 | - | - | 0 | - |
| - | - | 1669 | 238.1 | - | - | 0 | - |
| - | - | 2.14E+04 | 239.1 | - | - | 0 | - |
| - | - | 6297 | 240.1 | - | - | 0 | - |
| - | - | 5481 | 240.1 | - | - | 0 | - |
| - | - | 4452 | 241.1 | - | - | 0 | - |
| - | - | 854 | 241.1 | - | - | 0 | - |
| - | - | 989.7 | 241.2 | - | - | 0 | - |
| - | - | 874.7 | 243.1 | - | - | 0 | - |
| - | - | 4927 | 244.1 | - | - | 0 | - |
| - | - | 635.4 | 245.1 | - | - | 0 | - |
| - | - | 5564 | 247.1 | - | - | 0 | - |
| - | - | 1168 | 248.1 | - | - | 0 | - |
| - | - | 2022 | 248.1 | - | - | 0 | - |
| - | - | 632.4 | 249.1 | - | - | 0 | - |
| - | - | 3388 | 249.1 | - | - | 0 | - |
| - | - | 1387 | 249.1 | - | - | 0 | - |
| - | - | 870.3 | 250.1 | - | - | 0 | - |
| - | - | 4.283E+04 | 251.2 | - | - | 0 | - |
| - | - | 4904 | 252.2 | - | - | 0 | - |
| - | - | 3060 | 255.1 | - | - | 0 | - |
| - | - | 1181 | 257.1 | - | - | 0 | - |
| - | - | 1983 | 258.1 | - | - | 0 | - |
| - | - | 1242 | 261.1 | - | - | 0 | - |
| - | - | 534.2 | 261.1 | - | - | 0 | - |
| - | - | 670.5 | 262.1 | - | - | 0 | - |
| - | - | 6329 | 265.1 | - | - | 0 | - |
| - | - | 1605 | 266.1 | - | - | 0 | - |
| - | - | 1.034E+04 | 266.1 | - | - | 0 | - |
| - | - | 1092 | 267.1 | - | - | 0 | - |
| - | - | 927.3 | 268.2 | - | - | 0 | - |
| - | - | 834.2 | 269.2 | - | - | 0 | - |
| - | - | 1816 | 270.1 | - | - | 0 | - |
| - | - | 2965 | 272.2 | - | - | 0 | - |
| - | - | 1670 | 274.1 | - | - | 0 | - |
| - | - | 641.2 | 275.1 | - | - | 0 | - |
| - | - | 892.6 | 276.2 | - | - | 0 | - |
| - | - | 792.2 | 279.1 | - | - | 0 | - |
| - | - | 1521 | 280.1 | - | - | 0 | - |
| - | - | 1.351E+04 | 281.1 | - | - | 0 | - |
| - | - | 3387 | 282.2 | - | - | 0 | - |
| - | - | 1860 | 283 | - | - | 0 | - |
| - | - | 3.156E+04 | 283.1 | - | - | 0 | - |
| - | - | 1752 | 284.1 | - | - | 0 | - |
| - | - | 868.7 | 284.1 | - | - | 0 | - |
| - | - | 4212 | 284.1 | - | - | 0 | - |
| - | - | 1678 | 285 | - | - | 0 | - |
| - | - | 1628 | 286.1 | - | - | 0 | - |
| - | - | 674.8 | 287.2 | - | - | 0 | - |
| - | - | 1211 | 288.6 | - | - | 0 | - |
| 12 | y | 1.572E+04 | 293.1 | 0.0001747 | 0.5961 | +1 | 2 |
| - | - | 1047 | 293.1 | - | - | 0 | - |
| - | - | 3135 | 294.1 | - | - | 0 | - |
| - | - | 1799 | 295.1 | - | - | 0 | - |
| - | - | 1740 | 296.1 | - | - | 0 | - |
| 3 | b | 1169 | 296.2 | 0.0003365 | 1.136 | +1 | 3 |
| - | - | 2204 | 297.1 | - | - | 0 | - |
| - | - | 1096 | 297.1 | - | - | 0 | - |
| - | - | 5161 | 297.2 | - | - | 0 | - |
| - | - | 974.2 | 298.2 | - | - | 0 | - |
| - | - | 1.292E+05 | 299.1 | - | - | 0 | - |
| - | - | 3141 | 300.1 | - | - | 0 | - |
| - | - | 3839 | 300.2 | - | - | 0 | - |
| - | - | 1997 | 301.1 | - | - | 0 | - |
| - | - | 1.538E+04 | 301.2 | - | - | 0 | - |
| - | - | 1033 | 301.2 | - | - | 0 | - |
| - | - | 2831 | 302.1 | - | - | 0 | - |
| - | - | 1503 | 302.2 | - | - | 0 | - |
| - | - | 1220 | 305.6 | - | - | 0 | - |
| 9 | y | 1955 | 306.1 | 0.0006739 | 2.201 | +2 | 5 |
| - | - | 837.7 | 306.6 | - | - | 0 | - |
| 12 | y | 1996 | 311.1 | 0.0003802 | 1.222 | +1 | 2 |
| - | - | 827.1 | 311.1 | - | - | 0 | - |
| - | - | 696 | 314.2 | - | - | 0 | - |
| 3 | b | 2346 | 314.2 | 0.0001477 | 0.4702 | +1 | 3 |
| - | - | 8765 | 315.2 | - | - | 0 | - |
| - | - | 561.9 | 325.2 | - | - | 0 | - |
| - | - | 2061 | 326.7 | - | - | 0 | - |
| - | - | 981.7 | 327 | - | - | 0 | - |
| - | - | 885.7 | 327.2 | - | - | 0 | - |
| - | - | 828.2 | 332.2 | - | - | 0 | - |
| - | - | 1784 | 333.2 | - | - | 0 | - |
| - | - | 1694 | 341 | - | - | 0 | - |
| - | - | 741.1 | 342.2 | - | - | 0 | - |
| - | - | 1542 | 344.2 | - | - | 0 | - |
| - | - | 1.521E+04 | 345 | - | - | 0 | - |
| - | - | 1230 | 345.1 | - | - | 0 | - |
| - | - | 4893 | 346.1 | - | - | 0 | - |
| 6 | b | 2198 | 347.7 | 0.0006355 | 1.828 | +2 | 6 |
| - | - | 2919 | 350.2 | - | - | 0 | - |
| - | - | 910.9 | 351.2 | - | - | 0 | - |
| - | - | 566.5 | 351.7 | - | - | 0 | - |
| 8 | y | 989.9 | 353.7 | 0.0007285 | 2.06 | +2 | 6 |
| - | - | 647.3 | 353.7 | - | - | 0 | - |
| - | - | 2372 | 355.1 | - | - | 0 | - |
| - | - | 1.525E+04 | 359 | - | - | 0 | - |
| - | - | 1587 | 360.7 | - | - | 0 | - |
| - | - | 2025 | 362.2 | - | - | 0 | - |
| 8 | y | 4502 | 362.7 | 9.831E-05 | 0.2711 | +2 | 6 |
| - | - | 7246 | 363.1 | - | - | 0 | - |
| - | - | 3168 | 363.2 | - | - | 0 | - |
| - | - | 916.5 | 363.7 | - | - | 0 | - |
| - | - | 1304 | 364.1 | - | - | 0 | - |
| - | - | 715.5 | 364.1 | - | - | 0 | - |
| - | - | 883.1 | 366.2 | - | - | 0 | - |
| - | - | 565 | 367.2 | - | - | 0 | - |
| - | - | 1198 | 368.2 | - | - | 0 | - |
| - | - | 1279 | 369.1 | - | - | 0 | - |
| - | - | 1965 | 370.1 | - | - | 0 | - |
| - | - | 1525 | 371.1 | - | - | 0 | - |
| - | - | 1773 | 376.2 | - | - | 0 | - |
| - | - | 582.1 | 377.1 | - | - | 0 | - |
| - | - | 2436 | 378.2 | - | - | 0 | - |
| - | - | 1.106E+04 | 379.2 | - | - | 0 | - |
| - | - | 2030 | 380.2 | - | - | 0 | - |
| - | - | 1.938E+04 | 381.2 | - | - | 0 | - |
| - | - | 3253 | 382.2 | - | - | 0 | - |
| - | - | 802.9 | 382.2 | - | - | 0 | - |
| - | - | 1112 | 384.2 | - | - | 0 | - |
| - | - | 857.1 | 384.2 | - | - | 0 | - |
| - | - | 1504 | 385.2 | - | - | 0 | - |
| 5 | y | 877.2 | 385.2 | 0.002819 | 7.319 | +3 | 9 |
| - | - | 3628 | 394.2 | - | - | 0 | - |
| - | - | 1109 | 395.1 | - | - | 0 | - |
| - | - | 907.9 | 395.2 | - | - | 0 | - |
| - | - | 781.4 | 395.2 | - | - | 0 | - |
| - | - | 3783 | 396.2 | - | - | 0 | - |
| - | - | 1401 | 398.2 | - | - | 0 | - |
| - | - | 1107 | 402.2 | - | - | 0 | - |
| - | - | 1159 | 402.2 | - | - | 0 | - |
| - | - | 1154 | 403.2 | - | - | 0 | - |
| - | - | 600.6 | 405.3 | - | - | 0 | - |
| - | - | 634 | 407.2 | - | - | 0 | - |
| - | - | 1844 | 409.1 | - | - | 0 | - |
| - | - | 1323 | 411.2 | - | - | 0 | - |
| - | - | 1.102E+04 | 412.2 | - | - | 0 | - |
| - | - | 1003 | 413.1 | - | - | 0 | - |
| - | - | 1555 | 413.2 | - | - | 0 | - |
| - | - | 813.8 | 413.3 | - | - | 0 | - |
| - | - | 636.8 | 414.2 | - | - | 0 | - |
| - | - | 3024 | 414.2 | - | - | 0 | - |
| - | - | 2.899E+04 | 415 | - | - | 0 | - |
| - | - | 1198 | 415.2 | - | - | 0 | - |
| - | - | 888.9 | 415.2 | - | - | 0 | - |
| - | - | 2234 | 417.7 | - | - | 0 | - |
| - | - | 4601 | 418.2 | - | - | 0 | - |
| - | - | 1640 | 418.7 | - | - | 0 | - |
| - | - | 5275 | 423.3 | - | - | 0 | - |
| - | - | 1008 | 424.3 | - | - | 0 | - |
| - | - | 3.358E+04 | 426.7 | - | - | 0 | - |
| - | - | 1.607E+04 | 427.2 | - | - | 0 | - |
| - | - | 4834 | 427.7 | - | - | 0 | - |
| - | - | 1105 | 428.2 | - | - | 0 | - |
| - | - | 9225 | 429.2 | - | - | 0 | - |
| - | - | 2.566E+04 | 430.2 | - | - | 0 | - |
| - | - | 5254 | 431.2 | - | - | 0 | - |
| - | - | 806.5 | 431.6 | - | - | 0 | - |
| 7 | b | 4126 | 431.7 | 0.0004715 | 1.092 | +2 | 7 |
| 7 | b | 2631 | 432.2 | 0.005015 | 11.6 | +2 | 7 |
| - | - | 1080 | 432.7 | - | - | 0 | - |
| - | - | 1297 | 433 | - | - | 0 | - |
| 4 | b | 1616 | 433.3 | 7.321E-05 | 0.169 | +1 | 4 |
| 11 | y | 7476 | 439.2 | 0.0004209 | 0.9584 | +1 | 3 |
| - | - | 1842 | 440.2 | - | - | 0 | - |
| 7 | b | 1.369E+04 | 440.7 | 7.2E-05 | 0.1634 | +2 | 7 |
| - | - | 1065 | 440.9 | - | - | 0 | - |
| - | - | 1.002E+04 | 441.2 | - | - | 0 | - |
| - | - | 1970 | 441.7 | - | - | 0 | - |
| 7 | y | 819.9 | 447.2 | 0.002054 | 4.592 | +2 | 7 |
| - | - | 788.4 | 447.7 | - | - | 0 | - |
| - | - | 1646 | 448.2 | - | - | 0 | - |
| - | - | 906.6 | 450.3 | - | - | 0 | - |
| 4 | b | 7945 | 451.3 | 0.0004095 | 0.9074 | +1 | 4 |
| - | - | 2090 | 452.3 | - | - | 0 | - |
| 7 | y | 626.7 | 455.7 | 0.0002956 | 0.6485 | +2 | 7 |
| - | - | 970.5 | 459.2 | - | - | 0 | - |
| - | - | 1081 | 460.8 | - | - | 0 | - |
| - | - | 1110 | 461.2 | - | - | 0 | - |
| - | - | 2294 | 468.3 | - | - | 0 | - |
| - | - | 1565 | 474.3 | - | - | 0 | - |
| - | - | 2442 | 474.8 | - | - | 0 | - |
| - | - | 1168 | 475.3 | - | - | 0 | - |
| - | - | 2187 | 477.2 | - | - | 0 | - |
| 10 | y | 1458 | 478.2 | 0.001668 | 3.488 | +1 | 4 |
| - | - | 1015 | 478.3 | - | - | 0 | - |
| - | - | 6850 | 483.3 | - | - | 0 | - |
| - | - | 3210 | 483.8 | - | - | 0 | - |
| - | - | 860.6 | 484.3 | - | - | 0 | - |
| 8 | b | 688.5 | 488.3 | 0.0006759 | 1.384 | +2 | 8 |
| 8 | b | 1107 | 488.7 | 0.006257 | 12.8 | +2 | 8 |
| - | - | 1.915E+04 | 494.2 | - | - | 0 | - |
| - | - | 5694 | 495.2 | - | - | 0 | - |
| 10 | y | 2.401E+04 | 496.2 | 0.0002585 | 0.5209 | +1 | 4 |
| - | - | 5595 | 497.2 | - | - | 0 | - |
| - | - | 2177 | 497.8 | - | - | 0 | - |
| - | - | 832 | 498.2 | - | - | 0 | - |
| - | - | 629.8 | 499.3 | - | - | 0 | - |
| - | - | 2.897E+04 | 503.1 | - | - | 0 | - |
| - | - | 677.9 | 504.2 | - | - | 0 | - |
| - | - | 934 | 506.2 | - | - | 0 | - |
| - | - | 765.8 | 507.3 | - | - | 0 | - |
| - | - | 1889 | 511.3 | - | - | 0 | - |
| - | - | 793.8 | 512.3 | - | - | 0 | - |
| - | - | 1.052E+04 | 519.1 | - | - | 0 | - |
| - | - | 942.9 | 523.3 | - | - | 0 | - |
| - | - | 3494 | 525.3 | - | - | 0 | - |
| - | - | 797.7 | 526.3 | - | - | 0 | - |
| 0 | Precursor | 1025 | 529.3 | 0.002634 | 4.976 | +3 | -1 |
| 0 | Precursor | 2223 | 529.6 | 0.006538 | 12.34 | +3 | -1 |
| - | - | 696 | 532.2 | - | - | 0 | - |
| - | - | 750.5 | 534.6 | - | - | 0 | - |
| - | - | 985.1 | 534.6 | - | - | 0 | - |
| - | - | 765.1 | 535.2 | - | - | 0 | - |
| 0 | Precursor | 1351 | 535.3 | 0.001559 | 2.913 | +3 | -1 |
| - | - | 1771 | 535.8 | - | - | 0 | - |
| - | - | 1133 | 535.9 | - | - | 0 | - |
| - | - | 1865 | 536.1 | - | - | 0 | - |
| - | - | 6921 | 536.2 | - | - | 0 | - |
| - | - | 1033 | 536.3 | - | - | 0 | - |
| - | - | 4044 | 539.2 | - | - | 0 | - |
| - | - | 936.8 | 540.2 | - | - | 0 | - |
| - | - | 1769 | 542.3 | - | - | 0 | - |
| - | - | 5160 | 543.3 | - | - | 0 | - |
| - | - | 2192 | 544.3 | - | - | 0 | - |
| - | - | 1870 | 555.3 | - | - | 0 | - |
| - | - | 1188 | 555.8 | - | - | 0 | - |
| 5 | b | 1039 | 561.3 | 0.003025 | 5.39 | +1 | 5 |
| - | - | 757.2 | 563.3 | - | - | 0 | - |
| - | - | 1158 | 565.3 | - | - | 0 | - |
| - | - | 4457 | 567.2 | - | - | 0 | - |
| - | - | 2328 | 568.2 | - | - | 0 | - |
| - | - | 928.5 | 575.2 | - | - | 0 | - |
| - | - | 792.6 | 576.2 | - | - | 0 | - |
| 5 | b | 5330 | 579.3 | 0.0005203 | 0.8982 | +1 | 5 |
| - | - | 2464 | 580.3 | - | - | 0 | - |
| - | - | 1969 | 592.3 | - | - | 0 | - |
| 9 | y | 7788 | 593.3 | 0.0002812 | 0.4741 | +1 | 5 |
| - | - | 2671 | 593.3 | - | - | 0 | - |
| - | - | 2484 | 594.3 | - | - | 0 | - |
| - | - | 703.1 | 594.3 | - | - | 0 | - |
| - | - | 979.5 | 595.3 | - | - | 0 | - |
| - | - | 633.5 | 598.3 | - | - | 0 | - |
| - | - | 1158 | 602.8 | - | - | 0 | - |
| - | - | 1786 | 603.3 | - | - | 0 | - |
| - | - | 2.514E+04 | 610.3 | - | - | 0 | - |
| - | - | 1082 | 610.3 | - | - | 0 | - |
| 9 | y | 7.434E+04 | 611.3 | 0.0005676 | 0.9286 | +1 | 5 |
| - | - | 3741 | 611.8 | - | - | 0 | - |
| - | - | 2.146E+04 | 612.3 | - | - | 0 | - |
| - | - | 777.2 | 612.8 | - | - | 0 | - |
| - | - | 5070 | 613.3 | - | - | 0 | - |
| - | - | 1402 | 621.3 | - | - | 0 | - |
| - | - | 970.2 | 634.3 | - | - | 0 | - |
| - | - | 1392 | 635.3 | - | - | 0 | - |
| 4 | y | 1520 | 636.8 | 6.023E-05 | 0.09458 | +2 | 10 |
| 4 | y | 811.6 | 637.3 | 0.009458 | 14.84 | +2 | 10 |
| - | - | 2012 | 645.3 | - | - | 0 | - |
| 4 | y | 4765 | 645.8 | 0.002836 | 4.392 | +2 | 10 |
| - | - | 2262 | 646.3 | - | - | 0 | - |
| - | - | 1588 | 646.8 | - | - | 0 | - |
| - | - | 634.2 | 648.4 | - | - | 0 | - |
| - | - | 8095 | 652.3 | - | - | 0 | - |
| - | - | 2906 | 653.3 | - | - | 0 | - |
| - | - | 2577 | 660.9 | - | - | 0 | - |
| - | - | 1228 | 661.4 | - | - | 0 | - |
| - | - | 2457 | 662.3 | - | - | 0 | - |
| - | - | 1738 | 663.3 | - | - | 0 | - |
| 6 | b | 3489 | 676.3 | 0.001091 | 1.612 | +1 | 6 |
| 6 | b | 1933 | 677.3 | 0.006288 | 9.283 | +1 | 6 |
| - | - | 793.2 | 677.8 | - | - | 0 | - |
| - | - | 1056 | 679.3 | - | - | 0 | - |
| - | - | 1.335E+04 | 680.3 | - | - | 0 | - |
| - | - | 5199 | 681.3 | - | - | 0 | - |
| - | - | 1670 | 682.3 | - | - | 0 | - |
| - | - | 1162 | 684.3 | - | - | 0 | - |
| - | - | 625 | 684.8 | - | - | 0 | - |
| - | - | 702.7 | 685.3 | - | - | 0 | - |
| 3 | y | 4571 | 693.3 | 0.00283 | 4.081 | +2 | 11 |
| 3 | y | 3498 | 693.8 | 0.006183 | 8.912 | +2 | 11 |
| 6 | b | 4.204E+04 | 694.4 | 0.000852 | 1.227 | +1 | 6 |
| - | - | 745 | 694.8 | - | - | 0 | - |
| - | - | 1.501E+04 | 695.4 | - | - | 0 | - |
| - | - | 602.9 | 695.9 | - | - | 0 | - |
| - | - | 3528 | 696.4 | - | - | 0 | - |
| - | - | 937.8 | 697.4 | - | - | 0 | - |
| - | - | 1.048E+04 | 701.8 | - | - | 0 | - |
| 3 | y | 2.569E+04 | 702.3 | 0.001087 | 1.548 | +2 | 11 |
| 12 | b | 2.015E+04 | 702.8 | 0.01259 | 17.92 | +2 | 12 |
| 12 | b | 8142 | 703.3 | 0.003259 | 4.633 | +2 | 12 |
| - | - | 2033 | 703.8 | - | - | 0 | - |
| - | - | 1153 | 704.3 | - | - | 0 | - |
| 8 | y | 2186 | 706.3 | 0.001714 | 2.427 | +1 | 6 |
| - | - | 1112 | 707.4 | - | - | 0 | - |
| - | - | 1104 | 710.4 | - | - | 0 | - |
| - | - | 1549 | 710.9 | - | - | 0 | - |
| - | - | 1940 | 711.4 | - | - | 0 | - |
| 12 | b | 1841 | 711.9 | 0.0001294 | 0.1817 | +2 | 12 |
| - | - | 1796 | 712.4 | - | - | 0 | - |
| - | - | 925 | 712.9 | - | - | 0 | - |
| - | - | 2160 | 721.4 | - | - | 0 | - |
| - | - | 814.8 | 721.9 | - | - | 0 | - |
| - | - | 848.6 | 722.4 | - | - | 0 | - |
| - | - | 5455 | 723.4 | - | - | 0 | - |
| 8 | y | 2.185E+04 | 724.4 | 0.0003049 | 0.421 | +1 | 6 |
| - | - | 9138 | 725.4 | - | - | 0 | - |
| - | - | 2118 | 726.4 | - | - | 0 | - |
| - | - | 1209 | 729.4 | - | - | 0 | - |
| 2 | y | 1658 | 743.4 | 0.01018 | 13.7 | +2 | 12 |
| - | - | 781.4 | 747.4 | - | - | 0 | - |
| - | - | 5276 | 751.4 | - | - | 0 | - |
| 2 | y | 1747 | 751.9 | 0.001548 | 2.059 | +2 | 12 |
| - | - | 3427 | 752.4 | - | - | 0 | - |
| - | - | 820.2 | 752.9 | - | - | 0 | - |
| - | - | 918.7 | 753.4 | - | - | 0 | - |
| - | - | 655.6 | 755.4 | - | - | 0 | - |
| - | - | 2651 | 763.4 | - | - | 0 | - |
| - | - | 1468 | 764.4 | - | - | 0 | - |
| - | - | 1120 | 765.4 | - | - | 0 | - |
| - | - | 734.6 | 776.4 | - | - | 0 | - |
| - | - | 2220 | 779.4 | - | - | 0 | - |
| - | - | 766.2 | 780.4 | - | - | 0 | - |
| - | - | 1022 | 781.4 | - | - | 0 | - |
| - | - | 846.3 | 782.4 | - | - | 0 | - |
| 0 | Precursor | 6048 | 793.4 | 0.01241 | 15.64 | +2 | -1 |
| - | - | 1989 | 794.4 | - | - | 0 | - |
| - | - | 943.5 | 795.3 | - | - | 0 | - |
| - | - | 1085 | 807.4 | - | - | 0 | - |
| - | - | 798.8 | 826.4 | - | - | 0 | - |
| - | - | 879.9 | 834.4 | - | - | 0 | - |
| - | - | 1072 | 835.4 | - | - | 0 | - |
| - | - | 3927 | 836.5 | - | - | 0 | - |
| - | - | 1679 | 837.5 | - | - | 0 | - |
| - | - | 991.6 | 844.4 | - | - | 0 | - |
| - | - | 751.6 | 845.4 | - | - | 0 | - |
| - | - | 949.2 | 846.5 | - | - | 0 | - |
| - | - | 1102 | 847.5 | - | - | 0 | - |
| - | - | 1.086E+04 | 852.4 | - | - | 0 | - |
| - | - | 5234 | 853.4 | - | - | 0 | - |
| - | - | 1357 | 854.4 | - | - | 0 | - |
| 7 | b | 6175 | 862.4 | 0.002767 | 3.208 | +1 | 7 |
| 7 | b | 5173 | 863.4 | 0.01078 | 12.48 | +1 | 7 |
| - | - | 8969 | 864.5 | - | - | 0 | - |
| - | - | 3067 | 865.5 | - | - | 0 | - |
| - | - | 1061 | 866.5 | - | - | 0 | - |
| 7 | b | 4.791E+04 | 880.4 | 0.001613 | 1.832 | +1 | 7 |
| - | - | 2.657E+04 | 881.4 | - | - | 0 | - |
| - | - | 7354 | 882.4 | - | - | 0 | - |
| - | - | 1510 | 883.4 | - | - | 0 | - |
| - | - | 2568 | 908.4 | - | - | 0 | - |
| - | - | 2113 | 909.4 | - | - | 0 | - |
| 7 | y | 3354 | 910.4 | 0.00107 | 1.175 | +1 | 7 |
| - | - | 1647 | 911.4 | - | - | 0 | - |
| - | - | 1032 | 954.4 | - | - | 0 | - |
| - | - | 1570 | 965.5 | - | - | 0 | - |
| - | - | 1120 | 966.4 | - | - | 0 | - |
| 8 | b | 3508 | 993.5 | 0.003523 | 3.546 | +1 | 8 |
| - | - | 1787 | 994.5 | - | - | 0 | - |
| - | - | 800.4 | 995.5 | - | - | 0 | - |
| 6 | y | 1128 | 1025 | 0.001654 | 1.613 | +1 | 8 |
| - | - | 1203 | 1026 | - | - | 0 | - |
| - | - | 675.7 | 1093 | - | - | 0 | - |
| 5 | y | 1428 | 1136 | 0.002414 | 2.126 | +1 | 9 |
| 5 | y | 1727 | 1136 | 0.008932 | 7.859 | +1 | 9 |
| - | - | 1243 | 1137 | - | - | 0 | - |
| - | - | 1382 | 1153 | - | - | 0 | - |
| 5 | y | 3350 | 1154 | 0.0003274 | 0.2838 | +1 | 9 |
| - | - | 2293 | 1155 | - | - | 0 | - |
| - | - | 1051 | 1156 | - | - | 0 | - |
| - | - | 760 | 2655 | - | - | 0 | - |
| - | - | 802.3 | 3080 | - | - | 0 | - |
| - | - | 800.3 | 3204 | - | - | 0 | - |
| - | - | 724.3 | 3368 | - | - | 0 | - |

m/z Charge Intensity FragmentType MassShift Position
120.0810546875 0 5848.0205
121.08458709716797 0 639.31305
123.04434204101562 0 1072.562
125.10746765136719 0 481.6427
126.02745056152344 0 420.62164
127.05036163330078 0 1845.4167
128.08238220214844 0 376.95087
128.10728454589844 0 8347.589
129.0662078857422 0 1100.4103
129.1025390625 0 84621.44
130.06541442871094 0 36086.266
130.08653259277344 0 449.97284
130.10025024414062 0 847.21063
130.10585021972656 0 5628.7583
131.063720703125 0 584.5628
131.0688018798828 0 3658.3
131.1076202392578 0 402.24017
132.08106994628906 0 8494.931
132.10218811035156 0 1135.8739
133.08578491210938 0 546.8229
134.02740478515625 0 476.23346
136.0759735107422 0 12206.118
137.07925415039062 0 1561.2172
138.06649780273438 0 1019.4452
139.08682250976562 0 987.30865
140.67709350585938 0 551.2758
141.1025848388672 0 2598.1028
143.1180419921875 0 630.273
145.0609588623047 0 984.7412
146.06008911132812 0 1095.2192
148.9547119140625 0 1053.7222
149.0451202392578 0 14142.122
149.05186462402344 0 964.03784
150.0447998046875 0 796.1482
150.06613159179688 0 539.2614
151.0421600341797 0 676.65137
151.0870361328125 0 1540.4783
155.04542541503906 0 441.3177
155.09304809570312 0 4728.5156
155.1182098388672 0 3488.9014 a Water loss 1
157.0980682373047 0 439.88507
158.0836181640625 0 575.35205
159.0919189453125 0 62095.766
159.3428955078125 0 407.184
160.08865356445312 0 891.1705
160.09536743164062 0 6394.129
165.0548553466797 0 5351.252
166.061279296875 0 9399.677
166.08653259277344 0 3089.2607 y 9
167.05569458007812 0 22047.09
167.06398010253906 0 1282.3877
168.05557250976562 0 635.51245
168.08120727539062 0 755.99176
168.1016387939453 0 538.98
168.11329650878906 0 985.05945
169.0762939453125 0 797.1347
169.0972900390625 0 3026.6145
170.06015014648438 0 2876.7246
171.1497802734375 0 619.7874
172.07176208496094 0 1634.9734
173.0563507080078 0 897.2929
173.12876892089844 0 123529.92 a 1
173.4520721435547 0 1234.7085
174.13204956054688 0 12017.472
175.1349334716797 0 560.6246
177.1028289794922 0 793.0291
178.13404846191406 0 3473.0383
179.13746643066406 0 544.3279
182.08143615722656 0 11314.695 y 12
183.08477783203125 0 1506.8344
183.1129913330078 0 3816.6638 b Water loss 1
183.1495361328125 0 1181.215
185.1650848388672 0 943.92035
186.12396240234375 0 24460.068
187.08685302734375 0 4703.409
187.1077423095703 0 490.93613
187.12753295898438 0 1997.6222
187.14471435546875 0 1085.8771
188.0713653564453 0 649.98694
190.08224487304688 0 827.0318
190.13385009765625 0 1187.5607
195.11305236816406 0 5946.08
196.10836791992188 0 4022.641
197.11244201660156 0 578.38635
197.12811279296875 0 817.9613
198.0880584716797 0 575.6096
198.12759399414062 0 1855.0804
199.1446990966797 0 717.27686
201.12355041503906 0 33087.363 b 1
202.12696838378906 0 3325.7563
205.097412109375 0 2205.5688
206.10281372070312 0 508.6034
207.16021728515625 0 1204.8588
209.0916290283203 0 543.8373
210.12791442871094 0 1183.3557
211.10791015625 0 563.69684 y Water loss 10
211.1317901611328 0 494.29446
211.1439971923828 0 729.43555
212.13922119140625 0 931.7433
213.12359619140625 0 3618.3486
213.1602020263672 0 692.49005
215.1402587890625 0 572.125
216.09796142578125 0 671.1774
221.05943298339844 0 2037.3612
221.08445739746094 0 6474.101
221.10337829589844 0 854.0705
221.1403045654297 0 847.2358
222.08497619628906 0 1622.0482
222.1238555908203 0 663.4042
223.06396484375 0 4657.2764
223.08074951171875 0 1247.0298
223.1080322265625 0 2615.0054
223.15557861328125 0 10710.073
224.1021728515625 0 931.3262
224.1592559814453 0 1335.5697
225.0431365966797 0 9572.489
226.04344177246094 0 2807.1206
226.08236694335938 0 2206.3418
226.11859130859375 0 603.2291 b Water loss 5
227.02252197265625 0 1040.2587
227.04037475585938 0 1000.26154
227.06634521484375 0 1840.372
228.1347198486328 0 1156.0105
229.1181640625 0 1823.6887
229.1546173095703 0 805.6519
233.1396026611328 0 1213.1592
234.12396240234375 0 13252.4
235.12750244140625 0 1894.9812
237.1350860595703 0 935.8748
238.1190185546875 0 1669.4556
239.09519958496094 0 21396.557
240.09591674804688 0 6297.499
240.1344451904297 0 5481.4434
241.09246826171875 0 4451.6284
241.11972045898438 0 854.02496
241.19117736816406 0 989.71893
243.10850524902344 0 874.7034
244.09295654296875 0 4926.882
245.09689331054688 0 635.37244
247.10787963867188 0 5564.1
248.10177612304688 0 1168.0365
248.11422729492188 0 2021.9034
249.05279541015625 0 632.38794
249.0984344482422 0 3388.1133
249.13531494140625 0 1386.8126
250.1018524169922 0 870.2851
251.15049743652344 0 42829.395
252.15377807617188 0 4903.729
255.14761352539062 0 3059.719
257.1126708984375 0 1180.7722
258.1449279785156 0 1983.4469
261.119140625 0 1242.3682
261.13433837890625 0 534.2072
262.11761474609375 0 670.52905
265.12945556640625 0 6328.814
266.1130676269531 0 1605.4154
266.125 0 10344.876
267.1293029785156 0 1091.5012
268.1768493652344 0 927.3114
269.1863708496094 0 834.1684
270.1217956542969 0 1816.1783
272.17578125 0 2964.767
274.11834716796875 0 1669.5312
275.1024169921875 0 641.17554
276.1549072265625 0 892.60443
279.1458435058594 0 792.18494
280.12884521484375 0 1520.7086
281.0513000488281 0 13507.002
282.1565246582031 0 3387.3594
283.0305480957031 0 1859.7526
283.1402587890625 0 31560.775
284.103271484375 0 1752.4467
284.12548828125 0 868.6898
284.1433410644531 0 4212.184
285.0096435546875 0 1677.6145
286.14007568359375 0 1627.5103
287.1715393066406 0 674.81323
288.6197814941406 0 1210.9562
293.1133728027344 0 15718.984 y Water loss 11
293.13092041015625 0 1046.898
294.11761474609375 0 3135.218
295.10333251953125 0 1798.6349
296.1051025390625 0 1739.6392
296.19720458984375 0 1169.3348 b Water loss 2
297.08294677734375 0 2203.8667
297.1001892089844 0 1096.1328
297.1559753417969 0 5161.1084
298.1589050292969 0 974.18225
299.06195068359375 0 129220.73
300.0622863769531 0 3140.8882
300.1673278808594 0 3839.2183
301.0597229003906 0 1997.0527
301.1509704589844 0 15377.551
301.1700439453125 0 1032.9402
302.1138000488281 0 2831.0815
302.1545715332031 0 1503.2028
305.64471435546875 0 1220.2068
306.13787841796875 0 1955.0256 y 8
306.63916015625 0 837.6793
311.1233825683594 0 1995.8525 y 11
311.1392517089844 0 827.0924
314.1874694824219 0 695.96704
314.20758056640625 0 2345.9734 b 2
315.1664123535156 0 8764.866
325.1501770019531 0 561.8957
326.6634216308594 0 2061.1514
326.96624755859375 0 981.67053
327.16668701171875 0 885.653
332.2087097167969 0 828.2083
333.1766052246094 0 1783.7491
341.01861572265625 0 1693.8243
342.1817321777344 0 741.0916
344.1722412109375 0 1541.5197
344.9765625 0 15207.437
345.1303405761719 0 1229.6145
346.114501953125 0 4893.004
347.6802062988281 0 2198.2239 b 5
350.2185974121094 0 2918.9363
351.2215576171875 0 910.8533
351.6806945800781 0 566.4668
353.6746826171875 0 989.86005 y Water loss 7
353.70147705078125 0 647.3018
355.0704345703125 0 2372.2515
359.0284118652344 0 15245.411
360.6941223144531 0 1586.8456
362.1842346191406 0 2025.2358
362.67913818359375 0 4501.9277 y 7
363.1410827636719 0 7246.397
363.1813659667969 0 3168.335
363.6839294433594 0 916.4889
364.1238708496094 0 1304.1624
364.146240234375 0 715.5406
366.17877197265625 0 883.12085
367.2442321777344 0 564.97894
368.1935119628906 0 1198.1128
369.120849609375 0 1279.4012
370.1227722167969 0 1965.1494
371.1197509765625 0 1525.2661
376.19769287109375 0 1773.0897
377.14630126953125 0 582.13495
378.21343994140625 0 2436.4192
379.2088317871094 0 11056.782
380.21112060546875 0 2029.5781
381.15179443359375 0 19382.95
382.1541442871094 0 3253.3242
382.180908203125 0 802.8871
384.16357421875 0 1112.1207
384.1883850097656 0 857.132
385.1500549316406 0 1503.5953
385.1740417480469 0 877.1796 y 4
394.1720275878906 0 3628.339
395.1324768066406 0 1108.8339
395.1564025878906 0 907.8902
395.2409362792969 0 781.377
396.2242431640625 0 3782.6978
398.1785888671875 0 1401.0857
402.17498779296875 0 1106.5226
402.2006530761719 0 1158.5864
403.20159912109375 0 1154.4673
405.2624816894531 0 600.63324
407.2051086425781 0 634.00574
409.14727783203125 0 1844.023
411.1986999511719 0 1323.4082
412.1829833984375 0 11024.002
413.141845703125 0 1003.36957
413.1876525878906 0 1554.8103
413.25115966796875 0 813.8018
414.18896484375 0 636.78253
414.23486328125 0 3023.6533
415.036865234375 0 28991.615
415.1993713378906 0 1198.2694
415.2366027832031 0 888.856
417.717041015625 0 2233.863
418.2101745605469 0 4601.0015
418.71075439453125 0 1639.9105
423.27191162109375 0 5275.486
424.2727966308594 0 1008.0607
426.72186279296875 0 33583.125
427.2234191894531 0 16065.514
427.7245788574219 0 4833.863
428.2281188964844 0 1105.0048
429.20941162109375 0 9225.351
430.1936340332031 0 25664.277
431.197021484375 0 5254.1147
431.5850524902344 0 806.54596
431.71441650390625 0 4125.594 b Water loss 6
432.2109680175781 0 2630.719 b Ammonia loss 6
432.71075439453125 0 1080.4691
433.0470275878906 0 1297.1703
433.2557067871094 0 1616.2717 b Water loss 3
439.2191467285156 0 7475.891 y 10
440.22247314453125 0 1842.4575
440.71929931640625 0 13687.994 b 6
440.91552734375 0 1064.8497
441.2207336425781 0 10018.404
441.722412109375 0 1970.2501
447.2076721191406 0 819.87714 y Ammonia loss 6
447.7069091796875 0 788.3611
448.2041931152344 0 1646.3317
450.2822570800781 0 906.59564
451.2667541503906 0 7944.954 b 3
452.26898193359375 0 2089.8572
455.7185974121094 0 626.71344 y 6
459.1964111328125 0 970.4539
460.7545471191406 0 1081.328
461.20220947265625 0 1110.4495
468.2937927246094 0 2294.214
474.2580261230469 0 1564.9456
474.75335693359375 0 2442.462
475.2500915527344 0 1167.5403
477.2101135253906 0 2186.8538
478.2312927246094 0 1458.0804 y Water loss 9
478.2767333984375 0 1015.0379
483.2641296386719 0 6849.6694
483.764892578125 0 3209.8782
484.2668151855469 0 860.5553
488.25665283203125 0 688.4831 b Water loss 7
488.7542419433594 0 1107.0925 b Ammonia loss 7
494.2357177734375 0 19146.53
495.2392883300781 0 5694.1123
496.2404479980469 0 24014.102 y 9
497.2459411621094 0 5595.1934
497.7624206542969 0 2176.7737
498.2394104003906 0 832.02264
499.2556457519531 0 629.8081
503.107421875 0 28965.924
504.199462890625 0 677.9372
506.2268371582031 0 933.9639
507.25543212890625 0 765.82526
511.26239013671875 0 1889.403
512.2596435546875 0 793.7621
519.1387329101562 0 10524.795
523.260498046875 0 942.94586
525.2666625976562 0 3494.4678
526.2728881835938 0 797.729
529.2623291015625 0 1024.673 Precursor Water loss
529.59423828125 0 2223.1877 Precursor Ammonia loss
532.1921997070312 0 696.0351
534.5520629882812 0 750.5427
534.6338500976562 0 985.1288
535.2171630859375 0 765.07025
535.2616577148438 0 1350.9263 Precursor
535.80810546875 0 1771.2839
535.9298706054688 0 1132.547
536.059814453125 0 1864.7506
536.164794921875 0 6921.256
536.313720703125 0 1033.0963
539.2356567382812 0 4043.5408
540.2412719726562 0 936.7701
542.2938232421875 0 1769.3546
543.27734375 0 5159.7764
544.2798461914062 0 2192.039
555.2579345703125 0 1870.4354
555.7578735351562 0 1187.707
561.3173828125 0 1039.4929 b Water loss 4
563.2568359375 0 757.1508
565.26123046875 0 1157.6599
567.2305297851562 0 4457.0654
568.2351684570312 0 2327.9119
575.2454833984375 0 928.5256
576.2322998046875 0 792.6156
579.3244018554688 0 5329.7803 b 4
580.3271484375 0 2463.5186
592.2725219726562 0 1969.374
593.2562866210938 0 7788.2637 y Water loss 8
593.3034057617188 0 2671.0068
594.2529907226562 0 2484.1555
594.3090209960938 0 703.07544
595.2545166015625 0 979.4956
598.3026123046875 0 633.48346
602.79150390625 0 1157.564
603.2877807617188 0 1786.4833
610.28271484375 0 25139.277
610.331298828125 0 1082.2845
611.2677001953125 0 74344.67 y 8
611.7979125976562 0 3741.3418
612.2710571289062 0 21460.666
612.8028564453125 0 777.19855
613.27197265625 0 5069.816
621.251953125 0 1401.7202
634.3109130859375 0 970.21375
635.2947387695312 0 1392.1805
636.7857666015625 0 1519.9188 y Water loss 3
637.2872924804688 0 811.5932 y Ammonia loss 3
645.2969970703125 0 2012.3335
645.7939453125 0 4765.148 y 3
646.2919921875 0 2261.8784
646.7963256835938 0 1588.2561
648.3598022460938 0 634.182
652.3195190429688 0 8094.754
653.3221435546875 0 2905.7566
660.870849609375 0 2577.229
661.3762817382812 0 1228.3196
662.3040771484375 0 2457.4043
663.3009033203125 0 1737.7933
676.3402099609375 0 3488.8054 b Water loss 5
677.3316040039062 0 1933.1232 b Ammonia loss 5
677.8048095703125 0 793.1939
679.3312377929688 0 1055.7578
680.3151245117188 0 13350.087
681.317626953125 0 5198.955
682.3195190429688 0 1670.4646
684.3184204101562 0 1162.2467
684.81884765625 0 624.9958
685.3157348632812 0 702.7488
693.3306884765625 0 4571.275 y Water loss 2
693.8260498046875 0 3497.8 y Ammonia loss 2
694.3510131835938 0 42041.43 b 5
694.8316650390625 0 745.02795
695.35400390625 0 15006.558
695.8797607421875 0 602.8685
696.3569946289062 0 3528.1824
697.3544921875 0 937.8246
701.8402709960938 0 10483.558
702.334228515625 0 25691.531 y 2
702.8363647460938 0 20147.676 b Water loss 11
703.3377075195312 0 8142.244 b Ammonia loss 11
703.8382568359375 0 2033.4536
704.3344116210938 0 1153.4146
706.3423461914062 0 2185.7717 y Water loss 7
707.3501586914062 0 1112.4207
710.406982421875 0 1104.4355
710.9100952148438 0 1549.4529
711.3695068359375 0 1940.4453
711.8543701171875 0 1841.2476 b 11
712.3588256835938 0 1796.4657
712.8612060546875 0 925.00494
721.361572265625 0 2160.2227
721.8587036132812 0 814.81744
722.3639526367188 0 848.6252
723.3663940429688 0 5454.7617
724.3515014648438 0 21852.8 y 7
725.3548583984375 0 9137.968
726.357421875 0 2117.9438
729.3585205078125 0 1209.0753
743.3642578125 0 1658.2825 y Ammonia loss 1
747.3876953125 0 781.4257
751.3738403320312 0 5276.469
751.868896484375 0 1747.1283 y 1
752.372314453125 0 3427.0852
752.869140625 0 820.209
753.3748779296875 0 918.67535
755.4364624023438 0 655.58417
763.3640747070312 0 2651.4885
764.3666381835938 0 1468.3795
765.3988647460938 0 1120.4824
776.3823852539062 0 734.58026
779.3825073242188 0 2219.7793
780.37451171875 0 766.24207
781.3646850585938 0 1021.9192
782.3681030273438 0 846.3299
793.3983154296875 0 6047.5337 Precursor Water loss
794.3989868164062 0 1989.0613
795.3399047851562 0 943.48346
807.3756713867188 0 1085.4753
826.3871459960938 0 798.83777
834.4208374023438 0 879.8951
835.413818359375 0 1072.0936
836.4515991210938 0 3926.8738
837.4525756835938 0 1678.9983
844.3939208984375 0 991.5901
845.3975830078125 0 751.57666
846.4519653320312 0 949.1673
847.4518432617188 0 1102.2089
852.43505859375 0 10855.615
853.43798828125 0 5233.535
854.439453125 0 1357.0001
862.4178466796875 0 6174.59 b Water loss 6
863.4154052734375 0 5173.066 b Ammonia loss 6
864.455078125 0 8968.978
865.4557495117188 0 3066.5615
866.4707641601562 0 1061.4169
880.4295654296875 0 47908.94 b 6
881.432373046875 0 26571.797
882.4357299804688 0 7353.615
883.4351196289062 0 1509.8234
908.4259643554688 0 2568.3494
909.431640625 0 2113.3523
910.4315795898438 0 3354.361 y 6
911.4342041015625 0 1646.8716
954.4312133789062 0 1032.3628
965.4502563476562 0 1570.3599
966.4431762695312 0 1120.0242
993.51171875 0 3508.385 b 7
994.5159301757812 0 1787.1562
995.5191650390625 0 800.4415
1025.4591064453125 0 1128.3359 y 5
1026.4661865234375 0 1203.1036
1092.5556640625 0 675.6929
1135.5030517578125 0 1427.7336 y Water loss 4
1136.4984130859375 0 1726.8148 y Ammonia loss 4
1137.49609375 0 1243.0747
1152.52490234375 0 1381.52
1153.516357421875 0 3349.9446 y 4
1154.517578125 0 2293.0264
1155.5220947265625 0 1050.8689
2654.579833984375 0 759.9703
3080.156494140625 0 802.3475
3203.998046875 0 800.32886
3367.790771484375 0 724.338

Spectrum Details

|  |  |
| --- | --- |
| Matched peaks? Matched peaks The total absolute number of peaks matched. Additionally in brackets the total fraction of peaks matched and the total number of peaks is shown. | 62 (12.40% of 500) |
| FDR? FDR The false discovery rate estimated for this peptide. It is calculated by matching all theoretical fragments with a non-integer shift with the raw peaks for this spectrum. This is done with 40 different shifts. The resulting percentage is the average number of annotated peaks over the number of annotated peaks with the correct spectrum. | 0.81% |
| Satellite FDR? Satellite FDR See the FDR for details on its calculation. This satellite ion specific FDR only contains the satellite ions (d/w) for I/L/J positions. | ∞ |
| PSM Score? PSM Score The PSM Score as given by Hecklib to this annotated spectrum. It is shown with three significant figures. | 442 |

#### Spectrum 5785? Spectrum 5785 The raw spectrum of this peptide as annotated by Hecklib. The fragments are coloured according to ion type (see legend). Any peaks with a star '\*' as text can be hovered over to see the full details, first the ion type second the mass shift type. By hovering over the amino acids in the peptide or ions in the legend the corresponding peaks are highlighted. By toggling the 'Unassigned' label you can turn the background (unassigned) peaks on or off in the plot. By updating the slider in the Ion legend you can update the spectrum to only show the top X% of the peaks with labels. The top X% means any peak that is within X% of the highest intensity. By dragging in the spectrum you can zoom in to a specific part of the spectrum and use 'Zoom Out' to get back to the original zoom level. The annotation of the spectrum is based on the given sequence in the peptides file and is done with different software so inconsistencies are likely. The peaks are annotated based on the given sequence, with 20 ppm tolerance.

Copy Data

##### Spectrum 5785 (TSV)

###### Preview

```
Loading example...
```

*Click on the button to copy the data to your clipboard.*

Mz MinMz MaxIntensity Max

WidthHeightPeptide font sizePeptide stroke widthSpectrum font sizeSpectrum stroke widthCompact peptide

Ion legend

wxyz

abcd

OtherUnassignedIon^Charge^_Position_Show for top:%

TVLHQDWLDGKEY

01.36e+52.72e+54.08e+55.44e+5

Zoom Out

y^+3^_3_y^+3^_4_y^+1^_1_c^+1^_2_c^+2^_5_y^+1^_2_c^+3^_7_y^+1^_2_c^+1^_3_c^+2^_6_y^+2^_6_y^+2^_6_z^+1^_3_y^+3^_10_y^+1^_3_c^+2^_7_c^+1^_4_y^+2^_7_c^+1^_4_c^+3^_12_y^+1^_4_z^+2^_8_c^+2^_8_w^+1^_5_c^+1^_5_c^+2^_10_y^+1^_5_z^+1^_5_c^+1^_5_y^+1^_5_y^+2^_10_z^+2^_10_y^+2^_10_c^+2^_11_c^+2^_11_c^+2^_11_w^+1^_6_w^+2^_11_z^+2^_11_c^+1^_6_y^+2^_11_y^+2^_11_z^+2^_11_z^+1^_6_y^+2^_11_c^+1^_6_c^+2^_12_c^+2^_12_y^+1^_6_c^+2^_12_y^+2^_12_z^+2^_12_y^+2^_12_z^+1^_7_c^+1^_7_y^+1^_7_c^+1^_7_y^+1^_7_c^+1^_8_z^+1^_8_c^+1^_8_y^+1^_8_w^+1^_9_c^+1^_9_c^+1^_9_z^+1^_9_y^+1^_9_c^+1^_10_c^+1^_10_y^+1^_10_z^+1^_10_y^+1^_10_c^+1^_11_z^+1^_11_y^+1^_11_z^+1^_11_y^+1^_11_c^+1^_12_y^+1^_12_z^+1^_12_y^+1^_12_

0779155723363115

Fragment Matches Table

Show background peaks

| Position | Ion type | Intensity | mz Theoretical | mz Error (Th) | mz Error (ppm) | Charge | Series Number |
| --- | --- | --- | --- | --- | --- | --- | --- |
| - | - | 377 | 127.5 | - | - | 0 | - |
| - | - | 736.5 | 128.1 | - | - | 0 | - |
| - | - | 4923 | 129.1 | - | - | 0 | - |
| - | - | 827.5 | 130.1 | - | - | 0 | - |
| - | - | 385.7 | 130.4 | - | - | 0 | - |
| - | - | 1609 | 131.1 | - | - | 0 | - |
| - | - | 483.4 | 135 | - | - | 0 | - |
| - | - | 419.1 | 136 | - | - | 0 | - |
| - | - | 1431 | 136.1 | - | - | 0 | - |
| - | - | 2239 | 145.1 | - | - | 0 | - |
| - | - | 917 | 146.1 | - | - | 0 | - |
| 11 | y | 1973 | 147.1 | 0.0009468 | 6.438 | +3 | 3 |
| - | - | 1.08E+04 | 155.1 | - | - | 0 | - |
| - | - | 1489 | 157.1 | - | - | 0 | - |
| - | - | 555.5 | 160.1 | - | - | 0 | - |
| - | - | 490.9 | 163.5 | - | - | 0 | - |
| - | - | 514 | 165.1 | - | - | 0 | - |
| 10 | y | 3936 | 166.1 | 0.001573 | 9.469 | +3 | 4 |
| - | - | 732.1 | 170.1 | - | - | 0 | - |
| - | - | 1116 | 171.1 | - | - | 0 | - |
| - | - | 3.921E+04 | 173.1 | - | - | 0 | - |
| - | - | 3140 | 174.1 | - | - | 0 | - |
| - | - | 3007 | 175.1 | - | - | 0 | - |
| 13 | y | 6480 | 182.1 | 0.0003428 | 1.883 | +1 | 1 |
| - | - | 1603 | 183.1 | - | - | 0 | - |
| - | - | 719.6 | 184.1 | - | - | 0 | - |
| - | - | 848 | 185.1 | - | - | 0 | - |
| - | - | 2239 | 186.1 | - | - | 0 | - |
| - | - | 1255 | 188.1 | - | - | 0 | - |
| - | - | 501.3 | 188.7 | - | - | 0 | - |
| - | - | 2.829E+04 | 201.1 | - | - | 0 | - |
| - | - | 2995 | 201.1 | - | - | 0 | - |
| - | - | 2050 | 202.1 | - | - | 0 | - |
| - | - | 521.3 | 206.8 | - | - | 0 | - |
| - | - | 584.2 | 207.1 | - | - | 0 | - |
| - | - | 484.9 | 207.5 | - | - | 0 | - |
| - | - | 7862 | 215.1 | - | - | 0 | - |
| - | - | 853.9 | 216.1 | - | - | 0 | - |
| 2 | c | 2491 | 218.1 | 0.0004422 | 2.027 | +1 | 2 |
| - | - | 1256 | 223.2 | - | - | 0 | - |
| - | - | 1247 | 226.1 | - | - | 0 | - |
| - | - | 1748 | 228.1 | - | - | 0 | - |
| - | - | 2828 | 243.1 | - | - | 0 | - |
| - | - | 500.7 | 243.4 | - | - | 0 | - |
| - | - | 2937 | 245.1 | - | - | 0 | - |
| - | - | 5112 | 251.2 | - | - | 0 | - |
| - | - | 631.3 | 255.1 | - | - | 0 | - |
| - | - | 9323 | 258.1 | - | - | 0 | - |
| - | - | 1174 | 259.1 | - | - | 0 | - |
| - | - | 3509 | 260.2 | - | - | 0 | - |
| - | - | 6249 | 261.2 | - | - | 0 | - |
| - | - | 515.5 | 266 | - | - | 0 | - |
| - | - | 7213 | 268.2 | - | - | 0 | - |
| - | - | 646.4 | 272.2 | - | - | 0 | - |
| - | - | 730.1 | 283 | - | - | 0 | - |
| - | - | 1405 | 283.1 | - | - | 0 | - |
| - | - | 593.1 | 288.2 | - | - | 0 | - |
| 5 | c | 633 | 290.2 | 0.003471 | 11.96 | +2 | 5 |
| 12 | y | 4991 | 293.1 | 0.0004799 | 1.637 | +1 | 2 |
| - | - | 784.3 | 294.1 | - | - | 0 | - |
| - | - | 820.6 | 298.2 | - | - | 0 | - |
| - | - | 631.5 | 299.1 | - | - | 0 | - |
| 7 | c | 794.2 | 299.2 | 0.003416 | 11.42 | +3 | 7 |
| - | - | 848.8 | 300.1 | - | - | 0 | - |
| - | - | 793.8 | 300.2 | - | - | 0 | - |
| - | - | 906.6 | 301.1 | - | - | 0 | - |
| - | - | 1562 | 301.2 | - | - | 0 | - |
| - | - | 996.9 | 306.2 | - | - | 0 | - |
| - | - | 714.3 | 307.2 | - | - | 0 | - |
| 12 | y | 2749 | 311.1 | 0.001176 | 3.78 | +1 | 2 |
| - | - | 3983 | 312.2 | - | - | 0 | - |
| - | - | 2263 | 314.2 | - | - | 0 | - |
| - | - | 1786 | 315.2 | - | - | 0 | - |
| - | - | 1.683E+04 | 317.2 | - | - | 0 | - |
| - | - | 2223 | 318.2 | - | - | 0 | - |
| - | - | 688.7 | 319.2 | - | - | 0 | - |
| - | - | 2766 | 320.2 | - | - | 0 | - |
| - | - | 1320 | 322.2 | - | - | 0 | - |
| - | - | 1704 | 330.2 | - | - | 0 | - |
| 3 | c | 1.477E+04 | 331.2 | 0.0007288 | 2.2 | +1 | 3 |
| - | - | 580.5 | 332.2 | - | - | 0 | - |
| - | - | 2139 | 332.2 | - | - | 0 | - |
| - | - | 1491 | 339.2 | - | - | 0 | - |
| - | - | 1942 | 343.1 | - | - | 0 | - |
| 6 | c | 3449 | 347.7 | 0.0005439 | 1.564 | +2 | 6 |
| - | - | 1068 | 348.2 | - | - | 0 | - |
| 8 | y | 780.2 | 353.7 | 0.0001565 | 0.4426 | +2 | 6 |
| - | - | 2805 | 357.2 | - | - | 0 | - |
| - | - | 750.6 | 358.2 | - | - | 0 | - |
| - | - | 909 | 362.2 | - | - | 0 | - |
| 8 | y | 2227 | 362.7 | 0.0001288 | 0.3552 | +2 | 6 |
| - | - | 698.8 | 363.2 | - | - | 0 | - |
| - | - | 1957 | 364.2 | - | - | 0 | - |
| - | - | 861.3 | 365.2 | - | - | 0 | - |
| - | - | 832.6 | 366.1 | - | - | 0 | - |
| - | - | 2616 | 374.2 | - | - | 0 | - |
| - | - | 4446 | 376.2 | - | - | 0 | - |
| - | - | 809.1 | 377.2 | - | - | 0 | - |
| - | - | 1131 | 378.2 | - | - | 0 | - |
| - | - | 1249 | 379.2 | - | - | 0 | - |
| - | - | 1995 | 381.2 | - | - | 0 | - |
| - | - | 1022 | 381.2 | - | - | 0 | - |
| - | - | 693.7 | 382.2 | - | - | 0 | - |
| - | - | 790.8 | 387.2 | - | - | 0 | - |
| - | - | 910.7 | 389.2 | - | - | 0 | - |
| - | - | 2237 | 391.2 | - | - | 0 | - |
| - | - | 6393 | 392.2 | - | - | 0 | - |
| - | - | 2494 | 393.2 | - | - | 0 | - |
| - | - | 562.6 | 398.2 | - | - | 0 | - |
| - | - | 893.4 | 408.3 | - | - | 0 | - |
| - | - | 1244 | 409.2 | - | - | 0 | - |
| - | - | 2736 | 412.2 | - | - | 0 | - |
| - | - | 1278 | 412.7 | - | - | 0 | - |
| - | - | 2545 | 417.2 | - | - | 0 | - |
| - | - | 586.6 | 418.2 | - | - | 0 | - |
| 11 | z | 2639 | 423.2 | 0.001292 | 3.053 | +1 | 3 |
| - | - | 7255 | 423.3 | - | - | 0 | - |
| - | - | 996.4 | 424.2 | - | - | 0 | - |
| - | - | 1134 | 424.3 | - | - | 0 | - |
| - | - | 808.8 | 424.7 | - | - | 0 | - |
| - | - | 796.6 | 428.2 | - | - | 0 | - |
| - | - | 1695 | 429.2 | - | - | 0 | - |
| - | - | 1475 | 429.2 | - | - | 0 | - |
| 4 | y | 4659 | 430.2 | 0.004143 | 9.631 | +3 | 10 |
| - | - | 856.4 | 431.2 | - | - | 0 | - |
| - | - | 747.1 | 433.3 | - | - | 0 | - |
| - | - | 3435 | 434.7 | - | - | 0 | - |
| - | - | 2774 | 435.2 | - | - | 0 | - |
| 11 | y | 3060 | 439.2 | 0.0002989 | 0.6804 | +1 | 3 |
| - | - | 4633 | 439.7 | - | - | 0 | - |
| - | - | 2486 | 440.2 | - | - | 0 | - |
| - | - | 2459 | 442.3 | - | - | 0 | - |
| - | - | 1247 | 444.3 | - | - | 0 | - |
| - | - | 873 | 445.3 | - | - | 0 | - |
| - | - | 1043 | 446.3 | - | - | 0 | - |
| 7 | c | 7875 | 448.7 | 0.00321 | 7.155 | +2 | 7 |
| - | - | 3945 | 449.2 | - | - | 0 | - |
| - | - | 1619 | 449.7 | - | - | 0 | - |
| 4 | c | 924.9 | 450.3 | 0.0009351 | 2.077 | +1 | 4 |
| - | - | 1.675E+04 | 451.3 | - | - | 0 | - |
| - | - | 4093 | 452.3 | - | - | 0 | - |
| - | - | 7618 | 454.2 | - | - | 0 | - |
| 7 | y | 1.178E+04 | 454.7 | 0.005409 | 11.9 | +2 | 7 |
| - | - | 5918 | 455.2 | - | - | 0 | - |
| - | - | 2043 | 455.7 | - | - | 0 | - |
| - | - | 796.3 | 461.2 | - | - | 0 | - |
| - | - | 923.8 | 461.3 | - | - | 0 | - |
| - | - | 4938 | 463.2 | - | - | 0 | - |
| - | - | 1714 | 464.2 | - | - | 0 | - |
| 4 | c | 5.241E+04 | 468.3 | 0.0007769 | 1.659 | +1 | 4 |
| - | - | 899.5 | 468.8 | - | - | 0 | - |
| - | - | 1.267E+04 | 469.3 | - | - | 0 | - |
| - | - | 1075 | 470.3 | - | - | 0 | - |
| - | - | 627.9 | 471 | - | - | 0 | - |
| - | - | 889.8 | 473.7 | - | - | 0 | - |
| - | - | 857.6 | 476.2 | - | - | 0 | - |
| - | - | 2507 | 478.2 | - | - | 0 | - |
| - | - | 3725 | 479.2 | - | - | 0 | - |
| 12 | c | 8088 | 480.2 | 0.008019 | 16.7 | +3 | 12 |
| - | - | 5440 | 481.2 | - | - | 0 | - |
| - | - | 3892 | 482.3 | - | - | 0 | - |
| - | - | 2325 | 482.8 | - | - | 0 | - |
| - | - | 863.2 | 488.2 | - | - | 0 | - |
| - | - | 4027 | 488.8 | - | - | 0 | - |
| - | - | 4060 | 489.2 | - | - | 0 | - |
| - | - | 1787 | 489.3 | - | - | 0 | - |
| - | - | 854 | 489.8 | - | - | 0 | - |
| - | - | 3865 | 491.3 | - | - | 0 | - |
| - | - | 1625 | 491.8 | - | - | 0 | - |
| - | - | 6759 | 494.2 | - | - | 0 | - |
| - | - | 2971 | 494.3 | - | - | 0 | - |
| - | - | 4094 | 495.2 | - | - | 0 | - |
| - | - | 995 | 495.3 | - | - | 0 | - |
| 10 | y | 2.106E+04 | 496.2 | 0.002212 | 4.457 | +1 | 4 |
| - | - | 2214 | 496.8 | - | - | 0 | - |
| - | - | 5050 | 497.2 | - | - | 0 | - |
| - | - | 1844 | 498.2 | - | - | 0 | - |
| 6 | z | 2785 | 504.2 | 0.008807 | 17.47 | +2 | 8 |
| 8 | c | 3066 | 505.3 | 0.004117 | 8.148 | +2 | 8 |
| - | - | 1247 | 505.8 | - | - | 0 | - |
| - | - | 784.9 | 506.6 | - | - | 0 | - |
| - | - | 1062 | 507.1 | - | - | 0 | - |
| - | - | 9643 | 511.3 | - | - | 0 | - |
| - | - | 1881 | 512.3 | - | - | 0 | - |
| - | - | 4.88E+04 | 513.4 | - | - | 0 | - |
| - | - | 686.5 | 514.3 | - | - | 0 | - |
| - | - | 9467 | 514.4 | - | - | 0 | - |
| - | - | 805.1 | 520.2 | - | - | 0 | - |
| - | - | 952 | 523.2 | - | - | 0 | - |
| - | - | 7073 | 523.3 | - | - | 0 | - |
| - | - | 1436 | 524.2 | - | - | 0 | - |
| - | - | 769.6 | 524.2 | - | - | 0 | - |
| - | - | 1923 | 524.3 | - | - | 0 | - |
| - | - | 1327 | 524.3 | - | - | 0 | - |
| - | - | 1904 | 525.2 | - | - | 0 | - |
| - | - | 2542 | 525.6 | - | - | 0 | - |
| - | - | 2455 | 525.8 | - | - | 0 | - |
| - | - | 1809 | 526.3 | - | - | 0 | - |
| - | - | 772.4 | 527.3 | - | - | 0 | - |
| - | - | 2541 | 532.3 | - | - | 0 | - |
| - | - | 9983 | 533.3 | - | - | 0 | - |
| - | - | 2434 | 534.3 | - | - | 0 | - |
| - | - | 2317 | 534.6 | - | - | 0 | - |
| - | - | 2372 | 534.9 | - | - | 0 | - |
| - | - | 1608 | 535.3 | - | - | 0 | - |
| - | - | 1077 | 535.6 | - | - | 0 | - |
| - | - | 1096 | 535.9 | - | - | 0 | - |
| - | - | 862 | 536.2 | - | - | 0 | - |
| - | - | 2.37E+04 | 538.3 | - | - | 0 | - |
| - | - | 5425 | 539.3 | - | - | 0 | - |
| - | - | 6.043E+04 | 539.8 | - | - | 0 | - |
| - | - | 1862 | 540.3 | - | - | 0 | - |
| - | - | 4.136E+04 | 540.3 | - | - | 0 | - |
| - | - | 2663 | 540.6 | - | - | 0 | - |
| - | - | 1.541E+04 | 540.8 | - | - | 0 | - |
| - | - | 2311 | 540.9 | - | - | 0 | - |
| - | - | 5.386E+05 | 541.3 | - | - | 0 | - |
| - | - | 5653 | 541.6 | - | - | 0 | - |
| - | - | 815.6 | 541.8 | - | - | 0 | - |
| - | - | 6199 | 541.9 | - | - | 0 | - |
| - | - | 1.133E+05 | 542.3 | - | - | 0 | - |
| - | - | 718.4 | 543.3 | - | - | 0 | - |
| - | - | 5813 | 546.3 | - | - | 0 | - |
| - | - | 7125 | 546.8 | - | - | 0 | - |
| - | - | 2188 | 547.3 | - | - | 0 | - |
| - | - | 784.6 | 547.8 | - | - | 0 | - |
| 9 | w | 1.104E+04 | 550.3 | 0.0008937 | 1.624 | +1 | 5 |
| - | - | 874.3 | 550.3 | - | - | 0 | - |
| - | - | 3740 | 551.3 | - | - | 0 | - |
| - | - | 1026 | 552.3 | - | - | 0 | - |
| - | - | 3496 | 554.3 | - | - | 0 | - |
| - | - | 1338 | 554.8 | - | - | 0 | - |
| - | - | 857.8 | 555.3 | - | - | 0 | - |
| - | - | 1232 | 561.3 | - | - | 0 | - |
| - | - | 814.1 | 563.3 | - | - | 0 | - |
| - | - | 5379 | 564.3 | - | - | 0 | - |
| - | - | 890.8 | 565.3 | - | - | 0 | - |
| - | - | 1123 | 568.3 | - | - | 0 | - |
| - | - | 2307 | 571.3 | - | - | 0 | - |
| - | - | 1308 | 572.3 | - | - | 0 | - |
| - | - | 1892 | 578.3 | - | - | 0 | - |
| 5 | c | 4529 | 579.3 | 0.001802 | 3.111 | +1 | 5 |
| - | - | 884.3 | 580.3 | - | - | 0 | - |
| - | - | 1112 | 583.2 | - | - | 0 | - |
| - | - | 1129 | 588.8 | - | - | 0 | - |
| - | - | 2265 | 589.3 | - | - | 0 | - |
| - | - | 5961 | 589.8 | - | - | 0 | - |
| - | - | 1.051E+04 | 590.3 | - | - | 0 | - |
| 10 | c | 4581 | 590.8 | 0.005185 | 8.777 | +2 | 10 |
| - | - | 1131 | 591.3 | - | - | 0 | - |
| - | - | 3645 | 592.3 | - | - | 0 | - |
| 9 | y | 2712 | 593.3 | 0.001672 | 2.818 | +1 | 5 |
| - | - | 2371 | 594.3 | - | - | 0 | - |
| 9 | z | 1.122E+04 | 595.2 | 0.001714 | 2.879 | +1 | 5 |
| - | - | 1790 | 595.3 | - | - | 0 | - |
| - | - | 6701 | 596.3 | - | - | 0 | - |
| 5 | c | 5.822E+04 | 596.4 | 0.0007626 | 1.279 | +1 | 5 |
| - | - | 952.5 | 597.2 | - | - | 0 | - |
| - | - | 2119 | 597.3 | - | - | 0 | - |
| - | - | 1.992E+04 | 597.4 | - | - | 0 | - |
| - | - | 1.731E+04 | 597.8 | - | - | 0 | - |
| - | - | 7868 | 598.3 | - | - | 0 | - |
| - | - | 4870 | 598.4 | - | - | 0 | - |
| - | - | 2901 | 598.8 | - | - | 0 | - |
| - | - | 685.7 | 601.8 | - | - | 0 | - |
| - | - | 1.593E+04 | 610.3 | - | - | 0 | - |
| 9 | y | 2.998E+04 | 611.3 | 0.002338 | 3.824 | +1 | 5 |
| - | - | 8727 | 612.3 | - | - | 0 | - |
| - | - | 2007 | 613.3 | - | - | 0 | - |
| - | - | 1171 | 613.3 | - | - | 0 | - |
| - | - | 2161 | 614.3 | - | - | 0 | - |
| - | - | 1071 | 615.3 | - | - | 0 | - |
| - | - | 1237 | 619.8 | - | - | 0 | - |
| - | - | 1285 | 620.3 | - | - | 0 | - |
| - | - | 1052 | 621.3 | - | - | 0 | - |
| - | - | 1172 | 622.8 | - | - | 0 | - |
| - | - | 2076 | 623.3 | - | - | 0 | - |
| - | - | 885.4 | 624.3 | - | - | 0 | - |
| - | - | 1239 | 625.4 | - | - | 0 | - |
| - | - | 729.3 | 626.3 | - | - | 0 | - |
| - | - | 3730 | 626.4 | - | - | 0 | - |
| - | - | 1419 | 627.4 | - | - | 0 | - |
| - | - | 1464 | 631.8 | - | - | 0 | - |
| - | - | 1308 | 632.3 | - | - | 0 | - |
| - | - | 916.4 | 634.3 | - | - | 0 | - |
| - | - | 1.111E+04 | 636.3 | - | - | 0 | - |
| - | - | 4657 | 637.3 | - | - | 0 | - |
| - | - | 1.009E+04 | 639.3 | - | - | 0 | - |
| - | - | 6328 | 639.8 | - | - | 0 | - |
| - | - | 2400 | 640.3 | - | - | 0 | - |
| - | - | 811.9 | 641.8 | - | - | 0 | - |
| - | - | 2081 | 642.3 | - | - | 0 | - |
| - | - | 916.7 | 642.8 | - | - | 0 | - |
| 4 | y | 759.1 | 645.3 | 0.01196 | 18.53 | +2 | 10 |
| 4 | z | 1667 | 645.8 | 0.009752 | 15.1 | +2 | 10 |
| - | - | 1.705E+04 | 647.3 | - | - | 0 | - |
| - | - | 9741 | 647.8 | - | - | 0 | - |
| - | - | 8966 | 648.3 | - | - | 0 | - |
| - | - | 2598 | 649.3 | - | - | 0 | - |
| - | - | 2441 | 651.3 | - | - | 0 | - |
| - | - | 1.216E+04 | 652.3 | - | - | 0 | - |
| - | - | 1445 | 653.3 | - | - | 0 | - |
| - | - | 2603 | 653.3 | - | - | 0 | - |
| 4 | y | 3257 | 653.8 | 0.00631 | 9.652 | +2 | 10 |
| - | - | 1331 | 654.3 | - | - | 0 | - |
| - | - | 1165 | 654.3 | - | - | 0 | - |
| 11 | c | 683.6 | 654.8 | 0.004579 | 6.992 | +2 | 11 |
| 11 | c | 1966 | 655.3 | 0.01135 | 17.32 | +2 | 11 |
| - | - | 1078 | 657.8 | - | - | 0 | - |
| - | - | 2653 | 662.8 | - | - | 0 | - |
| - | - | 2.891E+04 | 663.4 | - | - | 0 | - |
| 11 | c | 7.116E+04 | 663.8 | 0.005644 | 8.502 | +2 | 11 |
| - | - | 5808 | 664.3 | - | - | 0 | - |
| - | - | 4.669E+04 | 664.3 | - | - | 0 | - |
| - | - | 1.818E+04 | 664.8 | - | - | 0 | - |
| 8 | w | 3001 | 665.3 | 0.0124 | 18.64 | +1 | 6 |
| - | - | 5525 | 665.4 | - | - | 0 | - |
| - | - | 1890 | 665.8 | - | - | 0 | - |
| - | - | 1401 | 666.3 | - | - | 0 | - |
| - | - | 1123 | 666.4 | - | - | 0 | - |
| - | - | 866.3 | 667.4 | - | - | 0 | - |
| - | - | 1009 | 668.3 | - | - | 0 | - |
| - | - | 1508 | 671.8 | - | - | 0 | - |
| - | - | 7908 | 674.3 | - | - | 0 | - |
| - | - | 4399 | 674.8 | - | - | 0 | - |
| - | - | 1546 | 675.3 | - | - | 0 | - |
| - | - | 3673 | 676.3 | - | - | 0 | - |
| - | - | 4129 | 676.9 | - | - | 0 | - |
| - | - | 4255 | 677.4 | - | - | 0 | - |
| - | - | 1320 | 677.9 | - | - | 0 | - |
| - | - | 899.5 | 679.3 | - | - | 0 | - |
| - | - | 1.937E+04 | 679.3 | - | - | 0 | - |
| - | - | 6600 | 680.3 | - | - | 0 | - |
| 3 | w | 9737 | 680.8 | 0.006338 | 9.31 | +2 | 11 |
| - | - | 4921 | 681.3 | - | - | 0 | - |
| - | - | 994.9 | 681.4 | - | - | 0 | - |
| - | - | 2053 | 681.8 | - | - | 0 | - |
| - | - | 999.3 | 682.3 | - | - | 0 | - |
| - | - | 730.3 | 692.3 | - | - | 0 | - |
| - | - | 843.8 | 692.8 | - | - | 0 | - |
| 3 | z | 1314 | 693.3 | 0.006755 | 9.743 | +2 | 11 |
| 6 | c | 5.332E+04 | 694.4 | 0.0006738 | 0.9704 | +1 | 6 |
| - | - | 1180 | 695.3 | - | - | 0 | - |
| - | - | 1.986E+04 | 695.4 | - | - | 0 | - |
| - | - | 923.7 | 695.8 | - | - | 0 | - |
| - | - | 3003 | 696.3 | - | - | 0 | - |
| - | - | 8465 | 696.4 | - | - | 0 | - |
| - | - | 1705 | 697.3 | - | - | 0 | - |
| - | - | 1886 | 697.4 | - | - | 0 | - |
| - | - | 1308 | 698.4 | - | - | 0 | - |
| - | - | 812.3 | 699.4 | - | - | 0 | - |
| - | - | 1109 | 700.4 | - | - | 0 | - |
| - | - | 2149 | 700.8 | - | - | 0 | - |
| 3 | y | 4381 | 701.3 | 0.005999 | 8.553 | +2 | 11 |
| 3 | y | 7275 | 701.8 | 0.01283 | 18.28 | +2 | 11 |
| 3 | z | 6008 | 702.3 | 0.007515 | 10.7 | +2 | 11 |
| - | - | 3417 | 702.8 | - | - | 0 | - |
| - | - | 998.8 | 703.3 | - | - | 0 | - |
| - | - | 5296 | 703.8 | - | - | 0 | - |
| - | - | 4834 | 704.3 | - | - | 0 | - |
| - | - | 768.4 | 704.4 | - | - | 0 | - |
| - | - | 1729 | 704.8 | - | - | 0 | - |
| - | - | 2970 | 706.4 | - | - | 0 | - |
| - | - | 1747 | 706.9 | - | - | 0 | - |
| - | - | 6509 | 707.4 | - | - | 0 | - |
| - | - | 1248 | 707.9 | - | - | 0 | - |
| 8 | z | 5808 | 708.3 | 0.01225 | 17.3 | +1 | 6 |
| - | - | 704.9 | 708.8 | - | - | 0 | - |
| - | - | 3505 | 709.3 | - | - | 0 | - |
| - | - | 1.13E+04 | 709.8 | - | - | 0 | - |
| 3 | y | 2.063E+04 | 710.3 | 0.006026 | 8.484 | +2 | 11 |
| - | - | 1.679E+04 | 710.8 | - | - | 0 | - |
| 6 | c | 1.112E+05 | 711.4 | 0.0007971 | 1.12 | +1 | 6 |
| - | - | 2.021E+04 | 711.9 | - | - | 0 | - |
| - | - | 4.323E+04 | 712.4 | - | - | 0 | - |
| - | - | 4947 | 712.9 | - | - | 0 | - |
| - | - | 1.038E+04 | 713.4 | - | - | 0 | - |
| - | - | 2177 | 714.4 | - | - | 0 | - |
| - | - | 3793 | 715.9 | - | - | 0 | - |
| - | - | 2169 | 716.4 | - | - | 0 | - |
| - | - | 930.4 | 716.9 | - | - | 0 | - |
| - | - | 943.8 | 717.9 | - | - | 0 | - |
| - | - | 1629 | 718.4 | - | - | 0 | - |
| - | - | 1099 | 718.9 | - | - | 0 | - |
| 12 | c | 2533 | 719.4 | 0.007025 | 9.765 | +2 | 12 |
| 12 | c | 1915 | 719.8 | 0.01038 | 14.42 | +2 | 12 |
| - | - | 2016 | 720.4 | - | - | 0 | - |
| - | - | 1888 | 720.9 | - | - | 0 | - |
| - | - | 3668 | 721.4 | - | - | 0 | - |
| - | - | 1102 | 721.9 | - | - | 0 | - |
| - | - | 1668 | 722.4 | - | - | 0 | - |
| - | - | 7837 | 723.4 | - | - | 0 | - |
| - | - | 1044 | 723.8 | - | - | 0 | - |
| 8 | y | 1.407E+04 | 724.4 | 0.002746 | 3.791 | +1 | 6 |
| - | - | 6752 | 725.4 | - | - | 0 | - |
| - | - | 1184 | 725.9 | - | - | 0 | - |
| - | - | 2497 | 726.4 | - | - | 0 | - |
| - | - | 1221 | 727.4 | - | - | 0 | - |
| - | - | 4.954E+04 | 727.9 | - | - | 0 | - |
| 12 | c | 8.624E+04 | 728.4 | 0.007724 | 10.6 | +2 | 12 |
| - | - | 5.093E+04 | 728.9 | - | - | 0 | - |
| - | - | 2.276E+04 | 729.4 | - | - | 0 | - |
| - | - | 1.364E+04 | 729.9 | - | - | 0 | - |
| - | - | 3871 | 730.4 | - | - | 0 | - |
| - | - | 2090 | 730.9 | - | - | 0 | - |
| - | - | 1753 | 734.4 | - | - | 0 | - |
| - | - | 1023 | 735.4 | - | - | 0 | - |
| - | - | 1921 | 735.9 | - | - | 0 | - |
| - | - | 3397 | 736.4 | - | - | 0 | - |
| - | - | 6641 | 736.9 | - | - | 0 | - |
| - | - | 1.476E+04 | 737.4 | - | - | 0 | - |
| - | - | 4.926E+04 | 737.9 | - | - | 0 | - |
| - | - | 3.055E+04 | 738.4 | - | - | 0 | - |
| - | - | 1.261E+04 | 738.9 | - | - | 0 | - |
| - | - | 4338 | 739.4 | - | - | 0 | - |
| - | - | 938.2 | 743.9 | - | - | 0 | - |
| - | - | 2435 | 744.4 | - | - | 0 | - |
| - | - | 1195 | 744.9 | - | - | 0 | - |
| - | - | 1015 | 745.3 | - | - | 0 | - |
| - | - | 3117 | 746.4 | - | - | 0 | - |
| - | - | 2262 | 746.4 | - | - | 0 | - |
| - | - | 2038 | 746.9 | - | - | 0 | - |
| - | - | 7806 | 747.4 | - | - | 0 | - |
| - | - | 4584 | 747.9 | - | - | 0 | - |
| - | - | 2858 | 748.4 | - | - | 0 | - |
| - | - | 4167 | 748.9 | - | - | 0 | - |
| - | - | 3373 | 749.4 | - | - | 0 | - |
| - | - | 2195 | 749.9 | - | - | 0 | - |
| - | - | 1083 | 750.4 | - | - | 0 | - |
| - | - | 1586 | 750.9 | - | - | 0 | - |
| 2 | y | 2769 | 751.3 | 0.01219 | 16.23 | +2 | 12 |
| 2 | z | 3862 | 751.9 | 0.007304 | 9.715 | +2 | 12 |
| - | - | 2852 | 752.4 | - | - | 0 | - |
| - | - | 1263 | 752.9 | - | - | 0 | - |
| - | - | 1204 | 753.4 | - | - | 0 | - |
| - | - | 2.841E+04 | 755.4 | - | - | 0 | - |
| - | - | 2.081E+04 | 755.9 | - | - | 0 | - |
| - | - | 7326 | 756.4 | - | - | 0 | - |
| - | - | 7804 | 756.9 | - | - | 0 | - |
| - | - | 7684 | 757.4 | - | - | 0 | - |
| - | - | 3892 | 757.9 | - | - | 0 | - |
| - | - | 1904 | 758.4 | - | - | 0 | - |
| - | - | 1643 | 758.9 | - | - | 0 | - |
| - | - | 2685 | 759.4 | - | - | 0 | - |
| 2 | y | 1760 | 759.9 | 0.005389 | 7.092 | +2 | 12 |
| - | - | 2835 | 760.4 | - | - | 0 | - |
| - | - | 1036 | 760.9 | - | - | 0 | - |
| - | - | 2048 | 763.4 | - | - | 0 | - |
| - | - | 2114 | 764.4 | - | - | 0 | - |
| - | - | 1319 | 765.9 | - | - | 0 | - |
| - | - | 1889 | 766.4 | - | - | 0 | - |
| - | - | 2427 | 766.9 | - | - | 0 | - |
| - | - | 1206 | 767.4 | - | - | 0 | - |
| - | - | 934.6 | 767.9 | - | - | 0 | - |
| - | - | 937.9 | 769.9 | - | - | 0 | - |
| - | - | 976.8 | 770.4 | - | - | 0 | - |
| - | - | 865.6 | 770.9 | - | - | 0 | - |
| - | - | 4759 | 771.4 | - | - | 0 | - |
| - | - | 9137 | 771.9 | - | - | 0 | - |
| - | - | 6186 | 772.4 | - | - | 0 | - |
| - | - | 4500 | 772.9 | - | - | 0 | - |
| - | - | 2414 | 773.4 | - | - | 0 | - |
| - | - | 2007 | 773.9 | - | - | 0 | - |
| - | - | 2279 | 774.4 | - | - | 0 | - |
| - | - | 1054 | 774.9 | - | - | 0 | - |
| - | - | 1584 | 775.4 | - | - | 0 | - |
| - | - | 1081 | 775.9 | - | - | 0 | - |
| - | - | 3.895E+04 | 777.4 | - | - | 0 | - |
| - | - | 8.951E+04 | 778.4 | - | - | 0 | - |
| - | - | 1071 | 778.9 | - | - | 0 | - |
| - | - | 3.684E+04 | 779.4 | - | - | 0 | - |
| - | - | 8519 | 779.9 | - | - | 0 | - |
| - | - | 1.946E+04 | 780.4 | - | - | 0 | - |
| - | - | 1.169E+04 | 780.9 | - | - | 0 | - |
| - | - | 1.126E+04 | 781.4 | - | - | 0 | - |
| - | - | 6593 | 781.9 | - | - | 0 | - |
| - | - | 9742 | 782.4 | - | - | 0 | - |
| - | - | 1483 | 782.5 | - | - | 0 | - |
| - | - | 7344 | 782.9 | - | - | 0 | - |
| - | - | 2757 | 783.4 | - | - | 0 | - |
| - | - | 929.1 | 783.5 | - | - | 0 | - |
| - | - | 1202 | 787.4 | - | - | 0 | - |
| - | - | 7068 | 787.9 | - | - | 0 | - |
| - | - | 6427 | 788.4 | - | - | 0 | - |
| - | - | 4844 | 788.9 | - | - | 0 | - |
| - | - | 1.562E+04 | 789.4 | - | - | 0 | - |
| - | - | 1.146E+04 | 789.9 | - | - | 0 | - |
| - | - | 4358 | 790.4 | - | - | 0 | - |
| - | - | 1.505E+04 | 790.9 | - | - | 0 | - |
| - | - | 1.077E+04 | 791.4 | - | - | 0 | - |
| - | - | 5343 | 791.9 | - | - | 0 | - |
| - | - | 980.3 | 792.4 | - | - | 0 | - |
| - | - | 3129 | 792.9 | - | - | 0 | - |
| - | - | 5111 | 793.4 | - | - | 0 | - |
| - | - | 4274 | 793.9 | - | - | 0 | - |
| - | - | 4887 | 794.4 | - | - | 0 | - |
| - | - | 4709 | 794.9 | - | - | 0 | - |
| - | - | 4647 | 795.4 | - | - | 0 | - |
| - | - | 9016 | 795.9 | - | - | 0 | - |
| - | - | 1631 | 796.4 | - | - | 0 | - |
| - | - | 741.3 | 796.9 | - | - | 0 | - |
| - | - | 7602 | 801.4 | - | - | 0 | - |
| - | - | 2.59E+04 | 801.9 | - | - | 0 | - |
| - | - | 3.698E+04 | 802.4 | - | - | 0 | - |
| - | - | 2.175E+04 | 802.9 | - | - | 0 | - |
| - | - | 1.386E+04 | 803.4 | - | - | 0 | - |
| - | - | 5.156E+04 | 803.9 | - | - | 0 | - |
| - | - | 3.394E+04 | 804.4 | - | - | 0 | - |
| - | - | 1.646E+04 | 804.9 | - | - | 0 | - |
| - | - | 1949 | 807.4 | - | - | 0 | - |
| - | - | 1822 | 808.4 | - | - | 0 | - |
| - | - | 2022 | 808.9 | - | - | 0 | - |
| - | - | 4264 | 809.4 | - | - | 0 | - |
| - | - | 4.865E+04 | 809.9 | - | - | 0 | - |
| - | - | 1.512E+05 | 810.4 | - | - | 0 | - |
| - | - | 1.9E+05 | 810.9 | - | - | 0 | - |
| - | - | 1.318E+05 | 811.4 | - | - | 0 | - |
| - | - | 6.483E+04 | 811.9 | - | - | 0 | - |
| - | - | 3.836E+04 | 812.4 | - | - | 0 | - |
| - | - | 1.842E+04 | 812.9 | - | - | 0 | - |
| - | - | 2368 | 813.4 | - | - | 0 | - |
| - | - | 3567 | 813.4 | - | - | 0 | - |
| - | - | 2527 | 823.5 | - | - | 0 | - |
| - | - | 2364 | 826.4 | - | - | 0 | - |
| - | - | 1652 | 827.4 | - | - | 0 | - |
| - | - | 1608 | 836.4 | - | - | 0 | - |
| - | - | 1923 | 840.5 | - | - | 0 | - |
| - | - | 1827 | 841.4 | - | - | 0 | - |
| - | - | 1286 | 842.4 | - | - | 0 | - |
| - | - | 3408 | 852.4 | - | - | 0 | - |
| - | - | 3100 | 857.5 | - | - | 0 | - |
| - | - | 5947 | 863.4 | - | - | 0 | - |
| - | - | 3092 | 864.4 | - | - | 0 | - |
| - | - | 1160 | 868.4 | - | - | 0 | - |
| - | - | 3843 | 878.4 | - | - | 0 | - |
| - | - | 2491 | 879.4 | - | - | 0 | - |
| - | - | 1109 | 890.4 | - | - | 0 | - |
| - | - | 3448 | 891.4 | - | - | 0 | - |
| 7 | z | 1957 | 892.4 | 0.01753 | 19.65 | +1 | 7 |
| - | - | 1250 | 893.4 | - | - | 0 | - |
| 7 | c | 1.417E+04 | 896.4 | 0.00558 | 6.224 | +1 | 7 |
| - | - | 878.5 | 896.5 | - | - | 0 | - |
| - | - | 7848 | 897.4 | - | - | 0 | - |
| - | - | 3115 | 898.4 | - | - | 0 | - |
| - | - | 1.167E+04 | 901.5 | - | - | 0 | - |
| - | - | 4541 | 902.5 | - | - | 0 | - |
| - | - | 926.2 | 903.5 | - | - | 0 | - |
| - | - | 2.754E+04 | 906.4 | - | - | 0 | - |
| - | - | 5.507E+04 | 907.4 | - | - | 0 | - |
| 7 | y | 3.018E+04 | 908.4 | 0.006376 | 7.019 | +1 | 7 |
| - | - | 1.55E+04 | 909.4 | - | - | 0 | - |
| - | - | 4925 | 910.4 | - | - | 0 | - |
| - | - | 1249 | 911.4 | - | - | 0 | - |
| - | - | 985.4 | 912.5 | - | - | 0 | - |
| 7 | c | 2.31E+04 | 913.4 | 0.004909 | 5.375 | +1 | 7 |
| - | - | 1.197E+04 | 914.5 | - | - | 0 | - |
| - | - | 3106 | 915.5 | - | - | 0 | - |
| - | - | 2000 | 924.4 | - | - | 0 | - |
| - | - | 2180 | 925.4 | - | - | 0 | - |
| 7 | y | 2110 | 926.4 | 0.01437 | 15.51 | +1 | 7 |
| - | - | 1066 | 932.5 | - | - | 0 | - |
| - | - | 1654 | 936.6 | - | - | 0 | - |
| - | - | 1242 | 937.6 | - | - | 0 | - |
| - | - | 7246 | 944.5 | - | - | 0 | - |
| - | - | 2768 | 945.5 | - | - | 0 | - |
| - | - | 1025 | 946.5 | - | - | 0 | - |
| - | - | 5052 | 953.6 | - | - | 0 | - |
| - | - | 1738 | 954.6 | - | - | 0 | - |
| - | - | 2919 | 963.5 | - | - | 0 | - |
| - | - | 1697 | 964.5 | - | - | 0 | - |
| - | - | 1214 | 965.5 | - | - | 0 | - |
| - | - | 6488 | 976.5 | - | - | 0 | - |
| - | - | 4176 | 977.5 | - | - | 0 | - |
| - | - | 1316 | 978.5 | - | - | 0 | - |
| - | - | 1242 | 980.5 | - | - | 0 | - |
| - | - | 1182 | 981.4 | - | - | 0 | - |
| - | - | 4235 | 982.5 | - | - | 0 | - |
| - | - | 2886 | 983.5 | - | - | 0 | - |
| - | - | 1319 | 987.5 | - | - | 0 | - |
| - | - | 3.001E+04 | 988.5 | - | - | 0 | - |
| - | - | 1.302E+04 | 989.5 | - | - | 0 | - |
| - | - | 3090 | 990.5 | - | - | 0 | - |
| - | - | 935.1 | 995.6 | - | - | 0 | - |
| - | - | 1541 | 1002 | - | - | 0 | - |
| - | - | 930.5 | 1003 | - | - | 0 | - |
| - | - | 1297 | 1008 | - | - | 0 | - |
| 8 | c | 2528 | 1010 | 0.00434 | 4.299 | +1 | 8 |
| - | - | 1689 | 1011 | - | - | 0 | - |
| - | - | 4302 | 1019 | - | - | 0 | - |
| - | - | 1.899E+04 | 1024 | - | - | 0 | - |
| 6 | z | 2.2E+04 | 1025 | 0.01426 | 13.91 | +1 | 8 |
| - | - | 6817 | 1026 | - | - | 0 | - |
| 8 | c | 4.906E+04 | 1027 | 0.006295 | 6.132 | +1 | 8 |
| - | - | 3762 | 1027 | - | - | 0 | - |
| - | - | 2.898E+04 | 1028 | - | - | 0 | - |
| - | - | 687.5 | 1028 | - | - | 0 | - |
| - | - | 1.051E+04 | 1029 | - | - | 0 | - |
| - | - | 3047 | 1030 | - | - | 0 | - |
| - | - | 4380 | 1040 | - | - | 0 | - |
| 6 | y | 4903 | 1041 | 0.01532 | 14.71 | +1 | 8 |
| - | - | 1265 | 1042 | - | - | 0 | - |
| - | - | 1.683E+04 | 1046 | - | - | 0 | - |
| - | - | 7311 | 1047 | - | - | 0 | - |
| - | - | 3438 | 1048 | - | - | 0 | - |
| - | - | 893.8 | 1049 | - | - | 0 | - |
| - | - | 2.044E+04 | 1052 | - | - | 0 | - |
| - | - | 1.577E+04 | 1053 | - | - | 0 | - |
| - | - | 717.6 | 1054 | - | - | 0 | - |
| - | - | 6164 | 1054 | - | - | 0 | - |
| - | - | 1454 | 1061 | - | - | 0 | - |
| - | - | 3911 | 1063 | - | - | 0 | - |
| - | - | 2125 | 1064 | - | - | 0 | - |
| - | - | 2041 | 1067 | - | - | 0 | - |
| - | - | 1028 | 1068 | - | - | 0 | - |
| - | - | 3497 | 1079 | - | - | 0 | - |
| - | - | 1.556E+04 | 1080 | - | - | 0 | - |
| - | - | 1.064E+04 | 1081 | - | - | 0 | - |
| - | - | 2472 | 1082 | - | - | 0 | - |
| - | - | 2264 | 1083 | - | - | 0 | - |
| - | - | 2.314E+04 | 1084 | - | - | 0 | - |
| - | - | 1941 | 1084 | - | - | 0 | - |
| - | - | 1.611E+04 | 1085 | - | - | 0 | - |
| - | - | 1206 | 1085 | - | - | 0 | - |
| - | - | 5338 | 1086 | - | - | 0 | - |
| - | - | 2156 | 1087 | - | - | 0 | - |
| - | - | 6148 | 1092 | - | - | 0 | - |
| - | - | 9850 | 1093 | - | - | 0 | - |
| - | - | 3885 | 1094 | - | - | 0 | - |
| 5 | w | 1514 | 1095 | 0.00939 | 8.572 | +1 | 9 |
| - | - | 938.9 | 1096 | - | - | 0 | - |
| - | - | 3979 | 1097 | - | - | 0 | - |
| - | - | 2824 | 1098 | - | - | 0 | - |
| - | - | 1659 | 1099 | - | - | 0 | - |
| - | - | 1244 | 1100 | - | - | 0 | - |
| - | - | 1027 | 1106 | - | - | 0 | - |
| - | - | 1969 | 1108 | - | - | 0 | - |
| - | - | 1165 | 1110 | - | - | 0 | - |
| 9 | c | 1281 | 1125 | 0.002727 | 2.425 | +1 | 9 |
| - | - | 1560 | 1126 | - | - | 0 | - |
| - | - | 1272 | 1127 | - | - | 0 | - |
| - | - | 2166 | 1140 | - | - | 0 | - |
| - | - | 9262 | 1141 | - | - | 0 | - |
| 9 | c | 1.067E+04 | 1142 | 0.0106 | 9.287 | +1 | 9 |
| - | - | 5288 | 1143 | - | - | 0 | - |
| - | - | 2203 | 1144 | - | - | 0 | - |
| - | - | 1.498E+04 | 1145 | - | - | 0 | - |
| - | - | 6193 | 1146 | - | - | 0 | - |
| - | - | 2719 | 1147 | - | - | 0 | - |
| - | - | 1.882E+04 | 1153 | - | - | 0 | - |
| 5 | z | 5.341E+04 | 1153 | 0.008788 | 7.618 | +1 | 9 |
| - | - | 3.148E+04 | 1154 | - | - | 0 | - |
| - | - | 1.256E+04 | 1155 | - | - | 0 | - |
| - | - | 3929 | 1157 | - | - | 0 | - |
| - | - | 1029 | 1158 | - | - | 0 | - |
| - | - | 2645 | 1169 | - | - | 0 | - |
| 5 | y | 1.002E+04 | 1170 | 0.009107 | 7.787 | +1 | 9 |
| - | - | 5531 | 1171 | - | - | 0 | - |
| - | - | 2339 | 1172 | - | - | 0 | - |
| - | - | 1225 | 1179 | - | - | 0 | - |
| - | - | 4396 | 1180 | - | - | 0 | - |
| 10 | c | 3137 | 1181 | 0.01252 | 10.61 | +1 | 10 |
| - | - | 1638 | 1182 | - | - | 0 | - |
| - | - | 1647 | 1188 | - | - | 0 | - |
| - | - | 1270 | 1189 | - | - | 0 | - |
| - | - | 2385 | 1195 | - | - | 0 | - |
| - | - | 912.1 | 1196 | - | - | 0 | - |
| - | - | 859.4 | 1197 | - | - | 0 | - |
| - | - | 3.008E+04 | 1198 | - | - | 0 | - |
| 10 | c | 6.654E+04 | 1199 | 0.01013 | 8.455 | +1 | 10 |
| - | - | 3.736E+04 | 1200 | - | - | 0 | - |
| - | - | 1.303E+04 | 1201 | - | - | 0 | - |
| - | - | 3573 | 1202 | - | - | 0 | - |
| - | - | 1024 | 1203 | - | - | 0 | - |
| - | - | 1319 | 1231 | - | - | 0 | - |
| - | - | 2.55E+04 | 1232 | - | - | 0 | - |
| - | - | 1.421E+04 | 1233 | - | - | 0 | - |
| - | - | 4257 | 1234 | - | - | 0 | - |
| - | - | 885.7 | 1240 | - | - | 0 | - |
| - | - | 918.9 | 1246 | - | - | 0 | - |
| - | - | 1055 | 1259 | - | - | 0 | - |
| - | - | 851.6 | 1262 | - | - | 0 | - |
| - | - | 962.5 | 1275 | - | - | 0 | - |
| - | - | 1251 | 1277 | - | - | 0 | - |
| - | - | 1225 | 1278 | - | - | 0 | - |
| - | - | 5748 | 1279 | - | - | 0 | - |
| - | - | 3482 | 1280 | - | - | 0 | - |
| - | - | 1944 | 1283 | - | - | 0 | - |
| - | - | 2965 | 1284 | - | - | 0 | - |
| - | - | 2199 | 1285 | - | - | 0 | - |
| 4 | y | 4594 | 1290 | 0.02472 | 19.17 | +1 | 10 |
| 4 | z | 2.355E+04 | 1291 | 0.00908 | 7.036 | +1 | 10 |
| - | - | 2.202E+04 | 1292 | - | - | 0 | - |
| - | - | 1.156E+04 | 1293 | - | - | 0 | - |
| - | - | 4198 | 1294 | - | - | 0 | - |
| - | - | 1159 | 1306 | - | - | 0 | - |
| 4 | y | 2572 | 1307 | 0.007934 | 6.072 | +1 | 10 |
| - | - | 2147 | 1308 | - | - | 0 | - |
| - | - | 1310 | 1311 | - | - | 0 | - |
| - | - | 1513 | 1312 | - | - | 0 | - |
| - | - | 1.671E+04 | 1319 | - | - | 0 | - |
| - | - | 9511 | 1320 | - | - | 0 | - |
| - | - | 3813 | 1321 | - | - | 0 | - |
| - | - | 3278 | 1325 | - | - | 0 | - |
| - | - | 1.237E+04 | 1326 | - | - | 0 | - |
| 11 | c | 2.734E+04 | 1327 | 0.008921 | 6.724 | +1 | 11 |
| - | - | 2.053E+04 | 1328 | - | - | 0 | - |
| - | - | 8950 | 1329 | - | - | 0 | - |
| - | - | 2946 | 1330 | - | - | 0 | - |
| - | - | 1612 | 1346 | - | - | 0 | - |
| - | - | 1243 | 1347 | - | - | 0 | - |
| 3 | z | 970.3 | 1387 | 0.0242 | 17.45 | +1 | 11 |
| - | - | 1735 | 1392 | - | - | 0 | - |
| 3 | y | 3387 | 1403 | 0.02696 | 19.22 | +1 | 11 |
| 3 | z | 1.23E+04 | 1404 | 0.01437 | 10.24 | +1 | 11 |
| - | - | 1.516E+04 | 1405 | - | - | 0 | - |
| - | - | 8514 | 1406 | - | - | 0 | - |
| - | - | 4471 | 1407 | - | - | 0 | - |
| - | - | 8920 | 1408 | - | - | 0 | - |
| - | - | 5497 | 1409 | - | - | 0 | - |
| - | - | 1600 | 1410 | - | - | 0 | - |
| - | - | 1531 | 1411 | - | - | 0 | - |
| - | - | 5004 | 1412 | - | - | 0 | - |
| - | - | 5205 | 1413 | - | - | 0 | - |
| - | - | 2886 | 1414 | - | - | 0 | - |
| - | - | 1637 | 1415 | - | - | 0 | - |
| - | - | 1332 | 1417 | - | - | 0 | - |
| 3 | y | 1762 | 1420 | 0.02067 | 14.56 | +1 | 11 |
| - | - | 1203 | 1421 | - | - | 0 | - |
| - | - | 1165 | 1422 | - | - | 0 | - |
| - | - | 1006 | 1427 | - | - | 0 | - |
| - | - | 3005 | 1432 | - | - | 0 | - |
| - | - | 1909 | 1433 | - | - | 0 | - |
| - | - | 1085 | 1438 | - | - | 0 | - |
| - | - | 3373 | 1439 | - | - | 0 | - |
| - | - | 4281 | 1440 | - | - | 0 | - |
| - | - | 2450 | 1441 | - | - | 0 | - |
| - | - | 1289 | 1442 | - | - | 0 | - |
| - | - | 3813 | 1455 | - | - | 0 | - |
| 12 | c | 1.282E+04 | 1456 | 0.01687 | 11.59 | +1 | 12 |
| - | - | 1.662E+04 | 1457 | - | - | 0 | - |
| - | - | 8193 | 1458 | - | - | 0 | - |
| - | - | 6111 | 1459 | - | - | 0 | - |
| - | - | 2398 | 1460 | - | - | 0 | - |
| - | - | 1429 | 1474 | - | - | 0 | - |
| - | - | 4160 | 1475 | - | - | 0 | - |
| - | - | 1.214E+04 | 1476 | - | - | 0 | - |
| - | - | 6033 | 1477 | - | - | 0 | - |
| - | - | 2448 | 1478 | - | - | 0 | - |
| - | - | 1660 | 1488 | - | - | 0 | - |
| - | - | 4437 | 1489 | - | - | 0 | - |
| - | - | 3051 | 1490 | - | - | 0 | - |
| - | - | 1110 | 1491 | - | - | 0 | - |
| - | - | 982.2 | 1494 | - | - | 0 | - |
| - | - | 6976 | 1495 | - | - | 0 | - |
| - | - | 4503 | 1496 | - | - | 0 | - |
| - | - | 1531 | 1497 | - | - | 0 | - |
| 2 | y | 1157 | 1502 | 0.02654 | 17.67 | +1 | 12 |
| 2 | z | 1589 | 1503 | 0.02066 | 13.75 | +1 | 12 |
| - | - | 4770 | 1504 | - | - | 0 | - |
| - | - | 2851 | 1505 | - | - | 0 | - |
| - | - | 1626 | 1506 | - | - | 0 | - |
| 2 | y | 969.6 | 1519 | 0.01061 | 6.985 | +1 | 12 |
| - | - | 2513 | 1523 | - | - | 0 | - |
| - | - | 1754 | 1524 | - | - | 0 | - |
| - | - | 879.7 | 1525 | - | - | 0 | - |
| - | - | 819.6 | 1545 | - | - | 0 | - |
| - | - | 865.2 | 1546 | - | - | 0 | - |
| - | - | 1238 | 1548 | - | - | 0 | - |
| - | - | 1986 | 1549 | - | - | 0 | - |
| - | - | 1040 | 1550 | - | - | 0 | - |
| - | - | 2098 | 1559 | - | - | 0 | - |
| - | - | 3819 | 1560 | - | - | 0 | - |
| - | - | 6789 | 1561 | - | - | 0 | - |
| - | - | 1.032E+04 | 1562 | - | - | 0 | - |
| - | - | 8705 | 1563 | - | - | 0 | - |
| - | - | 4446 | 1564 | - | - | 0 | - |
| - | - | 1.199E+04 | 1565 | - | - | 0 | - |
| - | - | 1.02E+04 | 1566 | - | - | 0 | - |
| - | - | 7435 | 1567 | - | - | 0 | - |
| - | - | 1880 | 1568 | - | - | 0 | - |
| - | - | 1611 | 1574 | - | - | 0 | - |
| - | - | 4220 | 1575 | - | - | 0 | - |
| - | - | 1.353E+04 | 1576 | - | - | 0 | - |
| - | - | 1.398E+04 | 1577 | - | - | 0 | - |
| - | - | 8618 | 1578 | - | - | 0 | - |
| - | - | 4509 | 1579 | - | - | 0 | - |
| - | - | 2968 | 1580 | - | - | 0 | - |
| - | - | 3010 | 1581 | - | - | 0 | - |
| - | - | 3938 | 1582 | - | - | 0 | - |
| - | - | 2354 | 1583 | - | - | 0 | - |
| - | - | 1184 | 1585 | - | - | 0 | - |
| - | - | 3266 | 1586 | - | - | 0 | - |
| - | - | 4400 | 1587 | - | - | 0 | - |
| - | - | 2758 | 1588 | - | - | 0 | - |
| - | - | 1854 | 1589 | - | - | 0 | - |
| - | - | 1374 | 1590 | - | - | 0 | - |
| - | - | 4234 | 1591 | - | - | 0 | - |
| - | - | 6739 | 1592 | - | - | 0 | - |
| - | - | 4881 | 1593 | - | - | 0 | - |
| - | - | 4845 | 1594 | - | - | 0 | - |
| - | - | 2961 | 1595 | - | - | 0 | - |
| - | - | 1258 | 1596 | - | - | 0 | - |
| - | - | 2792 | 1597 | - | - | 0 | - |
| - | - | 1964 | 1598 | - | - | 0 | - |
| - | - | 868.2 | 1599 | - | - | 0 | - |
| - | - | 2042 | 1602 | - | - | 0 | - |
| - | - | 1.286E+04 | 1603 | - | - | 0 | - |
| - | - | 4.705E+04 | 1604 | - | - | 0 | - |
| - | - | 6.237E+04 | 1605 | - | - | 0 | - |
| - | - | 3.989E+04 | 1606 | - | - | 0 | - |
| - | - | 2.358E+04 | 1607 | - | - | 0 | - |
| - | - | 3.455E+04 | 1608 | - | - | 0 | - |
| - | - | 2.658E+04 | 1609 | - | - | 0 | - |
| - | - | 1.007E+04 | 1610 | - | - | 0 | - |
| - | - | 5306 | 1619 | - | - | 0 | - |
| - | - | 2.25E+04 | 1620 | - | - | 0 | - |
| - | - | 5.395E+04 | 1621 | - | - | 0 | - |
| - | - | 7.915E+04 | 1622 | - | - | 0 | - |
| - | - | 5.864E+04 | 1623 | - | - | 0 | - |
| - | - | 3.186E+04 | 1624 | - | - | 0 | - |
| - | - | 3.651E+04 | 1625 | - | - | 0 | - |
| - | - | 2.298E+04 | 1626 | - | - | 0 | - |
| - | - | 1.092E+04 | 1627 | - | - | 0 | - |
| - | - | 946.6 | 3084 | - | - | 0 | - |

m/z Charge Intensity FragmentType MassShift Position
127.45966339111328 0 376.9658
128.0948486328125 0 736.48267
129.1025390625 0 4922.8633
130.065185546875 0 827.51276
130.38905334472656 0 385.74265
131.11817932128906 0 1608.7383
135.0414581298828 0 483.423
136.0350341796875 0 419.12183
136.0761260986328 0 1431.1962
145.0763397216797 0 2239.424
146.06057739257812 0 916.9523
147.07681274414062 0 1972.9562 y 10
155.09304809570312 0 10797.451
157.13389587402344 0 1488.6006
160.0733184814453 0 555.4532
163.46690368652344 0 490.88254
165.05499267578125 0 514.04663
166.08648681640625 0 3936.065 y 9
170.10574340820312 0 732.12305
171.11318969726562 0 1115.9493
173.1287841796875 0 39208.336
174.13226318359375 0 3139.8574
175.0869140625 0 3006.5808
182.08151245117188 0 6479.549 y 12
183.11300659179688 0 1603.2556
184.06036376953125 0 719.63965
185.07139587402344 0 848.00714
186.1241455078125 0 2239.0127
188.1398162841797 0 1255.2053
188.73655700683594 0 501.3266
201.12364196777344 0 28291.16
201.13502502441406 0 2995.3364
202.12652587890625 0 2050.2422
206.84107971191406 0 521.31146
207.11349487304688 0 584.1654
207.49038696289062 0 484.9056
215.1393585205078 0 7861.8633
216.14398193359375 0 853.89874
218.15036010742188 0 2490.6838 c 1
223.15538024902344 0 1255.5781
226.1080322265625 0 1247.0903
228.13462829589844 0 1747.9482
243.13414001464844 0 2828.0903
243.43267822265625 0 500.735
245.13731384277344 0 2937.025
251.1507568359375 0 5112.3784
255.08851623535156 0 631.2674
258.14520263671875 0 9323.019
259.1482238769531 0 1174.3076
260.1605529785156 0 3508.783
261.17108154296875 0 6248.944
266.01580810546875 0 515.5147
268.1770935058594 0 7212.7227
272.1600341796875 0 646.36206
283.04925537109375 0 730.0635
283.1394958496094 0 1405.0421
288.2287902832031 0 593.1258
290.1626281738281 0 632.9656 c Ammonia loss 4
293.1136779785156 0 4990.7344 y Water loss 11
294.11724853515625 0 784.2967
298.1566162109375 0 820.598
299.0602111816406 0 631.4549
299.14715576171875 0 794.2162 c Water loss 6
300.06353759765625 0 848.75934
300.16851806640625 0 793.8398
301.05950927734375 0 906.58386
301.15093994140625 0 1561.6569
306.16790771484375 0 996.91583
307.18951416015625 0 714.3219
311.12493896484375 0 2748.8362 y 11
312.156005859375 0 3983.264
314.2084045410156 0 2262.747
315.16693115234375 0 1786.1288
317.1587219238281 0 16825.902
318.16241455078125 0 2223.0747
319.1641845703125 0 688.7213
320.1979064941406 0 2766.4214
322.1884460449219 0 1320.3329
330.1668701171875 0 1704.2214
331.2347106933594 0 14767.019 c 2
332.1934509277344 0 580.4805
332.23846435546875 0 2138.6133
339.16619873046875 0 1490.7611
343.10205078125 0 1941.9849
347.68011474609375 0 3449.0984 c Ammonia loss 5
348.18310546875 0 1067.5403
353.6737976074219 0 780.1892 y Water loss 7
357.17767333984375 0 2804.7263
358.18084716796875 0 750.5734
362.18707275390625 0 908.995
362.6791076660156 0 2226.7063 y 7
363.1796875 0 698.8122
364.18646240234375 0 1957.4885
365.1897277832031 0 861.32263
366.14215087890625 0 832.56915
374.1801452636719 0 2616.0208
376.1981506347656 0 4445.964
377.2014465332031 0 809.12305
378.2036437988281 0 1131.2284
379.2095031738281 0 1249.407
381.1528015136719 0 1994.7615
381.22393798828125 0 1022.11664
382.1901550292969 0 693.73895
387.1871032714844 0 790.77576
389.1907958984375 0 910.67456
391.1848449707031 0 2237.2954
392.1930236816406 0 6393.2974
393.19854736328125 0 2493.5276
398.1988830566406 0 562.5868
408.2616882324219 0 893.4255
409.21905517578125 0 1244.332
412.24346923828125 0 2735.566
412.74542236328125 0 1277.9727
417.18780517578125 0 2544.7798
418.19061279296875 0 586.6182
423.2012939453125 0 2639.203 z 10
423.272216796875 0 7254.715
424.20550537109375 0 996.3784
424.2770690917969 0 1134.0012
424.7322692871094 0 808.79987
428.2273254394531 0 796.63605
429.20977783203125 0 1695.1617
429.2368469238281 0 1474.9974
430.19378662109375 0 4658.633 y Water loss 3
431.1974182128906 0 856.39075
433.2543640136719 0 747.0639
434.7196350097656 0 3435.0117
435.22430419921875 0 2774.3638
439.2190246582031 0 3059.585 y 10
439.7117919921875 0 4633.038
440.2136535644531 0 2485.69
442.27764892578125 0 2458.9177
444.255859375 0 1247.2001
445.2547912597656 0 873.0343
446.2632751464844 0 1042.9941
448.7174377441406 0 7874.533 c Ammonia loss 6
449.2192687988281 0 3944.9893
449.7203063964844 0 1618.5852
450.28326416015625 0 924.91766 c Water loss 3
451.26715087890625 0 16754.752
452.26971435546875 0 4092.8486
454.21942138671875 0 7618.401
454.7140197753906 0 11784.968 y Water loss 6
455.21478271484375 0 5917.5454
455.7180480957031 0 2043.1348
461.20623779296875 0 796.26447
461.2736511230469 0 923.78754
463.2305603027344 0 4937.7544
464.22967529296875 0 1714.195
468.2936706542969 0 52414.582 c 3
468.7835998535156 0 899.46924
469.2960510253906 0 12672.321
470.3003234863281 0 1074.7013
471.0030822753906 0 627.89655
473.7464904785156 0 889.78516
476.2181091308594 0 857.60126
478.2198791503906 0 2506.6982
479.2262268066406 0 3725.048
480.22723388671875 0 8088.3745 c Ammonia loss 11
481.23016357421875 0 5439.872
482.25677490234375 0 3892.4707
482.7557678222656 0 2324.8835
488.22430419921875 0 863.1809
488.75933837890625 0 4026.668
489.2077941894531 0 4059.7327
489.2608642578125 0 1787.4788
489.7603454589844 0 854.0142
491.262451171875 0 3865.0823
491.7629089355469 0 1624.6652
494.23590087890625 0 6759.4146
494.31005859375 0 2971.0085
495.2359924316406 0 4094.116
495.3107604980469 0 995.0179
496.2424011230469 0 21056.482 y 9
496.76055908203125 0 2214.2166
497.24530029296875 0 5050.418
498.2493896484375 0 1843.6006
504.2215270996094 0 2785.3328 z Water loss 5
505.2603759765625 0 3065.5784 c Ammonia loss 7
505.7619934082031 0 1246.6686
506.5566711425781 0 784.86615
507.1051330566406 0 1062.0754
511.33563232421875 0 9643.111
512.338134765625 0 1880.8828
513.3514404296875 0 48804.77
514.282958984375 0 686.4503
514.3543090820312 0 9466.673
520.2404174804688 0 805.0656
523.2296752929688 0 952.0209
523.3359375 0 7072.604
524.1760864257812 0 1435.9011
524.2352905273438 0 769.5524
524.27783203125 0 1922.5693
524.3379516601562 0 1327.2094
525.2313232421875 0 1904.4403
525.5631713867188 0 2541.659
525.8126220703125 0 2455.3052
526.3143310546875 0 1808.6464
527.3226928710938 0 772.40454
532.2858276367188 0 2541.363
533.293701171875 0 9983.064
534.2979736328125 0 2434.2236
534.5942993164062 0 2316.9934
534.9271850585938 0 2372.3716
535.258056640625 0 1608.4636
535.6041870117188 0 1077.1344
535.9412231445312 0 1096.4745
536.2361450195312 0 862.0203
538.2516479492188 0 23704.936
539.2547607421875 0 5424.903
539.8102416992188 0 60430.824
540.2620239257812 0 1862.0245
540.3115844726562 0 41358.43
540.5972290039062 0 2663.0725
540.8125 0 15407.217
540.9296875 0 2310.6802
541.3466186523438 0 538613.06
541.609375 0 5652.622
541.8184204101562 0 815.60516
541.9442138671875 0 6198.9546
542.3491821289062 0 113300.82
543.2756958007812 0 718.3954
546.2717895507812 0 5813.0825
546.7744140625 0 7124.7153
547.2757568359375 0 2188.4077
547.7789306640625 0 784.5664
550.2516479492188 0 11043.193 w 8
550.2925415039062 0 874.34015
551.2549438476562 0 3740.1367
552.25537109375 0 1026.1553
554.2806396484375 0 3495.7058
554.7831420898438 0 1337.942
555.2784423828125 0 857.7884
561.2899169921875 0 1232.3835
563.301025390625 0 814.1208
564.2785034179688 0 5378.569
565.2832641601562 0 890.76074
568.346435546875 0 1123.4246
571.31103515625 0 2306.9316
572.3073120117188 0 1307.6327
578.2958374023438 0 1891.7329
579.3231201171875 0 4528.7915 c Ammonia loss 4
580.3067626953125 0 884.295
583.225830078125 0 1112.1179
588.7802734375 0 1128.62
589.28369140625 0 2265.214
589.7877807617188 0 5961.0967
590.2922973632812 0 10505.493
590.7936401367188 0 4581.2246 c Water loss 9
591.2932739257812 0 1130.7386
592.3328857421875 0 3644.5059
593.2582397460938 0 2712.4832 y Water loss 8
594.2661743164062 0 2370.6636
595.2501220703125 0 11219.619 z 8
595.32080078125 0 1789.9373
596.25537109375 0 6701.2324
596.3522338867188 0 58219.203 c 4
597.2482299804688 0 952.4579
597.295654296875 0 2119.0527
597.3549194335938 0 19921.85
597.797119140625 0 17309.467
598.29833984375 0 7868.255
598.3573608398438 0 4870.469
598.79931640625 0 2900.6655
601.7832641601562 0 685.6601
610.2837524414062 0 15929.46
611.2694702148438 0 29984.174 y 8
612.2722778320312 0 8726.587
613.2741088867188 0 2006.6917
613.3333129882812 0 1170.659
614.325927734375 0 2161.0288
615.323974609375 0 1070.6356
619.7951049804688 0 1237.0779
620.30029296875 0 1284.9724
621.29345703125 0 1051.519
622.81884765625 0 1172.178
623.3191528320312 0 2076.0198
624.3252563476562 0 885.4053
625.3668212890625 0 1238.8417
626.2780151367188 0 729.2946
626.3734741210938 0 3730.4912
627.3768310546875 0 1419.2008
631.8178100585938 0 1463.8489
632.3228759765625 0 1308.1815
634.3179321289062 0 916.40643
636.3240966796875 0 11106.976
637.3280029296875 0 4656.506
639.322021484375 0 10094.799
639.8240966796875 0 6328.035
640.32568359375 0 2400.291
641.8427124023438 0 811.867
642.339111328125 0 2081.3813
642.84521484375 0 916.66223
645.2847900390625 0 759.0667 y Ammonia loss 3
645.7864990234375 0 1666.5648 z 3
647.3316040039062 0 17050.953
647.8330078125 0 9741.315
648.3246459960938 0 8965.931
649.3228759765625 0 2597.9226
651.3455810546875 0 2440.926
652.3426513671875 0 12156.9
653.2929077148438 0 1444.9044
653.3469848632812 0 2602.654
653.7924194335938 0 3257.34 y 3
654.2880249023438 0 1330.8844
654.34423828125 0 1165.2909
654.8405151367188 0 683.59314 c Water loss 10
655.3392944335938 0 1966.3273 c Ammonia loss 10
657.8347778320312 0 1078.4298
662.828857421875 0 2653.0593
663.3508911132812 0 28911.312
663.8468627929688 0 71164.63 c 10
664.2920532226562 0 5808.138
664.3475952148438 0 46687.926
664.8477783203125 0 18182.373
665.2901000976562 0 3000.5598 w 7
665.3504028320312 0 5525.431
665.847900390625 0 1889.9034
666.29296875 0 1400.956
666.353759765625 0 1122.6681
667.3614501953125 0 866.31226
668.31982421875 0 1009.22864
671.82861328125 0 1508.2628
674.3364868164062 0 7908.2925
674.8367919921875 0 4398.8784
675.3402099609375 0 1545.7639
676.3453369140625 0 3672.8545
676.8552856445312 0 4129.3896
677.3510131835938 0 4254.934
677.8551025390625 0 1319.7123
679.2753295898438 0 899.533
679.3423461914062 0 19369.805
680.3456420898438 0 6599.723
680.7977294921875 0 9737.207 w 2
681.2981567382812 0 4921.2114
681.3551635742188 0 994.9291
681.7989501953125 0 2052.9453
682.345703125 0 999.28394
692.3348388671875 0 730.28625
692.8181762695312 0 843.7753
693.3202514648438 0 1314.0009 z Water loss 2
694.3525390625 0 53315.5 c Ammonia loss 5
695.28076171875 0 1180.163
695.3554077148438 0 19864.492
695.8411865234375 0 923.6594
696.3081665039062 0 3002.6624
696.36328125 0 8464.85
697.3135986328125 0 1705.2925
697.3697509765625 0 1885.8235
698.365966796875 0 1308.0969
699.3648071289062 0 812.3485
700.364990234375 0 1109.13
700.833740234375 0 2149.4949
701.328857421875 0 4380.8115 y Water loss 2
701.8276977539062 0 7274.889 y Ammonia loss 2
702.3262939453125 0 6008.4766 z 2
702.8291625976562 0 3416.5452
703.3406982421875 0 998.7799
703.8431396484375 0 5295.6465
704.3450317382812 0 4833.874
704.404296875 0 768.4111
704.8446655273438 0 1729.3865
706.3563232421875 0 2969.6628
706.8577880859375 0 1746.718
707.3540649414062 0 6509.3887
707.8662719726562 0 1247.8845
708.3447265625 0 5807.705 z 7
708.8414916992188 0 704.9353
709.3369750976562 0 3504.7288
709.8387451171875 0 11299.151
710.3341674804688 0 20627.883 y 2
710.8348999023438 0 16793.045
711.3792114257812 0 111237.12 c 5
711.8512573242188 0 20208.574
712.3775634765625 0 43225.246
712.8546752929688 0 4946.772
713.3842163085938 0 10381.75
714.3854370117188 0 2176.885
715.875732421875 0 3793.2627
716.3738403320312 0 2169.3586
716.8801879882812 0 930.4122
717.8560180664062 0 943.78894
718.35400390625 0 1628.7457
718.8656616210938 0 1098.9442
719.3642578125 0 2532.7976 c Water loss 11
719.859619140625 0 1914.5226 c Ammonia loss 11
720.3594970703125 0 2016.003
720.8582763671875 0 1888.0043
721.3589477539062 0 3667.8035
721.8663940429688 0 1101.6766
722.3662719726562 0 1667.5718
723.3612060546875 0 7836.703
723.819580078125 0 1043.8733
724.3539428710938 0 14068.301 y 7
725.3572998046875 0 6751.802
725.865966796875 0 1184.4977
726.3629150390625 0 2497.0618
727.3650512695312 0 1220.6964
727.8735961914062 0 49539.5
728.3702392578125 0 86242.84 c 11
728.8700561523438 0 50928.355
729.3700561523438 0 22759.744
729.8710327148438 0 13642.49
730.3673095703125 0 3870.6738
730.8699340820312 0 2089.7158
734.3748168945312 0 1753.2073
735.3735961914062 0 1023.3703
735.8626708984375 0 1920.7283
736.3634033203125 0 3397.366
736.8659057617188 0 6640.7363
737.3618774414062 0 14761.36
737.8817749023438 0 49260.27
738.3841552734375 0 30552.734
738.8846435546875 0 12608.883
739.3705444335938 0 4337.604
743.8776245117188 0 938.2093
744.364013671875 0 2434.6309
744.8682861328125 0 1194.8809
745.340087890625 0 1014.673
746.3545532226562 0 3117.1863
746.4169921875 0 2261.9216
746.8626098632812 0 2038.0621
747.3580932617188 0 7806.4194
747.8611450195312 0 4583.866
748.3667602539062 0 2858.1438
748.8699340820312 0 4167.0576
749.3713989257812 0 3372.517
749.878173828125 0 2195.4673
750.3772583007812 0 1082.9325
750.8756713867188 0 1585.6233
751.3612670898438 0 2768.767 y Ammonia loss 1
751.8602905273438 0 3862.0461 z 1
752.3582763671875 0 2851.703
752.856201171875 0 1262.5498
753.36669921875 0 1203.5096
755.3687133789062 0 28412.658
755.8704833984375 0 20809.27
756.3717651367188 0 7326.1626
756.8648071289062 0 7804.1973
757.3659057617188 0 7683.9727
757.8707275390625 0 3892.479
758.3759765625 0 1903.5164
758.8790893554688 0 1643.4216
759.3834838867188 0 2684.596
759.8677368164062 0 1759.9281 y 1
760.366455078125 0 2834.7373
760.8617553710938 0 1036.1641
763.3785400390625 0 2048.1858
764.3741455078125 0 2113.6199
765.8807983398438 0 1318.9899
766.3677368164062 0 1888.663
766.865966796875 0 2427.273
767.3624267578125 0 1206.453
767.8770141601562 0 934.61584
769.8803100585938 0 937.8678
770.381103515625 0 976.84436
770.8790893554688 0 865.5957
771.3844604492188 0 4758.595
771.8809204101562 0 9137.095
772.3800048828125 0 6185.766
772.8818359375 0 4500.2793
773.3821411132812 0 2413.7693
773.8841552734375 0 2007.132
774.3834228515625 0 2279.4827
774.8932495117188 0 1054.3475
775.390869140625 0 1584.0928
775.8770141601562 0 1080.7659
777.378662109375 0 38948.23
778.3656616210938 0 89508.64
778.8817138671875 0 1070.7804
779.3690185546875 0 36842.992
779.8899536132812 0 8518.911
780.3756713867188 0 19461.48
780.882568359375 0 11685.787
781.3849487304688 0 11260.575
781.8983154296875 0 6593.379
782.4026489257812 0 9741.617
782.4767456054688 0 1482.9132
782.902587890625 0 7343.6904
783.4027099609375 0 2757.2825
783.4757080078125 0 929.10284
787.35302734375 0 1202.2815
787.8907470703125 0 7067.946
788.3876953125 0 6426.694
788.8869018554688 0 4843.768
789.383544921875 0 15624.326
789.8844604492188 0 11464.626
790.3915405273438 0 4357.7256
790.8995361328125 0 15046.882
791.4016723632812 0 10768.657
791.9027709960938 0 5343.4004
792.3862915039062 0 980.2683
792.8844604492188 0 3128.9148
793.3826904296875 0 5111.134
793.8818359375 0 4274.1455
794.3822021484375 0 4887.167
794.87939453125 0 4708.9375
795.3788452148438 0 4646.8525
795.8917236328125 0 9016.3955
796.3910522460938 0 1631.2894
796.8919067382812 0 741.26245
801.3941040039062 0 7601.686
801.8897705078125 0 25898.604
802.3869018554688 0 36980.67
802.8864135742188 0 21745.46
803.3928833007812 0 13858.169
803.9010009765625 0 51562.273
804.4030151367188 0 33942.27
804.9031982421875 0 16464.568
807.4231567382812 0 1949.247
808.431640625 0 1821.8658
808.8640747070312 0 2021.7249
809.3702392578125 0 4264.127
809.86181640625 0 48653.582
810.3909912109375 0 151239.81
810.893310546875 0 189951.02
811.393798828125 0 131829.42
811.8983764648438 0 64828.918
812.4059448242188 0 38357.336
812.9113159179688 0 18416.012
813.3540649414062 0 2367.5178
813.4153442382812 0 3567.1914
823.4780883789062 0 2527.4888
826.3991088867188 0 2363.5642
827.40234375 0 1652.2072
836.435791015625 0 1608.4989
840.505859375 0 1922.9272
841.4089965820312 0 1827.313
842.414306640625 0 1285.7943
852.436279296875 0 3408.1785
857.4625244140625 0 3099.5195
863.4187622070312 0 5947.287
864.42626953125 0 3092.2544
868.436279296875 0 1159.8126
878.4174194335938 0 3843.0647
879.4146728515625 0 2491.222
890.4312744140625 0 1109.4937
891.4137573242188 0 3447.8665
892.4087524414062 0 1956.6334 z Water loss 6
893.4041748046875 0 1249.7236
896.4267578125 0 14169.006 c Ammonia loss 6
896.5073852539062 0 878.5427
897.4302978515625 0 7847.845
898.4328002929688 0 3115.165
901.4749755859375 0 11674.702
902.477783203125 0 4541.1743
903.4846801757812 0 926.1847
906.423583984375 0 27542.566
907.41357421875 0 55068.223
908.4163208007812 0 30176.45 y Water loss 6
909.4207153320312 0 15501.91
910.4237670898438 0 4924.566
911.427490234375 0 1248.6859
912.5326538085938 0 985.41315
913.45263671875 0 23104.004 c 6
914.4560546875 0 11972.632
915.4577026367188 0 3105.5767
924.4324340820312 0 1999.8309
925.42626953125 0 2180.474
926.4348754882812 0 2109.6614 y 6
932.4937744140625 0 1066.3503
936.563720703125 0 1654.4594
937.5671997070312 0 1241.9366
944.4924926757812 0 7246.4575
945.4962158203125 0 2768.121
946.498779296875 0 1024.9498
953.5906372070312 0 5052.1577
954.5897216796875 0 1737.7041
963.524169921875 0 2919.4297
964.5265502929688 0 1697.2253
965.5184326171875 0 1214.4205
976.50830078125 0 6488.0044
977.5126953125 0 4176.1694
978.515625 0 1316.1705
980.4603881835938 0 1242.052
981.4437255859375 0 1182.2844
982.5243530273438 0 4234.818
983.5276489257812 0 2885.875
987.4967041015625 0 1319.49
988.5064697265625 0 30008.654
989.5089721679688 0 13016.221
990.5101318359375 0 3089.7979
995.5677490234375 0 935.0767
1001.517822265625 0 1541.1626
1002.5258178710938 0 930.4512
1007.5181274414062 0 1297.3489
1009.5095825195312 0 2527.593 c Ammonia loss 7
1010.52490234375 0 1688.8687
1018.5015258789062 0 4302.255
1024.4503173828125 0 18989.33
1025.4429931640625 0 22002.736 z 5
1026.4403076171875 0 6816.822
1026.5380859375 0 49064.23 c 7
1027.44140625 0 3762.3027
1027.5411376953125 0 28983.785
1028.4390869140625 0 687.5417
1028.5430908203125 0 10512.646
1029.5469970703125 0 3046.6697
1040.4720458984375 0 4380.3906
1041.4627685546875 0 4902.5747 y 5
1042.4600830078125 0 1265.3356
1045.5277099609375 0 16825.648
1046.531005859375 0 7310.9937
1047.539794921875 0 3437.6125
1048.548095703125 0 893.83966
1051.6248779296875 0 20444.805
1052.6282958984375 0 15773.459
1053.5374755859375 0 717.55566
1053.63134765625 0 6164.4756
1060.5535888671875 0 1453.7777
1062.5914306640625 0 3910.5479
1063.5843505859375 0 2125.483
1066.61181640625 0 2040.569
1067.625732421875 0 1027.5732
1078.601318359375 0 3497.3657
1079.6171875 0 15562.65
1080.6207275390625 0 10635.298
1081.6224365234375 0 2472.3352
1082.6275634765625 0 2264.3655
1083.533935546875 0 23143.287
1083.6358642578125 0 1941.0776
1084.5369873046875 0 16109.049
1084.634521484375 0 1205.7426
1085.5421142578125 0 5338.1855
1086.550537109375 0 2155.5183
1091.53466796875 0 6148.107
1092.540771484375 0 9849.728
1093.5421142578125 0 3885.1216
1095.4674072265625 0 1513.7369 w 4
1096.45654296875 0 938.8996
1096.5694580078125 0 3979.4834
1097.5655517578125 0 2824.1814
1098.572998046875 0 1658.7495
1099.56689453125 0 1244.1101
1105.5244140625 0 1027.3962
1107.5517578125 0 1969.0518
1109.5184326171875 0 1164.7725
1124.534912109375 0 1280.6946 c Ammonia loss 8
1125.535400390625 0 1559.8364
1126.5216064453125 0 1271.7316
1139.5703125 0 2165.9827
1140.5750732421875 0 9261.641
1141.5693359375 0 10672.889 c 8
1142.56982421875 0 5288.153
1143.580322265625 0 2202.6477
1144.5955810546875 0 14980.849
1145.5987548828125 0 6192.705
1146.60302734375 0 2718.7444
1152.5079345703125 0 18820.871
1153.49609375 0 53411.695 z 4
1154.4990234375 0 31484.186
1155.4993896484375 0 12560.34
1156.501220703125 0 3928.5728
1157.513916015625 0 1029.059
1168.523193359375 0 2645.0876
1169.51513671875 0 10019.047 y 4
1170.5159912109375 0 5530.609
1171.51708984375 0 2338.6094
1178.5693359375 0 1224.5427
1179.574462890625 0 4396.049
1180.5821533203125 0 3137.0266 c Water loss 9
1181.5782470703125 0 1638.2307
1187.6070556640625 0 1647.3727
1188.611572265625 0 1269.9755
1194.5877685546875 0 2385.29
1195.5828857421875 0 912.1387
1196.600830078125 0 859.3593
1197.60107421875 0 30079.152
1198.59033203125 0 66540.71 c 9
1199.5904541015625 0 37360.04
1200.5936279296875 0 13030.206
1201.5985107421875 0 3573.1868
1202.593017578125 0 1024.0586
1230.6187744140625 0 1319.3212
1231.6263427734375 0 25501.62
1232.6300048828125 0 14206.92
1233.632568359375 0 4257.138
1239.577392578125 0 885.7303
1245.6448974609375 0 918.8785
1258.6199951171875 0 1055.2946
1261.630126953125 0 851.6271
1274.6553955078125 0 962.49316
1276.6339111328125 0 1251.423
1277.6339111328125 0 1224.7352
1278.6412353515625 0 5747.7104
1279.6478271484375 0 3481.9465
1282.66796875 0 1944.3011
1283.6771240234375 0 2964.6404
1284.673583984375 0 2199.166
1289.5631103515625 0 4594.0327 y Ammonia loss 3
1290.5552978515625 0 23553.832 z 3
1291.558837890625 0 22015.615
1292.560791015625 0 11558.046
1293.5626220703125 0 4198.3213
1305.57470703125 0 1158.6239
1306.5728759765625 0 2571.5352 y 3
1307.5821533203125 0 2146.5273
1310.6697998046875 0 1310.3359
1311.6728515625 0 1513.065
1318.6588134765625 0 16707.02
1319.65966796875 0 9510.782
1320.6646728515625 0 3813.215
1324.6910400390625 0 3277.7073
1325.6939697265625 0 12373.677
1326.68408203125 0 27336.102 c 10
1327.687255859375 0 20528.43
1328.6890869140625 0 8949.508
1329.693359375 0 2945.5479
1345.656494140625 0 1612.1118
1346.6597900390625 0 1243.0955
1386.6279296875 0 970.3308 z Ammonia loss 2
1391.673095703125 0 1735.3868
1402.6494140625 0 3387.1985 y Ammonia loss 2
1403.6446533203125 0 12303.468 z 2
1404.6456298828125 0 15158.403
1405.6475830078125 0 8514.133
1406.6497802734375 0 4470.633
1407.6805419921875 0 8919.997
1408.6868896484375 0 5497.115
1409.6934814453125 0 1599.662
1410.719482421875 0 1530.8118
1411.7210693359375 0 5004.2446
1412.7200927734375 0 5204.677
1413.7232666015625 0 2885.9937
1414.7257080078125 0 1637.2634
1416.7183837890625 0 1332.3158
1419.669677734375 0 1762.2325 y 2
1420.66796875 0 1202.7451
1421.6871337890625 0 1164.707
1426.619384765625 0 1005.74066
1431.74755859375 0 3005.4531
1432.750732421875 0 1909.054
1437.7362060546875 0 1084.6108
1438.7203369140625 0 3372.8154
1439.7152099609375 0 4280.842
1440.7138671875 0 2450.057
1441.716064453125 0 1289.349
1454.736083984375 0 3813.351
1455.734619140625 0 12817.073 c 11
1456.7340087890625 0 16617.78
1457.735107421875 0 8192.825
1458.7393798828125 0 6110.826
1459.7432861328125 0 2398.2375
1473.7098388671875 0 1428.7245
1474.73876953125 0 4159.657
1475.7623291015625 0 12135.702
1476.7596435546875 0 6033.417
1477.763671875 0 2448.1848
1487.74853515625 0 1659.6234
1488.7354736328125 0 4436.5415
1489.73779296875 0 3051.179
1490.7467041015625 0 1109.5449
1493.706298828125 0 982.19116
1494.71484375 0 6976.1553
1495.7197265625 0 4502.5693
1496.717041015625 0 1530.9031
1501.7174072265625 0 1157.3331 y Ammonia loss 1
1502.7193603515625 0 1588.7825 z 1
1503.7158203125 0 4770.053
1504.71728515625 0 2850.9294
1505.71826171875 0 1625.5486
1518.72802734375 0 969.62805 y 1
1522.72705078125 0 2513.2375
1523.7333984375 0 1754.3054
1524.739013671875 0 879.6948
1544.753173828125 0 819.5824
1545.73095703125 0 865.19305
1547.7666015625 0 1237.7596
1548.7646484375 0 1986.176
1549.7725830078125 0 1039.7112
1558.762939453125 0 2097.7268
1559.774658203125 0 3818.9148
1560.77392578125 0 6789.1357
1561.7681884765625 0 10324.044
1562.76904296875 0 8705.064
1563.7679443359375 0 4445.8115
1564.780517578125 0 11988.562
1565.7803955078125 0 10197.741
1566.7818603515625 0 7435.2974
1567.77587890625 0 1879.6879
1573.72509765625 0 1611.2034
1574.7462158203125 0 4220.273
1575.7701416015625 0 13525.489
1576.7689208984375 0 13982.043
1577.7672119140625 0 8618.202
1578.769287109375 0 4509.46
1579.7916259765625 0 2968.3823
1580.8052978515625 0 3009.7983
1581.7989501953125 0 3938.1138
1582.79248046875 0 2354.3955
1584.6768798828125 0 1184.3895
1585.7254638671875 0 3265.8909
1586.75048828125 0 4400.2676
1587.7518310546875 0 2758.0483
1588.7640380859375 0 1854.0653
1589.787353515625 0 1374.1053
1590.7779541015625 0 4234.0615
1591.773681640625 0 6739.4062
1592.777587890625 0 4880.913
1593.7813720703125 0 4845.1274
1594.785400390625 0 2960.7625
1595.798095703125 0 1257.6403
1596.82958984375 0 2791.8315
1597.835205078125 0 1964.137
1598.826416015625 0 868.16095
1601.752197265625 0 2042.1075
1602.7716064453125 0 12863.658
1603.7701416015625 0 47047.023
1604.766845703125 0 62372.402
1605.7640380859375 0 39892.38
1606.7720947265625 0 23577.123
1607.794677734375 0 34553.145
1608.7994384765625 0 26577.637
1609.8004150390625 0 10070.873
1618.7528076171875 0 5305.7593
1619.751953125 0 22502.355
1620.7801513671875 0 53952.02
1621.7828369140625 0 79151.19
1622.78466796875 0 58642.625
1623.7919921875 0 31857.973
1624.815185546875 0 36512.27
1625.822265625 0 22975.314
1626.8248291015625 0 10922.585
3083.80029296875 0 946.604

Spectrum Details

|  |  |
| --- | --- |
| Matched peaks? Matched peaks The total absolute number of peaks matched. Additionally in brackets the total fraction of peaks matched and the total number of peaks is shown. | 81 (9.83% of 824) |
| FDR? FDR The false discovery rate estimated for this peptide. It is calculated by matching all theoretical fragments with a non-integer shift with the raw peaks for this spectrum. This is done with 40 different shifts. The resulting percentage is the average number of annotated peaks over the number of annotated peaks with the correct spectrum. | 5.35% |
| Satellite FDR? Satellite FDR See the FDR for details on its calculation. This satellite ion specific FDR only contains the satellite ions (d/w) for I/L/J positions. | 3.57% |
| PSM Score? PSM Score The PSM Score as given by Hecklib to this annotated spectrum. It is shown with three significant figures. | 664 |

#### Spectrum 8166? Spectrum 8166 The raw spectrum of this peptide as annotated by Hecklib. The fragments are coloured according to ion type (see legend). Any peaks with a star '\*' as text can be hovered over to see the full details, first the ion type second the mass shift type. By hovering over the amino acids in the peptide or ions in the legend the corresponding peaks are highlighted. By toggling the 'Unassigned' label you can turn the background (unassigned) peaks on or off in the plot. By updating the slider in the Ion legend you can update the spectrum to only show the top X% of the peaks with labels. The top X% means any peak that is within X% of the highest intensity. By dragging in the spectrum you can zoom in to a specific part of the spectrum and use 'Zoom Out' to get back to the original zoom level. The annotation of the spectrum is based on the given sequence in the peptides file and is done with different software so inconsistencies are likely. The peaks are annotated based on the given sequence, with 20 ppm tolerance.

Copy Data

##### Spectrum 8166 (TSV)

###### Preview

```
Loading example...
```

*Click on the button to copy the data to your clipboard.*

Mz MinMz MaxIntensity Max

WidthHeightPeptide font sizePeptide stroke widthSpectrum font sizeSpectrum stroke widthCompact peptide

Ion legend

wxyz

abcd

OtherUnassignedIon^Charge^_Position_Show for top:%

TVLHQDWLDGKEY

03.89e+47.77e+41.17e+51.55e+5

Zoom Out

a^+1^_2_y^+3^_4_a^+1^_2_y^+1^_1_b^+1^_2_b^+1^_2_y^+2^_3_y^+1^_2_y^+2^_5_y^+2^_5_y^+1^_2_b^+1^_3_b^+2^_6_y^+2^_6_b^+2^_7_b^+2^_7_b^+1^_4_y^+1^_3_b^+2^_7_b^+1^_4_y^+1^_4_b^+2^_8_b^+2^_8_y^+1^_4_y^+2^_8_\*\*b^+2^_9_b^+1^_5_b^+1^_5_y^+1^_5_y^+1^_5_y^+2^_10_y^+2^_10_b^+1^_6_b^+1^_6_y^+2^_11_y^+2^_11_b^+1^_6_y^+2^_11_b^+2^_12_b^+2^_12_y^+1^_6_b^+2^_12_y^+1^_6_y^+2^_12_\*b^+1^_7_b^+1^_7_b^+1^_7_y^+1^_7_b^+1^_8_y^+1^_8_y^+1^_9_y^+1^_9_y^+1^_9_

0777155523323110

Fragment Matches Table

Show background peaks

| Position | Ion type | Intensity | mz Theoretical | mz Error (Th) | mz Error (ppm) | Charge | Series Number |
| --- | --- | --- | --- | --- | --- | --- | --- |
| - | - | 5831 | 120.1 | - | - | 0 | - |
| - | - | 669.1 | 121.1 | - | - | 0 | - |
| - | - | 472.1 | 122.1 | - | - | 0 | - |
| - | - | 862.2 | 123 | - | - | 0 | - |
| - | - | 473.1 | 125.1 | - | - | 0 | - |
| - | - | 1862 | 127.1 | - | - | 0 | - |
| - | - | 6484 | 128.1 | - | - | 0 | - |
| - | - | 1375 | 129.1 | - | - | 0 | - |
| - | - | 6.959E+04 | 129.1 | - | - | 0 | - |
| - | - | 3.112E+04 | 130.1 | - | - | 0 | - |
| - | - | 4871 | 130.1 | - | - | 0 | - |
| - | - | 2883 | 131.1 | - | - | 0 | - |
| - | - | 442.3 | 131.5 | - | - | 0 | - |
| - | - | 8106 | 132.1 | - | - | 0 | - |
| - | - | 4274 | 132.1 | - | - | 0 | - |
| - | - | 885.7 | 133.1 | - | - | 0 | - |
| - | - | 1.164E+04 | 136.1 | - | - | 0 | - |
| - | - | 746.4 | 137.1 | - | - | 0 | - |
| - | - | 1476 | 138.1 | - | - | 0 | - |
| - | - | 643.5 | 138.1 | - | - | 0 | - |
| - | - | 569.1 | 139.1 | - | - | 0 | - |
| - | - | 1197 | 139.1 | - | - | 0 | - |
| - | - | 1719 | 141.1 | - | - | 0 | - |
| - | - | 699.5 | 142.1 | - | - | 0 | - |
| - | - | 759.1 | 145.1 | - | - | 0 | - |
| - | - | 876.8 | 146.1 | - | - | 0 | - |
| - | - | 663 | 147.1 | - | - | 0 | - |
| - | - | 1.886E+04 | 149 | - | - | 0 | - |
| - | - | 448.4 | 150.1 | - | - | 0 | - |
| - | - | 1586 | 151.1 | - | - | 0 | - |
| - | - | 415.2 | 151.3 | - | - | 0 | - |
| - | - | 566.9 | 153.1 | - | - | 0 | - |
| - | - | 431.6 | 153.9 | - | - | 0 | - |
| - | - | 524.1 | 155 | - | - | 0 | - |
| - | - | 562.9 | 155.1 | - | - | 0 | - |
| - | - | 4481 | 155.1 | - | - | 0 | - |
| 2 | a | 4013 | 155.1 | 0.0003203 | 2.065 | +1 | 2 |
| - | - | 500.2 | 156.1 | - | - | 0 | - |
| - | - | 618 | 157.1 | - | - | 0 | - |
| - | - | 841.8 | 157.1 | - | - | 0 | - |
| - | - | 416.7 | 158.1 | - | - | 0 | - |
| - | - | 5.246E+04 | 159.1 | - | - | 0 | - |
| - | - | 582.7 | 160.1 | - | - | 0 | - |
| - | - | 736.4 | 160.1 | - | - | 0 | - |
| - | - | 6427 | 160.1 | - | - | 0 | - |
| - | - | 478.6 | 161.1 | - | - | 0 | - |
| - | - | 5199 | 165.1 | - | - | 0 | - |
| - | - | 8762 | 166.1 | - | - | 0 | - |
| 10 | y | 2856 | 166.1 | 0.001557 | 9.377 | +3 | 4 |
| - | - | 727 | 166.1 | - | - | 0 | - |
| - | - | 2.606E+04 | 167.1 | - | - | 0 | - |
| - | - | 493.3 | 167.1 | - | - | 0 | - |
| - | - | 696.9 | 167.1 | - | - | 0 | - |
| - | - | 764.6 | 168.1 | - | - | 0 | - |
| - | - | 695.3 | 168.1 | - | - | 0 | - |
| - | - | 631.7 | 168.1 | - | - | 0 | - |
| - | - | 753 | 168.1 | - | - | 0 | - |
| - | - | 1207 | 169.1 | - | - | 0 | - |
| - | - | 3822 | 169.1 | - | - | 0 | - |
| - | - | 2591 | 170.1 | - | - | 0 | - |
| - | - | 886.6 | 171.1 | - | - | 0 | - |
| - | - | 1109 | 172.1 | - | - | 0 | - |
| - | - | 1329 | 173.1 | - | - | 0 | - |
| 2 | a | 1.033E+05 | 173.1 | 0.0003147 | 1.818 | +1 | 2 |
| - | - | 678.9 | 173.5 | - | - | 0 | - |
| - | - | 522.9 | 174.1 | - | - | 0 | - |
| - | - | 595.5 | 174.1 | - | - | 0 | - |
| - | - | 9357 | 174.1 | - | - | 0 | - |
| - | - | 773.3 | 175.1 | - | - | 0 | - |
| - | - | 436.8 | 177.1 | - | - | 0 | - |
| - | - | 3863 | 178.1 | - | - | 0 | - |
| - | - | 459.2 | 179.6 | - | - | 0 | - |
| 13 | y | 1.202E+04 | 182.1 | 0.000297 | 1.631 | +1 | 1 |
| - | - | 1070 | 183.1 | - | - | 0 | - |
| 2 | b | 2283 | 183.1 | 0.0003092 | 1.689 | +1 | 2 |
| - | - | 1152 | 183.1 | - | - | 0 | - |
| - | - | 987.6 | 185.2 | - | - | 0 | - |
| - | - | 1.967E+04 | 186.1 | - | - | 0 | - |
| - | - | 5047 | 187.1 | - | - | 0 | - |
| - | - | 495.5 | 187.1 | - | - | 0 | - |
| - | - | 1652 | 187.1 | - | - | 0 | - |
| - | - | 886.9 | 187.1 | - | - | 0 | - |
| - | - | 656 | 188.1 | - | - | 0 | - |
| - | - | 773.6 | 188.1 | - | - | 0 | - |
| - | - | 1162 | 190.1 | - | - | 0 | - |
| - | - | 623.9 | 190.1 | - | - | 0 | - |
| - | - | 581.4 | 194.1 | - | - | 0 | - |
| - | - | 5889 | 195.1 | - | - | 0 | - |
| - | - | 3982 | 196.1 | - | - | 0 | - |
| - | - | 563.3 | 196.1 | - | - | 0 | - |
| - | - | 1451 | 197.1 | - | - | 0 | - |
| - | - | 2070 | 198.1 | - | - | 0 | - |
| - | - | 638.7 | 199.1 | - | - | 0 | - |
| - | - | 1062 | 199.1 | - | - | 0 | - |
| - | - | 471.3 | 201.1 | - | - | 0 | - |
| 2 | b | 2.911E+04 | 201.1 | 0.0002426 | 1.206 | +1 | 2 |
| - | - | 3935 | 202.1 | - | - | 0 | - |
| - | - | 621.8 | 203.1 | - | - | 0 | - |
| - | - | 2642 | 205.1 | - | - | 0 | - |
| - | - | 641.8 | 206.1 | - | - | 0 | - |
| - | - | 1035 | 210.1 | - | - | 0 | - |
| 11 | y | 877.9 | 211.1 | 5.405E-05 | 0.256 | +2 | 3 |
| - | - | 1362 | 212.1 | - | - | 0 | - |
| - | - | 516.1 | 213.1 | - | - | 0 | - |
| - | - | 2221 | 213.1 | - | - | 0 | - |
| - | - | 564.3 | 215.1 | - | - | 0 | - |
| - | - | 586.1 | 216.1 | - | - | 0 | - |
| - | - | 535.7 | 216.7 | - | - | 0 | - |
| - | - | 2524 | 221.1 | - | - | 0 | - |
| - | - | 8201 | 221.1 | - | - | 0 | - |
| - | - | 585.6 | 221.1 | - | - | 0 | - |
| - | - | 2030 | 222.1 | - | - | 0 | - |
| - | - | 624.4 | 222.1 | - | - | 0 | - |
| - | - | 7519 | 223.1 | - | - | 0 | - |
| - | - | 1179 | 223.1 | - | - | 0 | - |
| - | - | 2321 | 223.1 | - | - | 0 | - |
| - | - | 9865 | 223.2 | - | - | 0 | - |
| - | - | 698.3 | 224.1 | - | - | 0 | - |
| - | - | 1215 | 224.2 | - | - | 0 | - |
| - | - | 1.124E+04 | 225 | - | - | 0 | - |
| - | - | 2611 | 226 | - | - | 0 | - |
| - | - | 3486 | 226.1 | - | - | 0 | - |
| - | - | 607 | 227 | - | - | 0 | - |
| - | - | 2336 | 227 | - | - | 0 | - |
| - | - | 1720 | 227.1 | - | - | 0 | - |
| - | - | 636.9 | 227.2 | - | - | 0 | - |
| - | - | 506.8 | 228 | - | - | 0 | - |
| - | - | 576.1 | 228.1 | - | - | 0 | - |
| - | - | 798.4 | 228.1 | - | - | 0 | - |
| - | - | 1440 | 229.1 | - | - | 0 | - |
| - | - | 2940 | 229.2 | - | - | 0 | - |
| - | - | 1432 | 233.1 | - | - | 0 | - |
| - | - | 2399 | 233.2 | - | - | 0 | - |
| - | - | 1.091E+04 | 234.1 | - | - | 0 | - |
| - | - | 902 | 235.1 | - | - | 0 | - |
| - | - | 824.5 | 237.1 | - | - | 0 | - |
| - | - | 1642 | 238.1 | - | - | 0 | - |
| - | - | 2.49E+04 | 239.1 | - | - | 0 | - |
| - | - | 6155 | 240.1 | - | - | 0 | - |
| - | - | 5148 | 240.1 | - | - | 0 | - |
| - | - | 4283 | 241.1 | - | - | 0 | - |
| - | - | 768.9 | 241.1 | - | - | 0 | - |
| - | - | 1024 | 241.1 | - | - | 0 | - |
| - | - | 672.3 | 241.2 | - | - | 0 | - |
| - | - | 458.2 | 243.1 | - | - | 0 | - |
| - | - | 4251 | 244.1 | - | - | 0 | - |
| - | - | 653.4 | 245.1 | - | - | 0 | - |
| - | - | 5558 | 247.1 | - | - | 0 | - |
| - | - | 2211 | 248.1 | - | - | 0 | - |
| - | - | 790.5 | 249.1 | - | - | 0 | - |
| - | - | 3089 | 249.1 | - | - | 0 | - |
| - | - | 968.9 | 249.1 | - | - | 0 | - |
| - | - | 3.684E+04 | 251.2 | - | - | 0 | - |
| - | - | 596.5 | 252.1 | - | - | 0 | - |
| - | - | 3422 | 252.2 | - | - | 0 | - |
| - | - | 1978 | 255.1 | - | - | 0 | - |
| - | - | 556.7 | 256.1 | - | - | 0 | - |
| - | - | 687.1 | 256.2 | - | - | 0 | - |
| - | - | 835.8 | 256.6 | - | - | 0 | - |
| - | - | 649.1 | 257.1 | - | - | 0 | - |
| - | - | 1835 | 258.1 | - | - | 0 | - |
| - | - | 935.1 | 261.1 | - | - | 0 | - |
| - | - | 638.4 | 261.2 | - | - | 0 | - |
| - | - | 738.8 | 262.1 | - | - | 0 | - |
| - | - | 550.5 | 264.3 | - | - | 0 | - |
| - | - | 484 | 264.3 | - | - | 0 | - |
| - | - | 4918 | 265.1 | - | - | 0 | - |
| - | - | 1108 | 266.1 | - | - | 0 | - |
| - | - | 9887 | 266.1 | - | - | 0 | - |
| - | - | 1102 | 267.1 | - | - | 0 | - |
| - | - | 1426 | 268.2 | - | - | 0 | - |
| - | - | 794.5 | 269.2 | - | - | 0 | - |
| - | - | 1011 | 270.1 | - | - | 0 | - |
| - | - | 662.4 | 270.6 | - | - | 0 | - |
| - | - | 1643 | 272.2 | - | - | 0 | - |
| - | - | 2184 | 274.1 | - | - | 0 | - |
| - | - | 712 | 275.1 | - | - | 0 | - |
| - | - | 1149 | 280.1 | - | - | 0 | - |
| - | - | 1.788E+04 | 281.1 | - | - | 0 | - |
| - | - | 2807 | 282.2 | - | - | 0 | - |
| - | - | 2949 | 283 | - | - | 0 | - |
| - | - | 479.5 | 283 | - | - | 0 | - |
| - | - | 2.672E+04 | 283.1 | - | - | 0 | - |
| - | - | 1312 | 284.1 | - | - | 0 | - |
| - | - | 3147 | 284.1 | - | - | 0 | - |
| - | - | 1367 | 285 | - | - | 0 | - |
| - | - | 1193 | 286.1 | - | - | 0 | - |
| - | - | 761.7 | 288.6 | - | - | 0 | - |
| 12 | y | 1.359E+04 | 293.1 | 0.0003273 | 1.117 | +1 | 2 |
| - | - | 956.6 | 293.1 | - | - | 0 | - |
| - | - | 2582 | 294.1 | - | - | 0 | - |
| - | - | 1634 | 295.1 | - | - | 0 | - |
| - | - | 2128 | 296.1 | - | - | 0 | - |
| - | - | 2939 | 297.1 | - | - | 0 | - |
| - | - | 1011 | 297.1 | - | - | 0 | - |
| 9 | y | 989.7 | 297.1 | 0.004949 | 16.66 | +2 | 5 |
| - | - | 4033 | 297.2 | - | - | 0 | - |
| - | - | 588.5 | 298.2 | - | - | 0 | - |
| - | - | 1.539E+05 | 299.1 | - | - | 0 | - |
| - | - | 3189 | 300.1 | - | - | 0 | - |
| - | - | 5492 | 300.2 | - | - | 0 | - |
| - | - | 2738 | 301.1 | - | - | 0 | - |
| - | - | 1.106E+04 | 301.2 | - | - | 0 | - |
| - | - | 1138 | 301.2 | - | - | 0 | - |
| - | - | 2856 | 302.1 | - | - | 0 | - |
| - | - | 1114 | 302.2 | - | - | 0 | - |
| - | - | 701.3 | 305.6 | - | - | 0 | - |
| 9 | y | 2479 | 306.1 | 0.001132 | 3.697 | +2 | 5 |
| 12 | y | 2105 | 311.1 | 0.0004743 | 1.524 | +1 | 2 |
| - | - | 617.7 | 313.5 | - | - | 0 | - |
| 3 | b | 1408 | 314.2 | 2.567E-05 | 0.0817 | +1 | 3 |
| - | - | 7051 | 315.2 | - | - | 0 | - |
| - | - | 1532 | 316.2 | - | - | 0 | - |
| - | - | 1604 | 326.7 | - | - | 0 | - |
| - | - | 1190 | 327 | - | - | 0 | - |
| - | - | 1346 | 327.2 | - | - | 0 | - |
| - | - | 2001 | 333.2 | - | - | 0 | - |
| - | - | 786.6 | 338.2 | - | - | 0 | - |
| - | - | 1604 | 341 | - | - | 0 | - |
| - | - | 609 | 343.2 | - | - | 0 | - |
| - | - | 1.934E+04 | 345 | - | - | 0 | - |
| - | - | 1375 | 345.1 | - | - | 0 | - |
| - | - | 4322 | 346.1 | - | - | 0 | - |
| - | - | 1509 | 347.1 | - | - | 0 | - |
| 6 | b | 1983 | 347.7 | 0.000666 | 1.915 | +2 | 6 |
| - | - | 3593 | 350.2 | - | - | 0 | - |
| - | - | 3066 | 355.1 | - | - | 0 | - |
| - | - | 1.727E+04 | 359 | - | - | 0 | - |
| - | - | 1550 | 361.2 | - | - | 0 | - |
| - | - | 2546 | 362.2 | - | - | 0 | - |
| 8 | y | 3971 | 362.7 | 0.0002204 | 0.6076 | +2 | 6 |
| - | - | 656.7 | 363.1 | - | - | 0 | - |
| - | - | 6755 | 363.1 | - | - | 0 | - |
| - | - | 3435 | 363.2 | - | - | 0 | - |
| - | - | 1066 | 364.1 | - | - | 0 | - |
| - | - | 737.2 | 364.1 | - | - | 0 | - |
| - | - | 702.7 | 366.2 | - | - | 0 | - |
| - | - | 1004 | 367.1 | - | - | 0 | - |
| - | - | 524.7 | 367.2 | - | - | 0 | - |
| - | - | 559.1 | 368.2 | - | - | 0 | - |
| - | - | 1123 | 369.1 | - | - | 0 | - |
| - | - | 2326 | 370.1 | - | - | 0 | - |
| - | - | 1020 | 371.1 | - | - | 0 | - |
| - | - | 1349 | 376.2 | - | - | 0 | - |
| - | - | 624.4 | 376.7 | - | - | 0 | - |
| - | - | 2244 | 378.2 | - | - | 0 | - |
| - | - | 8609 | 379.2 | - | - | 0 | - |
| - | - | 1245 | 380.2 | - | - | 0 | - |
| - | - | 751.4 | 381.1 | - | - | 0 | - |
| - | - | 1.765E+04 | 381.2 | - | - | 0 | - |
| - | - | 705.3 | 381.7 | - | - | 0 | - |
| - | - | 3404 | 382.2 | - | - | 0 | - |
| - | - | 1240 | 384.2 | - | - | 0 | - |
| - | - | 1258 | 385.1 | - | - | 0 | - |
| - | - | 3050 | 394.2 | - | - | 0 | - |
| - | - | 1357 | 395.1 | - | - | 0 | - |
| - | - | 802.8 | 395.2 | - | - | 0 | - |
| - | - | 2417 | 396.2 | - | - | 0 | - |
| - | - | 1865 | 398.2 | - | - | 0 | - |
| - | - | 841.7 | 402.2 | - | - | 0 | - |
| - | - | 1594 | 402.2 | - | - | 0 | - |
| - | - | 721.6 | 405.3 | - | - | 0 | - |
| - | - | 991 | 409.1 | - | - | 0 | - |
| - | - | 2033 | 411.2 | - | - | 0 | - |
| - | - | 1001 | 412.2 | - | - | 0 | - |
| - | - | 1.07E+04 | 412.2 | - | - | 0 | - |
| - | - | 1272 | 413.1 | - | - | 0 | - |
| - | - | 1140 | 413.2 | - | - | 0 | - |
| - | - | 2926 | 414.2 | - | - | 0 | - |
| - | - | 3.529E+04 | 415 | - | - | 0 | - |
| - | - | 875.5 | 415.2 | - | - | 0 | - |
| - | - | 1396 | 417.7 | - | - | 0 | - |
| - | - | 3099 | 418.2 | - | - | 0 | - |
| - | - | 1071 | 418.7 | - | - | 0 | - |
| - | - | 944.5 | 423.2 | - | - | 0 | - |
| - | - | 4115 | 423.3 | - | - | 0 | - |
| - | - | 959.3 | 424.3 | - | - | 0 | - |
| - | - | 2.789E+04 | 426.7 | - | - | 0 | - |
| - | - | 1.593E+04 | 427.2 | - | - | 0 | - |
| - | - | 4254 | 427.7 | - | - | 0 | - |
| - | - | 1428 | 429.1 | - | - | 0 | - |
| - | - | 8248 | 429.2 | - | - | 0 | - |
| - | - | 2.44E+04 | 430.2 | - | - | 0 | - |
| - | - | 661.4 | 431.1 | - | - | 0 | - |
| - | - | 4626 | 431.2 | - | - | 0 | - |
| 7 | b | 3239 | 431.7 | 0.0001358 | 0.3146 | +2 | 7 |
| 7 | b | 1846 | 432.2 | 0.000163 | 0.377 | +2 | 7 |
| - | - | 1686 | 433 | - | - | 0 | - |
| - | - | 619.7 | 433.2 | - | - | 0 | - |
| 4 | b | 661.7 | 433.3 | 0.00227 | 5.241 | +1 | 4 |
| - | - | 721 | 438.2 | - | - | 0 | - |
| - | - | 668.8 | 438.2 | - | - | 0 | - |
| 11 | y | 7507 | 439.2 | 0.0005125 | 1.167 | +1 | 3 |
| - | - | 1876 | 440.2 | - | - | 0 | - |
| 7 | b | 1.567E+04 | 440.7 | 0.0004687 | 1.064 | +2 | 7 |
| - | - | 1445 | 440.9 | - | - | 0 | - |
| - | - | 8672 | 441.2 | - | - | 0 | - |
| - | - | 695.8 | 441.3 | - | - | 0 | - |
| - | - | 1949 | 441.7 | - | - | 0 | - |
| - | - | 799.1 | 447.2 | - | - | 0 | - |
| - | - | 967.8 | 448.2 | - | - | 0 | - |
| - | - | 638.9 | 450.2 | - | - | 0 | - |
| 4 | b | 7621 | 451.3 | 0.0003484 | 0.7721 | +1 | 4 |
| - | - | 2227 | 452.3 | - | - | 0 | - |
| - | - | 1045 | 460.8 | - | - | 0 | - |
| - | - | 968.1 | 461.2 | - | - | 0 | - |
| - | - | 644.8 | 461.3 | - | - | 0 | - |
| - | - | 2448 | 468.3 | - | - | 0 | - |
| - | - | 1200 | 471.2 | - | - | 0 | - |
| - | - | 645 | 473.9 | - | - | 0 | - |
| - | - | 1479 | 474.3 | - | - | 0 | - |
| - | - | 1975 | 474.8 | - | - | 0 | - |
| - | - | 965.1 | 475.3 | - | - | 0 | - |
| - | - | 572.8 | 475.7 | - | - | 0 | - |
| - | - | 958.4 | 475.8 | - | - | 0 | - |
| - | - | 1602 | 477.2 | - | - | 0 | - |
| 10 | y | 885 | 478.2 | 0.005055 | 10.57 | +1 | 4 |
| - | - | 818 | 478.3 | - | - | 0 | - |
| - | - | 5112 | 483.3 | - | - | 0 | - |
| - | - | 2190 | 483.8 | - | - | 0 | - |
| - | - | 839.5 | 484.7 | - | - | 0 | - |
| 8 | b | 2428 | 488.3 | 0.001195 | 2.447 | +2 | 8 |
| 8 | b | 876.1 | 488.7 | 0.007295 | 14.93 | +2 | 8 |
| - | - | 656.3 | 492.3 | - | - | 0 | - |
| - | - | 1431 | 492.7 | - | - | 0 | - |
| - | - | 1.738E+04 | 494.2 | - | - | 0 | - |
| - | - | 4698 | 495.2 | - | - | 0 | - |
| 10 | y | 2.08E+04 | 496.2 | 0.0005942 | 1.197 | +1 | 4 |
| - | - | 5805 | 497.2 | - | - | 0 | - |
| - | - | 1295 | 497.8 | - | - | 0 | - |
| - | - | 1127 | 498.2 | - | - | 0 | - |
| - | - | 3.535E+04 | 503.1 | - | - | 0 | - |
| - | - | 1114 | 507.3 | - | - | 0 | - |
| - | - | 1347 | 511.3 | - | - | 0 | - |
| 6 | y | 682.7 | 513.2 | 0.009708 | 18.92 | +2 | 8 |
| - | - | 900.6 | 517.3 | - | - | 0 | - |
| - | - | 1.246E+04 | 519.1 | - | - | 0 | - |
| - | - | 667.9 | 521.1 | - | - | 0 | - |
| - | - | 2948 | 525.3 | - | - | 0 | - |
| - | - | 887.7 | 526.3 | - | - | 0 | - |
| 0 | Precursor | 853.7 | 529.3 | 0.0007233 | 1.367 | +3 | -1 |
| 0 | Precursor | 1006 | 529.6 | 0.004707 | 8.887 | +3 | -1 |
| - | - | 851.4 | 534.3 | - | - | 0 | - |
| - | - | 910.4 | 534.6 | - | - | 0 | - |
| - | - | 709.9 | 534.8 | - | - | 0 | - |
| - | - | 822.7 | 534.9 | - | - | 0 | - |
| - | - | 1002 | 535.6 | - | - | 0 | - |
| - | - | 765.5 | 535.9 | - | - | 0 | - |
| - | - | 818.7 | 536 | - | - | 0 | - |
| - | - | 9464 | 536.2 | - | - | 0 | - |
| - | - | 397.9 | 536.2 | - | - | 0 | - |
| - | - | 874.5 | 536.3 | - | - | 0 | - |
| - | - | 786.8 | 538.3 | - | - | 0 | - |
| - | - | 3149 | 539.2 | - | - | 0 | - |
| - | - | 663 | 540.2 | - | - | 0 | - |
| - | - | 1285 | 542.3 | - | - | 0 | - |
| - | - | 4057 | 543.3 | - | - | 0 | - |
| - | - | 887 | 544.3 | - | - | 0 | - |
| 9 | b | 1090 | 546.3 | 0.002423 | 4.436 | +2 | 9 |
| - | - | 1013 | 551.3 | - | - | 0 | - |
| - | - | 594.3 | 555.3 | - | - | 0 | - |
| - | - | 790.4 | 556.3 | - | - | 0 | - |
| 5 | b | 696.4 | 561.3 | 0.003758 | 6.695 | +1 | 5 |
| - | - | 834 | 563.3 | - | - | 0 | - |
| - | - | 1213 | 565.3 | - | - | 0 | - |
| - | - | 3827 | 567.2 | - | - | 0 | - |
| - | - | 1092 | 568.2 | - | - | 0 | - |
| - | - | 2206 | 575.2 | - | - | 0 | - |
| 5 | b | 4108 | 579.3 | 3.206E-05 | 0.05534 | +1 | 5 |
| - | - | 1549 | 580.3 | - | - | 0 | - |
| - | - | 753.1 | 581.3 | - | - | 0 | - |
| - | - | 712.1 | 584.2 | - | - | 0 | - |
| - | - | 1616 | 592.3 | - | - | 0 | - |
| - | - | 705.9 | 592.3 | - | - | 0 | - |
| 9 | y | 7223 | 593.3 | 0.0002681 | 0.4519 | +1 | 5 |
| - | - | 2401 | 593.3 | - | - | 0 | - |
| - | - | 2454 | 594.3 | - | - | 0 | - |
| - | - | 885.1 | 600.3 | - | - | 0 | - |
| - | - | 1195 | 602.8 | - | - | 0 | - |
| - | - | 775.3 | 603.8 | - | - | 0 | - |
| - | - | 1.919E+04 | 610.3 | - | - | 0 | - |
| - | - | 803.1 | 610.3 | - | - | 0 | - |
| 9 | y | 6.291E+04 | 611.3 | 0.0009949 | 1.628 | +1 | 5 |
| - | - | 3603 | 611.8 | - | - | 0 | - |
| - | - | 1.883E+04 | 612.3 | - | - | 0 | - |
| - | - | 808.9 | 612.8 | - | - | 0 | - |
| - | - | 3865 | 613.3 | - | - | 0 | - |
| - | - | 1252 | 621.3 | - | - | 0 | - |
| - | - | 758.8 | 623.2 | - | - | 0 | - |
| - | - | 1246 | 635.3 | - | - | 0 | - |
| - | - | 801.4 | 636.3 | - | - | 0 | - |
| 4 | y | 855 | 636.8 | 0.003968 | 6.231 | +2 | 10 |
| - | - | 963.4 | 640.3 | - | - | 0 | - |
| - | - | 2192 | 645.3 | - | - | 0 | - |
| 4 | y | 2791 | 645.8 | 0.004362 | 6.755 | +2 | 10 |
| - | - | 3239 | 646.3 | - | - | 0 | - |
| - | - | 7073 | 652.3 | - | - | 0 | - |
| - | - | 2770 | 653.3 | - | - | 0 | - |
| - | - | 1749 | 660.9 | - | - | 0 | - |
| - | - | 1594 | 661.4 | - | - | 0 | - |
| - | - | 2437 | 662.3 | - | - | 0 | - |
| - | - | 1842 | 663.3 | - | - | 0 | - |
| - | - | 1092 | 667.3 | - | - | 0 | - |
| 6 | b | 3530 | 676.3 | 0.0002361 | 0.3491 | +1 | 6 |
| 6 | b | 1843 | 677.3 | 0.0006701 | 0.9894 | +1 | 6 |
| - | - | 876 | 677.8 | - | - | 0 | - |
| - | - | 1756 | 678.3 | - | - | 0 | - |
| - | - | 1.299E+04 | 680.3 | - | - | 0 | - |
| - | - | 706.4 | 680.4 | - | - | 0 | - |
| - | - | 4532 | 681.3 | - | - | 0 | - |
| - | - | 1031 | 682.3 | - | - | 0 | - |
| - | - | 1127 | 684.3 | - | - | 0 | - |
| - | - | 777.7 | 684.8 | - | - | 0 | - |
| 3 | y | 4607 | 693.3 | 0.0008765 | 1.264 | +2 | 11 |
| 3 | y | 3138 | 693.8 | 0.007465 | 10.76 | +2 | 11 |
| 6 | b | 3.664E+04 | 694.4 | 0.0004858 | 0.6997 | +1 | 6 |
| - | - | 1.265E+04 | 695.4 | - | - | 0 | - |
| - | - | 3830 | 696.4 | - | - | 0 | - |
| - | - | 7145 | 701.8 | - | - | 0 | - |
| 3 | y | 2.453E+04 | 702.3 | 0.002613 | 3.721 | +2 | 11 |
| 12 | b | 1.646E+04 | 702.8 | 0.01284 | 18.27 | +2 | 12 |
| - | - | 902.6 | 702.9 | - | - | 0 | - |
| 12 | b | 6557 | 703.3 | 0.003686 | 5.241 | +2 | 12 |
| - | - | 2408 | 703.8 | - | - | 0 | - |
| - | - | 1004 | 704.3 | - | - | 0 | - |
| 8 | y | 2048 | 706.3 | 0.0006157 | 0.8717 | +1 | 6 |
| - | - | 858.8 | 707.3 | - | - | 0 | - |
| - | - | 2114 | 710.4 | - | - | 0 | - |
| - | - | 1157 | 710.9 | - | - | 0 | - |
| - | - | 2010 | 711.4 | - | - | 0 | - |
| 12 | b | 1758 | 711.9 | 0.003547 | 4.983 | +2 | 12 |
| - | - | 1444 | 712.4 | - | - | 0 | - |
| - | - | 1715 | 721.4 | - | - | 0 | - |
| - | - | 5374 | 723.4 | - | - | 0 | - |
| 8 | y | 1.822E+04 | 724.4 | 0.002136 | 2.949 | +1 | 6 |
| - | - | 8209 | 725.4 | - | - | 0 | - |
| - | - | 1582 | 726.4 | - | - | 0 | - |
| - | - | 1440 | 733.4 | - | - | 0 | - |
| - | - | 4043 | 751.4 | - | - | 0 | - |
| 2 | y | 2081 | 751.9 | 0.003196 | 4.251 | +2 | 12 |
| - | - | 4716 | 752.4 | - | - | 0 | - |
| - | - | 1172 | 752.9 | - | - | 0 | - |
| - | - | 804.9 | 755.4 | - | - | 0 | - |
| - | - | 1136 | 763.4 | - | - | 0 | - |
| - | - | 817.4 | 765.4 | - | - | 0 | - |
| - | - | 918.7 | 766.4 | - | - | 0 | - |
| - | - | 1974 | 779.4 | - | - | 0 | - |
| - | - | 1488 | 780.4 | - | - | 0 | - |
| - | - | 1481 | 781.4 | - | - | 0 | - |
| 0 | Precursor | 3935 | 793.4 | 0.01296 | 16.33 | +2 | -1 |
| - | - | 1953 | 794.4 | - | - | 0 | - |
| - | - | 961.9 | 807.4 | - | - | 0 | - |
| - | - | 903.4 | 812.3 | - | - | 0 | - |
| - | - | 2781 | 836.4 | - | - | 0 | - |
| - | - | 1243 | 837.5 | - | - | 0 | - |
| - | - | 867.3 | 846.4 | - | - | 0 | - |
| - | - | 8472 | 852.4 | - | - | 0 | - |
| - | - | 4776 | 853.4 | - | - | 0 | - |
| - | - | 959.9 | 854.4 | - | - | 0 | - |
| 7 | b | 5734 | 862.4 | 0.0001423 | 0.165 | +1 | 7 |
| 7 | b | 3576 | 863.4 | 0.01126 | 13.05 | +1 | 7 |
| - | - | 6375 | 864.5 | - | - | 0 | - |
| - | - | 3871 | 865.5 | - | - | 0 | - |
| - | - | 996.4 | 866.5 | - | - | 0 | - |
| - | - | 751.4 | 877.4 | - | - | 0 | - |
| 7 | b | 4.325E+04 | 880.4 | 0.0004531 | 0.5146 | +1 | 7 |
| - | - | 2.272E+04 | 881.4 | - | - | 0 | - |
| - | - | 6338 | 882.4 | - | - | 0 | - |
| - | - | 1124 | 883.4 | - | - | 0 | - |
| - | - | 1927 | 908.4 | - | - | 0 | - |
| - | - | 1737 | 909.4 | - | - | 0 | - |
| 7 | y | 3031 | 910.4 | 0.001986 | 2.181 | +1 | 7 |
| - | - | 1068 | 911.4 | - | - | 0 | - |
| 8 | b | 3297 | 993.5 | 0.001115 | 1.123 | +1 | 8 |
| - | - | 1077 | 994.5 | - | - | 0 | - |
| - | - | 994.4 | 1024 | - | - | 0 | - |
| 6 | y | 1374 | 1025 | 0.004339 | 4.232 | +1 | 8 |
| - | - | 1306 | 1094 | - | - | 0 | - |
| 5 | y | 2093 | 1136 | 0.001371 | 1.207 | +1 | 9 |
| 5 | y | 1198 | 1136 | 0.01613 | 14.2 | +1 | 9 |
| - | - | 720.6 | 1137 | - | - | 0 | - |
| - | - | 1417 | 1153 | - | - | 0 | - |
| 5 | y | 2776 | 1154 | 0.002403 | 2.083 | +1 | 9 |
| - | - | 1447 | 1155 | - | - | 0 | - |
| - | - | 942.2 | 1156 | - | - | 0 | - |
| - | - | 659 | 2000 | - | - | 0 | - |
| - | - | 698 | 2125 | - | - | 0 | - |
| - | - | 789 | 2567 | - | - | 0 | - |
| - | - | 643.5 | 2712 | - | - | 0 | - |
| - | - | 632 | 2902 | - | - | 0 | - |
| - | - | 732.6 | 3020 | - | - | 0 | - |
| - | - | 641.8 | 3020 | - | - | 0 | - |
| - | - | 685.6 | 3027 | - | - | 0 | - |
| - | - | 854.8 | 3079 | - | - | 0 | - |

m/z Charge Intensity FragmentType MassShift Position
120.0810546875 0 5831.4424
121.0843276977539 0 669.12634
122.07160949707031 0 472.10538
123.04430389404297 0 862.2303
125.10801696777344 0 473.0789
127.050537109375 0 1861.8196
128.10726928710938 0 6484.2095
129.06634521484375 0 1375.3588
129.1025390625 0 69592.484
130.06539916992188 0 31121.363
130.10585021972656 0 4871.328
131.06874084472656 0 2882.7017
131.5206298828125 0 442.32062
132.0810546875 0 8106.4546
132.10211181640625 0 4273.569
133.08531188964844 0 885.6774
136.075927734375 0 11636.026
137.0792236328125 0 746.3601
138.06651306152344 0 1475.9299
138.09130859375 0 643.50916
139.0505828857422 0 569.1252
139.08641052246094 0 1196.7207
141.10269165039062 0 1719.4183
142.0651092529297 0 699.4791
145.06117248535156 0 759.12115
146.06033325195312 0 876.8432
147.065673828125 0 663.0139
149.04513549804688 0 18860.885
150.06649780273438 0 448.3561
151.08702087402344 0 1585.7721
151.30137634277344 0 415.24835
153.10311889648438 0 566.94604
153.8965301513672 0 431.57407
155.04562377929688 0 524.061
155.08201599121094 0 562.9126
155.0929718017578 0 4480.762
155.1182098388672 0 4013.3801 a Water loss 1
156.12130737304688 0 500.24332
157.0974578857422 0 617.95764
157.1338348388672 0 841.76855
158.10154724121094 0 416.72382
159.09193420410156 0 52462.984
160.07565307617188 0 582.74786
160.08853149414062 0 736.42993
160.0953369140625 0 6426.5586
161.09799194335938 0 478.57153
165.054931640625 0 5199.2793
166.06134033203125 0 8762.066
166.0864715576172 0 2855.7197 y 9
166.0981903076172 0 726.9544
167.0557098388672 0 26064.572
167.0814971923828 0 493.3355
167.1178436279297 0 696.85645
168.05540466308594 0 764.6325
168.0806884765625 0 695.2536
168.10226440429688 0 631.65826
168.11346435546875 0 753.0038
169.07632446289062 0 1207.1198
169.09744262695312 0 3822.3083
170.06036376953125 0 2591.4888
171.14964294433594 0 886.5877
172.07171630859375 0 1109.1737
173.05601501464844 0 1329.4323
173.12876892089844 0 103331.766 a 1
173.45274353027344 0 678.87164
174.05502319335938 0 522.92535
174.12396240234375 0 595.48315
174.13209533691406 0 9356.944
175.08706665039062 0 773.25323
177.10301208496094 0 436.81668
178.13426208496094 0 3863.328
179.6286163330078 0 459.19434
182.0814666748047 0 12015.076 y 12
183.08502197265625 0 1070.2257
183.1131134033203 0 2282.8176 b Water loss 1
183.14981079101562 0 1151.8373
185.1647186279297 0 987.5575
186.12399291992188 0 19665.262
187.08692932128906 0 5046.6704
187.10899353027344 0 495.5281
187.1275634765625 0 1651.5227
187.14447021484375 0 886.85803
188.0709686279297 0 656.00354
188.09033203125 0 773.57666
190.0827178955078 0 1162.0267
190.13424682617188 0 623.94183
194.0917510986328 0 581.4439
195.11305236816406 0 5889.2026
196.10826110839844 0 3982.366
196.11676025390625 0 563.26337
197.1285858154297 0 1451.2212
198.1279296875 0 2070.1475
199.07151794433594 0 638.68005
199.14431762695312 0 1062.3292
201.08665466308594 0 471.29092
201.1236114501953 0 29106.984 b 1
202.12689208984375 0 3934.9192
203.10292053222656 0 621.77997
205.097412109375 0 2641.5706
206.1003875732422 0 641.82874
210.1269989013672 0 1034.7019
211.10777282714844 0 877.87915 y Water loss 10
212.13938903808594 0 1362.4429
213.06695556640625 0 516.14496
213.12364196777344 0 2220.9185
215.13938903808594 0 564.27264
216.09793090820312 0 586.121
216.72720336914062 0 535.686
221.05931091308594 0 2523.925
221.084716796875 0 8200.972
221.1399688720703 0 585.5583
222.08526611328125 0 2029.6843
222.12387084960938 0 624.37054
223.06394958496094 0 7519.3433
223.08212280273438 0 1179.4097
223.10780334472656 0 2320.765
223.1556396484375 0 9864.954
224.10214233398438 0 698.33014
224.1594696044922 0 1215.3748
225.043212890625 0 11236.315
226.04342651367188 0 2610.6592
226.08230590820312 0 3486.1948
227.02218627929688 0 606.95026
227.0400390625 0 2335.8997
227.06658935546875 0 1720.3049
227.15553283691406 0 636.9462
228.02304077148438 0 506.83105
228.09893798828125 0 576.14777
228.13426208496094 0 798.4231
229.11831665039062 0 1439.668
229.15489196777344 0 2940.482
233.13999938964844 0 1432.0186
233.1654510498047 0 2399.4067
234.12400817871094 0 10909.97
235.12713623046875 0 902.0088
237.13473510742188 0 824.4576
238.1190185546875 0 1642.4454
239.0952606201172 0 24902.076
240.09619140625 0 6155.0107
240.13462829589844 0 5147.9004
241.09275817871094 0 4283.1084
241.1188507080078 0 768.88434
241.13597106933594 0 1023.52246
241.19105529785156 0 672.3254
243.11032104492188 0 458.21072
244.09307861328125 0 4250.59
245.09654235839844 0 653.3602
247.1080780029297 0 5557.9873
248.1140594482422 0 2211.071
249.0541534423828 0 790.52734
249.09832763671875 0 3089.3228
249.13485717773438 0 968.90625
251.1505584716797 0 36836.1
252.13429260253906 0 596.51495
252.15370178222656 0 3422.2969
255.14724731445312 0 1977.5571
256.1083068847656 0 556.7281
256.1524353027344 0 687.1143
256.6099548339844 0 835.7622
257.1122741699219 0 649.1004
258.1449890136719 0 1834.6293
261.1187438964844 0 935.13916
261.1603698730469 0 638.44836
262.1185607910156 0 738.78296
264.27691650390625 0 550.53265
264.2956237792969 0 484.00375
265.12939453125 0 4917.9844
266.1142272949219 0 1107.6487
266.1250305175781 0 9887.292
267.1291198730469 0 1101.6115
268.1766662597656 0 1425.5837
269.18603515625 0 794.4626
270.12200927734375 0 1011.4022
270.6219787597656 0 662.3563
272.1759948730469 0 1642.5569
274.1185302734375 0 2183.9534
275.10284423828125 0 711.9947
280.1298828125 0 1149.4568
281.05157470703125 0 17878.77
282.1558837890625 0 2807.1707
283.0309143066406 0 2949.3442
283.04803466796875 0 479.5345
283.1403503417969 0 26724.47
284.1036071777344 0 1312.0565
284.14337158203125 0 3146.978
285.0097961425781 0 1366.6947
286.1383361816406 0 1192.918
288.6189880371094 0 761.6593
293.113525390625 0 13585.477 y Water loss 11
293.1309509277344 0 956.5508
294.11785888671875 0 2582.115
295.10357666015625 0 1633.8
296.1041564941406 0 2127.5845
297.0823059082031 0 2939.2512
297.10040283203125 0 1011.17505
297.1368713378906 0 989.6551 y Water loss 8
297.1564025878906 0 4033.4404
298.15887451171875 0 588.5172
299.06207275390625 0 153862.6
300.0633239746094 0 3188.7983
300.16748046875 0 5491.717
301.05987548828125 0 2737.6672
301.1510009765625 0 11064.69
301.1702880859375 0 1137.9537
302.1139221191406 0 2855.5498
302.1539611816406 0 1114.3138
305.6451110839844 0 701.30457
306.1383361816406 0 2478.6436 y 8
311.1242370605469 0 2105.1494 y 11
313.4514465332031 0 617.69855
314.20745849609375 0 1407.7874 b 2
315.1665344238281 0 7051.192
316.17041015625 0 1531.6385
326.66357421875 0 1604.0469
326.96612548828125 0 1190.2189
327.1657409667969 0 1346.3186
333.1772155761719 0 2001.1315
338.1839294433594 0 786.5819
341.01873779296875 0 1604.3064
343.1613464355469 0 608.9918
344.9767150878906 0 19336
345.1307678222656 0 1374.7269
346.11480712890625 0 4321.751
347.1178283691406 0 1508.725
347.68023681640625 0 1983.4989 b 5
350.2189025878906 0 3593.1384
355.0698547363281 0 3066.2227
359.0285949707031 0 17266.959
361.1981506347656 0 1549.702
362.1861877441406 0 2546.3657
362.67901611328125 0 3971.1597 y 7
363.0754089355469 0 656.73047
363.14129638671875 0 6755.089
363.1817321777344 0 3434.608
364.1240539550781 0 1065.5416
364.1457214355469 0 737.1784
366.1780090332031 0 702.6609
367.14007568359375 0 1004.4712
367.2440185546875 0 524.72815
368.1902160644531 0 559.0777
369.1228332519531 0 1123.167
370.122802734375 0 2325.982
371.12030029296875 0 1020.25977
376.1978454589844 0 1349.1782
376.69940185546875 0 624.35583
378.21417236328125 0 2244.269
379.2091064453125 0 8609.062
380.21380615234375 0 1244.73
381.0873718261719 0 751.4238
381.1518859863281 0 17653.234
381.7099304199219 0 705.2675
382.1551818847656 0 3404.463
384.1888122558594 0 1239.6794
385.14996337890625 0 1258.2472
394.17236328125 0 3049.6765
395.1348571777344 0 1357.1694
395.2398986816406 0 802.76685
396.2251281738281 0 2417.4155
398.1775207519531 0 1864.8553
402.1746520996094 0 841.7398
402.2003173828125 0 1593.8479
405.2613525390625 0 721.6164
409.1456604003906 0 990.9865
411.1985778808594 0 2033.2141
412.1606750488281 0 1001.453
412.18310546875 0 10698.808
413.14385986328125 0 1272.2231
413.18975830078125 0 1140.1696
414.23529052734375 0 2926.2778
415.0370788574219 0 35288.94
415.19879150390625 0 875.54736
417.7162780761719 0 1396.1963
418.2112731933594 0 3099.09
418.7097473144531 0 1070.6721
423.2024841308594 0 944.45264
423.2716064453125 0 4114.72
424.2745361328125 0 959.32733
426.72210693359375 0 27889.791
427.2236328125 0 15927.323
427.7248840332031 0 4253.64
429.089599609375 0 1427.7795
429.2098083496094 0 8248.435
430.1939392089844 0 24400.104
431.0679016113281 0 661.4093
431.1969299316406 0 4626.022
431.7140808105469 0 3238.896 b Water loss 6
432.20611572265625 0 1846.2805 b Ammonia loss 6
433.04827880859375 0 1686.1198
433.2151184082031 0 619.68945
433.2535095214844 0 661.67126 b Water loss 3
438.1742248535156 0 720.9974
438.2292175292969 0 668.84906
439.21923828125 0 7506.817 y 10
440.22308349609375 0 1875.9901
440.7196960449219 0 15671.318 b 6
440.9188537597656 0 1445.1306
441.22100830078125 0 8672.025
441.25146484375 0 695.79224
441.72113037109375 0 1948.7516
447.2169494628906 0 799.103
448.2026062011719 0 967.7912
450.2384338378906 0 638.913
451.2666931152344 0 7620.7563 b 3
452.2701110839844 0 2226.9243
460.7540283203125 0 1045.2743
461.2022705078125 0 968.07745
461.2535095214844 0 644.7846
468.2930603027344 0 2448.3013
471.17034912109375 0 1199.9946
473.9400329589844 0 645.0145
474.2599792480469 0 1479.225
474.7518005371094 0 1974.8525
475.2548828125 0 965.1142
475.6635437011719 0 572.84924
475.75433349609375 0 958.36707
477.2102966308594 0 1601.9148
478.23468017578125 0 884.96735 y Water loss 9
478.2747497558594 0 818.01465
483.26422119140625 0 5111.5303
483.76580810546875 0 2190.0566
484.7132873535156 0 839.5475
488.2571716308594 0 2427.9783 b Water loss 7
488.7552795410156 0 876.07684 b Ammonia loss 7
492.3273010253906 0 656.2707
492.6921691894531 0 1431.1814
494.236083984375 0 17376.924
495.239013671875 0 4697.5566
496.24078369140625 0 20798.104 y 9
497.24688720703125 0 5804.772
497.7625732421875 0 1295.4272
498.24420166015625 0 1126.9325
503.1077880859375 0 35347.348
507.2556457519531 0 1114.2814
511.262451171875 0 1347.0973
513.22265625 0 682.6515 y 5
517.283447265625 0 900.6441
519.1390991210938 0 12458.994
521.119873046875 0 667.8707
525.266357421875 0 2948.4155
526.2705688476562 0 887.6508
529.2589721679688 0 853.7323 Precursor Water loss
529.5924072265625 0 1005.832 Precursor Ammonia loss
534.2966918945312 0 851.4227
534.632568359375 0 910.38983
534.8311157226562 0 709.88983
534.8901977539062 0 822.6808
535.5989379882812 0 1002.0845
535.9326171875 0 765.4771
536.0489501953125 0 818.73804
536.1655883789062 0 9463.868
536.2064208984375 0 397.89276
536.2802734375 0 874.4527
538.2740478515625 0 786.81793
539.23583984375 0 3148.5378
540.2396850585938 0 662.97565
542.2940063476562 0 1284.7223
543.2777709960938 0 4057.379
544.2807006835938 0 886.98083
546.259033203125 0 1090.2699 b Ammonia loss 8
551.2572021484375 0 1012.797
555.261962890625 0 594.2929
556.2605590820312 0 790.3759
561.318115234375 0 696.4365 b Water loss 4
563.2567749023438 0 834.0289
565.2618408203125 0 1212.6222
567.2311401367188 0 3826.7961
568.2317504882812 0 1092.4377
575.2465209960938 0 2206.4336
579.3248901367188 0 4107.755 b 4
580.3258056640625 0 1549.0095
581.333740234375 0 753.1292
584.24609375 0 712.14716
592.2737426757812 0 1615.7396
592.3263549804688 0 705.8899
593.2568359375 0 7222.968 y Water loss 8
593.3052978515625 0 2400.7354
594.2546997070312 0 2453.944
600.3125610351562 0 885.05396
602.79052734375 0 1195.2161
603.7867431640625 0 775.3415
610.2833251953125 0 19185.006
610.3292846679688 0 803.13324
611.2681274414062 0 62909 y 8
611.7985229492188 0 3603.3542
612.271240234375 0 18833.863
612.8027954101562 0 808.87396
613.27294921875 0 3865.0706
621.2509765625 0 1252.3077
623.2298583984375 0 758.8296
635.2957763671875 0 1246.1765
636.32470703125 0 801.428
636.789794921875 0 854.97266 y Water loss 3
640.2789306640625 0 963.4055
645.299072265625 0 2191.572
645.7954711914062 0 2790.7832 y 3
646.2921752929688 0 3238.9858
652.320068359375 0 7073.3574
653.3236083984375 0 2769.7285
660.8723754882812 0 1748.8054
661.3719482421875 0 1593.5692
662.3046875 0 2437.411
663.2953491210938 0 1842.4531
667.278564453125 0 1092.4642
676.341064453125 0 3529.9436 b Water loss 5
677.3246459960938 0 1842.94 b Ammonia loss 5
677.8058471679688 0 875.99274
678.3125610351562 0 1756.3451
680.3153076171875 0 12993.564
680.3802490234375 0 706.43567
681.3181762695312 0 4532.3784
682.3201293945312 0 1031.3267
684.320556640625 0 1126.5554
684.8181762695312 0 777.6548
693.3287353515625 0 4606.8945 y Water loss 2
693.8273315429688 0 3138.3533 y Ammonia loss 2
694.3513793945312 0 36635.438 b 5
695.3546752929688 0 12646.746
696.3574829101562 0 3829.7417
701.8406982421875 0 7144.698
702.3357543945312 0 24527.326 y 2
702.8361206054688 0 16460.465 b Water loss 11
702.9061889648438 0 902.6154
703.3372802734375 0 6556.956 b Ammonia loss 11
703.8377075195312 0 2408.0286
704.3382568359375 0 1003.54895
706.3412475585938 0 2048.4785 y Water loss 7
707.3401489257812 0 858.814
710.4046020507812 0 2114.1145
710.9061889648438 0 1157.3673
711.3689575195312 0 2009.9039
711.8577880859375 0 1758.0981 b 11
712.3624267578125 0 1444.3665
721.362060546875 0 1714.5916
723.3670043945312 0 5374.4355
724.3533325195312 0 18220.494 y 7
725.3554077148438 0 8209.03
726.357666015625 0 1581.6694
733.3640747070312 0 1440.3253
751.3766479492188 0 4043.0625
751.8705444335938 0 2081.026 y 1
752.3729248046875 0 4715.9595
752.8753051757812 0 1171.8258
755.423583984375 0 804.9365
763.35888671875 0 1136.278
765.3939208984375 0 817.428
766.4087524414062 0 918.6881
779.3829956054688 0 1973.6772
780.3851928710938 0 1488.0835
781.366455078125 0 1481.394
793.3988647460938 0 3935.0576 Precursor Water loss
794.4015502929688 0 1953.1937
807.3739013671875 0 961.90607
812.2867431640625 0 903.38416
836.4466552734375 0 2781.3906
837.4705200195312 0 1243.3948
846.4478149414062 0 867.2603
852.4365234375 0 8472.45
853.4380493164062 0 4776.246
854.4404296875 0 959.92163
862.4204711914062 0 5733.8916 b Water loss 6
863.4158935546875 0 3575.731 b Ammonia loss 6
864.4556274414062 0 6375.299
865.4586791992188 0 3870.9353
866.4566040039062 0 996.4091
877.4417724609375 0 751.439
880.4307250976562 0 43249.355 b 6
881.4331665039062 0 22717.059
882.437255859375 0 6338.0464
883.4354858398438 0 1124.4354
908.4290161132812 0 1926.891
909.435791015625 0 1737.4114
910.4324951171875 0 3031.2668 y 6
911.4326171875 0 1068.3025
993.516357421875 0 3296.5557 b 7
994.5167236328125 0 1077.1072
1024.4696044921875 0 994.4147
1025.4617919921875 0 1373.6198 y 5
1093.5452880859375 0 1306.4365
1135.5068359375 0 2093.1343 y Water loss 4
1136.505615234375 0 1198.4016 y Ammonia loss 4
1137.48974609375 0 720.5681
1152.529052734375 0 1417.3812
1153.5184326171875 0 2776.1724 y 4
1154.5164794921875 0 1447.3126
1155.5352783203125 0 942.1897
1999.5606689453125 0 659.0243
2125.300537109375 0 698.0425
2567.07373046875 0 788.9979
2711.752197265625 0 643.4609
2901.924072265625 0 631.96844
3019.523193359375 0 732.63715
3020.13916015625 0 641.7797
3026.94091796875 0 685.6054
3078.85205078125 0 854.83484

Spectrum Details

|  |  |
| --- | --- |
| Matched peaks? Matched peaks The total absolute number of peaks matched. Additionally in brackets the total fraction of peaks matched and the total number of peaks is shown. | 56 (11.34% of 494) |
| FDR? FDR The false discovery rate estimated for this peptide. It is calculated by matching all theoretical fragments with a non-integer shift with the raw peaks for this spectrum. This is done with 40 different shifts. The resulting percentage is the average number of annotated peaks over the number of annotated peaks with the correct spectrum. | 1.49% |
| Satellite FDR? Satellite FDR See the FDR for details on its calculation. This satellite ion specific FDR only contains the satellite ions (d/w) for I/L/J positions. | ∞ |
| PSM Score? PSM Score The PSM Score as given by Hecklib to this annotated spectrum. It is shown with three significant figures. | 423 |

#### Spectrum 8472? Spectrum 8472 The raw spectrum of this peptide as annotated by Hecklib. The fragments are coloured according to ion type (see legend). Any peaks with a star '\*' as text can be hovered over to see the full details, first the ion type second the mass shift type. By hovering over the amino acids in the peptide or ions in the legend the corresponding peaks are highlighted. By toggling the 'Unassigned' label you can turn the background (unassigned) peaks on or off in the plot. By updating the slider in the Ion legend you can update the spectrum to only show the top X% of the peaks with labels. The top X% means any peak that is within X% of the highest intensity. By dragging in the spectrum you can zoom in to a specific part of the spectrum and use 'Zoom Out' to get back to the original zoom level. The annotation of the spectrum is based on the given sequence in the peptides file and is done with different software so inconsistencies are likely. The peaks are annotated based on the given sequence, with 20 ppm tolerance.

Copy Data

##### Spectrum 8472 (TSV)

###### Preview

```
Loading example...
```

*Click on the button to copy the data to your clipboard.*

Mz MinMz MaxIntensity Max

WidthHeightPeptide font sizePeptide stroke widthSpectrum font sizeSpectrum stroke widthCompact peptide

Ion legend

wxyz

abcd

OtherUnassignedIon^Charge^_Position_Show for top:%

TVLHQDWLDGKEY

03.60e+47.19e+41.08e+51.44e+5

Zoom Out

y^+3^_3_a^+1^_2_y^+3^_4_a^+1^_2_y^+1^_1_b^+1^_2_b^+1^_2_y^+2^_3_b^+2^_5_y^+1^_2_y^+2^_5_y^+1^_2_b^+1^_3_b^+2^_6_y^+2^_6_y^+2^_6_b^+2^_7_b^+2^_7_b^+1^_4_y^+1^_3_b^+2^_7_b^+1^_4_b^+3^_12_y^+1^_4_y^+1^_4_\*b^+1^_5_y^+1^_5_y^+1^_5_y^+2^_10_y^+2^_10_y^+2^_10_b^+1^_6_y^+2^_11_y^+2^_11_b^+1^_6_y^+2^_11_b^+2^_12_y^+1^_6_b^+2^_12_y^+1^_6_y^+2^_12_\*b^+1^_7_b^+1^_7_b^+1^_7_y^+1^_7_b^+1^_8_y^+1^_8_y^+1^_9_y^+1^_9_y^+1^_9_

0818163524533271

Fragment Matches Table

Show background peaks

| Position | Ion type | Intensity | mz Theoretical | mz Error (Th) | mz Error (ppm) | Charge | Series Number |
| --- | --- | --- | --- | --- | --- | --- | --- |
| - | - | 9425 | 120.1 | - | - | 0 | - |
| - | - | 881.4 | 121.1 | - | - | 0 | - |
| - | - | 365.9 | 121.7 | - | - | 0 | - |
| - | - | 517 | 122.1 | - | - | 0 | - |
| - | - | 593.9 | 123 | - | - | 0 | - |
| - | - | 365.8 | 125.5 | - | - | 0 | - |
| - | - | 1048 | 127.1 | - | - | 0 | - |
| - | - | 4738 | 128.1 | - | - | 0 | - |
| - | - | 1137 | 129.1 | - | - | 0 | - |
| - | - | 5.439E+04 | 129.1 | - | - | 0 | - |
| - | - | 2.532E+04 | 130.1 | - | - | 0 | - |
| - | - | 2851 | 130.1 | - | - | 0 | - |
| - | - | 2448 | 131.1 | - | - | 0 | - |
| - | - | 488.7 | 131.1 | - | - | 0 | - |
| - | - | 5275 | 132.1 | - | - | 0 | - |
| - | - | 1614 | 132.1 | - | - | 0 | - |
| - | - | 546.5 | 133.1 | - | - | 0 | - |
| - | - | 598.9 | 133.1 | - | - | 0 | - |
| - | - | 468.9 | 134 | - | - | 0 | - |
| - | - | 8491 | 136.1 | - | - | 0 | - |
| - | - | 1112 | 137.1 | - | - | 0 | - |
| - | - | 691 | 138.1 | - | - | 0 | - |
| - | - | 435.5 | 138.2 | - | - | 0 | - |
| - | - | 474.3 | 139.1 | - | - | 0 | - |
| - | - | 433.8 | 140.6 | - | - | 0 | - |
| - | - | 2548 | 141.1 | - | - | 0 | - |
| - | - | 651.7 | 142.1 | - | - | 0 | - |
| - | - | 1477 | 146.1 | - | - | 0 | - |
| 11 | y | 601.4 | 147.1 | 0.001054 | 7.164 | +3 | 3 |
| - | - | 687.1 | 148.9 | - | - | 0 | - |
| - | - | 1.614E+04 | 149 | - | - | 0 | - |
| - | - | 1277 | 151.1 | - | - | 0 | - |
| - | - | 3188 | 155.1 | - | - | 0 | - |
| 2 | a | 2315 | 155.1 | 0.0001677 | 1.081 | +1 | 2 |
| - | - | 922.6 | 157.1 | - | - | 0 | - |
| - | - | 2585 | 157.1 | - | - | 0 | - |
| - | - | 621.1 | 158.1 | - | - | 0 | - |
| - | - | 4.151E+04 | 159.1 | - | - | 0 | - |
| - | - | 5002 | 160.1 | - | - | 0 | - |
| - | - | 3950 | 165.1 | - | - | 0 | - |
| - | - | 6853 | 166.1 | - | - | 0 | - |
| 10 | y | 3807 | 166.1 | 0.001557 | 9.377 | +3 | 4 |
| - | - | 2.192E+04 | 167.1 | - | - | 0 | - |
| - | - | 1263 | 167.1 | - | - | 0 | - |
| - | - | 687 | 168.1 | - | - | 0 | - |
| - | - | 642.4 | 168.1 | - | - | 0 | - |
| - | - | 500.6 | 168.1 | - | - | 0 | - |
| - | - | 772.3 | 168.1 | - | - | 0 | - |
| - | - | 542.8 | 169.1 | - | - | 0 | - |
| - | - | 3927 | 169.1 | - | - | 0 | - |
| - | - | 1789 | 170.1 | - | - | 0 | - |
| - | - | 568.8 | 170.1 | - | - | 0 | - |
| - | - | 583.3 | 171.1 | - | - | 0 | - |
| - | - | 1630 | 172.1 | - | - | 0 | - |
| 2 | a | 7.686E+04 | 173.1 | 0.0002537 | 1.465 | +1 | 2 |
| - | - | 1240 | 174.1 | - | - | 0 | - |
| - | - | 7430 | 174.1 | - | - | 0 | - |
| - | - | 647.2 | 175.1 | - | - | 0 | - |
| - | - | 487.6 | 175.9 | - | - | 0 | - |
| - | - | 584.3 | 177.1 | - | - | 0 | - |
| - | - | 2109 | 178.1 | - | - | 0 | - |
| 13 | y | 9356 | 182.1 | 0.0001444 | 0.7931 | +1 | 1 |
| - | - | 1008 | 182.1 | - | - | 0 | - |
| - | - | 611 | 183.1 | - | - | 0 | - |
| 2 | b | 2391 | 183.1 | 0.0002482 | 1.356 | +1 | 2 |
| - | - | 2320 | 183.1 | - | - | 0 | - |
| - | - | 520.9 | 185.1 | - | - | 0 | - |
| - | - | 653 | 185.1 | - | - | 0 | - |
| - | - | 921.3 | 185.2 | - | - | 0 | - |
| - | - | 1.403E+04 | 186.1 | - | - | 0 | - |
| - | - | 3594 | 187.1 | - | - | 0 | - |
| - | - | 1747 | 187.1 | - | - | 0 | - |
| - | - | 752.8 | 187.1 | - | - | 0 | - |
| - | - | 1694 | 188.1 | - | - | 0 | - |
| - | - | 523.8 | 190.1 | - | - | 0 | - |
| - | - | 533.7 | 192.2 | - | - | 0 | - |
| - | - | 3211 | 195.1 | - | - | 0 | - |
| - | - | 2998 | 196.1 | - | - | 0 | - |
| - | - | 490.8 | 196.1 | - | - | 0 | - |
| - | - | 783.2 | 197.1 | - | - | 0 | - |
| - | - | 498.7 | 197.2 | - | - | 0 | - |
| - | - | 863.7 | 198.1 | - | - | 0 | - |
| - | - | 1961 | 198.1 | - | - | 0 | - |
| - | - | 469 | 199.1 | - | - | 0 | - |
| - | - | 704.9 | 199.1 | - | - | 0 | - |
| - | - | 537.9 | 200.7 | - | - | 0 | - |
| 2 | b | 1.969E+04 | 201.1 | 0.0001053 | 0.5234 | +1 | 2 |
| - | - | 1631 | 202.1 | - | - | 0 | - |
| - | - | 2444 | 205.1 | - | - | 0 | - |
| - | - | 587.7 | 205.1 | - | - | 0 | - |
| - | - | 746.1 | 207.2 | - | - | 0 | - |
| - | - | 692.7 | 209.1 | - | - | 0 | - |
| 11 | y | 623.5 | 211.1 | 0.0002066 | 0.9788 | +2 | 3 |
| - | - | 1937 | 211.1 | - | - | 0 | - |
| - | - | 2756 | 213.1 | - | - | 0 | - |
| - | - | 526.7 | 213.1 | - | - | 0 | - |
| - | - | 778.4 | 215.1 | - | - | 0 | - |
| - | - | 614.2 | 216.1 | - | - | 0 | - |
| - | - | 1028 | 216.1 | - | - | 0 | - |
| - | - | 1889 | 219.1 | - | - | 0 | - |
| - | - | 2178 | 221.1 | - | - | 0 | - |
| - | - | 8033 | 221.1 | - | - | 0 | - |
| - | - | 1173 | 221.1 | - | - | 0 | - |
| - | - | 1933 | 222.1 | - | - | 0 | - |
| - | - | 6075 | 223.1 | - | - | 0 | - |
| - | - | 1424 | 223.1 | - | - | 0 | - |
| - | - | 1890 | 223.1 | - | - | 0 | - |
| - | - | 7067 | 223.2 | - | - | 0 | - |
| - | - | 936.8 | 224.2 | - | - | 0 | - |
| - | - | 1.038E+04 | 225 | - | - | 0 | - |
| - | - | 686.5 | 225.1 | - | - | 0 | - |
| - | - | 2246 | 226 | - | - | 0 | - |
| - | - | 1594 | 226.1 | - | - | 0 | - |
| - | - | 825 | 227 | - | - | 0 | - |
| - | - | 1449 | 227 | - | - | 0 | - |
| - | - | 648.6 | 227.1 | - | - | 0 | - |
| - | - | 700.1 | 228.1 | - | - | 0 | - |
| - | - | 1053 | 229.1 | - | - | 0 | - |
| - | - | 1205 | 229.2 | - | - | 0 | - |
| - | - | 485.2 | 232.2 | - | - | 0 | - |
| - | - | 602.1 | 233.1 | - | - | 0 | - |
| - | - | 655 | 233.2 | - | - | 0 | - |
| - | - | 7783 | 234.1 | - | - | 0 | - |
| - | - | 1691 | 235.1 | - | - | 0 | - |
| - | - | 923.7 | 238.1 | - | - | 0 | - |
| - | - | 2.206E+04 | 239.1 | - | - | 0 | - |
| - | - | 5241 | 240.1 | - | - | 0 | - |
| - | - | 3731 | 240.1 | - | - | 0 | - |
| - | - | 3286 | 241.1 | - | - | 0 | - |
| - | - | 595.8 | 243.1 | - | - | 0 | - |
| - | - | 2838 | 244.1 | - | - | 0 | - |
| - | - | 530.3 | 244.1 | - | - | 0 | - |
| - | - | 3602 | 247.1 | - | - | 0 | - |
| - | - | 1251 | 247.1 | - | - | 0 | - |
| - | - | 1667 | 248.1 | - | - | 0 | - |
| - | - | 2696 | 249.1 | - | - | 0 | - |
| - | - | 766.8 | 249.1 | - | - | 0 | - |
| - | - | 516.5 | 250.3 | - | - | 0 | - |
| - | - | 2.529E+04 | 251.2 | - | - | 0 | - |
| - | - | 2490 | 252.2 | - | - | 0 | - |
| - | - | 585.9 | 253.2 | - | - | 0 | - |
| - | - | 1091 | 255.1 | - | - | 0 | - |
| - | - | 696 | 257.1 | - | - | 0 | - |
| - | - | 1225 | 258.1 | - | - | 0 | - |
| - | - | 779.1 | 261.1 | - | - | 0 | - |
| - | - | 509.1 | 262.1 | - | - | 0 | - |
| - | - | 3880 | 265.1 | - | - | 0 | - |
| - | - | 7171 | 266.1 | - | - | 0 | - |
| - | - | 499.1 | 267.1 | - | - | 0 | - |
| - | - | 934.8 | 267.1 | - | - | 0 | - |
| - | - | 655.3 | 268.1 | - | - | 0 | - |
| - | - | 745 | 268.2 | - | - | 0 | - |
| - | - | 1101 | 270.1 | - | - | 0 | - |
| - | - | 1439 | 272.2 | - | - | 0 | - |
| - | - | 1744 | 274.1 | - | - | 0 | - |
| - | - | 528.4 | 279.1 | - | - | 0 | - |
| - | - | 663.7 | 280.1 | - | - | 0 | - |
| - | - | 1.692E+04 | 281.1 | - | - | 0 | - |
| - | - | 842.3 | 281.2 | - | - | 0 | - |
| - | - | 546.1 | 282.1 | - | - | 0 | - |
| - | - | 1473 | 282.2 | - | - | 0 | - |
| - | - | 2510 | 283 | - | - | 0 | - |
| - | - | 2.239E+04 | 283.1 | - | - | 0 | - |
| - | - | 2737 | 284.1 | - | - | 0 | - |
| - | - | 1062 | 285 | - | - | 0 | - |
| - | - | 665.5 | 286.1 | - | - | 0 | - |
| - | - | 535.2 | 287.2 | - | - | 0 | - |
| - | - | 570 | 288.6 | - | - | 0 | - |
| 5 | b | 861.8 | 290.2 | 0.0008013 | 2.762 | +2 | 5 |
| - | - | 552.4 | 292.1 | - | - | 0 | - |
| 12 | y | 1.054E+04 | 293.1 | 0.0002052 | 0.7002 | +1 | 2 |
| - | - | 1725 | 294.1 | - | - | 0 | - |
| - | - | 2227 | 295.1 | - | - | 0 | - |
| - | - | 1773 | 296.1 | - | - | 0 | - |
| - | - | 1943 | 297.1 | - | - | 0 | - |
| - | - | 1004 | 297.1 | - | - | 0 | - |
| - | - | 3118 | 297.2 | - | - | 0 | - |
| - | - | 1.424E+05 | 299.1 | - | - | 0 | - |
| - | - | 3126 | 300.1 | - | - | 0 | - |
| - | - | 2906 | 300.2 | - | - | 0 | - |
| - | - | 1322 | 301.1 | - | - | 0 | - |
| - | - | 8145 | 301.2 | - | - | 0 | - |
| - | - | 870.4 | 301.2 | - | - | 0 | - |
| - | - | 1573 | 302.1 | - | - | 0 | - |
| - | - | 1104 | 302.2 | - | - | 0 | - |
| 9 | y | 1107 | 306.1 | 0.0001246 | 0.407 | +2 | 5 |
| - | - | 883.7 | 309.2 | - | - | 0 | - |
| 12 | y | 1703 | 311.1 | 0.0001055 | 0.3392 | +1 | 2 |
| - | - | 709.6 | 313.1 | - | - | 0 | - |
| - | - | 710.6 | 314.2 | - | - | 0 | - |
| 3 | b | 1012 | 314.2 | 9.64E-05 | 0.3068 | +1 | 3 |
| - | - | 5073 | 315.2 | - | - | 0 | - |
| - | - | 1214 | 316.2 | - | - | 0 | - |
| - | - | 788.9 | 325 | - | - | 0 | - |
| - | - | 655.6 | 326.7 | - | - | 0 | - |
| - | - | 784.9 | 327 | - | - | 0 | - |
| - | - | 647.3 | 329.2 | - | - | 0 | - |
| - | - | 1429 | 333.2 | - | - | 0 | - |
| - | - | 1494 | 341 | - | - | 0 | - |
| - | - | 608.5 | 343 | - | - | 0 | - |
| - | - | 1.909E+04 | 345 | - | - | 0 | - |
| - | - | 3764 | 346.1 | - | - | 0 | - |
| 6 | b | 2174 | 347.7 | 0.0003303 | 0.95 | +2 | 6 |
| - | - | 2645 | 350.2 | - | - | 0 | - |
| 8 | y | 584.7 | 353.7 | 0.001774 | 5.016 | +2 | 6 |
| - | - | 2878 | 355.1 | - | - | 0 | - |
| - | - | 1.678E+04 | 359 | - | - | 0 | - |
| - | - | 1409 | 361.2 | - | - | 0 | - |
| - | - | 1883 | 362.2 | - | - | 0 | - |
| 8 | y | 3505 | 362.7 | 3.727E-05 | 0.1028 | +2 | 6 |
| - | - | 5886 | 363.1 | - | - | 0 | - |
| - | - | 710.4 | 363.2 | - | - | 0 | - |
| - | - | 977.6 | 364.1 | - | - | 0 | - |
| - | - | 575.9 | 366.2 | - | - | 0 | - |
| - | - | 1518 | 369.1 | - | - | 0 | - |
| - | - | 2068 | 370.1 | - | - | 0 | - |
| - | - | 1548 | 371.1 | - | - | 0 | - |
| - | - | 535.6 | 371.2 | - | - | 0 | - |
| - | - | 636.6 | 376.2 | - | - | 0 | - |
| - | - | 1646 | 378.2 | - | - | 0 | - |
| - | - | 6554 | 379.2 | - | - | 0 | - |
| - | - | 949.2 | 380.2 | - | - | 0 | - |
| - | - | 1.257E+04 | 381.2 | - | - | 0 | - |
| - | - | 1939 | 382.2 | - | - | 0 | - |
| - | - | 867.6 | 384.2 | - | - | 0 | - |
| - | - | 2323 | 394.2 | - | - | 0 | - |
| - | - | 711.6 | 395.1 | - | - | 0 | - |
| - | - | 1062 | 395.2 | - | - | 0 | - |
| - | - | 1884 | 396.2 | - | - | 0 | - |
| - | - | 1702 | 398.2 | - | - | 0 | - |
| - | - | 1030 | 402.2 | - | - | 0 | - |
| - | - | 684.2 | 403.2 | - | - | 0 | - |
| - | - | 994.9 | 405.3 | - | - | 0 | - |
| - | - | 682.2 | 409.1 | - | - | 0 | - |
| - | - | 875.7 | 411.2 | - | - | 0 | - |
| - | - | 1129 | 412.2 | - | - | 0 | - |
| - | - | 7606 | 412.2 | - | - | 0 | - |
| - | - | 749 | 413.2 | - | - | 0 | - |
| - | - | 1150 | 414.2 | - | - | 0 | - |
| - | - | 3.434E+04 | 415 | - | - | 0 | - |
| - | - | 1414 | 417.7 | - | - | 0 | - |
| - | - | 1416 | 418.2 | - | - | 0 | - |
| - | - | 830.8 | 418.7 | - | - | 0 | - |
| - | - | 820.8 | 419.2 | - | - | 0 | - |
| - | - | 3659 | 423.3 | - | - | 0 | - |
| - | - | 1012 | 424.3 | - | - | 0 | - |
| - | - | 1114 | 426.7 | - | - | 0 | - |
| - | - | 2.162E+04 | 426.7 | - | - | 0 | - |
| - | - | 9681 | 427.2 | - | - | 0 | - |
| - | - | 3099 | 427.7 | - | - | 0 | - |
| - | - | 881.1 | 428.2 | - | - | 0 | - |
| - | - | 1575 | 429.1 | - | - | 0 | - |
| - | - | 6407 | 429.2 | - | - | 0 | - |
| - | - | 1.679E+04 | 430.2 | - | - | 0 | - |
| - | - | 4235 | 431.2 | - | - | 0 | - |
| 7 | b | 1647 | 431.7 | 0.001207 | 2.796 | +2 | 7 |
| 7 | b | 1733 | 432.2 | 0.00118 | 2.73 | +2 | 7 |
| - | - | 813.9 | 432.7 | - | - | 0 | - |
| - | - | 2112 | 433 | - | - | 0 | - |
| 4 | b | 741.6 | 433.3 | 0.0002868 | 0.662 | +1 | 4 |
| 11 | y | 7057 | 439.2 | 0.0004515 | 1.028 | +1 | 3 |
| - | - | 1288 | 440.2 | - | - | 0 | - |
| 7 | b | 1.066E+04 | 440.7 | 0.0003858 | 0.8753 | +2 | 7 |
| - | - | 766.6 | 440.9 | - | - | 0 | - |
| - | - | 4740 | 441.2 | - | - | 0 | - |
| - | - | 1093 | 441.3 | - | - | 0 | - |
| - | - | 1291 | 441.7 | - | - | 0 | - |
| - | - | 736.4 | 446.1 | - | - | 0 | - |
| - | - | 617.9 | 449.2 | - | - | 0 | - |
| 4 | b | 5916 | 451.3 | 0.0001043 | 0.2311 | +1 | 4 |
| - | - | 715.1 | 452.3 | - | - | 0 | - |
| - | - | 2034 | 452.9 | - | - | 0 | - |
| - | - | 882.2 | 453.2 | - | - | 0 | - |
| - | - | 1499 | 460.8 | - | - | 0 | - |
| - | - | 921.8 | 461.2 | - | - | 0 | - |
| - | - | 1305 | 468.3 | - | - | 0 | - |
| 12 | b | 568.3 | 468.9 | 0.006228 | 13.28 | +3 | 12 |
| - | - | 1068 | 471.2 | - | - | 0 | - |
| - | - | 1111 | 474.3 | - | - | 0 | - |
| - | - | 1436 | 474.8 | - | - | 0 | - |
| - | - | 908.8 | 475.3 | - | - | 0 | - |
| - | - | 777.8 | 476.2 | - | - | 0 | - |
| - | - | 1415 | 477.2 | - | - | 0 | - |
| - | - | 744.5 | 477.7 | - | - | 0 | - |
| 10 | y | 676 | 478.2 | 0.0007829 | 1.637 | +1 | 4 |
| - | - | 799.9 | 478.3 | - | - | 0 | - |
| - | - | 3354 | 483.3 | - | - | 0 | - |
| - | - | 2687 | 483.8 | - | - | 0 | - |
| - | - | 1330 | 488.1 | - | - | 0 | - |
| - | - | 1.225E+04 | 494.2 | - | - | 0 | - |
| - | - | 2430 | 495.2 | - | - | 0 | - |
| 10 | y | 1.572E+04 | 496.2 | 0.0004434 | 0.8936 | +1 | 4 |
| - | - | 3738 | 497.2 | - | - | 0 | - |
| - | - | 1872 | 497.8 | - | - | 0 | - |
| - | - | 3.331E+04 | 503.1 | - | - | 0 | - |
| - | - | 2062 | 504.1 | - | - | 0 | - |
| - | - | 2751 | 505.1 | - | - | 0 | - |
| - | - | 1209 | 511.3 | - | - | 0 | - |
| - | - | 620.6 | 515.3 | - | - | 0 | - |
| - | - | 805 | 516 | - | - | 0 | - |
| - | - | 924.2 | 517.3 | - | - | 0 | - |
| - | - | 1.109E+04 | 519.1 | - | - | 0 | - |
| - | - | 1064 | 521.1 | - | - | 0 | - |
| - | - | 2211 | 525.3 | - | - | 0 | - |
| - | - | 757 | 526.3 | - | - | 0 | - |
| 0 | Precursor | 1650 | 529.6 | 0.004829 | 9.118 | +3 | -1 |
| - | - | 999.7 | 534.3 | - | - | 0 | - |
| - | - | 1067 | 534.6 | - | - | 0 | - |
| - | - | 916.5 | 534.6 | - | - | 0 | - |
| - | - | 1027 | 535.2 | - | - | 0 | - |
| - | - | 7604 | 536.2 | - | - | 0 | - |
| - | - | 703.3 | 536.3 | - | - | 0 | - |
| - | - | 729.3 | 538.3 | - | - | 0 | - |
| - | - | 2351 | 539.2 | - | - | 0 | - |
| - | - | 583.9 | 540.2 | - | - | 0 | - |
| - | - | 1115 | 542.3 | - | - | 0 | - |
| - | - | 2750 | 543.3 | - | - | 0 | - |
| - | - | 1024 | 544.3 | - | - | 0 | - |
| - | - | 842.3 | 555.3 | - | - | 0 | - |
| - | - | 880.9 | 555.8 | - | - | 0 | - |
| - | - | 616.7 | 565.3 | - | - | 0 | - |
| - | - | 2409 | 567.2 | - | - | 0 | - |
| - | - | 965.2 | 573.8 | - | - | 0 | - |
| - | - | 926.9 | 574.3 | - | - | 0 | - |
| - | - | 987 | 575.2 | - | - | 0 | - |
| - | - | 862.7 | 576.3 | - | - | 0 | - |
| 5 | b | 3522 | 579.3 | 0.0006424 | 1.109 | +1 | 5 |
| - | - | 1205 | 580.3 | - | - | 0 | - |
| - | - | 766.4 | 587.3 | - | - | 0 | - |
| 9 | y | 4822 | 593.3 | 0.0008306 | 1.4 | +1 | 5 |
| - | - | 1715 | 593.3 | - | - | 0 | - |
| - | - | 1153 | 594.3 | - | - | 0 | - |
| - | - | 1291 | 596.3 | - | - | 0 | - |
| - | - | 911.6 | 596.7 | - | - | 0 | - |
| - | - | 1084 | 602.8 | - | - | 0 | - |
| - | - | 1.618E+04 | 610.3 | - | - | 0 | - |
| - | - | 547.8 | 610.3 | - | - | 0 | - |
| 9 | y | 4.415E+04 | 611.3 | 4.27E-05 | 0.06986 | +1 | 5 |
| - | - | 1436 | 611.8 | - | - | 0 | - |
| - | - | 1.389E+04 | 612.3 | - | - | 0 | - |
| - | - | 3063 | 613.3 | - | - | 0 | - |
| - | - | 1086 | 621.3 | - | - | 0 | - |
| - | - | 769.5 | 635.3 | - | - | 0 | - |
| 4 | y | 994.5 | 636.8 | 0.0009758 | 1.532 | +2 | 10 |
| 4 | y | 897.9 | 637.3 | 0.0072 | 11.3 | +2 | 10 |
| - | - | 1207 | 645.3 | - | - | 0 | - |
| 4 | y | 2476 | 645.8 | 0.00247 | 3.825 | +2 | 10 |
| - | - | 2303 | 646.3 | - | - | 0 | - |
| - | - | 4607 | 652.3 | - | - | 0 | - |
| - | - | 772.5 | 653 | - | - | 0 | - |
| - | - | 1576 | 653.3 | - | - | 0 | - |
| - | - | 689.3 | 653.7 | - | - | 0 | - |
| - | - | 2409 | 660.9 | - | - | 0 | - |
| - | - | 1404 | 661.4 | - | - | 0 | - |
| - | - | 1922 | 662.3 | - | - | 0 | - |
| 6 | b | 2370 | 676.3 | 0.001518 | 2.244 | +1 | 6 |
| - | - | 612.1 | 677.3 | - | - | 0 | - |
| - | - | 1551 | 678.9 | - | - | 0 | - |
| - | - | 1643 | 679.4 | - | - | 0 | - |
| - | - | 8506 | 680.3 | - | - | 0 | - |
| - | - | 2990 | 681.3 | - | - | 0 | - |
| - | - | 682.4 | 684.8 | - | - | 0 | - |
| 3 | y | 3660 | 693.3 | 0.0007104 | 1.025 | +2 | 11 |
| 3 | y | 2794 | 693.8 | 0.007892 | 11.37 | +2 | 11 |
| 6 | b | 2.582E+04 | 694.4 | 0.001646 | 2.37 | +1 | 6 |
| - | - | 9973 | 695.4 | - | - | 0 | - |
| - | - | 2044 | 696.4 | - | - | 0 | - |
| - | - | 5343 | 701.8 | - | - | 0 | - |
| 3 | y | 1.878E+04 | 702.3 | 0.001087 | 1.548 | +2 | 11 |
| - | - | 1.203E+04 | 702.8 | - | - | 0 | - |
| 12 | b | 6645 | 703.3 | 0.004174 | 5.935 | +2 | 12 |
| - | - | 1788 | 703.8 | - | - | 0 | - |
| - | - | 1454 | 704.3 | - | - | 0 | - |
| 8 | y | 1703 | 706.3 | 0.0007378 | 1.045 | +1 | 6 |
| - | - | 1560 | 710.4 | - | - | 0 | - |
| - | - | 1348 | 710.9 | - | - | 0 | - |
| - | - | 838.8 | 711.4 | - | - | 0 | - |
| 12 | b | 1490 | 711.9 | 5.373E-05 | 0.07548 | +2 | 12 |
| - | - | 1209 | 712.4 | - | - | 0 | - |
| - | - | 1316 | 721.4 | - | - | 0 | - |
| - | - | 4334 | 723.4 | - | - | 0 | - |
| 8 | y | 1.332E+04 | 724.4 | 0.0006716 | 0.9272 | +1 | 6 |
| - | - | 5912 | 725.4 | - | - | 0 | - |
| - | - | 1677 | 726.4 | - | - | 0 | - |
| - | - | 1111 | 734.4 | - | - | 0 | - |
| - | - | 2254 | 751.4 | - | - | 0 | - |
| 2 | y | 1094 | 751.9 | 0.002297 | 3.055 | +2 | 12 |
| - | - | 2133 | 752.4 | - | - | 0 | - |
| - | - | 1017 | 755.4 | - | - | 0 | - |
| - | - | 1701 | 763.4 | - | - | 0 | - |
| - | - | 912.1 | 764.4 | - | - | 0 | - |
| - | - | 1530 | 765.4 | - | - | 0 | - |
| - | - | 999.5 | 779.4 | - | - | 0 | - |
| - | - | 736.8 | 780.4 | - | - | 0 | - |
| 0 | Precursor | 2429 | 793.4 | 0.01131 | 14.26 | +2 | -1 |
| - | - | 1130 | 794.4 | - | - | 0 | - |
| - | - | 1250 | 807.4 | - | - | 0 | - |
| - | - | 619.5 | 835.4 | - | - | 0 | - |
| - | - | 1892 | 836.4 | - | - | 0 | - |
| - | - | 1169 | 837.5 | - | - | 0 | - |
| - | - | 906.3 | 847.4 | - | - | 0 | - |
| - | - | 6214 | 852.4 | - | - | 0 | - |
| - | - | 2955 | 853.4 | - | - | 0 | - |
| - | - | 579.9 | 859.4 | - | - | 0 | - |
| 7 | b | 3121 | 862.4 | 0.002645 | 3.067 | +1 | 7 |
| 7 | b | 2072 | 863.4 | 0.007236 | 8.381 | +1 | 7 |
| - | - | 5127 | 864.5 | - | - | 0 | - |
| - | - | 2305 | 865.5 | - | - | 0 | - |
| 7 | b | 3.043E+04 | 880.4 | 0.002528 | 2.872 | +1 | 7 |
| - | - | 1.727E+04 | 881.4 | - | - | 0 | - |
| - | - | 3989 | 882.4 | - | - | 0 | - |
| - | - | 1219 | 883.4 | - | - | 0 | - |
| - | - | 681.8 | 892.5 | - | - | 0 | - |
| - | - | 1534 | 908.4 | - | - | 0 | - |
| - | - | 1127 | 909.4 | - | - | 0 | - |
| 7 | y | 1829 | 910.4 | 0.00522 | 5.734 | +1 | 7 |
| - | - | 790.2 | 911.4 | - | - | 0 | - |
| - | - | 738.6 | 912.4 | - | - | 0 | - |
| - | - | 780.1 | 955.4 | - | - | 0 | - |
| 8 | b | 2117 | 993.5 | 0.002181 | 2.195 | +1 | 8 |
| - | - | 1534 | 994.5 | - | - | 0 | - |
| 6 | y | 1335 | 1025 | 0.001532 | 1.494 | +1 | 8 |
| - | - | 982.2 | 1077 | - | - | 0 | - |
| - | - | 920.6 | 1094 | - | - | 0 | - |
| 5 | y | 720.5 | 1136 | 0.001004 | 0.8845 | +1 | 9 |
| 5 | y | 891.9 | 1136 | 0.009176 | 8.074 | +1 | 9 |
| - | - | 633.6 | 1137 | - | - | 0 | - |
| 5 | y | 2721 | 1154 | 0.002358 | 2.044 | +1 | 9 |
| - | - | 1519 | 1155 | - | - | 0 | - |
| - | - | 1071 | 1156 | - | - | 0 | - |
| - | - | 932.7 | 3080 | - | - | 0 | - |
| - | - | 651.9 | 3189 | - | - | 0 | - |
| - | - | 777.2 | 3238 | - | - | 0 | - |

m/z Charge Intensity FragmentType MassShift Position
120.0810317993164 0 9424.517
121.08460998535156 0 881.44183
121.65116119384766 0 365.85413
122.07173919677734 0 516.95624
123.0445327758789 0 593.9325
125.46116638183594 0 365.75742
127.0503158569336 0 1048.3759
128.107177734375 0 4737.845
129.0660858154297 0 1137.1736
129.10250854492188 0 54390.18
130.06539916992188 0 25315.283
130.10595703125 0 2851.4275
131.06887817382812 0 2447.7134
131.11817932128906 0 488.671
132.08103942871094 0 5275.125
132.10214233398438 0 1613.7858
133.06114196777344 0 546.5447
133.08567810058594 0 598.9278
134.02755737304688 0 468.94144
136.07591247558594 0 8490.982
137.07928466796875 0 1111.8757
138.0667266845703 0 691.0235
138.24227905273438 0 435.46152
139.0867156982422 0 474.2954
140.59829711914062 0 433.78387
141.1025848388672 0 2547.6035
142.06546020507812 0 651.72144
146.0603485107422 0 1476.6617
147.0767059326172 0 601.4473 y 10
148.94735717773438 0 687.1333
149.0451202392578 0 16137.4375
151.08677673339844 0 1277.2091
155.09303283691406 0 3188.3425
155.11805725097656 0 2315.3735 a Water loss 1
157.09744262695312 0 922.5681
157.1336669921875 0 2584.5618
158.08380126953125 0 621.064
159.09190368652344 0 41511.49
160.09527587890625 0 5002.156
165.05487060546875 0 3950.0854
166.0612335205078 0 6852.916
166.0864715576172 0 3807.4512 y 9
167.0556640625 0 21923.715
167.0638885498047 0 1262.8048
168.05494689941406 0 687.04565
168.08062744140625 0 642.3945
168.10208129882812 0 500.56778
168.11361694335938 0 772.33057
169.0760955810547 0 542.8475
169.0972900390625 0 3927.1567
170.0601806640625 0 1789.3419
170.1012420654297 0 568.7762
171.14881896972656 0 583.32117
172.07174682617188 0 1629.9751
173.1287078857422 0 76864.4 a 1
174.0550994873047 0 1240.1223
174.132080078125 0 7430.4756
175.13404846191406 0 647.1982
175.8506317138672 0 487.60544
177.1124267578125 0 584.324
178.13401794433594 0 2109.4966
182.08131408691406 0 9355.766 y 12
182.090087890625 0 1008.23334
183.08518981933594 0 611.03546
183.11305236816406 0 2390.7104 b Water loss 1
183.14974975585938 0 2320.0454
185.0918731689453 0 520.89124
185.12803649902344 0 652.9905
185.1651611328125 0 921.2541
186.1239013671875 0 14027.886
187.08673095703125 0 3594.1335
187.12730407714844 0 1746.7478
187.1444091796875 0 752.8369
188.07083129882812 0 1694.0192
190.08270263671875 0 523.7565
192.1671142578125 0 533.71857
195.113037109375 0 3211.3555
196.10816955566406 0 2998.4526
196.11680603027344 0 490.80856
197.12852478027344 0 783.24915
197.1649932861328 0 498.66226
198.0878143310547 0 863.67706
198.12811279296875 0 1961.0059
199.07040405273438 0 469.03473
199.08616638183594 0 704.9003
200.73703002929688 0 537.8765
201.12347412109375 0 19686.621 b 1
202.12680053710938 0 1631.4496
205.0976104736328 0 2444.4463
205.14508056640625 0 587.7108
207.16046142578125 0 746.0673
209.09243774414062 0 692.6937
211.10792541503906 0 623.4569 y Water loss 10
211.1442413330078 0 1936.8405
213.12338256835938 0 2755.615
213.13356018066406 0 526.6994
215.13938903808594 0 778.35864
216.0975341796875 0 614.2485
216.13436889648438 0 1028.1476
219.14894104003906 0 1889.1787
221.05929565429688 0 2177.7163
221.0843505859375 0 8032.9043
221.10379028320312 0 1172.7083
222.08445739746094 0 1933.4825
223.0637664794922 0 6075.435
223.08172607421875 0 1424.0172
223.10791015625 0 1890.3289
223.1554412841797 0 7067.256
224.1596221923828 0 936.82196
225.04302978515625 0 10384.593
225.1239776611328 0 686.5325
226.04335021972656 0 2246.2205
226.08262634277344 0 1593.5905
227.02235412597656 0 825.031
227.03981018066406 0 1449.4594
227.06655883789062 0 648.60693
228.13414001464844 0 700.0699
229.11831665039062 0 1053.3268
229.15463256835938 0 1205.0176
232.15611267089844 0 485.18878
233.13925170898438 0 602.0835
233.16519165039062 0 655.0031
234.12384033203125 0 7782.625
235.12741088867188 0 1691.3632
238.1192626953125 0 923.71277
239.09506225585938 0 22064.28
240.0960693359375 0 5241.398
240.13430786132812 0 3731.316
241.09222412109375 0 3286.4756
243.109619140625 0 595.8067
244.0928192138672 0 2838.3674
244.10501098632812 0 530.30927
247.10775756835938 0 3602.4724
247.1441192626953 0 1250.7344
248.11419677734375 0 1666.5994
249.09849548339844 0 2695.9302
249.1347198486328 0 766.83276
250.33705139160156 0 516.46625
251.15036010742188 0 25288.047
252.15359497070312 0 2490.166
253.15554809570312 0 585.87885
255.14813232421875 0 1091.4587
257.1127014160156 0 695.9761
258.144287109375 0 1224.5284
261.1188049316406 0 779.1154
262.1199951171875 0 509.1098
265.12945556640625 0 3880.332
266.12481689453125 0 7171.255
267.1089172363281 0 499.12964
267.12841796875 0 934.8078
268.1279296875 0 655.27814
268.17657470703125 0 744.98114
270.1216735839844 0 1101.3486
272.17535400390625 0 1438.646
274.118408203125 0 1743.8782
279.1469421386719 0 528.3532
280.1292419433594 0 663.67255
281.0513610839844 0 16916.828
281.1500549316406 0 842.3098
282.0512390136719 0 546.11005
282.1564025878906 0 1473.2717
283.0306396484375 0 2510.333
283.1401062011719 0 22389.309
284.1432189941406 0 2737.3489
285.01019287109375 0 1061.6666
286.1398010253906 0 665.5259
287.16973876953125 0 535.2327
288.6195068359375 0 569.95386
290.1669006347656 0 861.7773 b 4
292.130859375 0 552.3593
293.1134033203125 0 10537.259 y Water loss 11
294.11767578125 0 1724.9719
295.1029968261719 0 2226.504
296.1042175292969 0 1773.1475
297.0824890136719 0 1942.5792
297.099365234375 0 1004.46027
297.1556091308594 0 3117.6675
299.06182861328125 0 142403.88
300.0625 0 3125.6611
300.16693115234375 0 2906.1167
301.0588684082031 0 1322.0974
301.15087890625 0 8144.99
301.1683349609375 0 870.41986
302.1138916015625 0 1573.2128
302.1533508300781 0 1104.3939
306.1373291015625 0 1106.546 y 8
309.2040710449219 0 883.7399
311.1236572265625 0 1703.2474 y 11
313.1139221191406 0 709.604
314.18780517578125 0 710.6146
314.20733642578125 0 1012.31744 b 2
315.16619873046875 0 5072.7373
316.16729736328125 0 1214.3822
324.9862365722656 0 788.8961
326.6639099121094 0 655.58655
326.9658508300781 0 784.9467
329.21710205078125 0 647.3479
333.1768798828125 0 1428.6104
341.0184020996094 0 1493.6914
342.99639892578125 0 608.4654
344.9764099121094 0 19093.664
346.1148986816406 0 3764.4468
347.6799011230469 0 2173.8154 b 5
350.2187805175781 0 2645.466
353.67218017578125 0 584.6982 y Water loss 7
355.0692443847656 0 2878.4924
359.0281066894531 0 16784.16
361.1968994140625 0 1409.3289
362.1855773925781 0 1883.1344
362.67919921875 0 3505.0396 y 7
363.14080810546875 0 5885.569
363.17999267578125 0 710.42334
364.1226501464844 0 977.5704
366.17755126953125 0 575.91486
369.1212463378906 0 1517.5533
370.1222839355469 0 2068.0535
371.1204833984375 0 1548.2067
371.2274169921875 0 535.6293
376.1977233886719 0 636.60675
378.21441650390625 0 1645.5752
379.2084045410156 0 6554.4927
380.2107849121094 0 949.1648
381.15142822265625 0 12568.784
382.15386962890625 0 1939.0614
384.1894226074219 0 867.59644
394.172119140625 0 2322.9397
395.1321105957031 0 711.5845
395.1561584472656 0 1061.8002
396.2248840332031 0 1883.7449
398.1762390136719 0 1701.829
402.2002258300781 0 1030.1204
403.20074462890625 0 684.15155
405.2615661621094 0 994.9266
409.1459655761719 0 682.2344
411.1988220214844 0 875.6978
412.1570129394531 0 1129.0736
412.1830139160156 0 7606.347
413.16619873046875 0 749.0372
414.2350769042969 0 1150.0135
415.03656005859375 0 34344.83
417.7172546386719 0 1414.087
418.2092590332031 0 1415.6989
418.7115173339844 0 830.82
419.1624450683594 0 820.797
423.2714538574219 0 3659.1082
424.2743225097656 0 1011.6675
426.68829345703125 0 1114.3169
426.72161865234375 0 21624.52
427.2230224609375 0 9681.086
427.72515869140625 0 3099.1973
428.22332763671875 0 881.06683
429.087890625 0 1575.1691
429.2091064453125 0 6406.556
430.1932678222656 0 16789.877
431.1965637207031 0 4235.3027
431.7127380371094 0 1647.4022 b Water loss 6
432.20477294921875 0 1733.2773 b Ammonia loss 6
432.71490478515625 0 813.8885
433.04638671875 0 2112.365
433.2554931640625 0 741.576 b Water loss 3
439.21917724609375 0 7057.431 y 10
440.219970703125 0 1287.6923
440.7188415527344 0 10657.144 b 6
440.9171447753906 0 766.5733
441.2207336425781 0 4739.5356
441.25225830078125 0 1092.7393
441.72174072265625 0 1291.3112
446.1112976074219 0 736.40717
449.21148681640625 0 617.86945
451.2664489746094 0 5915.98 b 3
452.2691955566406 0 715.13336
452.9130859375 0 2034.3959
453.24871826171875 0 882.22546
460.7523498535156 0 1498.8741
461.2023620605469 0 921.7556
468.2925720214844 0 1305.3344
468.907958984375 0 568.2618 b Water loss 11
471.1697692871094 0 1068.2452
474.2574157714844 0 1111.081
474.7544860839844 0 1436.2825
475.2559509277344 0 908.84644
476.2225341796875 0 777.7971
477.20947265625 0 1414.6589
477.6842346191406 0 744.53796
478.23040771484375 0 675.9565 y Water loss 9
478.2776184082031 0 799.94965
483.2628173828125 0 3353.9927
483.7650146484375 0 2686.7527
488.1219177246094 0 1330.0702
494.2354431152344 0 12247.065
495.23822021484375 0 2430.2095
496.23974609375 0 15721.561 y 9
497.244873046875 0 3738.4243
497.76275634765625 0 1872.1952
503.10699462890625 0 33306.734
504.1422119140625 0 2062.289
505.14642333984375 0 2750.9092
511.2642822265625 0 1209.2451
515.2861938476562 0 620.6228
516.0272827148438 0 804.98535
517.2578735351562 0 924.17725
519.13818359375 0 11092.299
521.1167602539062 0 1064.371
525.2655029296875 0 2210.5686
526.269775390625 0 756.9575
529.592529296875 0 1649.6069 Precursor Ammonia loss
534.2919921875 0 999.74664
534.552734375 0 1067.401
534.6300659179688 0 916.50446
535.2188110351562 0 1027.1632
536.16455078125 0 7603.735
536.2630615234375 0 703.30927
538.2803344726562 0 729.29395
539.2354736328125 0 2351.3262
540.2330932617188 0 583.8604
542.2947998046875 0 1115.1675
543.2769775390625 0 2749.6133
544.2828369140625 0 1023.7975
555.2554931640625 0 842.31647
555.7550048828125 0 880.9175
565.265625 0 616.7351
567.2301635742188 0 2408.9536
573.7977294921875 0 965.22864
574.2979736328125 0 926.9335
575.2430419921875 0 986.9704
576.3237915039062 0 862.67084
579.3242797851562 0 3521.883 b 4
580.3253173828125 0 1204.7212
587.3369140625 0 766.37
593.2557373046875 0 4821.723 y Water loss 8
593.3042602539062 0 1714.5991
594.2564086914062 0 1153.2594
596.3229370117188 0 1290.9287
596.65771484375 0 911.6436
602.7937622070312 0 1083.5698
610.2819213867188 0 16184.696
610.3334350585938 0 547.76587
611.26708984375 0 44154.195 y 8
611.7953491210938 0 1435.6687
612.2700805664062 0 13893.667
613.2724609375 0 3062.6777
621.2506103515625 0 1085.5306
635.297607421875 0 769.4904
636.7848510742188 0 994.54474 y Water loss 3
637.2850341796875 0 897.907 y Ammonia loss 3
645.294189453125 0 1207.492
645.7935791015625 0 2475.703 y 3
646.2945556640625 0 2303.0635
652.3192138671875 0 4607.242
653.0198974609375 0 772.54877
653.3240356445312 0 1576.1593
653.6900634765625 0 689.29376
660.87060546875 0 2408.7507
661.3712158203125 0 1404.26
662.3045654296875 0 1922.1088
676.3397827148438 0 2369.9316 b Water loss 5
677.303466796875 0 612.07794
678.8671875 0 1551.378
679.3668823242188 0 1642.6786
680.3142700195312 0 8505.765
681.3163452148438 0 2989.653
684.819091796875 0 682.4242
693.3271484375 0 3659.7136 y Water loss 2
693.8277587890625 0 2793.617 y Ammonia loss 2
694.3502197265625 0 25816.385 b 5
695.3534545898438 0 9973.426
696.3568115234375 0 2043.9487
701.8388061523438 0 5342.6694
702.334228515625 0 18783.26 y 2
702.8345947265625 0 12032.766
703.3367919921875 0 6644.9023 b Ammonia loss 11
703.8339233398438 0 1787.6385
704.3330078125 0 1453.5361
706.3413696289062 0 1702.7164 y Water loss 7
710.4050903320312 0 1560.4927
710.9068603515625 0 1347.7789
711.3629150390625 0 838.84094
711.8541870117188 0 1489.7283 b 11
712.3536376953125 0 1209.3878
721.3622436523438 0 1315.9099
723.3656005859375 0 4334.0054
724.3505249023438 0 13319.106 y 7
725.3543090820312 0 5912.049
726.3577270507812 0 1677.4407
734.3555297851562 0 1111.433
751.373046875 0 2253.9614
751.8650512695312 0 1093.7869 y 1
752.3707885742188 0 2133.006
755.4248657226562 0 1016.90283
763.3612670898438 0 1700.931
764.363037109375 0 912.1282
765.3994750976562 0 1529.7958
779.3827514648438 0 999.5198
780.380615234375 0 736.7664
793.397216796875 0 2428.818 Precursor Water loss
794.4026489257812 0 1130.1931
807.3731079101562 0 1249.9299
835.4185791015625 0 619.49396
836.4398803710938 0 1892.3534
837.4575805664062 0 1168.9271
847.4442138671875 0 906.25977
852.4344482421875 0 6214.067
853.4361572265625 0 2954.5903
859.4288940429688 0 579.8918
862.41796875 0 3121.4065 b Water loss 6
863.411865234375 0 2072.1997 b Ammonia loss 6
864.4519653320312 0 5127.1924
865.4580688476562 0 2304.5288
880.4286499023438 0 30433.508 b 6
881.4312133789062 0 17270.19
882.4342041015625 0 3989.4866
883.4353637695312 0 1218.9135
892.4598388671875 0 681.75116
908.4252319335938 0 1534.1819
909.4404907226562 0 1126.8925
910.4357299804688 0 1829.4489 y 6
911.4363403320312 0 790.17267
912.4263305664062 0 738.5527
955.4281616210938 0 780.1126
993.5130615234375 0 2116.6145 b 7
994.5155029296875 0 1533.8898
1025.458984375 0 1335.3027 y 5
1076.5379638671875 0 982.2032
1093.5374755859375 0 920.6046
1135.5064697265625 0 720.4703 y Water loss 4
1136.4986572265625 0 891.856 y Ammonia loss 4
1137.4893798828125 0 633.58545
1153.513671875 0 2721.1138 y 4
1154.519775390625 0 1518.966
1155.516845703125 0 1071.4163
3080.21142578125 0 932.7034
3188.883544921875 0 651.94507
3238.16259765625 0 777.1889

Spectrum Details

|  |  |
| --- | --- |
| Matched peaks? Matched peaks The total absolute number of peaks matched. Additionally in brackets the total fraction of peaks matched and the total number of peaks is shown. | 52 (12.01% of 433) |
| FDR? FDR The false discovery rate estimated for this peptide. It is calculated by matching all theoretical fragments with a non-integer shift with the raw peaks for this spectrum. This is done with 40 different shifts. The resulting percentage is the average number of annotated peaks over the number of annotated peaks with the correct spectrum. | 1.01% |
| Satellite FDR? Satellite FDR See the FDR for details on its calculation. This satellite ion specific FDR only contains the satellite ions (d/w) for I/L/J positions. | ∞ |
| PSM Score? PSM Score The PSM Score as given by Hecklib to this annotated spectrum. It is shown with three significant figures. | 386 |

#### Spectrum 7490? Spectrum 7490 The raw spectrum of this peptide as annotated by Hecklib. The fragments are coloured according to ion type (see legend). Any peaks with a star '\*' as text can be hovered over to see the full details, first the ion type second the mass shift type. By hovering over the amino acids in the peptide or ions in the legend the corresponding peaks are highlighted. By toggling the 'Unassigned' label you can turn the background (unassigned) peaks on or off in the plot. By updating the slider in the Ion legend you can update the spectrum to only show the top X% of the peaks with labels. The top X% means any peak that is within X% of the highest intensity. By dragging in the spectrum you can zoom in to a specific part of the spectrum and use 'Zoom Out' to get back to the original zoom level. The annotation of the spectrum is based on the given sequence in the peptides file and is done with different software so inconsistencies are likely. The peaks are annotated based on the given sequence, with 20 ppm tolerance.

Copy Data

##### Spectrum 7490 (TSV)

###### Preview

```
Loading example...
```

*Click on the button to copy the data to your clipboard.*

Mz MinMz MaxIntensity Max

WidthHeightPeptide font sizePeptide stroke widthSpectrum font sizeSpectrum stroke widthCompact peptide

Ion legend

wxyz

abcd

OtherUnassignedIon^Charge^_Position_Show for top:%

TVLHQDWLDGKEY

02.15e+44.29e+46.44e+48.58e+4

Zoom Out

y^+1^_2_y^+1^_4_c^+1^_5_c^+1^_5_y^+1^_5_y^+2^_10_y^+2^_10_c^+1^_6_y^+2^_11_c^+1^_6_c^+2^_12_c^+1^_7_y^+1^_7_c^+1^_7_y^+1^_7_c^+1^_8_c^+1^_8_z^+1^_8_c^+1^_8_y^+1^_8_c^+1^_9_c^+1^_9_y^+1^_9_z^+1^_9_y^+1^_9_c^+1^_10_c^+1^_10_z^+1^_10_y^+1^_10_c^+1^_11_c^+1^_11_z^+1^_11_y^+1^_11_c^+1^_12_z^+1^_12_

0778155623333111

Fragment Matches Table

Show background peaks

| Position | Ion type | Intensity | mz Theoretical | mz Error (Th) | mz Error (ppm) | Charge | Series Number |
| --- | --- | --- | --- | --- | --- | --- | --- |
| - | - | 913.3 | 129.1 | - | - | 0 | - |
| - | - | 467.1 | 148.9 | - | - | 0 | - |
| - | - | 532.7 | 148.9 | - | - | 0 | - |
| - | - | 610.1 | 148.9 | - | - | 0 | - |
| - | - | 705.9 | 148.9 | - | - | 0 | - |
| - | - | 1122 | 148.9 | - | - | 0 | - |
| - | - | 877.5 | 148.9 | - | - | 0 | - |
| - | - | 1443 | 148.9 | - | - | 0 | - |
| - | - | 3123 | 148.9 | - | - | 0 | - |
| - | - | 5263 | 148.9 | - | - | 0 | - |
| - | - | 3740 | 149 | - | - | 0 | - |
| - | - | 1689 | 149 | - | - | 0 | - |
| - | - | 1377 | 149 | - | - | 0 | - |
| - | - | 1116 | 149 | - | - | 0 | - |
| - | - | 1006 | 149 | - | - | 0 | - |
| - | - | 783.7 | 149 | - | - | 0 | - |
| - | - | 503.3 | 149 | - | - | 0 | - |
| - | - | 515.4 | 149 | - | - | 0 | - |
| - | - | 435.3 | 149 | - | - | 0 | - |
| - | - | 446.8 | 155.7 | - | - | 0 | - |
| - | - | 978.7 | 173.1 | - | - | 0 | - |
| - | - | 752.3 | 186.1 | - | - | 0 | - |
| - | - | 504.1 | 187.8 | - | - | 0 | - |
| - | - | 473.9 | 191.8 | - | - | 0 | - |
| - | - | 1515 | 201.1 | - | - | 0 | - |
| - | - | 436.3 | 209.1 | - | - | 0 | - |
| - | - | 500.5 | 212 | - | - | 0 | - |
| - | - | 468.9 | 251.1 | - | - | 0 | - |
| 12 | y | 1470 | 293.1 | 0.0004662 | 1.59 | +1 | 2 |
| - | - | 596.7 | 294.1 | - | - | 0 | - |
| - | - | 569.4 | 301.1 | - | - | 0 | - |
| - | - | 620.5 | 301.1 | - | - | 0 | - |
| - | - | 612.3 | 314.2 | - | - | 0 | - |
| - | - | 1069 | 451.3 | - | - | 0 | - |
| 10 | y | 2604 | 496.2 | 0.001451 | 2.923 | +1 | 4 |
| - | - | 794.8 | 497.2 | - | - | 0 | - |
| - | - | 674.7 | 555.8 | - | - | 0 | - |
| 5 | c | 728.1 | 579.3 | 0.00229 | 3.954 | +1 | 5 |
| 5 | c | 2060 | 596.4 | 0.001984 | 3.327 | +1 | 5 |
| - | - | 701.7 | 597.4 | - | - | 0 | - |
| - | - | 551.7 | 601.6 | - | - | 0 | - |
| - | - | 731.5 | 609.3 | - | - | 0 | - |
| - | - | 542.5 | 610.3 | - | - | 0 | - |
| 9 | y | 2440 | 611.3 | 0.001447 | 2.366 | +1 | 5 |
| - | - | 601.8 | 612.3 | - | - | 0 | - |
| 4 | y | 732.9 | 637.3 | 0.005246 | 8.233 | +2 | 10 |
| - | - | 875.7 | 638.8 | - | - | 0 | - |
| - | - | 561.6 | 645.3 | - | - | 0 | - |
| 4 | y | 1837 | 645.8 | 0.0007649 | 1.184 | +2 | 10 |
| - | - | 969.3 | 646.3 | - | - | 0 | - |
| - | - | 910.4 | 646.8 | - | - | 0 | - |
| - | - | 820.5 | 667.4 | - | - | 0 | - |
| 6 | c | 2257 | 694.4 | 0.002378 | 3.425 | +1 | 6 |
| - | - | 618.9 | 701.4 | - | - | 0 | - |
| 3 | y | 2575 | 702.3 | 0.003468 | 4.937 | +2 | 11 |
| - | - | 3130 | 702.8 | - | - | 0 | - |
| - | - | 1106 | 703.3 | - | - | 0 | - |
| 6 | c | 4023 | 711.4 | 0.007382 | 10.38 | +1 | 6 |
| 12 | c | 6603 | 711.9 | 0.0002514 | 0.3532 | +2 | 12 |
| - | - | 3585 | 712.4 | - | - | 0 | - |
| - | - | 1470 | 712.9 | - | - | 0 | - |
| - | - | 925.5 | 721.4 | - | - | 0 | - |
| - | - | 590.5 | 792.9 | - | - | 0 | - |
| - | - | 1016 | 793.4 | - | - | 0 | - |
| - | - | 1836 | 793.9 | - | - | 0 | - |
| - | - | 1179 | 794.4 | - | - | 0 | - |
| - | - | 4096 | 801.4 | - | - | 0 | - |
| - | - | 1.018E+04 | 801.9 | - | - | 0 | - |
| - | - | 4.513E+04 | 802.4 | - | - | 0 | - |
| - | - | 3.831E+04 | 802.9 | - | - | 0 | - |
| - | - | 2.088E+04 | 803.4 | - | - | 0 | - |
| - | - | 6844 | 803.9 | - | - | 0 | - |
| - | - | 1326 | 804.4 | - | - | 0 | - |
| - | - | 657.9 | 853.4 | - | - | 0 | - |
| - | - | 1349 | 854.4 | - | - | 0 | - |
| 7 | c | 1432 | 880.4 | 0.004787 | 5.437 | +1 | 7 |
| 7 | y | 939 | 893.4 | 0.01743 | 19.51 | +1 | 7 |
| - | - | 729.4 | 895.4 | - | - | 0 | - |
| - | - | 929.8 | 896.4 | - | - | 0 | - |
| 7 | c | 4780 | 897.5 | 0.00271 | 3.02 | +1 | 7 |
| - | - | 2402 | 898.5 | - | - | 0 | - |
| - | - | 694.7 | 909.4 | - | - | 0 | - |
| 7 | y | 1569 | 910.4 | 0.0005818 | 0.6391 | +1 | 7 |
| - | - | 1378 | 911.4 | - | - | 0 | - |
| - | - | 1460 | 917.4 | - | - | 0 | - |
| - | - | 762.1 | 932.5 | - | - | 0 | - |
| - | - | 976.1 | 939.4 | - | - | 0 | - |
| - | - | 927.2 | 966.5 | - | - | 0 | - |
| - | - | 1352 | 991.5 | - | - | 0 | - |
| 8 | c | 1630 | 992.5 | 0.01127 | 11.35 | +1 | 8 |
| 8 | c | 2580 | 993.5 | 0.0008101 | 0.8154 | +1 | 8 |
| - | - | 2607 | 994.5 | - | - | 0 | - |
| - | - | 729.1 | 995.5 | - | - | 0 | - |
| 6 | z | 1098 | 1009 | 0.00164 | 1.625 | +1 | 8 |
| - | - | 3844 | 1010 | - | - | 0 | - |
| - | - | 1663 | 1010 | - | - | 0 | - |
| 8 | c | 1.082E+04 | 1011 | 0.002118 | 2.096 | +1 | 8 |
| - | - | 751.7 | 1011 | - | - | 0 | - |
| - | - | 6082 | 1012 | - | - | 0 | - |
| - | - | 1986 | 1013 | - | - | 0 | - |
| - | - | 752.3 | 1018 | - | - | 0 | - |
| - | - | 800.5 | 1024 | - | - | 0 | - |
| 6 | y | 1337 | 1025 | 0.0006654 | 0.6489 | +1 | 8 |
| - | - | 736 | 1060 | - | - | 0 | - |
| - | - | 1741 | 1068 | - | - | 0 | - |
| - | - | 1454 | 1069 | - | - | 0 | - |
| - | - | 1334 | 1082 | - | - | 0 | - |
| - | - | 1645 | 1083 | - | - | 0 | - |
| 9 | c | 2416 | 1109 | 0.0004372 | 0.3943 | +1 | 9 |
| - | - | 1107 | 1110 | - | - | 0 | - |
| - | - | 890.7 | 1125 | - | - | 0 | - |
| 9 | c | 1343 | 1126 | 0.0002354 | 0.2092 | +1 | 9 |
| - | - | 812.8 | 1127 | - | - | 0 | - |
| 5 | y | 1715 | 1136 | 0.01479 | 13.02 | +1 | 9 |
| 5 | z | 7233 | 1137 | 0.00109 | 0.9584 | +1 | 9 |
| - | - | 4757 | 1139 | - | - | 0 | - |
| - | - | 2364 | 1140 | - | - | 0 | - |
| - | - | 925.2 | 1152 | - | - | 0 | - |
| - | - | 2528 | 1153 | - | - | 0 | - |
| 5 | y | 3767 | 1154 | 0.002158 | 1.871 | +1 | 9 |
| - | - | 1873 | 1155 | - | - | 0 | - |
| - | - | 1345 | 1164 | - | - | 0 | - |
| 10 | c | 962.4 | 1166 | 0.0004165 | 0.3573 | +1 | 10 |
| - | - | 6724 | 1182 | - | - | 0 | - |
| 10 | c | 1.265E+04 | 1183 | 0.000134 | 0.1133 | +1 | 10 |
| - | - | 7926 | 1184 | - | - | 0 | - |
| - | - | 2971 | 1185 | - | - | 0 | - |
| - | - | 1504 | 1202 | - | - | 0 | - |
| - | - | 1.007E+04 | 1218 | - | - | 0 | - |
| - | - | 5777 | 1219 | - | - | 0 | - |
| - | - | 3703 | 1220 | - | - | 0 | - |
| - | - | 986 | 1221 | - | - | 0 | - |
| - | - | 898.6 | 1242 | - | - | 0 | - |
| - | - | 908.3 | 1268 | - | - | 0 | - |
| 4 | z | 8236 | 1275 | 0.002751 | 2.158 | +1 | 10 |
| - | - | 6096 | 1276 | - | - | 0 | - |
| - | - | 2815 | 1277 | - | - | 0 | - |
| - | - | 779.9 | 1290 | - | - | 0 | - |
| 4 | y | 2296 | 1291 | 0.00524 | 4.06 | +1 | 10 |
| - | - | 1622 | 1292 | - | - | 0 | - |
| 11 | c | 1052 | 1294 | 0.001752 | 1.354 | +1 | 11 |
| - | - | 1790 | 1310 | - | - | 0 | - |
| 11 | c | 1.086E+04 | 1311 | 0.003276 | 2.5 | +1 | 11 |
| - | - | 7566 | 1312 | - | - | 0 | - |
| - | - | 3664 | 1313 | - | - | 0 | - |
| - | - | 974.6 | 1337 | - | - | 0 | - |
| 3 | z | 6144 | 1388 | 0.0008775 | 0.6324 | +1 | 11 |
| - | - | 5622 | 1389 | - | - | 0 | - |
| - | - | 1943 | 1390 | - | - | 0 | - |
| - | - | 977.4 | 1395 | - | - | 0 | - |
| - | - | 923.1 | 1396 | - | - | 0 | - |
| 3 | y | 2316 | 1404 | 0.003855 | 2.746 | +1 | 11 |
| - | - | 1535 | 1405 | - | - | 0 | - |
| - | - | 918.2 | 1406 | - | - | 0 | - |
| - | - | 9555 | 1439 | - | - | 0 | - |
| 12 | c | 2.394E+04 | 1440 | 0.00186 | 1.292 | +1 | 12 |
| - | - | 1.827E+04 | 1441 | - | - | 0 | - |
| - | - | 8745 | 1442 | - | - | 0 | - |
| - | - | 1799 | 1443 | - | - | 0 | - |
| - | - | 995.5 | 1459 | - | - | 0 | - |
| 2 | z | 849.1 | 1487 | 0.0179 | 12.04 | +1 | 12 |
| - | - | 787.2 | 1519 | - | - | 0 | - |
| - | - | 2697 | 1543 | - | - | 0 | - |
| - | - | 2463 | 1544 | - | - | 0 | - |
| - | - | 3670 | 1545 | - | - | 0 | - |
| - | - | 3629 | 1546 | - | - | 0 | - |
| - | - | 1714 | 1547 | - | - | 0 | - |
| - | - | 694.1 | 1549 | - | - | 0 | - |
| - | - | 4593 | 1559 | - | - | 0 | - |
| - | - | 7163 | 1560 | - | - | 0 | - |
| - | - | 8420 | 1561 | - | - | 0 | - |
| - | - | 3444 | 1562 | - | - | 0 | - |
| - | - | 2347 | 1563 | - | - | 0 | - |
| - | - | 2032 | 1576 | - | - | 0 | - |
| - | - | 6835 | 1577 | - | - | 0 | - |
| - | - | 6559 | 1578 | - | - | 0 | - |
| - | - | 3250 | 1579 | - | - | 0 | - |
| - | - | 981.7 | 1580 | - | - | 0 | - |
| - | - | 1571 | 1586 | - | - | 0 | - |
| - | - | 9513 | 1587 | - | - | 0 | - |
| - | - | 2.213E+04 | 1588 | - | - | 0 | - |
| - | - | 1.801E+04 | 1589 | - | - | 0 | - |
| - | - | 8162 | 1590 | - | - | 0 | - |
| - | - | 4112 | 1591 | - | - | 0 | - |
| - | - | 870.2 | 1592 | - | - | 0 | - |
| - | - | 1.302E+04 | 1603 | - | - | 0 | - |
| - | - | 5.21E+04 | 1604 | - | - | 0 | - |
| - | - | 8.496E+04 | 1605 | - | - | 0 | - |
| - | - | 5.847E+04 | 1606 | - | - | 0 | - |
| - | - | 3.184E+04 | 1607 | - | - | 0 | - |
| - | - | 9278 | 1608 | - | - | 0 | - |
| - | - | 1092 | 1609 | - | - | 0 | - |
| - | - | 715.5 | 1836 | - | - | 0 | - |
| - | - | 864.4 | 2404 | - | - | 0 | - |
| - | - | 942.6 | 2405 | - | - | 0 | - |
| - | - | 1395 | 2406 | - | - | 0 | - |
| - | - | 1543 | 2407 | - | - | 0 | - |
| - | - | 690.3 | 2410 | - | - | 0 | - |
| - | - | 779.3 | 3080 | - | - | 0 | - |

m/z Charge Intensity FragmentType MassShift Position
129.10203552246094 0 913.3234
148.8577880859375 0 467.10556
148.8645782470703 0 532.70264
148.9007568359375 0 610.06036
148.90890502929688 0 705.9446
148.915771484375 0 1121.7714
148.92361450195312 0 877.5246
148.93019104003906 0 1443.1426
148.9376678466797 0 3122.6675
148.94549560546875 0 5262.5225
148.962158203125 0 3739.5557
148.9698028564453 0 1688.6543
148.97702026367188 0 1376.5137
148.98440551757812 0 1115.556
148.991943359375 0 1005.92017
148.9991455078125 0 783.6625
149.00640869140625 0 503.28253
149.02069091796875 0 515.3882
149.0364227294922 0 435.33542
155.6680145263672 0 446.8227
173.12808227539062 0 978.6597
186.1236572265625 0 752.31024
187.83096313476562 0 504.13
191.83993530273438 0 473.87192
201.12271118164062 0 1514.8774
209.07522583007812 0 436.26352
211.9714813232422 0 500.49942
251.1497039794922 0 468.8592
293.11273193359375 0 1469.926 y Water loss 11
294.1152038574219 0 596.69415
301.1292724609375 0 569.4155
301.1494445800781 0 620.4636
314.207275390625 0 612.28595
451.2601318359375 0 1068.7094
496.2387390136719 0 2603.6152 y 9
497.24346923828125 0 794.8096
555.7552490234375 0 674.73944
579.3226318359375 0 728.08655 c Ammonia loss 4
596.3494873046875 0 2059.8408 c 4
597.3512573242188 0 701.6607
601.5804443359375 0 551.70294
609.2843627929688 0 731.478
610.2826538085938 0 542.45703
611.2656860351562 0 2439.6938 y 8
612.2646484375 0 601.7992
637.2830810546875 0 732.86395 y Ammonia loss 3
638.8289184570312 0 875.6736
645.2958374023438 0 561.57404
645.7903442382812 0 1836.6189 y 3
646.2904663085938 0 969.3189
646.7920532226562 0 910.36475
667.3624877929688 0 820.52655
694.3494873046875 0 2257.0574 c Ammonia loss 5
701.3709716796875 0 618.8638
702.3366088867188 0 2574.6665 y 2
702.8424072265625 0 3129.7717
703.333984375 0 1105.5204
711.3710327148438 0 4022.7068 c 5
711.8544921875 0 6602.766 c Ammonia loss 11
712.3576049804688 0 3585.1692
712.8558349609375 0 1469.5697
721.3637084960938 0 925.4732
792.8960571289062 0 590.5287
793.385986328125 0 1016.0929
793.8853759765625 0 1836.0743
794.3905029296875 0 1178.6848
801.406005859375 0 4095.5747
801.8961791992188 0 10175.728
802.3931884765625 0 45127.234
802.8929443359375 0 38310.45
803.3950805664062 0 20882.23
803.8958129882812 0 6843.9854
804.39697265625 0 1326.0038
853.4310302734375 0 657.89984
854.4459838867188 0 1348.6241
880.4263916015625 0 1431.7222 c Ammonia loss 6
893.42138671875 0 939.0493 y Ammonia loss 6
895.4202270507812 0 729.3608
896.4432983398438 0 929.82794
897.4550170898438 0 4779.802 c 6
898.4595336914062 0 2402.1929
909.4296264648438 0 694.6681
910.4310913085938 0 1568.5634 y 6
911.4318237304688 0 1378.1163
917.4402465820312 0 1460.0012
932.4819946289062 0 762.1391
939.3526000976562 0 976.08966
966.5322875976562 0 927.243
991.5223999023438 0 1352.0696
992.5199584960938 0 1630.4023 c Water loss 7
993.5160522460938 0 2579.723 c Ammonia loss 7
994.513671875 0 2607.283
995.5198974609375 0 729.1353
1009.4403686523438 0 1098.3076 z 5
1009.5360717773438 0 3843.7048
1010.4434204101562 0 1662.8184
1010.5396728515625 0 10815.447 c 7
1011.4494018554688 0 751.6953
1011.5425415039062 0 6081.9204
1012.5446166992188 0 1986.3693
1018.4989013671875 0 752.3146
1024.45458984375 0 800.4843
1025.456787109375 0 1336.5121 y 5
1059.536865234375 0 735.9965
1067.537109375 0 1741.4559
1068.5423583984375 0 1454.1788
1081.5552978515625 0 1334.4459
1082.5535888671875 0 1645.0006
1108.541748046875 0 2415.7793 c Ammonia loss 8
1109.5457763671875 0 1106.713
1124.5635986328125 0 890.6845
1125.5689697265625 0 1343.0526 c 8
1126.562744140625 0 812.7687
1136.5042724609375 0 1715.3901 y Ammonia loss 4
1137.4962158203125 0 7232.7656 z 4
1138.5001220703125 0 4756.5664
1139.5052490234375 0 2363.9675
1151.526611328125 0 925.2458
1152.5213623046875 0 2528.2046
1153.5181884765625 0 3766.836 y 4
1154.51318359375 0 1872.5862
1163.5321044921875 0 1345.3805
1165.563232421875 0 962.40686 c Ammonia loss 9
1181.5963134765625 0 6724.4824
1182.59033203125 0 12653.358 c 9
1183.5926513671875 0 7926.3745
1184.5970458984375 0 2971.1035
1201.538818359375 0 1504.0895
1217.5572509765625 0 10074.737
1218.559326171875 0 5777.456
1219.56103515625 0 3702.5737
1220.5579833984375 0 986.0385
1241.5201416015625 0 898.56976
1267.6590576171875 0 908.3097
1274.553466796875 0 8236.474 z 3
1275.5572509765625 0 6095.5854
1276.562255859375 0 2815.4233
1289.585693359375 0 779.9188
1290.5697021484375 0 2296.212 y 3
1291.5723876953125 0 1622.3167
1293.6568603515625 0 1052.1111 c Ammonia loss 10
1309.6944580078125 0 1789.7484
1310.681884765625 0 10857.262 c 10
1311.6846923828125 0 7566.119
1312.686279296875 0 3664.103
1336.7078857421875 0 974.585
1387.639404296875 0 6144.364 z 2
1388.6402587890625 0 5621.9873
1389.6439208984375 0 1942.9662
1394.7257080078125 0 977.3959
1395.7012939453125 0 923.1054
1403.6551513671875 0 2315.7505 y 2
1404.6536865234375 0 1535.1132
1405.6561279296875 0 918.1794
1438.7392578125 0 9555.186
1439.7296142578125 0 23935.5 c 11
1440.73046875 0 18274.826
1441.7322998046875 0 8745.273
1442.7196044921875 0 1799.237
1458.733154296875 0 995.5465
1486.6907958984375 0 849.14655 z 1
1518.721435546875 0 787.2344
1542.771240234375 0 2697.214
1543.7725830078125 0 2462.9695
1544.76025390625 0 3670.2234
1545.7645263671875 0 3629.2202
1546.763427734375 0 1714.1278
1548.70703125 0 694.057
1558.7740478515625 0 4592.89
1559.765869140625 0 7163.1074
1560.769775390625 0 8419.716
1561.7723388671875 0 3443.8342
1562.78271484375 0 2347.2573
1575.7919921875 0 2031.7296
1576.7860107421875 0 6835.1924
1577.79248046875 0 6558.5845
1578.7972412109375 0 3249.9329
1579.7874755859375 0 981.65826
1585.77294921875 0 1570.8613
1586.7659912109375 0 9512.715
1587.7601318359375 0 22131.273
1588.759033203125 0 18007.537
1589.75927734375 0 8161.5483
1590.7703857421875 0 4112.3965
1591.753173828125 0 870.22186
1602.782470703125 0 13021.677
1603.78076171875 0 52098.58
1604.7806396484375 0 84959.875
1605.78369140625 0 58471.926
1606.786376953125 0 31844.781
1607.7880859375 0 9278.048
1608.7919921875 0 1091.6062
1836.3603515625 0 715.5141
2404.195068359375 0 864.3682
2405.177490234375 0 942.6038
2406.195556640625 0 1394.8486
2407.201416015625 0 1542.6902
2410.224365234375 0 690.2957
3080.411376953125 0 779.2537

Spectrum Details

|  |  |
| --- | --- |
| Matched peaks? Matched peaks The total absolute number of peaks matched. Additionally in brackets the total fraction of peaks matched and the total number of peaks is shown. | 35 (17.59% of 199) |
| FDR? FDR The false discovery rate estimated for this peptide. It is calculated by matching all theoretical fragments with a non-integer shift with the raw peaks for this spectrum. This is done with 40 different shifts. The resulting percentage is the average number of annotated peaks over the number of annotated peaks with the correct spectrum. | 2.65% |
| Satellite FDR? Satellite FDR See the FDR for details on its calculation. This satellite ion specific FDR only contains the satellite ions (d/w) for I/L/J positions. | - |
| PSM Score? PSM Score The PSM Score as given by Hecklib to this annotated spectrum. It is shown with three significant figures. | 371 |

#### Spectrum 6890? Spectrum 6890 The raw spectrum of this peptide as annotated by Hecklib. The fragments are coloured according to ion type (see legend). Any peaks with a star '\*' as text can be hovered over to see the full details, first the ion type second the mass shift type. By hovering over the amino acids in the peptide or ions in the legend the corresponding peaks are highlighted. By toggling the 'Unassigned' label you can turn the background (unassigned) peaks on or off in the plot. By updating the slider in the Ion legend you can update the spectrum to only show the top X% of the peaks with labels. The top X% means any peak that is within X% of the highest intensity. By dragging in the spectrum you can zoom in to a specific part of the spectrum and use 'Zoom Out' to get back to the original zoom level. The annotation of the spectrum is based on the given sequence in the peptides file and is done with different software so inconsistencies are likely. The peaks are annotated based on the given sequence, with 20 ppm tolerance.

Copy Data

##### Spectrum 6890 (TSV)

###### Preview

```
Loading example...
```

*Click on the button to copy the data to your clipboard.*

Mz MinMz MaxIntensity Max

WidthHeightPeptide font sizePeptide stroke widthSpectrum font sizeSpectrum stroke widthCompact peptide

Ion legend

wxyz

abcd

OtherUnassignedIon^Charge^_Position_Show for top:%

TVLHQDWLDGKEY

03.05e+66.10e+69.15e+61.22e+7

Zoom Out

a^+1^_2_a^+1^_2_y^+1^_1_b^+1^_2_b^+1^_2_y^+2^_3_y^+1^_2_b^+1^_3_y^+1^_2_b^+1^_3_b^+2^_6_y^+1^_3_y^+1^_3_b^+2^_7_b^+1^_4_y^+1^_3_b^+2^_7_b^+1^_4_y^+1^_4_b^+2^_8_y^+1^_4_b^+2^_9_b^+2^_9_b^+1^_5_b^+2^_10_b^+1^_5_b^+2^_10_y^+1^_5_y^+1^_5_y^+2^_10_y^+2^_10_b^+2^_11_b^+2^_11_y^+2^_10_b^+2^_11_b^+1^_6_b^+1^_6_y^+2^_11_b^+1^_6_y^+2^_11_b^+2^_12_b^+2^_12_y^+1^_6_b^+2^_12_y^+1^_6_y^+2^_12_\*\*\*b^+1^_7_b^+1^_7_b^+1^_7_y^+1^_7_y^+1^_7_b^+1^_8_b^+1^_8_b^+1^_8_y^+1^_8_y^+1^_8_y^+1^_8_b^+1^_9_b^+1^_9_b^+1^_9_y^+1^_9_y^+1^_9_b^+1^_10_b^+1^_10_y^+1^_9_b^+1^_10_y^+1^_10_y^+1^_10_b^+1^_11_b^+1^_11_y^+1^_10_b^+1^_11_y^+1^_11_y^+1^_11_y^+1^_11_b^+1^_12_b^+1^_12_

0606121218182424

Fragment Matches Table

Show background peaks

| Position | Ion type | Intensity | mz Theoretical | mz Error (Th) | mz Error (ppm) | Charge | Series Number |
| --- | --- | --- | --- | --- | --- | --- | --- |
| - | - | 5.794E+04 | 122.1 | - | - | 0 | - |
| - | - | 8454 | 125.1 | - | - | 0 | - |
| - | - | 8.871E+04 | 127.1 | - | - | 0 | - |
| - | - | 1.657E+04 | 127.1 | - | - | 0 | - |
| - | - | 7832 | 127.1 | - | - | 0 | - |
| - | - | 5756 | 127.7 | - | - | 0 | - |
| - | - | 9638 | 128.1 | - | - | 0 | - |
| - | - | 1.362E+05 | 128.1 | - | - | 0 | - |
| - | - | 2.242E+04 | 129.1 | - | - | 0 | - |
| - | - | 4.636E+06 | 129.1 | - | - | 0 | - |
| - | - | 7621 | 130.1 | - | - | 0 | - |
| - | - | 8979 | 130.1 | - | - | 0 | - |
| - | - | 2.666E+05 | 130.1 | - | - | 0 | - |
| - | - | 3.95E+04 | 130.1 | - | - | 0 | - |
| - | - | 2.692E+05 | 130.1 | - | - | 0 | - |
| - | - | 2.937E+04 | 131.1 | - | - | 0 | - |
| - | - | 6896 | 131.1 | - | - | 0 | - |
| - | - | 4.785E+04 | 132.1 | - | - | 0 | - |
| - | - | 3.407E+05 | 136.1 | - | - | 0 | - |
| - | - | 2.201E+04 | 137.1 | - | - | 0 | - |
| - | - | 7.535E+04 | 138.1 | - | - | 0 | - |
| - | - | 7461 | 138.1 | - | - | 0 | - |
| - | - | 6.013E+04 | 139.1 | - | - | 0 | - |
| - | - | 8139 | 141.1 | - | - | 0 | - |
| - | - | 8.44E+04 | 141.1 | - | - | 0 | - |
| - | - | 2.34E+04 | 143 | - | - | 0 | - |
| - | - | 1.596E+04 | 144.1 | - | - | 0 | - |
| - | - | 4.79E+04 | 145.1 | - | - | 0 | - |
| - | - | 9434 | 146.1 | - | - | 0 | - |
| - | - | 1.477E+04 | 147 | - | - | 0 | - |
| - | - | 1.41E+04 | 149 | - | - | 0 | - |
| - | - | 9.287E+04 | 150.1 | - | - | 0 | - |
| - | - | 8646 | 150.1 | - | - | 0 | - |
| - | - | 1.127E+05 | 151.1 | - | - | 0 | - |
| - | - | 1.682E+04 | 152.1 | - | - | 0 | - |
| - | - | 1.003E+04 | 153.1 | - | - | 0 | - |
| - | - | 1.004E+04 | 154.1 | - | - | 0 | - |
| - | - | 1.793E+04 | 154.1 | - | - | 0 | - |
| - | - | 1.067E+04 | 155 | - | - | 0 | - |
| - | - | 4.939E+05 | 155.1 | - | - | 0 | - |
| 2 | a | 4.508E+04 | 155.1 | 0.0003966 | 2.557 | +1 | 2 |
| - | - | 7650 | 156.1 | - | - | 0 | - |
| - | - | 3.382E+04 | 156.1 | - | - | 0 | - |
| - | - | 3.525E+04 | 156.1 | - | - | 0 | - |
| - | - | 1.129E+04 | 157.1 | - | - | 0 | - |
| - | - | 7.038E+05 | 159.1 | - | - | 0 | - |
| - | - | 7.245E+04 | 160.1 | - | - | 0 | - |
| - | - | 1.728E+05 | 165.1 | - | - | 0 | - |
| - | - | 9661 | 165.1 | - | - | 0 | - |
| - | - | 3.458E+05 | 166.1 | - | - | 0 | - |
| - | - | 1.15E+05 | 166.1 | - | - | 0 | - |
| - | - | 2.305E+04 | 167.1 | - | - | 0 | - |
| - | - | 1.148E+04 | 167.1 | - | - | 0 | - |
| - | - | 1.892E+04 | 167.1 | - | - | 0 | - |
| - | - | 5.353E+04 | 168.1 | - | - | 0 | - |
| - | - | 4.063E+04 | 168.1 | - | - | 0 | - |
| - | - | 1.969E+04 | 169.1 | - | - | 0 | - |
| - | - | 1.281E+05 | 169.1 | - | - | 0 | - |
| - | - | 7435 | 169.2 | - | - | 0 | - |
| - | - | 4.474E+04 | 170.1 | - | - | 0 | - |
| - | - | 7881 | 170.1 | - | - | 0 | - |
| - | - | 1.221E+04 | 171.1 | - | - | 0 | - |
| - | - | 6.94E+04 | 172.1 | - | - | 0 | - |
| - | - | 2.865E+04 | 173.1 | - | - | 0 | - |
| 2 | a | 2.984E+06 | 173.1 | 0.0004825 | 2.787 | +1 | 2 |
| - | - | 2.483E+05 | 174.1 | - | - | 0 | - |
| - | - | 1.822E+04 | 176.1 | - | - | 0 | - |
| - | - | 7711 | 177.1 | - | - | 0 | - |
| - | - | 1.223E+04 | 177.1 | - | - | 0 | - |
| - | - | 1.371E+05 | 178.1 | - | - | 0 | - |
| - | - | 7634 | 179.1 | - | - | 0 | - |
| - | - | 1.871E+04 | 179.1 | - | - | 0 | - |
| - | - | 3.279E+04 | 181.1 | - | - | 0 | - |
| 13 | y | 4.629E+05 | 182.1 | 0.0004801 | 2.637 | +1 | 1 |
| - | - | 4.999E+04 | 183.1 | - | - | 0 | - |
| 2 | b | 4.57E+04 | 183.1 | 0.0003398 | 1.855 | +1 | 2 |
| - | - | 1.42E+04 | 185.2 | - | - | 0 | - |
| - | - | 1.267E+06 | 186.1 | - | - | 0 | - |
| - | - | 8.781E+04 | 187.1 | - | - | 0 | - |
| - | - | 9523 | 187.1 | - | - | 0 | - |
| - | - | 1.022E+05 | 187.1 | - | - | 0 | - |
| - | - | 1.116E+04 | 188.1 | - | - | 0 | - |
| - | - | 1.033E+04 | 189.1 | - | - | 0 | - |
| - | - | 4.794E+04 | 190.1 | - | - | 0 | - |
| - | - | 3.335E+04 | 190.1 | - | - | 0 | - |
| - | - | 1.104E+04 | 193.1 | - | - | 0 | - |
| - | - | 9.701E+04 | 194.1 | - | - | 0 | - |
| - | - | 3.094E+05 | 195.1 | - | - | 0 | - |
| - | - | 1.216E+05 | 196.1 | - | - | 0 | - |
| - | - | 2.191E+04 | 196.1 | - | - | 0 | - |
| - | - | 1.107E+04 | 197.1 | - | - | 0 | - |
| - | - | 1.677E+04 | 197.1 | - | - | 0 | - |
| - | - | 6.069E+04 | 198.1 | - | - | 0 | - |
| - | - | 2.235E+04 | 198.1 | - | - | 0 | - |
| - | - | 2.065E+04 | 199.1 | - | - | 0 | - |
| - | - | 1.921E+04 | 200.1 | - | - | 0 | - |
| 2 | b | 1.762E+06 | 201.1 | 0.0004257 | 2.117 | +1 | 2 |
| - | - | 3.751E+04 | 202.1 | - | - | 0 | - |
| - | - | 1.566E+05 | 202.1 | - | - | 0 | - |
| - | - | 1.422E+04 | 203.1 | - | - | 0 | - |
| - | - | 1.158E+05 | 205.1 | - | - | 0 | - |
| - | - | 7370 | 205.1 | - | - | 0 | - |
| - | - | 2.591E+04 | 205.1 | - | - | 0 | - |
| - | - | 1.375E+04 | 206.1 | - | - | 0 | - |
| - | - | 3.142E+04 | 206.1 | - | - | 0 | - |
| - | - | 9184 | 207.2 | - | - | 0 | - |
| - | - | 4.876E+04 | 207.2 | - | - | 0 | - |
| - | - | 8703 | 208.1 | - | - | 0 | - |
| - | - | 1.921E+04 | 210.1 | - | - | 0 | - |
| - | - | 8457 | 211.1 | - | - | 0 | - |
| 11 | y | 2.422E+04 | 211.1 | 0.0004813 | 2.28 | +2 | 3 |
| - | - | 9114 | 212.1 | - | - | 0 | - |
| - | - | 3.476E+04 | 212.1 | - | - | 0 | - |
| - | - | 1.321E+05 | 213.1 | - | - | 0 | - |
| - | - | 1.774E+04 | 213.2 | - | - | 0 | - |
| - | - | 1.844E+04 | 214.1 | - | - | 0 | - |
| - | - | 4.703E+04 | 216.1 | - | - | 0 | - |
| - | - | 1.297E+04 | 221.1 | - | - | 0 | - |
| - | - | 1.441E+04 | 221.1 | - | - | 0 | - |
| - | - | 6.454E+04 | 221.1 | - | - | 0 | - |
| - | - | 5.069E+04 | 221.1 | - | - | 0 | - |
| - | - | 5.951E+04 | 222.1 | - | - | 0 | - |
| - | - | 1.568E+04 | 223.1 | - | - | 0 | - |
| - | - | 5.684E+04 | 223.1 | - | - | 0 | - |
| - | - | 6.223E+05 | 223.2 | - | - | 0 | - |
| - | - | 1.562E+04 | 224.1 | - | - | 0 | - |
| - | - | 1.136E+04 | 224.1 | - | - | 0 | - |
| - | - | 7.522E+04 | 224.2 | - | - | 0 | - |
| - | - | 1.708E+05 | 226.1 | - | - | 0 | - |
| - | - | 9.618E+04 | 227.1 | - | - | 0 | - |
| - | - | 1.441E+04 | 227.1 | - | - | 0 | - |
| - | - | 1.883E+04 | 227.2 | - | - | 0 | - |
| - | - | 1.59E+04 | 228.1 | - | - | 0 | - |
| - | - | 4.034E+04 | 228.1 | - | - | 0 | - |
| - | - | 1.998E+04 | 229.1 | - | - | 0 | - |
| - | - | 2.091E+04 | 230.2 | - | - | 0 | - |
| - | - | 1.346E+04 | 230.2 | - | - | 0 | - |
| - | - | 6.911E+04 | 233.1 | - | - | 0 | - |
| - | - | 1.153E+04 | 233.2 | - | - | 0 | - |
| - | - | 4.094E+05 | 234.1 | - | - | 0 | - |
| - | - | 1.105E+04 | 235.1 | - | - | 0 | - |
| - | - | 4.015E+04 | 235.1 | - | - | 0 | - |
| - | - | 1.469E+04 | 235.2 | - | - | 0 | - |
| - | - | 2.513E+04 | 237.1 | - | - | 0 | - |
| - | - | 6.032E+04 | 238.1 | - | - | 0 | - |
| - | - | 3.251E+04 | 238.1 | - | - | 0 | - |
| - | - | 1.79E+04 | 239.1 | - | - | 0 | - |
| - | - | 3.227E+04 | 239.1 | - | - | 0 | - |
| - | - | 2.845E+05 | 240.1 | - | - | 0 | - |
| - | - | 1.696E+04 | 241.1 | - | - | 0 | - |
| - | - | 4.422E+04 | 241.1 | - | - | 0 | - |
| - | - | 2.497E+04 | 241.1 | - | - | 0 | - |
| - | - | 1.059E+04 | 241.2 | - | - | 0 | - |
| - | - | 1.853E+04 | 243.1 | - | - | 0 | - |
| - | - | 3.494E+05 | 244.1 | - | - | 0 | - |
| - | - | 4.09E+04 | 245.1 | - | - | 0 | - |
| - | - | 5.17E+05 | 247.1 | - | - | 0 | - |
| - | - | 1.639E+04 | 247.1 | - | - | 0 | - |
| - | - | 2.97E+04 | 247.2 | - | - | 0 | - |
| - | - | 5.487E+04 | 248.1 | - | - | 0 | - |
| - | - | 1.99E+05 | 248.1 | - | - | 0 | - |
| - | - | 3.304E+05 | 249.1 | - | - | 0 | - |
| - | - | 1.739E+04 | 249.1 | - | - | 0 | - |
| - | - | 1.426E+05 | 249.1 | - | - | 0 | - |
| - | - | 3.395E+04 | 250.1 | - | - | 0 | - |
| - | - | 1.639E+04 | 250.1 | - | - | 0 | - |
| - | - | 1.057E+04 | 250.1 | - | - | 0 | - |
| - | - | 1.059E+04 | 250.2 | - | - | 0 | - |
| - | - | 1.369E+06 | 251.2 | - | - | 0 | - |
| - | - | 1.491E+04 | 252.1 | - | - | 0 | - |
| - | - | 1.936E+05 | 252.2 | - | - | 0 | - |
| - | - | 1.586E+04 | 252.2 | - | - | 0 | - |
| - | - | 1.298E+04 | 253.2 | - | - | 0 | - |
| - | - | 1.63E+04 | 254.1 | - | - | 0 | - |
| - | - | 1.019E+05 | 255.1 | - | - | 0 | - |
| - | - | 1.376E+04 | 256.1 | - | - | 0 | - |
| - | - | 1.324E+04 | 256.2 | - | - | 0 | - |
| - | - | 4.398E+04 | 257.1 | - | - | 0 | - |
| - | - | 1.234E+04 | 258.1 | - | - | 0 | - |
| - | - | 1.117E+05 | 258.1 | - | - | 0 | - |
| - | - | 1.051E+04 | 259.1 | - | - | 0 | - |
| - | - | 1.044E+04 | 261.1 | - | - | 0 | - |
| - | - | 2.098E+04 | 261.1 | - | - | 0 | - |
| - | - | 1.099E+04 | 261.2 | - | - | 0 | - |
| - | - | 2.376E+04 | 262.1 | - | - | 0 | - |
| - | - | 2.802E+05 | 265.1 | - | - | 0 | - |
| - | - | 9296 | 265.2 | - | - | 0 | - |
| - | - | 8.849E+05 | 266.1 | - | - | 0 | - |
| - | - | 8.356E+04 | 267.1 | - | - | 0 | - |
| - | - | 3.392E+04 | 268.1 | - | - | 0 | - |
| - | - | 7.084E+04 | 268.2 | - | - | 0 | - |
| - | - | 1.67E+04 | 269.2 | - | - | 0 | - |
| - | - | 4.161E+04 | 269.2 | - | - | 0 | - |
| - | - | 2.329E+04 | 270.1 | - | - | 0 | - |
| - | - | 1.106E+04 | 270.1 | - | - | 0 | - |
| - | - | 1.077E+05 | 272.2 | - | - | 0 | - |
| - | - | 1.791E+04 | 273.2 | - | - | 0 | - |
| - | - | 1.415E+05 | 274.1 | - | - | 0 | - |
| - | - | 2.815E+04 | 274.1 | - | - | 0 | - |
| - | - | 4.303E+04 | 275.1 | - | - | 0 | - |
| - | - | 1.723E+04 | 275.1 | - | - | 0 | - |
| - | - | 1.992E+04 | 275.2 | - | - | 0 | - |
| - | - | 2.33E+04 | 276.1 | - | - | 0 | - |
| - | - | 4.567E+04 | 276.2 | - | - | 0 | - |
| - | - | 4.449E+04 | 279.1 | - | - | 0 | - |
| - | - | 5.493E+04 | 280.1 | - | - | 0 | - |
| - | - | 1.104E+05 | 282.2 | - | - | 0 | - |
| - | - | 2.907E+04 | 283.1 | - | - | 0 | - |
| - | - | 1.49E+06 | 283.1 | - | - | 0 | - |
| - | - | 1.675E+04 | 284.1 | - | - | 0 | - |
| - | - | 5.782E+04 | 284.1 | - | - | 0 | - |
| - | - | 1.943E+05 | 284.1 | - | - | 0 | - |
| - | - | 9394 | 285.1 | - | - | 0 | - |
| - | - | 1.678E+04 | 285.1 | - | - | 0 | - |
| - | - | 3.862E+04 | 285.2 | - | - | 0 | - |
| - | - | 8.631E+04 | 286.1 | - | - | 0 | - |
| - | - | 3.541E+04 | 287.2 | - | - | 0 | - |
| - | - | 2.068E+04 | 287.2 | - | - | 0 | - |
| 12 | y | 1.176E+06 | 293.1 | 0.000724 | 2.47 | +1 | 2 |
| - | - | 6.049E+04 | 293.2 | - | - | 0 | - |
| - | - | 2.324E+05 | 294.1 | - | - | 0 | - |
| - | - | 1.782E+04 | 295.1 | - | - | 0 | - |
| - | - | 1.98E+04 | 296.1 | - | - | 0 | - |
| 3 | b | 5.431E+04 | 296.2 | 0.0006416 | 2.166 | +1 | 3 |
| - | - | 2.515E+05 | 297.2 | - | - | 0 | - |
| - | - | 4.428E+04 | 298.1 | - | - | 0 | - |
| - | - | 3.326E+04 | 298.2 | - | - | 0 | - |
| - | - | 2.123E+05 | 300.2 | - | - | 0 | - |
| - | - | 2.323E+04 | 301.1 | - | - | 0 | - |
| - | - | 4.856E+05 | 301.2 | - | - | 0 | - |
| - | - | 2.74E+04 | 301.2 | - | - | 0 | - |
| - | - | 2.026E+05 | 302.1 | - | - | 0 | - |
| - | - | 4.615E+04 | 302.2 | - | - | 0 | - |
| - | - | 3.749E+04 | 303.1 | - | - | 0 | - |
| - | - | 2.399E+04 | 310.2 | - | - | 0 | - |
| 12 | y | 1.791E+05 | 311.1 | 0.000871 | 2.8 | +1 | 2 |
| - | - | 1.524E+04 | 311.2 | - | - | 0 | - |
| - | - | 3.22E+04 | 312.1 | - | - | 0 | - |
| - | - | 8921 | 314.2 | - | - | 0 | - |
| 3 | b | 2.227E+05 | 314.2 | 0.0008496 | 2.704 | +1 | 3 |
| - | - | 3.662E+05 | 315.2 | - | - | 0 | - |
| - | - | 3.915E+04 | 315.2 | - | - | 0 | - |
| - | - | 4.981E+04 | 316.2 | - | - | 0 | - |
| - | - | 1.048E+04 | 318.1 | - | - | 0 | - |
| - | - | 3.761E+04 | 319.2 | - | - | 0 | - |
| - | - | 1.595E+04 | 320.2 | - | - | 0 | - |
| - | - | 1.471E+04 | 322.2 | - | - | 0 | - |
| - | - | 1.101E+04 | 329.2 | - | - | 0 | - |
| - | - | 3.208E+04 | 332.2 | - | - | 0 | - |
| - | - | 8.266E+04 | 333.2 | - | - | 0 | - |
| - | - | 1.055E+04 | 333.2 | - | - | 0 | - |
| - | - | 3.245E+04 | 334.2 | - | - | 0 | - |
| - | - | 8843 | 334.2 | - | - | 0 | - |
| - | - | 1.602E+04 | 338.2 | - | - | 0 | - |
| 6 | b | 1.518E+04 | 339.2 | 0.003595 | 10.6 | +2 | 6 |
| - | - | 3.554E+04 | 344.2 | - | - | 0 | - |
| - | - | 7.597E+04 | 345.1 | - | - | 0 | - |
| - | - | 3.168E+05 | 346.1 | - | - | 0 | - |
| - | - | 2.032E+04 | 346.1 | - | - | 0 | - |
| - | - | 5.39E+04 | 347.1 | - | - | 0 | - |
| - | - | 9038 | 348.2 | - | - | 0 | - |
| - | - | 1.02E+05 | 350.2 | - | - | 0 | - |
| - | - | 1.692E+04 | 351.2 | - | - | 0 | - |
| - | - | 1.476E+04 | 352.2 | - | - | 0 | - |
| - | - | 1.546E+04 | 352.2 | - | - | 0 | - |
| - | - | 1.726E+04 | 360.2 | - | - | 0 | - |
| - | - | 4.46E+04 | 361.2 | - | - | 0 | - |
| - | - | 4.486E+04 | 362.2 | - | - | 0 | - |
| - | - | 4.075E+05 | 363.1 | - | - | 0 | - |
| - | - | 5.251E+04 | 364.1 | - | - | 0 | - |
| - | - | 4.693E+04 | 364.1 | - | - | 0 | - |
| - | - | 2.735E+04 | 366.2 | - | - | 0 | - |
| - | - | 3.263E+04 | 367.1 | - | - | 0 | - |
| - | - | 1.718E+04 | 367.2 | - | - | 0 | - |
| - | - | 1.082E+04 | 368.2 | - | - | 0 | - |
| - | - | 2.486E+04 | 376.2 | - | - | 0 | - |
| - | - | 1.602E+04 | 377.1 | - | - | 0 | - |
| - | - | 1.849E+04 | 377.1 | - | - | 0 | - |
| - | - | 2.469E+05 | 378.2 | - | - | 0 | - |
| - | - | 4.271E+05 | 379.2 | - | - | 0 | - |
| - | - | 2.362E+04 | 379.2 | - | - | 0 | - |
| - | - | 1.47E+04 | 380.2 | - | - | 0 | - |
| - | - | 7.569E+04 | 380.2 | - | - | 0 | - |
| - | - | 9.742E+05 | 381.2 | - | - | 0 | - |
| - | - | 1.606E+05 | 382.2 | - | - | 0 | - |
| - | - | 2.651E+04 | 383.2 | - | - | 0 | - |
| - | - | 4.747E+04 | 384.2 | - | - | 0 | - |
| - | - | 3.89E+04 | 384.2 | - | - | 0 | - |
| - | - | 8.841E+04 | 385.2 | - | - | 0 | - |
| - | - | 1.178E+04 | 386.2 | - | - | 0 | - |
| - | - | 1.488E+04 | 386.2 | - | - | 0 | - |
| - | - | 1.332E+04 | 389.2 | - | - | 0 | - |
| - | - | 4.331E+04 | 391.1 | - | - | 0 | - |
| - | - | 7.181E+04 | 394.2 | - | - | 0 | - |
| - | - | 6.407E+04 | 395.1 | - | - | 0 | - |
| - | - | 1.522E+04 | 395.2 | - | - | 0 | - |
| - | - | 4.058E+04 | 395.2 | - | - | 0 | - |
| - | - | 1.676E+04 | 396.2 | - | - | 0 | - |
| - | - | 2.199E+05 | 396.2 | - | - | 0 | - |
| - | - | 1.771E+04 | 397.2 | - | - | 0 | - |
| - | - | 3.996E+04 | 397.2 | - | - | 0 | - |
| - | - | 5.259E+04 | 398.2 | - | - | 0 | - |
| - | - | 4.68E+04 | 402.2 | - | - | 0 | - |
| - | - | 5.03E+04 | 402.2 | - | - | 0 | - |
| - | - | 1.425E+04 | 403.1 | - | - | 0 | - |
| - | - | 2.957E+04 | 403.2 | - | - | 0 | - |
| - | - | 1.167E+04 | 404.2 | - | - | 0 | - |
| - | - | 8592 | 405.2 | - | - | 0 | - |
| - | - | 3.74E+04 | 405.3 | - | - | 0 | - |
| - | - | 5.8E+04 | 406.2 | - | - | 0 | - |
| - | - | 2.866E+04 | 407.2 | - | - | 0 | - |
| - | - | 2.098E+04 | 407.2 | - | - | 0 | - |
| - | - | 5.677E+04 | 409.1 | - | - | 0 | - |
| - | - | 5.475E+04 | 411.2 | - | - | 0 | - |
| - | - | 5.725E+04 | 412.2 | - | - | 0 | - |
| - | - | 2.783E+05 | 412.2 | - | - | 0 | - |
| - | - | 1.17E+05 | 413.1 | - | - | 0 | - |
| - | - | 3.082E+04 | 413.2 | - | - | 0 | - |
| - | - | 2.28E+04 | 413.3 | - | - | 0 | - |
| - | - | 3.244E+04 | 414.1 | - | - | 0 | - |
| - | - | 1.358E+05 | 414.2 | - | - | 0 | - |
| - | - | 7.836E+04 | 415.2 | - | - | 0 | - |
| - | - | 2.244E+04 | 415.2 | - | - | 0 | - |
| - | - | 2.5E+04 | 416.2 | - | - | 0 | - |
| 11 | y | 2.444E+04 | 421.2 | 0.0006096 | 1.447 | +1 | 3 |
| 11 | y | 1.475E+04 | 422.2 | 0.0002061 | 0.4882 | +1 | 3 |
| - | - | 6.368E+05 | 423.3 | - | - | 0 | - |
| - | - | 1.551E+04 | 424.2 | - | - | 0 | - |
| - | - | 1.556E+05 | 424.3 | - | - | 0 | - |
| - | - | 1.846E+04 | 425.3 | - | - | 0 | - |
| - | - | 1.897E+04 | 426.2 | - | - | 0 | - |
| - | - | 5.931E+04 | 426.7 | - | - | 0 | - |
| - | - | 3.516E+04 | 427.2 | - | - | 0 | - |
| - | - | 1.581E+05 | 429.2 | - | - | 0 | - |
| - | - | 1.582E+05 | 430.2 | - | - | 0 | - |
| - | - | 5.078E+05 | 430.2 | - | - | 0 | - |
| - | - | 4.254E+04 | 431.2 | - | - | 0 | - |
| - | - | 7.699E+04 | 431.2 | - | - | 0 | - |
| 7 | b | 1.419E+04 | 431.7 | 0.001112 | 2.577 | +2 | 7 |
| - | - | 1.318E+04 | 433.2 | - | - | 0 | - |
| 4 | b | 6.276E+04 | 433.3 | 0.001147 | 2.649 | +1 | 4 |
| - | - | 1.886E+04 | 434.3 | - | - | 0 | - |
| - | - | 2.064E+04 | 438.2 | - | - | 0 | - |
| 11 | y | 5.23E+05 | 439.2 | 0.001123 | 2.556 | +1 | 3 |
| - | - | 1.206E+05 | 440.2 | - | - | 0 | - |
| 7 | b | 2.83E+04 | 440.7 | 0.00175 | 3.972 | +2 | 7 |
| - | - | 2.54E+04 | 441.2 | - | - | 0 | - |
| - | - | 1.022E+04 | 443.2 | - | - | 0 | - |
| - | - | 4.3E+04 | 448.2 | - | - | 0 | - |
| - | - | 1.641E+04 | 449.2 | - | - | 0 | - |
| - | - | 1.184E+04 | 450.2 | - | - | 0 | - |
| - | - | 1.688E+04 | 450.2 | - | - | 0 | - |
| - | - | 1.802E+04 | 450.3 | - | - | 0 | - |
| 4 | b | 1.116E+06 | 451.3 | 0.001264 | 2.801 | +1 | 4 |
| - | - | 2.411E+05 | 452.3 | - | - | 0 | - |
| - | - | 2.905E+04 | 453.3 | - | - | 0 | - |
| - | - | 1.064E+04 | 455.2 | - | - | 0 | - |
| - | - | 1.438E+04 | 455.3 | - | - | 0 | - |
| - | - | 2.8E+04 | 459.2 | - | - | 0 | - |
| - | - | 3.297E+04 | 461.2 | - | - | 0 | - |
| - | - | 1.804E+04 | 466.2 | - | - | 0 | - |
| - | - | 2.867E+05 | 468.3 | - | - | 0 | - |
| - | - | 6.365E+04 | 469.3 | - | - | 0 | - |
| - | - | 1.461E+04 | 471.2 | - | - | 0 | - |
| - | - | 1.374E+04 | 472.2 | - | - | 0 | - |
| - | - | 3.127E+04 | 476.2 | - | - | 0 | - |
| - | - | 5.519E+04 | 477.2 | - | - | 0 | - |
| 10 | y | 7.347E+04 | 478.2 | 0.000743 | 1.554 | +1 | 4 |
| - | - | 2.624E+04 | 478.3 | - | - | 0 | - |
| - | - | 1.383E+04 | 479.2 | - | - | 0 | - |
| - | - | 1.922E+04 | 480.2 | - | - | 0 | - |
| - | - | 2.347E+04 | 481.2 | - | - | 0 | - |
| - | - | 2.478E+04 | 481.7 | - | - | 0 | - |
| - | - | 1.147E+04 | 482.7 | - | - | 0 | - |
| - | - | 1.701E+04 | 484.2 | - | - | 0 | - |
| 8 | b | 1.52E+04 | 488.3 | 0.008885 | 18.2 | +2 | 8 |
| - | - | 1.847E+04 | 494.2 | - | - | 0 | - |
| - | - | 5.094E+05 | 494.2 | - | - | 0 | - |
| - | - | 1.299E+05 | 495.2 | - | - | 0 | - |
| 10 | y | 1.113E+06 | 496.2 | 0.001113 | 2.243 | +1 | 4 |
| - | - | 2.871E+05 | 497.2 | - | - | 0 | - |
| - | - | 1.426E+04 | 497.8 | - | - | 0 | - |
| - | - | 5.589E+04 | 498.2 | - | - | 0 | - |
| - | - | 7.072E+04 | 504.2 | - | - | 0 | - |
| - | - | 1.448E+04 | 505.2 | - | - | 0 | - |
| - | - | 1.166E+04 | 506.2 | - | - | 0 | - |
| - | - | 2.843E+04 | 506.3 | - | - | 0 | - |
| - | - | 1.176E+04 | 507.3 | - | - | 0 | - |
| - | - | 1.059E+04 | 511.3 | - | - | 0 | - |
| - | - | 1.067E+04 | 514.2 | - | - | 0 | - |
| - | - | 2.192E+04 | 515.3 | - | - | 0 | - |
| - | - | 1.499E+04 | 521.2 | - | - | 0 | - |
| - | - | 1.669E+04 | 522.2 | - | - | 0 | - |
| - | - | 7870 | 523.2 | - | - | 0 | - |
| - | - | 9948 | 523.3 | - | - | 0 | - |
| - | - | 3.224E+04 | 524.3 | - | - | 0 | - |
| - | - | 6.298E+04 | 525.3 | - | - | 0 | - |
| - | - | 5.68E+04 | 526.2 | - | - | 0 | - |
| - | - | 1.142E+04 | 526.3 | - | - | 0 | - |
| - | - | 1.611E+04 | 526.3 | - | - | 0 | - |
| - | - | 2.564E+04 | 527.2 | - | - | 0 | - |
| - | - | 1.087E+04 | 527.3 | - | - | 0 | - |
| - | - | 1.512E+04 | 528.7 | - | - | 0 | - |
| - | - | 1.2E+04 | 529.2 | - | - | 0 | - |
| - | - | 1.605E+04 | 530.2 | - | - | 0 | - |
| - | - | 1.265E+04 | 531.2 | - | - | 0 | - |
| - | - | 6.387E+04 | 532.2 | - | - | 0 | - |
| - | - | 2.838E+04 | 533.2 | - | - | 0 | - |
| - | - | 1.036E+04 | 533.2 | - | - | 0 | - |
| - | - | 1.671E+04 | 534.3 | - | - | 0 | - |
| - | - | 1.241E+04 | 535.3 | - | - | 0 | - |
| - | - | 3.019E+04 | 537.2 | - | - | 0 | - |
| - | - | 3.775E+04 | 537.7 | - | - | 0 | - |
| - | - | 1.856E+04 | 538.2 | - | - | 0 | - |
| - | - | 1.374E+04 | 538.8 | - | - | 0 | - |
| - | - | 2.263E+05 | 539.2 | - | - | 0 | - |
| - | - | 5.707E+04 | 540.2 | - | - | 0 | - |
| - | - | 2.883E+04 | 541.3 | - | - | 0 | - |
| - | - | 1.231E+04 | 541.8 | - | - | 0 | - |
| - | - | 5.978E+04 | 542.3 | - | - | 0 | - |
| - | - | 7.572E+04 | 543.3 | - | - | 0 | - |
| - | - | 1.507E+04 | 544.3 | - | - | 0 | - |
| 9 | b | 4.545E+04 | 545.8 | 0.01072 | 19.64 | +2 | 9 |
| 9 | b | 6.688E+04 | 546.3 | 0.008954 | 16.39 | +2 | 9 |
| - | - | 2.338E+04 | 546.8 | - | - | 0 | - |
| - | - | 1.251E+04 | 547.2 | - | - | 0 | - |
| - | - | 4.603E+04 | 549.2 | - | - | 0 | - |
| - | - | 2.106E+04 | 550.2 | - | - | 0 | - |
| - | - | 4.462E+04 | 551.3 | - | - | 0 | - |
| - | - | 6.246E+04 | 551.3 | - | - | 0 | - |
| - | - | 2.426E+04 | 552.3 | - | - | 0 | - |
| - | - | 9.34E+04 | 554.8 | - | - | 0 | - |
| - | - | 2.185E+05 | 555.3 | - | - | 0 | - |
| - | - | 9.825E+04 | 555.8 | - | - | 0 | - |
| - | - | 2.888E+04 | 556.3 | - | - | 0 | - |
| 5 | b | 7.717E+04 | 561.3 | 0.001316 | 2.345 | +1 | 5 |
| - | - | 2.575E+04 | 562.3 | - | - | 0 | - |
| - | - | 1.532E+04 | 563.3 | - | - | 0 | - |
| - | - | 1.078E+04 | 563.3 | - | - | 0 | - |
| - | - | 1.196E+04 | 563.8 | - | - | 0 | - |
| - | - | 3.248E+04 | 564.3 | - | - | 0 | - |
| - | - | 5.329E+04 | 565.3 | - | - | 0 | - |
| - | - | 2.627E+05 | 567.2 | - | - | 0 | - |
| - | - | 1.589E+04 | 567.3 | - | - | 0 | - |
| - | - | 9.053E+04 | 568.2 | - | - | 0 | - |
| - | - | 2.37E+04 | 569.2 | - | - | 0 | - |
| - | - | 1.022E+04 | 569.3 | - | - | 0 | - |
| 10 | b | 2.202E+04 | 574.3 | 0.001375 | 2.394 | +2 | 10 |
| - | - | 9896 | 574.8 | - | - | 0 | - |
| - | - | 2.757E+04 | 575.2 | - | - | 0 | - |
| - | - | 1.607E+04 | 576.2 | - | - | 0 | - |
| - | - | 1.252E+04 | 577.3 | - | - | 0 | - |
| - | - | 1.151E+04 | 578.3 | - | - | 0 | - |
| 5 | b | 7.35E+05 | 579.3 | 0.0008835 | 1.525 | +1 | 5 |
| - | - | 2.148E+05 | 580.3 | - | - | 0 | - |
| - | - | 3.983E+04 | 581.3 | - | - | 0 | - |
| - | - | 5.717E+04 | 582.3 | - | - | 0 | - |
| - | - | 1.222E+04 | 582.8 | - | - | 0 | - |
| 10 | b | 2.525E+04 | 583.3 | 0.01166 | 19.98 | +2 | 10 |
| - | - | 1.818E+04 | 586.3 | - | - | 0 | - |
| - | - | 1.087E+04 | 587.8 | - | - | 0 | - |
| - | - | 2.643E+04 | 592.3 | - | - | 0 | - |
| - | - | 1.424E+04 | 592.3 | - | - | 0 | - |
| 9 | y | 1.701E+05 | 593.3 | 0.0007564 | 1.275 | +1 | 5 |
| - | - | 4.576E+04 | 593.3 | - | - | 0 | - |
| - | - | 5.304E+04 | 594.3 | - | - | 0 | - |
| - | - | 1.65E+04 | 595.3 | - | - | 0 | - |
| - | - | 1.865E+04 | 596.4 | - | - | 0 | - |
| - | - | 2.477E+04 | 597.8 | - | - | 0 | - |
| - | - | 1.442E+04 | 598.3 | - | - | 0 | - |
| - | - | 1.549E+04 | 599.3 | - | - | 0 | - |
| - | - | 3.069E+04 | 600.3 | - | - | 0 | - |
| - | - | 1.535E+04 | 601.3 | - | - | 0 | - |
| - | - | 1.752E+04 | 602.3 | - | - | 0 | - |
| - | - | 4.894E+04 | 602.8 | - | - | 0 | - |
| - | - | 4.665E+04 | 603.3 | - | - | 0 | - |
| - | - | 2.013E+04 | 603.8 | - | - | 0 | - |
| - | - | 2.735E+05 | 610.3 | - | - | 0 | - |
| 9 | y | 1.321E+06 | 611.3 | 0.001788 | 2.926 | +1 | 5 |
| - | - | 1.443E+05 | 611.8 | - | - | 0 | - |
| - | - | 3.531E+05 | 612.3 | - | - | 0 | - |
| - | - | 1.479E+04 | 612.8 | - | - | 0 | - |
| - | - | 7.18E+04 | 613.3 | - | - | 0 | - |
| - | - | 1.394E+04 | 615.3 | - | - | 0 | - |
| - | - | 3.054E+04 | 617.3 | - | - | 0 | - |
| - | - | 1.707E+04 | 621.3 | - | - | 0 | - |
| - | - | 1.091E+04 | 622.3 | - | - | 0 | - |
| - | - | 1.063E+04 | 622.4 | - | - | 0 | - |
| - | - | 1.314E+04 | 623.4 | - | - | 0 | - |
| - | - | 1.107E+04 | 624.3 | - | - | 0 | - |
| - | - | 1.282E+04 | 624.8 | - | - | 0 | - |
| - | - | 1.215E+04 | 626.3 | - | - | 0 | - |
| - | - | 1.914E+04 | 628.3 | - | - | 0 | - |
| - | - | 1.128E+04 | 628.8 | - | - | 0 | - |
| - | - | 2.074E+04 | 629.3 | - | - | 0 | - |
| - | - | 2.216E+04 | 629.8 | - | - | 0 | - |
| - | - | 3.219E+04 | 634.3 | - | - | 0 | - |
| - | - | 3.887E+04 | 635.3 | - | - | 0 | - |
| - | - | 2.383E+04 | 636.3 | - | - | 0 | - |
| 4 | y | 5.391E+04 | 636.8 | 0.003541 | 5.561 | +2 | 10 |
| 4 | y | 5.822E+04 | 637.3 | 0.01257 | 19.73 | +2 | 10 |
| - | - | 1.13E+05 | 637.8 | - | - | 0 | - |
| 11 | b | 1.94E+05 | 638.3 | 0.004736 | 7.419 | +2 | 11 |
| 11 | b | 1.294E+05 | 638.8 | 0.01236 | 19.35 | +2 | 11 |
| - | - | 4.537E+04 | 639.3 | - | - | 0 | - |
| - | - | 2.249E+04 | 639.8 | - | - | 0 | - |
| - | - | 4.73E+04 | 640.3 | - | - | 0 | - |
| - | - | 1.169E+04 | 641.3 | - | - | 0 | - |
| - | - | 1.304E+04 | 644.3 | - | - | 0 | - |
| - | - | 2.532E+05 | 645.3 | - | - | 0 | - |
| 4 | y | 5.956E+05 | 645.8 | 0.00485 | 7.511 | +2 | 10 |
| - | - | 3.893E+05 | 646.3 | - | - | 0 | - |
| - | - | 9.747E+04 | 646.8 | - | - | 0 | - |
| - | - | 7.506E+04 | 646.8 | - | - | 0 | - |
| 11 | b | 1.731E+05 | 647.3 | 0.004275 | 6.604 | +2 | 11 |
| - | - | 9.68E+04 | 647.8 | - | - | 0 | - |
| - | - | 4.352E+04 | 648.3 | - | - | 0 | - |
| - | - | 1.556E+04 | 650.3 | - | - | 0 | - |
| - | - | 1.701E+04 | 650.3 | - | - | 0 | - |
| - | - | 2.41E+05 | 652.3 | - | - | 0 | - |
| - | - | 9.816E+04 | 653.3 | - | - | 0 | - |
| - | - | 2.846E+04 | 654.3 | - | - | 0 | - |
| - | - | 3.04E+04 | 658.3 | - | - | 0 | - |
| - | - | 1.497E+04 | 661.8 | - | - | 0 | - |
| - | - | 7.141E+04 | 662.3 | - | - | 0 | - |
| - | - | 5.374E+04 | 663.3 | - | - | 0 | - |
| - | - | 1.876E+04 | 664.3 | - | - | 0 | - |
| - | - | 1.328E+04 | 665.3 | - | - | 0 | - |
| - | - | 1.337E+04 | 668.3 | - | - | 0 | - |
| 6 | b | 1.058E+05 | 676.3 | 0.001656 | 2.448 | +1 | 6 |
| 6 | b | 9.966E+04 | 677.3 | 0.00879 | 12.98 | +1 | 6 |
| - | - | 3.544E+04 | 678.3 | - | - | 0 | - |
| - | - | 2.781E+04 | 679.3 | - | - | 0 | - |
| - | - | 5.024E+05 | 680.3 | - | - | 0 | - |
| - | - | 1.287E+04 | 680.8 | - | - | 0 | - |
| - | - | 2.131E+05 | 681.3 | - | - | 0 | - |
| - | - | 3.1E+04 | 682.3 | - | - | 0 | - |
| - | - | 1.337E+04 | 684.3 | - | - | 0 | - |
| - | - | 1.474E+04 | 685.3 | - | - | 0 | - |
| - | - | 2.55E+04 | 688.9 | - | - | 0 | - |
| - | - | 2.822E+04 | 689.3 | - | - | 0 | - |
| - | - | 2.564E+04 | 689.8 | - | - | 0 | - |
| - | - | 2.812E+04 | 692.8 | - | - | 0 | - |
| 3 | y | 7.687E+04 | 693.3 | 0.005271 | 7.603 | +2 | 11 |
| - | - | 8.987E+04 | 693.8 | - | - | 0 | - |
| 6 | b | 1.355E+06 | 694.4 | 0.0007959 | 1.146 | +1 | 6 |
| - | - | 2.059E+04 | 694.8 | - | - | 0 | - |
| - | - | 5.227E+05 | 695.4 | - | - | 0 | - |
| - | - | 9895 | 696.3 | - | - | 0 | - |
| - | - | 8.004E+04 | 696.4 | - | - | 0 | - |
| - | - | 2.772E+04 | 697.3 | - | - | 0 | - |
| - | - | 3.171E+04 | 697.4 | - | - | 0 | - |
| - | - | 9.558E+04 | 697.9 | - | - | 0 | - |
| - | - | 7.278E+04 | 698.4 | - | - | 0 | - |
| - | - | 2.845E+04 | 698.9 | - | - | 0 | - |
| - | - | 2.7E+05 | 701.8 | - | - | 0 | - |
| 3 | y | 7.547E+05 | 702.3 | 0.005604 | 7.979 | +2 | 11 |
| 12 | b | 7.149E+05 | 702.8 | 0.005575 | 7.931 | +2 | 12 |
| 12 | b | 3.503E+05 | 703.3 | 0.005836 | 8.297 | +2 | 12 |
| - | - | 8.687E+04 | 703.9 | - | - | 0 | - |
| - | - | 1.314E+04 | 704.3 | - | - | 0 | - |
| - | - | 1.371E+04 | 705.3 | - | - | 0 | - |
| 8 | y | 5.293E+04 | 706.3 | 0.003484 | 4.933 | +1 | 6 |
| - | - | 1.916E+04 | 707.3 | - | - | 0 | - |
| - | - | 9768 | 707.8 | - | - | 0 | - |
| - | - | 1.093E+04 | 708.3 | - | - | 0 | - |
| - | - | 2.761E+04 | 710.4 | - | - | 0 | - |
| - | - | 6.99E+05 | 711.4 | - | - | 0 | - |
| 12 | b | 1.416E+06 | 711.9 | 0.004707 | 6.612 | +2 | 12 |
| - | - | 9.809E+05 | 712.4 | - | - | 0 | - |
| - | - | 3.299E+05 | 712.9 | - | - | 0 | - |
| - | - | 1.83E+04 | 713.4 | - | - | 0 | - |
| - | - | 3.583E+04 | 720.4 | - | - | 0 | - |
| - | - | 8.636E+04 | 720.9 | - | - | 0 | - |
| - | - | 6.202E+04 | 721.4 | - | - | 0 | - |
| - | - | 3.158E+04 | 721.9 | - | - | 0 | - |
| - | - | 1.195E+05 | 723.4 | - | - | 0 | - |
| 8 | y | 2.943E+05 | 724.4 | 0.003906 | 5.392 | +1 | 6 |
| - | - | 1.239E+05 | 725.4 | - | - | 0 | - |
| - | - | 2.864E+04 | 726.4 | - | - | 0 | - |
| - | - | 5.07E+04 | 728.4 | - | - | 0 | - |
| - | - | 7.717E+04 | 729.4 | - | - | 0 | - |
| - | - | 2.826E+04 | 730.4 | - | - | 0 | - |
| - | - | 1.265E+04 | 733.4 | - | - | 0 | - |
| - | - | 1.704E+04 | 734.4 | - | - | 0 | - |
| - | - | 1.343E+04 | 735.4 | - | - | 0 | - |
| - | - | 2.557E+04 | 743.4 | - | - | 0 | - |
| - | - | 2.459E+04 | 748.4 | - | - | 0 | - |
| - | - | 2.517E+04 | 749.3 | - | - | 0 | - |
| - | - | 1.673E+04 | 750.3 | - | - | 0 | - |
| - | - | 6.696E+04 | 751.4 | - | - | 0 | - |
| 2 | y | 3.848E+04 | 751.9 | 0.00344 | 4.576 | +2 | 12 |
| - | - | 5.699E+04 | 752.4 | - | - | 0 | - |
| - | - | 1.362E+04 | 752.9 | - | - | 0 | - |
| - | - | 1.375E+04 | 753.4 | - | - | 0 | - |
| - | - | 1.352E+04 | 757.4 | - | - | 0 | - |
| - | - | 2.548E+04 | 760.3 | - | - | 0 | - |
| - | - | 4.494E+04 | 763.4 | - | - | 0 | - |
| - | - | 5.291E+04 | 765.4 | - | - | 0 | - |
| - | - | 1.42E+04 | 766.4 | - | - | 0 | - |
| - | - | 1.712E+04 | 767.3 | - | - | 0 | - |
| - | - | 1.312E+04 | 770.9 | - | - | 0 | - |
| - | - | 1.112E+04 | 771.4 | - | - | 0 | - |
| - | - | 1.715E+04 | 771.9 | - | - | 0 | - |
| - | - | 2.851E+04 | 775.4 | - | - | 0 | - |
| - | - | 2.913E+04 | 775.9 | - | - | 0 | - |
| - | - | 5.955E+04 | 776.4 | - | - | 0 | - |
| - | - | 1.493E+04 | 777.3 | - | - | 0 | - |
| - | - | 1.857E+04 | 778.3 | - | - | 0 | - |
| - | - | 2.77E+04 | 778.9 | - | - | 0 | - |
| - | - | 1.024E+05 | 779.4 | - | - | 0 | - |
| - | - | 7.249E+04 | 779.9 | - | - | 0 | - |
| - | - | 1.349E+05 | 780.4 | - | - | 0 | - |
| - | - | 8.709E+04 | 780.9 | - | - | 0 | - |
| - | - | 2.99E+04 | 781.4 | - | - | 0 | - |
| - | - | 1.783E+04 | 782.4 | - | - | 0 | - |
| - | - | 1.048E+04 | 783.9 | - | - | 0 | - |
| - | - | 3.351E+04 | 784.4 | - | - | 0 | - |
| - | - | 6.968E+04 | 784.9 | - | - | 0 | - |
| - | - | 3.806E+04 | 785.4 | - | - | 0 | - |
| - | - | 1.135E+04 | 786.4 | - | - | 0 | - |
| - | - | 2.014E+05 | 792.9 | - | - | 0 | - |
| 0 | Precursor | 1.003E+06 | 793.4 | 0.006124 | 7.719 | +2 | -1 |
| 0 | Precursor | 8.118E+05 | 793.9 | 0.01106 | 13.94 | +2 | -1 |
| - | - | 4.572E+05 | 794.4 | - | - | 0 | - |
| - | - | 9.685E+04 | 794.9 | - | - | 0 | - |
| - | - | 1.585E+05 | 795.3 | - | - | 0 | - |
| - | - | 7.761E+04 | 796.3 | - | - | 0 | - |
| - | - | 2.173E+04 | 797.3 | - | - | 0 | - |
| - | - | 4.596E+06 | 801.9 | - | - | 0 | - |
| 0 | Precursor | 1.208E+07 | 802.4 | 0.004931 | 6.145 | +2 | -1 |
| - | - | 9.579E+06 | 802.9 | - | - | 0 | - |
| - | - | 4.031E+06 | 803.4 | - | - | 0 | - |
| - | - | 4.275E+05 | 803.9 | - | - | 0 | - |
| - | - | 4.619E+04 | 807.4 | - | - | 0 | - |
| - | - | 3.502E+04 | 808.4 | - | - | 0 | - |
| - | - | 1.339E+04 | 818.4 | - | - | 0 | - |
| - | - | 1.952E+04 | 819.4 | - | - | 0 | - |
| - | - | 1.032E+04 | 823.4 | - | - | 0 | - |
| - | - | 1.106E+04 | 824.4 | - | - | 0 | - |
| - | - | 6.717E+04 | 825.4 | - | - | 0 | - |
| - | - | 7.092E+04 | 826.4 | - | - | 0 | - |
| - | - | 4.16E+04 | 827.4 | - | - | 0 | - |
| - | - | 2.28E+04 | 834.4 | - | - | 0 | - |
| - | - | 4.15E+04 | 834.4 | - | - | 0 | - |
| - | - | 6.549E+04 | 835.4 | - | - | 0 | - |
| - | - | 8.345E+04 | 836.4 | - | - | 0 | - |
| - | - | 2.638E+04 | 837.4 | - | - | 0 | - |
| - | - | 1.467E+04 | 842.4 | - | - | 0 | - |
| - | - | 8.686E+04 | 843.4 | - | - | 0 | - |
| - | - | 1.365E+05 | 844.4 | - | - | 0 | - |
| - | - | 6.309E+04 | 845.4 | - | - | 0 | - |
| - | - | 4.405E+04 | 851.4 | - | - | 0 | - |
| - | - | 6.022E+04 | 852.4 | - | - | 0 | - |
| - | - | 3.573E+05 | 852.4 | - | - | 0 | - |
| - | - | 2.586E+04 | 853.4 | - | - | 0 | - |
| - | - | 1.79E+05 | 853.4 | - | - | 0 | - |
| - | - | 4.492E+04 | 854.4 | - | - | 0 | - |
| 7 | b | 1.726E+05 | 862.4 | 0.001445 | 1.675 | +1 | 7 |
| 7 | b | 1.415E+05 | 863.4 | 0.01151 | 13.33 | +1 | 7 |
| - | - | 1.019E+05 | 864.5 | - | - | 0 | - |
| - | - | 5.236E+04 | 865.5 | - | - | 0 | - |
| - | - | 1.288E+04 | 868.4 | - | - | 0 | - |
| - | - | 3.278E+04 | 869.4 | - | - | 0 | - |
| - | - | 1.055E+04 | 873.4 | - | - | 0 | - |
| - | - | 3.214E+04 | 876.4 | - | - | 0 | - |
| - | - | 1.627E+04 | 877.5 | - | - | 0 | - |
| 7 | b | 1.059E+06 | 880.4 | 0.001134 | 1.288 | +1 | 7 |
| - | - | 4.973E+05 | 881.4 | - | - | 0 | - |
| - | - | 1.271E+05 | 882.4 | - | - | 0 | - |
| - | - | 1.446E+04 | 883.4 | - | - | 0 | - |
| - | - | 1.765E+04 | 891.4 | - | - | 0 | - |
| 7 | y | 5.655E+04 | 892.4 | 0.005836 | 6.54 | +1 | 7 |
| - | - | 2.693E+04 | 893.4 | - | - | 0 | - |
| - | - | 2.874E+04 | 894.4 | - | - | 0 | - |
| - | - | 1.378E+04 | 902.4 | - | - | 0 | - |
| - | - | 4.444E+04 | 907.4 | - | - | 0 | - |
| - | - | 1.115E+05 | 908.4 | - | - | 0 | - |
| - | - | 2.375E+05 | 909.4 | - | - | 0 | - |
| 7 | y | 4.379E+05 | 910.4 | 0.004732 | 5.198 | +1 | 7 |
| - | - | 1.931E+05 | 911.4 | - | - | 0 | - |
| - | - | 6.19E+04 | 912.4 | - | - | 0 | - |
| - | - | 1.615E+04 | 919.4 | - | - | 0 | - |
| - | - | 1.918E+04 | 927.4 | - | - | 0 | - |
| - | - | 1.323E+04 | 930.5 | - | - | 0 | - |
| - | - | 2.188E+04 | 935.4 | - | - | 0 | - |
| - | - | 5.432E+04 | 936.4 | - | - | 0 | - |
| - | - | 6.218E+04 | 937.4 | - | - | 0 | - |
| - | - | 2.655E+04 | 938.4 | - | - | 0 | - |
| - | - | 1.876E+04 | 944.4 | - | - | 0 | - |
| - | - | 1.379E+04 | 945.4 | - | - | 0 | - |
| - | - | 1.682E+04 | 946.4 | - | - | 0 | - |
| - | - | 1.727E+04 | 947.5 | - | - | 0 | - |
| - | - | 3.282E+04 | 948.5 | - | - | 0 | - |
| - | - | 2.886E+04 | 949.5 | - | - | 0 | - |
| - | - | 2.08E+04 | 950.5 | - | - | 0 | - |
| - | - | 3.186E+04 | 953.4 | - | - | 0 | - |
| - | - | 1.138E+05 | 954.4 | - | - | 0 | - |
| - | - | 1.046E+05 | 955.4 | - | - | 0 | - |
| - | - | 6.032E+04 | 956.4 | - | - | 0 | - |
| - | - | 2.365E+04 | 958.5 | - | - | 0 | - |
| - | - | 1.461E+04 | 961.5 | - | - | 0 | - |
| - | - | 6.029E+04 | 962.5 | - | - | 0 | - |
| - | - | 3.054E+04 | 963.5 | - | - | 0 | - |
| - | - | 4.248E+04 | 964.5 | - | - | 0 | - |
| - | - | 1.635E+05 | 965.5 | - | - | 0 | - |
| - | - | 8.893E+04 | 966.5 | - | - | 0 | - |
| - | - | 2.7E+04 | 967.5 | - | - | 0 | - |
| - | - | 5.831E+04 | 971.5 | - | - | 0 | - |
| - | - | 6.4E+04 | 972.5 | - | - | 0 | - |
| - | - | 4.321E+04 | 973.4 | - | - | 0 | - |
| 8 | b | 1.922E+05 | 975.5 | 0.00106 | 1.086 | +1 | 8 |
| 8 | b | 1.556E+05 | 976.5 | 0.01357 | 13.89 | +1 | 8 |
| - | - | 6.514E+04 | 977.5 | - | - | 0 | - |
| - | - | 1.851E+04 | 978.5 | - | - | 0 | - |
| - | - | 4.99E+04 | 979.5 | - | - | 0 | - |
| - | - | 7.779E+04 | 980.5 | - | - | 0 | - |
| - | - | 4.614E+04 | 981.5 | - | - | 0 | - |
| - | - | 3.194E+04 | 982.5 | - | - | 0 | - |
| 8 | b | 1.418E+06 | 993.5 | 0.001176 | 1.184 | +1 | 8 |
| - | - | 8.553E+05 | 994.5 | - | - | 0 | - |
| - | - | 2.242E+05 | 995.5 | - | - | 0 | - |
| - | - | 3.318E+04 | 996.5 | - | - | 0 | - |
| - | - | 2.28E+04 | 1006 | - | - | 0 | - |
| 6 | y | 5.894E+04 | 1007 | 0.003613 | 3.586 | +1 | 8 |
| 6 | y | 3.049E+04 | 1008 | 0.01825 | 18.1 | +1 | 8 |
| - | - | 3.704E+04 | 1011 | - | - | 0 | - |
| - | - | 5.493E+04 | 1012 | - | - | 0 | - |
| - | - | 2.271E+04 | 1013 | - | - | 0 | - |
| - | - | 1.316E+05 | 1024 | - | - | 0 | - |
| 6 | y | 2.664E+05 | 1025 | 0.005438 | 5.303 | +1 | 8 |
| - | - | 1.436E+05 | 1026 | - | - | 0 | - |
| - | - | 3.696E+04 | 1027 | - | - | 0 | - |
| - | - | 1.746E+04 | 1046 | - | - | 0 | - |
| - | - | 1.377E+04 | 1047 | - | - | 0 | - |
| - | - | 1.241E+04 | 1056 | - | - | 0 | - |
| - | - | 4.785E+04 | 1063 | - | - | 0 | - |
| - | - | 3.531E+04 | 1064 | - | - | 0 | - |
| - | - | 1.818E+04 | 1065 | - | - | 0 | - |
| - | - | 1.48E+04 | 1066 | - | - | 0 | - |
| - | - | 2.801E+04 | 1073 | - | - | 0 | - |
| - | - | 6.426E+04 | 1076 | - | - | 0 | - |
| - | - | 4.46E+04 | 1077 | - | - | 0 | - |
| - | - | 2.091E+04 | 1080 | - | - | 0 | - |
| - | - | 1.241E+04 | 1081 | - | - | 0 | - |
| - | - | 1.286E+04 | 1082 | - | - | 0 | - |
| - | - | 1.796E+04 | 1090 | - | - | 0 | - |
| 9 | b | 6.692E+04 | 1091 | 0.005742 | 5.265 | +1 | 9 |
| 9 | b | 1.141E+05 | 1092 | 0.008312 | 7.615 | +1 | 9 |
| - | - | 4.831E+04 | 1093 | - | - | 0 | - |
| - | - | 5.956E+04 | 1094 | - | - | 0 | - |
| - | - | 3.343E+04 | 1095 | - | - | 0 | - |
| - | - | 3.039E+05 | 1108 | - | - | 0 | - |
| 9 | b | 7.811E+05 | 1109 | 0.004446 | 4.01 | +1 | 9 |
| - | - | 4.007E+05 | 1110 | - | - | 0 | - |
| - | - | 1.197E+05 | 1111 | - | - | 0 | - |
| - | - | 3.513E+04 | 1117 | - | - | 0 | - |
| - | - | 6.217E+04 | 1118 | - | - | 0 | - |
| - | - | 2.876E+04 | 1119 | - | - | 0 | - |
| - | - | 4.799E+04 | 1135 | - | - | 0 | - |
| 5 | y | 1.885E+05 | 1136 | 0.002713 | 2.39 | +1 | 9 |
| 5 | y | 2.009E+05 | 1136 | 0.01027 | 9.041 | +1 | 9 |
| - | - | 8.225E+04 | 1137 | - | - | 0 | - |
| - | - | 2.832E+04 | 1139 | - | - | 0 | - |
| - | - | 1.06E+04 | 1140 | - | - | 0 | - |
| - | - | 2.195E+04 | 1147 | - | - | 0 | - |
| 10 | b | 8.442E+04 | 1148 | 0.006608 | 5.758 | +1 | 10 |
| 10 | b | 7.335E+04 | 1149 | 0.01759 | 15.31 | +1 | 10 |
| - | - | 3.602E+04 | 1150 | - | - | 0 | - |
| - | - | 2.01E+05 | 1153 | - | - | 0 | - |
| 5 | y | 3.693E+05 | 1154 | 0.005332 | 4.623 | +1 | 9 |
| - | - | 1.839E+05 | 1155 | - | - | 0 | - |
| - | - | 6.171E+04 | 1156 | - | - | 0 | - |
| - | - | 2.474E+05 | 1165 | - | - | 0 | - |
| 10 | b | 4.583E+05 | 1166 | 0.006786 | 5.822 | +1 | 10 |
| - | - | 2.665E+05 | 1167 | - | - | 0 | - |
| - | - | 6.85E+04 | 1168 | - | - | 0 | - |
| - | - | 5.283E+04 | 1182 | - | - | 0 | - |
| - | - | 8.245E+04 | 1183 | - | - | 0 | - |
| - | - | 4.566E+04 | 1184 | - | - | 0 | - |
| - | - | 1.573E+04 | 1185 | - | - | 0 | - |
| - | - | 1.37E+04 | 1204 | - | - | 0 | - |
| - | - | 4.074E+04 | 1205 | - | - | 0 | - |
| - | - | 2.975E+04 | 1206 | - | - | 0 | - |
| - | - | 1.108E+04 | 1207 | - | - | 0 | - |
| - | - | 2.092E+04 | 1222 | - | - | 0 | - |
| - | - | 3.171E+04 | 1223 | - | - | 0 | - |
| - | - | 2.189E+04 | 1224 | - | - | 0 | - |
| - | - | 1.337E+04 | 1256 | - | - | 0 | - |
| - | - | 1.847E+04 | 1272 | - | - | 0 | - |
| 4 | y | 7.752E+04 | 1273 | 0.001663 | 1.307 | +1 | 10 |
| 4 | y | 5.774E+04 | 1274 | 0.01875 | 14.72 | +1 | 10 |
| - | - | 2.099E+04 | 1275 | - | - | 0 | - |
| 11 | b | 5.92E+04 | 1276 | 0.001563 | 1.225 | +1 | 11 |
| 11 | b | 3.643E+04 | 1277 | 0.01357 | 10.63 | +1 | 11 |
| - | - | 2.328E+04 | 1278 | - | - | 0 | - |
| - | - | 1.811E+05 | 1290 | - | - | 0 | - |
| 4 | y | 4.417E+05 | 1291 | 0.005014 | 3.885 | +1 | 10 |
| - | - | 2.654E+05 | 1292 | - | - | 0 | - |
| - | - | 5.243E+04 | 1293 | - | - | 0 | - |
| - | - | 5.796E+04 | 1293 | - | - | 0 | - |
| 11 | b | 2.031E+05 | 1294 | 0.005084 | 3.93 | +1 | 11 |
| - | - | 1.338E+05 | 1295 | - | - | 0 | - |
| - | - | 4.217E+04 | 1296 | - | - | 0 | - |
| 3 | y | 4.939E+04 | 1386 | 0.001102 | 0.7955 | +1 | 11 |
| 3 | y | 3.654E+04 | 1387 | 0.01989 | 14.34 | +1 | 11 |
| - | - | 1.469E+04 | 1388 | - | - | 0 | - |
| - | - | 7.648E+04 | 1403 | - | - | 0 | - |
| 3 | y | 2.637E+05 | 1404 | 0.00469 | 3.342 | +1 | 11 |
| 12 | b | 1.859E+05 | 1405 | 0.02182 | 15.53 | +1 | 12 |
| 12 | b | 5.237E+04 | 1406 | 0.00449 | 3.194 | +1 | 12 |
| - | - | 1.285E+04 | 1407 | - | - | 0 | - |
| - | - | 1.061E+04 | 2400 | - | - | 0 | - |

m/z Charge Intensity FragmentType MassShift Position
122.0716781616211 0 57939.004
125.10750579833984 0 8453.792
127.05064392089844 0 88707.72
127.08689880371094 0 16573.092
127.12310791015625 0 7831.92
127.65061950683594 0 5755.7173
128.08238220214844 0 9637.534
128.10740661621094 0 136233.69
129.06625366210938 0 22415.957
129.10269165039062 0 4636078.5
130.05001831054688 0 7621.4688
130.0600128173828 0 8979.275
130.06552124023438 0 266604
130.09979248046875 0 39497.5
130.1060028076172 0 269214.3
131.0689239501953 0 29373.145
131.11753845214844 0 6895.792
132.08111572265625 0 47851.88
136.07611083984375 0 340726.38
137.07948303222656 0 22013.342
138.06663513183594 0 75347.08
138.13894653320312 0 7460.694
139.08700561523438 0 60127.12
141.09808349609375 0 8138.7427
141.10269165039062 0 84399.59
143.04556274414062 0 23399.916
144.07704162597656 0 15963.572
145.06118774414062 0 47903.26
146.09310913085938 0 9434.226
147.04428100585938 0 14769.718
148.9547119140625 0 14104.36
150.0666046142578 0 92874.57
150.0916748046875 0 8646.314
151.0870361328125 0 112689.61
152.09066772460938 0 16818.666
153.1030731201172 0 10025.46
154.06167602539062 0 10041.831
154.0978240966797 0 17933.914
155.04579162597656 0 10665.062
155.09317016601562 0 493896
155.1182861328125 0 45079.83 a Water loss 1
156.0714569091797 0 7649.701
156.077392578125 0 33818.367
156.0966033935547 0 35245.105
157.0979461669922 0 11294.788
159.09213256835938 0 703849.5
160.0955352783203 0 72451.07
165.0551300048828 0 172791.3
165.0775909423828 0 9660.615
166.0615234375 0 345823.47
166.09791564941406 0 115039.875
167.06478881835938 0 23052.133
167.10105895996094 0 11480.376
167.1187286376953 0 18919.258
168.10238647460938 0 53533.3
168.11355590820312 0 40628.297
169.0763702392578 0 19690.285
169.09765625 0 128141.734
169.202392578125 0 7435.272
170.06040954589844 0 44742.82
170.10093688964844 0 7880.5635
171.09217834472656 0 12207.586
172.07215881347656 0 69395.11
173.0562744140625 0 28650.682
173.12893676757812 0 2983527.8 a 1
174.13233947753906 0 248254.62
176.0820770263672 0 18223.504
177.08566284179688 0 7711.444
177.10252380371094 0 12225.07
178.13430786132812 0 137147.16
179.11944580078125 0 7633.899
179.1380615234375 0 18714.562
181.06130981445312 0 32794.574
182.08164978027344 0 462899.6 y 12
183.08547973632812 0 49990.766
183.11314392089844 0 45696.965 b Water loss 1
185.1654510498047 0 14197.251
186.12417602539062 0 1266832.4
187.08705139160156 0 87805.27
187.12059020996094 0 9522.58
187.1275634765625 0 102186.96
188.09056091308594 0 11159.643
189.09817504882812 0 10325.751
190.0825958251953 0 47935.613
190.13436889648438 0 33352.223
193.09825134277344 0 11040.888
194.09278869628906 0 97012.5
195.1132354736328 0 309427.9
196.10845947265625 0 121600.93
196.11685180664062 0 21909.549
197.11224365234375 0 11074.808
197.12884521484375 0 16772.158
198.08787536621094 0 60694.547
198.12791442871094 0 22345.87
199.07217407226562 0 20648.506
200.14364624023438 0 19207.852
201.12379455566406 0 1762239.2 b 1
202.08670043945312 0 37509.004
202.12713623046875 0 156556.56
203.1289825439453 0 14217.309
205.0976104736328 0 115810.9
205.10842895507812 0 7369.9775
205.14520263671875 0 25909.117
206.1013946533203 0 13754.104
206.12928771972656 0 31415.414
207.15086364746094 0 9183.676
207.1608428955078 0 48759.49
208.1078643798828 0 8703.347
210.12454223632812 0 19209.059
211.07273864746094 0 8456.709
211.1082000732422 0 24223.07 y Water loss 10
212.10313415527344 0 9114.028
212.1400604248047 0 34757.82
213.12384033203125 0 132096.39
213.16017150878906 0 17735.342
214.127197265625 0 18441.785
216.09835815429688 0 47033.793
221.07113647460938 0 12968.25
221.09228515625 0 14413.887
221.103759765625 0 64544.36
221.14024353027344 0 50686.15
222.12411499023438 0 59514.066
223.08729553222656 0 15684.143
223.10813903808594 0 56836.113
223.15586853027344 0 622342.6
224.10330200195312 0 15624.965
224.11875915527344 0 11360.337
224.15928649902344 0 75224.91
226.082763671875 0 170784.97
227.06675720214844 0 96181.38
227.08645629882812 0 14412.701
227.15431213378906 0 18827.715
228.0698699951172 0 15903.187
228.134765625 0 40343.137
229.11846923828125 0 19979.203
230.1502227783203 0 20907.252
230.18682861328125 0 13458.184
233.14010620117188 0 69105.98
233.16542053222656 0 11528.497
234.12428283691406 0 409350.03
235.10743713378906 0 11048.866
235.12771606445312 0 40149.25
235.15574645996094 0 14692.259
237.1353302001953 0 25130.217
238.11900329589844 0 60317.227
238.13087463378906 0 32513.836
239.1033172607422 0 17903.84
239.1146697998047 0 32269.482
240.1348114013672 0 284504.8
241.09768676757812 0 16958.896
241.11878967285156 0 44223.438
241.13804626464844 0 24967.525
241.1920623779297 0 10591.483
243.1101531982422 0 18530.768
244.0934295654297 0 349356.78
245.09640502929688 0 40904.535
247.10838317871094 0 516962.5
247.14356994628906 0 16390.191
247.15602111816406 0 29699.674
248.10308837890625 0 54873.215
248.11434936523438 0 199049.14
249.0987548828125 0 330416.47
249.11949157714844 0 17386.268
249.13525390625 0 142563
250.1022491455078 0 33954.062
250.11781311035156 0 16387.727
250.13906860351562 0 10571.039
250.16641235351562 0 10589.329
251.15089416503906 0 1369259.4
252.13662719726562 0 14911.78
252.15406799316406 0 193606.2
252.16946411132812 0 15863.262
253.15628051757812 0 12983.359
254.0770721435547 0 16297.776
255.1480255126953 0 101884.18
256.1097717285156 0 13758.372
256.1520080566406 0 13242.482
257.1135559082031 0 43975.77
258.1236572265625 0 12335.776
258.1454772949219 0 111668.34
259.1479187011719 0 10513.277
261.12042236328125 0 10437.614
261.13543701171875 0 20978.477
261.16021728515625 0 10989.712
262.1189880371094 0 23760.443
265.1299133300781 0 280152.06
265.16552734375 0 9295.615
266.12530517578125 0 884891.3
267.1285400390625 0 83562.86
268.1302490234375 0 33917.176
268.1773681640625 0 70839.07
269.1614685058594 0 16697.246
269.18603515625 0 41607.676
270.1224365234375 0 23292.408
270.1451416015625 0 11059.089
272.17626953125 0 107662.49
273.1791076660156 0 17908.92
274.1190185546875 0 141512.97
274.13116455078125 0 28154.035
275.10296630859375 0 43026.59
275.12054443359375 0 17234.61
275.1506042480469 0 19919.564
276.10931396484375 0 23297.387
276.1554870605469 0 45670.062
279.1457824707031 0 44493.008
280.1298522949219 0 54934.527
282.1569519042969 0 110430.2
283.1037292480469 0 29074.514
283.14068603515625 0 1490122.8
284.08782958984375 0 16750.934
284.1039733886719 0 57822.414
284.1438903808594 0 194342.22
285.0885009765625 0 9393.504
285.1061096191406 0 16782.996
285.1567077636719 0 38622.875
286.14044189453125 0 86305.12
287.171875 0 35407.938
287.18792724609375 0 20677.316
293.1139221191406 0 1176194.4 y Water loss 11
293.1617431640625 0 60491.72
294.1178894042969 0 232359.38
295.12005615234375 0 17820.645
296.08868408203125 0 19801.81
296.197509765625 0 54311.207 b Water loss 2
297.156494140625 0 251465.28
298.1404113769531 0 44282.777
298.1595153808594 0 33257.656
300.1683349609375 0 212272.84
301.1146545410156 0 23229.936
301.1513366699219 0 485645.06
301.1705322265625 0 27403.277
302.1143493652344 0 202605.56
302.1553649902344 0 46147.57
303.11724853515625 0 37486.34
310.15301513671875 0 23993.504
311.1246337890625 0 179070.53 y 11
311.1724548339844 0 15243.617
312.1280212402344 0 32201.404
314.1880798339844 0 8921.197
314.2082824707031 0 222692.08 b 2
315.1670227050781 0 366206
315.21099853515625 0 39149.83
316.17120361328125 0 49807.87
318.1204833984375 0 10484.454
319.1519775390625 0 37607.676
320.1729736328125 0 15949.293
322.2242431640625 0 14708.237
329.18359375 0 11008.58
332.2087097167969 0 32081.072
333.1778259277344 0 82662.54
333.2122497558594 0 10547.104
334.186279296875 0 32452.297
334.2239074707031 0 8843.477
338.18206787109375 0 16015.605
339.1698913574219 0 15181.864 b Ammonia loss 5
344.17230224609375 0 35540.277
345.1314392089844 0 75973.234
346.1153564453125 0 316790.94
346.1370849609375 0 20315.158
347.1188659667969 0 53899.227
348.167236328125 0 9037.892
350.21978759765625 0 101998.29
351.16619873046875 0 16918.357
352.1658020019531 0 14758.476
352.1997985839844 0 15460.356
360.2039489746094 0 17264.133
361.1985168457031 0 44601.188
362.1825866699219 0 44860.246
363.1417541503906 0 407453.94
364.12493896484375 0 52514.41
364.14569091796875 0 46930.33
366.1781311035156 0 27352.844
367.1405334472656 0 32633.203
367.1620178222656 0 17177.03
368.1958923339844 0 10816.107
376.1628723144531 0 24855.178
377.1241149902344 0 16016.592
377.14727783203125 0 18487.357
378.2143249511719 0 246850.86
379.2102355957031 0 427144.12
379.2441711425781 0 23623.889
380.165283203125 0 14704.703
380.212890625 0 75687.766
381.1524658203125 0 974175.25
382.1552429199219 0 160641.1
383.1575927734375 0 26505.742
384.1666259765625 0 47469.4
384.18975830078125 0 38896.062
385.15142822265625 0 88413.77
386.15557861328125 0 11784.012
386.1812438964844 0 14875.628
389.1933898925781 0 13317.567
391.1368713378906 0 43312.703
394.17340087890625 0 71809.93
395.1357421875 0 64065.39
395.1541748046875 0 15220.626
395.24139404296875 0 40580.484
396.1927795410156 0 16760.672
396.2251892089844 0 219918.27
397.1881408691406 0 17712.605
397.2275695800781 0 39963.92
398.176513671875 0 52588.57
402.1773681640625 0 46799.61
402.19989013671875 0 50303.254
403.1365966796875 0 14245.566
403.19927978515625 0 29570.545
404.1819763183594 0 11666.886
405.222412109375 0 8591.955
405.2623291015625 0 37401.508
406.2458190917969 0 58000.652
407.20458984375 0 28655.29
407.2439880371094 0 20981.271
409.1477966308594 0 56767.01
411.19964599609375 0 54752.72
412.1610107421875 0 57247.793
412.18389892578125 0 278342.34
413.1461181640625 0 117026.305
413.1878662109375 0 30818.086
413.25213623046875 0 22799.035
414.148681640625 0 32443.393
414.2357482910156 0 135813.61
415.19842529296875 0 78355.984
415.24041748046875 0 22443.494
416.202392578125 0 24997.031
421.2087707519531 0 24444.348 y Water loss 10
422.1923828125 0 14752.506 y Ammonia loss 10
423.2724304199219 0 636830.9
424.2203063964844 0 15505.867
424.27520751953125 0 155615.92
425.2763671875 0 18463.51
426.1993713378906 0 18973.293
426.72320556640625 0 59312.07
427.22369384765625 0 35161.543
429.21026611328125 0 158057.7
430.16937255859375 0 158234.1
430.1947326660156 0 507771.2
431.1739196777344 0 42538.508
431.19830322265625 0 76993.68
431.7150573730469 0 14193.62 b Water loss 6
433.2086181640625 0 13176.957
433.2569274902344 0 62760.594 b Water loss 3
434.2599182128906 0 18864.35
438.1744384765625 0 20639.152
439.2198486328125 0 523007.12 y 10
440.2230529785156 0 120583.82
440.7209777832031 0 28304.404 b 6
441.2236328125 0 25397.588
443.24237060546875 0 10217.08
448.20550537109375 0 42998.375
449.2115478515625 0 16412.139
450.1744384765625 0 11838.386
450.2359313964844 0 16876.19
450.2840576171875 0 18017.654
451.2676086425781 0 1116479.5 b 3
452.2704162597656 0 241118.8
453.2726135253906 0 29052.121
455.19769287109375 0 10635.873
455.2657470703125 0 14382.06
459.1998596191406 0 27998.002
461.2041015625 0 32967.727
466.2414245605469 0 18035.926
468.2938537597656 0 286725.66
469.2970886230469 0 63646.055
471.2340393066406 0 14608.41
472.2235412597656 0 13735.842
476.2271423339844 0 31272.28
477.2108154296875 0 55186.793
478.2288818359375 0 73472.73 y Water loss 9
478.2783203125 0 26239.84
479.23553466796875 0 13829.513
480.22369384765625 0 19223.748
481.23504638671875 0 23466.379
481.7300109863281 0 24777.662
482.7347106933594 0 11468.17
484.22052001953125 0 17012.045
488.2648620605469 0 15197.588 b Water loss 7
494.1947937011719 0 18473.16
494.23681640625 0 509374.38
495.239501953125 0 129855.586
496.2413024902344 0 1112814.4 y 9
497.2455139160156 0 287129.12
497.7645263671875 0 14258.547
498.2413024902344 0 55890.71
504.20086669921875 0 70722.91
505.197998046875 0 14483.3125
506.2349548339844 0 11658.639
506.2722473144531 0 28429.51
507.25701904296875 0 11755.584
511.266357421875 0 10589.628
514.1886596679688 0 10673.875
515.2623291015625 0 21917.88
521.2247314453125 0 14990.081
522.2084350585938 0 16688.098
523.2460327148438 0 7869.532
523.2666625976562 0 9948.463
524.2799072265625 0 32238.957
525.2672119140625 0 62975.125
526.2315673828125 0 56795.766
526.2594604492188 0 11422.161
526.2713012695312 0 16105.1045
527.23779296875 0 25635.928
527.2794799804688 0 10872.019
528.7366943359375 0 15121.858
529.2332763671875 0 11995.951
530.2255859375 0 16052.679
531.2171630859375 0 12651.498
532.1961059570312 0 63870.78
533.1991577148438 0 28384.787
533.2485961914062 0 10356.475
534.306640625 0 16712.299
535.301513671875 0 12405.668
537.2465209960938 0 30188.918
537.7404174804688 0 37745.45
538.2367553710938 0 18558.592
538.776123046875 0 13739.667
539.2369995117188 0 226290.34
540.2394409179688 0 57071.297
541.2538452148438 0 28832.61
541.76025390625 0 12308.271
542.2935180664062 0 59784.02
543.2588500976562 0 75721.67
544.2548828125 0 15073.586
545.7587280273438 0 45452.016 b Water loss 8
546.2525024414062 0 66875.61 b Ammonia loss 8
546.7512817382812 0 23378.357
547.2462158203125 0 12514.715
549.2220458984375 0 46026.332
550.2042846679688 0 21062.576
551.2579956054688 0 44618.277
551.3306884765625 0 62464.77
552.259765625 0 24264.709
554.7633056640625 0 93396.39
555.2573852539062 0 218480.9
555.7586669921875 0 98252.06
556.2596435546875 0 28877.234
561.315673828125 0 77165.86 b Water loss 4
562.3114013671875 0 25749.27
563.25927734375 0 15321.35
563.3007202148438 0 10780.494
563.7704467773438 0 11964.233
564.2627563476562 0 32480.963
565.266357421875 0 53285.387
567.2318725585938 0 262716.62
567.2816772460938 0 15892.447
568.2347412109375 0 90527.766
569.2366943359375 0 23696.56
569.2844848632812 0 10219.724
574.2815551757812 0 22019.357 b Water loss 9
574.7849731445312 0 9895.692
575.2469482421875 0 27573.676
576.23583984375 0 16073.665
577.2764282226562 0 12519.332
578.3431396484375 0 11505.239
579.3258056640625 0 734965.44 b 4
580.3289794921875 0 214771.12
581.3302612304688 0 39828.04
582.3046264648438 0 57174.38
582.7933349609375 0 12223.134
583.297119140625 0 25247.012 b 9
586.25830078125 0 18183.943
587.8092041015625 0 10874.682
592.2732543945312 0 26425.094
592.3240966796875 0 14243.168
593.25732421875 0 170051.69 y Water loss 8
593.3057861328125 0 45764.035
594.255126953125 0 53036.82
595.2545776367188 0 16498.854
596.3526000976562 0 18652.988
597.8037719726562 0 24769.982
598.298095703125 0 14419.716
599.3301391601562 0 15492.608
600.3172607421875 0 30694.918
601.3209838867188 0 15353.946
602.3020629882812 0 17521.986
602.7950439453125 0 48937.637
603.2928466796875 0 46652.316
603.7902221679688 0 20127.094
610.2841796875 0 273517.38
611.2689208984375 0 1320761.8 y 8
611.8004760742188 0 144254.22
612.2725830078125 0 353062.97
612.7966918945312 0 14787.061
613.2734985351562 0 71795.9
615.31591796875 0 13937.136
617.284912109375 0 30537.793
621.3060913085938 0 17069.031
622.3046264648438 0 10908.241
622.3671264648438 0 10631.043
623.3740844726562 0 13141.652
624.328857421875 0 11067.797
624.8284301757812 0 12816.712
626.295166015625 0 12149.505
628.277099609375 0 19140.764
628.7761840820312 0 11278.85
629.3248291015625 0 20742.217
629.8198852539062 0 22155.727
634.30908203125 0 32190.445
635.2947998046875 0 38865.83
636.2952880859375 0 23833.549
636.7893676757812 0 53914.8 y Water loss 3
637.2904052734375 0 58222.035 y Ammonia loss 3
637.8368530273438 0 113012.375
638.3323974609375 0 193993.67 b Water loss 10
638.83203125 0 129398.92 b Ammonia loss 10
639.33349609375 0 45367.723
639.83056640625 0 22485.121
640.2745361328125 0 47303.992
641.3096313476562 0 11691.879
644.2979736328125 0 13035.852
645.2989501953125 0 253213.55
645.7959594726562 0 595617.4 y 3
646.2952880859375 0 389264.78
646.79443359375 0 97471.766
646.8434448242188 0 75058.91
647.3372192382812 0 173111.27 b 10
647.8370971679688 0 96798.67
648.3396606445312 0 43515.406
650.268798828125 0 15559.705
650.3238525390625 0 17005.426
652.3215942382812 0 240955.06
653.32421875 0 98163.55
654.3262329101562 0 28457.203
658.2842407226562 0 30399.553
661.8367309570312 0 14967.553
662.3065795898438 0 71410
663.2951049804688 0 53735.543
664.3433837890625 0 18759.12
665.3427124023438 0 13284.394
668.2819213867188 0 13365.8125
676.3429565429688 0 105787.08 b Water loss 5
677.3341064453125 0 99658.42 b Ammonia loss 5
678.3367919921875 0 35442.17
679.3311767578125 0 27805.988
680.316650390625 0 502350.84
680.846923828125 0 12872.403
681.3198852539062 0 213143.83
682.3211059570312 0 30996.2
684.3314819335938 0 13366.712
685.328125 0 14737.977
688.8519287109375 0 25498.146
689.3493041992188 0 28224.117
689.8477172851562 0 25638.012
692.837158203125 0 28124.73
693.3331298828125 0 76873.555 y Water loss 2
693.8357543945312 0 89865.11
694.3526611328125 0 1355308 b 5
694.8355712890625 0 20587.693
695.3555908203125 0 522698.75
696.3035278320312 0 9894.838
696.3580932617188 0 80039.07
697.294677734375 0 27720.99
697.3569946289062 0 31707.04
697.8610229492188 0 95580.055
698.3565673828125 0 72776.055
698.8597412109375 0 28452.28
701.8421020507812 0 269975.38
702.3387451171875 0 754690.06 y 2
702.8433837890625 0 714920.2 b Water loss 11
703.3468017578125 0 350302.12 b Ammonia loss 11
703.8508911132812 0 86865.97
704.3473510742188 0 13138.517
705.348876953125 0 13713.9
706.3441162109375 0 52934.016 y Water loss 7
707.3429565429688 0 19162.238
707.8286743164062 0 9767.543
708.32373046875 0 10930.1045
710.363037109375 0 27607.434
711.3627319335938 0 698996.9
711.8589477539062 0 1416165 b 11
712.359130859375 0 980933.9
712.8596801757812 0 329902.97
713.3571166992188 0 18302.988
720.3701171875 0 35828.113
720.8634643554688 0 86360.81
721.3644409179688 0 62019.395
721.8638916015625 0 31581.988
723.3688354492188 0 119496.31
724.3551025390625 0 294326.88 y 7
725.357421875 0 123920.695
726.361572265625 0 28636.744
728.374267578125 0 50701.91
729.361572265625 0 77169.195
730.3634033203125 0 28260.238
733.3675537109375 0 12649.112
734.357666015625 0 17043.885
735.35888671875 0 13433.1
743.371337890625 0 25566.959
748.38037109375 0 24592.87
749.340576171875 0 25167.473
750.3356323242188 0 16728.21
751.3755493164062 0 66957.45
751.8707885742188 0 38475.414 y 1
752.3770141601562 0 56990.797
752.8723754882812 0 13617.649
753.3786010742188 0 13752.972
757.3666381835938 0 13518.768
760.3084106445312 0 25481.822
763.35888671875 0 44942.426
765.4058837890625 0 52908.66
766.4115600585938 0 14196.5205
767.3489990234375 0 17118.623
770.8845825195312 0 13117.627
771.3815307617188 0 11123.153
771.878662109375 0 17151.62
775.3857421875 0 28505.852
775.8706665039062 0 29126.953
776.3759765625 0 59548.723
777.3279418945312 0 14931.988
778.3263549804688 0 18568.355
778.8987426757812 0 27695.14
779.388916015625 0 102442.82
779.8909301757812 0 72494.695
780.3839721679688 0 134887.02
780.88232421875 0 87092.61
781.3817749023438 0 29899.38
782.3707885742188 0 17825.021
783.8897705078125 0 10483.974
784.382568359375 0 33514.336
784.8807983398438 0 69675.195
785.3836669921875 0 38061.254
786.373779296875 0 11348.746
792.8956298828125 0 201403.67
793.3920288085938 0 1003178.3 Precursor Water loss
793.8889770507812 0 811763.6 Precursor Ammonia loss
794.3898315429688 0 457200.16
794.88720703125 0 96845.21
795.34619140625 0 158506.05
796.347900390625 0 77611.52
797.3487548828125 0 21729.738
801.9007568359375 0 4596073
802.3961181640625 0 12078552 Precursor
802.8961181640625 0 9579024
803.396728515625 0 4030611.2
803.8971557617188 0 427525.3
807.3780517578125 0 46186.164
808.375244140625 0 35020.805
818.424072265625 0 13393.219
819.4381103515625 0 19524.037
823.3907470703125 0 10317.943
824.3742065429688 0 11057.517
825.3900756835938 0 67167.45
826.3816528320312 0 70915.81
827.3795166015625 0 41604.215
834.35400390625 0 22795.498
834.427490234375 0 41502.21
835.4154052734375 0 65492.99
836.4417724609375 0 83451.38
837.44775390625 0 26384.756
842.4171142578125 0 14669.748
843.4019165039062 0 86858.7
844.3925170898438 0 136515.73
845.39501953125 0 63094.492
851.3807983398438 0 44054.125
852.36279296875 0 60222.742
852.4380493164062 0 357255.25
853.364990234375 0 25857.045
853.4403686523438 0 179016.3
854.4420776367188 0 44923.434
862.4220581054688 0 172619.12 b Water loss 6
863.4161376953125 0 141545.73 b Ammonia loss 6
864.451904296875 0 101895.96
865.4584350585938 0 52356.355
868.4019775390625 0 12881.148
869.394775390625 0 32784.344
873.408935546875 0 10545.953
876.4468383789062 0 32142.615
877.4501953125 0 16265.343
880.4323120117188 0 1058946.5 b 6
881.4349365234375 0 497300.1
882.437744140625 0 127074.484
883.4393310546875 0 14459.965
891.4255981445312 0 17646.959
892.42578125 0 56552.445 y Water loss 6
893.4224853515625 0 26934.768
894.4439697265625 0 28743.717
902.4470825195312 0 13778.626
907.4425659179688 0 44440.758
908.4309692382812 0 111477.43
909.4442749023438 0 237456.72
910.4352416992188 0 437916.1 y 6
911.4375 0 193125.16
912.4373779296875 0 61903.12
919.3998413085938 0 16151.308
927.4226684570312 0 19176.752
930.4865112304688 0 13226.441
935.4376831054688 0 21883.85
936.4265747070312 0 54318.223
937.4155883789062 0 62184.55
938.418701171875 0 26548.385
944.44677734375 0 18756.05
945.442138671875 0 13789.697
946.4478149414062 0 16824.19
947.5175170898438 0 17271.457
948.50146484375 0 32818.64
949.5007934570312 0 28859.139
950.5014038085938 0 20804.65
953.4468383789062 0 31858.775
954.4354248046875 0 113825.25
955.4249877929688 0 104629.23
956.4220581054688 0 60315.613
958.4827880859375 0 23645.51
961.4656372070312 0 14608.561
962.4517822265625 0 60288.574
963.4515991210938 0 30539.914
964.4619750976562 0 42483.285
965.5209350585938 0 163487.39
966.5230712890625 0 88930.29
967.5248413085938 0 27003.568
971.4591674804688 0 58310.61
972.4501342773438 0 63998.98
973.4476318359375 0 43211.56
975.5057373046875 0 192237 b Water loss 7
976.5022583007812 0 155626.86 b Ammonia loss 7
977.5011596679688 0 65140.46
978.507080078125 0 18513.332
979.475830078125 0 49903.715
980.466796875 0 77789.43
981.4663696289062 0 46137.746
982.4727783203125 0 31943.65
993.5164184570312 0 1418142 b 7
994.51904296875 0 855274.9
995.5213012695312 0 224230.31
996.5206909179688 0 33177.48
1006.4669799804688 0 22798.178
1007.4505004882812 0 58937.92 y Water loss 5
1008.4491577148438 0 30486.973 y Ammonia loss 5
1010.5450439453125 0 37041.023
1011.531982421875 0 54926.75
1012.5311279296875 0 22706.78
1024.47509765625 0 131583.36
1025.462890625 0 266386.5 y 5
1026.46484375 0 143631.5
1027.4615478515625 0 36956.18
1045.5157470703125 0 17458.857
1046.52734375 0 13772.868
1056.4599609375 0 12408.907
1062.5361328125 0 47847.676
1063.5360107421875 0 35311.133
1064.5367431640625 0 18181.428
1065.505859375 0 14798.758
1073.490966796875 0 28008.865
1075.531982421875 0 64261.13
1076.5340576171875 0 44602.562
1079.5611572265625 0 20911.768
1080.5579833984375 0 12408.019
1081.503662109375 0 12863.978
1089.5458984375 0 17955.254
1090.52587890625 0 66923.945 b Water loss 8
1091.50732421875 0 114132.516 b Ammonia loss 8
1092.5133056640625 0 48307.293
1093.5423583984375 0 59556.242
1094.54736328125 0 33432.72
1107.5582275390625 0 303851.03
1108.546630859375 0 781108.5 b 8
1109.546142578125 0 400679.16
1110.5455322265625 0 119656.74
1117.494873046875 0 35133.734
1118.4891357421875 0 62168.008
1119.4893798828125 0 28759.9
1134.52392578125 0 47990.36
1135.5081787109375 0 188510.86 y Water loss 4
1136.499755859375 0 200904.17 y Ammonia loss 4
1137.4991455078125 0 82248.305
1138.576171875 0 28324.912
1139.5738525390625 0 10596.166
1146.5706787109375 0 21949.645
1147.5596923828125 0 84420.305 b Water loss 9
1148.5546875 0 73351.27 b Ammonia loss 9
1149.5531005859375 0 36020.125
1152.5325927734375 0 201021.66
1153.5213623046875 0 369299.7 y 4
1154.5230712890625 0 183932.34
1155.52197265625 0 61707.18
1164.5802001953125 0 247428.73
1165.5704345703125 0 458293.2 b 9
1166.5714111328125 0 266548.62
1167.5740966796875 0 68499.805
1181.608154296875 0 52828.754
1182.598388671875 0 82447.016
1183.5948486328125 0 45656.688
1184.5916748046875 0 15732.924
1203.59228515625 0 13702.867
1204.580322265625 0 40735.21
1205.580078125 0 29748.5
1206.5938720703125 0 11076.433
1221.602294921875 0 20921.738
1222.591552734375 0 31712.662
1223.58544921875 0 21885.309
1255.5518798828125 0 13373.385
1271.5836181640625 0 18469.447
1272.5660400390625 0 77515.984 y Water loss 3
1273.567138671875 0 57740.82 y Ammonia loss 3
1274.5667724609375 0 20993.438
1275.646484375 0 59199.35 b Water loss 10
1276.6456298828125 0 36429.17 b Ammonia loss 10
1277.6475830078125 0 23276.154
1289.589111328125 0 181133.17
1290.5799560546875 0 441747.66 y 3
1291.5804443359375 0 265373.22
1292.572998046875 0 52431.484
1292.6754150390625 0 57962.715
1293.6636962890625 0 203109.38 b 10
1294.664794921875 0 133822.34
1295.665283203125 0 42165.848
1385.6473388671875 0 49389.06 y Water loss 2
1386.65234375 0 36539.992 y Ammonia loss 2
1387.652099609375 0 14688.38
1402.67724609375 0 76477.19
1403.6636962890625 0 263692.7 y 2
1404.6688232421875 0 185939.14 b Water loss 11
1405.670166015625 0 52366.46 b Ammonia loss 11
1406.67578125 0 12850.555
2400.17919921875 0 10610.668

Spectrum Details

|  |  |
| --- | --- |
| Matched peaks? Matched peaks The total absolute number of peaks matched. Additionally in brackets the total fraction of peaks matched and the total number of peaks is shown. | 80 (9.84% of 813) |
| FDR? FDR The false discovery rate estimated for this peptide. It is calculated by matching all theoretical fragments with a non-integer shift with the raw peaks for this spectrum. This is done with 40 different shifts. The resulting percentage is the average number of annotated peaks over the number of annotated peaks with the correct spectrum. | 1.07% |
| Satellite FDR? Satellite FDR See the FDR for details on its calculation. This satellite ion specific FDR only contains the satellite ions (d/w) for I/L/J positions. | - |
| PSM Score? PSM Score The PSM Score as given by Hecklib to this annotated spectrum. It is shown with three significant figures. | 978 |

#### Spectrum 8727? Spectrum 8727 The raw spectrum of this peptide as annotated by Hecklib. The fragments are coloured according to ion type (see legend). Any peaks with a star '\*' as text can be hovered over to see the full details, first the ion type second the mass shift type. By hovering over the amino acids in the peptide or ions in the legend the corresponding peaks are highlighted. By toggling the 'Unassigned' label you can turn the background (unassigned) peaks on or off in the plot. By updating the slider in the Ion legend you can update the spectrum to only show the top X% of the peaks with labels. The top X% means any peak that is within X% of the highest intensity. By dragging in the spectrum you can zoom in to a specific part of the spectrum and use 'Zoom Out' to get back to the original zoom level. The annotation of the spectrum is based on the given sequence in the peptides file and is done with different software so inconsistencies are likely. The peaks are annotated based on the given sequence, with 20 ppm tolerance.

Copy Data

##### Spectrum 8727 (TSV)

###### Preview

```
Loading example...
```

*Click on the button to copy the data to your clipboard.*

Mz MinMz MaxIntensity Max

WidthHeightPeptide font sizePeptide stroke widthSpectrum font sizeSpectrum stroke widthCompact peptide

Ion legend

wxyz

abcd

OtherUnassignedIon^Charge^_Position_Show for top:%

TVLHQDWLDGKEY

02.72e+45.44e+48.16e+41.09e+5

Zoom Out

y^+3^_4_y^+1^_1_z^+3^_5_c^+1^_2_z^+1^_2_c^+1^_3_y^+1^_3_c^+2^_7_c^+1^_4_z^+1^_4_y^+1^_4_w^+1^_5_c^+1^_5_z^+1^_5_c^+1^_5_y^+1^_5_z^+2^_10_y^+2^_10_c^+2^_11_w^+1^_6_w^+2^_11_y^+2^_11_c^+1^_6_y^+2^_11_z^+1^_6_c^+1^_6_c^+2^_12_y^+1^_6_z^+2^_12_y^+2^_12_c^+1^_7_z^+1^_7_c^+1^_7_y^+1^_7_y^+1^_8_z^+1^_8_c^+1^_8_y^+1^_8_c^+1^_9_z^+1^_9_c^+1^_10_z^+1^_10_c^+1^_11_z^+1^_11_c^+1^_12_

0778155623343113

Fragment Matches Table

Show background peaks

| Position | Ion type | Intensity | mz Theoretical | mz Error (Th) | mz Error (ppm) | Charge | Series Number |
| --- | --- | --- | --- | --- | --- | --- | --- |
| - | - | 1249 | 129.1 | - | - | 0 | - |
| - | - | 1868 | 130.1 | - | - | 0 | - |
| - | - | 536.6 | 131.1 | - | - | 0 | - |
| - | - | 1688 | 132.1 | - | - | 0 | - |
| - | - | 406.3 | 135.4 | - | - | 0 | - |
| - | - | 385.1 | 140 | - | - | 0 | - |
| - | - | 1018 | 148.9 | - | - | 0 | - |
| - | - | 8445 | 149 | - | - | 0 | - |
| - | - | 445.2 | 152.4 | - | - | 0 | - |
| - | - | 413.9 | 153.1 | - | - | 0 | - |
| - | - | 3177 | 155.1 | - | - | 0 | - |
| - | - | 1509 | 159.1 | - | - | 0 | - |
| - | - | 403.1 | 165.9 | - | - | 0 | - |
| 10 | y | 893.1 | 166.1 | 0.001298 | 7.815 | +3 | 4 |
| - | - | 3216 | 167.1 | - | - | 0 | - |
| - | - | 469.5 | 169.1 | - | - | 0 | - |
| - | - | 635 | 169.1 | - | - | 0 | - |
| - | - | 413 | 169.9 | - | - | 0 | - |
| - | - | 5763 | 173.1 | - | - | 0 | - |
| - | - | 455.1 | 173.4 | - | - | 0 | - |
| - | - | 858.8 | 173.4 | - | - | 0 | - |
| - | - | 1513 | 173.4 | - | - | 0 | - |
| - | - | 413.4 | 173.9 | - | - | 0 | - |
| - | - | 433.4 | 174.1 | - | - | 0 | - |
| - | - | 519.3 | 174.2 | - | - | 0 | - |
| 13 | y | 1315 | 182.1 | 9.863E-05 | 0.5417 | +1 | 1 |
| - | - | 747.4 | 187.1 | - | - | 0 | - |
| - | - | 1282 | 189.1 | - | - | 0 | - |
| 9 | z | 491 | 199.1 | 0.0009228 | 4.635 | +3 | 5 |
| - | - | 4639 | 201.1 | - | - | 0 | - |
| - | - | 449 | 202.5 | - | - | 0 | - |
| - | - | 512 | 213.6 | - | - | 0 | - |
| - | - | 2319 | 215.1 | - | - | 0 | - |
| 2 | c | 1312 | 218.1 | 7.664E-05 | 0.3513 | +1 | 2 |
| - | - | 551.1 | 221.1 | - | - | 0 | - |
| - | - | 8688 | 221.1 | - | - | 0 | - |
| - | - | 1728 | 222.1 | - | - | 0 | - |
| - | - | 3471 | 223.1 | - | - | 0 | - |
| - | - | 1677 | 223.1 | - | - | 0 | - |
| - | - | 2634 | 225 | - | - | 0 | - |
| - | - | 1152 | 226 | - | - | 0 | - |
| - | - | 1.23E+04 | 239.1 | - | - | 0 | - |
| - | - | 3036 | 240.1 | - | - | 0 | - |
| - | - | 2464 | 241.1 | - | - | 0 | - |
| - | - | 618.2 | 243.1 | - | - | 0 | - |
| - | - | 582.5 | 245.1 | - | - | 0 | - |
| - | - | 935.3 | 251.2 | - | - | 0 | - |
| - | - | 2902 | 258.1 | - | - | 0 | - |
| - | - | 2705 | 268.2 | - | - | 0 | - |
| - | - | 2.547E+04 | 281.1 | - | - | 0 | - |
| - | - | 645.7 | 283 | - | - | 0 | - |
| 12 | z | 3320 | 295.1 | 0.001859 | 6.299 | +1 | 2 |
| - | - | 2656 | 296.1 | - | - | 0 | - |
| - | - | 3863 | 297.1 | - | - | 0 | - |
| - | - | 1692 | 297.1 | - | - | 0 | - |
| - | - | 1.077E+05 | 299.1 | - | - | 0 | - |
| - | - | 985.3 | 300.1 | - | - | 0 | - |
| - | - | 612.7 | 301.1 | - | - | 0 | - |
| - | - | 1110 | 313.1 | - | - | 0 | - |
| - | - | 685.1 | 314.1 | - | - | 0 | - |
| - | - | 743.4 | 314.2 | - | - | 0 | - |
| - | - | 3423 | 317.2 | - | - | 0 | - |
| - | - | 770.1 | 329.2 | - | - | 0 | - |
| 3 | c | 2037 | 331.2 | 6.469E-05 | 0.1953 | +1 | 3 |
| - | - | 1545 | 338.1 | - | - | 0 | - |
| - | - | 2746 | 341 | - | - | 0 | - |
| - | - | 836.2 | 345 | - | - | 0 | - |
| - | - | 1463 | 355.1 | - | - | 0 | - |
| - | - | 709.4 | 356.1 | - | - | 0 | - |
| - | - | 9660 | 359 | - | - | 0 | - |
| - | - | 616.9 | 359.8 | - | - | 0 | - |
| - | - | 548.7 | 364.2 | - | - | 0 | - |
| - | - | 2358 | 369.1 | - | - | 0 | - |
| - | - | 3139 | 370.1 | - | - | 0 | - |
| - | - | 2516 | 371.1 | - | - | 0 | - |
| - | - | 1548 | 390.2 | - | - | 0 | - |
| - | - | 1309 | 402.2 | - | - | 0 | - |
| - | - | 3760 | 410.7 | - | - | 0 | - |
| - | - | 1862 | 411.2 | - | - | 0 | - |
| - | - | 3.375E+04 | 415 | - | - | 0 | - |
| - | - | 1498 | 423.2 | - | - | 0 | - |
| - | - | 2439 | 423.3 | - | - | 0 | - |
| - | - | 658.4 | 424.3 | - | - | 0 | - |
| - | - | 2806 | 426.7 | - | - | 0 | - |
| - | - | 1918 | 427.2 | - | - | 0 | - |
| - | - | 1727 | 429.1 | - | - | 0 | - |
| 11 | y | 675 | 439.2 | 9.786E-05 | 0.2228 | +1 | 3 |
| 7 | c | 1829 | 440.7 | 4.148E-05 | 0.09412 | +2 | 7 |
| - | - | 2794 | 451.2 | - | - | 0 | - |
| - | - | 4591 | 451.3 | - | - | 0 | - |
| - | - | 850 | 452.2 | - | - | 0 | - |
| - | - | 762.5 | 452.3 | - | - | 0 | - |
| 4 | c | 1.587E+04 | 468.3 | 0.0002581 | 0.5511 | +1 | 4 |
| - | - | 4843 | 469.3 | - | - | 0 | - |
| - | - | 650.8 | 470.3 | - | - | 0 | - |
| 10 | z | 1562 | 480.2 | 0.0008551 | 1.781 | +1 | 4 |
| - | - | 761.2 | 489.2 | - | - | 0 | - |
| - | - | 1480 | 494.3 | - | - | 0 | - |
| 10 | y | 4569 | 496.2 | 0.0001364 | 0.2749 | +1 | 4 |
| - | - | 812.7 | 497.2 | - | - | 0 | - |
| - | - | 736.2 | 501.2 | - | - | 0 | - |
| - | - | 8.091E+04 | 503.1 | - | - | 0 | - |
| - | - | 3.133E+04 | 519.1 | - | - | 0 | - |
| - | - | 1475 | 519.2 | - | - | 0 | - |
| - | - | 866.8 | 522.3 | - | - | 0 | - |
| - | - | 1083 | 534.3 | - | - | 0 | - |
| - | - | 750.1 | 535.2 | - | - | 0 | - |
| - | - | 1.816E+04 | 536.2 | - | - | 0 | - |
| - | - | 1049 | 536.2 | - | - | 0 | - |
| - | - | 1622 | 538.2 | - | - | 0 | - |
| 9 | w | 597.7 | 550.3 | 0.0001003 | 0.1823 | +1 | 5 |
| - | - | 4689 | 550.3 | - | - | 0 | - |
| - | - | 1573 | 551.3 | - | - | 0 | - |
| - | - | 695 | 567.3 | - | - | 0 | - |
| 5 | c | 811.2 | 579.3 | 0.003084 | 5.323 | +1 | 5 |
| 9 | z | 1894 | 595.2 | 0.0004837 | 0.8126 | +1 | 5 |
| - | - | 633.7 | 596.3 | - | - | 0 | - |
| 5 | c | 1.038E+04 | 596.4 | 3.016E-05 | 0.05058 | +1 | 5 |
| - | - | 3315 | 597.4 | - | - | 0 | - |
| - | - | 927.4 | 598.4 | - | - | 0 | - |
| - | - | 844.7 | 602.3 | - | - | 0 | - |
| - | - | 2311 | 610.3 | - | - | 0 | - |
| 9 | y | 7543 | 611.3 | 0.001117 | 1.827 | +1 | 5 |
| - | - | 1757 | 612.3 | - | - | 0 | - |
| - | - | 722.1 | 617.4 | - | - | 0 | - |
| - | - | 655.8 | 618.3 | - | - | 0 | - |
| - | - | 558.1 | 625.3 | - | - | 0 | - |
| - | - | 690.3 | 633.8 | - | - | 0 | - |
| 4 | z | 575.7 | 637.8 | 0.0003574 | 0.5603 | +2 | 10 |
| 4 | y | 760.9 | 645.8 | 0.005827 | 9.023 | +2 | 10 |
| - | - | 4430 | 655.4 | - | - | 0 | - |
| 11 | c | 1.741E+04 | 655.8 | 0.001132 | 1.727 | +2 | 11 |
| - | - | 1.315E+04 | 656.3 | - | - | 0 | - |
| - | - | 5978 | 656.8 | - | - | 0 | - |
| - | - | 2214 | 657.3 | - | - | 0 | - |
| - | - | 3512 | 664.3 | - | - | 0 | - |
| - | - | 799.9 | 664.3 | - | - | 0 | - |
| 8 | w | 2680 | 665.3 | 0.00514 | 7.726 | +1 | 6 |
| - | - | 1178 | 666.3 | - | - | 0 | - |
| - | - | 885.1 | 667.4 | - | - | 0 | - |
| - | - | 1583 | 668.4 | - | - | 0 | - |
| - | - | 890.2 | 668.9 | - | - | 0 | - |
| - | - | 801 | 669.4 | - | - | 0 | - |
| - | - | 929.6 | 672.3 | - | - | 0 | - |
| 3 | w | 2409 | 672.8 | 0.002131 | 3.168 | +2 | 11 |
| - | - | 2230 | 673.3 | - | - | 0 | - |
| - | - | 873.9 | 673.8 | - | - | 0 | - |
| - | - | 1509 | 679.3 | - | - | 0 | - |
| - | - | 739.9 | 680.3 | - | - | 0 | - |
| 3 | y | 699.3 | 693.8 | 0.01137 | 16.39 | +2 | 11 |
| 6 | c | 1817 | 694.4 | 0.002927 | 4.216 | +1 | 6 |
| - | - | 936.4 | 694.8 | - | - | 0 | - |
| - | - | 707.3 | 695.3 | - | - | 0 | - |
| - | - | 898.7 | 701.8 | - | - | 0 | - |
| 3 | y | 3460 | 702.3 | 0.00127 | 1.809 | +2 | 11 |
| - | - | 2820 | 702.8 | - | - | 0 | - |
| - | - | 1757 | 703.3 | - | - | 0 | - |
| - | - | 2022 | 707.3 | - | - | 0 | - |
| 8 | z | 2339 | 708.3 | 0.004625 | 6.529 | +1 | 6 |
| - | - | 901.5 | 709.3 | - | - | 0 | - |
| 6 | c | 1.03E+04 | 711.4 | 0.0003015 | 0.4239 | +1 | 6 |
| - | - | 2851 | 712.4 | - | - | 0 | - |
| - | - | 924.7 | 713.4 | - | - | 0 | - |
| - | - | 996.9 | 715.4 | - | - | 0 | - |
| - | - | 1183 | 718.9 | - | - | 0 | - |
| - | - | 772.6 | 719.4 | - | - | 0 | - |
| - | - | 5955 | 719.9 | - | - | 0 | - |
| 12 | c | 1.282E+04 | 720.4 | 0.003823 | 5.306 | +2 | 12 |
| - | - | 8664 | 720.9 | - | - | 0 | - |
| - | - | 2958 | 721.4 | - | - | 0 | - |
| - | - | 1580 | 721.9 | - | - | 0 | - |
| - | - | 1078 | 723.4 | - | - | 0 | - |
| 8 | y | 2165 | 724.4 | 0.0006711 | 0.9265 | +1 | 6 |
| - | - | 1149 | 725.4 | - | - | 0 | - |
| - | - | 921.6 | 729.4 | - | - | 0 | - |
| - | - | 1743 | 736.9 | - | - | 0 | - |
| - | - | 3545 | 737.4 | - | - | 0 | - |
| - | - | 578.2 | 737.9 | - | - | 0 | - |
| 2 | z | 1195 | 743.9 | 0.008835 | 11.88 | +2 | 12 |
| - | - | 3478 | 748.9 | - | - | 0 | - |
| - | - | 2149 | 749.4 | - | - | 0 | - |
| - | - | 1743 | 749.9 | - | - | 0 | - |
| - | - | 882.6 | 750.9 | - | - | 0 | - |
| - | - | 757.7 | 751.4 | - | - | 0 | - |
| 2 | y | 755.9 | 751.9 | 0.009971 | 13.26 | +2 | 12 |
| - | - | 1092 | 771.9 | - | - | 0 | - |
| - | - | 1153 | 772.4 | - | - | 0 | - |
| - | - | 2848 | 772.9 | - | - | 0 | - |
| - | - | 2037 | 773.4 | - | - | 0 | - |
| - | - | 1719 | 779.9 | - | - | 0 | - |
| - | - | 2298 | 780.4 | - | - | 0 | - |
| - | - | 914.5 | 780.9 | - | - | 0 | - |
| - | - | 1205 | 785.9 | - | - | 0 | - |
| - | - | 1127 | 786.4 | - | - | 0 | - |
| - | - | 1211 | 786.9 | - | - | 0 | - |
| - | - | 988.2 | 787.4 | - | - | 0 | - |
| - | - | 692.8 | 793.4 | - | - | 0 | - |
| - | - | 3986 | 793.9 | - | - | 0 | - |
| - | - | 6645 | 794.4 | - | - | 0 | - |
| - | - | 6352 | 794.9 | - | - | 0 | - |
| - | - | 2565 | 795.4 | - | - | 0 | - |
| - | - | 2336 | 801.8 | - | - | 0 | - |
| - | - | 4306 | 801.9 | - | - | 0 | - |
| - | - | 3.491E+04 | 802.4 | - | - | 0 | - |
| - | - | 4.397E+04 | 802.9 | - | - | 0 | - |
| - | - | 2.992E+04 | 803.4 | - | - | 0 | - |
| - | - | 1.28E+04 | 803.9 | - | - | 0 | - |
| - | - | 3744 | 804.4 | - | - | 0 | - |
| - | - | 1576 | 804.9 | - | - | 0 | - |
| - | - | 825.3 | 817.4 | - | - | 0 | - |
| - | - | 870.2 | 837.4 | - | - | 0 | - |
| - | - | 1491 | 838.4 | - | - | 0 | - |
| - | - | 951.9 | 840.4 | - | - | 0 | - |
| 7 | c | 4524 | 880.4 | 0.0006972 | 0.7919 | +1 | 7 |
| - | - | 1908 | 881.4 | - | - | 0 | - |
| - | - | 1357 | 893.4 | - | - | 0 | - |
| 7 | z | 3989 | 894.4 | 0.003986 | 4.457 | +1 | 7 |
| - | - | 1663 | 895.4 | - | - | 0 | - |
| 7 | c | 1.385E+04 | 897.5 | 0.0021 | 2.34 | +1 | 7 |
| - | - | 7670 | 898.5 | - | - | 0 | - |
| - | - | 1801 | 899.5 | - | - | 0 | - |
| - | - | 1019 | 909.4 | - | - | 0 | - |
| 7 | y | 1754 | 910.4 | 0.01493 | 16.39 | +1 | 7 |
| - | - | 1465 | 932.5 | - | - | 0 | - |
| - | - | 1121 | 937.4 | - | - | 0 | - |
| - | - | 1568 | 954.4 | - | - | 0 | - |
| - | - | 857.1 | 955.5 | - | - | 0 | - |
| - | - | 1034 | 965.5 | - | - | 0 | - |
| 6 | y | 3291 | 1008 | 0.01978 | 19.61 | +1 | 8 |
| 6 | z | 5825 | 1009 | 0.008049 | 7.974 | +1 | 8 |
| - | - | 1963 | 1010 | - | - | 0 | - |
| 8 | c | 9208 | 1011 | 0.0002009 | 0.1988 | +1 | 8 |
| - | - | 723.4 | 1011 | - | - | 0 | - |
| - | - | 6017 | 1012 | - | - | 0 | - |
| - | - | 2099 | 1013 | - | - | 0 | - |
| - | - | 1023 | 1014 | - | - | 0 | - |
| - | - | 864.6 | 1024 | - | - | 0 | - |
| 6 | y | 1327 | 1025 | 0.008734 | 8.517 | +1 | 8 |
| - | - | 835.5 | 1026 | - | - | 0 | - |
| - | - | 3398 | 1027 | - | - | 0 | - |
| - | - | 1648 | 1042 | - | - | 0 | - |
| - | - | 957.6 | 1050 | - | - | 0 | - |
| - | - | 1.516E+04 | 1052 | - | - | 0 | - |
| - | - | 8454 | 1053 | - | - | 0 | - |
| - | - | 2291 | 1054 | - | - | 0 | - |
| - | - | 1379 | 1062 | - | - | 0 | - |
| - | - | 1407 | 1067 | - | - | 0 | - |
| - | - | 2577 | 1068 | - | - | 0 | - |
| - | - | 1564 | 1068 | - | - | 0 | - |
| - | - | 3954 | 1070 | - | - | 0 | - |
| - | - | 1884 | 1071 | - | - | 0 | - |
| - | - | 909.4 | 1072 | - | - | 0 | - |
| - | - | 1517 | 1072 | - | - | 0 | - |
| - | - | 1581 | 1082 | - | - | 0 | - |
| - | - | 1616 | 1125 | - | - | 0 | - |
| 9 | c | 2541 | 1126 | 0.003531 | 3.137 | +1 | 9 |
| - | - | 2014 | 1127 | - | - | 0 | - |
| - | - | 4188 | 1137 | - | - | 0 | - |
| 5 | z | 2.521E+04 | 1137 | 0.0006188 | 0.544 | +1 | 9 |
| - | - | 1.595E+04 | 1139 | - | - | 0 | - |
| - | - | 5119 | 1140 | - | - | 0 | - |
| - | - | 985.7 | 1140 | - | - | 0 | - |
| - | - | 1550 | 1149 | - | - | 0 | - |
| - | - | 772.7 | 1150 | - | - | 0 | - |
| - | - | 6340 | 1182 | - | - | 0 | - |
| 10 | c | 1.294E+04 | 1183 | 0.005505 | 4.655 | +1 | 10 |
| - | - | 7735 | 1184 | - | - | 0 | - |
| - | - | 2878 | 1185 | - | - | 0 | - |
| - | - | 873.5 | 1186 | - | - | 0 | - |
| - | - | 1305 | 1274 | - | - | 0 | - |
| 4 | z | 4854 | 1275 | 0.004329 | 3.397 | +1 | 10 |
| - | - | 4672 | 1276 | - | - | 0 | - |
| - | - | 2509 | 1277 | - | - | 0 | - |
| - | - | 1061 | 1278 | - | - | 0 | - |
| - | - | 971 | 1296 | - | - | 0 | - |
| - | - | 957.8 | 1297 | - | - | 0 | - |
| - | - | 2409 | 1310 | - | - | 0 | - |
| 11 | c | 7010 | 1311 | 0.002095 | 1.598 | +1 | 11 |
| - | - | 6011 | 1312 | - | - | 0 | - |
| - | - | 2635 | 1313 | - | - | 0 | - |
| - | - | 1022 | 1314 | - | - | 0 | - |
| 3 | z | 4062 | 1388 | 0.005592 | 4.03 | +1 | 11 |
| - | - | 5177 | 1389 | - | - | 0 | - |
| - | - | 3025 | 1390 | - | - | 0 | - |
| - | - | 1408 | 1391 | - | - | 0 | - |
| - | - | 1498 | 1396 | - | - | 0 | - |
| - | - | 1493 | 1397 | - | - | 0 | - |
| - | - | 1095 | 1398 | - | - | 0 | - |
| 12 | c | 2745 | 1440 | 0.01041 | 7.227 | +1 | 12 |
| - | - | 4548 | 1441 | - | - | 0 | - |
| - | - | 2253 | 1442 | - | - | 0 | - |
| - | - | 691.6 | 1488 | - | - | 0 | - |
| - | - | 1051 | 1489 | - | - | 0 | - |
| - | - | 879.4 | 1545 | - | - | 0 | - |
| - | - | 1967 | 1546 | - | - | 0 | - |
| - | - | 1376 | 1547 | - | - | 0 | - |
| - | - | 773.8 | 1548 | - | - | 0 | - |
| - | - | 1085 | 1559 | - | - | 0 | - |
| - | - | 3452 | 1560 | - | - | 0 | - |
| - | - | 2814 | 1561 | - | - | 0 | - |
| - | - | 1475 | 1562 | - | - | 0 | - |
| - | - | 734.4 | 1563 | - | - | 0 | - |
| - | - | 1517 | 1577 | - | - | 0 | - |
| - | - | 1636 | 1578 | - | - | 0 | - |
| - | - | 1008 | 1579 | - | - | 0 | - |
| - | - | 916 | 1587 | - | - | 0 | - |
| - | - | 7250 | 1588 | - | - | 0 | - |
| - | - | 1.176E+04 | 1589 | - | - | 0 | - |
| - | - | 9188 | 1590 | - | - | 0 | - |
| - | - | 4165 | 1591 | - | - | 0 | - |
| - | - | 1635 | 1592 | - | - | 0 | - |
| - | - | 1750 | 1593 | - | - | 0 | - |
| - | - | 2748 | 1604 | - | - | 0 | - |
| - | - | 1.316E+04 | 1605 | - | - | 0 | - |
| - | - | 2.163E+04 | 1606 | - | - | 0 | - |
| - | - | 1.622E+04 | 1607 | - | - | 0 | - |
| - | - | 8815 | 1608 | - | - | 0 | - |
| - | - | 3153 | 1609 | - | - | 0 | - |
| - | - | 2100 | 1610 | - | - | 0 | - |
| - | - | 847.7 | 3082 | - | - | 0 | - |

m/z Charge Intensity FragmentType MassShift Position
129.10238647460938 0 1248.5559
130.06532287597656 0 1868.2645
131.11790466308594 0 536.5946
132.1020965576172 0 1688.0804
135.43991088867188 0 406.29196
140.0190887451172 0 385.13763
148.94802856445312 0 1017.6405
149.044921875 0 8444.58
152.41969299316406 0 445.19174
153.08355712890625 0 413.8757
155.09278869628906 0 3177.202
159.0917205810547 0 1508.8097
165.8810272216797 0 403.09277
166.08621215820312 0 893.14404 y 9
167.0553436279297 0 3215.5437
169.08473205566406 0 469.48428
169.0969696044922 0 635.0171
169.8626708984375 0 412.9718
173.1285400390625 0 5762.6914
173.4178009033203 0 455.08127
173.4340057373047 0 858.81757
173.43946838378906 0 1512.6208
173.92364501953125 0 413.4414
174.13113403320312 0 433.3801
174.17112731933594 0 519.28204
182.08126831054688 0 1315.3768 y 12
187.1443634033203 0 747.3721
189.12342834472656 0 1281.699
199.08673095703125 0 490.9734 z 8
201.12338256835938 0 4638.513
202.46055603027344 0 448.98685
213.6340789794922 0 511.95972
215.13888549804688 0 2319.1758
218.14984130859375 0 1311.737 c 1
221.05844116210938 0 551.1405
221.08433532714844 0 8688.122
222.08489990234375 0 1728.405
223.0634307861328 0 3470.716
223.08169555664062 0 1676.5623
225.04269409179688 0 2634.0122
226.04315185546875 0 1152.397
239.094970703125 0 12304.455
240.09532165527344 0 3035.6482
241.09234619140625 0 2464.375
243.1332550048828 0 618.24335
245.13742065429688 0 582.46594
251.150390625 0 935.2956
258.1448974609375 0 2901.6753
268.1767578125 0 2705.4521
281.05120849609375 0 25469.156
283.0308532714844 0 645.6798
295.1031799316406 0 3320.4622 z 11
296.10357666015625 0 2656.3032
297.0825500488281 0 3862.9285
297.10028076171875 0 1692.03
299.0617980957031 0 107722.2
300.06268310546875 0 985.27673
301.0597229003906 0 612.6595
313.1129150390625 0 1110.2416
314.1131591796875 0 685.10913
314.2088623046875 0 743.4352
317.158203125 0 3423.4211
329.1503601074219 0 770.1432
331.2339172363281 0 2036.5466 c 2
338.1342468261719 0 1545.4507
341.01812744140625 0 2745.5674
344.9754943847656 0 836.18384
355.0699768066406 0 1463.3444
356.0733337402344 0 709.44434
359.0282897949219 0 9660.182
359.82904052734375 0 616.9188
364.1875 0 548.7263
369.12188720703125 0 2358.3213
370.12298583984375 0 3139.318
371.1200256347656 0 2516.3494
390.19775390625 0 1547.6339
402.1553955078125 0 1308.9241
410.66876220703125 0 3760.3096
411.1697998046875 0 1862.1544
415.0368347167969 0 33745.39
423.2238464355469 0 1498.2678
423.27105712890625 0 2439.45
424.27392578125 0 658.4083
426.7216796875 0 2805.6802
427.2241516113281 0 1917.9022
429.0884704589844 0 1726.6359
439.2186279296875 0 675.0094 y 10
440.7192687988281 0 1828.913 c Ammonia loss 6
451.21868896484375 0 2793.6167
451.2663269042969 0 4590.9375
452.2218322753906 0 849.98486
452.2679138183594 0 762.5127
468.29315185546875 0 15872.215 c 3
469.2962646484375 0 4843.3916
470.2984313964844 0 650.7831
480.2223205566406 0 1562.1571 z 9
489.2043151855469 0 761.23505
494.3084411621094 0 1479.5953
496.2403259277344 0 4568.6074 y 9
497.2432861328125 0 812.65985
501.2322998046875 0 736.1694
503.1075744628906 0 80907.234
519.1388549804688 0 31331.803
519.2395629882812 0 1475.1105
522.2911376953125 0 866.7831
534.3295288085938 0 1083.438
535.1998901367188 0 750.1244
536.1653442382812 0 18162.06
536.241455078125 0 1048.7572
538.249267578125 0 1622.1095
550.2508544921875 0 597.746 w 8
550.2875366210938 0 4688.7485
551.2894897460938 0 1572.9471
567.314208984375 0 694.95636
579.3218383789062 0 811.2189 c Ammonia loss 4
595.2479248046875 0 1893.6943 z 8
596.2506713867188 0 633.65045
596.3515014648438 0 10378.461 c 4
597.3540649414062 0 3314.939
598.3574829101562 0 927.41675
602.2901611328125 0 844.7009
610.2838745117188 0 2311.3013
611.2682495117188 0 7542.7896 y 8
612.2725219726562 0 1757.0475
617.359130859375 0 722.0657
618.3065185546875 0 655.7819
625.3256225585938 0 558.1259
633.841552734375 0 690.2955
637.7821044921875 0 575.6921 z 3
645.7969360351562 0 760.9119 y 3
655.35205078125 0 4429.637
655.8473510742188 0 17411.285 c 10
656.3485717773438 0 13146.148
656.849853515625 0 5978.114
657.3489379882812 0 2214.2795
664.2930297851562 0 3512.4167
664.3489379882812 0 799.93665
665.2828369140625 0 2680.3796 w 7
666.2827758789062 0 1177.5319
667.3629760742188 0 885.1415
668.3653564453125 0 1583.3562
668.85546875 0 890.20496
669.3660278320312 0 800.96423
672.3048706054688 0 929.565
672.7985229492188 0 2408.6133 w 2
673.2999877929688 0 2229.6047
673.7984008789062 0 873.89
679.3292236328125 0 1509.317
680.3265991210938 0 739.9273
693.8312377929688 0 699.2861 y Ammonia loss 2
694.3489379882812 0 1817.0223 c Ammonia loss 5
694.8251342773438 0 936.3954
695.3338623046875 0 707.3355
701.8340454101562 0 898.7418
702.3344116210938 0 3460.4587 y 2
702.8375854492188 0 2820.4539
703.338623046875 0 1756.5284
707.3489990234375 0 2022.3657
708.3370971679688 0 2339.1946 z 7
709.3392333984375 0 901.4983
711.3781127929688 0 10299.321 c 5
712.3814086914062 0 2851.4587
713.3846435546875 0 924.74835
715.372802734375 0 996.90955
718.92236328125 0 1182.8986
719.4244384765625 0 772.634
719.875 0 5955.0103
720.371337890625 0 12819 c 11
720.8714599609375 0 8663.506
721.3704223632812 0 2957.9565
721.8727416992188 0 1580.4058
723.3682250976562 0 1077.575
724.3518676757812 0 2164.8125 y 7
725.3567504882812 0 1148.5928
729.3565673828125 0 921.59454
736.8662719726562 0 1743.3928
737.3609008789062 0 3545.1074
737.8568115234375 0 578.2419
743.8668212890625 0 1195.32 z 1
748.86669921875 0 3477.881
749.367919921875 0 2148.6636
749.8682250976562 0 1743.3807
750.8743286132812 0 882.5602
751.367431640625 0 757.6525
751.8773193359375 0 755.8741 y 1
771.8851318359375 0 1091.8774
772.3926391601562 0 1152.519
772.8873291015625 0 2848.2585
773.3856811523438 0 2036.7441
779.8949584960938 0 1718.8497
780.392822265625 0 2297.685
780.894775390625 0 914.51
785.8819580078125 0 1205.0436
786.3817749023438 0 1126.745
786.8801879882812 0 1210.8259
787.3834228515625 0 988.2271
793.398193359375 0 692.83307
793.889404296875 0 3985.7214
794.3869018554688 0 6645.1357
794.886962890625 0 6351.7354
795.3882446289062 0 2565.0173
801.8291625976562 0 2336.2437
801.9008178710938 0 4305.7173
802.3961181640625 0 34909.56
802.8959350585938 0 43967.254
803.3971557617188 0 29918.455
803.8991088867188 0 12798.943
804.3999633789062 0 3744.2085
804.89501953125 0 1576.27
817.4412231445312 0 825.2634
837.359375 0 870.1726
838.3508911132812 0 1491.098
840.409423828125 0 951.87714
880.4304809570312 0 4523.9004 c Ammonia loss 6
881.4278564453125 0 1907.6423
893.421875 0 1356.5616
894.415771484375 0 3988.8884 z 6
895.421630859375 0 1663.4436
897.4556274414062 0 13847.077 c 6
898.4578247070312 0 7670.0024
899.4560546875 0 1800.867
909.446533203125 0 1019.2716
910.4454345703125 0 1754.0547 y 6
932.4684448242188 0 1464.7242
937.4223022460938 0 1120.5134
954.4489135742188 0 1568.4778
955.4564208984375 0 857.1163
965.4541625976562 0 1033.6537
1008.45068359375 0 3290.7664 y Ammonia loss 5
1009.44677734375 0 5824.9634 z 5
1010.439453125 0 1963.1971
1010.5419921875 0 9207.686 c 7
1011.4616088867188 0 723.4072
1011.5431518554688 0 6017.329
1012.5484619140625 0 2098.897
1013.5521240234375 0 1022.82495
1024.4736328125 0 864.6378
1025.4661865234375 0 1327.0763 y 5
1026.4595947265625 0 835.47205
1027.4178466796875 0 3398.3003
1041.5379638671875 0 1648.2961
1049.61181640625 0 957.58923
1052.448486328125 0 15161.012
1053.4512939453125 0 8454.492
1054.4554443359375 0 2290.7378
1061.5107421875 0 1378.755
1067.4306640625 0 1407.1418
1067.5428466796875 0 2577.3403
1068.4354248046875 0 1563.8895
1069.529052734375 0 3954.483
1070.537841796875 0 1883.5363
1071.60009765625 0 909.3861
1072.4285888671875 0 1516.792
1081.564697265625 0 1580.6542
1124.5797119140625 0 1616.1915
1125.572265625 0 2541.0833 c 8
1126.5714111328125 0 2014.2574
1136.5137939453125 0 4187.9717
1137.4979248046875 0 25210.78 z 4
1138.5008544921875 0 15954.096
1139.5013427734375 0 5118.9536
1140.4912109375 0 985.7358
1148.5399169921875 0 1549.9943
1149.5516357421875 0 772.7333
1181.6063232421875 0 6340.4385
1182.595703125 0 12935.712 c 9
1183.5963134765625 0 7735.0703
1184.59912109375 0 2877.6924
1185.594970703125 0 873.5379
1273.5787353515625 0 1305.2866
1274.560546875 0 4853.941 z 3
1275.56103515625 0 4671.597
1276.5706787109375 0 2508.6323
1277.5718994140625 0 1060.7319
1295.567138671875 0 971.047
1296.5751953125 0 957.7803
1309.693359375 0 2409.2878
1310.687255859375 0 7010.4478 c 10
1311.691162109375 0 6011.283
1312.690673828125 0 2635.1572
1313.705078125 0 1021.78314
1387.6458740234375 0 4062.388 z 2
1388.64501953125 0 5177.0254
1389.64697265625 0 3024.5056
1390.65576171875 0 1407.652
1395.7291259765625 0 1497.7341
1396.7359619140625 0 1493.2489
1397.733154296875 0 1095.1122
1439.7381591796875 0 2745.02 c 11
1440.73681640625 0 4548.0737
1441.743408203125 0 2252.6047
1487.714111328125 0 691.6471
1488.732177734375 0 1050.9054
1544.767822265625 0 879.4421
1545.76220703125 0 1966.5092
1546.7735595703125 0 1376.3292
1547.762939453125 0 773.7949
1558.7908935546875 0 1084.6821
1559.78564453125 0 3452.3013
1560.7769775390625 0 2813.5122
1561.777099609375 0 1474.5691
1562.7830810546875 0 734.40625
1576.80517578125 0 1517.3403
1577.797607421875 0 1635.9673
1578.8145751953125 0 1008.39954
1586.7698974609375 0 915.98474
1587.774169921875 0 7250.31
1588.7689208984375 0 11758.8
1589.7666015625 0 9188.37
1590.7720947265625 0 4164.672
1591.736083984375 0 1635.3486
1592.736083984375 0 1749.9176
1603.7916259765625 0 2748.3206
1604.7891845703125 0 13164.493
1605.791259765625 0 21631.963
1606.7933349609375 0 16219.4795
1607.79296875 0 8814.674
1608.791259765625 0 3153.2534
1609.7822265625 0 2099.5093
3081.7685546875 0 847.6879

Spectrum Details

|  |  |
| --- | --- |
| Matched peaks? Matched peaks The total absolute number of peaks matched. Additionally in brackets the total fraction of peaks matched and the total number of peaks is shown. | 45 (14.06% of 320) |
| FDR? FDR The false discovery rate estimated for this peptide. It is calculated by matching all theoretical fragments with a non-integer shift with the raw peaks for this spectrum. This is done with 40 different shifts. The resulting percentage is the average number of annotated peaks over the number of annotated peaks with the correct spectrum. | 3.76% |
| Satellite FDR? Satellite FDR See the FDR for details on its calculation. This satellite ion specific FDR only contains the satellite ions (d/w) for I/L/J positions. | 0.00% |
| PSM Score? PSM Score The PSM Score as given by Hecklib to this annotated spectrum. It is shown with three significant figures. | 371 |

#### Spectrum 6806? Spectrum 6806 The raw spectrum of this peptide as annotated by Hecklib. The fragments are coloured according to ion type (see legend). Any peaks with a star '\*' as text can be hovered over to see the full details, first the ion type second the mass shift type. By hovering over the amino acids in the peptide or ions in the legend the corresponding peaks are highlighted. By toggling the 'Unassigned' label you can turn the background (unassigned) peaks on or off in the plot. By updating the slider in the Ion legend you can update the spectrum to only show the top X% of the peaks with labels. The top X% means any peak that is within X% of the highest intensity. By dragging in the spectrum you can zoom in to a specific part of the spectrum and use 'Zoom Out' to get back to the original zoom level. The annotation of the spectrum is based on the given sequence in the peptides file and is done with different software so inconsistencies are likely. The peaks are annotated based on the given sequence, with 20 ppm tolerance.

Copy Data

##### Spectrum 6806 (TSV)

###### Preview

```
Loading example...
```

*Click on the button to copy the data to your clipboard.*

Mz MinMz MaxIntensity Max

WidthHeightPeptide font sizePeptide stroke widthSpectrum font sizeSpectrum stroke widthCompact peptide

Ion legend

wxyz

abcd

OtherUnassignedIon^Charge^_Position_Show for top:%

TVLHQDWLDGKEY

03.25e+46.50e+49.75e+41.30e+5

Zoom Out

y^+3^_3_a^+1^_2_y^+2^_2_d^+1^_2_y^+3^_4_a^+1^_2_y^+1^_1_b^+1^_2_b^+1^_2_y^+2^_3_b^+3^_6_y^+1^_2_b^+1^_3_y^+2^_5_y^+1^_2_b^+1^_3_b^+3^_8_y^+3^_8_b^+2^_6_y^+2^_6_y^+3^_10_b^+1^_4_y^+1^_3_b^+2^_7_b^+2^_7_b^+1^_4_y^+2^_7_y^+1^_4_b^+2^_8_b^+2^_8_\*\*\*b^+2^_9_b^+2^_9_b^+1^_5_y^+1^_5_y^+1^_5_y^+2^_10_y^+2^_10_b^+2^_11_y^+2^_10_b^+1^_6_b^+1^_6_b^+1^_6_y^+2^_11_y^+2^_11_y^+1^_6_y^+2^_11_b^+2^_12_b^+2^_12_b^+2^_12_y^+1^_6_y^+2^_12_\*b^+1^_7_b^+1^_7_y^+1^_7_y^+1^_7_b^+1^_8_y^+1^_8_y^+1^_9_y^+1^_9_y^+1^_9_

0859171825773436

Fragment Matches Table

Show background peaks

| Position | Ion type | Intensity | mz Theoretical | mz Error (Th) | mz Error (ppm) | Charge | Series Number |
| --- | --- | --- | --- | --- | --- | --- | --- |
| - | - | 7781 | 120.1 | - | - | 0 | - |
| - | - | 761.6 | 121.1 | - | - | 0 | - |
| - | - | 472.7 | 122.1 | - | - | 0 | - |
| - | - | 439.4 | 123 | - | - | 0 | - |
| - | - | 369.6 | 124.9 | - | - | 0 | - |
| - | - | 949.9 | 125.1 | - | - | 0 | - |
| - | - | 1204 | 126.1 | - | - | 0 | - |
| - | - | 702.1 | 127.1 | - | - | 0 | - |
| - | - | 7649 | 128.1 | - | - | 0 | - |
| - | - | 835.9 | 129.1 | - | - | 0 | - |
| - | - | 6.057E+04 | 129.1 | - | - | 0 | - |
| - | - | 7268 | 130.1 | - | - | 0 | - |
| - | - | 422.7 | 130.1 | - | - | 0 | - |
| - | - | 3730 | 130.1 | - | - | 0 | - |
| - | - | 593.1 | 131.1 | - | - | 0 | - |
| - | - | 2047 | 132 | - | - | 0 | - |
| - | - | 617.3 | 132.1 | - | - | 0 | - |
| - | - | 1144 | 132.1 | - | - | 0 | - |
| - | - | 1.176E+04 | 136.1 | - | - | 0 | - |
| - | - | 1513 | 137.1 | - | - | 0 | - |
| - | - | 506.5 | 138.1 | - | - | 0 | - |
| - | - | 1074 | 138.1 | - | - | 0 | - |
| - | - | 865.3 | 139.1 | - | - | 0 | - |
| - | - | 1220 | 141.1 | - | - | 0 | - |
| - | - | 451.2 | 144 | - | - | 0 | - |
| - | - | 4855 | 146.1 | - | - | 0 | - |
| - | - | 879.7 | 147 | - | - | 0 | - |
| 11 | y | 468.8 | 147.1 | 0.0009926 | 6.749 | +3 | 3 |
| - | - | 517.3 | 149 | - | - | 0 | - |
| - | - | 1719 | 151.1 | - | - | 0 | - |
| - | - | 677 | 154.1 | - | - | 0 | - |
| - | - | 600.6 | 154.1 | - | - | 0 | - |
| - | - | 4099 | 155.1 | - | - | 0 | - |
| 2 | a | 4187 | 155.1 | 0.0002288 | 1.475 | +1 | 2 |
| 12 | y | 448.3 | 156.1 | 0.0004899 | 3.139 | +2 | 2 |
| - | - | 3463 | 156.1 | - | - | 0 | - |
| - | - | 952.7 | 157.1 | - | - | 0 | - |
| - | - | 2484 | 157.1 | - | - | 0 | - |
| - | - | 2336 | 157.1 | - | - | 0 | - |
| - | - | 2806 | 157.1 | - | - | 0 | - |
| - | - | 755.8 | 157.1 | - | - | 0 | - |
| - | - | 5578 | 158.1 | - | - | 0 | - |
| - | - | 866.8 | 158.1 | - | - | 0 | - |
| - | - | 904.6 | 159.1 | - | - | 0 | - |
| - | - | 6603 | 159.1 | - | - | 0 | - |
| - | - | 590.5 | 159.1 | - | - | 0 | - |
| 2 | d | 1418 | 159.1 | 0.0004313 | 2.711 | +1 | 2 |
| - | - | 723.1 | 160.1 | - | - | 0 | - |
| - | - | 856.1 | 164 | - | - | 0 | - |
| - | - | 5372 | 165.1 | - | - | 0 | - |
| - | - | 8481 | 166.1 | - | - | 0 | - |
| 10 | y | 4125 | 166.1 | 0.001695 | 10.2 | +3 | 4 |
| - | - | 1001 | 167.1 | - | - | 0 | - |
| - | - | 512.5 | 167.1 | - | - | 0 | - |
| - | - | 772.7 | 167.1 | - | - | 0 | - |
| - | - | 559.9 | 167.1 | - | - | 0 | - |
| - | - | 566.5 | 167.1 | - | - | 0 | - |
| - | - | 651.9 | 168.1 | - | - | 0 | - |
| - | - | 576.9 | 168.1 | - | - | 0 | - |
| - | - | 409.2 | 168.8 | - | - | 0 | - |
| - | - | 579.8 | 169.1 | - | - | 0 | - |
| - | - | 2376 | 169.1 | - | - | 0 | - |
| - | - | 1460 | 169.1 | - | - | 0 | - |
| - | - | 1542 | 171.1 | - | - | 0 | - |
| - | - | 1021 | 172.1 | - | - | 0 | - |
| - | - | 743.9 | 173.1 | - | - | 0 | - |
| 2 | a | 1.287E+05 | 173.1 | 0.0004215 | 2.435 | +1 | 2 |
| - | - | 1.164E+04 | 174.1 | - | - | 0 | - |
| - | - | 2111 | 175.1 | - | - | 0 | - |
| - | - | 2.05E+04 | 175.1 | - | - | 0 | - |
| - | - | 1967 | 175.1 | - | - | 0 | - |
| - | - | 598.7 | 175.1 | - | - | 0 | - |
| - | - | 1859 | 176.1 | - | - | 0 | - |
| - | - | 595.2 | 177.1 | - | - | 0 | - |
| - | - | 454.4 | 177.1 | - | - | 0 | - |
| - | - | 3777 | 178.1 | - | - | 0 | - |
| - | - | 680.4 | 181.1 | - | - | 0 | - |
| 13 | y | 1.098E+04 | 182.1 | 0.0004496 | 2.469 | +1 | 1 |
| - | - | 596.1 | 183.1 | - | - | 0 | - |
| 2 | b | 1.377E+04 | 183.1 | 0.0003092 | 1.689 | +1 | 2 |
| - | - | 635.6 | 183.1 | - | - | 0 | - |
| - | - | 656.8 | 184.1 | - | - | 0 | - |
| - | - | 541.1 | 184.1 | - | - | 0 | - |
| - | - | 3108 | 185.1 | - | - | 0 | - |
| - | - | 923.3 | 185.1 | - | - | 0 | - |
| - | - | 835.7 | 185.2 | - | - | 0 | - |
| - | - | 845.7 | 186.1 | - | - | 0 | - |
| - | - | 1.94E+04 | 186.1 | - | - | 0 | - |
| - | - | 1572 | 187.1 | - | - | 0 | - |
| - | - | 1153 | 187.1 | - | - | 0 | - |
| - | - | 1231 | 187.1 | - | - | 0 | - |
| - | - | 1405 | 188.1 | - | - | 0 | - |
| - | - | 6310 | 189.1 | - | - | 0 | - |
| - | - | 728 | 190.1 | - | - | 0 | - |
| - | - | 888.9 | 190.1 | - | - | 0 | - |
| - | - | 634.7 | 194.1 | - | - | 0 | - |
| - | - | 4153 | 195.1 | - | - | 0 | - |
| - | - | 2701 | 196.1 | - | - | 0 | - |
| - | - | 886.4 | 196.1 | - | - | 0 | - |
| - | - | 1804 | 197.1 | - | - | 0 | - |
| - | - | 590 | 198.1 | - | - | 0 | - |
| - | - | 3086 | 199.1 | - | - | 0 | - |
| - | - | 706 | 199.1 | - | - | 0 | - |
| 2 | b | 4.927E+04 | 201.1 | 0.0003494 | 1.737 | +1 | 2 |
| - | - | 526.8 | 202.1 | - | - | 0 | - |
| - | - | 3849 | 202.1 | - | - | 0 | - |
| - | - | 1721 | 203.1 | - | - | 0 | - |
| - | - | 1231 | 203.1 | - | - | 0 | - |
| - | - | 2514 | 205.1 | - | - | 0 | - |
| - | - | 493.8 | 207.2 | - | - | 0 | - |
| - | - | 696.2 | 210.2 | - | - | 0 | - |
| 11 | y | 1971 | 211.1 | 0.0002677 | 1.268 | +2 | 3 |
| - | - | 7305 | 211.2 | - | - | 0 | - |
| - | - | 1157 | 212.1 | - | - | 0 | - |
| - | - | 2479 | 213.1 | - | - | 0 | - |
| - | - | 556.8 | 213.2 | - | - | 0 | - |
| - | - | 852.2 | 215.1 | - | - | 0 | - |
| - | - | 9957 | 217.1 | - | - | 0 | - |
| - | - | 811.7 | 218.1 | - | - | 0 | - |
| - | - | 483.1 | 218.9 | - | - | 0 | - |
| - | - | 783.4 | 222.1 | - | - | 0 | - |
| - | - | 863.4 | 223.1 | - | - | 0 | - |
| - | - | 2332 | 223.1 | - | - | 0 | - |
| - | - | 1.039E+04 | 223.2 | - | - | 0 | - |
| - | - | 1172 | 224.1 | - | - | 0 | - |
| - | - | 1076 | 224.2 | - | - | 0 | - |
| - | - | 1614 | 226.1 | - | - | 0 | - |
| 6 | b | 745.4 | 226.1 | 0.0004923 | 2.177 | +3 | 6 |
| - | - | 1471 | 227.1 | - | - | 0 | - |
| - | - | 2273 | 227.1 | - | - | 0 | - |
| - | - | 739 | 228.1 | - | - | 0 | - |
| - | - | 1657 | 228.2 | - | - | 0 | - |
| - | - | 3605 | 229.1 | - | - | 0 | - |
| - | - | 1467 | 233.1 | - | - | 0 | - |
| - | - | 1.035E+04 | 234.1 | - | - | 0 | - |
| - | - | 818.2 | 234.1 | - | - | 0 | - |
| - | - | 1166 | 235.1 | - | - | 0 | - |
| - | - | 768.1 | 237.1 | - | - | 0 | - |
| - | - | 791.1 | 238.1 | - | - | 0 | - |
| - | - | 4100 | 239.2 | - | - | 0 | - |
| - | - | 3111 | 240.1 | - | - | 0 | - |
| - | - | 549.7 | 241.1 | - | - | 0 | - |
| - | - | 759.8 | 241.1 | - | - | 0 | - |
| - | - | 623.8 | 241.2 | - | - | 0 | - |
| - | - | 2404 | 242.1 | - | - | 0 | - |
| - | - | 608.1 | 243.1 | - | - | 0 | - |
| - | - | 3112 | 244.1 | - | - | 0 | - |
| - | - | 6081 | 247.1 | - | - | 0 | - |
| - | - | 523.2 | 247.1 | - | - | 0 | - |
| - | - | 2066 | 248.1 | - | - | 0 | - |
| - | - | 3296 | 249.1 | - | - | 0 | - |
| - | - | 828.8 | 249.1 | - | - | 0 | - |
| - | - | 3.432E+04 | 251.2 | - | - | 0 | - |
| - | - | 4127 | 252.2 | - | - | 0 | - |
| - | - | 788.5 | 253.1 | - | - | 0 | - |
| - | - | 1763 | 255.1 | - | - | 0 | - |
| - | - | 1803 | 255.1 | - | - | 0 | - |
| - | - | 583.2 | 256.1 | - | - | 0 | - |
| - | - | 1701 | 256.2 | - | - | 0 | - |
| - | - | 910.9 | 258.1 | - | - | 0 | - |
| - | - | 538.1 | 261.1 | - | - | 0 | - |
| - | - | 704.2 | 262.1 | - | - | 0 | - |
| - | - | 2343 | 265.1 | - | - | 0 | - |
| - | - | 9228 | 266.1 | - | - | 0 | - |
| - | - | 966.3 | 267.1 | - | - | 0 | - |
| - | - | 779.7 | 268.2 | - | - | 0 | - |
| - | - | 5232 | 270.2 | - | - | 0 | - |
| - | - | 599.2 | 271.1 | - | - | 0 | - |
| - | - | 805.7 | 271.2 | - | - | 0 | - |
| - | - | 978.7 | 274.1 | - | - | 0 | - |
| - | - | 937.3 | 280.1 | - | - | 0 | - |
| - | - | 743.7 | 283 | - | - | 0 | - |
| - | - | 1.66E+04 | 283.1 | - | - | 0 | - |
| - | - | 720.2 | 284.1 | - | - | 0 | - |
| - | - | 2340 | 284.1 | - | - | 0 | - |
| - | - | 920.1 | 284.2 | - | - | 0 | - |
| - | - | 2315 | 286.1 | - | - | 0 | - |
| - | - | 751.3 | 287.2 | - | - | 0 | - |
| - | - | 574.1 | 288.1 | - | - | 0 | - |
| - | - | 674.1 | 288.2 | - | - | 0 | - |
| - | - | 1462 | 288.2 | - | - | 0 | - |
| 12 | y | 1.284E+04 | 293.1 | 0.0004799 | 1.637 | +1 | 2 |
| - | - | 2619 | 294.1 | - | - | 0 | - |
| 3 | b | 830 | 296.2 | 0.0008247 | 2.784 | +1 | 3 |
| - | - | 4270 | 297.2 | - | - | 0 | - |
| - | - | 1352 | 298.1 | - | - | 0 | - |
| - | - | 1927 | 298.2 | - | - | 0 | - |
| - | - | 2538 | 299.1 | - | - | 0 | - |
| - | - | 1351 | 299.1 | - | - | 0 | - |
| - | - | 3209 | 300.1 | - | - | 0 | - |
| - | - | 939.4 | 300.1 | - | - | 0 | - |
| - | - | 5074 | 301.1 | - | - | 0 | - |
| - | - | 9417 | 301.2 | - | - | 0 | - |
| - | - | 2608 | 302.1 | - | - | 0 | - |
| - | - | 1187 | 302.2 | - | - | 0 | - |
| - | - | 970.1 | 303.1 | - | - | 0 | - |
| - | - | 2594 | 305.2 | - | - | 0 | - |
| 9 | y | 1238 | 306.1 | 0.0007655 | 2.5 | +2 | 5 |
| - | - | 719.7 | 310.2 | - | - | 0 | - |
| 12 | y | 1731 | 311.1 | 0.0007489 | 2.407 | +1 | 2 |
| - | - | 577.9 | 311.1 | - | - | 0 | - |
| - | - | 1.049E+04 | 312.2 | - | - | 0 | - |
| - | - | 824.8 | 313.2 | - | - | 0 | - |
| - | - | 2308 | 313.2 | - | - | 0 | - |
| - | - | 2565 | 314.1 | - | - | 0 | - |
| 3 | b | 748.4 | 314.2 | 0.001002 | 3.19 | +1 | 3 |
| - | - | 5768 | 315.2 | - | - | 0 | - |
| - | - | 3086 | 316.2 | - | - | 0 | - |
| - | - | 1473 | 316.2 | - | - | 0 | - |
| - | - | 1484 | 318.1 | - | - | 0 | - |
| - | - | 1823 | 327.3 | - | - | 0 | - |
| - | - | 4718 | 330.2 | - | - | 0 | - |
| 8 | b | 655.8 | 331.2 | 0.0008803 | 2.658 | +3 | 8 |
| - | - | 696.4 | 332.2 | - | - | 0 | - |
| - | - | 930.4 | 333.2 | - | - | 0 | - |
| - | - | 550.7 | 337.2 | - | - | 0 | - |
| - | - | 1611 | 338.2 | - | - | 0 | - |
| - | - | 1462 | 341.2 | - | - | 0 | - |
| 6 | y | 2433 | 342.1 | 0.003154 | 9.218 | +3 | 8 |
| - | - | 1140 | 345.1 | - | - | 0 | - |
| - | - | 2965 | 346.1 | - | - | 0 | - |
| - | - | 607.4 | 347.1 | - | - | 0 | - |
| 6 | b | 1894 | 347.7 | 0.0001472 | 0.4233 | +2 | 6 |
| - | - | 667.8 | 348.7 | - | - | 0 | - |
| - | - | 957.8 | 349.2 | - | - | 0 | - |
| - | - | 3076 | 350.2 | - | - | 0 | - |
| - | - | 579.8 | 354.3 | - | - | 0 | - |
| - | - | 637.2 | 355.2 | - | - | 0 | - |
| - | - | 889.1 | 355.2 | - | - | 0 | - |
| - | - | 1514 | 359.2 | - | - | 0 | - |
| - | - | 611.1 | 361.2 | - | - | 0 | - |
| 8 | y | 2696 | 362.7 | 3.727E-05 | 0.1028 | +2 | 6 |
| - | - | 3955 | 363.1 | - | - | 0 | - |
| - | - | 1114 | 363.2 | - | - | 0 | - |
| - | - | 769.8 | 364.1 | - | - | 0 | - |
| - | - | 618.1 | 364.2 | - | - | 0 | - |
| - | - | 672.2 | 366.2 | - | - | 0 | - |
| - | - | 876.6 | 367.2 | - | - | 0 | - |
| - | - | 780.1 | 369.2 | - | - | 0 | - |
| - | - | 615.9 | 373.2 | - | - | 0 | - |
| - | - | 1340 | 378.2 | - | - | 0 | - |
| - | - | 7629 | 379.2 | - | - | 0 | - |
| - | - | 1463 | 380.2 | - | - | 0 | - |
| - | - | 1.009E+04 | 381.2 | - | - | 0 | - |
| - | - | 1701 | 382.2 | - | - | 0 | - |
| - | - | 1269 | 392.2 | - | - | 0 | - |
| - | - | 684.9 | 393.7 | - | - | 0 | - |
| - | - | 1664 | 394.2 | - | - | 0 | - |
| - | - | 2312 | 396.2 | - | - | 0 | - |
| - | - | 2320 | 397.3 | - | - | 0 | - |
| - | - | 730.7 | 398.2 | - | - | 0 | - |
| - | - | 743 | 399.2 | - | - | 0 | - |
| - | - | 2269 | 401.2 | - | - | 0 | - |
| - | - | 1257 | 402.2 | - | - | 0 | - |
| - | - | 1155 | 407.2 | - | - | 0 | - |
| - | - | 1119 | 409.2 | - | - | 0 | - |
| - | - | 1897 | 411.1 | - | - | 0 | - |
| - | - | 8103 | 412.2 | - | - | 0 | - |
| - | - | 2667 | 412.3 | - | - | 0 | - |
| - | - | 2785 | 413.2 | - | - | 0 | - |
| - | - | 2546 | 414.2 | - | - | 0 | - |
| - | - | 6286 | 414.3 | - | - | 0 | - |
| - | - | 1249 | 414.7 | - | - | 0 | - |
| - | - | 689.4 | 417.2 | - | - | 0 | - |
| - | - | 1043 | 418 | - | - | 0 | - |
| - | - | 1581 | 418.7 | - | - | 0 | - |
| - | - | 633.5 | 422.2 | - | - | 0 | - |
| - | - | 3389 | 423.2 | - | - | 0 | - |
| - | - | 3485 | 423.3 | - | - | 0 | - |
| - | - | 1220 | 423.7 | - | - | 0 | - |
| - | - | 4229 | 424.3 | - | - | 0 | - |
| - | - | 1530 | 425.3 | - | - | 0 | - |
| - | - | 1200 | 426.7 | - | - | 0 | - |
| - | - | 1373 | 427.2 | - | - | 0 | - |
| - | - | 2046 | 428.2 | - | - | 0 | - |
| - | - | 623 | 428.2 | - | - | 0 | - |
| - | - | 1964 | 429.1 | - | - | 0 | - |
| - | - | 1077 | 429.2 | - | - | 0 | - |
| - | - | 2007 | 429.2 | - | - | 0 | - |
| 4 | y | 1.733E+04 | 430.2 | 0.004357 | 10.13 | +3 | 10 |
| - | - | 635.6 | 430.7 | - | - | 0 | - |
| - | - | 4210 | 431.2 | - | - | 0 | - |
| - | - | 727.1 | 432.2 | - | - | 0 | - |
| 4 | b | 1007 | 433.3 | 0.0001709 | 0.3945 | +1 | 4 |
| - | - | 5780 | 434.7 | - | - | 0 | - |
| - | - | 2255 | 435.2 | - | - | 0 | - |
| 11 | y | 3269 | 439.2 | 0.001214 | 2.765 | +1 | 3 |
| 7 | b | 4673 | 439.7 | 0.003061 | 6.961 | +2 | 7 |
| - | - | 2924 | 440.2 | - | - | 0 | - |
| - | - | 926 | 440.7 | - | - | 0 | - |
| - | - | 1059 | 441.3 | - | - | 0 | - |
| - | - | 1.941E+04 | 442.3 | - | - | 0 | - |
| - | - | 2586 | 446.2 | - | - | 0 | - |
| - | - | 599.4 | 446.2 | - | - | 0 | - |
| - | - | 983.1 | 448.2 | - | - | 0 | - |
| 7 | b | 8535 | 448.7 | 0.003577 | 7.971 | +2 | 7 |
| - | - | 4175 | 449.2 | - | - | 0 | - |
| - | - | 814.8 | 449.7 | - | - | 0 | - |
| 4 | b | 6681 | 451.3 | 0.0006536 | 1.448 | +1 | 4 |
| - | - | 1849 | 452.3 | - | - | 0 | - |
| - | - | 1412 | 454.2 | - | - | 0 | - |
| 7 | y | 6121 | 454.7 | 0.003548 | 7.802 | +2 | 7 |
| - | - | 3617 | 455.2 | - | - | 0 | - |
| - | - | 1218 | 455.7 | - | - | 0 | - |
| - | - | 1202 | 458.2 | - | - | 0 | - |
| - | - | 838.9 | 461.2 | - | - | 0 | - |
| - | - | 2038 | 468.3 | - | - | 0 | - |
| - | - | 743.6 | 469.3 | - | - | 0 | - |
| - | - | 622.7 | 469.7 | - | - | 0 | - |
| - | - | 998.7 | 470.2 | - | - | 0 | - |
| - | - | 748.5 | 473.7 | - | - | 0 | - |
| - | - | 961.4 | 477.2 | - | - | 0 | - |
| 10 | y | 879.1 | 478.2 | 0.006429 | 13.44 | +1 | 4 |
| - | - | 1072 | 478.3 | - | - | 0 | - |
| - | - | 934.7 | 480.7 | - | - | 0 | - |
| - | - | 704.2 | 481.8 | - | - | 0 | - |
| - | - | 2586 | 482.3 | - | - | 0 | - |
| - | - | 1269 | 482.8 | - | - | 0 | - |
| - | - | 626.5 | 483.3 | - | - | 0 | - |
| - | - | 5419 | 483.3 | - | - | 0 | - |
| - | - | 1646 | 488.3 | - | - | 0 | - |
| - | - | 2448 | 491.3 | - | - | 0 | - |
| - | - | 1492 | 491.8 | - | - | 0 | - |
| - | - | 1.378E+04 | 494.2 | - | - | 0 | - |
| - | - | 3021 | 495.2 | - | - | 0 | - |
| - | - | 3709 | 495.3 | - | - | 0 | - |
| 8 | b | 1.935E+04 | 496.3 | 0.007447 | 15.01 | +2 | 8 |
| - | - | 5615 | 496.3 | - | - | 0 | - |
| - | - | 1.409E+04 | 496.8 | - | - | 0 | - |
| - | - | 2503 | 497.2 | - | - | 0 | - |
| - | - | 4455 | 497.3 | - | - | 0 | - |
| - | - | 2112 | 498.2 | - | - | 0 | - |
| - | - | 1364 | 503.9 | - | - | 0 | - |
| 8 | b | 2566 | 505.3 | 0.004605 | 9.114 | +2 | 8 |
| - | - | 2065 | 505.8 | - | - | 0 | - |
| - | - | 1439 | 506.1 | - | - | 0 | - |
| - | - | 963.1 | 507.1 | - | - | 0 | - |
| - | - | 655.9 | 507.3 | - | - | 0 | - |
| - | - | 886.9 | 510.2 | - | - | 0 | - |
| - | - | 1285 | 511.3 | - | - | 0 | - |
| - | - | 5225 | 511.3 | - | - | 0 | - |
| - | - | 8.308E+04 | 513.4 | - | - | 0 | - |
| - | - | 751.9 | 515.3 | - | - | 0 | - |
| - | - | 3165 | 523.3 | - | - | 0 | - |
| - | - | 3804 | 524.3 | - | - | 0 | - |
| - | - | 1950 | 524.8 | - | - | 0 | - |
| - | - | 2498 | 525.3 | - | - | 0 | - |
| - | - | 4257 | 525.8 | - | - | 0 | - |
| - | - | 1866 | 526.3 | - | - | 0 | - |
| - | - | 648.7 | 528.9 | - | - | 0 | - |
| 0 | Precursor | 1694 | 534.6 | 0.003871 | 7.242 | +3 | -1 |
| 0 | Precursor | 1128 | 534.9 | 0.005456 | 10.2 | +3 | -1 |
| - | - | 1221 | 535.3 | - | - | 0 | - |
| - | - | 1661 | 535.6 | - | - | 0 | - |
| - | - | 3798 | 535.9 | - | - | 0 | - |
| - | - | 925.6 | 539.2 | - | - | 0 | - |
| - | - | 1.405E+04 | 539.8 | - | - | 0 | - |
| - | - | 3765 | 540.3 | - | - | 0 | - |
| - | - | 7837 | 540.3 | - | - | 0 | - |
| - | - | 1207 | 540.4 | - | - | 0 | - |
| 0 | Precursor | 731 | 540.6 | 0.002791 | 5.163 | +3 | -1 |
| - | - | 3718 | 540.8 | - | - | 0 | - |
| - | - | 1332 | 540.9 | - | - | 0 | - |
| - | - | 1.254E+05 | 541.3 | - | - | 0 | - |
| - | - | 6784 | 541.6 | - | - | 0 | - |
| - | - | 7110 | 541.9 | - | - | 0 | - |
| - | - | 5397 | 543.3 | - | - | 0 | - |
| - | - | 1484 | 544.3 | - | - | 0 | - |
| - | - | 1425 | 548.2 | - | - | 0 | - |
| - | - | 660.8 | 551.3 | - | - | 0 | - |
| 9 | b | 1045 | 553.8 | 0.00136 | 2.455 | +2 | 9 |
| - | - | 856.6 | 554.2 | - | - | 0 | - |
| - | - | 6207 | 554.3 | - | - | 0 | - |
| - | - | 2936 | 554.8 | - | - | 0 | - |
| 9 | b | 1109 | 562.8 | 0.004927 | 8.756 | +2 | 9 |
| - | - | 2114 | 563.3 | - | - | 0 | - |
| - | - | 863.6 | 563.8 | - | - | 0 | - |
| - | - | 1080 | 564.3 | - | - | 0 | - |
| - | - | 1069 | 565.2 | - | - | 0 | - |
| - | - | 682.6 | 566.2 | - | - | 0 | - |
| - | - | 701.2 | 569.3 | - | - | 0 | - |
| - | - | 5845 | 573.8 | - | - | 0 | - |
| - | - | 2206 | 574.3 | - | - | 0 | - |
| 5 | b | 4726 | 579.3 | 0.0007614 | 1.314 | +1 | 5 |
| - | - | 1150 | 580.3 | - | - | 0 | - |
| - | - | 4575 | 583.2 | - | - | 0 | - |
| - | - | 1508 | 584.2 | - | - | 0 | - |
| - | - | 1108 | 587.3 | - | - | 0 | - |
| - | - | 1831 | 588.8 | - | - | 0 | - |
| - | - | 1382 | 589.3 | - | - | 0 | - |
| - | - | 1833 | 592.3 | - | - | 0 | - |
| 9 | y | 3345 | 593.3 | 0.001 | 1.686 | +1 | 5 |
| - | - | 1722 | 593.3 | - | - | 0 | - |
| - | - | 1000 | 594.3 | - | - | 0 | - |
| - | - | 1455 | 594.3 | - | - | 0 | - |
| - | - | 4.971E+04 | 597.8 | - | - | 0 | - |
| - | - | 2.796E+04 | 598.3 | - | - | 0 | - |
| - | - | 2989 | 610.3 | - | - | 0 | - |
| - | - | 958.7 | 610.8 | - | - | 0 | - |
| 9 | y | 4.209E+04 | 611.3 | 0.0009949 | 1.628 | +1 | 5 |
| - | - | 1.196E+04 | 612.3 | - | - | 0 | - |
| - | - | 2262 | 613.3 | - | - | 0 | - |
| - | - | 3347 | 619.8 | - | - | 0 | - |
| - | - | 3316 | 620.3 | - | - | 0 | - |
| - | - | 1590 | 620.8 | - | - | 0 | - |
| - | - | 945.5 | 621.3 | - | - | 0 | - |
| - | - | 856.9 | 624.3 | - | - | 0 | - |
| - | - | 3456 | 625.2 | - | - | 0 | - |
| - | - | 1727 | 627.3 | - | - | 0 | - |
| - | - | 735.6 | 628.3 | - | - | 0 | - |
| - | - | 2234 | 629.3 | - | - | 0 | - |
| - | - | 1293 | 629.8 | - | - | 0 | - |
| - | - | 8880 | 638.3 | - | - | 0 | - |
| - | - | 4164 | 638.8 | - | - | 0 | - |
| - | - | 769.5 | 639.3 | - | - | 0 | - |
| 4 | y | 939.4 | 644.8 | 0.00671 | 10.41 | +2 | 10 |
| 4 | y | 1153 | 645.3 | 0.005974 | 9.258 | +2 | 10 |
| 11 | b | 735.4 | 646.8 | 0.01272 | 19.67 | +2 | 11 |
| - | - | 4.406E+04 | 647.3 | - | - | 0 | - |
| - | - | 2.661E+04 | 647.8 | - | - | 0 | - |
| - | - | 851.9 | 648.3 | - | - | 0 | - |
| - | - | 2160 | 652.3 | - | - | 0 | - |
| - | - | 672.3 | 652.8 | - | - | 0 | - |
| 4 | y | 3962 | 653.8 | 0.003381 | 5.171 | +2 | 10 |
| - | - | 1757 | 654.3 | - | - | 0 | - |
| - | - | 820.4 | 654.8 | - | - | 0 | - |
| - | - | 902.6 | 656.3 | - | - | 0 | - |
| - | - | 2403 | 668.3 | - | - | 0 | - |
| - | - | 1339 | 669.3 | - | - | 0 | - |
| - | - | 2703 | 671.3 | - | - | 0 | - |
| - | - | 3412 | 672.8 | - | - | 0 | - |
| - | - | 999.4 | 673.3 | - | - | 0 | - |
| - | - | 1634 | 674.3 | - | - | 0 | - |
| 6 | b | 2200 | 676.3 | 0.0009236 | 1.366 | +1 | 6 |
| 6 | b | 891.7 | 677.3 | 0.0105 | 15.5 | +1 | 6 |
| - | - | 3170 | 678.3 | - | - | 0 | - |
| - | - | 1611 | 679.3 | - | - | 0 | - |
| - | - | 1096 | 680.3 | - | - | 0 | - |
| - | - | 1.075E+04 | 681.8 | - | - | 0 | - |
| - | - | 7315 | 682.3 | - | - | 0 | - |
| - | - | 1008 | 687.3 | - | - | 0 | - |
| - | - | 5158 | 689.4 | - | - | 0 | - |
| - | - | 1651 | 690.4 | - | - | 0 | - |
| - | - | 692.5 | 692.3 | - | - | 0 | - |
| 6 | b | 1.836E+04 | 694.4 | 0.0001856 | 0.2672 | +1 | 6 |
| - | - | 6973 | 695.4 | - | - | 0 | - |
| - | - | 1.23E+04 | 696.3 | - | - | 0 | - |
| - | - | 899.3 | 696.8 | - | - | 0 | - |
| - | - | 4037 | 697.3 | - | - | 0 | - |
| - | - | 1415 | 698.3 | - | - | 0 | - |
| - | - | 1928 | 699.3 | - | - | 0 | - |
| 3 | y | 3108 | 701.3 | 0.004961 | 7.074 | +2 | 11 |
| 3 | y | 5755 | 701.8 | 0.00984 | 14.02 | +2 | 11 |
| - | - | 2580 | 702.3 | - | - | 0 | - |
| - | - | 5962 | 702.8 | - | - | 0 | - |
| - | - | 5571 | 703.3 | - | - | 0 | - |
| 8 | y | 2236 | 706.3 | 0.005804 | 8.217 | +1 | 6 |
| - | - | 1148 | 709.3 | - | - | 0 | - |
| - | - | 2039 | 709.8 | - | - | 0 | - |
| 3 | y | 2.338E+04 | 710.3 | 0.003585 | 5.047 | +2 | 11 |
| 12 | b | 1.973E+04 | 710.8 | 0.0115 | 16.18 | +2 | 12 |
| 12 | b | 7718 | 711.3 | 0.001371 | 1.928 | +2 | 12 |
| - | - | 7.019E+04 | 711.9 | - | - | 0 | - |
| - | - | 4.526E+04 | 712.4 | - | - | 0 | - |
| - | - | 700.4 | 712.9 | - | - | 0 | - |
| 12 | b | 1112 | 719.8 | 0.003787 | 5.26 | +2 | 12 |
| - | - | 1068 | 720.4 | - | - | 0 | - |
| - | - | 706.4 | 720.9 | - | - | 0 | - |
| - | - | 810.7 | 721.4 | - | - | 0 | - |
| - | - | 720 | 722.4 | - | - | 0 | - |
| - | - | 882.4 | 723.4 | - | - | 0 | - |
| 8 | y | 1.449E+04 | 724.4 | 0.0008543 | 1.179 | +1 | 6 |
| - | - | 5430 | 725.4 | - | - | 0 | - |
| - | - | 2053 | 726.4 | - | - | 0 | - |
| - | - | 1380 | 728.4 | - | - | 0 | - |
| - | - | 1126 | 728.9 | - | - | 0 | - |
| - | - | 1773 | 737.4 | - | - | 0 | - |
| - | - | 796.2 | 737.9 | - | - | 0 | - |
| - | - | 1066 | 738.3 | - | - | 0 | - |
| - | - | 2648 | 739.4 | - | - | 0 | - |
| - | - | 1985 | 740.4 | - | - | 0 | - |
| - | - | 996.2 | 745.4 | - | - | 0 | - |
| - | - | 9959 | 746.4 | - | - | 0 | - |
| - | - | 7337 | 746.9 | - | - | 0 | - |
| - | - | 1755 | 751.4 | - | - | 0 | - |
| - | - | 765.2 | 753.4 | - | - | 0 | - |
| - | - | 1.222E+05 | 755.4 | - | - | 0 | - |
| - | - | 8.105E+04 | 755.9 | - | - | 0 | - |
| - | - | 871.4 | 756 | - | - | 0 | - |
| - | - | 2326 | 756.4 | - | - | 0 | - |
| 2 | y | 1920 | 759.9 | 0.0008721 | 1.148 | +2 | 12 |
| - | - | 5511 | 760.4 | - | - | 0 | - |
| - | - | 3093 | 760.9 | - | - | 0 | - |
| - | - | 5381 | 763.4 | - | - | 0 | - |
| - | - | 2196 | 764.4 | - | - | 0 | - |
| - | - | 944.6 | 770.4 | - | - | 0 | - |
| - | - | 1020 | 781.4 | - | - | 0 | - |
| - | - | 728.3 | 791.4 | - | - | 0 | - |
| - | - | 2039 | 795.4 | - | - | 0 | - |
| - | - | 719.5 | 797.4 | - | - | 0 | - |
| - | - | 2307 | 809.4 | - | - | 0 | - |
| 0 | Precursor | 2422 | 810.4 | 0.00783 | 9.663 | +2 | -1 |
| - | - | 797.2 | 811.3 | - | - | 0 | - |
| - | - | 774.3 | 812.3 | - | - | 0 | - |
| - | - | 935.5 | 818.4 | - | - | 0 | - |
| - | - | 757 | 824.4 | - | - | 0 | - |
| - | - | 1.088E+04 | 836.4 | - | - | 0 | - |
| - | - | 3682 | 837.4 | - | - | 0 | - |
| - | - | 1476 | 852.4 | - | - | 0 | - |
| - | - | 808.5 | 853.4 | - | - | 0 | - |
| - | - | 760.6 | 860.4 | - | - | 0 | - |
| - | - | 1333 | 861.4 | - | - | 0 | - |
| - | - | 1146 | 864.5 | - | - | 0 | - |
| - | - | 775.5 | 865.5 | - | - | 0 | - |
| - | - | 1166 | 868.4 | - | - | 0 | - |
| - | - | 897 | 869.4 | - | - | 0 | - |
| 7 | b | 4933 | 878.4 | 0.004853 | 5.525 | +1 | 7 |
| - | - | 2810 | 879.4 | - | - | 0 | - |
| - | - | 3857 | 880.4 | - | - | 0 | - |
| - | - | 684.2 | 882.4 | - | - | 0 | - |
| 7 | b | 1.696E+04 | 896.4 | 0.00503 | 5.612 | +1 | 7 |
| - | - | 9495 | 897.4 | - | - | 0 | - |
| - | - | 2812 | 898.4 | - | - | 0 | - |
| - | - | 819.7 | 906.4 | - | - | 0 | - |
| - | - | 1137 | 907.4 | - | - | 0 | - |
| 7 | y | 4174 | 908.4 | 0.007292 | 8.027 | +1 | 7 |
| - | - | 1943 | 909.4 | - | - | 0 | - |
| - | - | 902.6 | 916.4 | - | - | 0 | - |
| - | - | 3755 | 924.4 | - | - | 0 | - |
| - | - | 2187 | 925.4 | - | - | 0 | - |
| 7 | y | 1499 | 926.4 | 0.004845 | 5.229 | +1 | 7 |
| - | - | 998.9 | 971.4 | - | - | 0 | - |
| - | - | 1970 | 975.5 | - | - | 0 | - |
| - | - | 1252 | 976.5 | - | - | 0 | - |
| - | - | 1436 | 981.4 | - | - | 0 | - |
| - | - | 1436 | 986.5 | - | - | 0 | - |
| - | - | 1019 | 987.5 | - | - | 0 | - |
| - | - | 717.6 | 988.4 | - | - | 0 | - |
| - | - | 879.9 | 993.5 | - | - | 0 | - |
| 8 | b | 1089 | 1010 | 0.005927 | 5.871 | +1 | 8 |
| - | - | 1002 | 1012 | - | - | 0 | - |
| - | - | 1253 | 1014 | - | - | 0 | - |
| - | - | 988.4 | 1030 | - | - | 0 | - |
| 6 | y | 1229 | 1041 | 0.01568 | 15.06 | +1 | 8 |
| - | - | 796.5 | 1042 | - | - | 0 | - |
| - | - | 1199 | 1074 | - | - | 0 | - |
| - | - | 890.7 | 1092 | - | - | 0 | - |
| - | - | 718.4 | 1103 | - | - | 0 | - |
| - | - | 1106 | 1110 | - | - | 0 | - |
| 5 | y | 1739 | 1151 | 0.006244 | 5.422 | +1 | 9 |
| 5 | y | 1371 | 1152 | 0.002941 | 2.552 | +1 | 9 |
| - | - | 1064 | 1153 | - | - | 0 | - |
| - | - | 720.8 | 1161 | - | - | 0 | - |
| 5 | y | 3990 | 1170 | 0.004834 | 4.134 | +1 | 9 |
| - | - | 2534 | 1171 | - | - | 0 | - |
| - | - | 1137 | 1172 | - | - | 0 | - |
| - | - | 2941 | 1178 | - | - | 0 | - |
| - | - | 1890 | 1179 | - | - | 0 | - |
| - | - | 1162 | 1189 | - | - | 0 | - |
| - | - | 763.1 | 1277 | - | - | 0 | - |
| - | - | 728.5 | 1846 | - | - | 0 | - |
| - | - | 674.1 | 3402 | - | - | 0 | - |

m/z Charge Intensity FragmentType MassShift Position
120.0811538696289 0 7781.481
121.08441925048828 0 761.57404
122.07176971435547 0 472.73828
123.0444107055664 0 439.4446
124.93236541748047 0 369.58325
125.1077651977539 0 949.87286
126.05536651611328 0 1204.422
127.05076599121094 0 702.10913
128.1073455810547 0 7648.635
129.06642150878906 0 835.8776
129.1026153564453 0 60570.664
130.0654754638672 0 7267.5444
130.086669921875 0 422.6976
130.10594177246094 0 3729.949
131.0688018798828 0 593.05505
132.04476928710938 0 2046.9379
132.0811004638672 0 617.25574
132.1023712158203 0 1144.362
136.0760498046875 0 11758.912
137.0794677734375 0 1512.5135
138.0664825439453 0 506.51367
138.09178161621094 0 1074.204
139.0869140625 0 865.33905
141.102783203125 0 1219.9796
144.04531860351562 0 451.23145
146.0604248046875 0 4854.841
147.04444885253906 0 879.6803
147.07676696777344 0 468.79196 y 10
149.04476928710938 0 517.288
151.08700561523438 0 1718.9729
154.05014038085938 0 676.9557
154.08779907226562 0 600.56604
155.09300231933594 0 4098.6064
155.1181182861328 0 4187.4985 a Water loss 1
156.06600952148438 0 448.26947 y 11
156.07705688476562 0 3463.2087
157.06118774414062 0 952.65784
157.0764923095703 0 2484.4019
157.0974578857422 0 2336.1824
157.10865783691406 0 2805.77
157.13388061523438 0 755.7506
158.0604248046875 0 5577.7046
158.0930633544922 0 866.78375
159.0637969970703 0 904.5535
159.09210205078125 0 6603.4053
159.09869384765625 0 590.4677
159.1132354736328 0 1418.4412 d 1
160.09536743164062 0 723.1145
164.0474395751953 0 856.07745
165.05491638183594 0 5372.2183
166.0614013671875 0 8481.311
166.08660888671875 0 4124.9272 y 9
167.05557250976562 0 1001.1741
167.0645294189453 0 512.4987
167.0821990966797 0 772.6536
167.090087890625 0 559.9305
167.11837768554688 0 566.54724
168.10252380371094 0 651.8845
168.1142120361328 0 576.9447
168.83567810058594 0 409.18005
169.05264282226562 0 579.7629
169.0974578857422 0 2375.951
169.134033203125 0 1460.4861
171.0767822265625 0 1541.5813
172.07215881347656 0 1021.0677
173.05596923828125 0 743.8569
173.12887573242188 0 128692.89 a 1
174.1322021484375 0 11638.741
175.07176208496094 0 2111.4338
175.0869598388672 0 20497.707
175.11940002441406 0 1966.9075
175.13392639160156 0 598.6899
176.09022521972656 0 1858.8789
177.0928192138672 0 595.1744
177.10189819335938 0 454.40884
178.13417053222656 0 3777.4578
181.0611114501953 0 680.4464
182.0816192626953 0 10976.018 y 12
183.0848846435547 0 596.07684
183.1131134033203 0 13768.487 b Water loss 1
183.14952087402344 0 635.6429
184.1083984375 0 656.7574
184.11679077148438 0 541.0792
185.0714111328125 0 3107.572
185.092529296875 0 923.30853
185.165283203125 0 835.694
186.05543518066406 0 845.6977
186.12408447265625 0 19400.172
187.10801696777344 0 1572.024
187.1277313232422 0 1152.8284
187.1446990966797 0 1230.7808
188.07107543945312 0 1404.7029
189.08738708496094 0 6309.759
190.08242797851562 0 727.9523
190.13478088378906 0 888.92523
194.09246826171875 0 634.719
195.1132049560547 0 4153.0005
196.10830688476562 0 2701.1501
196.1172637939453 0 886.3977
197.12884521484375 0 1804.4883
198.08802795410156 0 589.96014
199.07167053222656 0 3086.2373
199.1441650390625 0 706.0079
201.12371826171875 0 49274.76 b 1
202.11842346191406 0 526.7694
202.12716674804688 0 3848.657
203.06629943847656 0 1721.2595
203.08204650878906 0 1231.0072
205.0976104736328 0 2514.3784
207.16046142578125 0 493.81915
210.23484802246094 0 696.2289
211.1079864501953 0 1970.9786 y Water loss 10
211.1555633544922 0 7304.6846
212.10328674316406 0 1156.5734
213.1237030029297 0 2479.1692
213.15908813476562 0 556.76746
215.1388397216797 0 852.1857
217.0822296142578 0 9957.083
218.08523559570312 0 811.7069
218.92202758789062 0 483.1444
222.1239471435547 0 783.35657
223.06369018554688 0 863.43
223.10838317871094 0 2332.4263
223.15574645996094 0 10388.215
224.10330200195312 0 1172.1038
224.15928649902344 0 1076.2308
226.08248901367188 0 1614.0459
226.11911010742188 0 745.44916 b Water loss 5
227.0668487548828 0 1471.0176
227.1143035888672 0 2273.1096
228.1339111328125 0 739.0254
228.1822509765625 0 1657.436
229.1185302734375 0 3604.6235
233.14035034179688 0 1466.5435
234.12420654296875 0 10354.409
234.13584899902344 0 818.23615
235.1277313232422 0 1165.9618
237.13522338867188 0 768.0593
238.11851501464844 0 791.1186
239.15069580078125 0 4099.986
240.1346435546875 0 3111.299
241.1182098388672 0 549.69525
241.13565063476562 0 759.7908
241.19052124023438 0 623.7744
242.11447143554688 0 2404.3125
243.14906311035156 0 608.1009
244.09329223632812 0 3112.4756
247.10826110839844 0 6080.7314
247.144775390625 0 523.20087
248.11358642578125 0 2065.8677
249.0985870361328 0 3295.7065
249.13583374023438 0 828.7987
251.15077209472656 0 34323.51
252.15396118164062 0 4126.5786
253.13365173339844 0 788.4688
255.1088409423828 0 1763.0981
255.14634704589844 0 1802.9751
256.1298522949219 0 583.1633
256.1769714355469 0 1700.7195
258.145263671875 0 910.87933
261.1338195800781 0 538.1354
262.119140625 0 704.2283
265.1297302246094 0 2343.2764
266.1251525878906 0 9228.087
267.12890625 0 966.3312
268.1770935058594 0 779.66315
270.1607360839844 0 5231.5503
271.1437683105469 0 599.2287
271.16357421875 0 805.7388
274.13067626953125 0 978.707
280.12939453125 0 937.31494
283.04888916015625 0 743.7292
283.1405029296875 0 16602.879
284.12579345703125 0 720.1766
284.143310546875 0 2339.6406
284.1595764160156 0 920.1333
286.1402893066406 0 2314.663
287.1719665527344 0 751.2521
288.1343688964844 0 574.07745
288.1546325683594 0 674.12976
288.1714782714844 0 1461.8553
293.1136779785156 0 12840.283 y Water loss 11
294.11749267578125 0 2619.0247
296.19769287109375 0 830.0401 b Water loss 2
297.1562805175781 0 4270.467
298.1397399902344 0 1352.4136
298.15704345703125 0 1927.4426
299.0621337890625 0 2537.8962
299.13543701171875 0 1351.4154
300.0626220703125 0 3208.8535
300.09783935546875 0 939.42834
301.0596008300781 0 5073.9453
301.15130615234375 0 9416.572
302.0599670410156 0 2608.1458
302.1541748046875 0 1187.334
303.0574951171875 0 970.1184
305.1614685058594 0 2594.4038
306.1379699707031 0 1237.7025 y 8
310.22393798828125 0 719.70996
311.12451171875 0 1730.5057 y 11
311.1379699707031 0 577.9309
312.1560363769531 0 10489.181
313.16009521484375 0 824.7582
313.1880798339844 0 2308.3008
314.1481018066406 0 2564.9778
314.20843505859375 0 748.3974 b 2
315.1670227050781 0 5767.875
316.150634765625 0 3086.1426
316.16845703125 0 1472.9465
318.10888671875 0 1484.1758
327.25048828125 0 1822.7646
330.16656494140625 0 4717.942
331.1706237792969 0 655.8113 b Water loss 7
332.15838623046875 0 696.36084
333.17767333984375 0 930.3698
337.1529541015625 0 550.70465
338.219482421875 0 1610.6442
341.18206787109375 0 1462.0315
342.1419982910156 0 2432.5088 y Ammonia loss 5
345.13134765625 0 1139.8427
346.1151123046875 0 2964.7605
347.119140625 0 607.3938
347.6797180175781 0 1894.3801 b 5
348.6576232910156 0 667.81995
349.1834716796875 0 957.80054
350.21905517578125 0 3075.88
354.26141357421875 0 579.7955
355.1687927246094 0 637.1688
355.24517822265625 0 889.0926
359.167724609375 0 1514.3495
361.1993408203125 0 611.0597
362.67919921875 0 2695.607 y 7
363.1415710449219 0 3955.188
363.1828918457031 0 1113.7538
364.14642333984375 0 769.75684
364.1728210449219 0 618.10754
366.1777038574219 0 672.1732
367.1627197265625 0 876.6336
369.2339782714844 0 780.05066
373.1833801269531 0 615.8867
378.2143249511719 0 1340.4932
379.20916748046875 0 7629.0005
380.2130432128906 0 1462.5956
381.1522521972656 0 10088.877
382.15594482421875 0 1700.5724
392.1923828125 0 1269.1091
393.7087707519531 0 684.89496
394.17236328125 0 1663.8324
396.2256164550781 0 2311.9329
397.2572326660156 0 2320.1128
398.1767578125 0 730.7003
399.1850280761719 0 743.002
401.2396240234375 0 2268.6025
402.1987609863281 0 1256.8326
407.2410888671875 0 1154.947
409.2205505371094 0 1118.9254
411.1304016113281 0 1897.3862
412.18328857421875 0 8102.647
412.2673034667969 0 2667.385
413.18511962890625 0 2785.3228
414.2342529296875 0 2545.832
414.2831115722656 0 6286.289
414.721923828125 0 1248.528
417.20172119140625 0 689.44495
418.0350341796875 0 1043.4032
418.7186279296875 0 1580.7448
422.20361328125 0 633.5248
423.23370361328125 0 3388.895
423.2724609375 0 3484.7466
423.7339172363281 0 1219.5408
424.267822265625 0 4228.9834
425.2506103515625 0 1529.8434
426.72467041015625 0 1200.3002
427.2216796875 0 1373.4231
428.1563720703125 0 2045.7324
428.21539306640625 0 622.9545
429.1404724121094 0 1964.2273
429.2079772949219 0 1077.4742
429.23651123046875 0 2007.3257
430.1940002441406 0 17328.81 y Water loss 3
430.7092590332031 0 635.61804
431.197509765625 0 4210.286
432.19708251953125 0 727.0757
433.2559509277344 0 1007.4433 b Water loss 3
434.7200622558594 0 5780.4365
435.2220764160156 0 2255.4575
439.2199401855469 0 3268.591 y 10
439.7120056152344 0 4673.1816 b Water loss 6
440.21612548828125 0 2924.2908
440.7166442871094 0 926.0096
441.2929382324219 0 1059.4186
442.2779235839844 0 19412.03
446.1673278808594 0 2585.7239
446.204345703125 0 599.4316
448.2063903808594 0 983.1461
448.7178039550781 0 8534.987 b 6
449.2183532714844 0 4175.3164
449.71875 0 814.8017
451.2669982910156 0 6680.885 b 3
452.26904296875 0 1849.3695
454.2073974609375 0 1412.4728
454.712158203125 0 6120.9316 y Water loss 6
455.2136535644531 0 3617.3313
455.7135009765625 0 1217.9116
458.2360534667969 0 1201.7368
461.2048645019531 0 838.9119
468.2938537597656 0 2038.4867
469.2906799316406 0 743.5693
469.6814880371094 0 622.73175
470.2031555175781 0 998.676
473.7407531738281 0 748.532
477.2127990722656 0 961.39465
478.2360534667969 0 879.14545 y Water loss 9
478.2767639160156 0 1071.866
480.7481994628906 0 934.7279
481.7632751464844 0 704.2379
482.2566833496094 0 2586.238
482.7554931640625 0 1269.4576
483.25982666015625 0 626.4577
483.3404541015625 0 5419.0464
488.2558288574219 0 1646.176
491.26263427734375 0 2448.4043
491.7638854980469 0 1492.3219
494.236328125 0 13781.255
495.23919677734375 0 3020.687
495.3413391113281 0 3708.995
496.2435302734375 0 19352.912 b Water loss 7
496.32513427734375 0 5614.874
496.7674560546875 0 14090.485
497.24072265625 0 2502.6675
497.2700500488281 0 4454.904
498.24774169921875 0 2112.0828
503.91448974609375 0 1364.4019
505.2608642578125 0 2565.961 b 7
505.7616271972656 0 2064.776
506.1063537597656 0 1438.9084
507.1053161621094 0 963.062
507.2559509277344 0 655.9467
510.2213439941406 0 886.9391
511.2642822265625 0 1285.038
511.3355407714844 0 5225.421
513.3514404296875 0 83082.73
515.2576293945312 0 751.9175
523.3355102539062 0 3164.7532
524.2630615234375 0 3804.2754
524.76513671875 0 1949.7506
525.2667236328125 0 2497.6064
525.8123779296875 0 4256.6567
526.3134765625 0 1866.0917
528.9130859375 0 648.7428
534.5935668945312 0 1694.1454 Precursor Water loss
534.9231567382812 0 1128.2734 Precursor Ammonia loss
535.2574462890625 0 1221.3091
535.604248046875 0 1661.3146
535.9407958984375 0 3797.685
539.2261962890625 0 925.5835
539.8099365234375 0 14054.522
540.2754516601562 0 3764.6978
540.3123168945312 0 7837.4517
540.3748168945312 0 1206.5684
540.5960083007812 0 730.96716 Precursor
540.8131103515625 0 3717.9617
540.9295654296875 0 1331.5421
541.3463134765625 0 125448.16
541.60986328125 0 6784.4937
541.9442138671875 0 7109.896
543.278564453125 0 5396.765
544.2804565429688 0 1483.501
548.1901245117188 0 1425.0469
551.2586059570312 0 660.8217
553.7658081054688 0 1045.2524 b Water loss 8
554.2393188476562 0 856.5817
554.2808837890625 0 6207.151
554.7835693359375 0 2936.2236
562.774658203125 0 1109.1482 b 8
563.2553100585938 0 2113.5432
563.7545166015625 0 863.61426
564.2573852539062 0 1079.6399
565.2157592773438 0 1069.4843
566.1990966796875 0 682.60986
569.2597045898438 0 701.174
573.7969360351562 0 5845.1094
574.2982788085938 0 2206.1028
579.32568359375 0 4725.6455 b 4
580.3285522460938 0 1150.1316
583.226806640625 0 4574.7603
584.229736328125 0 1508.353
587.2782592773438 0 1107.6567
588.7932739257812 0 1830.5743
589.2931518554688 0 1381.6455
592.306884765625 0 1832.7822
593.257568359375 0 3344.705 y Water loss 8
593.30615234375 0 1722.1849
594.2559814453125 0 1000.3158
594.3088989257812 0 1454.771
597.7971801757812 0 49710.027
598.2986450195312 0 27955.838
610.2849731445312 0 2988.5234
610.7864379882812 0 958.67426
611.2681274414062 0 42087.53 y 8
612.27099609375 0 11960.437
613.2752075195312 0 2261.678
619.7945556640625 0 3346.808
620.2965087890625 0 3315.7302
620.7980346679688 0 1590.4464
621.2994995117188 0 945.4682
624.2606201171875 0 856.8844
625.2473754882812 0 3456.0986
627.2626342773438 0 1726.6316
628.2618408203125 0 735.6304
629.3135375976562 0 2234.4258
629.8157348632812 0 1292.8141
638.3197021484375 0 8880.033
638.8216552734375 0 4163.606
639.31689453125 0 769.5049
644.7875366210938 0 939.3663 y Water loss 3
645.27880859375 0 1153.3309 y Ammonia loss 3
646.827392578125 0 735.40643 b Ammonia loss 10
647.3313598632812 0 44060.89
647.8329467773438 0 26606.271
648.338623046875 0 851.90906
652.3368530273438 0 2159.827
652.8475341796875 0 672.2792
653.7894897460938 0 3962.0542 y 3
654.288818359375 0 1756.7412
654.7877197265625 0 820.36615
656.2669677734375 0 902.62634
668.31494140625 0 2403.3247
669.3253784179688 0 1339.2068
671.3485107421875 0 2703.0518
672.8281860351562 0 3412.4019
673.33642578125 0 999.4196
674.3118896484375 0 1634.427
676.3422241210938 0 2200.46 b Water loss 5
677.3358154296875 0 891.71985 b Ammonia loss 5
678.3021240234375 0 3169.8486
679.2918090820312 0 1611.39
680.3096923828125 0 1096.0048
681.8343505859375 0 10752.625
682.334716796875 0 7314.7397
687.32861328125 0 1008.17865
689.3579711914062 0 5157.907
690.3611450195312 0 1650.9238
692.3217163085938 0 692.48584
694.35205078125 0 18356.947 b 5
695.3550415039062 0 6973.0903
696.310302734375 0 12297.983
696.8421630859375 0 899.3443
697.3128662109375 0 4037.2542
698.3170166015625 0 1414.9082
699.3390502929688 0 1927.7666
701.3278198242188 0 3108.4456 y Water loss 2
701.82470703125 0 5754.5913 y Ammonia loss 2
702.3248901367188 0 2580.2915
702.8463745117188 0 5961.7305
703.3474731445312 0 5570.976
706.346435546875 0 2235.9497 y Water loss 7
709.3280029296875 0 1148.0957
709.8326416015625 0 2039.4421
710.3317260742188 0 23377.879 y 2
710.8324584960938 0 19726.064 b Water loss 11
711.3345947265625 0 7717.982 b Ammonia loss 11
711.8521728515625 0 70187.52
712.354248046875 0 45263.76
712.8573608398438 0 700.37573
719.85302734375 0 1112.1276 b 11
720.3515014648438 0 1067.8274
720.8538208007812 0 706.4181
721.3651733398438 0 810.6617
722.3504028320312 0 720.04614
723.3672485351562 0 882.3735
724.35205078125 0 14489.678 y 7
725.3541259765625 0 5430.473
726.3565673828125 0 2052.711
728.3571166992188 0 1380.3735
728.8558349609375 0 1126.0284
737.3541870117188 0 1773.471
737.8584594726562 0 796.2037
738.3319091796875 0 1066.444
739.3740844726562 0 2648.323
740.36279296875 0 1985.0537
745.3516845703125 0 996.2034
746.3632202148438 0 9958.91
746.8646850585938 0 7337.439
751.3648071289062 0 1755.0007
753.365234375 0 765.2154
755.3684692382812 0 122227.164
755.8700561523438 0 81046.74
755.9530639648438 0 871.424
756.3731689453125 0 2325.7432
759.8632202148438 0 1920.3138 y 1
760.3627319335938 0 5510.628
760.8618774414062 0 3092.7058
763.3759765625 0 5380.9834
764.375 0 2196.2322
770.4150390625 0 944.56946
781.4034423828125 0 1020.15466
791.3818969726562 0 728.28925
795.374267578125 0 2039.2228
797.3733520507812 0 719.4703
809.3924560546875 0 2307.164
810.3783569335938 0 2421.7878 Precursor
811.3283081054688 0 797.1995
812.3409423828125 0 774.25073
818.4102172851562 0 935.4658
824.3606567382812 0 756.9933
836.427001953125 0 10875.362
837.43017578125 0 3682.0513
852.4332275390625 0 1475.9108
853.4331665039062 0 808.4908
860.39013671875 0 760.6155
861.3892211914062 0 1333.4426
864.4586791992188 0 1146.2626
865.4547729492188 0 775.50806
868.4374389648438 0 1166.2997
869.4354248046875 0 896.95685
878.4154663085938 0 4932.6284 b Water loss 6
879.4146728515625 0 2810.4543
880.4304809570312 0 3856.9392
882.4405517578125 0 684.18585
896.4262084960938 0 16964.771 b 6
897.4285278320312 0 9494.717
898.4287109375 0 2811.8687
906.414794921875 0 819.69275
907.4163208007812 0 1136.9755
908.417236328125 0 4173.877 y Water loss 6
909.4264526367188 0 1943.2878
916.4307250976562 0 902.5927
924.4210205078125 0 3755.474
925.4280395507812 0 2187.1472
926.4253540039062 0 1499.2734 y 6
971.4130249023438 0 998.9276
975.501220703125 0 1970.4734
976.4998168945312 0 1251.7863
981.4458618164062 0 1436.2566
986.4920043945312 0 1436.292
987.4913940429688 0 1019.13385
988.4401245117188 0 717.59705
993.5189819335938 0 879.9248
1009.5111694335938 0 1089.11 b 7
1012.484619140625 0 1002.26306
1014.4900512695312 0 1253.0973
1030.4822998046875 0 988.3597
1041.463134765625 0 1228.9088 y 5
1042.455322265625 0 796.4707
1073.5191650390625 0 1199.3292
1091.5233154296875 0 890.6771
1102.516357421875 0 718.3853
1109.5401611328125 0 1105.9795
1151.501708984375 0 1738.8339 y Water loss 4
1152.482421875 0 1371.0282 y Ammonia loss 4
1153.4915771484375 0 1063.7175
1160.5498046875 0 720.84033
1169.5108642578125 0 3989.6091 y 4
1170.5167236328125 0 2534.257
1171.5108642578125 0 1136.719
1177.561279296875 0 2940.7234
1178.5611572265625 0 1890.2444
1188.549072265625 0 1162.393
1276.6275634765625 0 763.085
1846.3431396484375 0 728.5081
3401.933837890625 0 674.1119

Spectrum Details

|  |  |
| --- | --- |
| Matched peaks? Matched peaks The total absolute number of peaks matched. Additionally in brackets the total fraction of peaks matched and the total number of peaks is shown. | 64 (11.39% of 562) |
| FDR? FDR The false discovery rate estimated for this peptide. It is calculated by matching all theoretical fragments with a non-integer shift with the raw peaks for this spectrum. This is done with 40 different shifts. The resulting percentage is the average number of annotated peaks over the number of annotated peaks with the correct spectrum. | 1.12% |
| Satellite FDR? Satellite FDR See the FDR for details on its calculation. This satellite ion specific FDR only contains the satellite ions (d/w) for I/L/J positions. | ∞ |
| PSM Score? PSM Score The PSM Score as given by Hecklib to this annotated spectrum. It is shown with three significant figures. | 423 |

#### Spectrum 6879? Spectrum 6879 The raw spectrum of this peptide as annotated by Hecklib. The fragments are coloured according to ion type (see legend). Any peaks with a star '\*' as text can be hovered over to see the full details, first the ion type second the mass shift type. By hovering over the amino acids in the peptide or ions in the legend the corresponding peaks are highlighted. By toggling the 'Unassigned' label you can turn the background (unassigned) peaks on or off in the plot. By updating the slider in the Ion legend you can update the spectrum to only show the top X% of the peaks with labels. The top X% means any peak that is within X% of the highest intensity. By dragging in the spectrum you can zoom in to a specific part of the spectrum and use 'Zoom Out' to get back to the original zoom level. The annotation of the spectrum is based on the given sequence in the peptides file and is done with different software so inconsistencies are likely. The peaks are annotated based on the given sequence, with 20 ppm tolerance.

Copy Data

##### Spectrum 6879 (TSV)

###### Preview

```
Loading example...
```

*Click on the button to copy the data to your clipboard.*

Mz MinMz MaxIntensity Max

WidthHeightPeptide font sizePeptide stroke widthSpectrum font sizeSpectrum stroke widthCompact peptide

Ion legend

wxyz

abcd

OtherUnassignedIon^Charge^_Position_Show for top:%

TVLHQDWLDGKEY

06.92e+61.38e+72.08e+72.77e+7

Zoom Out

y^+3^_3_a^+1^_2_a^+1^_2_y^+1^_1_b^+1^_2_b^+1^_2_y^+2^_3_y^+2^_4_a^+1^_3_b^+2^_5_y^+1^_2_b^+1^_3_y^+2^_5_y^+3^_7_y^+2^_5_y^+1^_2_b^+1^_3_b^+2^_6_b^+2^_6_b^+2^_6_y^+2^_6_y^+2^_6_b^+3^_9_y^+3^_9_b^+3^_10_y^+1^_3_y^+1^_3_b^+2^_7_b^+2^_7_b^+1^_4_y^+1^_3_b^+2^_7_y^+2^_7_y^+2^_7_b^+1^_4_y^+2^_7_y^+3^_11_b^+3^_12_y^+1^_4_b^+2^_8_b^+2^_8_y^+3^_12_y^+1^_4_\*\*\*b^+2^_9_b^+2^_9_b^+2^_9_b^+1^_5_b^+1^_5_b^+1^_5_y^+1^_5_y^+1^_5_y^+2^_10_y^+2^_10_b^+2^_11_y^+2^_10_b^+1^_6_b^+1^_6_y^+2^_11_y^+2^_11_b^+1^_6_y^+2^_11_b^+2^_12_b^+2^_12_y^+1^_6_b^+2^_12_y^+1^_6_y^+2^_12_y^+2^_12_b^+1^_7_b^+1^_7_b^+1^_7_y^+1^_7_b^+1^_8_b^+1^_8_y^+1^_8_y^+1^_8_y^+1^_8_b^+1^_9_b^+1^_9_y^+1^_9_y^+1^_9_y^+1^_9_b^+1^_10_y^+1^_10_

043987713161754

Fragment Matches Table

Show background peaks

| Position | Ion type | Intensity | mz Theoretical | mz Error (Th) | mz Error (ppm) | Charge | Series Number |
| --- | --- | --- | --- | --- | --- | --- | --- |
| - | - | 1.453E+04 | 121 | - | - | 0 | - |
| - | - | 1.24E+04 | 122.1 | - | - | 0 | - |
| - | - | 1.062E+05 | 122.1 | - | - | 0 | - |
| - | - | 1.007E+05 | 123 | - | - | 0 | - |
| - | - | 5.524E+04 | 123.1 | - | - | 0 | - |
| - | - | 1.39E+04 | 125.1 | - | - | 0 | - |
| - | - | 5.811E+04 | 125.1 | - | - | 0 | - |
| - | - | 8.929E+04 | 126.1 | - | - | 0 | - |
| - | - | 5.215E+05 | 127.1 | - | - | 0 | - |
| - | - | 2.237E+04 | 127.1 | - | - | 0 | - |
| - | - | 6.775E+04 | 127.1 | - | - | 0 | - |
| - | - | 3.463E+04 | 128.1 | - | - | 0 | - |
| - | - | 1.652E+06 | 128.1 | - | - | 0 | - |
| - | - | 1.034E+05 | 129.1 | - | - | 0 | - |
| - | - | 1.895E+07 | 129.1 | - | - | 0 | - |
| - | - | 8.965E+06 | 130.1 | - | - | 0 | - |
| - | - | 1.275E+05 | 130.1 | - | - | 0 | - |
| - | - | 1.266E+06 | 130.1 | - | - | 0 | - |
| - | - | 4.883E+04 | 131.1 | - | - | 0 | - |
| - | - | 9.473E+05 | 131.1 | - | - | 0 | - |
| - | - | 1.846E+04 | 131.1 | - | - | 0 | - |
| - | - | 2.824E+04 | 131.1 | - | - | 0 | - |
| - | - | 1.893E+06 | 132.1 | - | - | 0 | - |
| - | - | 3.001E+04 | 133.1 | - | - | 0 | - |
| - | - | 1.594E+05 | 133.1 | - | - | 0 | - |
| - | - | 1.926E+06 | 136.1 | - | - | 0 | - |
| - | - | 1.451E+05 | 137.1 | - | - | 0 | - |
| - | - | 2.323E+05 | 138.1 | - | - | 0 | - |
| - | - | 7.464E+04 | 138.1 | - | - | 0 | - |
| - | - | 3.565E+04 | 139.1 | - | - | 0 | - |
| - | - | 1.934E+05 | 139.1 | - | - | 0 | - |
| - | - | 1.698E+04 | 140.1 | - | - | 0 | - |
| - | - | 2.865E+05 | 141.1 | - | - | 0 | - |
| - | - | 2.52E+04 | 141.1 | - | - | 0 | - |
| - | - | 1.612E+05 | 142.1 | - | - | 0 | - |
| - | - | 2.679E+04 | 142.1 | - | - | 0 | - |
| - | - | 3.275E+04 | 142.1 | - | - | 0 | - |
| - | - | 1.039E+05 | 143 | - | - | 0 | - |
| - | - | 1.69E+04 | 143.1 | - | - | 0 | - |
| - | - | 2.129E+04 | 143.1 | - | - | 0 | - |
| - | - | 3.232E+04 | 144.1 | - | - | 0 | - |
| - | - | 9.491E+04 | 144.1 | - | - | 0 | - |
| - | - | 2.243E+05 | 145.1 | - | - | 0 | - |
| - | - | 3.49E+04 | 146.1 | - | - | 0 | - |
| - | - | 1.154E+05 | 147 | - | - | 0 | - |
| 11 | y | 1.668E+04 | 147.1 | 0.0005638 | 3.833 | +3 | 3 |
| - | - | 1.774E+04 | 148.1 | - | - | 0 | - |
| - | - | 2.325E+04 | 148.1 | - | - | 0 | - |
| - | - | 2.671E+04 | 149 | - | - | 0 | - |
| - | - | 1.324E+05 | 150.1 | - | - | 0 | - |
| - | - | 4.503E+05 | 151.1 | - | - | 0 | - |
| - | - | 1.624E+04 | 152.1 | - | - | 0 | - |
| - | - | 2.752E+04 | 152.1 | - | - | 0 | - |
| - | - | 3.625E+04 | 152.1 | - | - | 0 | - |
| - | - | 4.062E+04 | 153.1 | - | - | 0 | - |
| - | - | 6.237E+04 | 154.1 | - | - | 0 | - |
| - | - | 3.999E+04 | 154.1 | - | - | 0 | - |
| - | - | 4.776E+04 | 155 | - | - | 0 | - |
| - | - | 1.28E+06 | 155.1 | - | - | 0 | - |
| 2 | a | 7.502E+05 | 155.1 | 0.0007628 | 4.918 | +1 | 2 |
| - | - | 2.791E+04 | 155.1 | - | - | 0 | - |
| - | - | 6.583E+04 | 156.1 | - | - | 0 | - |
| - | - | 7.426E+04 | 156.1 | - | - | 0 | - |
| - | - | 3.77E+04 | 156.1 | - | - | 0 | - |
| - | - | 7.234E+04 | 156.1 | - | - | 0 | - |
| - | - | 7.184E+04 | 157.1 | - | - | 0 | - |
| - | - | 1.538E+05 | 158.1 | - | - | 0 | - |
| - | - | 1.566E+07 | 159.1 | - | - | 0 | - |
| - | - | 1.063E+05 | 160.1 | - | - | 0 | - |
| - | - | 8.325E+04 | 160.1 | - | - | 0 | - |
| - | - | 1.722E+06 | 160.1 | - | - | 0 | - |
| - | - | 8.072E+04 | 161.1 | - | - | 0 | - |
| - | - | 1.017E+06 | 165.1 | - | - | 0 | - |
| - | - | 5.996E+04 | 165.1 | - | - | 0 | - |
| - | - | 2.262E+06 | 166.1 | - | - | 0 | - |
| - | - | 1.909E+05 | 166.1 | - | - | 0 | - |
| - | - | 2.205E+04 | 167.1 | - | - | 0 | - |
| - | - | 1.663E+05 | 167.1 | - | - | 0 | - |
| - | - | 2.765E+04 | 167.1 | - | - | 0 | - |
| - | - | 2.064E+04 | 167.1 | - | - | 0 | - |
| - | - | 6.71E+04 | 167.1 | - | - | 0 | - |
| - | - | 2.652E+05 | 168.1 | - | - | 0 | - |
| - | - | 1.868E+05 | 168.1 | - | - | 0 | - |
| - | - | 1.517E+05 | 168.1 | - | - | 0 | - |
| - | - | 2.28E+05 | 169.1 | - | - | 0 | - |
| - | - | 2.81E+04 | 169.1 | - | - | 0 | - |
| - | - | 4.015E+05 | 169.1 | - | - | 0 | - |
| - | - | 1.744E+04 | 169.1 | - | - | 0 | - |
| - | - | 6.944E+05 | 170.1 | - | - | 0 | - |
| - | - | 8.314E+04 | 171.1 | - | - | 0 | - |
| - | - | 5.778E+04 | 171.1 | - | - | 0 | - |
| - | - | 3.221E+05 | 172.1 | - | - | 0 | - |
| - | - | 2.502E+04 | 172.1 | - | - | 0 | - |
| - | - | 3.157E+05 | 173.1 | - | - | 0 | - |
| 2 | a | 2.741E+07 | 173.1 | 0.0009098 | 5.255 | +1 | 2 |
| - | - | 2.493E+06 | 174.1 | - | - | 0 | - |
| - | - | 1.237E+05 | 175.1 | - | - | 0 | - |
| - | - | 3.953E+04 | 176.1 | - | - | 0 | - |
| - | - | 6.616E+04 | 177.1 | - | - | 0 | - |
| - | - | 8.635E+05 | 178.1 | - | - | 0 | - |
| - | - | 2.469E+04 | 179.1 | - | - | 0 | - |
| - | - | 9.531E+04 | 179.1 | - | - | 0 | - |
| - | - | 1.765E+05 | 181.1 | - | - | 0 | - |
| 13 | y | 2.284E+06 | 182.1 | 0.0008921 | 4.899 | +1 | 1 |
| - | - | 2.311E+05 | 183.1 | - | - | 0 | - |
| 2 | b | 7.035E+05 | 183.1 | 0.0009501 | 5.189 | +1 | 2 |
| - | - | 7.468E+04 | 184.1 | - | - | 0 | - |
| - | - | 1.881E+04 | 185.1 | - | - | 0 | - |
| - | - | 2.689E+04 | 185.1 | - | - | 0 | - |
| - | - | 1.846E+05 | 185.2 | - | - | 0 | - |
| - | - | 4.642E+06 | 186.1 | - | - | 0 | - |
| - | - | 1.239E+06 | 187.1 | - | - | 0 | - |
| - | - | 3.022E+04 | 187.1 | - | - | 0 | - |
| - | - | 3.564E+05 | 187.1 | - | - | 0 | - |
| - | - | 1.709E+04 | 187.1 | - | - | 0 | - |
| - | - | 1.064E+05 | 188.1 | - | - | 0 | - |
| - | - | 1.564E+05 | 188.1 | - | - | 0 | - |
| - | - | 2.726E+04 | 188.1 | - | - | 0 | - |
| - | - | 5.159E+04 | 189.1 | - | - | 0 | - |
| - | - | 4.399E+05 | 190.1 | - | - | 0 | - |
| - | - | 2.128E+05 | 190.1 | - | - | 0 | - |
| - | - | 2.627E+04 | 191.1 | - | - | 0 | - |
| - | - | 3.838E+04 | 193.1 | - | - | 0 | - |
| - | - | 1.54E+04 | 193.1 | - | - | 0 | - |
| - | - | 3.504E+04 | 194.1 | - | - | 0 | - |
| - | - | 1.925E+05 | 194.1 | - | - | 0 | - |
| - | - | 2.124E+04 | 194.1 | - | - | 0 | - |
| - | - | 1.38E+06 | 195.1 | - | - | 0 | - |
| - | - | 6.793E+05 | 196.1 | - | - | 0 | - |
| - | - | 9.924E+04 | 196.1 | - | - | 0 | - |
| - | - | 6.89E+04 | 197.1 | - | - | 0 | - |
| - | - | 1.11E+05 | 197.1 | - | - | 0 | - |
| - | - | 2.813E+04 | 197.2 | - | - | 0 | - |
| - | - | 1.653E+05 | 198.1 | - | - | 0 | - |
| - | - | 5.696E+05 | 198.1 | - | - | 0 | - |
| - | - | 1.377E+05 | 199.1 | - | - | 0 | - |
| - | - | 1.248E+05 | 199.1 | - | - | 0 | - |
| - | - | 1.039E+05 | 199.1 | - | - | 0 | - |
| - | - | 2.34E+04 | 200.1 | - | - | 0 | - |
| - | - | 1.145E+05 | 200.1 | - | - | 0 | - |
| 2 | b | 7.531E+06 | 201.1 | 0.0009598 | 4.772 | +1 | 2 |
| - | - | 6.483E+04 | 202.1 | - | - | 0 | - |
| - | - | 7.441E+05 | 202.1 | - | - | 0 | - |
| - | - | 6.25E+04 | 203.1 | - | - | 0 | - |
| - | - | 3.917E+04 | 203.2 | - | - | 0 | - |
| - | - | 1.9E+04 | 204.1 | - | - | 0 | - |
| - | - | 8.594E+04 | 204.1 | - | - | 0 | - |
| - | - | 4.401E+04 | 204.1 | - | - | 0 | - |
| - | - | 7.393E+05 | 205.1 | - | - | 0 | - |
| - | - | 8.188E+04 | 205.1 | - | - | 0 | - |
| - | - | 7.298E+04 | 206.1 | - | - | 0 | - |
| - | - | 1.215E+05 | 206.1 | - | - | 0 | - |
| - | - | 3.159E+04 | 207.2 | - | - | 0 | - |
| - | - | 2.444E+05 | 207.2 | - | - | 0 | - |
| - | - | 3.94E+04 | 208.1 | - | - | 0 | - |
| - | - | 3.329E+04 | 208.2 | - | - | 0 | - |
| - | - | 4.875E+04 | 209.1 | - | - | 0 | - |
| - | - | 2.624E+05 | 210.1 | - | - | 0 | - |
| 11 | y | 1.889E+05 | 211.1 | 0.001046 | 4.954 | +2 | 3 |
| - | - | 3.549E+04 | 211.1 | - | - | 0 | - |
| - | - | 2.862E+04 | 212.1 | - | - | 0 | - |
| - | - | 2.074E+05 | 212.1 | - | - | 0 | - |
| - | - | 6.202E+05 | 213.1 | - | - | 0 | - |
| - | - | 3.457E+04 | 213.1 | - | - | 0 | - |
| - | - | 6.358E+04 | 213.2 | - | - | 0 | - |
| - | - | 4.528E+04 | 214.1 | - | - | 0 | - |
| - | - | 4.466E+04 | 215.1 | - | - | 0 | - |
| - | - | 7.682E+04 | 215.1 | - | - | 0 | - |
| - | - | 1.723E+05 | 216.1 | - | - | 0 | - |
| - | - | 6.645E+04 | 221.1 | - | - | 0 | - |
| - | - | 7.053E+04 | 221.1 | - | - | 0 | - |
| - | - | 1.566E+05 | 221.1 | - | - | 0 | - |
| - | - | 1.065E+05 | 221.1 | - | - | 0 | - |
| - | - | 2.829E+05 | 222.1 | - | - | 0 | - |
| - | - | 5.6E+04 | 223.1 | - | - | 0 | - |
| - | - | 4.011E+05 | 223.1 | - | - | 0 | - |
| - | - | 5.169E+04 | 223.1 | - | - | 0 | - |
| - | - | 2.59E+06 | 223.2 | - | - | 0 | - |
| - | - | 8.319E+04 | 224.1 | - | - | 0 | - |
| - | - | 2.457E+04 | 224.1 | - | - | 0 | - |
| - | - | 3.066E+05 | 224.2 | - | - | 0 | - |
| - | - | 1.687E+04 | 225.1 | - | - | 0 | - |
| - | - | 2.997E+04 | 225.1 | - | - | 0 | - |
| - | - | 6.099E+04 | 225.2 | - | - | 0 | - |
| - | - | 1.653E+04 | 225.6 | - | - | 0 | - |
| - | - | 5.121E+05 | 226.1 | - | - | 0 | - |
| - | - | 2.506E+04 | 227.1 | - | - | 0 | - |
| - | - | 4.751E+05 | 227.1 | - | - | 0 | - |
| - | - | 4.148E+04 | 227.1 | - | - | 0 | - |
| - | - | 1.617E+05 | 227.2 | - | - | 0 | - |
| - | - | 5.23E+04 | 228.1 | - | - | 0 | - |
| - | - | 1.936E+05 | 228.1 | - | - | 0 | - |
| - | - | 1.622E+04 | 228.2 | - | - | 0 | - |
| - | - | 1.862E+05 | 229.1 | - | - | 0 | - |
| - | - | 4.537E+04 | 230.2 | - | - | 0 | - |
| - | - | 3.458E+04 | 230.2 | - | - | 0 | - |
| - | - | 4.246E+04 | 231.1 | - | - | 0 | - |
| - | - | 3.791E+05 | 233.1 | - | - | 0 | - |
| - | - | 2.668E+06 | 234.1 | - | - | 0 | - |
| - | - | 3.891E+04 | 234.7 | - | - | 0 | - |
| - | - | 2.492E+04 | 235.1 | - | - | 0 | - |
| - | - | 3.451E+05 | 235.1 | - | - | 0 | - |
| - | - | 7.957E+04 | 235.2 | - | - | 0 | - |
| - | - | 1.971E+04 | 235.6 | - | - | 0 | - |
| - | - | 4.056E+04 | 236.1 | - | - | 0 | - |
| - | - | 1.7E+05 | 237.1 | - | - | 0 | - |
| - | - | 2.459E+04 | 238.1 | - | - | 0 | - |
| - | - | 3.255E+05 | 238.1 | - | - | 0 | - |
| - | - | 7.426E+04 | 238.1 | - | - | 0 | - |
| - | - | 6.806E+04 | 239.1 | - | - | 0 | - |
| - | - | 4.662E+04 | 239.1 | - | - | 0 | - |
| - | - | 4.382E+04 | 239.1 | - | - | 0 | - |
| - | - | 1.176E+06 | 240.1 | - | - | 0 | - |
| - | - | 4.858E+04 | 241.1 | - | - | 0 | - |
| - | - | 1.645E+05 | 241.1 | - | - | 0 | - |
| - | - | 1.329E+05 | 241.1 | - | - | 0 | - |
| - | - | 1.55E+05 | 241.2 | - | - | 0 | - |
| - | - | 2.209E+05 | 243.1 | - | - | 0 | - |
| - | - | 1.195E+06 | 244.1 | - | - | 0 | - |
| - | - | 1.93E+04 | 244.6 | - | - | 0 | - |
| - | - | 1.319E+05 | 245.1 | - | - | 0 | - |
| - | - | 1.413E+06 | 247.1 | - | - | 0 | - |
| - | - | 8.395E+04 | 247.1 | - | - | 0 | - |
| - | - | 5.211E+04 | 247.2 | - | - | 0 | - |
| - | - | 1.924E+05 | 248.1 | - | - | 0 | - |
| - | - | 6.18E+05 | 248.1 | - | - | 0 | - |
| 10 | y | 3.846E+04 | 248.6 | 0.000794 | 3.194 | +2 | 4 |
| - | - | 9.043E+05 | 249.1 | - | - | 0 | - |
| - | - | 3.784E+04 | 249.1 | - | - | 0 | - |
| - | - | 2.67E+05 | 249.1 | - | - | 0 | - |
| - | - | 1.099E+05 | 250.1 | - | - | 0 | - |
| - | - | 5.586E+04 | 250.1 | - | - | 0 | - |
| - | - | 3.4E+04 | 250.2 | - | - | 0 | - |
| - | - | 9.764E+06 | 251.2 | - | - | 0 | - |
| - | - | 1.26E+06 | 252.2 | - | - | 0 | - |
| - | - | 4.319E+04 | 252.2 | - | - | 0 | - |
| - | - | 5.226E+04 | 252.6 | - | - | 0 | - |
| - | - | 2.976E+04 | 253.1 | - | - | 0 | - |
| - | - | 1.176E+05 | 253.2 | - | - | 0 | - |
| - | - | 1.784E+04 | 254.1 | - | - | 0 | - |
| - | - | 5.984E+05 | 255.1 | - | - | 0 | - |
| - | - | 2.161E+04 | 256.1 | - | - | 0 | - |
| - | - | 9.929E+04 | 256.1 | - | - | 0 | - |
| - | - | 3.022E+04 | 256.1 | - | - | 0 | - |
| - | - | 6.497E+04 | 256.2 | - | - | 0 | - |
| - | - | 7.41E+04 | 256.6 | - | - | 0 | - |
| - | - | 2.704E+05 | 257.1 | - | - | 0 | - |
| - | - | 6.863E+04 | 258.1 | - | - | 0 | - |
| - | - | 4.589E+05 | 258.1 | - | - | 0 | - |
| - | - | 5.602E+04 | 259.1 | - | - | 0 | - |
| - | - | 5.183E+04 | 259.1 | - | - | 0 | - |
| - | - | 2.732E+05 | 261.1 | - | - | 0 | - |
| - | - | 9.888E+04 | 261.1 | - | - | 0 | - |
| - | - | 2.509E+04 | 261.2 | - | - | 0 | - |
| - | - | 1.085E+05 | 262.1 | - | - | 0 | - |
| - | - | 1.463E+06 | 265.1 | - | - | 0 | - |
| - | - | 2.882E+06 | 266.1 | - | - | 0 | - |
| - | - | 3.732E+04 | 267.1 | - | - | 0 | - |
| - | - | 3.555E+05 | 267.1 | - | - | 0 | - |
| - | - | 3.468E+04 | 267.1 | - | - | 0 | - |
| - | - | 3.983E+04 | 267.6 | - | - | 0 | - |
| - | - | 2.016E+04 | 268.1 | - | - | 0 | - |
| - | - | 1.515E+05 | 268.1 | - | - | 0 | - |
| - | - | 2.334E+05 | 268.2 | - | - | 0 | - |
| 3 | a | 2.737E+04 | 268.2 | 0.001355 | 5.051 | +1 | 3 |
| - | - | 2.176E+04 | 269.1 | - | - | 0 | - |
| - | - | 9.273E+04 | 269.2 | - | - | 0 | - |
| - | - | 1.624E+05 | 269.2 | - | - | 0 | - |
| - | - | 3.739E+05 | 270.1 | - | - | 0 | - |
| - | - | 5.129E+04 | 270.1 | - | - | 0 | - |
| - | - | 3.939E+04 | 270.2 | - | - | 0 | - |
| - | - | 2.808E+04 | 270.2 | - | - | 0 | - |
| - | - | 1.383E+05 | 270.6 | - | - | 0 | - |
| - | - | 2.204E+04 | 271.1 | - | - | 0 | - |
| - | - | 6.611E+05 | 272.2 | - | - | 0 | - |
| - | - | 2.163E+04 | 273.2 | - | - | 0 | - |
| - | - | 1.028E+05 | 273.2 | - | - | 0 | - |
| - | - | 4.836E+05 | 274.1 | - | - | 0 | - |
| - | - | 1.65E+05 | 274.1 | - | - | 0 | - |
| - | - | 1.107E+05 | 275.1 | - | - | 0 | - |
| - | - | 7.542E+04 | 275.1 | - | - | 0 | - |
| - | - | 6.325E+04 | 275.2 | - | - | 0 | - |
| - | - | 1.277E+05 | 276.1 | - | - | 0 | - |
| - | - | 8.083E+04 | 276.1 | - | - | 0 | - |
| - | - | 2.142E+05 | 276.2 | - | - | 0 | - |
| - | - | 2.457E+04 | 277.1 | - | - | 0 | - |
| - | - | 3.931E+04 | 277.2 | - | - | 0 | - |
| - | - | 2.23E+04 | 278.2 | - | - | 0 | - |
| - | - | 2.19E+05 | 279.1 | - | - | 0 | - |
| - | - | 2.305E+04 | 280.1 | - | - | 0 | - |
| - | - | 2.032E+05 | 280.1 | - | - | 0 | - |
| - | - | 3.761E+04 | 280.1 | - | - | 0 | - |
| - | - | 2.54E+04 | 281.1 | - | - | 0 | - |
| - | - | 3.697E+04 | 281.1 | - | - | 0 | - |
| - | - | 1.823E+04 | 281.7 | - | - | 0 | - |
| - | - | 6.14E+05 | 282.2 | - | - | 0 | - |
| - | - | 4.507E+04 | 282.7 | - | - | 0 | - |
| - | - | 8.495E+06 | 283.1 | - | - | 0 | - |
| - | - | 1.575E+04 | 284.1 | - | - | 0 | - |
| - | - | 2.954E+05 | 284.1 | - | - | 0 | - |
| - | - | 1.334E+05 | 284.1 | - | - | 0 | - |
| - | - | 1.181E+06 | 284.1 | - | - | 0 | - |
| - | - | 2.468E+04 | 285.1 | - | - | 0 | - |
| - | - | 4.639E+04 | 285.1 | - | - | 0 | - |
| - | - | 2.341E+04 | 285.1 | - | - | 0 | - |
| - | - | 1.46E+05 | 285.2 | - | - | 0 | - |
| - | - | 4.281E+05 | 286.1 | - | - | 0 | - |
| - | - | 1.762E+04 | 286.2 | - | - | 0 | - |
| - | - | 7.44E+04 | 287.1 | - | - | 0 | - |
| - | - | 1.217E+05 | 287.2 | - | - | 0 | - |
| - | - | 4.14E+04 | 287.2 | - | - | 0 | - |
| - | - | 4.974E+04 | 288.1 | - | - | 0 | - |
| - | - | 1.2E+05 | 288.6 | - | - | 0 | - |
| 5 | b | 4.718E+04 | 290.2 | 0.001595 | 5.496 | +2 | 5 |
| - | - | 2.457E+04 | 291.2 | - | - | 0 | - |
| 12 | y | 3.534E+06 | 293.1 | 0.001548 | 5.281 | +1 | 2 |
| - | - | 1.034E+05 | 293.2 | - | - | 0 | - |
| - | - | 7.308E+05 | 294.1 | - | - | 0 | - |
| - | - | 7.694E+04 | 295.1 | - | - | 0 | - |
| - | - | 3.02E+04 | 296.1 | - | - | 0 | - |
| - | - | 4.506E+04 | 296.2 | - | - | 0 | - |
| 3 | b | 1.585E+05 | 296.2 | 0.001405 | 4.742 | +1 | 3 |
| - | - | 2.466E+04 | 296.6 | - | - | 0 | - |
| - | - | 5.576E+04 | 296.7 | - | - | 0 | - |
| 9 | y | 1.284E+05 | 297.1 | 0.002264 | 7.618 | +2 | 5 |
| - | - | 1.148E+06 | 297.2 | - | - | 0 | - |
| - | - | 4.02E+04 | 297.6 | - | - | 0 | - |
| 7 | y | 8.96E+04 | 298.1 | 0.004482 | 15.03 | +3 | 7 |
| - | - | 1.923E+05 | 298.2 | - | - | 0 | - |
| - | - | 4.557E+04 | 299.1 | - | - | 0 | - |
| - | - | 2.913E+04 | 300.1 | - | - | 0 | - |
| - | - | 9.752E+05 | 300.2 | - | - | 0 | - |
| - | - | 3.913E+06 | 301.2 | - | - | 0 | - |
| - | - | 1.518E+05 | 301.2 | - | - | 0 | - |
| - | - | 6.131E+05 | 302.1 | - | - | 0 | - |
| - | - | 5.2E+05 | 302.2 | - | - | 0 | - |
| - | - | 9.834E+04 | 303.1 | - | - | 0 | - |
| - | - | 8.004E+04 | 303.2 | - | - | 0 | - |
| - | - | 2.729E+04 | 304.2 | - | - | 0 | - |
| - | - | 1.239E+05 | 305.2 | - | - | 0 | - |
| - | - | 2.047E+04 | 305.2 | - | - | 0 | - |
| - | - | 2.092E+05 | 305.6 | - | - | 0 | - |
| - | - | 2.63E+04 | 305.7 | - | - | 0 | - |
| 9 | y | 9.821E+05 | 306.1 | 0.001773 | 5.79 | +2 | 5 |
| - | - | 5.733E+04 | 306.2 | - | - | 0 | - |
| - | - | 3.518E+05 | 306.6 | - | - | 0 | - |
| - | - | 7.875E+04 | 307.1 | - | - | 0 | - |
| - | - | 5.222E+04 | 309.2 | - | - | 0 | - |
| - | - | 7.834E+04 | 310.2 | - | - | 0 | - |
| - | - | 2.336E+04 | 310.2 | - | - | 0 | - |
| 12 | y | 4.753E+05 | 311.1 | 0.001939 | 6.233 | +1 | 2 |
| - | - | 8.882E+04 | 311.1 | - | - | 0 | - |
| - | - | 2.36E+04 | 311.2 | - | - | 0 | - |
| - | - | 1.047E+05 | 312.1 | - | - | 0 | - |
| - | - | 2.785E+04 | 313.1 | - | - | 0 | - |
| - | - | 1.786E+05 | 314.2 | - | - | 0 | - |
| 3 | b | 3.098E+05 | 314.2 | 0.001674 | 5.326 | +1 | 3 |
| - | - | 2.051E+06 | 315.2 | - | - | 0 | - |
| - | - | 6.926E+04 | 315.2 | - | - | 0 | - |
| - | - | 3.221E+05 | 316.2 | - | - | 0 | - |
| - | - | 2.933E+04 | 317.2 | - | - | 0 | - |
| - | - | 3.826E+04 | 317.7 | - | - | 0 | - |
| - | - | 5.83E+04 | 318.1 | - | - | 0 | - |
| - | - | 3.147E+04 | 318.1 | - | - | 0 | - |
| - | - | 2.808E+04 | 318.7 | - | - | 0 | - |
| - | - | 1.223E+05 | 319.2 | - | - | 0 | - |
| - | - | 2.86E+04 | 319.2 | - | - | 0 | - |
| - | - | 5.458E+04 | 321.2 | - | - | 0 | - |
| - | - | 3.569E+04 | 322.2 | - | - | 0 | - |
| - | - | 2.972E+04 | 324.2 | - | - | 0 | - |
| - | - | 2.808E+04 | 324.2 | - | - | 0 | - |
| - | - | 9.06E+04 | 325.2 | - | - | 0 | - |
| - | - | 5.081E+05 | 326.7 | - | - | 0 | - |
| - | - | 1.967E+05 | 327.2 | - | - | 0 | - |
| - | - | 6.26E+04 | 327.7 | - | - | 0 | - |
| - | - | 1.914E+04 | 330.2 | - | - | 0 | - |
| - | - | 4.407E+04 | 331.7 | - | - | 0 | - |
| - | - | 1.706E+05 | 332.2 | - | - | 0 | - |
| - | - | 6.877E+05 | 333.2 | - | - | 0 | - |
| - | - | 3.251E+04 | 333.2 | - | - | 0 | - |
| - | - | 3.945E+04 | 334.1 | - | - | 0 | - |
| - | - | 1.289E+05 | 334.2 | - | - | 0 | - |
| - | - | 4.62E+04 | 338.2 | - | - | 0 | - |
| 6 | b | 1.013E+05 | 338.7 | 0.001676 | 4.948 | +2 | 6 |
| 6 | b | 9.289E+04 | 339.2 | 0.005579 | 16.45 | +2 | 6 |
| - | - | 7.545E+04 | 339.7 | - | - | 0 | - |
| - | - | 1.717E+05 | 340.7 | - | - | 0 | - |
| - | - | 4.32E+04 | 341.2 | - | - | 0 | - |
| - | - | 2.544E+05 | 342.2 | - | - | 0 | - |
| - | - | 6.345E+04 | 343.2 | - | - | 0 | - |
| - | - | 3.615E+04 | 343.2 | - | - | 0 | - |
| - | - | 1.756E+05 | 344.2 | - | - | 0 | - |
| - | - | 2.978E+04 | 344.7 | - | - | 0 | - |
| - | - | 3.109E+05 | 345.1 | - | - | 0 | - |
| - | - | 2.58E+04 | 345.2 | - | - | 0 | - |
| - | - | 1.208E+06 | 346.1 | - | - | 0 | - |
| - | - | 6.35E+04 | 346.7 | - | - | 0 | - |
| - | - | 2.326E+05 | 347.1 | - | - | 0 | - |
| - | - | 2.964E+04 | 347.2 | - | - | 0 | - |
| 6 | b | 5.488E+05 | 347.7 | 0.001826 | 5.251 | +2 | 6 |
| - | - | 2.213E+04 | 348.1 | - | - | 0 | - |
| - | - | 1.937E+05 | 348.2 | - | - | 0 | - |
| - | - | 8.151E+04 | 348.7 | - | - | 0 | - |
| - | - | 3.323E+04 | 349.1 | - | - | 0 | - |
| - | - | 8.198E+04 | 349.2 | - | - | 0 | - |
| - | - | 8.174E+05 | 350.2 | - | - | 0 | - |
| - | - | 3.933E+04 | 351.1 | - | - | 0 | - |
| - | - | 4.061E+04 | 351.2 | - | - | 0 | - |
| - | - | 1.735E+05 | 351.2 | - | - | 0 | - |
| - | - | 3.541E+04 | 352.2 | - | - | 0 | - |
| - | - | 5.458E+04 | 352.7 | - | - | 0 | - |
| - | - | 3.234E+04 | 353.2 | - | - | 0 | - |
| 8 | y | 1.659E+05 | 353.7 | 0.00201 | 5.684 | +2 | 6 |
| - | - | 5.652E+04 | 354.2 | - | - | 0 | - |
| - | - | 2.271E+04 | 355.2 | - | - | 0 | - |
| - | - | 4.456E+04 | 356.2 | - | - | 0 | - |
| - | - | 6.278E+04 | 357.2 | - | - | 0 | - |
| - | - | 2.79E+04 | 358.2 | - | - | 0 | - |
| - | - | 5.785E+04 | 359.1 | - | - | 0 | - |
| - | - | 7.68E+04 | 360.2 | - | - | 0 | - |
| - | - | 2.436E+05 | 360.7 | - | - | 0 | - |
| - | - | 3.241E+05 | 361.2 | - | - | 0 | - |
| - | - | 8.234E+04 | 361.7 | - | - | 0 | - |
| - | - | 4.909E+05 | 362.2 | - | - | 0 | - |
| - | - | 2.641E+04 | 362.6 | - | - | 0 | - |
| - | - | 2.393E+04 | 362.7 | - | - | 0 | - |
| 8 | y | 1.752E+06 | 362.7 | 0.002007 | 5.535 | +2 | 6 |
| - | - | 1.799E+06 | 363.1 | - | - | 0 | - |
| - | - | 6.997E+05 | 363.2 | - | - | 0 | - |
| - | - | 1.753E+05 | 363.7 | - | - | 0 | - |
| - | - | 2.763E+05 | 364.1 | - | - | 0 | - |
| - | - | 2.543E+05 | 364.1 | - | - | 0 | - |
| 9 | b | 3.843E+04 | 364.2 | 0.003123 | 8.575 | +3 | 9 |
| - | - | 2.739E+04 | 364.7 | - | - | 0 | - |
| - | - | 5.606E+04 | 365.1 | - | - | 0 | - |
| - | - | 2.32E+04 | 365.2 | - | - | 0 | - |
| - | - | 2.93E+04 | 366.2 | - | - | 0 | - |
| - | - | 2.04E+05 | 366.2 | - | - | 0 | - |
| - | - | 1.138E+05 | 367.1 | - | - | 0 | - |
| - | - | 1.45E+05 | 367.2 | - | - | 0 | - |
| - | - | 1.215E+05 | 367.2 | - | - | 0 | - |
| - | - | 4.365E+04 | 367.7 | - | - | 0 | - |
| - | - | 3.159E+04 | 368.1 | - | - | 0 | - |
| - | - | 4.29E+04 | 368.2 | - | - | 0 | - |
| - | - | 8.396E+04 | 368.2 | - | - | 0 | - |
| - | - | 2.924E+04 | 368.2 | - | - | 0 | - |
| - | - | 2.9E+04 | 368.3 | - | - | 0 | - |
| - | - | 2.186E+04 | 369.2 | - | - | 0 | - |
| - | - | 3.064E+04 | 369.2 | - | - | 0 | - |
| - | - | 4.118E+04 | 369.2 | - | - | 0 | - |
| - | - | 2.183E+04 | 369.2 | - | - | 0 | - |
| - | - | 7.682E+04 | 373.2 | - | - | 0 | - |
| - | - | 5.858E+04 | 373.7 | - | - | 0 | - |
| - | - | 3.66E+04 | 374.7 | - | - | 0 | - |
| - | - | 2.42E+04 | 375.2 | - | - | 0 | - |
| - | - | 1.271E+05 | 376.2 | - | - | 0 | - |
| - | - | 2.198E+05 | 376.2 | - | - | 0 | - |
| - | - | 9.089E+04 | 376.7 | - | - | 0 | - |
| - | - | 3.664E+04 | 377.1 | - | - | 0 | - |
| - | - | 1.01E+05 | 377.1 | - | - | 0 | - |
| - | - | 1.73E+04 | 377.2 | - | - | 0 | - |
| - | - | 1.929E+04 | 377.2 | - | - | 0 | - |
| - | - | 3.235E+04 | 377.3 | - | - | 0 | - |
| - | - | 2.687E+04 | 378.2 | - | - | 0 | - |
| - | - | 6.347E+05 | 378.2 | - | - | 0 | - |
| - | - | 2.536E+06 | 379.2 | - | - | 0 | - |
| - | - | 5.712E+04 | 380.2 | - | - | 0 | - |
| - | - | 5.481E+05 | 380.2 | - | - | 0 | - |
| - | - | 4.853E+06 | 381.2 | - | - | 0 | - |
| - | - | 9.29E+04 | 381.2 | - | - | 0 | - |
| - | - | 1.771E+04 | 381.7 | - | - | 0 | - |
| - | - | 2.02E+05 | 381.7 | - | - | 0 | - |
| - | - | 8.753E+05 | 382.2 | - | - | 0 | - |
| - | - | 1.4E+05 | 382.2 | - | - | 0 | - |
| - | - | 7.779E+04 | 382.2 | - | - | 0 | - |
| - | - | 2.317E+04 | 382.3 | - | - | 0 | - |
| - | - | 5.06E+04 | 382.7 | - | - | 0 | - |
| - | - | 2.321E+04 | 382.7 | - | - | 0 | - |
| - | - | 1.679E+05 | 383.2 | - | - | 0 | - |
| - | - | 6.525E+04 | 383.2 | - | - | 0 | - |
| - | - | 2.177E+05 | 384.2 | - | - | 0 | - |
| - | - | 4.171E+05 | 384.2 | - | - | 0 | - |
| - | - | 2.811E+05 | 385.2 | - | - | 0 | - |
| 5 | y | 1.91E+05 | 385.2 | 0.002819 | 7.319 | +3 | 9 |
| - | - | 4.249E+04 | 385.2 | - | - | 0 | - |
| - | - | 5.685E+04 | 386.2 | - | - | 0 | - |
| - | - | 7.037E+04 | 386.2 | - | - | 0 | - |
| - | - | 4.509E+04 | 386.2 | - | - | 0 | - |
| - | - | 3.157E+04 | 387.2 | - | - | 0 | - |
| 10 | b | 7.616E+04 | 389.2 | 0.002426 | 6.233 | +3 | 10 |
| - | - | 1.751E+05 | 390.2 | - | - | 0 | - |
| - | - | 9.084E+04 | 390.7 | - | - | 0 | - |
| - | - | 1.977E+05 | 391.1 | - | - | 0 | - |
| - | - | 6.292E+04 | 391.2 | - | - | 0 | - |
| - | - | 4.367E+04 | 392.1 | - | - | 0 | - |
| - | - | 8.831E+04 | 392.2 | - | - | 0 | - |
| - | - | 3.95E+04 | 393.2 | - | - | 0 | - |
| - | - | 2.946E+04 | 393.2 | - | - | 0 | - |
| - | - | 8.274E+05 | 394.2 | - | - | 0 | - |
| - | - | 2.639E+05 | 395.1 | - | - | 0 | - |
| - | - | 2.807E+05 | 395.2 | - | - | 0 | - |
| - | - | 9.518E+04 | 395.2 | - | - | 0 | - |
| - | - | 3.976E+04 | 395.2 | - | - | 0 | - |
| - | - | 2.402E+04 | 395.2 | - | - | 0 | - |
| - | - | 1.414E+05 | 395.2 | - | - | 0 | - |
| - | - | 4.619E+04 | 396.1 | - | - | 0 | - |
| - | - | 5.627E+04 | 396.2 | - | - | 0 | - |
| - | - | 2.656E+04 | 396.2 | - | - | 0 | - |
| - | - | 6.781E+05 | 396.2 | - | - | 0 | - |
| - | - | 1.046E+05 | 397.2 | - | - | 0 | - |
| - | - | 1.376E+05 | 397.2 | - | - | 0 | - |
| - | - | 3.68E+04 | 397.7 | - | - | 0 | - |
| - | - | 3.432E+05 | 398.2 | - | - | 0 | - |
| - | - | 8.189E+04 | 399.2 | - | - | 0 | - |
| - | - | 2.402E+04 | 400.2 | - | - | 0 | - |
| - | - | 5.438E+04 | 401.2 | - | - | 0 | - |
| - | - | 7.692E+04 | 401.2 | - | - | 0 | - |
| - | - | 2.279E+05 | 402.2 | - | - | 0 | - |
| - | - | 5.681E+05 | 402.2 | - | - | 0 | - |
| - | - | 4.307E+04 | 403.1 | - | - | 0 | - |
| - | - | 2.332E+05 | 403.2 | - | - | 0 | - |
| - | - | 4.155E+04 | 404.2 | - | - | 0 | - |
| - | - | 2.238E+04 | 404.2 | - | - | 0 | - |
| - | - | 8.44E+04 | 404.7 | - | - | 0 | - |
| - | - | 3.217E+04 | 405.2 | - | - | 0 | - |
| - | - | 1.552E+05 | 405.3 | - | - | 0 | - |
| - | - | 3.256E+04 | 406.2 | - | - | 0 | - |
| - | - | 9.337E+04 | 406.2 | - | - | 0 | - |
| - | - | 8.493E+04 | 407.2 | - | - | 0 | - |
| - | - | 3.65E+04 | 407.2 | - | - | 0 | - |
| - | - | 3.548E+04 | 408.7 | - | - | 0 | - |
| - | - | 3.64E+05 | 409.1 | - | - | 0 | - |
| - | - | 3.227E+04 | 409.2 | - | - | 0 | - |
| - | - | 8.706E+04 | 409.2 | - | - | 0 | - |
| - | - | 3.795E+04 | 409.7 | - | - | 0 | - |
| - | - | 7.792E+04 | 410.2 | - | - | 0 | - |
| - | - | 3.755E+05 | 411.2 | - | - | 0 | - |
| - | - | 3.231E+06 | 412.2 | - | - | 0 | - |
| - | - | 1.481E+05 | 413.1 | - | - | 0 | - |
| - | - | 5.427E+05 | 413.2 | - | - | 0 | - |
| - | - | 4.141E+05 | 413.2 | - | - | 0 | - |
| - | - | 9.368E+04 | 413.3 | - | - | 0 | - |
| - | - | 4.842E+04 | 414.1 | - | - | 0 | - |
| - | - | 1.33E+05 | 414.2 | - | - | 0 | - |
| - | - | 5.434E+04 | 414.2 | - | - | 0 | - |
| - | - | 1.079E+06 | 414.2 | - | - | 0 | - |
| - | - | 2.94E+04 | 415.2 | - | - | 0 | - |
| - | - | 1.949E+05 | 415.2 | - | - | 0 | - |
| - | - | 2.389E+05 | 415.2 | - | - | 0 | - |
| - | - | 2.919E+04 | 416.2 | - | - | 0 | - |
| - | - | 5.543E+05 | 417.7 | - | - | 0 | - |
| - | - | 8.488E+05 | 418.2 | - | - | 0 | - |
| - | - | 3.714E+05 | 418.7 | - | - | 0 | - |
| - | - | 1.167E+05 | 419.2 | - | - | 0 | - |
| - | - | 2.643E+04 | 419.7 | - | - | 0 | - |
| - | - | 3.874E+04 | 420.2 | - | - | 0 | - |
| - | - | 2.81E+04 | 421.2 | - | - | 0 | - |
| 11 | y | 1.08E+05 | 421.2 | 0.002959 | 7.026 | +1 | 3 |
| 11 | y | 1.008E+05 | 422.2 | 0.002037 | 4.825 | +1 | 3 |
| - | - | 5.216E+04 | 422.3 | - | - | 0 | - |
| - | - | 9.25E+04 | 422.7 | - | - | 0 | - |
| - | - | 1.461E+05 | 423.2 | - | - | 0 | - |
| - | - | 1.277E+06 | 423.3 | - | - | 0 | - |
| - | - | 7.566E+04 | 423.7 | - | - | 0 | - |
| - | - | 5.145E+04 | 424.2 | - | - | 0 | - |
| - | - | 2.963E+05 | 424.3 | - | - | 0 | - |
| - | - | 8.633E+04 | 424.7 | - | - | 0 | - |
| - | - | 3.839E+04 | 425.2 | - | - | 0 | - |
| - | - | 5.496E+04 | 425.3 | - | - | 0 | - |
| - | - | 2.849E+04 | 426.2 | - | - | 0 | - |
| - | - | 8.674E+06 | 426.7 | - | - | 0 | - |
| - | - | 4.357E+06 | 427.2 | - | - | 0 | - |
| - | - | 1.371E+06 | 427.7 | - | - | 0 | - |
| - | - | 1.99E+05 | 428.2 | - | - | 0 | - |
| - | - | 2.497E+04 | 428.7 | - | - | 0 | - |
| - | - | 2.846E+04 | 429.2 | - | - | 0 | - |
| - | - | 1.858E+06 | 429.2 | - | - | 0 | - |
| - | - | 8.02E+06 | 430.2 | - | - | 0 | - |
| - | - | 1.621E+06 | 431.2 | - | - | 0 | - |
| 7 | b | 1.057E+06 | 431.7 | 0.002242 | 5.192 | +2 | 7 |
| 7 | b | 4.754E+05 | 432.2 | 0.007029 | 16.26 | +2 | 7 |
| - | - | 2.089E+05 | 432.7 | - | - | 0 | - |
| - | - | 2.828E+04 | 432.7 | - | - | 0 | - |
| - | - | 1.234E+05 | 433.2 | - | - | 0 | - |
| 4 | b | 2.896E+05 | 433.3 | 0.00191 | 4.409 | +1 | 4 |
| - | - | 2.28E+04 | 434.2 | - | - | 0 | - |
| - | - | 6.71E+04 | 434.3 | - | - | 0 | - |
| - | - | 3.695E+04 | 437.2 | - | - | 0 | - |
| - | - | 1.508E+05 | 438.2 | - | - | 0 | - |
| - | - | 9.88E+04 | 438.7 | - | - | 0 | - |
| 11 | y | 2.129E+06 | 439.2 | 0.002527 | 5.753 | +1 | 3 |
| - | - | 2.546E+04 | 439.7 | - | - | 0 | - |
| - | - | 1.105E+05 | 440.2 | - | - | 0 | - |
| - | - | 5.311E+05 | 440.2 | - | - | 0 | - |
| 7 | b | 4.589E+06 | 440.7 | 0.002391 | 5.426 | +2 | 7 |
| - | - | 2.2E+06 | 441.2 | - | - | 0 | - |
| - | - | 6.757E+05 | 441.7 | - | - | 0 | - |
| - | - | 1.198E+05 | 442.2 | - | - | 0 | - |
| - | - | 1.974E+04 | 442.3 | - | - | 0 | - |
| - | - | 5.731E+04 | 443.2 | - | - | 0 | - |
| - | - | 4.485E+04 | 444.2 | - | - | 0 | - |
| - | - | 1.99E+04 | 445.2 | - | - | 0 | - |
| - | - | 2.945E+04 | 446.2 | - | - | 0 | - |
| 7 | y | 4.679E+04 | 446.7 | 0.00072 | 1.612 | +2 | 7 |
| - | - | 3.759E+04 | 446.7 | - | - | 0 | - |
| 7 | y | 2.224E+05 | 447.2 | 0.004037 | 9.028 | +2 | 7 |
| - | - | 2.584E+04 | 447.3 | - | - | 0 | - |
| - | - | 7.56E+04 | 447.7 | - | - | 0 | - |
| - | - | 2.345E+04 | 448.2 | - | - | 0 | - |
| - | - | 4.476E+05 | 448.2 | - | - | 0 | - |
| - | - | 1.48E+05 | 449.2 | - | - | 0 | - |
| - | - | 9.232E+04 | 450.2 | - | - | 0 | - |
| - | - | 9.882E+04 | 450.2 | - | - | 0 | - |
| - | - | 1.139E+05 | 450.3 | - | - | 0 | - |
| 4 | b | 2.428E+06 | 451.3 | 0.002485 | 5.506 | +1 | 4 |
| - | - | 6.042E+05 | 452.3 | - | - | 0 | - |
| - | - | 2.956E+04 | 452.7 | - | - | 0 | - |
| - | - | 8.683E+04 | 453.3 | - | - | 0 | - |
| - | - | 3.732E+04 | 455.2 | - | - | 0 | - |
| - | - | 4.572E+04 | 455.2 | - | - | 0 | - |
| - | - | 1.568E+05 | 455.3 | - | - | 0 | - |
| 7 | y | 8.966E+04 | 455.7 | 0.005076 | 11.14 | +2 | 7 |
| - | - | 4.415E+04 | 456.2 | - | - | 0 | - |
| - | - | 4.592E+04 | 456.3 | - | - | 0 | - |
| - | - | 2.707E+04 | 456.7 | - | - | 0 | - |
| - | - | 3.257E+04 | 458.2 | - | - | 0 | - |
| - | - | 1.902E+05 | 459.2 | - | - | 0 | - |
| - | - | 1.827E+04 | 460.2 | - | - | 0 | - |
| - | - | 2.401E+04 | 460.2 | - | - | 0 | - |
| - | - | 3.068E+05 | 460.8 | - | - | 0 | - |
| - | - | 2.044E+05 | 461.2 | - | - | 0 | - |
| - | - | 3.953E+05 | 461.3 | - | - | 0 | - |
| - | - | 8.351E+04 | 461.8 | - | - | 0 | - |
| - | - | 4.334E+04 | 462.2 | - | - | 0 | - |
| - | - | 4.106E+04 | 462.3 | - | - | 0 | - |
| 3 | y | 2.699E+04 | 462.6 | 0.00237 | 5.124 | +3 | 11 |
| - | - | 2.683E+04 | 463.2 | - | - | 0 | - |
| - | - | 3.73E+04 | 465.3 | - | - | 0 | - |
| - | - | 8.575E+04 | 465.7 | - | - | 0 | - |
| - | - | 6.033E+04 | 466.2 | - | - | 0 | - |
| - | - | 1.153E+05 | 466.2 | - | - | 0 | - |
| - | - | 2.281E+04 | 468.2 | - | - | 0 | - |
| - | - | 5.839E+05 | 468.3 | - | - | 0 | - |
| - | - | 1.483E+05 | 469.3 | - | - | 0 | - |
| - | - | 2.345E+04 | 470.3 | - | - | 0 | - |
| - | - | 3.512E+04 | 472.2 | - | - | 0 | - |
| - | - | 4.971E+05 | 474.3 | - | - | 0 | - |
| - | - | 5.897E+05 | 474.8 | - | - | 0 | - |
| 12 | b | 2.587E+04 | 474.9 | 0.004659 | 9.811 | +3 | 12 |
| - | - | 3.667E+05 | 475.3 | - | - | 0 | - |
| - | - | 2.719E+04 | 475.6 | - | - | 0 | - |
| - | - | 8.01E+04 | 475.8 | - | - | 0 | - |
| - | - | 1.987E+05 | 476.2 | - | - | 0 | - |
| - | - | 4.035E+05 | 477.2 | - | - | 0 | - |
| - | - | 3.036E+04 | 478.2 | - | - | 0 | - |
| 10 | y | 2.924E+05 | 478.2 | 0.001332 | 2.786 | +1 | 4 |
| - | - | 2.495E+05 | 478.3 | - | - | 0 | - |
| - | - | 7.735E+04 | 479.2 | - | - | 0 | - |
| - | - | 7.734E+04 | 479.3 | - | - | 0 | - |
| - | - | 5.285E+04 | 479.7 | - | - | 0 | - |
| - | - | 6.353E+04 | 480.2 | - | - | 0 | - |
| - | - | 2.497E+04 | 480.3 | - | - | 0 | - |
| - | - | 5.128E+04 | 481.7 | - | - | 0 | - |
| - | - | 1.889E+06 | 483.3 | - | - | 0 | - |
| - | - | 1.032E+06 | 483.8 | - | - | 0 | - |
| - | - | 8.685E+04 | 484.2 | - | - | 0 | - |
| - | - | 3.118E+05 | 484.3 | - | - | 0 | - |
| - | - | 5.821E+04 | 484.8 | - | - | 0 | - |
| - | - | 2.107E+04 | 485.2 | - | - | 0 | - |
| - | - | 3.092E+04 | 487.2 | - | - | 0 | - |
| - | - | 3.168E+04 | 488.2 | - | - | 0 | - |
| 8 | b | 4.149E+05 | 488.3 | 0.003239 | 6.635 | +2 | 8 |
| 8 | b | 3.101E+05 | 488.7 | 0.009065 | 18.55 | +2 | 8 |
| - | - | 1.453E+05 | 489.3 | - | - | 0 | - |
| - | - | 2.55E+04 | 489.8 | - | - | 0 | - |
| - | - | 2.904E+04 | 490.2 | - | - | 0 | - |
| - | - | 4.61E+06 | 494.2 | - | - | 0 | - |
| - | - | 4.519E+04 | 495.2 | - | - | 0 | - |
| - | - | 1.289E+06 | 495.2 | - | - | 0 | - |
| 2 | y | 2.23E+04 | 495.9 | 0.007823 | 15.77 | +3 | 12 |
| 10 | y | 5.33E+06 | 496.2 | 0.0027 | 5.441 | +1 | 4 |
| - | - | 1.106E+06 | 497.2 | - | - | 0 | - |
| - | - | 6.805E+05 | 497.8 | - | - | 0 | - |
| - | - | 2.211E+05 | 498.2 | - | - | 0 | - |
| - | - | 3.097E+04 | 498.8 | - | - | 0 | - |
| - | - | 2.663E+04 | 499.2 | - | - | 0 | - |
| - | - | 5.347E+04 | 502.2 | - | - | 0 | - |
| - | - | 2.813E+04 | 503.2 | - | - | 0 | - |
| - | - | 2.211E+05 | 504.2 | - | - | 0 | - |
| - | - | 3.774E+04 | 505.2 | - | - | 0 | - |
| - | - | 3.112E+04 | 505.2 | - | - | 0 | - |
| - | - | 1.142E+05 | 506.2 | - | - | 0 | - |
| - | - | 6.502E+04 | 506.3 | - | - | 0 | - |
| - | - | 1.671E+05 | 507.3 | - | - | 0 | - |
| - | - | 2.214E+04 | 508.2 | - | - | 0 | - |
| - | - | 3.963E+04 | 508.3 | - | - | 0 | - |
| - | - | 4.292E+05 | 511.3 | - | - | 0 | - |
| - | - | 2.256E+04 | 512.2 | - | - | 0 | - |
| - | - | 1.299E+05 | 512.3 | - | - | 0 | - |
| - | - | 1.082E+05 | 513.2 | - | - | 0 | - |
| - | - | 1.977E+04 | 513.3 | - | - | 0 | - |
| - | - | 6.441E+04 | 514.2 | - | - | 0 | - |
| - | - | 2.638E+04 | 514.3 | - | - | 0 | - |
| - | - | 1.055E+05 | 515.3 | - | - | 0 | - |
| - | - | 2.746E+04 | 516.2 | - | - | 0 | - |
| - | - | 3.818E+04 | 516.3 | - | - | 0 | - |
| - | - | 2.477E+04 | 517.3 | - | - | 0 | - |
| - | - | 6.235E+04 | 519.3 | - | - | 0 | - |
| - | - | 5.509E+04 | 520.3 | - | - | 0 | - |
| - | - | 3.804E+04 | 521.3 | - | - | 0 | - |
| - | - | 7.626E+04 | 522.2 | - | - | 0 | - |
| - | - | 3.093E+04 | 522.9 | - | - | 0 | - |
| - | - | 1.772E+05 | 523.3 | - | - | 0 | - |
| - | - | 5.505E+04 | 523.6 | - | - | 0 | - |
| - | - | 2.98E+04 | 523.9 | - | - | 0 | - |
| - | - | 8.066E+04 | 524.3 | - | - | 0 | - |
| - | - | 8.479E+05 | 525.3 | - | - | 0 | - |
| - | - | 8.739E+04 | 526.2 | - | - | 0 | - |
| - | - | 2.535E+05 | 526.3 | - | - | 0 | - |
| - | - | 5.265E+04 | 527.3 | - | - | 0 | - |
| - | - | 5.678E+04 | 528.9 | - | - | 0 | - |
| 0 | Precursor | 3.331E+05 | 529.3 | 0.003854 | 7.282 | +3 | -1 |
| 0 | Precursor | 3.29E+05 | 529.6 | 0.008369 | 15.8 | +3 | -1 |
| - | - | 1.619E+05 | 529.9 | - | - | 0 | - |
| - | - | 1.014E+05 | 530.3 | - | - | 0 | - |
| - | - | 6.258E+04 | 531.2 | - | - | 0 | - |
| - | - | 1.694E+05 | 532.2 | - | - | 0 | - |
| - | - | 6.989E+04 | 533.2 | - | - | 0 | - |
| - | - | 3.34E+04 | 533.3 | - | - | 0 | - |
| - | - | 3.226E+04 | 534.2 | - | - | 0 | - |
| - | - | 5.275E+04 | 534.9 | - | - | 0 | - |
| 0 | Precursor | 1.835E+05 | 535.3 | 0.002896 | 5.411 | +3 | -1 |
| - | - | 3.838E+04 | 535.3 | - | - | 0 | - |
| - | - | 1.248E+05 | 535.6 | - | - | 0 | - |
| - | - | 9.025E+04 | 535.9 | - | - | 0 | - |
| - | - | 6.692E+04 | 537.2 | - | - | 0 | - |
| - | - | 6.277E+04 | 537.7 | - | - | 0 | - |
| - | - | 2.576E+04 | 538.2 | - | - | 0 | - |
| - | - | 8.024E+04 | 538.3 | - | - | 0 | - |
| - | - | 5.714E+04 | 538.8 | - | - | 0 | - |
| - | - | 9.66E+05 | 539.2 | - | - | 0 | - |
| - | - | 3.471E+04 | 539.3 | - | - | 0 | - |
| - | - | 2.866E+05 | 540.2 | - | - | 0 | - |
| - | - | 3.177E+04 | 540.8 | - | - | 0 | - |
| - | - | 6.437E+04 | 541.2 | - | - | 0 | - |
| - | - | 2.898E+05 | 542.3 | - | - | 0 | - |
| - | - | 1.06E+06 | 543.3 | - | - | 0 | - |
| - | - | 3.537E+05 | 544.3 | - | - | 0 | - |
| - | - | 8.101E+04 | 545.3 | - | - | 0 | - |
| 9 | b | 6.673E+04 | 545.8 | 0.002969 | 5.44 | +2 | 9 |
| 9 | b | 1.19E+05 | 546.3 | 0.008038 | 14.72 | +2 | 9 |
| - | - | 6.571E+04 | 546.8 | - | - | 0 | - |
| - | - | 1.004E+05 | 547.3 | - | - | 0 | - |
| - | - | 6.053E+04 | 547.8 | - | - | 0 | - |
| - | - | 4.854E+04 | 548.2 | - | - | 0 | - |
| - | - | 4.73E+04 | 548.3 | - | - | 0 | - |
| - | - | 2.176E+05 | 549.2 | - | - | 0 | - |
| - | - | 1.009E+05 | 550.2 | - | - | 0 | - |
| - | - | 3.809E+04 | 551.2 | - | - | 0 | - |
| - | - | 2.422E+05 | 551.3 | - | - | 0 | - |
| - | - | 1.173E+05 | 551.3 | - | - | 0 | - |
| - | - | 8.442E+04 | 552.3 | - | - | 0 | - |
| - | - | 3.634E+04 | 552.3 | - | - | 0 | - |
| - | - | 4.592E+04 | 553.3 | - | - | 0 | - |
| - | - | 3.537E+04 | 554.3 | - | - | 0 | - |
| 9 | b | 1.701E+05 | 554.8 | 0.007885 | 14.21 | +2 | 9 |
| - | - | 2.77E+05 | 555.3 | - | - | 0 | - |
| - | - | 1.805E+05 | 555.8 | - | - | 0 | - |
| - | - | 4.01E+04 | 556.3 | - | - | 0 | - |
| - | - | 3.322E+04 | 557.2 | - | - | 0 | - |
| - | - | 3.588E+04 | 558.2 | - | - | 0 | - |
| - | - | 3.994E+04 | 558.3 | - | - | 0 | - |
| - | - | 3.107E+04 | 559.3 | - | - | 0 | - |
| - | - | 4.845E+04 | 561.3 | - | - | 0 | - |
| 5 | b | 1.971E+05 | 561.3 | 0.002598 | 4.629 | +1 | 5 |
| 5 | b | 8.049E+04 | 562.3 | 0.006681 | 11.88 | +1 | 5 |
| - | - | 1.624E+05 | 563.3 | - | - | 0 | - |
| - | - | 2.772E+04 | 563.3 | - | - | 0 | - |
| - | - | 8.514E+04 | 564.3 | - | - | 0 | - |
| - | - | 6.21E+04 | 564.3 | - | - | 0 | - |
| - | - | 3.144E+04 | 564.8 | - | - | 0 | - |
| - | - | 4.591E+05 | 565.3 | - | - | 0 | - |
| - | - | 1.185E+05 | 566.3 | - | - | 0 | - |
| - | - | 1.039E+06 | 567.2 | - | - | 0 | - |
| - | - | 3.544E+04 | 567.8 | - | - | 0 | - |
| - | - | 3.015E+05 | 568.2 | - | - | 0 | - |
| - | - | 6.708E+04 | 568.8 | - | - | 0 | - |
| - | - | 6.976E+04 | 569.2 | - | - | 0 | - |
| - | - | 3.226E+04 | 569.3 | - | - | 0 | - |
| - | - | 3.564E+05 | 575.2 | - | - | 0 | - |
| - | - | 4.289E+04 | 575.3 | - | - | 0 | - |
| - | - | 1.836E+05 | 576.2 | - | - | 0 | - |
| - | - | 2.722E+04 | 576.3 | - | - | 0 | - |
| - | - | 6.617E+04 | 577.2 | - | - | 0 | - |
| - | - | 3.257E+04 | 577.3 | - | - | 0 | - |
| - | - | 3.845E+04 | 577.8 | - | - | 0 | - |
| - | - | 3.059E+04 | 578.3 | - | - | 0 | - |
| - | - | 4.115E+04 | 579.3 | - | - | 0 | - |
| 5 | b | 1.401E+06 | 579.3 | 0.003203 | 5.529 | +1 | 5 |
| - | - | 5.018E+05 | 580.3 | - | - | 0 | - |
| - | - | 3.966E+04 | 581.3 | - | - | 0 | - |
| - | - | 1.323E+05 | 581.3 | - | - | 0 | - |
| - | - | 5.8E+04 | 582.3 | - | - | 0 | - |
| - | - | 2.211E+04 | 582.8 | - | - | 0 | - |
| - | - | 2.771E+04 | 585.3 | - | - | 0 | - |
| - | - | 3.465E+04 | 586.3 | - | - | 0 | - |
| - | - | 3.751E+04 | 588.8 | - | - | 0 | - |
| - | - | 7.662E+04 | 589.3 | - | - | 0 | - |
| - | - | 2.875E+04 | 589.8 | - | - | 0 | - |
| - | - | 2.678E+05 | 592.3 | - | - | 0 | - |
| - | - | 1.755E+05 | 592.3 | - | - | 0 | - |
| 9 | y | 2.099E+06 | 593.3 | 0.003137 | 5.287 | +1 | 5 |
| - | - | 6.212E+05 | 593.3 | - | - | 0 | - |
| - | - | 6.551E+04 | 593.8 | - | - | 0 | - |
| - | - | 7.205E+05 | 594.3 | - | - | 0 | - |
| - | - | 1.71E+05 | 594.3 | - | - | 0 | - |
| - | - | 4.872E+04 | 594.8 | - | - | 0 | - |
| - | - | 1.942E+05 | 595.3 | - | - | 0 | - |
| - | - | 3.66E+04 | 596.3 | - | - | 0 | - |
| - | - | 6.126E+04 | 596.4 | - | - | 0 | - |
| - | - | 4.613E+04 | 597.3 | - | - | 0 | - |
| - | - | 3.997E+04 | 597.4 | - | - | 0 | - |
| - | - | 7.658E+04 | 597.8 | - | - | 0 | - |
| - | - | 4.126E+04 | 598.3 | - | - | 0 | - |
| - | - | 3.464E+04 | 599.3 | - | - | 0 | - |
| - | - | 3.177E+04 | 600.3 | - | - | 0 | - |
| - | - | 3.386E+04 | 601.3 | - | - | 0 | - |
| - | - | 1.146E+05 | 602.3 | - | - | 0 | - |
| - | - | 2.436E+05 | 602.8 | - | - | 0 | - |
| - | - | 2.639E+05 | 603.3 | - | - | 0 | - |
| - | - | 6.969E+04 | 603.8 | - | - | 0 | - |
| - | - | 4.008E+04 | 604.3 | - | - | 0 | - |
| - | - | 3.087E+04 | 605.3 | - | - | 0 | - |
| - | - | 2.651E+04 | 607.3 | - | - | 0 | - |
| - | - | 4.466E+04 | 609.4 | - | - | 0 | - |
| - | - | 4.158E+06 | 610.3 | - | - | 0 | - |
| 9 | y | 2.162E+07 | 611.3 | 0.003864 | 6.321 | +1 | 5 |
| - | - | 6.786E+05 | 611.8 | - | - | 0 | - |
| - | - | 7.132E+06 | 612.3 | - | - | 0 | - |
| - | - | 2.062E+05 | 612.8 | - | - | 0 | - |
| - | - | 1.597E+06 | 613.3 | - | - | 0 | - |
| - | - | 2.364E+05 | 614.3 | - | - | 0 | - |
| - | - | 3.455E+04 | 615.3 | - | - | 0 | - |
| - | - | 2.508E+04 | 616.3 | - | - | 0 | - |
| - | - | 1.027E+05 | 617.3 | - | - | 0 | - |
| - | - | 5.774E+04 | 618.3 | - | - | 0 | - |
| - | - | 3.61E+04 | 619.3 | - | - | 0 | - |
| - | - | 2.306E+04 | 620.3 | - | - | 0 | - |
| - | - | 7.813E+04 | 620.8 | - | - | 0 | - |
| - | - | 2.423E+05 | 621.3 | - | - | 0 | - |
| - | - | 1.057E+05 | 621.3 | - | - | 0 | - |
| - | - | 7.871E+04 | 622.3 | - | - | 0 | - |
| - | - | 3.506E+04 | 622.3 | - | - | 0 | - |
| - | - | 8.203E+04 | 622.4 | - | - | 0 | - |
| - | - | 2.789E+04 | 623.3 | - | - | 0 | - |
| - | - | 3.919E+04 | 623.4 | - | - | 0 | - |
| - | - | 4.902E+04 | 627.8 | - | - | 0 | - |
| - | - | 8.132E+04 | 628.3 | - | - | 0 | - |
| - | - | 5.036E+04 | 628.8 | - | - | 0 | - |
| - | - | 5.075E+04 | 630.3 | - | - | 0 | - |
| - | - | 2.311E+04 | 631.3 | - | - | 0 | - |
| - | - | 1.946E+05 | 634.3 | - | - | 0 | - |
| - | - | 2.744E+05 | 635.3 | - | - | 0 | - |
| - | - | 7.932E+04 | 636.3 | - | - | 0 | - |
| - | - | 1.281E+05 | 636.3 | - | - | 0 | - |
| 4 | y | 2.154E+05 | 636.8 | 0.004823 | 7.573 | +2 | 10 |
| 4 | y | 1.465E+05 | 637.3 | 0.01251 | 19.63 | +2 | 10 |
| - | - | 7.37E+04 | 637.3 | - | - | 0 | - |
| - | - | 7.199E+04 | 637.8 | - | - | 0 | - |
| 11 | b | 5.769E+04 | 638.3 | 0.01169 | 18.32 | +2 | 11 |
| - | - | 2.737E+04 | 638.8 | - | - | 0 | - |
| - | - | 1.159E+05 | 640.3 | - | - | 0 | - |
| - | - | 3.264E+04 | 641.3 | - | - | 0 | - |
| - | - | 3.157E+04 | 644.3 | - | - | 0 | - |
| - | - | 3.067E+05 | 645.3 | - | - | 0 | - |
| 4 | y | 8.888E+05 | 645.8 | 0.004972 | 7.7 | +2 | 10 |
| - | - | 8.065E+05 | 646.3 | - | - | 0 | - |
| - | - | 2.389E+05 | 646.8 | - | - | 0 | - |
| - | - | 7.659E+04 | 647.3 | - | - | 0 | - |
| - | - | 3.372E+04 | 647.4 | - | - | 0 | - |
| - | - | 5.754E+04 | 648.3 | - | - | 0 | - |
| - | - | 4.924E+04 | 649.3 | - | - | 0 | - |
| - | - | 9.262E+04 | 650.3 | - | - | 0 | - |
| - | - | 1.021E+05 | 650.3 | - | - | 0 | - |
| - | - | 4.257E+04 | 650.8 | - | - | 0 | - |
| - | - | 5.542E+04 | 651.3 | - | - | 0 | - |
| - | - | 1.93E+06 | 652.3 | - | - | 0 | - |
| - | - | 3.168E+04 | 652.8 | - | - | 0 | - |
| - | - | 7.649E+05 | 653.3 | - | - | 0 | - |
| - | - | 1.715E+05 | 654.3 | - | - | 0 | - |
| - | - | 6.48E+04 | 658.3 | - | - | 0 | - |
| - | - | 5.306E+04 | 658.3 | - | - | 0 | - |
| - | - | 1.063E+05 | 659.3 | - | - | 0 | - |
| - | - | 3.451E+04 | 660.3 | - | - | 0 | - |
| - | - | 5.096E+04 | 660.8 | - | - | 0 | - |
| - | - | 8.17E+04 | 661.3 | - | - | 0 | - |
| - | - | 3.008E+04 | 661.8 | - | - | 0 | - |
| - | - | 6.859E+05 | 662.3 | - | - | 0 | - |
| - | - | 4.248E+05 | 663.3 | - | - | 0 | - |
| - | - | 1.372E+05 | 664.3 | - | - | 0 | - |
| - | - | 1.103E+05 | 664.3 | - | - | 0 | - |
| - | - | 3.152E+04 | 665.3 | - | - | 0 | - |
| - | - | 5.976E+04 | 665.3 | - | - | 0 | - |
| - | - | 4.558E+04 | 666.4 | - | - | 0 | - |
| - | - | 5.114E+04 | 668.3 | - | - | 0 | - |
| - | - | 3.322E+04 | 669.3 | - | - | 0 | - |
| - | - | 2.972E+04 | 675.8 | - | - | 0 | - |
| 6 | b | 8.981E+05 | 676.3 | 0.003548 | 5.246 | +1 | 6 |
| 6 | b | 5.567E+05 | 677.3 | 0.01257 | 18.56 | +1 | 6 |
| - | - | 2.867E+05 | 678.3 | - | - | 0 | - |
| - | - | 3.157E+04 | 678.8 | - | - | 0 | - |
| - | - | 1.992E+05 | 679.3 | - | - | 0 | - |
| - | - | 4.129E+04 | 679.8 | - | - | 0 | - |
| - | - | 3.521E+06 | 680.3 | - | - | 0 | - |
| - | - | 1.386E+06 | 681.3 | - | - | 0 | - |
| - | - | 3.221E+05 | 682.3 | - | - | 0 | - |
| - | - | 2.372E+04 | 682.4 | - | - | 0 | - |
| - | - | 4.55E+04 | 683.3 | - | - | 0 | - |
| - | - | 1.34E+05 | 684.3 | - | - | 0 | - |
| - | - | 2.472E+05 | 684.8 | - | - | 0 | - |
| - | - | 1.744E+05 | 685.3 | - | - | 0 | - |
| - | - | 4.784E+04 | 685.8 | - | - | 0 | - |
| - | - | 3.341E+04 | 686.3 | - | - | 0 | - |
| - | - | 9.988E+04 | 688.3 | - | - | 0 | - |
| - | - | 4.219E+04 | 689.3 | - | - | 0 | - |
| - | - | 2.578E+04 | 691.3 | - | - | 0 | - |
| - | - | 2.442E+04 | 692.4 | - | - | 0 | - |
| - | - | 1.83E+05 | 692.8 | - | - | 0 | - |
| 3 | y | 9.758E+05 | 693.3 | 0.004783 | 6.898 | +2 | 11 |
| 3 | y | 8.61E+05 | 693.8 | 0.01186 | 17.09 | +2 | 11 |
| 6 | b | 9.863E+06 | 694.4 | 0.003176 | 4.574 | +1 | 6 |
| - | - | 2.07E+05 | 694.8 | - | - | 0 | - |
| - | - | 3.665E+06 | 695.4 | - | - | 0 | - |
| - | - | 8.797E+05 | 696.4 | - | - | 0 | - |
| - | - | 3.398E+04 | 697.3 | - | - | 0 | - |
| - | - | 1.908E+05 | 697.4 | - | - | 0 | - |
| - | - | 3.71E+04 | 697.9 | - | - | 0 | - |
| - | - | 6.789E+04 | 698.4 | - | - | 0 | - |
| - | - | 1.76E+06 | 701.8 | - | - | 0 | - |
| 3 | y | 5.735E+06 | 702.3 | 0.005482 | 7.805 | +2 | 11 |
| 12 | b | 4.582E+06 | 702.8 | 0.009237 | 13.14 | +2 | 12 |
| 12 | b | 2.17E+06 | 703.3 | 0.0004034 | 0.5736 | +2 | 12 |
| - | - | 5.511E+05 | 703.8 | - | - | 0 | - |
| - | - | 2.675E+05 | 704.3 | - | - | 0 | - |
| - | - | 1.334E+05 | 705.4 | - | - | 0 | - |
| 8 | y | 6.863E+05 | 706.3 | 0.004705 | 6.661 | +1 | 6 |
| - | - | 3.255E+05 | 707.3 | - | - | 0 | - |
| - | - | 8.348E+04 | 708.3 | - | - | 0 | - |
| - | - | 6.294E+04 | 708.3 | - | - | 0 | - |
| - | - | 3.5E+04 | 709.3 | - | - | 0 | - |
| - | - | 3.884E+05 | 711.4 | - | - | 0 | - |
| 12 | b | 4.529E+05 | 711.9 | 0.008186 | 11.5 | +2 | 12 |
| - | - | 3.139E+05 | 712.4 | - | - | 0 | - |
| - | - | 1.287E+05 | 712.9 | - | - | 0 | - |
| - | - | 4.086E+04 | 713.4 | - | - | 0 | - |
| - | - | 3.929E+04 | 715.3 | - | - | 0 | - |
| - | - | 2.378E+04 | 715.4 | - | - | 0 | - |
| - | - | 3.894E+04 | 716.3 | - | - | 0 | - |
| - | - | 2.933E+04 | 717.3 | - | - | 0 | - |
| - | - | 4.011E+04 | 718.4 | - | - | 0 | - |
| - | - | 8.538E+04 | 720.4 | - | - | 0 | - |
| - | - | 3.754E+04 | 720.9 | - | - | 0 | - |
| - | - | 4.548E+05 | 721.4 | - | - | 0 | - |
| - | - | 1.366E+05 | 722.4 | - | - | 0 | - |
| - | - | 1.226E+06 | 723.4 | - | - | 0 | - |
| 8 | y | 6.028E+06 | 724.4 | 0.004699 | 6.488 | +1 | 6 |
| - | - | 2.391E+06 | 725.4 | - | - | 0 | - |
| - | - | 6.649E+05 | 726.4 | - | - | 0 | - |
| - | - | 1.124E+05 | 727.4 | - | - | 0 | - |
| - | - | 7.872E+04 | 728.4 | - | - | 0 | - |
| - | - | 3.994E+04 | 728.9 | - | - | 0 | - |
| - | - | 1.535E+05 | 729.4 | - | - | 0 | - |
| - | - | 4.301E+04 | 729.9 | - | - | 0 | - |
| - | - | 7.518E+04 | 730.4 | - | - | 0 | - |
| - | - | 3.427E+04 | 732.3 | - | - | 0 | - |
| - | - | 3.075E+05 | 733.4 | - | - | 0 | - |
| - | - | 2.572E+05 | 734.4 | - | - | 0 | - |
| - | - | 9.422E+04 | 735.4 | - | - | 0 | - |
| - | - | 6.519E+04 | 737.9 | - | - | 0 | - |
| - | - | 6.046E+04 | 738.4 | - | - | 0 | - |
| - | - | 4.338E+04 | 739.4 | - | - | 0 | - |
| - | - | 2.541E+04 | 740.4 | - | - | 0 | - |
| 2 | y | 9.249E+04 | 742.9 | 0.004756 | 6.402 | +2 | 12 |
| - | - | 2.036E+05 | 743.4 | - | - | 0 | - |
| - | - | 6.637E+04 | 743.9 | - | - | 0 | - |
| - | - | 6.626E+04 | 744.4 | - | - | 0 | - |
| - | - | 3.968E+04 | 745.4 | - | - | 0 | - |
| - | - | 3.906E+04 | 746.3 | - | - | 0 | - |
| - | - | 9.72E+04 | 747.4 | - | - | 0 | - |
| - | - | 1.685E+05 | 748.4 | - | - | 0 | - |
| - | - | 7.449E+04 | 749.3 | - | - | 0 | - |
| - | - | 3.996E+04 | 749.4 | - | - | 0 | - |
| - | - | 3.725E+04 | 750.3 | - | - | 0 | - |
| - | - | 3.896E+04 | 750.4 | - | - | 0 | - |
| - | - | 1.025E+06 | 751.4 | - | - | 0 | - |
| 2 | y | 4.497E+05 | 751.9 | 0.006187 | 8.229 | +2 | 12 |
| - | - | 8.493E+05 | 752.4 | - | - | 0 | - |
| - | - | 1.708E+05 | 752.9 | - | - | 0 | - |
| - | - | 2.072E+05 | 753.4 | - | - | 0 | - |
| - | - | 3.381E+04 | 760.3 | - | - | 0 | - |
| - | - | 6.891E+04 | 761.4 | - | - | 0 | - |
| - | - | 1.544E+05 | 762.4 | - | - | 0 | - |
| - | - | 5.804E+05 | 763.4 | - | - | 0 | - |
| - | - | 2.878E+05 | 764.4 | - | - | 0 | - |
| - | - | 2.421E+05 | 765.4 | - | - | 0 | - |
| - | - | 1.393E+05 | 766.4 | - | - | 0 | - |
| - | - | 2.068E+05 | 767.4 | - | - | 0 | - |
| - | - | 2.93E+04 | 767.4 | - | - | 0 | - |
| - | - | 6.558E+04 | 768.4 | - | - | 0 | - |
| - | - | 5.27E+04 | 768.4 | - | - | 0 | - |
| - | - | 1.147E+05 | 769.4 | - | - | 0 | - |
| - | - | 4.959E+04 | 770.4 | - | - | 0 | - |
| - | - | 1.035E+05 | 775.4 | - | - | 0 | - |
| - | - | 1.229E+05 | 776.4 | - | - | 0 | - |
| - | - | 5.586E+04 | 777.4 | - | - | 0 | - |
| - | - | 2.595E+04 | 778.4 | - | - | 0 | - |
| - | - | 6.45E+05 | 779.4 | - | - | 0 | - |
| - | - | 2.527E+05 | 780.4 | - | - | 0 | - |
| - | - | 2.947E+05 | 781.4 | - | - | 0 | - |
| - | - | 1.399E+05 | 782.4 | - | - | 0 | - |
| - | - | 3.176E+04 | 783.4 | - | - | 0 | - |
| - | - | 4.043E+04 | 792.4 | - | - | 0 | - |
| - | - | 1.07E+06 | 793.4 | - | - | 0 | - |
| - | - | 4.911E+05 | 794.4 | - | - | 0 | - |
| - | - | 1.054E+05 | 795.3 | - | - | 0 | - |
| - | - | 1.028E+05 | 795.4 | - | - | 0 | - |
| - | - | 5.186E+04 | 796.3 | - | - | 0 | - |
| - | - | 3.637E+04 | 796.4 | - | - | 0 | - |
| - | - | 2.896E+04 | 799.4 | - | - | 0 | - |
| - | - | 1.886E+05 | 807.4 | - | - | 0 | - |
| - | - | 1.375E+05 | 807.4 | - | - | 0 | - |
| - | - | 9.831E+04 | 808.4 | - | - | 0 | - |
| - | - | 9.631E+04 | 808.4 | - | - | 0 | - |
| - | - | 5.73E+04 | 817.4 | - | - | 0 | - |
| - | - | 1.712E+05 | 818.4 | - | - | 0 | - |
| - | - | 8.982E+04 | 819.4 | - | - | 0 | - |
| - | - | 7.005E+04 | 820.4 | - | - | 0 | - |
| - | - | 8.218E+04 | 825.4 | - | - | 0 | - |
| - | - | 9.413E+04 | 826.4 | - | - | 0 | - |
| - | - | 5.558E+04 | 827.4 | - | - | 0 | - |
| - | - | 2.562E+04 | 829.4 | - | - | 0 | - |
| - | - | 1.568E+04 | 834.4 | - | - | 0 | - |
| - | - | 2.583E+05 | 834.4 | - | - | 0 | - |
| - | - | 3.19E+05 | 835.4 | - | - | 0 | - |
| - | - | 7.55E+05 | 836.5 | - | - | 0 | - |
| - | - | 3.497E+05 | 837.5 | - | - | 0 | - |
| - | - | 1.165E+05 | 838.5 | - | - | 0 | - |
| - | - | 1.003E+05 | 843.4 | - | - | 0 | - |
| - | - | 1.879E+05 | 844.4 | - | - | 0 | - |
| - | - | 1.694E+05 | 845.4 | - | - | 0 | - |
| - | - | 3.212E+05 | 846.5 | - | - | 0 | - |
| - | - | 2.207E+05 | 847.4 | - | - | 0 | - |
| - | - | 9.888E+04 | 848.4 | - | - | 0 | - |
| - | - | 7.48E+04 | 851.4 | - | - | 0 | - |
| - | - | 2.372E+06 | 852.4 | - | - | 0 | - |
| - | - | 1.179E+06 | 853.4 | - | - | 0 | - |
| - | - | 4.023E+05 | 854.4 | - | - | 0 | - |
| - | - | 7.115E+04 | 855.4 | - | - | 0 | - |
| - | - | 3.241E+04 | 856.4 | - | - | 0 | - |
| 7 | b | 1.306E+06 | 862.4 | 0.003642 | 4.223 | +1 | 7 |
| 7 | b | 1.044E+06 | 863.4 | 0.01657 | 19.2 | +1 | 7 |
| - | - | 1.87E+06 | 864.5 | - | - | 0 | - |
| - | - | 9.533E+05 | 865.5 | - | - | 0 | - |
| - | - | 2.764E+05 | 866.5 | - | - | 0 | - |
| - | - | 6.786E+04 | 867.5 | - | - | 0 | - |
| - | - | 2.699E+04 | 869.4 | - | - | 0 | - |
| - | - | 1.728E+05 | 876.4 | - | - | 0 | - |
| - | - | 1.033E+05 | 877.4 | - | - | 0 | - |
| - | - | 4.458E+04 | 878.4 | - | - | 0 | - |
| 7 | b | 1.138E+07 | 880.4 | 0.00388 | 4.407 | +1 | 7 |
| - | - | 6.107E+06 | 881.4 | - | - | 0 | - |
| - | - | 1.763E+06 | 882.4 | - | - | 0 | - |
| - | - | 3.405E+05 | 883.4 | - | - | 0 | - |
| - | - | 4.372E+04 | 884.4 | - | - | 0 | - |
| - | - | 6.021E+04 | 890.4 | - | - | 0 | - |
| - | - | 5.473E+04 | 891.4 | - | - | 0 | - |
| - | - | 2.127E+05 | 892.5 | - | - | 0 | - |
| - | - | 8.542E+04 | 893.5 | - | - | 0 | - |
| - | - | 1.006E+05 | 894.5 | - | - | 0 | - |
| - | - | 5.516E+04 | 895.5 | - | - | 0 | - |
| - | - | 1.445E+05 | 907.4 | - | - | 0 | - |
| - | - | 3.315E+05 | 908.4 | - | - | 0 | - |
| - | - | 3.162E+05 | 909.4 | - | - | 0 | - |
| 7 | y | 6.033E+05 | 910.4 | 0.006502 | 7.142 | +1 | 7 |
| - | - | 2.894E+05 | 911.4 | - | - | 0 | - |
| - | - | 7.451E+04 | 912.4 | - | - | 0 | - |
| - | - | 6.031E+04 | 936.4 | - | - | 0 | - |
| - | - | 8.64E+04 | 937.4 | - | - | 0 | - |
| - | - | 2.456E+04 | 938.4 | - | - | 0 | - |
| - | - | 6.729E+04 | 947.4 | - | - | 0 | - |
| - | - | 3.933E+04 | 948.4 | - | - | 0 | - |
| - | - | 1.119E+05 | 954.4 | - | - | 0 | - |
| - | - | 2.069E+05 | 955.4 | - | - | 0 | - |
| - | - | 8.43E+04 | 956.4 | - | - | 0 | - |
| - | - | 1.128E+05 | 964.5 | - | - | 0 | - |
| - | - | 1.988E+05 | 965.5 | - | - | 0 | - |
| - | - | 1.129E+05 | 966.5 | - | - | 0 | - |
| - | - | 8.666E+04 | 971.5 | - | - | 0 | - |
| - | - | 1.402E+05 | 972.5 | - | - | 0 | - |
| - | - | 5.761E+04 | 973.5 | - | - | 0 | - |
| - | - | 2.842E+04 | 974.5 | - | - | 0 | - |
| 8 | b | 1.087E+05 | 975.5 | 0.002403 | 2.463 | +1 | 8 |
| - | - | 9.501E+04 | 976.5 | - | - | 0 | - |
| - | - | 2.613E+04 | 977.5 | - | - | 0 | - |
| - | - | 5.922E+04 | 980.5 | - | - | 0 | - |
| - | - | 3.277E+04 | 981.5 | - | - | 0 | - |
| - | - | 3.081E+04 | 982.5 | - | - | 0 | - |
| 8 | b | 7.048E+05 | 993.5 | 0.004289 | 4.317 | +1 | 8 |
| - | - | 4.093E+05 | 994.5 | - | - | 0 | - |
| - | - | 1.598E+05 | 995.5 | - | - | 0 | - |
| - | - | 2.88E+04 | 996.5 | - | - | 0 | - |
| 6 | y | 7.45E+04 | 1007 | 0.01081 | 10.73 | +1 | 8 |
| 6 | y | 4.032E+04 | 1008 | 0.01233 | 12.23 | +1 | 8 |
| - | - | 8.49E+04 | 1024 | - | - | 0 | - |
| 6 | y | 3.739E+05 | 1025 | 0.007147 | 6.97 | +1 | 8 |
| - | - | 1.551E+05 | 1026 | - | - | 0 | - |
| - | - | 5.711E+04 | 1027 | - | - | 0 | - |
| - | - | 8.152E+04 | 1076 | - | - | 0 | - |
| - | - | 6.206E+04 | 1077 | - | - | 0 | - |
| 9 | b | 4.082E+04 | 1092 | 0.001842 | 1.688 | +1 | 9 |
| - | - | 3.901E+04 | 1093 | - | - | 0 | - |
| - | - | 1.489E+05 | 1094 | - | - | 0 | - |
| - | - | 8.274E+04 | 1095 | - | - | 0 | - |
| - | - | 3.321E+04 | 1096 | - | - | 0 | - |
| - | - | 3.737E+04 | 1108 | - | - | 0 | - |
| 9 | b | 3.923E+04 | 1109 | 0.01372 | 12.38 | +1 | 9 |
| - | - | 5.31E+04 | 1117 | - | - | 0 | - |
| - | - | 7.339E+04 | 1118 | - | - | 0 | - |
| - | - | 4.57E+04 | 1119 | - | - | 0 | - |
| - | - | 5.931E+04 | 1135 | - | - | 0 | - |
| 5 | y | 3.598E+05 | 1136 | 0.005521 | 4.862 | +1 | 9 |
| 5 | y | 3.736E+05 | 1136 | 0.01113 | 9.793 | +1 | 9 |
| - | - | 1.851E+05 | 1138 | - | - | 0 | - |
| - | - | 7.451E+04 | 1139 | - | - | 0 | - |
| - | - | 2.315E+05 | 1153 | - | - | 0 | - |
| 5 | y | 5.928E+05 | 1154 | 0.007896 | 6.845 | +1 | 9 |
| - | - | 4.008E+05 | 1155 | - | - | 0 | - |
| - | - | 1.406E+05 | 1156 | - | - | 0 | - |
| - | - | 3.241E+04 | 1157 | - | - | 0 | - |
| - | - | 2.645E+04 | 1165 | - | - | 0 | - |
| 10 | b | 6.75E+04 | 1166 | 0.008983 | 7.707 | +1 | 10 |
| 4 | y | 3.485E+04 | 1291 | 0.00524 | 4.06 | +1 | 10 |
| - | - | 2.266E+04 | 1737 | - | - | 0 | - |

m/z Charge Intensity FragmentType MassShift Position
121.04026794433594 0 14527.411
122.06812286376953 0 12397.871
122.07201385498047 0 106154.12
123.04472351074219 0 100671.84
123.05592346191406 0 55242.25
125.07140350341797 0 13901.272
125.10809326171875 0 58110.496
126.09199523925781 0 89286.48
127.05094909667969 0 521520.56
127.08758544921875 0 22368.95
127.1236572265625 0 67745.05
128.08255004882812 0 34626.35
128.1077423095703 0 1652295.9
129.0667266845703 0 103393
129.1029815673828 0 18950720
130.06582641601562 0 8965256
130.1001739501953 0 127542.47
130.10623168945312 0 1266204.1
131.0634307861328 0 48825.65
131.06912231445312 0 947341.5
131.1086883544922 0 18462.996
131.11827087402344 0 28243.729
132.08145141601562 0 1892993.1
133.06150817871094 0 30012.04
133.08485412597656 0 159374.86
136.076416015625 0 1925737
137.0797576904297 0 145059.56
138.06690979003906 0 232287.05
138.092041015625 0 74641.85
139.05096435546875 0 35654.02
139.0873565673828 0 193405.27
140.0908660888672 0 16981.219
141.10302734375 0 286482.66
141.1086883544922 0 25200.709
142.06590270996094 0 161164.08
142.1063995361328 0 26794.787
142.12355041503906 0 32754.65
143.04598999023438 0 103883.94
143.06912231445312 0 16898.17
143.11891174316406 0 21290.104
144.0763702392578 0 32317.246
144.0814971923828 0 94911.45
145.0615234375 0 224330.11
146.09317016601562 0 34895.98
147.04478454589844 0 115368.88
147.0783233642578 0 16676.23 y 10
148.05125427246094 0 17744.629
148.0612030029297 0 23245.404
148.95449829101562 0 26708.643
150.0669403076172 0 132401.8
151.08737182617188 0 450270.16
152.0717010498047 0 16241.592
152.0907745361328 0 27524.598
152.10781860351562 0 36251.582
153.10360717773438 0 40618.184
154.06202697753906 0 62373.855
154.0980682373047 0 39994.71
155.04603576660156 0 47762.92
155.09352111816406 0 1280055.4
155.11865234375 0 750187.06 a Water loss 1
155.12606811523438 0 27907.824
156.0775604248047 0 65833.54
156.0968017578125 0 74259.37
156.10317993164062 0 37702.58
156.12203979492188 0 72336.41
157.09805297851562 0 71838.414
158.0846405029297 0 153764.66
159.0924530029297 0 15662244
160.0763702392578 0 106268.36
160.08969116210938 0 83247.695
160.09579467773438 0 1722102
161.0990447998047 0 80717.48
165.055419921875 0 1017335.25
165.07789611816406 0 59962.67
166.06187438964844 0 2262333
166.09835815429688 0 190873.97
167.0599365234375 0 22046.307
167.06529235839844 0 166262.14
167.08248901367188 0 27654.473
167.10130310058594 0 20644.832
167.11866760253906 0 67101.414
168.08163452148438 0 265238.5
168.1027374267578 0 186829.84
168.11392211914062 0 151714.73
169.076904296875 0 228031.12
169.0846405029297 0 28096.373
169.09796142578125 0 401476.6
169.11769104003906 0 17436.51
170.0609588623047 0 694369.4
171.06431579589844 0 83135.14
171.09246826171875 0 57777.805
172.07269287109375 0 322147.16
172.10888671875 0 25019.39
173.056640625 0 315723
173.12936401367188 0 27407070 a 1
174.1326446533203 0 2492710.8
175.13473510742188 0 123671.73
176.0828857421875 0 39528.125
177.1030731201172 0 66157.6
178.13478088378906 0 863523.25
179.11892700195312 0 24693.37
179.13816833496094 0 95308.28
181.0616455078125 0 176528.78
182.08206176757812 0 2283961 y 12
183.0857391357422 0 231113.23
183.11375427246094 0 703466.8 b Water loss 1
184.11732482910156 0 74680.82
185.05650329589844 0 18807.367
185.07212829589844 0 26894.797
185.16583251953125 0 184631.4
186.12469482421875 0 4641670
187.08749389648438 0 1239421.6
187.12042236328125 0 30224.55
187.12811279296875 0 356398.9
187.14370727539062 0 17090.568
188.0716094970703 0 106441.16
188.0908966064453 0 156374.12
188.1294403076172 0 27255.074
189.09927368164062 0 51592.016
190.083251953125 0 439855.5
190.13491821289062 0 212811
191.13845825195312 0 26270.496
193.09828186035156 0 38380.81
193.10848999023438 0 15395.487
194.0828399658203 0 35041.984
194.0933380126953 0 192549.72
194.12960815429688 0 21237.916
195.1137237548828 0 1379723.2
196.1089630126953 0 679264.6
196.11727905273438 0 99236.875
197.10874938964844 0 68902.74
197.12936401367188 0 111003.7
197.1658172607422 0 28130.889
198.08822631835938 0 165302.97
198.12867736816406 0 569621.8
199.07217407226562 0 137724.56
199.08773803710938 0 124791.555
199.13201904296875 0 103867.71
200.10433959960938 0 23403.598
200.14381408691406 0 114535.41
201.12432861328125 0 7530558.5 b 1
202.08712768554688 0 64834.72
202.12754821777344 0 744052.3
203.12925720214844 0 62495.066
203.1506805419922 0 39173.168
204.07809448242188 0 18995.371
204.11399841308594 0 85943.31
204.13507080078125 0 44010.176
205.09808349609375 0 739275.5
205.1456756591797 0 81882.93
206.10183715820312 0 72979.336
206.1296844482422 0 121546.305
207.15077209472656 0 31587.76
207.1613311767578 0 244392.81
208.10946655273438 0 39404.93
208.1650390625 0 33286.16
209.05667114257812 0 48751.574
210.1271209716797 0 262391.72
211.1087646484375 0 188906.81 y Water loss 10
211.13160705566406 0 35485.63
212.11216735839844 0 28616.1
212.140380859375 0 207437.14
213.1243438720703 0 620186.25
213.14181518554688 0 34573.793
213.16090393066406 0 63580.168
214.1277313232422 0 45279.652
215.11917114257812 0 44656.926
215.14015197753906 0 76823.04
216.09896850585938 0 172275.6
221.07211303710938 0 66451.305
221.0929412841797 0 70532.68
221.10452270507812 0 156622.56
221.14085388183594 0 106515.016
222.12477111816406 0 282875.78
223.08778381347656 0 55997.41
223.10885620117188 0 401111.1
223.1295166015625 0 51687.562
223.1564178466797 0 2589957.8
224.10414123535156 0 83186.18
224.12083435058594 0 24565.32
224.15975952148438 0 306621.03
225.10336303710938 0 16872.385
225.12347412109375 0 29974.615
225.16116333007812 0 60992.734
225.62124633789062 0 16529.973
226.08335876464844 0 512121.6
227.05462646484375 0 25056.33
227.0673370361328 0 475052.3
227.08692932128906 0 41478.02
227.15548706054688 0 161747.39
228.07052612304688 0 52297.023
228.1354217529297 0 193613.67
228.1578826904297 0 16215.025
229.11936950683594 0 186209.78
230.15130615234375 0 45371.51
230.1873779296875 0 34580.57
231.08859252929688 0 42461.082
233.140869140625 0 379118.9
234.12490844726562 0 2667652.2
234.650634765625 0 38913.656
235.10830688476562 0 24915.43
235.12826538085938 0 345095.9
235.1561279296875 0 79566.66
235.6075439453125 0 19710.176
236.13059997558594 0 40558.46
237.13583374023438 0 170018.55
238.0995635986328 0 24593.248
238.11984252929688 0 325540.4
238.1315155029297 0 74259.68
239.08290100097656 0 68057.29
239.10354614257812 0 46617.875
239.11473083496094 0 43821.11
240.13540649414062 0 1176047.8
241.09857177734375 0 48578.023
241.11940002441406 0 164480.1
241.13877868652344 0 132894.23
241.19229125976562 0 155004.86
243.11044311523438 0 220902.12
244.09402465820312 0 1195131.8
244.61279296875 0 19296.072
245.0973663330078 0 131917.86
247.10894775390625 0 1413141.8
247.14540100097656 0 83953.8
247.15675354003906 0 52109.02
248.1036834716797 0 192370.83
248.11500549316406 0 618048.44
248.62452697753906 0 38458.277 y 9
249.09950256347656 0 904284.5
249.11964416503906 0 37842.58
249.13589477539062 0 267034
250.10316467285156 0 109889.29
250.11952209472656 0 55862.906
250.16744995117188 0 34003.406
251.15150451660156 0 9764276
252.1547088623047 0 1259977.2
252.1710968017578 0 43188.516
252.60421752929688 0 52257.758
253.12049865722656 0 29763.543
253.15701293945312 0 117594.66
254.0791015625 0 17835.934
255.14852905273438 0 598448.6
256.0959167480469 0 21606.574
256.1093444824219 0 99291.58
256.13018798828125 0 30215.959
256.152099609375 0 64966.06
256.61199951171875 0 74103.664
257.1142578125 0 270364.47
258.1239318847656 0 68633.18
258.1460876464844 0 458946.72
259.1297912597656 0 56017.934
259.1487731933594 0 51834.48
261.12030029296875 0 273245.84
261.1361389160156 0 98876.945
261.1601867675781 0 25093.23
262.12017822265625 0 108469.3
265.13055419921875 0 1462812.9
266.1261291503906 0 2881693
267.077392578125 0 37315.707
267.12933349609375 0 355472.84
267.1462097167969 0 34682.26
267.6138916015625 0 39830.938
268.09393310546875 0 20161.418
268.13055419921875 0 151535.27
268.1781311035156 0 233438.2
268.20330810546875 0 27369.518 a Water loss 2
269.1142272949219 0 21762.115
269.16192626953125 0 92728.75
269.1870422363281 0 162390.52
270.1229553222656 0 373896.25
270.1458435058594 0 51289.734
270.1605224609375 0 39391.184
270.18988037109375 0 28078.51
270.6247863769531 0 138270.64
271.12640380859375 0 22039.361
272.1770324707031 0 661076.8
273.1607971191406 0 21634.266
273.1803894042969 0 102808
274.1198425292969 0 483608.9
274.1322021484375 0 164957.97
275.1038818359375 0 110682.33
275.1227111816406 0 75422.89
275.1519470214844 0 63253.91
276.10986328125 0 127665.62
276.1278381347656 0 80832.88
276.15655517578125 0 214210.53
277.09515380859375 0 24574.234
277.15863037109375 0 39313.67
278.16192626953125 0 22302.541
279.1465759277344 0 218978.73
280.1151123046875 0 23051.861
280.13055419921875 0 203181.28
280.1477966308594 0 37608.36
281.0530700683594 0 25399.027
281.1342468261719 0 36967.52
281.6621398925781 0 18231.227
282.1575012207031 0 613953.5
282.6698913574219 0 45068.434
283.14154052734375 0 8494943
284.0884704589844 0 15745.447
284.10430908203125 0 295362.44
284.12469482421875 0 133361.19
284.1446228027344 0 1181169
285.091064453125 0 24676.723
285.1079406738281 0 46388.633
285.1304931640625 0 23409.521
285.1570739746094 0 146033.14
286.14117431640625 0 428127.56
286.1583251953125 0 17622.857
287.143798828125 0 74398.43
287.1728820800781 0 121743.15
287.1884765625 0 41402.043
288.12799072265625 0 49735.67
288.62030029296875 0 120006.77
290.1676940917969 0 47183.754 b 4
291.2183837890625 0 24567.97
293.11474609375 0 3534466.8 y Water loss 11
293.1627197265625 0 103403.914
294.1187744140625 0 730841.56
295.1214294433594 0 76942.43
296.08941650390625 0 30203.496
296.17840576171875 0 45063.27
296.1982727050781 0 158459.53 b Water loss 2
296.6430969238281 0 24656.676
296.6681823730469 0 55762.99
297.1341857910156 0 128440.66 y Water loss 8
297.1572570800781 0 1147994.4
297.63470458984375 0 40201.043
298.1403503417969 0 89595.47 y Water loss 6
298.1597900390625 0 192282.16
299.0630798339844 0 45573.348
300.11224365234375 0 29127.258
300.1688232421875 0 975164.56
301.1521911621094 0 3913027.2
301.1697692871094 0 151785.05
302.115234375 0 613060.8
302.1553955078125 0 520046.3
303.1189880371094 0 98339.484
303.1666259765625 0 80041.734
304.1713562011719 0 27293.305
305.18035888671875 0 123933.734
305.1980285644531 0 20471.473
305.6468811035156 0 209242.05
305.6828308105469 0 26297.777
306.13897705078125 0 982107.75 y 8
306.23016357421875 0 57327.016
306.6401062011719 0 351752.5
307.1413879394531 0 78746.5
309.1680908203125 0 52224.68
310.1542663574219 0 78335.64
310.173583984375 0 23361.72
311.1257019042969 0 475346.25 y 11
311.1395263671875 0 88816.58
311.1727294921875 0 23603.66
312.1290283203125 0 104696.94
313.1157531738281 0 27845.9
314.1876525878906 0 178609.2
314.2091064453125 0 309784.34 b 2
315.16790771484375 0 2050573.5
315.2121276855469 0 69257.234
316.17138671875 0 322091.16
317.1741943359375 0 29328.95
317.66058349609375 0 38264.004
318.1219482421875 0 58300.305
318.14990234375 0 31473.242
318.6531982421875 0 28077.146
319.1531677246094 0 122260.29
319.21368408203125 0 28603.66
321.1575622558594 0 54582.613
322.224365234375 0 35689.113
324.17327880859375 0 29715.676
324.1959533691406 0 28075.602
325.152099609375 0 90601.02
326.6654052734375 0 508148.28
327.16668701171875 0 196702.3
327.6688232421875 0 62597.27
330.1628723144531 0 19144.49
331.6563720703125 0 44072.453
332.20989990234375 0 170642.42
333.1785888671875 0 687706
333.2136535644531 0 32507.715
334.1198425292969 0 39445.684
334.1834411621094 0 128912.375
338.1840515136719 0 46195.645
338.67596435546875 0 101253.39 b Water loss 5
339.171875 0 92887 b Ammonia loss 5
339.6776428222656 0 75449.04
340.6629333496094 0 171658.33
341.164306640625 0 43199.13
342.18304443359375 0 254448.39
343.16314697265625 0 63450.293
343.18597412109375 0 36150.812
344.173095703125 0 175615.61
344.6700744628906 0 29782.297
345.13250732421875 0 310929.7
345.1744689941406 0 25802.924
346.1163635253906 0 1207620
346.69940185546875 0 63498.438
347.1194763183594 0 232560.48
347.20208740234375 0 29640.445
347.681396484375 0 548837.3 b 5
348.12030029296875 0 22127.229
348.1830749511719 0 193665.45
348.68359375 0 81505.51
349.12969970703125 0 33234.316
349.15283203125 0 81977.05
350.22039794921875 0 817439.8
351.14666748046875 0 39334.14
351.1676940917969 0 40610.465
351.22271728515625 0 173463
352.1669006347656 0 35406.66
352.6740417480469 0 54575.387
353.1806335449219 0 32337.855
353.67596435546875 0 165864.27 y Water loss 7
354.177978515625 0 56519.81
355.1625061035156 0 22709.516
356.19451904296875 0 44561.895
357.177734375 0 62779.52
358.1812438964844 0 27895.938
359.1376647949219 0 57848.547
360.20489501953125 0 76801.19
360.697021484375 0 243556.31
361.1986083984375 0 324050.7
361.6895751953125 0 82335.23
362.1866455078125 0 490898.5
362.64398193359375 0 26410.535
362.6502685546875 0 23931.582
362.6812438964844 0 1751842.9 y 7
363.1428527832031 0 1799274.5
363.18316650390625 0 699699.1
363.68408203125 0 175303.83
364.1263732910156 0 276311.1
364.1463928222656 0 254294.08
364.1851806640625 0 38425.145 b Water loss 8
364.6735534667969 0 27393.871
365.1286926269531 0 56058.215
365.1514892578125 0 23203.04
366.1576843261719 0 29296.262
366.1792297363281 0 204024.11
367.14111328125 0 113849.04
367.16326904296875 0 144985.17
367.24664306640625 0 121540.95
367.6857604980469 0 43646.816
368.146728515625 0 31588.89
368.1695861816406 0 42896.832
368.1940002441406 0 83956.81
368.22930908203125 0 29244.623
368.2527160644531 0 29004.08
369.15496826171875 0 21856.89
369.17474365234375 0 30643.79
369.1974792480469 0 41184.754
369.2354431152344 0 21829.979
373.1863708496094 0 76815.195
373.6877746582031 0 58578.695
374.69500732421875 0 36600.418
375.203857421875 0 24203.658
376.163818359375 0 127059.71
376.1994934082031 0 219801.31
376.70166015625 0 90885.16
377.12445068359375 0 36640.676
377.14739990234375 0 101017.31
377.19110107421875 0 17301.596
377.2039794921875 0 19285.361
377.26708984375 0 32352.803
378.15155029296875 0 26871.338
378.2154846191406 0 634732
379.2108459472656 0 2536459.8
380.1662292480469 0 57117.57
380.21380615234375 0 548069.8
381.1536865234375 0 4852914
381.2140808105469 0 92900.805
381.6909484863281 0 17709.348
381.712646484375 0 201983.19
382.1564025878906 0 875287.7
382.1904296875 0 139977.33
382.215087890625 0 77787.125
382.2714538574219 0 23171.457
382.6906433105469 0 50595.117
382.71746826171875 0 23206.213
383.1588439941406 0 167905.34
383.20770263671875 0 65245.863
384.16717529296875 0 217704.7
384.1901550292969 0 417146.03
385.15179443359375 0 281064.78
385.1740417480469 0 190965.66 y 4
385.1957092285156 0 42487.227
386.1559143066406 0 56849.168
386.1796875 0 70369.88
386.2060852050781 0 45087.305
387.1852111816406 0 31570.857
389.1951599121094 0 76158.94 b 9
390.19732666015625 0 175139.17
390.699462890625 0 90836.3
391.1379089355469 0 197669.19
391.188720703125 0 62923.03
392.1217346191406 0 43667.266
392.20941162109375 0 88312.58
393.1895446777344 0 39501.043
393.214111328125 0 29463.29
394.1741943359375 0 827430.2
395.1359558105469 0 263860.62
395.1585388183594 0 280691.62
395.17938232421875 0 95183.195
395.2060852050781 0 39763.188
395.2150573730469 0 24015.227
395.2422790527344 0 141365.5
396.138916015625 0 46189.08
396.1618957519531 0 56268.375
396.1922607421875 0 26563.357
396.2263488769531 0 678118.06
397.2051696777344 0 104632.09
397.2297058105469 0 137631.88
397.7063903808594 0 36800.35
398.1800537109375 0 343168.9
399.1844787597656 0 81887.875
400.2005615234375 0 24024.502
401.1845703125 0 54384.027
401.2171936035156 0 76919.71
402.17779541015625 0 227867.6
402.2008361816406 0 568104.3
403.13726806640625 0 43065.766
403.2020263671875 0 233234.17
404.1838073730469 0 41545.465
404.2183837890625 0 22378.678
404.7113952636719 0 84404.59
405.2304992675781 0 32170.053
405.2630920410156 0 155217.69
406.2164611816406 0 32558.496
406.24700927734375 0 93365.6
407.20587158203125 0 84925.26
407.2437744140625 0 36499.36
408.71331787109375 0 35475.598
409.1488037109375 0 363993.25
409.1778259277344 0 32266.049
409.20623779296875 0 87063.51
409.6988830566406 0 37953.133
410.1520080566406 0 77923.64
411.200927734375 0 375468.44
412.18511962890625 0 3231479
413.14495849609375 0 148132.58
413.1682434082031 0 542683.75
413.1902160644531 0 414060.38
413.25286865234375 0 93679.44
414.14910888671875 0 48417.195
414.17181396484375 0 132981.2
414.19873046875 0 54341.543
414.2370300292969 0 1078798.6
415.1720886230469 0 29401.89
415.2002258300781 0 194933.5
415.2409362792969 0 238852.08
416.20428466796875 0 29193.625
417.7188415527344 0 554349.6
418.2122497558594 0 848807.25
418.7130126953125 0 371374.78
419.2146911621094 0 116701.14
419.7130432128906 0 26428.975
420.1644592285156 0 38744.047
421.1507873535156 0 28103.775
421.21112060546875 0 107996.805 y Water loss 10
422.1942138671875 0 100819.89 y Ammonia loss 10
422.2539978027344 0 52161.527
422.7110290527344 0 92497.625
423.2044372558594 0 146133.7
423.2737121582031 0 1277478
423.7053527832031 0 75658.164
424.2247009277344 0 51446.902
424.2765197753906 0 296324.28
424.73516845703125 0 86327.914
425.2370300292969 0 38390.8
425.2796325683594 0 54964.625
426.19879150390625 0 28485.12
426.72418212890625 0 8674214
427.22564697265625 0 4357233.5
427.7268371582031 0 1371070.1
428.2279357910156 0 198954.58
428.72967529296875 0 24971.36
429.1839294433594 0 28459.53
429.21160888671875 0 1857527.1
430.1955871582031 0 8020369.5
431.19873046875 0 1621060
431.7161865234375 0 1056503 b Water loss 6
432.2129821777344 0 475422.8 b Ammonia loss 6
432.7144775390625 0 208914.16
432.7453918457031 0 28278.459
433.2116394042969 0 123434.19
433.2576904296875 0 289567.88 b Water loss 3
434.2099609375 0 22798.28
434.26123046875 0 67102.59
437.1944274902344 0 36946.277
438.17547607421875 0 150791.16
438.7301330566406 0 98799.36
439.22125244140625 0 2128698.2 y 10
439.7300109863281 0 25461.676
440.18017578125 0 110483.266
440.2243347167969 0 531051.75
440.72161865234375 0 4588738.5 b 6
441.2232971191406 0 2199738
441.72430419921875 0 675667.5
442.2262878417969 0 119791.72
442.2607421875 0 19742.094
443.2415771484375 0 57311.703
444.17864990234375 0 44851.504
445.18218994140625 0 19896.725
446.2054748535156 0 29452.99
446.712890625 0 46787.652 y Water loss 6
446.7428894042969 0 37589.863
447.20965576171875 0 222435.25 y Ammonia loss 6
447.27423095703125 0 25840.158
447.70977783203125 0 75604.484
448.1553649902344 0 23445.48
448.20660400390625 0 447620.25
449.21002197265625 0 148044.69
450.1751403808594 0 92320.305
450.23736572265625 0 98822.81
450.2849426269531 0 113922.57
451.2688293457031 0 2428422.5 b 3
452.271728515625 0 604210.1
452.7427978515625 0 29559.826
453.273681640625 0 86827.25
455.19903564453125 0 37320.098
455.22943115234375 0 45718.57
455.2679443359375 0 156797.05
455.7239685058594 0 89662.57 y 6
456.22515869140625 0 44146.777
456.27008056640625 0 45915.535
456.7247009277344 0 27069.965
458.18853759765625 0 32567.992
459.2013244628906 0 190236.36
460.20770263671875 0 18272.299
460.22576904296875 0 24014.082
460.7557373046875 0 306832.78
461.2062072753906 0 204391.45
461.2551574707031 0 395330.56
461.7581481933594 0 83514.75
462.2093200683594 0 43335.773
462.2541198730469 0 41056.043
462.55670166015625 0 26994.375 y Water loss 2
463.2323303222656 0 26834
465.25537109375 0 37300.133
465.749267578125 0 85748.82
466.1705017089844 0 60330.168
466.2453918457031 0 115281.56
468.1903381347656 0 22808.004
468.29534912109375 0 583851.5
469.298583984375 0 148347.75
470.2991943359375 0 23446.928
472.2237854003906 0 35119.996
474.26116943359375 0 497142.94
474.7563171386719 0 589677.8
474.909912109375 0 25871.56 b 11
475.25604248046875 0 366702.03
475.5755920410156 0 27185.03
475.7579650878906 0 80103.3
476.2271728515625 0 198679.02
477.21197509765625 0 403467.28
478.175537109375 0 30362.89
478.23095703125 0 292433.4 y Water loss 9
478.2800598144531 0 249508.62
479.2354736328125 0 77354.43
479.2836608886719 0 77340.36
479.74609375 0 52853.3
480.2247009277344 0 63530.402
480.2576599121094 0 24969.48
481.731201171875 0 51282.234
483.2664489746094 0 1888623.4
483.767822265625 0 1031560.25
484.22125244140625 0 86847.305
484.2694396972656 0 311804.38
484.7702331542969 0 58208.645
485.2230529785156 0 21072.768
487.1945495605469 0 30924.22
488.22198486328125 0 31679.203
488.25921630859375 0 414879.16 b Water loss 7
488.7570495605469 0 310110.75 b Ammonia loss 7
489.2566833496094 0 145270.73
489.75732421875 0 25498.373
490.2388610839844 0 29044.457
494.2385559082031 0 4610140.5
495.2027587890625 0 45192.06
495.2415466308594 0 1289304.1
495.9129638671875 0 22299.271 y Ammonia loss 1
496.2428894042969 0 5329913.5 y 9
497.2491760253906 0 1105961.9
497.76531982421875 0 680489.5
498.2478332519531 0 221054.05
498.76910400390625 0 30968.336
499.24566650390625 0 26631.576
502.23309326171875 0 53474.227
503.2156677246094 0 28130.648
504.2027893066406 0 221129.64
505.19940185546875 0 37741.016
505.2337341308594 0 31116.832
506.2295227050781 0 114191.98
506.274658203125 0 65015.098
507.259521484375 0 167145.78
508.22088623046875 0 22141.988
508.2632751464844 0 39629.473
511.2653503417969 0 429208.47
512.2252197265625 0 22555.613
512.2672729492188 0 129880.43
513.2122802734375 0 108245.52
513.271728515625 0 19771.441
514.1924438476562 0 64406.332
514.302734375 0 26380.426
515.28466796875 0 105460.38
516.2225341796875 0 27462.07
516.289306640625 0 38181.94
517.2536010742188 0 24773.133
519.3327026367188 0 62347.33
520.2681884765625 0 55086.55
521.2504272460938 0 38044.426
522.2304077148438 0 76264.82
522.929443359375 0 30934.057
523.260986328125 0 177158.39
523.5944213867188 0 55054.957
523.9237670898438 0 29795.314
524.2838745117188 0 80661.72
525.269287109375 0 847871.06
526.2327880859375 0 87385.96
526.2723999023438 0 253538.78
527.27392578125 0 52646.055
528.9349975585938 0 56779.324
529.2635498046875 0 333072 Precursor Water loss
529.5960693359375 0 328985.75 Precursor Ammonia loss
529.9309692382812 0 161918.27
530.2636108398438 0 101412.766
531.2183837890625 0 62578.05
532.1968383789062 0 169369.42
533.20068359375 0 69887.93
533.319580078125 0 33399.652
534.2349243164062 0 32263.717
534.9382934570312 0 52745.92
535.26611328125 0 183451.55 Precursor
535.3047485351562 0 38380.977
535.6002807617188 0 124816.5
535.9359741210938 0 90252.3
537.2474975585938 0 66922.86
537.7401733398438 0 62766.727
538.2369995117188 0 25762.715
538.2769775390625 0 80244.375
538.7753295898438 0 57136.58
539.2387084960938 0 965980.7
539.2792358398438 0 34705.11
540.24169921875 0 286606.56
540.7905883789062 0 31771.457
541.2488403320312 0 64369.227
542.296142578125 0 289810.78
543.27978515625 0 1060289.9
544.2828979492188 0 353672.44
545.2850341796875 0 81007.03
545.7664794921875 0 66726.94 b Water loss 8
546.25341796875 0 119020.97 b Ammonia loss 8
546.75341796875 0 65707.1
547.25341796875 0 100370.24
547.7800903320312 0 60526.848
548.2399291992188 0 48536.2
548.2988891601562 0 47304.8
549.2238159179688 0 217569.36
550.2107543945312 0 100857.055
551.2103271484375 0 38089.38
551.25927734375 0 242204.06
551.3331909179688 0 117288.164
552.2630615234375 0 84418.39
552.3327026367188 0 36339.08
553.2667236328125 0 45922.094
554.2822265625 0 35374.89
554.766845703125 0 170074.92 b 8
555.2591552734375 0 276970.6
555.7598876953125 0 180454.83
556.2608642578125 0 40103.766
557.2392578125 0 33222.81
558.2236938476562 0 35875.816
558.27197265625 0 39942.06
559.262451171875 0 31072.764
561.2821044921875 0 48446.324
561.3169555664062 0 197130.64 b Water loss 4
562.3050537109375 0 80494.67 b Ammonia loss 4
563.2598876953125 0 162364.7
563.2996826171875 0 27718.447
564.2662963867188 0 85135.13
564.333740234375 0 62095.145
564.7640380859375 0 31442.732
565.2653198242188 0 459091.84
566.2667846679688 0 118470.21
567.2338256835938 0 1038698.25
567.7686767578125 0 35438.234
568.2379150390625 0 301455.78
568.7608642578125 0 67075.97
569.2408447265625 0 69759.01
569.2881469726562 0 32264.527
575.2490844726562 0 356392.78
575.2970581054688 0 42889.074
576.2365112304688 0 183635.27
576.287841796875 0 27219.082
577.2244262695312 0 66170.14
577.273681640625 0 32573.873
577.7682495117188 0 38448.74
578.345458984375 0 30592.066
579.2505493164062 0 41146.59
579.328125 0 1400720.2 b 4
580.3306274414062 0 501835.97
581.2711181640625 0 39656.777
581.333251953125 0 132304.8
582.3079833984375 0 57997.65
582.7979125976562 0 22106.057
585.280029296875 0 27712.82
586.2651977539062 0 34654.906
588.7965087890625 0 37507.14
589.3087768554688 0 76624.484
589.797607421875 0 28746.635
592.2755737304688 0 267847.78
592.3275146484375 0 175475.55
593.2597045898438 0 2098847 y Water loss 8
593.3086547851562 0 621230.7
593.7886352539062 0 65507.617
594.2572021484375 0 720488.8
594.311767578125 0 170993.78
594.7870483398438 0 48722.203
595.2535400390625 0 194205.48
596.2545776367188 0 36596.188
596.3553466796875 0 61260.047
597.3074340820312 0 46129.125
597.3566284179688 0 39969.305
597.803466796875 0 76584.24
598.3049926757812 0 41263.05
599.3347778320312 0 34638.016
600.3184204101562 0 31765.088
601.31982421875 0 33858.523
602.3021850585938 0 114613.36
602.7955322265625 0 243565.36
603.293701171875 0 263917.44
603.7947998046875 0 69691.73
604.2867431640625 0 40084.62
605.3475952148438 0 30872.674
607.3200073242188 0 26513.055
609.3550415039062 0 44664.004
610.2862548828125 0 4158222.8
611.27099609375 0 21617316 y 8
611.8010864257812 0 678623.7
612.27392578125 0 7131722.5
612.8026733398438 0 206245.19
613.2755737304688 0 1596572.6
614.2789306640625 0 236356.6
615.3184204101562 0 34552.934
616.3007202148438 0 25083.625
617.28759765625 0 102693.73
618.3200073242188 0 57737.164
619.33056640625 0 36095.938
620.2781372070312 0 23063.93
620.8062744140625 0 78130.23
621.254150390625 0 242328.08
621.30517578125 0 105726.3
622.2581176757812 0 78713.07
622.3120727539062 0 35055.04
622.37158203125 0 82032.43
623.2555541992188 0 27888.498
623.3720703125 0 39194.5
627.78564453125 0 49020.25
628.2826538085938 0 81324.66
628.779541015625 0 50359.594
630.2926635742188 0 50750.484
631.322998046875 0 23112.955
634.3108520507812 0 194623.14
635.2991333007812 0 274369.66
636.300537109375 0 79316.34
636.330810546875 0 128105.04
636.7906494140625 0 215420.92 y Water loss 3
637.2903442382812 0 146529.67 y Ammonia loss 3
637.342529296875 0 73696.016
637.7935180664062 0 71986.61
638.33935546875 0 57688.637 b Water loss 10
638.8378295898438 0 27367.098
640.2769775390625 0 115864.89
641.308837890625 0 32640.523
644.3045654296875 0 31570.932
645.2998657226562 0 306723.38
645.7960815429688 0 888777.75 y 3
646.2962036132812 0 806499.94
646.7985229492188 0 238882.97
647.3011474609375 0 76589.95
647.390380859375 0 33719.01
648.3455200195312 0 57544.934
649.3444213867188 0 49241.227
650.2710571289062 0 92623.91
650.3314819335938 0 102140.12
650.7931518554688 0 42567.457
651.2811889648438 0 55415.746
652.3236083984375 0 1929697.6
652.82421875 0 31676.268
653.3262329101562 0 764908.8
654.328857421875 0 171526.48
658.2857666015625 0 64797.043
658.3344116210938 0 53062.695
659.31884765625 0 106286.05
660.3255004882812 0 34506.676
660.8430786132812 0 50957.484
661.33251953125 0 81701.3
661.8391723632812 0 30080.926
662.308837890625 0 685945.2
663.298828125 0 424841.2
664.2976684570312 0 137191.52
664.3455200195312 0 110291.695
665.2921142578125 0 31515.11
665.344482421875 0 59757.32
666.3547973632812 0 45583.625
668.2833862304688 0 51141.773
669.2879028320312 0 33224.67
675.8197631835938 0 29721.248
676.3448486328125 0 898122.7 b Water loss 5
677.337890625 0 556685.06 b Ammonia loss 5
678.3360595703125 0 286727
678.8411254882812 0 31574.434
679.33642578125 0 199173.25
679.831298828125 0 41290.918
680.3189697265625 0 3521104.8
681.3214721679688 0 1385698
682.324462890625 0 322131.16
682.3767700195312 0 23724.871
683.3276977539062 0 45498.54
684.3272094726562 0 133971.89
684.822021484375 0 247211.12
685.3209228515625 0 174437.55
685.8211669921875 0 47843.805
686.3263549804688 0 33406.04
688.334228515625 0 99883.14
689.32373046875 0 42194.418
691.3005981445312 0 25780.51
692.4130249023438 0 24422.646
692.8399047851562 0 183027.88
693.3326416015625 0 975844 y Water loss 2
693.8317260742188 0 861019.25 y Ammonia loss 2
694.3550415039062 0 9862975 b 5
694.8322143554688 0 206955.48
695.3583374023438 0 3664981
696.3607177734375 0 879731.7
697.2929077148438 0 33975.617
697.358154296875 0 190794.9
697.8610229492188 0 37103.11
698.3545532226562 0 67888.65
701.8447875976562 0 1759551.2
702.338623046875 0 5735229.5 y 2
702.8397216796875 0 4582264 b Water loss 11
703.3413696289062 0 2170129.5 b Ammonia loss 11
703.8421630859375 0 551067.1
704.341796875 0 267492.78
705.3561401367188 0 133439.45
706.3453369140625 0 686309.6 y Water loss 7
707.3477172851562 0 325525.78
708.3165893554688 0 83476.1
708.3479614257812 0 62942.05
709.31298828125 0 34996.92
711.3719482421875 0 388447.88
711.8624267578125 0 452942.62 b 11
712.363525390625 0 313878.72
712.8641967773438 0 128740.81
713.3643188476562 0 40855.85
715.3031005859375 0 39292.008
715.3685913085938 0 23779.37
716.3389282226562 0 38942.727
717.3385620117188 0 29327.059
718.3943481445312 0 40110.773
720.3826293945312 0 85383.984
720.8643798828125 0 37544.3
721.36669921875 0 454843.47
722.3674926757812 0 136641.7
723.3715209960938 0 1225900.6
724.3558959960938 0 6027693.5 y 7
725.35888671875 0 2390548.2
726.3612670898438 0 664921.44
727.3626098632812 0 112426.305
728.3741455078125 0 78717.41
728.85888671875 0 39940.508
729.3621215820312 0 153452.44
729.8624877929688 0 43011.27
730.3672485351562 0 75175.414
732.318115234375 0 34265.105
733.3668823242188 0 307549.38
734.3598022460938 0 257197.19
735.3629760742188 0 94224.16
737.8672485351562 0 65191.88
738.3653564453125 0 60460.973
739.3526000976562 0 43382.58
740.3585815429688 0 25405.412
742.8668212890625 0 92486.76 y Water loss 1
743.3721923828125 0 203623.12
743.865478515625 0 66366.01
744.3692626953125 0 66259.15
745.3512573242188 0 39680.56
746.3320922851562 0 39063.707
747.3956909179688 0 97204.7
748.3873291015625 0 168465.31
749.34130859375 0 74489.125
749.4072265625 0 39963.03
750.3297119140625 0 37246.695
750.3958740234375 0 38962.9
751.3797607421875 0 1024898.6
751.87353515625 0 449662.22 y 1
752.3780517578125 0 849335.8
752.8748168945312 0 170770.8
753.3803100585938 0 207228.1
760.313232421875 0 33813.855
761.3760375976562 0 68905.39
762.41796875 0 154404.92
763.3680419921875 0 580376.25
764.3704833984375 0 287762.9
765.4054565429688 0 242107.73
766.4096069335938 0 139313.06
767.350830078125 0 206783.78
767.4107055664062 0 29298.258
768.3510131835938 0 65580.3
768.4102172851562 0 52702.402
769.4049682617188 0 114667.82
770.4061889648438 0 49592.977
775.3958740234375 0 103526.03
776.3832397460938 0 122882.27
777.3853149414062 0 55864.027
778.4026489257812 0 25953.203
779.3878173828125 0 644962.94
780.3908081054688 0 252688.42
781.3706665039062 0 294683.1
782.3717041015625 0 139887.38
783.3710327148438 0 31758.246
792.4357299804688 0 40430.324
793.4032592773438 0 1070144
794.4059448242188 0 491105.88
795.3466796875 0 105398.18
795.409423828125 0 102773.53
796.3466186523438 0 51855.44
796.4202270507812 0 36366.867
799.3939208984375 0 28958.172
807.3817749023438 0 188557.66
807.4423828125 0 137474.86
808.3819580078125 0 98311.02
808.4389038085938 0 96311.51
817.4017944335938 0 57295.633
818.4385986328125 0 171212.31
819.442138671875 0 89816.74
820.436767578125 0 70050.04
825.3947143554688 0 82180.93
826.383056640625 0 94130.93
827.3863525390625 0 55575.58
829.4257202148438 0 25624.078
834.3607177734375 0 15684.945
834.4293823242188 0 258299.94
835.4190673828125 0 318981.12
836.4529418945312 0 755042.9
837.4566650390625 0 349654.66
838.4573364257812 0 116540.336
843.4036254882812 0 100270.49
844.3980102539062 0 187946.83
845.3991088867188 0 169354.33
846.4503784179688 0 321234.62
847.4485473632812 0 220707.94
848.449462890625 0 98878.55
851.3846435546875 0 74804.35
852.4401245117188 0 2371717
853.4430541992188 0 1179406
854.445556640625 0 402263.1
855.4492797851562 0 71146.836
856.4451904296875 0 32406.305
862.4242553710938 0 1306090.2 b Water loss 6
863.4212036132812 0 1043580 b Ammonia loss 6
864.4595336914062 0 1869771
865.4628295898438 0 953324.44
866.466552734375 0 276441.25
867.469482421875 0 67857.48
869.3905029296875 0 26988.26
876.443603515625 0 172786.89
877.4421997070312 0 103310.05
878.4435424804688 0 44584.258
880.43505859375 0 11379343 b 6
881.438232421875 0 6107007
882.4407348632812 0 1762546.9
883.4435424804688 0 340541.3
884.44677734375 0 43718.312
890.4201049804688 0 60209.273
891.4197387695312 0 54726.02
892.468505859375 0 212715.89
893.4733276367188 0 85420.05
894.454345703125 0 100565.266
895.456298828125 0 55160.53
907.44775390625 0 144506.48
908.43310546875 0 331517.78
909.4415283203125 0 316240.2
910.43701171875 0 603290.06 y 6
911.4392700195312 0 289391.5
912.4417724609375 0 74508.45
936.4306030273438 0 60310.797
937.424072265625 0 86397.414
938.4344482421875 0 24556.428
947.4410400390625 0 67286.89
948.42822265625 0 39330.08
954.4373779296875 0 111858.734
955.4280395507812 0 206941.25
956.4275512695312 0 84297.37
964.4656982421875 0 112760.77
965.455322265625 0 198751.73
966.456298828125 0 112893.64
971.4627075195312 0 86658.914
972.4508056640625 0 140246.92
973.4503173828125 0 57609.734
974.4542236328125 0 28417.094
975.507080078125 0 108656.625 b Water loss 7
976.5111694335938 0 95007.48
977.5081787109375 0 26126.656
980.4655151367188 0 59219.906
981.47216796875 0 32768.484
982.483154296875 0 30805.14
993.51953125 0 704837.9 b 7
994.5222778320312 0 409306.2
995.5240478515625 0 159758.25
996.52978515625 0 28796.8
1007.4577026367188 0 74503.9 y Water loss 5
1008.4432373046875 0 40315.9 y Ammonia loss 5
1024.47705078125 0 84901.52
1025.464599609375 0 373850.78 y 5
1026.467529296875 0 155130.8
1027.4674072265625 0 57110.02
1075.5341796875 0 81515.32
1076.536865234375 0 62060.805
1091.5137939453125 0 40823.41 b Ammonia loss 8
1092.558349609375 0 39012.74
1093.5494384765625 0 148903.95
1094.551513671875 0 82736.99
1095.5560302734375 0 33209.543
1107.5626220703125 0 37370.363
1108.555908203125 0 39230.344 b 8
1117.4957275390625 0 53101.574
1118.485107421875 0 73392.01
1119.4876708984375 0 45695.73
1134.5302734375 0 59308.465
1135.510986328125 0 359808.84 y Water loss 4
1136.5006103515625 0 373552.75 y Ammonia loss 4
1137.5020751953125 0 185113.61
1138.504150390625 0 74511.195
1152.536865234375 0 231488
1153.52392578125 0 592765.25 y 4
1154.5257568359375 0 400823.6
1155.5262451171875 0 140626.61
1156.530029296875 0 32407.14
1164.5780029296875 0 26448.848
1165.5726318359375 0 67503.37 b 9
1290.5697021484375 0 34846.633 y 3
1736.7259521484375 0 22657.254

Spectrum Details

|  |  |
| --- | --- |
| Matched peaks? Matched peaks The total absolute number of peaks matched. Additionally in brackets the total fraction of peaks matched and the total number of peaks is shown. | 87 (7.61% of 1143) |
| FDR? FDR The false discovery rate estimated for this peptide. It is calculated by matching all theoretical fragments with a non-integer shift with the raw peaks for this spectrum. This is done with 40 different shifts. The resulting percentage is the average number of annotated peaks over the number of annotated peaks with the correct spectrum. | 0.90% |
| Satellite FDR? Satellite FDR See the FDR for details on its calculation. This satellite ion specific FDR only contains the satellite ions (d/w) for I/L/J positions. | - |
| PSM Score? PSM Score The PSM Score as given by Hecklib to this annotated spectrum. It is shown with three significant figures. | 669 |

#### Spectrum 6124? Spectrum 6124 The raw spectrum of this peptide as annotated by Hecklib. The fragments are coloured according to ion type (see legend). Any peaks with a star '\*' as text can be hovered over to see the full details, first the ion type second the mass shift type. By hovering over the amino acids in the peptide or ions in the legend the corresponding peaks are highlighted. By toggling the 'Unassigned' label you can turn the background (unassigned) peaks on or off in the plot. By updating the slider in the Ion legend you can update the spectrum to only show the top X% of the peaks with labels. The top X% means any peak that is within X% of the highest intensity. By dragging in the spectrum you can zoom in to a specific part of the spectrum and use 'Zoom Out' to get back to the original zoom level. The annotation of the spectrum is based on the given sequence in the peptides file and is done with different software so inconsistencies are likely. The peaks are annotated based on the given sequence, with 20 ppm tolerance.

Copy Data

##### Spectrum 6124 (TSV)

###### Preview

```
Loading example...
```

*Click on the button to copy the data to your clipboard.*

Mz MinMz MaxIntensity Max

WidthHeightPeptide font sizePeptide stroke widthSpectrum font sizeSpectrum stroke widthCompact peptide

Ion legend

wxyz

abcd

OtherUnassignedIon^Charge^_Position_Show for top:%

TVLHQDWLDGKEY

05.88e+41.18e+51.76e+52.35e+5

Zoom Out

y^+3^_3_a^+1^_2_d^+1^_2_y^+3^_4_a^+1^_2_y^+1^_1_b^+1^_2_b^+1^_2_y^+2^_3_b^+3^_6_y^+1^_2_b^+1^_3_y^+2^_5_y^+1^_2_b^+1^_3_b^+3^_8_b^+2^_6_y^+3^_8_b^+2^_6_y^+2^_6_y^+2^_6_y^+1^_3_y^+3^_10_b^+1^_4_y^+1^_3_b^+2^_7_b^+2^_7_b^+1^_4_y^+2^_7_b^+2^_8_b^+2^_8_y^+2^_8_\*\*\*b^+1^_5_b^+2^_9_b^+1^_5_y^+1^_5_y^+1^_5_y^+2^_10_y^+2^_10_b^+2^_11_y^+2^_10_b^+2^_11_b^+1^_6_b^+1^_6_b^+1^_6_y^+2^_11_y^+2^_11_y^+1^_6_y^+2^_11_b^+2^_12_b^+2^_12_b^+2^_12_y^+1^_6_y^+2^_12_y^+2^_12_\*b^+1^_7_b^+1^_7_y^+1^_7_y^+1^_7_b^+1^_8_y^+1^_8_b^+1^_9_y^+1^_9_y^+1^_9_y^+1^_9_

0778155723353114

Fragment Matches Table

Show background peaks

| Position | Ion type | Intensity | mz Theoretical | mz Error (Th) | mz Error (ppm) | Charge | Series Number |
| --- | --- | --- | --- | --- | --- | --- | --- |
| - | - | 494.4 | 120 | - | - | 0 | - |
| - | - | 2.469E+04 | 120.1 | - | - | 0 | - |
| - | - | 2077 | 121.1 | - | - | 0 | - |
| - | - | 384.7 | 121.8 | - | - | 0 | - |
| - | - | 1010 | 122.1 | - | - | 0 | - |
| - | - | 961.2 | 123 | - | - | 0 | - |
| - | - | 462.8 | 123.1 | - | - | 0 | - |
| - | - | 1297 | 125.1 | - | - | 0 | - |
| - | - | 1494 | 126.1 | - | - | 0 | - |
| - | - | 1047 | 127.1 | - | - | 0 | - |
| - | - | 410.2 | 127.1 | - | - | 0 | - |
| - | - | 603.4 | 127.1 | - | - | 0 | - |
| - | - | 451.1 | 128.1 | - | - | 0 | - |
| - | - | 1.112E+04 | 128.1 | - | - | 0 | - |
| - | - | 1288 | 129.1 | - | - | 0 | - |
| - | - | 8.557E+04 | 129.1 | - | - | 0 | - |
| - | - | 430.3 | 130.1 | - | - | 0 | - |
| - | - | 1.118E+04 | 130.1 | - | - | 0 | - |
| - | - | 726.4 | 130.1 | - | - | 0 | - |
| - | - | 835.4 | 130.1 | - | - | 0 | - |
| - | - | 5166 | 130.1 | - | - | 0 | - |
| - | - | 668.9 | 131 | - | - | 0 | - |
| - | - | 874.7 | 131.1 | - | - | 0 | - |
| - | - | 3652 | 132 | - | - | 0 | - |
| - | - | 522.8 | 132.1 | - | - | 0 | - |
| - | - | 382.4 | 132.1 | - | - | 0 | - |
| - | - | 526 | 133.1 | - | - | 0 | - |
| - | - | 1.78E+04 | 136.1 | - | - | 0 | - |
| - | - | 1112 | 136.1 | - | - | 0 | - |
| - | - | 1187 | 137.1 | - | - | 0 | - |
| - | - | 1343 | 138.1 | - | - | 0 | - |
| - | - | 1407 | 138.1 | - | - | 0 | - |
| - | - | 1148 | 139.1 | - | - | 0 | - |
| - | - | 1089 | 140.1 | - | - | 0 | - |
| - | - | 2935 | 141.1 | - | - | 0 | - |
| - | - | 413.5 | 143 | - | - | 0 | - |
| - | - | 604.3 | 144.1 | - | - | 0 | - |
| - | - | 518.2 | 145.1 | - | - | 0 | - |
| - | - | 7655 | 146.1 | - | - | 0 | - |
| - | - | 471.5 | 146.5 | - | - | 0 | - |
| - | - | 870.5 | 147 | - | - | 0 | - |
| - | - | 718.5 | 147.1 | - | - | 0 | - |
| 11 | y | 1072 | 147.1 | 0.00113 | 7.683 | +3 | 3 |
| - | - | 692.4 | 149 | - | - | 0 | - |
| - | - | 1016 | 150.1 | - | - | 0 | - |
| - | - | 1789 | 151.1 | - | - | 0 | - |
| - | - | 1414 | 154.1 | - | - | 0 | - |
| - | - | 6963 | 155.1 | - | - | 0 | - |
| 2 | a | 7616 | 155.1 | 0.0003356 | 2.163 | +1 | 2 |
| - | - | 966 | 156.1 | - | - | 0 | - |
| - | - | 631.8 | 156.1 | - | - | 0 | - |
| - | - | 723.8 | 156.1 | - | - | 0 | - |
| - | - | 474.6 | 156.1 | - | - | 0 | - |
| - | - | 1838 | 157.1 | - | - | 0 | - |
| - | - | 7479 | 157.1 | - | - | 0 | - |
| - | - | 2318 | 157.1 | - | - | 0 | - |
| - | - | 4977 | 157.1 | - | - | 0 | - |
| - | - | 1256 | 157.1 | - | - | 0 | - |
| - | - | 1.013E+04 | 158.1 | - | - | 0 | - |
| - | - | 656 | 158.1 | - | - | 0 | - |
| - | - | 1600 | 158.1 | - | - | 0 | - |
| - | - | 1290 | 159.1 | - | - | 0 | - |
| - | - | 559.3 | 159.1 | - | - | 0 | - |
| - | - | 3142 | 159.1 | - | - | 0 | - |
| 2 | d | 2463 | 159.1 | 0.0004161 | 2.615 | +1 | 2 |
| - | - | 694.4 | 163.1 | - | - | 0 | - |
| - | - | 9378 | 165.1 | - | - | 0 | - |
| - | - | 1.295E+04 | 166.1 | - | - | 0 | - |
| 10 | y | 8979 | 166.1 | 0.001725 | 10.39 | +3 | 4 |
| - | - | 1184 | 166.1 | - | - | 0 | - |
| - | - | 477.6 | 167.1 | - | - | 0 | - |
| - | - | 900.3 | 167.1 | - | - | 0 | - |
| - | - | 985.9 | 167.1 | - | - | 0 | - |
| - | - | 542 | 167.1 | - | - | 0 | - |
| - | - | 617.5 | 168.1 | - | - | 0 | - |
| - | - | 1159 | 168.1 | - | - | 0 | - |
| - | - | 1069 | 168.1 | - | - | 0 | - |
| - | - | 2680 | 169.1 | - | - | 0 | - |
| - | - | 3246 | 169.1 | - | - | 0 | - |
| - | - | 519.6 | 170.1 | - | - | 0 | - |
| - | - | 1500 | 171.1 | - | - | 0 | - |
| - | - | 512.3 | 171.1 | - | - | 0 | - |
| - | - | 569.2 | 171.1 | - | - | 0 | - |
| - | - | 929.2 | 171.1 | - | - | 0 | - |
| - | - | 1569 | 172.1 | - | - | 0 | - |
| - | - | 456.1 | 172.1 | - | - | 0 | - |
| - | - | 911.8 | 173.1 | - | - | 0 | - |
| 2 | a | 1.86E+05 | 173.1 | 0.0004215 | 2.435 | +1 | 2 |
| - | - | 2159 | 173.4 | - | - | 0 | - |
| - | - | 4613 | 174.1 | - | - | 0 | - |
| - | - | 1.489E+04 | 174.1 | - | - | 0 | - |
| - | - | 2663 | 175.1 | - | - | 0 | - |
| - | - | 3.854E+04 | 175.1 | - | - | 0 | - |
| - | - | 2023 | 175.1 | - | - | 0 | - |
| - | - | 559.2 | 175.1 | - | - | 0 | - |
| - | - | 716.6 | 176.1 | - | - | 0 | - |
| - | - | 4479 | 176.1 | - | - | 0 | - |
| - | - | 699.8 | 177.1 | - | - | 0 | - |
| - | - | 4893 | 178.1 | - | - | 0 | - |
| - | - | 529.7 | 180.1 | - | - | 0 | - |
| - | - | 966.5 | 181.1 | - | - | 0 | - |
| 13 | y | 1.752E+04 | 182.1 | 0.0004191 | 2.302 | +1 | 1 |
| - | - | 972.8 | 183.1 | - | - | 0 | - |
| - | - | 1324 | 183.1 | - | - | 0 | - |
| 2 | b | 1.627E+04 | 183.1 | 0.0003703 | 2.022 | +1 | 2 |
| - | - | 4101 | 183.1 | - | - | 0 | - |
| - | - | 494.6 | 184.1 | - | - | 0 | - |
| - | - | 761.6 | 184.1 | - | - | 0 | - |
| - | - | 7601 | 185.1 | - | - | 0 | - |
| - | - | 1635 | 185.1 | - | - | 0 | - |
| - | - | 691.4 | 185.1 | - | - | 0 | - |
| - | - | 1015 | 185.1 | - | - | 0 | - |
| - | - | 1516 | 185.2 | - | - | 0 | - |
| - | - | 490.9 | 185.4 | - | - | 0 | - |
| - | - | 864.5 | 186.1 | - | - | 0 | - |
| - | - | 1077 | 186.1 | - | - | 0 | - |
| - | - | 2.562E+04 | 186.1 | - | - | 0 | - |
| - | - | 1028 | 187.1 | - | - | 0 | - |
| - | - | 1810 | 187.1 | - | - | 0 | - |
| - | - | 2155 | 187.1 | - | - | 0 | - |
| - | - | 1295 | 187.1 | - | - | 0 | - |
| - | - | 934.5 | 188.1 | - | - | 0 | - |
| - | - | 8168 | 189.1 | - | - | 0 | - |
| - | - | 1035 | 190.1 | - | - | 0 | - |
| - | - | 603 | 190.1 | - | - | 0 | - |
| - | - | 637.1 | 191.1 | - | - | 0 | - |
| - | - | 7035 | 195.1 | - | - | 0 | - |
| - | - | 3255 | 196.1 | - | - | 0 | - |
| - | - | 683 | 196.1 | - | - | 0 | - |
| - | - | 1249 | 196.1 | - | - | 0 | - |
| - | - | 2706 | 197.1 | - | - | 0 | - |
| - | - | 837.2 | 198.1 | - | - | 0 | - |
| - | - | 4292 | 199.1 | - | - | 0 | - |
| - | - | 1114 | 199.1 | - | - | 0 | - |
| 2 | b | 6.433E+04 | 201.1 | 0.0003342 | 1.661 | +1 | 2 |
| - | - | 822 | 202.1 | - | - | 0 | - |
| - | - | 5375 | 202.1 | - | - | 0 | - |
| - | - | 2397 | 203.1 | - | - | 0 | - |
| - | - | 2677 | 203.1 | - | - | 0 | - |
| - | - | 538.2 | 204.1 | - | - | 0 | - |
| - | - | 3530 | 205.1 | - | - | 0 | - |
| - | - | 1351 | 207.2 | - | - | 0 | - |
| - | - | 722.8 | 209.1 | - | - | 0 | - |
| - | - | 1314 | 209.1 | - | - | 0 | - |
| - | - | 571.8 | 210.1 | - | - | 0 | - |
| - | - | 552.8 | 211.1 | - | - | 0 | - |
| 11 | y | 1935 | 211.1 | 0.0003897 | 1.846 | +2 | 3 |
| - | - | 2168 | 211.1 | - | - | 0 | - |
| - | - | 1.267E+04 | 211.2 | - | - | 0 | - |
| - | - | 777.5 | 212.1 | - | - | 0 | - |
| - | - | 615.5 | 212.1 | - | - | 0 | - |
| - | - | 3014 | 213.1 | - | - | 0 | - |
| - | - | 682.6 | 214.2 | - | - | 0 | - |
| - | - | 510.1 | 215.1 | - | - | 0 | - |
| - | - | 1453 | 215.1 | - | - | 0 | - |
| - | - | 946.4 | 216.1 | - | - | 0 | - |
| - | - | 668 | 216.2 | - | - | 0 | - |
| - | - | 1.353E+04 | 217.1 | - | - | 0 | - |
| - | - | 763.7 | 218.1 | - | - | 0 | - |
| - | - | 2029 | 221.1 | - | - | 0 | - |
| - | - | 938.3 | 221.1 | - | - | 0 | - |
| - | - | 2238 | 222.1 | - | - | 0 | - |
| - | - | 755.3 | 223.1 | - | - | 0 | - |
| - | - | 1851 | 223.1 | - | - | 0 | - |
| - | - | 1.461E+04 | 223.2 | - | - | 0 | - |
| - | - | 1728 | 224.1 | - | - | 0 | - |
| - | - | 645.3 | 224.1 | - | - | 0 | - |
| - | - | 1280 | 224.2 | - | - | 0 | - |
| - | - | 520.3 | 225.1 | - | - | 0 | - |
| - | - | 2193 | 226.1 | - | - | 0 | - |
| 6 | b | 1546 | 226.1 | 0.0002939 | 1.3 | +3 | 6 |
| - | - | 3050 | 227.1 | - | - | 0 | - |
| - | - | 577.3 | 227.1 | - | - | 0 | - |
| - | - | 2123 | 227.1 | - | - | 0 | - |
| - | - | 2362 | 228.2 | - | - | 0 | - |
| - | - | 3157 | 229.1 | - | - | 0 | - |
| - | - | 549.4 | 229.1 | - | - | 0 | - |
| - | - | 872.4 | 230.1 | - | - | 0 | - |
| - | - | 2125 | 233.1 | - | - | 0 | - |
| - | - | 632 | 233.2 | - | - | 0 | - |
| - | - | 1.567E+04 | 234.1 | - | - | 0 | - |
| - | - | 731.2 | 234.1 | - | - | 0 | - |
| - | - | 2007 | 235.1 | - | - | 0 | - |
| - | - | 1221 | 237.1 | - | - | 0 | - |
| - | - | 1645 | 238.1 | - | - | 0 | - |
| - | - | 8171 | 239.2 | - | - | 0 | - |
| - | - | 617.2 | 240.1 | - | - | 0 | - |
| - | - | 5339 | 240.1 | - | - | 0 | - |
| - | - | 974.9 | 241.1 | - | - | 0 | - |
| - | - | 991 | 241.1 | - | - | 0 | - |
| - | - | 711 | 241.2 | - | - | 0 | - |
| - | - | 2353 | 242.1 | - | - | 0 | - |
| - | - | 650.9 | 243.1 | - | - | 0 | - |
| - | - | 500.3 | 243.1 | - | - | 0 | - |
| - | - | 1077 | 243.1 | - | - | 0 | - |
| - | - | 3956 | 244.1 | - | - | 0 | - |
| - | - | 556.8 | 245.1 | - | - | 0 | - |
| - | - | 9729 | 247.1 | - | - | 0 | - |
| - | - | 3316 | 248.1 | - | - | 0 | - |
| - | - | 3865 | 249.1 | - | - | 0 | - |
| - | - | 1141 | 249.1 | - | - | 0 | - |
| - | - | 5.48E+04 | 251.2 | - | - | 0 | - |
| - | - | 7580 | 252.2 | - | - | 0 | - |
| - | - | 688.3 | 253.1 | - | - | 0 | - |
| - | - | 748.2 | 254.1 | - | - | 0 | - |
| - | - | 2328 | 255.1 | - | - | 0 | - |
| - | - | 1375 | 255.1 | - | - | 0 | - |
| - | - | 3076 | 256.2 | - | - | 0 | - |
| - | - | 1210 | 257.1 | - | - | 0 | - |
| - | - | 2104 | 258.1 | - | - | 0 | - |
| - | - | 803.5 | 262.1 | - | - | 0 | - |
| - | - | 720.6 | 262.2 | - | - | 0 | - |
| - | - | 1069 | 263.1 | - | - | 0 | - |
| - | - | 691.2 | 263.1 | - | - | 0 | - |
| - | - | 5300 | 265.1 | - | - | 0 | - |
| - | - | 1046 | 265.2 | - | - | 0 | - |
| - | - | 983.2 | 266.1 | - | - | 0 | - |
| - | - | 1.247E+04 | 266.1 | - | - | 0 | - |
| - | - | 693.4 | 266.1 | - | - | 0 | - |
| - | - | 935 | 266.6 | - | - | 0 | - |
| - | - | 1712 | 267.1 | - | - | 0 | - |
| - | - | 771.6 | 268.1 | - | - | 0 | - |
| - | - | 900.5 | 268.2 | - | - | 0 | - |
| - | - | 671 | 269.1 | - | - | 0 | - |
| - | - | 1124 | 269.2 | - | - | 0 | - |
| - | - | 570.6 | 269.2 | - | - | 0 | - |
| - | - | 1254 | 270.1 | - | - | 0 | - |
| - | - | 1.45E+04 | 270.2 | - | - | 0 | - |
| - | - | 1863 | 271.2 | - | - | 0 | - |
| - | - | 1120 | 274.1 | - | - | 0 | - |
| - | - | 905.3 | 275.1 | - | - | 0 | - |
| - | - | 810.5 | 276.2 | - | - | 0 | - |
| - | - | 1313 | 278.1 | - | - | 0 | - |
| - | - | 1082 | 279.1 | - | - | 0 | - |
| - | - | 1281 | 280.1 | - | - | 0 | - |
| - | - | 1336 | 282.2 | - | - | 0 | - |
| - | - | 1091 | 282.2 | - | - | 0 | - |
| - | - | 3.298E+04 | 283.1 | - | - | 0 | - |
| - | - | 3750 | 284.1 | - | - | 0 | - |
| - | - | 1330 | 284.2 | - | - | 0 | - |
| - | - | 558.2 | 285.8 | - | - | 0 | - |
| - | - | 1353 | 286.1 | - | - | 0 | - |
| - | - | 1001 | 286.2 | - | - | 0 | - |
| - | - | 2609 | 288.2 | - | - | 0 | - |
| - | - | 1052 | 290.1 | - | - | 0 | - |
| - | - | 533.8 | 292.1 | - | - | 0 | - |
| 12 | y | 2.213E+04 | 293.1 | 0.0004799 | 1.637 | +1 | 2 |
| - | - | 576.6 | 293.2 | - | - | 0 | - |
| - | - | 3022 | 294.1 | - | - | 0 | - |
| - | - | 1451 | 294.2 | - | - | 0 | - |
| - | - | 763.2 | 295.2 | - | - | 0 | - |
| 3 | b | 1672 | 296.2 | 0.0005501 | 1.857 | +1 | 3 |
| - | - | 5841 | 297.2 | - | - | 0 | - |
| - | - | 1191 | 298.1 | - | - | 0 | - |
| - | - | 6135 | 298.2 | - | - | 0 | - |
| - | - | 2281 | 299.1 | - | - | 0 | - |
| - | - | 2574 | 300.1 | - | - | 0 | - |
| - | - | 1085 | 300.1 | - | - | 0 | - |
| - | - | 2149 | 300.2 | - | - | 0 | - |
| - | - | 4070 | 301.1 | - | - | 0 | - |
| - | - | 1035 | 301.1 | - | - | 0 | - |
| - | - | 1.098E+04 | 301.2 | - | - | 0 | - |
| - | - | 835.5 | 301.2 | - | - | 0 | - |
| - | - | 2780 | 302.1 | - | - | 0 | - |
| - | - | 1272 | 302.2 | - | - | 0 | - |
| - | - | 1141 | 303.1 | - | - | 0 | - |
| - | - | 3839 | 305.2 | - | - | 0 | - |
| - | - | 666.7 | 305.6 | - | - | 0 | - |
| 9 | y | 1723 | 306.1 | 0.0002162 | 0.7061 | +2 | 5 |
| - | - | 1611 | 310.2 | - | - | 0 | - |
| 12 | y | 2962 | 311.1 | 0.0003827 | 1.23 | +1 | 2 |
| - | - | 722.5 | 311.1 | - | - | 0 | - |
| - | - | 675.8 | 312.1 | - | - | 0 | - |
| - | - | 1.195E+04 | 312.2 | - | - | 0 | - |
| - | - | 876.5 | 313.2 | - | - | 0 | - |
| - | - | 3098 | 313.2 | - | - | 0 | - |
| - | - | 591.6 | 313.2 | - | - | 0 | - |
| - | - | 2947 | 314.1 | - | - | 0 | - |
| 3 | b | 816.2 | 314.2 | 0.001368 | 4.355 | +1 | 3 |
| - | - | 630.5 | 315.1 | - | - | 0 | - |
| - | - | 8205 | 315.2 | - | - | 0 | - |
| - | - | 593.6 | 315.2 | - | - | 0 | - |
| - | - | 3662 | 316.2 | - | - | 0 | - |
| - | - | 613.8 | 316.2 | - | - | 0 | - |
| - | - | 792 | 317.2 | - | - | 0 | - |
| - | - | 2524 | 318.1 | - | - | 0 | - |
| - | - | 907.4 | 322.2 | - | - | 0 | - |
| - | - | 954.2 | 323.2 | - | - | 0 | - |
| - | - | 2354 | 327.3 | - | - | 0 | - |
| - | - | 5116 | 330.2 | - | - | 0 | - |
| 8 | b | 815.7 | 331.2 | 0.0007888 | 2.382 | +3 | 8 |
| - | - | 564.3 | 332.1 | - | - | 0 | - |
| - | - | 1250 | 332.2 | - | - | 0 | - |
| - | - | 1267 | 333.2 | - | - | 0 | - |
| - | - | 788.6 | 334.1 | - | - | 0 | - |
| - | - | 590.1 | 337.2 | - | - | 0 | - |
| - | - | 2578 | 338.2 | - | - | 0 | - |
| 6 | b | 977.5 | 338.7 | 0.0006077 | 1.794 | +2 | 6 |
| - | - | 1996 | 341.2 | - | - | 0 | - |
| 6 | y | 3590 | 342.1 | 0.003886 | 11.36 | +3 | 8 |
| - | - | 860.2 | 343.2 | - | - | 0 | - |
| - | - | 2148 | 345.1 | - | - | 0 | - |
| - | - | 4737 | 346.1 | - | - | 0 | - |
| - | - | 988.4 | 347.1 | - | - | 0 | - |
| 6 | b | 4076 | 347.7 | 0.0003608 | 1.038 | +2 | 6 |
| - | - | 2141 | 348.2 | - | - | 0 | - |
| - | - | 1517 | 349.2 | - | - | 0 | - |
| - | - | 1266 | 350.2 | - | - | 0 | - |
| - | - | 3942 | 350.2 | - | - | 0 | - |
| - | - | 1049 | 351.2 | - | - | 0 | - |
| 8 | y | 878.5 | 353.7 | 0.001186 | 3.354 | +2 | 6 |
| - | - | 1070 | 355.2 | - | - | 0 | - |
| - | - | 1612 | 355.2 | - | - | 0 | - |
| - | - | 909.2 | 358.2 | - | - | 0 | - |
| - | - | 1839 | 359.2 | - | - | 0 | - |
| - | - | 1018 | 361.2 | - | - | 0 | - |
| - | - | 2289 | 362.2 | - | - | 0 | - |
| 8 | y | 3660 | 362.7 | 0.0007867 | 2.169 | +2 | 6 |
| - | - | 8223 | 363.1 | - | - | 0 | - |
| - | - | 2291 | 363.2 | - | - | 0 | - |
| - | - | 1465 | 364.1 | - | - | 0 | - |
| - | - | 1573 | 364.1 | - | - | 0 | - |
| - | - | 2584 | 364.2 | - | - | 0 | - |
| - | - | 774.1 | 364.7 | - | - | 0 | - |
| - | - | 1350 | 367.2 | - | - | 0 | - |
| - | - | 782.1 | 367.2 | - | - | 0 | - |
| - | - | 1942 | 373.2 | - | - | 0 | - |
| - | - | 2850 | 378.2 | - | - | 0 | - |
| - | - | 1.287E+04 | 379.2 | - | - | 0 | - |
| - | - | 2853 | 380.2 | - | - | 0 | - |
| - | - | 2.155E+04 | 381.2 | - | - | 0 | - |
| - | - | 4299 | 382.2 | - | - | 0 | - |
| - | - | 638.7 | 382.2 | - | - | 0 | - |
| - | - | 2101 | 383.2 | - | - | 0 | - |
| - | - | 1765 | 384.2 | - | - | 0 | - |
| - | - | 851.2 | 384.3 | - | - | 0 | - |
| - | - | 876.8 | 390.2 | - | - | 0 | - |
| - | - | 1132 | 391.1 | - | - | 0 | - |
| - | - | 920.7 | 392.2 | - | - | 0 | - |
| - | - | 2833 | 394.2 | - | - | 0 | - |
| - | - | 782.9 | 395.2 | - | - | 0 | - |
| - | - | 970.9 | 396.2 | - | - | 0 | - |
| - | - | 5556 | 396.2 | - | - | 0 | - |
| - | - | 1350 | 397.2 | - | - | 0 | - |
| - | - | 4257 | 397.3 | - | - | 0 | - |
| - | - | 1388 | 398.2 | - | - | 0 | - |
| - | - | 1556 | 398.3 | - | - | 0 | - |
| - | - | 940.4 | 400.2 | - | - | 0 | - |
| - | - | 1747 | 401.2 | - | - | 0 | - |
| - | - | 2140 | 402.2 | - | - | 0 | - |
| - | - | 721.8 | 405.3 | - | - | 0 | - |
| - | - | 1877 | 407.2 | - | - | 0 | - |
| - | - | 911.4 | 407.7 | - | - | 0 | - |
| - | - | 671.2 | 408.3 | - | - | 0 | - |
| - | - | 2159 | 409.1 | - | - | 0 | - |
| - | - | 1195 | 409.2 | - | - | 0 | - |
| - | - | 989.4 | 410.1 | - | - | 0 | - |
| - | - | 1587 | 411.1 | - | - | 0 | - |
| - | - | 820.2 | 411.2 | - | - | 0 | - |
| - | - | 1.175E+04 | 412.2 | - | - | 0 | - |
| - | - | 3649 | 412.3 | - | - | 0 | - |
| - | - | 5077 | 413.2 | - | - | 0 | - |
| - | - | 2885 | 414.2 | - | - | 0 | - |
| - | - | 1.239E+04 | 414.3 | - | - | 0 | - |
| - | - | 1896 | 414.7 | - | - | 0 | - |
| - | - | 791.7 | 417.2 | - | - | 0 | - |
| - | - | 747 | 418 | - | - | 0 | - |
| - | - | 1122 | 418.7 | - | - | 0 | - |
| - | - | 925.3 | 419 | - | - | 0 | - |
| - | - | 702.2 | 420.7 | - | - | 0 | - |
| 11 | y | 762.6 | 421.2 | 0.003936 | 9.345 | +1 | 3 |
| - | - | 708.1 | 422.2 | - | - | 0 | - |
| - | - | 4363 | 423.2 | - | - | 0 | - |
| - | - | 5463 | 423.3 | - | - | 0 | - |
| - | - | 1505 | 423.7 | - | - | 0 | - |
| - | - | 8006 | 424.3 | - | - | 0 | - |
| - | - | 934.8 | 424.7 | - | - | 0 | - |
| - | - | 3796 | 425.3 | - | - | 0 | - |
| - | - | 785.5 | 425.7 | - | - | 0 | - |
| - | - | 1461 | 426.2 | - | - | 0 | - |
| - | - | 3087 | 428.2 | - | - | 0 | - |
| - | - | 1132 | 428.2 | - | - | 0 | - |
| - | - | 683 | 429.1 | - | - | 0 | - |
| - | - | 4234 | 429.1 | - | - | 0 | - |
| - | - | 3235 | 429.2 | - | - | 0 | - |
| - | - | 2055 | 429.2 | - | - | 0 | - |
| - | - | 1163 | 430.1 | - | - | 0 | - |
| 4 | y | 2.39E+04 | 430.2 | 0.004479 | 10.41 | +3 | 10 |
| - | - | 1757 | 430.7 | - | - | 0 | - |
| - | - | 7231 | 431.2 | - | - | 0 | - |
| - | - | 1427 | 431.7 | - | - | 0 | - |
| - | - | 879.9 | 432.2 | - | - | 0 | - |
| 4 | b | 1251 | 433.3 | 0.0005371 | 1.24 | +1 | 4 |
| - | - | 9473 | 434.7 | - | - | 0 | - |
| - | - | 6050 | 435.2 | - | - | 0 | - |
| - | - | 1472 | 435.7 | - | - | 0 | - |
| 11 | y | 5960 | 439.2 | 0.001153 | 2.626 | +1 | 3 |
| 7 | b | 7053 | 439.7 | 0.003091 | 7.03 | +2 | 7 |
| - | - | 3918 | 440.2 | - | - | 0 | - |
| - | - | 1516 | 440.7 | - | - | 0 | - |
| - | - | 1483 | 441.3 | - | - | 0 | - |
| - | - | 3.731E+04 | 442.3 | - | - | 0 | - |
| - | - | 901.3 | 445.7 | - | - | 0 | - |
| - | - | 3945 | 446.2 | - | - | 0 | - |
| - | - | 1044 | 446.2 | - | - | 0 | - |
| - | - | 691.6 | 447.2 | - | - | 0 | - |
| - | - | 1112 | 448.2 | - | - | 0 | - |
| 7 | b | 9552 | 448.7 | 0.003119 | 6.951 | +2 | 7 |
| - | - | 5418 | 449.2 | - | - | 0 | - |
| - | - | 2471 | 449.7 | - | - | 0 | - |
| - | - | 695.5 | 451.2 | - | - | 0 | - |
| 4 | b | 9789 | 451.3 | 0.001081 | 2.395 | +1 | 4 |
| - | - | 1748 | 451.9 | - | - | 0 | - |
| - | - | 3808 | 452.3 | - | - | 0 | - |
| - | - | 835.4 | 452.6 | - | - | 0 | - |
| - | - | 3272 | 454.2 | - | - | 0 | - |
| 7 | y | 2.207E+04 | 454.7 | 0.003487 | 7.668 | +2 | 7 |
| - | - | 1.139E+04 | 455.2 | - | - | 0 | - |
| - | - | 1078 | 455.3 | - | - | 0 | - |
| - | - | 4026 | 455.7 | - | - | 0 | - |
| - | - | 760.5 | 456.2 | - | - | 0 | - |
| - | - | 892.1 | 456.7 | - | - | 0 | - |
| - | - | 2042 | 458.2 | - | - | 0 | - |
| - | - | 935.5 | 459.2 | - | - | 0 | - |
| - | - | 858.1 | 461.2 | - | - | 0 | - |
| - | - | 1401 | 461.3 | - | - | 0 | - |
| - | - | 1202 | 461.8 | - | - | 0 | - |
| - | - | 2403 | 468.3 | - | - | 0 | - |
| - | - | 1052 | 469.3 | - | - | 0 | - |
| - | - | 1902 | 470.2 | - | - | 0 | - |
| - | - | 917.5 | 470.2 | - | - | 0 | - |
| - | - | 679 | 470.3 | - | - | 0 | - |
| - | - | 1021 | 474.8 | - | - | 0 | - |
| - | - | 1691 | 474.9 | - | - | 0 | - |
| - | - | 753.3 | 475.2 | - | - | 0 | - |
| - | - | 1052 | 475.3 | - | - | 0 | - |
| - | - | 1618 | 477.2 | - | - | 0 | - |
| - | - | 1475 | 478.3 | - | - | 0 | - |
| - | - | 1076 | 478.3 | - | - | 0 | - |
| - | - | 864.8 | 479.2 | - | - | 0 | - |
| - | - | 6300 | 482.3 | - | - | 0 | - |
| - | - | 4577 | 482.8 | - | - | 0 | - |
| - | - | 1598 | 483.3 | - | - | 0 | - |
| - | - | 7726 | 483.3 | - | - | 0 | - |
| - | - | 775.4 | 486.8 | - | - | 0 | - |
| - | - | 1740 | 487.2 | - | - | 0 | - |
| - | - | 802.4 | 487.8 | - | - | 0 | - |
| - | - | 1255 | 488.3 | - | - | 0 | - |
| - | - | 3969 | 491.3 | - | - | 0 | - |
| - | - | 1979 | 491.8 | - | - | 0 | - |
| - | - | 2.701E+04 | 494.2 | - | - | 0 | - |
| - | - | 5298 | 495.2 | - | - | 0 | - |
| - | - | 6840 | 495.3 | - | - | 0 | - |
| 8 | b | 2.848E+04 | 496.3 | 0.007874 | 15.87 | +2 | 8 |
| - | - | 9219 | 496.3 | - | - | 0 | - |
| - | - | 1.737E+04 | 496.8 | - | - | 0 | - |
| - | - | 3889 | 497.2 | - | - | 0 | - |
| - | - | 5479 | 497.3 | - | - | 0 | - |
| - | - | 1651 | 498.2 | - | - | 0 | - |
| 8 | b | 4172 | 505.3 | 0.005124 | 10.14 | +2 | 8 |
| - | - | 2759 | 505.8 | - | - | 0 | - |
| - | - | 1317 | 506.1 | - | - | 0 | - |
| - | - | 1029 | 506.3 | - | - | 0 | - |
| - | - | 716 | 507.1 | - | - | 0 | - |
| - | - | 1267 | 507.3 | - | - | 0 | - |
| - | - | 2680 | 511.3 | - | - | 0 | - |
| - | - | 1.014E+04 | 511.3 | - | - | 0 | - |
| 6 | y | 735.7 | 512.2 | 0.002649 | 5.172 | +2 | 8 |
| - | - | 1.432E+05 | 513.4 | - | - | 0 | - |
| - | - | 1470 | 515.3 | - | - | 0 | - |
| - | - | 680.5 | 522.3 | - | - | 0 | - |
| - | - | 5532 | 523.3 | - | - | 0 | - |
| - | - | 4801 | 524.3 | - | - | 0 | - |
| - | - | 1505 | 524.8 | - | - | 0 | - |
| - | - | 4632 | 525.3 | - | - | 0 | - |
| - | - | 9641 | 525.8 | - | - | 0 | - |
| - | - | 2047 | 526.3 | - | - | 0 | - |
| - | - | 5060 | 526.3 | - | - | 0 | - |
| - | - | 1151 | 526.8 | - | - | 0 | - |
| - | - | 2193 | 526.8 | - | - | 0 | - |
| - | - | 1133 | 528.6 | - | - | 0 | - |
| - | - | 3164 | 532.3 | - | - | 0 | - |
| 0 | Precursor | 2306 | 534.6 | 0.002834 | 5.301 | +3 | -1 |
| 0 | Precursor | 2445 | 534.9 | 0.008325 | 15.56 | +3 | -1 |
| - | - | 3414 | 535.6 | - | - | 0 | - |
| - | - | 3083 | 535.9 | - | - | 0 | - |
| - | - | 1238 | 536.8 | - | - | 0 | - |
| - | - | 920.5 | 537.3 | - | - | 0 | - |
| - | - | 1141 | 539.3 | - | - | 0 | - |
| - | - | 3.51E+04 | 539.8 | - | - | 0 | - |
| - | - | 2.219E+04 | 540.3 | - | - | 0 | - |
| 0 | Precursor | 1183 | 540.6 | 0.003402 | 6.292 | +3 | -1 |
| - | - | 8170 | 540.8 | - | - | 0 | - |
| - | - | 1278 | 540.9 | - | - | 0 | - |
| - | - | 2.327E+05 | 541.3 | - | - | 0 | - |
| - | - | 7185 | 541.6 | - | - | 0 | - |
| - | - | 6659 | 541.9 | - | - | 0 | - |
| - | - | 1236 | 542.3 | - | - | 0 | - |
| - | - | 6463 | 543.3 | - | - | 0 | - |
| - | - | 2346 | 544.3 | - | - | 0 | - |
| - | - | 1503 | 548.2 | - | - | 0 | - |
| - | - | 1017 | 549.2 | - | - | 0 | - |
| - | - | 5248 | 554.3 | - | - | 0 | - |
| - | - | 3216 | 554.8 | - | - | 0 | - |
| - | - | 1816 | 555.2 | - | - | 0 | - |
| - | - | 1024 | 558.3 | - | - | 0 | - |
| - | - | 1074 | 562.2 | - | - | 0 | - |
| 5 | b | 779.3 | 562.3 | 0.003756 | 6.68 | +1 | 5 |
| 9 | b | 891.2 | 562.8 | 0.001631 | 2.899 | +2 | 9 |
| - | - | 4910 | 563.3 | - | - | 0 | - |
| - | - | 1722 | 563.8 | - | - | 0 | - |
| - | - | 1115 | 564.8 | - | - | 0 | - |
| - | - | 953.9 | 565.2 | - | - | 0 | - |
| - | - | 1388 | 565.3 | - | - | 0 | - |
| - | - | 996.4 | 569.3 | - | - | 0 | - |
| - | - | 760.8 | 570.3 | - | - | 0 | - |
| - | - | 1106 | 572.3 | - | - | 0 | - |
| - | - | 976.4 | 572.8 | - | - | 0 | - |
| - | - | 4256 | 573.8 | - | - | 0 | - |
| - | - | 1438 | 574.3 | - | - | 0 | - |
| - | - | 1503 | 575.2 | - | - | 0 | - |
| 5 | b | 6601 | 579.3 | 0.0009445 | 1.63 | +1 | 5 |
| - | - | 1877 | 580.3 | - | - | 0 | - |
| - | - | 5827 | 583.2 | - | - | 0 | - |
| - | - | 2471 | 584.2 | - | - | 0 | - |
| - | - | 740 | 584.3 | - | - | 0 | - |
| - | - | 1036 | 587.3 | - | - | 0 | - |
| - | - | 1666 | 588.8 | - | - | 0 | - |
| - | - | 1524 | 592.3 | - | - | 0 | - |
| 9 | y | 7076 | 593.3 | 0.0003291 | 0.5548 | +1 | 5 |
| - | - | 4455 | 593.3 | - | - | 0 | - |
| - | - | 2788 | 594.3 | - | - | 0 | - |
| - | - | 1384 | 594.3 | - | - | 0 | - |
| - | - | 683.1 | 595.3 | - | - | 0 | - |
| - | - | 1414 | 595.3 | - | - | 0 | - |
| - | - | 1111 | 596.4 | - | - | 0 | - |
| - | - | 5.565E+04 | 597.8 | - | - | 0 | - |
| - | - | 3.198E+04 | 598.3 | - | - | 0 | - |
| - | - | 855.5 | 598.7 | - | - | 0 | - |
| - | - | 1079 | 605.3 | - | - | 0 | - |
| - | - | 9510 | 610.3 | - | - | 0 | - |
| - | - | 960.7 | 610.3 | - | - | 0 | - |
| - | - | 2362 | 610.8 | - | - | 0 | - |
| 9 | y | 5.443E+04 | 611.3 | 0.001483 | 2.426 | +1 | 5 |
| - | - | 1.575E+04 | 612.3 | - | - | 0 | - |
| - | - | 3742 | 613.3 | - | - | 0 | - |
| - | - | 962 | 614.3 | - | - | 0 | - |
| - | - | 1357 | 616.3 | - | - | 0 | - |
| - | - | 1678 | 618.8 | - | - | 0 | - |
| - | - | 2469 | 619.3 | - | - | 0 | - |
| - | - | 5782 | 619.8 | - | - | 0 | - |
| - | - | 3782 | 620.3 | - | - | 0 | - |
| - | - | 1372 | 621.3 | - | - | 0 | - |
| - | - | 920.6 | 628.9 | - | - | 0 | - |
| - | - | 2465 | 629.3 | - | - | 0 | - |
| - | - | 1106 | 629.8 | - | - | 0 | - |
| - | - | 9291 | 638.3 | - | - | 0 | - |
| - | - | 4288 | 638.8 | - | - | 0 | - |
| - | - | 954.3 | 639.3 | - | - | 0 | - |
| 4 | y | 1500 | 644.8 | 0.002437 | 3.78 | +2 | 10 |
| 4 | y | 861.5 | 645.3 | 0.005913 | 9.163 | +2 | 10 |
| 11 | b | 950.7 | 646.3 | 0.004791 | 7.412 | +2 | 11 |
| - | - | 5.282E+04 | 647.3 | - | - | 0 | - |
| - | - | 3.145E+04 | 647.8 | - | - | 0 | - |
| - | - | 769.4 | 648.3 | - | - | 0 | - |
| - | - | 888.7 | 649 | - | - | 0 | - |
| - | - | 1152 | 650.3 | - | - | 0 | - |
| - | - | 1787 | 652.3 | - | - | 0 | - |
| - | - | 2121 | 652.8 | - | - | 0 | - |
| - | - | 1601 | 653.3 | - | - | 0 | - |
| - | - | 1040 | 653.3 | - | - | 0 | - |
| 4 | y | 4257 | 653.8 | 0.003808 | 5.824 | +2 | 10 |
| - | - | 2644 | 654.3 | - | - | 0 | - |
| - | - | 2160 | 654.8 | - | - | 0 | - |
| 11 | b | 758.2 | 655.3 | 0.01264 | 19.28 | +2 | 11 |
| - | - | 3746 | 657.4 | - | - | 0 | - |
| - | - | 1260 | 658.4 | - | - | 0 | - |
| - | - | 880.5 | 662.3 | - | - | 0 | - |
| - | - | 693.3 | 667.4 | - | - | 0 | - |
| - | - | 1174 | 667.8 | - | - | 0 | - |
| - | - | 5647 | 668.3 | - | - | 0 | - |
| - | - | 2726 | 669.3 | - | - | 0 | - |
| - | - | 3527 | 671.3 | - | - | 0 | - |
| - | - | 3038 | 672.8 | - | - | 0 | - |
| - | - | 1758 | 673.3 | - | - | 0 | - |
| - | - | 1938 | 674.3 | - | - | 0 | - |
| 6 | b | 1.009E+04 | 676.3 | 0.006218 | 9.193 | +1 | 6 |
| - | - | 4416 | 676.8 | - | - | 0 | - |
| 6 | b | 3402 | 677.3 | 0.003908 | 5.769 | +1 | 6 |
| - | - | 3283 | 677.4 | - | - | 0 | - |
| - | - | 3233 | 677.9 | - | - | 0 | - |
| - | - | 3885 | 678.3 | - | - | 0 | - |
| - | - | 1778 | 679.3 | - | - | 0 | - |
| - | - | 1.412E+04 | 681.8 | - | - | 0 | - |
| - | - | 7956 | 682.3 | - | - | 0 | - |
| - | - | 1024 | 684.3 | - | - | 0 | - |
| - | - | 952.7 | 686.8 | - | - | 0 | - |
| - | - | 7599 | 689.4 | - | - | 0 | - |
| - | - | 1831 | 690.4 | - | - | 0 | - |
| - | - | 925.3 | 692.3 | - | - | 0 | - |
| - | - | 1032 | 693.3 | - | - | 0 | - |
| 6 | b | 3.737E+04 | 694.4 | 0.0006128 | 0.8825 | +1 | 6 |
| - | - | 1.4E+04 | 695.4 | - | - | 0 | - |
| - | - | 1.326E+04 | 696.3 | - | - | 0 | - |
| - | - | 5412 | 697.3 | - | - | 0 | - |
| - | - | 2419 | 698.3 | - | - | 0 | - |
| - | - | 2440 | 699.3 | - | - | 0 | - |
| - | - | 2081 | 700.8 | - | - | 0 | - |
| 3 | y | 7993 | 701.3 | 0.002336 | 3.332 | +2 | 11 |
| 3 | y | 7223 | 701.8 | 0.01131 | 16.11 | +2 | 11 |
| - | - | 4527 | 702.3 | - | - | 0 | - |
| - | - | 9206 | 702.8 | - | - | 0 | - |
| - | - | 5267 | 703.3 | - | - | 0 | - |
| 8 | y | 3841 | 706.3 | 0.0008599 | 1.217 | +1 | 6 |
| - | - | 1518 | 707.3 | - | - | 0 | - |
| - | - | 1.25E+04 | 709.3 | - | - | 0 | - |
| - | - | 1.587E+04 | 709.8 | - | - | 0 | - |
| 3 | y | 3.545E+04 | 710.3 | 0.003585 | 5.047 | +2 | 11 |
| 12 | b | 2.665E+04 | 710.8 | 0.01065 | 14.98 | +2 | 12 |
| 12 | b | 9371 | 711.3 | 0.0001508 | 0.2119 | +2 | 12 |
| - | - | 8.39E+04 | 711.9 | - | - | 0 | - |
| - | - | 5.207E+04 | 712.4 | - | - | 0 | - |
| - | - | 1568 | 712.9 | - | - | 0 | - |
| - | - | 846.4 | 713.3 | - | - | 0 | - |
| - | - | 868.4 | 718.8 | - | - | 0 | - |
| 12 | b | 2779 | 719.8 | 0.005068 | 7.041 | +2 | 12 |
| - | - | 2109 | 720.4 | - | - | 0 | - |
| - | - | 900.5 | 720.9 | - | - | 0 | - |
| - | - | 1131 | 722.4 | - | - | 0 | - |
| - | - | 2073 | 723.4 | - | - | 0 | - |
| 8 | y | 2.109E+04 | 724.4 | 0.001037 | 1.432 | +1 | 6 |
| - | - | 7176 | 725.4 | - | - | 0 | - |
| - | - | 3271 | 725.9 | - | - | 0 | - |
| - | - | 5364 | 726.4 | - | - | 0 | - |
| - | - | 1413 | 726.9 | - | - | 0 | - |
| - | - | 1208 | 727.3 | - | - | 0 | - |
| - | - | 1101 | 737.4 | - | - | 0 | - |
| - | - | 3446 | 739.4 | - | - | 0 | - |
| - | - | 2249 | 740.4 | - | - | 0 | - |
| - | - | 850.8 | 741.4 | - | - | 0 | - |
| - | - | 1883 | 745.4 | - | - | 0 | - |
| - | - | 1.164E+04 | 746.4 | - | - | 0 | - |
| - | - | 9248 | 746.9 | - | - | 0 | - |
| - | - | 1733 | 747.4 | - | - | 0 | - |
| 2 | y | 1904 | 751.3 | 0.01097 | 14.6 | +2 | 12 |
| - | - | 1.457E+05 | 755.4 | - | - | 0 | - |
| - | - | 9.808E+04 | 755.9 | - | - | 0 | - |
| - | - | 2307 | 756.4 | - | - | 0 | - |
| - | - | 2271 | 758.9 | - | - | 0 | - |
| - | - | 2217 | 759.4 | - | - | 0 | - |
| 2 | y | 3176 | 759.9 | 0.002154 | 2.835 | +2 | 12 |
| - | - | 7660 | 760.4 | - | - | 0 | - |
| - | - | 4469 | 760.9 | - | - | 0 | - |
| - | - | 749.5 | 762.4 | - | - | 0 | - |
| - | - | 4699 | 763.4 | - | - | 0 | - |
| - | - | 1643 | 764.4 | - | - | 0 | - |
| - | - | 812.9 | 768.4 | - | - | 0 | - |
| - | - | 844 | 770.4 | - | - | 0 | - |
| - | - | 1548 | 779.3 | - | - | 0 | - |
| - | - | 2096 | 781.4 | - | - | 0 | - |
| - | - | 828.1 | 782.4 | - | - | 0 | - |
| - | - | 1044 | 786.4 | - | - | 0 | - |
| - | - | 961.3 | 787.4 | - | - | 0 | - |
| - | - | 992.9 | 791.4 | - | - | 0 | - |
| - | - | 1031 | 791.5 | - | - | 0 | - |
| - | - | 2137 | 795.4 | - | - | 0 | - |
| - | - | 1116 | 796.4 | - | - | 0 | - |
| - | - | 854.1 | 798.4 | - | - | 0 | - |
| - | - | 1501 | 804.4 | - | - | 0 | - |
| - | - | 779.4 | 806.4 | - | - | 0 | - |
| - | - | 2872 | 809.4 | - | - | 0 | - |
| 0 | Precursor | 3368 | 810.4 | 0.002154 | 2.658 | +2 | -1 |
| - | - | 1570 | 811.3 | - | - | 0 | - |
| - | - | 1580 | 813.4 | - | - | 0 | - |
| - | - | 968.9 | 819.5 | - | - | 0 | - |
| - | - | 977.4 | 823.4 | - | - | 0 | - |
| - | - | 1.405E+04 | 836.4 | - | - | 0 | - |
| - | - | 4601 | 837.4 | - | - | 0 | - |
| - | - | 1202 | 852.4 | - | - | 0 | - |
| - | - | 1703 | 860.4 | - | - | 0 | - |
| - | - | 1075 | 861.4 | - | - | 0 | - |
| - | - | 2071 | 868.4 | - | - | 0 | - |
| - | - | 1804 | 869.4 | - | - | 0 | - |
| 7 | b | 6710 | 878.4 | 0.004914 | 5.594 | +1 | 7 |
| - | - | 4861 | 879.4 | - | - | 0 | - |
| - | - | 965.2 | 880.4 | - | - | 0 | - |
| - | - | 781.1 | 885.4 | - | - | 0 | - |
| - | - | 974.9 | 890.4 | - | - | 0 | - |
| - | - | 823.7 | 892.4 | - | - | 0 | - |
| 7 | b | 2.86E+04 | 896.4 | 0.005274 | 5.884 | +1 | 7 |
| - | - | 1.674E+04 | 897.4 | - | - | 0 | - |
| - | - | 4062 | 898.4 | - | - | 0 | - |
| - | - | 813.7 | 899.4 | - | - | 0 | - |
| - | - | 810.6 | 906.4 | - | - | 0 | - |
| - | - | 1574 | 907.4 | - | - | 0 | - |
| 7 | y | 8640 | 908.4 | 0.005766 | 6.347 | +1 | 7 |
| - | - | 6110 | 909.4 | - | - | 0 | - |
| - | - | 1846 | 910.4 | - | - | 0 | - |
| - | - | 1205 | 912.4 | - | - | 0 | - |
| - | - | 4614 | 924.4 | - | - | 0 | - |
| - | - | 2224 | 925.4 | - | - | 0 | - |
| 7 | y | 1648 | 926.4 | 0.01003 | 10.83 | +1 | 7 |
| - | - | 789.9 | 927.4 | - | - | 0 | - |
| - | - | 834.8 | 953.4 | - | - | 0 | - |
| - | - | 845.1 | 968.5 | - | - | 0 | - |
| - | - | 1033 | 970.4 | - | - | 0 | - |
| - | - | 1711 | 971.4 | - | - | 0 | - |
| - | - | 1635 | 975.5 | - | - | 0 | - |
| - | - | 722.2 | 976.5 | - | - | 0 | - |
| - | - | 1658 | 981.4 | - | - | 0 | - |
| - | - | 2014 | 986.5 | - | - | 0 | - |
| - | - | 1059 | 987.5 | - | - | 0 | - |
| - | - | 1624 | 988.4 | - | - | 0 | - |
| - | - | 948.2 | 996.5 | - | - | 0 | - |
| 8 | b | 2449 | 1010 | 0.003181 | 3.151 | +1 | 8 |
| - | - | 870.7 | 1011 | - | - | 0 | - |
| - | - | 1261 | 1012 | - | - | 0 | - |
| - | - | 1270 | 1014 | - | - | 0 | - |
| - | - | 1285 | 1015 | - | - | 0 | - |
| - | - | 1296 | 1030 | - | - | 0 | - |
| - | - | 705.2 | 1031 | - | - | 0 | - |
| - | - | 1092 | 1039 | - | - | 0 | - |
| - | - | 786.9 | 1040 | - | - | 0 | - |
| 6 | y | 686.7 | 1041 | 0.002018 | 1.938 | +1 | 8 |
| - | - | 1153 | 1042 | - | - | 0 | - |
| - | - | 841.6 | 1073 | - | - | 0 | - |
| - | - | 1308 | 1091 | - | - | 0 | - |
| - | - | 1220 | 1092 | - | - | 0 | - |
| 9 | b | 904.5 | 1108 | 0.007303 | 6.594 | +1 | 9 |
| - | - | 1997 | 1110 | - | - | 0 | - |
| - | - | 949.8 | 1111 | - | - | 0 | - |
| - | - | 750.9 | 1112 | - | - | 0 | - |
| - | - | 1743 | 1150 | - | - | 0 | - |
| 5 | y | 2518 | 1151 | 0.002179 | 1.893 | +1 | 9 |
| 5 | y | 3470 | 1152 | 0.01088 | 9.437 | +1 | 9 |
| - | - | 1522 | 1153 | - | - | 0 | - |
| - | - | 946.3 | 1161 | - | - | 0 | - |
| - | - | 1430 | 1167 | - | - | 0 | - |
| - | - | 1742 | 1169 | - | - | 0 | - |
| 5 | y | 4593 | 1170 | 0.005689 | 4.864 | +1 | 9 |
| - | - | 2362 | 1171 | - | - | 0 | - |
| - | - | 3650 | 1178 | - | - | 0 | - |
| - | - | 1105 | 1179 | - | - | 0 | - |
| - | - | 661.8 | 1917 | - | - | 0 | - |
| - | - | 700.1 | 2728 | - | - | 0 | - |
| - | - | 1062 | 3083 | - | - | 0 | - |

m/z Charge Intensity FragmentType MassShift Position
120.04486083984375 0 494.44135
120.08113098144531 0 24686.06
121.08458709716797 0 2077.2937
121.77975463867188 0 384.69507
122.07180786132812 0 1010.399
123.04420471191406 0 961.2384
123.05587005615234 0 462.8252
125.10775756835938 0 1297.2733
126.05534362792969 0 1494.1068
127.05052947998047 0 1046.781
127.08671569824219 0 410.17404
127.12325286865234 0 603.3936
128.0954132080078 0 451.07657
128.10731506347656 0 11122.455
129.06622314453125 0 1287.5299
129.10260009765625 0 85568.8
130.05032348632812 0 430.3211
130.06546020507812 0 11176.218
130.08653259277344 0 726.39056
130.10012817382812 0 835.4134
130.1060028076172 0 5165.831
131.04981994628906 0 668.883
131.06890869140625 0 874.6985
132.04478454589844 0 3651.7952
132.08143615722656 0 522.8153
132.10272216796875 0 382.37518
133.06126403808594 0 525.95355
136.07606506347656 0 17803.326
136.081787109375 0 1111.5055
137.07952880859375 0 1186.6696
138.0668182373047 0 1342.9316
138.09169006347656 0 1406.7788
139.0869903564453 0 1148.3767
140.08218383789062 0 1089.0978
141.1026611328125 0 2934.9348
143.04612731933594 0 413.52597
144.06607055664062 0 604.3271
145.0610809326172 0 518.23334
146.06039428710938 0 7654.5913
146.53994750976562 0 471.52493
147.0442352294922 0 870.5268
147.0636749267578 0 718.4725
147.07662963867188 0 1071.9677 y 10
148.9539031982422 0 692.42584
150.066650390625 0 1016.41797
151.08697509765625 0 1788.7906
154.0502471923828 0 1413.7069
155.0930938720703 0 6963.366
155.11822509765625 0 7616.222 a Water loss 1
156.07716369628906 0 966.03955
156.09629821777344 0 631.8207
156.10256958007812 0 723.82745
156.12136840820312 0 474.5952
157.06106567382812 0 1838.3876
157.0763702392578 0 7479.1045
157.0973663330078 0 2318.077
157.10879516601562 0 4976.9688
157.13412475585938 0 1255.5795
158.06039428710938 0 10131.242
158.079833984375 0 656.0393
158.09286499023438 0 1599.7285
159.06385803222656 0 1290.2876
159.07650756835938 0 559.26825
159.09194946289062 0 3142.2327
159.11322021484375 0 2462.9585 d 1
163.08595275878906 0 694.4055
165.05496215820312 0 9378.102
166.06141662597656 0 12953.972
166.08663940429688 0 8979.431 y 9
166.09811401367188 0 1183.659
167.05711364746094 0 477.62146
167.06512451171875 0 900.27155
167.0818328857422 0 985.9413
167.08963012695312 0 541.9667
168.05551147460938 0 617.4521
168.10186767578125 0 1158.683
168.11341857910156 0 1069.1022
169.097412109375 0 2680.1433
169.13389587402344 0 3246.109
170.05221557617188 0 519.5613
171.07696533203125 0 1500.0597
171.09132385253906 0 512.33545
171.1131134033203 0 569.20544
171.14990234375 0 929.1614
172.0720977783203 0 1568.6282
172.08517456054688 0 456.09485
173.0563201904297 0 911.76245
173.12887573242188 0 186046.56 a 1
173.4392852783203 0 2159.1475
174.0552978515625 0 4612.5747
174.13223266601562 0 14892.146
175.0719451904297 0 2662.931
175.08694458007812 0 38544.45
175.11924743652344 0 2022.8834
175.13394165039062 0 559.24133
176.08309936523438 0 716.5968
176.09034729003906 0 4478.919
177.10272216796875 0 699.8385
178.13427734375 0 4893.2656
180.11351013183594 0 529.7388
181.0612030029297 0 966.45795
182.0815887451172 0 17517.523 y 12
183.0557403564453 0 972.8352
183.08514404296875 0 1323.9792
183.11317443847656 0 16266.49 b Water loss 1
183.14962768554688 0 4101.1816
184.0769500732422 0 494.63007
184.1169891357422 0 761.5579
185.0713653564453 0 7600.826
185.0925750732422 0 1635.301
185.104248046875 0 691.4122
185.12901306152344 0 1015.18555
185.1650848388672 0 1515.8064
185.44566345214844 0 490.89697
186.05523681640625 0 864.50793
186.0747833251953 0 1077.1338
186.1240997314453 0 25620.76
187.0869140625 0 1027.6128
187.1083526611328 0 1809.8522
187.12730407714844 0 2154.8892
187.14450073242188 0 1295.3702
188.0714874267578 0 934.5357
189.08738708496094 0 8168.0317
190.0825653076172 0 1034.654
190.13449096679688 0 602.95447
191.0821075439453 0 637.12585
195.1131591796875 0 7035.062
196.10829162597656 0 3254.7224
196.11705017089844 0 683.0482
196.14505004882812 0 1249.3322
197.12875366210938 0 2705.9548
198.08819580078125 0 837.23694
199.0716094970703 0 4291.9883
199.144287109375 0 1114.1182
201.1237030029297 0 64332.094 b 1
202.0500030517578 0 822.0251
202.1270751953125 0 5375.3765
203.06690979003906 0 2396.7056
203.08172607421875 0 2677.4868
204.13455200195312 0 538.2208
205.09756469726562 0 3530.1367
207.16017150878906 0 1351.0328
209.0561981201172 0 722.7886
209.09225463867188 0 1313.5175
210.08802795410156 0 571.8402
211.08726501464844 0 552.8191
211.1081085205078 0 1935.4855 y Water loss 10
211.14451599121094 0 2168.1377
211.1556854248047 0 12669.363
212.10292053222656 0 777.4705
212.13890075683594 0 615.5018
213.12364196777344 0 3014.0435
214.15524291992188 0 682.6498
215.08108520507812 0 510.10858
215.1392364501953 0 1452.8984
216.13426208496094 0 946.4286
216.17127990722656 0 668.00464
217.082275390625 0 13532.726
218.08522033691406 0 763.6756
221.0922088623047 0 2029.0846
221.1034698486328 0 938.28094
222.12417602539062 0 2237.7803
223.06349182128906 0 755.2864
223.10812377929688 0 1851.2952
223.15570068359375 0 14608.325
224.1032257080078 0 1728.1909
224.1402587890625 0 645.33704
224.15902709960938 0 1279.9495
225.06187438964844 0 520.3198
226.08274841308594 0 2193.4727
226.11891174316406 0 1546.2845 b Water loss 5
227.0668182373047 0 3050.06
227.08505249023438 0 577.32556
227.1142578125 0 2122.9429
228.18203735351562 0 2362.474
229.1188507080078 0 3156.9922
229.13153076171875 0 549.4302
230.11361694335938 0 872.35645
233.14010620117188 0 2124.8867
233.1653289794922 0 632.01117
234.12417602539062 0 15670.058
234.14393615722656 0 731.1918
235.12750244140625 0 2006.613
237.13465881347656 0 1221.3489
238.119384765625 0 1645.4249
239.15069580078125 0 8171.1177
240.09701538085938 0 617.2277
240.1347198486328 0 5338.631
241.11886596679688 0 974.9223
241.1326446533203 0 991.04694
241.19061279296875 0 710.9621
242.11399841308594 0 2352.852
243.10906982421875 0 650.9278
243.13526916503906 0 500.2741
243.14877319335938 0 1076.502
244.09317016601562 0 3955.947
245.12603759765625 0 556.76013
247.1082000732422 0 9728.722
248.1138458251953 0 3315.5583
249.09861755371094 0 3864.68
249.13426208496094 0 1140.58
251.15074157714844 0 54798.38
252.15386962890625 0 7579.6313
253.13430786132812 0 688.34595
254.1127471923828 0 748.24744
255.1091766357422 0 2327.635
255.14578247070312 0 1374.6184
256.1774597167969 0 3075.682
257.1121520996094 0 1210.474
258.1451721191406 0 2104.059
262.1198425292969 0 803.5461
262.15277099609375 0 720.6215
263.103271484375 0 1068.8359
263.13861083984375 0 691.162
265.1298828125 0 5299.853
265.1545715332031 0 1045.5059
266.11419677734375 0 983.15063
266.1252136230469 0 12469.602
266.1490783691406 0 693.35986
266.640869140625 0 934.9863
267.1282653808594 0 1712.4974
268.12811279296875 0 771.5798
268.1776123046875 0 900.48425
269.12908935546875 0 671.0052
269.1612548828125 0 1124.2102
269.18548583984375 0 570.5989
270.144287109375 0 1254.345
270.1604919433594 0 14500.511
271.1639099121094 0 1862.8484
274.1304016113281 0 1119.9587
275.1034851074219 0 905.3162
276.1551513671875 0 810.497
278.1136779785156 0 1312.572
279.14593505859375 0 1081.869
280.1290283203125 0 1281.1847
282.1564636230469 0 1335.7296
282.1817932128906 0 1090.8314
283.14044189453125 0 32984.6
284.1434020996094 0 3750.1736
284.16143798828125 0 1329.5607
285.7781982421875 0 558.1574
286.1398010253906 0 1353.0671
286.1554870605469 0 1000.76044
288.171142578125 0 2608.6372
290.114501953125 0 1052.0728
292.0513000488281 0 533.7531
293.1136779785156 0 22127.057 y Water loss 11
293.16107177734375 0 576.57886
294.1176452636719 0 3022.117
294.1562805175781 0 1451.3008
295.17718505859375 0 763.16943
296.1974182128906 0 1672.0635 b Water loss 2
297.15625 0 5840.953
298.13958740234375 0 1191.4531
298.1561279296875 0 6134.61
299.0624694824219 0 2280.8848
300.0625915527344 0 2574.0918
300.0987243652344 0 1085.0778
300.16693115234375 0 2148.616
301.0596008300781 0 4070.4138
301.0812072753906 0 1035.1896
301.15118408203125 0 10978.005
301.170654296875 0 835.50256
302.0599365234375 0 2779.7234
302.1553039550781 0 1272.2708
303.0579528808594 0 1140.9197
305.1612243652344 0 3839.3804
305.6446838378906 0 666.651
306.1374206542969 0 1723.3875 y 8
310.2238464355469 0 1611.0383
311.1241455078125 0 2962.3645 y 11
311.139892578125 0 722.4794
312.1259460449219 0 675.79297
312.156005859375 0 11950.375
313.1582336425781 0 876.4513
313.18756103515625 0 3097.546
313.22442626953125 0 591.6437
314.148193359375 0 2947.2842
314.20880126953125 0 816.2348 b 2
315.1352233886719 0 630.5424
315.1668701171875 0 8205.002
315.18365478515625 0 593.5698
316.15057373046875 0 3661.7483
316.1661071777344 0 613.7677
317.1677551269531 0 792.029
318.10919189453125 0 2524.4983
322.1868896484375 0 907.42487
323.1719055175781 0 954.21265
327.2512512207031 0 2353.6729
330.1667175292969 0 5115.6343
331.1705322265625 0 815.73584 b Water loss 7
332.1244812011719 0 564.28815
332.2088928222656 0 1250.1921
333.1772155761719 0 1267.3541
334.10205078125 0 788.62756
337.2354431152344 0 590.1001
338.2195129394531 0 2578.4277
338.6748962402344 0 977.4728 b Water loss 5
341.18206787109375 0 1995.5914
342.1412658691406 0 3590.1345 y Ammonia loss 5
343.1618957519531 0 860.1846
345.1312255859375 0 2147.916
346.115478515625 0 4736.782
347.1186828613281 0 988.3886
347.679931640625 0 4076.3945 b 5
348.1816711425781 0 2140.9807
349.1850891113281 0 1517.2822
350.1766357421875 0 1265.5735
350.21917724609375 0 3941.959
351.22174072265625 0 1048.9678
353.6751403808594 0 878.4895 y Water loss 7
355.1689453125 0 1069.5682
355.24554443359375 0 1612.1467
358.1773986816406 0 909.1986
359.1670227050781 0 1838.949
361.1986389160156 0 1017.533
362.1848449707031 0 2288.9226
362.6800231933594 0 3659.7654 y 7
363.1416015625 0 8223.454
363.18145751953125 0 2290.8127
364.1245422363281 0 1464.8425
364.1458435058594 0 1573.3884
364.1753234863281 0 2584.0698
364.677978515625 0 774.10944
367.1606750488281 0 1349.8472
367.24609375 0 782.12933
373.18505859375 0 1942.2119
378.21356201171875 0 2849.8506
379.2093200683594 0 12870.891
380.2118835449219 0 2853.4558
381.1521911621094 0 21554.432
382.15472412109375 0 4299.2695
382.1936340332031 0 638.7364
383.22808837890625 0 2101.4758
384.1902160644531 0 1764.6824
384.2723083496094 0 851.2483
390.17822265625 0 876.7983
391.1356506347656 0 1132.441
392.19110107421875 0 920.7022
394.1723937988281 0 2832.503
395.2396545410156 0 782.94794
396.1959228515625 0 970.91174
396.225341796875 0 5555.5415
397.226318359375 0 1350.4352
397.2565612792969 0 4257.3804
398.17706298828125 0 1387.6658
398.2884826660156 0 1555.9962
400.2232360839844 0 940.4139
401.2397766113281 0 1746.6829
402.1983337402344 0 2139.679
405.25927734375 0 721.76697
407.2383117675781 0 1876.8688
407.7290954589844 0 911.40247
408.2701721191406 0 671.19727
409.1465148925781 0 2158.5203
409.2212829589844 0 1195.1299
410.149169921875 0 989.411
411.13043212890625 0 1586.6448
411.199462890625 0 820.1836
412.1832580566406 0 11753.41
412.267333984375 0 3649.331
413.18341064453125 0 5076.9907
414.23388671875 0 2884.9907
414.2830505371094 0 12394.005
414.720947265625 0 1895.703
417.2021179199219 0 791.65
418.0347900390625 0 746.9664
418.7158508300781 0 1121.9644
419.0321350097656 0 925.2538
420.6589050292969 0 702.20844
421.21209716796875 0 762.6002 y Water loss 10
422.2073059082031 0 708.1203
423.23419189453125 0 4363.4272
423.2721862792969 0 5462.5083
423.7335205078125 0 1504.8518
424.2682189941406 0 8005.6177
424.73516845703125 0 934.8476
425.25164794921875 0 3796.43
425.7148132324219 0 785.454
426.20965576171875 0 1460.5332
428.1565246582031 0 3086.7463
428.2137756347656 0 1131.5776
429.08795166015625 0 683.0164
429.1408996582031 0 4233.5127
429.2087097167969 0 3234.7947
429.2357482910156 0 2055.213
430.14422607421875 0 1163.4438
430.1941223144531 0 23900.924 y Water loss 3
430.70758056640625 0 1756.7964
431.1978454589844 0 7231.028
431.7020263671875 0 1426.7825
432.1980285644531 0 879.8773
433.2563171386719 0 1251.3237 b Water loss 3
434.7196960449219 0 9473.079
435.2217712402344 0 6050.318
435.7230529785156 0 1471.6714
439.2198791503906 0 5960.218 y 10
439.7120361328125 0 7053.195 b Water loss 6
440.21343994140625 0 3918.2634
440.7113037109375 0 1516.3516
441.292724609375 0 1483.1743
442.2779235839844 0 37313.56
445.7049865722656 0 901.346
446.1677551269531 0 3944.7952
446.2041320800781 0 1043.7509
447.169921875 0 691.5609
448.2051086425781 0 1112.0544
448.71734619140625 0 9552.445 b 6
449.2182312011719 0 5418.2046
449.7204895019531 0 2470.692
451.2326354980469 0 695.5483
451.2674255371094 0 9788.81 b 3
451.9219970703125 0 1748.1038
452.2680969238281 0 3808.3882
452.5903625488281 0 835.4185
454.2192687988281 0 3271.8657
454.71209716796875 0 22074.414 y Water loss 6
455.2135925292969 0 11390.139
455.2697448730469 0 1077.6313
455.71405029296875 0 4026.1396
456.2135009765625 0 760.4788
456.7164611816406 0 892.08014
458.2369384765625 0 2042.4275
459.19964599609375 0 935.5245
461.2065734863281 0 858.05743
461.29248046875 0 1400.5103
461.79461669921875 0 1202.463
468.2935791015625 0 2403.1877
469.2784423828125 0 1052.1113
470.2044982910156 0 1901.6951
470.2401428222656 0 917.46265
470.27789306640625 0 679.03546
474.7979736328125 0 1021.2853
474.9131164550781 0 1690.5117
475.2469787597656 0 753.25574
475.29083251953125 0 1052.4854
477.2103576660156 0 1618.1962
478.2778015136719 0 1474.661
478.3136291503906 0 1076.2781
479.2308044433594 0 864.8058
482.2570495605469 0 6299.753
482.7571105957031 0 4577.1924
483.2590026855469 0 1598.1329
483.3409423828125 0 7725.824
486.7626037597656 0 775.4364
487.23162841796875 0 1739.7129
487.76171875 0 802.35486
488.25421142578125 0 1255.2992
491.2614440917969 0 3968.6655
491.7638244628906 0 1979.382
494.2364501953125 0 27009.988
495.2397155761719 0 5297.7256
495.3410339355469 0 6839.5815
496.24310302734375 0 28476.19 b Water loss 7
496.3245544433594 0 9218.647
496.7671813964844 0 17374.787
497.2407531738281 0 3889.1062
497.2717590332031 0 5478.801
498.24603271484375 0 1651.4064
505.2613830566406 0 4172.174 b 7
505.75982666015625 0 2759.4363
506.1058654785156 0 1316.6476
506.2709655761719 0 1028.9023
507.1043395996094 0 715.9977
507.2558898925781 0 1266.8416
511.26348876953125 0 2679.623
511.3359069824219 0 10135.316
512.2247314453125 0 735.6756 y Water loss 5
513.3514404296875 0 143225.5
515.25390625 0 1469.8651
522.283935546875 0 680.5128
523.335693359375 0 5532.209
524.26318359375 0 4801.1265
524.7637329101562 0 1504.7466
525.2669067382812 0 4632.1816
525.812744140625 0 9640.933
526.2704467773438 0 2047.4469
526.3143920898438 0 5059.7764
526.7695922851562 0 1151.0597
526.8148803710938 0 2192.6064
528.5914306640625 0 1132.5055
532.2730102539062 0 3164.4338
534.592529296875 0 2306.0757 Precursor Water loss
534.926025390625 0 2444.8972 Precursor Ammonia loss
535.6043701171875 0 3414.0059
535.9402465820312 0 3082.8772
536.7938232421875 0 1238.3147
537.29736328125 0 920.51587
539.27197265625 0 1141.1478
539.8099365234375 0 35096.73
540.311279296875 0 22186.512
540.5966186523438 0 1182.7595 Precursor
540.8124389648438 0 8169.567
540.930908203125 0 1277.8596
541.3463745117188 0 232749.6
541.6097412109375 0 7185.38
541.9442138671875 0 6659.18
542.29541015625 0 1236.0789
543.278564453125 0 6462.99
544.2791748046875 0 2345.659
548.190185546875 0 1502.6035
549.1929931640625 0 1017.3255
554.2808227539062 0 5247.665
554.7825927734375 0 3215.743
555.2313232421875 0 1815.6578
558.290283203125 0 1023.7547
562.2448120117188 0 1073.6287
562.2946166992188 0 779.32245 b Ammonia loss 4
562.7713623046875 0 891.23816 b 8
563.2546997070312 0 4909.5103
563.7532958984375 0 1722.2301
564.7915649414062 0 1114.7632
565.2151489257812 0 953.9279
565.2618408203125 0 1387.8245
569.2662963867188 0 996.4103
570.3132934570312 0 760.8423
572.3099365234375 0 1105.8097
572.8095092773438 0 976.3893
573.79736328125 0 4255.913
574.2990112304688 0 1438.2249
575.2437744140625 0 1503.2755
579.3258666992188 0 6600.6616 b 4
580.3274536132812 0 1877.2638
583.2276000976562 0 5827.0786
584.2301025390625 0 2471.1353
584.3176879882812 0 740.0314
587.2781372070312 0 1035.758
588.790771484375 0 1665.682
592.3073120117188 0 1524.1902
593.2568969726562 0 7075.509 y Water loss 8
593.3059692382812 0 4454.5854
594.2549438476562 0 2787.7612
594.3090209960938 0 1383.8014
595.2587280273438 0 683.0864
595.32080078125 0 1414.3604
596.3509521484375 0 1110.747
597.7973022460938 0 55648.504
598.2987060546875 0 31981.553
598.6671142578125 0 855.5188
605.3082275390625 0 1079.2467
610.283935546875 0 9509.913
610.3323974609375 0 960.6998
610.7887573242188 0 2362.3567
611.2686157226562 0 54434.39 y 8
612.2708740234375 0 15747.7295
613.2741088867188 0 3742.1948
614.2764282226562 0 962.03076
616.344970703125 0 1356.8129
618.7857055664062 0 1677.5729
619.2975463867188 0 2469.1167
619.7954711914062 0 5782.059
620.2999877929688 0 3782.0579
621.2511596679688 0 1372.0507
628.8519897460938 0 920.5588
629.315673828125 0 2465.1357
629.8136596679688 0 1106.2319
638.3198852539062 0 9291.226
638.8223266601562 0 4287.8613
639.3450927734375 0 954.3402
644.7832641601562 0 1499.7448 y Water loss 3
645.2787475585938 0 861.49884 y Ammonia loss 3
646.31787109375 0 950.6555 b Water loss 10
647.3314819335938 0 52817.062
647.8329467773438 0 31447.238
648.3464965820312 0 769.4286
649.0191040039062 0 888.68256
650.3046264648438 0 1152.3
652.3383178710938 0 1786.6748
652.7821044921875 0 2121.0747
653.2845458984375 0 1600.8566
653.34326171875 0 1040.3076
653.7899169921875 0 4257.488 y 3
654.2943725585938 0 2644.0935
654.7964477539062 0 2159.6587
655.3153076171875 0 758.2362 b 10
657.3568115234375 0 3745.5527
658.3605346679688 0 1260.4664
662.3320922851562 0 880.4635
667.3871459960938 0 693.3402
667.8185424804688 0 1174.3821
668.315185546875 0 5647.03
669.32177734375 0 2725.9312
671.3483276367188 0 3526.5278
672.8299560546875 0 3038.234
673.3345336914062 0 1757.852
674.307373046875 0 1938.3939
676.3350830078125 0 10091.269 b Water loss 5
676.8305053710938 0 4415.6006
677.3292236328125 0 3402.2173 b Ammonia loss 5
677.3799438476562 0 3282.689
677.8789672851562 0 3232.7112
678.3010864257812 0 3885.1191
679.2903442382812 0 1777.9435
681.83447265625 0 14124.079
682.3351440429688 0 7955.982
684.3081665039062 0 1024.2703
686.8219604492188 0 952.661
689.3585815429688 0 7599.263
690.3591918945312 0 1830.5043
692.3123779296875 0 925.2906
693.3099975585938 0 1031.9415
694.3524780273438 0 37370.14 b 5
695.35498046875 0 13998.277
696.3104858398438 0 13264.436
697.3118286132812 0 5412.19
698.3189086914062 0 2418.6165
699.3419799804688 0 2439.6492
700.8232421875 0 2080.8591
701.3251953125 0 7993.4844 y Water loss 2
701.826171875 0 7222.7637 y Ammonia loss 2
702.3255004882812 0 4526.8394
702.8458251953125 0 9205.912
703.3468017578125 0 5267.204
706.3414916992188 0 3840.8474 y Water loss 7
707.3438720703125 0 1518.149
709.3252563476562 0 12502.659
709.8286743164062 0 15873.912
710.3317260742188 0 35451.062 y 2
710.8333129882812 0 26652.18 b Water loss 11
711.3358154296875 0 9371.245 b Ammonia loss 11
711.8526000976562 0 83902.625
712.3546142578125 0 52068
712.8604125976562 0 1568.1669
713.2947387695312 0 846.4321
718.8470458984375 0 868.4435
719.8543090820312 0 2778.6987 b 11
720.3549194335938 0 2108.748
720.8571166992188 0 900.4889
722.3549194335938 0 1130.6593
723.36767578125 0 2073.4824
724.3522338867188 0 21085.223 y 7
725.3551025390625 0 7176.282
725.8617553710938 0 3270.9578
726.3616333007812 0 5363.5513
726.8667602539062 0 1413.0337
727.3460693359375 0 1208.1501
737.3555908203125 0 1101.2817
739.3740844726562 0 3445.9138
740.3670043945312 0 2249.1694
741.3564453125 0 850.79
745.3565673828125 0 1882.8927
746.3630981445312 0 11635.903
746.864501953125 0 9248.245
747.3552856445312 0 1733.0876
751.3600463867188 0 1904.1501 y Ammonia loss 1
755.36865234375 0 145700.11
755.8701171875 0 98076.98
756.3683471679688 0 2307.1697
758.8577270507812 0 2270.56
759.3648071289062 0 2217.4534
759.864501953125 0 3176.0352 y 1
760.3631591796875 0 7659.622
760.8632202148438 0 4469.111
762.413330078125 0 749.5074
763.3777465820312 0 4698.9443
764.3778686523438 0 1643.3613
768.376953125 0 812.89923
770.408935546875 0 843.97003
779.34814453125 0 1547.6545
781.3982543945312 0 2096.4097
782.3993530273438 0 828.1324
786.4105224609375 0 1043.7173
787.4100952148438 0 961.25104
791.38525390625 0 992.8781
791.4600830078125 0 1030.9586
795.3811645507812 0 2137.1948
796.3740234375 0 1116.2997
798.4148559570312 0 854.1186
804.4237060546875 0 1500.5835
806.4189453125 0 779.40247
809.3937377929688 0 2871.814
810.384033203125 0 3368.0635 Precursor
811.3352661132812 0 1569.6719
813.4428100585938 0 1579.9087
819.4583129882812 0 968.9157
823.3726806640625 0 977.4268
836.4265747070312 0 14048.844
837.4298706054688 0 4601.147
852.4202880859375 0 1202.3944
860.3920288085938 0 1703.0297
861.3967895507812 0 1074.7942
868.4295043945312 0 2070.659
869.4353637695312 0 1804.1389
878.41552734375 0 6709.7476 b Water loss 6
879.4141235351562 0 4861.0273
880.4191284179688 0 965.2126
885.4481201171875 0 781.10114
890.4212036132812 0 974.85815
892.4338989257812 0 823.68225
896.4264526367188 0 28600.93 b 6
897.4290161132812 0 16742.389
898.4320068359375 0 4062.161
899.4457397460938 0 813.7238
906.4077758789062 0 810.615
907.4124145507812 0 1574.4447
908.4157104492188 0 8639.74 y Water loss 6
909.420166015625 0 6109.9014
910.4288940429688 0 1845.6156
912.420166015625 0 1204.9514
924.4214477539062 0 4613.9272
925.4288330078125 0 2223.807
926.4305419921875 0 1647.5697 y 6
927.4354248046875 0 789.91095
953.4022827148438 0 834.7743
968.4861450195312 0 845.11566
970.424560546875 0 1032.5588
971.421630859375 0 1710.8539
975.5021362304688 0 1634.6642
976.498779296875 0 722.1595
981.442138671875 0 1657.9926
986.4906005859375 0 2013.7382
987.4944458007812 0 1058.6569
988.438232421875 0 1623.8882
996.4632568359375 0 948.192
1009.5084228515625 0 2449.0603 b 7
1010.5144653320312 0 870.72046
1012.478759765625 0 1260.9841
1014.4815673828125 0 1270.2153
1015.4786376953125 0 1284.8108
1030.4971923828125 0 1295.6309
1031.4866943359375 0 705.228
1039.43994140625 0 1091.701
1040.460693359375 0 786.885
1041.4454345703125 0 686.708 y 5
1042.458740234375 0 1152.9055
1072.5745849609375 0 841.59344
1090.5323486328125 0 1307.5858
1091.5250244140625 0 1219.7314
1107.512939453125 0 904.5428 b Ammonia loss 8
1109.5377197265625 0 1997.2125
1110.54296875 0 949.846
1111.5426025390625 0 750.8575
1150.4737548828125 0 1742.6693
1151.4932861328125 0 2518.2556 y Water loss 4
1152.4903564453125 0 3470.2341 y Ammonia loss 4
1153.491455078125 0 1521.7852
1160.5543212890625 0 946.2677
1167.4937744140625 0 1429.6952
1168.51025390625 0 1742.148
1169.51171875 0 4592.7754 y 4
1170.51708984375 0 2362.3699
1177.5618896484375 0 3649.6187
1178.55859375 0 1104.5232
1916.825927734375 0 661.84924
2728.41064453125 0 700.1244
3082.852294921875 0 1061.9934

Spectrum Details

|  |  |
| --- | --- |
| Matched peaks? Matched peaks The total absolute number of peaks matched. Additionally in brackets the total fraction of peaks matched and the total number of peaks is shown. | 69 (9.25% of 746) |
| FDR? FDR The false discovery rate estimated for this peptide. It is calculated by matching all theoretical fragments with a non-integer shift with the raw peaks for this spectrum. This is done with 40 different shifts. The resulting percentage is the average number of annotated peaks over the number of annotated peaks with the correct spectrum. | 1.69% |
| Satellite FDR? Satellite FDR See the FDR for details on its calculation. This satellite ion specific FDR only contains the satellite ions (d/w) for I/L/J positions. | ∞ |
| PSM Score? PSM Score The PSM Score as given by Hecklib to this annotated spectrum. It is shown with three significant figures. | 462 |

#### Spectrum 6412? Spectrum 6412 The raw spectrum of this peptide as annotated by Hecklib. The fragments are coloured according to ion type (see legend). Any peaks with a star '\*' as text can be hovered over to see the full details, first the ion type second the mass shift type. By hovering over the amino acids in the peptide or ions in the legend the corresponding peaks are highlighted. By toggling the 'Unassigned' label you can turn the background (unassigned) peaks on or off in the plot. By updating the slider in the Ion legend you can update the spectrum to only show the top X% of the peaks with labels. The top X% means any peak that is within X% of the highest intensity. By dragging in the spectrum you can zoom in to a specific part of the spectrum and use 'Zoom Out' to get back to the original zoom level. The annotation of the spectrum is based on the given sequence in the peptides file and is done with different software so inconsistencies are likely. The peaks are annotated based on the given sequence, with 20 ppm tolerance.

Copy Data

##### Spectrum 6412 (TSV)

###### Preview

```
Loading example...
```

*Click on the button to copy the data to your clipboard.*

Mz MinMz MaxIntensity Max

WidthHeightPeptide font sizePeptide stroke widthSpectrum font sizeSpectrum stroke widthCompact peptide

Ion legend

wxyz

abcd

OtherUnassignedIon^Charge^_Position_Show for top:%

TVLHQDWLDGKEY

06.27e+41.25e+51.88e+52.51e+5

Zoom Out

y^+1^_1_c^+1^_2_y^+1^_2_y^+1^_2_c^+1^_3_c^+2^_6_y^+1^_3_z^+1^_3_y^+3^_10_y^+1^_3_c^+2^_7_y^+2^_7_c^+1^_4_z^+1^_4_y^+1^_4_z^+2^_8_c^+2^_8_w^+1^_5_c^+1^_5_c^+2^_10_y^+1^_5_z^+1^_5_c^+1^_5_y^+1^_5_z^+2^_10_y^+2^_10_c^+2^_11_c^+2^_11_w^+1^_6_w^+2^_11_c^+1^_6_y^+2^_11_y^+2^_11_z^+2^_11_y^+2^_11_c^+1^_6_c^+2^_12_c^+2^_12_y^+1^_6_c^+2^_12_y^+2^_12_z^+2^_12_y^+2^_12_c^+1^_7_y^+1^_7_c^+1^_7_y^+1^_7_c^+1^_8_z^+1^_8_c^+1^_8_y^+1^_8_w^+1^_9_c^+1^_9_z^+1^_9_y^+1^_9_c^+1^_10_c^+1^_10_y^+1^_10_z^+1^_10_y^+1^_10_c^+1^_11_y^+1^_11_z^+1^_11_y^+1^_11_c^+1^_12_c^+1^_12_z^+1^_12_

0778155723353114

Fragment Matches Table

Show background peaks

| Position | Ion type | Intensity | mz Theoretical | mz Error (Th) | mz Error (ppm) | Charge | Series Number |
| --- | --- | --- | --- | --- | --- | --- | --- |
| - | - | 780.6 | 120.1 | - | - | 0 | - |
| - | - | 350.5 | 121.2 | - | - | 0 | - |
| - | - | 3543 | 129.1 | - | - | 0 | - |
| - | - | 811 | 131.1 | - | - | 0 | - |
| - | - | 727.4 | 136.1 | - | - | 0 | - |
| - | - | 363.2 | 140 | - | - | 0 | - |
| - | - | 508.3 | 146.1 | - | - | 0 | - |
| - | - | 500.9 | 147.6 | - | - | 0 | - |
| - | - | 1074 | 148.9 | - | - | 0 | - |
| - | - | 5804 | 155.1 | - | - | 0 | - |
| - | - | 830.4 | 157.1 | - | - | 0 | - |
| - | - | 414.6 | 170.9 | - | - | 0 | - |
| - | - | 1.939E+04 | 173.1 | - | - | 0 | - |
| - | - | 1654 | 174.1 | - | - | 0 | - |
| - | - | 457.4 | 179.4 | - | - | 0 | - |
| 13 | y | 3836 | 182.1 | 0.0001902 | 1.045 | +1 | 1 |
| - | - | 1536 | 186.1 | - | - | 0 | - |
| - | - | 873.9 | 188.1 | - | - | 0 | - |
| - | - | 1.326E+04 | 201.1 | - | - | 0 | - |
| - | - | 1299 | 201.1 | - | - | 0 | - |
| - | - | 1135 | 202.1 | - | - | 0 | - |
| - | - | 507.9 | 207.1 | - | - | 0 | - |
| - | - | 5578 | 215.1 | - | - | 0 | - |
| 2 | c | 977.5 | 218.1 | 0.0005642 | 2.586 | +1 | 2 |
| - | - | 568.6 | 223.1 | - | - | 0 | - |
| - | - | 920.4 | 223.2 | - | - | 0 | - |
| - | - | 872.7 | 226.1 | - | - | 0 | - |
| - | - | 561.6 | 226.5 | - | - | 0 | - |
| - | - | 633.5 | 240.1 | - | - | 0 | - |
| - | - | 885 | 243.1 | - | - | 0 | - |
| - | - | 1163 | 245.1 | - | - | 0 | - |
| - | - | 463.8 | 250.1 | - | - | 0 | - |
| - | - | 2348 | 251.2 | - | - | 0 | - |
| - | - | 5724 | 258.1 | - | - | 0 | - |
| - | - | 592.9 | 259.1 | - | - | 0 | - |
| - | - | 1244 | 260.2 | - | - | 0 | - |
| - | - | 4067 | 261.2 | - | - | 0 | - |
| - | - | 722.4 | 266.1 | - | - | 0 | - |
| - | - | 3170 | 268.2 | - | - | 0 | - |
| - | - | 508.9 | 283 | - | - | 0 | - |
| - | - | 912.7 | 283.1 | - | - | 0 | - |
| - | - | 1393 | 283.2 | - | - | 0 | - |
| - | - | 521.8 | 283.8 | - | - | 0 | - |
| 12 | y | 3357 | 293.1 | 0.0002358 | 0.8043 | +1 | 2 |
| - | - | 1340 | 299.1 | - | - | 0 | - |
| - | - | 1738 | 300.1 | - | - | 0 | - |
| - | - | 1013 | 300.2 | - | - | 0 | - |
| - | - | 1629 | 301.1 | - | - | 0 | - |
| - | - | 561 | 301.2 | - | - | 0 | - |
| - | - | 982.6 | 302.1 | - | - | 0 | - |
| - | - | 968.1 | 306.2 | - | - | 0 | - |
| - | - | 490.7 | 309.2 | - | - | 0 | - |
| 12 | y | 1412 | 311.1 | 0.0001055 | 0.3392 | +1 | 2 |
| - | - | 1324 | 312.2 | - | - | 0 | - |
| - | - | 1557 | 314.2 | - | - | 0 | - |
| - | - | 1180 | 315.2 | - | - | 0 | - |
| - | - | 8348 | 317.2 | - | - | 0 | - |
| - | - | 802.2 | 317.2 | - | - | 0 | - |
| - | - | 1508 | 318.2 | - | - | 0 | - |
| - | - | 1717 | 320.2 | - | - | 0 | - |
| - | - | 944.4 | 322.2 | - | - | 0 | - |
| - | - | 1440 | 330.2 | - | - | 0 | - |
| 3 | c | 4569 | 331.2 | 0.0001489 | 0.4496 | +1 | 3 |
| - | - | 1116 | 332.2 | - | - | 0 | - |
| - | - | 670.9 | 343.1 | - | - | 0 | - |
| 6 | c | 1027 | 347.7 | 0.0004937 | 1.42 | +2 | 6 |
| - | - | 766.4 | 357.2 | - | - | 0 | - |
| - | - | 575 | 361 | - | - | 0 | - |
| - | - | 864.6 | 364.2 | - | - | 0 | - |
| - | - | 1231 | 374.2 | - | - | 0 | - |
| - | - | 1835 | 376.2 | - | - | 0 | - |
| - | - | 541.3 | 376.2 | - | - | 0 | - |
| - | - | 825.3 | 378.2 | - | - | 0 | - |
| - | - | 698.7 | 379.2 | - | - | 0 | - |
| - | - | 708.1 | 381.2 | - | - | 0 | - |
| - | - | 991.8 | 391.2 | - | - | 0 | - |
| - | - | 4204 | 392.2 | - | - | 0 | - |
| - | - | 1281 | 393.2 | - | - | 0 | - |
| - | - | 856.5 | 393.7 | - | - | 0 | - |
| - | - | 584.8 | 399.2 | - | - | 0 | - |
| - | - | 700 | 409.2 | - | - | 0 | - |
| - | - | 627.3 | 415.5 | - | - | 0 | - |
| - | - | 1318 | 417.2 | - | - | 0 | - |
| - | - | 766.5 | 418 | - | - | 0 | - |
| 11 | y | 845.7 | 421.2 | 0.000274 | 0.6504 | +1 | 3 |
| 11 | z | 901.1 | 423.2 | 0.0007525 | 1.778 | +1 | 3 |
| - | - | 4237 | 423.3 | - | - | 0 | - |
| - | - | 1012 | 424.3 | - | - | 0 | - |
| - | - | 1358 | 429.2 | - | - | 0 | - |
| 4 | y | 1624 | 430.2 | 0.00573 | 13.32 | +3 | 10 |
| - | - | 2620 | 434.7 | - | - | 0 | - |
| - | - | 2148 | 435.2 | - | - | 0 | - |
| 11 | y | 1584 | 439.2 | 2.421E-05 | 0.05512 | +1 | 3 |
| - | - | 1561 | 439.7 | - | - | 0 | - |
| - | - | 1794 | 440.2 | - | - | 0 | - |
| - | - | 633.1 | 440.7 | - | - | 0 | - |
| - | - | 688.8 | 442.3 | - | - | 0 | - |
| - | - | 810.6 | 444.3 | - | - | 0 | - |
| - | - | 651.7 | 445.2 | - | - | 0 | - |
| - | - | 735.2 | 446.3 | - | - | 0 | - |
| 7 | c | 4341 | 448.7 | 0.002325 | 5.182 | +2 | 7 |
| - | - | 2286 | 449.2 | - | - | 0 | - |
| - | - | 9497 | 451.3 | - | - | 0 | - |
| - | - | 2295 | 452.3 | - | - | 0 | - |
| - | - | 690.5 | 453.3 | - | - | 0 | - |
| - | - | 1612 | 454.2 | - | - | 0 | - |
| 7 | y | 1107 | 454.7 | 0.00663 | 14.58 | +2 | 7 |
| - | - | 2474 | 463.2 | - | - | 0 | - |
| - | - | 756.8 | 464.2 | - | - | 0 | - |
| 4 | c | 2.376E+04 | 468.3 | 0.0002581 | 0.5511 | +1 | 4 |
| - | - | 6501 | 469.3 | - | - | 0 | - |
| - | - | 1143 | 470.3 | - | - | 0 | - |
| - | - | 925.8 | 478.2 | - | - | 0 | - |
| - | - | 886.4 | 479.2 | - | - | 0 | - |
| 10 | z | 4064 | 480.2 | 0.003663 | 7.627 | +1 | 4 |
| - | - | 2996 | 481.2 | - | - | 0 | - |
| - | - | 1402 | 482.3 | - | - | 0 | - |
| - | - | 1040 | 482.8 | - | - | 0 | - |
| - | - | 1734 | 488.2 | - | - | 0 | - |
| - | - | 1332 | 488.8 | - | - | 0 | - |
| - | - | 2179 | 489.2 | - | - | 0 | - |
| - | - | 1448 | 491.3 | - | - | 0 | - |
| - | - | 1861 | 494.2 | - | - | 0 | - |
| - | - | 7827 | 494.3 | - | - | 0 | - |
| - | - | 646.9 | 495.2 | - | - | 0 | - |
| - | - | 1658 | 495.3 | - | - | 0 | - |
| 10 | y | 1.575E+04 | 496.2 | 0.001052 | 2.12 | +1 | 4 |
| - | - | 624.5 | 496.3 | - | - | 0 | - |
| - | - | 969.9 | 496.8 | - | - | 0 | - |
| - | - | 3389 | 497.2 | - | - | 0 | - |
| 6 | z | 819.8 | 504.2 | 0.00719 | 14.26 | +2 | 8 |
| 8 | c | 2515 | 505.3 | 0.004544 | 8.993 | +2 | 8 |
| - | - | 1147 | 505.8 | - | - | 0 | - |
| - | - | 1840 | 506.1 | - | - | 0 | - |
| - | - | 2199 | 507.1 | - | - | 0 | - |
| - | - | 4098 | 511.3 | - | - | 0 | - |
| - | - | 2.378E+04 | 513.4 | - | - | 0 | - |
| - | - | 693.1 | 523.2 | - | - | 0 | - |
| - | - | 2996 | 523.3 | - | - | 0 | - |
| - | - | 1619 | 532.3 | - | - | 0 | - |
| - | - | 4914 | 533.3 | - | - | 0 | - |
| - | - | 947.6 | 534.3 | - | - | 0 | - |
| - | - | 2523 | 534.6 | - | - | 0 | - |
| - | - | 1084 | 534.9 | - | - | 0 | - |
| - | - | 777.8 | 535.3 | - | - | 0 | - |
| - | - | 933.4 | 535.6 | - | - | 0 | - |
| - | - | 5272 | 538.3 | - | - | 0 | - |
| - | - | 1115 | 539.2 | - | - | 0 | - |
| - | - | 1716 | 539.3 | - | - | 0 | - |
| - | - | 1.281E+04 | 539.8 | - | - | 0 | - |
| - | - | 768.5 | 540.3 | - | - | 0 | - |
| - | - | 7941 | 540.3 | - | - | 0 | - |
| - | - | 1988 | 540.6 | - | - | 0 | - |
| - | - | 2877 | 540.8 | - | - | 0 | - |
| - | - | 1049 | 540.9 | - | - | 0 | - |
| - | - | 2.485E+05 | 541.3 | - | - | 0 | - |
| - | - | 2452 | 541.6 | - | - | 0 | - |
| - | - | 1199 | 541.9 | - | - | 0 | - |
| - | - | 598.9 | 543.3 | - | - | 0 | - |
| - | - | 2853 | 546.3 | - | - | 0 | - |
| - | - | 2824 | 546.8 | - | - | 0 | - |
| - | - | 1105 | 547.3 | - | - | 0 | - |
| 9 | w | 8801 | 550.3 | 0.0004055 | 0.7369 | +1 | 5 |
| - | - | 3568 | 551.3 | - | - | 0 | - |
| - | - | 2242 | 554.3 | - | - | 0 | - |
| - | - | 736.6 | 559.7 | - | - | 0 | - |
| - | - | 822.1 | 560.3 | - | - | 0 | - |
| - | - | 1183 | 561.3 | - | - | 0 | - |
| - | - | 2201 | 564.3 | - | - | 0 | - |
| 5 | c | 4016 | 579.3 | 0.001924 | 3.321 | +1 | 5 |
| - | - | 1820 | 580.3 | - | - | 0 | - |
| - | - | 568.2 | 580.4 | - | - | 0 | - |
| - | - | 961.4 | 588.8 | - | - | 0 | - |
| - | - | 1214 | 589.3 | - | - | 0 | - |
| - | - | 3376 | 589.8 | - | - | 0 | - |
| - | - | 4069 | 590.3 | - | - | 0 | - |
| 10 | c | 1149 | 590.8 | 0.003842 | 6.504 | +2 | 10 |
| - | - | 2084 | 592.3 | - | - | 0 | - |
| 9 | y | 1382 | 593.3 | 0.0008174 | 1.378 | +1 | 5 |
| - | - | 3710 | 594.3 | - | - | 0 | - |
| 9 | z | 4682 | 595.2 | 0.002873 | 4.827 | +1 | 5 |
| - | - | 785.1 | 595.3 | - | - | 0 | - |
| - | - | 2941 | 596.3 | - | - | 0 | - |
| 5 | c | 7.015E+04 | 596.4 | 3.087E-05 | 0.05177 | +1 | 5 |
| - | - | 687 | 596.8 | - | - | 0 | - |
| - | - | 760.3 | 597.3 | - | - | 0 | - |
| - | - | 2.068E+04 | 597.4 | - | - | 0 | - |
| - | - | 8481 | 597.8 | - | - | 0 | - |
| - | - | 3430 | 598.3 | - | - | 0 | - |
| - | - | 3944 | 598.4 | - | - | 0 | - |
| - | - | 604.1 | 602.3 | - | - | 0 | - |
| - | - | 2085 | 603.3 | - | - | 0 | - |
| - | - | 635.2 | 605.3 | - | - | 0 | - |
| - | - | 1.203E+04 | 610.3 | - | - | 0 | - |
| - | - | 894.9 | 610.3 | - | - | 0 | - |
| 9 | y | 1.179E+04 | 611.3 | 0.002887 | 4.723 | +1 | 5 |
| - | - | 3574 | 612.3 | - | - | 0 | - |
| - | - | 803.7 | 613.3 | - | - | 0 | - |
| - | - | 820.3 | 614.3 | - | - | 0 | - |
| - | - | 1215 | 619.8 | - | - | 0 | - |
| - | - | 913.4 | 626.4 | - | - | 0 | - |
| - | - | 1014 | 630.3 | - | - | 0 | - |
| - | - | 681.3 | 634.3 | - | - | 0 | - |
| - | - | 6524 | 636.3 | - | - | 0 | - |
| - | - | 2410 | 637.3 | - | - | 0 | - |
| - | - | 5064 | 639.3 | - | - | 0 | - |
| - | - | 2977 | 639.8 | - | - | 0 | - |
| - | - | 1409 | 641.8 | - | - | 0 | - |
| - | - | 1781 | 645.3 | - | - | 0 | - |
| 4 | z | 1821 | 645.8 | 0.01195 | 18.5 | +2 | 10 |
| - | - | 1065 | 646.3 | - | - | 0 | - |
| - | - | 1448 | 646.8 | - | - | 0 | - |
| - | - | 7923 | 647.3 | - | - | 0 | - |
| - | - | 3735 | 647.8 | - | - | 0 | - |
| - | - | 5345 | 648.3 | - | - | 0 | - |
| - | - | 771 | 649.3 | - | - | 0 | - |
| - | - | 995.1 | 651.3 | - | - | 0 | - |
| - | - | 5391 | 652.3 | - | - | 0 | - |
| - | - | 1225 | 653.3 | - | - | 0 | - |
| - | - | 795 | 653.3 | - | - | 0 | - |
| 4 | y | 2305 | 653.8 | 0.005578 | 8.532 | +2 | 10 |
| - | - | 1159 | 654.3 | - | - | 0 | - |
| 11 | c | 1293 | 654.8 | 0.005067 | 7.738 | +2 | 11 |
| - | - | 2335 | 662.8 | - | - | 0 | - |
| - | - | 2.643E+04 | 663.4 | - | - | 0 | - |
| 11 | c | 3.451E+04 | 663.8 | 0.006804 | 10.25 | +2 | 11 |
| - | - | 3759 | 664.3 | - | - | 0 | - |
| - | - | 2.045E+04 | 664.3 | - | - | 0 | - |
| - | - | 9083 | 664.8 | - | - | 0 | - |
| 8 | w | 2864 | 665.3 | 0.01149 | 17.27 | +1 | 6 |
| - | - | 1544 | 665.4 | - | - | 0 | - |
| - | - | 931.2 | 666.4 | - | - | 0 | - |
| - | - | 740.1 | 667.4 | - | - | 0 | - |
| - | - | 751.5 | 670.4 | - | - | 0 | - |
| - | - | 3891 | 674.3 | - | - | 0 | - |
| - | - | 3808 | 674.8 | - | - | 0 | - |
| - | - | 3502 | 676.4 | - | - | 0 | - |
| - | - | 1993 | 676.9 | - | - | 0 | - |
| - | - | 1739 | 677.4 | - | - | 0 | - |
| - | - | 4910 | 679.3 | - | - | 0 | - |
| - | - | 757.4 | 679.8 | - | - | 0 | - |
| - | - | 2912 | 680.3 | - | - | 0 | - |
| 3 | w | 4632 | 680.8 | 0.009939 | 14.6 | +2 | 11 |
| - | - | 4098 | 681.3 | - | - | 0 | - |
| - | - | 790.3 | 685.3 | - | - | 0 | - |
| 6 | c | 1.626E+04 | 694.4 | 0.0003638 | 0.5239 | +1 | 6 |
| - | - | 7281 | 695.4 | - | - | 0 | - |
| - | - | 1949 | 696.3 | - | - | 0 | - |
| - | - | 3577 | 696.4 | - | - | 0 | - |
| - | - | 704.4 | 696.9 | - | - | 0 | - |
| - | - | 721 | 697.3 | - | - | 0 | - |
| - | - | 1355 | 697.4 | - | - | 0 | - |
| - | - | 1692 | 700.8 | - | - | 0 | - |
| 3 | y | 2286 | 701.3 | 0.006243 | 8.901 | +2 | 11 |
| 3 | y | 5662 | 701.8 | 0.01277 | 18.2 | +2 | 11 |
| 3 | z | 4756 | 702.3 | 0.004829 | 6.876 | +2 | 11 |
| - | - | 1345 | 702.4 | - | - | 0 | - |
| - | - | 2217 | 702.8 | - | - | 0 | - |
| - | - | 753.2 | 703.3 | - | - | 0 | - |
| - | - | 3918 | 703.8 | - | - | 0 | - |
| - | - | 1843 | 704.3 | - | - | 0 | - |
| - | - | 1365 | 705.9 | - | - | 0 | - |
| - | - | 2562 | 706.4 | - | - | 0 | - |
| - | - | 1373 | 706.9 | - | - | 0 | - |
| - | - | 3421 | 707.4 | - | - | 0 | - |
| - | - | 1384 | 707.9 | - | - | 0 | - |
| - | - | 3115 | 708.3 | - | - | 0 | - |
| - | - | 1623 | 709.3 | - | - | 0 | - |
| - | - | 7444 | 709.8 | - | - | 0 | - |
| 3 | y | 1.197E+04 | 710.3 | 0.00743 | 10.46 | +2 | 11 |
| - | - | 7517 | 710.8 | - | - | 0 | - |
| 6 | c | 4.384E+04 | 711.4 | 0.0007898 | 1.11 | +1 | 6 |
| - | - | 9840 | 711.9 | - | - | 0 | - |
| - | - | 1.763E+04 | 712.4 | - | - | 0 | - |
| - | - | 4014 | 713.4 | - | - | 0 | - |
| - | - | 1037 | 715.9 | - | - | 0 | - |
| - | - | 1125 | 717.9 | - | - | 0 | - |
| - | - | 994.2 | 718.4 | - | - | 0 | - |
| 12 | c | 2060 | 719.4 | 0.00025 | 0.3475 | +2 | 12 |
| 12 | c | 2339 | 719.8 | 0.0105 | 14.59 | +2 | 12 |
| - | - | 760.1 | 720.9 | - | - | 0 | - |
| - | - | 993.3 | 721.4 | - | - | 0 | - |
| - | - | 687 | 721.9 | - | - | 0 | - |
| - | - | 955.9 | 722.3 | - | - | 0 | - |
| - | - | 5587 | 723.4 | - | - | 0 | - |
| 8 | y | 5367 | 724.4 | 0.001343 | 1.853 | +1 | 6 |
| - | - | 2085 | 725.4 | - | - | 0 | - |
| - | - | 747.4 | 726.4 | - | - | 0 | - |
| - | - | 3.287E+04 | 727.9 | - | - | 0 | - |
| 12 | c | 4.722E+04 | 728.4 | 0.006564 | 9.012 | +2 | 12 |
| - | - | 3.159E+04 | 728.9 | - | - | 0 | - |
| - | - | 1.552E+04 | 729.4 | - | - | 0 | - |
| - | - | 7822 | 729.9 | - | - | 0 | - |
| - | - | 4142 | 730.4 | - | - | 0 | - |
| - | - | 803 | 730.9 | - | - | 0 | - |
| - | - | 676.5 | 734.4 | - | - | 0 | - |
| - | - | 928.1 | 735.4 | - | - | 0 | - |
| - | - | 729.6 | 735.9 | - | - | 0 | - |
| - | - | 2008 | 736.4 | - | - | 0 | - |
| - | - | 3127 | 736.9 | - | - | 0 | - |
| - | - | 4745 | 737.4 | - | - | 0 | - |
| - | - | 2.782E+04 | 737.9 | - | - | 0 | - |
| - | - | 1.603E+04 | 738.4 | - | - | 0 | - |
| - | - | 2255 | 739.4 | - | - | 0 | - |
| - | - | 1876 | 744.4 | - | - | 0 | - |
| - | - | 991.7 | 746.4 | - | - | 0 | - |
| - | - | 1063 | 746.9 | - | - | 0 | - |
| - | - | 4050 | 747.4 | - | - | 0 | - |
| - | - | 2744 | 747.9 | - | - | 0 | - |
| - | - | 835 | 748.9 | - | - | 0 | - |
| - | - | 1191 | 749.4 | - | - | 0 | - |
| - | - | 749.8 | 749.9 | - | - | 0 | - |
| - | - | 1485 | 750.4 | - | - | 0 | - |
| 2 | y | 1693 | 751.3 | 0.01433 | 19.07 | +2 | 12 |
| 2 | z | 1932 | 751.9 | 0.007243 | 9.634 | +2 | 12 |
| - | - | 2299 | 752.4 | - | - | 0 | - |
| - | - | 799.3 | 753.4 | - | - | 0 | - |
| - | - | 1.224E+04 | 755.4 | - | - | 0 | - |
| - | - | 9426 | 755.9 | - | - | 0 | - |
| - | - | 1108 | 756.4 | - | - | 0 | - |
| - | - | 2277 | 756.9 | - | - | 0 | - |
| - | - | 3206 | 757.4 | - | - | 0 | - |
| - | - | 1895 | 757.9 | - | - | 0 | - |
| - | - | 1483 | 758.4 | - | - | 0 | - |
| - | - | 965.1 | 759.4 | - | - | 0 | - |
| 2 | y | 2154 | 759.9 | 0.003191 | 4.2 | +2 | 12 |
| - | - | 1840 | 760.4 | - | - | 0 | - |
| - | - | 1287 | 760.9 | - | - | 0 | - |
| - | - | 1924 | 765.4 | - | - | 0 | - |
| - | - | 2231 | 765.9 | - | - | 0 | - |
| - | - | 1151 | 766.4 | - | - | 0 | - |
| - | - | 1346 | 766.9 | - | - | 0 | - |
| - | - | 1683 | 771.9 | - | - | 0 | - |
| - | - | 1069 | 772.4 | - | - | 0 | - |
| - | - | 1476 | 772.9 | - | - | 0 | - |
| - | - | 1204 | 773.4 | - | - | 0 | - |
| - | - | 959.1 | 773.9 | - | - | 0 | - |
| - | - | 1145 | 774.4 | - | - | 0 | - |
| - | - | 2.798E+04 | 777.4 | - | - | 0 | - |
| - | - | 2.342E+04 | 778.4 | - | - | 0 | - |
| - | - | 1205 | 778.9 | - | - | 0 | - |
| - | - | 9723 | 779.4 | - | - | 0 | - |
| - | - | 4863 | 779.9 | - | - | 0 | - |
| - | - | 9966 | 780.4 | - | - | 0 | - |
| - | - | 8075 | 780.9 | - | - | 0 | - |
| - | - | 7569 | 781.4 | - | - | 0 | - |
| - | - | 4157 | 781.9 | - | - | 0 | - |
| - | - | 5062 | 782.4 | - | - | 0 | - |
| - | - | 857.4 | 782.5 | - | - | 0 | - |
| - | - | 3893 | 782.9 | - | - | 0 | - |
| - | - | 731.3 | 783.4 | - | - | 0 | - |
| - | - | 845.4 | 786.4 | - | - | 0 | - |
| - | - | 1080 | 787.4 | - | - | 0 | - |
| - | - | 4135 | 787.9 | - | - | 0 | - |
| - | - | 4629 | 788.4 | - | - | 0 | - |
| - | - | 2210 | 788.9 | - | - | 0 | - |
| - | - | 9342 | 789.4 | - | - | 0 | - |
| - | - | 8156 | 789.9 | - | - | 0 | - |
| - | - | 6975 | 790.9 | - | - | 0 | - |
| - | - | 5332 | 791.4 | - | - | 0 | - |
| - | - | 1009 | 792.9 | - | - | 0 | - |
| - | - | 2148 | 793.4 | - | - | 0 | - |
| - | - | 2497 | 793.9 | - | - | 0 | - |
| - | - | 2989 | 794.4 | - | - | 0 | - |
| - | - | 2530 | 794.9 | - | - | 0 | - |
| - | - | 2725 | 795.4 | - | - | 0 | - |
| - | - | 1091 | 795.9 | - | - | 0 | - |
| - | - | 1319 | 796.4 | - | - | 0 | - |
| - | - | 3945 | 801.4 | - | - | 0 | - |
| - | - | 2.242E+04 | 801.9 | - | - | 0 | - |
| - | - | 2.999E+04 | 802.4 | - | - | 0 | - |
| - | - | 1.741E+04 | 802.9 | - | - | 0 | - |
| - | - | 8940 | 803.4 | - | - | 0 | - |
| - | - | 3.073E+04 | 803.9 | - | - | 0 | - |
| - | - | 2.088E+04 | 804.4 | - | - | 0 | - |
| - | - | 1871 | 809.4 | - | - | 0 | - |
| - | - | 2172 | 809.4 | - | - | 0 | - |
| - | - | 3.313E+04 | 809.9 | - | - | 0 | - |
| - | - | 9.843E+04 | 810.4 | - | - | 0 | - |
| - | - | 9.863E+04 | 810.9 | - | - | 0 | - |
| - | - | 6.251E+04 | 811.4 | - | - | 0 | - |
| - | - | 3.711E+04 | 811.9 | - | - | 0 | - |
| - | - | 2.485E+04 | 812.4 | - | - | 0 | - |
| - | - | 7186 | 812.9 | - | - | 0 | - |
| - | - | 1097 | 836.4 | - | - | 0 | - |
| - | - | 1098 | 841.4 | - | - | 0 | - |
| - | - | 840.9 | 852.4 | - | - | 0 | - |
| - | - | 1424 | 857.5 | - | - | 0 | - |
| - | - | 1445 | 863.4 | - | - | 0 | - |
| - | - | 1030 | 864.4 | - | - | 0 | - |
| - | - | 1499 | 878.4 | - | - | 0 | - |
| - | - | 1003 | 879.4 | - | - | 0 | - |
| 7 | c | 7558 | 896.4 | 0.003871 | 4.318 | +1 | 7 |
| - | - | 2827 | 897.4 | - | - | 0 | - |
| - | - | 989.2 | 898.4 | - | - | 0 | - |
| - | - | 5915 | 901.5 | - | - | 0 | - |
| - | - | 2383 | 902.5 | - | - | 0 | - |
| - | - | 8904 | 906.4 | - | - | 0 | - |
| - | - | 9534 | 907.4 | - | - | 0 | - |
| 7 | y | 6751 | 908.4 | 0.01315 | 14.48 | +1 | 7 |
| - | - | 3763 | 909.4 | - | - | 0 | - |
| - | - | 1665 | 910.4 | - | - | 0 | - |
| - | - | 1074 | 911.4 | - | - | 0 | - |
| 7 | c | 1.395E+04 | 913.4 | 0.003872 | 4.239 | +1 | 7 |
| - | - | 9266 | 914.5 | - | - | 0 | - |
| - | - | 1588 | 915.5 | - | - | 0 | - |
| - | - | 1960 | 917.5 | - | - | 0 | - |
| - | - | 1474 | 918.4 | - | - | 0 | - |
| - | - | 1197 | 919.4 | - | - | 0 | - |
| - | - | 830.5 | 924.4 | - | - | 0 | - |
| - | - | 3078 | 925.4 | - | - | 0 | - |
| 7 | y | 1137 | 926.4 | 0.005272 | 5.69 | +1 | 7 |
| - | - | 3097 | 944.5 | - | - | 0 | - |
| - | - | 1593 | 945.5 | - | - | 0 | - |
| - | - | 1622 | 946.5 | - | - | 0 | - |
| - | - | 768.8 | 947.5 | - | - | 0 | - |
| - | - | 775.1 | 964.5 | - | - | 0 | - |
| - | - | 4510 | 976.5 | - | - | 0 | - |
| - | - | 2003 | 977.5 | - | - | 0 | - |
| - | - | 1056 | 980.5 | - | - | 0 | - |
| - | - | 2528 | 981.4 | - | - | 0 | - |
| - | - | 3090 | 982.5 | - | - | 0 | - |
| - | - | 2460 | 983.5 | - | - | 0 | - |
| - | - | 1183 | 984.5 | - | - | 0 | - |
| - | - | 1082 | 987.5 | - | - | 0 | - |
| - | - | 1.653E+04 | 988.5 | - | - | 0 | - |
| - | - | 5887 | 989.5 | - | - | 0 | - |
| - | - | 938.7 | 1008 | - | - | 0 | - |
| 8 | c | 1041 | 1010 | 0.009467 | 9.378 | +1 | 8 |
| - | - | 974.7 | 1011 | - | - | 0 | - |
| - | - | 882.7 | 1017 | - | - | 0 | - |
| - | - | 3.927E+04 | 1024 | - | - | 0 | - |
| 6 | z | 4.04E+04 | 1025 | 0.0139 | 13.55 | +1 | 8 |
| - | - | 1.487E+04 | 1026 | - | - | 0 | - |
| 8 | c | 2.512E+04 | 1027 | 0.006051 | 5.894 | +1 | 8 |
| - | - | 3876 | 1027 | - | - | 0 | - |
| - | - | 1.602E+04 | 1028 | - | - | 0 | - |
| - | - | 872.6 | 1028 | - | - | 0 | - |
| - | - | 3649 | 1029 | - | - | 0 | - |
| - | - | 1347 | 1030 | - | - | 0 | - |
| - | - | 747.6 | 1036 | - | - | 0 | - |
| - | - | 8331 | 1040 | - | - | 0 | - |
| 6 | y | 7630 | 1041 | 0.01556 | 14.94 | +1 | 8 |
| - | - | 2834 | 1042 | - | - | 0 | - |
| - | - | 721.9 | 1043 | - | - | 0 | - |
| - | - | 9363 | 1046 | - | - | 0 | - |
| - | - | 3288 | 1047 | - | - | 0 | - |
| - | - | 697.7 | 1048 | - | - | 0 | - |
| - | - | 7217 | 1052 | - | - | 0 | - |
| - | - | 4086 | 1053 | - | - | 0 | - |
| - | - | 2633 | 1060 | - | - | 0 | - |
| - | - | 1994 | 1061 | - | - | 0 | - |
| - | - | 1111 | 1063 | - | - | 0 | - |
| - | - | 1048 | 1079 | - | - | 0 | - |
| - | - | 3976 | 1080 | - | - | 0 | - |
| - | - | 2620 | 1081 | - | - | 0 | - |
| - | - | 738.9 | 1081 | - | - | 0 | - |
| - | - | 5542 | 1084 | - | - | 0 | - |
| - | - | 4410 | 1085 | - | - | 0 | - |
| - | - | 1485 | 1086 | - | - | 0 | - |
| - | - | 3185 | 1092 | - | - | 0 | - |
| - | - | 5543 | 1093 | - | - | 0 | - |
| - | - | 1585 | 1094 | - | - | 0 | - |
| 5 | w | 1394 | 1095 | 0.01098 | 10.02 | +1 | 9 |
| - | - | 3242 | 1097 | - | - | 0 | - |
| - | - | 3954 | 1098 | - | - | 0 | - |
| - | - | 2322 | 1099 | - | - | 0 | - |
| - | - | 812.3 | 1101 | - | - | 0 | - |
| - | - | 1009 | 1108 | - | - | 0 | - |
| - | - | 831 | 1127 | - | - | 0 | - |
| - | - | 1055 | 1130 | - | - | 0 | - |
| - | - | 2051 | 1140 | - | - | 0 | - |
| - | - | 7798 | 1141 | - | - | 0 | - |
| 9 | c | 6754 | 1142 | 0.01378 | 12.07 | +1 | 9 |
| - | - | 2880 | 1143 | - | - | 0 | - |
| - | - | 907.1 | 1144 | - | - | 0 | - |
| - | - | 8442 | 1145 | - | - | 0 | - |
| - | - | 2953 | 1146 | - | - | 0 | - |
| - | - | 1.228E+04 | 1153 | - | - | 0 | - |
| 5 | z | 2.93E+04 | 1153 | 0.007933 | 6.878 | +1 | 9 |
| - | - | 1.814E+04 | 1154 | - | - | 0 | - |
| - | - | 6887 | 1155 | - | - | 0 | - |
| - | - | 1218 | 1157 | - | - | 0 | - |
| - | - | 3011 | 1169 | - | - | 0 | - |
| 5 | y | 5842 | 1170 | 0.005689 | 4.864 | +1 | 9 |
| - | - | 3064 | 1171 | - | - | 0 | - |
| - | - | 1357 | 1172 | - | - | 0 | - |
| - | - | 2736 | 1180 | - | - | 0 | - |
| 10 | c | 1650 | 1181 | 0.004952 | 4.194 | +1 | 10 |
| - | - | 1479 | 1195 | - | - | 0 | - |
| - | - | 2.621E+04 | 1198 | - | - | 0 | - |
| 10 | c | 3.491E+04 | 1199 | 0.01123 | 9.372 | +1 | 10 |
| - | - | 1.917E+04 | 1200 | - | - | 0 | - |
| - | - | 6538 | 1201 | - | - | 0 | - |
| - | - | 1212 | 1202 | - | - | 0 | - |
| - | - | 1.353E+04 | 1232 | - | - | 0 | - |
| - | - | 7048 | 1233 | - | - | 0 | - |
| - | - | 844.8 | 1240 | - | - | 0 | - |
| - | - | 809.3 | 1278 | - | - | 0 | - |
| - | - | 4002 | 1279 | - | - | 0 | - |
| - | - | 2211 | 1280 | - | - | 0 | - |
| - | - | 1087 | 1283 | - | - | 0 | - |
| - | - | 984.2 | 1284 | - | - | 0 | - |
| - | - | 1431 | 1286 | - | - | 0 | - |
| - | - | 1049 | 1287 | - | - | 0 | - |
| 4 | y | 4825 | 1290 | 0.02386 | 18.51 | +1 | 10 |
| 4 | z | 1.178E+04 | 1291 | 0.01347 | 10.44 | +1 | 10 |
| - | - | 1.087E+04 | 1292 | - | - | 0 | - |
| - | - | 4952 | 1293 | - | - | 0 | - |
| - | - | 819.3 | 1294 | - | - | 0 | - |
| 4 | y | 1421 | 1307 | 0.007324 | 5.605 | +1 | 10 |
| - | - | 883.4 | 1308 | - | - | 0 | - |
| - | - | 799.9 | 1310 | - | - | 0 | - |
| - | - | 863.1 | 1314 | - | - | 0 | - |
| - | - | 896.9 | 1315 | - | - | 0 | - |
| - | - | 9987 | 1319 | - | - | 0 | - |
| - | - | 5458 | 1320 | - | - | 0 | - |
| - | - | 1373 | 1324 | - | - | 0 | - |
| - | - | 2049 | 1325 | - | - | 0 | - |
| - | - | 1.206E+04 | 1326 | - | - | 0 | - |
| 11 | c | 1.853E+04 | 1327 | 0.01234 | 9.301 | +1 | 11 |
| - | - | 1.108E+04 | 1328 | - | - | 0 | - |
| - | - | 4824 | 1329 | - | - | 0 | - |
| - | - | 1707 | 1330 | - | - | 0 | - |
| - | - | 969.2 | 1346 | - | - | 0 | - |
| - | - | 868.7 | 1395 | - | - | 0 | - |
| - | - | 772.6 | 1398 | - | - | 0 | - |
| - | - | 2357 | 1402 | - | - | 0 | - |
| 3 | y | 1728 | 1403 | 0.02549 | 18.17 | +1 | 11 |
| 3 | z | 7334 | 1404 | 0.01742 | 12.41 | +1 | 11 |
| - | - | 7654 | 1405 | - | - | 0 | - |
| - | - | 3559 | 1406 | - | - | 0 | - |
| - | - | 2990 | 1407 | - | - | 0 | - |
| - | - | 6480 | 1408 | - | - | 0 | - |
| - | - | 4039 | 1409 | - | - | 0 | - |
| - | - | 1656 | 1411 | - | - | 0 | - |
| - | - | 3404 | 1412 | - | - | 0 | - |
| - | - | 3907 | 1413 | - | - | 0 | - |
| - | - | 2589 | 1414 | - | - | 0 | - |
| - | - | 1562 | 1415 | - | - | 0 | - |
| - | - | 1475 | 1416 | - | - | 0 | - |
| 3 | y | 1422 | 1420 | 0.01359 | 9.574 | +1 | 11 |
| - | - | 955.6 | 1421 | - | - | 0 | - |
| - | - | 926.4 | 1431 | - | - | 0 | - |
| - | - | 2095 | 1432 | - | - | 0 | - |
| - | - | 1570 | 1433 | - | - | 0 | - |
| 12 | c | 2216 | 1439 | 0.01766 | 12.27 | +1 | 12 |
| - | - | 4536 | 1440 | - | - | 0 | - |
| - | - | 2143 | 1441 | - | - | 0 | - |
| - | - | 774.2 | 1442 | - | - | 0 | - |
| - | - | 3331 | 1455 | - | - | 0 | - |
| 12 | c | 1.089E+04 | 1456 | 0.01638 | 11.25 | +1 | 12 |
| - | - | 1.222E+04 | 1457 | - | - | 0 | - |
| - | - | 5635 | 1458 | - | - | 0 | - |
| - | - | 3648 | 1459 | - | - | 0 | - |
| - | - | 1767 | 1460 | - | - | 0 | - |
| - | - | 761.6 | 1474 | - | - | 0 | - |
| - | - | 2673 | 1475 | - | - | 0 | - |
| - | - | 7614 | 1476 | - | - | 0 | - |
| - | - | 4431 | 1477 | - | - | 0 | - |
| - | - | 1033 | 1488 | - | - | 0 | - |
| - | - | 2440 | 1489 | - | - | 0 | - |
| - | - | 2287 | 1490 | - | - | 0 | - |
| - | - | 943.4 | 1491 | - | - | 0 | - |
| - | - | 4625 | 1495 | - | - | 0 | - |
| - | - | 2606 | 1496 | - | - | 0 | - |
| 2 | z | 2357 | 1503 | 0.02372 | 15.78 | +1 | 12 |
| - | - | 3510 | 1504 | - | - | 0 | - |
| - | - | 3021 | 1505 | - | - | 0 | - |
| - | - | 1053 | 1506 | - | - | 0 | - |
| - | - | 846.5 | 1523 | - | - | 0 | - |
| - | - | 1172 | 1539 | - | - | 0 | - |
| - | - | 801.3 | 1540 | - | - | 0 | - |
| - | - | 908.1 | 1545 | - | - | 0 | - |
| - | - | 1025 | 1546 | - | - | 0 | - |
| - | - | 1787 | 1559 | - | - | 0 | - |
| - | - | 1901 | 1560 | - | - | 0 | - |
| - | - | 4033 | 1561 | - | - | 0 | - |
| - | - | 5983 | 1562 | - | - | 0 | - |
| - | - | 4775 | 1563 | - | - | 0 | - |
| - | - | 2749 | 1564 | - | - | 0 | - |
| - | - | 7048 | 1565 | - | - | 0 | - |
| - | - | 8413 | 1566 | - | - | 0 | - |
| - | - | 2056 | 1567 | - | - | 0 | - |
| - | - | 2703 | 1575 | - | - | 0 | - |
| - | - | 9752 | 1576 | - | - | 0 | - |
| - | - | 1.048E+04 | 1577 | - | - | 0 | - |
| - | - | 5921 | 1578 | - | - | 0 | - |
| - | - | 3000 | 1579 | - | - | 0 | - |
| - | - | 1843 | 1580 | - | - | 0 | - |
| - | - | 1463 | 1581 | - | - | 0 | - |
| - | - | 1439 | 1582 | - | - | 0 | - |
| - | - | 2036 | 1586 | - | - | 0 | - |
| - | - | 1891 | 1587 | - | - | 0 | - |
| - | - | 2033 | 1588 | - | - | 0 | - |
| - | - | 884.1 | 1589 | - | - | 0 | - |
| - | - | 2633 | 1591 | - | - | 0 | - |
| - | - | 3654 | 1592 | - | - | 0 | - |
| - | - | 4761 | 1593 | - | - | 0 | - |
| - | - | 5240 | 1594 | - | - | 0 | - |
| - | - | 3151 | 1595 | - | - | 0 | - |
| - | - | 1480 | 1596 | - | - | 0 | - |
| - | - | 2400 | 1597 | - | - | 0 | - |
| - | - | 1062 | 1598 | - | - | 0 | - |
| - | - | 1559 | 1602 | - | - | 0 | - |
| - | - | 1.039E+04 | 1603 | - | - | 0 | - |
| - | - | 3.388E+04 | 1604 | - | - | 0 | - |
| - | - | 3.964E+04 | 1605 | - | - | 0 | - |
| - | - | 2.523E+04 | 1606 | - | - | 0 | - |
| - | - | 1.559E+04 | 1607 | - | - | 0 | - |
| - | - | 2.69E+04 | 1608 | - | - | 0 | - |
| - | - | 1.445E+04 | 1609 | - | - | 0 | - |
| - | - | 3851 | 1619 | - | - | 0 | - |
| - | - | 1.498E+04 | 1620 | - | - | 0 | - |
| - | - | 4.365E+04 | 1621 | - | - | 0 | - |
| - | - | 4.892E+04 | 1622 | - | - | 0 | - |
| - | - | 3.414E+04 | 1623 | - | - | 0 | - |
| - | - | 2.161E+04 | 1624 | - | - | 0 | - |
| - | - | 2.786E+04 | 1625 | - | - | 0 | - |
| - | - | 1.548E+04 | 1626 | - | - | 0 | - |
| - | - | 740.9 | 1718 | - | - | 0 | - |
| - | - | 812.1 | 2565 | - | - | 0 | - |
| - | - | 949.7 | 3083 | - | - | 0 | - |

m/z Charge Intensity FragmentType MassShift Position
120.08092498779297 0 780.6386
121.20516967773438 0 350.47107
129.10244750976562 0 3543.437
131.11813354492188 0 810.9955
136.07571411132812 0 727.3635
139.9889678955078 0 363.2058
146.06053161621094 0 508.26184
147.5709991455078 0 500.92078
148.94723510742188 0 1073.6857
155.09283447265625 0 5804.117
157.13381958007812 0 830.4219
170.92166137695312 0 414.59302
173.1286163330078 0 19391.61
174.13206481933594 0 1654.0215
179.41943359375 0 457.41278
182.08135986328125 0 3836.0515 y 12
186.12359619140625 0 1535.8599
188.13970947265625 0 873.9228
201.12344360351562 0 13263.476
201.13491821289062 0 1298.5856
202.1268310546875 0 1135.4606
207.11175537109375 0 507.90085
215.13926696777344 0 5577.872
218.15048217773438 0 977.4936 c 1
223.06370544433594 0 568.6258
223.1555633544922 0 920.4057
226.1076202392578 0 872.7368
226.5314483642578 0 561.5577
240.1342010498047 0 633.4814
243.13377380371094 0 884.97894
245.1369171142578 0 1162.5361
250.10595703125 0 463.75586
251.1504669189453 0 2347.7498
258.1449279785156 0 5723.6025
259.148681640625 0 592.8655
260.1607971191406 0 1243.8905
261.1708984375 0 4067.1155
266.1243896484375 0 722.419
268.1767272949219 0 3169.5544
283.0475158691406 0 508.8783
283.1373291015625 0 912.7334
283.1528625488281 0 1393.4248
283.82073974609375 0 521.77106
293.1134338378906 0 3356.5615 y Water loss 11
299.0618591308594 0 1340.0143
300.0614013671875 0 1737.5647
300.1676025390625 0 1013.1299
301.0596008300781 0 1628.9738
301.1514892578125 0 560.9926
302.0595703125 0 982.5714
306.1668395996094 0 968.10425
309.2048645019531 0 490.7356
311.1236572265625 0 1411.8623 y 11
312.1553955078125 0 1324.0977
314.2073669433594 0 1556.7756
315.1656188964844 0 1179.9514
317.15838623046875 0 8347.619
317.1812438964844 0 802.16296
318.1613464355469 0 1508.0306
320.1970520019531 0 1716.7438
322.1871643066406 0 944.36
330.1663818359375 0 1440.1261
331.234130859375 0 4569.411 c 2
332.2369689941406 0 1116.347
343.10015869140625 0 670.8844
347.6790771484375 0 1027.2645 c Ammonia loss 5
357.17742919921875 0 766.39703
361.0262756347656 0 574.9824
364.1865539550781 0 864.648
374.1788330078125 0 1230.561
376.1974792480469 0 1834.6094
376.2247314453125 0 541.29126
378.2081604003906 0 825.2577
379.2079162597656 0 698.70325
381.2220764160156 0 708.1341
391.1837463378906 0 991.7726
392.19287109375 0 4204.092
393.1995544433594 0 1281.1886
393.72088623046875 0 856.4604
399.17877197265625 0 584.77893
409.2231750488281 0 699.96967
415.5296325683594 0 627.3098
417.1870422363281 0 1317.5978
418.0355529785156 0 766.51044
421.20843505859375 0 845.70245 y Water loss 10
423.1992492675781 0 901.1139 z 10
423.2720642089844 0 4236.7007
424.2763977050781 0 1012.1794
429.20819091796875 0 1358.4923
430.19537353515625 0 1623.9678 y Water loss 3
434.7198791503906 0 2620.3027
435.2217102050781 0 2147.9346
439.21875 0 1583.5897 y 10
439.7123718261719 0 1560.7985
440.21435546875 0 1793.5359
440.7094421386719 0 633.1077
442.276611328125 0 688.81256
444.257080078125 0 810.6077
445.21881103515625 0 651.662
446.2604064941406 0 735.18304
448.716552734375 0 4341.0825 c Ammonia loss 6
449.2183837890625 0 2286.1873
451.2664489746094 0 9497.258
452.2693786621094 0 2294.9883
453.2727355957031 0 690.4997
454.2197570800781 0 1612.0621
454.7152404785156 0 1107.1613 y Water loss 6
463.2301025390625 0 2474.2017
464.2281799316406 0 756.79913
468.29315185546875 0 23755.672 c 3
469.2957458496094 0 6501
470.29644775390625 0 1142.56
478.2186584472656 0 925.76996
479.22406005859375 0 886.3986
480.2251281738281 0 4063.6748 z 9
481.22906494140625 0 2995.6895
482.259033203125 0 1402.2335
482.7550048828125 0 1040.2438
488.2209777832031 0 1733.8242
488.7577819824219 0 1331.6958
489.206787109375 0 2179.3208
491.26171875 0 1447.8065
494.2378234863281 0 1860.796
494.3087158203125 0 7827.053
495.2328796386719 0 646.94836
495.3105773925781 0 1657.643
496.2412414550781 0 15752.1 y 9
496.3099365234375 0 624.49286
496.75408935546875 0 969.8719
497.24322509765625 0 3388.8887
504.21990966796875 0 819.75226 z Water loss 5
505.26080322265625 0 2514.5105 c Ammonia loss 7
505.76019287109375 0 1147.1755
506.1063232421875 0 1839.9565
507.1036682128906 0 2199.4229
511.3348693847656 0 4098.3833
513.3510131835938 0 23778.54
523.1903686523438 0 693.13965
523.3351440429688 0 2996.4038
532.284423828125 0 1619.1304
533.2932739257812 0 4914.314
534.259765625 0 947.6233
534.5918579101562 0 2523.3923
534.9271240234375 0 1084.2852
535.2613525390625 0 777.78204
535.601318359375 0 933.4441
538.250732421875 0 5272.1885
539.162353515625 0 1114.651
539.2569580078125 0 1715.849
539.8094482421875 0 12813.848
540.269287109375 0 768.5458
540.3109741210938 0 7941.126
540.5980834960938 0 1987.5709
540.8123168945312 0 2877.3262
540.9341430664062 0 1049.4297
541.3458251953125 0 248458.7
541.6083374023438 0 2452.2722
541.9417114257812 0 1199.191
543.279541015625 0 598.9221
546.2708129882812 0 2853.1628
546.7725219726562 0 2823.9146
547.2769165039062 0 1104.5626
550.2511596679688 0 8801.271 w 8
551.2537841796875 0 3567.828
554.278564453125 0 2241.6157
559.7334594726562 0 736.60223
560.2781372070312 0 822.07306
561.2876586914062 0 1182.6028
564.278564453125 0 2201.0728
579.322998046875 0 4016.0305 c Ammonia loss 4
580.3281860351562 0 1819.9773
580.3892822265625 0 568.20135
588.7815551757812 0 961.3632
589.282958984375 0 1213.5547
589.78662109375 0 3376.0376
590.2906494140625 0 4069.0293
590.7922973632812 0 1149.4502 c Water loss 9
592.333984375 0 2084.311
593.2573852539062 0 1382.1116 y Water loss 8
594.2645874023438 0 3710.3489
595.2512817382812 0 4681.8076 z 8
595.3179321289062 0 785.10474
596.257568359375 0 2941.0376
596.3514404296875 0 70149.45 c 4
596.8283081054688 0 687.04364
597.3011474609375 0 760.3346
597.3543090820312 0 20682.844
597.7967529296875 0 8480.828
598.298828125 0 3430.4307
598.357177734375 0 3943.6409
602.28515625 0 604.05676
603.308349609375 0 2085.473
605.304931640625 0 635.23676
610.2830200195312 0 12029.775
610.3373413085938 0 894.8747
611.27001953125 0 11793.195 y 8
612.272216796875 0 3574.4375
613.27197265625 0 803.69507
614.3369140625 0 820.31573
619.7987670898438 0 1215.4304
626.3727416992188 0 913.3724
630.31103515625 0 1014.47284
634.3282470703125 0 681.2646
636.3220825195312 0 6523.976
637.3272094726562 0 2410.1204
639.3214721679688 0 5063.5947
639.8232421875 0 2977.1592
641.8368530273438 0 1409.469
645.2864379882812 0 1781.1787
645.7886962890625 0 1820.8562 z 3
646.2841186523438 0 1065.2211
646.82666015625 0 1447.8308
647.33056640625 0 7922.9663
647.8319091796875 0 3734.5676
648.3213500976562 0 5345.3984
649.3228759765625 0 770.9815
651.3397827148438 0 995.12585
652.3419189453125 0 5391.0312
653.2926025390625 0 1225.4093
653.3472290039062 0 794.9837
653.7916870117188 0 2304.5225 y 3
654.2885131835938 0 1158.816
654.8410034179688 0 1293.042 c Water loss 10
662.8458251953125 0 2334.568
663.3502807617188 0 26425.1
663.8480224609375 0 34511.152 c 10
664.291015625 0 3759.1726
664.3477783203125 0 20445.164
664.847900390625 0 9082.836
665.2891845703125 0 2863.753 w 7
665.350830078125 0 1544.2754
666.3560180664062 0 931.15045
667.366943359375 0 740.12054
670.4058837890625 0 751.46906
674.3359985351562 0 3891.3557
674.8373413085938 0 3808.1936
676.35595703125 0 3502.1458
676.85595703125 0 1993.3613
677.352783203125 0 1738.8044
679.3417358398438 0 4910.3657
679.8291625976562 0 757.3586
680.3017578125 0 2912.2131
680.8013305664062 0 4631.6113 w 2
681.30078125 0 4097.7437
685.3494873046875 0 790.2623
694.3515014648438 0 16261.931 c Ammonia loss 5
695.3544311523438 0 7280.537
696.307861328125 0 1949.0859
696.3638305664062 0 3577.293
696.8563842773438 0 704.3825
697.3089599609375 0 721.0486
697.3696899414062 0 1355.2617
700.831298828125 0 1692.2952
701.3291015625 0 2286.3838 y Water loss 2
701.82763671875 0 5662.3022 y Ammonia loss 2
702.3236083984375 0 4756.287 z 2
702.3799438476562 0 1344.5188
702.8291625976562 0 2216.5376
703.32861328125 0 753.18134
703.8413696289062 0 3918.4163
704.3453979492188 0 1843.4973
705.86572265625 0 1364.9822
706.360107421875 0 2561.5723
706.8645629882812 0 1373.3523
707.3502197265625 0 3420.9675
707.8697509765625 0 1383.9131
708.34814453125 0 3114.9685
709.3447875976562 0 1623.1252
709.8375854492188 0 7443.907
710.3355712890625 0 11973.761 y 2
710.8359985351562 0 7517.0093
711.3776245117188 0 43835.633 c 5
711.85107421875 0 9839.61
712.3765869140625 0 17634.062
713.3826904296875 0 4013.6345
715.8674926757812 0 1037.4044
717.8545532226562 0 1125.3755
718.3554077148438 0 994.214
719.3574829101562 0 2060.3499 c Water loss 11
719.8597412109375 0 2338.565 c Ammonia loss 11
720.857421875 0 760.1445
721.355712890625 0 993.31335
721.8690795898438 0 687.0218
722.34716796875 0 955.9458
723.359619140625 0 5587.1655
724.3525390625 0 5366.6895 y 7
725.3585815429688 0 2084.91
726.3527221679688 0 747.35114
727.8723754882812 0 32866.41
728.3690795898438 0 47223.832 c 11
728.869384765625 0 31585.98
729.369140625 0 15515.07
729.8624267578125 0 7822.4683
730.3595581054688 0 4142.474
730.8543090820312 0 803.0017
734.37060546875 0 676.45636
735.3782348632812 0 928.07135
735.8627319335938 0 729.581
736.36083984375 0 2008.0564
736.8651733398438 0 3126.528
737.3626708984375 0 4745.459
737.8821411132812 0 27816.158
738.3840942382812 0 16034.786
739.3712768554688 0 2255.12
744.3689575195312 0 1875.7991
746.3593139648438 0 991.68915
746.8590698242188 0 1062.9287
747.35595703125 0 4050.1997
747.8599243164062 0 2744.2693
748.8653564453125 0 835.0136
749.3681640625 0 1191.4205
749.8675537109375 0 749.8025
750.3711547851562 0 1484.6575
751.3634033203125 0 1692.7505 y Ammonia loss 1
751.8602294921875 0 1932.1777 z 1
752.3671875 0 2299.4443
753.3819580078125 0 799.2957
755.3677368164062 0 12241.959
755.8696899414062 0 9425.801
756.3674926757812 0 1107.5305
756.8648071289062 0 2276.8423
757.37060546875 0 3206.4026
757.872314453125 0 1894.5859
758.3787841796875 0 1483.2888
759.3690185546875 0 965.1052
759.8655395507812 0 2153.519 y 1
760.3637084960938 0 1840.4763
760.8641967773438 0 1287.4692
765.4201049804688 0 1923.5527
765.9173583984375 0 2231.3672
766.3641967773438 0 1150.7227
766.8634643554688 0 1346.0356
771.8814697265625 0 1683.1565
772.3868408203125 0 1068.9968
772.8858032226562 0 1476.3564
773.3697509765625 0 1204.2528
773.8677978515625 0 959.1279
774.3970336914062 0 1144.8059
777.3773803710938 0 27980.674
778.3687744140625 0 23422.53
778.8860473632812 0 1204.5396
779.37158203125 0 9723.358
779.8883666992188 0 4862.9907
780.378662109375 0 9966.0205
780.8804321289062 0 8074.864
781.3833618164062 0 7568.8804
781.892578125 0 4157.041
782.4009399414062 0 5061.538
782.4718017578125 0 857.355
782.9019775390625 0 3893.0754
783.380859375 0 731.3279
786.434326171875 0 845.3861
787.3977661132812 0 1079.5764
787.8914794921875 0 4135.1323
788.3905029296875 0 4628.8076
788.8897094726562 0 2209.8066
789.3861694335938 0 9341.515
789.88525390625 0 8155.799
790.8983764648438 0 6975.111
791.3992309570312 0 5331.788
792.887939453125 0 1009.26807
793.38525390625 0 2147.9548
793.8831787109375 0 2497.483
794.3809814453125 0 2988.6294
794.8800659179688 0 2530.2656
795.3738403320312 0 2724.7185
795.8720703125 0 1091.336
796.383544921875 0 1319.3926
801.3952026367188 0 3944.5325
801.8886108398438 0 22419.105
802.38525390625 0 29994.8
802.8851928710938 0 17411.775
803.3906860351562 0 8939.972
803.8994750976562 0 30725.854
804.4012451171875 0 20878.77
809.3851318359375 0 1870.7095
809.4249877929688 0 2171.71
809.8970336914062 0 33128.71
810.39599609375 0 98425.13
810.8953247070312 0 98632.914
811.3965454101562 0 62513.332
811.9012451171875 0 37114.79
812.4081420898438 0 24851.717
812.9098510742188 0 7186.385
836.4287719726562 0 1097.4431
841.4050903320312 0 1097.9377
852.4347534179688 0 840.87286
857.4618530273438 0 1424.3582
863.4074096679688 0 1445.0079
864.4239501953125 0 1029.844
878.412841796875 0 1498.9152
879.4147338867188 0 1003.15247
896.425048828125 0 7558.153 c Ammonia loss 6
897.4273071289062 0 2827.1672
898.4359741210938 0 989.16406
901.4733276367188 0 5914.716
902.4744873046875 0 2383.4521
906.4223022460938 0 8904.258
907.414794921875 0 9534.151
908.423095703125 0 6751.2437 y Water loss 6
909.4276733398438 0 3762.799
910.4256591796875 0 1665.1084
911.4337158203125 0 1074.4822
913.4515991210938 0 13946.992 c 6
914.4541015625 0 9266.031
915.4547119140625 0 1587.5227
917.4830932617188 0 1959.7609
918.4058837890625 0 1474.3871
919.3983764648438 0 1197.4423
924.4248657226562 0 830.4634
925.4327392578125 0 3077.9429
926.42578125 0 1136.7614 y 6
944.4896850585938 0 3097.4143
945.4931030273438 0 1592.9962
946.4971923828125 0 1622.3137
947.5044555664062 0 768.76044
964.5247192382812 0 775.1149
976.5072631835938 0 4510.1885
977.5106811523438 0 2002.5243
980.457763671875 0 1055.9492
981.4467163085938 0 2528.3545
982.5224609375 0 3090.402
983.52685546875 0 2459.8772
984.5245971679688 0 1182.9021
987.4993286132812 0 1081.934
988.504150390625 0 16525.758
989.5065307617188 0 5887.012
1007.51953125 0 938.7325
1009.5147094726562 0 1041.1383 c Ammonia loss 7
1010.5108032226562 0 974.73267
1016.5484619140625 0 882.6984
1024.4481201171875 0 39269.258
1025.442626953125 0 40396.562 z 5
1026.4384765625 0 14868.303
1026.537841796875 0 25121.506 c 7
1027.43896484375 0 3875.8855
1027.5394287109375 0 16023.265
1028.4351806640625 0 872.61505
1028.54150390625 0 3648.7585
1029.539794921875 0 1347.0553
1035.593505859375 0 747.64886
1040.468994140625 0 8330.558
1041.4630126953125 0 7629.615 y 5
1042.4615478515625 0 2834.4514
1043.457275390625 0 721.9369
1045.5267333984375 0 9362.947
1046.5299072265625 0 3287.963
1047.5545654296875 0 697.74414
1051.6220703125 0 7217.1807
1052.625 0 4085.9724
1059.5811767578125 0 2633.3542
1060.5810546875 0 1994.4229
1062.590087890625 0 1110.6919
1078.6229248046875 0 1048.3547
1079.6121826171875 0 3975.9878
1080.61572265625 0 2620.2844
1081.097900390625 0 738.86743
1083.5341796875 0 5541.87
1084.535888671875 0 4409.759
1085.5404052734375 0 1485.3298
1091.5330810546875 0 3184.81
1092.5389404296875 0 5542.8853
1093.546875 0 1584.5326
1095.468994140625 0 1393.5974 w 4
1096.5657958984375 0 3241.7666
1097.5615234375 0 3954.2512
1098.5643310546875 0 2321.7524
1100.5771484375 0 812.3456
1107.5458984375 0 1009.3831
1126.517822265625 0 830.96234
1129.5753173828125 0 1054.9344
1139.5662841796875 0 2051.459
1140.5738525390625 0 7797.997
1141.572509765625 0 6754.0415 c 8
1142.571044921875 0 2879.7996
1143.57861328125 0 907.0629
1144.59228515625 0 8441.708
1145.5972900390625 0 2952.9958
1152.5057373046875 0 12281.476
1153.4952392578125 0 29297.645 z 4
1154.496826171875 0 18143.648
1155.4984130859375 0 6887.353
1156.504150390625 0 1217.716
1168.5242919921875 0 3010.9426
1169.51171875 0 5842.3105 y 4
1170.5128173828125 0 3064.066
1171.5157470703125 0 1357.0713
1179.57275390625 0 2736.2273
1180.5745849609375 0 1649.704 c Water loss 9
1194.587158203125 0 1479.1039
1197.5986328125 0 26211.25
1198.5914306640625 0 34909.055 c 9
1199.59130859375 0 19169.975
1200.594482421875 0 6537.57
1201.6016845703125 0 1212.4298
1231.6236572265625 0 13534.87
1232.6285400390625 0 7047.8237
1239.589599609375 0 844.8412
1277.6312255859375 0 809.27466
1278.6376953125 0 4002.1372
1279.64208984375 0 2210.762
1282.67236328125 0 1087.037
1283.6641845703125 0 984.2482
1285.6546630859375 0 1431.0634
1286.6563720703125 0 1048.8898
1289.562255859375 0 4825.059 y Ammonia loss 3
1290.5596923828125 0 11781.284 z 3
1291.5604248046875 0 10866.135
1292.5625 0 4951.609
1293.5577392578125 0 819.29834
1306.572265625 0 1420.5482 y 3
1307.57861328125 0 883.3673
1309.67919921875 0 799.88806
1313.6788330078125 0 863.1032
1314.70361328125 0 896.87054
1318.6558837890625 0 9987.093
1319.6597900390625 0 5458.0776
1323.6964111328125 0 1372.8029
1324.690673828125 0 2049.2336
1325.691650390625 0 12060.95
1326.6875 0 18528.947 c 10
1327.6864013671875 0 11079.574
1328.68603515625 0 4824.0264
1329.691162109375 0 1707.0516
1345.6492919921875 0 969.1852
1394.7071533203125 0 868.71
1397.713134765625 0 772.5587
1401.728515625 0 2357.1206
1402.64794921875 0 1728.0562 y Ammonia loss 2
1403.647705078125 0 7333.71 z 2
1404.6451416015625 0 7654.4307
1405.6507568359375 0 3559.2175
1406.65625 0 2990.0684
1407.67919921875 0 6480.152
1408.6851806640625 0 4039.262
1410.7210693359375 0 1656.2509
1411.727783203125 0 3403.9048
1412.7216796875 0 3906.5237
1413.728515625 0 2588.7393
1414.73388671875 0 1562.1816
1415.750732421875 0 1474.5558
1419.66259765625 0 1422.4987 y 2
1420.6708984375 0 955.5669
1430.7413330078125 0 926.376
1431.749755859375 0 2095.486
1432.7359619140625 0 1569.7303
1438.7088623046875 0 2215.6792 c Ammonia loss 11
1439.7139892578125 0 4535.647
1440.7137451171875 0 2143.1863
1441.7174072265625 0 774.24054
1454.7333984375 0 3331.0435
1455.734130859375 0 10893.017 c 11
1456.732421875 0 12221.643
1457.7335205078125 0 5634.7163
1458.737060546875 0 3648.34
1459.731689453125 0 1766.9413
1473.72802734375 0 761.64374
1474.7540283203125 0 2672.7754
1475.760498046875 0 7613.8354
1476.7628173828125 0 4430.562
1487.7464599609375 0 1033.394
1488.738037109375 0 2439.5232
1489.7362060546875 0 2287.055
1490.73681640625 0 943.3776
1494.7139892578125 0 4625.2153
1495.72265625 0 2606.1128
1502.722412109375 0 2356.778 z 1
1503.7171630859375 0 3509.8997
1504.7183837890625 0 3021.1306
1505.739501953125 0 1052.6901
1522.7269287109375 0 846.54614
1538.742431640625 0 1172.2678
1539.7059326171875 0 801.2528
1544.7652587890625 0 908.0999
1545.743896484375 0 1025.0596
1558.767333984375 0 1787.4983
1559.7696533203125 0 1901.2648
1560.7696533203125 0 4032.547
1561.765380859375 0 5982.607
1562.767578125 0 4775.1123
1563.7647705078125 0 2749.0974
1564.7784423828125 0 7047.509
1565.77880859375 0 8412.691
1566.7843017578125 0 2055.7905
1574.78173828125 0 2703.195
1575.77685546875 0 9752.378
1576.7742919921875 0 10478.526
1577.7735595703125 0 5921.2695
1578.7750244140625 0 3000.0957
1579.795654296875 0 1843.3384
1580.7945556640625 0 1463.1578
1581.798828125 0 1438.8849
1585.7540283203125 0 2036.4292
1586.7529296875 0 1890.7225
1587.75634765625 0 2033.2722
1588.752197265625 0 884.12274
1590.774169921875 0 2633.256
1591.7845458984375 0 3654.081
1592.7982177734375 0 4760.541
1593.797119140625 0 5240.411
1594.80322265625 0 3150.5273
1595.8040771484375 0 1480.3375
1596.82763671875 0 2399.6042
1597.8203125 0 1061.5532
1601.7718505859375 0 1559.2866
1602.7763671875 0 10388.243
1603.770263671875 0 33880.613
1604.76611328125 0 39637.035
1605.76611328125 0 25228.52
1606.7762451171875 0 15586.966
1607.793701171875 0 26896.348
1608.798095703125 0 14449.5
1618.7874755859375 0 3851.4355
1619.787109375 0 14980.455
1620.7900390625 0 43651.18
1621.7891845703125 0 48924.188
1622.7908935546875 0 34140.637
1623.799560546875 0 21611.262
1624.8167724609375 0 27862.62
1625.82421875 0 15475.173
1717.617431640625 0 740.8928
2565.025146484375 0 812.1411
3082.90771484375 0 949.6976

Spectrum Details

|  |  |
| --- | --- |
| Matched peaks? Matched peaks The total absolute number of peaks matched. Additionally in brackets the total fraction of peaks matched and the total number of peaks is shown. | 67 (10.75% of 623) |
| FDR? FDR The false discovery rate estimated for this peptide. It is calculated by matching all theoretical fragments with a non-integer shift with the raw peaks for this spectrum. This is done with 40 different shifts. The resulting percentage is the average number of annotated peaks over the number of annotated peaks with the correct spectrum. | 5.15% |
| Satellite FDR? Satellite FDR See the FDR for details on its calculation. This satellite ion specific FDR only contains the satellite ions (d/w) for I/L/J positions. | 3.57% |
| PSM Score? PSM Score The PSM Score as given by Hecklib to this annotated spectrum. It is shown with three significant figures. | 567 |

#### Spectrum 8667? Spectrum 8667 The raw spectrum of this peptide as annotated by Hecklib. The fragments are coloured according to ion type (see legend). Any peaks with a star '\*' as text can be hovered over to see the full details, first the ion type second the mass shift type. By hovering over the amino acids in the peptide or ions in the legend the corresponding peaks are highlighted. By toggling the 'Unassigned' label you can turn the background (unassigned) peaks on or off in the plot. By updating the slider in the Ion legend you can update the spectrum to only show the top X% of the peaks with labels. The top X% means any peak that is within X% of the highest intensity. By dragging in the spectrum you can zoom in to a specific part of the spectrum and use 'Zoom Out' to get back to the original zoom level. The annotation of the spectrum is based on the given sequence in the peptides file and is done with different software so inconsistencies are likely. The peaks are annotated based on the given sequence, with 20 ppm tolerance.

Copy Data

##### Spectrum 8667 (TSV)

###### Preview

```
Loading example...
```

*Click on the button to copy the data to your clipboard.*

Mz MinMz MaxIntensity Max

WidthHeightPeptide font sizePeptide stroke widthSpectrum font sizeSpectrum stroke widthCompact peptide

Ion legend

wxyz

abcd

OtherUnassignedIon^Charge^_Position_Show for top:%

TVLHQDWLDGKEY

02.47e+44.95e+47.42e+49.90e+4

Zoom Out

y^+3^_4_y^+1^_1_c^+1^_2_z^+1^_2_y^+1^_2_c^+1^_3_y^+1^_3_c^+2^_7_c^+1^_4_z^+1^_4_y^+1^_4_w^+1^_5_c^+1^_5_z^+1^_5_c^+1^_5_y^+1^_5_y^+2^_10_c^+2^_11_w^+1^_6_w^+2^_11_y^+2^_11_c^+1^_6_y^+2^_11_z^+1^_6_c^+1^_6_c^+2^_12_y^+1^_6_z^+2^_12_c^+1^_7_z^+1^_7_c^+1^_7_y^+1^_7_w^+1^_8_c^+1^_8_z^+1^_8_c^+1^_8_y^+1^_8_w^+1^_9_c^+1^_9_z^+1^_9_y^+1^_9_c^+1^_10_y^+1^_10_z^+1^_10_c^+1^_11_z^+1^_11_c^+1^_12_z^+1^_12_

0778155623343112

Fragment Matches Table

Show background peaks

| Position | Ion type | Intensity | mz Theoretical | mz Error (Th) | mz Error (ppm) | Charge | Series Number |
| --- | --- | --- | --- | --- | --- | --- | --- |
| - | - | 1380 | 120.1 | - | - | 0 | - |
| - | - | 325.3 | 120.1 | - | - | 0 | - |
| - | - | 2716 | 120.1 | - | - | 0 | - |
| - | - | 340.2 | 122 | - | - | 0 | - |
| - | - | 1406 | 129.1 | - | - | 0 | - |
| - | - | 1626 | 130.1 | - | - | 0 | - |
| - | - | 3514 | 136.1 | - | - | 0 | - |
| - | - | 504.1 | 137 | - | - | 0 | - |
| - | - | 397.7 | 139.1 | - | - | 0 | - |
| - | - | 686.6 | 146.1 | - | - | 0 | - |
| - | - | 763.2 | 147 | - | - | 0 | - |
| - | - | 534.4 | 148.1 | - | - | 0 | - |
| - | - | 498.4 | 149 | - | - | 0 | - |
| - | - | 7032 | 149 | - | - | 0 | - |
| - | - | 901.5 | 150.1 | - | - | 0 | - |
| - | - | 3597 | 155.1 | - | - | 0 | - |
| - | - | 1325 | 159.1 | - | - | 0 | - |
| 10 | y | 3431 | 166.1 | 0.001359 | 8.183 | +3 | 4 |
| - | - | 2865 | 167.1 | - | - | 0 | - |
| - | - | 627.5 | 169.1 | - | - | 0 | - |
| - | - | 5610 | 173.1 | - | - | 0 | - |
| - | - | 3237 | 173.5 | - | - | 0 | - |
| - | - | 496.4 | 173.7 | - | - | 0 | - |
| 13 | y | 1443 | 182.1 | 0.0001302 | 0.7153 | +1 | 1 |
| - | - | 508.4 | 187.1 | - | - | 0 | - |
| - | - | 5360 | 187.1 | - | - | 0 | - |
| - | - | 518.5 | 195 | - | - | 0 | - |
| - | - | 5137 | 201.1 | - | - | 0 | - |
| - | - | 736.3 | 202.1 | - | - | 0 | - |
| - | - | 1941 | 203.1 | - | - | 0 | - |
| - | - | 3625 | 215.1 | - | - | 0 | - |
| 2 | c | 1912 | 218.1 | 0.0003208 | 1.47 | +1 | 2 |
| - | - | 675.5 | 219.1 | - | - | 0 | - |
| - | - | 7845 | 221.1 | - | - | 0 | - |
| - | - | 1723 | 222.1 | - | - | 0 | - |
| - | - | 3187 | 223.1 | - | - | 0 | - |
| - | - | 1622 | 223.1 | - | - | 0 | - |
| - | - | 1873 | 225 | - | - | 0 | - |
| - | - | 604.7 | 226 | - | - | 0 | - |
| - | - | 620.4 | 227 | - | - | 0 | - |
| - | - | 2613 | 231.1 | - | - | 0 | - |
| - | - | 1053 | 232.2 | - | - | 0 | - |
| - | - | 1110 | 235.1 | - | - | 0 | - |
| - | - | 1.029E+04 | 239.1 | - | - | 0 | - |
| - | - | 2890 | 240.1 | - | - | 0 | - |
| - | - | 2110 | 241.1 | - | - | 0 | - |
| - | - | 2505 | 258.1 | - | - | 0 | - |
| - | - | 2548 | 268.2 | - | - | 0 | - |
| - | - | 583.9 | 275.2 | - | - | 0 | - |
| - | - | 2.328E+04 | 281.1 | - | - | 0 | - |
| - | - | 1317 | 282.1 | - | - | 0 | - |
| - | - | 1192 | 283 | - | - | 0 | - |
| 12 | z | 1437 | 295.1 | 0.002439 | 8.264 | +1 | 2 |
| - | - | 2030 | 296.1 | - | - | 0 | - |
| - | - | 4099 | 297.1 | - | - | 0 | - |
| - | - | 1232 | 297.1 | - | - | 0 | - |
| - | - | 9.798E+04 | 299.1 | - | - | 0 | - |
| - | - | 4316 | 300.1 | - | - | 0 | - |
| - | - | 1055 | 301.1 | - | - | 0 | - |
| - | - | 825.7 | 301.2 | - | - | 0 | - |
| 12 | y | 712.9 | 311.1 | 0.0005938 | 1.909 | +1 | 2 |
| - | - | 3754 | 317.2 | - | - | 0 | - |
| - | - | 563.9 | 318.2 | - | - | 0 | - |
| - | - | 6748 | 329.1 | - | - | 0 | - |
| - | - | 1301 | 330.2 | - | - | 0 | - |
| 3 | c | 2076 | 331.2 | 0.0003015 | 0.9103 | +1 | 3 |
| - | - | 1055 | 334.1 | - | - | 0 | - |
| - | - | 1852 | 341 | - | - | 0 | - |
| - | - | 2044 | 355.1 | - | - | 0 | - |
| - | - | 8528 | 359 | - | - | 0 | - |
| - | - | 734.7 | 360 | - | - | 0 | - |
| - | - | 823.8 | 364.2 | - | - | 0 | - |
| - | - | 976.5 | 366.1 | - | - | 0 | - |
| - | - | 1638 | 369.1 | - | - | 0 | - |
| - | - | 3104 | 370.1 | - | - | 0 | - |
| - | - | 1776 | 371.1 | - | - | 0 | - |
| - | - | 683.1 | 374.2 | - | - | 0 | - |
| - | - | 566.2 | 376.2 | - | - | 0 | - |
| - | - | 581.3 | 390 | - | - | 0 | - |
| - | - | 2.917E+04 | 415 | - | - | 0 | - |
| - | - | 1507 | 416 | - | - | 0 | - |
| - | - | 2260 | 423.3 | - | - | 0 | - |
| - | - | 864.3 | 424.3 | - | - | 0 | - |
| - | - | 2832 | 426.7 | - | - | 0 | - |
| - | - | 1729 | 427.2 | - | - | 0 | - |
| - | - | 1319 | 429.1 | - | - | 0 | - |
| - | - | 1351 | 430.2 | - | - | 0 | - |
| - | - | 795.9 | 431.7 | - | - | 0 | - |
| 11 | y | 780.3 | 439.2 | 0.001367 | 3.112 | +1 | 3 |
| 7 | c | 1683 | 440.7 | 0.0005078 | 1.152 | +2 | 7 |
| - | - | 1024 | 441.2 | - | - | 0 | - |
| - | - | 567.3 | 442.7 | - | - | 0 | - |
| - | - | 584.1 | 445.2 | - | - | 0 | - |
| - | - | 3563 | 451.3 | - | - | 0 | - |
| - | - | 1047 | 452.1 | - | - | 0 | - |
| - | - | 1032 | 452.3 | - | - | 0 | - |
| - | - | 1075 | 454.3 | - | - | 0 | - |
| 4 | c | 1.52E+04 | 468.3 | 1.659E-05 | 0.03543 | +1 | 4 |
| - | - | 4964 | 469.3 | - | - | 0 | - |
| - | - | 669.6 | 470.3 | - | - | 0 | - |
| - | - | 2942 | 472.3 | - | - | 0 | - |
| - | - | 1990 | 476.2 | - | - | 0 | - |
| 10 | z | 1970 | 480.2 | 0.0005194 | 1.082 | +1 | 4 |
| - | - | 552.3 | 483.3 | - | - | 0 | - |
| - | - | 1595 | 494.3 | - | - | 0 | - |
| - | - | 711.2 | 495.2 | - | - | 0 | - |
| 10 | y | 5605 | 496.2 | 0.0001382 | 0.2786 | +1 | 4 |
| - | - | 1081 | 497.2 | - | - | 0 | - |
| - | - | 7.081E+04 | 503.1 | - | - | 0 | - |
| - | - | 6865 | 504.1 | - | - | 0 | - |
| - | - | 677.7 | 505.1 | - | - | 0 | - |
| - | - | 2.681E+04 | 519.1 | - | - | 0 | - |
| - | - | 1606 | 520.1 | - | - | 0 | - |
| - | - | 882 | 523.2 | - | - | 0 | - |
| - | - | 584.2 | 525.3 | - | - | 0 | - |
| - | - | 1.741E+04 | 536.2 | - | - | 0 | - |
| - | - | 1361 | 536.2 | - | - | 0 | - |
| - | - | 810.7 | 537.2 | - | - | 0 | - |
| - | - | 1625 | 538.3 | - | - | 0 | - |
| 9 | w | 2144 | 550.3 | 0.0005886 | 1.07 | +1 | 5 |
| - | - | 1181 | 551.3 | - | - | 0 | - |
| - | - | 1701 | 563.2 | - | - | 0 | - |
| 5 | c | 703.7 | 579.3 | 0.005156 | 8.9 | +1 | 5 |
| - | - | 906.3 | 582.3 | - | - | 0 | - |
| - | - | 1297 | 594.3 | - | - | 0 | - |
| 9 | z | 1543 | 595.2 | 0.000615 | 1.033 | +1 | 5 |
| - | - | 755 | 596.3 | - | - | 0 | - |
| 5 | c | 1.114E+04 | 596.4 | 0.000214 | 0.3588 | +1 | 5 |
| - | - | 3583 | 597.4 | - | - | 0 | - |
| - | - | 972.6 | 598.4 | - | - | 0 | - |
| - | - | 2020 | 600.3 | - | - | 0 | - |
| - | - | 607.4 | 601.3 | - | - | 0 | - |
| - | - | 1194 | 610.2 | - | - | 0 | - |
| - | - | 2928 | 610.3 | - | - | 0 | - |
| 9 | y | 8138 | 611.3 | 0.001544 | 2.526 | +1 | 5 |
| - | - | 1869 | 612.3 | - | - | 0 | - |
| - | - | 1149 | 617.2 | - | - | 0 | - |
| - | - | 3083 | 617.4 | - | - | 0 | - |
| - | - | 849.3 | 618.2 | - | - | 0 | - |
| - | - | 1337 | 618.4 | - | - | 0 | - |
| - | - | 1347 | 623.8 | - | - | 0 | - |
| - | - | 3448 | 631.3 | - | - | 0 | - |
| - | - | 1090 | 631.8 | - | - | 0 | - |
| - | - | 1048 | 632.3 | - | - | 0 | - |
| 4 | y | 905.5 | 645.8 | 0.004362 | 6.755 | +2 | 10 |
| - | - | 611.3 | 648.3 | - | - | 0 | - |
| - | - | 778.9 | 653.4 | - | - | 0 | - |
| - | - | 772.2 | 654.9 | - | - | 0 | - |
| - | - | 5659 | 655.4 | - | - | 0 | - |
| 11 | c | 1.971E+04 | 655.8 | 0.001132 | 1.727 | +2 | 11 |
| - | - | 1.399E+04 | 656.3 | - | - | 0 | - |
| - | - | 5061 | 656.9 | - | - | 0 | - |
| - | - | 1658 | 657.4 | - | - | 0 | - |
| - | - | 4312 | 664.3 | - | - | 0 | - |
| 8 | w | 3232 | 665.3 | 0.002881 | 4.331 | +1 | 6 |
| - | - | 1211 | 666.3 | - | - | 0 | - |
| - | - | 1877 | 667.4 | - | - | 0 | - |
| - | - | 950.6 | 668.4 | - | - | 0 | - |
| - | - | 1055 | 668.9 | - | - | 0 | - |
| - | - | 1764 | 671.4 | - | - | 0 | - |
| - | - | 1444 | 672.3 | - | - | 0 | - |
| 3 | w | 2178 | 672.8 | 0.005122 | 7.613 | +2 | 11 |
| - | - | 1453 | 673.3 | - | - | 0 | - |
| - | - | 5761 | 688.4 | - | - | 0 | - |
| - | - | 1395 | 689.4 | - | - | 0 | - |
| - | - | 561.8 | 692.3 | - | - | 0 | - |
| 3 | y | 1178 | 693.8 | 0.01216 | 17.53 | +2 | 11 |
| 6 | c | 2452 | 694.4 | 0.00488 | 7.029 | +1 | 6 |
| - | - | 676.6 | 697.3 | - | - | 0 | - |
| - | - | 1368 | 701.8 | - | - | 0 | - |
| 3 | y | 4155 | 702.3 | 0.00127 | 1.809 | +2 | 11 |
| - | - | 2723 | 702.8 | - | - | 0 | - |
| - | - | 1901 | 703.3 | - | - | 0 | - |
| - | - | 1667 | 707.3 | - | - | 0 | - |
| 8 | z | 2249 | 708.3 | 0.005296 | 7.477 | +1 | 6 |
| 6 | c | 9349 | 711.4 | 0.0007288 | 1.024 | +1 | 6 |
| - | - | 3711 | 712.4 | - | - | 0 | - |
| - | - | 680.6 | 712.9 | - | - | 0 | - |
| - | - | 2712 | 713.3 | - | - | 0 | - |
| - | - | 730.9 | 713.8 | - | - | 0 | - |
| - | - | 597.3 | 714.3 | - | - | 0 | - |
| - | - | 1426 | 718.9 | - | - | 0 | - |
| - | - | 6310 | 719.9 | - | - | 0 | - |
| 12 | c | 1.24E+04 | 720.4 | 0.003639 | 5.052 | +2 | 12 |
| - | - | 8945 | 720.9 | - | - | 0 | - |
| - | - | 994.4 | 721.3 | - | - | 0 | - |
| - | - | 4065 | 721.4 | - | - | 0 | - |
| - | - | 1366 | 721.8 | - | - | 0 | - |
| - | - | 1763 | 721.9 | - | - | 0 | - |
| - | - | 803.3 | 722.3 | - | - | 0 | - |
| - | - | 703.7 | 723.4 | - | - | 0 | - |
| 8 | y | 3365 | 724.4 | 0.0005496 | 0.7587 | +1 | 6 |
| - | - | 1118 | 725.4 | - | - | 0 | - |
| - | - | 620.1 | 730.4 | - | - | 0 | - |
| - | - | 1410 | 736.9 | - | - | 0 | - |
| - | - | 4799 | 737.4 | - | - | 0 | - |
| - | - | 3194 | 737.9 | - | - | 0 | - |
| 2 | z | 817.8 | 743.9 | 0.006089 | 8.185 | +2 | 12 |
| - | - | 3744 | 748.9 | - | - | 0 | - |
| - | - | 2034 | 749.4 | - | - | 0 | - |
| - | - | 1280 | 749.9 | - | - | 0 | - |
| - | - | 1076 | 750.9 | - | - | 0 | - |
| - | - | 956.5 | 751.4 | - | - | 0 | - |
| - | - | 663.4 | 755.2 | - | - | 0 | - |
| - | - | 936.3 | 758.9 | - | - | 0 | - |
| - | - | 602.8 | 767.4 | - | - | 0 | - |
| - | - | 686.6 | 771.9 | - | - | 0 | - |
| - | - | 2366 | 772.4 | - | - | 0 | - |
| - | - | 1679 | 772.9 | - | - | 0 | - |
| - | - | 1924 | 773.4 | - | - | 0 | - |
| - | - | 2841 | 779.9 | - | - | 0 | - |
| - | - | 1998 | 780.4 | - | - | 0 | - |
| - | - | 723.1 | 780.9 | - | - | 0 | - |
| - | - | 741.1 | 785.4 | - | - | 0 | - |
| - | - | 1821 | 785.9 | - | - | 0 | - |
| - | - | 736.3 | 786.4 | - | - | 0 | - |
| - | - | 708.5 | 786.9 | - | - | 0 | - |
| - | - | 1587 | 787.4 | - | - | 0 | - |
| - | - | 1799 | 793.3 | - | - | 0 | - |
| - | - | 831 | 793.4 | - | - | 0 | - |
| - | - | 3929 | 793.9 | - | - | 0 | - |
| - | - | 791.8 | 794.3 | - | - | 0 | - |
| - | - | 7713 | 794.4 | - | - | 0 | - |
| - | - | 4091 | 794.9 | - | - | 0 | - |
| - | - | 1867 | 795.4 | - | - | 0 | - |
| - | - | 1503 | 795.9 | - | - | 0 | - |
| - | - | 709.6 | 796.4 | - | - | 0 | - |
| - | - | 920.3 | 796.9 | - | - | 0 | - |
| - | - | 951.1 | 800.4 | - | - | 0 | - |
| - | - | 728.4 | 801.4 | - | - | 0 | - |
| - | - | 2498 | 801.8 | - | - | 0 | - |
| - | - | 4173 | 801.9 | - | - | 0 | - |
| - | - | 3.152E+04 | 802.4 | - | - | 0 | - |
| - | - | 4.771E+04 | 802.9 | - | - | 0 | - |
| - | - | 3.229E+04 | 803.4 | - | - | 0 | - |
| - | - | 1.396E+04 | 803.9 | - | - | 0 | - |
| - | - | 3953 | 804.4 | - | - | 0 | - |
| - | - | 2053 | 804.9 | - | - | 0 | - |
| - | - | 1731 | 807.3 | - | - | 0 | - |
| - | - | 621.6 | 808.3 | - | - | 0 | - |
| - | - | 2755 | 812.4 | - | - | 0 | - |
| - | - | 1195 | 813.4 | - | - | 0 | - |
| - | - | 5532 | 817.4 | - | - | 0 | - |
| - | - | 1901 | 818.4 | - | - | 0 | - |
| - | - | 920.3 | 826.3 | - | - | 0 | - |
| - | - | 687.7 | 837.4 | - | - | 0 | - |
| - | - | 1058 | 838.4 | - | - | 0 | - |
| - | - | 854.4 | 852.4 | - | - | 0 | - |
| - | - | 788.9 | 853.4 | - | - | 0 | - |
| - | - | 1211 | 864.5 | - | - | 0 | - |
| 7 | c | 4368 | 880.4 | 0.00027 | 0.3066 | +1 | 7 |
| - | - | 2316 | 881.4 | - | - | 0 | - |
| - | - | 720 | 884.5 | - | - | 0 | - |
| - | - | 1520 | 893.4 | - | - | 0 | - |
| 7 | z | 3306 | 894.4 | 0.002521 | 2.819 | +1 | 7 |
| - | - | 1326 | 895.4 | - | - | 0 | - |
| 7 | c | 1.404E+04 | 897.5 | 0.0003298 | 0.3675 | +1 | 7 |
| - | - | 5982 | 898.5 | - | - | 0 | - |
| - | - | 1588 | 899.5 | - | - | 0 | - |
| - | - | 1572 | 909.4 | - | - | 0 | - |
| 7 | y | 2081 | 910.4 | 0.004183 | 4.594 | +1 | 7 |
| - | - | 1168 | 911.4 | - | - | 0 | - |
| - | - | 2090 | 920.3 | - | - | 0 | - |
| - | - | 9426 | 932.5 | - | - | 0 | - |
| - | - | 3748 | 933.5 | - | - | 0 | - |
| - | - | 1214 | 934.5 | - | - | 0 | - |
| - | - | 1049 | 936.3 | - | - | 0 | - |
| - | - | 851 | 937.3 | - | - | 0 | - |
| 6 | w | 1308 | 964.4 | 0.01925 | 19.96 | +1 | 8 |
| - | - | 1529 | 965.5 | - | - | 0 | - |
| - | - | 733.8 | 966.5 | - | - | 0 | - |
| - | - | 1807 | 991.4 | - | - | 0 | - |
| - | - | 868.2 | 992.3 | - | - | 0 | - |
| 8 | c | 773.3 | 993.5 | 0.002641 | 2.658 | +1 | 8 |
| - | - | 1210 | 1007 | - | - | 0 | - |
| - | - | 4160 | 1008 | - | - | 0 | - |
| 6 | z | 6178 | 1009 | 0.00573 | 5.676 | +1 | 8 |
| - | - | 2467 | 1010 | - | - | 0 | - |
| 8 | c | 9356 | 1011 | 0.0007756 | 0.7676 | +1 | 8 |
| - | - | 698.2 | 1011 | - | - | 0 | - |
| - | - | 5808 | 1012 | - | - | 0 | - |
| - | - | 734.6 | 1012 | - | - | 0 | - |
| - | - | 1767 | 1013 | - | - | 0 | - |
| - | - | 686.6 | 1014 | - | - | 0 | - |
| - | - | 1803 | 1024 | - | - | 0 | - |
| 6 | y | 1806 | 1025 | 0.01093 | 10.66 | +1 | 8 |
| - | - | 999.7 | 1026 | - | - | 0 | - |
| - | - | 1772 | 1051 | - | - | 0 | - |
| - | - | 9233 | 1062 | - | - | 0 | - |
| - | - | 4610 | 1063 | - | - | 0 | - |
| - | - | 1556 | 1064 | - | - | 0 | - |
| - | - | 1502 | 1068 | - | - | 0 | - |
| - | - | 933.8 | 1069 | - | - | 0 | - |
| - | - | 704.4 | 1070 | - | - | 0 | - |
| - | - | 1542 | 1071 | - | - | 0 | - |
| - | - | 1396 | 1072 | - | - | 0 | - |
| - | - | 1796 | 1072 | - | - | 0 | - |
| - | - | 1275 | 1073 | - | - | 0 | - |
| - | - | 714.9 | 1073 | - | - | 0 | - |
| - | - | 705 | 1074 | - | - | 0 | - |
| 5 | w | 1059 | 1079 | 0.001831 | 1.697 | +1 | 9 |
| - | - | 1272 | 1081 | - | - | 0 | - |
| - | - | 1900 | 1082 | - | - | 0 | - |
| - | - | 1021 | 1083 | - | - | 0 | - |
| - | - | 2573 | 1119 | - | - | 0 | - |
| - | - | 1184 | 1120 | - | - | 0 | - |
| - | - | 1039 | 1125 | - | - | 0 | - |
| 9 | c | 2457 | 1126 | 0.00524 | 4.656 | +1 | 9 |
| - | - | 1072 | 1127 | - | - | 0 | - |
| - | - | 2677 | 1137 | - | - | 0 | - |
| 5 | z | 2.606E+04 | 1137 | 0.0006019 | 0.5291 | +1 | 9 |
| - | - | 1.615E+04 | 1139 | - | - | 0 | - |
| - | - | 6268 | 1140 | - | - | 0 | - |
| - | - | 1674 | 1140 | - | - | 0 | - |
| - | - | 9772 | 1149 | - | - | 0 | - |
| - | - | 5205 | 1150 | - | - | 0 | - |
| - | - | 1645 | 1151 | - | - | 0 | - |
| 5 | y | 846.6 | 1154 | 0.004112 | 3.564 | +1 | 9 |
| - | - | 6416 | 1182 | - | - | 0 | - |
| 10 | c | 1.419E+04 | 1183 | 0.003552 | 3.004 | +1 | 10 |
| - | - | 7844 | 1184 | - | - | 0 | - |
| - | - | 3304 | 1185 | - | - | 0 | - |
| - | - | 1083 | 1186 | - | - | 0 | - |
| - | - | 861.6 | 1187 | - | - | 0 | - |
| - | - | 640.2 | 1188 | - | - | 0 | - |
| - | - | 2337 | 1232 | - | - | 0 | - |
| - | - | 1444 | 1233 | - | - | 0 | - |
| - | - | 1842 | 1263 | - | - | 0 | - |
| - | - | 707.1 | 1264 | - | - | 0 | - |
| - | - | 690.5 | 1268 | - | - | 0 | - |
| 4 | y | 1040 | 1274 | 0.01569 | 12.32 | +1 | 10 |
| 4 | z | 5035 | 1275 | 0.001399 | 1.098 | +1 | 10 |
| - | - | 4879 | 1276 | - | - | 0 | - |
| - | - | 1454 | 1277 | - | - | 0 | - |
| - | - | 1132 | 1279 | - | - | 0 | - |
| - | - | 808.6 | 1280 | - | - | 0 | - |
| - | - | 1430 | 1281 | - | - | 0 | - |
| - | - | 1855 | 1290 | - | - | 0 | - |
| - | - | 1124 | 1291 | - | - | 0 | - |
| - | - | 577.6 | 1292 | - | - | 0 | - |
| - | - | 1.025E+04 | 1296 | - | - | 0 | - |
| - | - | 6173 | 1297 | - | - | 0 | - |
| - | - | 1907 | 1298 | - | - | 0 | - |
| - | - | 2609 | 1310 | - | - | 0 | - |
| 11 | c | 8823 | 1311 | 0.001851 | 1.412 | +1 | 11 |
| - | - | 5712 | 1312 | - | - | 0 | - |
| - | - | 2396 | 1313 | - | - | 0 | - |
| - | - | 761.6 | 1314 | - | - | 0 | - |
| - | - | 2879 | 1377 | - | - | 0 | - |
| - | - | 1773 | 1378 | - | - | 0 | - |
| - | - | 778.7 | 1379 | - | - | 0 | - |
| 3 | z | 4342 | 1388 | 0.005836 | 4.206 | +1 | 11 |
| - | - | 5086 | 1389 | - | - | 0 | - |
| - | - | 2815 | 1390 | - | - | 0 | - |
| - | - | 1230 | 1391 | - | - | 0 | - |
| - | - | 895.1 | 1393 | - | - | 0 | - |
| - | - | 1913 | 1396 | - | - | 0 | - |
| - | - | 1174 | 1397 | - | - | 0 | - |
| - | - | 813.4 | 1423 | - | - | 0 | - |
| - | - | 1128 | 1424 | - | - | 0 | - |
| - | - | 804.5 | 1425 | - | - | 0 | - |
| - | - | 869.7 | 1439 | - | - | 0 | - |
| 12 | c | 3463 | 1440 | 0.01224 | 8.499 | +1 | 12 |
| - | - | 4179 | 1441 | - | - | 0 | - |
| - | - | 2689 | 1442 | - | - | 0 | - |
| - | - | 686.9 | 1443 | - | - | 0 | - |
| - | - | 1878 | 1444 | - | - | 0 | - |
| - | - | 1560 | 1445 | - | - | 0 | - |
| - | - | 875.4 | 1446 | - | - | 0 | - |
| - | - | 1202 | 1460 | - | - | 0 | - |
| - | - | 797.3 | 1461 | - | - | 0 | - |
| 2 | z | 697 | 1487 | 0.0153 | 10.29 | +1 | 12 |
| - | - | 1183 | 1488 | - | - | 0 | - |
| - | - | 1072 | 1489 | - | - | 0 | - |
| - | - | 1394 | 1491 | - | - | 0 | - |
| - | - | 794.7 | 1517 | - | - | 0 | - |
| - | - | 1464 | 1545 | - | - | 0 | - |
| - | - | 1939 | 1546 | - | - | 0 | - |
| - | - | 1695 | 1547 | - | - | 0 | - |
| - | - | 958.8 | 1559 | - | - | 0 | - |
| - | - | 3241 | 1560 | - | - | 0 | - |
| - | - | 4225 | 1561 | - | - | 0 | - |
| - | - | 1538 | 1562 | - | - | 0 | - |
| - | - | 778.9 | 1563 | - | - | 0 | - |
| - | - | 2003 | 1564 | - | - | 0 | - |
| - | - | 680.2 | 1565 | - | - | 0 | - |
| - | - | 1083 | 1571 | - | - | 0 | - |
| - | - | 1452 | 1574 | - | - | 0 | - |
| - | - | 2483 | 1575 | - | - | 0 | - |
| - | - | 867.6 | 1577 | - | - | 0 | - |
| - | - | 2296 | 1578 | - | - | 0 | - |
| - | - | 1917 | 1579 | - | - | 0 | - |
| - | - | 1557 | 1587 | - | - | 0 | - |
| - | - | 7114 | 1588 | - | - | 0 | - |
| - | - | 1.334E+04 | 1589 | - | - | 0 | - |
| - | - | 1.008E+04 | 1590 | - | - | 0 | - |
| - | - | 5037 | 1591 | - | - | 0 | - |
| - | - | 7069 | 1592 | - | - | 0 | - |
| - | - | 5137 | 1593 | - | - | 0 | - |
| - | - | 2040 | 1594 | - | - | 0 | - |
| - | - | 2979 | 1604 | - | - | 0 | - |
| - | - | 1.364E+04 | 1605 | - | - | 0 | - |
| - | - | 2.237E+04 | 1606 | - | - | 0 | - |
| - | - | 1.624E+04 | 1607 | - | - | 0 | - |
| - | - | 7165 | 1608 | - | - | 0 | - |
| - | - | 3876 | 1609 | - | - | 0 | - |
| - | - | 2600 | 1610 | - | - | 0 | - |
| - | - | 1208 | 1611 | - | - | 0 | - |
| - | - | 721.6 | 1849 | - | - | 0 | - |
| - | - | 746 | 2193 | - | - | 0 | - |
| - | - | 752.9 | 3081 | - | - | 0 | - |

m/z Charge Intensity FragmentType MassShift Position
120.05699920654297 0 1380.1565
120.0768814086914 0 325.31873
120.08086395263672 0 2715.749
122.00196838378906 0 340.22592
129.1022491455078 0 1405.6173
130.06515502929688 0 1625.7948
136.07574462890625 0 3514.321
137.04763793945312 0 504.09866
139.0871124267578 0 397.65778
146.0601806640625 0 686.6456
147.04396057128906 0 763.20654
148.0518341064453 0 534.38245
149.03952026367188 0 498.38757
149.04483032226562 0 7031.6655
150.058349609375 0 901.5172
155.09266662597656 0 3597.2402
159.0917510986328 0 1324.6013
166.08627319335938 0 3430.6594 y 9
167.05532836914062 0 2865.3147
169.0972900390625 0 627.50073
173.1284942626953 0 5610.302
173.45066833496094 0 3236.725
173.69361877441406 0 496.40436
182.08103942871094 0 1443.3306 y 12
187.13694763183594 0 508.4404
187.1439971923828 0 5359.5806
194.9905242919922 0 518.4918
201.12327575683594 0 5136.9937
202.0536651611328 0 736.3455
203.09408569335938 0 1941.4874
215.13880920410156 0 3624.894
218.14959716796875 0 1912.3353 c 1
219.07937622070312 0 675.4663
221.08425903320312 0 7845.437
222.08493041992188 0 1723.1582
223.06353759765625 0 3186.561
223.08139038085938 0 1622.1578
225.04281616210938 0 1873.3411
226.0421600341797 0 604.67773
227.03956604003906 0 620.42334
231.08917236328125 0 2612.8274
232.16603088378906 0 1053.1788
235.07412719726562 0 1110.1655
239.0948944091797 0 10294.957
240.09591674804688 0 2889.9478
241.09169006347656 0 2110.4336
258.1448059082031 0 2504.6167
268.1763610839844 0 2547.7258
275.1703186035156 0 583.87256
281.0511169433594 0 23284.951
282.05181884765625 0 1317.4037
283.0309753417969 0 1191.6877
295.10260009765625 0 1436.7236 z 11
296.1046447753906 0 2029.7714
297.0823059082031 0 4099.2344
297.1006164550781 0 1231.9182
299.0616760253906 0 97975.91
300.0622863769531 0 4315.7295
301.05865478515625 0 1055.0449
301.15008544921875 0 825.65985
311.1231689453125 0 712.9279 y 11
317.15826416015625 0 3753.6284
318.16064453125 0 563.90424
329.14959716796875 0 6747.7207
330.15216064453125 0 1300.6162
331.2342834472656 0 2075.6067 c 2
334.1398010253906 0 1054.9431
341.01806640625 0 1852.1644
355.0695495605469 0 2044.0846
359.0283508300781 0 8527.544
360.02777099609375 0 734.696
364.18695068359375 0 823.8389
366.1336364746094 0 976.527
369.1213073730469 0 1637.5322
370.1224060058594 0 3103.7383
371.1202697753906 0 1775.9655
374.1786193847656 0 683.08405
376.2187194824219 0 566.1775
390.014404296875 0 581.2614
415.0367126464844 0 29167.895
416.03765869140625 0 1507.1941
423.2716369628906 0 2260.3542
424.2718200683594 0 864.2718
426.72186279296875 0 2832.469
427.2229919433594 0 1728.6088
429.08837890625 0 1319.1577
430.1925048828125 0 1350.5197
431.7120666503906 0 795.9131
439.2200927734375 0 780.27997 y 10
440.7187194824219 0 1682.5614 c Ammonia loss 6
441.2217712402344 0 1024.1764
442.71832275390625 0 567.3341
445.2161865234375 0 584.1
451.2657165527344 0 3563.0608
452.14849853515625 0 1047.3127
452.26824951171875 0 1032.1462
454.26568603515625 0 1074.8274
468.2928771972656 0 15203.901 c 3
469.2957763671875 0 4964.335
470.29656982421875 0 669.5528
472.2762145996094 0 2941.638
476.1849670410156 0 1989.9778
480.22198486328125 0 1970.0671 z 9
483.26715087890625 0 552.3157
494.30877685546875 0 1595.2373
495.2360534667969 0 711.22656
496.24005126953125 0 5605.139 y 9
497.2436218261719 0 1080.9393
503.1074523925781 0 70807.47
504.1080627441406 0 6865.3867
505.1417541503906 0 677.71246
519.1387939453125 0 26813.658
520.1387939453125 0 1606.2223
523.2260131835938 0 882.01227
525.2689819335938 0 584.2231
536.1650390625 0 17407.357
536.2471923828125 0 1361.0282
537.1653442382812 0 810.74274
538.2503051757812 0 1624.5531
550.2513427734375 0 2143.557 w 8
551.2560424804688 0 1180.6455
563.2161254882812 0 1701.314
579.330078125 0 703.7162 c Ammonia loss 4
582.3247680664062 0 906.2883
594.2636108398438 0 1297.21
595.2490234375 0 1542.8613 z 8
596.2565307617188 0 754.9577
596.3512573242188 0 11138.746 c 4
597.354736328125 0 3583.424
598.3584594726562 0 972.5701
600.3355712890625 0 2020.2909
601.3323974609375 0 607.3927
610.2022094726562 0 1193.9849
610.2833862304688 0 2928.1914
611.2686767578125 0 8137.7554 y 8
612.27197265625 0 1869.333
617.2295532226562 0 1149.2903
617.3623046875 0 3083.4028
618.2302856445312 0 849.3378
618.36474609375 0 1336.8184
623.7797241210938 0 1347.376
631.2617797851562 0 3448.155
631.7809448242188 0 1090.3102
632.2619018554688 0 1048.2477
645.7954711914062 0 905.4515 y 3
648.3324584960938 0 611.33
653.3609619140625 0 778.92523
654.8525390625 0 772.1786
655.3526611328125 0 5658.6655
655.8473510742188 0 19705.58 c 10
656.34765625 0 13986.555
656.8500366210938 0 5060.578
657.3521118164062 0 1658.0018
664.2935791015625 0 4311.8364
665.2805786132812 0 3231.7214 w 7
666.289306640625 0 1210.7916
667.3660888671875 0 1876.9033
668.3653564453125 0 950.589
668.857177734375 0 1055.1705
671.3710327148438 0 1763.5033
672.3043212890625 0 1444.4604
672.801513671875 0 2177.8157 w 2
673.2996215820312 0 1453.221
688.39892578125 0 5761.196
689.4016723632812 0 1395.0928
692.2565307617188 0 561.8315
693.83203125 0 1177.8129 y Ammonia loss 2
694.3469848632812 0 2452.2903 c Ammonia loss 5
697.32177734375 0 676.56744
701.8368530273438 0 1367.662
702.3344116210938 0 4154.5747 y 2
702.836669921875 0 2723.0403
703.336669921875 0 1900.5571
707.3486328125 0 1667.4327
708.3377685546875 0 2248.7817 z 7
711.377685546875 0 9349.208 c 5
712.3794555664062 0 3710.5024
712.8555908203125 0 680.5881
713.3125610351562 0 2711.974
713.8121948242188 0 730.9112
714.3220825195312 0 597.3407
718.9191284179688 0 1426.4061
719.8750610351562 0 6309.6094
720.3711547851562 0 12397.364 c 11
720.87109375 0 8945.319
721.3091430664062 0 994.42786
721.3724975585938 0 4064.5173
721.812744140625 0 1366.258
721.8753051757812 0 1763.353
722.3258666992188 0 803.28973
723.3651123046875 0 703.7024
724.3506469726562 0 3364.9824 y 7
725.3616943359375 0 1118.1635
730.3609008789062 0 620.0965
736.8658447265625 0 1409.8324
737.3613891601562 0 4798.7344
737.8605346679688 0 3194.163
743.8640747070312 0 817.8363 z 1
748.865234375 0 3743.7913
749.3667602539062 0 2033.6849
749.8704223632812 0 1280.057
750.8768310546875 0 1076.3846
751.380615234375 0 956.47125
755.23193359375 0 663.42926
758.8660278320312 0 936.29236
767.3547973632812 0 602.81647
771.8855590820312 0 686.56244
772.38818359375 0 2366.4673
772.8846435546875 0 1678.8463
773.3866577148438 0 1924.0363
779.892822265625 0 2840.5369
780.393310546875 0 1998.2526
780.891845703125 0 723.1241
785.3970947265625 0 741.09344
785.8848876953125 0 1820.728
786.3810424804688 0 736.25867
786.8712158203125 0 708.4879
787.3735961914062 0 1587.07
793.3164672851562 0 1798.9159
793.393798828125 0 831.0491
793.8898315429688 0 3929.105
794.3154907226562 0 791.83734
794.3862915039062 0 7713.003
794.885498046875 0 4090.9412
795.38671875 0 1866.6056
795.8514404296875 0 1502.5048
796.3894653320312 0 709.6256
796.8602905273438 0 920.28485
800.4160766601562 0 951.064
801.4298706054688 0 728.38495
801.8271484375 0 2498.107
801.90087890625 0 4173.1133
802.3956909179688 0 31515.18
802.8956298828125 0 47714.098
803.3963012695312 0 32288.309
803.897216796875 0 13957.175
804.3986206054688 0 3953.2212
804.8971557617188 0 2052.7961
807.2852172851562 0 1730.5142
808.2953491210938 0 621.6457
812.4148559570312 0 2755.1396
813.41357421875 0 1194.7506
817.4412231445312 0 5531.7456
818.4435424804688 0 1901.4917
826.2672119140625 0 920.2743
837.36865234375 0 687.68915
838.3511352539062 0 1058.3055
852.4318237304688 0 854.39105
853.43798828125 0 788.93634
864.4607543945312 0 1211.1863
880.430908203125 0 4368.4683 c Ammonia loss 6
881.4330444335938 0 2316.3352
884.5230712890625 0 720.0264
893.426025390625 0 1519.5184
894.414306640625 0 3306.0244 z 6
895.4180908203125 0 1326.0632
897.4573974609375 0 14044.844 c 6
898.4591064453125 0 5982.259
899.4637451171875 0 1588.2654
909.4488525390625 0 1572.4204
910.4346923828125 0 2080.983 y 6
911.4329223632812 0 1167.729
920.3101196289062 0 2090.331
932.467529296875 0 9425.515
933.47119140625 0 3747.9531
934.4718017578125 0 1214.3633
936.3311157226562 0 1048.8802
937.3329467773438 0 851.0346
964.4603271484375 0 1308.166 w 5
965.457763671875 0 1529.3528
966.4568481445312 0 733.7992
991.3505859375 0 1807.1993
992.3463745117188 0 868.23834
993.5178833007812 0 773.3036 c Ammonia loss 7
1007.3668212890625 0 1210.1866
1008.453857421875 0 4160.034
1009.4444580078125 0 6177.681 z 5
1010.443115234375 0 2467.2644
1010.541015625 0 9355.648 c 7
1011.4417114257812 0 698.19037
1011.5430908203125 0 5807.873
1012.4349975585938 0 734.6284
1012.5479125976562 0 1767.3147
1013.5491943359375 0 686.6315
1024.4764404296875 0 1802.6765
1025.4683837890625 0 1805.9353 y 5
1026.469970703125 0 999.68134
1050.6162109375 0 1772.3057
1061.509521484375 0 9232.924
1062.5130615234375 0 4609.897
1063.522216796875 0 1555.6895
1067.5369873046875 0 1501.6633
1068.5423583984375 0 933.8397
1070.4324951171875 0 704.35504
1070.5716552734375 0 1541.7698
1071.588623046875 0 1396.2175
1072.426025390625 0 1796.3896
1072.604248046875 0 1275.3406
1073.4267578125 0 714.8698
1073.580810546875 0 705.0114
1079.4698486328125 0 1058.7545 w 4
1080.5687255859375 0 1271.9182
1081.5618896484375 0 1899.8955
1082.5677490234375 0 1020.8589
1119.4056396484375 0 2573.3386
1120.4083251953125 0 1184.3646
1124.5828857421875 0 1038.8379
1125.573974609375 0 2457.0737 c 8
1126.5699462890625 0 1072.1499
1136.5145263671875 0 2676.7795
1137.4967041015625 0 26063.9 z 4
1138.500732421875 0 16150.227
1139.5028076171875 0 6267.5547
1140.498291015625 0 1673.7871
1148.5419921875 0 9771.953
1149.5443115234375 0 5204.5874
1150.5511474609375 0 1645.4604
1153.5201416015625 0 846.59283 y 4
1181.6041259765625 0 6415.7944
1182.59375 0 14192.159 c 9
1183.5958251953125 0 7844.1626
1184.5987548828125 0 3304.1003
1185.6141357421875 0 1082.8237
1186.6044921875 0 861.61786
1187.614990234375 0 640.1864
1232.490478515625 0 2336.5017
1233.4833984375 0 1444.4731
1262.5438232421875 0 1841.8438
1263.56298828125 0 707.1096
1267.6907958984375 0 690.5452
1273.5640869140625 0 1040.4692 y Ammonia loss 3
1274.5576171875 0 5035.45 z 3
1275.5604248046875 0 4879.266
1276.562255859375 0 1453.509
1278.5465087890625 0 1132.0927
1279.564697265625 0 808.5513
1280.5665283203125 0 1429.8503
1289.509765625 0 1855.4646
1290.5220947265625 0 1124.1082
1291.546142578125 0 577.5918
1295.5751953125 0 10254.193
1296.5791015625 0 6173.4375
1297.586669921875 0 1906.6309
1309.69873046875 0 2608.7783
1310.68701171875 0 8823.437 c 10
1311.691162109375 0 5712.4785
1312.688232421875 0 2396.1934
1313.6904296875 0 761.6239
1376.541748046875 0 2879.0474
1377.5401611328125 0 1772.8484
1378.5321044921875 0 778.737
1387.6461181640625 0 4341.782 z 2
1388.64404296875 0 5086.012
1389.650146484375 0 2814.92
1390.65185546875 0 1230.1376
1392.5675048828125 0 895.06464
1395.726318359375 0 1912.5801
1396.7362060546875 0 1174.2758
1422.7489013671875 0 813.43097
1423.7110595703125 0 1128.4502
1424.7152099609375 0 804.4635
1438.7498779296875 0 869.7333
1439.739990234375 0 3462.7388 c 11
1440.7347412109375 0 4179.1406
1441.73876953125 0 2688.9756
1442.7420654296875 0 686.87854
1443.6611328125 0 1877.5381
1444.6649169921875 0 1559.9081
1445.6785888671875 0 875.4032
1459.660400390625 0 1202.0458
1460.652587890625 0 797.3
1486.7239990234375 0 697.02094 z 1
1487.71533203125 0 1183.1942
1488.718505859375 0 1071.7378
1490.6383056640625 0 1394.0988
1516.6708984375 0 794.65607
1544.7637939453125 0 1463.8925
1545.7427978515625 0 1939.3207
1546.7432861328125 0 1695.1632
1558.7740478515625 0 958.8042
1559.7799072265625 0 3241.084
1560.7850341796875 0 4224.6787
1561.7796630859375 0 1537.9258
1562.78271484375 0 778.871
1563.6923828125 0 2003.4272
1564.6932373046875 0 680.24976
1570.765380859375 0 1082.8799
1573.701171875 0 1451.6619
1574.6744384765625 0 2483.095
1576.81884765625 0 867.64294
1577.8038330078125 0 2296.3687
1578.792724609375 0 1917.1815
1586.7821044921875 0 1557.4379
1587.771728515625 0 7113.8804
1588.7633056640625 0 13341.968
1589.7613525390625 0 10077.557
1590.743408203125 0 5037.264
1591.700927734375 0 7069.4277
1592.7022705078125 0 5136.803
1593.692138671875 0 2039.5767
1603.78173828125 0 2979.4136
1604.792236328125 0 13641.635
1605.79052734375 0 22366.035
1606.7882080078125 0 16235.569
1607.78466796875 0 7165.448
1608.757568359375 0 3875.8848
1609.763916015625 0 2599.8694
1610.7835693359375 0 1208.2538
1848.960693359375 0 721.5555
2192.75390625 0 746.026
3080.73779296875 0 752.91974

Spectrum Details

|  |  |
| --- | --- |
| Matched peaks? Matched peaks The total absolute number of peaks matched. Additionally in brackets the total fraction of peaks matched and the total number of peaks is shown. | 48 (11.68% of 411) |
| FDR? FDR The false discovery rate estimated for this peptide. It is calculated by matching all theoretical fragments with a non-integer shift with the raw peaks for this spectrum. This is done with 40 different shifts. The resulting percentage is the average number of annotated peaks over the number of annotated peaks with the correct spectrum. | 5.11% |
| Satellite FDR? Satellite FDR See the FDR for details on its calculation. This satellite ion specific FDR only contains the satellite ions (d/w) for I/L/J positions. | 1.19% |
| PSM Score? PSM Score The PSM Score as given by Hecklib to this annotated spectrum. It is shown with three significant figures. | 440 |

#### Spectrum 9201? Spectrum 9201 The raw spectrum of this peptide as annotated by Hecklib. The fragments are coloured according to ion type (see legend). Any peaks with a star '\*' as text can be hovered over to see the full details, first the ion type second the mass shift type. By hovering over the amino acids in the peptide or ions in the legend the corresponding peaks are highlighted. By toggling the 'Unassigned' label you can turn the background (unassigned) peaks on or off in the plot. By updating the slider in the Ion legend you can update the spectrum to only show the top X% of the peaks with labels. The top X% means any peak that is within X% of the highest intensity. By dragging in the spectrum you can zoom in to a specific part of the spectrum and use 'Zoom Out' to get back to the original zoom level. The annotation of the spectrum is based on the given sequence in the peptides file and is done with different software so inconsistencies are likely. The peaks are annotated based on the given sequence, with 20 ppm tolerance.

Copy Data

##### Spectrum 9201 (TSV)

###### Preview

```
Loading example...
```

*Click on the button to copy the data to your clipboard.*

Mz MinMz MaxIntensity Max

WidthHeightPeptide font sizePeptide stroke widthSpectrum font sizeSpectrum stroke widthCompact peptide

Ion legend

wxyz

abcd

OtherUnassignedIon^Charge^_Position_Show for top:%

TVLHQDWLDGKEY

02.50e+45.00e+47.50e+41.00e+5

Zoom Out

y^+3^_4_y^+1^_1_c^+1^_2_z^+1^_2_c^+1^_3_y^+1^_3_c^+2^_7_c^+1^_4_z^+1^_4_y^+1^_4_w^+1^_5_c^+1^_5_z^+1^_5_c^+1^_5_y^+1^_5_c^+2^_11_w^+1^_6_w^+2^_11_y^+2^_11_c^+1^_6_y^+2^_11_z^+1^_6_c^+1^_6_c^+2^_12_y^+1^_6_c^+1^_7_z^+1^_7_c^+1^_7_z^+1^_8_c^+1^_8_y^+1^_8_w^+1^_9_c^+1^_9_y^+1^_9_z^+1^_9_c^+1^_10_y^+1^_10_z^+1^_10_c^+1^_11_z^+1^_11_c^+1^_12_

0778155523333111

Fragment Matches Table

Show background peaks

| Position | Ion type | Intensity | mz Theoretical | mz Error (Th) | mz Error (ppm) | Charge | Series Number |
| --- | --- | --- | --- | --- | --- | --- | --- |
| - | - | 390.5 | 120.1 | - | - | 0 | - |
| - | - | 342.3 | 121.4 | - | - | 0 | - |
| - | - | 394.2 | 121.9 | - | - | 0 | - |
| - | - | 476.8 | 121.9 | - | - | 0 | - |
| - | - | 359.4 | 122.8 | - | - | 0 | - |
| - | - | 376.4 | 124.5 | - | - | 0 | - |
| - | - | 1124 | 129.1 | - | - | 0 | - |
| - | - | 1397 | 130.1 | - | - | 0 | - |
| - | - | 483.1 | 130.1 | - | - | 0 | - |
| - | - | 671 | 132.1 | - | - | 0 | - |
| - | - | 614.6 | 133.1 | - | - | 0 | - |
| - | - | 461 | 136.2 | - | - | 0 | - |
| - | - | 462.5 | 148.9 | - | - | 0 | - |
| - | - | 7927 | 149 | - | - | 0 | - |
| - | - | 428.7 | 150.4 | - | - | 0 | - |
| - | - | 2076 | 155.1 | - | - | 0 | - |
| - | - | 952.2 | 159.1 | - | - | 0 | - |
| 10 | y | 681 | 166.1 | 0.001206 | 7.264 | +3 | 4 |
| - | - | 2360 | 167.1 | - | - | 0 | - |
| - | - | 5463 | 173.1 | - | - | 0 | - |
| - | - | 613.6 | 173.4 | - | - | 0 | - |
| - | - | 946.4 | 173.4 | - | - | 0 | - |
| - | - | 1291 | 173.5 | - | - | 0 | - |
| - | - | 558.4 | 177.6 | - | - | 0 | - |
| 13 | y | 807.4 | 182.1 | 0.0001139 | 0.6255 | +1 | 1 |
| - | - | 3921 | 201.1 | - | - | 0 | - |
| - | - | 508.6 | 209.2 | - | - | 0 | - |
| - | - | 1696 | 215.1 | - | - | 0 | - |
| 2 | c | 936.7 | 218.1 | 0.0002292 | 1.051 | +1 | 2 |
| - | - | 9328 | 221.1 | - | - | 0 | - |
| - | - | 2692 | 222.1 | - | - | 0 | - |
| - | - | 3631 | 223.1 | - | - | 0 | - |
| - | - | 1113 | 223.1 | - | - | 0 | - |
| - | - | 2373 | 225 | - | - | 0 | - |
| - | - | 985.7 | 226 | - | - | 0 | - |
| - | - | 1.26E+04 | 239.1 | - | - | 0 | - |
| - | - | 3914 | 240.1 | - | - | 0 | - |
| - | - | 1634 | 241.1 | - | - | 0 | - |
| - | - | 712.6 | 243.1 | - | - | 0 | - |
| - | - | 1496 | 250.1 | - | - | 0 | - |
| - | - | 1773 | 258.1 | - | - | 0 | - |
| - | - | 494.4 | 267 | - | - | 0 | - |
| - | - | 1344 | 268.2 | - | - | 0 | - |
| - | - | 2.39E+04 | 281.1 | - | - | 0 | - |
| - | - | 844.3 | 283 | - | - | 0 | - |
| 12 | z | 2433 | 295.1 | 0.001798 | 6.092 | +1 | 2 |
| - | - | 3012 | 296.1 | - | - | 0 | - |
| - | - | 4279 | 297.1 | - | - | 0 | - |
| - | - | 2045 | 297.1 | - | - | 0 | - |
| - | - | 9.907E+04 | 299.1 | - | - | 0 | - |
| - | - | 1391 | 300.1 | - | - | 0 | - |
| - | - | 634.4 | 301.1 | - | - | 0 | - |
| - | - | 838.9 | 309.2 | - | - | 0 | - |
| - | - | 725.9 | 314.1 | - | - | 0 | - |
| - | - | 2809 | 317.2 | - | - | 0 | - |
| - | - | 672.4 | 327.6 | - | - | 0 | - |
| 3 | c | 1002 | 331.2 | 0.0009119 | 2.753 | +1 | 3 |
| - | - | 2485 | 341 | - | - | 0 | - |
| - | - | 1086 | 349.2 | - | - | 0 | - |
| - | - | 1172 | 355.1 | - | - | 0 | - |
| - | - | 9334 | 359 | - | - | 0 | - |
| - | - | 730.6 | 363.2 | - | - | 0 | - |
| - | - | 798.3 | 364.2 | - | - | 0 | - |
| - | - | 1356 | 369.1 | - | - | 0 | - |
| - | - | 3316 | 370.1 | - | - | 0 | - |
| - | - | 2678 | 371.1 | - | - | 0 | - |
| - | - | 574.9 | 397.3 | - | - | 0 | - |
| - | - | 3.217E+04 | 415 | - | - | 0 | - |
| - | - | 1428 | 423.3 | - | - | 0 | - |
| - | - | 2665 | 426.7 | - | - | 0 | - |
| - | - | 1346 | 427.2 | - | - | 0 | - |
| - | - | 673.4 | 427.7 | - | - | 0 | - |
| - | - | 1801 | 429.1 | - | - | 0 | - |
| 11 | y | 700 | 439.2 | 0.001258 | 2.863 | +1 | 3 |
| 7 | c | 1139 | 440.7 | 0.000813 | 1.845 | +2 | 7 |
| - | - | 967.5 | 441.2 | - | - | 0 | - |
| - | - | 3487 | 451.3 | - | - | 0 | - |
| - | - | 753.8 | 452.3 | - | - | 0 | - |
| 4 | c | 1.213E+04 | 468.3 | 0.0001692 | 0.3613 | +1 | 4 |
| - | - | 3035 | 469.3 | - | - | 0 | - |
| 10 | z | 1524 | 480.2 | 0.0009161 | 1.908 | +1 | 4 |
| - | - | 1166 | 483.3 | - | - | 0 | - |
| - | - | 855 | 494.3 | - | - | 0 | - |
| 10 | y | 4186 | 496.2 | 0.0002908 | 0.5861 | +1 | 4 |
| - | - | 764.7 | 497.2 | - | - | 0 | - |
| - | - | 7.409E+04 | 503.1 | - | - | 0 | - |
| - | - | 2.928E+04 | 519.1 | - | - | 0 | - |
| - | - | 902.1 | 523.2 | - | - | 0 | - |
| - | - | 655.4 | 534.3 | - | - | 0 | - |
| - | - | 1.739E+04 | 536.2 | - | - | 0 | - |
| - | - | 863.6 | 536.2 | - | - | 0 | - |
| - | - | 1456 | 538.2 | - | - | 0 | - |
| 9 | w | 1664 | 550.3 | 0.0006496 | 1.181 | +1 | 5 |
| - | - | 1210 | 551.3 | - | - | 0 | - |
| 5 | c | 644.1 | 579.3 | 0.002473 | 4.27 | +1 | 5 |
| 9 | z | 1471 | 595.2 | 0.001592 | 2.674 | +1 | 5 |
| - | - | 983 | 596.3 | - | - | 0 | - |
| 5 | c | 8066 | 596.4 | 0.000275 | 0.4612 | +1 | 5 |
| - | - | 2505 | 597.4 | - | - | 0 | - |
| - | - | 1776 | 610.3 | - | - | 0 | - |
| 9 | y | 4728 | 611.3 | 0.001117 | 1.827 | +1 | 5 |
| - | - | 1794 | 612.3 | - | - | 0 | - |
| - | - | 3139 | 655.4 | - | - | 0 | - |
| 11 | c | 1.403E+04 | 655.8 | 0.0008272 | 1.261 | +2 | 11 |
| - | - | 9504 | 656.3 | - | - | 0 | - |
| - | - | 3714 | 656.8 | - | - | 0 | - |
| - | - | 861.5 | 657.4 | - | - | 0 | - |
| - | - | 2348 | 664.3 | - | - | 0 | - |
| 8 | w | 2296 | 665.3 | 0.004468 | 6.716 | +1 | 6 |
| - | - | 880.8 | 666.3 | - | - | 0 | - |
| - | - | 614.2 | 667.4 | - | - | 0 | - |
| - | - | 689.8 | 671.9 | - | - | 0 | - |
| - | - | 1278 | 672.3 | - | - | 0 | - |
| 3 | w | 2181 | 672.8 | 0.006404 | 9.518 | +2 | 11 |
| - | - | 1078 | 673.3 | - | - | 0 | - |
| 3 | y | 1005 | 693.8 | 0.01039 | 14.98 | +2 | 11 |
| 6 | c | 1104 | 694.4 | 0.01123 | 16.17 | +1 | 6 |
| - | - | 670.1 | 694.8 | - | - | 0 | - |
| - | - | 951 | 701.8 | - | - | 0 | - |
| 3 | y | 3137 | 702.3 | 0.002796 | 3.981 | +2 | 11 |
| - | - | 2158 | 702.8 | - | - | 0 | - |
| - | - | 1240 | 707.3 | - | - | 0 | - |
| 8 | z | 2283 | 708.3 | 0.002366 | 3.341 | +1 | 6 |
| 6 | c | 6458 | 711.4 | 0.001156 | 1.625 | +1 | 6 |
| - | - | 2359 | 712.4 | - | - | 0 | - |
| - | - | 775.6 | 718.8 | - | - | 0 | - |
| - | - | 5084 | 719.9 | - | - | 0 | - |
| 12 | c | 9834 | 720.4 | 0.002785 | 3.866 | +2 | 12 |
| - | - | 6546 | 720.9 | - | - | 0 | - |
| - | - | 3659 | 721.4 | - | - | 0 | - |
| - | - | 715.3 | 721.9 | - | - | 0 | - |
| 8 | y | 2265 | 724.4 | 0.001159 | 1.601 | +1 | 6 |
| - | - | 857.6 | 729.9 | - | - | 0 | - |
| - | - | 865.3 | 730.4 | - | - | 0 | - |
| - | - | 1190 | 733.8 | - | - | 0 | - |
| - | - | 1492 | 734.3 | - | - | 0 | - |
| - | - | 1067 | 736.9 | - | - | 0 | - |
| - | - | 4208 | 737.4 | - | - | 0 | - |
| - | - | 973.4 | 737.9 | - | - | 0 | - |
| - | - | 1141 | 744.4 | - | - | 0 | - |
| - | - | 2194 | 748.9 | - | - | 0 | - |
| - | - | 1561 | 749.4 | - | - | 0 | - |
| - | - | 1075 | 749.9 | - | - | 0 | - |
| - | - | 1210 | 758.4 | - | - | 0 | - |
| - | - | 660.2 | 771.4 | - | - | 0 | - |
| - | - | 1372 | 772.4 | - | - | 0 | - |
| - | - | 1134 | 772.9 | - | - | 0 | - |
| - | - | 922.7 | 773.4 | - | - | 0 | - |
| - | - | 1646 | 779.9 | - | - | 0 | - |
| - | - | 1209 | 780.4 | - | - | 0 | - |
| - | - | 802.4 | 780.9 | - | - | 0 | - |
| - | - | 748.2 | 781.4 | - | - | 0 | - |
| - | - | 604.3 | 785.9 | - | - | 0 | - |
| - | - | 747.7 | 787.4 | - | - | 0 | - |
| - | - | 1082 | 787.9 | - | - | 0 | - |
| - | - | 695.6 | 793.3 | - | - | 0 | - |
| - | - | 3206 | 793.9 | - | - | 0 | - |
| - | - | 5461 | 794.4 | - | - | 0 | - |
| - | - | 3474 | 794.9 | - | - | 0 | - |
| - | - | 1701 | 795.4 | - | - | 0 | - |
| - | - | 1019 | 800.9 | - | - | 0 | - |
| - | - | 1364 | 801.4 | - | - | 0 | - |
| - | - | 2456 | 801.9 | - | - | 0 | - |
| - | - | 2.278E+04 | 802.4 | - | - | 0 | - |
| - | - | 2.899E+04 | 802.9 | - | - | 0 | - |
| - | - | 2.271E+04 | 803.4 | - | - | 0 | - |
| - | - | 8650 | 803.9 | - | - | 0 | - |
| - | - | 2869 | 804.4 | - | - | 0 | - |
| - | - | 1327 | 804.9 | - | - | 0 | - |
| - | - | 829.4 | 827.4 | - | - | 0 | - |
| - | - | 1113 | 838.4 | - | - | 0 | - |
| - | - | 729.3 | 839.3 | - | - | 0 | - |
| 7 | c | 3788 | 880.4 | 0.0007066 | 0.8025 | +1 | 7 |
| - | - | 1150 | 881.4 | - | - | 0 | - |
| - | - | 1443 | 893.4 | - | - | 0 | - |
| 7 | z | 2858 | 894.4 | 0.003559 | 3.979 | +1 | 7 |
| - | - | 860.4 | 895.4 | - | - | 0 | - |
| 7 | c | 9357 | 897.5 | 0.0008791 | 0.9796 | +1 | 7 |
| - | - | 4086 | 898.5 | - | - | 0 | - |
| - | - | 1771 | 899.5 | - | - | 0 | - |
| - | - | 892.4 | 904.5 | - | - | 0 | - |
| - | - | 680.4 | 905.5 | - | - | 0 | - |
| - | - | 590.8 | 907.1 | - | - | 0 | - |
| - | - | 1203 | 937.4 | - | - | 0 | - |
| - | - | 1407 | 938.4 | - | - | 0 | - |
| - | - | 1223 | 955.5 | - | - | 0 | - |
| - | - | 1449 | 965.5 | - | - | 0 | - |
| - | - | 2952 | 1008 | - | - | 0 | - |
| 6 | z | 3649 | 1009 | 0.003654 | 3.62 | +1 | 8 |
| - | - | 1474 | 1010 | - | - | 0 | - |
| 8 | c | 7141 | 1011 | 0.0005925 | 0.5864 | +1 | 8 |
| - | - | 4086 | 1012 | - | - | 0 | - |
| - | - | 1394 | 1013 | - | - | 0 | - |
| - | - | 1064 | 1024 | - | - | 0 | - |
| 6 | y | 895.4 | 1025 | 0.0124 | 12.09 | +1 | 8 |
| - | - | 2036 | 1068 | - | - | 0 | - |
| - | - | 846.9 | 1069 | - | - | 0 | - |
| - | - | 3401 | 1071 | - | - | 0 | - |
| - | - | 1558 | 1072 | - | - | 0 | - |
| 5 | w | 1150 | 1079 | 0.001099 | 1.018 | +1 | 9 |
| - | - | 932.8 | 1082 | - | - | 0 | - |
| - | - | 866.1 | 1125 | - | - | 0 | - |
| 9 | c | 2212 | 1126 | 0.001944 | 1.727 | +1 | 9 |
| - | - | 884.1 | 1127 | - | - | 0 | - |
| 5 | y | 3122 | 1136 | 0.02248 | 19.78 | +1 | 9 |
| 5 | z | 1.73E+04 | 1137 | 0.0001306 | 0.1148 | +1 | 9 |
| - | - | 9428 | 1139 | - | - | 0 | - |
| - | - | 3362 | 1140 | - | - | 0 | - |
| - | - | 813.4 | 1141 | - | - | 0 | - |
| - | - | 4185 | 1182 | - | - | 0 | - |
| 10 | c | 9813 | 1183 | 0.003064 | 2.591 | +1 | 10 |
| - | - | 5271 | 1184 | - | - | 0 | - |
| - | - | 2053 | 1185 | - | - | 0 | - |
| - | - | 1031 | 1186 | - | - | 0 | - |
| - | - | 894.6 | 1253 | - | - | 0 | - |
| - | - | 3214 | 1254 | - | - | 0 | - |
| - | - | 2677 | 1255 | - | - | 0 | - |
| - | - | 2211 | 1256 | - | - | 0 | - |
| 4 | y | 1054 | 1274 | 0.01691 | 13.28 | +1 | 10 |
| 4 | z | 3709 | 1275 | 0.00092 | 0.7218 | +1 | 10 |
| - | - | 3160 | 1276 | - | - | 0 | - |
| - | - | 1634 | 1277 | - | - | 0 | - |
| - | - | 1810 | 1310 | - | - | 0 | - |
| 11 | c | 5430 | 1311 | 0.0001417 | 0.1081 | +1 | 11 |
| - | - | 5346 | 1312 | - | - | 0 | - |
| - | - | 2262 | 1313 | - | - | 0 | - |
| 3 | z | 2780 | 1388 | 0.00254 | 1.831 | +1 | 11 |
| - | - | 3881 | 1389 | - | - | 0 | - |
| - | - | 2268 | 1390 | - | - | 0 | - |
| - | - | 1097 | 1391 | - | - | 0 | - |
| - | - | 913.1 | 1396 | - | - | 0 | - |
| - | - | 968.4 | 1397 | - | - | 0 | - |
| 12 | c | 2422 | 1440 | 0.007597 | 5.277 | +1 | 12 |
| - | - | 3466 | 1441 | - | - | 0 | - |
| - | - | 1932 | 1442 | - | - | 0 | - |
| - | - | 1198 | 1443 | - | - | 0 | - |
| - | - | 1222 | 1488 | - | - | 0 | - |
| - | - | 1071 | 1489 | - | - | 0 | - |
| - | - | 1278 | 1545 | - | - | 0 | - |
| - | - | 978.9 | 1546 | - | - | 0 | - |
| - | - | 818.1 | 1547 | - | - | 0 | - |
| - | - | 814 | 1559 | - | - | 0 | - |
| - | - | 1869 | 1560 | - | - | 0 | - |
| - | - | 2106 | 1561 | - | - | 0 | - |
| - | - | 1871 | 1577 | - | - | 0 | - |
| - | - | 1435 | 1578 | - | - | 0 | - |
| - | - | 770.5 | 1586 | - | - | 0 | - |
| - | - | 5501 | 1588 | - | - | 0 | - |
| - | - | 8440 | 1589 | - | - | 0 | - |
| - | - | 7046 | 1590 | - | - | 0 | - |
| - | - | 3307 | 1591 | - | - | 0 | - |
| - | - | 1754 | 1592 | - | - | 0 | - |
| - | - | 822.9 | 1593 | - | - | 0 | - |
| - | - | 1087 | 1602 | - | - | 0 | - |
| - | - | 987.3 | 1603 | - | - | 0 | - |
| - | - | 1815 | 1604 | - | - | 0 | - |
| - | - | 9814 | 1605 | - | - | 0 | - |
| - | - | 1.715E+04 | 1606 | - | - | 0 | - |
| - | - | 1.277E+04 | 1607 | - | - | 0 | - |
| - | - | 5878 | 1608 | - | - | 0 | - |
| - | - | 2871 | 1609 | - | - | 0 | - |
| - | - | 1435 | 1610 | - | - | 0 | - |
| - | - | 844.9 | 3080 | - | - | 0 | - |

m/z Charge Intensity FragmentType MassShift Position
120.08108520507812 0 390.47775
121.36935424804688 0 342.32413
121.93299865722656 0 394.21155
121.93827056884766 0 476.75165
122.83684539794922 0 359.37546
124.51727294921875 0 376.41217
129.1024169921875 0 1124.2379
130.06512451171875 0 1397.3291
130.0853271484375 0 483.14944
132.10189819335938 0 671.0178
133.0862274169922 0 614.5668
136.1622772216797 0 461.0206
148.9277801513672 0 462.47656
149.04481506347656 0 7926.545
150.3977508544922 0 428.65555
155.0926971435547 0 2076.0894
159.09214782714844 0 952.18256
166.08612060546875 0 681.0432 y 9
167.0552215576172 0 2360.0723
173.12841796875 0 5462.748
173.43222045898438 0 613.63715
173.43861389160156 0 946.3585
173.45263671875 0 1290.7856
177.55007934570312 0 558.37225
182.08128356933594 0 807.3732 y 12
201.12315368652344 0 3921.265
209.23611450195312 0 508.57834
215.13897705078125 0 1695.8682
218.14968872070312 0 936.6529 c 1
221.08384704589844 0 9327.625
222.08485412597656 0 2691.9294
223.06361389160156 0 3630.9395
223.08102416992188 0 1113.4077
225.04270935058594 0 2372.6814
226.04441833496094 0 985.6665
239.09474182128906 0 12601.435
240.09580993652344 0 3914.1875
241.09249877929688 0 1633.7338
243.133544921875 0 712.6031
250.12213134765625 0 1496.1796
258.14501953125 0 1772.7708
267.0061340332031 0 494.44302
268.1761169433594 0 1344.1908
281.05108642578125 0 23901.156
283.0310363769531 0 844.3237
295.1032409667969 0 2433.0015 z 11
296.1039733886719 0 3011.8328
297.0824279785156 0 4278.583
297.1010437011719 0 2044.5472
299.0616149902344 0 99073.484
300.062255859375 0 1390.8921
301.0582275390625 0 634.4438
309.2037658691406 0 838.92914
314.1142272949219 0 725.9299
317.15814208984375 0 2809.3271
327.6417541503906 0 672.3998
331.2348937988281 0 1002.2217 c 2
341.01800537109375 0 2484.7505
349.189453125 0 1086.1556
355.0703125 0 1172.0497
359.0280456542969 0 9334.266
363.17950439453125 0 730.5771
364.1898498535156 0 798.3375
369.1217956542969 0 1355.5769
370.1221008300781 0 3316.2185
371.11981201171875 0 2678.1006
397.3298645019531 0 574.85504
415.03662109375 0 32172.516
423.2714538574219 0 1427.5765
426.72174072265625 0 2665.1619
427.22265625 0 1345.8699
427.7265319824219 0 673.372
429.08770751953125 0 1800.7844
439.21746826171875 0 699.9958 y 10
440.7184143066406 0 1138.7804 c Ammonia loss 6
441.2202453613281 0 967.4619
451.2657775878906 0 3486.723
452.2680969238281 0 753.79877
468.292724609375 0 12126.489 c 3
469.295654296875 0 3035.4253
480.2223815917969 0 1524.3179 z 9
483.2633972167969 0 1165.8439
494.31005859375 0 855.00757
496.2398986816406 0 4185.7075 y 9
497.2450256347656 0 764.6841
503.1072692871094 0 74089.11
519.1385498046875 0 29283.22
523.2252197265625 0 902.1264
534.3384399414062 0 655.43933
536.1650390625 0 17391.576
536.2026977539062 0 863.62305
538.2491455078125 0 1455.9568
550.2514038085938 0 1663.9841 w 8
551.2576904296875 0 1209.8805
579.3224487304688 0 644.0814 c Ammonia loss 4
595.25 0 1470.769 z 8
596.25439453125 0 982.95245
596.3511962890625 0 8066.2314 c 4
597.3530883789062 0 2504.5635
610.2838134765625 0 1775.7703
611.2682495117188 0 4728.482 y 8
612.27392578125 0 1794.3972
655.3509521484375 0 3139.3408
655.8470458984375 0 14031.232 c 10
656.34765625 0 9504.361
656.8495483398438 0 3714.0293
657.3538208007812 0 861.4754
664.2923583984375 0 2347.7043
665.2821655273438 0 2296.117 w 7
666.284423828125 0 880.8289
667.367431640625 0 614.1596
671.8728637695312 0 689.7504
672.3058471679688 0 1277.7124
672.8027954101562 0 2180.7207 w 2
673.3031616210938 0 1078.4658
693.8302612304688 0 1004.5036 y Ammonia loss 2
694.3406372070312 0 1104.1599 c Ammonia loss 5
694.824951171875 0 670.1139
701.8405151367188 0 951.02997
702.3359375 0 3137.0938 y 2
702.8363647460938 0 2158.2466
707.347900390625 0 1240.1038
708.3348388671875 0 2283.2646 z 7
711.3772583007812 0 6458.1787 c 5
712.3789672851562 0 2359.4546
718.82080078125 0 775.6481
719.8746948242188 0 5083.945
720.3703002929688 0 9833.81 c 11
720.8711547851562 0 6545.9326
721.3696899414062 0 3658.7393
721.871826171875 0 715.32355
724.3523559570312 0 2264.8376 y 7
729.8517456054688 0 857.56067
730.3627319335938 0 865.3366
733.818603515625 0 1190.3483
734.3229370117188 0 1491.9823
736.8616333007812 0 1067.488
737.3623657226562 0 4208.0957
737.862548828125 0 973.38214
744.3639526367188 0 1140.9248
748.8658447265625 0 2193.9639
749.3679809570312 0 1561.3052
749.8699340820312 0 1075.3341
758.3594360351562 0 1210.1299
771.3884887695312 0 660.1638
772.3911743164062 0 1371.9967
772.8795166015625 0 1133.9558
773.389892578125 0 922.7091
779.893310546875 0 1646.1869
780.388916015625 0 1208.6437
780.9053955078125 0 802.44025
781.3825073242188 0 748.2235
785.8856811523438 0 604.3101
787.3731079101562 0 747.6623
787.8766479492188 0 1081.9128
793.3184204101562 0 695.5561
793.8905639648438 0 3206.4602
794.3864135742188 0 5461.3857
794.8840942382812 0 3474.449
795.3834228515625 0 1700.9528
800.937744140625 0 1019.3288
801.4420776367188 0 1364.0972
801.899658203125 0 2456.242
802.3950805664062 0 22782.732
802.8959350585938 0 28985.312
803.3961791992188 0 22712.217
803.8977661132812 0 8649.7705
804.4000244140625 0 2868.9463
804.8922119140625 0 1327.1187
827.412353515625 0 829.4303
838.3531494140625 0 1112.7186
839.3460083007812 0 729.3164
880.431884765625 0 3788.114 c Ammonia loss 6
881.435546875 0 1150.0791
893.4267578125 0 1443.3062
894.4153442382812 0 2858.0422 z 6
895.415283203125 0 860.4161
897.4568481445312 0 9356.676 c 6
898.4583740234375 0 4085.6716
899.4627685546875 0 1770.9122
904.47314453125 0 892.4241
905.4779052734375 0 680.40674
907.0798950195312 0 590.8301
937.4161987304688 0 1202.645
938.4266357421875 0 1407.2753
955.453857421875 0 1222.9388
965.4569091796875 0 1448.9558
1008.4560546875 0 2951.6997
1009.4423828125 0 3648.6987 z 5
1010.4390258789062 0 1474.325
1010.5411987304688 0 7141.2188 c 7
1011.544189453125 0 4085.8745
1012.549072265625 0 1394.1448
1024.484619140625 0 1063.8396
1025.4698486328125 0 895.3503 y 5
1067.5311279296875 0 2035.9092
1068.5380859375 0 846.8516
1070.529541015625 0 3401.2534
1071.5335693359375 0 1558.4243
1079.4691162109375 0 1149.9551 w 4
1081.5704345703125 0 932.8169
1124.5799560546875 0 866.0524
1125.5706787109375 0 2211.8706 c 8
1126.574951171875 0 884.135
1136.511962890625 0 3121.6655 y Ammonia loss 4
1137.4974365234375 0 17298.463 z 4
1138.5010986328125 0 9427.984
1139.502197265625 0 3362.4888
1140.5040283203125 0 813.4297
1181.6064453125 0 4184.9907
1182.59326171875 0 9813.013 c 9
1183.595947265625 0 5271.111
1184.59716796875 0 2053.2844
1185.596923828125 0 1031.1447
1252.6785888671875 0 894.55444
1253.6864013671875 0 3213.6362
1254.69384765625 0 2676.8933
1255.6866455078125 0 2211.338
1273.5653076171875 0 1053.8916 y Ammonia loss 3
1274.5552978515625 0 3709.3542 z 3
1275.5614013671875 0 3160.3533
1276.57080078125 0 1633.5193
1309.7021484375 0 1810.4852
1310.685302734375 0 5429.7925 c 10
1311.6898193359375 0 5345.513
1312.6893310546875 0 2261.7112
1387.642822265625 0 2780.058 z 2
1388.64306640625 0 3880.835
1389.646728515625 0 2268.2266
1390.6497802734375 0 1097.2616
1395.7401123046875 0 913.06134
1396.71875 0 968.42566
1439.7353515625 0 2421.5684 c 11
1440.7352294921875 0 3465.886
1441.740234375 0 1931.7006
1442.746826171875 0 1198.394
1487.724609375 0 1222.3578
1488.7255859375 0 1070.6107
1544.7957763671875 0 1277.641
1545.7786865234375 0 978.91364
1546.7496337890625 0 818.116
1558.7830810546875 0 814.0016
1559.7786865234375 0 1869.2104
1560.7764892578125 0 2105.9265
1576.791748046875 0 1870.9697
1577.7884521484375 0 1434.877
1585.678466796875 0 770.5066
1587.7734375 0 5501.49
1588.767822265625 0 8439.799
1589.7642822265625 0 7046.2974
1590.7603759765625 0 3307.4316
1591.7708740234375 0 1754.4475
1592.771728515625 0 822.861
1601.89404296875 0 1086.6487
1602.890380859375 0 987.3231
1603.7861328125 0 1814.5632
1604.7913818359375 0 9813.512
1605.791015625 0 17145.064
1606.7923583984375 0 12769.083
1607.7945556640625 0 5877.526
1608.80029296875 0 2871.262
1609.78857421875 0 1435.3584
3080.076904296875 0 844.8625

Spectrum Details

|  |  |
| --- | --- |
| Matched peaks? Matched peaks The total absolute number of peaks matched. Additionally in brackets the total fraction of peaks matched and the total number of peaks is shown. | 41 (15.59% of 263) |
| FDR? FDR The false discovery rate estimated for this peptide. It is calculated by matching all theoretical fragments with a non-integer shift with the raw peaks for this spectrum. This is done with 40 different shifts. The resulting percentage is the average number of annotated peaks over the number of annotated peaks with the correct spectrum. | 2.50% |
| Satellite FDR? Satellite FDR See the FDR for details on its calculation. This satellite ion specific FDR only contains the satellite ions (d/w) for I/L/J positions. | 0.00% |
| PSM Score? PSM Score The PSM Score as given by Hecklib to this annotated spectrum. It is shown with three significant figures. | 371 |

#### Spectrum 8963? Spectrum 8963 The raw spectrum of this peptide as annotated by Hecklib. The fragments are coloured according to ion type (see legend). Any peaks with a star '\*' as text can be hovered over to see the full details, first the ion type second the mass shift type. By hovering over the amino acids in the peptide or ions in the legend the corresponding peaks are highlighted. By toggling the 'Unassigned' label you can turn the background (unassigned) peaks on or off in the plot. By updating the slider in the Ion legend you can update the spectrum to only show the top X% of the peaks with labels. The top X% means any peak that is within X% of the highest intensity. By dragging in the spectrum you can zoom in to a specific part of the spectrum and use 'Zoom Out' to get back to the original zoom level. The annotation of the spectrum is based on the given sequence in the peptides file and is done with different software so inconsistencies are likely. The peaks are annotated based on the given sequence, with 20 ppm tolerance.

Copy Data

##### Spectrum 8963 (TSV)

###### Preview

```
Loading example...
```

*Click on the button to copy the data to your clipboard.*

Mz MinMz MaxIntensity Max

WidthHeightPeptide font sizePeptide stroke widthSpectrum font sizeSpectrum stroke widthCompact peptide

Ion legend

wxyz

abcd

OtherUnassignedIon^Charge^_Position_Show for top:%

TVLHQDWLDGKEY

02.72e+45.44e+48.16e+41.09e+5

Zoom Out

y^+3^_4_y^+1^_1_c^+1^_2_y^+1^_2_z^+1^_2_y^+1^_2_c^+1^_3_y^+2^_6_c^+2^_7_c^+1^_4_c^+1^_4_z^+1^_4_y^+1^_4_w^+1^_5_c^+1^_5_z^+1^_5_c^+1^_5_y^+1^_5_y^+2^_10_c^+2^_11_w^+1^_6_w^+2^_11_y^+2^_11_c^+1^_6_y^+2^_11_z^+1^_6_c^+1^_6_c^+2^_12_y^+1^_6_c^+1^_7_z^+1^_7_c^+1^_7_y^+1^_7_z^+1^_8_c^+1^_8_y^+1^_8_c^+1^_9_y^+1^_9_z^+1^_9_y^+1^_9_c^+1^_10_y^+1^_10_z^+1^_10_c^+1^_11_z^+1^_11_c^+1^_12_

0778155623343112

Fragment Matches Table

Show background peaks

| Position | Ion type | Intensity | mz Theoretical | mz Error (Th) | mz Error (ppm) | Charge | Series Number |
| --- | --- | --- | --- | --- | --- | --- | --- |
| - | - | 411.9 | 121.7 | - | - | 0 | - |
| - | - | 399.5 | 127.7 | - | - | 0 | - |
| - | - | 889.3 | 129.1 | - | - | 0 | - |
| - | - | 968.2 | 130.1 | - | - | 0 | - |
| - | - | 557.3 | 132.1 | - | - | 0 | - |
| - | - | 503.5 | 136.1 | - | - | 0 | - |
| - | - | 378.7 | 145.9 | - | - | 0 | - |
| - | - | 453.9 | 145.9 | - | - | 0 | - |
| - | - | 9386 | 149 | - | - | 0 | - |
| - | - | 450.5 | 150 | - | - | 0 | - |
| - | - | 2668 | 155.1 | - | - | 0 | - |
| - | - | 450.6 | 155.8 | - | - | 0 | - |
| - | - | 1267 | 159.1 | - | - | 0 | - |
| - | - | 482.3 | 162.7 | - | - | 0 | - |
| 10 | y | 576.7 | 166.1 | 0.001512 | 9.102 | +3 | 4 |
| - | - | 2815 | 167.1 | - | - | 0 | - |
| - | - | 6017 | 173.1 | - | - | 0 | - |
| - | - | 452.2 | 173.4 | - | - | 0 | - |
| - | - | 809.4 | 174.1 | - | - | 0 | - |
| - | - | 512.2 | 178.1 | - | - | 0 | - |
| 13 | y | 1459 | 182.1 | 8.178E-06 | 0.04492 | +1 | 1 |
| - | - | 715.6 | 186.1 | - | - | 0 | - |
| - | - | 567.5 | 200.8 | - | - | 0 | - |
| - | - | 3847 | 201.1 | - | - | 0 | - |
| - | - | 822.1 | 202.1 | - | - | 0 | - |
| - | - | 1951 | 215.1 | - | - | 0 | - |
| 2 | c | 1165 | 218.1 | 0.0001682 | 0.771 | +1 | 2 |
| - | - | 1.14E+04 | 221.1 | - | - | 0 | - |
| - | - | 2709 | 222.1 | - | - | 0 | - |
| - | - | 3793 | 223.1 | - | - | 0 | - |
| - | - | 1489 | 223.1 | - | - | 0 | - |
| - | - | 1936 | 225 | - | - | 0 | - |
| - | - | 1095 | 226 | - | - | 0 | - |
| - | - | 599.2 | 233.2 | - | - | 0 | - |
| - | - | 1.254E+04 | 239.1 | - | - | 0 | - |
| - | - | 4868 | 240.1 | - | - | 0 | - |
| - | - | 2010 | 241.1 | - | - | 0 | - |
| - | - | 542.1 | 244 | - | - | 0 | - |
| - | - | 1366 | 251.1 | - | - | 0 | - |
| - | - | 1627 | 258.1 | - | - | 0 | - |
| - | - | 493.8 | 261.1 | - | - | 0 | - |
| - | - | 1904 | 268.2 | - | - | 0 | - |
| - | - | 2.599E+04 | 281.1 | - | - | 0 | - |
| - | - | 1045 | 283 | - | - | 0 | - |
| - | - | 569.2 | 283 | - | - | 0 | - |
| 12 | y | 1349 | 293.1 | 0.000663 | 2.262 | +1 | 2 |
| 12 | z | 3659 | 295.1 | 0.001859 | 6.299 | +1 | 2 |
| - | - | 3179 | 296.1 | - | - | 0 | - |
| - | - | 4973 | 297.1 | - | - | 0 | - |
| - | - | 1073 | 297.1 | - | - | 0 | - |
| - | - | 1.078E+05 | 299.1 | - | - | 0 | - |
| - | - | 1127 | 300.1 | - | - | 0 | - |
| - | - | 751.4 | 301.1 | - | - | 0 | - |
| - | - | 831.6 | 309.2 | - | - | 0 | - |
| 12 | y | 636.2 | 311.1 | 0.002364 | 7.598 | +1 | 2 |
| - | - | 971.2 | 313.1 | - | - | 0 | - |
| - | - | 4266 | 317.2 | - | - | 0 | - |
| 3 | c | 1373 | 331.2 | 0.0004309 | 1.301 | +1 | 3 |
| - | - | 601.5 | 332.2 | - | - | 0 | - |
| - | - | 2697 | 341 | - | - | 0 | - |
| - | - | 2022 | 355.1 | - | - | 0 | - |
| - | - | 927 | 356.1 | - | - | 0 | - |
| - | - | 916.6 | 357.1 | - | - | 0 | - |
| - | - | 1.067E+04 | 359 | - | - | 0 | - |
| 8 | y | 536.9 | 362.7 | 0.002557 | 7.05 | +2 | 6 |
| - | - | 2643 | 369.1 | - | - | 0 | - |
| - | - | 3944 | 370.1 | - | - | 0 | - |
| - | - | 2117 | 371.1 | - | - | 0 | - |
| - | - | 608.3 | 372.1 | - | - | 0 | - |
| - | - | 941.5 | 374.2 | - | - | 0 | - |
| - | - | 549 | 386.9 | - | - | 0 | - |
| - | - | 2248 | 404.7 | - | - | 0 | - |
| - | - | 1317 | 405.2 | - | - | 0 | - |
| - | - | 3.159E+04 | 415 | - | - | 0 | - |
| - | - | 1508 | 423.3 | - | - | 0 | - |
| - | - | 804 | 424.3 | - | - | 0 | - |
| - | - | 564.2 | 424.7 | - | - | 0 | - |
| - | - | 2214 | 426.7 | - | - | 0 | - |
| - | - | 782.2 | 427.2 | - | - | 0 | - |
| - | - | 1403 | 429.1 | - | - | 0 | - |
| - | - | 769.3 | 429.2 | - | - | 0 | - |
| 7 | c | 930.2 | 440.7 | 0.001149 | 2.606 | +2 | 7 |
| - | - | 658.3 | 441.2 | - | - | 0 | - |
| 4 | c | 577.1 | 450.3 | 0.001362 | 3.025 | +1 | 4 |
| - | - | 3507 | 451.3 | - | - | 0 | - |
| - | - | 966.7 | 452.3 | - | - | 0 | - |
| 4 | c | 1.336E+04 | 468.3 | 0.0001387 | 0.2961 | +1 | 4 |
| - | - | 3483 | 469.3 | - | - | 0 | - |
| - | - | 630.1 | 473.2 | - | - | 0 | - |
| 10 | z | 994.5 | 480.2 | 0.0009149 | 1.905 | +1 | 4 |
| - | - | 1150 | 481.2 | - | - | 0 | - |
| - | - | 797.5 | 494.2 | - | - | 0 | - |
| - | - | 1015 | 494.3 | - | - | 0 | - |
| 10 | y | 4633 | 496.2 | 4.669E-05 | 0.09409 | +1 | 4 |
| - | - | 983 | 497.2 | - | - | 0 | - |
| - | - | 8.45E+04 | 503.1 | - | - | 0 | - |
| - | - | 3.035E+04 | 519.1 | - | - | 0 | - |
| - | - | 1188 | 534.2 | - | - | 0 | - |
| - | - | 662.6 | 534.3 | - | - | 0 | - |
| - | - | 854.4 | 535.3 | - | - | 0 | - |
| - | - | 632 | 535.3 | - | - | 0 | - |
| - | - | 1.881E+04 | 536.2 | - | - | 0 | - |
| - | - | 927 | 536.2 | - | - | 0 | - |
| - | - | 1292 | 536.3 | - | - | 0 | - |
| - | - | 1421 | 538.3 | - | - | 0 | - |
| 9 | w | 1847 | 550.3 | 0.0005101 | 0.927 | +1 | 5 |
| - | - | 1283 | 551.3 | - | - | 0 | - |
| 5 | c | 1198 | 579.3 | 0.0007645 | 1.32 | +1 | 5 |
| - | - | 1076 | 594.3 | - | - | 0 | - |
| 9 | z | 1387 | 595.2 | 0.0003616 | 0.6075 | +1 | 5 |
| 5 | c | 8937 | 596.4 | 0.0004581 | 0.7682 | +1 | 5 |
| - | - | 3046 | 597.4 | - | - | 0 | - |
| - | - | 740.8 | 598.4 | - | - | 0 | - |
| - | - | 1911 | 610.3 | - | - | 0 | - |
| 9 | y | 5965 | 611.3 | 0.0001404 | 0.2297 | +1 | 5 |
| - | - | 1429 | 612.3 | - | - | 0 | - |
| - | - | 566.8 | 628.4 | - | - | 0 | - |
| 4 | y | 683.2 | 645.8 | 0.00778 | 12.05 | +2 | 10 |
| - | - | 606.6 | 647.8 | - | - | 0 | - |
| - | - | 3921 | 655.4 | - | - | 0 | - |
| 11 | c | 1.57E+04 | 655.8 | 0.0008272 | 1.261 | +2 | 11 |
| - | - | 1.05E+04 | 656.3 | - | - | 0 | - |
| - | - | 5065 | 656.8 | - | - | 0 | - |
| - | - | 608.5 | 663.3 | - | - | 0 | - |
| - | - | 2294 | 664.3 | - | - | 0 | - |
| 8 | w | 2564 | 665.3 | 0.002088 | 3.138 | +1 | 6 |
| - | - | 1420 | 668.4 | - | - | 0 | - |
| - | - | 882.8 | 668.9 | - | - | 0 | - |
| - | - | 758 | 669.4 | - | - | 0 | - |
| - | - | 1939 | 672.3 | - | - | 0 | - |
| 3 | w | 1537 | 672.8 | 0.005733 | 8.52 | +2 | 11 |
| - | - | 1092 | 673.3 | - | - | 0 | - |
| 3 | y | 1168 | 693.8 | 0.004535 | 6.537 | +2 | 11 |
| 6 | c | 1796 | 694.4 | 0.009824 | 14.15 | +1 | 6 |
| - | - | 801 | 695.3 | - | - | 0 | - |
| - | - | 1041 | 701.8 | - | - | 0 | - |
| 3 | y | 2206 | 702.3 | 0.001454 | 2.07 | +2 | 11 |
| - | - | 2395 | 702.8 | - | - | 0 | - |
| - | - | 2654 | 707.3 | - | - | 0 | - |
| 8 | z | 1612 | 708.3 | 0.00493 | 6.96 | +1 | 6 |
| - | - | 983.1 | 709.3 | - | - | 0 | - |
| 6 | c | 9005 | 711.4 | 0.0007898 | 1.11 | +1 | 6 |
| - | - | 2542 | 712.4 | - | - | 0 | - |
| - | - | 897.1 | 713.4 | - | - | 0 | - |
| - | - | 1608 | 718.9 | - | - | 0 | - |
| - | - | 7419 | 719.9 | - | - | 0 | - |
| 12 | c | 1.086E+04 | 720.4 | 0.003639 | 5.052 | +2 | 12 |
| - | - | 7717 | 720.9 | - | - | 0 | - |
| - | - | 3423 | 721.4 | - | - | 0 | - |
| - | - | 1374 | 721.9 | - | - | 0 | - |
| - | - | 905.2 | 723.4 | - | - | 0 | - |
| 8 | y | 2205 | 724.4 | 0.002502 | 3.454 | +1 | 6 |
| - | - | 934.7 | 725.3 | - | - | 0 | - |
| - | - | 1421 | 736.9 | - | - | 0 | - |
| - | - | 4174 | 737.4 | - | - | 0 | - |
| - | - | 936.9 | 737.9 | - | - | 0 | - |
| - | - | 918.6 | 744.4 | - | - | 0 | - |
| - | - | 3608 | 748.9 | - | - | 0 | - |
| - | - | 2958 | 749.4 | - | - | 0 | - |
| - | - | 1121 | 749.9 | - | - | 0 | - |
| - | - | 1167 | 750.4 | - | - | 0 | - |
| - | - | 690 | 752.4 | - | - | 0 | - |
| - | - | 888.2 | 755.4 | - | - | 0 | - |
| - | - | 671.5 | 758.9 | - | - | 0 | - |
| - | - | 688.2 | 771.4 | - | - | 0 | - |
| - | - | 1466 | 772.4 | - | - | 0 | - |
| - | - | 1904 | 772.9 | - | - | 0 | - |
| - | - | 1975 | 773.4 | - | - | 0 | - |
| - | - | 846 | 773.9 | - | - | 0 | - |
| - | - | 2533 | 779.9 | - | - | 0 | - |
| - | - | 2080 | 780.4 | - | - | 0 | - |
| - | - | 961.9 | 780.9 | - | - | 0 | - |
| - | - | 908.8 | 785.9 | - | - | 0 | - |
| - | - | 1060 | 786.4 | - | - | 0 | - |
| - | - | 3354 | 793.9 | - | - | 0 | - |
| - | - | 6220 | 794.4 | - | - | 0 | - |
| - | - | 4114 | 794.9 | - | - | 0 | - |
| - | - | 1417 | 795.4 | - | - | 0 | - |
| - | - | 823.7 | 800.4 | - | - | 0 | - |
| - | - | 1351 | 801.8 | - | - | 0 | - |
| - | - | 5899 | 801.9 | - | - | 0 | - |
| - | - | 2.921E+04 | 802.4 | - | - | 0 | - |
| - | - | 3.56E+04 | 802.9 | - | - | 0 | - |
| - | - | 2.503E+04 | 803.4 | - | - | 0 | - |
| - | - | 1.11E+04 | 803.9 | - | - | 0 | - |
| - | - | 4969 | 804.4 | - | - | 0 | - |
| - | - | 1400 | 804.9 | - | - | 0 | - |
| - | - | 801.1 | 808.3 | - | - | 0 | - |
| - | - | 665.8 | 837.4 | - | - | 0 | - |
| - | - | 971.8 | 838.3 | - | - | 0 | - |
| 7 | c | 3973 | 880.4 | 0.0001479 | 0.168 | +1 | 7 |
| - | - | 1479 | 881.4 | - | - | 0 | - |
| - | - | 644.4 | 882.4 | - | - | 0 | - |
| - | - | 1443 | 893.4 | - | - | 0 | - |
| 7 | z | 2964 | 894.4 | 0.003193 | 3.57 | +1 | 7 |
| - | - | 1955 | 895.4 | - | - | 0 | - |
| 7 | c | 1.3E+04 | 897.5 | 0.001123 | 1.252 | +1 | 7 |
| - | - | 6178 | 898.5 | - | - | 0 | - |
| - | - | 1474 | 899.5 | - | - | 0 | - |
| - | - | 606 | 907.9 | - | - | 0 | - |
| - | - | 718.8 | 909.4 | - | - | 0 | - |
| 7 | y | 1335 | 910.4 | 0.005526 | 6.069 | +1 | 7 |
| - | - | 735.5 | 911.4 | - | - | 0 | - |
| - | - | 795.1 | 937.4 | - | - | 0 | - |
| - | - | 687.9 | 941.4 | - | - | 0 | - |
| - | - | 725.5 | 954.4 | - | - | 0 | - |
| - | - | 1308 | 964.5 | - | - | 0 | - |
| - | - | 1464 | 965.5 | - | - | 0 | - |
| - | - | 724.4 | 966.5 | - | - | 0 | - |
| - | - | 2176 | 1008 | - | - | 0 | - |
| 6 | z | 3965 | 1009 | 0.004204 | 4.164 | +1 | 8 |
| - | - | 1369 | 1010 | - | - | 0 | - |
| 8 | c | 8018 | 1011 | 7.885E-05 | 0.07803 | +1 | 8 |
| - | - | 660.2 | 1011 | - | - | 0 | - |
| - | - | 5972 | 1012 | - | - | 0 | - |
| - | - | 1652 | 1013 | - | - | 0 | - |
| - | - | 879.4 | 1014 | - | - | 0 | - |
| - | - | 1533 | 1024 | - | - | 0 | - |
| 6 | y | 1312 | 1025 | 0.005072 | 4.946 | +1 | 8 |
| - | - | 2338 | 1050 | - | - | 0 | - |
| - | - | 1571 | 1051 | - | - | 0 | - |
| - | - | 685.6 | 1052 | - | - | 0 | - |
| - | - | 1478 | 1068 | - | - | 0 | - |
| - | - | 672.4 | 1069 | - | - | 0 | - |
| - | - | 771.1 | 1071 | - | - | 0 | - |
| - | - | 1660 | 1071 | - | - | 0 | - |
| - | - | 820.1 | 1072 | - | - | 0 | - |
| - | - | 1545 | 1082 | - | - | 0 | - |
| - | - | 765.8 | 1083 | - | - | 0 | - |
| - | - | 1180 | 1125 | - | - | 0 | - |
| 9 | c | 2257 | 1126 | 0.00463 | 4.113 | +1 | 9 |
| - | - | 1411 | 1127 | - | - | 0 | - |
| 5 | y | 2876 | 1136 | 0.02224 | 19.57 | +1 | 9 |
| 5 | z | 2.018E+04 | 1137 | 0.0001136 | 0.09985 | +1 | 9 |
| - | - | 1.092E+04 | 1138 | - | - | 0 | - |
| - | - | 4630 | 1140 | - | - | 0 | - |
| 5 | y | 850 | 1154 | 0.006675 | 5.787 | +1 | 9 |
| - | - | 5753 | 1182 | - | - | 0 | - |
| 10 | c | 1.269E+04 | 1183 | 0.002697 | 2.281 | +1 | 10 |
| - | - | 7382 | 1184 | - | - | 0 | - |
| - | - | 2570 | 1185 | - | - | 0 | - |
| - | - | 1584 | 1186 | - | - | 0 | - |
| - | - | 866.4 | 1187 | - | - | 0 | - |
| - | - | 984.1 | 1267 | - | - | 0 | - |
| 4 | y | 1002 | 1274 | 0.01399 | 10.98 | +1 | 10 |
| 4 | z | 4786 | 1275 | 0.001888 | 1.481 | +1 | 10 |
| - | - | 4252 | 1276 | - | - | 0 | - |
| - | - | 1416 | 1277 | - | - | 0 | - |
| - | - | 2431 | 1310 | - | - | 0 | - |
| 11 | c | 6236 | 1311 | 0.002583 | 1.971 | +1 | 11 |
| - | - | 5065 | 1312 | - | - | 0 | - |
| - | - | 2961 | 1313 | - | - | 0 | - |
| 3 | z | 4242 | 1388 | 0.00193 | 1.391 | +1 | 11 |
| - | - | 4267 | 1389 | - | - | 0 | - |
| - | - | 2294 | 1390 | - | - | 0 | - |
| - | - | 1240 | 1391 | - | - | 0 | - |
| - | - | 1129 | 1396 | - | - | 0 | - |
| - | - | 1133 | 1397 | - | - | 0 | - |
| - | - | 868.9 | 1413 | - | - | 0 | - |
| - | - | 1152 | 1439 | - | - | 0 | - |
| 12 | c | 2391 | 1440 | 0.0137 | 9.516 | +1 | 12 |
| - | - | 3295 | 1441 | - | - | 0 | - |
| - | - | 2454 | 1442 | - | - | 0 | - |
| - | - | 717 | 1443 | - | - | 0 | - |
| - | - | 1327 | 1488 | - | - | 0 | - |
| - | - | 873.3 | 1489 | - | - | 0 | - |
| - | - | 742 | 1545 | - | - | 0 | - |
| - | - | 1497 | 1546 | - | - | 0 | - |
| - | - | 1121 | 1547 | - | - | 0 | - |
| - | - | 855.3 | 1548 | - | - | 0 | - |
| - | - | 2380 | 1560 | - | - | 0 | - |
| - | - | 2214 | 1561 | - | - | 0 | - |
| - | - | 1169 | 1562 | - | - | 0 | - |
| - | - | 857.7 | 1572 | - | - | 0 | - |
| - | - | 1520 | 1577 | - | - | 0 | - |
| - | - | 2156 | 1578 | - | - | 0 | - |
| - | - | 1510 | 1579 | - | - | 0 | - |
| - | - | 1016 | 1587 | - | - | 0 | - |
| - | - | 6753 | 1588 | - | - | 0 | - |
| - | - | 1.264E+04 | 1589 | - | - | 0 | - |
| - | - | 6990 | 1590 | - | - | 0 | - |
| - | - | 2992 | 1591 | - | - | 0 | - |
| - | - | 1637 | 1592 | - | - | 0 | - |
| - | - | 1195 | 1593 | - | - | 0 | - |
| - | - | 3177 | 1604 | - | - | 0 | - |
| - | - | 8883 | 1605 | - | - | 0 | - |
| - | - | 1.849E+04 | 1606 | - | - | 0 | - |
| - | - | 1.318E+04 | 1607 | - | - | 0 | - |
| - | - | 6267 | 1608 | - | - | 0 | - |
| - | - | 2726 | 1609 | - | - | 0 | - |
| - | - | 1802 | 1610 | - | - | 0 | - |
| - | - | 631.9 | 2037 | - | - | 0 | - |
| - | - | 716.4 | 2605 | - | - | 0 | - |
| - | - | 678.6 | 2700 | - | - | 0 | - |
| - | - | 769.4 | 3081 | - | - | 0 | - |

m/z Charge Intensity FragmentType MassShift Position
121.73789978027344 0 411.9075
127.7157974243164 0 399.5467
129.10206604003906 0 889.31494
130.06504821777344 0 968.22144
132.10183715820312 0 557.3489
136.07614135742188 0 503.4546
145.88633728027344 0 378.7304
145.89315795898438 0 453.85187
149.04483032226562 0 9386.482
150.04489135742188 0 450.49774
155.09263610839844 0 2667.517
155.75694274902344 0 450.63156
159.09149169921875 0 1266.5979
162.74900817871094 0 482.32983
166.08642578125 0 576.6522 y 9
167.0554656982422 0 2815.2131
173.12838745117188 0 6016.5146
173.43772888183594 0 452.23996
174.11216735839844 0 809.3992
178.0539093017578 0 512.21265
182.08116149902344 0 1459.1816 y 12
186.12335205078125 0 715.6114
200.7842559814453 0 567.49695
201.1232452392578 0 3846.7598
202.1263885498047 0 822.1458
215.1387481689453 0 1950.8507
218.14974975585938 0 1165.4695 c 1
221.08419799804688 0 11401.647
222.08482360839844 0 2708.9626
223.06349182128906 0 3793.0505
223.0819091796875 0 1488.979
225.04237365722656 0 1935.7417
226.0424346923828 0 1095.0195
233.16468811035156 0 599.22986
239.0947723388672 0 12536.418
240.0952911376953 0 4867.8477
241.09158325195312 0 2010.1825
243.9595489501953 0 542.09625
251.14981079101562 0 1366.4882
258.1446838378906 0 1627.3129
261.1430358886719 0 493.83557
268.1767883300781 0 1903.7051
281.0509338378906 0 25993.762
283.0307922363281 0 1045.095
283.0473937988281 0 569.238
293.1138610839844 0 1349.0527 y Water loss 11
295.1031799316406 0 3658.688 z 11
296.10382080078125 0 3178.8608
297.08233642578125 0 4972.551
297.09930419921875 0 1073.2816
299.06158447265625 0 107751.56
300.0633850097656 0 1126.8693
301.0605163574219 0 751.44025
309.2037353515625 0 831.5897
311.12139892578125 0 636.17694 y 11
313.1130676269531 0 971.2249
317.1578674316406 0 4265.6973
331.2335510253906 0 1373.4125 c 2
332.2361145019531 0 601.4651
341.0179138183594 0 2697.4119
355.06951904296875 0 2021.9209
356.07049560546875 0 926.9754
357.0677795410156 0 916.5584
359.0281677246094 0 10670.28
362.6817932128906 0 536.93945 y 7
369.1219177246094 0 2643.4587
370.1224365234375 0 3943.5686
371.11968994140625 0 2116.6506
372.1178283691406 0 608.3463
374.1791687011719 0 941.457
386.94708251953125 0 549.0447
404.66998291015625 0 2248.2278
405.1720275878906 0 1317.3835
415.03662109375 0 31587.61
423.2713928222656 0 1507.779
424.272705078125 0 803.966
424.7340087890625 0 564.2243
426.7210693359375 0 2214.1223
427.2234802246094 0 782.21765
429.0890808105469 0 1402.6392
429.2110290527344 0 769.3165
440.71807861328125 0 930.19855 c Ammonia loss 6
441.21881103515625 0 658.27545
450.28369140625 0 577.1285 c Water loss 3
451.2662048339844 0 3507.1118
452.27032470703125 0 966.73315
468.2927551269531 0 13360.705 c 3
469.29522705078125 0 3482.861
473.2008361816406 0 630.0834
480.2205505371094 0 994.532 z 9
481.2266540527344 0 1149.9117
494.2392578125 0 797.54016
494.3082275390625 0 1015.44604
496.2401428222656 0 4633.152 y 9
497.2413024902344 0 982.99493
503.10723876953125 0 84497.37
519.1386108398438 0 30352.863
534.2427978515625 0 1188.2927
534.29248046875 0 662.60626
535.250732421875 0 854.4331
535.2987060546875 0 632.018
536.1647338867188 0 18809.041
536.2037353515625 0 926.97833
536.317138671875 0 1292.0886
538.2525024414062 0 1421.1108
550.250244140625 0 1847.4 w 8
551.2567138671875 0 1282.9926
579.3241577148438 0 1198.462 c Ammonia loss 4
594.264892578125 0 1075.9406
595.248046875 0 1386.9581 z 8
596.3510131835938 0 8937.142 c 4
597.3539428710938 0 3046.0613
598.3567504882812 0 740.8173
610.2841186523438 0 1911.251
611.2672729492188 0 5965.3623 y 8
612.2699584960938 0 1429.481
628.3583984375 0 566.77405
645.7988891601562 0 683.16974 y 3
647.7567749023438 0 606.56244
655.352294921875 0 3920.6355
655.8470458984375 0 15704.17 c 10
656.34765625 0 10500.857
656.8490600585938 0 5064.9346
663.332763671875 0 608.4703
664.293701171875 0 2293.9502
665.27978515625 0 2564.4856 w 7
668.3611450195312 0 1420.2803
668.8549194335938 0 882.7754
669.36083984375 0 758.0178
672.3037109375 0 1938.7943
672.8021240234375 0 1536.9001 w 2
673.296875 0 1091.5908
693.8244018554688 0 1168.132 y Ammonia loss 2
694.342041015625 0 1795.6002 c Ammonia loss 5
695.3432006835938 0 800.9816
701.8373413085938 0 1041.4767
702.3345947265625 0 2206.0356 y 2
702.8369140625 0 2394.7285
707.3480224609375 0 2653.9033
708.33740234375 0 1611.597 z 7
709.3389892578125 0 983.13116
711.3776245117188 0 9004.942 c 5
712.3798217773438 0 2541.9524
713.3854370117188 0 897.0599
718.9169921875 0 1608.2023
719.8750610351562 0 7418.9683
720.3711547851562 0 10859.214 c 11
720.8707885742188 0 7716.8306
721.3717041015625 0 3423.3662
721.8756103515625 0 1374.3982
723.3658447265625 0 905.1553
724.3536987304688 0 2205.0913 y 7
725.3496704101562 0 934.6573
736.8678588867188 0 1420.6207
737.3602905273438 0 4173.7236
737.8575439453125 0 936.94904
744.3635864257812 0 918.6329
748.8661499023438 0 3608.2388
749.3665771484375 0 2957.7668
749.865234375 0 1121.3258
750.3726806640625 0 1166.6366
752.3646240234375 0 689.9777
755.4255981445312 0 888.18066
758.86865234375 0 671.4731
771.4019775390625 0 688.21826
772.3842163085938 0 1466.4768
772.8834838867188 0 1904.2589
773.3858032226562 0 1974.5345
773.8832397460938 0 845.98456
779.8897094726562 0 2533.2756
780.394775390625 0 2080.2847
780.8916625976562 0 961.8667
785.8780517578125 0 908.776
786.3848266601562 0 1060.3113
793.8864135742188 0 3354.174
794.385009765625 0 6220.202
794.8861083984375 0 4113.8657
795.385498046875 0 1417.0128
800.4006958007812 0 823.66144
801.8258666992188 0 1351.2256
801.8993530273438 0 5899.23
802.395263671875 0 29207.229
802.8961181640625 0 35596.72
803.3961791992188 0 25026.145
803.8978881835938 0 11099.752
804.397705078125 0 4969.192
804.8968505859375 0 1400.239
808.3338012695312 0 801.0518
837.367431640625 0 665.81573
838.3467407226562 0 971.8052
880.4310302734375 0 3972.7102 c Ammonia loss 6
881.4319458007812 0 1478.9154
882.4437866210938 0 644.4206
893.427001953125 0 1442.9309
894.4149780273438 0 2964.4446 z 6
895.41796875 0 1954.795
897.4566040039062 0 13000.113 c 6
898.458984375 0 6178.4556
899.4583740234375 0 1473.8787
907.9166870117188 0 605.98267
909.4426879882812 0 718.7757
910.43603515625 0 1335.0721 y 6
911.4431762695312 0 735.46533
937.4305419921875 0 795.07697
941.3635864257812 0 687.85535
954.4121704101562 0 725.46173
964.4632568359375 0 1307.9233
965.46044921875 0 1463.7491
966.5358276367188 0 724.41754
1008.4515991210938 0 2176.0708
1009.4429321289062 0 3964.6138 z 5
1010.4391479492188 0 1369.0305
1010.5418701171875 0 8017.6626 c 7
1011.4453125 0 660.20105
1011.5408325195312 0 5972.405
1012.5408935546875 0 1652.0067
1013.5504150390625 0 879.373
1024.475341796875 0 1533.4043
1025.4625244140625 0 1312.2744 y 5
1049.6158447265625 0 2337.792
1050.6180419921875 0 1570.5111
1051.627685546875 0 685.6369
1067.535888671875 0 1477.8534
1068.5416259765625 0 672.4263
1070.50537109375 0 771.0725
1071.4725341796875 0 1659.5023
1072.4434814453125 0 820.0751
1081.55810546875 0 1545.2494
1082.556396484375 0 765.7512
1124.582275390625 0 1179.7128
1125.5733642578125 0 2256.589 c 8
1126.5753173828125 0 1410.9431
1136.51171875 0 2875.9463 y Ammonia loss 4
1137.4971923828125 0 20175.877 z 4
1138.5 0 10920.058
1139.5037841796875 0 4629.8213
1153.522705078125 0 850.03503 y 4
1181.60546875 0 5753.3813
1182.5928955078125 0 12685.144 c 9
1183.59521484375 0 7382.335
1184.5927734375 0 2569.5935
1185.5992431640625 0 1584.4215
1186.6263427734375 0 866.3755
1266.668212890625 0 984.07697
1273.5623779296875 0 1002.4335 y Ammonia loss 3
1274.55810546875 0 4785.8257 z 3
1275.56298828125 0 4251.9204
1276.5650634765625 0 1415.6138
1309.698974609375 0 2430.9312
1310.687744140625 0 6236.3374 c 10
1311.6888427734375 0 5065.3433
1312.689453125 0 2961.0605
1387.6422119140625 0 4242.2383 z 2
1388.647216796875 0 4266.5938
1389.6495361328125 0 2294.3716
1390.6536865234375 0 1239.9115
1395.73095703125 0 1129.0162
1396.7279052734375 0 1132.782
1412.735107421875 0 868.92395
1438.7353515625 0 1152.2363
1439.741455078125 0 2390.8445 c 11
1440.73388671875 0 3295.4026
1441.73974609375 0 2453.698
1442.74267578125 0 716.98035
1487.7271728515625 0 1327.1913
1488.7137451171875 0 873.26117
1544.7830810546875 0 741.95966
1545.77099609375 0 1496.9563
1546.777587890625 0 1121.0576
1547.771484375 0 855.3381
1559.777099609375 0 2379.512
1560.7838134765625 0 2213.5164
1561.79248046875 0 1168.9512
1571.7601318359375 0 857.74384
1576.784912109375 0 1519.7368
1577.7911376953125 0 2155.783
1578.7996826171875 0 1509.7672
1586.7696533203125 0 1016.46857
1587.7728271484375 0 6752.649
1588.7674560546875 0 12635.271
1589.7679443359375 0 6989.8164
1590.7650146484375 0 2992.099
1591.7725830078125 0 1636.9999
1592.765869140625 0 1194.9761
1603.79150390625 0 3177.371
1604.788818359375 0 8882.914
1605.788330078125 0 18487.82
1606.78955078125 0 13175.129
1607.7940673828125 0 6266.5728
1608.796875 0 2725.5645
1609.7926025390625 0 1802.1357
2037.08203125 0 631.8769
2605.382568359375 0 716.4218
2700.047119140625 0 678.6147
3081.1416015625 0 769.41437

Spectrum Details

|  |  |
| --- | --- |
| Matched peaks? Matched peaks The total absolute number of peaks matched. Additionally in brackets the total fraction of peaks matched and the total number of peaks is shown. | 46 (15.59% of 295) |
| FDR? FDR The false discovery rate estimated for this peptide. It is calculated by matching all theoretical fragments with a non-integer shift with the raw peaks for this spectrum. This is done with 40 different shifts. The resulting percentage is the average number of annotated peaks over the number of annotated peaks with the correct spectrum. | 3.11% |
| Satellite FDR? Satellite FDR See the FDR for details on its calculation. This satellite ion specific FDR only contains the satellite ions (d/w) for I/L/J positions. | 0.00% |
| PSM Score? PSM Score The PSM Score as given by Hecklib to this annotated spectrum. It is shown with three significant figures. | 440 |

#### Reverse Lookup? Reverse Lookup All places where this read could be placed.

| Group | Segment | Template | Template Part | Read Part | Score | Unique |
| --- | --- | --- | --- | --- | --- | --- |
| Homo sapiens Heavy Chain | IGHC | IGHG1 | [189..202] | [0..13] | 99 | False |
| Homo sapiens Heavy Chain | IGHC | IGHG3 | [236..249] | [0..13] | 99 | False |
| Homo sapiens Heavy Chain | IGHC | IGHG2 | [185..198] | [0..13] | 90 | False |
| Homo sapiens Heavy Chain | IGHC | IGHG4 | [186..199] | [0..13] | 99 | False |

| Recombined | Template Part | Read Part | Score | Unique |
| --- | --- | --- | --- | --- |
| REC-0-1 | [314..327] | [0..13] | 104 | True |

#### Meta Information from Multiple reads

##### Number of combined reads

34

##### Intensity

0.779

##### TotalArea

1.1E+10

##### Changes to the peptide sequence

TVLHQDWLDGKEY

J→LSupport for Leucine based on side chain ions (1 for L 0 for I) (Position: 8)

J→LSupport for Leucine based on side chain ions (1 for L 0 for I) (Position: 3)

L→JNo support for either Leucine or Isoleucine based on side chain ions (Position: 8)

L→JNo support for either Leucine or Isoleucine based on side chain ions (Position: 3)

J→LSupport for Leucine based on side chain ions (1 for L 0 for I) (Position: 8)

J→LSupport for Leucine based on side chain ions (1 for L 0 for I) (Position: 3)

L→JNo support for either Leucine or Isoleucine based on side chain ions (Position: 8)

L→JNo support for either Leucine or Isoleucine based on side chain ions (Position: 3)

J→LSupport for Leucine based on side chain ions (1 for L 0 for I) (Position: 8)

J→LSupport for Leucine based on side chain ions (1 for L 0 for I) (Position: 3)

L→JNo support for either Leucine or Isoleucine based on side chain ions (Position: 8)

L→JNo support for either Leucine or Isoleucine based on side chain ions (Position: 3)

J→LSupport for Leucine based on side chain ions (1 for L 0 for I) (Position: 8)

J→LSupport for Leucine based on side chain ions (1 for L 0 for I) (Position: 3)

L→JNo support for either Leucine or Isoleucine based on side chain ions (Position: 8)

L→JNo support for either Leucine or Isoleucine based on side chain ions (Position: 3)

J→LSupport for Leucine based on side chain ions (1 for L 0 for I) (Position: 8)

J→LSupport for Leucine based on side chain ions (1 for L 0 for I) (Position: 3)

L→JNo support for either Leucine or Isoleucine based on side chain ions (Position: 8)

L→JNo support for either Leucine or Isoleucine based on side chain ions (Position: 3)

J→LSupport for Leucine based on side chain ions (1 for L 0 for I) (Position: 8)

J→LSupport for Leucine based on side chain ions (2 for L 0 for I) (Position: 3)

L→JNo support for either Leucine or Isoleucine based on side chain ions (Position: 3)

L→JNo support for either Leucine or Isoleucine based on side chain ions (Position: 8)

#### Positional Score

Copy Data

##### Positional Score (TSV)

###### Preview

```
Loading example...
```

*Click on the button to copy the data to your clipboard.*

100123456789101112

Label Value
"0" 0.886
"1" 0.899
"2" 0.933
"3" 0.916
"4" 0.901
"5" 0.925
"6" 0.927
"7" 0.939
"8" 0.936
"9" 0.909
"10" 0.899
"11" 0.929
"12" 0.938

#### Meta Information from PEAKS

##### Scan Identifier

F3:7243

##### Original sequence

T

V

L

H

Q

D

W

L

D

G

K

E

Y

##### Posttranslational Modifications

##### Source File

D:\separate\_stitch\_analyses\xle-disambiguation\raw\20210323\_F1\_UM1\_Peng0013\_SA\_F59\_ingel\_3ug\_chymo.raw

##### Fraction

3

##### Scan Feature

F3:14458

##### De Novo Score

99

##### ConfidenceScore

99

### m/z

802.3918

##### Mass

1602.7678

##### Charge

2

##### Retention Time

40.2

##### Predicted Retention Time

-

##### Area

1.044E+08

##### Parts Per Million

0.8

##### Fragmentation mode

ETHCD

##### Originating file

01 D:\separate\_stitch\_analyses\xle-disambiguation\20210325\_F59\_3ug\_DENOVO\_12.csv

#### Meta Information from PEAKS

##### Scan Identifier

F3:7331

##### Original sequence

T

V

L

H

Q

D

W

L

D

G

K

E

Y

##### Posttranslational Modifications

##### Source File

D:\separate\_stitch\_analyses\xle-disambiguation\raw\20210323\_F1\_UM1\_Peng0013\_SA\_F59\_ingel\_3ug\_chymo.raw

##### Fraction

3

##### Scan Feature

F3:5589

##### De Novo Score

99

##### ConfidenceScore

99

### m/z

535.2637

##### Mass

1602.7678

##### Charge

3

##### Retention Time

42.75

##### Predicted Retention Time

-

##### Area

1.543E+08

##### Parts Per Million

0.9

##### Fragmentation mode

HCD

##### Originating file

01 D:\separate\_stitch\_analyses\xle-disambiguation\20210325\_F59\_3ug\_DENOVO\_12.csv

#### Meta Information from PEAKS

##### Scan Identifier

F3:7274

##### Original sequence

T

V

L

H

Q

D

W

L

D

G

K

E

Y

##### Posttranslational Modifications

##### Source File

D:\separate\_stitch\_analyses\xle-disambiguation\raw\20210323\_F1\_UM1\_Peng0013\_SA\_F59\_ingel\_3ug\_chymo.raw

##### Fraction

3

##### Scan Feature

F3:5589

##### De Novo Score

98

##### ConfidenceScore

98

### m/z

535.2637

##### Mass

1602.7678

##### Charge

3

##### Retention Time

42.75

##### Predicted Retention Time

-

##### Area

1.543E+08

##### Parts Per Million

0.9

##### Fragmentation mode

HCD

##### Originating file

01 D:\separate\_stitch\_analyses\xle-disambiguation\20210325\_F59\_3ug\_DENOVO\_12.csv

#### Meta Information from PEAKS

##### Scan Identifier

F3:7394

##### Original sequence

T

V

L

H

Q

D

W

L

D

G

K

E

Y

##### Posttranslational Modifications

##### Source File

D:\separate\_stitch\_analyses\xle-disambiguation\raw\20210323\_F1\_UM1\_Peng0013\_SA\_F59\_ingel\_3ug\_chymo.raw

##### Fraction

3

##### Scan Feature

F3:5589

##### De Novo Score

98

##### ConfidenceScore

98

### m/z

535.2637

##### Mass

1602.7678

##### Charge

3

##### Retention Time

42.75

##### Predicted Retention Time

-

##### Area

1.543E+08

##### Parts Per Million

0.9

##### Fragmentation mode

HCD

##### Originating file

01 D:\separate\_stitch\_analyses\xle-disambiguation\20210325\_F59\_3ug\_DENOVO\_12.csv

#### Meta Information from PEAKS

##### Scan Identifier

F3:6887

##### Original sequence

T

V

L

H

Q

D

W

L

D

G

K

E

Y

##### Posttranslational Modifications

##### Source File

D:\separate\_stitch\_analyses\xle-disambiguation\raw\20210323\_F1\_UM1\_Peng0013\_SA\_F59\_ingel\_3ug\_chymo.raw

##### Fraction

3

##### Scan Feature

F3:5612

##### De Novo Score

98

##### ConfidenceScore

98

### m/z

535.2669

##### Mass

1602.7678

##### Charge

3

##### Retention Time

38.28

##### Predicted Retention Time

-

##### Area

1.498E+09

##### Parts Per Million

6.9

##### Fragmentation mode

ETHCD

##### Originating file

01 D:\separate\_stitch\_analyses\xle-disambiguation\20210325\_F59\_3ug\_DENOVO\_12.csv

#### Meta Information from PEAKS

##### Scan Identifier

F3:7458

##### Original sequence

T

V

L

H

Q

D

W

L

D

G

K

E

Y

##### Posttranslational Modifications

##### Source File

D:\separate\_stitch\_analyses\xle-disambiguation\raw\20210323\_F1\_UM1\_Peng0013\_SA\_F59\_ingel\_3ug\_chymo.raw

##### Fraction

3

##### Scan Feature

F3:5589

##### De Novo Score

98

##### ConfidenceScore

98

### m/z

535.2637

##### Mass

1602.7678

##### Charge

3

##### Retention Time

42.75

##### Predicted Retention Time

-

##### Area

1.543E+08

##### Parts Per Million

0.9

##### Fragmentation mode

HCD

##### Originating file

01 D:\separate\_stitch\_analyses\xle-disambiguation\20210325\_F59\_3ug\_DENOVO\_12.csv

#### Meta Information from PEAKS

##### Scan Identifier

F3:8106

##### Original sequence

T

V

L

H

Q

D

W

L

D

G

K

E

Y

##### Posttranslational Modifications

##### Source File

D:\separate\_stitch\_analyses\xle-disambiguation\raw\20210323\_F1\_UM1\_Peng0013\_SA\_F59\_ingel\_3ug\_chymo.raw

##### Fraction

3

##### Scan Feature

-

##### De Novo Score

98

##### ConfidenceScore

98

### m/z

535.2652

##### Mass

1602.7678

##### Charge

3

##### Retention Time

45.22

##### Predicted Retention Time

-

##### Area

0

##### Parts Per Million

3.7

##### Fragmentation mode

ETHCD

##### Originating file

01 D:\separate\_stitch\_analyses\xle-disambiguation\20210325\_F59\_3ug\_DENOVO\_12.csv

#### Meta Information from PEAKS

##### Scan Identifier

F3:7713

##### Original sequence

T

V

L

H

Q

D

W

L

D

G

K

E

Y

##### Posttranslational Modifications

##### Source File

D:\separate\_stitch\_analyses\xle-disambiguation\raw\20210323\_F1\_UM1\_Peng0013\_SA\_F59\_ingel\_3ug\_chymo.raw

##### Fraction

3

##### Scan Feature

F3:5589

##### De Novo Score

98

##### ConfidenceScore

98

### m/z

535.2637

##### Mass

1602.7678

##### Charge

3

##### Retention Time

42.75

##### Predicted Retention Time

-

##### Area

1.543E+08

##### Parts Per Million

0.9

##### Fragmentation mode

ETHCD

##### Originating file

01 D:\separate\_stitch\_analyses\xle-disambiguation\20210325\_F59\_3ug\_DENOVO\_12.csv

#### Meta Information from PEAKS

##### Scan Identifier

F3:7024

##### Original sequence

T

V

L

H

Q

D

W

L

D

G

K

E

Y

##### Posttranslational Modifications

##### Source File

D:\separate\_stitch\_analyses\xle-disambiguation\raw\20210323\_F1\_UM1\_Peng0013\_SA\_F59\_ingel\_3ug\_chymo.raw

##### Fraction

3

##### Scan Feature

F3:5612

##### De Novo Score

98

##### ConfidenceScore

98

### m/z

535.2669

##### Mass

1602.7678

##### Charge

3

##### Retention Time

38.28

##### Predicted Retention Time

-

##### Area

1.498E+09

##### Parts Per Million

6.9

##### Fragmentation mode

HCD

##### Originating file

01 D:\separate\_stitch\_analyses\xle-disambiguation\20210325\_F59\_3ug\_DENOVO\_12.csv

#### Meta Information from PEAKS

##### Scan Identifier

F3:7645

##### Original sequence

T

V

L

H

Q

D

W

L

D

G

K

E

Y

##### Posttranslational Modifications

##### Source File

D:\separate\_stitch\_analyses\xle-disambiguation\raw\20210323\_F1\_UM1\_Peng0013\_SA\_F59\_ingel\_3ug\_chymo.raw

##### Fraction

3

##### Scan Feature

F3:5589

##### De Novo Score

98

##### ConfidenceScore

98

### m/z

535.2637

##### Mass

1602.7678

##### Charge

3

##### Retention Time

42.75

##### Predicted Retention Time

-

##### Area

1.543E+08

##### Parts Per Million

0.9

##### Fragmentation mode

HCD

##### Originating file

01 D:\separate\_stitch\_analyses\xle-disambiguation\20210325\_F59\_3ug\_DENOVO\_12.csv

#### Meta Information from PEAKS

##### Scan Identifier

F3:7777

##### Original sequence

T

V

L

H

Q

D

W

L

D

G

K

E

Y

##### Posttranslational Modifications

##### Source File

D:\separate\_stitch\_analyses\xle-disambiguation\raw\20210323\_F1\_UM1\_Peng0013\_SA\_F59\_ingel\_3ug\_chymo.raw

##### Fraction

3

##### Scan Feature

F3:5589

##### De Novo Score

98

##### ConfidenceScore

98

### m/z

535.2637

##### Mass

1602.7678

##### Charge

3

##### Retention Time

42.75

##### Predicted Retention Time

-

##### Area

1.543E+08

##### Parts Per Million

0.9

##### Fragmentation mode

HCD

##### Originating file

01 D:\separate\_stitch\_analyses\xle-disambiguation\20210325\_F59\_3ug\_DENOVO\_12.csv

#### Meta Information from PEAKS

##### Scan Identifier

F3:6234

##### Original sequence

T

V

L

H

Q

D

W

+15.99

L

D

G

K

E

Y

##### Posttranslational Modifications

Oxidation (HW)

##### Source File

D:\separate\_stitch\_analyses\xle-disambiguation\raw\20210323\_F1\_UM1\_Peng0013\_SA\_F59\_ingel\_3ug\_chymo.raw

##### Fraction

3

##### Scan Feature

F3:5897

##### De Novo Score

98

##### ConfidenceScore

98

### m/z

540.5969

##### Mass

1618.7627

##### Charge

3

##### Retention Time

34.57

##### Predicted Retention Time

-

##### Area

2.856E+07

##### Parts Per Million

3.9

##### Fragmentation mode

HCD

##### Originating file

01 D:\separate\_stitch\_analyses\xle-disambiguation\20210325\_F59\_3ug\_DENOVO\_12.csv

#### Meta Information from PEAKS

##### Scan Identifier

F3:6988

##### Original sequence

T

V

L

H

Q

D

W

L

D

G

K

E

Y

##### Posttranslational Modifications

##### Source File

D:\separate\_stitch\_analyses\xle-disambiguation\raw\20210323\_F1\_UM1\_Peng0013\_SA\_F59\_ingel\_3ug\_chymo.raw

##### Fraction

3

##### Scan Feature

F3:5612

##### De Novo Score

98

##### ConfidenceScore

98

### m/z

535.2669

##### Mass

1602.7678

##### Charge

3

##### Retention Time

38.28

##### Predicted Retention Time

-

##### Area

1.498E+09

##### Parts Per Million

6.9

##### Fragmentation mode

HCD

##### Originating file

01 D:\separate\_stitch\_analyses\xle-disambiguation\20210325\_F59\_3ug\_DENOVO\_12.csv

#### Meta Information from PEAKS

##### Scan Identifier

F3:7091

##### Original sequence

T

V

L

H

Q

D

W

L

D

G

K

E

Y

##### Posttranslational Modifications

##### Source File

D:\separate\_stitch\_analyses\xle-disambiguation\raw\20210323\_F1\_UM1\_Peng0013\_SA\_F59\_ingel\_3ug\_chymo.raw

##### Fraction

3

##### Scan Feature

F3:5612

##### De Novo Score

98

##### ConfidenceScore

98

### m/z

535.2669

##### Mass

1602.7678

##### Charge

3

##### Retention Time

38.28

##### Predicted Retention Time

-

##### Area

1.498E+09

##### Parts Per Million

6.9

##### Fragmentation mode

HCD

##### Originating file

01 D:\separate\_stitch\_analyses\xle-disambiguation\20210325\_F59\_3ug\_DENOVO\_12.csv

#### Meta Information from PEAKS

##### Scan Identifier

F3:7158

##### Original sequence

T

V

L

H

Q

D

W

L

D

G

K

E

Y

##### Posttranslational Modifications

##### Source File

D:\separate\_stitch\_analyses\xle-disambiguation\raw\20210323\_F1\_UM1\_Peng0013\_SA\_F59\_ingel\_3ug\_chymo.raw

##### Fraction

3

##### Scan Feature

F3:5612

##### De Novo Score

97

##### ConfidenceScore

97

### m/z

535.2669

##### Mass

1602.7678

##### Charge

3

##### Retention Time

38.28

##### Predicted Retention Time

-

##### Area

1.498E+09

##### Parts Per Million

6.9

##### Fragmentation mode

HCD

##### Originating file

01 D:\separate\_stitch\_analyses\xle-disambiguation\20210325\_F59\_3ug\_DENOVO\_12.csv

#### Meta Information from PEAKS

##### Scan Identifier

F3:7841

##### Original sequence

T

V

L

H

Q

D

W

L

D

G

K

E

Y

##### Posttranslational Modifications

##### Source File

D:\separate\_stitch\_analyses\xle-disambiguation\raw\20210323\_F1\_UM1\_Peng0013\_SA\_F59\_ingel\_3ug\_chymo.raw

##### Fraction

3

##### Scan Feature

F3:5589

##### De Novo Score

97

##### ConfidenceScore

97

### m/z

535.2637

##### Mass

1602.7678

##### Charge

3

##### Retention Time

42.75

##### Predicted Retention Time

-

##### Area

1.543E+08

##### Parts Per Million

0.9

##### Fragmentation mode

HCD

##### Originating file

01 D:\separate\_stitch\_analyses\xle-disambiguation\20210325\_F59\_3ug\_DENOVO\_12.csv

#### Meta Information from PEAKS

##### Scan Identifier

F3:7971

##### Original sequence

T

V

L

H

Q

D

W

L

D

G

K

E

Y

##### Posttranslational Modifications

##### Source File

D:\separate\_stitch\_analyses\xle-disambiguation\raw\20210323\_F1\_UM1\_Peng0013\_SA\_F59\_ingel\_3ug\_chymo.raw

##### Fraction

3

##### Scan Feature

F3:5589

##### De Novo Score

97

##### ConfidenceScore

97

### m/z

535.2637

##### Mass

1602.7678

##### Charge

3

##### Retention Time

42.75

##### Predicted Retention Time

-

##### Area

1.543E+08

##### Parts Per Million

0.9

##### Fragmentation mode

ETHCD

##### Originating file

01 D:\separate\_stitch\_analyses\xle-disambiguation\20210325\_F59\_3ug\_DENOVO\_12.csv

#### Meta Information from PEAKS

##### Scan Identifier

F3:6245

##### Original sequence

T

V

L

H

Q

D

W

+15.99

L

D

G

K

E

Y

##### Posttranslational Modifications

Oxidation (HW)

##### Source File

D:\separate\_stitch\_analyses\xle-disambiguation\raw\20210323\_F1\_UM1\_Peng0013\_SA\_F59\_ingel\_3ug\_chymo.raw

##### Fraction

3

##### Scan Feature

F3:5894

##### De Novo Score

97

##### ConfidenceScore

97

### m/z

540.5956

##### Mass

1618.7627

##### Charge

3

##### Retention Time

31.45

##### Predicted Retention Time

-

##### Area

2.287E+07

##### Parts Per Million

1.4

##### Fragmentation mode

HCD

##### Originating file

01 D:\separate\_stitch\_analyses\xle-disambiguation\20210325\_F59\_3ug\_DENOVO\_12.csv

#### Meta Information from PEAKS

##### Scan Identifier

F3:7305

##### Original sequence

T

V

L

H

Q

D

W

L

D

G

K

E

Y

##### Posttranslational Modifications

##### Source File

D:\separate\_stitch\_analyses\xle-disambiguation\raw\20210323\_F1\_UM1\_Peng0013\_SA\_F59\_ingel\_3ug\_chymo.raw

##### Fraction

3

##### Scan Feature

F3:14458

##### De Novo Score

97

##### ConfidenceScore

97

### m/z

802.3918

##### Mass

1602.7678

##### Charge

2

##### Retention Time

40.2

##### Predicted Retention Time

-

##### Area

1.044E+08

##### Parts Per Million

0.8

##### Fragmentation mode

HCD

##### Originating file

01 D:\separate\_stitch\_analyses\xle-disambiguation\20210325\_F59\_3ug\_DENOVO\_12.csv

#### Meta Information from PEAKS

##### Scan Identifier

F3:7222

##### Original sequence

T

V

L

H

Q

D

W

L

D

G

K

E

Y

##### Posttranslational Modifications

##### Source File

D:\separate\_stitch\_analyses\xle-disambiguation\raw\20210323\_F1\_UM1\_Peng0013\_SA\_F59\_ingel\_3ug\_chymo.raw

##### Fraction

3

##### Scan Feature

-

##### De Novo Score

97

##### ConfidenceScore

97

### m/z

535.265

##### Mass

1602.7678

##### Charge

3

##### Retention Time

40.18

##### Predicted Retention Time

-

##### Area

0

##### Parts Per Million

3.4

##### Fragmentation mode

HCD

##### Originating file

01 D:\separate\_stitch\_analyses\xle-disambiguation\20210325\_F59\_3ug\_DENOVO\_12.csv

#### Meta Information from PEAKS

##### Scan Identifier

F3:8039

##### Original sequence

T

V

L

H

Q

D

W

L

D

G

K

E

Y

##### Posttranslational Modifications

##### Source File

D:\separate\_stitch\_analyses\xle-disambiguation\raw\20210323\_F1\_UM1\_Peng0013\_SA\_F59\_ingel\_3ug\_chymo.raw

##### Fraction

3

##### Scan Feature

F3:5589

##### De Novo Score

97

##### ConfidenceScore

97

### m/z

535.2637

##### Mass

1602.7678

##### Charge

3

##### Retention Time

42.75

##### Predicted Retention Time

-

##### Area

1.543E+08

##### Parts Per Million

0.9

##### Fragmentation mode

HCD

##### Originating file

01 D:\separate\_stitch\_analyses\xle-disambiguation\20210325\_F59\_3ug\_DENOVO\_12.csv

#### Meta Information from PEAKS

##### Scan Identifier

F3:5785

##### Original sequence

T

V

L

H

Q

D

W

+15.99

L

D

G

K

E

Y

##### Posttranslational Modifications

Oxidation (HW)

##### Source File

D:\separate\_stitch\_analyses\xle-disambiguation\raw\20210323\_F1\_UM1\_Peng0013\_SA\_F59\_ingel\_3ug\_chymo.raw

##### Fraction

3

##### Scan Feature

F3:5894

##### De Novo Score

97

##### ConfidenceScore

97

### m/z

540.5956

##### Mass

1618.7627

##### Charge

3

##### Retention Time

31.45

##### Predicted Retention Time

-

##### Area

2.287E+07

##### Parts Per Million

1.4

##### Fragmentation mode

ETHCD

##### Originating file

01 D:\separate\_stitch\_analyses\xle-disambiguation\20210325\_F59\_3ug\_DENOVO\_12.csv

#### Meta Information from PEAKS

##### Scan Identifier

F3:8166

##### Original sequence

T

V

L

H

Q

D

W

L

D

G

K

E

Y

##### Posttranslational Modifications

##### Source File

D:\separate\_stitch\_analyses\xle-disambiguation\raw\20210323\_F1\_UM1\_Peng0013\_SA\_F59\_ingel\_3ug\_chymo.raw

##### Fraction

3

##### Scan Feature

-

##### De Novo Score

97

##### ConfidenceScore

97

### m/z

535.2651

##### Mass

1602.7678

##### Charge

3

##### Retention Time

45.57

##### Predicted Retention Time

-

##### Area

0

##### Parts Per Million

3.6

##### Fragmentation mode

HCD

##### Originating file

01 D:\separate\_stitch\_analyses\xle-disambiguation\20210325\_F59\_3ug\_DENOVO\_12.csv

#### Meta Information from PEAKS

##### Scan Identifier

F3:8472

##### Original sequence

T

V

L

H

Q

D

W

L

D

G

K

E

Y

##### Posttranslational Modifications

##### Source File

D:\separate\_stitch\_analyses\xle-disambiguation\raw\20210323\_F1\_UM1\_Peng0013\_SA\_F59\_ingel\_3ug\_chymo.raw

##### Fraction

3

##### Scan Feature

F3:5600

##### De Novo Score

97

##### ConfidenceScore

97

### m/z

535.2659

##### Mass

1602.7678

##### Charge

3

##### Retention Time

48.27

##### Predicted Retention Time

-

##### Area

8.584E+06

##### Parts Per Million

5.1

##### Fragmentation mode

HCD

##### Originating file

01 D:\separate\_stitch\_analyses\xle-disambiguation\20210325\_F59\_3ug\_DENOVO\_12.csv

#### Meta Information from PEAKS

##### Scan Identifier

F3:7490

##### Original sequence

T

V

L

H

Q

D

W

L

D

G

K

E

Y

##### Posttranslational Modifications

##### Source File

D:\separate\_stitch\_analyses\xle-disambiguation\raw\20210323\_F1\_UM1\_Peng0013\_SA\_F59\_ingel\_3ug\_chymo.raw

##### Fraction

3

##### Scan Feature

F3:14458

##### De Novo Score

97

##### ConfidenceScore

97

### m/z

802.3918

##### Mass

1602.7678

##### Charge

2

##### Retention Time

40.2

##### Predicted Retention Time

-

##### Area

1.044E+08

##### Parts Per Million

0.8

##### Fragmentation mode

ETHCD

##### Originating file

01 D:\separate\_stitch\_analyses\xle-disambiguation\20210325\_F59\_3ug\_DENOVO\_12.csv

#### Meta Information from PEAKS

##### Scan Identifier

F3:6890

##### Original sequence

T

V

L

H

Q

D

W

L

D

G

K

E

Y

##### Posttranslational Modifications

##### Source File

D:\separate\_stitch\_analyses\xle-disambiguation\raw\20210323\_F1\_UM1\_Peng0013\_SA\_F59\_ingel\_3ug\_chymo.raw

##### Fraction

3

##### Scan Feature

-

##### De Novo Score

96

##### ConfidenceScore

96

### m/z

802.3925

##### Mass

1602.7678

##### Charge

2

##### Retention Time

38.29

##### Predicted Retention Time

-

##### Area

0

##### Parts Per Million

1.7

##### Fragmentation mode

HCD

##### Originating file

01 D:\separate\_stitch\_analyses\xle-disambiguation\20210325\_F59\_3ug\_DENOVO\_12.csv

#### Meta Information from PEAKS

##### Scan Identifier

F3:8727

##### Original sequence

T

V

L

H

Q

D

W

L

D

G

K

E

Y

##### Posttranslational Modifications

##### Source File

D:\separate\_stitch\_analyses\xle-disambiguation\raw\20210323\_F1\_UM1\_Peng0013\_SA\_F59\_ingel\_3ug\_chymo.raw

##### Fraction

3

##### Scan Feature

F3:5600

##### De Novo Score

96

##### ConfidenceScore

96

### m/z

535.2659

##### Mass

1602.7678

##### Charge

3

##### Retention Time

48.27

##### Predicted Retention Time

-

##### Area

8.584E+06

##### Parts Per Million

5.1

##### Fragmentation mode

ETHCD

##### Originating file

01 D:\separate\_stitch\_analyses\xle-disambiguation\20210325\_F59\_3ug\_DENOVO\_12.csv

#### Meta Information from PEAKS

##### Scan Identifier

F3:6806

##### Original sequence

T

V

L

H

Q

D

W

+15.99

L

D

G

K

E

Y

##### Posttranslational Modifications

Oxidation (HW)

##### Source File

D:\separate\_stitch\_analyses\xle-disambiguation\raw\20210323\_F1\_UM1\_Peng0013\_SA\_F59\_ingel\_3ug\_chymo.raw

##### Fraction

3

##### Scan Feature

F3:5895

##### De Novo Score

96

##### ConfidenceScore

96

### m/z

540.5959

##### Mass

1618.7627

##### Charge

3

##### Retention Time

37.81

##### Predicted Retention Time

-

##### Area

2.36E+06

##### Parts Per Million

2.1

##### Fragmentation mode

HCD

##### Originating file

01 D:\separate\_stitch\_analyses\xle-disambiguation\20210325\_F59\_3ug\_DENOVO\_12.csv

#### Meta Information from PEAKS

##### Scan Identifier

F3:6879

##### Original sequence

T

V

L

H

Q

D

W

L

D

G

K

E

Y

##### Posttranslational Modifications

##### Source File

D:\separate\_stitch\_analyses\xle-disambiguation\raw\20210323\_F1\_UM1\_Peng0013\_SA\_F59\_ingel\_3ug\_chymo.raw

##### Fraction

3

##### Scan Feature

F3:5612

##### De Novo Score

96

##### ConfidenceScore

96

### m/z

535.2669

##### Mass

1602.7678

##### Charge

3

##### Retention Time

38.28

##### Predicted Retention Time

-

##### Area

1.498E+09

##### Parts Per Million

6.9

##### Fragmentation mode

HCD

##### Originating file

01 D:\separate\_stitch\_analyses\xle-disambiguation\20210325\_F59\_3ug\_DENOVO\_12.csv

#### Meta Information from PEAKS

##### Scan Identifier

F3:6124

##### Original sequence

T

V

L

H

Q

D

W

+15.99

L

D

G

K

E

Y

##### Posttranslational Modifications

Oxidation (HW)

##### Source File

D:\separate\_stitch\_analyses\xle-disambiguation\raw\20210323\_F1\_UM1\_Peng0013\_SA\_F59\_ingel\_3ug\_chymo.raw

##### Fraction

3

##### Scan Feature

F3:5894

##### De Novo Score

96

##### ConfidenceScore

96

### m/z

540.5956

##### Mass

1618.7627

##### Charge

3

##### Retention Time

31.45

##### Predicted Retention Time

-

##### Area

2.287E+07

##### Parts Per Million

1.4

##### Fragmentation mode

HCD

##### Originating file

01 D:\separate\_stitch\_analyses\xle-disambiguation\20210325\_F59\_3ug\_DENOVO\_12.csv

#### Meta Information from PEAKS

##### Scan Identifier

F3:6412

##### Original sequence

T

V

L

H

Q

D

W

+15.99

L

D

G

K

E

Y

##### Posttranslational Modifications

Oxidation (HW)

##### Source File

D:\separate\_stitch\_analyses\xle-disambiguation\raw\20210323\_F1\_UM1\_Peng0013\_SA\_F59\_ingel\_3ug\_chymo.raw

##### Fraction

3

##### Scan Feature

F3:5894

##### De Novo Score

96

##### ConfidenceScore

96

### m/z

540.5956

##### Mass

1618.7627

##### Charge

3

##### Retention Time

31.45

##### Predicted Retention Time

-

##### Area

2.287E+07

##### Parts Per Million

1.4

##### Fragmentation mode

ETHCD

##### Originating file

01 D:\separate\_stitch\_analyses\xle-disambiguation\20210325\_F59\_3ug\_DENOVO\_12.csv

#### Meta Information from PEAKS

##### Scan Identifier

F3:8667

##### Original sequence

T

V

L

H

Q

D

W

L

D

G

K

E

Y

##### Posttranslational Modifications

##### Source File

D:\separate\_stitch\_analyses\xle-disambiguation\raw\20210323\_F1\_UM1\_Peng0013\_SA\_F59\_ingel\_3ug\_chymo.raw

##### Fraction

3

##### Scan Feature

F3:5600

##### De Novo Score

96

##### ConfidenceScore

96

### m/z

535.2659

##### Mass

1602.7678

##### Charge

3

##### Retention Time

48.27

##### Predicted Retention Time

-

##### Area

8.584E+06

##### Parts Per Million

5.1

##### Fragmentation mode

ETHCD

##### Originating file

01 D:\separate\_stitch\_analyses\xle-disambiguation\20210325\_F59\_3ug\_DENOVO\_12.csv

#### Meta Information from PEAKS

##### Scan Identifier

F3:9201

##### Original sequence

T

V

L

H

Q

D

W

L

D

G

K

E

Y

##### Posttranslational Modifications

##### Source File

D:\separate\_stitch\_analyses\xle-disambiguation\raw\20210323\_F1\_UM1\_Peng0013\_SA\_F59\_ingel\_3ug\_chymo.raw

##### Fraction

3

##### Scan Feature

F3:5599

##### De Novo Score

96

##### ConfidenceScore

96

### m/z

535.2659

##### Mass

1602.7678

##### Charge

3

##### Retention Time

50.75

##### Predicted Retention Time

-

##### Area

4.159E+06

##### Parts Per Million

5

##### Fragmentation mode

ETHCD

##### Originating file

01 D:\separate\_stitch\_analyses\xle-disambiguation\20210325\_F59\_3ug\_DENOVO\_12.csv

#### Meta Information from PEAKS

##### Scan Identifier

F3:8963

##### Original sequence

T

V

L

H

Q

D

W

L

D

G

K

E

Y

##### Posttranslational Modifications

##### Source File

D:\separate\_stitch\_analyses\xle-disambiguation\raw\20210323\_F1\_UM1\_Peng0013\_SA\_F59\_ingel\_3ug\_chymo.raw

##### Fraction

3

##### Scan Feature

F3:5599

##### De Novo Score

95

##### ConfidenceScore

95

### m/z

535.2659

##### Mass

1602.7678

##### Charge

3

##### Retention Time

50.75

##### Predicted Retention Time

-

##### Area

4.159E+06

##### Parts Per Million

5

##### Fragmentation mode

ETHCD

##### Originating file

01 D:\separate\_stitch\_analyses\xle-disambiguation\20210325\_F59\_3ug\_DENOVO\_12.csv
