## Supplementary material for "A handle on mass coincidence errors in *de novo* sequencing of antibodies by bottom-up proteomics": Combined_028.html

Details Combined\_028 | Stitch OverviewUndefined

### Read Combined\_028

#### Sequence (length=11)

JVKDYFPEPVT

#### Spectrum 7751? Spectrum 7751 The raw spectrum of this peptide as annotated by Hecklib. The fragments are coloured according to ion type (see legend). Any peaks with a star '\*' as text can be hovered over to see the full details, first the ion type second the mass shift type. By hovering over the amino acids in the peptide or ions in the legend the corresponding peaks are highlighted. By toggling the 'Unassigned' label you can turn the background (unassigned) peaks on or off in the plot. By updating the slider in the Ion legend you can update the spectrum to only show the top X% of the peaks with labels. The top X% means any peak that is within X% of the highest intensity. By dragging in the spectrum you can zoom in to a specific part of the spectrum and use 'Zoom Out' to get back to the original zoom level. The annotation of the spectrum is based on the given sequence in the peptides file and is done with different software so inconsistencies are likely. The peaks are annotated based on the given sequence, with 20 ppm tolerance.

Copy Data

##### Spectrum 7751 (TSV)

###### Preview

```
Loading example...
```

*Click on the button to copy the data to your clipboard.*

Mz MinMz MaxIntensity Max

WidthHeightPeptide font sizePeptide stroke widthSpectrum font sizeSpectrum stroke widthCompact peptide

Ion legend

wxyz

abcd

OtherUnassignedIon^Charge^_Position_Show for top:%

JVKDYFPEPVT

05.20e+51.04e+61.56e+62.08e+6

Zoom Out

y^+1^_1_y^+1^_2_y^+1^_2_y^+1^_3_y^+1^_3_c^+1^_3_c^+1^_3_c^+2^_6_y^+1^_4_c^+2^_7_y^+1^_4_c^+1^_4_c^+1^_4_c^+2^_8_y^+1^_5_z^+1^_5_y^+2^_9_y^+2^_9_y^+1^_5_c^+2^_9_c^+2^_10_c^+1^_5_c^+1^_5_y^+1^_6_c^+1^_6_y^+1^_7_z^+1^_7_y^+1^_7_c^+1^_7_c^+1^_7_z^+1^_8_y^+1^_8_c^+1^_8_y^+1^_9_z^+1^_9_y^+1^_9_c^+1^_9_z^+1^_10_c^+1^_10_y^+1^_10_c^+1^_10_

0790158123713161

Fragment Matches Table

Show background peaks

| Position | Ion type | Intensity | mz Theoretical | mz Error (Th) | mz Error (ppm) | Charge | Series Number |
| --- | --- | --- | --- | --- | --- | --- | --- |
| - | - | 1395 | 120.1 | - | - | 0 | - |
| - | - | 1829 | 120.1 | - | - | 0 | - |
| 11 | y | 2.872E+04 | 120.1 | 0.0002992 | 2.492 | +1 | 1 |
| - | - | 2718 | 120.1 | - | - | 0 | - |
| - | - | 2.331E+04 | 129.1 | - | - | 0 | - |
| - | - | 1422 | 130.1 | - | - | 0 | - |
| - | - | 1526 | 136.1 | - | - | 0 | - |
| - | - | 1361 | 162.8 | - | - | 0 | - |
| - | - | 1556 | 163.1 | - | - | 0 | - |
| - | - | 1276 | 163.2 | - | - | 0 | - |
| - | - | 4.777E+04 | 169.1 | - | - | 0 | - |
| - | - | 4200 | 170.1 | - | - | 0 | - |
| - | - | 1472 | 171.4 | - | - | 0 | - |
| - | - | 1.372E+04 | 173.5 | - | - | 0 | - |
| - | - | 1523 | 177 | - | - | 0 | - |
| - | - | 6.619E+04 | 185.2 | - | - | 0 | - |
| - | - | 7821 | 186.2 | - | - | 0 | - |
| - | - | 9.392E+04 | 197.1 | - | - | 0 | - |
| - | - | 9738 | 198.1 | - | - | 0 | - |
| - | - | 2703 | 199.1 | - | - | 0 | - |
| 10 | y | 2105 | 201.1 | 0.0001358 | 0.6752 | +1 | 2 |
| - | - | 5.179E+04 | 213.2 | - | - | 0 | - |
| - | - | 4509 | 214.2 | - | - | 0 | - |
| 10 | y | 7684 | 219.1 | 0.0002065 | 0.9423 | +1 | 2 |
| - | - | 2.599E+04 | 227.1 | - | - | 0 | - |
| - | - | 3054 | 228.1 | - | - | 0 | - |
| - | - | 2096 | 228.2 | - | - | 0 | - |
| - | - | 2960 | 244.1 | - | - | 0 | - |
| - | - | 1558 | 246.2 | - | - | 0 | - |
| - | - | 1588 | 247.4 | - | - | 0 | - |
| - | - | 1506 | 263.6 | - | - | 0 | - |
| - | - | 2530 | 279.1 | - | - | 0 | - |
| - | - | 3727 | 280.2 | - | - | 0 | - |
| 9 | y | 2.49E+04 | 298.2 | 0.000503 | 1.687 | +1 | 3 |
| - | - | 3984 | 299.2 | - | - | 0 | - |
| - | - | 5420 | 315.3 | - | - | 0 | - |
| 9 | y | 3.208E+05 | 316.2 | 0.0004974 | 1.573 | +1 | 3 |
| - | - | 5.075E+04 | 317.2 | - | - | 0 | - |
| - | - | 5350 | 318.2 | - | - | 0 | - |
| - | - | 5.616E+04 | 324.2 | - | - | 0 | - |
| - | - | 1.048E+04 | 325.2 | - | - | 0 | - |
| 3 | c | 6386 | 341.3 | 0.0001654 | 0.4848 | +1 | 3 |
| - | - | 4.532E+04 | 357.3 | - | - | 0 | - |
| 3 | c | 2.288E+04 | 358.3 | 0.001298 | 3.623 | +1 | 3 |
| - | - | 3262 | 359.3 | - | - | 0 | - |
| - | - | 4613 | 374.2 | - | - | 0 | - |
| 6 | c | 2178 | 383.7 | 0.0001041 | 0.2712 | +2 | 6 |
| - | - | 5721 | 395.2 | - | - | 0 | - |
| - | - | 8005 | 407.2 | - | - | 0 | - |
| - | - | 2617 | 418.2 | - | - | 0 | - |
| - | - | 3.021E+04 | 423.2 | - | - | 0 | - |
| - | - | 7581 | 424.2 | - | - | 0 | - |
| - | - | 1.086E+04 | 426.2 | - | - | 0 | - |
| - | - | 2750 | 427.2 | - | - | 0 | - |
| 8 | y | 2060 | 427.2 | 0.002435 | 5.7 | +1 | 4 |
| 7 | c | 9576 | 432.2 | 9.483E-05 | 0.2194 | +2 | 7 |
| - | - | 5360 | 432.7 | - | - | 0 | - |
| - | - | 2257 | 439.2 | - | - | 0 | - |
| 8 | y | 1.157E+04 | 445.2 | 0.0008121 | 1.824 | +1 | 4 |
| 4 | c | 1.387E+04 | 456.3 | 0.001024 | 2.244 | +1 | 4 |
| - | - | 2459 | 457.3 | - | - | 0 | - |
| - | - | 4201 | 471.2 | - | - | 0 | - |
| - | - | 9070 | 472.3 | - | - | 0 | - |
| 4 | c | 2.644E+04 | 473.3 | 0.0002622 | 0.5539 | +1 | 4 |
| - | - | 4557 | 474.3 | - | - | 0 | - |
| - | - | 7742 | 482.8 | - | - | 0 | - |
| - | - | 4432 | 483.3 | - | - | 0 | - |
| - | - | 2202 | 487.8 | - | - | 0 | - |
| - | - | 1983 | 488.7 | - | - | 0 | - |
| - | - | 2907 | 489.2 | - | - | 0 | - |
| 8 | c | 3.006E+04 | 496.8 | 0.0004963 | 0.999 | +2 | 8 |
| - | - | 1.358E+04 | 497.3 | - | - | 0 | - |
| - | - | 5097 | 497.8 | - | - | 0 | - |
| - | - | 1869 | 509.2 | - | - | 0 | - |
| - | - | 2357 | 511.3 | - | - | 0 | - |
| - | - | 1894 | 515.2 | - | - | 0 | - |
| - | - | 2155 | 521.3 | - | - | 0 | - |
| - | - | 2539 | 522.3 | - | - | 0 | - |
| - | - | 2324 | 523.3 | - | - | 0 | - |
| 7 | y | 8845 | 524.3 | 0.002314 | 4.414 | +1 | 5 |
| - | - | 2686 | 525.3 | - | - | 0 | - |
| 7 | z | 2015 | 526.3 | 0.004736 | 9 | +1 | 5 |
| - | - | 5216 | 531.3 | - | - | 0 | - |
| - | - | 4300 | 531.8 | - | - | 0 | - |
| - | - | 4272 | 534.3 | - | - | 0 | - |
| - | - | 3384 | 536.3 | - | - | 0 | - |
| - | - | 2536 | 536.8 | - | - | 0 | - |
| 3 | y | 5033 | 539.3 | 0.007052 | 13.08 | +2 | 9 |
| 3 | y | 2105 | 539.8 | 0.009729 | 18.03 | +2 | 9 |
| - | - | 3546 | 541.3 | - | - | 0 | - |
| 7 | y | 1.451E+05 | 542.3 | 0.0005995 | 1.105 | +1 | 5 |
| - | - | 4.618E+04 | 543.3 | - | - | 0 | - |
| - | - | 8639 | 544.3 | - | - | 0 | - |
| 9 | c | 9.886E+04 | 545.3 | 0.000451 | 0.8271 | +2 | 9 |
| - | - | 6.418E+04 | 545.8 | - | - | 0 | - |
| - | - | 2.332E+04 | 546.3 | - | - | 0 | - |
| - | - | 5299 | 546.8 | - | - | 0 | - |
| - | - | 1982 | 547.5 | - | - | 0 | - |
| - | - | 5.378E+04 | 554.3 | - | - | 0 | - |
| - | - | 1.99E+04 | 555.3 | - | - | 0 | - |
| - | - | 4501 | 556.3 | - | - | 0 | - |
| - | - | 2304 | 567.3 | - | - | 0 | - |
| - | - | 5323 | 570.3 | - | - | 0 | - |
| - | - | 1909 | 571.3 | - | - | 0 | - |
| - | - | 4424 | 571.8 | - | - | 0 | - |
| - | - | 4.027E+04 | 580.8 | - | - | 0 | - |
| - | - | 2.802E+04 | 581.3 | - | - | 0 | - |
| - | - | 9911 | 581.8 | - | - | 0 | - |
| - | - | 2446 | 585.8 | - | - | 0 | - |
| - | - | 5392 | 591.4 | - | - | 0 | - |
| - | - | 2221 | 592.4 | - | - | 0 | - |
| 10 | c | 2.984E+04 | 594.8 | 0.0005458 | 0.9176 | +2 | 10 |
| - | - | 2.256E+04 | 595.3 | - | - | 0 | - |
| - | - | 7125 | 595.8 | - | - | 0 | - |
| - | - | 1932 | 603.4 | - | - | 0 | - |
| 5 | c | 5.141E+04 | 619.3 | 0.000348 | 0.5618 | +1 | 5 |
| - | - | 2.038E+04 | 620.3 | - | - | 0 | - |
| - | - | 2895 | 621.4 | - | - | 0 | - |
| - | - | 1.045E+05 | 635.4 | - | - | 0 | - |
| 5 | c | 1.327E+05 | 636.4 | 0.0005053 | 0.7941 | +1 | 5 |
| - | - | 3.941E+04 | 637.4 | - | - | 0 | - |
| - | - | 2673 | 638.3 | - | - | 0 | - |
| - | - | 5721 | 638.4 | - | - | 0 | - |
| - | - | 1.461E+04 | 645.3 | - | - | 0 | - |
| - | - | 1.166E+04 | 645.8 | - | - | 0 | - |
| - | - | 5358 | 646.3 | - | - | 0 | - |
| - | - | 5034 | 651.3 | - | - | 0 | - |
| - | - | 3348 | 652.3 | - | - | 0 | - |
| - | - | 4732 | 653.3 | - | - | 0 | - |
| - | - | 2.368E+04 | 654.3 | - | - | 0 | - |
| - | - | 1.825E+04 | 654.9 | - | - | 0 | - |
| - | - | 8343 | 655.4 | - | - | 0 | - |
| - | - | 4456 | 681.3 | - | - | 0 | - |
| - | - | 8404 | 682.4 | - | - | 0 | - |
| - | - | 5067 | 688.3 | - | - | 0 | - |
| 6 | y | 7.672E+04 | 689.4 | 0.0003618 | 0.5249 | +1 | 6 |
| - | - | 3.026E+04 | 690.4 | - | - | 0 | - |
| - | - | 4676 | 691.4 | - | - | 0 | - |
| - | - | 1.282E+04 | 738.4 | - | - | 0 | - |
| - | - | 4.181E+04 | 739.4 | - | - | 0 | - |
| - | - | 6.934E+04 | 740.4 | - | - | 0 | - |
| - | - | 2.321E+04 | 741.4 | - | - | 0 | - |
| - | - | 5033 | 742.4 | - | - | 0 | - |
| - | - | 2253 | 744.4 | - | - | 0 | - |
| - | - | 6454 | 748.4 | - | - | 0 | - |
| - | - | 3780 | 749.4 | - | - | 0 | - |
| - | - | 3051 | 750.4 | - | - | 0 | - |
| 6 | c | 2.076E+05 | 766.4 | 0.0002324 | 0.3032 | +1 | 6 |
| - | - | 9.655E+04 | 767.4 | - | - | 0 | - |
| - | - | 2628 | 767.5 | - | - | 0 | - |
| - | - | 2.212E+04 | 768.4 | - | - | 0 | - |
| - | - | 4.622E+04 | 780.4 | - | - | 0 | - |
| - | - | 2.241E+04 | 781.4 | - | - | 0 | - |
| - | - | 6701 | 782.4 | - | - | 0 | - |
| - | - | 3139 | 823.4 | - | - | 0 | - |
| 5 | y | 2807 | 834.4 | 0.01657 | 19.86 | +1 | 7 |
| 5 | z | 5874 | 836.4 | 0.001534 | 1.834 | +1 | 7 |
| - | - | 3347 | 836.5 | - | - | 0 | - |
| - | - | 2824 | 848.4 | - | - | 0 | - |
| - | - | 1.407E+04 | 851.4 | - | - | 0 | - |
| 5 | y | 7.189E+04 | 852.4 | 0.0004667 | 0.5475 | +1 | 7 |
| - | - | 2.7E+04 | 853.4 | - | - | 0 | - |
| - | - | 8705 | 854.4 | - | - | 0 | - |
| 7 | c | 1.416E+04 | 863.5 | 0.000325 | 0.3764 | +1 | 7 |
| - | - | 8934 | 864.5 | - | - | 0 | - |
| - | - | 2.004E+04 | 877.4 | - | - | 0 | - |
| - | - | 1.026E+04 | 878.4 | - | - | 0 | - |
| - | - | 2297 | 878.5 | - | - | 0 | - |
| - | - | 1.96E+05 | 879.5 | - | - | 0 | - |
| 7 | c | 3.759E+05 | 880.5 | 0.001261 | 1.432 | +1 | 7 |
| - | - | 1.679E+05 | 881.5 | - | - | 0 | - |
| - | - | 3.795E+04 | 882.5 | - | - | 0 | - |
| - | - | 3883 | 883.5 | - | - | 0 | - |
| - | - | 6409 | 907.4 | - | - | 0 | - |
| - | - | 3279 | 934.5 | - | - | 0 | - |
| - | - | 2888 | 935.5 | - | - | 0 | - |
| - | - | 3502 | 948.4 | - | - | 0 | - |
| 4 | z | 5643 | 951.4 | 0.001361 | 1.431 | +1 | 8 |
| - | - | 9466 | 952.4 | - | - | 0 | - |
| - | - | 7544 | 953.4 | - | - | 0 | - |
| - | - | 4366 | 962.5 | - | - | 0 | - |
| - | - | 3725 | 963.5 | - | - | 0 | - |
| - | - | 1.829E+04 | 964.5 | - | - | 0 | - |
| - | - | 1.238E+04 | 965.5 | - | - | 0 | - |
| - | - | 2.172E+04 | 966.4 | - | - | 0 | - |
| - | - | 6398 | 966.5 | - | - | 0 | - |
| 4 | y | 4.424E+04 | 967.4 | 0.00147 | 1.519 | +1 | 8 |
| - | - | 3519 | 967.5 | - | - | 0 | - |
| - | - | 1.996E+04 | 968.4 | - | - | 0 | - |
| - | - | 6624 | 969.4 | - | - | 0 | - |
| - | - | 1.21E+04 | 974.5 | - | - | 0 | - |
| - | - | 5328 | 975.5 | - | - | 0 | - |
| - | - | 2.222E+04 | 976.5 | - | - | 0 | - |
| - | - | 1.692E+04 | 977.5 | - | - | 0 | - |
| - | - | 8509 | 978.5 | - | - | 0 | - |
| - | - | 4644 | 979.5 | - | - | 0 | - |
| - | - | 5480 | 990.6 | - | - | 0 | - |
| - | - | 4858 | 991.6 | - | - | 0 | - |
| 8 | c | 2.813E+05 | 992.5 | 2.926E-05 | 0.02948 | +1 | 8 |
| - | - | 1.699E+05 | 993.5 | - | - | 0 | - |
| - | - | 4.723E+04 | 994.5 | - | - | 0 | - |
| - | - | 3692 | 995.5 | - | - | 0 | - |
| - | - | 8408 | 1007 | - | - | 0 | - |
| - | - | 3092 | 1008 | - | - | 0 | - |
| - | - | 2.423E+04 | 1036 | - | - | 0 | - |
| - | - | 1.376E+04 | 1037 | - | - | 0 | - |
| - | - | 7120 | 1038 | - | - | 0 | - |
| - | - | 5634 | 1047 | - | - | 0 | - |
| - | - | 5361 | 1048 | - | - | 0 | - |
| - | - | 3012 | 1061 | - | - | 0 | - |
| - | - | 3058 | 1062 | - | - | 0 | - |
| - | - | 8408 | 1063 | - | - | 0 | - |
| - | - | 3712 | 1064 | - | - | 0 | - |
| - | - | 3208 | 1076 | - | - | 0 | - |
| 3 | y | 3891 | 1078 | 0.004647 | 4.313 | +1 | 9 |
| 3 | z | 7.845E+04 | 1080 | 1.099E-05 | 0.01018 | +1 | 9 |
| - | - | 4.988E+04 | 1081 | - | - | 0 | - |
| - | - | 2.041E+04 | 1082 | - | - | 0 | - |
| - | - | 5440 | 1083 | - | - | 0 | - |
| - | - | 8386 | 1090 | - | - | 0 | - |
| - | - | 5651 | 1091 | - | - | 0 | - |
| - | - | 2553 | 1092 | - | - | 0 | - |
| 3 | y | 6.678E+04 | 1096 | 0.0004245 | 0.3875 | +1 | 9 |
| - | - | 4.258E+04 | 1097 | - | - | 0 | - |
| - | - | 1.128E+04 | 1098 | - | - | 0 | - |
| - | - | 2464 | 1104 | - | - | 0 | - |
| 9 | c | 5.674E+04 | 1107 | 0.0009146 | 0.8265 | +1 | 9 |
| - | - | 3.755E+04 | 1108 | - | - | 0 | - |
| - | - | 1.043E+04 | 1109 | - | - | 0 | - |
| - | - | 5009 | 1118 | - | - | 0 | - |
| - | - | 3854 | 1119 | - | - | 0 | - |
| - | - | 5792 | 1125 | - | - | 0 | - |
| - | - | 5555 | 1126 | - | - | 0 | - |
| - | - | 1.76E+04 | 1147 | - | - | 0 | - |
| - | - | 1.809E+04 | 1148 | - | - | 0 | - |
| - | - | 1.155E+04 | 1149 | - | - | 0 | - |
| - | - | 2999 | 1161 | - | - | 0 | - |
| - | - | 1.063E+05 | 1162 | - | - | 0 | - |
| - | - | 7.97E+04 | 1163 | - | - | 0 | - |
| - | - | 5.116E+04 | 1164 | - | - | 0 | - |
| - | - | 1.817E+04 | 1165 | - | - | 0 | - |
| - | - | 4356 | 1166 | - | - | 0 | - |
| - | - | 4709 | 1173 | - | - | 0 | - |
| 2 | z | 3.831E+04 | 1179 | 0.001042 | 0.8842 | +1 | 10 |
| - | - | 2.631E+04 | 1180 | - | - | 0 | - |
| - | - | 8986 | 1181 | - | - | 0 | - |
| - | - | 2648 | 1182 | - | - | 0 | - |
| 10 | c | 2.349E+04 | 1189 | 0.001031 | 0.8677 | +1 | 10 |
| - | - | 1.956E+04 | 1190 | - | - | 0 | - |
| - | - | 4.387E+04 | 1191 | - | - | 0 | - |
| - | - | 3.105E+04 | 1192 | - | - | 0 | - |
| - | - | 1.17E+04 | 1193 | - | - | 0 | - |
| 2 | y | 6035 | 1195 | 0.006723 | 5.628 | +1 | 10 |
| - | - | 5889 | 1196 | - | - | 0 | - |
| - | - | 2651 | 1205 | - | - | 0 | - |
| 10 | c | 5.188E+05 | 1206 | 0.0004808 | 0.3988 | +1 | 10 |
| - | - | 3.583E+05 | 1207 | - | - | 0 | - |
| - | - | 1.972E+05 | 1208 | - | - | 0 | - |
| - | - | 6.971E+04 | 1209 | - | - | 0 | - |
| - | - | 1.829E+04 | 1210 | - | - | 0 | - |
| - | - | 2519 | 1211 | - | - | 0 | - |
| - | - | 4784 | 1217 | - | - | 0 | - |
| - | - | 1.159E+04 | 1220 | - | - | 0 | - |
| - | - | 8988 | 1221 | - | - | 0 | - |
| - | - | 4413 | 1230 | - | - | 0 | - |
| - | - | 5947 | 1231 | - | - | 0 | - |
| - | - | 4986 | 1232 | - | - | 0 | - |
| - | - | 4.227E+04 | 1237 | - | - | 0 | - |
| - | - | 3.258E+04 | 1238 | - | - | 0 | - |
| - | - | 1.174E+04 | 1239 | - | - | 0 | - |
| - | - | 2.8E+04 | 1245 | - | - | 0 | - |
| - | - | 2.411E+04 | 1246 | - | - | 0 | - |
| - | - | 2.354E+04 | 1247 | - | - | 0 | - |
| - | - | 1.916E+04 | 1248 | - | - | 0 | - |
| - | - | 1.445E+05 | 1249 | - | - | 0 | - |
| - | - | 1.118E+05 | 1250 | - | - | 0 | - |
| - | - | 4.203E+04 | 1251 | - | - | 0 | - |
| - | - | 7540 | 1252 | - | - | 0 | - |
| - | - | 2.323E+04 | 1253 | - | - | 0 | - |
| - | - | 1.417E+04 | 1254 | - | - | 0 | - |
| - | - | 6679 | 1255 | - | - | 0 | - |
| - | - | 3.483E+04 | 1263 | - | - | 0 | - |
| - | - | 2.928E+04 | 1264 | - | - | 0 | - |
| - | - | 5.969E+04 | 1265 | - | - | 0 | - |
| - | - | 3.941E+04 | 1266 | - | - | 0 | - |
| - | - | 1.43E+04 | 1267 | - | - | 0 | - |
| - | - | 3791 | 1273 | - | - | 0 | - |
| - | - | 2189 | 1274 | - | - | 0 | - |
| - | - | 2742 | 1275 | - | - | 0 | - |
| - | - | 1.014E+05 | 1281 | - | - | 0 | - |
| - | - | 7.281E+04 | 1282 | - | - | 0 | - |
| - | - | 3.399E+04 | 1283 | - | - | 0 | - |
| - | - | 4409 | 1284 | - | - | 0 | - |
| - | - | 5318 | 1290 | - | - | 0 | - |
| - | - | 3.859E+04 | 1291 | - | - | 0 | - |
| - | - | 5.435E+05 | 1292 | - | - | 0 | - |
| - | - | 4.191E+05 | 1293 | - | - | 0 | - |
| - | - | 1.767E+05 | 1294 | - | - | 0 | - |
| - | - | 2.807E+04 | 1295 | - | - | 0 | - |
| - | - | 4773 | 1306 | - | - | 0 | - |
| - | - | 1.506E+04 | 1307 | - | - | 0 | - |
| - | - | 4.311E+05 | 1308 | - | - | 0 | - |
| - | - | 2.06E+06 | 1309 | - | - | 0 | - |
| - | - | 1.498E+06 | 1310 | - | - | 0 | - |
| - | - | 5.659E+05 | 1311 | - | - | 0 | - |
| - | - | 8.046E+04 | 1312 | - | - | 0 | - |
| - | - | 5126 | 1324 | - | - | 0 | - |
| - | - | 2421 | 1325 | - | - | 0 | - |
| - | - | 5685 | 1341 | - | - | 0 | - |
| - | - | 3504 | 1342 | - | - | 0 | - |
| - | - | 2183 | 3130 | - | - | 0 | - |

m/z Charge Intensity FragmentType MassShift Position
120.05550384521484 0 1394.7797
120.06147003173828 0 1828.9865
120.0658187866211 0 28720.49 y 10
120.08122253417969 0 2718.4744
129.10250854492188 0 23313.232
130.106201171875 0 1422.1536
136.07627868652344 0 1525.5946
162.7814178466797 0 1360.624
163.09112548828125 0 1556.3225
163.20932006835938 0 1276.3848
169.13385009765625 0 47767.996
170.13748168945312 0 4199.8857
171.3583984375 0 1471.5151
173.4501953125 0 13715.871
177.00355529785156 0 1522.9626
185.1651153564453 0 66190.27
186.16847229003906 0 7821.1895
197.1287384033203 0 93924.11
198.1322784423828 0 9738.118
199.10838317871094 0 2703.4846
201.12350463867188 0 2104.9187 y Water loss 9
213.1600799560547 0 51794.69
214.1635284423828 0 4508.8354
219.13414001464844 0 7683.8066 y 9
227.1029815673828 0 25994.844
228.1061553955078 0 3053.7524
228.17105102539062 0 2095.7825
244.1297149658203 0 2959.5908
246.23353576660156 0 1558.1565
247.388671875 0 1588.4385
263.5632019042969 0 1505.8423
279.0967712402344 0 2529.73
280.1663513183594 0 3727.0383
298.1766357421875 0 24899.715 y Water loss 8
299.1804504394531 0 3983.5786
315.27557373046875 0 5420.3994
316.18719482421875 0 320782.44 y 8
317.1904296875 0 50745.31
318.1924743652344 0 5349.6157
324.156005859375 0 56156.67
325.1589660644531 0 10480.999
341.2548828125 0 6386.344 c Ammonia loss 2
357.2737731933594 0 45316.117
358.27996826171875 0 22883.48 c 2
359.2836608886719 0 3261.8655
374.1721496582031 0 4613.131
383.7102355957031 0 2178.0835 c Ammonia loss 5
395.2286376953125 0 5720.5566
407.1932067871094 0 8004.726
418.2406921386719 0 2616.869
423.2242126464844 0 30213.084
424.22674560546875 0 7581.3335
426.16632080078125 0 10860.673
427.1688537597656 0 2750.1982
427.2211608886719 0 2060.002 y Water loss 7
432.23681640625 0 9576.174 c Ammonia loss 6
432.7380676269531 0 5359.765
439.208251953125 0 2256.9922
445.2301025390625 0 11568.812 y 7
456.2826843261719 0 13869.803 c Ammonia loss 3
457.2843933105469 0 2459.244
471.2245788574219 0 4201.0654
472.3008728027344 0 9069.801
473.3084716796875 0 26438.404 c 3
474.3119812011719 0 4557.394
482.76214599609375 0 7742.2505
483.2630310058594 0 4431.7173
487.75421142578125 0 2201.866
488.74530029296875 0 1983.2256
489.2431335449219 0 2906.6252
496.7585144042969 0 30064.37 c Ammonia loss 7
497.25994873046875 0 13584.228
497.76116943359375 0 5097.1265
509.2431640625 0 1869.2385
511.2530517578125 0 2356.533
515.1605834960938 0 1894.406
521.2779541015625 0 2154.8423
522.284423828125 0 2538.988
523.2926025390625 0 2324.4114
524.2738037109375 0 8844.551 y Water loss 6
525.2750854492188 0 2685.964
526.26806640625 0 2015.1046 z 6
531.2901611328125 0 5216.225
531.7880249023438 0 4299.742
534.2578735351562 0 4271.863
536.3129272460938 0 3384.2944
536.7796630859375 0 2535.5046
539.2591552734375 0 5033.223 y Water loss 2
539.7679443359375 0 2105.2903 y Ammonia loss 2
541.2736206054688 0 3546.3005
542.2826538085938 0 145102.75 y 6
543.2854614257812 0 46178.25
544.2874145507812 0 8638.877
545.2848510742188 0 98856.74 c Ammonia loss 8
545.786376953125 0 64182.223
546.287841796875 0 23315.562
546.7906494140625 0 5299.1323
547.4794921875 0 1982.0153
554.26123046875 0 53775.04
555.264404296875 0 19902.06
556.2680053710938 0 4501.038
567.347900390625 0 2304.2405
570.291748046875 0 5322.563
571.2920532226562 0 1908.7125
571.8165283203125 0 4424.1353
580.82177734375 0 40269.496
581.3231811523438 0 28018.443
581.8253173828125 0 9911.37
585.8131103515625 0 2446.391
591.3504638671875 0 5392.372
592.3528442382812 0 2221.2698
594.8191528320312 0 29835.887 c Ammonia loss 9
595.3206787109375 0 22563.13
595.8221435546875 0 7125.4893
603.3500366210938 0 1931.6316
619.3453369140625 0 51406.88 c Ammonia loss 4
620.3487548828125 0 20384.195
621.3507690429688 0 2895.042
635.364990234375 0 104480.11
636.3710327148438 0 132738.84 c 4
637.3744506835938 0 39406.09
638.3218383789062 0 2672.7761
638.3763427734375 0 5720.961
645.3424682617188 0 14609.887
645.8433227539062 0 11661.38
646.34619140625 0 5357.8716
651.3140869140625 0 5033.954
652.2646484375 0 3348.0637
653.3274536132812 0 4731.879
654.3471069335938 0 23682.91
654.8502807617188 0 18246.715
655.3506469726562 0 8342.737
681.32275390625 0 4456.1084
682.3778076171875 0 8404.195
688.3414916992188 0 5066.5825
689.350830078125 0 76720.37 y 5
690.35400390625 0 30260.79
691.356689453125 0 4676.2295
738.418212890625 0 12822.567
739.426513671875 0 41808.406
740.4329833984375 0 69337.34
741.4368286132812 0 23205.602
742.4415893554688 0 5032.601
744.3547973632812 0 2252.6697
748.4019775390625 0 6453.6396
749.402587890625 0 3780.0725
750.3820190429688 0 3050.5518
766.4136352539062 0 207644.84 c Ammonia loss 5
767.4165649414062 0 96546.445
767.4993896484375 0 2628.057
768.4197387695312 0 22116.773
780.3567504882812 0 46218.113
781.3594360351562 0 22410.154
782.3633422851562 0 6700.845
823.4229125976562 0 3138.9055
834.3866577148438 0 2807.1902 y Water loss 4
836.3966064453125 0 5874.033 z 4
836.4801635742188 0 3346.8064
848.3775634765625 0 2823.6084
851.4063110351562 0 14070.174
852.413330078125 0 71893.1 y 4
853.4168090820312 0 26999.996
854.4186401367188 0 8705.122
863.4664916992188 0 14157.719 c Ammonia loss 6
864.4680786132812 0 8934.355
877.4088745117188 0 20039.25
878.4102783203125 0 10258.181
878.4948120117188 0 2297.1665
879.4849853515625 0 196040.7
880.491455078125 0 375885.12 c 6
881.4952392578125 0 167932.72
882.4984741210938 0 37945.492
883.4981689453125 0 3882.9011
907.4220581054688 0 6409.1675
934.4860229492188 0 3278.9402
935.482177734375 0 2887.5886
948.4155883789062 0 3501.8337
951.420654296875 0 5642.713 z 3
952.4288940429688 0 9466.351
953.4336547851562 0 7544.1025
962.5370483398438 0 4365.504
963.5339965820312 0 3725.0696
964.5142211914062 0 18293.06
965.518798828125 0 12384.824
966.4315185546875 0 21721.707
966.5302734375 0 6397.595
967.4392700195312 0 44241.715 y 3
967.5487060546875 0 3519.269
968.4434204101562 0 19958.848
969.4454956054688 0 6623.756
974.4963989257812 0 12101.471
975.4996948242188 0 5328.232
976.4788208007812 0 22220.314
977.48095703125 0 16920.273
978.485107421875 0 8508.607
979.4976196289062 0 4643.7354
990.5514526367188 0 5480.36
991.5576782226562 0 4858.101
992.5087890625 0 281339.25 c Ammonia loss 7
993.5119018554688 0 169941.23
994.5149536132812 0 47234.78
995.5125122070312 0 3692.1326
1007.4929809570312 0 8407.533
1008.4984741210938 0 3091.781
1035.5277099609375 0 24230.014
1036.5330810546875 0 13764.164
1037.5364990234375 0 7119.765
1046.5555419921875 0 5633.973
1047.5579833984375 0 5361.316
1060.5377197265625 0 3012.4104
1061.557861328125 0 3058.2144
1062.5712890625 0 8408.428
1063.577392578125 0 3711.8662
1075.55029296875 0 3208.1743
1077.52978515625 0 3890.743 y Water loss 2
1079.5169677734375 0 78452.35 z 2
1080.5194091796875 0 49875.93
1081.5228271484375 0 20408.275
1082.5308837890625 0 5440.325
1089.621826171875 0 8386.119
1090.578369140625 0 5651.391
1091.60693359375 0 2553.2417
1095.5352783203125 0 66775.09 y 2
1096.5380859375 0 42579.152
1097.54248046875 0 11281.832
1103.6343994140625 0 2464.154
1106.587158203125 0 56740.883 c 8
1107.5906982421875 0 37552.344
1108.5938720703125 0 10428.353
1117.6488037109375 0 5009.3247
1118.6552734375 0 3853.6868
1124.5345458984375 0 5791.8696
1125.542236328125 0 5554.6094
1146.63232421875 0 17595.008
1147.6417236328125 0 18090.639
1148.64697265625 0 11547.0205
1160.632080078125 0 2999.1514
1161.642333984375 0 106294.8
1162.6468505859375 0 79700.39
1163.6539306640625 0 51156.41
1164.660400390625 0 18170.506
1165.6661376953125 0 4356.4873
1172.631591796875 0 4708.521
1178.5843505859375 0 38313.18 z 1
1179.587158203125 0 26307.297
1180.5931396484375 0 8986.207
1181.58447265625 0 2647.7798
1188.62890625 0 23490.48 c Ammonia loss 9
1189.63232421875 0 19563.725
1190.6434326171875 0 43865.895
1191.64794921875 0 31053.678
1192.654052734375 0 11703.853
1194.61083984375 0 6034.886 y 1
1195.60302734375 0 5889.3267
1204.6422119140625 0 2650.9124
1205.656005859375 0 518795.34 c 9
1206.6591796875 0 358308.4
1207.664306640625 0 197190.42
1208.67138671875 0 69708.336
1209.6783447265625 0 18288.803
1210.6727294921875 0 2519.2993
1216.6287841796875 0 4784.491
1219.649658203125 0 11585.792
1220.6517333984375 0 8987.709
1229.6744384765625 0 4412.959
1230.6552734375 0 5946.869
1231.6575927734375 0 4986.188
1236.6739501953125 0 42265.508
1237.6719970703125 0 32580.88
1238.6700439453125 0 11737.481
1244.67724609375 0 28002.256
1245.6807861328125 0 24112.705
1246.6888427734375 0 23540.312
1247.68505859375 0 19162.896
1248.6534423828125 0 144538.2
1249.6590576171875 0 111778.055
1250.6640625 0 42031.84
1251.670166015625 0 7540.016
1252.6318359375 0 23233.287
1253.6346435546875 0 14166.92
1254.6357421875 0 6678.656
1262.6883544921875 0 34832.37
1263.691162109375 0 29284.95
1264.7020263671875 0 59690.605
1265.7049560546875 0 39410.574
1266.7012939453125 0 14299.179
1272.6678466796875 0 3790.9143
1273.6658935546875 0 2188.5874
1274.677490234375 0 2742.4685
1280.6990966796875 0 101449.66
1281.702392578125 0 72808.3
1282.7056884765625 0 33991.03
1283.701416015625 0 4409.4834
1289.658447265625 0 5317.9272
1290.6800537109375 0 38588.33
1291.66845703125 0 543479.8
1292.6717529296875 0 419100.53
1293.6734619140625 0 176746.77
1294.6748046875 0 28074.322
1305.6754150390625 0 4772.5312
1306.6767578125 0 15060.069
1307.6866455078125 0 431065.56
1308.69384765625 0 2060227.4
1309.697265625 0 1497746.6
1310.6998291015625 0 565861.25
1311.703125 0 80463.81
1323.6531982421875 0 5125.8604
1324.6605224609375 0 2420.7815
1340.68212890625 0 5685.444
1341.69677734375 0 3504.2087
3129.883544921875 0 2182.5596

Spectrum Details

|  |  |
| --- | --- |
| Matched peaks? Matched peaks The total absolute number of peaks matched. Additionally in brackets the total fraction of peaks matched and the total number of peaks is shown. | 41 (13.18% of 311) |
| FDR? FDR The false discovery rate estimated for this peptide. It is calculated by matching all theoretical fragments with a non-integer shift with the raw peaks for this spectrum. This is done with 40 different shifts. The resulting percentage is the average number of annotated peaks over the number of annotated peaks with the correct spectrum. | 2.85% |
| Satellite FDR? Satellite FDR See the FDR for details on its calculation. This satellite ion specific FDR only contains the satellite ions (d/w) for I/L/J positions. | - |
| PSM Score? PSM Score The PSM Score as given by Hecklib to this annotated spectrum. It is shown with three significant figures. | 487 |

#### Spectrum 7925? Spectrum 7925 The raw spectrum of this peptide as annotated by Hecklib. The fragments are coloured according to ion type (see legend). Any peaks with a star '\*' as text can be hovered over to see the full details, first the ion type second the mass shift type. By hovering over the amino acids in the peptide or ions in the legend the corresponding peaks are highlighted. By toggling the 'Unassigned' label you can turn the background (unassigned) peaks on or off in the plot. By updating the slider in the Ion legend you can update the spectrum to only show the top X% of the peaks with labels. The top X% means any peak that is within X% of the highest intensity. By dragging in the spectrum you can zoom in to a specific part of the spectrum and use 'Zoom Out' to get back to the original zoom level. The annotation of the spectrum is based on the given sequence in the peptides file and is done with different software so inconsistencies are likely. The peaks are annotated based on the given sequence, with 20 ppm tolerance.

Copy Data

##### Spectrum 7925 (TSV)

###### Preview

```
Loading example...
```

*Click on the button to copy the data to your clipboard.*

Mz MinMz MaxIntensity Max

WidthHeightPeptide font sizePeptide stroke widthSpectrum font sizeSpectrum stroke widthCompact peptide

Ion legend

wxyz

abcd

OtherUnassignedIon^Charge^_Position_Show for top:%

JVKDYFPEPVT

09.05e+41.81e+52.72e+53.62e+5

Zoom Out

y^+1^_1_y^+1^_2_y^+1^_3_y^+1^_3_c^+1^_3_y^+1^_4_c^+2^_7_y^+1^_4_c^+1^_4_c^+1^_4_c^+2^_8_y^+1^_5_y^+1^_5_c^+2^_9_c^+2^_10_c^+1^_5_c^+1^_5_y^+1^_6_c^+1^_6_z^+1^_7_y^+1^_7_c^+1^_7_c^+1^_7_z^+1^_8_y^+1^_8_c^+1^_8_y^+1^_9_z^+1^_9_y^+1^_9_c^+1^_9_z^+1^_10_c^+1^_10_y^+1^_10_c^+1^_10_

0721144221632884

Fragment Matches Table

Show background peaks

| Position | Ion type | Intensity | mz Theoretical | mz Error (Th) | mz Error (ppm) | Charge | Series Number |
| --- | --- | --- | --- | --- | --- | --- | --- |
| 11 | y | 4113 | 120.1 | 1.363E-05 | 0.1135 | +1 | 1 |
| - | - | 834.4 | 120.1 | - | - | 0 | - |
| - | - | 3265 | 129.1 | - | - | 0 | - |
| - | - | 1930 | 133.1 | - | - | 0 | - |
| - | - | 422.9 | 136.2 | - | - | 0 | - |
| - | - | 1041 | 149 | - | - | 0 | - |
| - | - | 466 | 162.7 | - | - | 0 | - |
| - | - | 499.7 | 167.9 | - | - | 0 | - |
| - | - | 7943 | 169.1 | - | - | 0 | - |
| - | - | 1403 | 173.4 | - | - | 0 | - |
| - | - | 1193 | 177.1 | - | - | 0 | - |
| - | - | 9659 | 185.2 | - | - | 0 | - |
| - | - | 1341 | 186.2 | - | - | 0 | - |
| - | - | 468.1 | 194.7 | - | - | 0 | - |
| - | - | 935.9 | 197.1 | - | - | 0 | - |
| - | - | 1.549E+04 | 197.1 | - | - | 0 | - |
| - | - | 1839 | 198.1 | - | - | 0 | - |
| - | - | 796.6 | 199.1 | - | - | 0 | - |
| - | - | 451.6 | 204.7 | - | - | 0 | - |
| - | - | 7354 | 213.2 | - | - | 0 | - |
| - | - | 808.2 | 214.2 | - | - | 0 | - |
| 10 | y | 973.7 | 219.1 | 8.441E-05 | 0.3852 | +1 | 2 |
| - | - | 4128 | 227.1 | - | - | 0 | - |
| - | - | 525.1 | 249.1 | - | - | 0 | - |
| - | - | 514.2 | 254.5 | - | - | 0 | - |
| - | - | 536.5 | 280.2 | - | - | 0 | - |
| - | - | 503.5 | 291.7 | - | - | 0 | - |
| 9 | y | 4352 | 298.2 | 1.477E-05 | 0.04952 | +1 | 3 |
| - | - | 641.7 | 299.1 | - | - | 0 | - |
| - | - | 495.5 | 309.9 | - | - | 0 | - |
| 9 | y | 4.731E+04 | 316.2 | 7.02E-05 | 0.222 | +1 | 3 |
| - | - | 8480 | 317.2 | - | - | 0 | - |
| - | - | 927.5 | 318.2 | - | - | 0 | - |
| - | - | 8872 | 324.2 | - | - | 0 | - |
| - | - | 1141 | 325.2 | - | - | 0 | - |
| - | - | 886.5 | 355.1 | - | - | 0 | - |
| - | - | 972.3 | 356.1 | - | - | 0 | - |
| - | - | 7553 | 357.3 | - | - | 0 | - |
| 3 | c | 3051 | 358.3 | 0.001359 | 3.794 | +1 | 3 |
| - | - | 589.5 | 373.2 | - | - | 0 | - |
| - | - | 803.3 | 395.2 | - | - | 0 | - |
| - | - | 729.3 | 407.2 | - | - | 0 | - |
| - | - | 618.8 | 418.2 | - | - | 0 | - |
| - | - | 3980 | 423.2 | - | - | 0 | - |
| - | - | 776.2 | 424.2 | - | - | 0 | - |
| - | - | 1322 | 426.2 | - | - | 0 | - |
| 8 | y | 571.7 | 427.2 | 0.0002505 | 0.5862 | +1 | 4 |
| 7 | c | 1748 | 432.2 | 0.0002714 | 0.6279 | +2 | 7 |
| - | - | 1203 | 432.7 | - | - | 0 | - |
| - | - | 674.8 | 438.3 | - | - | 0 | - |
| 8 | y | 2023 | 445.2 | 0.0003848 | 0.8643 | +1 | 4 |
| 4 | c | 2096 | 456.3 | 1.685E-05 | 0.03692 | +1 | 4 |
| - | - | 620 | 471.2 | - | - | 0 | - |
| - | - | 2451 | 472.3 | - | - | 0 | - |
| 4 | c | 4983 | 473.3 | 0.0001706 | 0.3605 | +1 | 4 |
| - | - | 1126 | 474.3 | - | - | 0 | - |
| - | - | 962.9 | 489.2 | - | - | 0 | - |
| - | - | 616.1 | 494.1 | - | - | 0 | - |
| 8 | c | 3793 | 496.8 | 0.0003132 | 0.6304 | +2 | 8 |
| - | - | 2320 | 497.3 | - | - | 0 | - |
| - | - | 781.3 | 497.8 | - | - | 0 | - |
| - | - | 585.5 | 522.3 | - | - | 0 | - |
| - | - | 1438 | 522.8 | - | - | 0 | - |
| 7 | y | 1134 | 524.3 | 0.004939 | 9.42 | +1 | 5 |
| - | - | 1136 | 531.3 | - | - | 0 | - |
| - | - | 853.5 | 531.8 | - | - | 0 | - |
| - | - | 671.9 | 532.3 | - | - | 0 | - |
| - | - | 895.4 | 539.3 | - | - | 0 | - |
| 7 | y | 2.018E+04 | 542.3 | 0.0004774 | 0.8804 | +1 | 5 |
| - | - | 4957 | 543.3 | - | - | 0 | - |
| - | - | 1060 | 544.3 | - | - | 0 | - |
| 9 | c | 1.593E+04 | 545.3 | 0.0005731 | 1.051 | +2 | 9 |
| - | - | 1.018E+04 | 545.8 | - | - | 0 | - |
| - | - | 2690 | 546.3 | - | - | 0 | - |
| - | - | 8980 | 554.3 | - | - | 0 | - |
| - | - | 2351 | 555.3 | - | - | 0 | - |
| - | - | 778.8 | 570.3 | - | - | 0 | - |
| - | - | 5245 | 580.8 | - | - | 0 | - |
| - | - | 3623 | 581.3 | - | - | 0 | - |
| - | - | 1559 | 581.8 | - | - | 0 | - |
| - | - | 1197 | 591.4 | - | - | 0 | - |
| 10 | c | 2546 | 594.8 | 0.001522 | 2.559 | +2 | 10 |
| - | - | 2866 | 595.3 | - | - | 0 | - |
| - | - | 1139 | 595.8 | - | - | 0 | - |
| - | - | 627.6 | 603.4 | - | - | 0 | - |
| 5 | c | 7164 | 619.3 | 0.000348 | 0.5618 | +1 | 5 |
| - | - | 2342 | 620.3 | - | - | 0 | - |
| - | - | 807.2 | 621.4 | - | - | 0 | - |
| - | - | 1.783E+04 | 635.4 | - | - | 0 | - |
| 5 | c | 2.3E+04 | 636.4 | 0.0004102 | 0.6446 | +1 | 5 |
| - | - | 7689 | 637.4 | - | - | 0 | - |
| - | - | 1372 | 638.4 | - | - | 0 | - |
| - | - | 2627 | 645.3 | - | - | 0 | - |
| - | - | 1451 | 645.8 | - | - | 0 | - |
| - | - | 892.5 | 652.3 | - | - | 0 | - |
| - | - | 2529 | 654.3 | - | - | 0 | - |
| - | - | 2847 | 654.8 | - | - | 0 | - |
| - | - | 1832 | 655.4 | - | - | 0 | - |
| - | - | 1651 | 681.3 | - | - | 0 | - |
| - | - | 793.6 | 682.4 | - | - | 0 | - |
| 6 | y | 1.275E+04 | 689.4 | 0.0005449 | 0.7905 | +1 | 6 |
| - | - | 4248 | 690.4 | - | - | 0 | - |
| - | - | 961.7 | 698.4 | - | - | 0 | - |
| - | - | 1293 | 705.4 | - | - | 0 | - |
| - | - | 798.3 | 706.4 | - | - | 0 | - |
| - | - | 668.8 | 716.4 | - | - | 0 | - |
| - | - | 1447 | 738.4 | - | - | 0 | - |
| - | - | 6540 | 739.4 | - | - | 0 | - |
| - | - | 1.122E+04 | 740.4 | - | - | 0 | - |
| - | - | 4242 | 741.4 | - | - | 0 | - |
| - | - | 698.2 | 742.4 | - | - | 0 | - |
| 6 | c | 2.871E+04 | 766.4 | 0.0008427 | 1.1 | +1 | 6 |
| - | - | 1.437E+04 | 767.4 | - | - | 0 | - |
| - | - | 3047 | 768.4 | - | - | 0 | - |
| - | - | 7451 | 780.4 | - | - | 0 | - |
| - | - | 2615 | 781.4 | - | - | 0 | - |
| - | - | 948.9 | 782.4 | - | - | 0 | - |
| 5 | z | 849.1 | 836.4 | 0.0009687 | 1.158 | +1 | 7 |
| - | - | 1072 | 837.4 | - | - | 0 | - |
| - | - | 1785 | 851.4 | - | - | 0 | - |
| 5 | y | 1.082E+04 | 852.4 | 0.0002047 | 0.2401 | +1 | 7 |
| - | - | 6811 | 853.4 | - | - | 0 | - |
| - | - | 1873 | 854.4 | - | - | 0 | - |
| - | - | 1096 | 859.4 | - | - | 0 | - |
| - | - | 1041 | 860.4 | - | - | 0 | - |
| 7 | c | 2866 | 863.5 | 0.002766 | 3.204 | +1 | 7 |
| - | - | 719 | 864.5 | - | - | 0 | - |
| - | - | 748.2 | 865.5 | - | - | 0 | - |
| - | - | 3483 | 877.4 | - | - | 0 | - |
| - | - | 1090 | 878.4 | - | - | 0 | - |
| - | - | 3.224E+04 | 879.5 | - | - | 0 | - |
| 7 | c | 6.422E+04 | 880.5 | 0.0004062 | 0.4614 | +1 | 7 |
| - | - | 2.935E+04 | 881.5 | - | - | 0 | - |
| - | - | 5759 | 882.5 | - | - | 0 | - |
| - | - | 654.2 | 904.5 | - | - | 0 | - |
| - | - | 793.4 | 906.5 | - | - | 0 | - |
| - | - | 775.2 | 908.4 | - | - | 0 | - |
| 4 | z | 849 | 951.4 | 0.001239 | 1.303 | +1 | 8 |
| - | - | 1771 | 952.4 | - | - | 0 | - |
| - | - | 981.6 | 953.4 | - | - | 0 | - |
| - | - | 1663 | 964.5 | - | - | 0 | - |
| - | - | 2472 | 965.5 | - | - | 0 | - |
| - | - | 3140 | 966.4 | - | - | 0 | - |
| - | - | 1033 | 966.5 | - | - | 0 | - |
| 4 | y | 6752 | 967.4 | 0.0002392 | 0.2472 | +1 | 8 |
| - | - | 3308 | 968.4 | - | - | 0 | - |
| - | - | 920.5 | 969.4 | - | - | 0 | - |
| - | - | 2017 | 974.5 | - | - | 0 | - |
| - | - | 1265 | 975.5 | - | - | 0 | - |
| - | - | 3775 | 976.5 | - | - | 0 | - |
| - | - | 1532 | 977.5 | - | - | 0 | - |
| - | - | 1210 | 978.5 | - | - | 0 | - |
| - | - | 787.9 | 991.6 | - | - | 0 | - |
| 8 | c | 4.151E+04 | 992.5 | 0.001311 | 1.321 | +1 | 8 |
| - | - | 2.338E+04 | 993.5 | - | - | 0 | - |
| - | - | 7005 | 994.5 | - | - | 0 | - |
| - | - | 1168 | 1007 | - | - | 0 | - |
| - | - | 900 | 1008 | - | - | 0 | - |
| - | - | 5126 | 1036 | - | - | 0 | - |
| - | - | 2634 | 1037 | - | - | 0 | - |
| - | - | 1326 | 1038 | - | - | 0 | - |
| - | - | 1878 | 1047 | - | - | 0 | - |
| - | - | 1629 | 1048 | - | - | 0 | - |
| - | - | 1876 | 1062 | - | - | 0 | - |
| - | - | 946.8 | 1063 | - | - | 0 | - |
| - | - | 811.4 | 1064 | - | - | 0 | - |
| 3 | y | 782.5 | 1078 | 0.00306 | 2.84 | +1 | 9 |
| - | - | 1005 | 1079 | - | - | 0 | - |
| 3 | z | 1.15E+04 | 1080 | 0.001332 | 1.234 | +1 | 9 |
| - | - | 8998 | 1081 | - | - | 0 | - |
| - | - | 2796 | 1082 | - | - | 0 | - |
| - | - | 1564 | 1090 | - | - | 0 | - |
| - | - | 993.2 | 1091 | - | - | 0 | - |
| 3 | y | 1.168E+04 | 1096 | 0.0009183 | 0.8382 | +1 | 9 |
| - | - | 6086 | 1097 | - | - | 0 | - |
| - | - | 1880 | 1098 | - | - | 0 | - |
| - | - | 1011 | 1104 | - | - | 0 | - |
| 9 | c | 8753 | 1107 | 0.0005503 | 0.4973 | +1 | 9 |
| - | - | 6677 | 1108 | - | - | 0 | - |
| - | - | 2286 | 1109 | - | - | 0 | - |
| - | - | 1166 | 1125 | - | - | 0 | - |
| - | - | 1033 | 1126 | - | - | 0 | - |
| - | - | 3211 | 1147 | - | - | 0 | - |
| - | - | 3785 | 1148 | - | - | 0 | - |
| - | - | 823.8 | 1149 | - | - | 0 | - |
| - | - | 1579 | 1150 | - | - | 0 | - |
| - | - | 650.6 | 1151 | - | - | 0 | - |
| - | - | 861 | 1157 | - | - | 0 | - |
| - | - | 743.2 | 1161 | - | - | 0 | - |
| - | - | 1.869E+04 | 1162 | - | - | 0 | - |
| - | - | 1.387E+04 | 1163 | - | - | 0 | - |
| - | - | 8449 | 1164 | - | - | 0 | - |
| - | - | 3735 | 1165 | - | - | 0 | - |
| 2 | z | 7698 | 1179 | 0.001766 | 1.498 | +1 | 10 |
| - | - | 4445 | 1180 | - | - | 0 | - |
| - | - | 1975 | 1181 | - | - | 0 | - |
| 10 | c | 2878 | 1189 | 0.0009218 | 0.7755 | +1 | 10 |
| - | - | 2211 | 1190 | - | - | 0 | - |
| - | - | 6065 | 1191 | - | - | 0 | - |
| - | - | 4602 | 1192 | - | - | 0 | - |
| - | - | 2011 | 1193 | - | - | 0 | - |
| 2 | y | 1017 | 1195 | 0.0008453 | 0.7076 | +1 | 10 |
| - | - | 897 | 1205 | - | - | 0 | - |
| 10 | c | 8.777E+04 | 1206 | 0.001594 | 1.322 | +1 | 10 |
| - | - | 6.144E+04 | 1207 | - | - | 0 | - |
| - | - | 3.378E+04 | 1208 | - | - | 0 | - |
| - | - | 1.038E+04 | 1209 | - | - | 0 | - |
| - | - | 3609 | 1210 | - | - | 0 | - |
| - | - | 2367 | 1220 | - | - | 0 | - |
| - | - | 1799 | 1221 | - | - | 0 | - |
| - | - | 8073 | 1237 | - | - | 0 | - |
| - | - | 5345 | 1238 | - | - | 0 | - |
| - | - | 1101 | 1239 | - | - | 0 | - |
| - | - | 4728 | 1245 | - | - | 0 | - |
| - | - | 5700 | 1246 | - | - | 0 | - |
| - | - | 3288 | 1247 | - | - | 0 | - |
| - | - | 3420 | 1248 | - | - | 0 | - |
| - | - | 2.581E+04 | 1249 | - | - | 0 | - |
| - | - | 1.805E+04 | 1250 | - | - | 0 | - |
| - | - | 8168 | 1251 | - | - | 0 | - |
| - | - | 1545 | 1252 | - | - | 0 | - |
| - | - | 4033 | 1253 | - | - | 0 | - |
| - | - | 2582 | 1254 | - | - | 0 | - |
| - | - | 1024 | 1255 | - | - | 0 | - |
| - | - | 5316 | 1263 | - | - | 0 | - |
| - | - | 3824 | 1264 | - | - | 0 | - |
| - | - | 1.108E+04 | 1265 | - | - | 0 | - |
| - | - | 7091 | 1266 | - | - | 0 | - |
| - | - | 2862 | 1267 | - | - | 0 | - |
| - | - | 1.89E+04 | 1281 | - | - | 0 | - |
| - | - | 1.255E+04 | 1282 | - | - | 0 | - |
| - | - | 4729 | 1283 | - | - | 0 | - |
| - | - | 763.2 | 1289 | - | - | 0 | - |
| - | - | 756.9 | 1289 | - | - | 0 | - |
| - | - | 1018 | 1290 | - | - | 0 | - |
| - | - | 7038 | 1291 | - | - | 0 | - |
| - | - | 9.687E+04 | 1292 | - | - | 0 | - |
| - | - | 7.272E+04 | 1293 | - | - | 0 | - |
| - | - | 3.195E+04 | 1294 | - | - | 0 | - |
| - | - | 4729 | 1295 | - | - | 0 | - |
| - | - | 714.5 | 1299 | - | - | 0 | - |
| - | - | 1180 | 1305 | - | - | 0 | - |
| - | - | 2525 | 1306 | - | - | 0 | - |
| - | - | 4339 | 1307 | - | - | 0 | - |
| - | - | 7.329E+04 | 1308 | - | - | 0 | - |
| - | - | 3.584E+05 | 1309 | - | - | 0 | - |
| - | - | 2.507E+05 | 1310 | - | - | 0 | - |
| - | - | 9.568E+04 | 1311 | - | - | 0 | - |
| - | - | 1.235E+04 | 1312 | - | - | 0 | - |
| - | - | 960.7 | 1325 | - | - | 0 | - |
| - | - | 911 | 1341 | - | - | 0 | - |
| - | - | 804.3 | 1342 | - | - | 0 | - |
| - | - | 726.9 | 1580 | - | - | 0 | - |
| - | - | 842.9 | 1860 | - | - | 0 | - |
| - | - | 709.8 | 1960 | - | - | 0 | - |
| - | - | 912.4 | 2638 | - | - | 0 | - |
| - | - | 724 | 2855 | - | - | 0 | - |

m/z Charge Intensity FragmentType MassShift Position
120.06550598144531 0 4113.2505 y 10
120.0807113647461 0 834.3656
129.1022491455078 0 3264.7546
133.0859375 0 1929.9515
136.20819091796875 0 422.93576
148.95469665527344 0 1041.3258
162.72544860839844 0 466.048
167.9169158935547 0 499.6954
169.13348388671875 0 7942.842
173.4387664794922 0 1402.6855
177.1122283935547 0 1193.0415
185.164794921875 0 9659.005
186.16812133789062 0 1341.309
194.69442749023438 0 468.10062
197.11912536621094 0 935.89404
197.12843322753906 0 15490.219
198.13194274902344 0 1839.3378
199.10757446289062 0 796.55206
204.66827392578125 0 451.63007
213.1596221923828 0 7354.078
214.16348266601562 0 808.1555
219.13401794433594 0 973.69604 y 9
227.1027069091797 0 4128.188
249.13156127929688 0 525.05414
254.50502014160156 0 514.2405
280.165283203125 0 536.45087
291.7145690917969 0 503.48184
298.1761474609375 0 4352.002 y Water loss 8
299.0634765625 0 641.71014
309.9388427734375 0 495.5144
316.186767578125 0 47312.816 y 8
317.1900634765625 0 8480.153
318.19183349609375 0 927.469
324.1555480957031 0 8871.83
325.15875244140625 0 1140.6841
355.0702819824219 0 886.498
356.0711669921875 0 972.3335
357.2730712890625 0 7553.2188
358.2799072265625 0 3050.5476 c 2
373.1866455078125 0 589.5148
395.2300109863281 0 803.3221
407.19085693359375 0 729.2684
418.2379150390625 0 618.8108
423.2243957519531 0 3979.9343
424.2266845703125 0 776.18854
426.166015625 0 1322.3639
427.2184753417969 0 571.69507 y Water loss 7
432.2364501953125 0 1748.2948 c Ammonia loss 6
432.7383728027344 0 1202.565
438.2707824707031 0 674.8261
445.22967529296875 0 2022.8579 y 7
456.28167724609375 0 2096.4497 c Ammonia loss 3
471.225830078125 0 620.0054
472.30029296875 0 2450.5325
473.3083801269531 0 4983.4453 c 3
474.3116149902344 0 1126.4631
489.2440490722656 0 962.9178
494.0513916015625 0 616.1391
496.7583312988281 0 3793.214 c Ammonia loss 7
497.260498046875 0 2320.1536
497.7602233886719 0 781.3075
522.2840576171875 0 585.48987
522.7863159179688 0 1438.1804
524.2764282226562 0 1133.8375 y Water loss 6
531.28759765625 0 1135.7201
531.7893676757812 0 853.4925
532.2962036132812 0 671.885
539.2522583007812 0 895.3874
542.2825317382812 0 20181.268 y 6
543.2854614257812 0 4956.8013
544.2860107421875 0 1060.3832
545.2849731445312 0 15931.149 c Ammonia loss 8
545.7864379882812 0 10183.295
546.2887573242188 0 2690.16
554.2611083984375 0 8980.362
555.2639770507812 0 2351.073
570.2910766601562 0 778.79156
580.8218383789062 0 5244.505
581.32275390625 0 3623.2798
581.8255615234375 0 1558.8749
591.3525390625 0 1196.816
594.8201293945312 0 2546.3105 c Ammonia loss 9
595.3209228515625 0 2865.5364
595.8225708007812 0 1139.0403
603.3592529296875 0 627.559
619.3453369140625 0 7163.7 c Ammonia loss 4
620.3496704101562 0 2341.545
621.351806640625 0 807.1747
635.3652954101562 0 17833.625
636.3719482421875 0 23000.602 c 4
637.3743896484375 0 7688.8423
638.377685546875 0 1372.3971
645.3428955078125 0 2627.446
645.8436889648438 0 1451.1299
652.2614135742188 0 892.51917
654.3472290039062 0 2528.9695
654.8497924804688 0 2846.7134
655.3506469726562 0 1831.8605
681.323486328125 0 1651.4387
682.3794555664062 0 793.5916
689.3510131835938 0 12753.021 y 5
690.3541259765625 0 4248.05
698.4419555664062 0 961.65564
705.3646850585938 0 1292.8279
706.3682250976562 0 798.275
716.393798828125 0 668.83795
738.4181518554688 0 1447.2662
739.4260864257812 0 6540.338
740.4334106445312 0 11224.517
741.4371337890625 0 4242.2397
742.4409790039062 0 698.20844
766.4142456054688 0 28712.37 c Ammonia loss 5
767.4169311523438 0 14371.614
768.4209594726562 0 3046.512
780.357177734375 0 7451.3633
781.3607788085938 0 2614.8594
782.3621826171875 0 948.92523
836.3941040039062 0 849.05145 z 4
837.3973999023438 0 1071.7388
851.4065551757812 0 1785.2155
852.4140014648438 0 10818.9 y 4
853.4173583984375 0 6811.2246
854.4193725585938 0 1873.4432
859.4373779296875 0 1095.6917
860.4443969726562 0 1040.5428
863.4689331054688 0 2866.429 c Ammonia loss 6
864.4768676757812 0 719.04535
865.48291015625 0 748.21655
877.4098510742188 0 3483.1797
878.4041137695312 0 1090.2919
879.4860229492188 0 32243.547
880.4923095703125 0 64218.383 c 6
881.4961547851562 0 29350.086
882.4996337890625 0 5759.225
904.47265625 0 654.1646
906.509765625 0 793.4256
908.4293823242188 0 775.1789
951.4207763671875 0 849.02277 z 3
952.4313354492188 0 1771.4663
953.4356079101562 0 981.5903
964.5187377929688 0 1662.8368
965.5211181640625 0 2471.8416
966.435302734375 0 3139.509
966.5302734375 0 1032.8273
967.4409790039062 0 6752.11 y 3
968.4458618164062 0 3307.9993
969.4459838867188 0 920.53986
974.5015869140625 0 2017.3429
975.5027465820312 0 1265.069
976.4837036132812 0 3774.7178
977.4814453125 0 1531.9836
978.4888305664062 0 1210.2113
991.5582885742188 0 787.9491
992.5100708007812 0 41505.31 c Ammonia loss 7
993.51318359375 0 23375.777
994.5169677734375 0 7004.759
1007.4954833984375 0 1168.1206
1008.49560546875 0 900.00836
1035.5283203125 0 5125.951
1036.5347900390625 0 2634.02
1037.5369873046875 0 1326.0156
1046.5523681640625 0 1877.6506
1047.5584716796875 0 1628.9507
1061.5494384765625 0 1876.1852
1062.5697021484375 0 946.77704
1063.5657958984375 0 811.39923
1077.5281982421875 0 782.46674 y Water loss 2
1078.53076171875 0 1004.7564
1079.518310546875 0 11500.559 z 2
1080.521728515625 0 8997.767
1081.523681640625 0 2796.2666
1089.617919921875 0 1564.2836
1090.6217041015625 0 993.1708
1095.53662109375 0 11680.427 y 2
1096.5401611328125 0 6085.82
1097.5450439453125 0 1879.8369
1103.64013671875 0 1010.6336
1106.588623046875 0 8753.3545 c 8
1107.593017578125 0 6676.8687
1108.595947265625 0 2285.6257
1124.5362548828125 0 1165.7908
1125.5401611328125 0 1032.9388
1146.6317138671875 0 3211.331
1147.6385498046875 0 3785.0864
1148.55712890625 0 823.7637
1149.583740234375 0 1578.7477
1150.5955810546875 0 650.56647
1156.7049560546875 0 860.99194
1160.628662109375 0 743.1768
1161.645263671875 0 18694.709
1162.649169921875 0 13872.992
1163.654541015625 0 8449.029
1164.6627197265625 0 3734.5247
1178.587158203125 0 7698.471 z 1
1179.5902099609375 0 4444.57
1180.5921630859375 0 1974.5931
1188.630859375 0 2878.0435 c Ammonia loss 9
1189.634765625 0 2210.7417
1190.645263671875 0 6065.474
1191.6502685546875 0 4602.1694
1192.652099609375 0 2010.8793
1194.603271484375 0 1016.69604 y 1
1204.6513671875 0 897.0211
1205.6580810546875 0 87768.42 c 9
1206.6610107421875 0 61439.062
1207.6663818359375 0 33777.566
1208.673583984375 0 10382.569
1209.6783447265625 0 3609.1406
1219.6513671875 0 2367.4006
1220.6512451171875 0 1799.0396
1236.67431640625 0 8072.8687
1237.6767578125 0 5345.3687
1238.6751708984375 0 1101.017
1244.6827392578125 0 4727.9336
1245.6845703125 0 5699.518
1246.6951904296875 0 3288.3376
1247.67724609375 0 3419.6772
1248.6571044921875 0 25814.25
1249.65771484375 0 18045.193
1250.6641845703125 0 8168.3574
1251.6710205078125 0 1544.6802
1252.6324462890625 0 4032.8323
1253.63623046875 0 2582.3137
1254.6290283203125 0 1023.54944
1262.689453125 0 5315.9595
1263.69384765625 0 3823.7195
1264.704833984375 0 11080.485
1265.7091064453125 0 7091.4004
1266.7108154296875 0 2862.062
1280.7025146484375 0 18902.674
1281.704833984375 0 12545.439
1282.7080078125 0 4728.7046
1288.68603515625 0 763.15594
1289.485107421875 0 756.8871
1289.6800537109375 0 1018.18567
1290.678955078125 0 7038.3154
1291.6715087890625 0 96871.17
1292.6734619140625 0 72715.96
1293.6756591796875 0 31951.912
1294.6748046875 0 4729.235
1299.1331787109375 0 714.45294
1304.718017578125 0 1179.7792
1305.69677734375 0 2525.3237
1306.6861572265625 0 4339.454
1307.6888427734375 0 73285.99
1308.69677734375 0 358445.3
1309.69970703125 0 250721.17
1310.7025146484375 0 95677.87
1311.7064208984375 0 12354.215
1324.65673828125 0 960.6788
1340.683837890625 0 910.9784
1341.68896484375 0 804.27313
1579.801513671875 0 726.88556
1859.9122314453125 0 842.9447
1959.9229736328125 0 709.7998
2637.969482421875 0 912.3915
2855.27783203125 0 723.98206

Spectrum Details

|  |  |
| --- | --- |
| Matched peaks? Matched peaks The total absolute number of peaks matched. Additionally in brackets the total fraction of peaks matched and the total number of peaks is shown. | 34 (13.23% of 257) |
| FDR? FDR The false discovery rate estimated for this peptide. It is calculated by matching all theoretical fragments with a non-integer shift with the raw peaks for this spectrum. This is done with 40 different shifts. The resulting percentage is the average number of annotated peaks over the number of annotated peaks with the correct spectrum. | 2.03% |
| Satellite FDR? Satellite FDR See the FDR for details on its calculation. This satellite ion specific FDR only contains the satellite ions (d/w) for I/L/J positions. | - |
| PSM Score? PSM Score The PSM Score as given by Hecklib to this annotated spectrum. It is shown with three significant figures. | 409 |

#### Spectrum 7808? Spectrum 7808 The raw spectrum of this peptide as annotated by Hecklib. The fragments are coloured according to ion type (see legend). Any peaks with a star '\*' as text can be hovered over to see the full details, first the ion type second the mass shift type. By hovering over the amino acids in the peptide or ions in the legend the corresponding peaks are highlighted. By toggling the 'Unassigned' label you can turn the background (unassigned) peaks on or off in the plot. By updating the slider in the Ion legend you can update the spectrum to only show the top X% of the peaks with labels. The top X% means any peak that is within X% of the highest intensity. By dragging in the spectrum you can zoom in to a specific part of the spectrum and use 'Zoom Out' to get back to the original zoom level. The annotation of the spectrum is based on the given sequence in the peptides file and is done with different software so inconsistencies are likely. The peaks are annotated based on the given sequence, with 20 ppm tolerance.

Copy Data

##### Spectrum 7808 (TSV)

###### Preview

```
Loading example...
```

*Click on the button to copy the data to your clipboard.*

Mz MinMz MaxIntensity Max

WidthHeightPeptide font sizePeptide stroke widthSpectrum font sizeSpectrum stroke widthCompact peptide

Ion legend

wxyz

abcd

OtherUnassignedIon^Charge^_Position_Show for top:%

JVKDYFPEPVT

09.26e+41.85e+52.78e+53.70e+5

Zoom Out

y^+1^_1_d^+1^_2_a^+1^_2_y^+1^_2_b^+1^_2_y^+1^_2_y^+1^_3_b^+2^_5_y^+1^_3_b^+1^_3_b^+2^_6_b^+2^_7_y^+1^_4_b^+2^_7_b^+1^_4_b^+1^_4_y^+1^_4_b^+1^_4_b^+2^_8_b^+2^_8_b^+2^_8_y^+1^_5_b^+2^_9_y^+1^_5_b^+2^_9_y^+2^_9_b^+2^_10_b^+2^_10_b^+1^_5_b^+1^_5_\*\*\*y^+1^_6_y^+1^_6_b^+1^_6_b^+1^_6_y^+1^_7_b^+1^_7_b^+1^_7_y^+1^_7_b^+1^_7_y^+1^_8_b^+1^_8_b^+1^_8_b^+1^_8_y^+1^_9_y^+1^_9_b^+1^_9_y^+1^_9_b^+1^_10_b^+1^_10_b^+1^_10_y^+1^_10_

0780156023393119

Fragment Matches Table

Show background peaks

| Position | Ion type | Intensity | mz Theoretical | mz Error (Th) | mz Error (ppm) | Charge | Series Number |
| --- | --- | --- | --- | --- | --- | --- | --- |
| 11 | y | 4.175E+04 | 120.1 | 0.0004518 | 3.763 | +1 | 1 |
| - | - | 5.52E+04 | 120.1 | - | - | 0 | - |
| - | - | 1523 | 121.1 | - | - | 0 | - |
| - | - | 4734 | 121.1 | - | - | 0 | - |
| - | - | 586.5 | 126.1 | - | - | 0 | - |
| - | - | 488.5 | 127.1 | - | - | 0 | - |
| - | - | 395.7 | 128.1 | - | - | 0 | - |
| - | - | 1.037E+05 | 129.1 | - | - | 0 | - |
| - | - | 1847 | 130.1 | - | - | 0 | - |
| - | - | 813.9 | 130.1 | - | - | 0 | - |
| - | - | 5473 | 130.1 | - | - | 0 | - |
| - | - | 438 | 134.1 | - | - | 0 | - |
| - | - | 2.418E+04 | 136.1 | - | - | 0 | - |
| - | - | 1606 | 137.1 | - | - | 0 | - |
| - | - | 750.4 | 138.1 | - | - | 0 | - |
| - | - | 2924 | 139.1 | - | - | 0 | - |
| - | - | 1040 | 140.1 | - | - | 0 | - |
| - | - | 714 | 146.1 | - | - | 0 | - |
| - | - | 685.6 | 146.1 | - | - | 0 | - |
| - | - | 678.6 | 151.1 | - | - | 0 | - |
| - | - | 1519 | 155.1 | - | - | 0 | - |
| - | - | 758.5 | 155.2 | - | - | 0 | - |
| - | - | 598.6 | 156.1 | - | - | 0 | - |
| - | - | 800.3 | 158 | - | - | 0 | - |
| - | - | 585 | 163.1 | - | - | 0 | - |
| - | - | 474.2 | 165.1 | - | - | 0 | - |
| - | - | 2212 | 165.1 | - | - | 0 | - |
| - | - | 1081 | 166.1 | - | - | 0 | - |
| - | - | 456.9 | 166.2 | - | - | 0 | - |
| - | - | 1418 | 167.1 | - | - | 0 | - |
| - | - | 2867 | 167.1 | - | - | 0 | - |
| - | - | 689.5 | 167.2 | - | - | 0 | - |
| - | - | 1890 | 168.1 | - | - | 0 | - |
| - | - | 435 | 168.3 | - | - | 0 | - |
| - | - | 3.11E+05 | 169.1 | - | - | 0 | - |
| - | - | 2.844E+04 | 170.1 | - | - | 0 | - |
| - | - | 928.9 | 171.1 | - | - | 0 | - |
| 2 | d | 2282 | 171.1 | 0.0005296 | 3.094 | +1 | 2 |
| - | - | 5439 | 172.1 | - | - | 0 | - |
| - | - | 507.8 | 172.1 | - | - | 0 | - |
| - | - | 713.5 | 172.2 | - | - | 0 | - |
| - | - | 796.2 | 173.1 | - | - | 0 | - |
| - | - | 1582 | 173.1 | - | - | 0 | - |
| - | - | 1891 | 173.4 | - | - | 0 | - |
| - | - | 2758 | 174.1 | - | - | 0 | - |
| - | - | 5109 | 181.1 | - | - | 0 | - |
| - | - | 6680 | 181.1 | - | - | 0 | - |
| - | - | 2656 | 182.1 | - | - | 0 | - |
| - | - | 500.5 | 182.1 | - | - | 0 | - |
| - | - | 739 | 182.1 | - | - | 0 | - |
| - | - | 2147 | 183.1 | - | - | 0 | - |
| - | - | 5325 | 183.1 | - | - | 0 | - |
| - | - | 810.1 | 184.1 | - | - | 0 | - |
| - | - | 608.5 | 185.1 | - | - | 0 | - |
| - | - | 1551 | 185.1 | - | - | 0 | - |
| 2 | a | 1.316E+05 | 185.2 | 0.000535 | 2.889 | +1 | 2 |
| - | - | 3567 | 186.1 | - | - | 0 | - |
| - | - | 1.292E+04 | 186.2 | - | - | 0 | - |
| - | - | 1005 | 188.1 | - | - | 0 | - |
| - | - | 4071 | 188.1 | - | - | 0 | - |
| - | - | 657.1 | 189.1 | - | - | 0 | - |
| - | - | 1528 | 190.1 | - | - | 0 | - |
| - | - | 1902 | 191.1 | - | - | 0 | - |
| - | - | 5585 | 195.1 | - | - | 0 | - |
| - | - | 2.462E+05 | 197.1 | - | - | 0 | - |
| - | - | 2.398E+04 | 198.1 | - | - | 0 | - |
| - | - | 3.814E+04 | 199.1 | - | - | 0 | - |
| - | - | 1176 | 199.1 | - | - | 0 | - |
| - | - | 745.2 | 199.1 | - | - | 0 | - |
| - | - | 2128 | 199.2 | - | - | 0 | - |
| - | - | 3469 | 200.1 | - | - | 0 | - |
| - | - | 1163 | 201.1 | - | - | 0 | - |
| 10 | y | 2.106E+04 | 201.1 | 0.000441 | 2.193 | +1 | 2 |
| - | - | 1127 | 202.1 | - | - | 0 | - |
| - | - | 999 | 203.1 | - | - | 0 | - |
| - | - | 1631 | 203.2 | - | - | 0 | - |
| - | - | 3062 | 204.1 | - | - | 0 | - |
| - | - | 513.8 | 206.2 | - | - | 0 | - |
| - | - | 712.4 | 207.1 | - | - | 0 | - |
| - | - | 1.453E+04 | 209.1 | - | - | 0 | - |
| - | - | 1777 | 210.1 | - | - | 0 | - |
| - | - | 911.5 | 210.2 | - | - | 0 | - |
| - | - | 546.8 | 211.1 | - | - | 0 | - |
| - | - | 2560 | 211.1 | - | - | 0 | - |
| - | - | 1619 | 212.1 | - | - | 0 | - |
| - | - | 1393 | 213.1 | - | - | 0 | - |
| - | - | 2700 | 213.1 | - | - | 0 | - |
| 2 | b | 6.225E+04 | 213.2 | 0.0005392 | 2.53 | +1 | 2 |
| - | - | 7179 | 214.2 | - | - | 0 | - |
| - | - | 4784 | 215.1 | - | - | 0 | - |
| - | - | 570.1 | 216.9 | - | - | 0 | - |
| - | - | 931.6 | 217.1 | - | - | 0 | - |
| - | - | 7179 | 217.1 | - | - | 0 | - |
| - | - | 650.6 | 218.1 | - | - | 0 | - |
| 10 | y | 3.432E+04 | 219.1 | 0.0005727 | 2.613 | +1 | 2 |
| - | - | 3075 | 220.1 | - | - | 0 | - |
| - | - | 623.9 | 223.1 | - | - | 0 | - |
| - | - | 1489 | 225.1 | - | - | 0 | - |
| - | - | 2.439E+04 | 226.1 | - | - | 0 | - |
| - | - | 1936 | 226.2 | - | - | 0 | - |
| - | - | 1.188E+05 | 227.1 | - | - | 0 | - |
| - | - | 1.157E+04 | 228.1 | - | - | 0 | - |
| - | - | 9904 | 228.2 | - | - | 0 | - |
| - | - | 927 | 229.1 | - | - | 0 | - |
| - | - | 941.1 | 229.1 | - | - | 0 | - |
| - | - | 1094 | 229.2 | - | - | 0 | - |
| - | - | 1759 | 230.2 | - | - | 0 | - |
| - | - | 1489 | 231.1 | - | - | 0 | - |
| - | - | 2.197E+04 | 231.1 | - | - | 0 | - |
| - | - | 3275 | 232.2 | - | - | 0 | - |
| - | - | 1298 | 233.1 | - | - | 0 | - |
| - | - | 3087 | 233.2 | - | - | 0 | - |
| - | - | 1096 | 237.1 | - | - | 0 | - |
| - | - | 2046 | 238.1 | - | - | 0 | - |
| - | - | 5102 | 238.2 | - | - | 0 | - |
| - | - | 1444 | 239.2 | - | - | 0 | - |
| - | - | 690.3 | 243.1 | - | - | 0 | - |
| - | - | 5.116E+04 | 244.1 | - | - | 0 | - |
| - | - | 1305 | 245.1 | - | - | 0 | - |
| - | - | 1.193E+04 | 245.1 | - | - | 0 | - |
| - | - | 1205 | 246.1 | - | - | 0 | - |
| - | - | 4103 | 247.1 | - | - | 0 | - |
| - | - | 1439 | 249.2 | - | - | 0 | - |
| - | - | 539.1 | 250.2 | - | - | 0 | - |
| - | - | 3.154E+04 | 251.1 | - | - | 0 | - |
| - | - | 603.2 | 251.2 | - | - | 0 | - |
| - | - | 3591 | 252.1 | - | - | 0 | - |
| - | - | 4420 | 254.1 | - | - | 0 | - |
| - | - | 2435 | 254.2 | - | - | 0 | - |
| - | - | 2122 | 254.2 | - | - | 0 | - |
| - | - | 2311 | 255.1 | - | - | 0 | - |
| - | - | 2082 | 256.2 | - | - | 0 | - |
| - | - | 1297 | 257.1 | - | - | 0 | - |
| - | - | 762.5 | 259.1 | - | - | 0 | - |
| - | - | 1032 | 261.1 | - | - | 0 | - |
| - | - | 4390 | 261.2 | - | - | 0 | - |
| - | - | 881.6 | 262.1 | - | - | 0 | - |
| - | - | 1385 | 263.1 | - | - | 0 | - |
| - | - | 1022 | 269.2 | - | - | 0 | - |
| - | - | 2596 | 270.2 | - | - | 0 | - |
| - | - | 7338 | 272.1 | - | - | 0 | - |
| - | - | 974.7 | 273.1 | - | - | 0 | - |
| - | - | 913.6 | 276.2 | - | - | 0 | - |
| - | - | 677.5 | 277.1 | - | - | 0 | - |
| - | - | 1470 | 277.2 | - | - | 0 | - |
| - | - | 662.5 | 277.2 | - | - | 0 | - |
| - | - | 1504 | 278.1 | - | - | 0 | - |
| - | - | 3.797E+04 | 279.1 | - | - | 0 | - |
| - | - | 5110 | 280.1 | - | - | 0 | - |
| - | - | 9315 | 280.2 | - | - | 0 | - |
| - | - | 976.2 | 281.1 | - | - | 0 | - |
| - | - | 1525 | 281.2 | - | - | 0 | - |
| - | - | 647.4 | 282.1 | - | - | 0 | - |
| - | - | 1.45E+04 | 283.1 | - | - | 0 | - |
| - | - | 739.9 | 283.2 | - | - | 0 | - |
| - | - | 2425 | 284.1 | - | - | 0 | - |
| - | - | 572.2 | 285.2 | - | - | 0 | - |
| - | - | 890.4 | 286.2 | - | - | 0 | - |
| - | - | 5334 | 287.2 | - | - | 0 | - |
| - | - | 1015 | 288.2 | - | - | 0 | - |
| - | - | 785.4 | 292.1 | - | - | 0 | - |
| - | - | 933.5 | 292.2 | - | - | 0 | - |
| - | - | 1108 | 294.2 | - | - | 0 | - |
| - | - | 670.1 | 295.2 | - | - | 0 | - |
| - | - | 910.7 | 296.1 | - | - | 0 | - |
| - | - | 810.6 | 296.2 | - | - | 0 | - |
| - | - | 702.4 | 296.2 | - | - | 0 | - |
| - | - | 933.3 | 296.2 | - | - | 0 | - |
| 9 | y | 5.971E+04 | 298.2 | 0.0007777 | 2.608 | +1 | 3 |
| - | - | 9907 | 299.2 | - | - | 0 | - |
| - | - | 1175 | 300.2 | - | - | 0 | - |
| - | - | 1924 | 301.2 | - | - | 0 | - |
| - | - | 4208 | 303.2 | - | - | 0 | - |
| - | - | 2942 | 304.2 | - | - | 0 | - |
| - | - | 3030 | 306.1 | - | - | 0 | - |
| - | - | 1045 | 306.2 | - | - | 0 | - |
| - | - | 600.1 | 307.1 | - | - | 0 | - |
| - | - | 2802 | 308.2 | - | - | 0 | - |
| - | - | 1506 | 310.1 | - | - | 0 | - |
| 5 | b | 2444 | 310.2 | 0.0006556 | 2.114 | +2 | 5 |
| - | - | 2791 | 310.2 | - | - | 0 | - |
| - | - | 1.237E+04 | 311.1 | - | - | 0 | - |
| - | - | 1962 | 312.1 | - | - | 0 | - |
| - | - | 1461 | 312.2 | - | - | 0 | - |
| - | - | 1251 | 313.2 | - | - | 0 | - |
| - | - | 2887 | 315.2 | - | - | 0 | - |
| 9 | y | 3.668E+05 | 316.2 | 0.0009857 | 3.118 | +1 | 3 |
| - | - | 5.701E+04 | 317.2 | - | - | 0 | - |
| - | - | 6589 | 318.2 | - | - | 0 | - |
| - | - | 667.6 | 319.1 | - | - | 0 | - |
| - | - | 678.8 | 319.2 | - | - | 0 | - |
| - | - | 633.5 | 322.2 | - | - | 0 | - |
| - | - | 2241 | 323.2 | - | - | 0 | - |
| - | - | 4199 | 323.2 | - | - | 0 | - |
| - | - | 1.059E+05 | 324.2 | - | - | 0 | - |
| - | - | 1.762E+04 | 325.2 | - | - | 0 | - |
| - | - | 4154 | 325.2 | - | - | 0 | - |
| - | - | 1446 | 325.2 | - | - | 0 | - |
| - | - | 2401 | 326.2 | - | - | 0 | - |
| - | - | 664.3 | 327.2 | - | - | 0 | - |
| - | - | 2446 | 328.2 | - | - | 0 | - |
| - | - | 1571 | 328.2 | - | - | 0 | - |
| - | - | 912.9 | 328.2 | - | - | 0 | - |
| - | - | 3628 | 329.2 | - | - | 0 | - |
| - | - | 549.4 | 330.2 | - | - | 0 | - |
| - | - | 2249 | 331.2 | - | - | 0 | - |
| - | - | 2472 | 335.2 | - | - | 0 | - |
| - | - | 611.7 | 335.2 | - | - | 0 | - |
| - | - | 3836 | 337.2 | - | - | 0 | - |
| - | - | 1670 | 337.2 | - | - | 0 | - |
| - | - | 1168 | 338.2 | - | - | 0 | - |
| - | - | 1392 | 339.2 | - | - | 0 | - |
| - | - | 5688 | 340.2 | - | - | 0 | - |
| - | - | 9390 | 341.2 | - | - | 0 | - |
| 3 | b | 9907 | 341.3 | 0.0008673 | 2.542 | +1 | 3 |
| - | - | 972.9 | 341.3 | - | - | 0 | - |
| - | - | 1656 | 342.1 | - | - | 0 | - |
| - | - | 1651 | 342.2 | - | - | 0 | - |
| - | - | 1450 | 342.3 | - | - | 0 | - |
| - | - | 1.185E+04 | 343.2 | - | - | 0 | - |
| - | - | 1297 | 344.2 | - | - | 0 | - |
| - | - | 2027 | 344.2 | - | - | 0 | - |
| - | - | 3111 | 344.7 | - | - | 0 | - |
| - | - | 633.6 | 345.1 | - | - | 0 | - |
| - | - | 1077 | 345.2 | - | - | 0 | - |
| - | - | 6490 | 346.2 | - | - | 0 | - |
| - | - | 859.7 | 347.2 | - | - | 0 | - |
| - | - | 5429 | 347.2 | - | - | 0 | - |
| - | - | 1439 | 347.7 | - | - | 0 | - |
| - | - | 1434 | 349.2 | - | - | 0 | - |
| - | - | 624.9 | 350.2 | - | - | 0 | - |
| - | - | 1089 | 351.2 | - | - | 0 | - |
| - | - | 963.3 | 351.2 | - | - | 0 | - |
| - | - | 6248 | 353.2 | - | - | 0 | - |
| - | - | 3881 | 353.2 | - | - | 0 | - |
| - | - | 1294 | 354.2 | - | - | 0 | - |
| - | - | 933 | 355.1 | - | - | 0 | - |
| - | - | 4772 | 355.2 | - | - | 0 | - |
| - | - | 3057 | 355.2 | - | - | 0 | - |
| - | - | 914.4 | 356.1 | - | - | 0 | - |
| - | - | 3392 | 356.2 | - | - | 0 | - |
| - | - | 842.5 | 356.2 | - | - | 0 | - |
| - | - | 646 | 357.1 | - | - | 0 | - |
| - | - | 941.8 | 357.2 | - | - | 0 | - |
| - | - | 1786 | 358.2 | - | - | 0 | - |
| - | - | 792.7 | 358.7 | - | - | 0 | - |
| - | - | 1022 | 359.1 | - | - | 0 | - |
| - | - | 793 | 360.2 | - | - | 0 | - |
| - | - | 1427 | 360.2 | - | - | 0 | - |
| - | - | 807.6 | 361.2 | - | - | 0 | - |
| - | - | 802.5 | 361.7 | - | - | 0 | - |
| - | - | 2519 | 362.2 | - | - | 0 | - |
| - | - | 888.3 | 363.2 | - | - | 0 | - |
| - | - | 1488 | 367.2 | - | - | 0 | - |
| - | - | 1247 | 369.2 | - | - | 0 | - |
| - | - | 1.319E+04 | 369.7 | - | - | 0 | - |
| - | - | 5630 | 370.2 | - | - | 0 | - |
| - | - | 2111 | 370.7 | - | - | 0 | - |
| - | - | 9697 | 371.2 | - | - | 0 | - |
| - | - | 1840 | 372.2 | - | - | 0 | - |
| - | - | 2170 | 372.2 | - | - | 0 | - |
| - | - | 1027 | 372.2 | - | - | 0 | - |
| - | - | 4674 | 373.2 | - | - | 0 | - |
| - | - | 1.462E+04 | 374.2 | - | - | 0 | - |
| - | - | 3784 | 375.2 | - | - | 0 | - |
| - | - | 888.2 | 375.2 | - | - | 0 | - |
| - | - | 724.5 | 375.7 | - | - | 0 | - |
| - | - | 1205 | 376.2 | - | - | 0 | - |
| - | - | 2562 | 377.2 | - | - | 0 | - |
| - | - | 941.3 | 378.2 | - | - | 0 | - |
| - | - | 9235 | 379.2 | - | - | 0 | - |
| - | - | 1129 | 380.2 | - | - | 0 | - |
| - | - | 1789 | 380.2 | - | - | 0 | - |
| - | - | 3643 | 381.1 | - | - | 0 | - |
| - | - | 1171 | 382.1 | - | - | 0 | - |
| - | - | 812 | 383.2 | - | - | 0 | - |
| - | - | 3256 | 383.2 | - | - | 0 | - |
| 6 | b | 3257 | 383.7 | 0.0005368 | 1.399 | +2 | 6 |
| - | - | 1186 | 384.2 | - | - | 0 | - |
| - | - | 821.4 | 384.7 | - | - | 0 | - |
| - | - | 784.5 | 385.2 | - | - | 0 | - |
| - | - | 739.8 | 386.2 | - | - | 0 | - |
| - | - | 9915 | 389.2 | - | - | 0 | - |
| - | - | 1767 | 390.2 | - | - | 0 | - |
| - | - | 1128 | 390.2 | - | - | 0 | - |
| - | - | 908.4 | 391.2 | - | - | 0 | - |
| - | - | 2.134E+04 | 391.2 | - | - | 0 | - |
| - | - | 4233 | 392.2 | - | - | 0 | - |
| - | - | 3723 | 392.2 | - | - | 0 | - |
| - | - | 1.109E+04 | 395.2 | - | - | 0 | - |
| - | - | 3065 | 396.2 | - | - | 0 | - |
| - | - | 1009 | 398.2 | - | - | 0 | - |
| - | - | 5353 | 403.2 | - | - | 0 | - |
| - | - | 1063 | 404.2 | - | - | 0 | - |
| - | - | 3854 | 405.2 | - | - | 0 | - |
| - | - | 2055 | 406.2 | - | - | 0 | - |
| - | - | 2.175E+04 | 407.2 | - | - | 0 | - |
| - | - | 679.1 | 407.3 | - | - | 0 | - |
| - | - | 1850 | 408.2 | - | - | 0 | - |
| - | - | 5836 | 408.2 | - | - | 0 | - |
| - | - | 1406 | 408.2 | - | - | 0 | - |
| - | - | 990.3 | 409.1 | - | - | 0 | - |
| - | - | 2340 | 409.2 | - | - | 0 | - |
| - | - | 1218 | 409.2 | - | - | 0 | - |
| - | - | 1444 | 409.3 | - | - | 0 | - |
| - | - | 1059 | 409.7 | - | - | 0 | - |
| - | - | 705.1 | 410.2 | - | - | 0 | - |
| - | - | 760.3 | 410.3 | - | - | 0 | - |
| - | - | 2279 | 411.3 | - | - | 0 | - |
| - | - | 757.7 | 412.3 | - | - | 0 | - |
| - | - | 1433 | 417.2 | - | - | 0 | - |
| - | - | 2.04E+04 | 418.2 | - | - | 0 | - |
| - | - | 8302 | 418.7 | - | - | 0 | - |
| - | - | 2118 | 419.2 | - | - | 0 | - |
| - | - | 2072 | 419.2 | - | - | 0 | - |
| - | - | 915.5 | 420.3 | - | - | 0 | - |
| - | - | 748.7 | 420.3 | - | - | 0 | - |
| - | - | 821.8 | 421.2 | - | - | 0 | - |
| 7 | b | 3.629E+04 | 423.2 | 0.006555 | 15.49 | +2 | 7 |
| - | - | 1.438E+04 | 424.2 | - | - | 0 | - |
| - | - | 2454 | 425.2 | - | - | 0 | - |
| - | - | 3.344E+04 | 426.2 | - | - | 0 | - |
| - | - | 8225 | 427.2 | - | - | 0 | - |
| 8 | y | 4866 | 427.2 | 0.00155 | 3.628 | +1 | 4 |
| - | - | 741 | 428.2 | - | - | 0 | - |
| - | - | 1596 | 428.2 | - | - | 0 | - |
| 7 | b | 1.58E+04 | 432.2 | 0.001102 | 2.549 | +2 | 7 |
| - | - | 8701 | 432.7 | - | - | 0 | - |
| - | - | 833.6 | 432.8 | - | - | 0 | - |
| - | - | 1797 | 433.2 | - | - | 0 | - |
| - | - | 1061 | 434.2 | - | - | 0 | - |
| - | - | 722.9 | 434.3 | - | - | 0 | - |
| - | - | 4832 | 435.2 | - | - | 0 | - |
| - | - | 835.6 | 436.2 | - | - | 0 | - |
| - | - | 3598 | 436.2 | - | - | 0 | - |
| - | - | 930.3 | 436.3 | - | - | 0 | - |
| - | - | 786 | 437.2 | - | - | 0 | - |
| 4 | b | 1.379E+04 | 438.3 | 0.001274 | 2.906 | +1 | 4 |
| - | - | 3340 | 438.3 | - | - | 0 | - |
| - | - | 2282 | 439.2 | - | - | 0 | - |
| 4 | b | 1683 | 439.3 | 0.0008694 | 1.979 | +1 | 4 |
| - | - | 1651 | 439.3 | - | - | 0 | - |
| - | - | 972.2 | 439.3 | - | - | 0 | - |
| - | - | 980.3 | 439.7 | - | - | 0 | - |
| - | - | 920.8 | 439.8 | - | - | 0 | - |
| - | - | 3741 | 440.3 | - | - | 0 | - |
| - | - | 756.1 | 441.3 | - | - | 0 | - |
| - | - | 982.5 | 443.3 | - | - | 0 | - |
| 8 | y | 1.021E+04 | 445.2 | 0.001575 | 3.538 | +1 | 4 |
| - | - | 2365 | 446.2 | - | - | 0 | - |
| - | - | 680.6 | 450.7 | - | - | 0 | - |
| - | - | 5757 | 452.2 | - | - | 0 | - |
| - | - | 1806 | 452.3 | - | - | 0 | - |
| - | - | 1524 | 453.2 | - | - | 0 | - |
| - | - | 701.4 | 454.2 | - | - | 0 | - |
| - | - | 2046 | 454.3 | - | - | 0 | - |
| - | - | 766.9 | 455.2 | - | - | 0 | - |
| 4 | b | 4.762E+04 | 456.3 | 0.001299 | 2.846 | +1 | 4 |
| - | - | 2526 | 457.2 | - | - | 0 | - |
| - | - | 1.157E+04 | 457.3 | - | - | 0 | - |
| - | - | 688.1 | 458.3 | - | - | 0 | - |
| - | - | 1128 | 458.3 | - | - | 0 | - |
| - | - | 1151 | 461.2 | - | - | 0 | - |
| - | - | 1857 | 464.3 | - | - | 0 | - |
| - | - | 1181 | 466.3 | - | - | 0 | - |
| - | - | 1789 | 467.8 | - | - | 0 | - |
| - | - | 1606 | 468.2 | - | - | 0 | - |
| - | - | 782.3 | 469.2 | - | - | 0 | - |
| - | - | 1.453E+04 | 470.2 | - | - | 0 | - |
| - | - | 1922 | 470.3 | - | - | 0 | - |
| - | - | 7149 | 471.2 | - | - | 0 | - |
| - | - | 1866 | 472.2 | - | - | 0 | - |
| - | - | 949.3 | 472.3 | - | - | 0 | - |
| - | - | 1267 | 473.2 | - | - | 0 | - |
| - | - | 1750 | 473.2 | - | - | 0 | - |
| - | - | 1356 | 473.3 | - | - | 0 | - |
| - | - | 893.1 | 473.8 | - | - | 0 | - |
| - | - | 2330 | 474.7 | - | - | 0 | - |
| - | - | 1877 | 476.3 | - | - | 0 | - |
| - | - | 3038 | 478.3 | - | - | 0 | - |
| - | - | 988.2 | 480.2 | - | - | 0 | - |
| - | - | 717.9 | 481.2 | - | - | 0 | - |
| - | - | 568.8 | 482.2 | - | - | 0 | - |
| - | - | 1.026E+04 | 482.3 | - | - | 0 | - |
| - | - | 7292 | 482.8 | - | - | 0 | - |
| - | - | 7046 | 483.3 | - | - | 0 | - |
| - | - | 1165 | 483.8 | - | - | 0 | - |
| - | - | 673.3 | 484.3 | - | - | 0 | - |
| - | - | 826.9 | 487.2 | - | - | 0 | - |
| 8 | b | 1514 | 487.8 | 0.0007737 | 1.586 | +2 | 8 |
| 8 | b | 3558 | 488.2 | 0.008705 | 17.83 | +2 | 8 |
| - | - | 4330 | 488.3 | - | - | 0 | - |
| - | - | 1896 | 488.7 | - | - | 0 | - |
| - | - | 2213 | 489.2 | - | - | 0 | - |
| - | - | 1489 | 489.3 | - | - | 0 | - |
| - | - | 705.8 | 489.9 | - | - | 0 | - |
| - | - | 894 | 490.3 | - | - | 0 | - |
| - | - | 911.5 | 491.2 | - | - | 0 | - |
| - | - | 1220 | 491.3 | - | - | 0 | - |
| - | - | 892.6 | 492.2 | - | - | 0 | - |
| - | - | 1348 | 495.2 | - | - | 0 | - |
| 8 | b | 2.469E+04 | 496.8 | 0.001076 | 2.166 | +2 | 8 |
| - | - | 1.529E+04 | 497.3 | - | - | 0 | - |
| - | - | 4041 | 497.8 | - | - | 0 | - |
| - | - | 1287 | 498.2 | - | - | 0 | - |
| - | - | 1568 | 500.3 | - | - | 0 | - |
| - | - | 2056 | 502.3 | - | - | 0 | - |
| - | - | 3258 | 504.2 | - | - | 0 | - |
| - | - | 1566 | 504.3 | - | - | 0 | - |
| - | - | 807.1 | 505.2 | - | - | 0 | - |
| - | - | 6033 | 506.3 | - | - | 0 | - |
| - | - | 1653 | 507.3 | - | - | 0 | - |
| - | - | 5590 | 508.3 | - | - | 0 | - |
| - | - | 7340 | 509.2 | - | - | 0 | - |
| - | - | 2516 | 510.2 | - | - | 0 | - |
| - | - | 820.7 | 510.3 | - | - | 0 | - |
| - | - | 1203 | 511.2 | - | - | 0 | - |
| - | - | 1822 | 516.2 | - | - | 0 | - |
| - | - | 2404 | 518.2 | - | - | 0 | - |
| - | - | 1961 | 518.3 | - | - | 0 | - |
| - | - | 1244 | 519.2 | - | - | 0 | - |
| - | - | 776.8 | 519.3 | - | - | 0 | - |
| - | - | 1424 | 520.3 | - | - | 0 | - |
| - | - | 3175 | 521.3 | - | - | 0 | - |
| - | - | 4601 | 522.3 | - | - | 0 | - |
| - | - | 3975 | 522.8 | - | - | 0 | - |
| - | - | 1888 | 523.2 | - | - | 0 | - |
| - | - | 4131 | 523.3 | - | - | 0 | - |
| - | - | 2372 | 523.8 | - | - | 0 | - |
| 7 | y | 1.296E+04 | 524.3 | 0.002131 | 4.065 | +1 | 5 |
| - | - | 4185 | 525.3 | - | - | 0 | - |
| - | - | 2.083E+04 | 526.3 | - | - | 0 | - |
| - | - | 4345 | 527.3 | - | - | 0 | - |
| - | - | 945.9 | 530.3 | - | - | 0 | - |
| - | - | 1273 | 530.3 | - | - | 0 | - |
| - | - | 6960 | 531.3 | - | - | 0 | - |
| - | - | 4514 | 531.8 | - | - | 0 | - |
| - | - | 1256 | 532.3 | - | - | 0 | - |
| - | - | 1.175E+04 | 534.3 | - | - | 0 | - |
| - | - | 2412 | 535.3 | - | - | 0 | - |
| - | - | 1090 | 535.3 | - | - | 0 | - |
| - | - | 1.305E+04 | 536.3 | - | - | 0 | - |
| - | - | 3590 | 536.3 | - | - | 0 | - |
| 9 | b | 2338 | 536.8 | 0.009697 | 18.07 | +2 | 9 |
| - | - | 4563 | 537.2 | - | - | 0 | - |
| - | - | 1257 | 537.3 | - | - | 0 | - |
| - | - | 1178 | 537.4 | - | - | 0 | - |
| - | - | 1830 | 538.2 | - | - | 0 | - |
| - | - | 7038 | 539.3 | - | - | 0 | - |
| - | - | 2299 | 539.8 | - | - | 0 | - |
| - | - | 3029 | 540.3 | - | - | 0 | - |
| - | - | 2097 | 540.3 | - | - | 0 | - |
| 7 | y | 1.318E+05 | 542.3 | 0.001332 | 2.456 | +1 | 5 |
| - | - | 3.431E+04 | 543.3 | - | - | 0 | - |
| - | - | 6671 | 544.3 | - | - | 0 | - |
| 9 | b | 7.769E+04 | 545.3 | 0.001 | 1.835 | +2 | 9 |
| - | - | 5.072E+04 | 545.8 | - | - | 0 | - |
| - | - | 1.834E+04 | 546.3 | - | - | 0 | - |
| - | - | 1576 | 546.8 | - | - | 0 | - |
| 3 | y | 1899 | 548.3 | 0.002619 | 4.777 | +2 | 9 |
| - | - | 2315 | 549.3 | - | - | 0 | - |
| - | - | 1538 | 551.3 | - | - | 0 | - |
| - | - | 6074 | 553.3 | - | - | 0 | - |
| - | - | 1.064E+05 | 554.3 | - | - | 0 | - |
| - | - | 3.328E+04 | 555.3 | - | - | 0 | - |
| - | - | 4860 | 556.3 | - | - | 0 | - |
| - | - | 770.8 | 556.3 | - | - | 0 | - |
| - | - | 6415 | 558.3 | - | - | 0 | - |
| - | - | 2950 | 559.3 | - | - | 0 | - |
| - | - | 1728 | 564.2 | - | - | 0 | - |
| - | - | 906.4 | 566.3 | - | - | 0 | - |
| - | - | 1039 | 567.3 | - | - | 0 | - |
| - | - | 9339 | 567.4 | - | - | 0 | - |
| - | - | 2743 | 568.4 | - | - | 0 | - |
| - | - | 1996 | 569.3 | - | - | 0 | - |
| - | - | 8138 | 570.3 | - | - | 0 | - |
| - | - | 2522 | 571.3 | - | - | 0 | - |
| - | - | 2094 | 571.8 | - | - | 0 | - |
| - | - | 927.8 | 572.3 | - | - | 0 | - |
| - | - | 1838 | 572.3 | - | - | 0 | - |
| - | - | 817 | 572.8 | - | - | 0 | - |
| - | - | 1250 | 573.3 | - | - | 0 | - |
| - | - | 2503 | 574.3 | - | - | 0 | - |
| - | - | 1487 | 575.3 | - | - | 0 | - |
| - | - | 1797 | 579.3 | - | - | 0 | - |
| - | - | 854.4 | 580.3 | - | - | 0 | - |
| - | - | 3.09E+04 | 580.8 | - | - | 0 | - |
| - | - | 2.16E+04 | 581.3 | - | - | 0 | - |
| - | - | 6756 | 581.8 | - | - | 0 | - |
| - | - | 5032 | 582.3 | - | - | 0 | - |
| - | - | 907.9 | 582.3 | - | - | 0 | - |
| - | - | 1735 | 583.3 | - | - | 0 | - |
| - | - | 728.2 | 584.3 | - | - | 0 | - |
| - | - | 2192 | 585.3 | - | - | 0 | - |
| 10 | b | 1939 | 585.8 | 0.003021 | 5.156 | +2 | 10 |
| - | - | 1716 | 586.3 | - | - | 0 | - |
| - | - | 746.3 | 590.3 | - | - | 0 | - |
| - | - | 1.651E+04 | 591.4 | - | - | 0 | - |
| - | - | 5593 | 592.4 | - | - | 0 | - |
| - | - | 972.6 | 593.4 | - | - | 0 | - |
| 10 | b | 1.947E+04 | 594.8 | 0.0009731 | 1.636 | +2 | 10 |
| - | - | 1.41E+04 | 595.3 | - | - | 0 | - |
| - | - | 5366 | 595.8 | - | - | 0 | - |
| - | - | 1276 | 596.3 | - | - | 0 | - |
| - | - | 3711 | 597.3 | - | - | 0 | - |
| - | - | 965.5 | 598.3 | - | - | 0 | - |
| - | - | 1238 | 599.3 | - | - | 0 | - |
| - | - | 786.5 | 600.3 | - | - | 0 | - |
| 5 | b | 5352 | 601.3 | 0.001208 | 2.009 | +1 | 5 |
| - | - | 1909 | 602.3 | - | - | 0 | - |
| - | - | 5792 | 603.4 | - | - | 0 | - |
| - | - | 2714 | 604.4 | - | - | 0 | - |
| - | - | 2070 | 605.3 | - | - | 0 | - |
| - | - | 1406 | 606.3 | - | - | 0 | - |
| - | - | 910.6 | 606.4 | - | - | 0 | - |
| - | - | 1036 | 607.3 | - | - | 0 | - |
| - | - | 1254 | 607.4 | - | - | 0 | - |
| - | - | 1600 | 608.3 | - | - | 0 | - |
| - | - | 1038 | 609.3 | - | - | 0 | - |
| - | - | 2276 | 615.3 | - | - | 0 | - |
| - | - | 790.2 | 616.3 | - | - | 0 | - |
| - | - | 1528 | 617.3 | - | - | 0 | - |
| - | - | 789.9 | 618.3 | - | - | 0 | - |
| 5 | b | 6.6E+04 | 619.3 | 0.001202 | 1.942 | +1 | 5 |
| - | - | 2.408E+04 | 620.3 | - | - | 0 | - |
| - | - | 4672 | 621.4 | - | - | 0 | - |
| - | - | 5105 | 623.3 | - | - | 0 | - |
| - | - | 855.5 | 624.3 | - | - | 0 | - |
| - | - | 1230 | 624.3 | - | - | 0 | - |
| - | - | 4165 | 625.3 | - | - | 0 | - |
| - | - | 1643 | 626.3 | - | - | 0 | - |
| - | - | 1421 | 629.3 | - | - | 0 | - |
| - | - | 866.1 | 630.3 | - | - | 0 | - |
| - | - | 1012 | 631.3 | - | - | 0 | - |
| - | - | 6472 | 633.3 | - | - | 0 | - |
| - | - | 1441 | 634.3 | - | - | 0 | - |
| - | - | 2742 | 635.3 | - | - | 0 | - |
| - | - | 7672 | 635.4 | - | - | 0 | - |
| - | - | 1795 | 636.3 | - | - | 0 | - |
| - | - | 3691 | 636.4 | - | - | 0 | - |
| - | - | 854.1 | 637.3 | - | - | 0 | - |
| - | - | 2931 | 638.3 | - | - | 0 | - |
| - | - | 681.8 | 639.4 | - | - | 0 | - |
| - | - | 932.9 | 645.3 | - | - | 0 | - |
| 0 | Precursor | 9717 | 645.3 | 0.001487 | 2.304 | +2 | -1 |
| 0 | Precursor | 7013 | 645.8 | 0.01027 | 15.91 | +2 | -1 |
| - | - | 3591 | 646.3 | - | - | 0 | - |
| - | - | 752.2 | 647.3 | - | - | 0 | - |
| - | - | 1368 | 649.3 | - | - | 0 | - |
| - | - | 1468 | 650.3 | - | - | 0 | - |
| - | - | 703.8 | 650.4 | - | - | 0 | - |
| - | - | 1.216E+04 | 651.3 | - | - | 0 | - |
| - | - | 3039 | 651.4 | - | - | 0 | - |
| - | - | 7462 | 652.3 | - | - | 0 | - |
| - | - | 3575 | 652.3 | - | - | 0 | - |
| - | - | 2926 | 652.4 | - | - | 0 | - |
| - | - | 1616 | 652.9 | - | - | 0 | - |
| - | - | 2717 | 653.3 | - | - | 0 | - |
| - | - | 1.388E+04 | 653.3 | - | - | 0 | - |
| 0 | Precursor | 1.495E+04 | 654.3 | 0.003002 | 4.588 | +2 | -1 |
| - | - | 9347 | 654.9 | - | - | 0 | - |
| - | - | 4433 | 655.3 | - | - | 0 | - |
| - | - | 4598 | 663.3 | - | - | 0 | - |
| - | - | 1436 | 664.3 | - | - | 0 | - |
| - | - | 2957 | 664.4 | - | - | 0 | - |
| - | - | 732.4 | 666.3 | - | - | 0 | - |
| - | - | 4615 | 667.3 | - | - | 0 | - |
| - | - | 2046 | 668.3 | - | - | 0 | - |
| - | - | 1007 | 669.4 | - | - | 0 | - |
| - | - | 894.6 | 670.4 | - | - | 0 | - |
| 6 | y | 2557 | 671.3 | 0.001954 | 2.911 | +1 | 6 |
| - | - | 1292 | 672.3 | - | - | 0 | - |
| - | - | 917.1 | 675.4 | - | - | 0 | - |
| - | - | 1.41E+04 | 681.3 | - | - | 0 | - |
| - | - | 3806 | 682.3 | - | - | 0 | - |
| - | - | 1.825E+04 | 682.4 | - | - | 0 | - |
| - | - | 1253 | 683.3 | - | - | 0 | - |
| - | - | 8271 | 683.4 | - | - | 0 | - |
| - | - | 1581 | 684.4 | - | - | 0 | - |
| - | - | 847.8 | 686.3 | - | - | 0 | - |
| - | - | 1757 | 688.4 | - | - | 0 | - |
| 6 | y | 4.441E+04 | 689.4 | 0.00146 | 2.119 | +1 | 6 |
| - | - | 1.865E+04 | 690.4 | - | - | 0 | - |
| - | - | 3505 | 691.4 | - | - | 0 | - |
| - | - | 874.8 | 692.4 | - | - | 0 | - |
| - | - | 1410 | 693.4 | - | - | 0 | - |
| - | - | 1262 | 694.4 | - | - | 0 | - |
| - | - | 1247 | 695.3 | - | - | 0 | - |
| - | - | 2045 | 703.4 | - | - | 0 | - |
| - | - | 1257 | 704.4 | - | - | 0 | - |
| - | - | 1093 | 714.3 | - | - | 0 | - |
| - | - | 1202 | 714.4 | - | - | 0 | - |
| - | - | 1015 | 715.3 | - | - | 0 | - |
| - | - | 716 | 715.4 | - | - | 0 | - |
| - | - | 8307 | 716.4 | - | - | 0 | - |
| - | - | 3566 | 717.4 | - | - | 0 | - |
| - | - | 2552 | 720.4 | - | - | 0 | - |
| - | - | 5240 | 721.4 | - | - | 0 | - |
| - | - | 3434 | 722.4 | - | - | 0 | - |
| - | - | 1149 | 723.4 | - | - | 0 | - |
| - | - | 1387 | 732.4 | - | - | 0 | - |
| - | - | 1573 | 733.4 | - | - | 0 | - |
| - | - | 1819 | 734.4 | - | - | 0 | - |
| - | - | 1118 | 735.4 | - | - | 0 | - |
| - | - | 2.662E+04 | 738.4 | - | - | 0 | - |
| - | - | 9637 | 739.4 | - | - | 0 | - |
| - | - | 2925 | 740.4 | - | - | 0 | - |
| - | - | 1618 | 744.3 | - | - | 0 | - |
| 6 | b | 1.07E+04 | 748.4 | 0.0004211 | 0.5627 | +1 | 6 |
| - | - | 696 | 749.3 | - | - | 0 | - |
| - | - | 6075 | 749.4 | - | - | 0 | - |
| - | - | 4128 | 750.4 | - | - | 0 | - |
| - | - | 2198 | 751.4 | - | - | 0 | - |
| - | - | 1.467E+04 | 752.4 | - | - | 0 | - |
| - | - | 5092 | 753.4 | - | - | 0 | - |
| - | - | 1600 | 754.4 | - | - | 0 | - |
| - | - | 836.8 | 760.4 | - | - | 0 | - |
| - | - | 6960 | 762.3 | - | - | 0 | - |
| - | - | 2757 | 763.3 | - | - | 0 | - |
| - | - | 1387 | 763.5 | - | - | 0 | - |
| - | - | 1510 | 764.4 | - | - | 0 | - |
| - | - | 1058 | 765.3 | - | - | 0 | - |
| 6 | b | 2.485E+05 | 766.4 | 0.001514 | 1.976 | +1 | 6 |
| - | - | 1.178E+05 | 767.4 | - | - | 0 | - |
| - | - | 2.575E+04 | 768.4 | - | - | 0 | - |
| - | - | 2793 | 769.4 | - | - | 0 | - |
| - | - | 689 | 778.4 | - | - | 0 | - |
| - | - | 8.628E+04 | 780.4 | - | - | 0 | - |
| - | - | 3.573E+04 | 781.4 | - | - | 0 | - |
| - | - | 9359 | 782.4 | - | - | 0 | - |
| - | - | 1032 | 782.4 | - | - | 0 | - |
| - | - | 2049 | 784.4 | - | - | 0 | - |
| - | - | 815.6 | 785.4 | - | - | 0 | - |
| - | - | 2442 | 796 | - | - | 0 | - |
| - | - | 2425 | 796.4 | - | - | 0 | - |
| - | - | 2575 | 796.7 | - | - | 0 | - |
| - | - | 1772 | 797.1 | - | - | 0 | - |
| - | - | 1691 | 811.4 | - | - | 0 | - |
| - | - | 972.4 | 812.4 | - | - | 0 | - |
| - | - | 1211 | 815.5 | - | - | 0 | - |
| - | - | 841 | 816.5 | - | - | 0 | - |
| - | - | 2491 | 817.4 | - | - | 0 | - |
| - | - | 1874 | 818.4 | - | - | 0 | - |
| - | - | 894.3 | 819.5 | - | - | 0 | - |
| - | - | 3554 | 820.5 | - | - | 0 | - |
| - | - | 1644 | 821.5 | - | - | 0 | - |
| - | - | 1401 | 822.4 | - | - | 0 | - |
| - | - | 2629 | 827.4 | - | - | 0 | - |
| - | - | 1238 | 828.4 | - | - | 0 | - |
| - | - | 2833 | 829.4 | - | - | 0 | - |
| - | - | 1682 | 830.4 | - | - | 0 | - |
| 5 | y | 816.2 | 834.4 | 0.01083 | 12.98 | +1 | 7 |
| - | - | 3426 | 835.5 | - | - | 0 | - |
| - | - | 1860 | 836.5 | - | - | 0 | - |
| 7 | b | 6805 | 845.5 | 0.01298 | 15.35 | +1 | 7 |
| 7 | b | 3376 | 846.4 | 0.00661 | 7.81 | +1 | 7 |
| - | - | 846.3 | 847.4 | - | - | 0 | - |
| - | - | 1968 | 848.4 | - | - | 0 | - |
| - | - | 5209 | 850.4 | - | - | 0 | - |
| - | - | 3637 | 851.4 | - | - | 0 | - |
| 5 | y | 2438 | 852.4 | 0.008628 | 10.12 | +1 | 7 |
| - | - | 1462 | 853.4 | - | - | 0 | - |
| - | - | 1668 | 861.4 | - | - | 0 | - |
| 7 | b | 1.879E+04 | 863.5 | 0.001912 | 2.214 | +1 | 7 |
| - | - | 9618 | 864.5 | - | - | 0 | - |
| - | - | 2652 | 865.5 | - | - | 0 | - |
| - | - | 2.998E+04 | 877.4 | - | - | 0 | - |
| - | - | 1.523E+04 | 878.4 | - | - | 0 | - |
| - | - | 1632 | 878.5 | - | - | 0 | - |
| - | - | 1.1E+04 | 879.4 | - | - | 0 | - |
| - | - | 1027 | 880 | - | - | 0 | - |
| - | - | 4095 | 880.4 | - | - | 0 | - |
| - | - | 887.1 | 881.4 | - | - | 0 | - |
| - | - | 2338 | 889.4 | - | - | 0 | - |
| - | - | 1469 | 890.4 | - | - | 0 | - |
| - | - | 2202 | 893.4 | - | - | 0 | - |
| - | - | 1312 | 894.4 | - | - | 0 | - |
| - | - | 2330 | 895.5 | - | - | 0 | - |
| - | - | 1054 | 896.5 | - | - | 0 | - |
| - | - | 1077 | 905.4 | - | - | 0 | - |
| - | - | 7779 | 907.4 | - | - | 0 | - |
| - | - | 4112 | 908.4 | - | - | 0 | - |
| - | - | 1506 | 909.4 | - | - | 0 | - |
| - | - | 1215 | 930.5 | - | - | 0 | - |
| - | - | 944.1 | 934.5 | - | - | 0 | - |
| - | - | 777.2 | 935.5 | - | - | 0 | - |
| - | - | 3338 | 946.5 | - | - | 0 | - |
| - | - | 3399 | 947.5 | - | - | 0 | - |
| - | - | 6186 | 948.5 | - | - | 0 | - |
| - | - | 2925 | 949.5 | - | - | 0 | - |
| - | - | 1775 | 956.5 | - | - | 0 | - |
| - | - | 1273 | 957.5 | - | - | 0 | - |
| - | - | 1695 | 958.5 | - | - | 0 | - |
| - | - | 940.8 | 959.5 | - | - | 0 | - |
| - | - | 5337 | 962.5 | - | - | 0 | - |
| - | - | 2818 | 963.5 | - | - | 0 | - |
| - | - | 2.724E+04 | 964.5 | - | - | 0 | - |
| - | - | 1.651E+04 | 965.5 | - | - | 0 | - |
| - | - | 4111 | 966.5 | - | - | 0 | - |
| 4 | y | 1757 | 967.4 | 0.0006153 | 0.636 | +1 | 8 |
| - | - | 934.1 | 968.4 | - | - | 0 | - |
| 8 | b | 1.332E+04 | 974.5 | 0.001256 | 1.288 | +1 | 8 |
| 8 | b | 8205 | 975.5 | 0.01944 | 19.93 | +1 | 8 |
| - | - | 2.992E+04 | 976.5 | - | - | 0 | - |
| - | - | 1.542E+04 | 977.5 | - | - | 0 | - |
| - | - | 4506 | 978.5 | - | - | 0 | - |
| - | - | 2712 | 990.5 | - | - | 0 | - |
| - | - | 2288 | 991.5 | - | - | 0 | - |
| 8 | b | 3.055E+05 | 992.5 | 0.001189 | 1.198 | +1 | 8 |
| - | - | 1.86E+05 | 993.5 | - | - | 0 | - |
| - | - | 5.731E+04 | 994.5 | - | - | 0 | - |
| - | - | 5592 | 995.5 | - | - | 0 | - |
| - | - | 987.4 | 1034 | - | - | 0 | - |
| - | - | 1672 | 1042 | - | - | 0 | - |
| - | - | 5912 | 1047 | - | - | 0 | - |
| - | - | 4635 | 1048 | - | - | 0 | - |
| - | - | 1303 | 1049 | - | - | 0 | - |
| - | - | 3252 | 1076 | - | - | 0 | - |
| - | - | 2109 | 1077 | - | - | 0 | - |
| 3 | y | 4065 | 1078 | 0.004525 | 4.199 | +1 | 9 |
| 3 | y | 2289 | 1079 | 0.021 | 19.47 | +1 | 9 |
| 9 | b | 8455 | 1090 | 0.0007322 | 0.672 | +1 | 9 |
| - | - | 4666 | 1091 | - | - | 0 | - |
| - | - | 2310 | 1092 | - | - | 0 | - |
| 3 | y | 6.936E+04 | 1096 | 0.0007962 | 0.7268 | +1 | 9 |
| - | - | 4.76E+04 | 1097 | - | - | 0 | - |
| - | - | 914.8 | 1097 | - | - | 0 | - |
| - | - | 1.418E+04 | 1098 | - | - | 0 | - |
| - | - | 1333 | 1099 | - | - | 0 | - |
| - | - | 1638 | 1161 | - | - | 0 | - |
| - | - | 1384 | 1162 | - | - | 0 | - |
| 10 | b | 2311 | 1171 | 0.0009647 | 0.8241 | +1 | 10 |
| 10 | b | 1651 | 1172 | 0.02234 | 19.07 | +1 | 10 |
| - | - | 935 | 1173 | - | - | 0 | - |
| 10 | b | 2.124E+04 | 1189 | 0.0004335 | 0.3647 | +1 | 10 |
| - | - | 1.627E+04 | 1190 | - | - | 0 | - |
| - | - | 6239 | 1191 | - | - | 0 | - |
| - | - | 843 | 1192 | - | - | 0 | - |
| 2 | y | 6596 | 1195 | 0.0004975 | 0.4165 | +1 | 10 |
| - | - | 3592 | 1196 | - | - | 0 | - |
| - | - | 1577 | 1197 | - | - | 0 | - |
| - | - | 772.7 | 1263 | - | - | 0 | - |
| - | - | 655.9 | 1659 | - | - | 0 | - |
| - | - | 858.9 | 3071 | - | - | 0 | - |
| - | - | 731.6 | 3088 | - | - | 0 | - |

m/z Charge Intensity FragmentType MassShift Position
120.06597137451172 0 41754.117 y 10
120.08121490478516 0 55196.867
121.06929779052734 0 1522.752
121.08454895019531 0 4734.162
126.05554962158203 0 586.51483
127.08720397949219 0 488.51785
128.07131958007812 0 395.72534
129.1027069091797 0 103708.484
130.05027770996094 0 1847.1521
130.1008758544922 0 813.92633
130.1060333251953 0 5473.1226
134.09706115722656 0 438.0067
136.07614135742188 0 24179.355
137.07972717285156 0 1606.4745
138.1281280517578 0 750.42444
139.08709716796875 0 2923.801
140.1438446044922 0 1039.6586
146.06048583984375 0 714.0284
146.1292724609375 0 685.5879
151.1235809326172 0 678.5554
155.11843872070312 0 1518.756
155.15464782714844 0 758.4884
156.1139678955078 0 598.56274
158.045654296875 0 800.25665
163.08718872070312 0 584.97314
165.0542755126953 0 474.23557
165.1025390625 0 2212.1438
166.08636474609375 0 1080.5864
166.2211151123047 0 456.93628
167.08206176757812 0 1417.6062
167.11827087402344 0 2866.7302
167.15444946289062 0 689.5426
168.10227966308594 0 1889.711
168.25637817382812 0 435.03134
169.1341094970703 0 311008.06
170.13739013671875 0 28437.92
171.14122009277344 0 928.87994
171.14971923828125 0 2281.5662 d 1
172.11248779296875 0 5439.3433
172.14501953125 0 507.823
172.15347290039062 0 713.50995
173.11605834960938 0 796.22107
173.12918090820312 0 1581.5135
173.4387969970703 0 1890.909
174.1282958984375 0 2757.5388
181.09759521484375 0 5109.327
181.1339874267578 0 6679.96
182.0816650390625 0 2656.0093
182.12940979003906 0 500.50122
182.13739013671875 0 738.99194
183.11343383789062 0 2147.1592
183.1497344970703 0 5324.602
184.1081085205078 0 810.1024
185.09249877929688 0 608.4687
185.12879943847656 0 1551.4196
185.16537475585938 0 131600.64 a 1
186.12814331054688 0 3566.9106
186.1687774658203 0 12924.098
188.1072540283203 0 1005.4266
188.1438446044922 0 4071.1558
189.14767456054688 0 657.1215
190.12301635742188 0 1528.4144
191.08203125 0 1901.6912
195.11318969726562 0 5585.0527
197.1289825439453 0 246210.44
198.1323699951172 0 23975.346
199.10818481445312 0 38142.47
199.13465881347656 0 1175.5227
199.14418029785156 0 745.1854
199.1811065673828 0 2128.1199
200.111328125 0 3468.837
201.113037109375 0 1162.8873
201.12380981445312 0 21063.36 y Water loss 9
202.1259307861328 0 1127.3649
203.10330200195312 0 999.0088
203.15455627441406 0 1630.8413
204.13882446289062 0 3062.3145
206.22438049316406 0 513.832
207.113037109375 0 712.35724
209.09262084960938 0 14530.167
210.09597778320312 0 1777.0906
210.16055297851562 0 911.5424
211.108642578125 0 546.7968
211.1444549560547 0 2559.7524
212.1398468017578 0 1619.1758
213.08775329589844 0 1392.8297
213.12400817871094 0 2699.846
213.16029357910156 0 62248.926 b 1
214.16378784179688 0 7178.5737
215.1395263671875 0 4784.3174
216.9047393798828 0 570.06537
217.08218383789062 0 931.5501
217.1340789794922 0 7178.9863
218.1373748779297 0 650.61084
219.13450622558594 0 34315.938 y 9
220.13795471191406 0 3075.1343
223.1083221435547 0 623.9135
225.0874786376953 0 1489.1031
226.1191864013672 0 24385.574
226.1553192138672 0 1936.2352
227.10325622558594 0 118847.266
228.10659790039062 0 11568.643
228.17123413085938 0 9904.359
229.10743713378906 0 926.9866
229.11883544921875 0 941.14685
229.1747589111328 0 1093.692
230.1867218017578 0 1758.8154
231.0988006591797 0 1488.589
231.1497344970703 0 21974.004
232.15316772460938 0 3275.066
233.09251403808594 0 1297.8962
233.16555786132812 0 3086.743
237.08726501464844 0 1096.2257
238.1232452392578 0 2046.1659
238.15554809570312 0 5101.584
239.2120361328125 0 1444.287
243.0978240966797 0 690.277
244.12982177734375 0 51161.383
245.11349487304688 0 1304.877
245.1302490234375 0 11933.504
246.13275146484375 0 1205.0238
247.14488220214844 0 4103.3447
249.1607666015625 0 1438.9744
250.16488647460938 0 539.14386
251.10324096679688 0 31535.232
251.21148681640625 0 603.1574
252.10630798339844 0 3590.7373
254.1142578125 0 4420.164
254.1506805419922 0 2434.5918
254.1869354248047 0 2122.1152
255.09829711914062 0 2311.2087
256.1659851074219 0 2082.4998
257.1278076171875 0 1297.3965
259.1446228027344 0 762.45996
261.0870666503906 0 1032.2914
261.1584777832031 0 4390.3623
262.0697326660156 0 881.589
263.13873291015625 0 1384.9487
269.1863708496094 0 1021.5948
270.1820373535156 0 2596.3782
272.12445068359375 0 7338.4805
273.1246032714844 0 974.6802
276.1715087890625 0 913.6135
277.1189270019531 0 677.49786
277.1554260253906 0 1469.5502
277.172607421875 0 662.5232
278.1498718261719 0 1503.5706
279.09820556640625 0 37969.617
280.1015319824219 0 5109.7783
280.1661682128906 0 9315.086
281.1037292480469 0 976.21564
281.1688232421875 0 1525.1497
282.1446533203125 0 647.4489
283.144775390625 0 14498.36
283.1788330078125 0 739.89374
284.1479797363281 0 2424.6895
285.1968688964844 0 572.1677
286.1557312011719 0 890.40436
287.21258544921875 0 5334.3384
288.2158508300781 0 1015.4102
292.1304016113281 0 785.4075
292.1665344238281 0 933.4517
294.1817626953125 0 1107.8973
295.17962646484375 0 670.05536
296.1243896484375 0 910.6533
296.16131591796875 0 810.6469
296.1796569824219 0 702.4102
296.1972961425781 0 933.3313
298.1769104003906 0 59707.945 y Water loss 8
299.179931640625 0 9906.859
300.18267822265625 0 1175.4315
301.1555480957031 0 1923.9294
303.2073974609375 0 4208.1045
304.1662902832031 0 2942.2627
306.14544677734375 0 3030.0151
306.21826171875 0 1045.1024
307.1481018066406 0 600.0994
308.1614685058594 0 2802.4558
310.1404113769531 0 1506.014
310.1767883300781 0 2444.3408 b 4
310.2135314941406 0 2791.2737
311.1400146484375 0 12365.921
312.1433410644531 0 1961.9558
312.1927185058594 0 1460.929
313.19140625 0 1251.2365
315.2076721191406 0 2886.9717
316.18768310546875 0 366760.06 y 8
317.19073486328125 0 57013.613
318.1925964355469 0 6589.187
319.1426086425781 0 667.5713
319.2144470214844 0 678.7512
322.17626953125 0 633.545
323.17230224609375 0 2240.7065
323.244873046875 0 4199.3496
324.15631103515625 0 105942.77
325.15948486328125 0 17617.883
325.18780517578125 0 4153.6333
325.223388671875 0 1446.3177
326.1709289550781 0 2401.3743
327.1693420410156 0 664.2753
328.15057373046875 0 2445.6492
328.1676025390625 0 1570.7202
328.190673828125 0 912.8633
329.1510314941406 0 3627.6548
330.1516418457031 0 549.37555
331.20269775390625 0 2249.142
335.1741027832031 0 2471.5293
335.2091064453125 0 611.6715
337.15185546875 0 3835.7336
337.18804931640625 0 1669.5398
338.1533203125 0 1168.1943
339.20330810546875 0 1392.3462
340.1875915527344 0 5688.491
341.18310546875 0 9389.565
341.2555847167969 0 9906.711 b 2
341.2771301269531 0 972.8698
342.1300354003906 0 1655.9668
342.1871337890625 0 1650.8342
342.2591247558594 0 1450.335
343.1983947753906 0 11849.426
344.16143798828125 0 1296.8053
344.200439453125 0 2027.2164
344.70574951171875 0 3111.259
345.1448669433594 0 633.6317
345.20904541015625 0 1077.0449
346.1769104003906 0 6490.421
347.1802062988281 0 859.7106
347.2030334472656 0 5428.636
347.7037353515625 0 1438.5481
349.1919860839844 0 1433.6154
350.19561767578125 0 624.8735
351.20672607421875 0 1088.882
351.23980712890625 0 963.3327
353.18292236328125 0 6248.389
353.218994140625 0 3880.6423
354.1857604980469 0 1294.3586
355.0718078613281 0 933.01556
355.1621398925781 0 4772.393
355.19854736328125 0 3057.325
356.07073974609375 0 914.3792
356.1626892089844 0 3392.2153
356.19976806640625 0 842.5364
357.0680847167969 0 645.98096
357.21392822265625 0 941.8242
358.2117919921875 0 1785.8579
358.70330810546875 0 792.69824
359.1246643066406 0 1022.0166
360.1556396484375 0 792.95667
360.2281494140625 0 1426.6964
361.18768310546875 0 807.5535
361.6973571777344 0 802.4807
362.17169189453125 0 2519.268
363.17242431640625 0 888.31885
367.1977233886719 0 1488.0491
369.17706298828125 0 1247.3495
369.71343994140625 0 13189.671
370.2150573730469 0 5630.1157
370.7168273925781 0 2111.027
371.19317626953125 0 9697.18
372.1563720703125 0 1839.5729
372.1958312988281 0 2170.4429
372.2261657714844 0 1027.4014
373.1877746582031 0 4674.2856
374.1719055175781 0 14619.424
375.17462158203125 0 3783.8247
375.2377624511719 0 888.196
375.69757080078125 0 724.4628
376.1512756347656 0 1204.723
377.2197265625 0 2562.0828
378.22271728515625 0 941.2859
379.1986389160156 0 9234.963
380.1615295410156 0 1129.4268
380.2015380859375 0 1788.7068
381.14532470703125 0 3642.7832
382.1475830078125 0 1171.2006
383.1561584472656 0 812.02924
383.19329833984375 0 3255.637
383.71087646484375 0 3256.6702 b 5
384.2141418457031 0 1186.3792
384.7118835449219 0 821.3635
385.2262268066406 0 784.53613
386.222412109375 0 739.8268
389.18292236328125 0 9914.503
390.1663513183594 0 1767.2527
390.1903991699219 0 1128.063
391.16851806640625 0 908.42847
391.1983947753906 0 21337.986
392.1816711425781 0 4232.7505
392.202392578125 0 3722.7874
395.2300109863281 0 11092.804
396.2323913574219 0 3064.5254
398.17327880859375 0 1008.5621
403.23486328125 0 5352.5615
404.23809814453125 0 1062.5138
405.214599609375 0 3853.8157
406.2132873535156 0 2054.743
407.1935119628906 0 21748.234
407.2656555175781 0 679.08923
408.1566467285156 0 1850.2699
408.196044921875 0 5835.9985
408.2259216308594 0 1406.476
409.138671875 0 990.2599
409.20709228515625 0 2340.0684
409.2355041503906 0 1218.0013
409.2818603515625 0 1443.6221
409.727294921875 0 1058.9219
410.2079772949219 0 705.1362
410.2822265625 0 760.29456
411.26116943359375 0 2278.5212
412.2622375488281 0 757.72736
417.17816162109375 0 1433.1265
418.240478515625 0 20396.676
418.741943359375 0 8302.371
419.1938171386719 0 2117.5996
419.241943359375 0 2071.9758
420.26153564453125 0 915.5475
420.2943115234375 0 748.6514
421.2448425292969 0 821.82184
423.2248840332031 0 36291.203 b Water loss 6
424.2253723144531 0 14378.034
425.2270812988281 0 2454.3452
426.1671142578125 0 33437.25
427.16998291015625 0 8225.188
427.22027587890625 0 4866.206 y Water loss 7
428.17376708984375 0 740.96027
428.221923828125 0 1595.9613
432.2378234863281 0 15797.861 b 6
432.739501953125 0 8701.035
432.7693786621094 0 833.61383
433.24261474609375 0 1796.8419
434.20556640625 0 1060.5574
434.2778625488281 0 722.869
435.1884765625 0 4832.163
436.15472412109375 0 835.6097
436.1889953613281 0 3597.8098
436.2549743652344 0 930.32544
437.24169921875 0 785.99036
438.2723693847656 0 13785.72 b Water loss 3
438.3083190917969 0 3340.4707
439.2066955566406 0 2281.638
439.2542419433594 0 1683.293 b Ammonia loss 3
439.2772216796875 0 1651.4296
439.3082275390625 0 972.20465
439.7109069824219 0 980.2783
439.7571716308594 0 920.8374
440.2532043457031 0 3741.2024
441.25408935546875 0 756.06525
443.29998779296875 0 982.48065
445.2308654785156 0 10207.407 y 7
446.2341613769531 0 2364.995
450.749755859375 0 680.647
452.2151184082031 0 5756.6523
452.2890625 0 1806.3555
453.21746826171875 0 1523.7068
454.2342529296875 0 701.3858
454.2688293457031 0 2045.848
455.2242431640625 0 766.9035
456.282958984375 0 47619.76 b 3
457.2461242675781 0 2526.3455
457.28564453125 0 11566.474
458.2503967285156 0 688.1359
458.2864990234375 0 1128.3785
461.24114990234375 0 1151.2788
464.2524719238281 0 1856.9678
466.2669982910156 0 1181.4869
467.7742614746094 0 1788.6665
468.2464599609375 0 1606.0404
469.2477111816406 0 782.2841
470.2257995605469 0 14527.931
470.3122863769531 0 1921.6781
471.2270202636719 0 7148.5547
472.2264404296875 0 1865.5234
472.2588195800781 0 949.3497
473.20318603515625 0 1266.5824
473.2404479980469 0 1750.4093
473.3101806640625 0 1356.0413
473.7555847167969 0 893.12067
474.74847412109375 0 2330.3748
476.2523193359375 0 1876.6636
478.2674865722656 0 3038.1694
480.2105407714844 0 988.23
481.24639892578125 0 717.93915
482.2235412597656 0 568.7994
482.26318359375 0 10261.327
482.76190185546875 0 7292.3833
483.2641906738281 0 7046.0205
483.763427734375 0 1165.2979
484.2708740234375 0 673.26337
487.2481384277344 0 826.93396
487.7535095214844 0 1513.7949 b Water loss 7
488.2534484863281 0 3557.806 b Ammonia loss 7
488.3245849609375 0 4330.333
488.74609375 0 1896.381
489.24658203125 0 2213.3677
489.3267822265625 0 1489.1294
489.91607666015625 0 705.8395
490.270263671875 0 893.97906
491.2151184082031 0 911.5414
491.2521667480469 0 1220.2487
492.21563720703125 0 892.64545
495.22320556640625 0 1347.5585
496.75909423828125 0 24694.72 b 7
497.26080322265625 0 15293
497.7618713378906 0 4041.204
498.222412109375 0 1286.7178
500.251220703125 0 1567.8647
502.2660827636719 0 2056.4592
504.2474670410156 0 3258.2917
504.3203125 0 1565.5034
505.195068359375 0 807.05634
506.2623596191406 0 6033.186
507.26177978515625 0 1653.3145
508.2560729980469 0 5589.9917
509.2420349121094 0 7339.9653
510.2442626953125 0 2516.1365
510.3283386230469 0 820.6808
511.2495422363281 0 1203.2908
516.247802734375 0 1821.5919
518.22607421875 0 2403.919
518.2618408203125 0 1961.0786
519.226318359375 0 1244.138
519.2665405273438 0 776.81464
520.2771606445312 0 1423.6399
521.2781982421875 0 3175.0273
522.2847900390625 0 4601.01
522.7855224609375 0 3974.8223
523.2197875976562 0 1887.7783
523.2888793945312 0 4130.927
523.7830200195312 0 2371.806
524.2736206054688 0 12955.9375 y Water loss 6
525.2750854492188 0 4185.17
526.2673950195312 0 20830.303
527.2701416015625 0 4345.0127
530.2590942382812 0 945.88617
530.2999877929688 0 1272.5419
531.2881469726562 0 6960.4263
531.7899780273438 0 4514.154
532.2879028320312 0 1255.6791
534.2570190429688 0 11750.78
535.26025390625 0 2411.7344
535.324951171875 0 1089.5023
536.250732421875 0 13046.157
536.3096923828125 0 3589.738
536.7808227539062 0 2338.2444 b Ammonia loss 8
537.2411499023438 0 4562.873
537.31298828125 0 1256.5381
537.3760986328125 0 1178.0892
538.2396240234375 0 1830.3379
539.2534790039062 0 7037.9297
539.7693481445312 0 2299.3396
540.2568969726562 0 3028.947
540.3085327148438 0 2097.4998
542.2833862304688 0 131752.83 y 6
543.286376953125 0 34306.953
544.2883911132812 0 6670.93
545.285400390625 0 77690.16 b 8
545.7869873046875 0 50720.492
546.2884521484375 0 18340.764
546.7901000976562 0 1575.5597
548.2741088867188 0 1899.0784 y 2
549.3403930664062 0 2314.6238
551.2849731445312 0 1537.9083
553.3355712890625 0 6073.879
554.2619018554688 0 106411.74
555.2647094726562 0 33282.957
556.2684326171875 0 4859.8447
556.312744140625 0 770.7648
558.3296508789062 0 6414.785
559.3330688476562 0 2949.9065
564.2468872070312 0 1728.2196
566.298828125 0 906.35675
567.3114624023438 0 1039.3635
567.3515014648438 0 9338.779
568.3538818359375 0 2742.8462
569.2958374023438 0 1996.2452
570.2931518554688 0 8138.0654
571.2948608398438 0 2522.3467
571.8173217773438 0 2093.7324
572.267578125 0 927.82935
572.31201171875 0 1838.2599
572.813232421875 0 817.0476
573.3450927734375 0 1250.3164
574.326416015625 0 2502.9797
575.3226928710938 0 1486.9094
579.2778930664062 0 1797.2996
580.3203735351562 0 854.4456
580.8223876953125 0 30901.594
581.3236083984375 0 21603.838
581.8250122070312 0 6756.415
582.2570190429688 0 5031.865
582.326416015625 0 907.9477
583.26171875 0 1735.4343
584.30908203125 0 728.24054
585.3388061523438 0 2192.2483
585.8163452148438 0 1939.3923 b Water loss 9
586.3182983398438 0 1716.3031
590.2945556640625 0 746.30334
591.351318359375 0 16506.488
592.3551635742188 0 5592.9106
593.3593139648438 0 972.6041
594.819580078125 0 19469.92 b 9
595.3211669921875 0 14102.171
595.8230590820312 0 5366.4766
596.320556640625 0 1276.1226
597.2879638671875 0 3711.2734
598.2941284179688 0 965.49976
599.2842407226562 0 1237.7495
600.2855224609375 0 786.4583
601.3356323242188 0 5352.206 b Water loss 4
602.3377075195312 0 1908.5159
603.35107421875 0 5791.5493
604.3550415039062 0 2713.9827
605.2969360351562 0 2069.8108
606.298095703125 0 1405.781
606.3582763671875 0 910.6142
607.3231811523438 0 1036.2936
607.3775024414062 0 1254.4031
608.3121948242188 0 1600.4852
609.3092651367188 0 1038.0874
615.2783813476562 0 2275.7422
616.282958984375 0 790.2057
617.293212890625 0 1528.2651
618.2932739257812 0 789.87805
619.34619140625 0 66002.85 b 4
620.3491821289062 0 24084.072
621.3512573242188 0 4671.522
623.3218383789062 0 5104.891
624.2662353515625 0 855.5087
624.322509765625 0 1229.9095
625.3358764648438 0 4164.8013
626.3385620117188 0 1643.1411
629.3292236328125 0 1420.9457
630.3275756835938 0 866.12134
631.3086547851562 0 1011.99695
633.291015625 0 6472.0034
634.2947387695312 0 1440.6573
635.3177490234375 0 2742.1372
635.3779907226562 0 7672.4644
636.312255859375 0 1795.1798
636.3789672851562 0 3691.177
637.3034057617188 0 854.1205
638.3194580078125 0 2931.2227
639.3515014648438 0 681.80994
645.2875366210938 0 932.86896
645.3439331054688 0 9717.211 Precursor Water loss
645.8447265625 0 7013.2085 Precursor Ammonia loss
646.3479614257812 0 3590.5867
647.3462524414062 0 752.1612
649.3311157226562 0 1368.0214
650.3241577148438 0 1467.7126
650.3921508789062 0 703.84326
651.3154907226562 0 12160.198
651.387451171875 0 3039.2405
652.26220703125 0 7462.2817
652.3201293945312 0 3574.5032
652.39892578125 0 2925.5618
652.8613891601562 0 1616.2784
653.2654418945312 0 2716.873
653.3304443359375 0 13882.241
654.3447265625 0 14945.287 Precursor
654.8501586914062 0 9346.697
655.3495483398438 0 4433.3525
663.3152465820312 0 4598.174
664.3141479492188 0 1435.6282
664.3692626953125 0 2956.519
666.3493041992188 0 732.36084
667.3482666015625 0 4614.616
668.3486328125 0 2046.0444
669.4014282226562 0 1006.8727
670.3997802734375 0 894.6475
671.3418579101562 0 2557.2317 y Water loss 5
672.3457641601562 0 1291.7233
675.3676147460938 0 917.10144
681.3258056640625 0 14101.374
682.3247680664062 0 3805.953
682.3790283203125 0 18253.127
683.3199462890625 0 1253.0978
683.3814086914062 0 8271.492
684.3851928710938 0 1581.0844
686.3455200195312 0 847.79004
688.4017333984375 0 1757.3452
689.3519287109375 0 44414.625 y 5
690.3548583984375 0 18652.254
691.3556518554688 0 3505.2847
692.3602294921875 0 874.8092
693.397705078125 0 1409.6428
694.4002075195312 0 1261.7122
695.3440551757812 0 1246.793
703.36962890625 0 2044.9875
704.369140625 0 1257.2982
714.3472900390625 0 1093.2191
714.4234008789062 0 1202.1924
715.3475952148438 0 1015.3761
715.420654296875 0 715.9969
716.3987426757812 0 8306.552
717.40234375 0 3566.0454
720.4088745117188 0 2551.6323
721.39453125 0 5239.7036
722.3934326171875 0 3433.6135
723.3932495117188 0 1148.7786
732.3644409179688 0 1386.8687
733.3576049804688 0 1573.3372
734.3546752929688 0 1819.1135
735.3512573242188 0 1117.6685
738.4197387695312 0 26623.424
739.4228515625 0 9636.516
740.426513671875 0 2924.9226
744.3358764648438 0 1617.628
748.4032592773438 0 10697.38 b Water loss 5
749.3240356445312 0 695.99713
749.4033813476562 0 6074.8955
750.3869018554688 0 4127.557
751.38525390625 0 2197.891
752.3636474609375 0 14673.113
753.3662109375 0 5091.6426
754.3656616210938 0 1600.1044
760.3562622070312 0 836.8487
762.3479614257812 0 6959.508
763.349365234375 0 2756.7334
763.4729614257812 0 1386.9202
764.4003295898438 0 1510.4977
765.3417358398438 0 1057.9193
766.4149169921875 0 248522.98 b 5
767.41796875 0 117791.836
768.4205932617188 0 25748.006
769.4224853515625 0 2793.3845
778.3814697265625 0 689.0473
780.3574829101562 0 86283.5
781.3604736328125 0 35732.523
782.3634643554688 0 9359.117
782.4387817382812 0 1032.1931
784.4274291992188 0 2049.1704
785.4252319335938 0 815.63745
796.0481567382812 0 2441.699
796.3834228515625 0 2425.2585
796.7156372070312 0 2574.7952
797.0523681640625 0 1772.4468
811.4400024414062 0 1691.0103
812.4417114257812 0 972.40686
815.4682006835938 0 1210.6565
816.4738159179688 0 840.96185
817.4472045898438 0 2490.912
818.4476318359375 0 1873.8608
819.4520874023438 0 894.3138
820.46044921875 0 3554.315
821.467041015625 0 1643.5093
822.4398193359375 0 1401.4595
827.4299926757812 0 2629.201
828.4347534179688 0 1238.108
829.4422607421875 0 2833.2986
830.4447631835938 0 1682.2925
834.4140625 0 816.21216 y Water loss 4
835.47216796875 0 3425.723
836.4766235351562 0 1860.2551
845.442626953125 0 6804.848 b Water loss 6
846.4462280273438 0 3376.2734 b Ammonia loss 6
847.4498901367188 0 846.3175
848.3864135742188 0 1967.6681
850.4348754882812 0 5209.387
851.435791015625 0 3637.0881
852.4224243164062 0 2438.0845 y 4
853.4227905273438 0 1461.6558
861.414794921875 0 1667.7473
863.4680786132812 0 18788.742 b 6
864.4703979492188 0 9617.851
865.4747314453125 0 2651.5818
877.4110717773438 0 29982.678
878.4137573242188 0 15227.169
878.4967651367188 0 1631.8262
879.4237060546875 0 10998.628
879.9883422851562 0 1027.1432
880.427978515625 0 4094.6294
881.4302978515625 0 887.0721
889.41064453125 0 2337.773
890.413818359375 0 1468.9106
893.4390869140625 0 2202.2363
894.4373168945312 0 1312.4497
895.4500122070312 0 2329.7314
896.462158203125 0 1053.703
905.4320068359375 0 1077.0557
907.421142578125 0 7778.762
908.4258422851562 0 4112.2905
909.424560546875 0 1506.4492
930.4774780273438 0 1215.3325
934.5399169921875 0 944.13464
935.5377197265625 0 777.2262
946.5044555664062 0 3337.7603
947.4990844726562 0 3399.2595
948.48876953125 0 6186.158
949.4906616210938 0 2925.397
956.4879150390625 0 1775.3217
957.49169921875 0 1273.4467
958.4703979492188 0 1695.0073
959.4747924804688 0 940.8043
962.5350341796875 0 5337.241
963.5394897460938 0 2817.5334
964.515380859375 0 27236.104
965.518310546875 0 16508.06
966.5201416015625 0 4110.6035
967.4401245117188 0 1756.8246 y 3
968.4473876953125 0 934.0629
974.4994506835938 0 13319.119 b Water loss 7
975.5016479492188 0 8205.481 b Ammonia loss 7
976.4803466796875 0 29915.793
977.4819946289062 0 15421.784
978.4868774414062 0 4505.7285
990.4954223632812 0 2712.1536
991.4981079101562 0 2288.277
992.5099487304688 0 305532.16 b 7
993.5131225585938 0 186000.17
994.5160522460938 0 57308.734
995.5186767578125 0 5592.434
1033.5394287109375 0 987.41144
1041.567626953125 0 1672.252
1046.5572509765625 0 5912.21
1047.5567626953125 0 4634.578
1048.5643310546875 0 1302.9032
1075.5477294921875 0 3251.729
1076.55224609375 0 2108.5479
1077.5296630859375 0 4064.689 y Water loss 2
1078.5301513671875 0 2288.8813 y Ammonia loss 2
1089.562255859375 0 8454.724 b 8
1090.564208984375 0 4666.0273
1091.5692138671875 0 2309.6582
1095.5364990234375 0 69359.695 y 2
1096.53955078125 0 47598.203
1096.6798095703125 0 914.83203
1097.542236328125 0 14181.738
1098.5457763671875 0 1333.1832
1160.6383056640625 0 1637.9738
1161.6412353515625 0 1384.0496
1170.618408203125 0 2310.9377 b Water loss 9
1171.625732421875 0 1651.4685 b Ammonia loss 9
1172.6160888671875 0 934.98724
1188.63037109375 0 21239.463 b 9
1189.6348876953125 0 16267.805
1190.634765625 0 6238.713
1191.6326904296875 0 843.0227
1194.6046142578125 0 6595.711 y 1
1195.6082763671875 0 3592.0952
1196.6142578125 0 1577.4436
1262.822509765625 0 772.65924
1659.474365234375 0 655.8865
3071.111328125 0 858.88617
3088.2919921875 0 731.6418

Spectrum Details

|  |  |
| --- | --- |
| Matched peaks? Matched peaks The total absolute number of peaks matched. Additionally in brackets the total fraction of peaks matched and the total number of peaks is shown. | 54 (7.25% of 745) |
| FDR? FDR The false discovery rate estimated for this peptide. It is calculated by matching all theoretical fragments with a non-integer shift with the raw peaks for this spectrum. This is done with 40 different shifts. The resulting percentage is the average number of annotated peaks over the number of annotated peaks with the correct spectrum. | 1.32% |
| Satellite FDR? Satellite FDR See the FDR for details on its calculation. This satellite ion specific FDR only contains the satellite ions (d/w) for I/L/J positions. | - |
| PSM Score? PSM Score The PSM Score as given by Hecklib to this annotated spectrum. It is shown with three significant figures. | 638 |

#### Spectrum 7692? Spectrum 7692 The raw spectrum of this peptide as annotated by Hecklib. The fragments are coloured according to ion type (see legend). Any peaks with a star '\*' as text can be hovered over to see the full details, first the ion type second the mass shift type. By hovering over the amino acids in the peptide or ions in the legend the corresponding peaks are highlighted. By toggling the 'Unassigned' label you can turn the background (unassigned) peaks on or off in the plot. By updating the slider in the Ion legend you can update the spectrum to only show the top X% of the peaks with labels. The top X% means any peak that is within X% of the highest intensity. By dragging in the spectrum you can zoom in to a specific part of the spectrum and use 'Zoom Out' to get back to the original zoom level. The annotation of the spectrum is based on the given sequence in the peptides file and is done with different software so inconsistencies are likely. The peaks are annotated based on the given sequence, with 20 ppm tolerance.

Copy Data

##### Spectrum 7692 (TSV)

###### Preview

```
Loading example...
```

*Click on the button to copy the data to your clipboard.*

Mz MinMz MaxIntensity Max

WidthHeightPeptide font sizePeptide stroke widthSpectrum font sizeSpectrum stroke widthCompact peptide

Ion legend

wxyz

abcd

OtherUnassignedIon^Charge^_Position_Show for top:%

JVKDYFPEPVT

03.56e+67.11e+61.07e+71.42e+7

Zoom Out

y^+1^_1_a^+1^_2_y^+1^_2_b^+1^_2_y^+1^_2_y^+1^_3_b^+2^_5_y^+1^_3_b^+1^_3_b^+2^_6_b^+2^_7_y^+1^_4_b^+2^_7_b^+1^_4_b^+1^_4_y^+1^_4_b^+1^_4_b^+2^_8_b^+2^_8_b^+2^_8_y^+1^_5_y^+2^_9_y^+1^_5_b^+2^_9_y^+2^_9_b^+2^_10_b^+2^_10_b^+1^_5_b^+1^_5_\*\*\*y^+1^_6_y^+1^_6_b^+1^_6_b^+1^_6_y^+1^_7_b^+1^_7_b^+1^_7_y^+1^_7_b^+1^_7_y^+1^_8_b^+1^_8_b^+1^_8_b^+1^_8_y^+1^_9_y^+1^_9_b^+1^_9_y^+1^_9_b^+1^_10_b^+1^_10_b^+1^_10_y^+1^_10_

03026049061209

Fragment Matches Table

Show background peaks

| Position | Ion type | Intensity | mz Theoretical | mz Error (Th) | mz Error (ppm) | Charge | Series Number |
| --- | --- | --- | --- | --- | --- | --- | --- |
| 11 | y | 1.538E+06 | 120.1 | 0.000406 | 3.381 | +1 | 1 |
| - | - | 2.033E+06 | 120.1 | - | - | 0 | - |
| - | - | 6.704E+04 | 121.1 | - | - | 0 | - |
| - | - | 1.615E+05 | 121.1 | - | - | 0 | - |
| - | - | 1.433E+04 | 124.4 | - | - | 0 | - |
| - | - | 3.435E+04 | 127.1 | - | - | 0 | - |
| - | - | 3.507E+04 | 128.1 | - | - | 0 | - |
| - | - | 3.931E+06 | 129.1 | - | - | 0 | - |
| - | - | 3.026E+04 | 130.1 | - | - | 0 | - |
| - | - | 3.379E+04 | 130.1 | - | - | 0 | - |
| - | - | 2.265E+05 | 130.1 | - | - | 0 | - |
| - | - | 2.89E+04 | 134.1 | - | - | 0 | - |
| - | - | 8.42E+05 | 136.1 | - | - | 0 | - |
| - | - | 7.563E+04 | 137.1 | - | - | 0 | - |
| - | - | 1.354E+05 | 139.1 | - | - | 0 | - |
| - | - | 5.884E+04 | 140.1 | - | - | 0 | - |
| - | - | 1.748E+04 | 141.1 | - | - | 0 | - |
| - | - | 1.966E+04 | 146.1 | - | - | 0 | - |
| - | - | 3.588E+04 | 146.1 | - | - | 0 | - |
| - | - | 4.496E+04 | 147 | - | - | 0 | - |
| - | - | 2.251E+04 | 147.1 | - | - | 0 | - |
| - | - | 4.388E+04 | 149 | - | - | 0 | - |
| - | - | 5.071E+04 | 155.1 | - | - | 0 | - |
| - | - | 2.217E+04 | 155.2 | - | - | 0 | - |
| - | - | 2.224E+04 | 156.1 | - | - | 0 | - |
| - | - | 5.354E+04 | 165.1 | - | - | 0 | - |
| - | - | 6.32E+04 | 167.1 | - | - | 0 | - |
| - | - | 1.041E+05 | 167.1 | - | - | 0 | - |
| - | - | 2.799E+04 | 167.2 | - | - | 0 | - |
| - | - | 1.157E+05 | 168.1 | - | - | 0 | - |
| - | - | 1.273E+07 | 169.1 | - | - | 0 | - |
| - | - | 1.135E+06 | 170.1 | - | - | 0 | - |
| - | - | 4.83E+04 | 171.1 | - | - | 0 | - |
| - | - | 1.908E+05 | 172.1 | - | - | 0 | - |
| - | - | 4.518E+04 | 172.1 | - | - | 0 | - |
| - | - | 3.164E+04 | 173.1 | - | - | 0 | - |
| - | - | 1.346E+05 | 173.4 | - | - | 0 | - |
| - | - | 7.494E+04 | 174.1 | - | - | 0 | - |
| - | - | 2.298E+05 | 181.1 | - | - | 0 | - |
| - | - | 2.039E+05 | 181.1 | - | - | 0 | - |
| - | - | 9.454E+04 | 182.1 | - | - | 0 | - |
| - | - | 2.485E+04 | 182.1 | - | - | 0 | - |
| - | - | 6.81E+04 | 183.1 | - | - | 0 | - |
| - | - | 2.038E+05 | 183.1 | - | - | 0 | - |
| - | - | 2.767E+04 | 184.1 | - | - | 0 | - |
| - | - | 7.736E+04 | 185.1 | - | - | 0 | - |
| 2 | a | 5.127E+06 | 185.2 | 0.0004893 | 2.642 | +1 | 2 |
| - | - | 1.588E+05 | 186.1 | - | - | 0 | - |
| - | - | 5.81E+05 | 186.2 | - | - | 0 | - |
| - | - | 2.916E+04 | 187.2 | - | - | 0 | - |
| - | - | 6.427E+04 | 188.1 | - | - | 0 | - |
| - | - | 1.292E+05 | 188.1 | - | - | 0 | - |
| - | - | 3.518E+04 | 189.1 | - | - | 0 | - |
| - | - | 5.684E+04 | 190.1 | - | - | 0 | - |
| - | - | 6.931E+04 | 191.1 | - | - | 0 | - |
| - | - | 2.657E+05 | 195.1 | - | - | 0 | - |
| - | - | 2.964E+04 | 196.1 | - | - | 0 | - |
| - | - | 9.737E+06 | 197.1 | - | - | 0 | - |
| - | - | 1.068E+06 | 198.1 | - | - | 0 | - |
| - | - | 2.553E+04 | 198.1 | - | - | 0 | - |
| - | - | 1.572E+06 | 199.1 | - | - | 0 | - |
| - | - | 5.147E+04 | 199.1 | - | - | 0 | - |
| - | - | 1.361E+05 | 200.1 | - | - | 0 | - |
| - | - | 4.189E+04 | 201.1 | - | - | 0 | - |
| 10 | y | 8.727E+05 | 201.1 | 0.0004105 | 2.041 | +1 | 2 |
| - | - | 7.866E+04 | 202.1 | - | - | 0 | - |
| - | - | 7.723E+04 | 203.2 | - | - | 0 | - |
| - | - | 1.93E+04 | 203.8 | - | - | 0 | - |
| - | - | 1.121E+05 | 204.1 | - | - | 0 | - |
| - | - | 2.622E+04 | 208.1 | - | - | 0 | - |
| - | - | 6.056E+05 | 209.1 | - | - | 0 | - |
| - | - | 5.179E+04 | 210.1 | - | - | 0 | - |
| - | - | 6.094E+04 | 210.2 | - | - | 0 | - |
| - | - | 2.312E+04 | 211.1 | - | - | 0 | - |
| - | - | 1.202E+05 | 211.1 | - | - | 0 | - |
| - | - | 7.295E+04 | 212.1 | - | - | 0 | - |
| - | - | 4.108E+04 | 213.1 | - | - | 0 | - |
| - | - | 1.211E+05 | 213.1 | - | - | 0 | - |
| 2 | b | 2.206E+06 | 213.2 | 0.0005087 | 2.387 | +1 | 2 |
| - | - | 2.318E+05 | 214.2 | - | - | 0 | - |
| - | - | 2.084E+05 | 215.1 | - | - | 0 | - |
| - | - | 2.908E+05 | 217.1 | - | - | 0 | - |
| - | - | 3.899E+04 | 218.1 | - | - | 0 | - |
| - | - | 1.998E+04 | 218.6 | - | - | 0 | - |
| 10 | y | 1.334E+06 | 219.1 | 0.0005117 | 2.335 | +1 | 2 |
| - | - | 1.18E+05 | 220.1 | - | - | 0 | - |
| - | - | 5.776E+04 | 225.1 | - | - | 0 | - |
| - | - | 1.11E+06 | 226.1 | - | - | 0 | - |
| - | - | 4.652E+06 | 227.1 | - | - | 0 | - |
| - | - | 4.722E+05 | 228.1 | - | - | 0 | - |
| - | - | 4.111E+05 | 228.2 | - | - | 0 | - |
| - | - | 4.299E+04 | 229.1 | - | - | 0 | - |
| - | - | 5.111E+04 | 229.2 | - | - | 0 | - |
| - | - | 5.389E+04 | 230.2 | - | - | 0 | - |
| - | - | 9.428E+05 | 231.1 | - | - | 0 | - |
| - | - | 2.202E+04 | 232.1 | - | - | 0 | - |
| - | - | 1.567E+05 | 232.2 | - | - | 0 | - |
| - | - | 4.854E+04 | 233.1 | - | - | 0 | - |
| - | - | 1.054E+05 | 233.2 | - | - | 0 | - |
| - | - | 2.996E+04 | 235.1 | - | - | 0 | - |
| - | - | 5.122E+04 | 237.1 | - | - | 0 | - |
| - | - | 1.035E+05 | 238.1 | - | - | 0 | - |
| - | - | 1.816E+05 | 238.2 | - | - | 0 | - |
| - | - | 3.021E+04 | 239.2 | - | - | 0 | - |
| - | - | 5.546E+04 | 239.2 | - | - | 0 | - |
| - | - | 4.658E+04 | 240.1 | - | - | 0 | - |
| - | - | 4.433E+04 | 243.1 | - | - | 0 | - |
| - | - | 2.192E+06 | 244.1 | - | - | 0 | - |
| - | - | 3.213E+04 | 245.1 | - | - | 0 | - |
| - | - | 4.634E+05 | 245.1 | - | - | 0 | - |
| - | - | 6.067E+04 | 246.1 | - | - | 0 | - |
| - | - | 1.861E+05 | 247.1 | - | - | 0 | - |
| - | - | 5.133E+04 | 249.2 | - | - | 0 | - |
| - | - | 1.188E+06 | 251.1 | - | - | 0 | - |
| - | - | 1.516E+05 | 252.1 | - | - | 0 | - |
| - | - | 2.891E+04 | 252.2 | - | - | 0 | - |
| - | - | 1.623E+05 | 254.1 | - | - | 0 | - |
| - | - | 8.817E+04 | 254.2 | - | - | 0 | - |
| - | - | 5.289E+04 | 254.2 | - | - | 0 | - |
| - | - | 9.546E+04 | 255.1 | - | - | 0 | - |
| - | - | 1.038E+05 | 256.2 | - | - | 0 | - |
| - | - | 5.71E+04 | 257.1 | - | - | 0 | - |
| - | - | 3.356E+04 | 259.1 | - | - | 0 | - |
| - | - | 3.352E+04 | 261.1 | - | - | 0 | - |
| - | - | 2.449E+05 | 261.2 | - | - | 0 | - |
| - | - | 2.801E+04 | 262.1 | - | - | 0 | - |
| - | - | 2.561E+04 | 262.2 | - | - | 0 | - |
| - | - | 3.302E+04 | 269.2 | - | - | 0 | - |
| - | - | 7.998E+04 | 270.2 | - | - | 0 | - |
| - | - | 3.337E+04 | 271.2 | - | - | 0 | - |
| - | - | 3.129E+05 | 272.1 | - | - | 0 | - |
| - | - | 3.93E+04 | 273.1 | - | - | 0 | - |
| - | - | 4.316E+04 | 276.2 | - | - | 0 | - |
| - | - | 6.133E+04 | 277.2 | - | - | 0 | - |
| - | - | 7.138E+04 | 278.2 | - | - | 0 | - |
| - | - | 1.547E+06 | 279.1 | - | - | 0 | - |
| - | - | 2.087E+05 | 280.1 | - | - | 0 | - |
| - | - | 3.727E+05 | 280.2 | - | - | 0 | - |
| - | - | 2.832E+04 | 281.2 | - | - | 0 | - |
| - | - | 5.723E+05 | 283.1 | - | - | 0 | - |
| - | - | 1.183E+05 | 284.1 | - | - | 0 | - |
| - | - | 2.405E+04 | 285.2 | - | - | 0 | - |
| - | - | 2.427E+05 | 287.2 | - | - | 0 | - |
| - | - | 2.352E+04 | 288.2 | - | - | 0 | - |
| - | - | 6.244E+04 | 288.2 | - | - | 0 | - |
| - | - | 3.027E+04 | 292.1 | - | - | 0 | - |
| - | - | 4.814E+04 | 292.2 | - | - | 0 | - |
| - | - | 2.8E+04 | 293.1 | - | - | 0 | - |
| - | - | 4.235E+04 | 294.2 | - | - | 0 | - |
| - | - | 3.762E+04 | 296.1 | - | - | 0 | - |
| - | - | 2.971E+04 | 296.2 | - | - | 0 | - |
| - | - | 4.327E+04 | 296.2 | - | - | 0 | - |
| 9 | y | 2.25E+06 | 298.2 | 0.0007472 | 2.506 | +1 | 3 |
| - | - | 3.362E+05 | 299.2 | - | - | 0 | - |
| - | - | 3.336E+04 | 300.2 | - | - | 0 | - |
| - | - | 5.095E+04 | 301.2 | - | - | 0 | - |
| - | - | 2.339E+04 | 302 | - | - | 0 | - |
| - | - | 1.553E+05 | 303.2 | - | - | 0 | - |
| - | - | 1.042E+05 | 304.2 | - | - | 0 | - |
| - | - | 3.085E+04 | 304.2 | - | - | 0 | - |
| - | - | 3.422E+04 | 305.2 | - | - | 0 | - |
| - | - | 2.431E+04 | 305.6 | - | - | 0 | - |
| - | - | 1.128E+05 | 306.1 | - | - | 0 | - |
| - | - | 3.848E+04 | 306.2 | - | - | 0 | - |
| - | - | 2.609E+04 | 308 | - | - | 0 | - |
| - | - | 1.019E+05 | 308.2 | - | - | 0 | - |
| - | - | 3.715E+04 | 309.2 | - | - | 0 | - |
| - | - | 5.172E+04 | 310.1 | - | - | 0 | - |
| 5 | b | 9.583E+04 | 310.2 | 0.0001674 | 0.5395 | +2 | 5 |
| - | - | 1.111E+05 | 310.2 | - | - | 0 | - |
| - | - | 4.757E+05 | 311.1 | - | - | 0 | - |
| - | - | 8.75E+04 | 312.1 | - | - | 0 | - |
| - | - | 1.023E+05 | 312.2 | - | - | 0 | - |
| - | - | 4.841E+04 | 313.2 | - | - | 0 | - |
| - | - | 1.194E+05 | 315.2 | - | - | 0 | - |
| 9 | y | 1.408E+07 | 316.2 | 0.0008942 | 2.828 | +1 | 3 |
| - | - | 2.109E+06 | 317.2 | - | - | 0 | - |
| - | - | 2.063E+05 | 318.2 | - | - | 0 | - |
| - | - | 2.45E+04 | 319.2 | - | - | 0 | - |
| - | - | 2.39E+04 | 321.1 | - | - | 0 | - |
| - | - | 8.799E+04 | 323.2 | - | - | 0 | - |
| - | - | 1.47E+05 | 323.2 | - | - | 0 | - |
| - | - | 4.068E+06 | 324.2 | - | - | 0 | - |
| - | - | 6.9E+05 | 325.2 | - | - | 0 | - |
| - | - | 1.634E+05 | 325.2 | - | - | 0 | - |
| - | - | 2.946E+04 | 325.2 | - | - | 0 | - |
| - | - | 9.9E+04 | 326.2 | - | - | 0 | - |
| - | - | 3.011E+04 | 328.1 | - | - | 0 | - |
| - | - | 8.939E+04 | 328.2 | - | - | 0 | - |
| - | - | 2.811E+04 | 328.2 | - | - | 0 | - |
| - | - | 1.239E+05 | 329.2 | - | - | 0 | - |
| - | - | 3.685E+04 | 329.2 | - | - | 0 | - |
| - | - | 3.173E+04 | 330.2 | - | - | 0 | - |
| - | - | 5.827E+04 | 331.2 | - | - | 0 | - |
| - | - | 7.112E+04 | 335.2 | - | - | 0 | - |
| - | - | 1.953E+05 | 337.2 | - | - | 0 | - |
| - | - | 4.922E+04 | 337.2 | - | - | 0 | - |
| - | - | 5.634E+04 | 338.2 | - | - | 0 | - |
| - | - | 3.633E+04 | 339.2 | - | - | 0 | - |
| - | - | 2.388E+05 | 340.2 | - | - | 0 | - |
| - | - | 3.997E+05 | 341.2 | - | - | 0 | - |
| 3 | b | 4.023E+05 | 341.3 | 0.0008063 | 2.363 | +1 | 3 |
| - | - | 5.134E+04 | 342.1 | - | - | 0 | - |
| - | - | 3.699E+04 | 342.2 | - | - | 0 | - |
| - | - | 7.91E+04 | 342.3 | - | - | 0 | - |
| - | - | 4.417E+05 | 343.2 | - | - | 0 | - |
| - | - | 4.508E+04 | 344.2 | - | - | 0 | - |
| - | - | 1.144E+05 | 344.2 | - | - | 0 | - |
| - | - | 9.652E+04 | 344.7 | - | - | 0 | - |
| - | - | 3.481E+04 | 345.2 | - | - | 0 | - |
| - | - | 2.777E+05 | 346.2 | - | - | 0 | - |
| - | - | 3.308E+04 | 347.2 | - | - | 0 | - |
| - | - | 2.06E+05 | 347.2 | - | - | 0 | - |
| - | - | 7.64E+04 | 347.7 | - | - | 0 | - |
| - | - | 2.596E+04 | 348.2 | - | - | 0 | - |
| - | - | 8.652E+04 | 349.2 | - | - | 0 | - |
| - | - | 2.352E+04 | 350.2 | - | - | 0 | - |
| - | - | 3.232E+04 | 351.2 | - | - | 0 | - |
| - | - | 2.96E+05 | 353.2 | - | - | 0 | - |
| - | - | 1.381E+05 | 353.2 | - | - | 0 | - |
| - | - | 7.975E+04 | 354.2 | - | - | 0 | - |
| - | - | 2.226E+05 | 355.2 | - | - | 0 | - |
| - | - | 1.618E+05 | 355.2 | - | - | 0 | - |
| - | - | 8.413E+04 | 356.2 | - | - | 0 | - |
| - | - | 3.071E+04 | 356.2 | - | - | 0 | - |
| - | - | 6.469E+04 | 357.2 | - | - | 0 | - |
| - | - | 3.778E+04 | 358.2 | - | - | 0 | - |
| - | - | 7.126E+04 | 358.7 | - | - | 0 | - |
| - | - | 3.377E+04 | 359.1 | - | - | 0 | - |
| - | - | 2.801E+04 | 359.2 | - | - | 0 | - |
| - | - | 5.47E+04 | 360.2 | - | - | 0 | - |
| - | - | 6.097E+04 | 361.2 | - | - | 0 | - |
| - | - | 9.361E+04 | 362.2 | - | - | 0 | - |
| - | - | 5.431E+04 | 363.1 | - | - | 0 | - |
| - | - | 3.001E+04 | 363.2 | - | - | 0 | - |
| - | - | 2.695E+04 | 363.2 | - | - | 0 | - |
| - | - | 3.053E+04 | 364.2 | - | - | 0 | - |
| - | - | 2.615E+04 | 365.2 | - | - | 0 | - |
| - | - | 6.979E+04 | 367.2 | - | - | 0 | - |
| - | - | 3.954E+04 | 369.2 | - | - | 0 | - |
| - | - | 4.128E+05 | 369.7 | - | - | 0 | - |
| - | - | 2.139E+05 | 370.2 | - | - | 0 | - |
| - | - | 4.669E+04 | 370.7 | - | - | 0 | - |
| - | - | 4.497E+05 | 371.2 | - | - | 0 | - |
| - | - | 9.042E+04 | 372.2 | - | - | 0 | - |
| - | - | 1.056E+05 | 372.2 | - | - | 0 | - |
| - | - | 3.619E+04 | 372.2 | - | - | 0 | - |
| - | - | 1.479E+05 | 373.2 | - | - | 0 | - |
| - | - | 6.729E+05 | 374.2 | - | - | 0 | - |
| - | - | 1.358E+05 | 375.2 | - | - | 0 | - |
| - | - | 4.694E+04 | 376.2 | - | - | 0 | - |
| - | - | 9.816E+04 | 377.2 | - | - | 0 | - |
| - | - | 3.077E+04 | 378.2 | - | - | 0 | - |
| - | - | 3.426E+05 | 379.2 | - | - | 0 | - |
| - | - | 3.34E+04 | 380.2 | - | - | 0 | - |
| - | - | 7.389E+04 | 380.2 | - | - | 0 | - |
| - | - | 1.724E+05 | 381.1 | - | - | 0 | - |
| - | - | 3.899E+04 | 383.2 | - | - | 0 | - |
| - | - | 1.274E+05 | 383.2 | - | - | 0 | - |
| 6 | b | 9.976E+04 | 383.7 | 0.0008725 | 2.274 | +2 | 6 |
| - | - | 3.659E+04 | 384.2 | - | - | 0 | - |
| - | - | 6.408E+04 | 385.2 | - | - | 0 | - |
| - | - | 2.765E+04 | 387.5 | - | - | 0 | - |
| - | - | 4.324E+05 | 389.2 | - | - | 0 | - |
| - | - | 7.02E+04 | 390.2 | - | - | 0 | - |
| - | - | 4.102E+04 | 390.2 | - | - | 0 | - |
| - | - | 2.636E+04 | 391.1 | - | - | 0 | - |
| - | - | 4.081E+04 | 391.2 | - | - | 0 | - |
| - | - | 8.941E+05 | 391.2 | - | - | 0 | - |
| - | - | 1.792E+05 | 392.2 | - | - | 0 | - |
| - | - | 1.003E+05 | 392.2 | - | - | 0 | - |
| - | - | 3.163E+04 | 393.3 | - | - | 0 | - |
| - | - | 2.62E+04 | 395 | - | - | 0 | - |
| - | - | 4.437E+05 | 395.2 | - | - | 0 | - |
| - | - | 5.207E+04 | 395.7 | - | - | 0 | - |
| - | - | 9.446E+04 | 396.2 | - | - | 0 | - |
| - | - | 2.552E+04 | 396.7 | - | - | 0 | - |
| - | - | 3.301E+04 | 398.2 | - | - | 0 | - |
| - | - | 1.89E+05 | 403.2 | - | - | 0 | - |
| - | - | 4.722E+04 | 404.2 | - | - | 0 | - |
| - | - | 1.383E+05 | 405.2 | - | - | 0 | - |
| - | - | 5.615E+04 | 406.2 | - | - | 0 | - |
| - | - | 9.368E+05 | 407.2 | - | - | 0 | - |
| - | - | 9.282E+04 | 408.2 | - | - | 0 | - |
| - | - | 2.485E+05 | 408.2 | - | - | 0 | - |
| - | - | 4.246E+04 | 408.2 | - | - | 0 | - |
| - | - | 4.352E+04 | 409.1 | - | - | 0 | - |
| - | - | 9.577E+04 | 409.2 | - | - | 0 | - |
| - | - | 4.541E+04 | 409.3 | - | - | 0 | - |
| - | - | 5.674E+04 | 411.3 | - | - | 0 | - |
| - | - | 4.918E+04 | 417.2 | - | - | 0 | - |
| - | - | 7.665E+05 | 418.2 | - | - | 0 | - |
| - | - | 3.999E+05 | 418.7 | - | - | 0 | - |
| - | - | 8.089E+04 | 419.2 | - | - | 0 | - |
| - | - | 6.445E+04 | 419.2 | - | - | 0 | - |
| - | - | 5.051E+04 | 420.3 | - | - | 0 | - |
| - | - | 4.599E+04 | 420.3 | - | - | 0 | - |
| - | - | 3.673E+04 | 422.2 | - | - | 0 | - |
| - | - | 2.84E+04 | 422.2 | - | - | 0 | - |
| 7 | b | 1.446E+06 | 423.2 | 0.006647 | 15.7 | +2 | 7 |
| - | - | 5.418E+05 | 424.2 | - | - | 0 | - |
| - | - | 6.849E+04 | 425.2 | - | - | 0 | - |
| - | - | 1.344E+06 | 426.2 | - | - | 0 | - |
| - | - | 3.236E+05 | 427.2 | - | - | 0 | - |
| 8 | y | 1.293E+05 | 427.2 | 0.001306 | 3.057 | +1 | 4 |
| - | - | 6.825E+04 | 428.2 | - | - | 0 | - |
| 7 | b | 6.405E+05 | 432.2 | 0.00101 | 2.338 | +2 | 7 |
| - | - | 3.405E+05 | 432.7 | - | - | 0 | - |
| - | - | 7.778E+04 | 433.2 | - | - | 0 | - |
| - | - | 7.91E+04 | 434.2 | - | - | 0 | - |
| - | - | 2.411E+05 | 435.2 | - | - | 0 | - |
| - | - | 3.477E+04 | 436.2 | - | - | 0 | - |
| - | - | 1.347E+05 | 436.2 | - | - | 0 | - |
| - | - | 3.591E+04 | 436.3 | - | - | 0 | - |
| - | - | 4.976E+04 | 437.2 | - | - | 0 | - |
| 4 | b | 5.232E+05 | 438.3 | 0.001121 | 2.558 | +1 | 4 |
| - | - | 1.326E+05 | 438.3 | - | - | 0 | - |
| - | - | 1.21E+05 | 439.2 | - | - | 0 | - |
| 4 | b | 8.683E+04 | 439.3 | 0.001206 | 2.745 | +1 | 4 |
| - | - | 4.071E+04 | 439.3 | - | - | 0 | - |
| - | - | 3.997E+04 | 440.2 | - | - | 0 | - |
| - | - | 1.72E+05 | 440.3 | - | - | 0 | - |
| - | - | 4.735E+04 | 443.2 | - | - | 0 | - |
| - | - | 4.335E+04 | 443.3 | - | - | 0 | - |
| - | - | 3.17E+04 | 444.2 | - | - | 0 | - |
| 8 | y | 3.894E+05 | 445.2 | 0.00185 | 4.154 | +1 | 4 |
| - | - | 9.036E+04 | 446.2 | - | - | 0 | - |
| - | - | 2.863E+04 | 447.2 | - | - | 0 | - |
| - | - | 3.68E+04 | 450.2 | - | - | 0 | - |
| - | - | 2.709E+05 | 452.2 | - | - | 0 | - |
| - | - | 7.508E+04 | 452.3 | - | - | 0 | - |
| - | - | 4.826E+04 | 453.2 | - | - | 0 | - |
| - | - | 2.7E+04 | 454.2 | - | - | 0 | - |
| - | - | 4.39E+04 | 454.2 | - | - | 0 | - |
| - | - | 7.928E+04 | 454.3 | - | - | 0 | - |
| - | - | 2.813E+04 | 455.2 | - | - | 0 | - |
| 4 | b | 1.963E+06 | 456.3 | 0.001115 | 2.445 | +1 | 4 |
| - | - | 1.021E+05 | 457.2 | - | - | 0 | - |
| - | - | 5.602E+05 | 457.3 | - | - | 0 | - |
| - | - | 3.189E+04 | 458.3 | - | - | 0 | - |
| - | - | 7.294E+04 | 458.3 | - | - | 0 | - |
| - | - | 3.955E+04 | 459.3 | - | - | 0 | - |
| - | - | 5.634E+04 | 461.2 | - | - | 0 | - |
| - | - | 4.268E+04 | 464.2 | - | - | 0 | - |
| - | - | 7.884E+04 | 464.3 | - | - | 0 | - |
| - | - | 5.295E+04 | 466.3 | - | - | 0 | - |
| - | - | 4.393E+04 | 466.8 | - | - | 0 | - |
| - | - | 6.745E+04 | 467.8 | - | - | 0 | - |
| - | - | 8.007E+04 | 468.2 | - | - | 0 | - |
| - | - | 6.082E+05 | 470.2 | - | - | 0 | - |
| - | - | 7.18E+04 | 470.3 | - | - | 0 | - |
| - | - | 3.391E+05 | 471.2 | - | - | 0 | - |
| - | - | 9.642E+04 | 472.2 | - | - | 0 | - |
| - | - | 4.964E+04 | 473.2 | - | - | 0 | - |
| - | - | 4.991E+04 | 473.2 | - | - | 0 | - |
| - | - | 5.724E+04 | 473.3 | - | - | 0 | - |
| - | - | 4.112E+04 | 473.8 | - | - | 0 | - |
| - | - | 2.737E+04 | 474.3 | - | - | 0 | - |
| - | - | 6.172E+04 | 475.3 | - | - | 0 | - |
| - | - | 7.173E+04 | 476.3 | - | - | 0 | - |
| - | - | 1.1E+05 | 478.3 | - | - | 0 | - |
| - | - | 3.091E+04 | 479.3 | - | - | 0 | - |
| - | - | 3.108E+04 | 480.2 | - | - | 0 | - |
| - | - | 4.539E+04 | 480.3 | - | - | 0 | - |
| - | - | 6.118E+04 | 481.2 | - | - | 0 | - |
| - | - | 3.35E+04 | 481.8 | - | - | 0 | - |
| - | - | 2.351E+04 | 482.2 | - | - | 0 | - |
| - | - | 3.999E+05 | 482.3 | - | - | 0 | - |
| - | - | 3.803E+05 | 482.8 | - | - | 0 | - |
| - | - | 3.047E+05 | 483.3 | - | - | 0 | - |
| - | - | 5.77E+04 | 483.8 | - | - | 0 | - |
| - | - | 5.011E+04 | 486.2 | - | - | 0 | - |
| - | - | 3.218E+04 | 487.3 | - | - | 0 | - |
| 8 | b | 5.854E+04 | 487.8 | 0.001048 | 2.149 | +2 | 8 |
| 8 | b | 1.735E+05 | 488.2 | 0.008491 | 17.39 | +2 | 8 |
| - | - | 1.905E+05 | 488.3 | - | - | 0 | - |
| - | - | 1.124E+05 | 488.7 | - | - | 0 | - |
| - | - | 6.88E+04 | 489.2 | - | - | 0 | - |
| - | - | 6.612E+04 | 489.3 | - | - | 0 | - |
| - | - | 6.746E+04 | 490.3 | - | - | 0 | - |
| - | - | 4.501E+04 | 491.2 | - | - | 0 | - |
| - | - | 3.946E+04 | 491.3 | - | - | 0 | - |
| - | - | 4.297E+04 | 492.3 | - | - | 0 | - |
| - | - | 5.323E+04 | 495.2 | - | - | 0 | - |
| - | - | 6.172E+04 | 495.8 | - | - | 0 | - |
| 8 | b | 8.175E+05 | 496.8 | 0.0008015 | 1.613 | +2 | 8 |
| - | - | 5.64E+05 | 497.3 | - | - | 0 | - |
| - | - | 1.719E+05 | 497.8 | - | - | 0 | - |
| - | - | 6.072E+04 | 498.2 | - | - | 0 | - |
| - | - | 5.902E+04 | 500.3 | - | - | 0 | - |
| - | - | 6.553E+04 | 502.3 | - | - | 0 | - |
| - | - | 1.377E+05 | 504.2 | - | - | 0 | - |
| - | - | 5.951E+04 | 504.3 | - | - | 0 | - |
| - | - | 5.623E+04 | 505.2 | - | - | 0 | - |
| - | - | 2.601E+05 | 506.3 | - | - | 0 | - |
| - | - | 5.175E+04 | 507.3 | - | - | 0 | - |
| - | - | 1.91E+05 | 508.3 | - | - | 0 | - |
| - | - | 3.162E+05 | 509.2 | - | - | 0 | - |
| - | - | 1.043E+05 | 510.2 | - | - | 0 | - |
| - | - | 5.681E+04 | 510.3 | - | - | 0 | - |
| - | - | 3.38E+04 | 511.3 | - | - | 0 | - |
| - | - | 4.157E+04 | 513.3 | - | - | 0 | - |
| - | - | 7.502E+04 | 516.2 | - | - | 0 | - |
| - | - | 8.632E+04 | 518.2 | - | - | 0 | - |
| - | - | 1.131E+05 | 518.3 | - | - | 0 | - |
| - | - | 2.919E+04 | 519.3 | - | - | 0 | - |
| - | - | 6.251E+04 | 520.3 | - | - | 0 | - |
| - | - | 1.32E+05 | 521.3 | - | - | 0 | - |
| - | - | 4.639E+04 | 521.8 | - | - | 0 | - |
| - | - | 1.163E+05 | 522.3 | - | - | 0 | - |
| - | - | 4.335E+04 | 522.8 | - | - | 0 | - |
| - | - | 1.17E+05 | 523.2 | - | - | 0 | - |
| - | - | 9.388E+04 | 523.3 | - | - | 0 | - |
| - | - | 8.525E+04 | 523.8 | - | - | 0 | - |
| 7 | y | 5.753E+05 | 524.3 | 0.00207 | 3.948 | +1 | 5 |
| - | - | 3.077E+04 | 524.8 | - | - | 0 | - |
| - | - | 3.454E+04 | 525.2 | - | - | 0 | - |
| - | - | 1.476E+05 | 525.3 | - | - | 0 | - |
| - | - | 6.829E+05 | 526.3 | - | - | 0 | - |
| - | - | 2.345E+05 | 527.3 | - | - | 0 | - |
| - | - | 2.946E+04 | 530.8 | - | - | 0 | - |
| - | - | 2.514E+05 | 531.3 | - | - | 0 | - |
| - | - | 1.489E+05 | 531.8 | - | - | 0 | - |
| - | - | 6.301E+04 | 532.3 | - | - | 0 | - |
| - | - | 4.619E+05 | 534.3 | - | - | 0 | - |
| - | - | 1.346E+05 | 535.3 | - | - | 0 | - |
| - | - | 3.479E+04 | 535.3 | - | - | 0 | - |
| - | - | 5.073E+05 | 536.3 | - | - | 0 | - |
| - | - | 1.409E+05 | 536.3 | - | - | 0 | - |
| - | - | 6.5E+04 | 536.8 | - | - | 0 | - |
| - | - | 1.772E+05 | 537.2 | - | - | 0 | - |
| - | - | 3.259E+04 | 538.2 | - | - | 0 | - |
| - | - | 2.946E+05 | 539.3 | - | - | 0 | - |
| 3 | y | 5.02E+04 | 539.8 | 0.01046 | 19.38 | +2 | 9 |
| - | - | 8.269E+04 | 540.3 | - | - | 0 | - |
| - | - | 7.506E+04 | 540.3 | - | - | 0 | - |
| - | - | 4.093E+04 | 541.3 | - | - | 0 | - |
| 7 | y | 4.631E+06 | 542.3 | 0.00121 | 2.231 | +1 | 5 |
| - | - | 1.299E+06 | 543.3 | - | - | 0 | - |
| - | - | 2.162E+05 | 544.3 | - | - | 0 | - |
| 9 | b | 2.982E+06 | 545.3 | 0.0008783 | 1.611 | +2 | 9 |
| - | - | 1.875E+06 | 545.8 | - | - | 0 | - |
| - | - | 5.559E+05 | 546.3 | - | - | 0 | - |
| - | - | 5.439E+04 | 546.8 | - | - | 0 | - |
| 3 | y | 6.923E+04 | 548.3 | 0.001521 | 2.773 | +2 | 9 |
| - | - | 7.455E+04 | 549.3 | - | - | 0 | - |
| - | - | 4.221E+04 | 551.3 | - | - | 0 | - |
| - | - | 4.389E+04 | 552.3 | - | - | 0 | - |
| - | - | 2.263E+05 | 553.3 | - | - | 0 | - |
| - | - | 4.052E+06 | 554.3 | - | - | 0 | - |
| - | - | 1.179E+06 | 555.3 | - | - | 0 | - |
| - | - | 2.133E+05 | 556.3 | - | - | 0 | - |
| - | - | 3.002E+05 | 558.3 | - | - | 0 | - |
| - | - | 7.581E+04 | 559.3 | - | - | 0 | - |
| - | - | 8.289E+04 | 564.2 | - | - | 0 | - |
| - | - | 3.51E+04 | 566.3 | - | - | 0 | - |
| - | - | 4.85E+04 | 567.3 | - | - | 0 | - |
| - | - | 3.404E+05 | 567.4 | - | - | 0 | - |
| - | - | 3.883E+04 | 568.3 | - | - | 0 | - |
| - | - | 1.18E+05 | 568.4 | - | - | 0 | - |
| - | - | 3.044E+05 | 570.3 | - | - | 0 | - |
| - | - | 1.099E+05 | 571.3 | - | - | 0 | - |
| - | - | 6.346E+04 | 571.8 | - | - | 0 | - |
| - | - | 4.825E+04 | 572.3 | - | - | 0 | - |
| - | - | 9.075E+04 | 572.3 | - | - | 0 | - |
| - | - | 6.097E+04 | 572.8 | - | - | 0 | - |
| - | - | 8.02E+04 | 573.3 | - | - | 0 | - |
| - | - | 6.002E+04 | 574.3 | - | - | 0 | - |
| - | - | 4.015E+04 | 575.3 | - | - | 0 | - |
| - | - | 3.61E+04 | 579.3 | - | - | 0 | - |
| - | - | 4.317E+04 | 580.3 | - | - | 0 | - |
| - | - | 1.001E+06 | 580.8 | - | - | 0 | - |
| - | - | 7.93E+05 | 581.3 | - | - | 0 | - |
| - | - | 2.615E+05 | 581.8 | - | - | 0 | - |
| - | - | 2.468E+05 | 582.3 | - | - | 0 | - |
| - | - | 4.287E+04 | 582.3 | - | - | 0 | - |
| - | - | 1.039E+05 | 583.3 | - | - | 0 | - |
| - | - | 7.62E+04 | 585.3 | - | - | 0 | - |
| 10 | b | 4.128E+04 | 585.8 | 0.001922 | 3.281 | +2 | 10 |
| - | - | 4.986E+04 | 586.3 | - | - | 0 | - |
| - | - | 2.417E+04 | 591.3 | - | - | 0 | - |
| - | - | 7.609E+05 | 591.4 | - | - | 0 | - |
| - | - | 1.96E+05 | 592.4 | - | - | 0 | - |
| - | - | 5.695E+04 | 593.4 | - | - | 0 | - |
| 10 | b | 8.156E+05 | 594.8 | 0.000851 | 1.431 | +2 | 10 |
| - | - | 5.433E+05 | 595.3 | - | - | 0 | - |
| - | - | 1.646E+05 | 595.8 | - | - | 0 | - |
| - | - | 4.317E+04 | 596.3 | - | - | 0 | - |
| - | - | 1.722E+05 | 597.3 | - | - | 0 | - |
| - | - | 6.267E+04 | 599.3 | - | - | 0 | - |
| 5 | b | 1.617E+05 | 601.3 | 0.001635 | 2.719 | +1 | 5 |
| - | - | 6.491E+04 | 602.3 | - | - | 0 | - |
| - | - | 2.994E+05 | 603.4 | - | - | 0 | - |
| - | - | 7.923E+04 | 604.4 | - | - | 0 | - |
| - | - | 9.097E+04 | 605.3 | - | - | 0 | - |
| - | - | 4.051E+04 | 606.3 | - | - | 0 | - |
| - | - | 3.114E+04 | 607.4 | - | - | 0 | - |
| - | - | 4.996E+04 | 608.3 | - | - | 0 | - |
| - | - | 4.159E+04 | 609.3 | - | - | 0 | - |
| - | - | 9.011E+04 | 615.3 | - | - | 0 | - |
| - | - | 6.72E+04 | 617.3 | - | - | 0 | - |
| 5 | b | 2.522E+06 | 619.3 | 0.001141 | 1.843 | +1 | 5 |
| - | - | 8.952E+05 | 620.3 | - | - | 0 | - |
| - | - | 1.889E+05 | 621.4 | - | - | 0 | - |
| - | - | 3.781E+04 | 622.3 | - | - | 0 | - |
| - | - | 1.777E+05 | 623.3 | - | - | 0 | - |
| - | - | 5.041E+04 | 624.3 | - | - | 0 | - |
| - | - | 7.93E+04 | 624.3 | - | - | 0 | - |
| - | - | 1.511E+05 | 625.3 | - | - | 0 | - |
| - | - | 5.332E+04 | 626.3 | - | - | 0 | - |
| - | - | 4.631E+04 | 629.3 | - | - | 0 | - |
| - | - | 6.383E+04 | 631.3 | - | - | 0 | - |
| - | - | 2.402E+05 | 633.3 | - | - | 0 | - |
| - | - | 8.447E+04 | 634.3 | - | - | 0 | - |
| - | - | 7.865E+04 | 635.3 | - | - | 0 | - |
| - | - | 3.232E+05 | 635.4 | - | - | 0 | - |
| - | - | 3.874E+04 | 636.3 | - | - | 0 | - |
| - | - | 1.335E+05 | 636.4 | - | - | 0 | - |
| - | - | 3.706E+04 | 637.4 | - | - | 0 | - |
| - | - | 1.339E+05 | 638.3 | - | - | 0 | - |
| - | - | 3.133E+04 | 640.4 | - | - | 0 | - |
| 0 | Precursor | 3.722E+05 | 645.3 | 0.0008765 | 1.358 | +2 | -1 |
| 0 | Precursor | 2.753E+05 | 645.8 | 0.01058 | 16.38 | +2 | -1 |
| - | - | 1.282E+05 | 646.3 | - | - | 0 | - |
| - | - | 4.059E+04 | 649.3 | - | - | 0 | - |
| - | - | 4.618E+04 | 650.3 | - | - | 0 | - |
| - | - | 3.963E+05 | 651.3 | - | - | 0 | - |
| - | - | 7.181E+04 | 651.4 | - | - | 0 | - |
| - | - | 3.407E+05 | 652.3 | - | - | 0 | - |
| - | - | 1.25E+05 | 652.3 | - | - | 0 | - |
| - | - | 6.409E+04 | 652.4 | - | - | 0 | - |
| - | - | 1.096E+05 | 653.3 | - | - | 0 | - |
| - | - | 5.155E+05 | 653.3 | - | - | 0 | - |
| 0 | Precursor | 5.547E+05 | 654.3 | 0.003124 | 4.774 | +2 | -1 |
| - | - | 3.099E+05 | 654.8 | - | - | 0 | - |
| - | - | 1.382E+05 | 655.4 | - | - | 0 | - |
| - | - | 1.599E+05 | 663.3 | - | - | 0 | - |
| - | - | 1.284E+05 | 664.4 | - | - | 0 | - |
| - | - | 5.305E+04 | 665.4 | - | - | 0 | - |
| - | - | 6.621E+04 | 666.3 | - | - | 0 | - |
| - | - | 2.033E+05 | 667.3 | - | - | 0 | - |
| - | - | 7.922E+04 | 668.3 | - | - | 0 | - |
| - | - | 4.713E+04 | 669.4 | - | - | 0 | - |
| 6 | y | 9.664E+04 | 671.3 | 0.001344 | 2.002 | +1 | 6 |
| - | - | 4.301E+04 | 672.3 | - | - | 0 | - |
| - | - | 5.123E+04 | 673.4 | - | - | 0 | - |
| - | - | 2.898E+04 | 679.3 | - | - | 0 | - |
| - | - | 5.918E+05 | 681.3 | - | - | 0 | - |
| - | - | 1.556E+05 | 682.3 | - | - | 0 | - |
| - | - | 7.622E+05 | 682.4 | - | - | 0 | - |
| - | - | 4.589E+04 | 683.3 | - | - | 0 | - |
| - | - | 2.873E+05 | 683.4 | - | - | 0 | - |
| - | - | 5.691E+04 | 684.4 | - | - | 0 | - |
| - | - | 5.833E+04 | 688.4 | - | - | 0 | - |
| 6 | y | 1.764E+06 | 689.4 | 0.0008501 | 1.233 | +1 | 6 |
| - | - | 6.228E+05 | 690.4 | - | - | 0 | - |
| - | - | 1.676E+05 | 691.4 | - | - | 0 | - |
| - | - | 5.884E+04 | 693.4 | - | - | 0 | - |
| - | - | 3.903E+04 | 694.4 | - | - | 0 | - |
| - | - | 4.439E+04 | 695.3 | - | - | 0 | - |
| - | - | 4.083E+04 | 700.4 | - | - | 0 | - |
| - | - | 3.39E+04 | 702.4 | - | - | 0 | - |
| - | - | 8.054E+04 | 703.4 | - | - | 0 | - |
| - | - | 4.627E+04 | 704.4 | - | - | 0 | - |
| - | - | 2.846E+04 | 714.4 | - | - | 0 | - |
| - | - | 7.215E+04 | 714.4 | - | - | 0 | - |
| - | - | 3.175E+05 | 716.4 | - | - | 0 | - |
| - | - | 1.566E+05 | 717.4 | - | - | 0 | - |
| - | - | 3.064E+04 | 718.4 | - | - | 0 | - |
| - | - | 1.226E+05 | 720.4 | - | - | 0 | - |
| - | - | 2.479E+05 | 721.4 | - | - | 0 | - |
| - | - | 1.477E+05 | 722.4 | - | - | 0 | - |
| - | - | 2.977E+04 | 723.4 | - | - | 0 | - |
| - | - | 3.704E+04 | 730.4 | - | - | 0 | - |
| - | - | 3.581E+04 | 731.4 | - | - | 0 | - |
| - | - | 7.007E+04 | 732.4 | - | - | 0 | - |
| - | - | 7.147E+04 | 734.4 | - | - | 0 | - |
| - | - | 8.067E+04 | 735.3 | - | - | 0 | - |
| - | - | 9.498E+05 | 738.4 | - | - | 0 | - |
| - | - | 4.526E+05 | 739.4 | - | - | 0 | - |
| - | - | 8.617E+04 | 740.4 | - | - | 0 | - |
| - | - | 3.107E+04 | 742.3 | - | - | 0 | - |
| - | - | 4.974E+04 | 744.3 | - | - | 0 | - |
| 6 | b | 3.818E+05 | 748.4 | 0.0003113 | 0.416 | +1 | 6 |
| - | - | 4.409E+04 | 749.3 | - | - | 0 | - |
| - | - | 2.164E+05 | 749.4 | - | - | 0 | - |
| - | - | 1.988E+05 | 750.4 | - | - | 0 | - |
| - | - | 1.005E+05 | 751.4 | - | - | 0 | - |
| - | - | 5.307E+05 | 752.4 | - | - | 0 | - |
| - | - | 2.095E+05 | 753.4 | - | - | 0 | - |
| - | - | 6.165E+04 | 754.4 | - | - | 0 | - |
| - | - | 4.257E+04 | 760.4 | - | - | 0 | - |
| - | - | 2.748E+05 | 762.3 | - | - | 0 | - |
| - | - | 1.365E+05 | 763.3 | - | - | 0 | - |
| - | - | 4.429E+04 | 763.5 | - | - | 0 | - |
| - | - | 7.505E+04 | 764.4 | - | - | 0 | - |
| - | - | 2.854E+04 | 764.5 | - | - | 0 | - |
| - | - | 3.928E+04 | 765.4 | - | - | 0 | - |
| 6 | b | 9.295E+06 | 766.4 | 0.00127 | 1.657 | +1 | 6 |
| - | - | 4.139E+06 | 767.4 | - | - | 0 | - |
| - | - | 8.788E+05 | 768.4 | - | - | 0 | - |
| - | - | 6.422E+04 | 769.4 | - | - | 0 | - |
| - | - | 5.932E+04 | 778.4 | - | - | 0 | - |
| - | - | 3.347E+06 | 780.4 | - | - | 0 | - |
| - | - | 1.553E+06 | 781.4 | - | - | 0 | - |
| - | - | 3.097E+05 | 782.4 | - | - | 0 | - |
| - | - | 4.724E+04 | 782.4 | - | - | 0 | - |
| - | - | 2.829E+04 | 783.4 | - | - | 0 | - |
| - | - | 7.206E+04 | 784.4 | - | - | 0 | - |
| - | - | 5.216E+04 | 785.4 | - | - | 0 | - |
| - | - | 4.306E+04 | 790.3 | - | - | 0 | - |
| - | - | 3.195E+04 | 799.4 | - | - | 0 | - |
| - | - | 4.567E+04 | 808.4 | - | - | 0 | - |
| - | - | 6.754E+04 | 811.4 | - | - | 0 | - |
| - | - | 6.344E+04 | 815.5 | - | - | 0 | - |
| - | - | 9.893E+04 | 817.4 | - | - | 0 | - |
| - | - | 6.319E+04 | 818.5 | - | - | 0 | - |
| - | - | 1.764E+05 | 820.5 | - | - | 0 | - |
| - | - | 8.759E+04 | 821.5 | - | - | 0 | - |
| - | - | 7.755E+04 | 822.4 | - | - | 0 | - |
| - | - | 3.44E+04 | 823.4 | - | - | 0 | - |
| - | - | 7.003E+04 | 827.4 | - | - | 0 | - |
| - | - | 1.192E+05 | 829.4 | - | - | 0 | - |
| - | - | 5.191E+04 | 830.5 | - | - | 0 | - |
| - | - | 4.327E+04 | 832.4 | - | - | 0 | - |
| 5 | y | 3.81E+04 | 834.4 | 0.008328 | 9.981 | +1 | 7 |
| - | - | 1.195E+05 | 835.5 | - | - | 0 | - |
| - | - | 6.494E+04 | 836.5 | - | - | 0 | - |
| 7 | b | 2.838E+05 | 845.5 | 0.01316 | 15.56 | +1 | 7 |
| 7 | b | 1.312E+05 | 846.4 | 0.005573 | 6.584 | +1 | 7 |
| - | - | 7.14E+04 | 847.5 | - | - | 0 | - |
| - | - | 8.355E+04 | 848.4 | - | - | 0 | - |
| - | - | 5.553E+04 | 849.4 | - | - | 0 | - |
| - | - | 2.06E+05 | 850.4 | - | - | 0 | - |
| - | - | 1.616E+05 | 851.4 | - | - | 0 | - |
| 5 | y | 8.448E+04 | 852.4 | 0.01082 | 12.7 | +1 | 7 |
| - | - | 3.895E+04 | 861.4 | - | - | 0 | - |
| 7 | b | 6.57E+05 | 863.5 | 0.001118 | 1.295 | +1 | 7 |
| - | - | 2.931E+05 | 864.5 | - | - | 0 | - |
| - | - | 1.095E+05 | 865.5 | - | - | 0 | - |
| - | - | 1.129E+06 | 877.4 | - | - | 0 | - |
| - | - | 5.203E+05 | 878.4 | - | - | 0 | - |
| - | - | 4.2E+04 | 878.5 | - | - | 0 | - |
| - | - | 4.983E+05 | 879.4 | - | - | 0 | - |
| - | - | 2.925E+04 | 879.5 | - | - | 0 | - |
| - | - | 1.966E+05 | 880.4 | - | - | 0 | - |
| - | - | 4.053E+04 | 881.4 | - | - | 0 | - |
| - | - | 1.043E+05 | 889.4 | - | - | 0 | - |
| - | - | 4.759E+04 | 890.4 | - | - | 0 | - |
| - | - | 7.49E+04 | 893.4 | - | - | 0 | - |
| - | - | 6.635E+04 | 894.4 | - | - | 0 | - |
| - | - | 6.216E+04 | 895.5 | - | - | 0 | - |
| - | - | 3.153E+05 | 907.4 | - | - | 0 | - |
| - | - | 1.41E+05 | 908.4 | - | - | 0 | - |
| - | - | 3.388E+04 | 909.4 | - | - | 0 | - |
| - | - | 4.444E+04 | 928.5 | - | - | 0 | - |
| - | - | 5.135E+04 | 929.5 | - | - | 0 | - |
| - | - | 4.449E+04 | 930.5 | - | - | 0 | - |
| - | - | 3.594E+04 | 931.5 | - | - | 0 | - |
| - | - | 4.201E+04 | 934.5 | - | - | 0 | - |
| - | - | 3.045E+04 | 942.5 | - | - | 0 | - |
| - | - | 3.878E+04 | 944.5 | - | - | 0 | - |
| - | - | 1.156E+05 | 946.5 | - | - | 0 | - |
| - | - | 9.368E+04 | 947.5 | - | - | 0 | - |
| - | - | 2.015E+05 | 948.5 | - | - | 0 | - |
| - | - | 1.154E+05 | 949.5 | - | - | 0 | - |
| - | - | 6.226E+04 | 956.5 | - | - | 0 | - |
| - | - | 4.549E+04 | 957.5 | - | - | 0 | - |
| - | - | 5.131E+04 | 958.5 | - | - | 0 | - |
| - | - | 5.118E+04 | 959.5 | - | - | 0 | - |
| - | - | 2.017E+05 | 962.5 | - | - | 0 | - |
| - | - | 9.818E+04 | 963.5 | - | - | 0 | - |
| - | - | 9.877E+05 | 964.5 | - | - | 0 | - |
| - | - | 6.327E+05 | 965.5 | - | - | 0 | - |
| - | - | 1.329E+05 | 966.5 | - | - | 0 | - |
| 4 | y | 9.53E+04 | 967.4 | 0.0001881 | 0.1944 | +1 | 8 |
| - | - | 4.947E+04 | 968.4 | - | - | 0 | - |
| 8 | b | 5.651E+05 | 974.5 | 0.001744 | 1.789 | +1 | 8 |
| 8 | b | 3.681E+05 | 975.5 | 0.01919 | 19.68 | +1 | 8 |
| - | - | 1.022E+06 | 976.5 | - | - | 0 | - |
| - | - | 5.945E+05 | 977.5 | - | - | 0 | - |
| - | - | 1.832E+05 | 978.5 | - | - | 0 | - |
| - | - | 1.001E+05 | 990.5 | - | - | 0 | - |
| - | - | 1.18E+05 | 991.5 | - | - | 0 | - |
| 8 | b | 1.089E+07 | 992.5 | 0.0008838 | 0.8904 | +1 | 8 |
| - | - | 6.363E+06 | 993.5 | - | - | 0 | - |
| - | - | 1.834E+06 | 994.5 | - | - | 0 | - |
| - | - | 1.97E+05 | 995.5 | - | - | 0 | - |
| - | - | 3.536E+04 | 1026 | - | - | 0 | - |
| - | - | 4.195E+04 | 1042 | - | - | 0 | - |
| - | - | 4.112E+04 | 1043 | - | - | 0 | - |
| - | - | 2.91E+05 | 1047 | - | - | 0 | - |
| - | - | 1.9E+05 | 1048 | - | - | 0 | - |
| - | - | 4.219E+04 | 1049 | - | - | 0 | - |
| - | - | 1.027E+05 | 1076 | - | - | 0 | - |
| - | - | 7.884E+04 | 1077 | - | - | 0 | - |
| 3 | y | 1.457E+05 | 1078 | 0.00306 | 2.84 | +1 | 9 |
| 3 | y | 1.001E+05 | 1079 | 0.02051 | 19.02 | +1 | 9 |
| - | - | 3.54E+04 | 1080 | - | - | 0 | - |
| 9 | b | 2.584E+05 | 1090 | 2.192E-07 | 0.0002012 | +1 | 9 |
| - | - | 1.626E+05 | 1091 | - | - | 0 | - |
| - | - | 7.265E+04 | 1092 | - | - | 0 | - |
| 3 | y | 2.43E+06 | 1096 | 0.0007962 | 0.7268 | +1 | 9 |
| - | - | 1.584E+06 | 1097 | - | - | 0 | - |
| - | - | 3.203E+04 | 1097 | - | - | 0 | - |
| - | - | 4.95E+05 | 1098 | - | - | 0 | - |
| - | - | 4.946E+04 | 1099 | - | - | 0 | - |
| - | - | 4.011E+04 | 1161 | - | - | 0 | - |
| - | - | 4.33E+04 | 1162 | - | - | 0 | - |
| 10 | b | 9.15E+04 | 1171 | 0.0009884 | 0.8444 | +1 | 10 |
| 10 | b | 6.224E+04 | 1172 | 0.009649 | 8.235 | +1 | 10 |
| - | - | 3.876E+04 | 1173 | - | - | 0 | - |
| 10 | b | 8.213E+05 | 1189 | 0.0001894 | 0.1593 | +1 | 10 |
| - | - | 5.912E+05 | 1190 | - | - | 0 | - |
| - | - | 2.097E+05 | 1191 | - | - | 0 | - |
| 2 | y | 2.082E+05 | 1195 | 0.0008453 | 0.7076 | +1 | 10 |
| - | - | 1.183E+05 | 1196 | - | - | 0 | - |
| - | - | 4.507E+04 | 1197 | - | - | 0 | - |

m/z Charge Intensity FragmentType MassShift Position
120.06592559814453 0 1538437.4 y 10
120.08119201660156 0 2033100.1
121.06925964355469 0 67037.38
121.08452606201172 0 161527.53
124.41740417480469 0 14326.688
127.08702087402344 0 34347.703
128.07125854492188 0 35067.34
129.10264587402344 0 3930805.5
130.05029296875 0 30260.115
130.10081481933594 0 33788.33
130.1060028076172 0 226483.97
134.09703063964844 0 28901.78
136.0760955810547 0 842049.75
137.0794219970703 0 75633.375
139.0869903564453 0 135371.98
140.1437530517578 0 58844.035
141.10308837890625 0 17484.488
146.06085205078125 0 19656.846
146.12925720214844 0 35884.11
147.04434204101562 0 44963.316
147.1131591796875 0 22508.766
148.95379638671875 0 43876.477
155.11830139160156 0 50708.094
155.15478515625 0 22170.193
156.11367797851562 0 22243.932
165.10296630859375 0 53542.547
167.08189392089844 0 63201.633
167.1183319091797 0 104120.76
167.15464782714844 0 27985.344
168.10243225097656 0 115681.37
169.13404846191406 0 12732121
170.13734436035156 0 1135079.2
171.14013671875 0 48301.695
172.1124725341797 0 190784.27
172.14468383789062 0 45180.914
173.11593627929688 0 31637.635
173.4398956298828 0 134639.22
174.12808227539062 0 74939.766
181.09768676757812 0 229796.3
181.134033203125 0 203873.14
182.08140563964844 0 94538.54
182.12918090820312 0 24853.967
183.1132049560547 0 68104.37
183.1497039794922 0 203834.58
184.1084442138672 0 27668.752
185.128662109375 0 77362.27
185.1653289794922 0 5127044.5 a 1
186.1282196044922 0 158801.44
186.16867065429688 0 581001.4
187.17274475097656 0 29161.36
188.1073455810547 0 64269.547
188.14390563964844 0 129181.34
189.147216796875 0 35175.723
190.12301635742188 0 56843.31
191.08212280273438 0 69313.24
195.11317443847656 0 265665.72
196.11679077148438 0 29635.678
197.12892150878906 0 9737033
198.13230895996094 0 1068001.6
198.1477508544922 0 25529.803
199.10813903808594 0 1571563.8
199.13507080078125 0 51474.727
200.1114501953125 0 136089.33
201.1129913330078 0 41885.953
201.123779296875 0 872675.44 y Water loss 9
202.12615966796875 0 78663.43
203.15475463867188 0 77225.61
203.77980041503906 0 19301.98
204.13868713378906 0 112060.7
208.10675048828125 0 26216.172
209.09254455566406 0 605605.25
210.09573364257812 0 51789.586
210.16065979003906 0 60935.832
211.10911560058594 0 23119.076
211.1444549560547 0 120232.38
212.13987731933594 0 72950.055
213.0875701904297 0 41077.457
213.1237335205078 0 121127.305
213.16026306152344 0 2205713 b 1
214.16354370117188 0 231753.55
215.1394805908203 0 208382.94
217.1341094970703 0 290768.72
218.13809204101562 0 38988.105
218.5554962158203 0 19976.684
219.1344451904297 0 1334204.4 y 9
220.13780212402344 0 117975.38
225.08731079101562 0 57762.473
226.11912536621094 0 1109883.6
227.1031951904297 0 4651964
228.10629272460938 0 472221.25
228.1712646484375 0 411085.94
229.1079559326172 0 42991.63
229.17425537109375 0 51108.29
230.18663024902344 0 53888.5
231.14974975585938 0 942761.4
232.13385009765625 0 22023.018
232.15325927734375 0 156690.53
233.0923309326172 0 48541.48
233.16546630859375 0 105368.55
235.14442443847656 0 29962.62
237.08712768554688 0 51219.688
238.12326049804688 0 103528.08
238.15548706054688 0 181633.64
239.15908813476562 0 30206.133
239.2127227783203 0 55463.14
240.134765625 0 46582.668
243.09796142578125 0 44328.56
244.12974548339844 0 2192105.5
245.09213256835938 0 32129.893
245.13015747070312 0 463440.7
246.13279724121094 0 60672.21
247.1446533203125 0 186093.83
249.16064453125 0 51331.4
251.10316467285156 0 1187629.6
252.10667419433594 0 151638.17
252.16941833496094 0 28905.725
254.11387634277344 0 162304.89
254.15066528320312 0 88172.58
254.18650817871094 0 52885.07
255.09814453125 0 95456.26
256.1658020019531 0 103836.66
257.1285400390625 0 57102.965
259.1444396972656 0 33556.44
261.08770751953125 0 33524.668
261.15863037109375 0 244873.47
262.0711975097656 0 28010.676
262.1585998535156 0 25611.373
269.1852111816406 0 33018.246
270.1815490722656 0 79983.34
271.18328857421875 0 33367.44
272.12469482421875 0 312890.22
273.1251525878906 0 39297.08
276.1707763671875 0 43164.758
277.1551208496094 0 61333.113
278.1507263183594 0 71382.8
279.09820556640625 0 1547409.9
280.1014709472656 0 208669.77
280.1661376953125 0 372666.16
281.1690979003906 0 28315.99
283.1447448730469 0 572346.2
284.14801025390625 0 118325.305
285.1968994140625 0 24052.902
287.21240234375 0 242723.67
288.16290283203125 0 23523.807
288.2158203125 0 62438.527
292.1293029785156 0 30267.773
292.16607666015625 0 48141.98
293.1297302246094 0 28003.035
294.1814880371094 0 42347.227
296.1248779296875 0 37619.08
296.1609191894531 0 29714.361
296.1797180175781 0 43266.184
298.1768798828125 0 2250068.5 y Water loss 8
299.1799621582031 0 336246.4
300.1830749511719 0 33359.94
301.1556396484375 0 50950.95
302.0123291015625 0 23387.6
303.2074890136719 0 155259.98
304.1662292480469 0 104164.89
304.2105712890625 0 30852.621
305.1697692871094 0 34224.47
305.5661926269531 0 24307.145
306.14501953125 0 112765.68
306.218505859375 0 38481.98
308.0395202636719 0 26093.113
308.16119384765625 0 101870.3
309.1578369140625 0 37153.348
310.1402587890625 0 51720.273
310.1763000488281 0 95832.06 b 4
310.21343994140625 0 111063.83
311.1398010253906 0 475725.2
312.14251708984375 0 87504.89
312.1923828125 0 102302.63
313.1924133300781 0 48406.105
315.207763671875 0 119423.27
316.1875915527344 0 14084603 y 8
317.1906433105469 0 2109168.5
318.19256591796875 0 206277.16
319.21356201171875 0 24500.53
321.1244812011719 0 23896.67
323.1725769042969 0 87992.99
323.244873046875 0 147017.64
324.1562194824219 0 4068311.2
325.1594543457031 0 689983.56
325.18743896484375 0 163417.31
325.22381591796875 0 29462.826
326.170654296875 0 98996.23
328.1478271484375 0 30106.93
328.16632080078125 0 89388.12
328.1933288574219 0 28108.906
329.1507873535156 0 123868.12
329.1885070800781 0 36848.266
330.2188415527344 0 31727.857
331.20245361328125 0 58267.258
335.1735534667969 0 71120.6
337.1517028808594 0 195346.33
337.1873779296875 0 49224.418
338.153076171875 0 56340.95
339.2022705078125 0 36332.22
340.18756103515625 0 238788.84
341.1831359863281 0 399707.53
341.2555236816406 0 402311.97 b 2
342.1300964355469 0 51337.1
342.1872863769531 0 36993.36
342.2586364746094 0 79096.68
343.1982116699219 0 441725.4
344.1617736816406 0 45084.156
344.2008361816406 0 114382.41
344.7059326171875 0 96519.95
345.1910095214844 0 34811.21
346.1769714355469 0 277703.03
347.18109130859375 0 33077.195
347.2028503417969 0 206026.73
347.7037658691406 0 76398.23
348.1559753417969 0 25962.65
349.1910400390625 0 86523.07
350.1944885253906 0 23519.861
351.2063293457031 0 32320.463
353.1826171875 0 296024.88
353.21917724609375 0 138111.56
354.1857604980469 0 79750.305
355.16192626953125 0 222573.02
355.19854736328125 0 161758.38
356.16259765625 0 84133.02
356.1997985839844 0 30710.838
357.2133483886719 0 64691.2
358.2140808105469 0 37782.72
358.70294189453125 0 71262.164
359.1249084472656 0 33772.31
359.2065124511719 0 28005.012
360.228271484375 0 54703.99
361.1877746582031 0 60968.695
362.17169189453125 0 93611.46
363.1354064941406 0 54310.625
363.173095703125 0 30011.543
363.2019958496094 0 26949.648
364.1888122558594 0 30525.354
365.182861328125 0 26151.154
367.19854736328125 0 69791.14
369.1767272949219 0 39542.773
369.7137451171875 0 412768.47
370.21490478515625 0 213868.2
370.7148742675781 0 46694.18
371.1929931640625 0 449662.72
372.15576171875 0 90421.54
372.19573974609375 0 105553.97
372.2228088378906 0 36192.773
373.18780517578125 0 147878.52
374.1717529296875 0 672943.3
375.175048828125 0 135815.11
376.1526794433594 0 46939.316
377.218505859375 0 98159.92
378.2235107421875 0 30767.8
379.19854736328125 0 342570.5
380.1625061035156 0 33396.598
380.2021789550781 0 73890.84
381.1455078125 0 172432.86
383.15655517578125 0 38988.8
383.1940002441406 0 127441.46
383.7112121582031 0 99762.984 b 5
384.2136535644531 0 36588.82
385.2252502441406 0 64077.684
387.5411682128906 0 27650.33
389.1827392578125 0 432424.7
390.1664123535156 0 70199.85
390.1899108886719 0 41016.85
391.1280517578125 0 26363.121
391.1698913574219 0 40808.08
391.1983642578125 0 894126.9
392.1817321777344 0 179232
392.2028503417969 0 100264.86
393.25048828125 0 31630.666
394.9561462402344 0 26202.78
395.2296142578125 0 443671.44
395.72979736328125 0 52070.234
396.2321472167969 0 94464.05
396.73638916015625 0 25518.176
398.1712646484375 0 33012.914
403.23492431640625 0 188982.22
404.2367248535156 0 47223.36
405.21392822265625 0 138267.28
406.2140197753906 0 56154.336
407.1933898925781 0 936806.75
408.1562194824219 0 92823.87
408.1954650878906 0 248469.9
408.22515869140625 0 42461.31
409.1385803222656 0 43519.863
409.2077331542969 0 95772.5
409.2818908691406 0 45405.246
411.26123046875 0 56735.598
417.17742919921875 0 49179.2
418.2403564453125 0 766457.9
418.7417297363281 0 399919.56
419.19317626953125 0 80891.625
419.2431335449219 0 64453.62
420.2633361816406 0 50514.215
420.296875 0 45986.324
422.2091369628906 0 36725.18
422.23992919921875 0 28399.041
423.22479248046875 0 1446282.1 b Water loss 6
424.2248840332031 0 541794.56
425.2276306152344 0 68491.99
426.16693115234375 0 1344283.6
427.1701965332031 0 323586.9
427.22003173828125 0 129312.39 y Water loss 7
428.22216796875 0 68249.055
432.23773193359375 0 640489.06 b 6
432.7392578125 0 340457.38
433.2409973144531 0 77781.37
434.20391845703125 0 79101.625
435.1881408691406 0 241070.19
436.1564025878906 0 34774.617
436.18963623046875 0 134738.05
436.2557678222656 0 35913.453
437.240234375 0 49758.96
438.272216796875 0 523237.16 b Water loss 3
438.3083190917969 0 132645.55
439.2083435058594 0 120980.18
439.2563171386719 0 86830.04 b Ammonia loss 3
439.2801208496094 0 40707.94
440.21435546875 0 39971.684
440.2529296875 0 172007.73
443.22894287109375 0 47346.812
443.3038330078125 0 43352.01
444.2322692871094 0 31695.691
445.23114013671875 0 389385.8 y 7
446.23553466796875 0 90360.63
447.2278137207031 0 28626.514
450.2353820800781 0 36797.418
452.2153015136719 0 270947.84
452.2882080078125 0 75078.01
453.2190246582031 0 48263.11
454.1614685058594 0 26997.355
454.231201171875 0 43903.8
454.2679138183594 0 79278.44
455.2165222167969 0 28128.357
456.28277587890625 0 1962715.8 b 3
457.2459411621094 0 102072.266
457.2853088378906 0 560182
458.25140380859375 0 31887.938
458.2876281738281 0 72937.24
459.2605895996094 0 39550
461.23992919921875 0 56335.445
464.2164611816406 0 42679.746
464.25225830078125 0 78837.086
466.26617431640625 0 52949.332
466.7640075683594 0 43930.234
467.77508544921875 0 67446.76
468.24505615234375 0 80067.44
470.2255859375 0 608168
470.3125915527344 0 71801.68
471.2264099121094 0 339107.72
472.22711181640625 0 96423.26
473.2041015625 0 49644.66
473.240234375 0 49909.43
473.30841064453125 0 57236.81
473.7564392089844 0 41119.38
474.31341552734375 0 27372.838
475.2539978027344 0 61724.63
476.2516174316406 0 71732.13
478.2674865722656 0 110037.73
479.2685546875 0 30912.99
480.20892333984375 0 31081.707
480.2816162109375 0 45385.17
481.246337890625 0 61184.168
481.77166748046875 0 33501.84
482.2247009277344 0 23512.299
482.2622375488281 0 399872.16
482.76153564453125 0 380262.06
483.263916015625 0 304688.47
483.7645568847656 0 57699.047
486.2362365722656 0 50108.562
487.2541809082031 0 32183.941
487.7537841796875 0 58540.46 b Water loss 7
488.25323486328125 0 173467.06 b Ammonia loss 7
488.32476806640625 0 190463.12
488.74420166015625 0 112367.06
489.244384765625 0 68799.26
489.3285217285156 0 66117.74
490.26678466796875 0 67462.01
491.21466064453125 0 45010.586
491.2532653808594 0 39461.996
492.28411865234375 0 42969.82
495.221435546875 0 53228.355
495.7515563964844 0 61715.53
496.7588195800781 0 817460.06 b 7
497.2606201171875 0 563967.2
497.7621154785156 0 171949.89
498.2205810546875 0 60717.68
500.25115966796875 0 59017.496
502.26531982421875 0 65532.59
504.2469482421875 0 137695.08
504.3206787109375 0 59512.7
505.1930847167969 0 56225.754
506.2624206542969 0 260142.11
507.2630310058594 0 51752.73
508.2561950683594 0 191007.92
509.2410583496094 0 316240.7
510.24310302734375 0 104270.414
510.3288879394531 0 56808.164
511.2535705566406 0 33799.832
513.2887573242188 0 41572.14
516.2471313476562 0 75018.67
518.2250366210938 0 86319.766
518.2623901367188 0 113088.82
519.265380859375 0 29186.977
520.2763671875 0 62514.48
521.2781982421875 0 132022.14
521.7853393554688 0 46393.008
522.2845458984375 0 116329.984
522.7843017578125 0 43353.41
523.21923828125 0 117006.13
523.2908325195312 0 93877.195
523.7828369140625 0 85253.25
524.2735595703125 0 575289.5 y Water loss 6
524.7836303710938 0 30767.227
525.2340698242188 0 34538.24
525.2755737304688 0 147619.66
526.2672119140625 0 682856.3
527.2705078125 0 234508.81
530.8018188476562 0 29455.584
531.2881469726562 0 251364.03
531.7882080078125 0 148909.77
532.2913208007812 0 63012.246
534.2567749023438 0 461920.34
535.259765625 0 134637.33
535.3245239257812 0 34788.99
536.2510986328125 0 507273.1
536.3094482421875 0 140851.47
536.783203125 0 64999.812
537.2422485351562 0 177190.61
538.2376098632812 0 32591.89
539.2537841796875 0 294551.62
539.7686767578125 0 50195.62 y Ammonia loss 2
540.2562255859375 0 82690.29
540.3093872070312 0 75060.88
541.3108520507812 0 40926.277
542.2832641601562 0 4630943 y 6
543.2862548828125 0 1299191.6
544.2883911132812 0 216173.81
545.2852783203125 0 2981549 b 8
545.7869873046875 0 1875380
546.2879638671875 0 555867.9
546.7902221679688 0 54392.348
548.2730102539062 0 69231.68 y 2
549.3406982421875 0 74550.93
551.2822875976562 0 42214.863
552.2823486328125 0 43894.54
553.3351440429688 0 226343.12
554.26171875 0 4051637.2
555.2647094726562 0 1178679.6
556.267578125 0 213337.53
558.3297119140625 0 300152.2
559.3331298828125 0 75806.71
564.2462768554688 0 82886.18
566.2972412109375 0 35096.824
567.3120727539062 0 48504.312
567.3519897460938 0 340356.44
568.3123779296875 0 38827.746
568.3553466796875 0 118045.4
570.2930297851562 0 304385.3
571.2952880859375 0 109881.07
571.8154296875 0 63458.1
572.2683715820312 0 48250.527
572.3128051757812 0 90751.69
572.8152465820312 0 60966.613
573.3397827148438 0 80201.414
574.324951171875 0 60018.625
575.3211059570312 0 40152.613
579.2762451171875 0 36100.305
580.316162109375 0 43166.418
580.8221435546875 0 1001431.5
581.3233642578125 0 793012.2
581.824951171875 0 261507.28
582.2572631835938 0 246777.78
582.3265991210938 0 42865.855
583.2587890625 0 103921.195
585.334716796875 0 76198.54
585.8152465820312 0 41279.53 b Water loss 9
586.3201904296875 0 49864.574
591.297119140625 0 24165.168
591.3512573242188 0 760884.25
592.3541259765625 0 196048.64
593.357421875 0 56953.254
594.8194580078125 0 815563.4 b 9
595.3212280273438 0 543285.56
595.8224487304688 0 164563.19
596.3197631835938 0 43170.734
597.2880249023438 0 172236.34
599.2844848632812 0 62665.594
601.3360595703125 0 161719.62 b Water loss 4
602.3330688476562 0 64909.348
603.3507080078125 0 299426.7
604.3539428710938 0 79227.55
605.296142578125 0 90972.516
606.2960815429688 0 40509.31
607.3809814453125 0 31143.492
608.3054809570312 0 49957.92
609.3121948242188 0 41593.56
615.278564453125 0 90107.734
617.2926635742188 0 67199.28
619.3461303710938 0 2522499.5 b 4
620.3489379882812 0 895156.44
621.3505249023438 0 188898.05
622.3232421875 0 37806.723
623.3203125 0 177672.39
624.2659912109375 0 50412.113
624.3224487304688 0 79304.08
625.3339233398438 0 151067.75
626.3385620117188 0 53323.67
629.330078125 0 46313.266
631.3106689453125 0 63834.332
633.290283203125 0 240175.44
634.2936401367188 0 84467.83
635.3175659179688 0 78651.984
635.3776245117188 0 323179.4
636.3009643554688 0 38735.55
636.3790893554688 0 133459.64
637.3820190429688 0 37063.203
638.3190307617188 0 133876.55
640.3577270507812 0 31331.922
645.3433227539062 0 372173.47 Precursor Water loss
645.8450317382812 0 275335.06 Precursor Ammonia loss
646.3491821289062 0 128244.945
649.3388671875 0 40591.312
650.3255004882812 0 46177.93
651.3155517578125 0 396267.62
651.386474609375 0 71813.94
652.2617797851562 0 340714.56
652.3189697265625 0 125043.375
652.399169921875 0 64090.133
653.265380859375 0 109639.164
653.330078125 0 515490.47
654.3446044921875 0 554660.44 Precursor
654.8492431640625 0 309896.3
655.3501586914062 0 138212.25
663.3133544921875 0 159911.53
664.3680419921875 0 128366.48
665.3690185546875 0 53047.285
666.3451538085938 0 66208.234
667.346435546875 0 203305.9
668.3479614257812 0 79218.06
669.4024658203125 0 47132.35
671.3412475585938 0 96642.47 y Water loss 5
672.3449096679688 0 43013.363
673.3950805664062 0 51232.26
679.31005859375 0 28979.502
681.3253173828125 0 591828.5
682.325439453125 0 155598.6
682.3787841796875 0 762202.3
683.3207397460938 0 45893.426
683.38134765625 0 287345.34
684.3828125 0 56913.395
688.4014892578125 0 58334.426
689.351318359375 0 1763587.4 y 5
690.354736328125 0 622765.7
691.3569946289062 0 167574.78
693.3975219726562 0 58838.12
694.40478515625 0 39029.42
695.3412475585938 0 44390.586
700.40478515625 0 40827.395
702.3839111328125 0 33904.508
703.3716430664062 0 80540.85
704.3704833984375 0 46274.215
714.3515014648438 0 28463.732
714.4199829101562 0 72148.945
716.3992309570312 0 317542.9
717.4028930664062 0 156632.28
718.4095458984375 0 30642.55
720.4063110351562 0 122583.24
721.3951416015625 0 247856.17
722.3953857421875 0 147673.83
723.396484375 0 29771.791
730.3812866210938 0 37035.164
731.3855590820312 0 35812.316
732.36767578125 0 70066.39
734.3528442382812 0 71474.78
735.3496704101562 0 80666.3
738.4194946289062 0 949787.25
739.4227905273438 0 452648.44
740.4243774414062 0 86171.12
742.343017578125 0 31072.047
744.3382568359375 0 49735.383
748.4025268554688 0 381832.78 b Water loss 5
749.317138671875 0 44094.926
749.4036865234375 0 216360.36
750.3880615234375 0 198759.55
751.3882446289062 0 100510.234
752.3628540039062 0 530702.6
753.3656616210938 0 209490.19
754.3667602539062 0 61653.195
760.3549194335938 0 42565.52
762.34716796875 0 274779.5
763.349853515625 0 136491.8
763.4729614257812 0 44288.527
764.4000244140625 0 75047.336
764.4732666015625 0 28536.078
765.4111938476562 0 39280.625
766.4146728515625 0 9295332 b 5
767.4175415039062 0 4139398.2
768.4201049804688 0 878811.94
769.4248657226562 0 64219.324
778.3789672851562 0 59320.496
780.357421875 0 3346747
781.3603515625 0 1552834.5
782.3631591796875 0 309656.84
782.4391479492188 0 47240.273
783.4439086914062 0 28292.746
784.4271850585938 0 72058.42
785.4324340820312 0 52159.883
790.3392333984375 0 43064.586
799.4393920898438 0 31954.846
808.35107421875 0 45674.42
811.43359375 0 67536.336
815.4672241210938 0 63438.26
817.445556640625 0 98925.06
818.4520874023438 0 63187.73
820.4623413085938 0 176373.86
821.4649047851562 0 87588.234
822.4437866210938 0 77551.9
823.4396362304688 0 34400.816
827.4306640625 0 70032.64
829.4432983398438 0 119186.875
830.4508056640625 0 51909.934
832.4252319335938 0 43273.65
834.4115600585938 0 38104.652 y Water loss 4
835.4713134765625 0 119503.85
836.4754638671875 0 64936.426
845.4424438476562 0 283819.3 b Water loss 6
846.4451904296875 0 131206.47 b Ammonia loss 6
847.4500122070312 0 71396.31
848.3851318359375 0 83554.79
849.3842163085938 0 55534.855
850.43603515625 0 205976.05
851.434326171875 0 161644.23
852.4246215820312 0 84484.08 y 4
861.4090576171875 0 38947.816
863.46728515625 0 657031.2 b 6
864.4703979492188 0 293109.94
865.4717407226562 0 109525.53
877.4108276367188 0 1128646.2
878.4134521484375 0 520255.56
878.4896850585938 0 41995.195
879.4230346679688 0 498325.2
879.5156860351562 0 29248.191
880.4293212890625 0 196575.88
881.42822265625 0 40527.848
889.411376953125 0 104310.68
890.4144897460938 0 47591.984
893.4401245117188 0 74895.555
894.447021484375 0 66353.984
895.450439453125 0 62162.164
907.4205322265625 0 315315.47
908.4223022460938 0 140966.48
909.4299926757812 0 33875.26
928.49072265625 0 44437.555
929.4817504882812 0 51349.98
930.474365234375 0 44493.83
931.4857788085938 0 35944.758
934.5457763671875 0 42011.938
942.4857788085938 0 30446.727
944.5052490234375 0 38781.293
946.5049438476562 0 115554.484
947.4938354492188 0 93681.87
948.489013671875 0 201535.77
949.4848022460938 0 115368.01
956.4888916015625 0 62264.25
957.4853515625 0 45485.965
958.4667358398438 0 51308.01
959.4701538085938 0 51176.49
962.5357055664062 0 201666.53
963.536865234375 0 98179.51
964.5145263671875 0 987682.1
965.5182495117188 0 632658.56
966.5203247070312 0 132909.7
967.4405517578125 0 95296.875 y 3
968.4425659179688 0 49469.527
974.4999389648438 0 565068.2 b Water loss 7
975.5014038085938 0 368095.88 b Ammonia loss 7
976.4808959960938 0 1021657.1
977.4818115234375 0 594533.06
978.4849243164062 0 183244.1
990.4956665039062 0 100109.875
991.4992065429688 0 118014.64
992.5096435546875 0 10888027 b 7
993.5126953125 0 6363270
994.5156860351562 0 1833726.2
995.51806640625 0 196958.08
1025.569091796875 0 35364.957
1041.5692138671875 0 41947.855
1042.5648193359375 0 41116.285
1046.556640625 0 291036.72
1047.5587158203125 0 190035.88
1048.564208984375 0 42194.31
1075.5477294921875 0 102747.625
1076.549072265625 0 78838.445
1077.5281982421875 0 145719.61 y Water loss 2
1078.5296630859375 0 100115.25 y Ammonia loss 2
1079.527587890625 0 35402.94
1089.5615234375 0 258369.02 b 8
1090.565185546875 0 162603.67
1091.56787109375 0 72650.99
1095.5364990234375 0 2430433.2 y 2
1096.5389404296875 0 1584071.1
1096.6815185546875 0 32025.85
1097.5421142578125 0 495046.03
1098.5467529296875 0 49461.56
1160.63427734375 0 40113.688
1161.6341552734375 0 43303.55
1170.620361328125 0 91502.37 b Water loss 9
1171.613037109375 0 62235.25 b Ammonia loss 9
1172.6279296875 0 38764.016
1188.630126953125 0 821271.75 b 9
1189.6324462890625 0 591201.56
1190.6348876953125 0 209651.81
1194.603271484375 0 208246.72 y 1
1195.6072998046875 0 118316.234
1196.615966796875 0 45070.68

Spectrum Details

|  |  |
| --- | --- |
| Matched peaks? Matched peaks The total absolute number of peaks matched. Additionally in brackets the total fraction of peaks matched and the total number of peaks is shown. | 53 (7.38% of 718) |
| FDR? FDR The false discovery rate estimated for this peptide. It is calculated by matching all theoretical fragments with a non-integer shift with the raw peaks for this spectrum. This is done with 40 different shifts. The resulting percentage is the average number of annotated peaks over the number of annotated peaks with the correct spectrum. | 1.08% |
| Satellite FDR? Satellite FDR See the FDR for details on its calculation. This satellite ion specific FDR only contains the satellite ions (d/w) for I/L/J positions. | - |
| PSM Score? PSM Score The PSM Score as given by Hecklib to this annotated spectrum. It is shown with three significant figures. | 638 |

#### Spectrum 7862? Spectrum 7862 The raw spectrum of this peptide as annotated by Hecklib. The fragments are coloured according to ion type (see legend). Any peaks with a star '\*' as text can be hovered over to see the full details, first the ion type second the mass shift type. By hovering over the amino acids in the peptide or ions in the legend the corresponding peaks are highlighted. By toggling the 'Unassigned' label you can turn the background (unassigned) peaks on or off in the plot. By updating the slider in the Ion legend you can update the spectrum to only show the top X% of the peaks with labels. The top X% means any peak that is within X% of the highest intensity. By dragging in the spectrum you can zoom in to a specific part of the spectrum and use 'Zoom Out' to get back to the original zoom level. The annotation of the spectrum is based on the given sequence in the peptides file and is done with different software so inconsistencies are likely. The peaks are annotated based on the given sequence, with 20 ppm tolerance.

Copy Data

##### Spectrum 7862 (TSV)

###### Preview

```
Loading example...
```

*Click on the button to copy the data to your clipboard.*

Mz MinMz MaxIntensity Max

WidthHeightPeptide font sizePeptide stroke widthSpectrum font sizeSpectrum stroke widthCompact peptide

Ion legend

wxyz

abcd

OtherUnassignedIon^Charge^_Position_Show for top:%

JVKDYFPEPVT

06.11e+41.22e+51.83e+52.44e+5

Zoom Out

y^+1^_1_a^+1^_2_y^+1^_2_b^+1^_2_y^+1^_2_y^+1^_3_b^+2^_5_y^+1^_3_b^+1^_3_b^+2^_6_b^+2^_7_y^+1^_4_b^+2^_7_b^+1^_4_b^+1^_4_y^+1^_4_b^+1^_4_b^+2^_8_b^+2^_8_b^+2^_8_y^+1^_5_y^+2^_9_y^+1^_5_b^+2^_9_y^+2^_9_b^+2^_10_b^+2^_10_b^+2^_10_y^+2^_10_b^+1^_5_b^+1^_5_\*\*\*y^+1^_6_y^+1^_6_b^+1^_6_b^+1^_6_y^+1^_7_b^+1^_7_b^+1^_7_y^+1^_7_b^+1^_7_y^+1^_8_b^+1^_8_b^+1^_8_b^+1^_8_y^+1^_9_y^+1^_9_b^+1^_9_y^+1^_9_b^+1^_10_b^+1^_10_b^+1^_10_y^+1^_10_

036272310851447

Fragment Matches Table

Show background peaks

| Position | Ion type | Intensity | mz Theoretical | mz Error (Th) | mz Error (ppm) | Charge | Series Number |
| --- | --- | --- | --- | --- | --- | --- | --- |
| 11 | y | 2.833E+04 | 120.1 | 0.0003526 | 2.937 | +1 | 1 |
| - | - | 3.843E+04 | 120.1 | - | - | 0 | - |
| - | - | 1127 | 121.1 | - | - | 0 | - |
| - | - | 3558 | 121.1 | - | - | 0 | - |
| - | - | 463.8 | 126.1 | - | - | 0 | - |
| - | - | 428.2 | 126.5 | - | - | 0 | - |
| - | - | 518 | 127.1 | - | - | 0 | - |
| - | - | 6.76E+04 | 129.1 | - | - | 0 | - |
| - | - | 1143 | 130.1 | - | - | 0 | - |
| - | - | 1026 | 130.1 | - | - | 0 | - |
| - | - | 4392 | 130.1 | - | - | 0 | - |
| - | - | 487 | 133.9 | - | - | 0 | - |
| - | - | 450.8 | 134.1 | - | - | 0 | - |
| - | - | 1.68E+04 | 136.1 | - | - | 0 | - |
| - | - | 418 | 136.9 | - | - | 0 | - |
| - | - | 1375 | 137.1 | - | - | 0 | - |
| - | - | 408.9 | 138.1 | - | - | 0 | - |
| - | - | 1931 | 139.1 | - | - | 0 | - |
| - | - | 708.8 | 140.1 | - | - | 0 | - |
| - | - | 560 | 146.1 | - | - | 0 | - |
| - | - | 762.1 | 147 | - | - | 0 | - |
| - | - | 665.8 | 148.9 | - | - | 0 | - |
| - | - | 481.7 | 152.1 | - | - | 0 | - |
| - | - | 1109 | 155.1 | - | - | 0 | - |
| - | - | 469.5 | 155.2 | - | - | 0 | - |
| - | - | 639.2 | 157.1 | - | - | 0 | - |
| - | - | 616.7 | 159.1 | - | - | 0 | - |
| - | - | 424.6 | 164.9 | - | - | 0 | - |
| - | - | 509.8 | 165.1 | - | - | 0 | - |
| - | - | 1549 | 165.1 | - | - | 0 | - |
| - | - | 472.3 | 166.1 | - | - | 0 | - |
| - | - | 1500 | 167.1 | - | - | 0 | - |
| - | - | 1660 | 167.1 | - | - | 0 | - |
| - | - | 423.9 | 167.3 | - | - | 0 | - |
| - | - | 1662 | 168.1 | - | - | 0 | - |
| - | - | 2.084E+05 | 169.1 | - | - | 0 | - |
| - | - | 1.936E+04 | 170.1 | - | - | 0 | - |
| - | - | 763.3 | 171.1 | - | - | 0 | - |
| - | - | 511.6 | 171.1 | - | - | 0 | - |
| - | - | 425.9 | 172 | - | - | 0 | - |
| - | - | 4287 | 172.1 | - | - | 0 | - |
| - | - | 676 | 172.1 | - | - | 0 | - |
| - | - | 4285 | 173.1 | - | - | 0 | - |
| - | - | 1037 | 173.1 | - | - | 0 | - |
| - | - | 1290 | 174.1 | - | - | 0 | - |
| - | - | 3725 | 181.1 | - | - | 0 | - |
| - | - | 3237 | 181.1 | - | - | 0 | - |
| - | - | 2416 | 182.1 | - | - | 0 | - |
| - | - | 595.9 | 182.1 | - | - | 0 | - |
| - | - | 1740 | 183.1 | - | - | 0 | - |
| - | - | 4005 | 183.1 | - | - | 0 | - |
| - | - | 1874 | 185.1 | - | - | 0 | - |
| 2 | a | 8.959E+04 | 185.2 | 0.0003977 | 2.148 | +1 | 2 |
| - | - | 2620 | 186.1 | - | - | 0 | - |
| - | - | 691.4 | 186.2 | - | - | 0 | - |
| - | - | 8595 | 186.2 | - | - | 0 | - |
| - | - | 643.8 | 187.1 | - | - | 0 | - |
| - | - | 577.7 | 187.1 | - | - | 0 | - |
| - | - | 1253 | 188.1 | - | - | 0 | - |
| - | - | 3127 | 188.1 | - | - | 0 | - |
| - | - | 1215 | 190.1 | - | - | 0 | - |
| - | - | 1262 | 191.1 | - | - | 0 | - |
| - | - | 508.1 | 192.6 | - | - | 0 | - |
| - | - | 4829 | 195.1 | - | - | 0 | - |
| - | - | 1.636E+05 | 197.1 | - | - | 0 | - |
| - | - | 1.691E+04 | 198.1 | - | - | 0 | - |
| - | - | 2.545E+04 | 199.1 | - | - | 0 | - |
| - | - | 320.8 | 199.1 | - | - | 0 | - |
| - | - | 1252 | 199.1 | - | - | 0 | - |
| - | - | 2219 | 199.2 | - | - | 0 | - |
| - | - | 1186 | 200.1 | - | - | 0 | - |
| 10 | y | 1.472E+04 | 201.1 | 0.0002731 | 1.358 | +1 | 2 |
| - | - | 1565 | 202.1 | - | - | 0 | - |
| - | - | 1439 | 203.2 | - | - | 0 | - |
| - | - | 1701 | 204.1 | - | - | 0 | - |
| - | - | 490.1 | 208.5 | - | - | 0 | - |
| - | - | 9036 | 209.1 | - | - | 0 | - |
| - | - | 529.9 | 210.1 | - | - | 0 | - |
| - | - | 706.4 | 210.2 | - | - | 0 | - |
| - | - | 1381 | 211.1 | - | - | 0 | - |
| - | - | 2170 | 212.1 | - | - | 0 | - |
| - | - | 734.1 | 213.1 | - | - | 0 | - |
| - | - | 2205 | 213.1 | - | - | 0 | - |
| 2 | b | 4.054E+04 | 213.2 | 0.0003866 | 1.814 | +1 | 2 |
| - | - | 3702 | 214.2 | - | - | 0 | - |
| - | - | 3832 | 215.1 | - | - | 0 | - |
| - | - | 822 | 217.1 | - | - | 0 | - |
| - | - | 761.9 | 217.1 | - | - | 0 | - |
| - | - | 5064 | 217.1 | - | - | 0 | - |
| - | - | 566.3 | 219.1 | - | - | 0 | - |
| 10 | y | 2.405E+04 | 219.1 | 0.0003133 | 1.43 | +1 | 2 |
| - | - | 2671 | 220.1 | - | - | 0 | - |
| - | - | 1391 | 225.1 | - | - | 0 | - |
| - | - | 1.933E+04 | 226.1 | - | - | 0 | - |
| - | - | 1012 | 226.2 | - | - | 0 | - |
| - | - | 7.718E+04 | 227.1 | - | - | 0 | - |
| - | - | 2540 | 227.1 | - | - | 0 | - |
| - | - | 697.6 | 227.2 | - | - | 0 | - |
| - | - | 6937 | 228.1 | - | - | 0 | - |
| - | - | 7963 | 228.2 | - | - | 0 | - |
| - | - | 668.7 | 229.1 | - | - | 0 | - |
| - | - | 1377 | 230.2 | - | - | 0 | - |
| - | - | 1.426E+04 | 231.1 | - | - | 0 | - |
| - | - | 2288 | 232.2 | - | - | 0 | - |
| - | - | 1019 | 233.1 | - | - | 0 | - |
| - | - | 1951 | 233.2 | - | - | 0 | - |
| - | - | 1152 | 237.1 | - | - | 0 | - |
| - | - | 1110 | 238.1 | - | - | 0 | - |
| - | - | 2696 | 238.2 | - | - | 0 | - |
| - | - | 567.6 | 240.1 | - | - | 0 | - |
| - | - | 3.196E+04 | 244.1 | - | - | 0 | - |
| - | - | 554.2 | 245.1 | - | - | 0 | - |
| - | - | 7590 | 245.1 | - | - | 0 | - |
| - | - | 943.1 | 246.1 | - | - | 0 | - |
| - | - | 3247 | 247.1 | - | - | 0 | - |
| - | - | 798.3 | 249.2 | - | - | 0 | - |
| - | - | 898.1 | 251.1 | - | - | 0 | - |
| - | - | 1.773E+04 | 251.1 | - | - | 0 | - |
| - | - | 2257 | 252.1 | - | - | 0 | - |
| - | - | 3215 | 254.1 | - | - | 0 | - |
| - | - | 1479 | 254.2 | - | - | 0 | - |
| - | - | 1305 | 254.2 | - | - | 0 | - |
| - | - | 1926 | 255.1 | - | - | 0 | - |
| - | - | 1284 | 256.2 | - | - | 0 | - |
| - | - | 534.4 | 257.9 | - | - | 0 | - |
| - | - | 929.5 | 261.1 | - | - | 0 | - |
| - | - | 3656 | 261.2 | - | - | 0 | - |
| - | - | 486.2 | 262.2 | - | - | 0 | - |
| - | - | 701.1 | 263.1 | - | - | 0 | - |
| - | - | 2094 | 263.1 | - | - | 0 | - |
| - | - | 591.6 | 266.1 | - | - | 0 | - |
| - | - | 1101 | 266.2 | - | - | 0 | - |
| - | - | 985.8 | 268.1 | - | - | 0 | - |
| - | - | 770.9 | 269.2 | - | - | 0 | - |
| - | - | 2112 | 270.2 | - | - | 0 | - |
| - | - | 4889 | 272.1 | - | - | 0 | - |
| - | - | 734.2 | 273.1 | - | - | 0 | - |
| - | - | 984.2 | 276.2 | - | - | 0 | - |
| - | - | 1405 | 277.2 | - | - | 0 | - |
| - | - | 1570 | 278.2 | - | - | 0 | - |
| - | - | 2.549E+04 | 279.1 | - | - | 0 | - |
| - | - | 3213 | 280.1 | - | - | 0 | - |
| - | - | 6210 | 280.2 | - | - | 0 | - |
| - | - | 1.096E+04 | 283.1 | - | - | 0 | - |
| - | - | 667.3 | 284.1 | - | - | 0 | - |
| - | - | 1703 | 284.1 | - | - | 0 | - |
| - | - | 4353 | 287.2 | - | - | 0 | - |
| - | - | 656.6 | 292.1 | - | - | 0 | - |
| - | - | 547.1 | 292.2 | - | - | 0 | - |
| - | - | 655.3 | 294.2 | - | - | 0 | - |
| - | - | 1061 | 296.1 | - | - | 0 | - |
| - | - | 924.3 | 296.2 | - | - | 0 | - |
| - | - | 1248 | 296.2 | - | - | 0 | - |
| 9 | y | 3.948E+04 | 298.2 | 0.000503 | 1.687 | +1 | 3 |
| - | - | 6464 | 299.2 | - | - | 0 | - |
| - | - | 913.4 | 301.2 | - | - | 0 | - |
| - | - | 2735 | 303.2 | - | - | 0 | - |
| - | - | 1594 | 304.2 | - | - | 0 | - |
| - | - | 1716 | 306.1 | - | - | 0 | - |
| - | - | 894.8 | 306.2 | - | - | 0 | - |
| - | - | 574.4 | 307.6 | - | - | 0 | - |
| - | - | 2087 | 308.2 | - | - | 0 | - |
| - | - | 904.1 | 310.1 | - | - | 0 | - |
| 5 | b | 1933 | 310.2 | 0.0003505 | 1.13 | +2 | 5 |
| - | - | 1839 | 310.2 | - | - | 0 | - |
| - | - | 7166 | 311.1 | - | - | 0 | - |
| - | - | 1830 | 312.1 | - | - | 0 | - |
| - | - | 968.6 | 312.2 | - | - | 0 | - |
| - | - | 1083 | 313.2 | - | - | 0 | - |
| - | - | 1899 | 315.2 | - | - | 0 | - |
| 9 | y | 2.421E+05 | 316.2 | 0.0005585 | 1.766 | +1 | 3 |
| - | - | 3.822E+04 | 317.2 | - | - | 0 | - |
| - | - | 3639 | 318.2 | - | - | 0 | - |
| - | - | 685.3 | 319.2 | - | - | 0 | - |
| - | - | 627.6 | 322.2 | - | - | 0 | - |
| - | - | 1692 | 323.2 | - | - | 0 | - |
| - | - | 2921 | 323.2 | - | - | 0 | - |
| - | - | 646.2 | 323.7 | - | - | 0 | - |
| - | - | 7.175E+04 | 324.2 | - | - | 0 | - |
| - | - | 1.063E+04 | 325.2 | - | - | 0 | - |
| - | - | 2382 | 325.2 | - | - | 0 | - |
| - | - | 1150 | 325.2 | - | - | 0 | - |
| - | - | 2194 | 326.2 | - | - | 0 | - |
| - | - | 994.5 | 328.1 | - | - | 0 | - |
| - | - | 1554 | 328.2 | - | - | 0 | - |
| - | - | 2453 | 329.1 | - | - | 0 | - |
| - | - | 1079 | 331.2 | - | - | 0 | - |
| - | - | 1399 | 335.2 | - | - | 0 | - |
| - | - | 803.4 | 335.2 | - | - | 0 | - |
| - | - | 2450 | 337.2 | - | - | 0 | - |
| - | - | 794.7 | 337.2 | - | - | 0 | - |
| - | - | 739.8 | 339.2 | - | - | 0 | - |
| - | - | 4219 | 340.2 | - | - | 0 | - |
| - | - | 6604 | 341.2 | - | - | 0 | - |
| 3 | b | 6132 | 341.3 | 0.0005317 | 1.558 | +1 | 3 |
| - | - | 844.4 | 342.1 | - | - | 0 | - |
| - | - | 1570 | 342.3 | - | - | 0 | - |
| - | - | 7413 | 343.2 | - | - | 0 | - |
| - | - | 1452 | 344.2 | - | - | 0 | - |
| - | - | 1872 | 344.7 | - | - | 0 | - |
| - | - | 1037 | 345.2 | - | - | 0 | - |
| - | - | 4656 | 346.2 | - | - | 0 | - |
| - | - | 1018 | 347.2 | - | - | 0 | - |
| - | - | 3879 | 347.2 | - | - | 0 | - |
| - | - | 1791 | 347.7 | - | - | 0 | - |
| - | - | 1822 | 349.2 | - | - | 0 | - |
| - | - | 717 | 351.2 | - | - | 0 | - |
| - | - | 4732 | 353.2 | - | - | 0 | - |
| - | - | 3065 | 353.2 | - | - | 0 | - |
| - | - | 852 | 354.2 | - | - | 0 | - |
| - | - | 833.4 | 354.2 | - | - | 0 | - |
| - | - | 1355 | 355.1 | - | - | 0 | - |
| - | - | 3676 | 355.2 | - | - | 0 | - |
| - | - | 2693 | 355.2 | - | - | 0 | - |
| - | - | 1642 | 356.2 | - | - | 0 | - |
| - | - | 882.3 | 357.2 | - | - | 0 | - |
| - | - | 671.6 | 358.3 | - | - | 0 | - |
| - | - | 780.6 | 359.2 | - | - | 0 | - |
| - | - | 1310 | 360.2 | - | - | 0 | - |
| - | - | 940.3 | 361.2 | - | - | 0 | - |
| - | - | 1703 | 362.2 | - | - | 0 | - |
| - | - | 1346 | 367.2 | - | - | 0 | - |
| - | - | 908.9 | 369.2 | - | - | 0 | - |
| - | - | 7585 | 369.7 | - | - | 0 | - |
| - | - | 2885 | 370.2 | - | - | 0 | - |
| - | - | 867.3 | 370.7 | - | - | 0 | - |
| - | - | 7055 | 371.2 | - | - | 0 | - |
| - | - | 733.4 | 372.2 | - | - | 0 | - |
| - | - | 1107 | 372.2 | - | - | 0 | - |
| - | - | 1853 | 373.2 | - | - | 0 | - |
| - | - | 1.116E+04 | 374.2 | - | - | 0 | - |
| - | - | 885.5 | 374.2 | - | - | 0 | - |
| - | - | 1839 | 375.2 | - | - | 0 | - |
| - | - | 785.1 | 375.2 | - | - | 0 | - |
| - | - | 808.5 | 376.2 | - | - | 0 | - |
| - | - | 1674 | 377.2 | - | - | 0 | - |
| - | - | 6438 | 379.2 | - | - | 0 | - |
| - | - | 564.4 | 380.2 | - | - | 0 | - |
| - | - | 1242 | 380.2 | - | - | 0 | - |
| - | - | 2523 | 381.1 | - | - | 0 | - |
| - | - | 2403 | 383.2 | - | - | 0 | - |
| 6 | b | 2175 | 383.7 | 0.0005978 | 1.558 | +2 | 6 |
| - | - | 7297 | 389.2 | - | - | 0 | - |
| - | - | 1179 | 390.2 | - | - | 0 | - |
| - | - | 1049 | 390.2 | - | - | 0 | - |
| - | - | 631.4 | 391.2 | - | - | 0 | - |
| - | - | 1.588E+04 | 391.2 | - | - | 0 | - |
| - | - | 3275 | 392.2 | - | - | 0 | - |
| - | - | 1773 | 392.2 | - | - | 0 | - |
| - | - | 808.1 | 393.3 | - | - | 0 | - |
| - | - | 727.1 | 394.2 | - | - | 0 | - |
| - | - | 7480 | 395.2 | - | - | 0 | - |
| - | - | 2164 | 396.2 | - | - | 0 | - |
| - | - | 704.3 | 400.7 | - | - | 0 | - |
| - | - | 3418 | 403.2 | - | - | 0 | - |
| - | - | 1003 | 404.2 | - | - | 0 | - |
| - | - | 2102 | 405.2 | - | - | 0 | - |
| - | - | 1149 | 406.2 | - | - | 0 | - |
| - | - | 1.645E+04 | 407.2 | - | - | 0 | - |
| - | - | 1835 | 408.2 | - | - | 0 | - |
| - | - | 3715 | 408.2 | - | - | 0 | - |
| - | - | 834.2 | 409.1 | - | - | 0 | - |
| - | - | 1579 | 409.2 | - | - | 0 | - |
| - | - | 1261 | 409.3 | - | - | 0 | - |
| - | - | 784.1 | 409.7 | - | - | 0 | - |
| - | - | 573.4 | 410.3 | - | - | 0 | - |
| - | - | 1053 | 411.3 | - | - | 0 | - |
| - | - | 1.365E+04 | 418.2 | - | - | 0 | - |
| - | - | 7420 | 418.7 | - | - | 0 | - |
| - | - | 1005 | 419.2 | - | - | 0 | - |
| - | - | 1637 | 419.2 | - | - | 0 | - |
| - | - | 659.5 | 420.3 | - | - | 0 | - |
| - | - | 655.4 | 422.2 | - | - | 0 | - |
| 7 | b | 2.499E+04 | 423.2 | 0.006921 | 16.35 | +2 | 7 |
| - | - | 9616 | 424.2 | - | - | 0 | - |
| - | - | 2235 | 425.2 | - | - | 0 | - |
| - | - | 2.331E+04 | 426.2 | - | - | 0 | - |
| - | - | 4957 | 427.2 | - | - | 0 | - |
| 8 | y | 3498 | 427.2 | 0.001153 | 2.7 | +1 | 4 |
| - | - | 974.9 | 428.2 | - | - | 0 | - |
| 7 | b | 8871 | 432.2 | 0.0005831 | 1.349 | +2 | 7 |
| - | - | 4934 | 432.7 | - | - | 0 | - |
| - | - | 1046 | 433.2 | - | - | 0 | - |
| - | - | 984.8 | 434.2 | - | - | 0 | - |
| - | - | 4458 | 435.2 | - | - | 0 | - |
| - | - | 1094 | 436.2 | - | - | 0 | - |
| - | - | 1783 | 436.2 | - | - | 0 | - |
| - | - | 1110 | 437.2 | - | - | 0 | - |
| 4 | b | 8026 | 438.3 | 0.0004497 | 1.026 | +1 | 4 |
| - | - | 1693 | 438.3 | - | - | 0 | - |
| - | - | 1538 | 439.2 | - | - | 0 | - |
| 4 | b | 2030 | 439.3 | 0.001297 | 2.954 | +1 | 4 |
| - | - | 682.1 | 439.3 | - | - | 0 | - |
| - | - | 2738 | 440.3 | - | - | 0 | - |
| - | - | 1232 | 443.2 | - | - | 0 | - |
| - | - | 1149 | 443.3 | - | - | 0 | - |
| 8 | y | 7772 | 445.2 | 0.0013 | 2.921 | +1 | 4 |
| - | - | 1018 | 446.2 | - | - | 0 | - |
| - | - | 3506 | 452.2 | - | - | 0 | - |
| - | - | 767.4 | 452.3 | - | - | 0 | - |
| - | - | 1103 | 453.2 | - | - | 0 | - |
| - | - | 1388 | 454.2 | - | - | 0 | - |
| - | - | 934.5 | 454.3 | - | - | 0 | - |
| 4 | b | 3.009E+04 | 456.3 | 0.0005356 | 1.174 | +1 | 4 |
| - | - | 1323 | 457.2 | - | - | 0 | - |
| - | - | 8392 | 457.3 | - | - | 0 | - |
| - | - | 1071 | 458.3 | - | - | 0 | - |
| - | - | 794.1 | 460.3 | - | - | 0 | - |
| - | - | 688.3 | 462.3 | - | - | 0 | - |
| - | - | 1106 | 464.3 | - | - | 0 | - |
| - | - | 819.5 | 466.3 | - | - | 0 | - |
| - | - | 947.1 | 468.2 | - | - | 0 | - |
| - | - | 602.1 | 470.2 | - | - | 0 | - |
| - | - | 1.143E+04 | 470.2 | - | - | 0 | - |
| - | - | 843.5 | 470.3 | - | - | 0 | - |
| - | - | 5850 | 471.2 | - | - | 0 | - |
| - | - | 1518 | 472.2 | - | - | 0 | - |
| - | - | 910.9 | 473.2 | - | - | 0 | - |
| - | - | 704.1 | 473.2 | - | - | 0 | - |
| - | - | 717.9 | 473.3 | - | - | 0 | - |
| - | - | 762.3 | 474.3 | - | - | 0 | - |
| - | - | 624.6 | 474.7 | - | - | 0 | - |
| - | - | 1029 | 475.2 | - | - | 0 | - |
| - | - | 1407 | 476.3 | - | - | 0 | - |
| - | - | 1867 | 478.3 | - | - | 0 | - |
| - | - | 609.1 | 480.2 | - | - | 0 | - |
| - | - | 724.5 | 480.8 | - | - | 0 | - |
| - | - | 5878 | 482.3 | - | - | 0 | - |
| - | - | 7258 | 482.8 | - | - | 0 | - |
| - | - | 4533 | 483.3 | - | - | 0 | - |
| - | - | 796 | 483.8 | - | - | 0 | - |
| - | - | 866.6 | 486.2 | - | - | 0 | - |
| 8 | b | 1817 | 487.8 | 0.001445 | 2.963 | +2 | 8 |
| 8 | b | 3315 | 488.2 | 0.006599 | 13.52 | +2 | 8 |
| - | - | 3537 | 488.3 | - | - | 0 | - |
| - | - | 2053 | 488.7 | - | - | 0 | - |
| - | - | 932.4 | 489.2 | - | - | 0 | - |
| - | - | 862.5 | 489.3 | - | - | 0 | - |
| - | - | 825.6 | 490.2 | - | - | 0 | - |
| - | - | 622.5 | 491.2 | - | - | 0 | - |
| - | - | 802.7 | 492.2 | - | - | 0 | - |
| - | - | 719.6 | 492.3 | - | - | 0 | - |
| - | - | 1181 | 495.7 | - | - | 0 | - |
| 8 | b | 1.758E+04 | 496.8 | 0.0003132 | 0.6304 | +2 | 8 |
| - | - | 9929 | 497.3 | - | - | 0 | - |
| - | - | 3502 | 497.8 | - | - | 0 | - |
| - | - | 910 | 498.2 | - | - | 0 | - |
| - | - | 950 | 500.2 | - | - | 0 | - |
| - | - | 1209 | 500.3 | - | - | 0 | - |
| - | - | 2140 | 504.2 | - | - | 0 | - |
| - | - | 1088 | 504.3 | - | - | 0 | - |
| - | - | 4777 | 506.3 | - | - | 0 | - |
| - | - | 2842 | 508.3 | - | - | 0 | - |
| - | - | 4686 | 509.2 | - | - | 0 | - |
| - | - | 1294 | 510.2 | - | - | 0 | - |
| - | - | 840.5 | 510.3 | - | - | 0 | - |
| - | - | 940.1 | 511.3 | - | - | 0 | - |
| - | - | 796.9 | 513.3 | - | - | 0 | - |
| - | - | 903.1 | 515.3 | - | - | 0 | - |
| - | - | 1375 | 518.2 | - | - | 0 | - |
| - | - | 1607 | 518.3 | - | - | 0 | - |
| - | - | 1036 | 520.3 | - | - | 0 | - |
| - | - | 1997 | 521.3 | - | - | 0 | - |
| - | - | 1250 | 521.8 | - | - | 0 | - |
| - | - | 3775 | 522.3 | - | - | 0 | - |
| - | - | 5635 | 522.8 | - | - | 0 | - |
| - | - | 1492 | 523.2 | - | - | 0 | - |
| - | - | 3546 | 523.3 | - | - | 0 | - |
| - | - | 2348 | 523.8 | - | - | 0 | - |
| 7 | y | 9797 | 524.3 | 0.001826 | 3.483 | +1 | 5 |
| - | - | 3057 | 525.3 | - | - | 0 | - |
| - | - | 1.307E+04 | 526.3 | - | - | 0 | - |
| - | - | 3616 | 527.3 | - | - | 0 | - |
| - | - | 719.4 | 528.3 | - | - | 0 | - |
| - | - | 797 | 530.8 | - | - | 0 | - |
| - | - | 3017 | 531.3 | - | - | 0 | - |
| - | - | 2724 | 531.8 | - | - | 0 | - |
| - | - | 906.2 | 532.2 | - | - | 0 | - |
| - | - | 790.8 | 532.3 | - | - | 0 | - |
| - | - | 1.055E+04 | 534.3 | - | - | 0 | - |
| - | - | 1917 | 535.3 | - | - | 0 | - |
| - | - | 8401 | 536.3 | - | - | 0 | - |
| - | - | 2717 | 536.3 | - | - | 0 | - |
| - | - | 1100 | 536.8 | - | - | 0 | - |
| - | - | 3137 | 537.2 | - | - | 0 | - |
| - | - | 5452 | 539.3 | - | - | 0 | - |
| 3 | y | 1140 | 539.8 | 0.008814 | 16.33 | +2 | 9 |
| - | - | 1475 | 540.3 | - | - | 0 | - |
| - | - | 1573 | 540.3 | - | - | 0 | - |
| - | - | 675.5 | 541.3 | - | - | 0 | - |
| 7 | y | 8.634E+04 | 542.3 | 0.0005995 | 1.105 | +1 | 5 |
| - | - | 2.34E+04 | 543.3 | - | - | 0 | - |
| - | - | 4059 | 544.3 | - | - | 0 | - |
| 9 | b | 5.259E+04 | 545.3 | 0.00039 | 0.7152 | +2 | 9 |
| - | - | 3.322E+04 | 545.8 | - | - | 0 | - |
| - | - | 1.289E+04 | 546.3 | - | - | 0 | - |
| - | - | 1911 | 546.8 | - | - | 0 | - |
| 3 | y | 1491 | 548.3 | 0.001887 | 3.441 | +2 | 9 |
| - | - | 1329 | 549.3 | - | - | 0 | - |
| - | - | 1096 | 551.3 | - | - | 0 | - |
| - | - | 2888 | 553.3 | - | - | 0 | - |
| - | - | 6.872E+04 | 554.3 | - | - | 0 | - |
| - | - | 2.356E+04 | 555.3 | - | - | 0 | - |
| - | - | 3867 | 556.3 | - | - | 0 | - |
| - | - | 4240 | 558.3 | - | - | 0 | - |
| - | - | 829.3 | 559.3 | - | - | 0 | - |
| - | - | 1047 | 564.2 | - | - | 0 | - |
| - | - | 6250 | 567.4 | - | - | 0 | - |
| - | - | 2604 | 568.4 | - | - | 0 | - |
| - | - | 784.3 | 569.3 | - | - | 0 | - |
| - | - | 5276 | 570.3 | - | - | 0 | - |
| - | - | 2040 | 571.3 | - | - | 0 | - |
| - | - | 1863 | 571.8 | - | - | 0 | - |
| - | - | 811.4 | 572.3 | - | - | 0 | - |
| - | - | 784.7 | 572.8 | - | - | 0 | - |
| - | - | 1243 | 574.3 | - | - | 0 | - |
| - | - | 919 | 579.3 | - | - | 0 | - |
| - | - | 1.809E+04 | 580.8 | - | - | 0 | - |
| - | - | 1.268E+04 | 581.3 | - | - | 0 | - |
| - | - | 5385 | 581.8 | - | - | 0 | - |
| - | - | 3608 | 582.3 | - | - | 0 | - |
| - | - | 1293 | 582.3 | - | - | 0 | - |
| - | - | 863.1 | 583.3 | - | - | 0 | - |
| - | - | 753.7 | 584.3 | - | - | 0 | - |
| - | - | 1904 | 585.3 | - | - | 0 | - |
| 10 | b | 1231 | 585.8 | 9.086E-05 | 0.1551 | +2 | 10 |
| 10 | b | 804.7 | 586.3 | 0.01168 | 19.93 | +2 | 10 |
| - | - | 788.7 | 586.8 | - | - | 0 | - |
| - | - | 1.195E+04 | 591.3 | - | - | 0 | - |
| - | - | 3261 | 592.4 | - | - | 0 | - |
| - | - | 733.2 | 593.4 | - | - | 0 | - |
| 10 | b | 1.351E+04 | 594.8 | 0.0001796 | 0.302 | +2 | 10 |
| - | - | 1.087E+04 | 595.3 | - | - | 0 | - |
| - | - | 4211 | 595.8 | - | - | 0 | - |
| - | - | 956.3 | 596.3 | - | - | 0 | - |
| - | - | 2602 | 597.3 | - | - | 0 | - |
| 2 | y | 873.1 | 597.8 | 0.001493 | 2.498 | +2 | 10 |
| - | - | 863.8 | 600.3 | - | - | 0 | - |
| 5 | b | 3233 | 601.3 | 0.0001094 | 0.182 | +1 | 5 |
| - | - | 1405 | 602.3 | - | - | 0 | - |
| - | - | 4311 | 603.4 | - | - | 0 | - |
| - | - | 1502 | 604.4 | - | - | 0 | - |
| - | - | 1118 | 605.3 | - | - | 0 | - |
| - | - | 1375 | 606.3 | - | - | 0 | - |
| - | - | 887.5 | 607.4 | - | - | 0 | - |
| - | - | 2015 | 608.3 | - | - | 0 | - |
| - | - | 643 | 609.3 | - | - | 0 | - |
| - | - | 1749 | 615.3 | - | - | 0 | - |
| - | - | 1958 | 617.3 | - | - | 0 | - |
| 5 | b | 4.415E+04 | 619.3 | 0.0001038 | 0.1677 | +1 | 5 |
| - | - | 1104 | 620.3 | - | - | 0 | - |
| - | - | 1.561E+04 | 620.3 | - | - | 0 | - |
| - | - | 2828 | 621.3 | - | - | 0 | - |
| - | - | 830.8 | 622.3 | - | - | 0 | - |
| - | - | 3186 | 623.3 | - | - | 0 | - |
| - | - | 1191 | 624.3 | - | - | 0 | - |
| - | - | 790.5 | 624.3 | - | - | 0 | - |
| - | - | 1807 | 625.3 | - | - | 0 | - |
| - | - | 1248 | 626.3 | - | - | 0 | - |
| - | - | 4398 | 633.3 | - | - | 0 | - |
| - | - | 917.9 | 634.3 | - | - | 0 | - |
| - | - | 1652 | 635.3 | - | - | 0 | - |
| - | - | 5374 | 635.4 | - | - | 0 | - |
| - | - | 1037 | 636.3 | - | - | 0 | - |
| - | - | 1680 | 636.4 | - | - | 0 | - |
| - | - | 2507 | 638.3 | - | - | 0 | - |
| - | - | 973.9 | 639.3 | - | - | 0 | - |
| - | - | 717.4 | 640 | - | - | 0 | - |
| - | - | 788.1 | 640.4 | - | - | 0 | - |
| - | - | 723.5 | 642.4 | - | - | 0 | - |
| 0 | Precursor | 7219 | 645.3 | 0.0001441 | 0.2233 | +2 | -1 |
| 0 | Precursor | 4575 | 645.8 | 0.007648 | 11.84 | +2 | -1 |
| - | - | 2911 | 646.3 | - | - | 0 | - |
| - | - | 7144 | 651.3 | - | - | 0 | - |
| - | - | 2313 | 651.4 | - | - | 0 | - |
| - | - | 5407 | 652.3 | - | - | 0 | - |
| - | - | 1963 | 652.3 | - | - | 0 | - |
| - | - | 1388 | 652.4 | - | - | 0 | - |
| - | - | 1206 | 652.9 | - | - | 0 | - |
| - | - | 1629 | 653.3 | - | - | 0 | - |
| - | - | 8552 | 653.3 | - | - | 0 | - |
| 0 | Precursor | 1.081E+04 | 654.3 | 0.003673 | 5.614 | +2 | -1 |
| - | - | 4881 | 654.8 | - | - | 0 | - |
| - | - | 3329 | 655.4 | - | - | 0 | - |
| - | - | 983.6 | 655.9 | - | - | 0 | - |
| - | - | 2858 | 663.3 | - | - | 0 | - |
| - | - | 1049 | 664.3 | - | - | 0 | - |
| - | - | 1547 | 664.4 | - | - | 0 | - |
| - | - | 689.5 | 666.3 | - | - | 0 | - |
| - | - | 2542 | 667.3 | - | - | 0 | - |
| - | - | 1127 | 668.3 | - | - | 0 | - |
| 6 | y | 1930 | 671.3 | 0.001769 | 2.635 | +1 | 6 |
| - | - | 924.1 | 672.3 | - | - | 0 | - |
| - | - | 9447 | 681.3 | - | - | 0 | - |
| - | - | 3085 | 682.3 | - | - | 0 | - |
| - | - | 1.368E+04 | 682.4 | - | - | 0 | - |
| - | - | 830.6 | 683.3 | - | - | 0 | - |
| - | - | 4639 | 683.4 | - | - | 0 | - |
| - | - | 1295 | 684.4 | - | - | 0 | - |
| - | - | 1400 | 688.4 | - | - | 0 | - |
| 6 | y | 3.289E+04 | 689.4 | 5.665E-05 | 0.08219 | +1 | 6 |
| - | - | 1.32E+04 | 690.4 | - | - | 0 | - |
| - | - | 2249 | 691.4 | - | - | 0 | - |
| - | - | 966.5 | 693.4 | - | - | 0 | - |
| - | - | 1343 | 695.3 | - | - | 0 | - |
| - | - | 1361 | 703.4 | - | - | 0 | - |
| - | - | 802.3 | 704.4 | - | - | 0 | - |
| - | - | 901.7 | 714.4 | - | - | 0 | - |
| - | - | 4953 | 716.4 | - | - | 0 | - |
| - | - | 1571 | 717.4 | - | - | 0 | - |
| - | - | 728.6 | 718.4 | - | - | 0 | - |
| - | - | 1375 | 720.4 | - | - | 0 | - |
| - | - | 3732 | 721.4 | - | - | 0 | - |
| - | - | 2120 | 722.4 | - | - | 0 | - |
| - | - | 1208 | 732.4 | - | - | 0 | - |
| - | - | 932.8 | 733.3 | - | - | 0 | - |
| - | - | 997 | 734.4 | - | - | 0 | - |
| - | - | 1.551E+04 | 738.4 | - | - | 0 | - |
| - | - | 6646 | 739.4 | - | - | 0 | - |
| - | - | 1385 | 740.4 | - | - | 0 | - |
| - | - | 722.1 | 744.3 | - | - | 0 | - |
| 6 | b | 7326 | 748.4 | 0.0007386 | 0.9868 | +1 | 6 |
| - | - | 3975 | 749.4 | - | - | 0 | - |
| - | - | 2931 | 750.4 | - | - | 0 | - |
| - | - | 1083 | 751.4 | - | - | 0 | - |
| - | - | 9558 | 752.4 | - | - | 0 | - |
| - | - | 3950 | 753.4 | - | - | 0 | - |
| - | - | 866.9 | 754.4 | - | - | 0 | - |
| - | - | 1008 | 760.4 | - | - | 0 | - |
| - | - | 5050 | 762.3 | - | - | 0 | - |
| - | - | 2408 | 763.3 | - | - | 0 | - |
| - | - | 780.2 | 763.5 | - | - | 0 | - |
| - | - | 974.9 | 765.3 | - | - | 0 | - |
| 6 | b | 1.625E+05 | 766.4 | 0.0002559 | 0.3339 | +1 | 6 |
| - | - | 7.496E+04 | 767.4 | - | - | 0 | - |
| - | - | 1.851E+04 | 768.4 | - | - | 0 | - |
| - | - | 1396 | 769.4 | - | - | 0 | - |
| - | - | 837.2 | 778.4 | - | - | 0 | - |
| - | - | 5.733E+04 | 780.4 | - | - | 0 | - |
| - | - | 2.506E+04 | 781.4 | - | - | 0 | - |
| - | - | 7173 | 782.4 | - | - | 0 | - |
| - | - | 790.5 | 783.4 | - | - | 0 | - |
| - | - | 919.6 | 784.4 | - | - | 0 | - |
| - | - | 4519 | 796 | - | - | 0 | - |
| - | - | 3190 | 796.4 | - | - | 0 | - |
| - | - | 3126 | 796.7 | - | - | 0 | - |
| - | - | 1091 | 808.3 | - | - | 0 | - |
| - | - | 1668 | 817.4 | - | - | 0 | - |
| - | - | 2504 | 820.5 | - | - | 0 | - |
| - | - | 2237 | 821.5 | - | - | 0 | - |
| - | - | 1648 | 827.4 | - | - | 0 | - |
| - | - | 2366 | 829.4 | - | - | 0 | - |
| - | - | 1394 | 830.4 | - | - | 0 | - |
| - | - | 886.8 | 833.4 | - | - | 0 | - |
| 5 | y | 752.4 | 834.4 | 0.004544 | 5.446 | +1 | 7 |
| - | - | 3041 | 835.5 | - | - | 0 | - |
| - | - | 1235 | 836.5 | - | - | 0 | - |
| 7 | b | 4743 | 845.5 | 0.01444 | 17.08 | +1 | 7 |
| 7 | b | 2652 | 846.4 | 0.003315 | 3.916 | +1 | 7 |
| - | - | 1253 | 848.4 | - | - | 0 | - |
| - | - | 3366 | 850.4 | - | - | 0 | - |
| - | - | 2757 | 851.4 | - | - | 0 | - |
| 5 | y | 1844 | 852.4 | 0.01131 | 13.27 | +1 | 7 |
| - | - | 682.5 | 853.4 | - | - | 0 | - |
| - | - | 824.7 | 861.4 | - | - | 0 | - |
| - | - | 772.5 | 862.4 | - | - | 0 | - |
| 7 | b | 1.17E+04 | 863.5 | 0.0002029 | 0.235 | +1 | 7 |
| - | - | 6601 | 864.5 | - | - | 0 | - |
| - | - | 2405 | 865.5 | - | - | 0 | - |
| - | - | 1.916E+04 | 877.4 | - | - | 0 | - |
| - | - | 9126 | 878.4 | - | - | 0 | - |
| - | - | 1029 | 878.5 | - | - | 0 | - |
| - | - | 7445 | 879.4 | - | - | 0 | - |
| - | - | 993.1 | 879.5 | - | - | 0 | - |
| - | - | 2569 | 880.4 | - | - | 0 | - |
| - | - | 1747 | 889.4 | - | - | 0 | - |
| - | - | 1011 | 890.4 | - | - | 0 | - |
| - | - | 1721 | 893.4 | - | - | 0 | - |
| - | - | 1258 | 894.4 | - | - | 0 | - |
| - | - | 1693 | 895.4 | - | - | 0 | - |
| - | - | 733.5 | 896.5 | - | - | 0 | - |
| - | - | 5726 | 907.4 | - | - | 0 | - |
| - | - | 2270 | 908.4 | - | - | 0 | - |
| - | - | 1146 | 910.5 | - | - | 0 | - |
| - | - | 776.2 | 919.5 | - | - | 0 | - |
| - | - | 2116 | 946.5 | - | - | 0 | - |
| - | - | 2415 | 947.5 | - | - | 0 | - |
| - | - | 4484 | 948.5 | - | - | 0 | - |
| - | - | 1555 | 949.5 | - | - | 0 | - |
| - | - | 1094 | 956.5 | - | - | 0 | - |
| - | - | 866.5 | 957.5 | - | - | 0 | - |
| - | - | 1224 | 958.5 | - | - | 0 | - |
| - | - | 3545 | 962.5 | - | - | 0 | - |
| - | - | 1462 | 963.5 | - | - | 0 | - |
| - | - | 1.787E+04 | 964.5 | - | - | 0 | - |
| - | - | 9233 | 965.5 | - | - | 0 | - |
| - | - | 2118 | 966.5 | - | - | 0 | - |
| 4 | y | 1592 | 967.4 | 0.0004932 | 0.5098 | +1 | 8 |
| 8 | b | 1.063E+04 | 974.5 | 0.0009417 | 0.9663 | +1 | 8 |
| 8 | b | 6087 | 975.5 | 0.01645 | 16.86 | +1 | 8 |
| - | - | 1.811E+04 | 976.5 | - | - | 0 | - |
| - | - | 1.157E+04 | 977.5 | - | - | 0 | - |
| - | - | 3272 | 978.5 | - | - | 0 | - |
| - | - | 1566 | 990.5 | - | - | 0 | - |
| - | - | 1470 | 991.5 | - | - | 0 | - |
| 8 | b | 1.983E+05 | 992.5 | 0.0008252 | 0.8315 | +1 | 8 |
| - | - | 1.194E+05 | 993.5 | - | - | 0 | - |
| - | - | 1174 | 993.6 | - | - | 0 | - |
| - | - | 3.724E+04 | 994.5 | - | - | 0 | - |
| - | - | 4123 | 995.5 | - | - | 0 | - |
| - | - | 825.3 | 1042 | - | - | 0 | - |
| - | - | 4119 | 1047 | - | - | 0 | - |
| - | - | 3045 | 1048 | - | - | 0 | - |
| - | - | 2078 | 1076 | - | - | 0 | - |
| - | - | 1471 | 1077 | - | - | 0 | - |
| 3 | y | 2474 | 1078 | 0.00306 | 2.84 | +1 | 9 |
| 3 | y | 1612 | 1079 | 0.01819 | 16.87 | +1 | 9 |
| 9 | b | 5075 | 1090 | 0.001587 | 1.457 | +1 | 9 |
| - | - | 3667 | 1091 | - | - | 0 | - |
| - | - | 1756 | 1092 | - | - | 0 | - |
| - | - | 731.9 | 1093 | - | - | 0 | - |
| 3 | y | 4.292E+04 | 1096 | 0.001035 | 0.9446 | +1 | 9 |
| - | - | 3.07E+04 | 1097 | - | - | 0 | - |
| - | - | 1.019E+04 | 1098 | - | - | 0 | - |
| - | - | 1063 | 1099 | - | - | 0 | - |
| 10 | b | 1720 | 1171 | 0.004016 | 3.431 | +1 | 10 |
| 10 | b | 902.4 | 1172 | 0.02271 | 19.38 | +1 | 10 |
| 10 | b | 1.399E+04 | 1189 | 0.002008 | 1.689 | +1 | 10 |
| - | - | 1.105E+04 | 1190 | - | - | 0 | - |
| - | - | 4841 | 1191 | - | - | 0 | - |
| 2 | y | 3549 | 1195 | 0.001944 | 1.627 | +1 | 10 |
| - | - | 2446 | 1196 | - | - | 0 | - |
| - | - | 1008 | 1197 | - | - | 0 | - |
| - | - | 709.7 | 1432 | - | - | 0 | - |

m/z Charge Intensity FragmentType MassShift Position
120.06587219238281 0 28334.87 y 10
120.08114624023438 0 38433.156
121.0693588256836 0 1126.8268
121.08444213867188 0 3557.786
126.05496978759766 0 463.79385
126.52264404296875 0 428.19147
127.08714294433594 0 518.0232
129.10260009765625 0 67595.44
130.05032348632812 0 1142.8364
130.06564331054688 0 1025.7383
130.10601806640625 0 4392.0806
133.85545349121094 0 487.02173
134.09690856933594 0 450.78992
136.0760498046875 0 16801.096
136.8589324951172 0 417.995
137.079345703125 0 1374.8109
138.1286163330078 0 408.9464
139.0869903564453 0 1931.2704
140.1436767578125 0 708.75446
146.12937927246094 0 560.0315
147.04469299316406 0 762.0648
148.94668579101562 0 665.79395
152.07102966308594 0 481.67273
155.11825561523438 0 1108.7366
155.15505981445312 0 469.45914
157.09793090820312 0 639.21265
159.0917205810547 0 616.6703
164.93984985351562 0 424.62396
165.05526733398438 0 509.80664
165.10276794433594 0 1549.0209
166.0972900390625 0 472.26617
167.08184814453125 0 1500.2831
167.1184539794922 0 1659.6587
167.2513427734375 0 423.94617
168.10206604003906 0 1662.0469
169.1339569091797 0 208415.6
170.13726806640625 0 19363.602
171.07716369628906 0 763.2759
171.14056396484375 0 511.6409
171.9581756591797 0 425.86426
172.11244201660156 0 4287.4653
172.1449737548828 0 676.0088
173.09231567382812 0 4285.433
173.1288604736328 0 1037.1554
174.1282501220703 0 1290.2278
181.0975341796875 0 3725.3801
181.1339569091797 0 3236.5007
182.08160400390625 0 2415.7834
182.12945556640625 0 595.9122
183.11314392089844 0 1739.5336
183.14967346191406 0 4004.629
185.12875366210938 0 1874.4524
185.1652374267578 0 89587.35 a 1
186.12811279296875 0 2620.0293
186.1608428955078 0 691.35046
186.1685791015625 0 8594.754
187.13137817382812 0 643.75684
187.1448974609375 0 577.71594
188.1073760986328 0 1253.2639
188.14366149902344 0 3126.5432
190.12271118164062 0 1215.3846
191.0814971923828 0 1262.0333
192.64849853515625 0 508.05646
195.11317443847656 0 4829.2227
197.1288299560547 0 163554.8
198.1322479248047 0 16914.75
199.1080322265625 0 25450.855
199.11708068847656 0 320.83923
199.13499450683594 0 1252.3655
199.18072509765625 0 2218.9104
200.11129760742188 0 1185.5682
201.12364196777344 0 14724.113 y Water loss 9
202.12599182128906 0 1564.955
203.1544952392578 0 1438.9705
204.1387939453125 0 1700.9856
208.53948974609375 0 490.05353
209.0924835205078 0 9036.405
210.096923828125 0 529.90063
210.1607666015625 0 706.4417
211.14443969726562 0 1381.391
212.13978576660156 0 2169.5984
213.0882110595703 0 734.0531
213.12347412109375 0 2205.082
213.16014099121094 0 40538.19 b 1
214.16354370117188 0 3701.665
215.1392364501953 0 3832.0593
217.0821990966797 0 821.96313
217.0970916748047 0 761.89264
217.1338348388672 0 5064.0103
219.07643127441406 0 566.32605
219.13424682617188 0 24049.434 y 9
220.13755798339844 0 2671.4312
225.08741760253906 0 1391.4728
226.1189727783203 0 19331.633
226.15577697753906 0 1011.835
227.10304260253906 0 77181.88
227.123046875 0 2540.3137
227.17547607421875 0 697.5775
228.10621643066406 0 6936.9185
228.17103576660156 0 7963.3906
229.1189727783203 0 668.65607
230.18695068359375 0 1376.7549
231.14955139160156 0 14260.215
232.15301513671875 0 2288.1792
233.09274291992188 0 1018.9915
233.16522216796875 0 1950.8865
237.08656311035156 0 1151.9462
238.12322998046875 0 1110.1206
238.1552734375 0 2695.7942
240.13522338867188 0 567.57544
244.1295928955078 0 31958.262
245.09251403808594 0 554.2367
245.12974548339844 0 7590.376
246.13363647460938 0 943.08685
247.14471435546875 0 3247.424
249.160888671875 0 798.3303
251.08815002441406 0 898.08185
251.10301208496094 0 17727.398
252.1064910888672 0 2256.8052
254.11422729492188 0 3215.0564
254.15016174316406 0 1479.0061
254.1864471435547 0 1305.2394
255.09812927246094 0 1926.1353
256.16558837890625 0 1284.3873
257.8708801269531 0 534.3825
261.087890625 0 929.5104
261.1583251953125 0 3656.4102
262.1607971191406 0 486.2427
263.1025695800781 0 701.0671
263.1390686035156 0 2094.472
266.11370849609375 0 591.6145
266.1506042480469 0 1100.7981
268.12921142578125 0 985.80475
269.1860656738281 0 770.8856
270.1814270019531 0 2111.972
272.1244812011719 0 4888.9546
273.1274719238281 0 734.2022
276.17047119140625 0 984.2242
277.15509033203125 0 1404.7178
278.15032958984375 0 1569.9148
279.097900390625 0 25491.44
280.10125732421875 0 3212.6418
280.1659851074219 0 6209.887
283.14447021484375 0 10956.636
284.12530517578125 0 667.32544
284.1474304199219 0 1703.1122
287.2120666503906 0 4352.6226
292.1279296875 0 656.627
292.16650390625 0 547.0571
294.181640625 0 655.3233
296.1242980957031 0 1061.1646
296.1785583496094 0 924.29144
296.1972351074219 0 1247.5978
298.1766357421875 0 39477.246 y Water loss 8
299.17987060546875 0 6464.371
301.1559753417969 0 913.4087
303.2071228027344 0 2735.3286
304.16583251953125 0 1594.1128
306.1451721191406 0 1715.6621
306.2182312011719 0 894.7596
307.6468811035156 0 574.3612
308.1613464355469 0 2087.217
310.1404724121094 0 904.07184
310.1764831542969 0 1932.588 b 4
310.212890625 0 1839.4827
311.1395263671875 0 7166.3823
312.1429443359375 0 1829.7206
312.19207763671875 0 968.6032
313.1907958984375 0 1082.7206
315.2069091796875 0 1899.3394
316.187255859375 0 242057.88 y 8
317.1903381347656 0 38217.055
318.1926574707031 0 3638.6406
319.21368408203125 0 685.26636
322.1771240234375 0 627.59125
323.1712951660156 0 1692.226
323.2447204589844 0 2920.8745
323.746337890625 0 646.2066
324.1559753417969 0 71753.27
325.1592102050781 0 10633.95
325.1877746582031 0 2381.8293
325.2232666015625 0 1149.721
326.17059326171875 0 2194.299
328.1484680175781 0 994.54425
328.1664733886719 0 1554.2601
329.1497497558594 0 2453.4976
331.2019958496094 0 1079.0184
335.1744384765625 0 1399.0654
335.207275390625 0 803.366
337.1517333984375 0 2449.8687
337.1869201660156 0 794.7144
339.20208740234375 0 739.7879
340.1872863769531 0 4219.1016
341.1827087402344 0 6604.351
341.2552490234375 0 6131.551 b 2
342.1286926269531 0 844.3565
342.25830078125 0 1570.3654
343.1979675292969 0 7412.581
344.2010192871094 0 1452.0186
344.70574951171875 0 1871.9557
345.20806884765625 0 1036.7057
346.1766052246094 0 4656.4155
347.1797790527344 0 1017.97485
347.2029113769531 0 3879.11
347.70361328125 0 1790.662
349.1915283203125 0 1822.2607
351.23931884765625 0 717.00555
353.1824035644531 0 4732.114
353.2190246582031 0 3064.544
354.18646240234375 0 851.9926
354.2214050292969 0 833.37897
355.0703430175781 0 1355.1959
355.1612854003906 0 3675.6753
355.1976623535156 0 2692.9182
356.16131591796875 0 1641.8126
357.21435546875 0 882.32477
358.2823791503906 0 671.6443
359.20709228515625 0 780.56384
360.2276916503906 0 1310.2415
361.18682861328125 0 940.2892
362.1709289550781 0 1703.2906
367.1975402832031 0 1346.469
369.17681884765625 0 908.92786
369.713134765625 0 7585.2915
370.2144470214844 0 2885.1377
370.71575927734375 0 867.28455
371.1928405761719 0 7055.194
372.1562194824219 0 733.4398
372.1951599121094 0 1106.7544
373.1883850097656 0 1852.8715
374.17108154296875 0 11157.572
374.1941223144531 0 885.53754
375.1745300292969 0 1839.3871
375.2396545410156 0 785.1311
376.1505432128906 0 808.48724
377.21820068359375 0 1673.9581
379.19818115234375 0 6437.581
380.1598205566406 0 564.36206
380.2012634277344 0 1241.8447
381.1448669433594 0 2523.3384
383.1932373046875 0 2403.1924
383.7109375 0 2174.8877 b 5
389.18267822265625 0 7296.8965
390.16375732421875 0 1179.3943
390.1878967285156 0 1049.1249
391.169921875 0 631.42883
391.19793701171875 0 15881.931
392.1815490722656 0 3275.499
392.20330810546875 0 1772.7725
393.2501525878906 0 808.06604
394.24139404296875 0 727.07965
395.2294921875 0 7479.548
396.2315673828125 0 2164.3342
400.7212219238281 0 704.3195
403.2341613769531 0 3418.0935
404.2374572753906 0 1003.16974
405.21368408203125 0 2101.9048
406.21441650390625 0 1148.8793
407.1930236816406 0 16449.887
408.1569519042969 0 1834.8795
408.1961669921875 0 3715.266
409.1379699707031 0 834.2262
409.2068786621094 0 1579.1641
409.2804870605469 0 1261.1488
409.73114013671875 0 784.0648
410.2780456542969 0 573.353
411.259033203125 0 1052.6837
418.23992919921875 0 13654.189
418.7409973144531 0 7419.908
419.1942443847656 0 1004.6927
419.24078369140625 0 1636.5712
420.29705810546875 0 659.4797
422.2423095703125 0 655.3615
423.2245178222656 0 24985.393 b Water loss 6
424.2247009277344 0 9615.656
425.2266845703125 0 2234.5088
426.1665344238281 0 23310.902
427.1695251464844 0 4956.8037
427.2198791503906 0 3497.6414 y Water loss 7
428.2223205566406 0 974.9327
432.2373046875 0 8870.573 b 6
432.7381286621094 0 4933.9224
433.2413024902344 0 1046.2739
434.20599365234375 0 984.8107
435.1883239746094 0 4457.968
436.1521301269531 0 1093.5798
436.18975830078125 0 1783.3328
437.2401428222656 0 1110.0795
438.27154541015625 0 8025.5522 b Water loss 3
438.3074035644531 0 1692.6277
439.2076721191406 0 1538.4438
439.25640869140625 0 2030.1078 b Ammonia loss 3
439.309326171875 0 682.1438
440.2532958984375 0 2737.8994
443.2286682128906 0 1232.2415
443.3020324707031 0 1148.8834
445.2305908203125 0 7772.3403 y 7
446.23846435546875 0 1017.9443
452.21453857421875 0 3506.0947
452.2886962890625 0 767.37
453.215576171875 0 1103.0509
454.2304992675781 0 1387.8262
454.26678466796875 0 934.4981
456.2821960449219 0 30094.479 b 3
457.2474365234375 0 1323.3424
457.28521728515625 0 8392.203
458.2850036621094 0 1071.3087
460.2544250488281 0 794.073
462.27276611328125 0 688.2688
464.2510986328125 0 1105.7894
466.26751708984375 0 819.5091
468.2453918457031 0 947.1141
470.18817138671875 0 602.13605
470.2251892089844 0 11427.446
470.3150939941406 0 843.47266
471.22625732421875 0 5849.7695
472.2251892089844 0 1518.463
473.2027587890625 0 910.93604
473.2378234863281 0 704.0831
473.3098449707031 0 717.85876
474.2539978027344 0 762.28467
474.74407958984375 0 624.61505
475.2498474121094 0 1028.88
476.2524719238281 0 1407.0968
478.2670593261719 0 1867.4827
480.2115783691406 0 609.09546
480.7650451660156 0 724.4731
482.2616882324219 0 5878.2314
482.76129150390625 0 7258.1836
483.26336669921875 0 4533.3096
483.7655029296875 0 796.01965
486.23553466796875 0 866.6365
487.7541809082031 0 1817.1658 b Water loss 7
488.2513427734375 0 3314.9395 b Ammonia loss 7
488.3236389160156 0 3537.3367
488.7431945800781 0 2052.667
489.2420349121094 0 932.35223
489.3294982910156 0 862.4864
490.2304382324219 0 825.5845
491.2218322753906 0 622.54736
492.2166442871094 0 802.6638
492.2804260253906 0 719.6115
495.7486267089844 0 1181.4491
496.7583312988281 0 17575.775 b 7
497.2601013183594 0 9928.885
497.7613830566406 0 3502.1462
498.2196960449219 0 909.9845
500.2123718261719 0 950.0091
500.25030517578125 0 1208.7689
504.24517822265625 0 2139.9883
504.31982421875 0 1088.2529
506.2608947753906 0 4777.037
508.25531005859375 0 2841.5427
509.240234375 0 4685.7026
510.2443542480469 0 1294.359
510.3290710449219 0 840.469
511.2515563964844 0 940.0573
513.2857666015625 0 796.8804
515.285888671875 0 903.05096
518.2244873046875 0 1374.8119
518.2627563476562 0 1606.9308
520.2752685546875 0 1035.7971
521.276611328125 0 1997.1323
521.7843017578125 0 1249.9315
522.2841796875 0 3774.5474
522.7852172851562 0 5635.404
523.2183227539062 0 1491.6074
523.2888793945312 0 3546.0112
523.7833862304688 0 2348.109
524.2733154296875 0 9796.662 y Water loss 6
525.2755126953125 0 3057.1582
526.266357421875 0 13065.467
527.27001953125 0 3616.219
528.2706298828125 0 719.3833
530.7981567382812 0 796.9528
531.2880859375 0 3016.8022
531.7881469726562 0 2724.1204
532.2396850585938 0 906.24347
532.2867431640625 0 790.772
534.2562255859375 0 10545.861
535.2588500976562 0 1916.5387
536.2503662109375 0 8400.518
536.30810546875 0 2717.3796
536.7822265625 0 1100.4698
537.2391967773438 0 3136.7292
539.2532348632812 0 5452.4214
539.7670288085938 0 1140.2933 y Ammonia loss 2
540.2553100585938 0 1474.8287
540.3089599609375 0 1572.5547
541.3080444335938 0 675.477
542.2826538085938 0 86335.51 y 6
543.2857666015625 0 23401.549
544.2879028320312 0 4058.6914
545.2847900390625 0 52587.703 b 8
545.7861328125 0 33221.18
546.2877197265625 0 12885.787
546.788818359375 0 1911.2382
548.2733764648438 0 1490.6624 y 2
549.3414306640625 0 1328.7972
551.2818603515625 0 1096.3278
553.3338012695312 0 2887.9277
554.2609252929688 0 68720.27
555.263916015625 0 23561.156
556.2672119140625 0 3866.952
558.3286743164062 0 4239.8296
559.3325805664062 0 829.27844
564.2444458007812 0 1047.4332
567.3505859375 0 6250.3774
568.3527221679688 0 2604.093
569.2919921875 0 784.28705
570.292236328125 0 5275.9824
571.2943725585938 0 2040.4177
571.8172607421875 0 1863.3976
572.3101806640625 0 811.4099
572.8154907226562 0 784.6592
574.32421875 0 1243.0028
579.2752075195312 0 919.02423
580.8216552734375 0 18087.238
581.3225708007812 0 12675.709
581.8240356445312 0 5385.021
582.2562866210938 0 3608.2856
582.321044921875 0 1293.3456
583.2566528320312 0 863.0569
584.3121948242188 0 753.7283
585.3375854492188 0 1903.9266
585.8134155273438 0 1231.2241 b Water loss 9
586.3170166015625 0 804.7172 b Ammonia loss 9
586.810302734375 0 788.65576
591.3499145507812 0 11946.623
592.3538818359375 0 3261.46
593.355712890625 0 733.2028
594.8187866210938 0 13512.205 b 9
595.3203735351562 0 10870.975
595.8211059570312 0 4211.219
596.3250122070312 0 956.3017
597.2864990234375 0 2601.626
597.8071899414062 0 873.0758 y 1
600.2840576171875 0 863.78625
601.3345336914062 0 3233.24 b Water loss 4
602.334716796875 0 1404.9989
603.3504638671875 0 4310.962
604.35302734375 0 1502.0608
605.296875 0 1118.3258
606.2944946289062 0 1374.7413
607.3738403320312 0 887.4955
608.3075561523438 0 2014.7137
609.3130493164062 0 643.0032
615.2777709960938 0 1748.8533
617.29150390625 0 1958.0109
619.3450927734375 0 44147.332 b 4
620.289306640625 0 1104.1188
620.348388671875 0 15609.624
621.3499755859375 0 2828.2493
622.3276977539062 0 830.8117
623.3177490234375 0 3186.4333
624.26708984375 0 1190.9535
624.3233032226562 0 790.52704
625.3363647460938 0 1806.5891
626.3378295898438 0 1248.1329
633.2891845703125 0 4398.277
634.290283203125 0 917.9447
635.3176879882812 0 1652.2417
635.3776245117188 0 5374.133
636.3109741210938 0 1036.5217
636.37744140625 0 1680.0292
638.3169555664062 0 2506.8882
639.3203735351562 0 973.88184
640.0042114257812 0 717.3684
640.3523559570312 0 788.1275
642.3529052734375 0 723.5258
645.3425903320312 0 7218.513 Precursor Water loss
645.8421020507812 0 4575.2866 Precursor Ammonia loss
646.348876953125 0 2911.2656
651.3135986328125 0 7144.0933
651.3847045898438 0 2313.1743
652.2605590820312 0 5406.719
652.319580078125 0 1962.787
652.395263671875 0 1387.9088
652.86376953125 0 1206.3159
653.26513671875 0 1629.3856
653.3289184570312 0 8552.426
654.3440551757812 0 10814.626 Precursor
654.8492431640625 0 4881.1353
655.3522338867188 0 3328.702
655.8533325195312 0 983.5968
663.3128662109375 0 2857.797
664.3104858398438 0 1049.3733
664.3678588867188 0 1547.4749
666.3417358398438 0 689.5146
667.3463134765625 0 2542.1365
668.3447265625 0 1127.1079
671.338134765625 0 1930.4647 y Water loss 5
672.3424072265625 0 924.06036
681.32470703125 0 9447.181
682.3240356445312 0 3085.0083
682.3778076171875 0 13684.827
683.3232421875 0 830.5802
683.3803100585938 0 4639.496
684.3814697265625 0 1294.8074
688.401123046875 0 1400.433
689.3505249023438 0 32894.504 y 5
690.3533935546875 0 13196.351
691.35302734375 0 2248.7358
693.399658203125 0 966.4777
695.3405151367188 0 1343.3971
703.3685302734375 0 1360.6539
704.3604736328125 0 802.26697
714.4148559570312 0 901.69574
716.3975219726562 0 4952.8696
717.4005126953125 0 1571.4803
718.4053955078125 0 728.6336
720.4072875976562 0 1374.6158
721.3925170898438 0 3732.3894
722.392578125 0 2119.7056
732.3638916015625 0 1207.6616
733.3497924804688 0 932.8302
734.353271484375 0 996.96277
738.4176635742188 0 15511.451
739.4216918945312 0 6646.4937
740.4234619140625 0 1385.0922
744.3477172851562 0 722.10284
748.402099609375 0 7326.4653 b Water loss 5
749.4019165039062 0 3975.1455
750.3856201171875 0 2930.8987
751.3843994140625 0 1083.2657
752.3615112304688 0 9558.16
753.3656616210938 0 3949.9668
754.370361328125 0 866.9379
760.3550415039062 0 1008.04846
762.3461303710938 0 5049.8184
763.3490600585938 0 2408.379
763.4674682617188 0 780.15466
765.3463134765625 0 974.9183
766.4131469726562 0 162528.3 b 5
767.4161376953125 0 74962.27
768.4189453125 0 18508.496
769.4227294921875 0 1395.5576
778.3764038085938 0 837.15857
780.3560791015625 0 57333.316
781.3592529296875 0 25064.557
782.3613891601562 0 7173.294
783.3646240234375 0 790.4831
784.4270629882812 0 919.5743
796.0452880859375 0 4519.0244
796.3810424804688 0 3190.3818
796.7135009765625 0 3125.9976
808.3479614257812 0 1091.2314
817.4464111328125 0 1667.7754
820.4600219726562 0 2504.2725
821.4654541015625 0 2237.4995
827.4310302734375 0 1648.1509
829.4395751953125 0 2366.4849
830.4444580078125 0 1394.264
833.425048828125 0 886.8409
834.4077758789062 0 752.4262 y Water loss 4
835.472900390625 0 3041.1838
836.4754028320312 0 1234.8652
845.441162109375 0 4742.7456 b Water loss 6
846.4429321289062 0 2652.2239 b Ammonia loss 6
848.3825073242188 0 1253.2192
850.4349365234375 0 3366.2268
851.4329223632812 0 2757.4346
852.4251098632812 0 1844.121 y 4
853.4255981445312 0 682.5466
861.409423828125 0 824.6509
862.4124755859375 0 772.4579
863.4663696289062 0 11698.352 b 6
864.46923828125 0 6601.4395
865.4692993164062 0 2404.781
877.4088134765625 0 19158.076
878.4119262695312 0 9125.535
878.4974365234375 0 1028.5864
879.4205322265625 0 7444.58
879.505615234375 0 993.07385
880.4251708984375 0 2569.287
889.40869140625 0 1746.9424
890.4096069335938 0 1010.96967
893.4386596679688 0 1721.3782
894.441162109375 0 1257.8254
895.4481201171875 0 1692.5347
896.468017578125 0 733.5021
907.420166015625 0 5726.3643
908.421630859375 0 2270.0386
910.541748046875 0 1146.2479
919.4944458007812 0 776.21014
946.5023193359375 0 2115.9495
947.4950561523438 0 2414.5337
948.4867553710938 0 4483.8667
949.49169921875 0 1554.9751
956.4822998046875 0 1093.678
957.4848022460938 0 866.46844
958.4678344726562 0 1223.6653
962.5339965820312 0 3544.6436
963.532470703125 0 1461.818
964.5132446289062 0 17866.377
965.5162353515625 0 9232.857
966.5195922851562 0 2117.8794
967.4402465820312 0 1592.0347 y 3
974.4972534179688 0 10634.079 b Water loss 7
975.4986572265625 0 6087.033 b Ammonia loss 7
976.47802734375 0 18112.262
977.4805908203125 0 11569.837
978.4805297851562 0 3272.2507
990.4898681640625 0 1566.4092
991.4935302734375 0 1470.1279
992.5079345703125 0 198323.52 b 7
993.5108032226562 0 119378.45
993.6057739257812 0 1174.3315
994.5135498046875 0 37237.035
995.515869140625 0 4122.964
1041.556884765625 0 825.3163
1046.554931640625 0 4118.995
1047.55419921875 0 3044.624
1075.54052734375 0 2077.5557
1076.5458984375 0 1470.5303
1077.5281982421875 0 2474.1663 y Water loss 2
1078.52734375 0 1612.1819 y Ammonia loss 2
1089.5599365234375 0 5075.082 b 8
1090.563232421875 0 3666.983
1091.56787109375 0 1756.1387
1092.566650390625 0 731.87
1095.53466796875 0 42921.277 y 2
1096.5367431640625 0 30702.424
1097.5404052734375 0 10186.484
1098.5478515625 0 1062.5928
1170.6153564453125 0 1720.1477 b Water loss 9
1171.6260986328125 0 902.4389 b Ammonia loss 9
1188.6279296875 0 13991.451 b 9
1189.6317138671875 0 11052.746
1190.6322021484375 0 4841.0703
1194.6021728515625 0 3549.1636 y 1
1195.603759765625 0 2445.5024
1196.6163330078125 0 1007.7445
1432.4486083984375 0 709.6979

Spectrum Details

|  |  |
| --- | --- |
| Matched peaks? Matched peaks The total absolute number of peaks matched. Additionally in brackets the total fraction of peaks matched and the total number of peaks is shown. | 55 (8.68% of 634) |
| FDR? FDR The false discovery rate estimated for this peptide. It is calculated by matching all theoretical fragments with a non-integer shift with the raw peaks for this spectrum. This is done with 40 different shifts. The resulting percentage is the average number of annotated peaks over the number of annotated peaks with the correct spectrum. | 1.08% |
| Satellite FDR? Satellite FDR See the FDR for details on its calculation. This satellite ion specific FDR only contains the satellite ions (d/w) for I/L/J positions. | - |
| PSM Score? PSM Score The PSM Score as given by Hecklib to this annotated spectrum. It is shown with three significant figures. | 638 |

#### Spectrum 7987? Spectrum 7987 The raw spectrum of this peptide as annotated by Hecklib. The fragments are coloured according to ion type (see legend). Any peaks with a star '\*' as text can be hovered over to see the full details, first the ion type second the mass shift type. By hovering over the amino acids in the peptide or ions in the legend the corresponding peaks are highlighted. By toggling the 'Unassigned' label you can turn the background (unassigned) peaks on or off in the plot. By updating the slider in the Ion legend you can update the spectrum to only show the top X% of the peaks with labels. The top X% means any peak that is within X% of the highest intensity. By dragging in the spectrum you can zoom in to a specific part of the spectrum and use 'Zoom Out' to get back to the original zoom level. The annotation of the spectrum is based on the given sequence in the peptides file and is done with different software so inconsistencies are likely. The peaks are annotated based on the given sequence, with 20 ppm tolerance.

Copy Data

##### Spectrum 7987 (TSV)

###### Preview

```
Loading example...
```

*Click on the button to copy the data to your clipboard.*

Mz MinMz MaxIntensity Max

WidthHeightPeptide font sizePeptide stroke widthSpectrum font sizeSpectrum stroke widthCompact peptide

Ion legend

wxyz

abcd

OtherUnassignedIon^Charge^_Position_Show for top:%

JVKDYFPEPVT

07.25e+41.45e+52.18e+52.90e+5

Zoom Out

y^+1^_1_y^+1^_2_y^+1^_3_y^+1^_3_c^+1^_3_c^+1^_3_c^+2^_7_y^+1^_4_c^+1^_4_c^+1^_4_c^+2^_8_y^+1^_5_y^+1^_5_c^+2^_9_c^+2^_10_c^+1^_5_c^+1^_5_y^+1^_6_c^+1^_6_z^+1^_7_y^+1^_7_c^+1^_7_c^+1^_7_y^+1^_8_c^+1^_8_c^+1^_8_z^+1^_9_c^+1^_9_y^+1^_9_c^+1^_9_z^+1^_10_c^+1^_10_y^+1^_10_c^+1^_10_

0661132119822642

Fragment Matches Table

Show background peaks

| Position | Ion type | Intensity | mz Theoretical | mz Error (Th) | mz Error (ppm) | Charge | Series Number |
| --- | --- | --- | --- | --- | --- | --- | --- |
| 11 | y | 3557 | 120.1 | 0.0002381 | 1.983 | +1 | 1 |
| - | - | 460.6 | 120.1 | - | - | 0 | - |
| - | - | 371.1 | 121.4 | - | - | 0 | - |
| - | - | 374.8 | 122.1 | - | - | 0 | - |
| - | - | 401.4 | 126.9 | - | - | 0 | - |
| - | - | 369.9 | 127.7 | - | - | 0 | - |
| - | - | 2711 | 129.1 | - | - | 0 | - |
| - | - | 458 | 131.2 | - | - | 0 | - |
| - | - | 436 | 136.4 | - | - | 0 | - |
| - | - | 425.6 | 145.3 | - | - | 0 | - |
| - | - | 411.4 | 150.9 | - | - | 0 | - |
| - | - | 457 | 158.3 | - | - | 0 | - |
| - | - | 498.8 | 165.8 | - | - | 0 | - |
| - | - | 6944 | 169.1 | - | - | 0 | - |
| - | - | 2016 | 173.4 | - | - | 0 | - |
| - | - | 896.3 | 185.1 | - | - | 0 | - |
| - | - | 6939 | 185.2 | - | - | 0 | - |
| - | - | 743.8 | 186.2 | - | - | 0 | - |
| - | - | 501.5 | 188.8 | - | - | 0 | - |
| - | - | 493.8 | 193.6 | - | - | 0 | - |
| - | - | 1.264E+04 | 197.1 | - | - | 0 | - |
| - | - | 1154 | 198.1 | - | - | 0 | - |
| - | - | 554.8 | 202 | - | - | 0 | - |
| - | - | 1.875E+04 | 203.1 | - | - | 0 | - |
| - | - | 714.2 | 204.1 | - | - | 0 | - |
| - | - | 475 | 209.1 | - | - | 0 | - |
| - | - | 6142 | 213.2 | - | - | 0 | - |
| - | - | 682.2 | 214.2 | - | - | 0 | - |
| 10 | y | 1177 | 219.1 | 0.0001454 | 0.6637 | +1 | 2 |
| - | - | 512.9 | 221.7 | - | - | 0 | - |
| - | - | 2592 | 227.1 | - | - | 0 | - |
| - | - | 634.7 | 242.1 | - | - | 0 | - |
| - | - | 481.6 | 244.6 | - | - | 0 | - |
| - | - | 547.9 | 257.8 | - | - | 0 | - |
| - | - | 3963 | 260.1 | - | - | 0 | - |
| 9 | y | 3597 | 298.2 | 0.0002589 | 0.8683 | +1 | 3 |
| - | - | 1810 | 299.1 | - | - | 0 | - |
| - | - | 667.4 | 315.3 | - | - | 0 | - |
| 9 | y | 3.804E+04 | 316.2 | 0.0003143 | 0.9941 | +1 | 3 |
| - | - | 5008 | 317.1 | - | - | 0 | - |
| - | - | 5707 | 317.2 | - | - | 0 | - |
| - | - | 811.5 | 318.2 | - | - | 0 | - |
| - | - | 7655 | 324.2 | - | - | 0 | - |
| - | - | 883.4 | 325.2 | - | - | 0 | - |
| 3 | c | 705.7 | 341.3 | 0.002546 | 7.46 | +1 | 3 |
| - | - | 1130 | 355.1 | - | - | 0 | - |
| - | - | 915.1 | 356.1 | - | - | 0 | - |
| - | - | 689.4 | 357.1 | - | - | 0 | - |
| - | - | 7735 | 357.3 | - | - | 0 | - |
| 3 | c | 2979 | 358.3 | 0.001115 | 3.112 | +1 | 3 |
| - | - | 679.8 | 359.3 | - | - | 0 | - |
| - | - | 807.1 | 407.2 | - | - | 0 | - |
| - | - | 518.6 | 413.1 | - | - | 0 | - |
| - | - | 612.8 | 415.1 | - | - | 0 | - |
| - | - | 737.4 | 418.2 | - | - | 0 | - |
| - | - | 3602 | 423.2 | - | - | 0 | - |
| - | - | 693.1 | 424.2 | - | - | 0 | - |
| - | - | 1032 | 426.2 | - | - | 0 | - |
| - | - | 1952 | 430.2 | - | - | 0 | - |
| 7 | c | 769.5 | 432.2 | 0.0002169 | 0.5018 | +2 | 7 |
| 8 | y | 1392 | 445.2 | 0.0005002 | 1.123 | +1 | 4 |
| 4 | c | 1411 | 456.3 | 0.0007798 | 1.709 | +1 | 4 |
| - | - | 3051 | 470.2 | - | - | 0 | - |
| - | - | 1138 | 472.3 | - | - | 0 | - |
| 4 | c | 3971 | 473.3 | 0.0003232 | 0.6829 | +1 | 4 |
| - | - | 1084 | 474.3 | - | - | 0 | - |
| - | - | 789.9 | 483.3 | - | - | 0 | - |
| - | - | 621 | 488.7 | - | - | 0 | - |
| 8 | c | 2680 | 496.8 | 0.0002521 | 0.5076 | +2 | 8 |
| - | - | 1136 | 497.3 | - | - | 0 | - |
| - | - | 1224 | 498.2 | - | - | 0 | - |
| - | - | 818.4 | 517.3 | - | - | 0 | - |
| - | - | 648.4 | 522.3 | - | - | 0 | - |
| - | - | 2007 | 522.8 | - | - | 0 | - |
| - | - | 775.3 | 523.8 | - | - | 0 | - |
| - | - | 1725 | 524.2 | - | - | 0 | - |
| 7 | y | 984.7 | 524.3 | 0.002558 | 4.88 | +1 | 5 |
| - | - | 990.6 | 531.3 | - | - | 0 | - |
| - | - | 799.5 | 534.3 | - | - | 0 | - |
| 7 | y | 1.7E+04 | 542.3 | 0.0005995 | 1.105 | +1 | 5 |
| - | - | 994.9 | 542.3 | - | - | 0 | - |
| - | - | 5073 | 543.3 | - | - | 0 | - |
| 9 | c | 1.162E+04 | 545.3 | 0.0002679 | 0.4913 | +2 | 9 |
| - | - | 7072 | 545.8 | - | - | 0 | - |
| - | - | 2107 | 546.3 | - | - | 0 | - |
| - | - | 6906 | 554.3 | - | - | 0 | - |
| - | - | 2037 | 555.3 | - | - | 0 | - |
| - | - | 2087 | 566.4 | - | - | 0 | - |
| - | - | 711.9 | 578.3 | - | - | 0 | - |
| - | - | 3309 | 580.8 | - | - | 0 | - |
| - | - | 3566 | 581.3 | - | - | 0 | - |
| - | - | 1336 | 581.8 | - | - | 0 | - |
| - | - | 640.8 | 591.4 | - | - | 0 | - |
| - | - | 691.6 | 594.3 | - | - | 0 | - |
| 10 | c | 3279 | 594.8 | 0.0002406 | 0.4046 | +2 | 10 |
| - | - | 2219 | 595.3 | - | - | 0 | - |
| - | - | 612.5 | 595.8 | - | - | 0 | - |
| 5 | c | 6479 | 619.3 | 0.0006532 | 1.055 | +1 | 5 |
| - | - | 2209 | 620.3 | - | - | 0 | - |
| - | - | 942.1 | 621.4 | - | - | 0 | - |
| - | - | 1462 | 622.3 | - | - | 0 | - |
| - | - | 1.45E+04 | 635.4 | - | - | 0 | - |
| 5 | c | 1.97E+04 | 636.4 | 0.0006884 | 1.082 | +1 | 5 |
| - | - | 2802 | 637.3 | - | - | 0 | - |
| - | - | 6374 | 637.4 | - | - | 0 | - |
| - | - | 2380 | 637.8 | - | - | 0 | - |
| - | - | 1.473E+04 | 638.3 | - | - | 0 | - |
| - | - | 870.3 | 638.4 | - | - | 0 | - |
| - | - | 2830 | 639.3 | - | - | 0 | - |
| - | - | 1283 | 645.3 | - | - | 0 | - |
| - | - | 1139 | 645.8 | - | - | 0 | - |
| - | - | 1043 | 646.8 | - | - | 0 | - |
| - | - | 1343 | 647.3 | - | - | 0 | - |
| - | - | 908.8 | 652.9 | - | - | 0 | - |
| - | - | 844.1 | 653.3 | - | - | 0 | - |
| - | - | 2100 | 654.3 | - | - | 0 | - |
| - | - | 1903 | 654.8 | - | - | 0 | - |
| - | - | 1337 | 655.4 | - | - | 0 | - |
| - | - | 941.8 | 681.3 | - | - | 0 | - |
| - | - | 1001 | 682.4 | - | - | 0 | - |
| - | - | 686.3 | 688.3 | - | - | 0 | - |
| 6 | y | 1.015E+04 | 689.4 | 0.0003618 | 0.5249 | +1 | 6 |
| - | - | 3489 | 690.4 | - | - | 0 | - |
| - | - | 646.6 | 697.4 | - | - | 0 | - |
| - | - | 1841 | 707.3 | - | - | 0 | - |
| - | - | 677.4 | 708.3 | - | - | 0 | - |
| - | - | 2796 | 723.3 | - | - | 0 | - |
| - | - | 1176 | 724.3 | - | - | 0 | - |
| - | - | 1896 | 738.4 | - | - | 0 | - |
| - | - | 5848 | 739.4 | - | - | 0 | - |
| - | - | 1.1E+04 | 740.4 | - | - | 0 | - |
| - | - | 2713 | 741.4 | - | - | 0 | - |
| - | - | 1152 | 742.4 | - | - | 0 | - |
| - | - | 1247 | 750.3 | - | - | 0 | - |
| - | - | 8555 | 766.3 | - | - | 0 | - |
| 6 | c | 2.538E+04 | 766.4 | 0.0003545 | 0.4625 | +1 | 6 |
| - | - | 5336 | 767.3 | - | - | 0 | - |
| - | - | 1.06E+04 | 767.4 | - | - | 0 | - |
| - | - | 2576 | 768.4 | - | - | 0 | - |
| - | - | 1011 | 769.4 | - | - | 0 | - |
| - | - | 6225 | 780.4 | - | - | 0 | - |
| - | - | 2271 | 781.4 | - | - | 0 | - |
| 5 | z | 2036 | 836.4 | 0.00721 | 8.62 | +1 | 7 |
| - | - | 778.5 | 837.4 | - | - | 0 | - |
| - | - | 1473 | 840.4 | - | - | 0 | - |
| - | - | 2766 | 851.4 | - | - | 0 | - |
| 5 | y | 8892 | 852.4 | 0.0002657 | 0.3117 | +1 | 7 |
| - | - | 4526 | 853.4 | - | - | 0 | - |
| - | - | 1172 | 854.4 | - | - | 0 | - |
| - | - | 3144 | 863.4 | - | - | 0 | - |
| 7 | c | 1452 | 863.5 | 0.004109 | 4.759 | +1 | 7 |
| - | - | 1208 | 876.4 | - | - | 0 | - |
| - | - | 2671 | 877.4 | - | - | 0 | - |
| - | - | 657.5 | 877.9 | - | - | 0 | - |
| - | - | 1008 | 878.4 | - | - | 0 | - |
| - | - | 2.783E+04 | 879.5 | - | - | 0 | - |
| 7 | c | 4.892E+04 | 880.5 | 0.0006504 | 0.7387 | +1 | 7 |
| - | - | 2.348E+04 | 881.5 | - | - | 0 | - |
| - | - | 4577 | 882.5 | - | - | 0 | - |
| - | - | 838.7 | 884.4 | - | - | 0 | - |
| - | - | 931.9 | 893.4 | - | - | 0 | - |
| - | - | 709.1 | 920.5 | - | - | 0 | - |
| - | - | 812.2 | 934.5 | - | - | 0 | - |
| - | - | 714 | 947.5 | - | - | 0 | - |
| - | - | 8729 | 949.5 | - | - | 0 | - |
| - | - | 3561 | 950.5 | - | - | 0 | - |
| - | - | 1493 | 952.4 | - | - | 0 | - |
| - | - | 819.4 | 953.4 | - | - | 0 | - |
| - | - | 1380 | 964.5 | - | - | 0 | - |
| - | - | 1905 | 965.5 | - | - | 0 | - |
| - | - | 2167 | 966.4 | - | - | 0 | - |
| - | - | 1116 | 966.5 | - | - | 0 | - |
| 4 | y | 4455 | 967.4 | 0.001531 | 1.582 | +1 | 8 |
| - | - | 2900 | 968.4 | - | - | 0 | - |
| - | - | 989.2 | 969.4 | - | - | 0 | - |
| - | - | 797.8 | 974.5 | - | - | 0 | - |
| - | - | 1.215E+04 | 976.5 | - | - | 0 | - |
| - | - | 3715 | 977.5 | - | - | 0 | - |
| - | - | 1331 | 978.5 | - | - | 0 | - |
| - | - | 2434 | 990.5 | - | - | 0 | - |
| 8 | c | 2840 | 991.5 | 0.0068 | 6.858 | +1 | 8 |
| 8 | c | 3.533E+04 | 992.5 | 0.0002759 | 0.278 | +1 | 8 |
| - | - | 2.593E+04 | 993.5 | - | - | 0 | - |
| - | - | 6519 | 994.5 | - | - | 0 | - |
| - | - | 728.5 | 995.5 | - | - | 0 | - |
| - | - | 1963 | 1007 | - | - | 0 | - |
| - | - | 3327 | 1008 | - | - | 0 | - |
| - | - | 1107 | 1009 | - | - | 0 | - |
| - | - | 5491 | 1034 | - | - | 0 | - |
| - | - | 2119 | 1035 | - | - | 0 | - |
| - | - | 3232 | 1036 | - | - | 0 | - |
| - | - | 1629 | 1037 | - | - | 0 | - |
| - | - | 1188 | 1038 | - | - | 0 | - |
| - | - | 2859 | 1050 | - | - | 0 | - |
| - | - | 1.323E+04 | 1051 | - | - | 0 | - |
| - | - | 4001 | 1052 | - | - | 0 | - |
| - | - | 761.2 | 1053 | - | - | 0 | - |
| - | - | 1158 | 1063 | - | - | 0 | - |
| - | - | 1027 | 1064 | - | - | 0 | - |
| - | - | 954.2 | 1065 | - | - | 0 | - |
| - | - | 701.3 | 1067 | - | - | 0 | - |
| - | - | 1066 | 1073 | - | - | 0 | - |
| 3 | z | 1.039E+04 | 1080 | 1.099E-05 | 0.01018 | +1 | 9 |
| - | - | 6609 | 1081 | - | - | 0 | - |
| - | - | 2429 | 1082 | - | - | 0 | - |
| 9 | c | 681.4 | 1090 | 0.005005 | 4.593 | +1 | 9 |
| - | - | 679.7 | 1090 | - | - | 0 | - |
| - | - | 1.44E+04 | 1091 | - | - | 0 | - |
| - | - | 4631 | 1092 | - | - | 0 | - |
| 3 | y | 7475 | 1096 | 0.0009183 | 0.8382 | +1 | 9 |
| - | - | 5997 | 1097 | - | - | 0 | - |
| - | - | 1850 | 1098 | - | - | 0 | - |
| - | - | 768.5 | 1106 | - | - | 0 | - |
| 9 | c | 7263 | 1107 | 0.001891 | 1.709 | +1 | 9 |
| - | - | 4383 | 1108 | - | - | 0 | - |
| - | - | 2066 | 1109 | - | - | 0 | - |
| - | - | 654.2 | 1119 | - | - | 0 | - |
| - | - | 1139 | 1125 | - | - | 0 | - |
| - | - | 736.7 | 1137 | - | - | 0 | - |
| - | - | 2203 | 1147 | - | - | 0 | - |
| - | - | 2490 | 1148 | - | - | 0 | - |
| - | - | 1101 | 1161 | - | - | 0 | - |
| - | - | 1.392E+04 | 1162 | - | - | 0 | - |
| - | - | 1.23E+04 | 1163 | - | - | 0 | - |
| - | - | 7481 | 1164 | - | - | 0 | - |
| - | - | 3701 | 1165 | - | - | 0 | - |
| 2 | z | 6617 | 1179 | 0.001408 | 1.195 | +1 | 10 |
| - | - | 2930 | 1180 | - | - | 0 | - |
| - | - | 1834 | 1181 | - | - | 0 | - |
| 10 | c | 2754 | 1189 | 0.0009093 | 0.765 | +1 | 10 |
| - | - | 2508 | 1190 | - | - | 0 | - |
| - | - | 5346 | 1191 | - | - | 0 | - |
| - | - | 3986 | 1192 | - | - | 0 | - |
| - | - | 1683 | 1193 | - | - | 0 | - |
| 2 | y | 728.5 | 1195 | 0.005136 | 4.299 | +1 | 10 |
| - | - | 8242 | 1203 | - | - | 0 | - |
| - | - | 3685 | 1204 | - | - | 0 | - |
| - | - | 9561 | 1205 | - | - | 0 | - |
| 10 | c | 6.827E+04 | 1206 | 0.000847 | 0.7025 | +1 | 10 |
| - | - | 5.028E+04 | 1207 | - | - | 0 | - |
| - | - | 2.78E+04 | 1208 | - | - | 0 | - |
| - | - | 1.059E+04 | 1209 | - | - | 0 | - |
| - | - | 1980 | 1210 | - | - | 0 | - |
| - | - | 1614 | 1220 | - | - | 0 | - |
| - | - | 1396 | 1221 | - | - | 0 | - |
| - | - | 831.1 | 1231 | - | - | 0 | - |
| - | - | 1030 | 1233 | - | - | 0 | - |
| - | - | 3162 | 1234 | - | - | 0 | - |
| - | - | 3.56E+04 | 1235 | - | - | 0 | - |
| - | - | 1.276E+04 | 1236 | - | - | 0 | - |
| - | - | 4794 | 1237 | - | - | 0 | - |
| - | - | 4267 | 1238 | - | - | 0 | - |
| - | - | 1814 | 1239 | - | - | 0 | - |
| - | - | 4660 | 1245 | - | - | 0 | - |
| - | - | 3428 | 1246 | - | - | 0 | - |
| - | - | 3287 | 1247 | - | - | 0 | - |
| - | - | 1603 | 1248 | - | - | 0 | - |
| - | - | 2.055E+04 | 1249 | - | - | 0 | - |
| - | - | 1.405E+04 | 1250 | - | - | 0 | - |
| - | - | 6506 | 1251 | - | - | 0 | - |
| - | - | 2477 | 1252 | - | - | 0 | - |
| - | - | 4162 | 1253 | - | - | 0 | - |
| - | - | 1934 | 1254 | - | - | 0 | - |
| - | - | 5183 | 1263 | - | - | 0 | - |
| - | - | 4402 | 1264 | - | - | 0 | - |
| - | - | 8290 | 1265 | - | - | 0 | - |
| - | - | 5855 | 1266 | - | - | 0 | - |
| - | - | 1034 | 1267 | - | - | 0 | - |
| - | - | 4615 | 1276 | - | - | 0 | - |
| - | - | 1772 | 1277 | - | - | 0 | - |
| - | - | 1.218E+04 | 1281 | - | - | 0 | - |
| - | - | 1.008E+04 | 1282 | - | - | 0 | - |
| - | - | 3178 | 1283 | - | - | 0 | - |
| - | - | 1522 | 1289 | - | - | 0 | - |
| - | - | 767.9 | 1290 | - | - | 0 | - |
| - | - | 6211 | 1291 | - | - | 0 | - |
| - | - | 7.545E+04 | 1292 | - | - | 0 | - |
| - | - | 6.197E+04 | 1293 | - | - | 0 | - |
| - | - | 8.197E+04 | 1294 | - | - | 0 | - |
| - | - | 3.06E+04 | 1295 | - | - | 0 | - |
| - | - | 1216 | 1305 | - | - | 0 | - |
| - | - | 2812 | 1306 | - | - | 0 | - |
| - | - | 4144 | 1307 | - | - | 0 | - |
| - | - | 5.896E+04 | 1308 | - | - | 0 | - |
| - | - | 2.871E+05 | 1309 | - | - | 0 | - |
| - | - | 2.04E+05 | 1310 | - | - | 0 | - |
| - | - | 7.678E+04 | 1311 | - | - | 0 | - |
| - | - | 1.079E+04 | 1312 | - | - | 0 | - |
| - | - | 893.9 | 1325 | - | - | 0 | - |
| - | - | 928 | 1754 | - | - | 0 | - |
| - | - | 1063 | 1859 | - | - | 0 | - |
| - | - | 650.9 | 2616 | - | - | 0 | - |

m/z Charge Intensity FragmentType MassShift Position
120.06575775146484 0 3557.1953 y 10
120.08100891113281 0 460.62518
121.37811279296875 0 371.0643
122.13825988769531 0 374.8025
126.894287109375 0 401.43265
127.71576690673828 0 369.93765
129.10235595703125 0 2710.548
131.1690673828125 0 457.9632
136.44114685058594 0 436.042
145.25466918945312 0 425.60284
150.92431640625 0 411.4493
158.3495330810547 0 457.0446
165.8311767578125 0 498.78775
169.1336669921875 0 6944.3413
173.43898010253906 0 2015.799
185.0922393798828 0 896.32
185.16500854492188 0 6938.978
186.1682891845703 0 743.81995
188.77542114257812 0 501.50644
193.6297149658203 0 493.7643
197.12860107421875 0 12636.645
198.13198852539062 0 1153.7444
201.99169921875 0 554.77386
203.10275268554688 0 18747.186
204.10618591308594 0 714.21423
209.09326171875 0 474.96323
213.1598663330078 0 6142.341
214.1626434326172 0 682.1979
219.1340789794922 0 1177.118 y 9
221.67637634277344 0 512.8552
227.10247802734375 0 2592.1172
242.1136474609375 0 634.73975
244.6398162841797 0 481.5944
257.8075866699219 0 547.877
260.1241149902344 0 3963.4683
298.1763916015625 0 3597.2554 y Water loss 8
299.1351013183594 0 1810.1694
315.27606201171875 0 667.39386
316.18701171875 0 38043.316 y 8
317.1458740234375 0 5008.158
317.1905822753906 0 5706.883
318.19244384765625 0 811.46265
324.1558532714844 0 7655.0127
325.1597595214844 0 883.39166
341.25726318359375 0 705.7123 c Ammonia loss 2
355.06884765625 0 1130.197
356.07171630859375 0 915.09674
357.0672912597656 0 689.44385
357.2734375 0 7735.172
358.2801513671875 0 2978.664 c 2
359.2850341796875 0 679.7869
407.1920166015625 0 807.0724
413.05877685546875 0 518.6245
415.0727233886719 0 612.8372
418.2393798828125 0 737.43524
423.2247619628906 0 3602.15
424.2284240722656 0 693.12866
426.1656494140625 0 1032.2505
430.2301025390625 0 1952.0825
432.2369384765625 0 769.52576 c Ammonia loss 6
445.2287902832031 0 1392.4521 y 7
456.2824401855469 0 1410.5858 c Ammonia loss 3
470.2073974609375 0 3050.652
472.3017272949219 0 1138.2858
473.30853271484375 0 3971.4644 c 3
474.3106994628906 0 1084.1577
483.2623291015625 0 789.9409
488.74212646484375 0 621.04663
496.7582702636719 0 2680.3647 c Ammonia loss 7
497.2601318359375 0 1135.7345
498.2381591796875 0 1223.5236
517.25537109375 0 818.4146
522.2835693359375 0 648.38983
522.7863159179688 0 2007.263
523.784423828125 0 775.29456
524.217529296875 0 1725.3871
524.2740478515625 0 984.6551 y Water loss 6
531.2894287109375 0 990.55914
534.2570190429688 0 799.4514
542.2826538085938 0 17002.984 y 6
542.3282470703125 0 994.86597
543.285888671875 0 5073.3657
545.28466796875 0 11624.326 c Ammonia loss 8
545.78564453125 0 7072.4136
546.2861328125 0 2106.8682
554.2608032226562 0 6906.319
555.2634887695312 0 2037.3171
566.4281005859375 0 2086.6096
578.2749633789062 0 711.8858
580.8212890625 0 3309.0107
581.3228149414062 0 3566.0999
581.826416015625 0 1335.614
591.3513793945312 0 640.8482
594.2855834960938 0 691.5735
594.81884765625 0 3279.1846 c Ammonia loss 9
595.3212280273438 0 2218.9248
595.828369140625 0 612.51447
619.3456420898438 0 6478.6533 c Ammonia loss 4
620.3486328125 0 2208.701
621.3560180664062 0 942.1457
622.277587890625 0 1462.4033
635.3648681640625 0 14504.389
636.370849609375 0 19704.549 c 4
637.2887573242188 0 2802.3264
637.373779296875 0 6373.5054
637.8088989257812 0 2380.0103
638.2982177734375 0 14732.733
638.3750610351562 0 870.339
639.3009033203125 0 2829.9429
645.3419799804688 0 1282.7258
645.8424072265625 0 1138.9515
646.8129272460938 0 1042.6105
647.3148803710938 0 1343.3838
652.8638916015625 0 908.81616
653.3358154296875 0 844.06366
654.3474731445312 0 2100.038
654.8485717773438 0 1903.3655
655.3505249023438 0 1336.9624
681.3285522460938 0 941.82623
682.3790283203125 0 1000.7519
688.3408813476562 0 686.2717
689.350830078125 0 10145.785 y 5
690.3533325195312 0 3489.4182
697.4392700195312 0 646.63947
707.3206176757812 0 1840.7703
708.3197631835938 0 677.36304
723.3245239257812 0 2795.9392
724.33056640625 0 1176.3433
738.4188232421875 0 1896.1672
739.4263305664062 0 5847.6123
740.4337158203125 0 10995.421
741.4375 0 2712.5703
742.443115234375 0 1152.3771
750.3131713867188 0 1247.4834
766.3317260742188 0 8555.106
766.4137573242188 0 25382.113 c Ammonia loss 5
767.336669921875 0 5336.2593
767.416748046875 0 10599.561
768.41552734375 0 2576.4097
769.4047241210938 0 1010.8528
780.3567504882812 0 6225.4844
781.359375 0 2270.6655
836.4022827148438 0 2036.0702 z 4
837.4049072265625 0 778.47455
840.443359375 0 1473.4625
851.4064331054688 0 2766.1248
852.4140625 0 8891.523 y 4
853.4164428710938 0 4525.5176
854.41748046875 0 1171.8163
863.395751953125 0 3144.0405
863.4702758789062 0 1452.0198 c Ammonia loss 6
876.3947143554688 0 1208.4845
877.4095458984375 0 2670.9746
877.9091796875 0 657.54297
878.4076538085938 0 1008.2462
879.485107421875 0 27833.143
880.4920654296875 0 48916.547 c 6
881.4953002929688 0 23483.166
882.4986572265625 0 4577.226
884.3892822265625 0 838.7312
893.4271850585938 0 931.85754
920.4718627929688 0 709.08575
934.4938354492188 0 812.1921
947.5009765625 0 713.9824
949.4944458007812 0 8728.965
950.4961547851562 0 3561.3523
952.4280395507812 0 1492.5138
953.4320678710938 0 819.42194
964.5144653320312 0 1380.4556
965.515869140625 0 1905.3313
966.4312133789062 0 2167.4653
966.5264892578125 0 1116.134
967.439208984375 0 4455.1675 y 3
968.444580078125 0 2900.0159
969.44921875 0 989.1797
974.4995727539062 0 797.75073
976.4813232421875 0 12154.536
977.4805908203125 0 3715.1243
978.4849243164062 0 1331.2866
990.5148315429688 0 2434.2925
991.5179443359375 0 2839.7678 c Water loss 7
992.5084838867188 0 35326.586 c Ammonia loss 7
993.5109252929688 0 25927.93
994.5134887695312 0 6519.348
995.5134887695312 0 728.53674
1006.5133666992188 0 1962.988
1007.5156860351562 0 3326.6086
1008.5205078125 0 1106.9526
1033.503173828125 0 5491.462
1034.5107421875 0 2119.2546
1035.5281982421875 0 3231.5623
1036.5308837890625 0 1629.3174
1037.5330810546875 0 1187.531
1049.5238037109375 0 2858.9565
1050.529052734375 0 13231.003
1051.5313720703125 0 4001.413
1052.5284423828125 0 761.2066
1062.550048828125 0 1158.2244
1063.556884765625 0 1027.3387
1064.5123291015625 0 954.17816
1066.5205078125 0 701.33826
1072.5211181640625 0 1066.2656
1079.5169677734375 0 10393.016 z 2
1080.518310546875 0 6608.5874
1081.5224609375 0 2428.9868
1089.5665283203125 0 681.3857 c Ammonia loss 8
1089.6187744140625 0 679.6511
1090.525146484375 0 14395.663
1091.52734375 0 4630.95
1095.53662109375 0 7475.1494 y 2
1096.5372314453125 0 5997.125
1097.5411376953125 0 1849.9829
1105.5687255859375 0 768.54565
1106.586181640625 0 7262.879 c 8
1107.590087890625 0 4383.443
1108.595458984375 0 2066.1177
1118.6536865234375 0 654.23016
1124.5345458984375 0 1139.0817
1136.58642578125 0 736.69617
1146.630859375 0 2202.7358
1147.6429443359375 0 2489.987
1160.6082763671875 0 1100.87
1161.642822265625 0 13916.4795
1162.646484375 0 12302.3125
1163.6522216796875 0 7481.1074
1164.6614990234375 0 3701.1428
1178.583984375 0 6617.304 z 1
1179.586669921875 0 2930.3613
1180.5869140625 0 1834.181
1188.6290283203125 0 2754.2273 c Ammonia loss 9
1189.635986328125 0 2508.3276
1190.6448974609375 0 5346.313
1191.641845703125 0 3985.6047
1192.6546630859375 0 1683.0619
1194.6092529296875 0 728.5181 y 1
1202.6414794921875 0 8242.306
1203.6435546875 0 3684.7388
1204.6033935546875 0 9560.866
1205.6556396484375 0 68267.55 c 9
1206.6595458984375 0 50277.316
1207.6649169921875 0 27802.705
1208.6702880859375 0 10587.225
1209.6793212890625 0 1979.7892
1219.644775390625 0 1614.3206
1220.642333984375 0 1395.9131
1230.654052734375 0 831.06964
1232.630615234375 0 1030.3815
1233.605224609375 0 3162.1826
1234.612548828125 0 35603.59
1235.616455078125 0 12760.961
1236.6744384765625 0 4794.4453
1237.672119140625 0 4267.2256
1238.681884765625 0 1814.1113
1244.6805419921875 0 4659.6475
1245.680419921875 0 3428.2058
1246.69091796875 0 3287.4243
1247.6871337890625 0 1603.1036
1248.6531982421875 0 20553.34
1249.653076171875 0 14050.457
1250.642578125 0 6505.55
1251.6265869140625 0 2476.6665
1252.63037109375 0 4162.054
1253.635986328125 0 1934.1385
1262.6868896484375 0 5182.851
1263.68994140625 0 4401.7856
1264.7012939453125 0 8290.36
1265.70556640625 0 5855.251
1266.701904296875 0 1034.2202
1275.6158447265625 0 4614.6406
1276.6212158203125 0 1771.9437
1280.700439453125 0 12183.403
1281.7027587890625 0 10077.126
1282.706787109375 0 3178.0122
1288.6959228515625 0 1522.4554
1289.6788330078125 0 767.8577
1290.680908203125 0 6210.9355
1291.668701171875 0 75449.17
1292.6646728515625 0 61966.04
1293.6343994140625 0 81972.86
1294.632080078125 0 30601.559
1304.7108154296875 0 1216.4358
1305.707275390625 0 2812.3196
1306.68359375 0 4143.9443
1307.6868896484375 0 58961.72
1308.694091796875 0 287140.5
1309.6973876953125 0 204012.77
1310.69970703125 0 76779.83
1311.7017822265625 0 10787.109
1324.6573486328125 0 893.8809
1753.820556640625 0 927.9818
1858.923095703125 0 1063.3247
2616.326171875 0 650.9091

Spectrum Details

|  |  |
| --- | --- |
| Matched peaks? Matched peaks The total absolute number of peaks matched. Additionally in brackets the total fraction of peaks matched and the total number of peaks is shown. | 34 (11.64% of 292) |
| FDR? FDR The false discovery rate estimated for this peptide. It is calculated by matching all theoretical fragments with a non-integer shift with the raw peaks for this spectrum. This is done with 40 different shifts. The resulting percentage is the average number of annotated peaks over the number of annotated peaks with the correct spectrum. | 2.31% |
| Satellite FDR? Satellite FDR See the FDR for details on its calculation. This satellite ion specific FDR only contains the satellite ions (d/w) for I/L/J positions. | - |
| PSM Score? PSM Score The PSM Score as given by Hecklib to this annotated spectrum. It is shown with three significant figures. | 409 |

#### Spectrum 8278? Spectrum 8278 The raw spectrum of this peptide as annotated by Hecklib. The fragments are coloured according to ion type (see legend). Any peaks with a star '\*' as text can be hovered over to see the full details, first the ion type second the mass shift type. By hovering over the amino acids in the peptide or ions in the legend the corresponding peaks are highlighted. By toggling the 'Unassigned' label you can turn the background (unassigned) peaks on or off in the plot. By updating the slider in the Ion legend you can update the spectrum to only show the top X% of the peaks with labels. The top X% means any peak that is within X% of the highest intensity. By dragging in the spectrum you can zoom in to a specific part of the spectrum and use 'Zoom Out' to get back to the original zoom level. The annotation of the spectrum is based on the given sequence in the peptides file and is done with different software so inconsistencies are likely. The peaks are annotated based on the given sequence, with 20 ppm tolerance.

Copy Data

##### Spectrum 8278 (TSV)

###### Preview

```
Loading example...
```

*Click on the button to copy the data to your clipboard.*

Mz MinMz MaxIntensity Max

WidthHeightPeptide font sizePeptide stroke widthSpectrum font sizeSpectrum stroke widthCompact peptide

Ion legend

wxyz

abcd

OtherUnassignedIon^Charge^_Position_Show for top:%

JVKDYFPEPVT

03.55e+47.10e+41.06e+51.42e+5

Zoom Out

y^+1^_1_y^+1^_2_y^+1^_3_y^+1^_3_c^+1^_3_c^+2^_7_c^+1^_4_c^+1^_4_c^+2^_8_y^+1^_5_c^+2^_9_c^+2^_10_c^+1^_5_c^+1^_5_y^+1^_6_c^+1^_6_y^+1^_7_c^+1^_7_c^+1^_7_y^+1^_8_c^+1^_8_z^+1^_9_y^+1^_9_c^+1^_9_z^+1^_10_c^+1^_10_c^+1^_10_

0881176326443525

Fragment Matches Table

Show background peaks

| Position | Ion type | Intensity | mz Theoretical | mz Error (Th) | mz Error (ppm) | Charge | Series Number |
| --- | --- | --- | --- | --- | --- | --- | --- |
| 11 | y | 1767 | 120.1 | 4.74E-05 | 0.3948 | +1 | 1 |
| - | - | 392.2 | 127.9 | - | - | 0 | - |
| - | - | 1004 | 129.1 | - | - | 0 | - |
| - | - | 1761 | 133.1 | - | - | 0 | - |
| - | - | 437.4 | 139.5 | - | - | 0 | - |
| - | - | 984.7 | 149 | - | - | 0 | - |
| - | - | 466.3 | 154 | - | - | 0 | - |
| - | - | 430.1 | 157.6 | - | - | 0 | - |
| - | - | 3232 | 169.1 | - | - | 0 | - |
| - | - | 1719 | 173.4 | - | - | 0 | - |
| - | - | 716 | 177.1 | - | - | 0 | - |
| - | - | 3781 | 185.2 | - | - | 0 | - |
| - | - | 569.3 | 186.2 | - | - | 0 | - |
| - | - | 490.8 | 196.9 | - | - | 0 | - |
| - | - | 5910 | 197.1 | - | - | 0 | - |
| - | - | 562.6 | 198.1 | - | - | 0 | - |
| - | - | 470.7 | 199.1 | - | - | 0 | - |
| - | - | 2888 | 213.2 | - | - | 0 | - |
| 10 | y | 520 | 219.1 | 0.000588 | 2.683 | +1 | 2 |
| - | - | 1097 | 227.1 | - | - | 0 | - |
| - | - | 648.3 | 262.6 | - | - | 0 | - |
| 9 | y | 1470 | 298.2 | 0.0003199 | 1.073 | +1 | 3 |
| - | - | 559 | 307.2 | - | - | 0 | - |
| 9 | y | 1.713E+04 | 316.2 | 0.0001923 | 0.6081 | +1 | 3 |
| - | - | 2348 | 317.2 | - | - | 0 | - |
| - | - | 3168 | 324.2 | - | - | 0 | - |
| - | - | 817.6 | 325.2 | - | - | 0 | - |
| - | - | 1403 | 355.1 | - | - | 0 | - |
| - | - | 931 | 356.1 | - | - | 0 | - |
| - | - | 3429 | 357.3 | - | - | 0 | - |
| 3 | c | 1441 | 358.3 | 0.000993 | 2.772 | +1 | 3 |
| - | - | 706.7 | 414.6 | - | - | 0 | - |
| - | - | 1588 | 423.2 | - | - | 0 | - |
| - | - | 849.1 | 426.2 | - | - | 0 | - |
| 7 | c | 723.7 | 432.2 | 0.00275 | 6.362 | +2 | 7 |
| - | - | 597.4 | 443.4 | - | - | 0 | - |
| 4 | c | 934.9 | 456.3 | 0.0004136 | 0.9064 | +1 | 4 |
| - | - | 925.7 | 472.3 | - | - | 0 | - |
| 4 | c | 2347 | 473.3 | 0.0009641 | 2.037 | +1 | 4 |
| 8 | c | 1816 | 496.8 | 0.0003887 | 0.7825 | +2 | 8 |
| - | - | 1092 | 497.3 | - | - | 0 | - |
| - | - | 657.1 | 497.8 | - | - | 0 | - |
| - | - | 656.2 | 521.9 | - | - | 0 | - |
| - | - | 1678 | 522.8 | - | - | 0 | - |
| - | - | 890.2 | 523.3 | - | - | 0 | - |
| 7 | y | 7622 | 542.3 | 0.0006605 | 1.218 | +1 | 5 |
| - | - | 2491 | 543.3 | - | - | 0 | - |
| 9 | c | 4998 | 545.3 | 0.0005121 | 0.9391 | +2 | 9 |
| - | - | 3663 | 545.8 | - | - | 0 | - |
| - | - | 1561 | 546.3 | - | - | 0 | - |
| - | - | 3104 | 554.3 | - | - | 0 | - |
| - | - | 1048 | 555.3 | - | - | 0 | - |
| - | - | 2217 | 580.8 | - | - | 0 | - |
| - | - | 1971 | 581.3 | - | - | 0 | - |
| - | - | 601.2 | 581.8 | - | - | 0 | - |
| 10 | c | 1211 | 594.8 | 0.002377 | 3.996 | +2 | 10 |
| - | - | 1469 | 595.3 | - | - | 0 | - |
| - | - | 554.5 | 606.4 | - | - | 0 | - |
| 5 | c | 3032 | 619.3 | 0.0007752 | 1.252 | +1 | 5 |
| - | - | 604.1 | 619.5 | - | - | 0 | - |
| - | - | 1291 | 620.3 | - | - | 0 | - |
| - | - | 7033 | 635.4 | - | - | 0 | - |
| 5 | c | 9161 | 636.4 | 0.002546 | 4.001 | +1 | 5 |
| - | - | 3639 | 637.4 | - | - | 0 | - |
| - | - | 1189 | 638.4 | - | - | 0 | - |
| - | - | 767.3 | 645.8 | - | - | 0 | - |
| - | - | 1284 | 653.4 | - | - | 0 | - |
| - | - | 1025 | 654.3 | - | - | 0 | - |
| - | - | 755.5 | 654.8 | - | - | 0 | - |
| - | - | 990.9 | 655.4 | - | - | 0 | - |
| - | - | 633.7 | 682.4 | - | - | 0 | - |
| 6 | y | 5084 | 689.4 | 0.0001265 | 0.1834 | +1 | 6 |
| - | - | 1627 | 690.4 | - | - | 0 | - |
| - | - | 839.9 | 738.4 | - | - | 0 | - |
| - | - | 3664 | 739.4 | - | - | 0 | - |
| - | - | 4210 | 740.4 | - | - | 0 | - |
| - | - | 1327 | 741.4 | - | - | 0 | - |
| 6 | c | 1.083E+04 | 766.4 | 0.0004155 | 0.5421 | +1 | 6 |
| - | - | 4412 | 767.4 | - | - | 0 | - |
| - | - | 1004 | 768.4 | - | - | 0 | - |
| - | - | 2528 | 780.4 | - | - | 0 | - |
| - | - | 1735 | 781.4 | - | - | 0 | - |
| - | - | 603.8 | 784.4 | - | - | 0 | - |
| - | - | 922.2 | 825.5 | - | - | 0 | - |
| - | - | 780.3 | 851.4 | - | - | 0 | - |
| 5 | y | 4038 | 852.4 | 0.0007719 | 0.9055 | +1 | 7 |
| - | - | 1714 | 853.4 | - | - | 0 | - |
| 7 | c | 1236 | 863.5 | 0.00352 | 4.077 | +1 | 7 |
| - | - | 474 | 879.4 | - | - | 0 | - |
| - | - | 1.445E+04 | 879.5 | - | - | 0 | - |
| 7 | c | 2.317E+04 | 880.5 | 0.001566 | 1.778 | +1 | 7 |
| - | - | 1.139E+04 | 881.5 | - | - | 0 | - |
| - | - | 3202 | 882.5 | - | - | 0 | - |
| - | - | 665.3 | 932.2 | - | - | 0 | - |
| - | - | 706.9 | 953.4 | - | - | 0 | - |
| - | - | 745 | 965.5 | - | - | 0 | - |
| - | - | 1234 | 966.4 | - | - | 0 | - |
| 4 | y | 2885 | 967.4 | 0.002019 | 2.087 | +1 | 8 |
| - | - | 1226 | 968.4 | - | - | 0 | - |
| - | - | 818 | 976.5 | - | - | 0 | - |
| - | - | 647.5 | 978.5 | - | - | 0 | - |
| 8 | c | 1.408E+04 | 992.5 | 0.0002149 | 0.2165 | +1 | 8 |
| - | - | 8605 | 993.5 | - | - | 0 | - |
| - | - | 2910 | 994.5 | - | - | 0 | - |
| - | - | 1439 | 1007 | - | - | 0 | - |
| - | - | 1540 | 1036 | - | - | 0 | - |
| - | - | 647.4 | 1037 | - | - | 0 | - |
| - | - | 678.3 | 1062 | - | - | 0 | - |
| 3 | z | 4932 | 1080 | 0.001354 | 1.254 | +1 | 9 |
| - | - | 3676 | 1081 | - | - | 0 | - |
| - | - | 1410 | 1082 | - | - | 0 | - |
| - | - | 649.1 | 1090 | - | - | 0 | - |
| 3 | y | 3822 | 1096 | 0.0005466 | 0.4989 | +1 | 9 |
| - | - | 2350 | 1097 | - | - | 0 | - |
| - | - | 1049 | 1098 | - | - | 0 | - |
| 9 | c | 4189 | 1107 | 0.001403 | 1.268 | +1 | 9 |
| - | - | 2032 | 1108 | - | - | 0 | - |
| - | - | 1199 | 1147 | - | - | 0 | - |
| - | - | 832.3 | 1148 | - | - | 0 | - |
| - | - | 961.9 | 1150 | - | - | 0 | - |
| - | - | 7744 | 1162 | - | - | 0 | - |
| - | - | 6184 | 1163 | - | - | 0 | - |
| - | - | 3187 | 1164 | - | - | 0 | - |
| - | - | 1418 | 1165 | - | - | 0 | - |
| 2 | z | 3057 | 1179 | 6.554E-05 | 0.0556 | +1 | 10 |
| - | - | 1970 | 1180 | - | - | 0 | - |
| 10 | c | 824.1 | 1189 | 0.001886 | 1.587 | +1 | 10 |
| - | - | 874.5 | 1190 | - | - | 0 | - |
| - | - | 3641 | 1191 | - | - | 0 | - |
| - | - | 2000 | 1192 | - | - | 0 | - |
| 10 | c | 3.325E+04 | 1206 | 0.0004808 | 0.3988 | +1 | 10 |
| - | - | 2.421E+04 | 1207 | - | - | 0 | - |
| - | - | 1.352E+04 | 1208 | - | - | 0 | - |
| - | - | 4243 | 1209 | - | - | 0 | - |
| - | - | 1204 | 1210 | - | - | 0 | - |
| - | - | 2915 | 1237 | - | - | 0 | - |
| - | - | 2626 | 1238 | - | - | 0 | - |
| - | - | 1276 | 1245 | - | - | 0 | - |
| - | - | 2013 | 1246 | - | - | 0 | - |
| - | - | 1623 | 1247 | - | - | 0 | - |
| - | - | 891.6 | 1248 | - | - | 0 | - |
| - | - | 9868 | 1249 | - | - | 0 | - |
| - | - | 7789 | 1250 | - | - | 0 | - |
| - | - | 3406 | 1251 | - | - | 0 | - |
| - | - | 2229 | 1253 | - | - | 0 | - |
| - | - | 1287 | 1254 | - | - | 0 | - |
| - | - | 1983 | 1263 | - | - | 0 | - |
| - | - | 1587 | 1264 | - | - | 0 | - |
| - | - | 4059 | 1265 | - | - | 0 | - |
| - | - | 2957 | 1266 | - | - | 0 | - |
| - | - | 6094 | 1281 | - | - | 0 | - |
| - | - | 5010 | 1282 | - | - | 0 | - |
| - | - | 1464 | 1283 | - | - | 0 | - |
| - | - | 1171 | 1289 | - | - | 0 | - |
| - | - | 2194 | 1291 | - | - | 0 | - |
| - | - | 3.57E+04 | 1292 | - | - | 0 | - |
| - | - | 2.984E+04 | 1293 | - | - | 0 | - |
| - | - | 1.086E+04 | 1294 | - | - | 0 | - |
| - | - | 1694 | 1295 | - | - | 0 | - |
| - | - | 1029 | 1305 | - | - | 0 | - |
| - | - | 2604 | 1306 | - | - | 0 | - |
| - | - | 3101 | 1307 | - | - | 0 | - |
| - | - | 2.758E+04 | 1308 | - | - | 0 | - |
| - | - | 1.405E+05 | 1309 | - | - | 0 | - |
| - | - | 1.015E+05 | 1310 | - | - | 0 | - |
| - | - | 4.031E+04 | 1311 | - | - | 0 | - |
| - | - | 4290 | 1312 | - | - | 0 | - |
| - | - | 636.6 | 1341 | - | - | 0 | - |
| - | - | 859.4 | 3490 | - | - | 0 | - |

m/z Charge Intensity FragmentType MassShift Position
120.06556701660156 0 1766.6411 y 10
127.91201782226562 0 392.1686
129.1028594970703 0 1003.5178
133.0861358642578 0 1760.748
139.52322387695312 0 437.3789
148.95376586914062 0 984.6541
153.96658325195312 0 466.3217
157.6160888671875 0 430.096
169.1336669921875 0 3232.004
173.4388427734375 0 1718.7711
177.11248779296875 0 715.9752
185.1649932861328 0 3781.0552
186.1686248779297 0 569.2596
196.8793487548828 0 490.84802
197.1285858154297 0 5910.0044
198.13230895996094 0 562.58203
199.12632751464844 0 470.68344
213.15989685058594 0 2888.2068
219.134521484375 0 519.96844 y 9
227.10231018066406 0 1097.2255
262.61407470703125 0 648.28064
298.17645263671875 0 1469.6213 y Water loss 8
307.17730712890625 0 559.0305
316.1868896484375 0 17128.229 y 8
317.1902770996094 0 2348.0125
324.1554870605469 0 3167.9067
325.158447265625 0 817.60547
355.0697021484375 0 1403.0813
356.06976318359375 0 930.957
357.27349853515625 0 3428.5981
358.2802734375 0 1440.9681 c 2
414.6413269042969 0 706.68207
423.224609375 0 1587.5669
426.1676025390625 0 849.1198
432.2394714355469 0 723.7464 c Ammonia loss 6
443.3859558105469 0 597.4137
456.2820739746094 0 934.9492 c Ammonia loss 3
472.3011169433594 0 925.7047
473.3091735839844 0 2346.5405 c 3
496.75762939453125 0 1815.6809 c Ammonia loss 7
497.2592468261719 0 1092.1039
497.76202392578125 0 657.09094
521.8908081054688 0 656.22797
522.7853393554688 0 1677.6462
523.28369140625 0 890.2062
542.28271484375 0 7621.9673 y 6
543.2855224609375 0 2490.5845
545.284912109375 0 4997.8276 c Ammonia loss 8
545.78662109375 0 3663.3376
546.2861938476562 0 1560.9661
554.2611694335938 0 3104.298
555.2633666992188 0 1048.4382
580.8217163085938 0 2217.1165
581.3216552734375 0 1971.173
581.826416015625 0 601.16907
594.8209838867188 0 1210.6859 c Ammonia loss 9
595.3193359375 0 1469.0992
606.3848266601562 0 554.4949
619.3457641601562 0 3031.576 c Ammonia loss 4
619.5298461914062 0 604.08997
620.3488159179688 0 1291.2393
635.3642578125 0 7033.29
636.3740844726562 0 9160.788 c 4
637.3768920898438 0 3639.2705
638.3779296875 0 1188.6176
645.84423828125 0 767.3052
653.41845703125 0 1284.0955
654.3438110351562 0 1025.3007
654.8453369140625 0 755.5229
655.354248046875 0 990.88086
682.3785400390625 0 633.7243
689.350341796875 0 5084.1494 y 5
690.3543090820312 0 1627.3281
738.4163208007812 0 839.8742
739.42431640625 0 3664.1772
740.4327392578125 0 4210.125
741.43798828125 0 1327.233
766.413818359375 0 10827.603 c Ammonia loss 5
767.41552734375 0 4411.615
768.41552734375 0 1004.16644
780.3566284179688 0 2528.2185
781.357421875 0 1734.8105
784.4302368164062 0 603.75226
825.5264892578125 0 922.1552
851.4061889648438 0 780.31805
852.4130249023438 0 4037.7769 y 4
853.4169921875 0 1713.5336
863.462646484375 0 1236.1998 c Ammonia loss 6
879.401611328125 0 473.97055
879.4844970703125 0 14453.701
880.4911499023438 0 23165.447 c 6
881.4947509765625 0 11392.794
882.4989624023438 0 3201.5923
932.1952514648438 0 665.31726
953.4339599609375 0 706.88525
965.5167236328125 0 744.9699
966.4269409179688 0 1234.1826
967.438720703125 0 2884.7249 y 3
968.4414672851562 0 1225.78
976.4758911132812 0 818.0132
978.4923706054688 0 647.4924
992.508544921875 0 14076.22 c Ammonia loss 7
993.5116577148438 0 8605.236
994.5136108398438 0 2909.6504
1007.4893188476562 0 1438.5005
1035.528564453125 0 1539.5542
1036.521240234375 0 647.40533
1061.54150390625 0 678.3059
1079.515625 0 4932.049 z 2
1080.5181884765625 0 3676.1094
1081.5299072265625 0 1409.5112
1089.6209716796875 0 649.0969
1095.53515625 0 3822.3445 y 2
1096.5380859375 0 2349.8406
1097.5423583984375 0 1049.3434
1106.586669921875 0 4189.004 c 8
1107.5916748046875 0 2031.5691
1146.625732421875 0 1198.5803
1147.6395263671875 0 832.3146
1149.577880859375 0 961.9233
1161.64208984375 0 7744.4062
1162.646728515625 0 6183.58
1163.6492919921875 0 3187.3936
1164.6663818359375 0 1417.6948
1178.5853271484375 0 3057.0208 z 1
1179.5855712890625 0 1969.772
1188.6280517578125 0 824.0882 c Ammonia loss 9
1189.6256103515625 0 874.48553
1190.64208984375 0 3640.951
1191.6453857421875 0 1999.8962
1205.656005859375 0 33248.855 c 9
1206.6590576171875 0 24205.244
1207.6640625 0 13517.986
1208.673583984375 0 4243.281
1209.6707763671875 0 1204.2383
1236.6759033203125 0 2914.5388
1237.6700439453125 0 2626.391
1244.681396484375 0 1276.3861
1245.677490234375 0 2013.333
1246.683837890625 0 1623.3661
1247.686279296875 0 891.62585
1248.652099609375 0 9868.332
1249.6551513671875 0 7789.2754
1250.66015625 0 3405.5388
1252.631103515625 0 2229.0176
1253.630859375 0 1287.3373
1262.6898193359375 0 1982.5493
1263.6807861328125 0 1586.7572
1264.7034912109375 0 4058.596
1265.70166015625 0 2956.554
1280.6986083984375 0 6094.088
1281.70263671875 0 5010.2764
1282.701171875 0 1463.6803
1288.70849609375 0 1171.3717
1290.6807861328125 0 2193.7483
1291.66796875 0 35696.895
1292.6707763671875 0 29836.383
1293.671875 0 10859.815
1294.6700439453125 0 1694.0814
1304.71044921875 0 1029.1696
1305.7095947265625 0 2603.73
1306.6923828125 0 3101.0122
1307.6856689453125 0 27581.658
1308.69287109375 0 140502.52
1309.6966552734375 0 101521.85
1310.6995849609375 0 40306.465
1311.6993408203125 0 4290.445
1340.6708984375 0 636.5621
3490.115478515625 0 859.3615

Spectrum Details

|  |  |
| --- | --- |
| Matched peaks? Matched peaks The total absolute number of peaks matched. Additionally in brackets the total fraction of peaks matched and the total number of peaks is shown. | 27 (15.98% of 169) |
| FDR? FDR The false discovery rate estimated for this peptide. It is calculated by matching all theoretical fragments with a non-integer shift with the raw peaks for this spectrum. This is done with 40 different shifts. The resulting percentage is the average number of annotated peaks over the number of annotated peaks with the correct spectrum. | 1.59% |
| Satellite FDR? Satellite FDR See the FDR for details on its calculation. This satellite ion specific FDR only contains the satellite ions (d/w) for I/L/J positions. | - |
| PSM Score? PSM Score The PSM Score as given by Hecklib to this annotated spectrum. It is shown with three significant figures. | 283 |

#### Spectrum 8165? Spectrum 8165 The raw spectrum of this peptide as annotated by Hecklib. The fragments are coloured according to ion type (see legend). Any peaks with a star '\*' as text can be hovered over to see the full details, first the ion type second the mass shift type. By hovering over the amino acids in the peptide or ions in the legend the corresponding peaks are highlighted. By toggling the 'Unassigned' label you can turn the background (unassigned) peaks on or off in the plot. By updating the slider in the Ion legend you can update the spectrum to only show the top X% of the peaks with labels. The top X% means any peak that is within X% of the highest intensity. By dragging in the spectrum you can zoom in to a specific part of the spectrum and use 'Zoom Out' to get back to the original zoom level. The annotation of the spectrum is based on the given sequence in the peptides file and is done with different software so inconsistencies are likely. The peaks are annotated based on the given sequence, with 20 ppm tolerance.

Copy Data

##### Spectrum 8165 (TSV)

###### Preview

```
Loading example...
```

*Click on the button to copy the data to your clipboard.*

Mz MinMz MaxIntensity Max

WidthHeightPeptide font sizePeptide stroke widthSpectrum font sizeSpectrum stroke widthCompact peptide

Ion legend

wxyz

abcd

OtherUnassignedIon^Charge^_Position_Show for top:%

JVKDYFPEPVT

04.46e+48.91e+41.34e+51.78e+5

Zoom Out

y^+1^_1_y^+1^_2_y^+1^_3_y^+1^_3_c^+1^_3_c^+1^_4_c^+1^_4_c^+2^_8_y^+2^_9_y^+1^_5_c^+2^_9_c^+2^_10_c^+1^_5_c^+1^_5_y^+1^_6_c^+1^_6_z^+1^_7_y^+1^_7_c^+1^_7_c^+1^_7_y^+1^_8_c^+1^_8_z^+1^_9_y^+1^_9_c^+1^_9_z^+1^_10_c^+1^_10_c^+1^_10_

0831166124923322

Fragment Matches Table

Show background peaks

| Position | Ion type | Intensity | mz Theoretical | mz Error (Th) | mz Error (ppm) | Charge | Series Number |
| --- | --- | --- | --- | --- | --- | --- | --- |
| 11 | y | 1657 | 120.1 | 4.74E-05 | 0.3948 | +1 | 1 |
| - | - | 1311 | 120.1 | - | - | 0 | - |
| - | - | 1727 | 129.1 | - | - | 0 | - |
| - | - | 451.2 | 132.2 | - | - | 0 | - |
| - | - | 401.8 | 139.5 | - | - | 0 | - |
| - | - | 503.9 | 139.8 | - | - | 0 | - |
| - | - | 400.7 | 148.8 | - | - | 0 | - |
| - | - | 1036 | 149 | - | - | 0 | - |
| - | - | 3704 | 169.1 | - | - | 0 | - |
| - | - | 2551 | 173.4 | - | - | 0 | - |
| - | - | 444.7 | 179.5 | - | - | 0 | - |
| - | - | 4807 | 185.2 | - | - | 0 | - |
| - | - | 472 | 193 | - | - | 0 | - |
| - | - | 9106 | 197.1 | - | - | 0 | - |
| - | - | 418 | 197.4 | - | - | 0 | - |
| - | - | 751.9 | 198.1 | - | - | 0 | - |
| - | - | 4159 | 213.2 | - | - | 0 | - |
| 10 | y | 535.9 | 219.1 | 0.0001597 | 0.7289 | +1 | 2 |
| - | - | 1528 | 227.1 | - | - | 0 | - |
| - | - | 709.6 | 239.1 | - | - | 0 | - |
| - | - | 604 | 239.2 | - | - | 0 | - |
| 9 | y | 1904 | 298.2 | 0.0002599 | 0.8716 | +1 | 3 |
| - | - | 1375 | 303.1 | - | - | 0 | - |
| 9 | y | 2.302E+04 | 316.2 | 0.0001923 | 0.6081 | +1 | 3 |
| - | - | 585.8 | 316.8 | - | - | 0 | - |
| - | - | 4562 | 317.2 | - | - | 0 | - |
| - | - | 3902 | 324.2 | - | - | 0 | - |
| - | - | 1299 | 355.1 | - | - | 0 | - |
| - | - | 789.7 | 356.1 | - | - | 0 | - |
| - | - | 3712 | 357.3 | - | - | 0 | - |
| 3 | c | 1214 | 358.3 | 0.002885 | 8.053 | +1 | 3 |
| - | - | 1858 | 423.2 | - | - | 0 | - |
| - | - | 633 | 426.2 | - | - | 0 | - |
| 4 | c | 1207 | 456.3 | 0.0008376 | 1.836 | +1 | 4 |
| - | - | 1164 | 472.3 | - | - | 0 | - |
| 4 | c | 2353 | 473.3 | 0.0004148 | 0.8763 | +1 | 4 |
| - | - | 637.3 | 488.7 | - | - | 0 | - |
| 8 | c | 2034 | 496.8 | 0.001182 | 2.38 | +2 | 8 |
| - | - | 1995 | 497.3 | - | - | 0 | - |
| - | - | 645.3 | 497.8 | - | - | 0 | - |
| - | - | 1680 | 522.8 | - | - | 0 | - |
| 3 | y | 670.3 | 539.3 | 0.00986 | 18.28 | +2 | 9 |
| 7 | y | 1.116E+04 | 542.3 | 0.0001722 | 0.3176 | +1 | 5 |
| - | - | 3029 | 543.3 | - | - | 0 | - |
| 9 | c | 7571 | 545.3 | 2.377E-05 | 0.0436 | +2 | 9 |
| - | - | 5954 | 545.8 | - | - | 0 | - |
| - | - | 1661 | 546.3 | - | - | 0 | - |
| - | - | 3564 | 554.3 | - | - | 0 | - |
| - | - | 1077 | 555.3 | - | - | 0 | - |
| - | - | 2710 | 580.8 | - | - | 0 | - |
| - | - | 1601 | 581.3 | - | - | 0 | - |
| - | - | 592.3 | 591.4 | - | - | 0 | - |
| 10 | c | 1904 | 594.8 | 0.001705 | 2.867 | +2 | 10 |
| - | - | 992.8 | 595.3 | - | - | 0 | - |
| 5 | c | 3315 | 619.3 | 7.927E-05 | 0.128 | +1 | 5 |
| - | - | 1606 | 620.3 | - | - | 0 | - |
| - | - | 1.068E+04 | 635.4 | - | - | 0 | - |
| 5 | c | 1.042E+04 | 636.4 | 0.001116 | 1.753 | +1 | 5 |
| - | - | 3831 | 637.4 | - | - | 0 | - |
| - | - | 623.7 | 638.4 | - | - | 0 | - |
| - | - | 978.7 | 645.8 | - | - | 0 | - |
| - | - | 882.9 | 652.9 | - | - | 0 | - |
| - | - | 1412 | 654.3 | - | - | 0 | - |
| - | - | 1447 | 654.8 | - | - | 0 | - |
| - | - | 840.3 | 655.4 | - | - | 0 | - |
| - | - | 1008 | 682.4 | - | - | 0 | - |
| 6 | y | 6040 | 689.4 | 0.0001875 | 0.272 | +1 | 6 |
| - | - | 2439 | 690.4 | - | - | 0 | - |
| - | - | 613.1 | 696.4 | - | - | 0 | - |
| - | - | 969.8 | 731.4 | - | - | 0 | - |
| - | - | 972.3 | 738.4 | - | - | 0 | - |
| - | - | 4181 | 739.4 | - | - | 0 | - |
| - | - | 5909 | 740.4 | - | - | 0 | - |
| - | - | 2424 | 741.4 | - | - | 0 | - |
| 6 | c | 1.363E+04 | 766.4 | 0.0001338 | 0.1746 | +1 | 6 |
| - | - | 7711 | 767.4 | - | - | 0 | - |
| - | - | 1511 | 768.4 | - | - | 0 | - |
| - | - | 4195 | 780.4 | - | - | 0 | - |
| - | - | 1560 | 781.4 | - | - | 0 | - |
| 5 | z | 986.5 | 836.4 | 0.002189 | 2.618 | +1 | 7 |
| - | - | 1382 | 851.4 | - | - | 0 | - |
| 5 | y | 6700 | 852.4 | 0.0001005 | 0.1179 | +1 | 7 |
| - | - | 2807 | 853.4 | - | - | 0 | - |
| 7 | c | 1348 | 863.5 | 0.003093 | 3.582 | +1 | 7 |
| - | - | 718.4 | 864.5 | - | - | 0 | - |
| - | - | 1653 | 877.4 | - | - | 0 | - |
| - | - | 660.8 | 878.4 | - | - | 0 | - |
| - | - | 1.771E+04 | 879.5 | - | - | 0 | - |
| 7 | c | 3.208E+04 | 880.5 | 0.0012 | 1.363 | +1 | 7 |
| - | - | 1.475E+04 | 881.5 | - | - | 0 | - |
| - | - | 2972 | 882.5 | - | - | 0 | - |
| - | - | 604.8 | 949.5 | - | - | 0 | - |
| - | - | 763.3 | 964.5 | - | - | 0 | - |
| - | - | 1722 | 965.5 | - | - | 0 | - |
| - | - | 1466 | 966.4 | - | - | 0 | - |
| 4 | y | 4507 | 967.4 | 0.0001881 | 0.1944 | +1 | 8 |
| - | - | 1728 | 968.4 | - | - | 0 | - |
| - | - | 817.3 | 974.5 | - | - | 0 | - |
| - | - | 1625 | 976.5 | - | - | 0 | - |
| - | - | 1137 | 977.5 | - | - | 0 | - |
| - | - | 708.5 | 978.5 | - | - | 0 | - |
| 8 | c | 1.902E+04 | 992.5 | 3.178E-05 | 0.03202 | +1 | 8 |
| - | - | 1.333E+04 | 993.5 | - | - | 0 | - |
| - | - | 3928 | 994.5 | - | - | 0 | - |
| - | - | 952 | 1008 | - | - | 0 | - |
| - | - | 1626 | 1036 | - | - | 0 | - |
| - | - | 1372 | 1037 | - | - | 0 | - |
| 3 | z | 6929 | 1080 | 0.0008655 | 0.8017 | +1 | 9 |
| - | - | 4896 | 1081 | - | - | 0 | - |
| - | - | 1567 | 1082 | - | - | 0 | - |
| 3 | y | 6316 | 1096 | 0.0005466 | 0.4989 | +1 | 9 |
| - | - | 2646 | 1097 | - | - | 0 | - |
| - | - | 2621 | 1098 | - | - | 0 | - |
| 9 | c | 5500 | 1107 | 0.0003042 | 0.2749 | +1 | 9 |
| - | - | 3506 | 1108 | - | - | 0 | - |
| - | - | 812.9 | 1109 | - | - | 0 | - |
| - | - | 2566 | 1114 | - | - | 0 | - |
| - | - | 1481 | 1115 | - | - | 0 | - |
| - | - | 744.4 | 1125 | - | - | 0 | - |
| - | - | 729 | 1146 | - | - | 0 | - |
| - | - | 1036 | 1147 | - | - | 0 | - |
| - | - | 1491 | 1148 | - | - | 0 | - |
| - | - | 831.1 | 1149 | - | - | 0 | - |
| - | - | 1123 | 1157 | - | - | 0 | - |
| - | - | 8899 | 1162 | - | - | 0 | - |
| - | - | 8074 | 1163 | - | - | 0 | - |
| - | - | 4170 | 1164 | - | - | 0 | - |
| - | - | 1347 | 1165 | - | - | 0 | - |
| 2 | z | 3764 | 1179 | 0.0003097 | 0.2628 | +1 | 10 |
| - | - | 2728 | 1180 | - | - | 0 | - |
| 10 | c | 1708 | 1189 | 0.003241 | 2.727 | +1 | 10 |
| - | - | 1056 | 1190 | - | - | 0 | - |
| - | - | 3655 | 1191 | - | - | 0 | - |
| - | - | 2272 | 1192 | - | - | 0 | - |
| - | - | 951.5 | 1193 | - | - | 0 | - |
| 10 | c | 4.473E+04 | 1206 | 7.466E-06 | 0.006192 | +1 | 10 |
| - | - | 3.299E+04 | 1207 | - | - | 0 | - |
| - | - | 1.638E+04 | 1208 | - | - | 0 | - |
| - | - | 6670 | 1209 | - | - | 0 | - |
| - | - | 1851 | 1210 | - | - | 0 | - |
| - | - | 1269 | 1220 | - | - | 0 | - |
| - | - | 4027 | 1237 | - | - | 0 | - |
| - | - | 2194 | 1238 | - | - | 0 | - |
| - | - | 730.5 | 1239 | - | - | 0 | - |
| - | - | 2220 | 1245 | - | - | 0 | - |
| - | - | 1963 | 1246 | - | - | 0 | - |
| - | - | 1780 | 1247 | - | - | 0 | - |
| - | - | 998.1 | 1248 | - | - | 0 | - |
| - | - | 1.277E+04 | 1249 | - | - | 0 | - |
| - | - | 9664 | 1250 | - | - | 0 | - |
| - | - | 3916 | 1251 | - | - | 0 | - |
| - | - | 748.7 | 1252 | - | - | 0 | - |
| - | - | 2288 | 1253 | - | - | 0 | - |
| - | - | 837.2 | 1254 | - | - | 0 | - |
| - | - | 917.4 | 1255 | - | - | 0 | - |
| - | - | 2696 | 1263 | - | - | 0 | - |
| - | - | 2465 | 1264 | - | - | 0 | - |
| - | - | 4234 | 1265 | - | - | 0 | - |
| - | - | 3391 | 1266 | - | - | 0 | - |
| - | - | 1431 | 1267 | - | - | 0 | - |
| - | - | 8196 | 1281 | - | - | 0 | - |
| - | - | 6707 | 1282 | - | - | 0 | - |
| - | - | 2344 | 1283 | - | - | 0 | - |
| - | - | 1189 | 1289 | - | - | 0 | - |
| - | - | 1134 | 1290 | - | - | 0 | - |
| - | - | 3899 | 1291 | - | - | 0 | - |
| - | - | 4.828E+04 | 1292 | - | - | 0 | - |
| - | - | 3.698E+04 | 1293 | - | - | 0 | - |
| - | - | 1.539E+04 | 1294 | - | - | 0 | - |
| - | - | 2891 | 1295 | - | - | 0 | - |
| - | - | 3730 | 1306 | - | - | 0 | - |
| - | - | 2354 | 1307 | - | - | 0 | - |
| - | - | 3.989E+04 | 1308 | - | - | 0 | - |
| - | - | 1.765E+05 | 1309 | - | - | 0 | - |
| - | - | 1.319E+05 | 1310 | - | - | 0 | - |
| - | - | 5.085E+04 | 1311 | - | - | 0 | - |
| - | - | 8750 | 1312 | - | - | 0 | - |
| - | - | 722 | 2847 | - | - | 0 | - |
| - | - | 777.7 | 3071 | - | - | 0 | - |
| - | - | 711.1 | 3289 | - | - | 0 | - |

m/z Charge Intensity FragmentType MassShift Position
120.06556701660156 0 1657.3303 y 10
120.0808334350586 0 1311.3484
129.10240173339844 0 1727.2961
132.21688842773438 0 451.2101
139.4886932373047 0 401.772
139.84432983398438 0 503.87656
148.78175354003906 0 400.73492
148.95474243164062 0 1035.8235
169.1336669921875 0 3703.8972
173.43954467773438 0 2550.8518
179.5242156982422 0 444.6991
185.16488647460938 0 4807.4897
192.99728393554688 0 471.99854
197.1285400390625 0 9106.452
197.4102325439453 0 418.04114
198.13218688964844 0 751.93365
213.159912109375 0 4159.1187
219.13377380371094 0 535.86383 y 9
227.1028289794922 0 1527.7416
239.09458923339844 0 709.5898
239.19326782226562 0 603.9793
298.1758728027344 0 1904.1353 y Water loss 8
303.0869445800781 0 1374.6614
316.1868896484375 0 23024.246 y 8
316.83026123046875 0 585.8066
317.1902770996094 0 4561.8516
324.156005859375 0 3901.5505
355.06982421875 0 1298.9177
356.07025146484375 0 789.6687
357.2731018066406 0 3711.5837
358.27838134765625 0 1214.3625 c 2
423.225341796875 0 1858.1215
426.1669921875 0 632.9892
456.28082275390625 0 1206.5398 c Ammonia loss 3
472.2996826171875 0 1164.1003
473.3086242675781 0 2353.2998 c 3
488.7425537109375 0 637.28314
496.7568359375 0 2034.196 c Ammonia loss 7
497.2580871582031 0 1995.4136
497.7608337402344 0 645.3195
522.7847290039062 0 1680.1378
539.25634765625 0 670.2576 y Water loss 2
542.2822265625 0 11164.532 y 6
543.2847900390625 0 3028.5415
545.284423828125 0 7570.627 c Ammonia loss 8
545.78662109375 0 5953.602
546.2885131835938 0 1660.625
554.2611083984375 0 3563.9058
555.2653198242188 0 1077.158
580.8208618164062 0 2709.5132
581.324462890625 0 1600.6588
591.3564453125 0 592.28546
594.8203125 0 1903.9279 c Ammonia loss 9
595.322998046875 0 992.7592
619.3449096679688 0 3315.0505 c Ammonia loss 4
620.3499145507812 0 1606.3876
635.364501953125 0 10682.382
636.3704223632812 0 10422.412 c 4
637.3746948242188 0 3831.4878
638.3800048828125 0 623.69824
645.8439331054688 0 978.7471
652.8619384765625 0 882.896
654.3482666015625 0 1411.7946
654.8488159179688 0 1446.8577
655.35693359375 0 840.3408
682.3776245117188 0 1007.938
689.3502807617188 0 6040.4253 y 5
690.3526000976562 0 2438.988
696.4307250976562 0 613.0927
731.4056396484375 0 969.76685
738.4168701171875 0 972.26196
739.4253540039062 0 4180.751
740.4323120117188 0 5909.3164
741.4374389648438 0 2424.1812
766.4132690429688 0 13629.291 c Ammonia loss 5
767.4171142578125 0 7710.788
768.421142578125 0 1510.995
780.356201171875 0 4195.244
781.3560791015625 0 1560.2164
836.3928833007812 0 986.5416 z 4
851.4061279296875 0 1381.543
852.4136962890625 0 6699.642 y 4
853.4174194335938 0 2807.3848
863.4630737304688 0 1347.6682 c Ammonia loss 6
864.4718627929688 0 718.43933
877.41015625 0 1653.4152
878.4047241210938 0 660.82654
879.4850463867188 0 17714.77
880.4915161132812 0 32083.855 c 6
881.4951171875 0 14751.191
882.498046875 0 2971.823
949.5051879882812 0 604.8296
964.5145874023438 0 763.289
965.5194091796875 0 1722.3177
966.4344482421875 0 1465.5011
967.4405517578125 0 4506.5347 y 3
968.4453125 0 1728.4979
974.5055541992188 0 817.2988
976.4822998046875 0 1624.7965
977.4728393554688 0 1137.4135
978.4947509765625 0 708.55
992.5087280273438 0 19015.805 c Ammonia loss 7
993.5122680664062 0 13332.511
994.5150756835938 0 3928.3855
1007.5014038085938 0 952.0433
1035.533447265625 0 1625.87
1036.5335693359375 0 1371.7185
1079.51611328125 0 6928.914 z 2
1080.51953125 0 4896.385
1081.525634765625 0 1567.4849
1095.53515625 0 6316.055 y 2
1096.5372314453125 0 2646.42
1097.540771484375 0 2620.7007
1106.5877685546875 0 5500.282 c 8
1107.590087890625 0 3505.6387
1108.59619140625 0 812.891
1113.5567626953125 0 2565.9905
1114.560302734375 0 1481.494
1124.5364990234375 0 744.39014
1145.623779296875 0 728.977
1146.64208984375 0 1035.7695
1147.64404296875 0 1490.9167
1148.6715087890625 0 831.0639
1156.6993408203125 0 1123.0099
1161.6429443359375 0 8898.841
1162.6468505859375 0 8073.914
1163.652099609375 0 4169.7627
1164.658935546875 0 1346.7435
1178.5850830078125 0 3764.3247 z 1
1179.58544921875 0 2727.621
1188.6331787109375 0 1707.7385 c Ammonia loss 9
1189.6376953125 0 1055.9996
1190.641357421875 0 3655.272
1191.649169921875 0 2271.7822
1192.6561279296875 0 951.5143
1205.656494140625 0 44732.605 c 9
1206.6593017578125 0 32993.81
1207.6650390625 0 16379.044
1208.6722412109375 0 6670.218
1209.671875 0 1850.7844
1219.65087890625 0 1268.7139
1236.6756591796875 0 4027.0542
1237.6761474609375 0 2194.042
1238.6839599609375 0 730.5348
1244.6793212890625 0 2219.627
1245.685302734375 0 1962.5892
1246.6900634765625 0 1779.7495
1247.6851806640625 0 998.138
1248.654296875 0 12771.4
1249.6495361328125 0 9664.192
1250.65869140625 0 3915.5542
1251.6494140625 0 748.6643
1252.6251220703125 0 2288.4727
1253.628662109375 0 837.2123
1254.6416015625 0 917.35736
1262.691162109375 0 2695.691
1263.6907958984375 0 2465.4258
1264.7052001953125 0 4233.8877
1265.706787109375 0 3390.626
1266.7039794921875 0 1430.7496
1280.6990966796875 0 8196.137
1281.7042236328125 0 6707.394
1282.700927734375 0 2343.8838
1288.6956787109375 0 1188.7177
1289.6839599609375 0 1133.8442
1290.6817626953125 0 3899.4893
1291.6695556640625 0 48280.83
1292.6712646484375 0 36984.86
1293.6719970703125 0 15387.052
1294.677734375 0 2890.8882
1305.7095947265625 0 3730.3545
1306.6871337890625 0 2354.3008
1307.686767578125 0 39891.957
1308.69384765625 0 176500.11
1309.6973876953125 0 131871.23
1310.7001953125 0 50851.574
1311.703125 0 8750.319
2847.094482421875 0 722.0181
3070.698486328125 0 777.72906
3289.130126953125 0 711.0879

Spectrum Details

|  |  |
| --- | --- |
| Matched peaks? Matched peaks The total absolute number of peaks matched. Additionally in brackets the total fraction of peaks matched and the total number of peaks is shown. | 28 (15.56% of 180) |
| FDR? FDR The false discovery rate estimated for this peptide. It is calculated by matching all theoretical fragments with a non-integer shift with the raw peaks for this spectrum. This is done with 40 different shifts. The resulting percentage is the average number of annotated peaks over the number of annotated peaks with the correct spectrum. | 1.45% |
| Satellite FDR? Satellite FDR See the FDR for details on its calculation. This satellite ion specific FDR only contains the satellite ions (d/w) for I/L/J positions. | - |
| PSM Score? PSM Score The PSM Score as given by Hecklib to this annotated spectrum. It is shown with three significant figures. | 300 |

#### Spectrum 8100? Spectrum 8100 The raw spectrum of this peptide as annotated by Hecklib. The fragments are coloured according to ion type (see legend). Any peaks with a star '\*' as text can be hovered over to see the full details, first the ion type second the mass shift type. By hovering over the amino acids in the peptide or ions in the legend the corresponding peaks are highlighted. By toggling the 'Unassigned' label you can turn the background (unassigned) peaks on or off in the plot. By updating the slider in the Ion legend you can update the spectrum to only show the top X% of the peaks with labels. The top X% means any peak that is within X% of the highest intensity. By dragging in the spectrum you can zoom in to a specific part of the spectrum and use 'Zoom Out' to get back to the original zoom level. The annotation of the spectrum is based on the given sequence in the peptides file and is done with different software so inconsistencies are likely. The peaks are annotated based on the given sequence, with 20 ppm tolerance.

Copy Data

##### Spectrum 8100 (TSV)

###### Preview

```
Loading example...
```

*Click on the button to copy the data to your clipboard.*

Mz MinMz MaxIntensity Max

WidthHeightPeptide font sizePeptide stroke widthSpectrum font sizeSpectrum stroke widthCompact peptide

Ion legend

wxyz

abcd

OtherUnassignedIon^Charge^_Position_Show for top:%

JVKDYFPEPVT

02.62e+45.24e+47.86e+41.05e+5

Zoom Out

y^+1^_1_a^+1^_2_y^+1^_2_b^+1^_2_y^+1^_2_y^+1^_3_b^+2^_5_y^+1^_3_b^+1^_3_b^+2^_6_b^+2^_7_y^+1^_4_b^+2^_7_b^+1^_4_b^+1^_4_y^+1^_4_b^+1^_4_b^+2^_8_y^+1^_5_y^+1^_5_b^+2^_9_b^+2^_10_b^+2^_10_b^+1^_5_b^+1^_5_\*\*\*y^+1^_6_b^+1^_6_b^+1^_6_b^+1^_7_b^+1^_7_y^+1^_7_b^+1^_7_y^+1^_8_b^+1^_8_b^+1^_8_b^+1^_8_y^+1^_9_b^+1^_9_y^+1^_9_b^+1^_10_y^+1^_10_

0703140521082810

Fragment Matches Table

Show background peaks

| Position | Ion type | Intensity | mz Theoretical | mz Error (Th) | mz Error (ppm) | Charge | Series Number |
| --- | --- | --- | --- | --- | --- | --- | --- |
| 11 | y | 1.232E+04 | 120.1 | 0.0002992 | 2.492 | +1 | 1 |
| - | - | 1063 | 120.1 | - | - | 0 | - |
| - | - | 1.828E+04 | 120.1 | - | - | 0 | - |
| - | - | 541.2 | 121.1 | - | - | 0 | - |
| - | - | 1835 | 121.1 | - | - | 0 | - |
| - | - | 363.2 | 122.7 | - | - | 0 | - |
| - | - | 382.7 | 123.2 | - | - | 0 | - |
| - | - | 795.5 | 128.1 | - | - | 0 | - |
| - | - | 3.144E+04 | 129.1 | - | - | 0 | - |
| - | - | 500.5 | 130 | - | - | 0 | - |
| - | - | 1727 | 130.1 | - | - | 0 | - |
| - | - | 558.5 | 131.1 | - | - | 0 | - |
| - | - | 3238 | 133.1 | - | - | 0 | - |
| - | - | 432.7 | 134.1 | - | - | 0 | - |
| - | - | 9107 | 136.1 | - | - | 0 | - |
| - | - | 1178 | 137.1 | - | - | 0 | - |
| - | - | 699.1 | 139.1 | - | - | 0 | - |
| - | - | 429.9 | 140.1 | - | - | 0 | - |
| - | - | 554.6 | 149 | - | - | 0 | - |
| - | - | 799.6 | 152.1 | - | - | 0 | - |
| - | - | 488.9 | 152.6 | - | - | 0 | - |
| - | - | 944.5 | 155.1 | - | - | 0 | - |
| - | - | 902.1 | 155.1 | - | - | 0 | - |
| - | - | 571.3 | 165.1 | - | - | 0 | - |
| - | - | 428.6 | 166 | - | - | 0 | - |
| - | - | 439.3 | 166.1 | - | - | 0 | - |
| - | - | 463.6 | 166.1 | - | - | 0 | - |
| - | - | 746.7 | 167.1 | - | - | 0 | - |
| - | - | 984.2 | 167.1 | - | - | 0 | - |
| - | - | 990 | 168.1 | - | - | 0 | - |
| - | - | 9.031E+04 | 169.1 | - | - | 0 | - |
| - | - | 550.2 | 170.1 | - | - | 0 | - |
| - | - | 8214 | 170.1 | - | - | 0 | - |
| - | - | 508.6 | 171.1 | - | - | 0 | - |
| - | - | 586.4 | 172.1 | - | - | 0 | - |
| - | - | 1413 | 172.1 | - | - | 0 | - |
| - | - | 1078 | 173.1 | - | - | 0 | - |
| - | - | 1942 | 173.4 | - | - | 0 | - |
| - | - | 893.2 | 174.1 | - | - | 0 | - |
| - | - | 618 | 175.1 | - | - | 0 | - |
| - | - | 478.4 | 175.1 | - | - | 0 | - |
| - | - | 1070 | 177.1 | - | - | 0 | - |
| - | - | 496.6 | 181.1 | - | - | 0 | - |
| - | - | 2202 | 181.1 | - | - | 0 | - |
| - | - | 1436 | 181.1 | - | - | 0 | - |
| - | - | 398.6 | 182.1 | - | - | 0 | - |
| - | - | 486.8 | 183.1 | - | - | 0 | - |
| - | - | 1424 | 183.1 | - | - | 0 | - |
| - | - | 1212 | 183.1 | - | - | 0 | - |
| - | - | 816.9 | 185.1 | - | - | 0 | - |
| 2 | a | 3.68E+04 | 185.2 | 0.0003214 | 1.736 | +1 | 2 |
| - | - | 1024 | 186.1 | - | - | 0 | - |
| - | - | 3184 | 186.2 | - | - | 0 | - |
| - | - | 1634 | 188.1 | - | - | 0 | - |
| - | - | 519.1 | 191.1 | - | - | 0 | - |
| - | - | 2022 | 195.1 | - | - | 0 | - |
| - | - | 7.374E+04 | 197.1 | - | - | 0 | - |
| - | - | 756 | 198.1 | - | - | 0 | - |
| - | - | 6946 | 198.1 | - | - | 0 | - |
| - | - | 1.199E+04 | 199.1 | - | - | 0 | - |
| - | - | 2397 | 199.2 | - | - | 0 | - |
| - | - | 701.4 | 200.1 | - | - | 0 | - |
| 10 | y | 7491 | 201.1 | 0.0002273 | 1.13 | +1 | 2 |
| - | - | 693.2 | 202.1 | - | - | 0 | - |
| - | - | 3103 | 203.1 | - | - | 0 | - |
| - | - | 508.1 | 203.2 | - | - | 0 | - |
| - | - | 510.8 | 203.8 | - | - | 0 | - |
| - | - | 809.1 | 204.1 | - | - | 0 | - |
| - | - | 974.6 | 205.1 | - | - | 0 | - |
| - | - | 569.3 | 205.2 | - | - | 0 | - |
| - | - | 5136 | 209.1 | - | - | 0 | - |
| - | - | 514.1 | 209.5 | - | - | 0 | - |
| - | - | 669.9 | 211.1 | - | - | 0 | - |
| - | - | 459.2 | 211.1 | - | - | 0 | - |
| - | - | 1193 | 212.1 | - | - | 0 | - |
| - | - | 801 | 213.1 | - | - | 0 | - |
| 2 | b | 1.56E+04 | 213.2 | 0.0003256 | 1.528 | +1 | 2 |
| - | - | 2073 | 214.2 | - | - | 0 | - |
| - | - | 1601 | 215.1 | - | - | 0 | - |
| - | - | 758.4 | 217.1 | - | - | 0 | - |
| - | - | 2486 | 217.1 | - | - | 0 | - |
| 10 | y | 9036 | 219.1 | 0.0003438 | 1.569 | +1 | 2 |
| - | - | 547.1 | 220.1 | - | - | 0 | - |
| - | - | 677.1 | 225.1 | - | - | 0 | - |
| - | - | 531.2 | 225.2 | - | - | 0 | - |
| - | - | 8954 | 226.1 | - | - | 0 | - |
| - | - | 1199 | 226.2 | - | - | 0 | - |
| - | - | 3.298E+04 | 227.1 | - | - | 0 | - |
| - | - | 1118 | 227.1 | - | - | 0 | - |
| - | - | 459.8 | 227.2 | - | - | 0 | - |
| - | - | 579.9 | 227.2 | - | - | 0 | - |
| - | - | 600.6 | 227.6 | - | - | 0 | - |
| - | - | 2938 | 228.1 | - | - | 0 | - |
| - | - | 2852 | 228.2 | - | - | 0 | - |
| - | - | 605.5 | 230.2 | - | - | 0 | - |
| - | - | 5551 | 231.1 | - | - | 0 | - |
| - | - | 747.6 | 232.2 | - | - | 0 | - |
| - | - | 519 | 233.1 | - | - | 0 | - |
| - | - | 1619 | 233.2 | - | - | 0 | - |
| - | - | 1457 | 238.2 | - | - | 0 | - |
| - | - | 828.7 | 239.1 | - | - | 0 | - |
| - | - | 1.435E+04 | 244.1 | - | - | 0 | - |
| - | - | 676.1 | 245.1 | - | - | 0 | - |
| - | - | 3365 | 245.1 | - | - | 0 | - |
| - | - | 554.8 | 247.1 | - | - | 0 | - |
| - | - | 1114 | 247.1 | - | - | 0 | - |
| - | - | 8917 | 251.1 | - | - | 0 | - |
| - | - | 739.4 | 252.1 | - | - | 0 | - |
| - | - | 513.6 | 253.9 | - | - | 0 | - |
| - | - | 1214 | 254.1 | - | - | 0 | - |
| - | - | 615.4 | 254.2 | - | - | 0 | - |
| - | - | 700.1 | 255.1 | - | - | 0 | - |
| - | - | 595.9 | 256 | - | - | 0 | - |
| - | - | 625.9 | 256.2 | - | - | 0 | - |
| - | - | 614.6 | 259.1 | - | - | 0 | - |
| - | - | 2148 | 261.2 | - | - | 0 | - |
| - | - | 573 | 262.1 | - | - | 0 | - |
| - | - | 789.3 | 263.1 | - | - | 0 | - |
| - | - | 1311 | 270.2 | - | - | 0 | - |
| - | - | 2431 | 272.1 | - | - | 0 | - |
| - | - | 839.3 | 277.2 | - | - | 0 | - |
| - | - | 1023 | 278.2 | - | - | 0 | - |
| - | - | 1.165E+04 | 279.1 | - | - | 0 | - |
| - | - | 1406 | 280.1 | - | - | 0 | - |
| - | - | 3289 | 280.2 | - | - | 0 | - |
| - | - | 4603 | 283.1 | - | - | 0 | - |
| - | - | 690 | 285 | - | - | 0 | - |
| - | - | 1691 | 287.2 | - | - | 0 | - |
| - | - | 1167 | 296.2 | - | - | 0 | - |
| - | - | 570 | 298.1 | - | - | 0 | - |
| 9 | y | 1.635E+04 | 298.2 | 0.0004115 | 1.38 | +1 | 3 |
| - | - | 902.3 | 299.1 | - | - | 0 | - |
| - | - | 3250 | 299.2 | - | - | 0 | - |
| - | - | 1391 | 303.2 | - | - | 0 | - |
| - | - | 670.3 | 304.2 | - | - | 0 | - |
| - | - | 792.9 | 306.1 | - | - | 0 | - |
| - | - | 1059 | 308.2 | - | - | 0 | - |
| - | - | 680.1 | 310.1 | - | - | 0 | - |
| 5 | b | 747.6 | 310.2 | 0.0001674 | 0.5395 | +2 | 5 |
| - | - | 1061 | 310.2 | - | - | 0 | - |
| - | - | 3757 | 311.1 | - | - | 0 | - |
| - | - | 705 | 315.2 | - | - | 0 | - |
| 9 | y | 1.038E+05 | 316.2 | 0.0004974 | 1.573 | +1 | 3 |
| - | - | 1.5E+04 | 317.2 | - | - | 0 | - |
| - | - | 668.7 | 318.2 | - | - | 0 | - |
| - | - | 683.9 | 323.2 | - | - | 0 | - |
| - | - | 1241 | 323.2 | - | - | 0 | - |
| - | - | 2.86E+04 | 324.2 | - | - | 0 | - |
| - | - | 5288 | 325.2 | - | - | 0 | - |
| - | - | 948 | 325.2 | - | - | 0 | - |
| - | - | 837 | 326.2 | - | - | 0 | - |
| - | - | 740.5 | 328.2 | - | - | 0 | - |
| - | - | 873.1 | 329.1 | - | - | 0 | - |
| - | - | 766 | 331.2 | - | - | 0 | - |
| - | - | 1096 | 337.2 | - | - | 0 | - |
| - | - | 1969 | 340.2 | - | - | 0 | - |
| - | - | 2597 | 341.2 | - | - | 0 | - |
| 3 | b | 2528 | 341.3 | 0.0008368 | 2.452 | +1 | 3 |
| - | - | 934.2 | 342.3 | - | - | 0 | - |
| - | - | 2227 | 343.2 | - | - | 0 | - |
| - | - | 854.1 | 344.2 | - | - | 0 | - |
| - | - | 598.4 | 344.7 | - | - | 0 | - |
| - | - | 1336 | 346.2 | - | - | 0 | - |
| - | - | 1492 | 347.2 | - | - | 0 | - |
| - | - | 2184 | 353.2 | - | - | 0 | - |
| - | - | 1113 | 353.2 | - | - | 0 | - |
| - | - | 720 | 355.1 | - | - | 0 | - |
| - | - | 1287 | 355.2 | - | - | 0 | - |
| - | - | 1144 | 355.2 | - | - | 0 | - |
| - | - | 759.6 | 356.1 | - | - | 0 | - |
| - | - | 859.5 | 356.2 | - | - | 0 | - |
| - | - | 663.8 | 357.1 | - | - | 0 | - |
| - | - | 818.1 | 357.2 | - | - | 0 | - |
| - | - | 992.9 | 358.2 | - | - | 0 | - |
| - | - | 871.3 | 362.2 | - | - | 0 | - |
| - | - | 651.4 | 367.2 | - | - | 0 | - |
| - | - | 2595 | 369.7 | - | - | 0 | - |
| - | - | 1549 | 370.2 | - | - | 0 | - |
| - | - | 3239 | 371.2 | - | - | 0 | - |
| - | - | 570.9 | 372.2 | - | - | 0 | - |
| - | - | 652.3 | 372.2 | - | - | 0 | - |
| - | - | 1137 | 373.2 | - | - | 0 | - |
| - | - | 4991 | 374.2 | - | - | 0 | - |
| - | - | 866.7 | 375.2 | - | - | 0 | - |
| - | - | 1315 | 377.2 | - | - | 0 | - |
| - | - | 2510 | 379.2 | - | - | 0 | - |
| - | - | 1349 | 381.1 | - | - | 0 | - |
| - | - | 1491 | 383.2 | - | - | 0 | - |
| 6 | b | 1295 | 383.7 | 0.0008365 | 2.18 | +2 | 6 |
| - | - | 2093 | 389.2 | - | - | 0 | - |
| - | - | 6329 | 391.2 | - | - | 0 | - |
| - | - | 1054 | 392.2 | - | - | 0 | - |
| - | - | 968.2 | 392.2 | - | - | 0 | - |
| - | - | 3111 | 395.2 | - | - | 0 | - |
| - | - | 1021 | 396.2 | - | - | 0 | - |
| - | - | 1339 | 403.2 | - | - | 0 | - |
| - | - | 1482 | 405.2 | - | - | 0 | - |
| - | - | 710.1 | 406.2 | - | - | 0 | - |
| - | - | 7246 | 407.2 | - | - | 0 | - |
| - | - | 1359 | 408.2 | - | - | 0 | - |
| - | - | 795.7 | 409.3 | - | - | 0 | - |
| - | - | 542.2 | 415.3 | - | - | 0 | - |
| - | - | 855 | 417.2 | - | - | 0 | - |
| - | - | 5929 | 418.2 | - | - | 0 | - |
| - | - | 1782 | 418.7 | - | - | 0 | - |
| - | - | 640 | 419.2 | - | - | 0 | - |
| - | - | 616.9 | 419.2 | - | - | 0 | - |
| - | - | 746.6 | 421.2 | - | - | 0 | - |
| 7 | b | 9507 | 423.2 | 0.006738 | 15.92 | +2 | 7 |
| - | - | 4401 | 424.2 | - | - | 0 | - |
| - | - | 893.4 | 425.2 | - | - | 0 | - |
| - | - | 8706 | 426.2 | - | - | 0 | - |
| - | - | 2226 | 427.2 | - | - | 0 | - |
| 8 | y | 1162 | 427.2 | 0.0002199 | 0.5148 | +1 | 4 |
| 7 | b | 4242 | 432.2 | 0.0004916 | 1.137 | +2 | 7 |
| - | - | 1755 | 432.7 | - | - | 0 | - |
| - | - | 1144 | 433.2 | - | - | 0 | - |
| - | - | 1780 | 435.2 | - | - | 0 | - |
| - | - | 1136 | 436.2 | - | - | 0 | - |
| 4 | b | 3610 | 438.3 | 0.0008159 | 1.862 | +1 | 4 |
| - | - | 810.1 | 438.3 | - | - | 0 | - |
| 4 | b | 778.5 | 439.3 | 0.0008999 | 2.049 | +1 | 4 |
| - | - | 1017 | 440.3 | - | - | 0 | - |
| 8 | y | 2406 | 445.2 | 0.001544 | 3.469 | +1 | 4 |
| - | - | 1440 | 452.2 | - | - | 0 | - |
| - | - | 741.2 | 453.2 | - | - | 0 | - |
| - | - | 601.8 | 454.2 | - | - | 0 | - |
| 4 | b | 1.285E+04 | 456.3 | 0.0007188 | 1.575 | +1 | 4 |
| - | - | 3458 | 457.3 | - | - | 0 | - |
| - | - | 4573 | 470.2 | - | - | 0 | - |
| - | - | 1130 | 470.3 | - | - | 0 | - |
| - | - | 3025 | 471.2 | - | - | 0 | - |
| - | - | 632.7 | 480.2 | - | - | 0 | - |
| - | - | 2064 | 482.3 | - | - | 0 | - |
| - | - | 2066 | 482.8 | - | - | 0 | - |
| - | - | 1920 | 483.3 | - | - | 0 | - |
| - | - | 1336 | 488.3 | - | - | 0 | - |
| - | - | 1197 | 488.7 | - | - | 0 | - |
| 8 | b | 8318 | 496.8 | 0.0005878 | 1.183 | +2 | 8 |
| - | - | 4541 | 497.3 | - | - | 0 | - |
| - | - | 1315 | 497.8 | - | - | 0 | - |
| - | - | 721 | 504.2 | - | - | 0 | - |
| - | - | 613.6 | 504.8 | - | - | 0 | - |
| - | - | 1743 | 506.3 | - | - | 0 | - |
| - | - | 1398 | 508.3 | - | - | 0 | - |
| - | - | 2788 | 509.2 | - | - | 0 | - |
| - | - | 715.3 | 518.3 | - | - | 0 | - |
| - | - | 969.1 | 521.8 | - | - | 0 | - |
| - | - | 2227 | 522.3 | - | - | 0 | - |
| - | - | 4290 | 522.8 | - | - | 0 | - |
| - | - | 662.3 | 523.2 | - | - | 0 | - |
| - | - | 1837 | 523.3 | - | - | 0 | - |
| 7 | y | 3393 | 524.3 | 0.002131 | 4.065 | +1 | 5 |
| - | - | 1054 | 525.3 | - | - | 0 | - |
| - | - | 5714 | 526.3 | - | - | 0 | - |
| - | - | 1883 | 527.3 | - | - | 0 | - |
| - | - | 1525 | 531.3 | - | - | 0 | - |
| - | - | 1891 | 531.8 | - | - | 0 | - |
| - | - | 3023 | 534.3 | - | - | 0 | - |
| - | - | 995.6 | 535.3 | - | - | 0 | - |
| - | - | 2872 | 536.3 | - | - | 0 | - |
| - | - | 1050 | 536.3 | - | - | 0 | - |
| - | - | 993.4 | 537.2 | - | - | 0 | - |
| - | - | 2296 | 539.3 | - | - | 0 | - |
| - | - | 913.3 | 540.3 | - | - | 0 | - |
| 7 | y | 3.558E+04 | 542.3 | 0.0005995 | 1.105 | +1 | 5 |
| - | - | 1.023E+04 | 543.3 | - | - | 0 | - |
| - | - | 1740 | 544.3 | - | - | 0 | - |
| 9 | b | 2.101E+04 | 545.3 | 0.00039 | 0.7152 | +2 | 9 |
| - | - | 1.357E+04 | 545.8 | - | - | 0 | - |
| - | - | 4773 | 546.3 | - | - | 0 | - |
| - | - | 730.5 | 549.3 | - | - | 0 | - |
| - | - | 792.3 | 551.3 | - | - | 0 | - |
| - | - | 1644 | 553.3 | - | - | 0 | - |
| - | - | 2.876E+04 | 554.3 | - | - | 0 | - |
| - | - | 1139 | 554.3 | - | - | 0 | - |
| - | - | 9169 | 555.3 | - | - | 0 | - |
| - | - | 1658 | 556.3 | - | - | 0 | - |
| - | - | 1288 | 558.3 | - | - | 0 | - |
| - | - | 696.4 | 559.3 | - | - | 0 | - |
| - | - | 1153 | 564.2 | - | - | 0 | - |
| - | - | 599.4 | 564.9 | - | - | 0 | - |
| - | - | 2781 | 567.4 | - | - | 0 | - |
| - | - | 834.9 | 568.4 | - | - | 0 | - |
| - | - | 1621 | 570.3 | - | - | 0 | - |
| - | - | 1253 | 571.3 | - | - | 0 | - |
| - | - | 687.6 | 571.8 | - | - | 0 | - |
| - | - | 755.8 | 572.3 | - | - | 0 | - |
| - | - | 666.3 | 574.3 | - | - | 0 | - |
| - | - | 8226 | 580.8 | - | - | 0 | - |
| - | - | 5263 | 581.3 | - | - | 0 | - |
| - | - | 1609 | 581.8 | - | - | 0 | - |
| - | - | 1364 | 582.3 | - | - | 0 | - |
| - | - | 857.2 | 583.3 | - | - | 0 | - |
| - | - | 724.2 | 585.3 | - | - | 0 | - |
| 10 | b | 853 | 585.8 | 3.121E-05 | 0.05328 | +2 | 10 |
| - | - | 4207 | 591.4 | - | - | 0 | - |
| - | - | 1813 | 592.4 | - | - | 0 | - |
| 10 | b | 5398 | 594.8 | 0.0005458 | 0.9176 | +2 | 10 |
| - | - | 5004 | 595.3 | - | - | 0 | - |
| - | - | 1251 | 595.8 | - | - | 0 | - |
| - | - | 735.9 | 597.3 | - | - | 0 | - |
| - | - | 542.5 | 600.5 | - | - | 0 | - |
| 5 | b | 950.6 | 601.3 | 0.0001094 | 0.182 | +1 | 5 |
| - | - | 1710 | 603.3 | - | - | 0 | - |
| - | - | 601.7 | 605.3 | - | - | 0 | - |
| - | - | 1185 | 615.3 | - | - | 0 | - |
| 5 | b | 1.814E+04 | 619.3 | 0.0001649 | 0.2662 | +1 | 5 |
| - | - | 7214 | 620.3 | - | - | 0 | - |
| - | - | 1604 | 621.4 | - | - | 0 | - |
| - | - | 848.4 | 623.3 | - | - | 0 | - |
| - | - | 672.1 | 625.3 | - | - | 0 | - |
| - | - | 845 | 631.3 | - | - | 0 | - |
| - | - | 1190 | 633.3 | - | - | 0 | - |
| - | - | 747.5 | 635.3 | - | - | 0 | - |
| - | - | 1918 | 635.4 | - | - | 0 | - |
| - | - | 732 | 636.3 | - | - | 0 | - |
| - | - | 1050 | 636.4 | - | - | 0 | - |
| - | - | 643.5 | 638.3 | - | - | 0 | - |
| 0 | Precursor | 3350 | 645.3 | 0.0004052 | 0.6279 | +2 | -1 |
| 0 | Precursor | 1672 | 645.8 | 0.008564 | 13.26 | +2 | -1 |
| - | - | 1084 | 646.3 | - | - | 0 | - |
| - | - | 2410 | 651.3 | - | - | 0 | - |
| - | - | 608.5 | 651.4 | - | - | 0 | - |
| - | - | 1318 | 652.3 | - | - | 0 | - |
| - | - | 905.7 | 652.3 | - | - | 0 | - |
| - | - | 639.1 | 652.4 | - | - | 0 | - |
| - | - | 1028 | 652.9 | - | - | 0 | - |
| - | - | 1187 | 653.3 | - | - | 0 | - |
| - | - | 3473 | 653.3 | - | - | 0 | - |
| - | - | 866.4 | 653.9 | - | - | 0 | - |
| 0 | Precursor | 4060 | 654.3 | 0.003551 | 5.427 | +2 | -1 |
| - | - | 2973 | 654.8 | - | - | 0 | - |
| - | - | 1730 | 655.4 | - | - | 0 | - |
| - | - | 1101 | 663.3 | - | - | 0 | - |
| - | - | 911.6 | 664.4 | - | - | 0 | - |
| - | - | 866.2 | 667.3 | - | - | 0 | - |
| - | - | 3693 | 681.3 | - | - | 0 | - |
| - | - | 1176 | 682.3 | - | - | 0 | - |
| - | - | 4384 | 682.4 | - | - | 0 | - |
| - | - | 2246 | 683.4 | - | - | 0 | - |
| - | - | 902.4 | 688.4 | - | - | 0 | - |
| 6 | y | 1.194E+04 | 689.4 | 0.0001787 | 0.2593 | +1 | 6 |
| - | - | 5666 | 690.4 | - | - | 0 | - |
| - | - | 1389 | 691.4 | - | - | 0 | - |
| - | - | 2993 | 716.4 | - | - | 0 | - |
| - | - | 650.2 | 720.4 | - | - | 0 | - |
| - | - | 1648 | 721.4 | - | - | 0 | - |
| - | - | 721.1 | 734.4 | - | - | 0 | - |
| - | - | 6716 | 738.4 | - | - | 0 | - |
| - | - | 2788 | 739.4 | - | - | 0 | - |
| 6 | b | 2448 | 748.4 | 0.0001282 | 0.1713 | +1 | 6 |
| - | - | 1318 | 749.4 | - | - | 0 | - |
| - | - | 1021 | 750.3 | - | - | 0 | - |
| - | - | 1860 | 750.4 | - | - | 0 | - |
| - | - | 1126 | 751.4 | - | - | 0 | - |
| - | - | 3511 | 752.4 | - | - | 0 | - |
| - | - | 1161 | 753.4 | - | - | 0 | - |
| - | - | 2454 | 762.3 | - | - | 0 | - |
| - | - | 932.7 | 763.3 | - | - | 0 | - |
| 6 | b | 6.566E+04 | 766.4 | 7.278E-05 | 0.09496 | +1 | 6 |
| - | - | 2.865E+04 | 767.4 | - | - | 0 | - |
| - | - | 6272 | 768.4 | - | - | 0 | - |
| - | - | 2.374E+04 | 780.4 | - | - | 0 | - |
| - | - | 9416 | 781.4 | - | - | 0 | - |
| - | - | 2916 | 782.4 | - | - | 0 | - |
| - | - | 606.5 | 784.4 | - | - | 0 | - |
| - | - | 3149 | 796 | - | - | 0 | - |
| - | - | 4501 | 796.4 | - | - | 0 | - |
| - | - | 3446 | 796.7 | - | - | 0 | - |
| - | - | 800.9 | 797 | - | - | 0 | - |
| - | - | 893.5 | 817.4 | - | - | 0 | - |
| - | - | 985 | 820.5 | - | - | 0 | - |
| - | - | 880.7 | 827.4 | - | - | 0 | - |
| - | - | 781.5 | 828.4 | - | - | 0 | - |
| - | - | 949.9 | 829.4 | - | - | 0 | - |
| - | - | 983.2 | 835.5 | - | - | 0 | - |
| 7 | b | 2387 | 845.5 | 0.01517 | 17.95 | +1 | 7 |
| 7 | b | 997.9 | 846.4 | 0.007709 | 9.108 | +1 | 7 |
| - | - | 1662 | 850.4 | - | - | 0 | - |
| - | - | 1353 | 851.4 | - | - | 0 | - |
| 5 | y | 854.2 | 852.4 | 0.01137 | 13.34 | +1 | 7 |
| 7 | b | 4090 | 863.5 | 0.0002854 | 0.3305 | +1 | 7 |
| - | - | 1877 | 864.5 | - | - | 0 | - |
| - | - | 756.1 | 865.5 | - | - | 0 | - |
| - | - | 8506 | 877.4 | - | - | 0 | - |
| - | - | 3730 | 878.4 | - | - | 0 | - |
| - | - | 3409 | 879.4 | - | - | 0 | - |
| - | - | 1215 | 880.4 | - | - | 0 | - |
| - | - | 833.8 | 895.5 | - | - | 0 | - |
| - | - | 2281 | 907.4 | - | - | 0 | - |
| - | - | 996.9 | 908.4 | - | - | 0 | - |
| - | - | 1104 | 946.5 | - | - | 0 | - |
| - | - | 1543 | 947.5 | - | - | 0 | - |
| - | - | 1675 | 948.5 | - | - | 0 | - |
| - | - | 1755 | 962.5 | - | - | 0 | - |
| - | - | 696.9 | 963.5 | - | - | 0 | - |
| - | - | 850.4 | 964.4 | - | - | 0 | - |
| - | - | 7271 | 964.5 | - | - | 0 | - |
| - | - | 4010 | 965.5 | - | - | 0 | - |
| - | - | 1000 | 966.5 | - | - | 0 | - |
| 4 | y | 687.6 | 967.4 | 0.00324 | 3.349 | +1 | 8 |
| 8 | b | 3302 | 974.5 | 0.002223 | 2.282 | +1 | 8 |
| 8 | b | 2521 | 975.5 | 0.01925 | 19.74 | +1 | 8 |
| - | - | 8946 | 976.5 | - | - | 0 | - |
| - | - | 4981 | 977.5 | - | - | 0 | - |
| - | - | 1408 | 978.5 | - | - | 0 | - |
| - | - | 805 | 990.5 | - | - | 0 | - |
| - | - | 841.2 | 991.5 | - | - | 0 | - |
| 8 | b | 8.222E+04 | 992.5 | 0.000398 | 0.401 | +1 | 8 |
| - | - | 4.628E+04 | 993.5 | - | - | 0 | - |
| - | - | 1.424E+04 | 994.5 | - | - | 0 | - |
| - | - | 2069 | 995.5 | - | - | 0 | - |
| - | - | 672.4 | 1015 | - | - | 0 | - |
| - | - | 1443 | 1045 | - | - | 0 | - |
| - | - | 1203 | 1047 | - | - | 0 | - |
| - | - | 970.3 | 1048 | - | - | 0 | - |
| 3 | y | 1087 | 1078 | 0.01673 | 15.53 | +1 | 9 |
| 9 | b | 2132 | 1090 | 0.003418 | 3.137 | +1 | 9 |
| - | - | 1535 | 1091 | - | - | 0 | - |
| - | - | 791.1 | 1092 | - | - | 0 | - |
| 3 | y | 1.662E+04 | 1096 | 0.0005466 | 0.4989 | +1 | 9 |
| - | - | 9596 | 1097 | - | - | 0 | - |
| - | - | 3587 | 1098 | - | - | 0 | - |
| 10 | b | 5511 | 1189 | 0.002252 | 1.895 | +1 | 10 |
| - | - | 4523 | 1190 | - | - | 0 | - |
| - | - | 1175 | 1191 | - | - | 0 | - |
| 2 | y | 1772 | 1195 | 0.003653 | 3.058 | +1 | 10 |
| - | - | 1174 | 1196 | - | - | 0 | - |
| - | - | 903.9 | 1197 | - | - | 0 | - |
| - | - | 674 | 1284 | - | - | 0 | - |
| - | - | 838.4 | 2782 | - | - | 0 | - |

m/z Charge Intensity FragmentType MassShift Position
120.0658187866211 0 12322.943 y 10
120.07624816894531 0 1063.3386
120.08108520507812 0 18282.426
121.06953430175781 0 541.21106
121.0843505859375 0 1834.6449
122.73821258544922 0 363.1716
123.23967742919922 0 382.66492
128.10763549804688 0 795.54144
129.1025390625 0 31442.06
130.0499725341797 0 500.509
130.10594177246094 0 1726.5715
131.07052612304688 0 558.46674
133.0862274169922 0 3237.729
134.08958435058594 0 432.74908
136.0760040283203 0 9107.159
137.0792999267578 0 1178.2723
139.08676147460938 0 699.1101
140.14366149902344 0 429.85977
148.95538330078125 0 554.55566
152.07058715820312 0 799.5719
152.64126586914062 0 488.8987
155.08203125 0 944.5023
155.11817932128906 0 902.114
165.10220336914062 0 571.3118
165.97198486328125 0 428.58835
166.05361938476562 0 439.3211
166.08560180664062 0 463.64822
167.08177185058594 0 746.66296
167.11830139160156 0 984.19226
168.10218811035156 0 990.0394
169.13388061523438 0 90310.2
170.12962341308594 0 550.1999
170.13719177246094 0 8213.831
171.12098693847656 0 508.62485
172.07154846191406 0 586.3913
172.11248779296875 0 1413.4524
173.1289520263672 0 1077.5923
173.43917846679688 0 1941.7166
174.12802124023438 0 893.17456
175.08697509765625 0 617.95575
175.09622192382812 0 478.4251
177.1126251220703 0 1070.4398
181.061279296875 0 496.6245
181.0974884033203 0 2201.7134
181.13394165039062 0 1435.7933
182.1280975341797 0 398.6323
183.08590698242188 0 486.82553
183.1134796142578 0 1424.2725
183.14947509765625 0 1211.6768
185.1285858154297 0 816.8922
185.1651611328125 0 36802.168 a 1
186.12826538085938 0 1023.5952
186.16845703125 0 3184.3047
188.1437530517578 0 1634.4454
191.0817413330078 0 519.0784
195.11318969726562 0 2022.4142
197.12875366210938 0 73742.39
198.12408447265625 0 756.00684
198.13209533691406 0 6946.2124
199.1079559326172 0 11989.88
199.18080139160156 0 2396.78
200.1115264892578 0 701.3529
201.12359619140625 0 7491.2837 y Water loss 9
202.12632751464844 0 693.15063
203.1027069091797 0 3102.7515
203.1545867919922 0 508.10062
203.75242614746094 0 510.81067
204.1389617919922 0 809.14655
205.10690307617188 0 974.551
205.20205688476562 0 569.34155
209.09228515625 0 5135.893
209.49769592285156 0 514.081
211.10748291015625 0 669.8599
211.14486694335938 0 459.18857
212.1396942138672 0 1192.8331
213.12384033203125 0 800.98517
213.1600799560547 0 15600.899 b 1
214.16360473632812 0 2072.7458
215.1394805908203 0 1601.0972
217.08154296875 0 758.352
217.13389587402344 0 2485.6362
219.13427734375 0 9035.689 y 9
220.1380615234375 0 547.05225
225.12347412109375 0 677.1023
225.16065979003906 0 531.17487
226.11888122558594 0 8953.928
226.15512084960938 0 1199.137
227.10299682617188 0 32981.223
227.12290954589844 0 1117.8633
227.15892028808594 0 459.76785
227.17572021484375 0 579.949
227.56851196289062 0 600.63135
228.10633850097656 0 2938.3254
228.17092895507812 0 2851.8872
230.1861114501953 0 605.4677
231.1496124267578 0 5551.344
232.15237426757812 0 747.57336
233.09359741210938 0 518.97186
233.16549682617188 0 1619.4703
238.15533447265625 0 1456.7886
239.09512329101562 0 828.7485
244.12954711914062 0 14345.543
245.1155242919922 0 676.1338
245.1299285888672 0 3364.5557
247.1072998046875 0 554.7651
247.14437866210938 0 1113.852
251.10293579101562 0 8917.252
252.10667419433594 0 739.36096
253.89134216308594 0 513.61084
254.1127166748047 0 1213.8528
254.15025329589844 0 615.3568
255.09817504882812 0 700.12634
256.0166320800781 0 595.9343
256.1664123535156 0 625.9225
259.14453125 0 614.6078
261.1586608886719 0 2148.2542
262.07073974609375 0 572.99524
263.1384582519531 0 789.30945
270.1813659667969 0 1310.763
272.12420654296875 0 2430.729
277.1557922363281 0 839.2772
278.1509704589844 0 1023.23016
279.09783935546875 0 11648.522
280.1009826660156 0 1405.8359
280.16595458984375 0 3288.9026
283.144287109375 0 4602.936
285.0105285644531 0 689.9864
287.2124938964844 0 1691.1675
296.1971740722656 0 1166.9635
298.13848876953125 0 570.02496
298.1765441894531 0 16352.815 y Water loss 8
299.0625 0 902.2777
299.1796569824219 0 3250.0002
303.2070617675781 0 1390.728
304.1645202636719 0 670.3358
306.1460876464844 0 792.9227
308.16082763671875 0 1058.605
310.1399841308594 0 680.0559
310.1763000488281 0 747.583 b 4
310.2120666503906 0 1060.7362
311.1395263671875 0 3757.3552
315.2081298828125 0 705.0107
316.18719482421875 0 103768.484 y 8
317.1905212402344 0 14995.155
318.1947021484375 0 668.7424
323.1719055175781 0 683.8608
323.2439880371094 0 1241.494
324.1559753417969 0 28595.518
325.15911865234375 0 5288.489
325.18511962890625 0 948.0329
326.17230224609375 0 837.0184
328.1680908203125 0 740.5456
329.1494445800781 0 873.0906
331.20355224609375 0 765.9898
337.1505432128906 0 1095.9436
340.1871337890625 0 1969.0177
341.1822509765625 0 2597.3647
341.25555419921875 0 2527.8904 b 2
342.2572937011719 0 934.2008
343.1980285644531 0 2226.9448
344.199951171875 0 854.08234
344.7074890136719 0 598.3505
346.1770324707031 0 1335.6926
347.202880859375 0 1491.9899
353.18170166015625 0 2184.1753
353.2182922363281 0 1113.34
355.0704040527344 0 719.9871
355.1611633300781 0 1287.1421
355.1973571777344 0 1143.6971
356.069580078125 0 759.6322
356.1628112792969 0 859.513
357.0668640136719 0 663.76904
357.213623046875 0 818.0657
358.2124328613281 0 992.90564
362.1718444824219 0 871.3174
367.19854736328125 0 651.403
369.7132263183594 0 2594.898
370.2132568359375 0 1548.9939
371.19244384765625 0 3238.5002
372.1560974121094 0 570.85645
372.1949157714844 0 652.2753
373.18756103515625 0 1137.4434
374.171142578125 0 4991.19
375.1741638183594 0 866.7321
377.21856689453125 0 1314.5696
379.19781494140625 0 2509.9417
381.1452331542969 0 1348.6492
383.19293212890625 0 1491.0184
383.7095031738281 0 1295.2448 b 5
389.18231201171875 0 2093.2915
391.1979675292969 0 6329.045
392.1794128417969 0 1053.8121
392.20404052734375 0 968.238
395.2292175292969 0 3111.4167
396.2303771972656 0 1020.89496
403.2349853515625 0 1339.0267
405.2140808105469 0 1482.0986
406.2143249511719 0 710.0536
407.1929626464844 0 7245.7666
408.1955871582031 0 1359.4171
409.2804260253906 0 795.7157
415.34033203125 0 542.23975
417.1773986816406 0 855.0365
418.2399597167969 0 5928.599
418.7427673339844 0 1781.984
419.1943054199219 0 639.97424
419.24005126953125 0 616.8709
421.2452392578125 0 746.5665
423.2247009277344 0 9507.382 b Water loss 6
424.223876953125 0 4401.2534
425.2274475097656 0 893.42773
426.1665344238281 0 8705.722
427.1690673828125 0 2225.6902
427.218505859375 0 1162.2538 y Water loss 7
432.2372131347656 0 4241.9434 b 6
432.7389221191406 0 1755.4354
433.2413330078125 0 1143.9352
435.18890380859375 0 1779.5417
436.1879577636719 0 1136.2465
438.27191162109375 0 3609.506 b Water loss 3
438.3044738769531 0 810.07513
439.25421142578125 0 778.4706 b Ammonia loss 3
440.25494384765625 0 1016.5035
445.2308349609375 0 2405.7227 y 7
452.21380615234375 0 1439.6241
453.2156982421875 0 741.23517
454.232177734375 0 601.80786
456.2823791503906 0 12848.999 b 3
457.2850036621094 0 3457.827
470.2251892089844 0 4572.9995
470.3153381347656 0 1130.0479
471.22625732421875 0 3024.5532
480.2107849121094 0 632.6676
482.2597961425781 0 2063.902
482.7620849609375 0 2066.4062
483.26361083984375 0 1919.5093
488.3227844238281 0 1336.2118
488.7449035644531 0 1196.6713
496.75860595703125 0 8318.152 b 7
497.2599792480469 0 4540.7163
497.7614440917969 0 1315.469
504.2469177246094 0 720.9843
504.7759094238281 0 613.6452
506.2613525390625 0 1742.5111
508.2568054199219 0 1397.506
509.2403869628906 0 2788.489
518.2603149414062 0 715.347
521.7874755859375 0 969.099
522.2859497070312 0 2226.8445
522.78515625 0 4289.8936
523.2197875976562 0 662.2982
523.28564453125 0 1837.4225
524.2736206054688 0 3392.853 y Water loss 6
525.2723999023438 0 1053.5076
526.26611328125 0 5713.8926
527.27099609375 0 1883.1624
531.2859497070312 0 1524.9928
531.7887573242188 0 1891.0059
534.2561645507812 0 3023.0132
535.2554321289062 0 995.56696
536.2506103515625 0 2872.4788
536.3093872070312 0 1050.3894
537.2372436523438 0 993.3655
539.25146484375 0 2295.5254
540.2554931640625 0 913.32196
542.2826538085938 0 35582.383 y 6
543.2857055664062 0 10226.418
544.2868041992188 0 1739.86
545.2847900390625 0 21013.193 b 8
545.7862548828125 0 13565.477
546.2884521484375 0 4772.6987
549.3370971679688 0 730.5261
551.2815551757812 0 792.3407
553.3339233398438 0 1644.2946
554.2610473632812 0 28759.738
554.3362426757812 0 1138.6973
555.2642822265625 0 9168.522
556.267822265625 0 1657.5194
558.3312377929688 0 1287.9097
559.329833984375 0 696.3916
564.24365234375 0 1153.2793
564.9459838867188 0 599.41943
567.3522338867188 0 2780.666
568.3516845703125 0 834.9278
570.2933349609375 0 1621.2292
571.29248046875 0 1253.4333
571.8163452148438 0 687.5948
572.3109741210938 0 755.7766
574.326416015625 0 666.2612
580.8215942382812 0 8225.728
581.3226318359375 0 5263.227
581.8252563476562 0 1608.9072
582.25634765625 0 1363.7552
583.2572631835938 0 857.2295
585.3428344726562 0 724.15985
585.8132934570312 0 852.9798 b Water loss 9
591.3505859375 0 4207.1924
592.3556518554688 0 1812.8617
594.8191528320312 0 5397.788 b 9
595.3209838867188 0 5004.0117
595.8226928710938 0 1250.8168
597.28759765625 0 735.8942
600.5264282226562 0 542.49713
601.3345336914062 0 950.5533 b Water loss 4
603.3495483398438 0 1710.065
605.2936401367188 0 601.7249
615.2760620117188 0 1184.9093
619.3451538085938 0 18143.51 b 4
620.3492431640625 0 7213.943
621.3523559570312 0 1604.3025
623.3175048828125 0 848.37994
625.3336791992188 0 672.0695
631.3102416992188 0 844.98627
633.2899780273438 0 1189.772
635.3208618164062 0 747.53735
635.377685546875 0 1918.0907
636.30908203125 0 731.9837
636.3786010742188 0 1049.7693
638.31982421875 0 643.4531
645.342041015625 0 3349.841 Precursor Water loss
645.843017578125 0 1672.2273 Precursor Ammonia loss
646.3461303710938 0 1084.4789
651.3151245117188 0 2410.0808
651.3804321289062 0 608.46027
652.2595825195312 0 1318.456
652.3162231445312 0 905.67535
652.4015502929688 0 639.0972
652.8624267578125 0 1027.6588
653.2661743164062 0 1187.3068
653.3302001953125 0 3473.2383
653.8639526367188 0 866.42535
654.3441772460938 0 4059.8794 Precursor
654.8499755859375 0 2972.7175
655.352294921875 0 1729.6395
663.3156127929688 0 1100.8138
664.3704223632812 0 911.6335
667.3471069335938 0 866.1636
681.3242797851562 0 3692.835
682.3246459960938 0 1176.4453
682.3785400390625 0 4383.548
683.3829345703125 0 2246.2468
688.400634765625 0 902.37134
689.3506469726562 0 11943.113 y 5
690.3538208007812 0 5665.763
691.3548583984375 0 1389.3613
716.3966674804688 0 2992.7888
720.407470703125 0 650.16693
721.3924560546875 0 1648.034
734.3529052734375 0 721.09894
738.418212890625 0 6716.232
739.4224243164062 0 2788.0388
748.4027099609375 0 2448.2834 b Water loss 5
749.4052124023438 0 1317.629
750.3132934570312 0 1020.6631
750.3851928710938 0 1859.9393
751.38134765625 0 1126.269
752.3621826171875 0 3511.3513
753.3621826171875 0 1161.3717
762.346435546875 0 2454.014
763.3497924804688 0 932.7461
766.413330078125 0 65660.66 b 5
767.416259765625 0 28647.412
768.4192504882812 0 6272.4985
780.3565673828125 0 23743.9
781.359375 0 9415.685
782.362548828125 0 2915.6963
784.4223022460938 0 606.5433
796.0457153320312 0 3149.3748
796.380126953125 0 4500.536
796.7134399414062 0 3445.6248
797.0493774414062 0 800.9457
817.4456176757812 0 893.46375
820.4603881835938 0 985.0489
827.4306030273438 0 880.7073
828.4314575195312 0 781.54193
829.4426879882812 0 949.8946
835.4739379882812 0 983.1795
845.4404296875 0 2386.9453 b Water loss 6
846.4473266601562 0 997.9022 b Ammonia loss 6
850.4376831054688 0 1661.5151
851.4326782226562 0 1353.3102
852.4251708984375 0 854.24567 y 4
863.4658813476562 0 4089.7314 b 6
864.4676513671875 0 1876.9036
865.4791259765625 0 756.09827
877.4095458984375 0 8506.448
878.411865234375 0 3729.7573
879.4224853515625 0 3409.0454
880.4215087890625 0 1215.2206
895.4537353515625 0 833.81036
907.4232788085938 0 2281.435
908.4297485351562 0 996.8883
946.5064697265625 0 1104.0703
947.4920043945312 0 1543.1049
948.4911499023438 0 1674.907
962.5321044921875 0 1755.1478
963.5253295898438 0 696.91064
964.4139404296875 0 850.40497
964.514404296875 0 7270.6245
965.515869140625 0 4010.0376
966.5156860351562 0 1000.42206
967.4375 0 687.58466 y 3
974.4959716796875 0 3302.078 b Water loss 7
975.50146484375 0 2521.4128 b Ammonia loss 7
976.4795532226562 0 8946.056
977.4796142578125 0 4981.2466
978.4820556640625 0 1408.0729
990.4967041015625 0 805.0109
991.51025390625 0 841.154
992.5083618164062 0 82217.2 b 7
993.5113525390625 0 46279.324
994.5136108398438 0 14241.028
995.5203247070312 0 2069.1372
1014.95751953125 0 672.3816
1044.566650390625 0 1443.3131
1046.5557861328125 0 1203.3827
1047.5643310546875 0 970.25275
1077.5418701171875 0 1086.8822 y Water loss 2
1089.55810546875 0 2131.764 b 8
1090.5460205078125 0 1534.906
1091.5621337890625 0 791.08093
1095.53515625 0 16616.71 y 2
1096.537841796875 0 9595.6045
1097.5426025390625 0 3586.645
1188.627685546875 0 5510.995 b 9
1189.6317138671875 0 4522.5435
1190.6348876953125 0 1175.3307
1194.6004638671875 0 1771.909 y 1
1195.6109619140625 0 1173.5569
1196.6102294921875 0 903.90643
1284.476806640625 0 674.0132
2782.347900390625 0 838.41016

Spectrum Details

|  |  |
| --- | --- |
| Matched peaks? Matched peaks The total absolute number of peaks matched. Additionally in brackets the total fraction of peaks matched and the total number of peaks is shown. | 44 (10.19% of 432) |
| FDR? FDR The false discovery rate estimated for this peptide. It is calculated by matching all theoretical fragments with a non-integer shift with the raw peaks for this spectrum. This is done with 40 different shifts. The resulting percentage is the average number of annotated peaks over the number of annotated peaks with the correct spectrum. | 0.76% |
| Satellite FDR? Satellite FDR See the FDR for details on its calculation. This satellite ion specific FDR only contains the satellite ions (d/w) for I/L/J positions. | - |
| PSM Score? PSM Score The PSM Score as given by Hecklib to this annotated spectrum. It is shown with three significant figures. | 522 |

#### Spectrum 8345? Spectrum 8345 The raw spectrum of this peptide as annotated by Hecklib. The fragments are coloured according to ion type (see legend). Any peaks with a star '\*' as text can be hovered over to see the full details, first the ion type second the mass shift type. By hovering over the amino acids in the peptide or ions in the legend the corresponding peaks are highlighted. By toggling the 'Unassigned' label you can turn the background (unassigned) peaks on or off in the plot. By updating the slider in the Ion legend you can update the spectrum to only show the top X% of the peaks with labels. The top X% means any peak that is within X% of the highest intensity. By dragging in the spectrum you can zoom in to a specific part of the spectrum and use 'Zoom Out' to get back to the original zoom level. The annotation of the spectrum is based on the given sequence in the peptides file and is done with different software so inconsistencies are likely. The peaks are annotated based on the given sequence, with 20 ppm tolerance.

Copy Data

##### Spectrum 8345 (TSV)

###### Preview

```
Loading example...
```

*Click on the button to copy the data to your clipboard.*

Mz MinMz MaxIntensity Max

WidthHeightPeptide font sizePeptide stroke widthSpectrum font sizeSpectrum stroke widthCompact peptide

Ion legend

wxyz

abcd

OtherUnassignedIon^Charge^_Position_Show for top:%

JVKDYFPEPVT

02.40e+44.80e+47.20e+49.60e+4

Zoom Out

y^+1^_1_y^+1^_3_y^+1^_3_c^+1^_3_c^+1^_4_c^+2^_8_y^+1^_5_c^+2^_9_c^+2^_10_c^+1^_5_c^+1^_5_y^+1^_6_c^+1^_6_y^+1^_7_c^+1^_7_c^+1^_7_y^+1^_8_c^+1^_8_z^+1^_9_y^+1^_9_c^+1^_9_z^+1^_10_c^+1^_10_c^+1^_10_

0787157523623149

Fragment Matches Table

Show background peaks

| Position | Ion type | Intensity | mz Theoretical | mz Error (Th) | mz Error (ppm) | Charge | Series Number |
| --- | --- | --- | --- | --- | --- | --- | --- |
| 11 | y | 1197 | 120.1 | 0.000139 | 1.157 | +1 | 1 |
| - | - | 364 | 120.9 | - | - | 0 | - |
| - | - | 413 | 125.7 | - | - | 0 | - |
| - | - | 552.4 | 129.1 | - | - | 0 | - |
| - | - | 392.8 | 132.2 | - | - | 0 | - |
| - | - | 424.4 | 132.8 | - | - | 0 | - |
| - | - | 438.8 | 133.1 | - | - | 0 | - |
| - | - | 392.6 | 139 | - | - | 0 | - |
| - | - | 364.2 | 143 | - | - | 0 | - |
| - | - | 410 | 144.7 | - | - | 0 | - |
| - | - | 483.9 | 148 | - | - | 0 | - |
| - | - | 404.1 | 150.8 | - | - | 0 | - |
| - | - | 417.8 | 155.7 | - | - | 0 | - |
| - | - | 393.3 | 159 | - | - | 0 | - |
| - | - | 458 | 161.1 | - | - | 0 | - |
| - | - | 3116 | 169.1 | - | - | 0 | - |
| - | - | 3499 | 173.5 | - | - | 0 | - |
| - | - | 493.4 | 181.5 | - | - | 0 | - |
| - | - | 2126 | 185.2 | - | - | 0 | - |
| - | - | 4593 | 197.1 | - | - | 0 | - |
| - | - | 473.1 | 202.2 | - | - | 0 | - |
| - | - | 517.7 | 211.4 | - | - | 0 | - |
| - | - | 2509 | 213.2 | - | - | 0 | - |
| - | - | 465.8 | 220.4 | - | - | 0 | - |
| - | - | 630.8 | 227.1 | - | - | 0 | - |
| 9 | y | 795.9 | 298.2 | 1.575E-05 | 0.05283 | +1 | 3 |
| - | - | 923.5 | 299.1 | - | - | 0 | - |
| 9 | y | 1.098E+04 | 316.2 | 0.0001129 | 0.3571 | +1 | 3 |
| - | - | 2092 | 317.2 | - | - | 0 | - |
| - | - | 2340 | 324.2 | - | - | 0 | - |
| - | - | 512.4 | 327.3 | - | - | 0 | - |
| - | - | 2985 | 355.1 | - | - | 0 | - |
| - | - | 1356 | 356.1 | - | - | 0 | - |
| - | - | 579.1 | 357.1 | - | - | 0 | - |
| - | - | 2121 | 357.3 | - | - | 0 | - |
| 3 | c | 881.8 | 358.3 | 0.005052 | 14.1 | +1 | 3 |
| - | - | 644.7 | 360 | - | - | 0 | - |
| - | - | 663.7 | 373.8 | - | - | 0 | - |
| - | - | 662.5 | 394.1 | - | - | 0 | - |
| - | - | 984.2 | 423.2 | - | - | 0 | - |
| 4 | c | 1842 | 473.3 | 0.0005923 | 1.251 | +1 | 4 |
| 8 | c | 683.6 | 496.8 | 0.0006939 | 1.397 | +2 | 8 |
| 7 | y | 4642 | 542.3 | 0.0001722 | 0.3176 | +1 | 5 |
| - | - | 1883 | 543.3 | - | - | 0 | - |
| 9 | c | 3250 | 545.3 | 0.0002069 | 0.3794 | +2 | 9 |
| - | - | 3939 | 545.8 | - | - | 0 | - |
| - | - | 930 | 546.3 | - | - | 0 | - |
| - | - | 1724 | 554.3 | - | - | 0 | - |
| - | - | 1631 | 580.8 | - | - | 0 | - |
| - | - | 1287 | 581.3 | - | - | 0 | - |
| 10 | c | 1344 | 594.8 | 0.0001186 | 0.1993 | +2 | 10 |
| - | - | 1039 | 595.3 | - | - | 0 | - |
| 5 | c | 1167 | 619.3 | 0.001178 | 1.902 | +1 | 5 |
| - | - | 1118 | 620.3 | - | - | 0 | - |
| - | - | 650.3 | 632.9 | - | - | 0 | - |
| - | - | 4892 | 635.4 | - | - | 0 | - |
| 5 | c | 5368 | 636.4 | 0.001665 | 2.616 | +1 | 5 |
| - | - | 1527 | 637.4 | - | - | 0 | - |
| - | - | 782.6 | 654.3 | - | - | 0 | - |
| - | - | 1117 | 655.4 | - | - | 0 | - |
| 6 | y | 2805 | 689.4 | 0.0006758 | 0.9803 | +1 | 6 |
| - | - | 1575 | 690.4 | - | - | 0 | - |
| - | - | 614.1 | 696.4 | - | - | 0 | - |
| - | - | 985.5 | 738.4 | - | - | 0 | - |
| - | - | 1897 | 739.4 | - | - | 0 | - |
| - | - | 2477 | 740.4 | - | - | 0 | - |
| - | - | 1315 | 741.4 | - | - | 0 | - |
| - | - | 664.6 | 749 | - | - | 0 | - |
| 6 | c | 7634 | 766.4 | 0.001355 | 1.767 | +1 | 6 |
| - | - | 3835 | 767.4 | - | - | 0 | - |
| - | - | 717.3 | 768.4 | - | - | 0 | - |
| - | - | 1459 | 780.4 | - | - | 0 | - |
| - | - | 720.1 | 851.4 | - | - | 0 | - |
| 5 | y | 2635 | 852.4 | 0.00303 | 3.555 | +1 | 7 |
| - | - | 1754 | 853.4 | - | - | 0 | - |
| 7 | c | 826.1 | 863.5 | 0.003865 | 4.476 | +1 | 7 |
| - | - | 1309 | 877.4 | - | - | 0 | - |
| - | - | 9104 | 879.5 | - | - | 0 | - |
| 7 | c | 1.527E+04 | 880.5 | 0.002542 | 2.888 | +1 | 7 |
| - | - | 7419 | 881.5 | - | - | 0 | - |
| - | - | 1728 | 882.5 | - | - | 0 | - |
| - | - | 843.9 | 934.5 | - | - | 0 | - |
| 4 | y | 2121 | 967.4 | 0.001836 | 1.898 | +1 | 8 |
| - | - | 578.3 | 974.5 | - | - | 0 | - |
| - | - | 1107 | 976.5 | - | - | 0 | - |
| 8 | c | 1.027E+04 | 992.5 | 0.001375 | 1.385 | +1 | 8 |
| - | - | 5787 | 993.5 | - | - | 0 | - |
| - | - | 1737 | 994.5 | - | - | 0 | - |
| - | - | 1245 | 1036 | - | - | 0 | - |
| - | - | 1071 | 1037 | - | - | 0 | - |
| 3 | z | 3761 | 1080 | 0.0003772 | 0.3494 | +1 | 9 |
| - | - | 2331 | 1081 | - | - | 0 | - |
| - | - | 735.8 | 1091 | - | - | 0 | - |
| 3 | y | 3184 | 1096 | 0.001157 | 1.056 | +1 | 9 |
| - | - | 986.5 | 1097 | - | - | 0 | - |
| 9 | c | 2966 | 1107 | 0.002746 | 2.481 | +1 | 9 |
| - | - | 1525 | 1108 | - | - | 0 | - |
| - | - | 878.3 | 1157 | - | - | 0 | - |
| - | - | 5451 | 1162 | - | - | 0 | - |
| - | - | 3832 | 1163 | - | - | 0 | - |
| - | - | 2884 | 1164 | - | - | 0 | - |
| 2 | z | 2316 | 1179 | 0.001643 | 1.394 | +1 | 10 |
| - | - | 1206 | 1180 | - | - | 0 | - |
| 10 | c | 900.8 | 1189 | 0.006769 | 5.694 | +1 | 10 |
| - | - | 2042 | 1191 | - | - | 0 | - |
| - | - | 1333 | 1192 | - | - | 0 | - |
| 10 | c | 2.212E+04 | 1206 | 0.0028 | 2.323 | +1 | 10 |
| - | - | 1.54E+04 | 1207 | - | - | 0 | - |
| - | - | 9008 | 1208 | - | - | 0 | - |
| - | - | 2581 | 1209 | - | - | 0 | - |
| - | - | 968.2 | 1210 | - | - | 0 | - |
| - | - | 2237 | 1237 | - | - | 0 | - |
| - | - | 1168 | 1238 | - | - | 0 | - |
| - | - | 1020 | 1245 | - | - | 0 | - |
| - | - | 746.4 | 1246 | - | - | 0 | - |
| - | - | 956.9 | 1247 | - | - | 0 | - |
| - | - | 920.9 | 1248 | - | - | 0 | - |
| - | - | 6899 | 1249 | - | - | 0 | - |
| - | - | 4392 | 1250 | - | - | 0 | - |
| - | - | 1946 | 1251 | - | - | 0 | - |
| - | - | 615.8 | 1253 | - | - | 0 | - |
| - | - | 762.1 | 1254 | - | - | 0 | - |
| - | - | 1982 | 1263 | - | - | 0 | - |
| - | - | 1380 | 1264 | - | - | 0 | - |
| - | - | 3084 | 1265 | - | - | 0 | - |
| - | - | 2190 | 1266 | - | - | 0 | - |
| - | - | 5089 | 1281 | - | - | 0 | - |
| - | - | 3449 | 1282 | - | - | 0 | - |
| - | - | 1448 | 1283 | - | - | 0 | - |
| - | - | 1354 | 1289 | - | - | 0 | - |
| - | - | 622.8 | 1290 | - | - | 0 | - |
| - | - | 2229 | 1291 | - | - | 0 | - |
| - | - | 2.565E+04 | 1292 | - | - | 0 | - |
| - | - | 1.746E+04 | 1293 | - | - | 0 | - |
| - | - | 6899 | 1294 | - | - | 0 | - |
| - | - | 1049 | 1295 | - | - | 0 | - |
| - | - | 890.1 | 1300 | - | - | 0 | - |
| - | - | 883.5 | 1305 | - | - | 0 | - |
| - | - | 2796 | 1306 | - | - | 0 | - |
| - | - | 3473 | 1307 | - | - | 0 | - |
| - | - | 2.253E+04 | 1308 | - | - | 0 | - |
| - | - | 9.509E+04 | 1309 | - | - | 0 | - |
| - | - | 6.942E+04 | 1310 | - | - | 0 | - |
| - | - | 2.679E+04 | 1311 | - | - | 0 | - |
| - | - | 3108 | 1312 | - | - | 0 | - |
| - | - | 661.7 | 1389 | - | - | 0 | - |
| - | - | 658.5 | 1526 | - | - | 0 | - |
| - | - | 588.7 | 2236 | - | - | 0 | - |
| - | - | 672.3 | 2471 | - | - | 0 | - |
| - | - | 771.2 | 3072 | - | - | 0 | - |
| - | - | 703 | 3118 | - | - | 0 | - |

m/z Charge Intensity FragmentType MassShift Position
120.06565856933594 0 1196.7666 y 10
120.90824890136719 0 364.04504
125.67642211914062 0 413.01428
129.10232543945312 0 552.4131
132.170654296875 0 392.82965
132.84454345703125 0 424.40222
133.0853729248047 0 438.75366
138.9875030517578 0 392.6089
143.04470825195312 0 364.18118
144.6851043701172 0 410.0042
148.00155639648438 0 483.92963
150.78810119628906 0 404.06912
155.718505859375 0 417.7932
159.01284790039062 0 393.3374
161.0941619873047 0 458.03687
169.1336212158203 0 3116.3413
173.45077514648438 0 3498.5085
181.50425720214844 0 493.38074
185.1648406982422 0 2126.257
197.12841796875 0 4592.6475
202.2443389892578 0 473.059
211.3523712158203 0 517.7373
213.1598358154297 0 2508.6116
220.38658142089844 0 465.8244
227.10260009765625 0 630.7638
298.1761169433594 0 795.8984 y Water loss 8
299.0622253417969 0 923.50146
316.18658447265625 0 10982.012 y 8
317.1902770996094 0 2092.291
324.1549377441406 0 2340.0298
327.32879638671875 0 512.38763
355.0693664550781 0 2984.5232
356.069580078125 0 1355.894
357.0666809082031 0 579.0771
357.27288818359375 0 2120.6787
358.2762145996094 0 881.84937 c 2
360.02777099609375 0 644.7213
373.7955017089844 0 663.6727
394.055419921875 0 662.4712
423.2241516113281 0 984.17
473.3076171875 0 1842.1656 c 3
496.75732421875 0 683.599 c Ammonia loss 7
542.2822265625 0 4642.296 y 6
543.2839965820312 0 1883.3417
545.2846069335938 0 3249.5981 c Ammonia loss 8
545.7850341796875 0 3938.9526
546.2880859375 0 929.96844
554.2607421875 0 1723.563
580.8216552734375 0 1630.8691
581.32177734375 0 1286.8347
594.8187255859375 0 1343.9833 c Ammonia loss 9
595.3203735351562 0 1038.6521
619.3438110351562 0 1167.2028 c Ammonia loss 4
620.3478393554688 0 1117.8734
632.8793334960938 0 650.32227
635.3634643554688 0 4892.268
636.369873046875 0 5368.198 c 4
637.374267578125 0 1527.175
654.3397216796875 0 782.56537
655.3543090820312 0 1117.0032
689.3497924804688 0 2804.6375 y 5
690.3511962890625 0 1575.3301
696.4290161132812 0 614.1385
738.4189453125 0 985.4971
739.4267578125 0 1897.4246
740.4310302734375 0 2477.1648
741.43701171875 0 1315.4315
748.9967041015625 0 664.6031
766.4120483398438 0 7634.247 c Ammonia loss 5
767.4143676757812 0 3835.4321
768.4129638671875 0 717.2846
780.3567504882812 0 1459.2037
851.4072875976562 0 720.0685
852.4107666015625 0 2634.5688 y 4
853.4137573242188 0 1754.3905
863.4700317382812 0 826.11066 c Ammonia loss 6
877.4120483398438 0 1308.7793
879.4834594726562 0 9104.395
880.4901733398438 0 15271.312 c 6
881.4938354492188 0 7418.8726
882.4981689453125 0 1727.5256
934.4777221679688 0 843.8994
967.4389038085938 0 2121.1934 y 3
974.4896850585938 0 578.33405
976.4810791015625 0 1106.8137
992.5073852539062 0 10267.501 c Ammonia loss 7
993.5112915039062 0 5787.1934
994.514892578125 0 1737.248
1035.51953125 0 1245.4067
1036.5291748046875 0 1070.5277
1079.5166015625 0 3761.4478 z 2
1080.521240234375 0 2331.3125
1090.6207275390625 0 735.8465
1095.5345458984375 0 3183.5088 y 2
1096.5350341796875 0 986.49066
1106.5853271484375 0 2966.0493 c 8
1107.587646484375 0 1524.9832
1156.703125 0 878.2954
1161.6414794921875 0 5451.4517
1162.6434326171875 0 3832.3372
1163.6500244140625 0 2884.403
1178.5870361328125 0 2316.099 z 1
1179.58935546875 0 1206.0789
1188.6231689453125 0 900.8456 c Ammonia loss 9
1190.64501953125 0 2042.4778
1191.647705078125 0 1333.4789
1205.6536865234375 0 22122.59 c 9
1206.6575927734375 0 15398.788
1207.6634521484375 0 9007.882
1208.6695556640625 0 2580.812
1209.6744384765625 0 968.22943
1236.6671142578125 0 2237.1387
1237.6806640625 0 1168.2402
1244.68701171875 0 1020.33374
1245.68896484375 0 746.35846
1246.683349609375 0 956.884
1247.6787109375 0 920.8579
1248.6474609375 0 6899.4185
1249.6552734375 0 4392.048
1250.6651611328125 0 1945.5048
1252.629638671875 0 615.796
1253.6453857421875 0 762.14264
1262.6851806640625 0 1981.5056
1263.6929931640625 0 1380.4557
1264.6995849609375 0 3084.3237
1265.704833984375 0 2189.6746
1280.696533203125 0 5088.819
1281.698486328125 0 3448.5327
1282.7064208984375 0 1448.3442
1288.6900634765625 0 1353.7639
1289.6845703125 0 622.7741
1290.680419921875 0 2228.9912
1291.666259765625 0 25654.889
1292.6685791015625 0 17464.574
1293.6680908203125 0 6898.979
1294.6754150390625 0 1048.8947
1300.1419677734375 0 890.0898
1304.6978759765625 0 883.5225
1305.7027587890625 0 2795.5183
1306.68310546875 0 3472.5466
1307.6839599609375 0 22530.656
1308.691162109375 0 95094.234
1309.6942138671875 0 69424.45
1310.69677734375 0 26794.33
1311.6993408203125 0 3107.6714
1388.8990478515625 0 661.6534
1526.337890625 0 658.51074
2235.70947265625 0 588.7134
2471.33544921875 0 672.281
3072.268798828125 0 771.2207
3118.2509765625 0 702.97296

Spectrum Details

|  |  |
| --- | --- |
| Matched peaks? Matched peaks The total absolute number of peaks matched. Additionally in brackets the total fraction of peaks matched and the total number of peaks is shown. | 24 (15.89% of 151) |
| FDR? FDR The false discovery rate estimated for this peptide. It is calculated by matching all theoretical fragments with a non-integer shift with the raw peaks for this spectrum. This is done with 40 different shifts. The resulting percentage is the average number of annotated peaks over the number of annotated peaks with the correct spectrum. | 1.29% |
| Satellite FDR? Satellite FDR See the FDR for details on its calculation. This satellite ion specific FDR only contains the satellite ions (d/w) for I/L/J positions. | - |
| PSM Score? PSM Score The PSM Score as given by Hecklib to this annotated spectrum. It is shown with three significant figures. | 249 |

#### Spectrum 8045? Spectrum 8045 The raw spectrum of this peptide as annotated by Hecklib. The fragments are coloured according to ion type (see legend). Any peaks with a star '\*' as text can be hovered over to see the full details, first the ion type second the mass shift type. By hovering over the amino acids in the peptide or ions in the legend the corresponding peaks are highlighted. By toggling the 'Unassigned' label you can turn the background (unassigned) peaks on or off in the plot. By updating the slider in the Ion legend you can update the spectrum to only show the top X% of the peaks with labels. The top X% means any peak that is within X% of the highest intensity. By dragging in the spectrum you can zoom in to a specific part of the spectrum and use 'Zoom Out' to get back to the original zoom level. The annotation of the spectrum is based on the given sequence in the peptides file and is done with different software so inconsistencies are likely. The peaks are annotated based on the given sequence, with 20 ppm tolerance.

Copy Data

##### Spectrum 8045 (TSV)

###### Preview

```
Loading example...
```

*Click on the button to copy the data to your clipboard.*

Mz MinMz MaxIntensity Max

WidthHeightPeptide font sizePeptide stroke widthSpectrum font sizeSpectrum stroke widthCompact peptide

Ion legend

wxyz

abcd

OtherUnassignedIon^Charge^_Position_Show for top:%

JVKDYFPEPVT

02.98e+45.97e+48.95e+41.19e+5

Zoom Out

y^+1^_1_a^+1^_2_y^+1^_2_b^+1^_2_y^+1^_2_y^+1^_3_y^+1^_3_b^+1^_3_b^+2^_6_b^+2^_7_y^+1^_4_b^+2^_7_b^+1^_4_y^+1^_4_b^+1^_4_b^+2^_8_b^+2^_8_b^+2^_8_y^+1^_5_y^+2^_9_y^+1^_5_b^+2^_9_b^+2^_10_b^+2^_10_b^+1^_5_b^+1^_5_\*\*\*y^+1^_6_y^+1^_6_b^+1^_6_b^+1^_6_b^+1^_7_b^+1^_7_b^+1^_7_y^+1^_8_b^+1^_8_b^+1^_8_b^+1^_8_y^+1^_9_b^+1^_9_y^+1^_9_b^+1^_10_b^+1^_10_b^+1^_10_y^+1^_10_

03026049061209

Fragment Matches Table

Show background peaks

| Position | Ion type | Intensity | mz Theoretical | mz Error (Th) | mz Error (ppm) | Charge | Series Number |
| --- | --- | --- | --- | --- | --- | --- | --- |
| 11 | y | 1.427E+04 | 120.1 | 0.0003449 | 2.873 | +1 | 1 |
| - | - | 2.028E+04 | 120.1 | - | - | 0 | - |
| - | - | 767 | 121.1 | - | - | 0 | - |
| - | - | 1170 | 121.1 | - | - | 0 | - |
| - | - | 352.7 | 124.6 | - | - | 0 | - |
| - | - | 833.8 | 127.1 | - | - | 0 | - |
| - | - | 470.1 | 128.1 | - | - | 0 | - |
| - | - | 799.6 | 128.1 | - | - | 0 | - |
| - | - | 499.9 | 129.1 | - | - | 0 | - |
| - | - | 3.764E+04 | 129.1 | - | - | 0 | - |
| - | - | 904.3 | 130.1 | - | - | 0 | - |
| - | - | 334 | 130.1 | - | - | 0 | - |
| - | - | 1921 | 130.1 | - | - | 0 | - |
| - | - | 634.5 | 131.1 | - | - | 0 | - |
| - | - | 893.2 | 133.1 | - | - | 0 | - |
| - | - | 1.071E+04 | 136.1 | - | - | 0 | - |
| - | - | 994.1 | 137.1 | - | - | 0 | - |
| - | - | 1253 | 139.1 | - | - | 0 | - |
| - | - | 1073 | 152.1 | - | - | 0 | - |
| - | - | 1151 | 155.1 | - | - | 0 | - |
| - | - | 1418 | 155.1 | - | - | 0 | - |
| - | - | 447.3 | 163.2 | - | - | 0 | - |
| - | - | 1027 | 165.1 | - | - | 0 | - |
| - | - | 530.2 | 166.1 | - | - | 0 | - |
| - | - | 1118 | 167.1 | - | - | 0 | - |
| - | - | 1120 | 167.1 | - | - | 0 | - |
| - | - | 553.4 | 167.2 | - | - | 0 | - |
| - | - | 1.028E+05 | 169.1 | - | - | 0 | - |
| - | - | 660.4 | 170.1 | - | - | 0 | - |
| - | - | 9589 | 170.1 | - | - | 0 | - |
| - | - | 1922 | 172.1 | - | - | 0 | - |
| - | - | 752.4 | 172.1 | - | - | 0 | - |
| - | - | 413.1 | 172.8 | - | - | 0 | - |
| - | - | 1118 | 173.1 | - | - | 0 | - |
| - | - | 2911 | 173.5 | - | - | 0 | - |
| - | - | 943 | 174.1 | - | - | 0 | - |
| - | - | 453.3 | 176.2 | - | - | 0 | - |
| - | - | 1473 | 181.1 | - | - | 0 | - |
| - | - | 1519 | 181.1 | - | - | 0 | - |
| - | - | 913.2 | 182.1 | - | - | 0 | - |
| - | - | 1427 | 183.1 | - | - | 0 | - |
| - | - | 2026 | 183.1 | - | - | 0 | - |
| - | - | 847.2 | 185.1 | - | - | 0 | - |
| 2 | a | 4.451E+04 | 185.2 | 0.0003672 | 1.983 | +1 | 2 |
| - | - | 481.9 | 186.1 | - | - | 0 | - |
| - | - | 732.9 | 186.1 | - | - | 0 | - |
| - | - | 4885 | 186.2 | - | - | 0 | - |
| - | - | 880.7 | 187.1 | - | - | 0 | - |
| - | - | 1766 | 188.1 | - | - | 0 | - |
| - | - | 894.9 | 190.1 | - | - | 0 | - |
| - | - | 537.6 | 191.1 | - | - | 0 | - |
| - | - | 2722 | 195.1 | - | - | 0 | - |
| - | - | 8.472E+04 | 197.1 | - | - | 0 | - |
| - | - | 528.7 | 198.1 | - | - | 0 | - |
| - | - | 9307 | 198.1 | - | - | 0 | - |
| - | - | 525.8 | 198.2 | - | - | 0 | - |
| - | - | 1.5E+04 | 199.1 | - | - | 0 | - |
| - | - | 1131 | 199.1 | - | - | 0 | - |
| - | - | 2806 | 199.2 | - | - | 0 | - |
| - | - | 806.9 | 200.2 | - | - | 0 | - |
| - | - | 498.8 | 200.9 | - | - | 0 | - |
| - | - | 625.8 | 201.1 | - | - | 0 | - |
| 10 | y | 8085 | 201.1 | 0.0003189 | 1.586 | +1 | 2 |
| - | - | 527.9 | 202.1 | - | - | 0 | - |
| - | - | 7024 | 203.1 | - | - | 0 | - |
| - | - | 1206 | 203.2 | - | - | 0 | - |
| - | - | 1157 | 204.1 | - | - | 0 | - |
| - | - | 610.3 | 208.1 | - | - | 0 | - |
| - | - | 4452 | 209.1 | - | - | 0 | - |
| - | - | 781.8 | 211.1 | - | - | 0 | - |
| - | - | 894.6 | 212.1 | - | - | 0 | - |
| - | - | 1299 | 213.1 | - | - | 0 | - |
| 2 | b | 1.921E+04 | 213.2 | 0.0003561 | 1.671 | +1 | 2 |
| - | - | 2731 | 214.2 | - | - | 0 | - |
| - | - | 1660 | 215.1 | - | - | 0 | - |
| - | - | 1193 | 217.1 | - | - | 0 | - |
| - | - | 2595 | 217.1 | - | - | 0 | - |
| 10 | y | 1.269E+04 | 219.1 | 0.0004048 | 1.847 | +1 | 2 |
| - | - | 761.9 | 220.1 | - | - | 0 | - |
| - | - | 622.3 | 225 | - | - | 0 | - |
| - | - | 701.1 | 225.1 | - | - | 0 | - |
| - | - | 9863 | 226.1 | - | - | 0 | - |
| - | - | 956.5 | 226.2 | - | - | 0 | - |
| - | - | 4.057E+04 | 227.1 | - | - | 0 | - |
| - | - | 579.7 | 227.2 | - | - | 0 | - |
| - | - | 484.1 | 227.4 | - | - | 0 | - |
| - | - | 5078 | 228.1 | - | - | 0 | - |
| - | - | 3284 | 228.2 | - | - | 0 | - |
| - | - | 648.4 | 229.1 | - | - | 0 | - |
| - | - | 587.3 | 230.2 | - | - | 0 | - |
| - | - | 7957 | 231.1 | - | - | 0 | - |
| - | - | 1036 | 232.2 | - | - | 0 | - |
| - | - | 669.2 | 233.1 | - | - | 0 | - |
| - | - | 1955 | 233.2 | - | - | 0 | - |
| - | - | 574.3 | 235.7 | - | - | 0 | - |
| - | - | 579.2 | 237.1 | - | - | 0 | - |
| - | - | 2182 | 238.2 | - | - | 0 | - |
| - | - | 1120 | 242.1 | - | - | 0 | - |
| - | - | 808.8 | 243.1 | - | - | 0 | - |
| - | - | 1.678E+04 | 244.1 | - | - | 0 | - |
| - | - | 4677 | 245.1 | - | - | 0 | - |
| - | - | 614 | 246.1 | - | - | 0 | - |
| - | - | 1808 | 247.1 | - | - | 0 | - |
| - | - | 1.072E+04 | 251.1 | - | - | 0 | - |
| - | - | 902.8 | 252.1 | - | - | 0 | - |
| - | - | 1707 | 254.1 | - | - | 0 | - |
| - | - | 1474 | 254.2 | - | - | 0 | - |
| - | - | 775.6 | 254.2 | - | - | 0 | - |
| - | - | 1428 | 255.1 | - | - | 0 | - |
| - | - | 918.4 | 256.2 | - | - | 0 | - |
| - | - | 1218 | 260.1 | - | - | 0 | - |
| - | - | 2109 | 261.2 | - | - | 0 | - |
| - | - | 1323 | 263.1 | - | - | 0 | - |
| - | - | 630.3 | 264 | - | - | 0 | - |
| - | - | 749.9 | 266.2 | - | - | 0 | - |
| - | - | 2682 | 272.1 | - | - | 0 | - |
| - | - | 840.8 | 278.1 | - | - | 0 | - |
| - | - | 1.174E+04 | 279.1 | - | - | 0 | - |
| - | - | 1983 | 280.1 | - | - | 0 | - |
| - | - | 2800 | 280.2 | - | - | 0 | - |
| - | - | 4851 | 283.1 | - | - | 0 | - |
| - | - | 701.8 | 284.1 | - | - | 0 | - |
| - | - | 2056 | 287.2 | - | - | 0 | - |
| - | - | 689.2 | 294.2 | - | - | 0 | - |
| - | - | 1052 | 296.2 | - | - | 0 | - |
| 9 | y | 1.941E+04 | 298.2 | 0.0005946 | 1.994 | +1 | 3 |
| - | - | 1091 | 299.1 | - | - | 0 | - |
| - | - | 2601 | 299.2 | - | - | 0 | - |
| - | - | 878 | 301.2 | - | - | 0 | - |
| - | - | 1662 | 303.2 | - | - | 0 | - |
| - | - | 1311 | 304.2 | - | - | 0 | - |
| - | - | 800.1 | 308.2 | - | - | 0 | - |
| - | - | 895.8 | 310.2 | - | - | 0 | - |
| - | - | 3705 | 311.1 | - | - | 0 | - |
| - | - | 533.7 | 312.2 | - | - | 0 | - |
| - | - | 1362 | 315.2 | - | - | 0 | - |
| 9 | y | 1.181E+05 | 316.2 | 0.000589 | 1.863 | +1 | 3 |
| - | - | 1093 | 317.1 | - | - | 0 | - |
| - | - | 1.808E+04 | 317.2 | - | - | 0 | - |
| - | - | 1884 | 318.2 | - | - | 0 | - |
| - | - | 856.3 | 323.2 | - | - | 0 | - |
| - | - | 838.7 | 323.2 | - | - | 0 | - |
| - | - | 3.413E+04 | 324.2 | - | - | 0 | - |
| - | - | 5736 | 325.2 | - | - | 0 | - |
| - | - | 1267 | 325.2 | - | - | 0 | - |
| - | - | 1133 | 326.2 | - | - | 0 | - |
| - | - | 1131 | 328.2 | - | - | 0 | - |
| - | - | 1057 | 329.2 | - | - | 0 | - |
| - | - | 926.4 | 335.2 | - | - | 0 | - |
| - | - | 1130 | 337.2 | - | - | 0 | - |
| - | - | 606.6 | 339.2 | - | - | 0 | - |
| - | - | 2501 | 340.2 | - | - | 0 | - |
| - | - | 2834 | 341.2 | - | - | 0 | - |
| 3 | b | 2133 | 341.3 | 0.0007453 | 2.184 | +1 | 3 |
| - | - | 726.2 | 342.2 | - | - | 0 | - |
| - | - | 3133 | 343.2 | - | - | 0 | - |
| - | - | 569.6 | 344.2 | - | - | 0 | - |
| - | - | 1291 | 344.7 | - | - | 0 | - |
| - | - | 1861 | 346.2 | - | - | 0 | - |
| - | - | 1410 | 347.2 | - | - | 0 | - |
| - | - | 761.8 | 347.7 | - | - | 0 | - |
| - | - | 2280 | 353.2 | - | - | 0 | - |
| - | - | 1737 | 353.2 | - | - | 0 | - |
| - | - | 1565 | 355.1 | - | - | 0 | - |
| - | - | 1494 | 355.2 | - | - | 0 | - |
| - | - | 1183 | 355.2 | - | - | 0 | - |
| - | - | 723.4 | 356.2 | - | - | 0 | - |
| - | - | 895.6 | 362.2 | - | - | 0 | - |
| - | - | 711.4 | 367.2 | - | - | 0 | - |
| - | - | 814.7 | 369.2 | - | - | 0 | - |
| - | - | 3934 | 369.7 | - | - | 0 | - |
| - | - | 1115 | 370.2 | - | - | 0 | - |
| - | - | 3897 | 371.2 | - | - | 0 | - |
| - | - | 742.8 | 372.2 | - | - | 0 | - |
| - | - | 1148 | 373.2 | - | - | 0 | - |
| - | - | 5460 | 374.2 | - | - | 0 | - |
| - | - | 1224 | 375.2 | - | - | 0 | - |
| - | - | 972.4 | 377.2 | - | - | 0 | - |
| - | - | 2057 | 379.2 | - | - | 0 | - |
| - | - | 988.1 | 380.2 | - | - | 0 | - |
| - | - | 1602 | 381.1 | - | - | 0 | - |
| 6 | b | 958.1 | 383.7 | 0.0005313 | 1.385 | +2 | 6 |
| - | - | 630.6 | 385.2 | - | - | 0 | - |
| - | - | 3744 | 389.2 | - | - | 0 | - |
| - | - | 886.4 | 390.2 | - | - | 0 | - |
| - | - | 7323 | 391.2 | - | - | 0 | - |
| - | - | 1436 | 392.2 | - | - | 0 | - |
| - | - | 748.5 | 392.2 | - | - | 0 | - |
| - | - | 571.4 | 393.2 | - | - | 0 | - |
| - | - | 3775 | 395.2 | - | - | 0 | - |
| - | - | 1007 | 396.2 | - | - | 0 | - |
| - | - | 1184 | 403.2 | - | - | 0 | - |
| - | - | 1739 | 405.2 | - | - | 0 | - |
| - | - | 872.7 | 406.2 | - | - | 0 | - |
| - | - | 7916 | 407.2 | - | - | 0 | - |
| - | - | 890 | 408.2 | - | - | 0 | - |
| - | - | 1642 | 408.2 | - | - | 0 | - |
| - | - | 852 | 409.2 | - | - | 0 | - |
| - | - | 612.1 | 417.2 | - | - | 0 | - |
| - | - | 5725 | 418.2 | - | - | 0 | - |
| - | - | 2340 | 418.7 | - | - | 0 | - |
| - | - | 918.7 | 421.2 | - | - | 0 | - |
| 7 | b | 1.228E+04 | 423.2 | 0.006738 | 15.92 | +2 | 7 |
| - | - | 5048 | 424.2 | - | - | 0 | - |
| - | - | 1.111E+04 | 426.2 | - | - | 0 | - |
| - | - | 3147 | 427.2 | - | - | 0 | - |
| 8 | y | 1320 | 427.2 | 0.0004515 | 1.057 | +1 | 4 |
| 7 | b | 4685 | 432.2 | 0.0008578 | 1.984 | +2 | 7 |
| - | - | 2617 | 432.7 | - | - | 0 | - |
| - | - | 786.7 | 433.2 | - | - | 0 | - |
| - | - | 703.3 | 434.2 | - | - | 0 | - |
| - | - | 1991 | 435.2 | - | - | 0 | - |
| - | - | 1265 | 436.2 | - | - | 0 | - |
| 4 | b | 4661 | 438.3 | 0.0008769 | 2.001 | +1 | 4 |
| - | - | 1270 | 438.3 | - | - | 0 | - |
| - | - | 983.5 | 439.2 | - | - | 0 | - |
| - | - | 908.7 | 439.3 | - | - | 0 | - |
| - | - | 695.4 | 439.3 | - | - | 0 | - |
| - | - | 1008 | 440.3 | - | - | 0 | - |
| 8 | y | 4093 | 445.2 | 0.001636 | 3.675 | +1 | 4 |
| - | - | 2988 | 452.2 | - | - | 0 | - |
| 4 | b | 1.55E+04 | 456.3 | 0.0007798 | 1.709 | +1 | 4 |
| - | - | 1226 | 457.2 | - | - | 0 | - |
| - | - | 4255 | 457.3 | - | - | 0 | - |
| - | - | 4939 | 470.2 | - | - | 0 | - |
| - | - | 594.8 | 471.2 | - | - | 0 | - |
| - | - | 2476 | 471.2 | - | - | 0 | - |
| - | - | 696.5 | 472.2 | - | - | 0 | - |
| - | - | 803.1 | 478.3 | - | - | 0 | - |
| - | - | 3348 | 482.3 | - | - | 0 | - |
| - | - | 2606 | 482.8 | - | - | 0 | - |
| - | - | 2298 | 483.3 | - | - | 0 | - |
| 8 | b | 934.2 | 487.8 | 0.002086 | 4.277 | +2 | 8 |
| 8 | b | 1612 | 488.2 | 0.005958 | 12.2 | +2 | 8 |
| - | - | 1804 | 488.3 | - | - | 0 | - |
| - | - | 1422 | 488.7 | - | - | 0 | - |
| - | - | 890.3 | 489.2 | - | - | 0 | - |
| 8 | b | 9012 | 496.8 | 0.0007099 | 1.429 | +2 | 8 |
| - | - | 5105 | 497.3 | - | - | 0 | - |
| - | - | 1665 | 497.8 | - | - | 0 | - |
| - | - | 765.1 | 498.2 | - | - | 0 | - |
| - | - | 2108 | 506.3 | - | - | 0 | - |
| - | - | 1683 | 508.3 | - | - | 0 | - |
| - | - | 2312 | 509.2 | - | - | 0 | - |
| - | - | 1234 | 510.2 | - | - | 0 | - |
| - | - | 1180 | 518.3 | - | - | 0 | - |
| - | - | 951.1 | 521.3 | - | - | 0 | - |
| - | - | 3289 | 522.3 | - | - | 0 | - |
| - | - | 5157 | 522.8 | - | - | 0 | - |
| - | - | 1229 | 523.2 | - | - | 0 | - |
| - | - | 2993 | 523.3 | - | - | 0 | - |
| - | - | 814.1 | 523.8 | - | - | 0 | - |
| - | - | 1555 | 524.2 | - | - | 0 | - |
| 7 | y | 4528 | 524.3 | 0.001093 | 2.085 | +1 | 5 |
| - | - | 1387 | 525.3 | - | - | 0 | - |
| - | - | 6895 | 526.3 | - | - | 0 | - |
| - | - | 1877 | 527.3 | - | - | 0 | - |
| - | - | 2617 | 531.3 | - | - | 0 | - |
| - | - | 1117 | 531.8 | - | - | 0 | - |
| - | - | 631.8 | 532.3 | - | - | 0 | - |
| - | - | 4462 | 534.3 | - | - | 0 | - |
| - | - | 1336 | 535.3 | - | - | 0 | - |
| - | - | 4400 | 536.3 | - | - | 0 | - |
| - | - | 1037 | 536.3 | - | - | 0 | - |
| - | - | 660.6 | 537 | - | - | 0 | - |
| - | - | 964.2 | 537.2 | - | - | 0 | - |
| - | - | 759.4 | 538.2 | - | - | 0 | - |
| - | - | 2092 | 539.3 | - | - | 0 | - |
| 3 | y | 784.2 | 539.8 | 0.007349 | 13.62 | +2 | 9 |
| - | - | 1033 | 540.3 | - | - | 0 | - |
| 7 | y | 3.957E+04 | 542.3 | 0.0008436 | 1.556 | +1 | 5 |
| - | - | 1.155E+04 | 543.3 | - | - | 0 | - |
| - | - | 1885 | 544.3 | - | - | 0 | - |
| 9 | b | 2.607E+04 | 545.3 | 0.0005121 | 0.9391 | +2 | 9 |
| - | - | 1.749E+04 | 545.8 | - | - | 0 | - |
| - | - | 6921 | 546.3 | - | - | 0 | - |
| - | - | 1107 | 546.8 | - | - | 0 | - |
| - | - | 741.9 | 549.3 | - | - | 0 | - |
| - | - | 1720 | 553.3 | - | - | 0 | - |
| - | - | 3.27E+04 | 554.3 | - | - | 0 | - |
| - | - | 1.116E+04 | 555.3 | - | - | 0 | - |
| - | - | 1509 | 556.3 | - | - | 0 | - |
| - | - | 2636 | 558.3 | - | - | 0 | - |
| - | - | 649.1 | 559.3 | - | - | 0 | - |
| - | - | 2932 | 567.4 | - | - | 0 | - |
| - | - | 2578 | 570.3 | - | - | 0 | - |
| - | - | 685.3 | 571.3 | - | - | 0 | - |
| - | - | 811.2 | 572.8 | - | - | 0 | - |
| - | - | 988.9 | 575.3 | - | - | 0 | - |
| - | - | 8912 | 580.8 | - | - | 0 | - |
| - | - | 8037 | 581.3 | - | - | 0 | - |
| - | - | 2632 | 581.8 | - | - | 0 | - |
| - | - | 1812 | 582.3 | - | - | 0 | - |
| - | - | 627.2 | 583.3 | - | - | 0 | - |
| 10 | b | 942 | 585.8 | 0.0005791 | 0.9886 | +2 | 10 |
| - | - | 6173 | 591.4 | - | - | 0 | - |
| - | - | 2456 | 592.4 | - | - | 0 | - |
| 10 | b | 5686 | 594.8 | 0.0006069 | 1.02 | +2 | 10 |
| - | - | 4407 | 595.3 | - | - | 0 | - |
| - | - | 1600 | 595.8 | - | - | 0 | - |
| - | - | 1727 | 597.3 | - | - | 0 | - |
| - | - | 613.9 | 598.9 | - | - | 0 | - |
| 5 | b | 1182 | 601.3 | 0.001147 | 1.907 | +1 | 5 |
| - | - | 1666 | 603.4 | - | - | 0 | - |
| - | - | 1278 | 606.4 | - | - | 0 | - |
| - | - | 1096 | 619.3 | - | - | 0 | - |
| 5 | b | 2.163E+04 | 619.3 | 0.000348 | 0.5618 | +1 | 5 |
| - | - | 7926 | 620.3 | - | - | 0 | - |
| - | - | 1521 | 621.4 | - | - | 0 | - |
| - | - | 828.5 | 622.8 | - | - | 0 | - |
| - | - | 1523 | 623.3 | - | - | 0 | - |
| - | - | 712.4 | 624.3 | - | - | 0 | - |
| - | - | 1416 | 625.3 | - | - | 0 | - |
| - | - | 761.3 | 626.3 | - | - | 0 | - |
| - | - | 1633 | 633.3 | - | - | 0 | - |
| - | - | 912 | 635.3 | - | - | 0 | - |
| - | - | 2210 | 635.4 | - | - | 0 | - |
| - | - | 1796 | 636.4 | - | - | 0 | - |
| 0 | Precursor | 3095 | 645.3 | 0.001077 | 1.668 | +2 | -1 |
| 0 | Precursor | 3207 | 645.8 | 0.008869 | 13.73 | +2 | -1 |
| - | - | 1462 | 646.3 | - | - | 0 | - |
| - | - | 3424 | 651.3 | - | - | 0 | - |
| - | - | 2249 | 652.3 | - | - | 0 | - |
| - | - | 1198 | 652.3 | - | - | 0 | - |
| - | - | 1093 | 652.4 | - | - | 0 | - |
| - | - | 1486 | 652.9 | - | - | 0 | - |
| - | - | 1351 | 653.3 | - | - | 0 | - |
| - | - | 4955 | 653.3 | - | - | 0 | - |
| - | - | 755.4 | 653.9 | - | - | 0 | - |
| 0 | Precursor | 4614 | 654.3 | 0.004345 | 6.64 | +2 | -1 |
| - | - | 2141 | 654.9 | - | - | 0 | - |
| - | - | 1864 | 655.3 | - | - | 0 | - |
| - | - | 1595 | 663.3 | - | - | 0 | - |
| - | - | 1114 | 667.3 | - | - | 0 | - |
| 6 | y | 774 | 671.3 | 0.0004895 | 0.7291 | +1 | 6 |
| - | - | 926.5 | 680 | - | - | 0 | - |
| - | - | 4691 | 681.3 | - | - | 0 | - |
| - | - | 1515 | 682.3 | - | - | 0 | - |
| - | - | 5146 | 682.4 | - | - | 0 | - |
| - | - | 2376 | 683.4 | - | - | 0 | - |
| - | - | 703.4 | 684.4 | - | - | 0 | - |
| - | - | 688.3 | 688.4 | - | - | 0 | - |
| 6 | y | 1.503E+04 | 689.4 | 0.0002398 | 0.3478 | +1 | 6 |
| - | - | 5652 | 690.4 | - | - | 0 | - |
| - | - | 1072 | 691.4 | - | - | 0 | - |
| - | - | 2959 | 716.4 | - | - | 0 | - |
| - | - | 1369 | 717.4 | - | - | 0 | - |
| - | - | 2091 | 721.4 | - | - | 0 | - |
| - | - | 1148 | 722.4 | - | - | 0 | - |
| - | - | 806.5 | 734.4 | - | - | 0 | - |
| - | - | 8008 | 738.4 | - | - | 0 | - |
| - | - | 3626 | 739.4 | - | - | 0 | - |
| 6 | b | 2972 | 748.4 | 0.001044 | 1.395 | +1 | 6 |
| - | - | 1270 | 749.4 | - | - | 0 | - |
| - | - | 849.5 | 750.3 | - | - | 0 | - |
| - | - | 1101 | 750.4 | - | - | 0 | - |
| - | - | 3759 | 752.4 | - | - | 0 | - |
| - | - | 1894 | 753.4 | - | - | 0 | - |
| - | - | 2437 | 762.3 | - | - | 0 | - |
| - | - | 1217 | 763.3 | - | - | 0 | - |
| 6 | b | 7.906E+04 | 766.4 | 0.0004155 | 0.5421 | +1 | 6 |
| - | - | 3.701E+04 | 767.4 | - | - | 0 | - |
| - | - | 8340 | 768.4 | - | - | 0 | - |
| - | - | 747.1 | 778.4 | - | - | 0 | - |
| - | - | 2.861E+04 | 780.4 | - | - | 0 | - |
| - | - | 1.149E+04 | 781.4 | - | - | 0 | - |
| - | - | 2843 | 782.4 | - | - | 0 | - |
| - | - | 952.6 | 784.4 | - | - | 0 | - |
| - | - | 2663 | 796 | - | - | 0 | - |
| - | - | 5832 | 796.4 | - | - | 0 | - |
| - | - | 3434 | 796.7 | - | - | 0 | - |
| - | - | 1340 | 797 | - | - | 0 | - |
| - | - | 670.8 | 817.5 | - | - | 0 | - |
| - | - | 1413 | 820.5 | - | - | 0 | - |
| - | - | 828.4 | 821.5 | - | - | 0 | - |
| - | - | 719.8 | 827.4 | - | - | 0 | - |
| - | - | 1270 | 829.5 | - | - | 0 | - |
| - | - | 720 | 831.4 | - | - | 0 | - |
| - | - | 1381 | 835.5 | - | - | 0 | - |
| 7 | b | 2362 | 845.5 | 0.01462 | 17.3 | +1 | 7 |
| 7 | b | 941.3 | 846.4 | 0.0008731 | 1.032 | +1 | 7 |
| - | - | 751.9 | 848.4 | - | - | 0 | - |
| - | - | 1105 | 850.4 | - | - | 0 | - |
| - | - | 1634 | 851.4 | - | - | 0 | - |
| - | - | 745.7 | 863.4 | - | - | 0 | - |
| 7 | b | 5308 | 863.5 | 0.001424 | 1.649 | +1 | 7 |
| - | - | 2911 | 864.5 | - | - | 0 | - |
| - | - | 9945 | 877.4 | - | - | 0 | - |
| - | - | 4616 | 878.4 | - | - | 0 | - |
| - | - | 4771 | 879.4 | - | - | 0 | - |
| - | - | 1258 | 880.4 | - | - | 0 | - |
| - | - | 940.3 | 893.4 | - | - | 0 | - |
| - | - | 2273 | 907.4 | - | - | 0 | - |
| - | - | 1089 | 908.4 | - | - | 0 | - |
| - | - | 1549 | 947.5 | - | - | 0 | - |
| - | - | 2194 | 948.5 | - | - | 0 | - |
| - | - | 1199 | 958.5 | - | - | 0 | - |
| - | - | 1464 | 962.5 | - | - | 0 | - |
| - | - | 700 | 963.5 | - | - | 0 | - |
| - | - | 9588 | 964.5 | - | - | 0 | - |
| - | - | 6062 | 965.5 | - | - | 0 | - |
| - | - | 1658 | 966.5 | - | - | 0 | - |
| 4 | y | 795.6 | 967.4 | 0.004084 | 4.222 | +1 | 8 |
| 8 | b | 4760 | 974.5 | 0.00204 | 2.094 | +1 | 8 |
| 8 | b | 2574 | 975.5 | 0.017 | 17.42 | +1 | 8 |
| - | - | 1.126E+04 | 976.5 | - | - | 0 | - |
| - | - | 3985 | 977.5 | - | - | 0 | - |
| - | - | 1696 | 978.5 | - | - | 0 | - |
| - | - | 940.9 | 990.5 | - | - | 0 | - |
| 8 | b | 9.889E+04 | 992.5 | 2.926E-05 | 0.02948 | +1 | 8 |
| - | - | 5.618E+04 | 993.5 | - | - | 0 | - |
| - | - | 1.73E+04 | 994.5 | - | - | 0 | - |
| - | - | 1510 | 995.5 | - | - | 0 | - |
| - | - | 914.8 | 1033 | - | - | 0 | - |
| - | - | 840.7 | 1044 | - | - | 0 | - |
| - | - | 838.6 | 1045 | - | - | 0 | - |
| - | - | 2330 | 1047 | - | - | 0 | - |
| - | - | 1350 | 1048 | - | - | 0 | - |
| 3 | y | 1221 | 1078 | 0.008187 | 7.598 | +1 | 9 |
| 9 | b | 2622 | 1090 | 0.00122 | 1.12 | +1 | 9 |
| - | - | 3420 | 1091 | - | - | 0 | - |
| - | - | 796.4 | 1092 | - | - | 0 | - |
| 3 | y | 2.126E+04 | 1096 | 0.0001804 | 0.1646 | +1 | 9 |
| - | - | 1.38E+04 | 1097 | - | - | 0 | - |
| - | - | 4749 | 1098 | - | - | 0 | - |
| - | - | 653.3 | 1161 | - | - | 0 | - |
| 10 | b | 990.4 | 1171 | 0.01158 | 9.896 | +1 | 10 |
| 10 | b | 709.6 | 1172 | 0.02173 | 18.55 | +1 | 10 |
| 10 | b | 6303 | 1189 | 0.002008 | 1.689 | +1 | 10 |
| - | - | 4456 | 1190 | - | - | 0 | - |
| - | - | 1651 | 1191 | - | - | 0 | - |
| 2 | y | 1664 | 1195 | 0.002695 | 2.256 | +1 | 10 |
| - | - | 783.6 | 1196 | - | - | 0 | - |
| - | - | 916.2 | 1197 | - | - | 0 | - |

m/z Charge Intensity FragmentType MassShift Position
120.06586456298828 0 14267.479 y 10
120.08113098144531 0 20279.684
121.06924438476562 0 767.02014
121.08451080322266 0 1170.1816
124.55619812011719 0 352.70486
127.08702850341797 0 833.7818
128.0710906982422 0 470.11212
128.10751342773438 0 799.641
129.06634521484375 0 499.90027
129.1025848388672 0 37635.11
130.0500030517578 0 904.34015
130.10018920898438 0 333.96753
130.10589599609375 0 1920.8606
131.07054138183594 0 634.48474
133.08644104003906 0 893.2044
136.07606506347656 0 10705.314
137.07931518554688 0 994.05585
139.08718872070312 0 1252.6398
152.07118225097656 0 1073.4707
155.08192443847656 0 1150.5557
155.11827087402344 0 1417.9641
163.22418212890625 0 447.34393
165.10267639160156 0 1027.221
166.05300903320312 0 530.2317
167.0819854736328 0 1117.906
167.1178436279297 0 1119.6265
167.15512084960938 0 553.35504
169.13394165039062 0 102784.64
170.13059997558594 0 660.41864
170.1372833251953 0 9588.796
172.11251831054688 0 1922.076
172.1449432373047 0 752.44086
172.82212829589844 0 413.11566
173.1288604736328 0 1118.2164
173.4518585205078 0 2911.4055
174.1277313232422 0 942.954
176.2356414794922 0 453.26465
181.09764099121094 0 1473.4384
181.1338653564453 0 1519.0541
182.0816650390625 0 913.1907
183.11314392089844 0 1427.4761
183.14947509765625 0 2025.6226
185.09243774414062 0 847.2293
185.1652069091797 0 44513.32 a 1
186.08839416503906 0 481.90106
186.12815856933594 0 732.9313
186.16859436035156 0 4885.138
187.14418029785156 0 880.69745
188.14361572265625 0 1766.167
190.12290954589844 0 894.8899
191.08200073242188 0 537.56775
195.11305236816406 0 2722.238
197.12879943847656 0 84723.46
198.1236572265625 0 528.68225
198.13226318359375 0 9307.3125
198.23802185058594 0 525.83026
199.10800170898438 0 14996.674
199.13485717773438 0 1130.9783
199.1809844970703 0 2806.0195
200.18389892578125 0 806.94037
200.89955139160156 0 498.82166
201.11338806152344 0 625.78064
201.12368774414062 0 8085.0825 y Water loss 9
202.12710571289062 0 527.9107
203.1029815673828 0 7024.31
203.15426635742188 0 1206.4883
204.1383056640625 0 1156.774
208.1081085205078 0 610.2582
209.09234619140625 0 4451.899
211.14512634277344 0 781.76953
212.13951110839844 0 894.60706
213.1241455078125 0 1299.306
213.1601104736328 0 19212.807 b 1
214.16351318359375 0 2730.6323
215.13931274414062 0 1660.3131
217.0819091796875 0 1193.1196
217.1339569091797 0 2594.6733
219.13433837890625 0 12693.15 y 9
220.1379852294922 0 761.94867
225.04360961914062 0 622.2773
225.0874786376953 0 701.1488
226.11903381347656 0 9862.865
226.1553192138672 0 956.4549
227.10305786132812 0 40574.33
227.1751251220703 0 579.70514
227.44772338867188 0 484.10764
228.10617065429688 0 5078.236
228.17103576660156 0 3284.3674
229.11935424804688 0 648.4464
230.18592834472656 0 587.3175
231.14964294433594 0 7956.757
232.15281677246094 0 1035.9972
233.0926055908203 0 669.24023
233.16519165039062 0 1955.1315
235.66671752929688 0 574.2944
237.08673095703125 0 579.1931
238.15528869628906 0 2181.7295
242.11471557617188 0 1119.8318
243.0977783203125 0 808.7696
244.12962341308594 0 16776.21
245.13003540039062 0 4677.223
246.1328582763672 0 613.9898
247.14439392089844 0 1807.9288
251.10302734375 0 10715.823
252.10659790039062 0 902.80475
254.11390686035156 0 1707.2388
254.15003967285156 0 1474.1337
254.18710327148438 0 775.564
255.09811401367188 0 1427.5016
256.1662292480469 0 918.3951
260.1239318847656 0 1218.0541
261.1581726074219 0 2108.798
263.1393737792969 0 1323.1608
264.0174560546875 0 630.27
266.15008544921875 0 749.88074
272.12445068359375 0 2681.757
278.14984130859375 0 840.8363
279.0980529785156 0 11737.747
280.1012878417969 0 1982.7971
280.1662902832031 0 2799.8596
283.14453125 0 4850.635
284.1472473144531 0 701.794
287.2123107910156 0 2055.7336
294.1813049316406 0 689.2275
296.1973571777344 0 1052.1708
298.1767272949219 0 19406.56 y Water loss 8
299.13702392578125 0 1091.4574
299.1795654296875 0 2600.7327
301.1551818847656 0 878.0011
303.207275390625 0 1661.5173
304.16619873046875 0 1310.8315
308.16064453125 0 800.14594
310.2134094238281 0 895.80646
311.14007568359375 0 3705.3083
312.1916809082031 0 533.6585
315.207275390625 0 1361.6592
316.1872863769531 0 118136.24 y 8
317.1461486816406 0 1092.8734
317.1903076171875 0 18075.957
318.1920166015625 0 1883.6752
323.1726989746094 0 856.3362
323.2445983886719 0 838.7138
324.1560974121094 0 34130.836
325.15911865234375 0 5735.7627
325.1876525878906 0 1266.9215
326.1715393066406 0 1132.5267
328.166748046875 0 1130.5173
329.15020751953125 0 1057.1155
335.173583984375 0 926.442
337.1517639160156 0 1130.1412
339.2030029296875 0 606.62683
340.18756103515625 0 2500.7837
341.1826477050781 0 2834.1248
341.2554626464844 0 2133.0115 b 2
342.1859130859375 0 726.23615
343.19805908203125 0 3133.041
344.2011413574219 0 569.55945
344.7051086425781 0 1290.829
346.1766357421875 0 1860.7831
347.2027893066406 0 1409.8535
347.70343017578125 0 761.81934
353.18194580078125 0 2280.144
353.2185363769531 0 1737.2642
355.07086181640625 0 1565.1233
355.161376953125 0 1493.9426
355.1981201171875 0 1182.9661
356.1640930175781 0 723.4011
362.17120361328125 0 895.58704
367.1966857910156 0 711.4297
369.17706298828125 0 814.7211
369.7127380371094 0 3933.6895
370.214599609375 0 1114.9624
371.1927795410156 0 3896.7004
372.1945495605469 0 742.79785
373.1871032714844 0 1147.6245
374.1714172363281 0 5459.9062
375.17462158203125 0 1223.7732
377.2198791503906 0 972.3915
379.19866943359375 0 2056.5085
380.2017822265625 0 988.12805
381.1455993652344 0 1602.3313
383.7098083496094 0 958.12897 b 5
385.2196960449219 0 630.5659
389.18212890625 0 3744.3552
390.1650695800781 0 886.3849
391.1978759765625 0 7323.445
392.18023681640625 0 1436.3243
392.2048645019531 0 748.5336
393.2497253417969 0 571.3526
395.22918701171875 0 3775.4854
396.2325744628906 0 1006.5065
403.2330627441406 0 1184.4015
405.2142639160156 0 1738.6443
406.21417236328125 0 872.6901
407.1933898925781 0 7915.9053
408.1567077636719 0 889.98773
408.1961364746094 0 1642.164
409.2088928222656 0 852.009
417.1773986816406 0 612.1175
418.23980712890625 0 5724.8623
418.7415466308594 0 2339.8003
421.245361328125 0 918.7364
423.2247009277344 0 12283.899 b Water loss 6
424.225341796875 0 5048.45
426.1667175292969 0 11105.65
427.169189453125 0 3146.814
427.21917724609375 0 1320.0032 y Water loss 7
432.2375793457031 0 4685.1113 b 6
432.7391052246094 0 2617.4124
433.2401123046875 0 786.73596
434.20477294921875 0 703.2705
435.18902587890625 0 1990.8049
436.18768310546875 0 1265.0317
438.27197265625 0 4660.9937 b Water loss 3
438.3074035644531 0 1270.1812
439.2071228027344 0 983.54205
439.27862548828125 0 908.6681
439.3114318847656 0 695.43115
440.2514343261719 0 1008.2786
445.2309265136719 0 4092.8176 y 7
452.214599609375 0 2988.3384
456.2824401855469 0 15499.206 b 3
457.2478332519531 0 1225.6183
457.28570556640625 0 4255.112
470.2255859375 0 4938.966
471.18878173828125 0 594.7959
471.22509765625 0 2476.101
472.22271728515625 0 696.5269
478.2680358886719 0 803.14185
482.2625427246094 0 3347.7107
482.76068115234375 0 2606.2869
483.26361083984375 0 2298.3806
487.75482177734375 0 934.19025 b Water loss 7
488.2507019042969 0 1611.8107 b Ammonia loss 7
488.3231506347656 0 1804.4941
488.7448425292969 0 1421.6815
489.2396545410156 0 890.3298
496.75872802734375 0 9012.078 b 7
497.26019287109375 0 5104.5566
497.7615661621094 0 1665.1844
498.21893310546875 0 765.1033
506.2621154785156 0 2107.742
508.2558288574219 0 1682.9366
509.24041748046875 0 2311.5193
510.24188232421875 0 1234.428
518.262451171875 0 1179.8702
521.2755126953125 0 951.11096
522.2854614257812 0 3289.3508
522.7852783203125 0 5156.882
523.2181396484375 0 1228.6239
523.2864379882812 0 2992.532
523.7854614257812 0 814.13025
524.2202758789062 0 1555.1677
524.2725830078125 0 4527.6284 y Water loss 6
525.27392578125 0 1387.3998
526.26708984375 0 6894.6343
527.269287109375 0 1876.6635
531.286865234375 0 2616.7144
531.7899169921875 0 1117.2997
532.2861328125 0 631.75146
534.2567138671875 0 4461.538
535.2581787109375 0 1335.8853
536.2515869140625 0 4399.936
536.3109130859375 0 1037.0878
537.040283203125 0 660.60114
537.2432861328125 0 964.20465
538.2391357421875 0 759.4205
539.25341796875 0 2092.1414
539.7655639648438 0 784.2118 y Ammonia loss 2
540.2548217773438 0 1033.1343
542.2828979492188 0 39566.13 y 6
543.2861328125 0 11546.279
544.2880249023438 0 1884.7238
545.284912109375 0 26067.9 b 8
545.7862548828125 0 17490.766
546.2882690429688 0 6921.3594
546.790283203125 0 1106.7189
549.341552734375 0 741.9491
553.33544921875 0 1719.6586
554.261474609375 0 32698
555.2646484375 0 11158.357
556.2665405273438 0 1508.9387
558.3291015625 0 2636.239
559.33154296875 0 649.0973
567.3502197265625 0 2932.2256
570.2925415039062 0 2577.6184
571.29345703125 0 685.34534
572.8102416992188 0 811.2222
575.3265991210938 0 988.9159
580.822021484375 0 8911.9
581.3228759765625 0 8036.683
581.8251342773438 0 2632.2927
582.2564697265625 0 1811.7992
583.2579345703125 0 627.1575
585.8139038085938 0 941.97375 b Water loss 9
591.3506469726562 0 6172.911
592.3546752929688 0 2456.231
594.8192138671875 0 5685.9517 b 9
595.3206176757812 0 4406.761
595.8226928710938 0 1600.1755
597.2883911132812 0 1726.6228
598.9346923828125 0 613.93585
601.3355712890625 0 1181.7285 b Water loss 4
603.3506469726562 0 1665.8665
606.3665161132812 0 1277.6992
619.286865234375 0 1095.832
619.3453369140625 0 21630.715 b 4
620.3482666015625 0 7926.212
621.3504638671875 0 1520.5088
622.8431396484375 0 828.53937
623.3204345703125 0 1523.3057
624.3177490234375 0 712.38306
625.3350830078125 0 1415.7495
626.338134765625 0 761.2776
633.2908325195312 0 1632.8021
635.3204345703125 0 912.03217
635.375732421875 0 2210.3813
636.3775024414062 0 1796.4877
645.3413696289062 0 3094.8523 Precursor Water loss
645.8433227539062 0 3206.8416 Precursor Ammonia loss
646.3447265625 0 1461.5995
651.3153686523438 0 3424.4976
652.2607421875 0 2249.0952
652.3182983398438 0 1197.5278
652.3989868164062 0 1093.1877
652.8619384765625 0 1486.108
653.263427734375 0 1351.1821
653.3300170898438 0 4955.297
653.8597412109375 0 755.392
654.3433837890625 0 4614.015 Precursor
654.8505249023438 0 2140.869
655.3492431640625 0 1864.4014
663.3128662109375 0 1594.6547
667.3455810546875 0 1114.0349
671.3403930664062 0 774.0304 y Water loss 5
679.986083984375 0 926.54224
681.3250122070312 0 4691.094
682.3252563476562 0 1515.193
682.3787841796875 0 5145.521
683.3807983398438 0 2375.8467
684.3836059570312 0 703.3918
688.4090576171875 0 688.2915
689.3507080078125 0 15033.609 y 5
690.3541870117188 0 5652.4976
691.35205078125 0 1071.9897
716.3978271484375 0 2958.5242
717.4028930664062 0 1368.7141
721.392822265625 0 2090.5437
722.3907470703125 0 1147.8373
734.3533325195312 0 806.4639
738.418212890625 0 8007.9697
739.4224243164062 0 3625.5242
748.4017944335938 0 2972.3 b Water loss 5
749.4053955078125 0 1270.3779
750.3112182617188 0 849.4635
750.385986328125 0 1101.0232
752.3634033203125 0 3758.93
753.3661499023438 0 1894.4244
762.3450317382812 0 2436.6477
763.3458251953125 0 1217.4498
766.413818359375 0 79060.33 b 5
767.4165649414062 0 37012.13
768.4199829101562 0 8340.384
778.3798828125 0 747.1242
780.35693359375 0 28612.047
781.35986328125 0 11491.678
782.3630981445312 0 2843.3035
784.4266967773438 0 952.58594
796.04638671875 0 2662.9639
796.380615234375 0 5832.3813
796.7132568359375 0 3434.1763
797.0471801757812 0 1339.9122
817.4520263671875 0 670.81055
820.4603271484375 0 1413.366
821.46142578125 0 828.38367
827.4296875 0 719.84784
829.4505615234375 0 1269.7045
831.427978515625 0 719.9928
835.4743041992188 0 1380.754
845.4409790039062 0 2362.3035 b Water loss 6
846.4404907226562 0 941.27014 b Ammonia loss 6
848.381591796875 0 751.9153
850.431884765625 0 1104.8132
851.4417724609375 0 1633.6511
863.4095458984375 0 745.7107
863.4675903320312 0 5307.6597 b 6
864.4696655273438 0 2910.707
877.408935546875 0 9944.873
878.4129638671875 0 4615.65
879.4203491210938 0 4771.251
880.4271850585938 0 1258.354
893.4419555664062 0 940.3169
907.420654296875 0 2272.6309
908.4182739257812 0 1088.6744
947.4971313476562 0 1549.214
948.4854125976562 0 2193.6675
958.4682006835938 0 1199.4376
962.5317993164062 0 1463.6803
963.5418701171875 0 700.0061
964.5140380859375 0 9587.661
965.5173950195312 0 6062.2197
966.5177001953125 0 1658.1024
967.44482421875 0 795.62915 y 3
974.4961547851562 0 4759.793 b Water loss 7
975.4992065429688 0 2574.4868 b Ammonia loss 7
976.4786987304688 0 11256.145
977.4832763671875 0 3985.4536
978.4818725585938 0 1696.4471
990.4977416992188 0 940.89514
992.5087890625 0 98889.59 b 7
993.5118408203125 0 56177.19
994.5146484375 0 17302.424
995.5166625976562 0 1510.3782
1033.499755859375 0 914.8317
1043.554931640625 0 840.7388
1044.55517578125 0 838.64386
1046.5560302734375 0 2329.5127
1047.5570068359375 0 1349.957
1077.5333251953125 0 1221.4598 y Water loss 2
1089.562744140625 0 2622.0745 b 8
1090.531494140625 0 3420.4312
1091.5291748046875 0 796.3915
1095.5355224609375 0 21258.213 y 2
1096.53759765625 0 13799.093
1097.544189453125 0 4749.1753
1160.6353759765625 0 653.2646
1170.6077880859375 0 990.4216 b Water loss 9
1171.6251220703125 0 709.58405 b Ammonia loss 9
1188.6279296875 0 6303.1914 b 9
1189.63232421875 0 4455.5483
1190.63134765625 0 1651.4403
1194.6068115234375 0 1664.1628 y 1
1195.60107421875 0 783.6272
1196.6190185546875 0 916.17865

Spectrum Details

|  |  |
| --- | --- |
| Matched peaks? Matched peaks The total absolute number of peaks matched. Additionally in brackets the total fraction of peaks matched and the total number of peaks is shown. | 47 (10.83% of 434) |
| FDR? FDR The false discovery rate estimated for this peptide. It is calculated by matching all theoretical fragments with a non-integer shift with the raw peaks for this spectrum. This is done with 40 different shifts. The resulting percentage is the average number of annotated peaks over the number of annotated peaks with the correct spectrum. | 0.86% |
| Satellite FDR? Satellite FDR See the FDR for details on its calculation. This satellite ion specific FDR only contains the satellite ions (d/w) for I/L/J positions. | - |
| PSM Score? PSM Score The PSM Score as given by Hecklib to this annotated spectrum. It is shown with three significant figures. | 544 |

#### Spectrum 8221? Spectrum 8221 The raw spectrum of this peptide as annotated by Hecklib. The fragments are coloured according to ion type (see legend). Any peaks with a star '\*' as text can be hovered over to see the full details, first the ion type second the mass shift type. By hovering over the amino acids in the peptide or ions in the legend the corresponding peaks are highlighted. By toggling the 'Unassigned' label you can turn the background (unassigned) peaks on or off in the plot. By updating the slider in the Ion legend you can update the spectrum to only show the top X% of the peaks with labels. The top X% means any peak that is within X% of the highest intensity. By dragging in the spectrum you can zoom in to a specific part of the spectrum and use 'Zoom Out' to get back to the original zoom level. The annotation of the spectrum is based on the given sequence in the peptides file and is done with different software so inconsistencies are likely. The peaks are annotated based on the given sequence, with 20 ppm tolerance.

Copy Data

##### Spectrum 8221 (TSV)

###### Preview

```
Loading example...
```

*Click on the button to copy the data to your clipboard.*

Mz MinMz MaxIntensity Max

WidthHeightPeptide font sizePeptide stroke widthSpectrum font sizeSpectrum stroke widthCompact peptide

Ion legend

wxyz

abcd

OtherUnassignedIon^Charge^_Position_Show for top:%

JVKDYFPEPVT

02.18e+44.36e+46.54e+48.72e+4

Zoom Out

y^+1^_1_a^+1^_2_y^+1^_2_b^+1^_2_y^+1^_2_y^+1^_3_b^+2^_5_y^+1^_3_b^+1^_3_b^+2^_7_b^+2^_7_b^+1^_4_b^+1^_4_y^+1^_4_b^+1^_4_b^+2^_8_y^+1^_5_y^+1^_5_b^+2^_9_y^+2^_9_b^+2^_10_b^+1^_5_\*\*\*y^+1^_6_b^+1^_6_b^+1^_6_b^+1^_6_b^+1^_7_y^+1^_8_b^+1^_8_b^+1^_8_b^+1^_8_y^+1^_9_b^+1^_9_y^+1^_9_b^+1^_10_b^+1^_10_y^+1^_10_

0531106315942125

Fragment Matches Table

Show background peaks

| Position | Ion type | Intensity | mz Theoretical | mz Error (Th) | mz Error (ppm) | Charge | Series Number |
| --- | --- | --- | --- | --- | --- | --- | --- |
| - | - | 878.4 | 120.1 | - | - | 0 | - |
| 11 | y | 8819 | 120.1 | 0.0002687 | 2.238 | +1 | 1 |
| - | - | 1.79E+04 | 120.1 | - | - | 0 | - |
| - | - | 972.9 | 120.1 | - | - | 0 | - |
| - | - | 1749 | 121.1 | - | - | 0 | - |
| - | - | 405.7 | 123.9 | - | - | 0 | - |
| - | - | 687.1 | 127.1 | - | - | 0 | - |
| - | - | 2.504E+04 | 129.1 | - | - | 0 | - |
| - | - | 422.3 | 129.7 | - | - | 0 | - |
| - | - | 529.5 | 130 | - | - | 0 | - |
| - | - | 1660 | 130.1 | - | - | 0 | - |
| - | - | 381.2 | 130.9 | - | - | 0 | - |
| - | - | 456.6 | 133.1 | - | - | 0 | - |
| - | - | 398.3 | 134.7 | - | - | 0 | - |
| - | - | 471.5 | 136.1 | - | - | 0 | - |
| - | - | 8681 | 136.1 | - | - | 0 | - |
| - | - | 531.4 | 137.1 | - | - | 0 | - |
| - | - | 582.5 | 140.1 | - | - | 0 | - |
| - | - | 449.3 | 142.7 | - | - | 0 | - |
| - | - | 394 | 145 | - | - | 0 | - |
| - | - | 448.9 | 153.5 | - | - | 0 | - |
| - | - | 479.7 | 155.1 | - | - | 0 | - |
| - | - | 809 | 167.1 | - | - | 0 | - |
| - | - | 570.9 | 167.1 | - | - | 0 | - |
| - | - | 463.7 | 167.1 | - | - | 0 | - |
| - | - | 464.2 | 167.9 | - | - | 0 | - |
| - | - | 6.893E+04 | 169.1 | - | - | 0 | - |
| - | - | 6928 | 170.1 | - | - | 0 | - |
| - | - | 581.7 | 171.1 | - | - | 0 | - |
| - | - | 1386 | 172.1 | - | - | 0 | - |
| - | - | 963.1 | 173.1 | - | - | 0 | - |
| - | - | 4322 | 173.4 | - | - | 0 | - |
| - | - | 719.9 | 175.1 | - | - | 0 | - |
| - | - | 477 | 180.3 | - | - | 0 | - |
| - | - | 1447 | 181.1 | - | - | 0 | - |
| - | - | 942.9 | 181.1 | - | - | 0 | - |
| - | - | 1246 | 182.1 | - | - | 0 | - |
| - | - | 479.1 | 182.1 | - | - | 0 | - |
| - | - | 1401 | 183.1 | - | - | 0 | - |
| - | - | 997.9 | 183.1 | - | - | 0 | - |
| - | - | 814.8 | 185.1 | - | - | 0 | - |
| 2 | a | 2.825E+04 | 185.2 | 0.0002604 | 1.406 | +1 | 2 |
| - | - | 2928 | 186.2 | - | - | 0 | - |
| - | - | 469.9 | 188.1 | - | - | 0 | - |
| - | - | 862 | 188.1 | - | - | 0 | - |
| - | - | 1689 | 195.1 | - | - | 0 | - |
| - | - | 5.605E+04 | 197.1 | - | - | 0 | - |
| - | - | 5627 | 198.1 | - | - | 0 | - |
| - | - | 8125 | 199.1 | - | - | 0 | - |
| - | - | 1682 | 199.2 | - | - | 0 | - |
| - | - | 751.6 | 200.1 | - | - | 0 | - |
| 10 | y | 5057 | 201.1 | 0.0002273 | 1.13 | +1 | 2 |
| - | - | 926.6 | 203.1 | - | - | 0 | - |
| - | - | 481 | 204.1 | - | - | 0 | - |
| - | - | 3434 | 209.1 | - | - | 0 | - |
| - | - | 697.1 | 209.1 | - | - | 0 | - |
| - | - | 526 | 209.5 | - | - | 0 | - |
| 2 | b | 1.377E+04 | 213.2 | 0.0002188 | 1.026 | +1 | 2 |
| - | - | 1830 | 214.2 | - | - | 0 | - |
| - | - | 958.6 | 215.1 | - | - | 0 | - |
| - | - | 903.3 | 217.1 | - | - | 0 | - |
| - | - | 1919 | 217.1 | - | - | 0 | - |
| 10 | y | 9177 | 219.1 | 0.000298 | 1.36 | +1 | 2 |
| - | - | 696.2 | 220.1 | - | - | 0 | - |
| - | - | 549.3 | 221.1 | - | - | 0 | - |
| - | - | 6265 | 226.1 | - | - | 0 | - |
| - | - | 482.6 | 226.2 | - | - | 0 | - |
| - | - | 2.808E+04 | 227.1 | - | - | 0 | - |
| - | - | 1044 | 227.2 | - | - | 0 | - |
| - | - | 2817 | 228.1 | - | - | 0 | - |
| - | - | 1961 | 228.2 | - | - | 0 | - |
| - | - | 739.8 | 229.1 | - | - | 0 | - |
| - | - | 626.4 | 230.2 | - | - | 0 | - |
| - | - | 5334 | 231.1 | - | - | 0 | - |
| - | - | 583.7 | 232.2 | - | - | 0 | - |
| - | - | 2587 | 233.2 | - | - | 0 | - |
| - | - | 684.7 | 235.1 | - | - | 0 | - |
| - | - | 735.5 | 239.1 | - | - | 0 | - |
| - | - | 602 | 240.1 | - | - | 0 | - |
| - | - | 1.208E+04 | 244.1 | - | - | 0 | - |
| - | - | 938.3 | 244.1 | - | - | 0 | - |
| - | - | 2230 | 245.1 | - | - | 0 | - |
| - | - | 988.8 | 247.1 | - | - | 0 | - |
| - | - | 6557 | 251.1 | - | - | 0 | - |
| - | - | 828 | 252.1 | - | - | 0 | - |
| - | - | 545.5 | 253.2 | - | - | 0 | - |
| - | - | 1590 | 254.1 | - | - | 0 | - |
| - | - | 651.7 | 255.1 | - | - | 0 | - |
| - | - | 619.8 | 256.1 | - | - | 0 | - |
| - | - | 662.5 | 256.2 | - | - | 0 | - |
| - | - | 1372 | 261.2 | - | - | 0 | - |
| - | - | 1044 | 263.1 | - | - | 0 | - |
| - | - | 658.5 | 270.2 | - | - | 0 | - |
| - | - | 576 | 270.7 | - | - | 0 | - |
| - | - | 1632 | 272.1 | - | - | 0 | - |
| - | - | 9838 | 279.1 | - | - | 0 | - |
| - | - | 949.4 | 280.1 | - | - | 0 | - |
| - | - | 2346 | 280.2 | - | - | 0 | - |
| - | - | 2953 | 283.1 | - | - | 0 | - |
| - | - | 807.8 | 287.2 | - | - | 0 | - |
| - | - | 1007 | 296.2 | - | - | 0 | - |
| 9 | y | 1.32E+04 | 298.2 | 0.0004115 | 1.38 | +1 | 3 |
| - | - | 1393 | 299.2 | - | - | 0 | - |
| - | - | 984.9 | 304.2 | - | - | 0 | - |
| - | - | 713.9 | 306.1 | - | - | 0 | - |
| - | - | 780.7 | 308.2 | - | - | 0 | - |
| 5 | b | 927.1 | 310.2 | 0.001144 | 3.688 | +2 | 5 |
| - | - | 2151 | 311.1 | - | - | 0 | - |
| 9 | y | 8.634E+04 | 316.2 | 0.0003449 | 1.091 | +1 | 3 |
| - | - | 1.204E+04 | 317.2 | - | - | 0 | - |
| - | - | 1239 | 318.2 | - | - | 0 | - |
| - | - | 870.7 | 323.2 | - | - | 0 | - |
| - | - | 2.379E+04 | 324.2 | - | - | 0 | - |
| - | - | 2719 | 325.2 | - | - | 0 | - |
| - | - | 868.8 | 325.2 | - | - | 0 | - |
| - | - | 824 | 326.2 | - | - | 0 | - |
| - | - | 599.1 | 328.2 | - | - | 0 | - |
| - | - | 1156 | 329.2 | - | - | 0 | - |
| - | - | 1369 | 340.2 | - | - | 0 | - |
| - | - | 2332 | 341.2 | - | - | 0 | - |
| 3 | b | 1803 | 341.3 | 0.001081 | 3.168 | +1 | 3 |
| - | - | 602.9 | 342.2 | - | - | 0 | - |
| - | - | 2436 | 343.2 | - | - | 0 | - |
| - | - | 528.5 | 344.2 | - | - | 0 | - |
| - | - | 516.4 | 344.2 | - | - | 0 | - |
| - | - | 933.5 | 344.7 | - | - | 0 | - |
| - | - | 1576 | 346.2 | - | - | 0 | - |
| - | - | 1745 | 353.2 | - | - | 0 | - |
| - | - | 1018 | 353.2 | - | - | 0 | - |
| - | - | 1377 | 355.1 | - | - | 0 | - |
| - | - | 891 | 355.2 | - | - | 0 | - |
| - | - | 621.7 | 355.2 | - | - | 0 | - |
| - | - | 687 | 356.1 | - | - | 0 | - |
| - | - | 749 | 356.2 | - | - | 0 | - |
| - | - | 772.6 | 357.1 | - | - | 0 | - |
| - | - | 2522 | 369.7 | - | - | 0 | - |
| - | - | 613.8 | 370.2 | - | - | 0 | - |
| - | - | 2208 | 371.2 | - | - | 0 | - |
| - | - | 844.5 | 373.2 | - | - | 0 | - |
| - | - | 3347 | 374.2 | - | - | 0 | - |
| - | - | 2048 | 379.2 | - | - | 0 | - |
| - | - | 673.4 | 380.2 | - | - | 0 | - |
| - | - | 1059 | 381.1 | - | - | 0 | - |
| - | - | 1872 | 389.2 | - | - | 0 | - |
| - | - | 4945 | 391.2 | - | - | 0 | - |
| - | - | 585 | 393.3 | - | - | 0 | - |
| - | - | 2378 | 395.2 | - | - | 0 | - |
| - | - | 800.7 | 403.2 | - | - | 0 | - |
| - | - | 969.7 | 405.2 | - | - | 0 | - |
| - | - | 5649 | 407.2 | - | - | 0 | - |
| - | - | 1025 | 408.2 | - | - | 0 | - |
| - | - | 616.1 | 409.2 | - | - | 0 | - |
| - | - | 687 | 411.3 | - | - | 0 | - |
| - | - | 5500 | 418.2 | - | - | 0 | - |
| - | - | 2501 | 418.7 | - | - | 0 | - |
| 7 | b | 7517 | 423.2 | 0.007013 | 16.57 | +2 | 7 |
| - | - | 2946 | 424.2 | - | - | 0 | - |
| - | - | 8109 | 426.2 | - | - | 0 | - |
| - | - | 1448 | 427.2 | - | - | 0 | - |
| 7 | b | 2874 | 432.2 | 0.000339 | 0.7842 | +2 | 7 |
| - | - | 2242 | 432.7 | - | - | 0 | - |
| - | - | 581.8 | 434.3 | - | - | 0 | - |
| - | - | 1615 | 435.2 | - | - | 0 | - |
| - | - | 784.6 | 436.2 | - | - | 0 | - |
| 4 | b | 2654 | 438.3 | 0.0008159 | 1.862 | +1 | 4 |
| - | - | 871.9 | 438.3 | - | - | 0 | - |
| 4 | b | 648.5 | 439.3 | 0.002548 | 5.8 | +1 | 4 |
| - | - | 1069 | 440.3 | - | - | 0 | - |
| 8 | y | 3259 | 445.2 | 0.000751 | 1.687 | +1 | 4 |
| - | - | 1355 | 452.2 | - | - | 0 | - |
| - | - | 577.2 | 454.3 | - | - | 0 | - |
| 4 | b | 9829 | 456.3 | 0.0003831 | 0.8395 | +1 | 4 |
| - | - | 2664 | 457.3 | - | - | 0 | - |
| - | - | 621.3 | 468.2 | - | - | 0 | - |
| - | - | 3736 | 470.2 | - | - | 0 | - |
| - | - | 2117 | 471.2 | - | - | 0 | - |
| - | - | 1670 | 482.3 | - | - | 0 | - |
| - | - | 708.8 | 482.8 | - | - | 0 | - |
| - | - | 1520 | 483.3 | - | - | 0 | - |
| - | - | 714.1 | 483.8 | - | - | 0 | - |
| - | - | 851.9 | 488.3 | - | - | 0 | - |
| - | - | 926.4 | 488.3 | - | - | 0 | - |
| - | - | 912.2 | 488.7 | - | - | 0 | - |
| 8 | b | 4715 | 496.8 | 0.0004658 | 0.9376 | +2 | 8 |
| - | - | 3795 | 497.3 | - | - | 0 | - |
| - | - | 742.8 | 498.2 | - | - | 0 | - |
| - | - | 2036 | 506.3 | - | - | 0 | - |
| - | - | 1052 | 508.3 | - | - | 0 | - |
| - | - | 1515 | 509.2 | - | - | 0 | - |
| - | - | 717.8 | 510.3 | - | - | 0 | - |
| - | - | 709.3 | 520.3 | - | - | 0 | - |
| - | - | 649.9 | 521.8 | - | - | 0 | - |
| - | - | 2243 | 522.3 | - | - | 0 | - |
| - | - | 4437 | 522.8 | - | - | 0 | - |
| - | - | 1577 | 523.3 | - | - | 0 | - |
| - | - | 867.2 | 523.8 | - | - | 0 | - |
| 7 | y | 3462 | 524.3 | 0.001154 | 2.202 | +1 | 5 |
| - | - | 1026 | 525.3 | - | - | 0 | - |
| - | - | 3478 | 526.3 | - | - | 0 | - |
| - | - | 832.6 | 527.3 | - | - | 0 | - |
| - | - | 1556 | 531.8 | - | - | 0 | - |
| - | - | 2888 | 534.3 | - | - | 0 | - |
| - | - | 2566 | 536.3 | - | - | 0 | - |
| - | - | 2077 | 539.3 | - | - | 0 | - |
| 7 | y | 2.688E+04 | 542.3 | 0.0005384 | 0.9929 | +1 | 5 |
| - | - | 7264 | 543.3 | - | - | 0 | - |
| - | - | 873.5 | 544.3 | - | - | 0 | - |
| 9 | b | 1.648E+04 | 545.3 | 0.00039 | 0.7152 | +2 | 9 |
| - | - | 8847 | 545.8 | - | - | 0 | - |
| - | - | 4225 | 546.3 | - | - | 0 | - |
| 3 | y | 865.6 | 548.3 | 0.002924 | 5.334 | +2 | 9 |
| - | - | 1087 | 553.3 | - | - | 0 | - |
| - | - | 2.034E+04 | 554.3 | - | - | 0 | - |
| - | - | 7250 | 555.3 | - | - | 0 | - |
| - | - | 1184 | 556.3 | - | - | 0 | - |
| - | - | 1600 | 558.3 | - | - | 0 | - |
| - | - | 1533 | 567.3 | - | - | 0 | - |
| - | - | 1011 | 568.4 | - | - | 0 | - |
| - | - | 2119 | 570.3 | - | - | 0 | - |
| - | - | 841.1 | 571.3 | - | - | 0 | - |
| - | - | 593.6 | 571.8 | - | - | 0 | - |
| - | - | 681.2 | 580.3 | - | - | 0 | - |
| - | - | 5726 | 580.8 | - | - | 0 | - |
| - | - | 4963 | 581.3 | - | - | 0 | - |
| - | - | 1064 | 581.8 | - | - | 0 | - |
| - | - | 1927 | 582.3 | - | - | 0 | - |
| - | - | 4167 | 591.4 | - | - | 0 | - |
| - | - | 1102 | 592.4 | - | - | 0 | - |
| 10 | b | 4809 | 594.8 | 0.0001866 | 0.3137 | +2 | 10 |
| - | - | 2233 | 595.3 | - | - | 0 | - |
| - | - | 1565 | 595.8 | - | - | 0 | - |
| - | - | 1463 | 603.3 | - | - | 0 | - |
| - | - | 608.5 | 606.3 | - | - | 0 | - |
| 5 | b | 1.442E+04 | 619.3 | 0.0001649 | 0.2662 | +1 | 5 |
| - | - | 3946 | 620.3 | - | - | 0 | - |
| - | - | 852.3 | 621.4 | - | - | 0 | - |
| - | - | 952.9 | 623.3 | - | - | 0 | - |
| - | - | 1143 | 625.3 | - | - | 0 | - |
| - | - | 1494 | 633.3 | - | - | 0 | - |
| - | - | 1219 | 635.4 | - | - | 0 | - |
| - | - | 789.1 | 638.3 | - | - | 0 | - |
| 0 | Precursor | 1920 | 645.3 | 0.0006494 | 1.006 | +2 | -1 |
| 0 | Precursor | 1885 | 645.8 | 0.01021 | 15.81 | +2 | -1 |
| - | - | 816.6 | 646.3 | - | - | 0 | - |
| - | - | 1985 | 651.3 | - | - | 0 | - |
| - | - | 1339 | 652.3 | - | - | 0 | - |
| - | - | 1755 | 652.9 | - | - | 0 | - |
| - | - | 1497 | 653.3 | - | - | 0 | - |
| - | - | 1593 | 653.8 | - | - | 0 | - |
| - | - | 1221 | 654.3 | - | - | 0 | - |
| 0 | Precursor | 3860 | 654.3 | 0.002941 | 4.495 | +2 | -1 |
| - | - | 1001 | 654.8 | - | - | 0 | - |
| - | - | 1840 | 654.8 | - | - | 0 | - |
| - | - | 1924 | 655.3 | - | - | 0 | - |
| - | - | 760 | 673.5 | - | - | 0 | - |
| - | - | 3307 | 681.3 | - | - | 0 | - |
| - | - | 918.7 | 682.3 | - | - | 0 | - |
| - | - | 3174 | 682.4 | - | - | 0 | - |
| - | - | 781.3 | 683.4 | - | - | 0 | - |
| 6 | y | 9098 | 689.4 | 0.0003096 | 0.4491 | +1 | 6 |
| - | - | 3864 | 690.4 | - | - | 0 | - |
| - | - | 1592 | 716.4 | - | - | 0 | - |
| - | - | 700.1 | 720.4 | - | - | 0 | - |
| - | - | 5161 | 738.4 | - | - | 0 | - |
| - | - | 2579 | 739.4 | - | - | 0 | - |
| - | - | 717.7 | 741 | - | - | 0 | - |
| 6 | b | 1607 | 748.4 | 0.0009827 | 1.313 | +1 | 6 |
| 6 | b | 1408 | 749.4 | 0.01494 | 19.94 | +1 | 6 |
| - | - | 3185 | 752.4 | - | - | 0 | - |
| - | - | 1416 | 753.4 | - | - | 0 | - |
| - | - | 1370 | 762.3 | - | - | 0 | - |
| - | - | 970.7 | 763.3 | - | - | 0 | - |
| 6 | b | 5.153E+04 | 766.4 | 0.0002559 | 0.3339 | +1 | 6 |
| - | - | 2.225E+04 | 767.4 | - | - | 0 | - |
| - | - | 5726 | 768.4 | - | - | 0 | - |
| - | - | 1.737E+04 | 780.4 | - | - | 0 | - |
| - | - | 9044 | 781.4 | - | - | 0 | - |
| - | - | 2442 | 782.4 | - | - | 0 | - |
| - | - | 1313 | 785.4 | - | - | 0 | - |
| - | - | 3034 | 796 | - | - | 0 | - |
| - | - | 2404 | 796.4 | - | - | 0 | - |
| - | - | 3085 | 796.7 | - | - | 0 | - |
| - | - | 1074 | 797 | - | - | 0 | - |
| - | - | 1189 | 835.5 | - | - | 0 | - |
| - | - | 1463 | 845.4 | - | - | 0 | - |
| - | - | 940.1 | 850.4 | - | - | 0 | - |
| - | - | 923.8 | 851.4 | - | - | 0 | - |
| 7 | b | 2778 | 863.5 | 0.0002029 | 0.235 | +1 | 7 |
| - | - | 2360 | 864.5 | - | - | 0 | - |
| - | - | 5228 | 877.4 | - | - | 0 | - |
| - | - | 861 | 877.5 | - | - | 0 | - |
| - | - | 3195 | 878.4 | - | - | 0 | - |
| - | - | 862.2 | 878.5 | - | - | 0 | - |
| - | - | 2936 | 879.4 | - | - | 0 | - |
| - | - | 1080 | 880.4 | - | - | 0 | - |
| - | - | 1152 | 893.4 | - | - | 0 | - |
| - | - | 1755 | 907.4 | - | - | 0 | - |
| - | - | 1023 | 908.4 | - | - | 0 | - |
| - | - | 865.8 | 946.5 | - | - | 0 | - |
| - | - | 1211 | 947.5 | - | - | 0 | - |
| - | - | 4306 | 964.5 | - | - | 0 | - |
| - | - | 3212 | 965.5 | - | - | 0 | - |
| - | - | 1140 | 966.5 | - | - | 0 | - |
| 4 | y | 851.1 | 967.4 | 0.003413 | 3.528 | +1 | 8 |
| 8 | b | 3378 | 974.5 | 0.0005755 | 0.5905 | +1 | 8 |
| 8 | b | 2101 | 975.5 | 0.01693 | 17.36 | +1 | 8 |
| - | - | 6268 | 976.5 | - | - | 0 | - |
| - | - | 2437 | 977.5 | - | - | 0 | - |
| - | - | 1115 | 978.5 | - | - | 0 | - |
| - | - | 797.1 | 990.5 | - | - | 0 | - |
| 8 | b | 6.3E+04 | 992.5 | 0.0006421 | 0.647 | +1 | 8 |
| - | - | 3.614E+04 | 993.5 | - | - | 0 | - |
| - | - | 1.187E+04 | 994.5 | - | - | 0 | - |
| - | - | 1546 | 995.5 | - | - | 0 | - |
| - | - | 1595 | 1047 | - | - | 0 | - |
| - | - | 748.7 | 1048 | - | - | 0 | - |
| 3 | y | 910.7 | 1078 | 0.008553 | 7.938 | +1 | 9 |
| 9 | b | 1772 | 1090 | 0.001099 | 1.009 | +1 | 9 |
| 3 | y | 1.334E+04 | 1096 | 0.001035 | 0.9446 | +1 | 9 |
| - | - | 1.016E+04 | 1097 | - | - | 0 | - |
| - | - | 2888 | 1098 | - | - | 0 | - |
| 10 | b | 711.4 | 1171 | 0.01063 | 9.082 | +1 | 10 |
| 10 | b | 4389 | 1189 | 0.0006651 | 0.5596 | +1 | 10 |
| - | - | 2960 | 1190 | - | - | 0 | - |
| - | - | 1355 | 1191 | - | - | 0 | - |
| 2 | y | 1281 | 1195 | 0.005728 | 4.795 | +1 | 10 |
| - | - | 1159 | 1196 | - | - | 0 | - |
| - | - | 649.1 | 1820 | - | - | 0 | - |
| - | - | 696.1 | 2104 | - | - | 0 | - |

m/z Charge Intensity FragmentType MassShift Position
120.06130981445312 0 878.4199
120.06578826904297 0 8818.537 y 10
120.0810546875 0 17900.318
120.08595275878906 0 972.9312
121.08451080322266 0 1749.1471
123.89981079101562 0 405.65292
127.08675384521484 0 687.12714
129.1024932861328 0 25035.248
129.7343292236328 0 422.31104
130.0499725341797 0 529.45825
130.1058807373047 0 1659.5768
130.9036865234375 0 381.24347
133.08612060546875 0 456.58545
134.69715881347656 0 398.2935
136.06207275390625 0 471.53943
136.0759735107422 0 8681.302
137.07936096191406 0 531.4369
140.14349365234375 0 582.49
142.72364807128906 0 449.27014
145.031982421875 0 393.99326
153.46473693847656 0 448.89954
155.11788940429688 0 479.67084
167.08212280273438 0 808.96265
167.0931396484375 0 570.91516
167.1177520751953 0 463.67755
167.9002685546875 0 464.19415
169.13381958007812 0 68928.01
170.13722229003906 0 6928.075
171.07618713378906 0 581.699
172.1124267578125 0 1385.792
173.1286163330078 0 963.0709
173.44058227539062 0 4322.2
175.0863800048828 0 719.85535
180.25450134277344 0 477.03403
181.0971221923828 0 1446.7689
181.13351440429688 0 942.87964
182.0815887451172 0 1246.2385
182.12942504882812 0 479.07614
183.113037109375 0 1400.5463
183.1496124267578 0 997.85754
185.12899780273438 0 814.80396
185.16510009765625 0 28247.55 a 1
186.1683807373047 0 2927.939
188.10667419433594 0 469.9353
188.14320373535156 0 861.9954
195.1129150390625 0 1688.556
197.12869262695312 0 56053.945
198.13209533691406 0 5626.557
199.10789489746094 0 8124.7163
199.18077087402344 0 1682.09
200.1117706298828 0 751.6268
201.12359619140625 0 5056.702 y Water loss 9
203.10252380371094 0 926.62915
204.13839721679688 0 480.98376
209.09225463867188 0 3434.145
209.10195922851562 0 697.05225
209.5316619873047 0 525.9845
213.15997314453125 0 13773.134 b 1
214.16346740722656 0 1830.3127
215.13919067382812 0 958.617
217.0824432373047 0 903.26276
217.13357543945312 0 1918.8477
219.1342315673828 0 9176.549 y 9
220.1382293701172 0 696.2217
221.08413696289062 0 549.3102
226.11878967285156 0 6264.8086
226.15492248535156 0 482.5974
227.10289001464844 0 28079.203
227.17518615722656 0 1043.5731
228.10621643066406 0 2816.5469
228.17088317871094 0 1960.9326
229.1182098388672 0 739.7598
230.18609619140625 0 626.3702
231.1494903564453 0 5334.3765
232.15272521972656 0 583.6991
233.1649627685547 0 2587.4314
235.1443328857422 0 684.7085
239.0949249267578 0 735.49677
240.13514709472656 0 602.0153
244.1294403076172 0 12077.393
244.1431884765625 0 938.3108
245.130126953125 0 2230.1716
247.1446075439453 0 988.8484
251.10292053222656 0 6556.6836
252.106689453125 0 828.0202
253.16734313964844 0 545.53485
254.11354064941406 0 1589.7045
255.09718322753906 0 651.6722
256.095947265625 0 619.77515
256.1666564941406 0 662.4816
261.15887451171875 0 1372.1172
263.1390075683594 0 1044.3877
270.1805419921875 0 658.549
270.6610412597656 0 576.0247
272.1242980957031 0 1632.366
279.0977783203125 0 9837.804
280.1014404296875 0 949.3974
280.1656188964844 0 2346.2383
283.14447021484375 0 2953.3367
287.2120056152344 0 807.8404
296.1964111328125 0 1007.3337
298.1765441894531 0 13195.131 y Water loss 8
299.1795349121094 0 1392.9948
304.16558837890625 0 984.90826
306.1442565917969 0 713.88354
308.1611633300781 0 780.7485
310.1772766113281 0 927.0506 b 4
311.139404296875 0 2150.7197
316.1870422363281 0 86339.055 y 8
317.1903381347656 0 12038.167
318.19256591796875 0 1239.4487
323.2444152832031 0 870.6637
324.1558532714844 0 23791.371
325.15924072265625 0 2718.8
325.1876220703125 0 868.7792
326.17071533203125 0 823.97656
328.1665954589844 0 599.0891
329.1501770019531 0 1155.9381
340.1875915527344 0 1368.9448
341.18231201171875 0 2331.7979
341.25579833984375 0 1803.224 b 2
342.18743896484375 0 602.92206
343.1981506347656 0 2436.4062
344.1589660644531 0 528.4856
344.20025634765625 0 516.3749
344.7069396972656 0 933.5171
346.17657470703125 0 1575.8958
353.1812744140625 0 1744.5134
353.218505859375 0 1018.0148
355.07000732421875 0 1377.4283
355.1606750488281 0 891.002
355.19732666015625 0 621.7153
356.0719299316406 0 687.0135
356.1629638671875 0 748.957
357.0678405761719 0 772.57947
369.7131652832031 0 2521.692
370.2156677246094 0 613.7505
371.1927795410156 0 2208.2905
373.18878173828125 0 844.45685
374.1712951660156 0 3346.9192
379.19866943359375 0 2047.5701
380.16326904296875 0 673.387
381.14385986328125 0 1058.8994
389.1820983886719 0 1872.1855
391.1978454589844 0 4945.0054
393.2503662109375 0 585.0455
395.2288513183594 0 2378.404
403.2340393066406 0 800.6968
405.2129821777344 0 969.71643
407.1927185058594 0 5649.1577
408.196533203125 0 1025.4716
409.2071228027344 0 616.07697
411.25927734375 0 687.00165
418.23968505859375 0 5499.899
418.7407531738281 0 2500.9075
423.22442626953125 0 7517.2593 b Water loss 6
424.2228698730469 0 2945.541
426.16644287109375 0 8108.5107
427.1693420410156 0 1448.0095
432.237060546875 0 2873.9626 b 6
432.7388000488281 0 2241.6702
434.27642822265625 0 581.818
435.1895446777344 0 1615.1865
436.189208984375 0 784.5791
438.27191162109375 0 2653.9612 b Water loss 3
438.30419921875 0 871.94385
439.2525634765625 0 648.5199 b Ammonia loss 3
440.2532958984375 0 1069.2551
445.23004150390625 0 3258.7969 y 7
452.2152404785156 0 1355.3882
454.2636413574219 0 577.1851
456.28204345703125 0 9828.885 b 3
457.284423828125 0 2664.131
468.2433776855469 0 621.3392
470.22528076171875 0 3735.554
471.2262878417969 0 2116.8972
482.2607421875 0 1670.0708
482.7611999511719 0 708.7808
483.2636413574219 0 1520.072
483.762939453125 0 714.12256
488.25457763671875 0 851.9133
488.3240051269531 0 926.3767
488.7449035644531 0 912.24023
496.75848388671875 0 4715.212 b 7
497.2590026855469 0 3794.801
498.21917724609375 0 742.81903
506.2604675292969 0 2035.9963
508.2548522949219 0 1052.4103
509.2406921386719 0 1514.8711
510.3279113769531 0 717.78754
520.2762451171875 0 709.33374
521.785888671875 0 649.9237
522.2844848632812 0 2242.8953
522.7859497070312 0 4437.24
523.2867431640625 0 1576.7759
523.782470703125 0 867.1745
524.2726440429688 0 3462.2715 y Water loss 6
525.277587890625 0 1026.0328
526.265380859375 0 3478.1423
527.26708984375 0 832.5817
531.7890625 0 1556.0448
534.2567749023438 0 2887.8977
536.2509155273438 0 2565.7341
539.253662109375 0 2076.7578
542.2825927734375 0 26875.02 y 6
543.2860717773438 0 7263.7446
544.2890625 0 873.4707
545.2847900390625 0 16481.725 b 8
545.7864379882812 0 8846.541
546.28759765625 0 4225.1704
548.2744140625 0 865.5754 y 2
553.3344116210938 0 1086.9443
554.260986328125 0 20338.771
555.2638549804688 0 7249.5786
556.2677612304688 0 1183.9962
558.3281860351562 0 1599.658
567.3499145507812 0 1533.2776
568.3551635742188 0 1011.40454
570.2931518554688 0 2119.1423
571.2919921875 0 841.1206
571.8154296875 0 593.64624
580.31494140625 0 681.24963
580.8216552734375 0 5725.652
581.3221435546875 0 4963.2817
581.8267211914062 0 1063.8395
582.2521362304688 0 1927.2062
591.3502197265625 0 4167.0786
592.3515014648438 0 1102.3281
594.8184204101562 0 4808.721 b 9
595.31884765625 0 2232.7322
595.822265625 0 1564.887
603.3485107421875 0 1462.626
606.2959594726562 0 608.505
619.3451538085938 0 14417.458 b 4
620.347412109375 0 3946.0237
621.3511962890625 0 852.2769
623.3211669921875 0 952.895
625.3314819335938 0 1142.6278
633.2905883789062 0 1493.7415
635.3775634765625 0 1218.5253
638.3178100585938 0 789.1482
645.341796875 0 1919.5524 Precursor Water loss
645.8446655273438 0 1884.9083 Precursor Ammonia loss
646.3455810546875 0 816.5553
651.3125610351562 0 1984.5286
652.259521484375 0 1339.1237
652.8606567382812 0 1755.2494
653.3219604492188 0 1496.6896
653.7685546875 0 1593.2885
654.26806640625 0 1221.0693
654.3447875976562 0 3859.5183 Precursor
654.7734375 0 1000.6225
654.8499755859375 0 1839.7166
655.3496704101562 0 1924.3286
673.521728515625 0 759.958
681.3245849609375 0 3307.1572
682.3203735351562 0 918.698
682.3782348632812 0 3174.1511
683.3795166015625 0 781.2982
689.3501586914062 0 9098.383 y 5
690.3532104492188 0 3864.4363
716.3969116210938 0 1592.0858
720.4093627929688 0 700.12866
738.4176025390625 0 5160.6
739.4221801757812 0 2579.0305
741.0490112304688 0 717.7046
748.40185546875 0 1606.619 b Water loss 5
749.4017944335938 0 1407.557 b Ammonia loss 5
752.3626708984375 0 3184.6873
753.3660888671875 0 1415.7341
762.3427734375 0 1370.2162
763.342529296875 0 970.7284
766.4131469726562 0 51531.66 b 5
767.4159545898438 0 22253.443
768.41943359375 0 5726.3696
780.3560791015625 0 17367.15
781.359375 0 9044.471
782.3639526367188 0 2442.29
785.4131469726562 0 1313.4247
796.0454711914062 0 3034.2063
796.3812866210938 0 2404.1748
796.7129516601562 0 3085.296
797.0474243164062 0 1073.9492
835.4712524414062 0 1189.4275
845.4371948242188 0 1463.221
850.4354858398438 0 940.0941
851.4358520507812 0 923.77936
863.4663696289062 0 2777.9956 b 6
864.4680786132812 0 2360.413
877.4085083007812 0 5227.7544
877.493408203125 0 860.968
878.4133911132812 0 3195.3787
878.4998779296875 0 862.20215
879.4207763671875 0 2936.2427
880.4240112304688 0 1079.6501
893.4356079101562 0 1152.0625
907.4190673828125 0 1755.4419
908.420654296875 0 1022.7901
946.5119018554688 0 865.8207
947.5016479492188 0 1210.7809
964.5139770507812 0 4305.7754
965.515869140625 0 3211.9255
966.5187377929688 0 1140.4358
967.4441528320312 0 851.0554 y 3
974.4976196289062 0 3378.0781 b Water loss 7
975.4991455078125 0 2100.6565 b Ammonia loss 7
976.4794311523438 0 6267.773
977.480224609375 0 2437.487
978.4827880859375 0 1114.7272
990.5012817382812 0 797.10516
992.5081176757812 0 62998.7 b 7
993.5113525390625 0 36143.895
994.5136108398438 0 11874.179
995.5164184570312 0 1546.1193
1046.5526123046875 0 1594.7554
1047.55517578125 0 748.697
1077.53369140625 0 910.7184 y Water loss 2
1089.5604248046875 0 1772.2122 b 8
1095.53466796875 0 13335.131 y 2
1096.5379638671875 0 10161.394
1097.5400390625 0 2888.0024
1170.6300048828125 0 711.4395 b Water loss 9
1188.6292724609375 0 4389.1284 b 9
1189.6329345703125 0 2960.2864
1190.631103515625 0 1354.6707
1194.598388671875 0 1281.0981 y 1
1195.610107421875 0 1158.5919
1819.9774169921875 0 649.0566
2104.280029296875 0 696.13257

Spectrum Details

|  |  |
| --- | --- |
| Matched peaks? Matched peaks The total absolute number of peaks matched. Additionally in brackets the total fraction of peaks matched and the total number of peaks is shown. | 40 (12.16% of 329) |
| FDR? FDR The false discovery rate estimated for this peptide. It is calculated by matching all theoretical fragments with a non-integer shift with the raw peaks for this spectrum. This is done with 40 different shifts. The resulting percentage is the average number of annotated peaks over the number of annotated peaks with the correct spectrum. | 0.77% |
| Satellite FDR? Satellite FDR See the FDR for details on its calculation. This satellite ion specific FDR only contains the satellite ions (d/w) for I/L/J positions. | - |
| PSM Score? PSM Score The PSM Score as given by Hecklib to this annotated spectrum. It is shown with three significant figures. | 457 |

#### Spectrum 8629? Spectrum 8629 The raw spectrum of this peptide as annotated by Hecklib. The fragments are coloured according to ion type (see legend). Any peaks with a star '\*' as text can be hovered over to see the full details, first the ion type second the mass shift type. By hovering over the amino acids in the peptide or ions in the legend the corresponding peaks are highlighted. By toggling the 'Unassigned' label you can turn the background (unassigned) peaks on or off in the plot. By updating the slider in the Ion legend you can update the spectrum to only show the top X% of the peaks with labels. The top X% means any peak that is within X% of the highest intensity. By dragging in the spectrum you can zoom in to a specific part of the spectrum and use 'Zoom Out' to get back to the original zoom level. The annotation of the spectrum is based on the given sequence in the peptides file and is done with different software so inconsistencies are likely. The peaks are annotated based on the given sequence, with 20 ppm tolerance.

Copy Data

##### Spectrum 8629 (TSV)

###### Preview

```
Loading example...
```

*Click on the button to copy the data to your clipboard.*

Mz MinMz MaxIntensity Max

WidthHeightPeptide font sizePeptide stroke widthSpectrum font sizeSpectrum stroke widthCompact peptide

Ion legend

wxyz

abcd

OtherUnassignedIon^Charge^_Position_Show for top:%

JVKDYFPEPVT

02.11e+44.21e+46.32e+48.43e+4

Zoom Out

y^+1^_1_y^+1^_2_y^+1^_3_y^+1^_3_c^+1^_3_c^+2^_7_c^+1^_4_c^+2^_8_y^+1^_5_c^+2^_9_c^+1^_5_c^+1^_5_y^+1^_6_c^+1^_6_y^+1^_7_c^+1^_7_y^+1^_8_c^+1^_8_z^+1^_9_y^+1^_9_c^+1^_9_y^+1^_10_z^+1^_10_c^+1^_10_

03316629941325

Fragment Matches Table

Show background peaks

| Position | Ion type | Intensity | mz Theoretical | mz Error (Th) | mz Error (ppm) | Charge | Series Number |
| --- | --- | --- | --- | --- | --- | --- | --- |
| 11 | y | 804.5 | 120.1 | 0.0001542 | 1.284 | +1 | 1 |
| - | - | 375.8 | 133.6 | - | - | 0 | - |
| - | - | 443.9 | 136.3 | - | - | 0 | - |
| - | - | 405.9 | 140.1 | - | - | 0 | - |
| - | - | 461.6 | 141.5 | - | - | 0 | - |
| - | - | 602.5 | 148.9 | - | - | 0 | - |
| - | - | 467.2 | 157.9 | - | - | 0 | - |
| - | - | 499.2 | 168 | - | - | 0 | - |
| - | - | 2235 | 169.1 | - | - | 0 | - |
| - | - | 2428 | 173.4 | - | - | 0 | - |
| - | - | 577.4 | 182.1 | - | - | 0 | - |
| - | - | 523.8 | 182.4 | - | - | 0 | - |
| - | - | 1857 | 185.2 | - | - | 0 | - |
| - | - | 496.8 | 186.2 | - | - | 0 | - |
| - | - | 3876 | 197.1 | - | - | 0 | - |
| 10 | y | 577.7 | 201.1 | 0.0002457 | 1.222 | +1 | 2 |
| - | - | 460.3 | 210.7 | - | - | 0 | - |
| - | - | 2029 | 213.2 | - | - | 0 | - |
| - | - | 943.4 | 227.1 | - | - | 0 | - |
| - | - | 590.5 | 235.2 | - | - | 0 | - |
| 9 | y | 942 | 298.2 | 0.000503 | 1.687 | +1 | 3 |
| 9 | y | 1.247E+04 | 316.2 | 0.0001923 | 0.6081 | +1 | 3 |
| - | - | 1635 | 317.2 | - | - | 0 | - |
| - | - | 2057 | 324.2 | - | - | 0 | - |
| - | - | 1856 | 355.1 | - | - | 0 | - |
| - | - | 1032 | 356.1 | - | - | 0 | - |
| - | - | 975.5 | 357.1 | - | - | 0 | - |
| - | - | 2253 | 357.3 | - | - | 0 | - |
| 3 | c | 697.1 | 358.3 | 0.001542 | 4.305 | +1 | 3 |
| - | - | 553.3 | 380.7 | - | - | 0 | - |
| - | - | 999.2 | 423.2 | - | - | 0 | - |
| - | - | 603.4 | 431.1 | - | - | 0 | - |
| 7 | c | 860 | 432.2 | 0.0005766 | 1.334 | +2 | 7 |
| 4 | c | 1093 | 473.3 | 0.0003177 | 0.6711 | +1 | 4 |
| 8 | c | 822.1 | 496.8 | 0.002098 | 4.223 | +2 | 8 |
| - | - | 746.9 | 497.3 | - | - | 0 | - |
| - | - | 988.3 | 522.8 | - | - | 0 | - |
| 7 | y | 4748 | 542.3 | 0.0001722 | 0.3176 | +1 | 5 |
| - | - | 1511 | 543.3 | - | - | 0 | - |
| 9 | c | 2815 | 545.3 | 0.0001593 | 0.2922 | +2 | 9 |
| - | - | 2327 | 545.8 | - | - | 0 | - |
| - | - | 632.5 | 546.8 | - | - | 0 | - |
| - | - | 1664 | 554.3 | - | - | 0 | - |
| - | - | 830.8 | 580.8 | - | - | 0 | - |
| - | - | 1082 | 581.3 | - | - | 0 | - |
| 5 | c | 2166 | 619.3 | 0.0001403 | 0.2265 | +1 | 5 |
| - | - | 718.7 | 620.3 | - | - | 0 | - |
| - | - | 4523 | 635.4 | - | - | 0 | - |
| 5 | c | 4915 | 636.4 | 0.0003492 | 0.5487 | +1 | 5 |
| - | - | 1474 | 637.4 | - | - | 0 | - |
| - | - | 890.5 | 655.4 | - | - | 0 | - |
| 6 | y | 2556 | 689.4 | 0.001103 | 1.6 | +1 | 6 |
| - | - | 1385 | 690.4 | - | - | 0 | - |
| - | - | 930.4 | 698.4 | - | - | 0 | - |
| - | - | 1743 | 739.4 | - | - | 0 | - |
| - | - | 3110 | 740.4 | - | - | 0 | - |
| - | - | 714.8 | 741.4 | - | - | 0 | - |
| - | - | 597.4 | 758.5 | - | - | 0 | - |
| 6 | c | 6293 | 766.4 | 0.000378 | 0.4931 | +1 | 6 |
| - | - | 2056 | 767.4 | - | - | 0 | - |
| - | - | 1553 | 768.4 | - | - | 0 | - |
| - | - | 1813 | 780.4 | - | - | 0 | - |
| - | - | 698.3 | 826.5 | - | - | 0 | - |
| 5 | y | 2157 | 852.4 | 0.002176 | 2.552 | +1 | 7 |
| - | - | 1054 | 853.4 | - | - | 0 | - |
| - | - | 1061 | 877.4 | - | - | 0 | - |
| - | - | 858.4 | 879.4 | - | - | 0 | - |
| - | - | 6783 | 879.5 | - | - | 0 | - |
| 7 | c | 1.498E+04 | 880.5 | 0.001749 | 1.986 | +1 | 7 |
| - | - | 5435 | 881.5 | - | - | 0 | - |
| - | - | 2081 | 882.5 | - | - | 0 | - |
| - | - | 739.9 | 947.5 | - | - | 0 | - |
| - | - | 695.2 | 952.4 | - | - | 0 | - |
| - | - | 1247 | 965.5 | - | - | 0 | - |
| - | - | 633.4 | 966.4 | - | - | 0 | - |
| 4 | y | 1562 | 967.4 | 0.001399 | 1.446 | +1 | 8 |
| - | - | 1271 | 968.4 | - | - | 0 | - |
| - | - | 796.5 | 976.5 | - | - | 0 | - |
| 8 | c | 1.028E+04 | 992.5 | 0.0009473 | 0.9545 | +1 | 8 |
| - | - | 5124 | 993.5 | - | - | 0 | - |
| - | - | 1555 | 994.5 | - | - | 0 | - |
| - | - | 660.7 | 1008 | - | - | 0 | - |
| - | - | 1582 | 1036 | - | - | 0 | - |
| 3 | z | 2861 | 1080 | 0.001476 | 1.367 | +1 | 9 |
| - | - | 3099 | 1081 | - | - | 0 | - |
| - | - | 625.3 | 1082 | - | - | 0 | - |
| 3 | y | 2517 | 1096 | 0.0006687 | 0.6103 | +1 | 9 |
| - | - | 1399 | 1097 | - | - | 0 | - |
| - | - | 794.9 | 1098 | - | - | 0 | - |
| 9 | c | 1650 | 1107 | 0.0009165 | 0.8282 | +1 | 9 |
| - | - | 1287 | 1108 | - | - | 0 | - |
| - | - | 888.9 | 1147 | - | - | 0 | - |
| - | - | 1463 | 1156 | - | - | 0 | - |
| - | - | 1072 | 1157 | - | - | 0 | - |
| - | - | 3904 | 1162 | - | - | 0 | - |
| - | - | 2919 | 1163 | - | - | 0 | - |
| - | - | 1643 | 1164 | - | - | 0 | - |
| 2 | y | 705.6 | 1177 | 0.02034 | 17.29 | +1 | 10 |
| 2 | z | 1730 | 1179 | 0.000911 | 0.773 | +1 | 10 |
| - | - | 1106 | 1180 | - | - | 0 | - |
| - | - | 1138 | 1191 | - | - | 0 | - |
| - | - | 1327 | 1192 | - | - | 0 | - |
| 10 | c | 2.135E+04 | 1206 | 0.002068 | 1.715 | +1 | 10 |
| - | - | 1.316E+04 | 1207 | - | - | 0 | - |
| - | - | 9429 | 1208 | - | - | 0 | - |
| - | - | 2454 | 1209 | - | - | 0 | - |
| - | - | 809.2 | 1210 | - | - | 0 | - |
| - | - | 925.8 | 1220 | - | - | 0 | - |
| - | - | 938.3 | 1237 | - | - | 0 | - |
| - | - | 1591 | 1237 | - | - | 0 | - |
| - | - | 1281 | 1238 | - | - | 0 | - |
| - | - | 777.7 | 1245 | - | - | 0 | - |
| - | - | 765.9 | 1246 | - | - | 0 | - |
| - | - | 893.4 | 1247 | - | - | 0 | - |
| - | - | 5109 | 1249 | - | - | 0 | - |
| - | - | 5368 | 1250 | - | - | 0 | - |
| - | - | 1589 | 1251 | - | - | 0 | - |
| - | - | 1237 | 1253 | - | - | 0 | - |
| - | - | 1141 | 1263 | - | - | 0 | - |
| - | - | 1176 | 1264 | - | - | 0 | - |
| - | - | 2808 | 1265 | - | - | 0 | - |
| - | - | 1523 | 1266 | - | - | 0 | - |
| - | - | 3913 | 1281 | - | - | 0 | - |
| - | - | 2070 | 1282 | - | - | 0 | - |
| - | - | 1006 | 1283 | - | - | 0 | - |
| - | - | 742.7 | 1290 | - | - | 0 | - |
| - | - | 1685 | 1291 | - | - | 0 | - |
| - | - | 2.187E+04 | 1292 | - | - | 0 | - |
| - | - | 1.625E+04 | 1293 | - | - | 0 | - |
| - | - | 575 | 1294 | - | - | 0 | - |
| - | - | 6316 | 1294 | - | - | 0 | - |
| - | - | 1120 | 1295 | - | - | 0 | - |
| - | - | 767.9 | 1305 | - | - | 0 | - |
| - | - | 2394 | 1306 | - | - | 0 | - |
| - | - | 1647 | 1307 | - | - | 0 | - |
| - | - | 857.9 | 1308 | - | - | 0 | - |
| - | - | 1.745E+04 | 1308 | - | - | 0 | - |
| - | - | 8.344E+04 | 1309 | - | - | 0 | - |
| - | - | 5.83E+04 | 1310 | - | - | 0 | - |
| - | - | 2.399E+04 | 1311 | - | - | 0 | - |
| - | - | 2993 | 1312 | - | - | 0 | - |

m/z Charge Intensity FragmentType MassShift Position
120.065673828125 0 804.5139 y 10
133.5923614501953 0 375.79935
136.26776123046875 0 443.92926
140.08839416503906 0 405.88403
141.47471618652344 0 461.64426
148.94729614257812 0 602.5369
157.94749450683594 0 467.15167
167.9989471435547 0 499.1716
169.1334991455078 0 2234.5823
173.4386749267578 0 2427.7205
182.08004760742188 0 577.3781
182.37710571289062 0 523.82526
185.16482543945312 0 1857.2024
186.16799926757812 0 496.8268
197.12843322753906 0 3876.098
201.1231231689453 0 577.72845 y Water loss 9
210.66665649414062 0 460.30692
213.15977478027344 0 2029.1083
227.1025848388672 0 943.39435
235.2018280029297 0 590.54846
298.1766357421875 0 941.9732 y Water loss 8
316.1868896484375 0 12466.557 y 8
317.1902160644531 0 1634.989
324.15594482421875 0 2056.7437
355.0699768066406 0 1856.2212
356.0704650878906 0 1031.5996
357.069091796875 0 975.4991
357.27362060546875 0 2253.0693
358.27972412109375 0 697.10724 c 2
380.6939697265625 0 553.333
423.224609375 0 999.17444
431.08880615234375 0 603.3886
432.23614501953125 0 860.0215 c Ammonia loss 6
473.3078918457031 0 1092.5455 c 3
496.75592041015625 0 822.1029 c Ammonia loss 7
497.2598876953125 0 746.9223
522.7889404296875 0 988.29663
542.2822265625 0 4747.6235 y 6
543.2852172851562 0 1511.4055
545.2842407226562 0 2814.9111 c Ammonia loss 8
545.7852172851562 0 2327.462
546.7916870117188 0 632.50446
554.2599487304688 0 1664.4492
580.8226318359375 0 830.7584
581.3225708007812 0 1082.0768
619.3448486328125 0 2165.743 c Ammonia loss 4
620.3453979492188 0 718.7263
635.3639526367188 0 4523.1934
636.3718872070312 0 4915.346 c 4
637.371826171875 0 1473.7124
655.3565673828125 0 890.4745
689.349365234375 0 2555.858 y 5
690.3529052734375 0 1384.9973
698.4451293945312 0 930.3629
739.421630859375 0 1743.4152
740.43359375 0 3110.035
741.43505859375 0 714.80084
758.4739990234375 0 597.4303
766.4130249023438 0 6292.8926 c Ammonia loss 5
767.4161987304688 0 2055.8118
768.4176635742188 0 1553.1809
780.3565673828125 0 1813.112
826.533203125 0 698.2555
852.41162109375 0 2156.8274 y 4
853.41845703125 0 1054.3855
877.4061889648438 0 1060.819
879.3993530273438 0 858.37103
879.4844360351562 0 6783.073
880.490966796875 0 14983.97 c 6
881.4947509765625 0 5435.216
882.4976196289062 0 2081.3193
947.5159912109375 0 739.90576
952.4291381835938 0 695.186
965.5174560546875 0 1246.712
966.4400634765625 0 633.4147
967.442138671875 0 1562.4467 y 3
968.441162109375 0 1271.2865
976.4776611328125 0 796.5074
992.5078125 0 10280.602 c Ammonia loss 7
993.5101928710938 0 5124.4053
994.516845703125 0 1555.3254
1008.49365234375 0 660.73755
1035.521728515625 0 1581.5084
1079.5155029296875 0 2860.9143 z 2
1080.5179443359375 0 3098.822
1081.5233154296875 0 625.3069
1095.5350341796875 0 2516.8723 y 2
1096.5377197265625 0 1399.4194
1097.5421142578125 0 794.91724
1106.5889892578125 0 1650.1779 c 8
1107.5865478515625 0 1286.9313
1146.63037109375 0 888.9493
1155.5255126953125 0 1462.7904
1156.5313720703125 0 1072.0455
1161.6424560546875 0 3904.0845
1162.648681640625 0 2919.37
1163.65087890625 0 1642.8904
1176.6138916015625 0 705.55164 y Water loss 1
1178.5863037109375 0 1729.7965 z 1
1179.5872802734375 0 1106.172
1190.6405029296875 0 1137.6621
1191.6435546875 0 1326.9098
1205.6544189453125 0 21348.1 c 9
1206.658203125 0 13164.727
1207.663330078125 0 9429.381
1208.669921875 0 2454.3564
1209.6669921875 0 809.1874
1219.6453857421875 0 925.80994
1236.5177001953125 0 938.3215
1236.67236328125 0 1591.2498
1237.6646728515625 0 1281.2262
1244.67578125 0 777.7109
1245.6881103515625 0 765.9182
1246.688232421875 0 893.38025
1248.650146484375 0 5108.684
1249.66015625 0 5368.301
1250.647216796875 0 1588.5049
1252.6375732421875 0 1237.4595
1262.6788330078125 0 1141.1813
1263.693359375 0 1175.6545
1264.698974609375 0 2807.7217
1265.7041015625 0 1523.1158
1280.69775390625 0 3913.1523
1281.7008056640625 0 2069.5884
1282.713623046875 0 1006.2944
1289.6693115234375 0 742.6753
1290.674560546875 0 1685.0677
1291.667724609375 0 21865.182
1292.6700439453125 0 16245.164
1293.5345458984375 0 575.0075
1293.673095703125 0 6315.652
1294.689453125 0 1119.866
1304.7115478515625 0 767.91125
1305.7078857421875 0 2393.7664
1306.6998291015625 0 1646.5074
1307.522705078125 0 857.8949
1307.6859130859375 0 17447.87
1308.692138671875 0 83442.75
1309.696044921875 0 58302.312
1310.698974609375 0 23990.45
1311.703369140625 0 2993.2317

Spectrum Details

|  |  |
| --- | --- |
| Matched peaks? Matched peaks The total absolute number of peaks matched. Additionally in brackets the total fraction of peaks matched and the total number of peaks is shown. | 24 (17.02% of 141) |
| FDR? FDR The false discovery rate estimated for this peptide. It is calculated by matching all theoretical fragments with a non-integer shift with the raw peaks for this spectrum. This is done with 40 different shifts. The resulting percentage is the average number of annotated peaks over the number of annotated peaks with the correct spectrum. | 1.29% |
| Satellite FDR? Satellite FDR See the FDR for details on its calculation. This satellite ion specific FDR only contains the satellite ions (d/w) for I/L/J positions. | - |
| PSM Score? PSM Score The PSM Score as given by Hecklib to this annotated spectrum. It is shown with three significant figures. | 249 |

#### Reverse Lookup? Reverse Lookup All places where this read could be placed.

| Group | Segment | Template | Template Part | Read Part | Score | Unique |
| --- | --- | --- | --- | --- | --- | --- |
| Homo sapiens Heavy Chain | IGHC | IGHG1 | [27..38] | [0..11] | 88 | False |
| Homo sapiens Heavy Chain | IGHC | IGHG3 | [27..38] | [0..11] | 88 | False |
| Homo sapiens Heavy Chain | IGHC | IGHG2 | [27..38] | [0..11] | 88 | False |
| Homo sapiens Heavy Chain | IGHC | IGHG4 | [27..38] | [0..11] | 88 | False |

| Recombined | Template Part | Read Part | Score | Unique |
| --- | --- | --- | --- | --- |
| REC-0-1 | [149..160] | [0..11] | 88 | True |

#### Meta Information from Multiple reads

##### Number of combined reads

13

##### Intensity

0.9873

##### TotalArea

3.162E+08

##### Changes to the peptide sequence

JVKDYFPEPVT

L→JNo support for either Leucine or Isoleucine based on side chain ions (Position: 1)

#### Positional Score

Copy Data

##### Positional Score (TSV)

###### Preview

```
Loading example...
```

*Click on the button to copy the data to your clipboard.*

10012345678910

Label Value
"0" 0.835
"1" 0.827
"2" 0.814
"3" 0.806
"4" 0.821
"5" 0.822
"6" 0.824
"7" 0.835
"8" 0.835
"9" 0.825
"10" 0.828

#### Meta Information from PEAKS

##### Scan Identifier

F2:7751

##### Original sequence

L

V

K

D

Y

F

P

E

P

V

T

##### Posttranslational Modifications

##### Source File

D:\separate\_stitch\_analyses\xle-disambiguation\raw\20210323\_F1\_UM1\_Peng0013\_SA\_F59\_ingel\_3ug\_TL.raw

##### Fraction

2

##### Scan Feature

-

##### De Novo Score

99

##### ConfidenceScore

99

### m/z

654.3489

##### Mass

1306.6809

##### Charge

2

##### Retention Time

43.48

##### Predicted Retention Time

-

##### Area

0

##### Parts Per Million

1.8

##### Fragmentation mode

ETHCD

##### Originating file

01 D:\separate\_stitch\_analyses\xle-disambiguation\20210325\_F59\_3ug\_DENOVO\_12.csv

#### Meta Information from PEAKS

##### Scan Identifier

F2:7925

##### Original sequence

L

V

K

D

Y

F

P

E

P

V

T

##### Posttranslational Modifications

##### Source File

D:\separate\_stitch\_analyses\xle-disambiguation\raw\20210323\_F1\_UM1\_Peng0013\_SA\_F59\_ingel\_3ug\_TL.raw

##### Fraction

2

##### Scan Feature

-

##### De Novo Score

99

##### ConfidenceScore

99

### m/z

654.3485

##### Mass

1306.6809

##### Charge

2

##### Retention Time

44.52

##### Predicted Retention Time

-

##### Area

0

##### Parts Per Million

1.2

##### Fragmentation mode

ETHCD

##### Originating file

01 D:\separate\_stitch\_analyses\xle-disambiguation\20210325\_F59\_3ug\_DENOVO\_12.csv

#### Meta Information from PEAKS

##### Scan Identifier

F2:7808

##### Original sequence

L

V

K

D

Y

F

P

E

P

V

T

##### Posttranslational Modifications

##### Source File

D:\separate\_stitch\_analyses\xle-disambiguation\raw\20210323\_F1\_UM1\_Peng0013\_SA\_F59\_ingel\_3ug\_TL.raw

##### Fraction

2

##### Scan Feature

-

##### De Novo Score

99

##### ConfidenceScore

99

### m/z

654.3486

##### Mass

1306.6809

##### Charge

2

##### Retention Time

43.83

##### Predicted Retention Time

-

##### Area

0

##### Parts Per Million

1.3

##### Fragmentation mode

HCD

##### Originating file

01 D:\separate\_stitch\_analyses\xle-disambiguation\20210325\_F59\_3ug\_DENOVO\_12.csv

#### Meta Information from PEAKS

##### Scan Identifier

F2:7692

##### Original sequence

L

V

K

D

Y

F

P

E

P

V

T

##### Posttranslational Modifications

##### Source File

D:\separate\_stitch\_analyses\xle-disambiguation\raw\20210323\_F1\_UM1\_Peng0013\_SA\_F59\_ingel\_3ug\_TL.raw

##### Fraction

2

##### Scan Feature

F2:10611

##### De Novo Score

98

##### ConfidenceScore

98

### m/z

654.3489

##### Mass

1306.6809

##### Charge

2

##### Retention Time

43.12

##### Predicted Retention Time

-

##### Area

3.162E+08

##### Parts Per Million

1.8

##### Fragmentation mode

HCD

##### Originating file

01 D:\separate\_stitch\_analyses\xle-disambiguation\20210325\_F59\_3ug\_DENOVO\_12.csv

#### Meta Information from PEAKS

##### Scan Identifier

F2:7862

##### Original sequence

L

V

K

D

Y

F

P

E

P

V

T

##### Posttranslational Modifications

##### Source File

D:\separate\_stitch\_analyses\xle-disambiguation\raw\20210323\_F1\_UM1\_Peng0013\_SA\_F59\_ingel\_3ug\_TL.raw

##### Fraction

2

##### Scan Feature

-

##### De Novo Score

98

##### ConfidenceScore

98

### m/z

654.3484

##### Mass

1306.6809

##### Charge

2

##### Retention Time

44.15

##### Predicted Retention Time

-

##### Area

0

##### Parts Per Million

1.1

##### Fragmentation mode

HCD

##### Originating file

01 D:\separate\_stitch\_analyses\xle-disambiguation\20210325\_F59\_3ug\_DENOVO\_12.csv

#### Meta Information from PEAKS

##### Scan Identifier

F2:7987

##### Original sequence

L

V

K

D

Y

F

P

E

P

V

T

##### Posttranslational Modifications

##### Source File

D:\separate\_stitch\_analyses\xle-disambiguation\raw\20210323\_F1\_UM1\_Peng0013\_SA\_F59\_ingel\_3ug\_TL.raw

##### Fraction

2

##### Scan Feature

-

##### De Novo Score

98

##### ConfidenceScore

98

### m/z

654.3485

##### Mass

1306.6809

##### Charge

2

##### Retention Time

44.9

##### Predicted Retention Time

-

##### Area

0

##### Parts Per Million

1.2

##### Fragmentation mode

ETHCD

##### Originating file

01 D:\separate\_stitch\_analyses\xle-disambiguation\20210325\_F59\_3ug\_DENOVO\_12.csv

#### Meta Information from PEAKS

##### Scan Identifier

F2:8278

##### Original sequence

L

V

K

D

Y

F

P

E

P

V

T

##### Posttranslational Modifications

##### Source File

D:\separate\_stitch\_analyses\xle-disambiguation\raw\20210323\_F1\_UM1\_Peng0013\_SA\_F59\_ingel\_3ug\_TL.raw

##### Fraction

2

##### Scan Feature

-

##### De Novo Score

97

##### ConfidenceScore

97

### m/z

654.3484

##### Mass

1306.6809

##### Charge

2

##### Retention Time

46.62

##### Predicted Retention Time

-

##### Area

0

##### Parts Per Million

1

##### Fragmentation mode

ETHCD

##### Originating file

01 D:\separate\_stitch\_analyses\xle-disambiguation\20210325\_F59\_3ug\_DENOVO\_12.csv

#### Meta Information from PEAKS

##### Scan Identifier

F2:8165

##### Original sequence

L

V

K

D

Y

F

P

E

P

V

T

##### Posttranslational Modifications

##### Source File

D:\separate\_stitch\_analyses\xle-disambiguation\raw\20210323\_F1\_UM1\_Peng0013\_SA\_F59\_ingel\_3ug\_TL.raw

##### Fraction

2

##### Scan Feature

-

##### De Novo Score

97

##### ConfidenceScore

97

### m/z

654.3492

##### Mass

1306.6809

##### Charge

2

##### Retention Time

45.97

##### Predicted Retention Time

-

##### Area

0

##### Parts Per Million

2.3

##### Fragmentation mode

ETHCD

##### Originating file

01 D:\separate\_stitch\_analyses\xle-disambiguation\20210325\_F59\_3ug\_DENOVO\_12.csv

#### Meta Information from PEAKS

##### Scan Identifier

F2:8100

##### Original sequence

L

V

K

D

Y

F

P

E

P

V

T

##### Posttranslational Modifications

##### Source File

D:\separate\_stitch\_analyses\xle-disambiguation\raw\20210323\_F1\_UM1\_Peng0013\_SA\_F59\_ingel\_3ug\_TL.raw

##### Fraction

2

##### Scan Feature

-

##### De Novo Score

97

##### ConfidenceScore

97

### m/z

654.3486

##### Mass

1306.6809

##### Charge

2

##### Retention Time

45.59

##### Predicted Retention Time

-

##### Area

0

##### Parts Per Million

1.3

##### Fragmentation mode

HCD

##### Originating file

01 D:\separate\_stitch\_analyses\xle-disambiguation\20210325\_F59\_3ug\_DENOVO\_12.csv

#### Meta Information from PEAKS

##### Scan Identifier

F2:8345

##### Original sequence

L

V

K

D

Y

F

P

E

P

V

T

##### Posttranslational Modifications

##### Source File

D:\separate\_stitch\_analyses\xle-disambiguation\raw\20210323\_F1\_UM1\_Peng0013\_SA\_F59\_ingel\_3ug\_TL.raw

##### Fraction

2

##### Scan Feature

-

##### De Novo Score

96

##### ConfidenceScore

96

### m/z

654.3483

##### Mass

1306.6809

##### Charge

2

##### Retention Time

47

##### Predicted Retention Time

-

##### Area

0

##### Parts Per Million

0.8

##### Fragmentation mode

ETHCD

##### Originating file

01 D:\separate\_stitch\_analyses\xle-disambiguation\20210325\_F59\_3ug\_DENOVO\_12.csv

#### Meta Information from PEAKS

##### Scan Identifier

F2:8045

##### Original sequence

L

V

K

D

Y

F

P

E

P

V

T

##### Posttranslational Modifications

##### Source File

D:\separate\_stitch\_analyses\xle-disambiguation\raw\20210323\_F1\_UM1\_Peng0013\_SA\_F59\_ingel\_3ug\_TL.raw

##### Fraction

2

##### Scan Feature

-

##### De Novo Score

96

##### ConfidenceScore

96

### m/z

654.3488

##### Mass

1306.6809

##### Charge

2

##### Retention Time

45.25

##### Predicted Retention Time

-

##### Area

0

##### Parts Per Million

1.6

##### Fragmentation mode

HCD

##### Originating file

01 D:\separate\_stitch\_analyses\xle-disambiguation\20210325\_F59\_3ug\_DENOVO\_12.csv

#### Meta Information from PEAKS

##### Scan Identifier

F2:8221

##### Original sequence

L

V

K

D

Y

F

P

E

P

V

T

##### Posttranslational Modifications

##### Source File

D:\separate\_stitch\_analyses\xle-disambiguation\raw\20210323\_F1\_UM1\_Peng0013\_SA\_F59\_ingel\_3ug\_TL.raw

##### Fraction

2

##### Scan Feature

-

##### De Novo Score

96

##### ConfidenceScore

96

### m/z

654.3482

##### Mass

1306.6809

##### Charge

2

##### Retention Time

46.31

##### Predicted Retention Time

-

##### Area

0

##### Parts Per Million

0.7

##### Fragmentation mode

HCD

##### Originating file

01 D:\separate\_stitch\_analyses\xle-disambiguation\20210325\_F59\_3ug\_DENOVO\_12.csv

#### Meta Information from PEAKS

##### Scan Identifier

F2:8629

##### Original sequence

L

V

K

D

Y

F

P

E

P

V

T

##### Posttranslational Modifications

##### Source File

D:\separate\_stitch\_analyses\xle-disambiguation\raw\20210323\_F1\_UM1\_Peng0013\_SA\_F59\_ingel\_3ug\_TL.raw

##### Fraction

2

##### Scan Feature

-

##### De Novo Score

95

##### ConfidenceScore

95

### m/z

654.3481

##### Mass

1306.6809

##### Charge

2

##### Retention Time

48.58

##### Predicted Retention Time

-

##### Area

0

##### Parts Per Million

0.5

##### Fragmentation mode

ETHCD

##### Originating file

01 D:\separate\_stitch\_analyses\xle-disambiguation\20210325\_F59\_3ug\_DENOVO\_12.csv
