## Supplementary material for "A handle on mass coincidence errors in *de novo* sequencing of antibodies by bottom-up proteomics": Combined_029.html

Details Combined\_029 | Stitch OverviewUndefined

### Read Combined\_029

#### Sequence (length=13)

VVVDVSHEDPEVK

#### Spectrum 4311? Spectrum 4311 The raw spectrum of this peptide as annotated by Hecklib. The fragments are coloured according to ion type (see legend). Any peaks with a star '\*' as text can be hovered over to see the full details, first the ion type second the mass shift type. By hovering over the amino acids in the peptide or ions in the legend the corresponding peaks are highlighted. By toggling the 'Unassigned' label you can turn the background (unassigned) peaks on or off in the plot. By updating the slider in the Ion legend you can update the spectrum to only show the top X% of the peaks with labels. The top X% means any peak that is within X% of the highest intensity. By dragging in the spectrum you can zoom in to a specific part of the spectrum and use 'Zoom Out' to get back to the original zoom level. The annotation of the spectrum is based on the given sequence in the peptides file and is done with different software so inconsistencies are likely. The peaks are annotated based on the given sequence, with 20 ppm tolerance.

Copy Data

##### Spectrum 4311 (TSV)

###### Preview

```
Loading example...
```

*Click on the button to copy the data to your clipboard.*

Mz MinMz MaxIntensity Max

WidthHeightPeptide font sizePeptide stroke widthSpectrum font sizeSpectrum stroke widthCompact peptide

Ion legend

wxyz

abcd

OtherUnassignedIon^Charge^_Position_Show for top:%

VVVDVSHEDPEVK

01.31e+42.62e+43.94e+45.25e+4

Zoom Out

y^+1^_1_y^+1^_1_a^+1^_2_y^+2^_3_b^+1^_2_y^+1^_2_b^+2^_6_y^+1^_3_y^+1^_3_y^+2^_7_y^+1^_4_y^+2^_8_y^+1^_4_y^+2^_9_y^+2^_10_y^+1^_5_y^+2^_11_y^+2^_11_y^+2^_11_y^+2^_12_y^+2^_12_y^+1^_6_y^+1^_7_y^+1^_8_y^+1^_9_y^+1^_10_

0779155923383118

Fragment Matches Table

Show background peaks

| Position | Ion type | Intensity | mz Theoretical | mz Error (Th) | mz Error (ppm) | Charge | Series Number |
| --- | --- | --- | --- | --- | --- | --- | --- |
| - | - | 3037 | 120.1 | - | - | 0 | - |
| - | - | 378.4 | 121 | - | - | 0 | - |
| - | - | 413.9 | 124.3 | - | - | 0 | - |
| - | - | 837.3 | 127.1 | - | - | 0 | - |
| - | - | 522.8 | 129.1 | - | - | 0 | - |
| - | - | 2.002E+04 | 129.1 | - | - | 0 | - |
| - | - | 1381 | 130.1 | - | - | 0 | - |
| - | - | 783.2 | 130.1 | - | - | 0 | - |
| 13 | y | 7552 | 130.1 | 0.0003081 | 2.368 | +1 | 1 |
| - | - | 1149 | 130.1 | - | - | 0 | - |
| - | - | 677.9 | 131.1 | - | - | 0 | - |
| - | - | 421.6 | 133.1 | - | - | 0 | - |
| - | - | 380.4 | 133.3 | - | - | 0 | - |
| - | - | 3673 | 136.1 | - | - | 0 | - |
| - | - | 514.5 | 139.1 | - | - | 0 | - |
| - | - | 498.5 | 140.1 | - | - | 0 | - |
| - | - | 568.6 | 141.1 | - | - | 0 | - |
| - | - | 515.1 | 141.1 | - | - | 0 | - |
| - | - | 540.8 | 142.1 | - | - | 0 | - |
| - | - | 613.8 | 142.1 | - | - | 0 | - |
| - | - | 1908 | 143.1 | - | - | 0 | - |
| - | - | 933.8 | 145.1 | - | - | 0 | - |
| - | - | 571.8 | 146.1 | - | - | 0 | - |
| - | - | 937.3 | 147.1 | - | - | 0 | - |
| 13 | y | 1.315E+04 | 147.1 | 0.0003092 | 2.102 | +1 | 1 |
| - | - | 468.5 | 152.1 | - | - | 0 | - |
| - | - | 561.1 | 152.1 | - | - | 0 | - |
| - | - | 715.3 | 153.1 | - | - | 0 | - |
| - | - | 738.2 | 155.1 | - | - | 0 | - |
| - | - | 633.6 | 157.1 | - | - | 0 | - |
| - | - | 467.2 | 158.1 | - | - | 0 | - |
| - | - | 663.4 | 158.1 | - | - | 0 | - |
| - | - | 476.2 | 159.1 | - | - | 0 | - |
| - | - | 858.4 | 159.1 | - | - | 0 | - |
| - | - | 5505 | 163.1 | - | - | 0 | - |
| - | - | 883.6 | 164.1 | - | - | 0 | - |
| - | - | 520.2 | 166.1 | - | - | 0 | - |
| - | - | 468.7 | 168.5 | - | - | 0 | - |
| - | - | 754.4 | 169.1 | - | - | 0 | - |
| - | - | 1615 | 169.1 | - | - | 0 | - |
| - | - | 574.1 | 169.1 | - | - | 0 | - |
| - | - | 445 | 170.1 | - | - | 0 | - |
| - | - | 488.5 | 171.1 | - | - | 0 | - |
| - | - | 2208 | 171.1 | - | - | 0 | - |
| 2 | a | 5.197E+04 | 171.1 | 0.000316 | 1.846 | +1 | 2 |
| - | - | 3783 | 172.2 | - | - | 0 | - |
| - | - | 516.8 | 172.4 | - | - | 0 | - |
| - | - | 599.2 | 173.1 | - | - | 0 | - |
| - | - | 1739 | 175.1 | - | - | 0 | - |
| - | - | 431.3 | 178.1 | - | - | 0 | - |
| - | - | 849 | 179.1 | - | - | 0 | - |
| - | - | 436 | 183 | - | - | 0 | - |
| - | - | 573.9 | 183.1 | - | - | 0 | - |
| - | - | 958.3 | 185.1 | - | - | 0 | - |
| - | - | 2.566E+04 | 185.2 | - | - | 0 | - |
| - | - | 3476 | 186.1 | - | - | 0 | - |
| - | - | 3056 | 186.2 | - | - | 0 | - |
| - | - | 6379 | 187.1 | - | - | 0 | - |
| - | - | 1571 | 187.1 | - | - | 0 | - |
| 11 | y | 585.9 | 188.1 | 0.0033 | 17.54 | +2 | 3 |
| - | - | 707.8 | 188.1 | - | - | 0 | - |
| - | - | 505.8 | 192.7 | - | - | 0 | - |
| - | - | 2049 | 197.1 | - | - | 0 | - |
| - | - | 606.3 | 197.1 | - | - | 0 | - |
| - | - | 3344 | 199.1 | - | - | 0 | - |
| 2 | b | 1.946E+04 | 199.1 | 0.0001828 | 0.918 | +1 | 2 |
| - | - | 1897 | 200.1 | - | - | 0 | - |
| - | - | 1240 | 201.1 | - | - | 0 | - |
| - | - | 875.4 | 202.1 | - | - | 0 | - |
| - | - | 1.747E+04 | 203.1 | - | - | 0 | - |
| - | - | 1363 | 204.1 | - | - | 0 | - |
| - | - | 1139 | 207.1 | - | - | 0 | - |
| - | - | 1033 | 208.1 | - | - | 0 | - |
| - | - | 932.1 | 210.1 | - | - | 0 | - |
| - | - | 512.1 | 211.1 | - | - | 0 | - |
| - | - | 1.498E+04 | 213.2 | - | - | 0 | - |
| - | - | 1981 | 214.2 | - | - | 0 | - |
| - | - | 6086 | 215.1 | - | - | 0 | - |
| - | - | 1053 | 215.1 | - | - | 0 | - |
| - | - | 628.8 | 216.1 | - | - | 0 | - |
| - | - | 1090 | 218.2 | - | - | 0 | - |
| - | - | 5579 | 225.1 | - | - | 0 | - |
| - | - | 9203 | 226.2 | - | - | 0 | - |
| - | - | 8984 | 227.1 | - | - | 0 | - |
| - | - | 895.8 | 227.2 | - | - | 0 | - |
| - | - | 1348 | 228.1 | - | - | 0 | - |
| - | - | 1051 | 228.1 | - | - | 0 | - |
| - | - | 1805 | 228.2 | - | - | 0 | - |
| - | - | 621.2 | 234.1 | - | - | 0 | - |
| - | - | 1010 | 239.1 | - | - | 0 | - |
| 12 | y | 1.628E+04 | 246.2 | 0.0002547 | 1.035 | +1 | 2 |
| - | - | 2080 | 247.2 | - | - | 0 | - |
| - | - | 885 | 249.1 | - | - | 0 | - |
| - | - | 1931 | 253.2 | - | - | 0 | - |
| - | - | 576.5 | 264.7 | - | - | 0 | - |
| - | - | 1040 | 266.1 | - | - | 0 | - |
| - | - | 2574 | 267.1 | - | - | 0 | - |
| - | - | 523 | 269.2 | - | - | 0 | - |
| - | - | 1592 | 270.2 | - | - | 0 | - |
| - | - | 888.5 | 270.2 | - | - | 0 | - |
| - | - | 560.6 | 271.2 | - | - | 0 | - |
| - | - | 563.5 | 272.9 | - | - | 0 | - |
| - | - | 1059 | 274.1 | - | - | 0 | - |
| - | - | 1529 | 279.2 | - | - | 0 | - |
| - | - | 901.6 | 283.1 | - | - | 0 | - |
| - | - | 2688 | 287.2 | - | - | 0 | - |
| - | - | 3919 | 297.2 | - | - | 0 | - |
| - | - | 683.2 | 298.2 | - | - | 0 | - |
| - | - | 6876 | 299.2 | - | - | 0 | - |
| 6 | b | 1511 | 300.2 | 0.001093 | 3.64 | +2 | 6 |
| - | - | 1245 | 302.1 | - | - | 0 | - |
| - | - | 568.5 | 302.9 | - | - | 0 | - |
| - | - | 653.1 | 306.2 | - | - | 0 | - |
| - | - | 1145 | 307.1 | - | - | 0 | - |
| - | - | 3405 | 314.2 | - | - | 0 | - |
| - | - | 1015 | 324.2 | - | - | 0 | - |
| - | - | 3231 | 326.2 | - | - | 0 | - |
| - | - | 856.9 | 327.1 | - | - | 0 | - |
| - | - | 1946 | 340.2 | - | - | 0 | - |
| - | - | 866.1 | 340.7 | - | - | 0 | - |
| - | - | 1164 | 342.1 | - | - | 0 | - |
| - | - | 568.4 | 344.2 | - | - | 0 | - |
| - | - | 648 | 346.1 | - | - | 0 | - |
| - | - | 1817 | 354.1 | - | - | 0 | - |
| 11 | y | 1324 | 357.2 | 0.0001629 | 0.4561 | +1 | 3 |
| - | - | 2135 | 364.1 | - | - | 0 | - |
| - | - | 841.6 | 365.1 | - | - | 0 | - |
| - | - | 8846 | 368.2 | - | - | 0 | - |
| - | - | 2513 | 369.2 | - | - | 0 | - |
| - | - | 1071 | 370.2 | - | - | 0 | - |
| - | - | 528.7 | 370.3 | - | - | 0 | - |
| 11 | y | 1107 | 375.2 | 0.001042 | 2.778 | +1 | 3 |
| - | - | 925.2 | 379.2 | - | - | 0 | - |
| - | - | 6544 | 382.1 | - | - | 0 | - |
| - | - | 1082 | 383.1 | - | - | 0 | - |
| - | - | 695.2 | 384.2 | - | - | 0 | - |
| - | - | 2071 | 398.2 | - | - | 0 | - |
| - | - | 1000 | 399.2 | - | - | 0 | - |
| - | - | 664.2 | 401.2 | - | - | 0 | - |
| - | - | 962.4 | 410.1 | - | - | 0 | - |
| - | - | 1072 | 411.2 | - | - | 0 | - |
| - | - | 577.2 | 422.5 | - | - | 0 | - |
| 7 | y | 2488 | 427.2 | 8.806E-05 | 0.2061 | +2 | 7 |
| - | - | 707.5 | 427.7 | - | - | 0 | - |
| - | - | 886.8 | 434.7 | - | - | 0 | - |
| - | - | 1503 | 436.3 | - | - | 0 | - |
| - | - | 1592 | 439.2 | - | - | 0 | - |
| - | - | 753.8 | 440.2 | - | - | 0 | - |
| - | - | 618.4 | 442.3 | - | - | 0 | - |
| - | - | 1164 | 451.2 | - | - | 0 | - |
| - | - | 753.9 | 453.2 | - | - | 0 | - |
| 10 | y | 747.3 | 454.3 | 0.001781 | 3.922 | +1 | 4 |
| - | - | 1439 | 457.2 | - | - | 0 | - |
| - | - | 682.4 | 464.3 | - | - | 0 | - |
| - | - | 832.6 | 465.3 | - | - | 0 | - |
| - | - | 5305 | 469.2 | - | - | 0 | - |
| - | - | 1785 | 470.2 | - | - | 0 | - |
| - | - | 655.3 | 470.3 | - | - | 0 | - |
| 6 | y | 5427 | 470.7 | 0.000118 | 0.2508 | +2 | 8 |
| - | - | 1742 | 471.2 | - | - | 0 | - |
| - | - | 841.4 | 471.7 | - | - | 0 | - |
| 10 | y | 2.766E+04 | 472.3 | 0.0002194 | 0.4646 | +1 | 4 |
| - | - | 7604 | 473.3 | - | - | 0 | - |
| - | - | 1159 | 474.3 | - | - | 0 | - |
| - | - | 711.8 | 475.3 | - | - | 0 | - |
| - | - | 688.3 | 475.8 | - | - | 0 | - |
| - | - | 2990 | 482.3 | - | - | 0 | - |
| - | - | 658.3 | 483.3 | - | - | 0 | - |
| - | - | 3269 | 484.3 | - | - | 0 | - |
| - | - | 1928 | 484.8 | - | - | 0 | - |
| - | - | 1785 | 485.3 | - | - | 0 | - |
| - | - | 1218 | 485.4 | - | - | 0 | - |
| - | - | 2401 | 510.3 | - | - | 0 | - |
| - | - | 1482 | 513.4 | - | - | 0 | - |
| - | - | 1527 | 514.7 | - | - | 0 | - |
| - | - | 1015 | 515.2 | - | - | 0 | - |
| 5 | y | 5224 | 520.3 | 0.0009998 | 1.922 | +2 | 9 |
| - | - | 2498 | 520.8 | - | - | 0 | - |
| - | - | 1138 | 521.3 | - | - | 0 | - |
| - | - | 729.3 | 525.3 | - | - | 0 | - |
| - | - | 991.2 | 536.3 | - | - | 0 | - |
| - | - | 1512 | 538.3 | - | - | 0 | - |
| - | - | 7568 | 553.3 | - | - | 0 | - |
| - | - | 2692 | 554.3 | - | - | 0 | - |
| - | - | 678.6 | 555.3 | - | - | 0 | - |
| - | - | 3866 | 568.2 | - | - | 0 | - |
| - | - | 1218 | 569.2 | - | - | 0 | - |
| - | - | 1.687E+04 | 570.3 | - | - | 0 | - |
| - | - | 968.2 | 571.3 | - | - | 0 | - |
| - | - | 4582 | 571.3 | - | - | 0 | - |
| - | - | 897.1 | 572.3 | - | - | 0 | - |
| 4 | y | 8158 | 577.8 | 0.0007384 | 1.278 | +2 | 10 |
| - | - | 5902 | 578.3 | - | - | 0 | - |
| - | - | 2592 | 578.8 | - | - | 0 | - |
| - | - | 904.9 | 580.3 | - | - | 0 | - |
| 9 | y | 1651 | 587.3 | 0.0006812 | 1.16 | +1 | 5 |
| - | - | 623.1 | 588.3 | - | - | 0 | - |
| - | - | 876.9 | 593.3 | - | - | 0 | - |
| - | - | 2628 | 605.3 | - | - | 0 | - |
| - | - | 1784 | 608.2 | - | - | 0 | - |
| 3 | y | 1024 | 618.3 | 0.001709 | 2.764 | +2 | 11 |
| 3 | y | 770 | 618.8 | 0.0108 | 17.45 | +2 | 11 |
| 3 | y | 4.533E+04 | 627.3 | 0.0005215 | 0.8314 | +2 | 11 |
| - | - | 2.742E+04 | 627.8 | - | - | 0 | - |
| - | - | 1.187E+04 | 628.3 | - | - | 0 | - |
| - | - | 2715 | 628.8 | - | - | 0 | - |
| - | - | 613.3 | 650.4 | - | - | 0 | - |
| - | - | 1802 | 666.4 | - | - | 0 | - |
| - | - | 2049 | 667.3 | - | - | 0 | - |
| 2 | y | 1109 | 668.3 | 0.01114 | 16.67 | +2 | 12 |
| - | - | 640.3 | 669.3 | - | - | 0 | - |
| 2 | y | 9543 | 676.8 | 0.000854 | 1.262 | +2 | 12 |
| - | - | 5356 | 677.3 | - | - | 0 | - |
| - | - | 3405 | 677.8 | - | - | 0 | - |
| - | - | 1624 | 678.3 | - | - | 0 | - |
| - | - | 4249 | 683.3 | - | - | 0 | - |
| - | - | 1862 | 684.3 | - | - | 0 | - |
| - | - | 1040 | 694.4 | - | - | 0 | - |
| - | - | 1365 | 695.3 | - | - | 0 | - |
| 8 | y | 4427 | 716.3 | 0.001629 | 2.274 | +1 | 6 |
| - | - | 1680 | 717.3 | - | - | 0 | - |
| - | - | 815.8 | 734.3 | - | - | 0 | - |
| - | - | 1374 | 738.4 | - | - | 0 | - |
| - | - | 1.546E+04 | 755.4 | - | - | 0 | - |
| - | - | 7463 | 756.4 | - | - | 0 | - |
| - | - | 1180 | 757.4 | - | - | 0 | - |
| - | - | 7540 | 765.5 | - | - | 0 | - |
| - | - | 3677 | 766.5 | - | - | 0 | - |
| - | - | 1294 | 767.5 | - | - | 0 | - |
| - | - | 1.573E+04 | 782.3 | - | - | 0 | - |
| - | - | 5724 | 783.3 | - | - | 0 | - |
| - | - | 1105 | 784.3 | - | - | 0 | - |
| - | - | 999.1 | 787.4 | - | - | 0 | - |
| - | - | 1557 | 794.3 | - | - | 0 | - |
| - | - | 1524 | 805.4 | - | - | 0 | - |
| - | - | 1178 | 806.4 | - | - | 0 | - |
| - | - | 637.5 | 807.4 | - | - | 0 | - |
| 7 | y | 4495 | 853.4 | 0.001031 | 1.208 | +1 | 7 |
| - | - | 1218 | 854.4 | - | - | 0 | - |
| - | - | 2483 | 868.5 | - | - | 0 | - |
| - | - | 1354 | 869.5 | - | - | 0 | - |
| - | - | 4221 | 881.4 | - | - | 0 | - |
| - | - | 1451 | 882.4 | - | - | 0 | - |
| - | - | 2182 | 909.4 | - | - | 0 | - |
| - | - | 1201 | 910.4 | - | - | 0 | - |
| 6 | y | 6218 | 940.4 | 0.002359 | 2.508 | +1 | 8 |
| - | - | 1740 | 941.4 | - | - | 0 | - |
| - | - | 998.5 | 942.4 | - | - | 0 | - |
| - | - | 846.7 | 980.4 | - | - | 0 | - |
| - | - | 6394 | 1008 | - | - | 0 | - |
| - | - | 3522 | 1009 | - | - | 0 | - |
| - | - | 1550 | 1010 | - | - | 0 | - |
| 5 | y | 1784 | 1040 | 0.005465 | 5.258 | +1 | 9 |
| - | - | 3084 | 1107 | - | - | 0 | - |
| - | - | 1780 | 1108 | - | - | 0 | - |
| 4 | y | 1645 | 1155 | 0.003478 | 3.012 | +1 | 10 |
| - | - | 1391 | 1156 | - | - | 0 | - |
| - | - | 620.6 | 1463 | - | - | 0 | - |
| - | - | 639.7 | 1653 | - | - | 0 | - |
| - | - | 670.1 | 2267 | - | - | 0 | - |
| - | - | 724.2 | 2304 | - | - | 0 | - |
| - | - | 711.6 | 2463 | - | - | 0 | - |
| - | - | 776.3 | 3074 | - | - | 0 | - |
| - | - | 705.9 | 3087 | - | - | 0 | - |

m/z Charge Intensity FragmentType MassShift Position
120.08106231689453 0 3036.5713
120.95418548583984 0 378.41727
124.34746551513672 0 413.91663
127.08712768554688 0 837.2859
129.06607055664062 0 522.8287
129.10252380371094 0 20018.48
130.050048828125 0 1381.1335
130.06532287597656 0 783.20496
130.08656311035156 0 7551.7803 y Ammonia loss 12
130.10574340820312 0 1148.6862
131.0899200439453 0 677.86847
133.06080627441406 0 421.64624
133.3066864013672 0 380.40466
136.07601928710938 0 3672.6494
139.0504608154297 0 514.4827
140.0820770263672 0 498.46997
141.06613159179688 0 568.6288
141.10244750976562 0 515.10065
142.08685302734375 0 540.77856
142.12254333496094 0 613.84564
143.11805725097656 0 1907.5082
145.06101989746094 0 933.75446
146.06063842773438 0 571.83484
147.07662963867188 0 937.3463
147.1131134033203 0 13153.129 y 12
152.07066345214844 0 468.49927
152.1068115234375 0 561.0872
153.0663604736328 0 715.26953
155.1179962158203 0 738.2256
157.09739685058594 0 633.5953
158.0598907470703 0 467.22977
158.09225463867188 0 663.4471
159.07662963867188 0 476.243
159.09190368652344 0 858.41473
163.0716094970703 0 5505.1216
164.1189422607422 0 883.569
166.06143188476562 0 520.24274
168.54615783691406 0 468.74344
169.06103515625 0 754.3924
169.09765625 0 1615.366
169.13377380371094 0 574.14813
170.1002655029297 0 445.03604
171.07669067382812 0 488.5129
171.11293029785156 0 2207.5486
171.14950561523438 0 51974.445 a 1
172.1527557373047 0 3783.33
172.42356872558594 0 516.7594
173.12896728515625 0 599.2185
175.08689880371094 0 1738.9558
178.1171112060547 0 431.25803
179.09315490722656 0 848.99603
182.955810546875 0 436.03806
183.1130828857422 0 573.8846
185.09251403808594 0 958.27856
185.16513061523438 0 25657.197
186.08767700195312 0 3476.1506
186.1685333251953 0 3056.1526
187.10800170898438 0 6378.8193
187.14447021484375 0 1570.517
188.11224365234375 0 585.87604 y 10
188.13916015625 0 707.76764
192.72891235351562 0 505.77335
197.10340881347656 0 2048.6443
197.1287078857422 0 606.3465
199.10787963867188 0 3343.817
199.144287109375 0 19462.664 b 1
200.1478271484375 0 1896.587
201.12342834472656 0 1240.268
202.10777282714844 0 875.3863
203.10284423828125 0 17468.814
204.10650634765625 0 1363.3015
207.087646484375 0 1138.6484
208.0722198486328 0 1033.2527
210.12374877929688 0 932.13684
211.1075897216797 0 512.1245
213.16006469726562 0 14976.543
214.16342163085938 0 1981.3179
215.10289001464844 0 6085.9136
215.13885498046875 0 1052.8793
216.10743713378906 0 628.82574
218.15066528320312 0 1090.1434
225.09840393066406 0 5578.8994
226.1552276611328 0 9202.508
227.10292053222656 0 8983.557
227.1584930419922 0 895.84503
228.1068572998047 0 1347.5873
228.13453674316406 0 1050.625
228.1710968017578 0 1805.4429
234.12351989746094 0 621.22614
239.1145477294922 0 1009.58295
246.1814727783203 0 16281.542 y 11
247.1846160888672 0 2079.863
249.0984344482422 0 885.0085
253.1664276123047 0 1931.1399
264.73455810546875 0 576.53577
266.12615966796875 0 1039.6846
267.1091003417969 0 2574.4568
269.19757080078125 0 523.0126
270.1810302734375 0 1592.374
270.1949768066406 0 888.5384
271.1766052246094 0 560.5816
272.944091796875 0 563.49316
274.1400451660156 0 1058.7545
279.181396484375 0 1528.6091
283.14190673828125 0 901.61176
287.2081298828125 0 2687.7532
297.1924133300781 0 3918.64
298.1963195800781 0 683.15094
299.1716613769531 0 6875.8003
300.1746826171875 0 1511.4884 b 5
302.134521484375 0 1244.6157
302.8784484863281 0 568.50256
306.1553039550781 0 653.1346
307.13983154296875 0 1144.9952
314.171142578125 0 3404.9504
324.1670227050781 0 1014.7097
326.1710205078125 0 3231.2283
327.1303405761719 0 856.86237
340.23480224609375 0 1945.7898
340.6899719238281 0 866.14026
342.1299133300781 0 1164.0863
344.1842956542969 0 568.43005
346.1145935058594 0 648.0388
354.1404724121094 0 1816.8257
357.2134094238281 0 1324.4985 y Water loss 10
364.12469482421875 0 2134.663
365.1284484863281 0 841.6299
368.2291564941406 0 8845.515
369.2325744628906 0 2512.8806
370.244873046875 0 1071.0023
370.27838134765625 0 528.6731
375.224853515625 0 1107.3539 y 10
379.2086486816406 0 925.2396
382.1354675292969 0 6543.853
383.1380920410156 0 1081.8048
384.1953430175781 0 695.1597
398.23944091796875 0 2071.0413
399.2425842285156 0 1000.2646
401.2024841308594 0 664.1501
410.1305847167969 0 962.3518
411.1962890625 0 1071.5549
422.5451965332031 0 577.2284
427.20623779296875 0 2487.8796 y 6
427.70849609375 0 707.5296
434.746337890625 0 886.7845
436.255615234375 0 1502.5525
439.19317626953125 0 1592.3745
440.19671630859375 0 753.7949
442.3108215332031 0 618.4329
451.15594482421875 0 1164.3674
453.2095642089844 0 753.85876
454.2677917480469 0 747.3457 y Water loss 9
457.2265930175781 0 1439.319
464.2616882324219 0 682.44934
465.2806396484375 0 832.62616
469.1677551269531 0 5304.5415
470.1708679199219 0 1784.531
470.3106689453125 0 655.292
470.7220458984375 0 5427.05 y 5
471.22283935546875 0 1742.1057
471.7237548828125 0 841.40344
472.27679443359375 0 27663.645 y 9
473.27935791015625 0 7603.858
474.2828674316406 0 1159.406
475.2762145996094 0 711.7542
475.7762145996094 0 688.2688
482.2725524902344 0 2989.899
483.2783508300781 0 658.32074
484.28253173828125 0 3269.2349
484.787353515625 0 1928.4858
485.28741455078125 0 1785.1875
485.3603210449219 0 1218.4487
510.2651062011719 0 2400.645
513.3510131835938 0 1481.6821
514.739501953125 0 1527.0017
515.2416381835938 0 1015.40063
520.25537109375 0 5224.183 y 4
520.7566528320312 0 2497.9634
521.25830078125 0 1137.7275
525.3153686523438 0 729.2902
536.2837524414062 0 991.17584
538.2615966796875 0 1511.9736
553.309326171875 0 7567.83
554.3114624023438 0 2691.667
555.3130493164062 0 678.64844
568.2360229492188 0 3866.1082
569.2380981445312 0 1218.1245
570.3243408203125 0 16870.918
571.28076171875 0 968.20013
571.327880859375 0 4582.4556
572.3304443359375 0 897.05005
577.7691040039062 0 8158.3374 y 3
578.27099609375 0 5902.104
578.7703857421875 0 2591.5076
580.3087768554688 0 904.9225
587.30419921875 0 1650.5333 y 8
588.3053588867188 0 623.0548
593.3071899414062 0 876.9033
605.3065185546875 0 2628.2449
608.229736328125 0 1784.4945
618.2970581054688 0 1024.3406 y Water loss 2
618.8015747070312 0 770.0429 y Ammonia loss 2
627.3035278320312 0 45326.63 y 2
627.8048095703125 0 27421.59
628.3065185546875 0 11872.678
628.8073120117188 0 2714.6223
650.366455078125 0 613.2955
666.3928833007812 0 1801.7601
667.3048706054688 0 2048.8887
668.3138427734375 0 1108.6576 y Ammonia loss 1
669.3150634765625 0 640.2831
676.83740234375 0 9542.836 y 1
677.3389892578125 0 5355.751
677.8403930664062 0 3405.0552
678.341796875 0 1623.9556
683.2626342773438 0 4249.2476
684.2647094726562 0 1862.4493
694.4251708984375 0 1040.0095
695.2625122070312 0 1365.4912
716.344482421875 0 4426.7227 y 7
717.3455810546875 0 1680.2615
734.349609375 0 815.80273
738.383056640625 0 1373.5399
755.4037475585938 0 15457.515
756.406005859375 0 7463.1274
757.4100952148438 0 1180.136
765.4605712890625 0 7540.194
766.4636840820312 0 3677.4185
767.4651489257812 0 1294.2424
782.3300170898438 0 15731.653
783.3335571289062 0 5724.4946
784.33935546875 0 1105.3568
787.3818969726562 0 999.114
794.3307495117188 0 1557.4666
805.38427734375 0 1523.7053
806.3901977539062 0 1177.627
807.3931884765625 0 637.52515
853.4039916992188 0 4495.411 y 6
854.401123046875 0 1218.3981
868.4866333007812 0 2483.0034
869.4911499023438 0 1354.0588
881.3955078125 0 4220.686
882.4013061523438 0 1450.8999
909.35595703125 0 2181.8625
910.3574829101562 0 1200.8499
940.4346923828125 0 6218.285 y 5
941.4385375976562 0 1739.5675
942.4446411132812 0 998.503
980.4408569335938 0 846.6953
1008.4240112304688 0 6394.3965
1009.4288330078125 0 3521.5913
1010.4298706054688 0 1549.8424
1039.5 0 1783.9817 y 4
1107.494384765625 0 3084.3503
1108.49267578125 0 1779.9159
1154.5289306640625 0 1644.7067 y 3
1155.53369140625 0 1391.4042
1462.6788330078125 0 620.5817
1652.672607421875 0 639.6997
2267.155029296875 0 670.1331
2304.08349609375 0 724.1703
2463.139892578125 0 711.5759
3073.68212890625 0 776.31805
3086.9111328125 0 705.9242

Spectrum Details

|  |  |
| --- | --- |
| Matched peaks? Matched peaks The total absolute number of peaks matched. Additionally in brackets the total fraction of peaks matched and the total number of peaks is shown. | 26 (9.85% of 264) |
| FDR? FDR The false discovery rate estimated for this peptide. It is calculated by matching all theoretical fragments with a non-integer shift with the raw peaks for this spectrum. This is done with 40 different shifts. The resulting percentage is the average number of annotated peaks over the number of annotated peaks with the correct spectrum. | 0.82% |
| Satellite FDR? Satellite FDR See the FDR for details on its calculation. This satellite ion specific FDR only contains the satellite ions (d/w) for I/L/J positions. | - |
| PSM Score? PSM Score The PSM Score as given by Hecklib to this annotated spectrum. It is shown with three significant figures. | 166 |

#### Spectrum 4253? Spectrum 4253 The raw spectrum of this peptide as annotated by Hecklib. The fragments are coloured according to ion type (see legend). Any peaks with a star '\*' as text can be hovered over to see the full details, first the ion type second the mass shift type. By hovering over the amino acids in the peptide or ions in the legend the corresponding peaks are highlighted. By toggling the 'Unassigned' label you can turn the background (unassigned) peaks on or off in the plot. By updating the slider in the Ion legend you can update the spectrum to only show the top X% of the peaks with labels. The top X% means any peak that is within X% of the highest intensity. By dragging in the spectrum you can zoom in to a specific part of the spectrum and use 'Zoom Out' to get back to the original zoom level. The annotation of the spectrum is based on the given sequence in the peptides file and is done with different software so inconsistencies are likely. The peaks are annotated based on the given sequence, with 20 ppm tolerance.

Copy Data

##### Spectrum 4253 (TSV)

###### Preview

```
Loading example...
```

*Click on the button to copy the data to your clipboard.*

Mz MinMz MaxIntensity Max

WidthHeightPeptide font sizePeptide stroke widthSpectrum font sizeSpectrum stroke widthCompact peptide

Ion legend

wxyz

abcd

OtherUnassignedIon^Charge^_Position_Show for top:%

VVVDVSHEDPEVK

03.02e+56.04e+59.06e+51.21e+6

Zoom Out

y^+1^_1_z^+1^_1_y^+1^_1_w^+1^_2_c^+1^_2_y^+1^_2_z^+1^_2_y^+1^_2_w^+1^_3_c^+1^_3_y^+1^_3_z^+1^_3_y^+1^_3_y^+3^_11_z^+2^_7_y^+2^_7_w^+1^_4_c^+1^_4_y^+3^_12_w^+2^_8_y^+1^_4_y^+1^_4_y^+2^_8_y^+2^_8_z^+2^_8_y^+2^_8_y^+1^_4_z^+2^_9_z^+2^_9_w^+2^_9_y^+2^_9_z^+2^_9_y^+2^_9_c^+1^_5_w^+2^_10_c^+2^_10_z^+1^_5_y^+2^_10_z^+2^_10_z^+1^_5_y^+2^_10_y^+1^_5_c^+1^_6_c^+2^_11_w^+2^_11_c^+2^_11_c^+1^_6_y^+2^_11_y^+2^_11_z^+2^_11_y^+2^_11_w^+1^_6_w^+2^_12_c^+2^_12_y^+2^_12_z^+2^_12_y^+2^_12_z^+1^_6_y^+1^_6_c^+1^_7_z^+1^_7_y^+1^_7_c^+1^_8_y^+1^_8_y^+1^_8_z^+1^_8_y^+1^_8_c^+1^_9_y^+1^_9_z^+1^_9_y^+1^_9_w^+1^_10_c^+1^_10_y^+1^_10_y^+1^_10_z^+1^_10_y^+1^_10_c^+1^_11_z^+1^_11_y^+1^_11_c^+1^_12_z^+1^_12_

036873611041472

Fragment Matches Table

Show background peaks

| Position | Ion type | Intensity | mz Theoretical | mz Error (Th) | mz Error (ppm) | Charge | Series Number |
| --- | --- | --- | --- | --- | --- | --- | --- |
| - | - | 418.3 | 122.1 | - | - | 0 | - |
| - | - | 402.6 | 125.9 | - | - | 0 | - |
| - | - | 786.4 | 126.1 | - | - | 0 | - |
| - | - | 506.3 | 127.1 | - | - | 0 | - |
| - | - | 561.9 | 127.1 | - | - | 0 | - |
| - | - | 442.1 | 129.1 | - | - | 0 | - |
| - | - | 1.646E+04 | 129.1 | - | - | 0 | - |
| 13 | y | 1.598E+04 | 130.1 | 0.0005064 | 3.893 | +1 | 1 |
| - | - | 997 | 130.1 | - | - | 0 | - |
| 13 | z | 2.118E+04 | 131.1 | 0.0005091 | 3.884 | +1 | 1 |
| - | - | 1709 | 131.1 | - | - | 0 | - |
| - | - | 688.5 | 132.1 | - | - | 0 | - |
| - | - | 4277 | 132.1 | - | - | 0 | - |
| - | - | 793.6 | 133.1 | - | - | 0 | - |
| - | - | 476.1 | 141.9 | - | - | 0 | - |
| - | - | 524.9 | 143.2 | - | - | 0 | - |
| - | - | 835.5 | 144.1 | - | - | 0 | - |
| 13 | y | 5.686E+04 | 147.1 | 0.0005381 | 3.658 | +1 | 1 |
| - | - | 4307 | 148.1 | - | - | 0 | - |
| - | - | 479.9 | 148.8 | - | - | 0 | - |
| - | - | 643.8 | 148.9 | - | - | 0 | - |
| - | - | 796.8 | 148.9 | - | - | 0 | - |
| - | - | 1226 | 148.9 | - | - | 0 | - |
| - | - | 1311 | 148.9 | - | - | 0 | - |
| - | - | 1920 | 148.9 | - | - | 0 | - |
| - | - | 3691 | 148.9 | - | - | 0 | - |
| - | - | 4294 | 149 | - | - | 0 | - |
| - | - | 2799 | 149 | - | - | 0 | - |
| - | - | 1118 | 149 | - | - | 0 | - |
| - | - | 1001 | 149 | - | - | 0 | - |
| - | - | 510.1 | 149 | - | - | 0 | - |
| - | - | 675.4 | 149 | - | - | 0 | - |
| - | - | 476.2 | 149 | - | - | 0 | - |
| - | - | 529.5 | 149 | - | - | 0 | - |
| - | - | 635.3 | 149 | - | - | 0 | - |
| - | - | 585.3 | 149.1 | - | - | 0 | - |
| - | - | 433.3 | 149.1 | - | - | 0 | - |
| - | - | 916.7 | 152.1 | - | - | 0 | - |
| - | - | 457.3 | 153.1 | - | - | 0 | - |
| - | - | 7725 | 154.1 | - | - | 0 | - |
| - | - | 2283 | 155.1 | - | - | 0 | - |
| - | - | 849.1 | 156.1 | - | - | 0 | - |
| - | - | 1699 | 163.1 | - | - | 0 | - |
| - | - | 773.7 | 169.1 | - | - | 0 | - |
| - | - | 1891 | 170.1 | - | - | 0 | - |
| - | - | 2282 | 171.1 | - | - | 0 | - |
| - | - | 3.391E+05 | 171.1 | - | - | 0 | - |
| - | - | 2079 | 172.1 | - | - | 0 | - |
| - | - | 3.02E+04 | 172.2 | - | - | 0 | - |
| - | - | 1244 | 173.2 | - | - | 0 | - |
| - | - | 630.2 | 173.4 | - | - | 0 | - |
| - | - | 1243 | 179.1 | - | - | 0 | - |
| - | - | 7204 | 185.2 | - | - | 0 | - |
| - | - | 6588 | 187.1 | - | - | 0 | - |
| - | - | 1617 | 189.1 | - | - | 0 | - |
| - | - | 1805 | 197.1 | - | - | 0 | - |
| - | - | 8418 | 198.1 | - | - | 0 | - |
| - | - | 2.806E+05 | 199.1 | - | - | 0 | - |
| - | - | 3.112E+04 | 200.1 | - | - | 0 | - |
| - | - | 655.7 | 201.1 | - | - | 0 | - |
| - | - | 1809 | 201.2 | - | - | 0 | - |
| - | - | 5202 | 203.1 | - | - | 0 | - |
| - | - | 592.3 | 204.1 | - | - | 0 | - |
| - | - | 776.3 | 211.1 | - | - | 0 | - |
| - | - | 6861 | 213.2 | - | - | 0 | - |
| - | - | 1.72E+04 | 215.1 | - | - | 0 | - |
| - | - | 943.4 | 215.1 | - | - | 0 | - |
| 12 | w | 4453 | 215.1 | 0.0007821 | 3.635 | +1 | 2 |
| - | - | 1236 | 216.1 | - | - | 0 | - |
| 2 | c | 2802 | 216.2 | 0.0007028 | 3.251 | +1 | 2 |
| - | - | 764.8 | 225.1 | - | - | 0 | - |
| - | - | 2996 | 227.1 | - | - | 0 | - |
| - | - | 2741 | 228.2 | - | - | 0 | - |
| 12 | y | 3346 | 229.2 | 0.0008944 | 3.903 | +1 | 2 |
| - | - | 856.5 | 229.2 | - | - | 0 | - |
| 12 | z | 6888 | 230.2 | 0.0006835 | 2.97 | +1 | 2 |
| - | - | 869.9 | 231.2 | - | - | 0 | - |
| - | - | 3.172E+04 | 232.1 | - | - | 0 | - |
| - | - | 3097 | 233.1 | - | - | 0 | - |
| - | - | 540.4 | 239.3 | - | - | 0 | - |
| 12 | y | 8.987E+04 | 246.2 | 0.0008498 | 3.452 | +1 | 2 |
| - | - | 9757 | 247.2 | - | - | 0 | - |
| - | - | 1119 | 248.2 | - | - | 0 | - |
| - | - | 7188 | 253.2 | - | - | 0 | - |
| - | - | 1209 | 254.2 | - | - | 0 | - |
| - | - | 790.9 | 270.2 | - | - | 0 | - |
| - | - | 2.132E+04 | 270.2 | - | - | 0 | - |
| - | - | 4441 | 271.2 | - | - | 0 | - |
| - | - | 889.2 | 272.2 | - | - | 0 | - |
| - | - | 3460 | 282.1 | - | - | 0 | - |
| - | - | 3716 | 283.1 | - | - | 0 | - |
| - | - | 1393 | 286.2 | - | - | 0 | - |
| - | - | 1331 | 287.2 | - | - | 0 | - |
| - | - | 4217 | 297.2 | - | - | 0 | - |
| - | - | 3.691E+04 | 298.2 | - | - | 0 | - |
| - | - | 632.5 | 299.2 | - | - | 0 | - |
| - | - | 6588 | 299.2 | - | - | 0 | - |
| 11 | w | 9.15E+04 | 300.2 | 0.001058 | 3.524 | +1 | 3 |
| - | - | 1.49E+04 | 301.2 | - | - | 0 | - |
| - | - | 870.4 | 302.1 | - | - | 0 | - |
| - | - | 1886 | 302.2 | - | - | 0 | - |
| - | - | 1.492E+04 | 314.2 | - | - | 0 | - |
| - | - | 3668 | 314.2 | - | - | 0 | - |
| - | - | 2159 | 315.2 | - | - | 0 | - |
| 3 | c | 8009 | 315.2 | 0.0007703 | 2.444 | +1 | 3 |
| - | - | 1652 | 316.2 | - | - | 0 | - |
| - | - | 3875 | 326.2 | - | - | 0 | - |
| - | - | 781.9 | 327.2 | - | - | 0 | - |
| - | - | 7013 | 331.2 | - | - | 0 | - |
| - | - | 1482 | 332.2 | - | - | 0 | - |
| - | - | 628.1 | 342.2 | - | - | 0 | - |
| - | - | 1474 | 343.2 | - | - | 0 | - |
| - | - | 610.9 | 348.6 | - | - | 0 | - |
| - | - | 3750 | 353.2 | - | - | 0 | - |
| - | - | 1.693E+04 | 354.2 | - | - | 0 | - |
| - | - | 4094 | 355.2 | - | - | 0 | - |
| 11 | y | 2250 | 357.2 | 0.001353 | 3.788 | +1 | 3 |
| 11 | z | 5000 | 359.2 | 0.000815 | 2.269 | +1 | 3 |
| - | - | 6230 | 368.2 | - | - | 0 | - |
| - | - | 1587 | 369.2 | - | - | 0 | - |
| - | - | 2496 | 371.2 | - | - | 0 | - |
| - | - | 1232 | 373.2 | - | - | 0 | - |
| 11 | y | 2259 | 375.2 | 0.0009508 | 2.534 | +1 | 3 |
| - | - | 2535 | 381.2 | - | - | 0 | - |
| - | - | 2608 | 382.1 | - | - | 0 | - |
| - | - | 2632 | 382.2 | - | - | 0 | - |
| - | - | 1819 | 383.2 | - | - | 0 | - |
| - | - | 7637 | 385.2 | - | - | 0 | - |
| - | - | 1.049E+04 | 386.3 | - | - | 0 | - |
| - | - | 2195 | 387.3 | - | - | 0 | - |
| - | - | 599.1 | 392.2 | - | - | 0 | - |
| - | - | 1447 | 395.2 | - | - | 0 | - |
| - | - | 911.7 | 398.2 | - | - | 0 | - |
| - | - | 2275 | 398.2 | - | - | 0 | - |
| - | - | 790.6 | 399.2 | - | - | 0 | - |
| - | - | 6040 | 401.2 | - | - | 0 | - |
| - | - | 2671 | 402.2 | - | - | 0 | - |
| - | - | 800.8 | 412.2 | - | - | 0 | - |
| - | - | 4.577E+04 | 413.2 | - | - | 0 | - |
| - | - | 1380 | 414.2 | - | - | 0 | - |
| - | - | 9195 | 414.2 | - | - | 0 | - |
| - | - | 832.4 | 415.2 | - | - | 0 | - |
| - | - | 1816 | 415.2 | - | - | 0 | - |
| - | - | 2087 | 416.2 | - | - | 0 | - |
| - | - | 1315 | 418.2 | - | - | 0 | - |
| 3 | y | 3183 | 418.5 | 0.001123 | 2.683 | +3 | 11 |
| - | - | 2462 | 418.9 | - | - | 0 | - |
| 7 | z | 1546 | 419.2 | 0.00411 | 9.803 | +2 | 7 |
| - | - | 644.4 | 419.5 | - | - | 0 | - |
| - | - | 712.6 | 426.2 | - | - | 0 | - |
| - | - | 786.8 | 426.3 | - | - | 0 | - |
| 7 | y | 1.118E+04 | 427.2 | 0.001614 | 3.778 | +2 | 7 |
| - | - | 4551 | 427.7 | - | - | 0 | - |
| - | - | 793.4 | 428.2 | - | - | 0 | - |
| 10 | w | 2.861E+04 | 429.2 | 0.001556 | 3.624 | +1 | 4 |
| - | - | 8380 | 430.2 | - | - | 0 | - |
| 4 | c | 5016 | 430.3 | 0.002666 | 6.197 | +1 | 4 |
| - | - | 1499 | 431.2 | - | - | 0 | - |
| - | - | 1559 | 431.3 | - | - | 0 | - |
| - | - | 816.5 | 431.8 | - | - | 0 | - |
| - | - | 1153 | 434.8 | - | - | 0 | - |
| - | - | 766.6 | 439.2 | - | - | 0 | - |
| - | - | 971.8 | 441.2 | - | - | 0 | - |
| - | - | 1074 | 441.3 | - | - | 0 | - |
| - | - | 2659 | 442.3 | - | - | 0 | - |
| - | - | 805.3 | 443.2 | - | - | 0 | - |
| - | - | 2961 | 444.3 | - | - | 0 | - |
| - | - | 1081 | 445.3 | - | - | 0 | - |
| 2 | y | 5146 | 451.6 | 0.001206 | 2.672 | +3 | 12 |
| - | - | 2507 | 451.9 | - | - | 0 | - |
| - | - | 774.5 | 452.2 | - | - | 0 | - |
| - | - | 727.5 | 453.2 | - | - | 0 | - |
| - | - | 690.5 | 453.3 | - | - | 0 | - |
| 6 | w | 833.7 | 454.2 | 0.001135 | 2.5 | +2 | 8 |
| 10 | y | 1038 | 454.3 | 0.004742 | 10.44 | +1 | 4 |
| - | - | 802.3 | 455.2 | - | - | 0 | - |
| 10 | y | 664.1 | 455.3 | 0.00143 | 3.141 | +1 | 4 |
| - | - | 8255 | 455.3 | - | - | 0 | - |
| - | - | 2837 | 456.2 | - | - | 0 | - |
| - | - | 1633 | 456.3 | - | - | 0 | - |
| - | - | 1114 | 457.3 | - | - | 0 | - |
| - | - | 1248 | 458.3 | - | - | 0 | - |
| 6 | y | 1574 | 461.7 | 0.005456 | 11.82 | +2 | 8 |
| 6 | y | 6572 | 462.2 | 0.002048 | 4.431 | +2 | 8 |
| 6 | z | 9056 | 462.7 | 0.00134 | 2.896 | +2 | 8 |
| - | - | 4342 | 463.2 | - | - | 0 | - |
| - | - | 1182 | 463.7 | - | - | 0 | - |
| - | - | 2.927E+04 | 468.3 | - | - | 0 | - |
| - | - | 3100 | 469.2 | - | - | 0 | - |
| - | - | 6225 | 469.3 | - | - | 0 | - |
| - | - | 4.367E+04 | 470.3 | - | - | 0 | - |
| - | - | 4630 | 470.3 | - | - | 0 | - |
| 6 | y | 1.799E+04 | 470.7 | 0.001835 | 3.898 | +2 | 8 |
| - | - | 8522 | 471.2 | - | - | 0 | - |
| - | - | 1.506E+04 | 471.3 | - | - | 0 | - |
| - | - | 1414 | 471.3 | - | - | 0 | - |
| - | - | 2756 | 471.7 | - | - | 0 | - |
| 10 | y | 1.313E+05 | 472.3 | 0.001684 | 3.566 | +1 | 4 |
| - | - | 3.416E+04 | 473.3 | - | - | 0 | - |
| - | - | 6221 | 474.3 | - | - | 0 | - |
| - | - | 3076 | 474.7 | - | - | 0 | - |
| - | - | 1137 | 475.2 | - | - | 0 | - |
| - | - | 1078 | 475.3 | - | - | 0 | - |
| - | - | 753.2 | 484.2 | - | - | 0 | - |
| - | - | 4796 | 484.3 | - | - | 0 | - |
| - | - | 7521 | 484.3 | - | - | 0 | - |
| - | - | 1.733E+04 | 484.6 | - | - | 0 | - |
| - | - | 1475 | 484.8 | - | - | 0 | - |
| - | - | 8709 | 484.9 | - | - | 0 | - |
| - | - | 1.918E+04 | 485.3 | - | - | 0 | - |
| - | - | 7410 | 485.3 | - | - | 0 | - |
| - | - | 962.7 | 485.4 | - | - | 0 | - |
| - | - | 1715 | 485.6 | - | - | 0 | - |
| - | - | 2427 | 486.3 | - | - | 0 | - |
| - | - | 2497 | 486.3 | - | - | 0 | - |
| - | - | 1768 | 494.3 | - | - | 0 | - |
| - | - | 920.4 | 496.3 | - | - | 0 | - |
| - | - | 1623 | 497.3 | - | - | 0 | - |
| - | - | 2.442E+04 | 499.3 | - | - | 0 | - |
| - | - | 1.169E+04 | 500.3 | - | - | 0 | - |
| - | - | 2.043E+04 | 501.3 | - | - | 0 | - |
| - | - | 5172 | 502.3 | - | - | 0 | - |
| 5 | z | 610.1 | 503.2 | 0.003628 | 7.21 | +2 | 9 |
| - | - | 1167 | 503.3 | - | - | 0 | - |
| 5 | z | 1157 | 503.7 | 0.009857 | 19.57 | +2 | 9 |
| 5 | w | 4921 | 504.7 | 0.0008737 | 1.731 | +2 | 9 |
| - | - | 1869 | 505.2 | - | - | 0 | - |
| - | - | 1282 | 505.7 | - | - | 0 | - |
| - | - | 1258 | 509.3 | - | - | 0 | - |
| - | - | 1822 | 510.3 | - | - | 0 | - |
| 5 | y | 2250 | 511.7 | 0.01005 | 19.63 | +2 | 9 |
| 5 | z | 1.67E+04 | 512.2 | 0.002259 | 4.409 | +2 | 9 |
| - | - | 6.485E+04 | 512.3 | - | - | 0 | - |
| - | - | 7330 | 512.8 | - | - | 0 | - |
| - | - | 2563 | 513.3 | - | - | 0 | - |
| - | - | 1.752E+04 | 513.3 | - | - | 0 | - |
| - | - | 3361 | 514.3 | - | - | 0 | - |
| 5 | y | 3.421E+04 | 520.3 | 0.001503 | 2.888 | +2 | 9 |
| - | - | 1.711E+04 | 520.8 | - | - | 0 | - |
| - | - | 5594 | 521.3 | - | - | 0 | - |
| - | - | 907.5 | 521.8 | - | - | 0 | - |
| - | - | 772.3 | 526.7 | - | - | 0 | - |
| - | - | 1.831E+05 | 527.3 | - | - | 0 | - |
| - | - | 5.242E+04 | 528.3 | - | - | 0 | - |
| - | - | 7935 | 529.3 | - | - | 0 | - |
| 5 | c | 1.668E+04 | 529.3 | 0.002673 | 5.05 | +1 | 5 |
| - | - | 687.9 | 530.3 | - | - | 0 | - |
| - | - | 5590 | 530.3 | - | - | 0 | - |
| - | - | 3813 | 533.8 | - | - | 0 | - |
| - | - | 1627 | 534.3 | - | - | 0 | - |
| - | - | 5658 | 538.3 | - | - | 0 | - |
| - | - | 729.1 | 538.8 | - | - | 0 | - |
| - | - | 3109 | 539.3 | - | - | 0 | - |
| - | - | 1754 | 540.3 | - | - | 0 | - |
| - | - | 4078 | 540.8 | - | - | 0 | - |
| - | - | 1604 | 541.3 | - | - | 0 | - |
| 4 | w | 945.1 | 547.3 | 0.0006759 | 1.235 | +2 | 10 |
| 10 | c | 1.224E+04 | 547.8 | 0.01038 | 18.95 | +2 | 10 |
| - | - | 8093 | 548.3 | - | - | 0 | - |
| - | - | 2718 | 548.8 | - | - | 0 | - |
| - | - | 1222 | 553.3 | - | - | 0 | - |
| 9 | z | 988.9 | 554.3 | 0.004217 | 7.608 | +1 | 5 |
| - | - | 2091 | 559.3 | - | - | 0 | - |
| - | - | 707.2 | 560.3 | - | - | 0 | - |
| - | - | 5568 | 568.2 | - | - | 0 | - |
| 4 | y | 1114 | 568.8 | 0.00375 | 6.594 | +2 | 10 |
| - | - | 1389 | 569.2 | - | - | 0 | - |
| 4 | z | 7940 | 569.8 | 0.00191 | 3.352 | +2 | 10 |
| - | - | 4637 | 570.3 | - | - | 0 | - |
| - | - | 7802 | 570.3 | - | - | 0 | - |
| - | - | 2060 | 570.8 | - | - | 0 | - |
| 9 | z | 3013 | 571.3 | 0.0007974 | 1.396 | +1 | 5 |
| - | - | 2179 | 571.3 | - | - | 0 | - |
| - | - | 1188 | 572.3 | - | - | 0 | - |
| 4 | y | 6.66E+04 | 577.8 | 0.001764 | 3.053 | +2 | 10 |
| - | - | 3.647E+04 | 578.3 | - | - | 0 | - |
| - | - | 1.424E+04 | 578.8 | - | - | 0 | - |
| - | - | 3543 | 579.3 | - | - | 0 | - |
| - | - | 3547 | 581.3 | - | - | 0 | - |
| - | - | 935.8 | 582.8 | - | - | 0 | - |
| 9 | y | 2.348E+04 | 587.3 | 0.001658 | 2.823 | +1 | 5 |
| - | - | 6780 | 588.3 | - | - | 0 | - |
| - | - | 876.5 | 589.3 | - | - | 0 | - |
| - | - | 1084 | 589.8 | - | - | 0 | - |
| - | - | 868.1 | 590.3 | - | - | 0 | - |
| - | - | 1073 | 590.8 | - | - | 0 | - |
| - | - | 1225 | 591.3 | - | - | 0 | - |
| - | - | 917.4 | 595.3 | - | - | 0 | - |
| - | - | 6690 | 597.3 | - | - | 0 | - |
| - | - | 6236 | 597.8 | - | - | 0 | - |
| - | - | 2365 | 598.3 | - | - | 0 | - |
| 6 | c | 955.3 | 598.4 | 0.003121 | 5.216 | +1 | 6 |
| - | - | 1.288E+04 | 599.3 | - | - | 0 | - |
| - | - | 4162 | 600.3 | - | - | 0 | - |
| - | - | 981 | 601.3 | - | - | 0 | - |
| 11 | c | 1536 | 603.3 | 0.008756 | 14.51 | +2 | 11 |
| - | - | 3615 | 603.8 | - | - | 0 | - |
| - | - | 2195 | 604.3 | - | - | 0 | - |
| - | - | 1628 | 604.8 | - | - | 0 | - |
| - | - | 1337 | 605.3 | - | - | 0 | - |
| - | - | 1215 | 605.8 | - | - | 0 | - |
| - | - | 1102 | 606.3 | - | - | 0 | - |
| 3 | w | 7335 | 611.8 | 0.00239 | 3.906 | +2 | 11 |
| 11 | c | 4268 | 612.3 | 0.0116 | 18.94 | +2 | 11 |
| - | - | 1694 | 612.8 | - | - | 0 | - |
| - | - | 855.3 | 613.3 | - | - | 0 | - |
| 6 | c | 4.276E+04 | 616.4 | 0.001956 | 3.173 | +1 | 6 |
| - | - | 1.535E+04 | 617.4 | - | - | 0 | - |
| 3 | y | 7641 | 618.3 | 0.002747 | 4.442 | +2 | 11 |
| - | - | 3340 | 618.4 | - | - | 0 | - |
| 3 | y | 4156 | 618.8 | 0.01074 | 17.35 | +2 | 11 |
| 3 | z | 4.476E+04 | 619.3 | 0.002554 | 4.124 | +2 | 11 |
| - | - | 2.65E+04 | 619.8 | - | - | 0 | - |
| - | - | 1.052E+04 | 620.3 | - | - | 0 | - |
| - | - | 2545 | 620.8 | - | - | 0 | - |
| - | - | 976.9 | 624.3 | - | - | 0 | - |
| - | - | 3897 | 626.3 | - | - | 0 | - |
| - | - | 2472 | 626.8 | - | - | 0 | - |
| 3 | y | 4.305E+05 | 627.3 | 0.002103 | 3.352 | +2 | 11 |
| - | - | 2.661E+05 | 627.8 | - | - | 0 | - |
| - | - | 1.138E+05 | 628.3 | - | - | 0 | - |
| - | - | 3.087E+04 | 628.8 | - | - | 0 | - |
| - | - | 5582 | 629.3 | - | - | 0 | - |
| - | - | 1645 | 632.3 | - | - | 0 | - |
| - | - | 1773 | 639.3 | - | - | 0 | - |
| - | - | 3538 | 640.4 | - | - | 0 | - |
| - | - | 2683 | 640.8 | - | - | 0 | - |
| 8 | w | 7.844E+04 | 641.3 | 0.001957 | 3.052 | +1 | 6 |
| - | - | 1191 | 641.8 | - | - | 0 | - |
| - | - | 2.63E+04 | 642.3 | - | - | 0 | - |
| - | - | 5359 | 643.3 | - | - | 0 | - |
| - | - | 1007 | 644.3 | - | - | 0 | - |
| - | - | 2502 | 646 | - | - | 0 | - |
| - | - | 1954 | 646.3 | - | - | 0 | - |
| - | - | 1577 | 646.6 | - | - | 0 | - |
| - | - | 6946 | 646.8 | - | - | 0 | - |
| - | - | 949.1 | 647 | - | - | 0 | - |
| - | - | 5183 | 647.3 | - | - | 0 | - |
| - | - | 1690 | 647.8 | - | - | 0 | - |
| - | - | 1389 | 647.8 | - | - | 0 | - |
| - | - | 2232 | 648.3 | - | - | 0 | - |
| - | - | 1502 | 648.8 | - | - | 0 | - |
| - | - | 5740 | 653.3 | - | - | 0 | - |
| - | - | 5811 | 653.8 | - | - | 0 | - |
| - | - | 3800 | 654.3 | - | - | 0 | - |
| - | - | 9584 | 656.3 | - | - | 0 | - |
| - | - | 3365 | 657.3 | - | - | 0 | - |
| - | - | 864.1 | 658.3 | - | - | 0 | - |
| - | - | 2415 | 660.9 | - | - | 0 | - |
| 2 | w | 1.276E+04 | 661.3 | 0.003339 | 5.049 | +2 | 12 |
| 12 | c | 1.418E+04 | 661.8 | 0.00607 | 9.171 | +2 | 12 |
| - | - | 8944 | 662.3 | - | - | 0 | - |
| - | - | 3091 | 662.8 | - | - | 0 | - |
| - | - | 995.6 | 665.4 | - | - | 0 | - |
| - | - | 6111 | 667.3 | - | - | 0 | - |
| - | - | 3184 | 667.4 | - | - | 0 | - |
| - | - | 5371 | 667.9 | - | - | 0 | - |
| 2 | y | 3925 | 668.3 | 0.004303 | 6.439 | +2 | 12 |
| 2 | z | 2.141E+04 | 668.8 | 0.00216 | 3.23 | +2 | 12 |
| - | - | 1.402E+04 | 669.3 | - | - | 0 | - |
| - | - | 5561 | 669.8 | - | - | 0 | - |
| - | - | 1176 | 670.3 | - | - | 0 | - |
| - | - | 1489 | 674.4 | - | - | 0 | - |
| - | - | 1729 | 675.4 | - | - | 0 | - |
| - | - | 2298 | 675.8 | - | - | 0 | - |
| - | - | 2191 | 676.3 | - | - | 0 | - |
| 2 | y | 8.959E+04 | 676.8 | 0.00232 | 3.427 | +2 | 12 |
| - | - | 5.804E+04 | 677.3 | - | - | 0 | - |
| - | - | 3.03E+04 | 677.8 | - | - | 0 | - |
| - | - | 9266 | 678.3 | - | - | 0 | - |
| - | - | 1476 | 678.8 | - | - | 0 | - |
| - | - | 1099 | 681.8 | - | - | 0 | - |
| - | - | 3610 | 682.4 | - | - | 0 | - |
| - | - | 2172 | 682.9 | - | - | 0 | - |
| - | - | 8264 | 683.3 | - | - | 0 | - |
| - | - | 1824 | 683.4 | - | - | 0 | - |
| - | - | 954.8 | 683.9 | - | - | 0 | - |
| - | - | 2796 | 684.3 | - | - | 0 | - |
| - | - | 1053 | 685.3 | - | - | 0 | - |
| - | - | 1372 | 685.9 | - | - | 0 | - |
| - | - | 1578 | 686.4 | - | - | 0 | - |
| - | - | 1610 | 686.9 | - | - | 0 | - |
| - | - | 1479 | 687.9 | - | - | 0 | - |
| - | - | 1437 | 688.4 | - | - | 0 | - |
| - | - | 2084 | 688.9 | - | - | 0 | - |
| - | - | 2128 | 689.4 | - | - | 0 | - |
| - | - | 3415 | 689.9 | - | - | 0 | - |
| - | - | 2391 | 690.4 | - | - | 0 | - |
| - | - | 1.184E+04 | 690.9 | - | - | 0 | - |
| - | - | 4827 | 691.4 | - | - | 0 | - |
| - | - | 2958 | 691.9 | - | - | 0 | - |
| - | - | 1313 | 693.4 | - | - | 0 | - |
| - | - | 1251 | 695.3 | - | - | 0 | - |
| - | - | 3067 | 695.9 | - | - | 0 | - |
| - | - | 8999 | 696.4 | - | - | 0 | - |
| - | - | 1.67E+05 | 696.9 | - | - | 0 | - |
| - | - | 1.489E+05 | 697.4 | - | - | 0 | - |
| - | - | 7.614E+04 | 697.9 | - | - | 0 | - |
| - | - | 2.784E+04 | 698.4 | - | - | 0 | - |
| - | - | 6459 | 698.9 | - | - | 0 | - |
| 8 | z | 5.426E+04 | 700.3 | 0.002447 | 3.494 | +1 | 6 |
| - | - | 2.095E+04 | 701.3 | - | - | 0 | - |
| - | - | 6059 | 702.3 | - | - | 0 | - |
| - | - | 1684 | 703.3 | - | - | 0 | - |
| - | - | 1.386E+04 | 703.9 | - | - | 0 | - |
| - | - | 1.063E+04 | 704.4 | - | - | 0 | - |
| - | - | 2.754E+04 | 704.9 | - | - | 0 | - |
| - | - | 2.129E+04 | 705.4 | - | - | 0 | - |
| - | - | 8818 | 705.9 | - | - | 0 | - |
| - | - | 1485 | 706.4 | - | - | 0 | - |
| - | - | 1529 | 707.4 | - | - | 0 | - |
| - | - | 1662 | 708.4 | - | - | 0 | - |
| - | - | 2.67E+04 | 709.4 | - | - | 0 | - |
| - | - | 4050 | 709.9 | - | - | 0 | - |
| - | - | 837.7 | 710.4 | - | - | 0 | - |
| - | - | 1.072E+04 | 710.4 | - | - | 0 | - |
| - | - | 1.177E+05 | 710.9 | - | - | 0 | - |
| - | - | 7.954E+04 | 711.4 | - | - | 0 | - |
| - | - | 4.046E+04 | 711.9 | - | - | 0 | - |
| - | - | 1.166E+04 | 712.4 | - | - | 0 | - |
| - | - | 3029 | 712.9 | - | - | 0 | - |
| 8 | y | 4.926E+04 | 716.3 | 0.002644 | 3.691 | +1 | 6 |
| - | - | 1.864E+04 | 717.4 | - | - | 0 | - |
| - | - | 1.168E+04 | 717.9 | - | - | 0 | - |
| - | - | 3.254E+05 | 718.4 | - | - | 0 | - |
| - | - | 2.214E+05 | 718.9 | - | - | 0 | - |
| - | - | 1.133E+05 | 719.4 | - | - | 0 | - |
| - | - | 3.61E+04 | 719.9 | - | - | 0 | - |
| - | - | 9734 | 720.4 | - | - | 0 | - |
| - | - | 6247 | 725.4 | - | - | 0 | - |
| - | - | 1.446E+04 | 725.9 | - | - | 0 | - |
| - | - | 6.81E+05 | 726.4 | - | - | 0 | - |
| - | - | 1.196E+06 | 726.9 | - | - | 0 | - |
| - | - | 7.423E+05 | 727.4 | - | - | 0 | - |
| - | - | 3.234E+05 | 727.9 | - | - | 0 | - |
| - | - | 9.165E+04 | 728.4 | - | - | 0 | - |
| - | - | 1.768E+04 | 728.9 | - | - | 0 | - |
| - | - | 2268 | 734.4 | - | - | 0 | - |
| - | - | 1282 | 736.4 | - | - | 0 | - |
| 7 | c | 1.71E+05 | 753.4 | 0.002736 | 3.631 | +1 | 7 |
| - | - | 6.385E+04 | 754.4 | - | - | 0 | - |
| - | - | 1.958E+04 | 755.4 | - | - | 0 | - |
| - | - | 5485 | 756.4 | - | - | 0 | - |
| - | - | 1540 | 764.3 | - | - | 0 | - |
| - | - | 6780 | 765.4 | - | - | 0 | - |
| - | - | 4832 | 765.5 | - | - | 0 | - |
| - | - | 3422 | 766.4 | - | - | 0 | - |
| - | - | 1431 | 766.5 | - | - | 0 | - |
| - | - | 3103 | 778.4 | - | - | 0 | - |
| - | - | 9229 | 779.4 | - | - | 0 | - |
| - | - | 3777 | 780.4 | - | - | 0 | - |
| - | - | 1962 | 781.4 | - | - | 0 | - |
| - | - | 7.2E+04 | 782.3 | - | - | 0 | - |
| - | - | 3.07E+04 | 783.3 | - | - | 0 | - |
| - | - | 7927 | 784.3 | - | - | 0 | - |
| - | - | 1038 | 785.3 | - | - | 0 | - |
| - | - | 1534 | 793.4 | - | - | 0 | - |
| - | - | 2659 | 794.3 | - | - | 0 | - |
| - | - | 1711 | 795.3 | - | - | 0 | - |
| - | - | 1466 | 805.4 | - | - | 0 | - |
| 7 | z | 6.703E+04 | 837.4 | 0.0028 | 3.344 | +1 | 7 |
| - | - | 2.99E+04 | 838.4 | - | - | 0 | - |
| - | - | 4397 | 838.5 | - | - | 0 | - |
| - | - | 7203 | 839.4 | - | - | 0 | - |
| - | - | 2345 | 839.5 | - | - | 0 | - |
| - | - | 1283 | 840.4 | - | - | 0 | - |
| - | - | 3490 | 852.4 | - | - | 0 | - |
| 7 | y | 1.868E+04 | 853.4 | 0.00202 | 2.368 | +1 | 7 |
| - | - | 7586 | 854.4 | - | - | 0 | - |
| - | - | 1713 | 855.4 | - | - | 0 | - |
| - | - | 1080 | 879.4 | - | - | 0 | - |
| - | - | 2045 | 879.5 | - | - | 0 | - |
| - | - | 3238 | 880.4 | - | - | 0 | - |
| - | - | 855.7 | 880.5 | - | - | 0 | - |
| - | - | 1.511E+04 | 881.4 | - | - | 0 | - |
| 8 | c | 3.487E+05 | 882.5 | 0.00299 | 3.388 | +1 | 8 |
| - | - | 1.611E+05 | 883.5 | - | - | 0 | - |
| - | - | 4.807E+04 | 884.5 | - | - | 0 | - |
| - | - | 8976 | 885.5 | - | - | 0 | - |
| - | - | 1391 | 886.5 | - | - | 0 | - |
| - | - | 3323 | 909.4 | - | - | 0 | - |
| - | - | 957.7 | 910.4 | - | - | 0 | - |
| - | - | 912.7 | 921.4 | - | - | 0 | - |
| 6 | y | 5095 | 922.4 | 0.01688 | 18.3 | +1 | 8 |
| 6 | y | 1.22E+04 | 923.4 | 0.002401 | 2.6 | +1 | 8 |
| 6 | z | 1.157E+05 | 924.4 | 0.002754 | 2.979 | +1 | 8 |
| - | - | 5.582E+04 | 925.4 | - | - | 0 | - |
| - | - | 1.824E+04 | 926.4 | - | - | 0 | - |
| - | - | 4281 | 927.4 | - | - | 0 | - |
| - | - | 1992 | 939.4 | - | - | 0 | - |
| 6 | y | 7.961E+04 | 940.4 | 0.003317 | 3.527 | +1 | 8 |
| - | - | 3.562E+04 | 941.4 | - | - | 0 | - |
| - | - | 1.151E+04 | 942.4 | - | - | 0 | - |
| - | - | 2111 | 943.5 | - | - | 0 | - |
| - | - | 1847 | 948.4 | - | - | 0 | - |
| - | - | 2369 | 949.4 | - | - | 0 | - |
| - | - | 4435 | 950.4 | - | - | 0 | - |
| - | - | 2373 | 951.4 | - | - | 0 | - |
| - | - | 2060 | 951.5 | - | - | 0 | - |
| - | - | 3.171E+04 | 952.5 | - | - | 0 | - |
| - | - | 2.184E+04 | 953.5 | - | - | 0 | - |
| - | - | 9141 | 954.5 | - | - | 0 | - |
| - | - | 2802 | 955.5 | - | - | 0 | - |
| - | - | 1832 | 964.5 | - | - | 0 | - |
| - | - | 1855 | 965.4 | - | - | 0 | - |
| - | - | 1808 | 967.4 | - | - | 0 | - |
| - | - | 2445 | 967.6 | - | - | 0 | - |
| - | - | 3222 | 968.4 | - | - | 0 | - |
| - | - | 6711 | 968.6 | - | - | 0 | - |
| - | - | 1841 | 969 | - | - | 0 | - |
| - | - | 4883 | 969.5 | - | - | 0 | - |
| - | - | 3081 | 969.6 | - | - | 0 | - |
| - | - | 1614 | 970 | - | - | 0 | - |
| - | - | 1261 | 970.5 | - | - | 0 | - |
| 9 | c | 1006 | 979.5 | 0.01304 | 13.31 | +1 | 9 |
| - | - | 7095 | 980.5 | - | - | 0 | - |
| - | - | 6692 | 981.5 | - | - | 0 | - |
| - | - | 3238 | 982.5 | - | - | 0 | - |
| - | - | 3.039E+04 | 1008 | - | - | 0 | - |
| - | - | 1.511E+04 | 1009 | - | - | 0 | - |
| - | - | 4060 | 1010 | - | - | 0 | - |
| - | - | 1260 | 1011 | - | - | 0 | - |
| 5 | y | 1040 | 1022 | 0.0137 | 13.4 | +1 | 9 |
| 5 | z | 1.073E+05 | 1023 | 0.004348 | 4.248 | +1 | 9 |
| - | - | 6.074E+04 | 1024 | - | - | 0 | - |
| - | - | 2.318E+04 | 1025 | - | - | 0 | - |
| - | - | 4993 | 1027 | - | - | 0 | - |
| - | - | 2052 | 1039 | - | - | 0 | - |
| 5 | y | 8.488E+04 | 1040 | 0.00308 | 2.963 | +1 | 9 |
| - | - | 4.41E+04 | 1041 | - | - | 0 | - |
| - | - | 1.554E+04 | 1042 | - | - | 0 | - |
| - | - | 4010 | 1043 | - | - | 0 | - |
| - | - | 2987 | 1051 | - | - | 0 | - |
| - | - | 2092 | 1052 | - | - | 0 | - |
| - | - | 763.6 | 1065 | - | - | 0 | - |
| - | - | 1849 | 1066 | - | - | 0 | - |
| - | - | 1659 | 1068 | - | - | 0 | - |
| - | - | 2091 | 1080 | - | - | 0 | - |
| - | - | 4075 | 1081 | - | - | 0 | - |
| - | - | 1776 | 1082 | - | - | 0 | - |
| - | - | 918.5 | 1093 | - | - | 0 | - |
| 4 | w | 1591 | 1094 | 0.01327 | 12.13 | +1 | 10 |
| 10 | c | 1.729E+05 | 1095 | 0.007503 | 6.855 | +1 | 10 |
| - | - | 9.452E+04 | 1096 | - | - | 0 | - |
| - | - | 3.664E+04 | 1097 | - | - | 0 | - |
| - | - | 7660 | 1098 | - | - | 0 | - |
| - | - | 1056 | 1099 | - | - | 0 | - |
| - | - | 1.474E+04 | 1107 | - | - | 0 | - |
| - | - | 9524 | 1109 | - | - | 0 | - |
| - | - | 2508 | 1110 | - | - | 0 | - |
| - | - | 2.23E+04 | 1121 | - | - | 0 | - |
| - | - | 1.284E+04 | 1122 | - | - | 0 | - |
| - | - | 4779 | 1123 | - | - | 0 | - |
| - | - | 985.1 | 1124 | - | - | 0 | - |
| - | - | 1153 | 1126 | - | - | 0 | - |
| - | - | 1054 | 1127 | - | - | 0 | - |
| 4 | y | 1020 | 1137 | 0.004876 | 4.29 | +1 | 10 |
| 4 | y | 1266 | 1138 | 0.0127 | 11.16 | +1 | 10 |
| 4 | z | 9.324E+04 | 1139 | 0.003283 | 2.884 | +1 | 10 |
| - | - | 8.191E+04 | 1140 | - | - | 0 | - |
| - | - | 3.93E+04 | 1141 | - | - | 0 | - |
| - | - | 1.136E+04 | 1142 | - | - | 0 | - |
| - | - | 2270 | 1143 | - | - | 0 | - |
| - | - | 1109 | 1154 | - | - | 0 | - |
| 4 | y | 1.333E+04 | 1155 | 0.003358 | 2.909 | +1 | 10 |
| - | - | 9322 | 1156 | - | - | 0 | - |
| - | - | 3197 | 1157 | - | - | 0 | - |
| - | - | 915.5 | 1158 | - | - | 0 | - |
| - | - | 1119 | 1163 | - | - | 0 | - |
| - | - | 917.8 | 1164 | - | - | 0 | - |
| - | - | 1234 | 1179 | - | - | 0 | - |
| - | - | 7.044E+04 | 1180 | - | - | 0 | - |
| - | - | 4.784E+04 | 1181 | - | - | 0 | - |
| - | - | 1.966E+04 | 1182 | - | - | 0 | - |
| - | - | 5592 | 1183 | - | - | 0 | - |
| - | - | 1264 | 1184 | - | - | 0 | - |
| - | - | 6322 | 1207 | - | - | 0 | - |
| - | - | 5728 | 1208 | - | - | 0 | - |
| - | - | 2236 | 1209 | - | - | 0 | - |
| - | - | 1923 | 1210 | - | - | 0 | - |
| - | - | 1912 | 1212 | - | - | 0 | - |
| - | - | 2309 | 1220 | - | - | 0 | - |
| - | - | 1556 | 1221 | - | - | 0 | - |
| - | - | 1161 | 1223 | - | - | 0 | - |
| 11 | c | 7.295E+04 | 1224 | 0.002516 | 2.056 | +1 | 11 |
| - | - | 4.288E+04 | 1225 | - | - | 0 | - |
| - | - | 1.733E+04 | 1226 | - | - | 0 | - |
| - | - | 4666 | 1227 | - | - | 0 | - |
| - | - | 1073 | 1235 | - | - | 0 | - |
| 3 | z | 1.427E+04 | 1238 | 0.003351 | 2.708 | +1 | 11 |
| - | - | 1.708E+04 | 1239 | - | - | 0 | - |
| - | - | 7454 | 1240 | - | - | 0 | - |
| - | - | 2504 | 1241 | - | - | 0 | - |
| 3 | y | 1594 | 1254 | 0.000374 | 0.2983 | +1 | 11 |
| - | - | 2507 | 1255 | - | - | 0 | - |
| - | - | 1203 | 1263 | - | - | 0 | - |
| - | - | 1155 | 1278 | - | - | 0 | - |
| - | - | 2.361E+04 | 1279 | - | - | 0 | - |
| - | - | 1.955E+04 | 1280 | - | - | 0 | - |
| - | - | 1.012E+04 | 1281 | - | - | 0 | - |
| - | - | 2891 | 1282 | - | - | 0 | - |
| - | - | 1148 | 1307 | - | - | 0 | - |
| - | - | 1699 | 1308 | - | - | 0 | - |
| - | - | 5753 | 1322 | - | - | 0 | - |
| 12 | c | 9.734E+04 | 1323 | 0.002583 | 1.953 | +1 | 12 |
| - | - | 6.148E+04 | 1324 | - | - | 0 | - |
| - | - | 1196 | 1324 | - | - | 0 | - |
| - | - | 2.697E+04 | 1325 | - | - | 0 | - |
| - | - | 8104 | 1326 | - | - | 0 | - |
| - | - | 1189 | 1327 | - | - | 0 | - |
| 2 | z | 2250 | 1337 | 0.005616 | 4.201 | +1 | 12 |
| - | - | 1.114E+04 | 1338 | - | - | 0 | - |
| - | - | 7870 | 1339 | - | - | 0 | - |
| - | - | 3511 | 1340 | - | - | 0 | - |
| - | - | 1307 | 1371 | - | - | 0 | - |
| - | - | 2453 | 1377 | - | - | 0 | - |
| - | - | 972 | 1378 | - | - | 0 | - |
| - | - | 6292 | 1393 | - | - | 0 | - |
| - | - | 3.085E+04 | 1394 | - | - | 0 | - |
| - | - | 2.079E+04 | 1395 | - | - | 0 | - |
| - | - | 1.12E+04 | 1396 | - | - | 0 | - |
| - | - | 2990 | 1397 | - | - | 0 | - |
| - | - | 830.7 | 1407 | - | - | 0 | - |
| - | - | 3089 | 1408 | - | - | 0 | - |
| - | - | 6138 | 1409 | - | - | 0 | - |
| - | - | 3912 | 1410 | - | - | 0 | - |
| - | - | 1142 | 1411 | - | - | 0 | - |
| - | - | 962.3 | 1419 | - | - | 0 | - |
| - | - | 1693 | 1425 | - | - | 0 | - |
| - | - | 4715 | 1426 | - | - | 0 | - |
| - | - | 2865 | 1427 | - | - | 0 | - |
| - | - | 1075 | 1428 | - | - | 0 | - |
| - | - | 1275 | 1435 | - | - | 0 | - |
| - | - | 1.536E+04 | 1436 | - | - | 0 | - |
| - | - | 9.033E+04 | 1437 | - | - | 0 | - |
| - | - | 6.113E+04 | 1438 | - | - | 0 | - |
| - | - | 2.882E+04 | 1439 | - | - | 0 | - |
| - | - | 7735 | 1440 | - | - | 0 | - |
| - | - | 2040 | 1441 | - | - | 0 | - |
| - | - | 1.626E+04 | 1452 | - | - | 0 | - |
| - | - | 6.227E+04 | 1453 | - | - | 0 | - |
| - | - | 1.483E+05 | 1454 | - | - | 0 | - |
| - | - | 9.778E+04 | 1455 | - | - | 0 | - |
| - | - | 4.363E+04 | 1456 | - | - | 0 | - |
| - | - | 1.318E+04 | 1457 | - | - | 0 | - |
| - | - | 2721 | 1458 | - | - | 0 | - |

m/z Charge Intensity FragmentType MassShift Position
122.0514144897461 0 418.26147
125.94807434082031 0 402.56357
126.0921859741211 0 786.4246
127.0870361328125 0 506.28378
127.13605499267578 0 561.9194
129.06068420410156 0 442.07834
129.10272216796875 0 16463.045
130.08676147460938 0 15981.598 y Ammonia loss 12
130.106201171875 0 997.0351
131.09458923339844 0 21179.205 z 12
131.11830139160156 0 1708.7297
132.09877014160156 0 688.5007
132.10240173339844 0 4276.6226
133.06130981445312 0 793.5795
141.94569396972656 0 476.0785
143.15492248535156 0 524.9243
144.1262969970703 0 835.49023
147.11334228515625 0 56863.996 y 12
148.11669921875 0 4307.266
148.84828186035156 0 479.91675
148.90597534179688 0 643.8087
148.91319274902344 0 796.7705
148.9203643798828 0 1225.7164
148.92771911621094 0 1311.3933
148.9346160888672 0 1919.6245
148.9423370361328 0 3690.6614
148.95895385742188 0 4294.0913
148.966796875 0 2798.7505
148.9739532470703 0 1118.1545
148.98092651367188 0 1000.7671
148.9882049560547 0 510.1493
149.0029296875 0 675.4101
149.01011657714844 0 476.2116
149.03945922851562 0 529.5097
149.04635620117188 0 635.2685
149.06146240234375 0 585.2678
149.11138916015625 0 433.3457
152.10731506347656 0 916.7326
153.07861328125 0 457.32053
154.1231231689453 0 7725.382
155.13107299804688 0 2282.9824
156.13916015625 0 849.1267
163.07203674316406 0 1699.128
169.13429260253906 0 773.65155
170.14212036132812 0 1890.5779
171.0896453857422 0 2281.842
171.14981079101562 0 339146.28
172.09735107421875 0 2078.755
172.1531524658203 0 30204.64
173.15623474121094 0 1244.4288
173.44007873535156 0 630.15247
179.1185760498047 0 1243.0444
185.1654815673828 0 7204.311
187.1083221435547 0 6588.3525
189.12408447265625 0 1617.3127
197.1292266845703 0 1805.1968
198.13693237304688 0 8417.911
199.14471435546875 0 280646.25
200.1481170654297 0 31115.686
201.12432861328125 0 655.7494
201.15028381347656 0 1808.645
203.1031494140625 0 5202.1943
204.1070556640625 0 592.3253
211.14488220214844 0 776.2593
213.16050720214844 0 6861.123
215.103271484375 0 17203.21
215.11500549316406 0 943.41766
215.13980102539062 0 4453.462 w 11
216.107421875 0 1235.5173
216.17135620117188 0 2802.497 c 1
225.098876953125 0 764.84406
227.10345458984375 0 2996.2747
228.17153930664062 0 2741.1733
229.1555633544922 0 3345.7842 y Ammonia loss 11
229.17552185058594 0 856.4763
230.16317749023438 0 6887.7593 z 11
231.16668701171875 0 869.89514
232.1299591064453 0 31717.629
233.13330078125 0 3097.3484
239.2740478515625 0 540.44
246.18206787109375 0 89870.58 y 11
247.1852569580078 0 9757.248
248.189208984375 0 1119.2894
253.1918487548828 0 7187.9272
254.19622802734375 0 1208.8241
270.1824035644531 0 790.8585
270.2182922363281 0 21320.162
271.2224426269531 0 4440.907
272.17431640625 0 889.1916
282.1221923828125 0 3460.265
283.1296081542969 0 3716.4702
286.1766357421875 0 1393.3743
287.2080993652344 0 1331.4814
297.2058410644531 0 4217.479
298.2135925292969 0 36905.426
299.1727294921875 0 632.4786
299.2167663574219 0 6587.5054
300.1928405761719 0 91500.69 w 10
301.1960144042969 0 14896.073
302.13580322265625 0 870.3685
302.1983337402344 0 1886.1447
314.1722412109375 0 14924.375
314.2317810058594 0 3668.2012
315.1749267578125 0 2159.2856
315.2398376464844 0 8009.46 c 2
316.243896484375 0 1651.7255
326.1720886230469 0 3875.16
327.1750183105469 0 781.8815
331.19879150390625 0 7013.3887
332.2016906738281 0 1482.4608
342.202880859375 0 628.1076
343.1990966796875 0 1474.1074
348.5955810546875 0 610.92035
353.1961364746094 0 3749.5005
354.2032470703125 0 16931.104
355.2073059082031 0 4093.86
357.214599609375 0 2249.7427 y Water loss 10
359.2059020996094 0 4999.591 z 10
368.2428894042969 0 6230.4717
369.246337890625 0 1587.3455
371.2387390136719 0 2496.1804
373.2100830078125 0 1232.0905
375.2247619628906 0 2258.945 y 10
381.1903381347656 0 2534.6392
382.1375427246094 0 2607.9736
382.1976318359375 0 2631.8955
383.1921691894531 0 1818.6212
385.24591064453125 0 7637.0806
386.2532043457031 0 10494.796
387.2581787109375 0 2194.7166
392.1732177734375 0 599.07434
395.23114013671875 0 1446.9327
398.1951904296875 0 911.7369
398.2357177734375 0 2275.2168
399.2427978515625 0 790.5637
401.2046203613281 0 6040.021
402.2110595703125 0 2670.6255
412.23126220703125 0 800.84247
413.2408142089844 0 45768.01
414.1769714355469 0 1380.1718
414.244140625 0 9194.627
415.17657470703125 0 832.38556
415.24951171875 0 1816.127
416.2287902832031 0 2087.302
418.2318420410156 0 1315.3076
418.5395812988281 0 3183.1497 y 2
418.8745422363281 0 2461.544
419.2008972167969 0 1545.8188 z 6
419.5440979003906 0 644.415
426.2371826171875 0 712.6242
426.3109436035156 0 786.83923
427.207763671875 0 11177.8125 y 6
427.7094421386719 0 4551.236
428.21002197265625 0 793.4161
429.2359313964844 0 28606.092 w 9
430.2402648925781 0 8379.813
430.2686767578125 0 5015.657 c 3
431.2447814941406 0 1498.8127
431.2713928222656 0 1558.9216
431.7627868652344 0 816.47015
434.7514343261719 0 1153.1277
439.1927795410156 0 766.628
441.2331237792969 0 971.75116
441.33056640625 0 1073.6536
442.26788330078125 0 2659.3164
443.2459716796875 0 805.2648
444.2582702636719 0 2961.1187
445.2602844238281 0 1080.6385
451.5624694824219 0 5146.376 y 1
451.8966064453125 0 2506.5203
452.23297119140625 0 774.52563
453.2093200683594 0 727.4898
453.25677490234375 0 690.5276
454.2102966308594 0 833.7253 w 5
454.270751953125 0 1038.3236 y Water loss 9
455.2144470214844 0 802.2836
455.24859619140625 0 664.1126 y Ammonia loss 9
455.2754821777344 0 8255.473
456.2235412597656 0 2836.959
456.27777099609375 0 1633.3419
457.2554931640625 0 1113.6487
458.2701416015625 0 1248.318
461.71142578125 0 1573.796 y Water loss 5
462.2109375 0 6572.0107 y Ammonia loss 5
462.7141418457031 0 9056.1455 z 5
463.21527099609375 0 4341.637
463.71612548828125 0 1182.422
468.2835998535156 0 29265.006
469.16998291015625 0 3100.214
469.28985595703125 0 6224.657
470.2627868652344 0 43671.56
470.3089599609375 0 4629.9707
470.7239990234375 0 17994.383 y 5
471.2252502441406 0 8521.785
471.267333984375 0 15060.101
471.30938720703125 0 1414.4587
471.7262878417969 0 2756.1494
472.27825927734375 0 131285.12 y 9
473.2815856933594 0 34160.555
474.28448486328125 0 6220.7065
474.7086486816406 0 3075.9846
475.209228515625 0 1136.7465
475.2867431640625 0 1077.9026
484.2209777832031 0 753.2074
484.2851257324219 0 4796.163
484.315185546875 0 7520.547
484.5857849121094 0 17329.977
484.7872314453125 0 1475.2476
484.91986083984375 0 8709.453
485.2518005371094 0 19176.09
485.3210754394531 0 7410.2437
485.3601989746094 0 962.72845
485.589111328125 0 1714.9692
486.25177001953125 0 2426.7166
486.3260803222656 0 2496.9788
494.3001403808594 0 1768.3251
496.26751708984375 0 920.4272
497.2984619140625 0 1623.4834
499.26544189453125 0 24420.166
500.2712097167969 0 11693.604
501.28021240234375 0 20434.037
502.28411865234375 0 5172.3306
503.23809814453125 0 610.08417 z Water loss 4
503.2835388183594 0 1167.0378
503.74359130859375 0 1157.4781 z Ammonia loss 4
504.73614501953125 0 4920.6826 w 4
505.2341003417969 0 1868.999
505.740234375 0 1282.0469
509.28643798828125 0 1257.676
510.26861572265625 0 1821.7432
511.7531433105469 0 2250.1858 y Ammonia loss 4
512.249267578125 0 16696.121 z 4
512.309814453125 0 64847.3
512.7507934570312 0 7329.921
513.250244140625 0 2563.465
513.3129272460938 0 17524.803
514.3158569335938 0 3361.117
520.2578735351562 0 34207.547 y 4
520.7599487304688 0 17113.42
521.2598876953125 0 5593.539
521.7628784179688 0 907.5366
526.7415161132812 0 772.267
527.296875 0 183054.25
528.2999267578125 0 52420.812
529.3001708984375 0 7934.719
529.3370971679688 0 16682.012 c 4
530.3026123046875 0 687.91626
530.3401489257812 0 5590.036
533.7511596679688 0 3812.5508
534.2518920898438 0 1627.4028
538.2640380859375 0 5657.5454
538.7603149414062 0 729.08246
539.2660522460938 0 3109.285
540.2588500976562 0 1754.2136
540.7587890625 0 4078.074
541.2606811523438 0 1604.3513
547.2623291015625 0 945.1277 w 3
547.76708984375 0 12235.398 c 9
548.2686157226562 0 8093.098
548.7715454101562 0 2718.049
553.31005859375 0 1222.1055
554.2540283203125 0 988.947 z Ammonia loss 8
559.2852783203125 0 2090.7058
560.2960815429688 0 707.16754
568.2396850585938 0 5567.874
568.768310546875 0 1114.2129 y Water loss 3
569.2389526367188 0 1388.8091
569.7623901367188 0 7939.722 z 3
570.2650756835938 0 4636.5684
570.326171875 0 7801.871
570.764892578125 0 2060.414
571.2839965820312 0 3012.7312 z 8
571.3338012695312 0 2178.7258
572.2855834960938 0 1187.8562
577.7716064453125 0 66600.6 y 3
578.27294921875 0 36471.918
578.7747192382812 0 14235.77
579.2772827148438 0 3542.5503
581.3306274414062 0 3546.6006
582.7705688476562 0 935.7628
587.30517578125 0 23483.27 y 8
588.308349609375 0 6780.4297
589.31005859375 0 876.54333
589.7880859375 0 1084.1372
590.2935180664062 0 868.1113
590.7978515625 0 1073.2833
591.2650756835938 0 1225.0708
595.3322143554688 0 917.4234
597.3019409179688 0 6690.477
597.8028564453125 0 6236.4395
598.3028564453125 0 2364.8833
598.3590087890625 0 955.2982 c Water loss 5
599.3419799804688 0 12883.647
600.3456420898438 0 4161.9883
601.3428344726562 0 981.0335
603.2847290039062 0 1535.5809 c Water loss 10
603.78759765625 0 3615.2078
604.2886962890625 0 2194.6973
604.795654296875 0 1627.857
605.2745361328125 0 1336.595
605.7739868164062 0 1214.9164
606.2733154296875 0 1102.0272
611.7853393554688 0 7334.952 w 2
612.2871704101562 0 4267.8247 c 10
612.7885131835938 0 1693.8136
613.2880859375 0 855.32666
616.368408203125 0 42756.91 c 5
617.3715209960938 0 15354.896
618.301513671875 0 7640.522 y Water loss 2
618.374267578125 0 3339.8926
618.801513671875 0 4155.9243 y Ammonia loss 2
619.2972412109375 0 44755.645 z 2
619.7982177734375 0 26499.297
620.2997436523438 0 10521.377
620.79931640625 0 2544.7546
624.3353271484375 0 976.8651
626.298583984375 0 3897.2502
626.7980346679688 0 2471.9832
627.30615234375 0 430456.88 y 2
627.8077392578125 0 266094.94
628.3089599609375 0 113778.25
628.8098754882812 0 30873.037
629.310791015625 0 5582.3105
632.301513671875 0 1645.2037
639.3187866210938 0 1773.488
640.41552734375 0 3537.9702
640.7986450195312 0 2683.3599
641.3160400390625 0 78444.27 w 7
641.8021850585938 0 1190.7877
642.3191528320312 0 26296.127
643.3218383789062 0 5359.419
644.326904296875 0 1006.5819
645.9723510742188 0 2502.2842
646.3182983398438 0 1953.7108
646.6454467773438 0 1576.6903
646.8350830078125 0 6945.5825
646.975341796875 0 949.12585
647.3356323242188 0 5183.168
647.8052978515625 0 1690.2268
647.8441162109375 0 1389.1394
648.312255859375 0 2232.11
648.80859375 0 1501.5719
653.3218994140625 0 5739.84
653.8236083984375 0 5811.475
654.3265380859375 0 3799.9463
656.3392944335938 0 9583.723
657.3403930664062 0 3364.9268
658.3479614257812 0 864.1252
660.8566284179688 0 2414.962
661.3204956054688 0 12763.607 w 1
661.826904296875 0 14179.948 c 11
662.332763671875 0 8943.928
662.8370361328125 0 3091.2463
665.4220581054688 0 995.58636
667.3053588867188 0 6111.09
667.3643798828125 0 3183.5469
667.861083984375 0 5371.465
668.3206787109375 0 3924.7922 y Ammonia loss 1
668.8310546875 0 21407.092 z 1
669.3321533203125 0 14023.908
669.8328857421875 0 5561.122
670.3370971679688 0 1175.8965
674.368408203125 0 1489.3076
675.3743896484375 0 1728.9038
675.8326416015625 0 2297.8567
676.3394775390625 0 2191.4065
676.840576171875 0 89591.92 y 1
677.3419799804688 0 58043.613
677.8430786132812 0 30297.717
678.344970703125 0 9266.315
678.8475341796875 0 1476.0105
681.8272094726562 0 1098.8469
682.3568115234375 0 3609.993
682.8555908203125 0 2171.7944
683.2651977539062 0 8264.278
683.3580322265625 0 1823.7596
683.8545532226562 0 954.75165
684.2696533203125 0 2796.464
685.2714233398438 0 1053.0881
685.8543701171875 0 1372.252
686.3526611328125 0 1577.8351
686.858642578125 0 1609.5868
687.8648681640625 0 1479.1538
688.367431640625 0 1437.4088
688.8622436523438 0 2083.9043
689.3540649414062 0 2127.886
689.8600463867188 0 3415.4705
690.3567504882812 0 2390.9792
690.8671875 0 11841.09
691.3673706054688 0 4827.0957
691.8707885742188 0 2958.3232
693.3988037109375 0 1313.1924
695.2614135742188 0 1251.1758
695.8685913085938 0 3066.5706
696.3690795898438 0 8998.784
696.8681640625 0 166980.62
697.3698120117188 0 148865.05
697.8711547851562 0 76141.805
698.3727416992188 0 27840.725
698.8742065429688 0 6458.5674
700.329833984375 0 54255.555 z 7
701.3328247070312 0 20945.232
702.3365478515625 0 6058.715
703.3466796875 0 1683.6411
703.8748168945312 0 13863.484
704.3775024414062 0 10626.483
704.88232421875 0 27537.979
705.3845825195312 0 21291.168
705.884521484375 0 8817.826
706.3944091796875 0 1485.4012
707.3743896484375 0 1528.6503
708.398193359375 0 1661.6873
709.4144287109375 0 26699.02
709.860595703125 0 4049.7869
710.3551025390625 0 837.71027
710.416748046875 0 10722.327
710.85400390625 0 117656.96
711.35546875 0 79537.46
711.856689453125 0 40464.06
712.3585205078125 0 11660.868
712.859375 0 3028.807
716.3487548828125 0 49257.97 y 7
717.3531494140625 0 18638.248
717.8699951171875 0 11680.96
718.3657836914062 0 325425.97
718.8673095703125 0 221419.27
719.368408203125 0 113263.01
719.8697509765625 0 36104.508
720.3707885742188 0 9733.946
725.3668823242188 0 6246.8984
725.8700561523438 0 14462.333
726.375244140625 0 680963
726.878173828125 0 1195803.6
727.379638671875 0 742333.56
727.880859375 0 323386.97
728.3822021484375 0 91646.86
728.8841552734375 0 17684.564
734.3561401367188 0 2267.9285
736.3988037109375 0 1282.3077
753.4281005859375 0 170965.16 c 6
754.4306640625 0 63851.46
755.425537109375 0 19581.574
756.4207763671875 0 5485.311
764.3263549804688 0 1540.1326
765.3687133789062 0 6779.57
765.4644165039062 0 4832.189
766.373046875 0 3422.3242
766.4681396484375 0 1430.7141
778.3758544921875 0 3103.0876
779.4429931640625 0 9228.662
780.44677734375 0 3777.0874
781.447265625 0 1962.3989
782.3344116210938 0 71999.984
783.3374633789062 0 30699.977
784.3399658203125 0 7926.9546
785.3439331054688 0 1038.0586
793.4004516601562 0 1533.5405
794.3320922851562 0 2658.6812
795.3369750976562 0 1711.0735
805.3922119140625 0 1465.615
837.3890991210938 0 67032.766 z 6
838.3912963867188 0 29904.13
838.4581298828125 0 4397.251
839.3925170898438 0 7202.6846
839.4652099609375 0 2345.494
840.3941040039062 0 1283.2068
852.3991088867188 0 3490.2969
853.4070434570312 0 18677.104 y 6
854.4098510742188 0 7586.3613
855.4137573242188 0 1712.9985
879.3930053710938 0 1080.3055
879.546142578125 0 2044.9509
880.4414672851562 0 3238.1362
880.5372314453125 0 855.70917
881.4038696289062 0 15111.779
882.470947265625 0 348735.6 c 7
883.4735107421875 0 161104.27
884.4759521484375 0 48070.5
885.4779663085938 0 8975.858
886.4833374023438 0 1391.3914
909.3653564453125 0 3323.3435
910.3712768554688 0 957.7482
921.44482421875 0 912.66705
922.4096069335938 0 5095.1675 y Water loss 5
923.4129028320312 0 12198.967 y Ammonia loss 5
924.4210815429688 0 115743 z 5
925.4244384765625 0 55821.42
926.4266967773438 0 18238.078
927.4310913085938 0 4280.745
939.43115234375 0 1992.488
940.4403686523438 0 79611.66 y 5
941.4426879882812 0 35619.438
942.4459838867188 0 11514.874
943.4505004882812 0 2111.11
948.4081420898438 0 1846.5157
949.4183349609375 0 2368.6375
950.4246826171875 0 4434.5493
951.4497680664062 0 2372.7837
951.5455932617188 0 2059.5151
952.4761962890625 0 31710.668
953.480712890625 0 21837.043
954.4845581054688 0 9141.283
955.4874267578125 0 2802.3472
964.4578857421875 0 1831.857
965.4388427734375 0 1854.9293
967.4381103515625 0 1808.279
967.5606689453125 0 2444.7744
968.4468383789062 0 3221.7114
968.5670776367188 0 6710.7466
968.96240234375 0 1840.7535
969.459228515625 0 4883.473
969.5730590820312 0 3081.4731
969.9644775390625 0 1614.2375
970.468994140625 0 1261.1432
979.4973754882812 0 1006.31964 c Water loss 8
980.46875 0 7095.0063
981.4760131835938 0 6692.151
982.48046875 0 3238.3079
1008.4309692382812 0 30389.629
1009.4334716796875 0 15107.675
1010.4364624023438 0 4059.6592
1011.43701171875 0 1259.8153
1022.4926147460938 0 1039.5399 y Ammonia loss 4
1023.4910888671875 0 107349.086 z 4
1024.494873046875 0 60739.953
1025.4981689453125 0 23184.748
1026.5006103515625 0 4992.8306
1038.5067138671875 0 2052.1458
1039.508544921875 0 84876.73 y 4
1040.5118408203125 0 44098.426
1041.514404296875 0 15540.797
1042.5147705078125 0 4010.483
1050.53369140625 0 2986.5984
1051.5340576171875 0 2091.8672
1064.501953125 0 763.6148
1066.4952392578125 0 1848.6698
1067.506591796875 0 1659.1112
1079.51025390625 0 2091.0647
1080.5111083984375 0 4074.6406
1081.51513671875 0 1775.8129
1092.507568359375 0 918.51953
1093.529296875 0 1590.5984 w 3
1094.5401611328125 0 172898.9 c 9
1095.54296875 0 94515.77
1096.5474853515625 0 36638.957
1097.5458984375 0 7660.0083
1098.5565185546875 0 1056.3883
1107.4979248046875 0 14737.296
1108.5023193359375 0 9524.19
1109.5069580078125 0 2507.91
1120.5675048828125 0 22303.951
1121.56884765625 0 12837.815
1122.5743408203125 0 4779.473
1123.565673828125 0 985.14844
1125.505126953125 0 1152.5729
1126.5167236328125 0 1054.2335
1136.5169677734375 0 1020.04736 y Water loss 3
1137.4931640625 0 1265.8522 y Ammonia loss 3
1138.5169677734375 0 93240.086 z 3
1139.5218505859375 0 81908.85
1140.524658203125 0 39296.344
1141.529296875 0 11361.639
1142.5323486328125 0 2269.5535
1153.519287109375 0 1109.1528
1154.5357666015625 0 13333.116 y 3
1155.5380859375 0 9322.032
1156.5408935546875 0 3196.8464
1157.5452880859375 0 915.5459
1162.564208984375 0 1119.1621
1163.57373046875 0 917.7611
1178.5653076171875 0 1234.2935
1179.5802001953125 0 70441.82
1180.5836181640625 0 47835.1
1181.5867919921875 0 19656.496
1182.5867919921875 0 5591.787
1183.602783203125 0 1264.3147
1206.5804443359375 0 6322.259
1207.5833740234375 0 5727.932
1208.585205078125 0 2236.3208
1209.579833984375 0 1923.3264
1211.560546875 0 1911.8816
1219.636962890625 0 2308.7556
1220.6439208984375 0 1555.515
1222.5885009765625 0 1160.5818
1223.5927734375 0 72951.05 c 10
1224.5963134765625 0 42881.105
1225.59912109375 0 17325.662
1226.602783203125 0 4666.412
1234.659912109375 0 1072.7877
1237.58544921875 0 14268.827 z 2
1238.5902099609375 0 17081.709
1239.59326171875 0 7453.73
1240.5948486328125 0 2503.661
1253.6011962890625 0 1593.9612 y 2
1254.6048583984375 0 2507.4568
1262.645751953125 0 1203.1986
1277.630615234375 0 1154.977
1278.647705078125 0 23610.477
1279.651123046875 0 19548.215
1280.655029296875 0 10122.243
1281.6595458984375 0 2891.49
1306.6424560546875 0 1148.3363
1307.645263671875 0 1699.3517
1321.6546630859375 0 5752.6504
1322.6612548828125 0 97335.45 c 11
1323.6639404296875 0 61484.004
1323.8541259765625 0 1196.4762
1324.6676025390625 0 26973.797
1325.66943359375 0 8103.705
1326.6627197265625 0 1188.6547
1336.6561279296875 0 2250.3564 z 1
1337.659912109375 0 11140.022
1338.66455078125 0 7870.0527
1339.6666259765625 0 3511.1094
1370.6954345703125 0 1307.4888
1376.7086181640625 0 2453.1694
1377.69189453125 0 971.9698
1392.7265625 0 6292.0493
1393.73388671875 0 30847.314
1394.737060546875 0 20786.432
1395.7396240234375 0 11198.747
1396.74560546875 0 2990.497
1406.733642578125 0 830.74695
1407.7425537109375 0 3088.7395
1408.7506103515625 0 6137.719
1409.75439453125 0 3912.255
1410.7392578125 0 1141.5023
1418.7181396484375 0 962.26447
1424.7489013671875 0 1692.786
1425.7598876953125 0 4715.1294
1426.7581787109375 0 2865.1772
1427.7691650390625 0 1075.1176
1434.730712890625 0 1275.4712
1435.72314453125 0 15363.94
1436.72900390625 0 90333.336
1437.7314453125 0 61133.086
1438.7354736328125 0 28819.91
1439.7376708984375 0 7734.936
1440.7340087890625 0 2040.0674
1451.7398681640625 0 16255.481
1452.74658203125 0 62271.688
1453.7532958984375 0 148281.03
1454.7569580078125 0 97779.586
1455.760498046875 0 43628.176
1456.7623291015625 0 13182.325
1457.7689208984375 0 2720.9597

Spectrum Details

|  |  |
| --- | --- |
| Matched peaks? Matched peaks The total absolute number of peaks matched. Additionally in brackets the total fraction of peaks matched and the total number of peaks is shown. | 82 (12.69% of 646) |
| FDR? FDR The false discovery rate estimated for this peptide. It is calculated by matching all theoretical fragments with a non-integer shift with the raw peaks for this spectrum. This is done with 40 different shifts. The resulting percentage is the average number of annotated peaks over the number of annotated peaks with the correct spectrum. | 1.05% |
| Satellite FDR? Satellite FDR See the FDR for details on its calculation. This satellite ion specific FDR only contains the satellite ions (d/w) for I/L/J positions. | - |
| PSM Score? PSM Score The PSM Score as given by Hecklib to this annotated spectrum. It is shown with three significant figures. | 648 |

#### Spectrum 4197? Spectrum 4197 The raw spectrum of this peptide as annotated by Hecklib. The fragments are coloured according to ion type (see legend). Any peaks with a star '\*' as text can be hovered over to see the full details, first the ion type second the mass shift type. By hovering over the amino acids in the peptide or ions in the legend the corresponding peaks are highlighted. By toggling the 'Unassigned' label you can turn the background (unassigned) peaks on or off in the plot. By updating the slider in the Ion legend you can update the spectrum to only show the top X% of the peaks with labels. The top X% means any peak that is within X% of the highest intensity. By dragging in the spectrum you can zoom in to a specific part of the spectrum and use 'Zoom Out' to get back to the original zoom level. The annotation of the spectrum is based on the given sequence in the peptides file and is done with different software so inconsistencies are likely. The peaks are annotated based on the given sequence, with 20 ppm tolerance.

Copy Data

##### Spectrum 4197 (TSV)

###### Preview

```
Loading example...
```

*Click on the button to copy the data to your clipboard.*

Mz MinMz MaxIntensity Max

WidthHeightPeptide font sizePeptide stroke widthSpectrum font sizeSpectrum stroke widthCompact peptide

Ion legend

wxyz

abcd

OtherUnassignedIon^Charge^_Position_Show for top:%

VVVDVSHEDPEVK

01.51e+43.03e+44.54e+46.05e+4

Zoom Out

y^+1^_1_z^+1^_1_y^+1^_1_y^+1^_2_w^+1^_3_c^+1^_3_w^+1^_4_y^+1^_4_z^+2^_9_y^+2^_9_c^+1^_5_z^+2^_10_y^+2^_10_y^+1^_5_c^+1^_6_z^+2^_11_y^+2^_11_w^+1^_6_w^+2^_12_y^+2^_12_z^+1^_6_y^+1^_6_c^+1^_7_z^+1^_7_y^+1^_7_c^+1^_8_z^+1^_8_y^+1^_8_z^+1^_9_y^+1^_9_c^+1^_10_z^+1^_10_y^+1^_10_c^+1^_11_z^+1^_11_c^+1^_12_

0778155623333111

Fragment Matches Table

Show background peaks

| Position | Ion type | Intensity | mz Theoretical | mz Error (Th) | mz Error (ppm) | Charge | Series Number |
| --- | --- | --- | --- | --- | --- | --- | --- |
| - | - | 356.1 | 120.2 | - | - | 0 | - |
| - | - | 1068 | 129.1 | - | - | 0 | - |
| - | - | 354.3 | 129.4 | - | - | 0 | - |
| 13 | y | 1326 | 130.1 | 0.0004854 | 3.731 | +1 | 1 |
| - | - | 454.4 | 130.8 | - | - | 0 | - |
| 13 | z | 964.8 | 131.1 | 0.0003148 | 2.402 | +1 | 1 |
| - | - | 2340 | 133.1 | - | - | 0 | - |
| 13 | y | 3620 | 147.1 | 0.0004995 | 3.395 | +1 | 1 |
| - | - | 849.7 | 148.9 | - | - | 0 | - |
| - | - | 539.6 | 154.1 | - | - | 0 | - |
| - | - | 1.364E+04 | 171.1 | - | - | 0 | - |
| - | - | 1148 | 172.2 | - | - | 0 | - |
| - | - | 4543 | 173.4 | - | - | 0 | - |
| - | - | 448.8 | 185.4 | - | - | 0 | - |
| - | - | 853.2 | 199.1 | - | - | 0 | - |
| - | - | 1.199E+04 | 199.1 | - | - | 0 | - |
| - | - | 1341 | 200.1 | - | - | 0 | - |
| - | - | 708.9 | 215.1 | - | - | 0 | - |
| - | - | 504.5 | 231.2 | - | - | 0 | - |
| - | - | 1856 | 232.1 | - | - | 0 | - |
| 12 | y | 2638 | 246.2 | 0.0005693 | 2.312 | +1 | 2 |
| - | - | 689.4 | 247.2 | - | - | 0 | - |
| - | - | 981.6 | 270.2 | - | - | 0 | - |
| - | - | 520.7 | 283 | - | - | 0 | - |
| - | - | 1975 | 298.2 | - | - | 0 | - |
| - | - | 662.6 | 299.2 | - | - | 0 | - |
| 11 | w | 8228 | 300.2 | 0.0008038 | 2.677 | +1 | 3 |
| - | - | 889 | 301.2 | - | - | 0 | - |
| - | - | 754.7 | 314.2 | - | - | 0 | - |
| 3 | c | 714.5 | 315.2 | 0.001411 | 4.477 | +1 | 3 |
| - | - | 549.5 | 325.4 | - | - | 0 | - |
| - | - | 589.1 | 344.4 | - | - | 0 | - |
| - | - | 1714 | 354.2 | - | - | 0 | - |
| - | - | 512.4 | 361.1 | - | - | 0 | - |
| - | - | 642.7 | 386.2 | - | - | 0 | - |
| - | - | 3221 | 413.2 | - | - | 0 | - |
| - | - | 530.6 | 414.2 | - | - | 0 | - |
| 10 | w | 2101 | 429.2 | 0.0007638 | 1.779 | +1 | 4 |
| - | - | 3491 | 468.3 | - | - | 0 | - |
| - | - | 4232 | 470.3 | - | - | 0 | - |
| - | - | 1218 | 471.3 | - | - | 0 | - |
| 10 | y | 4128 | 472.3 | 0.001337 | 2.831 | +1 | 4 |
| - | - | 1471 | 473.3 | - | - | 0 | - |
| - | - | 1105 | 484.3 | - | - | 0 | - |
| - | - | 1104 | 485.2 | - | - | 0 | - |
| - | - | 2127 | 499.3 | - | - | 0 | - |
| - | - | 691.1 | 500.3 | - | - | 0 | - |
| - | - | 755.8 | 501.3 | - | - | 0 | - |
| - | - | 904.6 | 507.3 | - | - | 0 | - |
| 5 | z | 1475 | 512.2 | 0.0004268 | 0.8333 | +2 | 9 |
| - | - | 5394 | 512.3 | - | - | 0 | - |
| - | - | 1076 | 513.3 | - | - | 0 | - |
| 5 | y | 788.7 | 520.3 | 0.001793 | 3.447 | +2 | 9 |
| - | - | 1.326E+04 | 527.3 | - | - | 0 | - |
| - | - | 2994 | 528.3 | - | - | 0 | - |
| - | - | 824.9 | 529.3 | - | - | 0 | - |
| 5 | c | 1117 | 529.3 | 0.002429 | 4.588 | +1 | 5 |
| - | - | 1456 | 547.8 | - | - | 0 | - |
| 4 | z | 905.2 | 569.8 | 0.001386 | 2.433 | +2 | 10 |
| 4 | y | 2507 | 577.8 | 0.001532 | 2.651 | +2 | 10 |
| - | - | 612.7 | 578.3 | - | - | 0 | - |
| 9 | y | 1314 | 587.3 | 0.001455 | 2.478 | +1 | 5 |
| - | - | 704.2 | 588.3 | - | - | 0 | - |
| - | - | 1295 | 599.3 | - | - | 0 | - |
| - | - | 918.5 | 608.3 | - | - | 0 | - |
| 6 | c | 3516 | 616.4 | 0.001279 | 2.076 | +1 | 6 |
| - | - | 1157 | 617.4 | - | - | 0 | - |
| 3 | z | 2604 | 619.3 | 0.0008641 | 1.395 | +2 | 11 |
| - | - | 1702 | 619.8 | - | - | 0 | - |
| 3 | y | 7735 | 627.3 | 0.001559 | 2.485 | +2 | 11 |
| - | - | 4492 | 627.8 | - | - | 0 | - |
| - | - | 2816 | 628.3 | - | - | 0 | - |
| 8 | w | 6004 | 641.3 | 0.001033 | 1.611 | +1 | 6 |
| - | - | 1952 | 642.3 | - | - | 0 | - |
| - | - | 631.2 | 648.3 | - | - | 0 | - |
| - | - | 814.2 | 656.3 | - | - | 0 | - |
| 2 | w | 1058 | 661.3 | 0.00401 | 6.064 | +2 | 12 |
| 2 | y | 1762 | 676.8 | 0.0004878 | 0.7207 | +2 | 12 |
| - | - | 1675 | 677.3 | - | - | 0 | - |
| - | - | 777.9 | 684.3 | - | - | 0 | - |
| - | - | 745.5 | 690.9 | - | - | 0 | - |
| - | - | 8964 | 696.9 | - | - | 0 | - |
| - | - | 8684 | 697.4 | - | - | 0 | - |
| - | - | 4356 | 697.9 | - | - | 0 | - |
| - | - | 1413 | 698.4 | - | - | 0 | - |
| 8 | z | 4187 | 700.3 | 0.000849 | 1.212 | +1 | 6 |
| - | - | 2104 | 701.3 | - | - | 0 | - |
| - | - | 688.8 | 703.9 | - | - | 0 | - |
| - | - | 1231 | 704.9 | - | - | 0 | - |
| - | - | 546 | 705.4 | - | - | 0 | - |
| - | - | 2814 | 709.4 | - | - | 0 | - |
| - | - | 1483 | 710.4 | - | - | 0 | - |
| - | - | 5355 | 710.9 | - | - | 0 | - |
| - | - | 5119 | 711.4 | - | - | 0 | - |
| - | - | 3711 | 711.9 | - | - | 0 | - |
| - | - | 1001 | 712.4 | - | - | 0 | - |
| 8 | y | 3634 | 716.3 | 0.002483 | 3.467 | +1 | 6 |
| - | - | 1227 | 717.3 | - | - | 0 | - |
| - | - | 757 | 717.9 | - | - | 0 | - |
| - | - | 1.657E+04 | 718.4 | - | - | 0 | - |
| - | - | 1.165E+04 | 718.9 | - | - | 0 | - |
| - | - | 7678 | 719.4 | - | - | 0 | - |
| - | - | 2529 | 719.9 | - | - | 0 | - |
| - | - | 3.557E+04 | 726.4 | - | - | 0 | - |
| - | - | 5.993E+04 | 726.9 | - | - | 0 | - |
| - | - | 4.208E+04 | 727.4 | - | - | 0 | - |
| - | - | 1063 | 727.5 | - | - | 0 | - |
| - | - | 1.913E+04 | 727.9 | - | - | 0 | - |
| - | - | 4555 | 728.4 | - | - | 0 | - |
| - | - | 777.6 | 728.9 | - | - | 0 | - |
| 7 | c | 1.318E+04 | 753.4 | 0.001781 | 2.363 | +1 | 7 |
| - | - | 4984 | 754.4 | - | - | 0 | - |
| - | - | 1649 | 755.4 | - | - | 0 | - |
| - | - | 1589 | 782.3 | - | - | 0 | - |
| 7 | z | 5408 | 837.4 | 0.001533 | 1.831 | +1 | 7 |
| - | - | 2597 | 838.4 | - | - | 0 | - |
| - | - | 870.8 | 839.4 | - | - | 0 | - |
| 7 | y | 1523 | 853.4 | 0.008246 | 9.663 | +1 | 7 |
| - | - | 1069 | 854.4 | - | - | 0 | - |
| 8 | c | 3.022E+04 | 882.5 | 0.001344 | 1.523 | +1 | 8 |
| - | - | 1.441E+04 | 883.5 | - | - | 0 | - |
| - | - | 3931 | 884.5 | - | - | 0 | - |
| - | - | 997.1 | 885.5 | - | - | 0 | - |
| 6 | z | 8537 | 924.4 | 0.000969 | 1.048 | +1 | 8 |
| - | - | 4581 | 925.4 | - | - | 0 | - |
| - | - | 1597 | 926.4 | - | - | 0 | - |
| 6 | y | 4171 | 940.4 | 0.002054 | 2.184 | +1 | 8 |
| - | - | 2124 | 941.4 | - | - | 0 | - |
| - | - | 944.7 | 942.4 | - | - | 0 | - |
| - | - | 1185 | 951.5 | - | - | 0 | - |
| - | - | 2164 | 952.5 | - | - | 0 | - |
| - | - | 1641 | 953.5 | - | - | 0 | - |
| - | - | 785.4 | 968.4 | - | - | 0 | - |
| - | - | 1205 | 969.5 | - | - | 0 | - |
| - | - | 663.5 | 970.4 | - | - | 0 | - |
| - | - | 747.4 | 970.5 | - | - | 0 | - |
| 5 | z | 1.052E+04 | 1023 | 0.001085 | 1.06 | +1 | 9 |
| - | - | 5777 | 1024 | - | - | 0 | - |
| - | - | 1888 | 1025 | - | - | 0 | - |
| 5 | y | 5773 | 1040 | 9.425E-05 | 0.09067 | +1 | 9 |
| - | - | 3878 | 1041 | - | - | 0 | - |
| - | - | 1370 | 1042 | - | - | 0 | - |
| 10 | c | 1.385E+04 | 1095 | 0.01312 | 11.99 | +1 | 10 |
| - | - | 8106 | 1096 | - | - | 0 | - |
| - | - | 3647 | 1097 | - | - | 0 | - |
| - | - | 608.3 | 1098 | - | - | 0 | - |
| - | - | 1498 | 1121 | - | - | 0 | - |
| - | - | 826.8 | 1122 | - | - | 0 | - |
| 4 | z | 8632 | 1139 | 0.0007449 | 0.6542 | +1 | 10 |
| - | - | 8825 | 1140 | - | - | 0 | - |
| - | - | 4598 | 1141 | - | - | 0 | - |
| - | - | 976.8 | 1142 | - | - | 0 | - |
| 4 | y | 1532 | 1155 | 0.00287 | 2.486 | +1 | 10 |
| - | - | 915.4 | 1156 | - | - | 0 | - |
| - | - | 828.3 | 1157 | - | - | 0 | - |
| - | - | 6220 | 1180 | - | - | 0 | - |
| - | - | 4717 | 1181 | - | - | 0 | - |
| - | - | 1342 | 1182 | - | - | 0 | - |
| 11 | c | 6648 | 1224 | 0.002123 | 1.735 | +1 | 11 |
| - | - | 3429 | 1225 | - | - | 0 | - |
| - | - | 2694 | 1226 | - | - | 0 | - |
| 3 | z | 1368 | 1238 | 0.00202 | 1.632 | +1 | 11 |
| - | - | 2088 | 1239 | - | - | 0 | - |
| - | - | 1007 | 1240 | - | - | 0 | - |
| - | - | 1494 | 1279 | - | - | 0 | - |
| - | - | 2658 | 1280 | - | - | 0 | - |
| - | - | 1038 | 1281 | - | - | 0 | - |
| 12 | c | 9151 | 1323 | 0.003764 | 2.846 | +1 | 12 |
| - | - | 6581 | 1324 | - | - | 0 | - |
| - | - | 2549 | 1325 | - | - | 0 | - |
| - | - | 1065 | 1326 | - | - | 0 | - |
| - | - | 1836 | 1338 | - | - | 0 | - |
| - | - | 1351 | 1339 | - | - | 0 | - |
| - | - | 1088 | 1340 | - | - | 0 | - |
| - | - | 1585 | 1393 | - | - | 0 | - |
| - | - | 7180 | 1394 | - | - | 0 | - |
| - | - | 5625 | 1395 | - | - | 0 | - |
| - | - | 2509 | 1396 | - | - | 0 | - |
| - | - | 1084 | 1408 | - | - | 0 | - |
| - | - | 1668 | 1409 | - | - | 0 | - |
| - | - | 765 | 1410 | - | - | 0 | - |
| - | - | 1198 | 1426 | - | - | 0 | - |
| - | - | 3332 | 1436 | - | - | 0 | - |
| - | - | 2.05E+04 | 1437 | - | - | 0 | - |
| - | - | 1.383E+04 | 1438 | - | - | 0 | - |
| - | - | 9156 | 1439 | - | - | 0 | - |
| - | - | 2529 | 1440 | - | - | 0 | - |
| - | - | 2338 | 1452 | - | - | 0 | - |
| - | - | 1.615E+04 | 1453 | - | - | 0 | - |
| - | - | 3.38E+04 | 1454 | - | - | 0 | - |
| - | - | 2.51E+04 | 1455 | - | - | 0 | - |
| - | - | 1.245E+04 | 1456 | - | - | 0 | - |
| - | - | 4017 | 1457 | - | - | 0 | - |
| - | - | 756.2 | 1458 | - | - | 0 | - |
| - | - | 1035 | 3080 | - | - | 0 | - |

m/z Charge Intensity FragmentType MassShift Position
120.1557846069336 0 356.11868
129.10215759277344 0 1067.9237
129.36334228515625 0 354.25143
130.0857696533203 0 1325.8374 y Ammonia loss 12
130.79681396484375 0 454.3706
131.09376525878906 0 964.76605 z 12
133.0604705810547 0 2339.9905
147.1123046875 0 3619.713 y 12
148.94703674316406 0 849.687
154.12249755859375 0 539.639
171.1486053466797 0 13639.368
172.15213012695312 0 1148.2196
173.4403533935547 0 4542.6514
185.4085693359375 0 448.79825
199.13436889648438 0 853.1903
199.1434326171875 0 11992.411
200.14707946777344 0 1341.2548
215.10215759277344 0 708.9057
231.16131591796875 0 504.53326
232.12840270996094 0 1856.484
246.18064880371094 0 2638.3806 y 11
247.18421936035156 0 689.3887
270.216796875 0 981.61115
282.9604187011719 0 520.6965
298.2119445800781 0 1975.0343
299.21575927734375 0 662.55115
300.19097900390625 0 8227.7 w 10
301.1937255859375 0 889.019
314.170654296875 0 754.73114
315.240478515625 0 714.47375 c 2
325.4361572265625 0 549.5155
344.40045166015625 0 589.08105
354.20111083984375 0 1714.4543
361.0732116699219 0 512.41345
386.24969482421875 0 642.65704
413.238525390625 0 3220.8481
414.2437744140625 0 530.62604
429.2336120605469 0 2101.086 w 9
468.2804260253906 0 3490.9485
470.25970458984375 0 4231.5786
471.2646484375 0 1217.9972
472.2752380371094 0 4127.974 y 9
473.2789611816406 0 1470.7533
484.3117370605469 0 1104.9567
485.2442932128906 0 1104.2856
499.2621765136719 0 2126.68
500.26531982421875 0 691.12555
501.2723693847656 0 755.7832
507.29083251953125 0 904.5506
512.24658203125 0 1474.5342 z 4
512.306884765625 0 5394.385
513.3109130859375 0 1076.2811
520.2545776367188 0 788.70306 y 4
527.2936401367188 0 13263.396
528.2965087890625 0 2993.9458
529.2965087890625 0 824.9167
529.3368530273438 0 1116.6807 c 4
547.7630615234375 0 1456.3715
569.7590942382812 0 905.174 z 3
577.768310546875 0 2507.1377 y 3
578.269775390625 0 612.747
587.3020629882812 0 1313.7704 y 8
588.3065185546875 0 704.1963
599.3392944335938 0 1295.4924
608.3388671875 0 918.5356
616.3651733398438 0 3515.6646 c 5
617.3682861328125 0 1156.8716
619.2938232421875 0 2604.4736 z 2
619.7965087890625 0 1701.8833
627.302490234375 0 7735.049 y 2
627.8033447265625 0 4492.009
628.304443359375 0 2816.0034
641.3130493164062 0 6004.043 w 7
642.31640625 0 1952.0695
648.3182983398438 0 631.2067
656.3361206054688 0 814.20135
661.3211669921875 0 1058.4044 w 1
676.8377685546875 0 1762.11 y 1
677.3385009765625 0 1674.8857
684.26220703125 0 777.9389
690.8624267578125 0 745.54266
696.8641967773438 0 8964.429
697.3656616210938 0 8684.178
697.867431640625 0 4355.8896
698.36767578125 0 1412.5913
700.3265380859375 0 4186.534 z 7
701.3280639648438 0 2104.3735
703.8692016601562 0 688.81396
704.8821411132812 0 1230.77
705.377197265625 0 546.03265
709.4088745117188 0 2814.011
710.41259765625 0 1482.6025
710.8505249023438 0 5354.943
711.350341796875 0 5119.0703
711.8513793945312 0 3710.5442
712.3524780273438 0 1000.5577
716.3436279296875 0 3634.4482 y 7
717.3463745117188 0 1227.1433
717.8666381835938 0 756.9625
718.3610229492188 0 16566.246
718.86328125 0 11651.3
719.3635864257812 0 7677.592
719.8658447265625 0 2529.3372
726.3709716796875 0 35571.62
726.873779296875 0 59926.11
727.375244140625 0 42079.727
727.4521484375 0 1062.6555
727.8770751953125 0 19134.07
728.3788452148438 0 4554.664
728.882080078125 0 777.61316
753.423583984375 0 13178.014 c 6
754.4261474609375 0 4983.9775
755.4299926757812 0 1649.2301
782.3306274414062 0 1588.601
837.384765625 0 5407.8247 z 6
838.3861694335938 0 2597.3462
839.3890380859375 0 870.82794
853.4132690429688 0 1523.3895 y 6
854.4195556640625 0 1068.608
882.4666137695312 0 30222.084 c 7
883.4690551757812 0 14412.987
884.4729614257812 0 3930.9084
885.4739379882812 0 997.05493
924.4173583984375 0 8536.549 z 5
925.4198608398438 0 4580.8613
926.4237670898438 0 1597.0979
940.4349975585938 0 4171.074 y 5
941.4370727539062 0 2124.1006
942.4428100585938 0 944.6629
951.455078125 0 1184.5168
952.4686889648438 0 2163.8665
953.4677124023438 0 1640.7819
968.4435424804688 0 785.43604
969.452880859375 0 1205.3544
970.3533935546875 0 663.4506
970.4637451171875 0 747.36505
1023.4856567382812 0 10518.5 z 4
1024.490478515625 0 5777.308
1025.4937744140625 0 1887.6669
1039.50537109375 0 5773.273 y 4
1040.50537109375 0 3877.9292
1041.5133056640625 0 1370.4003
1094.5345458984375 0 13852.15 c 9
1095.5389404296875 0 8105.503
1096.5384521484375 0 3647.223
1097.54931640625 0 608.2984
1120.55810546875 0 1498.3319
1121.5611572265625 0 826.83234
1138.512939453125 0 8632.237 z 3
1139.5159912109375 0 8825.238
1140.5205078125 0 4597.9805
1141.5330810546875 0 976.8116
1154.5352783203125 0 1532.4191 y 3
1155.5306396484375 0 915.4324
1156.5369873046875 0 828.3086
1179.5755615234375 0 6220.469
1180.5777587890625 0 4717.4424
1181.581787109375 0 1341.532
1223.588134765625 0 6647.544 c 10
1224.5909423828125 0 3429.3462
1225.59375 0 2693.5095
1237.580078125 0 1367.699 z 2
1238.582763671875 0 2088.4722
1239.575927734375 0 1007.12384
1278.640869140625 0 1493.5769
1279.648681640625 0 2658.277
1280.6512451171875 0 1038.3252
1322.6549072265625 0 9151.258 c 11
1323.657958984375 0 6580.997
1324.657958984375 0 2549.2957
1325.6575927734375 0 1064.535
1337.6536865234375 0 1835.8635
1338.6541748046875 0 1350.6045
1339.6571044921875 0 1087.6603
1392.720703125 0 1584.5485
1393.7276611328125 0 7179.548
1394.731689453125 0 5625.1655
1395.7373046875 0 2509.4458
1407.743408203125 0 1084.3052
1408.741943359375 0 1667.8403
1409.748046875 0 765.0481
1425.764404296875 0 1198.1779
1435.7183837890625 0 3332.3801
1436.723388671875 0 20498.324
1437.7254638671875 0 13826.717
1438.7288818359375 0 9156.063
1439.728271484375 0 2528.8857
1451.7298583984375 0 2338.0864
1452.740966796875 0 16150.784
1453.7486572265625 0 33799.91
1454.7518310546875 0 25104.973
1455.7550048828125 0 12451.009
1456.7578125 0 4017.2292
1457.765380859375 0 756.2255
3080.369384765625 0 1035.4404

Spectrum Details

|  |  |
| --- | --- |
| Matched peaks? Matched peaks The total absolute number of peaks matched. Additionally in brackets the total fraction of peaks matched and the total number of peaks is shown. | 36 (18.46% of 195) |
| FDR? FDR The false discovery rate estimated for this peptide. It is calculated by matching all theoretical fragments with a non-integer shift with the raw peaks for this spectrum. This is done with 40 different shifts. The resulting percentage is the average number of annotated peaks over the number of annotated peaks with the correct spectrum. | 0.40% |
| Satellite FDR? Satellite FDR See the FDR for details on its calculation. This satellite ion specific FDR only contains the satellite ions (d/w) for I/L/J positions. | - |
| PSM Score? PSM Score The PSM Score as given by Hecklib to this annotated spectrum. It is shown with three significant figures. | 285 |

#### Reverse Lookup? Reverse Lookup All places where this read could be placed.

| Group | Segment | Template | Template Part | Read Part | Score | Unique |
| --- | --- | --- | --- | --- | --- | --- |
| Homo sapiens Heavy Chain | IGHC | IGHG1 | [144..157] | [0..13] | 104 | False |
| Homo sapiens Heavy Chain | IGHC | IGHG3 | [191..204] | [0..13] | 95 | False |
| Homo sapiens Heavy Chain | IGHC | IGHG2 | [140..153] | [0..13] | 95 | False |

| Recombined | Template Part | Read Part | Score | Unique |
| --- | --- | --- | --- | --- |
| REC-0-1 | [267..280] | [0..13] | 104 | True |

#### Meta Information from Multiple reads

##### Number of combined reads

3

##### Intensity

0.7224

##### TotalArea

1.48E+07

#### Positional Score

Copy Data

##### Positional Score (TSV)

###### Preview

```
Loading example...
```

*Click on the button to copy the data to your clipboard.*

100123456789101112

Label Value
"0" 0.333
"1" 0.333
"2" 0.333
"3" 0.33
"4" 0.327
"5" 0.33
"6" 0.33
"7" 0.327
"8" 0.297
"9" 0.277
"10" 0.313
"11" 0.33
"12" 0.333

#### Meta Information from PEAKS

##### Scan Identifier

F2:4311

##### Original sequence

V

V

V

D

V

S

H

E

D

P

E

V

K

##### Posttranslational Modifications

##### Source File

D:\separate\_stitch\_analyses\xle-disambiguation\raw\20210323\_F1\_UM1\_Peng0013\_SA\_F59\_ingel\_3ug\_TL.raw

##### Fraction

2

##### Scan Feature

-

##### De Novo Score

99

##### ConfidenceScore

99

### m/z

484.5848

##### Mass

1450.7302

##### Charge

3

##### Retention Time

23

##### Predicted Retention Time

-

##### Area

0

##### Parts Per Million

1.6

##### Fragmentation mode

HCD

##### Originating file

01 D:\separate\_stitch\_analyses\xle-disambiguation\20210325\_F59\_3ug\_DENOVO\_12.csv

#### Meta Information from PEAKS

##### Scan Identifier

F2:4253

##### Original sequence

V

V

V

D

V

S

H

E

D

P

E

V

K

##### Posttranslational Modifications

##### Source File

D:\separate\_stitch\_analyses\xle-disambiguation\raw\20210323\_F1\_UM1\_Peng0013\_SA\_F59\_ingel\_3ug\_TL.raw

##### Fraction

2

##### Scan Feature

F2:3488

##### De Novo Score

99

##### ConfidenceScore

99

### m/z

484.5848

##### Mass

1450.7302

##### Charge

3

##### Retention Time

22.67

##### Predicted Retention Time

-

##### Area

8.124E+06

##### Parts Per Million

1.6

##### Fragmentation mode

ETHCD

##### Originating file

01 D:\separate\_stitch\_analyses\xle-disambiguation\20210325\_F59\_3ug\_DENOVO\_12.csv

#### Meta Information from PEAKS

##### Scan Identifier

F4:4197

##### Original sequence

V

V

V

D

V

S

H

E

D

P

E

V

K

##### Posttranslational Modifications

##### Source File

D:\separate\_stitch\_analyses\xle-disambiguation\raw\20210323\_F1\_UM1\_Peng0013\_SA\_F59\_ingel\_3ug\_tryp.raw

##### Fraction

4

##### Scan Feature

F4:2922

##### De Novo Score

94

##### ConfidenceScore

96

### m/z

484.5849

##### Mass

1450.7302

##### Charge

3

##### Retention Time

22.6

##### Predicted Retention Time

25.38

##### Area

6.681E+06

##### Parts Per Million

1.8

##### Fragmentation mode

ETHCD

##### Originating file

01 D:\separate\_stitch\_analyses\xle-disambiguation\20210325\_F59\_3ug\_DENOVO\_12.csv
