## Supplementary material for "A handle on mass coincidence errors in *de novo* sequencing of antibodies by bottom-up proteomics": Combined_030.html

Details Combined\_030 | Stitch OverviewUndefined

### Read Combined\_030

#### Sequence (length=17)

VTJFPPSSEEJQANKAT

#### Spectrum 8009? Spectrum 8009 The raw spectrum of this peptide as annotated by Hecklib. The fragments are coloured according to ion type (see legend). Any peaks with a star '\*' as text can be hovered over to see the full details, first the ion type second the mass shift type. By hovering over the amino acids in the peptide or ions in the legend the corresponding peaks are highlighted. By toggling the 'Unassigned' label you can turn the background (unassigned) peaks on or off in the plot. By updating the slider in the Ion legend you can update the spectrum to only show the top X% of the peaks with labels. The top X% means any peak that is within X% of the highest intensity. By dragging in the spectrum you can zoom in to a specific part of the spectrum and use 'Zoom Out' to get back to the original zoom level. The annotation of the spectrum is based on the given sequence in the peptides file and is done with different software so inconsistencies are likely. The peaks are annotated based on the given sequence, with 20 ppm tolerance.

Copy Data

##### Spectrum 8009 (TSV)

###### Preview

```
Loading example...
```

*Click on the button to copy the data to your clipboard.*

Mz MinMz MaxIntensity Max

WidthHeightPeptide font sizePeptide stroke widthSpectrum font sizeSpectrum stroke widthCompact peptide

Ion legend

wxyz

abcd

OtherUnassignedIon^Charge^_Position_Show for top:%

VTJFPPSSEEJQANKAT

02.05e+44.09e+46.14e+48.19e+4

Zoom Out

y^+1^_1_y^+1^_2_y^+1^_2_c^+1^_2_c^+1^_2_y^+3^_7_z^+1^_3_y^+1^_3_c^+1^_3_y^+2^_7_y^+1^_4_z^+1^_4_y^+1^_4_y^+1^_5_c^+2^_9_y^+2^_9_y^+1^_5_z^+3^_15_w^+1^_6_y^+2^_11_y^+2^_11_y^+1^_6_z^+1^_6_y^+2^_12_y^+2^_12_y^+1^_6_y^+2^_12_c^+2^_12_c^+1^_6_y^+2^_13_y^+2^_13_z^+2^_13_y^+2^_13_z^+1^_7_z^+2^_14_y^+1^_7_y^+2^_14_y^+2^_14_z^+2^_14_c^+1^_7_y^+2^_14_w^+1^_8_z^+2^_15_y^+2^_15_y^+2^_15_y^+2^_15_c^+2^_15_z^+1^_8_z^+1^_8_c^+1^_8_z^+2^_16_z^+2^_16_c^+2^_16_y^+1^_8_y^+2^_16_y^+2^_16_z^+2^_16_c^+2^_16_y^+1^_8_w^+1^_9_c^+1^_9_z^+1^_9_y^+1^_9_z^+1^_10_z^+1^_10_y^+1^_10_z^+1^_10_y^+1^_10_c^+1^_10_y^+1^_11_y^+1^_11_z^+1^_11_y^+1^_11_c^+1^_11_c^+1^_11_y^+1^_12_c^+1^_12_c^+1^_12_y^+1^_13_z^+1^_13_y^+1^_13_c^+1^_13_y^+1^_14_z^+1^_14_c^+1^_14_y^+1^_14_c^+1^_14_z^+1^_15_c^+1^_15_z^+1^_16_y^+1^_16_z^+1^_16_c^+1^_16_

046492813921856

Fragment Matches Table

Show background peaks

| Position | Ion type | Intensity | mz Theoretical | mz Error (Th) | mz Error (ppm) | Charge | Series Number |
| --- | --- | --- | --- | --- | --- | --- | --- |
| - | - | 696.7 | 120.1 | - | - | 0 | - |
| 17 | y | 1.112E+04 | 120.1 | 8.555E-05 | 0.7125 | +1 | 1 |
| - | - | 1810 | 120.1 | - | - | 0 | - |
| - | - | 547.5 | 121.1 | - | - | 0 | - |
| - | - | 371.4 | 124.6 | - | - | 0 | - |
| - | - | 739.2 | 126.1 | - | - | 0 | - |
| - | - | 960.5 | 128.1 | - | - | 0 | - |
| - | - | 408.5 | 128.1 | - | - | 0 | - |
| - | - | 4562 | 129.1 | - | - | 0 | - |
| - | - | 396.3 | 129.2 | - | - | 0 | - |
| - | - | 746.6 | 131.1 | - | - | 0 | - |
| - | - | 474.5 | 140.1 | - | - | 0 | - |
| - | - | 589.4 | 148.9 | - | - | 0 | - |
| - | - | 1105 | 149 | - | - | 0 | - |
| - | - | 454.2 | 151.3 | - | - | 0 | - |
| - | - | 708.5 | 155.1 | - | - | 0 | - |
| - | - | 565.4 | 155.1 | - | - | 0 | - |
| - | - | 534.1 | 158.1 | - | - | 0 | - |
| - | - | 2070 | 169.1 | - | - | 0 | - |
| - | - | 657.1 | 171.1 | - | - | 0 | - |
| 16 | y | 1295 | 173.1 | 7.911E-05 | 0.4571 | +1 | 2 |
| - | - | 2900 | 173.1 | - | - | 0 | - |
| - | - | 495.3 | 181.1 | - | - | 0 | - |
| - | - | 3672 | 183.1 | - | - | 0 | - |
| - | - | 548.6 | 185.1 | - | - | 0 | - |
| 16 | y | 3994 | 191.1 | 0.0001345 | 0.704 | +1 | 2 |
| - | - | 1702 | 197.1 | - | - | 0 | - |
| 2 | c | 626.5 | 200.1 | 0.0002189 | 1.094 | +1 | 2 |
| - | - | 7029 | 201.1 | - | - | 0 | - |
| - | - | 552.6 | 211.1 | - | - | 0 | - |
| - | - | 8754 | 212.1 | - | - | 0 | - |
| - | - | 795.8 | 213.1 | - | - | 0 | - |
| - | - | 1416 | 215.1 | - | - | 0 | - |
| 2 | c | 4587 | 218.1 | 1.492E-05 | 0.06838 | +1 | 2 |
| - | - | 4220 | 221.1 | - | - | 0 | - |
| - | - | 749.8 | 222.1 | - | - | 0 | - |
| - | - | 763.3 | 223.1 | - | - | 0 | - |
| - | - | 478.8 | 225.8 | - | - | 0 | - |
| - | - | 862.4 | 226.1 | - | - | 0 | - |
| - | - | 988.1 | 227.1 | - | - | 0 | - |
| - | - | 1642 | 227.1 | - | - | 0 | - |
| - | - | 706.2 | 227.1 | - | - | 0 | - |
| - | - | 1176 | 233.2 | - | - | 0 | - |
| - | - | 1014 | 235.1 | - | - | 0 | - |
| - | - | 5012 | 239.1 | - | - | 0 | - |
| - | - | 1903 | 240.1 | - | - | 0 | - |
| - | - | 727.1 | 242.2 | - | - | 0 | - |
| 11 | y | 1364 | 243.1 | 0.003444 | 14.17 | +3 | 7 |
| - | - | 658.9 | 251.2 | - | - | 0 | - |
| - | - | 1394 | 254.1 | - | - | 0 | - |
| - | - | 1005 | 261.2 | - | - | 0 | - |
| - | - | 2040 | 263.1 | - | - | 0 | - |
| - | - | 806 | 272.2 | - | - | 0 | - |
| - | - | 698.4 | 280.2 | - | - | 0 | - |
| - | - | 1261 | 281.1 | - | - | 0 | - |
| - | - | 655.5 | 281.1 | - | - | 0 | - |
| - | - | 851 | 282.1 | - | - | 0 | - |
| - | - | 835.7 | 296.2 | - | - | 0 | - |
| - | - | 1.083E+04 | 296.2 | - | - | 0 | - |
| - | - | 631.1 | 297.1 | - | - | 0 | - |
| - | - | 1001 | 297.2 | - | - | 0 | - |
| - | - | 1634 | 297.2 | - | - | 0 | - |
| - | - | 1104 | 298.2 | - | - | 0 | - |
| - | - | 5175 | 299.1 | - | - | 0 | - |
| - | - | 1.318E+04 | 299.2 | - | - | 0 | - |
| - | - | 1785 | 300.1 | - | - | 0 | - |
| - | - | 1104 | 300.2 | - | - | 0 | - |
| - | - | 657.3 | 301.1 | - | - | 0 | - |
| 15 | z | 6849 | 303.2 | 0.0001004 | 0.3311 | +1 | 3 |
| - | - | 1270 | 304.2 | - | - | 0 | - |
| - | - | 982.8 | 314.2 | - | - | 0 | - |
| - | - | 4472 | 314.2 | - | - | 0 | - |
| - | - | 935.4 | 315.2 | - | - | 0 | - |
| - | - | 5855 | 316.2 | - | - | 0 | - |
| - | - | 911.1 | 317.2 | - | - | 0 | - |
| 15 | y | 4222 | 319.2 | 6.593E-05 | 0.2066 | +1 | 3 |
| - | - | 740.8 | 320.2 | - | - | 0 | - |
| - | - | 1443 | 325.2 | - | - | 0 | - |
| 3 | c | 2.554E+04 | 331.2 | 8.79E-05 | 0.2654 | +1 | 3 |
| - | - | 4461 | 332.2 | - | - | 0 | - |
| - | - | 741.9 | 344.2 | - | - | 0 | - |
| - | - | 986.3 | 345.2 | - | - | 0 | - |
| - | - | 607.7 | 346.1 | - | - | 0 | - |
| - | - | 551.8 | 346.2 | - | - | 0 | - |
| - | - | 653.5 | 347.2 | - | - | 0 | - |
| - | - | 540.6 | 348.2 | - | - | 0 | - |
| - | - | 809 | 350.1 | - | - | 0 | - |
| - | - | 3.835E+04 | 355.1 | - | - | 0 | - |
| - | - | 1.684E+04 | 356.1 | - | - | 0 | - |
| - | - | 9743 | 357.1 | - | - | 0 | - |
| - | - | 1037 | 358.1 | - | - | 0 | - |
| - | - | 1414 | 362.2 | - | - | 0 | - |
| - | - | 894.1 | 370.2 | - | - | 0 | - |
| - | - | 1854 | 371.1 | - | - | 0 | - |
| - | - | 610.8 | 371.2 | - | - | 0 | - |
| - | - | 1293 | 372.1 | - | - | 0 | - |
| - | - | 916.8 | 373.1 | - | - | 0 | - |
| 11 | y | 3372 | 373.2 | 0.004092 | 10.96 | +2 | 7 |
| - | - | 763.6 | 373.7 | - | - | 0 | - |
| - | - | 639.8 | 374.1 | - | - | 0 | - |
| - | - | 1406 | 385.2 | - | - | 0 | - |
| - | - | 699.3 | 385.4 | - | - | 0 | - |
| - | - | 1754 | 386.2 | - | - | 0 | - |
| - | - | 880.6 | 415 | - | - | 0 | - |
| 14 | y | 859.5 | 415.2 | 0.0001008 | 0.2427 | +1 | 4 |
| - | - | 1176 | 416 | - | - | 0 | - |
| - | - | 1075 | 417 | - | - | 0 | - |
| 14 | z | 2280 | 417.2 | 0.0006123 | 1.468 | +1 | 4 |
| - | - | 7195 | 418.2 | - | - | 0 | - |
| - | - | 1167 | 419.2 | - | - | 0 | - |
| 14 | y | 2724 | 433.2 | 0.0007481 | 1.727 | +1 | 4 |
| - | - | 8121 | 433.3 | - | - | 0 | - |
| - | - | 4886 | 434.3 | - | - | 0 | - |
| - | - | 1830 | 435.3 | - | - | 0 | - |
| - | - | 729.2 | 442.2 | - | - | 0 | - |
| - | - | 5846 | 444.2 | - | - | 0 | - |
| - | - | 3144 | 445.3 | - | - | 0 | - |
| - | - | 2588 | 446.3 | - | - | 0 | - |
| - | - | 1514 | 460.3 | - | - | 0 | - |
| - | - | 1.34E+04 | 461.3 | - | - | 0 | - |
| - | - | 3151 | 462.3 | - | - | 0 | - |
| - | - | 743.3 | 463.3 | - | - | 0 | - |
| - | - | 799.1 | 480.7 | - | - | 0 | - |
| - | - | 1450 | 480.8 | - | - | 0 | - |
| - | - | 1270 | 481.2 | - | - | 0 | - |
| 13 | y | 689.8 | 486.3 | 0.001268 | 2.608 | +1 | 5 |
| 9 | c | 9198 | 488.3 | 0.001861 | 3.812 | +2 | 9 |
| - | - | 1116 | 489.1 | - | - | 0 | - |
| - | - | 7165 | 489.3 | - | - | 0 | - |
| - | - | 627.2 | 490.1 | - | - | 0 | - |
| - | - | 1420 | 490.3 | - | - | 0 | - |
| - | - | 2480 | 493.7 | - | - | 0 | - |
| - | - | 1086 | 494.2 | - | - | 0 | - |
| - | - | 731.7 | 497.3 | - | - | 0 | - |
| - | - | 696.4 | 498.2 | - | - | 0 | - |
| 9 | y | 1959 | 502.3 | 0.008705 | 17.33 | +2 | 9 |
| - | - | 3364 | 502.7 | - | - | 0 | - |
| - | - | 2320 | 503.2 | - | - | 0 | - |
| - | - | 740.8 | 503.7 | - | - | 0 | - |
| 13 | y | 9221 | 504.3 | 0.0001028 | 0.2039 | +1 | 5 |
| - | - | 2058 | 505.3 | - | - | 0 | - |
| - | - | 789.6 | 515.3 | - | - | 0 | - |
| - | - | 4717 | 538.3 | - | - | 0 | - |
| - | - | 3371 | 538.8 | - | - | 0 | - |
| 3 | z | 769 | 539.3 | 0.001137 | 2.108 | +3 | 15 |
| - | - | 2266 | 543.3 | - | - | 0 | - |
| - | - | 928.3 | 543.8 | - | - | 0 | - |
| - | - | 6683 | 552.3 | - | - | 0 | - |
| - | - | 4851 | 552.8 | - | - | 0 | - |
| - | - | 1449 | 553.3 | - | - | 0 | - |
| - | - | 894.5 | 554.3 | - | - | 0 | - |
| 12 | w | 1.616E+04 | 558.3 | 0.0002995 | 0.5365 | +1 | 6 |
| - | - | 1775 | 558.3 | - | - | 0 | - |
| - | - | 3962 | 559.3 | - | - | 0 | - |
| - | - | 1247 | 559.3 | - | - | 0 | - |
| - | - | 814.3 | 560.3 | - | - | 0 | - |
| - | - | 1.093E+04 | 566.4 | - | - | 0 | - |
| - | - | 4071 | 567.4 | - | - | 0 | - |
| - | - | 767.7 | 572.3 | - | - | 0 | - |
| - | - | 679.6 | 577.1 | - | - | 0 | - |
| - | - | 1336 | 578.1 | - | - | 0 | - |
| - | - | 1188 | 579.1 | - | - | 0 | - |
| 7 | y | 704.8 | 580.3 | 0.002416 | 4.164 | +2 | 11 |
| 7 | y | 1297 | 580.8 | 0.006736 | 11.6 | +2 | 11 |
| - | - | 1306 | 583.3 | - | - | 0 | - |
| - | - | 727.8 | 584.3 | - | - | 0 | - |
| - | - | 998.4 | 591.3 | - | - | 0 | - |
| - | - | 690.6 | 594.4 | - | - | 0 | - |
| - | - | 680 | 599.3 | - | - | 0 | - |
| - | - | 1991 | 599.8 | - | - | 0 | - |
| - | - | 3473 | 602.8 | - | - | 0 | - |
| - | - | 3200 | 603.3 | - | - | 0 | - |
| - | - | 1043 | 603.8 | - | - | 0 | - |
| - | - | 8593 | 610.2 | - | - | 0 | - |
| - | - | 8211 | 611.2 | - | - | 0 | - |
| - | - | 746.6 | 611.3 | - | - | 0 | - |
| - | - | 7656 | 611.8 | - | - | 0 | - |
| - | - | 6954 | 612.2 | - | - | 0 | - |
| - | - | 720.8 | 612.2 | - | - | 0 | - |
| - | - | 731.9 | 612.2 | - | - | 0 | - |
| - | - | 5423 | 612.3 | - | - | 0 | - |
| - | - | 847 | 612.4 | - | - | 0 | - |
| - | - | 1752 | 612.8 | - | - | 0 | - |
| - | - | 767.1 | 613.2 | - | - | 0 | - |
| 12 | y | 7142 | 614.3 | 0.0109 | 17.74 | +1 | 6 |
| - | - | 753.5 | 614.8 | - | - | 0 | - |
| - | - | 9537 | 615.3 | - | - | 0 | - |
| 12 | z | 3974 | 616.3 | 0.0005355 | 0.8689 | +1 | 6 |
| - | - | 1401 | 616.4 | - | - | 0 | - |
| - | - | 1177 | 617.3 | - | - | 0 | - |
| - | - | 1141 | 617.8 | - | - | 0 | - |
| - | - | 630.7 | 618.3 | - | - | 0 | - |
| - | - | 1754 | 626.3 | - | - | 0 | - |
| - | - | 5045 | 626.8 | - | - | 0 | - |
| - | - | 3215 | 627.3 | - | - | 0 | - |
| - | - | 1921 | 627.3 | - | - | 0 | - |
| - | - | 1146 | 627.4 | - | - | 0 | - |
| - | - | 900.1 | 627.8 | - | - | 0 | - |
| - | - | 1171 | 628.3 | - | - | 0 | - |
| - | - | 2938 | 628.4 | - | - | 0 | - |
| 6 | y | 586.8 | 628.8 | 0.001393 | 2.216 | +2 | 12 |
| 6 | y | 905.5 | 629.3 | 0.005744 | 9.128 | +2 | 12 |
| - | - | 1203 | 629.4 | - | - | 0 | - |
| 12 | y | 5575 | 632.3 | 3.353E-05 | 0.05302 | +1 | 6 |
| - | - | 2243 | 633.3 | - | - | 0 | - |
| - | - | 1616 | 635.3 | - | - | 0 | - |
| - | - | 1175 | 635.8 | - | - | 0 | - |
| - | - | 695.5 | 636.3 | - | - | 0 | - |
| 6 | y | 563.9 | 637.8 | 0.00193 | 3.026 | +2 | 12 |
| - | - | 1049 | 638.3 | - | - | 0 | - |
| - | - | 1540 | 655.3 | - | - | 0 | - |
| - | - | 1208 | 655.8 | - | - | 0 | - |
| - | - | 820.9 | 656.3 | - | - | 0 | - |
| - | - | 1419 | 658.4 | - | - | 0 | - |
| 12 | c | 1151 | 664.8 | 0.005246 | 7.89 | +2 | 12 |
| - | - | 781.6 | 665.3 | - | - | 0 | - |
| - | - | 811 | 668.3 | - | - | 0 | - |
| - | - | 636.4 | 668.8 | - | - | 0 | - |
| - | - | 2542 | 671.4 | - | - | 0 | - |
| 6 | c | 3280 | 672.4 | 0.001744 | 2.594 | +1 | 6 |
| - | - | 1999 | 673.4 | - | - | 0 | - |
| - | - | 907.2 | 674.4 | - | - | 0 | - |
| - | - | 1631 | 676.8 | - | - | 0 | - |
| 5 | y | 7903 | 677.3 | 0.0004926 | 0.7273 | +2 | 13 |
| 5 | y | 6494 | 677.8 | 0.007866 | 11.6 | +2 | 13 |
| 5 | z | 3724 | 678.3 | 0.002977 | 4.388 | +2 | 13 |
| - | - | 1824 | 678.8 | - | - | 0 | - |
| - | - | 5147 | 685.8 | - | - | 0 | - |
| 5 | y | 5.207E+04 | 686.3 | 0.00277 | 4.036 | +2 | 13 |
| - | - | 1.948E+04 | 686.8 | - | - | 0 | - |
| - | - | 1.53E+04 | 687.3 | - | - | 0 | - |
| - | - | 2725 | 687.8 | - | - | 0 | - |
| - | - | 2074 | 688.4 | - | - | 0 | - |
| - | - | 1169 | 697.4 | - | - | 0 | - |
| - | - | 1013 | 716.4 | - | - | 0 | - |
| - | - | 1925 | 722.3 | - | - | 0 | - |
| - | - | 786.5 | 723.4 | - | - | 0 | - |
| - | - | 593.8 | 724.9 | - | - | 0 | - |
| 11 | z | 1.17E+04 | 729.4 | 0.0006151 | 0.8433 | +1 | 7 |
| - | - | 5276 | 730.4 | - | - | 0 | - |
| - | - | 1827 | 731.4 | - | - | 0 | - |
| - | - | 1353 | 740.3 | - | - | 0 | - |
| - | - | 1442 | 742.4 | - | - | 0 | - |
| 4 | z | 875.7 | 742.9 | 0.0002362 | 0.3179 | +2 | 14 |
| 11 | y | 7051 | 745.4 | 0.0009245 | 1.24 | +1 | 7 |
| - | - | 2718 | 746.4 | - | - | 0 | - |
| - | - | 835.6 | 747.4 | - | - | 0 | - |
| - | - | 748.8 | 750.4 | - | - | 0 | - |
| 4 | y | 2043 | 750.9 | 0.00125 | 1.665 | +2 | 14 |
| 4 | y | 1714 | 751.4 | 0.009486 | 12.63 | +2 | 14 |
| 4 | z | 763.6 | 751.9 | 0.007161 | 9.524 | +2 | 14 |
| - | - | 3003 | 757.4 | - | - | 0 | - |
| - | - | 928.3 | 758.4 | - | - | 0 | - |
| - | - | 1963 | 759.4 | - | - | 0 | - |
| 7 | c | 3.302E+04 | 759.4 | 0.0001115 | 0.1468 | +1 | 7 |
| 4 | y | 1.367E+04 | 759.9 | 0.000187 | 0.2461 | +2 | 14 |
| - | - | 7818 | 760.4 | - | - | 0 | - |
| - | - | 1.413E+04 | 760.4 | - | - | 0 | - |
| - | - | 4441 | 760.9 | - | - | 0 | - |
| - | - | 1352 | 761.4 | - | - | 0 | - |
| - | - | 4009 | 761.4 | - | - | 0 | - |
| - | - | 1020 | 762.5 | - | - | 0 | - |
| - | - | 6256 | 764.4 | - | - | 0 | - |
| - | - | 3320 | 765.4 | - | - | 0 | - |
| - | - | 1361 | 766.4 | - | - | 0 | - |
| - | - | 1004 | 790.4 | - | - | 0 | - |
| - | - | 791 | 798.4 | - | - | 0 | - |
| 10 | w | 1.203E+04 | 799.4 | 0.0001187 | 0.1485 | +1 | 8 |
| 3 | z | 997.9 | 799.9 | 0.01155 | 14.43 | +2 | 15 |
| - | - | 4880 | 800.4 | - | - | 0 | - |
| - | - | 1148 | 801.4 | - | - | 0 | - |
| - | - | 5037 | 802.4 | - | - | 0 | - |
| - | - | 2009 | 803.4 | - | - | 0 | - |
| 3 | y | 996.3 | 807.4 | 0.006826 | 8.454 | +2 | 15 |
| 3 | y | 1712 | 807.9 | 0.009141 | 11.32 | +2 | 15 |
| - | - | 858.9 | 808.4 | - | - | 0 | - |
| - | - | 1053 | 813.9 | - | - | 0 | - |
| - | - | 1197 | 815.4 | - | - | 0 | - |
| 3 | y | 3014 | 816.4 | 0.003584 | 4.39 | +2 | 15 |
| - | - | 2507 | 816.9 | - | - | 0 | - |
| - | - | 1123 | 817.4 | - | - | 0 | - |
| - | - | 863.2 | 821.9 | - | - | 0 | - |
| - | - | 4166 | 822.4 | - | - | 0 | - |
| - | - | 4511 | 823.4 | - | - | 0 | - |
| - | - | 1905 | 824.4 | - | - | 0 | - |
| - | - | 1086 | 829.4 | - | - | 0 | - |
| 15 | c | 3.334E+04 | 829.9 | 0.0003095 | 0.3729 | +2 | 15 |
| - | - | 2.889E+04 | 830.4 | - | - | 0 | - |
| - | - | 1.476E+04 | 830.9 | - | - | 0 | - |
| - | - | 6720 | 831.4 | - | - | 0 | - |
| - | - | 1716 | 831.9 | - | - | 0 | - |
| - | - | 745.3 | 835.9 | - | - | 0 | - |
| 10 | z | 688.6 | 840.4 | 0.001088 | 1.295 | +1 | 8 |
| - | - | 813.2 | 840.9 | - | - | 0 | - |
| 10 | z | 1099 | 841.4 | 0.0149 | 17.7 | +1 | 8 |
| - | - | 972.4 | 842.4 | - | - | 0 | - |
| - | - | 1518 | 843.4 | - | - | 0 | - |
| - | - | 1932 | 844 | - | - | 0 | - |
| - | - | 1753 | 844.5 | - | - | 0 | - |
| - | - | 792.9 | 845 | - | - | 0 | - |
| - | - | 875.8 | 845.5 | - | - | 0 | - |
| 8 | c | 2.795E+04 | 846.5 | 6.553E-05 | 0.07742 | +1 | 8 |
| - | - | 1.392E+04 | 847.5 | - | - | 0 | - |
| - | - | 4162 | 848.5 | - | - | 0 | - |
| - | - | 1968 | 848.9 | - | - | 0 | - |
| - | - | 1748 | 849.4 | - | - | 0 | - |
| 2 | z | 3225 | 849.9 | 0.0005881 | 0.6919 | +2 | 16 |
| 2 | z | 2805 | 850.4 | 0.009191 | 10.81 | +2 | 16 |
| - | - | 1163 | 850.9 | - | - | 0 | - |
| 16 | c | 3285 | 856.9 | 0.0004039 | 0.4713 | +2 | 16 |
| 10 | y | 3962 | 857.4 | 0.007891 | 9.203 | +1 | 8 |
| 2 | y | 4690 | 857.9 | 0.01265 | 14.74 | +2 | 16 |
| 2 | y | 2.412E+04 | 858.4 | 0.01674 | 19.5 | +2 | 16 |
| 2 | z | 2513 | 858.9 | 0.01667 | 19.41 | +2 | 16 |
| - | - | 1.034E+04 | 859.4 | - | - | 0 | - |
| - | - | 1197 | 859.9 | - | - | 0 | - |
| - | - | 4199 | 860.4 | - | - | 0 | - |
| - | - | 1123 | 864.5 | - | - | 0 | - |
| - | - | 3451 | 865 | - | - | 0 | - |
| 16 | c | 4.452E+04 | 865.5 | 0.0002986 | 0.3451 | +2 | 16 |
| - | - | 3.841E+04 | 866 | - | - | 0 | - |
| - | - | 2.65E+04 | 866.5 | - | - | 0 | - |
| - | - | 1.142E+04 | 867 | - | - | 0 | - |
| - | - | 2790 | 867.5 | - | - | 0 | - |
| - | - | 1523 | 867.9 | - | - | 0 | - |
| - | - | 1870 | 868.4 | - | - | 0 | - |
| - | - | 1014 | 869.4 | - | - | 0 | - |
| - | - | 766.5 | 870.9 | - | - | 0 | - |
| - | - | 1166 | 871.4 | - | - | 0 | - |
| - | - | 851.9 | 872.5 | - | - | 0 | - |
| - | - | 1529 | 872.9 | - | - | 0 | - |
| - | - | 2848 | 873.5 | - | - | 0 | - |
| 10 | y | 5332 | 874.5 | 1.838E-05 | 0.02102 | +1 | 8 |
| - | - | 1478 | 875.5 | - | - | 0 | - |
| - | - | 2462 | 880.4 | - | - | 0 | - |
| - | - | 6164 | 881 | - | - | 0 | - |
| - | - | 7958 | 881.5 | - | - | 0 | - |
| - | - | 2568 | 882 | - | - | 0 | - |
| - | - | 1386 | 882.5 | - | - | 0 | - |
| - | - | 3672 | 885.5 | - | - | 0 | - |
| - | - | 4602 | 886 | - | - | 0 | - |
| - | - | 2885 | 886.5 | - | - | 0 | - |
| - | - | 818.9 | 887 | - | - | 0 | - |
| - | - | 1548 | 887.5 | - | - | 0 | - |
| - | - | 1700 | 888 | - | - | 0 | - |
| - | - | 1118 | 888.5 | - | - | 0 | - |
| - | - | 1502 | 889 | - | - | 0 | - |
| - | - | 1878 | 889.4 | - | - | 0 | - |
| - | - | 893.3 | 894 | - | - | 0 | - |
| - | - | 2.028E+04 | 894.5 | - | - | 0 | - |
| - | - | 2.024E+04 | 895 | - | - | 0 | - |
| - | - | 1.07E+04 | 895.5 | - | - | 0 | - |
| - | - | 5898 | 896 | - | - | 0 | - |
| - | - | 1542 | 896.5 | - | - | 0 | - |
| - | - | 1473 | 899.5 | - | - | 0 | - |
| - | - | 2466 | 900 | - | - | 0 | - |
| - | - | 852.5 | 902.5 | - | - | 0 | - |
| - | - | 7494 | 907.4 | - | - | 0 | - |
| - | - | 4622 | 908 | - | - | 0 | - |
| - | - | 2081 | 908.4 | - | - | 0 | - |
| - | - | 4386 | 908.5 | - | - | 0 | - |
| - | - | 3640 | 909 | - | - | 0 | - |
| - | - | 1088 | 909.5 | - | - | 0 | - |
| - | - | 877.6 | 915.5 | - | - | 0 | - |
| - | - | 1927 | 916 | - | - | 0 | - |
| - | - | 4.978E+04 | 916.5 | - | - | 0 | - |
| - | - | 8.108E+04 | 917 | - | - | 0 | - |
| - | - | 5.426E+04 | 917.5 | - | - | 0 | - |
| - | - | 3.286E+04 | 918 | - | - | 0 | - |
| - | - | 1.032E+04 | 918.5 | - | - | 0 | - |
| - | - | 3389 | 919 | - | - | 0 | - |
| 9 | w | 5379 | 928.5 | 0.0005365 | 0.5779 | +1 | 9 |
| - | - | 5492 | 929.5 | - | - | 0 | - |
| - | - | 2484 | 930.5 | - | - | 0 | - |
| - | - | 1021 | 931.5 | - | - | 0 | - |
| - | - | 1037 | 939.4 | - | - | 0 | - |
| - | - | 689.8 | 940.5 | - | - | 0 | - |
| - | - | 2549 | 944.5 | - | - | 0 | - |
| - | - | 3704 | 945.5 | - | - | 0 | - |
| - | - | 1598 | 946.5 | - | - | 0 | - |
| - | - | 938.7 | 956.5 | - | - | 0 | - |
| - | - | 1237 | 958.5 | - | - | 0 | - |
| - | - | 1692 | 959.5 | - | - | 0 | - |
| - | - | 5403 | 960.5 | - | - | 0 | - |
| - | - | 2708 | 961.5 | - | - | 0 | - |
| - | - | 1.484E+04 | 973.5 | - | - | 0 | - |
| - | - | 8162 | 974.5 | - | - | 0 | - |
| 9 | c | 4.301E+04 | 975.5 | 0.0005354 | 0.5488 | +1 | 9 |
| - | - | 2.24E+04 | 976.5 | - | - | 0 | - |
| - | - | 7890 | 977.5 | - | - | 0 | - |
| - | - | 1447 | 978.5 | - | - | 0 | - |
| 9 | z | 1.922E+04 | 987.5 | 4.694E-05 | 0.04753 | +1 | 9 |
| - | - | 1.065E+04 | 988.5 | - | - | 0 | - |
| - | - | 3756 | 989.5 | - | - | 0 | - |
| - | - | 938.5 | 990.5 | - | - | 0 | - |
| - | - | 1357 | 1002 | - | - | 0 | - |
| - | - | 1081 | 1003 | - | - | 0 | - |
| 9 | y | 4198 | 1004 | 0.00748 | 7.453 | +1 | 9 |
| - | - | 3682 | 1004 | - | - | 0 | - |
| - | - | 866.9 | 1013 | - | - | 0 | - |
| - | - | 1156 | 1014 | - | - | 0 | - |
| - | - | 1192 | 1018 | - | - | 0 | - |
| - | - | 840.7 | 1019 | - | - | 0 | - |
| - | - | 883.9 | 1032 | - | - | 0 | - |
| - | - | 3277 | 1055 | - | - | 0 | - |
| 8 | z | 2735 | 1057 | 0.003444 | 3.26 | +1 | 10 |
| 8 | z | 1721 | 1057 | 0.01828 | 17.28 | +1 | 10 |
| - | - | 1145 | 1059 | - | - | 0 | - |
| - | - | 1618 | 1060 | - | - | 0 | - |
| - | - | 2802 | 1061 | - | - | 0 | - |
| - | - | 1680 | 1062 | - | - | 0 | - |
| - | - | 1438 | 1073 | - | - | 0 | - |
| 8 | y | 1467 | 1074 | 0.001384 | 1.29 | +1 | 10 |
| 8 | z | 2.094E+04 | 1075 | 0.0008253 | 0.7681 | +1 | 10 |
| - | - | 1.502E+04 | 1076 | - | - | 0 | - |
| - | - | 5781 | 1077 | - | - | 0 | - |
| - | - | 2097 | 1078 | - | - | 0 | - |
| - | - | 4240 | 1088 | - | - | 0 | - |
| - | - | 2232 | 1089 | - | - | 0 | - |
| - | - | 1847 | 1090 | - | - | 0 | - |
| 8 | y | 1.169E+04 | 1091 | 0.001727 | 1.584 | +1 | 10 |
| - | - | 8110 | 1092 | - | - | 0 | - |
| - | - | 2525 | 1093 | - | - | 0 | - |
| - | - | 2747 | 1099 | - | - | 0 | - |
| - | - | 1623 | 1100 | - | - | 0 | - |
| - | - | 3134 | 1104 | - | - | 0 | - |
| 10 | c | 4.582E+04 | 1105 | 0.001136 | 1.029 | +1 | 10 |
| - | - | 2.955E+04 | 1106 | - | - | 0 | - |
| - | - | 1.027E+04 | 1107 | - | - | 0 | - |
| - | - | 3052 | 1108 | - | - | 0 | - |
| - | - | 808.1 | 1109 | - | - | 0 | - |
| - | - | 753.6 | 1119 | - | - | 0 | - |
| - | - | 7800 | 1121 | - | - | 0 | - |
| - | - | 4613 | 1122 | - | - | 0 | - |
| - | - | 1806 | 1123 | - | - | 0 | - |
| 7 | y | 1089 | 1160 | 0.01269 | 10.95 | +1 | 11 |
| 7 | y | 937.4 | 1161 | 0.009639 | 8.306 | +1 | 11 |
| 7 | z | 8957 | 1162 | 0.0004714 | 0.4059 | +1 | 11 |
| - | - | 6635 | 1163 | - | - | 0 | - |
| - | - | 3747 | 1164 | - | - | 0 | - |
| - | - | 1572 | 1165 | - | - | 0 | - |
| - | - | 875 | 1166 | - | - | 0 | - |
| 7 | y | 6011 | 1178 | 0.0004304 | 0.3655 | +1 | 11 |
| - | - | 5934 | 1179 | - | - | 0 | - |
| - | - | 2311 | 1180 | - | - | 0 | - |
| - | - | 1133 | 1181 | - | - | 0 | - |
| - | - | 1605 | 1182 | - | - | 0 | - |
| - | - | 1293 | 1199 | - | - | 0 | - |
| 11 | c | 1510 | 1200 | 0.02349 | 19.58 | +1 | 11 |
| - | - | 1907 | 1201 | - | - | 0 | - |
| - | - | 1137 | 1202 | - | - | 0 | - |
| - | - | 1051 | 1206 | - | - | 0 | - |
| - | - | 1.08E+04 | 1207 | - | - | 0 | - |
| - | - | 8034 | 1208 | - | - | 0 | - |
| - | - | 2563 | 1209 | - | - | 0 | - |
| 11 | c | 2.664E+04 | 1218 | 0.001338 | 1.099 | +1 | 11 |
| - | - | 2.093E+04 | 1219 | - | - | 0 | - |
| - | - | 8206 | 1220 | - | - | 0 | - |
| - | - | 1480 | 1221 | - | - | 0 | - |
| - | - | 5851 | 1223 | - | - | 0 | - |
| - | - | 1.051E+04 | 1224 | - | - | 0 | - |
| - | - | 5454 | 1225 | - | - | 0 | - |
| - | - | 2300 | 1226 | - | - | 0 | - |
| - | - | 829.4 | 1245 | - | - | 0 | - |
| - | - | 2230 | 1253 | - | - | 0 | - |
| - | - | 1350 | 1254 | - | - | 0 | - |
| - | - | 1139 | 1270 | - | - | 0 | - |
| - | - | 1025 | 1271 | - | - | 0 | - |
| 6 | y | 2144 | 1275 | 0.003324 | 2.608 | +1 | 12 |
| - | - | 2258 | 1276 | - | - | 0 | - |
| - | - | 1665 | 1302 | - | - | 0 | - |
| - | - | 1697 | 1303 | - | - | 0 | - |
| - | - | 893.4 | 1328 | - | - | 0 | - |
| 12 | c | 2459 | 1329 | 0.01615 | 12.16 | +1 | 12 |
| - | - | 2422 | 1330 | - | - | 0 | - |
| - | - | 1805 | 1331 | - | - | 0 | - |
| - | - | 749.9 | 1332 | - | - | 0 | - |
| - | - | 3324 | 1345 | - | - | 0 | - |
| 12 | c | 3.581E+04 | 1346 | 0.00242 | 1.799 | +1 | 12 |
| - | - | 2.443E+04 | 1347 | - | - | 0 | - |
| - | - | 1.173E+04 | 1348 | - | - | 0 | - |
| - | - | 3958 | 1349 | - | - | 0 | - |
| - | - | 978.2 | 1350 | - | - | 0 | - |
| - | - | 770.2 | 1353 | - | - | 0 | - |
| 5 | y | 910.4 | 1355 | 0.005308 | 3.918 | +1 | 13 |
| 5 | z | 1063 | 1356 | 0.02177 | 16.06 | +1 | 13 |
| - | - | 2399 | 1371 | - | - | 0 | - |
| 5 | y | 9870 | 1372 | 0.003053 | 2.226 | +1 | 13 |
| - | - | 7254 | 1373 | - | - | 0 | - |
| - | - | 3877 | 1374 | - | - | 0 | - |
| - | - | 2027 | 1375 | - | - | 0 | - |
| - | - | 840.6 | 1398 | - | - | 0 | - |
| - | - | 1003 | 1399 | - | - | 0 | - |
| - | - | 2713 | 1400 | - | - | 0 | - |
| - | - | 4327 | 1401 | - | - | 0 | - |
| - | - | 4202 | 1402 | - | - | 0 | - |
| - | - | 3129 | 1403 | - | - | 0 | - |
| - | - | 1270 | 1404 | - | - | 0 | - |
| - | - | 4411 | 1416 | - | - | 0 | - |
| 13 | c | 2.469E+04 | 1417 | 0.003523 | 2.487 | +1 | 13 |
| - | - | 1.68E+04 | 1418 | - | - | 0 | - |
| - | - | 8412 | 1419 | - | - | 0 | - |
| - | - | 1906 | 1420 | - | - | 0 | - |
| - | - | 2403 | 1431 | - | - | 0 | - |
| - | - | 2075 | 1432 | - | - | 0 | - |
| - | - | 1244 | 1433 | - | - | 0 | - |
| - | - | 933.4 | 1476 | - | - | 0 | - |
| - | - | 1953 | 1487 | - | - | 0 | - |
| - | - | 1692 | 1488 | - | - | 0 | - |
| - | - | 1380 | 1489 | - | - | 0 | - |
| 4 | y | 1572 | 1502 | 0.007695 | 5.124 | +1 | 14 |
| 4 | z | 2.037E+04 | 1503 | 0.002083 | 1.386 | +1 | 14 |
| - | - | 3.356E+04 | 1504 | - | - | 0 | - |
| - | - | 2.315E+04 | 1505 | - | - | 0 | - |
| - | - | 1055 | 1505 | - | - | 0 | - |
| - | - | 9786 | 1506 | - | - | 0 | - |
| - | - | 3315 | 1507 | - | - | 0 | - |
| 14 | c | 1009 | 1514 | 0.003902 | 2.578 | +1 | 14 |
| 4 | y | 2859 | 1519 | 0.005793 | 3.814 | +1 | 14 |
| - | - | 2638 | 1520 | - | - | 0 | - |
| - | - | 2011 | 1521 | - | - | 0 | - |
| - | - | 738 | 1522 | - | - | 0 | - |
| 14 | c | 4.032E+04 | 1531 | 0.00275 | 1.796 | +1 | 14 |
| - | - | 3.334E+04 | 1532 | - | - | 0 | - |
| - | - | 1039 | 1532 | - | - | 0 | - |
| - | - | 1.603E+04 | 1533 | - | - | 0 | - |
| - | - | 6444 | 1534 | - | - | 0 | - |
| - | - | 1583 | 1535 | - | - | 0 | - |
| - | - | 885.5 | 1544 | - | - | 0 | - |
| - | - | 1120 | 1545 | - | - | 0 | - |
| - | - | 2752 | 1615 | - | - | 0 | - |
| 3 | z | 5132 | 1616 | 0.01993 | 12.34 | +1 | 15 |
| - | - | 7393 | 1617 | - | - | 0 | - |
| - | - | 5110 | 1618 | - | - | 0 | - |
| - | - | 2889 | 1619 | - | - | 0 | - |
| - | - | 808.9 | 1633 | - | - | 0 | - |
| - | - | 1893 | 1643 | - | - | 0 | - |
| - | - | 1521 | 1644 | - | - | 0 | - |
| - | - | 1338 | 1645 | - | - | 0 | - |
| 15 | c | 3696 | 1659 | 0.000798 | 0.4811 | +1 | 15 |
| - | - | 1.182E+04 | 1660 | - | - | 0 | - |
| - | - | 9544 | 1661 | - | - | 0 | - |
| - | - | 4313 | 1662 | - | - | 0 | - |
| - | - | 1172 | 1663 | - | - | 0 | - |
| - | - | 1016 | 1673 | - | - | 0 | - |
| - | - | 2732 | 1686 | - | - | 0 | - |
| - | - | 3459 | 1687 | - | - | 0 | - |
| - | - | 2763 | 1688 | - | - | 0 | - |
| - | - | 1009 | 1689 | - | - | 0 | - |
| - | - | 1240 | 1690 | - | - | 0 | - |
| 2 | z | 1425 | 1700 | 0.01516 | 8.918 | +1 | 16 |
| - | - | 1222 | 1701 | - | - | 0 | - |
| - | - | 4904 | 1714 | - | - | 0 | - |
| 2 | y | 3557 | 1715 | 0.0305 | 17.79 | +1 | 16 |
| - | - | 3573 | 1716 | - | - | 0 | - |
| 2 | z | 3913 | 1717 | 0.008142 | 4.742 | +1 | 16 |
| - | - | 9331 | 1718 | - | - | 0 | - |
| - | - | 7767 | 1719 | - | - | 0 | - |
| - | - | 3572 | 1720 | - | - | 0 | - |
| - | - | 1377 | 1721 | - | - | 0 | - |
| 16 | c | 5666 | 1730 | 0.00238 | 1.376 | +1 | 16 |
| - | - | 1.259E+04 | 1731 | - | - | 0 | - |
| - | - | 1.254E+04 | 1732 | - | - | 0 | - |
| - | - | 6199 | 1733 | - | - | 0 | - |
| - | - | 2912 | 1734 | - | - | 0 | - |
| - | - | 1147 | 1761 | - | - | 0 | - |
| - | - | 2811 | 1771 | - | - | 0 | - |
| - | - | 4364 | 1772 | - | - | 0 | - |
| - | - | 3849 | 1773 | - | - | 0 | - |
| - | - | 1007 | 1774 | - | - | 0 | - |
| - | - | 1203 | 1775 | - | - | 0 | - |
| - | - | 5462 | 1788 | - | - | 0 | - |
| - | - | 2.204E+04 | 1789 | - | - | 0 | - |
| - | - | 1.98E+04 | 1790 | - | - | 0 | - |
| - | - | 1.111E+04 | 1791 | - | - | 0 | - |
| - | - | 3739 | 1792 | - | - | 0 | - |
| - | - | 1065 | 1793 | - | - | 0 | - |
| - | - | 1150 | 1799 | - | - | 0 | - |
| - | - | 1663 | 1800 | - | - | 0 | - |
| - | - | 1472 | 1805 | - | - | 0 | - |
| - | - | 4754 | 1806 | - | - | 0 | - |
| - | - | 3916 | 1807 | - | - | 0 | - |
| - | - | 2373 | 1808 | - | - | 0 | - |
| - | - | 842.7 | 1809 | - | - | 0 | - |
| - | - | 991.8 | 1815 | - | - | 0 | - |
| - | - | 5588 | 1816 | - | - | 0 | - |
| - | - | 3.342E+04 | 1817 | - | - | 0 | - |
| - | - | 2.996E+04 | 1818 | - | - | 0 | - |
| - | - | 1.845E+04 | 1819 | - | - | 0 | - |
| - | - | 6392 | 1820 | - | - | 0 | - |
| - | - | 1409 | 1821 | - | - | 0 | - |
| - | - | 3907 | 1832 | - | - | 0 | - |
| - | - | 1.949E+04 | 1833 | - | - | 0 | - |
| - | - | 5.763E+04 | 1834 | - | - | 0 | - |
| - | - | 5.279E+04 | 1835 | - | - | 0 | - |
| - | - | 2.649E+04 | 1836 | - | - | 0 | - |
| - | - | 1.061E+04 | 1837 | - | - | 0 | - |
| - | - | 3358 | 1838 | - | - | 0 | - |

m/z Charge Intensity FragmentType MassShift Position
120.06143188476562 0 696.68396
120.06560516357422 0 11118.019 y 16
120.08100128173828 0 1809.5883
121.0692367553711 0 547.46875
124.55908966064453 0 371.43878
126.05525207519531 0 739.15735
128.09442138671875 0 960.52716
128.10696411132812 0 408.47168
129.10231018066406 0 4562.2944
129.1953125 0 396.3229
131.11807250976562 0 746.56866
140.10699462890625 0 474.4931
148.94754028320312 0 589.4428
149.0448455810547 0 1105.2852
151.33648681640625 0 454.15222
155.09420776367188 0 708.4674
155.1181182861328 0 565.4288
158.14134216308594 0 534.0505
169.1336212158203 0 2069.8303
171.0767822265625 0 657.1482
173.09214782714844 0 1294.6191 y Water loss 15
173.12843322753906 0 2899.6445
181.0966033935547 0 495.31393
183.1128387451172 0 3672.1968
185.1281280517578 0 548.62115
191.10276794433594 0 3994.2212 y 15
197.12840270996094 0 1701.5481
200.1395721435547 0 626.5208 c Water loss 1
201.12327575683594 0 7028.981
211.10830688476562 0 552.59735
212.13937377929688 0 8753.775
213.1427001953125 0 795.75684
215.1392822265625 0 1416.1409
218.14993286132812 0 4586.633 c 1
221.08424377441406 0 4220.372
222.08499145507812 0 749.8085
223.08212280273438 0 763.26935
225.80661010742188 0 478.75153
226.1187286376953 0 862.42975
227.09054565429688 0 988.1465
227.10255432128906 0 1642.3906
227.12693786621094 0 706.22107
233.16494750976562 0 1176.1436
235.10739135742188 0 1013.6435
239.09487915039062 0 5011.5386
240.09573364257812 0 1903.0446
242.1501007080078 0 727.1105
243.14486694335938 0 1363.7866 y Water loss 10
251.1764373779297 0 658.9261
254.14907836914062 0 1394.1986
261.15966796875 0 1005.4822
263.10235595703125 0 2039.936
272.1849670410156 0 805.99274
280.1778259277344 0 698.4395
281.05072021484375 0 1260.9365
281.13665771484375 0 655.50183
282.14544677734375 0 850.9801
296.1772766113281 0 835.7221
296.1968688964844 0 10828.605
297.0820617675781 0 631.09845
297.1562805175781 0 1001.0995
297.2003173828125 0 1633.8605
298.16241455078125 0 1104.414
299.06201171875 0 5174.771
299.1715087890625 0 13179.412
300.0614013671875 0 1784.9369
300.1752014160156 0 1103.9227
301.0579528808594 0 657.2571
303.17877197265625 0 6848.7134 z 14
304.1832275390625 0 1270.0298
314.1842346191406 0 982.837
314.207763671875 0 4471.845
315.21075439453125 0 935.43945
316.186767578125 0 5854.699
317.1905212402344 0 911.1123
319.1976623535156 0 4222.1284 y 14
320.2003479003906 0 740.7932
325.1875 0 1443.2091
331.23406982421875 0 25535.438 c 2
332.2375183105469 0 4461.1816
344.172607421875 0 741.8729
345.1795959472656 0 986.3441
346.1246337890625 0 607.65393
346.1844482421875 0 551.76715
347.19598388671875 0 653.45
348.1918029785156 0 540.60626
350.13433837890625 0 808.9845
355.0697021484375 0 38351.93
356.0704345703125 0 16836.96
357.06756591796875 0 9743.368
358.0696105957031 0 1037.3506
362.2078552246094 0 1413.5902
370.2080383300781 0 894.0711
371.1010437011719 0 1853.8386
371.1926574707031 0 610.83875
372.1015625 0 1292.6399
373.0789489746094 0 916.8136
373.2096862792969 0 3371.7341 y 10
373.71478271484375 0 763.63684
374.08074951171875 0 639.8141
385.2192077636719 0 1406.2267
385.41058349609375 0 699.2544
386.2024230957031 0 1753.8217
415.0357971191406 0 880.58704
415.2298583984375 0 859.51666 y Water loss 13
416.03570556640625 0 1175.9215
417.03411865234375 0 1075.0479
417.222412109375 0 2279.971 z 13
418.2294006347656 0 7195.1724
419.2333068847656 0 1166.9637
433.24127197265625 0 2723.7634 y 13
433.2810974121094 0 8120.567
434.2864990234375 0 4885.598
435.2946472167969 0 1829.8246
442.2394104003906 0 729.2164
444.24578857421875 0 5845.6177
445.2520751953125 0 3143.6953
446.2651672363281 0 2588.2551
460.26812744140625 0 1514.0149
461.276123046875 0 13395.945
462.27911376953125 0 3151.3176
463.2819519042969 0 743.3332
480.716552734375 0 799.12177
480.750732421875 0 1449.7213
481.2488098144531 0 1270.2938
486.2683410644531 0 689.80835 y Water loss 12
488.2590637207031 0 9197.543 c 8
489.05584716796875 0 1115.8505
489.26513671875 0 7165.095
490.0542297363281 0 627.1861
490.2680358886719 0 1419.8853
493.72802734375 0 2480.212
494.2323913574219 0 1086.4792
497.2613525390625 0 731.74744
498.2216491699219 0 696.4088
502.26507568359375 0 1958.8936 y 8
502.7331848144531 0 3364.4785
503.2338562011719 0 2320.2095
503.73516845703125 0 740.7586
504.2777404785156 0 9220.98 y 12
505.27984619140625 0 2057.6526
515.2930908203125 0 789.5569
538.2696533203125 0 4717.083
538.7706909179688 0 3370.6687
539.2733154296875 0 768.95526 z 2
543.2604370117188 0 2265.7397
543.760498046875 0 928.30115
552.2672119140625 0 6683.28
552.7693481445312 0 4851.3193
553.2711791992188 0 1448.6636
554.3196411132812 0 894.46124
558.2879028320312 0 16157.418 w 11
558.3289794921875 0 1774.9677
559.290283203125 0 3961.6
559.3318481445312 0 1247.353
560.2948608398438 0 814.3374
566.4273681640625 0 10933.563
567.4308471679688 0 4070.914
572.3045654296875 0 767.74884
577.1273803710938 0 679.64355
578.1268310546875 0 1335.6788
579.1235961914062 0 1188.3715
580.2807006835938 0 704.8489 y Water loss 6
580.7818603515625 0 1297.1383 y Ammonia loss 6
583.3348388671875 0 1306.02
584.3356323242188 0 727.7889
591.2978515625 0 998.39343
594.3878173828125 0 690.6424
599.2669067382812 0 680.00574
599.8055419921875 0 1990.7046
602.7907104492188 0 3473.1714
603.29296875 0 3200.3223
603.7950439453125 0 1043.2185
610.184326171875 0 8593.225
611.1849975585938 0 8211.243
611.32666015625 0 746.64514
611.7965087890625 0 7655.914
612.18310546875 0 6954.3027
612.2317504882812 0 720.8184
612.2421875 0 731.8558
612.2977294921875 0 5423.461
612.3963623046875 0 846.95337
612.8012084960938 0 1751.8846
613.18359375 0 767.1128
614.3365478515625 0 7141.6445 y Water loss 11
614.82177734375 0 753.47974
615.3448486328125 0 9537.43
616.3169555664062 0 3974.1753 z 11
616.3577880859375 0 1401.0656
617.3201293945312 0 1176.6735
617.8060913085938 0 1140.7832
618.3114013671875 0 630.7427
626.3431396484375 0 1753.6548
626.81201171875 0 5044.642
627.261474609375 0 3215.4858
627.3121337890625 0 1920.9808
627.357177734375 0 1146.1622
627.8157958984375 0 900.11487
628.264892578125 0 1171.4436
628.392578125 0 2938.293
628.80810546875 0 586.82605 y Water loss 5
629.3072509765625 0 905.5447 y Ammonia loss 5
629.3973999023438 0 1202.694
632.336181640625 0 5574.67 y 11
633.3370361328125 0 2243.4092
635.3232421875 0 1615.7711
635.8255615234375 0 1175.4073
636.332275390625 0 695.5097
637.8167114257812 0 563.8762 y 5
638.3157958984375 0 1048.5276
655.3425903320312 0 1539.5638
655.8419799804688 0 1207.7747
656.3387451171875 0 820.90436
658.36669921875 0 1418.888
664.8350219726562 0 1151.0624 c Ammonia loss 11
665.3370361328125 0 781.5896
668.3349609375 0 810.96643
668.8260498046875 0 636.41016
671.3739013671875 0 2541.9683
672.40966796875 0 3279.8433 c 5
673.4097900390625 0 1998.8588
674.4227905273438 0 907.19745
676.8320922851562 0 1631.1814
677.3353881835938 0 7903.17 y Water loss 4
677.8357543945312 0 6494.48 y Ammonia loss 4
678.3347778320312 0 3724.4158 z 4
678.8335571289062 0 1823.7065
685.8372802734375 0 5146.709
686.3439331054688 0 52071.87 y 4
686.8425903320312 0 19479.47
687.3485717773438 0 15303.316
687.8472290039062 0 2725.2693
688.3751831054688 0 2074.4546
697.416748046875 0 1169.472
716.4360961914062 0 1013.40045
722.3480834960938 0 1924.701
723.3538818359375 0 786.51807
724.869140625 0 593.78876
729.4009399414062 0 11698.823 z 10
730.4037475585938 0 5275.701
731.406494140625 0 1826.7053
740.3458862304688 0 1352.7936
742.4135131835938 0 1442.0316
742.8609619140625 0 875.74194 z Water loss 3
745.4212036132812 0 7050.812 y 10
746.4226684570312 0 2717.7275
747.4231567382812 0 835.58514
750.3651123046875 0 748.82983
750.871337890625 0 2043.3398 y Water loss 3
751.37158203125 0 1714.0638 y Ammonia loss 3
751.8731689453125 0 763.6055 z 3
757.4198608398438 0 3002.6448
758.4309692382812 0 928.25476
759.3731689453125 0 1963.4966
759.4400634765625 0 33023.85 c 6
759.8751831054688 0 13674.993 y 3
760.3750610351562 0 7818.0005
760.4435424804688 0 14132.394
760.8778686523438 0 4440.882
761.37744140625 0 1352.3866
761.447509765625 0 4008.7842
762.451904296875 0 1020.4848
764.36083984375 0 6255.659
765.3614501953125 0 3320.217
766.3661499023438 0 1361.0808
790.3980712890625 0 1003.5643
798.38916015625 0 790.9712
799.4307250976562 0 12026.912 w 9
799.9063110351562 0 997.94104 z Ammonia loss 2
800.4310302734375 0 4880.4355
801.43212890625 0 1147.6182
802.3812866210938 0 5037.455
803.385009765625 0 2009.324
807.4189453125 0 996.305 y Water loss 2
807.9132690429688 0 1711.7184 y Ammonia loss 2
808.366455078125 0 858.9135
813.9266357421875 0 1053.2239
815.39306640625 0 1197.2548
816.413818359375 0 3013.6267 y 2
816.9186401367188 0 2507.112
817.4231567382812 0 1122.8007
821.9346313476562 0 863.1866
822.3797607421875 0 4165.635
823.3849487304688 0 4511.0283
824.3896484375 0 1905.2839
829.443359375 0 1085.9658
829.9407348632812 0 33341.22 c 14
830.4422607421875 0 28890.94
830.942626953125 0 14755.6045
831.4439086914062 0 6720.021
831.9419555664062 0 1715.8325
835.93994140625 0 745.3289
840.4324951171875 0 688.5819 z Water loss 9
840.9248046875 0 813.20184
841.4324951171875 0 1099.2399 z Ammonia loss 9
842.4461059570312 0 972.4243
843.4486083984375 0 1517.7545
843.9559936523438 0 1931.9935
844.45458984375 0 1752.7014
844.9552612304688 0 792.8647
845.457763671875 0 875.75793
846.4720458984375 0 27949.777 c 7
847.4746704101562 0 13923.5
848.4769287109375 0 4162.043
848.9317016601562 0 1968.0967
849.433349609375 0 1747.7991
849.9271850585938 0 3225.3901 z Water loss 1
850.4277954101562 0 2804.8455 z Ammonia loss 1
850.9312133789062 0 1163.031
856.9459228515625 0 3285.3052 c Ammonia loss 15
857.4442138671875 0 3962.0574 y Ammonia loss 9
857.9486083984375 0 4690.0225 y Water loss 1
858.4447021484375 0 24116.197 y Ammonia loss 1
858.9485473632812 0 2513.4424 z 1
859.4448852539062 0 10344.589
859.9380493164062 0 1197.047
860.44580078125 0 4199.0405
864.4573974609375 0 1122.9949
864.9605712890625 0 3451.3564
865.4598999023438 0 44515.3 c 15
865.9608154296875 0 38407.523
866.4603881835938 0 26497.152
866.959716796875 0 11420.18
867.456787109375 0 2789.9536
867.94970703125 0 1522.54
868.4034423828125 0 1869.667
869.4031372070312 0 1013.86285
870.94677734375 0 766.5229
871.44677734375 0 1165.7356
872.4583129882812 0 851.9393
872.9473266601562 0 1529.0188
873.4569091796875 0 2848.0378
874.462890625 0 5331.9233 y 9
875.4650268554688 0 1477.8989
880.4489135742188 0 2462.1816
880.9639282226562 0 6163.988
881.4638061523438 0 7957.777
881.966796875 0 2567.7585
882.4686279296875 0 1385.6455
885.466796875 0 3671.5188
885.96923828125 0 4602.333
886.4645385742188 0 2885.0732
886.9652099609375 0 818.8678
887.463134765625 0 1547.6096
887.9620971679688 0 1700.3796
888.4641723632812 0 1118.0537
888.9545288085938 0 1502.4143
889.44921875 0 1878.0476
893.9700927734375 0 893.2771
894.4691162109375 0 20283.959
894.9697265625 0 20238.596
895.4710693359375 0 10701.106
895.9690551757812 0 5898.279
896.466064453125 0 1541.7344
899.4642333984375 0 1473.436
899.9608764648438 0 2466.0515
902.4679565429688 0 852.4573
907.404541015625 0 7494.463
907.9733276367188 0 4622.4233
908.396728515625 0 2080.7876
908.472412109375 0 4386.4683
908.9716186523438 0 3640.4739
909.4736328125 0 1087.5881
915.4640502929688 0 877.59265
915.9666748046875 0 1926.8352
916.4752807617188 0 49775.938
916.977294921875 0 81076.89
917.47900390625 0 54260.26
917.981201171875 0 32860.605
918.4822998046875 0 10321.611
918.9830932617188 0 3389.2366
928.472900390625 0 5378.949 w 8
929.4765625 0 5491.97
930.4811401367188 0 2483.9417
931.4921875 0 1020.5977
939.4413452148438 0 1036.9401
940.475830078125 0 689.7747
944.4716186523438 0 2549.3345
945.4774169921875 0 3703.884
946.4844360351562 0 1598.346
956.462158203125 0 938.65234
958.4854125976562 0 1237.0774
959.4853515625 0 1692.3762
960.4896850585938 0 5403.0137
961.4915771484375 0 2708.2512
973.4948120117188 0 14840.024
974.4968872070312 0 8161.5186
975.5140380859375 0 43008.477 c 8
976.5172729492188 0 22401.248
977.5198974609375 0 7890.1167
978.5191040039062 0 1446.509
987.4866943359375 0 19222.06 z 8
988.4886474609375 0 10647.476
989.4916381835938 0 3756.3428
990.501953125 0 938.49133
1001.5269775390625 0 1357.0725
1002.5226440429688 0 1080.7354
1003.4979858398438 0 4197.589 y 8
1004.4979248046875 0 3682.383
1012.5249633789062 0 866.91
1013.5242309570312 0 1155.7948
1018.4523315429688 0 1191.5348
1019.4691772460938 0 840.7484
1031.5076904296875 0 883.9204
1055.497314453125 0 3277.4722
1056.5047607421875 0 2735.1116 z Water loss 7
1057.510498046875 0 1720.977 z Ammonia loss 7
1058.513427734375 0 1144.5432
1059.512939453125 0 1617.8539
1060.532470703125 0 2802.3347
1061.5338134765625 0 1680.3995
1072.504638671875 0 1438.3601
1073.5123291015625 0 1466.718 y Ammonia loss 7
1074.5179443359375 0 20942.018 z 7
1075.5225830078125 0 15017.986
1076.52587890625 0 5780.8394
1077.5289306640625 0 2097.136
1087.5299072265625 0 4240.1157
1088.5308837890625 0 2232.2097
1089.5301513671875 0 1846.8885
1090.5357666015625 0 11692.886 y 7
1091.5394287109375 0 8110.4272
1092.5401611328125 0 2524.7764
1098.50634765625 0 2746.5854
1099.52001953125 0 1623.3922
1103.5262451171875 0 3133.964
1104.5560302734375 0 45822.855 c 9
1105.5587158203125 0 29552.39
1106.561279296875 0 10266.072
1107.564697265625 0 3051.7712
1108.57763671875 0 808.082
1118.5396728515625 0 753.6067
1120.5523681640625 0 7799.969
1121.556640625 0 4612.5356
1122.558349609375 0 1806.272
1159.5462646484375 0 1088.6387 y Water loss 6
1160.5526123046875 0 937.36926 y Ammonia loss 6
1161.55126953125 0 8957.04 z 6
1162.5531005859375 0 6634.722
1163.5631103515625 0 3746.934
1164.5628662109375 0 1571.7963
1165.56884765625 0 874.98553
1177.569091796875 0 6010.8447 y 6
1178.5733642578125 0 5934.189
1179.5728759765625 0 2311.0344
1180.5802001953125 0 1133.1832
1181.5748291015625 0 1604.6415
1198.608154296875 0 1292.9309
1199.607177734375 0 1509.7904 c Water loss 10
1200.6121826171875 0 1906.9375
1201.615234375 0 1136.5193
1205.5826416015625 0 1051.1027
1206.568603515625 0 10797.521
1207.568603515625 0 8033.9336
1208.5714111328125 0 2562.5044
1217.639892578125 0 26644.934 c 10
1218.6422119140625 0 20927.285
1219.6466064453125 0 8206.347
1220.6395263671875 0 1480.136
1222.5826416015625 0 5851.207
1223.5885009765625 0 10505.1045
1224.59326171875 0 5454.478
1225.59326171875 0 2300.1836
1244.65380859375 0 829.366
1252.6126708984375 0 2229.875
1253.6160888671875 0 1349.8323
1269.6409912109375 0 1139.2095
1270.644287109375 0 1025.1066
1274.6256103515625 0 2144.4827 y 5
1275.6240234375 0 2258.1118
1301.6719970703125 0 1664.5652
1302.6787109375 0 1697.4569
1327.65234375 0 893.44
1328.6571044921875 0 2459.2017 c Ammonia loss 11
1329.658203125 0 2422.3674
1330.6552734375 0 1805.2306
1331.654541015625 0 749.8601
1344.686279296875 0 3324.2021
1345.6973876953125 0 35807.234 c 11
1346.70166015625 0 24433.438
1347.70458984375 0 11729.617
1348.7020263671875 0 3957.9968
1349.700927734375 0 978.22815
1352.6556396484375 0 770.2027
1354.65380859375 0 910.3952 y Ammonia loss 4
1355.6781005859375 0 1062.984 z 4
1370.6683349609375 0 2399.26
1371.6719970703125 0 9870.104 y 4
1372.6785888671875 0 7253.833
1373.685546875 0 3876.5378
1374.6888427734375 0 2026.9241
1397.708984375 0 840.5546
1398.673828125 0 1003.2554
1399.6724853515625 0 2713.214
1400.68359375 0 4326.669
1401.6944580078125 0 4201.5317
1402.694580078125 0 3128.67
1403.6993408203125 0 1269.753
1415.726318359375 0 4411.0522
1416.7333984375 0 24688.152 c 12
1417.7371826171875 0 16798.627
1418.7391357421875 0 8412.495
1419.7396240234375 0 1906.093
1430.7005615234375 0 2403.0144
1431.6988525390625 0 2075.4336
1432.702392578125 0 1244.489
1475.72705078125 0 933.3515
1486.7481689453125 0 1952.5719
1487.769287109375 0 1692.3373
1488.7635498046875 0 1379.8442
1501.724609375 0 1572.2651 y Ammonia loss 3
1502.72265625 0 20371.217 z 3
1503.728759765625 0 33563.965
1504.731201171875 0 23153.047
1504.9486083984375 0 1054.8438
1505.734619140625 0 9786.046
1506.734130859375 0 3315.313
1513.7572021484375 0 1008.67316 c Ammonia loss 13
1518.7376708984375 0 2858.8147 y 3
1519.74267578125 0 2638.0732
1520.7567138671875 0 2010.982
1521.7525634765625 0 737.9565
1530.777099609375 0 40316.656 c 13
1531.7796630859375 0 33343.63
1532.015869140625 0 1039.2485
1532.78271484375 0 16027.735
1533.7867431640625 0 6444.3965
1534.789306640625 0 1582.9382
1543.7750244140625 0 885.49817
1544.7880859375 0 1120.215
1614.8431396484375 0 2751.9893
1615.8287353515625 0 5131.703 z 2
1616.8173828125 0 7392.6465
1617.82470703125 0 5110.1772
1618.82421875 0 2889.4268
1632.8323974609375 0 808.9215
1642.8624267578125 0 1892.9862
1643.859375 0 1521.0769
1644.8660888671875 0 1337.8071
1658.8756103515625 0 3695.659 c 14
1659.8798828125 0 11820.969
1660.8829345703125 0 9544.221
1661.883056640625 0 4313.3115
1662.876708984375 0 1172.1746
1672.847412109375 0 1015.721
1685.8939208984375 0 2732.4727
1686.9053955078125 0 3458.5154
1687.902099609375 0 2763.0125
1688.913818359375 0 1008.73773
1689.9083251953125 0 1239.6312
1699.8450927734375 0 1424.7391 z Ammonia loss 1
1700.8638916015625 0 1221.6735
1713.887939453125 0 4904.2285
1714.8951416015625 0 3557.0032 y Water loss 1
1715.8848876953125 0 3573.3254
1716.8646240234375 0 3912.731 z 1
1717.860595703125 0 9331.384
1718.865478515625 0 7767.077
1719.866455078125 0 3572.08
1720.8802490234375 0 1376.9133
1729.9095458984375 0 5666.0176 c 15
1730.9180908203125 0 12594.802
1731.9212646484375 0 12540.045
1732.9212646484375 0 6198.667
1733.9263916015625 0 2911.5894
1760.9420166015625 0 1147.1172
1770.9217529296875 0 2811.0913
1771.920654296875 0 4364.4917
1772.916748046875 0 3848.8948
1773.9117431640625 0 1006.76685
1774.92626953125 0 1202.5145
1787.93505859375 0 5461.5156
1788.936279296875 0 22040.844
1789.93896484375 0 19804.004
1790.9403076171875 0 11111.916
1791.9427490234375 0 3739.0825
1792.9471435546875 0 1064.81
1798.908447265625 0 1149.8037
1799.9197998046875 0 1663.2184
1804.9483642578125 0 1471.9006
1805.9635009765625 0 4753.7266
1806.962158203125 0 3915.7092
1807.9637451171875 0 2373.2976
1808.964111328125 0 842.7259
1814.9329833984375 0 991.80426
1815.9305419921875 0 5588.0615
1816.93115234375 0 33422.438
1817.93408203125 0 29962.709
1818.9368896484375 0 18452.557
1819.9351806640625 0 6391.639
1820.943359375 0 1409.1987
1831.9415283203125 0 3907.4263
1832.9476318359375 0 19489.297
1833.955322265625 0 57630.895
1834.9598388671875 0 52789.152
1835.9615478515625 0 26490.283
1836.9637451171875 0 10608.607
1837.964111328125 0 3357.6519

Spectrum Details

|  |  |
| --- | --- |
| Matched peaks? Matched peaks The total absolute number of peaks matched. Additionally in brackets the total fraction of peaks matched and the total number of peaks is shown. | 93 (15.55% of 598) |
| FDR? FDR The false discovery rate estimated for this peptide. It is calculated by matching all theoretical fragments with a non-integer shift with the raw peaks for this spectrum. This is done with 40 different shifts. The resulting percentage is the average number of annotated peaks over the number of annotated peaks with the correct spectrum. | 1.05% |
| Satellite FDR? Satellite FDR See the FDR for details on its calculation. This satellite ion specific FDR only contains the satellite ions (d/w) for I/L/J positions. | - |
| PSM Score? PSM Score The PSM Score as given by Hecklib to this annotated spectrum. It is shown with three significant figures. | 833 |

#### Spectrum 7949? Spectrum 7949 The raw spectrum of this peptide as annotated by Hecklib. The fragments are coloured according to ion type (see legend). Any peaks with a star '\*' as text can be hovered over to see the full details, first the ion type second the mass shift type. By hovering over the amino acids in the peptide or ions in the legend the corresponding peaks are highlighted. By toggling the 'Unassigned' label you can turn the background (unassigned) peaks on or off in the plot. By updating the slider in the Ion legend you can update the spectrum to only show the top X% of the peaks with labels. The top X% means any peak that is within X% of the highest intensity. By dragging in the spectrum you can zoom in to a specific part of the spectrum and use 'Zoom Out' to get back to the original zoom level. The annotation of the spectrum is based on the given sequence in the peptides file and is done with different software so inconsistencies are likely. The peaks are annotated based on the given sequence, with 20 ppm tolerance.

Copy Data

##### Spectrum 7949 (TSV)

###### Preview

```
Loading example...
```

*Click on the button to copy the data to your clipboard.*

Mz MinMz MaxIntensity Max

WidthHeightPeptide font sizePeptide stroke widthSpectrum font sizeSpectrum stroke widthCompact peptide

Ion legend

wxyz

abcd

OtherUnassignedIon^Charge^_Position_Show for top:%

VTJFPPSSEEJQANKAT

02.35e+44.70e+47.04e+49.39e+4

Zoom Out

y^+1^_1_y^+1^_2_y^+1^_2_c^+1^_2_c^+1^_2_y^+3^_7_z^+3^_8_z^+1^_3_y^+2^_6_y^+2^_6_y^+1^_3_c^+1^_3_y^+2^_7_y^+1^_4_z^+1^_4_y^+1^_4_y^+3^_13_y^+1^_5_c^+2^_9_y^+2^_9_y^+1^_5_w^+2^_10_z^+3^_15_w^+1^_6_y^+2^_11_c^+2^_11_y^+1^_6_z^+1^_6_y^+2^_12_y^+1^_6_y^+2^_12_c^+2^_12_c^+1^_6_c^+2^_12_y^+2^_13_y^+2^_13_z^+2^_13_y^+2^_13_c^+2^_13_z^+1^_7_z^+2^_14_y^+1^_7_y^+2^_14_y^+2^_14_z^+2^_14_c^+1^_7_y^+2^_14_w^+1^_8_z^+2^_15_y^+2^_15_y^+2^_15_z^+2^_15_y^+2^_15_c^+2^_15_z^+1^_8_z^+1^_8_c^+1^_8_z^+2^_16_z^+2^_16_c^+2^_16_y^+1^_8_y^+2^_16_y^+2^_16_z^+2^_16_c^+2^_16_y^+1^_8_w^+1^_9_z^+1^_9_c^+1^_9_z^+1^_9_y^+1^_9_z^+1^_10_z^+1^_10_y^+1^_10_y^+1^_10_z^+1^_10_y^+1^_10_c^+1^_10_z^+1^_11_y^+1^_11_y^+1^_11_z^+1^_11_y^+1^_11_c^+1^_11_c^+1^_11_y^+1^_12_c^+1^_12_c^+1^_12_y^+1^_13_y^+1^_13_c^+1^_13_y^+1^_14_z^+1^_14_y^+1^_14_c^+1^_14_z^+1^_15_c^+1^_15_z^+1^_16_y^+1^_16_z^+1^_16_c^+1^_16_

0775155123263102

Fragment Matches Table

Show background peaks

| Position | Ion type | Intensity | mz Theoretical | mz Error (Th) | mz Error (ppm) | Charge | Series Number |
| --- | --- | --- | --- | --- | --- | --- | --- |
| 17 | y | 1.152E+04 | 120.1 | 0.0001008 | 0.8396 | +1 | 1 |
| - | - | 2237 | 120.1 | - | - | 0 | - |
| - | - | 363.4 | 121.1 | - | - | 0 | - |
| - | - | 405.9 | 125.9 | - | - | 0 | - |
| - | - | 1777 | 126.1 | - | - | 0 | - |
| - | - | 338 | 126.6 | - | - | 0 | - |
| - | - | 807.9 | 128.1 | - | - | 0 | - |
| - | - | 956.6 | 128.1 | - | - | 0 | - |
| - | - | 5174 | 129.1 | - | - | 0 | - |
| - | - | 436.6 | 130.3 | - | - | 0 | - |
| - | - | 1084 | 131.1 | - | - | 0 | - |
| - | - | 391 | 132.6 | - | - | 0 | - |
| - | - | 379.5 | 136.6 | - | - | 0 | - |
| - | - | 407.2 | 137.9 | - | - | 0 | - |
| - | - | 1009 | 140.1 | - | - | 0 | - |
| - | - | 417.4 | 143.7 | - | - | 0 | - |
| - | - | 1019 | 149 | - | - | 0 | - |
| - | - | 642.7 | 149 | - | - | 0 | - |
| - | - | 861.7 | 155.1 | - | - | 0 | - |
| - | - | 625 | 157.1 | - | - | 0 | - |
| - | - | 537 | 158.1 | - | - | 0 | - |
| - | - | 2864 | 169.1 | - | - | 0 | - |
| - | - | 1249 | 171.1 | - | - | 0 | - |
| 16 | y | 1365 | 173.1 | 0.000308 | 1.779 | +1 | 2 |
| - | - | 3363 | 173.1 | - | - | 0 | - |
| - | - | 1460 | 173.4 | - | - | 0 | - |
| - | - | 2508 | 183.1 | - | - | 0 | - |
| 16 | y | 2857 | 191.1 | 0.000104 | 0.5443 | +1 | 2 |
| - | - | 2298 | 197.1 | - | - | 0 | - |
| 2 | c | 689.8 | 200.1 | 0.0004372 | 2.185 | +1 | 2 |
| - | - | 7348 | 201.1 | - | - | 0 | - |
| - | - | 630.8 | 211.1 | - | - | 0 | - |
| - | - | 9954 | 212.1 | - | - | 0 | - |
| - | - | 1236 | 213.1 | - | - | 0 | - |
| - | - | 750.9 | 215.1 | - | - | 0 | - |
| 2 | c | 4409 | 218.1 | 3.018E-05 | 0.1383 | +1 | 2 |
| - | - | 3218 | 221.1 | - | - | 0 | - |
| - | - | 1058 | 222.1 | - | - | 0 | - |
| - | - | 573.6 | 223.1 | - | - | 0 | - |
| - | - | 1350 | 227.1 | - | - | 0 | - |
| - | - | 1873 | 227.1 | - | - | 0 | - |
| - | - | 1507 | 233.2 | - | - | 0 | - |
| - | - | 863.4 | 235.1 | - | - | 0 | - |
| - | - | 4216 | 239.1 | - | - | 0 | - |
| - | - | 1336 | 240.1 | - | - | 0 | - |
| - | - | 566.8 | 241.1 | - | - | 0 | - |
| - | - | 808.2 | 242.2 | - | - | 0 | - |
| 11 | y | 1134 | 243.1 | 0.004314 | 17.74 | +3 | 7 |
| - | - | 1105 | 254.1 | - | - | 0 | - |
| - | - | 598.4 | 256.1 | - | - | 0 | - |
| - | - | 1251 | 261.2 | - | - | 0 | - |
| - | - | 3035 | 263.1 | - | - | 0 | - |
| - | - | 754.1 | 264.1 | - | - | 0 | - |
| - | - | 777.8 | 280.2 | - | - | 0 | - |
| - | - | 1243 | 281.1 | - | - | 0 | - |
| 10 | z | 801.8 | 281.1 | 0.005165 | 18.37 | +3 | 8 |
| - | - | 624.4 | 281.2 | - | - | 0 | - |
| - | - | 630 | 282.1 | - | - | 0 | - |
| - | - | 965.1 | 282.1 | - | - | 0 | - |
| - | - | 1.181E+04 | 296.2 | - | - | 0 | - |
| - | - | 1136 | 297.2 | - | - | 0 | - |
| - | - | 2066 | 297.2 | - | - | 0 | - |
| - | - | 585.5 | 298.1 | - | - | 0 | - |
| - | - | 846.4 | 298.2 | - | - | 0 | - |
| - | - | 4584 | 299.1 | - | - | 0 | - |
| - | - | 1.493E+04 | 299.2 | - | - | 0 | - |
| - | - | 2242 | 300.1 | - | - | 0 | - |
| - | - | 562.5 | 300.2 | - | - | 0 | - |
| - | - | 2249 | 300.2 | - | - | 0 | - |
| - | - | 1064 | 301.1 | - | - | 0 | - |
| 15 | z | 7922 | 303.2 | 3.934E-05 | 0.1298 | +1 | 3 |
| - | - | 1647 | 304.2 | - | - | 0 | - |
| - | - | 729.8 | 307.2 | - | - | 0 | - |
| 12 | y | 798.4 | 307.7 | 9.922E-06 | 0.03225 | +2 | 6 |
| - | - | 713.4 | 314.2 | - | - | 0 | - |
| - | - | 4762 | 314.2 | - | - | 0 | - |
| - | - | 871.4 | 315.2 | - | - | 0 | - |
| - | - | 7705 | 316.2 | - | - | 0 | - |
| 12 | y | 1149 | 316.7 | 3.764E-05 | 0.1189 | +2 | 6 |
| - | - | 523.8 | 317.1 | - | - | 0 | - |
| - | - | 871.9 | 317.2 | - | - | 0 | - |
| 15 | y | 5298 | 319.2 | 2.562E-05 | 0.08026 | +1 | 3 |
| - | - | 946.9 | 325.2 | - | - | 0 | - |
| 3 | c | 2.766E+04 | 331.2 | 8.79E-05 | 0.2654 | +1 | 3 |
| - | - | 5090 | 332.2 | - | - | 0 | - |
| - | - | 669 | 333.2 | - | - | 0 | - |
| - | - | 593.2 | 344.2 | - | - | 0 | - |
| - | - | 867.7 | 345.2 | - | - | 0 | - |
| - | - | 658.5 | 346.2 | - | - | 0 | - |
| - | - | 1185 | 347.2 | - | - | 0 | - |
| - | - | 895 | 350.1 | - | - | 0 | - |
| - | - | 3.708E+04 | 355.1 | - | - | 0 | - |
| - | - | 1.617E+04 | 356.1 | - | - | 0 | - |
| - | - | 8737 | 357.1 | - | - | 0 | - |
| - | - | 882.5 | 358.1 | - | - | 0 | - |
| - | - | 1692 | 362.2 | - | - | 0 | - |
| - | - | 811.2 | 369.2 | - | - | 0 | - |
| - | - | 1106 | 371.1 | - | - | 0 | - |
| - | - | 989.6 | 372.1 | - | - | 0 | - |
| - | - | 1045 | 373.1 | - | - | 0 | - |
| 11 | y | 4325 | 373.2 | 0.004732 | 12.68 | +2 | 7 |
| - | - | 1329 | 378.2 | - | - | 0 | - |
| - | - | 781.5 | 385.2 | - | - | 0 | - |
| - | - | 1325 | 386.2 | - | - | 0 | - |
| - | - | 1792 | 415 | - | - | 0 | - |
| 14 | y | 905.4 | 415.2 | 0.002249 | 5.416 | +1 | 4 |
| - | - | 1112 | 416 | - | - | 0 | - |
| - | - | 911.9 | 417 | - | - | 0 | - |
| 14 | z | 2614 | 417.2 | 0.00107 | 2.565 | +1 | 4 |
| - | - | 7721 | 418.2 | - | - | 0 | - |
| - | - | 2174 | 419.2 | - | - | 0 | - |
| - | - | 813.1 | 426.2 | - | - | 0 | - |
| 14 | y | 2721 | 433.2 | 0.001755 | 4.051 | +1 | 4 |
| - | - | 8883 | 433.3 | - | - | 0 | - |
| - | - | 598.7 | 434.3 | - | - | 0 | - |
| - | - | 4960 | 434.3 | - | - | 0 | - |
| - | - | 678.7 | 434.7 | - | - | 0 | - |
| - | - | 2119 | 435.3 | - | - | 0 | - |
| - | - | 6883 | 444.2 | - | - | 0 | - |
| - | - | 611 | 445.2 | - | - | 0 | - |
| - | - | 2406 | 445.3 | - | - | 0 | - |
| - | - | 875.6 | 446.2 | - | - | 0 | - |
| - | - | 1780 | 446.3 | - | - | 0 | - |
| 5 | y | 861.6 | 451.9 | 0.001335 | 2.955 | +3 | 13 |
| - | - | 723.3 | 453.2 | - | - | 0 | - |
| - | - | 1503 | 460.3 | - | - | 0 | - |
| - | - | 1.288E+04 | 461.3 | - | - | 0 | - |
| - | - | 3667 | 462.3 | - | - | 0 | - |
| - | - | 605.1 | 472.2 | - | - | 0 | - |
| - | - | 698 | 473.2 | - | - | 0 | - |
| 13 | y | 760.3 | 486.3 | 0.0005052 | 1.039 | +1 | 5 |
| 9 | c | 1.127E+04 | 488.3 | 0.001739 | 3.562 | +2 | 9 |
| - | - | 1125 | 488.7 | - | - | 0 | - |
| - | - | 861.6 | 489.1 | - | - | 0 | - |
| - | - | 7632 | 489.3 | - | - | 0 | - |
| - | - | 1230 | 490.1 | - | - | 0 | - |
| - | - | 1492 | 490.3 | - | - | 0 | - |
| - | - | 1757 | 493.7 | - | - | 0 | - |
| - | - | 1422 | 494.2 | - | - | 0 | - |
| - | - | 713.9 | 496.2 | - | - | 0 | - |
| - | - | 979 | 497.3 | - | - | 0 | - |
| - | - | 1175 | 498.2 | - | - | 0 | - |
| 9 | y | 1891 | 502.3 | 0.008674 | 17.27 | +2 | 9 |
| - | - | 2414 | 502.7 | - | - | 0 | - |
| - | - | 1187 | 503.2 | - | - | 0 | - |
| - | - | 1033 | 503.3 | - | - | 0 | - |
| - | - | 589.4 | 503.7 | - | - | 0 | - |
| 13 | y | 8432 | 504.3 | 0.0002859 | 0.567 | +1 | 5 |
| - | - | 1866 | 505.3 | - | - | 0 | - |
| - | - | 998.7 | 515.3 | - | - | 0 | - |
| 8 | w | 795.7 | 529.3 | 0.002202 | 4.16 | +2 | 10 |
| - | - | 765.3 | 529.8 | - | - | 0 | - |
| - | - | 7050 | 538.3 | - | - | 0 | - |
| - | - | 2664 | 538.8 | - | - | 0 | - |
| 3 | z | 2239 | 539.3 | 0.001625 | 3.013 | +3 | 15 |
| - | - | 725.5 | 540.3 | - | - | 0 | - |
| - | - | 1732 | 543.3 | - | - | 0 | - |
| - | - | 1348 | 543.8 | - | - | 0 | - |
| - | - | 752.2 | 544.3 | - | - | 0 | - |
| - | - | 921.5 | 544.8 | - | - | 0 | - |
| - | - | 5422 | 552.3 | - | - | 0 | - |
| - | - | 3865 | 552.8 | - | - | 0 | - |
| - | - | 693.1 | 554.3 | - | - | 0 | - |
| - | - | 1009 | 557.3 | - | - | 0 | - |
| 12 | w | 1.564E+04 | 558.3 | 0.0001774 | 0.3178 | +1 | 6 |
| - | - | 2311 | 558.3 | - | - | 0 | - |
| - | - | 3847 | 559.3 | - | - | 0 | - |
| - | - | 1542 | 559.3 | - | - | 0 | - |
| - | - | 737.7 | 560.3 | - | - | 0 | - |
| - | - | 849.8 | 572.3 | - | - | 0 | - |
| - | - | 1069 | 577.1 | - | - | 0 | - |
| - | - | 701.2 | 578.1 | - | - | 0 | - |
| - | - | 649 | 579.1 | - | - | 0 | - |
| 7 | y | 594.3 | 580.8 | 0.002829 | 4.872 | +2 | 11 |
| - | - | 865.1 | 582.3 | - | - | 0 | - |
| - | - | 876.6 | 582.8 | - | - | 0 | - |
| - | - | 1557 | 583.3 | - | - | 0 | - |
| - | - | 708.4 | 599.3 | - | - | 0 | - |
| - | - | 2208 | 599.8 | - | - | 0 | - |
| 11 | c | 1204 | 600.3 | 0.01013 | 16.88 | +2 | 11 |
| - | - | 747.9 | 600.8 | - | - | 0 | - |
| - | - | 3201 | 602.8 | - | - | 0 | - |
| - | - | 2786 | 603.3 | - | - | 0 | - |
| - | - | 8862 | 610.2 | - | - | 0 | - |
| - | - | 551 | 610.4 | - | - | 0 | - |
| - | - | 9045 | 611.2 | - | - | 0 | - |
| - | - | 9002 | 611.8 | - | - | 0 | - |
| - | - | 6785 | 612.2 | - | - | 0 | - |
| - | - | 1050 | 612.2 | - | - | 0 | - |
| - | - | 5931 | 612.3 | - | - | 0 | - |
| - | - | 1640 | 612.8 | - | - | 0 | - |
| - | - | 998 | 613.2 | - | - | 0 | - |
| 12 | y | 8225 | 614.3 | 0.01169 | 19.03 | +1 | 6 |
| - | - | 1.036E+04 | 615.3 | - | - | 0 | - |
| 12 | z | 3458 | 616.3 | 0.001878 | 3.048 | +1 | 6 |
| - | - | 1696 | 616.4 | - | - | 0 | - |
| - | - | 898.5 | 617.3 | - | - | 0 | - |
| - | - | 1093 | 617.8 | - | - | 0 | - |
| - | - | 1605 | 618.3 | - | - | 0 | - |
| - | - | 929.3 | 618.8 | - | - | 0 | - |
| - | - | 3144 | 626.3 | - | - | 0 | - |
| - | - | 5457 | 626.8 | - | - | 0 | - |
| - | - | 5097 | 627.3 | - | - | 0 | - |
| - | - | 2242 | 627.3 | - | - | 0 | - |
| - | - | 1631 | 628.3 | - | - | 0 | - |
| - | - | 3145 | 628.4 | - | - | 0 | - |
| 6 | y | 814.3 | 628.8 | 0.0007831 | 1.245 | +2 | 12 |
| - | - | 1906 | 629.4 | - | - | 0 | - |
| - | - | 705.4 | 630.4 | - | - | 0 | - |
| - | - | 862.4 | 630.4 | - | - | 0 | - |
| 12 | y | 6222 | 632.3 | 2.751E-05 | 0.0435 | +1 | 6 |
| - | - | 1917 | 633.3 | - | - | 0 | - |
| - | - | 809.1 | 634.3 | - | - | 0 | - |
| - | - | 974.5 | 635.3 | - | - | 0 | - |
| - | - | 1559 | 635.8 | - | - | 0 | - |
| 6 | y | 924.2 | 637.8 | 0.004845 | 7.596 | +2 | 12 |
| - | - | 2887 | 655.3 | - | - | 0 | - |
| - | - | 2754 | 655.8 | - | - | 0 | - |
| - | - | 763.5 | 656.3 | - | - | 0 | - |
| - | - | 928.7 | 658.4 | - | - | 0 | - |
| 12 | c | 1332 | 664.8 | 0.003842 | 5.779 | +2 | 12 |
| - | - | 1194 | 667.4 | - | - | 0 | - |
| - | - | 921.3 | 668.8 | - | - | 0 | - |
| - | - | 2319 | 671.4 | - | - | 0 | - |
| 6 | c | 3716 | 672.4 | 2.56E-05 | 0.03807 | +1 | 6 |
| 12 | c | 627 | 673.4 | 0.01175 | 17.44 | +2 | 12 |
| - | - | 2140 | 673.4 | - | - | 0 | - |
| - | - | 2184 | 676.8 | - | - | 0 | - |
| 5 | y | 7750 | 677.3 | 5.668E-05 | 0.08369 | +2 | 13 |
| 5 | y | 7599 | 677.8 | 0.007927 | 11.69 | +2 | 13 |
| 5 | z | 2774 | 678.3 | 0.004075 | 6.008 | +2 | 13 |
| - | - | 2158 | 678.8 | - | - | 0 | - |
| - | - | 810.6 | 685.3 | - | - | 0 | - |
| - | - | 4574 | 685.8 | - | - | 0 | - |
| 5 | y | 5.648E+04 | 686.3 | 0.002526 | 3.68 | +2 | 13 |
| - | - | 2.152E+04 | 686.8 | - | - | 0 | - |
| - | - | 1.862E+04 | 687.3 | - | - | 0 | - |
| - | - | 2560 | 687.8 | - | - | 0 | - |
| - | - | 1845 | 688.4 | - | - | 0 | - |
| - | - | 1586 | 697.4 | - | - | 0 | - |
| - | - | 1218 | 698.4 | - | - | 0 | - |
| 13 | c | 1051 | 700.4 | 0.009459 | 13.51 | +2 | 13 |
| - | - | 1451 | 716.4 | - | - | 0 | - |
| - | - | 2016 | 722.3 | - | - | 0 | - |
| - | - | 1046 | 723.3 | - | - | 0 | - |
| 11 | z | 1.258E+04 | 729.4 | 0.0006762 | 0.927 | +1 | 7 |
| - | - | 5008 | 730.4 | - | - | 0 | - |
| - | - | 785.3 | 731.4 | - | - | 0 | - |
| - | - | 702.8 | 738.4 | - | - | 0 | - |
| - | - | 1697 | 740.3 | - | - | 0 | - |
| - | - | 1774 | 742.4 | - | - | 0 | - |
| 4 | z | 784.7 | 742.9 | 0.0009076 | 1.222 | +2 | 14 |
| - | - | 1186 | 743.4 | - | - | 0 | - |
| 11 | y | 6625 | 745.4 | 0.0004362 | 0.5852 | +1 | 7 |
| - | - | 3063 | 746.4 | - | - | 0 | - |
| 4 | y | 3444 | 750.9 | 0.000764 | 1.018 | +2 | 14 |
| 4 | y | 2867 | 751.4 | 0.008144 | 10.84 | +2 | 14 |
| 4 | z | 1473 | 751.9 | 0.000447 | 0.5945 | +2 | 14 |
| - | - | 846.1 | 752.4 | - | - | 0 | - |
| - | - | 2162 | 757.4 | - | - | 0 | - |
| - | - | 1823 | 758.4 | - | - | 0 | - |
| - | - | 2632 | 759.4 | - | - | 0 | - |
| 7 | c | 3.355E+04 | 759.4 | 5.048E-05 | 0.06648 | +1 | 7 |
| 4 | y | 1.155E+04 | 759.9 | 0.0006143 | 0.8084 | +2 | 14 |
| - | - | 1.039E+04 | 760.4 | - | - | 0 | - |
| - | - | 1.35E+04 | 760.4 | - | - | 0 | - |
| - | - | 3554 | 760.9 | - | - | 0 | - |
| - | - | 1545 | 761.4 | - | - | 0 | - |
| - | - | 3375 | 761.4 | - | - | 0 | - |
| - | - | 5616 | 764.4 | - | - | 0 | - |
| - | - | 3534 | 765.4 | - | - | 0 | - |
| - | - | 1098 | 766.4 | - | - | 0 | - |
| - | - | 1203 | 798.9 | - | - | 0 | - |
| 10 | w | 1.321E+04 | 799.4 | 0.0003018 | 0.3776 | +1 | 8 |
| 3 | z | 1093 | 799.9 | 0.011 | 13.75 | +2 | 15 |
| - | - | 7236 | 800.4 | - | - | 0 | - |
| - | - | 1547 | 801.4 | - | - | 0 | - |
| - | - | 6079 | 802.4 | - | - | 0 | - |
| - | - | 2965 | 803.4 | - | - | 0 | - |
| 3 | y | 1721 | 807.4 | 0.003347 | 4.145 | +2 | 15 |
| 3 | y | 2075 | 807.9 | 0.009447 | 11.69 | +2 | 15 |
| 3 | z | 1211 | 808.4 | 0.01292 | 15.98 | +2 | 15 |
| - | - | 1236 | 808.9 | - | - | 0 | - |
| - | - | 714.8 | 812.9 | - | - | 0 | - |
| - | - | 1634 | 813.4 | - | - | 0 | - |
| - | - | 953.5 | 813.9 | - | - | 0 | - |
| - | - | 843 | 814.4 | - | - | 0 | - |
| - | - | 1232 | 815.4 | - | - | 0 | - |
| 3 | y | 2878 | 816.4 | 0.003584 | 4.39 | +2 | 15 |
| - | - | 2215 | 816.9 | - | - | 0 | - |
| - | - | 965.9 | 817.4 | - | - | 0 | - |
| - | - | 3968 | 822.4 | - | - | 0 | - |
| - | - | 5093 | 823.4 | - | - | 0 | - |
| - | - | 1920 | 824.4 | - | - | 0 | - |
| - | - | 1434 | 829.4 | - | - | 0 | - |
| 15 | c | 3.702E+04 | 829.9 | 0.0001264 | 0.1523 | +2 | 15 |
| - | - | 3.4E+04 | 830.4 | - | - | 0 | - |
| - | - | 1.616E+04 | 830.9 | - | - | 0 | - |
| - | - | 5920 | 831.4 | - | - | 0 | - |
| - | - | 1006 | 832 | - | - | 0 | - |
| - | - | 896 | 834.9 | - | - | 0 | - |
| - | - | 1318 | 836.4 | - | - | 0 | - |
| - | - | 670 | 836.9 | - | - | 0 | - |
| 10 | z | 1227 | 840.4 | 0.0006206 | 0.7384 | +1 | 8 |
| 10 | z | 992.4 | 841.4 | 0.009159 | 10.88 | +1 | 8 |
| - | - | 1332 | 842.4 | - | - | 0 | - |
| - | - | 3204 | 843.5 | - | - | 0 | - |
| - | - | 2530 | 844 | - | - | 0 | - |
| - | - | 2207 | 844.5 | - | - | 0 | - |
| - | - | 919.4 | 845 | - | - | 0 | - |
| 8 | c | 2.926E+04 | 846.5 | 0.0001876 | 0.2216 | +1 | 8 |
| - | - | 1.419E+04 | 847.5 | - | - | 0 | - |
| - | - | 4388 | 848.5 | - | - | 0 | - |
| - | - | 1783 | 848.9 | - | - | 0 | - |
| - | - | 2082 | 849.4 | - | - | 0 | - |
| 2 | z | 3479 | 849.9 | 0.001809 | 2.128 | +2 | 16 |
| 2 | z | 2209 | 850.4 | 0.004003 | 4.707 | +2 | 16 |
| - | - | 1968 | 850.9 | - | - | 0 | - |
| - | - | 877.1 | 852.4 | - | - | 0 | - |
| 16 | c | 2652 | 856.9 | 3.766E-05 | 0.04395 | +2 | 16 |
| 10 | y | 4822 | 857.4 | 0.007891 | 9.203 | +1 | 8 |
| 2 | y | 3648 | 857.9 | 0.01051 | 12.25 | +2 | 16 |
| 2 | y | 2.368E+04 | 858.4 | 0.01643 | 19.14 | +2 | 16 |
| 2 | z | 4011 | 858.9 | 0.008367 | 9.741 | +2 | 16 |
| - | - | 1.081E+04 | 859.4 | - | - | 0 | - |
| - | - | 1346 | 859.9 | - | - | 0 | - |
| - | - | 2935 | 860.4 | - | - | 0 | - |
| - | - | 1139 | 864 | - | - | 0 | - |
| - | - | 4760 | 865 | - | - | 0 | - |
| 16 | c | 4.939E+04 | 865.5 | 0.0002376 | 0.2745 | +2 | 16 |
| - | - | 5.132E+04 | 866 | - | - | 0 | - |
| - | - | 2.704E+04 | 866.5 | - | - | 0 | - |
| - | - | 1.186E+04 | 867 | - | - | 0 | - |
| - | - | 2998 | 867.5 | - | - | 0 | - |
| - | - | 2023 | 868 | - | - | 0 | - |
| - | - | 2352 | 868.4 | - | - | 0 | - |
| - | - | 1270 | 869.4 | - | - | 0 | - |
| - | - | 967.7 | 871.4 | - | - | 0 | - |
| - | - | 1093 | 872.5 | - | - | 0 | - |
| - | - | 1601 | 872.9 | - | - | 0 | - |
| - | - | 3510 | 873.5 | - | - | 0 | - |
| 10 | y | 4794 | 874.5 | 0.00108 | 1.235 | +1 | 8 |
| - | - | 1862 | 875.5 | - | - | 0 | - |
| - | - | 818.5 | 878.5 | - | - | 0 | - |
| - | - | 4399 | 880.5 | - | - | 0 | - |
| - | - | 7555 | 881 | - | - | 0 | - |
| - | - | 7463 | 881.5 | - | - | 0 | - |
| - | - | 3712 | 882 | - | - | 0 | - |
| - | - | 2176 | 882.5 | - | - | 0 | - |
| - | - | 758.3 | 883 | - | - | 0 | - |
| - | - | 3354 | 885.5 | - | - | 0 | - |
| - | - | 4561 | 886 | - | - | 0 | - |
| - | - | 2867 | 886.5 | - | - | 0 | - |
| - | - | 1097 | 887 | - | - | 0 | - |
| - | - | 2454 | 887.5 | - | - | 0 | - |
| - | - | 1464 | 888 | - | - | 0 | - |
| - | - | 1029 | 888.5 | - | - | 0 | - |
| - | - | 2218 | 888.9 | - | - | 0 | - |
| - | - | 1512 | 889.5 | - | - | 0 | - |
| - | - | 1186 | 894 | - | - | 0 | - |
| - | - | 2.239E+04 | 894.5 | - | - | 0 | - |
| - | - | 2.012E+04 | 895 | - | - | 0 | - |
| - | - | 1.249E+04 | 895.5 | - | - | 0 | - |
| - | - | 6082 | 896 | - | - | 0 | - |
| - | - | 1266 | 896.5 | - | - | 0 | - |
| - | - | 2114 | 899.5 | - | - | 0 | - |
| - | - | 2253 | 900 | - | - | 0 | - |
| - | - | 1569 | 900.5 | - | - | 0 | - |
| - | - | 892 | 902.5 | - | - | 0 | - |
| - | - | 9515 | 907.4 | - | - | 0 | - |
| - | - | 4270 | 908 | - | - | 0 | - |
| - | - | 2894 | 908.4 | - | - | 0 | - |
| - | - | 3897 | 908.5 | - | - | 0 | - |
| - | - | 5410 | 909 | - | - | 0 | - |
| - | - | 832.3 | 910 | - | - | 0 | - |
| - | - | 1453 | 915.5 | - | - | 0 | - |
| - | - | 5.516E+04 | 916.5 | - | - | 0 | - |
| - | - | 9.299E+04 | 917 | - | - | 0 | - |
| - | - | 6.96E+04 | 917.5 | - | - | 0 | - |
| - | - | 3.16E+04 | 918 | - | - | 0 | - |
| - | - | 1.217E+04 | 918.5 | - | - | 0 | - |
| - | - | 3528 | 919 | - | - | 0 | - |
| 9 | w | 6013 | 928.5 | 0.0006586 | 0.7093 | +1 | 9 |
| - | - | 6385 | 929.5 | - | - | 0 | - |
| - | - | 2366 | 930.5 | - | - | 0 | - |
| - | - | 1007 | 931.5 | - | - | 0 | - |
| - | - | 3096 | 944.5 | - | - | 0 | - |
| - | - | 3597 | 945.5 | - | - | 0 | - |
| - | - | 1772 | 946.5 | - | - | 0 | - |
| - | - | 905.9 | 957.5 | - | - | 0 | - |
| - | - | 1474 | 958.5 | - | - | 0 | - |
| - | - | 1468 | 959.5 | - | - | 0 | - |
| - | - | 5485 | 960.5 | - | - | 0 | - |
| - | - | 3101 | 961.5 | - | - | 0 | - |
| 9 | z | 1148 | 969.5 | 0.0131 | 13.52 | +1 | 9 |
| - | - | 1.963E+04 | 973.5 | - | - | 0 | - |
| - | - | 9534 | 974.5 | - | - | 0 | - |
| 9 | c | 4.414E+04 | 975.5 | 0.0007795 | 0.7991 | +1 | 9 |
| - | - | 2.4E+04 | 976.5 | - | - | 0 | - |
| - | - | 7543 | 977.5 | - | - | 0 | - |
| - | - | 1539 | 978.5 | - | - | 0 | - |
| 9 | z | 2.04E+04 | 987.5 | 0.0005352 | 0.542 | +1 | 9 |
| - | - | 1.169E+04 | 988.5 | - | - | 0 | - |
| - | - | 3876 | 989.5 | - | - | 0 | - |
| - | - | 1154 | 990.5 | - | - | 0 | - |
| - | - | 1036 | 1002 | - | - | 0 | - |
| - | - | 909.6 | 1003 | - | - | 0 | - |
| 9 | y | 5558 | 1004 | 0.00986 | 9.825 | +1 | 9 |
| - | - | 3150 | 1005 | - | - | 0 | - |
| - | - | 1237 | 1006 | - | - | 0 | - |
| - | - | 1505 | 1013 | - | - | 0 | - |
| - | - | 923.4 | 1015 | - | - | 0 | - |
| - | - | 1107 | 1017 | - | - | 0 | - |
| - | - | 1346 | 1018 | - | - | 0 | - |
| - | - | 758.9 | 1019 | - | - | 0 | - |
| - | - | 973.3 | 1031 | - | - | 0 | - |
| - | - | 1001 | 1032 | - | - | 0 | - |
| - | - | 826.3 | 1053 | - | - | 0 | - |
| - | - | 4543 | 1056 | - | - | 0 | - |
| 8 | z | 3704 | 1057 | 2.628E-05 | 0.02488 | +1 | 10 |
| 8 | z | 1698 | 1057 | 0.01828 | 17.28 | +1 | 10 |
| - | - | 1169 | 1059 | - | - | 0 | - |
| - | - | 2867 | 1061 | - | - | 0 | - |
| - | - | 1971 | 1062 | - | - | 0 | - |
| - | - | 860 | 1071 | - | - | 0 | - |
| - | - | 1249 | 1072 | - | - | 0 | - |
| 8 | y | 1593 | 1073 | 0.02107 | 19.64 | +1 | 10 |
| 8 | y | 1730 | 1074 | 0.000652 | 0.6074 | +1 | 10 |
| 8 | z | 2.043E+04 | 1075 | 0.0005812 | 0.5409 | +1 | 10 |
| - | - | 1.568E+04 | 1076 | - | - | 0 | - |
| - | - | 6975 | 1077 | - | - | 0 | - |
| - | - | 2245 | 1078 | - | - | 0 | - |
| - | - | 4040 | 1088 | - | - | 0 | - |
| - | - | 2561 | 1089 | - | - | 0 | - |
| - | - | 1506 | 1090 | - | - | 0 | - |
| 8 | y | 1.37E+04 | 1091 | 0.001361 | 1.248 | +1 | 10 |
| - | - | 7676 | 1092 | - | - | 0 | - |
| - | - | 2734 | 1093 | - | - | 0 | - |
| - | - | 817.7 | 1094 | - | - | 0 | - |
| - | - | 1543 | 1099 | - | - | 0 | - |
| - | - | 1488 | 1100 | - | - | 0 | - |
| - | - | 2791 | 1104 | - | - | 0 | - |
| 10 | c | 4.665E+04 | 1105 | 0.00138 | 1.25 | +1 | 10 |
| - | - | 3.146E+04 | 1106 | - | - | 0 | - |
| - | - | 1.143E+04 | 1107 | - | - | 0 | - |
| - | - | 2952 | 1108 | - | - | 0 | - |
| - | - | 882.1 | 1118 | - | - | 0 | - |
| - | - | 904.2 | 1119 | - | - | 0 | - |
| - | - | 8278 | 1121 | - | - | 0 | - |
| - | - | 5892 | 1122 | - | - | 0 | - |
| - | - | 1952 | 1123 | - | - | 0 | - |
| - | - | 1138 | 1134 | - | - | 0 | - |
| 7 | z | 1069 | 1144 | 4.98E-05 | 0.04355 | +1 | 11 |
| 7 | y | 1214 | 1160 | 0.01806 | 15.58 | +1 | 11 |
| 7 | y | 887.4 | 1161 | 0.0008588 | 0.74 | +1 | 11 |
| 7 | z | 1.002E+04 | 1162 | 0.0006272 | 0.54 | +1 | 11 |
| - | - | 6896 | 1163 | - | - | 0 | - |
| - | - | 3804 | 1164 | - | - | 0 | - |
| - | - | 2255 | 1165 | - | - | 0 | - |
| 7 | y | 6402 | 1178 | 0.0003021 | 0.2565 | +1 | 11 |
| - | - | 4414 | 1179 | - | - | 0 | - |
| - | - | 1446 | 1180 | - | - | 0 | - |
| - | - | 1420 | 1181 | - | - | 0 | - |
| - | - | 1905 | 1182 | - | - | 0 | - |
| - | - | 1200 | 1183 | - | - | 0 | - |
| - | - | 918.5 | 1186 | - | - | 0 | - |
| - | - | 2232 | 1199 | - | - | 0 | - |
| 11 | c | 1634 | 1200 | 0.02251 | 18.77 | +1 | 11 |
| - | - | 1632 | 1201 | - | - | 0 | - |
| - | - | 1248 | 1202 | - | - | 0 | - |
| - | - | 1167 | 1206 | - | - | 0 | - |
| - | - | 1.164E+04 | 1207 | - | - | 0 | - |
| - | - | 8976 | 1208 | - | - | 0 | - |
| - | - | 2427 | 1209 | - | - | 0 | - |
| 11 | c | 3.038E+04 | 1218 | 0.001704 | 1.4 | +1 | 11 |
| - | - | 1.967E+04 | 1219 | - | - | 0 | - |
| - | - | 8611 | 1220 | - | - | 0 | - |
| - | - | 2767 | 1221 | - | - | 0 | - |
| - | - | 7141 | 1223 | - | - | 0 | - |
| - | - | 1.14E+04 | 1224 | - | - | 0 | - |
| - | - | 7047 | 1225 | - | - | 0 | - |
| - | - | 1483 | 1226 | - | - | 0 | - |
| - | - | 971.4 | 1236 | - | - | 0 | - |
| - | - | 2736 | 1253 | - | - | 0 | - |
| - | - | 1475 | 1254 | - | - | 0 | - |
| - | - | 1415 | 1270 | - | - | 0 | - |
| - | - | 1191 | 1271 | - | - | 0 | - |
| 6 | y | 3085 | 1275 | 0.0005819 | 0.4565 | +1 | 12 |
| - | - | 2079 | 1276 | - | - | 0 | - |
| - | - | 1063 | 1277 | - | - | 0 | - |
| - | - | 1142 | 1302 | - | - | 0 | - |
| - | - | 1521 | 1303 | - | - | 0 | - |
| - | - | 781.3 | 1304 | - | - | 0 | - |
| 12 | c | 2720 | 1329 | 0.02079 | 15.65 | +1 | 12 |
| - | - | 2908 | 1330 | - | - | 0 | - |
| - | - | 1692 | 1331 | - | - | 0 | - |
| - | - | 4216 | 1345 | - | - | 0 | - |
| 12 | c | 3.969E+04 | 1346 | 0.002176 | 1.617 | +1 | 12 |
| - | - | 3.08E+04 | 1347 | - | - | 0 | - |
| - | - | 1.279E+04 | 1348 | - | - | 0 | - |
| - | - | 3850 | 1349 | - | - | 0 | - |
| 5 | y | 1143 | 1355 | 0.01495 | 11.04 | +1 | 13 |
| - | - | 753.5 | 1358 | - | - | 0 | - |
| - | - | 2117 | 1371 | - | - | 0 | - |
| 5 | y | 1.181E+04 | 1372 | 0.003053 | 2.226 | +1 | 13 |
| - | - | 8972 | 1373 | - | - | 0 | - |
| - | - | 4721 | 1374 | - | - | 0 | - |
| - | - | 1428 | 1375 | - | - | 0 | - |
| - | - | 1555 | 1399 | - | - | 0 | - |
| - | - | 3684 | 1400 | - | - | 0 | - |
| - | - | 5318 | 1401 | - | - | 0 | - |
| - | - | 5105 | 1402 | - | - | 0 | - |
| - | - | 2702 | 1403 | - | - | 0 | - |
| - | - | 953 | 1404 | - | - | 0 | - |
| - | - | 5353 | 1416 | - | - | 0 | - |
| 13 | c | 2.715E+04 | 1417 | 0.003401 | 2.401 | +1 | 13 |
| - | - | 2.122E+04 | 1418 | - | - | 0 | - |
| - | - | 7748 | 1419 | - | - | 0 | - |
| - | - | 3134 | 1420 | - | - | 0 | - |
| - | - | 2321 | 1431 | - | - | 0 | - |
| - | - | 2906 | 1432 | - | - | 0 | - |
| - | - | 1320 | 1433 | - | - | 0 | - |
| - | - | 818.9 | 1444 | - | - | 0 | - |
| - | - | 930.4 | 1486 | - | - | 0 | - |
| - | - | 2174 | 1487 | - | - | 0 | - |
| - | - | 2048 | 1488 | - | - | 0 | - |
| - | - | 1035 | 1489 | - | - | 0 | - |
| 4 | y | 898.6 | 1502 | 0.007206 | 4.799 | +1 | 14 |
| 4 | z | 2.391E+04 | 1503 | 0.002938 | 1.955 | +1 | 14 |
| - | - | 3.587E+04 | 1504 | - | - | 0 | - |
| - | - | 2.491E+04 | 1505 | - | - | 0 | - |
| - | - | 1.077E+04 | 1506 | - | - | 0 | - |
| - | - | 3230 | 1507 | - | - | 0 | - |
| - | - | 772 | 1508 | - | - | 0 | - |
| 4 | y | 3203 | 1519 | 0.0004327 | 0.2849 | +1 | 14 |
| - | - | 2574 | 1520 | - | - | 0 | - |
| - | - | 2078 | 1521 | - | - | 0 | - |
| - | - | 881 | 1522 | - | - | 0 | - |
| - | - | 949.3 | 1529 | - | - | 0 | - |
| - | - | 1214 | 1530 | - | - | 0 | - |
| 14 | c | 4.52E+04 | 1531 | 0.002628 | 1.717 | +1 | 14 |
| - | - | 4.18E+04 | 1532 | - | - | 0 | - |
| - | - | 1.909E+04 | 1533 | - | - | 0 | - |
| - | - | 7935 | 1534 | - | - | 0 | - |
| - | - | 1906 | 1535 | - | - | 0 | - |
| - | - | 1137 | 1544 | - | - | 0 | - |
| - | - | 815.6 | 1545 | - | - | 0 | - |
| - | - | 3222 | 1615 | - | - | 0 | - |
| 3 | z | 5212 | 1616 | 0.01554 | 9.616 | +1 | 15 |
| - | - | 9227 | 1617 | - | - | 0 | - |
| - | - | 6891 | 1618 | - | - | 0 | - |
| - | - | 2141 | 1619 | - | - | 0 | - |
| - | - | 698.2 | 1633 | - | - | 0 | - |
| - | - | 2100 | 1643 | - | - | 0 | - |
| - | - | 1970 | 1644 | - | - | 0 | - |
| - | - | 918 | 1645 | - | - | 0 | - |
| 15 | c | 4788 | 1659 | 0.0009109 | 0.5491 | +1 | 15 |
| - | - | 1.283E+04 | 1660 | - | - | 0 | - |
| - | - | 9232 | 1661 | - | - | 0 | - |
| - | - | 4350 | 1662 | - | - | 0 | - |
| - | - | 2094 | 1663 | - | - | 0 | - |
| - | - | 1501 | 1673 | - | - | 0 | - |
| - | - | 3300 | 1686 | - | - | 0 | - |
| - | - | 3624 | 1687 | - | - | 0 | - |
| - | - | 3130 | 1688 | - | - | 0 | - |
| - | - | 1906 | 1689 | - | - | 0 | - |
| 2 | z | 1599 | 1700 | 0.02261 | 13.3 | +1 | 16 |
| - | - | 1618 | 1701 | - | - | 0 | - |
| - | - | 3876 | 1714 | - | - | 0 | - |
| 2 | y | 4095 | 1715 | 0.03258 | 19 | +1 | 16 |
| - | - | 3250 | 1716 | - | - | 0 | - |
| 2 | z | 3708 | 1717 | 0.00924 | 5.382 | +1 | 16 |
| - | - | 1.082E+04 | 1718 | - | - | 0 | - |
| - | - | 8672 | 1719 | - | - | 0 | - |
| - | - | 4608 | 1720 | - | - | 0 | - |
| - | - | 1881 | 1721 | - | - | 0 | - |
| - | - | 1020 | 1722 | - | - | 0 | - |
| 16 | c | 4579 | 1730 | 0.004211 | 2.434 | +1 | 16 |
| - | - | 1.49E+04 | 1731 | - | - | 0 | - |
| - | - | 1.428E+04 | 1732 | - | - | 0 | - |
| - | - | 7508 | 1733 | - | - | 0 | - |
| - | - | 2937 | 1734 | - | - | 0 | - |
| - | - | 1190 | 1761 | - | - | 0 | - |
| - | - | 1572 | 1762 | - | - | 0 | - |
| - | - | 1033 | 1763 | - | - | 0 | - |
| - | - | 3069 | 1771 | - | - | 0 | - |
| - | - | 3556 | 1772 | - | - | 0 | - |
| - | - | 4200 | 1773 | - | - | 0 | - |
| - | - | 1348 | 1774 | - | - | 0 | - |
| - | - | 1567 | 1775 | - | - | 0 | - |
| - | - | 845.1 | 1777 | - | - | 0 | - |
| - | - | 863.4 | 1787 | - | - | 0 | - |
| - | - | 6278 | 1788 | - | - | 0 | - |
| - | - | 2.573E+04 | 1789 | - | - | 0 | - |
| - | - | 2.071E+04 | 1790 | - | - | 0 | - |
| - | - | 1.233E+04 | 1791 | - | - | 0 | - |
| - | - | 4196 | 1792 | - | - | 0 | - |
| - | - | 1705 | 1793 | - | - | 0 | - |
| - | - | 907.4 | 1800 | - | - | 0 | - |
| - | - | 883.7 | 1801 | - | - | 0 | - |
| - | - | 1207 | 1805 | - | - | 0 | - |
| - | - | 5033 | 1806 | - | - | 0 | - |
| - | - | 4494 | 1807 | - | - | 0 | - |
| - | - | 2967 | 1808 | - | - | 0 | - |
| - | - | 1335 | 1815 | - | - | 0 | - |
| - | - | 8253 | 1816 | - | - | 0 | - |
| - | - | 3.579E+04 | 1817 | - | - | 0 | - |
| - | - | 3.484E+04 | 1818 | - | - | 0 | - |
| - | - | 1.816E+04 | 1819 | - | - | 0 | - |
| - | - | 6248 | 1820 | - | - | 0 | - |
| - | - | 2377 | 1821 | - | - | 0 | - |
| - | - | 4353 | 1832 | - | - | 0 | - |
| - | - | 2.103E+04 | 1833 | - | - | 0 | - |
| - | - | 6.362E+04 | 1834 | - | - | 0 | - |
| - | - | 5.726E+04 | 1835 | - | - | 0 | - |
| - | - | 2.771E+04 | 1836 | - | - | 0 | - |
| - | - | 1.162E+04 | 1837 | - | - | 0 | - |
| - | - | 3427 | 1838 | - | - | 0 | - |
| - | - | 874 | 1839 | - | - | 0 | - |
| - | - | 675.7 | 2325 | - | - | 0 | - |
| - | - | 659 | 3071 | - | - | 0 | - |

m/z Charge Intensity FragmentType MassShift Position
120.06562042236328 0 11523.408 y 16
120.080810546875 0 2237.279
121.06889343261719 0 363.37018
125.88548278808594 0 405.85184
126.0550765991211 0 1776.8569
126.6019515991211 0 338.03076
128.0944061279297 0 807.917
128.10702514648438 0 956.6497
129.10226440429688 0 5174.2876
130.34983825683594 0 436.60165
131.1181182861328 0 1084.4545
132.62591552734375 0 390.97568
136.6066131591797 0 379.48633
137.87278747558594 0 407.2008
140.10711669921875 0 1009.37067
143.68194580078125 0 417.38028
148.95382690429688 0 1018.8542
149.04478454589844 0 642.68207
155.11798095703125 0 861.6861
157.13330078125 0 624.953
158.141845703125 0 537.0474
169.1337890625 0 2863.6948
171.07652282714844 0 1249.0331
173.09237670898438 0 1365.4236 y Water loss 15
173.12843322753906 0 3363.037
173.43881225585938 0 1460.0138
183.11280822753906 0 2507.8835
191.1027374267578 0 2856.581 y 15
197.12855529785156 0 2298.1797
200.138916015625 0 689.7674 c Water loss 1
201.12332153320312 0 7347.5913
211.10816955566406 0 630.7899
212.13931274414062 0 9954.045
213.14273071289062 0 1235.5834
215.1396026611328 0 750.93536
218.1499481201172 0 4408.948 c 1
221.0843505859375 0 3218.033
222.08448791503906 0 1058.3654
223.08203125 0 573.56616
227.09002685546875 0 1349.775
227.1023712158203 0 1873.1945
233.16490173339844 0 1506.709
235.10745239257812 0 863.4167
239.09519958496094 0 4216.048
240.09664916992188 0 1336.2607
241.09242248535156 0 566.75635
242.1501007080078 0 808.23206
243.14573669433594 0 1133.5236 y Water loss 10
254.14984130859375 0 1105.4111
256.1419677734375 0 598.3834
261.1598205566406 0 1251.4275
263.1023864746094 0 3034.7847
264.1073913574219 0 754.0732
280.1782531738281 0 777.8479
281.051025390625 0 1243.0227
281.1388854980469 0 801.837 z Ammonia loss 9
281.18505859375 0 624.41376
282.0514221191406 0 630.04944
282.1445007324219 0 965.1153
296.1969299316406 0 11807.428
297.1551513671875 0 1136.3287
297.2002868652344 0 2066.1892
298.1270751953125 0 585.5158
298.1617736816406 0 846.431
299.0617370605469 0 4584.023
299.1715087890625 0 14933.883
300.0617980957031 0 2241.6523
300.1559753417969 0 562.4711
300.1745300292969 0 2248.5486
301.0603332519531 0 1064.2556
303.1788330078125 0 7922.1494 z 14
304.1830749511719 0 1647.3577
307.1756286621094 0 729.79504
307.6664733886719 0 798.41296 y Water loss 11
314.1866149902344 0 713.3759
314.2074279785156 0 4762.06
315.21142578125 0 871.3882
316.1867370605469 0 7705.005
316.6717834472656 0 1148.611 y 11
317.0541687011719 0 523.75
317.1902160644531 0 871.9004
319.19757080078125 0 5297.6177 y 14
325.18701171875 0 946.92645
331.23406982421875 0 27655.201 c 2
332.2374267578125 0 5089.832
333.2393798828125 0 668.9756
344.17291259765625 0 593.22656
345.1802062988281 0 867.72754
346.18890380859375 0 658.54846
347.1947326660156 0 1185.4178
350.1361389160156 0 895.0099
355.06976318359375 0 37080.09
356.0704040527344 0 16167.544
357.0673828125 0 8737.248
358.0686950683594 0 882.52167
362.2068176269531 0 1692.3315
369.2002868652344 0 811.18304
371.1004638671875 0 1106.4883
372.10076904296875 0 989.6184
373.07928466796875 0 1044.5762
373.20904541015625 0 4324.663 y 10
378.2265930175781 0 1329.1509
385.21844482421875 0 781.4554
386.2028503417969 0 1324.592
415.03704833984375 0 1792.2623
415.2322082519531 0 905.4295 y Water loss 13
416.0376892089844 0 1111.9762
417.0358581542969 0 911.86395
417.2228698730469 0 2613.7942 z 13
418.22955322265625 0 7720.6016
419.23272705078125 0 2173.7317
426.23394775390625 0 813.13916
433.2422790527344 0 2721.0276 y 13
433.28118896484375 0 8883.417
434.2555236816406 0 598.7478
434.28662109375 0 4960.3496
434.7042541503906 0 678.73206
435.29443359375 0 2118.8057
444.2456970214844 0 6882.824
445.20306396484375 0 610.995
445.2531433105469 0 2406.219
446.2352600097656 0 875.6187
446.2648010253906 0 1780.2716
451.89434814453125 0 861.5997 y Water loss 4
453.218505859375 0 723.3415
460.2674560546875 0 1502.754
461.2761535644531 0 12883.7
462.2792663574219 0 3666.8882
472.24554443359375 0 605.0841
473.23382568359375 0 698.0319
486.267578125 0 760.345 y Water loss 12
488.2591857910156 0 11267.053 c 8
488.7345275878906 0 1124.6962
489.05596923828125 0 861.6082
489.26513671875 0 7632.4414
490.0565185546875 0 1229.5596
490.26849365234375 0 1491.8195
493.7278137207031 0 1757.4233
494.22955322265625 0 1421.8601
496.24822998046875 0 713.855
497.2576904296875 0 978.95886
498.21923828125 0 1174.9822
502.2650451660156 0 1891.0902 y 8
502.7322998046875 0 2413.551
503.2325744628906 0 1186.6223
503.2698974609375 0 1032.6888
503.73724365234375 0 589.4461
504.2779235839844 0 8432.016 y 12
505.28033447265625 0 1866.4769
515.2942504882812 0 998.72125
529.2638549804688 0 795.68384 w 7
529.7650756835938 0 765.30383
538.2694091796875 0 7049.682
538.7708740234375 0 2663.786
539.2728271484375 0 2239.2231 z 2
540.31787109375 0 725.5495
543.2610473632812 0 1732.4749
543.76513671875 0 1347.7756
544.2660522460938 0 752.1627
544.7706298828125 0 921.4524
552.267578125 0 5421.577
552.7693481445312 0 3865.3125
554.32080078125 0 693.0858
557.3220825195312 0 1008.8082
558.2880249023438 0 15641.981 w 11
558.329833984375 0 2310.9812
559.2900390625 0 3846.6501
559.3341674804688 0 1542.4291
560.2945556640625 0 737.6785
572.3045654296875 0 849.83124
577.1263427734375 0 1068.8077
578.1263427734375 0 701.15686
579.1215209960938 0 648.95526
580.7779541015625 0 594.289 y Ammonia loss 6
582.281982421875 0 865.1007
582.7792358398438 0 876.6496
583.3366088867188 0 1556.7338
599.2660522460938 0 708.4492
599.8078002929688 0 2207.7295
600.308837890625 0 1204.3954 c Water loss 10
600.8093872070312 0 747.9042
602.7908325195312 0 3201.3562
603.2919921875 0 2785.5063
610.1842651367188 0 8862.088
610.3616333007812 0 551.00494
611.1846313476562 0 9045.362
611.7958984375 0 9002.332
612.182373046875 0 6784.7837
612.2408447265625 0 1049.5511
612.2982177734375 0 5931.0073
612.8023681640625 0 1640.2288
613.1807250976562 0 998.01013
614.3373413085938 0 8225.104 y Water loss 11
615.3448486328125 0 10362.112
616.3156127929688 0 3458.1807 z 11
616.353759765625 0 1696.2216
617.3182983398438 0 898.5086
617.8045043945312 0 1092.6915
618.3072509765625 0 1604.5782
618.805908203125 0 929.3029
626.3419189453125 0 3144.013
626.8125 0 5456.916
627.2615356445312 0 5096.9204
627.31494140625 0 2241.8936
628.2669677734375 0 1630.8917
628.3923950195312 0 3144.5437
628.8087158203125 0 814.2768 y Water loss 5
629.396728515625 0 1905.7325
630.3577270507812 0 705.43646
630.4095458984375 0 862.42334
632.3362426757812 0 6222.092 y 11
633.3388061523438 0 1916.7163
634.341064453125 0 809.11035
635.3252563476562 0 974.53534
635.8245849609375 0 1558.878
637.8099365234375 0 924.1999 y 5
655.3401489257812 0 2886.7563
655.8421630859375 0 2754.1414
656.3426513671875 0 763.52515
658.3619384765625 0 928.6791
664.83642578125 0 1332.0271 c Ammonia loss 11
667.3981323242188 0 1193.5032
668.8251953125 0 921.2698
671.3714599609375 0 2318.6438
672.4078979492188 0 3715.9263 c 5
673.341796875 0 626.982 c 11
673.410400390625 0 2140.017
676.8301391601562 0 2184.2188
677.3359375 0 7749.844 y Water loss 4
677.8358154296875 0 7598.947 y Ammonia loss 4
678.3358764648438 0 2773.6182 z 4
678.832763671875 0 2157.886
685.3242797851562 0 810.6029
685.837890625 0 4573.6377
686.3436889648438 0 56477.61 y 4
686.8426513671875 0 21515.86
687.3477172851562 0 18624.803
687.8453979492188 0 2559.7788
688.3534545898438 0 1844.7242
697.4151611328125 0 1586.0878
698.421630859375 0 1217.9404
700.349365234375 0 1050.9651 c Ammonia loss 12
716.4329833984375 0 1451.1783
722.3475952148438 0 2015.6172
723.3497924804688 0 1045.8584
729.40087890625 0 12579.541 z 10
730.4031372070312 0 5007.9883
731.4085083007812 0 785.31866
738.3751220703125 0 702.83875
740.3443603515625 0 1697.271
742.4156494140625 0 1774.3164
742.8616333007812 0 784.715 z Water loss 3
743.421142578125 0 1186.0101
745.4207153320312 0 6624.703 y 10
746.4223022460938 0 3062.6396
750.8693237304688 0 3443.5618 y Water loss 3
751.3702392578125 0 2867.119 y Ammonia loss 3
751.866455078125 0 1472.9985 z 3
752.3658447265625 0 846.10504
757.4197998046875 0 2161.5515
758.42578125 0 1822.9799
759.3712768554688 0 2632.3425
759.4400024414062 0 33546.32 c 6
759.874755859375 0 11548.593 y 3
760.3744506835938 0 10394.089
760.4436645507812 0 13499.963
760.8782958984375 0 3554.2664
761.3779296875 0 1544.8152
761.4463500976562 0 3375.3774
764.3607177734375 0 5615.999
765.3632202148438 0 3534.355
766.3618774414062 0 1098.3237
798.9033203125 0 1203.4395
799.4305419921875 0 13205.909 w 9
799.90576171875 0 1092.538 z Ammonia loss 2
800.4306640625 0 7235.6655
801.4319458007812 0 1546.5696
802.3821411132812 0 6078.992
803.3837280273438 0 2964.581
807.4154663085938 0 1720.8522 y Water loss 2
807.91357421875 0 2075.2683 y Ammonia loss 2
808.4209594726562 0 1210.6318 z 2
808.9296875 0 1236.193
812.9239501953125 0 714.7635
813.4188842773438 0 1634.1877
813.9207153320312 0 953.46594
814.4282836914062 0 842.9552
815.3845825195312 0 1231.6317
816.413818359375 0 2878.0974 y 2
816.9196166992188 0 2215.4087
817.4208984375 0 965.9111
822.3797607421875 0 3968.2307
823.3820190429688 0 5093.151
824.3878784179688 0 1920.4755
829.443603515625 0 1434.453
829.94091796875 0 37016.63 c 14
830.4420776367188 0 34002.74
830.943359375 0 16164.45
831.44384765625 0 5920.4033
831.9521484375 0 1006.16986
834.9363403320312 0 896.02423
836.4320678710938 0 1318.1403
836.9276733398438 0 670.0089
840.4342041015625 0 1227.3944 z Water loss 9
841.4267578125 0 992.4445 z Ammonia loss 9
842.4421997070312 0 1332.4336
843.451416015625 0 3203.742
843.9549560546875 0 2530.0217
844.4555053710938 0 2207.134
844.9580078125 0 919.3941
846.47216796875 0 29257.805 c 7
847.4744262695312 0 14192.109
848.4765014648438 0 4388.31
848.93115234375 0 1783.3513
849.4306640625 0 2082.488
849.9284057617188 0 3478.9963 z Water loss 1
850.422607421875 0 2208.9363 z Ammonia loss 1
850.9298095703125 0 1967.7782
852.4224853515625 0 877.09314
856.9462890625 0 2651.5728 c Ammonia loss 15
857.4442138671875 0 4821.707 y Ammonia loss 9
857.9464721679688 0 3648.39 y Water loss 1
858.4443969726562 0 23678.434 y Ammonia loss 1
858.9402465820312 0 4011.3735 z 1
859.4447021484375 0 10812.551
859.936767578125 0 1345.8552
860.44580078125 0 2935.4243
863.96435546875 0 1138.5984
864.9608154296875 0 4760.3003
865.4598388671875 0 49390.516 c 15
865.9607543945312 0 51316.996
866.461181640625 0 27036.473
866.961669921875 0 11859.071
867.4600219726562 0 2998.058
867.9603881835938 0 2022.9735
868.4032592773438 0 2351.87
869.4083862304688 0 1270.123
871.4444580078125 0 967.6648
872.4586791992188 0 1093.4417
872.9473266601562 0 1601.1686
873.4536743164062 0 3510.402
874.4617919921875 0 4793.542 y 9
875.462890625 0 1862.271
878.4534912109375 0 818.4717
880.4520874023438 0 4399.358
880.9620971679688 0 7555.026
881.4641723632812 0 7463.0024
881.9686889648438 0 3712.4148
882.4647827148438 0 2175.6177
882.972900390625 0 758.31415
885.462158203125 0 3354.3264
885.9660034179688 0 4560.551
886.46435546875 0 2866.6133
886.9654541015625 0 1097.4773
887.4654541015625 0 2454.1943
887.9666137695312 0 1463.7759
888.4718017578125 0 1029.3972
888.94775390625 0 2217.847
889.4530639648438 0 1512.43
893.9710083007812 0 1185.5878
894.4685668945312 0 22392.785
894.9700317382812 0 20123.775
895.4705200195312 0 12490.665
895.969482421875 0 6082.2744
896.47265625 0 1265.5068
899.4635620117188 0 2114.4744
899.96435546875 0 2253.222
900.4577026367188 0 1569.2872
902.45458984375 0 891.9958
907.4059448242188 0 9515.151
907.9733276367188 0 4270.273
908.4024658203125 0 2893.7507
908.4746704101562 0 3897.1272
908.9707641601562 0 5410.3696
909.9712524414062 0 832.3214
915.472412109375 0 1452.5981
916.4754638671875 0 55157.98
916.9774169921875 0 92994.125
917.4793701171875 0 69599.24
917.98095703125 0 31602.498
918.4822387695312 0 12169.22
918.9822387695312 0 3528.0488
928.4727783203125 0 6013.4453 w 8
929.4794311523438 0 6385.1343
930.4796752929688 0 2365.88
931.4974975585938 0 1006.52295
944.4664916992188 0 3095.8027
945.4762573242188 0 3597.2817
946.4796142578125 0 1772.306
957.4613037109375 0 905.9197
958.4868774414062 0 1474.3646
959.4840087890625 0 1467.6338
960.4892578125 0 5484.9146
961.48974609375 0 3100.5352
969.4630737304688 0 1148.4338 z Water loss 8
973.4945068359375 0 19626.125
974.4976196289062 0 9534.05
975.5137939453125 0 44142.13 c 8
976.5170288085938 0 24004.225
977.5202026367188 0 7542.5317
978.5270385742188 0 1539.4749
987.4862060546875 0 20395.17 z 8
988.489990234375 0 11689.588
989.4929809570312 0 3875.7446
990.4891357421875 0 1154.264
1001.5220947265625 0 1035.969
1002.5254516601562 0 909.61084
1003.49560546875 0 5557.9043 y 8
1004.5005493164062 0 3150.1077
1005.5020141601562 0 1236.9374
1012.5198364257812 0 1505.0969
1015.4895629882812 0 923.38135
1016.5089721679688 0 1106.5411
1018.4569091796875 0 1345.87
1019.4678344726562 0 758.9433
1030.5068359375 0 973.25195
1031.5079345703125 0 1000.68744
1053.4810791015625 0 826.33984
1055.5015869140625 0 4542.736
1056.5081787109375 0 3703.5208 z Water loss 7
1057.510498046875 0 1697.5778 z Ammonia loss 7
1058.519775390625 0 1169.076
1060.532470703125 0 2866.6348
1061.54052734375 0 1971.232
1070.5201416015625 0 859.9943
1071.51171875 0 1249.2789
1072.505859375 0 1593.173 y Water loss 7
1073.5115966796875 0 1729.5599 y Ammonia loss 7
1074.5181884765625 0 20431.914 z 7
1075.523193359375 0 15682.32
1076.5255126953125 0 6975.3276
1077.528564453125 0 2244.6626
1087.529541015625 0 4039.704
1088.532470703125 0 2561.0234
1089.5321044921875 0 1506.44
1090.5361328125 0 13696.504 y 7
1091.538818359375 0 7676.056
1092.541015625 0 2734.1785
1093.5494384765625 0 817.72455
1098.508056640625 0 1542.9812
1099.5133056640625 0 1487.8507
1103.5263671875 0 2790.9102
1104.5557861328125 0 46645.46 c 9
1105.5584716796875 0 31457.764
1106.562744140625 0 11427.803
1107.5635986328125 0 2952.472
1117.5260009765625 0 882.0583
1118.5283203125 0 904.16705
1120.552734375 0 8278.067
1121.5562744140625 0 5891.99
1122.5552978515625 0 1951.6302
1133.5570068359375 0 1138.4916
1143.540283203125 0 1069.4191 z Water loss 6
1159.5408935546875 0 1213.8992 y Water loss 6
1160.5421142578125 0 887.44116 y Ammonia loss 6
1161.5501708984375 0 10021.674 z 6
1162.5538330078125 0 6895.5156
1163.56103515625 0 3804.4587
1164.5628662109375 0 2254.7014
1177.56982421875 0 6402.3066 y 6
1178.57177734375 0 4414.309
1179.5823974609375 0 1446.3154
1180.575439453125 0 1419.6041
1181.5767822265625 0 1905.399
1182.5809326171875 0 1200.1257
1185.53173828125 0 918.5385
1198.610595703125 0 2231.5142
1199.608154296875 0 1633.9797 c Water loss 10
1200.609375 0 1632.4164
1201.6162109375 0 1248.3048
1205.5819091796875 0 1167.0376
1206.565185546875 0 11642.349
1207.56884765625 0 8976.186
1208.5673828125 0 2426.5508
1217.6395263671875 0 30378.148 c 10
1218.6424560546875 0 19673.451
1219.64599609375 0 8611.385
1220.6385498046875 0 2767.2847
1222.582275390625 0 7141.3633
1223.58984375 0 11398.304
1224.5916748046875 0 7047.205
1225.59375 0 1483.0035
1235.609375 0 971.40686
1252.6114501953125 0 2736.1016
1253.6141357421875 0 1474.9187
1269.6395263671875 0 1415.4154
1270.6470947265625 0 1190.9015
1274.6217041015625 0 3084.9724 y 5
1275.626708984375 0 2079.446
1276.62646484375 0 1062.6677
1301.662841796875 0 1142.426
1302.6744384765625 0 1521.414
1303.6400146484375 0 781.3091
1328.6524658203125 0 2719.521 c Ammonia loss 11
1329.6575927734375 0 2908.3247
1330.6539306640625 0 1691.9569
1344.6895751953125 0 4216.0312
1345.6976318359375 0 39688.523 c 11
1346.7001953125 0 30795.797
1347.70361328125 0 12794.064
1348.7078857421875 0 3850.1997
1354.6634521484375 0 1142.5585 y Ammonia loss 4
1357.676025390625 0 753.45404
1370.663330078125 0 2116.5564
1371.6719970703125 0 11806.766 y 4
1372.6776123046875 0 8971.951
1373.6871337890625 0 4721.2715
1374.6961669921875 0 1428.3541
1398.6732177734375 0 1554.8483
1399.6759033203125 0 3683.6719
1400.6783447265625 0 5317.9385
1401.6944580078125 0 5105.491
1402.69775390625 0 2702.3235
1403.6971435546875 0 952.9784
1415.7275390625 0 5353.2373
1416.7335205078125 0 27152.072 c 12
1417.7364501953125 0 21223.031
1418.7392578125 0 7748.3037
1419.7420654296875 0 3133.6511
1430.699951171875 0 2320.6714
1431.6932373046875 0 2905.8303
1432.697998046875 0 1319.7758
1443.736083984375 0 818.9274
1485.7332763671875 0 930.3671
1486.7537841796875 0 2173.7893
1487.7637939453125 0 2047.9148
1488.77490234375 0 1035.4661
1501.72412109375 0 898.5785 y Ammonia loss 3
1502.7218017578125 0 23913.855 z 3
1503.7275390625 0 35872.285
1504.7315673828125 0 24905.24
1505.735595703125 0 10769.822
1506.7332763671875 0 3229.8206
1507.7464599609375 0 772.0301
1518.743896484375 0 3202.8005 y 3
1519.74560546875 0 2574.0269
1520.7454833984375 0 2078.0186
1521.7520751953125 0 880.9992
1528.7642822265625 0 949.3073
1529.775634765625 0 1213.791
1530.7772216796875 0 45204.883 c 13
1531.7801513671875 0 41804.582
1532.7822265625 0 19092.973
1533.7867431640625 0 7935.082
1534.7813720703125 0 1905.6893
1543.781005859375 0 1136.5703
1544.792236328125 0 815.5941
1614.8466796875 0 3221.9397
1615.8243408203125 0 5211.7275 z 2
1616.8206787109375 0 9226.758
1617.8236083984375 0 6891.4707
1618.81884765625 0 2140.8213
1632.870361328125 0 698.18506
1642.8538818359375 0 2100.326
1643.85302734375 0 1969.8948
1644.850341796875 0 917.9799
1658.8739013671875 0 4787.645 c 14
1659.877685546875 0 12830.791
1660.88134765625 0 9231.601
1661.8779296875 0 4350.0376
1662.891357421875 0 2093.7134
1672.8533935546875 0 1501.407
1685.8961181640625 0 3300.199
1686.9000244140625 0 3624.305
1687.8992919921875 0 3129.6309
1688.911865234375 0 1905.5542
1699.8525390625 0 1599.0616 z Ammonia loss 1
1700.8460693359375 0 1617.583
1713.889404296875 0 3876.4385
1714.897216796875 0 4095.3025 y Water loss 1
1715.885009765625 0 3249.5652
1716.86572265625 0 3708.4534 z 1
1717.862548828125 0 10819.652
1718.8665771484375 0 8671.636
1719.8675537109375 0 4608.0854
1720.867919921875 0 1881.1091
1721.8583984375 0 1020.4126
1729.90771484375 0 4578.978 c 15
1730.9154052734375 0 14896.259
1731.9190673828125 0 14280.0205
1732.920166015625 0 7507.999
1733.9140625 0 2936.9385
1760.9495849609375 0 1189.5121
1761.9261474609375 0 1572.0165
1762.923583984375 0 1032.6564
1770.9326171875 0 3068.6792
1771.919189453125 0 3555.8066
1772.918212890625 0 4200.1704
1773.9183349609375 0 1348.2195
1774.914306640625 0 1567.3446
1776.89892578125 0 845.0797
1786.929443359375 0 863.41016
1787.937744140625 0 6277.7017
1788.9373779296875 0 25733.305
1789.9395751953125 0 20706.906
1790.9412841796875 0 12326.269
1791.943603515625 0 4195.6934
1792.93310546875 0 1704.5919
1799.92041015625 0 907.44574
1800.9229736328125 0 883.6949
1804.955810546875 0 1206.7693
1805.95947265625 0 5032.9946
1806.9649658203125 0 4493.5044
1807.9727783203125 0 2967.212
1814.9337158203125 0 1335.353
1815.9302978515625 0 8252.96
1816.9310302734375 0 35794.023
1817.9342041015625 0 34841.6
1818.9361572265625 0 18155.686
1819.9371337890625 0 6248.153
1820.9454345703125 0 2376.5925
1831.942626953125 0 4352.6245
1832.94775390625 0 21026.969
1833.9552001953125 0 63615.38
1834.9593505859375 0 57260.24
1835.96142578125 0 27710.47
1836.9639892578125 0 11615.544
1837.962890625 0 3426.842
1838.958984375 0 873.98895
2325.19921875 0 675.73737
3070.9375 0 658.969

Spectrum Details

|  |  |
| --- | --- |
| Matched peaks? Matched peaks The total absolute number of peaks matched. Additionally in brackets the total fraction of peaks matched and the total number of peaks is shown. | 101 (16.26% of 621) |
| FDR? FDR The false discovery rate estimated for this peptide. It is calculated by matching all theoretical fragments with a non-integer shift with the raw peaks for this spectrum. This is done with 40 different shifts. The resulting percentage is the average number of annotated peaks over the number of annotated peaks with the correct spectrum. | 0.78% |
| Satellite FDR? Satellite FDR See the FDR for details on its calculation. This satellite ion specific FDR only contains the satellite ions (d/w) for I/L/J positions. | - |
| PSM Score? PSM Score The PSM Score as given by Hecklib to this annotated spectrum. It is shown with three significant figures. | 852 |

#### Spectrum 7896? Spectrum 7896 The raw spectrum of this peptide as annotated by Hecklib. The fragments are coloured according to ion type (see legend). Any peaks with a star '\*' as text can be hovered over to see the full details, first the ion type second the mass shift type. By hovering over the amino acids in the peptide or ions in the legend the corresponding peaks are highlighted. By toggling the 'Unassigned' label you can turn the background (unassigned) peaks on or off in the plot. By updating the slider in the Ion legend you can update the spectrum to only show the top X% of the peaks with labels. The top X% means any peak that is within X% of the highest intensity. By dragging in the spectrum you can zoom in to a specific part of the spectrum and use 'Zoom Out' to get back to the original zoom level. The annotation of the spectrum is based on the given sequence in the peptides file and is done with different software so inconsistencies are likely. The peaks are annotated based on the given sequence, with 20 ppm tolerance.

Copy Data

##### Spectrum 7896 (TSV)

###### Preview

```
Loading example...
```

*Click on the button to copy the data to your clipboard.*

Mz MinMz MaxIntensity Max

WidthHeightPeptide font sizePeptide stroke widthSpectrum font sizeSpectrum stroke widthCompact peptide

Ion legend

wxyz

abcd

OtherUnassignedIon^Charge^_Position_Show for top:%

VTJFPPSSEEJQANKAT

01.75e+43.50e+45.25e+47.00e+4

Zoom Out

y^+1^_1_y^+1^_2_y^+1^_2_c^+1^_2_y^+3^_7_z^+1^_3_y^+2^_6_y^+2^_6_y^+1^_3_c^+1^_3_y^+2^_7_y^+1^_4_z^+1^_4_y^+1^_4_c^+2^_9_y^+2^_9_y^+1^_5_w^+2^_10_z^+3^_15_w^+1^_6_y^+3^_16_y^+2^_11_c^+2^_11_y^+1^_6_z^+1^_6_y^+1^_6_y^+2^_12_c^+1^_6_c^+2^_12_c^+1^_6_y^+2^_13_y^+2^_13_z^+2^_13_y^+2^_13_z^+1^_7_y^+1^_7_y^+2^_14_y^+2^_14_z^+2^_14_c^+1^_7_y^+2^_14_w^+1^_8_y^+2^_15_y^+2^_15_c^+2^_15_z^+1^_8_c^+1^_8_z^+2^_16_z^+2^_16_c^+2^_16_y^+1^_8_y^+2^_16_y^+2^_16_z^+2^_16_c^+2^_16_y^+1^_8_w^+1^_9_c^+1^_9_z^+1^_9_y^+1^_9_z^+1^_10_z^+1^_10_y^+1^_10_y^+1^_10_z^+1^_10_c^+1^_10_y^+1^_10_c^+1^_10_y^+1^_11_z^+1^_11_y^+1^_11_c^+1^_11_y^+1^_12_y^+1^_12_c^+1^_12_c^+1^_12_y^+1^_13_c^+1^_13_y^+1^_14_z^+1^_14_c^+1^_14_y^+1^_14_c^+1^_14_y^+1^_15_z^+1^_15_y^+1^_15_c^+1^_15_y^+1^_16_y^+1^_16_z^+1^_16_c^+1^_16_

0775155023253100

Fragment Matches Table

Show background peaks

| Position | Ion type | Intensity | mz Theoretical | mz Error (Th) | mz Error (ppm) | Charge | Series Number |
| --- | --- | --- | --- | --- | --- | --- | --- |
| - | - | 695.7 | 120.1 | - | - | 0 | - |
| 17 | y | 1.064E+04 | 120.1 | 7.029E-05 | 0.5854 | +1 | 1 |
| - | - | 2015 | 120.1 | - | - | 0 | - |
| - | - | 1158 | 126.1 | - | - | 0 | - |
| - | - | 830.4 | 128.1 | - | - | 0 | - |
| - | - | 4639 | 129.1 | - | - | 0 | - |
| - | - | 394.2 | 129.2 | - | - | 0 | - |
| - | - | 929.7 | 131.1 | - | - | 0 | - |
| - | - | 1187 | 140.1 | - | - | 0 | - |
| - | - | 450.4 | 146.7 | - | - | 0 | - |
| - | - | 1075 | 149 | - | - | 0 | - |
| - | - | 439 | 153.8 | - | - | 0 | - |
| - | - | 575.5 | 155.1 | - | - | 0 | - |
| - | - | 491.9 | 160.6 | - | - | 0 | - |
| - | - | 2654 | 169.1 | - | - | 0 | - |
| - | - | 727.5 | 171.1 | - | - | 0 | - |
| 16 | y | 1382 | 173.1 | 0.0002927 | 1.691 | +1 | 2 |
| - | - | 2950 | 173.1 | - | - | 0 | - |
| - | - | 544.9 | 177.7 | - | - | 0 | - |
| - | - | 516.9 | 181.1 | - | - | 0 | - |
| - | - | 2398 | 183.1 | - | - | 0 | - |
| - | - | 628.4 | 187.1 | - | - | 0 | - |
| 16 | y | 2414 | 191.1 | 6.382E-05 | 0.334 | +1 | 2 |
| - | - | 2236 | 197.1 | - | - | 0 | - |
| - | - | 5983 | 201.1 | - | - | 0 | - |
| - | - | 502.8 | 207.4 | - | - | 0 | - |
| - | - | 527.6 | 211.1 | - | - | 0 | - |
| - | - | 8244 | 212.1 | - | - | 0 | - |
| - | - | 815 | 213.1 | - | - | 0 | - |
| - | - | 871 | 215.1 | - | - | 0 | - |
| - | - | 618.9 | 217.1 | - | - | 0 | - |
| 2 | c | 3148 | 218.1 | 0.000214 | 0.9808 | +1 | 2 |
| - | - | 3301 | 221.1 | - | - | 0 | - |
| - | - | 717.2 | 226.1 | - | - | 0 | - |
| - | - | 1348 | 227.1 | - | - | 0 | - |
| - | - | 1744 | 227.1 | - | - | 0 | - |
| - | - | 1259 | 233.2 | - | - | 0 | - |
| - | - | 1139 | 235.1 | - | - | 0 | - |
| - | - | 600 | 238.2 | - | - | 0 | - |
| - | - | 5430 | 239.1 | - | - | 0 | - |
| - | - | 1594 | 240.1 | - | - | 0 | - |
| - | - | 729.4 | 241.1 | - | - | 0 | - |
| 11 | y | 898.3 | 243.1 | 0.003368 | 13.85 | +3 | 7 |
| - | - | 1014 | 254.2 | - | - | 0 | - |
| - | - | 2808 | 263.1 | - | - | 0 | - |
| - | - | 1051 | 281.1 | - | - | 0 | - |
| - | - | 1198 | 282.1 | - | - | 0 | - |
| - | - | 738.6 | 295.1 | - | - | 0 | - |
| - | - | 1.136E+04 | 296.2 | - | - | 0 | - |
| - | - | 698.4 | 297.1 | - | - | 0 | - |
| - | - | 863.4 | 297.2 | - | - | 0 | - |
| - | - | 1572 | 297.2 | - | - | 0 | - |
| - | - | 785.2 | 298.2 | - | - | 0 | - |
| - | - | 6149 | 299.1 | - | - | 0 | - |
| - | - | 1.431E+04 | 299.2 | - | - | 0 | - |
| - | - | 1232 | 300.1 | - | - | 0 | - |
| - | - | 2019 | 300.2 | - | - | 0 | - |
| - | - | 773 | 301.1 | - | - | 0 | - |
| 15 | z | 6998 | 303.2 | 2.169E-05 | 0.07155 | +1 | 3 |
| - | - | 1125 | 304.2 | - | - | 0 | - |
| 12 | y | 710.8 | 307.7 | 0.0006813 | 2.214 | +2 | 6 |
| - | - | 1105 | 314.2 | - | - | 0 | - |
| - | - | 4226 | 314.2 | - | - | 0 | - |
| - | - | 620 | 315.2 | - | - | 0 | - |
| - | - | 7792 | 316.2 | - | - | 0 | - |
| 12 | y | 1256 | 316.7 | 0.0004344 | 1.372 | +2 | 6 |
| - | - | 664.1 | 317.2 | - | - | 0 | - |
| 15 | y | 3264 | 319.2 | 0.0003711 | 1.163 | +1 | 3 |
| 3 | c | 2.425E+04 | 331.2 | 8.79E-05 | 0.2654 | +1 | 3 |
| - | - | 4363 | 332.2 | - | - | 0 | - |
| - | - | 735.1 | 343.4 | - | - | 0 | - |
| - | - | 783.3 | 345.2 | - | - | 0 | - |
| - | - | 583.5 | 347.2 | - | - | 0 | - |
| - | - | 713.9 | 350.1 | - | - | 0 | - |
| - | - | 3.817E+04 | 355.1 | - | - | 0 | - |
| - | - | 1.663E+04 | 356.1 | - | - | 0 | - |
| - | - | 1.009E+04 | 357.1 | - | - | 0 | - |
| - | - | 1091 | 358.1 | - | - | 0 | - |
| - | - | 1013 | 362.2 | - | - | 0 | - |
| - | - | 615.8 | 369.2 | - | - | 0 | - |
| - | - | 1687 | 371.1 | - | - | 0 | - |
| - | - | 1024 | 372.1 | - | - | 0 | - |
| - | - | 840.7 | 373.1 | - | - | 0 | - |
| 11 | y | 3453 | 373.2 | 0.004763 | 12.76 | +2 | 7 |
| - | - | 855.9 | 378.2 | - | - | 0 | - |
| - | - | 924.3 | 385.2 | - | - | 0 | - |
| - | - | 1711 | 386.2 | - | - | 0 | - |
| - | - | 667.3 | 407.2 | - | - | 0 | - |
| - | - | 2670 | 415 | - | - | 0 | - |
| 14 | y | 752.5 | 415.2 | 0.0001008 | 0.2427 | +1 | 4 |
| 14 | z | 2014 | 417.2 | 0.00165 | 3.955 | +1 | 4 |
| - | - | 5839 | 418.2 | - | - | 0 | - |
| - | - | 578.8 | 419 | - | - | 0 | - |
| - | - | 1533 | 419.2 | - | - | 0 | - |
| 14 | y | 2337 | 433.2 | 0.000626 | 1.445 | +1 | 4 |
| - | - | 7895 | 433.3 | - | - | 0 | - |
| - | - | 845 | 434.3 | - | - | 0 | - |
| - | - | 4359 | 434.3 | - | - | 0 | - |
| - | - | 2693 | 435.3 | - | - | 0 | - |
| - | - | 816 | 442.2 | - | - | 0 | - |
| - | - | 5423 | 444.2 | - | - | 0 | - |
| - | - | 576 | 445.2 | - | - | 0 | - |
| - | - | 2234 | 445.3 | - | - | 0 | - |
| - | - | 1639 | 446.3 | - | - | 0 | - |
| - | - | 1249 | 460.3 | - | - | 0 | - |
| - | - | 1.117E+04 | 461.3 | - | - | 0 | - |
| - | - | 2550 | 462.3 | - | - | 0 | - |
| - | - | 785.6 | 463.3 | - | - | 0 | - |
| - | - | 653.6 | 480.2 | - | - | 0 | - |
| - | - | 1476 | 480.7 | - | - | 0 | - |
| 9 | c | 9223 | 488.3 | 0.001922 | 3.937 | +2 | 9 |
| - | - | 1449 | 489.1 | - | - | 0 | - |
| - | - | 6158 | 489.3 | - | - | 0 | - |
| - | - | 912.8 | 490.1 | - | - | 0 | - |
| - | - | 1844 | 490.3 | - | - | 0 | - |
| - | - | 1904 | 493.7 | - | - | 0 | - |
| - | - | 1163 | 494.2 | - | - | 0 | - |
| - | - | 1044 | 498.2 | - | - | 0 | - |
| 9 | y | 2048 | 502.3 | 0.008278 | 16.48 | +2 | 9 |
| - | - | 3179 | 502.7 | - | - | 0 | - |
| - | - | 2229 | 503.2 | - | - | 0 | - |
| - | - | 828.6 | 503.3 | - | - | 0 | - |
| 13 | y | 8076 | 504.3 | 0.0001333 | 0.2644 | +1 | 5 |
| - | - | 1532 | 505.3 | - | - | 0 | - |
| - | - | 740.2 | 518.2 | - | - | 0 | - |
| 8 | w | 623.6 | 529.3 | 0.004643 | 8.773 | +2 | 10 |
| - | - | 638.2 | 536.3 | - | - | 0 | - |
| - | - | 6252 | 538.3 | - | - | 0 | - |
| - | - | 3655 | 538.8 | - | - | 0 | - |
| 3 | z | 973.2 | 539.3 | 0.002174 | 4.032 | +3 | 15 |
| - | - | 2101 | 543.3 | - | - | 0 | - |
| - | - | 636 | 543.3 | - | - | 0 | - |
| - | - | 1123 | 543.8 | - | - | 0 | - |
| - | - | 1444 | 544.3 | - | - | 0 | - |
| - | - | 6640 | 552.3 | - | - | 0 | - |
| - | - | 812.2 | 552.3 | - | - | 0 | - |
| - | - | 5595 | 552.8 | - | - | 0 | - |
| - | - | 1613 | 553.3 | - | - | 0 | - |
| - | - | 1272 | 554.3 | - | - | 0 | - |
| 12 | w | 1.511E+04 | 558.3 | 0.0004826 | 0.8645 | +1 | 6 |
| - | - | 2710 | 558.3 | - | - | 0 | - |
| - | - | 4718 | 559.3 | - | - | 0 | - |
| - | - | 595.3 | 559.3 | - | - | 0 | - |
| - | - | 1420 | 560.3 | - | - | 0 | - |
| 2 | y | 1470 | 572.3 | 0.01022 | 17.86 | +3 | 16 |
| - | - | 1179 | 577.1 | - | - | 0 | - |
| - | - | 702.9 | 578.1 | - | - | 0 | - |
| - | - | 775.5 | 579.1 | - | - | 0 | - |
| 7 | y | 942.3 | 580.3 | 0.003138 | 5.408 | +2 | 11 |
| - | - | 842.5 | 583.3 | - | - | 0 | - |
| - | - | 800.6 | 591.3 | - | - | 0 | - |
| - | - | 1842 | 599.8 | - | - | 0 | - |
| 11 | c | 1266 | 600.3 | 0.01135 | 18.91 | +2 | 11 |
| - | - | 2963 | 602.8 | - | - | 0 | - |
| - | - | 1736 | 603.3 | - | - | 0 | - |
| - | - | 8401 | 610.2 | - | - | 0 | - |
| - | - | 8783 | 611.2 | - | - | 0 | - |
| - | - | 7301 | 611.8 | - | - | 0 | - |
| - | - | 6641 | 612.2 | - | - | 0 | - |
| - | - | 4486 | 612.3 | - | - | 0 | - |
| - | - | 2311 | 612.8 | - | - | 0 | - |
| - | - | 712.8 | 613.2 | - | - | 0 | - |
| 12 | y | 8255 | 614.3 | 0.01114 | 18.14 | +1 | 6 |
| - | - | 1.106E+04 | 615.3 | - | - | 0 | - |
| 12 | z | 4304 | 616.3 | 0.001512 | 2.453 | +1 | 6 |
| - | - | 1540 | 616.4 | - | - | 0 | - |
| - | - | 1237 | 617.3 | - | - | 0 | - |
| - | - | 1531 | 617.8 | - | - | 0 | - |
| - | - | 2541 | 626.3 | - | - | 0 | - |
| - | - | 3252 | 626.8 | - | - | 0 | - |
| - | - | 3599 | 627.3 | - | - | 0 | - |
| - | - | 1729 | 627.3 | - | - | 0 | - |
| - | - | 1494 | 627.4 | - | - | 0 | - |
| - | - | 910 | 627.8 | - | - | 0 | - |
| - | - | 1423 | 628.3 | - | - | 0 | - |
| - | - | 3443 | 628.4 | - | - | 0 | - |
| - | - | 1330 | 629.4 | - | - | 0 | - |
| - | - | 1251 | 630.4 | - | - | 0 | - |
| 12 | y | 5802 | 632.3 | 0.0001496 | 0.2366 | +1 | 6 |
| - | - | 1226 | 633.3 | - | - | 0 | - |
| - | - | 1192 | 635.3 | - | - | 0 | - |
| 6 | y | 553.3 | 637.8 | 0.00132 | 2.069 | +2 | 12 |
| 6 | c | 789.2 | 654.4 | 0.01176 | 17.97 | +1 | 6 |
| - | - | 2042 | 655.3 | - | - | 0 | - |
| - | - | 1808 | 655.8 | - | - | 0 | - |
| - | - | 937.2 | 656.3 | - | - | 0 | - |
| - | - | 675.8 | 658.2 | - | - | 0 | - |
| - | - | 1023 | 658.4 | - | - | 0 | - |
| 12 | c | 767.3 | 664.8 | 0.005734 | 8.625 | +2 | 12 |
| - | - | 990.7 | 668.3 | - | - | 0 | - |
| - | - | 2738 | 671.4 | - | - | 0 | - |
| 6 | c | 3850 | 672.4 | 0.001002 | 1.49 | +1 | 6 |
| - | - | 1208 | 673.4 | - | - | 0 | - |
| - | - | 1848 | 676.8 | - | - | 0 | - |
| 5 | y | 7562 | 677.3 | 0.0005537 | 0.8174 | +2 | 13 |
| 5 | y | 5782 | 677.8 | 0.008659 | 12.77 | +2 | 13 |
| 5 | z | 3493 | 678.3 | 0.002305 | 3.399 | +2 | 13 |
| - | - | 4529 | 685.8 | - | - | 0 | - |
| 5 | y | 4.995E+04 | 686.3 | 0.002465 | 3.591 | +2 | 13 |
| - | - | 2.066E+04 | 686.8 | - | - | 0 | - |
| - | - | 1.375E+04 | 687.3 | - | - | 0 | - |
| - | - | 3332 | 687.8 | - | - | 0 | - |
| - | - | 3031 | 688.4 | - | - | 0 | - |
| - | - | 1399 | 697.4 | - | - | 0 | - |
| - | - | 1021 | 698.4 | - | - | 0 | - |
| - | - | 852.2 | 713.4 | - | - | 0 | - |
| - | - | 870.1 | 716.4 | - | - | 0 | - |
| - | - | 1796 | 722.3 | - | - | 0 | - |
| - | - | 988.8 | 723.4 | - | - | 0 | - |
| 11 | z | 1.134E+04 | 729.4 | 0.0008593 | 1.178 | +1 | 7 |
| - | - | 3920 | 730.4 | - | - | 0 | - |
| - | - | 1428 | 731.4 | - | - | 0 | - |
| - | - | 1470 | 740.3 | - | - | 0 | - |
| - | - | 2223 | 742.4 | - | - | 0 | - |
| 11 | y | 6890 | 745.4 | 0.0005583 | 0.7489 | +1 | 7 |
| - | - | 2029 | 746.4 | - | - | 0 | - |
| 4 | y | 1039 | 750.9 | 0.002961 | 3.944 | +2 | 14 |
| 4 | y | 1786 | 751.4 | 0.005092 | 6.777 | +2 | 14 |
| 4 | z | 939 | 751.9 | 0.001201 | 1.597 | +2 | 14 |
| - | - | 2292 | 757.4 | - | - | 0 | - |
| - | - | 2065 | 759.4 | - | - | 0 | - |
| 7 | c | 2.977E+04 | 759.4 | 0.0001937 | 0.255 | +1 | 7 |
| 4 | y | 8736 | 759.9 | 0.0004312 | 0.5674 | +2 | 14 |
| - | - | 6643 | 760.4 | - | - | 0 | - |
| - | - | 1.332E+04 | 760.4 | - | - | 0 | - |
| - | - | 2805 | 760.9 | - | - | 0 | - |
| - | - | 2071 | 761.4 | - | - | 0 | - |
| - | - | 4425 | 761.4 | - | - | 0 | - |
| - | - | 5880 | 764.4 | - | - | 0 | - |
| - | - | 3327 | 765.4 | - | - | 0 | - |
| - | - | 1260 | 766.4 | - | - | 0 | - |
| - | - | 988.7 | 798.4 | - | - | 0 | - |
| - | - | 1183 | 798.9 | - | - | 0 | - |
| 10 | w | 1.098E+04 | 799.4 | 0.001645 | 2.057 | +1 | 8 |
| - | - | 5272 | 800.4 | - | - | 0 | - |
| - | - | 1614 | 801.4 | - | - | 0 | - |
| - | - | 4429 | 802.4 | - | - | 0 | - |
| - | - | 2203 | 803.4 | - | - | 0 | - |
| - | - | 848.2 | 803.5 | - | - | 0 | - |
| 3 | y | 1862 | 807.4 | 0.009328 | 11.55 | +2 | 15 |
| - | - | 2240 | 807.9 | - | - | 0 | - |
| - | - | 1050 | 808.4 | - | - | 0 | - |
| - | - | 1195 | 813.9 | - | - | 0 | - |
| - | - | 1897 | 814.4 | - | - | 0 | - |
| - | - | 1228 | 815.4 | - | - | 0 | - |
| 3 | y | 2861 | 816.4 | 0.0001047 | 0.1283 | +2 | 15 |
| - | - | 1605 | 816.9 | - | - | 0 | - |
| - | - | 1026 | 817.4 | - | - | 0 | - |
| - | - | 750.8 | 817.9 | - | - | 0 | - |
| - | - | 3665 | 822.4 | - | - | 0 | - |
| - | - | 5501 | 823.4 | - | - | 0 | - |
| - | - | 2258 | 824.4 | - | - | 0 | - |
| - | - | 858.5 | 829.4 | - | - | 0 | - |
| 15 | c | 2.653E+04 | 829.9 | 0.0005537 | 0.6671 | +2 | 15 |
| - | - | 3.067E+04 | 830.4 | - | - | 0 | - |
| - | - | 1.598E+04 | 830.9 | - | - | 0 | - |
| - | - | 5393 | 831.4 | - | - | 0 | - |
| - | - | 935.7 | 832 | - | - | 0 | - |
| - | - | 1016 | 835.9 | - | - | 0 | - |
| 10 | z | 1660 | 841.4 | 0.01319 | 15.67 | +1 | 8 |
| - | - | 1031 | 842.4 | - | - | 0 | - |
| - | - | 2444 | 843.5 | - | - | 0 | - |
| - | - | 2800 | 844 | - | - | 0 | - |
| - | - | 2321 | 844.5 | - | - | 0 | - |
| 8 | c | 2.461E+04 | 846.5 | 0.0003007 | 0.3552 | +1 | 8 |
| - | - | 1.135E+04 | 847.5 | - | - | 0 | - |
| - | - | 897.8 | 847.9 | - | - | 0 | - |
| - | - | 3519 | 848.5 | - | - | 0 | - |
| - | - | 2773 | 848.9 | - | - | 0 | - |
| - | - | 848.4 | 849.4 | - | - | 0 | - |
| 2 | z | 2621 | 849.9 | 0.002785 | 3.277 | +2 | 16 |
| 2 | z | 3091 | 850.4 | 0.01102 | 12.96 | +2 | 16 |
| - | - | 859.6 | 852.9 | - | - | 0 | - |
| 16 | c | 3473 | 856.9 | 0.002906 | 3.391 | +2 | 16 |
| 10 | y | 4373 | 857.4 | 0.008928 | 10.41 | +1 | 8 |
| 2 | y | 3210 | 857.9 | 0.008316 | 9.693 | +2 | 16 |
| 2 | y | 2.361E+04 | 858.4 | 0.0171 | 19.92 | +2 | 16 |
| 2 | z | 2705 | 858.9 | 0.01063 | 12.37 | +2 | 16 |
| - | - | 1.027E+04 | 859.4 | - | - | 0 | - |
| - | - | 1188 | 859.9 | - | - | 0 | - |
| - | - | 2007 | 860.4 | - | - | 0 | - |
| - | - | 2383 | 865 | - | - | 0 | - |
| 16 | c | 4.247E+04 | 865.5 | 6.547E-06 | 0.007565 | +2 | 16 |
| - | - | 3.878E+04 | 866 | - | - | 0 | - |
| - | - | 2.631E+04 | 866.5 | - | - | 0 | - |
| - | - | 1.146E+04 | 867 | - | - | 0 | - |
| - | - | 5007 | 867.5 | - | - | 0 | - |
| - | - | 2010 | 868.4 | - | - | 0 | - |
| - | - | 776 | 869.4 | - | - | 0 | - |
| - | - | 917 | 871 | - | - | 0 | - |
| - | - | 1184 | 871.5 | - | - | 0 | - |
| - | - | 867.8 | 872 | - | - | 0 | - |
| - | - | 1094 | 872.9 | - | - | 0 | - |
| - | - | 2686 | 873.5 | - | - | 0 | - |
| - | - | 848.3 | 873.9 | - | - | 0 | - |
| 10 | y | 4696 | 874.5 | 0.001385 | 1.584 | +1 | 8 |
| - | - | 1751 | 875.5 | - | - | 0 | - |
| - | - | 2413 | 880.5 | - | - | 0 | - |
| - | - | 6732 | 881 | - | - | 0 | - |
| - | - | 6680 | 881.5 | - | - | 0 | - |
| - | - | 3973 | 882 | - | - | 0 | - |
| - | - | 1883 | 882.5 | - | - | 0 | - |
| - | - | 2007 | 885.5 | - | - | 0 | - |
| - | - | 3629 | 886 | - | - | 0 | - |
| - | - | 3384 | 886.5 | - | - | 0 | - |
| - | - | 1484 | 887 | - | - | 0 | - |
| - | - | 1409 | 888 | - | - | 0 | - |
| - | - | 927.9 | 888.5 | - | - | 0 | - |
| - | - | 1055 | 889 | - | - | 0 | - |
| - | - | 2614 | 889.4 | - | - | 0 | - |
| - | - | 2.154E+04 | 894.5 | - | - | 0 | - |
| - | - | 1.899E+04 | 895 | - | - | 0 | - |
| - | - | 1.038E+04 | 895.5 | - | - | 0 | - |
| - | - | 3521 | 896 | - | - | 0 | - |
| - | - | 1143 | 896.5 | - | - | 0 | - |
| - | - | 980.2 | 899.5 | - | - | 0 | - |
| - | - | 1733 | 900 | - | - | 0 | - |
| - | - | 1390 | 900.5 | - | - | 0 | - |
| - | - | 831.5 | 902 | - | - | 0 | - |
| - | - | 8636 | 907.4 | - | - | 0 | - |
| - | - | 3925 | 908 | - | - | 0 | - |
| - | - | 2581 | 908.4 | - | - | 0 | - |
| - | - | 5058 | 908.5 | - | - | 0 | - |
| - | - | 3538 | 909 | - | - | 0 | - |
| - | - | 1927 | 909.5 | - | - | 0 | - |
| - | - | 1223 | 910 | - | - | 0 | - |
| - | - | 1531 | 916 | - | - | 0 | - |
| - | - | 4.853E+04 | 916.5 | - | - | 0 | - |
| - | - | 6.93E+04 | 917 | - | - | 0 | - |
| - | - | 5.297E+04 | 917.5 | - | - | 0 | - |
| - | - | 2.917E+04 | 918 | - | - | 0 | - |
| - | - | 1.058E+04 | 918.5 | - | - | 0 | - |
| - | - | 2778 | 919 | - | - | 0 | - |
| 9 | w | 5386 | 928.5 | 0.0003534 | 0.3807 | +1 | 9 |
| - | - | 5279 | 929.5 | - | - | 0 | - |
| - | - | 2299 | 930.5 | - | - | 0 | - |
| - | - | 3039 | 944.5 | - | - | 0 | - |
| - | - | 5080 | 945.5 | - | - | 0 | - |
| - | - | 1803 | 946.5 | - | - | 0 | - |
| - | - | 1781 | 959.5 | - | - | 0 | - |
| - | - | 6777 | 960.5 | - | - | 0 | - |
| - | - | 2337 | 961.5 | - | - | 0 | - |
| - | - | 828.8 | 962.5 | - | - | 0 | - |
| - | - | 1.225E+04 | 973.5 | - | - | 0 | - |
| - | - | 7510 | 974.5 | - | - | 0 | - |
| 9 | c | 4.003E+04 | 975.5 | 0.0009016 | 0.9242 | +1 | 9 |
| - | - | 2.006E+04 | 976.5 | - | - | 0 | - |
| - | - | 7939 | 977.5 | - | - | 0 | - |
| - | - | 1994 | 978.5 | - | - | 0 | - |
| 9 | z | 1.888E+04 | 987.5 | 0.0004742 | 0.4802 | +1 | 9 |
| - | - | 9858 | 988.5 | - | - | 0 | - |
| - | - | 3963 | 989.5 | - | - | 0 | - |
| - | - | 887.9 | 990.5 | - | - | 0 | - |
| - | - | 1526 | 1002 | - | - | 0 | - |
| 9 | y | 4518 | 1004 | 0.005404 | 5.385 | +1 | 9 |
| - | - | 2223 | 1005 | - | - | 0 | - |
| - | - | 1303 | 1005 | - | - | 0 | - |
| - | - | 1499 | 1013 | - | - | 0 | - |
| - | - | 970.9 | 1014 | - | - | 0 | - |
| - | - | 984.1 | 1018 | - | - | 0 | - |
| - | - | 840.8 | 1019 | - | - | 0 | - |
| - | - | 823.2 | 1021 | - | - | 0 | - |
| - | - | 772.7 | 1032 | - | - | 0 | - |
| - | - | 4089 | 1055 | - | - | 0 | - |
| 8 | z | 3380 | 1057 | 0.003688 | 3.491 | +1 | 10 |
| 8 | z | 1507 | 1057 | 0.01962 | 18.55 | +1 | 10 |
| - | - | 892.4 | 1059 | - | - | 0 | - |
| - | - | 1062 | 1060 | - | - | 0 | - |
| - | - | 2570 | 1061 | - | - | 0 | - |
| - | - | 1510 | 1062 | - | - | 0 | - |
| 8 | y | 1878 | 1073 | 0.01301 | 12.13 | +1 | 10 |
| 8 | y | 1552 | 1074 | 0.001751 | 1.631 | +1 | 10 |
| 8 | z | 1.997E+04 | 1075 | 0.0005812 | 0.5409 | +1 | 10 |
| - | - | 1.246E+04 | 1076 | - | - | 0 | - |
| - | - | 6937 | 1077 | - | - | 0 | - |
| - | - | 1467 | 1078 | - | - | 0 | - |
| 10 | c | 729.6 | 1087 | 0.01804 | 16.6 | +1 | 10 |
| - | - | 3653 | 1088 | - | - | 0 | - |
| - | - | 1864 | 1089 | - | - | 0 | - |
| - | - | 1822 | 1090 | - | - | 0 | - |
| 8 | y | 1.324E+04 | 1091 | 0.0009947 | 0.9121 | +1 | 10 |
| - | - | 6197 | 1092 | - | - | 0 | - |
| - | - | 2398 | 1093 | - | - | 0 | - |
| - | - | 1137 | 1094 | - | - | 0 | - |
| - | - | 1739 | 1099 | - | - | 0 | - |
| - | - | 815.3 | 1100 | - | - | 0 | - |
| - | - | 842.9 | 1103 | - | - | 0 | - |
| - | - | 2621 | 1104 | - | - | 0 | - |
| 10 | c | 4.416E+04 | 1105 | 0.001747 | 1.581 | +1 | 10 |
| - | - | 2.622E+04 | 1106 | - | - | 0 | - |
| - | - | 8579 | 1107 | - | - | 0 | - |
| - | - | 3009 | 1108 | - | - | 0 | - |
| - | - | 8517 | 1121 | - | - | 0 | - |
| - | - | 5003 | 1122 | - | - | 0 | - |
| - | - | 2068 | 1123 | - | - | 0 | - |
| - | - | 904.8 | 1134 | - | - | 0 | - |
| - | - | 1111 | 1143 | - | - | 0 | - |
| 7 | y | 1016 | 1161 | 0.008419 | 7.254 | +1 | 11 |
| 7 | z | 9591 | 1162 | 1.684E-05 | 0.0145 | +1 | 11 |
| - | - | 6247 | 1163 | - | - | 0 | - |
| - | - | 3986 | 1164 | - | - | 0 | - |
| - | - | 2241 | 1165 | - | - | 0 | - |
| 7 | y | 5162 | 1178 | 0.0006745 | 0.5728 | +1 | 11 |
| - | - | 4960 | 1179 | - | - | 0 | - |
| - | - | 2029 | 1180 | - | - | 0 | - |
| - | - | 1447 | 1181 | - | - | 0 | - |
| - | - | 1192 | 1182 | - | - | 0 | - |
| - | - | 1404 | 1183 | - | - | 0 | - |
| - | - | 3277 | 1199 | - | - | 0 | - |
| - | - | 1497 | 1200 | - | - | 0 | - |
| - | - | 1821 | 1201 | - | - | 0 | - |
| - | - | 1.292E+04 | 1207 | - | - | 0 | - |
| - | - | 9053 | 1208 | - | - | 0 | - |
| - | - | 2310 | 1209 | - | - | 0 | - |
| 11 | c | 2.551E+04 | 1218 | 0.001826 | 1.5 | +1 | 11 |
| - | - | 1.846E+04 | 1219 | - | - | 0 | - |
| - | - | 7426 | 1220 | - | - | 0 | - |
| - | - | 1853 | 1221 | - | - | 0 | - |
| - | - | 7799 | 1223 | - | - | 0 | - |
| - | - | 1.211E+04 | 1224 | - | - | 0 | - |
| - | - | 7131 | 1225 | - | - | 0 | - |
| - | - | 1458 | 1226 | - | - | 0 | - |
| - | - | 2334 | 1253 | - | - | 0 | - |
| - | - | 1507 | 1254 | - | - | 0 | - |
| 6 | y | 901.4 | 1258 | 0.01742 | 13.85 | +1 | 12 |
| - | - | 1486 | 1270 | - | - | 0 | - |
| - | - | 816.7 | 1271 | - | - | 0 | - |
| 6 | y | 2348 | 1275 | 0.0003947 | 0.3096 | +1 | 12 |
| - | - | 1855 | 1276 | - | - | 0 | - |
| - | - | 1010 | 1277 | - | - | 0 | - |
| - | - | 1548 | 1302 | - | - | 0 | - |
| - | - | 784.5 | 1303 | - | - | 0 | - |
| - | - | 852.7 | 1328 | - | - | 0 | - |
| 12 | c | 1449 | 1329 | 0.02067 | 15.56 | +1 | 12 |
| - | - | 2507 | 1330 | - | - | 0 | - |
| - | - | 2399 | 1331 | - | - | 0 | - |
| - | - | 678.6 | 1332 | - | - | 0 | - |
| - | - | 789.8 | 1344 | - | - | 0 | - |
| - | - | 3240 | 1345 | - | - | 0 | - |
| 12 | c | 3.56E+04 | 1346 | 0.002664 | 1.98 | +1 | 12 |
| - | - | 2.999E+04 | 1347 | - | - | 0 | - |
| - | - | 1.009E+04 | 1348 | - | - | 0 | - |
| - | - | 3497 | 1349 | - | - | 0 | - |
| - | - | 2708 | 1371 | - | - | 0 | - |
| 5 | y | 9397 | 1372 | 0.002809 | 2.048 | +1 | 13 |
| - | - | 7692 | 1373 | - | - | 0 | - |
| - | - | 3862 | 1374 | - | - | 0 | - |
| - | - | 1576 | 1375 | - | - | 0 | - |
| - | - | 1375 | 1399 | - | - | 0 | - |
| - | - | 2788 | 1400 | - | - | 0 | - |
| - | - | 4796 | 1401 | - | - | 0 | - |
| - | - | 4672 | 1402 | - | - | 0 | - |
| - | - | 2363 | 1403 | - | - | 0 | - |
| - | - | 1096 | 1404 | - | - | 0 | - |
| - | - | 4567 | 1416 | - | - | 0 | - |
| 13 | c | 2.155E+04 | 1417 | 0.004134 | 2.918 | +1 | 13 |
| - | - | 1.558E+04 | 1418 | - | - | 0 | - |
| - | - | 7139 | 1419 | - | - | 0 | - |
| - | - | 1796 | 1420 | - | - | 0 | - |
| - | - | 2341 | 1431 | - | - | 0 | - |
| - | - | 1728 | 1432 | - | - | 0 | - |
| - | - | 924.7 | 1433 | - | - | 0 | - |
| - | - | 1001 | 1443 | - | - | 0 | - |
| - | - | 966.2 | 1459 | - | - | 0 | - |
| - | - | 1231 | 1487 | - | - | 0 | - |
| - | - | 1595 | 1488 | - | - | 0 | - |
| 4 | y | 1535 | 1502 | 0.005253 | 3.498 | +1 | 14 |
| 4 | z | 2.007E+04 | 1503 | 0.002816 | 1.874 | +1 | 14 |
| - | - | 3.141E+04 | 1504 | - | - | 0 | - |
| - | - | 1.952E+04 | 1505 | - | - | 0 | - |
| - | - | 1.012E+04 | 1506 | - | - | 0 | - |
| - | - | 3259 | 1507 | - | - | 0 | - |
| 14 | c | 1056 | 1514 | 0.01147 | 7.577 | +1 | 14 |
| 4 | y | 3262 | 1519 | 0.0001886 | 0.1242 | +1 | 14 |
| - | - | 2611 | 1520 | - | - | 0 | - |
| - | - | 1667 | 1521 | - | - | 0 | - |
| - | - | 1096 | 1522 | - | - | 0 | - |
| - | - | 914.5 | 1530 | - | - | 0 | - |
| 14 | c | 3.828E+04 | 1531 | 0.002872 | 1.876 | +1 | 14 |
| - | - | 3.434E+04 | 1532 | - | - | 0 | - |
| - | - | 1.606E+04 | 1533 | - | - | 0 | - |
| - | - | 5950 | 1534 | - | - | 0 | - |
| - | - | 1395 | 1535 | - | - | 0 | - |
| 3 | y | 1605 | 1615 | 0.03044 | 18.85 | +1 | 15 |
| 3 | z | 4508 | 1616 | 0.02066 | 12.79 | +1 | 15 |
| - | - | 6757 | 1617 | - | - | 0 | - |
| - | - | 5261 | 1618 | - | - | 0 | - |
| - | - | 1529 | 1619 | - | - | 0 | - |
| 3 | y | 912.7 | 1632 | 0.03038 | 18.62 | +1 | 15 |
| - | - | 902.2 | 1643 | - | - | 0 | - |
| - | - | 1514 | 1644 | - | - | 0 | - |
| - | - | 1115 | 1645 | - | - | 0 | - |
| 15 | c | 4158 | 1659 | 0.00262 | 1.579 | +1 | 15 |
| - | - | 1.043E+04 | 1660 | - | - | 0 | - |
| - | - | 8741 | 1661 | - | - | 0 | - |
| - | - | 5457 | 1662 | - | - | 0 | - |
| - | - | 832.1 | 1663 | - | - | 0 | - |
| - | - | 1571 | 1673 | - | - | 0 | - |
| - | - | 802.9 | 1674 | - | - | 0 | - |
| - | - | 2429 | 1686 | - | - | 0 | - |
| - | - | 3207 | 1687 | - | - | 0 | - |
| - | - | 2108 | 1688 | - | - | 0 | - |
| - | - | 915.1 | 1689 | - | - | 0 | - |
| - | - | 1782 | 1701 | - | - | 0 | - |
| - | - | 2932 | 1714 | - | - | 0 | - |
| 2 | y | 3103 | 1715 | 0.0272 | 15.86 | +1 | 16 |
| 2 | y | 2509 | 1716 | 0.02805 | 16.35 | +1 | 16 |
| 2 | z | 2689 | 1717 | 0.01339 | 7.8 | +1 | 16 |
| - | - | 9204 | 1718 | - | - | 0 | - |
| - | - | 6645 | 1719 | - | - | 0 | - |
| - | - | 3192 | 1720 | - | - | 0 | - |
| 16 | c | 4992 | 1730 | 0.005676 | 3.281 | +1 | 16 |
| - | - | 1.4E+04 | 1731 | - | - | 0 | - |
| - | - | 1.247E+04 | 1732 | - | - | 0 | - |
| - | - | 6436 | 1733 | - | - | 0 | - |
| - | - | 2313 | 1734 | - | - | 0 | - |
| - | - | 970.5 | 1735 | - | - | 0 | - |
| - | - | 1083 | 1762 | - | - | 0 | - |
| - | - | 2466 | 1771 | - | - | 0 | - |
| - | - | 4948 | 1772 | - | - | 0 | - |
| - | - | 3158 | 1773 | - | - | 0 | - |
| - | - | 2541 | 1774 | - | - | 0 | - |
| - | - | 1120 | 1775 | - | - | 0 | - |
| - | - | 6549 | 1788 | - | - | 0 | - |
| - | - | 1.934E+04 | 1789 | - | - | 0 | - |
| - | - | 1.951E+04 | 1790 | - | - | 0 | - |
| - | - | 9028 | 1791 | - | - | 0 | - |
| - | - | 3622 | 1792 | - | - | 0 | - |
| - | - | 1389 | 1800 | - | - | 0 | - |
| - | - | 1263 | 1805 | - | - | 0 | - |
| - | - | 5799 | 1806 | - | - | 0 | - |
| - | - | 4170 | 1807 | - | - | 0 | - |
| - | - | 2658 | 1808 | - | - | 0 | - |
| - | - | 983.6 | 1815 | - | - | 0 | - |
| - | - | 7257 | 1816 | - | - | 0 | - |
| - | - | 3.144E+04 | 1817 | - | - | 0 | - |
| - | - | 2.7E+04 | 1818 | - | - | 0 | - |
| - | - | 1.578E+04 | 1819 | - | - | 0 | - |
| - | - | 5965 | 1820 | - | - | 0 | - |
| - | - | 1534 | 1821 | - | - | 0 | - |
| - | - | 3793 | 1832 | - | - | 0 | - |
| - | - | 2.102E+04 | 1833 | - | - | 0 | - |
| - | - | 5.354E+04 | 1834 | - | - | 0 | - |
| - | - | 4.809E+04 | 1835 | - | - | 0 | - |
| - | - | 2.588E+04 | 1836 | - | - | 0 | - |
| - | - | 9787 | 1837 | - | - | 0 | - |
| - | - | 2609 | 1838 | - | - | 0 | - |
| - | - | 779.6 | 1839 | - | - | 0 | - |
| - | - | 825.3 | 3069 | - | - | 0 | - |

m/z Charge Intensity FragmentType MassShift Position
120.06145477294922 0 695.6927
120.06558990478516 0 10644.592 y 16
120.0809326171875 0 2014.8103
126.0550308227539 0 1157.7925
128.0946502685547 0 830.38763
129.10227966308594 0 4638.9907
129.22695922851562 0 394.20547
131.11801147460938 0 929.6794
140.10690307617188 0 1186.6373
146.6717987060547 0 450.35645
149.0447998046875 0 1075.2426
153.8125762939453 0 439.04633
155.11795043945312 0 575.4869
160.5535888671875 0 491.85083
169.1337432861328 0 2654.1638
171.07699584960938 0 727.4811
173.0923614501953 0 1382.1315 y Water loss 15
173.1284942626953 0 2949.7917
177.67529296875 0 544.94916
181.0967559814453 0 516.9248
183.1127471923828 0 2397.856
187.14402770996094 0 628.4204
191.10256958007812 0 2413.902 y 15
197.1284637451172 0 2236.384
201.12330627441406 0 5982.756
207.37342834472656 0 502.82608
211.10739135742188 0 527.6333
212.13934326171875 0 8243.809
213.14260864257812 0 815.0349
215.1387176513672 0 871.0363
217.13307189941406 0 618.8862
218.1497039794922 0 3147.7217 c 1
221.08433532714844 0 3300.861
226.1187744140625 0 717.24963
227.08999633789062 0 1347.6384
227.10296630859375 0 1744.1807
233.16497802734375 0 1258.8066
235.10702514648438 0 1139.1215
238.15493774414062 0 599.9901
239.09495544433594 0 5430.1455
240.09571838378906 0 1594.1486
241.0916290283203 0 729.372
243.14479064941406 0 898.2774 y Water loss 10
254.15069580078125 0 1013.51337
263.1021423339844 0 2807.7314
281.05145263671875 0 1051.4863
282.1448059082031 0 1197.9187
295.1037902832031 0 738.5546
296.19683837890625 0 11361.847
297.08270263671875 0 698.36523
297.155517578125 0 863.44366
297.2002258300781 0 1572.0522
298.1618347167969 0 785.1759
299.0616760253906 0 6149.4663
299.1715087890625 0 14314.773
300.06329345703125 0 1231.8602
300.1744079589844 0 2018.9429
301.0594787597656 0 772.9765
303.17889404296875 0 6997.7744 z 14
304.1827087402344 0 1124.5336
307.6671447753906 0 710.82446 y Water loss 11
314.18426513671875 0 1104.9292
314.20745849609375 0 4225.806
315.20953369140625 0 620.0324
316.18670654296875 0 7792.422
316.67218017578125 0 1256.1443 y 11
317.1889343261719 0 664.0662
319.1979675292969 0 3263.6194 y 14
331.23406982421875 0 24248.996 c 2
332.2375183105469 0 4362.5054
343.3712463378906 0 735.0915
345.1793518066406 0 783.26654
347.1959228515625 0 583.472
350.13427734375 0 713.8748
355.06976318359375 0 38171.684
356.0702819824219 0 16629.2
357.067626953125 0 10089.685
358.0687561035156 0 1090.9176
362.2061462402344 0 1012.63245
369.2012023925781 0 615.7883
371.1013488769531 0 1687.2555
372.0997314453125 0 1024.017
373.07965087890625 0 840.66833
373.2090148925781 0 3453.2139 y 10
378.2270812988281 0 855.87897
385.2192687988281 0 924.34467
386.2035217285156 0 1711.2217
407.2237548828125 0 667.3139
415.0370178222656 0 2670.3718
415.2298583984375 0 752.4796 y Water loss 13
417.22344970703125 0 2013.876 z 13
418.2296447753906 0 5839.215
418.99517822265625 0 578.83484
419.23309326171875 0 1533.1431
433.24114990234375 0 2337.379 y 13
433.2809753417969 0 7895.476
434.25091552734375 0 844.97076
434.28619384765625 0 4358.6665
435.29486083984375 0 2692.9788
442.23785400390625 0 815.9972
444.2458801269531 0 5423.2163
445.2001647949219 0 576.0258
445.252685546875 0 2233.5051
446.2649841308594 0 1638.6981
460.2676086425781 0 1249.2542
461.27587890625 0 11167.735
462.2790832519531 0 2550.2788
463.2818298339844 0 785.558
480.208984375 0 653.5853
480.7205505371094 0 1476.2212
488.2590026855469 0 9222.817 c 8
489.057373046875 0 1449.2042
489.26470947265625 0 6157.7393
490.05889892578125 0 912.81165
490.26788330078125 0 1843.8018
493.7291564941406 0 1904.1252
494.2294616699219 0 1163.0884
498.2203369140625 0 1044.1583
502.2646484375 0 2047.9257 y 8
502.73211669921875 0 3178.7947
503.2337646484375 0 2228.5188
503.2702941894531 0 828.61273
504.27777099609375 0 8076.141 y 12
505.28106689453125 0 1531.6573
518.2000122070312 0 740.2337
529.2662963867188 0 623.57684 w 7
536.2800903320312 0 638.18286
538.2693481445312 0 6252.1255
538.7711791992188 0 3654.597
539.2722778320312 0 973.1751 z 2
543.2611694335938 0 2100.7974
543.3170776367188 0 635.98914
543.7638549804688 0 1123.4258
544.26513671875 0 1443.6423
552.2669067382812 0 6640.4624
552.3169555664062 0 812.1667
552.7689208984375 0 5595.1978
553.271728515625 0 1613.1914
554.3172607421875 0 1272.0215
558.2877197265625 0 15105.327 w 11
558.3301391601562 0 2709.75
559.2915649414062 0 4718.3794
559.3310546875 0 595.2604
560.29345703125 0 1420.2976
572.3032836914062 0 1470.4111 y Water loss 1
577.1273193359375 0 1179.459
578.1296997070312 0 702.93634
579.1221313476562 0 775.5068
580.2862548828125 0 942.2793 y Water loss 6
583.3380737304688 0 842.5252
591.2913818359375 0 800.5753
599.8067626953125 0 1841.8367
600.3076171875 0 1265.642 c Water loss 10
602.7908325195312 0 2962.5632
603.2918701171875 0 1736.3708
610.1841430664062 0 8400.849
611.1854248046875 0 8783.417
611.7964477539062 0 7300.8433
612.1828002929688 0 6640.5674
612.2966918945312 0 4486.1753
612.8007202148438 0 2310.901
613.184326171875 0 712.81976
614.3367919921875 0 8255.054 y Water loss 11
615.3448486328125 0 11058.466
616.3159790039062 0 4304.299 z 11
616.3578491210938 0 1539.6042
617.3225708007812 0 1236.7311
617.805908203125 0 1530.8658
626.3426513671875 0 2540.573
626.8116455078125 0 3251.9316
627.260498046875 0 3599.2756
627.310791015625 0 1729.4713
627.3552856445312 0 1494.3419
627.8158569335938 0 909.9957
628.2631225585938 0 1423.1256
628.3927612304688 0 3443.3093
629.3950805664062 0 1330.3497
630.3549194335938 0 1251.0865
632.3363647460938 0 5802.429 y 11
633.33642578125 0 1225.5581
635.3238525390625 0 1191.7262
637.8161010742188 0 553.30304 y 5
654.4091186523438 0 789.1812 c Water loss 5
655.341064453125 0 2042.1375
655.842529296875 0 1808.119
656.3364868164062 0 937.24316
658.2335205078125 0 675.798
658.3690795898438 0 1022.5538
664.8345336914062 0 767.34467 c Ammonia loss 11
668.33056640625 0 990.7204
671.3721923828125 0 2737.9312
672.4069213867188 0 3850.367 c 5
673.4155883789062 0 1208.2761
676.8328247070312 0 1847.9686
677.3353271484375 0 7561.794 y Water loss 4
677.8365478515625 0 5781.937 y Ammonia loss 4
678.3341064453125 0 3492.9517 z 4
685.8370361328125 0 4528.8184
686.3436279296875 0 49952.11 y 4
686.8423461914062 0 20659.107
687.3482666015625 0 13747.774
687.8458251953125 0 3332.3396
688.3546142578125 0 3031.3533
697.415283203125 0 1399.1997
698.4212646484375 0 1020.93036
713.4056396484375 0 852.2203
716.44287109375 0 870.06335
722.3484497070312 0 1796.1183
723.3519287109375 0 988.77814
729.4006958007812 0 11339.072 z 10
730.4041748046875 0 3920.1584
731.40966796875 0 1427.9559
740.344970703125 0 1470.0236
742.4138793945312 0 2223.1965
745.4208374023438 0 6890.0083 y 10
746.423095703125 0 2029.4661
750.8671264648438 0 1039.1274 y Water loss 3
751.3671875 0 1786.1438 y Ammonia loss 3
751.8648071289062 0 939.04803 z 3
757.42138671875 0 2292.425
759.3709106445312 0 2065.1646
759.4397583007812 0 29769.883 c 6
759.8749389648438 0 8735.98 y 3
760.3749389648438 0 6642.873
760.4432983398438 0 13319.851
760.87646484375 0 2805.4617
761.376953125 0 2071.1401
761.4464721679688 0 4424.555
764.3612060546875 0 5879.807
765.3626098632812 0 3327.3193
766.3600463867188 0 1259.9114
798.3911743164062 0 988.6802
798.890869140625 0 1183.2502
799.42919921875 0 10977.121 w 9
800.4308471679688 0 5272.498
801.4359741210938 0 1613.5872
802.3816528320312 0 4428.97
803.3822021484375 0 2202.7886
803.461181640625 0 848.22003
807.4214477539062 0 1861.9136 y Water loss 2
807.92578125 0 2239.5984
808.433349609375 0 1049.9572
813.9231567382812 0 1195.0376
814.4254150390625 0 1896.7983
815.384521484375 0 1228.0444
816.4172973632812 0 2861.342 y 2
816.91650390625 0 1604.8901
817.4193115234375 0 1026.1016
817.923095703125 0 750.84766
822.3775024414062 0 3665.0728
823.3850708007812 0 5500.9487
824.3883666992188 0 2258.1655
829.4432373046875 0 858.46484
829.9404907226562 0 26525.72 c 14
830.4417724609375 0 30668.963
830.943115234375 0 15978.563
831.4432373046875 0 5393.1875
831.950439453125 0 935.6799
835.9376220703125 0 1015.61127
841.4307861328125 0 1660.489 z Ammonia loss 9
842.4362182617188 0 1031.3224
843.4506225585938 0 2443.66
843.9523315429688 0 2799.858
844.45654296875 0 2321.3325
846.4716796875 0 24607.709 c 7
847.4745483398438 0 11346.335
847.9415893554688 0 897.775
848.4763793945312 0 3519.4766
848.9320068359375 0 2773.2358
849.4285278320312 0 848.44055
849.9293823242188 0 2621.1333 z Water loss 1
850.4296264648438 0 3091.1287 z Ammonia loss 1
852.928466796875 0 859.5739
856.9434204101562 0 3473.1838 c Ammonia loss 15
857.4452514648438 0 4372.979 y Ammonia loss 9
857.9442749023438 0 3209.5483 y Water loss 1
858.445068359375 0 23614.49 y Ammonia loss 1
858.9425048828125 0 2704.9177 z 1
859.4454345703125 0 10269.244
859.938720703125 0 1188.1891
860.4494018554688 0 2006.6498
864.9580688476562 0 2383.023
865.4595947265625 0 42467.555 c 15
865.9606323242188 0 38777.11
866.4609375 0 26310.896
866.9593505859375 0 11463.912
867.460693359375 0 5006.5366
868.4017333984375 0 2009.5785
869.4105224609375 0 775.96216
870.9500122070312 0 916.99115
871.4534301757812 0 1184.1
871.956298828125 0 867.83636
872.94287109375 0 1094.4783
873.456298828125 0 2685.9307
873.94091796875 0 848.34955
874.4614868164062 0 4695.9556 y 9
875.46533203125 0 1751.0615
880.45361328125 0 2413.1506
880.963134765625 0 6732.333
881.4605712890625 0 6679.5854
881.9657592773438 0 3972.614
882.4666748046875 0 1883.0986
885.4651489257812 0 2006.6549
885.9649658203125 0 3628.7217
886.4657592773438 0 3383.6677
886.9609375 0 1484.265
887.971435546875 0 1408.6293
888.4717407226562 0 927.8738
888.955078125 0 1055.4893
889.4464111328125 0 2613.8345
894.4688720703125 0 21540.191
894.9697875976562 0 18987.896
895.470947265625 0 10381.562
895.9699096679688 0 3520.964
896.4710693359375 0 1142.889
899.4659423828125 0 980.2487
899.959228515625 0 1732.556
900.4571533203125 0 1389.9124
901.9580078125 0 831.5362
907.4049072265625 0 8635.783
907.9724731445312 0 3924.8188
908.3978881835938 0 2581.178
908.4724731445312 0 5058.3726
908.9710083007812 0 3537.5378
909.4739990234375 0 1927.1152
909.9667358398438 0 1222.6113
915.97265625 0 1531.4542
916.4747314453125 0 48533.098
916.9771118164062 0 69304.984
917.4788208007812 0 52971.855
917.980712890625 0 29169.494
918.4818115234375 0 10576.805
918.9826049804688 0 2777.7056
928.4730834960938 0 5386.094 w 8
929.4779663085938 0 5278.8154
930.4819946289062 0 2299.1604
944.4738159179688 0 3039.131
945.4771728515625 0 5080
946.479248046875 0 1802.9259
959.4896240234375 0 1781.0271
960.4893798828125 0 6777.312
961.4915161132812 0 2337.4211
962.4867553710938 0 828.75287
973.49462890625 0 12248.538
974.4985961914062 0 7509.5337
975.513671875 0 40033.016 c 8
976.5166625976562 0 20061.965
977.5196533203125 0 7939.252
978.52197265625 0 1993.9608
987.4862670898438 0 18876.648 z 8
988.4871215820312 0 9857.757
989.49267578125 0 3963.1887
990.4965209960938 0 887.86176
1001.5252685546875 0 1526.3319
1003.5000610351562 0 4517.7 y 8
1004.5022583007812 0 2222.6445
1005.4910278320312 0 1302.6964
1012.5177001953125 0 1498.6368
1013.519287109375 0 970.93353
1018.4541625976562 0 984.142
1019.4592895507812 0 840.7939
1021.4791259765625 0 823.24963
1031.5018310546875 0 772.7209
1055.49951171875 0 4088.525
1056.5045166015625 0 3379.5837 z Water loss 7
1057.5118408203125 0 1507.2471 z Ammonia loss 7
1058.5040283203125 0 892.44244
1059.5118408203125 0 1061.8564
1060.532470703125 0 2570.3528
1061.534912109375 0 1509.8735
1072.513916015625 0 1878.4806 y Water loss 7
1073.5126953125 0 1551.5817 y Ammonia loss 7
1074.5181884765625 0 19974.027 z 7
1075.5228271484375 0 12458.48
1076.5269775390625 0 6936.5015
1077.5355224609375 0 1467.3867
1086.528564453125 0 729.6356 c Water loss 9
1087.529296875 0 3653.1301
1088.5284423828125 0 1864.1333
1089.53173828125 0 1822.419
1090.5364990234375 0 13238.733 y 7
1091.5384521484375 0 6196.623
1092.5411376953125 0 2397.9597
1093.5419921875 0 1137.0056
1098.5018310546875 0 1738.7721
1099.516845703125 0 815.25934
1102.5452880859375 0 842.86835
1103.5277099609375 0 2621.138
1104.555419921875 0 44162.41 c 9
1105.557861328125 0 26218.74
1106.5606689453125 0 8579.054
1107.564208984375 0 3009.4385
1120.5528564453125 0 8517.204
1121.55615234375 0 5003.027
1122.5589599609375 0 2067.6487
1133.5570068359375 0 904.75854
1142.5301513671875 0 1111.1588
1160.5513916015625 0 1015.9002 y Ammonia loss 6
1161.55078125 0 9591.195 z 6
1162.5533447265625 0 6247.12
1163.55908203125 0 3986.3525
1164.556884765625 0 2241.2505
1177.56884765625 0 5162.0425 y 6
1178.5733642578125 0 4959.6704
1179.5740966796875 0 2028.7639
1180.5709228515625 0 1447.3413
1181.572998046875 0 1192.0359
1182.58984375 0 1404.4271
1198.6094970703125 0 3277.1147
1199.60498046875 0 1496.8423
1200.608642578125 0 1821.3893
1206.5665283203125 0 12917.076
1207.5677490234375 0 9053.001
1208.5709228515625 0 2309.643
1217.639404296875 0 25514.26 c 10
1218.64208984375 0 18463.926
1219.6441650390625 0 7426.428
1220.6474609375 0 1853.3901
1222.5828857421875 0 7799.4
1223.587646484375 0 12111.51
1224.5927734375 0 7131.4937
1225.598388671875 0 1458.4199
1252.6146240234375 0 2333.598
1253.619873046875 0 1506.7709
1257.6131591796875 0 901.3734 y Ammonia loss 5
1269.6400146484375 0 1486.2504
1270.640625 0 816.66724
1274.6226806640625 0 2348.034 y 5
1275.6251220703125 0 1855.0304
1276.628662109375 0 1010.21844
1301.67431640625 0 1547.97
1302.6533203125 0 784.4756
1327.65869140625 0 852.7251
1328.652587890625 0 1449.1246 c Ammonia loss 11
1329.6424560546875 0 2507.054
1330.659423828125 0 2398.5693
1331.6517333984375 0 678.5872
1343.6756591796875 0 789.7853
1344.685302734375 0 3239.734
1345.6971435546875 0 35599.953 c 11
1346.700439453125 0 29992.482
1347.70458984375 0 10088.227
1348.70361328125 0 3496.8345
1370.666015625 0 2708.1414
1371.6722412109375 0 9396.957 y 4
1372.6800537109375 0 7691.503
1373.682861328125 0 3862.1091
1374.69970703125 0 1575.542
1398.6973876953125 0 1375.1616
1399.674072265625 0 2788.1433
1400.6781005859375 0 4795.9062
1401.68994140625 0 4671.9565
1402.6976318359375 0 2363.1792
1403.7003173828125 0 1096.2229
1415.72705078125 0 4566.756
1416.7327880859375 0 21547.156 c 12
1417.73583984375 0 15577.481
1418.73876953125 0 7138.7915
1419.7445068359375 0 1795.6353
1430.697509765625 0 2340.9055
1431.69970703125 0 1728.4412
1432.709228515625 0 924.6582
1442.7374267578125 0 1000.9458
1458.7154541015625 0 966.1849
1486.7545166015625 0 1230.7655
1487.759765625 0 1595.2644
1501.72216796875 0 1535.0818 y Ammonia loss 3
1502.721923828125 0 20070.727 z 3
1503.7276611328125 0 31413.826
1504.7314453125 0 19516.582
1505.7335205078125 0 10116.03
1506.7381591796875 0 3258.882
1513.7647705078125 0 1056.1193 c Ammonia loss 13
1518.74365234375 0 3262.3882 y 3
1519.7447509765625 0 2611.2903
1520.749267578125 0 1667.2034
1521.751708984375 0 1096.3734
1529.76318359375 0 914.5304
1530.7769775390625 0 38280.09 c 13
1531.7794189453125 0 34342.3
1532.78271484375 0 16056.64
1533.78564453125 0 5950.0737
1534.783447265625 0 1395.2397
1614.8314208984375 0 1605.29 y Ammonia loss 2
1615.8294677734375 0 4507.908 z 2
1616.818115234375 0 6756.9834
1617.8228759765625 0 5261.3286
1618.8328857421875 0 1529.0969
1631.85791015625 0 912.69934 y 2
1642.8414306640625 0 902.1647
1643.862060546875 0 1514.3328
1644.85107421875 0 1114.548
1658.8721923828125 0 4157.9463 c 14
1659.879150390625 0 10433.043
1660.882568359375 0 8741.136
1661.88232421875 0 5457.1143
1662.8839111328125 0 832.13055
1672.8487548828125 0 1570.5674
1673.8651123046875 0 802.856
1685.8909912109375 0 2428.805
1686.895751953125 0 3207.414
1687.9046630859375 0 2108.1106
1688.912109375 0 915.139
1700.8529052734375 0 1782.1567
1713.8873291015625 0 2931.8977
1714.891845703125 0 3103.1633 y Water loss 1
1715.876708984375 0 2508.976 y Ammonia loss 1
1716.869873046875 0 2689.307 z 1
1717.86376953125 0 9204.25
1718.8656005859375 0 6644.9937
1719.87451171875 0 3192.129
1729.90625 0 4992.152 c 15
1730.91552734375 0 14000.845
1731.91748046875 0 12474.106
1732.9180908203125 0 6435.847
1733.9212646484375 0 2313.411
1734.894287109375 0 970.45404
1761.92529296875 0 1083.1332
1770.9200439453125 0 2466.3347
1771.9180908203125 0 4948.4077
1772.914794921875 0 3158.4998
1773.9161376953125 0 2540.7158
1774.908447265625 0 1120.3378
1787.9344482421875 0 6549.036
1788.935791015625 0 19339.375
1789.9393310546875 0 19505.484
1790.943115234375 0 9027.509
1791.943115234375 0 3622.0925
1799.9193115234375 0 1389.2721
1804.9547119140625 0 1262.9215
1805.9573974609375 0 5799.4316
1806.9599609375 0 4170.1353
1807.9625244140625 0 2658.253
1814.932373046875 0 983.5895
1815.9287109375 0 7257.4736
1816.9296875 0 31439.072
1817.9322509765625 0 27004.832
1818.9354248046875 0 15775.033
1819.937255859375 0 5965.0664
1820.9412841796875 0 1533.6272
1831.9393310546875 0 3793.2544
1832.946533203125 0 21015.86
1833.95458984375 0 53544.67
1834.95849609375 0 48093.08
1835.9605712890625 0 25883.123
1836.96435546875 0 9786.894
1837.9627685546875 0 2609.494
1838.9517822265625 0 779.57544
3069.306396484375 0 825.2526

Spectrum Details

|  |  |
| --- | --- |
| Matched peaks? Matched peaks The total absolute number of peaks matched. Additionally in brackets the total fraction of peaks matched and the total number of peaks is shown. | 91 (16.58% of 549) |
| FDR? FDR The false discovery rate estimated for this peptide. It is calculated by matching all theoretical fragments with a non-integer shift with the raw peaks for this spectrum. This is done with 40 different shifts. The resulting percentage is the average number of annotated peaks over the number of annotated peaks with the correct spectrum. | 0.84% |
| Satellite FDR? Satellite FDR See the FDR for details on its calculation. This satellite ion specific FDR only contains the satellite ions (d/w) for I/L/J positions. | - |
| PSM Score? PSM Score The PSM Score as given by Hecklib to this annotated spectrum. It is shown with three significant figures. | 813 |

#### Spectrum 7881? Spectrum 7881 The raw spectrum of this peptide as annotated by Hecklib. The fragments are coloured according to ion type (see legend). Any peaks with a star '\*' as text can be hovered over to see the full details, first the ion type second the mass shift type. By hovering over the amino acids in the peptide or ions in the legend the corresponding peaks are highlighted. By toggling the 'Unassigned' label you can turn the background (unassigned) peaks on or off in the plot. By updating the slider in the Ion legend you can update the spectrum to only show the top X% of the peaks with labels. The top X% means any peak that is within X% of the highest intensity. By dragging in the spectrum you can zoom in to a specific part of the spectrum and use 'Zoom Out' to get back to the original zoom level. The annotation of the spectrum is based on the given sequence in the peptides file and is done with different software so inconsistencies are likely. The peaks are annotated based on the given sequence, with 20 ppm tolerance.

Copy Data

##### Spectrum 7881 (TSV)

###### Preview

```
Loading example...
```

*Click on the button to copy the data to your clipboard.*

Mz MinMz MaxIntensity Max

WidthHeightPeptide font sizePeptide stroke widthSpectrum font sizeSpectrum stroke widthCompact peptide

Ion legend

wxyz

abcd

OtherUnassignedIon^Charge^_Position_Show for top:%

VTJFPPSSEEJQANKAT

04.32e+58.63e+51.30e+61.73e+6

Zoom Out

y^+1^_3_y^+1^_4_y^+1^_5_y^+1^_5_y^+1^_6_z^+1^_6_y^+1^_6_y^+2^_12_z^+2^_13_y^+2^_13_y^+2^_13_z^+2^_13_y^+2^_13_c^+2^_13_z^+1^_7_y^+1^_7_y^+2^_14_y^+2^_14_z^+2^_14_y^+2^_14_y^+2^_15_y^+2^_15_c^+2^_16_y^+1^_8_y^+2^_16_y^+2^_16_y^+2^_16_y^+1^_8_c^+1^_9_z^+1^_9_y^+1^_9_y^+1^_10_y^+1^_10_z^+1^_10_y^+1^_10_c^+1^_10_y^+1^_11_z^+1^_11_y^+1^_11_c^+1^_11_y^+1^_12_y^+1^_12_y^+1^_12_c^+1^_12_c^+1^_12_c^+1^_12_y^+1^_13_y^+1^_13_z^+1^_13_y^+1^_13_c^+1^_13_z^+1^_14_c^+1^_14_y^+1^_14_c^+1^_14_y^+1^_15_z^+1^_15_y^+1^_15_c^+1^_15_z^+1^_16_z^+1^_16_y^+1^_16_y^+1^_16_z^+1^_16_c^+1^_16_

0797159323903187

Fragment Matches Table

Show background peaks

| Position | Ion type | Intensity | mz Theoretical | mz Error (Th) | mz Error (ppm) | Charge | Series Number |
| --- | --- | --- | --- | --- | --- | --- | --- |
| - | - | 4521 | 120.1 | - | - | 0 | - |
| - | - | 8931 | 129.1 | - | - | 0 | - |
| - | - | 2352 | 148.9 | - | - | 0 | - |
| - | - | 2033 | 169.1 | - | - | 0 | - |
| - | - | 7042 | 173.1 | - | - | 0 | - |
| - | - | 1554 | 173.5 | - | - | 0 | - |
| - | - | 1376 | 177 | - | - | 0 | - |
| - | - | 9279 | 183.1 | - | - | 0 | - |
| - | - | 1282 | 187.1 | - | - | 0 | - |
| - | - | 2.486E+04 | 201.1 | - | - | 0 | - |
| - | - | 3916 | 215.1 | - | - | 0 | - |
| - | - | 2912 | 217.1 | - | - | 0 | - |
| - | - | 2001 | 226.1 | - | - | 0 | - |
| - | - | 1.373E+04 | 233.2 | - | - | 0 | - |
| - | - | 4665 | 243.1 | - | - | 0 | - |
| - | - | 5172 | 245.1 | - | - | 0 | - |
| - | - | 1306 | 246 | - | - | 0 | - |
| - | - | 3684 | 247.1 | - | - | 0 | - |
| - | - | 5845 | 251.2 | - | - | 0 | - |
| - | - | 1.207E+04 | 261.2 | - | - | 0 | - |
| - | - | 3550 | 269.2 | - | - | 0 | - |
| - | - | 1200 | 271.2 | - | - | 0 | - |
| - | - | 2766 | 280.2 | - | - | 0 | - |
| - | - | 2254 | 282.1 | - | - | 0 | - |
| - | - | 2.521E+05 | 296.2 | - | - | 0 | - |
| - | - | 1746 | 297.2 | - | - | 0 | - |
| - | - | 3.973E+04 | 297.2 | - | - | 0 | - |
| - | - | 2902 | 298.2 | - | - | 0 | - |
| - | - | 1827 | 298.2 | - | - | 0 | - |
| - | - | 4659 | 303.2 | - | - | 0 | - |
| - | - | 1395 | 304.2 | - | - | 0 | - |
| - | - | 1145 | 307.4 | - | - | 0 | - |
| - | - | 4302 | 314.2 | - | - | 0 | - |
| - | - | 1.27E+05 | 314.2 | - | - | 0 | - |
| - | - | 1.887E+04 | 315.2 | - | - | 0 | - |
| - | - | 1534 | 317.2 | - | - | 0 | - |
| 15 | y | 1.665E+04 | 319.2 | 0.000188 | 0.589 | +1 | 3 |
| - | - | 4733 | 330.2 | - | - | 0 | - |
| - | - | 1816 | 344.2 | - | - | 0 | - |
| - | - | 1.156E+04 | 346.2 | - | - | 0 | - |
| - | - | 1658 | 347.2 | - | - | 0 | - |
| - | - | 1.099E+04 | 348.2 | - | - | 0 | - |
| - | - | 2136 | 349.2 | - | - | 0 | - |
| - | - | 1655 | 360.2 | - | - | 0 | - |
| - | - | 3.821E+04 | 362.2 | - | - | 0 | - |
| - | - | 6922 | 363.2 | - | - | 0 | - |
| - | - | 1663 | 385.2 | - | - | 0 | - |
| - | - | 1991 | 390.2 | - | - | 0 | - |
| - | - | 2070 | 393.3 | - | - | 0 | - |
| - | - | 2836 | 398.2 | - | - | 0 | - |
| - | - | 7586 | 411.3 | - | - | 0 | - |
| - | - | 3026 | 415.3 | - | - | 0 | - |
| - | - | 1.419E+04 | 416.3 | - | - | 0 | - |
| - | - | 4922 | 417.3 | - | - | 0 | - |
| - | - | 1540 | 425.2 | - | - | 0 | - |
| - | - | 1758 | 431.3 | - | - | 0 | - |
| 14 | y | 1.759E+04 | 433.2 | 0.002366 | 5.46 | +1 | 4 |
| - | - | 1.859E+05 | 433.3 | - | - | 0 | - |
| - | - | 3267 | 434.2 | - | - | 0 | - |
| - | - | 4.336E+04 | 434.3 | - | - | 0 | - |
| - | - | 5216 | 435.3 | - | - | 0 | - |
| - | - | 1465 | 441.9 | - | - | 0 | - |
| - | - | 1564 | 442.2 | - | - | 0 | - |
| - | - | 2520 | 443.2 | - | - | 0 | - |
| - | - | 1.91E+04 | 443.3 | - | - | 0 | - |
| - | - | 4826 | 444.3 | - | - | 0 | - |
| - | - | 2229 | 459.3 | - | - | 0 | - |
| - | - | 1.878E+05 | 461.3 | - | - | 0 | - |
| - | - | 4.525E+04 | 462.3 | - | - | 0 | - |
| - | - | 5589 | 463.3 | - | - | 0 | - |
| 13 | y | 1858 | 487.3 | 0.001994 | 4.092 | +1 | 5 |
| - | - | 2617 | 498.2 | - | - | 0 | - |
| 13 | y | 3.087E+04 | 504.3 | 0.0005301 | 1.051 | +1 | 5 |
| - | - | 5227 | 505.3 | - | - | 0 | - |
| - | - | 1672 | 513.3 | - | - | 0 | - |
| - | - | 7905 | 530.3 | - | - | 0 | - |
| - | - | 2362 | 531.3 | - | - | 0 | - |
| - | - | 6979 | 540.3 | - | - | 0 | - |
| - | - | 3356 | 541.3 | - | - | 0 | - |
| - | - | 2.808E+04 | 558.3 | - | - | 0 | - |
| - | - | 6734 | 559.3 | - | - | 0 | - |
| - | - | 2009 | 582.8 | - | - | 0 | - |
| - | - | 4308 | 591.3 | - | - | 0 | - |
| - | - | 1772 | 591.8 | - | - | 0 | - |
| 12 | y | 4238 | 615.3 | 0.0007587 | 1.233 | +1 | 6 |
| 12 | z | 7537 | 616.3 | 0.0009628 | 1.562 | +1 | 6 |
| - | - | 5329 | 617.3 | - | - | 0 | - |
| - | - | 8180 | 617.8 | - | - | 0 | - |
| - | - | 3988 | 618.3 | - | - | 0 | - |
| - | - | 1873 | 618.8 | - | - | 0 | - |
| - | - | 2.712E+04 | 626.8 | - | - | 0 | - |
| - | - | 3235 | 627.3 | - | - | 0 | - |
| - | - | 1.99E+04 | 627.3 | - | - | 0 | - |
| - | - | 4508 | 627.8 | - | - | 0 | - |
| 12 | y | 1.943E+04 | 632.3 | 0.0006379 | 1.009 | +1 | 6 |
| - | - | 5796 | 633.3 | - | - | 0 | - |
| 6 | y | 2440 | 637.8 | 0.001198 | 1.878 | +2 | 12 |
| - | - | 3052 | 638.3 | - | - | 0 | - |
| - | - | 1471 | 654.4 | - | - | 0 | - |
| - | - | 2013 | 655.3 | - | - | 0 | - |
| - | - | 2372 | 663.3 | - | - | 0 | - |
| - | - | 1480 | 663.8 | - | - | 0 | - |
| 5 | z | 1625 | 669.3 | 0.001972 | 2.947 | +2 | 13 |
| 5 | y | 1.531E+04 | 677.3 | 0.000606 | 0.8947 | +2 | 13 |
| 5 | y | 2.05E+04 | 677.8 | 0.004875 | 7.192 | +2 | 13 |
| 5 | z | 1.182E+04 | 678.3 | 0.002244 | 3.309 | +2 | 13 |
| - | - | 3833 | 678.8 | - | - | 0 | - |
| - | - | 4433 | 685.3 | - | - | 0 | - |
| - | - | 1726 | 685.8 | - | - | 0 | - |
| 5 | y | 6.872E+05 | 686.3 | 0.0006948 | 1.012 | +2 | 13 |
| - | - | 4.947E+05 | 686.8 | - | - | 0 | - |
| - | - | 1.558E+05 | 687.3 | - | - | 0 | - |
| - | - | 1.749E+04 | 687.8 | - | - | 0 | - |
| - | - | 1338 | 692.6 | - | - | 0 | - |
| 13 | c | 1.131E+04 | 700.4 | 0.01086 | 15.51 | +2 | 13 |
| - | - | 6617 | 700.9 | - | - | 0 | - |
| - | - | 2264 | 724.4 | - | - | 0 | - |
| 11 | z | 1.119E+04 | 729.4 | 0.0001269 | 0.1739 | +1 | 7 |
| - | - | 9680 | 730.4 | - | - | 0 | - |
| - | - | 2839 | 731.4 | - | - | 0 | - |
| - | - | 4708 | 740.3 | - | - | 0 | - |
| - | - | 1732 | 741.3 | - | - | 0 | - |
| - | - | 5859 | 742.4 | - | - | 0 | - |
| - | - | 3328 | 743.4 | - | - | 0 | - |
| 11 | y | 9730 | 745.4 | 0.001474 | 1.977 | +1 | 7 |
| - | - | 3719 | 746.4 | - | - | 0 | - |
| 4 | y | 4013 | 750.9 | 0.002898 | 3.86 | +2 | 14 |
| 4 | y | 5507 | 751.4 | 0.009181 | 12.22 | +2 | 14 |
| 4 | z | 2886 | 751.9 | 0.002095 | 2.786 | +2 | 14 |
| - | - | 2140 | 757.4 | - | - | 0 | - |
| 4 | y | 9.491E+04 | 759.9 | 0.0006064 | 0.7981 | +2 | 14 |
| - | - | 8.77E+04 | 760.4 | - | - | 0 | - |
| - | - | 3.444E+04 | 760.9 | - | - | 0 | - |
| - | - | 5063 | 761.4 | - | - | 0 | - |
| - | - | 1367 | 774.3 | - | - | 0 | - |
| 3 | y | 2925 | 807.9 | 0.00731 | 9.049 | +2 | 15 |
| - | - | 2572 | 813.4 | - | - | 0 | - |
| 3 | y | 2.107E+04 | 816.4 | 0.0008718 | 1.068 | +2 | 15 |
| - | - | 2.125E+04 | 816.9 | - | - | 0 | - |
| - | - | 1.023E+04 | 817.4 | - | - | 0 | - |
| - | - | 1571 | 821.9 | - | - | 0 | - |
| - | - | 5307 | 829.4 | - | - | 0 | - |
| - | - | 1758 | 830.5 | - | - | 0 | - |
| - | - | 2186 | 839.4 | - | - | 0 | - |
| - | - | 4221 | 847.9 | - | - | 0 | - |
| - | - | 7244 | 848.4 | - | - | 0 | - |
| - | - | 4647 | 848.9 | - | - | 0 | - |
| - | - | 2267 | 849.4 | - | - | 0 | - |
| 16 | c | 2.04E+04 | 856.9 | 0.001305 | 1.523 | +2 | 16 |
| 10 | y | 2.491E+04 | 857.4 | 0.01192 | 13.9 | +1 | 8 |
| 2 | y | 1.052E+04 | 857.9 | 0.01326 | 15.46 | +2 | 16 |
| 2 | y | 3.044E+04 | 858.4 | 0.0171 | 19.92 | +2 | 16 |
| - | - | 2.616E+04 | 859.5 | - | - | 0 | - |
| - | - | 7610 | 860.5 | - | - | 0 | - |
| 2 | y | 5315 | 866.9 | 0.0003844 | 0.4434 | +2 | 16 |
| - | - | 6973 | 867.4 | - | - | 0 | - |
| - | - | 3325 | 867.9 | - | - | 0 | - |
| - | - | 1.023E+04 | 868.4 | - | - | 0 | - |
| - | - | 4152 | 869.4 | - | - | 0 | - |
| 10 | y | 9591 | 874.5 | 0.002155 | 2.464 | +1 | 8 |
| - | - | 2712 | 875.5 | - | - | 0 | - |
| - | - | 3175 | 886.5 | - | - | 0 | - |
| - | - | 2382 | 898.5 | - | - | 0 | - |
| - | - | 4284 | 899 | - | - | 0 | - |
| - | - | 1.209E+04 | 907.5 | - | - | 0 | - |
| - | - | 2.152E+04 | 908 | - | - | 0 | - |
| - | - | 9939 | 908.5 | - | - | 0 | - |
| - | - | 2767 | 909 | - | - | 0 | - |
| - | - | 2229 | 916 | - | - | 0 | - |
| - | - | 3070 | 916.4 | - | - | 0 | - |
| - | - | 1.863E+05 | 916.5 | - | - | 0 | - |
| - | - | 1.877E+05 | 917 | - | - | 0 | - |
| - | - | 9.062E+04 | 917.5 | - | - | 0 | - |
| - | - | 1.416E+04 | 918 | - | - | 0 | - |
| - | - | 6878 | 939.4 | - | - | 0 | - |
| - | - | 2883 | 940.5 | - | - | 0 | - |
| - | - | 3873 | 958.5 | - | - | 0 | - |
| - | - | 1530 | 971.5 | - | - | 0 | - |
| - | - | 6295 | 973.5 | - | - | 0 | - |
| - | - | 3359 | 974.5 | - | - | 0 | - |
| 9 | c | 2313 | 975.5 | 0.01177 | 12.06 | +1 | 9 |
| 9 | z | 4.973E+04 | 987.5 | 0.0005634 | 0.5706 | +1 | 9 |
| - | - | 6.914E+04 | 988.5 | - | - | 0 | - |
| - | - | 2.883E+04 | 989.5 | - | - | 0 | - |
| - | - | 6204 | 990.5 | - | - | 0 | - |
| - | - | 6275 | 1002 | - | - | 0 | - |
| 9 | y | 1.392E+04 | 1004 | 0.002475 | 2.466 | +1 | 9 |
| - | - | 5719 | 1005 | - | - | 0 | - |
| - | - | 1844 | 1015 | - | - | 0 | - |
| - | - | 1921 | 1032 | - | - | 0 | - |
| - | - | 6962 | 1053 | - | - | 0 | - |
| - | - | 3356 | 1054 | - | - | 0 | - |
| 8 | y | 3315 | 1073 | 0.01655 | 15.43 | +1 | 10 |
| 8 | y | 5772 | 1074 | 0.001911 | 1.781 | +1 | 10 |
| 8 | z | 5.076E+04 | 1075 | 0.0002733 | 0.2543 | +1 | 10 |
| - | - | 1.393E+05 | 1076 | - | - | 0 | - |
| - | - | 6.388E+04 | 1077 | - | - | 0 | - |
| - | - | 1.031E+04 | 1078 | - | - | 0 | - |
| - | - | 4834 | 1088 | - | - | 0 | - |
| - | - | 4233 | 1089 | - | - | 0 | - |
| - | - | 1.774E+04 | 1090 | - | - | 0 | - |
| 8 | y | 6.52E+04 | 1091 | 0.0001402 | 0.1286 | +1 | 10 |
| - | - | 2.908E+04 | 1092 | - | - | 0 | - |
| - | - | 9096 | 1093 | - | - | 0 | - |
| - | - | 2806 | 1104 | - | - | 0 | - |
| 10 | c | 1.737E+04 | 1105 | 0.0007701 | 0.6972 | +1 | 10 |
| - | - | 9354 | 1106 | - | - | 0 | - |
| - | - | 2277 | 1107 | - | - | 0 | - |
| - | - | 4661 | 1117 | - | - | 0 | - |
| - | - | 8254 | 1118 | - | - | 0 | - |
| - | - | 4559 | 1119 | - | - | 0 | - |
| - | - | 1852 | 1156 | - | - | 0 | - |
| - | - | 2897 | 1157 | - | - | 0 | - |
| - | - | 3116 | 1160 | - | - | 0 | - |
| 7 | y | 4910 | 1161 | 0.00317 | 2.731 | +1 | 11 |
| 7 | z | 2.607E+04 | 1162 | 0.0008377 | 0.7211 | +1 | 11 |
| - | - | 5.288E+04 | 1163 | - | - | 0 | - |
| - | - | 2.58E+04 | 1164 | - | - | 0 | - |
| - | - | 9787 | 1165 | - | - | 0 | - |
| - | - | 3077 | 1166 | - | - | 0 | - |
| - | - | 2228 | 1175 | - | - | 0 | - |
| - | - | 6291 | 1177 | - | - | 0 | - |
| 7 | y | 3.982E+04 | 1178 | 0.0009124 | 0.7748 | +1 | 11 |
| - | - | 1.799E+04 | 1179 | - | - | 0 | - |
| - | - | 5431 | 1180 | - | - | 0 | - |
| - | - | 1.802E+04 | 1182 | - | - | 0 | - |
| - | - | 1.296E+04 | 1183 | - | - | 0 | - |
| - | - | 3989 | 1184 | - | - | 0 | - |
| - | - | 1.097E+04 | 1201 | - | - | 0 | - |
| - | - | 6864 | 1202 | - | - | 0 | - |
| - | - | 2765 | 1203 | - | - | 0 | - |
| - | - | 2.015E+04 | 1217 | - | - | 0 | - |
| 11 | c | 4.307E+04 | 1218 | 0.001826 | 1.5 | +1 | 11 |
| - | - | 2.472E+04 | 1219 | - | - | 0 | - |
| - | - | 4282 | 1220 | - | - | 0 | - |
| - | - | 3513 | 1235 | - | - | 0 | - |
| - | - | 4607 | 1236 | - | - | 0 | - |
| - | - | 2.059E+04 | 1253 | - | - | 0 | - |
| - | - | 1.481E+04 | 1254 | - | - | 0 | - |
| - | - | 4125 | 1255 | - | - | 0 | - |
| 6 | y | 2391 | 1257 | 0.009671 | 7.696 | +1 | 12 |
| 6 | y | 2776 | 1258 | 0.007358 | 5.851 | +1 | 12 |
| 6 | y | 8.717E+04 | 1275 | 0.0001505 | 0.1181 | +1 | 12 |
| - | - | 5.54E+04 | 1276 | - | - | 0 | - |
| - | - | 1.524E+04 | 1277 | - | - | 0 | - |
| - | - | 1952 | 1278 | - | - | 0 | - |
| - | - | 3746 | 1300 | - | - | 0 | - |
| - | - | 2372 | 1302 | - | - | 0 | - |
| - | - | 4726 | 1303 | - | - | 0 | - |
| - | - | 3905 | 1311 | - | - | 0 | - |
| - | - | 2539 | 1312 | - | - | 0 | - |
| - | - | 3407 | 1326 | - | - | 0 | - |
| - | - | 4864 | 1327 | - | - | 0 | - |
| 12 | c | 2352 | 1328 | 0.003941 | 2.968 | +1 | 12 |
| 12 | c | 1.915E+04 | 1329 | 0.002361 | 1.777 | +1 | 12 |
| - | - | 1.288E+04 | 1330 | - | - | 0 | - |
| - | - | 5167 | 1331 | - | - | 0 | - |
| - | - | 2304 | 1344 | - | - | 0 | - |
| - | - | 2.602E+04 | 1345 | - | - | 0 | - |
| 12 | c | 1.115E+05 | 1346 | 0.0007114 | 0.5286 | +1 | 12 |
| - | - | 7.612E+04 | 1347 | - | - | 0 | - |
| - | - | 2.593E+04 | 1348 | - | - | 0 | - |
| - | - | 2681 | 1349 | - | - | 0 | - |
| 5 | y | 1.119E+04 | 1354 | 0.002995 | 2.213 | +1 | 13 |
| 5 | y | 2.059E+04 | 1355 | 0.01153 | 8.514 | +1 | 13 |
| 5 | z | 1.211E+04 | 1356 | 0.0005346 | 0.3943 | +1 | 13 |
| - | - | 2523 | 1357 | - | - | 0 | - |
| - | - | 3982 | 1358 | - | - | 0 | - |
| - | - | 1892 | 1359 | - | - | 0 | - |
| - | - | 5234 | 1360 | - | - | 0 | - |
| - | - | 4422 | 1370 | - | - | 0 | - |
| - | - | 3.934E+04 | 1371 | - | - | 0 | - |
| 5 | y | 7.925E+05 | 1372 | 1.037E-06 | 0.0007563 | +1 | 13 |
| - | - | 5.216E+05 | 1373 | - | - | 0 | - |
| - | - | 1.835E+05 | 1374 | - | - | 0 | - |
| - | - | 1.95E+04 | 1375 | - | - | 0 | - |
| - | - | 1807 | 1382 | - | - | 0 | - |
| - | - | 4152 | 1398 | - | - | 0 | - |
| - | - | 1.636E+04 | 1399 | - | - | 0 | - |
| - | - | 3.218E+04 | 1400 | - | - | 0 | - |
| - | - | 2.156E+04 | 1401 | - | - | 0 | - |
| - | - | 1.518E+05 | 1402 | - | - | 0 | - |
| - | - | 1.238E+05 | 1403 | - | - | 0 | - |
| - | - | 4.211E+04 | 1404 | - | - | 0 | - |
| - | - | 4416 | 1405 | - | - | 0 | - |
| - | - | 8.79E+04 | 1416 | - | - | 0 | - |
| 13 | c | 1.27E+05 | 1417 | 0.003645 | 2.573 | +1 | 13 |
| - | - | 6.505E+04 | 1418 | - | - | 0 | - |
| - | - | 1.729E+04 | 1419 | - | - | 0 | - |
| - | - | 6414 | 1431 | - | - | 0 | - |
| - | - | 6756 | 1432 | - | - | 0 | - |
| - | - | 2726 | 1433 | - | - | 0 | - |
| - | - | 9455 | 1487 | - | - | 0 | - |
| - | - | 9353 | 1488 | - | - | 0 | - |
| - | - | 3524 | 1489 | - | - | 0 | - |
| 4 | z | 6.625E+04 | 1503 | 0.0002524 | 0.168 | +1 | 14 |
| - | - | 6.285E+04 | 1504 | - | - | 0 | - |
| - | - | 2.533E+04 | 1505 | - | - | 0 | - |
| - | - | 2895 | 1506 | - | - | 0 | - |
| 14 | c | 6277 | 1514 | 0.0006061 | 0.4004 | +1 | 14 |
| - | - | 6035 | 1515 | - | - | 0 | - |
| - | - | 2504 | 1516 | - | - | 0 | - |
| - | - | 2647 | 1518 | - | - | 0 | - |
| 4 | y | 9.016E+04 | 1519 | 0.0004327 | 0.2849 | +1 | 14 |
| - | - | 6.945E+04 | 1520 | - | - | 0 | - |
| - | - | 2.739E+04 | 1521 | - | - | 0 | - |
| - | - | 2944 | 1522 | - | - | 0 | - |
| - | - | 2355 | 1529 | - | - | 0 | - |
| - | - | 1.239E+04 | 1530 | - | - | 0 | - |
| 14 | c | 2.109E+05 | 1531 | 0.0007966 | 0.5204 | +1 | 14 |
| - | - | 1.687E+05 | 1532 | - | - | 0 | - |
| - | - | 6.864E+04 | 1533 | - | - | 0 | - |
| - | - | 7169 | 1534 | - | - | 0 | - |
| - | - | 2821 | 1544 | - | - | 0 | - |
| - | - | 3236 | 1545 | - | - | 0 | - |
| - | - | 3504 | 1571 | - | - | 0 | - |
| - | - | 5417 | 1572 | - | - | 0 | - |
| - | - | 3183 | 1573 | - | - | 0 | - |
| - | - | 2277 | 1574 | - | - | 0 | - |
| 3 | y | 3807 | 1614 | 0.008232 | 5.101 | +1 | 15 |
| - | - | 2.929E+04 | 1615 | - | - | 0 | - |
| 3 | z | 4.57E+04 | 1616 | 0.02872 | 17.77 | +1 | 15 |
| - | - | 3.433E+04 | 1617 | - | - | 0 | - |
| - | - | 1.195E+04 | 1618 | - | - | 0 | - |
| 3 | y | 3.287E+04 | 1632 | 0.001356 | 0.8309 | +1 | 15 |
| - | - | 2.845E+04 | 1633 | - | - | 0 | - |
| - | - | 1.173E+04 | 1634 | - | - | 0 | - |
| - | - | 2112 | 1645 | - | - | 0 | - |
| - | - | 3649 | 1646 | - | - | 0 | - |
| - | - | 2492 | 1647 | - | - | 0 | - |
| - | - | 3692 | 1655 | - | - | 0 | - |
| - | - | 5370 | 1656 | - | - | 0 | - |
| - | - | 3435 | 1657 | - | - | 0 | - |
| - | - | 2550 | 1658 | - | - | 0 | - |
| 15 | c | 8.572E+04 | 1659 | 0.001888 | 1.138 | +1 | 15 |
| - | - | 9.41E+04 | 1660 | - | - | 0 | - |
| - | - | 4.289E+04 | 1661 | - | - | 0 | - |
| - | - | 1.205E+04 | 1662 | - | - | 0 | - |
| - | - | 5918 | 1663 | - | - | 0 | - |
| - | - | 1.391E+04 | 1672 | - | - | 0 | - |
| - | - | 1.642E+04 | 1673 | - | - | 0 | - |
| - | - | 8620 | 1674 | - | - | 0 | - |
| - | - | 2828 | 1675 | - | - | 0 | - |
| - | - | 2.489E+04 | 1686 | - | - | 0 | - |
| - | - | 2.998E+04 | 1687 | - | - | 0 | - |
| - | - | 1.807E+04 | 1688 | - | - | 0 | - |
| - | - | 7545 | 1689 | - | - | 0 | - |
| - | - | 2963 | 1690 | - | - | 0 | - |
| 2 | z | 7243 | 1699 | 0.006011 | 3.538 | +1 | 16 |
| 2 | z | 7523 | 1700 | 0.02139 | 12.58 | +1 | 16 |
| 2 | y | 3.91E+04 | 1715 | 1.753E-05 | 0.01022 | +1 | 16 |
| 2 | y | 6.367E+04 | 1716 | 0.01426 | 8.31 | +1 | 16 |
| 2 | z | 2.014E+05 | 1717 | 0.00155 | 0.9028 | +1 | 16 |
| - | - | 2.018E+05 | 1718 | - | - | 0 | - |
| - | - | 9.803E+04 | 1719 | - | - | 0 | - |
| - | - | 1.75E+04 | 1720 | - | - | 0 | - |
| 16 | c | 1.468E+05 | 1730 | 0.001038 | 0.5999 | +1 | 16 |
| - | - | 1.395E+05 | 1731 | - | - | 0 | - |
| - | - | 9.673E+04 | 1732 | - | - | 0 | - |
| - | - | 3.859E+04 | 1733 | - | - | 0 | - |
| - | - | 1.492E+04 | 1734 | - | - | 0 | - |
| - | - | 2555 | 1735 | - | - | 0 | - |
| - | - | 8178 | 1744 | - | - | 0 | - |
| - | - | 1.862E+04 | 1745 | - | - | 0 | - |
| - | - | 1.416E+04 | 1746 | - | - | 0 | - |
| - | - | 5745 | 1747 | - | - | 0 | - |
| - | - | 1.027E+04 | 1760 | - | - | 0 | - |
| - | - | 2.242E+04 | 1761 | - | - | 0 | - |
| - | - | 3.437E+04 | 1762 | - | - | 0 | - |
| - | - | 2.015E+04 | 1763 | - | - | 0 | - |
| - | - | 7662 | 1764 | - | - | 0 | - |
| - | - | 1.435E+04 | 1769 | - | - | 0 | - |
| - | - | 1.936E+04 | 1770 | - | - | 0 | - |
| - | - | 1.358E+04 | 1771 | - | - | 0 | - |
| - | - | 7285 | 1772 | - | - | 0 | - |
| - | - | 4858 | 1773 | - | - | 0 | - |
| - | - | 1.603E+04 | 1774 | - | - | 0 | - |
| - | - | 1.701E+04 | 1775 | - | - | 0 | - |
| - | - | 1.052E+04 | 1776 | - | - | 0 | - |
| - | - | 2.297E+04 | 1777 | - | - | 0 | - |
| - | - | 2.014E+04 | 1778 | - | - | 0 | - |
| - | - | 9161 | 1779 | - | - | 0 | - |
| - | - | 3.884E+04 | 1787 | - | - | 0 | - |
| - | - | 2.331E+05 | 1788 | - | - | 0 | - |
| - | - | 2.275E+05 | 1789 | - | - | 0 | - |
| - | - | 1.234E+05 | 1790 | - | - | 0 | - |
| - | - | 2.125E+04 | 1791 | - | - | 0 | - |
| - | - | 2616 | 1792 | - | - | 0 | - |
| - | - | 2407 | 1797 | - | - | 0 | - |
| - | - | 6806 | 1798 | - | - | 0 | - |
| - | - | 4505 | 1799 | - | - | 0 | - |
| - | - | 3483 | 1800 | - | - | 0 | - |
| - | - | 3397 | 1801 | - | - | 0 | - |
| - | - | 3525 | 1802 | - | - | 0 | - |
| - | - | 1.817E+04 | 1805 | - | - | 0 | - |
| - | - | 1.775E+04 | 1806 | - | - | 0 | - |
| - | - | 1.107E+04 | 1807 | - | - | 0 | - |
| - | - | 5210 | 1814 | - | - | 0 | - |
| - | - | 5.442E+04 | 1815 | - | - | 0 | - |
| - | - | 3.547E+05 | 1816 | - | - | 0 | - |
| - | - | 3.416E+05 | 1817 | - | - | 0 | - |
| - | - | 1.735E+05 | 1818 | - | - | 0 | - |
| - | - | 2.587E+04 | 1819 | - | - | 0 | - |
| - | - | 7577 | 1830 | - | - | 0 | - |
| - | - | 2.674E+04 | 1831 | - | - | 0 | - |
| - | - | 6.528E+05 | 1832 | - | - | 0 | - |
| - | - | 1.71E+06 | 1833 | - | - | 0 | - |
| - | - | 1.457E+06 | 1834 | - | - | 0 | - |
| - | - | 6.161E+05 | 1835 | - | - | 0 | - |
| - | - | 8.24E+04 | 1836 | - | - | 0 | - |
| - | - | 2967 | 1865 | - | - | 0 | - |
| - | - | 3525 | 1866 | - | - | 0 | - |
| - | - | 1852 | 3155 | - | - | 0 | - |

m/z Charge Intensity FragmentType MassShift Position
120.0809555053711 0 4521.0728
129.10238647460938 0 8930.654
148.9471435546875 0 2352.3416
169.1337432861328 0 2032.6084
173.12863159179688 0 7041.5557
173.4530792236328 0 1554.0836
177.02857971191406 0 1376.1522
183.11297607421875 0 9278.826
187.14515686035156 0 1281.551
201.12347412109375 0 24855.508
215.13897705078125 0 3916.376
217.13412475585938 0 2911.8364
226.11837768554688 0 2001.41
233.16477966308594 0 13727.597
243.14512634277344 0 4665.0806
245.1287841796875 0 5172.43
246.04254150390625 0 1305.6047
247.14410400390625 0 3684.3115
251.17523193359375 0 5844.555
261.15985107421875 0 12073.432
269.1859436035156 0 3550.1055
271.204833984375 0 1199.5994
280.16607666015625 0 2765.5823
282.14483642578125 0 2254.1284
296.19720458984375 0 252138.9
297.1566162109375 0 1746.291
297.2004699707031 0 39730.277
298.1769104003906 0 2901.9785
298.2037048339844 0 1827.4575
303.17010498046875 0 4659.2666
304.1934814453125 0 1394.5188
307.3913879394531 0 1145.0864
314.1861877441406 0 4301.7783
314.20782470703125 0 127033.984
315.2109069824219 0 18866.723
317.1874084472656 0 1533.735
319.1977844238281 0 16654.951 y 14
330.182373046875 0 4732.505
344.1961364746094 0 1815.9423
346.1766357421875 0 11564.764
347.17779541015625 0 1657.7942
348.1917419433594 0 10992.1
349.1940002441406 0 2136.123
360.22808837890625 0 1654.7795
362.20758056640625 0 38214.723
363.210693359375 0 6921.7236
385.21929931640625 0 1663.1241
390.2022399902344 0 1990.8085
393.251220703125 0 2069.7576
398.2447204589844 0 2835.9546
411.26043701171875 0 7586.256
415.2715759277344 0 3025.586
416.2548522949219 0 14191.055
417.25604248046875 0 4921.5107
425.21502685546875 0 1540.4767
431.2659606933594 0 1757.7015
433.2428894042969 0 17588.057 y 13
433.2816467285156 0 185900.31
434.24884033203125 0 3266.685
434.2848205566406 0 43361.875
435.287353515625 0 5216.461
441.93597412109375 0 1464.8583
442.2415771484375 0 1563.6412
443.23187255859375 0 2519.9963
443.2658996582031 0 19097.549
444.26947021484375 0 4825.856
459.2609558105469 0 2229.0024
461.27655029296875 0 187778.06
462.2796325683594 0 45251.438
463.281494140625 0 5588.5913
487.2530822753906 0 1857.7228 y Ammonia loss 12
498.21893310546875 0 2617.465
504.2781677246094 0 30866.035 y 12
505.281982421875 0 5226.5337
513.2783203125 0 1671.6414
530.3349609375 0 7905.4434
531.3358764648438 0 2362.1445
540.31884765625 0 6978.7637
541.3230590820312 0 3355.773
558.3292236328125 0 28075.98
559.332275390625 0 6734.278
582.778076171875 0 2009.0297
591.2940063476562 0 4308.0728
591.7947387695312 0 1771.8381
615.3104248046875 0 4238.458 y Ammonia loss 11
616.3165283203125 0 7537.1577 z 11
617.3250732421875 0 5329.3765
617.8079223632812 0 8179.696
618.3102416992188 0 3988.2717
618.8040771484375 0 1873.0693
626.8123779296875 0 27115.418
627.2610473632812 0 3234.6353
627.314208984375 0 19896.994
627.81298828125 0 4507.6494
632.3368530273438 0 19434.568 y 11
633.3370971679688 0 5795.671
637.8159790039062 0 2439.8516 y 5
638.3167724609375 0 3051.6138
654.3616333007812 0 1471.1613
655.33837890625 0 2013.3085
663.3391723632812 0 2371.932
663.8378295898438 0 1480.4413
669.3284912109375 0 1624.656 z Water loss 4
677.3364868164062 0 15307.108 y Water loss 4
677.832763671875 0 20496.156 y Ammonia loss 4
678.3340454101562 0 11815.452 z 4
678.8355102539062 0 3832.5068
685.3328247070312 0 4432.6533
685.8308715820312 0 1725.9215
686.3418579101562 0 687205.5 y 4
686.8433227539062 0 494723.6
687.3442993164062 0 155783.77
687.8458251953125 0 17485.75
692.5582885742188 0 1337.7174
700.3479614257812 0 11305.038 c Ammonia loss 12
700.8502807617188 0 6616.8193
724.3993530273438 0 2264.2495
729.4014282226562 0 11194.838 z 10
730.4088134765625 0 9680.48
731.41162109375 0 2839.1853
740.34716796875 0 4707.815
741.3493041992188 0 1731.7667
742.414306640625 0 5858.8203
743.4151000976562 0 3328.1672
745.4217529296875 0 9729.959 y 10
746.4227905273438 0 3718.5244
750.8729858398438 0 4013.1536 y Water loss 3
751.3712768554688 0 5506.684 y Ammonia loss 3
751.8681030273438 0 2886.366 z 3
757.4202270507812 0 2140.403
759.8759765625 0 94906.87 y 3
760.3775024414062 0 87696.45
760.8786010742188 0 34441.133
761.3806762695312 0 5063.499
774.342041015625 0 1366.9186
807.9114379882812 0 2925.0254 y Ammonia loss 2
813.4141845703125 0 2572.2822
816.4182739257812 0 21072.674 y 2
816.9194946289062 0 21253.072
817.4222412109375 0 10225.719
821.9262084960938 0 1570.5948
829.4481201171875 0 5307.363
830.454833984375 0 1757.9484
839.4212646484375 0 2186.329
847.9406127929688 0 4221.2065
848.439697265625 0 7243.6104
848.9395751953125 0 4646.617
849.4320678710938 0 2266.5518
856.9476318359375 0 20398.125 c Ammonia loss 15
857.4482421875 0 24905.104 y Ammonia loss 9
857.94921875 0 10515.634 y Water loss 1
858.445068359375 0 30437.396 y Ammonia loss 1
859.4505004882812 0 26164.613
860.455810546875 0 7609.5503
866.9408569335938 0 5314.703 y 1
867.442138671875 0 6973.193
867.9468994140625 0 3325.4756
868.4055786132812 0 10225.428
869.406494140625 0 4152.428
874.4650268554688 0 9591.054 y 9
875.4696655273438 0 2712.2456
886.46728515625 0 3174.9482
898.4616088867188 0 2381.5754
898.96435546875 0 4284.064
907.469482421875 0 12089.281
907.9705810546875 0 21519.57
908.4717407226562 0 9939.203
908.9624633789062 0 2766.8938
915.9724731445312 0 2229.055
916.36572265625 0 3070.292
916.4762573242188 0 186298.53
916.9773559570312 0 187718.8
917.4786376953125 0 90623.414
917.979736328125 0 14161.729
939.4418334960938 0 6878.314
940.4547729492188 0 2882.987
958.4911499023438 0 3872.7043
971.4800415039062 0 1529.796
973.49560546875 0 6294.8486
974.498046875 0 3359.3958
975.5028076171875 0 2312.6025 c 8
987.4873046875 0 49725.3 z 8
988.4942626953125 0 69136.62
989.4979248046875 0 28831.062
990.50341796875 0 6203.684
1002.4972534179688 0 6274.769
1003.5029907226562 0 13920.838 y 8
1004.5073852539062 0 5719.241
1015.4851684570312 0 1843.6805
1031.5045166015625 0 1921.2036
1053.487548828125 0 6961.6704
1054.4864501953125 0 3355.9893
1072.5103759765625 0 3314.9685 y Water loss 7
1073.509033203125 0 5771.5137 y Ammonia loss 7
1074.51904296875 0 50763.598 z 7
1075.52685546875 0 139261.5
1076.5306396484375 0 63882.47
1077.533935546875 0 10305.188
1087.5306396484375 0 4834.336
1088.5382080078125 0 4233.024
1089.5313720703125 0 17738.613
1090.537353515625 0 65199.34 y 7
1091.5404052734375 0 29080.541
1092.5440673828125 0 9096.323
1103.549560546875 0 2805.9492
1104.556396484375 0 17369.12 c 9
1105.5606689453125 0 9354.114
1106.560546875 0 2277.4636
1116.532958984375 0 4660.8213
1117.534423828125 0 8254.296
1118.538818359375 0 4558.564
1155.581787109375 0 1851.6211
1156.57568359375 0 2896.758
1159.5286865234375 0 3115.8455
1160.546142578125 0 4910.367 y Ammonia loss 6
1161.5516357421875 0 26072.975 z 6
1162.5582275390625 0 52878.92
1163.5621337890625 0 25799.889
1164.5592041015625 0 9787.424
1165.552734375 0 3077.3965
1174.6292724609375 0 2227.96
1176.560302734375 0 6290.8413
1177.5704345703125 0 39815.53 y 6
1178.5712890625 0 17991.793
1179.5789794921875 0 5430.7734
1181.582275390625 0 18021.717
1182.587890625 0 12962.516
1183.59130859375 0 3988.583
1200.615234375 0 10967.332
1201.6151123046875 0 6863.697
1202.617431640625 0 2765.462
1216.6337890625 0 20145.125
1217.639404296875 0 43074.59 c 10
1218.6436767578125 0 24724.17
1219.645751953125 0 4282.2065
1234.6041259765625 0 3513.1726
1235.6134033203125 0 4607.245
1252.616455078125 0 20593.076
1253.6217041015625 0 14811.981
1254.6163330078125 0 4124.522
1256.60205078125 0 2390.884 y Water loss 5
1257.58837890625 0 2775.6333 y Ammonia loss 5
1274.6224365234375 0 87166.5 y 5
1275.6253662109375 0 55395.246
1276.629638671875 0 15239.666
1277.6246337890625 0 1951.7666
1299.6392822265625 0 3746.2246
1301.6807861328125 0 2372.1482
1302.6915283203125 0 4726.089
1310.6724853515625 0 3905.0693
1311.6431884765625 0 2538.9907
1325.6695556640625 0 3406.5884
1326.6728515625 0 4864.01
1327.685302734375 0 2352.3567 c Water loss 11
1328.6708984375 0 19145.646 c Ammonia loss 11
1329.6719970703125 0 12880.017
1330.679443359375 0 5166.589
1343.6959228515625 0 2304.409
1344.692138671875 0 26017.412
1345.6990966796875 0 111498.96 c 11
1346.7021484375 0 76118.13
1347.7056884765625 0 25929.174
1348.7161865234375 0 2680.8796
1353.66748046875 0 11186.8545 y Water loss 4
1354.6600341796875 0 20594.023 y Ammonia loss 4
1355.6568603515625 0 12109.579 z 4
1356.650146484375 0 2522.7883
1357.6827392578125 0 3981.6194
1358.6885986328125 0 1892.449
1359.669921875 0 5233.636
1369.6541748046875 0 4421.801
1370.6680908203125 0 39335.617
1371.675048828125 0 792488.44 y 4
1372.6785888671875 0 521642.6
1373.6812744140625 0 183480.69
1374.6900634765625 0 19503.385
1381.6981201171875 0 1807.335
1397.7120361328125 0 4151.609
1398.6690673828125 0 16360.39
1399.682373046875 0 32176.908
1400.689208984375 0 21559.787
1401.701416015625 0 151772.61
1402.704345703125 0 123787.48
1403.707275390625 0 42106.32
1404.70849609375 0 4416.175
1415.7286376953125 0 87895.12
1416.7332763671875 0 126970.69 c 12
1417.7373046875 0 65051.355
1418.743408203125 0 17290.557
1430.69580078125 0 6413.6675
1431.6939697265625 0 6756.3887
1432.686767578125 0 2726.2449
1486.7659912109375 0 9455.032
1487.7684326171875 0 9352.516
1488.773193359375 0 3523.6736
1502.7244873046875 0 66251.92 z 3
1503.727294921875 0 62848.71
1504.7314453125 0 25328.979
1505.7392578125 0 2894.6287
1513.75390625 0 6276.6343 c Ammonia loss 13
1514.7557373046875 0 6034.613
1515.7730712890625 0 2503.8567
1517.757080078125 0 2646.8333
1518.743896484375 0 90163.586 y 3
1519.7457275390625 0 69454.6
1520.7481689453125 0 27394.021
1521.742919921875 0 2943.898
1528.7757568359375 0 2354.6445
1529.7724609375 0 12389.563
1530.779052734375 0 210914.77 c 13
1531.7818603515625 0 168665
1532.7855224609375 0 68640.87
1533.7821044921875 0 7168.8633
1543.7926025390625 0 2821.0178
1544.7916259765625 0 3236.107
1570.8172607421875 0 3503.7107
1571.8221435546875 0 5416.9136
1572.8094482421875 0 3182.9263
1573.773681640625 0 2276.593
1613.8251953125 0 3806.7258 y Water loss 2
1614.8599853515625 0 29290.441
1615.8375244140625 0 45699.31 z 2
1616.831298828125 0 34325.707
1617.8216552734375 0 11951.854
1631.826171875 0 32873.316 y 2
1632.8302001953125 0 28454.637
1633.8309326171875 0 11728.193
1644.8455810546875 0 2112.199
1645.8382568359375 0 3649.0437
1646.8421630859375 0 2491.804
1654.853759765625 0 3692.252
1655.8509521484375 0 5370.3857
1656.85498046875 0 3435.427
1657.8641357421875 0 2550.246
1658.8729248046875 0 85719.49 c 14
1659.8765869140625 0 94098.14
1660.8701171875 0 42887.387
1661.818359375 0 12052.823
1662.8074951171875 0 5918.354
1671.8455810546875 0 13911.451
1672.84521484375 0 16423.443
1673.8460693359375 0 8619.829
1674.845703125 0 2827.7703
1685.8961181640625 0 24891.115
1686.9022216796875 0 29979.057
1687.9085693359375 0 18073.662
1688.912109375 0 7544.5815
1689.899169921875 0 2962.7666
1698.8519287109375 0 7242.7725 z Water loss 1
1699.851318359375 0 7522.8496 z Ammonia loss 1
1714.8646240234375 0 39099.707 y Water loss 1
1715.8629150390625 0 63672.543 y Ammonia loss 1
1716.8580322265625 0 201426.42 z 1
1717.8597412109375 0 201770.14
1718.86474609375 0 98033.93
1719.8670654296875 0 17503.924
1729.9129638671875 0 146759.72 c 15
1730.9154052734375 0 139530.42
1731.9189453125 0 96732.266
1732.9251708984375 0 38589.926
1733.9248046875 0 14920.5
1734.92822265625 0 2555.4546
1743.8956298828125 0 8177.897
1744.8953857421875 0 18623.283
1745.89892578125 0 14159.807
1746.89453125 0 5745.484
1759.8895263671875 0 10272.98
1760.912109375 0 22415.404
1761.9168701171875 0 34369.594
1762.914306640625 0 20147.44
1763.9122314453125 0 7662.5
1768.936279296875 0 14346.72
1769.9349365234375 0 19356.934
1770.9375 0 13581.811
1771.9354248046875 0 7284.5356
1772.9276123046875 0 4858.0747
1773.9189453125 0 16031.204
1774.919677734375 0 17005.598
1775.9207763671875 0 10524.383
1776.89501953125 0 22970.797
1777.8916015625 0 20141.676
1778.8951416015625 0 9160.739
1786.9449462890625 0 38838.715
1787.93310546875 0 233107.05
1788.935302734375 0 227548.77
1789.9359130859375 0 123419.97
1790.92724609375 0 21249.531
1791.8896484375 0 2616.0054
1796.91064453125 0 2406.615
1797.91259765625 0 6805.858
1798.9132080078125 0 4504.6577
1799.9136962890625 0 3483.4395
1800.8892822265625 0 3396.8643
1801.9063720703125 0 3525.023
1804.957763671875 0 18171.258
1805.9578857421875 0 17753.908
1806.9576416015625 0 11072.519
1813.904052734375 0 5210.277
1814.9381103515625 0 54424.277
1815.9288330078125 0 354702.38
1816.9296875 0 341577.62
1817.931396484375 0 173522.69
1818.9327392578125 0 25865.889
1829.9263916015625 0 7576.755
1830.93603515625 0 26744.457
1831.9447021484375 0 652777.3
1832.95166015625 0 1709681.2
1833.95458984375 0 1457438.8
1834.9580078125 0 616064.7
1835.9588623046875 0 82403.195
1864.9334716796875 0 2966.5625
1865.931640625 0 3524.9658
3155.2294921875 0 1852.0929

Spectrum Details

|  |  |
| --- | --- |
| Matched peaks? Matched peaks The total absolute number of peaks matched. Additionally in brackets the total fraction of peaks matched and the total number of peaks is shown. | 65 (15.74% of 413) |
| FDR? FDR The false discovery rate estimated for this peptide. It is calculated by matching all theoretical fragments with a non-integer shift with the raw peaks for this spectrum. This is done with 40 different shifts. The resulting percentage is the average number of annotated peaks over the number of annotated peaks with the correct spectrum. | 0.48% |
| Satellite FDR? Satellite FDR See the FDR for details on its calculation. This satellite ion specific FDR only contains the satellite ions (d/w) for I/L/J positions. | - |
| PSM Score? PSM Score The PSM Score as given by Hecklib to this annotated spectrum. It is shown with three significant figures. | 627 |

#### Spectrum 8046? Spectrum 8046 The raw spectrum of this peptide as annotated by Hecklib. The fragments are coloured according to ion type (see legend). Any peaks with a star '\*' as text can be hovered over to see the full details, first the ion type second the mass shift type. By hovering over the amino acids in the peptide or ions in the legend the corresponding peaks are highlighted. By toggling the 'Unassigned' label you can turn the background (unassigned) peaks on or off in the plot. By updating the slider in the Ion legend you can update the spectrum to only show the top X% of the peaks with labels. The top X% means any peak that is within X% of the highest intensity. By dragging in the spectrum you can zoom in to a specific part of the spectrum and use 'Zoom Out' to get back to the original zoom level. The annotation of the spectrum is based on the given sequence in the peptides file and is done with different software so inconsistencies are likely. The peaks are annotated based on the given sequence, with 20 ppm tolerance.

Copy Data

##### Spectrum 8046 (TSV)

###### Preview

```
Loading example...
```

*Click on the button to copy the data to your clipboard.*

Mz MinMz MaxIntensity Max

WidthHeightPeptide font sizePeptide stroke widthSpectrum font sizeSpectrum stroke widthCompact peptide

Ion legend

wxyz

abcd

OtherUnassignedIon^Charge^_Position_Show for top:%

VTJFPPSSEEJQANKAT

04.60e+59.19e+51.38e+61.84e+6

Zoom Out

y^+1^_3_y^+1^_4_y^+1^_5_y^+1^_6_z^+1^_6_y^+1^_6_y^+2^_12_c^+2^_12_y^+2^_13_y^+2^_13_z^+2^_13_y^+2^_13_c^+2^_13_z^+1^_7_y^+1^_7_y^+2^_14_y^+2^_14_z^+2^_14_y^+2^_14_y^+2^_15_y^+2^_15_z^+2^_15_y^+2^_15_c^+2^_16_y^+1^_8_y^+2^_16_y^+2^_16_y^+2^_16_y^+1^_8_w^+1^_9_z^+1^_9_y^+1^_9_y^+1^_10_z^+1^_10_y^+1^_10_c^+1^_10_y^+1^_11_y^+1^_11_z^+1^_11_y^+1^_11_c^+1^_11_y^+1^_12_c^+1^_12_c^+1^_12_c^+1^_12_y^+1^_13_y^+1^_13_z^+1^_13_y^+1^_13_c^+1^_13_y^+1^_14_z^+1^_14_c^+1^_14_y^+1^_14_c^+1^_14_y^+1^_15_z^+1^_15_y^+1^_15_c^+1^_15_z^+1^_16_z^+1^_16_y^+1^_16_y^+1^_16_z^+1^_16_c^+1^_16_

047194214131885

Fragment Matches Table

Show background peaks

| Position | Ion type | Intensity | mz Theoretical | mz Error (Th) | mz Error (ppm) | Charge | Series Number |
| --- | --- | --- | --- | --- | --- | --- | --- |
| - | - | 5537 | 120.1 | - | - | 0 | - |
| - | - | 1068 | 122.3 | - | - | 0 | - |
| - | - | 8410 | 129.1 | - | - | 0 | - |
| - | - | 1048 | 143.1 | - | - | 0 | - |
| - | - | 1250 | 165.4 | - | - | 0 | - |
| - | - | 4929 | 173.1 | - | - | 0 | - |
| - | - | 1.65E+04 | 173.5 | - | - | 0 | - |
| - | - | 1660 | 183.1 | - | - | 0 | - |
| - | - | 1.061E+04 | 183.1 | - | - | 0 | - |
| - | - | 2.006E+04 | 201.1 | - | - | 0 | - |
| - | - | 5638 | 215.1 | - | - | 0 | - |
| - | - | 2527 | 217.1 | - | - | 0 | - |
| - | - | 2107 | 224.6 | - | - | 0 | - |
| - | - | 2588 | 226.1 | - | - | 0 | - |
| - | - | 1441 | 229 | - | - | 0 | - |
| - | - | 1.351E+04 | 233.2 | - | - | 0 | - |
| - | - | 1744 | 233.6 | - | - | 0 | - |
| - | - | 2007 | 234.2 | - | - | 0 | - |
| - | - | 5060 | 243.1 | - | - | 0 | - |
| - | - | 4520 | 245.1 | - | - | 0 | - |
| - | - | 4506 | 251.2 | - | - | 0 | - |
| - | - | 1614 | 253.4 | - | - | 0 | - |
| - | - | 1.194E+04 | 261.2 | - | - | 0 | - |
| - | - | 2340 | 280.2 | - | - | 0 | - |
| - | - | 2024 | 282.1 | - | - | 0 | - |
| - | - | 1745 | 294.2 | - | - | 0 | - |
| - | - | 2.647E+05 | 296.2 | - | - | 0 | - |
| - | - | 3542 | 297.2 | - | - | 0 | - |
| - | - | 3.919E+04 | 297.2 | - | - | 0 | - |
| - | - | 4765 | 298.2 | - | - | 0 | - |
| - | - | 2775 | 298.2 | - | - | 0 | - |
| - | - | 1561 | 301.1 | - | - | 0 | - |
| - | - | 4420 | 303.2 | - | - | 0 | - |
| - | - | 4855 | 314.2 | - | - | 0 | - |
| - | - | 1.284E+05 | 314.2 | - | - | 0 | - |
| - | - | 1.925E+04 | 315.2 | - | - | 0 | - |
| - | - | 2291 | 317.8 | - | - | 0 | - |
| 15 | y | 1.245E+04 | 319.2 | 0.0003711 | 1.163 | +1 | 3 |
| - | - | 6308 | 330.2 | - | - | 0 | - |
| - | - | 2453 | 331.2 | - | - | 0 | - |
| - | - | 1645 | 345.7 | - | - | 0 | - |
| - | - | 7839 | 346.2 | - | - | 0 | - |
| - | - | 2697 | 347.2 | - | - | 0 | - |
| - | - | 1.203E+04 | 348.2 | - | - | 0 | - |
| - | - | 2460 | 349.2 | - | - | 0 | - |
| - | - | 1645 | 360 | - | - | 0 | - |
| - | - | 2407 | 360.2 | - | - | 0 | - |
| - | - | 3.487E+04 | 362.2 | - | - | 0 | - |
| - | - | 4390 | 363.2 | - | - | 0 | - |
| - | - | 2223 | 385.2 | - | - | 0 | - |
| - | - | 2225 | 390.2 | - | - | 0 | - |
| - | - | 3350 | 398.2 | - | - | 0 | - |
| - | - | 1.144E+04 | 411.3 | - | - | 0 | - |
| - | - | 2282 | 412.3 | - | - | 0 | - |
| - | - | 3695 | 415.3 | - | - | 0 | - |
| - | - | 1.564E+04 | 416.3 | - | - | 0 | - |
| - | - | 5305 | 417.3 | - | - | 0 | - |
| 14 | y | 1.708E+04 | 433.2 | 0.002121 | 4.897 | +1 | 4 |
| - | - | 1.797E+05 | 433.3 | - | - | 0 | - |
| - | - | 3698 | 434.2 | - | - | 0 | - |
| - | - | 4.254E+04 | 434.3 | - | - | 0 | - |
| - | - | 5550 | 435.3 | - | - | 0 | - |
| - | - | 2166 | 442.2 | - | - | 0 | - |
| - | - | 3552 | 443.2 | - | - | 0 | - |
| - | - | 1.823E+04 | 443.3 | - | - | 0 | - |
| - | - | 4458 | 444.3 | - | - | 0 | - |
| - | - | 1935 | 459.3 | - | - | 0 | - |
| - | - | 1.812E+05 | 461.3 | - | - | 0 | - |
| - | - | 4.869E+04 | 462.3 | - | - | 0 | - |
| - | - | 4852 | 463.3 | - | - | 0 | - |
| - | - | 1701 | 469.2 | - | - | 0 | - |
| - | - | 1868 | 498.2 | - | - | 0 | - |
| 13 | y | 3.106E+04 | 504.3 | 0.0001944 | 0.3855 | +1 | 5 |
| - | - | 7204 | 505.3 | - | - | 0 | - |
| - | - | 7468 | 530.3 | - | - | 0 | - |
| - | - | 6836 | 540.3 | - | - | 0 | - |
| - | - | 2.487E+04 | 558.3 | - | - | 0 | - |
| - | - | 9412 | 559.3 | - | - | 0 | - |
| - | - | 1591 | 582.3 | - | - | 0 | - |
| - | - | 2760 | 582.8 | - | - | 0 | - |
| - | - | 2000 | 583.3 | - | - | 0 | - |
| - | - | 3902 | 591.3 | - | - | 0 | - |
| 12 | y | 2108 | 615.3 | 0.003139 | 5.102 | +1 | 6 |
| 12 | z | 5255 | 616.3 | 4.725E-05 | 0.07666 | +1 | 6 |
| - | - | 6963 | 617.3 | - | - | 0 | - |
| - | - | 4181 | 617.8 | - | - | 0 | - |
| - | - | 6266 | 618.3 | - | - | 0 | - |
| - | - | 3.028E+04 | 626.8 | - | - | 0 | - |
| - | - | 4171 | 627.3 | - | - | 0 | - |
| - | - | 2.259E+04 | 627.3 | - | - | 0 | - |
| - | - | 4166 | 627.8 | - | - | 0 | - |
| - | - | 1960 | 628.7 | - | - | 0 | - |
| 12 | y | 1.84E+04 | 632.3 | 0.0003327 | 0.5261 | +1 | 6 |
| - | - | 6667 | 633.3 | - | - | 0 | - |
| 6 | y | 2085 | 637.8 | 0.0007705 | 1.208 | +2 | 12 |
| - | - | 2982 | 663.3 | - | - | 0 | - |
| - | - | 3230 | 663.8 | - | - | 0 | - |
| 12 | c | 2057 | 664.3 | 0.007989 | 12.03 | +2 | 12 |
| - | - | 2221 | 668.8 | - | - | 0 | - |
| 5 | y | 1.924E+04 | 677.3 | 0.0004229 | 0.6244 | +2 | 13 |
| 5 | y | 2.098E+04 | 677.8 | 0.004753 | 7.012 | +2 | 13 |
| 5 | z | 1.317E+04 | 678.3 | 0.0004743 | 0.6992 | +2 | 13 |
| - | - | 2220 | 685.3 | - | - | 0 | - |
| 5 | y | 6.538E+05 | 686.3 | 0.0004506 | 0.6565 | +2 | 13 |
| - | - | 4.895E+05 | 686.8 | - | - | 0 | - |
| - | - | 1.551E+05 | 687.3 | - | - | 0 | - |
| - | - | 1.427E+04 | 687.8 | - | - | 0 | - |
| 13 | c | 8270 | 700.4 | 0.01086 | 15.51 | +2 | 13 |
| - | - | 4666 | 700.8 | - | - | 0 | - |
| - | - | 3194 | 701.3 | - | - | 0 | - |
| 11 | z | 1.439E+04 | 729.4 | 4.785E-06 | 0.00656 | +1 | 7 |
| - | - | 1.085E+04 | 730.4 | - | - | 0 | - |
| - | - | 5728 | 740.3 | - | - | 0 | - |
| - | - | 2134 | 741.3 | - | - | 0 | - |
| - | - | 5405 | 742.4 | - | - | 0 | - |
| - | - | 2691 | 743.4 | - | - | 0 | - |
| 11 | y | 1.254E+04 | 745.4 | 0.0005583 | 0.7489 | +1 | 7 |
| - | - | 3826 | 746.4 | - | - | 0 | - |
| 4 | y | 4940 | 750.9 | 0.001983 | 2.64 | +2 | 14 |
| 4 | y | 6558 | 751.4 | 0.007289 | 9.701 | +2 | 14 |
| 4 | z | 3353 | 751.9 | 0.0001418 | 0.1886 | +2 | 14 |
| 4 | y | 1.009E+05 | 759.9 | 0.0002402 | 0.3161 | +2 | 14 |
| - | - | 7.373E+04 | 760.4 | - | - | 0 | - |
| - | - | 3.581E+04 | 760.9 | - | - | 0 | - |
| 3 | y | 1971 | 807.4 | 0.001393 | 1.726 | +2 | 15 |
| 3 | y | 1954 | 807.9 | 0.005601 | 6.933 | +2 | 15 |
| 3 | z | 1827 | 808.4 | 0.003499 | 4.328 | +2 | 15 |
| - | - | 1899 | 810.3 | - | - | 0 | - |
| - | - | 2535 | 812.9 | - | - | 0 | - |
| 3 | y | 2.165E+04 | 816.4 | 0.00136 | 1.666 | +2 | 15 |
| - | - | 1.867E+04 | 816.9 | - | - | 0 | - |
| - | - | 8542 | 817.4 | - | - | 0 | - |
| - | - | 3711 | 829.4 | - | - | 0 | - |
| - | - | 2802 | 842.9 | - | - | 0 | - |
| - | - | 5139 | 847.9 | - | - | 0 | - |
| - | - | 4900 | 848.4 | - | - | 0 | - |
| - | - | 4405 | 848.9 | - | - | 0 | - |
| 16 | c | 1.753E+04 | 856.9 | 3.766E-05 | 0.04395 | +2 | 16 |
| 10 | y | 2.071E+04 | 857.4 | 0.01003 | 11.69 | +1 | 8 |
| 2 | y | 1.467E+04 | 857.9 | 0.01369 | 15.95 | +2 | 16 |
| 2 | y | 3.351E+04 | 858.4 | 0.016 | 18.64 | +2 | 16 |
| - | - | 2.622E+04 | 859.5 | - | - | 0 | - |
| - | - | 6364 | 860.5 | - | - | 0 | - |
| 2 | y | 3059 | 866.9 | 0.002911 | 3.358 | +2 | 16 |
| - | - | 6831 | 867.4 | - | - | 0 | - |
| - | - | 4290 | 867.9 | - | - | 0 | - |
| - | - | 8631 | 868.4 | - | - | 0 | - |
| - | - | 3468 | 869.4 | - | - | 0 | - |
| 10 | y | 7934 | 874.5 | 0.001117 | 1.277 | +1 | 8 |
| - | - | 3487 | 875.5 | - | - | 0 | - |
| - | - | 3282 | 886.5 | - | - | 0 | - |
| - | - | 2324 | 899.5 | - | - | 0 | - |
| - | - | 1.463E+04 | 907.5 | - | - | 0 | - |
| - | - | 1.536E+04 | 908 | - | - | 0 | - |
| - | - | 1.347E+04 | 908.5 | - | - | 0 | - |
| - | - | 4001 | 909 | - | - | 0 | - |
| - | - | 3187 | 916 | - | - | 0 | - |
| - | - | 1.63E+05 | 916.5 | - | - | 0 | - |
| - | - | 1.8E+05 | 917 | - | - | 0 | - |
| - | - | 1.043E+05 | 917.5 | - | - | 0 | - |
| - | - | 1.348E+04 | 918 | - | - | 0 | - |
| 9 | w | 2043 | 928.5 | 0.0009638 | 1.038 | +1 | 9 |
| - | - | 2079 | 929.5 | - | - | 0 | - |
| - | - | 5151 | 939.4 | - | - | 0 | - |
| - | - | 1861 | 940.4 | - | - | 0 | - |
| - | - | 4717 | 958.5 | - | - | 0 | - |
| - | - | 2791 | 959.5 | - | - | 0 | - |
| - | - | 5883 | 973.5 | - | - | 0 | - |
| - | - | 3665 | 974.5 | - | - | 0 | - |
| - | - | 1933 | 975.3 | - | - | 0 | - |
| 9 | z | 5.159E+04 | 987.5 | 0.0007465 | 0.756 | +1 | 9 |
| - | - | 8.349E+04 | 988.5 | - | - | 0 | - |
| - | - | 2.552E+04 | 989.5 | - | - | 0 | - |
| - | - | 6829 | 990.5 | - | - | 0 | - |
| - | - | 3507 | 1003 | - | - | 0 | - |
| 9 | y | 1.632E+04 | 1004 | 0.004184 | 4.169 | +1 | 9 |
| - | - | 6460 | 1005 | - | - | 0 | - |
| - | - | 2090 | 1032 | - | - | 0 | - |
| - | - | 7153 | 1053 | - | - | 0 | - |
| - | - | 3817 | 1054 | - | - | 0 | - |
| - | - | 2342 | 1073 | - | - | 0 | - |
| 8 | y | 7487 | 1074 | 0.00114 | 1.062 | +1 | 10 |
| 8 | z | 5.046E+04 | 1075 | 2.915E-05 | 0.02713 | +1 | 10 |
| - | - | 1.313E+05 | 1076 | - | - | 0 | - |
| - | - | 7.014E+04 | 1077 | - | - | 0 | - |
| - | - | 1.544E+04 | 1078 | - | - | 0 | - |
| - | - | 5526 | 1088 | - | - | 0 | - |
| - | - | 4483 | 1089 | - | - | 0 | - |
| - | - | 2.183E+04 | 1090 | - | - | 0 | - |
| 8 | y | 6.465E+04 | 1091 | 0.0009947 | 0.9121 | +1 | 10 |
| - | - | 3.215E+04 | 1092 | - | - | 0 | - |
| - | - | 6585 | 1093 | - | - | 0 | - |
| - | - | 4236 | 1104 | - | - | 0 | - |
| 10 | c | 1.468E+04 | 1105 | 0.002235 | 2.023 | +1 | 10 |
| - | - | 9404 | 1106 | - | - | 0 | - |
| - | - | 4537 | 1117 | - | - | 0 | - |
| - | - | 1.001E+04 | 1118 | - | - | 0 | - |
| - | - | 4499 | 1119 | - | - | 0 | - |
| - | - | 2147 | 1157 | - | - | 0 | - |
| 7 | y | 2728 | 1160 | 0.02014 | 17.37 | +1 | 11 |
| 7 | y | 3545 | 1161 | 0.002324 | 2.002 | +1 | 11 |
| 7 | z | 2.392E+04 | 1162 | 0.0009934 | 0.8552 | +1 | 11 |
| - | - | 5.402E+04 | 1163 | - | - | 0 | - |
| - | - | 2.807E+04 | 1164 | - | - | 0 | - |
| - | - | 9814 | 1165 | - | - | 0 | - |
| - | - | 3097 | 1166 | - | - | 0 | - |
| - | - | 4557 | 1177 | - | - | 0 | - |
| 7 | y | 3.42E+04 | 1178 | 5.792E-05 | 0.04918 | +1 | 11 |
| - | - | 2.018E+04 | 1179 | - | - | 0 | - |
| - | - | 2646 | 1180 | - | - | 0 | - |
| - | - | 1.437E+04 | 1182 | - | - | 0 | - |
| - | - | 1.119E+04 | 1183 | - | - | 0 | - |
| - | - | 3624 | 1184 | - | - | 0 | - |
| - | - | 1.055E+04 | 1201 | - | - | 0 | - |
| - | - | 5260 | 1202 | - | - | 0 | - |
| - | - | 2.243E+04 | 1217 | - | - | 0 | - |
| 11 | c | 4.545E+04 | 1218 | 0.002681 | 2.202 | +1 | 11 |
| - | - | 2.666E+04 | 1219 | - | - | 0 | - |
| - | - | 7102 | 1220 | - | - | 0 | - |
| - | - | 2598 | 1231 | - | - | 0 | - |
| - | - | 5242 | 1235 | - | - | 0 | - |
| - | - | 3480 | 1236 | - | - | 0 | - |
| - | - | 2.131E+04 | 1253 | - | - | 0 | - |
| - | - | 1.432E+04 | 1254 | - | - | 0 | - |
| - | - | 3624 | 1255 | - | - | 0 | - |
| 6 | y | 8.972E+04 | 1275 | 0.0002157 | 0.1692 | +1 | 12 |
| - | - | 5.602E+04 | 1276 | - | - | 0 | - |
| - | - | 1.59E+04 | 1277 | - | - | 0 | - |
| - | - | 3030 | 1278 | - | - | 0 | - |
| - | - | 2926 | 1300 | - | - | 0 | - |
| - | - | 3192 | 1301 | - | - | 0 | - |
| - | - | 2165 | 1302 | - | - | 0 | - |
| - | - | 4716 | 1303 | - | - | 0 | - |
| - | - | 3418 | 1311 | - | - | 0 | - |
| - | - | 2935 | 1312 | - | - | 0 | - |
| - | - | 2834 | 1326 | - | - | 0 | - |
| - | - | 4189 | 1327 | - | - | 0 | - |
| 12 | c | 3171 | 1328 | 0.008579 | 6.462 | +1 | 12 |
| 12 | c | 1.437E+04 | 1329 | 0.004436 | 3.338 | +1 | 12 |
| - | - | 1.232E+04 | 1330 | - | - | 0 | - |
| - | - | 4027 | 1331 | - | - | 0 | - |
| - | - | 3.195E+04 | 1345 | - | - | 0 | - |
| 12 | c | 1.199E+05 | 1346 | 0.001444 | 1.073 | +1 | 12 |
| - | - | 7.993E+04 | 1347 | - | - | 0 | - |
| - | - | 2.162E+04 | 1348 | - | - | 0 | - |
| - | - | 2714 | 1349 | - | - | 0 | - |
| 5 | y | 1.145E+04 | 1354 | 0.001033 | 0.7631 | +1 | 13 |
| 5 | y | 1.918E+04 | 1355 | 0.01056 | 7.793 | +1 | 13 |
| 5 | z | 8028 | 1356 | 0.004959 | 3.658 | +1 | 13 |
| - | - | 3535 | 1357 | - | - | 0 | - |
| - | - | 3659 | 1358 | - | - | 0 | - |
| - | - | 2790 | 1359 | - | - | 0 | - |
| - | - | 2891 | 1360 | - | - | 0 | - |
| - | - | 2797 | 1361 | - | - | 0 | - |
| - | - | 2608 | 1370 | - | - | 0 | - |
| - | - | 4.479E+04 | 1371 | - | - | 0 | - |
| 5 | y | 8.064E+05 | 1372 | 0.0008555 | 0.6237 | +1 | 13 |
| - | - | 5.66E+05 | 1373 | - | - | 0 | - |
| - | - | 1.82E+05 | 1374 | - | - | 0 | - |
| - | - | 1.58E+04 | 1375 | - | - | 0 | - |
| - | - | 3257 | 1398 | - | - | 0 | - |
| - | - | 1.565E+04 | 1399 | - | - | 0 | - |
| - | - | 3.146E+04 | 1400 | - | - | 0 | - |
| - | - | 1.707E+04 | 1401 | - | - | 0 | - |
| - | - | 1.622E+05 | 1402 | - | - | 0 | - |
| - | - | 1.219E+05 | 1403 | - | - | 0 | - |
| - | - | 3.865E+04 | 1404 | - | - | 0 | - |
| - | - | 5126 | 1405 | - | - | 0 | - |
| - | - | 9.526E+04 | 1416 | - | - | 0 | - |
| 13 | c | 1.269E+05 | 1417 | 0.0045 | 3.176 | +1 | 13 |
| - | - | 6.594E+04 | 1418 | - | - | 0 | - |
| - | - | 2.026E+04 | 1419 | - | - | 0 | - |
| - | - | 5027 | 1431 | - | - | 0 | - |
| - | - | 5862 | 1432 | - | - | 0 | - |
| - | - | 2978 | 1433 | - | - | 0 | - |
| - | - | 9555 | 1487 | - | - | 0 | - |
| - | - | 1.221E+04 | 1488 | - | - | 0 | - |
| - | - | 4011 | 1489 | - | - | 0 | - |
| 4 | y | 2366 | 1501 | 0.01329 | 8.859 | +1 | 14 |
| 4 | z | 6.658E+04 | 1503 | 0.001839 | 1.224 | +1 | 14 |
| - | - | 6.522E+04 | 1504 | - | - | 0 | - |
| - | - | 3.008E+04 | 1505 | - | - | 0 | - |
| 14 | c | 5692 | 1514 | 0.0006061 | 0.4004 | +1 | 14 |
| - | - | 7555 | 1515 | - | - | 0 | - |
| - | - | 4258 | 1516 | - | - | 0 | - |
| - | - | 3080 | 1518 | - | - | 0 | - |
| 4 | y | 8.841E+04 | 1519 | 5.558E-05 | 0.0366 | +1 | 14 |
| - | - | 7.745E+04 | 1520 | - | - | 0 | - |
| - | - | 2.952E+04 | 1521 | - | - | 0 | - |
| - | - | 1.071E+04 | 1530 | - | - | 0 | - |
| 14 | c | 2.044E+05 | 1531 | 0.001651 | 1.079 | +1 | 14 |
| - | - | 1.828E+05 | 1532 | - | - | 0 | - |
| - | - | 2819 | 1532 | - | - | 0 | - |
| - | - | 6.517E+04 | 1533 | - | - | 0 | - |
| - | - | 1.009E+04 | 1534 | - | - | 0 | - |
| - | - | 2630 | 1544 | - | - | 0 | - |
| - | - | 4976 | 1545 | - | - | 0 | - |
| - | - | 3111 | 1571 | - | - | 0 | - |
| - | - | 4898 | 1572 | - | - | 0 | - |
| - | - | 3440 | 1573 | - | - | 0 | - |
| 3 | y | 3109 | 1614 | 0.00312 | 1.933 | +1 | 15 |
| - | - | 3.021E+04 | 1615 | - | - | 0 | - |
| 3 | z | 4.268E+04 | 1616 | 0.02604 | 16.11 | +1 | 15 |
| - | - | 2.851E+04 | 1617 | - | - | 0 | - |
| - | - | 1.005E+04 | 1618 | - | - | 0 | - |
| - | - | 2774 | 1619 | - | - | 0 | - |
| 3 | y | 2.628E+04 | 1632 | 0.0003793 | 0.2325 | +1 | 15 |
| - | - | 2.791E+04 | 1633 | - | - | 0 | - |
| - | - | 1.334E+04 | 1634 | - | - | 0 | - |
| - | - | 2361 | 1643 | - | - | 0 | - |
| - | - | 2627 | 1644 | - | - | 0 | - |
| - | - | 2981 | 1645 | - | - | 0 | - |
| - | - | 4929 | 1646 | - | - | 0 | - |
| - | - | 3310 | 1647 | - | - | 0 | - |
| - | - | 3990 | 1655 | - | - | 0 | - |
| - | - | 5662 | 1656 | - | - | 0 | - |
| - | - | 3639 | 1657 | - | - | 0 | - |
| 15 | c | 1.033E+05 | 1659 | 0.001888 | 1.138 | +1 | 15 |
| - | - | 9.189E+04 | 1660 | - | - | 0 | - |
| - | - | 4.667E+04 | 1661 | - | - | 0 | - |
| - | - | 1.276E+04 | 1662 | - | - | 0 | - |
| - | - | 4887 | 1663 | - | - | 0 | - |
| - | - | 1.173E+04 | 1672 | - | - | 0 | - |
| - | - | 1.71E+04 | 1673 | - | - | 0 | - |
| - | - | 7474 | 1674 | - | - | 0 | - |
| - | - | 3891 | 1675 | - | - | 0 | - |
| - | - | 3.173E+04 | 1686 | - | - | 0 | - |
| - | - | 3.204E+04 | 1687 | - | - | 0 | - |
| - | - | 2.229E+04 | 1688 | - | - | 0 | - |
| - | - | 9580 | 1689 | - | - | 0 | - |
| 2 | z | 5507 | 1699 | 0.01504 | 8.856 | +1 | 16 |
| 2 | z | 5471 | 1700 | 0.01662 | 9.78 | +1 | 16 |
| - | - | 3159 | 1701 | - | - | 0 | - |
| 2 | y | 3.802E+04 | 1715 | 0.001691 | 0.9864 | +1 | 16 |
| 2 | y | 6.46E+04 | 1716 | 0.01121 | 6.531 | +1 | 16 |
| 2 | z | 2.25E+05 | 1717 | 0.0002073 | 0.1207 | +1 | 16 |
| - | - | 2.246E+05 | 1718 | - | - | 0 | - |
| - | - | 2885 | 1719 | - | - | 0 | - |
| - | - | 9.441E+04 | 1719 | - | - | 0 | - |
| - | - | 1.899E+04 | 1720 | - | - | 0 | - |
| 16 | c | 1.572E+05 | 1730 | 0.0001829 | 0.1057 | +1 | 16 |
| - | - | 1.575E+05 | 1731 | - | - | 0 | - |
| - | - | 1.072E+05 | 1732 | - | - | 0 | - |
| - | - | 4.416E+04 | 1733 | - | - | 0 | - |
| - | - | 2.103E+04 | 1734 | - | - | 0 | - |
| - | - | 3103 | 1743 | - | - | 0 | - |
| - | - | 1.181E+04 | 1744 | - | - | 0 | - |
| - | - | 2.299E+04 | 1745 | - | - | 0 | - |
| - | - | 1.565E+04 | 1746 | - | - | 0 | - |
| - | - | 7300 | 1747 | - | - | 0 | - |
| - | - | 2821 | 1759 | - | - | 0 | - |
| - | - | 1.28E+04 | 1760 | - | - | 0 | - |
| - | - | 2.678E+04 | 1761 | - | - | 0 | - |
| - | - | 3.123E+04 | 1762 | - | - | 0 | - |
| - | - | 2.201E+04 | 1763 | - | - | 0 | - |
| - | - | 9400 | 1764 | - | - | 0 | - |
| - | - | 1.92E+04 | 1769 | - | - | 0 | - |
| - | - | 2.483E+04 | 1770 | - | - | 0 | - |
| - | - | 1.601E+04 | 1771 | - | - | 0 | - |
| - | - | 9402 | 1772 | - | - | 0 | - |
| - | - | 4448 | 1773 | - | - | 0 | - |
| - | - | 1.548E+04 | 1774 | - | - | 0 | - |
| - | - | 1.98E+04 | 1775 | - | - | 0 | - |
| - | - | 9388 | 1776 | - | - | 0 | - |
| - | - | 2.083E+04 | 1777 | - | - | 0 | - |
| - | - | 2.455E+04 | 1778 | - | - | 0 | - |
| - | - | 8891 | 1779 | - | - | 0 | - |
| - | - | 3177 | 1786 | - | - | 0 | - |
| - | - | 4.218E+04 | 1787 | - | - | 0 | - |
| - | - | 2.663E+05 | 1788 | - | - | 0 | - |
| - | - | 2.554E+05 | 1789 | - | - | 0 | - |
| - | - | 1.253E+05 | 1790 | - | - | 0 | - |
| - | - | 2.222E+04 | 1791 | - | - | 0 | - |
| - | - | 5387 | 1797 | - | - | 0 | - |
| - | - | 8289 | 1798 | - | - | 0 | - |
| - | - | 6913 | 1799 | - | - | 0 | - |
| - | - | 4382 | 1800 | - | - | 0 | - |
| - | - | 3956 | 1801 | - | - | 0 | - |
| - | - | 2550 | 1803 | - | - | 0 | - |
| - | - | 1.872E+04 | 1805 | - | - | 0 | - |
| - | - | 2.315E+04 | 1806 | - | - | 0 | - |
| - | - | 1.127E+04 | 1807 | - | - | 0 | - |
| - | - | 4164 | 1814 | - | - | 0 | - |
| - | - | 6.072E+04 | 1815 | - | - | 0 | - |
| - | - | 4.009E+05 | 1816 | - | - | 0 | - |
| - | - | 3.731E+05 | 1817 | - | - | 0 | - |
| - | - | 1.965E+05 | 1818 | - | - | 0 | - |
| - | - | 2963 | 1818 | - | - | 0 | - |
| - | - | 3.043E+04 | 1819 | - | - | 0 | - |
| - | - | 3451 | 1820 | - | - | 0 | - |
| - | - | 6246 | 1830 | - | - | 0 | - |
| - | - | 2.609E+04 | 1831 | - | - | 0 | - |
| - | - | 6.786E+05 | 1832 | - | - | 0 | - |
| - | - | 1.82E+06 | 1833 | - | - | 0 | - |
| - | - | 1.572E+06 | 1834 | - | - | 0 | - |
| - | - | 6.62E+05 | 1835 | - | - | 0 | - |
| - | - | 9.563E+04 | 1836 | - | - | 0 | - |
| - | - | 5579 | 1866 | - | - | 0 | - |

m/z Charge Intensity FragmentType MassShift Position
120.08097839355469 0 5536.8774
122.33557891845703 0 1067.5835
129.1023712158203 0 8410.003
143.12428283691406 0 1047.7318
165.4038848876953 0 1249.9261
173.1286163330078 0 4929.395
173.4501953125 0 16499.486
183.10523986816406 0 1659.5571
183.11297607421875 0 10610.647
201.12335205078125 0 20055.432
215.13912963867188 0 5637.737
217.1337890625 0 2526.752
224.60670471191406 0 2107.0093
226.11875915527344 0 2587.7578
229.0153045654297 0 1440.8817
233.16502380371094 0 13512.206
233.60089111328125 0 1743.7103
234.1679229736328 0 2007.4443
243.1453094482422 0 5060.2075
245.12835693359375 0 4519.5933
251.1758270263672 0 4506.033
253.39984130859375 0 1613.523
261.1598815917969 0 11938.391
280.164306640625 0 2339.7998
282.1460266113281 0 2023.6187
294.1814880371094 0 1745.4854
296.1971435546875 0 264691.56
297.1565246582031 0 3542.2383
297.2004089355469 0 39190.51
298.1767883300781 0 4764.852
298.201904296875 0 2775.2996
301.08428955078125 0 1560.9652
303.1708679199219 0 4419.997
314.1861572265625 0 4854.794
314.2077331542969 0 128420.52
315.2109680175781 0 19251.762
317.8280944824219 0 2290.988
319.1979675292969 0 12454.946 y 14
330.1822814941406 0 6308.478
331.1854553222656 0 2453.3079
345.7443542480469 0 1644.9579
346.1758728027344 0 7838.8784
347.1797790527344 0 2696.6267
348.1916198730469 0 12026.337
349.1960754394531 0 2460.0376
360.0343933105469 0 1644.9562
360.22802734375 0 2406.9622
362.2074279785156 0 34867.734
363.2109375 0 4390.421
385.2208557128906 0 2223.406
390.20355224609375 0 2225.4539
398.245849609375 0 3350.4573
411.2607727050781 0 11438.161
412.26470947265625 0 2282.254
415.2695617675781 0 3695.1416
416.25469970703125 0 15644.145
417.255859375 0 5304.749
433.2426452636719 0 17077.066 y 13
433.2814636230469 0 179701.23
434.24853515625 0 3698.133
434.2843933105469 0 42538.52
435.2874755859375 0 5550.2935
442.24700927734375 0 2166.1216
443.2301330566406 0 3551.7283
443.26568603515625 0 18225.738
444.2690734863281 0 4457.9077
459.2615966796875 0 1935.1113
461.2764587402344 0 181210.62
462.2794189453125 0 48685.023
463.2828674316406 0 4852.3306
469.24432373046875 0 1700.544
498.21826171875 0 1868.3374
504.27783203125 0 31060.414 y 12
505.2801818847656 0 7204.231
530.3342895507812 0 7468.375
540.3175048828125 0 6836.3438
558.3291625976562 0 24867.979
559.3319702148438 0 9411.829
582.2835693359375 0 1590.9889
582.7810668945312 0 2760.1133
583.2803955078125 0 2000.0458
591.2950439453125 0 3901.7893
615.3128051757812 0 2107.588 y Ammonia loss 11
616.3174438476562 0 5255.0747 z 11
617.3250122070312 0 6962.742
617.810302734375 0 4180.994
618.3080444335938 0 6266.0054
626.8126220703125 0 30283.225
627.26025390625 0 4170.8555
627.3140869140625 0 22589.74
627.8171997070312 0 4166.0483
628.731689453125 0 1960.114
632.3365478515625 0 18402.014 y 11
633.33984375 0 6666.5674
637.8155517578125 0 2085.3152 y 5
663.3402099609375 0 2982.3125
663.8370971679688 0 3230.0044
664.3402709960938 0 2056.7202 c Water loss 11
668.826171875 0 2221.3833
677.3363037109375 0 19241.467 y Water loss 4
677.8326416015625 0 20979.865 y Ammonia loss 4
678.332275390625 0 13174.197 z 4
685.3318481445312 0 2219.5732
686.3416137695312 0 653821 y 4
686.8429565429688 0 489535.12
687.3438720703125 0 155066.64
687.8455810546875 0 14270.105
700.3479614257812 0 8269.562 c Ammonia loss 12
700.849853515625 0 4666.366
701.346923828125 0 3194.143
729.4015502929688 0 14393.622 z 10
730.4068603515625 0 10846.628
740.3464965820312 0 5728.129
741.3467407226562 0 2133.552
742.4148559570312 0 5405.313
743.4163208007812 0 2691.0315
745.4208374023438 0 12537.878 y 10
746.4171142578125 0 3825.8337
750.8720703125 0 4940.125 y Water loss 3
751.369384765625 0 6557.617 y Ammonia loss 3
751.8661499023438 0 3352.603 z 3
759.8756103515625 0 100913.83 y 3
760.376953125 0 73725.25
760.8781127929688 0 35814.91
807.4135131835938 0 1971.0668 y Water loss 2
807.9097290039062 0 1954.0134 y Ammonia loss 2
808.404541015625 0 1827.1187 z 2
810.2609252929688 0 1899.1115
812.9097900390625 0 2534.9695
816.4187622070312 0 21654.78 y 2
816.9195556640625 0 18672.877
817.4204711914062 0 8541.786
829.4388427734375 0 3711.2524
842.9432373046875 0 2801.9956
847.9402465820312 0 5138.891
848.4385986328125 0 4900.131
848.9385986328125 0 4404.714
856.9462890625 0 17525.508 c Ammonia loss 15
857.4463500976562 0 20712.533 y Ammonia loss 9
857.9496459960938 0 14665.226 y Water loss 1
858.4439697265625 0 33509.32 y Ammonia loss 1
859.4500122070312 0 26221.09
860.4528198242188 0 6364.25
866.9441528320312 0 3059.0068 y 1
867.4407958984375 0 6831.2095
867.9445190429688 0 4289.812
868.4036865234375 0 8631.29
869.4034423828125 0 3468.4727
874.4639892578125 0 7934.3276 y 9
875.4623413085938 0 3487.19
886.4605102539062 0 3281.9734
899.4602661132812 0 2324.2551
907.470703125 0 14627.937
907.96875 0 15356.42
908.4700927734375 0 13469.71
908.9693603515625 0 4000.7883
915.964599609375 0 3187.1604
916.4758911132812 0 163024.83
916.9766845703125 0 180019.62
917.4783935546875 0 104312.414
917.9784545898438 0 13477.065
928.4724731445312 0 2042.9987 w 8
929.48291015625 0 2078.891
939.440185546875 0 5150.5967
940.4459228515625 0 1860.6488
958.4898071289062 0 4717.454
959.48974609375 0 2791.3784
973.489501953125 0 5883.363
974.5005493164062 0 3665.4197
975.3219604492188 0 1932.7743
987.4874877929688 0 51586.19 z 8
988.4937744140625 0 83489.09
989.4971313476562 0 25523.646
990.5 0 6829.382
1002.5005493164062 0 3507.1113
1003.5012817382812 0 16321.071 y 8
1004.5103149414062 0 6460.4727
1031.502685546875 0 2090.256
1053.4857177734375 0 7153.248
1054.49560546875 0 3817.373
1072.5006103515625 0 2342.2874
1073.5120849609375 0 7487.3926 y Ammonia loss 7
1074.518798828125 0 50463.562 z 7
1075.526123046875 0 131348.77
1076.5294189453125 0 70142.84
1077.53271484375 0 15438.128
1087.53173828125 0 5526.456
1088.5333251953125 0 4483.3677
1089.530517578125 0 21834.543
1090.5364990234375 0 64645.13 y 7
1091.5396728515625 0 32152.96
1092.5418701171875 0 6585.0205
1103.5462646484375 0 4236.412
1104.554931640625 0 14680.751 c 9
1105.5596923828125 0 9403.877
1116.53369140625 0 4537.4
1117.5369873046875 0 10010.215
1118.5421142578125 0 4499.273
1156.567626953125 0 2147.2432
1159.538818359375 0 2727.5093 y Water loss 6
1160.5406494140625 0 3545.22 y Ammonia loss 6
1161.5498046875 0 23924.31 z 6
1162.5567626953125 0 54017.977
1163.5615234375 0 28073.36
1164.557373046875 0 9814.338
1165.556884765625 0 3097.3416
1176.5604248046875 0 4557.0337
1177.569580078125 0 34204.51 y 6
1178.5697021484375 0 20183.148
1179.574951171875 0 2646.2405
1181.5819091796875 0 14374.038
1182.5845947265625 0 11191.569
1183.5972900390625 0 3624.0278
1200.6119384765625 0 10553.452
1201.613037109375 0 5260.1924
1216.634521484375 0 22433.596
1217.6385498046875 0 45448.07 c 10
1218.6431884765625 0 26660.518
1219.6456298828125 0 7102.267
1230.603759765625 0 2598.314
1234.603759765625 0 5242.294
1235.602783203125 0 3480.3853
1252.6170654296875 0 21311.316
1253.6195068359375 0 14321.442
1254.609130859375 0 3623.6912
1274.6220703125 0 89716.33 y 5
1275.6251220703125 0 56021.344
1276.6268310546875 0 15903.209
1277.6268310546875 0 3029.5789
1299.6357421875 0 2925.771
1300.638427734375 0 3191.5168
1301.6680908203125 0 2165.337
1302.675048828125 0 4716.083
1310.661376953125 0 3417.986
1311.6632080078125 0 2934.5562
1325.669921875 0 2833.9263
1326.6729736328125 0 4189.0225
1327.6806640625 0 3170.6213 c Water loss 11
1328.6688232421875 0 14366.873 c Ammonia loss 11
1329.6685791015625 0 12317.791
1330.6798095703125 0 4027.1685
1344.69091796875 0 31945.322
1345.6983642578125 0 119948.336 c 11
1346.70166015625 0 79928.56
1347.703857421875 0 21624
1348.7127685546875 0 2714.156
1353.6634521484375 0 11447.957 y Water loss 4
1354.6590576171875 0 19181.959 y Ammonia loss 4
1355.6513671875 0 8028.2817 z 4
1356.6497802734375 0 3535.056
1357.6739501953125 0 3658.57
1358.675537109375 0 2789.7905
1359.6712646484375 0 2891.1575
1360.6749267578125 0 2797.3806
1369.671630859375 0 2607.8833
1370.6666259765625 0 44785.31
1371.6741943359375 0 806432.56 y 4
1372.6773681640625 0 565997.4
1373.680419921875 0 182015.62
1374.6876220703125 0 15803.857
1397.720458984375 0 3257.4458
1398.6676025390625 0 15654.261
1399.680419921875 0 31464.113
1400.6837158203125 0 17069.873
1401.7005615234375 0 162160.36
1402.7032470703125 0 121921.67
1403.7069091796875 0 38646.973
1404.7081298828125 0 5126.1953
1415.728271484375 0 95257.18
1416.732421875 0 126938.2 c 12
1417.7364501953125 0 65936.35
1418.7415771484375 0 20257.81
1430.7010498046875 0 5026.6763
1431.697021484375 0 5862.034
1432.7001953125 0 2978.2393
1486.7635498046875 0 9554.889
1487.767822265625 0 12211.611
1488.7733154296875 0 4010.6938
1500.7196044921875 0 2366.185 y Water loss 3
1502.722900390625 0 66576.52 z 3
1503.7252197265625 0 65218.027
1504.7283935546875 0 30075.615
1513.75390625 0 5692.182 c Ammonia loss 13
1514.7525634765625 0 7555.3833
1515.7642822265625 0 4258.0444
1517.7508544921875 0 3079.7163
1518.743408203125 0 88405.164 y 3
1519.744140625 0 77454.76
1520.7457275390625 0 29520.568
1529.767578125 0 10706.896
1530.7781982421875 0 204373.77 c 13
1531.78076171875 0 182753.36
1532.0167236328125 0 2819.47
1532.7845458984375 0 65170.977
1533.7862548828125 0 10086.607
1543.7803955078125 0 2630.0925
1544.7916259765625 0 4975.6694
1570.8238525390625 0 3110.973
1571.80712890625 0 4898.4004
1572.8153076171875 0 3440.3464
1613.8138427734375 0 3109.1565 y Water loss 2
1614.85498046875 0 30206.645
1615.8348388671875 0 42677.32 z 2
1616.828125 0 28511.348
1617.8173828125 0 10051.303
1618.8277587890625 0 2773.7917
1631.8271484375 0 26284.635 y 2
1632.828125 0 27911.762
1633.8331298828125 0 13339.054
1642.86865234375 0 2360.5398
1643.82421875 0 2626.737
1644.84765625 0 2980.6592
1645.825927734375 0 4929.323
1646.8275146484375 0 3310.289
1654.849609375 0 3990.1565
1655.8505859375 0 5661.6343
1656.8489990234375 0 3638.7288
1658.8729248046875 0 103346.37 c 14
1659.8743896484375 0 91894.44
1660.865234375 0 46668.824
1661.8138427734375 0 12758.486
1662.804443359375 0 4887.4204
1671.8453369140625 0 11725.871
1672.8443603515625 0 17100.652
1673.83642578125 0 7473.798
1674.8214111328125 0 3891.3938
1685.896484375 0 31733.93
1686.900634765625 0 32039.438
1687.906982421875 0 22294.658
1688.9053955078125 0 9580.019
1698.8609619140625 0 5506.699 z Water loss 1
1699.8465576171875 0 5470.661 z Ammonia loss 1
1700.8468017578125 0 3159.2952
1714.8663330078125 0 38021.418 y Water loss 1
1715.85986328125 0 64597.883 y Ammonia loss 1
1716.856689453125 0 225034.52 z 1
1717.85888671875 0 224608.12
1718.585693359375 0 2885.127
1718.8631591796875 0 94405.82
1719.8660888671875 0 18990.238
1729.9117431640625 0 157214.23 c 15
1730.9132080078125 0 157543.39
1731.9176025390625 0 107227.38
1732.924072265625 0 44155.117
1733.9267578125 0 21034.023
1742.8968505859375 0 3103.0981
1743.90185546875 0 11807.759
1744.8896484375 0 22991.28
1745.886962890625 0 15653.821
1746.8975830078125 0 7300.183
1758.9049072265625 0 2821.0696
1759.88525390625 0 12797.362
1760.912841796875 0 26779.547
1761.9154052734375 0 31231.264
1762.91162109375 0 22008.764
1763.9093017578125 0 9400.474
1768.9371337890625 0 19196.611
1769.9346923828125 0 24830.637
1770.9368896484375 0 16014.409
1771.9403076171875 0 9402.392
1772.9058837890625 0 4448.4316
1773.9180908203125 0 15476.642
1774.91748046875 0 19797.668
1775.9180908203125 0 9388.227
1776.8917236328125 0 20830.314
1777.889892578125 0 24554.459
1778.8961181640625 0 8890.531
1785.9105224609375 0 3176.739
1786.9425048828125 0 42180.8
1787.9312744140625 0 266268.78
1788.9334716796875 0 255437.61
1789.9342041015625 0 125341.12
1790.9271240234375 0 22217.496
1796.938232421875 0 5387.2563
1797.9207763671875 0 8289.117
1798.918212890625 0 6912.9253
1799.930908203125 0 4381.516
1800.911376953125 0 3956.168
1802.93603515625 0 2549.7405
1804.9549560546875 0 18716.443
1805.9560546875 0 23150.736
1806.9554443359375 0 11272.854
1813.91943359375 0 4163.7183
1814.9375 0 60724.74
1815.927001953125 0 400854.12
1816.92822265625 0 373142
1817.9293212890625 0 196481.84
1818.233154296875 0 2963.345
1818.9307861328125 0 30427.498
1819.92822265625 0 3451.0256
1829.912353515625 0 6246.2114
1830.9337158203125 0 26093.506
1831.943359375 0 678603.1
1832.9503173828125 0 1820112.1
1833.9530029296875 0 1572178.1
1834.9560546875 0 662022.5
1835.9564208984375 0 95631.8
1865.942626953125 0 5578.991

Spectrum Details

|  |  |
| --- | --- |
| Matched peaks? Matched peaks The total absolute number of peaks matched. Additionally in brackets the total fraction of peaks matched and the total number of peaks is shown. | 65 (16.33% of 398) |
| FDR? FDR The false discovery rate estimated for this peptide. It is calculated by matching all theoretical fragments with a non-integer shift with the raw peaks for this spectrum. This is done with 40 different shifts. The resulting percentage is the average number of annotated peaks over the number of annotated peaks with the correct spectrum. | 0.33% |
| Satellite FDR? Satellite FDR See the FDR for details on its calculation. This satellite ion specific FDR only contains the satellite ions (d/w) for I/L/J positions. | - |
| PSM Score? PSM Score The PSM Score as given by Hecklib to this annotated spectrum. It is shown with three significant figures. | 574 |

#### Spectrum 8281? Spectrum 8281 The raw spectrum of this peptide as annotated by Hecklib. The fragments are coloured according to ion type (see legend). Any peaks with a star '\*' as text can be hovered over to see the full details, first the ion type second the mass shift type. By hovering over the amino acids in the peptide or ions in the legend the corresponding peaks are highlighted. By toggling the 'Unassigned' label you can turn the background (unassigned) peaks on or off in the plot. By updating the slider in the Ion legend you can update the spectrum to only show the top X% of the peaks with labels. The top X% means any peak that is within X% of the highest intensity. By dragging in the spectrum you can zoom in to a specific part of the spectrum and use 'Zoom Out' to get back to the original zoom level. The annotation of the spectrum is based on the given sequence in the peptides file and is done with different software so inconsistencies are likely. The peaks are annotated based on the given sequence, with 20 ppm tolerance.

Copy Data

##### Spectrum 8281 (TSV)

###### Preview

```
Loading example...
```

*Click on the button to copy the data to your clipboard.*

Mz MinMz MaxIntensity Max

WidthHeightPeptide font sizePeptide stroke widthSpectrum font sizeSpectrum stroke widthCompact peptide

Ion legend

wxyz

abcd

OtherUnassignedIon^Charge^_Position_Show for top:%

VTJFPPSSEEJQANKAT

01.33e+52.66e+53.98e+55.31e+5

Zoom Out

y^+1^_3_y^+1^_4_y^+1^_5_y^+1^_5_y^+1^_6_z^+1^_6_y^+1^_6_y^+2^_13_y^+2^_13_z^+2^_13_y^+2^_13_z^+1^_7_y^+1^_7_y^+2^_14_c^+2^_14_y^+2^_14_y^+2^_15_y^+2^_15_y^+2^_15_c^+2^_16_y^+1^_8_y^+2^_16_z^+1^_8_y^+2^_16_y^+1^_8_z^+1^_9_y^+1^_9_y^+1^_10_y^+1^_10_z^+1^_10_y^+1^_10_c^+1^_10_y^+1^_11_z^+1^_11_y^+1^_11_c^+1^_11_y^+1^_12_c^+1^_12_c^+1^_12_y^+1^_13_y^+1^_13_z^+1^_13_y^+1^_13_c^+1^_13_c^+1^_13_z^+1^_14_c^+1^_14_y^+1^_14_c^+1^_14_y^+1^_15_z^+1^_15_y^+1^_15_c^+1^_15_z^+1^_16_z^+1^_16_y^+1^_16_y^+1^_16_z^+1^_16_c^+1^_16_

046492713911854

Fragment Matches Table

Show background peaks

| Position | Ion type | Intensity | mz Theoretical | mz Error (Th) | mz Error (ppm) | Charge | Series Number |
| --- | --- | --- | --- | --- | --- | --- | --- |
| - | - | 1173 | 120.1 | - | - | 0 | - |
| - | - | 348.4 | 128.5 | - | - | 0 | - |
| - | - | 2004 | 129.1 | - | - | 0 | - |
| - | - | 399.5 | 133.2 | - | - | 0 | - |
| - | - | 415.6 | 137.5 | - | - | 0 | - |
| - | - | 407.7 | 144.3 | - | - | 0 | - |
| - | - | 505.7 | 148.8 | - | - | 0 | - |
| - | - | 445.5 | 148.9 | - | - | 0 | - |
| - | - | 821 | 148.9 | - | - | 0 | - |
| - | - | 615 | 148.9 | - | - | 0 | - |
| - | - | 704.6 | 148.9 | - | - | 0 | - |
| - | - | 761.4 | 148.9 | - | - | 0 | - |
| - | - | 946.6 | 148.9 | - | - | 0 | - |
| - | - | 1224 | 148.9 | - | - | 0 | - |
| - | - | 1321 | 148.9 | - | - | 0 | - |
| - | - | 3119 | 148.9 | - | - | 0 | - |
| - | - | 5299 | 148.9 | - | - | 0 | - |
| - | - | 3960 | 149 | - | - | 0 | - |
| - | - | 1749 | 149 | - | - | 0 | - |
| - | - | 1401 | 149 | - | - | 0 | - |
| - | - | 932.9 | 149 | - | - | 0 | - |
| - | - | 629 | 149 | - | - | 0 | - |
| - | - | 508.8 | 149 | - | - | 0 | - |
| - | - | 392.8 | 149 | - | - | 0 | - |
| - | - | 430.3 | 149 | - | - | 0 | - |
| - | - | 562 | 149 | - | - | 0 | - |
| - | - | 446.4 | 151.7 | - | - | 0 | - |
| - | - | 552.3 | 160.6 | - | - | 0 | - |
| - | - | 1461 | 173.1 | - | - | 0 | - |
| - | - | 435.1 | 174.3 | - | - | 0 | - |
| - | - | 2884 | 183.1 | - | - | 0 | - |
| - | - | 6664 | 201.1 | - | - | 0 | - |
| - | - | 469 | 201.1 | - | - | 0 | - |
| - | - | 1448 | 215.1 | - | - | 0 | - |
| - | - | 765.4 | 217.1 | - | - | 0 | - |
| - | - | 4182 | 233.2 | - | - | 0 | - |
| - | - | 544.4 | 234.2 | - | - | 0 | - |
| - | - | 1470 | 243.1 | - | - | 0 | - |
| - | - | 988.2 | 245.1 | - | - | 0 | - |
| - | - | 1169 | 251.2 | - | - | 0 | - |
| - | - | 2939 | 261.2 | - | - | 0 | - |
| - | - | 811.6 | 262.2 | - | - | 0 | - |
| - | - | 713.8 | 280.2 | - | - | 0 | - |
| - | - | 869.1 | 282.1 | - | - | 0 | - |
| - | - | 6.28E+04 | 296.2 | - | - | 0 | - |
| - | - | 1.007E+04 | 297.2 | - | - | 0 | - |
| - | - | 667.6 | 298.2 | - | - | 0 | - |
| - | - | 1047 | 298.2 | - | - | 0 | - |
| - | - | 1306 | 303.2 | - | - | 0 | - |
| - | - | 3.567E+04 | 314.2 | - | - | 0 | - |
| - | - | 4601 | 315.2 | - | - | 0 | - |
| 15 | y | 3562 | 319.2 | 5.614E-05 | 0.1759 | +1 | 3 |
| - | - | 1146 | 330.2 | - | - | 0 | - |
| - | - | 2946 | 346.2 | - | - | 0 | - |
| - | - | 2538 | 348.2 | - | - | 0 | - |
| - | - | 1005 | 349.2 | - | - | 0 | - |
| - | - | 783.3 | 360.2 | - | - | 0 | - |
| - | - | 9908 | 362.2 | - | - | 0 | - |
| - | - | 2031 | 363.2 | - | - | 0 | - |
| - | - | 622.9 | 389.3 | - | - | 0 | - |
| - | - | 726.3 | 390.2 | - | - | 0 | - |
| - | - | 2427 | 411.3 | - | - | 0 | - |
| - | - | 1328 | 415.3 | - | - | 0 | - |
| - | - | 4651 | 416.3 | - | - | 0 | - |
| - | - | 976.4 | 417.3 | - | - | 0 | - |
| 14 | y | 4710 | 433.2 | 0.002549 | 5.883 | +1 | 4 |
| - | - | 4.867E+04 | 433.3 | - | - | 0 | - |
| - | - | 925.5 | 434.2 | - | - | 0 | - |
| - | - | 1.275E+04 | 434.3 | - | - | 0 | - |
| - | - | 813.4 | 435.3 | - | - | 0 | - |
| - | - | 687.9 | 441.2 | - | - | 0 | - |
| - | - | 976.7 | 443.2 | - | - | 0 | - |
| - | - | 5882 | 443.3 | - | - | 0 | - |
| - | - | 1358 | 444.3 | - | - | 0 | - |
| - | - | 5.003E+04 | 461.3 | - | - | 0 | - |
| - | - | 1.313E+04 | 462.3 | - | - | 0 | - |
| - | - | 1438 | 463.3 | - | - | 0 | - |
| 13 | y | 648.7 | 487.3 | 0.003581 | 7.349 | +1 | 5 |
| - | - | 846.3 | 498.2 | - | - | 0 | - |
| 13 | y | 8042 | 504.3 | 0.0002554 | 0.5065 | +1 | 5 |
| - | - | 1474 | 505.3 | - | - | 0 | - |
| - | - | 1965 | 530.3 | - | - | 0 | - |
| - | - | 673.4 | 531.3 | - | - | 0 | - |
| - | - | 1534 | 540.3 | - | - | 0 | - |
| - | - | 671 | 541.3 | - | - | 0 | - |
| - | - | 6040 | 558.3 | - | - | 0 | - |
| - | - | 2768 | 559.3 | - | - | 0 | - |
| - | - | 1332 | 582.8 | - | - | 0 | - |
| - | - | 1182 | 583.3 | - | - | 0 | - |
| - | - | 904.1 | 591.3 | - | - | 0 | - |
| 12 | y | 1204 | 615.3 | 0.003444 | 5.598 | +1 | 6 |
| 12 | z | 1204 | 616.3 | 0.001234 | 2.003 | +1 | 6 |
| - | - | 1569 | 617.3 | - | - | 0 | - |
| - | - | 683.6 | 617.8 | - | - | 0 | - |
| - | - | 1790 | 618.3 | - | - | 0 | - |
| - | - | 860.2 | 618.8 | - | - | 0 | - |
| - | - | 647.1 | 624.2 | - | - | 0 | - |
| - | - | 7896 | 626.8 | - | - | 0 | - |
| - | - | 732.6 | 627.3 | - | - | 0 | - |
| - | - | 4137 | 627.3 | - | - | 0 | - |
| - | - | 1098 | 627.8 | - | - | 0 | - |
| 12 | y | 4779 | 632.3 | 0.0007599 | 1.202 | +1 | 6 |
| - | - | 1195 | 633.3 | - | - | 0 | - |
| - | - | 682.8 | 638.3 | - | - | 0 | - |
| 5 | y | 4932 | 677.3 | 0.001583 | 2.336 | +2 | 13 |
| 5 | y | 6842 | 677.8 | 0.004204 | 6.202 | +2 | 13 |
| 5 | z | 2278 | 678.3 | 0.0001971 | 0.2906 | +2 | 13 |
| - | - | 843.3 | 678.8 | - | - | 0 | - |
| - | - | 818.1 | 679.3 | - | - | 0 | - |
| - | - | 1336 | 685.8 | - | - | 0 | - |
| 5 | y | 1.749E+05 | 686.3 | 0.0003896 | 0.5676 | +2 | 13 |
| - | - | 1.321E+05 | 686.8 | - | - | 0 | - |
| - | - | 3.905E+04 | 687.3 | - | - | 0 | - |
| - | - | 4504 | 687.8 | - | - | 0 | - |
| - | - | 917.5 | 700.3 | - | - | 0 | - |
| - | - | 2316 | 700.8 | - | - | 0 | - |
| - | - | 631.8 | 726.2 | - | - | 0 | - |
| 11 | z | 2690 | 729.4 | 0.001216 | 1.667 | +1 | 7 |
| - | - | 2322 | 730.4 | - | - | 0 | - |
| - | - | 1222 | 740.3 | - | - | 0 | - |
| - | - | 1839 | 742.4 | - | - | 0 | - |
| - | - | 614.2 | 743.4 | - | - | 0 | - |
| 11 | y | 3160 | 745.4 | 0.000192 | 0.2576 | +1 | 7 |
| 4 | y | 746.4 | 751.4 | 0.001125 | 1.497 | +2 | 14 |
| 14 | c | 680.5 | 756.9 | 0.00564 | 7.452 | +2 | 14 |
| 4 | y | 2.482E+04 | 759.9 | 0.0003623 | 0.4768 | +2 | 14 |
| - | - | 2.203E+04 | 760.4 | - | - | 0 | - |
| - | - | 1.05E+04 | 760.9 | - | - | 0 | - |
| - | - | 1327 | 761.4 | - | - | 0 | - |
| 3 | y | 857 | 807.4 | 0.001271 | 1.575 | +2 | 15 |
| 3 | y | 924.2 | 807.9 | 0.002245 | 2.778 | +2 | 15 |
| - | - | 963.6 | 812.9 | - | - | 0 | - |
| - | - | 780.6 | 813.4 | - | - | 0 | - |
| 3 | y | 5467 | 816.4 | 0.0008718 | 1.068 | +2 | 15 |
| - | - | 5920 | 816.9 | - | - | 0 | - |
| - | - | 1422 | 817.4 | - | - | 0 | - |
| - | - | 769.6 | 829.4 | - | - | 0 | - |
| - | - | 637.8 | 842.4 | - | - | 0 | - |
| - | - | 670.5 | 843.5 | - | - | 0 | - |
| - | - | 1383 | 847.9 | - | - | 0 | - |
| - | - | 884.8 | 848.4 | - | - | 0 | - |
| 16 | c | 5287 | 856.9 | 0.0004039 | 0.4713 | +2 | 16 |
| 10 | y | 5636 | 857.4 | 0.01143 | 13.33 | +1 | 8 |
| 2 | y | 3811 | 857.9 | 0.01198 | 13.96 | +2 | 16 |
| 10 | z | 8815 | 858.4 | 0.001286 | 1.499 | +1 | 8 |
| - | - | 6692 | 859.4 | - | - | 0 | - |
| - | - | 1921 | 860.4 | - | - | 0 | - |
| 2 | y | 966 | 866.9 | 0.002179 | 2.514 | +2 | 16 |
| - | - | 1483 | 867.4 | - | - | 0 | - |
| - | - | 2662 | 868.4 | - | - | 0 | - |
| - | - | 954.1 | 869.4 | - | - | 0 | - |
| 10 | y | 2507 | 874.5 | 0.0002868 | 0.328 | +1 | 8 |
| - | - | 1064 | 875.5 | - | - | 0 | - |
| - | - | 672.6 | 886.5 | - | - | 0 | - |
| - | - | 4173 | 907.5 | - | - | 0 | - |
| - | - | 4392 | 908 | - | - | 0 | - |
| - | - | 3810 | 908.5 | - | - | 0 | - |
| - | - | 924.2 | 909 | - | - | 0 | - |
| - | - | 1188 | 916 | - | - | 0 | - |
| - | - | 4.186E+04 | 916.5 | - | - | 0 | - |
| - | - | 5.352E+04 | 917 | - | - | 0 | - |
| - | - | 2.806E+04 | 917.5 | - | - | 0 | - |
| - | - | 3801 | 918 | - | - | 0 | - |
| - | - | 2307 | 939.4 | - | - | 0 | - |
| - | - | 752.3 | 958.5 | - | - | 0 | - |
| - | - | 1368 | 973.5 | - | - | 0 | - |
| - | - | 690.8 | 974.5 | - | - | 0 | - |
| 9 | z | 1.339E+04 | 987.5 | 0.0002582 | 0.2615 | +1 | 9 |
| - | - | 1.801E+04 | 988.5 | - | - | 0 | - |
| - | - | 7555 | 989.5 | - | - | 0 | - |
| - | - | 1447 | 990.5 | - | - | 0 | - |
| - | - | 975.9 | 1002 | - | - | 0 | - |
| 9 | y | 4120 | 1004 | 0.003634 | 3.622 | +1 | 9 |
| - | - | 2668 | 1005 | - | - | 0 | - |
| - | - | 1004 | 1053 | - | - | 0 | - |
| - | - | 1226 | 1054 | - | - | 0 | - |
| 8 | y | 1521 | 1073 | 0.01509 | 14.07 | +1 | 10 |
| 8 | y | 2222 | 1074 | 0.00114 | 1.062 | +1 | 10 |
| 8 | z | 1.369E+04 | 1075 | 0.000215 | 0.2001 | +1 | 10 |
| - | - | 3.778E+04 | 1076 | - | - | 0 | - |
| - | - | 1.793E+04 | 1077 | - | - | 0 | - |
| - | - | 4737 | 1078 | - | - | 0 | - |
| - | - | 1672 | 1088 | - | - | 0 | - |
| - | - | 1122 | 1089 | - | - | 0 | - |
| - | - | 4260 | 1090 | - | - | 0 | - |
| 8 | y | 1.53E+04 | 1091 | 0.001239 | 1.136 | +1 | 10 |
| - | - | 7008 | 1092 | - | - | 0 | - |
| - | - | 1821 | 1093 | - | - | 0 | - |
| - | - | 1058 | 1104 | - | - | 0 | - |
| 10 | c | 4143 | 1105 | 0.003456 | 3.129 | +1 | 10 |
| - | - | 2175 | 1106 | - | - | 0 | - |
| - | - | 1373 | 1117 | - | - | 0 | - |
| - | - | 2212 | 1118 | - | - | 0 | - |
| - | - | 1666 | 1119 | - | - | 0 | - |
| 7 | y | 1191 | 1160 | 0.01379 | 11.89 | +1 | 11 |
| 7 | z | 6702 | 1162 | 0.0004714 | 0.4059 | +1 | 11 |
| - | - | 1.535E+04 | 1163 | - | - | 0 | - |
| - | - | 7402 | 1164 | - | - | 0 | - |
| - | - | 2945 | 1165 | - | - | 0 | - |
| - | - | 938.9 | 1175 | - | - | 0 | - |
| - | - | 1617 | 1177 | - | - | 0 | - |
| 7 | y | 9041 | 1178 | 0.0006745 | 0.5728 | +1 | 11 |
| - | - | 4829 | 1179 | - | - | 0 | - |
| - | - | 919.3 | 1180 | - | - | 0 | - |
| - | - | 3573 | 1182 | - | - | 0 | - |
| - | - | 2744 | 1183 | - | - | 0 | - |
| - | - | 3317 | 1201 | - | - | 0 | - |
| - | - | 2126 | 1202 | - | - | 0 | - |
| - | - | 5502 | 1217 | - | - | 0 | - |
| 11 | c | 1.26E+04 | 1218 | 0.002315 | 1.901 | +1 | 11 |
| - | - | 6445 | 1219 | - | - | 0 | - |
| - | - | 1428 | 1220 | - | - | 0 | - |
| - | - | 868.7 | 1235 | - | - | 0 | - |
| - | - | 5646 | 1253 | - | - | 0 | - |
| - | - | 3935 | 1254 | - | - | 0 | - |
| - | - | 1351 | 1255 | - | - | 0 | - |
| 6 | y | 2.08E+04 | 1275 | 0.0002157 | 0.1692 | +1 | 12 |
| - | - | 1.467E+04 | 1276 | - | - | 0 | - |
| - | - | 4553 | 1277 | - | - | 0 | - |
| - | - | 1177 | 1301 | - | - | 0 | - |
| - | - | 1059 | 1302 | - | - | 0 | - |
| - | - | 1491 | 1311 | - | - | 0 | - |
| - | - | 1313 | 1327 | - | - | 0 | - |
| 12 | c | 5886 | 1329 | 0.006755 | 5.084 | +1 | 12 |
| - | - | 3858 | 1330 | - | - | 0 | - |
| - | - | 8531 | 1345 | - | - | 0 | - |
| 12 | c | 3.131E+04 | 1346 | 0.001688 | 1.254 | +1 | 12 |
| - | - | 2.025E+04 | 1347 | - | - | 0 | - |
| - | - | 6913 | 1348 | - | - | 0 | - |
| - | - | 1168 | 1349 | - | - | 0 | - |
| 5 | y | 2871 | 1354 | 0.002864 | 2.116 | +1 | 13 |
| 5 | y | 4070 | 1355 | 0.006162 | 4.549 | +1 | 13 |
| 5 | z | 2809 | 1356 | 0.001633 | 1.205 | +1 | 13 |
| - | - | 1224 | 1357 | - | - | 0 | - |
| - | - | 1100 | 1358 | - | - | 0 | - |
| - | - | 1622 | 1360 | - | - | 0 | - |
| - | - | 1158 | 1361 | - | - | 0 | - |
| - | - | 1073 | 1370 | - | - | 0 | - |
| - | - | 1.253E+04 | 1371 | - | - | 0 | - |
| 5 | y | 2.132E+05 | 1372 | 0.0011 | 0.8017 | +1 | 13 |
| - | - | 1.546E+05 | 1373 | - | - | 0 | - |
| - | - | 5.562E+04 | 1374 | - | - | 0 | - |
| - | - | 5589 | 1375 | - | - | 0 | - |
| - | - | 990.3 | 1398 | - | - | 0 | - |
| - | - | 4628 | 1399 | - | - | 0 | - |
| 13 | c | 7172 | 1400 | 0.02727 | 19.48 | +1 | 13 |
| - | - | 5254 | 1401 | - | - | 0 | - |
| - | - | 3.884E+04 | 1402 | - | - | 0 | - |
| - | - | 2.91E+04 | 1403 | - | - | 0 | - |
| - | - | 8795 | 1404 | - | - | 0 | - |
| - | - | 1226 | 1405 | - | - | 0 | - |
| - | - | 2.526E+04 | 1416 | - | - | 0 | - |
| 13 | c | 3.631E+04 | 1417 | 0.00389 | 2.745 | +1 | 13 |
| - | - | 1.923E+04 | 1418 | - | - | 0 | - |
| - | - | 5443 | 1419 | - | - | 0 | - |
| - | - | 1606 | 1431 | - | - | 0 | - |
| - | - | 1212 | 1432 | - | - | 0 | - |
| - | - | 946.9 | 1474 | - | - | 0 | - |
| - | - | 772 | 1475 | - | - | 0 | - |
| - | - | 3454 | 1487 | - | - | 0 | - |
| - | - | 3080 | 1488 | - | - | 0 | - |
| - | - | 1226 | 1489 | - | - | 0 | - |
| 4 | z | 1.618E+04 | 1503 | 0.0009848 | 0.6554 | +1 | 14 |
| - | - | 1.6E+04 | 1504 | - | - | 0 | - |
| - | - | 7453 | 1505 | - | - | 0 | - |
| - | - | 813 | 1506 | - | - | 0 | - |
| 14 | c | 1694 | 1514 | 0.007206 | 4.761 | +1 | 14 |
| - | - | 1996 | 1515 | - | - | 0 | - |
| - | - | 1292 | 1516 | - | - | 0 | - |
| 4 | y | 2.495E+04 | 1519 | 0.0006659 | 0.4385 | +1 | 14 |
| - | - | 1.943E+04 | 1520 | - | - | 0 | - |
| - | - | 8755 | 1521 | - | - | 0 | - |
| - | - | 903.9 | 1522 | - | - | 0 | - |
| - | - | 2481 | 1530 | - | - | 0 | - |
| 14 | c | 6.02E+04 | 1531 | 0.001895 | 1.238 | +1 | 14 |
| - | - | 5.318E+04 | 1532 | - | - | 0 | - |
| - | - | 1180 | 1532 | - | - | 0 | - |
| - | - | 2.049E+04 | 1533 | - | - | 0 | - |
| - | - | 2454 | 1534 | - | - | 0 | - |
| - | - | 1241 | 1544 | - | - | 0 | - |
| - | - | 1266 | 1545 | - | - | 0 | - |
| - | - | 1624 | 1572 | - | - | 0 | - |
| - | - | 1713 | 1573 | - | - | 0 | - |
| - | - | 828.1 | 1574 | - | - | 0 | - |
| 3 | y | 1584 | 1614 | 0.002266 | 1.404 | +1 | 15 |
| - | - | 5838 | 1615 | - | - | 0 | - |
| 3 | z | 1.276E+04 | 1616 | 0.02567 | 15.89 | +1 | 15 |
| - | - | 8493 | 1617 | - | - | 0 | - |
| - | - | 3485 | 1618 | - | - | 0 | - |
| 3 | y | 7577 | 1632 | 0.003919 | 2.402 | +1 | 15 |
| - | - | 6340 | 1633 | - | - | 0 | - |
| - | - | 2994 | 1634 | - | - | 0 | - |
| - | - | 911.7 | 1645 | - | - | 0 | - |
| - | - | 1520 | 1646 | - | - | 0 | - |
| - | - | 1375 | 1647 | - | - | 0 | - |
| - | - | 1500 | 1655 | - | - | 0 | - |
| - | - | 1590 | 1656 | - | - | 0 | - |
| 15 | c | 2.865E+04 | 1659 | 0.002986 | 1.8 | +1 | 15 |
| - | - | 2.877E+04 | 1660 | - | - | 0 | - |
| - | - | 1.344E+04 | 1661 | - | - | 0 | - |
| - | - | 3736 | 1662 | - | - | 0 | - |
| - | - | 2237 | 1663 | - | - | 0 | - |
| - | - | 911.3 | 1671 | - | - | 0 | - |
| - | - | 3313 | 1672 | - | - | 0 | - |
| - | - | 4342 | 1673 | - | - | 0 | - |
| - | - | 2034 | 1674 | - | - | 0 | - |
| - | - | 8619 | 1686 | - | - | 0 | - |
| - | - | 1.052E+04 | 1687 | - | - | 0 | - |
| - | - | 6526 | 1688 | - | - | 0 | - |
| - | - | 2134 | 1689 | - | - | 0 | - |
| - | - | 1120 | 1690 | - | - | 0 | - |
| - | - | 899.9 | 1698 | - | - | 0 | - |
| 2 | z | 1519 | 1699 | 0.009551 | 5.622 | +1 | 16 |
| 2 | z | 1538 | 1700 | 0.02651 | 15.6 | +1 | 16 |
| 2 | y | 1.051E+04 | 1715 | 0.002459 | 1.434 | +1 | 16 |
| 2 | y | 1.708E+04 | 1716 | 0.01536 | 8.95 | +1 | 16 |
| 2 | z | 6.763E+04 | 1717 | 0.0004514 | 0.2629 | +1 | 16 |
| - | - | 6.178E+04 | 1718 | - | - | 0 | - |
| - | - | 3.052E+04 | 1719 | - | - | 0 | - |
| - | - | 5840 | 1720 | - | - | 0 | - |
| - | - | 1118 | 1727 | - | - | 0 | - |
| - | - | 1185 | 1728 | - | - | 0 | - |
| - | - | 1052 | 1729 | - | - | 0 | - |
| 16 | c | 4.429E+04 | 1730 | 0.0006712 | 0.388 | +1 | 16 |
| - | - | 4.433E+04 | 1731 | - | - | 0 | - |
| - | - | 3.094E+04 | 1732 | - | - | 0 | - |
| - | - | 1.232E+04 | 1733 | - | - | 0 | - |
| - | - | 5110 | 1734 | - | - | 0 | - |
| - | - | 1444 | 1735 | - | - | 0 | - |
| - | - | 1058 | 1742 | - | - | 0 | - |
| - | - | 1176 | 1743 | - | - | 0 | - |
| - | - | 2857 | 1744 | - | - | 0 | - |
| - | - | 5191 | 1745 | - | - | 0 | - |
| - | - | 5771 | 1746 | - | - | 0 | - |
| - | - | 1763 | 1747 | - | - | 0 | - |
| - | - | 877.4 | 1754 | - | - | 0 | - |
| - | - | 1267 | 1759 | - | - | 0 | - |
| - | - | 4049 | 1760 | - | - | 0 | - |
| - | - | 7082 | 1761 | - | - | 0 | - |
| - | - | 9982 | 1762 | - | - | 0 | - |
| - | - | 6237 | 1763 | - | - | 0 | - |
| - | - | 2010 | 1764 | - | - | 0 | - |
| - | - | 5767 | 1769 | - | - | 0 | - |
| - | - | 7566 | 1770 | - | - | 0 | - |
| - | - | 5678 | 1771 | - | - | 0 | - |
| - | - | 2592 | 1772 | - | - | 0 | - |
| - | - | 1734 | 1773 | - | - | 0 | - |
| - | - | 4366 | 1774 | - | - | 0 | - |
| - | - | 5018 | 1775 | - | - | 0 | - |
| - | - | 2013 | 1776 | - | - | 0 | - |
| - | - | 7521 | 1777 | - | - | 0 | - |
| - | - | 5835 | 1778 | - | - | 0 | - |
| - | - | 2941 | 1779 | - | - | 0 | - |
| - | - | 1631 | 1786 | - | - | 0 | - |
| - | - | 1.314E+04 | 1787 | - | - | 0 | - |
| - | - | 8.317E+04 | 1788 | - | - | 0 | - |
| - | - | 7.241E+04 | 1789 | - | - | 0 | - |
| - | - | 3.9E+04 | 1790 | - | - | 0 | - |
| - | - | 7288 | 1791 | - | - | 0 | - |
| - | - | 973 | 1792 | - | - | 0 | - |
| - | - | 2736 | 1798 | - | - | 0 | - |
| - | - | 2590 | 1799 | - | - | 0 | - |
| - | - | 1049 | 1800 | - | - | 0 | - |
| - | - | 995.1 | 1803 | - | - | 0 | - |
| - | - | 6010 | 1805 | - | - | 0 | - |
| - | - | 5569 | 1806 | - | - | 0 | - |
| - | - | 3837 | 1807 | - | - | 0 | - |
| - | - | 2041 | 1814 | - | - | 0 | - |
| - | - | 1.556E+04 | 1815 | - | - | 0 | - |
| - | - | 1.133E+05 | 1816 | - | - | 0 | - |
| - | - | 1.161E+05 | 1817 | - | - | 0 | - |
| - | - | 5.682E+04 | 1818 | - | - | 0 | - |
| - | - | 9360 | 1819 | - | - | 0 | - |
| - | - | 2044 | 1830 | - | - | 0 | - |
| - | - | 7494 | 1831 | - | - | 0 | - |
| - | - | 1.916E+05 | 1832 | - | - | 0 | - |
| - | - | 5.258E+05 | 1833 | - | - | 0 | - |
| - | - | 4.417E+05 | 1834 | - | - | 0 | - |
| - | - | 2.031E+05 | 1835 | - | - | 0 | - |
| - | - | 2.835E+04 | 1836 | - | - | 0 | - |

m/z Charge Intensity FragmentType MassShift Position
120.08103942871094 0 1173.1749
128.50204467773438 0 348.41776
129.1024627685547 0 2003.5388
133.2472381591797 0 399.48773
137.53005981445312 0 415.55002
144.31802368164062 0 407.74838
148.8440399169922 0 505.65167
148.8804473876953 0 445.4587
148.88751220703125 0 820.9501
148.89492797851562 0 615.0221
148.9020538330078 0 704.6363
148.90882873535156 0 761.4099
148.91639709472656 0 946.62415
148.9235076904297 0 1224.1974
148.9306640625 0 1320.6179
148.93804931640625 0 3118.6348
148.94580078125 0 5299.3794
148.9622802734375 0 3960.2458
148.97000122070312 0 1749.0684
148.97702026367188 0 1400.8641
148.98428344726562 0 932.9145
148.99163818359375 0 628.9628
148.99876403808594 0 508.8083
149.00621032714844 0 392.7972
149.02740478515625 0 430.28592
149.0486602783203 0 562.0084
151.73284912109375 0 446.39694
160.6261749267578 0 552.3387
173.12855529785156 0 1460.713
174.2852325439453 0 435.06256
183.1129913330078 0 2883.8618
201.1233367919922 0 6664.064
201.1490936279297 0 468.962
215.13938903808594 0 1447.6464
217.13414001464844 0 765.38885
233.164794921875 0 4181.551
234.16940307617188 0 544.3715
243.14529418945312 0 1469.9634
245.1285400390625 0 988.2372
251.17498779296875 0 1169.3917
261.1595458984375 0 2939.069
262.1641540527344 0 811.64075
280.1654968261719 0 713.757
282.1439514160156 0 869.0917
296.1971130371094 0 62800.32
297.2003479003906 0 10070.6
298.1754150390625 0 667.5708
298.203369140625 0 1047.1204
303.1702880859375 0 1305.828
314.20770263671875 0 35669.72
315.210693359375 0 4601.3657
319.1975402832031 0 3561.9395 y 14
330.1809387207031 0 1146.1915
346.17620849609375 0 2946.4802
348.1919860839844 0 2537.94
349.1971435546875 0 1005.376
360.2276306152344 0 783.30396
362.2074890136719 0 9908.15
363.2110595703125 0 2031.1508
389.2547912597656 0 622.9059
390.2016906738281 0 726.3138
411.2601318359375 0 2427.3608
415.2701110839844 0 1328.0017
416.2552795410156 0 4651.096
417.2539367675781 0 976.44617
433.2430725097656 0 4709.792 y 13
433.281494140625 0 48670.93
434.2484436035156 0 925.49786
434.28436279296875 0 12751.508
435.2884826660156 0 813.4176
441.2475891113281 0 687.88477
443.23199462890625 0 976.7081
443.2660217285156 0 5882.1396
444.2674560546875 0 1358.2944
461.2764587402344 0 50025.82
462.27978515625 0 13130.8955
463.28216552734375 0 1437.5559
487.2546691894531 0 648.7444 y Ammonia loss 12
498.2213439941406 0 846.2757
504.27789306640625 0 8042.2534 y 12
505.2795715332031 0 1473.7656
530.3340454101562 0 1965.2592
531.3367919921875 0 673.3984
540.318359375 0 1533.7211
541.3240356445312 0 670.9956
558.3289184570312 0 6039.702
559.3325805664062 0 2767.8003
582.7825927734375 0 1331.927
583.2838745117188 0 1182.3583
591.2950439453125 0 904.1444
615.3131103515625 0 1203.9851 y Ammonia loss 11
616.3187255859375 0 1203.8351 z 11
617.3209838867188 0 1569.0746
617.8088989257812 0 683.6445
618.3034057617188 0 1789.8231
618.8060913085938 0 860.16327
624.1806640625 0 647.0506
626.8126220703125 0 7896.4365
627.2627563476562 0 732.5745
627.3131713867188 0 4136.828
627.8189086914062 0 1098.3597
632.3369750976562 0 4778.5547 y 11
633.3360595703125 0 1194.9957
638.3176879882812 0 682.7536
677.3374633789062 0 4931.5083 y Water loss 4
677.8320922851562 0 6841.8096 y Ammonia loss 4
678.3316040039062 0 2278.269 z 4
678.8262939453125 0 843.31824
679.3326416015625 0 818.0688
685.8348388671875 0 1336.4338
686.341552734375 0 174873.19 y 4
686.8427124023438 0 132054.31
687.3436889648438 0 39047.2
687.8453979492188 0 4503.552
700.341796875 0 917.4844
700.847900390625 0 2315.9858
726.2018432617188 0 631.82825
729.4027709960938 0 2690.3403 z 10
730.4055786132812 0 2321.7134
740.345947265625 0 1221.7162
742.4141845703125 0 1839.4009
743.4266967773438 0 614.17706
745.4204711914062 0 3160.1433 y 10
751.3632202148438 0 746.408 y Ammonia loss 3
756.8939208984375 0 680.5047 c Water loss 13
759.875732421875 0 24824.135 y 3
760.376708984375 0 22025.72
760.8779907226562 0 10496.774
761.3809204101562 0 1327.0935
807.4133911132812 0 856.9885 y Water loss 2
807.9063720703125 0 924.212 y Ammonia loss 2
812.9066162109375 0 963.6238
813.4092407226562 0 780.6206
816.4182739257812 0 5467.206 y 2
816.9180908203125 0 5920.475
817.4180297851562 0 1422.4723
829.4469604492188 0 769.5786
842.3838500976562 0 637.7583
843.4528198242188 0 670.47876
847.938720703125 0 1382.6917
848.438720703125 0 884.77167
856.9459228515625 0 5286.6094 c Ammonia loss 15
857.44775390625 0 5635.787 y Ammonia loss 9
857.9479370117188 0 3810.7117 y Water loss 1
858.4454345703125 0 8814.888 z 9
859.4493408203125 0 6691.59
860.448974609375 0 1920.9443
866.9434204101562 0 966.00085 y 1
867.4412231445312 0 1482.547
868.4036254882812 0 2661.527
869.4061889648438 0 954.099
874.4625854492188 0 2507.1235 y 9
875.4637451171875 0 1064.1387
886.4584350585938 0 672.598
907.4702758789062 0 4172.7046
907.9693603515625 0 4391.8555
908.4714965820312 0 3809.541
908.9714965820312 0 924.1878
915.967529296875 0 1188.3463
916.4755249023438 0 41858.062
916.9766235351562 0 53523.72
917.4774169921875 0 28063.314
917.977783203125 0 3800.969
939.4418334960938 0 2307.116
958.4860229492188 0 752.26117
973.49267578125 0 1367.7882
974.4971923828125 0 690.7649
987.4869995117188 0 13385.599 z 8
988.4931640625 0 18010.994
989.496337890625 0 7555.472
990.5030517578125 0 1446.5286
1002.4930419921875 0 975.89813
1003.5018310546875 0 4119.6265 y 8
1004.5029296875 0 2667.9983
1053.48193359375 0 1004.46265
1054.48193359375 0 1225.9421
1072.5118408203125 0 1520.8933 y Water loss 7
1073.5120849609375 0 2221.7114 y Ammonia loss 7
1074.5185546875 0 13690.953 z 7
1075.5260009765625 0 37781.08
1076.5286865234375 0 17931.67
1077.531982421875 0 4736.974
1087.5247802734375 0 1672.1361
1088.5380859375 0 1121.7858
1089.5318603515625 0 4260.4126
1090.5362548828125 0 15301.0625 y 7
1091.5391845703125 0 7007.585
1092.543701171875 0 1821.4319
1103.5570068359375 0 1057.8566
1104.5537109375 0 4142.931 c 9
1105.5645751953125 0 2174.6602
1116.534912109375 0 1372.6525
1117.5345458984375 0 2211.5322
1118.5360107421875 0 1666.4504
1159.545166015625 0 1191.3713 y Water loss 6
1161.55126953125 0 6702.196 z 6
1162.5567626953125 0 15345.081
1163.56005859375 0 7402.302
1164.558837890625 0 2944.9253
1174.64404296875 0 938.8995
1176.5616455078125 0 1616.9663
1177.56884765625 0 9041.163 y 6
1178.5728759765625 0 4828.9893
1179.5733642578125 0 919.3293
1181.5811767578125 0 3573.4473
1182.5797119140625 0 2744.0847
1200.612548828125 0 3316.7402
1201.62158203125 0 2126.2007
1216.63232421875 0 5502.4966
1217.638916015625 0 12598.07 c 10
1218.6441650390625 0 6444.8784
1219.6387939453125 0 1427.9846
1234.5989990234375 0 868.683
1252.6143798828125 0 5645.713
1253.619140625 0 3935.1467
1254.6142578125 0 1350.757
1274.6220703125 0 20804.207 y 5
1275.62353515625 0 14672.432
1276.6275634765625 0 4553.432
1300.645263671875 0 1177.4052
1301.6815185546875 0 1059.333
1310.657958984375 0 1491.153
1326.660888671875 0 1312.6725
1328.66650390625 0 5886.0933 c Ammonia loss 11
1329.6702880859375 0 3857.7227
1344.6910400390625 0 8531.476
1345.6981201171875 0 31309.768 c 11
1346.700927734375 0 20249.21
1347.7052001953125 0 6913.3926
1348.7027587890625 0 1168.1542
1353.66162109375 0 2871.2773 y Water loss 4
1354.6546630859375 0 4070.2036 y Ammonia loss 4
1355.657958984375 0 2809.0422 z 4
1356.65234375 0 1223.9213
1357.678466796875 0 1099.7977
1359.67431640625 0 1622.151
1360.677001953125 0 1157.6929
1369.6683349609375 0 1072.871
1370.6658935546875 0 12532.973
1371.6739501953125 0 213213.8 y 4
1372.6767578125 0 154566.06
1373.6793212890625 0 55618.25
1374.683349609375 0 5588.761
1397.7078857421875 0 990.2998
1398.6673583984375 0 4628.3438
1399.68310546875 0 7172.42 c Ammonia loss 12
1400.685546875 0 5253.66
1401.7003173828125 0 38836.04
1402.702880859375 0 29103.426
1403.7061767578125 0 8794.851
1404.7015380859375 0 1225.7474
1415.7279052734375 0 25258.438
1416.7330322265625 0 36311.883 c 12
1417.736572265625 0 19233.176
1418.7412109375 0 5442.641
1430.6954345703125 0 1606.4103
1431.697998046875 0 1212.4272
1473.7379150390625 0 946.9007
1474.7286376953125 0 772.0498
1486.7640380859375 0 3453.7827
1487.7672119140625 0 3079.959
1488.7608642578125 0 1226.2584
1502.7237548828125 0 16184.89 z 3
1503.7249755859375 0 16000.2295
1504.72509765625 0 7453.118
1505.7403564453125 0 812.98627
1513.74609375 0 1694.0989 c Ammonia loss 13
1514.74853515625 0 1995.9083
1515.762451171875 0 1291.8132
1518.7427978515625 0 24947.582 y 3
1519.7454833984375 0 19434.965
1520.7452392578125 0 8754.95
1521.740478515625 0 903.9313
1529.7703857421875 0 2480.9707
1530.7779541015625 0 60198.03 c 13
1531.77978515625 0 53178.383
1532.0155029296875 0 1180.245
1532.78369140625 0 20487.48
1533.7822265625 0 2453.5273
1543.7818603515625 0 1241.1144
1544.7894287109375 0 1266.1455
1571.816162109375 0 1624.1198
1572.8082275390625 0 1712.5027
1573.7852783203125 0 828.0782
1613.814697265625 0 1584.1075 y Water loss 2
1614.850830078125 0 5837.7026
1615.83447265625 0 12755.254 z 2
1616.82666015625 0 8493.114
1617.8255615234375 0 3484.6228
1631.8236083984375 0 7577.071 y 2
1632.828857421875 0 6339.6094
1633.8388671875 0 2994.3726
1644.832275390625 0 911.72205
1645.845703125 0 1520.1056
1646.83056640625 0 1374.8208
1654.834228515625 0 1500.4138
1655.8636474609375 0 1590.2681
1658.871826171875 0 28646.652 c 14
1659.8746337890625 0 28767.848
1660.8629150390625 0 13436.354
1661.810546875 0 3735.6677
1662.79296875 0 2237.3308
1670.878662109375 0 911.3019
1671.8465576171875 0 3313.01
1672.840576171875 0 4341.784
1673.84130859375 0 2034.2279
1685.8978271484375 0 8618.526
1686.899658203125 0 10519.215
1687.90380859375 0 6525.84
1688.91162109375 0 2134.176
1689.9161376953125 0 1119.9773
1697.8375244140625 0 899.93964
1698.85546875 0 1518.81 z Water loss 1
1699.8564453125 0 1537.5518 z Ammonia loss 1
1714.8621826171875 0 10508.114 y Water loss 1
1715.864013671875 0 17075.617 y Ammonia loss 1
1716.85693359375 0 67633.914 z 1
1717.8582763671875 0 61780
1718.863037109375 0 30522.33
1719.865966796875 0 5839.7466
1726.9068603515625 0 1117.8395
1727.937255859375 0 1185.3096
1728.9039306640625 0 1051.6825
1729.9112548828125 0 44291.12 c 15
1730.912841796875 0 44331.734
1731.9180908203125 0 30935.285
1732.92236328125 0 12317.654
1733.928955078125 0 5110.4
1734.922607421875 0 1444.3655
1741.8746337890625 0 1057.804
1742.875244140625 0 1176.2382
1743.9053955078125 0 2857.0342
1744.8968505859375 0 5190.8633
1745.8897705078125 0 5771.256
1746.899658203125 0 1763.1917
1753.937744140625 0 877.3529
1758.9190673828125 0 1266.7766
1759.8890380859375 0 4048.8425
1760.9150390625 0 7082.457
1761.91552734375 0 9982.237
1762.916748046875 0 6237.128
1763.9007568359375 0 2009.6387
1768.9342041015625 0 5767.029
1769.9356689453125 0 7565.561
1770.93359375 0 5678.398
1771.9354248046875 0 2591.5461
1772.9190673828125 0 1734.3997
1773.9112548828125 0 4366.428
1774.9195556640625 0 5018.023
1775.90625 0 2013.3594
1776.888427734375 0 7521.255
1777.8890380859375 0 5834.645
1778.8941650390625 0 2941.468
1785.9158935546875 0 1631.3887
1786.9423828125 0 13137.733
1787.931640625 0 83167.414
1788.933837890625 0 72408.84
1789.9337158203125 0 38995.566
1790.9276123046875 0 7288.181
1791.9075927734375 0 973.0036
1797.914794921875 0 2736.1782
1798.9149169921875 0 2590.01
1799.8936767578125 0 1048.9102
1802.947998046875 0 995.0648
1804.955322265625 0 6010.151
1805.95654296875 0 5568.5166
1806.9586181640625 0 3836.7844
1813.905517578125 0 2040.5425
1814.9344482421875 0 15560.187
1815.9268798828125 0 113342.02
1816.9281005859375 0 116116.59
1817.9283447265625 0 56823.062
1818.929443359375 0 9360.454
1829.9276123046875 0 2043.959
1830.9296875 0 7493.6367
1831.9432373046875 0 191630.16
1832.949462890625 0 525769.4
1833.952392578125 0 441694.66
1834.9552001953125 0 203091.42
1835.9554443359375 0 28350.594

Spectrum Details

|  |  |
| --- | --- |
| Matched peaks? Matched peaks The total absolute number of peaks matched. Additionally in brackets the total fraction of peaks matched and the total number of peaks is shown. | 59 (15.53% of 380) |
| FDR? FDR The false discovery rate estimated for this peptide. It is calculated by matching all theoretical fragments with a non-integer shift with the raw peaks for this spectrum. This is done with 40 different shifts. The resulting percentage is the average number of annotated peaks over the number of annotated peaks with the correct spectrum. | 0.56% |
| Satellite FDR? Satellite FDR See the FDR for details on its calculation. This satellite ion specific FDR only contains the satellite ions (d/w) for I/L/J positions. | - |
| PSM Score? PSM Score The PSM Score as given by Hecklib to this annotated spectrum. It is shown with three significant figures. | 574 |

#### Spectrum 7814? Spectrum 7814 The raw spectrum of this peptide as annotated by Hecklib. The fragments are coloured according to ion type (see legend). Any peaks with a star '\*' as text can be hovered over to see the full details, first the ion type second the mass shift type. By hovering over the amino acids in the peptide or ions in the legend the corresponding peaks are highlighted. By toggling the 'Unassigned' label you can turn the background (unassigned) peaks on or off in the plot. By updating the slider in the Ion legend you can update the spectrum to only show the top X% of the peaks with labels. The top X% means any peak that is within X% of the highest intensity. By dragging in the spectrum you can zoom in to a specific part of the spectrum and use 'Zoom Out' to get back to the original zoom level. The annotation of the spectrum is based on the given sequence in the peptides file and is done with different software so inconsistencies are likely. The peaks are annotated based on the given sequence, with 20 ppm tolerance.

Copy Data

##### Spectrum 7814 (TSV)

###### Preview

```
Loading example...
```

*Click on the button to copy the data to your clipboard.*

Mz MinMz MaxIntensity Max

WidthHeightPeptide font sizePeptide stroke widthSpectrum font sizeSpectrum stroke widthCompact peptide

Ion legend

wxyz

abcd

OtherUnassignedIon^Charge^_Position_Show for top:%

VTJFPPSSEEJQANKAT

02.88e+55.77e+58.65e+51.15e+6

Zoom Out

y^+1^_3_y^+1^_4_y^+1^_5_y^+1^_5_y^+1^_6_y^+1^_6_z^+1^_6_y^+1^_6_y^+2^_12_c^+2^_12_y^+2^_13_y^+2^_13_z^+2^_13_y^+2^_13_c^+2^_13_z^+1^_7_y^+1^_7_y^+2^_14_y^+2^_14_z^+2^_14_c^+2^_14_y^+2^_14_z^+2^_15_y^+2^_15_y^+2^_15_c^+2^_16_y^+1^_8_y^+2^_16_y^+2^_16_y^+2^_16_y^+1^_8_c^+1^_9_z^+1^_9_y^+1^_9_y^+1^_10_y^+1^_10_z^+1^_10_y^+1^_10_c^+1^_10_y^+1^_11_z^+1^_11_y^+1^_11_c^+1^_11_y^+1^_12_y^+1^_12_y^+1^_12_c^+1^_12_c^+1^_12_c^+1^_12_y^+1^_13_y^+1^_13_z^+1^_13_y^+1^_13_c^+1^_13_y^+1^_14_y^+1^_14_z^+1^_14_c^+1^_14_y^+1^_14_c^+1^_14_y^+1^_15_z^+1^_15_y^+1^_15_c^+1^_15_z^+1^_16_z^+1^_16_y^+1^_16_y^+1^_16_z^+1^_16_c^+1^_16_

047194214131885

Fragment Matches Table

Show background peaks

| Position | Ion type | Intensity | mz Theoretical | mz Error (Th) | mz Error (ppm) | Charge | Series Number |
| --- | --- | --- | --- | --- | --- | --- | --- |
| - | - | 2824 | 120.1 | - | - | 0 | - |
| - | - | 466.3 | 120.1 | - | - | 0 | - |
| - | - | 580.5 | 125.9 | - | - | 0 | - |
| - | - | 5050 | 129.1 | - | - | 0 | - |
| - | - | 542.4 | 129.8 | - | - | 0 | - |
| - | - | 1127 | 148.9 | - | - | 0 | - |
| - | - | 751.4 | 148.9 | - | - | 0 | - |
| - | - | 842.1 | 148.9 | - | - | 0 | - |
| - | - | 1058 | 148.9 | - | - | 0 | - |
| - | - | 1418 | 148.9 | - | - | 0 | - |
| - | - | 1742 | 148.9 | - | - | 0 | - |
| - | - | 3121 | 148.9 | - | - | 0 | - |
| - | - | 5427 | 148.9 | - | - | 0 | - |
| - | - | 7283 | 149 | - | - | 0 | - |
| - | - | 4789 | 149 | - | - | 0 | - |
| - | - | 2203 | 149 | - | - | 0 | - |
| - | - | 1756 | 149 | - | - | 0 | - |
| - | - | 1901 | 149 | - | - | 0 | - |
| - | - | 1221 | 149 | - | - | 0 | - |
| - | - | 1026 | 149 | - | - | 0 | - |
| - | - | 687 | 149 | - | - | 0 | - |
| - | - | 1019 | 149 | - | - | 0 | - |
| - | - | 603.7 | 149 | - | - | 0 | - |
| - | - | 773.3 | 149.1 | - | - | 0 | - |
| - | - | 3117 | 173.1 | - | - | 0 | - |
| - | - | 5571 | 183.1 | - | - | 0 | - |
| - | - | 818.8 | 185.3 | - | - | 0 | - |
| - | - | 651.7 | 193.2 | - | - | 0 | - |
| - | - | 1.481E+04 | 201.1 | - | - | 0 | - |
| - | - | 2880 | 215.1 | - | - | 0 | - |
| - | - | 2189 | 217.1 | - | - | 0 | - |
| - | - | 1084 | 219.1 | - | - | 0 | - |
| - | - | 1239 | 226.1 | - | - | 0 | - |
| - | - | 9342 | 233.2 | - | - | 0 | - |
| - | - | 2739 | 243.1 | - | - | 0 | - |
| - | - | 2543 | 245.1 | - | - | 0 | - |
| - | - | 1903 | 247.1 | - | - | 0 | - |
| - | - | 4450 | 251.2 | - | - | 0 | - |
| - | - | 961.9 | 252.2 | - | - | 0 | - |
| - | - | 863.2 | 253.2 | - | - | 0 | - |
| - | - | 6935 | 261.2 | - | - | 0 | - |
| - | - | 1320 | 262.2 | - | - | 0 | - |
| - | - | 954.1 | 269.2 | - | - | 0 | - |
| - | - | 919.8 | 273.1 | - | - | 0 | - |
| - | - | 1940 | 280.2 | - | - | 0 | - |
| - | - | 2427 | 282.1 | - | - | 0 | - |
| - | - | 1.52E+05 | 296.2 | - | - | 0 | - |
| - | - | 1817 | 297.2 | - | - | 0 | - |
| - | - | 2.459E+04 | 297.2 | - | - | 0 | - |
| - | - | 2541 | 298.2 | - | - | 0 | - |
| - | - | 2127 | 298.2 | - | - | 0 | - |
| - | - | 2575 | 303.2 | - | - | 0 | - |
| - | - | 941.5 | 303.7 | - | - | 0 | - |
| - | - | 2994 | 314.2 | - | - | 0 | - |
| - | - | 7.951E+04 | 314.2 | - | - | 0 | - |
| - | - | 1311 | 315.2 | - | - | 0 | - |
| - | - | 1.339E+04 | 315.2 | - | - | 0 | - |
| - | - | 1461 | 317.2 | - | - | 0 | - |
| 15 | y | 9216 | 319.2 | 0.0004321 | 1.354 | +1 | 3 |
| - | - | 3527 | 330.2 | - | - | 0 | - |
| - | - | 1634 | 344.2 | - | - | 0 | - |
| - | - | 6724 | 346.2 | - | - | 0 | - |
| - | - | 1848 | 347.2 | - | - | 0 | - |
| - | - | 5726 | 348.2 | - | - | 0 | - |
| - | - | 2071 | 349.2 | - | - | 0 | - |
| - | - | 2171 | 360.2 | - | - | 0 | - |
| - | - | 2.289E+04 | 362.2 | - | - | 0 | - |
| - | - | 3721 | 363.2 | - | - | 0 | - |
| - | - | 1391 | 385.2 | - | - | 0 | - |
| - | - | 1388 | 390.2 | - | - | 0 | - |
| - | - | 1288 | 393.3 | - | - | 0 | - |
| - | - | 2124 | 398.2 | - | - | 0 | - |
| - | - | 7002 | 411.3 | - | - | 0 | - |
| - | - | 1157 | 412.3 | - | - | 0 | - |
| - | - | 2935 | 415.3 | - | - | 0 | - |
| - | - | 8979 | 416.3 | - | - | 0 | - |
| - | - | 2223 | 417.3 | - | - | 0 | - |
| - | - | 894.4 | 425.1 | - | - | 0 | - |
| 14 | y | 1.261E+04 | 433.2 | 0.002366 | 5.46 | +1 | 4 |
| - | - | 1.178E+05 | 433.3 | - | - | 0 | - |
| - | - | 1804 | 434.2 | - | - | 0 | - |
| - | - | 3.192E+04 | 434.3 | - | - | 0 | - |
| - | - | 2930 | 435.3 | - | - | 0 | - |
| - | - | 970.1 | 442.2 | - | - | 0 | - |
| - | - | 2358 | 443.2 | - | - | 0 | - |
| - | - | 1.437E+04 | 443.3 | - | - | 0 | - |
| - | - | 3252 | 444.3 | - | - | 0 | - |
| - | - | 1045 | 445.2 | - | - | 0 | - |
| - | - | 1.271E+05 | 461.3 | - | - | 0 | - |
| - | - | 2.892E+04 | 462.3 | - | - | 0 | - |
| - | - | 3014 | 463.3 | - | - | 0 | - |
| 13 | y | 1074 | 487.3 | 0.003733 | 7.662 | +1 | 5 |
| - | - | 1574 | 498.2 | - | - | 0 | - |
| 13 | y | 2.059E+04 | 504.3 | 0.0005301 | 1.051 | +1 | 5 |
| - | - | 3863 | 505.3 | - | - | 0 | - |
| - | - | 5855 | 530.3 | - | - | 0 | - |
| - | - | 2467 | 531.3 | - | - | 0 | - |
| - | - | 4696 | 540.3 | - | - | 0 | - |
| - | - | 992.1 | 541.3 | - | - | 0 | - |
| - | - | 1.787E+04 | 558.3 | - | - | 0 | - |
| - | - | 5223 | 559.3 | - | - | 0 | - |
| - | - | 1266 | 560.3 | - | - | 0 | - |
| - | - | 1524 | 582.8 | - | - | 0 | - |
| - | - | 1972 | 591.3 | - | - | 0 | - |
| - | - | 3335 | 591.8 | - | - | 0 | - |
| 12 | y | 1356 | 614.3 | 0.001681 | 2.736 | +1 | 6 |
| 12 | y | 2948 | 615.3 | 0.001674 | 2.721 | +1 | 6 |
| 12 | z | 3894 | 616.3 | 0.0005631 | 0.9137 | +1 | 6 |
| - | - | 2364 | 617.3 | - | - | 0 | - |
| - | - | 3953 | 617.8 | - | - | 0 | - |
| - | - | 2976 | 618.3 | - | - | 0 | - |
| - | - | 1323 | 618.8 | - | - | 0 | - |
| - | - | 2.158E+04 | 626.8 | - | - | 0 | - |
| - | - | 2196 | 627.3 | - | - | 0 | - |
| - | - | 1.233E+04 | 627.3 | - | - | 0 | - |
| - | - | 2096 | 627.8 | - | - | 0 | - |
| 12 | y | 1.246E+04 | 632.3 | 0.0005768 | 0.9122 | +1 | 6 |
| - | - | 4472 | 633.3 | - | - | 0 | - |
| - | - | 1084 | 635.8 | - | - | 0 | - |
| 6 | y | 3254 | 637.8 | 0.0006485 | 1.017 | +2 | 12 |
| - | - | 2640 | 638.3 | - | - | 0 | - |
| - | - | 1457 | 655.3 | - | - | 0 | - |
| - | - | 1816 | 655.4 | - | - | 0 | - |
| - | - | 876.2 | 656.3 | - | - | 0 | - |
| - | - | 3320 | 663.3 | - | - | 0 | - |
| 12 | c | 1377 | 664.8 | 0.005978 | 8.992 | +2 | 12 |
| 5 | y | 1.333E+04 | 677.3 | 0.000545 | 0.8046 | +2 | 13 |
| 5 | y | 1.196E+04 | 677.8 | 0.00695 | 10.25 | +2 | 13 |
| 5 | z | 9259 | 678.3 | 0.001878 | 2.769 | +2 | 13 |
| - | - | 2779 | 678.8 | - | - | 0 | - |
| - | - | 1446 | 685.3 | - | - | 0 | - |
| - | - | 1715 | 685.8 | - | - | 0 | - |
| 5 | y | 4.473E+05 | 686.3 | 0.0007558 | 1.101 | +2 | 13 |
| - | - | 3.233E+05 | 686.8 | - | - | 0 | - |
| - | - | 1.075E+05 | 687.3 | - | - | 0 | - |
| - | - | 1.012E+04 | 687.8 | - | - | 0 | - |
| 13 | c | 5885 | 700.4 | 0.01056 | 15.08 | +2 | 13 |
| - | - | 3789 | 700.8 | - | - | 0 | - |
| - | - | 1176 | 701.3 | - | - | 0 | - |
| - | - | 1132 | 724.4 | - | - | 0 | - |
| 11 | z | 7129 | 729.4 | 0.0008497 | 1.165 | +1 | 7 |
| - | - | 6115 | 730.4 | - | - | 0 | - |
| - | - | 2402 | 731.4 | - | - | 0 | - |
| - | - | 3667 | 740.3 | - | - | 0 | - |
| - | - | 4095 | 742.4 | - | - | 0 | - |
| - | - | 1798 | 743.4 | - | - | 0 | - |
| 11 | y | 7136 | 745.4 | 0.0003751 | 0.5033 | +1 | 7 |
| - | - | 2687 | 746.4 | - | - | 0 | - |
| 4 | y | 3309 | 750.9 | 0.0001537 | 0.2047 | +2 | 14 |
| 4 | y | 5294 | 751.4 | 0.007106 | 9.458 | +2 | 14 |
| 4 | z | 2732 | 751.9 | 0.002095 | 2.786 | +2 | 14 |
| - | - | 1084 | 752.4 | - | - | 0 | - |
| 14 | c | 1569 | 757.4 | 0.009238 | 12.2 | +2 | 14 |
| 4 | y | 7.123E+04 | 759.9 | 0.0006675 | 0.8784 | +2 | 14 |
| - | - | 5.599E+04 | 760.4 | - | - | 0 | - |
| - | - | 2.423E+04 | 760.9 | - | - | 0 | - |
| - | - | 1608 | 761.4 | - | - | 0 | - |
| 3 | z | 822.3 | 799.4 | 0.004137 | 5.175 | +2 | 15 |
| 3 | y | 2566 | 807.9 | 0.00908 | 11.24 | +2 | 15 |
| - | - | 2357 | 812.9 | - | - | 0 | - |
| - | - | 1926 | 813.4 | - | - | 0 | - |
| 3 | y | 1.56E+04 | 816.4 | 0.001116 | 1.367 | +2 | 15 |
| - | - | 1.269E+04 | 816.9 | - | - | 0 | - |
| - | - | 6413 | 817.4 | - | - | 0 | - |
| - | - | 2686 | 829.4 | - | - | 0 | - |
| - | - | 1519 | 830.4 | - | - | 0 | - |
| - | - | 1448 | 842.9 | - | - | 0 | - |
| - | - | 1334 | 847.9 | - | - | 0 | - |
| - | - | 4496 | 848.4 | - | - | 0 | - |
| - | - | 2628 | 848.9 | - | - | 0 | - |
| 16 | c | 1.34E+04 | 856.9 | 0.0009389 | 1.096 | +2 | 16 |
| 10 | y | 1.321E+04 | 857.4 | 0.01192 | 13.9 | +1 | 8 |
| 2 | y | 6550 | 857.9 | 0.01448 | 16.88 | +2 | 16 |
| 2 | y | 2.074E+04 | 858.4 | 0.01667 | 19.42 | +2 | 16 |
| - | - | 1.792E+04 | 859.5 | - | - | 0 | - |
| - | - | 5360 | 860.5 | - | - | 0 | - |
| 2 | y | 5654 | 866.9 | 0.0008363 | 0.9646 | +2 | 16 |
| - | - | 3894 | 867.4 | - | - | 0 | - |
| - | - | 1445 | 867.9 | - | - | 0 | - |
| - | - | 3858 | 868.4 | - | - | 0 | - |
| - | - | 2513 | 869.4 | - | - | 0 | - |
| 10 | y | 8884 | 874.5 | 0.001117 | 1.277 | +1 | 8 |
| - | - | 2794 | 875.5 | - | - | 0 | - |
| - | - | 1051 | 886 | - | - | 0 | - |
| - | - | 1266 | 886.5 | - | - | 0 | - |
| - | - | 1933 | 899 | - | - | 0 | - |
| - | - | 9803 | 907.5 | - | - | 0 | - |
| - | - | 1.451E+04 | 908 | - | - | 0 | - |
| - | - | 9415 | 908.5 | - | - | 0 | - |
| - | - | 2937 | 909 | - | - | 0 | - |
| - | - | 1.262E+05 | 916.5 | - | - | 0 | - |
| - | - | 1.216E+05 | 917 | - | - | 0 | - |
| - | - | 1729 | 917.1 | - | - | 0 | - |
| - | - | 6.928E+04 | 917.5 | - | - | 0 | - |
| - | - | 9051 | 918 | - | - | 0 | - |
| - | - | 3689 | 939.4 | - | - | 0 | - |
| - | - | 1657 | 940.5 | - | - | 0 | - |
| - | - | 2993 | 958.5 | - | - | 0 | - |
| - | - | 2187 | 959.5 | - | - | 0 | - |
| - | - | 4077 | 973.5 | - | - | 0 | - |
| - | - | 2144 | 974.5 | - | - | 0 | - |
| 9 | c | 1693 | 975.5 | 0.002489 | 2.551 | +1 | 9 |
| 9 | z | 3.453E+04 | 987.5 | 0.001174 | 1.189 | +1 | 9 |
| - | - | 4.439E+04 | 988.5 | - | - | 0 | - |
| - | - | 1.958E+04 | 989.5 | - | - | 0 | - |
| - | - | 4389 | 990.5 | - | - | 0 | - |
| - | - | 3764 | 1002 | - | - | 0 | - |
| 9 | y | 1.153E+04 | 1004 | 0.002719 | 2.709 | +1 | 9 |
| - | - | 4309 | 1005 | - | - | 0 | - |
| - | - | 1226 | 1032 | - | - | 0 | - |
| - | - | 4846 | 1053 | - | - | 0 | - |
| - | - | 2210 | 1054 | - | - | 0 | - |
| - | - | 1424 | 1059 | - | - | 0 | - |
| - | - | 1183 | 1062 | - | - | 0 | - |
| 8 | y | 1583 | 1073 | 0.01057 | 9.857 | +1 | 10 |
| 8 | y | 4622 | 1074 | 4.168E-05 | 0.03882 | +1 | 10 |
| 8 | z | 3.179E+04 | 1075 | 0.000215 | 0.2001 | +1 | 10 |
| - | - | 8.946E+04 | 1076 | - | - | 0 | - |
| - | - | 4.369E+04 | 1077 | - | - | 0 | - |
| - | - | 1498 | 1077 | - | - | 0 | - |
| - | - | 1.005E+04 | 1078 | - | - | 0 | - |
| - | - | 1172 | 1085 | - | - | 0 | - |
| - | - | 1001 | 1087 | - | - | 0 | - |
| - | - | 3612 | 1088 | - | - | 0 | - |
| - | - | 1977 | 1089 | - | - | 0 | - |
| - | - | 1.449E+04 | 1090 | - | - | 0 | - |
| 8 | y | 4.354E+04 | 1091 | 0.0002623 | 0.2405 | +1 | 10 |
| - | - | 1.881E+04 | 1092 | - | - | 0 | - |
| - | - | 4390 | 1093 | - | - | 0 | - |
| - | - | 2904 | 1104 | - | - | 0 | - |
| 10 | c | 1.172E+04 | 1105 | 0.0008168 | 0.7395 | +1 | 10 |
| - | - | 4818 | 1106 | - | - | 0 | - |
| - | - | 1252 | 1107 | - | - | 0 | - |
| - | - | 3398 | 1117 | - | - | 0 | - |
| - | - | 5377 | 1118 | - | - | 0 | - |
| - | - | 3158 | 1119 | - | - | 0 | - |
| - | - | 1977 | 1160 | - | - | 0 | - |
| 7 | y | 3162 | 1161 | 0.00269 | 2.318 | +1 | 11 |
| 7 | z | 1.787E+04 | 1162 | 0.0002273 | 0.1957 | +1 | 11 |
| - | - | 3.823E+04 | 1163 | - | - | 0 | - |
| - | - | 1.995E+04 | 1164 | - | - | 0 | - |
| - | - | 7580 | 1165 | - | - | 0 | - |
| - | - | 2354 | 1166 | - | - | 0 | - |
| - | - | 1347 | 1174 | - | - | 0 | - |
| - | - | 1343 | 1175 | - | - | 0 | - |
| - | - | 3770 | 1177 | - | - | 0 | - |
| 7 | y | 2.386E+04 | 1178 | 0.0004241 | 0.3602 | +1 | 11 |
| - | - | 1.27E+04 | 1179 | - | - | 0 | - |
| - | - | 2645 | 1180 | - | - | 0 | - |
| - | - | 1.125E+04 | 1182 | - | - | 0 | - |
| - | - | 6653 | 1183 | - | - | 0 | - |
| - | - | 2864 | 1184 | - | - | 0 | - |
| - | - | 7776 | 1201 | - | - | 0 | - |
| - | - | 4191 | 1202 | - | - | 0 | - |
| - | - | 1796 | 1203 | - | - | 0 | - |
| - | - | 1.401E+04 | 1217 | - | - | 0 | - |
| 11 | c | 2.556E+04 | 1218 | 0.001826 | 1.5 | +1 | 11 |
| - | - | 1.523E+04 | 1219 | - | - | 0 | - |
| - | - | 2998 | 1220 | - | - | 0 | - |
| - | - | 3109 | 1235 | - | - | 0 | - |
| - | - | 2190 | 1236 | - | - | 0 | - |
| - | - | 1055 | 1237 | - | - | 0 | - |
| - | - | 1.497E+04 | 1253 | - | - | 0 | - |
| - | - | 9127 | 1254 | - | - | 0 | - |
| - | - | 2943 | 1255 | - | - | 0 | - |
| 6 | y | 1406 | 1257 | 0.0002171 | 0.1728 | +1 | 12 |
| 6 | y | 2069 | 1258 | 0.0112 | 8.903 | +1 | 12 |
| - | - | 1387 | 1274 | - | - | 0 | - |
| 6 | y | 5.255E+04 | 1275 | 0.0002726 | 0.2139 | +1 | 12 |
| - | - | 3.839E+04 | 1276 | - | - | 0 | - |
| - | - | 1.287E+04 | 1277 | - | - | 0 | - |
| - | - | 3524 | 1302 | - | - | 0 | - |
| - | - | 2501 | 1303 | - | - | 0 | - |
| - | - | 1682 | 1304 | - | - | 0 | - |
| - | - | 1671 | 1311 | - | - | 0 | - |
| - | - | 1798 | 1326 | - | - | 0 | - |
| - | - | 1415 | 1327 | - | - | 0 | - |
| 12 | c | 2060 | 1328 | 0.01676 | 12.62 | +1 | 12 |
| 12 | c | 1.345E+04 | 1329 | 0.002849 | 2.144 | +1 | 12 |
| - | - | 7150 | 1330 | - | - | 0 | - |
| - | - | 3813 | 1331 | - | - | 0 | - |
| - | - | 2.116E+04 | 1345 | - | - | 0 | - |
| 12 | c | 7.825E+04 | 1346 | 0.0007114 | 0.5286 | +1 | 12 |
| - | - | 4.948E+04 | 1347 | - | - | 0 | - |
| - | - | 1.603E+04 | 1348 | - | - | 0 | - |
| - | - | 2150 | 1349 | - | - | 0 | - |
| 5 | y | 9617 | 1354 | 0.00323 | 2.386 | +1 | 13 |
| 5 | y | 1.187E+04 | 1355 | 0.01092 | 8.063 | +1 | 13 |
| 5 | z | 7630 | 1356 | 0.005173 | 3.816 | +1 | 13 |
| - | - | 2361 | 1357 | - | - | 0 | - |
| - | - | 3642 | 1358 | - | - | 0 | - |
| - | - | 1226 | 1359 | - | - | 0 | - |
| - | - | 2188 | 1360 | - | - | 0 | - |
| - | - | 1780 | 1361 | - | - | 0 | - |
| - | - | 2438 | 1370 | - | - | 0 | - |
| - | - | 2.668E+04 | 1371 | - | - | 0 | - |
| 5 | y | 5.131E+05 | 1372 | 0.0001231 | 0.08975 | +1 | 13 |
| - | - | 3.616E+05 | 1373 | - | - | 0 | - |
| - | - | 1.18E+05 | 1374 | - | - | 0 | - |
| - | - | 1.171E+04 | 1375 | - | - | 0 | - |
| - | - | 2534 | 1398 | - | - | 0 | - |
| - | - | 1.02E+04 | 1399 | - | - | 0 | - |
| - | - | 1.895E+04 | 1400 | - | - | 0 | - |
| - | - | 1.197E+04 | 1401 | - | - | 0 | - |
| - | - | 1.016E+05 | 1402 | - | - | 0 | - |
| - | - | 7.398E+04 | 1403 | - | - | 0 | - |
| - | - | 2.828E+04 | 1404 | - | - | 0 | - |
| - | - | 4379 | 1405 | - | - | 0 | - |
| - | - | 5.609E+04 | 1416 | - | - | 0 | - |
| 13 | c | 8.014E+04 | 1417 | 0.003645 | 2.573 | +1 | 13 |
| - | - | 4.547E+04 | 1418 | - | - | 0 | - |
| - | - | 1.074E+04 | 1419 | - | - | 0 | - |
| - | - | 1230 | 1420 | - | - | 0 | - |
| - | - | 3106 | 1431 | - | - | 0 | - |
| - | - | 3699 | 1432 | - | - | 0 | - |
| - | - | 1063 | 1433 | - | - | 0 | - |
| - | - | 1266 | 1434 | - | - | 0 | - |
| - | - | 5903 | 1487 | - | - | 0 | - |
| - | - | 4868 | 1488 | - | - | 0 | - |
| - | - | 3084 | 1489 | - | - | 0 | - |
| 4 | y | 1497 | 1501 | 0.01759 | 11.72 | +1 | 14 |
| 4 | y | 1076 | 1502 | 0.02222 | 14.8 | +1 | 14 |
| 4 | z | 4.512E+04 | 1503 | 0.0004965 | 0.3304 | +1 | 14 |
| - | - | 4.019E+04 | 1504 | - | - | 0 | - |
| - | - | 1.609E+04 | 1505 | - | - | 0 | - |
| 14 | c | 4932 | 1514 | 0.001094 | 0.7229 | +1 | 14 |
| - | - | 4523 | 1515 | - | - | 0 | - |
| - | - | 2857 | 1516 | - | - | 0 | - |
| - | - | 2753 | 1518 | - | - | 0 | - |
| 4 | y | 5.355E+04 | 1519 | 0.0001776 | 0.117 | +1 | 14 |
| - | - | 4.653E+04 | 1520 | - | - | 0 | - |
| - | - | 1.644E+04 | 1521 | - | - | 0 | - |
| - | - | 1822 | 1522 | - | - | 0 | - |
| - | - | 8451 | 1530 | - | - | 0 | - |
| 14 | c | 1.286E+05 | 1531 | 0.0007966 | 0.5204 | +1 | 14 |
| - | - | 1.131E+05 | 1532 | - | - | 0 | - |
| - | - | 4.349E+04 | 1533 | - | - | 0 | - |
| - | - | 6439 | 1534 | - | - | 0 | - |
| - | - | 3292 | 1544 | - | - | 0 | - |
| - | - | 1969 | 1545 | - | - | 0 | - |
| - | - | 1500 | 1558 | - | - | 0 | - |
| - | - | 1421 | 1571 | - | - | 0 | - |
| - | - | 3036 | 1572 | - | - | 0 | - |
| - | - | 1715 | 1573 | - | - | 0 | - |
| 3 | y | 2287 | 1614 | 0.0004348 | 0.2694 | +1 | 15 |
| - | - | 1.962E+04 | 1615 | - | - | 0 | - |
| 3 | z | 3.084E+04 | 1616 | 0.02787 | 17.25 | +1 | 15 |
| - | - | 2.122E+04 | 1617 | - | - | 0 | - |
| - | - | 7866 | 1618 | - | - | 0 | - |
| 3 | y | 2.091E+04 | 1632 | 0.0005014 | 0.3073 | +1 | 15 |
| - | - | 1.826E+04 | 1633 | - | - | 0 | - |
| - | - | 7786 | 1634 | - | - | 0 | - |
| - | - | 1386 | 1643 | - | - | 0 | - |
| - | - | 1846 | 1645 | - | - | 0 | - |
| - | - | 1688 | 1646 | - | - | 0 | - |
| - | - | 1496 | 1647 | - | - | 0 | - |
| - | - | 3948 | 1655 | - | - | 0 | - |
| - | - | 2733 | 1656 | - | - | 0 | - |
| - | - | 2007 | 1658 | - | - | 0 | - |
| 15 | c | 6.057E+04 | 1659 | 0.002132 | 1.285 | +1 | 15 |
| - | - | 6.187E+04 | 1660 | - | - | 0 | - |
| - | - | 2.724E+04 | 1661 | - | - | 0 | - |
| - | - | 7415 | 1662 | - | - | 0 | - |
| - | - | 3323 | 1663 | - | - | 0 | - |
| - | - | 9519 | 1672 | - | - | 0 | - |
| - | - | 1.275E+04 | 1673 | - | - | 0 | - |
| - | - | 5998 | 1674 | - | - | 0 | - |
| - | - | 1720 | 1675 | - | - | 0 | - |
| - | - | 1.82E+04 | 1686 | - | - | 0 | - |
| - | - | 1.729E+04 | 1687 | - | - | 0 | - |
| - | - | 1.462E+04 | 1688 | - | - | 0 | - |
| - | - | 6096 | 1689 | - | - | 0 | - |
| - | - | 2904 | 1690 | - | - | 0 | - |
| - | - | 1419 | 1698 | - | - | 0 | - |
| 2 | z | 3793 | 1699 | 0.007598 | 4.473 | +1 | 16 |
| 2 | z | 2970 | 1700 | 0.009667 | 5.687 | +1 | 16 |
| - | - | 1729 | 1701 | - | - | 0 | - |
| 2 | y | 2.117E+04 | 1715 | 0.002337 | 1.363 | +1 | 16 |
| 2 | y | 3.879E+04 | 1716 | 0.01414 | 8.238 | +1 | 16 |
| 2 | z | 1.37E+05 | 1717 | 0.002038 | 1.187 | +1 | 16 |
| - | - | 1.27E+05 | 1718 | - | - | 0 | - |
| - | - | 6.402E+04 | 1719 | - | - | 0 | - |
| - | - | 1.133E+04 | 1720 | - | - | 0 | - |
| - | - | 2025 | 1729 | - | - | 0 | - |
| 16 | c | 9.034E+04 | 1730 | 0.0007936 | 0.4588 | +1 | 16 |
| - | - | 9.679E+04 | 1731 | - | - | 0 | - |
| - | - | 6.707E+04 | 1732 | - | - | 0 | - |
| - | - | 2.61E+04 | 1733 | - | - | 0 | - |
| - | - | 1.134E+04 | 1734 | - | - | 0 | - |
| - | - | 2022 | 1735 | - | - | 0 | - |
| - | - | 5688 | 1744 | - | - | 0 | - |
| - | - | 1.028E+04 | 1745 | - | - | 0 | - |
| - | - | 8506 | 1746 | - | - | 0 | - |
| - | - | 4965 | 1747 | - | - | 0 | - |
| - | - | 1304 | 1748 | - | - | 0 | - |
| - | - | 7384 | 1760 | - | - | 0 | - |
| - | - | 1.532E+04 | 1761 | - | - | 0 | - |
| - | - | 1.879E+04 | 1762 | - | - | 0 | - |
| - | - | 1.418E+04 | 1763 | - | - | 0 | - |
| - | - | 4040 | 1764 | - | - | 0 | - |
| - | - | 1.031E+04 | 1769 | - | - | 0 | - |
| - | - | 1.268E+04 | 1770 | - | - | 0 | - |
| - | - | 7629 | 1771 | - | - | 0 | - |
| - | - | 4924 | 1772 | - | - | 0 | - |
| - | - | 3100 | 1773 | - | - | 0 | - |
| - | - | 1.162E+04 | 1774 | - | - | 0 | - |
| - | - | 1.277E+04 | 1775 | - | - | 0 | - |
| - | - | 4886 | 1776 | - | - | 0 | - |
| - | - | 1.39E+04 | 1777 | - | - | 0 | - |
| - | - | 1.563E+04 | 1778 | - | - | 0 | - |
| - | - | 6899 | 1779 | - | - | 0 | - |
| - | - | 1652 | 1780 | - | - | 0 | - |
| - | - | 2577 | 1786 | - | - | 0 | - |
| - | - | 2.449E+04 | 1787 | - | - | 0 | - |
| - | - | 1.567E+05 | 1788 | - | - | 0 | - |
| - | - | 1.516E+05 | 1789 | - | - | 0 | - |
| - | - | 7.489E+04 | 1790 | - | - | 0 | - |
| - | - | 1.332E+04 | 1791 | - | - | 0 | - |
| - | - | 2095 | 1792 | - | - | 0 | - |
| - | - | 1803 | 1797 | - | - | 0 | - |
| - | - | 5167 | 1798 | - | - | 0 | - |
| - | - | 5018 | 1799 | - | - | 0 | - |
| - | - | 1484 | 1802 | - | - | 0 | - |
| - | - | 1213 | 1803 | - | - | 0 | - |
| - | - | 1.402E+04 | 1805 | - | - | 0 | - |
| - | - | 1.349E+04 | 1806 | - | - | 0 | - |
| - | - | 8109 | 1807 | - | - | 0 | - |
| - | - | 1457 | 1813 | - | - | 0 | - |
| - | - | 3502 | 1814 | - | - | 0 | - |
| - | - | 3.696E+04 | 1815 | - | - | 0 | - |
| - | - | 2.303E+05 | 1816 | - | - | 0 | - |
| - | - | 2.24E+05 | 1817 | - | - | 0 | - |
| - | - | 1.171E+05 | 1818 | - | - | 0 | - |
| - | - | 1.875E+04 | 1819 | - | - | 0 | - |
| - | - | 1420 | 1820 | - | - | 0 | - |
| - | - | 4624 | 1830 | - | - | 0 | - |
| - | - | 1.566E+04 | 1831 | - | - | 0 | - |
| - | - | 4.355E+05 | 1832 | - | - | 0 | - |
| - | - | 1.142E+06 | 1833 | - | - | 0 | - |
| - | - | 9.657E+05 | 1834 | - | - | 0 | - |
| - | - | 4.07E+05 | 1835 | - | - | 0 | - |
| - | - | 5.35E+04 | 1836 | - | - | 0 | - |
| - | - | 2023 | 1866 | - | - | 0 | - |

m/z Charge Intensity FragmentType MassShift Position
120.08102416992188 0 2824.1958
120.13362884521484 0 466.28857
125.88520050048828 0 580.45734
129.10240173339844 0 5050.359
129.82728576660156 0 542.41254
148.8828887939453 0 1127.4854
148.89064025878906 0 751.44446
148.905029296875 0 842.12067
148.91200256347656 0 1058.4781
148.9197998046875 0 1418.2476
148.9264678955078 0 1741.9448
148.93357849121094 0 3120.6853
148.94137573242188 0 5426.7075
148.95790100097656 0 7282.9067
148.9657440185547 0 4788.657
148.97299194335938 0 2203.3447
148.98019409179688 0 1755.555
148.98736572265625 0 1901.2825
148.9945068359375 0 1221.3206
149.00173950195312 0 1025.5151
149.0086212158203 0 686.97375
149.0161895751953 0 1018.807
149.02294921875 0 603.7326
149.05264282226562 0 773.2812
173.12876892089844 0 3116.7744
183.1129608154297 0 5571.359
185.32168579101562 0 818.8198
193.15292358398438 0 651.7317
201.12350463867188 0 14812.793
215.13916015625 0 2879.5063
217.13375854492188 0 2189.2698
219.1492462158203 0 1083.5284
226.1190185546875 0 1239.0282
233.1650848388672 0 9341.76
243.14535522460938 0 2739.2952
245.12872314453125 0 2542.6606
247.14474487304688 0 1902.6436
251.17562866210938 0 4449.979
252.1782684326172 0 961.86707
253.2028045654297 0 863.2349
261.16009521484375 0 6935.378
262.1637878417969 0 1320.0171
269.185302734375 0 954.10516
273.07135009765625 0 919.7587
280.165771484375 0 1940.4592
282.14508056640625 0 2426.916
296.1972351074219 0 152024.62
297.1571350097656 0 1816.5426
297.2004699707031 0 24586.02
298.1764831542969 0 2540.805
298.2035827636719 0 2126.6467
303.1701354980469 0 2575.0325
303.67816162109375 0 941.4764
314.18487548828125 0 2994.3657
314.2078857421875 0 79507.29
315.1895446777344 0 1311.497
315.21099853515625 0 13393.435
317.1865234375 0 1460.8081
319.1980285644531 0 9216.291 y 14
330.1814880371094 0 3527.0198
344.1973571777344 0 1634.2585
346.17657470703125 0 6724
347.18035888671875 0 1847.7015
348.1921691894531 0 5726.0586
349.19427490234375 0 2071.3384
360.2284851074219 0 2170.8174
362.2075500488281 0 22890.477
363.2098693847656 0 3720.9377
385.2196350097656 0 1391.212
390.20196533203125 0 1388.0365
393.251708984375 0 1287.8495
398.2431640625 0 2123.6667
411.2608947753906 0 7001.875
412.2650146484375 0 1157.2756
415.271240234375 0 2934.8503
416.2550048828125 0 8978.513
417.256591796875 0 2223.4739
425.11993408203125 0 894.4486
433.2428894042969 0 12606.858 y 13
433.2816467285156 0 117809.195
434.24859619140625 0 1803.6328
434.28472900390625 0 31917.535
435.28814697265625 0 2930.4023
442.2415466308594 0 970.1318
443.2296142578125 0 2357.6987
443.2660827636719 0 14369.436
444.269775390625 0 3252.1978
445.244140625 0 1045.0656
461.2766418457031 0 127114.02
462.27972412109375 0 28915.213
463.28289794921875 0 3013.8284
487.25482177734375 0 1073.6461 y Ammonia loss 12
498.2198181152344 0 1573.5953
504.2781677246094 0 20593.223 y 12
505.28094482421875 0 3863.4607
530.3341674804688 0 5855.18
531.3380126953125 0 2467.1858
540.3189697265625 0 4695.641
541.3184814453125 0 992.1016
558.3294677734375 0 17873.438
559.3319091796875 0 5223.2227
560.33154296875 0 1265.7242
582.7817993164062 0 1524.2084
591.2907104492188 0 1971.7505
591.7942504882812 0 3334.7493
614.3273315429688 0 1355.715 y Water loss 11
615.3113403320312 0 2947.784 y Ammonia loss 11
616.3180541992188 0 3894.1128 z 11
617.322998046875 0 2363.9438
617.807373046875 0 3953.1184
618.3095092773438 0 2976.463
618.8021240234375 0 1322.7424
626.8125610351562 0 21582.662
627.26171875 0 2195.6035
627.3141479492188 0 12326.07
627.8141479492188 0 2096.4648
632.3367919921875 0 12462.841 y 11
633.3394775390625 0 4471.5347
635.8165893554688 0 1083.7593
637.8154296875 0 3254.3713 y 5
638.317138671875 0 2640.0146
655.3348999023438 0 1456.66
655.3876953125 0 1815.5117
656.3273315429688 0 876.17053
663.3366088867188 0 3320
664.8342895507812 0 1377.2267 c Ammonia loss 11
677.33642578125 0 13329.244 y Water loss 4
677.8348388671875 0 11959.391 y Ammonia loss 4
678.3336791992188 0 9258.57 z 4
678.83447265625 0 2778.9595
685.33447265625 0 1446.2798
685.8331909179688 0 1714.8784
686.3419189453125 0 447346 y 4
686.84326171875 0 323330.84
687.344482421875 0 107541.69
687.8460083007812 0 10119.557
700.3482666015625 0 5884.6855 c Ammonia loss 12
700.8491821289062 0 3788.6062
701.3485717773438 0 1176.0186
724.4043579101562 0 1131.9808
729.4024047851562 0 7128.511 z 10
730.4071044921875 0 6115.345
731.4111328125 0 2401.769
740.3457641601562 0 3666.71
742.4132690429688 0 4095.3354
743.4215698242188 0 1797.5161
745.420654296875 0 7136.182 y 10
746.4237060546875 0 2687.1504
750.8699340820312 0 3309.4307 y Water loss 3
751.3692016601562 0 5294.211 y Ammonia loss 3
751.8681030273438 0 2732.0989 z 3
752.3577270507812 0 1083.7555
757.3895263671875 0 1569.0568 c Ammonia loss 13
759.8760375976562 0 71227.69 y 3
760.3772583007812 0 55993.832
760.8790283203125 0 24233.17
761.3807983398438 0 1608.4207
799.3986206054688 0 822.34033 z Water loss 2
807.9132080078125 0 2565.887 y Ammonia loss 2
812.9169311523438 0 2357.0989
813.411376953125 0 1926.3951
816.4185180664062 0 15598.975 y 2
816.9198608398438 0 12687.332
817.4225463867188 0 6412.929
829.4454956054688 0 2686.156
830.44921875 0 1518.9426
842.944580078125 0 1448.1332
847.9427490234375 0 1333.8284
848.4404907226562 0 4496.1753
848.938232421875 0 2628.498
856.947265625 0 13400.695 c Ammonia loss 15
857.4482421875 0 13210.876 y Ammonia loss 9
857.950439453125 0 6549.623 y Water loss 1
858.4446411132812 0 20741.158 y Ammonia loss 1
859.450439453125 0 17917.643
860.4539184570312 0 5360.1226
866.9420776367188 0 5653.697 y 1
867.4436645507812 0 3893.7668
867.9415893554688 0 1444.502
868.4061889648438 0 3858.2793
869.409912109375 0 2512.8025
874.4639892578125 0 8884.034 y 9
875.466064453125 0 2794.0266
885.9661254882812 0 1051.0239
886.473388671875 0 1266.4681
898.9617919921875 0 1932.9434
907.4706420898438 0 9803.241
907.9696655273438 0 14507.36
908.4712524414062 0 9414.541
908.9708251953125 0 2937.076
916.476318359375 0 126247.734
916.9771728515625 0 121617.45
917.083984375 0 1728.993
917.4789428710938 0 69280.695
917.9798583984375 0 9050.557
939.4411010742188 0 3688.8765
940.4511108398438 0 1657.0293
958.48974609375 0 2992.5413
959.4893188476562 0 2186.6592
973.497314453125 0 4077.0083
974.4984130859375 0 2143.784
975.5120849609375 0 1692.7515 c 8
987.4879150390625 0 34529.402 z 8
988.494140625 0 44392.37
989.4990234375 0 19577.283
990.4990844726562 0 4389.1187
1002.49755859375 0 3763.893
1003.5027465820312 0 11526.496 y 8
1004.5054931640625 0 4308.5664
1031.50390625 0 1225.6006
1053.4853515625 0 4846.045
1054.4921875 0 2210.0667
1058.5072021484375 0 1423.6538
1061.5396728515625 0 1182.8221
1072.516357421875 0 1582.6918 y Water loss 7
1073.510986328125 0 4621.8174 y Ammonia loss 7
1074.5185546875 0 31785.553 z 7
1075.5272216796875 0 89463.38
1076.5301513671875 0 43690.863
1076.6619873046875 0 1498.4283
1077.53271484375 0 10048.968
1084.5277099609375 0 1171.6709
1086.5074462890625 0 1000.6635
1087.531494140625 0 3612.0085
1088.5465087890625 0 1977.1328
1089.5316162109375 0 14494.946
1090.5372314453125 0 43543.152 y 7
1091.5404052734375 0 18810.322
1092.54296875 0 4389.9624
1103.5501708984375 0 2904.371
1104.5579833984375 0 11719.268 c 9
1105.56005859375 0 4818.078
1106.56787109375 0 1252.3168
1116.529296875 0 3398.398
1117.535400390625 0 5377.3467
1118.5396728515625 0 3157.7683
1159.5291748046875 0 1977.3054
1160.540283203125 0 3161.554 y Ammonia loss 6
1161.551025390625 0 17871.559 z 6
1162.5576171875 0 38230.93
1163.56298828125 0 19950.5
1164.5601806640625 0 7579.521
1165.558349609375 0 2354.3086
1173.6199951171875 0 1347.1736
1174.63916015625 0 1343.3717
1176.556884765625 0 3770.3123
1177.5699462890625 0 23862.098 y 6
1178.5721435546875 0 12704.943
1179.573486328125 0 2644.5662
1181.581298828125 0 11252.078
1182.585205078125 0 6652.6206
1183.5975341796875 0 2863.5212
1200.61474609375 0 7776.126
1201.6207275390625 0 4190.9224
1202.624755859375 0 1796.1407
1216.63330078125 0 14008.538
1217.639404296875 0 25558.91 c 10
1218.642822265625 0 15225.29
1219.6424560546875 0 2997.879
1234.610107421875 0 3108.6968
1235.60302734375 0 2189.6301
1236.5792236328125 0 1055.4335
1252.6163330078125 0 14970.19
1253.6229248046875 0 9127.081
1254.615966796875 0 2942.6765
1256.6119384765625 0 1405.798 y Water loss 5
1257.60693359375 0 2069.175 y Ammonia loss 5
1273.613037109375 0 1387.2454
1274.62255859375 0 52546.883 y 5
1275.6259765625 0 38389.598
1276.6279296875 0 12874.609
1301.6715087890625 0 3523.5896
1302.69482421875 0 2501.1653
1303.6990966796875 0 1682.2299
1310.6717529296875 0 1671.0018
1325.6785888671875 0 1797.8789
1326.67236328125 0 1415.4788
1327.6724853515625 0 2059.6624 c Water loss 11
1328.67041015625 0 13451.278 c Ammonia loss 11
1329.6722412109375 0 7149.635
1330.6832275390625 0 3813.3674
1344.69140625 0 21158.697
1345.6990966796875 0 78249.65 c 11
1346.703125 0 49483.7
1347.7049560546875 0 16030.146
1348.713134765625 0 2150.2493
1353.6612548828125 0 9617.003 y Water loss 4
1354.659423828125 0 11873.238 y Ammonia loss 4
1355.6614990234375 0 7630.332 z 4
1356.6558837890625 0 2360.8816
1357.6834716796875 0 3642.443
1358.68701171875 0 1226.0819
1359.672607421875 0 2187.5505
1360.6630859375 0 1779.5479
1369.65673828125 0 2437.8035
1370.66650390625 0 26684.098
1371.6749267578125 0 513080.22 y 4
1372.678466796875 0 361641.66
1373.6812744140625 0 118002.69
1374.6895751953125 0 11709.388
1397.7293701171875 0 2534.374
1398.671630859375 0 10197.337
1399.6807861328125 0 18949.623
1400.6878662109375 0 11969.009
1401.7010498046875 0 101606.77
1402.7041015625 0 73983.91
1403.7073974609375 0 28280.082
1404.7027587890625 0 4378.7207
1415.7288818359375 0 56085.367
1416.7332763671875 0 80139.9 c 12
1417.7376708984375 0 45467.21
1418.7420654296875 0 10735.827
1419.751708984375 0 1230.083
1430.6988525390625 0 3105.868
1431.7069091796875 0 3699.0652
1432.694580078125 0 1063.2341
1433.6907958984375 0 1265.9858
1486.76416015625 0 5903.4756
1487.7667236328125 0 4868.113
1488.772216796875 0 3084.262
1500.75048828125 0 1497.0674 y Water loss 3
1501.7391357421875 0 1075.9984 y Ammonia loss 3
1502.7242431640625 0 45117.17 z 3
1503.7281494140625 0 40186.945
1504.732177734375 0 16087.765
1513.75439453125 0 4931.522 c Ammonia loss 13
1514.7550048828125 0 4523.2104
1515.7674560546875 0 2856.7527
1517.7327880859375 0 2753.1064
1518.7432861328125 0 53548.64 y 3
1519.745361328125 0 46531.35
1520.7481689453125 0 16437.354
1521.7490234375 0 1821.5035
1529.770751953125 0 8451.211
1530.779052734375 0 128615.79 c 13
1531.781982421875 0 113101.22
1532.78564453125 0 43492.863
1533.791015625 0 6439.4873
1543.783935546875 0 3292.2905
1544.787109375 0 1969.0074
1557.789794921875 0 1499.9158
1570.8187255859375 0 1421.0421
1571.8182373046875 0 3036.0618
1572.7921142578125 0 1715.2147
1613.8165283203125 0 2286.9946 y Water loss 2
1614.857421875 0 19621.72
1615.836669921875 0 30843.432 z 2
1616.829345703125 0 21218.617
1617.8209228515625 0 7865.7075
1631.8270263671875 0 20914.934 y 2
1632.82763671875 0 18258.299
1633.8345947265625 0 7786.417
1642.8621826171875 0 1386.3269
1644.84423828125 0 1846.0281
1645.8380126953125 0 1687.9381
1646.834716796875 0 1496.1249
1654.8531494140625 0 3947.7527
1655.858154296875 0 2732.9763
1657.849609375 0 2007.0406
1658.8726806640625 0 60566.93 c 14
1659.8763427734375 0 61865.586
1660.8681640625 0 27237.004
1661.815673828125 0 7415.0713
1662.7930908203125 0 3323.0422
1671.8436279296875 0 9519.385
1672.8431396484375 0 12748.447
1673.8443603515625 0 5997.7573
1674.8382568359375 0 1720.0752
1685.8968505859375 0 18196.475
1686.90234375 0 17286.818
1687.9068603515625 0 14615.792
1688.9080810546875 0 6095.78
1689.9078369140625 0 2904.3257
1697.849609375 0 1419.0265
1698.853515625 0 3793.3455 z Water loss 1
1699.839599609375 0 2969.7454 z Ammonia loss 1
1700.85205078125 0 1729.319
1714.8623046875 0 21168.13 y Water loss 1
1715.86279296875 0 38787.38 y Ammonia loss 1
1716.8585205078125 0 136969.72 z 1
1717.85986328125 0 127005.47
1718.8648681640625 0 64019.445
1719.8690185546875 0 11333.577
1728.9056396484375 0 2024.502
1729.9127197265625 0 90344.22 c 15
1730.9156494140625 0 96792.836
1731.9183349609375 0 67066.555
1732.923828125 0 26097.848
1733.924072265625 0 11338.733
1734.932373046875 0 2021.5789
1743.90625 0 5687.526
1744.8939208984375 0 10276.234
1745.8919677734375 0 8506.176
1746.9041748046875 0 4965.3105
1747.8988037109375 0 1304.072
1759.8887939453125 0 7383.626
1760.9141845703125 0 15320.523
1761.91748046875 0 18793.088
1762.9178466796875 0 14176.8545
1763.9110107421875 0 4040.1277
1768.93408203125 0 10310.414
1769.9339599609375 0 12681.885
1770.9395751953125 0 7628.7495
1771.94140625 0 4923.8394
1772.92333984375 0 3100.4636
1773.9146728515625 0 11624.366
1774.91748046875 0 12773.02
1775.9163818359375 0 4886.3584
1776.8941650390625 0 13895.866
1777.89208984375 0 15628.303
1778.896484375 0 6898.879
1779.8880615234375 0 1651.8073
1785.917236328125 0 2577.2512
1786.94677734375 0 24486.578
1787.93310546875 0 156664.69
1788.93505859375 0 151627.62
1789.9359130859375 0 74887.66
1790.9307861328125 0 13318.206
1791.893798828125 0 2094.8184
1796.9412841796875 0 1802.5502
1797.912353515625 0 5167.384
1798.924560546875 0 5017.9434
1801.9068603515625 0 1484.0692
1802.939453125 0 1212.7926
1804.95751953125 0 14022.818
1805.9581298828125 0 13493.773
1806.9595947265625 0 8109.056
1812.900634765625 0 1456.8407
1813.9173583984375 0 3502.0964
1814.937744140625 0 36960.86
1815.9278564453125 0 230267.69
1816.929443359375 0 224044.3
1817.9306640625 0 117149.59
1818.930908203125 0 18748.678
1819.918701171875 0 1420.2955
1829.927978515625 0 4623.6953
1830.9354248046875 0 15661.489
1831.9444580078125 0 435492.3
1832.951416015625 0 1142439.5
1833.954345703125 0 965719.25
1834.9578857421875 0 407006.16
1835.9581298828125 0 53503.03
1865.9400634765625 0 2022.7128

Spectrum Details

|  |  |
| --- | --- |
| Matched peaks? Matched peaks The total absolute number of peaks matched. Additionally in brackets the total fraction of peaks matched and the total number of peaks is shown. | 70 (15.80% of 443) |
| FDR? FDR The false discovery rate estimated for this peptide. It is calculated by matching all theoretical fragments with a non-integer shift with the raw peaks for this spectrum. This is done with 40 different shifts. The resulting percentage is the average number of annotated peaks over the number of annotated peaks with the correct spectrum. | 0.51% |
| Satellite FDR? Satellite FDR See the FDR for details on its calculation. This satellite ion specific FDR only contains the satellite ions (d/w) for I/L/J positions. | - |
| PSM Score? PSM Score The PSM Score as given by Hecklib to this annotated spectrum. It is shown with three significant figures. | 681 |

#### Spectrum 8106? Spectrum 8106 The raw spectrum of this peptide as annotated by Hecklib. The fragments are coloured according to ion type (see legend). Any peaks with a star '\*' as text can be hovered over to see the full details, first the ion type second the mass shift type. By hovering over the amino acids in the peptide or ions in the legend the corresponding peaks are highlighted. By toggling the 'Unassigned' label you can turn the background (unassigned) peaks on or off in the plot. By updating the slider in the Ion legend you can update the spectrum to only show the top X% of the peaks with labels. The top X% means any peak that is within X% of the highest intensity. By dragging in the spectrum you can zoom in to a specific part of the spectrum and use 'Zoom Out' to get back to the original zoom level. The annotation of the spectrum is based on the given sequence in the peptides file and is done with different software so inconsistencies are likely. The peaks are annotated based on the given sequence, with 20 ppm tolerance.

Copy Data

##### Spectrum 8106 (TSV)

###### Preview

```
Loading example...
```

*Click on the button to copy the data to your clipboard.*

Mz MinMz MaxIntensity Max

WidthHeightPeptide font sizePeptide stroke widthSpectrum font sizeSpectrum stroke widthCompact peptide

Ion legend

wxyz

abcd

OtherUnassignedIon^Charge^_Position_Show for top:%

VTJFPPSSEEJQANKAT

03.36e+56.73e+51.01e+61.35e+6

Zoom Out

y^+1^_1_a^+1^_2_y^+1^_2_a^+1^_2_b^+1^_2_y^+1^_2_b^+1^_2_y^+2^_5_a^+1^_3_a^+1^_3_b^+1^_3_y^+1^_3_y^+1^_3_y^+2^_6_b^+1^_3_y^+1^_3_b^+2^_8_y^+1^_4_b^+1^_4_b^+1^_4_y^+1^_5_y^+1^_5_y^+1^_5_b^+2^_10_b^+1^_5_b^+1^_5_b^+2^_11_y^+1^_6_y^+1^_6_y^+2^_12_y^+1^_6_y^+2^_12_b^+2^_12_b^+2^_12_y^+2^_13_y^+2^_13_y^+2^_13_b^+2^_13_b^+2^_13_y^+1^_7_b^+1^_7_y^+1^_7_y^+2^_14_y^+2^_14_b^+2^_14_y^+2^_14_y^+2^_15_b^+2^_15_y^+2^_15_b^+1^_8_b^+2^_16_b^+2^_16_y^+1^_8_b^+2^_16_y^+1^_8_y^+2^_16_y^+2^_16_y^+1^_8_\*\*\*b^+1^_9_y^+1^_9_y^+1^_9_b^+1^_10_y^+1^_10_y^+1^_10_b^+1^_10_y^+1^_10_y^+1^_11_y^+1^_11_y^+1^_11_b^+1^_11_b^+1^_11_y^+1^_12_y^+1^_12_y^+1^_12_b^+1^_12_b^+1^_12_b^+1^_12_y^+1^_13_y^+1^_13_y^+1^_13_b^+1^_13_b^+1^_13_b^+1^_13_b^+1^_14_y^+1^_14_y^+1^_14_b^+1^_14_y^+1^_14_y^+1^_15_

0861172125823442

Fragment Matches Table

Show background peaks

| Position | Ion type | Intensity | mz Theoretical | mz Error (Th) | mz Error (ppm) | Charge | Series Number |
| --- | --- | --- | --- | --- | --- | --- | --- |
| 17 | y | 8626 | 120.1 | 0.0003297 | 2.746 | +1 | 1 |
| - | - | 1.181E+05 | 120.1 | - | - | 0 | - |
| - | - | 8964 | 121.1 | - | - | 0 | - |
| - | - | 1051 | 121.2 | - | - | 0 | - |
| - | - | 2.45E+04 | 126.1 | - | - | 0 | - |
| - | - | 2336 | 127.1 | - | - | 0 | - |
| - | - | 3.172E+04 | 128.1 | - | - | 0 | - |
| - | - | 6639 | 129.1 | - | - | 0 | - |
| - | - | 2.114E+05 | 129.1 | - | - | 0 | - |
| - | - | 5069 | 130.1 | - | - | 0 | - |
| - | - | 1.127E+04 | 130.1 | - | - | 0 | - |
| - | - | 8415 | 131.1 | - | - | 0 | - |
| - | - | 2472 | 132.1 | - | - | 0 | - |
| - | - | 1231 | 133.1 | - | - | 0 | - |
| - | - | 5764 | 138.1 | - | - | 0 | - |
| - | - | 1.814E+04 | 138.1 | - | - | 0 | - |
| - | - | 1305 | 139.1 | - | - | 0 | - |
| - | - | 955.1 | 139.1 | - | - | 0 | - |
| - | - | 9755 | 139.1 | - | - | 0 | - |
| - | - | 1391 | 139.1 | - | - | 0 | - |
| - | - | 1121 | 139.1 | - | - | 0 | - |
| - | - | 1868 | 139.1 | - | - | 0 | - |
| - | - | 2225 | 141.1 | - | - | 0 | - |
| - | - | 2175 | 141.1 | - | - | 0 | - |
| - | - | 1514 | 143.1 | - | - | 0 | - |
| - | - | 1234 | 144.1 | - | - | 0 | - |
| - | - | 6400 | 147.1 | - | - | 0 | - |
| - | - | 3667 | 153.1 | - | - | 0 | - |
| - | - | 7826 | 154.1 | - | - | 0 | - |
| - | - | 3104 | 154.1 | - | - | 0 | - |
| - | - | 6652 | 155.1 | - | - | 0 | - |
| 2 | a | 3.283E+04 | 155.1 | 0.000305 | 1.967 | +1 | 2 |
| - | - | 2034 | 156.1 | - | - | 0 | - |
| - | - | 7187 | 157.1 | - | - | 0 | - |
| - | - | 2031 | 157.1 | - | - | 0 | - |
| - | - | 2.787E+04 | 157.1 | - | - | 0 | - |
| - | - | 3400 | 165.1 | - | - | 0 | - |
| - | - | 4188 | 166.1 | - | - | 0 | - |
| - | - | 1.832E+04 | 167.1 | - | - | 0 | - |
| - | - | 1.362E+04 | 167.1 | - | - | 0 | - |
| - | - | 2108 | 168.1 | - | - | 0 | - |
| - | - | 2269 | 168.1 | - | - | 0 | - |
| - | - | 3251 | 169.1 | - | - | 0 | - |
| - | - | 1367 | 169.1 | - | - | 0 | - |
| - | - | 3.189E+04 | 169.1 | - | - | 0 | - |
| - | - | 1693 | 170.1 | - | - | 0 | - |
| - | - | 2779 | 171.1 | - | - | 0 | - |
| - | - | 1584 | 171.1 | - | - | 0 | - |
| - | - | 4295 | 172.1 | - | - | 0 | - |
| 16 | y | 9248 | 173.1 | 0.0003385 | 1.956 | +1 | 2 |
| 2 | a | 1.038E+05 | 173.1 | 0.0003147 | 1.818 | +1 | 2 |
| - | - | 6444 | 174.1 | - | - | 0 | - |
| - | - | 1.074E+04 | 175.1 | - | - | 0 | - |
| - | - | 2334 | 176.1 | - | - | 0 | - |
| - | - | 3233 | 180.1 | - | - | 0 | - |
| - | - | 9416 | 181.1 | - | - | 0 | - |
| - | - | 6175 | 182.1 | - | - | 0 | - |
| - | - | 1.119E+04 | 182.1 | - | - | 0 | - |
| - | - | 1.061E+04 | 183.1 | - | - | 0 | - |
| 2 | b | 3.875E+05 | 183.1 | 0.000355 | 1.939 | +1 | 2 |
| - | - | 1.02E+04 | 184.1 | - | - | 0 | - |
| - | - | 3.245E+04 | 184.1 | - | - | 0 | - |
| - | - | 3.66E+04 | 185.1 | - | - | 0 | - |
| - | - | 3320 | 185.1 | - | - | 0 | - |
| - | - | 7065 | 186.1 | - | - | 0 | - |
| - | - | 2394 | 186.1 | - | - | 0 | - |
| - | - | 5.643E+04 | 187.1 | - | - | 0 | - |
| - | - | 3785 | 188.1 | - | - | 0 | - |
| - | - | 7728 | 189.1 | - | - | 0 | - |
| - | - | 2812 | 191.1 | - | - | 0 | - |
| 16 | y | 1.017E+04 | 191.1 | 0.0003176 | 1.662 | +1 | 2 |
| - | - | 2339 | 195.1 | - | - | 0 | - |
| - | - | 3.148E+04 | 195.1 | - | - | 0 | - |
| - | - | 3026 | 196.1 | - | - | 0 | - |
| - | - | 1225 | 196.1 | - | - | 0 | - |
| - | - | 1111 | 197.1 | - | - | 0 | - |
| - | - | 2924 | 197.1 | - | - | 0 | - |
| - | - | 4.094E+04 | 197.1 | - | - | 0 | - |
| - | - | 6671 | 198.1 | - | - | 0 | - |
| - | - | 4156 | 198.1 | - | - | 0 | - |
| - | - | 1.491E+04 | 199.1 | - | - | 0 | - |
| - | - | 6288 | 199.1 | - | - | 0 | - |
| - | - | 2992 | 199.1 | - | - | 0 | - |
| - | - | 1.106E+04 | 200.1 | - | - | 0 | - |
| - | - | 1.436E+04 | 200.1 | - | - | 0 | - |
| 2 | b | 1.932E+05 | 201.1 | 0.0002731 | 1.358 | +1 | 2 |
| - | - | 1.606E+04 | 202.1 | - | - | 0 | - |
| - | - | 1797 | 203.1 | - | - | 0 | - |
| - | - | 1233 | 207.1 | - | - | 0 | - |
| - | - | 1880 | 208.1 | - | - | 0 | - |
| - | - | 1.965E+04 | 208.1 | - | - | 0 | - |
| - | - | 5.051E+04 | 209.1 | - | - | 0 | - |
| - | - | 3097 | 210.1 | - | - | 0 | - |
| - | - | 2442 | 210.1 | - | - | 0 | - |
| - | - | 2707 | 211.1 | - | - | 0 | - |
| - | - | 5905 | 212.1 | - | - | 0 | - |
| - | - | 4270 | 213.1 | - | - | 0 | - |
| - | - | 3259 | 213.1 | - | - | 0 | - |
| - | - | 1800 | 214.2 | - | - | 0 | - |
| - | - | 7.431E+04 | 215.1 | - | - | 0 | - |
| - | - | 7950 | 216.1 | - | - | 0 | - |
| - | - | 3.669E+04 | 217.1 | - | - | 0 | - |
| - | - | 4.41E+04 | 217.1 | - | - | 0 | - |
| - | - | 2494 | 218.1 | - | - | 0 | - |
| - | - | 1372 | 218.1 | - | - | 0 | - |
| - | - | 4548 | 218.1 | - | - | 0 | - |
| - | - | 3079 | 218.1 | - | - | 0 | - |
| - | - | 1.577E+04 | 219.1 | - | - | 0 | - |
| - | - | 1620 | 220.2 | - | - | 0 | - |
| - | - | 2243 | 224.2 | - | - | 0 | - |
| - | - | 1.702E+04 | 225.1 | - | - | 0 | - |
| - | - | 9485 | 225.1 | - | - | 0 | - |
| - | - | 3752 | 225.2 | - | - | 0 | - |
| - | - | 1.717E+05 | 226.1 | - | - | 0 | - |
| - | - | 4667 | 227.1 | - | - | 0 | - |
| - | - | 1.434E+04 | 227.1 | - | - | 0 | - |
| - | - | 2119 | 228.1 | - | - | 0 | - |
| - | - | 1355 | 230.6 | - | - | 0 | - |
| - | - | 4467 | 231.1 | - | - | 0 | - |
| - | - | 9.625E+04 | 233.2 | - | - | 0 | - |
| - | - | 1.52E+04 | 234.2 | - | - | 0 | - |
| - | - | 1.05E+04 | 236.1 | - | - | 0 | - |
| - | - | 1544 | 236.1 | - | - | 0 | - |
| - | - | 2212 | 236.1 | - | - | 0 | - |
| - | - | 1669 | 237.1 | - | - | 0 | - |
| - | - | 1.016E+04 | 241.1 | - | - | 0 | - |
| - | - | 1398 | 241.2 | - | - | 0 | - |
| - | - | 3153 | 241.2 | - | - | 0 | - |
| - | - | 1337 | 242.1 | - | - | 0 | - |
| - | - | 3.269E+04 | 242.2 | - | - | 0 | - |
| - | - | 5112 | 243.1 | - | - | 0 | - |
| - | - | 5.69E+04 | 243.1 | - | - | 0 | - |
| 13 | y | 7882 | 244.1 | 0.0009445 | 3.869 | +2 | 5 |
| - | - | 5652 | 244.1 | - | - | 0 | - |
| - | - | 2.818E+04 | 245.1 | - | - | 0 | - |
| - | - | 3556 | 246.1 | - | - | 0 | - |
| - | - | 1.738E+04 | 247.1 | - | - | 0 | - |
| - | - | 3083 | 248.1 | - | - | 0 | - |
| - | - | 5.177E+04 | 251.2 | - | - | 0 | - |
| - | - | 3347 | 252.1 | - | - | 0 | - |
| - | - | 5807 | 252.1 | - | - | 0 | - |
| - | - | 1716 | 252.2 | - | - | 0 | - |
| - | - | 4962 | 252.2 | - | - | 0 | - |
| - | - | 2674 | 253.1 | - | - | 0 | - |
| - | - | 4.331E+04 | 254.1 | - | - | 0 | - |
| - | - | 1.18E+04 | 254.2 | - | - | 0 | - |
| - | - | 2773 | 255.1 | - | - | 0 | - |
| - | - | 1737 | 255.2 | - | - | 0 | - |
| - | - | 2047 | 257.2 | - | - | 0 | - |
| - | - | 2418 | 258.1 | - | - | 0 | - |
| - | - | 2.962E+04 | 259.1 | - | - | 0 | - |
| - | - | 2595 | 260.1 | - | - | 0 | - |
| - | - | 3.466E+04 | 261.2 | - | - | 0 | - |
| - | - | 5873 | 262.1 | - | - | 0 | - |
| - | - | 4168 | 262.2 | - | - | 0 | - |
| - | - | 3008 | 263.1 | - | - | 0 | - |
| - | - | 8031 | 264.1 | - | - | 0 | - |
| - | - | 4604 | 268.1 | - | - | 0 | - |
| - | - | 3213 | 268.2 | - | - | 0 | - |
| 3 | a | 1.032E+04 | 268.2 | 0.000378 | 1.409 | +1 | 3 |
| - | - | 1626 | 269.1 | - | - | 0 | - |
| - | - | 1863 | 269.2 | - | - | 0 | - |
| - | - | 1.421E+04 | 269.2 | - | - | 0 | - |
| - | - | 1677 | 270.1 | - | - | 0 | - |
| - | - | 6164 | 270.2 | - | - | 0 | - |
| - | - | 1463 | 271.2 | - | - | 0 | - |
| - | - | 2.142E+04 | 272.1 | - | - | 0 | - |
| - | - | 3181 | 273.1 | - | - | 0 | - |
| - | - | 2.208E+04 | 275.2 | - | - | 0 | - |
| - | - | 3410 | 278.1 | - | - | 0 | - |
| - | - | 1.959E+04 | 279.1 | - | - | 0 | - |
| - | - | 8387 | 280.1 | - | - | 0 | - |
| - | - | 2086 | 280.1 | - | - | 0 | - |
| - | - | 3.311E+04 | 280.2 | - | - | 0 | - |
| - | - | 3239 | 281.2 | - | - | 0 | - |
| - | - | 2488 | 282.1 | - | - | 0 | - |
| - | - | 6.463E+04 | 282.1 | - | - | 0 | - |
| - | - | 7340 | 283.1 | - | - | 0 | - |
| - | - | 2.562E+04 | 285.2 | - | - | 0 | - |
| - | - | 4107 | 285.2 | - | - | 0 | - |
| - | - | 2.021E+04 | 286.1 | - | - | 0 | - |
| - | - | 3763 | 286.2 | - | - | 0 | - |
| 3 | a | 3279 | 286.2 | 6.725E-05 | 0.235 | +1 | 3 |
| - | - | 2327 | 287.1 | - | - | 0 | - |
| - | - | 2019 | 289.2 | - | - | 0 | - |
| - | - | 1606 | 292.1 | - | - | 0 | - |
| - | - | 1779 | 294.1 | - | - | 0 | - |
| - | - | 6469 | 294.2 | - | - | 0 | - |
| - | - | 1481 | 295.2 | - | - | 0 | - |
| 3 | b | 9.893E+05 | 296.2 | 0.0005806 | 1.96 | +1 | 3 |
| - | - | 8488 | 297.1 | - | - | 0 | - |
| - | - | 5.502E+04 | 297.2 | - | - | 0 | - |
| - | - | 1.389E+05 | 297.2 | - | - | 0 | - |
| - | - | 2001 | 298.1 | - | - | 0 | - |
| - | - | 6502 | 298.2 | - | - | 0 | - |
| - | - | 3.402E+04 | 298.2 | - | - | 0 | - |
| - | - | 7704 | 298.2 | - | - | 0 | - |
| - | - | 2533 | 299.1 | - | - | 0 | - |
| - | - | 7492 | 299.2 | - | - | 0 | - |
| - | - | 2527 | 300.1 | - | - | 0 | - |
| - | - | 3041 | 300.2 | - | - | 0 | - |
| - | - | 2829 | 300.2 | - | - | 0 | - |
| - | - | 2487 | 301.2 | - | - | 0 | - |
| 15 | y | 6110 | 301.2 | 0.0004377 | 1.453 | +1 | 3 |
| 15 | y | 3004 | 302.2 | 0.0002173 | 0.7192 | +1 | 3 |
| - | - | 3.975E+04 | 303.2 | - | - | 0 | - |
| - | - | 1.143E+04 | 304.1 | - | - | 0 | - |
| - | - | 6198 | 304.2 | - | - | 0 | - |
| - | - | 4209 | 306.1 | - | - | 0 | - |
| - | - | 1913 | 307.1 | - | - | 0 | - |
| 12 | y | 4582 | 308.2 | 0.002845 | 9.231 | +2 | 6 |
| - | - | 4968 | 310.1 | - | - | 0 | - |
| - | - | 1623 | 311.2 | - | - | 0 | - |
| - | - | 1587 | 312.2 | - | - | 0 | - |
| - | - | 5768 | 313.2 | - | - | 0 | - |
| - | - | 2209 | 314.1 | - | - | 0 | - |
| - | - | 7286 | 314.1 | - | - | 0 | - |
| - | - | 3.916E+04 | 314.2 | - | - | 0 | - |
| 3 | b | 2.457E+05 | 314.2 | 0.0006055 | 1.927 | +1 | 3 |
| - | - | 1989 | 315.2 | - | - | 0 | - |
| - | - | 5743 | 315.2 | - | - | 0 | - |
| - | - | 4.03E+04 | 315.2 | - | - | 0 | - |
| - | - | 2357 | 316.2 | - | - | 0 | - |
| - | - | 8610 | 317.2 | - | - | 0 | - |
| - | - | 1834 | 318.1 | - | - | 0 | - |
| 15 | y | 5.921E+04 | 319.2 | 0.0004627 | 1.449 | +1 | 3 |
| - | - | 9680 | 320.2 | - | - | 0 | - |
| - | - | 1840 | 321.2 | - | - | 0 | - |
| - | - | 2112 | 322.2 | - | - | 0 | - |
| - | - | 4939 | 323.2 | - | - | 0 | - |
| - | - | 1907 | 324.2 | - | - | 0 | - |
| - | - | 2637 | 325.2 | - | - | 0 | - |
| - | - | 3421 | 326.2 | - | - | 0 | - |
| - | - | 1.808E+04 | 328.1 | - | - | 0 | - |
| - | - | 2080 | 328.2 | - | - | 0 | - |
| - | - | 1985 | 329.1 | - | - | 0 | - |
| - | - | 2183 | 329.2 | - | - | 0 | - |
| - | - | 2.545E+04 | 330.2 | - | - | 0 | - |
| - | - | 3040 | 331.2 | - | - | 0 | - |
| - | - | 7575 | 333.2 | - | - | 0 | - |
| - | - | 8452 | 334.2 | - | - | 0 | - |
| - | - | 2139 | 335.2 | - | - | 0 | - |
| - | - | 4307 | 336.2 | - | - | 0 | - |
| - | - | 8318 | 337.2 | - | - | 0 | - |
| - | - | 3715 | 338.1 | - | - | 0 | - |
| - | - | 5844 | 339.2 | - | - | 0 | - |
| - | - | 1756 | 339.2 | - | - | 0 | - |
| - | - | 2455 | 340.2 | - | - | 0 | - |
| - | - | 1.86E+04 | 341.2 | - | - | 0 | - |
| - | - | 6638 | 342.2 | - | - | 0 | - |
| - | - | 9519 | 344.2 | - | - | 0 | - |
| - | - | 4.701E+04 | 346.1 | - | - | 0 | - |
| - | - | 3.709E+04 | 346.2 | - | - | 0 | - |
| - | - | 7445 | 347.1 | - | - | 0 | - |
| - | - | 7004 | 347.2 | - | - | 0 | - |
| - | - | 2.885E+04 | 348.2 | - | - | 0 | - |
| - | - | 1680 | 349.2 | - | - | 0 | - |
| - | - | 5392 | 349.2 | - | - | 0 | - |
| - | - | 6862 | 350.2 | - | - | 0 | - |
| - | - | 2.845E+04 | 351.2 | - | - | 0 | - |
| - | - | 1450 | 351.2 | - | - | 0 | - |
| - | - | 5331 | 352.2 | - | - | 0 | - |
| - | - | 1.55E+04 | 353.2 | - | - | 0 | - |
| - | - | 1.196E+04 | 354.2 | - | - | 0 | - |
| - | - | 2495 | 354.2 | - | - | 0 | - |
| - | - | 9435 | 355.2 | - | - | 0 | - |
| - | - | 2804 | 356.1 | - | - | 0 | - |
| - | - | 1908 | 356.2 | - | - | 0 | - |
| - | - | 6615 | 360.2 | - | - | 0 | - |
| - | - | 8.354E+04 | 362.2 | - | - | 0 | - |
| - | - | 2077 | 363.2 | - | - | 0 | - |
| - | - | 1.251E+04 | 363.2 | - | - | 0 | - |
| - | - | 1.658E+04 | 365.1 | - | - | 0 | - |
| - | - | 6047 | 365.3 | - | - | 0 | - |
| - | - | 3185 | 366.1 | - | - | 0 | - |
| - | - | 1.031E+04 | 367.2 | - | - | 0 | - |
| - | - | 1713 | 368.2 | - | - | 0 | - |
| - | - | 7968 | 368.2 | - | - | 0 | - |
| - | - | 2126 | 369.1 | - | - | 0 | - |
| - | - | 4.43E+04 | 369.2 | - | - | 0 | - |
| - | - | 2181 | 370.1 | - | - | 0 | - |
| - | - | 7227 | 370.2 | - | - | 0 | - |
| - | - | 5714 | 370.7 | - | - | 0 | - |
| - | - | 1772 | 371.2 | - | - | 0 | - |
| - | - | 1.024E+04 | 371.2 | - | - | 0 | - |
| - | - | 1.028E+04 | 372.2 | - | - | 0 | - |
| - | - | 1.491E+04 | 373.2 | - | - | 0 | - |
| - | - | 2493 | 374.2 | - | - | 0 | - |
| - | - | 2138 | 375.2 | - | - | 0 | - |
| - | - | 1965 | 377.2 | - | - | 0 | - |
| - | - | 2194 | 379.2 | - | - | 0 | - |
| - | - | 1736 | 380.2 | - | - | 0 | - |
| - | - | 5171 | 381.2 | - | - | 0 | - |
| - | - | 2.727E+04 | 383.2 | - | - | 0 | - |
| - | - | 7055 | 383.3 | - | - | 0 | - |
| - | - | 4887 | 384.2 | - | - | 0 | - |
| - | - | 2.553E+04 | 385.2 | - | - | 0 | - |
| - | - | 2787 | 386.2 | - | - | 0 | - |
| - | - | 2514 | 386.2 | - | - | 0 | - |
| - | - | 2276 | 387.2 | - | - | 0 | - |
| - | - | 1798 | 388.2 | - | - | 0 | - |
| - | - | 2048 | 390.2 | - | - | 0 | - |
| - | - | 4640 | 390.2 | - | - | 0 | - |
| - | - | 5122 | 393.2 | - | - | 0 | - |
| - | - | 1.27E+04 | 393.3 | - | - | 0 | - |
| - | - | 3914 | 394.3 | - | - | 0 | - |
| - | - | 2202 | 395.2 | - | - | 0 | - |
| - | - | 6861 | 397.1 | - | - | 0 | - |
| - | - | 1895 | 398.1 | - | - | 0 | - |
| - | - | 2706 | 398.2 | - | - | 0 | - |
| - | - | 1.905E+04 | 398.2 | - | - | 0 | - |
| - | - | 5004 | 399.2 | - | - | 0 | - |
| - | - | 4527 | 399.2 | - | - | 0 | - |
| - | - | 4178 | 400.2 | - | - | 0 | - |
| - | - | 5.214E+04 | 401.2 | - | - | 0 | - |
| - | - | 2739 | 401.2 | - | - | 0 | - |
| - | - | 9303 | 402.2 | - | - | 0 | - |
| - | - | 3418 | 405.2 | - | - | 0 | - |
| - | - | 1840 | 406.2 | - | - | 0 | - |
| - | - | 2171 | 407.2 | - | - | 0 | - |
| - | - | 9933 | 407.2 | - | - | 0 | - |
| - | - | 1.804E+04 | 408.2 | - | - | 0 | - |
| - | - | 2208 | 409.2 | - | - | 0 | - |
| - | - | 4356 | 410.2 | - | - | 0 | - |
| - | - | 1763 | 411.2 | - | - | 0 | - |
| - | - | 2.811E+04 | 411.3 | - | - | 0 | - |
| - | - | 4798 | 412.3 | - | - | 0 | - |
| - | - | 2550 | 413.2 | - | - | 0 | - |
| - | - | 1.526E+04 | 415.1 | - | - | 0 | - |
| 8 | b | 5623 | 415.2 | 0.00442 | 10.64 | +2 | 8 |
| - | - | 1.136E+04 | 415.3 | - | - | 0 | - |
| - | - | 3065 | 416.1 | - | - | 0 | - |
| - | - | 2432 | 416.2 | - | - | 0 | - |
| - | - | 3.847E+04 | 416.3 | - | - | 0 | - |
| - | - | 1.084E+04 | 417.3 | - | - | 0 | - |
| - | - | 2636 | 418.3 | - | - | 0 | - |
| - | - | 2153 | 419.2 | - | - | 0 | - |
| - | - | 2576 | 422.2 | - | - | 0 | - |
| - | - | 2979 | 423.2 | - | - | 0 | - |
| - | - | 7521 | 424.2 | - | - | 0 | - |
| - | - | 4835 | 425.1 | - | - | 0 | - |
| - | - | 1.674E+04 | 425.2 | - | - | 0 | - |
| - | - | 2977 | 426.2 | - | - | 0 | - |
| - | - | 9128 | 427.2 | - | - | 0 | - |
| - | - | 2138 | 428.2 | - | - | 0 | - |
| - | - | 1903 | 429.2 | - | - | 0 | - |
| - | - | 2443 | 429.2 | - | - | 0 | - |
| - | - | 2492 | 431.3 | - | - | 0 | - |
| - | - | 2.345E+04 | 433.2 | - | - | 0 | - |
| 14 | y | 4.91E+04 | 433.2 | 0.001908 | 4.403 | +1 | 4 |
| - | - | 3.102E+05 | 433.3 | - | - | 0 | - |
| - | - | 3468 | 434.2 | - | - | 0 | - |
| - | - | 7540 | 434.2 | - | - | 0 | - |
| - | - | 6843 | 434.2 | - | - | 0 | - |
| - | - | 7.942E+04 | 434.3 | - | - | 0 | - |
| - | - | 5921 | 434.7 | - | - | 0 | - |
| - | - | 6153 | 435.2 | - | - | 0 | - |
| - | - | 9966 | 435.3 | - | - | 0 | - |
| - | - | 4768 | 440.2 | - | - | 0 | - |
| - | - | 8551 | 441.2 | - | - | 0 | - |
| - | - | 2917 | 441.3 | - | - | 0 | - |
| - | - | 1.383E+04 | 442.2 | - | - | 0 | - |
| - | - | 6034 | 443.1 | - | - | 0 | - |
| - | - | 3307 | 443.2 | - | - | 0 | - |
| - | - | 1.583E+04 | 443.2 | - | - | 0 | - |
| 4 | b | 3.545E+04 | 443.3 | 0.0009533 | 2.151 | +1 | 4 |
| - | - | 3176 | 444.2 | - | - | 0 | - |
| - | - | 2512 | 444.2 | - | - | 0 | - |
| - | - | 7687 | 444.3 | - | - | 0 | - |
| - | - | 4796 | 445.2 | - | - | 0 | - |
| - | - | 2000 | 446.3 | - | - | 0 | - |
| - | - | 2411 | 448.2 | - | - | 0 | - |
| - | - | 3357 | 451.2 | - | - | 0 | - |
| - | - | 9078 | 452.2 | - | - | 0 | - |
| - | - | 2580 | 453.2 | - | - | 0 | - |
| - | - | 2045 | 459.2 | - | - | 0 | - |
| - | - | 1.272E+04 | 459.2 | - | - | 0 | - |
| - | - | 4978 | 459.3 | - | - | 0 | - |
| - | - | 3708 | 460.2 | - | - | 0 | - |
| - | - | 1922 | 460.3 | - | - | 0 | - |
| 4 | b | 2.756E+05 | 461.3 | 0.0008561 | 1.856 | +1 | 4 |
| - | - | 1.641E+04 | 462.2 | - | - | 0 | - |
| - | - | 6.843E+04 | 462.3 | - | - | 0 | - |
| - | - | 2977 | 463.2 | - | - | 0 | - |
| - | - | 7709 | 463.3 | - | - | 0 | - |
| - | - | 1993 | 464.2 | - | - | 0 | - |
| - | - | 5556 | 466.2 | - | - | 0 | - |
| - | - | 3700 | 468.2 | - | - | 0 | - |
| - | - | 1750 | 468.3 | - | - | 0 | - |
| - | - | 6784 | 469.2 | - | - | 0 | - |
| - | - | 3.746E+04 | 470.2 | - | - | 0 | - |
| - | - | 1.035E+04 | 471.2 | - | - | 0 | - |
| - | - | 4088 | 476.2 | - | - | 0 | - |
| - | - | 3775 | 477.3 | - | - | 0 | - |
| - | - | 8133 | 478.2 | - | - | 0 | - |
| - | - | 4642 | 479.2 | - | - | 0 | - |
| - | - | 2.941E+04 | 480.2 | - | - | 0 | - |
| - | - | 8298 | 481.2 | - | - | 0 | - |
| - | - | 5667 | 482.2 | - | - | 0 | - |
| - | - | 1971 | 483.2 | - | - | 0 | - |
| - | - | 1.264E+04 | 484.2 | - | - | 0 | - |
| 13 | y | 5955 | 486.3 | 0.0004746 | 0.9761 | +1 | 5 |
| - | - | 1950 | 486.3 | - | - | 0 | - |
| 13 | y | 7954 | 487.3 | 0.00178 | 3.653 | +1 | 5 |
| - | - | 2929 | 492.2 | - | - | 0 | - |
| - | - | 1.267E+04 | 494.2 | - | - | 0 | - |
| - | - | 2462 | 495.2 | - | - | 0 | - |
| - | - | 6204 | 495.3 | - | - | 0 | - |
| - | - | 1.819E+04 | 496.3 | - | - | 0 | - |
| - | - | 2897 | 497.3 | - | - | 0 | - |
| - | - | 1.005E+05 | 498.2 | - | - | 0 | - |
| - | - | 2.383E+04 | 499.2 | - | - | 0 | - |
| - | - | 7696 | 500.2 | - | - | 0 | - |
| - | - | 1789 | 501.2 | - | - | 0 | - |
| - | - | 2.251E+04 | 502.2 | - | - | 0 | - |
| - | - | 4368 | 503.2 | - | - | 0 | - |
| - | - | 2059 | 503.3 | - | - | 0 | - |
| 13 | y | 6.937E+04 | 504.3 | 0.0006216 | 1.233 | +1 | 5 |
| - | - | 1.474E+04 | 505.3 | - | - | 0 | - |
| - | - | 1785 | 506.2 | - | - | 0 | - |
| - | - | 1790 | 506.3 | - | - | 0 | - |
| - | - | 5300 | 510.2 | - | - | 0 | - |
| - | - | 2.185E+04 | 512.2 | - | - | 0 | - |
| - | - | 2420 | 512.3 | - | - | 0 | - |
| - | - | 5376 | 513.2 | - | - | 0 | - |
| - | - | 6786 | 513.3 | - | - | 0 | - |
| - | - | 3168 | 515.2 | - | - | 0 | - |
| - | - | 1965 | 516.2 | - | - | 0 | - |
| - | - | 4896 | 518.2 | - | - | 0 | - |
| - | - | 1787 | 521.3 | - | - | 0 | - |
| - | - | 3655 | 524.2 | - | - | 0 | - |
| - | - | 2053 | 527.2 | - | - | 0 | - |
| - | - | 6356 | 528.2 | - | - | 0 | - |
| - | - | 7.272E+04 | 530.2 | - | - | 0 | - |
| - | - | 2.163E+04 | 530.3 | - | - | 0 | - |
| - | - | 1.836E+04 | 531.2 | - | - | 0 | - |
| - | - | 5550 | 531.3 | - | - | 0 | - |
| - | - | 2727 | 532.2 | - | - | 0 | - |
| 10 | b | 1918 | 535.3 | 0.008904 | 16.63 | +2 | 10 |
| - | - | 1867 | 536.2 | - | - | 0 | - |
| - | - | 1685 | 538.3 | - | - | 0 | - |
| - | - | 7403 | 538.3 | - | - | 0 | - |
| - | - | 1669 | 539.3 | - | - | 0 | - |
| 5 | b | 1.456E+04 | 540.3 | 0.0009238 | 1.71 | +1 | 5 |
| - | - | 2461 | 541.3 | - | - | 0 | - |
| - | - | 8307 | 546.2 | - | - | 0 | - |
| - | - | 2467 | 547.2 | - | - | 0 | - |
| - | - | 2652 | 553.3 | - | - | 0 | - |
| - | - | 5869 | 555.3 | - | - | 0 | - |
| - | - | 4098 | 556.2 | - | - | 0 | - |
| - | - | 2182 | 556.3 | - | - | 0 | - |
| 5 | b | 4.109E+04 | 558.3 | 0.000552 | 0.9887 | +1 | 5 |
| - | - | 1.058E+04 | 559.3 | - | - | 0 | - |
| - | - | 3873 | 563.2 | - | - | 0 | - |
| - | - | 2104 | 564.2 | - | - | 0 | - |
| - | - | 9419 | 569.3 | - | - | 0 | - |
| - | - | 9634 | 569.8 | - | - | 0 | - |
| - | - | 3353 | 570.3 | - | - | 0 | - |
| - | - | 4892 | 571.3 | - | - | 0 | - |
| - | - | 2918 | 573.8 | - | - | 0 | - |
| - | - | 3139 | 574.3 | - | - | 0 | - |
| - | - | 1.66E+04 | 578.3 | - | - | 0 | - |
| - | - | 7591 | 578.8 | - | - | 0 | - |
| - | - | 3814 | 579.3 | - | - | 0 | - |
| - | - | 1.622E+04 | 581.3 | - | - | 0 | - |
| - | - | 7752 | 582.3 | - | - | 0 | - |
| - | - | 2.353E+04 | 582.8 | - | - | 0 | - |
| - | - | 1.226E+04 | 583.3 | - | - | 0 | - |
| - | - | 1738 | 583.8 | - | - | 0 | - |
| - | - | 4422 | 587.3 | - | - | 0 | - |
| - | - | 5680 | 591.2 | - | - | 0 | - |
| - | - | 7271 | 591.3 | - | - | 0 | - |
| 11 | b | 5858 | 591.8 | 0.009798 | 16.56 | +2 | 11 |
| - | - | 3000 | 592.3 | - | - | 0 | - |
| - | - | 1.077E+04 | 597.3 | - | - | 0 | - |
| - | - | 4455 | 598.3 | - | - | 0 | - |
| - | - | 7.077E+04 | 599.3 | - | - | 0 | - |
| - | - | 1.876E+04 | 600.3 | - | - | 0 | - |
| - | - | 1952 | 601.3 | - | - | 0 | - |
| - | - | 3653 | 604.3 | - | - | 0 | - |
| - | - | 3469 | 604.8 | - | - | 0 | - |
| - | - | 4337 | 607.3 | - | - | 0 | - |
| - | - | 1637 | 608.3 | - | - | 0 | - |
| - | - | 4448 | 608.4 | - | - | 0 | - |
| - | - | 3856 | 608.8 | - | - | 0 | - |
| - | - | 2.071E+04 | 609.3 | - | - | 0 | - |
| - | - | 4191 | 609.3 | - | - | 0 | - |
| - | - | 2339 | 609.3 | - | - | 0 | - |
| - | - | 6115 | 609.8 | - | - | 0 | - |
| - | - | 5291 | 610.3 | - | - | 0 | - |
| - | - | 1892 | 610.3 | - | - | 0 | - |
| - | - | 1.399E+04 | 612.8 | - | - | 0 | - |
| - | - | 1.22E+04 | 613.3 | - | - | 0 | - |
| - | - | 2904 | 613.8 | - | - | 0 | - |
| 12 | y | 9832 | 614.3 | 8.896E-05 | 0.1448 | +1 | 6 |
| 12 | y | 3.692E+04 | 615.3 | 0.005406 | 8.786 | +1 | 6 |
| - | - | 8894 | 616.3 | - | - | 0 | - |
| - | - | 2586 | 617.3 | - | - | 0 | - |
| - | - | 6.536E+04 | 617.8 | - | - | 0 | - |
| - | - | 5.337E+04 | 618.3 | - | - | 0 | - |
| - | - | 2.043E+04 | 618.8 | - | - | 0 | - |
| - | - | 5241 | 619.3 | - | - | 0 | - |
| - | - | 1802 | 624.3 | - | - | 0 | - |
| - | - | 1.084E+04 | 625.3 | - | - | 0 | - |
| - | - | 5625 | 626.3 | - | - | 0 | - |
| - | - | 9951 | 626.4 | - | - | 0 | - |
| - | - | 1.452E+05 | 626.8 | - | - | 0 | - |
| - | - | 1.6E+05 | 627.3 | - | - | 0 | - |
| - | - | 9.555E+04 | 627.3 | - | - | 0 | - |
| - | - | 3.407E+04 | 627.8 | - | - | 0 | - |
| - | - | 4.19E+04 | 628.3 | - | - | 0 | - |
| - | - | 2351 | 628.3 | - | - | 0 | - |
| 6 | y | 3161 | 628.8 | 0.002431 | 3.866 | +2 | 12 |
| - | - | 6637 | 629.3 | - | - | 0 | - |
| - | - | 2720 | 629.3 | - | - | 0 | - |
| - | - | 3241 | 629.8 | - | - | 0 | - |
| 12 | y | 4.827E+04 | 632.3 | 0.0005768 | 0.9122 | +1 | 6 |
| - | - | 1.285E+04 | 633.3 | - | - | 0 | - |
| - | - | 5841 | 635.8 | - | - | 0 | - |
| - | - | 5005 | 636.3 | - | - | 0 | - |
| - | - | 2448 | 636.8 | - | - | 0 | - |
| 6 | y | 2.046E+04 | 637.8 | 0.0002822 | 0.4425 | +2 | 12 |
| - | - | 1.268E+04 | 638.3 | - | - | 0 | - |
| - | - | 3599 | 638.8 | - | - | 0 | - |
| - | - | 4299 | 640.3 | - | - | 0 | - |
| - | - | 4698 | 642.3 | - | - | 0 | - |
| - | - | 3.263E+04 | 643.3 | - | - | 0 | - |
| - | - | 1.071E+04 | 644.3 | - | - | 0 | - |
| - | - | 6474 | 645.3 | - | - | 0 | - |
| - | - | 1885 | 646.3 | - | - | 0 | - |
| - | - | 3070 | 653.3 | - | - | 0 | - |
| - | - | 9762 | 655.3 | - | - | 0 | - |
| 12 | b | 5200 | 655.8 | 0.006873 | 10.48 | +2 | 12 |
| 12 | b | 2193 | 656.3 | 0.01273 | 19.39 | +2 | 12 |
| - | - | 2672 | 656.8 | - | - | 0 | - |
| - | - | 2553 | 657.3 | - | - | 0 | - |
| - | - | 4915 | 658.3 | - | - | 0 | - |
| - | - | 2174 | 659.3 | - | - | 0 | - |
| - | - | 1.202E+04 | 660.3 | - | - | 0 | - |
| - | - | 4005 | 661.3 | - | - | 0 | - |
| - | - | 5314 | 663.3 | - | - | 0 | - |
| - | - | 4059 | 663.8 | - | - | 0 | - |
| - | - | 4239 | 664.3 | - | - | 0 | - |
| - | - | 1906 | 667.3 | - | - | 0 | - |
| - | - | 2643 | 668.3 | - | - | 0 | - |
| - | - | 1.21E+04 | 668.8 | - | - | 0 | - |
| - | - | 7449 | 669.3 | - | - | 0 | - |
| - | - | 4442 | 674.3 | - | - | 0 | - |
| 5 | y | 4.854E+04 | 677.3 | 0.000545 | 0.8046 | +2 | 13 |
| 5 | y | 6.635E+04 | 677.8 | 0.004448 | 6.562 | +2 | 13 |
| - | - | 3.479E+04 | 678.3 | - | - | 0 | - |
| - | - | 1.212E+04 | 678.8 | - | - | 0 | - |
| - | - | 2352 | 679.3 | - | - | 0 | - |
| - | - | 2086 | 684.4 | - | - | 0 | - |
| - | - | 4874 | 685.3 | - | - | 0 | - |
| - | - | 3352 | 685.8 | - | - | 0 | - |
| 5 | y | 1.124E+06 | 686.3 | 0.0007558 | 1.101 | +2 | 13 |
| - | - | 8.527E+05 | 686.8 | - | - | 0 | - |
| - | - | 2.748E+05 | 687.3 | - | - | 0 | - |
| - | - | 2.351E+04 | 687.8 | - | - | 0 | - |
| 13 | b | 2545 | 691.8 | 0.005218 | 7.542 | +2 | 13 |
| - | - | 1.094E+04 | 694.3 | - | - | 0 | - |
| - | - | 3817 | 695.3 | - | - | 0 | - |
| - | - | 2697 | 698.3 | - | - | 0 | - |
| 13 | b | 1.357E+04 | 700.4 | 0.01214 | 17.34 | +2 | 13 |
| - | - | 1.261E+04 | 700.8 | - | - | 0 | - |
| - | - | 3348 | 701.3 | - | - | 0 | - |
| - | - | 4635 | 704.3 | - | - | 0 | - |
| - | - | 2384 | 705.3 | - | - | 0 | - |
| - | - | 2086 | 709.3 | - | - | 0 | - |
| - | - | 8.951E+04 | 712.4 | - | - | 0 | - |
| - | - | 2.921E+04 | 713.4 | - | - | 0 | - |
| - | - | 4367 | 714.4 | - | - | 0 | - |
| - | - | 2357 | 721.4 | - | - | 0 | - |
| - | - | 2180 | 721.7 | - | - | 0 | - |
| - | - | 1.208E+04 | 722.3 | - | - | 0 | - |
| - | - | 4703 | 723.3 | - | - | 0 | - |
| - | - | 2001 | 725.3 | - | - | 0 | - |
| - | - | 2577 | 726.3 | - | - | 0 | - |
| - | - | 2979 | 727.3 | - | - | 0 | - |
| - | - | 2184 | 728.3 | - | - | 0 | - |
| 11 | y | 2700 | 728.4 | 0.002205 | 3.027 | +1 | 7 |
| - | - | 1814 | 731.4 | - | - | 0 | - |
| - | - | 2391 | 735.3 | - | - | 0 | - |
| - | - | 2042 | 737.4 | - | - | 0 | - |
| - | - | 4529 | 738.3 | - | - | 0 | - |
| - | - | 5057 | 738.4 | - | - | 0 | - |
| - | - | 6849 | 739.4 | - | - | 0 | - |
| - | - | 1.055E+05 | 740.3 | - | - | 0 | - |
| - | - | 3.995E+04 | 741.3 | - | - | 0 | - |
| - | - | 6995 | 742.4 | - | - | 0 | - |
| 7 | b | 7146 | 742.4 | 0.001941 | 2.615 | +1 | 7 |
| - | - | 2124 | 743.4 | - | - | 0 | - |
| - | - | 4228 | 743.4 | - | - | 0 | - |
| - | - | 5850 | 745.3 | - | - | 0 | - |
| 11 | y | 3.776E+04 | 745.4 | 0.0006803 | 0.9127 | +1 | 7 |
| - | - | 2245 | 746.3 | - | - | 0 | - |
| - | - | 1.389E+04 | 746.4 | - | - | 0 | - |
| 4 | y | 6999 | 750.9 | 9.045E-05 | 0.1205 | +2 | 14 |
| 4 | y | 4795 | 751.4 | 0.005824 | 7.752 | +2 | 14 |
| - | - | 4826 | 751.9 | - | - | 0 | - |
| - | - | 7475 | 753.3 | - | - | 0 | - |
| - | - | 6073 | 754.3 | - | - | 0 | - |
| - | - | 4768 | 755.3 | - | - | 0 | - |
| - | - | 6563 | 755.4 | - | - | 0 | - |
| - | - | 2690 | 756.4 | - | - | 0 | - |
| - | - | 2953 | 756.9 | - | - | 0 | - |
| 14 | b | 5.045E+04 | 757.4 | 0.006143 | 8.111 | +2 | 14 |
| - | - | 1.735E+04 | 758.4 | - | - | 0 | - |
| - | - | 4129 | 759.4 | - | - | 0 | - |
| 4 | y | 1.495E+05 | 759.9 | 0.0005454 | 0.7178 | +2 | 14 |
| - | - | 1.35E+05 | 760.4 | - | - | 0 | - |
| - | - | 5.439E+04 | 760.9 | - | - | 0 | - |
| - | - | 8869 | 761.4 | - | - | 0 | - |
| - | - | 1760 | 766.4 | - | - | 0 | - |
| - | - | 2.677E+04 | 771.4 | - | - | 0 | - |
| - | - | 1.18E+04 | 772.4 | - | - | 0 | - |
| - | - | 4857 | 774.3 | - | - | 0 | - |
| - | - | 1993 | 775.3 | - | - | 0 | - |
| - | - | 2181 | 797.4 | - | - | 0 | - |
| 3 | y | 2795 | 807.9 | 0.01268 | 15.7 | +2 | 15 |
| - | - | 2769 | 809.4 | - | - | 0 | - |
| - | - | 3373 | 809.5 | - | - | 0 | - |
| - | - | 2874 | 811.4 | - | - | 0 | - |
| 15 | b | 4266 | 812.9 | 0.0001886 | 0.2321 | +2 | 15 |
| - | - | 4253 | 813.4 | - | - | 0 | - |
| - | - | 3749 | 814.4 | - | - | 0 | - |
| 3 | y | 3.647E+04 | 816.4 | 0.0008718 | 1.068 | +2 | 15 |
| - | - | 2.88E+04 | 816.9 | - | - | 0 | - |
| - | - | 1.673E+04 | 817.4 | - | - | 0 | - |
| - | - | 2597 | 821.9 | - | - | 0 | - |
| - | - | 3001 | 822.4 | - | - | 0 | - |
| - | - | 4081 | 823.4 | - | - | 0 | - |
| - | - | 7777 | 824.4 | - | - | 0 | - |
| - | - | 3443 | 825.4 | - | - | 0 | - |
| - | - | 3918 | 828.4 | - | - | 0 | - |
| 8 | b | 8079 | 829.4 | 0.004208 | 5.073 | +1 | 8 |
| - | - | 2701 | 830.4 | - | - | 0 | - |
| - | - | 3701 | 832.4 | - | - | 0 | - |
| - | - | 2130 | 839.4 | - | - | 0 | - |
| - | - | 1.554E+04 | 840.4 | - | - | 0 | - |
| - | - | 7776 | 841.4 | - | - | 0 | - |
| - | - | 2.12E+04 | 842.4 | - | - | 0 | - |
| - | - | 2268 | 843 | - | - | 0 | - |
| - | - | 8021 | 843.4 | - | - | 0 | - |
| 16 | b | 2923 | 847.9 | 0.001591 | 1.877 | +2 | 16 |
| 16 | b | 8985 | 848.4 | 0.004509 | 5.314 | +2 | 16 |
| - | - | 6480 | 848.9 | - | - | 0 | - |
| - | - | 3862 | 849.4 | - | - | 0 | - |
| - | - | 1.708E+04 | 850.4 | - | - | 0 | - |
| - | - | 1.024E+04 | 851.4 | - | - | 0 | - |
| - | - | 4545 | 852.4 | - | - | 0 | - |
| 10 | y | 3318 | 856.5 | 0.005578 | 6.513 | +1 | 8 |
| 16 | b | 2.801E+04 | 856.9 | 0.0002675 | 0.3122 | +2 | 16 |
| 10 | y | 3.907E+04 | 857.4 | 0.011 | 12.83 | +1 | 8 |
| 2 | y | 1.406E+04 | 857.9 | 0.01106 | 12.89 | +2 | 16 |
| - | - | 6036 | 858.4 | - | - | 0 | - |
| - | - | 1898 | 859.4 | - | - | 0 | - |
| - | - | 1996 | 860.4 | - | - | 0 | - |
| - | - | 2743 | 866.4 | - | - | 0 | - |
| 2 | y | 1.069E+04 | 866.9 | 0.00108 | 1.246 | +2 | 16 |
| - | - | 1.587E+04 | 867.4 | - | - | 0 | - |
| - | - | 3818 | 867.9 | - | - | 0 | - |
| - | - | 1.341E+05 | 868.4 | - | - | 0 | - |
| - | - | 5.69E+04 | 869.4 | - | - | 0 | - |
| - | - | 1.067E+04 | 870.4 | - | - | 0 | - |
| 10 | y | 2.575E+04 | 874.5 | 0.0001405 | 0.1606 | +1 | 8 |
| - | - | 1.036E+04 | 875.5 | - | - | 0 | - |
| - | - | 4525 | 884.4 | - | - | 0 | - |
| - | - | 7001 | 885.4 | - | - | 0 | - |
| - | - | 1991 | 886 | - | - | 0 | - |
| - | - | 2140 | 886.4 | - | - | 0 | - |
| - | - | 8797 | 887.4 | - | - | 0 | - |
| - | - | 2985 | 888.4 | - | - | 0 | - |
| - | - | 3342 | 893.4 | - | - | 0 | - |
| - | - | 8823 | 894.4 | - | - | 0 | - |
| - | - | 5427 | 895.4 | - | - | 0 | - |
| - | - | 2978 | 899 | - | - | 0 | - |
| - | - | 3121 | 899.5 | - | - | 0 | - |
| - | - | 3755 | 900.4 | - | - | 0 | - |
| - | - | 2048 | 904.4 | - | - | 0 | - |
| 0 | Precursor | 2.251E+04 | 907.5 | 0.0009644 | 1.063 | +2 | -1 |
| 0 | Precursor | 4.151E+04 | 908 | 0.008346 | 9.192 | +2 | -1 |
| - | - | 2.078E+04 | 908.5 | - | - | 0 | - |
| - | - | 4853 | 909 | - | - | 0 | - |
| - | - | 1996 | 909.5 | - | - | 0 | - |
| - | - | 2.261E+04 | 911.4 | - | - | 0 | - |
| - | - | 9881 | 912.4 | - | - | 0 | - |
| - | - | 2147 | 913.4 | - | - | 0 | - |
| - | - | 2362 | 915.5 | - | - | 0 | - |
| - | - | 2577 | 916 | - | - | 0 | - |
| - | - | 3430 | 916.4 | - | - | 0 | - |
| 0 | Precursor | 2.463E+05 | 916.5 | 0.0005038 | 0.5498 | +2 | -1 |
| - | - | 2.816E+05 | 917 | - | - | 0 | - |
| - | - | 1.337E+05 | 917.5 | - | - | 0 | - |
| - | - | 2.014E+04 | 918 | - | - | 0 | - |
| - | - | 1.225E+04 | 921.4 | - | - | 0 | - |
| - | - | 1.307E+04 | 922.4 | - | - | 0 | - |
| - | - | 6385 | 923.4 | - | - | 0 | - |
| - | - | 2468 | 936.4 | - | - | 0 | - |
| - | - | 2091 | 938.4 | - | - | 0 | - |
| - | - | 1.011E+05 | 939.4 | - | - | 0 | - |
| - | - | 5.118E+04 | 940.4 | - | - | 0 | - |
| - | - | 1.042E+04 | 941.4 | - | - | 0 | - |
| - | - | 4823 | 953.5 | - | - | 0 | - |
| - | - | 5237 | 954.5 | - | - | 0 | - |
| - | - | 1.562E+04 | 956.5 | - | - | 0 | - |
| - | - | 8013 | 957.4 | - | - | 0 | - |
| 9 | b | 5926 | 958.5 | 0.005541 | 5.781 | +1 | 9 |
| - | - | 3435 | 970.4 | - | - | 0 | - |
| - | - | 9609 | 971.5 | - | - | 0 | - |
| - | - | 2580 | 972.5 | - | - | 0 | - |
| - | - | 1934 | 974.4 | - | - | 0 | - |
| 9 | y | 2648 | 986.5 | 0.0008811 | 0.8932 | +1 | 9 |
| - | - | 3176 | 987.5 | - | - | 0 | - |
| - | - | 3403 | 988.5 | - | - | 0 | - |
| 9 | y | 1.705E+04 | 1004 | 0.0007046 | 0.7021 | +1 | 9 |
| - | - | 9698 | 1005 | - | - | 0 | - |
| - | - | 2659 | 1006 | - | - | 0 | - |
| - | - | 1.06E+04 | 1008 | - | - | 0 | - |
| - | - | 3200 | 1009 | - | - | 0 | - |
| - | - | 8395 | 1015 | - | - | 0 | - |
| - | - | 3126 | 1016 | - | - | 0 | - |
| - | - | 3274 | 1018 | - | - | 0 | - |
| - | - | 2685 | 1019 | - | - | 0 | - |
| - | - | 8724 | 1025 | - | - | 0 | - |
| - | - | 3603 | 1026 | - | - | 0 | - |
| - | - | 5816 | 1035 | - | - | 0 | - |
| - | - | 1.828E+04 | 1036 | - | - | 0 | - |
| - | - | 8170 | 1037 | - | - | 0 | - |
| - | - | 2399 | 1038 | - | - | 0 | - |
| - | - | 8076 | 1041 | - | - | 0 | - |
| - | - | 9084 | 1041 | - | - | 0 | - |
| - | - | 3082 | 1042 | - | - | 0 | - |
| - | - | 7.659E+04 | 1053 | - | - | 0 | - |
| - | - | 4.527E+04 | 1054 | - | - | 0 | - |
| - | - | 1.211E+04 | 1055 | - | - | 0 | - |
| - | - | 9914 | 1059 | - | - | 0 | - |
| - | - | 9891 | 1060 | - | - | 0 | - |
| - | - | 1889 | 1067 | - | - | 0 | - |
| - | - | 3012 | 1068 | - | - | 0 | - |
| - | - | 4893 | 1069 | - | - | 0 | - |
| 10 | b | 3227 | 1070 | 0.01688 | 15.78 | +1 | 10 |
| - | - | 2.629E+04 | 1071 | - | - | 0 | - |
| - | - | 1.469E+04 | 1072 | - | - | 0 | - |
| 8 | y | 6185 | 1073 | 0.006421 | 5.987 | +1 | 10 |
| 8 | y | 5048 | 1074 | 0.009319 | 8.681 | +1 | 10 |
| - | - | 1.728E+04 | 1085 | - | - | 0 | - |
| - | - | 1.039E+04 | 1086 | - | - | 0 | - |
| - | - | 1.063E+04 | 1087 | - | - | 0 | - |
| 10 | b | 1.037E+04 | 1088 | 0.001321 | 1.214 | +1 | 10 |
| - | - | 5134 | 1089 | - | - | 0 | - |
| 8 | y | 3.072E+04 | 1091 | 0.0004701 | 0.4311 | +1 | 10 |
| - | - | 1.544E+04 | 1092 | - | - | 0 | - |
| - | - | 6038 | 1093 | - | - | 0 | - |
| - | - | 4104 | 1121 | - | - | 0 | - |
| - | - | 2474 | 1122 | - | - | 0 | - |
| - | - | 2527 | 1128 | - | - | 0 | - |
| - | - | 4318 | 1129 | - | - | 0 | - |
| - | - | 1.235E+04 | 1138 | - | - | 0 | - |
| - | - | 1.461E+04 | 1139 | - | - | 0 | - |
| - | - | 7500 | 1140 | - | - | 0 | - |
| - | - | 1990 | 1141 | - | - | 0 | - |
| - | - | 2338 | 1147 | - | - | 0 | - |
| - | - | 2.607E+04 | 1156 | - | - | 0 | - |
| - | - | 2.012E+04 | 1157 | - | - | 0 | - |
| - | - | 3470 | 1158 | - | - | 0 | - |
| 7 | y | 6764 | 1160 | 0.001951 | 1.682 | +1 | 11 |
| 7 | y | 7081 | 1161 | 0.007198 | 6.202 | +1 | 11 |
| - | - | 2706 | 1162 | - | - | 0 | - |
| - | - | 5187 | 1164 | - | - | 0 | - |
| - | - | 1.797E+04 | 1165 | - | - | 0 | - |
| - | - | 1.028E+04 | 1166 | - | - | 0 | - |
| - | - | 2875 | 1167 | - | - | 0 | - |
| 7 | y | 4.019E+04 | 1178 | 6.415E-05 | 0.05448 | +1 | 11 |
| - | - | 2.203E+04 | 1179 | - | - | 0 | - |
| - | - | 7730 | 1180 | - | - | 0 | - |
| - | - | 8.936E+04 | 1182 | - | - | 0 | - |
| 11 | b | 6.131E+04 | 1183 | 0.02123 | 17.95 | +1 | 11 |
| - | - | 1.844E+04 | 1184 | - | - | 0 | - |
| - | - | 1908 | 1185 | - | - | 0 | - |
| 11 | b | 1.631E+04 | 1201 | 0.006893 | 5.742 | +1 | 11 |
| - | - | 1.186E+04 | 1202 | - | - | 0 | - |
| - | - | 3405 | 1203 | - | - | 0 | - |
| - | - | 2618 | 1208 | - | - | 0 | - |
| - | - | 2574 | 1209 | - | - | 0 | - |
| - | - | 5607 | 1218 | - | - | 0 | - |
| - | - | 3384 | 1219 | - | - | 0 | - |
| - | - | 6287 | 1225 | - | - | 0 | - |
| - | - | 4017 | 1226 | - | - | 0 | - |
| - | - | 4.132E+04 | 1235 | - | - | 0 | - |
| - | - | 3.526E+04 | 1236 | - | - | 0 | - |
| - | - | 1.489E+04 | 1237 | - | - | 0 | - |
| - | - | 4184 | 1238 | - | - | 0 | - |
| - | - | 2799 | 1240 | - | - | 0 | - |
| - | - | 8.661E+04 | 1253 | - | - | 0 | - |
| - | - | 6.199E+04 | 1254 | - | - | 0 | - |
| - | - | 1.574E+04 | 1255 | - | - | 0 | - |
| - | - | 2289 | 1256 | - | - | 0 | - |
| 6 | y | 9117 | 1257 | 0.001004 | 0.7986 | +1 | 12 |
| 6 | y | 2.044E+04 | 1258 | 0.002041 | 1.623 | +1 | 12 |
| - | - | 1.218E+04 | 1259 | - | - | 0 | - |
| - | - | 5498 | 1271 | - | - | 0 | - |
| - | - | 2806 | 1272 | - | - | 0 | - |
| - | - | 2250 | 1273 | - | - | 0 | - |
| 6 | y | 2.253E+05 | 1275 | 0.0009481 | 0.7438 | +1 | 12 |
| - | - | 1.474E+05 | 1276 | - | - | 0 | - |
| - | - | 5.086E+04 | 1277 | - | - | 0 | - |
| - | - | 5490 | 1278 | - | - | 0 | - |
| - | - | 2792 | 1293 | - | - | 0 | - |
| - | - | 2051 | 1294 | - | - | 0 | - |
| - | - | 3427 | 1301 | - | - | 0 | - |
| - | - | 1.274E+04 | 1310 | - | - | 0 | - |
| 12 | b | 1.433E+04 | 1311 | 0.006007 | 4.583 | +1 | 12 |
| 12 | b | 7320 | 1312 | 0.01747 | 13.32 | +1 | 12 |
| - | - | 4008 | 1313 | - | - | 0 | - |
| - | - | 6407 | 1326 | - | - | 0 | - |
| - | - | 6234 | 1327 | - | - | 0 | - |
| - | - | 6828 | 1328 | - | - | 0 | - |
| 12 | b | 3.332E+04 | 1329 | 0.006023 | 4.533 | +1 | 12 |
| - | - | 2.367E+04 | 1330 | - | - | 0 | - |
| - | - | 7327 | 1331 | - | - | 0 | - |
| - | - | 3857 | 1336 | - | - | 0 | - |
| - | - | 7988 | 1337 | - | - | 0 | - |
| - | - | 7289 | 1338 | - | - | 0 | - |
| - | - | 3422 | 1339 | - | - | 0 | - |
| 5 | y | 4.504E+04 | 1354 | 0.001888 | 1.394 | +1 | 13 |
| 5 | y | 7.577E+04 | 1355 | 0.004942 | 3.648 | +1 | 13 |
| - | - | 4.027E+04 | 1356 | - | - | 0 | - |
| - | - | 1.412E+04 | 1357 | - | - | 0 | - |
| - | - | 5875 | 1370 | - | - | 0 | - |
| 5 | y | 1.333E+06 | 1372 | 0.001588 | 1.158 | +1 | 13 |
| - | - | 9.439E+05 | 1373 | - | - | 0 | - |
| - | - | 3.273E+05 | 1374 | - | - | 0 | - |
| - | - | 3.305E+04 | 1375 | - | - | 0 | - |
| 13 | b | 5031 | 1382 | 0.005716 | 4.137 | +1 | 13 |
| 13 | b | 5869 | 1383 | 0.003648 | 2.638 | +1 | 13 |
| 13 | b | 2.184E+04 | 1400 | 0.006515 | 4.655 | +1 | 13 |
| - | - | 1.655E+04 | 1401 | - | - | 0 | - |
| - | - | 5776 | 1402 | - | - | 0 | - |
| 14 | b | 2599 | 1497 | 0.003851 | 2.573 | +1 | 14 |
| 4 | y | 2877 | 1501 | 0.005482 | 3.653 | +1 | 14 |
| 4 | y | 3800 | 1502 | 0.0002399 | 0.1597 | +1 | 14 |
| 14 | b | 1.106E+04 | 1514 | 0.0009723 | 0.6423 | +1 | 14 |
| - | - | 9070 | 1515 | - | - | 0 | - |
| - | - | 4307 | 1516 | - | - | 0 | - |
| 4 | y | 7.147E+04 | 1519 | 0.002253 | 1.483 | +1 | 14 |
| - | - | 6.024E+04 | 1520 | - | - | 0 | - |
| - | - | 1305 | 1520 | - | - | 0 | - |
| - | - | 2.393E+04 | 1521 | - | - | 0 | - |
| 3 | y | 1.336E+04 | 1632 | 0.005628 | 3.449 | +1 | 15 |
| - | - | 1.146E+04 | 1633 | - | - | 0 | - |
| - | - | 4863 | 1634 | - | - | 0 | - |
| - | - | 2063 | 2235 | - | - | 0 | - |
| - | - | 1998 | 3072 | - | - | 0 | - |
| - | - | 1937 | 3408 | - | - | 0 | - |

m/z Charge Intensity FragmentType MassShift Position
120.06584930419922 0 8626.144 y 16
120.08109283447266 0 118124.445
121.08441162109375 0 8963.634
121.23117065429688 0 1050.6869
126.0552749633789 0 24496.156
127.05858612060547 0 2336.182
128.10728454589844 0 31716.725
129.06619262695312 0 6638.7295
129.10255432128906 0 211360.02
130.0501708984375 0 5069.45
130.1058807373047 0 11270.714
131.1181182861328 0 8415.057
132.07693481445312 0 2472.2598
133.06121826171875 0 1230.605
138.0916748046875 0 5763.8164
138.12802124023438 0 18142.87
139.05038452148438 0 1304.9515
139.08274841308594 0 955.1086
139.08688354492188 0 9754.693
139.0927734375 0 1391.031
139.09490966796875 0 1120.9705
139.13148498535156 0 1868.1005
141.06637573242188 0 2225.004
141.10264587402344 0 2174.9111
143.1175537109375 0 1513.724
144.06578063964844 0 1233.5667
147.07667541503906 0 6399.501
153.10244750976562 0 3666.7576
154.0865936279297 0 7825.639
154.09791564941406 0 3104.1646
155.0818634033203 0 6651.9834
155.11819458007812 0 32832.457 a Water loss 1
156.1218719482422 0 2034.1859
157.06101989746094 0 7186.785
157.0908203125 0 2030.6165
157.09747314453125 0 27866.828
165.10238647460938 0 3400.4878
166.0867919921875 0 4188.2773
167.08180236816406 0 18322.11
167.11822509765625 0 13616.513
168.0849151611328 0 2107.9373
168.12123107910156 0 2268.733
169.0611114501953 0 3251.415
169.09756469726562 0 1367.1417
169.13385009765625 0 31893.9
170.1378936767578 0 1693.0033
171.07679748535156 0 2778.5447
171.11402893066406 0 1584.0377
172.10816955566406 0 4294.937
173.0924072265625 0 9248.095 y Water loss 15
173.12876892089844 0 103846.266 a 1
174.12818908691406 0 6443.8335
175.0716094970703 0 10736.217
176.10757446289062 0 2333.8296
180.11349487304688 0 3233.2334
181.0974578857422 0 9416.167
182.0927734375 0 6175.156
182.12916564941406 0 11193.107
183.07691955566406 0 10606.1875
183.1131591796875 0 387544.12 b Water loss 1
184.10826110839844 0 10196.731
184.11663818359375 0 32452.516
185.0923614501953 0 36601.523
185.12852478027344 0 3320.177
186.08758544921875 0 7064.692
186.09552001953125 0 2394.2969
187.1444549560547 0 56433.305
188.1481475830078 0 3785.1357
189.08731079101562 0 7728.4883
191.08154296875 0 2812.1848
191.1029510498047 0 10168.666 y 15
195.0770263671875 0 2339.3062
195.11309814453125 0 31481.123
196.11672973632812 0 3026.3022
196.134033203125 0 1225.2286
197.0919647216797 0 1111.4817
197.1188201904297 0 2923.8691
197.12875366210938 0 40942.305
198.1239013671875 0 6670.578
198.13241577148438 0 4155.672
199.07162475585938 0 14906.437
199.10789489746094 0 6287.8174
199.11839294433594 0 2992.3918
200.10336303710938 0 11064.118
200.1395263671875 0 14364.264
201.12364196777344 0 193235.61 b 1
202.12713623046875 0 16064.59
203.103271484375 0 1796.6782
207.11228942871094 0 1232.5061
208.09751892089844 0 1879.6228
208.1083526611328 0 19651.648
209.09239196777344 0 50512.66
210.0962677001953 0 3096.5981
210.12388610839844 0 2441.7947
211.14442443847656 0 2706.9373
212.13954162597656 0 5905.236
213.0873565673828 0 4270.3896
213.12356567382812 0 3258.847
214.1544952392578 0 1799.6261
215.13934326171875 0 74308.41
216.1427459716797 0 7949.672
217.0822296142578 0 36685.6
217.13381958007812 0 44096.305
218.08627319335938 0 2493.8384
218.11544799804688 0 1371.7942
218.13720703125 0 4547.5117
218.14952087402344 0 3079.279
219.14968872070312 0 15773.753
220.15286254882812 0 1620.4172
224.17616271972656 0 2242.7053
225.12351989746094 0 17017.68
225.13514709472656 0 9485.072
225.16012573242188 0 3752.257
226.11895751953125 0 171668.3
227.1029052734375 0 4666.767
227.12246704101562 0 14342.865
228.13482666015625 0 2119.2244
230.5900421142578 0 1355.1584
231.09779357910156 0 4466.7534
233.1652374267578 0 96254.04
234.16859436035156 0 15196.113
236.10345458984375 0 10495.324
236.1204071044922 0 1543.6854
236.13999938964844 0 2211.6536
237.08743286132812 0 1668.5786
241.082275390625 0 10162.575
241.17849731445312 0 1398.004
241.19100952148438 0 3153.3801
242.11260986328125 0 1336.7883
242.15032958984375 0 32687.648
243.1338348388672 0 5112.4683
243.14572143554688 0 56897.598
244.130126953125 0 7882.0024 y Ammonia loss 12
244.1486053466797 0 5652.2544
245.12892150878906 0 28178.38
246.13223266601562 0 3556.0564
247.14451599121094 0 17381.338
248.14842224121094 0 3083.2627
251.17576599121094 0 51773.832
252.0985107421875 0 3347.211
252.13482666015625 0 5806.948
252.16796875 0 1716.3625
252.17938232421875 0 4961.8325
253.1306915283203 0 2674.4458
254.11387634277344 0 43305.27
254.150390625 0 11796.678
255.11819458007812 0 2772.735
255.15419006347656 0 1737.2098
257.1647033691406 0 2047.386
258.1094665527344 0 2418.2324
259.0927429199219 0 29622.15
260.0964660644531 0 2594.958
261.1600341796875 0 34660.79
262.1189880371094 0 5873.3564
262.1640319824219 0 4168.4927
263.1387634277344 0 3008.2217
264.13458251953125 0 8030.9863
268.0933532714844 0 4603.7495
268.16619873046875 0 3212.6152
268.20233154296875 0 10322.043 a Water loss 2
269.075927734375 0 1626.0404
269.160400390625 0 1862.6008
269.186279296875 0 14206.202
270.1445007324219 0 1676.7155
270.18194580078125 0 6163.895
271.18170166015625 0 1463.0936
272.1242370605469 0 21423.148
273.12786865234375 0 3180.7085
275.1756591796875 0 22075.727
278.1497802734375 0 3410.0374
279.14556884765625 0 19589.848
280.1294250488281 0 8386.634
280.1496887207031 0 2085.9204
280.1659240722656 0 33106.45
281.1701354980469 0 3239.3748
282.10821533203125 0 2487.6797
282.14520263671875 0 64633.79
283.1484069824219 0 7339.6094
285.16021728515625 0 25617.691
285.1974182128906 0 4106.7334
286.1037902832031 0 20208.082
286.16302490234375 0 3762.6294
286.21258544921875 0 3278.5315 a 2
287.1075439453125 0 2327.3088
289.1512145996094 0 2018.8519
292.13006591796875 0 1605.7728
294.1476135253906 0 1778.8508
294.1816711425781 0 6469.211
295.1839904785156 0 1480.6649
296.19744873046875 0 989319.3 b Water loss 2
297.1199951171875 0 8488.037
297.1563415527344 0 55021.63
297.2005310058594 0 138876.08
298.1037902832031 0 2000.5469
298.1588439941406 0 6502.187
298.1766357421875 0 34023.49
298.2033996582031 0 7704.078
299.08795166015625 0 2533.0933
299.176513671875 0 7492.2876
300.1194763183594 0 2526.925
300.1551818847656 0 3040.784
300.170166015625 0 2828.6963
301.1553039550781 0 2486.997
301.1874694824219 0 6110.131 y Water loss 14
302.1712646484375 0 3004.4934 y Ammonia loss 14
303.1708679199219 0 39754.926
304.1147155761719 0 11429.408
304.1722412109375 0 6197.9287
306.1452941894531 0 4209.1
307.1405944824219 0 1913.482
308.16131591796875 0 4581.6865 y Ammonia loss 11
310.1038818359375 0 4968.069
311.171875 0 1623.2433
312.1911926269531 0 1586.9425
313.1884460449219 0 5768.1216
314.0992126464844 0 2209.4624
314.1466979980469 0 7285.6333
314.1835021972656 0 39163.31
314.2080383300781 0 245683.06 b 2
315.1676330566406 0 1988.931
315.1863098144531 0 5743.0654
315.211181640625 0 40301.465
316.2146911621094 0 2357.4146
317.1864013671875 0 8610.215
318.130615234375 0 1834.0238
319.19805908203125 0 59205.24 y 14
320.201171875 0 9679.948
321.1553955078125 0 1840.2998
322.1770324707031 0 2111.7397
323.1718444824219 0 4939.184
324.1542663574219 0 1906.7379
325.1512756347656 0 2637.0178
326.17041015625 0 3421.3801
328.11444091796875 0 18084.035
328.1647644042969 0 2079.569
329.1163024902344 0 1984.7103
329.1828918457031 0 2183.3264
330.1817321777344 0 25453.871
331.185302734375 0 3040.4785
333.1564025878906 0 7574.9937
334.2129211425781 0 8451.5205
335.2156677246094 0 2138.7815
336.1557922363281 0 4307.154
337.1512145996094 0 8317.984
338.1349792480469 0 3714.9102
339.1667785644531 0 5844.1055
339.20355224609375 0 1756.0839
340.1985778808594 0 2455.4307
341.1841735839844 0 18597.096
342.181396484375 0 6637.8687
344.1974792480469 0 9518.771
346.1250915527344 0 47012.484
346.17657470703125 0 37087.336
347.1298522949219 0 7445.345
347.1791687011719 0 7003.904
348.1921081542969 0 28849.496
349.1529541015625 0 1679.6124
349.19561767578125 0 5391.683
350.1829833984375 0 6861.623
351.1667175292969 0 28448.428
351.18511962890625 0 1450.0144
352.1700439453125 0 5330.975
353.182373046875 0 15499.556
354.1659240722656 0 11959.628
354.18377685546875 0 2494.5042
355.1625061035156 0 9434.701
356.14404296875 0 2803.7407
356.16552734375 0 1908.3995
360.2284240722656 0 6614.969
362.2078552246094 0 83542.45
363.1669921875 0 2076.8718
363.21051025390625 0 12509.826
365.1458740234375 0 16579.041
365.2549133300781 0 6046.785
366.14947509765625 0 3184.9963
367.20843505859375 0 10311.696
368.1659851074219 0 1713.4437
368.19305419921875 0 7967.931
369.1387939453125 0 2125.9197
369.17724609375 0 44300.11
370.12384033203125 0 2180.9185
370.17987060546875 0 7226.79
370.67718505859375 0 5714.0044
371.1544189453125 0 1772.1981
371.19256591796875 0 10236.915
372.1767272949219 0 10281.331
373.17266845703125 0 14908.181
374.17462158203125 0 2493.1543
375.1676940917969 0 2138.2617
377.21893310546875 0 1964.569
379.1959533691406 0 2193.602
380.1948547363281 0 1736.1238
381.177001953125 0 5171.4463
383.1567077636719 0 27272.91
383.2659606933594 0 7055.0137
384.1594543457031 0 4887.4116
385.219970703125 0 25525.709
386.2018737792969 0 2787.4941
386.2259826660156 0 2514.2239
387.1509094238281 0 2275.849
388.22406005859375 0 1798.279
390.1763610839844 0 2048.1873
390.2030029296875 0 4639.824
393.17742919921875 0 5122.233
393.25067138671875 0 12701.738
394.25262451171875 0 3914.3308
395.22991943359375 0 2202.4172
397.1356201171875 0 6860.6323
398.1178894042969 0 1895.4402
398.2041320800781 0 2706.0012
398.2442932128906 0 19050.209
399.1882629394531 0 5004.2925
399.2465515136719 0 4527.1235
400.2237548828125 0 4177.7188
401.1673583984375 0 52143.27
401.2169189453125 0 2739.0278
402.1701354980469 0 9303.209
405.1617126464844 0 3418.1484
406.1630859375 0 1840.0952
407.15838623046875 0 2171.2542
407.2036437988281 0 9932.809
408.18853759765625 0 18036.434
409.1936340332031 0 2208.4827
410.2031555175781 0 4355.969
411.1539611816406 0 1763.3027
411.2608947753906 0 28108.79
412.2633361816406 0 4798.1875
413.2064514160156 0 2549.647
415.14666748046875 0 15264.728
415.23077392578125 0 5623.003 b 7
415.2711486816406 0 11360.645
416.14862060546875 0 3065.2852
416.18963623046875 0 2431.7358
416.25518798828125 0 38472.96
417.2561340332031 0 10840.436
418.2572021484375 0 2635.7969
419.1770324707031 0 2153.262
422.2049865722656 0 2576.0732
423.18695068359375 0 2978.5488
424.2290344238281 0 7521.263
425.1306457519531 0 4835.248
425.2150573730469 0 16742.162
426.21636962890625 0 2977.0764
427.232666015625 0 9128.095
428.23529052734375 0 2138.1777
429.16790771484375 0 1903.4174
429.2126159667969 0 2443.4285
431.2646484375 0 2492.1436
433.15740966796875 0 23453
433.242431640625 0 49100.4 y 13
433.2817687988281 0 310156.78
434.16082763671875 0 3467.9106
434.20428466796875 0 7539.8213
434.2455139160156 0 6843.3647
434.2848815917969 0 79418.87
434.7059020996094 0 5920.7373
435.2080993652344 0 6153.233
435.2872009277344 0 9965.899
440.2152404785156 0 4768.444
441.19903564453125 0 8550.544
441.2515563964844 0 2917.415
442.2403259277344 0 13831.38
443.14056396484375 0 6034.167
443.1751403808594 0 3306.8076
443.22991943359375 0 15830.36
443.2662353515625 0 35446.523 b Water loss 3
444.1873474121094 0 3176.142
444.23602294921875 0 2512.2998
444.2690734863281 0 7686.866
445.2444763183594 0 4795.627
446.2500305175781 0 2000.4272
448.18377685546875 0 2410.6682
451.23138427734375 0 3356.6233
452.2147521972656 0 9077.618
453.21881103515625 0 2579.5881
459.1733703613281 0 2045.3134
459.2090759277344 0 12723.18
459.26104736328125 0 4978.4194
460.21343994140625 0 3708.3572
460.2642822265625 0 1921.6537
461.2767028808594 0 275617.97 b 3
462.1993103027344 0 16410.338
462.27972412109375 0 68434.94
463.2033996582031 0 2977.1582
463.2834167480469 0 7708.5044
464.2144775390625 0 1992.9375
466.19366455078125 0 5556.0356
468.2096862792969 0 3699.9497
468.2564392089844 0 1750.3538
469.2447204589844 0 6784.205
470.2254638671875 0 37459.164
471.22833251953125 0 10353.129
476.1792297363281 0 4087.7205
477.25750732421875 0 3774.9917
478.2408142089844 0 8132.657
479.2262878417969 0 4642.2046
480.20977783203125 0 29410.629
481.21270751953125 0 8298.273
482.2243347167969 0 5666.542
483.2117614746094 0 1971.079
484.20452880859375 0 12639.59
486.2675476074219 0 5955.088 y Water loss 12
486.3069152832031 0 1950.013
487.25286865234375 0 7954.264 y Ammonia loss 12
492.2086181640625 0 2929.1946
494.1888122558594 0 12671.087
495.19378662109375 0 2462.2578
495.2677307128906 0 6203.8765
496.25164794921875 0 18190.43
497.25244140625 0 2896.5264
498.2203063964844 0 100533.555
499.22314453125 0 23827.797
500.2352600097656 0 7696.3896
501.21673583984375 0 1788.5823
502.2151794433594 0 22511.166
503.21771240234375 0 4367.6934
503.2608337402344 0 2058.758
504.27825927734375 0 69367.414 y 12
505.28076171875 0 14737.171
506.2354431152344 0 1785.2914
506.2845153808594 0 1789.8573
510.2218322753906 0 5300.176
512.199462890625 0 21848.8
512.3237915039062 0 2419.7478
513.2033081054688 0 5375.984
513.2783203125 0 6786.351
515.2487182617188 0 3168.1257
516.2418212890625 0 1964.5072
518.2457885742188 0 4895.758
521.2727661132812 0 1787.3345
524.236083984375 0 3654.529
527.2464599609375 0 2052.9023
528.2301025390625 0 6355.999
530.2098388671875 0 72721.55
530.334228515625 0 21632.814
531.213134765625 0 18356.416
531.3375244140625 0 5549.819
532.216064453125 0 2726.6492
535.2547607421875 0 1917.846 b Water loss 9
536.2356567382812 0 1866.9993
538.2564086914062 0 1684.8048
538.298828125 0 7403.0234
539.3037719726562 0 1669.4674
540.3189697265625 0 14556.581 b Water loss 4
541.3235473632812 0 2461.3513
546.2407836914062 0 8307.216
547.2396240234375 0 2467.4458
553.262939453125 0 2652.0645
555.3247680664062 0 5869.4126
556.224365234375 0 4097.7925
556.2688598632812 0 2182.337
558.3291625976562 0 41094.24 b 4
559.3333129882812 0 10583.441
563.246337890625 0 3872.8496
564.24072265625 0 2103.7744
569.2797241210938 0 9419.177
569.7793579101562 0 9633.659
570.2738037109375 0 3352.5168
571.2731323242188 0 4892.003
573.776611328125 0 2917.9858
574.2700805664062 0 3139.089
578.2860717773438 0 16596.342
578.7870483398438 0 7590.6807
579.28466796875 0 3814.4407
581.2576904296875 0 16220.452
582.2891235351562 0 7751.8135
582.7826538085938 0 23532.875
583.2823486328125 0 12256.09
583.7877197265625 0 1737.8044
587.267578125 0 4422.173
591.2427368164062 0 5680.2607
591.2950439453125 0 7271.377
591.7958984375 0 5857.936 b Water loss 10
592.2997436523438 0 3000.2239
597.2916259765625 0 10767.249
598.28466796875 0 4454.9126
599.2677612304688 0 70773.62
600.2719116210938 0 18762.637
601.279052734375 0 1951.8745
604.3062744140625 0 3653.207
604.8043212890625 0 3468.5413
607.272216796875 0 4337.1377
608.2716674804688 0 1637.2823
608.3511352539062 0 4447.665
608.8012084960938 0 3855.7378
609.2518310546875 0 20706.393
609.2952270507812 0 4190.83
609.341064453125 0 2339.3228
609.7969360351562 0 6114.6025
610.2540283203125 0 5291.423
610.3499755859375 0 1892.056
612.8150024414062 0 13991.744
613.31640625 0 12198.701
613.8180541992188 0 2903.989
614.3255615234375 0 9832.129 y Water loss 11
615.3042602539062 0 36923.023 y Ammonia loss 11
616.3067626953125 0 8894.123
617.3024291992188 0 2586.4495
617.8071899414062 0 65356.863
618.3062744140625 0 53368.94
618.8047485351562 0 20427.145
619.3031616210938 0 5241.3564
624.2979736328125 0 1802.1177
625.2846069335938 0 10836.838
626.2847900390625 0 5624.9414
626.3624267578125 0 9951.194
626.8126220703125 0 145229.33
627.26220703125 0 160031.05
627.314453125 0 95546.94
627.8153076171875 0 34065.34
628.2658081054688 0 41898.184
628.3231811523438 0 2350.638
628.8070678710938 0 3160.607 y Water loss 5
629.267333984375 0 6636.915
629.3157348632812 0 2719.7468
629.8067626953125 0 3240.7893
632.3367919921875 0 48274.516 y 11
633.339599609375 0 12845.93
635.8190307617188 0 5840.6626
636.3184814453125 0 5004.5117
636.8209228515625 0 2448.419
637.8150634765625 0 20462.4 y 5
638.3162231445312 0 12683.514
638.82080078125 0 3598.552
640.2935180664062 0 4299.2964
642.3110961914062 0 4697.672
643.294189453125 0 32628.736
644.2937622070312 0 10706.72
645.2908935546875 0 6473.8247
646.2871704101562 0 1884.8015
653.277099609375 0 3070.0493
655.3411865234375 0 9762.205
655.8418579101562 0 5199.7505 b Water loss 11
656.3397216796875 0 2193.4348 b Ammonia loss 11
656.815673828125 0 2671.572
657.3218383789062 0 2553.0886
658.3045043945312 0 4915.0254
659.3135375976562 0 2174.0427
660.3204345703125 0 12023.236
661.3235473632812 0 4005.032
663.3380126953125 0 5313.748
663.8383178710938 0 4059.3445
664.3359375 0 4239.378
667.3380737304688 0 1905.6729
668.3311157226562 0 2643.415
668.8257446289062 0 12104.124
669.3225708007812 0 7449.15
674.3002319335938 0 4441.5034
677.33642578125 0 48535.62 y Water loss 4
677.8323364257812 0 66347.375 y Ammonia loss 4
678.3327026367188 0 34786.21
678.8319091796875 0 12124.107
679.2590942382812 0 2351.625
684.3719482421875 0 2085.815
685.3330078125 0 4874.258
685.8375244140625 0 3351.778
686.3419189453125 0 1124319 y 4
686.8432006835938 0 852732.1
687.3440551757812 0 274775.34
687.8448486328125 0 23507.22
691.84033203125 0 2545.0474 b Ammonia loss 12
694.341796875 0 10943.746
695.34521484375 0 3817.014
698.298583984375 0 2697.4536
700.3466796875 0 13568.349 b 12
700.8492431640625 0 12610.087
701.3482055664062 0 3347.947
704.3263549804688 0 4634.9834
705.3344116210938 0 2384.227
709.321533203125 0 2085.62
712.3517456054688 0 89509.47
713.35498046875 0 29214.309
714.3526611328125 0 4367.0757
721.3633422851562 0 2357.224
721.6515502929688 0 2179.5127
722.3372192382812 0 12078.133
723.3394165039062 0 4702.762
725.3451538085938 0 2000.6805
726.3342895507812 0 2576.6736
727.3269653320312 0 2979.0027
728.3284301757812 0 2183.6833
728.3959350585938 0 2700.002 y Ammonia loss 10
731.3630981445312 0 1814.0385
735.3328247070312 0 2391.0396
737.3926391601562 0 2042.4672
738.2987060546875 0 4529.196
738.3778076171875 0 5056.597
739.3622436523438 0 6848.911
740.346923828125 0 105487.53
741.3499145507812 0 39949.77
742.351806640625 0 6994.9414
742.4153442382812 0 7145.6646 b 6
743.3502197265625 0 2123.6306
743.4179077148438 0 4227.676
745.337890625 0 5849.5586
745.4209594726562 0 37761.32 y 10
746.3496704101562 0 2245.4436
746.4238891601562 0 13885.515
750.8701782226562 0 6999.312 y Water loss 3
751.367919921875 0 4795.0503 y Ammonia loss 3
751.8687744140625 0 4825.8994
753.3407592773438 0 7474.643
754.3383178710938 0 6072.8213
755.3237915039062 0 4767.8745
755.4052734375 0 6563.3896
756.3980712890625 0 2689.7922
756.89013671875 0 2953.1135
757.3741455078125 0 50449.043 b 13
758.3761596679688 0 17348.41
759.371337890625 0 4128.589
759.8759155273438 0 149463.44 y 3
760.3771362304688 0 135003.28
760.8784790039062 0 54390.125
761.3787231445312 0 8868.644
766.36328125 0 1759.7998
771.3521728515625 0 26769.78
772.3541259765625 0 11802.027
774.3338623046875 0 4857.446
775.335205078125 0 1993.1874
797.3700561523438 0 2181.3816
807.9168090820312 0 2794.7437 y Ammonia loss 2
809.3740844726562 0 2769.053
809.4627075195312 0 3372.9158
811.3795776367188 0 2874.3523
812.914306640625 0 4266.4688 b Ammonia loss 14
813.4175415039062 0 4253.1836
814.3965454101562 0 3748.7988
816.4182739257812 0 36472.656 y 2
816.9197387695312 0 28800.158
817.4197998046875 0 16726.809
821.93017578125 0 2596.553
822.4271850585938 0 3000.7246
823.394287109375 0 4081.3213
824.380859375 0 7776.8613
825.3695678710938 0 3443.2314
828.4038696289062 0 3918.4714
829.4412231445312 0 8079.2085 b 7
830.4497680664062 0 2700.8542
832.3834838867188 0 3700.8293
839.42431640625 0 2129.7363
840.4109497070312 0 15535.653
841.4107055664062 0 7775.6587
842.390625 0 21200.443
842.9562377929688 0 2268.023
843.3906860351562 0 8021.1523
847.939453125 0 2923.3389 b Water loss 15
848.4375610351562 0 8985.161 b Ammonia loss 15
848.9415283203125 0 6479.5356
849.4304809570312 0 3862.0454
850.3952026367188 0 17075.293
851.3858032226562 0 10242.379
852.3934936523438 0 4545.4
856.4578857421875 0 3317.567 y Water loss 9
856.9465942382812 0 28005.422 b 15
857.4473266601562 0 39069.277 y Ammonia loss 9
857.947021484375 0 14063.858 y Water loss 1
858.4453125 0 6035.6826
859.4236450195312 0 1897.6755
860.42236328125 0 1996.3501
866.4338989257812 0 2742.7378
866.9423217773438 0 10690.912 y 1
867.4390869140625 0 15865.673
867.939208984375 0 3817.5417
868.4056396484375 0 134062.4
869.408447265625 0 56903.63
870.410400390625 0 10665.203
874.4630126953125 0 25754.29 y 9
875.4636840820312 0 10355.289
884.4420166015625 0 4525.351
885.4462280273438 0 7001.2026
885.9690551757812 0 1990.5734
886.4366455078125 0 2139.695
887.413818359375 0 8796.595
888.4193725585938 0 2985.4026
893.4375610351562 0 3342.243
894.42333984375 0 8822.953
895.4296875 0 5427.4224
898.955810546875 0 2978.0588
899.4645385742188 0 3120.9556
900.4463500976562 0 3754.5237
904.4218139648438 0 2047.5786
907.4711303710938 0 22505.41 Precursor Water loss
907.9705200195312 0 41509.035 Precursor Ammonia loss
908.469970703125 0 20778.008
908.9682006835938 0 4852.5044
909.4558715820312 0 1996.1438
911.4454956054688 0 22608.076
912.4489135742188 0 9881.138
913.4483642578125 0 2147.0586
915.46728515625 0 2362.2944
915.9724731445312 0 2577.418
916.366455078125 0 3430.2131
916.4759521484375 0 246325.31 Precursor
916.9768676757812 0 281562.1
917.4782104492188 0 133667.66
917.9785766601562 0 20140.908
921.4300537109375 0 12246.98
922.4199829101562 0 13072.41
923.4210205078125 0 6384.6206
936.44677734375 0 2467.9404
938.4476928710938 0 2091.3213
939.44140625 0 101115.05
940.444580078125 0 51179.26
941.446044921875 0 10422.039
953.4644775390625 0 4823.212
954.4606323242188 0 5237.467
956.452392578125 0 15624.93
957.4472045898438 0 8013.084
958.4824829101562 0 5925.8394 b 8
970.4459838867188 0 3434.9546
971.478271484375 0 9608.877
972.4808349609375 0 2579.6516
974.44287109375 0 1934.3472
986.4797973632812 0 2647.8452 y Ammonia loss 8
987.4758911132812 0 3176.2046
988.47705078125 0 3403.4148
1003.5047607421875 0 17053.963 y 8
1004.5078735351562 0 9698.284
1005.5025634765625 0 2659.3416
1008.462646484375 0 10601.975
1009.4703979492188 0 3199.962
1015.4736328125 0 8394.882
1016.4750366210938 0 3125.501
1018.4525146484375 0 3274.037
1019.4398193359375 0 2685.1726
1025.4888916015625 0 8723.592
1026.495361328125 0 3603.3938
1035.471923828125 0 5816
1036.4608154296875 0 18278.104
1037.465576171875 0 8170.0596
1038.485595703125 0 2399.1667
1040.50048828125 0 8076.134
1041.4945068359375 0 9084.471
1042.49462890625 0 3081.6167
1053.4853515625 0 76593.94
1054.4859619140625 0 45269.582
1055.4886474609375 0 12114.125
1058.5115966796875 0 9913.576
1059.5145263671875 0 9890.925
1066.5174560546875 0 1889.2747
1067.508056640625 0 3011.581
1068.502197265625 0 4892.584
1069.503173828125 0 3227.2297 b Water loss 9
1070.5115966796875 0 26288.236
1071.5113525390625 0 14687.971
1072.5205078125 0 6185.2466 y Water loss 7
1073.520263671875 0 5048.025 y Ammonia loss 7
1084.52734375 0 17278.502
1085.5284423828125 0 10385.88
1086.51708984375 0 10629.359
1087.529296875 0 10367.901 b 9
1088.5302734375 0 5134.1284
1090.5379638671875 0 30718.344 y 7
1091.5401611328125 0 15437.539
1092.5411376953125 0 6037.898
1120.53564453125 0 4104.287
1121.531005859375 0 2473.8674
1127.5673828125 0 2527.2888
1128.55712890625 0 4318.1045
1137.5533447265625 0 12354.592
1138.544677734375 0 14606.573
1139.5447998046875 0 7500.4575
1140.546142578125 0 1989.6139
1146.5350341796875 0 2338.0938
1155.56396484375 0 26071.902
1156.5657958984375 0 20116.78
1157.5675048828125 0 3470.3164
1159.5570068359375 0 6764.216 y Water loss 6
1160.5501708984375 0 7080.9473 y Ammonia loss 6
1161.552978515625 0 2706.389
1163.571533203125 0 5187.2153
1164.5557861328125 0 17973.115
1165.55810546875 0 10278.785
1166.5587158203125 0 2874.7734
1177.5694580078125 0 40188.996 y 6
1178.5704345703125 0 22026.803
1179.57177734375 0 7729.5093
1181.5791015625 0 89359.2
1182.5828857421875 0 61306.543 b Water loss 10
1183.583251953125 0 18442.945
1184.566162109375 0 1908.2509
1200.6077880859375 0 16309.157 b 10
1201.6134033203125 0 11861.729
1202.6165771484375 0 3404.8528
1207.6064453125 0 2618.395
1208.603271484375 0 2574.4915
1217.591552734375 0 5606.536
1218.5877685546875 0 3383.9702
1224.6190185546875 0 6287.327
1225.6220703125 0 4017.1985
1234.604736328125 0 41324.266
1235.6005859375 0 35261.645
1236.5977783203125 0 14893.377
1237.592529296875 0 4183.775
1239.589111328125 0 2798.8926
1252.615478515625 0 86611.055
1253.6182861328125 0 61985.504
1254.618896484375 0 15735.148
1255.617919921875 0 2288.5632
1256.6107177734375 0 9117.122 y Water loss 5
1257.5977783203125 0 20435.758 y Ammonia loss 5
1258.5994873046875 0 12177.842
1270.628173828125 0 5498.3955
1271.62939453125 0 2806.4443
1272.603759765625 0 2250.0586
1274.621337890625 0 225298.92 y 5
1275.6236572265625 0 147396.22
1276.6263427734375 0 50856.71
1277.6246337890625 0 5490.3296
1292.6527099609375 0 2792.0637
1293.6461181640625 0 2051.048
1300.6492919921875 0 3427.1682
1309.66748046875 0 12743.6455
1310.668701171875 0 14331.545 b Water loss 11
1311.6641845703125 0 7319.684 b Ammonia loss 11
1312.6463623046875 0 4008.2944
1325.664794921875 0 6406.5493
1326.66552734375 0 6234.392
1327.65966796875 0 6827.74
1328.667236328125 0 33319.53 b 11
1329.6702880859375 0 23669.96
1330.67724609375 0 7326.559
1335.6463623046875 0 3856.747
1336.6422119140625 0 7988.3657
1337.63916015625 0 7289.0146
1338.637451171875 0 3421.9011
1353.66259765625 0 45036.074 y Water loss 4
1354.6534423828125 0 75766.875 y Ammonia loss 4
1355.6546630859375 0 40267.215
1356.6556396484375 0 14121.418
1369.6510009765625 0 5875.027
1371.6734619140625 0 1332500.4 y 4
1372.6763916015625 0 943923.6
1373.678466796875 0 327258.16
1374.6795654296875 0 33046.45
1381.694091796875 0 5031.142 b Water loss 12
1382.68017578125 0 5869.4517 b Ammonia loss 12
1399.703857421875 0 21839.44 b 12
1400.7060546875 0 16546.97
1401.707763671875 0 5775.707
1496.722900390625 0 2598.615 b Ammonia loss 13
1500.7274169921875 0 2877.4373 y Water loss 3
1501.7166748046875 0 3800.4397 y Ammonia loss 3
1513.7542724609375 0 11062.202 b 13
1514.751220703125 0 9070.497
1515.746337890625 0 4307.4634
1518.7412109375 0 71473.02 y 3
1519.7432861328125 0 60237.37
1519.959228515625 0 1305.1681
1520.746826171875 0 23926.393
1631.8218994140625 0 13362.671 y 2
1632.8271484375 0 11458.376
1633.8189697265625 0 4863.4487
2234.514404296875 0 2063.2102
3072.246337890625 0 1997.6465
3408.21044921875 0 1937.1259

Spectrum Details

|  |  |
| --- | --- |
| Matched peaks? Matched peaks The total absolute number of peaks matched. Additionally in brackets the total fraction of peaks matched and the total number of peaks is shown. | 92 (10.75% of 856) |
| FDR? FDR The false discovery rate estimated for this peptide. It is calculated by matching all theoretical fragments with a non-integer shift with the raw peaks for this spectrum. This is done with 40 different shifts. The resulting percentage is the average number of annotated peaks over the number of annotated peaks with the correct spectrum. | 0.65% |
| Satellite FDR? Satellite FDR See the FDR for details on its calculation. This satellite ion specific FDR only contains the satellite ions (d/w) for I/L/J positions. | - |
| PSM Score? PSM Score The PSM Score as given by Hecklib to this annotated spectrum. It is shown with three significant figures. | 1.07E+03 |

#### Spectrum 8067? Spectrum 8067 The raw spectrum of this peptide as annotated by Hecklib. The fragments are coloured according to ion type (see legend). Any peaks with a star '\*' as text can be hovered over to see the full details, first the ion type second the mass shift type. By hovering over the amino acids in the peptide or ions in the legend the corresponding peaks are highlighted. By toggling the 'Unassigned' label you can turn the background (unassigned) peaks on or off in the plot. By updating the slider in the Ion legend you can update the spectrum to only show the top X% of the peaks with labels. The top X% means any peak that is within X% of the highest intensity. By dragging in the spectrum you can zoom in to a specific part of the spectrum and use 'Zoom Out' to get back to the original zoom level. The annotation of the spectrum is based on the given sequence in the peptides file and is done with different software so inconsistencies are likely. The peaks are annotated based on the given sequence, with 20 ppm tolerance.

Copy Data

##### Spectrum 8067 (TSV)

###### Preview

```
Loading example...
```

*Click on the button to copy the data to your clipboard.*

Mz MinMz MaxIntensity Max

WidthHeightPeptide font sizePeptide stroke widthSpectrum font sizeSpectrum stroke widthCompact peptide

Ion legend

wxyz

abcd

OtherUnassignedIon^Charge^_Position_Show for top:%

VTJFPPSSEEJQANKAT

02.79e+45.57e+48.36e+41.11e+5

Zoom Out

y^+1^_1_a^+1^_2_y^+1^_2_a^+1^_2_b^+1^_2_y^+1^_2_b^+1^_2_b^+3^_6_y^+3^_7_y^+2^_5_a^+1^_3_b^+2^_5_b^+1^_3_y^+1^_3_y^+2^_6_b^+1^_3_y^+2^_6_y^+1^_3_b^+2^_6_y^+2^_7_b^+2^_8_b^+2^_8_y^+1^_4_y^+2^_8_y^+1^_4_b^+1^_4_y^+3^_13_y^+3^_13_b^+1^_4_b^+2^_9_y^+1^_5_y^+1^_5_y^+1^_5_y^+2^_10_y^+3^_15_b^+1^_5_b^+2^_10_y^+2^_10_b^+3^_15_b^+1^_5_y^+2^_11_y^+2^_11_b^+2^_11_b^+2^_11_\*y^+1^_6_y^+1^_6_y^+2^_12_y^+2^_12_y^+1^_6_y^+2^_12_b^+2^_12_b^+2^_12_b^+2^_12_y^+2^_13_y^+2^_13_y^+2^_13_b^+2^_13_b^+2^_13_b^+1^_7_y^+1^_7_y^+1^_7_b^+1^_7_y^+1^_7_y^+2^_14_y^+2^_14_b^+2^_14_y^+2^_14_y^+2^_15_y^+2^_15_b^+1^_8_b^+2^_15_y^+2^_15_b^+1^_8_b^+2^_16_y^+1^_8_b^+2^_16_y^+1^_8_y^+2^_16_y^+2^_16_y^+2^_16_y^+1^_8_b^+1^_9_y^+1^_9_y^+1^_9_b^+1^_10_y^+1^_10_b^+1^_10_y^+1^_10_y^+1^_11_y^+1^_11_y^+1^_11_b^+1^_11_b^+1^_11_y^+1^_12_y^+1^_12_y^+1^_12_b^+1^_12_y^+1^_13_y^+1^_13_b^+1^_13_b^+1^_14_

0857171425713428

Fragment Matches Table

Show background peaks

| Position | Ion type | Intensity | mz Theoretical | mz Error (Th) | mz Error (ppm) | Charge | Series Number |
| --- | --- | --- | --- | --- | --- | --- | --- |
| 17 | y | 4.802E+04 | 120.1 | 0.0002763 | 2.301 | +1 | 1 |
| - | - | 3.921E+04 | 120.1 | - | - | 0 | - |
| - | - | 278.4 | 121.1 | - | - | 0 | - |
| - | - | 277.8 | 121.1 | - | - | 0 | - |
| - | - | 1833 | 121.1 | - | - | 0 | - |
| - | - | 3676 | 121.1 | - | - | 0 | - |
| - | - | 4205 | 126.1 | - | - | 0 | - |
| - | - | 756.4 | 127.1 | - | - | 0 | - |
| - | - | 1.393E+04 | 128.1 | - | - | 0 | - |
| - | - | 2757 | 129.1 | - | - | 0 | - |
| - | - | 477.7 | 129.1 | - | - | 0 | - |
| - | - | 6.79E+04 | 129.1 | - | - | 0 | - |
| - | - | 3850 | 130.1 | - | - | 0 | - |
| - | - | 1208 | 130.1 | - | - | 0 | - |
| - | - | 399.3 | 130.1 | - | - | 0 | - |
| - | - | 629.3 | 130.1 | - | - | 0 | - |
| - | - | 3581 | 130.1 | - | - | 0 | - |
| - | - | 3105 | 131.1 | - | - | 0 | - |
| - | - | 747 | 132.1 | - | - | 0 | - |
| - | - | 517 | 133.1 | - | - | 0 | - |
| - | - | 375 | 134 | - | - | 0 | - |
| - | - | 1649 | 136.1 | - | - | 0 | - |
| - | - | 2852 | 138.1 | - | - | 0 | - |
| - | - | 1482 | 138.1 | - | - | 0 | - |
| - | - | 1910 | 139.1 | - | - | 0 | - |
| - | - | 1543 | 141.1 | - | - | 0 | - |
| - | - | 519.7 | 142.1 | - | - | 0 | - |
| - | - | 1670 | 147.1 | - | - | 0 | - |
| - | - | 668.2 | 148.1 | - | - | 0 | - |
| - | - | 1077 | 149 | - | - | 0 | - |
| - | - | 2672 | 149 | - | - | 0 | - |
| - | - | 485.2 | 152.1 | - | - | 0 | - |
| - | - | 1705 | 154.1 | - | - | 0 | - |
| - | - | 2753 | 154.1 | - | - | 0 | - |
| - | - | 4131 | 155.1 | - | - | 0 | - |
| 2 | a | 9715 | 155.1 | 0.0002288 | 1.475 | +1 | 2 |
| - | - | 1630 | 157.1 | - | - | 0 | - |
| - | - | 6237 | 157.1 | - | - | 0 | - |
| - | - | 5027 | 159.1 | - | - | 0 | - |
| - | - | 1543 | 165.1 | - | - | 0 | - |
| - | - | 888.9 | 166.1 | - | - | 0 | - |
| - | - | 3173 | 167.1 | - | - | 0 | - |
| - | - | 3347 | 167.1 | - | - | 0 | - |
| - | - | 2537 | 167.1 | - | - | 0 | - |
| - | - | 512.6 | 168.1 | - | - | 0 | - |
| - | - | 711.5 | 169.1 | - | - | 0 | - |
| - | - | 1.012E+04 | 169.1 | - | - | 0 | - |
| - | - | 468.6 | 170.1 | - | - | 0 | - |
| - | - | 753.4 | 170.1 | - | - | 0 | - |
| - | - | 1848 | 171.1 | - | - | 0 | - |
| - | - | 2891 | 172.1 | - | - | 0 | - |
| - | - | 476.5 | 172.1 | - | - | 0 | - |
| 16 | y | 7662 | 173.1 | 0.0001707 | 0.986 | +1 | 2 |
| 2 | a | 3.486E+04 | 173.1 | 0.0002384 | 1.377 | +1 | 2 |
| - | - | 2169 | 173.4 | - | - | 0 | - |
| - | - | 2637 | 174.1 | - | - | 0 | - |
| - | - | 1811 | 175.1 | - | - | 0 | - |
| - | - | 885.3 | 180.1 | - | - | 0 | - |
| - | - | 1629 | 181.1 | - | - | 0 | - |
| - | - | 3456 | 182.1 | - | - | 0 | - |
| - | - | 3276 | 182.1 | - | - | 0 | - |
| - | - | 3726 | 183.1 | - | - | 0 | - |
| 2 | b | 4.431E+04 | 183.1 | 0.0002024 | 1.106 | +1 | 2 |
| - | - | 547.1 | 184.1 | - | - | 0 | - |
| - | - | 2218 | 184.1 | - | - | 0 | - |
| - | - | 4256 | 184.1 | - | - | 0 | - |
| - | - | 6874 | 185.1 | - | - | 0 | - |
| - | - | 658.4 | 185.1 | - | - | 0 | - |
| - | - | 2740 | 186.1 | - | - | 0 | - |
| - | - | 1.464E+04 | 187.1 | - | - | 0 | - |
| - | - | 637.2 | 188.1 | - | - | 0 | - |
| - | - | 3537 | 189.1 | - | - | 0 | - |
| - | - | 860.2 | 191.1 | - | - | 0 | - |
| 16 | y | 1.092E+04 | 191.1 | 0.0002566 | 1.343 | +1 | 2 |
| - | - | 856.8 | 191.1 | - | - | 0 | - |
| - | - | 1144 | 192.1 | - | - | 0 | - |
| - | - | 585 | 195.1 | - | - | 0 | - |
| - | - | 6813 | 195.1 | - | - | 0 | - |
| - | - | 508.9 | 197.1 | - | - | 0 | - |
| - | - | 1.426E+04 | 197.1 | - | - | 0 | - |
| - | - | 2720 | 198.1 | - | - | 0 | - |
| - | - | 680.9 | 198.1 | - | - | 0 | - |
| - | - | 4523 | 199.1 | - | - | 0 | - |
| - | - | 693.5 | 199.1 | - | - | 0 | - |
| - | - | 1674 | 199.1 | - | - | 0 | - |
| - | - | 1067 | 199.1 | - | - | 0 | - |
| - | - | 4561 | 200.1 | - | - | 0 | - |
| - | - | 3384 | 200.1 | - | - | 0 | - |
| - | - | 979.2 | 201.1 | - | - | 0 | - |
| 2 | b | 4.332E+04 | 201.1 | 0.0001663 | 0.8269 | +1 | 2 |
| - | - | 4231 | 202.1 | - | - | 0 | - |
| - | - | 464.5 | 203.1 | - | - | 0 | - |
| - | - | 735.4 | 208.1 | - | - | 0 | - |
| - | - | 5199 | 208.1 | - | - | 0 | - |
| - | - | 1.175E+04 | 209.1 | - | - | 0 | - |
| - | - | 1503 | 210.1 | - | - | 0 | - |
| - | - | 564.6 | 210.1 | - | - | 0 | - |
| - | - | 728.3 | 211.1 | - | - | 0 | - |
| - | - | 1710 | 212.1 | - | - | 0 | - |
| - | - | 1031 | 213.1 | - | - | 0 | - |
| - | - | 708.6 | 213.1 | - | - | 0 | - |
| - | - | 648.7 | 214.1 | - | - | 0 | - |
| - | - | 650 | 214.2 | - | - | 0 | - |
| - | - | 1.647E+04 | 215.1 | - | - | 0 | - |
| - | - | 1812 | 216.1 | - | - | 0 | - |
| - | - | 8437 | 217.1 | - | - | 0 | - |
| - | - | 4937 | 217.1 | - | - | 0 | - |
| - | - | 1.154E+04 | 217.1 | - | - | 0 | - |
| - | - | 832.3 | 218.1 | - | - | 0 | - |
| - | - | 1544 | 218.1 | - | - | 0 | - |
| - | - | 1040 | 218.1 | - | - | 0 | - |
| 6 | b | 802.7 | 219.1 | 0.001645 | 7.509 | +3 | 6 |
| - | - | 1240 | 219.1 | - | - | 0 | - |
| - | - | 4525 | 221.1 | - | - | 0 | - |
| - | - | 718.4 | 222.1 | - | - | 0 | - |
| - | - | 754.5 | 223.1 | - | - | 0 | - |
| - | - | 791.8 | 223.1 | - | - | 0 | - |
| - | - | 570.8 | 224.1 | - | - | 0 | - |
| - | - | 1852 | 225 | - | - | 0 | - |
| - | - | 5349 | 225.1 | - | - | 0 | - |
| - | - | 2810 | 225.1 | - | - | 0 | - |
| - | - | 4.843E+04 | 226.1 | - | - | 0 | - |
| - | - | 753.4 | 227 | - | - | 0 | - |
| - | - | 1341 | 227.1 | - | - | 0 | - |
| - | - | 4925 | 227.1 | - | - | 0 | - |
| - | - | 525.3 | 228.1 | - | - | 0 | - |
| - | - | 747.3 | 228.1 | - | - | 0 | - |
| - | - | 1729 | 231.1 | - | - | 0 | - |
| - | - | 1.931E+04 | 233.2 | - | - | 0 | - |
| - | - | 652.9 | 234.1 | - | - | 0 | - |
| - | - | 2401 | 234.1 | - | - | 0 | - |
| - | - | 518 | 234.1 | - | - | 0 | - |
| - | - | 2178 | 234.2 | - | - | 0 | - |
| - | - | 5248 | 235.1 | - | - | 0 | - |
| - | - | 2684 | 236.1 | - | - | 0 | - |
| - | - | 722.4 | 236.1 | - | - | 0 | - |
| - | - | 888.9 | 238.1 | - | - | 0 | - |
| - | - | 1.08E+04 | 239.1 | - | - | 0 | - |
| - | - | 545.9 | 239.6 | - | - | 0 | - |
| - | - | 3468 | 240.1 | - | - | 0 | - |
| - | - | 3139 | 241.1 | - | - | 0 | - |
| - | - | 1.001E+04 | 242.2 | - | - | 0 | - |
| - | - | 1356 | 243.1 | - | - | 0 | - |
| 11 | y | 1.824E+04 | 243.1 | 0.004177 | 17.18 | +3 | 7 |
| 13 | y | 1865 | 244.1 | 0.0004562 | 1.869 | +2 | 5 |
| - | - | 1959 | 244.1 | - | - | 0 | - |
| - | - | 6678 | 245.1 | - | - | 0 | - |
| - | - | 1648 | 246.1 | - | - | 0 | - |
| - | - | 1758 | 247.1 | - | - | 0 | - |
| - | - | 691.3 | 248.1 | - | - | 0 | - |
| - | - | 530.7 | 251.2 | - | - | 0 | - |
| - | - | 6753 | 251.2 | - | - | 0 | - |
| - | - | 1068 | 252.1 | - | - | 0 | - |
| - | - | 1144 | 252.1 | - | - | 0 | - |
| - | - | 737.7 | 252.2 | - | - | 0 | - |
| - | - | 1120 | 253.1 | - | - | 0 | - |
| - | - | 6874 | 254.1 | - | - | 0 | - |
| - | - | 1588 | 254.2 | - | - | 0 | - |
| - | - | 564.6 | 254.3 | - | - | 0 | - |
| - | - | 1263 | 257.2 | - | - | 0 | - |
| - | - | 554.5 | 258.1 | - | - | 0 | - |
| - | - | 6276 | 259.1 | - | - | 0 | - |
| - | - | 8844 | 261.2 | - | - | 0 | - |
| - | - | 1979 | 262.1 | - | - | 0 | - |
| - | - | 791.5 | 262.2 | - | - | 0 | - |
| - | - | 8688 | 263.1 | - | - | 0 | - |
| - | - | 529.7 | 264.1 | - | - | 0 | - |
| - | - | 872.3 | 264.1 | - | - | 0 | - |
| - | - | 7435 | 264.1 | - | - | 0 | - |
| - | - | 1265 | 265.1 | - | - | 0 | - |
| - | - | 639.8 | 266.1 | - | - | 0 | - |
| - | - | 1160 | 268.1 | - | - | 0 | - |
| 3 | a | 1261 | 268.2 | 0.0002323 | 0.8662 | +1 | 3 |
| - | - | 1550 | 269.2 | - | - | 0 | - |
| - | - | 632.2 | 270.2 | - | - | 0 | - |
| - | - | 2677 | 272.1 | - | - | 0 | - |
| - | - | 1885 | 274.1 | - | - | 0 | - |
| - | - | 1706 | 275.2 | - | - | 0 | - |
| - | - | 624.9 | 276.1 | - | - | 0 | - |
| - | - | 548.2 | 277.1 | - | - | 0 | - |
| - | - | 4945 | 279.1 | - | - | 0 | - |
| 5 | b | 752.4 | 279.7 | 0.0001168 | 0.4176 | +2 | 5 |
| - | - | 937 | 280.1 | - | - | 0 | - |
| - | - | 2286 | 280.1 | - | - | 0 | - |
| - | - | 1003 | 280.1 | - | - | 0 | - |
| - | - | 4050 | 280.2 | - | - | 0 | - |
| - | - | 1615 | 281.1 | - | - | 0 | - |
| - | - | 1.722E+04 | 282.1 | - | - | 0 | - |
| - | - | 1909 | 283.1 | - | - | 0 | - |
| - | - | 1.195E+04 | 285 | - | - | 0 | - |
| - | - | 3221 | 285.2 | - | - | 0 | - |
| - | - | 3914 | 286 | - | - | 0 | - |
| - | - | 3742 | 286.1 | - | - | 0 | - |
| - | - | 936.8 | 287 | - | - | 0 | - |
| - | - | 613.7 | 294.2 | - | - | 0 | - |
| - | - | 3272 | 296.1 | - | - | 0 | - |
| - | - | 5090 | 296.2 | - | - | 0 | - |
| 3 | b | 1.104E+05 | 296.2 | 0.0002449 | 0.8268 | +1 | 3 |
| - | - | 2781 | 297.1 | - | - | 0 | - |
| - | - | 1.705E+04 | 297.2 | - | - | 0 | - |
| - | - | 1253 | 297.2 | - | - | 0 | - |
| - | - | 2.075E+04 | 297.2 | - | - | 0 | - |
| - | - | 2530 | 298.2 | - | - | 0 | - |
| - | - | 4656 | 298.2 | - | - | 0 | - |
| - | - | 1629 | 298.2 | - | - | 0 | - |
| - | - | 1.423E+04 | 299.1 | - | - | 0 | - |
| - | - | 915.7 | 299.2 | - | - | 0 | - |
| - | - | 763 | 299.2 | - | - | 0 | - |
| - | - | 4156 | 300.1 | - | - | 0 | - |
| - | - | 1507 | 300.2 | - | - | 0 | - |
| - | - | 1720 | 301.1 | - | - | 0 | - |
| 15 | y | 2295 | 301.2 | 0.0003767 | 1.251 | +1 | 3 |
| - | - | 4124 | 303.2 | - | - | 0 | - |
| - | - | 1742 | 304.1 | - | - | 0 | - |
| - | - | 1057 | 306.1 | - | - | 0 | - |
| - | - | 748.3 | 307.1 | - | - | 0 | - |
| 12 | y | 7216 | 307.7 | 0.0002541 | 0.8258 | +2 | 6 |
| - | - | 1938 | 308.2 | - | - | 0 | - |
| - | - | 1397 | 310.1 | - | - | 0 | - |
| - | - | 671.7 | 313.2 | - | - | 0 | - |
| - | - | 2321 | 313.2 | - | - | 0 | - |
| - | - | 2857 | 314.1 | - | - | 0 | - |
| - | - | 1.621E+04 | 314.2 | - | - | 0 | - |
| 3 | b | 2.24E+04 | 314.2 | 0.0003003 | 0.9558 | +1 | 3 |
| - | - | 2312 | 315.2 | - | - | 0 | - |
| - | - | 4505 | 315.2 | - | - | 0 | - |
| - | - | 9103 | 316.2 | - | - | 0 | - |
| 12 | y | 3040 | 316.7 | 8.443E-05 | 0.2666 | +2 | 6 |
| - | - | 2781 | 317.2 | - | - | 0 | - |
| 15 | y | 1.559E+04 | 319.2 | 0.0003101 | 0.9714 | +1 | 3 |
| - | - | 1569 | 320.2 | - | - | 0 | - |
| - | - | 968.1 | 323.2 | - | - | 0 | - |
| - | - | 749.1 | 324.2 | - | - | 0 | - |
| - | - | 1127 | 325.1 | - | - | 0 | - |
| - | - | 2918 | 328.1 | - | - | 0 | - |
| 6 | b | 775.4 | 328.2 | 0.0005598 | 1.706 | +2 | 6 |
| - | - | 1145 | 329.2 | - | - | 0 | - |
| - | - | 3285 | 330.2 | - | - | 0 | - |
| - | - | 536.1 | 331.1 | - | - | 0 | - |
| - | - | 617.3 | 332.9 | - | - | 0 | - |
| - | - | 3746 | 333.2 | - | - | 0 | - |
| - | - | 952.9 | 334.2 | - | - | 0 | - |
| - | - | 1035 | 337.2 | - | - | 0 | - |
| - | - | 1456 | 339.2 | - | - | 0 | - |
| - | - | 3383 | 341.2 | - | - | 0 | - |
| - | - | 2428 | 342.2 | - | - | 0 | - |
| - | - | 1809 | 344.2 | - | - | 0 | - |
| - | - | 1405 | 345 | - | - | 0 | - |
| - | - | 7252 | 346.1 | - | - | 0 | - |
| - | - | 4813 | 346.2 | - | - | 0 | - |
| - | - | 946.9 | 347.1 | - | - | 0 | - |
| - | - | 3056 | 348.2 | - | - | 0 | - |
| - | - | 1029 | 349.2 | - | - | 0 | - |
| - | - | 1780 | 350.1 | - | - | 0 | - |
| - | - | 3123 | 350.2 | - | - | 0 | - |
| - | - | 1.033E+04 | 351.2 | - | - | 0 | - |
| - | - | 954.6 | 351.2 | - | - | 0 | - |
| - | - | 2317 | 352.2 | - | - | 0 | - |
| - | - | 5311 | 353.2 | - | - | 0 | - |
| - | - | 1440 | 354.2 | - | - | 0 | - |
| - | - | 1125 | 354.2 | - | - | 0 | - |
| - | - | 4.117E+04 | 355.1 | - | - | 0 | - |
| - | - | 1795 | 355.2 | - | - | 0 | - |
| - | - | 1.698E+04 | 356.1 | - | - | 0 | - |
| - | - | 9382 | 357.1 | - | - | 0 | - |
| - | - | 998 | 359 | - | - | 0 | - |
| - | - | 798.7 | 360.2 | - | - | 0 | - |
| - | - | 9193 | 362.2 | - | - | 0 | - |
| - | - | 835.3 | 363.2 | - | - | 0 | - |
| - | - | 2602 | 363.2 | - | - | 0 | - |
| - | - | 2304 | 365.1 | - | - | 0 | - |
| - | - | 4409 | 367.2 | - | - | 0 | - |
| - | - | 3023 | 368.2 | - | - | 0 | - |
| - | - | 1.131E+04 | 369.2 | - | - | 0 | - |
| - | - | 896.9 | 369.2 | - | - | 0 | - |
| - | - | 2029 | 370.2 | - | - | 0 | - |
| - | - | 1343 | 371.1 | - | - | 0 | - |
| - | - | 2629 | 371.2 | - | - | 0 | - |
| - | - | 706 | 371.2 | - | - | 0 | - |
| - | - | 688 | 372.2 | - | - | 0 | - |
| - | - | 804.5 | 372.2 | - | - | 0 | - |
| - | - | 3776 | 373.1 | - | - | 0 | - |
| - | - | 3185 | 373.2 | - | - | 0 | - |
| 11 | y | 4457 | 373.2 | 0.0003946 | 1.057 | +2 | 7 |
| - | - | 1926 | 373.7 | - | - | 0 | - |
| - | - | 2113 | 374.1 | - | - | 0 | - |
| - | - | 1018 | 374.2 | - | - | 0 | - |
| - | - | 713.3 | 381.2 | - | - | 0 | - |
| - | - | 971.8 | 382.2 | - | - | 0 | - |
| - | - | 3638 | 383.2 | - | - | 0 | - |
| - | - | 1031 | 383.3 | - | - | 0 | - |
| - | - | 983.6 | 384.2 | - | - | 0 | - |
| - | - | 725 | 384.3 | - | - | 0 | - |
| - | - | 505.9 | 385.2 | - | - | 0 | - |
| - | - | 1.489E+04 | 385.2 | - | - | 0 | - |
| - | - | 893.7 | 386.2 | - | - | 0 | - |
| - | - | 2650 | 386.2 | - | - | 0 | - |
| - | - | 1447 | 393.2 | - | - | 0 | - |
| - | - | 1164 | 397.1 | - | - | 0 | - |
| - | - | 1371 | 397.2 | - | - | 0 | - |
| - | - | 824.5 | 397.7 | - | - | 0 | - |
| - | - | 794.9 | 398.2 | - | - | 0 | - |
| - | - | 1454 | 398.2 | - | - | 0 | - |
| - | - | 794.2 | 399.2 | - | - | 0 | - |
| - | - | 1236 | 400.2 | - | - | 0 | - |
| - | - | 948.2 | 401 | - | - | 0 | - |
| - | - | 8401 | 401.2 | - | - | 0 | - |
| - | - | 2163 | 402.2 | - | - | 0 | - |
| 8 | b | 1456 | 406.2 | 6.321E-05 | 0.1556 | +2 | 8 |
| - | - | 798.6 | 406.7 | - | - | 0 | - |
| - | - | 2259 | 407.2 | - | - | 0 | - |
| - | - | 758.7 | 407.7 | - | - | 0 | - |
| - | - | 7576 | 408.2 | - | - | 0 | - |
| - | - | 918.9 | 409.2 | - | - | 0 | - |
| - | - | 719.4 | 410.2 | - | - | 0 | - |
| - | - | 747.1 | 411.2 | - | - | 0 | - |
| - | - | 3362 | 411.3 | - | - | 0 | - |
| - | - | 1318 | 412.3 | - | - | 0 | - |
| - | - | 644.8 | 413.2 | - | - | 0 | - |
| - | - | 1968 | 415 | - | - | 0 | - |
| - | - | 1615 | 415.1 | - | - | 0 | - |
| 8 | b | 1716 | 415.2 | 0.00442 | 10.64 | +2 | 8 |
| - | - | 1380 | 415.3 | - | - | 0 | - |
| - | - | 1461 | 416 | - | - | 0 | - |
| 14 | y | 1166 | 416.2 | 0.004287 | 10.3 | +1 | 4 |
| - | - | 4071 | 416.3 | - | - | 0 | - |
| - | - | 1298 | 417.3 | - | - | 0 | - |
| - | - | 4142 | 419 | - | - | 0 | - |
| - | - | 1852 | 420 | - | - | 0 | - |
| - | - | 852.8 | 421 | - | - | 0 | - |
| - | - | 4349 | 424.2 | - | - | 0 | - |
| - | - | 774.4 | 425.1 | - | - | 0 | - |
| - | - | 6617 | 425.2 | - | - | 0 | - |
| - | - | 864.9 | 426.2 | - | - | 0 | - |
| - | - | 1698 | 427.2 | - | - | 0 | - |
| 10 | y | 755.1 | 429.2 | 0.006315 | 14.71 | +2 | 8 |
| - | - | 2619 | 433.2 | - | - | 0 | - |
| 14 | y | 1.549E+04 | 433.2 | 0.000626 | 1.445 | +1 | 4 |
| - | - | 3.319E+04 | 433.3 | - | - | 0 | - |
| - | - | 1020 | 434.2 | - | - | 0 | - |
| - | - | 2873 | 434.2 | - | - | 0 | - |
| - | - | 7944 | 434.3 | - | - | 0 | - |
| - | - | 994.1 | 434.7 | - | - | 0 | - |
| - | - | 1857 | 435.2 | - | - | 0 | - |
| - | - | 1280 | 435.3 | - | - | 0 | - |
| - | - | 727.3 | 438.2 | - | - | 0 | - |
| - | - | 1687 | 440.2 | - | - | 0 | - |
| - | - | 919.5 | 441.2 | - | - | 0 | - |
| - | - | 934.5 | 441.3 | - | - | 0 | - |
| - | - | 5349 | 442.2 | - | - | 0 | - |
| - | - | 1001 | 443.1 | - | - | 0 | - |
| - | - | 2173 | 443.2 | - | - | 0 | - |
| 4 | b | 3110 | 443.3 | 0.0007397 | 1.669 | +1 | 4 |
| - | - | 895 | 444.3 | - | - | 0 | - |
| - | - | 1136 | 445.2 | - | - | 0 | - |
| - | - | 667.4 | 445.2 | - | - | 0 | - |
| - | - | 1680 | 451.2 | - | - | 0 | - |
| 5 | y | 2745 | 451.9 | 0.001519 | 3.36 | +3 | 13 |
| 5 | y | 4094 | 452.2 | 0.003824 | 8.457 | +3 | 13 |
| - | - | 1328 | 452.6 | - | - | 0 | - |
| - | - | 1007 | 453.2 | - | - | 0 | - |
| - | - | 905.5 | 454.2 | - | - | 0 | - |
| - | - | 588.4 | 458.3 | - | - | 0 | - |
| - | - | 1717 | 459.2 | - | - | 0 | - |
| - | - | 1455 | 459.3 | - | - | 0 | - |
| - | - | 964.6 | 461.2 | - | - | 0 | - |
| 4 | b | 3.222E+04 | 461.3 | 0.0001848 | 0.4005 | +1 | 4 |
| - | - | 3147 | 462.2 | - | - | 0 | - |
| - | - | 9370 | 462.3 | - | - | 0 | - |
| - | - | 1013 | 463.3 | - | - | 0 | - |
| - | - | 768.1 | 464.2 | - | - | 0 | - |
| - | - | 718.3 | 464.3 | - | - | 0 | - |
| - | - | 602.2 | 465.2 | - | - | 0 | - |
| - | - | 620.3 | 466.2 | - | - | 0 | - |
| - | - | 1245 | 468.2 | - | - | 0 | - |
| - | - | 1994 | 468.3 | - | - | 0 | - |
| - | - | 2268 | 469.2 | - | - | 0 | - |
| - | - | 9865 | 470.2 | - | - | 0 | - |
| - | - | 1077 | 470.7 | - | - | 0 | - |
| - | - | 2384 | 471.2 | - | - | 0 | - |
| - | - | 1045 | 472.2 | - | - | 0 | - |
| - | - | 737.3 | 476.2 | - | - | 0 | - |
| - | - | 2684 | 477.3 | - | - | 0 | - |
| - | - | 5616 | 478.2 | - | - | 0 | - |
| - | - | 3983 | 479.2 | - | - | 0 | - |
| 9 | b | 961.8 | 479.7 | 0.0009763 | 2.035 | +2 | 9 |
| - | - | 1.024E+04 | 480.2 | - | - | 0 | - |
| - | - | 1273 | 480.2 | - | - | 0 | - |
| - | - | 1026 | 480.7 | - | - | 0 | - |
| - | - | 1541 | 480.8 | - | - | 0 | - |
| - | - | 3482 | 481.2 | - | - | 0 | - |
| - | - | 891.5 | 482.2 | - | - | 0 | - |
| - | - | 1567 | 484.2 | - | - | 0 | - |
| 13 | y | 5002 | 486.3 | 0.0004136 | 0.8506 | +1 | 5 |
| 13 | y | 2840 | 487.3 | 0.001211 | 2.485 | +1 | 5 |
| - | - | 1087 | 488.3 | - | - | 0 | - |
| - | - | 799.2 | 488.7 | - | - | 0 | - |
| - | - | 833.3 | 489.1 | - | - | 0 | - |
| - | - | 684.4 | 490.1 | - | - | 0 | - |
| - | - | 889.1 | 490.2 | - | - | 0 | - |
| - | - | 4217 | 493.7 | - | - | 0 | - |
| - | - | 1083 | 494.2 | - | - | 0 | - |
| - | - | 1494 | 494.2 | - | - | 0 | - |
| - | - | 626.7 | 494.7 | - | - | 0 | - |
| - | - | 3404 | 495.3 | - | - | 0 | - |
| - | - | 1.31E+04 | 496.3 | - | - | 0 | - |
| - | - | 2812 | 497.3 | - | - | 0 | - |
| - | - | 3.079E+04 | 498.2 | - | - | 0 | - |
| - | - | 7211 | 499.2 | - | - | 0 | - |
| - | - | 1628 | 500.2 | - | - | 0 | - |
| - | - | 3199 | 502.2 | - | - | 0 | - |
| - | - | 4602 | 502.7 | - | - | 0 | - |
| - | - | 1677 | 503.2 | - | - | 0 | - |
| 13 | y | 3.604E+04 | 504.3 | 1.127E-05 | 0.02235 | +1 | 5 |
| - | - | 8484 | 505.3 | - | - | 0 | - |
| - | - | 1455 | 506.3 | - | - | 0 | - |
| - | - | 1028 | 508.2 | - | - | 0 | - |
| - | - | 817.8 | 510.2 | - | - | 0 | - |
| - | - | 2292 | 512.2 | - | - | 0 | - |
| - | - | 4770 | 513.3 | - | - | 0 | - |
| - | - | 741.4 | 514.3 | - | - | 0 | - |
| - | - | 1059 | 515.2 | - | - | 0 | - |
| - | - | 870.3 | 518.2 | - | - | 0 | - |
| - | - | 718.6 | 526.3 | - | - | 0 | - |
| - | - | 703.3 | 528.2 | - | - | 0 | - |
| - | - | 937.8 | 529.3 | - | - | 0 | - |
| - | - | 1746 | 529.8 | - | - | 0 | - |
| - | - | 1.062E+04 | 530.2 | - | - | 0 | - |
| - | - | 2686 | 530.3 | - | - | 0 | - |
| - | - | 2685 | 531.2 | - | - | 0 | - |
| - | - | 1274 | 531.3 | - | - | 0 | - |
| - | - | 638.4 | 534.8 | - | - | 0 | - |
| - | - | 693.3 | 535.9 | - | - | 0 | - |
| - | - | 748.1 | 536.3 | - | - | 0 | - |
| 8 | y | 800.5 | 536.8 | 0.002088 | 3.89 | +2 | 10 |
| - | - | 5403 | 538.3 | - | - | 0 | - |
| - | - | 3367 | 538.8 | - | - | 0 | - |
| 3 | y | 609.5 | 538.9 | 0.003567 | 6.619 | +3 | 15 |
| 5 | b | 3375 | 540.3 | 0.0006186 | 1.145 | +1 | 5 |
| - | - | 665 | 541.3 | - | - | 0 | - |
| - | - | 716.8 | 542.8 | - | - | 0 | - |
| - | - | 2542 | 543.3 | - | - | 0 | - |
| - | - | 1113 | 543.8 | - | - | 0 | - |
| 10 | b | 2517 | 544.3 | 0.001857 | 3.412 | +2 | 10 |
| - | - | 878.7 | 544.8 | - | - | 0 | - |
| 8 | y | 707.9 | 545.8 | 0.00438 | 8.025 | +2 | 10 |
| - | - | 829 | 546.2 | - | - | 0 | - |
| 15 | b | 899.4 | 548 | 0.001294 | 2.362 | +3 | 15 |
| - | - | 6234 | 552.3 | - | - | 0 | - |
| - | - | 4305 | 552.8 | - | - | 0 | - |
| - | - | 1577 | 553.3 | - | - | 0 | - |
| - | - | 774.5 | 553.3 | - | - | 0 | - |
| - | - | 2184 | 555.3 | - | - | 0 | - |
| - | - | 1369 | 556.3 | - | - | 0 | - |
| - | - | 1103 | 556.3 | - | - | 0 | - |
| 5 | b | 7845 | 558.3 | 0.0004246 | 0.7604 | +1 | 5 |
| - | - | 2323 | 559.3 | - | - | 0 | - |
| - | - | 988.7 | 560.8 | - | - | 0 | - |
| - | - | 800.8 | 561.3 | - | - | 0 | - |
| - | - | 800.8 | 563.2 | - | - | 0 | - |
| - | - | 757.4 | 566.4 | - | - | 0 | - |
| - | - | 1177 | 569.3 | - | - | 0 | - |
| - | - | 2289 | 569.8 | - | - | 0 | - |
| - | - | 707.1 | 570.3 | - | - | 0 | - |
| - | - | 602.6 | 571.3 | - | - | 0 | - |
| - | - | 737 | 573.8 | - | - | 0 | - |
| - | - | 1023 | 574.3 | - | - | 0 | - |
| - | - | 833.7 | 578.1 | - | - | 0 | - |
| - | - | 1984 | 578.3 | - | - | 0 | - |
| - | - | 1190 | 578.8 | - | - | 0 | - |
| - | - | 1905 | 579.3 | - | - | 0 | - |
| 7 | y | 1698 | 580.3 | 0.005102 | 8.792 | +2 | 11 |
| 7 | y | 1198 | 580.8 | 0.008078 | 13.91 | +2 | 11 |
| - | - | 2587 | 581.3 | - | - | 0 | - |
| - | - | 1961 | 582.3 | - | - | 0 | - |
| - | - | 3904 | 582.8 | - | - | 0 | - |
| - | - | 2398 | 583.3 | - | - | 0 | - |
| - | - | 1464 | 583.8 | - | - | 0 | - |
| - | - | 660.6 | 586.3 | - | - | 0 | - |
| - | - | 2202 | 591.2 | - | - | 0 | - |
| - | - | 2285 | 591.3 | - | - | 0 | - |
| 11 | b | 1801 | 591.8 | 0.007479 | 12.64 | +2 | 11 |
| - | - | 1033 | 592.2 | - | - | 0 | - |
| - | - | 3515 | 597.3 | - | - | 0 | - |
| - | - | 2188 | 598.3 | - | - | 0 | - |
| - | - | 1.375E+04 | 599.3 | - | - | 0 | - |
| - | - | 1541 | 599.6 | - | - | 0 | - |
| - | - | 904.4 | 600 | - | - | 0 | - |
| - | - | 4538 | 600.3 | - | - | 0 | - |
| 11 | b | 844.7 | 600.8 | 0.001775 | 2.954 | +2 | 11 |
| - | - | 829.9 | 601.3 | - | - | 0 | - |
| - | - | 719.2 | 602.1 | - | - | 0 | - |
| - | - | 2850 | 602.8 | - | - | 0 | - |
| - | - | 3076 | 603.3 | - | - | 0 | - |
| - | - | 970.5 | 604.3 | - | - | 0 | - |
| - | - | 826.1 | 606.8 | - | - | 0 | - |
| - | - | 1224 | 608.4 | - | - | 0 | - |
| - | - | 1516 | 608.8 | - | - | 0 | - |
| - | - | 6847 | 609.3 | - | - | 0 | - |
| - | - | 877.6 | 609.3 | - | - | 0 | - |
| - | - | 1005 | 609.3 | - | - | 0 | - |
| - | - | 1438 | 609.8 | - | - | 0 | - |
| - | - | 4307 | 610.2 | - | - | 0 | - |
| - | - | 1634 | 610.3 | - | - | 0 | - |
| - | - | 4694 | 611.2 | - | - | 0 | - |
| 0 | Precursor | 882.8 | 611.3 | 0.002386 | 3.904 | +3 | -1 |
| - | - | 5312 | 611.8 | - | - | 0 | - |
| - | - | 4354 | 612.2 | - | - | 0 | - |
| - | - | 866.9 | 612.2 | - | - | 0 | - |
| - | - | 3582 | 612.3 | - | - | 0 | - |
| - | - | 3833 | 612.8 | - | - | 0 | - |
| - | - | 2349 | 613.3 | - | - | 0 | - |
| - | - | 743.4 | 613.8 | - | - | 0 | - |
| 12 | y | 1.019E+04 | 614.3 | 0.001188 | 1.933 | +1 | 6 |
| 12 | y | 2.077E+04 | 615.3 | 3.472E-05 | 0.05642 | +1 | 6 |
| - | - | 6552 | 616.3 | - | - | 0 | - |
| - | - | 3649 | 617.3 | - | - | 0 | - |
| - | - | 1.501E+04 | 617.8 | - | - | 0 | - |
| - | - | 1.181E+04 | 618.3 | - | - | 0 | - |
| - | - | 7848 | 618.8 | - | - | 0 | - |
| - | - | 2186 | 619.3 | - | - | 0 | - |
| - | - | 938.6 | 620.3 | - | - | 0 | - |
| - | - | 977.4 | 620.8 | - | - | 0 | - |
| - | - | 1096 | 625.3 | - | - | 0 | - |
| - | - | 4540 | 626.4 | - | - | 0 | - |
| - | - | 3.176E+04 | 626.8 | - | - | 0 | - |
| - | - | 6.148E+04 | 627.3 | - | - | 0 | - |
| - | - | 2.039E+04 | 627.3 | - | - | 0 | - |
| - | - | 9294 | 627.8 | - | - | 0 | - |
| - | - | 1.939E+04 | 628.3 | - | - | 0 | - |
| - | - | 1681 | 628.3 | - | - | 0 | - |
| 6 | y | 3519 | 628.8 | 0.0009259 | 1.472 | +2 | 12 |
| - | - | 2623 | 629.3 | - | - | 0 | - |
| 6 | y | 1728 | 629.3 | 0.009834 | 15.63 | +2 | 12 |
| - | - | 1972 | 629.8 | - | - | 0 | - |
| 12 | y | 2.983E+04 | 632.3 | 0.0003387 | 0.5356 | +1 | 6 |
| - | - | 9630 | 633.3 | - | - | 0 | - |
| - | - | 1577 | 634.3 | - | - | 0 | - |
| - | - | 683.8 | 635.8 | - | - | 0 | - |
| 6 | y | 5294 | 637.8 | 0.0003891 | 0.6101 | +2 | 12 |
| - | - | 3257 | 638.3 | - | - | 0 | - |
| - | - | 1646 | 638.8 | - | - | 0 | - |
| - | - | 4005 | 643.3 | - | - | 0 | - |
| - | - | 1946 | 644.3 | - | - | 0 | - |
| - | - | 2912 | 645.3 | - | - | 0 | - |
| - | - | 825.9 | 646.3 | - | - | 0 | - |
| - | - | 958 | 646.8 | - | - | 0 | - |
| - | - | 610.3 | 647.8 | - | - | 0 | - |
| - | - | 1.101E+04 | 655.3 | - | - | 0 | - |
| 12 | b | 8796 | 655.8 | 0.006323 | 9.642 | +2 | 12 |
| 12 | b | 4352 | 656.3 | 0.01181 | 18 | +2 | 12 |
| - | - | 1097 | 656.8 | - | - | 0 | - |
| - | - | 828.6 | 658.3 | - | - | 0 | - |
| - | - | 794.2 | 659.8 | - | - | 0 | - |
| - | - | 1601 | 660.3 | - | - | 0 | - |
| 12 | b | 746 | 664.8 | 0.01104 | 16.61 | +2 | 12 |
| - | - | 1086 | 667.3 | - | - | 0 | - |
| - | - | 3291 | 668.3 | - | - | 0 | - |
| - | - | 4606 | 668.8 | - | - | 0 | - |
| - | - | 2371 | 669.3 | - | - | 0 | - |
| - | - | 954.5 | 669.8 | - | - | 0 | - |
| - | - | 1017 | 676.3 | - | - | 0 | - |
| 5 | y | 3.591E+04 | 677.3 | 0.0006757 | 0.9976 | +2 | 13 |
| 5 | y | 3.172E+04 | 677.8 | 0.00634 | 9.353 | +2 | 13 |
| - | - | 1.662E+04 | 678.3 | - | - | 0 | - |
| - | - | 5757 | 678.8 | - | - | 0 | - |
| - | - | 1522 | 679.3 | - | - | 0 | - |
| - | - | 1395 | 684.4 | - | - | 0 | - |
| 5 | y | 7.457E+04 | 686.3 | 0.000587 | 0.8552 | +2 | 13 |
| - | - | 5.669E+04 | 686.8 | - | - | 0 | - |
| - | - | 522.3 | 686.9 | - | - | 0 | - |
| - | - | 2.207E+04 | 687.3 | - | - | 0 | - |
| - | - | 6778 | 687.8 | - | - | 0 | - |
| - | - | 894.7 | 688.3 | - | - | 0 | - |
| 13 | b | 886.4 | 691.4 | 0.009365 | 13.55 | +2 | 13 |
| - | - | 2292 | 694.3 | - | - | 0 | - |
| - | - | 1223 | 695.3 | - | - | 0 | - |
| - | - | 972.4 | 698.3 | - | - | 0 | - |
| 13 | b | 3987 | 700.4 | 0.01257 | 17.95 | +2 | 13 |
| - | - | 3897 | 700.8 | - | - | 0 | - |
| - | - | 1404 | 701.4 | - | - | 0 | - |
| - | - | 1644 | 704.3 | - | - | 0 | - |
| - | - | 1342 | 710.4 | - | - | 0 | - |
| - | - | 921.6 | 711.4 | - | - | 0 | - |
| - | - | 9541 | 712.4 | - | - | 0 | - |
| - | - | 3694 | 713.4 | - | - | 0 | - |
| - | - | 826.2 | 714.4 | - | - | 0 | - |
| - | - | 2976 | 722.3 | - | - | 0 | - |
| - | - | 927.4 | 723.3 | - | - | 0 | - |
| 7 | b | 1891 | 724.4 | 0.0007996 | 1.104 | +1 | 7 |
| - | - | 1289 | 726.3 | - | - | 0 | - |
| 11 | y | 1469 | 727.4 | 4.649E-05 | 0.06392 | +1 | 7 |
| 11 | y | 2206 | 728.4 | 0.001839 | 2.524 | +1 | 7 |
| - | - | 2326 | 728.9 | - | - | 0 | - |
| - | - | 1888 | 729.4 | - | - | 0 | - |
| - | - | 1132 | 730.4 | - | - | 0 | - |
| - | - | 797.8 | 731.4 | - | - | 0 | - |
| - | - | 1666 | 737.4 | - | - | 0 | - |
| - | - | 1394 | 738.4 | - | - | 0 | - |
| - | - | 1075 | 739.4 | - | - | 0 | - |
| - | - | 2.227E+04 | 740.3 | - | - | 0 | - |
| - | - | 8870 | 741.3 | - | - | 0 | - |
| - | - | 2958 | 742.3 | - | - | 0 | - |
| 7 | b | 3886 | 742.4 | 0.0008427 | 1.135 | +1 | 7 |
| - | - | 737.8 | 742.9 | - | - | 0 | - |
| - | - | 1429 | 743.4 | - | - | 0 | - |
| - | - | 860.1 | 745.3 | - | - | 0 | - |
| 11 | y | 1.332E+04 | 745.4 | 0.0007235 | 0.9706 | +1 | 7 |
| - | - | 5299 | 746.4 | - | - | 0 | - |
| - | - | 1272 | 747.4 | - | - | 0 | - |
| - | - | 1088 | 749.3 | - | - | 0 | - |
| 4 | y | 9339 | 750.9 | 0.0008861 | 1.18 | +2 | 14 |
| 4 | y | 8337 | 751.4 | 0.007228 | 9.62 | +2 | 14 |
| - | - | 4192 | 751.9 | - | - | 0 | - |
| - | - | 1785 | 752.4 | - | - | 0 | - |
| - | - | 813.9 | 752.9 | - | - | 0 | - |
| - | - | 1540 | 753.3 | - | - | 0 | - |
| - | - | 2770 | 755.4 | - | - | 0 | - |
| - | - | 1295 | 756.4 | - | - | 0 | - |
| 14 | b | 7694 | 757.4 | 0.006753 | 8.916 | +2 | 14 |
| - | - | 3624 | 758.4 | - | - | 0 | - |
| - | - | 1659 | 759.4 | - | - | 0 | - |
| 4 | y | 8639 | 759.9 | 0.0006143 | 0.8084 | +2 | 14 |
| - | - | 9520 | 760.4 | - | - | 0 | - |
| - | - | 4001 | 760.9 | - | - | 0 | - |
| - | - | 920.7 | 761.4 | - | - | 0 | - |
| - | - | 762.5 | 761.9 | - | - | 0 | - |
| - | - | 4536 | 771.4 | - | - | 0 | - |
| - | - | 2217 | 772.4 | - | - | 0 | - |
| - | - | 775.3 | 773.4 | - | - | 0 | - |
| - | - | 4241 | 774.3 | - | - | 0 | - |
| - | - | 2222 | 775.3 | - | - | 0 | - |
| - | - | 1475 | 793.4 | - | - | 0 | - |
| - | - | 773 | 797.4 | - | - | 0 | - |
| - | - | 1114 | 806.4 | - | - | 0 | - |
| 3 | y | 1925 | 807.4 | 0.00233 | 2.885 | +2 | 15 |
| 3 | y | 3446 | 807.9 | 0.006761 | 8.369 | +2 | 15 |
| - | - | 2065 | 808.4 | - | - | 0 | - |
| - | - | 1359 | 808.9 | - | - | 0 | - |
| 8 | b | 3428 | 811.4 | 0.001578 | 1.945 | +1 | 8 |
| 15 | b | 2031 | 812.4 | 0.01239 | 15.25 | +2 | 15 |
| - | - | 2229 | 813.4 | - | - | 0 | - |
| 3 | y | 1837 | 816.4 | 0.001081 | 1.324 | +2 | 15 |
| - | - | 1380 | 816.9 | - | - | 0 | - |
| - | - | 681.5 | 822.4 | - | - | 0 | - |
| 8 | b | 3261 | 829.4 | 0.001767 | 2.13 | +1 | 8 |
| - | - | 1628 | 830.4 | - | - | 0 | - |
| - | - | 1009 | 832.4 | - | - | 0 | - |
| - | - | 1800 | 839.4 | - | - | 0 | - |
| - | - | 2734 | 840.4 | - | - | 0 | - |
| - | - | 2407 | 841.4 | - | - | 0 | - |
| - | - | 1752 | 842.4 | - | - | 0 | - |
| - | - | 1178 | 842.9 | - | - | 0 | - |
| - | - | 897.9 | 843.4 | - | - | 0 | - |
| - | - | 811.7 | 845.4 | - | - | 0 | - |
| 16 | b | 1834 | 848.4 | 0.002022 | 2.383 | +2 | 16 |
| - | - | 829.9 | 848.9 | - | - | 0 | - |
| - | - | 3014 | 850.4 | - | - | 0 | - |
| - | - | 1982 | 851.4 | - | - | 0 | - |
| - | - | 884.9 | 852.4 | - | - | 0 | - |
| 10 | y | 1744 | 856.5 | 0.004798 | 5.602 | +1 | 8 |
| 16 | b | 2151 | 856.9 | 0.002967 | 3.463 | +2 | 16 |
| 10 | y | 3994 | 857.4 | 0.006914 | 8.064 | +1 | 8 |
| 2 | y | 2505 | 857.9 | 0.004166 | 4.855 | +2 | 16 |
| 2 | y | 2116 | 858.4 | 0.008679 | 10.11 | +2 | 16 |
| - | - | 1991 | 859.4 | - | - | 0 | - |
| - | - | 1056 | 861.4 | - | - | 0 | - |
| - | - | 2693 | 863.4 | - | - | 0 | - |
| 2 | y | 899.7 | 866.9 | 7.924E-05 | 0.0914 | +2 | 16 |
| - | - | 1470 | 867.4 | - | - | 0 | - |
| - | - | 2.585E+04 | 868.4 | - | - | 0 | - |
| - | - | 1.377E+04 | 869.4 | - | - | 0 | - |
| - | - | 3235 | 870.4 | - | - | 0 | - |
| - | - | 977.4 | 871.4 | - | - | 0 | - |
| - | - | 3671 | 873.5 | - | - | 0 | - |
| 10 | y | 6906 | 874.5 | 0.001263 | 1.445 | +1 | 8 |
| - | - | 1561 | 875.5 | - | - | 0 | - |
| - | - | 4443 | 879.4 | - | - | 0 | - |
| - | - | 1921 | 880.4 | - | - | 0 | - |
| - | - | 1103 | 885.4 | - | - | 0 | - |
| - | - | 4135 | 887.4 | - | - | 0 | - |
| - | - | 1908 | 888.4 | - | - | 0 | - |
| - | - | 3934 | 889.4 | - | - | 0 | - |
| - | - | 1487 | 890.4 | - | - | 0 | - |
| - | - | 803.3 | 895.4 | - | - | 0 | - |
| - | - | 1.627E+04 | 907.4 | - | - | 0 | - |
| - | - | 8028 | 908.4 | - | - | 0 | - |
| - | - | 979.3 | 909.4 | - | - | 0 | - |
| - | - | 2673 | 911.4 | - | - | 0 | - |
| - | - | 1136 | 912.4 | - | - | 0 | - |
| - | - | 1564 | 921.4 | - | - | 0 | - |
| - | - | 2676 | 922.4 | - | - | 0 | - |
| - | - | 1.279E+04 | 939.4 | - | - | 0 | - |
| - | - | 7953 | 940.4 | - | - | 0 | - |
| - | - | 3566 | 941.5 | - | - | 0 | - |
| - | - | 837 | 942.5 | - | - | 0 | - |
| - | - | 867 | 953.5 | - | - | 0 | - |
| - | - | 855.5 | 954.5 | - | - | 0 | - |
| - | - | 3698 | 956.4 | - | - | 0 | - |
| - | - | 1987 | 957.5 | - | - | 0 | - |
| 9 | b | 8360 | 958.5 | 0.001696 | 1.77 | +1 | 9 |
| - | - | 7277 | 959.5 | - | - | 0 | - |
| - | - | 1.098E+04 | 960.5 | - | - | 0 | - |
| - | - | 4553 | 961.5 | - | - | 0 | - |
| - | - | 2590 | 971.5 | - | - | 0 | - |
| 9 | y | 1055 | 986.5 | 0.0008201 | 0.8313 | +1 | 9 |
| - | - | 1039 | 987.5 | - | - | 0 | - |
| - | - | 2197 | 988.5 | - | - | 0 | - |
| - | - | 1567 | 989.5 | - | - | 0 | - |
| - | - | 1619 | 1000 | - | - | 0 | - |
| 9 | y | 1847 | 1004 | 8.886E-05 | 0.08855 | +1 | 9 |
| - | - | 1007 | 1005 | - | - | 0 | - |
| - | - | 841.9 | 1008 | - | - | 0 | - |
| - | - | 1749 | 1015 | - | - | 0 | - |
| - | - | 1614 | 1016 | - | - | 0 | - |
| - | - | 1268 | 1025 | - | - | 0 | - |
| - | - | 2653 | 1036 | - | - | 0 | - |
| - | - | 939.8 | 1037 | - | - | 0 | - |
| - | - | 917.9 | 1041 | - | - | 0 | - |
| - | - | 1640 | 1041 | - | - | 0 | - |
| - | - | 924.3 | 1053 | - | - | 0 | - |
| - | - | 1.024E+04 | 1053 | - | - | 0 | - |
| - | - | 6235 | 1054 | - | - | 0 | - |
| - | - | 2258 | 1055 | - | - | 0 | - |
| - | - | 873.8 | 1056 | - | - | 0 | - |
| - | - | 3160 | 1059 | - | - | 0 | - |
| - | - | 2583 | 1060 | - | - | 0 | - |
| - | - | 878 | 1061 | - | - | 0 | - |
| - | - | 1099 | 1067 | - | - | 0 | - |
| 10 | b | 3246 | 1070 | 0.005893 | 5.51 | +1 | 10 |
| - | - | 5571 | 1071 | - | - | 0 | - |
| - | - | 3675 | 1072 | - | - | 0 | - |
| 8 | y | 1644 | 1073 | 0.01265 | 11.79 | +1 | 10 |
| - | - | 4120 | 1076 | - | - | 0 | - |
| - | - | 2229 | 1077 | - | - | 0 | - |
| - | - | 1621 | 1085 | - | - | 0 | - |
| - | - | 1831 | 1086 | - | - | 0 | - |
| - | - | 2497 | 1087 | - | - | 0 | - |
| 10 | b | 2.434E+04 | 1088 | 0.003518 | 3.235 | +1 | 10 |
| - | - | 1.325E+04 | 1089 | - | - | 0 | - |
| - | - | 5818 | 1090 | - | - | 0 | - |
| 8 | y | 2884 | 1091 | 0.003314 | 3.039 | +1 | 10 |
| - | - | 1541 | 1102 | - | - | 0 | - |
| - | - | 5292 | 1104 | - | - | 0 | - |
| - | - | 2729 | 1105 | - | - | 0 | - |
| - | - | 814 | 1106 | - | - | 0 | - |
| - | - | 751.1 | 1128 | - | - | 0 | - |
| - | - | 1059 | 1130 | - | - | 0 | - |
| - | - | 1593 | 1138 | - | - | 0 | - |
| - | - | 2492 | 1139 | - | - | 0 | - |
| - | - | 1058 | 1140 | - | - | 0 | - |
| - | - | 4346 | 1156 | - | - | 0 | - |
| - | - | 2540 | 1157 | - | - | 0 | - |
| - | - | 1124 | 1158 | - | - | 0 | - |
| 7 | y | 921.9 | 1160 | 0.004031 | 3.476 | +1 | 11 |
| 7 | y | 801.9 | 1161 | 0.007442 | 6.412 | +1 | 11 |
| - | - | 3166 | 1165 | - | - | 0 | - |
| - | - | 2011 | 1166 | - | - | 0 | - |
| 7 | y | 1680 | 1178 | 0.0003021 | 0.2565 | +1 | 11 |
| - | - | 1068 | 1179 | - | - | 0 | - |
| - | - | 9552 | 1182 | - | - | 0 | - |
| 11 | b | 5708 | 1183 | 0.02331 | 19.71 | +1 | 11 |
| - | - | 2834 | 1184 | - | - | 0 | - |
| 11 | b | 3910 | 1201 | 0.009457 | 7.877 | +1 | 11 |
| - | - | 2025 | 1202 | - | - | 0 | - |
| - | - | 783.3 | 1203 | - | - | 0 | - |
| - | - | 1382 | 1213 | - | - | 0 | - |
| - | - | 1362 | 1218 | - | - | 0 | - |
| - | - | 5886 | 1235 | - | - | 0 | - |
| - | - | 4675 | 1236 | - | - | 0 | - |
| - | - | 2053 | 1237 | - | - | 0 | - |
| - | - | 1.461E+04 | 1253 | - | - | 0 | - |
| - | - | 1.009E+04 | 1254 | - | - | 0 | - |
| - | - | 4775 | 1255 | - | - | 0 | - |
| - | - | 1009 | 1256 | - | - | 0 | - |
| 6 | y | 2513 | 1257 | 0.0007054 | 0.5614 | +1 | 12 |
| 6 | y | 1985 | 1258 | 0.002408 | 1.914 | +1 | 12 |
| - | - | 1492 | 1259 | - | - | 0 | - |
| 6 | y | 8009 | 1275 | 0.004366 | 3.425 | +1 | 12 |
| - | - | 4616 | 1276 | - | - | 0 | - |
| - | - | 1086 | 1277 | - | - | 0 | - |
| 12 | b | 1733 | 1329 | 0.01005 | 7.565 | +1 | 12 |
| - | - | 1101 | 1330 | - | - | 0 | - |
| 5 | y | 904.4 | 1354 | 0.01031 | 7.617 | +1 | 13 |
| 5 | y | 2684 | 1372 | 0.005006 | 3.649 | +1 | 13 |
| - | - | 2421 | 1373 | - | - | 0 | - |
| - | - | 894.1 | 1374 | - | - | 0 | - |
| 13 | b | 1835 | 1400 | 0.01225 | 8.754 | +1 | 13 |
| - | - | 1716 | 1401 | - | - | 0 | - |
| 14 | b | 1231 | 1514 | 0.001347 | 0.8899 | +1 | 14 |
| - | - | 1032 | 1515 | - | - | 0 | - |
| - | - | 612.8 | 2226 | - | - | 0 | - |
| - | - | 839.7 | 3070 | - | - | 0 | - |
| - | - | 676.3 | 3394 | - | - | 0 | - |

m/z Charge Intensity FragmentType MassShift Position
120.0657958984375 0 48024.516 y 16
120.08103942871094 0 39209.15
121.06279754638672 0 278.37137
121.06446075439453 0 277.82547
121.0692138671875 0 1832.9965
121.08441162109375 0 3675.7322
126.05525970458984 0 4204.773
127.08673095703125 0 756.4404
128.1072540283203 0 13933.934
129.06607055664062 0 2756.5444
129.07064819335938 0 477.6919
129.10250854492188 0 67897.15
130.05014038085938 0 3850.486
130.0655517578125 0 1208.2766
130.0820770263672 0 399.25488
130.10073852539062 0 629.2786
130.10586547851562 0 3581.4673
131.1180877685547 0 3105.4004
132.0769500732422 0 747.02795
133.06101989746094 0 516.9579
133.9902801513672 0 375.0168
136.07583618164062 0 1649.4329
138.0916290283203 0 2852.0662
138.12811279296875 0 1482.4999
139.08706665039062 0 1909.7589
141.066162109375 0 1542.5334
142.06101989746094 0 519.6732
147.07664489746094 0 1669.8123
148.0606231689453 0 668.16327
148.95445251464844 0 1077.4479
149.0449676513672 0 2672.1077
152.07052612304688 0 485.18164
154.08653259277344 0 1705.263
154.0977325439453 0 2753.2502
155.0817413330078 0 4131.2
155.1181182861328 0 9715.455 a Water loss 1
157.06112670898438 0 1630.2992
157.09739685058594 0 6236.781
159.09188842773438 0 5026.732
165.10247802734375 0 1542.8506
166.08644104003906 0 888.87976
167.0556640625 0 3172.5674
167.08177185058594 0 3346.9631
167.1180419921875 0 2536.587
168.05577087402344 0 512.6298
169.06048583984375 0 711.49133
169.13375854492188 0 10117.613
170.1185302734375 0 468.57315
170.1376190185547 0 753.4204
171.07687377929688 0 1848.4126
172.10850524902344 0 2890.6692
172.14340209960938 0 476.50848
173.0922393798828 0 7661.835 y Water loss 15
173.12869262695312 0 34856.57 a 1
173.4391632080078 0 2169.2195
174.1318359375 0 2637.0679
175.07174682617188 0 1811.0322
180.1133270263672 0 885.31195
181.09716796875 0 1628.6167
182.0926055908203 0 3455.7783
182.12905883789062 0 3275.5166
183.07666015625 0 3726.3354
183.11300659179688 0 44311.043 b Water loss 1
184.0797882080078 0 547.13043
184.10818481445312 0 2218.2515
184.11643981933594 0 4255.614
185.09230041503906 0 6874.01
185.12814331054688 0 658.3997
186.0874481201172 0 2739.7876
187.14439392089844 0 14636.527
188.1485595703125 0 637.2087
189.0872344970703 0 3537.4778
191.0820770263672 0 860.21484
191.10289001464844 0 10920.467 y 15
191.1179962158203 0 856.82806
192.1060028076172 0 1144.0942
195.07614135742188 0 584.9567
195.1129913330078 0 6812.7935
197.10386657714844 0 508.88766
197.12864685058594 0 14263.762
198.123779296875 0 2719.6038
198.13259887695312 0 680.8679
199.07156372070312 0 4522.5425
199.0811004638672 0 693.4666
199.10780334472656 0 1674.445
199.1186065673828 0 1066.716
200.10308837890625 0 4561.3193
200.1395263671875 0 3383.7651
201.0868682861328 0 979.1575
201.12353515625 0 43321.914 b 1
202.12692260742188 0 4230.5034
203.11343383789062 0 464.53824
208.09825134277344 0 735.3702
208.1083221435547 0 5198.933
209.09228515625 0 11752.357
210.0955810546875 0 1502.7673
210.12368774414062 0 564.60016
211.14486694335938 0 728.3157
212.13975524902344 0 1709.7249
213.08750915527344 0 1030.5618
213.1234130859375 0 708.618
214.1298065185547 0 648.6729
214.1558837890625 0 650.0023
215.13922119140625 0 16468.547
216.14231872558594 0 1811.7446
217.0821075439453 0 8436.835
217.09710693359375 0 4937.434
217.1336669921875 0 11536.149
218.0848388671875 0 832.3103
218.13723754882812 0 1544.0406
218.14930725097656 0 1040.3608
219.1336212158203 0 802.67975 b 5
219.1498565673828 0 1240.1613
221.08448791503906 0 4525.036
222.08677673339844 0 718.4202
223.063232421875 0 754.53186
223.08163452148438 0 791.7627
224.13987731933594 0 570.7931
225.0431671142578 0 1852.469
225.12342834472656 0 5349.114
225.1351776123047 0 2810.1516
226.1188201904297 0 48425.16
227.0400390625 0 753.356
227.10281372070312 0 1341.156
227.12222290039062 0 4924.818
228.0972137451172 0 525.2783
228.1342315673828 0 747.26074
231.09805297851562 0 1729.4808
233.16500854492188 0 19312.95
234.0880126953125 0 652.9052
234.12387084960938 0 2401.2246
234.1391143798828 0 517.9609
234.16868591308594 0 2177.9102
235.10775756835938 0 5247.9575
236.10301208496094 0 2683.7197
236.1400604248047 0 722.4394
238.1182861328125 0 888.8644
239.09518432617188 0 10796.418
239.625244140625 0 545.8529
240.0963592529297 0 3467.903
241.0820770263672 0 3139.4492
242.15011596679688 0 10013.415
243.13375854492188 0 1355.8356
243.14559936523438 0 18236.588 y Water loss 10
244.129638671875 0 1864.8458 y Ammonia loss 12
244.14862060546875 0 1959.1309
245.1287078857422 0 6678.214
246.12379455566406 0 1648.4781
247.14422607421875 0 1758.0353
248.13754272460938 0 691.2504
251.15170288085938 0 530.6731
251.1756134033203 0 6753.405
252.09762573242188 0 1067.6371
252.1343231201172 0 1143.6858
252.18051147460938 0 737.74664
253.1296844482422 0 1120.0886
254.11349487304688 0 6873.5645
254.1503143310547 0 1588.1411
254.25306701660156 0 564.57886
257.19769287109375 0 1262.916
258.1089172363281 0 554.48474
259.0925598144531 0 6276.25
261.15972900390625 0 8844.467
262.1188049316406 0 1979.4249
262.16339111328125 0 791.49713
263.1027526855469 0 8687.501
264.0671081542969 0 529.6774
264.1059265136719 0 872.259
264.1344299316406 0 7434.5737
265.1369323730469 0 1265.3438
266.1489562988281 0 639.82
268.0927734375 0 1159.8506
268.20172119140625 0 1260.9089 a Water loss 2
269.1861877441406 0 1549.7689
270.18023681640625 0 632.17145
272.1241455078125 0 2677.1987
274.11907958984375 0 1884.9441
275.1753234863281 0 1705.7295
276.1201171875 0 624.9443
277.11944580078125 0 548.2325
279.1454162597656 0 4945.0015
279.6680603027344 0 752.4024 b 4
280.0930480957031 0 936.97925
280.12957763671875 0 2286.1365
280.1495361328125 0 1002.5733
280.1663818359375 0 4049.563
281.0510559082031 0 1614.9362
282.14501953125 0 17219.732
283.1478576660156 0 1909.0294
285.00982666015625 0 11950.774
285.1599426269531 0 3220.8884
286.01019287109375 0 3914.0513
286.1034851074219 0 3742.0667
287.00750732421875 0 936.7593
294.1813659667969 0 613.74554
296.13555908203125 0 3272.0227
296.1731262207031 0 5089.798
296.1971130371094 0 110384.4 b Water loss 2
297.1195983886719 0 2780.8098
297.156005859375 0 17051.398
297.17572021484375 0 1253.3801
297.2002868652344 0 20748.316
298.1584777832031 0 2530.4927
298.176513671875 0 4655.5693
298.2025451660156 0 1628.731
299.0619201660156 0 14234.566
299.1537780761719 0 915.66394
299.1730651855469 0 762.9898
300.06231689453125 0 4155.792
300.15582275390625 0 1507.3674
301.0592346191406 0 1719.8116
301.1874084472656 0 2295.487 y Water loss 14
303.17041015625 0 4123.571
304.1142883300781 0 1741.771
306.1455993652344 0 1056.8099
307.1404113769531 0 748.2577
307.6667175292969 0 7216.367 y Water loss 11
308.1668395996094 0 1937.7676
310.1033935546875 0 1396.587
313.16436767578125 0 671.6675
313.1881103515625 0 2321.3916
314.1466979980469 0 2857.1343
314.1827392578125 0 16209.371
314.2077331542969 0 22396.014 b 2
315.1856689453125 0 2311.8992
315.2106628417969 0 4504.966
316.1868591308594 0 9103.295
316.6716613769531 0 3040.3496 y 11
317.1883850097656 0 2781.42
319.1979064941406 0 15591.029 y 14
320.200927734375 0 1568.5281
323.1712646484375 0 968.0824
324.15350341796875 0 749.092
325.1499938964844 0 1126.6285
328.1141357421875 0 2917.6194
328.19488525390625 0 775.4498 b 5
329.18231201171875 0 1145.4987
330.18194580078125 0 3284.8704
331.13873291015625 0 536.0911
332.85784912109375 0 617.28815
333.1556396484375 0 3745.763
334.2125244140625 0 952.87225
337.1531066894531 0 1035.4984
339.1661071777344 0 1456.4655
341.1835021972656 0 3383.0525
342.1788635253906 0 2427.659
344.1989440917969 0 1809.109
344.9773254394531 0 1404.741
346.1245422363281 0 7251.6694
346.1763916015625 0 4812.752
347.12933349609375 0 946.8884
348.19207763671875 0 3055.7178
349.1957702636719 0 1028.5277
350.1346130371094 0 1780.1908
350.1822814941406 0 3123.23
351.16632080078125 0 10330.47
351.18731689453125 0 954.6348
352.16998291015625 0 2317.421
353.18182373046875 0 5311.0156
354.16595458984375 0 1440.2141
354.1866455078125 0 1124.9917
355.0699462890625 0 41173.51
355.1628723144531 0 1794.6003
356.07073974609375 0 16978.295
357.06817626953125 0 9382.257
359.02752685546875 0 998.02185
360.2293701171875 0 798.7301
362.2074890136719 0 9193.145
363.1665344238281 0 835.2689
363.2092590332031 0 2601.868
365.145751953125 0 2303.5796
367.2083740234375 0 4408.8726
368.1926574707031 0 3023.3103
369.1768798828125 0 11314.1
369.20050048828125 0 896.8711
370.1803283691406 0 2029.0508
371.10137939453125 0 1343.2532
371.19171142578125 0 2629.4004
371.2425537109375 0 706.0242
372.1747741699219 0 687.9941
372.1993103027344 0 804.5128
373.08056640625 0 3775.5398
373.1717834472656 0 3184.8691
373.21417236328125 0 4456.8325 y 10
373.7151184082031 0 1926.2533
374.080078125 0 2113.0566
374.21673583984375 0 1017.62604
381.1759033203125 0 713.29175
382.20849609375 0 971.75726
383.15631103515625 0 3637.9573
383.2655944824219 0 1031.2002
384.15802001953125 0 983.59863
384.2664489746094 0 725.00464
385.19488525390625 0 505.88504
385.2194519042969 0 14894.95
386.198974609375 0 893.68274
386.2230529785156 0 2650.3987
393.25 0 1446.8503
397.1355285644531 0 1164.3792
397.2165832519531 0 1370.8147
397.71795654296875 0 824.5032
398.20257568359375 0 794.9391
398.2447204589844 0 1454.2769
399.1867980957031 0 794.24255
400.22296142578125 0 1236.0046
400.9850158691406 0 948.19214
401.166748046875 0 8400.959
402.1705017089844 0 2162.6257
406.22100830078125 0 1455.6565 b Water loss 7
406.7228698730469 0 798.6044
407.2039794921875 0 2259.3157
407.7218322753906 0 758.6749
408.1877746582031 0 7575.907
409.1900634765625 0 918.92487
410.20123291015625 0 719.4287
411.20623779296875 0 747.1253
411.2598876953125 0 3361.9727
412.2641906738281 0 1317.9573
413.21136474609375 0 644.8197
415.036376953125 0 1968.1492
415.1452941894531 0 1615.4526
415.23077392578125 0 1716.2404 b 7
415.26959228515625 0 1379.8008
416.0377197265625 0 1461.3948
416.21826171875 0 1166.3794 y Ammonia loss 13
416.25445556640625 0 4071.2783
417.2555236816406 0 1298.1886
418.9956359863281 0 4142.048
419.9960021972656 0 1851.7594
420.99371337890625 0 852.8156
424.2290954589844 0 4348.9565
425.1341552734375 0 774.35175
425.2143249511719 0 6616.8667
426.2189025878906 0 864.92676
427.2303771972656 0 1698.2725
429.2154846191406 0 755.11896 y Ammonia loss 9
433.1560363769531 0 2618.775
433.24114990234375 0 15494.303 y 13
433.28118896484375 0 33186.72
434.2056884765625 0 1019.94525
434.2440490722656 0 2872.9849
434.283935546875 0 7943.6724
434.7067565917969 0 994.072
435.207275390625 0 1856.7645
435.2862854003906 0 1280.1307
438.2362365722656 0 727.2543
440.2110900878906 0 1687.3329
441.1971130371094 0 919.5009
441.2513427734375 0 934.5265
442.24053955078125 0 5348.5093
443.1422119140625 0 1001.27203
443.2316589355469 0 2173.1138
443.2660217285156 0 3110.0984 b Water loss 3
444.26739501953125 0 895.04047
445.2021484375 0 1135.9623
445.2417907714844 0 667.41174
451.2305603027344 0 1679.8165
451.89453125 0 2744.78 y Water loss 4
452.2171936035156 0 4094.2852 y Ammonia loss 4
452.56060791015625 0 1327.7206
453.2145690917969 0 1006.7681
454.2044372558594 0 905.4873
458.2717590332031 0 588.3773
459.2099304199219 0 1716.6965
459.26055908203125 0 1455.4662
461.2178039550781 0 964.59485
461.2760314941406 0 32221.896 b 3
462.1990661621094 0 3146.7324
462.2790832519531 0 9369.82
463.28204345703125 0 1012.72864
464.21856689453125 0 768.0994
464.25341796875 0 718.349
465.2444152832031 0 602.21344
466.194091796875 0 620.27234
468.20733642578125 0 1245.3232
468.2559814453125 0 1993.593
469.2444763183594 0 2267.5142
470.2250061035156 0 9864.717
470.7320556640625 0 1076.9933
471.2281494140625 0 2383.7603
472.2433166503906 0 1045.104
476.17779541015625 0 737.33594
477.25634765625 0 2684.137
478.24151611328125 0 5615.8506
479.2272644042969 0 3982.6465
479.7486267089844 0 961.8219 b 8
480.20916748046875 0 10242.91
480.25 0 1272.7227
480.71734619140625 0 1026.2009
480.7504577636719 0 1540.6195
481.21380615234375 0 3481.743
482.2216491699219 0 891.5295
484.2033996582031 0 1567.2208
486.2674865722656 0 5002.3774 y Water loss 12
487.2498779296875 0 2840.1108 y Ammonia loss 12
488.2512512207031 0 1087.4738
488.73614501953125 0 799.19025
489.0557556152344 0 833.33124
490.05438232421875 0 684.4365
490.20758056640625 0 889.1347
493.7272644042969 0 4216.7607
494.1895446777344 0 1083.3036
494.2291564941406 0 1493.8466
494.7300720214844 0 626.6862
495.2669677734375 0 3403.9011
496.25146484375 0 13102.471
497.2531433105469 0 2812.2258
498.2197265625 0 30793.086
499.2225036621094 0 7211.073
500.2315368652344 0 1628.0963
502.2145690917969 0 3199.3298
502.7328186035156 0 4601.5713
503.2301025390625 0 1676.9421
504.27764892578125 0 36039.914 y 12
505.2799377441406 0 8484.002
506.2847595214844 0 1454.712
508.2190856933594 0 1027.8705
510.2195739746094 0 817.7798
512.1988525390625 0 2291.7227
513.2774047851562 0 4769.533
514.2630004882812 0 741.42865
515.2481689453125 0 1059.2914
518.2052612304688 0 870.2771
526.25537109375 0 718.5944
528.232177734375 0 703.2901
529.2637939453125 0 937.78345
529.7642211914062 0 1745.6252
530.208984375 0 10623.659
530.3327026367188 0 2685.9814
531.2120361328125 0 2684.9763
531.3370971679688 0 1274.4637
534.7532958984375 0 638.4446
535.9453735351562 0 693.2618
536.2757568359375 0 748.132
536.7650146484375 0 800.50116 y Water loss 7
538.2691650390625 0 5402.889
538.7705688476562 0 3367.4
538.9420776367188 0 609.48615 y Ammonia loss 2
540.3186645507812 0 3375.0837 b Water loss 4
541.3221435546875 0 665.0074
542.7689208984375 0 716.8098
543.2625732421875 0 2541.6145
543.766357421875 0 1112.7965
544.26708984375 0 2516.9465 b 9
544.771728515625 0 878.6639
545.7680053710938 0 707.88684 y 7
546.2387084960938 0 828.9821
547.95556640625 0 899.43866 b 14
552.2669677734375 0 6233.998
552.7674560546875 0 4304.7217
553.265380859375 0 1576.6024
553.3099365234375 0 774.45074
555.324951171875 0 2183.5366
556.276123046875 0 1369.417
556.3275146484375 0 1102.9636
558.3281860351562 0 7845.359 b 4
559.3309326171875 0 2323.2783
560.7757568359375 0 988.7459
561.2837524414062 0 800.8181
563.2450561523438 0 800.75073
566.42919921875 0 757.3908
569.2914428710938 0 1176.8818
569.7769165039062 0 2288.5146
570.2781372070312 0 707.07074
571.2710571289062 0 602.6495
573.7738037109375 0 736.97864
574.2744140625 0 1022.736
578.1233520507812 0 833.69934
578.2847900390625 0 1984.4685
578.7872924804688 0 1189.8284
579.2868041992188 0 1905.0498
580.2780151367188 0 1697.8768 y Water loss 6
580.783203125 0 1197.8932 y Ammonia loss 6
581.257568359375 0 2586.5408
582.28662109375 0 1961.1456
582.7826538085938 0 3904.232
583.2823486328125 0 2397.6748
583.7860717773438 0 1464.1161
586.3348388671875 0 660.5515
591.2413330078125 0 2202.2104
591.2936401367188 0 2285.0962
591.7982177734375 0 1801.0977 b Water loss 10
592.2401123046875 0 1032.7224
597.2951049804688 0 3514.8975
598.2892456054688 0 2187.627
599.2672119140625 0 13746.321
599.6446533203125 0 1540.9509
599.979736328125 0 904.39154
600.27001953125 0 4537.7944
600.8092041015625 0 844.6509 b 10
601.2734375 0 829.89667
602.0714111328125 0 719.1532
602.7910766601562 0 2850.2983
603.29296875 0 3075.852
604.2990112304688 0 970.53595
606.8123168945312 0 826.1294
608.3538818359375 0 1224.444
608.8018798828125 0 1516.3348
609.2509155273438 0 6846.5537
609.2940673828125 0 877.6154
609.3405151367188 0 1005.2865
609.8005981445312 0 1438.1439
610.1839599609375 0 4306.665
610.2571411132812 0 1633.9833
611.1843872070312 0 4693.7446
611.32177734375 0 882.79614 Precursor
611.796875 0 5311.7266
612.182373046875 0 4354.1743
612.244140625 0 866.8823
612.2994384765625 0 3581.6997
612.8114624023438 0 3832.9268
613.3143920898438 0 2349.3398
613.8119506835938 0 743.395
614.324462890625 0 10192.999 y Water loss 11
615.3096313476562 0 20767.424 y Ammonia loss 11
616.31103515625 0 6551.596
617.3130493164062 0 3648.6265
617.8065795898438 0 15006.078
618.3063354492188 0 11813.079
618.8051147460938 0 7848.389
619.3040161132812 0 2185.8755
620.301025390625 0 938.64215
620.798095703125 0 977.41895
625.2828369140625 0 1095.6151
626.3626708984375 0 4539.564
626.8116455078125 0 31761.191
627.261474609375 0 61479.883
627.3135375976562 0 20385.654
627.813232421875 0 9293.98
628.2647705078125 0 19391.66
628.3153686523438 0 1681.3538
628.8104248046875 0 3518.767 y Water loss 5
629.2657470703125 0 2622.642
629.3113403320312 0 1727.8452 y Ammonia loss 5
629.8050537109375 0 1972.025
632.3358764648438 0 29830.764 y 11
633.337158203125 0 9629.734
634.341064453125 0 1577.0768
635.814453125 0 683.8328
637.8143920898438 0 5294.2964 y 5
638.3168334960938 0 3257.4058
638.814697265625 0 1645.5092
643.2942504882812 0 4004.7725
644.29248046875 0 1945.9271
645.300048828125 0 2912.4028
646.2863159179688 0 825.88464
646.8289794921875 0 958.0437
647.8173217773438 0 610.30316
655.3394165039062 0 11007.565
655.84130859375 0 8795.566 b Water loss 11
656.3388061523438 0 4352.294 b Ammonia loss 11
656.8427124023438 0 1097.4958
658.3032836914062 0 828.55005
659.8196411132812 0 794.1873
660.3192138671875 0 1600.8224
664.8292236328125 0 745.98206 b 11
667.33837890625 0 1085.5024
668.3313598632812 0 3291.2178
668.8245849609375 0 4606.0835
669.32421875 0 2370.5317
669.825439453125 0 954.5444
676.3274536132812 0 1016.71277
677.335205078125 0 35908.875 y Water loss 4
677.834228515625 0 31719.002 y Ammonia loss 4
678.3342895507812 0 16624.52
678.835693359375 0 5757.2495
679.3306884765625 0 1521.5404
684.3668823242188 0 1395.2474
686.340576171875 0 74572.82 y 4
686.8416748046875 0 56692.395
686.895751953125 0 522.27997
687.343505859375 0 22071.639
687.844482421875 0 6777.715
688.3380737304688 0 894.6647
691.3441772460938 0 886.43933 b Water loss 12
694.3387451171875 0 2292.0352
695.34375 0 1223.2955
698.3385009765625 0 972.38196
700.3462524414062 0 3986.628 b 12
700.8489990234375 0 3897.2266
701.3502807617188 0 1404.4237
704.3276977539062 0 1644.1556
710.3814086914062 0 1342.0621
711.3844604492188 0 921.5645
712.3504638671875 0 9540.654
713.3530883789062 0 3694.257
714.35888671875 0 826.2331
722.3343505859375 0 2976.2773
723.3384399414062 0 927.43317
724.4020385742188 0 1891.0887 b Water loss 6
726.338134765625 0 1288.6146
727.40966796875 0 1469.1597 y Water loss 10
728.3955688476562 0 2206.2935 y Ammonia loss 10
728.8744506835938 0 2325.5933
729.3818359375 0 1888.2736
730.3862915039062 0 1132.0793
731.3528442382812 0 797.8408
737.3929443359375 0 1666.2661
738.3797607421875 0 1394.1151
739.3697509765625 0 1074.6405
740.3458862304688 0 22266.139
741.3492431640625 0 8870.434
742.34814453125 0 2957.8733
742.4142456054688 0 3885.9424 b 6
742.8621826171875 0 737.76886
743.4171142578125 0 1428.8707
745.3451538085938 0 860.11395
745.4195556640625 0 13315.387 y 10
746.4220581054688 0 5298.935
747.4239501953125 0 1271.5322
749.3342895507812 0 1088.3918
750.8692016601562 0 9338.551 y Water loss 3
751.3693237304688 0 8336.686 y Ammonia loss 3
751.8697509765625 0 4191.669
752.3745727539062 0 1784.6344
752.8701782226562 0 813.9464
753.34033203125 0 1540.4843
755.4030151367188 0 2769.6023
756.406982421875 0 1295.2798
757.37353515625 0 7693.6846 b 13
758.3738403320312 0 3623.9238
759.3801879882812 0 1659.3208
759.874755859375 0 8638.542 y 3
760.37548828125 0 9519.977
760.8768920898438 0 4001.4004
761.379638671875 0 920.6645
761.8939819335938 0 762.49243
771.3508911132812 0 4535.797
772.3507080078125 0 2217.1257
773.3548583984375 0 775.2922
774.329833984375 0 4240.932
775.333740234375 0 2222.0415
793.4176025390625 0 1475.1763
797.376220703125 0 772.9552
806.3594970703125 0 1114.0935
807.4097900390625 0 1925.3583 y Water loss 2
807.910888671875 0 3445.5427 y Ammonia loss 2
808.4117431640625 0 2065.3193
808.9130859375 0 1359.1165
811.4332885742188 0 3427.6306 b Water loss 7
812.4348754882812 0 2031.3916 b Water loss 14
813.4318237304688 0 2228.835
816.4163208007812 0 1837.1328 y 2
816.9192504882812 0 1379.7336
822.4022216796875 0 681.4703
829.4436645507812 0 3261.34 b 7
830.444580078125 0 1628.4991
832.3862915039062 0 1008.55743
839.423583984375 0 1800.3898
840.4061279296875 0 2734.216
841.418212890625 0 2407.454
842.400146484375 0 1751.7473
842.9468994140625 0 1177.7698
843.38623046875 0 897.89233
845.396240234375 0 811.6579
848.4310302734375 0 1834.4622 b Ammonia loss 15
848.9434204101562 0 829.9213
850.39599609375 0 3013.8293
851.3904418945312 0 1982.0801
852.382080078125 0 884.8851
856.447509765625 0 1743.9052 y Water loss 9
856.943359375 0 2150.809 b 15
857.4432373046875 0 3994.4768 y Ammonia loss 9
857.9401245117188 0 2504.602 y Water loss 1
858.4366455078125 0 2116.2546 y Ammonia loss 1
859.4262084960938 0 1990.7994
861.396728515625 0 1056.3738
863.381103515625 0 2692.522
866.941162109375 0 899.7058 y 1
867.4284057617188 0 1470.4459
868.4033203125 0 25849.266
869.4058837890625 0 13768.805
870.406982421875 0 3235.4565
871.4083251953125 0 977.38556
873.4566650390625 0 3670.869
874.4616088867188 0 6906.4756 y 9
875.4594116210938 0 1560.7815
879.409912109375 0 4442.7256
880.4113159179688 0 1921.1744
885.4341430664062 0 1103.0339
887.4134521484375 0 4134.826
888.4132690429688 0 1907.5148
889.3965454101562 0 3934.2031
890.3973388671875 0 1486.987
895.4334716796875 0 803.25714
907.404541015625 0 16266.876
908.4059448242188 0 8028.3784
909.4131469726562 0 979.251
911.444580078125 0 2673.089
912.44921875 0 1135.797
921.4280395507812 0 1564.3285
922.4273681640625 0 2676.3816
939.4403686523438 0 12789.003
940.4481201171875 0 7953.124
941.4580688476562 0 3566.1594
942.475341796875 0 836.9649
953.4698486328125 0 867.0352
954.4716186523438 0 855.4793
956.4478149414062 0 3697.981
957.4607543945312 0 1987.2032
958.486328125 0 8360.069 b 8
959.4898681640625 0 7277.1377
960.4888916015625 0 10982.342
961.4907836914062 0 4553.045
971.4707641601562 0 2589.8455
986.479736328125 0 1055.2332 y Ammonia loss 8
987.4743041992188 0 1038.7485
988.46533203125 0 2197.0813
989.470458984375 0 1566.9176
1000.4906616210938 0 1619.4943
1003.5055541992188 0 1846.8287 y 8
1004.5040893554688 0 1006.85077
1008.4617919921875 0 841.89276
1015.4712524414062 0 1748.7062
1016.4765014648438 0 1614.2306
1025.488525390625 0 1267.728
1036.463134765625 0 2652.8625
1037.4669189453125 0 939.7913
1040.5001220703125 0 917.94507
1041.49365234375 0 1639.6752
1052.5001220703125 0 924.3046
1053.482421875 0 10243.741
1054.48388671875 0 6234.7456
1055.4871826171875 0 2257.839
1056.48388671875 0 873.83777
1058.5107421875 0 3160.4248
1059.519287109375 0 2583.1719
1060.52392578125 0 878.0395
1067.4949951171875 0 1098.6205
1069.51416015625 0 3245.98 b Water loss 9
1070.5123291015625 0 5571.354
1071.511474609375 0 3674.7256
1072.5142822265625 0 1643.921 y Water loss 7
1075.5203857421875 0 4120.004
1076.5205078125 0 2229.093
1084.5250244140625 0 1620.9835
1085.525390625 0 1830.542
1086.509033203125 0 2496.8032
1087.527099609375 0 24335.816 b 9
1088.5306396484375 0 13254.964
1089.5328369140625 0 5817.6147
1090.5341796875 0 2884.1643 y 7
1101.541748046875 0 1540.7356
1103.52490234375 0 5291.6445
1104.5244140625 0 2728.5146
1105.5367431640625 0 814.02496
1127.56396484375 0 751.06744
1129.5518798828125 0 1059.3918
1137.5517578125 0 1592.7036
1138.5479736328125 0 2491.625
1139.5604248046875 0 1058.068
1155.5621337890625 0 4345.692
1156.5616455078125 0 2540.4868
1157.5576171875 0 1123.7386
1159.56298828125 0 921.92145 y Water loss 6
1160.5504150390625 0 801.9225 y Ammonia loss 6
1164.55517578125 0 3165.5671
1165.5570068359375 0 2010.9062
1177.56982421875 0 1680.2083 y 6
1178.5662841796875 0 1067.8573
1181.5772705078125 0 9552.443
1182.580810546875 0 5708.3003 b Water loss 10
1183.58056640625 0 2834.1577
1200.605224609375 0 3910.2686 b 10
1201.6092529296875 0 2024.8105
1202.6046142578125 0 783.30347
1212.6219482421875 0 1382.3577
1217.581787109375 0 1362.2386
1234.601318359375 0 5886.266
1235.6007080078125 0 4675.2393
1236.6002197265625 0 2053.128
1252.6114501953125 0 14610.117
1253.6143798828125 0 10093.895
1254.617431640625 0 4775.21
1255.629638671875 0 1008.7074
1256.6124267578125 0 2513.253 y Water loss 5
1257.59814453125 0 1985.3677 y Ammonia loss 5
1258.6041259765625 0 1491.6366
1274.617919921875 0 8008.908 y 5
1275.6209716796875 0 4616.4624
1276.6246337890625 0 1086.0095
1328.6632080078125 0 1732.5989 b 11
1329.66064453125 0 1101.3218
1353.6541748046875 0 904.3754 y Water loss 4
1371.6700439453125 0 2683.8257 y 4
1372.672607421875 0 2420.9001
1373.661376953125 0 894.0872
1399.6981201171875 0 1835.3975 b 12
1400.6959228515625 0 1715.6418
1513.751953125 0 1230.7931 b 13
1514.7613525390625 0 1032.2653
2225.93115234375 0 612.80585
3070.04736328125 0 839.68494
3394.049072265625 0 676.2764

Spectrum Details

|  |  |
| --- | --- |
| Matched peaks? Matched peaks The total absolute number of peaks matched. Additionally in brackets the total fraction of peaks matched and the total number of peaks is shown. | 102 (12.85% of 794) |
| FDR? FDR The false discovery rate estimated for this peptide. It is calculated by matching all theoretical fragments with a non-integer shift with the raw peaks for this spectrum. This is done with 40 different shifts. The resulting percentage is the average number of annotated peaks over the number of annotated peaks with the correct spectrum. | 0.84% |
| Satellite FDR? Satellite FDR See the FDR for details on its calculation. This satellite ion specific FDR only contains the satellite ions (d/w) for I/L/J positions. | ∞ |
| PSM Score? PSM Score The PSM Score as given by Hecklib to this annotated spectrum. It is shown with three significant figures. | 859 |

#### Spectrum 8383? Spectrum 8383 The raw spectrum of this peptide as annotated by Hecklib. The fragments are coloured according to ion type (see legend). Any peaks with a star '\*' as text can be hovered over to see the full details, first the ion type second the mass shift type. By hovering over the amino acids in the peptide or ions in the legend the corresponding peaks are highlighted. By toggling the 'Unassigned' label you can turn the background (unassigned) peaks on or off in the plot. By updating the slider in the Ion legend you can update the spectrum to only show the top X% of the peaks with labels. The top X% means any peak that is within X% of the highest intensity. By dragging in the spectrum you can zoom in to a specific part of the spectrum and use 'Zoom Out' to get back to the original zoom level. The annotation of the spectrum is based on the given sequence in the peptides file and is done with different software so inconsistencies are likely. The peaks are annotated based on the given sequence, with 20 ppm tolerance.

Copy Data

##### Spectrum 8383 (TSV)

###### Preview

```
Loading example...
```

*Click on the button to copy the data to your clipboard.*

Mz MinMz MaxIntensity Max

WidthHeightPeptide font sizePeptide stroke widthSpectrum font sizeSpectrum stroke widthCompact peptide

Ion legend

wxyz

abcd

OtherUnassignedIon^Charge^_Position_Show for top:%

VTJFPPSSEEJQANKAT

06.41e+41.28e+51.92e+52.56e+5

Zoom Out

y^+1^_3_y^+1^_4_y^+1^_5_y^+1^_6_z^+1^_6_y^+1^_6_y^+2^_12_y^+2^_13_y^+2^_13_z^+2^_13_y^+2^_13_c^+2^_13_z^+1^_7_y^+1^_7_y^+2^_14_y^+2^_14_y^+2^_15_c^+2^_16_y^+1^_8_y^+2^_16_y^+2^_16_y^+2^_16_y^+1^_8_z^+1^_9_y^+1^_9_z^+1^_10_y^+1^_10_z^+1^_10_y^+1^_10_c^+1^_10_z^+1^_11_y^+1^_11_c^+1^_11_y^+1^_12_c^+1^_12_c^+1^_12_y^+1^_13_y^+1^_13_z^+1^_13_y^+1^_13_c^+1^_13_z^+1^_14_c^+1^_14_y^+1^_14_c^+1^_14_z^+1^_15_y^+1^_15_c^+1^_15_z^+1^_16_y^+1^_16_y^+1^_16_z^+1^_16_c^+1^_16_

046492813911855

Fragment Matches Table

Show background peaks

| Position | Ion type | Intensity | mz Theoretical | mz Error (Th) | mz Error (ppm) | Charge | Series Number |
| --- | --- | --- | --- | --- | --- | --- | --- |
| - | - | 667.7 | 120.1 | - | - | 0 | - |
| - | - | 413.1 | 127.1 | - | - | 0 | - |
| - | - | 1060 | 129.1 | - | - | 0 | - |
| - | - | 390 | 131.8 | - | - | 0 | - |
| - | - | 445.1 | 148.8 | - | - | 0 | - |
| - | - | 465.6 | 148.9 | - | - | 0 | - |
| - | - | 447.5 | 148.9 | - | - | 0 | - |
| - | - | 675.9 | 148.9 | - | - | 0 | - |
| - | - | 668.8 | 148.9 | - | - | 0 | - |
| - | - | 676.3 | 148.9 | - | - | 0 | - |
| - | - | 789.1 | 148.9 | - | - | 0 | - |
| - | - | 1157 | 148.9 | - | - | 0 | - |
| - | - | 1482 | 148.9 | - | - | 0 | - |
| - | - | 3225 | 148.9 | - | - | 0 | - |
| - | - | 5124 | 148.9 | - | - | 0 | - |
| - | - | 3883 | 149 | - | - | 0 | - |
| - | - | 2015 | 149 | - | - | 0 | - |
| - | - | 1325 | 149 | - | - | 0 | - |
| - | - | 1027 | 149 | - | - | 0 | - |
| - | - | 602.6 | 149 | - | - | 0 | - |
| - | - | 439.7 | 149 | - | - | 0 | - |
| - | - | 627.5 | 149 | - | - | 0 | - |
| - | - | 431.7 | 161.6 | - | - | 0 | - |
| - | - | 466.8 | 167.7 | - | - | 0 | - |
| - | - | 569.3 | 173.1 | - | - | 0 | - |
| - | - | 805.9 | 173.4 | - | - | 0 | - |
| - | - | 532.2 | 174.1 | - | - | 0 | - |
| - | - | 466.7 | 180.2 | - | - | 0 | - |
| - | - | 933.8 | 183.1 | - | - | 0 | - |
| - | - | 483.4 | 186.2 | - | - | 0 | - |
| - | - | 432.8 | 194.6 | - | - | 0 | - |
| - | - | 3510 | 201.1 | - | - | 0 | - |
| - | - | 596.4 | 217.1 | - | - | 0 | - |
| - | - | 1861 | 233.2 | - | - | 0 | - |
| - | - | 610.7 | 243.1 | - | - | 0 | - |
| - | - | 481.8 | 245 | - | - | 0 | - |
| - | - | 1025 | 251.2 | - | - | 0 | - |
| - | - | 933.4 | 261.2 | - | - | 0 | - |
| - | - | 625.2 | 291.3 | - | - | 0 | - |
| - | - | 3.151E+04 | 296.2 | - | - | 0 | - |
| - | - | 4906 | 297.2 | - | - | 0 | - |
| - | - | 797.7 | 298.2 | - | - | 0 | - |
| - | - | 553.5 | 301.2 | - | - | 0 | - |
| - | - | 1.642E+04 | 314.2 | - | - | 0 | - |
| - | - | 2745 | 315.2 | - | - | 0 | - |
| - | - | 700.8 | 316.2 | - | - | 0 | - |
| 15 | y | 1820 | 319.2 | 8.665E-05 | 0.2715 | +1 | 3 |
| - | - | 568.2 | 320.5 | - | - | 0 | - |
| - | - | 787.9 | 330.2 | - | - | 0 | - |
| - | - | 1129 | 346.2 | - | - | 0 | - |
| - | - | 1556 | 348.2 | - | - | 0 | - |
| - | - | 5030 | 362.2 | - | - | 0 | - |
| - | - | 759.5 | 363.2 | - | - | 0 | - |
| - | - | 1477 | 411.3 | - | - | 0 | - |
| - | - | 565 | 415.6 | - | - | 0 | - |
| - | - | 3233 | 416.3 | - | - | 0 | - |
| 14 | y | 1922 | 433.2 | 0.00206 | 4.756 | +1 | 4 |
| - | - | 2.302E+04 | 433.3 | - | - | 0 | - |
| - | - | 7060 | 434.3 | - | - | 0 | - |
| - | - | 791.8 | 443.2 | - | - | 0 | - |
| - | - | 2103 | 443.3 | - | - | 0 | - |
| - | - | 2.475E+04 | 461.3 | - | - | 0 | - |
| - | - | 7326 | 462.3 | - | - | 0 | - |
| - | - | 710.1 | 463.3 | - | - | 0 | - |
| 13 | y | 3262 | 504.3 | 0.0005075 | 1.006 | +1 | 5 |
| - | - | 868.2 | 505.3 | - | - | 0 | - |
| - | - | 654.2 | 505.9 | - | - | 0 | - |
| - | - | 622.3 | 530.3 | - | - | 0 | - |
| - | - | 3113 | 558.3 | - | - | 0 | - |
| - | - | 1075 | 559.3 | - | - | 0 | - |
| - | - | 655.1 | 569.3 | - | - | 0 | - |
| - | - | 619.5 | 570.3 | - | - | 0 | - |
| 12 | y | 624.8 | 615.3 | 0.003575 | 5.81 | +1 | 6 |
| 12 | z | 855.5 | 616.3 | 0.00316 | 5.127 | +1 | 6 |
| - | - | 818.5 | 618.3 | - | - | 0 | - |
| - | - | 3723 | 626.8 | - | - | 0 | - |
| - | - | 840.2 | 627.3 | - | - | 0 | - |
| - | - | 2749 | 627.3 | - | - | 0 | - |
| - | - | 1849 | 627.8 | - | - | 0 | - |
| 12 | y | 3083 | 632.3 | 0.001065 | 1.684 | +1 | 6 |
| - | - | 667.7 | 633.3 | - | - | 0 | - |
| 6 | y | 639.4 | 637.8 | 0.0003433 | 0.5382 | +2 | 12 |
| - | - | 769.1 | 641.4 | - | - | 0 | - |
| - | - | 1092 | 668.4 | - | - | 0 | - |
| - | - | 836.8 | 669.4 | - | - | 0 | - |
| 5 | y | 1557 | 677.3 | 0.0001875 | 0.2768 | +2 | 13 |
| 5 | y | 3019 | 677.8 | 0.004936 | 7.282 | +2 | 13 |
| 5 | z | 1661 | 678.3 | 0.0002301 | 0.3393 | +2 | 13 |
| - | - | 702.1 | 685.3 | - | - | 0 | - |
| 5 | y | 8.169E+04 | 686.3 | 0.0001454 | 0.2119 | +2 | 13 |
| - | - | 6.797E+04 | 686.8 | - | - | 0 | - |
| - | - | 2.769E+04 | 687.3 | - | - | 0 | - |
| - | - | 6303 | 687.8 | - | - | 0 | - |
| 13 | c | 1346 | 700.4 | 0.01312 | 18.74 | +2 | 13 |
| - | - | 641.1 | 700.9 | - | - | 0 | - |
| 11 | z | 855 | 729.4 | 0.0001879 | 0.2576 | +1 | 7 |
| - | - | 1436 | 730.4 | - | - | 0 | - |
| - | - | 980.7 | 740.4 | - | - | 0 | - |
| 11 | y | 1535 | 745.4 | 0.0006624 | 0.8887 | +1 | 7 |
| 4 | y | 853.5 | 750.9 | 0.00186 | 2.478 | +2 | 14 |
| 4 | y | 9582 | 759.9 | 0.0006064 | 0.7981 | +2 | 14 |
| - | - | 1.198E+04 | 760.4 | - | - | 0 | - |
| - | - | 5365 | 760.9 | - | - | 0 | - |
| - | - | 1398 | 761.4 | - | - | 0 | - |
| 3 | y | 2280 | 816.4 | 0.0004099 | 0.5021 | +2 | 15 |
| - | - | 3021 | 816.9 | - | - | 0 | - |
| - | - | 775.7 | 817.4 | - | - | 0 | - |
| - | - | 1009 | 848.4 | - | - | 0 | - |
| 16 | c | 2158 | 856.9 | 0.0002818 | 0.3288 | +2 | 16 |
| 10 | y | 2287 | 857.4 | 0.01045 | 12.19 | +1 | 8 |
| 2 | y | 1652 | 857.9 | 0.006973 | 8.128 | +2 | 16 |
| 2 | y | 4314 | 858.4 | 0.0157 | 18.29 | +2 | 16 |
| - | - | 3918 | 859.4 | - | - | 0 | - |
| - | - | 761.5 | 860.4 | - | - | 0 | - |
| 2 | y | 692 | 866.9 | 0.00462 | 5.33 | +2 | 16 |
| - | - | 684.9 | 867.4 | - | - | 0 | - |
| - | - | 1297 | 868.4 | - | - | 0 | - |
| 10 | y | 1240 | 874.5 | 0.003986 | 4.558 | +1 | 8 |
| - | - | 1543 | 907.5 | - | - | 0 | - |
| - | - | 2877 | 908 | - | - | 0 | - |
| - | - | 1198 | 908.5 | - | - | 0 | - |
| - | - | 1.644E+04 | 916.5 | - | - | 0 | - |
| - | - | 2.27E+04 | 917 | - | - | 0 | - |
| - | - | 1.396E+04 | 917.5 | - | - | 0 | - |
| - | - | 4702 | 918 | - | - | 0 | - |
| - | - | 777.9 | 939.4 | - | - | 0 | - |
| 9 | z | 6181 | 987.5 | 0.0006855 | 0.6942 | +1 | 9 |
| - | - | 9622 | 988.5 | - | - | 0 | - |
| - | - | 4734 | 989.5 | - | - | 0 | - |
| - | - | 1449 | 990.5 | - | - | 0 | - |
| - | - | 831.7 | 1002 | - | - | 0 | - |
| 9 | y | 2007 | 1004 | 8.886E-05 | 0.08855 | +1 | 9 |
| - | - | 888.5 | 1005 | - | - | 0 | - |
| - | - | 3900 | 1040 | - | - | 0 | - |
| - | - | 2014 | 1041 | - | - | 0 | - |
| - | - | 960.8 | 1053 | - | - | 0 | - |
| - | - | 792.4 | 1054 | - | - | 0 | - |
| 8 | z | 1050 | 1057 | 0.02102 | 19.9 | +1 | 10 |
| - | - | 764.9 | 1072 | - | - | 0 | - |
| 8 | y | 1264 | 1074 | 4.168E-05 | 0.03882 | +1 | 10 |
| 8 | z | 7927 | 1075 | 0.0007033 | 0.6545 | +1 | 10 |
| - | - | 2.006E+04 | 1076 | - | - | 0 | - |
| - | - | 9284 | 1077 | - | - | 0 | - |
| - | - | 2866 | 1078 | - | - | 0 | - |
| - | - | 3083 | 1090 | - | - | 0 | - |
| 8 | y | 7535 | 1091 | 0.0004701 | 0.4311 | +1 | 10 |
| - | - | 3959 | 1092 | - | - | 0 | - |
| - | - | 1848 | 1093 | - | - | 0 | - |
| 10 | c | 1753 | 1105 | 0.00216 | 1.955 | +1 | 10 |
| - | - | 840.3 | 1106 | - | - | 0 | - |
| - | - | 1217 | 1118 | - | - | 0 | - |
| - | - | 1158 | 1119 | - | - | 0 | - |
| 7 | z | 3229 | 1162 | 0.001482 | 1.276 | +1 | 11 |
| - | - | 6933 | 1163 | - | - | 0 | - |
| - | - | 4673 | 1164 | - | - | 0 | - |
| - | - | 1828 | 1165 | - | - | 0 | - |
| - | - | 964 | 1175 | - | - | 0 | - |
| 7 | y | 4773 | 1178 | 0.003116 | 2.646 | +1 | 11 |
| - | - | 3042 | 1179 | - | - | 0 | - |
| - | - | 1458 | 1182 | - | - | 0 | - |
| - | - | 1339 | 1183 | - | - | 0 | - |
| - | - | 1147 | 1202 | - | - | 0 | - |
| - | - | 2871 | 1217 | - | - | 0 | - |
| 11 | c | 5891 | 1218 | 0.0009718 | 0.7981 | +1 | 11 |
| - | - | 4308 | 1219 | - | - | 0 | - |
| - | - | 1053 | 1220 | - | - | 0 | - |
| - | - | 2734 | 1253 | - | - | 0 | - |
| - | - | 1776 | 1254 | - | - | 0 | - |
| - | - | 1704 | 1255 | - | - | 0 | - |
| 6 | y | 1.136E+04 | 1275 | 0.0009481 | 0.7438 | +1 | 12 |
| - | - | 6147 | 1276 | - | - | 0 | - |
| - | - | 3192 | 1277 | - | - | 0 | - |
| - | - | 721.6 | 1303 | - | - | 0 | - |
| 12 | c | 1919 | 1329 | 0.01103 | 8.3 | +1 | 12 |
| - | - | 1896 | 1330 | - | - | 0 | - |
| - | - | 3731 | 1345 | - | - | 0 | - |
| 12 | c | 1.511E+04 | 1346 | 0.001566 | 1.164 | +1 | 12 |
| - | - | 1.203E+04 | 1347 | - | - | 0 | - |
| - | - | 4969 | 1348 | - | - | 0 | - |
| - | - | 1471 | 1349 | - | - | 0 | - |
| 5 | y | 1552 | 1354 | 0.002742 | 2.026 | +1 | 13 |
| 5 | y | 2833 | 1355 | 0.006406 | 4.729 | +1 | 13 |
| 5 | z | 1436 | 1356 | 0.007644 | 5.639 | +1 | 13 |
| - | - | 6931 | 1371 | - | - | 0 | - |
| 5 | y | 1.025E+05 | 1372 | 0.001222 | 0.8907 | +1 | 13 |
| - | - | 8.115E+04 | 1373 | - | - | 0 | - |
| - | - | 3.563E+04 | 1374 | - | - | 0 | - |
| - | - | 7279 | 1375 | - | - | 0 | - |
| - | - | 1644 | 1399 | - | - | 0 | - |
| - | - | 3279 | 1400 | - | - | 0 | - |
| - | - | 1888 | 1401 | - | - | 0 | - |
| - | - | 1.839E+04 | 1402 | - | - | 0 | - |
| - | - | 1.573E+04 | 1403 | - | - | 0 | - |
| - | - | 7488 | 1404 | - | - | 0 | - |
| - | - | 1840 | 1405 | - | - | 0 | - |
| - | - | 1.051E+04 | 1416 | - | - | 0 | - |
| 13 | c | 1.875E+04 | 1417 | 0.004378 | 3.09 | +1 | 13 |
| - | - | 1.232E+04 | 1418 | - | - | 0 | - |
| - | - | 3595 | 1419 | - | - | 0 | - |
| - | - | 991.7 | 1420 | - | - | 0 | - |
| - | - | 883.3 | 1431 | - | - | 0 | - |
| - | - | 811.2 | 1432 | - | - | 0 | - |
| - | - | 947.4 | 1462 | - | - | 0 | - |
| - | - | 1158 | 1487 | - | - | 0 | - |
| - | - | 1827 | 1488 | - | - | 0 | - |
| - | - | 777 | 1489 | - | - | 0 | - |
| 4 | z | 6414 | 1503 | 0.001107 | 0.7366 | +1 | 14 |
| - | - | 8062 | 1504 | - | - | 0 | - |
| - | - | 4638 | 1505 | - | - | 0 | - |
| - | - | 1104 | 1506 | - | - | 0 | - |
| 14 | c | 944.8 | 1514 | 0.01135 | 7.497 | +1 | 14 |
| - | - | 824.6 | 1515 | - | - | 0 | - |
| 4 | y | 1.176E+04 | 1519 | 0.001154 | 0.76 | +1 | 14 |
| - | - | 1.15E+04 | 1520 | - | - | 0 | - |
| - | - | 5177 | 1521 | - | - | 0 | - |
| - | - | 1084 | 1522 | - | - | 0 | - |
| - | - | 1545 | 1530 | - | - | 0 | - |
| 14 | c | 2.538E+04 | 1531 | 0.001529 | 0.9988 | +1 | 14 |
| - | - | 2.693E+04 | 1532 | - | - | 0 | - |
| - | - | 1.339E+04 | 1533 | - | - | 0 | - |
| - | - | 3190 | 1534 | - | - | 0 | - |
| - | - | 1082 | 1572 | - | - | 0 | - |
| - | - | 2766 | 1615 | - | - | 0 | - |
| 3 | z | 6241 | 1616 | 0.03141 | 19.44 | +1 | 15 |
| - | - | 4109 | 1617 | - | - | 0 | - |
| - | - | 2569 | 1618 | - | - | 0 | - |
| 3 | y | 2626 | 1632 | 0.004041 | 2.477 | +1 | 15 |
| - | - | 3206 | 1633 | - | - | 0 | - |
| - | - | 2805 | 1634 | - | - | 0 | - |
| - | - | 1448 | 1635 | - | - | 0 | - |
| - | - | 791.6 | 1645 | - | - | 0 | - |
| - | - | 991.6 | 1646 | - | - | 0 | - |
| - | - | 949.4 | 1655 | - | - | 0 | - |
| 15 | c | 1.427E+04 | 1659 | 0.001888 | 1.138 | +1 | 15 |
| - | - | 1.348E+04 | 1660 | - | - | 0 | - |
| - | - | 8387 | 1661 | - | - | 0 | - |
| - | - | 3144 | 1662 | - | - | 0 | - |
| - | - | 1599 | 1663 | - | - | 0 | - |
| - | - | 1846 | 1672 | - | - | 0 | - |
| - | - | 2467 | 1673 | - | - | 0 | - |
| - | - | 1430 | 1674 | - | - | 0 | - |
| - | - | 3318 | 1686 | - | - | 0 | - |
| - | - | 4333 | 1687 | - | - | 0 | - |
| - | - | 4433 | 1688 | - | - | 0 | - |
| - | - | 1527 | 1689 | - | - | 0 | - |
| 2 | z | 1446 | 1699 | 0.001861 | 1.095 | +1 | 16 |
| - | - | 871.7 | 1701 | - | - | 0 | - |
| 2 | y | 4469 | 1715 | 0.004046 | 2.359 | +1 | 16 |
| 2 | y | 9816 | 1716 | 0.01267 | 7.385 | +1 | 16 |
| 2 | z | 3.273E+04 | 1717 | 0.0004514 | 0.2629 | +1 | 16 |
| - | - | 3.431E+04 | 1718 | - | - | 0 | - |
| - | - | 1.974E+04 | 1719 | - | - | 0 | - |
| - | - | 6680 | 1720 | - | - | 0 | - |
| - | - | 1480 | 1721 | - | - | 0 | - |
| - | - | 4429 | 1726 | - | - | 0 | - |
| - | - | 4715 | 1727 | - | - | 0 | - |
| - | - | 4357 | 1728 | - | - | 0 | - |
| - | - | 1690 | 1729 | - | - | 0 | - |
| 16 | c | 2.167E+04 | 1730 | 0.0001829 | 0.1057 | +1 | 16 |
| - | - | 2.261E+04 | 1731 | - | - | 0 | - |
| - | - | 1.888E+04 | 1732 | - | - | 0 | - |
| - | - | 1.069E+04 | 1733 | - | - | 0 | - |
| - | - | 3021 | 1734 | - | - | 0 | - |
| - | - | 1466 | 1735 | - | - | 0 | - |
| - | - | 1128 | 1741 | - | - | 0 | - |
| - | - | 2892 | 1742 | - | - | 0 | - |
| - | - | 2652 | 1743 | - | - | 0 | - |
| - | - | 3790 | 1744 | - | - | 0 | - |
| - | - | 3331 | 1745 | - | - | 0 | - |
| - | - | 2606 | 1746 | - | - | 0 | - |
| - | - | 1163 | 1747 | - | - | 0 | - |
| - | - | 955.2 | 1748 | - | - | 0 | - |
| - | - | 2446 | 1758 | - | - | 0 | - |
| - | - | 2577 | 1759 | - | - | 0 | - |
| - | - | 2523 | 1760 | - | - | 0 | - |
| - | - | 4329 | 1761 | - | - | 0 | - |
| - | - | 6114 | 1762 | - | - | 0 | - |
| - | - | 4558 | 1763 | - | - | 0 | - |
| - | - | 1479 | 1764 | - | - | 0 | - |
| - | - | 2496 | 1769 | - | - | 0 | - |
| - | - | 4262 | 1770 | - | - | 0 | - |
| - | - | 3368 | 1771 | - | - | 0 | - |
| - | - | 2317 | 1772 | - | - | 0 | - |
| - | - | 1978 | 1774 | - | - | 0 | - |
| - | - | 4080 | 1775 | - | - | 0 | - |
| - | - | 2099 | 1776 | - | - | 0 | - |
| - | - | 3667 | 1777 | - | - | 0 | - |
| - | - | 3640 | 1778 | - | - | 0 | - |
| - | - | 2162 | 1779 | - | - | 0 | - |
| - | - | 6742 | 1787 | - | - | 0 | - |
| - | - | 3.675E+04 | 1788 | - | - | 0 | - |
| - | - | 3.854E+04 | 1789 | - | - | 0 | - |
| - | - | 2.32E+04 | 1790 | - | - | 0 | - |
| - | - | 8539 | 1791 | - | - | 0 | - |
| - | - | 2294 | 1798 | - | - | 0 | - |
| - | - | 2714 | 1799 | - | - | 0 | - |
| - | - | 1704 | 1800 | - | - | 0 | - |
| - | - | 1197 | 1801 | - | - | 0 | - |
| - | - | 3149 | 1805 | - | - | 0 | - |
| - | - | 3266 | 1806 | - | - | 0 | - |
| - | - | 1934 | 1807 | - | - | 0 | - |
| - | - | 1839 | 1814 | - | - | 0 | - |
| - | - | 1.121E+04 | 1815 | - | - | 0 | - |
| - | - | 5.639E+04 | 1816 | - | - | 0 | - |
| - | - | 6.296E+04 | 1817 | - | - | 0 | - |
| - | - | 3.679E+04 | 1818 | - | - | 0 | - |
| - | - | 1.288E+04 | 1819 | - | - | 0 | - |
| - | - | 2195 | 1820 | - | - | 0 | - |
| - | - | 3145 | 1831 | - | - | 0 | - |
| - | - | 9.185E+04 | 1832 | - | - | 0 | - |
| - | - | 2.537E+05 | 1833 | - | - | 0 | - |
| - | - | 2.396E+05 | 1834 | - | - | 0 | - |
| - | - | 1.319E+05 | 1835 | - | - | 0 | - |
| - | - | 4.786E+04 | 1836 | - | - | 0 | - |
| - | - | 4865 | 1837 | - | - | 0 | - |

m/z Charge Intensity FragmentType MassShift Position
120.08097076416016 0 667.71234
127.0871353149414 0 413.1162
129.1024169921875 0 1059.9595
131.82553100585938 0 389.99207
148.8448486328125 0 445.13947
148.8520050048828 0 465.6042
148.85890197753906 0 447.5025
148.89501953125 0 675.8668
148.9020538330078 0 668.7721
148.90921020507812 0 676.34656
148.91615295410156 0 789.1273
148.92344665527344 0 1156.6019
148.93045043945312 0 1482.1593
148.9376678466797 0 3225.2527
148.9453125 0 5124.0864
148.9617156982422 0 3882.6335
148.96945190429688 0 2015.0428
148.9763946533203 0 1324.6584
148.98353576660156 0 1026.9263
148.99057006835938 0 602.5571
148.99755859375 0 439.71127
149.00506591796875 0 627.50287
161.58840942382812 0 431.65112
167.6685333251953 0 466.84845
173.12893676757812 0 569.32764
173.44149780273438 0 805.886
174.13807678222656 0 532.2284
180.15841674804688 0 466.70416
183.11297607421875 0 933.7723
186.19393920898438 0 483.38892
194.58059692382812 0 432.75607
201.12342834472656 0 3510.348
217.13363647460938 0 596.4215
233.1645965576172 0 1861.2104
243.14523315429688 0 610.6841
245.04498291015625 0 481.82315
251.17572021484375 0 1024.8513
261.15948486328125 0 933.3799
291.2903137207031 0 625.25
296.1969299316406 0 31514.521
297.1999206542969 0 4906.154
298.1763610839844 0 797.67786
301.1933288574219 0 553.5096
314.20751953125 0 16422.523
315.2107849121094 0 2745.2588
316.2144470214844 0 700.78723
319.197509765625 0 1819.5946 y 14
320.5184326171875 0 568.2479
330.18194580078125 0 787.9223
346.1772155761719 0 1129.1699
348.1919860839844 0 1556.0625
362.2073059082031 0 5030.139
363.2106018066406 0 759.507
411.261474609375 0 1476.9054
415.5987548828125 0 564.9816
416.2547912597656 0 3232.9358
433.2425842285156 0 1921.5834 y 13
433.2813720703125 0 23024.523
434.284423828125 0 7060.1045
443.2320861816406 0 791.8146
443.26605224609375 0 2103.081
461.2763977050781 0 24750.223
462.2795104980469 0 7325.652
463.28204345703125 0 710.08936
504.2771301269531 0 3262.4377 y 12
505.2794494628906 0 868.18994
505.8778076171875 0 654.1551
530.336669921875 0 622.32227
558.3284912109375 0 3112.5117
559.33203125 0 1074.7208
569.3310546875 0 655.0619
570.331787109375 0 619.4642
615.3060913085938 0 624.78485 y Ammonia loss 11
616.3143310546875 0 855.52625 z 11
618.3099365234375 0 818.4523
626.8123168945312 0 3723.291
627.26171875 0 840.23224
627.3140869140625 0 2748.9458
627.8145751953125 0 1848.6426
632.3372802734375 0 3082.5544 y 11
633.34228515625 0 667.66846
637.8151245117188 0 639.3556 y 5
641.3526000976562 0 769.0891
668.3970336914062 0 1091.9493
669.4005126953125 0 836.80096
677.335693359375 0 1557.4272 y Water loss 4
677.8328247070312 0 3019.0928 y Ammonia loss 4
678.33203125 0 1661.2261 z 4
685.3305053710938 0 702.1178
686.34130859375 0 81686.414 y 4
686.8424682617188 0 67974.94
687.343505859375 0 27685.729
687.8438720703125 0 6303.2886
700.345703125 0 1345.7404 c Ammonia loss 12
700.8534545898438 0 641.1246
729.4013671875 0 854.9726 z 10
730.4075317382812 0 1436.3605
740.4173583984375 0 980.65015
745.4196166992188 0 1535.4133 y 10
750.8719482421875 0 853.4789 y Water loss 3
759.8759765625 0 9581.903 y 3
760.3765869140625 0 11976.905
760.8773803710938 0 5364.85
761.3807983398438 0 1397.6581
816.4169921875 0 2279.6323 y 2
816.9183349609375 0 3021.1455
817.4168090820312 0 775.7099
848.434814453125 0 1008.89844
856.946044921875 0 2158.0732 c Ammonia loss 15
857.44677734375 0 2287.2107 y Ammonia loss 9
857.9429321289062 0 1651.9926 y Water loss 1
858.4436645507812 0 4313.579 y Ammonia loss 1
859.4479370117188 0 3917.8184
860.445068359375 0 761.4696
866.9458618164062 0 691.9818 y 1
867.4463500976562 0 684.9235
868.4068603515625 0 1297.3943
874.4668579101562 0 1239.9381 y 9
907.470703125 0 1543.4188
907.970458984375 0 2877.0942
908.4754638671875 0 1198.2911
916.4755859375 0 16435.398
916.976318359375 0 22700.316
917.477783203125 0 13958.24
917.9789428710938 0 4701.599
939.4368896484375 0 777.8783
987.4874267578125 0 6180.6626 z 8
988.49267578125 0 9622.441
989.49462890625 0 4733.684
990.4990844726562 0 1449.4968
1002.4889526367188 0 831.7115
1003.5055541992188 0 2006.7759 y 8
1004.5043334960938 0 888.5277
1040.4893798828125 0 3899.6982
1041.4908447265625 0 2013.9495
1053.48583984375 0 960.7782
1054.486328125 0 792.38617
1056.4871826171875 0 1049.9988 z Water loss 7
1072.4908447265625 0 764.8964
1073.510986328125 0 1263.8367 y Ammonia loss 7
1074.51806640625 0 7927.4707 z 7
1075.525390625 0 20057.521
1076.5279541015625 0 9284.403
1077.527587890625 0 2865.8335
1089.5301513671875 0 3083.2097
1090.5379638671875 0 7534.8037 y 7
1091.53857421875 0 3959.0095
1092.5438232421875 0 1848.4615
1104.559326171875 0 1752.5149 c 9
1105.5640869140625 0 840.2675
1117.5338134765625 0 1216.6278
1118.540283203125 0 1158.4039
1161.54931640625 0 3228.5522 z 6
1162.5550537109375 0 6933.211
1163.5621337890625 0 4672.6543
1164.5582275390625 0 1827.8828
1174.6549072265625 0 964.0466
1177.56640625 0 4772.832 y 6
1178.5701904296875 0 3041.637
1181.5830078125 0 1458.1377
1182.575927734375 0 1338.6873
1201.6142578125 0 1147.0732
1216.6317138671875 0 2871.3662
1217.6402587890625 0 5891.3325 c 10
1218.644287109375 0 4307.9707
1219.650146484375 0 1053.0923
1252.617919921875 0 2733.6846
1253.62060546875 0 1776.1008
1254.6185302734375 0 1703.8934
1274.621337890625 0 11363.669 y 5
1275.625 0 6147.15
1276.6270751953125 0 3191.7932
1302.6826171875 0 721.55835
1328.6622314453125 0 1919.1925 c Ammonia loss 11
1329.67724609375 0 1895.559
1344.690673828125 0 3731.1707
1345.6982421875 0 15109.761 c 11
1346.70166015625 0 12025.325
1347.7034912109375 0 4969.487
1348.716552734375 0 1470.768
1353.6617431640625 0 1552.2509 y Water loss 4
1354.6549072265625 0 2833.2708 y Ammonia loss 4
1355.648681640625 0 1435.8679 z 4
1370.666259765625 0 6930.526
1371.673828125 0 102453.04 y 4
1372.676513671875 0 81147.62
1373.67919921875 0 35626.12
1374.681640625 0 7279.2764
1398.6778564453125 0 1644.1631
1399.6810302734375 0 3279.022
1400.677490234375 0 1888.4287
1401.7001953125 0 18392.316
1402.7034912109375 0 15733.546
1403.707275390625 0 7487.714
1404.714111328125 0 1839.8615
1415.72802734375 0 10507.894
1416.7325439453125 0 18752.232 c 12
1417.737548828125 0 12322.883
1418.7413330078125 0 3595.484
1419.7529296875 0 991.6754
1430.7103271484375 0 883.3453
1431.704345703125 0 811.23474
1461.7845458984375 0 947.42444
1486.7635498046875 0 1157.6483
1487.7684326171875 0 1826.6055
1488.7703857421875 0 777.014
1502.7236328125 0 6413.5757 z 3
1503.72216796875 0 8062.4004
1504.7239990234375 0 4637.8345
1505.728759765625 0 1104.4985
1513.7646484375 0 944.8239 c Ammonia loss 13
1514.751953125 0 824.5858
1518.7423095703125 0 11756.067 y 3
1519.74365234375 0 11497.741
1520.74609375 0 5177.3477
1521.748291015625 0 1084.1787
1529.774169921875 0 1544.72
1530.7783203125 0 25378.06 c 13
1531.78076171875 0 26931.945
1532.78369140625 0 13388.544
1533.7864990234375 0 3190.4539
1571.8270263671875 0 1081.9517
1614.8621826171875 0 2765.8774
1615.8402099609375 0 6241.088 z 2
1616.8280029296875 0 4108.9736
1617.82470703125 0 2569.2944
1631.823486328125 0 2626.0542 y 2
1632.8310546875 0 3206.4326
1633.8375244140625 0 2804.7485
1634.8388671875 0 1448.3435
1644.8486328125 0 791.6089
1645.82177734375 0 991.5971
1654.8621826171875 0 949.4041
1658.8729248046875 0 14267.054 c 14
1659.8740234375 0 13481.379
1660.866943359375 0 8386.937
1661.8406982421875 0 3144.4978
1662.8033447265625 0 1598.935
1671.838623046875 0 1845.8171
1672.83984375 0 2467.4653
1673.8604736328125 0 1430.3198
1685.8958740234375 0 3317.6792
1686.8984375 0 4332.8203
1687.9046630859375 0 4432.821
1688.9168701171875 0 1526.9867
1698.8477783203125 0 1446.0273 z Water loss 1
1700.8427734375 0 871.69476
1714.860595703125 0 4469.2163 y Water loss 1
1715.861328125 0 9816.003 y Ammonia loss 1
1716.85693359375 0 32727.121 z 1
1717.8585205078125 0 34311.47
1718.8623046875 0 19739.67
1719.8616943359375 0 6679.7793
1720.870361328125 0 1480.267
1725.88671875 0 4428.772
1726.8929443359375 0 4714.5693
1727.8974609375 0 4356.8003
1728.8968505859375 0 1689.9285
1729.9117431640625 0 21670.36 c 15
1730.9134521484375 0 22612.588
1731.918212890625 0 18882.006
1732.9210205078125 0 10693.654
1733.9246826171875 0 3021.183
1734.9228515625 0 1466.0907
1740.8720703125 0 1127.8113
1741.882568359375 0 2891.7432
1742.88623046875 0 2651.7766
1743.90234375 0 3790.4294
1744.8885498046875 0 3330.5583
1745.8974609375 0 2605.6233
1746.9080810546875 0 1163.0903
1747.909423828125 0 955.22406
1757.880126953125 0 2445.7517
1758.87646484375 0 2577.3843
1759.889404296875 0 2522.7502
1760.914306640625 0 4328.6187
1761.9156494140625 0 6114.166
1762.9093017578125 0 4557.5977
1763.9151611328125 0 1478.7728
1768.9263916015625 0 2495.998
1769.934326171875 0 4262.011
1770.9332275390625 0 3368.3774
1771.9267578125 0 2317.047
1773.90234375 0 1978.2987
1774.909912109375 0 4079.995
1775.9210205078125 0 2098.9631
1776.89697265625 0 3666.692
1777.88916015625 0 3640.4163
1778.8839111328125 0 2161.7761
1786.939208984375 0 6742.2397
1787.9327392578125 0 36748.562
1788.934326171875 0 38536.266
1789.934814453125 0 23196.705
1790.9312744140625 0 8538.942
1797.9058837890625 0 2293.531
1798.900634765625 0 2714.3474
1799.9061279296875 0 1704.1903
1800.9134521484375 0 1197.2156
1804.95361328125 0 3149.0125
1805.9571533203125 0 3266.2979
1806.96240234375 0 1933.9846
1813.89404296875 0 1839.1711
1814.93310546875 0 11209.316
1815.9271240234375 0 56391.8
1816.9281005859375 0 62955.45
1817.9293212890625 0 36790.902
1818.9300537109375 0 12875.078
1819.928955078125 0 2195.2314
1830.9178466796875 0 3145.0935
1831.943359375 0 91849.664
1832.9488525390625 0 253700.22
1833.951904296875 0 239632.19
1834.95458984375 0 131905.7
1835.9560546875 0 47855.93
1836.95849609375 0 4865.314

Spectrum Details

|  |  |
| --- | --- |
| Matched peaks? Matched peaks The total absolute number of peaks matched. Additionally in brackets the total fraction of peaks matched and the total number of peaks is shown. | 53 (16.83% of 315) |
| FDR? FDR The false discovery rate estimated for this peptide. It is calculated by matching all theoretical fragments with a non-integer shift with the raw peaks for this spectrum. This is done with 40 different shifts. The resulting percentage is the average number of annotated peaks over the number of annotated peaks with the correct spectrum. | 0.63% |
| Satellite FDR? Satellite FDR See the FDR for details on its calculation. This satellite ion specific FDR only contains the satellite ions (d/w) for I/L/J positions. | - |
| PSM Score? PSM Score The PSM Score as given by Hecklib to this annotated spectrum. It is shown with three significant figures. | 488 |

#### Spectrum 8223? Spectrum 8223 The raw spectrum of this peptide as annotated by Hecklib. The fragments are coloured according to ion type (see legend). Any peaks with a star '\*' as text can be hovered over to see the full details, first the ion type second the mass shift type. By hovering over the amino acids in the peptide or ions in the legend the corresponding peaks are highlighted. By toggling the 'Unassigned' label you can turn the background (unassigned) peaks on or off in the plot. By updating the slider in the Ion legend you can update the spectrum to only show the top X% of the peaks with labels. The top X% means any peak that is within X% of the highest intensity. By dragging in the spectrum you can zoom in to a specific part of the spectrum and use 'Zoom Out' to get back to the original zoom level. The annotation of the spectrum is based on the given sequence in the peptides file and is done with different software so inconsistencies are likely. The peaks are annotated based on the given sequence, with 20 ppm tolerance.

Copy Data

##### Spectrum 8223 (TSV)

###### Preview

```
Loading example...
```

*Click on the button to copy the data to your clipboard.*

Mz MinMz MaxIntensity Max

WidthHeightPeptide font sizePeptide stroke widthSpectrum font sizeSpectrum stroke widthCompact peptide

Ion legend

wxyz

abcd

OtherUnassignedIon^Charge^_Position_Show for top:%

VTJFPPSSEEJQANKAT

01.62e+53.23e+54.85e+56.46e+5

Zoom Out

y^+1^_1_a^+1^_2_d^+1^_2_y^+1^_2_a^+1^_2_b^+1^_2_y^+1^_2_b^+1^_2_y^+2^_5_a^+1^_3_a^+1^_3_b^+1^_3_y^+1^_3_y^+1^_3_y^+2^_6_b^+1^_3_y^+1^_3_b^+2^_8_y^+1^_4_y^+2^_8_y^+1^_4_b^+1^_4_b^+1^_4_y^+1^_5_y^+1^_5_y^+1^_5_b^+1^_5_b^+1^_5_b^+2^_11_y^+1^_6_y^+1^_6_y^+2^_12_y^+1^_6_y^+2^_12_b^+2^_12_b^+2^_12_y^+2^_13_y^+2^_13_y^+2^_13_b^+2^_13_b^+2^_13_b^+2^_13_b^+1^_7_y^+1^_7_y^+1^_7_b^+1^_7_y^+1^_7_y^+2^_14_y^+2^_14_b^+2^_14_y^+2^_14_y^+2^_15_y^+2^_15_b^+2^_15_y^+2^_15_b^+2^_15_b^+1^_8_b^+2^_16_b^+2^_16_y^+1^_8_b^+2^_16_y^+1^_8_y^+2^_16_y^+2^_16_y^+1^_8_\*\*\*b^+1^_9_y^+1^_9_y^+1^_9_b^+1^_10_y^+1^_10_y^+1^_10_b^+1^_10_y^+1^_10_y^+1^_11_y^+1^_11_y^+1^_11_b^+1^_11_b^+1^_11_y^+1^_12_y^+1^_12_y^+1^_12_b^+1^_12_b^+1^_12_b^+1^_12_y^+1^_13_y^+1^_13_y^+1^_13_b^+1^_13_b^+1^_13_b^+1^_13_b^+1^_14_y^+1^_14_y^+1^_14_b^+1^_14_y^+1^_14_y^+1^_15_

0776155223283104

Fragment Matches Table

Show background peaks

| Position | Ion type | Intensity | mz Theoretical | mz Error (Th) | mz Error (ppm) | Charge | Series Number |
| --- | --- | --- | --- | --- | --- | --- | --- |
| 17 | y | 3398 | 120.1 | 0.0002992 | 2.492 | +1 | 1 |
| - | - | 5.826E+04 | 120.1 | - | - | 0 | - |
| - | - | 3846 | 121.1 | - | - | 0 | - |
| - | - | 474 | 125.9 | - | - | 0 | - |
| - | - | 1.148E+04 | 126.1 | - | - | 0 | - |
| - | - | 679.6 | 127.1 | - | - | 0 | - |
| - | - | 861.4 | 127.1 | - | - | 0 | - |
| - | - | 1.549E+04 | 128.1 | - | - | 0 | - |
| - | - | 3248 | 129.1 | - | - | 0 | - |
| - | - | 1.032E+05 | 129.1 | - | - | 0 | - |
| - | - | 2588 | 130.1 | - | - | 0 | - |
| - | - | 915.5 | 130.1 | - | - | 0 | - |
| - | - | 6903 | 130.1 | - | - | 0 | - |
| - | - | 2193 | 131.1 | - | - | 0 | - |
| - | - | 1693 | 132.1 | - | - | 0 | - |
| - | - | 476.8 | 132.1 | - | - | 0 | - |
| - | - | 1045 | 136.1 | - | - | 0 | - |
| - | - | 503.3 | 137.1 | - | - | 0 | - |
| - | - | 3180 | 138.1 | - | - | 0 | - |
| - | - | 8457 | 138.1 | - | - | 0 | - |
| - | - | 4474 | 139.1 | - | - | 0 | - |
| - | - | 652.5 | 139.1 | - | - | 0 | - |
| - | - | 691.1 | 139.1 | - | - | 0 | - |
| - | - | 602.9 | 140.1 | - | - | 0 | - |
| - | - | 523 | 140.1 | - | - | 0 | - |
| - | - | 610.3 | 140.1 | - | - | 0 | - |
| - | - | 1573 | 141.1 | - | - | 0 | - |
| - | - | 1532 | 141.1 | - | - | 0 | - |
| - | - | 656 | 142.1 | - | - | 0 | - |
| - | - | 3082 | 147.1 | - | - | 0 | - |
| - | - | 678.8 | 151.1 | - | - | 0 | - |
| - | - | 554.6 | 152.1 | - | - | 0 | - |
| - | - | 1295 | 153.1 | - | - | 0 | - |
| - | - | 558.8 | 153.5 | - | - | 0 | - |
| - | - | 2287 | 154.1 | - | - | 0 | - |
| - | - | 895.6 | 154.1 | - | - | 0 | - |
| - | - | 2816 | 155.1 | - | - | 0 | - |
| 2 | a | 1.484E+04 | 155.1 | 0.0003356 | 2.163 | +1 | 2 |
| - | - | 851.4 | 156.1 | - | - | 0 | - |
| - | - | 4204 | 157.1 | - | - | 0 | - |
| - | - | 1.234E+04 | 157.1 | - | - | 0 | - |
| 2 | d | 745.1 | 157.1 | 0.0007988 | 5.083 | +1 | 2 |
| - | - | 2533 | 165.1 | - | - | 0 | - |
| - | - | 1811 | 166.1 | - | - | 0 | - |
| - | - | 9349 | 167.1 | - | - | 0 | - |
| - | - | 7071 | 167.1 | - | - | 0 | - |
| - | - | 628.7 | 168.1 | - | - | 0 | - |
| - | - | 1452 | 169.1 | - | - | 0 | - |
| - | - | 869.6 | 169.1 | - | - | 0 | - |
| - | - | 1.429E+04 | 169.1 | - | - | 0 | - |
| - | - | 1680 | 170.1 | - | - | 0 | - |
| - | - | 1014 | 171.1 | - | - | 0 | - |
| - | - | 1753 | 172.1 | - | - | 0 | - |
| - | - | 563.6 | 172.1 | - | - | 0 | - |
| 16 | y | 3991 | 173.1 | 0.0003233 | 1.868 | +1 | 2 |
| 2 | a | 4.48E+04 | 173.1 | 0.0003605 | 2.082 | +1 | 2 |
| - | - | 5780 | 173.4 | - | - | 0 | - |
| - | - | 4451 | 174.1 | - | - | 0 | - |
| - | - | 4412 | 175.1 | - | - | 0 | - |
| - | - | 693.1 | 175.1 | - | - | 0 | - |
| - | - | 713.6 | 176.1 | - | - | 0 | - |
| - | - | 609 | 177.1 | - | - | 0 | - |
| - | - | 1416 | 180.1 | - | - | 0 | - |
| - | - | 4899 | 181.1 | - | - | 0 | - |
| - | - | 547.7 | 182.1 | - | - | 0 | - |
| - | - | 3007 | 182.1 | - | - | 0 | - |
| - | - | 5312 | 182.1 | - | - | 0 | - |
| - | - | 3793 | 183.1 | - | - | 0 | - |
| 2 | b | 1.826E+05 | 183.1 | 0.000355 | 1.939 | +1 | 2 |
| - | - | 4519 | 184.1 | - | - | 0 | - |
| - | - | 1.229E+04 | 184.1 | - | - | 0 | - |
| - | - | 965.5 | 185.1 | - | - | 0 | - |
| - | - | 1.878E+04 | 185.1 | - | - | 0 | - |
| - | - | 549.4 | 185.1 | - | - | 0 | - |
| - | - | 3755 | 186.1 | - | - | 0 | - |
| - | - | 959 | 186.1 | - | - | 0 | - |
| - | - | 2.87E+04 | 187.1 | - | - | 0 | - |
| - | - | 2550 | 188.1 | - | - | 0 | - |
| - | - | 5514 | 189.1 | - | - | 0 | - |
| - | - | 1164 | 191.1 | - | - | 0 | - |
| 16 | y | 4778 | 191.1 | 0.0004397 | 2.301 | +1 | 2 |
| - | - | 1389 | 195.1 | - | - | 0 | - |
| - | - | 1.286E+04 | 195.1 | - | - | 0 | - |
| - | - | 1141 | 196.1 | - | - | 0 | - |
| - | - | 2.009E+04 | 197.1 | - | - | 0 | - |
| - | - | 4622 | 198.1 | - | - | 0 | - |
| - | - | 1323 | 198.1 | - | - | 0 | - |
| - | - | 5928 | 199.1 | - | - | 0 | - |
| - | - | 2504 | 199.1 | - | - | 0 | - |
| - | - | 1283 | 199.1 | - | - | 0 | - |
| - | - | 6566 | 200.1 | - | - | 0 | - |
| - | - | 6826 | 200.1 | - | - | 0 | - |
| - | - | 1759 | 201.1 | - | - | 0 | - |
| 2 | b | 9.281E+04 | 201.1 | 0.0003036 | 1.51 | +1 | 2 |
| - | - | 1080 | 202.1 | - | - | 0 | - |
| - | - | 6458 | 202.1 | - | - | 0 | - |
| - | - | 1846 | 203.1 | - | - | 0 | - |
| - | - | 1113 | 203.1 | - | - | 0 | - |
| - | - | 711.3 | 208.1 | - | - | 0 | - |
| - | - | 7912 | 208.1 | - | - | 0 | - |
| - | - | 2.241E+04 | 209.1 | - | - | 0 | - |
| - | - | 832.6 | 209.1 | - | - | 0 | - |
| - | - | 1638 | 210.1 | - | - | 0 | - |
| - | - | 1007 | 210.1 | - | - | 0 | - |
| - | - | 1080 | 211.1 | - | - | 0 | - |
| - | - | 2208 | 212.1 | - | - | 0 | - |
| - | - | 1499 | 213.1 | - | - | 0 | - |
| - | - | 2468 | 213.1 | - | - | 0 | - |
| - | - | 890.3 | 214.2 | - | - | 0 | - |
| - | - | 3.493E+04 | 215.1 | - | - | 0 | - |
| - | - | 3335 | 216.1 | - | - | 0 | - |
| - | - | 1.575E+04 | 217.1 | - | - | 0 | - |
| - | - | 2.168E+04 | 217.1 | - | - | 0 | - |
| - | - | 1377 | 218.1 | - | - | 0 | - |
| - | - | 625.6 | 218.1 | - | - | 0 | - |
| - | - | 2201 | 218.1 | - | - | 0 | - |
| - | - | 1129 | 218.1 | - | - | 0 | - |
| - | - | 6913 | 219.1 | - | - | 0 | - |
| - | - | 1511 | 220.2 | - | - | 0 | - |
| - | - | 736.2 | 224.1 | - | - | 0 | - |
| - | - | 1082 | 224.2 | - | - | 0 | - |
| - | - | 8325 | 225.1 | - | - | 0 | - |
| - | - | 4728 | 225.1 | - | - | 0 | - |
| - | - | 1713 | 225.2 | - | - | 0 | - |
| - | - | 8.301E+04 | 226.1 | - | - | 0 | - |
| - | - | 2413 | 227.1 | - | - | 0 | - |
| - | - | 6145 | 227.1 | - | - | 0 | - |
| - | - | 714.4 | 228.1 | - | - | 0 | - |
| - | - | 1285 | 228.1 | - | - | 0 | - |
| - | - | 951.3 | 231.1 | - | - | 0 | - |
| - | - | 5.027E+04 | 233.2 | - | - | 0 | - |
| - | - | 6112 | 234.2 | - | - | 0 | - |
| - | - | 1357 | 235.1 | - | - | 0 | - |
| - | - | 6018 | 236.1 | - | - | 0 | - |
| - | - | 1215 | 236.1 | - | - | 0 | - |
| - | - | 1669 | 237.1 | - | - | 0 | - |
| - | - | 714 | 237.1 | - | - | 0 | - |
| - | - | 606.6 | 240.1 | - | - | 0 | - |
| - | - | 5592 | 241.1 | - | - | 0 | - |
| - | - | 1274 | 241.2 | - | - | 0 | - |
| - | - | 713.5 | 242.1 | - | - | 0 | - |
| - | - | 1.446E+04 | 242.2 | - | - | 0 | - |
| - | - | 763.2 | 242.2 | - | - | 0 | - |
| - | - | 3121 | 243.1 | - | - | 0 | - |
| - | - | 2.391E+04 | 243.1 | - | - | 0 | - |
| 13 | y | 4622 | 244.1 | 0.0008376 | 3.431 | +2 | 5 |
| - | - | 1638 | 244.1 | - | - | 0 | - |
| - | - | 1.544E+04 | 245.1 | - | - | 0 | - |
| - | - | 1673 | 246.1 | - | - | 0 | - |
| - | - | 8130 | 247.1 | - | - | 0 | - |
| - | - | 2.521E+04 | 251.2 | - | - | 0 | - |
| - | - | 703 | 252.1 | - | - | 0 | - |
| - | - | 2533 | 252.1 | - | - | 0 | - |
| - | - | 843.6 | 252.2 | - | - | 0 | - |
| - | - | 2326 | 252.2 | - | - | 0 | - |
| - | - | 775 | 253.1 | - | - | 0 | - |
| - | - | 1300 | 253.1 | - | - | 0 | - |
| - | - | 857.8 | 253.2 | - | - | 0 | - |
| - | - | 1.762E+04 | 254.1 | - | - | 0 | - |
| - | - | 4795 | 254.2 | - | - | 0 | - |
| - | - | 1597 | 255.1 | - | - | 0 | - |
| - | - | 642.5 | 257.2 | - | - | 0 | - |
| - | - | 615.4 | 257.2 | - | - | 0 | - |
| - | - | 982.6 | 258.1 | - | - | 0 | - |
| - | - | 1.467E+04 | 259.1 | - | - | 0 | - |
| - | - | 748.4 | 259.1 | - | - | 0 | - |
| - | - | 742 | 259.1 | - | - | 0 | - |
| - | - | 1109 | 260.1 | - | - | 0 | - |
| - | - | 978.2 | 260.2 | - | - | 0 | - |
| - | - | 604.9 | 261.1 | - | - | 0 | - |
| - | - | 1.53E+04 | 261.2 | - | - | 0 | - |
| - | - | 3151 | 262.1 | - | - | 0 | - |
| - | - | 2439 | 262.2 | - | - | 0 | - |
| - | - | 1045 | 263.1 | - | - | 0 | - |
| - | - | 1055 | 264.1 | - | - | 0 | - |
| - | - | 4788 | 264.1 | - | - | 0 | - |
| - | - | 592.1 | 265.1 | - | - | 0 | - |
| - | - | 651 | 266.1 | - | - | 0 | - |
| - | - | 825.8 | 266.1 | - | - | 0 | - |
| - | - | 2304 | 268.1 | - | - | 0 | - |
| 3 | a | 5469 | 268.2 | 0.0004085 | 1.523 | +1 | 3 |
| - | - | 929.7 | 269.1 | - | - | 0 | - |
| - | - | 1386 | 269.2 | - | - | 0 | - |
| - | - | 5696 | 269.2 | - | - | 0 | - |
| - | - | 995.3 | 269.2 | - | - | 0 | - |
| - | - | 1287 | 270.1 | - | - | 0 | - |
| - | - | 3018 | 270.2 | - | - | 0 | - |
| - | - | 915.2 | 271.1 | - | - | 0 | - |
| - | - | 624.6 | 271.2 | - | - | 0 | - |
| - | - | 1.102E+04 | 272.1 | - | - | 0 | - |
| - | - | 1151 | 273.1 | - | - | 0 | - |
| - | - | 9315 | 275.2 | - | - | 0 | - |
| - | - | 748.8 | 276.1 | - | - | 0 | - |
| - | - | 1426 | 276.2 | - | - | 0 | - |
| - | - | 1201 | 277.1 | - | - | 0 | - |
| - | - | 904.4 | 278.1 | - | - | 0 | - |
| - | - | 1314 | 278.1 | - | - | 0 | - |
| - | - | 8637 | 279.1 | - | - | 0 | - |
| - | - | 3921 | 280.1 | - | - | 0 | - |
| - | - | 1155 | 280.1 | - | - | 0 | - |
| - | - | 1.397E+04 | 280.2 | - | - | 0 | - |
| - | - | 2097 | 281.2 | - | - | 0 | - |
| - | - | 1172 | 282.1 | - | - | 0 | - |
| - | - | 2.79E+04 | 282.1 | - | - | 0 | - |
| - | - | 4019 | 283.1 | - | - | 0 | - |
| - | - | 937.7 | 284.2 | - | - | 0 | - |
| - | - | 1.205E+04 | 285.2 | - | - | 0 | - |
| - | - | 1182 | 285.2 | - | - | 0 | - |
| - | - | 1.013E+04 | 286.1 | - | - | 0 | - |
| - | - | 1819 | 286.2 | - | - | 0 | - |
| 3 | a | 1180 | 286.2 | 0.0006471 | 2.261 | +1 | 3 |
| - | - | 1463 | 287.1 | - | - | 0 | - |
| - | - | 2290 | 294.2 | - | - | 0 | - |
| - | - | 2321 | 296.1 | - | - | 0 | - |
| 3 | b | 4.715E+05 | 296.2 | 0.0005806 | 1.96 | +1 | 3 |
| - | - | 4331 | 297.1 | - | - | 0 | - |
| - | - | 2.642E+04 | 297.2 | - | - | 0 | - |
| - | - | 6.909E+04 | 297.2 | - | - | 0 | - |
| - | - | 3211 | 298.2 | - | - | 0 | - |
| - | - | 1.74E+04 | 298.2 | - | - | 0 | - |
| - | - | 4867 | 298.2 | - | - | 0 | - |
| - | - | 1276 | 299.1 | - | - | 0 | - |
| - | - | 2754 | 299.2 | - | - | 0 | - |
| - | - | 1467 | 300.1 | - | - | 0 | - |
| - | - | 1711 | 300.2 | - | - | 0 | - |
| - | - | 1864 | 300.2 | - | - | 0 | - |
| - | - | 1359 | 301.2 | - | - | 0 | - |
| 15 | y | 2297 | 301.2 | 0.000804 | 2.669 | +1 | 3 |
| 15 | y | 880.4 | 302.2 | 0.001163 | 3.85 | +1 | 3 |
| - | - | 1.941E+04 | 303.2 | - | - | 0 | - |
| - | - | 5385 | 304.1 | - | - | 0 | - |
| - | - | 2767 | 304.2 | - | - | 0 | - |
| - | - | 2576 | 306.1 | - | - | 0 | - |
| - | - | 589.3 | 307.1 | - | - | 0 | - |
| - | - | 1781 | 307.1 | - | - | 0 | - |
| 12 | y | 1769 | 308.2 | 0.002723 | 8.835 | +2 | 6 |
| - | - | 2087 | 310.1 | - | - | 0 | - |
| - | - | 803.5 | 310.1 | - | - | 0 | - |
| - | - | 761.8 | 311.1 | - | - | 0 | - |
| - | - | 702.2 | 311.2 | - | - | 0 | - |
| - | - | 864.9 | 312.2 | - | - | 0 | - |
| - | - | 714.6 | 313.2 | - | - | 0 | - |
| - | - | 4489 | 313.2 | - | - | 0 | - |
| - | - | 992.9 | 314.1 | - | - | 0 | - |
| - | - | 4537 | 314.1 | - | - | 0 | - |
| - | - | 1.778E+04 | 314.2 | - | - | 0 | - |
| 3 | b | 1.186E+05 | 314.2 | 0.0006055 | 1.927 | +1 | 3 |
| - | - | 792.1 | 315.1 | - | - | 0 | - |
| - | - | 1592 | 315.2 | - | - | 0 | - |
| - | - | 2254 | 315.2 | - | - | 0 | - |
| - | - | 1.821E+04 | 315.2 | - | - | 0 | - |
| - | - | 697.6 | 316.2 | - | - | 0 | - |
| - | - | 716.4 | 316.2 | - | - | 0 | - |
| - | - | 5094 | 317.2 | - | - | 0 | - |
| - | - | 798.4 | 318.1 | - | - | 0 | - |
| 15 | y | 3.146E+04 | 319.2 | 0.0005237 | 1.641 | +1 | 3 |
| - | - | 4234 | 320.2 | - | - | 0 | - |
| - | - | 800.3 | 321.2 | - | - | 0 | - |
| - | - | 2439 | 323.2 | - | - | 0 | - |
| - | - | 1234 | 324.2 | - | - | 0 | - |
| - | - | 1794 | 325.2 | - | - | 0 | - |
| - | - | 1636 | 326.2 | - | - | 0 | - |
| - | - | 7579 | 328.1 | - | - | 0 | - |
| - | - | 919 | 328.2 | - | - | 0 | - |
| - | - | 917.5 | 329.1 | - | - | 0 | - |
| - | - | 1201 | 329.2 | - | - | 0 | - |
| - | - | 1.281E+04 | 330.2 | - | - | 0 | - |
| - | - | 2459 | 331.2 | - | - | 0 | - |
| - | - | 3067 | 333.2 | - | - | 0 | - |
| - | - | 3643 | 334.2 | - | - | 0 | - |
| - | - | 909.1 | 335.2 | - | - | 0 | - |
| - | - | 1948 | 336.2 | - | - | 0 | - |
| - | - | 3936 | 337.2 | - | - | 0 | - |
| - | - | 1145 | 338.1 | - | - | 0 | - |
| - | - | 751.5 | 338.2 | - | - | 0 | - |
| - | - | 2631 | 339.2 | - | - | 0 | - |
| - | - | 831.1 | 339.2 | - | - | 0 | - |
| - | - | 8230 | 341.2 | - | - | 0 | - |
| - | - | 3059 | 342.2 | - | - | 0 | - |
| - | - | 3336 | 344.2 | - | - | 0 | - |
| - | - | 752.9 | 345.2 | - | - | 0 | - |
| - | - | 2.192E+04 | 346.1 | - | - | 0 | - |
| - | - | 1.754E+04 | 346.2 | - | - | 0 | - |
| - | - | 3660 | 347.1 | - | - | 0 | - |
| - | - | 2888 | 347.2 | - | - | 0 | - |
| - | - | 1.601E+04 | 348.2 | - | - | 0 | - |
| - | - | 3085 | 349.2 | - | - | 0 | - |
| - | - | 4708 | 350.2 | - | - | 0 | - |
| - | - | 1.461E+04 | 351.2 | - | - | 0 | - |
| - | - | 1887 | 352.2 | - | - | 0 | - |
| - | - | 8982 | 353.2 | - | - | 0 | - |
| - | - | 5296 | 354.2 | - | - | 0 | - |
| - | - | 1086 | 354.2 | - | - | 0 | - |
| - | - | 4992 | 355.2 | - | - | 0 | - |
| - | - | 1488 | 356.1 | - | - | 0 | - |
| - | - | 861.1 | 358.2 | - | - | 0 | - |
| - | - | 2410 | 360.2 | - | - | 0 | - |
| - | - | 916.5 | 361.2 | - | - | 0 | - |
| - | - | 3.708E+04 | 362.2 | - | - | 0 | - |
| - | - | 1375 | 363.2 | - | - | 0 | - |
| - | - | 7250 | 363.2 | - | - | 0 | - |
| - | - | 8529 | 365.1 | - | - | 0 | - |
| - | - | 963.4 | 365.2 | - | - | 0 | - |
| - | - | 849.1 | 365.2 | - | - | 0 | - |
| - | - | 2407 | 365.3 | - | - | 0 | - |
| - | - | 846.8 | 366.3 | - | - | 0 | - |
| - | - | 712.8 | 367.2 | - | - | 0 | - |
| - | - | 5322 | 367.2 | - | - | 0 | - |
| - | - | 1982 | 367.2 | - | - | 0 | - |
| - | - | 3293 | 368.2 | - | - | 0 | - |
| - | - | 671.9 | 368.2 | - | - | 0 | - |
| - | - | 1209 | 369.1 | - | - | 0 | - |
| - | - | 2.283E+04 | 369.2 | - | - | 0 | - |
| - | - | 3157 | 370.2 | - | - | 0 | - |
| - | - | 2716 | 370.7 | - | - | 0 | - |
| - | - | 1121 | 371.2 | - | - | 0 | - |
| - | - | 5593 | 371.2 | - | - | 0 | - |
| - | - | 4500 | 372.2 | - | - | 0 | - |
| - | - | 8003 | 373.2 | - | - | 0 | - |
| - | - | 1701 | 374.2 | - | - | 0 | - |
| - | - | 1117 | 375.2 | - | - | 0 | - |
| - | - | 1191 | 379.1 | - | - | 0 | - |
| - | - | 692.9 | 379.2 | - | - | 0 | - |
| - | - | 1121 | 379.2 | - | - | 0 | - |
| - | - | 1998 | 381.2 | - | - | 0 | - |
| - | - | 1.401E+04 | 383.2 | - | - | 0 | - |
| - | - | 3348 | 383.3 | - | - | 0 | - |
| - | - | 2520 | 384.2 | - | - | 0 | - |
| - | - | 1282 | 384.3 | - | - | 0 | - |
| - | - | 1.108E+04 | 385.2 | - | - | 0 | - |
| - | - | 2037 | 386.2 | - | - | 0 | - |
| - | - | 1153 | 386.2 | - | - | 0 | - |
| - | - | 1074 | 387.2 | - | - | 0 | - |
| - | - | 1583 | 388.1 | - | - | 0 | - |
| - | - | 1109 | 389.3 | - | - | 0 | - |
| - | - | 1141 | 390.2 | - | - | 0 | - |
| - | - | 777 | 392.2 | - | - | 0 | - |
| - | - | 2533 | 393.2 | - | - | 0 | - |
| - | - | 7108 | 393.3 | - | - | 0 | - |
| - | - | 1815 | 394.3 | - | - | 0 | - |
| - | - | 1117 | 395.2 | - | - | 0 | - |
| - | - | 2868 | 397.1 | - | - | 0 | - |
| - | - | 1384 | 398.1 | - | - | 0 | - |
| - | - | 1708 | 398.2 | - | - | 0 | - |
| - | - | 7691 | 398.2 | - | - | 0 | - |
| - | - | 2014 | 399.2 | - | - | 0 | - |
| - | - | 2591 | 399.2 | - | - | 0 | - |
| - | - | 2415 | 400.2 | - | - | 0 | - |
| - | - | 2.156E+04 | 401.2 | - | - | 0 | - |
| - | - | 3725 | 402.2 | - | - | 0 | - |
| - | - | 1913 | 405.2 | - | - | 0 | - |
| - | - | 750.4 | 405.2 | - | - | 0 | - |
| - | - | 3803 | 407.2 | - | - | 0 | - |
| - | - | 7064 | 408.2 | - | - | 0 | - |
| - | - | 1277 | 410.2 | - | - | 0 | - |
| - | - | 985.9 | 411.2 | - | - | 0 | - |
| - | - | 1.13E+04 | 411.3 | - | - | 0 | - |
| - | - | 1682 | 412.3 | - | - | 0 | - |
| - | - | 1267 | 413.2 | - | - | 0 | - |
| - | - | 7063 | 415.1 | - | - | 0 | - |
| 8 | b | 2390 | 415.2 | 0.005366 | 12.92 | +2 | 8 |
| - | - | 5722 | 415.3 | - | - | 0 | - |
| - | - | 1530 | 416.2 | - | - | 0 | - |
| 14 | y | 1942 | 416.2 | 0.002212 | 5.314 | +1 | 4 |
| - | - | 1.653E+04 | 416.3 | - | - | 0 | - |
| - | - | 5413 | 417.3 | - | - | 0 | - |
| - | - | 984.9 | 422.2 | - | - | 0 | - |
| - | - | 3980 | 424.2 | - | - | 0 | - |
| - | - | 2237 | 425.1 | - | - | 0 | - |
| - | - | 1123 | 425.2 | - | - | 0 | - |
| - | - | 7755 | 425.2 | - | - | 0 | - |
| - | - | 719.9 | 426.2 | - | - | 0 | - |
| - | - | 605.5 | 426.2 | - | - | 0 | - |
| - | - | 752 | 427.2 | - | - | 0 | - |
| - | - | 4753 | 427.2 | - | - | 0 | - |
| 10 | y | 1578 | 429.2 | 0.008268 | 19.26 | +2 | 8 |
| - | - | 1269 | 431.2 | - | - | 0 | - |
| - | - | 1140 | 431.3 | - | - | 0 | - |
| - | - | 1.135E+04 | 433.2 | - | - | 0 | - |
| 14 | y | 2.32E+04 | 433.2 | 0.001999 | 4.615 | +1 | 4 |
| - | - | 1.572E+05 | 433.3 | - | - | 0 | - |
| - | - | 1393 | 434.2 | - | - | 0 | - |
| - | - | 3260 | 434.2 | - | - | 0 | - |
| - | - | 3727 | 434.2 | - | - | 0 | - |
| - | - | 3.492E+04 | 434.3 | - | - | 0 | - |
| - | - | 2086 | 434.7 | - | - | 0 | - |
| - | - | 2027 | 435.2 | - | - | 0 | - |
| - | - | 3639 | 435.3 | - | - | 0 | - |
| - | - | 1554 | 440.2 | - | - | 0 | - |
| - | - | 3380 | 441.2 | - | - | 0 | - |
| - | - | 1646 | 441.3 | - | - | 0 | - |
| - | - | 4448 | 442.2 | - | - | 0 | - |
| - | - | 3221 | 443.1 | - | - | 0 | - |
| - | - | 1006 | 443.2 | - | - | 0 | - |
| - | - | 6558 | 443.2 | - | - | 0 | - |
| 4 | b | 1.619E+04 | 443.3 | 0.0008312 | 1.875 | +1 | 4 |
| - | - | 809.5 | 444.1 | - | - | 0 | - |
| - | - | 1404 | 444.2 | - | - | 0 | - |
| - | - | 1183 | 444.2 | - | - | 0 | - |
| - | - | 3796 | 444.3 | - | - | 0 | - |
| - | - | 1517 | 445.2 | - | - | 0 | - |
| - | - | 1690 | 451.2 | - | - | 0 | - |
| - | - | 4082 | 452.2 | - | - | 0 | - |
| - | - | 1261 | 453.2 | - | - | 0 | - |
| - | - | 833 | 454.2 | - | - | 0 | - |
| - | - | 1198 | 459.2 | - | - | 0 | - |
| - | - | 5948 | 459.2 | - | - | 0 | - |
| - | - | 1659 | 459.3 | - | - | 0 | - |
| - | - | 711.6 | 460.2 | - | - | 0 | - |
| 4 | b | 1.285E+05 | 461.3 | 0.0008561 | 1.856 | +1 | 4 |
| - | - | 6679 | 462.2 | - | - | 0 | - |
| - | - | 3.462E+04 | 462.3 | - | - | 0 | - |
| - | - | 1082 | 463.2 | - | - | 0 | - |
| - | - | 4923 | 463.3 | - | - | 0 | - |
| - | - | 1470 | 464.2 | - | - | 0 | - |
| - | - | 892.9 | 464.2 | - | - | 0 | - |
| - | - | 2926 | 466.2 | - | - | 0 | - |
| - | - | 2347 | 468.2 | - | - | 0 | - |
| - | - | 1644 | 468.3 | - | - | 0 | - |
| - | - | 3213 | 469.2 | - | - | 0 | - |
| - | - | 2.015E+04 | 470.2 | - | - | 0 | - |
| - | - | 4411 | 471.2 | - | - | 0 | - |
| - | - | 1673 | 476.2 | - | - | 0 | - |
| - | - | 1645 | 477.3 | - | - | 0 | - |
| - | - | 3220 | 478.2 | - | - | 0 | - |
| - | - | 3154 | 479.2 | - | - | 0 | - |
| - | - | 1.695E+04 | 480.2 | - | - | 0 | - |
| - | - | 4335 | 481.2 | - | - | 0 | - |
| - | - | 3066 | 482.2 | - | - | 0 | - |
| - | - | 739 | 483.2 | - | - | 0 | - |
| - | - | 6493 | 484.2 | - | - | 0 | - |
| - | - | 925.4 | 485.3 | - | - | 0 | - |
| 13 | y | 2535 | 486.3 | 0.001512 | 3.11 | +1 | 5 |
| 13 | y | 2646 | 487.3 | 0.00178 | 3.653 | +1 | 5 |
| - | - | 857.1 | 488.3 | - | - | 0 | - |
| - | - | 924.1 | 492.2 | - | - | 0 | - |
| - | - | 7452 | 494.2 | - | - | 0 | - |
| - | - | 1021 | 495.2 | - | - | 0 | - |
| - | - | 3175 | 495.3 | - | - | 0 | - |
| - | - | 7027 | 496.3 | - | - | 0 | - |
| - | - | 1266 | 497.3 | - | - | 0 | - |
| - | - | 5.006E+04 | 498.2 | - | - | 0 | - |
| - | - | 1.126E+04 | 499.2 | - | - | 0 | - |
| - | - | 2873 | 500.2 | - | - | 0 | - |
| - | - | 742.7 | 501.2 | - | - | 0 | - |
| - | - | 1.101E+04 | 502.2 | - | - | 0 | - |
| - | - | 2524 | 503.2 | - | - | 0 | - |
| - | - | 839.9 | 503.3 | - | - | 0 | - |
| 13 | y | 3.291E+04 | 504.3 | 0.0006521 | 1.293 | +1 | 5 |
| - | - | 6076 | 505.3 | - | - | 0 | - |
| - | - | 2620 | 510.2 | - | - | 0 | - |
| - | - | 1.107E+04 | 512.2 | - | - | 0 | - |
| - | - | 1697 | 512.3 | - | - | 0 | - |
| - | - | 2869 | 513.2 | - | - | 0 | - |
| - | - | 3644 | 513.3 | - | - | 0 | - |
| - | - | 1645 | 515.2 | - | - | 0 | - |
| - | - | 855.6 | 516.2 | - | - | 0 | - |
| - | - | 1866 | 518.2 | - | - | 0 | - |
| - | - | 1005 | 521.3 | - | - | 0 | - |
| - | - | 911.6 | 524.2 | - | - | 0 | - |
| - | - | 1021 | 527.2 | - | - | 0 | - |
| - | - | 3072 | 528.2 | - | - | 0 | - |
| - | - | 1485 | 529.2 | - | - | 0 | - |
| - | - | 3.253E+04 | 530.2 | - | - | 0 | - |
| - | - | 9589 | 530.3 | - | - | 0 | - |
| - | - | 7832 | 531.2 | - | - | 0 | - |
| - | - | 2924 | 531.3 | - | - | 0 | - |
| - | - | 1028 | 532.2 | - | - | 0 | - |
| - | - | 971.1 | 534.3 | - | - | 0 | - |
| - | - | 2832 | 538.3 | - | - | 0 | - |
| 5 | b | 6268 | 540.3 | 0.0006186 | 1.145 | +1 | 5 |
| - | - | 1688 | 541.3 | - | - | 0 | - |
| - | - | 950.4 | 542.8 | - | - | 0 | - |
| - | - | 887.8 | 545.2 | - | - | 0 | - |
| - | - | 3745 | 546.2 | - | - | 0 | - |
| - | - | 1439 | 547.2 | - | - | 0 | - |
| - | - | 808.6 | 551.2 | - | - | 0 | - |
| - | - | 2294 | 553.3 | - | - | 0 | - |
| - | - | 2442 | 555.3 | - | - | 0 | - |
| - | - | 2922 | 556.2 | - | - | 0 | - |
| - | - | 2211 | 556.3 | - | - | 0 | - |
| - | - | 920 | 556.3 | - | - | 0 | - |
| - | - | 1039 | 558.2 | - | - | 0 | - |
| 5 | b | 1.642E+04 | 558.3 | 0.000491 | 0.8794 | +1 | 5 |
| - | - | 5550 | 559.3 | - | - | 0 | - |
| - | - | 2214 | 563.2 | - | - | 0 | - |
| - | - | 712.3 | 564.2 | - | - | 0 | - |
| - | - | 3001 | 569.3 | - | - | 0 | - |
| - | - | 2754 | 569.8 | - | - | 0 | - |
| - | - | 698.8 | 570.3 | - | - | 0 | - |
| - | - | 1809 | 571.3 | - | - | 0 | - |
| - | - | 1013 | 573.2 | - | - | 0 | - |
| - | - | 1029 | 574.3 | - | - | 0 | - |
| - | - | 4872 | 578.3 | - | - | 0 | - |
| - | - | 4425 | 578.8 | - | - | 0 | - |
| - | - | 2330 | 579.3 | - | - | 0 | - |
| - | - | 6292 | 581.3 | - | - | 0 | - |
| - | - | 4544 | 582.3 | - | - | 0 | - |
| - | - | 1.106E+04 | 582.8 | - | - | 0 | - |
| - | - | 1236 | 583.2 | - | - | 0 | - |
| - | - | 7587 | 583.3 | - | - | 0 | - |
| - | - | 962.7 | 583.8 | - | - | 0 | - |
| - | - | 3069 | 587.3 | - | - | 0 | - |
| - | - | 1119 | 589.3 | - | - | 0 | - |
| - | - | 4185 | 591.2 | - | - | 0 | - |
| - | - | 3593 | 591.3 | - | - | 0 | - |
| 11 | b | 3821 | 591.8 | 0.01041 | 17.59 | +2 | 11 |
| - | - | 930 | 592.2 | - | - | 0 | - |
| - | - | 907.3 | 596.3 | - | - | 0 | - |
| - | - | 4731 | 597.3 | - | - | 0 | - |
| - | - | 1710 | 598.3 | - | - | 0 | - |
| - | - | 3.224E+04 | 599.3 | - | - | 0 | - |
| - | - | 8591 | 600.3 | - | - | 0 | - |
| - | - | 1258 | 601.3 | - | - | 0 | - |
| - | - | 2719 | 604.3 | - | - | 0 | - |
| - | - | 1030 | 604.8 | - | - | 0 | - |
| - | - | 1700 | 607.3 | - | - | 0 | - |
| - | - | 1310 | 608.3 | - | - | 0 | - |
| - | - | 2403 | 608.4 | - | - | 0 | - |
| - | - | 867.4 | 608.8 | - | - | 0 | - |
| - | - | 9341 | 609.3 | - | - | 0 | - |
| - | - | 2997 | 609.3 | - | - | 0 | - |
| - | - | 1678 | 609.3 | - | - | 0 | - |
| - | - | 2608 | 609.8 | - | - | 0 | - |
| - | - | 2863 | 610.3 | - | - | 0 | - |
| - | - | 7559 | 612.8 | - | - | 0 | - |
| - | - | 3651 | 613.3 | - | - | 0 | - |
| - | - | 1244 | 613.8 | - | - | 0 | - |
| 12 | y | 4437 | 614.3 | 0.0004552 | 0.7409 | +1 | 6 |
| 12 | y | 1.828E+04 | 615.3 | 0.004551 | 7.397 | +1 | 6 |
| - | - | 5997 | 616.3 | - | - | 0 | - |
| - | - | 1817 | 617.3 | - | - | 0 | - |
| - | - | 2.692E+04 | 617.8 | - | - | 0 | - |
| - | - | 3.044E+04 | 618.3 | - | - | 0 | - |
| - | - | 1.088E+04 | 618.8 | - | - | 0 | - |
| - | - | 2773 | 619.3 | - | - | 0 | - |
| - | - | 807.7 | 619.8 | - | - | 0 | - |
| - | - | 944.6 | 620.3 | - | - | 0 | - |
| - | - | 823.8 | 624.3 | - | - | 0 | - |
| - | - | 7153 | 625.3 | - | - | 0 | - |
| - | - | 2035 | 626.3 | - | - | 0 | - |
| - | - | 4483 | 626.4 | - | - | 0 | - |
| - | - | 907.5 | 626.7 | - | - | 0 | - |
| - | - | 6.643E+04 | 626.8 | - | - | 0 | - |
| - | - | 7.659E+04 | 627.3 | - | - | 0 | - |
| - | - | 4.716E+04 | 627.3 | - | - | 0 | - |
| - | - | 1.557E+04 | 627.8 | - | - | 0 | - |
| - | - | 2.28E+04 | 628.3 | - | - | 0 | - |
| - | - | 1295 | 628.3 | - | - | 0 | - |
| 6 | y | 1320 | 628.8 | 0.001292 | 2.055 | +2 | 12 |
| - | - | 2749 | 629.3 | - | - | 0 | - |
| - | - | 1434 | 629.3 | - | - | 0 | - |
| - | - | 1565 | 629.8 | - | - | 0 | - |
| - | - | 930.7 | 630.3 | - | - | 0 | - |
| 12 | y | 2.198E+04 | 632.3 | 0.0005768 | 0.9122 | +1 | 6 |
| - | - | 5264 | 633.3 | - | - | 0 | - |
| - | - | 2391 | 635.8 | - | - | 0 | - |
| - | - | 2692 | 636.3 | - | - | 0 | - |
| 6 | y | 1.039E+04 | 637.8 | 0.0005874 | 0.921 | +2 | 12 |
| - | - | 5724 | 638.3 | - | - | 0 | - |
| - | - | 1019 | 638.8 | - | - | 0 | - |
| - | - | 942.6 | 639.3 | - | - | 0 | - |
| - | - | 1177 | 642.3 | - | - | 0 | - |
| - | - | 1.504E+04 | 643.3 | - | - | 0 | - |
| - | - | 6795 | 644.3 | - | - | 0 | - |
| - | - | 3072 | 645.3 | - | - | 0 | - |
| - | - | 1505 | 646.8 | - | - | 0 | - |
| - | - | 975.2 | 649.3 | - | - | 0 | - |
| - | - | 1233 | 653.3 | - | - | 0 | - |
| - | - | 5149 | 655.3 | - | - | 0 | - |
| 12 | b | 3926 | 655.8 | 0.006018 | 9.176 | +2 | 12 |
| - | - | 863.4 | 656.3 | - | - | 0 | - |
| - | - | 941.2 | 656.3 | - | - | 0 | - |
| - | - | 2228 | 658.3 | - | - | 0 | - |
| - | - | 1034 | 659.3 | - | - | 0 | - |
| - | - | 876.5 | 659.8 | - | - | 0 | - |
| - | - | 6397 | 660.3 | - | - | 0 | - |
| - | - | 2292 | 661.3 | - | - | 0 | - |
| - | - | 2517 | 663.3 | - | - | 0 | - |
| - | - | 2598 | 663.8 | - | - | 0 | - |
| - | - | 1928 | 664.3 | - | - | 0 | - |
| 12 | b | 1304 | 664.8 | 0.008725 | 13.12 | +2 | 12 |
| - | - | 1088 | 665.3 | - | - | 0 | - |
| - | - | 1498 | 667.3 | - | - | 0 | - |
| - | - | 2532 | 668.3 | - | - | 0 | - |
| - | - | 5101 | 668.8 | - | - | 0 | - |
| - | - | 2697 | 669.3 | - | - | 0 | - |
| - | - | 1295 | 669.8 | - | - | 0 | - |
| - | - | 894.2 | 670.3 | - | - | 0 | - |
| - | - | 1583 | 674.3 | - | - | 0 | - |
| - | - | 1098 | 676.3 | - | - | 0 | - |
| 5 | y | 2.108E+04 | 677.3 | 0.001033 | 1.525 | +2 | 13 |
| 5 | y | 3.051E+04 | 677.8 | 0.004631 | 6.832 | +2 | 13 |
| - | - | 1.757E+04 | 678.3 | - | - | 0 | - |
| - | - | 4711 | 678.8 | - | - | 0 | - |
| - | - | 1190 | 679.3 | - | - | 0 | - |
| - | - | 785.9 | 679.3 | - | - | 0 | - |
| - | - | 916.3 | 684.4 | - | - | 0 | - |
| - | - | 2286 | 685.3 | - | - | 0 | - |
| - | - | 1740 | 685.8 | - | - | 0 | - |
| 5 | y | 5.462E+05 | 686.3 | 0.0006948 | 1.012 | +2 | 13 |
| - | - | 3.92E+05 | 686.8 | - | - | 0 | - |
| - | - | 1.356E+05 | 687.3 | - | - | 0 | - |
| - | - | 1.489E+04 | 687.8 | - | - | 0 | - |
| 13 | b | 1325 | 691.4 | 0.008938 | 12.93 | +2 | 13 |
| 13 | b | 1075 | 691.8 | 0.001007 | 1.455 | +2 | 13 |
| - | - | 4266 | 694.3 | - | - | 0 | - |
| - | - | 2299 | 695.3 | - | - | 0 | - |
| - | - | 983 | 696.4 | - | - | 0 | - |
| - | - | 953.6 | 698.3 | - | - | 0 | - |
| 13 | b | 6324 | 700.4 | 0.01263 | 18.04 | +2 | 13 |
| - | - | 4866 | 700.8 | - | - | 0 | - |
| - | - | 2226 | 701.4 | - | - | 0 | - |
| - | - | 2572 | 704.3 | - | - | 0 | - |
| - | - | 1173 | 705.3 | - | - | 0 | - |
| - | - | 841.1 | 709.3 | - | - | 0 | - |
| - | - | 4.484E+04 | 712.4 | - | - | 0 | - |
| - | - | 1.423E+04 | 713.4 | - | - | 0 | - |
| - | - | 2198 | 714.4 | - | - | 0 | - |
| - | - | 793.5 | 720.3 | - | - | 0 | - |
| - | - | 1103 | 720.4 | - | - | 0 | - |
| - | - | 6817 | 722.3 | - | - | 0 | - |
| - | - | 2398 | 723.3 | - | - | 0 | - |
| 7 | b | 1621 | 724.4 | 0.001959 | 2.705 | +1 | 7 |
| - | - | 1014 | 726.3 | - | - | 0 | - |
| - | - | 1090 | 727.3 | - | - | 0 | - |
| 11 | y | 886.9 | 727.4 | 0.001113 | 1.53 | +1 | 7 |
| 11 | y | 1877 | 728.4 | 0.002998 | 4.117 | +1 | 7 |
| - | - | 889.5 | 735.3 | - | - | 0 | - |
| - | - | 884 | 735.5 | - | - | 0 | - |
| - | - | 1333 | 737.3 | - | - | 0 | - |
| - | - | 1829 | 737.4 | - | - | 0 | - |
| - | - | 2514 | 738.3 | - | - | 0 | - |
| - | - | 1926 | 738.4 | - | - | 0 | - |
| - | - | 934 | 739.3 | - | - | 0 | - |
| - | - | 2854 | 739.4 | - | - | 0 | - |
| - | - | 5.35E+04 | 740.3 | - | - | 0 | - |
| - | - | 1.651E+04 | 741.4 | - | - | 0 | - |
| - | - | 4490 | 742.4 | - | - | 0 | - |
| 7 | b | 3747 | 742.4 | 0.002552 | 3.437 | +1 | 7 |
| - | - | 1546 | 743.4 | - | - | 0 | - |
| - | - | 1711 | 743.4 | - | - | 0 | - |
| - | - | 2782 | 745.3 | - | - | 0 | - |
| 11 | y | 1.598E+04 | 745.4 | 0.0009855 | 1.322 | +1 | 7 |
| - | - | 2033 | 746.3 | - | - | 0 | - |
| - | - | 7459 | 746.4 | - | - | 0 | - |
| - | - | 1207 | 747.4 | - | - | 0 | - |
| 4 | y | 2325 | 750.9 | 0.003081 | 4.103 | +2 | 14 |
| 4 | y | 4278 | 751.4 | 0.005092 | 6.777 | +2 | 14 |
| - | - | 2238 | 751.9 | - | - | 0 | - |
| - | - | 1119 | 752.4 | - | - | 0 | - |
| - | - | 3146 | 753.3 | - | - | 0 | - |
| - | - | 2511 | 754.3 | - | - | 0 | - |
| - | - | 1585 | 755.3 | - | - | 0 | - |
| - | - | 2631 | 755.4 | - | - | 0 | - |
| - | - | 1322 | 756.3 | - | - | 0 | - |
| - | - | 1120 | 756.4 | - | - | 0 | - |
| - | - | 1604 | 756.9 | - | - | 0 | - |
| 14 | b | 2.713E+04 | 757.4 | 0.006021 | 7.949 | +2 | 14 |
| - | - | 791.2 | 757.9 | - | - | 0 | - |
| - | - | 9361 | 758.4 | - | - | 0 | - |
| - | - | 2397 | 759.4 | - | - | 0 | - |
| 4 | y | 7.034E+04 | 759.9 | 0.0006064 | 0.7981 | +2 | 14 |
| - | - | 6.253E+04 | 760.4 | - | - | 0 | - |
| - | - | 2.603E+04 | 760.9 | - | - | 0 | - |
| - | - | 2105 | 761.4 | - | - | 0 | - |
| - | - | 927.5 | 766.4 | - | - | 0 | - |
| - | - | 1.284E+04 | 771.4 | - | - | 0 | - |
| - | - | 5372 | 772.4 | - | - | 0 | - |
| - | - | 1195 | 773.4 | - | - | 0 | - |
| - | - | 3132 | 774.3 | - | - | 0 | - |
| - | - | 967.5 | 775.3 | - | - | 0 | - |
| - | - | 1087 | 796.4 | - | - | 0 | - |
| - | - | 1350 | 797.4 | - | - | 0 | - |
| 3 | y | 1431 | 807.4 | 0.005605 | 6.942 | +2 | 15 |
| 3 | y | 1652 | 807.9 | 0.00609 | 7.538 | +2 | 15 |
| - | - | 1025 | 808.4 | - | - | 0 | - |
| - | - | 2079 | 811.4 | - | - | 0 | - |
| 15 | b | 1790 | 812.9 | 0.00384 | 4.723 | +2 | 15 |
| - | - | 1312 | 813.4 | - | - | 0 | - |
| - | - | 993.1 | 813.9 | - | - | 0 | - |
| - | - | 1900 | 814.4 | - | - | 0 | - |
| - | - | 1275 | 815.4 | - | - | 0 | - |
| 3 | y | 1.384E+04 | 816.4 | 0.0001394 | 0.1707 | +2 | 15 |
| - | - | 1.341E+04 | 816.9 | - | - | 0 | - |
| - | - | 6761 | 817.4 | - | - | 0 | - |
| 15 | b | 1083 | 821.4 | 0.008153 | 9.926 | +2 | 15 |
| - | - | 2383 | 821.9 | - | - | 0 | - |
| - | - | 1768 | 822.4 | - | - | 0 | - |
| - | - | 2050 | 823.4 | - | - | 0 | - |
| - | - | 2802 | 824.4 | - | - | 0 | - |
| - | - | 1666 | 825.4 | - | - | 0 | - |
| - | - | 1308 | 828.4 | - | - | 0 | - |
| 8 | b | 3903 | 829.4 | 0.002865 | 3.455 | +1 | 8 |
| - | - | 2037 | 830.5 | - | - | 0 | - |
| - | - | 1639 | 832.4 | - | - | 0 | - |
| - | - | 1231 | 833.4 | - | - | 0 | - |
| - | - | 1877 | 839.4 | - | - | 0 | - |
| - | - | 7516 | 840.4 | - | - | 0 | - |
| - | - | 3044 | 841.4 | - | - | 0 | - |
| - | - | 9104 | 842.4 | - | - | 0 | - |
| - | - | 1469 | 843 | - | - | 0 | - |
| - | - | 3680 | 843.4 | - | - | 0 | - |
| - | - | 776.1 | 844.4 | - | - | 0 | - |
| 16 | b | 3188 | 847.9 | 0.0009112 | 1.075 | +2 | 16 |
| 16 | b | 5554 | 848.4 | 0.003715 | 4.379 | +2 | 16 |
| - | - | 3635 | 848.9 | - | - | 0 | - |
| - | - | 1023 | 849.4 | - | - | 0 | - |
| - | - | 8176 | 850.4 | - | - | 0 | - |
| - | - | 6575 | 851.4 | - | - | 0 | - |
| - | - | 2076 | 852.4 | - | - | 0 | - |
| 10 | y | 1554 | 856.5 | 0.003686 | 4.304 | +1 | 8 |
| 16 | b | 1.44E+04 | 856.9 | 0.0002065 | 0.2409 | +2 | 16 |
| 10 | y | 1.768E+04 | 857.4 | 0.009966 | 11.62 | +1 | 8 |
| 2 | y | 7575 | 857.9 | 0.01161 | 13.53 | +2 | 16 |
| - | - | 1734 | 858.4 | - | - | 0 | - |
| - | - | 2588 | 859.4 | - | - | 0 | - |
| - | - | 1249 | 866.4 | - | - | 0 | - |
| 2 | y | 4574 | 866.9 | 0.0005922 | 0.683 | +2 | 16 |
| - | - | 6911 | 867.4 | - | - | 0 | - |
| - | - | 994.5 | 867.9 | - | - | 0 | - |
| - | - | 6.493E+04 | 868.4 | - | - | 0 | - |
| - | - | 2.886E+04 | 869.4 | - | - | 0 | - |
| - | - | 5788 | 870.4 | - | - | 0 | - |
| 10 | y | 1.274E+04 | 874.5 | 0.0001037 | 0.1186 | +1 | 8 |
| - | - | 5062 | 875.5 | - | - | 0 | - |
| - | - | 989.2 | 876.5 | - | - | 0 | - |
| - | - | 1820 | 884.4 | - | - | 0 | - |
| - | - | 1858 | 885.4 | - | - | 0 | - |
| - | - | 999.8 | 886.4 | - | - | 0 | - |
| - | - | 4423 | 887.4 | - | - | 0 | - |
| - | - | 1720 | 888.4 | - | - | 0 | - |
| - | - | 936.2 | 893.4 | - | - | 0 | - |
| - | - | 5282 | 894.4 | - | - | 0 | - |
| - | - | 2568 | 895.4 | - | - | 0 | - |
| - | - | 1581 | 899 | - | - | 0 | - |
| - | - | 1815 | 899.5 | - | - | 0 | - |
| - | - | 1253 | 900.4 | - | - | 0 | - |
| - | - | 1842 | 904.4 | - | - | 0 | - |
| - | - | 809.8 | 905.4 | - | - | 0 | - |
| 0 | Precursor | 1.085E+04 | 907.5 | 0.0005615 | 0.6187 | +2 | -1 |
| 0 | Precursor | 1.774E+04 | 908 | 0.007309 | 8.05 | +2 | -1 |
| - | - | 9129 | 908.5 | - | - | 0 | - |
| - | - | 2598 | 909 | - | - | 0 | - |
| - | - | 1.265E+04 | 911.4 | - | - | 0 | - |
| - | - | 4670 | 912.4 | - | - | 0 | - |
| - | - | 1473 | 913.4 | - | - | 0 | - |
| - | - | 1691 | 915.5 | - | - | 0 | - |
| - | - | 1784 | 916 | - | - | 0 | - |
| - | - | 962.2 | 916.4 | - | - | 0 | - |
| 0 | Precursor | 1.168E+05 | 916.5 | 0.0003818 | 0.4166 | +2 | -1 |
| - | - | 1.276E+05 | 917 | - | - | 0 | - |
| - | - | 6.3E+04 | 917.5 | - | - | 0 | - |
| - | - | 9276 | 918 | - | - | 0 | - |
| - | - | 6390 | 921.4 | - | - | 0 | - |
| - | - | 8294 | 922.4 | - | - | 0 | - |
| - | - | 2628 | 923.4 | - | - | 0 | - |
| - | - | 875.2 | 926.5 | - | - | 0 | - |
| - | - | 963.3 | 938.4 | - | - | 0 | - |
| - | - | 4.776E+04 | 939.4 | - | - | 0 | - |
| - | - | 2.205E+04 | 940.4 | - | - | 0 | - |
| - | - | 5403 | 941.4 | - | - | 0 | - |
| - | - | 1168 | 942.4 | - | - | 0 | - |
| - | - | 2119 | 953.5 | - | - | 0 | - |
| - | - | 2478 | 954.5 | - | - | 0 | - |
| - | - | 1014 | 955.5 | - | - | 0 | - |
| - | - | 7396 | 956.4 | - | - | 0 | - |
| - | - | 3091 | 957.5 | - | - | 0 | - |
| 9 | b | 3308 | 958.5 | 0.004748 | 4.954 | +1 | 9 |
| - | - | 1733 | 959.5 | - | - | 0 | - |
| - | - | 1985 | 970.4 | - | - | 0 | - |
| - | - | 4056 | 971.5 | - | - | 0 | - |
| - | - | 1393 | 972.5 | - | - | 0 | - |
| - | - | 1026 | 973.5 | - | - | 0 | - |
| 9 | y | 1229 | 986.5 | 0.007778 | 7.885 | +1 | 9 |
| - | - | 1790 | 987.5 | - | - | 0 | - |
| - | - | 1045 | 988.5 | - | - | 0 | - |
| - | - | 977.8 | 998.4 | - | - | 0 | - |
| 9 | y | 8446 | 1004 | 0.0002774 | 0.2764 | +1 | 9 |
| - | - | 4684 | 1005 | - | - | 0 | - |
| - | - | 1755 | 1006 | - | - | 0 | - |
| - | - | 3740 | 1008 | - | - | 0 | - |
| - | - | 1939 | 1009 | - | - | 0 | - |
| - | - | 960.1 | 1011 | - | - | 0 | - |
| - | - | 3436 | 1015 | - | - | 0 | - |
| - | - | 2013 | 1016 | - | - | 0 | - |
| - | - | 1318 | 1017 | - | - | 0 | - |
| - | - | 1903 | 1018 | - | - | 0 | - |
| - | - | 960.3 | 1022 | - | - | 0 | - |
| - | - | 1042 | 1023 | - | - | 0 | - |
| - | - | 3705 | 1025 | - | - | 0 | - |
| - | - | 1684 | 1026 | - | - | 0 | - |
| - | - | 3008 | 1035 | - | - | 0 | - |
| - | - | 8634 | 1036 | - | - | 0 | - |
| - | - | 5005 | 1037 | - | - | 0 | - |
| - | - | 935.8 | 1038 | - | - | 0 | - |
| - | - | 4275 | 1041 | - | - | 0 | - |
| - | - | 4958 | 1041 | - | - | 0 | - |
| - | - | 1838 | 1042 | - | - | 0 | - |
| - | - | 1270 | 1052 | - | - | 0 | - |
| - | - | 3.833E+04 | 1053 | - | - | 0 | - |
| - | - | 2.414E+04 | 1054 | - | - | 0 | - |
| - | - | 7532 | 1055 | - | - | 0 | - |
| - | - | 1046 | 1056 | - | - | 0 | - |
| - | - | 6130 | 1059 | - | - | 0 | - |
| - | - | 3793 | 1060 | - | - | 0 | - |
| - | - | 2051 | 1068 | - | - | 0 | - |
| - | - | 1928 | 1069 | - | - | 0 | - |
| 10 | b | 1834 | 1070 | 0.01834 | 17.15 | +1 | 10 |
| - | - | 1.034E+04 | 1071 | - | - | 0 | - |
| - | - | 6477 | 1072 | - | - | 0 | - |
| 8 | y | 3087 | 1073 | 0.01472 | 13.73 | +1 | 10 |
| 8 | y | 2277 | 1074 | 0.004924 | 4.587 | +1 | 10 |
| - | - | 8309 | 1085 | - | - | 0 | - |
| - | - | 4941 | 1086 | - | - | 0 | - |
| - | - | 3785 | 1087 | - | - | 0 | - |
| 10 | b | 6889 | 1088 | 0.004128 | 3.796 | +1 | 10 |
| - | - | 3055 | 1089 | - | - | 0 | - |
| 8 | y | 1.544E+04 | 1091 | 1.816E-05 | 0.01666 | +1 | 10 |
| - | - | 8042 | 1092 | - | - | 0 | - |
| - | - | 2463 | 1093 | - | - | 0 | - |
| - | - | 996.8 | 1102 | - | - | 0 | - |
| - | - | 1035 | 1111 | - | - | 0 | - |
| - | - | 1543 | 1121 | - | - | 0 | - |
| - | - | 900.1 | 1122 | - | - | 0 | - |
| - | - | 1839 | 1129 | - | - | 0 | - |
| - | - | 1252 | 1130 | - | - | 0 | - |
| - | - | 7141 | 1138 | - | - | 0 | - |
| - | - | 7912 | 1139 | - | - | 0 | - |
| - | - | 3974 | 1140 | - | - | 0 | - |
| - | - | 1088 | 1143 | - | - | 0 | - |
| - | - | 1015 | 1147 | - | - | 0 | - |
| - | - | 1.29E+04 | 1156 | - | - | 0 | - |
| - | - | 8686 | 1157 | - | - | 0 | - |
| - | - | 3207 | 1158 | - | - | 0 | - |
| 7 | y | 3163 | 1160 | 0.002439 | 2.103 | +1 | 11 |
| 7 | y | 4020 | 1161 | 0.006221 | 5.361 | +1 | 11 |
| - | - | 1703 | 1162 | - | - | 0 | - |
| - | - | 1884 | 1164 | - | - | 0 | - |
| - | - | 7304 | 1165 | - | - | 0 | - |
| - | - | 4528 | 1166 | - | - | 0 | - |
| - | - | 1604 | 1167 | - | - | 0 | - |
| - | - | 1137 | 1173 | - | - | 0 | - |
| - | - | 907.6 | 1174 | - | - | 0 | - |
| 7 | y | 1.876E+04 | 1178 | 6.415E-05 | 0.05448 | +1 | 11 |
| - | - | 1.206E+04 | 1179 | - | - | 0 | - |
| - | - | 3576 | 1180 | - | - | 0 | - |
| - | - | 4.512E+04 | 1182 | - | - | 0 | - |
| 11 | b | 3.406E+04 | 1183 | 0.02074 | 17.54 | +1 | 11 |
| - | - | 8980 | 1184 | - | - | 0 | - |
| - | - | 1119 | 1185 | - | - | 0 | - |
| - | - | 1663 | 1200 | - | - | 0 | - |
| 11 | b | 7938 | 1201 | 0.009945 | 8.283 | +1 | 11 |
| - | - | 5812 | 1202 | - | - | 0 | - |
| - | - | 1397 | 1203 | - | - | 0 | - |
| - | - | 1643 | 1208 | - | - | 0 | - |
| - | - | 1396 | 1209 | - | - | 0 | - |
| - | - | 1823 | 1213 | - | - | 0 | - |
| - | - | 1176 | 1217 | - | - | 0 | - |
| - | - | 2678 | 1218 | - | - | 0 | - |
| - | - | 2615 | 1219 | - | - | 0 | - |
| - | - | 3700 | 1225 | - | - | 0 | - |
| - | - | 2893 | 1226 | - | - | 0 | - |
| - | - | 854.5 | 1231 | - | - | 0 | - |
| - | - | 960.3 | 1232 | - | - | 0 | - |
| - | - | 1.842E+04 | 1235 | - | - | 0 | - |
| - | - | 1.745E+04 | 1236 | - | - | 0 | - |
| - | - | 9632 | 1237 | - | - | 0 | - |
| - | - | 1505 | 1238 | - | - | 0 | - |
| - | - | 1158 | 1240 | - | - | 0 | - |
| - | - | 4.086E+04 | 1253 | - | - | 0 | - |
| - | - | 2.947E+04 | 1254 | - | - | 0 | - |
| - | - | 8120 | 1255 | - | - | 0 | - |
| 6 | y | 4818 | 1257 | 0.0001491 | 0.1186 | +1 | 12 |
| 6 | y | 9617 | 1258 | 0.003506 | 2.788 | +1 | 12 |
| - | - | 6691 | 1259 | - | - | 0 | - |
| - | - | 1516 | 1260 | - | - | 0 | - |
| - | - | 2141 | 1271 | - | - | 0 | - |
| - | - | 1186 | 1272 | - | - | 0 | - |
| 6 | y | 1.085E+05 | 1275 | 0.001436 | 1.127 | +1 | 12 |
| - | - | 7.096E+04 | 1276 | - | - | 0 | - |
| - | - | 1.951E+04 | 1277 | - | - | 0 | - |
| - | - | 2273 | 1278 | - | - | 0 | - |
| - | - | 1324 | 1293 | - | - | 0 | - |
| - | - | 1406 | 1301 | - | - | 0 | - |
| - | - | 832.1 | 1309 | - | - | 0 | - |
| - | - | 6076 | 1310 | - | - | 0 | - |
| 12 | b | 5735 | 1311 | 0.00735 | 5.608 | +1 | 12 |
| 12 | b | 3188 | 1312 | 0.01198 | 9.135 | +1 | 12 |
| - | - | 2523 | 1326 | - | - | 0 | - |
| - | - | 2392 | 1327 | - | - | 0 | - |
| - | - | 2503 | 1328 | - | - | 0 | - |
| 12 | b | 1.435E+04 | 1329 | 0.006755 | 5.084 | +1 | 12 |
| - | - | 9758 | 1330 | - | - | 0 | - |
| - | - | 3029 | 1331 | - | - | 0 | - |
| - | - | 2584 | 1336 | - | - | 0 | - |
| - | - | 3914 | 1337 | - | - | 0 | - |
| - | - | 2796 | 1338 | - | - | 0 | - |
| - | - | 950 | 1353 | - | - | 0 | - |
| 5 | y | 2.132E+04 | 1354 | 0.002498 | 1.845 | +1 | 13 |
| 5 | y | 3.775E+04 | 1355 | 0.004942 | 3.648 | +1 | 13 |
| - | - | 2.163E+04 | 1356 | - | - | 0 | - |
| - | - | 7177 | 1357 | - | - | 0 | - |
| - | - | 962.6 | 1358 | - | - | 0 | - |
| - | - | 3118 | 1370 | - | - | 0 | - |
| - | - | 3252 | 1371 | - | - | 0 | - |
| 5 | y | 6.398E+05 | 1372 | 0.001588 | 1.158 | +1 | 13 |
| - | - | 4.618E+05 | 1373 | - | - | 0 | - |
| - | - | 1.643E+05 | 1374 | - | - | 0 | - |
| - | - | 1.59E+04 | 1375 | - | - | 0 | - |
| 13 | b | 2436 | 1382 | 0.007669 | 5.551 | +1 | 13 |
| 13 | b | 2430 | 1383 | 0.007583 | 5.484 | +1 | 13 |
| - | - | 908.6 | 1384 | - | - | 0 | - |
| 13 | b | 1.079E+04 | 1400 | 0.00798 | 5.701 | +1 | 13 |
| - | - | 8134 | 1401 | - | - | 0 | - |
| - | - | 3038 | 1402 | - | - | 0 | - |
| 14 | b | 1291 | 1497 | 0.004206 | 2.81 | +1 | 14 |
| - | - | 1123 | 1498 | - | - | 0 | - |
| 4 | y | 1984 | 1501 | 0.004261 | 2.84 | +1 | 14 |
| 4 | y | 1983 | 1502 | 0.00378 | 2.517 | +1 | 14 |
| - | - | 1747 | 1503 | - | - | 0 | - |
| 14 | b | 4854 | 1514 | 0.003544 | 2.341 | +1 | 14 |
| - | - | 5111 | 1515 | - | - | 0 | - |
| - | - | 2328 | 1516 | - | - | 0 | - |
| 4 | y | 3.24E+04 | 1519 | 0.002619 | 1.724 | +1 | 14 |
| - | - | 3.157E+04 | 1520 | - | - | 0 | - |
| - | - | 1.128E+04 | 1521 | - | - | 0 | - |
| - | - | 1376 | 1522 | - | - | 0 | - |
| 3 | y | 6758 | 1632 | 0.006727 | 4.122 | +1 | 15 |
| - | - | 6241 | 1633 | - | - | 0 | - |
| - | - | 2542 | 1634 | - | - | 0 | - |
| - | - | 911.5 | 3073 | - | - | 0 | - |

m/z Charge Intensity FragmentType MassShift Position
120.0658187866211 0 3397.846 y 16
120.08110809326172 0 58256.918
121.08444213867188 0 3845.5957
125.86992645263672 0 474.02066
126.0552749633789 0 11476.65
127.05860900878906 0 679.6393
127.08674621582031 0 861.42346
128.1072998046875 0 15488.092
129.0661163330078 0 3248.0337
129.10256958007812 0 103184.77
130.05015563964844 0 2587.6165
130.1002655029297 0 915.48206
130.1059112548828 0 6903.153
131.1182098388672 0 2193.4358
132.07705688476562 0 1692.8088
132.10260009765625 0 476.8071
136.0762939453125 0 1044.8718
137.10792541503906 0 503.27295
138.09152221679688 0 3180.282
138.12802124023438 0 8456.944
139.08688354492188 0 4473.805
139.123291015625 0 652.5096
139.13111877441406 0 691.0612
140.07113647460938 0 602.8659
140.09043884277344 0 523.01385
140.14385986328125 0 610.30444
141.06626892089844 0 1572.6852
141.10272216796875 0 1531.9364
142.1230926513672 0 655.9808
147.07676696777344 0 3082.32
151.12318420410156 0 678.83905
152.1072998046875 0 554.6272
153.1024627685547 0 1294.809
153.45082092285156 0 558.7834
154.08651733398438 0 2287.067
154.0977783203125 0 895.64197
155.081787109375 0 2815.93
155.11822509765625 0 14835.414 a Water loss 1
156.12152099609375 0 851.4358
157.0610809326172 0 4204.435
157.0974578857422 0 12335.015
157.13433837890625 0 745.07935 d 1
165.1024627685547 0 2533.3677
166.08651733398438 0 1810.6333
167.0818634033203 0 9349.083
167.11814880371094 0 7071.16
168.12127685546875 0 628.73566
169.0612030029297 0 1452.2716
169.09744262695312 0 869.6491
169.13385009765625 0 14291.544
170.13726806640625 0 1680.2535
171.07681274414062 0 1014.3965
172.10809326171875 0 1752.7438
172.1452178955078 0 563.5757
173.09239196777344 0 3991.4917 y Water loss 15
173.12881469726562 0 44796.9 a 1
173.4404296875 0 5779.817
174.1282196044922 0 4450.6187
175.07164001464844 0 4411.9277
175.13218688964844 0 693.0785
176.10777282714844 0 713.6344
177.10276794433594 0 609.04425
180.1133270263672 0 1415.6487
181.09747314453125 0 4898.784
182.08511352539062 0 547.7235
182.09291076660156 0 3007.153
182.12925720214844 0 5312.158
183.07688903808594 0 3792.9126
183.1131591796875 0 182583.02 b Water loss 1
184.10842895507812 0 4519.105
184.1166229248047 0 12293.237
185.08282470703125 0 965.4756
185.09237670898438 0 18775.635
185.11862182617188 0 549.39014
186.08767700195312 0 3755.2527
186.09547424316406 0 958.9684
187.1444549560547 0 28702.768
188.1473846435547 0 2550.022
189.08738708496094 0 5513.7954
191.0818328857422 0 1163.7164
191.1030731201172 0 4778.1274 y 15
195.07638549804688 0 1389.0569
195.1131134033203 0 12855.981
196.11680603027344 0 1141.2273
197.12875366210938 0 20091.588
198.12408447265625 0 4621.8164
198.1324462890625 0 1322.7941
199.07168579101562 0 5927.743
199.10789489746094 0 2504.4302
199.11839294433594 0 1283.4546
200.10333251953125 0 6565.79
200.1396484375 0 6825.759
201.0875244140625 0 1758.9299
201.12367248535156 0 92810.25 b 1
202.1179962158203 0 1080.1475
202.12713623046875 0 6458.4956
203.10284423828125 0 1846.1311
203.1138916015625 0 1113.0382
208.09767150878906 0 711.312
208.1083984375 0 7912.1396
209.09239196777344 0 22410.686
209.12867736816406 0 832.57874
210.09596252441406 0 1638.0891
210.12461853027344 0 1006.94336
211.14474487304688 0 1080.1049
212.1398468017578 0 2207.693
213.08743286132812 0 1498.6237
213.1236114501953 0 2468.2832
214.1552276611328 0 890.3274
215.1393585205078 0 34934.082
216.1425018310547 0 3334.5825
217.08216857910156 0 15751.733
217.13381958007812 0 21676.047
218.08560180664062 0 1377.2864
218.11473083496094 0 625.6149
218.13719177246094 0 2201.076
218.14968872070312 0 1129.3331
219.1496124267578 0 6913.3843
220.1528778076172 0 1511.1406
224.1041717529297 0 736.20807
224.17562866210938 0 1081.6082
225.12356567382812 0 8324.956
225.13526916503906 0 4728.048
225.16026306152344 0 1713.262
226.11898803710938 0 83012.516
227.10287475585938 0 2413.346
227.12222290039062 0 6145.3745
228.09825134277344 0 714.3811
228.13462829589844 0 1285.0538
231.09812927246094 0 951.2978
233.16525268554688 0 50270.42
234.1686553955078 0 6112.362
235.1440887451172 0 1356.8458
236.10340881347656 0 6018.17
236.1397705078125 0 1215.0443
237.08709716796875 0 1668.5148
237.1241455078125 0 713.96716
240.09877014160156 0 606.6186
241.08230590820312 0 5591.842
241.19140625 0 1274.1926
242.1133270263672 0 713.5167
242.1503448486328 0 14456.379
242.1643524169922 0 763.2052
243.13381958007812 0 3120.5967
243.145751953125 0 23907.957
244.13002014160156 0 4622.09 y Ammonia loss 12
244.14886474609375 0 1637.713
245.12899780273438 0 15443.418
246.13241577148438 0 1672.8892
247.1444549560547 0 8130.412
251.17578125 0 25206.79
252.0985565185547 0 703.0073
252.13470458984375 0 2532.7366
252.16831970214844 0 843.62976
252.17919921875 0 2326.0063
253.1173858642578 0 775.00574
253.1303253173828 0 1299.7046
253.1532745361328 0 857.8356
254.1138916015625 0 17616.068
254.15029907226562 0 4794.617
255.11727905273438 0 1597.407
257.1646728515625 0 642.51483
257.19610595703125 0 615.4098
258.1085510253906 0 982.61237
259.0928039550781 0 14674.293
259.1082763671875 0 748.4291
259.1446228027344 0 741.9981
260.0964050292969 0 1109.0231
260.1609802246094 0 978.225
261.1364440917969 0 604.93805
261.1600341796875 0 15299.1
262.1187744140625 0 3151.3152
262.163818359375 0 2438.6704
263.1396179199219 0 1045.15
264.0990295410156 0 1055.4943
264.1346130371094 0 4787.9307
265.13885498046875 0 592.1411
266.1150817871094 0 650.98846
266.1498718261719 0 825.8294
268.09326171875 0 2303.7227
268.2023620605469 0 5469.311 a Water loss 2
269.12591552734375 0 929.6964
269.16119384765625 0 1385.8093
269.1861267089844 0 5695.577
269.2030029296875 0 995.2788
270.14544677734375 0 1286.9231
270.1817932128906 0 3018.308
271.1422424316406 0 915.22723
271.18267822265625 0 624.55133
272.1243591308594 0 11015.259
273.1264343261719 0 1151.2229
275.1756896972656 0 9314.54
276.1190185546875 0 748.8371
276.17889404296875 0 1426.1302
277.1187438964844 0 1200.5258
278.1121520996094 0 904.447
278.1498718261719 0 1314.0928
279.1458435058594 0 8637.286
280.1296691894531 0 3920.651
280.14862060546875 0 1155.1754
280.166015625 0 13972.891
281.168212890625 0 2097.1316
282.1092529296875 0 1171.7374
282.1452331542969 0 27895.936
283.1480407714844 0 4019.4045
284.1614685058594 0 937.74176
285.1601867675781 0 12048.912
285.1972961425781 0 1181.5415
286.1038513183594 0 10133.81
286.1637268066406 0 1818.934
286.2131652832031 0 1180.115 a 2
287.1064147949219 0 1463.2189
294.1820068359375 0 2290.2432
296.08831787109375 0 2321.4531
296.19744873046875 0 471524.62 b Water loss 2
297.1199645996094 0 4331.2373
297.1563415527344 0 26415.893
297.2005920410156 0 69091.28
298.1589660644531 0 3210.9045
298.1766662597656 0 17404.62
298.20306396484375 0 4866.788
299.088134765625 0 1276.1241
299.17352294921875 0 2754.0747
300.1189880371094 0 1467.3291
300.15557861328125 0 1710.8406
300.1717834472656 0 1863.9309
301.1561584472656 0 1358.7172
301.1878356933594 0 2296.5457 y Water loss 14
302.1722106933594 0 880.37866 y Ammonia loss 14
303.1708984375 0 19414.805
304.1144714355469 0 5385.009
304.17254638671875 0 2767.3704
306.1455383300781 0 2576.0803
307.11083984375 0 589.3171
307.14093017578125 0 1780.675
308.16119384765625 0 1768.7646 y Ammonia loss 11
310.1035461425781 0 2087.376
310.1384582519531 0 803.50116
311.0882873535156 0 761.836
311.17254638671875 0 702.191
312.19207763671875 0 864.9121
313.1697692871094 0 714.55457
313.1883544921875 0 4489.1436
314.099853515625 0 992.85425
314.14697265625 0 4536.859
314.1836242675781 0 17782.494
314.2080383300781 0 118602.24 b 2
315.1487121582031 0 792.088
315.16680908203125 0 1591.7101
315.18609619140625 0 2254.024
315.21099853515625 0 18214.295
316.2017517089844 0 697.6113
316.2138977050781 0 716.41235
317.18658447265625 0 5094.3594
318.13031005859375 0 798.3618
319.1981201171875 0 31463.57 y 14
320.20135498046875 0 4234.4243
321.1563720703125 0 800.34235
323.1717224121094 0 2438.5408
324.15423583984375 0 1234.1787
325.1510925292969 0 1793.9872
326.17047119140625 0 1636.0603
328.11431884765625 0 7579.059
328.1669616699219 0 918.9835
329.119384765625 0 917.482
329.18341064453125 0 1201.4191
330.1817626953125 0 12808.011
331.1852111816406 0 2459.0361
333.156005859375 0 3067.4976
334.2129211425781 0 3643.4934
335.1715393066406 0 909.0949
336.1558837890625 0 1948.427
337.15179443359375 0 3935.7576
338.1372375488281 0 1144.9061
338.15478515625 0 751.5005
339.1671447753906 0 2630.7654
339.2385559082031 0 831.0613
341.1842956542969 0 8229.646
342.182861328125 0 3059.4607
344.19744873046875 0 3335.7917
345.201416015625 0 752.8956
346.125 0 21916.836
346.1766357421875 0 17544.275
347.1294250488281 0 3660.1074
347.1798095703125 0 2887.7493
348.1921081542969 0 16008.062
349.195068359375 0 3084.8071
350.1828918457031 0 4708.169
351.1666564941406 0 14607.9795
352.1699523925781 0 1887.4058
353.1823425292969 0 8981.742
354.16650390625 0 5295.926
354.1882629394531 0 1086.2172
355.1622009277344 0 4991.655
356.1443786621094 0 1488.3683
358.211181640625 0 861.1489
360.2284240722656 0 2409.5444
361.2308349609375 0 916.48
362.2079162597656 0 37081.402
363.1668701171875 0 1374.5822
363.2110595703125 0 7250.314
365.1459655761719 0 8528.855
365.172607421875 0 963.3984
365.1815490722656 0 849.1498
365.25494384765625 0 2406.8499
366.2602233886719 0 846.7856
367.1622314453125 0 712.7774
367.2087707519531 0 5321.7144
367.23486328125 0 1981.6696
368.1934509277344 0 3293.042
368.2364807128906 0 671.8908
369.13946533203125 0 1209.0269
369.1772155761719 0 22831.615
370.1798400878906 0 3156.8264
370.6768493652344 0 2715.9888
371.1557312011719 0 1121.3623
371.1926574707031 0 5593.3467
372.1764221191406 0 4500.032
373.17303466796875 0 8002.816
374.1751403808594 0 1701.448
375.1662902832031 0 1116.8447
379.1265869140625 0 1191.2618
379.1641540527344 0 692.9372
379.1936950683594 0 1120.5977
381.17755126953125 0 1997.5045
383.156494140625 0 14010.305
383.26593017578125 0 3348.4214
384.1595458984375 0 2519.566
384.26690673828125 0 1281.8094
385.2197265625 0 11083.201
386.2044982910156 0 2036.9974
386.2269287109375 0 1152.5325
387.1502990722656 0 1073.9655
388.13470458984375 0 1582.5905
389.2550964355469 0 1108.9213
390.2040100097656 0 1140.5052
392.19305419921875 0 777.04553
393.17742919921875 0 2532.876
393.25054931640625 0 7107.8696
394.2535095214844 0 1814.5724
395.1939697265625 0 1117.4402
397.1360168457031 0 2868.352
398.11981201171875 0 1384.1927
398.2041931152344 0 1708.4521
398.2440490722656 0 7691.2114
399.187255859375 0 2013.9478
399.24615478515625 0 2590.7231
400.22412109375 0 2415.4836
401.16729736328125 0 21558.39
402.1709899902344 0 3724.9902
405.1627502441406 0 1912.9655
405.21270751953125 0 750.4247
407.2039489746094 0 3803.3743
408.18829345703125 0 7063.727
410.2033996582031 0 1277.0527
411.1515808105469 0 985.8988
411.2608642578125 0 11299.313
412.2627258300781 0 1681.6372
413.2062683105469 0 1267.2788
415.1466979980469 0 7062.735
415.2317199707031 0 2390.016 b 7
415.27117919921875 0 5722.316
416.1500244140625 0 1529.6344
416.2161865234375 0 1942.1335 y Ammonia loss 13
416.2550354003906 0 16531.113
417.2559814453125 0 5413.214
422.2019348144531 0 984.9231
424.22857666015625 0 3979.9712
425.1307067871094 0 2237.2825
425.1658630371094 0 1123.3417
425.2153015136719 0 7754.8994
426.166015625 0 719.9461
426.21246337890625 0 605.48773
427.1801452636719 0 752.0459
427.2328796386719 0 4753.065
429.2135314941406 0 1577.6239 y Ammonia loss 9
431.2150573730469 0 1269.3026
431.267578125 0 1140.2878
433.1572570800781 0 11346.749
433.2425231933594 0 23199.34 y 13
433.2817687988281 0 157163.95
434.16046142578125 0 1392.5024
434.2049255371094 0 3260.157
434.2455749511719 0 3727.2148
434.2847900390625 0 34917.21
434.7063903808594 0 2085.62
435.2077331542969 0 2026.5978
435.28790283203125 0 3639.4807
440.2151184082031 0 1554.2803
441.19866943359375 0 3379.921
441.2505798339844 0 1646.2513
442.2394104003906 0 4447.8154
443.1412353515625 0 3221.443
443.1747741699219 0 1006.3008
443.2301025390625 0 6558.2896
443.26611328125 0 16194.638 b Water loss 3
444.1423645019531 0 809.5005
444.18505859375 0 1403.7401
444.23583984375 0 1183.4587
444.2691650390625 0 3796.1902
445.2453308105469 0 1517.0922
451.22900390625 0 1689.925
452.21490478515625 0 4081.8315
453.21728515625 0 1261.1744
454.1971740722656 0 832.95355
459.1738586425781 0 1197.5123
459.2094421386719 0 5947.5913
459.2599182128906 0 1658.8501
460.2180480957031 0 711.58026
461.2767028808594 0 128481.125 b 3
462.1992492675781 0 6678.7764
462.2798156738281 0 34616.93
463.2010192871094 0 1082.1079
463.28253173828125 0 4923.013
464.2131652832031 0 1470.1195
464.2496337890625 0 892.9127
466.1936950683594 0 2926.3245
468.2099914550781 0 2346.7217
468.25274658203125 0 1644.4005
469.2427673339844 0 3213.4714
470.2252502441406 0 20149.135
471.2283935546875 0 4411.3687
476.1772155761719 0 1673.1116
477.25787353515625 0 1645.3868
478.2393798828125 0 3220.4368
479.2264404296875 0 3154.4722
480.2098388671875 0 16945.535
481.21240234375 0 4335.216
482.22515869140625 0 3066.0513
483.20965576171875 0 738.96674
484.20501708984375 0 6493.4834
485.28485107421875 0 925.4241
486.2685852050781 0 2534.7397 y Water loss 12
487.25286865234375 0 2645.7476 y Ammonia loss 12
488.2520446777344 0 857.12335
492.2086486816406 0 924.1041
494.18878173828125 0 7452.487
495.193359375 0 1021.33167
495.2681884765625 0 3174.7256
496.25128173828125 0 7027.209
497.2550048828125 0 1266.0052
498.2203369140625 0 50059.18
499.22320556640625 0 11264.656
500.2322998046875 0 2872.937
501.23516845703125 0 742.7246
502.21527099609375 0 11005.019
503.21856689453125 0 2523.689
503.2602233886719 0 839.94916
504.2782897949219 0 32914.613 y 12
505.2812805175781 0 6075.874
510.2202453613281 0 2619.987
512.1994018554688 0 11069.471
512.3222045898438 0 1697.4775
513.20263671875 0 2869.1704
513.2783203125 0 3643.8696
515.2479248046875 0 1644.739
516.2396850585938 0 855.642
518.24658203125 0 1865.9166
521.2733154296875 0 1004.7309
524.2362670898438 0 911.61176
527.2468872070312 0 1021.445
528.2301025390625 0 3071.5383
529.2273559570312 0 1484.8848
530.2100219726562 0 32529.104
530.334228515625 0 9589.372
531.2127685546875 0 7832.1895
531.337158203125 0 2923.6033
532.2147827148438 0 1028.4407
534.2576293945312 0 971.0577
538.2971801757812 0 2832.065
540.3186645507812 0 6268.23 b Water loss 4
541.3216552734375 0 1688.3303
542.7656860351562 0 950.42737
545.2380981445312 0 887.77826
546.2402954101562 0 3745.3132
547.2415161132812 0 1438.7573
551.2474365234375 0 808.5722
553.2616577148438 0 2293.5225
555.3253173828125 0 2441.8762
556.224609375 0 2921.8032
556.2726440429688 0 2210.783
556.3222045898438 0 919.9596
558.2078857421875 0 1039.0193
558.3291015625 0 16422.014 b 4
559.331787109375 0 5549.956
563.2454833984375 0 2214.056
564.2354125976562 0 712.2814
569.281494140625 0 3000.9512
569.7787475585938 0 2753.9968
570.2904052734375 0 698.8081
571.2743530273438 0 1809.4563
573.2318725585938 0 1013.0715
574.2745361328125 0 1029.2812
578.2860717773438 0 4872.21
578.7876586914062 0 4425.3735
579.2855224609375 0 2330.0476
581.2572021484375 0 6292.152
582.2880249023438 0 4544.417
582.7826538085938 0 11061.05
583.2369384765625 0 1236.1509
583.2833862304688 0 7587.319
583.7848510742188 0 962.7054
587.2667846679688 0 3068.5718
589.2620849609375 0 1119.3802
591.24072265625 0 4185.323
591.2946166992188 0 3592.566
591.7952880859375 0 3821.076 b Water loss 10
592.2416381835938 0 930.02374
596.3126220703125 0 907.31323
597.2921752929688 0 4730.6816
598.2876586914062 0 1709.7192
599.267822265625 0 32237.81
600.2711181640625 0 8590.587
601.27490234375 0 1258.4642
604.3032836914062 0 2718.9795
604.8038330078125 0 1029.5245
607.273681640625 0 1700.0894
608.2702026367188 0 1309.7921
608.3523559570312 0 2402.7612
608.7962646484375 0 867.4279
609.2517700195312 0 9341.276
609.2977294921875 0 2997.1194
609.3417358398438 0 1677.816
609.7955322265625 0 2607.9849
610.254638671875 0 2862.9045
612.8146362304688 0 7558.725
613.316162109375 0 3651.1433
613.8175659179688 0 1244.2213
614.3251953125 0 4436.8267 y Water loss 11
615.3051147460938 0 18281.916 y Ammonia loss 11
616.3058471679688 0 5997.4526
617.2994384765625 0 1816.5261
617.8073120117188 0 26916.523
618.3056030273438 0 30435.889
618.8048706054688 0 10882.493
619.302978515625 0 2773.0012
619.805908203125 0 807.66095
620.3043823242188 0 944.5902
624.3050537109375 0 823.8249
625.2836303710938 0 7153.1606
626.2855834960938 0 2035.2552
626.3622436523438 0 4482.761
626.7492065429688 0 907.5226
626.8126831054688 0 66426.21
627.26220703125 0 76586.266
627.3145751953125 0 47161.47
627.814208984375 0 15570.778
628.265625 0 22802.133
628.3136596679688 0 1295.1964
628.810791015625 0 1320.0776 y Water loss 5
629.2673950195312 0 2749.3525
629.3162231445312 0 1433.9409
629.8096313476562 0 1564.9744
630.3071899414062 0 930.6953
632.3367919921875 0 21983.861 y 11
633.3389892578125 0 5263.955
635.8185424804688 0 2390.6826
636.31884765625 0 2691.8555
637.8153686523438 0 10387.771 y 5
638.316162109375 0 5723.993
638.8168334960938 0 1018.7497
639.2586669921875 0 942.55316
642.3167114257812 0 1177.416
643.2943725585938 0 15041.211
644.294189453125 0 6794.6807
645.2899780273438 0 3071.8184
646.8272094726562 0 1505.448
649.3319702148438 0 975.2356
653.282958984375 0 1232.8635
655.3404541015625 0 5149.489
655.8410034179688 0 3925.8914 b Water loss 11
656.2885131835938 0 863.3793
656.34521484375 0 941.2394
658.3074951171875 0 2227.814
659.3143920898438 0 1033.6117
659.8126220703125 0 876.5331
660.3192749023438 0 6396.558
661.3228759765625 0 2292.1973
663.3363647460938 0 2517.2625
663.8401489257812 0 2598.2134
664.3358154296875 0 1928.4946
664.83154296875 0 1304.0175 b 11
665.3352661132812 0 1088.0507
667.337646484375 0 1498.307
668.3304443359375 0 2532.1072
668.8241577148438 0 5101.059
669.323486328125 0 2697.1387
669.8272094726562 0 1295.3959
670.313720703125 0 894.22485
674.2982177734375 0 1583.1611
676.334716796875 0 1098.2266
677.3369140625 0 21081.99 y Water loss 4
677.83251953125 0 30506.13 y Ammonia loss 4
678.33203125 0 17566.863
678.8322143554688 0 4711.418
679.2589111328125 0 1190.0343
679.3270874023438 0 785.85834
684.3728637695312 0 916.255
685.3338623046875 0 2286.1401
685.8330688476562 0 1739.5657
686.3418579101562 0 546249.25 y 4
686.843017578125 0 392037.97
687.3439331054688 0 135570.62
687.8451538085938 0 14894.434
691.3446044921875 0 1324.6158 b Water loss 12
691.8445434570312 0 1075.0269 b Ammonia loss 12
694.3408203125 0 4265.5293
695.3407592773438 0 2299.1592
696.3545532226562 0 983.0323
698.29736328125 0 953.5812
700.34619140625 0 6323.9956 b 12
700.8472290039062 0 4865.804
701.35107421875 0 2226.2432
704.324462890625 0 2572.119
705.3297729492188 0 1172.5132
709.30810546875 0 841.08014
712.3519287109375 0 44844.12
713.3548583984375 0 14232.63
714.3515014648438 0 2197.7202
720.2904052734375 0 793.5203
720.3682250976562 0 1102.9539
722.3364868164062 0 6816.7256
723.3392333984375 0 2398.1858
724.40087890625 0 1620.5367 b Water loss 6
726.3339233398438 0 1014.30695
727.329345703125 0 1090.1964
727.4108276367188 0 886.91516 y Water loss 10
728.396728515625 0 1876.794 y Ammonia loss 10
735.33544921875 0 889.5297
735.5277709960938 0 884.03705
737.3181762695312 0 1333.1571
737.390625 0 1828.9731
738.2987670898438 0 2514.0098
738.3834838867188 0 1926.278
739.29638671875 0 933.99054
739.3641967773438 0 2853.6362
740.3468627929688 0 53495.785
741.3500366210938 0 16509.775
742.3529663085938 0 4489.5625
742.4159545898438 0 3747.3591 b 6
743.3555297851562 0 1546.0085
743.4192504882812 0 1710.5685
745.3392333984375 0 2781.9678
745.4212646484375 0 15977.298 y 10
746.3384399414062 0 2032.668
746.4227294921875 0 7459.1704
747.4288330078125 0 1207.4841
750.8731689453125 0 2324.98 y Water loss 3
751.3671875 0 4277.559 y Ammonia loss 3
751.8668212890625 0 2237.768
752.365234375 0 1119.223
753.3424682617188 0 3146.4944
754.3351440429688 0 2511.3752
755.3294677734375 0 1584.7789
755.4050903320312 0 2631.4712
756.3262329101562 0 1322.4789
756.4037475585938 0 1119.9146
756.8881225585938 0 1603.8306
757.374267578125 0 27133.553 b 13
757.8906860351562 0 791.2165
758.3759765625 0 9361.21
759.3756713867188 0 2397.2522
759.8759765625 0 70335.266 y 3
760.3767700195312 0 62533.586
760.8781127929688 0 26033.943
761.3779907226562 0 2105.305
766.36279296875 0 927.49475
771.3523559570312 0 12843.111
772.3529052734375 0 5371.609
773.3525390625 0 1194.898
774.33154296875 0 3131.9485
775.342041015625 0 967.49316
796.3869018554688 0 1087.3256
797.37060546875 0 1350.4832
807.417724609375 0 1430.7808 y Water loss 2
807.9102172851562 0 1652.3448 y Ammonia loss 2
808.4175415039062 0 1025.2196
811.3837280273438 0 2079.1838
812.9183349609375 0 1789.8267 b Ammonia loss 14
813.4197387695312 0 1311.5006
813.919189453125 0 993.1296
814.3934936523438 0 1899.9421
815.39013671875 0 1275.3407
816.4175415039062 0 13835.041 y 2
816.919677734375 0 13407.727
817.42041015625 0 6760.759
821.4196166992188 0 1083.0916 b 14
821.9292602539062 0 2382.7168
822.4037475585938 0 1767.8265
823.3915405273438 0 2049.8452
824.3793334960938 0 2801.7502
825.372802734375 0 1665.8577
828.4130249023438 0 1307.7135
829.4425659179688 0 3903.2622 b 7
830.4500732421875 0 2037.0161
832.3847045898438 0 1638.6506
833.3782348632812 0 1231.2784
839.4230346679688 0 1877.4152
840.4108276367188 0 7515.8564
841.4077758789062 0 3043.809
842.3898315429688 0 9104.157
842.9529418945312 0 1469.2119
843.3897094726562 0 3680.3496
844.3972778320312 0 776.11475
847.9419555664062 0 3188.1265 b Water loss 15
848.436767578125 0 5553.9624 b Ammonia loss 15
848.9382934570312 0 3635.3284
849.4321899414062 0 1023.1735
850.3950805664062 0 8175.782
851.3909301757812 0 6575.448
852.39404296875 0 2076.0105
856.4559936523438 0 1554.1399 y Water loss 9
856.946533203125 0 14400.561 b 15
857.4462890625 0 17682.35 y Ammonia loss 9
857.9475708007812 0 7575.2104 y Water loss 1
858.4459228515625 0 1734.0093
859.414306640625 0 2588.1604
866.443115234375 0 1249.0858
866.9418334960938 0 4574.0444 y 1
867.4382934570312 0 6911.2695
867.9456787109375 0 994.5245
868.4052734375 0 64932.2
869.4080810546875 0 28860.049
870.4100341796875 0 5788.1226
874.4627685546875 0 12739.272 y 9
875.4650268554688 0 5061.8066
876.4738159179688 0 989.2319
884.4423217773438 0 1819.8417
885.4451904296875 0 1858.0323
886.4495239257812 0 999.75354
887.4152221679688 0 4422.916
888.4195556640625 0 1720.345
893.4298706054688 0 936.1671
894.4223022460938 0 5281.5396
895.43408203125 0 2567.7773
898.9613647460938 0 1581.4192
899.4620971679688 0 1815.1073
900.4409790039062 0 1252.649
904.4237670898438 0 1841.9744
905.420166015625 0 809.7622
907.4696044921875 0 10849.69 Precursor Water loss
907.969482421875 0 17736.48 Precursor Ammonia loss
908.4697875976562 0 9129.142
908.9692993164062 0 2597.838
911.4459228515625 0 12651.62
912.4476318359375 0 4670.423
913.449462890625 0 1472.6926
915.470458984375 0 1691.001
915.96630859375 0 1783.9458
916.368408203125 0 962.1822
916.475830078125 0 116772.52 Precursor
916.976806640625 0 127603.59
917.4777221679688 0 63000.703
917.9794311523438 0 9276.133
921.429443359375 0 6390.096
922.4213256835938 0 8294.48
923.421875 0 2628.2517
926.4526977539062 0 875.19135
938.4397583007812 0 963.3314
939.4413452148438 0 47762.848
940.4447631835938 0 22048.73
941.4485473632812 0 5402.925
942.4445190429688 0 1168.0115
953.4667358398438 0 2118.7537
954.4590454101562 0 2478.361
955.46484375 0 1013.94073
956.4479370117188 0 7395.7524
957.4551391601562 0 3091.2297
958.4832763671875 0 3307.9153 b 8
959.4896850585938 0 1733.042
970.4489135742188 0 1985.3396
971.4734497070312 0 4055.583
972.4833984375 0 1392.6677
973.459716796875 0 1025.6919
986.4866943359375 0 1228.755 y Ammonia loss 8
987.4766235351562 0 1789.9092
988.4758911132812 0 1045.1074
998.4464721679688 0 977.75104
1003.5051879882812 0 8445.988 y 8
1004.50390625 0 4684.3213
1005.5113525390625 0 1755.1394
1008.4628295898438 0 3739.634
1009.4658203125 0 1939.2965
1011.4823608398438 0 960.1111
1015.4749145507812 0 3436.3386
1016.4763793945312 0 2012.5134
1017.4717407226562 0 1317.858
1018.447998046875 0 1902.6115
1022.4901733398438 0 960.3198
1023.4762573242188 0 1042.257
1025.489013671875 0 3705.2485
1026.4888916015625 0 1684.3772
1035.4786376953125 0 3007.7092
1036.46142578125 0 8634.112
1037.4652099609375 0 5004.845
1038.479248046875 0 935.7847
1040.5013427734375 0 4274.504
1041.491943359375 0 4957.557
1042.4932861328125 0 1838.2942
1052.4951171875 0 1269.938
1053.4849853515625 0 38329.93
1054.4854736328125 0 24138.244
1055.4874267578125 0 7532.251
1056.49169921875 0 1045.9652
1058.51220703125 0 6129.756
1059.517578125 0 3792.882
1067.50390625 0 2051.4583
1068.5068359375 0 1928.2085
1069.501708984375 0 1833.5453 b Water loss 9
1070.5111083984375 0 10343.734
1071.513671875 0 6477.207
1072.51220703125 0 3086.66 y Water loss 7
1073.515869140625 0 2276.934 y Ammonia loss 7
1084.5269775390625 0 8308.666
1085.5284423828125 0 4940.9663
1086.5135498046875 0 3785.4973
1087.5264892578125 0 6889.0083 b 9
1088.5277099609375 0 3054.9187
1090.5374755859375 0 15436.393 y 7
1091.5377197265625 0 8041.934
1092.54248046875 0 2462.9026
1101.5335693359375 0 996.8443
1110.541748046875 0 1035.248
1120.5264892578125 0 1543.2578
1121.5335693359375 0 900.13464
1128.55517578125 0 1839.1604
1129.570068359375 0 1251.5336
1137.5537109375 0 7141.0176
1138.5478515625 0 7912.263
1139.5489501953125 0 3973.7722
1142.5362548828125 0 1088.192
1146.5576171875 0 1014.9385
1155.5635986328125 0 12900.559
1156.5638427734375 0 8686.371
1157.56689453125 0 3207.3745
1159.5565185546875 0 3163.374 y Water loss 6
1160.5491943359375 0 4020.3237 y Ammonia loss 6
1161.5565185546875 0 1703.4979
1163.573974609375 0 1884.1605
1164.5565185546875 0 7303.666
1165.55908203125 0 4527.863
1166.5572509765625 0 1604.06
1172.6044921875 0 1136.5541
1173.5701904296875 0 907.57153
1177.5694580078125 0 18761.715 y 6
1178.57080078125 0 12058.903
1179.5743408203125 0 3576.3406
1181.57958984375 0 45116.938
1182.5833740234375 0 34062.777 b Water loss 10
1183.585205078125 0 8980.208
1184.5675048828125 0 1119.2107
1199.5966796875 0 1662.6444
1200.604736328125 0 7938.004 b 10
1201.610595703125 0 5811.6514
1202.6181640625 0 1396.6628
1207.601318359375 0 1643.1116
1208.599609375 0 1395.9152
1212.623046875 0 1822.8839
1216.5926513671875 0 1175.9026
1217.5906982421875 0 2677.5593
1218.5794677734375 0 2615.1597
1224.61962890625 0 3699.6904
1225.619873046875 0 2892.5525
1230.593994140625 0 854.4824
1231.60400390625 0 960.29346
1234.6053466796875 0 18424.541
1235.600341796875 0 17452.385
1236.59912109375 0 9632.133
1237.591796875 0 1504.7645
1239.594482421875 0 1158.3783
1252.6153564453125 0 40858.406
1253.617919921875 0 29470.504
1254.6195068359375 0 8120.189
1256.611572265625 0 4817.8276 y Water loss 5
1257.5992431640625 0 9616.768 y Ammonia loss 5
1258.60107421875 0 6690.959
1259.603515625 0 1516.0779
1270.6328125 0 2140.9702
1271.623779296875 0 1186.2719
1274.620849609375 0 108466.586 y 5
1275.6231689453125 0 70958.555
1276.6248779296875 0 19508.344
1277.6273193359375 0 2273.3762
1292.64453125 0 1324.1569
1300.6617431640625 0 1405.514
1308.6318359375 0 832.1099
1309.671142578125 0 6075.714
1310.6700439453125 0 5734.972 b Water loss 11
1311.65869140625 0 3187.7375 b Ammonia loss 11
1325.6678466796875 0 2522.507
1326.6669921875 0 2392.396
1327.659423828125 0 2502.9958
1328.66650390625 0 14348.351 b 11
1329.6702880859375 0 9757.715
1330.67333984375 0 3028.6099
1335.65185546875 0 2584.1465
1336.643798828125 0 3913.7563
1337.64013671875 0 2796.376
1353.478759765625 0 949.95917
1353.6619873046875 0 21321.371 y Water loss 4
1354.6534423828125 0 37746.496 y Ammonia loss 4
1355.654052734375 0 21628.715
1356.65576171875 0 7176.898
1357.6524658203125 0 962.6494
1369.6551513671875 0 3117.5808
1370.659423828125 0 3251.587
1371.6734619140625 0 639806.3 y 4
1372.6759033203125 0 461819.72
1373.677734375 0 164295.78
1374.67919921875 0 15900.118
1381.692138671875 0 2435.77 b Water loss 12
1382.69140625 0 2429.9353 b Ammonia loss 12
1383.692138671875 0 908.57513
1399.702392578125 0 10792.041 b 12
1400.7034912109375 0 8133.8525
1401.70751953125 0 3038.0393
1496.73095703125 0 1291.3665 b Ammonia loss 13
1497.7264404296875 0 1122.8275
1500.7286376953125 0 1983.727 y Water loss 3
1501.713134765625 0 1983.2649 y Ammonia loss 3
1502.7132568359375 0 1746.8607
1513.749755859375 0 4853.842 b 13
1514.7523193359375 0 5111.159
1515.7523193359375 0 2328.3564
1518.7408447265625 0 32400.861 y 3
1519.7430419921875 0 31567.717
1520.74462890625 0 11282.125
1521.7451171875 0 1375.9692
1631.82080078125 0 6758.007 y 2
1632.8255615234375 0 6241.1675
1633.8251953125 0 2541.729
3073.2890625 0 911.53217

Spectrum Details

|  |  |
| --- | --- |
| Matched peaks? Matched peaks The total absolute number of peaks matched. Additionally in brackets the total fraction of peaks matched and the total number of peaks is shown. | 99 (10.62% of 932) |
| FDR? FDR The false discovery rate estimated for this peptide. It is calculated by matching all theoretical fragments with a non-integer shift with the raw peaks for this spectrum. This is done with 40 different shifts. The resulting percentage is the average number of annotated peaks over the number of annotated peaks with the correct spectrum. | 0.82% |
| Satellite FDR? Satellite FDR See the FDR for details on its calculation. This satellite ion specific FDR only contains the satellite ions (d/w) for I/L/J positions. | - |
| PSM Score? PSM Score The PSM Score as given by Hecklib to this annotated spectrum. It is shown with three significant figures. | 1.14E+03 |

#### Spectrum 8167? Spectrum 8167 The raw spectrum of this peptide as annotated by Hecklib. The fragments are coloured according to ion type (see legend). Any peaks with a star '\*' as text can be hovered over to see the full details, first the ion type second the mass shift type. By hovering over the amino acids in the peptide or ions in the legend the corresponding peaks are highlighted. By toggling the 'Unassigned' label you can turn the background (unassigned) peaks on or off in the plot. By updating the slider in the Ion legend you can update the spectrum to only show the top X% of the peaks with labels. The top X% means any peak that is within X% of the highest intensity. By dragging in the spectrum you can zoom in to a specific part of the spectrum and use 'Zoom Out' to get back to the original zoom level. The annotation of the spectrum is based on the given sequence in the peptides file and is done with different software so inconsistencies are likely. The peaks are annotated based on the given sequence, with 20 ppm tolerance.

Copy Data

##### Spectrum 8167 (TSV)

###### Preview

```
Loading example...
```

*Click on the button to copy the data to your clipboard.*

Mz MinMz MaxIntensity Max

WidthHeightPeptide font sizePeptide stroke widthSpectrum font sizeSpectrum stroke widthCompact peptide

Ion legend

wxyz

abcd

OtherUnassignedIon^Charge^_Position_Show for top:%

VTJFPPSSEEJQANKAT

02.15e+54.29e+56.44e+58.58e+5

Zoom Out

y^+1^_1_a^+1^_2_y^+1^_2_a^+1^_2_b^+1^_2_y^+1^_2_b^+1^_2_y^+2^_5_a^+1^_3_a^+1^_3_b^+1^_3_y^+1^_3_y^+1^_3_y^+2^_6_b^+1^_3_y^+1^_3_b^+2^_8_y^+1^_4_y^+2^_8_y^+1^_4_b^+1^_4_b^+1^_4_y^+1^_5_y^+1^_5_y^+1^_5_b^+1^_5_b^+1^_5_b^+2^_11_y^+1^_6_y^+1^_6_y^+2^_12_y^+1^_6_y^+2^_12_b^+2^_12_b^+2^_12_y^+2^_13_y^+2^_13_y^+2^_13_b^+2^_13_b^+2^_13_y^+1^_7_b^+1^_7_y^+1^_7_y^+2^_14_y^+2^_14_b^+2^_14_y^+2^_14_y^+2^_15_b^+2^_15_y^+2^_15_b^+1^_8_b^+2^_16_b^+2^_16_y^+1^_8_b^+2^_16_y^+1^_8_y^+2^_16_y^+2^_16_y^+1^_8_\*\*\*b^+1^_9_y^+1^_9_y^+1^_9_b^+1^_10_y^+1^_10_y^+1^_10_b^+1^_10_y^+1^_10_y^+1^_11_y^+1^_11_y^+1^_11_b^+1^_11_b^+1^_11_y^+1^_12_y^+1^_12_y^+1^_12_b^+1^_12_b^+1^_12_b^+1^_12_y^+1^_13_y^+1^_13_y^+1^_13_b^+1^_13_b^+1^_13_b^+1^_13_y^+1^_14_b^+1^_14_y^+1^_14_y^+1^_15_

041382612381651

Fragment Matches Table

Show background peaks

| Position | Ion type | Intensity | mz Theoretical | mz Error (Th) | mz Error (ppm) | Charge | Series Number |
| --- | --- | --- | --- | --- | --- | --- | --- |
| 17 | y | 6628 | 120.1 | 0.0004289 | 3.572 | +1 | 1 |
| - | - | 9.429E+04 | 120.1 | - | - | 0 | - |
| - | - | 7392 | 121.1 | - | - | 0 | - |
| - | - | 1234 | 126.1 | - | - | 0 | - |
| - | - | 2.054E+04 | 126.1 | - | - | 0 | - |
| - | - | 1275 | 127.1 | - | - | 0 | - |
| - | - | 625.1 | 127.1 | - | - | 0 | - |
| - | - | 2.473E+04 | 128.1 | - | - | 0 | - |
| - | - | 4625 | 129.1 | - | - | 0 | - |
| - | - | 1.628E+05 | 129.1 | - | - | 0 | - |
| - | - | 4087 | 130.1 | - | - | 0 | - |
| - | - | 1232 | 130.1 | - | - | 0 | - |
| - | - | 1.13E+04 | 130.1 | - | - | 0 | - |
| - | - | 3650 | 131.1 | - | - | 0 | - |
| - | - | 2458 | 132.1 | - | - | 0 | - |
| - | - | 844.6 | 132.1 | - | - | 0 | - |
| - | - | 1065 | 133.1 | - | - | 0 | - |
| - | - | 724.3 | 134 | - | - | 0 | - |
| - | - | 688.1 | 134.5 | - | - | 0 | - |
| - | - | 1092 | 136.1 | - | - | 0 | - |
| - | - | 955.3 | 137.1 | - | - | 0 | - |
| - | - | 4923 | 138.1 | - | - | 0 | - |
| - | - | 1.197E+04 | 138.1 | - | - | 0 | - |
| - | - | 6378 | 139.1 | - | - | 0 | - |
| - | - | 1128 | 139.1 | - | - | 0 | - |
| - | - | 1020 | 140.1 | - | - | 0 | - |
| - | - | 2695 | 141.1 | - | - | 0 | - |
| - | - | 980.1 | 141.1 | - | - | 0 | - |
| - | - | 807.1 | 142.1 | - | - | 0 | - |
| - | - | 1482 | 143.1 | - | - | 0 | - |
| - | - | 4585 | 147.1 | - | - | 0 | - |
| - | - | 2226 | 153.1 | - | - | 0 | - |
| - | - | 5070 | 154.1 | - | - | 0 | - |
| - | - | 2673 | 154.1 | - | - | 0 | - |
| - | - | 7864 | 155.1 | - | - | 0 | - |
| 2 | a | 2.213E+04 | 155.1 | 0.0004119 | 2.655 | +1 | 2 |
| - | - | 1831 | 155.1 | - | - | 0 | - |
| - | - | 2342 | 156.1 | - | - | 0 | - |
| - | - | 7260 | 157.1 | - | - | 0 | - |
| - | - | 1.872E+04 | 157.1 | - | - | 0 | - |
| - | - | 775.1 | 159.6 | - | - | 0 | - |
| - | - | 1022 | 163.1 | - | - | 0 | - |
| - | - | 4579 | 165.1 | - | - | 0 | - |
| - | - | 2569 | 166.1 | - | - | 0 | - |
| - | - | 1.36E+04 | 167.1 | - | - | 0 | - |
| - | - | 1376 | 167.1 | - | - | 0 | - |
| - | - | 1.528E+04 | 167.1 | - | - | 0 | - |
| - | - | 1739 | 168.1 | - | - | 0 | - |
| - | - | 871.8 | 168.1 | - | - | 0 | - |
| - | - | 2332 | 169.1 | - | - | 0 | - |
| - | - | 2424 | 169.1 | - | - | 0 | - |
| - | - | 2.303E+04 | 169.1 | - | - | 0 | - |
| - | - | 2274 | 170.1 | - | - | 0 | - |
| - | - | 1929 | 171.1 | - | - | 0 | - |
| - | - | 2162 | 172.1 | - | - | 0 | - |
| 16 | y | 7330 | 173.1 | 0.0005369 | 3.102 | +1 | 2 |
| 2 | a | 7.289E+04 | 173.1 | 0.0004673 | 2.699 | +1 | 2 |
| - | - | 1.002E+04 | 173.5 | - | - | 0 | - |
| - | - | 4709 | 174.1 | - | - | 0 | - |
| - | - | 4030 | 174.1 | - | - | 0 | - |
| - | - | 7303 | 175.1 | - | - | 0 | - |
| - | - | 1062 | 175.1 | - | - | 0 | - |
| - | - | 1588 | 176.1 | - | - | 0 | - |
| - | - | 1734 | 180.1 | - | - | 0 | - |
| - | - | 7216 | 181.1 | - | - | 0 | - |
| - | - | 5549 | 182.1 | - | - | 0 | - |
| - | - | 1352 | 182.1 | - | - | 0 | - |
| - | - | 7652 | 182.1 | - | - | 0 | - |
| - | - | 6735 | 183.1 | - | - | 0 | - |
| 2 | b | 2.848E+05 | 183.1 | 0.0005381 | 2.939 | +1 | 2 |
| - | - | 6762 | 184.1 | - | - | 0 | - |
| - | - | 2.377E+04 | 184.1 | - | - | 0 | - |
| - | - | 2.74E+04 | 185.1 | - | - | 0 | - |
| - | - | 1233 | 185.1 | - | - | 0 | - |
| - | - | 1428 | 185.1 | - | - | 0 | - |
| - | - | 950.2 | 185.8 | - | - | 0 | - |
| - | - | 5711 | 186.1 | - | - | 0 | - |
| - | - | 1558 | 186.1 | - | - | 0 | - |
| - | - | 1237 | 187.1 | - | - | 0 | - |
| - | - | 4.786E+04 | 187.1 | - | - | 0 | - |
| - | - | 904.1 | 187.4 | - | - | 0 | - |
| - | - | 3422 | 188.1 | - | - | 0 | - |
| - | - | 8591 | 189.1 | - | - | 0 | - |
| - | - | 1561 | 191.1 | - | - | 0 | - |
| 16 | y | 6584 | 191.1 | 0.0006076 | 3.179 | +1 | 2 |
| - | - | 1383 | 193.1 | - | - | 0 | - |
| - | - | 1880 | 195.1 | - | - | 0 | - |
| - | - | 2.254E+04 | 195.1 | - | - | 0 | - |
| - | - | 2237 | 196.1 | - | - | 0 | - |
| - | - | 2.624E+04 | 197.1 | - | - | 0 | - |
| - | - | 7142 | 198.1 | - | - | 0 | - |
| - | - | 2824 | 198.1 | - | - | 0 | - |
| - | - | 1.183E+04 | 199.1 | - | - | 0 | - |
| - | - | 4071 | 199.1 | - | - | 0 | - |
| - | - | 1121 | 199.1 | - | - | 0 | - |
| - | - | 1302 | 200.1 | - | - | 0 | - |
| - | - | 1.197E+04 | 200.1 | - | - | 0 | - |
| - | - | 9184 | 200.1 | - | - | 0 | - |
| 2 | b | 1.52E+05 | 201.1 | 0.000441 | 2.193 | +1 | 2 |
| - | - | 1.344E+04 | 202.1 | - | - | 0 | - |
| - | - | 2365 | 203.1 | - | - | 0 | - |
| - | - | 1063 | 207.1 | - | - | 0 | - |
| - | - | 1209 | 208.1 | - | - | 0 | - |
| - | - | 1.313E+04 | 208.1 | - | - | 0 | - |
| - | - | 3.814E+04 | 209.1 | - | - | 0 | - |
| - | - | 1750 | 209.1 | - | - | 0 | - |
| - | - | 3444 | 210.1 | - | - | 0 | - |
| - | - | 2788 | 210.1 | - | - | 0 | - |
| - | - | 2185 | 211.1 | - | - | 0 | - |
| - | - | 3141 | 212.1 | - | - | 0 | - |
| - | - | 2614 | 213.1 | - | - | 0 | - |
| - | - | 2813 | 213.1 | - | - | 0 | - |
| - | - | 1667 | 214.2 | - | - | 0 | - |
| - | - | 5.879E+04 | 215.1 | - | - | 0 | - |
| - | - | 4909 | 216.1 | - | - | 0 | - |
| - | - | 2.649E+04 | 217.1 | - | - | 0 | - |
| - | - | 3.187E+04 | 217.1 | - | - | 0 | - |
| - | - | 808.2 | 217.1 | - | - | 0 | - |
| - | - | 2668 | 218.1 | - | - | 0 | - |
| - | - | 4090 | 218.1 | - | - | 0 | - |
| - | - | 1752 | 218.1 | - | - | 0 | - |
| - | - | 1.261E+04 | 219.1 | - | - | 0 | - |
| - | - | 1167 | 219.2 | - | - | 0 | - |
| - | - | 1169 | 220.2 | - | - | 0 | - |
| - | - | 1.318E+04 | 225.1 | - | - | 0 | - |
| - | - | 6282 | 225.1 | - | - | 0 | - |
| - | - | 1978 | 225.2 | - | - | 0 | - |
| - | - | 1.317E+05 | 226.1 | - | - | 0 | - |
| - | - | 3991 | 227.1 | - | - | 0 | - |
| - | - | 1.377E+04 | 227.1 | - | - | 0 | - |
| - | - | 3412 | 228.1 | - | - | 0 | - |
| - | - | 2572 | 231.1 | - | - | 0 | - |
| - | - | 7.588E+04 | 233.2 | - | - | 0 | - |
| - | - | 1107 | 234.1 | - | - | 0 | - |
| - | - | 2311 | 234.1 | - | - | 0 | - |
| - | - | 1.126E+04 | 234.2 | - | - | 0 | - |
| - | - | 7569 | 236.1 | - | - | 0 | - |
| - | - | 2210 | 236.1 | - | - | 0 | - |
| - | - | 2781 | 237.1 | - | - | 0 | - |
| - | - | 1113 | 237.1 | - | - | 0 | - |
| - | - | 1297 | 240.1 | - | - | 0 | - |
| - | - | 8955 | 241.1 | - | - | 0 | - |
| - | - | 1984 | 241.2 | - | - | 0 | - |
| - | - | 1205 | 242.1 | - | - | 0 | - |
| - | - | 2.288E+04 | 242.2 | - | - | 0 | - |
| - | - | 4807 | 243.1 | - | - | 0 | - |
| - | - | 3.737E+04 | 243.1 | - | - | 0 | - |
| 13 | y | 6578 | 244.1 | 0.0006545 | 2.681 | +2 | 5 |
| - | - | 4796 | 244.1 | - | - | 0 | - |
| - | - | 2.175E+04 | 245.1 | - | - | 0 | - |
| - | - | 2173 | 246.1 | - | - | 0 | - |
| - | - | 1.227E+04 | 247.1 | - | - | 0 | - |
| - | - | 2212 | 248.1 | - | - | 0 | - |
| - | - | 1243 | 250.1 | - | - | 0 | - |
| - | - | 1692 | 251.2 | - | - | 0 | - |
| - | - | 4.165E+04 | 251.2 | - | - | 0 | - |
| - | - | 1944 | 252.1 | - | - | 0 | - |
| - | - | 3888 | 252.1 | - | - | 0 | - |
| - | - | 4672 | 252.2 | - | - | 0 | - |
| - | - | 1355 | 253.1 | - | - | 0 | - |
| - | - | 3.237E+04 | 254.1 | - | - | 0 | - |
| - | - | 5815 | 254.2 | - | - | 0 | - |
| - | - | 3481 | 255.1 | - | - | 0 | - |
| - | - | 1517 | 258.1 | - | - | 0 | - |
| - | - | 2.496E+04 | 259.1 | - | - | 0 | - |
| - | - | 1191 | 259.1 | - | - | 0 | - |
| - | - | 2015 | 260.1 | - | - | 0 | - |
| - | - | 2064 | 260.2 | - | - | 0 | - |
| - | - | 2.412E+04 | 261.2 | - | - | 0 | - |
| - | - | 4125 | 262.1 | - | - | 0 | - |
| - | - | 4098 | 262.2 | - | - | 0 | - |
| - | - | 1516 | 263.1 | - | - | 0 | - |
| - | - | 1331 | 264.1 | - | - | 0 | - |
| - | - | 7416 | 264.1 | - | - | 0 | - |
| - | - | 1814 | 266.2 | - | - | 0 | - |
| - | - | 3734 | 268.1 | - | - | 0 | - |
| - | - | 1951 | 268.2 | - | - | 0 | - |
| 3 | a | 7758 | 268.2 | 0.0007442 | 2.775 | +1 | 3 |
| - | - | 1334 | 269.1 | - | - | 0 | - |
| - | - | 2414 | 269.2 | - | - | 0 | - |
| - | - | 1312 | 269.2 | - | - | 0 | - |
| - | - | 1.125E+04 | 269.2 | - | - | 0 | - |
| - | - | 1628 | 269.2 | - | - | 0 | - |
| - | - | 3188 | 270.2 | - | - | 0 | - |
| - | - | 1.759E+04 | 272.1 | - | - | 0 | - |
| - | - | 2035 | 273.1 | - | - | 0 | - |
| - | - | 1.606E+04 | 275.2 | - | - | 0 | - |
| - | - | 2235 | 276.2 | - | - | 0 | - |
| - | - | 2209 | 278.1 | - | - | 0 | - |
| - | - | 1.372E+04 | 279.1 | - | - | 0 | - |
| - | - | 7748 | 280.1 | - | - | 0 | - |
| - | - | 2020 | 280.1 | - | - | 0 | - |
| - | - | 2.63E+04 | 280.2 | - | - | 0 | - |
| - | - | 4178 | 281.2 | - | - | 0 | - |
| - | - | 1424 | 282.1 | - | - | 0 | - |
| - | - | 4.575E+04 | 282.1 | - | - | 0 | - |
| - | - | 7070 | 283.1 | - | - | 0 | - |
| - | - | 2.008E+04 | 285.2 | - | - | 0 | - |
| - | - | 2325 | 285.2 | - | - | 0 | - |
| - | - | 1.613E+04 | 286.1 | - | - | 0 | - |
| - | - | 2917 | 286.2 | - | - | 0 | - |
| 3 | a | 2517 | 286.2 | 0.0008912 | 3.114 | +1 | 3 |
| - | - | 1821 | 287.1 | - | - | 0 | - |
| - | - | 1475 | 289.2 | - | - | 0 | - |
| - | - | 3255 | 294.2 | - | - | 0 | - |
| - | - | 3556 | 296.1 | - | - | 0 | - |
| 3 | b | 7.312E+05 | 296.2 | 0.0008553 | 2.887 | +1 | 3 |
| - | - | 7435 | 297.1 | - | - | 0 | - |
| - | - | 4.455E+04 | 297.2 | - | - | 0 | - |
| - | - | 1.303E+05 | 297.2 | - | - | 0 | - |
| - | - | 4846 | 298.2 | - | - | 0 | - |
| - | - | 2.574E+04 | 298.2 | - | - | 0 | - |
| - | - | 1.173E+04 | 298.2 | - | - | 0 | - |
| - | - | 3530 | 299.2 | - | - | 0 | - |
| - | - | 1747 | 300.1 | - | - | 0 | - |
| - | - | 2350 | 300.2 | - | - | 0 | - |
| - | - | 1647 | 300.2 | - | - | 0 | - |
| 15 | y | 2523 | 301.2 | 0.0007429 | 2.467 | +1 | 3 |
| 15 | y | 1993 | 302.2 | 0.0009498 | 3.143 | +1 | 3 |
| - | - | 2.859E+04 | 303.2 | - | - | 0 | - |
| - | - | 8734 | 304.1 | - | - | 0 | - |
| - | - | 4571 | 304.2 | - | - | 0 | - |
| - | - | 5559 | 306.1 | - | - | 0 | - |
| 12 | y | 2957 | 308.2 | 0.004004 | 12.99 | +2 | 6 |
| - | - | 2261 | 310.1 | - | - | 0 | - |
| - | - | 1588 | 311.2 | - | - | 0 | - |
| - | - | 1239 | 312.2 | - | - | 0 | - |
| - | - | 1407 | 313.2 | - | - | 0 | - |
| - | - | 7012 | 313.2 | - | - | 0 | - |
| - | - | 1442 | 314.1 | - | - | 0 | - |
| - | - | 7429 | 314.1 | - | - | 0 | - |
| - | - | 3.032E+04 | 314.2 | - | - | 0 | - |
| 3 | b | 1.79E+05 | 314.2 | 0.0009107 | 2.898 | +1 | 3 |
| - | - | 1774 | 315.2 | - | - | 0 | - |
| - | - | 3913 | 315.2 | - | - | 0 | - |
| - | - | 3.115E+04 | 315.2 | - | - | 0 | - |
| - | - | 858.7 | 316.2 | - | - | 0 | - |
| - | - | 1587 | 316.2 | - | - | 0 | - |
| - | - | 6599 | 317.2 | - | - | 0 | - |
| - | - | 1478 | 318.2 | - | - | 0 | - |
| 15 | y | 4.481E+04 | 319.2 | 0.0007678 | 2.406 | +1 | 3 |
| - | - | 7368 | 320.2 | - | - | 0 | - |
| - | - | 1333 | 321.2 | - | - | 0 | - |
| - | - | 3400 | 323.2 | - | - | 0 | - |
| - | - | 1777 | 324.2 | - | - | 0 | - |
| - | - | 2073 | 325.2 | - | - | 0 | - |
| - | - | 1274 | 325.2 | - | - | 0 | - |
| - | - | 2358 | 326.2 | - | - | 0 | - |
| - | - | 1607 | 326.2 | - | - | 0 | - |
| - | - | 1.25E+04 | 328.1 | - | - | 0 | - |
| - | - | 2573 | 329.1 | - | - | 0 | - |
| - | - | 1365 | 329.2 | - | - | 0 | - |
| - | - | 1.807E+04 | 330.2 | - | - | 0 | - |
| - | - | 3804 | 331.2 | - | - | 0 | - |
| - | - | 4920 | 333.2 | - | - | 0 | - |
| - | - | 5418 | 334.2 | - | - | 0 | - |
| - | - | 2265 | 335.2 | - | - | 0 | - |
| - | - | 3098 | 336.2 | - | - | 0 | - |
| - | - | 1086 | 337.1 | - | - | 0 | - |
| - | - | 6458 | 337.2 | - | - | 0 | - |
| - | - | 3296 | 338.1 | - | - | 0 | - |
| - | - | 2020 | 338.2 | - | - | 0 | - |
| - | - | 4972 | 339.2 | - | - | 0 | - |
| - | - | 1186 | 340.2 | - | - | 0 | - |
| - | - | 1.296E+04 | 341.2 | - | - | 0 | - |
| - | - | 4084 | 342.2 | - | - | 0 | - |
| - | - | 7068 | 344.2 | - | - | 0 | - |
| - | - | 1144 | 345.2 | - | - | 0 | - |
| - | - | 3.455E+04 | 346.1 | - | - | 0 | - |
| - | - | 2.704E+04 | 346.2 | - | - | 0 | - |
| - | - | 6252 | 347.1 | - | - | 0 | - |
| - | - | 4762 | 347.2 | - | - | 0 | - |
| - | - | 2.134E+04 | 348.2 | - | - | 0 | - |
| - | - | 5298 | 349.2 | - | - | 0 | - |
| - | - | 5203 | 350.2 | - | - | 0 | - |
| - | - | 1184 | 351.1 | - | - | 0 | - |
| - | - | 2.229E+04 | 351.2 | - | - | 0 | - |
| - | - | 5347 | 352.2 | - | - | 0 | - |
| - | - | 1.224E+04 | 353.2 | - | - | 0 | - |
| - | - | 7243 | 354.2 | - | - | 0 | - |
| - | - | 2160 | 354.2 | - | - | 0 | - |
| - | - | 7271 | 355.2 | - | - | 0 | - |
| - | - | 2685 | 356.1 | - | - | 0 | - |
| - | - | 1218 | 356.2 | - | - | 0 | - |
| - | - | 1367 | 358.2 | - | - | 0 | - |
| - | - | 3863 | 360.2 | - | - | 0 | - |
| - | - | 6.166E+04 | 362.2 | - | - | 0 | - |
| - | - | 1884 | 363.2 | - | - | 0 | - |
| - | - | 1.35E+04 | 363.2 | - | - | 0 | - |
| - | - | 1.065E+04 | 365.1 | - | - | 0 | - |
| - | - | 1672 | 365.2 | - | - | 0 | - |
| - | - | 3206 | 365.3 | - | - | 0 | - |
| - | - | 2345 | 366.1 | - | - | 0 | - |
| - | - | 7280 | 367.2 | - | - | 0 | - |
| - | - | 1745 | 367.2 | - | - | 0 | - |
| - | - | 7251 | 368.2 | - | - | 0 | - |
| - | - | 3.211E+04 | 369.2 | - | - | 0 | - |
| - | - | 1077 | 370.1 | - | - | 0 | - |
| - | - | 5764 | 370.2 | - | - | 0 | - |
| - | - | 4147 | 370.7 | - | - | 0 | - |
| - | - | 1357 | 371.2 | - | - | 0 | - |
| - | - | 7268 | 371.2 | - | - | 0 | - |
| - | - | 7251 | 372.2 | - | - | 0 | - |
| - | - | 1.325E+04 | 373.2 | - | - | 0 | - |
| - | - | 2309 | 374.2 | - | - | 0 | - |
| - | - | 1669 | 375.2 | - | - | 0 | - |
| - | - | 1329 | 379.1 | - | - | 0 | - |
| - | - | 1666 | 379.2 | - | - | 0 | - |
| - | - | 1853 | 380.2 | - | - | 0 | - |
| - | - | 1159 | 381.1 | - | - | 0 | - |
| - | - | 2838 | 381.2 | - | - | 0 | - |
| - | - | 2.271E+04 | 383.2 | - | - | 0 | - |
| - | - | 4638 | 383.3 | - | - | 0 | - |
| - | - | 4031 | 384.2 | - | - | 0 | - |
| - | - | 1.751E+04 | 385.2 | - | - | 0 | - |
| - | - | 3573 | 386.2 | - | - | 0 | - |
| - | - | 1509 | 389.3 | - | - | 0 | - |
| - | - | 2984 | 390.2 | - | - | 0 | - |
| - | - | 2781 | 393.2 | - | - | 0 | - |
| - | - | 9797 | 393.3 | - | - | 0 | - |
| - | - | 2847 | 394.3 | - | - | 0 | - |
| - | - | 1828 | 395.2 | - | - | 0 | - |
| - | - | 5408 | 397.1 | - | - | 0 | - |
| - | - | 1874 | 398.1 | - | - | 0 | - |
| - | - | 1812 | 398.2 | - | - | 0 | - |
| - | - | 1.473E+04 | 398.2 | - | - | 0 | - |
| - | - | 1996 | 399.2 | - | - | 0 | - |
| - | - | 3109 | 399.2 | - | - | 0 | - |
| - | - | 1149 | 400.2 | - | - | 0 | - |
| - | - | 2921 | 400.2 | - | - | 0 | - |
| - | - | 3.818E+04 | 401.2 | - | - | 0 | - |
| - | - | 7334 | 402.2 | - | - | 0 | - |
| - | - | 3568 | 405.2 | - | - | 0 | - |
| - | - | 5589 | 407.2 | - | - | 0 | - |
| - | - | 1.315E+04 | 408.2 | - | - | 0 | - |
| - | - | 2071 | 409.2 | - | - | 0 | - |
| - | - | 3556 | 410.2 | - | - | 0 | - |
| - | - | 1609 | 411.2 | - | - | 0 | - |
| - | - | 1.73E+04 | 411.3 | - | - | 0 | - |
| - | - | 3563 | 412.3 | - | - | 0 | - |
| - | - | 2353 | 413.2 | - | - | 0 | - |
| - | - | 1.109E+04 | 415.1 | - | - | 0 | - |
| 8 | b | 4073 | 415.2 | 0.006404 | 15.42 | +2 | 8 |
| - | - | 7515 | 415.3 | - | - | 0 | - |
| - | - | 1798 | 416.2 | - | - | 0 | - |
| 14 | y | 1983 | 416.2 | 0.003616 | 8.687 | +1 | 4 |
| - | - | 2.84E+04 | 416.3 | - | - | 0 | - |
| - | - | 9883 | 417.3 | - | - | 0 | - |
| - | - | 1330 | 418.3 | - | - | 0 | - |
| - | - | 1837 | 423.2 | - | - | 0 | - |
| - | - | 5344 | 424.2 | - | - | 0 | - |
| - | - | 4007 | 425.1 | - | - | 0 | - |
| - | - | 1123 | 425.2 | - | - | 0 | - |
| - | - | 1.2E+04 | 425.2 | - | - | 0 | - |
| - | - | 1454 | 426.2 | - | - | 0 | - |
| - | - | 6448 | 427.2 | - | - | 0 | - |
| - | - | 1838 | 428.2 | - | - | 0 | - |
| 10 | y | 1633 | 429.2 | 0.007658 | 17.84 | +2 | 8 |
| - | - | 1469 | 431.2 | - | - | 0 | - |
| - | - | 1.464E+04 | 433.2 | - | - | 0 | - |
| 14 | y | 3.345E+04 | 433.2 | 0.002518 | 5.812 | +1 | 4 |
| - | - | 2.302E+05 | 433.3 | - | - | 0 | - |
| - | - | 3632 | 434.2 | - | - | 0 | - |
| - | - | 5957 | 434.2 | - | - | 0 | - |
| - | - | 6959 | 434.2 | - | - | 0 | - |
| - | - | 6.348E+04 | 434.3 | - | - | 0 | - |
| - | - | 5325 | 434.7 | - | - | 0 | - |
| - | - | 5030 | 435.2 | - | - | 0 | - |
| - | - | 1225 | 435.2 | - | - | 0 | - |
| - | - | 1352 | 435.3 | - | - | 0 | - |
| - | - | 1.036E+04 | 435.3 | - | - | 0 | - |
| - | - | 2309 | 440.2 | - | - | 0 | - |
| - | - | 4430 | 441.2 | - | - | 0 | - |
| - | - | 2296 | 441.3 | - | - | 0 | - |
| - | - | 1568 | 442.2 | - | - | 0 | - |
| - | - | 9415 | 442.2 | - | - | 0 | - |
| - | - | 4544 | 443.1 | - | - | 0 | - |
| - | - | 1955 | 443.2 | - | - | 0 | - |
| - | - | 1.174E+04 | 443.2 | - | - | 0 | - |
| 4 | b | 2.388E+04 | 443.3 | 0.001533 | 3.459 | +1 | 4 |
| - | - | 1477 | 444.2 | - | - | 0 | - |
| - | - | 1851 | 444.2 | - | - | 0 | - |
| - | - | 7167 | 444.3 | - | - | 0 | - |
| - | - | 3778 | 445.2 | - | - | 0 | - |
| - | - | 1166 | 450.2 | - | - | 0 | - |
| - | - | 3153 | 451.2 | - | - | 0 | - |
| - | - | 7159 | 452.2 | - | - | 0 | - |
| - | - | 1553 | 458.2 | - | - | 0 | - |
| - | - | 1404 | 459.2 | - | - | 0 | - |
| - | - | 6392 | 459.2 | - | - | 0 | - |
| - | - | 1847 | 459.3 | - | - | 0 | - |
| - | - | 2069 | 460.2 | - | - | 0 | - |
| 4 | b | 2.008E+05 | 461.3 | 0.001344 | 2.915 | +1 | 4 |
| - | - | 1.043E+04 | 462.2 | - | - | 0 | - |
| - | - | 5.55E+04 | 462.3 | - | - | 0 | - |
| - | - | 2918 | 463.2 | - | - | 0 | - |
| - | - | 9407 | 463.3 | - | - | 0 | - |
| - | - | 2788 | 464.2 | - | - | 0 | - |
| - | - | 3823 | 466.2 | - | - | 0 | - |
| - | - | 4196 | 468.2 | - | - | 0 | - |
| - | - | 1473 | 468.2 | - | - | 0 | - |
| - | - | 5306 | 469.2 | - | - | 0 | - |
| - | - | 2.87E+04 | 470.2 | - | - | 0 | - |
| - | - | 7647 | 471.2 | - | - | 0 | - |
| - | - | 3519 | 476.2 | - | - | 0 | - |
| - | - | 2318 | 477.3 | - | - | 0 | - |
| - | - | 5089 | 478.2 | - | - | 0 | - |
| - | - | 3584 | 479.2 | - | - | 0 | - |
| - | - | 2.066E+04 | 480.2 | - | - | 0 | - |
| - | - | 4849 | 481.2 | - | - | 0 | - |
| - | - | 5689 | 482.2 | - | - | 0 | - |
| - | - | 1563 | 483.2 | - | - | 0 | - |
| - | - | 1.091E+04 | 484.2 | - | - | 0 | - |
| - | - | 1730 | 485.2 | - | - | 0 | - |
| 13 | y | 3387 | 486.3 | 0.001146 | 2.357 | +1 | 5 |
| 13 | y | 5214 | 487.3 | 0.002299 | 4.718 | +1 | 5 |
| - | - | 2257 | 488.3 | - | - | 0 | - |
| - | - | 2095 | 492.2 | - | - | 0 | - |
| - | - | 8093 | 494.2 | - | - | 0 | - |
| - | - | 1974 | 495.2 | - | - | 0 | - |
| - | - | 4365 | 495.3 | - | - | 0 | - |
| - | - | 1.136E+04 | 496.3 | - | - | 0 | - |
| - | - | 2806 | 497.3 | - | - | 0 | - |
| - | - | 7.402E+04 | 498.2 | - | - | 0 | - |
| - | - | 1.779E+04 | 499.2 | - | - | 0 | - |
| - | - | 5394 | 500.2 | - | - | 0 | - |
| - | - | 1.622E+04 | 502.2 | - | - | 0 | - |
| - | - | 3666 | 503.2 | - | - | 0 | - |
| 13 | y | 5.145E+04 | 504.3 | 0.001171 | 2.322 | +1 | 5 |
| - | - | 1.122E+04 | 505.3 | - | - | 0 | - |
| - | - | 4335 | 510.2 | - | - | 0 | - |
| - | - | 1.58E+04 | 512.2 | - | - | 0 | - |
| - | - | 2102 | 512.3 | - | - | 0 | - |
| - | - | 4880 | 513.2 | - | - | 0 | - |
| - | - | 6115 | 513.3 | - | - | 0 | - |
| - | - | 2221 | 514.3 | - | - | 0 | - |
| - | - | 2611 | 515.2 | - | - | 0 | - |
| - | - | 1896 | 516.2 | - | - | 0 | - |
| - | - | 2536 | 518.2 | - | - | 0 | - |
| - | - | 1390 | 520.3 | - | - | 0 | - |
| - | - | 1804 | 521.3 | - | - | 0 | - |
| - | - | 1676 | 524.2 | - | - | 0 | - |
| - | - | 1346 | 527.2 | - | - | 0 | - |
| - | - | 4994 | 528.2 | - | - | 0 | - |
| - | - | 4.784E+04 | 530.2 | - | - | 0 | - |
| - | - | 1.441E+04 | 530.3 | - | - | 0 | - |
| - | - | 1.471E+04 | 531.2 | - | - | 0 | - |
| - | - | 4972 | 531.3 | - | - | 0 | - |
| - | - | 3106 | 532.2 | - | - | 0 | - |
| - | - | 1567 | 535.3 | - | - | 0 | - |
| - | - | 4436 | 538.3 | - | - | 0 | - |
| - | - | 1596 | 539.2 | - | - | 0 | - |
| - | - | 1852 | 539.3 | - | - | 0 | - |
| 5 | b | 1.012E+04 | 540.3 | 0.001595 | 2.952 | +1 | 5 |
| - | - | 2209 | 541.3 | - | - | 0 | - |
| - | - | 5418 | 546.2 | - | - | 0 | - |
| - | - | 2134 | 547.2 | - | - | 0 | - |
| - | - | 2907 | 553.3 | - | - | 0 | - |
| - | - | 1317 | 554.3 | - | - | 0 | - |
| - | - | 3892 | 555.3 | - | - | 0 | - |
| - | - | 2385 | 556.2 | - | - | 0 | - |
| - | - | 1393 | 556.3 | - | - | 0 | - |
| - | - | 2285 | 556.3 | - | - | 0 | - |
| 5 | b | 3.085E+04 | 558.3 | 0.001162 | 2.082 | +1 | 5 |
| - | - | 1.021E+04 | 559.3 | - | - | 0 | - |
| - | - | 1701 | 560.3 | - | - | 0 | - |
| - | - | 1887 | 560.8 | - | - | 0 | - |
| - | - | 2849 | 563.2 | - | - | 0 | - |
| - | - | 3386 | 569.3 | - | - | 0 | - |
| - | - | 3621 | 569.3 | - | - | 0 | - |
| - | - | 7151 | 569.8 | - | - | 0 | - |
| - | - | 2296 | 570.3 | - | - | 0 | - |
| - | - | 2628 | 571.3 | - | - | 0 | - |
| - | - | 1362 | 572.3 | - | - | 0 | - |
| - | - | 1649 | 573.2 | - | - | 0 | - |
| - | - | 1.204E+04 | 578.3 | - | - | 0 | - |
| - | - | 4533 | 578.8 | - | - | 0 | - |
| - | - | 3029 | 579.3 | - | - | 0 | - |
| - | - | 9662 | 581.3 | - | - | 0 | - |
| - | - | 4186 | 582.3 | - | - | 0 | - |
| - | - | 1.671E+04 | 582.8 | - | - | 0 | - |
| - | - | 8518 | 583.3 | - | - | 0 | - |
| - | - | 2295 | 583.8 | - | - | 0 | - |
| - | - | 2863 | 587.3 | - | - | 0 | - |
| - | - | 3976 | 591.2 | - | - | 0 | - |
| - | - | 5546 | 591.3 | - | - | 0 | - |
| 11 | b | 5230 | 591.8 | 0.009737 | 16.45 | +2 | 11 |
| - | - | 1463 | 592.3 | - | - | 0 | - |
| - | - | 6086 | 597.3 | - | - | 0 | - |
| - | - | 3572 | 598.3 | - | - | 0 | - |
| - | - | 4.429E+04 | 599.3 | - | - | 0 | - |
| - | - | 1.842E+04 | 600.3 | - | - | 0 | - |
| - | - | 2762 | 601.3 | - | - | 0 | - |
| - | - | 3236 | 604.3 | - | - | 0 | - |
| - | - | 3135 | 604.8 | - | - | 0 | - |
| - | - | 3658 | 607.3 | - | - | 0 | - |
| - | - | 1425 | 608.3 | - | - | 0 | - |
| - | - | 3297 | 608.4 | - | - | 0 | - |
| - | - | 1.635E+04 | 609.3 | - | - | 0 | - |
| - | - | 3405 | 609.3 | - | - | 0 | - |
| - | - | 2029 | 609.3 | - | - | 0 | - |
| - | - | 4527 | 609.8 | - | - | 0 | - |
| - | - | 5040 | 610.3 | - | - | 0 | - |
| - | - | 7666 | 612.8 | - | - | 0 | - |
| - | - | 7738 | 613.3 | - | - | 0 | - |
| - | - | 3237 | 613.8 | - | - | 0 | - |
| 12 | y | 9620 | 614.3 | 0.0008876 | 1.445 | +1 | 6 |
| 12 | y | 3.183E+04 | 615.3 | 0.004429 | 7.198 | +1 | 6 |
| - | - | 8408 | 616.3 | - | - | 0 | - |
| - | - | 2125 | 617.3 | - | - | 0 | - |
| - | - | 3.881E+04 | 617.8 | - | - | 0 | - |
| - | - | 4.264E+04 | 618.3 | - | - | 0 | - |
| - | - | 1.901E+04 | 618.8 | - | - | 0 | - |
| - | - | 6218 | 619.3 | - | - | 0 | - |
| - | - | 1335 | 619.8 | - | - | 0 | - |
| - | - | 9203 | 625.3 | - | - | 0 | - |
| - | - | 4127 | 626.3 | - | - | 0 | - |
| - | - | 6665 | 626.4 | - | - | 0 | - |
| - | - | 1.088E+05 | 626.8 | - | - | 0 | - |
| - | - | 1.126E+05 | 627.3 | - | - | 0 | - |
| - | - | 6.952E+04 | 627.3 | - | - | 0 | - |
| - | - | 2.823E+04 | 627.8 | - | - | 0 | - |
| - | - | 3.723E+04 | 628.3 | - | - | 0 | - |
| - | - | 5326 | 628.3 | - | - | 0 | - |
| 6 | y | 2156 | 628.8 | 0.000722 | 1.148 | +2 | 12 |
| - | - | 7544 | 629.3 | - | - | 0 | - |
| - | - | 2987 | 629.8 | - | - | 0 | - |
| 12 | y | 3.144E+04 | 632.3 | 0.001431 | 2.264 | +1 | 6 |
| - | - | 1.118E+04 | 633.3 | - | - | 0 | - |
| - | - | 2157 | 634.3 | - | - | 0 | - |
| - | - | 6007 | 635.8 | - | - | 0 | - |
| - | - | 3024 | 636.3 | - | - | 0 | - |
| 6 | y | 1.573E+04 | 637.8 | 0.001869 | 2.931 | +2 | 12 |
| - | - | 8741 | 638.3 | - | - | 0 | - |
| - | - | 5554 | 638.8 | - | - | 0 | - |
| - | - | 2750 | 642.3 | - | - | 0 | - |
| - | - | 2.428E+04 | 643.3 | - | - | 0 | - |
| - | - | 1.166E+04 | 644.3 | - | - | 0 | - |
| - | - | 4113 | 645.3 | - | - | 0 | - |
| - | - | 1491 | 646.3 | - | - | 0 | - |
| - | - | 2098 | 653.3 | - | - | 0 | - |
| - | - | 5963 | 655.3 | - | - | 0 | - |
| - | - | 1637 | 655.4 | - | - | 0 | - |
| 12 | b | 5579 | 655.8 | 0.006628 | 10.11 | +2 | 12 |
| 12 | b | 2660 | 656.3 | 0.01242 | 18.93 | +2 | 12 |
| - | - | 1896 | 658.3 | - | - | 0 | - |
| - | - | 1279 | 659.3 | - | - | 0 | - |
| - | - | 8785 | 660.3 | - | - | 0 | - |
| - | - | 4642 | 661.3 | - | - | 0 | - |
| - | - | 1260 | 662.3 | - | - | 0 | - |
| - | - | 2653 | 663.3 | - | - | 0 | - |
| - | - | 2859 | 663.8 | - | - | 0 | - |
| - | - | 1591 | 664.3 | - | - | 0 | - |
| - | - | 1460 | 667.3 | - | - | 0 | - |
| - | - | 5259 | 668.3 | - | - | 0 | - |
| - | - | 5593 | 668.8 | - | - | 0 | - |
| - | - | 4950 | 669.3 | - | - | 0 | - |
| - | - | 2565 | 669.8 | - | - | 0 | - |
| - | - | 1394 | 670.3 | - | - | 0 | - |
| - | - | 2438 | 674.3 | - | - | 0 | - |
| 5 | y | 3.483E+04 | 677.3 | 0.001338 | 1.976 | +2 | 13 |
| 5 | y | 4.651E+04 | 677.8 | 0.00695 | 10.25 | +2 | 13 |
| - | - | 2.932E+04 | 678.3 | - | - | 0 | - |
| - | - | 1.225E+04 | 678.8 | - | - | 0 | - |
| - | - | 1400 | 679.3 | - | - | 0 | - |
| - | - | 2462 | 679.3 | - | - | 0 | - |
| - | - | 2236 | 685.3 | - | - | 0 | - |
| - | - | 2915 | 685.8 | - | - | 0 | - |
| 5 | y | 7.671E+05 | 686.3 | 0.001549 | 2.257 | +2 | 13 |
| - | - | 6.319E+05 | 686.8 | - | - | 0 | - |
| - | - | 2.688E+05 | 687.3 | - | - | 0 | - |
| - | - | 6.313E+04 | 687.8 | - | - | 0 | - |
| - | - | 7294 | 688.3 | - | - | 0 | - |
| 13 | b | 1721 | 691.8 | 0.005645 | 8.16 | +2 | 13 |
| - | - | 1624 | 692.3 | - | - | 0 | - |
| - | - | 6274 | 694.3 | - | - | 0 | - |
| - | - | 3647 | 695.3 | - | - | 0 | - |
| - | - | 2053 | 696.4 | - | - | 0 | - |
| - | - | 1565 | 698.3 | - | - | 0 | - |
| 13 | b | 8544 | 700.4 | 0.01123 | 16.03 | +2 | 13 |
| - | - | 7382 | 700.8 | - | - | 0 | - |
| - | - | 4183 | 701.3 | - | - | 0 | - |
| - | - | 2788 | 704.3 | - | - | 0 | - |
| - | - | 2602 | 710.3 | - | - | 0 | - |
| - | - | 6.515E+04 | 712.4 | - | - | 0 | - |
| - | - | 2.373E+04 | 713.4 | - | - | 0 | - |
| - | - | 5837 | 714.4 | - | - | 0 | - |
| - | - | 1880 | 720.4 | - | - | 0 | - |
| - | - | 1887 | 721.4 | - | - | 0 | - |
| - | - | 8988 | 722.3 | - | - | 0 | - |
| - | - | 3480 | 723.3 | - | - | 0 | - |
| - | - | 1612 | 724.3 | - | - | 0 | - |
| - | - | 1618 | 725.3 | - | - | 0 | - |
| - | - | 2926 | 727.3 | - | - | 0 | - |
| 11 | y | 2364 | 728.4 | 0.004768 | 6.547 | +1 | 7 |
| - | - | 1884 | 731.4 | - | - | 0 | - |
| - | - | 2304 | 737.3 | - | - | 0 | - |
| - | - | 2621 | 737.4 | - | - | 0 | - |
| - | - | 3142 | 738.3 | - | - | 0 | - |
| - | - | 3420 | 738.4 | - | - | 0 | - |
| - | - | 4928 | 739.4 | - | - | 0 | - |
| - | - | 7.751E+04 | 740.3 | - | - | 0 | - |
| - | - | 3.434E+04 | 741.4 | - | - | 0 | - |
| - | - | 6510 | 742.4 | - | - | 0 | - |
| 7 | b | 5718 | 742.4 | 0.004505 | 6.068 | +1 | 7 |
| - | - | 2141 | 743.4 | - | - | 0 | - |
| - | - | 3506 | 743.4 | - | - | 0 | - |
| - | - | 4230 | 745.3 | - | - | 0 | - |
| 11 | y | 2.677E+04 | 745.4 | 0.001962 | 2.632 | +1 | 7 |
| - | - | 3153 | 746.3 | - | - | 0 | - |
| - | - | 1.165E+04 | 746.4 | - | - | 0 | - |
| - | - | 2462 | 747.4 | - | - | 0 | - |
| 4 | y | 4042 | 750.9 | 0.00418 | 5.567 | +2 | 14 |
| 4 | y | 6471 | 751.4 | 0.006679 | 8.889 | +2 | 14 |
| - | - | 3242 | 751.9 | - | - | 0 | - |
| - | - | 1904 | 752.4 | - | - | 0 | - |
| - | - | 5885 | 753.3 | - | - | 0 | - |
| - | - | 3543 | 754.3 | - | - | 0 | - |
| - | - | 2446 | 755.3 | - | - | 0 | - |
| - | - | 4310 | 755.4 | - | - | 0 | - |
| - | - | 2602 | 756.3 | - | - | 0 | - |
| - | - | 1923 | 756.4 | - | - | 0 | - |
| - | - | 2261 | 756.9 | - | - | 0 | - |
| 14 | b | 3.768E+04 | 757.4 | 0.005227 | 6.902 | +2 | 14 |
| - | - | 1.472E+04 | 758.4 | - | - | 0 | - |
| - | - | 2827 | 759.4 | - | - | 0 | - |
| - | - | 827.2 | 759.8 | - | - | 0 | - |
| 4 | y | 9.723E+04 | 759.9 | 0.001766 | 2.324 | +2 | 14 |
| - | - | 1.026E+05 | 760.4 | - | - | 0 | - |
| - | - | 4.371E+04 | 760.9 | - | - | 0 | - |
| - | - | 1.572E+04 | 761.4 | - | - | 0 | - |
| - | - | 1688 | 766.4 | - | - | 0 | - |
| - | - | 1.883E+04 | 771.4 | - | - | 0 | - |
| - | - | 9680 | 772.4 | - | - | 0 | - |
| - | - | 1758 | 773.4 | - | - | 0 | - |
| - | - | 2901 | 774.3 | - | - | 0 | - |
| - | - | 2567 | 775.3 | - | - | 0 | - |
| - | - | 1363 | 775.5 | - | - | 0 | - |
| - | - | 1758 | 783.4 | - | - | 0 | - |
| - | - | 1801 | 797.4 | - | - | 0 | - |
| - | - | 1597 | 805.4 | - | - | 0 | - |
| 3 | y | 1731 | 807.9 | 0.006456 | 7.991 | +2 | 15 |
| - | - | 1468 | 811.4 | - | - | 0 | - |
| 15 | b | 2203 | 812.9 | 0.005427 | 6.675 | +2 | 15 |
| - | - | 2216 | 813.4 | - | - | 0 | - |
| - | - | 1903 | 813.9 | - | - | 0 | - |
| - | - | 2909 | 814.4 | - | - | 0 | - |
| 3 | y | 2.127E+04 | 816.4 | 0.002031 | 2.488 | +2 | 15 |
| - | - | 2.56E+04 | 816.9 | - | - | 0 | - |
| - | - | 1.174E+04 | 817.4 | - | - | 0 | - |
| - | - | 3209 | 817.9 | - | - | 0 | - |
| - | - | 2943 | 822.4 | - | - | 0 | - |
| - | - | 2131 | 823.4 | - | - | 0 | - |
| - | - | 5325 | 824.4 | - | - | 0 | - |
| - | - | 2441 | 825.4 | - | - | 0 | - |
| - | - | 1481 | 826.4 | - | - | 0 | - |
| - | - | 1627 | 827.4 | - | - | 0 | - |
| 8 | b | 4819 | 829.4 | 0.0005526 | 0.6663 | +1 | 8 |
| - | - | 2879 | 830.4 | - | - | 0 | - |
| - | - | 1745 | 832.4 | - | - | 0 | - |
| - | - | 2198 | 833.4 | - | - | 0 | - |
| - | - | 2321 | 839.4 | - | - | 0 | - |
| - | - | 1.059E+04 | 840.4 | - | - | 0 | - |
| - | - | 6427 | 841.4 | - | - | 0 | - |
| - | - | 1.577E+04 | 842.4 | - | - | 0 | - |
| - | - | 6752 | 843.4 | - | - | 0 | - |
| 16 | b | 3589 | 847.9 | 0.003841 | 4.53 | +2 | 16 |
| 16 | b | 6378 | 848.4 | 0.004082 | 4.811 | +2 | 16 |
| - | - | 5305 | 848.9 | - | - | 0 | - |
| - | - | 3142 | 849.4 | - | - | 0 | - |
| - | - | 1.281E+04 | 850.4 | - | - | 0 | - |
| - | - | 1.004E+04 | 851.4 | - | - | 0 | - |
| - | - | 2950 | 852.4 | - | - | 0 | - |
| 10 | y | 4075 | 856.5 | 0.003821 | 4.462 | +1 | 8 |
| 16 | b | 1.708E+04 | 856.9 | 0.001915 | 2.235 | +2 | 16 |
| 10 | y | 2.371E+04 | 857.4 | 0.01277 | 14.9 | +1 | 8 |
| 2 | y | 1.478E+04 | 857.9 | 0.01515 | 17.66 | +2 | 16 |
| - | - | 4743 | 858.4 | - | - | 0 | - |
| - | - | 2274 | 859.4 | - | - | 0 | - |
| 2 | y | 5776 | 866.9 | 0.003522 | 4.062 | +2 | 16 |
| - | - | 9364 | 867.4 | - | - | 0 | - |
| - | - | 4903 | 867.9 | - | - | 0 | - |
| - | - | 9.649E+04 | 868.4 | - | - | 0 | - |
| - | - | 4.558E+04 | 869.4 | - | - | 0 | - |
| - | - | 1.44E+04 | 870.4 | - | - | 0 | - |
| 10 | y | 1.781E+04 | 874.5 | 0.001361 | 1.557 | +1 | 8 |
| - | - | 8876 | 875.5 | - | - | 0 | - |
| - | - | 2064 | 876.5 | - | - | 0 | - |
| - | - | 3117 | 884.4 | - | - | 0 | - |
| - | - | 4514 | 885.4 | - | - | 0 | - |
| - | - | 5884 | 887.4 | - | - | 0 | - |
| - | - | 4517 | 888.4 | - | - | 0 | - |
| - | - | 1773 | 893.4 | - | - | 0 | - |
| - | - | 5718 | 894.4 | - | - | 0 | - |
| - | - | 3662 | 895.4 | - | - | 0 | - |
| - | - | 2375 | 898.5 | - | - | 0 | - |
| - | - | 2299 | 899.5 | - | - | 0 | - |
| - | - | 2391 | 900.4 | - | - | 0 | - |
| - | - | 1921 | 903.4 | - | - | 0 | - |
| - | - | 1868 | 904.4 | - | - | 0 | - |
| 0 | Precursor | 1.573E+04 | 907.5 | 0.002307 | 2.542 | +2 | -1 |
| 0 | Precursor | 2.252E+04 | 908 | 0.009506 | 10.47 | +2 | -1 |
| - | - | 1.623E+04 | 908.5 | - | - | 0 | - |
| - | - | 5348 | 909 | - | - | 0 | - |
| - | - | 1.848E+04 | 911.4 | - | - | 0 | - |
| - | - | 9427 | 912.5 | - | - | 0 | - |
| - | - | 2364 | 913.4 | - | - | 0 | - |
| - | - | 2888 | 916 | - | - | 0 | - |
| 0 | Precursor | 1.522E+05 | 916.5 | 0.001786 | 1.948 | +2 | -1 |
| - | - | 1.67E+05 | 917 | - | - | 0 | - |
| - | - | 1.081E+05 | 917.5 | - | - | 0 | - |
| - | - | 3.816E+04 | 918 | - | - | 0 | - |
| - | - | 4888 | 918.5 | - | - | 0 | - |
| - | - | 9092 | 921.4 | - | - | 0 | - |
| - | - | 1.181E+04 | 922.4 | - | - | 0 | - |
| - | - | 4139 | 923.4 | - | - | 0 | - |
| - | - | 1662 | 924.4 | - | - | 0 | - |
| - | - | 6.586E+04 | 939.4 | - | - | 0 | - |
| - | - | 3.223E+04 | 940.4 | - | - | 0 | - |
| - | - | 1.063E+04 | 941.4 | - | - | 0 | - |
| - | - | 1964 | 942.4 | - | - | 0 | - |
| - | - | 2817 | 953.5 | - | - | 0 | - |
| - | - | 3911 | 954.5 | - | - | 0 | - |
| - | - | 2166 | 955.5 | - | - | 0 | - |
| - | - | 1.139E+04 | 956.5 | - | - | 0 | - |
| - | - | 6134 | 957.5 | - | - | 0 | - |
| 9 | b | 6509 | 958.5 | 0.001391 | 1.451 | +1 | 9 |
| - | - | 2013 | 959.5 | - | - | 0 | - |
| - | - | 1906 | 967.5 | - | - | 0 | - |
| - | - | 3151 | 970.5 | - | - | 0 | - |
| - | - | 5497 | 971.5 | - | - | 0 | - |
| - | - | 3210 | 972.5 | - | - | 0 | - |
| - | - | 1486 | 973.5 | - | - | 0 | - |
| - | - | 1339 | 982.5 | - | - | 0 | - |
| 9 | y | 2291 | 985.5 | 0.0003328 | 0.3377 | +1 | 9 |
| - | - | 3338 | 987.5 | - | - | 0 | - |
| - | - | 1916 | 988.5 | - | - | 0 | - |
| - | - | 1692 | 989.5 | - | - | 0 | - |
| 9 | y | 1.177E+04 | 1004 | 0.002835 | 2.826 | +1 | 9 |
| - | - | 8655 | 1005 | - | - | 0 | - |
| - | - | 1782 | 1006 | - | - | 0 | - |
| - | - | 6166 | 1008 | - | - | 0 | - |
| - | - | 3949 | 1009 | - | - | 0 | - |
| - | - | 1753 | 1010 | - | - | 0 | - |
| - | - | 5300 | 1015 | - | - | 0 | - |
| - | - | 3913 | 1016 | - | - | 0 | - |
| - | - | 1865 | 1017 | - | - | 0 | - |
| - | - | 2058 | 1018 | - | - | 0 | - |
| - | - | 2293 | 1019 | - | - | 0 | - |
| - | - | 1741 | 1023 | - | - | 0 | - |
| - | - | 2243 | 1024 | - | - | 0 | - |
| - | - | 3408 | 1025 | - | - | 0 | - |
| - | - | 2261 | 1026 | - | - | 0 | - |
| - | - | 3944 | 1035 | - | - | 0 | - |
| - | - | 1.437E+04 | 1036 | - | - | 0 | - |
| - | - | 6716 | 1037 | - | - | 0 | - |
| - | - | 3783 | 1038 | - | - | 0 | - |
| - | - | 5124 | 1041 | - | - | 0 | - |
| - | - | 6218 | 1041 | - | - | 0 | - |
| - | - | 3412 | 1042 | - | - | 0 | - |
| - | - | 1872 | 1052 | - | - | 0 | - |
| - | - | 5.385E+04 | 1053 | - | - | 0 | - |
| - | - | 3.468E+04 | 1054 | - | - | 0 | - |
| - | - | 1.269E+04 | 1055 | - | - | 0 | - |
| - | - | 3131 | 1056 | - | - | 0 | - |
| - | - | 9026 | 1059 | - | - | 0 | - |
| - | - | 4383 | 1060 | - | - | 0 | - |
| - | - | 1367 | 1067 | - | - | 0 | - |
| - | - | 4123 | 1068 | - | - | 0 | - |
| - | - | 3851 | 1069 | - | - | 0 | - |
| 10 | b | 2929 | 1070 | 0.009189 | 8.591 | +1 | 10 |
| - | - | 1.797E+04 | 1071 | - | - | 0 | - |
| - | - | 1.182E+04 | 1072 | - | - | 0 | - |
| 8 | y | 5270 | 1073 | 0.006299 | 5.873 | +1 | 10 |
| 8 | y | 3483 | 1074 | 0.007488 | 6.975 | +1 | 10 |
| - | - | 1794 | 1075 | - | - | 0 | - |
| - | - | 1357 | 1077 | - | - | 0 | - |
| - | - | 1.071E+04 | 1085 | - | - | 0 | - |
| - | - | 7932 | 1086 | - | - | 0 | - |
| - | - | 6760 | 1087 | - | - | 0 | - |
| 10 | b | 1.083E+04 | 1088 | 0.005349 | 4.918 | +1 | 10 |
| - | - | 3045 | 1089 | - | - | 0 | - |
| - | - | 1700 | 1090 | - | - | 0 | - |
| 8 | y | 2.155E+04 | 1091 | 0.001203 | 1.103 | +1 | 10 |
| - | - | 1.37E+04 | 1092 | - | - | 0 | - |
| - | - | 4313 | 1093 | - | - | 0 | - |
| - | - | 1305 | 1094 | - | - | 0 | - |
| - | - | 1559 | 1103 | - | - | 0 | - |
| - | - | 2541 | 1121 | - | - | 0 | - |
| - | - | 1933 | 1122 | - | - | 0 | - |
| - | - | 1899 | 1128 | - | - | 0 | - |
| - | - | 1959 | 1129 | - | - | 0 | - |
| - | - | 9175 | 1138 | - | - | 0 | - |
| - | - | 1.174E+04 | 1139 | - | - | 0 | - |
| - | - | 6481 | 1140 | - | - | 0 | - |
| - | - | 1771 | 1141 | - | - | 0 | - |
| - | - | 2167 | 1147 | - | - | 0 | - |
| - | - | 1553 | 1148 | - | - | 0 | - |
| - | - | 1.9E+04 | 1156 | - | - | 0 | - |
| - | - | 1.301E+04 | 1157 | - | - | 0 | - |
| - | - | 5479 | 1158 | - | - | 0 | - |
| 7 | y | 5354 | 1160 | 0.0004858 | 0.4189 | +1 | 11 |
| 7 | y | 6007 | 1161 | 0.009883 | 8.516 | +1 | 11 |
| - | - | 3368 | 1162 | - | - | 0 | - |
| - | - | 2765 | 1164 | - | - | 0 | - |
| - | - | 1.294E+04 | 1165 | - | - | 0 | - |
| - | - | 5855 | 1166 | - | - | 0 | - |
| - | - | 3287 | 1167 | - | - | 0 | - |
| 7 | y | 2.647E+04 | 1178 | 0.002255 | 1.915 | +1 | 11 |
| - | - | 1.98E+04 | 1179 | - | - | 0 | - |
| - | - | 6695 | 1180 | - | - | 0 | - |
| - | - | 6.259E+04 | 1182 | - | - | 0 | - |
| 11 | b | 4.822E+04 | 1183 | 0.01842 | 15.58 | +1 | 11 |
| - | - | 1.901E+04 | 1184 | - | - | 0 | - |
| - | - | 4057 | 1185 | - | - | 0 | - |
| - | - | 1750 | 1199 | - | - | 0 | - |
| 11 | b | 1.068E+04 | 1201 | 0.003231 | 2.691 | +1 | 11 |
| - | - | 7983 | 1202 | - | - | 0 | - |
| - | - | 3932 | 1203 | - | - | 0 | - |
| - | - | 2306 | 1208 | - | - | 0 | - |
| - | - | 1733 | 1209 | - | - | 0 | - |
| - | - | 2888 | 1213 | - | - | 0 | - |
| - | - | 4220 | 1218 | - | - | 0 | - |
| - | - | 2733 | 1219 | - | - | 0 | - |
| - | - | 5429 | 1225 | - | - | 0 | - |
| - | - | 3812 | 1226 | - | - | 0 | - |
| - | - | 1466 | 1227 | - | - | 0 | - |
| - | - | 2.27E+04 | 1235 | - | - | 0 | - |
| - | - | 2.615E+04 | 1236 | - | - | 0 | - |
| - | - | 1.474E+04 | 1237 | - | - | 0 | - |
| - | - | 3727 | 1238 | - | - | 0 | - |
| - | - | 2681 | 1239 | - | - | 0 | - |
| - | - | 2479 | 1240 | - | - | 0 | - |
| - | - | 1420 | 1241 | - | - | 0 | - |
| - | - | 5.765E+04 | 1253 | - | - | 0 | - |
| - | - | 4.271E+04 | 1254 | - | - | 0 | - |
| - | - | 1.647E+04 | 1255 | - | - | 0 | - |
| - | - | 4515 | 1256 | - | - | 0 | - |
| 6 | y | 8150 | 1257 | 0.0001491 | 0.1186 | +1 | 12 |
| 6 | y | 1.394E+04 | 1258 | 0.007168 | 5.7 | +1 | 12 |
| - | - | 8487 | 1259 | - | - | 0 | - |
| - | - | 3000 | 1260 | - | - | 0 | - |
| - | - | 3093 | 1271 | - | - | 0 | - |
| 6 | y | 1.49E+05 | 1275 | 0.0007609 | 0.5969 | +1 | 12 |
| - | - | 1.097E+05 | 1276 | - | - | 0 | - |
| - | - | 4.724E+04 | 1277 | - | - | 0 | - |
| - | - | 9664 | 1278 | - | - | 0 | - |
| - | - | 6604 | 1310 | - | - | 0 | - |
| 12 | b | 8723 | 1311 | 0.007228 | 5.514 | +1 | 12 |
| 12 | b | 5518 | 1312 | 0.0237 | 18.07 | +1 | 12 |
| - | - | 3065 | 1313 | - | - | 0 | - |
| - | - | 3238 | 1326 | - | - | 0 | - |
| - | - | 2598 | 1327 | - | - | 0 | - |
| - | - | 3556 | 1328 | - | - | 0 | - |
| 12 | b | 1.728E+04 | 1329 | 0.003459 | 2.603 | +1 | 12 |
| - | - | 1.369E+04 | 1330 | - | - | 0 | - |
| - | - | 7233 | 1331 | - | - | 0 | - |
| - | - | 2788 | 1336 | - | - | 0 | - |
| - | - | 4821 | 1337 | - | - | 0 | - |
| - | - | 5404 | 1338 | - | - | 0 | - |
| - | - | 1753 | 1342 | - | - | 0 | - |
| 5 | y | 2.677E+04 | 1354 | 0.0005539 | 0.4092 | +1 | 13 |
| 5 | y | 5.19E+04 | 1355 | 0.008238 | 6.081 | +1 | 13 |
| - | - | 3.6E+04 | 1356 | - | - | 0 | - |
| - | - | 1.286E+04 | 1357 | - | - | 0 | - |
| - | - | 3532 | 1358 | - | - | 0 | - |
| - | - | 2299 | 1370 | - | - | 0 | - |
| - | - | 4171 | 1371 | - | - | 0 | - |
| 5 | y | 8.5E+05 | 1372 | 0.0009755 | 0.7112 | +1 | 13 |
| - | - | 7.078E+05 | 1373 | - | - | 0 | - |
| - | - | 2.977E+05 | 1374 | - | - | 0 | - |
| - | - | 7.45E+04 | 1375 | - | - | 0 | - |
| - | - | 5207 | 1376 | - | - | 0 | - |
| 13 | b | 3086 | 1382 | 0.008036 | 5.816 | +1 | 13 |
| 13 | b | 2289 | 1383 | 0.003188 | 2.306 | +1 | 13 |
| 13 | b | 1.386E+04 | 1400 | 0.003586 | 2.562 | +1 | 13 |
| - | - | 1.14E+04 | 1401 | - | - | 0 | - |
| - | - | 5958 | 1402 | - | - | 0 | - |
| - | - | 1580 | 1403 | - | - | 0 | - |
| 4 | y | 2722 | 1502 | 0.008793 | 5.856 | +1 | 14 |
| - | - | 2123 | 1503 | - | - | 0 | - |
| 14 | b | 5477 | 1514 | 0.008174 | 5.4 | +1 | 14 |
| - | - | 6611 | 1515 | - | - | 0 | - |
| - | - | 3300 | 1516 | - | - | 0 | - |
| 4 | y | 4.982E+04 | 1519 | 0.000921 | 0.6064 | +1 | 14 |
| - | - | 4.438E+04 | 1520 | - | - | 0 | - |
| - | - | 2.004E+04 | 1521 | - | - | 0 | - |
| - | - | 7693 | 1522 | - | - | 0 | - |
| 3 | y | 8727 | 1632 | 0.003187 | 1.953 | +1 | 15 |
| - | - | 9845 | 1633 | - | - | 0 | - |
| - | - | 5352 | 1634 | - | - | 0 | - |
| - | - | 1656 | 1635 | - | - | 0 | - |

m/z Charge Intensity FragmentType MassShift Position
120.06594848632812 0 6627.9263 y 16
120.08119201660156 0 94285.98
121.08456420898438 0 7392.227
126.05013275146484 0 1234.1139
126.05536651611328 0 20540.053
127.0588150024414 0 1275.2206
127.08663177490234 0 625.0641
128.10739135742188 0 24729.426
129.06629943847656 0 4624.7007
129.1026611328125 0 162751.31
130.05032348632812 0 4086.5386
130.10035705566406 0 1231.9663
130.10597229003906 0 11298.465
131.1182403564453 0 3650.091
132.07701110839844 0 2457.574
132.10264587402344 0 844.6372
133.0610809326172 0 1065.2349
133.9544677734375 0 724.25494
134.50140380859375 0 688.10376
136.0760955810547 0 1092.2761
137.10769653320312 0 955.3106
138.0917205810547 0 4922.6924
138.128173828125 0 11965.46
139.08702087402344 0 6377.806
139.1315460205078 0 1127.5282
140.07102966308594 0 1019.75543
141.06619262695312 0 2695.4111
141.10281372070312 0 980.0877
142.12355041503906 0 807.0692
143.1184539794922 0 1482.1401
147.07684326171875 0 4585.254
153.10305786132812 0 2225.7146
154.086669921875 0 5069.708
154.09786987304688 0 2672.8372
155.08197021484375 0 7863.768
155.11830139160156 0 22128.5 a Water loss 1
155.12513732910156 0 1831.4369
156.12144470214844 0 2342.4392
157.0611572265625 0 7259.7134
157.0976104736328 0 18719.535
159.59678649902344 0 775.14966
163.08734130859375 0 1021.81445
165.10276794433594 0 4578.594
166.08651733398438 0 2569.4714
167.08197021484375 0 13599.13
167.08863830566406 0 1376.3684
167.11830139160156 0 15279.045
168.08511352539062 0 1739.4846
168.12196350097656 0 871.8357
169.0609588623047 0 2331.5532
169.09765625 0 2423.7283
169.1339874267578 0 23032.34
170.13726806640625 0 2273.561
171.07691955566406 0 1929.4374
172.10873413085938 0 2161.8198
173.0926055908203 0 7329.5977 y Water loss 15
173.12892150878906 0 72886.164 a 1
173.4505157470703 0 10021.4795
174.12811279296875 0 4708.8403
174.13209533691406 0 4029.6729
175.07186889648438 0 7302.677
175.13168334960938 0 1061.9088
176.10777282714844 0 1588.4004
180.11354064941406 0 1734.3857
181.0977325439453 0 7215.617
182.09278869628906 0 5548.8403
182.100830078125 0 1351.6368
182.1292266845703 0 7651.679
183.07688903808594 0 6734.6733
183.11334228515625 0 284818 b Water loss 1
184.10865783691406 0 6761.773
184.1167449951172 0 23766.559
185.092529296875 0 27395.379
185.119384765625 0 1232.5548
185.12838745117188 0 1428.2959
185.80615234375 0 950.2463
186.08786010742188 0 5711.299
186.0956268310547 0 1557.7452
187.07164001464844 0 1236.7306
187.14462280273438 0 47864.902
187.416259765625 0 904.1494
188.1476287841797 0 3421.74
189.0874786376953 0 8590.573
191.08181762695312 0 1560.7455
191.10324096679688 0 6583.5356 y 15
193.09800720214844 0 1382.9747
195.07696533203125 0 1879.5996
195.11328125 0 22539.91
196.1168670654297 0 2236.6406
197.12892150878906 0 26240.361
198.1240997314453 0 7141.5254
198.13284301757812 0 2824.2983
199.07171630859375 0 11828.897
199.1080322265625 0 4071.262
199.1178436279297 0 1120.9106
200.07513427734375 0 1301.6754
200.1034698486328 0 11970.649
200.13973999023438 0 9184.219
201.12380981445312 0 151977.47 b 1
202.1271514892578 0 13444.117
203.10342407226562 0 2365.4004
207.11349487304688 0 1062.5032
208.09793090820312 0 1208.7721
208.10850524902344 0 13130.045
209.09254455566406 0 38136.543
209.1107635498047 0 1750.3766
210.0961151123047 0 3443.799
210.1243896484375 0 2787.603
211.1443328857422 0 2185.383
212.14010620117188 0 3140.5732
213.0868377685547 0 2614.385
213.1240997314453 0 2812.631
214.15480041503906 0 1667.1636
215.1395263671875 0 58787.43
216.14306640625 0 4909.1704
217.0823974609375 0 26492.676
217.1339569091797 0 31872.62
217.14273071289062 0 808.22174
218.0855255126953 0 2668.3584
218.13735961914062 0 4089.6519
218.14979553222656 0 1752.4421
219.1497344970703 0 12607.236
219.18594360351562 0 1166.7426
220.15264892578125 0 1169.2252
225.1237335205078 0 13179.488
225.13531494140625 0 6282.403
225.16024780273438 0 1978.2449
226.1191864013672 0 131703.9
227.10292053222656 0 3991.0227
227.12245178222656 0 13765.049
228.13467407226562 0 3412.471
231.0988006591797 0 2571.7407
233.16546630859375 0 75882.914
234.0870361328125 0 1106.663
234.1240692138672 0 2310.5332
234.16864013671875 0 11257.895
236.1033935546875 0 7568.9404
236.13998413085938 0 2210.2454
237.0875244140625 0 2781.304
237.12423706054688 0 1112.7664
240.09909057617188 0 1297.2611
241.08253479003906 0 8955.24
241.19117736816406 0 1983.9177
242.11343383789062 0 1204.985
242.15057373046875 0 22880.371
243.1338653564453 0 4807.1494
243.1460418701172 0 37369.234
244.1298370361328 0 6577.764 y Ammonia loss 12
244.1493682861328 0 4796.0156
245.129150390625 0 21748.191
246.1324005126953 0 2173.3271
247.144775390625 0 12274.676
248.14871215820312 0 2211.5015
250.1180419921875 0 1243.4868
251.1525421142578 0 1691.6213
251.176025390625 0 41645.87
252.09812927246094 0 1943.8073
252.13511657714844 0 3888.4077
252.1788787841797 0 4672.149
253.11843872070312 0 1354.5587
254.11412048339844 0 32374.326
254.15048217773438 0 5815.4775
255.11764526367188 0 3481.1023
258.10986328125 0 1517.106
259.09307861328125 0 24960.225
259.14483642578125 0 1190.919
260.0964050292969 0 2014.8284
260.1615295410156 0 2064.1404
261.16033935546875 0 24119.258
262.1193542480469 0 4124.802
262.16357421875 0 4097.715
263.138427734375 0 1516.43
264.09954833984375 0 1331.4545
264.134765625 0 7415.947
266.15032958984375 0 1813.7915
268.0930480957031 0 3734.4297
268.1658020019531 0 1951.107
268.20269775390625 0 7758.445 a Water loss 2
269.1253356933594 0 1333.6624
269.16217041015625 0 2414.2385
269.1702880859375 0 1311.7689
269.1864929199219 0 11253.549
269.2029113769531 0 1627.769
270.1821594238281 0 3187.996
272.1245422363281 0 17591.275
273.12640380859375 0 2035.2528
275.1760559082031 0 16061.819
276.17852783203125 0 2234.8982
278.1498107910156 0 2209.0552
279.14593505859375 0 13721.717
280.1299133300781 0 7748.358
280.14910888671875 0 2019.9083
280.16619873046875 0 26304.105
281.1687316894531 0 4177.7583
282.1081237792969 0 1424.3837
282.1454772949219 0 45754.496
283.1485290527344 0 7069.7407
285.16046142578125 0 20078.582
285.1964111328125 0 2325.111
286.10418701171875 0 16131.141
286.1639099121094 0 2916.8708
286.2134094238281 0 2517.07 a 2
287.10748291015625 0 1820.557
289.1916809082031 0 1475.2559
294.18280029296875 0 3254.8694
296.0891418457031 0 3556.0327
296.1977233886719 0 731193.4 b Water loss 2
297.1203308105469 0 7434.9946
297.1565856933594 0 44548.516
297.2008361816406 0 130262.85
298.15899658203125 0 4846.1836
298.17694091796875 0 25737.959
298.2030029296875 0 11727.091
299.1775207519531 0 3529.9678
300.12078857421875 0 1747.2631
300.1559753417969 0 2350.0044
300.1728515625 0 1646.5269
301.1877746582031 0 2522.6301 y Water loss 14
302.1719970703125 0 1993.0201 y Ammonia loss 14
303.171142578125 0 28591.916
304.11474609375 0 8733.733
304.1744079589844 0 4570.9883
306.1459045410156 0 5559.3022
308.1624755859375 0 2957.1323 y Ammonia loss 11
310.1051330566406 0 2260.5952
311.1717529296875 0 1588.1388
312.1917724609375 0 1238.6997
313.16705322265625 0 1407.4126
313.18865966796875 0 7011.6714
314.1001281738281 0 1442.3883
314.1474304199219 0 7428.7827
314.1839599609375 0 30320.006
314.2083435058594 0 178984.02 b 2
315.16766357421875 0 1774.2737
315.18646240234375 0 3912.5178
315.21124267578125 0 31151.184
316.20477294921875 0 858.6563
316.2153015136719 0 1586.709
317.1868591308594 0 6599.0337
318.1853942871094 0 1478.0017
319.1983642578125 0 44805.234 y 14
320.2014465332031 0 7367.616
321.1560974121094 0 1333.226
323.171875 0 3400.443
324.1546936035156 0 1776.7601
325.15106201171875 0 2072.579
325.1878967285156 0 1273.9249
326.17047119140625 0 2358.399
326.1897277832031 0 1606.6708
328.11492919921875 0 12501.301
329.117919921875 0 2573.2102
329.184326171875 0 1365.2107
330.1822509765625 0 18071.861
331.1853942871094 0 3803.9692
333.1568908691406 0 4919.9224
334.2135009765625 0 5417.971
335.1725158691406 0 2264.5454
336.15667724609375 0 3097.7544
337.1207580566406 0 1086.416
337.1518249511719 0 6457.5957
338.13519287109375 0 3296.3123
338.15496826171875 0 2020.2052
339.1670227050781 0 4972.394
340.16790771484375 0 1186.49
341.18426513671875 0 12961.892
342.18316650390625 0 4083.5347
344.19775390625 0 7068.1885
345.2034606933594 0 1143.8353
346.1253967285156 0 34549.508
346.177001953125 0 27044.14
347.1302490234375 0 6251.591
347.1792907714844 0 4762.008
348.1926574707031 0 21343.918
349.1959228515625 0 5298.0312
350.1830749511719 0 5202.5913
351.1296691894531 0 1183.8243
351.1670837402344 0 22288.094
352.1704406738281 0 5346.513
353.1827697753906 0 12241.252
354.1668395996094 0 7243.234
354.1885681152344 0 2159.5676
355.1626281738281 0 7271.345
356.1443786621094 0 2685.1416
356.1661071777344 0 1217.593
358.213623046875 0 1367.1593
360.2290954589844 0 3862.6453
362.20819091796875 0 61663.383
363.1672058105469 0 1883.6614
363.21136474609375 0 13500.783
365.1462097167969 0 10648.117
365.1831970214844 0 1671.7245
365.25579833984375 0 3205.6006
366.14862060546875 0 2344.7537
367.2089538574219 0 7280.32
367.2337951660156 0 1744.7383
368.19342041015625 0 7250.5225
369.1776428222656 0 32113.842
370.1242370605469 0 1077.2383
370.1809997558594 0 5763.5015
370.67742919921875 0 4146.52
371.1559143066406 0 1357.0137
371.1932373046875 0 7268.3335
372.17755126953125 0 7251.2324
373.1729431152344 0 13248.602
374.1748962402344 0 2309.3652
375.1679382324219 0 1668.7798
379.125 0 1329.4332
379.19439697265625 0 1666.0753
380.1936950683594 0 1853.2723
381.1399841308594 0 1159.4827
381.17840576171875 0 2837.695
383.1569519042969 0 22712.434
383.2666320800781 0 4638.244
384.1598815917969 0 4031.1697
385.22015380859375 0 17514.363
386.2228088378906 0 3573.0493
389.25421142578125 0 1508.6703
390.2031555175781 0 2984.2053
393.17681884765625 0 2780.6787
393.25091552734375 0 9796.831
394.2533874511719 0 2847.3728
395.1929626464844 0 1828.1736
397.1360778808594 0 5408.351
398.1188659667969 0 1874.1859
398.20379638671875 0 1812.0554
398.2445983886719 0 14734.552
399.1873474121094 0 1996.154
399.2484436035156 0 3108.8953
400.1945495605469 0 1148.8778
400.22503662109375 0 2921.3455
401.1678466796875 0 38175.543
402.1711730957031 0 7334.2104
405.1622314453125 0 3567.7332
407.2040100097656 0 5588.9
408.1888427734375 0 13145.65
409.1893310546875 0 2070.7317
410.2043762207031 0 3556.281
411.15179443359375 0 1609.4784
411.26129150390625 0 17302.355
412.2629089355469 0 3562.63
413.20526123046875 0 2353.4639
415.1468200683594 0 11087.965
415.2327575683594 0 4072.7644 b 7
415.2718811035156 0 7514.999
416.1513977050781 0 1798.2922
416.21759033203125 0 1983.0773 y Ammonia loss 13
416.25555419921875 0 28396.451
417.25726318359375 0 9882.908
418.257080078125 0 1330.0756
423.1880187988281 0 1837.3279
424.2273254394531 0 5344.3857
425.1311950683594 0 4007.4678
425.1664123535156 0 1123.383
425.2154235839844 0 11999.1045
426.2198791503906 0 1454.218
427.2332763671875 0 6447.6465
428.2373962402344 0 1838.3702
429.2141418457031 0 1632.6581 y Ammonia loss 9
431.2160949707031 0 1469.2996
433.15771484375 0 14643.657
433.2430419921875 0 33447.57 y 13
433.2821960449219 0 230179.61
434.16082763671875 0 3631.9597
434.2036437988281 0 5956.86
434.2458190917969 0 6958.7646
434.285400390625 0 63484.83
434.70721435546875 0 5324.93
435.20782470703125 0 5030.03
435.2397766113281 0 1225.402
435.2532958984375 0 1351.5037
435.2876281738281 0 10361.95
440.2153015136719 0 2309.471
441.19903564453125 0 4430.4927
441.25177001953125 0 2296.2937
442.2039794921875 0 1568.4553
442.24090576171875 0 9414.765
443.1417236328125 0 4543.5005
443.1754455566406 0 1954.894
443.2304382324219 0 11738.938
443.2668151855469 0 23882.697 b Water loss 3
444.18609619140625 0 1476.5297
444.2358093261719 0 1851.2299
444.26995849609375 0 7166.5317
445.2452087402344 0 3777.6506
450.2356262207031 0 1165.6005
451.2312316894531 0 3152.878
452.2152404785156 0 7159.036
458.22430419921875 0 1552.6847
459.17718505859375 0 1403.7057
459.2102355957031 0 6392.0386
459.2583312988281 0 1846.8204
460.2176818847656 0 2069.103
461.2771911621094 0 200804.88 b 3
462.20037841796875 0 10427.267
462.28033447265625 0 55497.945
463.2021484375 0 2917.5618
463.2831726074219 0 9406.762
464.21539306640625 0 2788.153
466.1957702636719 0 3822.628
468.2091369628906 0 4196.3066
468.2481689453125 0 1473.0115
469.24322509765625 0 5306.0664
470.22613525390625 0 28697.83
471.2285461425781 0 7646.995
476.1795654296875 0 3519.1567
477.25958251953125 0 2317.987
478.2415771484375 0 5089.0444
479.2288513183594 0 3584.4026
480.2106018066406 0 20660.457
481.2128601074219 0 4849.071
482.22442626953125 0 5689.286
483.21002197265625 0 1562.7474
484.20489501953125 0 10909.866
485.2115478515625 0 1730.2579
486.2682189941406 0 3387.0215 y Water loss 12
487.2533874511719 0 5213.742 y Ammonia loss 12
488.2574768066406 0 2256.5286
492.209228515625 0 2094.7625
494.1895446777344 0 8093.484
495.1925354003906 0 1974.2097
495.2684020996094 0 4365.335
496.25225830078125 0 11359.689
497.25689697265625 0 2806.0593
498.2207946777344 0 74020.54
499.2239990234375 0 17793.254
500.23272705078125 0 5393.5166
502.2156982421875 0 16215.327
503.2190856933594 0 3666.0305
504.27880859375 0 51452.008 y 12
505.2820129394531 0 11224.598
510.2207946777344 0 4335.4365
512.1998901367188 0 15799.966
512.3211059570312 0 2101.93
513.202880859375 0 4880.3525
513.2788696289062 0 6114.9575
514.3046875 0 2220.7349
515.2477416992188 0 2611.119
516.242431640625 0 1895.9586
518.2464599609375 0 2535.8635
520.2842407226562 0 1390.1354
521.26953125 0 1804.27
524.2374877929688 0 1676.1522
527.2478637695312 0 1346.052
528.231689453125 0 4993.8555
530.21044921875 0 47838.023
530.3348999023438 0 14406.438
531.2138671875 0 14713.807
531.3377075195312 0 4972.335
532.2152709960938 0 3105.601
535.2525634765625 0 1566.6274
538.3007202148438 0 4436.3745
539.25 0 1595.6921
539.3045654296875 0 1851.8833
540.3196411132812 0 10122.23 b Water loss 4
541.3235473632812 0 2209.262
546.2412719726562 0 5417.5347
547.2386474609375 0 2133.9795
553.2627563476562 0 2907.4397
554.2540893554688 0 1316.6613
555.326171875 0 3892.0337
556.22509765625 0 2385.0024
556.2724609375 0 1392.5525
556.3260498046875 0 2285.1016
558.3297729492188 0 30853.14 b 4
559.3330688476562 0 10206.946
560.3343505859375 0 1700.891
560.7713012695312 0 1886.6991
563.2470703125 0 2848.555
569.26318359375 0 3385.5017
569.2801513671875 0 3621.3296
569.7798461914062 0 7151.4463
570.2791137695312 0 2295.6365
571.2720336914062 0 2627.808
572.2674560546875 0 1362.3169
573.2295532226562 0 1648.7571
578.2872314453125 0 12040.268
578.787353515625 0 4533.4897
579.2876586914062 0 3029.361
581.2581176757812 0 9662.465
582.2883911132812 0 4186.0684
582.7841796875 0 16712.969
583.2831420898438 0 8517.677
583.786376953125 0 2295.0554
587.2695922851562 0 2863.333
591.2418823242188 0 3976.1392
591.2944946289062 0 5545.6997
591.7959594726562 0 5229.9004 b Water loss 10
592.29541015625 0 1463.2736
597.2926635742188 0 6085.9214
598.2899780273438 0 3572.3286
599.2687377929688 0 44286.95
600.271728515625 0 18422.691
601.277099609375 0 2761.7212
604.3049926757812 0 3235.8862
604.8038940429688 0 3134.9565
607.275390625 0 3658.2861
608.2726440429688 0 1425.2169
608.3529663085938 0 3296.5654
609.2530517578125 0 16351.584
609.296142578125 0 3405.461
609.3419189453125 0 2028.6354
609.7975463867188 0 4526.501
610.2550048828125 0 5039.736
612.8161010742188 0 7665.678
613.3165893554688 0 7738.4844
613.820068359375 0 3236.6807
614.3265380859375 0 9620.193 y Water loss 11
615.3052368164062 0 31829.307 y Ammonia loss 11
616.3074340820312 0 8407.652
617.309326171875 0 2124.7725
617.80810546875 0 38810.047
618.30615234375 0 42637.695
618.8065185546875 0 19010.607
619.3077392578125 0 6217.9277
619.8045654296875 0 1334.961
625.2845458984375 0 9203.005
626.2853393554688 0 4126.5063
626.3629150390625 0 6665.134
626.8135375976562 0 108787.17
627.262939453125 0 112571.914
627.3153076171875 0 69517.91
627.8157348632812 0 28234.346
628.2666015625 0 37228.92
628.3173217773438 0 5326.2715
628.8087768554688 0 2155.775 y Water loss 5
629.2698974609375 0 7543.643
629.805908203125 0 2987.1633
632.337646484375 0 31438.377 y 11
633.3405151367188 0 11176.414
634.3443603515625 0 2156.7405
635.8206176757812 0 6007.149
636.3206176757812 0 3023.953
637.816650390625 0 15734.955 y 5
638.3168334960938 0 8740.555
638.8179931640625 0 5554.0273
642.3132934570312 0 2749.5342
643.2954711914062 0 24284.424
644.294677734375 0 11661.742
645.2922973632812 0 4113.314
646.295654296875 0 1491.0769
653.2804565429688 0 2098.1936
655.3406372070312 0 5962.52
655.39453125 0 1636.8308
655.8416137695312 0 5579.335 b Water loss 11
656.3394165039062 0 2659.5967 b Ammonia loss 11
658.3080444335938 0 1896.4883
659.3084716796875 0 1279.3152
660.3209228515625 0 8785.352
661.3233032226562 0 4641.699
662.3351440429688 0 1260.2063
663.337158203125 0 2652.8467
663.8403930664062 0 2859.09
664.341552734375 0 1591.4131
667.3380737304688 0 1460.3171
668.3319091796875 0 5259.027
668.8258056640625 0 5592.853
669.3258666992188 0 4950.3423
669.8290405273438 0 2565.11
670.3203125 0 1393.795
674.2998046875 0 2437.5466
677.3372192382812 0 34834.88 y Water loss 4
677.8348388671875 0 46510.734 y Ammonia loss 4
678.3336791992188 0 29321.088
678.8336791992188 0 12245.789
679.2603759765625 0 1399.5012
679.3275756835938 0 2461.5134
685.3292236328125 0 2236.3752
685.838623046875 0 2915.1423
686.3427124023438 0 767077.4 y 4
686.8439331054688 0 631878.3
687.3450927734375 0 268812.2
687.8462524414062 0 63134.3
688.3446655273438 0 7293.822
691.8399047851562 0 1720.9299 b Ammonia loss 12
692.3366088867188 0 1623.5422
694.3425903320312 0 6273.8833
695.3397216796875 0 3646.7349
696.35693359375 0 2052.6672
698.2938842773438 0 1564.991
700.3475952148438 0 8543.909 b 12
700.84912109375 0 7382.1206
701.3494873046875 0 4182.671
704.3286743164062 0 2787.6406
710.3009643554688 0 2601.6748
712.3529663085938 0 65149.348
713.3560791015625 0 23728.096
714.35693359375 0 5836.9404
720.3689575195312 0 1880.0405
721.3677368164062 0 1886.7957
722.3381958007812 0 8987.5625
723.342041015625 0 3479.9275
724.3479614257812 0 1611.6776
725.3456420898438 0 1617.995
727.3305053710938 0 2926.1465
728.3984985351562 0 2363.7834 y Ammonia loss 10
731.3552856445312 0 1884.0707
737.3140258789062 0 2303.858
737.396728515625 0 2621.052
738.2993774414062 0 3142.1477
738.38037109375 0 3420.0198
739.369384765625 0 4928.383
740.34814453125 0 77509.31
741.3512573242188 0 34336.73
742.3515625 0 6509.655
742.4179077148438 0 5718.434 b 6
743.3524169921875 0 2141.2778
743.4181518554688 0 3506.3022
745.3411254882812 0 4229.58
745.4222412109375 0 26766.66 y 10
746.3404541015625 0 3152.9167
746.4242553710938 0 11648.591
747.42626953125 0 2461.8586
750.874267578125 0 4041.8875 y Water loss 3
751.3687744140625 0 6471.0786 y Ammonia loss 3
751.8701171875 0 3241.7188
752.3668823242188 0 1903.5204
753.3430786132812 0 5885.4316
754.3368530273438 0 3543.3425
755.3253784179688 0 2445.8284
755.4043579101562 0 4310.044
756.3228149414062 0 2601.5442
756.4061889648438 0 1923.355
756.8828735351562 0 2261.0032
757.3750610351562 0 37684.3 b 13
758.376953125 0 14722.762
759.3768920898438 0 2827.1091
759.8162231445312 0 827.2106
759.8771362304688 0 97225.9 y 3
760.378173828125 0 102595.36
760.8793334960938 0 43709.457
761.3797607421875 0 15720.635
766.35888671875 0 1688.4557
771.3541870117188 0 18834.924
772.3563842773438 0 9679.856
773.3565063476562 0 1758.1705
774.3375244140625 0 2901.3596
775.3359985351562 0 2566.779
775.4539184570312 0 1362.7867
783.4033203125 0 1758.2312
797.3703002929688 0 1800.8737
805.3740844726562 0 1597.3864
807.9105834960938 0 1730.8257 y Ammonia loss 2
811.3753662109375 0 1467.97
812.919921875 0 2202.6782 b Ammonia loss 14
813.4193725585938 0 2216.3135
813.9189453125 0 1903.3439
814.4022216796875 0 2909.1602
816.41943359375 0 21274.75 y 2
816.9208374023438 0 25598.506
817.420654296875 0 11737.3125
817.9251098632812 0 3208.7937
822.4103393554688 0 2942.6436
823.3862915039062 0 2130.7844
824.3800659179688 0 5325.1665
825.3636474609375 0 2441.319
826.37255859375 0 1481.4225
827.3597412109375 0 1627.4038
829.4459838867188 0 4818.5986 b 7
830.4459838867188 0 2878.9539
832.3876953125 0 1745.4055
833.384765625 0 2198.079
839.4244384765625 0 2321.4695
840.4134521484375 0 10587.494
841.4114990234375 0 6427.2324
842.3914184570312 0 15767.533
843.392578125 0 6752.221
847.9448852539062 0 3589.0085 b Water loss 15
848.4371337890625 0 6377.8335 b Ammonia loss 15
848.9395751953125 0 5304.506
849.4359741210938 0 3142.093
850.3973999023438 0 12809.947
851.3914184570312 0 10040.3
852.4012451171875 0 2950.024
856.448486328125 0 4074.7031 y Water loss 9
856.9482421875 0 17075.63 b 15
857.4490966796875 0 23709.514 y Ammonia loss 9
857.9511108398438 0 14781.705 y Water loss 1
858.447998046875 0 4743.4565
859.4194946289062 0 2273.541
866.9447631835938 0 5776.0625 y 1
867.4401245117188 0 9363.691
867.9447021484375 0 4903.26
868.4067993164062 0 96488.13
869.409423828125 0 45581.17
870.4105834960938 0 14403.819
874.4642333984375 0 17807.602 y 9
875.4677124023438 0 8875.778
876.4700317382812 0 2063.7896
884.4420166015625 0 3117.4414
885.4423828125 0 4513.658
887.4153442382812 0 5884.217
888.4177856445312 0 4516.7886
893.4306030273438 0 1773.1691
894.4259643554688 0 5718.454
895.4396362304688 0 3662.4429
898.4692993164062 0 2375.4673
899.459716796875 0 2299.4592
900.44921875 0 2390.787
903.421875 0 1920.9836
904.419677734375 0 1868.1876
907.4724731445312 0 15728.487 Precursor Water loss
907.9716796875 0 22522.703 Precursor Ammonia loss
908.4706420898438 0 16231.274
908.9707641601562 0 5348.3633
911.4468383789062 0 18478.59
912.4502563476562 0 9426.916
913.4449462890625 0 2364.085
915.9705810546875 0 2887.9807
916.4772338867188 0 152161.1 Precursor
916.9779052734375 0 166956.48
917.4799194335938 0 108084
917.9811401367188 0 38158.414
918.4828491210938 0 4887.6113
921.42919921875 0 9092.387
922.424072265625 0 11811.495
923.4203491210938 0 4139.239
924.4252319335938 0 1662.2595
939.4427490234375 0 65858.51
940.4463500976562 0 32231.73
941.4486694335938 0 10633.943
942.4462890625 0 1964.4972
953.46630859375 0 2816.751
954.4617309570312 0 3911.2598
955.4581298828125 0 2165.8774
956.45068359375 0 11385.052
957.45458984375 0 6133.845
958.4866333007812 0 6508.658 b 8
959.4972534179688 0 2012.5061
967.4732055664062 0 1905.588
970.4517211914062 0 3151.1838
971.4754638671875 0 5496.5796
972.4794921875 0 3210.2222
973.4671020507812 0 1486.2961
982.4777221679688 0 1339.3209
985.4945678710938 0 2291.3809 y Water loss 8
987.4829711914062 0 3338.2861
988.4779663085938 0 1915.5392
989.484375 0 1691.7542
1003.50830078125 0 11769.942 y 8
1004.5068359375 0 8655.38
1005.5093994140625 0 1781.5892
1008.4642944335938 0 6165.6445
1009.4695434570312 0 3949.1677
1010.4671630859375 0 1753.2534
1015.4761352539062 0 5300.3535
1016.4783325195312 0 3913.4968
1017.4768676757812 0 1864.5734
1018.4529418945312 0 2057.5327
1019.447021484375 0 2292.9402
1023.483154296875 0 1740.5907
1024.48193359375 0 2243.4688
1025.49072265625 0 3408.149
1026.4888916015625 0 2261.4646
1035.47802734375 0 3944.2546
1036.465087890625 0 14371.176
1037.4666748046875 0 6716.1235
1038.471435546875 0 3783.2983
1040.5023193359375 0 5124.0356
1041.4991455078125 0 6217.545
1042.49755859375 0 3411.5789
1052.496337890625 0 1872.4934
1053.4871826171875 0 53852.594
1054.48828125 0 34678.684
1055.4892578125 0 12690.477
1056.4971923828125 0 3131.0698
1058.5140380859375 0 9026.293
1059.5152587890625 0 4383.4194
1066.51806640625 0 1367.3822
1067.5047607421875 0 4122.5415
1068.51025390625 0 3850.782
1069.5108642578125 0 2929.3535 b Water loss 9
1070.513427734375 0 17968.125
1071.5137939453125 0 11815.843
1072.5206298828125 0 5270.348 y Water loss 7
1073.5184326171875 0 3483.3667 y Ammonia loss 7
1074.5162353515625 0 1793.9382
1076.5169677734375 0 1357.1842
1084.5272216796875 0 10714.524
1085.5283203125 0 7932.2964
1086.5162353515625 0 6760.16
1087.5252685546875 0 10831.526 b 9
1088.5301513671875 0 3045.2842
1089.5289306640625 0 1699.7618
1090.5386962890625 0 21547.91 y 7
1091.540771484375 0 13699.845
1092.5435791015625 0 4313.1597
1093.537353515625 0 1305.1434
1102.5511474609375 0 1559.1141
1120.5347900390625 0 2541.1467
1121.5386962890625 0 1932.6329
1127.568115234375 0 1899.3856
1128.557373046875 0 1958.7716
1137.554443359375 0 9175.1455
1138.5484619140625 0 11744.708
1139.5467529296875 0 6480.9805
1140.54931640625 0 1771.2104
1146.54931640625 0 2167.098
1147.5574951171875 0 1552.6437
1155.5655517578125 0 19003.94
1156.56787109375 0 13013.317
1157.5697021484375 0 5479.048
1159.5584716796875 0 5353.937 y Water loss 6
1160.5528564453125 0 6007.3345 y Ammonia loss 6
1161.5526123046875 0 3368.4321
1163.580078125 0 2764.5803
1164.5577392578125 0 12944.557
1165.5621337890625 0 5855.481
1166.5648193359375 0 3287.353
1177.57177734375 0 26474.55 y 6
1178.5732421875 0 19801.086
1179.5770263671875 0 6695.147
1181.58203125 0 62590.594
1182.585693359375 0 48220.445 b Water loss 10
1183.5865478515625 0 19010.826
1184.590576171875 0 4056.5186
1198.592529296875 0 1750.3948
1200.6114501953125 0 10681.428 b 10
1201.607177734375 0 7982.9473
1202.614990234375 0 3931.908
1207.598876953125 0 2306.2454
1208.60986328125 0 1732.8342
1212.6241455078125 0 2888.115
1217.5892333984375 0 4219.532
1218.582275390625 0 2732.9072
1224.6290283203125 0 5428.7314
1225.6280517578125 0 3811.7515
1226.6143798828125 0 1465.5363
1234.6072998046875 0 22702.895
1235.6031494140625 0 26145.703
1236.6038818359375 0 14739.828
1237.607421875 0 3726.6025
1238.5927734375 0 2681.287
1239.5960693359375 0 2479.4897
1240.6021728515625 0 1419.641
1252.617919921875 0 57651.67
1253.6201171875 0 42711.406
1254.621826171875 0 16469.602
1255.621337890625 0 4514.559
1256.611572265625 0 8150.3286 y Water loss 5
1257.6029052734375 0 13935.752 y Ammonia loss 5
1258.606201171875 0 8486.686
1259.6068115234375 0 3000.044
1270.6280517578125 0 3093.1917
1274.623046875 0 148959.58 y 5
1275.62548828125 0 109745.51
1276.6275634765625 0 47240.375
1277.6309814453125 0 9664.473
1309.66943359375 0 6604.4863
1310.669921875 0 8722.915 b Water loss 11
1311.67041015625 0 5517.5234 b Ammonia loss 11
1312.6558837890625 0 3065.1057
1325.6707763671875 0 3237.5017
1326.6695556640625 0 2598.232
1327.653564453125 0 3556.3225
1328.6697998046875 0 17276.963 b 11
1329.6741943359375 0 13687.572
1330.67333984375 0 7232.622
1335.655029296875 0 2788.151
1336.6473388671875 0 4820.944
1337.640625 0 5403.874
1341.65673828125 0 1752.6296
1353.6650390625 0 26769.418 y Water loss 4
1354.65673828125 0 51904.52 y Ammonia loss 4
1355.6575927734375 0 36000.61
1356.6580810546875 0 12864.129
1357.6611328125 0 3531.939
1369.6650390625 0 2299.4722
1370.661376953125 0 4171.2383
1371.676025390625 0 849995 y 4
1372.678466796875 0 707775.9
1373.6805419921875 0 297723.53
1374.682861328125 0 74502.555
1375.6865234375 0 5206.6875
1381.6917724609375 0 3085.9094 b Water loss 12
1382.68701171875 0 2289.4773 b Ammonia loss 12
1399.706787109375 0 13857.637 b 12
1400.7069091796875 0 11396.528
1401.7113037109375 0 5958.4824
1402.71484375 0 1580.3068
1501.7257080078125 0 2722.4854 y Ammonia loss 3
1502.723876953125 0 2123.0508
1513.761474609375 0 5477.137 b 13
1514.7591552734375 0 6611.472
1515.7659912109375 0 3299.6836
1518.744384765625 0 49817.293 y 3
1519.7454833984375 0 44380.664
1520.7490234375 0 20044.076
1521.7508544921875 0 7693.4985
1631.8243408203125 0 8727.084 y 2
1632.830810546875 0 9845.349
1633.82861328125 0 5352.154
1634.8323974609375 0 1656.1906

Spectrum Details

|  |  |
| --- | --- |
| Matched peaks? Matched peaks The total absolute number of peaks matched. Additionally in brackets the total fraction of peaks matched and the total number of peaks is shown. | 91 (10.20% of 892) |
| FDR? FDR The false discovery rate estimated for this peptide. It is calculated by matching all theoretical fragments with a non-integer shift with the raw peaks for this spectrum. This is done with 40 different shifts. The resulting percentage is the average number of annotated peaks over the number of annotated peaks with the correct spectrum. | 0.71% |
| Satellite FDR? Satellite FDR See the FDR for details on its calculation. This satellite ion specific FDR only contains the satellite ions (d/w) for I/L/J positions. | - |
| PSM Score? PSM Score The PSM Score as given by Hecklib to this annotated spectrum. It is shown with three significant figures. | 1.04E+03 |

#### Spectrum 7834? Spectrum 7834 The raw spectrum of this peptide as annotated by Hecklib. The fragments are coloured according to ion type (see legend). Any peaks with a star '\*' as text can be hovered over to see the full details, first the ion type second the mass shift type. By hovering over the amino acids in the peptide or ions in the legend the corresponding peaks are highlighted. By toggling the 'Unassigned' label you can turn the background (unassigned) peaks on or off in the plot. By updating the slider in the Ion legend you can update the spectrum to only show the top X% of the peaks with labels. The top X% means any peak that is within X% of the highest intensity. By dragging in the spectrum you can zoom in to a specific part of the spectrum and use 'Zoom Out' to get back to the original zoom level. The annotation of the spectrum is based on the given sequence in the peptides file and is done with different software so inconsistencies are likely. The peaks are annotated based on the given sequence, with 20 ppm tolerance.

Copy Data

##### Spectrum 7834 (TSV)

###### Preview

```
Loading example...
```

*Click on the button to copy the data to your clipboard.*

Mz MinMz MaxIntensity Max

WidthHeightPeptide font sizePeptide stroke widthSpectrum font sizeSpectrum stroke widthCompact peptide

Ion legend

wxyz

abcd

OtherUnassignedIon^Charge^_Position_Show for top:%

VTJFPPSSEEJQANKAT

02.37e+44.73e+47.10e+49.46e+4

Zoom Out

y^+1^_1_a^+1^_2_y^+1^_2_a^+1^_2_b^+1^_2_y^+1^_2_b^+1^_2_y^+3^_7_y^+2^_5_a^+1^_3_b^+2^_5_b^+1^_3_y^+1^_3_y^+2^_6_y^+2^_6_b^+1^_3_y^+2^_6_y^+1^_3_y^+3^_9_y^+2^_7_y^+2^_7_b^+2^_8_b^+2^_8_y^+1^_4_y^+2^_8_y^+1^_4_y^+2^_8_b^+1^_4_y^+3^_13_y^+3^_13_b^+1^_4_y^+1^_5_y^+1^_5_y^+3^_14_y^+1^_5_b^+1^_5_b^+2^_10_b^+1^_5_y^+3^_16_y^+2^_11_y^+2^_11_b^+2^_11_b^+2^_11_y^+1^_6_y^+1^_6_y^+2^_12_y^+1^_6_y^+2^_12_b^+2^_12_b^+2^_12_y^+2^_13_y^+2^_13_y^+2^_13_b^+2^_13_b^+2^_13_b^+2^_13_b^+1^_7_y^+1^_7_y^+1^_7_b^+1^_7_y^+1^_7_y^+2^_14_y^+2^_14_b^+2^_14_y^+2^_14_y^+2^_15_y^+2^_15_b^+1^_8_b^+2^_15_y^+2^_15_b^+1^_8_b^+2^_16_b^+2^_16_y^+1^_8_b^+2^_16_y^+1^_8_y^+2^_16_y^+2^_16_y^+1^_8_b^+1^_9_y^+1^_9_b^+1^_10_y^+1^_10_y^+1^_10_b^+1^_10_y^+1^_10_y^+1^_11_b^+1^_11_b^+1^_11_y^+1^_12_y^+1^_12_y^+1^_12_b^+1^_12_b^+1^_12_y^+1^_13_y^+1^_13_y^+1^_13_b^+1^_13_b^+1^_14_

0775154923243099

Fragment Matches Table

Show background peaks

| Position | Ion type | Intensity | mz Theoretical | mz Error (Th) | mz Error (ppm) | Charge | Series Number |
| --- | --- | --- | --- | --- | --- | --- | --- |
| 17 | y | 4.344E+04 | 120.1 | 0.0003678 | 3.064 | +1 | 1 |
| - | - | 4.054E+04 | 120.1 | - | - | 0 | - |
| - | - | 1830 | 121.1 | - | - | 0 | - |
| - | - | 3580 | 121.1 | - | - | 0 | - |
| - | - | 446.8 | 122.1 | - | - | 0 | - |
| - | - | 356.1 | 125.1 | - | - | 0 | - |
| - | - | 3566 | 126.1 | - | - | 0 | - |
| - | - | 675.1 | 127.1 | - | - | 0 | - |
| - | - | 1.056E+04 | 128.1 | - | - | 0 | - |
| - | - | 1552 | 129.1 | - | - | 0 | - |
| - | - | 5.667E+04 | 129.1 | - | - | 0 | - |
| - | - | 3612 | 130.1 | - | - | 0 | - |
| - | - | 1429 | 130.1 | - | - | 0 | - |
| - | - | 904.9 | 130.1 | - | - | 0 | - |
| - | - | 4354 | 130.1 | - | - | 0 | - |
| - | - | 3414 | 131.1 | - | - | 0 | - |
| - | - | 1194 | 132.1 | - | - | 0 | - |
| - | - | 594.4 | 132.1 | - | - | 0 | - |
| - | - | 2757 | 136.1 | - | - | 0 | - |
| - | - | 2052 | 138.1 | - | - | 0 | - |
| - | - | 1480 | 138.1 | - | - | 0 | - |
| - | - | 1544 | 139.1 | - | - | 0 | - |
| - | - | 775 | 141.1 | - | - | 0 | - |
| - | - | 561.6 | 145.1 | - | - | 0 | - |
| - | - | 1747 | 147.1 | - | - | 0 | - |
| - | - | 512.4 | 148.1 | - | - | 0 | - |
| - | - | 432.2 | 148.2 | - | - | 0 | - |
| - | - | 2840 | 149 | - | - | 0 | - |
| - | - | 536 | 153.1 | - | - | 0 | - |
| - | - | 684.5 | 153.1 | - | - | 0 | - |
| - | - | 1030 | 154.1 | - | - | 0 | - |
| - | - | 1665 | 154.1 | - | - | 0 | - |
| - | - | 3610 | 155.1 | - | - | 0 | - |
| 2 | a | 6940 | 155.1 | 0.0003813 | 2.458 | +1 | 2 |
| - | - | 718.6 | 156.1 | - | - | 0 | - |
| - | - | 1692 | 157.1 | - | - | 0 | - |
| - | - | 3783 | 157.1 | - | - | 0 | - |
| - | - | 5173 | 159.1 | - | - | 0 | - |
| - | - | 1067 | 165.1 | - | - | 0 | - |
| - | - | 4205 | 167.1 | - | - | 0 | - |
| - | - | 2357 | 167.1 | - | - | 0 | - |
| - | - | 2548 | 167.1 | - | - | 0 | - |
| - | - | 439.3 | 168.1 | - | - | 0 | - |
| - | - | 1008 | 169.1 | - | - | 0 | - |
| - | - | 1.206E+04 | 169.1 | - | - | 0 | - |
| - | - | 754.6 | 169.1 | - | - | 0 | - |
| - | - | 1068 | 170.1 | - | - | 0 | - |
| - | - | 1129 | 171.1 | - | - | 0 | - |
| - | - | 1559 | 172.1 | - | - | 0 | - |
| - | - | 583.6 | 172.1 | - | - | 0 | - |
| - | - | 400.1 | 172.5 | - | - | 0 | - |
| 16 | y | 6198 | 173.1 | 0.000369 | 2.132 | +1 | 2 |
| 2 | a | 2.9E+04 | 173.1 | 0.000391 | 2.258 | +1 | 2 |
| - | - | 2050 | 174.1 | - | - | 0 | - |
| - | - | 1593 | 175.1 | - | - | 0 | - |
| - | - | 444.3 | 177.1 | - | - | 0 | - |
| - | - | 1314 | 181.1 | - | - | 0 | - |
| - | - | 3742 | 182.1 | - | - | 0 | - |
| - | - | 2416 | 182.1 | - | - | 0 | - |
| - | - | 473.5 | 182.8 | - | - | 0 | - |
| - | - | 2814 | 183.1 | - | - | 0 | - |
| 2 | b | 3.937E+04 | 183.1 | 0.0004313 | 2.355 | +1 | 2 |
| - | - | 519.6 | 183.1 | - | - | 0 | - |
| - | - | 2945 | 184.1 | - | - | 0 | - |
| - | - | 3452 | 184.1 | - | - | 0 | - |
| - | - | 5229 | 185.1 | - | - | 0 | - |
| - | - | 2816 | 186.1 | - | - | 0 | - |
| - | - | 909.7 | 186.1 | - | - | 0 | - |
| - | - | 550.9 | 187.1 | - | - | 0 | - |
| - | - | 1054 | 187.1 | - | - | 0 | - |
| - | - | 1.151E+04 | 187.1 | - | - | 0 | - |
| - | - | 3710 | 189.1 | - | - | 0 | - |
| 16 | y | 8825 | 191.1 | 0.0003939 | 2.061 | +1 | 2 |
| - | - | 1085 | 191.1 | - | - | 0 | - |
| - | - | 789.6 | 195.1 | - | - | 0 | - |
| - | - | 5997 | 195.1 | - | - | 0 | - |
| - | - | 835.4 | 196.1 | - | - | 0 | - |
| - | - | 941.2 | 197.1 | - | - | 0 | - |
| - | - | 1.493E+04 | 197.1 | - | - | 0 | - |
| - | - | 1845 | 198.1 | - | - | 0 | - |
| - | - | 1077 | 198.1 | - | - | 0 | - |
| - | - | 4544 | 199.1 | - | - | 0 | - |
| - | - | 1037 | 199.1 | - | - | 0 | - |
| - | - | 612.8 | 199.1 | - | - | 0 | - |
| - | - | 3548 | 200.1 | - | - | 0 | - |
| - | - | 3919 | 200.1 | - | - | 0 | - |
| 2 | b | 3.559E+04 | 201.1 | 0.0003647 | 1.813 | +1 | 2 |
| - | - | 3557 | 202.1 | - | - | 0 | - |
| - | - | 663.2 | 203.1 | - | - | 0 | - |
| - | - | 787 | 208.1 | - | - | 0 | - |
| - | - | 4537 | 208.1 | - | - | 0 | - |
| - | - | 1.014E+04 | 209.1 | - | - | 0 | - |
| - | - | 1249 | 210.1 | - | - | 0 | - |
| - | - | 995.8 | 210.1 | - | - | 0 | - |
| - | - | 514 | 211.1 | - | - | 0 | - |
| - | - | 1292 | 212.1 | - | - | 0 | - |
| - | - | 865.4 | 213.1 | - | - | 0 | - |
| - | - | 586.2 | 214.2 | - | - | 0 | - |
| - | - | 1.468E+04 | 215.1 | - | - | 0 | - |
| - | - | 1529 | 216.1 | - | - | 0 | - |
| - | - | 7381 | 217.1 | - | - | 0 | - |
| - | - | 4202 | 217.1 | - | - | 0 | - |
| - | - | 8519 | 217.1 | - | - | 0 | - |
| - | - | 1150 | 218.1 | - | - | 0 | - |
| - | - | 582.4 | 218.2 | - | - | 0 | - |
| - | - | 1515 | 219.1 | - | - | 0 | - |
| - | - | 5719 | 221.1 | - | - | 0 | - |
| - | - | 1017 | 222.1 | - | - | 0 | - |
| - | - | 511.9 | 223.7 | - | - | 0 | - |
| - | - | 2075 | 225 | - | - | 0 | - |
| - | - | 4916 | 225.1 | - | - | 0 | - |
| - | - | 1984 | 225.1 | - | - | 0 | - |
| - | - | 4.117E+04 | 226.1 | - | - | 0 | - |
| - | - | 1063 | 227.1 | - | - | 0 | - |
| - | - | 4172 | 227.1 | - | - | 0 | - |
| - | - | 892.8 | 228.1 | - | - | 0 | - |
| - | - | 683.4 | 229.4 | - | - | 0 | - |
| - | - | 908.5 | 231.1 | - | - | 0 | - |
| - | - | 1.652E+04 | 233.2 | - | - | 0 | - |
| - | - | 2136 | 234.2 | - | - | 0 | - |
| - | - | 6884 | 235.1 | - | - | 0 | - |
| - | - | 2187 | 236.1 | - | - | 0 | - |
| - | - | 752.9 | 236.1 | - | - | 0 | - |
| - | - | 1.233E+04 | 239.1 | - | - | 0 | - |
| - | - | 3126 | 240.1 | - | - | 0 | - |
| - | - | 3032 | 241.1 | - | - | 0 | - |
| - | - | 9422 | 242.2 | - | - | 0 | - |
| - | - | 1040 | 243.1 | - | - | 0 | - |
| 11 | y | 1.678E+04 | 243.1 | 0.004299 | 17.68 | +3 | 7 |
| - | - | 821.6 | 243.2 | - | - | 0 | - |
| 13 | y | 1583 | 244.1 | 0.0006545 | 2.681 | +2 | 5 |
| - | - | 1279 | 244.1 | - | - | 0 | - |
| - | - | 6076 | 245.1 | - | - | 0 | - |
| - | - | 1794 | 246.1 | - | - | 0 | - |
| - | - | 1153 | 247.1 | - | - | 0 | - |
| - | - | 738.6 | 251.2 | - | - | 0 | - |
| - | - | 5288 | 251.2 | - | - | 0 | - |
| - | - | 933.1 | 252.1 | - | - | 0 | - |
| - | - | 1126 | 252.1 | - | - | 0 | - |
| - | - | 780.5 | 252.2 | - | - | 0 | - |
| - | - | 593.4 | 254.1 | - | - | 0 | - |
| - | - | 610.2 | 254.1 | - | - | 0 | - |
| - | - | 5686 | 254.1 | - | - | 0 | - |
| - | - | 1486 | 254.1 | - | - | 0 | - |
| - | - | 657 | 255.1 | - | - | 0 | - |
| - | - | 5288 | 259.1 | - | - | 0 | - |
| - | - | 7123 | 261.2 | - | - | 0 | - |
| - | - | 1123 | 262.1 | - | - | 0 | - |
| - | - | 1091 | 262.2 | - | - | 0 | - |
| - | - | 1.025E+04 | 263.1 | - | - | 0 | - |
| - | - | 1117 | 264.1 | - | - | 0 | - |
| - | - | 4660 | 264.1 | - | - | 0 | - |
| - | - | 613 | 265.1 | - | - | 0 | - |
| - | - | 572.3 | 268.2 | - | - | 0 | - |
| 3 | a | 1125 | 268.2 | 0.0007137 | 2.661 | +1 | 3 |
| - | - | 1002 | 269 | - | - | 0 | - |
| - | - | 692.1 | 269.2 | - | - | 0 | - |
| - | - | 1235 | 269.2 | - | - | 0 | - |
| - | - | 2901 | 272.1 | - | - | 0 | - |
| - | - | 1378 | 274.1 | - | - | 0 | - |
| - | - | 1150 | 275.2 | - | - | 0 | - |
| - | - | 5451 | 279.1 | - | - | 0 | - |
| 5 | b | 1232 | 279.7 | 0.0008797 | 3.146 | +2 | 5 |
| - | - | 1790 | 280.1 | - | - | 0 | - |
| - | - | 1150 | 280.1 | - | - | 0 | - |
| - | - | 3478 | 280.2 | - | - | 0 | - |
| - | - | 2018 | 281.1 | - | - | 0 | - |
| - | - | 1.428E+04 | 282.1 | - | - | 0 | - |
| - | - | 2109 | 283.1 | - | - | 0 | - |
| - | - | 761.4 | 284.2 | - | - | 0 | - |
| - | - | 1.071E+04 | 285 | - | - | 0 | - |
| - | - | 1720 | 285.2 | - | - | 0 | - |
| - | - | 3008 | 286 | - | - | 0 | - |
| - | - | 2603 | 286.1 | - | - | 0 | - |
| - | - | 1061 | 287 | - | - | 0 | - |
| - | - | 714.5 | 294.2 | - | - | 0 | - |
| - | - | 2411 | 296.1 | - | - | 0 | - |
| - | - | 5754 | 296.2 | - | - | 0 | - |
| 3 | b | 9.368E+04 | 296.2 | 0.0006111 | 2.063 | +1 | 3 |
| - | - | 2372 | 297.1 | - | - | 0 | - |
| - | - | 1.59E+04 | 297.2 | - | - | 0 | - |
| - | - | 965.5 | 297.2 | - | - | 0 | - |
| - | - | 1.487E+04 | 297.2 | - | - | 0 | - |
| - | - | 2433 | 298.2 | - | - | 0 | - |
| - | - | 5087 | 298.2 | - | - | 0 | - |
| - | - | 1443 | 298.2 | - | - | 0 | - |
| - | - | 1.223E+04 | 299.1 | - | - | 0 | - |
| - | - | 1077 | 299.2 | - | - | 0 | - |
| - | - | 3218 | 300.1 | - | - | 0 | - |
| - | - | 823.2 | 300.2 | - | - | 0 | - |
| - | - | 1295 | 301.1 | - | - | 0 | - |
| 15 | y | 1554 | 301.2 | 0.001018 | 3.379 | +1 | 3 |
| - | - | 1098 | 303 | - | - | 0 | - |
| - | - | 3454 | 303.2 | - | - | 0 | - |
| - | - | 1585 | 304.1 | - | - | 0 | - |
| 12 | y | 5094 | 307.7 | 0.0004067 | 1.322 | +2 | 6 |
| 12 | y | 1538 | 308.2 | 0.003028 | 9.825 | +2 | 6 |
| - | - | 740.6 | 313.2 | - | - | 0 | - |
| - | - | 1996 | 313.2 | - | - | 0 | - |
| - | - | 2130 | 314.1 | - | - | 0 | - |
| - | - | 1.347E+04 | 314.2 | - | - | 0 | - |
| 3 | b | 2.002E+04 | 314.2 | 0.0006665 | 2.121 | +1 | 3 |
| - | - | 634.8 | 315.2 | - | - | 0 | - |
| - | - | 2294 | 315.2 | - | - | 0 | - |
| - | - | 3646 | 315.2 | - | - | 0 | - |
| - | - | 1.049E+04 | 316.2 | - | - | 0 | - |
| - | - | 690.6 | 316.2 | - | - | 0 | - |
| 12 | y | 4549 | 316.7 | 0.0008311 | 2.624 | +2 | 6 |
| - | - | 1164 | 317.2 | - | - | 0 | - |
| - | - | 2170 | 317.2 | - | - | 0 | - |
| 15 | y | 1.326E+04 | 319.2 | 0.0006153 | 1.927 | +1 | 3 |
| - | - | 1176 | 320.2 | - | - | 0 | - |
| - | - | 838.6 | 324.2 | - | - | 0 | - |
| - | - | 875.8 | 325.2 | - | - | 0 | - |
| - | - | 848.4 | 326.2 | - | - | 0 | - |
| - | - | 2168 | 328.1 | - | - | 0 | - |
| - | - | 2465 | 330.2 | - | - | 0 | - |
| - | - | 706.9 | 332.1 | - | - | 0 | - |
| - | - | 2893 | 333.2 | - | - | 0 | - |
| - | - | 1174 | 334.2 | - | - | 0 | - |
| 9 | y | 861 | 335.2 | 0.00177 | 5.28 | +3 | 9 |
| - | - | 1178 | 336.2 | - | - | 0 | - |
| - | - | 1156 | 339.2 | - | - | 0 | - |
| - | - | 850.3 | 340.2 | - | - | 0 | - |
| - | - | 2063 | 341.2 | - | - | 0 | - |
| - | - | 1001 | 342.2 | - | - | 0 | - |
| - | - | 1883 | 344.2 | - | - | 0 | - |
| - | - | 1828 | 345 | - | - | 0 | - |
| - | - | 6015 | 346.1 | - | - | 0 | - |
| - | - | 3546 | 346.2 | - | - | 0 | - |
| - | - | 2712 | 348.2 | - | - | 0 | - |
| - | - | 763.9 | 349.2 | - | - | 0 | - |
| - | - | 2287 | 350.1 | - | - | 0 | - |
| - | - | 2072 | 350.2 | - | - | 0 | - |
| - | - | 9568 | 351.2 | - | - | 0 | - |
| - | - | 683.9 | 351.2 | - | - | 0 | - |
| - | - | 1778 | 352.2 | - | - | 0 | - |
| - | - | 4336 | 353.2 | - | - | 0 | - |
| - | - | 1586 | 354.2 | - | - | 0 | - |
| - | - | 1150 | 354.2 | - | - | 0 | - |
| - | - | 4.197E+04 | 355.1 | - | - | 0 | - |
| - | - | 1715 | 355.2 | - | - | 0 | - |
| - | - | 605 | 355.2 | - | - | 0 | - |
| - | - | 1.719E+04 | 356.1 | - | - | 0 | - |
| - | - | 8969 | 357.1 | - | - | 0 | - |
| - | - | 918.1 | 358.2 | - | - | 0 | - |
| - | - | 8259 | 362.2 | - | - | 0 | - |
| - | - | 2337 | 363.2 | - | - | 0 | - |
| 11 | y | 907.8 | 364.7 | 0.0008212 | 2.252 | +2 | 7 |
| - | - | 1713 | 365.1 | - | - | 0 | - |
| - | - | 5934 | 367.2 | - | - | 0 | - |
| - | - | 2735 | 368.2 | - | - | 0 | - |
| - | - | 7897 | 369.2 | - | - | 0 | - |
| - | - | 1600 | 370.2 | - | - | 0 | - |
| - | - | 816.4 | 370.7 | - | - | 0 | - |
| - | - | 1370 | 371.1 | - | - | 0 | - |
| - | - | 2500 | 371.2 | - | - | 0 | - |
| - | - | 674.7 | 371.7 | - | - | 0 | - |
| - | - | 3364 | 373.1 | - | - | 0 | - |
| - | - | 2130 | 373.2 | - | - | 0 | - |
| 11 | y | 5062 | 373.2 | 0.000303 | 0.8119 | +2 | 7 |
| - | - | 2309 | 373.7 | - | - | 0 | - |
| - | - | 1264 | 374.1 | - | - | 0 | - |
| - | - | 839 | 375.1 | - | - | 0 | - |
| - | - | 620.9 | 379.2 | - | - | 0 | - |
| - | - | 962.1 | 381.2 | - | - | 0 | - |
| - | - | 3039 | 383.2 | - | - | 0 | - |
| - | - | 798.5 | 383.2 | - | - | 0 | - |
| - | - | 659.8 | 384.2 | - | - | 0 | - |
| - | - | 1.32E+04 | 385.2 | - | - | 0 | - |
| - | - | 1754 | 386.2 | - | - | 0 | - |
| - | - | 1538 | 393.2 | - | - | 0 | - |
| - | - | 887.2 | 397.1 | - | - | 0 | - |
| - | - | 1293 | 397.2 | - | - | 0 | - |
| - | - | 843.3 | 398.2 | - | - | 0 | - |
| - | - | 1412 | 398.2 | - | - | 0 | - |
| - | - | 5853 | 401.2 | - | - | 0 | - |
| - | - | 1100 | 402.2 | - | - | 0 | - |
| 8 | b | 1194 | 406.2 | 0.000521 | 1.283 | +2 | 8 |
| - | - | 2717 | 407.2 | - | - | 0 | - |
| - | - | 1233 | 407.7 | - | - | 0 | - |
| - | - | 4466 | 408.2 | - | - | 0 | - |
| - | - | 1417 | 409.2 | - | - | 0 | - |
| - | - | 3048 | 411.3 | - | - | 0 | - |
| - | - | 878.4 | 412.3 | - | - | 0 | - |
| - | - | 738.3 | 413.2 | - | - | 0 | - |
| - | - | 2286 | 415 | - | - | 0 | - |
| - | - | 1817 | 415.1 | - | - | 0 | - |
| 8 | b | 1756 | 415.2 | 0.004115 | 9.91 | +2 | 8 |
| - | - | 1423 | 415.3 | - | - | 0 | - |
| - | - | 1465 | 416 | - | - | 0 | - |
| 14 | y | 1093 | 416.2 | 0.0006859 | 1.648 | +1 | 4 |
| - | - | 2696 | 416.3 | - | - | 0 | - |
| - | - | 868.8 | 417 | - | - | 0 | - |
| - | - | 4221 | 419 | - | - | 0 | - |
| - | - | 1492 | 420 | - | - | 0 | - |
| - | - | 1458 | 421 | - | - | 0 | - |
| - | - | 803 | 423.2 | - | - | 0 | - |
| - | - | 754.1 | 423.7 | - | - | 0 | - |
| - | - | 3664 | 424.2 | - | - | 0 | - |
| - | - | 5693 | 425.2 | - | - | 0 | - |
| - | - | 1141 | 426.2 | - | - | 0 | - |
| - | - | 1752 | 427.2 | - | - | 0 | - |
| 10 | y | 667.3 | 429.2 | 0.007353 | 17.13 | +2 | 8 |
| - | - | 2532 | 433.2 | - | - | 0 | - |
| 14 | y | 1.221E+04 | 433.2 | 0.001236 | 2.854 | +1 | 4 |
| - | - | 2.596E+04 | 433.3 | - | - | 0 | - |
| - | - | 1608 | 434.2 | - | - | 0 | - |
| - | - | 1985 | 434.2 | - | - | 0 | - |
| - | - | 7908 | 434.3 | - | - | 0 | - |
| - | - | 1384 | 434.7 | - | - | 0 | - |
| - | - | 1280 | 435.2 | - | - | 0 | - |
| - | - | 1070 | 435.3 | - | - | 0 | - |
| 10 | y | 675.1 | 437.7 | 0.003476 | 7.942 | +2 | 8 |
| - | - | 886.8 | 440.2 | - | - | 0 | - |
| - | - | 622.6 | 441.2 | - | - | 0 | - |
| - | - | 871.5 | 441.3 | - | - | 0 | - |
| - | - | 3486 | 442.2 | - | - | 0 | - |
| - | - | 877.6 | 442.3 | - | - | 0 | - |
| - | - | 861.4 | 443.1 | - | - | 0 | - |
| - | - | 1705 | 443.2 | - | - | 0 | - |
| 4 | b | 3182 | 443.3 | 0.002052 | 4.629 | +1 | 4 |
| - | - | 1054 | 445.2 | - | - | 0 | - |
| - | - | 609.5 | 450.2 | - | - | 0 | - |
| - | - | 1165 | 451.2 | - | - | 0 | - |
| 5 | y | 2973 | 451.9 | 0.0006641 | 1.47 | +3 | 13 |
| 5 | y | 2429 | 452.2 | 0.00181 | 4.003 | +3 | 13 |
| - | - | 974.7 | 453.2 | - | - | 0 | - |
| - | - | 1839 | 454.2 | - | - | 0 | - |
| - | - | 1045 | 454.7 | - | - | 0 | - |
| - | - | 1329 | 459.2 | - | - | 0 | - |
| 4 | b | 2.456E+04 | 461.3 | 0.0008867 | 1.922 | +1 | 4 |
| - | - | 2471 | 462.2 | - | - | 0 | - |
| - | - | 7968 | 462.3 | - | - | 0 | - |
| - | - | 1463 | 463.3 | - | - | 0 | - |
| - | - | 668.6 | 467.2 | - | - | 0 | - |
| - | - | 1156 | 468.2 | - | - | 0 | - |
| - | - | 1979 | 468.3 | - | - | 0 | - |
| - | - | 2145 | 469.2 | - | - | 0 | - |
| - | - | 8827 | 470.2 | - | - | 0 | - |
| - | - | 802.5 | 470.7 | - | - | 0 | - |
| - | - | 2169 | 471.2 | - | - | 0 | - |
| - | - | 1035 | 471.7 | - | - | 0 | - |
| - | - | 648.2 | 473.7 | - | - | 0 | - |
| - | - | 817.7 | 476.2 | - | - | 0 | - |
| - | - | 1597 | 477.3 | - | - | 0 | - |
| - | - | 5157 | 478.2 | - | - | 0 | - |
| - | - | 2065 | 479.2 | - | - | 0 | - |
| - | - | 9434 | 480.2 | - | - | 0 | - |
| - | - | 1582 | 480.7 | - | - | 0 | - |
| - | - | 2740 | 481.2 | - | - | 0 | - |
| - | - | 816.4 | 482.2 | - | - | 0 | - |
| - | - | 1096 | 484.2 | - | - | 0 | - |
| 13 | y | 4618 | 486.3 | 0.002153 | 4.428 | +1 | 5 |
| 13 | y | 2245 | 487.3 | 0.001109 | 2.275 | +1 | 5 |
| - | - | 713.6 | 488.2 | - | - | 0 | - |
| - | - | 751.6 | 488.7 | - | - | 0 | - |
| - | - | 1191 | 489.1 | - | - | 0 | - |
| - | - | 591.9 | 489.3 | - | - | 0 | - |
| - | - | 893.3 | 490.1 | - | - | 0 | - |
| - | - | 807.5 | 490.2 | - | - | 0 | - |
| - | - | 3668 | 493.7 | - | - | 0 | - |
| - | - | 887.8 | 494.2 | - | - | 0 | - |
| - | - | 2159 | 494.2 | - | - | 0 | - |
| - | - | 753.3 | 494.7 | - | - | 0 | - |
| - | - | 2426 | 495.3 | - | - | 0 | - |
| - | - | 1.166E+04 | 496.3 | - | - | 0 | - |
| - | - | 2311 | 497.3 | - | - | 0 | - |
| - | - | 2.408E+04 | 498.2 | - | - | 0 | - |
| - | - | 6876 | 499.2 | - | - | 0 | - |
| - | - | 1636 | 500.2 | - | - | 0 | - |
| 4 | y | 798.9 | 501.2 | 0.004774 | 9.524 | +3 | 14 |
| - | - | 1861 | 502.2 | - | - | 0 | - |
| - | - | 5491 | 502.7 | - | - | 0 | - |
| - | - | 3194 | 503.2 | - | - | 0 | - |
| - | - | 1040 | 503.7 | - | - | 0 | - |
| 13 | y | 2.856E+04 | 504.3 | 0.0005911 | 1.172 | +1 | 5 |
| - | - | 7390 | 505.3 | - | - | 0 | - |
| - | - | 622.6 | 506.3 | - | - | 0 | - |
| - | - | 970.1 | 508.2 | - | - | 0 | - |
| - | - | 964.2 | 510.2 | - | - | 0 | - |
| - | - | 1950 | 512.2 | - | - | 0 | - |
| - | - | 4366 | 513.3 | - | - | 0 | - |
| - | - | 756 | 515.2 | - | - | 0 | - |
| - | - | 916.9 | 518.2 | - | - | 0 | - |
| - | - | 651 | 521.3 | - | - | 0 | - |
| - | - | 1034 | 529.3 | - | - | 0 | - |
| - | - | 1231 | 529.8 | - | - | 0 | - |
| - | - | 8057 | 530.2 | - | - | 0 | - |
| - | - | 837.6 | 530.3 | - | - | 0 | - |
| - | - | 741 | 530.3 | - | - | 0 | - |
| - | - | 2409 | 530.3 | - | - | 0 | - |
| - | - | 2429 | 531.2 | - | - | 0 | - |
| - | - | 1310 | 536.3 | - | - | 0 | - |
| - | - | 8784 | 538.3 | - | - | 0 | - |
| - | - | 4911 | 538.8 | - | - | 0 | - |
| 5 | b | 2460 | 540.3 | 0.002694 | 4.986 | +1 | 5 |
| - | - | 801.3 | 542.6 | - | - | 0 | - |
| - | - | 3323 | 543.3 | - | - | 0 | - |
| - | - | 1616 | 543.8 | - | - | 0 | - |
| 10 | b | 2595 | 544.3 | 0.0008195 | 1.506 | +2 | 10 |
| - | - | 717.1 | 545.3 | - | - | 0 | - |
| - | - | 1374 | 546.2 | - | - | 0 | - |
| - | - | 844.9 | 551.3 | - | - | 0 | - |
| - | - | 7388 | 552.3 | - | - | 0 | - |
| - | - | 4437 | 552.8 | - | - | 0 | - |
| - | - | 1628 | 553.3 | - | - | 0 | - |
| - | - | 1942 | 555.3 | - | - | 0 | - |
| - | - | 754.5 | 555.8 | - | - | 0 | - |
| - | - | 998.4 | 556.3 | - | - | 0 | - |
| - | - | 686.4 | 556.3 | - | - | 0 | - |
| - | - | 1113 | 558.3 | - | - | 0 | - |
| 5 | b | 7157 | 558.3 | 0.001162 | 2.082 | +1 | 5 |
| - | - | 1963 | 559.3 | - | - | 0 | - |
| - | - | 704.2 | 569.8 | - | - | 0 | - |
| - | - | 791.4 | 571.3 | - | - | 0 | - |
| - | - | 603.7 | 573.2 | - | - | 0 | - |
| - | - | 769.3 | 573.3 | - | - | 0 | - |
| - | - | 757.5 | 574.3 | - | - | 0 | - |
| 2 | y | 1769 | 578.3 | 0.008378 | 14.49 | +3 | 16 |
| - | - | 1658 | 578.8 | - | - | 0 | - |
| - | - | 1021 | 579.3 | - | - | 0 | - |
| 7 | y | 1340 | 580.3 | 0.0003303 | 0.5692 | +2 | 11 |
| 7 | y | 846.8 | 580.8 | 0.01028 | 17.69 | +2 | 11 |
| - | - | 1850 | 581.3 | - | - | 0 | - |
| - | - | 1828 | 582.3 | - | - | 0 | - |
| - | - | 3711 | 582.8 | - | - | 0 | - |
| - | - | 2114 | 583.3 | - | - | 0 | - |
| - | - | 1505 | 583.8 | - | - | 0 | - |
| - | - | 754.6 | 586.3 | - | - | 0 | - |
| - | - | 1026 | 587.3 | - | - | 0 | - |
| - | - | 1173 | 591.2 | - | - | 0 | - |
| - | - | 1759 | 591.3 | - | - | 0 | - |
| 11 | b | 1670 | 591.8 | 0.01059 | 17.9 | +2 | 11 |
| - | - | 999.1 | 592.3 | - | - | 0 | - |
| - | - | 3104 | 597.3 | - | - | 0 | - |
| - | - | 2280 | 598.3 | - | - | 0 | - |
| - | - | 1.183E+04 | 599.3 | - | - | 0 | - |
| - | - | 1377 | 599.3 | - | - | 0 | - |
| - | - | 3374 | 600.3 | - | - | 0 | - |
| 11 | b | 1638 | 600.8 | 0.001887 | 3.141 | +2 | 11 |
| - | - | 870.4 | 601.3 | - | - | 0 | - |
| - | - | 870.2 | 602.3 | - | - | 0 | - |
| - | - | 2394 | 602.8 | - | - | 0 | - |
| - | - | 2013 | 603.3 | - | - | 0 | - |
| - | - | 1454 | 608.4 | - | - | 0 | - |
| - | - | 1236 | 608.8 | - | - | 0 | - |
| - | - | 5625 | 609.3 | - | - | 0 | - |
| - | - | 1675 | 609.3 | - | - | 0 | - |
| - | - | 722 | 609.3 | - | - | 0 | - |
| - | - | 5101 | 610.2 | - | - | 0 | - |
| - | - | 1861 | 610.3 | - | - | 0 | - |
| - | - | 4920 | 611.2 | - | - | 0 | - |
| - | - | 7979 | 611.8 | - | - | 0 | - |
| - | - | 3965 | 612.2 | - | - | 0 | - |
| - | - | 5275 | 612.3 | - | - | 0 | - |
| - | - | 2701 | 612.8 | - | - | 0 | - |
| - | - | 2722 | 613.3 | - | - | 0 | - |
| 12 | y | 9491 | 614.3 | 0.001132 | 1.842 | +1 | 6 |
| 12 | y | 1.745E+04 | 615.3 | 0.0008808 | 1.431 | +1 | 6 |
| - | - | 5174 | 616.3 | - | - | 0 | - |
| - | - | 3763 | 617.3 | - | - | 0 | - |
| - | - | 1.093E+04 | 617.8 | - | - | 0 | - |
| - | - | 1.085E+04 | 618.3 | - | - | 0 | - |
| - | - | 4325 | 618.8 | - | - | 0 | - |
| - | - | 758.4 | 619.3 | - | - | 0 | - |
| - | - | 844.9 | 619.8 | - | - | 0 | - |
| - | - | 1226 | 625.3 | - | - | 0 | - |
| - | - | 3643 | 626.4 | - | - | 0 | - |
| - | - | 2.4E+04 | 626.8 | - | - | 0 | - |
| - | - | 4.814E+04 | 627.3 | - | - | 0 | - |
| - | - | 1.645E+04 | 627.3 | - | - | 0 | - |
| - | - | 6385 | 627.8 | - | - | 0 | - |
| - | - | 1.36E+04 | 628.3 | - | - | 0 | - |
| - | - | 1662 | 628.3 | - | - | 0 | - |
| 6 | y | 1928 | 628.8 | 0.0008649 | 1.375 | +2 | 12 |
| - | - | 2529 | 629.3 | - | - | 0 | - |
| - | - | 1465 | 629.3 | - | - | 0 | - |
| - | - | 1114 | 629.8 | - | - | 0 | - |
| 12 | y | 2.279E+04 | 632.3 | 0.0004548 | 0.7192 | +1 | 6 |
| - | - | 7296 | 633.3 | - | - | 0 | - |
| - | - | 1042 | 634.3 | - | - | 0 | - |
| - | - | 886.9 | 635.8 | - | - | 0 | - |
| - | - | 1064 | 636.3 | - | - | 0 | - |
| 6 | y | 3826 | 637.8 | 0.0005264 | 0.8253 | +2 | 12 |
| - | - | 2551 | 638.3 | - | - | 0 | - |
| - | - | 1598 | 638.8 | - | - | 0 | - |
| - | - | 789.6 | 639.3 | - | - | 0 | - |
| - | - | 3943 | 643.3 | - | - | 0 | - |
| - | - | 1943 | 644.3 | - | - | 0 | - |
| - | - | 3856 | 645.3 | - | - | 0 | - |
| - | - | 1249 | 646.8 | - | - | 0 | - |
| - | - | 677.5 | 650.3 | - | - | 0 | - |
| - | - | 1.076E+04 | 655.3 | - | - | 0 | - |
| 12 | b | 8085 | 655.8 | 0.007239 | 11.04 | +2 | 12 |
| - | - | 3274 | 656.3 | - | - | 0 | - |
| - | - | 921.1 | 656.8 | - | - | 0 | - |
| - | - | 964.7 | 657.3 | - | - | 0 | - |
| - | - | 967.4 | 658.3 | - | - | 0 | - |
| - | - | 838.2 | 659.3 | - | - | 0 | - |
| - | - | 967.9 | 659.8 | - | - | 0 | - |
| - | - | 1534 | 660.3 | - | - | 0 | - |
| 12 | b | 997 | 664.8 | 0.001651 | 2.484 | +2 | 12 |
| - | - | 657.3 | 665.3 | - | - | 0 | - |
| - | - | 685.6 | 666.4 | - | - | 0 | - |
| - | - | 2984 | 668.3 | - | - | 0 | - |
| - | - | 3308 | 668.8 | - | - | 0 | - |
| - | - | 2230 | 669.3 | - | - | 0 | - |
| - | - | 983.1 | 669.8 | - | - | 0 | - |
| 5 | y | 2.843E+04 | 677.3 | 0.0008501 | 1.255 | +2 | 13 |
| 5 | y | 2.832E+04 | 677.8 | 0.007927 | 11.69 | +2 | 13 |
| - | - | 1.549E+04 | 678.3 | - | - | 0 | - |
| - | - | 4900 | 678.8 | - | - | 0 | - |
| - | - | 879.2 | 679.3 | - | - | 0 | - |
| - | - | 828 | 680.3 | - | - | 0 | - |
| - | - | 1572 | 684.4 | - | - | 0 | - |
| 5 | y | 6.293E+04 | 686.3 | 0.0005116 | 0.7455 | +2 | 13 |
| - | - | 4.654E+04 | 686.8 | - | - | 0 | - |
| - | - | 1.745E+04 | 687.3 | - | - | 0 | - |
| - | - | 5891 | 687.8 | - | - | 0 | - |
| - | - | 1788 | 688.4 | - | - | 0 | - |
| 13 | b | 737.5 | 691.4 | 0.008938 | 12.93 | +2 | 13 |
| 13 | b | 949.6 | 691.8 | 0.005767 | 8.336 | +2 | 13 |
| - | - | 1150 | 694.3 | - | - | 0 | - |
| - | - | 659.9 | 698.3 | - | - | 0 | - |
| - | - | 712.9 | 699.3 | - | - | 0 | - |
| 13 | b | 3834 | 700.4 | 0.01153 | 16.47 | +2 | 13 |
| - | - | 1452 | 700.9 | - | - | 0 | - |
| - | - | 1482 | 701.4 | - | - | 0 | - |
| - | - | 826.7 | 704.3 | - | - | 0 | - |
| - | - | 2076 | 710.4 | - | - | 0 | - |
| - | - | 8553 | 712.4 | - | - | 0 | - |
| - | - | 701.1 | 712.9 | - | - | 0 | - |
| - | - | 3722 | 713.4 | - | - | 0 | - |
| - | - | 931.7 | 714.4 | - | - | 0 | - |
| - | - | 1338 | 720.4 | - | - | 0 | - |
| - | - | 887.7 | 721.4 | - | - | 0 | - |
| - | - | 2779 | 722.3 | - | - | 0 | - |
| - | - | 803.1 | 723.3 | - | - | 0 | - |
| 7 | b | 1998 | 724.4 | 0.0006165 | 0.851 | +1 | 7 |
| - | - | 853.5 | 725.4 | - | - | 0 | - |
| 11 | y | 1167 | 727.4 | 0.003037 | 4.175 | +1 | 7 |
| 11 | y | 1778 | 728.4 | 0.001351 | 1.854 | +1 | 7 |
| - | - | 1134 | 728.9 | - | - | 0 | - |
| - | - | 1851 | 729.4 | - | - | 0 | - |
| - | - | 798 | 730.4 | - | - | 0 | - |
| - | - | 862.9 | 732.3 | - | - | 0 | - |
| - | - | 728.9 | 737.3 | - | - | 0 | - |
| - | - | 1681 | 737.4 | - | - | 0 | - |
| - | - | 804.5 | 738.3 | - | - | 0 | - |
| - | - | 961 | 738.4 | - | - | 0 | - |
| - | - | 809.4 | 739.4 | - | - | 0 | - |
| - | - | 1.634E+04 | 740.3 | - | - | 0 | - |
| - | - | 7874 | 741.4 | - | - | 0 | - |
| - | - | 2296 | 742.4 | - | - | 0 | - |
| 7 | b | 3524 | 742.4 | 0.002308 | 3.108 | +1 | 7 |
| - | - | 821.6 | 743.4 | - | - | 0 | - |
| - | - | 913.5 | 743.4 | - | - | 0 | - |
| - | - | 895.9 | 745.4 | - | - | 0 | - |
| 11 | y | 1.088E+04 | 745.4 | 0.001108 | 1.486 | +1 | 7 |
| - | - | 4973 | 746.4 | - | - | 0 | - |
| - | - | 1047 | 747.4 | - | - | 0 | - |
| - | - | 1381 | 749.3 | - | - | 0 | - |
| 4 | y | 6120 | 750.9 | 0.0008839 | 1.177 | +2 | 14 |
| 4 | y | 7425 | 751.4 | 0.008693 | 11.57 | +2 | 14 |
| - | - | 3737 | 751.9 | - | - | 0 | - |
| - | - | 1482 | 752.4 | - | - | 0 | - |
| - | - | 1939 | 755.4 | - | - | 0 | - |
| - | - | 1468 | 756.4 | - | - | 0 | - |
| 14 | b | 5325 | 757.4 | 0.005899 | 7.788 | +2 | 14 |
| - | - | 3042 | 758.4 | - | - | 0 | - |
| - | - | 1986 | 759.4 | - | - | 0 | - |
| 4 | y | 8451 | 759.9 | 0.0006064 | 0.7981 | +2 | 14 |
| - | - | 6547 | 760.4 | - | - | 0 | - |
| - | - | 3422 | 760.9 | - | - | 0 | - |
| - | - | 3142 | 771.4 | - | - | 0 | - |
| - | - | 1360 | 772.4 | - | - | 0 | - |
| - | - | 3697 | 774.3 | - | - | 0 | - |
| - | - | 918.7 | 774.5 | - | - | 0 | - |
| - | - | 1586 | 775.3 | - | - | 0 | - |
| - | - | 854.4 | 793.4 | - | - | 0 | - |
| - | - | 1083 | 806.4 | - | - | 0 | - |
| 3 | y | 2655 | 807.4 | 1.039E-05 | 0.01287 | +2 | 15 |
| 3 | y | 1769 | 807.9 | 0.006151 | 7.613 | +2 | 15 |
| - | - | 885.3 | 808.4 | - | - | 0 | - |
| 8 | b | 2955 | 811.4 | 0.002371 | 2.923 | +1 | 8 |
| 15 | b | 1983 | 812.4 | 0.008299 | 10.21 | +2 | 15 |
| - | - | 2252 | 813.4 | - | - | 0 | - |
| 3 | y | 997.2 | 816.4 | 0.001142 | 1.399 | +2 | 15 |
| - | - | 1344 | 816.9 | - | - | 0 | - |
| - | - | 715.7 | 817.4 | - | - | 0 | - |
| 8 | b | 2813 | 829.4 | 0.001163 | 1.402 | +1 | 8 |
| - | - | 1210 | 830.5 | - | - | 0 | - |
| - | - | 986.7 | 835.4 | - | - | 0 | - |
| - | - | 1074 | 839.4 | - | - | 0 | - |
| - | - | 1166 | 840.4 | - | - | 0 | - |
| - | - | 2711 | 841.4 | - | - | 0 | - |
| - | - | 1076 | 842.4 | - | - | 0 | - |
| 16 | b | 1060 | 847.9 | 0.002193 | 2.586 | +2 | 16 |
| 16 | b | 876.9 | 848.4 | 0.003715 | 4.379 | +2 | 16 |
| - | - | 771 | 849.4 | - | - | 0 | - |
| - | - | 2180 | 850.4 | - | - | 0 | - |
| - | - | 1879 | 851.4 | - | - | 0 | - |
| 10 | y | 1481 | 856.5 | 0.003503 | 4.09 | +1 | 8 |
| 16 | b | 1279 | 856.9 | 0.001319 | 1.54 | +2 | 16 |
| 10 | y | 4158 | 857.4 | 0.0107 | 12.48 | +1 | 8 |
| 2 | y | 997 | 857.9 | 0.007095 | 8.27 | +2 | 16 |
| 2 | y | 2212 | 858.4 | 0.01063 | 12.39 | +2 | 16 |
| - | - | 1126 | 858.9 | - | - | 0 | - |
| - | - | 1375 | 859.4 | - | - | 0 | - |
| - | - | 3003 | 863.4 | - | - | 0 | - |
| - | - | 901.4 | 864.4 | - | - | 0 | - |
| - | - | 722.9 | 867.4 | - | - | 0 | - |
| - | - | 1.96E+04 | 868.4 | - | - | 0 | - |
| - | - | 1.01E+04 | 869.4 | - | - | 0 | - |
| - | - | 2282 | 870.4 | - | - | 0 | - |
| - | - | 3587 | 873.5 | - | - | 0 | - |
| 10 | y | 5557 | 874.5 | 0.0001405 | 0.1606 | +1 | 8 |
| - | - | 1936 | 875.5 | - | - | 0 | - |
| - | - | 5241 | 879.4 | - | - | 0 | - |
| - | - | 1755 | 880.4 | - | - | 0 | - |
| - | - | 3948 | 887.4 | - | - | 0 | - |
| - | - | 2634 | 888.4 | - | - | 0 | - |
| - | - | 3858 | 889.4 | - | - | 0 | - |
| - | - | 2213 | 890.4 | - | - | 0 | - |
| - | - | 1558 | 894.4 | - | - | 0 | - |
| - | - | 2.079E+04 | 907.4 | - | - | 0 | - |
| - | - | 9096 | 908.4 | - | - | 0 | - |
| - | - | 1159 | 909.4 | - | - | 0 | - |
| - | - | 1964 | 911.4 | - | - | 0 | - |
| - | - | 1202 | 912.5 | - | - | 0 | - |
| - | - | 861.8 | 917.4 | - | - | 0 | - |
| - | - | 1438 | 921.4 | - | - | 0 | - |
| - | - | 1781 | 922.4 | - | - | 0 | - |
| - | - | 1.104E+04 | 939.4 | - | - | 0 | - |
| - | - | 6471 | 940.5 | - | - | 0 | - |
| - | - | 2169 | 941.5 | - | - | 0 | - |
| - | - | 1586 | 942.5 | - | - | 0 | - |
| - | - | 914.9 | 954.4 | - | - | 0 | - |
| - | - | 2499 | 956.5 | - | - | 0 | - |
| - | - | 1234 | 957.4 | - | - | 0 | - |
| 9 | b | 6134 | 958.5 | 0.0009028 | 0.9419 | +1 | 9 |
| - | - | 2940 | 959.5 | - | - | 0 | - |
| - | - | 1.118E+04 | 960.5 | - | - | 0 | - |
| - | - | 5651 | 961.5 | - | - | 0 | - |
| - | - | 1090 | 962.5 | - | - | 0 | - |
| - | - | 1188 | 971.5 | - | - | 0 | - |
| - | - | 755.9 | 987.5 | - | - | 0 | - |
| - | - | 1450 | 988.5 | - | - | 0 | - |
| - | - | 1114 | 1000 | - | - | 0 | - |
| - | - | 667.9 | 1001 | - | - | 0 | - |
| 9 | y | 1179 | 1004 | 0.006381 | 6.359 | +1 | 9 |
| - | - | 963.8 | 1005 | - | - | 0 | - |
| - | - | 1152 | 1015 | - | - | 0 | - |
| - | - | 784.7 | 1016 | - | - | 0 | - |
| - | - | 795.4 | 1035 | - | - | 0 | - |
| - | - | 2282 | 1036 | - | - | 0 | - |
| - | - | 1562 | 1037 | - | - | 0 | - |
| - | - | 1086 | 1040 | - | - | 0 | - |
| - | - | 1598 | 1041 | - | - | 0 | - |
| - | - | 9083 | 1053 | - | - | 0 | - |
| - | - | 4924 | 1054 | - | - | 0 | - |
| - | - | 1712 | 1055 | - | - | 0 | - |
| - | - | 1214 | 1058 | - | - | 0 | - |
| - | - | 2557 | 1059 | - | - | 0 | - |
| - | - | 1650 | 1060 | - | - | 0 | - |
| - | - | 813.1 | 1061 | - | - | 0 | - |
| - | - | 891.9 | 1069 | - | - | 0 | - |
| 10 | b | 2218 | 1070 | 0.007113 | 6.651 | +1 | 10 |
| - | - | 5696 | 1071 | - | - | 0 | - |
| - | - | 2689 | 1072 | - | - | 0 | - |
| 8 | y | 892.9 | 1073 | 0.01118 | 10.43 | +1 | 10 |
| 8 | y | 1273 | 1074 | 0.009563 | 8.908 | +1 | 10 |
| - | - | 4618 | 1076 | - | - | 0 | - |
| - | - | 3243 | 1077 | - | - | 0 | - |
| - | - | 1889 | 1085 | - | - | 0 | - |
| - | - | 1488 | 1086 | - | - | 0 | - |
| - | - | 2313 | 1087 | - | - | 0 | - |
| 10 | b | 1.858E+04 | 1088 | 0.0007102 | 0.6531 | +1 | 10 |
| - | - | 1.084E+04 | 1089 | - | - | 0 | - |
| - | - | 4035 | 1090 | - | - | 0 | - |
| 8 | y | 2000 | 1091 | 0.001203 | 1.103 | +1 | 10 |
| - | - | 833.5 | 1092 | - | - | 0 | - |
| - | - | 933 | 1102 | - | - | 0 | - |
| - | - | 6810 | 1104 | - | - | 0 | - |
| - | - | 3593 | 1105 | - | - | 0 | - |
| - | - | 884.7 | 1106 | - | - | 0 | - |
| - | - | 2407 | 1138 | - | - | 0 | - |
| - | - | 1417 | 1139 | - | - | 0 | - |
| - | - | 1103 | 1140 | - | - | 0 | - |
| - | - | 3953 | 1156 | - | - | 0 | - |
| - | - | 2549 | 1157 | - | - | 0 | - |
| - | - | 1013 | 1158 | - | - | 0 | - |
| - | - | 2476 | 1165 | - | - | 0 | - |
| - | - | 1815 | 1166 | - | - | 0 | - |
| 7 | y | 1506 | 1178 | 0.003604 | 3.061 | +1 | 11 |
| - | - | 1204 | 1179 | - | - | 0 | - |
| - | - | 6945 | 1182 | - | - | 0 | - |
| 11 | b | 5758 | 1183 | 0.01891 | 15.99 | +1 | 11 |
| - | - | 2057 | 1184 | - | - | 0 | - |
| 11 | b | 3430 | 1201 | 0.005306 | 4.42 | +1 | 11 |
| - | - | 2344 | 1202 | - | - | 0 | - |
| - | - | 907.6 | 1214 | - | - | 0 | - |
| - | - | 4402 | 1235 | - | - | 0 | - |
| - | - | 4033 | 1236 | - | - | 0 | - |
| - | - | 2519 | 1237 | - | - | 0 | - |
| - | - | 1.265E+04 | 1253 | - | - | 0 | - |
| - | - | 8966 | 1254 | - | - | 0 | - |
| - | - | 3220 | 1255 | - | - | 0 | - |
| - | - | 1295 | 1256 | - | - | 0 | - |
| 6 | y | 1389 | 1257 | 0.003269 | 2.601 | +1 | 12 |
| 6 | y | 1501 | 1258 | 0.009976 | 7.933 | +1 | 12 |
| - | - | 887.3 | 1259 | - | - | 0 | - |
| 6 | y | 7368 | 1275 | 0.002291 | 1.797 | +1 | 12 |
| - | - | 4806 | 1276 | - | - | 0 | - |
| - | - | 1936 | 1277 | - | - | 0 | - |
| 12 | b | 797.9 | 1311 | 0.008204 | 6.26 | +1 | 12 |
| 12 | b | 1693 | 1329 | 0.009441 | 7.105 | +1 | 12 |
| - | - | 971 | 1330 | - | - | 0 | - |
| 5 | y | 1565 | 1354 | 0.003606 | 2.664 | +1 | 13 |
| 5 | y | 1194 | 1355 | 0.003477 | 2.567 | +1 | 13 |
| 5 | y | 2194 | 1372 | 0.00818 | 5.963 | +1 | 13 |
| - | - | 1442 | 1373 | - | - | 0 | - |
| 13 | b | 1851 | 1400 | 0.01933 | 13.81 | +1 | 13 |
| - | - | 1673 | 1401 | - | - | 0 | - |
| 14 | b | 1158 | 1514 | 0.004512 | 2.981 | +1 | 14 |
| - | - | 931.2 | 1515 | - | - | 0 | - |
| - | - | 716.2 | 1537 | - | - | 0 | - |
| - | - | 767.7 | 3068 | - | - | 0 | - |

m/z Charge Intensity FragmentType MassShift Position
120.06588745117188 0 43440.066 y 16
120.0811538696289 0 40538.812
121.06924438476562 0 1829.5543
121.08446502685547 0 3580.371
122.07038116455078 0 446.83835
125.0711669921875 0 356.05927
126.0552749633789 0 3566.1472
127.08712005615234 0 675.1086
128.10736083984375 0 10558.544
129.06617736816406 0 1552.056
129.1026153564453 0 56672.555
130.05027770996094 0 3612.259
130.06556701660156 0 1428.5835
130.10072326660156 0 904.8747
130.10598754882812 0 4353.8916
131.11830139160156 0 3414.0051
132.07676696777344 0 1194.2189
132.08177185058594 0 594.4353
136.07606506347656 0 2756.7234
138.09164428710938 0 2051.7441
138.12808227539062 0 1480.3141
139.0871124267578 0 1543.6781
141.06617736816406 0 774.9843
145.09764099121094 0 561.64594
147.07687377929688 0 1747.1085
148.06015014648438 0 512.3605
148.1520538330078 0 432.1705
149.04510498046875 0 2839.9375
153.0660858154297 0 535.9988
153.10250854492188 0 684.4911
154.08673095703125 0 1029.8123
154.09791564941406 0 1665.4094
155.0819091796875 0 3609.9683
155.11827087402344 0 6940.423 a Water loss 1
156.12181091308594 0 718.6178
157.0611572265625 0 1692.4857
157.09759521484375 0 3783.2231
159.09210205078125 0 5172.7026
165.10256958007812 0 1066.8263
167.055908203125 0 4205.4683
167.08177185058594 0 2356.5723
167.11834716796875 0 2548.1865
168.0858154296875 0 439.32257
169.06088256835938 0 1008.2976
169.13389587402344 0 12058.066
169.14208984375 0 754.59174
170.1378173828125 0 1068.0647
171.07675170898438 0 1129.1361
172.10812377929688 0 1558.9902
172.14486694335938 0 583.6112
172.51646423339844 0 400.13794
173.09243774414062 0 6197.696 y Water loss 15
173.12884521484375 0 28997.178 a 1
174.13185119628906 0 2049.5977
175.07176208496094 0 1593.3328
177.10195922851562 0 444.25412
181.0975341796875 0 1313.7289
182.09286499023438 0 3742.1025
182.12905883789062 0 2416.2458
182.78858947753906 0 473.534
183.07679748535156 0 2813.5466
183.1132354736328 0 39366.938 b Water loss 1
183.14967346191406 0 519.63116
184.10862731933594 0 2945.107
184.11683654785156 0 3451.9875
185.09243774414062 0 5229.253
186.08787536621094 0 2815.7966
186.09616088867188 0 909.7124
187.07122802734375 0 550.92255
187.0872344970703 0 1053.9099
187.14454650878906 0 11505.606
189.08729553222656 0 3709.8945
191.10302734375 0 8825.154 y 15
191.1180419921875 0 1084.6691
195.0762939453125 0 789.6183
195.1131591796875 0 5996.9585
196.1163330078125 0 835.44165
197.10369873046875 0 941.2482
197.1288299560547 0 14931.008
198.12393188476562 0 1844.9695
198.13299560546875 0 1076.5636
199.07164001464844 0 4543.594
199.10841369628906 0 1037.3542
199.11915588378906 0 612.761
200.1033935546875 0 3548.3142
200.1395721435547 0 3918.657
201.1237335205078 0 35590.605 b 1
202.1270751953125 0 3556.9282
203.11473083496094 0 663.1828
208.09811401367188 0 786.9814
208.10842895507812 0 4536.5737
209.09246826171875 0 10139.823
210.09622192382812 0 1249.4011
210.12387084960938 0 995.80023
211.1087188720703 0 514.03375
212.1396484375 0 1292.3354
213.08740234375 0 865.41595
214.15530395507812 0 586.1559
215.139404296875 0 14681.733
216.14303588867188 0 1529.429
217.08230590820312 0 7380.9907
217.097412109375 0 4201.741
217.13389587402344 0 8519.094
218.1376953125 0 1150.1014
218.15000915527344 0 582.37646
219.14981079101562 0 1514.9961
221.08474731445312 0 5719.2896
222.08470153808594 0 1016.6874
223.73362731933594 0 511.9499
225.04306030273438 0 2074.98
225.12368774414062 0 4916.2876
225.1352996826172 0 1983.9786
226.11898803710938 0 41169.133
227.10305786132812 0 1062.8304
227.12237548828125 0 4172.488
228.13475036621094 0 892.75104
229.351806640625 0 683.4248
231.09799194335938 0 908.53326
233.16531372070312 0 16517.855
234.16856384277344 0 2136.073
235.10813903808594 0 6884.0703
236.1035919189453 0 2187.2805
236.13986206054688 0 752.9007
239.09544372558594 0 12332.257
240.09634399414062 0 3125.8186
241.0821990966797 0 3031.9358
242.15042114257812 0 9422.055
243.1342315673828 0 1039.9684
243.14572143554688 0 16777.018 y Water loss 10
243.15890502929688 0 821.5597
244.1298370361328 0 1583.4537 y Ammonia loss 12
244.1490478515625 0 1279.3085
245.12904357910156 0 6076.411
246.12423706054688 0 1793.6696
247.1444549560547 0 1153.4283
251.15237426757812 0 738.635
251.17588806152344 0 5288.3584
252.09796142578125 0 933.1099
252.13467407226562 0 1126.284
252.1800994873047 0 780.5278
254.09210205078125 0 593.41364
254.09954833984375 0 610.1883
254.11386108398438 0 5685.743
254.1498260498047 0 1485.8522
255.11790466308594 0 656.9792
259.0929260253906 0 5287.5815
261.15997314453125 0 7123.0244
262.1188659667969 0 1123.2184
262.1650390625 0 1090.9387
263.1029357910156 0 10245.511
264.10723876953125 0 1116.5693
264.13470458984375 0 4659.5645
265.1387939453125 0 613.0473
268.1649169921875 0 572.29315
268.2026672363281 0 1124.7721 a Water loss 2
268.978515625 0 1001.81683
269.1609802246094 0 692.14606
269.1859130859375 0 1235.144
272.1244812011719 0 2900.6387
274.1185607910156 0 1377.8749
275.17535400390625 0 1149.8396
279.14581298828125 0 5450.512
279.6688232421875 0 1231.6716 b 4
280.1297912597656 0 1789.6589
280.1494445800781 0 1150.4255
280.1664733886719 0 3478.4622
281.0517578125 0 2018.1713
282.1452941894531 0 14278.306
283.1482238769531 0 2108.897
284.1618957519531 0 761.3518
285.01019287109375 0 10707.77
285.1606750488281 0 1719.8203
286.0104064941406 0 3008.0273
286.1041564941406 0 2603.0286
287.0075378417969 0 1060.766
294.1812744140625 0 714.455
296.1358947753906 0 2411.2026
296.17352294921875 0 5754.173
296.1974792480469 0 93676.26 b Water loss 2
297.1197509765625 0 2371.9941
297.1564636230469 0 15896.175
297.1764221191406 0 965.45245
297.2005920410156 0 14865.322
298.15899658203125 0 2432.8972
298.17681884765625 0 5087.1255
298.203125 0 1442.6896
299.0622253417969 0 12232.388
299.15350341796875 0 1076.589
300.0627746582031 0 3217.9846
300.1561279296875 0 823.2013
301.058349609375 0 1294.5562
301.18804931640625 0 1554.1471 y Water loss 14
303.02142333984375 0 1098.474
303.1708984375 0 3453.9104
304.114501953125 0 1585.2162
307.6668701171875 0 5094.2217 y Water loss 11
308.1614990234375 0 1537.9576 y Ammonia loss 11
313.16400146484375 0 740.5952
313.1880798339844 0 1995.9962
314.1462707519531 0 2130.3213
314.1831970214844 0 13469.201
314.2080993652344 0 20018.164 b 2
315.1676330566406 0 634.7505
315.1866455078125 0 2294.3303
315.2109069824219 0 3645.8818
316.1873779296875 0 10486.182
316.2056579589844 0 690.5543
316.6725769042969 0 4548.9385 y 11
317.1716003417969 0 1163.9008
317.1896057128906 0 2169.7483
319.1982116699219 0 13255.974 y 14
320.2005615234375 0 1175.9497
324.1553955078125 0 838.622
325.1517333984375 0 875.81
326.17059326171875 0 848.4207
328.11468505859375 0 2167.6062
330.18157958984375 0 2465.1504
332.12451171875 0 706.9379
333.15673828125 0 2893.0095
334.2125244140625 0 1173.9995
335.17156982421875 0 860.953 y 8
336.15557861328125 0 1178.0956
339.16650390625 0 1155.8875
340.1988830566406 0 850.3094
341.18377685546875 0 2062.9136
342.18023681640625 0 1000.81067
344.201171875 0 1883.1748
344.9767150878906 0 1828.3226
346.1251525878906 0 6015.0967
346.1763916015625 0 3545.7346
348.19207763671875 0 2711.622
349.1984558105469 0 763.9041
350.134765625 0 2287.1584
350.1822204589844 0 2071.6099
351.16680908203125 0 9568.086
351.187744140625 0 683.934
352.1693420410156 0 1777.7336
353.1824035644531 0 4336.1226
354.1661682128906 0 1586.208
354.1872253417969 0 1150.2875
355.0705261230469 0 41969.812
355.1629333496094 0 1714.8245
355.19818115234375 0 604.966
356.0710754394531 0 17193.016
357.06817626953125 0 8968.706
358.21160888671875 0 918.1492
362.2080993652344 0 8259.078
363.2104797363281 0 2337.0876
364.7013244628906 0 907.77094 y Ammonia loss 10
365.1462707519531 0 1712.8417
367.2091064453125 0 5934.074
368.1924743652344 0 2735.1611
369.1772766113281 0 7896.6714
370.1804504394531 0 1600.453
370.67681884765625 0 816.3803
371.1023254394531 0 1369.5753
371.19207763671875 0 2500.455
371.68463134765625 0 674.6668
373.08087158203125 0 3363.7886
373.1725158691406 0 2129.9885
373.2140808105469 0 5062.4575 y 10
373.7156677246094 0 2309.46
374.0819091796875 0 1264.1893
375.0804748535156 0 838.9894
379.1961669921875 0 620.9024
381.1763610839844 0 962.10236
383.1567077636719 0 3039.4258
383.1817626953125 0 798.4752
384.15997314453125 0 659.79535
385.2198181152344 0 13199.473
386.22314453125 0 1754.4788
393.24993896484375 0 1538.3362
397.13616943359375 0 887.19556
397.21636962890625 0 1293.4188
398.2040710449219 0 843.33234
398.2449035644531 0 1411.7097
401.1673583984375 0 5853.159
402.17230224609375 0 1100.102
406.2205505371094 0 1193.6937 b Water loss 7
407.20318603515625 0 2717.17
407.7245788574219 0 1233.3433
408.1885070800781 0 4466.0225
409.19195556640625 0 1416.8738
411.2613525390625 0 3047.9807
412.26336669921875 0 878.36505
413.21343994140625 0 738.2647
415.0368347167969 0 2286.092
415.14697265625 0 1816.6736
415.23046875 0 1756.4761 b 7
415.2696228027344 0 1423.0369
416.0381164550781 0 1464.6887
416.21466064453125 0 1093.4321 y Ammonia loss 13
416.255126953125 0 2695.545
417.0346984863281 0 868.7961
418.9965515136719 0 4221.4614
419.9969482421875 0 1491.9363
420.9954528808594 0 1458.4622
423.1856994628906 0 802.9645
423.7003479003906 0 754.0823
424.2293701171875 0 3663.707
425.2156066894531 0 5692.527
426.2184753417969 0 1141.421
427.2317810058594 0 1751.8407
429.2144470214844 0 667.2869 y Ammonia loss 9
433.1567077636719 0 2532.3066
433.24176025390625 0 12205.7705 y 13
433.2817687988281 0 25961.008
434.2048645019531 0 1608.3291
434.2439270019531 0 1985.4064
434.2850341796875 0 7908.1035
434.70660400390625 0 1383.7351
435.2072448730469 0 1280.2041
435.2893981933594 0 1069.5428
437.7315979003906 0 675.1152 y 9
440.2109069824219 0 886.8256
441.2009582519531 0 622.5873
441.25054931640625 0 871.50183
442.2411804199219 0 3486.3667
442.2738342285156 0 877.6341
443.1415710449219 0 861.4214
443.2315979003906 0 1704.8138
443.267333984375 0 3181.5688 b Water loss 3
445.1998291015625 0 1053.5769
450.2405700683594 0 609.51086
451.23150634765625 0 1164.5745
451.8936767578125 0 2973.4014 y Water loss 4
452.2192077636719 0 2428.9858 y Ammonia loss 4
453.215576171875 0 974.6858
454.2074890136719 0 1839.4723
454.70831298828125 0 1044.7249
459.21160888671875 0 1329.1155
461.2767333984375 0 24556.219 b 3
462.19921875 0 2470.8213
462.2797546386719 0 7967.6963
463.2833557128906 0 1463.0833
467.2247314453125 0 668.6028
468.2092590332031 0 1155.9
468.25653076171875 0 1979.2539
469.2430725097656 0 2144.9648
470.2255859375 0 8826.737
470.7261047363281 0 802.4518
471.2290954589844 0 2168.7974
471.71466064453125 0 1035.3651
473.73638916015625 0 648.165
476.1793518066406 0 817.65564
477.2575378417969 0 1597.253
478.24188232421875 0 5157.289
479.2271423339844 0 2064.7458
480.2102966308594 0 9433.596
480.7204284667969 0 1581.7769
481.2156982421875 0 2740.382
482.2220764160156 0 816.4025
484.20458984375 0 1096.0018
486.26922607421875 0 4618.481 y Water loss 12
487.252197265625 0 2245.1592 y Ammonia loss 12
488.25 0 713.58215
488.7353515625 0 751.5877
489.057373046875 0 1191.3373
489.25067138671875 0 591.8755
490.0581970214844 0 893.3212
490.2103271484375 0 807.5417
493.7289733886719 0 3668.4517
494.19085693359375 0 887.8493
494.2304992675781 0 2159.173
494.7361755371094 0 753.29193
495.26904296875 0 2426.4993
496.2524108886719 0 11663.972
497.25482177734375 0 2311.1055
498.22039794921875 0 24081.908
499.2233581542969 0 6875.855
500.2313537597656 0 1635.9036
501.24859619140625 0 798.93604 y Ammonia loss 3
502.2151184082031 0 1860.604
502.7334899902344 0 5491.1187
503.234130859375 0 3193.715
503.7362060546875 0 1040.03
504.2782287597656 0 28556.166 y 12
505.2817077636719 0 7390.1304
506.2830505371094 0 622.599
508.2196960449219 0 970.0815
510.2206726074219 0 964.1986
512.199951171875 0 1950.4929
513.2785034179688 0 4366.1816
515.2488403320312 0 756.0017
518.2031860351562 0 916.89716
521.2739868164062 0 651.0379
529.2681884765625 0 1034.0413
529.7582397460938 0 1231.3756
530.2100830078125 0 8057.211
530.2567138671875 0 837.56854
530.27490234375 0 740.9947
530.3340454101562 0 2408.7612
531.2141723632812 0 2428.6528
536.2816162109375 0 1309.5734
538.2701416015625 0 8783.927
538.7717895507812 0 4911.352
540.3207397460938 0 2459.5989 b Water loss 4
542.61376953125 0 801.27924
543.262451171875 0 3323.3984
543.7640991210938 0 1615.9586
544.2681274414062 0 2594.8474 b 9
545.2655029296875 0 717.066
546.24365234375 0 1373.8466
551.2831420898438 0 844.9335
552.2681884765625 0 7388.0757
552.7683715820312 0 4436.9087
553.267578125 0 1628.0377
555.3244018554688 0 1941.9589
555.7831420898438 0 754.47644
556.2708129882812 0 998.3677
556.3233032226562 0 686.36084
558.2816162109375 0 1113.4064
558.3297729492188 0 7157.319 b 4
559.3329467773438 0 1962.5333
569.7747192382812 0 704.1919
571.2805786132812 0 791.4038
573.2340087890625 0 603.6671
573.2837524414062 0 769.293
574.273193359375 0 757.48096
578.2882080078125 0 1769.3063 y 1
578.7877197265625 0 1657.6104
579.2845458984375 0 1020.9942
580.283447265625 0 1339.7773 y Water loss 6
580.785400390625 0 846.7854 y Ammonia loss 6
581.2579956054688 0 1850.4321
582.2882690429688 0 1827.5105
582.7833862304688 0 3710.5215
583.2824096679688 0 2113.936
583.7824096679688 0 1505.2258
586.3346557617188 0 754.60516
587.2675170898438 0 1026.2368
591.2426147460938 0 1173.0894
591.293212890625 0 1758.908
591.7951049804688 0 1670.2468 b Water loss 10
592.2974853515625 0 999.06415
597.29541015625 0 3104.1436
598.2892456054688 0 2279.602
599.2676391601562 0 11834.905
599.31298828125 0 1376.7871
600.2727661132812 0 3373.8196
600.8128662109375 0 1638.2422 b 10
601.2703857421875 0 870.37646
602.2777709960938 0 870.2082
602.7905883789062 0 2393.915
603.294189453125 0 2012.906
608.3513793945312 0 1453.6653
608.8018798828125 0 1236.0049
609.2520141601562 0 5625.1978
609.298095703125 0 1674.8606
609.3440551757812 0 722.0446
610.1843872070312 0 5101.348
610.2551879882812 0 1861.0527
611.1852416992188 0 4920.088
611.79736328125 0 7978.644
612.1835327148438 0 3965.4001
612.2980346679688 0 5274.667
612.8108520507812 0 2700.8706
613.3192138671875 0 2721.792
614.3267822265625 0 9491.067 y Water loss 11
615.310546875 0 17454.273 y Ammonia loss 11
616.3129272460938 0 5174.134
617.314453125 0 3763.1743
617.8074951171875 0 10927.755
618.3065185546875 0 10848.064
618.8057861328125 0 4324.607
619.3074951171875 0 758.4018
619.803955078125 0 844.8856
625.2842407226562 0 1225.612
626.3624267578125 0 3643.1797
626.8126220703125 0 23995.723
627.2625122070312 0 48138.492
627.31396484375 0 16448.617
627.8157348632812 0 6385.377
628.2655639648438 0 13601.857
628.316162109375 0 1661.935
628.8103637695312 0 1928.3523 y Water loss 5
629.2677612304688 0 2529.144
629.31689453125 0 1464.6514
629.812255859375 0 1113.6017
632.336669921875 0 22794.76 y 11
633.3399047851562 0 7296.4663
634.3427734375 0 1041.9409
635.8185424804688 0 886.85516
636.3214111328125 0 1063.8918
637.8153076171875 0 3826.4097 y 5
638.3167724609375 0 2551.4888
638.81787109375 0 1598.46
639.3160400390625 0 789.55426
643.2941284179688 0 3943.295
644.2916870117188 0 1942.9629
645.3055419921875 0 3855.5764
646.8301391601562 0 1249.3633
650.32177734375 0 677.5024
655.34130859375 0 10764.844
655.8422241210938 0 8084.824 b Water loss 11
656.342041015625 0 3273.963
656.8447265625 0 921.0869
657.333740234375 0 964.6766
658.306396484375 0 967.3815
659.3260498046875 0 838.1727
659.8218994140625 0 967.86255
660.3204956054688 0 1533.7345
664.8419189453125 0 996.98627 b 11
665.344482421875 0 657.2532
666.3604736328125 0 685.618
668.3326416015625 0 2984.067
668.825439453125 0 3307.609
669.3285522460938 0 2229.897
669.8231201171875 0 983.09875
677.3367309570312 0 28425.629 y Water loss 4
677.8358154296875 0 28323.955 y Ammonia loss 4
678.3362426757812 0 15492.77
678.8365478515625 0 4900.3613
679.3360595703125 0 879.1956
680.328125 0 827.9963
684.3655395507812 0 1572.2836
686.3416748046875 0 62931.656 y 4
686.84326171875 0 46541.645
687.3447265625 0 17448.15
687.84521484375 0 5891.2925
688.3806762695312 0 1788.2533
691.3446044921875 0 737.5495 b Water loss 12
691.8397827148438 0 949.55634 b Ammonia loss 12
694.3428344726562 0 1150.2776
698.33935546875 0 659.8861
699.342529296875 0 712.9492
700.3472900390625 0 3833.5098 b 12
700.8540649414062 0 1452.2355
701.3511352539062 0 1481.7472
704.3335571289062 0 826.7093
710.3844604492188 0 2076.3105
712.353515625 0 8553.479
712.8590087890625 0 701.12085
713.3547973632812 0 3721.6216
714.3666381835938 0 931.7087
720.3697509765625 0 1337.864
721.3603515625 0 887.71423
722.3368530273438 0 2779.3008
723.3421630859375 0 803.09985
724.4022216796875 0 1998.3573 b Water loss 6
725.4056396484375 0 853.4763
727.4066772460938 0 1167.1588 y Water loss 10
728.3950805664062 0 1777.875 y Ammonia loss 10
728.8729248046875 0 1133.7356
729.3836059570312 0 1851.39
730.3778076171875 0 798.01276
732.3424072265625 0 862.8905
737.3184814453125 0 728.8608
737.3969116210938 0 1680.6027
738.3056030273438 0 804.50775
738.3849487304688 0 961.0105
739.360595703125 0 809.3617
740.3475341796875 0 16343.249
741.3500366210938 0 7873.6455
742.3506469726562 0 2295.918
742.4157104492188 0 3523.9604 b 6
743.3546752929688 0 821.5897
743.4235229492188 0 913.45953
745.3500366210938 0 895.9098
745.42138671875 0 10877.546 y 10
746.4234619140625 0 4973.0186
747.4260864257812 0 1046.6191
749.342529296875 0 1381.0157
750.8709716796875 0 6120.409 y Water loss 3
751.3707885742188 0 7425.32 y Ammonia loss 3
751.8723754882812 0 3736.5588
752.3707275390625 0 1481.9335
755.4061279296875 0 1938.6609
756.4081420898438 0 1467.8967
757.3743896484375 0 5325.135 b 13
758.3748168945312 0 3041.622
759.3822631835938 0 1986.1805
759.8759765625 0 8450.717 y 3
760.37744140625 0 6547.4507
760.87744140625 0 3422.0408
771.3512573242188 0 3141.9067
772.3505859375 0 1360.3877
774.3314819335938 0 3697.3784
774.4534912109375 0 918.72205
775.3327026367188 0 1585.8131
793.4196166992188 0 854.37463
806.35693359375 0 1083.0198
807.412109375 0 2655.076 y Water loss 2
807.9102783203125 0 1768.802 y Ammonia loss 2
808.4149169921875 0 885.3157
811.4324951171875 0 2954.6746 b Water loss 7
812.4307861328125 0 1982.9325 b Water loss 14
813.4346923828125 0 2252.2607
816.416259765625 0 997.22314 y 2
816.9194946289062 0 1343.9702
817.4121704101562 0 715.68304
829.4465942382812 0 2812.9812 b 7
830.4500732421875 0 1209.7281
835.3829956054688 0 986.6501
839.4190673828125 0 1073.553
840.41796875 0 1165.8522
841.4258422851562 0 2711.3906
842.3853759765625 0 1075.9379
847.9432373046875 0 1060.4403 b Water loss 15
848.436767578125 0 876.9174 b Ammonia loss 15
849.4235229492188 0 770.98645
850.3963623046875 0 2180.1047
851.3935546875 0 1879.3599
856.455810546875 0 1481.3225 y Water loss 9
856.9450073242188 0 1279.3168 b 15
857.447021484375 0 4158.1797 y Ammonia loss 9
857.9430541992188 0 997.0093 y Water loss 1
858.4385986328125 0 2212.2317 y Ammonia loss 1
858.9398803710938 0 1125.7855
859.4275512695312 0 1375.2335
863.3844604492188 0 3002.5017
864.3880615234375 0 901.3803
867.4351806640625 0 722.9199
868.4050903320312 0 19596.35
869.4075317382812 0 10099.798
870.4093627929688 0 2282.049
873.4575805664062 0 3587.01
874.4630126953125 0 5557.3926 y 9
875.464599609375 0 1936.375
879.411376953125 0 5241.4106
880.4168701171875 0 1755.021
887.4146118164062 0 3948.3794
888.4202270507812 0 2634.051
889.39794921875 0 3858.0574
890.394775390625 0 2213.4067
894.4202880859375 0 1558.1981
907.406005859375 0 20792.389
908.409912109375 0 9096.086
909.4155883789062 0 1159.45
911.445068359375 0 1964.0758
912.4556274414062 0 1202.4272
917.4494018554688 0 861.83386
921.4290161132812 0 1437.9084
922.4351806640625 0 1780.8551
939.4425048828125 0 11043.805
940.4534301757812 0 6470.65
941.4595336914062 0 2168.5535
942.4738159179688 0 1585.6497
954.44970703125 0 914.908
956.45361328125 0 2498.8967
957.4451904296875 0 1234.3617
958.4871215820312 0 6133.5894 b 8
959.4910278320312 0 2940.137
960.4905395507812 0 11180.792
961.4900512695312 0 5650.8516
962.4950561523438 0 1089.7505
971.4775390625 0 1188.3248
987.4855346679688 0 755.9002
988.466796875 0 1450.1898
1000.4954833984375 0 1113.6042
1001.1045532226562 0 667.8807
1003.4990844726562 0 1179.2606 y 8
1004.5051879882812 0 963.752
1015.4708251953125 0 1151.69
1016.48046875 0 784.723
1035.4783935546875 0 795.3637
1036.4671630859375 0 2282.2708
1037.45703125 0 1561.7113
1040.498291015625 0 1086.2338
1041.49853515625 0 1597.656
1053.486083984375 0 9083.467
1054.488525390625 0 4924.0073
1055.491943359375 0 1712.0226
1057.5184326171875 0 1214.1035
1058.5145263671875 0 2557.2544
1059.52587890625 0 1649.8948
1060.520751953125 0 813.05334
1068.509765625 0 891.9178
1069.512939453125 0 2217.743 b Water loss 9
1070.5159912109375 0 5696.486
1071.5152587890625 0 2689.0889
1072.5157470703125 0 892.8788 y Water loss 7
1073.5205078125 0 1272.5066 y Ammonia loss 7
1075.5216064453125 0 4617.739
1076.525634765625 0 3242.9512
1084.5234375 0 1888.6562
1085.5196533203125 0 1487.7545
1086.513427734375 0 2312.8367
1087.5299072265625 0 18576.727 b 9
1088.5323486328125 0 10836.916
1089.53857421875 0 4035.3713
1090.5386962890625 0 2000.047 y 7
1091.546142578125 0 833.5483
1101.55126953125 0 932.95013
1103.5267333984375 0 6809.822
1104.530517578125 0 3593.439
1105.5386962890625 0 884.69977
1137.554931640625 0 2406.9697
1138.5499267578125 0 1416.8064
1139.5550537109375 0 1102.5594
1155.5616455078125 0 3952.9436
1156.56640625 0 2548.5325
1157.574951171875 0 1012.6541
1164.5614013671875 0 2475.947
1165.558837890625 0 1815.0582
1177.56591796875 0 1505.5645 y 6
1178.570556640625 0 1203.8177
1181.5782470703125 0 6945.181
1182.585205078125 0 5758.3447 b Water loss 10
1183.58837890625 0 2057.0078
1200.609375 0 3429.863 b 10
1201.6129150390625 0 2343.7207
1213.6212158203125 0 907.5668
1234.6041259765625 0 4402.135
1235.6053466796875 0 4032.865
1236.614013671875 0 2518.793
1252.6143798828125 0 12647.106
1253.617919921875 0 8966.168
1254.623046875 0 3220.065
1255.6298828125 0 1295.4719
1256.614990234375 0 1388.9608 y Water loss 5
1257.605712890625 0 1501.4507 y Ammonia loss 5
1258.596923828125 0 887.29895
1274.6199951171875 0 7367.7856 y 5
1275.6234130859375 0 4806.326
1276.628173828125 0 1936.1915
1310.6708984375 0 797.90436 b Water loss 11
1328.663818359375 0 1692.8704 b 11
1329.6680908203125 0 970.96454
1353.6680908203125 0 1564.5381 y Water loss 4
1354.6519775390625 0 1194.0922 y Ammonia loss 4
1371.6668701171875 0 2194.2183 y 4
1372.6845703125 0 1442.4038
1399.6910400390625 0 1851.0444 b 12
1400.70068359375 0 1672.6365
1513.7578125 0 1157.6577 b 13
1514.75146484375 0 931.19507
1536.61279296875 0 716.2447
3067.984130859375 0 767.6979

Spectrum Details

|  |  |
| --- | --- |
| Matched peaks? Matched peaks The total absolute number of peaks matched. Additionally in brackets the total fraction of peaks matched and the total number of peaks is shown. | 99 (13.58% of 729) |
| FDR? FDR The false discovery rate estimated for this peptide. It is calculated by matching all theoretical fragments with a non-integer shift with the raw peaks for this spectrum. This is done with 40 different shifts. The resulting percentage is the average number of annotated peaks over the number of annotated peaks with the correct spectrum. | 0.82% |
| Satellite FDR? Satellite FDR See the FDR for details on its calculation. This satellite ion specific FDR only contains the satellite ions (d/w) for I/L/J positions. | - |
| PSM Score? PSM Score The PSM Score as given by Hecklib to this annotated spectrum. It is shown with three significant figures. | 859 |

#### Spectrum 8439? Spectrum 8439 The raw spectrum of this peptide as annotated by Hecklib. The fragments are coloured according to ion type (see legend). Any peaks with a star '\*' as text can be hovered over to see the full details, first the ion type second the mass shift type. By hovering over the amino acids in the peptide or ions in the legend the corresponding peaks are highlighted. By toggling the 'Unassigned' label you can turn the background (unassigned) peaks on or off in the plot. By updating the slider in the Ion legend you can update the spectrum to only show the top X% of the peaks with labels. The top X% means any peak that is within X% of the highest intensity. By dragging in the spectrum you can zoom in to a specific part of the spectrum and use 'Zoom Out' to get back to the original zoom level. The annotation of the spectrum is based on the given sequence in the peptides file and is done with different software so inconsistencies are likely. The peaks are annotated based on the given sequence, with 20 ppm tolerance.

Copy Data

##### Spectrum 8439 (TSV)

###### Preview

```
Loading example...
```

*Click on the button to copy the data to your clipboard.*

Mz MinMz MaxIntensity Max

WidthHeightPeptide font sizePeptide stroke widthSpectrum font sizeSpectrum stroke widthCompact peptide

Ion legend

wxyz

abcd

OtherUnassignedIon^Charge^_Position_Show for top:%

VTJFPPSSEEJQANKAT

03.66e+47.33e+41.10e+51.47e+5

Zoom Out

y^+1^_1_a^+1^_2_y^+1^_2_a^+1^_2_b^+1^_2_y^+1^_2_b^+1^_2_y^+2^_5_a^+1^_3_b^+1^_3_y^+2^_6_b^+1^_3_y^+1^_3_y^+1^_4_b^+1^_4_b^+1^_4_y^+1^_5_y^+1^_5_b^+1^_5_b^+1^_5_b^+2^_11_y^+1^_6_y^+1^_6_y^+1^_6_y^+2^_12_b^+2^_12_y^+2^_13_y^+2^_13_y^+2^_13_b^+2^_13_b^+1^_7_y^+1^_7_y^+2^_14_b^+2^_14_y^+2^_14_y^+2^_15_b^+1^_8_b^+2^_16_b^+2^_16_y^+1^_8_y^+2^_16_y^+2^_16_y^+1^_8_\*\*\*b^+1^_9_y^+1^_9_y^+1^_10_b^+1^_10_y^+1^_10_y^+1^_11_y^+1^_11_y^+1^_11_b^+1^_11_b^+1^_11_y^+1^_12_y^+1^_12_y^+1^_12_b^+1^_12_b^+1^_12_b^+1^_12_y^+1^_13_y^+1^_13_y^+1^_13_b^+1^_13_b^+1^_13_y^+1^_14_y^+1^_15_

041282512371649

Fragment Matches Table

Show background peaks

| Position | Ion type | Intensity | mz Theoretical | mz Error (Th) | mz Error (ppm) | Charge | Series Number |
| --- | --- | --- | --- | --- | --- | --- | --- |
| 17 | y | 847.9 | 120.1 | 0.0003907 | 3.254 | +1 | 1 |
| - | - | 1.599E+04 | 120.1 | - | - | 0 | - |
| - | - | 960.5 | 121.1 | - | - | 0 | - |
| - | - | 386.5 | 126 | - | - | 0 | - |
| - | - | 3822 | 126.1 | - | - | 0 | - |
| - | - | 354.8 | 126.7 | - | - | 0 | - |
| - | - | 457.4 | 127.1 | - | - | 0 | - |
| - | - | 517.6 | 128.1 | - | - | 0 | - |
| - | - | 3952 | 128.1 | - | - | 0 | - |
| - | - | 820.5 | 129.1 | - | - | 0 | - |
| - | - | 2.867E+04 | 129.1 | - | - | 0 | - |
| - | - | 802.3 | 130.1 | - | - | 0 | - |
| - | - | 1459 | 130.1 | - | - | 0 | - |
| - | - | 662.9 | 131.1 | - | - | 0 | - |
| - | - | 402.6 | 133.1 | - | - | 0 | - |
| - | - | 1894 | 136.1 | - | - | 0 | - |
| - | - | 738.3 | 138.1 | - | - | 0 | - |
| - | - | 803.9 | 138.1 | - | - | 0 | - |
| - | - | 1907 | 138.1 | - | - | 0 | - |
| - | - | 837.5 | 139.1 | - | - | 0 | - |
| - | - | 691.6 | 141.1 | - | - | 0 | - |
| - | - | 441.7 | 141.1 | - | - | 0 | - |
| - | - | 1220 | 147.1 | - | - | 0 | - |
| - | - | 438.1 | 148.2 | - | - | 0 | - |
| - | - | 571.8 | 148.8 | - | - | 0 | - |
| - | - | 406.1 | 148.9 | - | - | 0 | - |
| - | - | 544.3 | 148.9 | - | - | 0 | - |
| - | - | 643.8 | 148.9 | - | - | 0 | - |
| - | - | 793.1 | 148.9 | - | - | 0 | - |
| - | - | 841.5 | 148.9 | - | - | 0 | - |
| - | - | 930.9 | 148.9 | - | - | 0 | - |
| - | - | 1369 | 148.9 | - | - | 0 | - |
| - | - | 1288 | 148.9 | - | - | 0 | - |
| - | - | 3309 | 148.9 | - | - | 0 | - |
| - | - | 6216 | 149 | - | - | 0 | - |
| - | - | 3692 | 149 | - | - | 0 | - |
| - | - | 1711 | 149 | - | - | 0 | - |
| - | - | 1249 | 149 | - | - | 0 | - |
| - | - | 1264 | 149 | - | - | 0 | - |
| - | - | 711.1 | 149 | - | - | 0 | - |
| - | - | 674.7 | 149 | - | - | 0 | - |
| - | - | 803.3 | 149 | - | - | 0 | - |
| - | - | 650 | 149 | - | - | 0 | - |
| - | - | 594.6 | 149 | - | - | 0 | - |
| - | - | 438.6 | 149 | - | - | 0 | - |
| - | - | 1380 | 155.1 | - | - | 0 | - |
| 2 | a | 3740 | 155.1 | 0.0002898 | 1.868 | +1 | 2 |
| - | - | 1329 | 157.1 | - | - | 0 | - |
| - | - | 3860 | 157.1 | - | - | 0 | - |
| - | - | 435.5 | 165.1 | - | - | 0 | - |
| - | - | 708.2 | 165.1 | - | - | 0 | - |
| - | - | 2452 | 167.1 | - | - | 0 | - |
| - | - | 1661 | 167.1 | - | - | 0 | - |
| - | - | 3902 | 169.1 | - | - | 0 | - |
| - | - | 489.7 | 170.1 | - | - | 0 | - |
| - | - | 583.8 | 172.1 | - | - | 0 | - |
| - | - | 460.7 | 173.1 | - | - | 0 | - |
| 16 | y | 777.7 | 173.1 | 0.0004301 | 2.485 | +1 | 2 |
| 2 | a | 1.278E+04 | 173.1 | 0.0002994 | 1.73 | +1 | 2 |
| - | - | 1067 | 174.1 | - | - | 0 | - |
| - | - | 1124 | 175.1 | - | - | 0 | - |
| - | - | 763.3 | 181.1 | - | - | 0 | - |
| - | - | 1162 | 182.1 | - | - | 0 | - |
| - | - | 1630 | 182.1 | - | - | 0 | - |
| - | - | 1261 | 183.1 | - | - | 0 | - |
| 2 | b | 4.852E+04 | 183.1 | 0.0002787 | 1.522 | +1 | 2 |
| - | - | 1507 | 184.1 | - | - | 0 | - |
| - | - | 5517 | 184.1 | - | - | 0 | - |
| - | - | 4617 | 185.1 | - | - | 0 | - |
| - | - | 1175 | 186.1 | - | - | 0 | - |
| - | - | 8119 | 187.1 | - | - | 0 | - |
| - | - | 1259 | 189.1 | - | - | 0 | - |
| - | - | 804 | 191.1 | - | - | 0 | - |
| 16 | y | 1097 | 191.1 | 0.0003482 | 1.822 | +1 | 2 |
| - | - | 4062 | 195.1 | - | - | 0 | - |
| - | - | 5040 | 197.1 | - | - | 0 | - |
| - | - | 1235 | 198.1 | - | - | 0 | - |
| - | - | 2226 | 199.1 | - | - | 0 | - |
| - | - | 522.3 | 199.1 | - | - | 0 | - |
| - | - | 1541 | 200.1 | - | - | 0 | - |
| - | - | 1588 | 200.1 | - | - | 0 | - |
| 2 | b | 2.527E+04 | 201.1 | 0.0002426 | 1.206 | +1 | 2 |
| - | - | 2213 | 202.1 | - | - | 0 | - |
| - | - | 666.1 | 204.1 | - | - | 0 | - |
| - | - | 2092 | 208.1 | - | - | 0 | - |
| - | - | 5966 | 209.1 | - | - | 0 | - |
| - | - | 702.9 | 213.1 | - | - | 0 | - |
| - | - | 678.8 | 214.2 | - | - | 0 | - |
| - | - | 8764 | 215.1 | - | - | 0 | - |
| - | - | 1192 | 216.1 | - | - | 0 | - |
| - | - | 4089 | 217.1 | - | - | 0 | - |
| - | - | 5049 | 217.1 | - | - | 0 | - |
| - | - | 766.8 | 218.1 | - | - | 0 | - |
| - | - | 658.2 | 218.1 | - | - | 0 | - |
| - | - | 2099 | 219.1 | - | - | 0 | - |
| - | - | 509.6 | 220.2 | - | - | 0 | - |
| - | - | 3025 | 225.1 | - | - | 0 | - |
| - | - | 893 | 225.1 | - | - | 0 | - |
| - | - | 2.105E+04 | 226.1 | - | - | 0 | - |
| - | - | 722.4 | 227.1 | - | - | 0 | - |
| - | - | 1912 | 227.1 | - | - | 0 | - |
| - | - | 1.226E+04 | 233.2 | - | - | 0 | - |
| - | - | 1422 | 234.2 | - | - | 0 | - |
| - | - | 1255 | 236.1 | - | - | 0 | - |
| - | - | 1179 | 241.1 | - | - | 0 | - |
| - | - | 3457 | 242.2 | - | - | 0 | - |
| - | - | 1033 | 243.1 | - | - | 0 | - |
| - | - | 6844 | 243.1 | - | - | 0 | - |
| 13 | y | 1615 | 244.1 | 0.000624 | 2.556 | +2 | 5 |
| - | - | 586 | 244.1 | - | - | 0 | - |
| - | - | 4135 | 245.1 | - | - | 0 | - |
| - | - | 2345 | 247.1 | - | - | 0 | - |
| - | - | 5849 | 251.2 | - | - | 0 | - |
| - | - | 1228 | 252.1 | - | - | 0 | - |
| - | - | 5283 | 254.1 | - | - | 0 | - |
| - | - | 1019 | 254.1 | - | - | 0 | - |
| - | - | 709.4 | 255.1 | - | - | 0 | - |
| - | - | 692.3 | 256.1 | - | - | 0 | - |
| - | - | 2476 | 259.1 | - | - | 0 | - |
| - | - | 3755 | 261.2 | - | - | 0 | - |
| - | - | 730.2 | 262.2 | - | - | 0 | - |
| - | - | 1523 | 264.1 | - | - | 0 | - |
| 3 | a | 1224 | 268.2 | 0.0007511 | 2.801 | +1 | 3 |
| - | - | 730.6 | 269.1 | - | - | 0 | - |
| - | - | 1775 | 269.2 | - | - | 0 | - |
| - | - | 1268 | 270.1 | - | - | 0 | - |
| - | - | 712.5 | 270.2 | - | - | 0 | - |
| - | - | 2904 | 272.1 | - | - | 0 | - |
| - | - | 622.6 | 274.1 | - | - | 0 | - |
| - | - | 2369 | 275.2 | - | - | 0 | - |
| - | - | 2525 | 279.1 | - | - | 0 | - |
| - | - | 963.5 | 280.1 | - | - | 0 | - |
| - | - | 4338 | 280.2 | - | - | 0 | - |
| - | - | 5434 | 282.1 | - | - | 0 | - |
| - | - | 991.4 | 283.1 | - | - | 0 | - |
| - | - | 2528 | 285.2 | - | - | 0 | - |
| - | - | 2307 | 286.1 | - | - | 0 | - |
| - | - | 860.9 | 294.2 | - | - | 0 | - |
| - | - | 721.6 | 296.1 | - | - | 0 | - |
| 3 | b | 1.227E+05 | 296.2 | 0.0004585 | 1.548 | +1 | 3 |
| - | - | 811.2 | 297.1 | - | - | 0 | - |
| - | - | 6823 | 297.2 | - | - | 0 | - |
| - | - | 2.018E+04 | 297.2 | - | - | 0 | - |
| - | - | 914 | 298.2 | - | - | 0 | - |
| - | - | 4189 | 298.2 | - | - | 0 | - |
| - | - | 1653 | 298.2 | - | - | 0 | - |
| - | - | 4004 | 303.2 | - | - | 0 | - |
| - | - | 1416 | 304.1 | - | - | 0 | - |
| - | - | 1225 | 304.2 | - | - | 0 | - |
| 12 | y | 780 | 308.2 | 0.002204 | 7.151 | +2 | 6 |
| - | - | 595.6 | 312.1 | - | - | 0 | - |
| - | - | 636 | 313.2 | - | - | 0 | - |
| - | - | 762.4 | 314.1 | - | - | 0 | - |
| - | - | 5406 | 314.2 | - | - | 0 | - |
| 3 | b | 2.973E+04 | 314.2 | 0.000514 | 1.636 | +1 | 3 |
| - | - | 828.3 | 315.2 | - | - | 0 | - |
| - | - | 5566 | 315.2 | - | - | 0 | - |
| - | - | 1762 | 317.2 | - | - | 0 | - |
| 15 | y | 7427 | 319.2 | 0.0005237 | 1.641 | +1 | 3 |
| - | - | 1437 | 320.2 | - | - | 0 | - |
| - | - | 2282 | 328.1 | - | - | 0 | - |
| - | - | 3625 | 330.2 | - | - | 0 | - |
| - | - | 682 | 333.2 | - | - | 0 | - |
| - | - | 939 | 334.2 | - | - | 0 | - |
| - | - | 1007 | 337.2 | - | - | 0 | - |
| - | - | 1375 | 339.2 | - | - | 0 | - |
| - | - | 2312 | 341.2 | - | - | 0 | - |
| - | - | 961.2 | 342.2 | - | - | 0 | - |
| - | - | 1494 | 344.2 | - | - | 0 | - |
| - | - | 4785 | 346.1 | - | - | 0 | - |
| - | - | 4697 | 346.2 | - | - | 0 | - |
| - | - | 634.9 | 347.1 | - | - | 0 | - |
| - | - | 4669 | 348.2 | - | - | 0 | - |
| - | - | 787.5 | 349.2 | - | - | 0 | - |
| - | - | 938.6 | 350.2 | - | - | 0 | - |
| - | - | 3816 | 351.2 | - | - | 0 | - |
| - | - | 931.4 | 352.2 | - | - | 0 | - |
| - | - | 1991 | 353.2 | - | - | 0 | - |
| - | - | 1261 | 354.2 | - | - | 0 | - |
| - | - | 1118 | 355.2 | - | - | 0 | - |
| - | - | 697.5 | 360.2 | - | - | 0 | - |
| - | - | 9875 | 362.2 | - | - | 0 | - |
| - | - | 1779 | 363.2 | - | - | 0 | - |
| - | - | 1777 | 365.1 | - | - | 0 | - |
| - | - | 928.8 | 367.2 | - | - | 0 | - |
| - | - | 1113 | 368.2 | - | - | 0 | - |
| - | - | 4820 | 369.2 | - | - | 0 | - |
| - | - | 1162 | 369.2 | - | - | 0 | - |
| - | - | 816.9 | 370.2 | - | - | 0 | - |
| - | - | 1343 | 371.2 | - | - | 0 | - |
| - | - | 1418 | 372.2 | - | - | 0 | - |
| - | - | 1633 | 373.2 | - | - | 0 | - |
| - | - | 697.9 | 381.2 | - | - | 0 | - |
| - | - | 3508 | 383.2 | - | - | 0 | - |
| - | - | 537.1 | 383.3 | - | - | 0 | - |
| - | - | 3560 | 385.2 | - | - | 0 | - |
| - | - | 875.6 | 393.2 | - | - | 0 | - |
| - | - | 1733 | 393.3 | - | - | 0 | - |
| - | - | 684.9 | 395.2 | - | - | 0 | - |
| - | - | 782.9 | 397.1 | - | - | 0 | - |
| - | - | 2405 | 398.2 | - | - | 0 | - |
| - | - | 635.6 | 399.2 | - | - | 0 | - |
| - | - | 694.5 | 400.2 | - | - | 0 | - |
| - | - | 5399 | 401.2 | - | - | 0 | - |
| - | - | 967 | 402.2 | - | - | 0 | - |
| - | - | 954.4 | 407.2 | - | - | 0 | - |
| - | - | 1462 | 408.2 | - | - | 0 | - |
| - | - | 2610 | 411.3 | - | - | 0 | - |
| - | - | 688.3 | 412.3 | - | - | 0 | - |
| - | - | 2059 | 415.1 | - | - | 0 | - |
| - | - | 1196 | 415.3 | - | - | 0 | - |
| - | - | 629.5 | 416.2 | - | - | 0 | - |
| - | - | 4448 | 416.3 | - | - | 0 | - |
| - | - | 1105 | 417.3 | - | - | 0 | - |
| - | - | 656 | 423.2 | - | - | 0 | - |
| - | - | 866.7 | 424.2 | - | - | 0 | - |
| - | - | 728.9 | 425.1 | - | - | 0 | - |
| - | - | 1757 | 425.2 | - | - | 0 | - |
| - | - | 839.2 | 427.2 | - | - | 0 | - |
| - | - | 2346 | 433.2 | - | - | 0 | - |
| 14 | y | 5497 | 433.2 | 0.001816 | 4.192 | +1 | 4 |
| - | - | 3.817E+04 | 433.3 | - | - | 0 | - |
| - | - | 672.7 | 434.2 | - | - | 0 | - |
| - | - | 1092 | 434.2 | - | - | 0 | - |
| - | - | 1.181E+04 | 434.3 | - | - | 0 | - |
| - | - | 684.6 | 435.3 | - | - | 0 | - |
| - | - | 605.1 | 440.2 | - | - | 0 | - |
| - | - | 1132 | 441.2 | - | - | 0 | - |
| - | - | 1806 | 442.2 | - | - | 0 | - |
| - | - | 861 | 443.1 | - | - | 0 | - |
| - | - | 1962 | 443.2 | - | - | 0 | - |
| 4 | b | 4003 | 443.3 | 0.0007397 | 1.669 | +1 | 4 |
| - | - | 606.4 | 444.2 | - | - | 0 | - |
| - | - | 1429 | 444.3 | - | - | 0 | - |
| - | - | 832.8 | 452.2 | - | - | 0 | - |
| - | - | 1428 | 459.2 | - | - | 0 | - |
| 4 | b | 2.934E+04 | 461.3 | 0.000673 | 1.459 | +1 | 4 |
| - | - | 1383 | 462.2 | - | - | 0 | - |
| - | - | 9741 | 462.3 | - | - | 0 | - |
| - | - | 865.4 | 463.2 | - | - | 0 | - |
| - | - | 1437 | 463.3 | - | - | 0 | - |
| - | - | 955.9 | 464.2 | - | - | 0 | - |
| - | - | 890.4 | 466.2 | - | - | 0 | - |
| - | - | 938 | 469.2 | - | - | 0 | - |
| - | - | 4526 | 470.2 | - | - | 0 | - |
| - | - | 839 | 471.2 | - | - | 0 | - |
| - | - | 670 | 477.3 | - | - | 0 | - |
| - | - | 703.3 | 478.2 | - | - | 0 | - |
| - | - | 1021 | 479.2 | - | - | 0 | - |
| - | - | 3929 | 480.2 | - | - | 0 | - |
| - | - | 983.7 | 481.2 | - | - | 0 | - |
| - | - | 1444 | 484.2 | - | - | 0 | - |
| 13 | y | 1216 | 487.3 | 0.002482 | 5.094 | +1 | 5 |
| - | - | 1225 | 494.2 | - | - | 0 | - |
| - | - | 1172 | 495.3 | - | - | 0 | - |
| - | - | 1142 | 496.2 | - | - | 0 | - |
| - | - | 1.198E+04 | 498.2 | - | - | 0 | - |
| - | - | 3291 | 499.2 | - | - | 0 | - |
| - | - | 1156 | 500.2 | - | - | 0 | - |
| - | - | 2999 | 502.2 | - | - | 0 | - |
| - | - | 712 | 503.2 | - | - | 0 | - |
| 13 | y | 8142 | 504.3 | 0.0005606 | 1.112 | +1 | 5 |
| - | - | 1866 | 505.3 | - | - | 0 | - |
| - | - | 1844 | 512.2 | - | - | 0 | - |
| - | - | 1234 | 513.3 | - | - | 0 | - |
| - | - | 859.5 | 529.2 | - | - | 0 | - |
| - | - | 8942 | 530.2 | - | - | 0 | - |
| - | - | 2535 | 530.3 | - | - | 0 | - |
| - | - | 2366 | 531.2 | - | - | 0 | - |
| - | - | 1390 | 531.3 | - | - | 0 | - |
| - | - | 1465 | 538.3 | - | - | 0 | - |
| - | - | 930.8 | 540.2 | - | - | 0 | - |
| 5 | b | 1436 | 540.3 | 0.001168 | 2.162 | +1 | 5 |
| - | - | 699.1 | 541.3 | - | - | 0 | - |
| - | - | 828.8 | 546.2 | - | - | 0 | - |
| - | - | 741.2 | 551.3 | - | - | 0 | - |
| - | - | 626.8 | 555.3 | - | - | 0 | - |
| 5 | b | 3836 | 558.3 | 6.372E-05 | 0.1141 | +1 | 5 |
| - | - | 2165 | 559.3 | - | - | 0 | - |
| - | - | 693.8 | 563.2 | - | - | 0 | - |
| - | - | 814.8 | 569.3 | - | - | 0 | - |
| - | - | 889.9 | 569.3 | - | - | 0 | - |
| - | - | 737.4 | 569.8 | - | - | 0 | - |
| - | - | 1739 | 578.3 | - | - | 0 | - |
| - | - | 1109 | 578.8 | - | - | 0 | - |
| - | - | 614.2 | 579.3 | - | - | 0 | - |
| - | - | 1156 | 581.3 | - | - | 0 | - |
| - | - | 799.8 | 582.3 | - | - | 0 | - |
| - | - | 2528 | 582.8 | - | - | 0 | - |
| - | - | 865.7 | 583.3 | - | - | 0 | - |
| - | - | 1048 | 591.2 | - | - | 0 | - |
| - | - | 1191 | 591.3 | - | - | 0 | - |
| 11 | b | 1580 | 591.8 | 0.01181 | 19.96 | +2 | 11 |
| - | - | 1721 | 597.3 | - | - | 0 | - |
| - | - | 616 | 598.3 | - | - | 0 | - |
| - | - | 8506 | 599.3 | - | - | 0 | - |
| - | - | 2302 | 600.3 | - | - | 0 | - |
| - | - | 743.1 | 608.4 | - | - | 0 | - |
| - | - | 746.4 | 608.8 | - | - | 0 | - |
| - | - | 2357 | 609.3 | - | - | 0 | - |
| - | - | 900.4 | 610.3 | - | - | 0 | - |
| - | - | 1986 | 612.8 | - | - | 0 | - |
| - | - | 1069 | 613.3 | - | - | 0 | - |
| 12 | y | 1819 | 614.3 | 0.0005162 | 0.8403 | +1 | 6 |
| 12 | y | 4267 | 615.3 | 0.005711 | 9.281 | +1 | 6 |
| - | - | 2322 | 616.3 | - | - | 0 | - |
| - | - | 7155 | 617.8 | - | - | 0 | - |
| - | - | 7089 | 618.3 | - | - | 0 | - |
| - | - | 3821 | 618.8 | - | - | 0 | - |
| - | - | 1442 | 619.3 | - | - | 0 | - |
| - | - | 894.9 | 625.3 | - | - | 0 | - |
| - | - | 985.5 | 626.3 | - | - | 0 | - |
| - | - | 903.3 | 626.4 | - | - | 0 | - |
| - | - | 1.845E+04 | 626.8 | - | - | 0 | - |
| - | - | 1.823E+04 | 627.3 | - | - | 0 | - |
| - | - | 1.171E+04 | 627.3 | - | - | 0 | - |
| - | - | 6008 | 627.8 | - | - | 0 | - |
| - | - | 7038 | 628.3 | - | - | 0 | - |
| - | - | 1875 | 629.3 | - | - | 0 | - |
| - | - | 668.3 | 630.3 | - | - | 0 | - |
| 12 | y | 4577 | 632.3 | 0.0003387 | 0.5356 | +1 | 6 |
| - | - | 2116 | 633.3 | - | - | 0 | - |
| - | - | 733.7 | 634.3 | - | - | 0 | - |
| - | - | 1122 | 635.8 | - | - | 0 | - |
| - | - | 697.5 | 636.3 | - | - | 0 | - |
| 6 | y | 3030 | 637.8 | 0.0002822 | 0.4425 | +2 | 12 |
| - | - | 1847 | 638.3 | - | - | 0 | - |
| - | - | 774.2 | 640.3 | - | - | 0 | - |
| - | - | 3667 | 643.3 | - | - | 0 | - |
| - | - | 2425 | 644.3 | - | - | 0 | - |
| - | - | 1001 | 655.3 | - | - | 0 | - |
| 12 | b | 792.9 | 655.8 | 0.007971 | 12.15 | +2 | 12 |
| - | - | 1966 | 660.3 | - | - | 0 | - |
| - | - | 617.8 | 661.3 | - | - | 0 | - |
| - | - | 1094 | 663.3 | - | - | 0 | - |
| - | - | 881.6 | 663.8 | - | - | 0 | - |
| - | - | 981.5 | 668.3 | - | - | 0 | - |
| - | - | 1230 | 668.4 | - | - | 0 | - |
| - | - | 1111 | 668.8 | - | - | 0 | - |
| - | - | 650.6 | 669.3 | - | - | 0 | - |
| - | - | 838 | 669.4 | - | - | 0 | - |
| - | - | 680.6 | 669.8 | - | - | 0 | - |
| 5 | y | 4587 | 677.3 | 0.0003706 | 0.5471 | +2 | 13 |
| 5 | y | 7715 | 677.8 | 0.005058 | 7.462 | +2 | 13 |
| - | - | 6348 | 678.3 | - | - | 0 | - |
| - | - | 2250 | 678.8 | - | - | 0 | - |
| - | - | 632.3 | 679.3 | - | - | 0 | - |
| - | - | 912.2 | 685.3 | - | - | 0 | - |
| 5 | y | 1.237E+05 | 686.3 | 0.0002065 | 0.3008 | +2 | 13 |
| - | - | 1.023E+05 | 686.8 | - | - | 0 | - |
| - | - | 4.803E+04 | 687.3 | - | - | 0 | - |
| - | - | 1.151E+04 | 687.8 | - | - | 0 | - |
| - | - | 993.3 | 694.3 | - | - | 0 | - |
| - | - | 736.6 | 695.3 | - | - | 0 | - |
| 13 | b | 2133 | 700.4 | 0.01196 | 17.08 | +2 | 13 |
| - | - | 1043 | 700.8 | - | - | 0 | - |
| - | - | 772.4 | 704.3 | - | - | 0 | - |
| - | - | 1.057E+04 | 712.4 | - | - | 0 | - |
| - | - | 4068 | 713.4 | - | - | 0 | - |
| - | - | 1631 | 722.3 | - | - | 0 | - |
| - | - | 1.305E+04 | 740.3 | - | - | 0 | - |
| - | - | 5498 | 741.3 | - | - | 0 | - |
| - | - | 1631 | 742.3 | - | - | 0 | - |
| 7 | b | 1396 | 742.4 | 0.001392 | 1.875 | +1 | 7 |
| 11 | y | 3911 | 745.4 | 0.0003751 | 0.5033 | +1 | 7 |
| - | - | 1945 | 746.4 | - | - | 0 | - |
| 4 | y | 1014 | 751.4 | 0.008937 | 11.89 | +2 | 14 |
| - | - | 961.4 | 755.3 | - | - | 0 | - |
| 14 | b | 6355 | 757.4 | 0.006936 | 9.158 | +2 | 14 |
| - | - | 2477 | 758.4 | - | - | 0 | - |
| - | - | 1142 | 759.4 | - | - | 0 | - |
| 4 | y | 1.541E+04 | 759.9 | 3.906E-06 | 0.005141 | +2 | 14 |
| - | - | 1.683E+04 | 760.4 | - | - | 0 | - |
| - | - | 6930 | 760.9 | - | - | 0 | - |
| - | - | 1846 | 761.4 | - | - | 0 | - |
| - | - | 2241 | 771.4 | - | - | 0 | - |
| - | - | 1767 | 772.4 | - | - | 0 | - |
| 3 | y | 2545 | 816.4 | 0.0006541 | 0.8011 | +2 | 15 |
| - | - | 3648 | 816.9 | - | - | 0 | - |
| - | - | 1741 | 817.4 | - | - | 0 | - |
| - | - | 743.1 | 817.9 | - | - | 0 | - |
| - | - | 802.8 | 824.4 | - | - | 0 | - |
| - | - | 884.9 | 826.9 | - | - | 0 | - |
| 8 | b | 1057 | 829.4 | 0.003842 | 4.632 | +1 | 8 |
| - | - | 1258 | 840.4 | - | - | 0 | - |
| - | - | 1653 | 841.4 | - | - | 0 | - |
| - | - | 1922 | 842.4 | - | - | 0 | - |
| - | - | 1033 | 843.4 | - | - | 0 | - |
| 16 | b | 1176 | 848.4 | 0.005546 | 6.537 | +2 | 16 |
| - | - | 1903 | 850.4 | - | - | 0 | - |
| - | - | 1550 | 851.4 | - | - | 0 | - |
| 16 | b | 3437 | 856.9 | 0.001244 | 1.452 | +2 | 16 |
| 10 | y | 4196 | 857.4 | 0.009905 | 11.55 | +1 | 8 |
| 2 | y | 1972 | 857.9 | 0.01064 | 12.4 | +2 | 16 |
| - | - | 989.3 | 858.4 | - | - | 0 | - |
| - | - | 847 | 859.4 | - | - | 0 | - |
| 2 | y | 1333 | 866.9 | 0.0003234 | 0.373 | +2 | 16 |
| - | - | 1048 | 867.4 | - | - | 0 | - |
| - | - | 847.6 | 867.9 | - | - | 0 | - |
| - | - | 1.494E+04 | 868.4 | - | - | 0 | - |
| - | - | 8395 | 869.4 | - | - | 0 | - |
| - | - | 2260 | 870.4 | - | - | 0 | - |
| 10 | y | 2436 | 874.5 | 0.001691 | 1.933 | +1 | 8 |
| - | - | 1954 | 875.5 | - | - | 0 | - |
| 0 | Precursor | 1586 | 907.5 | 0.006787 | 7.479 | +2 | -1 |
| 0 | Precursor | 3565 | 908 | 0.007492 | 8.251 | +2 | -1 |
| - | - | 2808 | 908.5 | - | - | 0 | - |
| - | - | 1022 | 909 | - | - | 0 | - |
| - | - | 2618 | 911.4 | - | - | 0 | - |
| - | - | 1812 | 912.4 | - | - | 0 | - |
| - | - | 751 | 913.4 | - | - | 0 | - |
| 0 | Precursor | 2.033E+04 | 916.5 | 0.0007169 | 0.7822 | +2 | -1 |
| - | - | 2.732E+04 | 917 | - | - | 0 | - |
| - | - | 1.627E+04 | 917.5 | - | - | 0 | - |
| - | - | 7530 | 918 | - | - | 0 | - |
| - | - | 918.7 | 921.4 | - | - | 0 | - |
| - | - | 2105 | 922.4 | - | - | 0 | - |
| - | - | 1092 | 923.4 | - | - | 0 | - |
| - | - | 1287 | 927.3 | - | - | 0 | - |
| - | - | 749.6 | 937.4 | - | - | 0 | - |
| - | - | 1.236E+04 | 939.4 | - | - | 0 | - |
| - | - | 7640 | 940.4 | - | - | 0 | - |
| - | - | 2412 | 941.4 | - | - | 0 | - |
| - | - | 744.2 | 954.5 | - | - | 0 | - |
| - | - | 1612 | 956.4 | - | - | 0 | - |
| - | - | 1272 | 957.4 | - | - | 0 | - |
| 9 | b | 757.6 | 958.5 | 0.008837 | 9.22 | +1 | 9 |
| - | - | 964 | 971.5 | - | - | 0 | - |
| 9 | y | 2525 | 1004 | 0.0001499 | 0.1494 | +1 | 9 |
| - | - | 1251 | 1005 | - | - | 0 | - |
| - | - | 980.9 | 1005 | - | - | 0 | - |
| - | - | 1120 | 1015 | - | - | 0 | - |
| - | - | 855.9 | 1024 | - | - | 0 | - |
| - | - | 861.2 | 1025 | - | - | 0 | - |
| - | - | 833 | 1035 | - | - | 0 | - |
| - | - | 1928 | 1036 | - | - | 0 | - |
| - | - | 1749 | 1037 | - | - | 0 | - |
| - | - | 4043 | 1040 | - | - | 0 | - |
| - | - | 2689 | 1041 | - | - | 0 | - |
| - | - | 853.4 | 1042 | - | - | 0 | - |
| - | - | 1.063E+04 | 1053 | - | - | 0 | - |
| - | - | 5322 | 1054 | - | - | 0 | - |
| - | - | 2969 | 1055 | - | - | 0 | - |
| - | - | 1520 | 1056 | - | - | 0 | - |
| - | - | 1577 | 1059 | - | - | 0 | - |
| - | - | 875.8 | 1068 | - | - | 0 | - |
| - | - | 689.6 | 1069 | - | - | 0 | - |
| - | - | 2749 | 1071 | - | - | 0 | - |
| - | - | 1986 | 1072 | - | - | 0 | - |
| 8 | y | 1408 | 1073 | 0.01826 | 17.03 | +1 | 10 |
| - | - | 1581 | 1085 | - | - | 0 | - |
| - | - | 1410 | 1086 | - | - | 0 | - |
| - | - | 1197 | 1087 | - | - | 0 | - |
| 10 | b | 1822 | 1088 | 0.003762 | 3.459 | +1 | 10 |
| 8 | y | 2559 | 1091 | 0.002338 | 2.143 | +1 | 10 |
| - | - | 2480 | 1092 | - | - | 0 | - |
| - | - | 1467 | 1093 | - | - | 0 | - |
| - | - | 1495 | 1138 | - | - | 0 | - |
| - | - | 2574 | 1139 | - | - | 0 | - |
| - | - | 1455 | 1140 | - | - | 0 | - |
| - | - | 3507 | 1156 | - | - | 0 | - |
| - | - | 2916 | 1157 | - | - | 0 | - |
| - | - | 901.2 | 1158 | - | - | 0 | - |
| 7 | y | 1167 | 1160 | 0.001101 | 0.9496 | +1 | 11 |
| 7 | y | 1153 | 1161 | 0.004756 | 4.098 | +1 | 11 |
| - | - | 2307 | 1165 | - | - | 0 | - |
| - | - | 746.8 | 1166 | - | - | 0 | - |
| 7 | y | 5403 | 1178 | 0.004215 | 3.579 | +1 | 11 |
| - | - | 3189 | 1179 | - | - | 0 | - |
| - | - | 1138 | 1180 | - | - | 0 | - |
| - | - | 1.008E+04 | 1182 | - | - | 0 | - |
| 11 | b | 8829 | 1183 | 0.02221 | 18.78 | +1 | 11 |
| - | - | 3425 | 1184 | - | - | 0 | - |
| 11 | b | 1496 | 1201 | 0.004696 | 3.911 | +1 | 11 |
| - | - | 900.2 | 1218 | - | - | 0 | - |
| - | - | 917.5 | 1225 | - | - | 0 | - |
| - | - | 3133 | 1235 | - | - | 0 | - |
| - | - | 5470 | 1236 | - | - | 0 | - |
| - | - | 1884 | 1237 | - | - | 0 | - |
| - | - | 9724 | 1253 | - | - | 0 | - |
| - | - | 8055 | 1254 | - | - | 0 | - |
| - | - | 3317 | 1255 | - | - | 0 | - |
| 6 | y | 1151 | 1257 | 0.003757 | 2.99 | +1 | 12 |
| 6 | y | 2358 | 1258 | 0.005215 | 4.147 | +1 | 12 |
| - | - | 2036 | 1259 | - | - | 0 | - |
| 6 | y | 2.424E+04 | 1275 | 0.002291 | 1.797 | +1 | 12 |
| - | - | 1.955E+04 | 1276 | - | - | 0 | - |
| - | - | 7270 | 1277 | - | - | 0 | - |
| - | - | 1418 | 1278 | - | - | 0 | - |
| - | - | 758.7 | 1310 | - | - | 0 | - |
| 12 | b | 1196 | 1311 | 0.002223 | 1.696 | +1 | 12 |
| 12 | b | 1038 | 1312 | 0.02553 | 19.47 | +1 | 12 |
| 12 | b | 3183 | 1329 | 0.004436 | 3.338 | +1 | 12 |
| - | - | 2572 | 1330 | - | - | 0 | - |
| - | - | 1220 | 1331 | - | - | 0 | - |
| - | - | 1113 | 1337 | - | - | 0 | - |
| 5 | y | 3764 | 1354 | 0.001399 | 1.034 | +1 | 13 |
| 5 | y | 8419 | 1355 | 0.004209 | 3.107 | +1 | 13 |
| - | - | 5489 | 1356 | - | - | 0 | - |
| - | - | 2310 | 1357 | - | - | 0 | - |
| 5 | y | 1.451E+05 | 1372 | 0.002931 | 2.137 | +1 | 13 |
| - | - | 1.154E+05 | 1373 | - | - | 0 | - |
| - | - | 5.499E+04 | 1374 | - | - | 0 | - |
| - | - | 1.323E+04 | 1375 | - | - | 0 | - |
| - | - | 784.5 | 1376 | - | - | 0 | - |
| 13 | b | 743.5 | 1383 | 0.004775 | 3.453 | +1 | 13 |
| 13 | b | 2920 | 1400 | 0.01189 | 8.492 | +1 | 13 |
| - | - | 1844 | 1401 | - | - | 0 | - |
| 4 | y | 7331 | 1519 | 0.003718 | 2.448 | +1 | 14 |
| - | - | 8400 | 1520 | - | - | 0 | - |
| - | - | 3099 | 1521 | - | - | 0 | - |
| - | - | 835.1 | 1522 | - | - | 0 | - |
| 3 | y | 1368 | 1632 | 0.00255 | 1.563 | +1 | 15 |
| - | - | 1411 | 1633 | - | - | 0 | - |

m/z Charge Intensity FragmentType MassShift Position
120.06591033935547 0 847.87177 y 16
120.08106231689453 0 15990.141
121.08441162109375 0 960.46686
125.9787368774414 0 386.50055
126.05526733398438 0 3821.5325
126.6500473022461 0 354.8049
127.0864486694336 0 457.42297
128.08193969726562 0 517.5595
128.10726928710938 0 3951.7712
129.0659942626953 0 820.5157
129.10252380371094 0 28670.617
130.05007934570312 0 802.3422
130.10577392578125 0 1458.5027
131.11813354492188 0 662.8549
133.06094360351562 0 402.5875
136.07611083984375 0 1893.7458
138.05508422851562 0 738.3449
138.09185791015625 0 803.8626
138.1280975341797 0 1906.9012
139.08682250976562 0 837.4682
141.06597900390625 0 691.613
141.1026153564453 0 441.66125
147.07676696777344 0 1220.4224
148.2191619873047 0 438.0574
148.80274963378906 0 571.76917
148.87440490722656 0 406.10803
148.88162231445312 0 544.2552
148.89617919921875 0 643.7661
148.903076171875 0 793.11017
148.9105682373047 0 841.4539
148.91786193847656 0 930.9458
148.9248809814453 0 1368.6783
148.93199157714844 0 1287.906
148.93948364257812 0 3308.7168
148.95584106445312 0 6215.519
148.963623046875 0 3691.6357
148.97120666503906 0 1711.0353
148.97825622558594 0 1248.9752
148.9854278564453 0 1263.9229
148.99267578125 0 711.1305
149.00021362304688 0 674.66925
149.00709533691406 0 803.31744
149.0140380859375 0 649.9568
149.02114868164062 0 594.57764
149.0282745361328 0 438.58456
155.08180236816406 0 1379.8408
155.11817932128906 0 3739.5474 a Water loss 1
157.0608673095703 0 1329.3878
157.09744262695312 0 3860.3418
165.0653076171875 0 435.50922
165.10243225097656 0 708.1829
167.08184814453125 0 2452.2554
167.1179656982422 0 1661.3398
169.13377380371094 0 3901.673
170.1376953125 0 489.695
172.10818481445312 0 583.80396
173.0562286376953 0 460.68927
173.09249877929688 0 777.7351 y Water loss 15
173.12875366210938 0 12778.308 a 1
174.13229370117188 0 1066.6414
175.07138061523438 0 1124.3904
181.09744262695312 0 763.2866
182.0930633544922 0 1162.4402
182.1291961669922 0 1630.423
183.0768585205078 0 1261.3431
183.1130828857422 0 48518.88 b Water loss 1
184.10836791992188 0 1506.6523
184.11666870117188 0 5516.7974
185.09230041503906 0 4616.804
186.08741760253906 0 1174.8588
187.1444549560547 0 8118.962
189.0868682861328 0 1259.3561
191.08203125 0 804.0046
191.1029815673828 0 1096.797 y 15
195.1129913330078 0 4061.5015
197.12869262695312 0 5040.0713
198.12379455566406 0 1235.3818
199.07186889648438 0 2226.0771
199.10838317871094 0 522.3294
200.1033477783203 0 1540.5789
200.13963317871094 0 1588.2842
201.1236114501953 0 25269.244 b 1
202.1271209716797 0 2212.8362
204.08685302734375 0 666.1205
208.10829162597656 0 2091.8833
209.09230041503906 0 5965.923
213.08653259277344 0 702.922
214.15541076660156 0 678.848
215.13926696777344 0 8763.949
216.14283752441406 0 1191.7449
217.08221435546875 0 4088.676
217.1337890625 0 5049.0596
218.1372528076172 0 766.84814
218.14968872070312 0 658.15594
219.1494598388672 0 2099.2937
220.22666931152344 0 509.6156
225.1237030029297 0 3025.2908
225.13458251953125 0 892.9619
226.11886596679688 0 21047.525
227.10304260253906 0 722.4247
227.1226806640625 0 1912.1178
233.1651153564453 0 12257.697
234.16844177246094 0 1421.6035
236.10353088378906 0 1255.4066
241.08261108398438 0 1178.5604
242.1502685546875 0 3457.4783
243.13327026367188 0 1032.5403
243.14564514160156 0 6843.886
244.1298065185547 0 1615.3208 y Ammonia loss 12
244.14883422851562 0 585.97815
245.12881469726562 0 4134.765
247.14451599121094 0 2345.4348
251.1757354736328 0 5848.6543
252.13389587402344 0 1228.4652
254.11387634277344 0 5283.399
254.14988708496094 0 1018.66785
255.11737060546875 0 709.39294
256.1295471191406 0 692.3469
259.0927734375 0 2475.6208
261.1600341796875 0 3754.8474
262.1642150878906 0 730.2162
264.1346130371094 0 1523.0469
268.2012023925781 0 1224.2527 a Water loss 2
269.1249694824219 0 730.63916
269.1856689453125 0 1775.3857
270.1448669433594 0 1268.2301
270.1800537109375 0 712.5432
272.124267578125 0 2904.1995
274.13983154296875 0 622.6242
275.17547607421875 0 2369.374
279.1450500488281 0 2525.016
280.13018798828125 0 963.4795
280.16583251953125 0 4337.9043
282.1451416015625 0 5433.993
283.1473693847656 0 991.4176
285.159912109375 0 2528.3608
286.1034851074219 0 2306.729
294.1823425292969 0 860.9131
296.08807373046875 0 721.6321
296.19732666015625 0 122696.37 b Water loss 2
297.1200256347656 0 811.2256
297.15625 0 6823.4585
297.2005310058594 0 20177.96
298.1593017578125 0 914.0329
298.1767883300781 0 4189.361
298.2031555175781 0 1653.3531
303.1705627441406 0 4004.4683
304.1144714355469 0 1416.3845
304.1728210449219 0 1225.3344
308.1606750488281 0 780.0128 y Ammonia loss 11
312.1169738769531 0 595.56
313.18804931640625 0 635.97217
314.1477355957031 0 762.4117
314.1834716796875 0 5405.713
314.20794677734375 0 29726.312 b 2
315.1909484863281 0 828.2578
315.2109375 0 5566.1006
317.1857604980469 0 1762.3953
319.1981201171875 0 7426.6025 y 14
320.2007751464844 0 1437.0925
328.11444091796875 0 2282.4868
330.1814270019531 0 3625.1345
333.15679931640625 0 681.98413
334.2127380371094 0 939.04736
337.1510314941406 0 1006.81415
339.1676330566406 0 1374.9056
341.18408203125 0 2311.705
342.18231201171875 0 961.22284
344.19775390625 0 1493.6371
346.1246643066406 0 4785.233
346.17645263671875 0 4697.203
347.1303405761719 0 634.9313
348.1920166015625 0 4668.576
349.19549560546875 0 787.4723
350.181640625 0 938.5768
351.16668701171875 0 3816.2043
352.1683654785156 0 931.43665
353.18243408203125 0 1990.6499
354.1657409667969 0 1261.2916
355.1607360839844 0 1117.8485
360.2283020019531 0 697.51685
362.20770263671875 0 9874.563
363.21002197265625 0 1779.359
365.14501953125 0 1777.0306
367.2099609375 0 928.7899
368.1929626464844 0 1112.5681
369.1769104003906 0 4819.904
369.215087890625 0 1161.6716
370.17755126953125 0 816.91614
371.19403076171875 0 1342.6306
372.1763610839844 0 1417.9392
373.1722412109375 0 1633.3873
381.1748962402344 0 697.9258
383.1564636230469 0 3507.866
383.26654052734375 0 537.0858
385.2196960449219 0 3559.6704
393.1775817871094 0 875.617
393.2502746582031 0 1732.8663
395.1937255859375 0 684.94696
397.1364440917969 0 782.925
398.2434387207031 0 2405.0957
399.2467956542969 0 635.6458
400.2221374511719 0 694.5021
401.1671142578125 0 5399.2856
402.169921875 0 966.9539
407.20281982421875 0 954.3863
408.1885070800781 0 1461.8352
411.2613525390625 0 2610.1716
412.2640686035156 0 688.2645
415.1464538574219 0 2059.3516
415.2713928222656 0 1195.6998
416.1518249511719 0 629.5145
416.2549743652344 0 4447.6357
417.2578125 0 1105.0381
423.1854553222656 0 656.00635
424.2300109863281 0 866.7102
425.1318054199219 0 728.92377
425.2141418457031 0 1757.3148
427.2332763671875 0 839.2239
433.15692138671875 0 2345.5679
433.2423400878906 0 5497.032 y 13
433.28155517578125 0 38174.848
434.1585998535156 0 672.71747
434.2473449707031 0 1092.3082
434.28466796875 0 11814.941
435.2870788574219 0 684.6191
440.2149658203125 0 605.06494
441.1976623535156 0 1132.0458
442.2386474609375 0 1805.7972
443.1432800292969 0 860.9696
443.2286682128906 0 1961.6459
443.2660217285156 0 4003.1597 b Water loss 3
444.1903076171875 0 606.37537
444.268798828125 0 1428.5212
452.2154235839844 0 832.76794
459.2085266113281 0 1428.2223
461.2765197753906 0 29339.916 b 3
462.1982116699219 0 1383.2162
462.27978515625 0 9740.703
463.2001953125 0 865.3509
463.2816467285156 0 1436.709
464.2161560058594 0 955.88495
466.19287109375 0 890.4378
469.2415466308594 0 938.0245
470.2257080078125 0 4525.957
471.2285461425781 0 838.9985
477.2555847167969 0 669.99506
478.2374267578125 0 703.2628
479.2267150878906 0 1021.0273
480.2098693847656 0 3929.497
481.2135925292969 0 983.68585
484.2039794921875 0 1444.175
487.2535705566406 0 1216.466 y Ammonia loss 12
494.18890380859375 0 1224.5027
495.2669982910156 0 1172.4875
496.24725341796875 0 1142.0566
498.2201843261719 0 11978.676
499.2228088378906 0 3291.4097
500.2328796386719 0 1156.0444
502.2149353027344 0 2999.2625
503.21673583984375 0 711.966
504.2781982421875 0 8141.9233 y 12
505.2796936035156 0 1866.0492
512.1993408203125 0 1844.4308
513.2779541015625 0 1234.3207
529.2276611328125 0 859.5278
530.2100219726562 0 8942.208
530.3345336914062 0 2534.6323
531.2127685546875 0 2366.4778
531.33984375 0 1390.3705
538.2990112304688 0 1465.4491
540.1965942382812 0 930.754
540.3192138671875 0 1435.5624 b Water loss 4
541.3213500976562 0 699.05005
546.239501953125 0 828.7912
551.3141479492188 0 741.207
555.3182373046875 0 626.8391
558.3286743164062 0 3836.166 b 4
559.3321533203125 0 2165.4578
563.2496948242188 0 693.7959
569.281982421875 0 814.8001
569.330810546875 0 889.90314
569.7785034179688 0 737.4458
578.28515625 0 1738.7479
578.7896728515625 0 1109.3148
579.2867431640625 0 614.1767
581.2535400390625 0 1155.6582
582.2909545898438 0 799.7987
582.7825927734375 0 2527.8152
583.2817993164062 0 865.70746
591.2406616210938 0 1047.8945
591.293701171875 0 1191.3875
591.7938842773438 0 1579.7617 b Water loss 10
597.2913818359375 0 1720.848
598.2821655273438 0 615.9633
599.267578125 0 8505.994
600.2721557617188 0 2301.717
608.3507690429688 0 743.07367
608.8017578125 0 746.3812
609.2509765625 0 2357.4983
610.251953125 0 900.36285
612.8157958984375 0 1986.3733
613.3119506835938 0 1069.3197
614.3251342773438 0 1818.8794 y Water loss 11
615.303955078125 0 4266.6514 y Ammonia loss 11
616.3045654296875 0 2321.5925
617.806396484375 0 7154.871
618.3046875 0 7089.199
618.8035888671875 0 3820.595
619.3052978515625 0 1441.516
625.2861328125 0 894.8751
626.2803955078125 0 985.5189
626.365234375 0 903.3089
626.8123168945312 0 18446.707
627.2617797851562 0 18231.195
627.314208984375 0 11708.195
627.8134765625 0 6008.0166
628.265625 0 7038.099
629.2672119140625 0 1875.2368
630.3023681640625 0 668.32275
632.3358764648438 0 4576.8164 y 11
633.3390502929688 0 2115.8582
634.3418579101562 0 733.7409
635.8175659179688 0 1122.07
636.320556640625 0 697.503
637.8150634765625 0 3030.17 y 5
638.3168334960938 0 1847.0387
640.292236328125 0 774.1855
643.2935180664062 0 3666.7185
644.2953491210938 0 2425.3567
655.33935546875 0 1000.8281
655.8429565429688 0 792.9127 b Water loss 11
660.3187866210938 0 1966.3145
661.3209228515625 0 617.7936
663.3350219726562 0 1093.811
663.8433227539062 0 881.58636
668.3240966796875 0 981.46716
668.3945922851562 0 1230.4917
668.8264770507812 0 1110.5027
669.3280029296875 0 650.5872
669.3934326171875 0 837.9788
669.8279418945312 0 680.5753
677.3355102539062 0 4586.6597 y Water loss 4
677.8329467773438 0 7715.032 y Ammonia loss 4
678.3297729492188 0 6348.023
678.8321533203125 0 2250.2551
679.3287963867188 0 632.3129
685.324462890625 0 912.1708
686.3413696289062 0 123724.055 y 4
686.8424072265625 0 102312.02
687.343017578125 0 48030.086
687.8451538085938 0 11507.616
694.3374633789062 0 993.2918
695.3389282226562 0 736.557
700.3468627929688 0 2132.8958 b 12
700.84521484375 0 1043.15
704.3264770507812 0 772.4065
712.3511962890625 0 10573.566
713.3538818359375 0 4068.011
722.3369140625 0 1631.262
740.346435546875 0 13053.653
741.3486938476562 0 5497.974
742.3480834960938 0 1631.1653
742.414794921875 0 1396.1738 b 6
745.420654296875 0 3910.5056 y 10
746.4193725585938 0 1945.0034
751.3710327148438 0 1014.12335 y Ammonia loss 3
755.3278198242188 0 961.41327
757.3733520507812 0 6354.9854 b 13
758.3746948242188 0 2477.2861
759.3750610351562 0 1141.8589
759.8753662109375 0 15412.339 y 3
760.3760986328125 0 16833.164
760.8775024414062 0 6929.6274
761.3762817382812 0 1846.1594
771.3506469726562 0 2240.7305
772.35205078125 0 1766.7825
816.416748046875 0 2544.7466 y 2
816.9185180664062 0 3648.3384
817.4205322265625 0 1740.768
817.9127197265625 0 743.1348
824.3848876953125 0 802.8149
826.8512573242188 0 884.9264
829.4415893554688 0 1056.7152 b 7
840.4144287109375 0 1258.1871
841.4047241210938 0 1653.0747
842.39111328125 0 1921.729
843.3892822265625 0 1032.8601
848.4385986328125 0 1176.0116 b Ammonia loss 15
850.39794921875 0 1902.9329
851.3923950195312 0 1549.8353
856.9475708007812 0 3436.5674 b 15
857.4462280273438 0 4196.4053 y Ammonia loss 9
857.9465942382812 0 1972.1329 y Water loss 1
858.4461059570312 0 989.2707
859.4188842773438 0 847.02155
866.94091796875 0 1332.7004 y 1
867.4312744140625 0 1047.7572
867.9411010742188 0 847.6339
868.4043579101562 0 14941.658
869.4070434570312 0 8394.638
870.4071655273438 0 2259.949
874.461181640625 0 2435.5298 y 9
875.4597778320312 0 1954.0609
907.46337890625 0 1585.7019 Precursor Water loss
907.9696655273438 0 3565.0193 Precursor Ammonia loss
908.4667358398438 0 2808.0896
908.9632568359375 0 1022.1594
911.4452514648438 0 2617.9893
912.4456176757812 0 1812.0359
913.431640625 0 751.038
916.4747314453125 0 20325.17 Precursor
916.97607421875 0 27323.008
917.4766235351562 0 16274.863
917.9771118164062 0 7529.8994
921.422607421875 0 918.7356
922.4254760742188 0 2105.2393
923.4219970703125 0 1092.1163
927.25146484375 0 1287.1519
937.4364013671875 0 749.5523
939.4410400390625 0 12357.871
940.443359375 0 7639.702
941.4470825195312 0 2411.8098
954.4589233398438 0 744.2088
956.44384765625 0 1612.3992
957.4402465820312 0 1272.2567
958.4791870117188 0 757.58594 b 8
971.47021484375 0 963.9903
1003.505615234375 0 2525.4048 y 8
1004.50341796875 0 1250.7621
1005.4946899414062 0 980.88947
1015.4689331054688 0 1120.1069
1024.47705078125 0 855.8584
1025.491943359375 0 861.16003
1035.468017578125 0 833.0239
1036.4615478515625 0 1928.1691
1037.4669189453125 0 1748.7903
1040.4903564453125 0 4042.884
1041.49267578125 0 2689.0376
1042.490966796875 0 853.42664
1053.48388671875 0 10630.587
1054.4842529296875 0 5321.6724
1055.4808349609375 0 2969.068
1056.4879150390625 0 1519.5809
1058.511962890625 0 1577.0557
1067.5008544921875 0 875.7697
1068.516845703125 0 689.62225
1070.5155029296875 0 2748.6318
1071.5098876953125 0 1986.0623
1072.5086669921875 0 1408.3511 y Water loss 7
1084.5269775390625 0 1581.3108
1085.53125 0 1410.4591
1086.5128173828125 0 1196.925
1087.52685546875 0 1822.0199 b 9
1090.53515625 0 2558.8203 y 7
1091.5389404296875 0 2479.801
1092.535400390625 0 1467.397
1137.5560302734375 0 1494.8752
1138.5465087890625 0 2574.114
1139.5487060546875 0 1454.8944
1155.5615234375 0 3506.9143
1156.5670166015625 0 2916.349
1157.55859375 0 901.2402
1159.56005859375 0 1167.4954 y Water loss 6
1160.5477294921875 0 1152.5859 y Ammonia loss 6
1164.5560302734375 0 2306.775
1165.5625 0 746.80865
1177.5653076171875 0 5402.615 y 6
1178.5701904296875 0 3188.8381
1179.5673828125 0 1138.057
1181.578369140625 0 10079.028
1182.5819091796875 0 8828.634 b Water loss 10
1183.5845947265625 0 3425.0864
1200.6099853515625 0 1496.2894 b 10
1217.5880126953125 0 900.2184
1224.6190185546875 0 917.49384
1234.6055908203125 0 3133.1345
1235.5980224609375 0 5469.98
1236.6005859375 0 1883.9939
1252.6134033203125 0 9724.474
1253.6151123046875 0 8055.001
1254.6214599609375 0 3316.8103
1256.615478515625 0 1151.2529 y Water loss 5
1257.6009521484375 0 2358.3118 y Ammonia loss 5
1258.5928955078125 0 2035.6927
1274.6199951171875 0 24237.297 y 5
1275.621337890625 0 19550.582
1276.6234130859375 0 7270.3394
1277.63525390625 0 1418.4172
1309.6796875 0 758.6604
1310.6649169921875 0 1195.8563 b Water loss 11
1311.6722412109375 0 1038.2944 b Ammonia loss 11
1328.6688232421875 0 3183.4426 b 11
1329.6707763671875 0 2571.8008
1330.676025390625 0 1220.3296
1336.6361083984375 0 1113.4852
1353.6630859375 0 3764.2585 y Water loss 4
1354.6527099609375 0 8419.12 y Ammonia loss 4
1355.65478515625 0 5489.214
1356.6500244140625 0 2310.3987
1371.672119140625 0 145108.12 y 4
1372.674072265625 0 115388.695
1373.6761474609375 0 54991.273
1374.677490234375 0 13233.489
1375.6834716796875 0 784.4869
1382.6885986328125 0 743.53925 b Ammonia loss 12
1399.698486328125 0 2920.1934 b 12
1400.7021484375 0 1843.5844
1518.73974609375 0 7331.112 y 3
1519.7413330078125 0 8400.023
1520.742431640625 0 3099.1187
1521.753173828125 0 835.0565
1631.830078125 0 1368.3357 y 2
1632.8245849609375 0 1411.3242

Spectrum Details

|  |  |
| --- | --- |
| Matched peaks? Matched peaks The total absolute number of peaks matched. Additionally in brackets the total fraction of peaks matched and the total number of peaks is shown. | 69 (13.42% of 514) |
| FDR? FDR The false discovery rate estimated for this peptide. It is calculated by matching all theoretical fragments with a non-integer shift with the raw peaks for this spectrum. This is done with 40 different shifts. The resulting percentage is the average number of annotated peaks over the number of annotated peaks with the correct spectrum. | 0.66% |
| Satellite FDR? Satellite FDR See the FDR for details on its calculation. This satellite ion specific FDR only contains the satellite ions (d/w) for I/L/J positions. | - |
| PSM Score? PSM Score The PSM Score as given by Hecklib to this annotated spectrum. It is shown with three significant figures. | 730 |

#### Spectrum 8129? Spectrum 8129 The raw spectrum of this peptide as annotated by Hecklib. The fragments are coloured according to ion type (see legend). Any peaks with a star '\*' as text can be hovered over to see the full details, first the ion type second the mass shift type. By hovering over the amino acids in the peptide or ions in the legend the corresponding peaks are highlighted. By toggling the 'Unassigned' label you can turn the background (unassigned) peaks on or off in the plot. By updating the slider in the Ion legend you can update the spectrum to only show the top X% of the peaks with labels. The top X% means any peak that is within X% of the highest intensity. By dragging in the spectrum you can zoom in to a specific part of the spectrum and use 'Zoom Out' to get back to the original zoom level. The annotation of the spectrum is based on the given sequence in the peptides file and is done with different software so inconsistencies are likely. The peaks are annotated based on the given sequence, with 20 ppm tolerance.

Copy Data

##### Spectrum 8129 (TSV)

###### Preview

```
Loading example...
```

*Click on the button to copy the data to your clipboard.*

Mz MinMz MaxIntensity Max

WidthHeightPeptide font sizePeptide stroke widthSpectrum font sizeSpectrum stroke widthCompact peptide

Ion legend

wxyz

abcd

OtherUnassignedIon^Charge^_Position_Show for top:%

VTJFPPSSEEJQANKAT

02.00e+44.00e+45.99e+47.99e+4

Zoom Out

y^+1^_1_a^+1^_2_y^+1^_2_a^+1^_2_b^+1^_2_y^+1^_2_b^+1^_2_b^+3^_6_y^+3^_7_y^+2^_5_a^+1^_3_b^+2^_5_b^+1^_3_y^+1^_3_y^+2^_6_y^+2^_6_b^+1^_3_y^+2^_6_y^+1^_3_y^+2^_7_y^+2^_7_b^+2^_8_b^+2^_8_y^+1^_4_y^+2^_8_y^+1^_4_b^+1^_4_y^+3^_13_y^+3^_13_b^+1^_4_b^+2^_9_y^+1^_5_y^+1^_5_y^+3^_14_y^+3^_14_y^+1^_5_b^+1^_5_b^+2^_10_b^+3^_15_b^+1^_5_y^+2^_11_b^+2^_11_y^+1^_6_y^+1^_6_y^+2^_12_y^+1^_6_y^+2^_12_b^+2^_12_y^+2^_13_y^+2^_13_y^+2^_13_b^+2^_13_b^+2^_13_b^+1^_7_y^+1^_7_y^+1^_7_b^+1^_7_y^+1^_7_y^+2^_14_y^+2^_14_b^+2^_14_y^+2^_14_y^+2^_15_y^+2^_15_b^+1^_8_y^+2^_15_b^+1^_8_b^+2^_16_y^+1^_8_b^+2^_16_y^+1^_8_y^+2^_16_y^+1^_8_b^+1^_9_y^+1^_9_y^+1^_9_b^+1^_10_y^+1^_10_b^+1^_10_y^+1^_10_y^+1^_11_b^+1^_11_b^+1^_11_y^+1^_12_y^+1^_12_y^+1^_12_y^+1^_13_b^+1^_13_b^+1^_14_

0775155123263102

Fragment Matches Table

Show background peaks

| Position | Ion type | Intensity | mz Theoretical | mz Error (Th) | mz Error (ppm) | Charge | Series Number |
| --- | --- | --- | --- | --- | --- | --- | --- |
| 17 | y | 3.926E+04 | 120.1 | 0.0003678 | 3.064 | +1 | 1 |
| - | - | 3.265E+04 | 120.1 | - | - | 0 | - |
| - | - | 327.9 | 120.1 | - | - | 0 | - |
| - | - | 1302 | 121.1 | - | - | 0 | - |
| - | - | 2243 | 121.1 | - | - | 0 | - |
| - | - | 400.9 | 123.1 | - | - | 0 | - |
| - | - | 4058 | 126.1 | - | - | 0 | - |
| - | - | 1.168E+04 | 128.1 | - | - | 0 | - |
| - | - | 4132 | 129.1 | - | - | 0 | - |
| - | - | 2074 | 129.1 | - | - | 0 | - |
| - | - | 4.796E+04 | 129.1 | - | - | 0 | - |
| - | - | 3519 | 130.1 | - | - | 0 | - |
| - | - | 1177 | 130.1 | - | - | 0 | - |
| - | - | 555.2 | 130.1 | - | - | 0 | - |
| - | - | 3802 | 130.1 | - | - | 0 | - |
| - | - | 2837 | 131.1 | - | - | 0 | - |
| - | - | 911.1 | 132.1 | - | - | 0 | - |
| - | - | 408 | 134.7 | - | - | 0 | - |
| - | - | 388.5 | 135 | - | - | 0 | - |
| - | - | 2492 | 136.1 | - | - | 0 | - |
| - | - | 1951 | 138.1 | - | - | 0 | - |
| - | - | 1086 | 138.1 | - | - | 0 | - |
| - | - | 1410 | 139.1 | - | - | 0 | - |
| - | - | 1657 | 141.1 | - | - | 0 | - |
| - | - | 479.4 | 147.1 | - | - | 0 | - |
| - | - | 1075 | 147.1 | - | - | 0 | - |
| - | - | 2536 | 149 | - | - | 0 | - |
| - | - | 754.6 | 152.1 | - | - | 0 | - |
| - | - | 3920 | 153 | - | - | 0 | - |
| - | - | 523.6 | 153.1 | - | - | 0 | - |
| - | - | 836 | 153.1 | - | - | 0 | - |
| - | - | 1279 | 154.1 | - | - | 0 | - |
| - | - | 1880 | 154.1 | - | - | 0 | - |
| - | - | 3186 | 155.1 | - | - | 0 | - |
| 2 | a | 6679 | 155.1 | 0.0003508 | 2.262 | +1 | 2 |
| - | - | 1272 | 157.1 | - | - | 0 | - |
| - | - | 3497 | 157.1 | - | - | 0 | - |
| - | - | 4033 | 159.1 | - | - | 0 | - |
| - | - | 1527 | 165.1 | - | - | 0 | - |
| - | - | 4113 | 167.1 | - | - | 0 | - |
| - | - | 2374 | 167.1 | - | - | 0 | - |
| - | - | 1667 | 167.1 | - | - | 0 | - |
| - | - | 520.5 | 169.1 | - | - | 0 | - |
| - | - | 8314 | 169.1 | - | - | 0 | - |
| - | - | 1784 | 171 | - | - | 0 | - |
| - | - | 531.8 | 171.1 | - | - | 0 | - |
| - | - | 1747 | 172.1 | - | - | 0 | - |
| 16 | y | 6108 | 173.1 | 0.0003843 | 2.22 | +1 | 2 |
| 2 | a | 2.519E+04 | 173.1 | 0.0003605 | 2.082 | +1 | 2 |
| - | - | 1417 | 173.5 | - | - | 0 | - |
| - | - | 1759 | 174.1 | - | - | 0 | - |
| - | - | 1935 | 175.1 | - | - | 0 | - |
| - | - | 468.8 | 176.1 | - | - | 0 | - |
| - | - | 4371 | 177.1 | - | - | 0 | - |
| - | - | 428.9 | 177.8 | - | - | 0 | - |
| - | - | 1656 | 179 | - | - | 0 | - |
| - | - | 473 | 179.1 | - | - | 0 | - |
| - | - | 476.1 | 179.8 | - | - | 0 | - |
| - | - | 1201 | 181.1 | - | - | 0 | - |
| - | - | 2750 | 182.1 | - | - | 0 | - |
| - | - | 2262 | 182.1 | - | - | 0 | - |
| - | - | 551.8 | 182.2 | - | - | 0 | - |
| - | - | 2865 | 183.1 | - | - | 0 | - |
| 2 | b | 3.398E+04 | 183.1 | 0.000355 | 1.939 | +1 | 2 |
| - | - | 2469 | 184.1 | - | - | 0 | - |
| - | - | 3505 | 184.1 | - | - | 0 | - |
| - | - | 5627 | 185.1 | - | - | 0 | - |
| - | - | 2092 | 186.1 | - | - | 0 | - |
| - | - | 736.4 | 187.1 | - | - | 0 | - |
| - | - | 1.14E+04 | 187.1 | - | - | 0 | - |
| - | - | 3034 | 189.1 | - | - | 0 | - |
| - | - | 602 | 191.1 | - | - | 0 | - |
| 16 | y | 6792 | 191.1 | 0.0003787 | 1.982 | +1 | 2 |
| - | - | 1277 | 191.1 | - | - | 0 | - |
| - | - | 3530 | 195 | - | - | 0 | - |
| - | - | 541.3 | 195.1 | - | - | 0 | - |
| - | - | 4954 | 195.1 | - | - | 0 | - |
| - | - | 777.6 | 197.1 | - | - | 0 | - |
| - | - | 1.259E+04 | 197.1 | - | - | 0 | - |
| - | - | 2568 | 198.1 | - | - | 0 | - |
| - | - | 1230 | 198.1 | - | - | 0 | - |
| - | - | 3413 | 199.1 | - | - | 0 | - |
| - | - | 1194 | 199.1 | - | - | 0 | - |
| - | - | 3160 | 200.1 | - | - | 0 | - |
| - | - | 2746 | 200.1 | - | - | 0 | - |
| - | - | 723.1 | 201.1 | - | - | 0 | - |
| 2 | b | 3.232E+04 | 201.1 | 0.0002884 | 1.434 | +1 | 2 |
| - | - | 2345 | 202.1 | - | - | 0 | - |
| - | - | 772.3 | 203.1 | - | - | 0 | - |
| - | - | 540.6 | 203.1 | - | - | 0 | - |
| - | - | 573 | 207.4 | - | - | 0 | - |
| - | - | 726.6 | 208.1 | - | - | 0 | - |
| - | - | 3770 | 208.1 | - | - | 0 | - |
| - | - | 9514 | 209.1 | - | - | 0 | - |
| - | - | 970 | 210.1 | - | - | 0 | - |
| - | - | 745.2 | 210.1 | - | - | 0 | - |
| - | - | 792.5 | 212.1 | - | - | 0 | - |
| - | - | 994.2 | 213.1 | - | - | 0 | - |
| - | - | 765.5 | 213.1 | - | - | 0 | - |
| - | - | 1.346E+04 | 215.1 | - | - | 0 | - |
| - | - | 654.6 | 216.1 | - | - | 0 | - |
| - | - | 7778 | 217.1 | - | - | 0 | - |
| - | - | 3348 | 217.1 | - | - | 0 | - |
| - | - | 7410 | 217.1 | - | - | 0 | - |
| - | - | 766.2 | 217.1 | - | - | 0 | - |
| - | - | 894.3 | 218.1 | - | - | 0 | - |
| - | - | 667.4 | 218.1 | - | - | 0 | - |
| - | - | 1953 | 219 | - | - | 0 | - |
| 6 | b | 878.4 | 219.1 | 0.001081 | 4.932 | +3 | 6 |
| - | - | 1285 | 219.1 | - | - | 0 | - |
| - | - | 4371 | 221.1 | - | - | 0 | - |
| - | - | 811.7 | 222.1 | - | - | 0 | - |
| - | - | 788.2 | 223.1 | - | - | 0 | - |
| - | - | 1918 | 225 | - | - | 0 | - |
| - | - | 4464 | 225.1 | - | - | 0 | - |
| - | - | 2069 | 225.1 | - | - | 0 | - |
| - | - | 541.6 | 226 | - | - | 0 | - |
| - | - | 3.829E+04 | 226.1 | - | - | 0 | - |
| - | - | 944.2 | 227.1 | - | - | 0 | - |
| - | - | 3546 | 227.1 | - | - | 0 | - |
| - | - | 1345 | 228.1 | - | - | 0 | - |
| - | - | 1469 | 231.1 | - | - | 0 | - |
| - | - | 1.347E+04 | 233.2 | - | - | 0 | - |
| - | - | 526.5 | 234.1 | - | - | 0 | - |
| - | - | 1795 | 234.2 | - | - | 0 | - |
| - | - | 5757 | 235.1 | - | - | 0 | - |
| - | - | 518.9 | 235.3 | - | - | 0 | - |
| - | - | 1667 | 236.1 | - | - | 0 | - |
| - | - | 723.2 | 236.1 | - | - | 0 | - |
| - | - | 1.256E+04 | 239.1 | - | - | 0 | - |
| - | - | 3159 | 240.1 | - | - | 0 | - |
| - | - | 3004 | 241.1 | - | - | 0 | - |
| - | - | 1106 | 241.1 | - | - | 0 | - |
| - | - | 8731 | 242.2 | - | - | 0 | - |
| - | - | 716.9 | 243.1 | - | - | 0 | - |
| 11 | y | 1.458E+04 | 243.1 | 0.004268 | 17.56 | +3 | 7 |
| 13 | y | 1039 | 244.1 | 0.000151 | 0.6185 | +2 | 5 |
| - | - | 1193 | 244.1 | - | - | 0 | - |
| - | - | 1397 | 245 | - | - | 0 | - |
| - | - | 6368 | 245.1 | - | - | 0 | - |
| - | - | 1399 | 246.1 | - | - | 0 | - |
| - | - | 1122 | 247.1 | - | - | 0 | - |
| - | - | 5270 | 251.2 | - | - | 0 | - |
| - | - | 676.4 | 252.1 | - | - | 0 | - |
| - | - | 891.8 | 252.1 | - | - | 0 | - |
| - | - | 5068 | 254.1 | - | - | 0 | - |
| - | - | 1278 | 254.2 | - | - | 0 | - |
| - | - | 878.5 | 257.2 | - | - | 0 | - |
| - | - | 4122 | 259.1 | - | - | 0 | - |
| - | - | 7208 | 261.2 | - | - | 0 | - |
| - | - | 845.6 | 262.1 | - | - | 0 | - |
| - | - | 1333 | 262.2 | - | - | 0 | - |
| - | - | 4499 | 263.1 | - | - | 0 | - |
| - | - | 7623 | 263.1 | - | - | 0 | - |
| - | - | 755.3 | 264.1 | - | - | 0 | - |
| - | - | 1061 | 264.1 | - | - | 0 | - |
| - | - | 4647 | 264.1 | - | - | 0 | - |
| - | - | 718.8 | 265.1 | - | - | 0 | - |
| - | - | 601.8 | 268.1 | - | - | 0 | - |
| 3 | a | 839.6 | 268.2 | 0.0005985 | 2.232 | +1 | 3 |
| - | - | 1179 | 269 | - | - | 0 | - |
| - | - | 1322 | 269.2 | - | - | 0 | - |
| - | - | 665.6 | 270.2 | - | - | 0 | - |
| - | - | 2722 | 272.1 | - | - | 0 | - |
| - | - | 1504 | 274.1 | - | - | 0 | - |
| - | - | 1133 | 275.2 | - | - | 0 | - |
| - | - | 3591 | 279.1 | - | - | 0 | - |
| 5 | b | 1483 | 279.7 | 0.0002083 | 0.7449 | +2 | 5 |
| - | - | 1580 | 280.1 | - | - | 0 | - |
| - | - | 683.7 | 280.1 | - | - | 0 | - |
| - | - | 2787 | 280.2 | - | - | 0 | - |
| - | - | 1267 | 281.1 | - | - | 0 | - |
| - | - | 2452 | 281.1 | - | - | 0 | - |
| - | - | 1.196E+04 | 282.1 | - | - | 0 | - |
| - | - | 1622 | 283.1 | - | - | 0 | - |
| - | - | 1.045E+04 | 285 | - | - | 0 | - |
| - | - | 803.9 | 285.1 | - | - | 0 | - |
| - | - | 1994 | 285.2 | - | - | 0 | - |
| - | - | 3676 | 286 | - | - | 0 | - |
| - | - | 2528 | 286.1 | - | - | 0 | - |
| - | - | 1324 | 287 | - | - | 0 | - |
| - | - | 701 | 294.2 | - | - | 0 | - |
| - | - | 2497 | 296.1 | - | - | 0 | - |
| - | - | 4058 | 296.2 | - | - | 0 | - |
| 3 | b | 7.912E+04 | 296.2 | 0.0004585 | 1.548 | +1 | 3 |
| - | - | 561.4 | 297.1 | - | - | 0 | - |
| - | - | 3187 | 297.1 | - | - | 0 | - |
| - | - | 863.6 | 297.1 | - | - | 0 | - |
| - | - | 510.4 | 297.1 | - | - | 0 | - |
| - | - | 1.262E+04 | 297.2 | - | - | 0 | - |
| - | - | 693.9 | 297.2 | - | - | 0 | - |
| - | - | 1.428E+04 | 297.2 | - | - | 0 | - |
| - | - | 1611 | 298.2 | - | - | 0 | - |
| - | - | 3956 | 298.2 | - | - | 0 | - |
| - | - | 1036 | 298.2 | - | - | 0 | - |
| - | - | 834.8 | 298.7 | - | - | 0 | - |
| - | - | 1.153E+04 | 299.1 | - | - | 0 | - |
| - | - | 660.8 | 299.2 | - | - | 0 | - |
| - | - | 649.6 | 299.2 | - | - | 0 | - |
| - | - | 4369 | 300.1 | - | - | 0 | - |
| - | - | 752.1 | 300.2 | - | - | 0 | - |
| - | - | 1544 | 301.1 | - | - | 0 | - |
| 15 | y | 1989 | 301.2 | 0.001109 | 3.683 | +1 | 3 |
| - | - | 1403 | 303 | - | - | 0 | - |
| - | - | 7.538E+04 | 303.1 | - | - | 0 | - |
| - | - | 3267 | 303.2 | - | - | 0 | - |
| - | - | 1.266E+04 | 304.1 | - | - | 0 | - |
| - | - | 738.5 | 304.1 | - | - | 0 | - |
| - | - | 617.6 | 305.1 | - | - | 0 | - |
| 12 | y | 4227 | 307.7 | 0.0004677 | 1.52 | +2 | 6 |
| 12 | y | 1080 | 308.2 | 0.002967 | 9.627 | +2 | 6 |
| - | - | 665 | 310.1 | - | - | 0 | - |
| - | - | 2109 | 313.2 | - | - | 0 | - |
| - | - | 2815 | 314.1 | - | - | 0 | - |
| - | - | 1.27E+04 | 314.2 | - | - | 0 | - |
| 3 | b | 1.711E+04 | 314.2 | 0.0004834 | 1.539 | +1 | 3 |
| - | - | 642.2 | 315.1 | - | - | 0 | - |
| - | - | 593.1 | 315.2 | - | - | 0 | - |
| - | - | 1968 | 315.2 | - | - | 0 | - |
| - | - | 2926 | 315.2 | - | - | 0 | - |
| - | - | 7445 | 316.2 | - | - | 0 | - |
| 12 | y | 2547 | 316.7 | 0.0001902 | 0.6007 | +2 | 6 |
| - | - | 1240 | 317.2 | - | - | 0 | - |
| 15 | y | 1.092E+04 | 319.2 | 0.0003711 | 1.163 | +1 | 3 |
| - | - | 1939 | 320.2 | - | - | 0 | - |
| - | - | 888.7 | 325.2 | - | - | 0 | - |
| - | - | 684 | 326.2 | - | - | 0 | - |
| - | - | 1069 | 327.1 | - | - | 0 | - |
| - | - | 2593 | 328.1 | - | - | 0 | - |
| - | - | 864.6 | 329.2 | - | - | 0 | - |
| - | - | 1828 | 330.2 | - | - | 0 | - |
| - | - | 2758 | 333.2 | - | - | 0 | - |
| - | - | 614.6 | 333.7 | - | - | 0 | - |
| - | - | 1189 | 334.2 | - | - | 0 | - |
| - | - | 687.3 | 336.2 | - | - | 0 | - |
| - | - | 1395 | 339.2 | - | - | 0 | - |
| - | - | 2259 | 341.2 | - | - | 0 | - |
| - | - | 1594 | 342.2 | - | - | 0 | - |
| - | - | 844.3 | 344.2 | - | - | 0 | - |
| - | - | 1477 | 345 | - | - | 0 | - |
| - | - | 3318 | 345.1 | - | - | 0 | - |
| - | - | 797.9 | 346 | - | - | 0 | - |
| - | - | 1118 | 346.1 | - | - | 0 | - |
| - | - | 5539 | 346.1 | - | - | 0 | - |
| - | - | 2702 | 346.2 | - | - | 0 | - |
| - | - | 1244 | 347.1 | - | - | 0 | - |
| - | - | 2222 | 348.2 | - | - | 0 | - |
| - | - | 1061 | 349.2 | - | - | 0 | - |
| - | - | 1944 | 350.1 | - | - | 0 | - |
| - | - | 1759 | 350.2 | - | - | 0 | - |
| - | - | 8597 | 351.2 | - | - | 0 | - |
| - | - | 823.9 | 351.2 | - | - | 0 | - |
| - | - | 737.2 | 352.2 | - | - | 0 | - |
| - | - | 3972 | 353.2 | - | - | 0 | - |
| - | - | 829.8 | 354.2 | - | - | 0 | - |
| - | - | 953.4 | 354.2 | - | - | 0 | - |
| - | - | 4.336E+04 | 355.1 | - | - | 0 | - |
| - | - | 1.652E+04 | 356.1 | - | - | 0 | - |
| - | - | 9101 | 357.1 | - | - | 0 | - |
| - | - | 8003 | 362.2 | - | - | 0 | - |
| - | - | 1933 | 363.2 | - | - | 0 | - |
| 11 | y | 658.9 | 364.2 | 0.0007636 | 2.097 | +2 | 7 |
| - | - | 1565 | 365.1 | - | - | 0 | - |
| - | - | 3327 | 367.2 | - | - | 0 | - |
| - | - | 841.3 | 367.2 | - | - | 0 | - |
| - | - | 3183 | 368.2 | - | - | 0 | - |
| - | - | 2274 | 369.1 | - | - | 0 | - |
| - | - | 6870 | 369.2 | - | - | 0 | - |
| - | - | 1308 | 370.2 | - | - | 0 | - |
| - | - | 876.3 | 370.7 | - | - | 0 | - |
| - | - | 1031 | 371.1 | - | - | 0 | - |
| - | - | 1727 | 371.2 | - | - | 0 | - |
| - | - | 4159 | 373.1 | - | - | 0 | - |
| - | - | 510.5 | 373.1 | - | - | 0 | - |
| - | - | 2076 | 373.2 | - | - | 0 | - |
| 11 | y | 2528 | 373.2 | 0.0004861 | 1.303 | +2 | 7 |
| - | - | 1828 | 373.7 | - | - | 0 | - |
| - | - | 1743 | 374.1 | - | - | 0 | - |
| - | - | 827.6 | 375.1 | - | - | 0 | - |
| - | - | 1057 | 381.2 | - | - | 0 | - |
| - | - | 2353 | 383.2 | - | - | 0 | - |
| - | - | 712.4 | 383.3 | - | - | 0 | - |
| - | - | 1.061E+04 | 385.2 | - | - | 0 | - |
| - | - | 2158 | 386.2 | - | - | 0 | - |
| - | - | 575.4 | 394.2 | - | - | 0 | - |
| - | - | 1380 | 395.1 | - | - | 0 | - |
| - | - | 1254 | 397.2 | - | - | 0 | - |
| - | - | 764.9 | 398.2 | - | - | 0 | - |
| - | - | 1451 | 398.2 | - | - | 0 | - |
| - | - | 615.4 | 399.2 | - | - | 0 | - |
| - | - | 655 | 399.2 | - | - | 0 | - |
| - | - | 656.2 | 401 | - | - | 0 | - |
| - | - | 3831 | 401.2 | - | - | 0 | - |
| - | - | 856 | 402.2 | - | - | 0 | - |
| 8 | b | 703 | 406.2 | 0.0002768 | 0.6815 | +2 | 8 |
| - | - | 1927 | 407.2 | - | - | 0 | - |
| - | - | 4964 | 408.2 | - | - | 0 | - |
| - | - | 1203 | 409.2 | - | - | 0 | - |
| - | - | 887.8 | 410.2 | - | - | 0 | - |
| - | - | 720.9 | 411.1 | - | - | 0 | - |
| - | - | 2357 | 411.3 | - | - | 0 | - |
| - | - | 4679 | 413.1 | - | - | 0 | - |
| - | - | 1552 | 414.1 | - | - | 0 | - |
| - | - | 2963 | 415 | - | - | 0 | - |
| - | - | 1200 | 415.1 | - | - | 0 | - |
| 8 | b | 1422 | 415.2 | 0.004145 | 9.983 | +2 | 8 |
| - | - | 1021 | 415.3 | - | - | 0 | - |
| - | - | 1446 | 416 | - | - | 0 | - |
| 14 | y | 1650 | 416.2 | 0.0007469 | 1.795 | +1 | 4 |
| - | - | 2721 | 416.3 | - | - | 0 | - |
| - | - | 630.9 | 417 | - | - | 0 | - |
| - | - | 894.7 | 417.3 | - | - | 0 | - |
| - | - | 4929 | 419 | - | - | 0 | - |
| - | - | 2014 | 420 | - | - | 0 | - |
| - | - | 1276 | 421 | - | - | 0 | - |
| - | - | 3333 | 424.2 | - | - | 0 | - |
| - | - | 4081 | 425.2 | - | - | 0 | - |
| - | - | 988 | 426.2 | - | - | 0 | - |
| - | - | 1883 | 427.2 | - | - | 0 | - |
| - | - | 705.9 | 429.1 | - | - | 0 | - |
| 10 | y | 898.3 | 429.2 | 0.008299 | 19.33 | +2 | 8 |
| - | - | 1728 | 431.1 | - | - | 0 | - |
| - | - | 651 | 432.1 | - | - | 0 | - |
| - | - | 3476 | 433.2 | - | - | 0 | - |
| 14 | y | 1.264E+04 | 433.2 | 0.0006566 | 1.515 | +1 | 4 |
| - | - | 2.526E+04 | 433.3 | - | - | 0 | - |
| - | - | 1469 | 434.2 | - | - | 0 | - |
| - | - | 2781 | 434.2 | - | - | 0 | - |
| - | - | 6167 | 434.3 | - | - | 0 | - |
| - | - | 1039 | 434.7 | - | - | 0 | - |
| - | - | 1097 | 435.2 | - | - | 0 | - |
| - | - | 827.7 | 435.3 | - | - | 0 | - |
| - | - | 720.5 | 440.2 | - | - | 0 | - |
| - | - | 654.8 | 441.2 | - | - | 0 | - |
| - | - | 3613 | 442.2 | - | - | 0 | - |
| - | - | 1778 | 443.2 | - | - | 0 | - |
| 4 | b | 3946 | 443.3 | 0.0005261 | 1.187 | +1 | 4 |
| - | - | 945.2 | 444.2 | - | - | 0 | - |
| - | - | 1306 | 444.3 | - | - | 0 | - |
| - | - | 2201 | 449.1 | - | - | 0 | - |
| - | - | 998.4 | 450.2 | - | - | 0 | - |
| - | - | 845.4 | 451.2 | - | - | 0 | - |
| 5 | y | 3056 | 451.9 | 0.0002063 | 0.4565 | +3 | 13 |
| 5 | y | 2916 | 452.2 | 0.00123 | 2.721 | +3 | 13 |
| - | - | 760.9 | 454.7 | - | - | 0 | - |
| - | - | 914.7 | 459.2 | - | - | 0 | - |
| 4 | b | 2.196E+04 | 461.3 | 0.0004289 | 0.9298 | +1 | 4 |
| - | - | 1997 | 462.2 | - | - | 0 | - |
| - | - | 6723 | 462.3 | - | - | 0 | - |
| - | - | 931 | 463.3 | - | - | 0 | - |
| - | - | 1931 | 465.1 | - | - | 0 | - |
| - | - | 837.3 | 466.2 | - | - | 0 | - |
| - | - | 669.4 | 466.7 | - | - | 0 | - |
| - | - | 1301 | 468.3 | - | - | 0 | - |
| - | - | 1534 | 469.2 | - | - | 0 | - |
| - | - | 7766 | 470.2 | - | - | 0 | - |
| 9 | b | 928.8 | 470.7 | 0.002469 | 5.246 | +2 | 9 |
| - | - | 1617 | 471.2 | - | - | 0 | - |
| - | - | 1279 | 471.7 | - | - | 0 | - |
| - | - | 1738 | 477.3 | - | - | 0 | - |
| - | - | 4037 | 478.2 | - | - | 0 | - |
| - | - | 2090 | 479.2 | - | - | 0 | - |
| - | - | 7095 | 480.2 | - | - | 0 | - |
| - | - | 2022 | 480.7 | - | - | 0 | - |
| - | - | 3096 | 481.2 | - | - | 0 | - |
| - | - | 1199 | 484.2 | - | - | 0 | - |
| - | - | 637.4 | 484.7 | - | - | 0 | - |
| 13 | y | 4234 | 486.3 | 0.001207 | 2.482 | +1 | 5 |
| 13 | y | 1905 | 487.3 | 0.001302 | 2.672 | +1 | 5 |
| - | - | 1353 | 488.7 | - | - | 0 | - |
| - | - | 1083 | 489.1 | - | - | 0 | - |
| - | - | 1080 | 490.2 | - | - | 0 | - |
| - | - | 2458 | 493.7 | - | - | 0 | - |
| - | - | 921.1 | 494.2 | - | - | 0 | - |
| - | - | 2093 | 494.2 | - | - | 0 | - |
| - | - | 595 | 494.7 | - | - | 0 | - |
| - | - | 2852 | 495.3 | - | - | 0 | - |
| - | - | 9517 | 496.3 | - | - | 0 | - |
| - | - | 2001 | 497.3 | - | - | 0 | - |
| - | - | 2.062E+04 | 498.2 | - | - | 0 | - |
| - | - | 6060 | 499.2 | - | - | 0 | - |
| - | - | 1508 | 500.2 | - | - | 0 | - |
| 4 | y | 904.1 | 500.9 | 0.00148 | 2.955 | +3 | 14 |
| 4 | y | 598.6 | 501.2 | 0.007703 | 15.37 | +3 | 14 |
| - | - | 1608 | 502.2 | - | - | 0 | - |
| - | - | 3672 | 502.7 | - | - | 0 | - |
| - | - | 1242 | 503.2 | - | - | 0 | - |
| 13 | y | 2.833E+04 | 504.3 | 0.0002554 | 0.5065 | +1 | 5 |
| - | - | 7094 | 505.3 | - | - | 0 | - |
| - | - | 673.2 | 506.3 | - | - | 0 | - |
| - | - | 1074 | 512.2 | - | - | 0 | - |
| - | - | 3816 | 513.3 | - | - | 0 | - |
| - | - | 729.6 | 515.2 | - | - | 0 | - |
| - | - | 1101 | 521.3 | - | - | 0 | - |
| - | - | 839.4 | 529.3 | - | - | 0 | - |
| - | - | 8083 | 530.2 | - | - | 0 | - |
| - | - | 2112 | 530.3 | - | - | 0 | - |
| - | - | 2081 | 531.2 | - | - | 0 | - |
| - | - | 756.5 | 536.6 | - | - | 0 | - |
| - | - | 6044 | 538.3 | - | - | 0 | - |
| - | - | 4607 | 538.8 | - | - | 0 | - |
| - | - | 807.1 | 539.3 | - | - | 0 | - |
| 5 | b | 3008 | 540.3 | 0.001961 | 3.63 | +1 | 5 |
| - | - | 2338 | 543.3 | - | - | 0 | - |
| - | - | 1400 | 543.8 | - | - | 0 | - |
| 10 | b | 2564 | 544.3 | 0.002651 | 4.87 | +2 | 10 |
| - | - | 662.5 | 544.8 | - | - | 0 | - |
| - | - | 624.6 | 545.3 | - | - | 0 | - |
| - | - | 1108 | 546.2 | - | - | 0 | - |
| 15 | b | 695.3 | 548 | 0.002856 | 5.212 | +3 | 15 |
| - | - | 960.2 | 548.3 | - | - | 0 | - |
| - | - | 682.4 | 551.3 | - | - | 0 | - |
| - | - | 4609 | 552.3 | - | - | 0 | - |
| - | - | 916.2 | 552.3 | - | - | 0 | - |
| - | - | 3795 | 552.8 | - | - | 0 | - |
| - | - | 750.3 | 553.3 | - | - | 0 | - |
| - | - | 3059 | 555.3 | - | - | 0 | - |
| - | - | 571.9 | 556.3 | - | - | 0 | - |
| - | - | 875.4 | 556.3 | - | - | 0 | - |
| 5 | b | 4985 | 558.3 | 0.0002415 | 0.4325 | +1 | 5 |
| - | - | 2423 | 559.3 | - | - | 0 | - |
| - | - | 630 | 560.8 | - | - | 0 | - |
| - | - | 729.4 | 561.3 | - | - | 0 | - |
| - | - | 740.1 | 567.3 | - | - | 0 | - |
| - | - | 698 | 569.8 | - | - | 0 | - |
| - | - | 689.8 | 570.3 | - | - | 0 | - |
| - | - | 785.4 | 571.3 | - | - | 0 | - |
| - | - | 757.4 | 573.2 | - | - | 0 | - |
| - | - | 886.6 | 578.1 | - | - | 0 | - |
| - | - | 2861 | 578.3 | - | - | 0 | - |
| - | - | 1357 | 578.8 | - | - | 0 | - |
| - | - | 1287 | 579.3 | - | - | 0 | - |
| 7 | y | 1718 | 580.3 | 0.001623 | 2.797 | +2 | 11 |
| - | - | 1616 | 581.3 | - | - | 0 | - |
| - | - | 1086 | 582.3 | - | - | 0 | - |
| - | - | 3750 | 582.8 | - | - | 0 | - |
| - | - | 2773 | 583.3 | - | - | 0 | - |
| - | - | 1104 | 591.2 | - | - | 0 | - |
| - | - | 2405 | 591.3 | - | - | 0 | - |
| 11 | b | 1532 | 591.8 | 0.01145 | 19.34 | +2 | 11 |
| - | - | 853.2 | 592.3 | - | - | 0 | - |
| - | - | 830.1 | 595.3 | - | - | 0 | - |
| - | - | 2771 | 597.3 | - | - | 0 | - |
| - | - | 1989 | 598.3 | - | - | 0 | - |
| - | - | 1.046E+04 | 599.3 | - | - | 0 | - |
| - | - | 736.3 | 599.3 | - | - | 0 | - |
| - | - | 2862 | 600.3 | - | - | 0 | - |
| - | - | 1007 | 601.3 | - | - | 0 | - |
| - | - | 2773 | 602.8 | - | - | 0 | - |
| - | - | 2020 | 603.3 | - | - | 0 | - |
| - | - | 617.4 | 608.4 | - | - | 0 | - |
| - | - | 4079 | 609.3 | - | - | 0 | - |
| - | - | 828.6 | 609.3 | - | - | 0 | - |
| - | - | 1036 | 609.8 | - | - | 0 | - |
| - | - | 5425 | 610.2 | - | - | 0 | - |
| - | - | 1158 | 610.3 | - | - | 0 | - |
| - | - | 5787 | 611.2 | - | - | 0 | - |
| - | - | 5436 | 611.8 | - | - | 0 | - |
| - | - | 4216 | 612.2 | - | - | 0 | - |
| - | - | 3844 | 612.3 | - | - | 0 | - |
| - | - | 878.3 | 612.4 | - | - | 0 | - |
| - | - | 2445 | 612.8 | - | - | 0 | - |
| 12 | y | 6761 | 614.3 | 0.0001552 | 0.2526 | +1 | 6 |
| 12 | y | 1.573E+04 | 615.3 | 8.735E-05 | 0.142 | +1 | 6 |
| - | - | 5355 | 616.3 | - | - | 0 | - |
| - | - | 2538 | 617.3 | - | - | 0 | - |
| - | - | 8990 | 617.8 | - | - | 0 | - |
| - | - | 1.126E+04 | 618.3 | - | - | 0 | - |
| - | - | 4716 | 618.8 | - | - | 0 | - |
| - | - | 764.5 | 619.3 | - | - | 0 | - |
| - | - | 738 | 620.3 | - | - | 0 | - |
| - | - | 720.7 | 625.3 | - | - | 0 | - |
| - | - | 757 | 626.3 | - | - | 0 | - |
| - | - | 2886 | 626.4 | - | - | 0 | - |
| - | - | 2.584E+04 | 626.8 | - | - | 0 | - |
| - | - | 4.169E+04 | 627.3 | - | - | 0 | - |
| - | - | 1.478E+04 | 627.3 | - | - | 0 | - |
| - | - | 6847 | 627.8 | - | - | 0 | - |
| - | - | 1.418E+04 | 628.3 | - | - | 0 | - |
| 6 | y | 1016 | 628.8 | 0.0002948 | 0.4688 | +2 | 12 |
| - | - | 2637 | 629.3 | - | - | 0 | - |
| - | - | 888.8 | 629.8 | - | - | 0 | - |
| 12 | y | 1.991E+04 | 632.3 | 8.854E-05 | 0.14 | +1 | 6 |
| - | - | 7396 | 633.3 | - | - | 0 | - |
| - | - | 1539 | 634.3 | - | - | 0 | - |
| - | - | 951.8 | 636.3 | - | - | 0 | - |
| 6 | y | 4062 | 637.8 | 0.0008926 | 1.399 | +2 | 12 |
| - | - | 3938 | 638.3 | - | - | 0 | - |
| - | - | 1106 | 638.8 | - | - | 0 | - |
| - | - | 2721 | 643.3 | - | - | 0 | - |
| - | - | 1757 | 644.3 | - | - | 0 | - |
| - | - | 2341 | 645.3 | - | - | 0 | - |
| - | - | 1.039E+04 | 655.3 | - | - | 0 | - |
| 12 | b | 7709 | 655.8 | 0.006445 | 9.828 | +2 | 12 |
| - | - | 2108 | 656.3 | - | - | 0 | - |
| - | - | 786.5 | 659.8 | - | - | 0 | - |
| - | - | 1874 | 660.3 | - | - | 0 | - |
| - | - | 645 | 663.3 | - | - | 0 | - |
| - | - | 871.9 | 667.3 | - | - | 0 | - |
| - | - | 2039 | 668.3 | - | - | 0 | - |
| - | - | 4534 | 668.8 | - | - | 0 | - |
| - | - | 2010 | 669.3 | - | - | 0 | - |
| - | - | 801.4 | 669.8 | - | - | 0 | - |
| - | - | 1066 | 676.3 | - | - | 0 | - |
| 5 | y | 2.559E+04 | 677.3 | 0.0003706 | 0.5471 | +2 | 13 |
| 5 | y | 2.682E+04 | 677.8 | 0.007194 | 10.61 | +2 | 13 |
| - | - | 1.394E+04 | 678.3 | - | - | 0 | - |
| - | - | 4178 | 678.8 | - | - | 0 | - |
| - | - | 1089 | 679.3 | - | - | 0 | - |
| - | - | 1146 | 684.4 | - | - | 0 | - |
| 5 | y | 5.541E+04 | 686.3 | 0.0001597 | 0.2327 | +2 | 13 |
| - | - | 4.078E+04 | 686.8 | - | - | 0 | - |
| - | - | 1.655E+04 | 687.3 | - | - | 0 | - |
| - | - | 3839 | 687.8 | - | - | 0 | - |
| 13 | b | 1380 | 691.4 | 0.0104 | 15.05 | +2 | 13 |
| - | - | 1952 | 694.3 | - | - | 0 | - |
| - | - | 769.8 | 695.3 | - | - | 0 | - |
| - | - | 952.5 | 698.3 | - | - | 0 | - |
| 13 | b | 1738 | 700.4 | 0.01086 | 15.51 | +2 | 13 |
| - | - | 2300 | 700.8 | - | - | 0 | - |
| - | - | 1446 | 701.3 | - | - | 0 | - |
| - | - | 1922 | 710.4 | - | - | 0 | - |
| - | - | 6401 | 712.4 | - | - | 0 | - |
| - | - | 3399 | 713.4 | - | - | 0 | - |
| - | - | 1183 | 714.4 | - | - | 0 | - |
| - | - | 755.9 | 720.4 | - | - | 0 | - |
| - | - | 2894 | 722.3 | - | - | 0 | - |
| 7 | b | 1831 | 724.4 | 0.001825 | 2.519 | +1 | 7 |
| - | - | 1188 | 725.4 | - | - | 0 | - |
| - | - | 790.2 | 726.3 | - | - | 0 | - |
| - | - | 678.2 | 726.4 | - | - | 0 | - |
| 11 | y | 832.4 | 727.4 | 0.001328 | 1.826 | +1 | 7 |
| 11 | y | 1469 | 728.4 | 0.002144 | 2.943 | +1 | 7 |
| - | - | 1580 | 728.9 | - | - | 0 | - |
| - | - | 722.7 | 729.4 | - | - | 0 | - |
| - | - | 860.1 | 729.9 | - | - | 0 | - |
| - | - | 664.5 | 732.3 | - | - | 0 | - |
| - | - | 1960 | 737.4 | - | - | 0 | - |
| - | - | 870.1 | 738.3 | - | - | 0 | - |
| - | - | 1200 | 738.4 | - | - | 0 | - |
| - | - | 845.6 | 739.4 | - | - | 0 | - |
| - | - | 1.749E+04 | 740.3 | - | - | 0 | - |
| - | - | 6139 | 741.3 | - | - | 0 | - |
| - | - | 2103 | 742.4 | - | - | 0 | - |
| 7 | b | 3258 | 742.4 | 0.001697 | 2.286 | +1 | 7 |
| - | - | 1632 | 743.4 | - | - | 0 | - |
| - | - | 789 | 745.3 | - | - | 0 | - |
| 11 | y | 9168 | 745.4 | 0.0001131 | 0.1518 | +1 | 7 |
| - | - | 3403 | 746.4 | - | - | 0 | - |
| - | - | 1457 | 747.4 | - | - | 0 | - |
| - | - | 747.3 | 749.3 | - | - | 0 | - |
| 4 | y | 6377 | 750.9 | 0.0001515 | 0.2017 | +2 | 14 |
| 4 | y | 5993 | 751.4 | 0.007777 | 10.35 | +2 | 14 |
| - | - | 2118 | 751.9 | - | - | 0 | - |
| - | - | 1446 | 752.4 | - | - | 0 | - |
| - | - | 1663 | 755.4 | - | - | 0 | - |
| 14 | b | 4898 | 757.4 | 0.005471 | 7.224 | +2 | 14 |
| - | - | 1831 | 758.4 | - | - | 0 | - |
| - | - | 1371 | 759.4 | - | - | 0 | - |
| 4 | y | 5783 | 759.9 | 0.0007363 | 0.969 | +2 | 14 |
| - | - | 5373 | 760.4 | - | - | 0 | - |
| - | - | 2251 | 760.9 | - | - | 0 | - |
| - | - | 1162 | 761.4 | - | - | 0 | - |
| - | - | 3180 | 771.4 | - | - | 0 | - |
| - | - | 1525 | 772.4 | - | - | 0 | - |
| - | - | 2707 | 774.3 | - | - | 0 | - |
| - | - | 1263 | 775.3 | - | - | 0 | - |
| 3 | y | 1720 | 807.4 | 0.000661 | 0.8187 | +2 | 15 |
| 3 | y | 1794 | 807.9 | 0.008775 | 10.86 | +2 | 15 |
| - | - | 1742 | 808.4 | - | - | 0 | - |
| 8 | b | 1995 | 811.4 | 8.909E-06 | 0.01098 | +1 | 8 |
| - | - | 1097 | 812.4 | - | - | 0 | - |
| - | - | 1790 | 813.4 | - | - | 0 | - |
| 3 | y | 860.4 | 816.4 | 0.0009329 | 1.143 | +2 | 15 |
| - | - | 778.2 | 824.4 | - | - | 0 | - |
| 8 | b | 1885 | 829.4 | 0.0006681 | 0.8055 | +1 | 8 |
| - | - | 1678 | 830.4 | - | - | 0 | - |
| - | - | 876.5 | 839.4 | - | - | 0 | - |
| - | - | 2029 | 840.4 | - | - | 0 | - |
| - | - | 2543 | 841.4 | - | - | 0 | - |
| - | - | 1024 | 842.4 | - | - | 0 | - |
| 16 | b | 952.1 | 848.4 | 0.0004195 | 0.4945 | +2 | 16 |
| - | - | 1218 | 848.9 | - | - | 0 | - |
| - | - | 1100 | 849.4 | - | - | 0 | - |
| - | - | 2478 | 850.4 | - | - | 0 | - |
| - | - | 1566 | 851.4 | - | - | 0 | - |
| 10 | y | 1554 | 856.5 | 0.002051 | 2.395 | +1 | 8 |
| 16 | b | 1630 | 856.9 | 0.001197 | 1.397 | +2 | 16 |
| 10 | y | 3226 | 857.4 | 0.003984 | 4.647 | +1 | 8 |
| 2 | y | 1235 | 858.4 | 0.008923 | 10.39 | +2 | 16 |
| - | - | 1373 | 859.4 | - | - | 0 | - |
| - | - | 952.9 | 861.4 | - | - | 0 | - |
| - | - | 2432 | 863.4 | - | - | 0 | - |
| - | - | 1.999E+04 | 868.4 | - | - | 0 | - |
| - | - | 1.002E+04 | 869.4 | - | - | 0 | - |
| - | - | 2743 | 870.4 | - | - | 0 | - |
| - | - | 2903 | 873.5 | - | - | 0 | - |
| 10 | y | 4958 | 874.5 | 0.00163 | 1.864 | +1 | 8 |
| - | - | 2182 | 875.5 | - | - | 0 | - |
| - | - | 3602 | 879.4 | - | - | 0 | - |
| - | - | 1356 | 880.4 | - | - | 0 | - |
| - | - | 4308 | 887.4 | - | - | 0 | - |
| - | - | 1633 | 888.4 | - | - | 0 | - |
| - | - | 2301 | 889.4 | - | - | 0 | - |
| - | - | 1690 | 890.4 | - | - | 0 | - |
| - | - | 1.271E+04 | 907.4 | - | - | 0 | - |
| - | - | 6708 | 908.4 | - | - | 0 | - |
| - | - | 1175 | 909.4 | - | - | 0 | - |
| - | - | 1091 | 911.4 | - | - | 0 | - |
| - | - | 1706 | 921.4 | - | - | 0 | - |
| - | - | 1831 | 922.4 | - | - | 0 | - |
| - | - | 1.017E+04 | 939.4 | - | - | 0 | - |
| - | - | 6639 | 940.4 | - | - | 0 | - |
| - | - | 2251 | 941.5 | - | - | 0 | - |
| - | - | 2358 | 956.5 | - | - | 0 | - |
| - | - | 993.3 | 957.5 | - | - | 0 | - |
| 9 | b | 4596 | 958.5 | 0.001818 | 1.897 | +1 | 9 |
| - | - | 3708 | 959.5 | - | - | 0 | - |
| - | - | 8496 | 960.5 | - | - | 0 | - |
| - | - | 3066 | 961.5 | - | - | 0 | - |
| - | - | 848.3 | 962.5 | - | - | 0 | - |
| - | - | 979.4 | 971.5 | - | - | 0 | - |
| 9 | y | 822.1 | 986.5 | 0.01108 | 11.23 | +1 | 9 |
| - | - | 1477 | 988.5 | - | - | 0 | - |
| - | - | 783 | 1001 | - | - | 0 | - |
| 9 | y | 1392 | 1004 | 0.002347 | 2.339 | +1 | 9 |
| - | - | 776.3 | 1008 | - | - | 0 | - |
| - | - | 1516 | 1015 | - | - | 0 | - |
| - | - | 1041 | 1016 | - | - | 0 | - |
| - | - | 1024 | 1025 | - | - | 0 | - |
| - | - | 801.5 | 1035 | - | - | 0 | - |
| - | - | 2248 | 1036 | - | - | 0 | - |
| - | - | 998.9 | 1037 | - | - | 0 | - |
| - | - | 1310 | 1040 | - | - | 0 | - |
| - | - | 7372 | 1053 | - | - | 0 | - |
| - | - | 5114 | 1054 | - | - | 0 | - |
| - | - | 1854 | 1055 | - | - | 0 | - |
| - | - | 2025 | 1059 | - | - | 0 | - |
| - | - | 1657 | 1060 | - | - | 0 | - |
| 10 | b | 2977 | 1070 | 0.003817 | 3.569 | +1 | 10 |
| - | - | 3357 | 1071 | - | - | 0 | - |
| - | - | 1788 | 1072 | - | - | 0 | - |
| 8 | y | 883.1 | 1073 | 0.0005619 | 0.5239 | +1 | 10 |
| - | - | 4196 | 1076 | - | - | 0 | - |
| - | - | 2885 | 1077 | - | - | 0 | - |
| - | - | 1218 | 1085 | - | - | 0 | - |
| - | - | 1299 | 1086 | - | - | 0 | - |
| - | - | 1359 | 1087 | - | - | 0 | - |
| 10 | b | 1.607E+04 | 1088 | 0.00303 | 2.786 | +1 | 10 |
| - | - | 8847 | 1089 | - | - | 0 | - |
| - | - | 4192 | 1090 | - | - | 0 | - |
| 8 | y | 2759 | 1091 | 0.0008727 | 0.8002 | +1 | 10 |
| - | - | 1786 | 1092 | - | - | 0 | - |
| - | - | 4844 | 1104 | - | - | 0 | - |
| - | - | 2184 | 1105 | - | - | 0 | - |
| - | - | 2232 | 1139 | - | - | 0 | - |
| - | - | 1255 | 1140 | - | - | 0 | - |
| - | - | 3222 | 1156 | - | - | 0 | - |
| - | - | 1775 | 1157 | - | - | 0 | - |
| - | - | 2114 | 1165 | - | - | 0 | - |
| - | - | 1380 | 1166 | - | - | 0 | - |
| 7 | y | 994.9 | 1178 | 0.001163 | 0.9874 | +1 | 11 |
| - | - | 1115 | 1179 | - | - | 0 | - |
| - | - | 6180 | 1182 | - | - | 0 | - |
| 11 | b | 3848 | 1183 | 0.02013 | 17.02 | +1 | 11 |
| - | - | 1809 | 1184 | - | - | 0 | - |
| 11 | b | 3026 | 1201 | 0.009701 | 8.08 | +1 | 11 |
| - | - | 2601 | 1202 | - | - | 0 | - |
| - | - | 1758 | 1213 | - | - | 0 | - |
| - | - | 1008 | 1214 | - | - | 0 | - |
| - | - | 700.5 | 1225 | - | - | 0 | - |
| - | - | 4573 | 1235 | - | - | 0 | - |
| - | - | 4011 | 1236 | - | - | 0 | - |
| - | - | 1928 | 1237 | - | - | 0 | - |
| - | - | 1.159E+04 | 1253 | - | - | 0 | - |
| - | - | 7627 | 1254 | - | - | 0 | - |
| - | - | 2943 | 1255 | - | - | 0 | - |
| 6 | y | 1396 | 1257 | 0.001492 | 1.187 | +1 | 12 |
| 6 | y | 1791 | 1258 | 0.004727 | 3.759 | +1 | 12 |
| 6 | y | 6124 | 1275 | 0.005343 | 4.192 | +1 | 12 |
| - | - | 3227 | 1276 | - | - | 0 | - |
| - | - | 1982 | 1277 | - | - | 0 | - |
| - | - | 732.5 | 1310 | - | - | 0 | - |
| - | - | 1032 | 1330 | - | - | 0 | - |
| 5 | y | 1755 | 1372 | 0.007814 | 5.696 | +1 | 13 |
| - | - | 1219 | 1373 | - | - | 0 | - |
| 13 | b | 1444 | 1400 | 0.02568 | 18.35 | +1 | 13 |
| 14 | b | 1037 | 1514 | 0.001094 | 0.7229 | +1 | 14 |
| - | - | 698.6 | 2581 | - | - | 0 | - |
| - | - | 852.6 | 3071 | - | - | 0 | - |

m/z Charge Intensity FragmentType MassShift Position
120.06588745117188 0 39260.492 y 16
120.08114624023438 0 32649.3
120.13687896728516 0 327.86673
121.069091796875 0 1301.8446
121.08452606201172 0 2242.533
123.09138488769531 0 400.85513
126.05535125732422 0 4057.943
128.10736083984375 0 11680.273
129.05496215820312 0 4131.9253
129.0662384033203 0 2073.8562
129.1025848388672 0 47962.812
130.05023193359375 0 3518.7344
130.06544494628906 0 1177.1027
130.10081481933594 0 555.2489
130.10594177246094 0 3801.921
131.1182403564453 0 2836.7224
132.0769500732422 0 911.10675
134.69137573242188 0 407.98358
135.0051727294922 0 388.48975
136.07603454589844 0 2492.3604
138.09169006347656 0 1950.8882
138.12818908691406 0 1085.807
139.08700561523438 0 1409.9508
141.06629943847656 0 1657.4823
147.06578063964844 0 479.38458
147.0770263671875 0 1074.5074
149.04518127441406 0 2536.2837
152.0708465576172 0 754.62836
153.01869201660156 0 3919.541
153.0550079345703 0 523.5548
153.10301208496094 0 835.95544
154.08648681640625 0 1278.6685
154.09783935546875 0 1880.1711
155.08184814453125 0 3185.9067
155.1182403564453 0 6679.413 a Water loss 1
157.06141662597656 0 1271.6539
157.0974884033203 0 3497.081
159.0919647216797 0 4033.4565
165.10272216796875 0 1527.0771
167.0558624267578 0 4112.527
167.081787109375 0 2373.8452
167.11801147460938 0 1666.7246
169.0612335205078 0 520.46686
169.1338653564453 0 8314.135
171.0289764404297 0 1784.0537
171.07675170898438 0 531.78815
172.10858154296875 0 1746.6272
173.0924530029297 0 6107.5005 y Water loss 15
173.12881469726562 0 25191.115 a 1
173.45187377929688 0 1416.7655
174.1319580078125 0 1758.8834
175.07162475585938 0 1935.0757
176.0660858154297 0 468.7913
177.05503845214844 0 4370.8545
177.80865478515625 0 428.86646
179.0341339111328 0 1655.5621
179.08218383789062 0 473.0265
179.78501892089844 0 476.14627
181.0974884033203 0 1201.0446
182.09286499023438 0 2749.6477
182.12928771972656 0 2261.932
182.1988525390625 0 551.84546
183.07662963867188 0 2864.5193
183.1131591796875 0 33982.297 b Water loss 1
184.10826110839844 0 2469.176
184.11668395996094 0 3505.4355
185.09242248535156 0 5627.256
186.08779907226562 0 2092.4187
187.0869903564453 0 736.4025
187.14447021484375 0 11399.862
189.0876007080078 0 3033.5332
191.0819091796875 0 601.9701
191.10301208496094 0 6791.5845 y 15
191.1180877685547 0 1277.2085
195.02911376953125 0 3529.835
195.07696533203125 0 541.25275
195.1131134033203 0 4953.82
197.1033935546875 0 777.6367
197.12875366210938 0 12588.114
198.12399291992188 0 2568.3972
198.1330108642578 0 1230.1302
199.0715789794922 0 3413.4236
199.10826110839844 0 1193.7738
200.10321044921875 0 3159.8928
200.13963317871094 0 2745.5144
201.08758544921875 0 723.05975
201.1236572265625 0 32318.762 b 1
202.12698364257812 0 2345.2686
203.1030731201172 0 772.3389
203.11378479003906 0 540.58844
207.40310668945312 0 572.9716
208.09869384765625 0 726.55475
208.10838317871094 0 3770.2808
209.09243774414062 0 9514.14
210.09642028808594 0 970.0032
210.12417602539062 0 745.2334
212.13914489746094 0 792.5003
213.08700561523438 0 994.1728
213.12342834472656 0 765.53754
215.1393585205078 0 13456.914
216.1425323486328 0 654.6061
217.08213806152344 0 7778.158
217.09747314453125 0 3348.1653
217.13375854492188 0 7409.809
217.1446990966797 0 766.1664
218.13807678222656 0 894.3387
218.14947509765625 0 667.42944
219.02923583984375 0 1953.4467
219.133056640625 0 878.4241 b 5
219.14974975585938 0 1284.7501
221.0847930908203 0 4371.21
222.0854949951172 0 811.6913
223.0638427734375 0 788.17664
225.04347229003906 0 1917.5538
225.12364196777344 0 4464.3325
225.13511657714844 0 2069.202
226.04290771484375 0 541.62195
226.11892700195312 0 38287.297
227.1027374267578 0 944.18225
227.1222381591797 0 3545.9336
228.1348419189453 0 1344.8412
231.09815979003906 0 1469.4955
233.16517639160156 0 13467.761
234.1250457763672 0 526.53827
234.168701171875 0 1794.8474
235.1080322265625 0 5757.447
235.26344299316406 0 518.9009
236.1033172607422 0 1667.3888
236.1400604248047 0 723.1543
239.0953369140625 0 12561.221
240.096435546875 0 3159.455
241.0820770263672 0 3003.649
241.0937957763672 0 1106.1847
242.15037536621094 0 8730.721
243.13406372070312 0 716.9219
243.14569091796875 0 14575.43 y Water loss 10
244.12933349609375 0 1039.419 y Ammonia loss 12
244.14947509765625 0 1193.0806
245.04495239257812 0 1397.1637
245.12889099121094 0 6367.871
246.12338256835938 0 1398.5424
247.14483642578125 0 1122.0396
251.17575073242188 0 5270.359
252.09942626953125 0 676.4086
252.13401794433594 0 891.82806
254.1138153076172 0 5068.2803
254.15032958984375 0 1278.1991
257.1971740722656 0 878.48517
259.0927429199219 0 4121.5513
261.1599426269531 0 7208.007
262.11920166015625 0 845.6258
262.1638488769531 0 1332.7886
263.05523681640625 0 4498.912
263.1029052734375 0 7622.9165
264.0589599609375 0 755.2684
264.1083068847656 0 1060.7329
264.1344299316406 0 4647.354
265.1391296386719 0 718.8166
268.0932922363281 0 601.8053
268.20135498046875 0 839.5784 a Water loss 2
268.9786682128906 0 1179.2968
269.18658447265625 0 1321.9731
270.1820068359375 0 665.60565
272.1241455078125 0 2722.0103
274.1192932128906 0 1503.8923
275.17529296875 0 1132.5615
279.1453552246094 0 3590.6272
279.66815185546875 0 1482.8936 b 4
280.1295471191406 0 1579.7106
280.14947509765625 0 683.70135
280.1661682128906 0 2787
281.0509948730469 0 1267.3322
281.06610107421875 0 2451.7822
282.14520263671875 0 11964.459
283.1483154296875 0 1622.1587
285.01019287109375 0 10447.927
285.07598876953125 0 803.8659
285.16009521484375 0 1993.5948
286.01019287109375 0 3675.709
286.1037292480469 0 2528.0989
287.00732421875 0 1323.5099
294.1819763183594 0 700.96375
296.1361083984375 0 2496.997
296.17333984375 0 4058.4312
296.19732666015625 0 79120.71 b Water loss 2
297.08380126953125 0 561.38544
297.11944580078125 0 3187.3164
297.13623046875 0 863.6089
297.14044189453125 0 510.424
297.15631103515625 0 12621.956
297.1763000488281 0 693.8564
297.20050048828125 0 14278.41
298.158935546875 0 1611.4775
298.1768798828125 0 3955.8008
298.2033386230469 0 1035.5294
298.6623840332031 0 834.79395
299.06201171875 0 11525.554
299.15283203125 0 660.8004
299.1801452636719 0 649.5849
300.06256103515625 0 4369.307
300.1556701660156 0 752.08997
301.0583190917969 0 1543.5228
301.1881408691406 0 1988.9497 y Water loss 14
303.021240234375 0 1402.9218
303.08673095703125 0 75384.555
303.1708984375 0 3267.4053
304.09014892578125 0 12659.826
304.11163330078125 0 738.5414
305.11480712890625 0 617.6407
307.66693115234375 0 4227.4346 y Water loss 11
308.16143798828125 0 1079.6964 y Ammonia loss 11
310.1035461425781 0 664.98584
313.18841552734375 0 2108.9836
314.14691162109375 0 2815.4182
314.18292236328125 0 12696.317
314.2079162597656 0 17114.523 b 2
315.0866394042969 0 642.17786
315.16796875 0 593.11743
315.1864929199219 0 1968.1177
315.21087646484375 0 2925.8516
316.18707275390625 0 7444.694
316.67193603515625 0 2546.94 y 11
317.190185546875 0 1240.2689
319.1979675292969 0 10920.387 y 14
320.2007751464844 0 1938.5619
325.15228271484375 0 888.73334
326.1708068847656 0 684.0459
327.087158203125 0 1068.7166
328.1148986816406 0 2593.148
329.1812744140625 0 864.64923
330.18243408203125 0 1827.8297
333.15582275390625 0 2758.0159
333.6827697753906 0 614.64886
334.2129821777344 0 1188.7101
336.155029296875 0 687.26227
339.16796875 0 1394.7919
341.183349609375 0 2259.278
342.1795349121094 0 1594.3796
344.1980895996094 0 844.26843
344.9778747558594 0 1477.0212
345.0972595214844 0 3318.0576
345.9786071777344 0 797.9085
346.1004943847656 0 1117.7097
346.1250915527344 0 5539.376
346.1766052246094 0 2702.4646
347.1292419433594 0 1243.9235
348.192138671875 0 2222.2224
349.197509765625 0 1060.7269
350.13446044921875 0 1943.6477
350.1830139160156 0 1758.5629
351.1667785644531 0 8597.206
351.18853759765625 0 823.9429
352.16943359375 0 737.16724
353.1820068359375 0 3972.1206
354.164794921875 0 829.76556
354.1876525878906 0 953.44965
355.0700378417969 0 43362.09
356.0706787109375 0 16520.951
357.0682678222656 0 9101.159
362.20770263671875 0 8002.5625
363.2099304199219 0 1933.2911
364.2092590332031 0 658.86804 y Water loss 10
365.145751953125 0 1565.1721
367.2087097167969 0 3327.4277
367.233642578125 0 841.26086
368.1929626464844 0 3182.6975
369.0968322753906 0 2274.4377
369.17718505859375 0 6869.7383
370.1802673339844 0 1308.3992
370.68060302734375 0 876.32135
371.1005554199219 0 1031.2166
371.1929016113281 0 1726.9783
373.0805358886719 0 4158.532
373.10064697265625 0 510.45093
373.1717224121094 0 2076.1084
373.2142639160156 0 2527.9912 y 10
373.7149658203125 0 1827.549
374.0815734863281 0 1743.3219
375.0812683105469 0 827.6006
381.1772766113281 0 1057.2208
383.15625 0 2353.498
383.263427734375 0 712.40857
385.21954345703125 0 10611.676
386.22259521484375 0 2158.1345
394.1767883300781 0 575.4418
395.1124267578125 0 1380.3704
397.2164306640625 0 1253.6396
398.20184326171875 0 764.9433
398.2447509765625 0 1450.9005
399.18701171875 0 615.38324
399.2439270019531 0 655.023
400.9855041503906 0 656.20087
401.1673583984375 0 3830.5916
402.1700134277344 0 855.9649
406.2207946777344 0 703.0341 b Water loss 7
407.2049255371094 0 1927.0115
408.1883544921875 0 4964.3613
409.1918029785156 0 1202.5732
410.203857421875 0 887.8274
411.1104431152344 0 720.85864
411.2611083984375 0 2356.5867
413.1232604980469 0 4678.6875
414.12744140625 0 1551.5735
415.0367431640625 0 2962.9062
415.1483459472656 0 1200.3627
415.2304992675781 0 1421.6389 b 7
415.26904296875 0 1020.7834
416.0384826660156 0 1446.1915
416.2147216796875 0 1650.4 y Ammonia loss 13
416.25537109375 0 2721.2295
417.0367126464844 0 630.9024
417.2585144042969 0 894.7205
418.995849609375 0 4928.9824
419.9957580566406 0 2013.9158
420.99468994140625 0 1275.8486
424.2295227050781 0 3332.5967
425.2152404785156 0 4080.7417
426.2196044921875 0 988.0034
427.2308044433594 0 1882.792
429.11614990234375 0 705.8765
429.2135009765625 0 898.2784 y Ammonia loss 9
431.1348571777344 0 1727.5514
432.1373291015625 0 651.02454
433.1566467285156 0 3475.6643
433.2411804199219 0 12635.893 y 13
433.2813720703125 0 25263.209
434.205322265625 0 1469.2162
434.2439880371094 0 2780.754
434.2847595214844 0 6166.7256
434.7069091796875 0 1039.4458
435.2059631347656 0 1097.1711
435.2857666015625 0 827.65
440.2115478515625 0 720.4521
441.24920654296875 0 654.75085
442.2406311035156 0 3612.8923
443.2295227050781 0 1777.9238
443.26580810546875 0 3946.3967 b Water loss 3
444.210205078125 0 945.17053
444.26763916015625 0 1305.8063
449.14447021484375 0 2201.1484
450.2367858886719 0 998.421
451.2319641113281 0 845.43945
451.8932189941406 0 3055.6423 y Water loss 4
452.21978759765625 0 2915.7666 y Ammonia loss 4
454.7071533203125 0 760.919
459.211181640625 0 914.7112
461.2762756347656 0 21959.307 b 3
462.19903564453125 0 1997.3696
462.279052734375 0 6722.705
463.283203125 0 930.9566
465.1399841308594 0 1931.029
466.1935729980469 0 837.2951
466.7247619628906 0 669.4052
468.25714111328125 0 1301.0769
469.2446594238281 0 1533.7773
470.2252197265625 0 7766.4395
470.7398986816406 0 928.7554 b Water loss 8
471.22637939453125 0 1616.51
471.7446594238281 0 1278.6799
477.2561950683594 0 1737.5767
478.2415466308594 0 4036.5352
479.226318359375 0 2089.5186
480.20989990234375 0 7094.799
480.72015380859375 0 2021.6089
481.2158508300781 0 3095.524
484.2043762207031 0 1198.7695
484.7169494628906 0 637.3708
486.2682800292969 0 4234.273 y Water loss 12
487.2497863769531 0 1905.4218 y Ammonia loss 12
488.736083984375 0 1352.5802
489.05438232421875 0 1082.8182
490.20733642578125 0 1080.0197
493.7279968261719 0 2457.8699
494.18841552734375 0 921.09406
494.230712890625 0 2093.1538
494.73675537109375 0 594.9533
495.2674560546875 0 2852.4604
496.2514953613281 0 9517.448
497.2532043457031 0 2001.3655
498.2200012207031 0 20621.918
499.22283935546875 0 6060.437
500.23065185546875 0 1508.2335
500.91729736328125 0 904.0957 y Water loss 3
501.25152587890625 0 598.6496 y Ammonia loss 3
502.2145690917969 0 1608.1857
502.7334899902344 0 3672.2158
503.2274475097656 0 1242.4666
504.27789306640625 0 28334.568 y 12
505.28057861328125 0 7093.783
506.27679443359375 0 673.2175
512.1975708007812 0 1074.3695
513.2779541015625 0 3816.4172
515.244384765625 0 729.5712
521.2679443359375 0 1100.6499
529.2660522460938 0 839.427
530.2091674804688 0 8082.8594
530.333251953125 0 2111.9185
531.2132568359375 0 2081.3633
536.6073608398438 0 756.4691
538.2689819335938 0 6044.314
538.7711791992188 0 4606.9004
539.2700805664062 0 807.0818
540.3200073242188 0 3008.3914 b Water loss 4
543.2614135742188 0 2337.9382
543.76416015625 0 1399.8071
544.2662963867188 0 2564.4985 b 9
544.7667236328125 0 662.4728
545.269287109375 0 624.6035
546.2416381835938 0 1107.9795
547.951416015625 0 695.30096 b 14
548.2863159179688 0 960.1568
551.281494140625 0 682.3574
552.267578125 0 4608.545
552.3191528320312 0 916.235
552.7691650390625 0 3794.6206
553.2669677734375 0 750.30884
555.3242797851562 0 3059.104
556.2685546875 0 571.9386
556.3264770507812 0 875.4159
558.328369140625 0 4985.3647 b 4
559.3316650390625 0 2422.6074
560.7689208984375 0 629.95953
561.2765502929688 0 729.3699
567.284912109375 0 740.1104
569.7771606445312 0 697.9847
570.2823486328125 0 689.8344
571.2711791992188 0 785.3908
573.2266235351562 0 757.3739
578.1270751953125 0 886.64044
578.2849731445312 0 2860.8809
578.7845458984375 0 1356.8567
579.2865600585938 0 1287.1102
580.281494140625 0 1717.6732 y Water loss 6
581.2598876953125 0 1616.1049
582.2896118164062 0 1085.8342
582.7838745117188 0 3749.626
583.282958984375 0 2772.6296
591.2427368164062 0 1104.3882
591.294189453125 0 2404.598
591.7942504882812 0 1531.7858 b Water loss 10
592.2971801757812 0 853.24054
595.3428955078125 0 830.08844
597.2974853515625 0 2770.9404
598.2871704101562 0 1988.9451
599.2671508789062 0 10456.855
599.3175659179688 0 736.28827
600.2705078125 0 2862.035
601.2740478515625 0 1006.74927
602.7910766601562 0 2773.1582
603.2916870117188 0 2019.5083
608.3521728515625 0 617.4081
609.2510375976562 0 4079.4683
609.296630859375 0 828.62744
609.7918090820312 0 1035.7605
610.1845092773438 0 5424.8926
610.2505493164062 0 1157.8896
611.1846313476562 0 5787.1426
611.7964477539062 0 5436.083
612.1827392578125 0 4216.0273
612.297607421875 0 3844.1401
612.4335327148438 0 878.31396
612.8091430664062 0 2444.8315
614.3258056640625 0 6761.488 y Water loss 11
615.3097534179688 0 15725.916 y Ammonia loss 11
616.3123168945312 0 5354.8125
617.3135375976562 0 2538.4136
617.8067016601562 0 8990.409
618.3060302734375 0 11263.858
618.8054809570312 0 4716.109
619.3114624023438 0 764.5415
620.289306640625 0 738.0291
625.2828979492188 0 720.7281
626.2860107421875 0 756.97894
626.3617553710938 0 2886.4255
626.81201171875 0 25844.469
627.2617797851562 0 41691.543
627.3136596679688 0 14780.164
627.813720703125 0 6847.4165
628.2650146484375 0 14183.378
628.8092041015625 0 1015.91644 y Water loss 5
629.2672729492188 0 2636.7795
629.80419921875 0 888.7752
632.3363037109375 0 19910.312 y 11
633.3381958007812 0 7396.0586
634.3379516601562 0 1538.902
636.3153076171875 0 951.8023
637.815673828125 0 4061.9941 y 5
638.3167114257812 0 3938.2668
638.81640625 0 1105.8978
643.29345703125 0 2721.4602
644.2930297851562 0 1756.94
645.3048095703125 0 2341.1501
655.3403930664062 0 10391.919
655.8414306640625 0 7709.2227 b Water loss 11
656.3414916992188 0 2108.0637
659.8253784179688 0 786.5291
660.31787109375 0 1873.6107
663.3328247070312 0 644.98
667.337890625 0 871.8583
668.332275390625 0 2038.6512
668.8257446289062 0 4534.183
669.3241577148438 0 2009.7543
669.8233642578125 0 801.4162
676.3310546875 0 1066.197
677.3355102539062 0 25593.787 y Water loss 4
677.8350830078125 0 26820.771 y Ammonia loss 4
678.3339233398438 0 13938.986
678.8348999023438 0 4177.538
679.3374633789062 0 1089.4414
684.3682250976562 0 1146.1968
686.3410034179688 0 55414.66 y 4
686.8422241210938 0 40778.465
687.3439331054688 0 16548.37
687.8450927734375 0 3838.866
691.3431396484375 0 1379.7781 b Water loss 12
694.3384399414062 0 1952.2606
695.343017578125 0 769.828
698.3400268554688 0 952.4606
700.3479614257812 0 1737.6752 b 12
700.846435546875 0 2300.1943
701.3489990234375 0 1445.6804
710.3840942382812 0 1922.2189
712.35107421875 0 6401.054
713.354736328125 0 3399.1284
714.3575439453125 0 1182.547
720.3659057617188 0 755.9009
722.3362426757812 0 2894.2334
724.4046630859375 0 1830.5989 b Water loss 6
725.4080200195312 0 1187.5627
726.337646484375 0 790.1938
726.4102172851562 0 678.2072
727.4083862304688 0 832.4156 y Water loss 10
728.3958740234375 0 1468.7114 y Ammonia loss 10
728.87646484375 0 1580.3229
729.3773193359375 0 722.67004
729.879638671875 0 860.09753
732.3458862304688 0 664.4912
737.3941040039062 0 1959.8768
738.307373046875 0 870.06586
738.3807373046875 0 1200.3499
739.3673095703125 0 845.5945
740.3460693359375 0 17485.348
741.3486328125 0 6139.4624
742.350830078125 0 2102.978
742.4151000976562 0 3258.402 b 6
743.4163208007812 0 1632.2498
745.3452758789062 0 789.01733
745.420166015625 0 9168.401 y 10
746.4216918945312 0 3402.5212
747.4195556640625 0 1456.9797
749.3390502929688 0 747.3224
750.8702392578125 0 6377.1323 y Water loss 3
751.369873046875 0 5993.023 y Ammonia loss 3
751.867431640625 0 2117.518
752.3702392578125 0 1445.7529
755.4047241210938 0 1663.4336
757.3748168945312 0 4898.1284 b 13
758.375732421875 0 1830.7267
759.3853759765625 0 1370.8787
759.8746337890625 0 5782.7744 y 3
760.3761596679688 0 5373.4336
760.8782348632812 0 2250.5125
761.38525390625 0 1161.9064
771.3508911132812 0 3180.0857
772.3549194335938 0 1524.828
774.33056640625 0 2707.483
775.3351440429688 0 1263.1309
807.4127807617188 0 1719.8041 y Water loss 2
807.9129028320312 0 1793.7285 y Ammonia loss 2
808.410400390625 0 1741.9982
811.4348754882812 0 1995.2067 b Water loss 7
812.4390258789062 0 1096.5804
813.43359375 0 1790.3179
816.4183349609375 0 860.4033 y 2
824.3978881835938 0 778.185
829.4447631835938 0 1885.0939 b 7
830.4442749023438 0 1678.3966
839.4122314453125 0 876.51294
840.4168090820312 0 2029.2384
841.4254150390625 0 2543.2112
842.3844604492188 0 1023.7526
848.4334716796875 0 952.0649 b Ammonia loss 15
848.93505859375 0 1218.4299
849.4220581054688 0 1100.1249
850.392578125 0 2477.5706
851.3941040039062 0 1565.6912
856.4502563476562 0 1553.5165 y Water loss 9
856.9451293945312 0 1629.5221 b 15
857.4403076171875 0 3225.6606 y Ammonia loss 9
858.4368896484375 0 1235.0802 y Ammonia loss 1
859.4267578125 0 1372.7094
861.4085083007812 0 952.919
863.3807983398438 0 2432.3608
868.4041137695312 0 19991.994
869.40576171875 0 10024.039
870.408203125 0 2742.6797
873.4566040039062 0 2902.6418
874.4612426757812 0 4957.91 y 9
875.4600830078125 0 2182.2668
879.4111328125 0 3602.0598
880.4127197265625 0 1356.4688
887.41357421875 0 4307.746
888.4166870117188 0 1633.4365
889.3955078125 0 2300.986
890.402587890625 0 1690.3835
907.4046020507812 0 12709.602
908.4063720703125 0 6707.708
909.408935546875 0 1175.2352
911.448974609375 0 1091.2091
921.4314575195312 0 1706.4866
922.4258422851562 0 1830.745
939.4402465820312 0 10170.124
940.4486083984375 0 6639.1436
941.4560546875 0 2251.2253
956.455078125 0 2358.1797
957.4581909179688 0 993.2575
958.4862060546875 0 4596.034 b 8
959.4904174804688 0 3707.7783
960.4881591796875 0 8495.662
961.4898681640625 0 3065.5725
962.4660034179688 0 848.261
971.4755249023438 0 979.42676
986.4678344726562 0 822.06384 y Ammonia loss 8
988.4653930664062 0 1476.7915
1001.4990844726562 0 783.036
1003.5078125 0 1392.2335 y 8
1008.4714965820312 0 776.3034
1015.4686279296875 0 1516.4314
1016.475341796875 0 1041.2806
1025.49072265625 0 1024.1246
1035.4794921875 0 801.53723
1036.4603271484375 0 2247.9868
1037.4696044921875 0 998.87067
1040.4964599609375 0 1309.984
1053.4847412109375 0 7372.3228
1054.4857177734375 0 5113.923
1055.4853515625 0 1853.9443
1058.509521484375 0 2025.2471
1059.5213623046875 0 1657.174
1069.5162353515625 0 2977.416 b Water loss 9
1070.5135498046875 0 3356.9482
1071.5106201171875 0 1788.444
1072.5263671875 0 883.1371 y Water loss 7
1075.5220947265625 0 4196.11
1076.522216796875 0 2885.2446
1084.5242919921875 0 1218.4863
1085.52685546875 0 1298.6897
1086.51611328125 0 1358.9037
1087.527587890625 0 16069.56 b 9
1088.53076171875 0 8847.197
1089.5318603515625 0 4191.757
1090.53662109375 0 2759.3074 y 7
1091.540771484375 0 1786.0334
1103.525634765625 0 4843.5264
1104.526123046875 0 2183.8413
1138.5484619140625 0 2231.8809
1139.5426025390625 0 1255.0251
1155.55908203125 0 3221.8413
1156.564208984375 0 1775.4635
1164.5545654296875 0 2114.3308
1165.5631103515625 0 1380.4403
1177.568359375 0 994.85834 y 6
1178.569580078125 0 1115.286
1181.5765380859375 0 6179.8623
1182.583984375 0 3847.9712 b Water loss 10
1183.5836181640625 0 1809.2902
1200.60498046875 0 3026.0684 b 10
1201.609619140625 0 2600.9023
1212.61328125 0 1758.3325
1213.6180419921875 0 1008.0522
1224.6171875 0 700.46906
1234.6025390625 0 4572.61
1235.6041259765625 0 4011.4114
1236.6036376953125 0 1928.3883
1252.6119384765625 0 11586.65
1253.6153564453125 0 7626.5825
1254.6163330078125 0 2942.8618
1256.6102294921875 0 1396.2438 y Water loss 5
1257.6004638671875 0 1791.313 y Ammonia loss 5
1274.616943359375 0 6124.295 y 5
1275.6185302734375 0 3227.295
1276.6260986328125 0 1981.5697
1309.6710205078125 0 732.53394
1329.66943359375 0 1032.2059
1371.667236328125 0 1754.7019 y 4
1372.68017578125 0 1219.3353
1399.6846923828125 0 1443.744 b 12
1513.75439453125 0 1036.8684 b 13
2580.81591796875 0 698.597
3071.2314453125 0 852.6034

Spectrum Details

|  |  |
| --- | --- |
| Matched peaks? Matched peaks The total absolute number of peaks matched. Additionally in brackets the total fraction of peaks matched and the total number of peaks is shown. | 89 (12.90% of 690) |
| FDR? FDR The false discovery rate estimated for this peptide. It is calculated by matching all theoretical fragments with a non-integer shift with the raw peaks for this spectrum. This is done with 40 different shifts. The resulting percentage is the average number of annotated peaks over the number of annotated peaks with the correct spectrum. | 1.34% |
| Satellite FDR? Satellite FDR See the FDR for details on its calculation. This satellite ion specific FDR only contains the satellite ions (d/w) for I/L/J positions. | ∞ |
| PSM Score? PSM Score The PSM Score as given by Hecklib to this annotated spectrum. It is shown with three significant figures. | 772 |

#### Spectrum 8190? Spectrum 8190 The raw spectrum of this peptide as annotated by Hecklib. The fragments are coloured according to ion type (see legend). Any peaks with a star '\*' as text can be hovered over to see the full details, first the ion type second the mass shift type. By hovering over the amino acids in the peptide or ions in the legend the corresponding peaks are highlighted. By toggling the 'Unassigned' label you can turn the background (unassigned) peaks on or off in the plot. By updating the slider in the Ion legend you can update the spectrum to only show the top X% of the peaks with labels. The top X% means any peak that is within X% of the highest intensity. By dragging in the spectrum you can zoom in to a specific part of the spectrum and use 'Zoom Out' to get back to the original zoom level. The annotation of the spectrum is based on the given sequence in the peptides file and is done with different software so inconsistencies are likely. The peaks are annotated based on the given sequence, with 20 ppm tolerance.

Copy Data

##### Spectrum 8190 (TSV)

###### Preview

```
Loading example...
```

*Click on the button to copy the data to your clipboard.*

Mz MinMz MaxIntensity Max

WidthHeightPeptide font sizePeptide stroke widthSpectrum font sizeSpectrum stroke widthCompact peptide

Ion legend

wxyz

abcd

OtherUnassignedIon^Charge^_Position_Show for top:%

VTJFPPSSEEJQANKAT

01.92e+43.85e+45.77e+47.69e+4

Zoom Out

y^+1^_1_a^+1^_2_y^+1^_2_a^+1^_2_b^+1^_2_y^+1^_2_b^+1^_2_y^+3^_7_y^+2^_5_a^+1^_3_b^+2^_5_b^+1^_3_y^+1^_3_y^+2^_6_b^+1^_3_y^+2^_6_y^+1^_3_y^+2^_7_y^+2^_7_b^+2^_8_y^+1^_4_y^+1^_4_b^+1^_4_y^+3^_13_y^+3^_13_b^+1^_4_y^+1^_5_y^+1^_5_y^+1^_5_b^+1^_5_b^+2^_10_b^+3^_15_b^+1^_5_y^+3^_16_y^+2^_11_b^+2^_11_y^+1^_6_y^+1^_6_y^+2^_12_y^+1^_6_y^+2^_12_b^+2^_12_y^+2^_13_y^+2^_13_y^+2^_13_b^+2^_13_y^+1^_7_b^+1^_7_y^+1^_7_y^+2^_14_y^+2^_14_b^+2^_14_y^+2^_14_y^+2^_15_y^+2^_15_b^+1^_8_b^+2^_15_b^+1^_8_b^+2^_16_y^+1^_8_y^+2^_16_y^+2^_16_y^+1^_8_b^+1^_9_y^+1^_9_b^+1^_10_y^+1^_10_b^+1^_10_y^+1^_10_y^+1^_11_y^+1^_11_b^+1^_11_y^+1^_12_y^+1^_12_y^+1^_12_y^+1^_13_

0776155223283104

Fragment Matches Table

Show background peaks

| Position | Ion type | Intensity | mz Theoretical | mz Error (Th) | mz Error (ppm) | Charge | Series Number |
| --- | --- | --- | --- | --- | --- | --- | --- |
| 17 | y | 2.86E+04 | 120.1 | 0.0002992 | 2.492 | +1 | 1 |
| - | - | 4.107E+04 | 120.1 | - | - | 0 | - |
| - | - | 1428 | 121.1 | - | - | 0 | - |
| - | - | 3904 | 121.1 | - | - | 0 | - |
| - | - | 352.9 | 121.4 | - | - | 0 | - |
| - | - | 377.3 | 122.1 | - | - | 0 | - |
| - | - | 652 | 123 | - | - | 0 | - |
| - | - | 2751 | 126.1 | - | - | 0 | - |
| - | - | 8057 | 128.1 | - | - | 0 | - |
| - | - | 4142 | 129.1 | - | - | 0 | - |
| - | - | 1290 | 129.1 | - | - | 0 | - |
| - | - | 3.738E+04 | 129.1 | - | - | 0 | - |
| - | - | 1951 | 130.1 | - | - | 0 | - |
| - | - | 1279 | 130.1 | - | - | 0 | - |
| - | - | 2488 | 130.1 | - | - | 0 | - |
| - | - | 1851 | 131.1 | - | - | 0 | - |
| - | - | 1376 | 134.1 | - | - | 0 | - |
| - | - | 523 | 135 | - | - | 0 | - |
| - | - | 7512 | 136.1 | - | - | 0 | - |
| - | - | 951.5 | 137.1 | - | - | 0 | - |
| - | - | 1837 | 138.1 | - | - | 0 | - |
| - | - | 1003 | 138.1 | - | - | 0 | - |
| - | - | 1072 | 139.1 | - | - | 0 | - |
| - | - | 1060 | 141.1 | - | - | 0 | - |
| - | - | 502.8 | 145 | - | - | 0 | - |
| - | - | 509.5 | 145.1 | - | - | 0 | - |
| - | - | 457 | 147 | - | - | 0 | - |
| - | - | 696.7 | 147.1 | - | - | 0 | - |
| - | - | 932.7 | 147.1 | - | - | 0 | - |
| - | - | 2342 | 149 | - | - | 0 | - |
| - | - | 2942 | 152.1 | - | - | 0 | - |
| - | - | 4234 | 153 | - | - | 0 | - |
| - | - | 527.6 | 154 | - | - | 0 | - |
| - | - | 757.5 | 154.1 | - | - | 0 | - |
| - | - | 1344 | 154.1 | - | - | 0 | - |
| - | - | 2012 | 155.1 | - | - | 0 | - |
| 2 | a | 5736 | 155.1 | 0.000305 | 1.967 | +1 | 2 |
| - | - | 642.1 | 156.1 | - | - | 0 | - |
| - | - | 885.5 | 157.1 | - | - | 0 | - |
| - | - | 2577 | 157.1 | - | - | 0 | - |
| - | - | 6051 | 159.1 | - | - | 0 | - |
| - | - | 708.5 | 160.1 | - | - | 0 | - |
| - | - | 1270 | 161.1 | - | - | 0 | - |
| - | - | 656 | 162.1 | - | - | 0 | - |
| - | - | 1101 | 164.1 | - | - | 0 | - |
| - | - | 1407 | 165.1 | - | - | 0 | - |
| - | - | 1903 | 165.1 | - | - | 0 | - |
| - | - | 494.3 | 166.1 | - | - | 0 | - |
| - | - | 3558 | 167.1 | - | - | 0 | - |
| - | - | 2460 | 167.1 | - | - | 0 | - |
| - | - | 1707 | 167.1 | - | - | 0 | - |
| - | - | 452.1 | 167.3 | - | - | 0 | - |
| - | - | 641 | 169.1 | - | - | 0 | - |
| - | - | 517.4 | 169.1 | - | - | 0 | - |
| - | - | 8708 | 169.1 | - | - | 0 | - |
| - | - | 796.3 | 170.1 | - | - | 0 | - |
| - | - | 2416 | 171 | - | - | 0 | - |
| - | - | 1242 | 171.1 | - | - | 0 | - |
| - | - | 498.4 | 171.1 | - | - | 0 | - |
| - | - | 909.2 | 172.1 | - | - | 0 | - |
| 16 | y | 4274 | 173.1 | 0.0002012 | 1.162 | +1 | 2 |
| 2 | a | 1.964E+04 | 173.1 | 0.0002842 | 1.641 | +1 | 2 |
| - | - | 1064 | 174.1 | - | - | 0 | - |
| - | - | 1312 | 175.1 | - | - | 0 | - |
| - | - | 4526 | 177.1 | - | - | 0 | - |
| - | - | 1896 | 179 | - | - | 0 | - |
| - | - | 609.9 | 181.1 | - | - | 0 | - |
| - | - | 963.4 | 181.1 | - | - | 0 | - |
| - | - | 3195 | 182.1 | - | - | 0 | - |
| - | - | 2143 | 182.1 | - | - | 0 | - |
| - | - | 2020 | 182.1 | - | - | 0 | - |
| - | - | 1974 | 183.1 | - | - | 0 | - |
| 2 | b | 2.515E+04 | 183.1 | 0.0002787 | 1.522 | +1 | 2 |
| - | - | 2434 | 184.1 | - | - | 0 | - |
| - | - | 2610 | 184.1 | - | - | 0 | - |
| - | - | 4005 | 185.1 | - | - | 0 | - |
| - | - | 668.9 | 185.1 | - | - | 0 | - |
| - | - | 1422 | 186.1 | - | - | 0 | - |
| - | - | 753.3 | 187.1 | - | - | 0 | - |
| - | - | 8285 | 187.1 | - | - | 0 | - |
| - | - | 1166 | 188.1 | - | - | 0 | - |
| - | - | 2440 | 189.1 | - | - | 0 | - |
| - | - | 684.6 | 191.1 | - | - | 0 | - |
| 16 | y | 6482 | 191.1 | 0.0002871 | 1.502 | +1 | 2 |
| - | - | 1815 | 191.1 | - | - | 0 | - |
| - | - | 640 | 192.1 | - | - | 0 | - |
| - | - | 564.5 | 195 | - | - | 0 | - |
| - | - | 3556 | 195 | - | - | 0 | - |
| - | - | 3672 | 195.1 | - | - | 0 | - |
| - | - | 1579 | 196.1 | - | - | 0 | - |
| - | - | 1.131E+04 | 197.1 | - | - | 0 | - |
| - | - | 2547 | 198.1 | - | - | 0 | - |
| - | - | 1637 | 198.1 | - | - | 0 | - |
| - | - | 770.9 | 198.1 | - | - | 0 | - |
| - | - | 2999 | 199.1 | - | - | 0 | - |
| - | - | 2577 | 200.1 | - | - | 0 | - |
| - | - | 2473 | 200.1 | - | - | 0 | - |
| - | - | 856 | 201.1 | - | - | 0 | - |
| 2 | b | 2.473E+04 | 201.1 | 0.0002121 | 1.055 | +1 | 2 |
| - | - | 2129 | 202.1 | - | - | 0 | - |
| - | - | 752.8 | 203.1 | - | - | 0 | - |
| - | - | 539.4 | 206.2 | - | - | 0 | - |
| - | - | 3060 | 208.1 | - | - | 0 | - |
| - | - | 7710 | 209.1 | - | - | 0 | - |
| - | - | 926.4 | 210.1 | - | - | 0 | - |
| - | - | 794.9 | 212.1 | - | - | 0 | - |
| - | - | 795.8 | 213.1 | - | - | 0 | - |
| - | - | 678.7 | 213.1 | - | - | 0 | - |
| - | - | 910.3 | 214.1 | - | - | 0 | - |
| - | - | 825.2 | 214.1 | - | - | 0 | - |
| - | - | 7536 | 215.1 | - | - | 0 | - |
| - | - | 1444 | 216.1 | - | - | 0 | - |
| - | - | 4910 | 217.1 | - | - | 0 | - |
| - | - | 2523 | 217.1 | - | - | 0 | - |
| - | - | 6703 | 217.1 | - | - | 0 | - |
| - | - | 938.7 | 218.1 | - | - | 0 | - |
| - | - | 1306 | 219 | - | - | 0 | - |
| - | - | 1490 | 219.2 | - | - | 0 | - |
| - | - | 4247 | 221.1 | - | - | 0 | - |
| - | - | 1349 | 222.1 | - | - | 0 | - |
| - | - | 1860 | 225 | - | - | 0 | - |
| - | - | 2452 | 225.1 | - | - | 0 | - |
| - | - | 1397 | 225.1 | - | - | 0 | - |
| - | - | 2.82E+04 | 226.1 | - | - | 0 | - |
| - | - | 720.7 | 227 | - | - | 0 | - |
| - | - | 2165 | 227.1 | - | - | 0 | - |
| - | - | 1525 | 228.1 | - | - | 0 | - |
| - | - | 1068 | 229.1 | - | - | 0 | - |
| - | - | 552.4 | 230.5 | - | - | 0 | - |
| - | - | 726.4 | 231.1 | - | - | 0 | - |
| - | - | 543.2 | 232.2 | - | - | 0 | - |
| - | - | 1.032E+04 | 233.2 | - | - | 0 | - |
| - | - | 1753 | 234.2 | - | - | 0 | - |
| - | - | 6084 | 235.1 | - | - | 0 | - |
| - | - | 1127 | 236.1 | - | - | 0 | - |
| - | - | 886.5 | 236.1 | - | - | 0 | - |
| - | - | 516.3 | 237.1 | - | - | 0 | - |
| - | - | 1.169E+04 | 239.1 | - | - | 0 | - |
| - | - | 2413 | 240.1 | - | - | 0 | - |
| - | - | 1476 | 241.1 | - | - | 0 | - |
| - | - | 1375 | 241.1 | - | - | 0 | - |
| - | - | 5955 | 242.2 | - | - | 0 | - |
| - | - | 939.8 | 243.1 | - | - | 0 | - |
| 11 | y | 1.1E+04 | 243.1 | 0.004146 | 17.05 | +3 | 7 |
| 13 | y | 1095 | 244.1 | 0.0004257 | 1.744 | +2 | 5 |
| - | - | 843 | 244.1 | - | - | 0 | - |
| - | - | 1381 | 245 | - | - | 0 | - |
| - | - | 3487 | 245.1 | - | - | 0 | - |
| - | - | 1357 | 246.1 | - | - | 0 | - |
| - | - | 1008 | 247.1 | - | - | 0 | - |
| - | - | 546.3 | 248.7 | - | - | 0 | - |
| - | - | 3778 | 251.2 | - | - | 0 | - |
| - | - | 768 | 252.1 | - | - | 0 | - |
| - | - | 1061 | 252.1 | - | - | 0 | - |
| - | - | 626 | 253.1 | - | - | 0 | - |
| - | - | 4070 | 253.2 | - | - | 0 | - |
| - | - | 4195 | 254.1 | - | - | 0 | - |
| - | - | 1244 | 254.1 | - | - | 0 | - |
| - | - | 583.8 | 254.2 | - | - | 0 | - |
| - | - | 631.7 | 257.2 | - | - | 0 | - |
| - | - | 4133 | 259.1 | - | - | 0 | - |
| - | - | 5474 | 261.2 | - | - | 0 | - |
| - | - | 1113 | 262.1 | - | - | 0 | - |
| - | - | 704.3 | 262.2 | - | - | 0 | - |
| - | - | 4212 | 263.1 | - | - | 0 | - |
| - | - | 8330 | 263.1 | - | - | 0 | - |
| - | - | 545.3 | 263.1 | - | - | 0 | - |
| - | - | 654.9 | 264.1 | - | - | 0 | - |
| - | - | 3503 | 264.1 | - | - | 0 | - |
| - | - | 657 | 267 | - | - | 0 | - |
| 3 | a | 684.8 | 268.2 | 0.0006222 | 2.32 | +1 | 3 |
| - | - | 615.2 | 269 | - | - | 0 | - |
| - | - | 1927 | 270.2 | - | - | 0 | - |
| - | - | 1735 | 272.1 | - | - | 0 | - |
| - | - | 604.7 | 274.1 | - | - | 0 | - |
| - | - | 963 | 275.2 | - | - | 0 | - |
| - | - | 3465 | 279.1 | - | - | 0 | - |
| 5 | b | 1358 | 279.7 | 0.0005241 | 1.874 | +2 | 5 |
| - | - | 996.3 | 280.1 | - | - | 0 | - |
| - | - | 800.6 | 280.1 | - | - | 0 | - |
| - | - | 2683 | 280.2 | - | - | 0 | - |
| - | - | 1123 | 281.1 | - | - | 0 | - |
| - | - | 1667 | 281.1 | - | - | 0 | - |
| - | - | 9898 | 282.1 | - | - | 0 | - |
| - | - | 859.4 | 283.1 | - | - | 0 | - |
| - | - | 1.038E+04 | 285 | - | - | 0 | - |
| - | - | 985 | 285.2 | - | - | 0 | - |
| - | - | 3619 | 286 | - | - | 0 | - |
| - | - | 2152 | 286.1 | - | - | 0 | - |
| - | - | 1238 | 287 | - | - | 0 | - |
| - | - | 1003 | 289.2 | - | - | 0 | - |
| - | - | 1489 | 296.1 | - | - | 0 | - |
| - | - | 3094 | 296.2 | - | - | 0 | - |
| 3 | b | 5.9E+04 | 296.2 | 0.0003059 | 1.033 | +1 | 3 |
| - | - | 1731 | 297.1 | - | - | 0 | - |
| - | - | 1.096E+04 | 297.2 | - | - | 0 | - |
| - | - | 382.8 | 297.2 | - | - | 0 | - |
| - | - | 650.6 | 297.2 | - | - | 0 | - |
| - | - | 1.135E+04 | 297.2 | - | - | 0 | - |
| - | - | 1564 | 298.2 | - | - | 0 | - |
| - | - | 4205 | 298.2 | - | - | 0 | - |
| - | - | 1248 | 298.2 | - | - | 0 | - |
| - | - | 1.274E+04 | 299.1 | - | - | 0 | - |
| - | - | 1604 | 299.2 | - | - | 0 | - |
| - | - | 610.8 | 299.2 | - | - | 0 | - |
| - | - | 584.5 | 299.4 | - | - | 0 | - |
| - | - | 577.3 | 299.7 | - | - | 0 | - |
| - | - | 2988 | 300.1 | - | - | 0 | - |
| - | - | 2004 | 301.1 | - | - | 0 | - |
| 15 | y | 1546 | 301.2 | 0.000926 | 3.075 | +1 | 3 |
| - | - | 7.618E+04 | 303.1 | - | - | 0 | - |
| - | - | 2116 | 303.2 | - | - | 0 | - |
| - | - | 1.376E+04 | 304.1 | - | - | 0 | - |
| - | - | 1086 | 304.1 | - | - | 0 | - |
| 12 | y | 2582 | 307.7 | 2.06E-05 | 0.06694 | +2 | 6 |
| - | - | 822.5 | 308.2 | - | - | 0 | - |
| - | - | 2084 | 313.2 | - | - | 0 | - |
| - | - | 1147 | 314.1 | - | - | 0 | - |
| - | - | 8385 | 314.2 | - | - | 0 | - |
| 3 | b | 1.246E+04 | 314.2 | 0.0002698 | 0.8587 | +1 | 3 |
| - | - | 1189 | 315.2 | - | - | 0 | - |
| - | - | 2289 | 315.2 | - | - | 0 | - |
| - | - | 7871 | 316.2 | - | - | 0 | - |
| 12 | y | 1487 | 316.7 | 0.0007395 | 2.335 | +2 | 6 |
| - | - | 2124 | 317.2 | - | - | 0 | - |
| 15 | y | 7178 | 319.2 | 0.0004627 | 1.449 | +1 | 3 |
| - | - | 1673 | 320.2 | - | - | 0 | - |
| - | - | 639.2 | 322.2 | - | - | 0 | - |
| - | - | 644.3 | 323.2 | - | - | 0 | - |
| - | - | 714.3 | 325.2 | - | - | 0 | - |
| - | - | 668.7 | 327.1 | - | - | 0 | - |
| - | - | 1491 | 328.1 | - | - | 0 | - |
| - | - | 581.8 | 329.1 | - | - | 0 | - |
| - | - | 1461 | 330.2 | - | - | 0 | - |
| - | - | 798.9 | 332.1 | - | - | 0 | - |
| - | - | 1917 | 333.2 | - | - | 0 | - |
| - | - | 685.5 | 337.2 | - | - | 0 | - |
| - | - | 512.3 | 340.2 | - | - | 0 | - |
| - | - | 1923 | 341.2 | - | - | 0 | - |
| - | - | 910.9 | 342.2 | - | - | 0 | - |
| - | - | 989.6 | 344.2 | - | - | 0 | - |
| - | - | 848.5 | 345 | - | - | 0 | - |
| - | - | 4084 | 345.1 | - | - | 0 | - |
| - | - | 1254 | 346 | - | - | 0 | - |
| - | - | 879.4 | 346.1 | - | - | 0 | - |
| - | - | 3243 | 346.1 | - | - | 0 | - |
| - | - | 2496 | 346.2 | - | - | 0 | - |
| - | - | 1043 | 348.2 | - | - | 0 | - |
| - | - | 2689 | 350.1 | - | - | 0 | - |
| - | - | 1647 | 350.2 | - | - | 0 | - |
| - | - | 5222 | 351.2 | - | - | 0 | - |
| - | - | 673.4 | 351.2 | - | - | 0 | - |
| - | - | 1111 | 352.2 | - | - | 0 | - |
| - | - | 3375 | 353.2 | - | - | 0 | - |
| - | - | 1153 | 354.2 | - | - | 0 | - |
| - | - | 3.776E+04 | 355.1 | - | - | 0 | - |
| - | - | 970.4 | 355.2 | - | - | 0 | - |
| - | - | 1.792E+04 | 356.1 | - | - | 0 | - |
| - | - | 8937 | 357.1 | - | - | 0 | - |
| - | - | 789.5 | 358.1 | - | - | 0 | - |
| - | - | 728 | 360.2 | - | - | 0 | - |
| - | - | 820.5 | 361.1 | - | - | 0 | - |
| - | - | 6428 | 362.2 | - | - | 0 | - |
| - | - | 707 | 363.2 | - | - | 0 | - |
| 11 | y | 1222 | 364.7 | 0.0002414 | 0.6619 | +2 | 7 |
| - | - | 1228 | 365.1 | - | - | 0 | - |
| - | - | 3442 | 367.2 | - | - | 0 | - |
| - | - | 3089 | 368.2 | - | - | 0 | - |
| - | - | 2528 | 369.1 | - | - | 0 | - |
| - | - | 5858 | 369.2 | - | - | 0 | - |
| - | - | 591.7 | 370.1 | - | - | 0 | - |
| - | - | 1254 | 370.2 | - | - | 0 | - |
| - | - | 846.3 | 370.7 | - | - | 0 | - |
| - | - | 1090 | 371.1 | - | - | 0 | - |
| - | - | 1494 | 371.2 | - | - | 0 | - |
| - | - | 636.4 | 371.7 | - | - | 0 | - |
| - | - | 658.6 | 372.1 | - | - | 0 | - |
| - | - | 3528 | 373.1 | - | - | 0 | - |
| - | - | 1668 | 373.2 | - | - | 0 | - |
| 11 | y | 2357 | 373.2 | 0.0001548 | 0.4146 | +2 | 7 |
| - | - | 1265 | 373.7 | - | - | 0 | - |
| - | - | 1628 | 374.1 | - | - | 0 | - |
| - | - | 1206 | 375.1 | - | - | 0 | - |
| - | - | 2104 | 383.2 | - | - | 0 | - |
| - | - | 541.2 | 383.2 | - | - | 0 | - |
| - | - | 9823 | 385.2 | - | - | 0 | - |
| - | - | 900.3 | 386.2 | - | - | 0 | - |
| - | - | 1214 | 386.2 | - | - | 0 | - |
| - | - | 1052 | 393.2 | - | - | 0 | - |
| - | - | 910.4 | 395.1 | - | - | 0 | - |
| - | - | 988.8 | 397.1 | - | - | 0 | - |
| - | - | 661.4 | 397.2 | - | - | 0 | - |
| - | - | 927.7 | 398.2 | - | - | 0 | - |
| - | - | 706.1 | 399.2 | - | - | 0 | - |
| - | - | 765.3 | 401 | - | - | 0 | - |
| - | - | 4531 | 401.2 | - | - | 0 | - |
| - | - | 854.6 | 402.2 | - | - | 0 | - |
| - | - | 2132 | 407.2 | - | - | 0 | - |
| - | - | 4102 | 408.2 | - | - | 0 | - |
| - | - | 1002 | 409.2 | - | - | 0 | - |
| - | - | 1891 | 411.3 | - | - | 0 | - |
| - | - | 739.5 | 412.3 | - | - | 0 | - |
| - | - | 3926 | 413.1 | - | - | 0 | - |
| - | - | 1280 | 414.1 | - | - | 0 | - |
| - | - | 2369 | 415 | - | - | 0 | - |
| - | - | 714.2 | 415.1 | - | - | 0 | - |
| 8 | b | 1282 | 415.2 | 0.003382 | 8.146 | +2 | 8 |
| - | - | 1143 | 415.3 | - | - | 0 | - |
| - | - | 1402 | 416 | - | - | 0 | - |
| 14 | y | 1213 | 416.2 | 0.0008385 | 2.015 | +1 | 4 |
| - | - | 2850 | 416.3 | - | - | 0 | - |
| - | - | 3800 | 419 | - | - | 0 | - |
| - | - | 1666 | 420 | - | - | 0 | - |
| - | - | 839.7 | 421.2 | - | - | 0 | - |
| - | - | 2062 | 424.2 | - | - | 0 | - |
| - | - | 3086 | 425.2 | - | - | 0 | - |
| - | - | 813.4 | 426.2 | - | - | 0 | - |
| - | - | 1119 | 427.2 | - | - | 0 | - |
| - | - | 709.5 | 429.2 | - | - | 0 | - |
| - | - | 1862 | 431.1 | - | - | 0 | - |
| - | - | 653.2 | 433.1 | - | - | 0 | - |
| - | - | 1401 | 433.2 | - | - | 0 | - |
| 14 | y | 8289 | 433.2 | 0.0007786 | 1.797 | +1 | 4 |
| - | - | 1.747E+04 | 433.3 | - | - | 0 | - |
| - | - | 727.8 | 434.2 | - | - | 0 | - |
| - | - | 1733 | 434.2 | - | - | 0 | - |
| - | - | 4848 | 434.3 | - | - | 0 | - |
| - | - | 926 | 435.3 | - | - | 0 | - |
| - | - | 893.2 | 440.2 | - | - | 0 | - |
| - | - | 2135 | 442.2 | - | - | 0 | - |
| - | - | 1933 | 443.2 | - | - | 0 | - |
| 4 | b | 2191 | 443.3 | 0.0002209 | 0.4983 | +1 | 4 |
| - | - | 650.9 | 444.3 | - | - | 0 | - |
| - | - | 1005 | 445.2 | - | - | 0 | - |
| - | - | 2139 | 449.1 | - | - | 0 | - |
| - | - | 916 | 451.2 | - | - | 0 | - |
| 5 | y | 1734 | 451.9 | 0.001244 | 2.753 | +3 | 13 |
| 5 | y | 2813 | 452.2 | 0.003 | 6.635 | +3 | 13 |
| - | - | 1789 | 454.2 | - | - | 0 | - |
| - | - | 873.6 | 459.2 | - | - | 0 | - |
| - | - | 990.5 | 461.2 | - | - | 0 | - |
| 4 | b | 1.691E+04 | 461.3 | 0.0003068 | 0.6652 | +1 | 4 |
| - | - | 727.6 | 461.7 | - | - | 0 | - |
| - | - | 1929 | 462.2 | - | - | 0 | - |
| - | - | 4515 | 462.3 | - | - | 0 | - |
| - | - | 911.4 | 463.3 | - | - | 0 | - |
| - | - | 1986 | 465.1 | - | - | 0 | - |
| - | - | 896.2 | 468.2 | - | - | 0 | - |
| - | - | 1114 | 468.3 | - | - | 0 | - |
| - | - | 1016 | 469.2 | - | - | 0 | - |
| - | - | 6120 | 470.2 | - | - | 0 | - |
| - | - | 1726 | 471.2 | - | - | 0 | - |
| - | - | 1604 | 477.3 | - | - | 0 | - |
| - | - | 2703 | 478.2 | - | - | 0 | - |
| - | - | 648.1 | 478.7 | - | - | 0 | - |
| - | - | 973.2 | 479.2 | - | - | 0 | - |
| - | - | 6120 | 480.2 | - | - | 0 | - |
| - | - | 1107 | 480.7 | - | - | 0 | - |
| - | - | 858.4 | 480.8 | - | - | 0 | - |
| - | - | 1758 | 481.2 | - | - | 0 | - |
| - | - | 1028 | 482.2 | - | - | 0 | - |
| - | - | 991.7 | 484.2 | - | - | 0 | - |
| - | - | 627.4 | 484.7 | - | - | 0 | - |
| - | - | 656.1 | 485.2 | - | - | 0 | - |
| 13 | y | 1927 | 486.3 | 0.001634 | 3.361 | +1 | 5 |
| 13 | y | 2541 | 487.3 | 0.0001425 | 0.2925 | +1 | 5 |
| - | - | 824.4 | 488.2 | - | - | 0 | - |
| - | - | 1005 | 490.2 | - | - | 0 | - |
| - | - | 2467 | 493.7 | - | - | 0 | - |
| - | - | 1002 | 494.2 | - | - | 0 | - |
| - | - | 2181 | 494.2 | - | - | 0 | - |
| - | - | 1458 | 495.3 | - | - | 0 | - |
| - | - | 7093 | 496.3 | - | - | 0 | - |
| - | - | 1545 | 497.3 | - | - | 0 | - |
| - | - | 1.593E+04 | 498.2 | - | - | 0 | - |
| - | - | 2454 | 498.3 | - | - | 0 | - |
| - | - | 3770 | 499.2 | - | - | 0 | - |
| - | - | 783.2 | 499.3 | - | - | 0 | - |
| - | - | 1712 | 502.2 | - | - | 0 | - |
| - | - | 4347 | 502.7 | - | - | 0 | - |
| - | - | 1885 | 503.2 | - | - | 0 | - |
| - | - | 979.9 | 503.7 | - | - | 0 | - |
| 13 | y | 1.848E+04 | 504.3 | 4.976E-05 | 0.09868 | +1 | 5 |
| - | - | 5097 | 505.3 | - | - | 0 | - |
| - | - | 612.4 | 506.3 | - | - | 0 | - |
| - | - | 701.5 | 511.3 | - | - | 0 | - |
| - | - | 1641 | 512.2 | - | - | 0 | - |
| - | - | 2270 | 513.3 | - | - | 0 | - |
| - | - | 804.8 | 517.3 | - | - | 0 | - |
| - | - | 1490 | 529.3 | - | - | 0 | - |
| - | - | 4530 | 530.2 | - | - | 0 | - |
| - | - | 630.2 | 530.3 | - | - | 0 | - |
| - | - | 1706 | 530.3 | - | - | 0 | - |
| - | - | 1546 | 531.2 | - | - | 0 | - |
| - | - | 772.9 | 536.3 | - | - | 0 | - |
| - | - | 6685 | 538.3 | - | - | 0 | - |
| - | - | 4982 | 538.8 | - | - | 0 | - |
| - | - | 1450 | 539.3 | - | - | 0 | - |
| 5 | b | 2015 | 540.3 | 0.0008462 | 1.566 | +1 | 5 |
| - | - | 1548 | 543.3 | - | - | 0 | - |
| - | - | 1685 | 543.8 | - | - | 0 | - |
| 10 | b | 1225 | 544.3 | 0.002528 | 4.646 | +2 | 10 |
| 15 | b | 789.8 | 548 | 0.0005977 | 1.091 | +3 | 15 |
| - | - | 3660 | 552.3 | - | - | 0 | - |
| - | - | 3265 | 552.8 | - | - | 0 | - |
| - | - | 791.9 | 553.3 | - | - | 0 | - |
| - | - | 1182 | 555.3 | - | - | 0 | - |
| - | - | 999.4 | 556.3 | - | - | 0 | - |
| - | - | 846.3 | 556.3 | - | - | 0 | - |
| 5 | b | 4062 | 558.3 | 0.0003079 | 0.5514 | +1 | 5 |
| - | - | 1041 | 559.3 | - | - | 0 | - |
| - | - | 859.1 | 569.3 | - | - | 0 | - |
| - | - | 1259 | 569.8 | - | - | 0 | - |
| 2 | y | 1999 | 578.3 | 0.009721 | 16.81 | +3 | 16 |
| - | - | 1405 | 578.8 | - | - | 0 | - |
| - | - | 713.4 | 579.1 | - | - | 0 | - |
| - | - | 886.3 | 579.3 | - | - | 0 | - |
| 7 | y | 680.5 | 580.3 | 0.0005744 | 0.9899 | +2 | 11 |
| - | - | 900.1 | 581.3 | - | - | 0 | - |
| - | - | 2600 | 582.8 | - | - | 0 | - |
| - | - | 1786 | 583.3 | - | - | 0 | - |
| - | - | 1540 | 591.2 | - | - | 0 | - |
| - | - | 1013 | 591.3 | - | - | 0 | - |
| 11 | b | 1259 | 591.8 | 0.01096 | 18.52 | +2 | 11 |
| - | - | 1728 | 597.3 | - | - | 0 | - |
| - | - | 1028 | 598.3 | - | - | 0 | - |
| - | - | 7885 | 599.3 | - | - | 0 | - |
| - | - | 2647 | 600.3 | - | - | 0 | - |
| - | - | 896.9 | 601.3 | - | - | 0 | - |
| - | - | 2557 | 602.8 | - | - | 0 | - |
| - | - | 3079 | 603.3 | - | - | 0 | - |
| - | - | 904.6 | 603.8 | - | - | 0 | - |
| - | - | 935.9 | 604.3 | - | - | 0 | - |
| - | - | 1392 | 608.3 | - | - | 0 | - |
| - | - | 725.4 | 608.8 | - | - | 0 | - |
| - | - | 4211 | 609.3 | - | - | 0 | - |
| - | - | 1037 | 609.3 | - | - | 0 | - |
| - | - | 1539 | 609.8 | - | - | 0 | - |
| - | - | 3998 | 610.2 | - | - | 0 | - |
| - | - | 1088 | 610.3 | - | - | 0 | - |
| - | - | 4687 | 611.2 | - | - | 0 | - |
| - | - | 4199 | 611.8 | - | - | 0 | - |
| - | - | 3975 | 612.2 | - | - | 0 | - |
| - | - | 3887 | 612.3 | - | - | 0 | - |
| - | - | 1841 | 612.8 | - | - | 0 | - |
| - | - | 705 | 613.2 | - | - | 0 | - |
| - | - | 6728 | 613.3 | - | - | 0 | - |
| 12 | y | 7760 | 614.3 | 0.002108 | 3.432 | +1 | 6 |
| 12 | y | 1.214E+04 | 615.3 | 0.0003925 | 0.6379 | +1 | 6 |
| - | - | 3286 | 616.3 | - | - | 0 | - |
| - | - | 3307 | 617.3 | - | - | 0 | - |
| - | - | 6334 | 617.8 | - | - | 0 | - |
| - | - | 6395 | 618.3 | - | - | 0 | - |
| - | - | 2721 | 618.8 | - | - | 0 | - |
| - | - | 692.8 | 624.1 | - | - | 0 | - |
| - | - | 649.7 | 625.4 | - | - | 0 | - |
| - | - | 1907 | 626.4 | - | - | 0 | - |
| - | - | 1.643E+04 | 626.8 | - | - | 0 | - |
| - | - | 3.134E+04 | 627.3 | - | - | 0 | - |
| - | - | 1.004E+04 | 627.3 | - | - | 0 | - |
| - | - | 4053 | 627.8 | - | - | 0 | - |
| - | - | 1.072E+04 | 628.3 | - | - | 0 | - |
| - | - | 1225 | 628.3 | - | - | 0 | - |
| 6 | y | 1103 | 628.8 | 0.0003766 | 0.5989 | +2 | 12 |
| - | - | 2021 | 629.3 | - | - | 0 | - |
| - | - | 1024 | 629.8 | - | - | 0 | - |
| 12 | y | 1.54E+04 | 632.3 | 0.0003387 | 0.5356 | +1 | 6 |
| - | - | 4963 | 633.3 | - | - | 0 | - |
| - | - | 806 | 636.8 | - | - | 0 | - |
| 6 | y | 2998 | 637.8 | 0.0001602 | 0.2511 | +2 | 12 |
| - | - | 1130 | 638.3 | - | - | 0 | - |
| - | - | 3124 | 643.3 | - | - | 0 | - |
| - | - | 948 | 644.3 | - | - | 0 | - |
| - | - | 887.4 | 645.3 | - | - | 0 | - |
| - | - | 854.3 | 646.3 | - | - | 0 | - |
| - | - | 6156 | 655.3 | - | - | 0 | - |
| 12 | b | 5170 | 655.8 | 0.005713 | 8.711 | +2 | 12 |
| - | - | 2181 | 656.3 | - | - | 0 | - |
| - | - | 1153 | 656.8 | - | - | 0 | - |
| - | - | 1166 | 659.8 | - | - | 0 | - |
| - | - | 1531 | 660.3 | - | - | 0 | - |
| - | - | 1742 | 668.3 | - | - | 0 | - |
| - | - | 2901 | 668.8 | - | - | 0 | - |
| - | - | 1827 | 669.3 | - | - | 0 | - |
| - | - | 1564 | 669.8 | - | - | 0 | - |
| - | - | 652.3 | 670.4 | - | - | 0 | - |
| 5 | y | 1.877E+04 | 677.3 | 0.0005537 | 0.8174 | +2 | 13 |
| 5 | y | 2.192E+04 | 677.8 | 0.006889 | 10.16 | +2 | 13 |
| - | - | 9382 | 678.3 | - | - | 0 | - |
| - | - | 3897 | 678.8 | - | - | 0 | - |
| - | - | 1358 | 679.3 | - | - | 0 | - |
| - | - | 694.3 | 680.5 | - | - | 0 | - |
| - | - | 764.4 | 684.4 | - | - | 0 | - |
| - | - | 804.6 | 685.7 | - | - | 0 | - |
| 5 | y | 3.78E+04 | 686.3 | 0.000648 | 0.9442 | +2 | 13 |
| - | - | 3.001E+04 | 686.8 | - | - | 0 | - |
| - | - | 1.373E+04 | 687.3 | - | - | 0 | - |
| - | - | 4549 | 687.8 | - | - | 0 | - |
| - | - | 964.9 | 688.4 | - | - | 0 | - |
| - | - | 1495 | 694.3 | - | - | 0 | - |
| - | - | 1367 | 696.4 | - | - | 0 | - |
| - | - | 983.1 | 697.4 | - | - | 0 | - |
| - | - | 926.6 | 698.3 | - | - | 0 | - |
| 13 | b | 1809 | 700.4 | 0.01269 | 18.13 | +2 | 13 |
| - | - | 1775 | 700.8 | - | - | 0 | - |
| - | - | 750.3 | 701.3 | - | - | 0 | - |
| - | - | 5913 | 712.4 | - | - | 0 | - |
| - | - | 2346 | 713.4 | - | - | 0 | - |
| - | - | 3490 | 714.4 | - | - | 0 | - |
| - | - | 1995 | 715.4 | - | - | 0 | - |
| - | - | 722.4 | 720.4 | - | - | 0 | - |
| - | - | 2378 | 722.3 | - | - | 0 | - |
| - | - | 942.9 | 727.3 | - | - | 0 | - |
| 11 | y | 823.6 | 727.4 | 0.004075 | 5.602 | +1 | 7 |
| - | - | 1711 | 728.9 | - | - | 0 | - |
| - | - | 976.7 | 729.4 | - | - | 0 | - |
| - | - | 602.4 | 731.3 | - | - | 0 | - |
| - | - | 938.2 | 739.4 | - | - | 0 | - |
| - | - | 1.044E+04 | 740.3 | - | - | 0 | - |
| - | - | 4074 | 741.4 | - | - | 0 | - |
| - | - | 1921 | 742.3 | - | - | 0 | - |
| 7 | b | 2117 | 742.4 | 0.001599 | 2.153 | +1 | 7 |
| - | - | 874.6 | 743.4 | - | - | 0 | - |
| 11 | y | 7466 | 745.4 | 0.0009676 | 1.298 | +1 | 7 |
| - | - | 835.2 | 746.3 | - | - | 0 | - |
| - | - | 2946 | 746.4 | - | - | 0 | - |
| - | - | 681.3 | 747.4 | - | - | 0 | - |
| - | - | 1749 | 749.3 | - | - | 0 | - |
| 4 | y | 4075 | 750.9 | 0.0003368 | 0.4485 | +2 | 14 |
| 4 | y | 4916 | 751.4 | 0.007472 | 9.945 | +2 | 14 |
| - | - | 1085 | 751.9 | - | - | 0 | - |
| - | - | 805.1 | 755.3 | - | - | 0 | - |
| - | - | 1844 | 755.4 | - | - | 0 | - |
| - | - | 841.6 | 756.4 | - | - | 0 | - |
| 14 | b | 2079 | 757.4 | 0.01182 | 15.61 | +2 | 14 |
| - | - | 2770 | 757.4 | - | - | 0 | - |
| - | - | 1112 | 758.4 | - | - | 0 | - |
| - | - | 1570 | 758.4 | - | - | 0 | - |
| - | - | 846.8 | 759.4 | - | - | 0 | - |
| 4 | y | 4787 | 759.9 | 0.0009194 | 1.21 | +2 | 14 |
| - | - | 2890 | 760.4 | - | - | 0 | - |
| - | - | 2410 | 760.9 | - | - | 0 | - |
| - | - | 2941 | 771.4 | - | - | 0 | - |
| - | - | 1039 | 772.3 | - | - | 0 | - |
| - | - | 2791 | 774.3 | - | - | 0 | - |
| - | - | 815.7 | 775.3 | - | - | 0 | - |
| - | - | 9417 | 785.4 | - | - | 0 | - |
| - | - | 7673 | 786.4 | - | - | 0 | - |
| - | - | 1503 | 787.4 | - | - | 0 | - |
| - | - | 768.6 | 798.4 | - | - | 0 | - |
| - | - | 748.9 | 798.9 | - | - | 0 | - |
| - | - | 1043 | 802.4 | - | - | 0 | - |
| - | - | 915 | 806.4 | - | - | 0 | - |
| 3 | y | 811.5 | 807.4 | 0.00353 | 4.372 | +2 | 15 |
| 3 | y | 1339 | 807.9 | 0.009019 | 11.16 | +2 | 15 |
| 8 | b | 1552 | 811.4 | 0.002616 | 3.223 | +1 | 8 |
| 15 | b | 963.5 | 812.4 | 0.01343 | 16.53 | +2 | 15 |
| - | - | 1998 | 813.4 | - | - | 0 | - |
| - | - | 941 | 817.4 | - | - | 0 | - |
| - | - | 760.9 | 822.4 | - | - | 0 | - |
| 8 | b | 2258 | 829.4 | 0.00195 | 2.351 | +1 | 8 |
| - | - | 1037 | 830.4 | - | - | 0 | - |
| - | - | 1244 | 835.4 | - | - | 0 | - |
| - | - | 761.8 | 840.4 | - | - | 0 | - |
| - | - | 2472 | 841.4 | - | - | 0 | - |
| - | - | 1087 | 842.4 | - | - | 0 | - |
| - | - | 729.3 | 849.4 | - | - | 0 | - |
| - | - | 2701 | 850.4 | - | - | 0 | - |
| - | - | 910.7 | 851.4 | - | - | 0 | - |
| 16 | b | 1148 | 856.9 | 0.005592 | 6.525 | +2 | 16 |
| 10 | y | 2204 | 857.4 | 0.007158 | 8.348 | +1 | 8 |
| 2 | y | 1065 | 857.9 | 0.005417 | 6.314 | +2 | 16 |
| 2 | y | 1162 | 858.4 | 0.004284 | 4.991 | +2 | 16 |
| - | - | 812.6 | 859.4 | - | - | 0 | - |
| - | - | 3126 | 863.4 | - | - | 0 | - |
| - | - | 1166 | 864.4 | - | - | 0 | - |
| - | - | 1.397E+04 | 868.4 | - | - | 0 | - |
| - | - | 6949 | 869.4 | - | - | 0 | - |
| - | - | 1630 | 870.4 | - | - | 0 | - |
| - | - | 3621 | 873.5 | - | - | 0 | - |
| 10 | y | 3634 | 874.5 | 0.001507 | 1.724 | +1 | 8 |
| - | - | 1817 | 875.5 | - | - | 0 | - |
| - | - | 3761 | 879.4 | - | - | 0 | - |
| - | - | 1383 | 880.4 | - | - | 0 | - |
| - | - | 968.3 | 884.4 | - | - | 0 | - |
| - | - | 2448 | 887.4 | - | - | 0 | - |
| - | - | 1133 | 888.4 | - | - | 0 | - |
| - | - | 3695 | 889.4 | - | - | 0 | - |
| - | - | 1745 | 890.4 | - | - | 0 | - |
| - | - | 705.9 | 900.4 | - | - | 0 | - |
| - | - | 841.1 | 905.5 | - | - | 0 | - |
| - | - | 1.485E+04 | 907.4 | - | - | 0 | - |
| - | - | 8111 | 908.4 | - | - | 0 | - |
| - | - | 1820 | 909.4 | - | - | 0 | - |
| - | - | 1169 | 911.5 | - | - | 0 | - |
| - | - | 1135 | 921.4 | - | - | 0 | - |
| - | - | 861.4 | 922.4 | - | - | 0 | - |
| - | - | 1572 | 932.5 | - | - | 0 | - |
| - | - | 777.5 | 933.5 | - | - | 0 | - |
| - | - | 678.8 | 934.5 | - | - | 0 | - |
| - | - | 5385 | 939.4 | - | - | 0 | - |
| - | - | 4664 | 940.5 | - | - | 0 | - |
| - | - | 1281 | 941.5 | - | - | 0 | - |
| - | - | 676.1 | 942.5 | - | - | 0 | - |
| - | - | 762.2 | 948.5 | - | - | 0 | - |
| - | - | 723.4 | 949.5 | - | - | 0 | - |
| - | - | 2246 | 956.5 | - | - | 0 | - |
| - | - | 951.1 | 957.5 | - | - | 0 | - |
| 9 | b | 4024 | 958.5 | 0.002123 | 2.215 | +1 | 9 |
| - | - | 2229 | 959.5 | - | - | 0 | - |
| - | - | 7793 | 960.5 | - | - | 0 | - |
| - | - | 4502 | 961.5 | - | - | 0 | - |
| - | - | 1649 | 962.5 | - | - | 0 | - |
| - | - | 1051 | 971.5 | - | - | 0 | - |
| - | - | 683.5 | 988.5 | - | - | 0 | - |
| 9 | y | 758.7 | 1004 | 0.002047 | 2.04 | +1 | 9 |
| - | - | 784.2 | 1015 | - | - | 0 | - |
| - | - | 850.3 | 1036 | - | - | 0 | - |
| - | - | 926.4 | 1037 | - | - | 0 | - |
| - | - | 1417 | 1040 | - | - | 0 | - |
| - | - | 698.2 | 1041 | - | - | 0 | - |
| - | - | 5592 | 1053 | - | - | 0 | - |
| - | - | 3512 | 1054 | - | - | 0 | - |
| - | - | 1239 | 1055 | - | - | 0 | - |
| - | - | 1854 | 1059 | - | - | 0 | - |
| - | - | 1618 | 1060 | - | - | 0 | - |
| - | - | 902.6 | 1068 | - | - | 0 | - |
| 10 | b | 1418 | 1070 | 0.006991 | 6.537 | +1 | 10 |
| - | - | 3292 | 1071 | - | - | 0 | - |
| - | - | 1332 | 1072 | - | - | 0 | - |
| 8 | y | 686.1 | 1073 | 0.003125 | 2.914 | +1 | 10 |
| - | - | 4428 | 1076 | - | - | 0 | - |
| - | - | 2076 | 1077 | - | - | 0 | - |
| - | - | 1012 | 1085 | - | - | 0 | - |
| - | - | 776.3 | 1086 | - | - | 0 | - |
| - | - | 1352 | 1087 | - | - | 0 | - |
| 10 | b | 1.248E+04 | 1088 | 0.003884 | 3.571 | +1 | 10 |
| - | - | 6072 | 1089 | - | - | 0 | - |
| - | - | 2460 | 1090 | - | - | 0 | - |
| 8 | y | 1625 | 1091 | 0.001117 | 1.024 | +1 | 10 |
| - | - | 689.8 | 1092 | - | - | 0 | - |
| - | - | 3924 | 1104 | - | - | 0 | - |
| - | - | 3028 | 1105 | - | - | 0 | - |
| - | - | 893.4 | 1106 | - | - | 0 | - |
| - | - | 1262 | 1138 | - | - | 0 | - |
| - | - | 1658 | 1139 | - | - | 0 | - |
| - | - | 1920 | 1156 | - | - | 0 | - |
| - | - | 1282 | 1157 | - | - | 0 | - |
| 7 | y | 989.2 | 1161 | 0.01684 | 14.51 | +1 | 11 |
| - | - | 1377 | 1165 | - | - | 0 | - |
| - | - | 893.3 | 1166 | - | - | 0 | - |
| 7 | y | 1394 | 1178 | 0.007144 | 6.067 | +1 | 11 |
| - | - | 1074 | 1179 | - | - | 0 | - |
| - | - | 4751 | 1182 | - | - | 0 | - |
| - | - | 3718 | 1183 | - | - | 0 | - |
| - | - | 1888 | 1184 | - | - | 0 | - |
| 11 | b | 2035 | 1201 | 0.005917 | 4.928 | +1 | 11 |
| - | - | 1154 | 1202 | - | - | 0 | - |
| - | - | 940.7 | 1213 | - | - | 0 | - |
| - | - | 3093 | 1235 | - | - | 0 | - |
| - | - | 2302 | 1236 | - | - | 0 | - |
| - | - | 1121 | 1237 | - | - | 0 | - |
| - | - | 8871 | 1253 | - | - | 0 | - |
| - | - | 5437 | 1254 | - | - | 0 | - |
| - | - | 2420 | 1255 | - | - | 0 | - |
| 6 | y | 1026 | 1257 | 0.008152 | 6.487 | +1 | 12 |
| 6 | y | 1592 | 1258 | 0.008999 | 7.156 | +1 | 12 |
| - | - | 743 | 1259 | - | - | 0 | - |
| 6 | y | 3622 | 1275 | 0.00693 | 5.437 | +1 | 12 |
| - | - | 2771 | 1276 | - | - | 0 | - |
| - | - | 1032 | 1277 | - | - | 0 | - |
| - | - | 709.8 | 1356 | - | - | 0 | - |
| 5 | y | 1615 | 1372 | 0.006593 | 4.806 | +1 | 13 |
| - | - | 907 | 1373 | - | - | 0 | - |
| - | - | 894.6 | 1400 | - | - | 0 | - |
| - | - | 745.8 | 1401 | - | - | 0 | - |
| - | - | 870 | 1515 | - | - | 0 | - |
| - | - | 669.1 | 2466 | - | - | 0 | - |
| - | - | 804.4 | 3073 | - | - | 0 | - |

m/z Charge Intensity FragmentType MassShift Position
120.0658187866211 0 28598.426 y 16
120.08109283447266 0 41074.516
121.0693359375 0 1428.4729
121.08439636230469 0 3904.4565
121.35698699951172 0 352.94623
122.08751678466797 0 377.2603
123.04418182373047 0 652.0231
126.0552978515625 0 2750.6196
128.10728454589844 0 8057.212
129.05494689941406 0 4141.781
129.06625366210938 0 1290.3468
129.1025390625 0 37378.445
130.0501708984375 0 1951.1497
130.06549072265625 0 1278.8304
130.10598754882812 0 2488.2664
131.1182098388672 0 1851.0529
134.0604248046875 0 1376.0634
135.04461669921875 0 523.00586
136.07598876953125 0 7512.2534
137.07936096191406 0 951.54443
138.0917510986328 0 1837.4662
138.1282501220703 0 1002.7996
139.08676147460938 0 1072.408
141.06614685058594 0 1059.8264
145.04995727539062 0 502.81653
145.0978240966797 0 509.5081
147.04441833496094 0 457.01218
147.06590270996094 0 696.7313
147.07669067382812 0 932.7097
149.04507446289062 0 2341.5398
152.0708770751953 0 2942.3225
153.0186004638672 0 4233.8843
154.0499267578125 0 527.5736
154.0865936279297 0 757.4826
154.09771728515625 0 1344.2618
155.081787109375 0 2011.78
155.11819458007812 0 5736.303 a Water loss 1
156.12135314941406 0 642.1246
157.0611572265625 0 885.4956
157.09744262695312 0 2576.9246
159.09194946289062 0 6050.566
160.09530639648438 0 708.46436
161.07115173339844 0 1269.8506
162.055419921875 0 655.9684
164.07077026367188 0 1100.6726
165.05519104003906 0 1406.5839
165.10247802734375 0 1903.069
166.0858917236328 0 494.2881
167.05569458007812 0 3558.409
167.0817413330078 0 2459.6797
167.11801147460938 0 1707.2739
167.32550048828125 0 452.08234
169.06149291992188 0 640.96375
169.0976104736328 0 517.35834
169.1337890625 0 8707.889
170.13735961914062 0 796.252
171.02899169921875 0 2415.9878
171.07672119140625 0 1242.2648
171.11300659179688 0 498.3675
172.1077117919922 0 909.1977
173.09226989746094 0 4274.1025 y Water loss 15
173.1287384033203 0 19642.266 a 1
174.13246154785156 0 1063.9004
175.07174682617188 0 1311.9393
177.05482482910156 0 4526.2188
179.0339813232422 0 1896.2971
181.05010986328125 0 609.91724
181.0975799560547 0 963.35205
182.08152770996094 0 3195.2407
182.09251403808594 0 2142.7556
182.12911987304688 0 2019.9573
183.07647705078125 0 1973.8848
183.1130828857422 0 25149.049 b Water loss 1
184.10818481445312 0 2434.4087
184.11672973632812 0 2610.3225
185.09228515625 0 4004.9302
185.1009521484375 0 668.9238
186.087646484375 0 1422.2063
187.08633422851562 0 753.2695
187.14443969726562 0 8285.303
188.14805603027344 0 1166.431
189.0872802734375 0 2440.1863
191.08203125 0 684.5734
191.10292053222656 0 6481.676 y 15
191.1182403564453 0 1814.7836
192.106201171875 0 639.9523
195.02041625976562 0 564.48456
195.02899169921875 0 3555.9016
195.11306762695312 0 3672.4927
196.06057739257812 0 1579.1465
197.12875366210938 0 11314.165
198.07640075683594 0 2546.612
198.12387084960938 0 1637.2961
198.13291931152344 0 770.94684
199.07144165039062 0 2998.6619
200.103271484375 0 2577.0894
200.1396942138672 0 2472.7107
201.08731079101562 0 855.9508
201.1235809326172 0 24728.557 b 1
202.12709045410156 0 2128.7075
203.1029510498047 0 752.81506
206.2314910888672 0 539.3934
208.10826110839844 0 3060.0264
209.09234619140625 0 7710.0693
210.09622192382812 0 926.43823
212.1391143798828 0 794.8642
213.08689880371094 0 795.83594
213.1239776611328 0 678.66003
214.07142639160156 0 910.2647
214.13058471679688 0 825.16296
215.13934326171875 0 7535.899
216.1426544189453 0 1444.1941
217.0821075439453 0 4910.1094
217.09725952148438 0 2523.158
217.13369750976562 0 6703.0825
218.13717651367188 0 938.66644
219.0288543701172 0 1306.4331
219.15025329589844 0 1489.9626
221.08448791503906 0 4247.333
222.08441162109375 0 1348.8379
225.04287719726562 0 1860.2223
225.12350463867188 0 2452.4517
225.13507080078125 0 1397.1207
226.11883544921875 0 28198.018
227.04010009765625 0 720.7257
227.1223602294922 0 2165.2249
228.1345977783203 0 1525.434
229.1187286376953 0 1068.0865
230.5080108642578 0 552.3579
231.09776306152344 0 726.38257
232.1577911376953 0 543.2189
233.1650848388672 0 10315.033
234.16835021972656 0 1753.2625
235.1080780029297 0 6083.9165
236.10275268554688 0 1126.5991
236.13946533203125 0 886.49963
237.0867462158203 0 516.3467
239.0952606201172 0 11694.871
240.0962371826172 0 2412.7158
241.08184814453125 0 1475.7102
241.0934295654297 0 1374.7893
242.1503143310547 0 5955.096
243.13340759277344 0 939.79486
243.14556884765625 0 11001.466 y Water loss 10
244.12960815429688 0 1094.8712 y Ammonia loss 12
244.1484375 0 842.9889
245.0445098876953 0 1380.741
245.12881469726562 0 3486.5642
246.12330627441406 0 1356.5869
247.14443969726562 0 1008.0035
248.6970977783203 0 546.2593
251.1756134033203 0 3778.4746
252.0990753173828 0 768.0154
252.13511657714844 0 1061.1954
253.1308135986328 0 626.049
253.16612243652344 0 4070.1333
254.11373901367188 0 4194.5576
254.14959716796875 0 1243.8032
254.1701202392578 0 583.7651
257.1960144042969 0 631.7252
259.0926818847656 0 4132.779
261.15972900390625 0 5474.213
262.118896484375 0 1112.7227
262.1650085449219 0 704.3332
263.0552062988281 0 4212.436
263.10284423828125 0 8330.476
263.13946533203125 0 545.3368
264.0596618652344 0 654.85376
264.13433837890625 0 3502.9807
266.9985656738281 0 657.03735
268.20257568359375 0 684.8007 a Water loss 2
268.97894287109375 0 615.2386
270.1923522949219 0 1926.9292
272.12445068359375 0 1735.1769
274.117431640625 0 604.6664
275.1759338378906 0 963.01056
279.1453552246094 0 3465.453
279.66741943359375 0 1358.2462 b 4
280.129150390625 0 996.3241
280.1491394042969 0 800.634
280.166015625 0 2683.2383
281.0506286621094 0 1123.1399
281.06597900390625 0 1666.7554
282.1450500488281 0 9897.755
283.148681640625 0 859.38696
285.00994873046875 0 10375.266
285.1604919433594 0 984.9727
286.0101623535156 0 3618.9753
286.1039733886719 0 2152.3335
287.00640869140625 0 1238.369
289.1549987792969 0 1003.4994
296.13555908203125 0 1489.123
296.173583984375 0 3094.4192
296.1971740722656 0 58996.617 b Water loss 2
297.1196594238281 0 1731.3885
297.1559143066406 0 10957.461
297.1717529296875 0 382.82376
297.17584228515625 0 650.5732
297.20025634765625 0 11350.691
298.1590881347656 0 1563.8933
298.1766052246094 0 4205.428
298.2039794921875 0 1248.2863
299.06201171875 0 12739.141
299.15374755859375 0 1603.9772
299.1742248535156 0 610.82086
299.4413757324219 0 584.45074
299.6567077636719 0 577.27954
300.0625 0 2988.1233
301.0593566894531 0 2003.9187
301.1879577636719 0 1546.1198 y Water loss 14
303.08660888671875 0 76181.1
303.1708984375 0 2116.4924
304.0899963378906 0 13755.002
304.1101379394531 0 1085.6648
307.66644287109375 0 2582.426 y Water loss 11
308.16766357421875 0 822.463
313.1883239746094 0 2083.8389
314.1483154296875 0 1146.5118
314.18280029296875 0 8385.44
314.20770263671875 0 12464.706 b 2
315.1868896484375 0 1189.2128
315.2101745605469 0 2288.991
316.1869812011719 0 7871.3955
316.6724853515625 0 1487.4934 y 11
317.1885681152344 0 2123.5986
319.19805908203125 0 7177.812 y 14
320.2008972167969 0 1672.8984
322.177001953125 0 639.21814
323.1729736328125 0 644.3334
325.1504211425781 0 714.263
327.08648681640625 0 668.7179
328.1153869628906 0 1491.4265
329.148193359375 0 581.8381
330.1819763183594 0 1461.4304
332.1253662109375 0 798.9349
333.1565246582031 0 1916.8081
337.15057373046875 0 685.4991
340.1968688964844 0 512.28845
341.1830139160156 0 1923.2927
342.1811828613281 0 910.86945
344.1988220214844 0 989.6276
344.9765319824219 0 848.4835
345.09716796875 0 4084.1956
345.9785461425781 0 1253.6738
346.102783203125 0 879.3913
346.1248474121094 0 3243.4849
346.1767578125 0 2496.4854
348.190673828125 0 1042.8705
350.1347351074219 0 2689.1833
350.18206787109375 0 1647.4641
351.1665344238281 0 5221.8696
351.1860046386719 0 673.4403
352.1700134277344 0 1110.8048
353.1823425292969 0 3375.1304
354.16473388671875 0 1152.5719
355.0699462890625 0 37759.062
355.16162109375 0 970.3564
356.0706481933594 0 17917.852
357.0679626464844 0 8936.787
358.0679016113281 0 789.52985
360.1541748046875 0 728.0484
361.13897705078125 0 820.5336
362.2073669433594 0 6428.135
363.2089538574219 0 707.0016
364.70074462890625 0 1221.6223 y Ammonia loss 10
365.1452331542969 0 1228.3323
367.2091064453125 0 3442.188
368.19281005859375 0 3088.7231
369.0968322753906 0 2527.9912
369.1767883300781 0 5857.9136
370.0984802246094 0 591.722
370.17987060546875 0 1254.3492
370.6768798828125 0 846.339
371.0999450683594 0 1089.8972
371.19219970703125 0 1494.0859
371.6827697753906 0 636.4063
372.1029968261719 0 658.61
373.0809326171875 0 3527.8752
373.17132568359375 0 1667.7944
373.213623046875 0 2357.2214 y 10
373.7149353027344 0 1265.4613
374.0802001953125 0 1628.454
375.0783996582031 0 1206.2383
383.156005859375 0 2104.2886
383.1844787597656 0 541.2249
385.2191467285156 0 9822.712
386.2001647949219 0 900.3205
386.22454833984375 0 1213.693
393.2484436035156 0 1052.2875
395.11297607421875 0 910.3735
397.13555908203125 0 988.8366
397.2183837890625 0 661.3974
398.2445373535156 0 927.708
399.18658447265625 0 706.1212
400.98431396484375 0 765.3374
401.16656494140625 0 4530.5283
402.1681213378906 0 854.6316
407.202880859375 0 2132.0881
408.1882019042969 0 4101.668
409.1911926269531 0 1002.4078
411.26080322265625 0 1891.2025
412.26470947265625 0 739.5242
413.1229553222656 0 3925.9958
414.1270446777344 0 1280.0638
415.0367736816406 0 2369.4211
415.1463317871094 0 714.2237
415.229736328125 0 1282.2965 b 7
415.2724304199219 0 1143.3644
416.0382385253906 0 1402.0188
416.2148132324219 0 1212.735 y Ammonia loss 13
416.2544860839844 0 2850.3042
418.9953308105469 0 3799.7134
419.9961853027344 0 1666.3483
421.2198791503906 0 839.73535
424.228515625 0 2062.1072
425.21405029296875 0 3086.2017
426.2161560058594 0 813.43756
427.2308349609375 0 1119.2163
429.2115783691406 0 709.5373
431.1337585449219 0 1861.6714
433.1245422363281 0 653.2177
433.1560363769531 0 1400.8865
433.2413024902344 0 8288.577 y 13
433.28143310546875 0 17472.06
434.20709228515625 0 727.8021
434.2449645996094 0 1733.3279
434.2843322753906 0 4848.3086
435.28509521484375 0 925.9968
440.2125549316406 0 893.1823
442.2412109375 0 2135.181
443.2287292480469 0 1932.538
443.2655029296875 0 2191.345 b Water loss 3
444.2691955566406 0 650.9181
445.1996154785156 0 1004.6929
449.1449279785156 0 2138.5464
451.2304992675781 0 915.9838
451.8942565917969 0 1733.9794 y Water loss 4
452.218017578125 0 2813.0105 y Ammonia loss 4
454.205810546875 0 1788.6316
459.2070007324219 0 873.5579
461.21661376953125 0 990.5188
461.2761535644531 0 16908.932 b 3
461.73870849609375 0 727.61456
462.1997375488281 0 1929.4846
462.27911376953125 0 4514.5625
463.2820739746094 0 911.4305
465.1392822265625 0 1985.8577
468.2100524902344 0 896.20386
468.25579833984375 0 1113.6178
469.24383544921875 0 1016.4573
470.22479248046875 0 6119.927
471.228271484375 0 1726.1158
477.2575378417969 0 1603.576
478.2414855957031 0 2702.6938
478.7342529296875 0 648.0808
479.2210998535156 0 973.19727
480.2096862792969 0 6119.756
480.7177734375 0 1106.5714
480.75250244140625 0 858.4167
481.2161560058594 0 1757.5011
482.225341796875 0 1027.5594
484.2038879394531 0 991.6692
484.72393798828125 0 627.36365
485.2207336425781 0 656.08295
486.2687072753906 0 1926.9514 y Water loss 12
487.2509460449219 0 2541.453 y Ammonia loss 12
488.2498779296875 0 824.3621
490.2098083496094 0 1004.7562
493.7278137207031 0 2466.7827
494.18963623046875 0 1001.8805
494.2288513183594 0 2181.1323
495.2662048339844 0 1457.7134
496.2516784667969 0 7092.8633
497.25347900390625 0 1544.92
498.2197265625 0 15933.79
498.3038635253906 0 2453.647
499.2230529785156 0 3769.5247
499.3072509765625 0 783.21893
502.21514892578125 0 1712.1315
502.7330017089844 0 4347.289
503.2314147949219 0 1885.3304
503.7378845214844 0 979.9401
504.277587890625 0 18476.402 y 12
505.28045654296875 0 5096.718
506.2821044921875 0 612.3961
511.2508239746094 0 701.5407
512.198486328125 0 1641.0002
513.27734375 0 2269.652
517.252685546875 0 804.75256
529.262939453125 0 1490.0139
530.208740234375 0 4530.4805
530.2743530273438 0 630.164
530.3330688476562 0 1705.5872
531.2105712890625 0 1546.0789
536.2752075195312 0 772.938
538.2691650390625 0 6684.8955
538.7715454101562 0 4981.756
539.2723388671875 0 1449.8862
540.3171997070312 0 2015.4875 b Water loss 4
543.2607421875 0 1548.4954
543.7625122070312 0 1684.5034
544.2664184570312 0 1225.0693 b 9
547.9536743164062 0 789.8479 b 14
552.2670288085938 0 3660.383
552.7680053710938 0 3265.145
553.2664794921875 0 791.9035
555.3247680664062 0 1181.6726
556.2736206054688 0 999.3741
556.3269653320312 0 846.25653
558.3289184570312 0 4061.656 b 4
559.3308715820312 0 1040.6912
569.2574462890625 0 859.1238
569.77685546875 0 1258.7982
578.286865234375 0 1998.7144 y 1
578.7865600585938 0 1405.4336
579.1238403320312 0 713.4359
579.2789306640625 0 886.28595
580.28369140625 0 680.5446 y Water loss 6
581.253662109375 0 900.1199
582.7808227539062 0 2600.2693
583.2830200195312 0 1785.6095
591.2383422851562 0 1539.7844
591.2877197265625 0 1012.628
591.7947387695312 0 1259.339 b Water loss 10
597.2972412109375 0 1728.0823
598.2857055664062 0 1027.8008
599.266845703125 0 7884.876
600.2701416015625 0 2647.0977
601.2743530273438 0 896.90753
602.790283203125 0 2556.774
603.2914428710938 0 3078.7615
603.7931518554688 0 904.5545
604.3063354492188 0 935.92523
608.3487548828125 0 1391.9683
608.7976684570312 0 725.4065
609.2507934570312 0 4211.275
609.2987670898438 0 1037.2006
609.793701171875 0 1538.902
610.1841430664062 0 3998.4524
610.2523803710938 0 1088.0684
611.1844482421875 0 4686.515
611.7957763671875 0 4199.2534
612.1820068359375 0 3975.2678
612.2975463867188 0 3887.476
612.81005859375 0 1841.4575
613.1810913085938 0 704.96277
613.3283081054688 0 6727.698
614.3277587890625 0 7759.6724 y Water loss 11
615.31005859375 0 12140.747 y Ammonia loss 11
616.3125 0 3286.1096
617.31298828125 0 3307.391
617.8067016601562 0 6334.3477
618.305419921875 0 6394.731
618.8038330078125 0 2720.9014
624.132568359375 0 692.835
625.3778076171875 0 649.6682
626.3617553710938 0 1906.9214
626.8117065429688 0 16434.19
627.2613525390625 0 31337.521
627.3135986328125 0 10043.443
627.8131713867188 0 4053.0044
628.2644653320312 0 10722.881
628.314453125 0 1225.0002
628.8098754882812 0 1102.5206 y Water loss 5
629.2677001953125 0 2021.277
629.8048706054688 0 1023.72095
632.3358764648438 0 15402.28 y 11
633.3380737304688 0 4963.052
636.8233642578125 0 806.02856
637.81494140625 0 2997.927 y 5
638.3143920898438 0 1130.3229
643.2952270507812 0 3123.9414
644.29345703125 0 948.01324
645.3154296875 0 887.3501
646.3111572265625 0 854.26965
655.3394165039062 0 6156.4834
655.8406982421875 0 5169.837 b Water loss 11
656.3422241210938 0 2181.3037
656.8417358398438 0 1153.0936
659.818115234375 0 1165.6703
660.317626953125 0 1531.274
668.3290405273438 0 1742.1451
668.8268432617188 0 2901.1794
669.3212280273438 0 1827.3036
669.822509765625 0 1564.418
670.3509521484375 0 652.2504
677.3353271484375 0 18766.838 y Water loss 4
677.8347778320312 0 21916.4 y Ammonia loss 4
678.3353881835938 0 9382.348
678.8348388671875 0 3897.3982
679.3402709960938 0 1357.6237
680.5264282226562 0 694.30676
684.366943359375 0 764.4291
685.7063598632812 0 804.6122
686.3405151367188 0 37799.18 y 4
686.841796875 0 30010.713
687.3434448242188 0 13725.678
687.8449096679688 0 4549.108
688.3883056640625 0 964.8793
694.3432006835938 0 1494.8375
696.362548828125 0 1367.0048
697.3650512695312 0 983.1189
698.3362426757812 0 926.5663
700.3461303710938 0 1808.661 b 12
700.84619140625 0 1775.4753
701.3490600585938 0 750.3155
712.3510131835938 0 5913.037
713.3565673828125 0 2346.2913
714.3748779296875 0 3489.7869
715.3792724609375 0 1995.1407
720.3717041015625 0 722.43353
722.3357543945312 0 2378.4133
727.3334350585938 0 942.8954
727.4056396484375 0 823.6319 y Water loss 10
728.870361328125 0 1711.0875
729.379638671875 0 976.7303
731.3366088867188 0 602.4342
739.3670654296875 0 938.1647
740.3453979492188 0 10441.744
741.3504638671875 0 4073.8425
742.3421020507812 0 1921.2384
742.4118041992188 0 2116.5242 b 6
743.4200439453125 0 874.5691
745.4193115234375 0 7466.18 y 10
746.3484497070312 0 835.2187
746.4221801757812 0 2946.4307
747.4279174804688 0 681.29144
749.3370971679688 0 1748.7727
750.8697509765625 0 4075.0588 y Water loss 3
751.3695678710938 0 4916.2754 y Ammonia loss 3
751.8697509765625 0 1085.374
755.3273315429688 0 805.1216
755.4050903320312 0 1844.1444
756.4022216796875 0 841.61066
757.3684692382812 0 2078.911 b 13
757.4207153320312 0 2769.8308
758.3645629882812 0 1111.9231
758.4304809570312 0 1569.8419
759.3754272460938 0 846.7867
759.8744506835938 0 4787.359 y 3
760.3748779296875 0 2889.8247
760.8775634765625 0 2410.195
771.3509521484375 0 2941.1584
772.348876953125 0 1039.0845
774.3307495117188 0 2791.1562
775.33251953125 0 815.72034
785.4144287109375 0 9416.564
786.4164428710938 0 7672.799
787.4164428710938 0 1503.3293
798.3770751953125 0 768.6358
798.9058837890625 0 748.9429
802.445068359375 0 1042.5814
806.3570556640625 0 915.0198
807.4156494140625 0 811.54724 y Water loss 2
807.9131469726562 0 1338.8942 y Ammonia loss 2
811.4322509765625 0 1552.2445 b Water loss 7
812.4359130859375 0 963.5398 b Water loss 14
813.432861328125 0 1998.4623
817.4066772460938 0 941.01886
822.4110107421875 0 760.92126
829.4434814453125 0 2258.479 b 7
830.4469604492188 0 1036.5212
835.3819580078125 0 1243.9033
840.4070434570312 0 761.8014
841.4271850585938 0 2472.2244
842.4215087890625 0 1087.0345
849.4154663085938 0 729.3477
850.3947143554688 0 2700.8635
851.3936767578125 0 910.703
856.9407348632812 0 1148.3723 b 15
857.4434814453125 0 2203.6072 y Ammonia loss 9
857.9305419921875 0 1065.3389 y Water loss 1
858.4322509765625 0 1161.9601 y Ammonia loss 1
859.4238891601562 0 812.62854
863.3799438476562 0 3125.896
864.3837280273438 0 1166.2498
868.40283203125 0 13969.573
869.4053344726562 0 6948.9146
870.40869140625 0 1629.5966
873.4575805664062 0 3620.8403
874.4613647460938 0 3633.647 y 9
875.4656982421875 0 1816.7
879.4087524414062 0 3761.1123
880.4132690429688 0 1382.8939
884.4429931640625 0 968.3145
887.412841796875 0 2448.242
888.4170532226562 0 1133.4216
889.3967895507812 0 3695.1943
890.39599609375 0 1744.7946
900.441650390625 0 705.89276
905.4882202148438 0 841.116
907.4036254882812 0 14854.045
908.4058837890625 0 8110.868
909.4088745117188 0 1820.2516
911.4505004882812 0 1169.3302
921.4183349609375 0 1135.0292
922.4174194335938 0 861.3992
932.4832763671875 0 1571.7302
933.4865112304688 0 777.47906
934.4826049804688 0 678.7739
939.440185546875 0 5384.5776
940.4512939453125 0 4664.2354
941.46044921875 0 1280.6644
942.46142578125 0 676.0747
948.4716186523438 0 762.15784
949.4849853515625 0 723.3774
956.4530639648438 0 2245.5166
957.4578247070312 0 951.1005
958.4859008789062 0 4023.7307 b 8
959.4868774414062 0 2228.6384
960.48779296875 0 7793.0986
961.4881591796875 0 4501.5835
962.4805908203125 0 1649.1948
971.474365234375 0 1050.7433
988.4586791992188 0 683.52997
1003.50341796875 0 758.6578 y 8
1015.4697265625 0 784.16595
1036.4586181640625 0 850.3338
1037.4659423828125 0 926.43665
1040.4979248046875 0 1416.721
1041.48193359375 0 698.1551
1053.483642578125 0 5592.1226
1054.48583984375 0 3511.7432
1055.4967041015625 0 1238.9395
1058.50732421875 0 1853.5487
1059.5211181640625 0 1617.8638
1067.5025634765625 0 902.5872
1069.5130615234375 0 1418.201 b Water loss 9
1070.51025390625 0 3292.13
1071.51171875 0 1332.3958
1072.5238037109375 0 686.1254 y Water loss 7
1075.51953125 0 4428.2627
1076.522216796875 0 2075.728
1084.5302734375 0 1011.9271
1085.5069580078125 0 776.2767
1086.509521484375 0 1351.6825
1087.5267333984375 0 12478.321 b 9
1088.5302734375 0 6072.186
1089.531982421875 0 2459.7534
1090.536376953125 0 1625.1138 y 7
1091.5404052734375 0 689.81464
1103.523681640625 0 3924.0708
1104.5289306640625 0 3028.1233
1105.5323486328125 0 893.4241
1137.5465087890625 0 1261.5424
1138.548828125 0 1657.5787
1155.5623779296875 0 1919.8234
1156.561279296875 0 1281.655
1160.559814453125 0 989.2278 y Ammonia loss 6
1164.5496826171875 0 1376.9209
1165.5521240234375 0 893.2869
1177.5623779296875 0 1394.3595 y 6
1178.570068359375 0 1074.4646
1181.5762939453125 0 4750.8975
1182.5797119140625 0 3718.1196
1183.58251953125 0 1888.0553
1200.6087646484375 0 2035.2349 b 10
1201.607666015625 0 1153.7983
1212.6153564453125 0 940.6727
1234.6033935546875 0 3092.504
1235.600341796875 0 2302.0862
1236.6080322265625 0 1121.3604
1252.6121826171875 0 8870.516
1253.612548828125 0 5437.4385
1254.6187744140625 0 2419.5686
1256.619873046875 0 1026.0197 y Water loss 5
1257.604736328125 0 1591.6232 y Ammonia loss 5
1258.601806640625 0 743.0432
1274.6153564453125 0 3622.3916 y 5
1275.620361328125 0 2771.4736
1276.6239013671875 0 1031.7498
1355.6627197265625 0 709.7741
1371.66845703125 0 1614.5576 y 4
1372.674560546875 0 906.9811
1399.680419921875 0 894.64825
1400.6923828125 0 745.81244
1514.765869140625 0 869.9987
2466.253662109375 0 669.0714
3072.822998046875 0 804.3903

Spectrum Details

|  |  |
| --- | --- |
| Matched peaks? Matched peaks The total absolute number of peaks matched. Additionally in brackets the total fraction of peaks matched and the total number of peaks is shown. | 76 (11.19% of 679) |
| FDR? FDR The false discovery rate estimated for this peptide. It is calculated by matching all theoretical fragments with a non-integer shift with the raw peaks for this spectrum. This is done with 40 different shifts. The resulting percentage is the average number of annotated peaks over the number of annotated peaks with the correct spectrum. | 1.47% |
| Satellite FDR? Satellite FDR See the FDR for details on its calculation. This satellite ion specific FDR only contains the satellite ions (d/w) for I/L/J positions. | - |
| PSM Score? PSM Score The PSM Score as given by Hecklib to this annotated spectrum. It is shown with three significant figures. | 648 |

#### Reverse Lookup? Reverse Lookup All places where this read could be placed.

| Group | Segment | Template | Template Part | Read Part | Score | Unique |
| --- | --- | --- | --- | --- | --- | --- |
| Homo sapiens Light Chain | IGLC | IGLC2 | [8..25] | [0..17] | 136 | False |
| Homo sapiens Light Chain | IGLC | IGLC3 | [6..23] | [0..17] | 136 | False |
| Homo sapiens Light Chain | IGLC | IGLC6 | [8..25] | [0..17] | 136 | False |
| Homo sapiens Light Chain | IGLC | IGLC7 | [8..25] | [0..17] | 136 | False |

| Recombined | Template Part | Read Part | Score | Unique |
| --- | --- | --- | --- | --- |
| REC-0-1\_002 | [119..136] | [0..17] | 136 | True |

#### Meta Information from Multiple reads

##### Number of combined reads

16

##### Intensity

0.7887

##### TotalArea

4.373E+09

##### Changes to the peptide sequence

VTJFPPSSEEJQANKAT

L→JNo support for either Leucine or Isoleucine based on side chain ions (Position: 11)

L→JNo support for either Leucine or Isoleucine based on side chain ions (Position: 3)

#### Positional Score

Copy Data

##### Positional Score (TSV)

###### Preview

```
Loading example...
```

*Click on the button to copy the data to your clipboard.*

10012345678910111213141516

Label Value
"0" 0.815
"1" 0.834
"2" 0.869
"3" 0.867
"4" 0.868
"5" 0.869
"6" 0.868
"7" 0.861
"8" 0.869
"9" 0.874
"10" 0.874
"11" 0.853
"12" 0.854
"13" 0.842
"14" 0.809
"15" 0.821
"16" 0.849

#### Meta Information from PEAKS

##### Scan Identifier

F2:8009

##### Original sequence

V

T

L

F

P

P

S

S

E

E

L

Q

A

N

K

A

T

##### Posttranslational Modifications

##### Source File

D:\separate\_stitch\_analyses\xle-disambiguation\raw\20210323\_F1\_UM1\_Peng0013\_SA\_F59\_ingel\_3ug\_TL.raw

##### Fraction

2

##### Scan Feature

F2:8672

##### De Novo Score

99

##### ConfidenceScore

99

### m/z

611.3208

##### Mass

1830.9363

##### Charge

3

##### Retention Time

44.66

##### Predicted Retention Time

-

##### Area

4.031E+07

##### Parts Per Million

2.3

##### Fragmentation mode

ETHCD

##### Originating file

01 D:\separate\_stitch\_analyses\xle-disambiguation\20210325\_F59\_3ug\_DENOVO\_12.csv

#### Meta Information from PEAKS

##### Scan Identifier

F2:7949

##### Original sequence

V

T

L

F

P

P

S

S

E

E

L

Q

A

N

K

A

T

##### Posttranslational Modifications

##### Source File

D:\separate\_stitch\_analyses\xle-disambiguation\raw\20210323\_F1\_UM1\_Peng0013\_SA\_F59\_ingel\_3ug\_TL.raw

##### Fraction

2

##### Scan Feature

F2:8672

##### De Novo Score

99

##### ConfidenceScore

99

### m/z

611.3208

##### Mass

1830.9363

##### Charge

3

##### Retention Time

44.66

##### Predicted Retention Time

-

##### Area

4.031E+07

##### Parts Per Million

2.3

##### Fragmentation mode

ETHCD

##### Originating file

01 D:\separate\_stitch\_analyses\xle-disambiguation\20210325\_F59\_3ug\_DENOVO\_12.csv

#### Meta Information from PEAKS

##### Scan Identifier

F2:7896

##### Original sequence

V

T

L

F

P

P

S

S

E

E

L

Q

A

N

K

A

T

##### Posttranslational Modifications

##### Source File

D:\separate\_stitch\_analyses\xle-disambiguation\raw\20210323\_F1\_UM1\_Peng0013\_SA\_F59\_ingel\_3ug\_TL.raw

##### Fraction

2

##### Scan Feature

F2:8672

##### De Novo Score

98

##### ConfidenceScore

98

### m/z

611.3208

##### Mass

1830.9363

##### Charge

3

##### Retention Time

44.66

##### Predicted Retention Time

-

##### Area

4.031E+07

##### Parts Per Million

2.3

##### Fragmentation mode

ETHCD

##### Originating file

01 D:\separate\_stitch\_analyses\xle-disambiguation\20210325\_F59\_3ug\_DENOVO\_12.csv

#### Meta Information from PEAKS

##### Scan Identifier

F2:7881

##### Original sequence

V

T

L

F

P

P

S

S

E

E

L

Q

A

N

K

A

T

##### Posttranslational Modifications

##### Source File

D:\separate\_stitch\_analyses\xle-disambiguation\raw\20210323\_F1\_UM1\_Peng0013\_SA\_F59\_ingel\_3ug\_TL.raw

##### Fraction

2

##### Scan Feature

F2:17534

##### De Novo Score

98

##### ConfidenceScore

98

### m/z

916.4772

##### Mass

1830.9363

##### Charge

2

##### Retention Time

44.66

##### Predicted Retention Time

-

##### Area

4.545E+08

##### Parts Per Million

1.9

##### Fragmentation mode

ETHCD

##### Originating file

01 D:\separate\_stitch\_analyses\xle-disambiguation\20210325\_F59\_3ug\_DENOVO\_12.csv

#### Meta Information from PEAKS

##### Scan Identifier

F2:8046

##### Original sequence

V

T

L

F

P

P

S

S

E

E

L

Q

A

N

K

A

T

##### Posttranslational Modifications

##### Source File

D:\separate\_stitch\_analyses\xle-disambiguation\raw\20210323\_F1\_UM1\_Peng0013\_SA\_F59\_ingel\_3ug\_TL.raw

##### Fraction

2

##### Scan Feature

F2:17534

##### De Novo Score

98

##### ConfidenceScore

98

### m/z

916.4772

##### Mass

1830.9363

##### Charge

2

##### Retention Time

44.66

##### Predicted Retention Time

-

##### Area

4.545E+08

##### Parts Per Million

1.9

##### Fragmentation mode

ETHCD

##### Originating file

01 D:\separate\_stitch\_analyses\xle-disambiguation\20210325\_F59\_3ug\_DENOVO\_12.csv

#### Meta Information from PEAKS

##### Scan Identifier

F2:8281

##### Original sequence

V

T

L

F

P

P

S

S

E

E

L

Q

A

N

K

A

T

##### Posttranslational Modifications

##### Source File

D:\separate\_stitch\_analyses\xle-disambiguation\raw\20210323\_F1\_UM1\_Peng0013\_SA\_F59\_ingel\_3ug\_TL.raw

##### Fraction

2

##### Scan Feature

F2:17534

##### De Novo Score

98

##### ConfidenceScore

98

### m/z

916.4772

##### Mass

1830.9363

##### Charge

2

##### Retention Time

44.66

##### Predicted Retention Time

-

##### Area

4.545E+08

##### Parts Per Million

1.9

##### Fragmentation mode

ETHCD

##### Originating file

01 D:\separate\_stitch\_analyses\xle-disambiguation\20210325\_F59\_3ug\_DENOVO\_12.csv

#### Meta Information from PEAKS

##### Scan Identifier

F2:7814

##### Original sequence

V

T

L

F

P

P

S

S

E

E

L

Q

A

N

K

A

T

##### Posttranslational Modifications

##### Source File

D:\separate\_stitch\_analyses\xle-disambiguation\raw\20210323\_F1\_UM1\_Peng0013\_SA\_F59\_ingel\_3ug\_TL.raw

##### Fraction

2

##### Scan Feature

F2:17534

##### De Novo Score

98

##### ConfidenceScore

98

### m/z

916.4772

##### Mass

1830.9363

##### Charge

2

##### Retention Time

44.66

##### Predicted Retention Time

-

##### Area

4.545E+08

##### Parts Per Million

1.9

##### Fragmentation mode

ETHCD

##### Originating file

01 D:\separate\_stitch\_analyses\xle-disambiguation\20210325\_F59\_3ug\_DENOVO\_12.csv

#### Meta Information from PEAKS

##### Scan Identifier

F2:8106

##### Original sequence

V

T

L

F

P

P

S

S

E

E

L

Q

A

N

K

A

T

##### Posttranslational Modifications

##### Source File

D:\separate\_stitch\_analyses\xle-disambiguation\raw\20210323\_F1\_UM1\_Peng0013\_SA\_F59\_ingel\_3ug\_TL.raw

##### Fraction

2

##### Scan Feature

F2:17534

##### De Novo Score

98

##### ConfidenceScore

98

### m/z

916.4772

##### Mass

1830.9363

##### Charge

2

##### Retention Time

44.66

##### Predicted Retention Time

-

##### Area

4.545E+08

##### Parts Per Million

1.9

##### Fragmentation mode

HCD

##### Originating file

01 D:\separate\_stitch\_analyses\xle-disambiguation\20210325\_F59\_3ug\_DENOVO\_12.csv

#### Meta Information from PEAKS

##### Scan Identifier

F2:8067

##### Original sequence

V

T

L

F

P

P

S

S

E

E

L

Q

A

N

K

A

T

##### Posttranslational Modifications

##### Source File

D:\separate\_stitch\_analyses\xle-disambiguation\raw\20210323\_F1\_UM1\_Peng0013\_SA\_F59\_ingel\_3ug\_TL.raw

##### Fraction

2

##### Scan Feature

F2:8672

##### De Novo Score

97

##### ConfidenceScore

97

### m/z

611.3208

##### Mass

1830.9363

##### Charge

3

##### Retention Time

44.66

##### Predicted Retention Time

-

##### Area

4.031E+07

##### Parts Per Million

2.3

##### Fragmentation mode

HCD

##### Originating file

01 D:\separate\_stitch\_analyses\xle-disambiguation\20210325\_F59\_3ug\_DENOVO\_12.csv

#### Meta Information from PEAKS

##### Scan Identifier

F2:8383

##### Original sequence

V

T

L

F

P

P

S

S

E

E

L

Q

A

N

K

A

T

##### Posttranslational Modifications

##### Source File

D:\separate\_stitch\_analyses\xle-disambiguation\raw\20210323\_F1\_UM1\_Peng0013\_SA\_F59\_ingel\_3ug\_TL.raw

##### Fraction

2

##### Scan Feature

F2:17534

##### De Novo Score

97

##### ConfidenceScore

97

### m/z

916.4772

##### Mass

1830.9363

##### Charge

2

##### Retention Time

44.66

##### Predicted Retention Time

-

##### Area

4.545E+08

##### Parts Per Million

1.9

##### Fragmentation mode

ETHCD

##### Originating file

01 D:\separate\_stitch\_analyses\xle-disambiguation\20210325\_F59\_3ug\_DENOVO\_12.csv

#### Meta Information from PEAKS

##### Scan Identifier

F2:8223

##### Original sequence

V

T

L

F

P

P

S

S

E

E

L

Q

A

N

K

A

T

##### Posttranslational Modifications

##### Source File

D:\separate\_stitch\_analyses\xle-disambiguation\raw\20210323\_F1\_UM1\_Peng0013\_SA\_F59\_ingel\_3ug\_TL.raw

##### Fraction

2

##### Scan Feature

F2:17534

##### De Novo Score

97

##### ConfidenceScore

97

### m/z

916.4772

##### Mass

1830.9363

##### Charge

2

##### Retention Time

44.66

##### Predicted Retention Time

-

##### Area

4.545E+08

##### Parts Per Million

1.9

##### Fragmentation mode

HCD

##### Originating file

01 D:\separate\_stitch\_analyses\xle-disambiguation\20210325\_F59\_3ug\_DENOVO\_12.csv

#### Meta Information from PEAKS

##### Scan Identifier

F2:8167

##### Original sequence

V

T

L

F

P

P

S

S

E

E

L

Q

A

N

K

A

T

##### Posttranslational Modifications

##### Source File

D:\separate\_stitch\_analyses\xle-disambiguation\raw\20210323\_F1\_UM1\_Peng0013\_SA\_F59\_ingel\_3ug\_TL.raw

##### Fraction

2

##### Scan Feature

F2:17534

##### De Novo Score

97

##### ConfidenceScore

97

### m/z

916.4772

##### Mass

1830.9363

##### Charge

2

##### Retention Time

44.66

##### Predicted Retention Time

-

##### Area

4.545E+08

##### Parts Per Million

1.9

##### Fragmentation mode

HCD

##### Originating file

01 D:\separate\_stitch\_analyses\xle-disambiguation\20210325\_F59\_3ug\_DENOVO\_12.csv

#### Meta Information from PEAKS

##### Scan Identifier

F2:7834

##### Original sequence

V

T

L

F

P

P

S

S

E

E

L

Q

A

N

K

A

T

##### Posttranslational Modifications

##### Source File

D:\separate\_stitch\_analyses\xle-disambiguation\raw\20210323\_F1\_UM1\_Peng0013\_SA\_F59\_ingel\_3ug\_TL.raw

##### Fraction

2

##### Scan Feature

F2:8672

##### De Novo Score

96

##### ConfidenceScore

96

### m/z

611.3208

##### Mass

1830.9363

##### Charge

3

##### Retention Time

44.66

##### Predicted Retention Time

-

##### Area

4.031E+07

##### Parts Per Million

2.3

##### Fragmentation mode

HCD

##### Originating file

01 D:\separate\_stitch\_analyses\xle-disambiguation\20210325\_F59\_3ug\_DENOVO\_12.csv

#### Meta Information from PEAKS

##### Scan Identifier

F2:8439

##### Original sequence

V

T

L

F

P

P

S

S

E

E

L

Q

A

N

K

A

T

##### Posttranslational Modifications

##### Source File

D:\separate\_stitch\_analyses\xle-disambiguation\raw\20210323\_F1\_UM1\_Peng0013\_SA\_F59\_ingel\_3ug\_TL.raw

##### Fraction

2

##### Scan Feature

F2:17534

##### De Novo Score

96

##### ConfidenceScore

96

### m/z

916.4772

##### Mass

1830.9363

##### Charge

2

##### Retention Time

44.66

##### Predicted Retention Time

-

##### Area

4.545E+08

##### Parts Per Million

1.9

##### Fragmentation mode

HCD

##### Originating file

01 D:\separate\_stitch\_analyses\xle-disambiguation\20210325\_F59\_3ug\_DENOVO\_12.csv

#### Meta Information from PEAKS

##### Scan Identifier

F2:8129

##### Original sequence

V

T

L

F

P

P

S

S

E

E

L

Q

A

N

K

A

T

##### Posttranslational Modifications

##### Source File

D:\separate\_stitch\_analyses\xle-disambiguation\raw\20210323\_F1\_UM1\_Peng0013\_SA\_F59\_ingel\_3ug\_TL.raw

##### Fraction

2

##### Scan Feature

F2:8672

##### De Novo Score

95

##### ConfidenceScore

95

### m/z

611.3208

##### Mass

1830.9363

##### Charge

3

##### Retention Time

44.66

##### Predicted Retention Time

-

##### Area

4.031E+07

##### Parts Per Million

2.3

##### Fragmentation mode

HCD

##### Originating file

01 D:\separate\_stitch\_analyses\xle-disambiguation\20210325\_F59\_3ug\_DENOVO\_12.csv

#### Meta Information from PEAKS

##### Scan Identifier

F2:8190

##### Original sequence

V

T

L

F

P

P

S

S

E

E

L

Q

A

N

K

A

T

##### Posttranslational Modifications

##### Source File

D:\separate\_stitch\_analyses\xle-disambiguation\raw\20210323\_F1\_UM1\_Peng0013\_SA\_F59\_ingel\_3ug\_TL.raw

##### Fraction

2

##### Scan Feature

F2:8672

##### De Novo Score

95

##### ConfidenceScore

95

### m/z

611.3208

##### Mass

1830.9363

##### Charge

3

##### Retention Time

44.66

##### Predicted Retention Time

-

##### Area

4.031E+07

##### Parts Per Million

2.3

##### Fragmentation mode

HCD

##### Originating file

01 D:\separate\_stitch\_analyses\xle-disambiguation\20210325\_F59\_3ug\_DENOVO\_12.csv
