## Supplementary material for "A handle on mass coincidence errors in *de novo* sequencing of antibodies by bottom-up proteomics": Combined_031.html

Details Combined\_031 | Stitch OverviewUndefined

### Read Combined\_031

#### Sequence (length=8)

FDDYAMHW

#### Spectrum 5490? Spectrum 5490 The raw spectrum of this peptide as annotated by Hecklib. The fragments are coloured according to ion type (see legend). Any peaks with a star '\*' as text can be hovered over to see the full details, first the ion type second the mass shift type. By hovering over the amino acids in the peptide or ions in the legend the corresponding peaks are highlighted. By toggling the 'Unassigned' label you can turn the background (unassigned) peaks on or off in the plot. By updating the slider in the Ion legend you can update the spectrum to only show the top X% of the peaks with labels. The top X% means any peak that is within X% of the highest intensity. By dragging in the spectrum you can zoom in to a specific part of the spectrum and use 'Zoom Out' to get back to the original zoom level. The annotation of the spectrum is based on the given sequence in the peptides file and is done with different software so inconsistencies are likely. The peaks are annotated based on the given sequence, with 20 ppm tolerance.

Copy Data

##### Spectrum 5490 (TSV)

###### Preview

```
Loading example...
```

*Click on the button to copy the data to your clipboard.*

Mz MinMz MaxIntensity Max

WidthHeightPeptide font sizePeptide stroke widthSpectrum font sizeSpectrum stroke widthCompact peptide

Ion legend

wxyz

abcd

OtherUnassignedIon^Charge^_Position_Show for top:%

FDDYAMHW

01.64e+43.27e+44.91e+46.55e+4

Zoom Out

d^+1^_2_a^+1^_2_a^+1^_2_b^+1^_2_a^+1^_3_y^+1^_2_y^+2^_5_b^+1^_3_y^+2^_6_b^+1^_4_\*\*y^+1^_4_b^+1^_5_y^+1^_5_y^+1^_6_y^+1^_6_b^+1^_7_y^+1^_7_y^+1^_7_

045691113671822

Fragment Matches Table

Show background peaks

| Position | Ion type | Intensity | mz Theoretical | mz Error (Th) | mz Error (ppm) | Charge | Series Number |
| --- | --- | --- | --- | --- | --- | --- | --- |
| - | - | 6.482E+04 | 120.1 | - | - | 0 | - |
| - | - | 365.9 | 121.1 | - | - | 0 | - |
| - | - | 4989 | 121.1 | - | - | 0 | - |
| - | - | 362.5 | 122.1 | - | - | 0 | - |
| - | - | 391.3 | 123.4 | - | - | 0 | - |
| - | - | 373.6 | 124.6 | - | - | 0 | - |
| - | - | 357.7 | 127.7 | - | - | 0 | - |
| - | - | 600.6 | 128.1 | - | - | 0 | - |
| - | - | 4318 | 129.1 | - | - | 0 | - |
| - | - | 1295 | 130.1 | - | - | 0 | - |
| - | - | 386.9 | 132.3 | - | - | 0 | - |
| - | - | 478.4 | 133.1 | - | - | 0 | - |
| - | - | 475.5 | 133.1 | - | - | 0 | - |
| - | - | 9400 | 136.1 | - | - | 0 | - |
| - | - | 423.7 | 136.6 | - | - | 0 | - |
| - | - | 488.4 | 138.1 | - | - | 0 | - |
| - | - | 672.5 | 141.1 | - | - | 0 | - |
| - | - | 688.4 | 142.1 | - | - | 0 | - |
| - | - | 989.1 | 146.1 | - | - | 0 | - |
| - | - | 645.9 | 146.1 | - | - | 0 | - |
| - | - | 403.2 | 147 | - | - | 0 | - |
| - | - | 652.7 | 147.1 | - | - | 0 | - |
| - | - | 389.9 | 148.5 | - | - | 0 | - |
| - | - | 583.5 | 148.9 | - | - | 0 | - |
| - | - | 438.9 | 150.5 | - | - | 0 | - |
| - | - | 398 | 150.7 | - | - | 0 | - |
| - | - | 950.1 | 152.1 | - | - | 0 | - |
| - | - | 543.6 | 155.1 | - | - | 0 | - |
| - | - | 2404 | 156.1 | - | - | 0 | - |
| - | - | 534.5 | 156.1 | - | - | 0 | - |
| - | - | 536.7 | 157.1 | - | - | 0 | - |
| - | - | 538.6 | 159.1 | - | - | 0 | - |
| - | - | 1031 | 163.1 | - | - | 0 | - |
| - | - | 630 | 165.1 | - | - | 0 | - |
| - | - | 1571 | 173.1 | - | - | 0 | - |
| - | - | 1343 | 173.1 | - | - | 0 | - |
| - | - | 813.5 | 174.1 | - | - | 0 | - |
| - | - | 443.1 | 174.1 | - | - | 0 | - |
| - | - | 3788 | 175.1 | - | - | 0 | - |
| - | - | 601.8 | 175.1 | - | - | 0 | - |
| - | - | 514.5 | 176.1 | - | - | 0 | - |
| - | - | 732 | 177.1 | - | - | 0 | - |
| - | - | 562 | 183.1 | - | - | 0 | - |
| - | - | 445.8 | 183.7 | - | - | 0 | - |
| - | - | 574.8 | 185.1 | - | - | 0 | - |
| - | - | 649 | 185.1 | - | - | 0 | - |
| - | - | 4143 | 187.1 | - | - | 0 | - |
| - | - | 1358 | 187.1 | - | - | 0 | - |
| - | - | 1193 | 190.1 | - | - | 0 | - |
| - | - | 665.6 | 191.1 | - | - | 0 | - |
| 2 | d | 2393 | 191.1 | 0.0001982 | 1.037 | +1 | 2 |
| - | - | 818.8 | 193.1 | - | - | 0 | - |
| - | - | 774.1 | 195.2 | - | - | 0 | - |
| - | - | 614.1 | 197.1 | - | - | 0 | - |
| - | - | 978.8 | 201.1 | - | - | 0 | - |
| - | - | 782.3 | 203.1 | - | - | 0 | - |
| - | - | 702.2 | 203.1 | - | - | 0 | - |
| - | - | 749.7 | 204.1 | - | - | 0 | - |
| - | - | 1859 | 215.1 | - | - | 0 | - |
| 2 | a | 8361 | 217.1 | 0.0001359 | 0.6262 | +1 | 2 |
| - | - | 5702 | 221.1 | - | - | 0 | - |
| - | - | 3274 | 221.1 | - | - | 0 | - |
| - | - | 905.8 | 222.1 | - | - | 0 | - |
| - | - | 1319 | 226.2 | - | - | 0 | - |
| - | - | 785 | 227.1 | - | - | 0 | - |
| - | - | 1070 | 231.1 | - | - | 0 | - |
| - | - | 906.2 | 233.2 | - | - | 0 | - |
| 2 | a | 1.453E+04 | 235.1 | 0.0001914 | 0.814 | +1 | 2 |
| - | - | 1890 | 236.1 | - | - | 0 | - |
| - | - | 930 | 239.1 | - | - | 0 | - |
| - | - | 516.4 | 242.1 | - | - | 0 | - |
| - | - | 688.1 | 244.1 | - | - | 0 | - |
| - | - | 3532 | 251.1 | - | - | 0 | - |
| - | - | 597.4 | 261.2 | - | - | 0 | - |
| 2 | b | 1.142E+04 | 263.1 | 2.786E-06 | 0.01059 | +1 | 2 |
| - | - | 1468 | 264.1 | - | - | 0 | - |
| - | - | 508.4 | 265.2 | - | - | 0 | - |
| - | - | 539.2 | 278.1 | - | - | 0 | - |
| - | - | 1960 | 279.1 | - | - | 0 | - |
| - | - | 724.4 | 280.1 | - | - | 0 | - |
| - | - | 1624 | 285.1 | - | - | 0 | - |
| - | - | 1054 | 292.1 | - | - | 0 | - |
| - | - | 596.6 | 312.1 | - | - | 0 | - |
| - | - | 2145 | 340.1 | - | - | 0 | - |
| 3 | a | 1282 | 350.1 | 0.0002259 | 0.6451 | +1 | 3 |
| - | - | 1100 | 356.1 | - | - | 0 | - |
| 7 | y | 1.454E+04 | 358.1 | 0.005118 | 14.29 | +1 | 2 |
| - | - | 2719 | 359.2 | - | - | 0 | - |
| 4 | y | 560.9 | 370.1 | 0.005021 | 13.57 | +2 | 5 |
| 3 | b | 9055 | 378.1 | 0.0002148 | 0.5681 | +1 | 3 |
| - | - | 1449 | 379.1 | - | - | 0 | - |
| - | - | 1230 | 394.1 | - | - | 0 | - |
| - | - | 565.2 | 411.7 | - | - | 0 | - |
| 3 | y | 724.6 | 427.7 | 0.001132 | 2.647 | +2 | 6 |
| - | - | 2606 | 441.2 | - | - | 0 | - |
| - | - | 648.6 | 455.2 | - | - | 0 | - |
| - | - | 1320 | 465.2 | - | - | 0 | - |
| - | - | 802.1 | 477.2 | - | - | 0 | - |
| - | - | 698.4 | 491.2 | - | - | 0 | - |
| - | - | 1703 | 495.2 | - | - | 0 | - |
| - | - | 763.7 | 496.2 | - | - | 0 | - |
| - | - | 3578 | 505.2 | - | - | 0 | - |
| - | - | 648.5 | 506.2 | - | - | 0 | - |
| - | - | 2140 | 512.2 | - | - | 0 | - |
| - | - | 1741 | 513.2 | - | - | 0 | - |
| - | - | 677 | 524.3 | - | - | 0 | - |
| - | - | 1019 | 526.7 | - | - | 0 | - |
| 4 | b | 601.8 | 541.2 | 0.005154 | 9.524 | +1 | 4 |
| 0 | Precursor | 1126 | 549.7 | 0.004695 | 8.542 | +2 | -1 |
| - | - | 857.4 | 557.3 | - | - | 0 | - |
| 0 | Precursor | 3167 | 558.7 | 0.005028 | 9 | +2 | -1 |
| - | - | 2739 | 559.2 | - | - | 0 | - |
| - | - | 532.3 | 559.3 | - | - | 0 | - |
| - | - | 855.6 | 559.7 | - | - | 0 | - |
| - | - | 758.3 | 560 | - | - | 0 | - |
| - | - | 1311 | 570.2 | - | - | 0 | - |
| 5 | y | 8276 | 576.2 | 0.01009 | 17.51 | +1 | 4 |
| - | - | 2673 | 577.2 | - | - | 0 | - |
| - | - | 694.2 | 578.2 | - | - | 0 | - |
| - | - | 796.7 | 606.2 | - | - | 0 | - |
| 5 | b | 599.3 | 612.2 | 0.002358 | 3.851 | +1 | 5 |
| - | - | 815.5 | 634.2 | - | - | 0 | - |
| - | - | 1442 | 675.3 | - | - | 0 | - |
| - | - | 826.4 | 721.3 | - | - | 0 | - |
| 4 | y | 5451 | 739.3 | 0.009384 | 12.69 | +1 | 5 |
| - | - | 2711 | 740.3 | - | - | 0 | - |
| - | - | 793.7 | 749.3 | - | - | 0 | - |
| - | - | 4295 | 790.3 | - | - | 0 | - |
| - | - | 1593 | 791.3 | - | - | 0 | - |
| 3 | y | 968 | 836.3 | 0.008448 | 10.1 | +1 | 6 |
| 3 | y | 1.186E+04 | 854.3 | 0.01033 | 12.1 | +1 | 6 |
| - | - | 4919 | 855.3 | - | - | 0 | - |
| - | - | 2416 | 856.3 | - | - | 0 | - |
| 7 | b | 642.3 | 896.3 | 0.0007142 | 0.7968 | +1 | 7 |
| - | - | 3090 | 905.3 | - | - | 0 | - |
| - | - | 1365 | 906.3 | - | - | 0 | - |
| 2 | y | 882.1 | 951.3 | 0.01513 | 15.91 | +1 | 7 |
| - | - | 844.2 | 952.3 | - | - | 0 | - |
| 2 | y | 8425 | 969.3 | 0.009209 | 9.5 | +1 | 7 |
| - | - | 3707 | 970.3 | - | - | 0 | - |
| - | - | 1764 | 971.3 | - | - | 0 | - |
| - | - | 824.6 | 979.3 | - | - | 0 | - |
| - | - | 713.2 | 1004 | - | - | 0 | - |
| - | - | 602.1 | 1804 | - | - | 0 | - |

m/z Charge Intensity FragmentType MassShift Position
120.08100891113281 0 64819.938
121.08077239990234 0 365.90186
121.0843276977539 0 4988.877
122.0715560913086 0 362.48105
123.35560607910156 0 391.31857
124.57332611083984 0 373.61014
127.6724624633789 0 357.70834
128.107177734375 0 600.58734
129.1024169921875 0 4318.1206
130.06529235839844 0 1294.7273
132.29061889648438 0 386.85837
133.0612030029297 0 478.40472
133.08615112304688 0 475.47858
136.07589721679688 0 9399.667
136.55291748046875 0 423.6748
138.06614685058594 0 488.38983
141.10243225097656 0 672.52435
142.1227264404297 0 688.43555
146.06031799316406 0 989.0525
146.0970001220703 0 645.93195
147.04434204101562 0 403.17636
147.0765838623047 0 652.65564
148.48199462890625 0 389.87473
148.9478302001953 0 583.45044
150.46116638183594 0 438.94077
150.7327117919922 0 397.96368
152.0709686279297 0 950.1102
155.08154296875 0 543.55536
156.076904296875 0 2404.3167
156.0835723876953 0 534.46655
157.1333465576172 0 536.67267
159.0921630859375 0 538.63696
163.07167053222656 0 1031.4313
165.1023712158203 0 629.96497
173.09214782714844 0 1570.9604
173.12879943847656 0 1343.338
174.0552520751953 0 813.4581
174.0671844482422 0 443.08798
175.08680725097656 0 3787.6133
175.11936950683594 0 601.7976
176.09010314941406 0 514.4526
177.06781005859375 0 732.03204
183.1134796142578 0 562.0331
183.69688415527344 0 445.81033
185.05572509765625 0 574.8084
185.0917205810547 0 649.0249
187.10800170898438 0 4142.8467
187.1444854736328 0 1358.3499
190.0863800048828 0 1192.7772
191.08143615722656 0 665.59485
191.1180877685547 0 2393.2065 d 1
193.1087188720703 0 818.84467
195.185546875 0 774.14105
197.12881469726562 0 614.0664
201.12376403808594 0 978.8284
203.06651306152344 0 782.29944
203.08169555664062 0 702.1735
204.07650756835938 0 749.6504
215.13894653320312 0 1859.1455
217.0972900390625 0 8361.443 a Water loss 1
221.0922393798828 0 5701.6875
221.1036834716797 0 3274.4817
222.0951690673828 0 905.77734
226.15493774414062 0 1318.9236
227.10260009765625 0 785.00104
231.0623016357422 0 1069.6411
233.1649932861328 0 906.17957
235.10791015625 0 14525.679 a 1
236.1117401123047 0 1889.5154
239.1135711669922 0 930.03076
242.1488800048828 0 516.4271
244.12965393066406 0 688.0833
251.10263061523438 0 3531.7766
261.1603698730469 0 597.4391
263.1026306152344 0 11423.1045 b 1
264.10540771484375 0 1467.5824
265.15350341796875 0 508.38608
278.1148376464844 0 539.1986
279.0972595214844 0 1960.1575
280.0994567871094 0 724.42316
285.10150146484375 0 1623.929
292.1419982910156 0 1053.6477
312.14617919921875 0 596.60016
340.14093017578125 0 2144.9788
350.1348876953125 0 1282.365 a 2
356.1385803222656 0 1099.8435
358.15118408203125 0 14544.689 y 6
359.15399169921875 0 2718.7
370.14715576171875 0 560.8753 y 3
378.1297912597656 0 9055.081 b 2
379.1333923339844 0 1449.4656
394.1228332519531 0 1229.527
411.6748352050781 0 565.2172
427.65673828125 0 724.6246 y 2
441.18939208984375 0 2606.1428
455.2048645019531 0 648.6141
465.1615295410156 0 1320.4366
477.18096923828125 0 802.08405
491.2071533203125 0 698.4132
495.1880798339844 0 1702.5626
496.18878173828125 0 763.69916
505.1871643066406 0 3578.126
506.18646240234375 0 648.5261
512.2258911132812 0 2140.1406
513.1983642578125 0 1740.8522
524.2642822265625 0 677.0342
526.7085571289062 0 1019.3288
541.1980590820312 0 601.7529 b 3
549.7026977539062 0 1125.5564 Precursor Water loss
557.3001708984375 0 857.4174
558.7083129882812 0 3167.2334 Precursor
559.2108154296875 0 2739.4846
559.256591796875 0 532.33026
559.7053833007812 0 855.5785
559.9511108398438 0 758.3449
570.2310791015625 0 1310.6982
576.2237548828125 0 8276.357 y 4
577.2255859375 0 2672.9958
578.224609375 0 694.23724
606.235595703125 0 796.6882
612.2276611328125 0 599.3327 b 4
634.227783203125 0 815.5435
675.2894897460938 0 1441.7811
721.2643432617188 0 826.4439
739.286376953125 0 5451.11 y 3
740.2906494140625 0 2710.868
749.2639770507812 0 793.67554
790.3148193359375 0 4295.085
791.3179321289062 0 1592.7731
836.3018188476562 0 967.95605 y Water loss 2
854.3142700195312 0 11861.93 y 2
855.316162109375 0 4919.431
856.3162841796875 0 2416.3372
896.3201293945312 0 642.33075 b 6
905.3421630859375 0 3089.952
906.3496704101562 0 1364.989
951.33544921875 0 882.09344 y Water loss 1
952.3331298828125 0 844.2194
969.340087890625 0 8425.188 y 1
970.3421630859375 0 3707.379
971.3438110351562 0 1763.9553
979.328857421875 0 824.586
1003.7855834960938 0 713.20087
1804.0595703125 0 602.1271

Spectrum Details

|  |  |
| --- | --- |
| Matched peaks? Matched peaks The total absolute number of peaks matched. Additionally in brackets the total fraction of peaks matched and the total number of peaks is shown. | 20 (13.89% of 144) |
| FDR? FDR The false discovery rate estimated for this peptide. It is calculated by matching all theoretical fragments with a non-integer shift with the raw peaks for this spectrum. This is done with 40 different shifts. The resulting percentage is the average number of annotated peaks over the number of annotated peaks with the correct spectrum. | 0.24% |
| Satellite FDR? Satellite FDR See the FDR for details on its calculation. This satellite ion specific FDR only contains the satellite ions (d/w) for I/L/J positions. | - |
| PSM Score? PSM Score The PSM Score as given by Hecklib to this annotated spectrum. It is shown with three significant figures. | 206 |

#### Spectrum 6681? Spectrum 6681 The raw spectrum of this peptide as annotated by Hecklib. The fragments are coloured according to ion type (see legend). Any peaks with a star '\*' as text can be hovered over to see the full details, first the ion type second the mass shift type. By hovering over the amino acids in the peptide or ions in the legend the corresponding peaks are highlighted. By toggling the 'Unassigned' label you can turn the background (unassigned) peaks on or off in the plot. By updating the slider in the Ion legend you can update the spectrum to only show the top X% of the peaks with labels. The top X% means any peak that is within X% of the highest intensity. By dragging in the spectrum you can zoom in to a specific part of the spectrum and use 'Zoom Out' to get back to the original zoom level. The annotation of the spectrum is based on the given sequence in the peptides file and is done with different software so inconsistencies are likely. The peaks are annotated based on the given sequence, with 20 ppm tolerance.

Copy Data

##### Spectrum 6681 (TSV)

###### Preview

```
Loading example...
```

*Click on the button to copy the data to your clipboard.*

Mz MinMz MaxIntensity Max

WidthHeightPeptide font sizePeptide stroke widthSpectrum font sizeSpectrum stroke widthCompact peptide

Ion legend

wxyz

abcd

OtherUnassignedIon^Charge^_Position_Show for top:%

FDDYAMHW

06.96e+41.39e+52.09e+52.78e+5

Zoom Out

a^+2^_3_d^+1^_2_y^+1^_1_a^+1^_2_a^+1^_2_a^+1^_2_b^+1^_2_b^+1^_2_b^+1^_2_d^+1^_3_a^+1^_3_y^+1^_2_a^+1^_3_b^+1^_3_y^+2^_5_b^+1^_3_b^+2^_6_y^+2^_6_b^+2^_7_y^+2^_7_y^+1^_3_b^+1^_4_b^+1^_4_\*\*y^+1^_4_b^+1^_5_y^+1^_5_y^+1^_6_y^+1^_6_b^+1^_7_y^+1^_7_y^+1^_7_

0762152322853046

Fragment Matches Table

Show background peaks

| Position | Ion type | Intensity | mz Theoretical | mz Error (Th) | mz Error (ppm) | Charge | Series Number |
| --- | --- | --- | --- | --- | --- | --- | --- |
| - | - | 2.757E+05 | 120.1 | - | - | 0 | - |
| - | - | 2.36E+04 | 121.1 | - | - | 0 | - |
| - | - | 575.3 | 122.1 | - | - | 0 | - |
| - | - | 849.4 | 127.1 | - | - | 0 | - |
| - | - | 781.2 | 128.1 | - | - | 0 | - |
| - | - | 447.2 | 129.1 | - | - | 0 | - |
| - | - | 3394 | 129.1 | - | - | 0 | - |
| - | - | 3846 | 130.1 | - | - | 0 | - |
| - | - | 543.1 | 130.1 | - | - | 0 | - |
| - | - | 456.5 | 131.1 | - | - | 0 | - |
| - | - | 766.4 | 131.1 | - | - | 0 | - |
| - | - | 550.1 | 132.1 | - | - | 0 | - |
| - | - | 501.5 | 133.1 | - | - | 0 | - |
| - | - | 1148 | 133.1 | - | - | 0 | - |
| - | - | 2.598E+04 | 136.1 | - | - | 0 | - |
| - | - | 1494 | 137.1 | - | - | 0 | - |
| - | - | 2622 | 138.1 | - | - | 0 | - |
| - | - | 2466 | 138.1 | - | - | 0 | - |
| - | - | 582.3 | 139.1 | - | - | 0 | - |
| - | - | 2205 | 146.1 | - | - | 0 | - |
| - | - | 3009 | 146.1 | - | - | 0 | - |
| - | - | 769 | 148.1 | - | - | 0 | - |
| - | - | 832 | 149 | - | - | 0 | - |
| - | - | 3264 | 149 | - | - | 0 | - |
| - | - | 1643 | 155.1 | - | - | 0 | - |
| - | - | 1116 | 155.1 | - | - | 0 | - |
| - | - | 7170 | 156.1 | - | - | 0 | - |
| - | - | 535.4 | 157.1 | - | - | 0 | - |
| - | - | 933.5 | 158.1 | - | - | 0 | - |
| - | - | 473.6 | 159.1 | - | - | 0 | - |
| - | - | 8623 | 159.1 | - | - | 0 | - |
| - | - | 1131 | 159.1 | - | - | 0 | - |
| - | - | 1488 | 160.1 | - | - | 0 | - |
| - | - | 625 | 162.1 | - | - | 0 | - |
| - | - | 495.9 | 165.1 | - | - | 0 | - |
| - | - | 1166 | 165.1 | - | - | 0 | - |
| - | - | 704.8 | 166.1 | - | - | 0 | - |
| - | - | 1233 | 166.1 | - | - | 0 | - |
| 3 | a | 4654 | 167.1 | 0.002107 | 12.61 | +2 | 3 |
| - | - | 1403 | 167.1 | - | - | 0 | - |
| - | - | 489.6 | 171.1 | - | - | 0 | - |
| - | - | 1767 | 172.1 | - | - | 0 | - |
| - | - | 538.9 | 172.1 | - | - | 0 | - |
| - | - | 519.3 | 173.1 | - | - | 0 | - |
| - | - | 646 | 174.1 | - | - | 0 | - |
| - | - | 2831 | 175.1 | - | - | 0 | - |
| - | - | 422.9 | 175.1 | - | - | 0 | - |
| - | - | 1578 | 176.1 | - | - | 0 | - |
| - | - | 514.4 | 176.1 | - | - | 0 | - |
| - | - | 1186 | 177.1 | - | - | 0 | - |
| - | - | 1099 | 180.1 | - | - | 0 | - |
| - | - | 1044 | 181.1 | - | - | 0 | - |
| - | - | 450.4 | 181.3 | - | - | 0 | - |
| - | - | 2134 | 185.1 | - | - | 0 | - |
| - | - | 611.5 | 185.1 | - | - | 0 | - |
| - | - | 487.6 | 185.2 | - | - | 0 | - |
| - | - | 721.1 | 187.1 | - | - | 0 | - |
| - | - | 1424 | 187.1 | - | - | 0 | - |
| - | - | 519.9 | 187.1 | - | - | 0 | - |
| - | - | 9304 | 188.1 | - | - | 0 | - |
| - | - | 1006 | 189.1 | - | - | 0 | - |
| - | - | 550.1 | 189.1 | - | - | 0 | - |
| - | - | 518.7 | 189.1 | - | - | 0 | - |
| - | - | 459.7 | 189.8 | - | - | 0 | - |
| - | - | 2811 | 190.1 | - | - | 0 | - |
| - | - | 671.4 | 191.1 | - | - | 0 | - |
| - | - | 839.2 | 191.1 | - | - | 0 | - |
| 2 | d | 1.06E+04 | 191.1 | 0.0001525 | 0.7977 | +1 | 2 |
| - | - | 930 | 192.1 | - | - | 0 | - |
| - | - | 4737 | 193.1 | - | - | 0 | - |
| - | - | 500.8 | 194.1 | - | - | 0 | - |
| - | - | 1914 | 195.1 | - | - | 0 | - |
| - | - | 576 | 195.1 | - | - | 0 | - |
| - | - | 467.3 | 195.1 | - | - | 0 | - |
| - | - | 661.1 | 197.1 | - | - | 0 | - |
| - | - | 754 | 199.1 | - | - | 0 | - |
| - | - | 896.4 | 199.2 | - | - | 0 | - |
| - | - | 574.1 | 200.1 | - | - | 0 | - |
| - | - | 802.8 | 201.1 | - | - | 0 | - |
| - | - | 1010 | 202.1 | - | - | 0 | - |
| - | - | 1793 | 203.1 | - | - | 0 | - |
| - | - | 520.7 | 203.1 | - | - | 0 | - |
| - | - | 2935 | 204.1 | - | - | 0 | - |
| 8 | y | 5668 | 205.1 | 0.0001359 | 0.6628 | +1 | 1 |
| - | - | 914.2 | 205.1 | - | - | 0 | - |
| - | - | 980.6 | 206.1 | - | - | 0 | - |
| - | - | 2288 | 207.1 | - | - | 0 | - |
| - | - | 1050 | 208.1 | - | - | 0 | - |
| - | - | 1215 | 209.1 | - | - | 0 | - |
| - | - | 513.3 | 212.1 | - | - | 0 | - |
| - | - | 543.1 | 213.1 | - | - | 0 | - |
| - | - | 792.5 | 215.1 | - | - | 0 | - |
| 2 | a | 3.692E+04 | 217.1 | 0.0001359 | 0.6262 | +1 | 2 |
| 2 | a | 891 | 218.1 | 0.0005259 | 2.411 | +1 | 2 |
| - | - | 3479 | 218.1 | - | - | 0 | - |
| - | - | 1180 | 219.1 | - | - | 0 | - |
| - | - | 808.4 | 219.1 | - | - | 0 | - |
| - | - | 486.3 | 220.1 | - | - | 0 | - |
| - | - | 6504 | 221.1 | - | - | 0 | - |
| - | - | 1.451E+04 | 221.1 | - | - | 0 | - |
| - | - | 1233 | 222.1 | - | - | 0 | - |
| - | - | 2017 | 222.1 | - | - | 0 | - |
| - | - | 853.1 | 223.1 | - | - | 0 | - |
| - | - | 9175 | 225 | - | - | 0 | - |
| - | - | 801.5 | 226 | - | - | 0 | - |
| - | - | 529.1 | 226.1 | - | - | 0 | - |
| - | - | 786.2 | 226.2 | - | - | 0 | - |
| - | - | 1095 | 227 | - | - | 0 | - |
| - | - | 755.7 | 227.1 | - | - | 0 | - |
| - | - | 1214 | 228.1 | - | - | 0 | - |
| - | - | 745.7 | 229.1 | - | - | 0 | - |
| - | - | 4022 | 231.1 | - | - | 0 | - |
| - | - | 808.3 | 233.1 | - | - | 0 | - |
| 2 | a | 5.363E+04 | 235.1 | 0.0001761 | 0.7491 | +1 | 2 |
| - | - | 6663 | 236.1 | - | - | 0 | - |
| - | - | 1.937E+04 | 239.1 | - | - | 0 | - |
| - | - | 1293 | 239.1 | - | - | 0 | - |
| - | - | 3678 | 240.1 | - | - | 0 | - |
| - | - | 670.6 | 241.1 | - | - | 0 | - |
| - | - | 1096 | 242.2 | - | - | 0 | - |
| - | - | 559.9 | 245.1 | - | - | 0 | - |
| 2 | b | 1463 | 245.1 | 0.0002775 | 1.132 | +1 | 2 |
| 2 | b | 763.5 | 246.1 | 0.0009878 | 4.014 | +1 | 2 |
| - | - | 799.1 | 246.1 | - | - | 0 | - |
| - | - | 592.3 | 248.2 | - | - | 0 | - |
| - | - | 1.433E+04 | 251.1 | - | - | 0 | - |
| - | - | 1963 | 252.1 | - | - | 0 | - |
| - | - | 2683 | 257.1 | - | - | 0 | - |
| - | - | 962 | 261.1 | - | - | 0 | - |
| 2 | b | 4.83E+04 | 263.1 | 2.773E-05 | 0.1054 | +1 | 2 |
| - | - | 6914 | 264.1 | - | - | 0 | - |
| - | - | 1149 | 264.1 | - | - | 0 | - |
| - | - | 676.7 | 265.1 | - | - | 0 | - |
| - | - | 662.3 | 267.1 | - | - | 0 | - |
| - | - | 556.5 | 269.1 | - | - | 0 | - |
| - | - | 3405 | 274.1 | - | - | 0 | - |
| - | - | 8652 | 279.1 | - | - | 0 | - |
| - | - | 1039 | 281.1 | - | - | 0 | - |
| - | - | 7103 | 285.1 | - | - | 0 | - |
| - | - | 809.2 | 286.1 | - | - | 0 | - |
| - | - | 559.6 | 287.1 | - | - | 0 | - |
| - | - | 5331 | 292.1 | - | - | 0 | - |
| - | - | 811.9 | 293.1 | - | - | 0 | - |
| - | - | 3795 | 295.1 | - | - | 0 | - |
| - | - | 1599 | 296.1 | - | - | 0 | - |
| - | - | 4047 | 296.2 | - | - | 0 | - |
| - | - | 644.9 | 297.2 | - | - | 0 | - |
| - | - | 9889 | 299.1 | - | - | 0 | - |
| - | - | 1138 | 300.1 | - | - | 0 | - |
| - | - | 976.8 | 303.1 | - | - | 0 | - |
| 3 | d | 1772 | 306.1 | 6.489E-05 | 0.212 | +1 | 3 |
| - | - | 619.9 | 310.2 | - | - | 0 | - |
| - | - | 1427 | 313.1 | - | - | 0 | - |
| - | - | 1040 | 318.1 | - | - | 0 | - |
| - | - | 1075 | 323.1 | - | - | 0 | - |
| - | - | 9886 | 324.1 | - | - | 0 | - |
| - | - | 1987 | 325.1 | - | - | 0 | - |
| 3 | a | 888.3 | 332.1 | 1.265E-05 | 0.03809 | +1 | 3 |
| - | - | 731.1 | 334.1 | - | - | 0 | - |
| - | - | 2036 | 338.1 | - | - | 0 | - |
| 7 | y | 5.533E+04 | 342.2 | 0.000184 | 0.5379 | +1 | 2 |
| - | - | 805.1 | 343.1 | - | - | 0 | - |
| - | - | 1.03E+04 | 343.2 | - | - | 0 | - |
| - | - | 657 | 344.2 | - | - | 0 | - |
| - | - | 2496 | 348.1 | - | - | 0 | - |
| 3 | a | 6021 | 350.1 | 0.0001038 | 0.2965 | +1 | 3 |
| - | - | 863 | 351.1 | - | - | 0 | - |
| - | - | 986.8 | 352.1 | - | - | 0 | - |
| - | - | 980.3 | 355.1 | - | - | 0 | - |
| - | - | 4899 | 356.1 | - | - | 0 | - |
| - | - | 777 | 357.2 | - | - | 0 | - |
| - | - | 1102 | 359 | - | - | 0 | - |
| - | - | 531.6 | 360.1 | - | - | 0 | - |
| 3 | b | 2783 | 360.1 | 6.808E-06 | 0.0189 | +1 | 3 |
| - | - | 876.8 | 361.2 | - | - | 0 | - |
| 4 | y | 1040 | 362.1 | 0.002829 | 7.811 | +2 | 5 |
| - | - | 1071 | 366.1 | - | - | 0 | - |
| - | - | 5126 | 369.1 | - | - | 0 | - |
| - | - | 1556 | 370.1 | - | - | 0 | - |
| - | - | 1004 | 374.1 | - | - | 0 | - |
| - | - | 1114 | 376.1 | - | - | 0 | - |
| 3 | b | 3.294E+04 | 378.1 | 2.931E-05 | 0.07752 | +1 | 3 |
| - | - | 6429 | 379.1 | - | - | 0 | - |
| 6 | b | 770.6 | 380.1 | 0.0017 | 4.471 | +2 | 6 |
| - | - | 1771 | 382.1 | - | - | 0 | - |
| - | - | 876.9 | 390.2 | - | - | 0 | - |
| - | - | 6481 | 394.1 | - | - | 0 | - |
| - | - | 1982 | 395.1 | - | - | 0 | - |
| - | - | 1074 | 404.1 | - | - | 0 | - |
| - | - | 669.7 | 407.2 | - | - | 0 | - |
| - | - | 1419 | 408.2 | - | - | 0 | - |
| - | - | 945.4 | 415 | - | - | 0 | - |
| - | - | 633 | 419 | - | - | 0 | - |
| 3 | y | 678.9 | 419.7 | 0.002205 | 5.254 | +2 | 6 |
| - | - | 888.4 | 420.2 | - | - | 0 | - |
| - | - | 7992 | 425.2 | - | - | 0 | - |
| - | - | 1493 | 426.2 | - | - | 0 | - |
| - | - | 1915 | 427.2 | - | - | 0 | - |
| - | - | 889.3 | 434.7 | - | - | 0 | - |
| - | - | 682.8 | 442.2 | - | - | 0 | - |
| - | - | 1228 | 447.2 | - | - | 0 | - |
| 7 | b | 580.4 | 448.7 | 0.00209 | 4.658 | +2 | 7 |
| - | - | 676.7 | 449.2 | - | - | 0 | - |
| - | - | 1926 | 455.2 | - | - | 0 | - |
| - | - | 1346 | 461.2 | - | - | 0 | - |
| - | - | 5575 | 465.2 | - | - | 0 | - |
| - | - | 783.6 | 466.2 | - | - | 0 | - |
| - | - | 896.2 | 469.2 | - | - | 0 | - |
| - | - | 951 | 471.2 | - | - | 0 | - |
| - | - | 887.8 | 473.2 | - | - | 0 | - |
| 2 | y | 2009 | 477.2 | 0.003016 | 6.32 | +2 | 7 |
| - | - | 746.7 | 477.7 | - | - | 0 | - |
| - | - | 719.1 | 482.2 | - | - | 0 | - |
| - | - | 1138 | 486.2 | - | - | 0 | - |
| 6 | y | 1.466E+04 | 489.2 | 0.004886 | 9.988 | +1 | 3 |
| - | - | 2850 | 490.2 | - | - | 0 | - |
| - | - | 2216 | 491.2 | - | - | 0 | - |
| - | - | 907.2 | 492.2 | - | - | 0 | - |
| - | - | 638.3 | 494.9 | - | - | 0 | - |
| - | - | 5985 | 495.2 | - | - | 0 | - |
| - | - | 1697 | 496.2 | - | - | 0 | - |
| - | - | 8799 | 496.2 | - | - | 0 | - |
| - | - | 1449 | 497.2 | - | - | 0 | - |
| - | - | 3042 | 497.2 | - | - | 0 | - |
| - | - | 797.4 | 498.2 | - | - | 0 | - |
| - | - | 725.8 | 509.7 | - | - | 0 | - |
| - | - | 802.2 | 510.2 | - | - | 0 | - |
| - | - | 7753 | 513.2 | - | - | 0 | - |
| - | - | 1478 | 514.2 | - | - | 0 | - |
| - | - | 3532 | 518.7 | - | - | 0 | - |
| - | - | 3397 | 519.2 | - | - | 0 | - |
| - | - | 804.9 | 519.7 | - | - | 0 | - |
| - | - | 929 | 520.2 | - | - | 0 | - |
| 4 | b | 1007 | 523.2 | 0.001925 | 3.679 | +1 | 4 |
| - | - | 1256 | 527.7 | - | - | 0 | - |
| - | - | 636.5 | 528.2 | - | - | 0 | - |
| 4 | b | 1764 | 541.2 | 0.0008273 | 1.529 | +1 | 4 |
| 0 | Precursor | 4671 | 541.7 | 0.002564 | 4.733 | +2 | -1 |
| - | - | 2285 | 542.2 | - | - | 0 | - |
| - | - | 1267 | 542.2 | - | - | 0 | - |
| - | - | 761.2 | 542.7 | - | - | 0 | - |
| - | - | 730 | 543.2 | - | - | 0 | - |
| - | - | 957.3 | 548.2 | - | - | 0 | - |
| - | - | 996.1 | 550 | - | - | 0 | - |
| - | - | 1397 | 550.3 | - | - | 0 | - |
| 0 | Precursor | 2.222E+04 | 550.7 | 0.002348 | 4.263 | +2 | -1 |
| - | - | 1.297E+04 | 551.2 | - | - | 0 | - |
| - | - | 513.1 | 551.3 | - | - | 0 | - |
| - | - | 5694 | 551.7 | - | - | 0 | - |
| - | - | 1440 | 552.2 | - | - | 0 | - |
| 5 | y | 2.995E+04 | 560.2 | 0.004607 | 8.224 | +1 | 4 |
| - | - | 1.024E+04 | 561.2 | - | - | 0 | - |
| - | - | 2394 | 562.2 | - | - | 0 | - |
| - | - | 3414 | 570.2 | - | - | 0 | - |
| - | - | 714.4 | 571.2 | - | - | 0 | - |
| - | - | 912.6 | 584.2 | - | - | 0 | - |
| - | - | 665.1 | 597.5 | - | - | 0 | - |
| - | - | 5204 | 606.2 | - | - | 0 | - |
| - | - | 1591 | 607.2 | - | - | 0 | - |
| 5 | b | 1083 | 612.2 | 0.006248 | 10.21 | +1 | 5 |
| - | - | 4724 | 634.2 | - | - | 0 | - |
| - | - | 1618 | 635.2 | - | - | 0 | - |
| - | - | 742.2 | 638.9 | - | - | 0 | - |
| - | - | 934.7 | 652.2 | - | - | 0 | - |
| - | - | 2704 | 657.3 | - | - | 0 | - |
| - | - | 8222 | 659.3 | - | - | 0 | - |
| - | - | 2588 | 660.3 | - | - | 0 | - |
| - | - | 931.3 | 661.3 | - | - | 0 | - |
| - | - | 674.6 | 677.3 | - | - | 0 | - |
| - | - | 891.8 | 681.8 | - | - | 0 | - |
| - | - | 652.2 | 682.3 | - | - | 0 | - |
| - | - | 2664 | 685.3 | - | - | 0 | - |
| - | - | 827.8 | 686.3 | - | - | 0 | - |
| - | - | 1049 | 703.3 | - | - | 0 | - |
| - | - | 675.4 | 709.3 | - | - | 0 | - |
| - | - | 706.1 | 710.4 | - | - | 0 | - |
| - | - | 2424 | 721.3 | - | - | 0 | - |
| - | - | 1130 | 722.3 | - | - | 0 | - |
| 4 | y | 2.216E+04 | 723.3 | 0.004267 | 5.899 | +1 | 5 |
| - | - | 9529 | 724.3 | - | - | 0 | - |
| - | - | 2739 | 725.3 | - | - | 0 | - |
| - | - | 891.2 | 728.3 | - | - | 0 | - |
| - | - | 2985 | 749.3 | - | - | 0 | - |
| - | - | 1342 | 750.3 | - | - | 0 | - |
| - | - | 636.9 | 755.3 | - | - | 0 | - |
| - | - | 2319 | 756.3 | - | - | 0 | - |
| - | - | 769.8 | 757.3 | - | - | 0 | - |
| - | - | 779.1 | 767.3 | - | - | 0 | - |
| - | - | 1.797E+04 | 774.3 | - | - | 0 | - |
| - | - | 7916 | 775.3 | - | - | 0 | - |
| - | - | 1567 | 776.3 | - | - | 0 | - |
| - | - | 865.4 | 792.3 | - | - | 0 | - |
| 3 | y | 2355 | 820.3 | 0.005039 | 6.143 | +1 | 6 |
| - | - | 1141 | 821.3 | - | - | 0 | - |
| - | - | 745.1 | 822.3 | - | - | 0 | - |
| - | - | 1161 | 832.3 | - | - | 0 | - |
| 3 | y | 5.324E+04 | 838.3 | 0.003996 | 4.767 | +1 | 6 |
| - | - | 2.648E+04 | 839.3 | - | - | 0 | - |
| - | - | 8613 | 840.3 | - | - | 0 | - |
| - | - | 1172 | 841.3 | - | - | 0 | - |
| - | - | 1002 | 868.3 | - | - | 0 | - |
| - | - | 2685 | 871.3 | - | - | 0 | - |
| - | - | 1084 | 872.3 | - | - | 0 | - |
| - | - | 1.605E+04 | 889.3 | - | - | 0 | - |
| - | - | 7592 | 890.3 | - | - | 0 | - |
| - | - | 1997 | 891.4 | - | - | 0 | - |
| 7 | b | 2312 | 896.3 | 0.001813 | 2.022 | +1 | 7 |
| - | - | 793.1 | 897.3 | - | - | 0 | - |
| - | - | 801.3 | 917.3 | - | - | 0 | - |
| 2 | y | 3795 | 935.3 | 0.003487 | 3.728 | +1 | 7 |
| - | - | 2623 | 936.3 | - | - | 0 | - |
| - | - | 997.2 | 937.3 | - | - | 0 | - |
| 2 | y | 3.912E+04 | 953.3 | 0.003665 | 3.844 | +1 | 7 |
| - | - | 1029 | 954.2 | - | - | 0 | - |
| - | - | 1.907E+04 | 954.3 | - | - | 0 | - |
| - | - | 7417 | 955.3 | - | - | 0 | - |
| - | - | 1068 | 956.4 | - | - | 0 | - |
| - | - | 2932 | 963.3 | - | - | 0 | - |
| - | - | 736.4 | 964.3 | - | - | 0 | - |
| - | - | 643.6 | 1155 | - | - | 0 | - |
| - | - | 676 | 1530 | - | - | 0 | - |
| - | - | 611.3 | 3016 | - | - | 0 | - |

m/z Charge Intensity FragmentType MassShift Position
120.08106231689453 0 275699.44
121.08432006835938 0 23597.533
122.08778381347656 0 575.3123
127.08716583251953 0 849.3749
128.10736083984375 0 781.1737
129.06556701660156 0 447.18573
129.10250854492188 0 3394.3608
130.06536865234375 0 3846.157
130.0867156982422 0 543.0805
131.0693359375 0 456.5294
131.11813354492188 0 766.38104
132.08079528808594 0 550.07166
133.06092834472656 0 501.54742
133.08592224121094 0 1147.8334
136.0758819580078 0 25979.523
137.07908630371094 0 1494.2203
138.05520629882812 0 2621.582
138.06626892089844 0 2465.9963
139.0863494873047 0 582.25543
146.06019592285156 0 2205.096
146.09664916992188 0 3008.5627
148.0870819091797 0 768.95953
148.95474243164062 0 832.01715
149.04502868652344 0 3263.5393
155.0816192626953 0 1642.5327
155.09303283691406 0 1115.8928
156.07684326171875 0 7169.8926
157.06089782714844 0 535.38104
158.09226989746094 0 933.45605
159.07652282714844 0 473.5725
159.09176635742188 0 8622.994
159.11268615722656 0 1131.2842
160.0950927734375 0 1488.0576
162.0915069580078 0 625.0415
165.05540466308594 0 495.9192
165.0771942138672 0 1165.8528
166.05361938476562 0 704.77155
166.06106567382812 0 1232.8224
167.0555877685547 0 4653.8223 a Ammonia loss 2
167.09286499023438 0 1402.8868
171.14962768554688 0 489.57025
172.0757598876953 0 1767.2926
172.108642578125 0 538.89453
173.07962036132812 0 519.29456
174.0660858154297 0 645.9906
175.0867919921875 0 2831.0747
175.09552001953125 0 422.88925
176.08224487304688 0 1578.2058
176.09005737304688 0 514.4347
177.11306762695312 0 1185.704
180.07733154296875 0 1099.0747
181.09764099121094 0 1043.7006
181.2983856201172 0 450.3904
185.05592346191406 0 2134.0066
185.12889099121094 0 611.45123
185.1649627685547 0 487.55255
187.063232421875 0 721.08105
187.10787963867188 0 1423.7697
187.14456176757812 0 519.91113
188.07077026367188 0 9303.515
189.0744171142578 0 1005.96344
189.08697509765625 0 550.14246
189.1022491455078 0 518.7074
189.8035430908203 0 459.68027
190.08644104003906 0 2811.2046
191.08230590820312 0 671.3632
191.09213256835938 0 839.15875
191.1180419921875 0 10603.223 d 1
192.1212615966797 0 929.9583
193.10845947265625 0 4737.293
194.08006286621094 0 500.838
195.0875244140625 0 1914.0376
195.11282348632812 0 575.9507
195.12351989746094 0 467.31555
197.09291076660156 0 661.123
199.10777282714844 0 753.9691
199.180419921875 0 896.4162
200.13925170898438 0 574.13965
201.12359619140625 0 802.7932
202.0535125732422 0 1010.3765
203.06629943847656 0 1793.2339
203.07815551757812 0 520.6566
204.0768280029297 0 2934.8645
205.0972900390625 0 5667.5845 y 7
205.10784912109375 0 914.214
206.1007080078125 0 980.6125
207.11265563964844 0 2287.6658
208.0721435546875 0 1050.3203
209.0921173095703 0 1215.2533
212.1158447265625 0 513.3001
213.09896850585938 0 543.0994
215.1396484375 0 792.5339
217.0972900390625 0 36924.215 a Water loss 1
218.08169555664062 0 890.95514 a Ammonia loss 1
218.10079956054688 0 3479.1985
219.0796356201172 0 1179.5187
219.11338806152344 0 808.4495
220.11825561523438 0 486.25003
221.0844268798828 0 6504.2173
221.10340881347656 0 14513.575
222.08486938476562 0 1233.4683
222.1069793701172 0 2017.4305
223.0636749267578 0 853.14545
225.0430908203125 0 9175.407
226.04315185546875 0 801.5217
226.11874389648438 0 529.08765
226.15457153320312 0 786.19977
227.0223388671875 0 1094.7343
227.1031036376953 0 755.66064
228.1131134033203 0 1213.9305
229.1075897216797 0 745.73364
231.06129455566406 0 4022.436
233.09201049804688 0 808.2886
235.10789489746094 0 53627.984 a 1
236.1110382080078 0 6663.134
239.0950927734375 0 19370.883
239.11354064941406 0 1293.3977
240.09559631347656 0 3677.8564
241.09271240234375 0 670.59985
242.15029907226562 0 1095.5621
245.07852172851562 0 559.8603
245.09234619140625 0 1463.1809 b Water loss 1
246.0770721435547 0 763.4769 b Ammonia loss 1
246.13536071777344 0 799.05444
248.15086364746094 0 592.34216
251.10269165039062 0 14325.112
252.10586547851562 0 1963.4584
257.10626220703125 0 2683.1755
261.0869140625 0 961.98535
263.1026611328125 0 48300.234 b 1
264.1060791015625 0 6914.1245
264.1455993652344 0 1148.5725
265.10675048828125 0 676.65094
267.09039306640625 0 662.2544
269.0762634277344 0 556.48944
274.1297302246094 0 3405.4775
279.0976257324219 0 8651.674
281.05133056640625 0 1039.3883
285.101806640625 0 7102.75
286.10504150390625 0 809.16986
287.09967041015625 0 559.63007
292.1405029296875 0 5330.872
293.1427307128906 0 811.857
295.1029968261719 0 3794.6445
296.1029052734375 0 1599.3025
296.15087890625 0 4047.109
297.1531677246094 0 644.91925
299.0618896484375 0 9889.075
300.06298828125 0 1137.8254
303.11212158203125 0 976.7653
306.1448974609375 0 1771.9963 d 2
310.1521911621094 0 619.8531
313.1141357421875 0 1426.7795
318.14605712890625 0 1040.1952
323.0985412597656 0 1074.9318
324.145751953125 0 9885.774
325.1491394042969 0 1986.8234
332.12408447265625 0 888.315 a Water loss 2
334.1293640136719 0 731.0801
338.1335144042969 0 2035.8121
342.15625 0 55325.418 y 6
343.1375732421875 0 805.14813
343.15948486328125 0 10302.583
344.1630859375 0 657.0216
348.1191711425781 0 2496.0774
350.134765625 0 6020.9126 a 2
351.1361389160156 0 862.9504
352.1407775878906 0 986.7713
355.069091796875 0 980.3143
356.1390380859375 0 4899.137
357.2492980957031 0 777.02344
359.02923583984375 0 1101.7227
360.0954284667969 0 531.60547
360.1190185546875 0 2783.402 b Water loss 2
361.17205810546875 0 876.8067
362.14996337890625 0 1039.6542 y 3
366.1292419433594 0 1071.373
369.1219482421875 0 5126.037
370.12237548828125 0 1556.413
374.1484375 0 1003.9199
376.1128234863281 0 1113.5043
378.1295471191406 0 32942.562 b 2
379.13238525390625 0 6428.599
380.1355895996094 0 770.60504 b 5
382.1435241699219 0 1771.0344
390.15478515625 0 876.9319
394.12469482421875 0 6480.5015
395.1283874511719 0 1982.2557
404.1460876464844 0 1073.8461
407.1812438964844 0 669.7402
408.1659851074219 0 1418.7776
415.0362854003906 0 945.44946
418.9947204589844 0 633.01337
419.6628112792969 0 678.87445 y 2
420.16595458984375 0 888.39526
425.1932067871094 0 7992.033
426.19573974609375 0 1493.3695
427.20928955078125 0 1915.042
434.6702575683594 0 889.2557
442.1708068847656 0 682.8177
447.15252685546875 0 1227.7118
448.6654357910156 0 580.4057 b 6
449.1697082519531 0 676.67065
455.2032775878906 0 1925.6833
461.1942138671875 0 1346.2823
465.16217041015625 0 5574.72
466.1663818359375 0 783.5776
469.2060546875 0 896.21045
471.1803894042969 0 950.95825
473.2121887207031 0 887.7602
477.1770935058594 0 2008.7124 y 1
477.6787414550781 0 746.7195
482.169921875 0 719.1042
486.1600036621094 0 1137.965
489.1914367675781 0 14660.668 y 5
490.1940002441406 0 2849.7473
491.20330810546875 0 2216.3406
492.2085876464844 0 907.1528
494.9266052246094 0 638.3098
495.1875305175781 0 5984.782
496.1892395019531 0 1696.543
496.23040771484375 0 8799.46
497.1706237792969 0 1448.7676
497.23345947265625 0 3041.5095
498.23651123046875 0 797.4102
509.7073669433594 0 725.7958
510.2063293457031 0 802.1787
513.1976928710938 0 7752.9585
514.2003784179688 0 1477.7361
518.7098388671875 0 3531.9805
519.2061767578125 0 3397.2544
519.71044921875 0 804.89197
520.20458984375 0 928.97656
523.1842651367188 0 1006.61426 b Water loss 3
527.7076416015625 0 1255.7571
528.2092895507812 0 636.53625
541.1920776367188 0 1764.0192 b 3
541.70556640625 0 4671.3 Precursor Water loss
542.2066650390625 0 2284.5796
542.2413940429688 0 1266.8652
542.7039794921875 0 761.17993
543.2128295898438 0 730.04956
548.1970825195312 0 957.26483
549.959716796875 0 996.1496
550.2931518554688 0 1396.9398
550.7106323242188 0 22224.99 Precursor
551.212158203125 0 12970.53
551.26123046875 0 513.05347
551.7122192382812 0 5693.8623
552.2107543945312 0 1439.793
560.228271484375 0 29946.348 y 4
561.2321166992188 0 10240.399
562.229736328125 0 2394.382
570.2306518554688 0 3413.71
571.237060546875 0 714.356
584.2369995117188 0 912.5707
597.4656372070312 0 665.0638
606.2335205078125 0 5203.978
607.2376708984375 0 1590.535
612.2362670898438 0 1083.1359 b 4
634.2285766601562 0 4724.0576
635.2294311523438 0 1617.6882
638.8514404296875 0 742.1937
652.238037109375 0 934.72565
657.2609252929688 0 2704.1877
659.2931518554688 0 8222.278
660.2972412109375 0 2587.6956
661.3016357421875 0 931.3146
677.2803955078125 0 674.64246
681.8349609375 0 891.76337
682.3435668945312 0 652.24915
685.2579956054688 0 2663.804
686.2630615234375 0 827.8045
703.263671875 0 1049.3602
709.3306274414062 0 675.38257
710.4485473632812 0 706.0724
721.2603759765625 0 2423.803
722.2640380859375 0 1130.0287
723.291259765625 0 22164.268 y 3
724.2943115234375 0 9528.552
725.294921875 0 2738.7803
728.3167724609375 0 891.17883
749.2552490234375 0 2985.2756
750.2608642578125 0 1341.5156
755.3255615234375 0 636.8791
756.310546875 0 2318.9753
757.3096313476562 0 769.7915
767.2646484375 0 779.1043
774.3198852539062 0 17971.086
775.3233032226562 0 7916.1416
776.3238525390625 0 1567.3782
792.3089599609375 0 865.3991
820.3084106445312 0 2355.1245 y Water loss 2
821.3071899414062 0 1140.9663
822.3026123046875 0 745.0577
832.3281860351562 0 1160.7954
838.3179321289062 0 53236.492 y 2
839.3208618164062 0 26483.65
840.3204345703125 0 8612.743
841.3243408203125 0 1171.522
868.3291625976562 0 1001.9503
871.3372192382812 0 2685.3823
872.3388671875 0 1084.4697
889.34619140625 0 16046.429
890.348876953125 0 7591.751
891.35205078125 0 1996.6792
896.3212280273438 0 2311.6099 b 6
897.3279418945312 0 793.0741
917.3275146484375 0 801.2741
935.3338012695312 0 3795.4922 y Water loss 1
936.3306274414062 0 2623.4617
937.3265991210938 0 997.218
953.3445434570312 0 39120.047 y 1
954.2376708984375 0 1029.0851
954.3478393554688 0 19071.225
955.3466186523438 0 7416.959
956.350830078125 0 1067.557
963.3298950195312 0 2932.423
964.3335571289062 0 736.35803
1155.1805419921875 0 643.60156
1529.529541015625 0 676.0032
3016.069580078125 0 611.2949

Spectrum Details

|  |  |
| --- | --- |
| Matched peaks? Matched peaks The total absolute number of peaks matched. Additionally in brackets the total fraction of peaks matched and the total number of peaks is shown. | 33 (10.25% of 322) |
| FDR? FDR The false discovery rate estimated for this peptide. It is calculated by matching all theoretical fragments with a non-integer shift with the raw peaks for this spectrum. This is done with 40 different shifts. The resulting percentage is the average number of annotated peaks over the number of annotated peaks with the correct spectrum. | 0.51% |
| Satellite FDR? Satellite FDR See the FDR for details on its calculation. This satellite ion specific FDR only contains the satellite ions (d/w) for I/L/J positions. | - |
| PSM Score? PSM Score The PSM Score as given by Hecklib to this annotated spectrum. It is shown with three significant figures. | 371 |

#### Spectrum 6738? Spectrum 6738 The raw spectrum of this peptide as annotated by Hecklib. The fragments are coloured according to ion type (see legend). Any peaks with a star '\*' as text can be hovered over to see the full details, first the ion type second the mass shift type. By hovering over the amino acids in the peptide or ions in the legend the corresponding peaks are highlighted. By toggling the 'Unassigned' label you can turn the background (unassigned) peaks on or off in the plot. By updating the slider in the Ion legend you can update the spectrum to only show the top X% of the peaks with labels. The top X% means any peak that is within X% of the highest intensity. By dragging in the spectrum you can zoom in to a specific part of the spectrum and use 'Zoom Out' to get back to the original zoom level. The annotation of the spectrum is based on the given sequence in the peptides file and is done with different software so inconsistencies are likely. The peaks are annotated based on the given sequence, with 20 ppm tolerance.

Copy Data

##### Spectrum 6738 (TSV)

###### Preview

```
Loading example...
```

*Click on the button to copy the data to your clipboard.*

Mz MinMz MaxIntensity Max

WidthHeightPeptide font sizePeptide stroke widthSpectrum font sizeSpectrum stroke widthCompact peptide

Ion legend

wxyz

abcd

OtherUnassignedIon^Charge^_Position_Show for top:%

FDDYAMHW

04.17e+48.33e+41.25e+51.67e+5

Zoom Out

a^+2^_3_d^+1^_2_y^+1^_1_a^+1^_2_a^+1^_2_b^+1^_2_b^+1^_2_d^+1^_3_a^+1^_3_y^+1^_2_a^+1^_3_b^+1^_3_y^+2^_5_b^+1^_3_y^+2^_6_y^+2^_7_y^+1^_3_b^+1^_4_\*\*y^+1^_4_y^+1^_5_y^+1^_6_y^+1^_6_b^+1^_7_y^+1^_7_y^+1^_7_

0846169225383384

Fragment Matches Table

Show background peaks

| Position | Ion type | Intensity | mz Theoretical | mz Error (Th) | mz Error (ppm) | Charge | Series Number |
| --- | --- | --- | --- | --- | --- | --- | --- |
| - | - | 1.65E+05 | 120.1 | - | - | 0 | - |
| - | - | 587.8 | 121.1 | - | - | 0 | - |
| - | - | 760.1 | 121.1 | - | - | 0 | - |
| - | - | 1.417E+04 | 121.1 | - | - | 0 | - |
| - | - | 485.7 | 127.1 | - | - | 0 | - |
| - | - | 1274 | 128.1 | - | - | 0 | - |
| - | - | 404.6 | 128.4 | - | - | 0 | - |
| - | - | 4270 | 129.1 | - | - | 0 | - |
| - | - | 2009 | 130.1 | - | - | 0 | - |
| - | - | 807.5 | 131.1 | - | - | 0 | - |
| - | - | 509.7 | 132.1 | - | - | 0 | - |
| - | - | 5003 | 133.1 | - | - | 0 | - |
| - | - | 918.2 | 134.1 | - | - | 0 | - |
| - | - | 1.889E+04 | 136.1 | - | - | 0 | - |
| - | - | 1118 | 137.1 | - | - | 0 | - |
| - | - | 1695 | 138.1 | - | - | 0 | - |
| - | - | 2014 | 138.1 | - | - | 0 | - |
| - | - | 635.9 | 141.1 | - | - | 0 | - |
| - | - | 1120 | 146.1 | - | - | 0 | - |
| - | - | 2476 | 146.1 | - | - | 0 | - |
| - | - | 483.4 | 146.4 | - | - | 0 | - |
| - | - | 561.4 | 148.1 | - | - | 0 | - |
| - | - | 3256 | 149 | - | - | 0 | - |
| - | - | 576 | 153.1 | - | - | 0 | - |
| - | - | 882 | 155.1 | - | - | 0 | - |
| - | - | 3687 | 156.1 | - | - | 0 | - |
| - | - | 748.8 | 159.1 | - | - | 0 | - |
| - | - | 5426 | 159.1 | - | - | 0 | - |
| - | - | 566.2 | 160.5 | - | - | 0 | - |
| - | - | 477 | 162 | - | - | 0 | - |
| - | - | 510.5 | 162.1 | - | - | 0 | - |
| - | - | 517.1 | 163 | - | - | 0 | - |
| - | - | 748.7 | 163.1 | - | - | 0 | - |
| - | - | 1110 | 165.1 | - | - | 0 | - |
| - | - | 718 | 165.1 | - | - | 0 | - |
| - | - | 468.7 | 165.9 | - | - | 0 | - |
| - | - | 918.1 | 166.1 | - | - | 0 | - |
| 3 | a | 4715 | 167.1 | 0.002229 | 13.34 | +2 | 3 |
| - | - | 688.8 | 167.1 | - | - | 0 | - |
| - | - | 454 | 171.3 | - | - | 0 | - |
| - | - | 881.7 | 172.1 | - | - | 0 | - |
| - | - | 1582 | 173.1 | - | - | 0 | - |
| - | - | 1245 | 175.1 | - | - | 0 | - |
| - | - | 711.8 | 175.1 | - | - | 0 | - |
| - | - | 495.9 | 175.1 | - | - | 0 | - |
| - | - | 1249 | 176.1 | - | - | 0 | - |
| - | - | 547.5 | 176.1 | - | - | 0 | - |
| - | - | 2694 | 177.1 | - | - | 0 | - |
| - | - | 697.2 | 178.1 | - | - | 0 | - |
| - | - | 959.8 | 181.1 | - | - | 0 | - |
| - | - | 1040 | 182.1 | - | - | 0 | - |
| - | - | 982.4 | 183.1 | - | - | 0 | - |
| - | - | 1446 | 185.1 | - | - | 0 | - |
| - | - | 479.2 | 186.1 | - | - | 0 | - |
| - | - | 2273 | 187.1 | - | - | 0 | - |
| - | - | 546.1 | 187.1 | - | - | 0 | - |
| - | - | 5942 | 188.1 | - | - | 0 | - |
| - | - | 635.2 | 189.1 | - | - | 0 | - |
| - | - | 1365 | 190.1 | - | - | 0 | - |
| - | - | 492.8 | 190.1 | - | - | 0 | - |
| - | - | 501.4 | 191.1 | - | - | 0 | - |
| 2 | d | 6131 | 191.1 | 0.0002593 | 1.357 | +1 | 2 |
| - | - | 3702 | 193.1 | - | - | 0 | - |
| - | - | 496.9 | 194.1 | - | - | 0 | - |
| - | - | 681.7 | 199.1 | - | - | 0 | - |
| - | - | 629.1 | 200.1 | - | - | 0 | - |
| - | - | 1699 | 201.1 | - | - | 0 | - |
| - | - | 1319 | 203.1 | - | - | 0 | - |
| - | - | 1139 | 204.1 | - | - | 0 | - |
| 8 | y | 3380 | 205.1 | 1.383E-06 | 0.006741 | +1 | 1 |
| - | - | 904.2 | 205.1 | - | - | 0 | - |
| - | - | 1173 | 207.1 | - | - | 0 | - |
| - | - | 556.9 | 208.1 | - | - | 0 | - |
| - | - | 1594 | 209.1 | - | - | 0 | - |
| 2 | a | 2.187E+04 | 217.1 | 0.0001359 | 0.6262 | +1 | 2 |
| - | - | 3317 | 218.1 | - | - | 0 | - |
| - | - | 704.5 | 219.1 | - | - | 0 | - |
| - | - | 9594 | 221.1 | - | - | 0 | - |
| - | - | 7724 | 221.1 | - | - | 0 | - |
| - | - | 1354 | 221.1 | - | - | 0 | - |
| - | - | 688.8 | 221.1 | - | - | 0 | - |
| - | - | 464.7 | 222.1 | - | - | 0 | - |
| - | - | 722.1 | 223.1 | - | - | 0 | - |
| - | - | 688 | 223.1 | - | - | 0 | - |
| - | - | 1.033E+04 | 225 | - | - | 0 | - |
| - | - | 697.9 | 225.1 | - | - | 0 | - |
| - | - | 1576 | 226 | - | - | 0 | - |
| - | - | 959.1 | 226.2 | - | - | 0 | - |
| - | - | 864.8 | 227 | - | - | 0 | - |
| - | - | 883.7 | 229.1 | - | - | 0 | - |
| - | - | 2494 | 231.1 | - | - | 0 | - |
| - | - | 548.2 | 231.1 | - | - | 0 | - |
| 2 | a | 3.264E+04 | 235.1 | 6.931E-05 | 0.2948 | +1 | 2 |
| - | - | 4727 | 236.1 | - | - | 0 | - |
| - | - | 2.398E+04 | 239.1 | - | - | 0 | - |
| - | - | 984.5 | 239.1 | - | - | 0 | - |
| - | - | 3247 | 240.1 | - | - | 0 | - |
| - | - | 601.7 | 241.1 | - | - | 0 | - |
| - | - | 1051 | 242.1 | - | - | 0 | - |
| 2 | b | 1112 | 245.1 | 0.000104 | 0.4243 | +1 | 2 |
| - | - | 739.4 | 249.1 | - | - | 0 | - |
| - | - | 9058 | 251.1 | - | - | 0 | - |
| - | - | 717.8 | 252.1 | - | - | 0 | - |
| - | - | 614 | 253.1 | - | - | 0 | - |
| - | - | 612.6 | 256.1 | - | - | 0 | - |
| - | - | 2190 | 257.1 | - | - | 0 | - |
| - | - | 615.3 | 261.1 | - | - | 0 | - |
| 2 | b | 2.86E+04 | 263.1 | 3.33E-05 | 0.1266 | +1 | 2 |
| - | - | 3771 | 264.1 | - | - | 0 | - |
| - | - | 699.8 | 264.1 | - | - | 0 | - |
| - | - | 649.2 | 266.1 | - | - | 0 | - |
| - | - | 557.5 | 269.4 | - | - | 0 | - |
| - | - | 2317 | 274.1 | - | - | 0 | - |
| - | - | 5219 | 279.1 | - | - | 0 | - |
| - | - | 670.2 | 280.1 | - | - | 0 | - |
| - | - | 835.8 | 281.1 | - | - | 0 | - |
| - | - | 786.6 | 283 | - | - | 0 | - |
| - | - | 744.7 | 285 | - | - | 0 | - |
| - | - | 3496 | 285.1 | - | - | 0 | - |
| - | - | 871.1 | 285.1 | - | - | 0 | - |
| - | - | 2601 | 292.1 | - | - | 0 | - |
| - | - | 6469 | 295.1 | - | - | 0 | - |
| - | - | 2228 | 296.1 | - | - | 0 | - |
| - | - | 3303 | 296.2 | - | - | 0 | - |
| - | - | 9988 | 299.1 | - | - | 0 | - |
| - | - | 1349 | 300.1 | - | - | 0 | - |
| 3 | d | 983.1 | 306.1 | 0.00156 | 5.096 | +1 | 3 |
| - | - | 622.8 | 308.4 | - | - | 0 | - |
| - | - | 720.6 | 310.2 | - | - | 0 | - |
| - | - | 1409 | 313.1 | - | - | 0 | - |
| - | - | 693 | 313.2 | - | - | 0 | - |
| - | - | 585.8 | 318.1 | - | - | 0 | - |
| - | - | 5424 | 324.1 | - | - | 0 | - |
| - | - | 919.8 | 325.1 | - | - | 0 | - |
| - | - | 646.3 | 327.1 | - | - | 0 | - |
| - | - | 580.4 | 328.1 | - | - | 0 | - |
| - | - | 516.6 | 329.1 | - | - | 0 | - |
| - | - | 665.3 | 330.1 | - | - | 0 | - |
| 3 | a | 628.8 | 332.1 | 0.001691 | 5.092 | +1 | 3 |
| - | - | 550.1 | 335.1 | - | - | 0 | - |
| - | - | 516.9 | 337.1 | - | - | 0 | - |
| - | - | 1929 | 338.1 | - | - | 0 | - |
| - | - | 565.1 | 338.9 | - | - | 0 | - |
| - | - | 688.7 | 341.1 | - | - | 0 | - |
| 7 | y | 3.155E+04 | 342.2 | 3.146E-05 | 0.09194 | +1 | 2 |
| - | - | 6891 | 343.2 | - | - | 0 | - |
| - | - | 920.8 | 344.2 | - | - | 0 | - |
| - | - | 1118 | 348.1 | - | - | 0 | - |
| 3 | a | 3468 | 350.1 | 0.0001403 | 0.4008 | +1 | 3 |
| - | - | 605.3 | 351.1 | - | - | 0 | - |
| - | - | 1104 | 355.1 | - | - | 0 | - |
| - | - | 3067 | 356.1 | - | - | 0 | - |
| - | - | 695.6 | 359 | - | - | 0 | - |
| 3 | b | 853.6 | 360.1 | 0.000573 | 1.591 | +1 | 3 |
| 4 | y | 743.7 | 362.1 | 0.001883 | 5.199 | +2 | 5 |
| - | - | 6005 | 369.1 | - | - | 0 | - |
| - | - | 2227 | 370.1 | - | - | 0 | - |
| - | - | 892.1 | 374.1 | - | - | 0 | - |
| - | - | 823.2 | 376.1 | - | - | 0 | - |
| 3 | b | 1.968E+04 | 378.1 | 0.0001514 | 0.4004 | +1 | 3 |
| - | - | 3553 | 379.1 | - | - | 0 | - |
| - | - | 791.2 | 381.1 | - | - | 0 | - |
| - | - | 937.7 | 382.1 | - | - | 0 | - |
| - | - | 614.6 | 384.1 | - | - | 0 | - |
| - | - | 3698 | 394.1 | - | - | 0 | - |
| - | - | 749.3 | 407.2 | - | - | 0 | - |
| - | - | 1418 | 408.2 | - | - | 0 | - |
| - | - | 770 | 415 | - | - | 0 | - |
| - | - | 913.6 | 419.2 | - | - | 0 | - |
| 3 | y | 1061 | 419.7 | 0.001351 | 3.218 | +2 | 6 |
| - | - | 4384 | 425.2 | - | - | 0 | - |
| - | - | 990.8 | 426.2 | - | - | 0 | - |
| - | - | 626.5 | 442.2 | - | - | 0 | - |
| - | - | 627.3 | 449.2 | - | - | 0 | - |
| - | - | 1095 | 453.3 | - | - | 0 | - |
| - | - | 591.4 | 454.2 | - | - | 0 | - |
| - | - | 754.6 | 455.2 | - | - | 0 | - |
| - | - | 2293 | 465.2 | - | - | 0 | - |
| - | - | 961 | 471.2 | - | - | 0 | - |
| 2 | y | 1825 | 477.2 | 0.001948 | 4.082 | +2 | 7 |
| 6 | y | 8864 | 489.2 | 0.004551 | 9.302 | +1 | 3 |
| - | - | 2779 | 490.2 | - | - | 0 | - |
| - | - | 711.1 | 491.2 | - | - | 0 | - |
| - | - | 4316 | 495.2 | - | - | 0 | - |
| - | - | 965.3 | 496.2 | - | - | 0 | - |
| - | - | 5090 | 496.2 | - | - | 0 | - |
| - | - | 630.1 | 496.8 | - | - | 0 | - |
| - | - | 1147 | 497.2 | - | - | 0 | - |
| - | - | 943.3 | 497.2 | - | - | 0 | - |
| - | - | 2743 | 503.3 | - | - | 0 | - |
| - | - | 1656 | 504.3 | - | - | 0 | - |
| - | - | 573 | 509.9 | - | - | 0 | - |
| - | - | 3964 | 513.2 | - | - | 0 | - |
| - | - | 1137 | 518.7 | - | - | 0 | - |
| - | - | 1736 | 519.2 | - | - | 0 | - |
| - | - | 751.2 | 520.2 | - | - | 0 | - |
| - | - | 1047 | 532.3 | - | - | 0 | - |
| - | - | 843.6 | 533.2 | - | - | 0 | - |
| 4 | b | 1119 | 541.2 | 0.0004001 | 0.7393 | +1 | 4 |
| 0 | Precursor | 1666 | 541.7 | 0.001832 | 3.381 | +2 | -1 |
| - | - | 1078 | 542.2 | - | - | 0 | - |
| - | - | 686.7 | 548.2 | - | - | 0 | - |
| - | - | 2540 | 550.3 | - | - | 0 | - |
| - | - | 685.3 | 550.6 | - | - | 0 | - |
| 0 | Precursor | 1.106E+04 | 550.7 | 0.001981 | 3.598 | +2 | -1 |
| - | - | 7420 | 551.2 | - | - | 0 | - |
| - | - | 4073 | 551.7 | - | - | 0 | - |
| 5 | y | 1.653E+04 | 560.2 | 0.004546 | 8.115 | +1 | 4 |
| - | - | 5452 | 561.2 | - | - | 0 | - |
| - | - | 1385 | 562.2 | - | - | 0 | - |
| - | - | 1889 | 570.2 | - | - | 0 | - |
| - | - | 653.8 | 573.8 | - | - | 0 | - |
| - | - | 2426 | 606.2 | - | - | 0 | - |
| - | - | 813.8 | 607.2 | - | - | 0 | - |
| - | - | 2617 | 634.2 | - | - | 0 | - |
| - | - | 947.3 | 635.2 | - | - | 0 | - |
| - | - | 735.7 | 638.9 | - | - | 0 | - |
| - | - | 1108 | 657.3 | - | - | 0 | - |
| - | - | 5075 | 659.3 | - | - | 0 | - |
| - | - | 1421 | 660.3 | - | - | 0 | - |
| - | - | 811.9 | 678.3 | - | - | 0 | - |
| - | - | 1533 | 681.8 | - | - | 0 | - |
| - | - | 826.4 | 682.3 | - | - | 0 | - |
| - | - | 1479 | 685.3 | - | - | 0 | - |
| - | - | 746.3 | 686.3 | - | - | 0 | - |
| - | - | 1038 | 705.3 | - | - | 0 | - |
| - | - | 651.2 | 706.4 | - | - | 0 | - |
| - | - | 736.2 | 710.5 | - | - | 0 | - |
| - | - | 1472 | 721.3 | - | - | 0 | - |
| 4 | y | 1.173E+04 | 723.3 | 0.004023 | 5.562 | +1 | 5 |
| - | - | 4980 | 724.3 | - | - | 0 | - |
| - | - | 1694 | 725.3 | - | - | 0 | - |
| - | - | 1913 | 749.3 | - | - | 0 | - |
| - | - | 727.4 | 756.3 | - | - | 0 | - |
| - | - | 1.084E+04 | 774.3 | - | - | 0 | - |
| - | - | 4472 | 775.3 | - | - | 0 | - |
| - | - | 835.8 | 776.3 | - | - | 0 | - |
| - | - | 774.1 | 782.4 | - | - | 0 | - |
| 3 | y | 1771 | 820.3 | 0.004063 | 4.953 | +1 | 6 |
| 3 | y | 3.092E+04 | 838.3 | 0.00363 | 4.33 | +1 | 6 |
| - | - | 1.539E+04 | 839.3 | - | - | 0 | - |
| - | - | 4704 | 840.3 | - | - | 0 | - |
| - | - | 631 | 868.3 | - | - | 0 | - |
| - | - | 2087 | 871.3 | - | - | 0 | - |
| - | - | 9441 | 889.3 | - | - | 0 | - |
| - | - | 3857 | 890.3 | - | - | 0 | - |
| - | - | 1080 | 891.4 | - | - | 0 | - |
| 7 | b | 1826 | 896.3 | 0.002667 | 2.976 | +1 | 7 |
| - | - | 690.1 | 900.3 | - | - | 0 | - |
| 2 | y | 2078 | 935.3 | 0.0005413 | 0.5787 | +1 | 7 |
| - | - | 1703 | 936.3 | - | - | 0 | - |
| 2 | y | 2.242E+04 | 953.3 | 0.003298 | 3.46 | +1 | 7 |
| - | - | 1.124E+04 | 954.3 | - | - | 0 | - |
| - | - | 3886 | 955.3 | - | - | 0 | - |
| - | - | 1474 | 963.3 | - | - | 0 | - |
| - | - | 720.9 | 964.3 | - | - | 0 | - |
| - | - | 718.5 | 1908 | - | - | 0 | - |
| - | - | 666.7 | 2848 | - | - | 0 | - |
| - | - | 777 | 3072 | - | - | 0 | - |
| - | - | 749.3 | 3350 | - | - | 0 | - |

m/z Charge Intensity FragmentType MassShift Position
120.08102416992188 0 164955.03
121.07737731933594 0 587.8444
121.07896423339844 0 760.0549
121.08431243896484 0 14172.737
127.08654022216797 0 485.6955
128.107177734375 0 1273.6887
128.40733337402344 0 404.6182
129.10244750976562 0 4269.61
130.0652618408203 0 2008.8347
131.11819458007812 0 807.5023
132.08094787597656 0 509.7361
133.08607482910156 0 5003.132
134.08963012695312 0 918.23944
136.07586669921875 0 18892.174
137.0791778564453 0 1117.6254
138.05517578125 0 1694.7566
138.06626892089844 0 2013.6582
141.10256958007812 0 635.94116
146.06031799316406 0 1119.944
146.09664916992188 0 2475.7031
146.36068725585938 0 483.42764
148.0872039794922 0 561.4164
149.04498291015625 0 3255.6433
153.06625366210938 0 575.97473
155.08169555664062 0 881.9736
156.07684326171875 0 3686.709
159.0764923095703 0 748.84314
159.0916748046875 0 5425.9663
160.48423767089844 0 566.2291
162.04026794433594 0 476.95956
162.0546112060547 0 510.48834
163.027587890625 0 517.06036
163.0718231201172 0 748.69916
165.0547637939453 0 1109.5881
165.0772247314453 0 718.0286
165.90408325195312 0 468.68933
166.06118774414062 0 918.1486
167.0554656982422 0 4715.211 a Ammonia loss 2
167.0933380126953 0 688.7943
171.30039978027344 0 453.98444
172.0761260986328 0 881.68665
173.12864685058594 0 1581.6877
175.08706665039062 0 1245.0328
175.0965576171875 0 711.77783
175.1191864013672 0 495.91064
176.0819549560547 0 1248.8097
176.10690307617188 0 547.47174
177.11231994628906 0 2694.115
178.08631896972656 0 697.19507
181.0972137451172 0 959.7611
182.0821533203125 0 1039.794
183.11317443847656 0 982.37573
185.05581665039062 0 1445.7719
186.08937072753906 0 479.19934
187.1079864501953 0 2273.1345
187.1449737548828 0 546.11
188.07066345214844 0 5942.2344
189.0864715576172 0 635.15753
190.08624267578125 0 1365.163
190.10154724121094 0 492.76996
191.0918731689453 0 501.4313
191.11814880371094 0 6130.7275 d 1
193.10842895507812 0 3702.4937
194.1125030517578 0 496.92517
199.07159423828125 0 681.7019
200.10308837890625 0 629.1206
201.1233673095703 0 1699.2455
203.06658935546875 0 1319.4031
204.077392578125 0 1138.8867
205.09715270996094 0 3379.6013 y 7
205.10736083984375 0 904.22504
207.11306762695312 0 1172.5372
208.0714874267578 0 556.92035
209.09193420410156 0 1593.8562
217.0972900390625 0 21869.807 a Water loss 1
218.10073852539062 0 3317.2253
219.07948303222656 0 704.5141
221.08432006835938 0 9594.068
221.10340881347656 0 7724.233
221.12841796875 0 1354.2858
221.13934326171875 0 688.80853
222.0843505859375 0 464.69968
223.0635223388672 0 722.1206
223.11912536621094 0 687.9878
225.04296875 0 10325.34
225.1237335205078 0 697.93964
226.0437774658203 0 1575.8353
226.1549530029297 0 959.1069
227.02313232421875 0 864.8412
229.10801696777344 0 883.6724
231.06126403808594 0 2493.6948
231.11212158203125 0 548.1543
235.1077880859375 0 32639.021 a 1
236.11105346679688 0 4727.4
239.0950164794922 0 23976.324
239.11480712890625 0 984.5007
240.09588623046875 0 3247.1587
241.0947265625 0 601.693
242.14984130859375 0 1050.6836
245.0919647216797 0 1111.9429 b Water loss 1
249.12257385253906 0 739.44293
251.1027069091797 0 9057.719
252.1056671142578 0 717.76025
253.10870361328125 0 614.0063
256.1279296875 0 612.57135
257.1064453125 0 2190.3342
261.0871276855469 0 615.3226
263.10260009765625 0 28596.639 b 1
264.1060485839844 0 3770.8413
264.14263916015625 0 699.7669
266.1240234375 0 649.2313
269.3560485839844 0 557.476
274.1302490234375 0 2316.5232
279.097412109375 0 5219.106
280.10003662109375 0 670.1874
281.05206298828125 0 835.81476
283.0289306640625 0 786.5744
285.0094299316406 0 744.70264
285.1017150878906 0 3495.8186
285.11798095703125 0 871.0691
292.1402587890625 0 2601.3052
295.103271484375 0 6468.621
296.1036071777344 0 2227.7761
296.150634765625 0 3303.3386
299.0617980957031 0 9987.74
300.0624084472656 0 1349.0089
306.1463928222656 0 983.05536 d 2
308.3695068359375 0 622.80804
310.15283203125 0 720.6466
313.1141662597656 0 1409.4276
313.1867370605469 0 692.98083
318.14422607421875 0 585.8414
324.1456604003906 0 5424.1416
325.1499938964844 0 919.76337
327.1446838378906 0 646.3019
328.14483642578125 0 580.3782
329.12060546875 0 516.5514
330.1070251464844 0 665.2773
332.1224060058594 0 628.8205 a Water loss 2
335.0843811035156 0 550.1059
337.0700378417969 0 516.8885
338.132568359375 0 1928.64
338.859375 0 565.0699
341.1484680175781 0 688.7134
342.1560974121094 0 31550.941 y 6
343.15948486328125 0 6891.3545
344.1623840332031 0 920.77954
348.1188659667969 0 1118.03
350.134521484375 0 3468.3645 a 2
351.1391296386719 0 605.2725
355.07000732421875 0 1104.2882
356.1387634277344 0 3066.5674
359.029296875 0 695.62604
360.1184387207031 0 853.5918 b Water loss 2
362.1490173339844 0 743.6778 y 3
369.1219482421875 0 6004.6123
370.1220703125 0 2226.6895
374.148193359375 0 892.1374
376.1143493652344 0 823.1883
378.1294250488281 0 19677.182 b 2
379.1320495605469 0 3553.3005
381.142333984375 0 791.2358
382.14398193359375 0 937.66077
384.13555908203125 0 614.62085
394.1242370605469 0 3697.7869
407.1834411621094 0 749.25604
408.1663513183594 0 1417.5168
415.0353088378906 0 770.0041
419.15765380859375 0 913.6007
419.6619567871094 0 1060.6097 y 2
425.1930236816406 0 4383.9795
426.1967468261719 0 990.80066
442.17236328125 0 626.48663
449.1671142578125 0 627.2576
453.3441467285156 0 1095.2704
454.1898498535156 0 591.4147
455.20166015625 0 754.6058
465.1620788574219 0 2292.823
471.1829833984375 0 960.9914
477.176025390625 0 1824.91 y 1
489.19110107421875 0 8864.225 y 5
490.1955261230469 0 2779.4292
491.2085266113281 0 711.1425
495.1869812011719 0 4315.615
496.1923828125 0 965.278
496.2304992675781 0 5089.7407
496.81396484375 0 630.0553
497.172607421875 0 1147.4027
497.2340393066406 0 943.3362
503.3059997558594 0 2742.857
504.3085632324219 0 1655.7947
509.9465026855469 0 573.0499
513.1979370117188 0 3964.4956
518.7109985351562 0 1137.0562
519.2037963867188 0 1735.6807
520.2041015625 0 751.2042
532.3032836914062 0 1047.1693
533.2009887695312 0 843.6262
541.1925048828125 0 1118.9672 b 3
541.704833984375 0 1665.7063 Precursor Water loss
542.2001342773438 0 1078.1932
548.1994018554688 0 686.65186
550.2944946289062 0 2539.7095
550.633544921875 0 685.33856
550.7102661132812 0 11064.181 Precursor
551.2116088867188 0 7420.3135
551.7119140625 0 4073.234
560.2282104492188 0 16530.654 y 4
561.2312622070312 0 5451.9644
562.2323608398438 0 1385.0354
570.2279052734375 0 1888.9583
573.7990112304688 0 653.78815
606.2340087890625 0 2426.1099
607.2349853515625 0 813.8141
634.2294921875 0 2616.7512
635.232666015625 0 947.32294
638.8533935546875 0 735.72534
657.2616577148438 0 1108.0178
659.2927856445312 0 5074.5923
660.2959594726562 0 1420.5469
678.30078125 0 811.87103
681.8357543945312 0 1533.2372
682.3339233398438 0 826.40924
685.2553100585938 0 1478.619
686.2611694335938 0 746.2769
705.2803955078125 0 1037.8961
706.3523559570312 0 651.20465
710.457763671875 0 736.1579
721.259033203125 0 1471.8942
723.291015625 0 11729.259 y 3
724.2943725585938 0 4980.3926
725.2935791015625 0 1694.0564
749.255859375 0 1912.8474
756.3126831054688 0 727.3755
774.3197631835938 0 10835.042
775.3233032226562 0 4471.7505
776.3284912109375 0 835.80566
782.4450073242188 0 774.1126
820.3074340820312 0 1771.0829 y Water loss 2
838.3175659179688 0 30920.225 y 2
839.3204956054688 0 15394.593
840.3200073242188 0 4704.305
868.3250122070312 0 630.99603
871.338623046875 0 2086.826
889.3463134765625 0 9440.741
890.34912109375 0 3857.2957
891.3516845703125 0 1079.8204
896.3220825195312 0 1825.6769 b 6
900.3225708007812 0 690.109
935.3297729492188 0 2077.8804 y Water loss 1
936.3324584960938 0 1703.0144
953.3441772460938 0 22424.838 y 1
954.3471069335938 0 11239.082
955.3452758789062 0 3885.5947
963.3291015625 0 1474.3933
964.3284301757812 0 720.93896
1907.6851806640625 0 718.4664
2848.13720703125 0 666.73303
3071.807861328125 0 776.9541
3350.30810546875 0 749.26227

Spectrum Details

|  |  |
| --- | --- |
| Matched peaks? Matched peaks The total absolute number of peaks matched. Additionally in brackets the total fraction of peaks matched and the total number of peaks is shown. | 27 (10.38% of 260) |
| FDR? FDR The false discovery rate estimated for this peptide. It is calculated by matching all theoretical fragments with a non-integer shift with the raw peaks for this spectrum. This is done with 40 different shifts. The resulting percentage is the average number of annotated peaks over the number of annotated peaks with the correct spectrum. | 0.35% |
| Satellite FDR? Satellite FDR See the FDR for details on its calculation. This satellite ion specific FDR only contains the satellite ions (d/w) for I/L/J positions. | - |
| PSM Score? PSM Score The PSM Score as given by Hecklib to this annotated spectrum. It is shown with three significant figures. | 285 |

#### Spectrum 6797? Spectrum 6797 The raw spectrum of this peptide as annotated by Hecklib. The fragments are coloured according to ion type (see legend). Any peaks with a star '\*' as text can be hovered over to see the full details, first the ion type second the mass shift type. By hovering over the amino acids in the peptide or ions in the legend the corresponding peaks are highlighted. By toggling the 'Unassigned' label you can turn the background (unassigned) peaks on or off in the plot. By updating the slider in the Ion legend you can update the spectrum to only show the top X% of the peaks with labels. The top X% means any peak that is within X% of the highest intensity. By dragging in the spectrum you can zoom in to a specific part of the spectrum and use 'Zoom Out' to get back to the original zoom level. The annotation of the spectrum is based on the given sequence in the peptides file and is done with different software so inconsistencies are likely. The peaks are annotated based on the given sequence, with 20 ppm tolerance.

Copy Data

##### Spectrum 6797 (TSV)

###### Preview

```
Loading example...
```

*Click on the button to copy the data to your clipboard.*

Mz MinMz MaxIntensity Max

WidthHeightPeptide font sizePeptide stroke widthSpectrum font sizeSpectrum stroke widthCompact peptide

Ion legend

wxyz

abcd

OtherUnassignedIon^Charge^_Position_Show for top:%

FDDYAMHW

02.75e+45.50e+48.25e+41.10e+5

Zoom Out

a^+2^_3_d^+1^_2_y^+1^_1_a^+1^_2_a^+1^_2_b^+1^_2_d^+1^_3_y^+1^_2_a^+1^_3_b^+1^_3_b^+1^_3_y^+2^_7_y^+1^_3_\*\*y^+1^_4_y^+1^_5_y^+1^_6_y^+1^_6_b^+1^_7_y^+1^_7_y^+1^_7_

0738147722152954

Fragment Matches Table

Show background peaks

| Position | Ion type | Intensity | mz Theoretical | mz Error (Th) | mz Error (ppm) | Charge | Series Number |
| --- | --- | --- | --- | --- | --- | --- | --- |
| - | - | 1.089E+05 | 120.1 | - | - | 0 | - |
| - | - | 284.8 | 121 | - | - | 0 | - |
| - | - | 8581 | 121.1 | - | - | 0 | - |
| - | - | 395.1 | 126.1 | - | - | 0 | - |
| - | - | 815.9 | 128.1 | - | - | 0 | - |
| - | - | 3949 | 129.1 | - | - | 0 | - |
| - | - | 1321 | 130.1 | - | - | 0 | - |
| - | - | 554.8 | 131.1 | - | - | 0 | - |
| - | - | 529.9 | 133.1 | - | - | 0 | - |
| - | - | 1196 | 133.1 | - | - | 0 | - |
| - | - | 1.262E+04 | 136.1 | - | - | 0 | - |
| - | - | 827.6 | 137.1 | - | - | 0 | - |
| - | - | 488.5 | 137.9 | - | - | 0 | - |
| - | - | 553.4 | 138.1 | - | - | 0 | - |
| - | - | 1506 | 138.1 | - | - | 0 | - |
| - | - | 434 | 141.1 | - | - | 0 | - |
| - | - | 525.1 | 143.1 | - | - | 0 | - |
| - | - | 458.4 | 144.1 | - | - | 0 | - |
| - | - | 1041 | 146.1 | - | - | 0 | - |
| - | - | 1040 | 146.1 | - | - | 0 | - |
| - | - | 470.4 | 148.9 | - | - | 0 | - |
| - | - | 3948 | 149 | - | - | 0 | - |
| - | - | 583.9 | 154.1 | - | - | 0 | - |
| - | - | 615.6 | 155.1 | - | - | 0 | - |
| - | - | 2440 | 156.1 | - | - | 0 | - |
| - | - | 503.3 | 157.1 | - | - | 0 | - |
| - | - | 722.1 | 159.1 | - | - | 0 | - |
| - | - | 4652 | 159.1 | - | - | 0 | - |
| - | - | 966.8 | 159.1 | - | - | 0 | - |
| - | - | 668 | 163.1 | - | - | 0 | - |
| - | - | 621.8 | 165.1 | - | - | 0 | - |
| - | - | 651.5 | 165.1 | - | - | 0 | - |
| - | - | 882.1 | 166.1 | - | - | 0 | - |
| 3 | a | 5291 | 167.1 | 0.002122 | 12.7 | +2 | 3 |
| - | - | 671.9 | 171.1 | - | - | 0 | - |
| - | - | 644.4 | 172.1 | - | - | 0 | - |
| - | - | 549.4 | 173.1 | - | - | 0 | - |
| - | - | 684.6 | 173.1 | - | - | 0 | - |
| - | - | 2381 | 173.5 | - | - | 0 | - |
| - | - | 490.1 | 174.1 | - | - | 0 | - |
| - | - | 1355 | 175.1 | - | - | 0 | - |
| - | - | 559.8 | 175.1 | - | - | 0 | - |
| - | - | 884.3 | 176.1 | - | - | 0 | - |
| - | - | 617.7 | 177.1 | - | - | 0 | - |
| - | - | 986.4 | 181.1 | - | - | 0 | - |
| - | - | 1055 | 182.1 | - | - | 0 | - |
| - | - | 652.1 | 183.1 | - | - | 0 | - |
| - | - | 514.9 | 184.1 | - | - | 0 | - |
| - | - | 951.1 | 185.1 | - | - | 0 | - |
| - | - | 779.3 | 185.1 | - | - | 0 | - |
| - | - | 542 | 186.1 | - | - | 0 | - |
| - | - | 2181 | 187.1 | - | - | 0 | - |
| - | - | 3179 | 188.1 | - | - | 0 | - |
| - | - | 573.2 | 189.1 | - | - | 0 | - |
| - | - | 875.1 | 190.1 | - | - | 0 | - |
| 2 | d | 4545 | 191.1 | 0.0002288 | 1.197 | +1 | 2 |
| - | - | 2898 | 193.1 | - | - | 0 | - |
| - | - | 582.3 | 194.5 | - | - | 0 | - |
| - | - | 736 | 195.1 | - | - | 0 | - |
| - | - | 705.4 | 199.1 | - | - | 0 | - |
| - | - | 669.1 | 199.2 | - | - | 0 | - |
| - | - | 680 | 201.1 | - | - | 0 | - |
| - | - | 868.6 | 203.1 | - | - | 0 | - |
| - | - | 958.5 | 204.1 | - | - | 0 | - |
| 8 | y | 1946 | 205.1 | 0.0002733 | 1.332 | +1 | 1 |
| - | - | 1104 | 207.1 | - | - | 0 | - |
| - | - | 1343 | 209.1 | - | - | 0 | - |
| - | - | 807.1 | 211.1 | - | - | 0 | - |
| 2 | a | 1.561E+04 | 217.1 | 0.0001359 | 0.6262 | +1 | 2 |
| - | - | 455.4 | 217.1 | - | - | 0 | - |
| - | - | 1439 | 218.1 | - | - | 0 | - |
| - | - | 727.9 | 219.1 | - | - | 0 | - |
| - | - | 8420 | 221.1 | - | - | 0 | - |
| - | - | 6404 | 221.1 | - | - | 0 | - |
| - | - | 1542 | 222.1 | - | - | 0 | - |
| - | - | 591.6 | 222.1 | - | - | 0 | - |
| - | - | 1328 | 223.1 | - | - | 0 | - |
| - | - | 1.04E+04 | 225 | - | - | 0 | - |
| - | - | 1632 | 226 | - | - | 0 | - |
| - | - | 1748 | 226.2 | - | - | 0 | - |
| - | - | 963.8 | 227 | - | - | 0 | - |
| - | - | 694.1 | 228.1 | - | - | 0 | - |
| - | - | 1322 | 231.1 | - | - | 0 | - |
| - | - | 513.4 | 233.1 | - | - | 0 | - |
| 2 | a | 2.079E+04 | 235.1 | 0.0001914 | 0.814 | +1 | 2 |
| - | - | 2557 | 236.1 | - | - | 0 | - |
| - | - | 2.518E+04 | 239.1 | - | - | 0 | - |
| - | - | 3551 | 240.1 | - | - | 0 | - |
| - | - | 1500 | 241.1 | - | - | 0 | - |
| - | - | 816.6 | 242.1 | - | - | 0 | - |
| - | - | 5570 | 251.1 | - | - | 0 | - |
| - | - | 613.4 | 252.1 | - | - | 0 | - |
| - | - | 574.7 | 253.9 | - | - | 0 | - |
| - | - | 965.8 | 257.1 | - | - | 0 | - |
| 2 | b | 1.833E+04 | 263.1 | 3.33E-05 | 0.1266 | +1 | 2 |
| - | - | 2561 | 264.1 | - | - | 0 | - |
| - | - | 588.7 | 265.1 | - | - | 0 | - |
| - | - | 1404 | 274.1 | - | - | 0 | - |
| - | - | 3231 | 279.1 | - | - | 0 | - |
| - | - | 638.2 | 282.1 | - | - | 0 | - |
| - | - | 801.7 | 285 | - | - | 0 | - |
| - | - | 2668 | 285.1 | - | - | 0 | - |
| - | - | 2034 | 292.1 | - | - | 0 | - |
| - | - | 6207 | 295.1 | - | - | 0 | - |
| - | - | 1121 | 296.1 | - | - | 0 | - |
| - | - | 1808 | 296.2 | - | - | 0 | - |
| - | - | 9658 | 299.1 | - | - | 0 | - |
| - | - | 1510 | 300.1 | - | - | 0 | - |
| 3 | d | 875.7 | 306.1 | 9.541E-05 | 0.3117 | +1 | 3 |
| - | - | 566.4 | 307 | - | - | 0 | - |
| - | - | 541.2 | 310.1 | - | - | 0 | - |
| - | - | 551.3 | 313.1 | - | - | 0 | - |
| - | - | 1056 | 313.1 | - | - | 0 | - |
| - | - | 3978 | 324.1 | - | - | 0 | - |
| - | - | 1083 | 325.1 | - | - | 0 | - |
| - | - | 1307 | 338.1 | - | - | 0 | - |
| - | - | 569.5 | 341.1 | - | - | 0 | - |
| 7 | y | 2.156E+04 | 342.2 | 0.0001535 | 0.4487 | +1 | 2 |
| - | - | 4694 | 343.2 | - | - | 0 | - |
| - | - | 803.3 | 344.2 | - | - | 0 | - |
| - | - | 843 | 345 | - | - | 0 | - |
| - | - | 718.2 | 348.1 | - | - | 0 | - |
| 3 | a | 2185 | 350.1 | 4.877E-05 | 0.1393 | +1 | 3 |
| - | - | 680.3 | 351.1 | - | - | 0 | - |
| - | - | 1468 | 355.1 | - | - | 0 | - |
| - | - | 1223 | 356.1 | - | - | 0 | - |
| - | - | 676.7 | 357.1 | - | - | 0 | - |
| - | - | 700.6 | 357.2 | - | - | 0 | - |
| 3 | b | 718.8 | 360.1 | 0.0004204 | 1.167 | +1 | 3 |
| - | - | 725.4 | 361.2 | - | - | 0 | - |
| - | - | 7720 | 369.1 | - | - | 0 | - |
| - | - | 2281 | 370.1 | - | - | 0 | - |
| 3 | b | 1.359E+04 | 378.1 | 3.172E-05 | 0.08389 | +1 | 3 |
| - | - | 2254 | 379.1 | - | - | 0 | - |
| - | - | 2134 | 394.1 | - | - | 0 | - |
| - | - | 616.4 | 407.2 | - | - | 0 | - |
| - | - | 984.6 | 408.2 | - | - | 0 | - |
| - | - | 697.7 | 409.2 | - | - | 0 | - |
| - | - | 2569 | 425.2 | - | - | 0 | - |
| - | - | 749.6 | 427.2 | - | - | 0 | - |
| - | - | 769.4 | 453.3 | - | - | 0 | - |
| - | - | 911.4 | 455.2 | - | - | 0 | - |
| - | - | 1207 | 465.2 | - | - | 0 | - |
| - | - | 708.6 | 471.2 | - | - | 0 | - |
| 2 | y | 805.6 | 477.2 | 0.001581 | 3.314 | +2 | 7 |
| 6 | y | 6042 | 489.2 | 0.005466 | 11.17 | +1 | 3 |
| - | - | 1208 | 490.2 | - | - | 0 | - |
| - | - | 1886 | 495.2 | - | - | 0 | - |
| - | - | 989.1 | 496.2 | - | - | 0 | - |
| - | - | 3283 | 496.2 | - | - | 0 | - |
| - | - | 1174 | 497.2 | - | - | 0 | - |
| - | - | 590.1 | 508.3 | - | - | 0 | - |
| - | - | 2337 | 513.2 | - | - | 0 | - |
| - | - | 961 | 514.2 | - | - | 0 | - |
| - | - | 1200 | 518.7 | - | - | 0 | - |
| - | - | 1498 | 519.2 | - | - | 0 | - |
| 0 | Precursor | 1897 | 541.7 | 0.002137 | 3.945 | +2 | -1 |
| - | - | 1151 | 542.2 | - | - | 0 | - |
| - | - | 961.2 | 542.7 | - | - | 0 | - |
| - | - | 558.7 | 543.2 | - | - | 0 | - |
| - | - | 1801 | 550 | - | - | 0 | - |
| - | - | 2024 | 550.3 | - | - | 0 | - |
| 0 | Precursor | 8975 | 550.7 | 0.002165 | 3.93 | +2 | -1 |
| - | - | 4249 | 551.2 | - | - | 0 | - |
| - | - | 2506 | 551.7 | - | - | 0 | - |
| 5 | y | 1.32E+04 | 560.2 | 0.004668 | 8.333 | +1 | 4 |
| - | - | 3265 | 561.2 | - | - | 0 | - |
| - | - | 761.1 | 562.2 | - | - | 0 | - |
| - | - | 675.5 | 570.2 | - | - | 0 | - |
| - | - | 830.7 | 573.8 | - | - | 0 | - |
| - | - | 2184 | 606.2 | - | - | 0 | - |
| - | - | 677 | 607.2 | - | - | 0 | - |
| - | - | 1541 | 634.2 | - | - | 0 | - |
| - | - | 3872 | 659.3 | - | - | 0 | - |
| - | - | 905 | 681.8 | - | - | 0 | - |
| - | - | 1482 | 682.3 | - | - | 0 | - |
| - | - | 1833 | 685.3 | - | - | 0 | - |
| - | - | 723.3 | 689.3 | - | - | 0 | - |
| - | - | 752.2 | 710.5 | - | - | 0 | - |
| - | - | 1269 | 721.3 | - | - | 0 | - |
| 4 | y | 9428 | 723.3 | 0.003718 | 5.14 | +1 | 5 |
| - | - | 3678 | 724.3 | - | - | 0 | - |
| - | - | 730.8 | 725.3 | - | - | 0 | - |
| - | - | 681.6 | 731.4 | - | - | 0 | - |
| - | - | 1537 | 749.3 | - | - | 0 | - |
| - | - | 670.8 | 756.3 | - | - | 0 | - |
| - | - | 6447 | 774.3 | - | - | 0 | - |
| - | - | 2619 | 775.3 | - | - | 0 | - |
| - | - | 1075 | 775.4 | - | - | 0 | - |
| - | - | 912.1 | 776.3 | - | - | 0 | - |
| 3 | y | 1029 | 820.3 | 0.00742 | 9.045 | +1 | 6 |
| 3 | y | 2.216E+04 | 838.3 | 0.00363 | 4.33 | +1 | 6 |
| - | - | 8953 | 839.3 | - | - | 0 | - |
| - | - | 3995 | 840.3 | - | - | 0 | - |
| - | - | 1068 | 871.3 | - | - | 0 | - |
| - | - | 631.4 | 888.7 | - | - | 0 | - |
| - | - | 4876 | 889.3 | - | - | 0 | - |
| - | - | 3479 | 890.3 | - | - | 0 | - |
| 7 | b | 1172 | 896.3 | 0.004926 | 5.495 | +1 | 7 |
| 2 | y | 1874 | 935.3 | 0.005257 | 5.62 | +1 | 7 |
| - | - | 1382 | 936.3 | - | - | 0 | - |
| 2 | y | 1.527E+04 | 953.3 | 0.003787 | 3.972 | +1 | 7 |
| - | - | 7201 | 954.3 | - | - | 0 | - |
| - | - | 2652 | 955.3 | - | - | 0 | - |
| - | - | 832 | 964.3 | - | - | 0 | - |
| - | - | 687 | 2690 | - | - | 0 | - |
| - | - | 727.4 | 2924 | - | - | 0 | - |

m/z Charge Intensity FragmentType MassShift Position
120.08103942871094 0 108865.164
121.03753662109375 0 284.81693
121.08434295654297 0 8581.349
126.05519104003906 0 395.06705
128.10704040527344 0 815.8828
129.10240173339844 0 3948.5742
130.06533813476562 0 1321.2856
131.11810302734375 0 554.7817
133.0607147216797 0 529.89526
133.0861053466797 0 1195.5352
136.07589721679688 0 12624.319
137.07907104492188 0 827.6134
137.9129180908203 0 488.50964
138.05506896972656 0 553.3532
138.06626892089844 0 1505.9435
141.06658935546875 0 433.98312
143.11814880371094 0 525.0838
144.08116149902344 0 458.35922
146.0602264404297 0 1040.6732
146.0972900390625 0 1039.6742
148.94650268554688 0 470.4454
149.04505920410156 0 3947.9143
154.06170654296875 0 583.93256
155.08212280273438 0 615.6401
156.07699584960938 0 2440.4104
157.09690856933594 0 503.28027
159.0768585205078 0 722.1442
159.0917205810547 0 4651.5654
159.11264038085938 0 966.80634
163.0713348388672 0 668.0043
165.07740783691406 0 621.8235
165.10272216796875 0 651.5228
166.0616912841797 0 882.0614
167.05557250976562 0 5290.58 a Ammonia loss 2
171.14920043945312 0 671.8759
172.07630920410156 0 644.3768
173.09237670898438 0 549.4295
173.12905883789062 0 684.57513
173.45118713378906 0 2380.5334
174.0663299560547 0 490.13684
175.0868682861328 0 1354.7277
175.09617614746094 0 559.8311
176.08209228515625 0 884.2657
177.11297607421875 0 617.69495
181.0975341796875 0 986.4281
182.0816650390625 0 1055.3127
183.11241149902344 0 652.0939
184.07176208496094 0 514.89197
185.05581665039062 0 951.14154
185.09222412109375 0 779.2618
186.12359619140625 0 541.9579
187.1078643798828 0 2181.2024
188.07080078125 0 3178.8083
189.0865478515625 0 573.1858
190.0861053466797 0 875.14734
191.1181182861328 0 4544.8403 d 1
193.10862731933594 0 2897.9722
194.54359436035156 0 582.3093
195.08827209472656 0 736.0002
199.1439208984375 0 705.3596
199.18128967285156 0 669.14026
201.12322998046875 0 679.954
203.0662841796875 0 868.6174
204.07664489746094 0 958.53784
205.09742736816406 0 1946.3896 y 7
207.11300659179688 0 1104.2428
209.09213256835938 0 1343.0426
211.14402770996094 0 807.1119
217.0972900390625 0 15612.976 a Water loss 1
217.13319396972656 0 455.3528
218.10057067871094 0 1439.1672
219.07968139648438 0 727.93256
221.0845489501953 0 8419.658
221.10337829589844 0 6403.6377
222.0855255126953 0 1542.01
222.10763549804688 0 591.55536
223.0635986328125 0 1327.7687
225.0430450439453 0 10401.91
226.04315185546875 0 1631.5237
226.15530395507812 0 1747.9565
227.02200317382812 0 963.80676
228.11407470703125 0 694.0507
231.06126403808594 0 1321.974
233.0914306640625 0 513.3719
235.10791015625 0 20793.559 a 1
236.11087036132812 0 2556.655
239.09506225585938 0 25184.389
240.09573364257812 0 3551.302
241.09271240234375 0 1499.748
242.1486358642578 0 816.55225
251.10284423828125 0 5570.031
252.1056671142578 0 613.407
253.90777587890625 0 574.7222
257.1061096191406 0 965.7734
263.10260009765625 0 18331.959 b 1
264.1059265136719 0 2560.944
265.119140625 0 588.74866
274.13055419921875 0 1404.3477
279.09735107421875 0 3231.3433
282.05218505859375 0 638.19006
285.00958251953125 0 801.7204
285.10174560546875 0 2668.002
292.1400451660156 0 2034.4178
295.1031799316406 0 6207.109
296.1039733886719 0 1120.5743
296.1508483886719 0 1808.1544
299.06182861328125 0 9658.068
300.06243896484375 0 1510.3884
306.1449279785156 0 875.6934 d 2
306.983154296875 0 566.4152
310.1379089355469 0 541.20026
313.0787658691406 0 551.2529
313.11419677734375 0 1056.306
324.14544677734375 0 3977.848
325.1499938964844 0 1082.7084
338.1345520019531 0 1306.88
341.1492004394531 0 569.49603
342.1562194824219 0 21556.582 y 6
343.15985107421875 0 4694.3516
344.1641540527344 0 803.319
344.97698974609375 0 843.0244
348.11944580078125 0 718.24207
350.1346130371094 0 2184.6985 a 2
351.1379699707031 0 680.2718
355.0696105957031 0 1467.7653
356.1379089355469 0 1223.0337
357.1419372558594 0 676.6593
357.2493896484375 0 700.6135
360.11859130859375 0 718.8376 b Water loss 2
361.1715393066406 0 725.388
369.1219482421875 0 7720.0806
370.1230773925781 0 2281.0037
378.1296081542969 0 13586.956 b 2
379.1319885253906 0 2253.6692
394.12506103515625 0 2133.8296
407.181396484375 0 616.39685
408.1666259765625 0 984.59467
409.1723327636719 0 697.6536
425.19415283203125 0 2568.7883
427.2087097167969 0 749.621
453.34674072265625 0 769.3831
455.2038879394531 0 911.4473
465.1623229980469 0 1206.7288
471.1800842285156 0 708.6276
477.1756591796875 0 805.5707 y 1
489.1920166015625 0 6042.187 y 5
490.19476318359375 0 1207.6627
495.1874694824219 0 1886.1736
496.19305419921875 0 989.1011
496.23138427734375 0 3283.248
497.23291015625 0 1173.5339
508.250732421875 0 590.0557
513.1986083984375 0 2337.3132
514.1967163085938 0 960.96234
518.7113647460938 0 1200.0382
519.2047729492188 0 1498.4728
541.7051391601562 0 1896.6915 Precursor Water loss
542.2019653320312 0 1150.65
542.7084350585938 0 961.22327
543.21142578125 0 558.66644
549.9627075195312 0 1800.5962
550.2958984375 0 2024.0205
550.71044921875 0 8975.028 Precursor
551.211669921875 0 4248.9385
551.7132568359375 0 2505.5254
560.2283325195312 0 13195.996 y 4
561.2323608398438 0 3264.6282
562.2297973632812 0 761.1208
570.2310180664062 0 675.5451
573.7979736328125 0 830.651
606.233154296875 0 2183.7864
607.2365112304688 0 677.032
634.2286376953125 0 1541.3914
659.2931518554688 0 3871.9585
681.8377685546875 0 904.9603
682.3348388671875 0 1481.5343
685.2560424804688 0 1833.0035
689.3404541015625 0 723.3265
710.4588012695312 0 752.21027
721.2603759765625 0 1268.5182
723.2907104492188 0 9427.774 y 3
724.2938842773438 0 3677.6475
725.2882080078125 0 730.82684
731.39013671875 0 681.5502
749.2536010742188 0 1537.3301
756.3156127929688 0 670.80176
774.3196411132812 0 6447.258
775.3235473632812 0 2619.3594
775.4017944335938 0 1074.7589
776.3225708007812 0 912.06116
820.310791015625 0 1029.421 y Water loss 2
838.3175659179688 0 22163.473 y 2
839.3203125 0 8953.364
840.3211669921875 0 3995.0332
871.3387451171875 0 1067.5026
888.6915893554688 0 631.3847
889.3460083007812 0 4875.976
890.3488159179688 0 3479.2576
896.3243408203125 0 1171.8126 b 6
935.3355712890625 0 1874.0326 y Water loss 1
936.3380126953125 0 1382.4736
953.3446655273438 0 15265.37 y 1
954.3477783203125 0 7200.565
955.3475952148438 0 2651.6284
964.3290405273438 0 831.994
2690.343505859375 0 686.9632
2924.322021484375 0 727.4473

Spectrum Details

|  |  |
| --- | --- |
| Matched peaks? Matched peaks The total absolute number of peaks matched. Additionally in brackets the total fraction of peaks matched and the total number of peaks is shown. | 22 (10.63% of 207) |
| FDR? FDR The false discovery rate estimated for this peptide. It is calculated by matching all theoretical fragments with a non-integer shift with the raw peaks for this spectrum. This is done with 40 different shifts. The resulting percentage is the average number of annotated peaks over the number of annotated peaks with the correct spectrum. | 0.22% |
| Satellite FDR? Satellite FDR See the FDR for details on its calculation. This satellite ion specific FDR only contains the satellite ions (d/w) for I/L/J positions. | - |
| PSM Score? PSM Score The PSM Score as given by Hecklib to this annotated spectrum. It is shown with three significant figures. | 225 |

#### Spectrum 5428? Spectrum 5428 The raw spectrum of this peptide as annotated by Hecklib. The fragments are coloured according to ion type (see legend). Any peaks with a star '\*' as text can be hovered over to see the full details, first the ion type second the mass shift type. By hovering over the amino acids in the peptide or ions in the legend the corresponding peaks are highlighted. By toggling the 'Unassigned' label you can turn the background (unassigned) peaks on or off in the plot. By updating the slider in the Ion legend you can update the spectrum to only show the top X% of the peaks with labels. The top X% means any peak that is within X% of the highest intensity. By dragging in the spectrum you can zoom in to a specific part of the spectrum and use 'Zoom Out' to get back to the original zoom level. The annotation of the spectrum is based on the given sequence in the peptides file and is done with different software so inconsistencies are likely. The peaks are annotated based on the given sequence, with 20 ppm tolerance.

Copy Data

##### Spectrum 5428 (TSV)

###### Preview

```
Loading example...
```

*Click on the button to copy the data to your clipboard.*

Mz MinMz MaxIntensity Max

WidthHeightPeptide font sizePeptide stroke widthSpectrum font sizeSpectrum stroke widthCompact peptide

Ion legend

wxyz

abcd

OtherUnassignedIon^Charge^_Position_Show for top:%

FDDYAMHW

01.66e+43.32e+44.97e+46.63e+4

Zoom Out

d^+1^_2_a^+1^_2_a^+1^_2_b^+1^_2_a^+1^_3_y^+1^_2_b^+1^_3_y^+1^_3_\*\*y^+1^_4_y^+1^_5_y^+1^_6_y^+1^_7_y^+1^_7_

0812162324353247

Fragment Matches Table

Show background peaks

| Position | Ion type | Intensity | mz Theoretical | mz Error (Th) | mz Error (ppm) | Charge | Series Number |
| --- | --- | --- | --- | --- | --- | --- | --- |
| - | - | 6.566E+04 | 120.1 | - | - | 0 | - |
| - | - | 5345 | 121.1 | - | - | 0 | - |
| - | - | 445.5 | 125.1 | - | - | 0 | - |
| - | - | 733.2 | 128.1 | - | - | 0 | - |
| - | - | 5332 | 129.1 | - | - | 0 | - |
| - | - | 426.7 | 130.1 | - | - | 0 | - |
| - | - | 1390 | 130.1 | - | - | 0 | - |
| - | - | 736.6 | 130.1 | - | - | 0 | - |
| - | - | 465.6 | 131.1 | - | - | 0 | - |
| - | - | 359.5 | 131.3 | - | - | 0 | - |
| - | - | 623.2 | 133.1 | - | - | 0 | - |
| - | - | 946.3 | 133.1 | - | - | 0 | - |
| - | - | 9202 | 136.1 | - | - | 0 | - |
| - | - | 374.9 | 136.4 | - | - | 0 | - |
| - | - | 1044 | 137.1 | - | - | 0 | - |
| - | - | 385.7 | 138.1 | - | - | 0 | - |
| - | - | 486 | 140.1 | - | - | 0 | - |
| - | - | 762.9 | 141.1 | - | - | 0 | - |
| - | - | 581.1 | 142.1 | - | - | 0 | - |
| - | - | 567.7 | 143.1 | - | - | 0 | - |
| - | - | 504 | 146.1 | - | - | 0 | - |
| - | - | 568.2 | 147 | - | - | 0 | - |
| - | - | 497.9 | 152.3 | - | - | 0 | - |
| - | - | 452.2 | 153.1 | - | - | 0 | - |
| - | - | 484.5 | 155.1 | - | - | 0 | - |
| - | - | 507.9 | 155.1 | - | - | 0 | - |
| - | - | 2012 | 156.1 | - | - | 0 | - |
| - | - | 6815 | 158.1 | - | - | 0 | - |
| - | - | 855.2 | 159.1 | - | - | 0 | - |
| - | - | 1021 | 159.1 | - | - | 0 | - |
| - | - | 494 | 165 | - | - | 0 | - |
| - | - | 862.8 | 165.1 | - | - | 0 | - |
| - | - | 782.7 | 166.1 | - | - | 0 | - |
| - | - | 889.4 | 166.1 | - | - | 0 | - |
| - | - | 566.4 | 169.1 | - | - | 0 | - |
| - | - | 810.1 | 170.1 | - | - | 0 | - |
| - | - | 541.6 | 171.1 | - | - | 0 | - |
| - | - | 793.1 | 171.1 | - | - | 0 | - |
| - | - | 1641 | 173.1 | - | - | 0 | - |
| - | - | 826.8 | 173.1 | - | - | 0 | - |
| - | - | 1569 | 173.5 | - | - | 0 | - |
| - | - | 3618 | 175.1 | - | - | 0 | - |
| - | - | 745.9 | 175.1 | - | - | 0 | - |
| - | - | 584.3 | 176.1 | - | - | 0 | - |
| - | - | 1717 | 177.1 | - | - | 0 | - |
| - | - | 616.1 | 177.1 | - | - | 0 | - |
| - | - | 583.4 | 182.1 | - | - | 0 | - |
| - | - | 782.6 | 183.1 | - | - | 0 | - |
| - | - | 485.5 | 185.1 | - | - | 0 | - |
| - | - | 1764 | 185.1 | - | - | 0 | - |
| - | - | 645.7 | 185.2 | - | - | 0 | - |
| - | - | 9376 | 187.1 | - | - | 0 | - |
| - | - | 1730 | 187.1 | - | - | 0 | - |
| - | - | 753.7 | 188.1 | - | - | 0 | - |
| - | - | 487.3 | 189.1 | - | - | 0 | - |
| - | - | 549.1 | 190.1 | - | - | 0 | - |
| 2 | d | 2315 | 191.1 | 1.305E-07 | 0.0006827 | +1 | 2 |
| - | - | 1329 | 193.1 | - | - | 0 | - |
| - | - | 760.8 | 195.1 | - | - | 0 | - |
| - | - | 1550 | 199.2 | - | - | 0 | - |
| - | - | 1464 | 201.1 | - | - | 0 | - |
| - | - | 999.8 | 203.1 | - | - | 0 | - |
| - | - | 585.7 | 204.1 | - | - | 0 | - |
| - | - | 655.1 | 207.1 | - | - | 0 | - |
| - | - | 514.6 | 212.1 | - | - | 0 | - |
| - | - | 576.5 | 213.2 | - | - | 0 | - |
| - | - | 1213 | 215.1 | - | - | 0 | - |
| - | - | 900.9 | 217.1 | - | - | 0 | - |
| 2 | a | 8390 | 217.1 | 6.242E-05 | 0.2875 | +1 | 2 |
| - | - | 885.6 | 218.1 | - | - | 0 | - |
| - | - | 7216 | 221.1 | - | - | 0 | - |
| - | - | 3585 | 221.1 | - | - | 0 | - |
| - | - | 529.5 | 225.1 | - | - | 0 | - |
| - | - | 702.1 | 225.2 | - | - | 0 | - |
| - | - | 1573 | 226.2 | - | - | 0 | - |
| - | - | 937.3 | 227.1 | - | - | 0 | - |
| - | - | 840.5 | 227.2 | - | - | 0 | - |
| - | - | 683.2 | 231.1 | - | - | 0 | - |
| - | - | 739.1 | 233.2 | - | - | 0 | - |
| 2 | a | 1.159E+04 | 235.1 | 6.987E-06 | 0.02972 | +1 | 2 |
| - | - | 940.6 | 235.1 | - | - | 0 | - |
| - | - | 1175 | 236.1 | - | - | 0 | - |
| - | - | 637.8 | 239.1 | - | - | 0 | - |
| - | - | 818.6 | 240.1 | - | - | 0 | - |
| - | - | 489.7 | 240.3 | - | - | 0 | - |
| - | - | 830.4 | 241.2 | - | - | 0 | - |
| - | - | 508.2 | 249.1 | - | - | 0 | - |
| - | - | 2964 | 251.1 | - | - | 0 | - |
| - | - | 635.6 | 257.1 | - | - | 0 | - |
| - | - | 710.6 | 258.1 | - | - | 0 | - |
| - | - | 489.2 | 261.2 | - | - | 0 | - |
| 2 | b | 1.064E+04 | 263.1 | 0.0001859 | 0.7065 | +1 | 2 |
| - | - | 1099 | 264.1 | - | - | 0 | - |
| - | - | 735.8 | 274.1 | - | - | 0 | - |
| - | - | 1215 | 279.1 | - | - | 0 | - |
| - | - | 1684 | 285.1 | - | - | 0 | - |
| - | - | 986.9 | 287.1 | - | - | 0 | - |
| - | - | 939.9 | 292.1 | - | - | 0 | - |
| - | - | 515.2 | 313.5 | - | - | 0 | - |
| - | - | 531.4 | 328.6 | - | - | 0 | - |
| - | - | 2838 | 340.1 | - | - | 0 | - |
| 3 | a | 1688 | 350.1 | 0.0005921 | 1.691 | +1 | 3 |
| - | - | 814.8 | 353.2 | - | - | 0 | - |
| - | - | 656 | 356.1 | - | - | 0 | - |
| 7 | y | 1.325E+04 | 358.1 | 0.004691 | 13.1 | +1 | 2 |
| - | - | 3265 | 359.2 | - | - | 0 | - |
| - | - | 532.3 | 362.4 | - | - | 0 | - |
| 3 | b | 8629 | 378.1 | 0.0002453 | 0.6488 | +1 | 3 |
| - | - | 1547 | 379.1 | - | - | 0 | - |
| - | - | 717.1 | 394.1 | - | - | 0 | - |
| - | - | 1961 | 441.2 | - | - | 0 | - |
| - | - | 1169 | 465.2 | - | - | 0 | - |
| - | - | 1502 | 473.3 | - | - | 0 | - |
| - | - | 758.5 | 477.2 | - | - | 0 | - |
| - | - | 737.4 | 478.8 | - | - | 0 | - |
| - | - | 671.3 | 487.2 | - | - | 0 | - |
| - | - | 1410 | 495.2 | - | - | 0 | - |
| - | - | 615.3 | 496.2 | - | - | 0 | - |
| 6 | y | 3852 | 505.2 | 0.00924 | 18.29 | +1 | 3 |
| - | - | 1243 | 506.2 | - | - | 0 | - |
| - | - | 2584 | 512.2 | - | - | 0 | - |
| - | - | 1301 | 513.2 | - | - | 0 | - |
| - | - | 650.8 | 514.2 | - | - | 0 | - |
| - | - | 761.2 | 521.8 | - | - | 0 | - |
| - | - | 822.4 | 533.4 | - | - | 0 | - |
| 0 | Precursor | 1681 | 549.7 | 0.005001 | 9.097 | +2 | -1 |
| - | - | 1016 | 557.6 | - | - | 0 | - |
| - | - | 858 | 558 | - | - | 0 | - |
| - | - | 1645 | 558.3 | - | - | 0 | - |
| 0 | Precursor | 2676 | 558.7 | 0.004906 | 8.782 | +2 | -1 |
| - | - | 1161 | 559.1 | - | - | 0 | - |
| - | - | 1755 | 559.2 | - | - | 0 | - |
| - | - | 598.2 | 559.3 | - | - | 0 | - |
| - | - | 995.7 | 559.7 | - | - | 0 | - |
| - | - | 801.5 | 560.2 | - | - | 0 | - |
| 5 | y | 7192 | 576.2 | 0.009846 | 17.09 | +1 | 4 |
| - | - | 1812 | 577.2 | - | - | 0 | - |
| - | - | 900.4 | 606.2 | - | - | 0 | - |
| - | - | 979.3 | 634.2 | - | - | 0 | - |
| - | - | 660.5 | 638.3 | - | - | 0 | - |
| - | - | 1140 | 657.3 | - | - | 0 | - |
| - | - | 1951 | 675.3 | - | - | 0 | - |
| - | - | 1162 | 681.8 | - | - | 0 | - |
| - | - | 1686 | 682.3 | - | - | 0 | - |
| - | - | 840.2 | 682.8 | - | - | 0 | - |
| - | - | 717.5 | 685.3 | - | - | 0 | - |
| - | - | 629.8 | 696.3 | - | - | 0 | - |
| - | - | 1792 | 738.9 | - | - | 0 | - |
| 4 | y | 4620 | 739.3 | 0.009018 | 12.2 | +1 | 5 |
| - | - | 2265 | 740.3 | - | - | 0 | - |
| - | - | 764.3 | 741.3 | - | - | 0 | - |
| - | - | 655 | 749.3 | - | - | 0 | - |
| - | - | 4078 | 790.3 | - | - | 0 | - |
| - | - | 1680 | 791.3 | - | - | 0 | - |
| 3 | y | 1.077E+04 | 854.3 | 0.009113 | 10.67 | +1 | 6 |
| - | - | 5345 | 855.3 | - | - | 0 | - |
| - | - | 1926 | 856.3 | - | - | 0 | - |
| - | - | 3691 | 905.3 | - | - | 0 | - |
| - | - | 1332 | 906.3 | - | - | 0 | - |
| 2 | y | 881.4 | 951.3 | 0.008726 | 9.173 | +1 | 7 |
| 2 | y | 7348 | 969.3 | 0.00866 | 8.934 | +1 | 7 |
| - | - | 4438 | 970.3 | - | - | 0 | - |
| - | - | 1967 | 971.3 | - | - | 0 | - |
| - | - | 583.1 | 1229 | - | - | 0 | - |
| - | - | 738.5 | 1650 | - | - | 0 | - |
| - | - | 634 | 1826 | - | - | 0 | - |
| - | - | 656.4 | 2008 | - | - | 0 | - |
| - | - | 681.6 | 3199 | - | - | 0 | - |
| - | - | 792.2 | 3214 | - | - | 0 | - |

m/z Charge Intensity FragmentType MassShift Position
120.08090209960938 0 65657.34
121.08423614501953 0 5345.4653
125.10746765136719 0 445.48383
128.1072235107422 0 733.1608
129.10227966308594 0 5331.9663
130.05001831054688 0 426.68124
130.0652313232422 0 1390.4496
130.08624267578125 0 736.618
131.1184539794922 0 465.55057
131.30282592773438 0 359.4838
133.06076049804688 0 623.24396
133.08583068847656 0 946.3348
136.0757598876953 0 9202.181
136.38595581054688 0 374.9336
137.07911682128906 0 1043.9626
138.06605529785156 0 385.6866
140.0816650390625 0 486.00818
141.1024932861328 0 762.9268
142.1230926513672 0 581.0752
143.1182861328125 0 567.716
146.09619140625 0 503.98358
147.04396057128906 0 568.24475
152.33311462402344 0 497.88522
153.06614685058594 0 452.19922
155.0817413330078 0 484.51724
155.11709594726562 0 507.8896
156.0767822265625 0 2012.3586
158.1175994873047 0 6814.769
159.09165954589844 0 855.1586
159.1128387451172 0 1021.2924
164.9977569580078 0 493.96686
165.1023406982422 0 862.8406
166.0535430908203 0 782.68677
166.1339874267578 0 889.4169
169.13345336914062 0 566.3709
170.09190368652344 0 810.06067
171.07672119140625 0 541.61414
171.11289978027344 0 793.144
173.0922393798828 0 1641.0344
173.12823486328125 0 826.79486
173.45172119140625 0 1569.0222
175.08670043945312 0 3617.6836
175.11878967285156 0 745.8564
176.0814666748047 0 584.3398
177.0673828125 0 1717.1378
177.1127471923828 0 616.1241
182.08192443847656 0 583.4434
183.1130828857422 0 782.581
185.0555419921875 0 485.47305
185.09217834472656 0 1763.5068
185.16488647460938 0 645.6939
187.1077880859375 0 9376.471
187.1442413330078 0 1729.9044
188.10279846191406 0 753.72986
189.1022186279297 0 487.29285
190.08627319335938 0 549.0781
191.11788940429688 0 2315.4756 d 1
193.1080322265625 0 1328.8256
195.11279296875 0 760.7532
199.1804656982422 0 1549.6016
201.12338256835938 0 1463.6804
203.0818328857422 0 999.8326
204.07659912109375 0 585.7041
207.11231994628906 0 655.1413
212.1387481689453 0 514.5502
213.1595458984375 0 576.5114
215.13868713378906 0 1212.6122
217.08473205566406 0 900.94727
217.0970916748047 0 8389.602 a Water loss 1
218.10101318359375 0 885.5952
221.0919952392578 0 7215.692
221.10353088378906 0 3584.7227
225.1228790283203 0 529.47327
225.19630432128906 0 702.111
226.15524291992188 0 1572.6635
227.10202026367188 0 937.2916
227.17481994628906 0 840.50726
231.06130981445312 0 683.1819
233.1649932861328 0 739.1153
235.1077117919922 0 11585.563 a 1
235.12001037597656 0 940.59753
236.11109924316406 0 1175.086
239.11436462402344 0 637.81146
240.1341094970703 0 818.618
240.2982177734375 0 489.66876
241.1547393798828 0 830.4475
249.09814453125 0 508.19336
251.1020965576172 0 2963.6785
257.10638427734375 0 635.62286
258.1451721191406 0 710.64044
261.15924072265625 0 489.22174
263.1024475097656 0 10643.913 b 1
264.10589599609375 0 1099.181
274.13055419921875 0 735.7872
279.0967102050781 0 1215.0684
285.1016845703125 0 1684.0874
287.08074951171875 0 986.90533
292.14117431640625 0 939.9134
313.51165771484375 0 515.1839
328.620849609375 0 531.4426
340.1402587890625 0 2837.9978
350.13525390625 0 1688.303 a 2
353.1928405761719 0 814.81915
356.1385803222656 0 656.0115
358.1507568359375 0 13249.274 y 6
359.1540222167969 0 3264.7505
362.3736572265625 0 532.3324
378.12982177734375 0 8629.035 b 2
379.1328125 0 1547.3245
394.1249694824219 0 717.0872
441.1880798339844 0 1961.4218
465.16070556640625 0 1169.358
473.2601318359375 0 1502.2554
477.1856689453125 0 758.5441
478.7695617675781 0 737.4057
487.1772766113281 0 671.2981
495.1878356933594 0 1409.607
496.1824035644531 0 615.25446
505.185791015625 0 3852.1824 y 5
506.1888427734375 0 1242.5763
512.2247924804688 0 2584.0317
513.1952514648438 0 1300.6366
514.1956176757812 0 650.843
521.7815551757812 0 761.18317
533.4451904296875 0 822.3526
549.7030029296875 0 1681.0197 Precursor Water loss
557.628173828125 0 1016.0358
557.9579467773438 0 857.9948
558.3216552734375 0 1645.1235
558.7081909179688 0 2675.9656 Precursor
559.1386108398438 0 1160.993
559.2106323242188 0 1754.9147
559.316650390625 0 598.1744
559.71337890625 0 995.74005
560.210205078125 0 801.54285
576.2235107421875 0 7191.5166 y 4
577.2254028320312 0 1812.2551
606.2340087890625 0 900.3631
634.2313842773438 0 979.34296
638.3197631835938 0 660.5137
657.2615356445312 0 1140.1443
675.2883911132812 0 1950.9185
681.8323364257812 0 1161.7783
682.3357543945312 0 1686.2904
682.8378295898438 0 840.19055
685.2542114257812 0 717.54846
696.3320922851562 0 629.7542
738.87451171875 0 1792.44
739.2860107421875 0 4620.2896 y 3
740.2891845703125 0 2264.5078
741.2838134765625 0 764.3289
749.2610473632812 0 655.0101
790.3157958984375 0 4077.6187
791.3154907226562 0 1679.6934
854.3130493164062 0 10765.343 y 2
855.3161010742188 0 5344.99
856.3153686523438 0 1926.4319
905.342041015625 0 3691.4067
906.3433227539062 0 1331.9644
951.3290405273438 0 881.42786 y Water loss 1
969.3395385742188 0 7348.2905 y 1
970.3425903320312 0 4437.9434
971.343017578125 0 1966.5715
1228.517333984375 0 583.11224
1650.2099609375 0 738.52026
1826.3094482421875 0 634.04865
2008.4052734375 0 656.413
3199.4462890625 0 681.6169
3214.39404296875 0 792.1625

Spectrum Details

|  |  |
| --- | --- |
| Matched peaks? Matched peaks The total absolute number of peaks matched. Additionally in brackets the total fraction of peaks matched and the total number of peaks is shown. | 15 (8.88% of 169) |
| FDR? FDR The false discovery rate estimated for this peptide. It is calculated by matching all theoretical fragments with a non-integer shift with the raw peaks for this spectrum. This is done with 40 different shifts. The resulting percentage is the average number of annotated peaks over the number of annotated peaks with the correct spectrum. | 0.16% |
| Satellite FDR? Satellite FDR See the FDR for details on its calculation. This satellite ion specific FDR only contains the satellite ions (d/w) for I/L/J positions. | - |
| PSM Score? PSM Score The PSM Score as given by Hecklib to this annotated spectrum. It is shown with three significant figures. | 152 |

#### Reverse Lookup? Reverse Lookup All places where this read could be placed.

| Group | Segment | Template | Template Part | Read Part | Score | Unique |
| --- | --- | --- | --- | --- | --- | --- |
| Homo sapiens Heavy Chain | IGHV | IGHV3-9 | [28..36] | [0..8] | 64 | True |

| Recombined | Template Part | Read Part | Score | Unique |
| --- | --- | --- | --- | --- |
| REC-0-1 | [28..36] | [0..8] | 64 | True |

#### Meta Information from Multiple reads

##### Number of combined reads

5

##### Intensity

0.5832

##### TotalArea

5.639E+07

#### Positional Score

Copy Data

##### Positional Score (TSV)

###### Preview

```
Loading example...
```

*Click on the button to copy the data to your clipboard.*

1001234567

Label Value
"0" 0.598
"1" 0.6
"2" 0.6
"3" 0.596
"4" 0.59
"5" 0.588
"6" 0.562
"7" 0.576

#### Meta Information from PEAKS

##### Scan Identifier

F2:5490

##### Original sequence

F

D

D

Y

A

M

+15.99

H

W

+15.99

##### Posttranslational Modifications

Oxidation (M); Oxidation (HW)

##### Source File

D:\separate\_stitch\_analyses\xle-disambiguation\raw\20210323\_F1\_UM1\_Peng0013\_SA\_F59\_ingel\_3ug\_TL.raw

##### Fraction

2

##### Scan Feature

F2:6626

##### De Novo Score

99

##### ConfidenceScore

99

### m/z

558.7093

##### Mass

1115.4019

##### Charge

2

##### Retention Time

29.64

##### Predicted Retention Time

-

##### Area

1.981E+06

##### Parts Per Million

1.9

##### Fragmentation mode

HCD

##### Originating file

01 D:\separate\_stitch\_analyses\xle-disambiguation\20210325\_F59\_3ug\_DENOVO\_12.csv

#### Meta Information from PEAKS

##### Scan Identifier

F2:6681

##### Original sequence

F

D

D

Y

A

M

+15.99

H

W

##### Posttranslational Modifications

Oxidation (M)

##### Source File

D:\separate\_stitch\_analyses\xle-disambiguation\raw\20210323\_F1\_UM1\_Peng0013\_SA\_F59\_ingel\_3ug\_TL.raw

##### Fraction

2

##### Scan Feature

F2:6291

##### De Novo Score

99

##### ConfidenceScore

99

### m/z

550.7115

##### Mass

1099.407

##### Charge

2

##### Retention Time

36.67

##### Predicted Retention Time

-

##### Area

1.748E+07

##### Parts Per Million

1.4

##### Fragmentation mode

HCD

##### Originating file

01 D:\separate\_stitch\_analyses\xle-disambiguation\20210325\_F59\_3ug\_DENOVO\_12.csv

#### Meta Information from PEAKS

##### Scan Identifier

F2:6738

##### Original sequence

F

D

D

Y

A

M

+15.99

H

W

##### Posttranslational Modifications

Oxidation (M)

##### Source File

D:\separate\_stitch\_analyses\xle-disambiguation\raw\20210323\_F1\_UM1\_Peng0013\_SA\_F59\_ingel\_3ug\_TL.raw

##### Fraction

2

##### Scan Feature

F2:6291

##### De Novo Score

98

##### ConfidenceScore

98

### m/z

550.7115

##### Mass

1099.407

##### Charge

2

##### Retention Time

36.67

##### Predicted Retention Time

-

##### Area

1.748E+07

##### Parts Per Million

1.4

##### Fragmentation mode

HCD

##### Originating file

01 D:\separate\_stitch\_analyses\xle-disambiguation\20210325\_F59\_3ug\_DENOVO\_12.csv

#### Meta Information from PEAKS

##### Scan Identifier

F2:6797

##### Original sequence

F

D

D

Y

A

M

+15.99

H

W

##### Posttranslational Modifications

Oxidation (M)

##### Source File

D:\separate\_stitch\_analyses\xle-disambiguation\raw\20210323\_F1\_UM1\_Peng0013\_SA\_F59\_ingel\_3ug\_TL.raw

##### Fraction

2

##### Scan Feature

F2:6291

##### De Novo Score

98

##### ConfidenceScore

98

### m/z

550.7115

##### Mass

1099.407

##### Charge

2

##### Retention Time

36.67

##### Predicted Retention Time

-

##### Area

1.748E+07

##### Parts Per Million

1.4

##### Fragmentation mode

HCD

##### Originating file

01 D:\separate\_stitch\_analyses\xle-disambiguation\20210325\_F59\_3ug\_DENOVO\_12.csv

#### Meta Information from PEAKS

##### Scan Identifier

F2:5428

##### Original sequence

F

D

D

Y

A

M

+15.99

H

W

+15.99

##### Posttranslational Modifications

Oxidation (M); Oxidation (HW)

##### Source File

D:\separate\_stitch\_analyses\xle-disambiguation\raw\20210323\_F1\_UM1\_Peng0013\_SA\_F59\_ingel\_3ug\_TL.raw

##### Fraction

2

##### Scan Feature

F2:6626

##### De Novo Score

97

##### ConfidenceScore

97

### m/z

558.7093

##### Mass

1115.4019

##### Charge

2

##### Retention Time

29.64

##### Predicted Retention Time

-

##### Area

1.981E+06

##### Parts Per Million

1.9

##### Fragmentation mode

HCD

##### Originating file

01 D:\separate\_stitch\_analyses\xle-disambiguation\20210325\_F59\_3ug\_DENOVO\_12.csv
