## Supplementary material for "A handle on mass coincidence errors in *de novo* sequencing of antibodies by bottom-up proteomics": Combined_032.html

Details Combined\_032 | Stitch OverviewUndefined

### Read Combined\_032

#### Sequence (length=13)

SCSVMHEAJHNHY

#### Spectrum 2789? Spectrum 2789 The raw spectrum of this peptide as annotated by Hecklib. The fragments are coloured according to ion type (see legend). Any peaks with a star '\*' as text can be hovered over to see the full details, first the ion type second the mass shift type. By hovering over the amino acids in the peptide or ions in the legend the corresponding peaks are highlighted. By toggling the 'Unassigned' label you can turn the background (unassigned) peaks on or off in the plot. By updating the slider in the Ion legend you can update the spectrum to only show the top X% of the peaks with labels. The top X% means any peak that is within X% of the highest intensity. By dragging in the spectrum you can zoom in to a specific part of the spectrum and use 'Zoom Out' to get back to the original zoom level. The annotation of the spectrum is based on the given sequence in the peptides file and is done with different software so inconsistencies are likely. The peaks are annotated based on the given sequence, with 20 ppm tolerance.

Copy Data

##### Spectrum 2789 (TSV)

###### Preview

```
Loading example...
```

*Click on the button to copy the data to your clipboard.*

Mz MinMz MaxIntensity Max

WidthHeightPeptide font sizePeptide stroke widthSpectrum font sizeSpectrum stroke widthCompact peptide

Ion legend

wxyz

abcd

OtherUnassignedIon^Charge^_Position_Show for top:%

SCSVMHEAJHNHY

09.05e+51.81e+62.72e+63.62e+6

Zoom Out

y^+1^_1_a^+1^_2_a^+1^_2_b^+1^_2_b^+1^_2_y^+2^_4_y^+2^_4_b^+1^_3_y^+1^_2_y^+3^_8_b^+1^_3_y^+3^_8_y^+2^_5_b^+2^_6_y^+2^_6_b^+2^_7_y^+1^_3_b^+1^_4_y^+3^_10_b^+2^_7_y^+1^_3_b^+1^_4_y^+2^_7_y^+3^_11_y^+3^_11_y^+3^_11_b^+2^_8_b^+3^_12_y^+3^_12_y^+3^_12_y^+2^_8_y^+2^_8_y^+3^_12_y^+2^_8_\*\*\*y^+1^_4_y^+1^_4_y^+2^_9_y^+2^_9_b^+2^_10_y^+2^_9_y^+2^_10_y^+2^_10_y^+2^_10_b^+2^_11_y^+1^_5_y^+2^_11_y^+2^_11_y^+2^_11_y^+1^_5_b^+2^_12_b^+1^_6_y^+1^_6_y^+2^_12_y^+2^_12_y^+1^_6_y^+2^_12_b^+1^_7_y^+1^_7_y^+1^_7_y^+1^_7_b^+1^_8_y^+1^_8_y^+1^_8_y^+1^_8_b^+1^_9_b^+1^_10_y^+1^_9_b^+1^_10_b^+1^_11_

0649129719462595

Fragment Matches Table

Show background peaks

| Position | Ion type | Intensity | mz Theoretical | mz Error (Th) | mz Error (ppm) | Charge | Series Number |
| --- | --- | --- | --- | --- | --- | --- | --- |
| - | - | 7599 | 121 | - | - | 0 | - |
| - | - | 1.184E+04 | 122.1 | - | - | 0 | - |
| - | - | 1.623E+04 | 123 | - | - | 0 | - |
| - | - | 1.382E+04 | 124.1 | - | - | 0 | - |
| - | - | 2.738E+04 | 127.1 | - | - | 0 | - |
| - | - | 1.265E+05 | 129.1 | - | - | 0 | - |
| - | - | 3.91E+04 | 129.1 | - | - | 0 | - |
| - | - | 4957 | 129.3 | - | - | 0 | - |
| - | - | 3.826E+04 | 130 | - | - | 0 | - |
| - | - | 7331 | 130.1 | - | - | 0 | - |
| - | - | 5974 | 130.6 | - | - | 0 | - |
| - | - | 3.82E+04 | 131.1 | - | - | 0 | - |
| - | - | 5096 | 132.2 | - | - | 0 | - |
| - | - | 1.058E+04 | 133 | - | - | 0 | - |
| - | - | 6.453E+05 | 134 | - | - | 0 | - |
| - | - | 2.864E+04 | 135 | - | - | 0 | - |
| - | - | 2.68E+04 | 136 | - | - | 0 | - |
| - | - | 7240 | 136.1 | - | - | 0 | - |
| - | - | 4.647E+05 | 136.1 | - | - | 0 | - |
| - | - | 5175 | 136.2 | - | - | 0 | - |
| - | - | 6155 | 137.1 | - | - | 0 | - |
| - | - | 4.299E+04 | 137.1 | - | - | 0 | - |
| - | - | 3.969E+04 | 138.1 | - | - | 0 | - |
| - | - | 5342 | 139.1 | - | - | 0 | - |
| - | - | 2.642E+04 | 141.1 | - | - | 0 | - |
| - | - | 8749 | 143 | - | - | 0 | - |
| - | - | 5741 | 143.8 | - | - | 0 | - |
| - | - | 5289 | 144 | - | - | 0 | - |
| - | - | 1.206E+04 | 147 | - | - | 0 | - |
| - | - | 1.713E+04 | 148.1 | - | - | 0 | - |
| - | - | 6229 | 148.9 | - | - | 0 | - |
| - | - | 8407 | 148.9 | - | - | 0 | - |
| - | - | 7995 | 148.9 | - | - | 0 | - |
| - | - | 7751 | 148.9 | - | - | 0 | - |
| - | - | 7022 | 148.9 | - | - | 0 | - |
| - | - | 1.45E+04 | 148.9 | - | - | 0 | - |
| - | - | 1.668E+04 | 148.9 | - | - | 0 | - |
| - | - | 1.976E+04 | 148.9 | - | - | 0 | - |
| - | - | 4.514E+04 | 148.9 | - | - | 0 | - |
| - | - | 6.431E+04 | 149 | - | - | 0 | - |
| - | - | 4.424E+04 | 149 | - | - | 0 | - |
| - | - | 1.735E+04 | 149 | - | - | 0 | - |
| - | - | 1.671E+04 | 149 | - | - | 0 | - |
| - | - | 1.178E+04 | 149 | - | - | 0 | - |
| - | - | 9658 | 149 | - | - | 0 | - |
| - | - | 7085 | 149 | - | - | 0 | - |
| - | - | 6703 | 149 | - | - | 0 | - |
| - | - | 5694 | 149 | - | - | 0 | - |
| - | - | 5813 | 149 | - | - | 0 | - |
| - | - | 1.789E+04 | 149 | - | - | 0 | - |
| - | - | 2.705E+04 | 152.1 | - | - | 0 | - |
| - | - | 7529 | 154.1 | - | - | 0 | - |
| - | - | 6289 | 155.1 | - | - | 0 | - |
| - | - | 3.734E+04 | 155.1 | - | - | 0 | - |
| - | - | 1.753E+04 | 155.1 | - | - | 0 | - |
| - | - | 1.079E+05 | 156.1 | - | - | 0 | - |
| - | - | 6682 | 156.1 | - | - | 0 | - |
| - | - | 1.223E+05 | 157.1 | - | - | 0 | - |
| - | - | 1.033E+04 | 157.1 | - | - | 0 | - |
| - | - | 5934 | 157.1 | - | - | 0 | - |
| - | - | 3.594E+05 | 158 | - | - | 0 | - |
| - | - | 6087 | 158.1 | - | - | 0 | - |
| - | - | 1.445E+04 | 159 | - | - | 0 | - |
| - | - | 8091 | 159.1 | - | - | 0 | - |
| - | - | 4.578E+05 | 159.1 | - | - | 0 | - |
| - | - | 1.105E+04 | 160 | - | - | 0 | - |
| - | - | 4.218E+04 | 160.1 | - | - | 0 | - |
| - | - | 1.884E+05 | 161 | - | - | 0 | - |
| - | - | 1.344E+04 | 162 | - | - | 0 | - |
| - | - | 1.064E+04 | 162 | - | - | 0 | - |
| - | - | 6250 | 163.1 | - | - | 0 | - |
| - | - | 5969 | 164.1 | - | - | 0 | - |
| - | - | 1.496E+05 | 165.1 | - | - | 0 | - |
| - | - | 3.583E+04 | 165.1 | - | - | 0 | - |
| - | - | 5.68E+04 | 166.1 | - | - | 0 | - |
| - | - | 1.018E+05 | 166.1 | - | - | 0 | - |
| - | - | 2.274E+04 | 167.1 | - | - | 0 | - |
| - | - | 5.19E+04 | 168 | - | - | 0 | - |
| - | - | 7715 | 169.1 | - | - | 0 | - |
| - | - | 4823 | 170.9 | - | - | 0 | - |
| - | - | 8922 | 171.1 | - | - | 0 | - |
| - | - | 7727 | 173.1 | - | - | 0 | - |
| - | - | 6702 | 174 | - | - | 0 | - |
| - | - | 1.213E+04 | 174.1 | - | - | 0 | - |
| - | - | 8624 | 175.1 | - | - | 0 | - |
| - | - | 4.084E+05 | 176 | - | - | 0 | - |
| - | - | 6.125E+04 | 176.1 | - | - | 0 | - |
| - | - | 2.89E+04 | 177 | - | - | 0 | - |
| - | - | 1.091E+04 | 177.1 | - | - | 0 | - |
| - | - | 1.659E+04 | 178 | - | - | 0 | - |
| - | - | 1.439E+04 | 179.1 | - | - | 0 | - |
| - | - | 1.433E+04 | 180.1 | - | - | 0 | - |
| - | - | 1.262E+04 | 181.1 | - | - | 0 | - |
| 13 | y | 2.969E+05 | 182.1 | 0.0005106 | 2.804 | +1 | 1 |
| - | - | 7627 | 183.1 | - | - | 0 | - |
| - | - | 2.674E+04 | 183.1 | - | - | 0 | - |
| - | - | 2.523E+04 | 183.1 | - | - | 0 | - |
| - | - | 6347 | 184 | - | - | 0 | - |
| - | - | 2.409E+04 | 185 | - | - | 0 | - |
| - | - | 7280 | 185.1 | - | - | 0 | - |
| - | - | 3.318E+05 | 186 | - | - | 0 | - |
| - | - | 8972 | 186.1 | - | - | 0 | - |
| - | - | 1.751E+04 | 187 | - | - | 0 | - |
| - | - | 2.163E+05 | 187.1 | - | - | 0 | - |
| - | - | 1.586E+04 | 188 | - | - | 0 | - |
| - | - | 1.459E+04 | 188.1 | - | - | 0 | - |
| - | - | 2.708E+04 | 189.1 | - | - | 0 | - |
| - | - | 5.54E+04 | 190.1 | - | - | 0 | - |
| - | - | 8311 | 191.1 | - | - | 0 | - |
| - | - | 3.06E+04 | 191.1 | - | - | 0 | - |
| - | - | 1.291E+04 | 193.1 | - | - | 0 | - |
| - | - | 1.12E+05 | 193.1 | - | - | 0 | - |
| - | - | 1.081E+04 | 194.1 | - | - | 0 | - |
| - | - | 1.174E+04 | 194.1 | - | - | 0 | - |
| - | - | 7269 | 194.1 | - | - | 0 | - |
| - | - | 1.185E+04 | 195.1 | - | - | 0 | - |
| - | - | 1.434E+04 | 196.1 | - | - | 0 | - |
| - | - | 8559 | 198.1 | - | - | 0 | - |
| - | - | 3.665E+04 | 199.1 | - | - | 0 | - |
| - | - | 1.112E+04 | 200.1 | - | - | 0 | - |
| - | - | 2.129E+04 | 201.1 | - | - | 0 | - |
| - | - | 1.467E+04 | 202.1 | - | - | 0 | - |
| 2 | a | 7.998E+04 | 203.1 | 0.004029 | 19.84 | +1 | 2 |
| - | - | 1.306E+04 | 203.1 | - | - | 0 | - |
| - | - | 6473 | 203.1 | - | - | 0 | - |
| - | - | 7.762E+04 | 204.1 | - | - | 0 | - |
| - | - | 1.006E+04 | 205.1 | - | - | 0 | - |
| - | - | 7968 | 205.1 | - | - | 0 | - |
| - | - | 1.204E+04 | 206.1 | - | - | 0 | - |
| - | - | 1.338E+05 | 207.1 | - | - | 0 | - |
| - | - | 6838 | 208.1 | - | - | 0 | - |
| - | - | 1.374E+04 | 210.1 | - | - | 0 | - |
| - | - | 1.939E+04 | 211.1 | - | - | 0 | - |
| - | - | 8526 | 213 | - | - | 0 | - |
| - | - | 2.12E+04 | 216.1 | - | - | 0 | - |
| - | - | 8982 | 216.1 | - | - | 0 | - |
| - | - | 2.798E+04 | 217.1 | - | - | 0 | - |
| - | - | 3.558E+04 | 218.1 | - | - | 0 | - |
| - | - | 9950 | 219.1 | - | - | 0 | - |
| 2 | a | 3.585E+06 | 221.1 | 0.003943 | 17.84 | +1 | 2 |
| - | - | 2.931E+05 | 221.1 | - | - | 0 | - |
| - | - | 2.628E+05 | 222.1 | - | - | 0 | - |
| - | - | 7782 | 222.1 | - | - | 0 | - |
| - | - | 3.537E+04 | 222.1 | - | - | 0 | - |
| - | - | 1.232E+05 | 223.1 | - | - | 0 | - |
| - | - | 1.795E+04 | 223.1 | - | - | 0 | - |
| - | - | 2.399E+04 | 223.2 | - | - | 0 | - |
| - | - | 7974 | 224.1 | - | - | 0 | - |
| - | - | 1.312E+05 | 224.1 | - | - | 0 | - |
| - | - | 8261 | 225 | - | - | 0 | - |
| - | - | 1.283E+04 | 225.1 | - | - | 0 | - |
| - | - | 7662 | 226.1 | - | - | 0 | - |
| - | - | 1.141E+05 | 226.1 | - | - | 0 | - |
| - | - | 9299 | 226.1 | - | - | 0 | - |
| - | - | 1.831E+04 | 227.1 | - | - | 0 | - |
| - | - | 9434 | 228.1 | - | - | 0 | - |
| - | - | 1.533E+04 | 230.1 | - | - | 0 | - |
| 2 | b | 1.118E+05 | 231 | 0.003949 | 17.09 | +1 | 2 |
| - | - | 1.036E+04 | 232 | - | - | 0 | - |
| - | - | 2.309E+04 | 233.1 | - | - | 0 | - |
| - | - | 2.737E+04 | 234.1 | - | - | 0 | - |
| - | - | 2.589E+05 | 235.1 | - | - | 0 | - |
| - | - | 1.5E+04 | 236.1 | - | - | 0 | - |
| - | - | 3.134E+04 | 236.1 | - | - | 0 | - |
| - | - | 1.267E+04 | 238.1 | - | - | 0 | - |
| - | - | 1.329E+04 | 239.1 | - | - | 0 | - |
| - | - | 7.697E+04 | 239.1 | - | - | 0 | - |
| - | - | 2.667E+05 | 244.1 | - | - | 0 | - |
| - | - | 6393 | 244.3 | - | - | 0 | - |
| - | - | 2.998E+04 | 245.1 | - | - | 0 | - |
| - | - | 2.738E+04 | 247.1 | - | - | 0 | - |
| 2 | b | 9.624E+05 | 249.1 | 0.003924 | 15.75 | +1 | 2 |
| - | - | 7.053E+04 | 249.1 | - | - | 0 | - |
| - | - | 9390 | 249.1 | - | - | 0 | - |
| - | - | 7.386E+04 | 250.1 | - | - | 0 | - |
| - | - | 3.247E+04 | 251.1 | - | - | 0 | - |
| - | - | 7.415E+04 | 251.2 | - | - | 0 | - |
| - | - | 5.229E+05 | 252.1 | - | - | 0 | - |
| - | - | 2.569E+04 | 252.1 | - | - | 0 | - |
| - | - | 1.304E+04 | 252.2 | - | - | 0 | - |
| - | - | 5.194E+04 | 253.1 | - | - | 0 | - |
| - | - | 5.71E+04 | 253.1 | - | - | 0 | - |
| - | - | 8492 | 254.1 | - | - | 0 | - |
| - | - | 4.967E+04 | 257.1 | - | - | 0 | - |
| - | - | 1.091E+04 | 262.1 | - | - | 0 | - |
| - | - | 1.389E+04 | 267.1 | - | - | 0 | - |
| - | - | 1.902E+05 | 267.1 | - | - | 0 | - |
| - | - | 2.528E+04 | 268.1 | - | - | 0 | - |
| - | - | 1.155E+04 | 268.2 | - | - | 0 | - |
| - | - | 5.014E+04 | 270.1 | - | - | 0 | - |
| - | - | 3.28E+04 | 270.1 | - | - | 0 | - |
| - | - | 6.203E+04 | 273.1 | - | - | 0 | - |
| - | - | 7960 | 274.1 | - | - | 0 | - |
| - | - | 9122 | 275.1 | - | - | 0 | - |
| - | - | 5.407E+04 | 275.1 | - | - | 0 | - |
| - | - | 9080 | 277.1 | - | - | 0 | - |
| 10 | y | 1.634E+04 | 277.1 | 0.0003397 | 1.226 | +2 | 4 |
| - | - | 1.584E+04 | 279.1 | - | - | 0 | - |
| - | - | 2.875E+04 | 280.1 | - | - | 0 | - |
| - | - | 1.066E+04 | 281.1 | - | - | 0 | - |
| - | - | 8318 | 284.1 | - | - | 0 | - |
| - | - | 1.253E+05 | 285.1 | - | - | 0 | - |
| 10 | y | 1.064E+05 | 285.6 | 0.0005845 | 2.046 | +2 | 4 |
| - | - | 1.693E+04 | 286.1 | - | - | 0 | - |
| - | - | 3.742E+04 | 286.1 | - | - | 0 | - |
| - | - | 1.011E+04 | 287.2 | - | - | 0 | - |
| - | - | 8786 | 292.2 | - | - | 0 | - |
| - | - | 1.694E+04 | 295.1 | - | - | 0 | - |
| - | - | 6587 | 296.7 | - | - | 0 | - |
| - | - | 7692 | 299.1 | - | - | 0 | - |
| - | - | 1.333E+05 | 301.1 | - | - | 0 | - |
| - | - | 1.171E+04 | 302.1 | - | - | 0 | - |
| - | - | 1.242E+04 | 302.2 | - | - | 0 | - |
| - | - | 1.122E+04 | 304.1 | - | - | 0 | - |
| - | - | 2.557E+04 | 304.2 | - | - | 0 | - |
| - | - | 1.458E+04 | 309.1 | - | - | 0 | - |
| - | - | 1.127E+04 | 313.1 | - | - | 0 | - |
| - | - | 1.508E+04 | 316.1 | - | - | 0 | - |
| 3 | b | 1.444E+05 | 318.1 | 0.003751 | 11.79 | +1 | 3 |
| 12 | y | 5.954E+05 | 319.1 | 0.0007876 | 2.468 | +1 | 2 |
| - | - | 1.287E+05 | 320.1 | - | - | 0 | - |
| - | - | 3.194E+04 | 320.2 | - | - | 0 | - |
| - | - | 2.033E+04 | 321.1 | - | - | 0 | - |
| - | - | 8232 | 322.1 | - | - | 0 | - |
| - | - | 6.898E+04 | 322.2 | - | - | 0 | - |
| - | - | 7549 | 323.2 | - | - | 0 | - |
| - | - | 1.457E+04 | 323.2 | - | - | 0 | - |
| - | - | 2.127E+04 | 326.1 | - | - | 0 | - |
| - | - | 5.56E+04 | 329.1 | - | - | 0 | - |
| - | - | 6.571E+04 | 330.1 | - | - | 0 | - |
| - | - | 8799 | 331.1 | - | - | 0 | - |
| - | - | 2.648E+04 | 332.1 | - | - | 0 | - |
| - | - | 1.01E+05 | 334.1 | - | - | 0 | - |
| 6 | y | 1.851E+04 | 335.2 | 0.004293 | 12.81 | +3 | 8 |
| 3 | b | 1.11E+05 | 336.1 | 0.003756 | 11.18 | +1 | 3 |
| - | - | 1.875E+04 | 337.1 | - | - | 0 | - |
| - | - | 2.146E+04 | 337.1 | - | - | 0 | - |
| - | - | 1.22E+05 | 338.1 | - | - | 0 | - |
| - | - | 1.581E+04 | 339.1 | - | - | 0 | - |
| - | - | 2.099E+04 | 339.2 | - | - | 0 | - |
| 6 | y | 8501 | 340.8 | 0.001312 | 3.85 | +3 | 8 |
| 9 | y | 2.984E+04 | 342.2 | 0.001002 | 2.93 | +2 | 5 |
| - | - | 1.202E+04 | 342.7 | - | - | 0 | - |
| - | - | 1.329E+04 | 344.1 | - | - | 0 | - |
| - | - | 3.22E+04 | 344.1 | - | - | 0 | - |
| - | - | 8731 | 345.2 | - | - | 0 | - |
| - | - | 8271 | 347.2 | - | - | 0 | - |
| - | - | 4.234E+04 | 348.1 | - | - | 0 | - |
| - | - | 1.946E+04 | 348.2 | - | - | 0 | - |
| - | - | 1.098E+05 | 350.1 | - | - | 0 | - |
| - | - | 2.575E+04 | 351.1 | - | - | 0 | - |
| - | - | 9662 | 351.7 | - | - | 0 | - |
| - | - | 1.273E+04 | 353.1 | - | - | 0 | - |
| - | - | 1.134E+04 | 353.2 | - | - | 0 | - |
| - | - | 2.299E+04 | 354.1 | - | - | 0 | - |
| - | - | 1.133E+04 | 355.1 | - | - | 0 | - |
| - | - | 8287 | 355.2 | - | - | 0 | - |
| - | - | 6312 | 358.8 | - | - | 0 | - |
| 6 | b | 1.57E+04 | 360.1 | 0.0006767 | 1.879 | +2 | 6 |
| - | - | 3.094E+04 | 361.2 | - | - | 0 | - |
| - | - | 5.302E+04 | 365.2 | - | - | 0 | - |
| - | - | 4.943E+04 | 366.1 | - | - | 0 | - |
| - | - | 7577 | 368.8 | - | - | 0 | - |
| - | - | 1.235E+05 | 371.2 | - | - | 0 | - |
| - | - | 7.871E+04 | 372.1 | - | - | 0 | - |
| - | - | 1.651E+04 | 372.2 | - | - | 0 | - |
| - | - | 1.929E+04 | 373.1 | - | - | 0 | - |
| - | - | 1.046E+04 | 375.2 | - | - | 0 | - |
| - | - | 8639 | 376.1 | - | - | 0 | - |
| 8 | y | 3.641E+04 | 377.7 | 0.0008781 | 2.325 | +2 | 6 |
| - | - | 2.328E+04 | 378.2 | - | - | 0 | - |
| - | - | 4.437E+04 | 379.2 | - | - | 0 | - |
| - | - | 1.207E+04 | 380.2 | - | - | 0 | - |
| - | - | 7.215E+04 | 381.1 | - | - | 0 | - |
| - | - | 1.923E+04 | 382.1 | - | - | 0 | - |
| - | - | 8621 | 384.2 | - | - | 0 | - |
| - | - | 1.009E+04 | 386.1 | - | - | 0 | - |
| - | - | 8321 | 387.2 | - | - | 0 | - |
| - | - | 1.593E+04 | 389.1 | - | - | 0 | - |
| - | - | 2.396E+04 | 389.2 | - | - | 0 | - |
| - | - | 1.34E+04 | 389.2 | - | - | 0 | - |
| - | - | 4.733E+04 | 390.2 | - | - | 0 | - |
| - | - | 8643 | 393.2 | - | - | 0 | - |
| - | - | 2.823E+04 | 393.2 | - | - | 0 | - |
| - | - | 7393 | 393.5 | - | - | 0 | - |
| - | - | 9178 | 395.2 | - | - | 0 | - |
| - | - | 1.104E+04 | 396.1 | - | - | 0 | - |
| - | - | 1.53E+04 | 397.7 | - | - | 0 | - |
| - | - | 3.327E+04 | 398.1 | - | - | 0 | - |
| - | - | 8.359E+04 | 399.1 | - | - | 0 | - |
| - | - | 2.207E+04 | 400.1 | - | - | 0 | - |
| - | - | 1.054E+04 | 402.1 | - | - | 0 | - |
| - | - | 9159 | 404.2 | - | - | 0 | - |
| - | - | 1.006E+04 | 405.2 | - | - | 0 | - |
| - | - | 3.413E+04 | 405.2 | - | - | 0 | - |
| - | - | 4.516E+04 | 406.2 | - | - | 0 | - |
| - | - | 2.092E+04 | 406.7 | - | - | 0 | - |
| - | - | 2.349E+04 | 407.2 | - | - | 0 | - |
| - | - | 5.55E+04 | 407.2 | - | - | 0 | - |
| - | - | 1.697E+04 | 408.2 | - | - | 0 | - |
| - | - | 1.658E+04 | 410.7 | - | - | 0 | - |
| - | - | 1.188E+04 | 412.2 | - | - | 0 | - |
| - | - | 9.431E+04 | 414.1 | - | - | 0 | - |
| - | - | 2.448E+04 | 415.2 | - | - | 0 | - |
| - | - | 9060 | 415.2 | - | - | 0 | - |
| 7 | b | 7428 | 415.6 | 1.285E-05 | 0.03091 | +2 | 7 |
| 11 | y | 1.701E+05 | 416.2 | 0.0009498 | 2.282 | +1 | 3 |
| 4 | b | 4.013E+04 | 417.1 | 0.002676 | 6.415 | +1 | 4 |
| - | - | 2.848E+04 | 417.2 | - | - | 0 | - |
| - | - | 2.274E+04 | 418.1 | - | - | 0 | - |
| - | - | 3.097E+04 | 420.2 | - | - | 0 | - |
| - | - | 2.435E+04 | 420.7 | - | - | 0 | - |
| - | - | 1.071E+05 | 421.2 | - | - | 0 | - |
| - | - | 1.016E+04 | 421.2 | - | - | 0 | - |
| - | - | 1.792E+04 | 422.2 | - | - | 0 | - |
| 4 | y | 1.945E+04 | 422.9 | 0.002434 | 5.757 | +3 | 10 |
| - | - | 1.378E+04 | 423.2 | - | - | 0 | - |
| - | - | 5.894E+04 | 423.2 | - | - | 0 | - |
| - | - | 1.346E+04 | 424.2 | - | - | 0 | - |
| - | - | 2.633E+04 | 424.5 | - | - | 0 | - |
| 7 | b | 1.732E+04 | 424.6 | 0.001475 | 3.473 | +2 | 7 |
| - | - | 7879 | 424.7 | - | - | 0 | - |
| - | - | 2.158E+04 | 424.9 | - | - | 0 | - |
| - | - | 9599 | 425.2 | - | - | 0 | - |
| - | - | 1.377E+04 | 425.2 | - | - | 0 | - |
| - | - | 1.036E+04 | 425.6 | - | - | 0 | - |
| - | - | 1.064E+04 | 426.2 | - | - | 0 | - |
| - | - | 2.441E+04 | 429.2 | - | - | 0 | - |
| - | - | 1.128E+04 | 429.7 | - | - | 0 | - |
| - | - | 1.09E+04 | 430.5 | - | - | 0 | - |
| - | - | 1.089E+04 | 431.2 | - | - | 0 | - |
| - | - | 9370 | 431.2 | - | - | 0 | - |
| 11 | y | 2.928E+05 | 433.2 | 0.001043 | 2.407 | +1 | 3 |
| - | - | 2.861E+04 | 433.2 | - | - | 0 | - |
| - | - | 1.354E+04 | 433.7 | - | - | 0 | - |
| - | - | 6.317E+04 | 434.2 | - | - | 0 | - |
| - | - | 8858 | 434.5 | - | - | 0 | - |
| 4 | b | 1.755E+04 | 435.2 | 0.003536 | 8.126 | +1 | 4 |
| - | - | 1.457E+04 | 435.2 | - | - | 0 | - |
| - | - | 7.818E+04 | 436.2 | - | - | 0 | - |
| - | - | 1.145E+04 | 437.2 | - | - | 0 | - |
| - | - | 2.158E+04 | 439.2 | - | - | 0 | - |
| - | - | 1.765E+04 | 440.2 | - | - | 0 | - |
| - | - | 9931 | 440.9 | - | - | 0 | - |
| - | - | 2.285E+04 | 440.9 | - | - | 0 | - |
| - | - | 1.425E+04 | 441.3 | - | - | 0 | - |
| - | - | 1.253E+04 | 441.6 | - | - | 0 | - |
| 7 | y | 2.397E+04 | 442.2 | 0.0007346 | 1.661 | +2 | 7 |
| - | - | 1.897E+04 | 442.7 | - | - | 0 | - |
| - | - | 1.548E+04 | 443.2 | - | - | 0 | - |
| - | - | 5.835E+04 | 443.2 | - | - | 0 | - |
| - | - | 9907 | 444.2 | - | - | 0 | - |
| - | - | 1.618E+04 | 444.2 | - | - | 0 | - |
| 3 | y | 1.373E+05 | 445.9 | 0.002695 | 6.045 | +3 | 11 |
| - | - | 2.074E+04 | 446.2 | - | - | 0 | - |
| 3 | y | 8.204E+04 | 446.2 | 0.008705 | 19.51 | +3 | 11 |
| - | - | 6.571E+04 | 446.5 | - | - | 0 | - |
| - | - | 1.992E+04 | 446.7 | - | - | 0 | - |
| - | - | 1.025E+04 | 446.9 | - | - | 0 | - |
| - | - | 1.653E+04 | 447.2 | - | - | 0 | - |
| - | - | 2.848E+04 | 447.7 | - | - | 0 | - |
| - | - | 1.939E+04 | 448.2 | - | - | 0 | - |
| - | - | 8689 | 449.2 | - | - | 0 | - |
| - | - | 1.694E+04 | 449.2 | - | - | 0 | - |
| - | - | 1.357E+05 | 451.2 | - | - | 0 | - |
| 3 | y | 1.499E+05 | 451.9 | 0.002836 | 6.276 | +3 | 11 |
| - | - | 8.922E+04 | 452.2 | - | - | 0 | - |
| - | - | 1.951E+04 | 452.2 | - | - | 0 | - |
| - | - | 3.003E+04 | 452.5 | - | - | 0 | - |
| - | - | 9197 | 453.2 | - | - | 0 | - |
| - | - | 8585 | 453.7 | - | - | 0 | - |
| - | - | 1.131E+04 | 457.2 | - | - | 0 | - |
| 8 | b | 2.971E+04 | 460.2 | 0.00129 | 2.802 | +2 | 8 |
| - | - | 2.958E+04 | 461.2 | - | - | 0 | - |
| - | - | 4.385E+04 | 461.7 | - | - | 0 | - |
| - | - | 2.187E+04 | 462.2 | - | - | 0 | - |
| - | - | 3.327E+04 | 464.9 | - | - | 0 | - |
| - | - | 2.272E+04 | 465.2 | - | - | 0 | - |
| - | - | 1.315E+04 | 465.5 | - | - | 0 | - |
| - | - | 9593 | 468.3 | - | - | 0 | - |
| - | - | 1.905E+04 | 468.9 | - | - | 0 | - |
| - | - | 2.673E+04 | 469.2 | - | - | 0 | - |
| - | - | 1.136E+04 | 469.5 | - | - | 0 | - |
| - | - | 8582 | 469.9 | - | - | 0 | - |
| - | - | 2.927E+04 | 470.7 | - | - | 0 | - |
| - | - | 6.494E+04 | 471.2 | - | - | 0 | - |
| - | - | 1.654E+04 | 472.2 | - | - | 0 | - |
| - | - | 1.192E+04 | 473.9 | - | - | 0 | - |
| 12 | b | 2.489E+04 | 474.2 | 0.0009458 | 1.995 | +3 | 12 |
| - | - | 2.084E+04 | 474.3 | - | - | 0 | - |
| - | - | 2.814E+04 | 474.5 | - | - | 0 | - |
| - | - | 1.391E+04 | 474.9 | - | - | 0 | - |
| - | - | 2.844E+04 | 476.6 | - | - | 0 | - |
| - | - | 2.141E+04 | 476.9 | - | - | 0 | - |
| - | - | 7652 | 477.2 | - | - | 0 | - |
| - | - | 2.012E+04 | 478.2 | - | - | 0 | - |
| - | - | 1.604E+04 | 478.5 | - | - | 0 | - |
| - | - | 1.263E+04 | 478.9 | - | - | 0 | - |
| - | - | 1.536E+04 | 479.2 | - | - | 0 | - |
| - | - | 5.452E+04 | 479.7 | - | - | 0 | - |
| - | - | 1.5E+04 | 480.2 | - | - | 0 | - |
| - | - | 1.308E+04 | 480.5 | - | - | 0 | - |
| - | - | 1.082E+04 | 480.7 | - | - | 0 | - |
| - | - | 1.281E+04 | 483.2 | - | - | 0 | - |
| - | - | 2.023E+04 | 484.2 | - | - | 0 | - |
| - | - | 1.654E+04 | 484.5 | - | - | 0 | - |
| - | - | 5.634E+04 | 485.2 | - | - | 0 | - |
| - | - | 1.436E+04 | 485.2 | - | - | 0 | - |
| - | - | 1.986E+04 | 486.2 | - | - | 0 | - |
| - | - | 1.073E+04 | 486.7 | - | - | 0 | - |
| - | - | 9630 | 487.7 | - | - | 0 | - |
| - | - | 1.224E+04 | 489.2 | - | - | 0 | - |
| - | - | 1.06E+04 | 489.2 | - | - | 0 | - |
| - | - | 1.28E+04 | 490.2 | - | - | 0 | - |
| - | - | 1.384E+04 | 490.2 | - | - | 0 | - |
| - | - | 2.04E+04 | 491.9 | - | - | 0 | - |
| - | - | 1.385E+04 | 492.2 | - | - | 0 | - |
| - | - | 2.191E+04 | 492.5 | - | - | 0 | - |
| - | - | 2.225E+04 | 493.2 | - | - | 0 | - |
| - | - | 5.409E+04 | 493.7 | - | - | 0 | - |
| - | - | 9236 | 493.9 | - | - | 0 | - |
| - | - | 3.417E+04 | 494.2 | - | - | 0 | - |
| - | - | 1.879E+04 | 494.7 | - | - | 0 | - |
| - | - | 1.623E+04 | 495.2 | - | - | 0 | - |
| - | - | 1.021E+04 | 495.2 | - | - | 0 | - |
| - | - | 1.109E+04 | 495.5 | - | - | 0 | - |
| - | - | 9817 | 496.2 | - | - | 0 | - |
| - | - | 1.433E+05 | 497.9 | - | - | 0 | - |
| - | - | 1.562E+05 | 498.2 | - | - | 0 | - |
| - | - | 6.446E+04 | 498.6 | - | - | 0 | - |
| - | - | 2.327E+04 | 498.9 | - | - | 0 | - |
| - | - | 8413 | 499.2 | - | - | 0 | - |
| 2 | y | 4.826E+04 | 499.5 | 0.001204 | 2.41 | +3 | 12 |
| 2 | y | 7.114E+04 | 499.9 | 0.005443 | 10.89 | +3 | 12 |
| - | - | 3.124E+04 | 500.2 | - | - | 0 | - |
| - | - | 1.789E+04 | 500.5 | - | - | 0 | - |
| 6 | y | 2.287E+04 | 501.7 | 0.0001163 | 0.2317 | +2 | 8 |
| - | - | 7235 | 501.9 | - | - | 0 | - |
| 6 | y | 5.285E+04 | 502.2 | 0.002188 | 4.357 | +2 | 8 |
| - | - | 1.708E+04 | 502.3 | - | - | 0 | - |
| - | - | 6.208E+04 | 502.7 | - | - | 0 | - |
| - | - | 2.784E+04 | 503.2 | - | - | 0 | - |
| - | - | 1.387E+04 | 503.7 | - | - | 0 | - |
| - | - | 9461 | 504.2 | - | - | 0 | - |
| 2 | y | 1.759E+05 | 505.5 | 0.0009781 | 1.935 | +3 | 12 |
| - | - | 1.316E+05 | 505.9 | - | - | 0 | - |
| - | - | 7.156E+04 | 506.2 | - | - | 0 | - |
| - | - | 1.029E+04 | 506.3 | - | - | 0 | - |
| - | - | 4.667E+04 | 506.5 | - | - | 0 | - |
| - | - | 9403 | 506.9 | - | - | 0 | - |
| - | - | 2.796E+04 | 507.2 | - | - | 0 | - |
| - | - | 2.467E+04 | 507.6 | - | - | 0 | - |
| - | - | 2.102E+04 | 507.9 | - | - | 0 | - |
| - | - | 2.288E+04 | 508.2 | - | - | 0 | - |
| - | - | 2.914E+04 | 508.3 | - | - | 0 | - |
| - | - | 7594 | 508.9 | - | - | 0 | - |
| 6 | y | 2.705E+05 | 510.7 | 0.0008764 | 1.716 | +2 | 8 |
| - | - | 1.716E+05 | 511.2 | - | - | 0 | - |
| - | - | 1.384E+04 | 511.5 | - | - | 0 | - |
| - | - | 6.66E+04 | 511.7 | - | - | 0 | - |
| - | - | 1.024E+04 | 512.2 | - | - | 0 | - |
| - | - | 2.102E+05 | 513.2 | - | - | 0 | - |
| - | - | 1.956E+05 | 513.6 | - | - | 0 | - |
| - | - | 9.972E+04 | 513.9 | - | - | 0 | - |
| - | - | 2.718E+04 | 514.2 | - | - | 0 | - |
| - | - | 1.316E+04 | 514.6 | - | - | 0 | - |
| - | - | 2.146E+04 | 516.5 | - | - | 0 | - |
| - | - | 3.625E+04 | 516.9 | - | - | 0 | - |
| - | - | 2.663E+04 | 517.2 | - | - | 0 | - |
| - | - | 1.16E+04 | 517.5 | - | - | 0 | - |
| - | - | 6.151E+04 | 518.2 | - | - | 0 | - |
| - | - | 2.7E+04 | 518.7 | - | - | 0 | - |
| - | - | 1.067E+04 | 519.2 | - | - | 0 | - |
| - | - | 5.139E+04 | 520.3 | - | - | 0 | - |
| - | - | 1.29E+04 | 522.2 | - | - | 0 | - |
| - | - | 3.031E+04 | 522.9 | - | - | 0 | - |
| - | - | 1.192E+04 | 523.2 | - | - | 0 | - |
| - | - | 1.204E+04 | 523.5 | - | - | 0 | - |
| - | - | 8742 | 523.7 | - | - | 0 | - |
| - | - | 1.274E+04 | 524.2 | - | - | 0 | - |
| - | - | 1.099E+04 | 528.3 | - | - | 0 | - |
| 0 | Precursor | 9.997E+04 | 528.6 | 0.001453 | 2.749 | +3 | -1 |
| - | - | 1.059E+04 | 528.6 | - | - | 0 | - |
| 0 | Precursor | 1.634E+05 | 528.9 | 0.005357 | 10.13 | +3 | -1 |
| - | - | 1.304E+05 | 529.2 | - | - | 0 | - |
| - | - | 2.74E+04 | 529.3 | - | - | 0 | - |
| - | - | 8.629E+04 | 529.5 | - | - | 0 | - |
| - | - | 8913 | 529.8 | - | - | 0 | - |
| - | - | 2.634E+04 | 529.9 | - | - | 0 | - |
| - | - | 1.576E+04 | 530.2 | - | - | 0 | - |
| - | - | 1.065E+04 | 531.2 | - | - | 0 | - |
| - | - | 1.669E+04 | 531.8 | - | - | 0 | - |
| - | - | 1.818E+04 | 532.3 | - | - | 0 | - |
| - | - | 9189 | 532.8 | - | - | 0 | - |
| - | - | 4.829E+04 | 534.3 | - | - | 0 | - |
| - | - | 1.115E+04 | 534.3 | - | - | 0 | - |
| 0 | Precursor | 1.707E+06 | 534.6 | 0.001044 | 1.953 | +3 | -1 |
| - | - | 1.918E+04 | 534.7 | - | - | 0 | - |
| - | - | 1.563E+06 | 534.9 | - | - | 0 | - |
| - | - | 9.098E+05 | 535.2 | - | - | 0 | - |
| - | - | 1.407E+04 | 535.3 | - | - | 0 | - |
| - | - | 4.012E+05 | 535.6 | - | - | 0 | - |
| - | - | 1.183E+04 | 535.7 | - | - | 0 | - |
| - | - | 1.378E+05 | 535.9 | - | - | 0 | - |
| - | - | 2.233E+04 | 536.2 | - | - | 0 | - |
| - | - | 9.127E+04 | 536.2 | - | - | 0 | - |
| - | - | 1.808E+04 | 537.2 | - | - | 0 | - |
| - | - | 1.174E+04 | 538.2 | - | - | 0 | - |
| - | - | 1.282E+04 | 538.7 | - | - | 0 | - |
| - | - | 2.453E+04 | 539.2 | - | - | 0 | - |
| - | - | 1.501E+04 | 540.2 | - | - | 0 | - |
| - | - | 5.736E+04 | 540.8 | - | - | 0 | - |
| - | - | 8359 | 541.2 | - | - | 0 | - |
| - | - | 4.133E+04 | 541.3 | - | - | 0 | - |
| - | - | 4.48E+04 | 543.3 | - | - | 0 | - |
| - | - | 8.125E+04 | 543.7 | - | - | 0 | - |
| - | - | 4.883E+04 | 544.2 | - | - | 0 | - |
| - | - | 1.256E+04 | 544.7 | - | - | 0 | - |
| - | - | 4.749E+04 | 545.3 | - | - | 0 | - |
| - | - | 3.243E+04 | 547.3 | - | - | 0 | - |
| - | - | 8763 | 548.3 | - | - | 0 | - |
| - | - | 2.432E+05 | 552.3 | - | - | 0 | - |
| - | - | 1.338E+05 | 552.8 | - | - | 0 | - |
| 10 | y | 1.213E+05 | 553.2 | 0.00173 | 3.128 | +1 | 4 |
| - | - | 3.691E+04 | 553.3 | - | - | 0 | - |
| - | - | 3.346E+04 | 554.2 | - | - | 0 | - |
| - | - | 4.405E+04 | 554.8 | - | - | 0 | - |
| - | - | 5.302E+04 | 555.3 | - | - | 0 | - |
| - | - | 2.106E+04 | 555.8 | - | - | 0 | - |
| - | - | 9519 | 556.3 | - | - | 0 | - |
| - | - | 9887 | 557.7 | - | - | 0 | - |
| - | - | 1.275E+04 | 558.2 | - | - | 0 | - |
| - | - | 3.019E+04 | 560.3 | - | - | 0 | - |
| - | - | 1.073E+04 | 561.7 | - | - | 0 | - |
| - | - | 5.849E+04 | 563.8 | - | - | 0 | - |
| - | - | 3.25E+04 | 564.3 | - | - | 0 | - |
| - | - | 2.78E+04 | 564.8 | - | - | 0 | - |
| - | - | 7.293E+04 | 565.3 | - | - | 0 | - |
| - | - | 2.76E+04 | 566.3 | - | - | 0 | - |
| - | - | 2.006E+04 | 566.7 | - | - | 0 | - |
| - | - | 2.616E+04 | 568.2 | - | - | 0 | - |
| 10 | y | 3.28E+05 | 570.2 | 0.000816 | 1.431 | +1 | 4 |
| - | - | 1.129E+05 | 571.2 | - | - | 0 | - |
| - | - | 7.367E+04 | 572.3 | - | - | 0 | - |
| - | - | 1.269E+05 | 572.8 | - | - | 0 | - |
| - | - | 1.074E+05 | 573.3 | - | - | 0 | - |
| - | - | 3.546E+04 | 573.8 | - | - | 0 | - |
| - | - | 3.488E+04 | 574.3 | - | - | 0 | - |
| 5 | y | 2.26E+04 | 575.2 | 0.003062 | 5.324 | +2 | 9 |
| 5 | y | 7.749E+04 | 575.7 | 0.005073 | 8.812 | +2 | 9 |
| 10 | b | 4.683E+04 | 576.2 | 0.01089 | 18.9 | +2 | 10 |
| - | - | 2.007E+04 | 576.7 | - | - | 0 | - |
| - | - | 1.943E+04 | 577.8 | - | - | 0 | - |
| - | - | 1.298E+04 | 578.3 | - | - | 0 | - |
| - | - | 2.246E+04 | 578.8 | - | - | 0 | - |
| - | - | 1.192E+04 | 581.3 | - | - | 0 | - |
| - | - | 1.399E+04 | 582.2 | - | - | 0 | - |
| 5 | y | 4.659E+05 | 584.3 | 0.003334 | 5.707 | +2 | 9 |
| - | - | 2.653E+05 | 584.8 | - | - | 0 | - |
| - | - | 1.117E+05 | 585.3 | - | - | 0 | - |
| - | - | 3.667E+04 | 585.8 | - | - | 0 | - |
| - | - | 1.37E+05 | 586.8 | - | - | 0 | - |
| - | - | 1.163E+05 | 587.3 | - | - | 0 | - |
| - | - | 6.26E+04 | 587.8 | - | - | 0 | - |
| - | - | 4.91E+04 | 588.3 | - | - | 0 | - |
| - | - | 3.787E+04 | 589.3 | - | - | 0 | - |
| - | - | 2.444E+04 | 591.3 | - | - | 0 | - |
| - | - | 2.203E+04 | 593.3 | - | - | 0 | - |
| - | - | 1.354E+04 | 593.8 | - | - | 0 | - |
| - | - | 1.152E+05 | 595.8 | - | - | 0 | - |
| - | - | 6.152E+04 | 596.3 | - | - | 0 | - |
| - | - | 2.108E+04 | 596.8 | - | - | 0 | - |
| - | - | 4.331E+04 | 598.3 | - | - | 0 | - |
| - | - | 1.218E+05 | 600.2 | - | - | 0 | - |
| - | - | 1.298E+04 | 600.3 | - | - | 0 | - |
| - | - | 4.9E+04 | 601.2 | - | - | 0 | - |
| - | - | 9.357E+04 | 601.8 | - | - | 0 | - |
| - | - | 8399 | 602.2 | - | - | 0 | - |
| - | - | 9.01E+04 | 602.3 | - | - | 0 | - |
| - | - | 3.431E+04 | 602.8 | - | - | 0 | - |
| - | - | 1.141E+04 | 603.3 | - | - | 0 | - |
| - | - | 1.926E+04 | 604.2 | - | - | 0 | - |
| - | - | 9441 | 604.8 | - | - | 0 | - |
| - | - | 1.413E+04 | 605.4 | - | - | 0 | - |
| - | - | 9.635E+04 | 607.3 | - | - | 0 | - |
| - | - | 2.542E+04 | 608.3 | - | - | 0 | - |
| - | - | 1.08E+04 | 610.8 | - | - | 0 | - |
| - | - | 1.167E+04 | 612.2 | - | - | 0 | - |
| - | - | 1.418E+04 | 616.3 | - | - | 0 | - |
| - | - | 1.524E+04 | 621.3 | - | - | 0 | - |
| - | - | 1.194E+04 | 622.3 | - | - | 0 | - |
| 4 | y | 1.214E+04 | 624.8 | 0.003401 | 5.444 | +2 | 10 |
| 4 | y | 3.982E+04 | 625.3 | 0.004497 | 7.191 | +2 | 10 |
| - | - | 2.506E+04 | 625.8 | - | - | 0 | - |
| - | - | 1.872E+04 | 626.3 | - | - | 0 | - |
| - | - | 1.486E+04 | 627.8 | - | - | 0 | - |
| - | - | 1.133E+04 | 628.8 | - | - | 0 | - |
| - | - | 2.034E+04 | 632.2 | - | - | 0 | - |
| 4 | y | 2.563E+05 | 633.8 | 0.003246 | 5.122 | +2 | 10 |
| - | - | 1.878E+05 | 634.3 | - | - | 0 | - |
| - | - | 7.538E+04 | 634.8 | - | - | 0 | - |
| - | - | 3.93E+04 | 635.3 | - | - | 0 | - |
| - | - | 1.004E+04 | 635.8 | - | - | 0 | - |
| - | - | 3.646E+04 | 636.3 | - | - | 0 | - |
| - | - | 9.704E+04 | 636.8 | - | - | 0 | - |
| - | - | 4.964E+04 | 637.3 | - | - | 0 | - |
| - | - | 3.039E+04 | 637.8 | - | - | 0 | - |
| - | - | 1.055E+04 | 639.3 | - | - | 0 | - |
| 11 | b | 4.4E+04 | 642.3 | 0.001844 | 2.871 | +2 | 11 |
| - | - | 1.884E+04 | 642.8 | - | - | 0 | - |
| - | - | 1.258E+04 | 643.3 | - | - | 0 | - |
| - | - | 1.423E+04 | 643.3 | - | - | 0 | - |
| - | - | 1.579E+04 | 644.3 | - | - | 0 | - |
| - | - | 1.612E+04 | 644.8 | - | - | 0 | - |
| - | - | 5.589E+05 | 645.3 | - | - | 0 | - |
| - | - | 4.101E+05 | 645.8 | - | - | 0 | - |
| - | - | 1.424E+05 | 646.3 | - | - | 0 | - |
| - | - | 2.688E+04 | 646.8 | - | - | 0 | - |
| - | - | 1.133E+04 | 648.3 | - | - | 0 | - |
| - | - | 1.037E+04 | 650.8 | - | - | 0 | - |
| - | - | 5.177E+04 | 653.3 | - | - | 0 | - |
| - | - | 2.594E+04 | 653.8 | - | - | 0 | - |
| - | - | 2.256E+04 | 654.3 | - | - | 0 | - |
| - | - | 1.506E+04 | 654.8 | - | - | 0 | - |
| - | - | 2.651E+04 | 657.3 | - | - | 0 | - |
| - | - | 4.656E+04 | 659.8 | - | - | 0 | - |
| - | - | 1.73E+04 | 660.3 | - | - | 0 | - |
| - | - | 1.518E+04 | 660.8 | - | - | 0 | - |
| - | - | 7.361E+04 | 660.9 | - | - | 0 | - |
| - | - | 1.088E+04 | 661.3 | - | - | 0 | - |
| - | - | 5.84E+04 | 661.4 | - | - | 0 | - |
| - | - | 2.922E+04 | 661.9 | - | - | 0 | - |
| - | - | 1.279E+04 | 664.8 | - | - | 0 | - |
| - | - | 1.637E+04 | 665.3 | - | - | 0 | - |
| 9 | y | 4.382E+04 | 666.3 | 0.002566 | 3.851 | +1 | 5 |
| - | - | 2.717E+04 | 667.3 | - | - | 0 | - |
| - | - | 2.326E+04 | 667.8 | - | - | 0 | - |
| 3 | y | 1.365E+05 | 668.3 | 0.002585 | 3.868 | +2 | 11 |
| 3 | y | 2.459E+05 | 668.8 | 0.00661 | 9.883 | +2 | 11 |
| - | - | 1.565E+05 | 669.3 | - | - | 0 | - |
| - | - | 1.009E+05 | 669.8 | - | - | 0 | - |
| - | - | 3.411E+04 | 670.3 | - | - | 0 | - |
| - | - | 1.139E+04 | 670.8 | - | - | 0 | - |
| - | - | 1.079E+05 | 671.3 | - | - | 0 | - |
| - | - | 4.27E+04 | 672.3 | - | - | 0 | - |
| - | - | 1.78E+04 | 673.3 | - | - | 0 | - |
| - | - | 4.404E+04 | 674.3 | - | - | 0 | - |
| - | - | 2.056E+04 | 675.3 | - | - | 0 | - |
| - | - | 1.132E+04 | 676.3 | - | - | 0 | - |
| - | - | 1.361E+04 | 676.8 | - | - | 0 | - |
| 3 | y | 1.953E+06 | 677.3 | 0.003589 | 5.299 | +2 | 11 |
| - | - | 1.43E+06 | 677.8 | - | - | 0 | - |
| - | - | 7.311E+05 | 678.3 | - | - | 0 | - |
| - | - | 2.171E+05 | 678.8 | - | - | 0 | - |
| - | - | 5.864E+04 | 679.3 | - | - | 0 | - |
| - | - | 3.858E+04 | 682.3 | - | - | 0 | - |
| - | - | 1.063E+04 | 682.4 | - | - | 0 | - |
| - | - | 3.405E+04 | 682.8 | - | - | 0 | - |
| 9 | y | 1.802E+05 | 683.3 | 0.0006143 | 0.899 | +1 | 5 |
| - | - | 9.902E+04 | 684.3 | - | - | 0 | - |
| - | - | 3.252E+04 | 685.3 | - | - | 0 | - |
| - | - | 1.452E+04 | 686.3 | - | - | 0 | - |
| - | - | 1.492E+04 | 687.8 | - | - | 0 | - |
| - | - | 1.486E+04 | 688.3 | - | - | 0 | - |
| - | - | 1.073E+04 | 688.8 | - | - | 0 | - |
| - | - | 1.875E+04 | 691.3 | - | - | 0 | - |
| - | - | 1.873E+04 | 694.3 | - | - | 0 | - |
| - | - | 1.434E+04 | 696.4 | - | - | 0 | - |
| - | - | 7.275E+04 | 696.8 | - | - | 0 | - |
| - | - | 6.999E+04 | 697.3 | - | - | 0 | - |
| - | - | 2.455E+04 | 697.8 | - | - | 0 | - |
| - | - | 1.075E+04 | 698.3 | - | - | 0 | - |
| - | - | 9.851E+04 | 702.3 | - | - | 0 | - |
| - | - | 2.495E+04 | 702.8 | - | - | 0 | - |
| - | - | 5.233E+04 | 703.3 | - | - | 0 | - |
| - | - | 1.982E+04 | 704.3 | - | - | 0 | - |
| - | - | 9693 | 708.3 | - | - | 0 | - |
| - | - | 5.631E+04 | 710.4 | - | - | 0 | - |
| 12 | b | 1.219E+05 | 710.8 | 0.001441 | 2.027 | +2 | 12 |
| - | - | 3.859E+04 | 710.9 | - | - | 0 | - |
| - | - | 9.278E+04 | 711.3 | - | - | 0 | - |
| - | - | 3.075E+04 | 711.4 | - | - | 0 | - |
| - | - | 4.912E+04 | 711.8 | - | - | 0 | - |
| - | - | 1.603E+04 | 712.3 | - | - | 0 | - |
| - | - | 2.024E+04 | 716.8 | - | - | 0 | - |
| - | - | 2.915E+04 | 717.3 | - | - | 0 | - |
| - | - | 2.448E+04 | 717.8 | - | - | 0 | - |
| 6 | b | 2.079E+04 | 719.2 | 0.0002813 | 0.391 | +1 | 6 |
| - | - | 5.537E+04 | 719.8 | - | - | 0 | - |
| - | - | 3.301E+04 | 720.3 | - | - | 0 | - |
| - | - | 7.061E+04 | 720.4 | - | - | 0 | - |
| - | - | 1.626E+04 | 720.8 | - | - | 0 | - |
| - | - | 2.721E+04 | 721.4 | - | - | 0 | - |
| - | - | 9532 | 722.4 | - | - | 0 | - |
| - | - | 9.042E+04 | 725.8 | - | - | 0 | - |
| - | - | 6.651E+04 | 726.3 | - | - | 0 | - |
| - | - | 4.672E+04 | 726.8 | - | - | 0 | - |
| - | - | 2.445E+04 | 727.3 | - | - | 0 | - |
| - | - | 3.235E+04 | 735.3 | - | - | 0 | - |
| 8 | y | 8.322E+04 | 737.3 | 0.00189 | 2.564 | +1 | 6 |
| - | - | 2.955E+04 | 738.3 | - | - | 0 | - |
| - | - | 1.382E+04 | 739.4 | - | - | 0 | - |
| - | - | 1.096E+04 | 743.2 | - | - | 0 | - |
| 2 | y | 3.576E+04 | 748.8 | 0.0005986 | 0.7994 | +2 | 12 |
| 2 | y | 6.773E+04 | 749.3 | 0.007943 | 10.6 | +2 | 12 |
| - | - | 4.738E+04 | 749.8 | - | - | 0 | - |
| - | - | 1.242E+04 | 750.3 | - | - | 0 | - |
| 8 | y | 2.839E+05 | 754.4 | 0.001037 | 1.375 | +1 | 6 |
| - | - | 1.197E+05 | 755.4 | - | - | 0 | - |
| - | - | 4.786E+04 | 756.4 | - | - | 0 | - |
| - | - | 1.232E+04 | 757.4 | - | - | 0 | - |
| 2 | y | 3.329E+05 | 757.8 | 0.001321 | 1.743 | +2 | 12 |
| - | - | 2.605E+05 | 758.3 | - | - | 0 | - |
| - | - | 1.652E+05 | 758.8 | - | - | 0 | - |
| - | - | 3.874E+04 | 759.3 | - | - | 0 | - |
| - | - | 2.41E+04 | 759.8 | - | - | 0 | - |
| - | - | 2.738E+04 | 761.3 | - | - | 0 | - |
| - | - | 1.428E+04 | 766.4 | - | - | 0 | - |
| - | - | 1.099E+04 | 767.4 | - | - | 0 | - |
| - | - | 2.95E+04 | 768.3 | - | - | 0 | - |
| - | - | 1.46E+04 | 769.3 | - | - | 0 | - |
| - | - | 1.694E+04 | 784.3 | - | - | 0 | - |
| - | - | 1.093E+05 | 784.4 | - | - | 0 | - |
| - | - | 1.317E+04 | 785.3 | - | - | 0 | - |
| - | - | 7.942E+04 | 785.4 | - | - | 0 | - |
| - | - | 3.623E+04 | 786.4 | - | - | 0 | - |
| - | - | 1.064E+04 | 793.4 | - | - | 0 | - |
| - | - | 1.012E+04 | 794.4 | - | - | 0 | - |
| - | - | 1.303E+04 | 808.4 | - | - | 0 | - |
| - | - | 3.534E+04 | 811.4 | - | - | 0 | - |
| - | - | 2.012E+04 | 812.4 | - | - | 0 | - |
| - | - | 1.126E+04 | 814.3 | - | - | 0 | - |
| - | - | 1.023E+04 | 821.4 | - | - | 0 | - |
| - | - | 1.708E+04 | 829.4 | - | - | 0 | - |
| - | - | 1.269E+04 | 830.4 | - | - | 0 | - |
| - | - | 2.646E+04 | 832.3 | - | - | 0 | - |
| - | - | 1.935E+04 | 833.3 | - | - | 0 | - |
| - | - | 3.661E+04 | 839.4 | - | - | 0 | - |
| - | - | 3.114E+04 | 840.4 | - | - | 0 | - |
| - | - | 1.631E+04 | 841.4 | - | - | 0 | - |
| 7 | b | 3.813E+04 | 848.3 | 0.001437 | 1.694 | +1 | 7 |
| - | - | 2.542E+04 | 848.4 | - | - | 0 | - |
| - | - | 1.643E+04 | 849.3 | - | - | 0 | - |
| - | - | 4.819E+04 | 849.4 | - | - | 0 | - |
| - | - | 2.391E+04 | 850.4 | - | - | 0 | - |
| - | - | 1.032E+04 | 851.4 | - | - | 0 | - |
| - | - | 1.371E+04 | 855.3 | - | - | 0 | - |
| - | - | 1.057E+04 | 856.3 | - | - | 0 | - |
| - | - | 4.317E+04 | 857.4 | - | - | 0 | - |
| - | - | 1.927E+04 | 858.4 | - | - | 0 | - |
| - | - | 1.038E+04 | 863.4 | - | - | 0 | - |
| - | - | 9944 | 864.5 | - | - | 0 | - |
| 7 | y | 6.474E+04 | 865.4 | 0.0001685 | 0.1947 | +1 | 7 |
| 7 | y | 9.32E+04 | 866.4 | 0.00898 | 10.36 | +1 | 7 |
| - | - | 4.843E+04 | 867.4 | - | - | 0 | - |
| - | - | 1.301E+04 | 868.4 | - | - | 0 | - |
| - | - | 1.202E+04 | 874.5 | - | - | 0 | - |
| - | - | 1.798E+04 | 881.4 | - | - | 0 | - |
| - | - | 1.354E+04 | 882.4 | - | - | 0 | - |
| 7 | y | 2.443E+05 | 883.4 | 0.0009855 | 1.116 | +1 | 7 |
| - | - | 1.143E+05 | 884.4 | - | - | 0 | - |
| - | - | 2.821E+04 | 885.4 | - | - | 0 | - |
| - | - | 1.043E+04 | 893.2 | - | - | 0 | - |
| - | - | 1.641E+04 | 893.4 | - | - | 0 | - |
| - | - | 1.379E+04 | 894.4 | - | - | 0 | - |
| 8 | b | 8.226E+04 | 919.3 | 0.0008834 | 0.961 | +1 | 8 |
| - | - | 3.457E+04 | 920.3 | - | - | 0 | - |
| - | - | 1.658E+04 | 921.3 | - | - | 0 | - |
| - | - | 4.68E+04 | 921.4 | - | - | 0 | - |
| - | - | 4.144E+04 | 922.4 | - | - | 0 | - |
| - | - | 1.905E+04 | 923.4 | - | - | 0 | - |
| - | - | 1.062E+04 | 940.4 | - | - | 0 | - |
| - | - | 1.146E+04 | 944.5 | - | - | 0 | - |
| - | - | 1.775E+04 | 945.4 | - | - | 0 | - |
| - | - | 1.529E+04 | 946.4 | - | - | 0 | - |
| - | - | 2.234E+04 | 948.4 | - | - | 0 | - |
| - | - | 1.14E+04 | 953.5 | - | - | 0 | - |
| - | - | 2.193E+04 | 968.4 | - | - | 0 | - |
| - | - | 7.605E+04 | 971.5 | - | - | 0 | - |
| - | - | 3.589E+04 | 972.5 | - | - | 0 | - |
| - | - | 1.446E+04 | 986.4 | - | - | 0 | - |
| 6 | y | 1.261E+04 | 1002 | 0.001829 | 1.825 | +1 | 8 |
| 6 | y | 5.287E+04 | 1003 | 0.003108 | 3.097 | +1 | 8 |
| - | - | 4.167E+04 | 1004 | - | - | 0 | - |
| - | - | 1.798E+04 | 1005 | - | - | 0 | - |
| - | - | 1.119E+04 | 1008 | - | - | 0 | - |
| - | - | 1.314E+04 | 1017 | - | - | 0 | - |
| - | - | 1.918E+04 | 1018 | - | - | 0 | - |
| - | - | 1.097E+04 | 1019 | - | - | 0 | - |
| 6 | y | 2.627E+05 | 1020 | 0.0004233 | 0.4148 | +1 | 8 |
| - | - | 1.538E+05 | 1021 | - | - | 0 | - |
| - | - | 3.762E+04 | 1022 | - | - | 0 | - |
| 9 | b | 4.868E+04 | 1032 | 0.002086 | 2.02 | +1 | 9 |
| - | - | 2.51E+04 | 1033 | - | - | 0 | - |
| - | - | 1.036E+05 | 1035 | - | - | 0 | - |
| - | - | 6.207E+04 | 1036 | - | - | 0 | - |
| - | - | 2.876E+04 | 1037 | - | - | 0 | - |
| - | - | 1.032E+04 | 1038 | - | - | 0 | - |
| - | - | 1.306E+04 | 1082 | - | - | 0 | - |
| - | - | 1.594E+04 | 1086 | - | - | 0 | - |
| - | - | 1.442E+04 | 1087 | - | - | 0 | - |
| - | - | 5.843E+04 | 1104 | - | - | 0 | - |
| - | - | 3.552E+04 | 1104 | - | - | 0 | - |
| - | - | 1.294E+04 | 1106 | - | - | 0 | - |
| - | - | 2.791E+04 | 1132 | - | - | 0 | - |
| 10 | b | 1.082E+04 | 1151 | 0.01673 | 14.53 | +1 | 10 |
| 5 | y | 3.789E+04 | 1167 | 0.005888 | 5.044 | +1 | 9 |
| - | - | 2.794E+04 | 1168 | - | - | 0 | - |
| 10 | b | 2.111E+04 | 1169 | 0.01825 | 15.6 | +1 | 10 |
| - | - | 2.609E+04 | 1196 | - | - | 0 | - |
| - | - | 1.428E+04 | 1197 | - | - | 0 | - |
| 11 | b | 2.307E+04 | 1284 | 0.001853 | 1.444 | +1 | 11 |
| - | - | 1.031E+04 | 1285 | - | - | 0 | - |
| - | - | 8022 | 2282 | - | - | 0 | - |
| - | - | 8719 | 2569 | - | - | 0 | - |

m/z Charge Intensity FragmentType MassShift Position
121.04000091552734 0 7598.82
122.0717544555664 0 11840.102
123.0445785522461 0 16230.343
124.0762710571289 0 13822.943
127.0506591796875 0 27384.498
129.0662841796875 0 126471.69
129.1026611328125 0 39102.785
129.29397583007812 0 4956.989
130.03253173828125 0 38256.887
130.06919860839844 0 7330.558
130.62399291992188 0 5973.8916
131.08197021484375 0 38201.3
132.19989013671875 0 5096.2866
133.04356384277344 0 10576.415
134.02748107910156 0 645308.94
135.0308380126953 0 28636.58
136.02320861816406 0 26797.533
136.05128479003906 0 7239.65
136.07611083984375 0 464658.84
136.16983032226562 0 5175.4717
137.074462890625 0 6155.0605
137.07948303222656 0 42989.676
138.0665740966797 0 39685.348
139.0508270263672 0 5342.1772
141.1027069091797 0 26416.902
143.027587890625 0 8748.824
143.81459045410156 0 5741.2812
144.01182556152344 0 5288.735
147.04458618164062 0 12063.554
148.08753967285156 0 17125.621
148.86077880859375 0 6229.39
148.87571716308594 0 8407.489
148.89724731445312 0 7995.317
148.9041748046875 0 7751.0664
148.91168212890625 0 7022.02
148.91908264160156 0 14503.478
148.92628479003906 0 16677.207
148.93304443359375 0 19757.04
148.94070434570312 0 45142.91
148.95738220214844 0 64311.31
148.96514892578125 0 44237.484
148.9726104736328 0 17354.531
148.9799041748047 0 16708.312
148.98696899414062 0 11782.508
148.99423217773438 0 9658.181
149.00140380859375 0 7085.1064
149.008544921875 0 6703.2363
149.01551818847656 0 5694.064
149.03775024414062 0 5813.3125
149.04507446289062 0 17894.85
152.08226013183594 0 27050.953
154.06153869628906 0 7529.204
155.08251953125 0 6288.923
155.09304809570312 0 37336.97
155.1181182861328 0 17531.72
156.07717895507812 0 107890.92
156.09622192382812 0 6681.616
157.0611572265625 0 122257.28
157.08035278320312 0 10334.003
157.13404846191406 0 5933.5317
158.02743530273438 0 359351.38
158.0644989013672 0 6086.9106
159.03103637695312 0 14445.589
159.0559539794922 0 8091.3506
159.1132354736328 0 457779.8
160.0233154296875 0 11052.12
160.1165008544922 0 42176.676
161.038330078125 0 188393.28
162.0225067138672 0 13439.543
162.041748046875 0 10643.527
163.0973663330078 0 6249.8286
164.08241271972656 0 5968.795
165.05503845214844 0 149648.17
165.0774383544922 0 35828.766
166.06124877929688 0 56800.727
166.0866241455078 0 101806.195
167.09304809570312 0 22740.26
168.01177978515625 0 51904.5
169.09764099121094 0 7715.131
170.94224548339844 0 4823.1997
171.1129150390625 0 8922.024
173.0924530029297 0 7727.1294
174.02197265625 0 6702.147
174.06654357910156 0 12132.667
175.09873962402344 0 8623.965
176.03807067871094 0 408402.12
176.08224487304688 0 61252.156
177.04141235351562 0 28895.56
177.113525390625 0 10906.991
178.03384399414062 0 16594.838
179.09312438964844 0 14389.24
180.07730102539062 0 14326.335
181.0901641845703 0 12615.202
182.08168029785156 0 296905.34 y 12
183.07725524902344 0 7626.9185
183.0852508544922 0 26735.031
183.1134490966797 0 25233.973
183.97454833984375 0 6346.903
185.0384063720703 0 24094.14
185.12942504882812 0 7279.581
186.0224609375 0 331766.66
186.124267578125 0 8972.427
187.025634765625 0 17509.527
187.10818481445312 0 216300.84
188.0182342529297 0 15857.886
188.11155700683594 0 14589.5625
189.07772827148438 0 27084.316
190.0615997314453 0 55398.207
191.06472778320312 0 8310.741
191.09335327148438 0 30597.113
193.07276916503906 0 12905.308
193.10885620117188 0 112028.336
194.0809326171875 0 10814.738
194.09292602539062 0 11742.503
194.1129913330078 0 7269.0503
195.088134765625 0 11846.175
196.07203674316406 0 14340.691
198.08753967285156 0 8558.56
199.07183837890625 0 36645.23
200.13970947265625 0 11124.539
201.08729553222656 0 21285.21
202.1090545654297 0 14666.184
203.04898071289062 0 79977.86 a Water loss 1
203.09353637695312 0 13055.2
203.1040496826172 0 6473.2236
204.07723999023438 0 77615.29
205.08071899414062 0 10058.522
205.1079864501953 0 7967.5073
206.1036834716797 0 12041.084
207.08810424804688 0 133753.75
208.0730438232422 0 6838.4136
210.12461853027344 0 13739.593
211.1079559326172 0 19387.846
213.0334930419922 0 8526.391
216.0984344482422 0 21204.352
216.13482666015625 0 8982.158
217.07225036621094 0 27980.375
218.05661010742188 0 35582.727
219.0885009765625 0 9950.042
221.05963134765625 0 3585256.5 a 1
221.10377502441406 0 293129.9
222.0626678466797 0 262803.88
222.08767700195312 0 7781.71
222.10714721679688 0 35374.965
223.0553436279297 0 123208.484
223.0642852783203 0 17950.598
223.15576171875 0 23989.129
224.0589141845703 0 7974.453
224.11468505859375 0 131227.48
225.0432891845703 0 8261.3955
225.11770629882812 0 12827.755
226.06405639648438 0 7661.8823
226.082763671875 0 114140.53
226.11895751953125 0 9298.853
227.0859832763672 0 18312.096
228.11407470703125 0 9433.548
230.0852813720703 0 15333.725
231.04397583007812 0 111757.03 b Water loss 1
232.04832458496094 0 10360.605
233.14039611816406 0 23092.84
234.099365234375 0 27366.025
235.0831298828125 0 258854.56
236.0670928955078 0 14996.978
236.08653259277344 0 31342.38
238.1188507080078 0 12670.626
239.09552001953125 0 13287.274
239.114501953125 0 76966.87
244.09339904785156 0 266681.38
244.32369995117188 0 6393.3403
245.09698486328125 0 29977.89
247.11199951171875 0 27376.021
249.0545654296875 0 962378.25 b 1
249.09860229492188 0 70525.99
249.13478088378906 0 9390.312
250.05764770507812 0 73864.984
251.0501708984375 0 32465.53
251.1508331298828 0 74146.54
252.1096954345703 0 522898.25
252.13453674316406 0 25694.744
252.15420532226562 0 13039.692
253.09361267089844 0 51943.3
253.112548828125 0 57095.77
254.06015014648438 0 8491.86
257.1071472167969 0 49666.05
262.0935974121094 0 10906.469
267.0927429199219 0 13887.939
267.1092834472656 0 190219.25
268.1122131347656 0 25277.453
268.1768493652344 0 11546.29
270.1201171875 0 50137.438
270.14556884765625 0 32796.254
273.13531494140625 0 62029.477
274.1390075683594 0 7960.0073
275.0681457519531 0 9122.491
275.1067810058594 0 54068.62
277.09454345703125 0 9080.1
277.1116638183594 0 16340.868 y Ammonia loss 9
279.1463623046875 0 15836.894
280.10455322265625 0 28754.293
281.1497497558594 0 10662.346
284.104736328125 0 8318.188
285.1021728515625 0 125292.85
285.62518310546875 0 106375.36 y 9
286.1056213378906 0 16929.982
286.1268615722656 0 37416.05
287.17108154296875 0 10113.376
292.1772155761719 0 8786.263
295.1040954589844 0 16939.537
296.6982727050781 0 6586.686
299.0608215332031 0 7692.367
301.1302795410156 0 133342.62
302.132568359375 0 11710.037
302.1611633300781 0 12416.4375
304.14093017578125 0 11220.199
304.17779541015625 0 25566.344
309.1361389160156 0 14577.78
313.0986022949219 0 11265.016
316.1334228515625 0 15075.219
318.0762023925781 0 144411.62 b Water loss 2
319.140869140625 0 595385.94 y 11
320.1390075683594 0 128693.12
320.173095703125 0 31941.512
321.14141845703125 0 20334.924
322.11614990234375 0 8232.419
322.18804931640625 0 68984.65
323.15631103515625 0 7549.151
323.19122314453125 0 14570.465
326.1373596191406 0 21270.852
329.1253356933594 0 55600.31
330.11279296875 0 65709.41
331.11590576171875 0 8799.165
332.13623046875 0 26483.31
334.1440124511719 0 100984.39
335.1465759277344 0 18512.49 y Ammonia loss 5
336.0867614746094 0 111026.55 b 2
337.09051513671875 0 18745.521
337.13031005859375 0 21460.89
338.14666748046875 0 122021.1
339.1489562988281 0 15811.156
339.2138671875 0 20993.477
340.82769775390625 0 8500.985 y 5
342.1676330566406 0 29840.422 y 8
342.66827392578125 0 12018.608
344.126220703125 0 13294.358
344.1474609375 0 32197.055
345.1507263183594 0 8731.022
347.1828308105469 0 8270.652
348.1241455078125 0 42338.12
348.1674499511719 0 19455.184
350.1466979980469 0 109828.68
351.1499328613281 0 25749.379
351.66815185546875 0 9661.832
353.1255798339844 0 12734.921
353.1944885253906 0 11335.937
354.1309814453125 0 22988.94
355.1397399902344 0 11331.877
355.1632080078125 0 8287.006
358.8482971191406 0 6312.372
360.12847900390625 0 15703.103 b 5
361.1737060546875 0 30943.916
365.1936950683594 0 53024.484
366.1413879394531 0 49430.78
368.8397216796875 0 7577.243
371.15802001953125 0 123471.27
372.1418762207031 0 78707.13
372.1627502441406 0 16507.996
373.14599609375 0 19291.832
375.1768493652344 0 10462.534
376.12451171875 0 8639.178
377.6860656738281 0 36414.016 y 7
378.1866455078125 0 23277.6
379.2098083496094 0 44368.332
380.212158203125 0 12072.989
381.1201477050781 0 72145.29
382.1243896484375 0 19234.064
384.1699523925781 0 8620.856
386.1495056152344 0 10086.35
387.2142333984375 0 8321.404
389.1467590332031 0 15930.158
389.1695251464844 0 23959.03
389.1940002441406 0 13404.1455
390.1524658203125 0 47330.383
393.1562805175781 0 8643.45
393.18896484375 0 28234.396
393.5105285644531 0 7392.747
395.1722412109375 0 9178.229
396.1351318359375 0 11038.468
397.68927001953125 0 15297.448
398.1475524902344 0 33265.75
399.130615234375 0 83591.32
400.13482666015625 0 22072.963
402.14208984375 0 10538.992
404.1578369140625 0 9158.969
405.1648864746094 0 10055.819
405.2259216308594 0 34134.66
406.2028503417969 0 45161.793
406.70440673828125 0 20917.941
407.1786804199219 0 23490.562
407.205078125 0 55504.883
408.2371520996094 0 16974.744
410.6526184082031 0 16584.8
412.1666564941406 0 11883.543
414.1451110839844 0 94310.68
415.1734313964844 0 24478.398
415.2186584472656 0 9060.126
415.6438293457031 0 7427.691 b Water loss 6
416.15740966796875 0 170073.38 y Ammonia loss 10
417.14569091796875 0 40134.09 b Water loss 3
417.16229248046875 0 28478.64
418.1476745605469 0 22742.738
420.2002258300781 0 30965.05
420.69781494140625 0 24347.861
421.18450927734375 0 107145.96
421.2130432128906 0 10157.857
422.18804931640625 0 17924.15
422.8617858886719 0 19450.197 y 3
423.19232177734375 0 13776.46
423.2357177734375 0 58937.363
424.2391662597656 0 13457.276
424.53631591796875 0 26331.375
424.65057373046875 0 17319.992 b 6
424.6911926269531 0 7878.5825
424.8695983886719 0 21582.375
425.1542663574219 0 9598.53
425.1897277832031 0 13769.067
425.64801025390625 0 10363.428
426.18389892578125 0 10639.225
429.2060241699219 0 24413.371
429.7091064453125 0 11279.286
430.53778076171875 0 10895.955
431.16448974609375 0 10885.922
431.20428466796875 0 9369.859
433.1840515136719 0 292786.25 y 10
433.21826171875 0 28606.283
433.7026672363281 0 13537.964
434.1876220703125 0 63165.875
434.52508544921875 0 8858.29
435.1553955078125 0 17549.432 b 3
435.1894836425781 0 14566.003
436.2315979003906 0 78182.48
437.2347412109375 0 11451.357
439.2088623046875 0 21580.516
440.1952209472656 0 17654.713
440.86309814453125 0 9931.144
440.9202880859375 0 22849.64
441.2540283203125 0 14246.783
441.5854187011719 0 12531.341
442.20574951171875 0 23971.205 y 6
442.710205078125 0 18969.166
443.1709289550781 0 15478.45
443.2082824707031 0 58345.766
444.1519470214844 0 9906.531
444.20770263671875 0 16183.986
445.86920166015625 0 137328.73 y Water loss 2
446.1695556640625 0 20735.137
446.2032165527344 0 82035.805 y Ammonia loss 2
446.5362854003906 0 65713.7
446.67352294921875 0 19924.3
446.86981201171875 0 10246.465
447.2193908691406 0 16534.1
447.7206726074219 0 28480.326
448.223876953125 0 19389.742
449.1807556152344 0 8688.633
449.2161560058594 0 16937.96
451.2306213378906 0 135729.64
451.87286376953125 0 149856.55 y 2
452.2066345214844 0 89222.42
452.23614501953125 0 19510.773
452.5398254394531 0 30032.9
453.20501708984375 0 9197.171
453.7020263671875 0 8584.871
457.229736328125 0 11314.222
460.1689453125 0 29709.55 b 7
461.215576171875 0 29579.76
461.7181396484375 0 43852.582
462.2154846191406 0 21870.889
464.8654479980469 0 33268.402
465.19854736328125 0 22718.193
465.5333557128906 0 13145.5
468.2560119628906 0 9593.229
468.8747863769531 0 19052.87
469.20794677734375 0 26733.87
469.5433349609375 0 11362.1455
469.8731689453125 0 8582.212
470.7220458984375 0 29267.338
471.20440673828125 0 64940.277
472.20343017578125 0 16544.576
473.9412536621094 0 11922.501
474.1967468261719 0 24890.947 b 11
474.2597351074219 0 20836.184
474.52996826171875 0 28144.049
474.8667297363281 0 13912.189
476.5546569824219 0 28435.762
476.8902893066406 0 21412.467
477.2152404785156 0 7651.723
478.2091369628906 0 20117.785
478.5392761230469 0 16041.764
478.8766784667969 0 12633.423
479.22454833984375 0 15363.626
479.72003173828125 0 54522.625
480.22137451171875 0 15000.232
480.5331115722656 0 13082.733
480.7193908691406 0 10816.291
483.22882080078125 0 12805.291
484.21051025390625 0 20228.1
484.5469970703125 0 16536.21
485.18206787109375 0 56344.785
485.216064453125 0 14364.322
486.23944091796875 0 19855.393
486.74114990234375 0 10729.683
487.7303161621094 0 9629.865
489.2039794921875 0 12244.585
489.2443542480469 0 10599.213
490.2046813964844 0 12796.13
490.24298095703125 0 13842.251
491.88446044921875 0 20398.25
492.2132873535156 0 13852.589
492.5465087890625 0 21905.955
493.2236022949219 0 22250.906
493.71820068359375 0 54093.52
493.8662414550781 0 9235.752
494.2147521972656 0 34166.29
494.7137451171875 0 18786.752
495.1598205566406 0 16234.519
495.2073059082031 0 10213.116
495.540771484375 0 11088.038
496.1605529785156 0 9817.347
497.88677978515625 0 143325.05
498.2207946777344 0 156216.89
498.5545654296875 0 64459.48
498.88824462890625 0 23272.02
499.22637939453125 0 8413.478
499.540771484375 0 48257.523 y Water loss 1
499.8730163574219 0 71143.28 y Ammonia loss 1
500.2058410644531 0 31237.396
500.53948974609375 0 17889.531
501.73077392578125 0 22871.295 y Water loss 5
501.88299560546875 0 7234.8423
502.224853515625 0 52848.445 y Ammonia loss 5
502.25811767578125 0 17078.48
502.72296142578125 0 62082.062
503.22772216796875 0 27843.38
503.7174987792969 0 13871.217
504.2124328613281 0 9461.129
505.5440673828125 0 175907.12 y 1
505.8785095214844 0 131563.69
506.21124267578125 0 71560.15
506.27142333984375 0 10290.95
506.54510498046875 0 46665.18
506.8818359375 0 9402.793
507.21478271484375 0 27959.965
507.554443359375 0 24671.877
507.88555908203125 0 21024.465
508.2162170410156 0 22884.273
508.2530212402344 0 29142.254
508.8616943359375 0 7593.6064
510.73681640625 0 270460.16 y 5
511.2391052246094 0 171631.55
511.5321350097656 0 13835.594
511.74017333984375 0 66597.664
512.243896484375 0 10239.29
513.220947265625 0 210229.8
513.5563354492188 0 195622.95
513.890380859375 0 99724.13
514.2202758789062 0 27182.959
514.5569458007812 0 13156.682
516.5442504882812 0 21459.627
516.8753662109375 0 36248.85
517.2068481445312 0 26626.371
517.5443725585938 0 11602.946
518.2377319335938 0 61506.438
518.7395629882812 0 26997.498
519.2357788085938 0 10669.586
520.2557983398438 0 51392.688
522.2133178710938 0 12898.0205
522.8759765625 0 30313.908
523.2056274414062 0 11915.185
523.5444946289062 0 12041.864
523.7281494140625 0 8742.358
524.2365112304688 0 12738.731
528.2671508789062 0 10990.666
528.5516967773438 0 99969.1 Precursor Water loss
528.6220092773438 0 10589.591
528.8836059570312 0 163396.39 Precursor Ammonia loss
529.2172241210938 0 130439.555
529.2564086914062 0 27403.021
529.5491333007812 0 86288.35
529.7531127929688 0 8913.365
529.8812255859375 0 26339.648
530.2362670898438 0 15758.537
531.2373657226562 0 10648.925
531.7643432617188 0 16686.506
532.2547607421875 0 18181.248
532.7630004882812 0 9189.337
534.2677001953125 0 48285.29
534.3072509765625 0 11151.503
534.5548095703125 0 1706971.6 Precursor
534.7378540039062 0 19184.299
534.8890380859375 0 1563155.6
535.2221069335938 0 909791.8
535.2633666992188 0 14069.876
535.5558471679688 0 401164.38
535.7354736328125 0 11831.966
535.89013671875 0 137822.8
536.2068481445312 0 22328.822
536.2474365234375 0 91273.305
537.245849609375 0 18075.291
538.2447509765625 0 11743.779
538.7412109375 0 12821.48
539.2451171875 0 24530.217
540.2213134765625 0 15010.342
540.7714233398438 0 57361.39
541.2164916992188 0 8358.545
541.271728515625 0 41333.883
543.2510986328125 0 44797.6
543.7438354492188 0 81251.57
544.2447509765625 0 48828.08
544.7488403320312 0 12563.756
545.294921875 0 47488.67
547.2628173828125 0 32430.191
548.260498046875 0 8763.074
552.2551879882812 0 243201.6
552.756103515625 0 133785.2
553.2171020507812 0 121331.39 y Ammonia loss 9
553.2572631835938 0 36910.457
554.2186279296875 0 33458.74
554.767822265625 0 44045.18
555.267822265625 0 53016.145
555.7660522460938 0 21058.81
556.2631225585938 0 9518.651
557.7403564453125 0 9887.155
558.242431640625 0 12751.188
560.2957153320312 0 30191.31
561.7484741210938 0 10727.594
563.7734985351562 0 58490.504
564.2706909179688 0 32498.193
564.7682495117188 0 27804.514
565.2730102539062 0 72928.76
566.2767944335938 0 27597.477
566.7413940429688 0 20062.969
568.2206420898438 0 26161.488
570.2427368164062 0 328017.66 y 9
571.2451171875 0 112947.375
572.2500610351562 0 73673.39
572.7700805664062 0 126936.92
573.2731323242188 0 107439.28
573.7711181640625 0 35459.527
574.2742309570312 0 34875.73
575.2489624023438 0 22600.584 y Water loss 4
575.7429809570312 0 77488.09 y Ammonia loss 4
576.2447509765625 0 46825.594 b Water loss 9
576.7447509765625 0 20066.1
577.7622680664062 0 19426.95
578.2554931640625 0 12984.338
578.761962890625 0 22464.191
581.273681640625 0 11916.429
582.2376708984375 0 13986.7
584.2545166015625 0 465902.28 y 4
584.7560424804688 0 265340.38
585.2537231445312 0 111663.2
585.7540893554688 0 36669.324
586.767578125 0 136956.88
587.2670288085938 0 116322.63
587.7670288085938 0 62603.55
588.2880249023438 0 49102.766
589.2767944335938 0 37869.906
591.3001098632812 0 24438.576
593.2807006835938 0 22030.344
593.7786865234375 0 13537.186
595.7733764648438 0 115158.01
596.2740478515625 0 61520.125
596.7758178710938 0 21079.72
598.2672729492188 0 43313.55
600.2455444335938 0 121835.12
600.2958984375 0 12975.461
601.2488403320312 0 49001.363
601.7898559570312 0 93569.69
602.2429809570312 0 8398.737
602.291015625 0 90095.27
602.79248046875 0 34309.855
603.2946166992188 0 11414.551
604.2227172851562 0 19261.746
604.8325805664062 0 9440.552
605.358154296875 0 14128.041
607.2836303710938 0 96345.63
608.2869262695312 0 25420.7
610.7692260742188 0 10801.217
612.2406005859375 0 11667.914
616.275390625 0 14179.693
621.2811889648438 0 15241.672
622.3009033203125 0 11944.973
624.7835083007812 0 12139.572 y Water loss 3
625.276611328125 0 39822.168 y Ammonia loss 3
625.7828369140625 0 25064.139
626.2759399414062 0 18721.973
627.7884521484375 0 14860.337
628.7816162109375 0 11333.723
632.2178955078125 0 20337.637
633.7886352539062 0 256347.81 y 3
634.2900390625 0 187757.62
634.7904663085938 0 75383.62
635.2871704101562 0 39301.426
635.778076171875 0 10036.109
636.298828125 0 36459.25
636.794921875 0 97041.48
637.297607421875 0 49637.67
637.7978515625 0 30385.738
639.2982177734375 0 10553.058
642.262451171875 0 44000.68 b 10
642.7631225585938 0 18837.006
643.2730102539062 0 12578.392
643.3339233398438 0 14227.574
644.271728515625 0 15786.877
644.7759399414062 0 16115.234
645.3054809570312 0 558946.25
645.8069458007812 0 410113.56
646.3081665039062 0 142375.92
646.810791015625 0 26881.045
648.2920532226562 0 11331.525
650.791259765625 0 10371.666
653.27490234375 0 51773.887
653.7801513671875 0 25938.945
654.2909545898438 0 22558.297
654.7975463867188 0 15056.507
657.3133544921875 0 26505.742
659.7865600585938 0 46561.773
660.2885131835938 0 17297.312
660.7857055664062 0 15176.036
660.8736572265625 0 73607.66
661.2901000976562 0 10879.127
661.3740844726562 0 58400.715
661.8759765625 0 29219.422
664.7953491210938 0 12790.555
665.3035278320312 0 16370.373
666.302001953125 0 43817.043 y Ammonia loss 8
667.2786865234375 0 27170.45
667.77587890625 0 23259.879
668.2987060546875 0 136499.06 y Water loss 2
668.7947387695312 0 245922.33 y Ammonia loss 2
669.2950439453125 0 156461.75
669.7944946289062 0 100865.08
670.2919921875 0 34111.93
670.7974243164062 0 11393.505
671.283203125 0 107914.78
672.2852783203125 0 42699.25
673.2846069335938 0 17797.916
674.337890625 0 44039.586
675.3403930664062 0 20556.764
676.2806396484375 0 11323.827
676.7807006835938 0 13610.801
677.3049926757812 0 1952663.2 y 2
677.806396484375 0 1429877.1
678.3067016601562 0 731067.1
678.8063354492188 0 217078.64
679.3064575195312 0 58643.598
682.295166015625 0 38576.848
682.35888671875 0 10625.723
682.798828125 0 34046.074
683.3265991210938 0 180205.84 y 8
684.327392578125 0 99018.27
685.3123168945312 0 32520.615
686.313232421875 0 14517.186
687.7960815429688 0 14917.936
688.2945556640625 0 14858.234
688.7898559570312 0 10732.271
691.25390625 0 18753.854
694.301025390625 0 18734.09
696.411376953125 0 14343.376
696.7941284179688 0 72752.51
697.2927856445312 0 69985.64
697.7969970703125 0 24551.383
698.2870483398438 0 10748.667
702.3340454101562 0 98512.69
702.8065185546875 0 24946.373
703.3255004882812 0 52334.28
704.3261108398438 0 19820.188
708.2935180664062 0 9692.829
710.4077758789062 0 56305.832
710.79150390625 0 121886.14 b 11
710.9075317382812 0 38590.574
711.2921142578125 0 92781.05
711.4105834960938 0 30745.125
711.793701171875 0 49123.207
712.298828125 0 16033.931
716.8084106445312 0 20238.463
717.3050537109375 0 29152.225
717.8031616210938 0 24477.729
719.248046875 0 20790.523 b 5
719.795654296875 0 55372.418
720.2960815429688 0 33013.01
720.36572265625 0 70613.26
720.794677734375 0 16258.612
721.36962890625 0 27213.887
722.3714599609375 0 9531.96
725.8131103515625 0 90417.67
726.3145141601562 0 66508.66
726.813720703125 0 46721.48
727.3180541992188 0 24449.922
735.3216552734375 0 32346.883
737.3384399414062 0 83221.05 y Ammonia loss 7
738.3422241210938 0 29553.984
739.3544311523438 0 13824.4
743.244873046875 0 10958.653
748.8051147460938 0 35763.246 y Water loss 1
749.3056640625 0 67725.59 y Ammonia loss 1
749.8042602539062 0 47380.273
750.299560546875 0 12417.32
754.3641357421875 0 283887.78 y 7
755.367431640625 0 119701.16
756.3707275390625 0 47861.81
757.3741455078125 0 12322.831
757.8123168945312 0 332922.4 y 1
758.3133544921875 0 260490.14
758.8132934570312 0 165209.25
759.31494140625 0 38744.18
759.8148193359375 0 24096.191
761.25830078125 0 27378.252
766.3505859375 0 14282.903
767.3613891601562 0 10988.929
768.2986450195312 0 29502.266
769.299072265625 0 14600.446
784.2935180664062 0 16941.635
784.3665771484375 0 109342.33
785.2952880859375 0 13172.188
785.3707885742188 0 79422.72
786.369140625 0 36229.87
793.3866577148438 0 10639.149
794.3716430664062 0 10121.939
808.3795776367188 0 13034.904
811.3971557617188 0 35344.445
812.3949584960938 0 20121.443
814.2843627929688 0 11264.002
821.378173828125 0 10226.618
829.4290161132812 0 17082.164
830.4327392578125 0 12693.258
832.2946166992188 0 26456.31
833.2974853515625 0 19349.49
839.3964233398438 0 36611.82
840.391845703125 0 31140.076
841.3838500976562 0 16310.236
848.2923583984375 0 38126.773 b 6
848.3722534179688 0 25420.686
849.29150390625 0 16434.266
849.3715209960938 0 48185.297
850.373291015625 0 23908.922
851.3724975585938 0 10315.891
855.3279418945312 0 13708.954
856.3370971679688 0 10573.775
857.4171752929688 0 43166.473
858.4190673828125 0 19273.842
863.3696899414062 0 10382.152
864.4688720703125 0 9943.953
865.3949584960938 0 64736.992 y Water loss 6
866.3881225585938 0 93196.555 y Ammonia loss 6
867.3897094726562 0 48429.457
868.3892211914062 0 13008.705
874.4530639648438 0 12016.554
881.3876342773438 0 17981.475
882.3856201171875 0 13536.764
883.4066772460938 0 244301.92 y 6
884.4114379882812 0 114326.43
885.418212890625 0 28210.277
893.1585083007812 0 10427.189
893.4203491210938 0 16406.664
894.4313354492188 0 13791.332
919.3289184570312 0 82257.61 b 7
920.3309326171875 0 34568.367
921.33203125 0 16576.79
921.4263916015625 0 46799.664
922.4281616210938 0 41437.535
923.4197998046875 0 19051.4
940.4403686523438 0 10617.64
944.4798583984375 0 11461.875
945.3844604492188 0 17748.064
946.3809814453125 0 15294.138
948.4359741210938 0 22337.611
953.4638061523438 0 11397.31
968.413818359375 0 21927.254
971.4714965820312 0 76045.195
972.4686889648438 0 35889.03
986.428466796875 0 14458.747
1002.4522094726562 0 12614.858 y Water loss 5
1003.441162109375 0 52873.473 y Ammonia loss 5
1004.4425048828125 0 41669.28
1005.4384765625 0 17979.428
1008.4774169921875 0 11190.487
1017.4586791992188 0 13143.879
1018.44677734375 0 19177.686
1019.4508056640625 0 10974.314
1020.4650268554688 0 262650.03 y 5
1021.4683837890625 0 153848.73
1022.4725341796875 0 37621.496
1032.4141845703125 0 48682.926 b 8
1033.4158935546875 0 25100.05
1035.4678955078125 0 103607.35
1036.4700927734375 0 62071.836
1037.470703125 0 28762.92
1038.4747314453125 0 10324.763
1082.4410400390625 0 13055.162
1086.474365234375 0 15941.475
1087.47509765625 0 14418.44
1103.501708984375 0 58425.562
1104.4996337890625 0 35523.414
1105.505859375 0 12941.095
1132.4853515625 0 27906.611
1151.4771728515625 0 10824.336 b Water loss 9
1167.5009765625 0 37889.246 y 4
1168.4967041015625 0 27944.174
1169.4892578125 0 21112.191 b 9
1196.48193359375 0 26087.383
1197.4892578125 0 14281.92
1283.5120849609375 0 23069.992 b 10
1284.514892578125 0 10310.66
2282.2294921875 0 8021.5923
2569.19482421875 0 8718.885

Spectrum Details

|  |  |
| --- | --- |
| Matched peaks? Matched peaks The total absolute number of peaks matched. Additionally in brackets the total fraction of peaks matched and the total number of peaks is shown. | 72 (8.83% of 815) |
| FDR? FDR The false discovery rate estimated for this peptide. It is calculated by matching all theoretical fragments with a non-integer shift with the raw peaks for this spectrum. This is done with 40 different shifts. The resulting percentage is the average number of annotated peaks over the number of annotated peaks with the correct spectrum. | 1.59% |
| Satellite FDR? Satellite FDR See the FDR for details on its calculation. This satellite ion specific FDR only contains the satellite ions (d/w) for I/L/J positions. | - |
| PSM Score? PSM Score The PSM Score as given by Hecklib to this annotated spectrum. It is shown with three significant figures. | 474 |

#### Spectrum 2941? Spectrum 2941 The raw spectrum of this peptide as annotated by Hecklib. The fragments are coloured according to ion type (see legend). Any peaks with a star '\*' as text can be hovered over to see the full details, first the ion type second the mass shift type. By hovering over the amino acids in the peptide or ions in the legend the corresponding peaks are highlighted. By toggling the 'Unassigned' label you can turn the background (unassigned) peaks on or off in the plot. By updating the slider in the Ion legend you can update the spectrum to only show the top X% of the peaks with labels. The top X% means any peak that is within X% of the highest intensity. By dragging in the spectrum you can zoom in to a specific part of the spectrum and use 'Zoom Out' to get back to the original zoom level. The annotation of the spectrum is based on the given sequence in the peptides file and is done with different software so inconsistencies are likely. The peaks are annotated based on the given sequence, with 20 ppm tolerance.

Copy Data

##### Spectrum 2941 (TSV)

###### Preview

```
Loading example...
```

*Click on the button to copy the data to your clipboard.*

Mz MinMz MaxIntensity Max

WidthHeightPeptide font sizePeptide stroke widthSpectrum font sizeSpectrum stroke widthCompact peptide

Ion legend

wxyz

abcd

OtherUnassignedIon^Charge^_Position_Show for top:%

SCSVMHEAJHNHY

04.72e+49.43e+41.42e+51.89e+5

Zoom Out

y^+1^_1_a^+1^_2_b^+1^_2_b^+1^_2_y^+2^_4_y^+2^_4_b^+1^_3_y^+1^_2_b^+1^_3_y^+2^_5_b^+2^_6_y^+2^_6_b^+2^_7_y^+1^_3_b^+1^_4_y^+3^_10_y^+3^_10_b^+2^_7_y^+1^_3_b^+1^_4_y^+2^_7_y^+3^_11_y^+3^_11_y^+3^_11_b^+2^_8_b^+3^_12_y^+3^_12_y^+3^_12_y^+2^_8_y^+2^_8_y^+3^_12_y^+2^_8_\*\*\*y^+1^_4_y^+1^_4_y^+2^_9_b^+2^_10_y^+2^_9_y^+2^_10_y^+2^_10_b^+2^_11_y^+1^_5_y^+2^_11_y^+2^_11_y^+2^_11_y^+1^_5_b^+2^_12_b^+1^_6_y^+1^_6_y^+2^_12_y^+2^_12_y^+1^_6_y^+2^_12_b^+1^_7_y^+1^_7_y^+1^_7_y^+1^_7_b^+1^_8_y^+1^_8_y^+1^_8_y^+1^_8_b^+1^_9_y^+1^_9_b^+1^_11_

0778155623343112

Fragment Matches Table

Show background peaks

| Position | Ion type | Intensity | mz Theoretical | mz Error (Th) | mz Error (ppm) | Charge | Series Number |
| --- | --- | --- | --- | --- | --- | --- | --- |
| - | - | 1663 | 123 | - | - | 0 | - |
| - | - | 624.1 | 124.1 | - | - | 0 | - |
| - | - | 1831 | 127.1 | - | - | 0 | - |
| - | - | 7717 | 129.1 | - | - | 0 | - |
| - | - | 1.506E+04 | 129.1 | - | - | 0 | - |
| - | - | 2163 | 130 | - | - | 0 | - |
| - | - | 582.1 | 130.1 | - | - | 0 | - |
| - | - | 2147 | 131.1 | - | - | 0 | - |
| - | - | 1211 | 132.1 | - | - | 0 | - |
| - | - | 395.8 | 133 | - | - | 0 | - |
| - | - | 905.8 | 133 | - | - | 0 | - |
| - | - | 3.55E+04 | 134 | - | - | 0 | - |
| - | - | 1632 | 135 | - | - | 0 | - |
| - | - | 1102 | 136 | - | - | 0 | - |
| - | - | 495.6 | 136.1 | - | - | 0 | - |
| - | - | 3.46E+04 | 136.1 | - | - | 0 | - |
| - | - | 2156 | 137.1 | - | - | 0 | - |
| - | - | 2237 | 138.1 | - | - | 0 | - |
| - | - | 449.7 | 139.1 | - | - | 0 | - |
| - | - | 367.3 | 141.1 | - | - | 0 | - |
| - | - | 2156 | 141.1 | - | - | 0 | - |
| - | - | 875.9 | 144 | - | - | 0 | - |
| - | - | 422.2 | 144 | - | - | 0 | - |
| - | - | 1372 | 147 | - | - | 0 | - |
| - | - | 1460 | 148.1 | - | - | 0 | - |
| - | - | 5487 | 149 | - | - | 0 | - |
| - | - | 478.5 | 151.6 | - | - | 0 | - |
| - | - | 693.2 | 152.1 | - | - | 0 | - |
| - | - | 494.9 | 152.4 | - | - | 0 | - |
| - | - | 438.4 | 153.1 | - | - | 0 | - |
| - | - | 493.2 | 154.1 | - | - | 0 | - |
| - | - | 999.9 | 155.1 | - | - | 0 | - |
| - | - | 2544 | 155.1 | - | - | 0 | - |
| - | - | 1853 | 155.1 | - | - | 0 | - |
| - | - | 7632 | 156.1 | - | - | 0 | - |
| - | - | 6303 | 157.1 | - | - | 0 | - |
| - | - | 620.6 | 157.1 | - | - | 0 | - |
| - | - | 666.6 | 157.1 | - | - | 0 | - |
| - | - | 645.2 | 157.1 | - | - | 0 | - |
| - | - | 1.955E+04 | 158 | - | - | 0 | - |
| - | - | 779.4 | 159 | - | - | 0 | - |
| - | - | 2.575E+04 | 159.1 | - | - | 0 | - |
| - | - | 951.2 | 160 | - | - | 0 | - |
| - | - | 442.7 | 160.1 | - | - | 0 | - |
| - | - | 1945 | 160.1 | - | - | 0 | - |
| - | - | 1.173E+04 | 161 | - | - | 0 | - |
| - | - | 507.3 | 162.1 | - | - | 0 | - |
| - | - | 1.672E+04 | 165.1 | - | - | 0 | - |
| - | - | 1307 | 165.1 | - | - | 0 | - |
| - | - | 593.9 | 165.1 | - | - | 0 | - |
| - | - | 2556 | 166.1 | - | - | 0 | - |
| - | - | 446.6 | 166.1 | - | - | 0 | - |
| - | - | 4757 | 166.1 | - | - | 0 | - |
| - | - | 6419 | 167.1 | - | - | 0 | - |
| - | - | 1371 | 167.1 | - | - | 0 | - |
| - | - | 426 | 168 | - | - | 0 | - |
| - | - | 437.1 | 168 | - | - | 0 | - |
| - | - | 3348 | 168 | - | - | 0 | - |
| - | - | 1073 | 169.1 | - | - | 0 | - |
| - | - | 901.6 | 171.1 | - | - | 0 | - |
| - | - | 2482 | 173.1 | - | - | 0 | - |
| - | - | 3130 | 173.5 | - | - | 0 | - |
| - | - | 535.2 | 174 | - | - | 0 | - |
| - | - | 684.9 | 174.1 | - | - | 0 | - |
| - | - | 516.9 | 175.1 | - | - | 0 | - |
| - | - | 2.118E+04 | 176 | - | - | 0 | - |
| - | - | 2718 | 176.1 | - | - | 0 | - |
| - | - | 661.4 | 177 | - | - | 0 | - |
| - | - | 876.8 | 178 | - | - | 0 | - |
| - | - | 999.6 | 179.1 | - | - | 0 | - |
| - | - | 1294 | 180.1 | - | - | 0 | - |
| - | - | 493.1 | 180.1 | - | - | 0 | - |
| - | - | 894.2 | 181.1 | - | - | 0 | - |
| 13 | y | 2.86E+04 | 182.1 | 0.0004191 | 2.302 | +1 | 1 |
| - | - | 1052 | 182.1 | - | - | 0 | - |
| - | - | 3012 | 183.1 | - | - | 0 | - |
| - | - | 1684 | 183.1 | - | - | 0 | - |
| - | - | 1934 | 185 | - | - | 0 | - |
| - | - | 618.6 | 185.1 | - | - | 0 | - |
| - | - | 1.862E+04 | 186 | - | - | 0 | - |
| - | - | 557.2 | 186.1 | - | - | 0 | - |
| - | - | 801.3 | 186.1 | - | - | 0 | - |
| - | - | 1256 | 187 | - | - | 0 | - |
| - | - | 1.084E+04 | 187.1 | - | - | 0 | - |
| - | - | 948 | 188 | - | - | 0 | - |
| - | - | 891.5 | 188.1 | - | - | 0 | - |
| - | - | 1334 | 189.1 | - | - | 0 | - |
| - | - | 3521 | 190.1 | - | - | 0 | - |
| - | - | 2211 | 191.1 | - | - | 0 | - |
| - | - | 640.8 | 193.1 | - | - | 0 | - |
| - | - | 6339 | 193.1 | - | - | 0 | - |
| - | - | 1010 | 194.1 | - | - | 0 | - |
| - | - | 751.8 | 194.1 | - | - | 0 | - |
| - | - | 490 | 195.1 | - | - | 0 | - |
| - | - | 967.6 | 197.1 | - | - | 0 | - |
| - | - | 497.2 | 198.1 | - | - | 0 | - |
| - | - | 1724 | 199.1 | - | - | 0 | - |
| - | - | 832.3 | 199.1 | - | - | 0 | - |
| - | - | 520.9 | 200.1 | - | - | 0 | - |
| - | - | 1603 | 200.1 | - | - | 0 | - |
| - | - | 519.3 | 201.1 | - | - | 0 | - |
| - | - | 1119 | 201.1 | - | - | 0 | - |
| - | - | 1183 | 201.1 | - | - | 0 | - |
| - | - | 842.8 | 202.1 | - | - | 0 | - |
| - | - | 5066 | 203 | - | - | 0 | - |
| - | - | 1212 | 203.1 | - | - | 0 | - |
| - | - | 846.7 | 204 | - | - | 0 | - |
| - | - | 4140 | 204.1 | - | - | 0 | - |
| - | - | 714.3 | 205.1 | - | - | 0 | - |
| - | - | 595.1 | 205.1 | - | - | 0 | - |
| - | - | 830.5 | 206.1 | - | - | 0 | - |
| - | - | 8371 | 207.1 | - | - | 0 | - |
| - | - | 511.2 | 208.1 | - | - | 0 | - |
| - | - | 517.7 | 208.1 | - | - | 0 | - |
| - | - | 1033 | 210.1 | - | - | 0 | - |
| - | - | 767.3 | 211.1 | - | - | 0 | - |
| - | - | 559.2 | 212.1 | - | - | 0 | - |
| - | - | 658.5 | 214.1 | - | - | 0 | - |
| - | - | 880.6 | 216.1 | - | - | 0 | - |
| - | - | 4442 | 216.1 | - | - | 0 | - |
| - | - | 1955 | 217.1 | - | - | 0 | - |
| - | - | 2344 | 218.1 | - | - | 0 | - |
| 2 | a | 1.868E+05 | 221.1 | 0.00405 | 18.32 | +1 | 2 |
| - | - | 9714 | 221.1 | - | - | 0 | - |
| - | - | 1.725E+04 | 221.1 | - | - | 0 | - |
| - | - | 1.524E+04 | 222.1 | - | - | 0 | - |
| - | - | 1878 | 222.1 | - | - | 0 | - |
| - | - | 1584 | 222.1 | - | - | 0 | - |
| - | - | 6601 | 223.1 | - | - | 0 | - |
| - | - | 1449 | 223.1 | - | - | 0 | - |
| - | - | 968.9 | 223.1 | - | - | 0 | - |
| - | - | 1053 | 223.2 | - | - | 0 | - |
| - | - | 7688 | 224.1 | - | - | 0 | - |
| - | - | 1.275E+04 | 225 | - | - | 0 | - |
| - | - | 1046 | 225.1 | - | - | 0 | - |
| - | - | 631.2 | 225.1 | - | - | 0 | - |
| - | - | 2512 | 226 | - | - | 0 | - |
| - | - | 7595 | 226.1 | - | - | 0 | - |
| - | - | 1415 | 226.1 | - | - | 0 | - |
| - | - | 1654 | 226.2 | - | - | 0 | - |
| - | - | 1599 | 227 | - | - | 0 | - |
| - | - | 725.4 | 227 | - | - | 0 | - |
| - | - | 586.8 | 227.1 | - | - | 0 | - |
| - | - | 571.7 | 228.1 | - | - | 0 | - |
| - | - | 680.6 | 228.1 | - | - | 0 | - |
| - | - | 554.5 | 230.1 | - | - | 0 | - |
| 2 | b | 5873 | 231 | 0.004178 | 18.08 | +1 | 2 |
| - | - | 1061 | 233.1 | - | - | 0 | - |
| - | - | 1548 | 234.1 | - | - | 0 | - |
| - | - | 1008 | 234.1 | - | - | 0 | - |
| - | - | 1.394E+04 | 235.1 | - | - | 0 | - |
| - | - | 632.9 | 236.1 | - | - | 0 | - |
| - | - | 1257 | 236.1 | - | - | 0 | - |
| - | - | 876.4 | 238.1 | - | - | 0 | - |
| - | - | 2.186E+04 | 239.1 | - | - | 0 | - |
| - | - | 3661 | 239.1 | - | - | 0 | - |
| - | - | 4323 | 240.1 | - | - | 0 | - |
| - | - | 2101 | 241.1 | - | - | 0 | - |
| - | - | 1.497E+04 | 244.1 | - | - | 0 | - |
| - | - | 1520 | 245.1 | - | - | 0 | - |
| - | - | 1797 | 247.1 | - | - | 0 | - |
| 2 | b | 5.479E+04 | 249.1 | 0.004076 | 16.37 | +1 | 2 |
| - | - | 3613 | 249.1 | - | - | 0 | - |
| - | - | 960.8 | 249.1 | - | - | 0 | - |
| - | - | 4983 | 250.1 | - | - | 0 | - |
| - | - | 708 | 250.1 | - | - | 0 | - |
| - | - | 1856 | 251.1 | - | - | 0 | - |
| - | - | 4730 | 251.2 | - | - | 0 | - |
| - | - | 2.924E+04 | 252.1 | - | - | 0 | - |
| - | - | 1816 | 252.1 | - | - | 0 | - |
| - | - | 3766 | 253.1 | - | - | 0 | - |
| - | - | 2751 | 253.1 | - | - | 0 | - |
| - | - | 1216 | 254.2 | - | - | 0 | - |
| - | - | 2139 | 257.1 | - | - | 0 | - |
| - | - | 714.1 | 258.1 | - | - | 0 | - |
| - | - | 606.2 | 266.1 | - | - | 0 | - |
| - | - | 1007 | 267.1 | - | - | 0 | - |
| - | - | 9829 | 267.1 | - | - | 0 | - |
| - | - | 800.4 | 268.1 | - | - | 0 | - |
| - | - | 612.8 | 268.1 | - | - | 0 | - |
| - | - | 711.4 | 269.1 | - | - | 0 | - |
| - | - | 915.2 | 269.2 | - | - | 0 | - |
| - | - | 549.7 | 270.1 | - | - | 0 | - |
| - | - | 2944 | 270.1 | - | - | 0 | - |
| - | - | 1968 | 270.1 | - | - | 0 | - |
| - | - | 2692 | 273.1 | - | - | 0 | - |
| - | - | 2673 | 275.1 | - | - | 0 | - |
| - | - | 570.3 | 275.1 | - | - | 0 | - |
| - | - | 905 | 277.1 | - | - | 0 | - |
| 10 | y | 1394 | 277.1 | 4.024E-06 | 0.01452 | +2 | 4 |
| - | - | 1066 | 279.1 | - | - | 0 | - |
| - | - | 1537 | 280.1 | - | - | 0 | - |
| - | - | 908.2 | 281.1 | - | - | 0 | - |
| - | - | 4311 | 281.2 | - | - | 0 | - |
| - | - | 689.6 | 282.1 | - | - | 0 | - |
| - | - | 6969 | 285.1 | - | - | 0 | - |
| 10 | y | 6077 | 285.6 | 0.0003708 | 1.298 | +2 | 4 |
| - | - | 790.4 | 286.1 | - | - | 0 | - |
| - | - | 1983 | 286.1 | - | - | 0 | - |
| - | - | 827.7 | 287.2 | - | - | 0 | - |
| - | - | 4923 | 295.1 | - | - | 0 | - |
| - | - | 1819 | 296.1 | - | - | 0 | - |
| - | - | 799.7 | 297.2 | - | - | 0 | - |
| - | - | 1.063E+04 | 299.1 | - | - | 0 | - |
| - | - | 2812 | 300.1 | - | - | 0 | - |
| - | - | 1302 | 301.1 | - | - | 0 | - |
| - | - | 6566 | 301.1 | - | - | 0 | - |
| - | - | 911.4 | 302.1 | - | - | 0 | - |
| - | - | 1312 | 302.2 | - | - | 0 | - |
| - | - | 1694 | 304.2 | - | - | 0 | - |
| - | - | 585.4 | 305.1 | - | - | 0 | - |
| - | - | 1124 | 313.1 | - | - | 0 | - |
| - | - | 652.9 | 314.1 | - | - | 0 | - |
| - | - | 817.9 | 316.1 | - | - | 0 | - |
| 3 | b | 6400 | 318.1 | 0.004147 | 13.04 | +1 | 3 |
| - | - | 936.7 | 319.1 | - | - | 0 | - |
| 12 | y | 3.277E+04 | 319.1 | 0.000574 | 1.799 | +1 | 2 |
| - | - | 551.8 | 319.6 | - | - | 0 | - |
| - | - | 7144 | 320.1 | - | - | 0 | - |
| - | - | 1675 | 320.2 | - | - | 0 | - |
| - | - | 878 | 321.1 | - | - | 0 | - |
| - | - | 4033 | 322.2 | - | - | 0 | - |
| - | - | 711.2 | 323.2 | - | - | 0 | - |
| - | - | 1142 | 326.1 | - | - | 0 | - |
| - | - | 1123 | 328.2 | - | - | 0 | - |
| - | - | 2677 | 329.1 | - | - | 0 | - |
| - | - | 1078 | 329.2 | - | - | 0 | - |
| - | - | 3486 | 330.1 | - | - | 0 | - |
| - | - | 1312 | 332.1 | - | - | 0 | - |
| - | - | 617.7 | 333.1 | - | - | 0 | - |
| - | - | 4153 | 334.1 | - | - | 0 | - |
| 3 | b | 4423 | 336.1 | 0.004061 | 12.08 | +1 | 3 |
| - | - | 682.8 | 337.1 | - | - | 0 | - |
| - | - | 1269 | 337.1 | - | - | 0 | - |
| - | - | 713.3 | 337.2 | - | - | 0 | - |
| - | - | 5685 | 338.1 | - | - | 0 | - |
| - | - | 834.5 | 339.1 | - | - | 0 | - |
| 9 | y | 1308 | 342.2 | 0.0008193 | 2.394 | +2 | 5 |
| - | - | 1085 | 342.7 | - | - | 0 | - |
| - | - | 576.9 | 343.2 | - | - | 0 | - |
| - | - | 1408 | 344.1 | - | - | 0 | - |
| - | - | 900.7 | 345.7 | - | - | 0 | - |
| - | - | 1546 | 348.1 | - | - | 0 | - |
| - | - | 1497 | 348.2 | - | - | 0 | - |
| - | - | 7054 | 350.1 | - | - | 0 | - |
| - | - | 1622 | 351.1 | - | - | 0 | - |
| - | - | 670 | 353.2 | - | - | 0 | - |
| - | - | 939.7 | 354.1 | - | - | 0 | - |
| - | - | 1143 | 355.1 | - | - | 0 | - |
| - | - | 632.7 | 355.1 | - | - | 0 | - |
| - | - | 656 | 357.2 | - | - | 0 | - |
| 6 | b | 850.7 | 360.1 | 0.001775 | 4.93 | +2 | 6 |
| - | - | 1212 | 361.2 | - | - | 0 | - |
| - | - | 682.8 | 361.2 | - | - | 0 | - |
| - | - | 600.4 | 362.2 | - | - | 0 | - |
| - | - | 623.3 | 362.8 | - | - | 0 | - |
| - | - | 3692 | 365.2 | - | - | 0 | - |
| - | - | 2410 | 366.1 | - | - | 0 | - |
| - | - | 828.8 | 366.2 | - | - | 0 | - |
| - | - | 555 | 366.7 | - | - | 0 | - |
| - | - | 614 | 367.2 | - | - | 0 | - |
| - | - | 3291 | 369.1 | - | - | 0 | - |
| - | - | 1523 | 370.1 | - | - | 0 | - |
| - | - | 841.2 | 370.1 | - | - | 0 | - |
| - | - | 696.7 | 370.7 | - | - | 0 | - |
| - | - | 930.7 | 371.1 | - | - | 0 | - |
| - | - | 6699 | 371.2 | - | - | 0 | - |
| - | - | 4745 | 372.1 | - | - | 0 | - |
| - | - | 1056 | 372.2 | - | - | 0 | - |
| - | - | 592.7 | 373.1 | - | - | 0 | - |
| - | - | 675.4 | 374.2 | - | - | 0 | - |
| 8 | y | 2179 | 377.7 | 5.416E-05 | 0.1434 | +2 | 6 |
| - | - | 1671 | 379.2 | - | - | 0 | - |
| - | - | 3547 | 381.1 | - | - | 0 | - |
| - | - | 768.5 | 382.1 | - | - | 0 | - |
| - | - | 1089 | 382.2 | - | - | 0 | - |
| - | - | 939.3 | 383.2 | - | - | 0 | - |
| - | - | 1799 | 384.2 | - | - | 0 | - |
| - | - | 905.6 | 389.1 | - | - | 0 | - |
| - | - | 1728 | 389.2 | - | - | 0 | - |
| - | - | 2470 | 390.2 | - | - | 0 | - |
| - | - | 677 | 390.7 | - | - | 0 | - |
| - | - | 839.2 | 391.2 | - | - | 0 | - |
| - | - | 1214 | 393.2 | - | - | 0 | - |
| - | - | 546.1 | 394.2 | - | - | 0 | - |
| - | - | 604.5 | 396.1 | - | - | 0 | - |
| - | - | 2187 | 398.1 | - | - | 0 | - |
| - | - | 3962 | 399.1 | - | - | 0 | - |
| - | - | 856.7 | 399.7 | - | - | 0 | - |
| - | - | 1485 | 400.1 | - | - | 0 | - |
| - | - | 1668 | 400.3 | - | - | 0 | - |
| - | - | 625.1 | 401.3 | - | - | 0 | - |
| - | - | 919.7 | 402.7 | - | - | 0 | - |
| - | - | 628.1 | 403.2 | - | - | 0 | - |
| - | - | 884.3 | 403.9 | - | - | 0 | - |
| - | - | 717.3 | 404.2 | - | - | 0 | - |
| - | - | 605.1 | 405.2 | - | - | 0 | - |
| - | - | 2200 | 405.2 | - | - | 0 | - |
| - | - | 2617 | 406.2 | - | - | 0 | - |
| - | - | 1566 | 406.7 | - | - | 0 | - |
| - | - | 778.2 | 407.2 | - | - | 0 | - |
| - | - | 941.7 | 407.2 | - | - | 0 | - |
| - | - | 2821 | 407.2 | - | - | 0 | - |
| - | - | 1637 | 408.2 | - | - | 0 | - |
| - | - | 717.3 | 408.2 | - | - | 0 | - |
| - | - | 1308 | 410.7 | - | - | 0 | - |
| - | - | 656.4 | 411.2 | - | - | 0 | - |
| - | - | 674.7 | 411.7 | - | - | 0 | - |
| - | - | 1154 | 412.2 | - | - | 0 | - |
| - | - | 2984 | 414.1 | - | - | 0 | - |
| - | - | 1282 | 414.2 | - | - | 0 | - |
| - | - | 1381 | 414.7 | - | - | 0 | - |
| - | - | 850.9 | 415 | - | - | 0 | - |
| - | - | 799.7 | 415.1 | - | - | 0 | - |
| - | - | 1259 | 415.2 | - | - | 0 | - |
| - | - | 587 | 415.2 | - | - | 0 | - |
| 7 | b | 638.2 | 415.6 | 0.0005365 | 1.291 | +2 | 7 |
| - | - | 612.7 | 415.8 | - | - | 0 | - |
| - | - | 687.4 | 415.9 | - | - | 0 | - |
| 11 | y | 9281 | 416.2 | 0.000431 | 1.036 | +1 | 3 |
| - | - | 1975 | 416.7 | - | - | 0 | - |
| 4 | b | 1207 | 417.1 | 0.006735 | 16.15 | +1 | 4 |
| - | - | 1424 | 417.2 | - | - | 0 | - |
| 4 | y | 862.1 | 417.2 | 0.005252 | 12.59 | +3 | 10 |
| - | - | 2466 | 420.2 | - | - | 0 | - |
| - | - | 1030 | 420.7 | - | - | 0 | - |
| - | - | 5109 | 421.2 | - | - | 0 | - |
| - | - | 686.4 | 422.2 | - | - | 0 | - |
| - | - | 1161 | 422.2 | - | - | 0 | - |
| - | - | 710.4 | 422.2 | - | - | 0 | - |
| 4 | y | 792.4 | 422.9 | 0.002892 | 6.839 | +3 | 10 |
| - | - | 718.7 | 423.2 | - | - | 0 | - |
| - | - | 3428 | 423.2 | - | - | 0 | - |
| - | - | 843.7 | 423.5 | - | - | 0 | - |
| - | - | 2394 | 424.5 | - | - | 0 | - |
| 7 | b | 1362 | 424.6 | 0.0006814 | 1.605 | +2 | 7 |
| - | - | 1249 | 424.9 | - | - | 0 | - |
| - | - | 904.5 | 425.7 | - | - | 0 | - |
| - | - | 913 | 426.2 | - | - | 0 | - |
| - | - | 1318 | 429.2 | - | - | 0 | - |
| - | - | 1897 | 429.2 | - | - | 0 | - |
| - | - | 949.6 | 429.7 | - | - | 0 | - |
| - | - | 725.8 | 430.5 | - | - | 0 | - |
| - | - | 820.2 | 430.7 | - | - | 0 | - |
| - | - | 626 | 431.2 | - | - | 0 | - |
| - | - | 2018 | 431.6 | - | - | 0 | - |
| - | - | 819 | 431.7 | - | - | 0 | - |
| - | - | 2497 | 431.9 | - | - | 0 | - |
| 11 | y | 1.686E+04 | 433.2 | 0.0006153 | 1.42 | +1 | 3 |
| - | - | 1434 | 433.2 | - | - | 0 | - |
| - | - | 3127 | 434.2 | - | - | 0 | - |
| 4 | b | 907.3 | 435.2 | 0.004513 | 10.37 | +1 | 4 |
| - | - | 4685 | 436.2 | - | - | 0 | - |
| - | - | 807.3 | 440.2 | - | - | 0 | - |
| - | - | 6854 | 440.9 | - | - | 0 | - |
| - | - | 5870 | 441.3 | - | - | 0 | - |
| - | - | 2709 | 441.6 | - | - | 0 | - |
| 7 | y | 1054 | 442.2 | 0.001493 | 3.377 | +2 | 7 |
| - | - | 714.7 | 443.2 | - | - | 0 | - |
| - | - | 3612 | 443.2 | - | - | 0 | - |
| - | - | 674.5 | 443.9 | - | - | 0 | - |
| - | - | 654.6 | 444.2 | - | - | 0 | - |
| 3 | y | 5122 | 445.9 | 0.002512 | 5.635 | +3 | 11 |
| - | - | 1271 | 446.2 | - | - | 0 | - |
| 3 | y | 8728 | 446.2 | 0.008705 | 19.51 | +3 | 11 |
| - | - | 3181 | 446.5 | - | - | 0 | - |
| - | - | 1522 | 446.9 | - | - | 0 | - |
| - | - | 1101 | 447.2 | - | - | 0 | - |
| - | - | 2241 | 447.7 | - | - | 0 | - |
| - | - | 1109 | 448.2 | - | - | 0 | - |
| - | - | 1090 | 449.2 | - | - | 0 | - |
| - | - | 6450 | 451.2 | - | - | 0 | - |
| 3 | y | 7106 | 451.9 | 0.002073 | 4.588 | +3 | 11 |
| - | - | 6897 | 452.2 | - | - | 0 | - |
| - | - | 929.3 | 452.2 | - | - | 0 | - |
| - | - | 2512 | 452.5 | - | - | 0 | - |
| - | - | 1142 | 452.7 | - | - | 0 | - |
| - | - | 1551 | 452.9 | - | - | 0 | - |
| - | - | 1274 | 455.2 | - | - | 0 | - |
| - | - | 764 | 456.2 | - | - | 0 | - |
| - | - | 1004 | 458.7 | - | - | 0 | - |
| - | - | 694.8 | 459.3 | - | - | 0 | - |
| - | - | 839.7 | 459.5 | - | - | 0 | - |
| 8 | b | 1577 | 460.2 | 0.000374 | 0.8129 | +2 | 8 |
| - | - | 6465 | 461.2 | - | - | 0 | - |
| - | - | 3379 | 461.7 | - | - | 0 | - |
| - | - | 1628 | 462.2 | - | - | 0 | - |
| - | - | 940.6 | 462.7 | - | - | 0 | - |
| - | - | 1155 | 463.6 | - | - | 0 | - |
| - | - | 1660 | 463.9 | - | - | 0 | - |
| - | - | 1027 | 464.3 | - | - | 0 | - |
| - | - | 893.8 | 464.6 | - | - | 0 | - |
| - | - | 1772 | 464.9 | - | - | 0 | - |
| - | - | 1239 | 465.2 | - | - | 0 | - |
| - | - | 985 | 465.5 | - | - | 0 | - |
| - | - | 3358 | 467.2 | - | - | 0 | - |
| - | - | 1725 | 467.7 | - | - | 0 | - |
| - | - | 1040 | 468.3 | - | - | 0 | - |
| - | - | 1785 | 470.7 | - | - | 0 | - |
| - | - | 3568 | 471.2 | - | - | 0 | - |
| - | - | 6961 | 471.6 | - | - | 0 | - |
| - | - | 5099 | 471.9 | - | - | 0 | - |
| - | - | 1116 | 472.3 | - | - | 0 | - |
| - | - | 1440 | 473.2 | - | - | 0 | - |
| - | - | 992.2 | 473.7 | - | - | 0 | - |
| - | - | 3479 | 473.9 | - | - | 0 | - |
| 12 | b | 1601 | 474.2 | 0.002167 | 4.569 | +3 | 12 |
| - | - | 2295 | 474.3 | - | - | 0 | - |
| - | - | 1865 | 474.5 | - | - | 0 | - |
| - | - | 967.4 | 474.9 | - | - | 0 | - |
| - | - | 1.166E+04 | 476.2 | - | - | 0 | - |
| - | - | 1745 | 476.6 | - | - | 0 | - |
| - | - | 3449 | 476.7 | - | - | 0 | - |
| - | - | 1564 | 476.9 | - | - | 0 | - |
| - | - | 775 | 477.2 | - | - | 0 | - |
| - | - | 1038 | 478.2 | - | - | 0 | - |
| - | - | 866.1 | 478.5 | - | - | 0 | - |
| - | - | 763.5 | 479.2 | - | - | 0 | - |
| - | - | 2662 | 479.7 | - | - | 0 | - |
| - | - | 714.8 | 479.8 | - | - | 0 | - |
| - | - | 1340 | 480.2 | - | - | 0 | - |
| - | - | 790 | 480.7 | - | - | 0 | - |
| - | - | 3625 | 482.2 | - | - | 0 | - |
| - | - | 2037 | 482.7 | - | - | 0 | - |
| - | - | 748.4 | 483.2 | - | - | 0 | - |
| - | - | 2581 | 484.2 | - | - | 0 | - |
| - | - | 2593 | 485.2 | - | - | 0 | - |
| - | - | 1188 | 485.2 | - | - | 0 | - |
| - | - | 783.8 | 486.2 | - | - | 0 | - |
| - | - | 937.6 | 486.2 | - | - | 0 | - |
| - | - | 762.6 | 486.7 | - | - | 0 | - |
| - | - | 758.8 | 489.2 | - | - | 0 | - |
| - | - | 3.068E+04 | 491.2 | - | - | 0 | - |
| - | - | 1.219E+04 | 491.7 | - | - | 0 | - |
| - | - | 1368 | 492.2 | - | - | 0 | - |
| - | - | 1134 | 492.5 | - | - | 0 | - |
| - | - | 861.9 | 492.7 | - | - | 0 | - |
| - | - | 961.2 | 492.9 | - | - | 0 | - |
| - | - | 2880 | 493.7 | - | - | 0 | - |
| - | - | 1943 | 494.2 | - | - | 0 | - |
| - | - | 1057 | 494.5 | - | - | 0 | - |
| - | - | 3635 | 495.2 | - | - | 0 | - |
| - | - | 2469 | 495.7 | - | - | 0 | - |
| - | - | 1255 | 497.6 | - | - | 0 | - |
| - | - | 7518 | 497.9 | - | - | 0 | - |
| - | - | 8373 | 498.2 | - | - | 0 | - |
| - | - | 3465 | 498.6 | - | - | 0 | - |
| - | - | 1044 | 498.9 | - | - | 0 | - |
| - | - | 1266 | 499.2 | - | - | 0 | - |
| 2 | y | 3055 | 499.5 | 0.001234 | 2.471 | +3 | 12 |
| 2 | y | 1743 | 499.9 | 0.005108 | 10.22 | +3 | 12 |
| - | - | 771.1 | 500.5 | - | - | 0 | - |
| 6 | y | 840.5 | 501.7 | 0.001349 | 2.688 | +2 | 8 |
| - | - | 948.3 | 501.9 | - | - | 0 | - |
| 6 | y | 2179 | 502.2 | 0.001364 | 2.716 | +2 | 8 |
| - | - | 4139 | 502.7 | - | - | 0 | - |
| - | - | 844.8 | 503.1 | - | - | 0 | - |
| - | - | 1886 | 503.2 | - | - | 0 | - |
| - | - | 714.8 | 503.9 | - | - | 0 | - |
| - | - | 2151 | 504.2 | - | - | 0 | - |
| - | - | 1057 | 504.7 | - | - | 0 | - |
| 2 | y | 1.115E+04 | 505.5 | 0.0004593 | 0.9085 | +3 | 12 |
| - | - | 8139 | 505.9 | - | - | 0 | - |
| - | - | 5053 | 506.2 | - | - | 0 | - |
| - | - | 993.4 | 506.3 | - | - | 0 | - |
| - | - | 2269 | 506.5 | - | - | 0 | - |
| - | - | 1048 | 506.9 | - | - | 0 | - |
| - | - | 2726 | 507.2 | - | - | 0 | - |
| - | - | 1996 | 507.6 | - | - | 0 | - |
| - | - | 873.2 | 507.9 | - | - | 0 | - |
| - | - | 910.1 | 508.2 | - | - | 0 | - |
| - | - | 1772 | 508.3 | - | - | 0 | - |
| - | - | 695.6 | 508.5 | - | - | 0 | - |
| 6 | y | 1.661E+04 | 510.7 | 8.293E-05 | 0.1624 | +2 | 8 |
| - | - | 575.8 | 510.8 | - | - | 0 | - |
| - | - | 6943 | 511.2 | - | - | 0 | - |
| - | - | 2472 | 511.7 | - | - | 0 | - |
| - | - | 2.947E+04 | 513.2 | - | - | 0 | - |
| - | - | 8702 | 513.6 | - | - | 0 | - |
| - | - | 1.059E+04 | 513.7 | - | - | 0 | - |
| - | - | 6805 | 513.9 | - | - | 0 | - |
| - | - | 2310 | 516.5 | - | - | 0 | - |
| - | - | 2129 | 516.9 | - | - | 0 | - |
| - | - | 1516 | 517.2 | - | - | 0 | - |
| - | - | 1175 | 517.2 | - | - | 0 | - |
| - | - | 1573 | 517.5 | - | - | 0 | - |
| - | - | 1086 | 517.9 | - | - | 0 | - |
| - | - | 3439 | 518.2 | - | - | 0 | - |
| - | - | 1149 | 518.7 | - | - | 0 | - |
| - | - | 2515 | 520.3 | - | - | 0 | - |
| - | - | 1619 | 523.2 | - | - | 0 | - |
| - | - | 725.5 | 526.2 | - | - | 0 | - |
| - | - | 883.9 | 528.2 | - | - | 0 | - |
| - | - | 1165 | 528.3 | - | - | 0 | - |
| 0 | Precursor | 6587 | 528.6 | 0.0004392 | 0.8309 | +3 | -1 |
| - | - | 1229 | 528.6 | - | - | 0 | - |
| 0 | Precursor | 1.17E+04 | 528.9 | 0.004563 | 8.629 | +3 | -1 |
| - | - | 7525 | 529.2 | - | - | 0 | - |
| - | - | 1183 | 529.3 | - | - | 0 | - |
| - | - | 5597 | 529.5 | - | - | 0 | - |
| - | - | 856.2 | 529.9 | - | - | 0 | - |
| - | - | 901 | 531.2 | - | - | 0 | - |
| - | - | 671.1 | 532.3 | - | - | 0 | - |
| - | - | 783.2 | 532.8 | - | - | 0 | - |
| - | - | 967.9 | 533.3 | - | - | 0 | - |
| - | - | 2822 | 533.3 | - | - | 0 | - |
| - | - | 1286 | 533.8 | - | - | 0 | - |
| - | - | 880.2 | 533.9 | - | - | 0 | - |
| - | - | 1254 | 534.3 | - | - | 0 | - |
| - | - | 9549 | 534.3 | - | - | 0 | - |
| 0 | Precursor | 8.061E+04 | 534.6 | 0.0004948 | 0.9257 | +3 | -1 |
| - | - | 7041 | 534.6 | - | - | 0 | - |
| - | - | 7.067E+04 | 534.9 | - | - | 0 | - |
| - | - | 3870 | 535 | - | - | 0 | - |
| - | - | 3.841E+04 | 535.2 | - | - | 0 | - |
| - | - | 1439 | 535.3 | - | - | 0 | - |
| - | - | 1.69E+04 | 535.6 | - | - | 0 | - |
| - | - | 1054 | 535.7 | - | - | 0 | - |
| - | - | 7319 | 535.8 | - | - | 0 | - |
| - | - | 5066 | 535.9 | - | - | 0 | - |
| - | - | 1101 | 536.2 | - | - | 0 | - |
| - | - | 4310 | 536.2 | - | - | 0 | - |
| - | - | 1180 | 537.2 | - | - | 0 | - |
| - | - | 933 | 538.7 | - | - | 0 | - |
| - | - | 1121 | 539.2 | - | - | 0 | - |
| - | - | 742.9 | 540.3 | - | - | 0 | - |
| - | - | 2283 | 540.8 | - | - | 0 | - |
| - | - | 1055 | 541.2 | - | - | 0 | - |
| - | - | 1946 | 541.3 | - | - | 0 | - |
| - | - | 2515 | 543.2 | - | - | 0 | - |
| - | - | 5594 | 543.7 | - | - | 0 | - |
| - | - | 1829 | 544.2 | - | - | 0 | - |
| - | - | 2382 | 545.3 | - | - | 0 | - |
| - | - | 1827 | 547.3 | - | - | 0 | - |
| - | - | 1304 | 548.3 | - | - | 0 | - |
| - | - | 1131 | 548.8 | - | - | 0 | - |
| - | - | 1.078E+04 | 552.3 | - | - | 0 | - |
| - | - | 7550 | 552.8 | - | - | 0 | - |
| 10 | y | 8094 | 553.2 | 0.0004486 | 0.8108 | +1 | 4 |
| - | - | 3026 | 553.3 | - | - | 0 | - |
| - | - | 715.2 | 553.3 | - | - | 0 | - |
| - | - | 1072 | 553.8 | - | - | 0 | - |
| - | - | 1989 | 554.2 | - | - | 0 | - |
| - | - | 2378 | 554.8 | - | - | 0 | - |
| - | - | 2316 | 555.3 | - | - | 0 | - |
| - | - | 963.4 | 555.8 | - | - | 0 | - |
| - | - | 1049 | 556.3 | - | - | 0 | - |
| - | - | 1899 | 560.3 | - | - | 0 | - |
| - | - | 1716 | 560.8 | - | - | 0 | - |
| - | - | 1094 | 561.3 | - | - | 0 | - |
| - | - | 2670 | 563.8 | - | - | 0 | - |
| - | - | 1795 | 564.3 | - | - | 0 | - |
| - | - | 1110 | 564.8 | - | - | 0 | - |
| - | - | 4011 | 565.3 | - | - | 0 | - |
| - | - | 815.8 | 566.7 | - | - | 0 | - |
| - | - | 776.1 | 568.2 | - | - | 0 | - |
| 10 | y | 1.811E+04 | 570.2 | 0.0002056 | 0.3606 | +1 | 4 |
| - | - | 963.2 | 570.3 | - | - | 0 | - |
| - | - | 5919 | 571.2 | - | - | 0 | - |
| - | - | 3446 | 572.2 | - | - | 0 | - |
| - | - | 6821 | 572.8 | - | - | 0 | - |
| - | - | 3306 | 573.3 | - | - | 0 | - |
| - | - | 2269 | 573.8 | - | - | 0 | - |
| - | - | 1029 | 574.3 | - | - | 0 | - |
| 5 | y | 3258 | 575.7 | 0.004951 | 8.6 | +2 | 9 |
| 10 | b | 1347 | 576.2 | 0.01119 | 19.43 | +2 | 10 |
| - | - | 1518 | 576.7 | - | - | 0 | - |
| - | - | 1821 | 577.8 | - | - | 0 | - |
| - | - | 1132 | 578.3 | - | - | 0 | - |
| - | - | 1472 | 580.3 | - | - | 0 | - |
| - | - | 828.4 | 581.3 | - | - | 0 | - |
| - | - | 950.9 | 582.2 | - | - | 0 | - |
| 5 | y | 2.146E+04 | 584.3 | 0.002602 | 4.453 | +2 | 9 |
| - | - | 1.339E+04 | 584.8 | - | - | 0 | - |
| - | - | 7440 | 585.3 | - | - | 0 | - |
| - | - | 3172 | 585.8 | - | - | 0 | - |
| - | - | 8841 | 586.8 | - | - | 0 | - |
| - | - | 6586 | 587.3 | - | - | 0 | - |
| - | - | 2762 | 587.8 | - | - | 0 | - |
| - | - | 2750 | 588.3 | - | - | 0 | - |
| - | - | 1022 | 589.3 | - | - | 0 | - |
| - | - | 1224 | 591.3 | - | - | 0 | - |
| - | - | 893 | 593.3 | - | - | 0 | - |
| - | - | 1557 | 593.8 | - | - | 0 | - |
| - | - | 705.6 | 594.3 | - | - | 0 | - |
| - | - | 907.6 | 595.3 | - | - | 0 | - |
| - | - | 5303 | 595.8 | - | - | 0 | - |
| - | - | 3251 | 596.3 | - | - | 0 | - |
| - | - | 849.9 | 596.8 | - | - | 0 | - |
| - | - | 1411 | 596.8 | - | - | 0 | - |
| - | - | 1712 | 598.3 | - | - | 0 | - |
| - | - | 6418 | 600.2 | - | - | 0 | - |
| - | - | 3120 | 601.2 | - | - | 0 | - |
| - | - | 5225 | 601.8 | - | - | 0 | - |
| - | - | 928.4 | 602.2 | - | - | 0 | - |
| - | - | 3511 | 602.3 | - | - | 0 | - |
| - | - | 1958 | 602.8 | - | - | 0 | - |
| - | - | 944.5 | 604.2 | - | - | 0 | - |
| - | - | 3916 | 604.3 | - | - | 0 | - |
| - | - | 2196 | 604.8 | - | - | 0 | - |
| - | - | 2785 | 605.4 | - | - | 0 | - |
| - | - | 1484 | 605.9 | - | - | 0 | - |
| - | - | 4458 | 607.3 | - | - | 0 | - |
| - | - | 1001 | 608.3 | - | - | 0 | - |
| - | - | 707.2 | 610.3 | - | - | 0 | - |
| - | - | 844 | 616.3 | - | - | 0 | - |
| - | - | 1162 | 621.3 | - | - | 0 | - |
| 4 | y | 2310 | 625.3 | 0.004313 | 6.899 | +2 | 10 |
| - | - | 895.4 | 625.8 | - | - | 0 | - |
| - | - | 878 | 631.8 | - | - | 0 | - |
| - | - | 827.9 | 631.9 | - | - | 0 | - |
| - | - | 765 | 632.2 | - | - | 0 | - |
| 4 | y | 1.422E+04 | 633.8 | 0.002636 | 4.159 | +2 | 10 |
| - | - | 8125 | 634.3 | - | - | 0 | - |
| - | - | 4069 | 634.8 | - | - | 0 | - |
| - | - | 2691 | 635.3 | - | - | 0 | - |
| - | - | 2045 | 635.8 | - | - | 0 | - |
| - | - | 2312 | 636.3 | - | - | 0 | - |
| - | - | 4433 | 636.8 | - | - | 0 | - |
| - | - | 2769 | 637.3 | - | - | 0 | - |
| - | - | 835.7 | 639.9 | - | - | 0 | - |
| 11 | b | 835.9 | 642.3 | 0.004626 | 7.203 | +2 | 11 |
| - | - | 1663 | 642.8 | - | - | 0 | - |
| - | - | 2.994E+04 | 645.3 | - | - | 0 | - |
| - | - | 2.344E+04 | 645.8 | - | - | 0 | - |
| - | - | 1.116E+04 | 646.3 | - | - | 0 | - |
| - | - | 2318 | 646.8 | - | - | 0 | - |
| - | - | 1277 | 646.9 | - | - | 0 | - |
| - | - | 1051 | 648.3 | - | - | 0 | - |
| - | - | 830.5 | 649.3 | - | - | 0 | - |
| - | - | 1158 | 651.9 | - | - | 0 | - |
| - | - | 925 | 652.4 | - | - | 0 | - |
| - | - | 2195 | 653.3 | - | - | 0 | - |
| - | - | 1020 | 653.8 | - | - | 0 | - |
| - | - | 1590 | 653.9 | - | - | 0 | - |
| - | - | 1430 | 654.3 | - | - | 0 | - |
| - | - | 1681 | 654.4 | - | - | 0 | - |
| - | - | 1235 | 654.8 | - | - | 0 | - |
| - | - | 1503 | 657.3 | - | - | 0 | - |
| - | - | 1167 | 659.8 | - | - | 0 | - |
| - | - | 2.599E+04 | 660.9 | - | - | 0 | - |
| - | - | 1.608E+04 | 661.4 | - | - | 0 | - |
| - | - | 6194 | 661.9 | - | - | 0 | - |
| - | - | 2926 | 662.4 | - | - | 0 | - |
| - | - | 1507 | 664.8 | - | - | 0 | - |
| 9 | y | 2682 | 666.3 | 0.002932 | 4.401 | +1 | 5 |
| - | - | 1848 | 667.3 | - | - | 0 | - |
| - | - | 2164 | 667.8 | - | - | 0 | - |
| 3 | y | 5780 | 668.3 | 0.00112 | 1.676 | +2 | 11 |
| 3 | y | 1.341E+04 | 668.8 | 0.005267 | 7.876 | +2 | 11 |
| - | - | 1.007E+04 | 669.3 | - | - | 0 | - |
| - | - | 4455 | 669.8 | - | - | 0 | - |
| - | - | 887.4 | 670.3 | - | - | 0 | - |
| - | - | 6600 | 671.3 | - | - | 0 | - |
| - | - | 2651 | 672.3 | - | - | 0 | - |
| - | - | 2542 | 674.3 | - | - | 0 | - |
| - | - | 1987 | 676.3 | - | - | 0 | - |
| 3 | y | 1.098E+05 | 677.3 | 0.002735 | 4.038 | +2 | 11 |
| - | - | 8.273E+04 | 677.8 | - | - | 0 | - |
| - | - | 3.849E+04 | 678.3 | - | - | 0 | - |
| - | - | 1.195E+04 | 678.8 | - | - | 0 | - |
| - | - | 2589 | 679.3 | - | - | 0 | - |
| - | - | 729.8 | 680.3 | - | - | 0 | - |
| - | - | 1970 | 682.3 | - | - | 0 | - |
| - | - | 1131 | 682.4 | - | - | 0 | - |
| - | - | 1617 | 682.8 | - | - | 0 | - |
| 9 | y | 7925 | 683.3 | 0.0014 | 2.049 | +1 | 5 |
| - | - | 4048 | 684.3 | - | - | 0 | - |
| - | - | 1674 | 685.3 | - | - | 0 | - |
| - | - | 1255 | 687.8 | - | - | 0 | - |
| - | - | 1526 | 692.4 | - | - | 0 | - |
| - | - | 1087 | 695.3 | - | - | 0 | - |
| - | - | 4364 | 696.4 | - | - | 0 | - |
| - | - | 3918 | 696.8 | - | - | 0 | - |
| - | - | 2716 | 696.9 | - | - | 0 | - |
| - | - | 1916 | 697.3 | - | - | 0 | - |
| - | - | 1863 | 697.4 | - | - | 0 | - |
| - | - | 2804 | 697.8 | - | - | 0 | - |
| - | - | 747 | 698.3 | - | - | 0 | - |
| - | - | 1388 | 701.4 | - | - | 0 | - |
| - | - | 1788 | 701.9 | - | - | 0 | - |
| - | - | 5479 | 702.3 | - | - | 0 | - |
| - | - | 1578 | 702.4 | - | - | 0 | - |
| - | - | 1496 | 702.8 | - | - | 0 | - |
| - | - | 770.3 | 702.9 | - | - | 0 | - |
| - | - | 2888 | 703.3 | - | - | 0 | - |
| - | - | 1028 | 704.3 | - | - | 0 | - |
| - | - | 1.792E+04 | 710.4 | - | - | 0 | - |
| 12 | b | 6206 | 710.8 | 0.000464 | 0.6528 | +2 | 12 |
| - | - | 1.488E+04 | 710.9 | - | - | 0 | - |
| - | - | 4901 | 711.3 | - | - | 0 | - |
| - | - | 5965 | 711.4 | - | - | 0 | - |
| - | - | 1957 | 711.8 | - | - | 0 | - |
| - | - | 1906 | 711.9 | - | - | 0 | - |
| - | - | 1084 | 716.8 | - | - | 0 | - |
| - | - | 1483 | 717.3 | - | - | 0 | - |
| - | - | 1495 | 717.8 | - | - | 0 | - |
| 6 | b | 856.5 | 719.2 | 0.0004033 | 0.5608 | +1 | 6 |
| - | - | 951 | 719.3 | - | - | 0 | - |
| - | - | 3640 | 719.8 | - | - | 0 | - |
| - | - | 1426 | 720.3 | - | - | 0 | - |
| - | - | 3380 | 720.4 | - | - | 0 | - |
| - | - | 1715 | 720.8 | - | - | 0 | - |
| - | - | 1184 | 721.4 | - | - | 0 | - |
| - | - | 7417 | 725.8 | - | - | 0 | - |
| - | - | 4257 | 726.3 | - | - | 0 | - |
| - | - | 1370 | 726.8 | - | - | 0 | - |
| - | - | 916.1 | 727.3 | - | - | 0 | - |
| - | - | 1935 | 735.3 | - | - | 0 | - |
| - | - | 792.7 | 736.3 | - | - | 0 | - |
| 8 | y | 3948 | 737.3 | 0.0003645 | 0.4944 | +1 | 6 |
| - | - | 1431 | 738.3 | - | - | 0 | - |
| - | - | 957.6 | 739.3 | - | - | 0 | - |
| - | - | 1011 | 740.3 | - | - | 0 | - |
| - | - | 2754 | 744.4 | - | - | 0 | - |
| - | - | 2416 | 744.9 | - | - | 0 | - |
| - | - | 850 | 745.4 | - | - | 0 | - |
| 2 | y | 2907 | 748.8 | 0.00127 | 1.696 | +2 | 12 |
| 2 | y | 3081 | 749.3 | 0.005685 | 7.587 | +2 | 12 |
| - | - | 3117 | 749.8 | - | - | 0 | - |
| - | - | 1902 | 750.3 | - | - | 0 | - |
| 8 | y | 1.624E+04 | 754.4 | 0.0001216 | 0.1612 | +1 | 6 |
| - | - | 6791 | 755.4 | - | - | 0 | - |
| - | - | 2971 | 756.4 | - | - | 0 | - |
| 2 | y | 1.508E+04 | 757.8 | 0.0002657 | 0.3507 | +2 | 12 |
| - | - | 1.372E+04 | 758.3 | - | - | 0 | - |
| - | - | 8384 | 758.8 | - | - | 0 | - |
| - | - | 3390 | 759.3 | - | - | 0 | - |
| - | - | 1901 | 761.3 | - | - | 0 | - |
| - | - | 1868 | 765.4 | - | - | 0 | - |
| - | - | 831.6 | 766.4 | - | - | 0 | - |
| - | - | 1233 | 768.3 | - | - | 0 | - |
| - | - | 1042 | 784.3 | - | - | 0 | - |
| - | - | 5426 | 784.4 | - | - | 0 | - |
| - | - | 4811 | 785.4 | - | - | 0 | - |
| - | - | 1505 | 786.4 | - | - | 0 | - |
| - | - | 2252 | 793.4 | - | - | 0 | - |
| - | - | 1091 | 808.4 | - | - | 0 | - |
| - | - | 1080 | 811.4 | - | - | 0 | - |
| - | - | 821.4 | 812.4 | - | - | 0 | - |
| - | - | 975 | 821.4 | - | - | 0 | - |
| - | - | 1152 | 829.4 | - | - | 0 | - |
| - | - | 1111 | 832.3 | - | - | 0 | - |
| - | - | 1045 | 833.3 | - | - | 0 | - |
| - | - | 1466 | 839.4 | - | - | 0 | - |
| - | - | 817.8 | 840.4 | - | - | 0 | - |
| 7 | b | 1989 | 848.3 | 0.0008211 | 0.968 | +1 | 7 |
| - | - | 1351 | 848.4 | - | - | 0 | - |
| - | - | 863.8 | 849.3 | - | - | 0 | - |
| - | - | 2309 | 849.4 | - | - | 0 | - |
| - | - | 1554 | 850.4 | - | - | 0 | - |
| - | - | 1096 | 855.3 | - | - | 0 | - |
| - | - | 2101 | 857.4 | - | - | 0 | - |
| - | - | 1467 | 864.5 | - | - | 0 | - |
| 7 | y | 3768 | 865.4 | 0.0001075 | 0.1242 | +1 | 7 |
| 7 | y | 3601 | 866.4 | 0.005501 | 6.349 | +1 | 7 |
| - | - | 2478 | 867.4 | - | - | 0 | - |
| - | - | 1054 | 881.4 | - | - | 0 | - |
| - | - | 1442 | 881.5 | - | - | 0 | - |
| 7 | y | 1.32E+04 | 883.4 | 0.001334 | 1.51 | +1 | 7 |
| - | - | 6953 | 884.4 | - | - | 0 | - |
| - | - | 2113 | 885.4 | - | - | 0 | - |
| - | - | 1029 | 888.5 | - | - | 0 | - |
| - | - | 1183 | 893.4 | - | - | 0 | - |
| 8 | b | 3716 | 919.3 | 0.0002731 | 0.297 | +1 | 8 |
| - | - | 1788 | 920.3 | - | - | 0 | - |
| - | - | 2403 | 921.4 | - | - | 0 | - |
| - | - | 924.9 | 921.5 | - | - | 0 | - |
| - | - | 2690 | 922.4 | - | - | 0 | - |
| - | - | 944.1 | 938.4 | - | - | 0 | - |
| - | - | 843.3 | 941.4 | - | - | 0 | - |
| - | - | 2261 | 964.5 | - | - | 0 | - |
| - | - | 3301 | 971.5 | - | - | 0 | - |
| - | - | 1753 | 972.5 | - | - | 0 | - |
| - | - | 762.7 | 987.5 | - | - | 0 | - |
| - | - | 786.9 | 991.4 | - | - | 0 | - |
| - | - | 1620 | 992.5 | - | - | 0 | - |
| 6 | y | 833 | 1002 | 0.00482 | 4.808 | +1 | 8 |
| 6 | y | 2668 | 1003 | 0.0003611 | 0.3598 | +1 | 8 |
| - | - | 1997 | 1004 | - | - | 0 | - |
| - | - | 841.3 | 1005 | - | - | 0 | - |
| - | - | 911.8 | 1018 | - | - | 0 | - |
| 6 | y | 1.084E+04 | 1020 | 0.001408 | 1.38 | +1 | 8 |
| - | - | 771.1 | 1021 | - | - | 0 | - |
| - | - | 6648 | 1021 | - | - | 0 | - |
| - | - | 2596 | 1022 | - | - | 0 | - |
| - | - | 9008 | 1032 | - | - | 0 | - |
| 9 | b | 2585 | 1032 | 0.0006207 | 0.6012 | +1 | 9 |
| - | - | 4079 | 1033 | - | - | 0 | - |
| - | - | 1774 | 1033 | - | - | 0 | - |
| - | - | 5790 | 1035 | - | - | 0 | - |
| - | - | 2261 | 1036 | - | - | 0 | - |
| - | - | 1854 | 1037 | - | - | 0 | - |
| - | - | 1078 | 1082 | - | - | 0 | - |
| - | - | 1064 | 1086 | - | - | 0 | - |
| - | - | 3361 | 1103 | - | - | 0 | - |
| - | - | 1795 | 1105 | - | - | 0 | - |
| - | - | 1096 | 1132 | - | - | 0 | - |
| 5 | y | 2180 | 1167 | 0.002959 | 2.534 | +1 | 9 |
| - | - | 1170 | 1168 | - | - | 0 | - |
| - | - | 717.4 | 1220 | - | - | 0 | - |
| 11 | b | 1232 | 1284 | 0.0007547 | 0.588 | +1 | 11 |
| - | - | 860.5 | 3081 | - | - | 0 | - |

m/z Charge Intensity FragmentType MassShift Position
123.04436492919922 0 1662.9977
124.0757827758789 0 624.1047
127.05052185058594 0 1830.5292
129.0662384033203 0 7717.4478
129.1026153564453 0 15056.308
130.03248596191406 0 2163.2832
130.10580444335938 0 582.14026
131.08184814453125 0 2147.101
132.1024627685547 0 1211.1041
132.9528045654297 0 395.8167
133.04342651367188 0 905.7753
134.02743530273438 0 35504.59
135.03076171875 0 1632.0455
136.0232696533203 0 1101.6877
136.0509490966797 0 495.6004
136.07608032226562 0 34600.277
137.07937622070312 0 2156.3945
138.0665740966797 0 2237.4065
139.0868682861328 0 449.68234
141.06735229492188 0 367.2965
141.10263061523438 0 2155.9302
144.0118408203125 0 875.9024
144.03549194335938 0 422.158
147.0446014404297 0 1372.3066
148.08724975585938 0 1460.1729
149.0453338623047 0 5487.184
151.61676025390625 0 478.49908
152.0819549560547 0 693.19
152.42080688476562 0 494.8633
153.1021728515625 0 438.40735
154.06153869628906 0 493.2197
155.08218383789062 0 999.90027
155.093017578125 0 2544.3726
155.1182861328125 0 1853.104
156.07713317871094 0 7631.531
157.06118774414062 0 6302.7085
157.080322265625 0 620.60443
157.0975341796875 0 666.56805
157.1338653564453 0 645.1736
158.0274200439453 0 19548.896
159.03091430664062 0 779.407
159.11318969726562 0 25748.852
160.02320861816406 0 951.1764
160.0870819091797 0 442.6699
160.11639404296875 0 1945.1877
161.0382843017578 0 11727.299
162.06654357910156 0 507.3088
165.05499267578125 0 16716.934
165.07778930664062 0 1307.1722
165.1028594970703 0 593.92267
166.06088256835938 0 2556.1172
166.08119201660156 0 446.56808
166.08665466308594 0 4757.13
167.0558319091797 0 6418.5103
167.09329223632812 0 1371.4215
167.99319458007812 0 426.0251
168.0050506591797 0 437.0922
168.01174926757812 0 3347.9043
169.09739685058594 0 1073.0554
171.1130828857422 0 901.6173
173.12884521484375 0 2482.0784
173.4505157470703 0 3130.246
174.0220184326172 0 535.17255
174.06680297851562 0 684.9244
175.0538787841797 0 516.88043
176.0380096435547 0 21182.668
176.08230590820312 0 2718.3257
177.04226684570312 0 661.37726
178.03427124023438 0 876.7686
179.09283447265625 0 999.61395
180.0773162841797 0 1294.3801
180.11318969726562 0 493.056
181.09805297851562 0 894.19635
182.0815887451172 0 28601.342 y 12
182.1288604736328 0 1052.3619
183.08493041992188 0 3011.6917
183.11302185058594 0 1683.9653
185.03834533691406 0 1933.6489
185.09303283691406 0 618.64874
186.02236938476562 0 18619.889
186.08798217773438 0 557.184
186.1240997314453 0 801.2776
187.02574157714844 0 1256.0209
187.10804748535156 0 10842.6045
188.01829528808594 0 948.0188
188.11148071289062 0 891.4789
189.07754516601562 0 1333.6342
190.0615997314453 0 3520.7078
191.0932159423828 0 2211.1174
193.07272338867188 0 640.8271
193.1086883544922 0 6338.7266
194.0924530029297 0 1009.6247
194.11305236816406 0 751.84814
195.1135711669922 0 490.0404
197.1033935546875 0 967.5586
198.08815002441406 0 497.22818
199.07180786132812 0 1723.9066
199.1075897216797 0 832.28125
200.1046142578125 0 520.92725
200.13951110839844 0 1603.2594
201.0696563720703 0 519.2671
201.08750915527344 0 1119.4921
201.12380981445312 0 1182.9384
202.10888671875 0 842.8278
203.04879760742188 0 5066.41
203.09231567382812 0 1211.626
204.0327606201172 0 846.6868
204.07720947265625 0 4139.889
205.08103942871094 0 714.2844
205.1090850830078 0 595.07996
206.10372924804688 0 830.4523
207.08799743652344 0 8371.042
208.07223510742188 0 511.2069
208.09202575683594 0 517.7091
210.12416076660156 0 1033.044
211.1087188720703 0 767.3469
212.13987731933594 0 559.19867
214.130126953125 0 658.5379
216.0980987548828 0 880.6363
216.13453674316406 0 4442.0527
217.07264709472656 0 1954.9281
218.05653381347656 0 2344.2458
221.0595245361328 0 186799.2 a 1
221.08493041992188 0 9713.626
221.10374450683594 0 17249.865
222.0625762939453 0 15243.112
222.0864715576172 0 1878.1965
222.10739135742188 0 1583.5674
223.05514526367188 0 6600.586
223.0644989013672 0 1449.3973
223.08216857910156 0 968.8638
223.15586853027344 0 1052.9174
224.114501953125 0 7688.052
225.04331970214844 0 12753.03
225.0987091064453 0 1045.9138
225.1181640625 0 631.16626
226.04371643066406 0 2512.307
226.08261108398438 0 7594.8516
226.11935424804688 0 1414.9585
226.15562438964844 0 1654.4963
227.02294921875 0 1599.0055
227.03948974609375 0 725.43616
227.08644104003906 0 586.79364
228.113525390625 0 571.70026
228.13475036621094 0 680.6214
230.0840606689453 0 554.45874
231.0437469482422 0 5872.749 b Water loss 1
233.1397705078125 0 1060.6482
234.0988006591797 0 1548.3304
234.12380981445312 0 1007.76135
235.0830078125 0 13940.813
236.06716918945312 0 632.87317
236.08644104003906 0 1256.5856
238.11912536621094 0 876.366
239.09542846679688 0 21864.068
239.11441040039062 0 3661.0247
240.09609985351562 0 4323.3003
241.0926055908203 0 2101.3267
244.09317016601562 0 14968.763
245.09642028808594 0 1519.7354
247.11170959472656 0 1797.0786
249.05441284179688 0 54786.625 b 1
249.0986328125 0 3613.2156
249.13514709472656 0 960.7773
250.05772399902344 0 4982.7354
250.0820770263672 0 707.9763
251.05052185058594 0 1855.7437
251.1507110595703 0 4730.0444
252.1095733642578 0 29235.883
252.1345977783203 0 1815.6672
253.09375 0 3765.7874
253.11265563964844 0 2751.0366
254.1608428955078 0 1215.7776
257.1072692871094 0 2139.1511
258.1104431152344 0 714.07434
266.1137390136719 0 606.1518
267.09295654296875 0 1007.20483
267.1091613769531 0 9829.031
268.1123046875 0 800.4426
268.12872314453125 0 612.82776
269.13702392578125 0 711.40106
269.1613464355469 0 915.23035
270.0815734863281 0 549.731
270.1200866699219 0 2943.7556
270.1448059082031 0 1968.427
273.13494873046875 0 2692.3433
275.1064758300781 0 2673.1404
275.1249084472656 0 570.32153
277.0941162109375 0 904.9663
277.111328125 0 1393.5432 y Ammonia loss 9
279.1468811035156 0 1066.3076
280.10455322265625 0 1536.6111
281.05224609375 0 908.24365
281.1500244140625 0 4310.7944
282.0520324707031 0 689.6166
285.1020812988281 0 6969.3853
285.6249694824219 0 6077.3154 y 9
286.10504150390625 0 790.4023
286.12652587890625 0 1982.5247
287.1717529296875 0 827.69275
295.1036682128906 0 4923.0044
296.1044616699219 0 1818.6685
297.1555480957031 0 799.6721
299.06219482421875 0 10630.596
300.0627136230469 0 2811.714
301.05877685546875 0 1302.0095
301.1300048828125 0 6565.812
302.13360595703125 0 911.42365
302.16241455078125 0 1311.7583
304.1770324707031 0 1693.7806
305.1255187988281 0 585.43713
313.11395263671875 0 1124.2681
314.1174011230469 0 652.9498
316.1336669921875 0 817.85364
318.0758056640625 0 6400.4272 b Water loss 2
319.07928466796875 0 936.7415
319.1406555175781 0 32770.84 y 11
319.6399230957031 0 551.82825
320.1386413574219 0 7143.755
320.1729736328125 0 1675.39
321.1412048339844 0 878.03033
322.18768310546875 0 4032.8418
323.1925964355469 0 711.1997
326.1366882324219 0 1142.3474
328.15252685546875 0 1122.6863
329.125244140625 0 2677.108
329.21734619140625 0 1078.3718
330.1123352050781 0 3485.8975
332.1347961425781 0 1312.4594
333.11907958984375 0 617.6657
334.1436462402344 0 4152.685
336.0864562988281 0 4423.1836 b 2
337.09014892578125 0 682.7794
337.12939453125 0 1269.3722
337.1577453613281 0 713.2741
338.14654541015625 0 5684.979
339.14990234375 0 834.47955
342.1674499511719 0 1308.3187 y 8
342.66778564453125 0 1085.32
343.1632995605469 0 576.8778
344.1477355957031 0 1408.1841
345.67205810546875 0 900.7058
348.12408447265625 0 1545.695
348.1675109863281 0 1497.2795
350.14642333984375 0 7054.1104
351.14935302734375 0 1622.2385
353.1911315917969 0 670.0188
354.1324768066406 0 939.6592
355.06964111328125 0 1143.4326
355.1361083984375 0 632.69135
357.21875 0 655.96576
360.12957763671875 0 850.711 b 5
361.1729431152344 0 1211.5729
361.201171875 0 682.7584
362.1772155761719 0 600.3817
362.832275390625 0 623.31104
365.19329833984375 0 3691.6646
366.1410827636719 0 2409.8757
366.1775207519531 0 828.7992
366.6549072265625 0 554.9689
367.1751403808594 0 613.9602
369.1225280761719 0 3290.8047
370.122314453125 0 1523.2352
370.1495056152344 0 841.20197
370.6598815917969 0 696.66614
371.12109375 0 930.73047
371.1575927734375 0 6698.539
372.1412353515625 0 4745.131
372.16357421875 0 1056.3025
373.1443176269531 0 592.70746
374.1830749511719 0 675.4407
377.68524169921875 0 2179.204 y 7
379.2099914550781 0 1670.7098
381.1197814941406 0 3547.1724
382.1231994628906 0 768.484
382.2449035644531 0 1088.6926
383.20318603515625 0 939.31586
384.17034912109375 0 1799.1799
389.1454772949219 0 905.59814
389.1690673828125 0 1728.0149
390.1521301269531 0 2470.3018
390.65899658203125 0 676.989
391.208740234375 0 839.2238
393.188720703125 0 1214.3529
394.18096923828125 0 546.1221
396.1334533691406 0 604.48083
398.1458435058594 0 2187.2075
399.1306457519531 0 3961.5554
399.6722412109375 0 856.7286
400.13433837890625 0 1485.3182
400.2566833496094 0 1668.4551
401.2575378417969 0 625.07947
402.689697265625 0 919.73047
403.17828369140625 0 628.13446
403.9064636230469 0 884.3105
404.2413635253906 0 717.29254
405.1708679199219 0 605.14044
405.2257080078125 0 2200.158
406.20257568359375 0 2616.784
406.7039489746094 0 1566.1907
407.1558532714844 0 778.188
407.18023681640625 0 941.6711
407.20465087890625 0 2821.19
408.1787414550781 0 1637.4913
408.2366943359375 0 717.32684
410.6517333984375 0 1307.9185
411.19427490234375 0 656.4276
411.68988037109375 0 674.6646
412.1634521484375 0 1154.0099
414.1443786621094 0 2984.1812
414.1732482910156 0 1282.1788
414.6731872558594 0 1381.0605
415.0382995605469 0 850.8871
415.1455993652344 0 799.713
415.1739196777344 0 1259.2656
415.21881103515625 0 586.95605
415.6432800292969 0 638.18097 b Water loss 6
415.81072998046875 0 612.7144
415.88818359375 0 687.4425
416.1568908691406 0 9280.614 y Ammonia loss 10
416.69024658203125 0 1974.7756
417.1416320800781 0 1206.666 b Water loss 3
417.1642150878906 0 1424.3704
417.1890869140625 0 862.1483 y Ammonia loss 3
420.2015380859375 0 2465.7905
420.69677734375 0 1029.9917
421.1836242675781 0 5109.4404
422.15338134765625 0 686.42755
422.1859130859375 0 1161.0277
422.2167053222656 0 710.4365
422.86224365234375 0 792.3821 y 3
423.18890380859375 0 718.68677
423.23486328125 0 3427.6726
423.52752685546875 0 843.7284
424.5355529785156 0 2393.96
424.6497802734375 0 1361.515 b 6
424.8714294433594 0 1249.045
425.69195556640625 0 904.4721
426.1842346191406 0 913.03394
429.1766357421875 0 1317.717
429.205810546875 0 1896.6025
429.7070617675781 0 949.6154
430.5389709472656 0 725.79315
430.6961975097656 0 820.2225
431.1640930175781 0 626.0372
431.58563232421875 0 2018.1458
431.69818115234375 0 819.04364
431.92034912109375 0 2497.022
433.1836242675781 0 16857.537 y 10
433.218017578125 0 1434.3263
434.1866760253906 0 3126.7805
435.1544189453125 0 907.2716 b 3
436.23089599609375 0 4685.327
440.1923522949219 0 807.27905
440.9182434082031 0 6854.388
441.252197265625 0 5869.8374
441.58709716796875 0 2709.136
442.2079772949219 0 1053.5668 y 6
443.17425537109375 0 714.68555
443.2074890136719 0 3611.63
443.86407470703125 0 674.53827
444.2087707519531 0 654.64856
445.8690185546875 0 5121.8804 y Water loss 2
446.1711120605469 0 1271.4476
446.2032165527344 0 8728.348 y Ammonia loss 2
446.5370178222656 0 3181.2112
446.8702087402344 0 1522.421
447.21875 0 1101.2588
447.7196350097656 0 2240.6682
448.22344970703125 0 1109.3965
449.2141418457031 0 1089.6449
451.2304382324219 0 6449.6675
451.8721008300781 0 7105.969 y 2
452.20635986328125 0 6897.4443
452.234130859375 0 929.2793
452.5408935546875 0 2511.7617
452.7026672363281 0 1142.1757
452.8756408691406 0 1550.7053
455.2010498046875 0 1273.7013
456.24542236328125 0 763.9588
458.70196533203125 0 1004.1913
459.2682189941406 0 694.8066
459.5321044921875 0 839.67676
460.16802978515625 0 1577.12 b 7
461.21490478515625 0 6464.7407
461.7165832519531 0 3379.0396
462.21502685546875 0 1627.7495
462.7165832519531 0 940.5909
463.59869384765625 0 1155.3561
463.9328918457031 0 1659.9857
464.2624206542969 0 1026.8718
464.6063537597656 0 893.7791
464.8641662597656 0 1772.2719
465.19635009765625 0 1239.2723
465.5365295410156 0 985.0252
467.21490478515625 0 3358.4973
467.7127990722656 0 1724.8307
468.2566833496094 0 1039.5468
470.72027587890625 0 1785.0975
471.20294189453125 0 3567.7896
471.58489990234375 0 6960.9155
471.91900634765625 0 5098.576
472.2550048828125 0 1115.8417
473.2152404785156 0 1439.5797
473.7071838378906 0 992.17755
473.9405517578125 0 3478.6826
474.1979675292969 0 1601.2828 b 11
474.2735900878906 0 2294.6274
474.5312805175781 0 1864.8047
474.86248779296875 0 967.42035
476.22003173828125 0 11658.158
476.55419921875 0 1744.9143
476.72149658203125 0 3448.5042
476.8883361816406 0 1563.9701
477.2203063964844 0 774.95984
478.20684814453125 0 1037.8557
478.54351806640625 0 866.11505
479.2223815917969 0 763.5201
479.7201232910156 0 2661.5962
479.75689697265625 0 714.84296
480.2237243652344 0 1340.2223
480.72186279296875 0 789.9743
482.2197570800781 0 3625.2188
482.7169189453125 0 2036.7957
483.21881103515625 0 748.3833
484.21063232421875 0 2580.5984
485.1818542480469 0 2593.29
485.21844482421875 0 1187.8253
486.1834716796875 0 783.75287
486.2388916015625 0 937.5909
486.70068359375 0 762.6282
489.23529052734375 0 758.78455
491.2255859375 0 30682.064
491.7268371582031 0 12193.177
492.218505859375 0 1368.4912
492.5466003417969 0 1134.1705
492.7225036621094 0 861.9196
492.87640380859375 0 961.24646
493.71722412109375 0 2879.7212
494.20989990234375 0 1943.1161
494.5346984863281 0 1057.373
495.2132263183594 0 3635.0085
495.7179870605469 0 2468.8337
497.5786437988281 0 1254.9321
497.8861083984375 0 7517.551
498.2229309082031 0 8373.313
498.5531005859375 0 3465.2131
498.8849182128906 0 1044.1432
499.220458984375 0 1266.351
499.5408020019531 0 3054.8535 y Water loss 1
499.8726806640625 0 1742.9377 y Ammonia loss 1
500.54266357421875 0 771.11115
501.72930908203125 0 840.4976 y Water loss 5
501.8776550292969 0 948.29315
502.2240295410156 0 2178.6775 y Ammonia loss 5
502.7230224609375 0 4139.4565
503.10675048828125 0 844.7649
503.2273254394531 0 1886.1082
503.8919372558594 0 714.78217
504.2316589355469 0 2150.85
504.7296142578125 0 1056.6112
505.5435485839844 0 11146.761 y 1
505.8780517578125 0 8138.581
506.210693359375 0 5052.8438
506.2716064453125 0 993.44293
506.54412841796875 0 2269.2485
506.87945556640625 0 1047.589
507.2148132324219 0 2725.7644
507.550048828125 0 1995.5823
507.8871154785156 0 873.2096
508.21295166015625 0 910.0993
508.2523498535156 0 1771.7448
508.5475769042969 0 695.5776
510.73602294921875 0 16608.379 y 5
510.7734680175781 0 575.80145
511.2376403808594 0 6943.0034
511.74072265625 0 2471.8254
513.2363891601562 0 29471.4
513.55615234375 0 8702.452
513.7398681640625 0 10588.122
513.8903198242188 0 6804.606
516.5442504882812 0 2310.058
516.8746948242188 0 2128.566
517.2056274414062 0 1516.4095
517.246826171875 0 1175.4052
517.5410766601562 0 1572.9552
517.8764038085938 0 1085.7938
518.237548828125 0 3438.8562
518.7378540039062 0 1148.8162
520.2562255859375 0 2515.1501
523.209716796875 0 1619.4048
526.2447509765625 0 725.53796
528.2265014648438 0 883.93146
528.26708984375 0 1164.8695
528.5498046875 0 6587.055 Precursor Water loss
528.6239013671875 0 1229.4047
528.8828125 0 11697.486 Precursor Ammonia loss
529.216064453125 0 7524.698
529.2933959960938 0 1182.9495
529.5493774414062 0 5597.136
529.8814697265625 0 856.1652
531.23583984375 0 900.9598
532.2605590820312 0 671.101
532.7664184570312 0 783.1731
533.2636108398438 0 967.9467
533.3065185546875 0 2822.1992
533.807373046875 0 1285.831
533.8949584960938 0 880.16095
534.2662353515625 0 1254.089
534.3014526367188 0 9549.274
534.5542602539062 0 80607.54 Precursor
534.6322631835938 0 7041.2393
534.8884887695312 0 70668.01
534.9669799804688 0 3869.6943
535.22119140625 0 38405.3
535.3104248046875 0 1439.2444
535.5557250976562 0 16900.834
535.6967163085938 0 1054.1912
535.7530517578125 0 7318.9946
535.8881225585938 0 5065.596
536.2053833007812 0 1101.0709
536.2477416992188 0 4309.9136
537.2471313476562 0 1180.3625
538.7421264648438 0 932.99884
539.2448120117188 0 1120.7378
540.2926635742188 0 742.9461
540.7696533203125 0 2282.8962
541.2086181640625 0 1055.4808
541.27197265625 0 1945.7302
543.2493896484375 0 2515.3752
543.7440185546875 0 5594.218
544.2470092773438 0 1828.7994
545.29345703125 0 2382.4436
547.2605590820312 0 1827.0051
548.3170776367188 0 1304.2751
548.8165283203125 0 1130.5961
552.2548217773438 0 10779.676
552.756103515625 0 7549.822
553.2158203125 0 8094 y Ammonia loss 9
553.2586059570312 0 3025.7495
553.3125 0 715.2047
553.7590942382812 0 1071.8713
554.220458984375 0 1988.8964
554.768798828125 0 2377.6616
555.268798828125 0 2315.7366
555.7671508789062 0 963.42255
556.2628173828125 0 1048.8131
560.2929077148438 0 1898.7996
560.8150634765625 0 1716.3998
561.3154907226562 0 1094.0261
563.7726440429688 0 2669.8374
564.2717895507812 0 1794.5612
564.770263671875 0 1109.7848
565.2721557617188 0 4011.1147
566.7412109375 0 815.8262
568.2186889648438 0 776.11096
570.2421264648438 0 18105.63 y 9
570.2886352539062 0 963.23065
571.2446899414062 0 5919.16
572.2493896484375 0 3446.3896
572.7694702148438 0 6820.963
573.2689819335938 0 3305.7622
573.7703857421875 0 2269.4143
574.2747802734375 0 1029.3356
575.7428588867188 0 3257.5344 y Ammonia loss 4
576.2450561523438 0 1347.4069 b Water loss 9
576.7442626953125 0 1517.7877
577.7612915039062 0 1820.8688
578.258056640625 0 1131.8276
580.3182373046875 0 1471.7506
581.3206787109375 0 828.3564
582.2325439453125 0 950.9402
584.2537841796875 0 21455.648 y 4
584.7555541992188 0 13390.533
585.2547607421875 0 7440.161
585.7536010742188 0 3171.9421
586.7667846679688 0 8841.055
587.2666015625 0 6585.642
587.7675170898438 0 2761.517
588.2879638671875 0 2749.6301
589.2791137695312 0 1022.3865
591.2960815429688 0 1224.3302
593.2745361328125 0 893.00165
593.779296875 0 1556.6296
594.2562866210938 0 705.6447
595.3294677734375 0 907.5689
595.7725219726562 0 5302.9746
596.274169921875 0 3250.694
596.7743530273438 0 849.9248
596.8466186523438 0 1411.0742
598.266357421875 0 1711.758
600.2451782226562 0 6418.1743
601.2467041015625 0 3120.0403
601.7890014648438 0 5224.873
602.241943359375 0 928.4116
602.2908325195312 0 3511.4133
602.7921142578125 0 1958.1709
604.2203979492188 0 944.4981
604.3306884765625 0 3915.9077
604.832275390625 0 2195.6475
605.3553466796875 0 2784.72
605.85791015625 0 1484.1755
607.2841796875 0 4457.6167
608.3197631835938 0 1001.271
610.34765625 0 707.20734
616.2808837890625 0 843.95355
621.2759399414062 0 1162.0725
625.2764282226562 0 2310.1704 y Ammonia loss 3
625.7766723632812 0 895.4087
631.794677734375 0 877.966
631.8523559570312 0 827.8653
632.21435546875 0 765.04535
633.7880249023438 0 14222.67 y 3
634.2890014648438 0 8124.632
634.78955078125 0 4068.8457
635.2846069335938 0 2690.7346
635.7767333984375 0 2045.1196
636.2977905273438 0 2311.9187
636.7921752929688 0 4432.8755
637.2941284179688 0 2768.7354
639.8699951171875 0 835.72485
642.2559814453125 0 835.87744 b 10
642.764404296875 0 1663.4751
645.3047485351562 0 29939.69
645.8060302734375 0 23442.686
646.3074951171875 0 11156.298
646.80810546875 0 2318.4744
646.8699340820312 0 1277.0552
648.2964477539062 0 1050.9316
649.326416015625 0 830.531
651.8687133789062 0 1158.127
652.3651733398438 0 925.03925
653.273193359375 0 2195.3499
653.7780151367188 0 1019.5105
653.8670043945312 0 1590.0974
654.3015747070312 0 1430.2908
654.3659057617188 0 1681.3477
654.8001098632812 0 1235.4158
657.31103515625 0 1503.4478
659.7859497070312 0 1166.5667
660.8729858398438 0 25990.459
661.3743286132812 0 16076.122
661.875732421875 0 6193.5957
662.3756713867188 0 2926.006
664.7935180664062 0 1507.4062
666.3023681640625 0 2682.4604 y Ammonia loss 8
667.2802124023438 0 1847.5922
667.7747802734375 0 2163.5454
668.2972412109375 0 5779.5923 y Water loss 2
668.7933959960938 0 13414.151 y Ammonia loss 2
669.2921752929688 0 10074.403
669.7932739257812 0 4455.22
670.2933349609375 0 887.4395
671.281982421875 0 6600.431
672.2855834960938 0 2650.984
674.3370971679688 0 2541.5837
676.2803955078125 0 1987.2965
677.3041381835938 0 109798.18 y 2
677.805419921875 0 82730.87
678.3059692382812 0 38485.043
678.806396484375 0 11947.713
679.3057861328125 0 2588.727
680.3079833984375 0 729.7956
682.2958984375 0 1970.2701
682.3634643554688 0 1131.3683
682.7979125976562 0 1617.3276
683.3245849609375 0 7925.17 y 8
684.32763671875 0 4047.5771
685.3168334960938 0 1674.4767
687.7926635742188 0 1255.1984
692.3706665039062 0 1525.555
695.2919311523438 0 1086.8351
696.4093627929688 0 4363.6064
696.793212890625 0 3918.495
696.9113159179688 0 2716.4036
697.2890625 0 1916.124
697.4111328125 0 1862.8704
697.7936401367188 0 2804.4563
698.2864379882812 0 746.9754
701.3895874023438 0 1388.3087
701.8942260742188 0 1788.1348
702.3333129882812 0 5479.4087
702.3978271484375 0 1577.9796
702.810302734375 0 1495.7831
702.9031372070312 0 770.2649
703.3262329101562 0 2888.1074
704.3189697265625 0 1028.1436
710.4072875976562 0 17921.629
710.79052734375 0 6205.5796 b 11
710.908447265625 0 14876.948
711.2910766601562 0 4901.359
711.4083862304688 0 5964.8804
711.7926635742188 0 1957.3131
711.909912109375 0 1906.2692
716.8038330078125 0 1083.965
717.3030395507812 0 1483.3143
717.80615234375 0 1495.0923
719.2479248046875 0 856.5059 b 5
719.3217163085938 0 951.0007
719.7957763671875 0 3640.204
720.296875 0 1426.3815
720.36328125 0 3380.4119
720.798828125 0 1714.9243
721.36669921875 0 1183.8884
725.8117065429688 0 7417.31
726.3139038085938 0 4257.367
726.814208984375 0 1370.2068
727.3109130859375 0 916.103
735.3232421875 0 1934.7201
736.3370971679688 0 792.67755
737.3369140625 0 3947.6147 y Ammonia loss 7
738.3414306640625 0 1430.7706
739.3466186523438 0 957.637
740.2935180664062 0 1011.13776
744.4017333984375 0 2754.0518
744.9025268554688 0 2415.533
745.4015502929688 0 850.0469
748.804443359375 0 2907.092 y Water loss 1
749.3034057617188 0 3081.3635 y Ammonia loss 1
749.803466796875 0 3116.5454
750.298583984375 0 1902.1136
754.3632202148438 0 16237.475 y 7
755.366455078125 0 6790.8584
756.369384765625 0 2971.4167
757.8107299804688 0 15081.589 y 1
758.3125610351562 0 13716.69
758.8124389648438 0 8384.265
759.3116455078125 0 3390.095
761.260498046875 0 1900.9945
765.401123046875 0 1868.0925
766.3527221679688 0 831.56226
768.2959594726562 0 1232.648
784.2946166992188 0 1042.4282
784.3652954101562 0 5426.217
785.3695068359375 0 4811.162
786.3676147460938 0 1505.2463
793.3970336914062 0 2252.3389
808.3705444335938 0 1091.3547
811.3952026367188 0 1079.54
812.3921508789062 0 821.38403
821.3699951171875 0 974.9744
829.4307861328125 0 1151.9939
832.289306640625 0 1110.5297
833.2952270507812 0 1045.1805
839.3939819335938 0 1466.2458
840.3807983398438 0 817.7846
848.2901000976562 0 1989.3342 b 6
848.3726806640625 0 1351.4175
849.2908935546875 0 863.7859
849.3699951171875 0 2308.728
850.3751220703125 0 1554.4519
855.3338012695312 0 1096.1892
857.4231567382812 0 2101.2053
864.469970703125 0 1466.8682
865.39501953125 0 3768.311 y Water loss 6
866.3846435546875 0 3601.1333 y Ammonia loss 6
867.3833618164062 0 2478.0706
881.3861694335938 0 1053.7958
881.49853515625 0 1441.5314
883.4043579101562 0 13195.679 y 6
884.40966796875 0 6952.535
885.4141845703125 0 2113.425
888.4584350585938 0 1029.014
893.4232177734375 0 1183.2404
919.3283081054688 0 3716.3489 b 7
920.3333129882812 0 1787.6711
921.4203491210938 0 2403.1653
921.5077514648438 0 924.9129
922.4239501953125 0 2690.2236
938.447265625 0 944.0555
941.4212036132812 0 843.33215
964.5305786132812 0 2260.8127
971.4689331054688 0 3300.5938
972.4697265625 0 1753.1251
987.5240478515625 0 762.7217
991.4474487304688 0 786.87476
992.5310668945312 0 1620.1185
1002.44921875 0 833.0154 y Water loss 5
1003.4384155273438 0 2667.7048 y Ammonia loss 5
1004.4353637695312 0 1996.5612
1005.4401245117188 0 841.30676
1018.4396362304688 0 911.8207
1020.4631958007812 0 10842.605 y 5
1020.5765991210938 0 771.13916
1021.46484375 0 6648.453
1022.4682006835938 0 2596.3018
1031.5458984375 0 9008.075
1032.4127197265625 0 2584.6365 b 8
1032.5506591796875 0 4079.3608
1033.41455078125 0 1774.2036
1035.4661865234375 0 5789.9453
1036.4671630859375 0 2261.3367
1037.465087890625 0 1854.1917
1082.4288330078125 0 1078.0912
1086.46875 0 1064.1876
1103.4991455078125 0 3360.6575
1104.5025634765625 0 1794.8091
1132.4757080078125 0 1096.0579
1167.498046875 0 2180.0312 y 4
1168.499755859375 0 1170.087
1219.5032958984375 0 717.43
1283.51318359375 0 1231.9867 b 10
3081.3525390625 0 860.4912

Spectrum Details

|  |  |
| --- | --- |
| Matched peaks? Matched peaks The total absolute number of peaks matched. Additionally in brackets the total fraction of peaks matched and the total number of peaks is shown. | 66 (8.22% of 803) |
| FDR? FDR The false discovery rate estimated for this peptide. It is calculated by matching all theoretical fragments with a non-integer shift with the raw peaks for this spectrum. This is done with 40 different shifts. The resulting percentage is the average number of annotated peaks over the number of annotated peaks with the correct spectrum. | 2.45% |
| Satellite FDR? Satellite FDR See the FDR for details on its calculation. This satellite ion specific FDR only contains the satellite ions (d/w) for I/L/J positions. | - |
| PSM Score? PSM Score The PSM Score as given by Hecklib to this annotated spectrum. It is shown with three significant figures. | 414 |

#### Spectrum 2840? Spectrum 2840 The raw spectrum of this peptide as annotated by Hecklib. The fragments are coloured according to ion type (see legend). Any peaks with a star '\*' as text can be hovered over to see the full details, first the ion type second the mass shift type. By hovering over the amino acids in the peptide or ions in the legend the corresponding peaks are highlighted. By toggling the 'Unassigned' label you can turn the background (unassigned) peaks on or off in the plot. By updating the slider in the Ion legend you can update the spectrum to only show the top X% of the peaks with labels. The top X% means any peak that is within X% of the highest intensity. By dragging in the spectrum you can zoom in to a specific part of the spectrum and use 'Zoom Out' to get back to the original zoom level. The annotation of the spectrum is based on the given sequence in the peptides file and is done with different software so inconsistencies are likely. The peaks are annotated based on the given sequence, with 20 ppm tolerance.

Copy Data

##### Spectrum 2840 (TSV)

###### Preview

```
Loading example...
```

*Click on the button to copy the data to your clipboard.*

Mz MinMz MaxIntensity Max

WidthHeightPeptide font sizePeptide stroke widthSpectrum font sizeSpectrum stroke widthCompact peptide

Ion legend

wxyz

abcd

OtherUnassignedIon^Charge^_Position_Show for top:%

SCSVMHEAJHNHY

08.18e+51.64e+62.45e+63.27e+6

Zoom Out

y^+1^_1_a^+1^_2_a^+1^_2_b^+4^_8_b^+1^_2_b^+1^_2_y^+2^_4_y^+2^_4_b^+2^_5_b^+4^_10_b^+1^_3_y^+1^_2_y^+2^_5_y^+4^_11_y^+3^_8_b^+1^_3_y^+4^_11_y^+3^_8_y^+2^_5_b^+2^_6_y^+2^_6_y^+4^_12_y^+4^_12_y^+2^_6_y^+4^_12_b^+3^_10_y^+3^_9_\*\*\*y^+1^_3_y^+3^_10_b^+1^_4_y^+3^_10_y^+3^_10_b^+2^_7_y^+1^_3_b^+1^_4_y^+2^_7_y^+3^_11_y^+3^_11_y^+3^_11_b^+3^_12_y^+3^_12_y^+3^_12_y^+2^_8_y^+2^_8_y^+3^_12_y^+2^_8_y^+1^_4_y^+1^_4_y^+2^_9_y^+2^_9_b^+2^_10_y^+2^_9_y^+2^_10_y^+2^_10_y^+2^_10_b^+2^_11_y^+1^_5_y^+2^_11_y^+2^_11_y^+2^_11_y^+1^_5_b^+1^_6_y^+1^_6_y^+1^_6_b^+1^_7_y^+1^_7_y^+1^_7_y^+1^_7_b^+1^_8_y^+1^_8_y^+1^_8_y^+1^_8_b^+1^_9_y^+1^_9_

0778155723353113

Fragment Matches Table

Show background peaks

| Position | Ion type | Intensity | mz Theoretical | mz Error (Th) | mz Error (ppm) | Charge | Series Number |
| --- | --- | --- | --- | --- | --- | --- | --- |
| - | - | 3320 | 120 | - | - | 0 | - |
| - | - | 4463 | 120.1 | - | - | 0 | - |
| - | - | 1.566E+04 | 120.1 | - | - | 0 | - |
| - | - | 2803 | 120.8 | - | - | 0 | - |
| - | - | 2.22E+04 | 121 | - | - | 0 | - |
| - | - | 2.404E+04 | 122.1 | - | - | 0 | - |
| - | - | 8.13E+04 | 123 | - | - | 0 | - |
| - | - | 1.823E+04 | 123.1 | - | - | 0 | - |
| - | - | 8985 | 124 | - | - | 0 | - |
| - | - | 5561 | 124 | - | - | 0 | - |
| - | - | 3.419E+04 | 124.1 | - | - | 0 | - |
| - | - | 1.364E+04 | 126 | - | - | 0 | - |
| - | - | 6861 | 126.1 | - | - | 0 | - |
| - | - | 3.753E+04 | 127.1 | - | - | 0 | - |
| - | - | 7814 | 127.1 | - | - | 0 | - |
| - | - | 2836 | 128.1 | - | - | 0 | - |
| - | - | 1.068E+05 | 129.1 | - | - | 0 | - |
| - | - | 2.828E+04 | 129.1 | - | - | 0 | - |
| - | - | 6.409E+04 | 130 | - | - | 0 | - |
| - | - | 3413 | 130.1 | - | - | 0 | - |
| - | - | 5030 | 130.1 | - | - | 0 | - |
| - | - | 4795 | 131 | - | - | 0 | - |
| - | - | 3.835E+04 | 131.1 | - | - | 0 | - |
| - | - | 6374 | 132 | - | - | 0 | - |
| - | - | 4309 | 132.1 | - | - | 0 | - |
| - | - | 1.151E+04 | 133 | - | - | 0 | - |
| - | - | 3327 | 133.1 | - | - | 0 | - |
| - | - | 1.364E+06 | 134 | - | - | 0 | - |
| - | - | 5011 | 134.1 | - | - | 0 | - |
| - | - | 7120 | 135 | - | - | 0 | - |
| - | - | 5.806E+04 | 135 | - | - | 0 | - |
| - | - | 5.828E+04 | 136 | - | - | 0 | - |
| - | - | 5263 | 136 | - | - | 0 | - |
| - | - | 4720 | 136 | - | - | 0 | - |
| - | - | 1.444E+04 | 136.1 | - | - | 0 | - |
| - | - | 1.353E+06 | 136.1 | - | - | 0 | - |
| - | - | 5477 | 137.1 | - | - | 0 | - |
| - | - | 1.154E+05 | 137.1 | - | - | 0 | - |
| - | - | 6.99E+04 | 138.1 | - | - | 0 | - |
| - | - | 5151 | 138.1 | - | - | 0 | - |
| - | - | 1.92E+04 | 139.1 | - | - | 0 | - |
| - | - | 1.449E+04 | 140 | - | - | 0 | - |
| - | - | 3483 | 141.1 | - | - | 0 | - |
| - | - | 4.899E+04 | 141.1 | - | - | 0 | - |
| - | - | 6211 | 142.1 | - | - | 0 | - |
| - | - | 5411 | 142.1 | - | - | 0 | - |
| - | - | 1.315E+04 | 143 | - | - | 0 | - |
| - | - | 1.496E+04 | 144 | - | - | 0 | - |
| - | - | 5402 | 144 | - | - | 0 | - |
| - | - | 7800 | 146 | - | - | 0 | - |
| - | - | 7.15E+04 | 147 | - | - | 0 | - |
| - | - | 2.604E+04 | 148 | - | - | 0 | - |
| - | - | 7731 | 148 | - | - | 0 | - |
| - | - | 5.571E+04 | 148.1 | - | - | 0 | - |
| - | - | 5006 | 149 | - | - | 0 | - |
| - | - | 3509 | 149 | - | - | 0 | - |
| - | - | 3784 | 149.1 | - | - | 0 | - |
| - | - | 4358 | 149.1 | - | - | 0 | - |
| - | - | 4559 | 149.1 | - | - | 0 | - |
| - | - | 7820 | 150.1 | - | - | 0 | - |
| - | - | 3.225E+04 | 152.1 | - | - | 0 | - |
| - | - | 4326 | 153.1 | - | - | 0 | - |
| - | - | 1.769E+04 | 154.1 | - | - | 0 | - |
| - | - | 2.142E+04 | 155.1 | - | - | 0 | - |
| - | - | 5.93E+04 | 155.1 | - | - | 0 | - |
| - | - | 3.765E+04 | 155.1 | - | - | 0 | - |
| - | - | 9266 | 156 | - | - | 0 | - |
| - | - | 1.453E+05 | 156.1 | - | - | 0 | - |
| - | - | 5275 | 156.1 | - | - | 0 | - |
| - | - | 9940 | 157 | - | - | 0 | - |
| - | - | 5.546E+04 | 157.1 | - | - | 0 | - |
| - | - | 9407 | 157.1 | - | - | 0 | - |
| - | - | 3.382E+04 | 157.1 | - | - | 0 | - |
| - | - | 5.037E+05 | 158 | - | - | 0 | - |
| - | - | 2.734E+04 | 159 | - | - | 0 | - |
| - | - | 1.227E+04 | 159.1 | - | - | 0 | - |
| - | - | 1.918E+06 | 159.1 | - | - | 0 | - |
| - | - | 2.357E+04 | 160 | - | - | 0 | - |
| - | - | 7394 | 160 | - | - | 0 | - |
| - | - | 4519 | 160.1 | - | - | 0 | - |
| - | - | 9501 | 160.1 | - | - | 0 | - |
| - | - | 1.504E+05 | 160.1 | - | - | 0 | - |
| - | - | 2.207E+05 | 161 | - | - | 0 | - |
| - | - | 8234 | 161.1 | - | - | 0 | - |
| - | - | 5.621E+04 | 162 | - | - | 0 | - |
| - | - | 4772 | 162 | - | - | 0 | - |
| - | - | 2.498E+04 | 162.1 | - | - | 0 | - |
| - | - | 7444 | 163 | - | - | 0 | - |
| - | - | 4444 | 163 | - | - | 0 | - |
| - | - | 8912 | 163 | - | - | 0 | - |
| - | - | 8936 | 163.1 | - | - | 0 | - |
| - | - | 8066 | 164.1 | - | - | 0 | - |
| - | - | 5377 | 164.1 | - | - | 0 | - |
| - | - | 4.91E+05 | 165.1 | - | - | 0 | - |
| - | - | 8.358E+04 | 165.1 | - | - | 0 | - |
| - | - | 9296 | 165.1 | - | - | 0 | - |
| - | - | 1.331E+05 | 166.1 | - | - | 0 | - |
| - | - | 1.7E+05 | 166.1 | - | - | 0 | - |
| - | - | 1.082E+04 | 167.1 | - | - | 0 | - |
| - | - | 2.226E+04 | 167.1 | - | - | 0 | - |
| - | - | 4930 | 167.1 | - | - | 0 | - |
| - | - | 6.527E+04 | 168 | - | - | 0 | - |
| - | - | 3524 | 169 | - | - | 0 | - |
| - | - | 2.509E+04 | 169.1 | - | - | 0 | - |
| - | - | 1.088E+04 | 171.1 | - | - | 0 | - |
| - | - | 1.033E+04 | 172.1 | - | - | 0 | - |
| - | - | 5639 | 172.6 | - | - | 0 | - |
| - | - | 7155 | 173 | - | - | 0 | - |
| - | - | 3307 | 173.1 | - | - | 0 | - |
| - | - | 3.41E+04 | 173.1 | - | - | 0 | - |
| - | - | 4603 | 173.5 | - | - | 0 | - |
| - | - | 2.119E+04 | 174 | - | - | 0 | - |
| - | - | 3.233E+04 | 174.1 | - | - | 0 | - |
| - | - | 9792 | 175.1 | - | - | 0 | - |
| - | - | 1.307E+04 | 175.1 | - | - | 0 | - |
| - | - | 6.678E+05 | 176 | - | - | 0 | - |
| - | - | 1.412E+05 | 176.1 | - | - | 0 | - |
| - | - | 3.825E+04 | 177 | - | - | 0 | - |
| - | - | 4703 | 177.1 | - | - | 0 | - |
| - | - | 1.393E+04 | 177.1 | - | - | 0 | - |
| - | - | 2.42E+04 | 177.1 | - | - | 0 | - |
| - | - | 2.732E+04 | 178 | - | - | 0 | - |
| - | - | 7927 | 178 | - | - | 0 | - |
| - | - | 5837 | 178.1 | - | - | 0 | - |
| - | - | 6531 | 178.1 | - | - | 0 | - |
| - | - | 1.394E+04 | 178.1 | - | - | 0 | - |
| - | - | 3.748E+04 | 179.1 | - | - | 0 | - |
| - | - | 1.171E+04 | 180 | - | - | 0 | - |
| - | - | 2.516E+04 | 180.1 | - | - | 0 | - |
| - | - | 5376 | 180.1 | - | - | 0 | - |
| - | - | 4491 | 180.1 | - | - | 0 | - |
| - | - | 5.257E+04 | 181.1 | - | - | 0 | - |
| - | - | 6095 | 181.6 | - | - | 0 | - |
| 13 | y | 5.897E+05 | 182.1 | 0.0005259 | 2.888 | +1 | 1 |
| - | - | 5005 | 182.1 | - | - | 0 | - |
| - | - | 1.58E+04 | 183.1 | - | - | 0 | - |
| - | - | 5.707E+04 | 183.1 | - | - | 0 | - |
| - | - | 9.183E+04 | 183.1 | - | - | 0 | - |
| - | - | 4345 | 184.1 | - | - | 0 | - |
| - | - | 7617 | 184.1 | - | - | 0 | - |
| - | - | 5.959E+04 | 185 | - | - | 0 | - |
| - | - | 2.416E+04 | 185.1 | - | - | 0 | - |
| - | - | 3.468E+05 | 186 | - | - | 0 | - |
| - | - | 2.674E+04 | 187 | - | - | 0 | - |
| - | - | 3.5E+05 | 187.1 | - | - | 0 | - |
| - | - | 1.586E+04 | 188 | - | - | 0 | - |
| - | - | 2.782E+04 | 188.1 | - | - | 0 | - |
| - | - | 4.917E+04 | 189.1 | - | - | 0 | - |
| - | - | 1.375E+05 | 190.1 | - | - | 0 | - |
| - | - | 5725 | 190.1 | - | - | 0 | - |
| - | - | 9111 | 190.1 | - | - | 0 | - |
| - | - | 3671 | 191 | - | - | 0 | - |
| - | - | 4655 | 191.1 | - | - | 0 | - |
| - | - | 5.622E+04 | 191.1 | - | - | 0 | - |
| - | - | 9849 | 192.1 | - | - | 0 | - |
| - | - | 4521 | 192.1 | - | - | 0 | - |
| - | - | 2.863E+04 | 193.1 | - | - | 0 | - |
| - | - | 2.431E+05 | 193.1 | - | - | 0 | - |
| - | - | 2.282E+04 | 194.1 | - | - | 0 | - |
| - | - | 4.86E+04 | 194.1 | - | - | 0 | - |
| - | - | 1.745E+04 | 194.1 | - | - | 0 | - |
| - | - | 9154 | 195.1 | - | - | 0 | - |
| - | - | 3.102E+04 | 195.1 | - | - | 0 | - |
| - | - | 7154 | 195.1 | - | - | 0 | - |
| - | - | 5390 | 196.1 | - | - | 0 | - |
| - | - | 5517 | 197.1 | - | - | 0 | - |
| - | - | 4690 | 198.1 | - | - | 0 | - |
| - | - | 7727 | 199.1 | - | - | 0 | - |
| - | - | 4538 | 199.1 | - | - | 0 | - |
| - | - | 1.042E+04 | 200.1 | - | - | 0 | - |
| - | - | 5883 | 201.1 | - | - | 0 | - |
| - | - | 4.564E+04 | 201.1 | - | - | 0 | - |
| - | - | 5551 | 201.1 | - | - | 0 | - |
| - | - | 3640 | 202.1 | - | - | 0 | - |
| - | - | 2.008E+04 | 202.1 | - | - | 0 | - |
| 2 | a | 1.628E+05 | 203.1 | 0.004045 | 19.92 | +1 | 2 |
| - | - | 2.056E+04 | 203.1 | - | - | 0 | - |
| - | - | 3.001E+04 | 204 | - | - | 0 | - |
| - | - | 1.137E+04 | 204.1 | - | - | 0 | - |
| - | - | 1.567E+05 | 204.1 | - | - | 0 | - |
| - | - | 5543 | 204.1 | - | - | 0 | - |
| - | - | 4158 | 204.1 | - | - | 0 | - |
| - | - | 9677 | 205.1 | - | - | 0 | - |
| - | - | 1.391E+04 | 205.1 | - | - | 0 | - |
| - | - | 5365 | 205.1 | - | - | 0 | - |
| - | - | 5853 | 206.1 | - | - | 0 | - |
| - | - | 2.093E+04 | 206.1 | - | - | 0 | - |
| - | - | 4343 | 206.1 | - | - | 0 | - |
| - | - | 2.724E+05 | 207.1 | - | - | 0 | - |
| - | - | 7756 | 207.2 | - | - | 0 | - |
| - | - | 9521 | 208.1 | - | - | 0 | - |
| - | - | 2.836E+04 | 208.1 | - | - | 0 | - |
| - | - | 6381 | 208.1 | - | - | 0 | - |
| - | - | 4193 | 209.1 | - | - | 0 | - |
| - | - | 2.909E+04 | 210.1 | - | - | 0 | - |
| - | - | 2.527E+04 | 211.1 | - | - | 0 | - |
| - | - | 5011 | 211.1 | - | - | 0 | - |
| - | - | 1.68E+04 | 213 | - | - | 0 | - |
| - | - | 6016 | 213.1 | - | - | 0 | - |
| - | - | 1.922E+04 | 214 | - | - | 0 | - |
| - | - | 4656 | 215.6 | - | - | 0 | - |
| - | - | 5199 | 216.1 | - | - | 0 | - |
| - | - | 6774 | 216.1 | - | - | 0 | - |
| - | - | 5.151E+04 | 217.1 | - | - | 0 | - |
| - | - | 5.781E+04 | 218.1 | - | - | 0 | - |
| - | - | 8722 | 218.1 | - | - | 0 | - |
| - | - | 5430 | 219.1 | - | - | 0 | - |
| - | - | 2.088E+04 | 219.1 | - | - | 0 | - |
| - | - | 3823 | 220.1 | - | - | 0 | - |
| 2 | a | 3.239E+06 | 221.1 | 0.003928 | 17.77 | +1 | 2 |
| - | - | 6.479E+05 | 221.1 | - | - | 0 | - |
| - | - | 2.316E+05 | 222.1 | - | - | 0 | - |
| - | - | 1.682E+04 | 222.1 | - | - | 0 | - |
| - | - | 5.709E+04 | 222.1 | - | - | 0 | - |
| - | - | 4386 | 222.1 | - | - | 0 | - |
| - | - | 3165 | 222.1 | - | - | 0 | - |
| - | - | 1.366E+05 | 223.1 | - | - | 0 | - |
| - | - | 1.588E+04 | 223.1 | - | - | 0 | - |
| - | - | 1.435E+04 | 223.1 | - | - | 0 | - |
| - | - | 3.993E+04 | 223.2 | - | - | 0 | - |
| - | - | 1.314E+04 | 224.1 | - | - | 0 | - |
| - | - | 2.181E+05 | 224.1 | - | - | 0 | - |
| - | - | 4016 | 224.2 | - | - | 0 | - |
| - | - | 1.702E+04 | 225.1 | - | - | 0 | - |
| - | - | 1.804E+04 | 225.1 | - | - | 0 | - |
| - | - | 9867 | 226.1 | - | - | 0 | - |
| 8 | b | 5.072E+04 | 226.1 | 0.001985 | 8.78 | +4 | 8 |
| - | - | 3828 | 226.2 | - | - | 0 | - |
| - | - | 1.374E+04 | 228.1 | - | - | 0 | - |
| - | - | 4218 | 229.1 | - | - | 0 | - |
| - | - | 4085 | 230.1 | - | - | 0 | - |
| - | - | 1.163E+04 | 230.1 | - | - | 0 | - |
| - | - | 8858 | 230.1 | - | - | 0 | - |
| 2 | b | 1.874E+05 | 231 | 0.003918 | 16.96 | +1 | 2 |
| - | - | 1.791E+04 | 231.1 | - | - | 0 | - |
| - | - | 9465 | 232 | - | - | 0 | - |
| - | - | 1.334E+04 | 232 | - | - | 0 | - |
| - | - | 2.68E+04 | 233.1 | - | - | 0 | - |
| - | - | 5.544E+04 | 234.1 | - | - | 0 | - |
| - | - | 2.844E+04 | 234.1 | - | - | 0 | - |
| - | - | 4.676E+05 | 235.1 | - | - | 0 | - |
| - | - | 5556 | 236 | - | - | 0 | - |
| - | - | 1.724E+04 | 236.1 | - | - | 0 | - |
| - | - | 5.705E+04 | 236.1 | - | - | 0 | - |
| - | - | 7928 | 236.1 | - | - | 0 | - |
| - | - | 7451 | 237.1 | - | - | 0 | - |
| - | - | 6174 | 237.1 | - | - | 0 | - |
| - | - | 9169 | 237.6 | - | - | 0 | - |
| - | - | 1.853E+04 | 238.1 | - | - | 0 | - |
| - | - | 8269 | 238.1 | - | - | 0 | - |
| - | - | 6701 | 239.1 | - | - | 0 | - |
| - | - | 1.322E+05 | 239.1 | - | - | 0 | - |
| - | - | 6666 | 240.1 | - | - | 0 | - |
| - | - | 1.596E+04 | 240.1 | - | - | 0 | - |
| - | - | 8661 | 243 | - | - | 0 | - |
| - | - | 6610 | 244.1 | - | - | 0 | - |
| - | - | 7.736E+04 | 244.1 | - | - | 0 | - |
| - | - | 7645 | 245.1 | - | - | 0 | - |
| - | - | 3.93E+04 | 247.1 | - | - | 0 | - |
| - | - | 4893 | 248.1 | - | - | 0 | - |
| 2 | b | 6.062E+05 | 249.1 | 0.003878 | 15.57 | +1 | 2 |
| - | - | 1.243E+05 | 249.1 | - | - | 0 | - |
| - | - | 6622 | 249.1 | - | - | 0 | - |
| - | - | 5517 | 249.2 | - | - | 0 | - |
| - | - | 5.209E+04 | 250.1 | - | - | 0 | - |
| - | - | 1.553E+04 | 250.1 | - | - | 0 | - |
| - | - | 8127 | 250.1 | - | - | 0 | - |
| - | - | 1.458E+04 | 250.2 | - | - | 0 | - |
| - | - | 1.9E+04 | 251.1 | - | - | 0 | - |
| - | - | 1.112E+05 | 251.2 | - | - | 0 | - |
| - | - | 7.434E+05 | 252.1 | - | - | 0 | - |
| - | - | 2.457E+04 | 252.1 | - | - | 0 | - |
| - | - | 1.076E+04 | 252.2 | - | - | 0 | - |
| - | - | 8.856E+04 | 253.1 | - | - | 0 | - |
| - | - | 8.68E+04 | 253.1 | - | - | 0 | - |
| - | - | 4493 | 254.1 | - | - | 0 | - |
| - | - | 4896 | 254.1 | - | - | 0 | - |
| - | - | 7189 | 254.1 | - | - | 0 | - |
| - | - | 7879 | 256.1 | - | - | 0 | - |
| - | - | 7436 | 257.1 | - | - | 0 | - |
| - | - | 7.582E+04 | 257.1 | - | - | 0 | - |
| - | - | 6435 | 257.1 | - | - | 0 | - |
| - | - | 9061 | 258.1 | - | - | 0 | - |
| - | - | 4305 | 259.1 | - | - | 0 | - |
| - | - | 9788 | 259.2 | - | - | 0 | - |
| - | - | 1.058E+04 | 261.1 | - | - | 0 | - |
| - | - | 1.907E+04 | 262.1 | - | - | 0 | - |
| - | - | 1.095E+04 | 263.1 | - | - | 0 | - |
| - | - | 4324 | 263.1 | - | - | 0 | - |
| - | - | 4120 | 264.1 | - | - | 0 | - |
| - | - | 3.69E+04 | 267.1 | - | - | 0 | - |
| - | - | 3.047E+05 | 267.1 | - | - | 0 | - |
| - | - | 1.758E+04 | 268.1 | - | - | 0 | - |
| - | - | 4558 | 268.1 | - | - | 0 | - |
| - | - | 3.283E+04 | 268.1 | - | - | 0 | - |
| - | - | 1.213E+04 | 268.1 | - | - | 0 | - |
| - | - | 1.045E+04 | 268.2 | - | - | 0 | - |
| - | - | 2.009E+04 | 269.1 | - | - | 0 | - |
| - | - | 7528 | 269.2 | - | - | 0 | - |
| - | - | 4898 | 270.1 | - | - | 0 | - |
| - | - | 7.129E+04 | 270.1 | - | - | 0 | - |
| - | - | 2.695E+04 | 270.1 | - | - | 0 | - |
| - | - | 7196 | 271.1 | - | - | 0 | - |
| - | - | 4842 | 271.2 | - | - | 0 | - |
| - | - | 8959 | 272.1 | - | - | 0 | - |
| - | - | 7.06E+04 | 273.1 | - | - | 0 | - |
| - | - | 3854 | 273.2 | - | - | 0 | - |
| - | - | 1.468E+04 | 274.1 | - | - | 0 | - |
| - | - | 5994 | 274.2 | - | - | 0 | - |
| - | - | 3.483E+04 | 275.1 | - | - | 0 | - |
| - | - | 5462 | 275.2 | - | - | 0 | - |
| - | - | 9162 | 276.1 | - | - | 0 | - |
| - | - | 1.604E+04 | 276.2 | - | - | 0 | - |
| - | - | 6025 | 276.6 | - | - | 0 | - |
| - | - | 3.861E+04 | 277.1 | - | - | 0 | - |
| 10 | y | 3.854E+04 | 277.1 | 0.0006754 | 2.437 | +2 | 4 |
| - | - | 1.07E+04 | 277.6 | - | - | 0 | - |
| - | - | 4266 | 278.1 | - | - | 0 | - |
| - | - | 5488 | 278.2 | - | - | 0 | - |
| - | - | 6912 | 278.6 | - | - | 0 | - |
| - | - | 1.776E+04 | 279.1 | - | - | 0 | - |
| - | - | 4.176E+04 | 280.1 | - | - | 0 | - |
| - | - | 6607 | 280.1 | - | - | 0 | - |
| - | - | 1.724E+04 | 280.7 | - | - | 0 | - |
| - | - | 5363 | 281.1 | - | - | 0 | - |
| - | - | 5020 | 281.2 | - | - | 0 | - |
| - | - | 2.162E+04 | 284.1 | - | - | 0 | - |
| - | - | 6111 | 284.1 | - | - | 0 | - |
| - | - | 2.324E+05 | 285.1 | - | - | 0 | - |
| 10 | y | 2.912E+05 | 285.6 | 0.000615 | 2.153 | +2 | 4 |
| - | - | 2.592E+04 | 286.1 | - | - | 0 | - |
| - | - | 8.825E+04 | 286.1 | - | - | 0 | - |
| - | - | 3.543E+04 | 286.6 | - | - | 0 | - |
| - | - | 1.132E+04 | 287.1 | - | - | 0 | - |
| - | - | 4853 | 287.1 | - | - | 0 | - |
| - | - | 8545 | 287.2 | - | - | 0 | - |
| - | - | 4604 | 287.2 | - | - | 0 | - |
| - | - | 1.144E+04 | 290.1 | - | - | 0 | - |
| 5 | b | 5828 | 291.6 | 0.00224 | 7.68 | +2 | 5 |
| - | - | 1.104E+04 | 292.1 | - | - | 0 | - |
| - | - | 1.34E+04 | 292.2 | - | - | 0 | - |
| 10 | b | 1.657E+04 | 293.1 | 0.002065 | 7.043 | +4 | 10 |
| - | - | 6517 | 293.1 | - | - | 0 | - |
| - | - | 1.153E+04 | 294.2 | - | - | 0 | - |
| - | - | 2.245E+04 | 294.6 | - | - | 0 | - |
| - | - | 3.842E+04 | 295.1 | - | - | 0 | - |
| - | - | 5483 | 296.1 | - | - | 0 | - |
| - | - | 4423 | 296.2 | - | - | 0 | - |
| - | - | 1.389E+04 | 297.2 | - | - | 0 | - |
| - | - | 1.05E+04 | 298.8 | - | - | 0 | - |
| - | - | 1.236E+04 | 300.1 | - | - | 0 | - |
| - | - | 9760 | 300.6 | - | - | 0 | - |
| - | - | 1.336E+05 | 301.1 | - | - | 0 | - |
| - | - | 1.752E+04 | 302.1 | - | - | 0 | - |
| - | - | 1.222E+04 | 302.1 | - | - | 0 | - |
| - | - | 3.291E+04 | 302.2 | - | - | 0 | - |
| - | - | 6332 | 302.6 | - | - | 0 | - |
| - | - | 6403 | 303.1 | - | - | 0 | - |
| - | - | 1.489E+04 | 304.1 | - | - | 0 | - |
| - | - | 1.676E+04 | 304.1 | - | - | 0 | - |
| - | - | 4.377E+04 | 304.2 | - | - | 0 | - |
| - | - | 1.34E+04 | 305.1 | - | - | 0 | - |
| - | - | 2.255E+04 | 305.1 | - | - | 0 | - |
| - | - | 6240 | 305.2 | - | - | 0 | - |
| - | - | 5295 | 308.1 | - | - | 0 | - |
| - | - | 2.23E+04 | 309.1 | - | - | 0 | - |
| - | - | 5577 | 310.1 | - | - | 0 | - |
| - | - | 5115 | 311.5 | - | - | 0 | - |
| - | - | 2.481E+04 | 312.1 | - | - | 0 | - |
| - | - | 9033 | 312.1 | - | - | 0 | - |
| - | - | 1.033E+04 | 312.6 | - | - | 0 | - |
| - | - | 9289 | 313.1 | - | - | 0 | - |
| - | - | 7240 | 314.2 | - | - | 0 | - |
| - | - | 4844 | 315.1 | - | - | 0 | - |
| - | - | 1.948E+04 | 316.1 | - | - | 0 | - |
| - | - | 4771 | 316.6 | - | - | 0 | - |
| - | - | 5066 | 317.6 | - | - | 0 | - |
| 3 | b | 5.674E+04 | 318.1 | 0.00372 | 11.7 | +1 | 3 |
| 12 | y | 5.947E+05 | 319.1 | 0.0007876 | 2.468 | +1 | 2 |
| - | - | 4512 | 319.7 | - | - | 0 | - |
| - | - | 2.01E+05 | 320.1 | - | - | 0 | - |
| - | - | 5.846E+04 | 320.2 | - | - | 0 | - |
| - | - | 5434 | 320.5 | - | - | 0 | - |
| - | - | 3.388E+04 | 321.1 | - | - | 0 | - |
| - | - | 8063 | 321.2 | - | - | 0 | - |
| - | - | 1.86E+04 | 322.1 | - | - | 0 | - |
| - | - | 4.787E+04 | 322.1 | - | - | 0 | - |
| - | - | 9.648E+04 | 322.2 | - | - | 0 | - |
| - | - | 1.362E+04 | 322.6 | - | - | 0 | - |
| - | - | 4067 | 322.7 | - | - | 0 | - |
| - | - | 6041 | 323.1 | - | - | 0 | - |
| - | - | 1.653E+04 | 323.2 | - | - | 0 | - |
| - | - | 7941 | 324.1 | - | - | 0 | - |
| - | - | 5165 | 324.6 | - | - | 0 | - |
| - | - | 1.367E+04 | 325.2 | - | - | 0 | - |
| - | - | 4.658E+04 | 326.1 | - | - | 0 | - |
| - | - | 1.175E+04 | 327.1 | - | - | 0 | - |
| - | - | 1.124E+04 | 328.2 | - | - | 0 | - |
| - | - | 3.025E+04 | 328.7 | - | - | 0 | - |
| - | - | 6.323E+04 | 329.1 | - | - | 0 | - |
| - | - | 1.014E+05 | 330.1 | - | - | 0 | - |
| - | - | 1.509E+04 | 330.2 | - | - | 0 | - |
| - | - | 7439 | 331.1 | - | - | 0 | - |
| - | - | 8421 | 331.1 | - | - | 0 | - |
| - | - | 4663 | 331.4 | - | - | 0 | - |
| - | - | 5050 | 332.1 | - | - | 0 | - |
| - | - | 6.578E+04 | 332.1 | - | - | 0 | - |
| - | - | 6851 | 332.2 | - | - | 0 | - |
| - | - | 4354 | 332.8 | - | - | 0 | - |
| - | - | 2.068E+04 | 333.1 | - | - | 0 | - |
| - | - | 7103 | 333.1 | - | - | 0 | - |
| 9 | y | 1.88E+04 | 333.7 | 0.003016 | 9.039 | +2 | 5 |
| - | - | 7554 | 334.1 | - | - | 0 | - |
| - | - | 5.613E+04 | 334.1 | - | - | 0 | - |
| 3 | y | 1.121E+04 | 334.7 | 0.003392 | 10.13 | +4 | 11 |
| - | - | 4824 | 334.9 | - | - | 0 | - |
| - | - | 5521 | 335.1 | - | - | 0 | - |
| 6 | y | 1.232E+04 | 335.2 | 0.0001624 | 0.4845 | +3 | 8 |
| - | - | 8671 | 335.5 | - | - | 0 | - |
| - | - | 2.495E+04 | 335.9 | - | - | 0 | - |
| 3 | b | 2.573E+04 | 336.1 | 0.003665 | 10.9 | +1 | 3 |
| - | - | 1.256E+04 | 336.1 | - | - | 0 | - |
| - | - | 1.451E+04 | 336.2 | - | - | 0 | - |
| - | - | 9018 | 336.6 | - | - | 0 | - |
| - | - | 3.784E+04 | 337.1 | - | - | 0 | - |
| - | - | 5001 | 337.2 | - | - | 0 | - |
| - | - | 5896 | 337.6 | - | - | 0 | - |
| - | - | 1.223E+04 | 337.7 | - | - | 0 | - |
| - | - | 1.608E+05 | 338.1 | - | - | 0 | - |
| 3 | y | 2.924E+04 | 339.2 | 0.004346 | 12.81 | +4 | 11 |
| - | - | 1.581E+04 | 339.2 | - | - | 0 | - |
| - | - | 4567 | 339.4 | - | - | 0 | - |
| - | - | 9307 | 340.2 | - | - | 0 | - |
| - | - | 8572 | 340.5 | - | - | 0 | - |
| 6 | y | 8.497E+04 | 340.8 | 0.000824 | 2.418 | +3 | 8 |
| - | - | 4.52E+04 | 341.2 | - | - | 0 | - |
| - | - | 1.843E+04 | 341.5 | - | - | 0 | - |
| 9 | y | 8.573E+04 | 342.2 | 0.0007583 | 2.216 | +2 | 5 |
| - | - | 3.061E+04 | 342.7 | - | - | 0 | - |
| - | - | 1.997E+04 | 343.2 | - | - | 0 | - |
| - | - | 8647 | 344.1 | - | - | 0 | - |
| - | - | 6.204E+04 | 344.1 | - | - | 0 | - |
| - | - | 4551 | 344.5 | - | - | 0 | - |
| - | - | 9610 | 344.9 | - | - | 0 | - |
| - | - | 6415 | 345.2 | - | - | 0 | - |
| - | - | 5845 | 345.4 | - | - | 0 | - |
| - | - | 6613 | 346.4 | - | - | 0 | - |
| - | - | 5662 | 346.6 | - | - | 0 | - |
| - | - | 1.236E+04 | 347.2 | - | - | 0 | - |
| - | - | 1.067E+04 | 347.7 | - | - | 0 | - |
| - | - | 5858 | 347.9 | - | - | 0 | - |
| - | - | 5.052E+04 | 348.1 | - | - | 0 | - |
| - | - | 1.358E+04 | 348.2 | - | - | 0 | - |
| - | - | 6192 | 348.7 | - | - | 0 | - |
| - | - | 7476 | 348.9 | - | - | 0 | - |
| - | - | 1.165E+04 | 349.1 | - | - | 0 | - |
| - | - | 7604 | 349.6 | - | - | 0 | - |
| - | - | 2.412E+05 | 350.1 | - | - | 0 | - |
| - | - | 8850 | 350.7 | - | - | 0 | - |
| - | - | 3.861E+04 | 351.1 | - | - | 0 | - |
| - | - | 3.817E+04 | 351.7 | - | - | 0 | - |
| - | - | 2.026E+04 | 351.9 | - | - | 0 | - |
| - | - | 1.064E+04 | 352.2 | - | - | 0 | - |
| - | - | 1.49E+04 | 352.2 | - | - | 0 | - |
| - | - | 1.052E+04 | 352.4 | - | - | 0 | - |
| - | - | 1.863E+04 | 353.1 | - | - | 0 | - |
| - | - | 5238 | 353.2 | - | - | 0 | - |
| - | - | 6990 | 353.4 | - | - | 0 | - |
| - | - | 4.426E+04 | 354.1 | - | - | 0 | - |
| - | - | 6672 | 354.2 | - | - | 0 | - |
| - | - | 5644 | 354.4 | - | - | 0 | - |
| - | - | 7878 | 354.8 | - | - | 0 | - |
| - | - | 9038 | 354.8 | - | - | 0 | - |
| - | - | 1.586E+04 | 355.1 | - | - | 0 | - |
| - | - | 8014 | 355.2 | - | - | 0 | - |
| - | - | 4384 | 356.8 | - | - | 0 | - |
| - | - | 6100 | 357.1 | - | - | 0 | - |
| - | - | 7.461E+04 | 357.7 | - | - | 0 | - |
| - | - | 4.499E+04 | 357.9 | - | - | 0 | - |
| - | - | 5522 | 358.1 | - | - | 0 | - |
| - | - | 2.574E+04 | 358.2 | - | - | 0 | - |
| - | - | 1.379E+04 | 358.7 | - | - | 0 | - |
| - | - | 2.531E+04 | 358.9 | - | - | 0 | - |
| - | - | 2.017E+04 | 359.2 | - | - | 0 | - |
| - | - | 1.062E+04 | 359.4 | - | - | 0 | - |
| - | - | 1.663E+04 | 359.5 | - | - | 0 | - |
| - | - | 1.954E+04 | 359.8 | - | - | 0 | - |
| 6 | b | 7611 | 360.1 | 0.001104 | 3.065 | +2 | 6 |
| - | - | 8598 | 360.2 | - | - | 0 | - |
| - | - | 5590 | 360.7 | - | - | 0 | - |
| - | - | 5673 | 360.9 | - | - | 0 | - |
| - | - | 4.553E+04 | 361.2 | - | - | 0 | - |
| - | - | 1.037E+04 | 361.2 | - | - | 0 | - |
| - | - | 7589 | 361.9 | - | - | 0 | - |
| - | - | 1.381E+04 | 362.2 | - | - | 0 | - |
| - | - | 6698 | 362.2 | - | - | 0 | - |
| - | - | 3.634E+04 | 362.8 | - | - | 0 | - |
| - | - | 2.484E+04 | 363.2 | - | - | 0 | - |
| - | - | 5764 | 363.4 | - | - | 0 | - |
| - | - | 1.267E+04 | 363.5 | - | - | 0 | - |
| - | - | 1.167E+04 | 363.7 | - | - | 0 | - |
| - | - | 7174 | 363.9 | - | - | 0 | - |
| - | - | 9035 | 364.1 | - | - | 0 | - |
| - | - | 6045 | 364.4 | - | - | 0 | - |
| - | - | 7338 | 364.9 | - | - | 0 | - |
| - | - | 1.145E+04 | 365.2 | - | - | 0 | - |
| - | - | 7.37E+04 | 365.2 | - | - | 0 | - |
| - | - | 7.053E+04 | 366.1 | - | - | 0 | - |
| - | - | 1.393E+04 | 366.2 | - | - | 0 | - |
| - | - | 5058 | 366.4 | - | - | 0 | - |
| - | - | 1.437E+04 | 367.1 | - | - | 0 | - |
| - | - | 6766 | 368.1 | - | - | 0 | - |
| - | - | 1.201E+04 | 368.2 | - | - | 0 | - |
| - | - | 1.64E+05 | 368.5 | - | - | 0 | - |
| - | - | 7976 | 368.7 | - | - | 0 | - |
| - | - | 9.653E+04 | 368.8 | - | - | 0 | - |
| - | - | 2.201E+04 | 368.9 | - | - | 0 | - |
| - | - | 1.541E+04 | 369.2 | - | - | 0 | - |
| 8 | y | 2.733E+04 | 369.2 | 0.001915 | 5.188 | +2 | 6 |
| - | - | 2.293E+04 | 369.4 | - | - | 0 | - |
| - | - | 1.206E+04 | 369.7 | - | - | 0 | - |
| - | - | 1.919E+04 | 370.2 | - | - | 0 | - |
| - | - | 7984 | 370.2 | - | - | 0 | - |
| - | - | 6400 | 370.4 | - | - | 0 | - |
| - | - | 8322 | 370.7 | - | - | 0 | - |
| - | - | 9910 | 370.9 | - | - | 0 | - |
| - | - | 2.322E+05 | 371.2 | - | - | 0 | - |
| - | - | 1.104E+04 | 371.9 | - | - | 0 | - |
| - | - | 1.302E+05 | 372.1 | - | - | 0 | - |
| - | - | 2.234E+04 | 372.2 | - | - | 0 | - |
| - | - | 5844 | 372.4 | - | - | 0 | - |
| - | - | 4813 | 372.6 | - | - | 0 | - |
| - | - | 2.615E+04 | 373.1 | - | - | 0 | - |
| - | - | 1.291E+04 | 373.4 | - | - | 0 | - |
| - | - | 1.041E+05 | 373.7 | - | - | 0 | - |
| - | - | 6.067E+04 | 373.9 | - | - | 0 | - |
| - | - | 7125 | 374.1 | - | - | 0 | - |
| - | - | 5.068E+04 | 374.2 | - | - | 0 | - |
| - | - | 1.27E+04 | 374.4 | - | - | 0 | - |
| - | - | 4.237E+04 | 374.7 | - | - | 0 | - |
| 2 | y | 5.635E+04 | 374.9 | 0.0001228 | 0.3277 | +4 | 12 |
| 2 | y | 3.576E+04 | 375.2 | 0.003843 | 10.24 | +4 | 12 |
| - | - | 1.106E+04 | 375.2 | - | - | 0 | - |
| - | - | 1.597E+04 | 375.4 | - | - | 0 | - |
| - | - | 6019 | 375.7 | - | - | 0 | - |
| - | - | 9858 | 376.1 | - | - | 0 | - |
| - | - | 6.614E+04 | 376.2 | - | - | 0 | - |
| - | - | 3.707E+04 | 376.4 | - | - | 0 | - |
| - | - | 8424 | 376.5 | - | - | 0 | - |
| - | - | 2.402E+04 | 376.7 | - | - | 0 | - |
| - | - | 1.179E+04 | 376.9 | - | - | 0 | - |
| - | - | 6454 | 377.2 | - | - | 0 | - |
| 8 | y | 8.486E+04 | 377.7 | 0.0007256 | 1.921 | +2 | 6 |
| - | - | 8753 | 377.9 | - | - | 0 | - |
| - | - | 7107 | 378.1 | - | - | 0 | - |
| - | - | 3.548E+04 | 378.2 | - | - | 0 | - |
| - | - | 1.456E+04 | 378.4 | - | - | 0 | - |
| - | - | 2.816E+04 | 378.7 | - | - | 0 | - |
| - | - | 3.08E+04 | 379.2 | - | - | 0 | - |
| 2 | y | 6763 | 379.4 | 0.0003183 | 0.8388 | +4 | 12 |
| - | - | 1.005E+04 | 379.9 | - | - | 0 | - |
| - | - | 6456 | 380.4 | - | - | 0 | - |
| - | - | 1.866E+04 | 380.7 | - | - | 0 | - |
| - | - | 2.782E+04 | 380.9 | - | - | 0 | - |
| - | - | 9.528E+04 | 381.1 | - | - | 0 | - |
| - | - | 1.357E+04 | 381.2 | - | - | 0 | - |
| - | - | 2.445E+04 | 382.1 | - | - | 0 | - |
| - | - | 4.176E+04 | 382.2 | - | - | 0 | - |
| - | - | 1.674E+04 | 382.5 | - | - | 0 | - |
| - | - | 8150 | 382.9 | - | - | 0 | - |
| - | - | 6129 | 383.2 | - | - | 0 | - |
| - | - | 5953 | 383.4 | - | - | 0 | - |
| - | - | 7072 | 384.1 | - | - | 0 | - |
| - | - | 3.783E+04 | 384.2 | - | - | 0 | - |
| 10 | b | 9606 | 384.5 | 0.005465 | 14.21 | +3 | 10 |
| - | - | 8155 | 384.7 | - | - | 0 | - |
| - | - | 2.019E+04 | 384.9 | - | - | 0 | - |
| - | - | 9.578E+04 | 385.2 | - | - | 0 | - |
| - | - | 5.777E+04 | 385.4 | - | - | 0 | - |
| - | - | 2.609E+04 | 385.7 | - | - | 0 | - |
| - | - | 6359 | 385.8 | - | - | 0 | - |
| - | - | 1.354E+04 | 385.9 | - | - | 0 | - |
| - | - | 2.03E+04 | 386.1 | - | - | 0 | - |
| - | - | 9934 | 387.2 | - | - | 0 | - |
| - | - | 1.222E+04 | 387.2 | - | - | 0 | - |
| - | - | 8293 | 387.7 | - | - | 0 | - |
| - | - | 1.121E+04 | 387.9 | - | - | 0 | - |
| - | - | 8616 | 388.7 | - | - | 0 | - |
| - | - | 6935 | 389.1 | - | - | 0 | - |
| - | - | 6.387E+04 | 389.2 | - | - | 0 | - |
| - | - | 1.652E+04 | 389.2 | - | - | 0 | - |
| - | - | 1.446E+04 | 389.4 | - | - | 0 | - |
| - | - | 2.373E+04 | 389.7 | - | - | 0 | - |
| 5 | y | 1.455E+05 | 389.8 | 0.002625 | 6.734 | +3 | 9 |
| - | - | 4.895E+04 | 390.2 | - | - | 0 | - |
| - | - | 8.164E+04 | 390.2 | - | - | 0 | - |
| - | - | 3.21E+04 | 390.5 | - | - | 0 | - |
| - | - | 8817 | 390.8 | - | - | 0 | - |
| - | - | 1.062E+04 | 391.2 | - | - | 0 | - |
| - | - | 7926 | 391.2 | - | - | 0 | - |
| - | - | 2.425E+04 | 391.5 | - | - | 0 | - |
| - | - | 1.961E+04 | 391.8 | - | - | 0 | - |
| - | - | 2.795E+04 | 392.2 | - | - | 0 | - |
| - | - | 4.373E+04 | 392.4 | - | - | 0 | - |
| - | - | 2.496E+04 | 392.7 | - | - | 0 | - |
| - | - | 1.287E+04 | 393.2 | - | - | 0 | - |
| - | - | 4.956E+04 | 393.2 | - | - | 0 | - |
| - | - | 6289 | 393.7 | - | - | 0 | - |
| - | - | 6859 | 394.2 | - | - | 0 | - |
| - | - | 8326 | 394.2 | - | - | 0 | - |
| - | - | 1.062E+04 | 395.9 | - | - | 0 | - |
| - | - | 1.979E+04 | 396.1 | - | - | 0 | - |
| 0 | Precursor | 3.512E+04 | 396.7 | 0.0006591 | 1.662 | +4 | -1 |
| 0 | Precursor | 3.061E+04 | 396.9 | 0.005082 | 12.81 | +4 | -1 |
| - | - | 2.164E+04 | 397.2 | - | - | 0 | - |
| - | - | 8800 | 397.2 | - | - | 0 | - |
| - | - | 7470 | 397.4 | - | - | 0 | - |
| - | - | 6937 | 397.5 | - | - | 0 | - |
| - | - | 4.193E+04 | 397.7 | - | - | 0 | - |
| - | - | 4.853E+04 | 398.1 | - | - | 0 | - |
| - | - | 1.671E+04 | 398.2 | - | - | 0 | - |
| - | - | 9.715E+04 | 399.1 | - | - | 0 | - |
| - | - | 3.334E+04 | 400.1 | - | - | 0 | - |
| - | - | 7989 | 400.2 | - | - | 0 | - |
| - | - | 8294 | 401.1 | - | - | 0 | - |
| 0 | Precursor | 7.89E+04 | 401.2 | 0.001039 | 2.59 | +4 | -1 |
| - | - | 8058 | 401.2 | - | - | 0 | - |
| - | - | 7.934E+04 | 401.4 | - | - | 0 | - |
| - | - | 9.863E+04 | 401.5 | - | - | 0 | - |
| - | - | 5.008E+04 | 401.7 | - | - | 0 | - |
| - | - | 6.005E+04 | 401.9 | - | - | 0 | - |
| - | - | 1.609E+04 | 401.9 | - | - | 0 | - |
| - | - | 1.594E+04 | 402.1 | - | - | 0 | - |
| - | - | 5918 | 402.2 | - | - | 0 | - |
| - | - | 2.243E+04 | 402.2 | - | - | 0 | - |
| - | - | 1.064E+04 | 402.7 | - | - | 0 | - |
| - | - | 6981 | 402.7 | - | - | 0 | - |
| - | - | 6102 | 402.8 | - | - | 0 | - |
| - | - | 1.529E+04 | 403.2 | - | - | 0 | - |
| - | - | 8085 | 403.7 | - | - | 0 | - |
| - | - | 1.096E+04 | 403.9 | - | - | 0 | - |
| - | - | 3.306E+04 | 404.2 | - | - | 0 | - |
| - | - | 7653 | 404.2 | - | - | 0 | - |
| - | - | 4.894E+04 | 404.7 | - | - | 0 | - |
| - | - | 6469 | 404.9 | - | - | 0 | - |
| - | - | 1.27E+04 | 405.2 | - | - | 0 | - |
| - | - | 2.002E+04 | 405.2 | - | - | 0 | - |
| - | - | 6.154E+04 | 405.2 | - | - | 0 | - |
| - | - | 5069 | 405.7 | - | - | 0 | - |
| - | - | 8957 | 405.7 | - | - | 0 | - |
| - | - | 1.371E+05 | 406.2 | - | - | 0 | - |
| - | - | 1.05E+04 | 406.2 | - | - | 0 | - |
| - | - | 5.677E+04 | 406.7 | - | - | 0 | - |
| - | - | 3.671E+04 | 407.2 | - | - | 0 | - |
| - | - | 4.518E+04 | 407.2 | - | - | 0 | - |
| - | - | 8715 | 407.6 | - | - | 0 | - |
| - | - | 9858 | 408.2 | - | - | 0 | - |
| - | - | 1.376E+04 | 408.2 | - | - | 0 | - |
| - | - | 1.068E+04 | 408.5 | - | - | 0 | - |
| - | - | 1.813E+04 | 408.8 | - | - | 0 | - |
| - | - | 7573 | 409.2 | - | - | 0 | - |
| - | - | 5226 | 409.5 | - | - | 0 | - |
| - | - | 2.484E+04 | 411.2 | - | - | 0 | - |
| - | - | 5651 | 411.7 | - | - | 0 | - |
| - | - | 1.952E+04 | 411.7 | - | - | 0 | - |
| - | - | 1.358E+04 | 412.2 | - | - | 0 | - |
| - | - | 1.268E+04 | 413.2 | - | - | 0 | - |
| - | - | 1.498E+05 | 414.1 | - | - | 0 | - |
| - | - | 7553 | 414.5 | - | - | 0 | - |
| - | - | 2.651E+04 | 415.1 | - | - | 0 | - |
| - | - | 3.122E+04 | 415.2 | - | - | 0 | - |
| - | - | 2.217E+04 | 415.2 | - | - | 0 | - |
| 11 | y | 2.542E+05 | 416.2 | 0.0008887 | 2.136 | +1 | 3 |
| - | - | 1.22E+04 | 416.7 | - | - | 0 | - |
| - | - | 7066 | 416.7 | - | - | 0 | - |
| 4 | y | 6511 | 416.9 | 0.001958 | 4.697 | +3 | 10 |
| 4 | b | 2.896E+04 | 417.1 | 0.001028 | 2.465 | +1 | 4 |
| - | - | 4.511E+04 | 417.2 | - | - | 0 | - |
| 4 | y | 2.122E+04 | 417.2 | 0.006472 | 15.51 | +3 | 10 |
| - | - | 9475 | 417.5 | - | - | 0 | - |
| - | - | 1.103E+04 | 417.9 | - | - | 0 | - |
| - | - | 7804 | 418.1 | - | - | 0 | - |
| - | - | 9530 | 418.9 | - | - | 0 | - |
| - | - | 1.438E+04 | 419.2 | - | - | 0 | - |
| - | - | 9500 | 419.5 | - | - | 0 | - |
| - | - | 9.894E+04 | 420.2 | - | - | 0 | - |
| - | - | 5.24E+04 | 420.7 | - | - | 0 | - |
| - | - | 1.32E+05 | 421.2 | - | - | 0 | - |
| - | - | 1.223E+04 | 421.5 | - | - | 0 | - |
| - | - | 7568 | 421.9 | - | - | 0 | - |
| - | - | 7522 | 422.2 | - | - | 0 | - |
| - | - | 2.986E+04 | 422.2 | - | - | 0 | - |
| 4 | y | 1.842E+05 | 422.9 | 0.002709 | 6.406 | +3 | 10 |
| - | - | 1.541E+05 | 423.2 | - | - | 0 | - |
| - | - | 7.662E+04 | 423.2 | - | - | 0 | - |
| - | - | 6.834E+04 | 423.5 | - | - | 0 | - |
| - | - | 2.476E+04 | 423.9 | - | - | 0 | - |
| - | - | 9515 | 424.2 | - | - | 0 | - |
| - | - | 1.712E+04 | 424.2 | - | - | 0 | - |
| - | - | 4.876E+04 | 424.5 | - | - | 0 | - |
| 7 | b | 8082 | 424.6 | 0.001231 | 2.898 | +2 | 7 |
| - | - | 1.693E+04 | 424.7 | - | - | 0 | - |
| - | - | 5.534E+04 | 424.9 | - | - | 0 | - |
| - | - | 6.406E+04 | 425.2 | - | - | 0 | - |
| - | - | 1.482E+04 | 425.5 | - | - | 0 | - |
| - | - | 2.487E+04 | 425.7 | - | - | 0 | - |
| - | - | 5739 | 426.1 | - | - | 0 | - |
| - | - | 1.494E+04 | 426.2 | - | - | 0 | - |
| - | - | 2.32E+04 | 428.2 | - | - | 0 | - |
| - | - | 1.341E+04 | 428.5 | - | - | 0 | - |
| - | - | 6.938E+04 | 429.2 | - | - | 0 | - |
| - | - | 3.44E+04 | 429.7 | - | - | 0 | - |
| - | - | 2.42E+04 | 429.9 | - | - | 0 | - |
| - | - | 2.316E+04 | 430.2 | - | - | 0 | - |
| - | - | 6292 | 430.2 | - | - | 0 | - |
| - | - | 3.964E+05 | 430.5 | - | - | 0 | - |
| - | - | 1.075E+04 | 430.7 | - | - | 0 | - |
| - | - | 2.87E+05 | 430.9 | - | - | 0 | - |
| - | - | 1.318E+04 | 431.2 | - | - | 0 | - |
| - | - | 1.173E+05 | 431.2 | - | - | 0 | - |
| - | - | 3.041E+04 | 431.5 | - | - | 0 | - |
| - | - | 8391 | 431.9 | - | - | 0 | - |
| - | - | 6081 | 432.9 | - | - | 0 | - |
| 11 | y | 3.705E+05 | 433.2 | 0.001409 | 3.252 | +1 | 3 |
| - | - | 4.539E+04 | 433.7 | - | - | 0 | - |
| - | - | 1.753E+04 | 433.9 | - | - | 0 | - |
| - | - | 9.642E+04 | 434.2 | - | - | 0 | - |
| - | - | 5906 | 434.7 | - | - | 0 | - |
| 4 | b | 2.211E+04 | 435.2 | 0.003872 | 8.898 | +1 | 4 |
| - | - | 1.619E+04 | 435.2 | - | - | 0 | - |
| - | - | 3.204E+04 | 435.9 | - | - | 0 | - |
| - | - | 6192 | 436.2 | - | - | 0 | - |
| - | - | 1.657E+04 | 436.2 | - | - | 0 | - |
| - | - | 8.617E+04 | 436.2 | - | - | 0 | - |
| - | - | 5784 | 436.9 | - | - | 0 | - |
| - | - | 1.183E+04 | 437.2 | - | - | 0 | - |
| - | - | 7061 | 437.5 | - | - | 0 | - |
| - | - | 5754 | 437.9 | - | - | 0 | - |
| - | - | 1.606E+04 | 438.7 | - | - | 0 | - |
| - | - | 5596 | 439.2 | - | - | 0 | - |
| - | - | 4.987E+04 | 439.2 | - | - | 0 | - |
| - | - | 6929 | 439.5 | - | - | 0 | - |
| - | - | 1.083E+04 | 439.7 | - | - | 0 | - |
| - | - | 9009 | 440.2 | - | - | 0 | - |
| - | - | 1.184E+04 | 440.5 | - | - | 0 | - |
| - | - | 1.29E+04 | 440.9 | - | - | 0 | - |
| - | - | 1.1E+04 | 440.9 | - | - | 0 | - |
| - | - | 8918 | 441.2 | - | - | 0 | - |
| - | - | 2.055E+04 | 441.9 | - | - | 0 | - |
| - | - | 1.172E+04 | 442.1 | - | - | 0 | - |
| 7 | y | 1.148E+05 | 442.2 | 0.0007303 | 1.651 | +2 | 7 |
| - | - | 1.035E+04 | 442.5 | - | - | 0 | - |
| - | - | 5.152E+04 | 442.7 | - | - | 0 | - |
| - | - | 1.559E+04 | 443.2 | - | - | 0 | - |
| - | - | 6.77E+04 | 443.2 | - | - | 0 | - |
| - | - | 6724 | 443.5 | - | - | 0 | - |
| - | - | 5765 | 443.9 | - | - | 0 | - |
| - | - | 1.495E+04 | 444.2 | - | - | 0 | - |
| - | - | 7888 | 444.2 | - | - | 0 | - |
| - | - | 8121 | 444.7 | - | - | 0 | - |
| - | - | 1.703E+04 | 445.2 | - | - | 0 | - |
| - | - | 1.415E+04 | 445.5 | - | - | 0 | - |
| 3 | y | 8.434E+04 | 445.9 | 0.002451 | 5.498 | +3 | 11 |
| 3 | y | 1.096E+05 | 446.2 | 0.006325 | 14.17 | +3 | 11 |
| - | - | 6.369E+04 | 446.5 | - | - | 0 | - |
| - | - | 3.151E+04 | 446.9 | - | - | 0 | - |
| - | - | 1.471E+04 | 447.2 | - | - | 0 | - |
| - | - | 1.48E+04 | 447.2 | - | - | 0 | - |
| - | - | 4.792E+04 | 447.5 | - | - | 0 | - |
| - | - | 1.008E+05 | 447.7 | - | - | 0 | - |
| - | - | 5.122E+04 | 447.9 | - | - | 0 | - |
| - | - | 4.072E+04 | 448.2 | - | - | 0 | - |
| - | - | 3.863E+04 | 448.5 | - | - | 0 | - |
| - | - | 1.38E+04 | 448.7 | - | - | 0 | - |
| - | - | 1.194E+04 | 448.9 | - | - | 0 | - |
| - | - | 1.168E+04 | 449.2 | - | - | 0 | - |
| - | - | 4.742E+04 | 449.2 | - | - | 0 | - |
| - | - | 6775 | 449.9 | - | - | 0 | - |
| - | - | 7838 | 450.1 | - | - | 0 | - |
| - | - | 6805 | 450.2 | - | - | 0 | - |
| - | - | 1.244E+04 | 450.2 | - | - | 0 | - |
| - | - | 7210 | 451.2 | - | - | 0 | - |
| - | - | 4190 | 451.2 | - | - | 0 | - |
| - | - | 1.692E+05 | 451.2 | - | - | 0 | - |
| 3 | y | 9.698E+05 | 451.9 | 0.002775 | 6.141 | +3 | 11 |
| - | - | 6.621E+05 | 452.2 | - | - | 0 | - |
| - | - | 3.128E+05 | 452.5 | - | - | 0 | - |
| - | - | 1.234E+05 | 452.9 | - | - | 0 | - |
| - | - | 4.745E+04 | 453.2 | - | - | 0 | - |
| - | - | 9055 | 453.2 | - | - | 0 | - |
| - | - | 2.043E+04 | 453.7 | - | - | 0 | - |
| - | - | 7847 | 453.9 | - | - | 0 | - |
| - | - | 1.662E+04 | 454.2 | - | - | 0 | - |
| - | - | 2.682E+04 | 455.2 | - | - | 0 | - |
| - | - | 1.396E+04 | 455.5 | - | - | 0 | - |
| - | - | 1.235E+04 | 455.9 | - | - | 0 | - |
| - | - | 1.022E+04 | 458.9 | - | - | 0 | - |
| - | - | 2.308E+04 | 459.2 | - | - | 0 | - |
| - | - | 5.288E+04 | 461.2 | - | - | 0 | - |
| - | - | 2E+04 | 461.3 | - | - | 0 | - |
| - | - | 9.231E+04 | 461.7 | - | - | 0 | - |
| - | - | 7901 | 461.8 | - | - | 0 | - |
| - | - | 6.533E+04 | 462.2 | - | - | 0 | - |
| - | - | 2.286E+04 | 462.7 | - | - | 0 | - |
| - | - | 8497 | 462.9 | - | - | 0 | - |
| - | - | 3.703E+04 | 463.2 | - | - | 0 | - |
| - | - | 1.786E+04 | 463.5 | - | - | 0 | - |
| - | - | 2.212E+04 | 463.9 | - | - | 0 | - |
| - | - | 1.339E+04 | 464.2 | - | - | 0 | - |
| - | - | 1.545E+04 | 464.9 | - | - | 0 | - |
| - | - | 1.34E+04 | 465.2 | - | - | 0 | - |
| - | - | 8875 | 466.5 | - | - | 0 | - |
| - | - | 8976 | 467.2 | - | - | 0 | - |
| - | - | 6265 | 467.2 | - | - | 0 | - |
| - | - | 1.513E+04 | 468.3 | - | - | 0 | - |
| - | - | 9.601E+04 | 468.9 | - | - | 0 | - |
| - | - | 8.817E+04 | 469.2 | - | - | 0 | - |
| - | - | 5.925E+04 | 469.5 | - | - | 0 | - |
| - | - | 5.117E+04 | 469.9 | - | - | 0 | - |
| - | - | 4.11E+04 | 470.2 | - | - | 0 | - |
| - | - | 1.659E+04 | 470.5 | - | - | 0 | - |
| - | - | 6.345E+04 | 470.7 | - | - | 0 | - |
| - | - | 6070 | 470.9 | - | - | 0 | - |
| - | - | 5.226E+04 | 471.2 | - | - | 0 | - |
| - | - | 8723 | 471.7 | - | - | 0 | - |
| - | - | 3.01E+04 | 472.2 | - | - | 0 | - |
| - | - | 8387 | 472.2 | - | - | 0 | - |
| - | - | 3.787E+04 | 472.5 | - | - | 0 | - |
| - | - | 5507 | 472.7 | - | - | 0 | - |
| - | - | 2.795E+04 | 472.9 | - | - | 0 | - |
| - | - | 2.226E+04 | 473.2 | - | - | 0 | - |
| - | - | 1.134E+04 | 473.5 | - | - | 0 | - |
| 12 | b | 1.401E+04 | 474.2 | 0.0001834 | 0.3867 | +3 | 12 |
| - | - | 3.527E+04 | 474.3 | - | - | 0 | - |
| - | - | 7887 | 474.7 | - | - | 0 | - |
| - | - | 5780 | 475.2 | - | - | 0 | - |
| - | - | 8772 | 475.3 | - | - | 0 | - |
| - | - | 1.112E+04 | 477.1 | - | - | 0 | - |
| - | - | 1.817E+05 | 478.2 | - | - | 0 | - |
| - | - | 2.155E+05 | 478.5 | - | - | 0 | - |
| - | - | 6109 | 478.7 | - | - | 0 | - |
| - | - | 1.282E+05 | 478.9 | - | - | 0 | - |
| - | - | 2.912E+04 | 479.2 | - | - | 0 | - |
| - | - | 2.184E+04 | 479.2 | - | - | 0 | - |
| - | - | 1.62E+04 | 479.5 | - | - | 0 | - |
| - | - | 1.247E+05 | 479.7 | - | - | 0 | - |
| - | - | 6.665E+04 | 480.2 | - | - | 0 | - |
| - | - | 9099 | 480.5 | - | - | 0 | - |
| - | - | 2.448E+04 | 480.7 | - | - | 0 | - |
| - | - | 1.535E+05 | 484.2 | - | - | 0 | - |
| - | - | 1.2E+05 | 484.5 | - | - | 0 | - |
| - | - | 1.413E+04 | 484.7 | - | - | 0 | - |
| - | - | 5.601E+04 | 484.9 | - | - | 0 | - |
| - | - | 1.165E+05 | 485.2 | - | - | 0 | - |
| - | - | 3.674E+04 | 485.2 | - | - | 0 | - |
| - | - | 1.212E+04 | 485.7 | - | - | 0 | - |
| - | - | 2.418E+04 | 486.2 | - | - | 0 | - |
| - | - | 2.188E+04 | 486.2 | - | - | 0 | - |
| - | - | 6855 | 486.7 | - | - | 0 | - |
| - | - | 7694 | 487.7 | - | - | 0 | - |
| - | - | 1.342E+04 | 488.2 | - | - | 0 | - |
| - | - | 7313 | 488.3 | - | - | 0 | - |
| - | - | 6083 | 488.7 | - | - | 0 | - |
| - | - | 7099 | 489.2 | - | - | 0 | - |
| - | - | 1.709E+04 | 489.2 | - | - | 0 | - |
| - | - | 8197 | 490.2 | - | - | 0 | - |
| - | - | 2.262E+04 | 490.2 | - | - | 0 | - |
| - | - | 2.196E+04 | 492.7 | - | - | 0 | - |
| - | - | 2.092E+04 | 493.2 | - | - | 0 | - |
| - | - | 3.236E+04 | 493.5 | - | - | 0 | - |
| - | - | 9.745E+04 | 493.7 | - | - | 0 | - |
| - | - | 5.06E+04 | 493.9 | - | - | 0 | - |
| - | - | 5.058E+04 | 494.2 | - | - | 0 | - |
| - | - | 2.854E+04 | 494.5 | - | - | 0 | - |
| - | - | 1.889E+04 | 494.7 | - | - | 0 | - |
| - | - | 6876 | 494.9 | - | - | 0 | - |
| - | - | 1.542E+04 | 495.2 | - | - | 0 | - |
| - | - | 1.295E+04 | 495.2 | - | - | 0 | - |
| - | - | 6874 | 495.7 | - | - | 0 | - |
| - | - | 7130 | 496.2 | - | - | 0 | - |
| - | - | 8847 | 496.8 | - | - | 0 | - |
| - | - | 2.147E+04 | 497.3 | - | - | 0 | - |
| - | - | 1.87E+04 | 497.8 | - | - | 0 | - |
| 2 | y | 1.943E+05 | 499.5 | 0.001326 | 2.654 | +3 | 12 |
| 2 | y | 2.879E+05 | 499.9 | 0.005047 | 10.1 | +3 | 12 |
| - | - | 1.82E+05 | 500.2 | - | - | 0 | - |
| - | - | 9.559E+04 | 500.5 | - | - | 0 | - |
| - | - | 8368 | 500.7 | - | - | 0 | - |
| - | - | 4.057E+04 | 500.9 | - | - | 0 | - |
| - | - | 1.408E+04 | 501.2 | - | - | 0 | - |
| 6 | y | 3.615E+04 | 501.7 | 0.0008487 | 1.691 | +2 | 8 |
| 6 | y | 9.268E+04 | 502.2 | 0.001913 | 3.81 | +2 | 8 |
| - | - | 1.413E+05 | 502.7 | - | - | 0 | - |
| - | - | 5.904E+04 | 503.2 | - | - | 0 | - |
| - | - | 8350 | 503.7 | - | - | 0 | - |
| - | - | 1.066E+04 | 504.2 | - | - | 0 | - |
| - | - | 6619 | 505.2 | - | - | 0 | - |
| - | - | 7381 | 505.2 | - | - | 0 | - |
| 2 | y | 2.947E+05 | 505.5 | 0.001222 | 2.418 | +3 | 12 |
| - | - | 2.566E+05 | 505.9 | - | - | 0 | - |
| - | - | 1.307E+05 | 506.2 | - | - | 0 | - |
| - | - | 2.071E+04 | 506.3 | - | - | 0 | - |
| - | - | 4.784E+04 | 506.5 | - | - | 0 | - |
| - | - | 9601 | 506.9 | - | - | 0 | - |
| - | - | 2.87E+04 | 507.2 | - | - | 0 | - |
| - | - | 1.097E+04 | 508.2 | - | - | 0 | - |
| - | - | 2.524E+04 | 508.3 | - | - | 0 | - |
| - | - | 6127 | 509.2 | - | - | 0 | - |
| - | - | 1.327E+04 | 509.7 | - | - | 0 | - |
| 6 | y | 6.413E+05 | 510.7 | 0.0009679 | 1.895 | +2 | 8 |
| - | - | 3.825E+05 | 511.2 | - | - | 0 | - |
| - | - | 1.139E+05 | 511.7 | - | - | 0 | - |
| - | - | 3.606E+04 | 512.2 | - | - | 0 | - |
| - | - | 6040 | 513.2 | - | - | 0 | - |
| - | - | 4.181E+04 | 513.2 | - | - | 0 | - |
| - | - | 1.509E+04 | 514.2 | - | - | 0 | - |
| - | - | 2.356E+04 | 515.7 | - | - | 0 | - |
| - | - | 1.22E+04 | 516.2 | - | - | 0 | - |
| - | - | 7559 | 516.7 | - | - | 0 | - |
| - | - | 2.645E+04 | 517.2 | - | - | 0 | - |
| - | - | 7.998E+04 | 518.2 | - | - | 0 | - |
| - | - | 3.038E+04 | 518.7 | - | - | 0 | - |
| - | - | 1.399E+04 | 519.2 | - | - | 0 | - |
| - | - | 8.264E+04 | 520.3 | - | - | 0 | - |
| - | - | 6826 | 520.7 | - | - | 0 | - |
| - | - | 2.183E+04 | 521.3 | - | - | 0 | - |
| - | - | 6323 | 521.7 | - | - | 0 | - |
| - | - | 2.626E+04 | 522.2 | - | - | 0 | - |
| - | - | 3.904E+04 | 523.2 | - | - | 0 | - |
| - | - | 9242 | 524.2 | - | - | 0 | - |
| - | - | 9937 | 524.2 | - | - | 0 | - |
| - | - | 7039 | 524.3 | - | - | 0 | - |
| - | - | 6510 | 525.7 | - | - | 0 | - |
| - | - | 1.182E+04 | 526.2 | - | - | 0 | - |
| - | - | 9018 | 527.7 | - | - | 0 | - |
| - | - | 2.689E+04 | 528.3 | - | - | 0 | - |
| - | - | 4.952E+04 | 529.3 | - | - | 0 | - |
| - | - | 8253 | 529.3 | - | - | 0 | - |
| - | - | 2.624E+04 | 529.8 | - | - | 0 | - |
| - | - | 2.361E+04 | 530.2 | - | - | 0 | - |
| - | - | 6798 | 530.7 | - | - | 0 | - |
| - | - | 1.441E+04 | 531.8 | - | - | 0 | - |
| - | - | 1.946E+04 | 532.3 | - | - | 0 | - |
| - | - | 9525 | 532.7 | - | - | 0 | - |
| - | - | 8416 | 532.8 | - | - | 0 | - |
| - | - | 1.109E+04 | 533.2 | - | - | 0 | - |
| - | - | 5936 | 533.3 | - | - | 0 | - |
| - | - | 7816 | 534.2 | - | - | 0 | - |
| - | - | 8.742E+04 | 534.3 | - | - | 0 | - |
| - | - | 5.126E+04 | 534.7 | - | - | 0 | - |
| - | - | 1.002E+05 | 535.2 | - | - | 0 | - |
| - | - | 1.08E+04 | 535.3 | - | - | 0 | - |
| - | - | 1.821E+04 | 535.7 | - | - | 0 | - |
| - | - | 2.736E+04 | 536.2 | - | - | 0 | - |
| - | - | 7.179E+04 | 536.2 | - | - | 0 | - |
| - | - | 7539 | 537.2 | - | - | 0 | - |
| - | - | 2.246E+04 | 537.2 | - | - | 0 | - |
| - | - | 8416 | 537.3 | - | - | 0 | - |
| - | - | 2.63E+04 | 538.2 | - | - | 0 | - |
| - | - | 3.004E+04 | 538.7 | - | - | 0 | - |
| - | - | 4.587E+04 | 539.2 | - | - | 0 | - |
| - | - | 2.184E+04 | 539.7 | - | - | 0 | - |
| - | - | 1.359E+04 | 540.2 | - | - | 0 | - |
| - | - | 6.746E+04 | 540.8 | - | - | 0 | - |
| - | - | 6120 | 541.2 | - | - | 0 | - |
| - | - | 5769 | 541.2 | - | - | 0 | - |
| - | - | 4.091E+04 | 541.3 | - | - | 0 | - |
| - | - | 1.606E+04 | 541.7 | - | - | 0 | - |
| - | - | 1.31E+04 | 541.8 | - | - | 0 | - |
| - | - | 1.722E+04 | 542.7 | - | - | 0 | - |
| - | - | 6.77E+04 | 543.3 | - | - | 0 | - |
| - | - | 1.687E+05 | 543.7 | - | - | 0 | - |
| - | - | 9.909E+04 | 544.2 | - | - | 0 | - |
| - | - | 2.677E+04 | 544.7 | - | - | 0 | - |
| - | - | 1.058E+04 | 545.2 | - | - | 0 | - |
| - | - | 5.442E+04 | 545.3 | - | - | 0 | - |
| - | - | 9282 | 545.8 | - | - | 0 | - |
| - | - | 1.272E+04 | 546.3 | - | - | 0 | - |
| - | - | 4.907E+04 | 547.3 | - | - | 0 | - |
| - | - | 2.049E+04 | 548.3 | - | - | 0 | - |
| - | - | 1.985E+04 | 550.2 | - | - | 0 | - |
| - | - | 1.862E+04 | 551.2 | - | - | 0 | - |
| - | - | 7120 | 551.7 | - | - | 0 | - |
| - | - | 6.728E+05 | 552.3 | - | - | 0 | - |
| - | - | 4.367E+05 | 552.8 | - | - | 0 | - |
| 10 | y | 1.754E+05 | 553.2 | 0.0008148 | 1.473 | +1 | 4 |
| - | - | 1.156E+05 | 553.3 | - | - | 0 | - |
| - | - | 3.261E+04 | 553.8 | - | - | 0 | - |
| - | - | 5.2E+04 | 554.2 | - | - | 0 | - |
| - | - | 6846 | 554.3 | - | - | 0 | - |
| - | - | 4.699E+04 | 554.8 | - | - | 0 | - |
| - | - | 8253 | 555.2 | - | - | 0 | - |
| - | - | 3.887E+04 | 555.3 | - | - | 0 | - |
| - | - | 1.057E+04 | 555.8 | - | - | 0 | - |
| - | - | 1.858E+04 | 556.3 | - | - | 0 | - |
| - | - | 1.378E+04 | 557.2 | - | - | 0 | - |
| - | - | 1.051E+04 | 557.7 | - | - | 0 | - |
| - | - | 1.148E+04 | 558.2 | - | - | 0 | - |
| - | - | 5366 | 558.7 | - | - | 0 | - |
| - | - | 5621 | 560.3 | - | - | 0 | - |
| - | - | 3.687E+04 | 560.3 | - | - | 0 | - |
| - | - | 1.428E+04 | 561.3 | - | - | 0 | - |
| - | - | 1.19E+04 | 561.3 | - | - | 0 | - |
| - | - | 1.178E+04 | 561.7 | - | - | 0 | - |
| - | - | 1.199E+04 | 562.3 | - | - | 0 | - |
| - | - | 5724 | 562.3 | - | - | 0 | - |
| - | - | 7271 | 563.2 | - | - | 0 | - |
| - | - | 4.333E+04 | 563.8 | - | - | 0 | - |
| - | - | 4.087E+04 | 564.3 | - | - | 0 | - |
| - | - | 1.954E+04 | 564.8 | - | - | 0 | - |
| - | - | 9.784E+04 | 565.3 | - | - | 0 | - |
| - | - | 2.059E+04 | 566.3 | - | - | 0 | - |
| - | - | 2.986E+04 | 566.7 | - | - | 0 | - |
| - | - | 2.245E+04 | 567.2 | - | - | 0 | - |
| - | - | 1.175E+04 | 567.7 | - | - | 0 | - |
| - | - | 1.887E+04 | 568.2 | - | - | 0 | - |
| 10 | y | 4.62E+05 | 570.2 | 0.0007549 | 1.324 | +1 | 4 |
| - | - | 1.319E+05 | 571.2 | - | - | 0 | - |
| - | - | 6168 | 571.7 | - | - | 0 | - |
| - | - | 6.187E+04 | 572.2 | - | - | 0 | - |
| - | - | 1.569E+05 | 572.8 | - | - | 0 | - |
| - | - | 1.073E+05 | 573.3 | - | - | 0 | - |
| - | - | 3.398E+04 | 573.8 | - | - | 0 | - |
| - | - | 3.363E+04 | 574.3 | - | - | 0 | - |
| 5 | y | 3.958E+04 | 575.2 | 0.004955 | 8.613 | +2 | 9 |
| 5 | y | 1.164E+05 | 575.7 | 0.005439 | 9.448 | +2 | 9 |
| 10 | b | 7.927E+04 | 576.2 | 0.01022 | 17.73 | +2 | 10 |
| - | - | 4.006E+04 | 576.7 | - | - | 0 | - |
| - | - | 6545 | 577.2 | - | - | 0 | - |
| - | - | 1.856E+04 | 577.8 | - | - | 0 | - |
| - | - | 1.544E+04 | 578.3 | - | - | 0 | - |
| - | - | 9256 | 578.8 | - | - | 0 | - |
| - | - | 1.163E+04 | 579.3 | - | - | 0 | - |
| - | - | 8150 | 580.2 | - | - | 0 | - |
| - | - | 5397 | 581.3 | - | - | 0 | - |
| - | - | 1.941E+04 | 582.2 | - | - | 0 | - |
| - | - | 8196 | 583.2 | - | - | 0 | - |
| 5 | y | 7.713E+05 | 584.3 | 0.003395 | 5.811 | +2 | 9 |
| - | - | 4.703E+05 | 584.8 | - | - | 0 | - |
| - | - | 2.193E+05 | 585.3 | - | - | 0 | - |
| - | - | 7.31E+04 | 585.8 | - | - | 0 | - |
| - | - | 2.529E+04 | 586.3 | - | - | 0 | - |
| - | - | 9.937E+04 | 586.8 | - | - | 0 | - |
| - | - | 1.308E+04 | 587.2 | - | - | 0 | - |
| - | - | 7.993E+04 | 587.3 | - | - | 0 | - |
| - | - | 3.506E+04 | 587.8 | - | - | 0 | - |
| - | - | 5.521E+04 | 588.3 | - | - | 0 | - |
| - | - | 3.92E+04 | 589.3 | - | - | 0 | - |
| - | - | 1.26E+04 | 589.7 | - | - | 0 | - |
| - | - | 9078 | 590.2 | - | - | 0 | - |
| - | - | 5687 | 591.2 | - | - | 0 | - |
| - | - | 2.305E+04 | 591.3 | - | - | 0 | - |
| - | - | 8457 | 592.3 | - | - | 0 | - |
| - | - | 1.221E+04 | 592.8 | - | - | 0 | - |
| - | - | 2.759E+04 | 593.3 | - | - | 0 | - |
| - | - | 1.324E+04 | 593.8 | - | - | 0 | - |
| - | - | 1.118E+04 | 594.3 | - | - | 0 | - |
| - | - | 5.571E+04 | 595.8 | - | - | 0 | - |
| - | - | 3.345E+04 | 596.3 | - | - | 0 | - |
| - | - | 1.83E+04 | 596.8 | - | - | 0 | - |
| - | - | 7528 | 596.8 | - | - | 0 | - |
| - | - | 8.797E+04 | 598.3 | - | - | 0 | - |
| - | - | 2.357E+04 | 598.7 | - | - | 0 | - |
| - | - | 2.462E+04 | 599.3 | - | - | 0 | - |
| - | - | 7982 | 599.7 | - | - | 0 | - |
| - | - | 1.063E+05 | 600.2 | - | - | 0 | - |
| - | - | 3.225E+04 | 601.2 | - | - | 0 | - |
| - | - | 1.492E+05 | 601.8 | - | - | 0 | - |
| - | - | 6529 | 602.2 | - | - | 0 | - |
| - | - | 1.06E+05 | 602.3 | - | - | 0 | - |
| - | - | 2.877E+04 | 602.8 | - | - | 0 | - |
| - | - | 1.508E+04 | 603.3 | - | - | 0 | - |
| - | - | 5794 | 603.8 | - | - | 0 | - |
| - | - | 1.694E+04 | 604.2 | - | - | 0 | - |
| - | - | 7009 | 604.8 | - | - | 0 | - |
| - | - | 6457 | 605.2 | - | - | 0 | - |
| - | - | 1.237E+04 | 605.3 | - | - | 0 | - |
| - | - | 9.596E+04 | 607.3 | - | - | 0 | - |
| - | - | 3.248E+04 | 608.3 | - | - | 0 | - |
| - | - | 8822 | 610.2 | - | - | 0 | - |
| - | - | 1.816E+04 | 612.3 | - | - | 0 | - |
| - | - | 2.408E+04 | 612.8 | - | - | 0 | - |
| - | - | 1.666E+04 | 613.3 | - | - | 0 | - |
| - | - | 1.395E+04 | 614.2 | - | - | 0 | - |
| - | - | 5366 | 614.3 | - | - | 0 | - |
| - | - | 8043 | 615.2 | - | - | 0 | - |
| - | - | 7225 | 615.8 | - | - | 0 | - |
| - | - | 7985 | 616.3 | - | - | 0 | - |
| - | - | 1.288E+04 | 617.3 | - | - | 0 | - |
| - | - | 7293 | 617.8 | - | - | 0 | - |
| - | - | 7662 | 618.3 | - | - | 0 | - |
| - | - | 1.676E+04 | 621.3 | - | - | 0 | - |
| - | - | 1.683E+04 | 621.8 | - | - | 0 | - |
| - | - | 5983 | 622.3 | - | - | 0 | - |
| 4 | y | 1.169E+04 | 624.8 | 0.002181 | 3.49 | +2 | 10 |
| 4 | y | 3.897E+04 | 625.3 | 0.00468 | 7.484 | +2 | 10 |
| - | - | 1.964E+04 | 625.8 | - | - | 0 | - |
| - | - | 2.163E+04 | 626.3 | - | - | 0 | - |
| - | - | 1.929E+04 | 626.8 | - | - | 0 | - |
| - | - | 1.04E+04 | 627.3 | - | - | 0 | - |
| - | - | 7466 | 627.8 | - | - | 0 | - |
| - | - | 2.073E+04 | 632.2 | - | - | 0 | - |
| - | - | 6093 | 633.2 | - | - | 0 | - |
| - | - | 1.971E+04 | 633.3 | - | - | 0 | - |
| 4 | y | 1.868E+05 | 633.8 | 0.003368 | 5.314 | +2 | 10 |
| - | - | 7763 | 634.2 | - | - | 0 | - |
| - | - | 1.16E+05 | 634.3 | - | - | 0 | - |
| - | - | 9343 | 634.3 | - | - | 0 | - |
| - | - | 6.41E+04 | 634.8 | - | - | 0 | - |
| - | - | 3.265E+04 | 635.3 | - | - | 0 | - |
| - | - | 2.094E+04 | 635.8 | - | - | 0 | - |
| - | - | 7375 | 636.3 | - | - | 0 | - |
| - | - | 6923 | 636.3 | - | - | 0 | - |
| - | - | 2.204E+04 | 636.8 | - | - | 0 | - |
| - | - | 1.533E+04 | 637.3 | - | - | 0 | - |
| - | - | 1.5E+04 | 638.8 | - | - | 0 | - |
| - | - | 1.537E+04 | 639.3 | - | - | 0 | - |
| 11 | b | 1.48E+04 | 642.3 | 0.004163 | 6.482 | +2 | 11 |
| - | - | 7840 | 643.3 | - | - | 0 | - |
| - | - | 1.995E+04 | 643.3 | - | - | 0 | - |
| - | - | 2.295E+04 | 644.3 | - | - | 0 | - |
| - | - | 7921 | 644.3 | - | - | 0 | - |
| - | - | 4.073E+04 | 644.8 | - | - | 0 | - |
| - | - | 9.349E+04 | 645.3 | - | - | 0 | - |
| - | - | 6.527E+04 | 645.8 | - | - | 0 | - |
| - | - | 2.105E+04 | 646.3 | - | - | 0 | - |
| - | - | 1.363E+04 | 648.3 | - | - | 0 | - |
| - | - | 1.274E+04 | 650.3 | - | - | 0 | - |
| - | - | 1.444E+04 | 651.3 | - | - | 0 | - |
| - | - | 1.068E+04 | 652.2 | - | - | 0 | - |
| - | - | 6.688E+04 | 653.3 | - | - | 0 | - |
| - | - | 2.278E+04 | 653.8 | - | - | 0 | - |
| - | - | 9021 | 654.3 | - | - | 0 | - |
| - | - | 2.099E+04 | 656.3 | - | - | 0 | - |
| - | - | 4.539E+04 | 657.3 | - | - | 0 | - |
| - | - | 7906 | 658.3 | - | - | 0 | - |
| - | - | 6109 | 658.3 | - | - | 0 | - |
| - | - | 2.043E+04 | 658.8 | - | - | 0 | - |
| - | - | 1.267E+04 | 659.3 | - | - | 0 | - |
| - | - | 7874 | 659.8 | - | - | 0 | - |
| - | - | 2.423E+04 | 660.9 | - | - | 0 | - |
| - | - | 1.779E+04 | 661.4 | - | - | 0 | - |
| - | - | 1.147E+04 | 665.3 | - | - | 0 | - |
| 9 | y | 5.173E+04 | 666.3 | 0.003421 | 5.134 | +1 | 5 |
| - | - | 3.718E+04 | 667.3 | - | - | 0 | - |
| - | - | 3.467E+04 | 667.8 | - | - | 0 | - |
| 3 | y | 2.999E+04 | 668.3 | 0.002237 | 3.347 | +2 | 11 |
| 3 | y | 3.215E+04 | 668.8 | 0.004535 | 6.78 | +2 | 11 |
| - | - | 1.316E+04 | 669.3 | - | - | 0 | - |
| - | - | 8715 | 669.8 | - | - | 0 | - |
| - | - | 1.547E+05 | 671.3 | - | - | 0 | - |
| - | - | 2.178E+04 | 671.3 | - | - | 0 | - |
| - | - | 5.303E+04 | 672.3 | - | - | 0 | - |
| - | - | 8166 | 672.3 | - | - | 0 | - |
| - | - | 1.428E+04 | 673.3 | - | - | 0 | - |
| - | - | 7.353E+04 | 674.3 | - | - | 0 | - |
| - | - | 2.49E+04 | 675.3 | - | - | 0 | - |
| - | - | 2.429E+04 | 676.3 | - | - | 0 | - |
| - | - | 1.677E+04 | 676.8 | - | - | 0 | - |
| 3 | y | 1.438E+05 | 677.3 | 0.002857 | 4.218 | +2 | 11 |
| - | - | 8.352E+04 | 677.8 | - | - | 0 | - |
| - | - | 2.903E+04 | 678.3 | - | - | 0 | - |
| - | - | 1.49E+04 | 678.8 | - | - | 0 | - |
| - | - | 3.415E+04 | 679.3 | - | - | 0 | - |
| - | - | 3.552E+04 | 680.2 | - | - | 0 | - |
| - | - | 8124 | 681.2 | - | - | 0 | - |
| - | - | 1.169E+04 | 682.3 | - | - | 0 | - |
| - | - | 7830 | 682.4 | - | - | 0 | - |
| - | - | 1.599E+04 | 682.8 | - | - | 0 | - |
| 9 | y | 1.634E+05 | 683.3 | 0.0008584 | 1.256 | +1 | 5 |
| - | - | 9.285E+04 | 684.3 | - | - | 0 | - |
| - | - | 3.787E+04 | 685.3 | - | - | 0 | - |
| - | - | 7423 | 686.3 | - | - | 0 | - |
| - | - | 9996 | 688.4 | - | - | 0 | - |
| - | - | 7531 | 691.3 | - | - | 0 | - |
| - | - | 1.158E+04 | 692.4 | - | - | 0 | - |
| - | - | 1.125E+04 | 694.3 | - | - | 0 | - |
| - | - | 2.412E+04 | 697.3 | - | - | 0 | - |
| - | - | 2.196E+04 | 697.3 | - | - | 0 | - |
| - | - | 8162 | 698.3 | - | - | 0 | - |
| - | - | 1.547E+05 | 702.3 | - | - | 0 | - |
| - | - | 7.718E+04 | 703.3 | - | - | 0 | - |
| - | - | 1.704E+04 | 704.3 | - | - | 0 | - |
| - | - | 1.948E+04 | 707.3 | - | - | 0 | - |
| - | - | 1.081E+04 | 708.3 | - | - | 0 | - |
| 6 | b | 8505 | 719.2 | 0.003442 | 4.785 | +1 | 6 |
| - | - | 1.734E+04 | 719.3 | - | - | 0 | - |
| - | - | 6.022E+04 | 720.4 | - | - | 0 | - |
| - | - | 2.017E+04 | 721.4 | - | - | 0 | - |
| - | - | 1.214E+04 | 722.3 | - | - | 0 | - |
| - | - | 8084 | 723.3 | - | - | 0 | - |
| - | - | 1.117E+04 | 733.3 | - | - | 0 | - |
| - | - | 3.518E+04 | 735.3 | - | - | 0 | - |
| - | - | 1.603E+04 | 736.3 | - | - | 0 | - |
| 8 | y | 8.567E+04 | 737.3 | 0.0006697 | 0.9083 | +1 | 6 |
| - | - | 3.601E+04 | 738.3 | - | - | 0 | - |
| - | - | 9099 | 739.3 | - | - | 0 | - |
| - | - | 9178 | 740.4 | - | - | 0 | - |
| - | - | 2.779E+04 | 743.3 | - | - | 0 | - |
| - | - | 1.992E+04 | 744.2 | - | - | 0 | - |
| - | - | 8692 | 745.2 | - | - | 0 | - |
| - | - | 3.014E+04 | 750.3 | - | - | 0 | - |
| - | - | 4.006E+04 | 751.3 | - | - | 0 | - |
| - | - | 9728 | 752.3 | - | - | 0 | - |
| - | - | 1.369E+04 | 752.3 | - | - | 0 | - |
| 8 | y | 2.826E+05 | 754.4 | 0.000793 | 1.051 | +1 | 6 |
| - | - | 1.048E+05 | 755.4 | - | - | 0 | - |
| - | - | 3.887E+04 | 756.4 | - | - | 0 | - |
| - | - | 1.922E+04 | 757.4 | - | - | 0 | - |
| - | - | 7483 | 758.4 | - | - | 0 | - |
| - | - | 5553 | 759.4 | - | - | 0 | - |
| - | - | 3.052E+04 | 761.3 | - | - | 0 | - |
| - | - | 1.695E+04 | 766.4 | - | - | 0 | - |
| - | - | 1.338E+04 | 767.4 | - | - | 0 | - |
| - | - | 1.309E+04 | 768.3 | - | - | 0 | - |
| - | - | 8840 | 768.4 | - | - | 0 | - |
| - | - | 8935 | 770.4 | - | - | 0 | - |
| - | - | 8129 | 771.4 | - | - | 0 | - |
| - | - | 4612 | 784.3 | - | - | 0 | - |
| - | - | 7.922E+04 | 784.4 | - | - | 0 | - |
| - | - | 1.082E+05 | 785.4 | - | - | 0 | - |
| - | - | 4.252E+04 | 786.4 | - | - | 0 | - |
| - | - | 9485 | 787.4 | - | - | 0 | - |
| - | - | 1.517E+04 | 794.4 | - | - | 0 | - |
| - | - | 1.259E+04 | 808.4 | - | - | 0 | - |
| - | - | 2.965E+04 | 811.4 | - | - | 0 | - |
| - | - | 1.604E+04 | 812.4 | - | - | 0 | - |
| - | - | 3.886E+04 | 814.3 | - | - | 0 | - |
| - | - | 3.248E+04 | 815.3 | - | - | 0 | - |
| - | - | 1.492E+04 | 816.3 | - | - | 0 | - |
| - | - | 1.115E+04 | 821.4 | - | - | 0 | - |
| - | - | 1.236E+04 | 822.4 | - | - | 0 | - |
| - | - | 8202 | 829.4 | - | - | 0 | - |
| - | - | 8623 | 831.4 | - | - | 0 | - |
| - | - | 4.199E+04 | 832.3 | - | - | 0 | - |
| - | - | 1.719E+04 | 833.3 | - | - | 0 | - |
| - | - | 1.1E+04 | 834.3 | - | - | 0 | - |
| - | - | 8514 | 834.4 | - | - | 0 | - |
| - | - | 8846 | 835.4 | - | - | 0 | - |
| - | - | 7994 | 836.4 | - | - | 0 | - |
| - | - | 8733 | 837.4 | - | - | 0 | - |
| - | - | 7817 | 838.4 | - | - | 0 | - |
| - | - | 2.442E+04 | 839.4 | - | - | 0 | - |
| - | - | 2.028E+04 | 840.4 | - | - | 0 | - |
| - | - | 6618 | 841.4 | - | - | 0 | - |
| 7 | b | 7534 | 848.3 | 0.002225 | 2.623 | +1 | 7 |
| - | - | 4.813E+04 | 848.4 | - | - | 0 | - |
| - | - | 8.178E+04 | 849.4 | - | - | 0 | - |
| - | - | 3.597E+04 | 850.4 | - | - | 0 | - |
| - | - | 8888 | 851.4 | - | - | 0 | - |
| - | - | 6346 | 856.3 | - | - | 0 | - |
| - | - | 2.794E+04 | 857.4 | - | - | 0 | - |
| - | - | 1.197E+04 | 858.4 | - | - | 0 | - |
| - | - | 1.41E+04 | 863.4 | - | - | 0 | - |
| - | - | 1.5E+04 | 864.4 | - | - | 0 | - |
| 7 | y | 7.76E+04 | 865.4 | 0.0005958 | 0.6884 | +1 | 7 |
| 7 | y | 1.032E+05 | 866.4 | 0.005989 | 6.913 | +1 | 7 |
| - | - | 5.191E+04 | 867.4 | - | - | 0 | - |
| - | - | 1.739E+04 | 868.4 | - | - | 0 | - |
| - | - | 5686 | 874.5 | - | - | 0 | - |
| - | - | 2.578E+04 | 881.4 | - | - | 0 | - |
| - | - | 7848 | 882.4 | - | - | 0 | - |
| 7 | y | 2.909E+05 | 883.4 | 0.0005583 | 0.632 | +1 | 7 |
| - | - | 1.311E+05 | 884.4 | - | - | 0 | - |
| - | - | 3.699E+04 | 885.4 | - | - | 0 | - |
| - | - | 1.002E+04 | 886.4 | - | - | 0 | - |
| - | - | 5801 | 893.4 | - | - | 0 | - |
| - | - | 1.321E+04 | 894.4 | - | - | 0 | - |
| - | - | 6171 | 917.4 | - | - | 0 | - |
| 8 | b | 1.597E+04 | 919.3 | 0.0003952 | 0.4298 | +1 | 8 |
| - | - | 8348 | 921.3 | - | - | 0 | - |
| - | - | 2.311E+04 | 921.4 | - | - | 0 | - |
| - | - | 2.006E+04 | 922.4 | - | - | 0 | - |
| - | - | 1.212E+04 | 923.4 | - | - | 0 | - |
| - | - | 1.216E+04 | 927.4 | - | - | 0 | - |
| - | - | 1.585E+04 | 928.4 | - | - | 0 | - |
| - | - | 6931 | 929.4 | - | - | 0 | - |
| - | - | 8049 | 940.4 | - | - | 0 | - |
| - | - | 7790 | 943.5 | - | - | 0 | - |
| - | - | 3.032E+04 | 945.4 | - | - | 0 | - |
| - | - | 1.725E+04 | 946.4 | - | - | 0 | - |
| - | - | 1.48E+04 | 948.4 | - | - | 0 | - |
| - | - | 8383 | 949.4 | - | - | 0 | - |
| - | - | 7491 | 953.5 | - | - | 0 | - |
| - | - | 8039 | 954.5 | - | - | 0 | - |
| - | - | 3.529E+04 | 971.5 | - | - | 0 | - |
| - | - | 1.643E+04 | 972.5 | - | - | 0 | - |
| - | - | 6874 | 973.5 | - | - | 0 | - |
| - | - | 6264 | 986.4 | - | - | 0 | - |
| - | - | 6237 | 987.4 | - | - | 0 | - |
| 6 | y | 9597 | 1002 | 0.002016 | 2.011 | +1 | 8 |
| 6 | y | 2.051E+04 | 1003 | 0.002436 | 2.428 | +1 | 8 |
| - | - | 1.678E+04 | 1004 | - | - | 0 | - |
| - | - | 1.017E+04 | 1005 | - | - | 0 | - |
| - | - | 9466 | 1007 | - | - | 0 | - |
| - | - | 8681 | 1017 | - | - | 0 | - |
| - | - | 1.105E+04 | 1018 | - | - | 0 | - |
| - | - | 5931 | 1019 | - | - | 0 | - |
| 6 | y | 1.045E+05 | 1020 | 0.000126 | 0.1235 | +1 | 8 |
| - | - | 5.279E+04 | 1021 | - | - | 0 | - |
| - | - | 1.683E+04 | 1022 | - | - | 0 | - |
| - | - | 6340 | 1030 | - | - | 0 | - |
| 9 | b | 8687 | 1032 | 0.00294 | 2.848 | +1 | 9 |
| - | - | 5.127E+04 | 1035 | - | - | 0 | - |
| - | - | 2.661E+04 | 1036 | - | - | 0 | - |
| - | - | 1.1E+04 | 1037 | - | - | 0 | - |
| - | - | 6262 | 1086 | - | - | 0 | - |
| - | - | 5507 | 1087 | - | - | 0 | - |
| - | - | 9446 | 1104 | - | - | 0 | - |
| - | - | 7066 | 1105 | - | - | 0 | - |
| - | - | 1.102E+04 | 1114 | - | - | 0 | - |
| - | - | 7954 | 1115 | - | - | 0 | - |
| - | - | 7885 | 1132 | - | - | 0 | - |
| - | - | 6974 | 1133 | - | - | 0 | - |
| 5 | y | 5244 | 1167 | 0.01047 | 8.967 | +1 | 9 |
| - | - | 7727 | 1178 | - | - | 0 | - |
| - | - | 5828 | 1179 | - | - | 0 | - |
| - | - | 1.602E+04 | 1196 | - | - | 0 | - |
| - | - | 1.088E+04 | 1197 | - | - | 0 | - |
| - | - | 4427 | 1565 | - | - | 0 | - |
| - | - | 5725 | 1971 | - | - | 0 | - |
| - | - | 4828 | 3082 | - | - | 0 | - |

m/z Charge Intensity FragmentType MassShift Position
120.04825592041016 0 3320.047
120.05213165283203 0 4462.752
120.05622863769531 0 15659.779
120.78741455078125 0 2802.8123
121.0400390625 0 22196.043
122.07171630859375 0 24041.64
123.04450988769531 0 81303.516
123.05575561523438 0 18230.26
124.0397720336914 0 8985.103
124.0481185913086 0 5561.0317
124.07611083984375 0 34188.47
126.00130462646484 0 13637.718
126.09174346923828 0 6861.025
127.05064392089844 0 37534.24
127.08706665039062 0 7813.553
128.07150268554688 0 2836.4944
129.06629943847656 0 106825.52
129.10264587402344 0 28277.09
130.03253173828125 0 64094.53
130.05026245117188 0 3412.512
130.0694580078125 0 5030.376
131.0360107421875 0 4795.0674
131.08192443847656 0 38348.15
132.01162719726562 0 6373.892
132.08555603027344 0 4309.405
133.0435791015625 0 11505.047
133.0764923095703 0 3326.905
134.0275115966797 0 1364038
134.0713653564453 0 5010.8237
135.02618408203125 0 7120.09
135.0308837890625 0 58058.8
136.02328491210938 0 58283.09
136.02926635742188 0 5262.5557
136.03201293945312 0 4720.2188
136.0508575439453 0 14444.77
136.07614135742188 0 1352598
137.07371520996094 0 5476.9097
137.07948303222656 0 115414.54
138.06668090820312 0 69904.21
138.09188842773438 0 5151.201
139.05056762695312 0 19204.729
140.0167694091797 0 14492.538
141.06716918945312 0 3483.364
141.1026611328125 0 48985.613
142.0502166748047 0 6210.8926
142.08688354492188 0 5410.6733
143.02777099609375 0 13151.471
144.01187133789062 0 14963.132
144.03565979003906 0 5401.881
146.02757263183594 0 7800.438
147.04452514648438 0 71497.305
148.04298400878906 0 26042.828
148.0493621826172 0 7730.5034
148.08741760253906 0 55708.062
148.95433044433594 0 5006.346
149.0275115966797 0 3508.769
149.07150268554688 0 3783.8745
149.082763671875 0 4358.4824
149.0904998779297 0 4558.9854
150.06642150878906 0 7819.9233
152.08229064941406 0 32254.916
153.10292053222656 0 4325.5054
154.06153869628906 0 17693.264
155.0819091796875 0 21417.654
155.09323120117188 0 59301.71
155.11831665039062 0 37645.906
156.01185607910156 0 9266.461
156.07720947265625 0 145259.44
156.09657287597656 0 5274.8887
157.0433807373047 0 9940.311
157.06118774414062 0 55459.285
157.08074951171875 0 9406.971
157.13394165039062 0 33820.19
158.02748107910156 0 503747.7
159.03091430664062 0 27338.283
159.0557403564453 0 12273.75
159.11325073242188 0 1918044.1
160.02317810058594 0 23574.518
160.04293823242188 0 7393.73
160.0871124267578 0 4518.64
160.1103973388672 0 9500.8125
160.11663818359375 0 150378.75
161.03836059570312 0 220711.06
161.11802673339844 0 8233.557
162.0223846435547 0 56208.293
162.0419464111328 0 4771.6904
162.06674194335938 0 24979.615
163.00656127929688 0 7443.746
163.02651977539062 0 4443.8687
163.03392028808594 0 8912.3545
163.0984649658203 0 8935.8125
164.08267211914062 0 8066.1987
164.11892700195312 0 5376.634
165.0550994873047 0 491029.78
165.07752990722656 0 83584.09
165.10263061523438 0 9296.285
166.06118774414062 0 133085.33
166.0865936279297 0 169966.72
167.06503295898438 0 10820.545
167.09271240234375 0 22256.477
167.11785888671875 0 4929.813
168.01182556152344 0 65271.684
169.0152587890625 0 3524.2634
169.0976104736328 0 25091.617
171.1133270263672 0 10883.188
172.05096435546875 0 10334.305
172.577392578125 0 5639.2026
173.03819274902344 0 7154.781
173.0792694091797 0 3307.4824
173.0924072265625 0 34103.715
173.45281982421875 0 4602.803
174.0225372314453 0 21186.068
174.066650390625 0 32329.785
175.05404663085938 0 9792.221
175.09817504882812 0 13073.859
176.03810119628906 0 667764.9
176.08233642578125 0 141154.39
177.0413055419922 0 38247.086
177.07757568359375 0 4702.834
177.0857391357422 0 13929.362
177.11395263671875 0 24203.38
178.0338134765625 0 27316.766
178.04208374023438 0 7926.988
178.06178283691406 0 5837.071
178.09805297851562 0 6531.404
178.1342315673828 0 13940.847
179.09326171875 0 37481.574
180.03274536132812 0 11709.643
180.07723999023438 0 25155.082
180.0963897705078 0 5375.828
180.11416625976562 0 4491.4214
181.09066772460938 0 52570.805
181.59234619140625 0 6095.3667
182.08169555664062 0 589739.25 y 12
182.1300048828125 0 5005.33
183.0769805908203 0 15795.478
183.08514404296875 0 57067.57
183.11325073242188 0 91829.97
184.08706665039062 0 4345.302
184.11656188964844 0 7616.964
185.03843688964844 0 59585.05
185.12896728515625 0 24157.295
186.02249145507812 0 346802
187.02566528320312 0 26735.666
187.10821533203125 0 349993.4
188.01812744140625 0 15857.368
188.11172485351562 0 27816.969
189.07760620117188 0 49166.902
190.06163024902344 0 137549.8
190.0811309814453 0 5724.637
190.13397216796875 0 9110.916
191.04916381835938 0 3670.7195
191.0652313232422 0 4654.679
191.09326171875 0 56222.27
192.0775604248047 0 9848.632
192.09703063964844 0 4520.658
193.07254028320312 0 28631.55
193.10885620117188 0 243067.27
194.08018493652344 0 22815.049
194.0928497314453 0 48601.97
194.11221313476562 0 17445.9
195.07867431640625 0 9153.886
195.08824157714844 0 31024.338
195.11367797851562 0 7153.672
196.07177734375 0 5390.384
197.09262084960938 0 5517.0063
198.08775329589844 0 4689.6357
199.0714874267578 0 7727.446
199.1086883544922 0 4537.7954
200.13992309570312 0 10418.932
201.07748413085938 0 5882.7437
201.08749389648438 0 45638.676
201.10577392578125 0 5551.3936
202.091064453125 0 3640.0952
202.1091766357422 0 20078.986
203.04896545410156 0 162774.88 a Water loss 1
203.09291076660156 0 20559.475
204.03292846679688 0 30013.793
204.05230712890625 0 11371.069
204.07725524902344 0 156742.12
204.09353637695312 0 5542.524
204.1055908203125 0 4158.0693
205.08111572265625 0 9677.306
205.10879516601562 0 13908.857
205.14549255371094 0 5365.055
206.09451293945312 0 5853.416
206.10423278808594 0 20931.652
206.1283416748047 0 4342.716
207.088134765625 0 272400.44
207.16062927246094 0 7756.065
208.07278442382812 0 9521.001
208.0915069580078 0 28359.277
208.11976623535156 0 6380.8306
209.09230041503906 0 4193.4365
210.12428283691406 0 29087.771
211.1083526611328 0 25271.049
211.1284637451172 0 5010.7344
213.03358459472656 0 16797.738
213.1236572265625 0 6016.374
214.0174102783203 0 19224.998
215.614501953125 0 4656.274
216.09852600097656 0 5198.558
216.1341552734375 0 6774.027
217.0725555419922 0 51513.58
218.0565643310547 0 57810.992
218.07579040527344 0 8722.407
219.06097412109375 0 5430.274
219.0882568359375 0 20884.82
220.09156799316406 0 3822.9424
221.0596466064453 0 3238852.2 a 1
221.10377502441406 0 647923.5
222.06275939941406 0 231618.98
222.08792114257812 0 16815.451
222.1072540283203 0 57088.258
222.12049865722656 0 4385.7925
222.122802734375 0 3165.3037
223.0553436279297 0 136611.73
223.0644989013672 0 15880.296
223.1077880859375 0 14345.88
223.15582275390625 0 39933.938
224.0588836669922 0 13142.934
224.1147003173828 0 218050.86
224.15809631347656 0 4015.8362
225.09901428222656 0 17018.045
225.11814880371094 0 18040.734
226.0648193359375 0 9866.922
226.0828399658203 0 50723.652 b Water loss 7
226.15530395507812 0 3827.9795
228.11367797851562 0 13738.664
229.1185302734375 0 4217.5166
230.0592041015625 0 4085.0627
230.08558654785156 0 11629.1045
230.10394287109375 0 8857.632
231.04400634765625 0 187392.03 b Water loss 1
231.08822631835938 0 17913.893
232.02806091308594 0 9465.306
232.046875 0 13335.099
233.14024353027344 0 26804.982
234.09918212890625 0 55435.816
234.12440490722656 0 28437.96
235.08316040039062 0 467641.56
236.04859924316406 0 5556.0825
236.0667724609375 0 17238.678
236.08648681640625 0 57054.31
236.11488342285156 0 7927.741
237.0999298095703 0 7451.415
237.13502502441406 0 6173.957
237.63368225097656 0 9169.224
238.11883544921875 0 18526.574
238.13099670410156 0 8268.92
239.10052490234375 0 6701.0767
239.114501953125 0 132238.1
240.08078002929688 0 6666.088
240.11781311035156 0 15964.535
243.04371643066406 0 8660.706
244.0803680419922 0 6609.815
244.0934295654297 0 77357.36
245.0972442626953 0 7644.7964
247.11181640625 0 39301.945
248.1155242919922 0 4892.9863
249.0546112060547 0 606153.4 b 1
249.09889221191406 0 124258.21
249.1354217529297 0 6621.7505
249.17083740234375 0 5516.9185
250.0579376220703 0 52088.23
250.08290100097656 0 15525.43
250.10299682617188 0 8127.074
250.16680908203125 0 14576.4795
251.05027770996094 0 18999.098
251.15090942382812 0 111237.01
252.10975646972656 0 743379.06
252.13461303710938 0 24570.916
252.15328979492188 0 10760.56
253.09368896484375 0 88562.805
253.11277770996094 0 86797.05
254.0602569580078 0 4493.4233
254.09674072265625 0 4895.8623
254.11419677734375 0 7188.915
256.1087341308594 0 7878.594
257.0955505371094 0 7436.3394
257.107421875 0 75818.336
257.13995361328125 0 6435.3506
258.111083984375 0 9061.379
259.10284423828125 0 4305.3306
259.15582275390625 0 9788.265
261.1349182128906 0 10581.943
262.0940246582031 0 19069.08
263.07806396484375 0 10952.403
263.0968017578125 0 4324.438
264.13848876953125 0 4120.1587
267.0918273925781 0 36902.37
267.1093444824219 0 304699.75
268.07501220703125 0 17576.629
268.09429931640625 0 4558.0103
268.1119384765625 0 32826.312
268.1297912597656 0 12130.574
268.1775817871094 0 10453.782
269.1361083984375 0 20091.941
269.1615905761719 0 7528.1714
270.0787048339844 0 4898.041
270.1202087402344 0 71291.586
270.1454162597656 0 26946.477
271.0988464355469 0 7195.601
271.1508483886719 0 4841.5527
272.07061767578125 0 8959.234
273.1351318359375 0 70602.54
273.1508483886719 0 3854.0002
274.1306457519531 0 14678.133
274.1670837402344 0 5994.277
275.1067199707031 0 34826.312
275.1507873535156 0 5462.359
276.11029052734375 0 9162.296
276.1826477050781 0 16039.579
276.61956787109375 0 6024.962
277.0938720703125 0 38612.96
277.11199951171875 0 38541.65 y Ammonia loss 9
277.61322021484375 0 10701.373
278.0971984863281 0 4266.2334
278.162841796875 0 5487.7344
278.6315002441406 0 6912.4854
279.145751953125 0 17758.852
280.1046142578125 0 41756.77
280.13201904296875 0 6607.337
280.6513671875 0 17241.215
281.08758544921875 0 5363.141
281.15252685546875 0 5019.8804
284.1060485839844 0 21620.605
284.13726806640625 0 6111.3403
285.1022033691406 0 232420.47
285.6252136230469 0 291206.72 y 9
286.1054382324219 0 25919.451
286.1268310546875 0 88254.99
286.6283874511719 0 35428.26
287.0987854003906 0 11319.609
287.11517333984375 0 4852.7104
287.1509704589844 0 8544.81
287.1712646484375 0 4604.283
290.14862060546875 0 11444.953
291.6005859375 0 5827.8096 b 4
292.1406555175781 0 11036.865
292.1776123046875 0 13400.307
293.1252746582031 0 16574.201 b 9
293.1416931152344 0 6516.9355
294.1934509277344 0 11534.21
294.64898681640625 0 22452.846
295.1044921875 0 38421.71
296.1080017089844 0 5482.995
296.1585693359375 0 4423.475
297.1567687988281 0 13888.915
298.8163146972656 0 10500.602
300.0655212402344 0 12356.859
300.6260986328125 0 9759.934
301.1300964355469 0 133594.05
302.11859130859375 0 17521.207
302.1333923339844 0 12222.173
302.1618347167969 0 32906.043
302.615478515625 0 6331.6523
303.10595703125 0 6402.7725
304.1048583984375 0 14893.568
304.14239501953125 0 16757.436
304.17742919921875 0 43771.33
305.0888977050781 0 13403.981
305.12567138671875 0 22552.588
305.17987060546875 0 6240.2153
308.1498107910156 0 5294.7793
309.1355285644531 0 22299.26
310.14007568359375 0 5577.277
311.48052978515625 0 5115.4946
312.1021728515625 0 24811.377
312.1204833984375 0 9032.743
312.6296081542969 0 10325.787
313.0994873046875 0 9289.422
314.1718444824219 0 7240.489
315.1279602050781 0 4843.8545
316.13385009765625 0 19475.643
316.6138916015625 0 4770.963
317.6191101074219 0 5066.257
318.07623291015625 0 56739.57 b Water loss 2
319.140869140625 0 594730 y 11
319.6620178222656 0 4511.5874
320.1376647949219 0 201035.9
320.1727600097656 0 58461.066
320.48382568359375 0 5433.5967
321.1400146484375 0 33882.543
321.17486572265625 0 8062.55
322.11529541015625 0 18596.582
322.14898681640625 0 47869.92
322.18829345703125 0 96479.68
322.6491394042969 0 13617.3955
322.670654296875 0 4066.8079
323.0820617675781 0 6040.551
323.1915283203125 0 16532.533
324.1320495605469 0 7940.935
324.64886474609375 0 5165.074
325.1512756347656 0 13665.365
326.1366271972656 0 46582.88
327.1404113769531 0 11745.921
328.15216064453125 0 11243.777
328.6673583984375 0 30248.67
329.12518310546875 0 63225.26
330.11285400390625 0 101414.98
330.1568603515625 0 15087.969
331.09844970703125 0 7438.5083
331.1181945800781 0 8421.052
331.4080505371094 0 4663.0415
332.11126708984375 0 5049.8413
332.1362609863281 0 65780.72
332.1728820800781 0 6851.236
332.827880859375 0 4353.869
333.1199035644531 0 20682.684
333.13836669921875 0 7103.061
333.6563720703125 0 18804.719 y Ammonia loss 8
334.12237548828125 0 7554.079
334.14453125 0 56127.145
334.65509033203125 0 11207.068 y Water loss 2
334.9045715332031 0 4823.8228
335.11590576171875 0 5520.7266
335.1510314941406 0 12315.524 y Ammonia loss 5
335.48687744140625 0 8671.084
335.90960693359375 0 24953.121
336.08685302734375 0 25728.32 b 2
336.1449890136719 0 12557.101
336.1638488769531 0 14508.304
336.64593505859375 0 9017.603
337.1302490234375 0 37838.164
337.1998291015625 0 5000.9175
337.64935302734375 0 5895.9395
337.6729431152344 0 12228.973
338.14678955078125 0 160780.27
339.1499938964844 0 29243.95 y 2
339.2152099609375 0 15807.113
339.4070129394531 0 4566.7944
340.15411376953125 0 9307.248
340.48956298828125 0 8571.765
340.82720947265625 0 84969.75 y 5
341.16162109375 0 45200.37
341.4956359863281 0 18428.156
342.1673889160156 0 85731.17 y 8
342.6680603027344 0 30607.441
343.1643981933594 0 19970.475
344.127685546875 0 8646.784
344.1477966308594 0 62035.594
344.4931335449219 0 4550.623
344.9100646972656 0 9610.278
345.152587890625 0 6414.5786
345.4100646972656 0 5845.148
346.4127197265625 0 6612.642
346.6428527832031 0 5662.158
347.1834411621094 0 12356.285
347.6524658203125 0 10671.259
347.90338134765625 0 5857.888
348.1278076171875 0 50522.87
348.1670227050781 0 13578.311
348.65484619140625 0 6192.3354
348.90325927734375 0 7475.7466
349.13134765625 0 11646.42
349.6170349121094 0 7603.6313
350.14666748046875 0 241164.19
350.6612243652344 0 8850.028
351.14990234375 0 38606.836
351.67041015625 0 38169.414
351.90875244140625 0 20260.469
352.1552429199219 0 10642.235
352.1722106933594 0 14904.161
352.40972900390625 0 10517.507
353.1256408691406 0 18633.846
353.1710205078125 0 5238.416
353.4125671386719 0 6990.411
354.1314697265625 0 44255.363
354.17071533203125 0 6672.3545
354.4041442871094 0 5643.5474
354.8263854980469 0 7877.523
354.8480529785156 0 9038.224
355.13916015625 0 15857.998
355.1759948730469 0 8013.8916
356.82867431640625 0 4383.5195
357.1199951171875 0 6099.7583
357.6676330566406 0 74607.89
357.9184875488281 0 44990.117
358.13018798828125 0 5521.6445
358.1688232421875 0 25744
358.6558532714844 0 13789.569
358.90625 0 25306.545
359.160888671875 0 20171.312
359.40606689453125 0 10619.706
359.4976806640625 0 16626.941
359.83331298828125 0 19536.041
360.12890625 0 7611.208 b 5
360.1669616699219 0 8598.182
360.6772766113281 0 5590.4517
360.8504333496094 0 5673.116
361.1740417480469 0 45531.414
361.199462890625 0 10372.876
361.9092102050781 0 7588.9624
362.1586608886719 0 13813.327
362.1805419921875 0 6697.597
362.8309326171875 0 36339.023
363.16534423828125 0 24842.621
363.4088134765625 0 5763.8633
363.4994812011719 0 12673.146
363.6567687988281 0 11667.25
363.9013671875 0 7173.8823
364.1252136230469 0 9034.686
364.4029846191406 0 6045.195
364.9082336425781 0 7338.1157
365.15869140625 0 11445.059
365.1938781738281 0 73702.24
366.141357421875 0 70530.22
366.19720458984375 0 13928.38
366.4112243652344 0 5057.6626
367.1422424316406 0 14372.368
368.1415100097656 0 6766.2188
368.16680908203125 0 12006.986
368.5061340332031 0 163952.98
368.6592712402344 0 7976
368.8403625488281 0 96527.29
368.9087219238281 0 22008.979
369.16143798828125 0 15410.41
369.173828125 0 27330.297 y Ammonia loss 7
369.4107666015625 0 22925.803
369.6583557128906 0 12062.708
370.15167236328125 0 19185.357
370.17864990234375 0 7983.8726
370.4030456542969 0 6399.8945
370.65570068359375 0 8322.193
370.904541015625 0 9910.354
371.1580810546875 0 232212.75
371.90673828125 0 11042.393
372.1419372558594 0 130172.34
372.162109375 0 22342.879
372.40997314453125 0 5843.847
372.63031005859375 0 4813.3677
373.1453552246094 0 26152.701
373.4089050292969 0 12910.292
373.6663513183594 0 104143.29
373.91748046875 0 60674.883
374.1457214355469 0 7125.16
374.1676940917969 0 50678.656
374.4174499511719 0 12695.451
374.6561584472656 0 42372.344
374.9063720703125 0 56353.004 y Water loss 1
375.1563415527344 0 35761.168 y Ammonia loss 1
375.1789855957031 0 11056.951
375.40789794921875 0 15971.626
375.653076171875 0 6018.797
376.1250915527344 0 9857.587
376.1625671386719 0 66144.99
376.41229248046875 0 37070.65
376.5115966796875 0 8424.427
376.6622009277344 0 24024.113
376.9127197265625 0 11786.202
377.1660461425781 0 6453.807
377.6859130859375 0 84863.875 y 7
377.85418701171875 0 8752.959
378.1416931152344 0 7106.9463
378.1872253417969 0 35482.492
378.41229248046875 0 14558.902
378.689208984375 0 28160.42
379.2099914550781 0 30801.361
379.4094543457031 0 6763.4653 y 1
379.91156005859375 0 10054.842
380.4076843261719 0 6455.975
380.6649475097656 0 18660.525
380.9163818359375 0 27815.322
381.1206359863281 0 95276.52
381.1639404296875 0 13574.198
382.12353515625 0 24453.475
382.18267822265625 0 41756.906
382.5157165527344 0 16742.135
382.85101318359375 0 8149.8003
383.1598815917969 0 6129.248
383.4110107421875 0 5953.2314
384.13677978515625 0 7072.3457
384.1687927246094 0 37827.105
384.49713134765625 0 9605.862 b Water loss 9
384.6545715332031 0 8155.0024
384.90728759765625 0 20189.107
385.16796875 0 95781.984
385.4185791015625 0 57774.61
385.66943359375 0 26086.865
385.8440856933594 0 6359.3633
385.91864013671875 0 13541.717
386.1490783691406 0 20302.576
387.18218994140625 0 9933.86
387.2144775390625 0 12216.338
387.6595153808594 0 8292.726
387.9078674316406 0 11214.338
388.685791015625 0 8616.022
389.1444396972656 0 6935.3384
389.1689758300781 0 63873.18
389.1943054199219 0 16523.734
389.4089050292969 0 14459.11
389.6587829589844 0 23726.28
389.83917236328125 0 145506.36 y 4
390.1517639160156 0 48953.742
390.1739807128906 0 81638.55
390.50628662109375 0 32103.527
390.8406982421875 0 8817.468
391.1554260253906 0 10618.946
391.20928955078125 0 7926.3584
391.5146484375 0 24245.598
391.8483581542969 0 19614.062
392.1623840332031 0 27946.252
392.4124755859375 0 43732.156
392.66156005859375 0 24960.09
393.1540832519531 0 12871.944
393.1888732910156 0 49564.516
393.692626953125 0 6288.6387
394.157470703125 0 6858.853
394.19256591796875 0 8325.534
395.8557434082031 0 10620.943
396.13446044921875 0 19785.838
396.6651611328125 0 35115.69 Precursor Water loss
396.91558837890625 0 30607.463 Precursor Ammonia loss
397.16473388671875 0 21637.793
397.19403076171875 0 8799.855
397.4171447753906 0 7470.2656
397.5198974609375 0 6936.86
397.6895751953125 0 41926.89
398.14678955078125 0 48528.07
398.19219970703125 0 16706.906
399.1308898925781 0 97146.336
400.1344299316406 0 33338.1
400.1849670410156 0 7988.992
401.1378173828125 0 8293.873
401.1681823730469 0 78903.35 Precursor
401.1956481933594 0 8057.598
401.4190368652344 0 79335.86
401.52923583984375 0 98628.23
401.6690673828125 0 50084.887
401.8634948730469 0 60052.27
401.91900634765625 0 16087.577
402.14154052734375 0 15943.375
402.1690673828125 0 5918.374
402.1971130371094 0 22429.594
402.65374755859375 0 10635.693
402.6824645996094 0 6980.8447
402.84674072265625 0 6102.442
403.1737976074219 0 15291.302
403.70361328125 0 8085.0024
403.9075622558594 0 10957.359
404.1574401855469 0 33059.29
404.24224853515625 0 7652.656
404.69171142578125 0 48942.207
404.85723876953125 0 6469.473
405.1628723144531 0 12704.579
405.1929931640625 0 20020.633
405.22564697265625 0 61539.91
405.6969299316406 0 5069.185
405.73443603515625 0 8956.952
406.2026672363281 0 137120.88
406.22979736328125 0 10502.701
406.7030944824219 0 56774.562
407.1793212890625 0 36710.867
407.20477294921875 0 45178.25
407.6474609375 0 8715.485
408.18133544921875 0 9858.065
408.2362060546875 0 13760.04
408.5187683105469 0 10679.309
408.8489685058594 0 18125.998
409.17962646484375 0 7572.5024
409.5166931152344 0 5226.324
411.1922912597656 0 24836.588
411.6569519042969 0 5651.103
411.6901550292969 0 19519.258
412.16650390625 0 13580.451
413.1901550292969 0 12681.327
414.1451721191406 0 149815.23
414.5215148925781 0 7552.8667
415.1468200683594 0 26506.91
415.17413330078125 0 31223.23
415.2105712890625 0 22172.26
416.1573486328125 0 254194.2 y Ammonia loss 10
416.6511535644531 0 12201.41
416.6788330078125 0 7066.346
416.8577880859375 0 6510.761 y Water loss 3
417.1473388671875 0 28964.322 b Water loss 3
417.1603698730469 0 45105.953
417.1903076171875 0 21223.777 y Ammonia loss 3
417.52239990234375 0 9475.409
417.85235595703125 0 11029.435
418.1453857421875 0 7803.9614
418.8617248535156 0 9530.259
419.1972351074219 0 14375.142
419.5300598144531 0 9500.111
420.20001220703125 0 98937.625
420.6974792480469 0 52404.562
421.1844177246094 0 132020.45
421.53143310546875 0 12231.941
421.8632507324219 0 7567.9194
422.1553649902344 0 7521.7446
422.1885986328125 0 29860.375
422.862060546875 0 184160.64 y 3
423.1964111328125 0 154133.44
423.2359619140625 0 76620.164
423.5298156738281 0 68335.266
423.8623046875 0 24757.025
424.1996765136719 0 9514.562
424.23773193359375 0 17115.824
424.5364685058594 0 48760.15
424.65032958984375 0 8081.7197 b 6
424.6899108886719 0 16931.215
424.86798095703125 0 55336.168
425.19354248046875 0 64064.254
425.53289794921875 0 14823.336
425.6903076171875 0 24871.81
426.145751953125 0 5739.0425
426.1864929199219 0 14944.325
428.1969909667969 0 23197.715
428.5307922363281 0 13409.299
429.2056884765625 0 69382.09
429.708251953125 0 34400.062
429.8516540527344 0 24195.2
430.1827697753906 0 23157.703
430.2102966308594 0 6292.34
430.5400390625 0 396426.78
430.69940185546875 0 10754.745
430.87408447265625 0 286972
431.1697082519531 0 13180.113
431.20794677734375 0 117345.47
431.54205322265625 0 30407.348
431.92169189453125 0 8391
432.8693542480469 0 6081.0127
433.1844177246094 0 370543.47 y 10
433.70269775390625 0 45386.19
433.8682861328125 0 17525.504
434.1888427734375 0 96418.086
434.7023620605469 0 5906.0215
435.1550598144531 0 22114.213 b 3
435.1893615722656 0 16191.248
435.85504150390625 0 32038.838
436.1595153808594 0 6192.48
436.19110107421875 0 16569.154
436.2314147949219 0 86169.57
436.87005615234375 0 5783.886
437.2347412109375 0 11833.646
437.53558349609375 0 7061.4424
437.8666076660156 0 5753.866
438.7122802734375 0 16058.57
439.1809387207031 0 5595.7246
439.2089538574219 0 49865.77
439.5169677734375 0 6928.8887
439.7110900878906 0 10827.451
440.19775390625 0 9008.606
440.5281982421875 0 11841.737
440.8612060546875 0 12898.034
440.9174499511719 0 10996.983
441.194580078125 0 8918.476
441.86859130859375 0 20552.957
442.1398010253906 0 11718.774
442.20721435546875 0 114808.62 y 6
442.53887939453125 0 10350.757
442.7099609375 0 51518.35
443.16845703125 0 15589.459
443.2084655761719 0 67700.82
443.5327453613281 0 6723.7344
443.8691101074219 0 5765.2593
444.15191650390625 0 14954.284
444.2063903808594 0 7888.4487
444.6963806152344 0 8121.274
445.1859436035156 0 17031.752
445.521240234375 0 14145.973
445.86895751953125 0 84337.06 y Water loss 2
446.2008361816406 0 109609.73 y Ammonia loss 2
446.5335388183594 0 63692.387
446.8655090332031 0 31506.256
447.1949462890625 0 14705.349
447.2201843261719 0 14800.528
447.5428466796875 0 47919.15
447.7208251953125 0 100751.8
447.8755798339844 0 51217.555
448.2198181152344 0 40720.72
448.5433654785156 0 38634.457
448.7196960449219 0 13802.861
448.8765563964844 0 11944.433
449.180419921875 0 11684.875
449.21514892578125 0 47416.668
449.8594970703125 0 6774.977
450.1435852050781 0 7838.4487
450.1875915527344 0 6804.5938
450.2469482421875 0 12438.878
451.1948547363281 0 7210.4624
451.2031555175781 0 4190.0146
451.2311096191406 0 169205.97
451.872802734375 0 969761.5 y 2
452.2068786621094 0 662137.7
452.5408935546875 0 312839.8
452.87457275390625 0 123403.93
453.20599365234375 0 47449.54
453.2393798828125 0 9054.672
453.7049865722656 0 20427.604
453.8709411621094 0 7847.1035
454.202880859375 0 16620.842
455.20159912109375 0 26819.445
455.5355224609375 0 13955.217
455.8695983886719 0 12346.894
458.8625183105469 0 10217.064
459.20025634765625 0 23075.02
461.21533203125 0 52878.344
461.2508544921875 0 19997.723
461.7179260253906 0 92309.59
461.7509460449219 0 7900.847
462.2154541015625 0 65332.68
462.71697998046875 0 22860.625
462.8726806640625 0 8496.659
463.20245361328125 0 37028.707
463.5373229980469 0 17858.08
463.86993408203125 0 22121.803
464.20196533203125 0 13387.762
464.8652648925781 0 15447.325
465.19781494140625 0 13397.453
466.5314025878906 0 8874.685
467.1695861816406 0 8976.32
467.2085876464844 0 6265.112
468.2581481933594 0 15130.328
468.8763427734375 0 96011.65
469.2087097167969 0 88174.3
469.54254150390625 0 59252.117
469.875732421875 0 51170.754
470.21075439453125 0 41101.21
470.54498291015625 0 16586.045
470.7223815917969 0 63445.227
470.87969970703125 0 6069.6025
471.2051696777344 0 52256.125
471.7222900390625 0 8722.775
472.20501708984375 0 30099.691
472.2385559082031 0 8387.341
472.53497314453125 0 37870.727
472.7243957519531 0 5507.1196
472.8675842285156 0 27949.246
473.198486328125 0 22257.803
473.5303649902344 0 11344.102
474.19561767578125 0 14008.372 b 11
474.2575988769531 0 35270.008
474.7224426269531 0 7887.0415
475.2287292480469 0 5779.631
475.2627258300781 0 8772.39
477.148193359375 0 11120.415
478.20806884765625 0 181650.36
478.5400695800781 0 215459.19
478.7257995605469 0 6109.486
478.8728942871094 0 128162.63
479.20526123046875 0 29118.785
479.223388671875 0 21844.238
479.5424499511719 0 16198.552
479.7203369140625 0 124682.805
480.22100830078125 0 66650.63
480.5330810546875 0 9099.048
480.72125244140625 0 24482.865
484.2115173339844 0 153511.39
484.5450439453125 0 119957.26
484.713134765625 0 14132.518
484.8793029785156 0 56008.773
485.18218994140625 0 116522.03
485.21783447265625 0 36736.742
485.7073059082031 0 12120.51
486.18499755859375 0 24177.082
486.2388000488281 0 21875.27
486.744140625 0 6855.3433
487.7344665527344 0 7694.1406
488.2330627441406 0 13416.902
488.2704162597656 0 7313.4536
488.7429504394531 0 6083.492
489.2041015625 0 7099.1714
489.245849609375 0 17089.104
490.2056884765625 0 8196.733
490.24346923828125 0 22623.334
492.7259216308594 0 21960.709
493.22216796875 0 20921.047
493.5371398925781 0 32357.441
493.7181701660156 0 97451.76
493.8681945800781 0 50601.96
494.2149658203125 0 50582.203
494.53338623046875 0 28537.973
494.71722412109375 0 18889.557
494.86444091796875 0 6876.015
495.1598815917969 0 15424.151
495.2065124511719 0 12947.929
495.73028564453125 0 6873.6997
496.2007141113281 0 7130.2803
496.78558349609375 0 8847.084
497.25518798828125 0 21467.742
497.7576904296875 0 18697.107
499.5408935546875 0 194322.62 y Water loss 1
499.87261962890625 0 287915.12 y Ammonia loss 1
500.2059326171875 0 181994.64
500.53863525390625 0 95594.25
500.7218933105469 0 8367.892
500.8716125488281 0 40568.39
501.2065734863281 0 14079.188
501.73150634765625 0 36145.82 y Water loss 5
502.2245788574219 0 92676.08 y Ammonia loss 5
502.7247619628906 0 141258.22
503.227294921875 0 59035.35
503.7286071777344 0 8349.959
504.239990234375 0 10660.101
505.18798828125 0 6618.702
505.24517822265625 0 7381.32
505.5443115234375 0 294698.7 y 1
505.8784484863281 0 256606.1
506.21209716796875 0 130700.305
506.274658203125 0 20706.566
506.54541015625 0 47838.5
506.87811279296875 0 9601.303
507.212646484375 0 28697.104
508.2131652832031 0 10967.034
508.2525939941406 0 25239.834
509.2307434082031 0 6127.122
509.7280578613281 0 13269.846
510.7369079589844 0 641268.94 y 5
511.2386169433594 0 382497.34
511.74072265625 0 113869.43
512.2413330078125 0 36062.44
513.1758422851562 0 6039.8584
513.2147827148438 0 41809.582
514.2172241210938 0 15086.24
515.7294921875 0 23557.22
516.2279052734375 0 12202.023
516.7328491210938 0 7558.6235
517.2423706054688 0 26449.824
518.2386474609375 0 79977.78
518.7399291992188 0 30380.197
519.2386474609375 0 13987.947
520.255615234375 0 82638.17
520.7413330078125 0 6826.0312
521.2568969726562 0 21830.879
521.714599609375 0 6322.6465
522.2144165039062 0 26264.363
523.1990966796875 0 39039.973
524.19775390625 0 9241.845
524.2351684570312 0 9937.022
524.2740478515625 0 7038.7383
525.7291259765625 0 6509.9624
526.2256469726562 0 11819.674
527.7297973632812 0 9018.045
528.2683715820312 0 26890.98
529.2542114257812 0 49515.723
529.2996826171875 0 8253.276
529.7549438476562 0 26238.664
530.2355346679688 0 23614.69
530.7349243164062 0 6797.5996
531.76513671875 0 14410.655
532.2604370117188 0 19459.846
532.71875 0 9525.322
532.7650146484375 0 8415.807
533.2172241210938 0 11091.674
533.266357421875 0 5935.567
534.225341796875 0 7816.317
534.268310546875 0 87416.68
534.7374267578125 0 51259.605
535.2058715820312 0 100246.44
535.273193359375 0 10795.153
535.7361450195312 0 18211.264
536.20703125 0 27356.37
536.2470703125 0 71785.35
537.208740234375 0 7539.332
537.2491455078125 0 22455.277
537.2857055664062 0 8415.82
538.2437744140625 0 26297.402
538.7432861328125 0 30041.975
539.2457885742188 0 45865.65
539.7478637695312 0 21835.406
540.2259521484375 0 13591.899
540.7708740234375 0 67462.28
541.206298828125 0 6119.95
541.2227172851562 0 5768.994
541.2714233398438 0 40908.453
541.7239990234375 0 16059.073
541.7747802734375 0 13101.123
542.7342529296875 0 17219.826
543.25048828125 0 67699.87
543.74462890625 0 168658.5
544.2446899414062 0 99089.95
544.74560546875 0 26766.746
545.24609375 0 10582.209
545.29541015625 0 54423.79
545.76025390625 0 9281.846
546.2991333007812 0 12722.655
547.2635498046875 0 49071.188
548.2509155273438 0 20490.66
550.2079467773438 0 19849.705
551.1965942382812 0 18620.08
551.7434692382812 0 7120.1753
552.25537109375 0 672831.94
552.7568969726562 0 436694.62
553.2161865234375 0 175368.31 y Ammonia loss 9
553.2589721679688 0 115648.664
553.7588500976562 0 32605.955
554.2192993164062 0 51997.766
554.2630615234375 0 6846.1875
554.7686157226562 0 46989.684
555.220703125 0 8252.913
555.268798828125 0 38871.72
555.7659912109375 0 10567.55
556.2628784179688 0 18579.877
557.2498779296875 0 13784.842
557.7357177734375 0 10513.707
558.236328125 0 11479.445
558.7327880859375 0 5366.46
560.2540893554688 0 5621.412
560.294677734375 0 36865.7
561.2510986328125 0 14281.269
561.2953491210938 0 11897.995
561.7494506835938 0 11781.283
562.2567138671875 0 11987.567
562.30419921875 0 5724.2593
563.1976928710938 0 7270.799
563.7710571289062 0 43328.312
564.2660522460938 0 40867.67
564.7606811523438 0 19537.041
565.273193359375 0 97844.86
566.2754516601562 0 20585.557
566.7405395507812 0 29856.828
567.2389526367188 0 22446.363
567.7398071289062 0 11747.132
568.2220458984375 0 18870.832
570.24267578125 0 461952.28 y 9
571.2454223632812 0 131858.48
571.7452392578125 0 6168.4536
572.2498168945312 0 61865.26
572.7703857421875 0 156913.86
573.2754516601562 0 107322.66
573.7716064453125 0 33982.38
574.2756958007812 0 33629.574
575.2508544921875 0 39577.438 y Water loss 4
575.7433471679688 0 116391.39 y Ammonia loss 4
576.2440795898438 0 79272.234 b Water loss 9
576.7435913085938 0 40056.383
577.2423706054688 0 6545.117
577.7601928710938 0 18560.066
578.256591796875 0 15442.042
578.7579345703125 0 9256.14
579.2882080078125 0 11629.791
580.22314453125 0 8149.938
581.2691040039062 0 5396.897
582.2342529296875 0 19411.389
583.2403564453125 0 8196.046
584.2545776367188 0 771319.2 y 4
584.7560424804688 0 470305.75
585.2561645507812 0 219331.97
585.7567749023438 0 73101.836
586.255126953125 0 25289.088
586.7673950195312 0 99372.18
587.1986083984375 0 13077.606
587.266845703125 0 79932.945
587.76708984375 0 35055.703
588.287841796875 0 55211.49
589.2780151367188 0 39195.29
589.7411499023438 0 12603.693
590.23681640625 0 9077.89
591.2489624023438 0 5687.254
591.30126953125 0 23045.414
592.3054809570312 0 8457.172
592.7841796875 0 12210.107
593.2744750976562 0 27589.547
593.7777099609375 0 13243.203
594.2805786132812 0 11175.13
595.7726440429688 0 55705.86
596.2747192382812 0 33451.023
596.7732543945312 0 18304.518
596.846435546875 0 7528.278
598.267822265625 0 87966.13
598.7471923828125 0 23568.822
599.266845703125 0 24618.824
599.7489624023438 0 7981.7036
600.24609375 0 106326.38
601.2489013671875 0 32248.914
601.7894897460938 0 149155.08
602.2433471679688 0 6528.5527
602.2916259765625 0 105977.95
602.79248046875 0 28766.82
603.2915649414062 0 15078.526
603.7630004882812 0 5794.2656
604.2235107421875 0 16936.053
604.757080078125 0 7009.49
605.2220458984375 0 6457.427
605.31640625 0 12373.293
607.2843017578125 0 95961.664
608.2879638671875 0 32480.045
610.226318359375 0 8822.136
612.2745361328125 0 18158.875
612.7693481445312 0 24081.338
613.2716064453125 0 16657.236
614.2058715820312 0 13953.012
614.2677001953125 0 5366.1626
615.1951904296875 0 8042.6133
615.7886962890625 0 7224.707
616.2789916992188 0 7985.078
617.2697143554688 0 12883.673
617.7617797851562 0 7293.3774
618.2617797851562 0 7661.604
621.2801513671875 0 16764.021
621.7793579101562 0 16825.363
622.2831420898438 0 5982.718
624.7822875976562 0 11690.139 y Water loss 3
625.2767944335938 0 38972.27 y Ammonia loss 3
625.7786254882812 0 19637.502
626.2708740234375 0 21625.697
626.7671508789062 0 19294.875
627.2615966796875 0 10404.452
627.7832641601562 0 7465.8564
632.2171630859375 0 20734.256
633.2210083007812 0 6093.4478
633.3374633789062 0 19708.906
633.7887573242188 0 186821.62 y 3
634.2295532226562 0 7762.7593
634.2899780273438 0 115982.914
634.33935546875 0 9343.457
634.7904663085938 0 64101.113
635.2830200195312 0 32645.191
635.7764282226562 0 20944.623
636.2743530273438 0 7374.7944
636.3034057617188 0 6923.088
636.7949829101562 0 22040.19
637.2958374023438 0 15330.967
638.7808227539062 0 14999.705
639.3009033203125 0 15373.441
642.2647705078125 0 14802.603 b 10
643.2803344726562 0 7839.8955
643.332275390625 0 19952.473
644.27880859375 0 22948.254
644.3380737304688 0 7921.2915
644.7706298828125 0 40726.066
645.3052368164062 0 93492.13
645.8063354492188 0 65272.797
646.3082885742188 0 21051.314
648.292236328125 0 13634.999
650.296142578125 0 12737.226
651.2572021484375 0 14443.669
652.2418823242188 0 10682.439
653.275390625 0 66875.44
653.779541015625 0 22782.896
654.2871704101562 0 9021.083
656.3260498046875 0 20991.053
657.3143310546875 0 45389.586
658.2655029296875 0 7905.87
658.3174438476562 0 6108.546
658.7675170898438 0 20428.617
659.2730102539062 0 12673.314
659.7830200195312 0 7873.569
660.8737182617188 0 24229.133
661.3736572265625 0 17793.53
665.3189086914062 0 11471.459
666.3028564453125 0 51725.58 y Ammonia loss 8
667.2804565429688 0 37184.676
667.7750854492188 0 34672.285
668.2938842773438 0 29986.32 y Water loss 2
668.7926635742188 0 32150.438 y Ammonia loss 2
669.2890625 0 13164.972
669.7979736328125 0 8715.136
671.2822875976562 0 154741.56
671.3262329101562 0 21781.436
672.2855834960938 0 53034.45
672.3406372070312 0 8165.923
673.2852783203125 0 14280.091
674.338623046875 0 73528.91
675.34130859375 0 24899.523
676.2816772460938 0 24289.26
676.782470703125 0 16771.371
677.3042602539062 0 143802.84 y 2
677.8058471679688 0 83515.48
678.3059692382812 0 29025.674
678.8084716796875 0 14896.936
679.2517700195312 0 34153.484
680.23828125 0 35515.51
681.241943359375 0 8124.1167
682.2926635742188 0 11694.28
682.3657836914062 0 7829.9683
682.7989501953125 0 15992.15
683.3268432617188 0 163411.5 y 8
684.3275146484375 0 92848.16
685.3192749023438 0 37874.027
686.3164672851562 0 7422.612
688.3527221679688 0 9995.844
691.3350219726562 0 7530.6226
692.3735961914062 0 11582.499
694.3001708984375 0 11246.484
697.261474609375 0 24124.69
697.3363647460938 0 21956.9
698.3357543945312 0 8162.3896
702.3335571289062 0 154691.61
703.3313598632812 0 77182.33
704.3349609375 0 17038.717
707.3310546875 0 19483.895
708.343017578125 0 10812.036
719.2517700195312 0 8504.892 b 5
719.3278198242188 0 17341.781
720.359619140625 0 60219.125
721.3607177734375 0 20174.705
722.2921752929688 0 12144.839
723.287353515625 0 8083.607
733.2676391601562 0 11165.501
735.3245849609375 0 35176.723
736.3368530273438 0 16031.844
737.3372192382812 0 85671.16 y Ammonia loss 7
738.341552734375 0 36012.46
739.33984375 0 9098.973
740.3572998046875 0 9177.907
743.2507934570312 0 27794.336
744.240234375 0 19922.441
745.2430419921875 0 8692.13
750.2882690429688 0 30140.156
751.2776489257812 0 40059.69
752.2786865234375 0 9728.061
752.3489379882812 0 13686.622
754.3638916015625 0 282626.9 y 7
755.3668212890625 0 104834.13
756.369873046875 0 38866.51
757.3727416992188 0 19221.25
758.3754272460938 0 7482.761
759.3629150390625 0 5553.1953
761.2598876953125 0 30524.508
766.3552856445312 0 16946.973
767.3575439453125 0 13382.757
768.2948608398438 0 13090.476
768.3578491210938 0 8840.424
770.3989868164062 0 8935.447
771.3873291015625 0 8128.923
784.2943725585938 0 4612.4688
784.36669921875 0 79221.17
785.3692626953125 0 108224.48
786.3703002929688 0 42524.36
787.3685302734375 0 9484.903
794.3652954101562 0 15168.65
808.3751220703125 0 12591.262
811.399658203125 0 29653.186
812.3951416015625 0 16037.459
814.2862548828125 0 38859.168
815.275390625 0 32483.7
816.27880859375 0 14924.298
821.378173828125 0 11149.135
822.3721313476562 0 12364.09
829.43408203125 0 8202.092
831.35791015625 0 8622.853
832.2958984375 0 41994.113
833.2999267578125 0 17193.965
834.2972412109375 0 11002.661
834.3890991210938 0 8513.551
835.3864135742188 0 8846.058
836.3731079101562 0 7993.589
837.4044189453125 0 8733.455
838.3850708007812 0 7816.6035
839.3964233398438 0 24417.783
840.3893432617188 0 20275.877
841.3870849609375 0 6617.78
848.2886962890625 0 7533.6953 b 6
848.3702392578125 0 48128.867
849.3688354492188 0 81777.14
850.370361328125 0 35973.246
851.3680419921875 0 8888.222
856.3347778320312 0 6346.245
857.4183959960938 0 27936.055
858.4168090820312 0 11965.323
863.37353515625 0 14099.098
864.365478515625 0 15000.05
865.39453125 0 77604.336 y Water loss 6
866.3851318359375 0 103228.97 y Ammonia loss 6
867.3869018554688 0 51908.88
868.3870849609375 0 17387.938
874.461669921875 0 5686.258
881.3833618164062 0 25778.676
882.3849487304688 0 7847.6016
883.40625 0 290870.75 y 6
884.4114379882812 0 131133.98
885.4140014648438 0 36986.934
886.425048828125 0 10024.007
893.3958129882812 0 5801.4375
894.4224853515625 0 13212.846
917.387451171875 0 6171.375
919.3284301757812 0 15972.597 b 7
921.333251953125 0 8347.684
921.4262084960938 0 23109.11
922.4268188476562 0 20055.379
923.4227905273438 0 12116.503
927.3706665039062 0 12163.157
928.3613891601562 0 15847.435
929.3655395507812 0 6931.323
940.4347534179688 0 8049.4043
943.482421875 0 7789.7856
945.3809814453125 0 30320.23
946.3836669921875 0 17250.738
948.4338989257812 0 14795.02
949.4429931640625 0 8383.379
953.4617919921875 0 7490.5527
954.454345703125 0 8038.9917
971.46875 0 35294.21
972.468505859375 0 16434.62
973.46533203125 0 6873.6973
986.424072265625 0 6264.055
987.4239501953125 0 6236.656
1002.4560546875 0 9597.228 y Water loss 5
1003.4404907226562 0 20508.87 y Ammonia loss 5
1004.4371948242188 0 16781.594
1005.4446411132812 0 10172.802
1007.4722900390625 0 9465.601
1017.455322265625 0 8681.277
1018.4425659179688 0 11047.465
1019.4595336914062 0 5931.343
1020.4644775390625 0 104488.15 y 5
1021.4671630859375 0 52789.55
1022.4725952148438 0 16828.354
1030.4530029296875 0 6340.25
1032.4150390625 0 8687.182 b 8
1035.46728515625 0 51269.336
1036.47021484375 0 26612.05
1037.469482421875 0 10998.388
1086.489501953125 0 6262.1665
1087.4796142578125 0 5507.173
1103.50634765625 0 9446.345
1104.508056640625 0 7066.008
1114.477294921875 0 11020.892
1115.470947265625 0 7954.178
1132.4854736328125 0 7885.2886
1133.48974609375 0 6974.308
1167.484619140625 0 5243.802 y 4
1178.472900390625 0 7726.8413
1179.4566650390625 0 5828.356
1196.4835205078125 0 16023.225
1197.4835205078125 0 10883.059
1564.74072265625 0 4426.885
1971.1353759765625 0 5724.671
3082.21142578125 0 4828.0435

Spectrum Details

|  |  |
| --- | --- |
| Matched peaks? Matched peaks The total absolute number of peaks matched. Additionally in brackets the total fraction of peaks matched and the total number of peaks is shown. | 77 (5.82% of 1322) |
| FDR? FDR The false discovery rate estimated for this peptide. It is calculated by matching all theoretical fragments with a non-integer shift with the raw peaks for this spectrum. This is done with 40 different shifts. The resulting percentage is the average number of annotated peaks over the number of annotated peaks with the correct spectrum. | 2.35% |
| Satellite FDR? Satellite FDR See the FDR for details on its calculation. This satellite ion specific FDR only contains the satellite ions (d/w) for I/L/J positions. | - |
| PSM Score? PSM Score The PSM Score as given by Hecklib to this annotated spectrum. It is shown with three significant figures. | 414 |

#### Reverse Lookup? Reverse Lookup All places where this read could be placed.

| Group | Segment | Template | Template Part | Read Part | Score | Unique |
| --- | --- | --- | --- | --- | --- | --- |
| Homo sapiens Heavy Chain | IGHC | IGHG1 | [306..319] | [0..13] | 104 | False |
| Homo sapiens Heavy Chain | IGHC | IGHG2 | [302..315] | [0..13] | 104 | False |
| Homo sapiens Heavy Chain | IGHC | IGHG4 | [303..316] | [0..13] | 104 | False |

| Recombined | Template Part | Read Part | Score | Unique |
| --- | --- | --- | --- | --- |
| REC-0-1 | [431..444] | [0..13] | 104 | True |

#### Meta Information from Multiple reads

##### Number of combined reads

3

##### Intensity

0.7881

##### TotalArea

4.732E+08

##### Changes to the peptide sequence

SCSVMHEAJHNHY

L→JNo support for either Leucine or Isoleucine based on side chain ions (Position: 9)

#### Positional Score

Copy Data

##### Positional Score (TSV)

###### Preview

```
Loading example...
```

*Click on the button to copy the data to your clipboard.*

100123456789101112

Label Value
"0" 0.317
"1" 0.33
"2" 0.33
"3" 0.323
"4" 0.33
"5" 0.33
"6" 0.33
"7" 0.327
"8" 0.333
"9" 0.33
"10" 0.33
"11" 0.33
"12" 0.333

#### Meta Information from PEAKS

##### Scan Identifier

F3:2789

##### Original sequence

S

C

+58.01

S

V

M

+15.99

H

E

A

L

H

N

H

Y

##### Posttranslational Modifications

Carboxymethyl; Oxidation (M)

##### Source File

D:\separate\_stitch\_analyses\xle-disambiguation\raw\20210323\_F1\_UM1\_Peng0013\_SA\_F59\_ingel\_3ug\_chymo.raw

##### Fraction

3

##### Scan Feature

F3:5548

##### De Novo Score

99

##### ConfidenceScore

99

### m/z

534.5554

##### Mass

1600.6399

##### Charge

3

##### Retention Time

15.16

##### Predicted Retention Time

-

##### Area

1.728E+08

##### Parts Per Million

2.8

##### Fragmentation mode

HCD

##### Originating file

01 D:\separate\_stitch\_analyses\xle-disambiguation\20210325\_F59\_3ug\_DENOVO\_12.csv

#### Meta Information from PEAKS

##### Scan Identifier

F3:2941

##### Original sequence

S

C

+58.01

S

V

M

+15.99

H

E

A

L

H

N

H

Y

##### Posttranslational Modifications

Carboxymethyl; Oxidation (M)

##### Source File

D:\separate\_stitch\_analyses\xle-disambiguation\raw\20210323\_F1\_UM1\_Peng0013\_SA\_F59\_ingel\_3ug\_chymo.raw

##### Fraction

3

##### Scan Feature

F3:5548

##### De Novo Score

99

##### ConfidenceScore

99

### m/z

534.5554

##### Mass

1600.6399

##### Charge

3

##### Retention Time

15.16

##### Predicted Retention Time

-

##### Area

1.728E+08

##### Parts Per Million

2.8

##### Fragmentation mode

HCD

##### Originating file

01 D:\separate\_stitch\_analyses\xle-disambiguation\20210325\_F59\_3ug\_DENOVO\_12.csv

#### Meta Information from PEAKS

##### Scan Identifier

F3:2840

##### Original sequence

S

C

+58.01

S

V

M

+15.99

H

E

A

L

H

N

H

Y

##### Posttranslational Modifications

Carboxymethyl; Oxidation (M)

##### Source File

D:\separate\_stitch\_analyses\xle-disambiguation\raw\20210323\_F1\_UM1\_Peng0013\_SA\_F59\_ingel\_3ug\_chymo.raw

##### Fraction

3

##### Scan Feature

F3:753

##### De Novo Score

98

##### ConfidenceScore

98

### m/z

401.1684

##### Mass

1600.6399

##### Charge

4

##### Retention Time

15.16

##### Predicted Retention Time

-

##### Area

1.276E+08

##### Parts Per Million

2.9

##### Fragmentation mode

HCD

##### Originating file

01 D:\separate\_stitch\_analyses\xle-disambiguation\20210325\_F59\_3ug\_DENOVO\_12.csv
