## Supplementary material for "A handle on mass coincidence errors in *de novo* sequencing of antibodies by bottom-up proteomics": Combined_033.html

Details Combined\_033 | Stitch OverviewUndefined

### Read Combined\_033

#### Sequence (length=12)

VLGQPKAAPSVT

#### Spectrum 4087? Spectrum 4087 The raw spectrum of this peptide as annotated by Hecklib. The fragments are coloured according to ion type (see legend). Any peaks with a star '\*' as text can be hovered over to see the full details, first the ion type second the mass shift type. By hovering over the amino acids in the peptide or ions in the legend the corresponding peaks are highlighted. By toggling the 'Unassigned' label you can turn the background (unassigned) peaks on or off in the plot. By updating the slider in the Ion legend you can update the spectrum to only show the top X% of the peaks with labels. The top X% means any peak that is within X% of the highest intensity. By dragging in the spectrum you can zoom in to a specific part of the spectrum and use 'Zoom Out' to get back to the original zoom level. The annotation of the spectrum is based on the given sequence in the peptides file and is done with different software so inconsistencies are likely. The peaks are annotated based on the given sequence, with 20 ppm tolerance.

Copy Data

##### Spectrum 4087 (TSV)

###### Preview

```
Loading example...
```

*Click on the button to copy the data to your clipboard.*

Mz MinMz MaxIntensity Max

WidthHeightPeptide font sizePeptide stroke widthSpectrum font sizeSpectrum stroke widthCompact peptide

Ion legend

wxyz

abcd

OtherUnassignedIon^Charge^_Position_Show for top:%

VLGQPKAAPSVT

09.63e+61.93e+72.89e+73.85e+7

Zoom Out

y^+1^_1_y^+1^_2_y^+1^_2_c^+1^_3_y^+1^_3_z^+2^_8_c^+2^_8_y^+1^_4_y^+2^_8_c^+1^_4_y^+1^_4_c^+2^_9_y^+2^_9_y^+2^_9_y^+2^_9_y^+2^_10_y^+2^_10_y^+1^_5_c^+2^_10_c^+2^_10_y^+2^_10_c^+1^_5_c^+1^_5_c^+2^_11_c^+2^_11_z^+2^_11_y^+2^_11_y^+1^_6_c^+1^_6_c^+1^_6_z^+1^_7_y^+1^_7_c^+1^_7_c^+1^_7_z^+1^_8_y^+1^_8_z^+1^_8_c^+1^_8_y^+1^_8_w^+1^_9_c^+1^_9_c^+1^_9_y^+1^_9_z^+1^_9_y^+1^_9_z^+1^_10_y^+1^_10_y^+1^_10_z^+1^_10_c^+1^_10_y^+1^_10_c^+1^_10_w^+1^_11_c^+1^_11_z^+1^_11_c^+1^_11_y^+1^_11_

0769153823063075

Fragment Matches Table

Show background peaks

| Position | Ion type | Intensity | mz Theoretical | mz Error (Th) | mz Error (ppm) | Charge | Series Number |
| --- | --- | --- | --- | --- | --- | --- | --- |
| 12 | y | 4.847E+06 | 120.1 | 0.0003068 | 2.555 | +1 | 1 |
| - | - | 2.161E+05 | 121.1 | - | - | 0 | - |
| - | - | 3.623E+04 | 157.1 | - | - | 0 | - |
| - | - | 6.07E+04 | 158.1 | - | - | 0 | - |
| - | - | 2.387E+04 | 159.1 | - | - | 0 | - |
| - | - | 1.928E+05 | 167.1 | - | - | 0 | - |
| - | - | 1.825E+04 | 167.6 | - | - | 0 | - |
| - | - | 8.45E+04 | 169.1 | - | - | 0 | - |
| - | - | 2.297E+04 | 169.1 | - | - | 0 | - |
| - | - | 6.988E+04 | 171.1 | - | - | 0 | - |
| - | - | 5.608E+04 | 173.5 | - | - | 0 | - |
| - | - | 1.56E+04 | 176.4 | - | - | 0 | - |
| - | - | 3.883E+04 | 184.1 | - | - | 0 | - |
| - | - | 2.339E+05 | 185.1 | - | - | 0 | - |
| - | - | 5.972E+06 | 185.2 | - | - | 0 | - |
| - | - | 6.705E+05 | 186.1 | - | - | 0 | - |
| - | - | 3.347E+04 | 186.1 | - | - | 0 | - |
| - | - | 7.058E+05 | 186.2 | - | - | 0 | - |
| - | - | 3.991E+04 | 187.1 | - | - | 0 | - |
| - | - | 1.027E+05 | 187.1 | - | - | 0 | - |
| - | - | 2.077E+04 | 187.2 | - | - | 0 | - |
| - | - | 6.195E+04 | 188.1 | - | - | 0 | - |
| - | - | 2.267E+04 | 196.8 | - | - | 0 | - |
| 11 | y | 3.075E+05 | 201.1 | 0.0003036 | 1.51 | +1 | 2 |
| - | - | 5.008E+04 | 202.1 | - | - | 0 | - |
| - | - | 2.883E+04 | 202.1 | - | - | 0 | - |
| - | - | 7.504E+06 | 213.2 | - | - | 0 | - |
| - | - | 8.726E+05 | 214.2 | - | - | 0 | - |
| - | - | 2.737E+04 | 215.2 | - | - | 0 | - |
| 11 | y | 2.734E+06 | 219.1 | 0.000298 | 1.36 | +1 | 2 |
| - | - | 2.591E+05 | 226.2 | - | - | 0 | - |
| - | - | 2.921E+04 | 227.2 | - | - | 0 | - |
| - | - | 2.413E+04 | 238.2 | - | - | 0 | - |
| - | - | 2.079E+04 | 245.4 | - | - | 0 | - |
| - | - | 1.783E+05 | 256.2 | - | - | 0 | - |
| - | - | 5.397E+05 | 266.2 | - | - | 0 | - |
| - | - | 7.062E+04 | 267.2 | - | - | 0 | - |
| - | - | 2.453E+04 | 267.2 | - | - | 0 | - |
| - | - | 8.701E+05 | 270.2 | - | - | 0 | - |
| - | - | 1.126E+05 | 271.2 | - | - | 0 | - |
| - | - | 6.286E+05 | 284.2 | - | - | 0 | - |
| - | - | 2.412E+05 | 285.2 | - | - | 0 | - |
| - | - | 3.969E+04 | 286.2 | - | - | 0 | - |
| 3 | c | 1.582E+06 | 287.2 | 0.0004237 | 1.475 | +1 | 3 |
| - | - | 1.976E+05 | 288.2 | - | - | 0 | - |
| - | - | 2.945E+04 | 297.2 | - | - | 0 | - |
| - | - | 9.096E+05 | 299.2 | - | - | 0 | - |
| - | - | 1.7E+05 | 300.2 | - | - | 0 | - |
| 10 | y | 1.166E+05 | 306.2 | 0.0003283 | 1.072 | +1 | 3 |
| - | - | 2.994E+05 | 327.2 | - | - | 0 | - |
| - | - | 2.212E+04 | 327.2 | - | - | 0 | - |
| - | - | 2.775E+04 | 328.2 | - | - | 0 | - |
| - | - | 4.29E+04 | 337.2 | - | - | 0 | - |
| - | - | 2.635E+04 | 341.2 | - | - | 0 | - |
| - | - | 6.913E+04 | 353.2 | - | - | 0 | - |
| - | - | 3.606E+04 | 355.2 | - | - | 0 | - |
| - | - | 5.061E+04 | 360.2 | - | - | 0 | - |
| - | - | 2.618E+04 | 365.2 | - | - | 0 | - |
| - | - | 7.347E+05 | 368.2 | - | - | 0 | - |
| 5 | z | 3.393E+04 | 369.2 | 0.001844 | 4.995 | +2 | 8 |
| - | - | 1.589E+05 | 369.2 | - | - | 0 | - |
| - | - | 8.456E+05 | 370.2 | - | - | 0 | - |
| - | - | 1.318E+05 | 371.2 | - | - | 0 | - |
| - | - | 2.518E+04 | 372.3 | - | - | 0 | - |
| - | - | 2.684E+04 | 372.7 | - | - | 0 | - |
| - | - | 4.579E+04 | 375.7 | - | - | 0 | - |
| - | - | 3.645E+04 | 380.2 | - | - | 0 | - |
| - | - | 6.986E+04 | 381.2 | - | - | 0 | - |
| - | - | 2.947E+04 | 382.2 | - | - | 0 | - |
| - | - | 2.989E+04 | 382.7 | - | - | 0 | - |
| 8 | c | 1.306E+05 | 383.2 | 0.0008707 | 2.272 | +2 | 8 |
| - | - | 6.008E+04 | 383.7 | - | - | 0 | - |
| 9 | y | 2.603E+05 | 385.2 | 0.000335 | 0.8696 | +1 | 4 |
| 5 | y | 9.336E+04 | 385.7 | 0.0002038 | 0.5283 | +2 | 8 |
| - | - | 2.56E+04 | 394.2 | - | - | 0 | - |
| - | - | 4.687E+04 | 395.7 | - | - | 0 | - |
| - | - | 2.509E+04 | 396.2 | - | - | 0 | - |
| - | - | 2.52E+04 | 396.7 | - | - | 0 | - |
| 4 | c | 4.576E+06 | 398.2 | 0.0005914 | 1.485 | +1 | 4 |
| - | - | 1.081E+06 | 399.2 | - | - | 0 | - |
| - | - | 9.481E+04 | 400.2 | - | - | 0 | - |
| - | - | 2.407E+04 | 401.2 | - | - | 0 | - |
| 9 | y | 3.979E+06 | 403.2 | 0.0005125 | 1.271 | +1 | 4 |
| - | - | 8.219E+05 | 404.2 | - | - | 0 | - |
| - | - | 1.159E+05 | 404.7 | - | - | 0 | - |
| - | - | 7.046E+04 | 405.2 | - | - | 0 | - |
| - | - | 2.21E+04 | 405.7 | - | - | 0 | - |
| - | - | 5.571E+04 | 408.2 | - | - | 0 | - |
| - | - | 2.189E+05 | 409.7 | - | - | 0 | - |
| - | - | 1.385E+05 | 410.2 | - | - | 0 | - |
| - | - | 3.145E+04 | 410.8 | - | - | 0 | - |
| - | - | 5.762E+04 | 411.2 | - | - | 0 | - |
| - | - | 2.823E+04 | 412.3 | - | - | 0 | - |
| - | - | 9.606E+04 | 416.7 | - | - | 0 | - |
| - | - | 2.844E+04 | 417.2 | - | - | 0 | - |
| - | - | 1.266E+05 | 417.8 | - | - | 0 | - |
| - | - | 5.695E+04 | 418.3 | - | - | 0 | - |
| - | - | 1.411E+05 | 418.7 | - | - | 0 | - |
| - | - | 3.291E+04 | 418.8 | - | - | 0 | - |
| - | - | 4.561E+04 | 419.2 | - | - | 0 | - |
| - | - | 2.774E+04 | 419.8 | - | - | 0 | - |
| - | - | 2.387E+04 | 423.2 | - | - | 0 | - |
| - | - | 2.722E+04 | 423.3 | - | - | 0 | - |
| - | - | 3.055E+04 | 423.8 | - | - | 0 | - |
| - | - | 4.781E+04 | 425.7 | - | - | 0 | - |
| - | - | 5.674E+04 | 426.2 | - | - | 0 | - |
| - | - | 3.891E+04 | 430.8 | - | - | 0 | - |
| 9 | c | 8.252E+05 | 431.8 | 0.0004879 | 1.13 | +2 | 9 |
| - | - | 4.743E+05 | 432.3 | - | - | 0 | - |
| - | - | 1.354E+05 | 432.8 | - | - | 0 | - |
| - | - | 4.428E+04 | 439.8 | - | - | 0 | - |
| - | - | 2.855E+04 | 440.3 | - | - | 0 | - |
| 4 | y | 1.747E+05 | 440.7 | 0.001405 | 3.188 | +2 | 9 |
| 4 | y | 9.004E+04 | 441.2 | 0.007322 | 16.59 | +2 | 9 |
| - | - | 7.45E+04 | 444.8 | - | - | 0 | - |
| - | - | 4.165E+04 | 445.3 | - | - | 0 | - |
| - | - | 2.873E+04 | 448.8 | - | - | 0 | - |
| 4 | y | 6.018E+04 | 449.8 | 0.00125 | 2.778 | +2 | 9 |
| - | - | 2.278E+04 | 450.3 | - | - | 0 | - |
| - | - | 3.474E+04 | 450.8 | - | - | 0 | - |
| - | - | 3.901E+04 | 451.8 | - | - | 0 | - |
| - | - | 9.994E+04 | 452.3 | - | - | 0 | - |
| - | - | 1.845E+05 | 452.8 | - | - | 0 | - |
| - | - | 1.024E+05 | 453.3 | - | - | 0 | - |
| - | - | 9.556E+04 | 453.8 | - | - | 0 | - |
| - | - | 5.9E+04 | 454.3 | - | - | 0 | - |
| - | - | 2.194E+04 | 457.3 | - | - | 0 | - |
| - | - | 9.047E+04 | 457.8 | - | - | 0 | - |
| - | - | 6.064E+04 | 458.3 | - | - | 0 | - |
| - | - | 3.954E+04 | 458.8 | - | - | 0 | - |
| - | - | 1.499E+05 | 459.3 | - | - | 0 | - |
| - | - | 6.9E+04 | 459.8 | - | - | 0 | - |
| - | - | 2.901E+05 | 460.3 | - | - | 0 | - |
| - | - | 1.748E+05 | 460.8 | - | - | 0 | - |
| - | - | 3.81E+05 | 461.3 | - | - | 0 | - |
| - | - | 2.066E+05 | 461.8 | - | - | 0 | - |
| - | - | 6.161E+04 | 462.3 | - | - | 0 | - |
| - | - | 1.234E+05 | 465.3 | - | - | 0 | - |
| - | - | 3.953E+06 | 466.3 | - | - | 0 | - |
| - | - | 2.188E+06 | 466.8 | - | - | 0 | - |
| - | - | 7.328E+05 | 467.3 | - | - | 0 | - |
| - | - | 2.438E+04 | 467.3 | - | - | 0 | - |
| - | - | 9.282E+04 | 467.8 | - | - | 0 | - |
| - | - | 1.094E+05 | 468.3 | - | - | 0 | - |
| - | - | 7.05E+04 | 468.8 | - | - | 0 | - |
| 3 | y | 1.952E+05 | 469.3 | 0.001171 | 2.496 | +2 | 10 |
| - | - | 3.195E+05 | 469.3 | - | - | 0 | - |
| 3 | y | 1.169E+05 | 469.8 | 0.00895 | 19.05 | +2 | 10 |
| - | - | 2.02E+05 | 469.8 | - | - | 0 | - |
| - | - | 5.204E+04 | 470.3 | - | - | 0 | - |
| 8 | y | 3.412E+05 | 474.3 | 0.0005996 | 1.264 | +1 | 5 |
| 10 | c | 4.396E+04 | 474.8 | 0.0009871 | 2.079 | +2 | 10 |
| 10 | c | 3.048E+06 | 475.3 | 0.0006785 | 1.428 | +2 | 10 |
| - | - | 1.61E+06 | 475.8 | - | - | 0 | - |
| - | - | 4.621E+05 | 476.3 | - | - | 0 | - |
| - | - | 4.285E+04 | 476.8 | - | - | 0 | - |
| 3 | y | 1.877E+05 | 478.3 | 0.001199 | 2.507 | +2 | 10 |
| - | - | 5.079E+04 | 478.3 | - | - | 0 | - |
| - | - | 1.148E+05 | 478.8 | - | - | 0 | - |
| - | - | 1.028E+05 | 480.3 | - | - | 0 | - |
| - | - | 5.671E+04 | 480.8 | - | - | 0 | - |
| - | - | 1.226E+05 | 482.3 | - | - | 0 | - |
| - | - | 3.786E+04 | 483.3 | - | - | 0 | - |
| - | - | 1.3E+05 | 489.3 | - | - | 0 | - |
| - | - | 5.284E+04 | 489.8 | - | - | 0 | - |
| - | - | 2.826E+04 | 490.3 | - | - | 0 | - |
| - | - | 2.974E+04 | 491.3 | - | - | 0 | - |
| - | - | 7.179E+04 | 493.3 | - | - | 0 | - |
| - | - | 2.66E+04 | 493.8 | - | - | 0 | - |
| - | - | 2.659E+04 | 494.3 | - | - | 0 | - |
| 5 | c | 1.059E+05 | 495.3 | 0.0002262 | 0.4567 | +1 | 5 |
| - | - | 1.187E+05 | 496.3 | - | - | 0 | - |
| - | - | 4.035E+04 | 497.3 | - | - | 0 | - |
| - | - | 4.252E+04 | 498.3 | - | - | 0 | - |
| - | - | 1.375E+06 | 501.8 | - | - | 0 | - |
| - | - | 9.429E+05 | 502.3 | - | - | 0 | - |
| - | - | 2.257E+05 | 502.8 | - | - | 0 | - |
| - | - | 9.711E+04 | 506.8 | - | - | 0 | - |
| - | - | 2.578E+05 | 507.3 | - | - | 0 | - |
| - | - | 1.079E+05 | 507.8 | - | - | 0 | - |
| - | - | 5.636E+04 | 508.3 | - | - | 0 | - |
| - | - | 2.495E+04 | 509.3 | - | - | 0 | - |
| - | - | 3.955E+04 | 509.8 | - | - | 0 | - |
| - | - | 2.689E+06 | 510.8 | - | - | 0 | - |
| - | - | 1.661E+06 | 511.3 | - | - | 0 | - |
| - | - | 5.569E+05 | 511.8 | - | - | 0 | - |
| 5 | c | 8.581E+04 | 512.3 | 0.003401 | 6.639 | +1 | 5 |
| - | - | 3.09E+04 | 513.3 | - | - | 0 | - |
| - | - | 4.694E+06 | 515.8 | - | - | 0 | - |
| - | - | 3.307E+06 | 516.3 | - | - | 0 | - |
| - | - | 4.099E+04 | 516.4 | - | - | 0 | - |
| - | - | 1.042E+06 | 516.8 | - | - | 0 | - |
| - | - | 7.542E+04 | 517.3 | - | - | 0 | - |
| 11 | c | 4.194E+04 | 524.3 | 0.002425 | 4.624 | +2 | 11 |
| 11 | c | 2.91E+06 | 524.8 | 0.0005292 | 1.008 | +2 | 11 |
| - | - | 1.896E+06 | 525.3 | - | - | 0 | - |
| - | - | 6.015E+05 | 525.8 | - | - | 0 | - |
| - | - | 1.254E+05 | 526.3 | - | - | 0 | - |
| 2 | z | 4.805E+04 | 526.8 | 0.007958 | 15.11 | +2 | 11 |
| - | - | 2.363E+05 | 534.3 | - | - | 0 | - |
| 2 | y | 3.216E+05 | 534.8 | 0.001098 | 2.053 | +2 | 11 |
| - | - | 2.449E+05 | 535.3 | - | - | 0 | - |
| - | - | 5.976E+04 | 535.8 | - | - | 0 | - |
| - | - | 4.194E+04 | 536.3 | - | - | 0 | - |
| - | - | 4.607E+04 | 539.8 | - | - | 0 | - |
| - | - | 7.454E+04 | 540.3 | - | - | 0 | - |
| 7 | y | 7.833E+04 | 545.3 | 0.0005037 | 0.9237 | +1 | 6 |
| - | - | 3.163E+04 | 551.3 | - | - | 0 | - |
| - | - | 7.258E+05 | 552.3 | - | - | 0 | - |
| - | - | 7.839E+05 | 553.3 | - | - | 0 | - |
| - | - | 2.237E+05 | 554.3 | - | - | 0 | - |
| - | - | 9.243E+05 | 556.3 | - | - | 0 | - |
| - | - | 2.871E+05 | 557.3 | - | - | 0 | - |
| - | - | 3.165E+04 | 558.4 | - | - | 0 | - |
| - | - | 4.485E+04 | 561.3 | - | - | 0 | - |
| - | - | 3.331E+04 | 562.3 | - | - | 0 | - |
| - | - | 1.321E+05 | 566.3 | - | - | 0 | - |
| - | - | 1.947E+05 | 566.8 | - | - | 0 | - |
| - | - | 6.795E+04 | 567.3 | - | - | 0 | - |
| - | - | 4.399E+04 | 568.4 | - | - | 0 | - |
| - | - | 3.256E+06 | 575.3 | - | - | 0 | - |
| - | - | 2.173E+06 | 575.8 | - | - | 0 | - |
| - | - | 7.167E+05 | 576.3 | - | - | 0 | - |
| - | - | 7.434E+04 | 576.8 | - | - | 0 | - |
| - | - | 3.394E+04 | 578.4 | - | - | 0 | - |
| - | - | 7.278E+04 | 579.4 | - | - | 0 | - |
| - | - | 3.567E+04 | 580.3 | - | - | 0 | - |
| - | - | 4.147E+04 | 580.4 | - | - | 0 | - |
| - | - | 5.426E+04 | 582.3 | - | - | 0 | - |
| - | - | 5.392E+04 | 583.3 | - | - | 0 | - |
| - | - | 3.04E+06 | 584.3 | - | - | 0 | - |
| - | - | 2.042E+06 | 584.8 | - | - | 0 | - |
| - | - | 2.298E+04 | 585.2 | - | - | 0 | - |
| - | - | 7.314E+05 | 585.3 | - | - | 0 | - |
| - | - | 4.925E+04 | 585.8 | - | - | 0 | - |
| - | - | 7.498E+04 | 595.4 | - | - | 0 | - |
| - | - | 2.267E+04 | 596.4 | - | - | 0 | - |
| - | - | 1.839E+05 | 596.4 | - | - | 0 | - |
| - | - | 5.922E+04 | 597.4 | - | - | 0 | - |
| - | - | 9.293E+04 | 605.4 | - | - | 0 | - |
| - | - | 3.999E+04 | 606.4 | - | - | 0 | - |
| - | - | 7.441E+04 | 608.4 | - | - | 0 | - |
| - | - | 2.559E+04 | 609.4 | - | - | 0 | - |
| 6 | c | 6.532E+05 | 623.4 | 0.0006001 | 0.9626 | +1 | 6 |
| - | - | 2.192E+05 | 624.4 | - | - | 0 | - |
| - | - | 3.707E+04 | 625.4 | - | - | 0 | - |
| - | - | 7.81E+05 | 633.4 | - | - | 0 | - |
| - | - | 2.787E+05 | 634.4 | - | - | 0 | - |
| - | - | 2.757E+04 | 635.4 | - | - | 0 | - |
| - | - | 1.658E+06 | 639.4 | - | - | 0 | - |
| 6 | c | 8.686E+06 | 640.4 | 0.0002961 | 0.4624 | +1 | 6 |
| - | - | 3.071E+06 | 641.4 | - | - | 0 | - |
| - | - | 5.287E+05 | 642.4 | - | - | 0 | - |
| - | - | 1.312E+05 | 650.4 | - | - | 0 | - |
| - | - | 2.121E+06 | 651.4 | - | - | 0 | - |
| - | - | 8.108E+05 | 652.4 | - | - | 0 | - |
| - | - | 1.869E+05 | 653.4 | - | - | 0 | - |
| 6 | z | 1.699E+06 | 657.4 | 0.0004366 | 0.6641 | +1 | 7 |
| - | - | 1.957E+06 | 658.4 | - | - | 0 | - |
| - | - | 5.814E+05 | 659.4 | - | - | 0 | - |
| - | - | 8.612E+04 | 660.4 | - | - | 0 | - |
| - | - | 2.508E+04 | 662.4 | - | - | 0 | - |
| - | - | 1.46E+05 | 663.4 | - | - | 0 | - |
| - | - | 6.464E+04 | 664.4 | - | - | 0 | - |
| - | - | 1.622E+05 | 666.4 | - | - | 0 | - |
| - | - | 5.203E+05 | 667.4 | - | - | 0 | - |
| - | - | 2.091E+05 | 668.4 | - | - | 0 | - |
| - | - | 3.154E+04 | 669.4 | - | - | 0 | - |
| - | - | 4.202E+04 | 669.4 | - | - | 0 | - |
| - | - | 2.01E+05 | 672.4 | - | - | 0 | - |
| 6 | y | 2.09E+06 | 673.4 | 0.0004503 | 0.6687 | +1 | 7 |
| - | - | 7.521E+05 | 674.4 | - | - | 0 | - |
| - | - | 1.594E+05 | 675.4 | - | - | 0 | - |
| - | - | 3.161E+04 | 676.4 | - | - | 0 | - |
| - | - | 2.64E+04 | 677.4 | - | - | 0 | - |
| - | - | 2.776E+04 | 680.4 | - | - | 0 | - |
| - | - | 7.496E+04 | 681.4 | - | - | 0 | - |
| 7 | c | 4.678E+05 | 694.4 | 0.0002905 | 0.4184 | +1 | 7 |
| - | - | 1.976E+05 | 695.4 | - | - | 0 | - |
| - | - | 3.298E+04 | 704.4 | - | - | 0 | - |
| - | - | 3.883E+04 | 708.4 | - | - | 0 | - |
| - | - | 7.989E+05 | 710.4 | - | - | 0 | - |
| 7 | c | 5.867E+06 | 711.5 | 0.0002917 | 0.41 | +1 | 7 |
| - | - | 2.365E+06 | 712.5 | - | - | 0 | - |
| - | - | 4.168E+05 | 713.5 | - | - | 0 | - |
| - | - | 3.169E+05 | 719.4 | - | - | 0 | - |
| - | - | 1.098E+05 | 720.4 | - | - | 0 | - |
| - | - | 4.458E+04 | 721.4 | - | - | 0 | - |
| - | - | 3.472E+04 | 722.5 | - | - | 0 | - |
| - | - | 7.141E+04 | 732.4 | - | - | 0 | - |
| - | - | 5.197E+04 | 733.4 | - | - | 0 | - |
| - | - | 3.23E+04 | 734.4 | - | - | 0 | - |
| - | - | 3.868E+04 | 736.4 | - | - | 0 | - |
| 5 | z | 4.169E+05 | 737.4 | 0.00112 | 1.519 | +1 | 8 |
| - | - | 8.823E+04 | 737.5 | - | - | 0 | - |
| - | - | 1.874E+05 | 738.4 | - | - | 0 | - |
| - | - | 2.068E+06 | 738.5 | - | - | 0 | - |
| - | - | 1.85E+06 | 739.5 | - | - | 0 | - |
| - | - | 4.841E+05 | 740.5 | - | - | 0 | - |
| - | - | 1.076E+05 | 741.5 | - | - | 0 | - |
| - | - | 3.868E+04 | 748.4 | - | - | 0 | - |
| - | - | 5.393E+04 | 749.4 | - | - | 0 | - |
| - | - | 1.575E+05 | 750.5 | - | - | 0 | - |
| - | - | 5.622E+04 | 751.5 | - | - | 0 | - |
| 5 | y | 5.177E+05 | 752.4 | 0.001525 | 2.027 | +1 | 8 |
| - | - | 2.114E+05 | 753.4 | - | - | 0 | - |
| 5 | z | 5.299E+04 | 754.4 | 0.01243 | 16.48 | +1 | 8 |
| - | - | 3.682E+04 | 761.4 | - | - | 0 | - |
| - | - | 6.762E+04 | 762.4 | - | - | 0 | - |
| - | - | 2.903E+04 | 763.4 | - | - | 0 | - |
| - | - | 7.397E+04 | 764.5 | - | - | 0 | - |
| 8 | c | 5.503E+06 | 765.5 | 0.0005913 | 0.7724 | +1 | 8 |
| - | - | 2.436E+06 | 766.5 | - | - | 0 | - |
| - | - | 5.286E+05 | 767.5 | - | - | 0 | - |
| - | - | 3.189E+04 | 768.4 | - | - | 0 | - |
| - | - | 5.647E+05 | 769.4 | - | - | 0 | - |
| 5 | y | 1.289E+07 | 770.4 | 0.000787 | 1.022 | +1 | 8 |
| - | - | 5.371E+06 | 771.4 | - | - | 0 | - |
| - | - | 1.154E+06 | 772.4 | - | - | 0 | - |
| - | - | 6.051E+04 | 773.4 | - | - | 0 | - |
| - | - | 7.79E+04 | 779.4 | - | - | 0 | - |
| - | - | 4.796E+04 | 780.5 | - | - | 0 | - |
| - | - | 1.75E+05 | 781.5 | - | - | 0 | - |
| - | - | 6.053E+04 | 782.5 | - | - | 0 | - |
| - | - | 6.884E+04 | 790.5 | - | - | 0 | - |
| - | - | 8.057E+04 | 791.5 | - | - | 0 | - |
| - | - | 5.309E+04 | 793.5 | - | - | 0 | - |
| - | - | 9.381E+04 | 794.5 | - | - | 0 | - |
| - | - | 3.938E+04 | 795.5 | - | - | 0 | - |
| - | - | 6.547E+04 | 797.4 | - | - | 0 | - |
| - | - | 4.178E+05 | 798.4 | - | - | 0 | - |
| - | - | 1.402E+05 | 799.4 | - | - | 0 | - |
| - | - | 3.894E+04 | 800.4 | - | - | 0 | - |
| - | - | 4.373E+04 | 801.4 | - | - | 0 | - |
| - | - | 2.474E+04 | 802.4 | - | - | 0 | - |
| - | - | 4.097E+04 | 806.5 | - | - | 0 | - |
| - | - | 5.237E+05 | 807.5 | - | - | 0 | - |
| - | - | 4.615E+05 | 808.5 | - | - | 0 | - |
| - | - | 1.889E+05 | 809.5 | - | - | 0 | - |
| - | - | 2.845E+04 | 810.5 | - | - | 0 | - |
| - | - | 3.032E+04 | 817.4 | - | - | 0 | - |
| - | - | 4.993E+05 | 818.5 | - | - | 0 | - |
| - | - | 2.475E+05 | 819.5 | - | - | 0 | - |
| - | - | 5.049E+04 | 820.4 | - | - | 0 | - |
| - | - | 9.581E+04 | 820.5 | - | - | 0 | - |
| - | - | 6.587E+04 | 821.5 | - | - | 0 | - |
| - | - | 6.334E+05 | 822.5 | - | - | 0 | - |
| - | - | 3.305E+05 | 823.5 | - | - | 0 | - |
| 4 | w | 3.081E+06 | 824.5 | 0.0008425 | 1.022 | +1 | 9 |
| - | - | 1.452E+06 | 825.5 | - | - | 0 | - |
| - | - | 3.937E+05 | 826.5 | - | - | 0 | - |
| - | - | 5.593E+04 | 827.5 | - | - | 0 | - |
| - | - | 1.397E+05 | 832.5 | - | - | 0 | - |
| - | - | 5.554E+04 | 833.5 | - | - | 0 | - |
| - | - | 5.319E+04 | 834.5 | - | - | 0 | - |
| - | - | 4.725E+05 | 835.5 | - | - | 0 | - |
| - | - | 1.145E+06 | 836.5 | - | - | 0 | - |
| - | - | 1.788E+05 | 836.5 | - | - | 0 | - |
| - | - | 5.754E+05 | 837.5 | - | - | 0 | - |
| - | - | 5.754E+04 | 837.5 | - | - | 0 | - |
| - | - | 2.173E+05 | 838.5 | - | - | 0 | - |
| - | - | 5.552E+04 | 839.5 | - | - | 0 | - |
| - | - | 3.48E+04 | 840.4 | - | - | 0 | - |
| - | - | 3.897E+04 | 845.5 | - | - | 0 | - |
| - | - | 1.682E+05 | 850.5 | - | - | 0 | - |
| - | - | 8.559E+04 | 851.5 | - | - | 0 | - |
| - | - | 4.884E+04 | 852.5 | - | - | 0 | - |
| 9 | c | 1.967E+05 | 862.5 | 0.0006589 | 0.7639 | +1 | 9 |
| - | - | 3.792E+05 | 863.5 | - | - | 0 | - |
| - | - | 3.818E+05 | 864.5 | - | - | 0 | - |
| - | - | 1.552E+05 | 865.5 | - | - | 0 | - |
| - | - | 3.809E+04 | 866.5 | - | - | 0 | - |
| - | - | 3.072E+04 | 868.5 | - | - | 0 | - |
| - | - | 1.988E+05 | 877.5 | - | - | 0 | - |
| - | - | 9.723E+06 | 878.5 | - | - | 0 | - |
| 9 | c | 1.226E+07 | 879.5 | 0.001756 | 1.997 | +1 | 9 |
| - | - | 5.139E+06 | 880.5 | - | - | 0 | - |
| 4 | y | 5.241E+05 | 881.5 | 0.001029 | 1.167 | +1 | 9 |
| - | - | 1.069E+06 | 881.6 | - | - | 0 | - |
| 4 | z | 5.1E+06 | 882.5 | 0.000126 | 0.1427 | +1 | 9 |
| - | - | 2.554E+06 | 883.5 | - | - | 0 | - |
| - | - | 6.205E+05 | 884.5 | - | - | 0 | - |
| - | - | 3.435E+04 | 893.5 | - | - | 0 | - |
| - | - | 2.795E+04 | 896.5 | - | - | 0 | - |
| - | - | 9.373E+05 | 897.5 | - | - | 0 | - |
| 4 | y | 6.197E+06 | 898.5 | 9.828E-06 | 0.01094 | +1 | 9 |
| - | - | 3.048E+06 | 899.5 | - | - | 0 | - |
| - | - | 3.875E+04 | 900.4 | - | - | 0 | - |
| - | - | 7.707E+05 | 900.5 | - | - | 0 | - |
| - | - | 5.706E+04 | 901.5 | - | - | 0 | - |
| - | - | 4.919E+04 | 902.5 | - | - | 0 | - |
| - | - | 3.607E+04 | 903.5 | - | - | 0 | - |
| - | - | 6.033E+04 | 906.5 | - | - | 0 | - |
| - | - | 9.931E+04 | 907.5 | - | - | 0 | - |
| - | - | 1.047E+05 | 914.5 | - | - | 0 | - |
| - | - | 5.806E+04 | 915.5 | - | - | 0 | - |
| - | - | 7.878E+04 | 916.5 | - | - | 0 | - |
| - | - | 1.035E+05 | 917.5 | - | - | 0 | - |
| - | - | 4.653E+04 | 918.5 | - | - | 0 | - |
| - | - | 3.807E+04 | 919.5 | - | - | 0 | - |
| - | - | 3.672E+04 | 920.5 | - | - | 0 | - |
| - | - | 7.483E+04 | 920.6 | - | - | 0 | - |
| 3 | z | 4.417E+04 | 921.5 | 0.01414 | 15.34 | +1 | 10 |
| - | - | 8.709E+04 | 921.6 | - | - | 0 | - |
| - | - | 5.657E+06 | 931.5 | - | - | 0 | - |
| - | - | 3.048E+06 | 932.5 | - | - | 0 | - |
| - | - | 1.125E+06 | 933.5 | - | - | 0 | - |
| - | - | 2.431E+05 | 934.6 | - | - | 0 | - |
| - | - | 1.043E+05 | 935.5 | - | - | 0 | - |
| - | - | 3.953E+04 | 936.5 | - | - | 0 | - |
| 3 | y | 3.697E+05 | 937.5 | 0.0004633 | 0.4942 | +1 | 10 |
| 3 | y | 2.787E+05 | 938.5 | 0.009795 | 10.44 | +1 | 10 |
| 3 | z | 1.429E+06 | 939.5 | 0.000444 | 0.4726 | +1 | 10 |
| - | - | 7.501E+05 | 940.5 | - | - | 0 | - |
| - | - | 2.316E+05 | 941.5 | - | - | 0 | - |
| 10 | c | 8.919E+06 | 949.5 | 0.0006989 | 0.7361 | +1 | 10 |
| - | - | 4.854E+06 | 950.5 | - | - | 0 | - |
| - | - | 1.41E+06 | 951.6 | - | - | 0 | - |
| - | - | 1.758E+05 | 952.6 | - | - | 0 | - |
| 3 | y | 7.976E+06 | 955.5 | 0.0002136 | 0.2235 | +1 | 10 |
| - | - | 4.402E+06 | 956.5 | - | - | 0 | - |
| - | - | 1.178E+06 | 957.5 | - | - | 0 | - |
| - | - | 5.544E+04 | 958.5 | - | - | 0 | - |
| 10 | c | 6.978E+06 | 966.6 | 8.976E-05 | 0.09286 | +1 | 10 |
| - | - | 3.929E+06 | 967.6 | - | - | 0 | - |
| - | - | 1.162E+06 | 968.6 | - | - | 0 | - |
| - | - | 5.022E+04 | 969.6 | - | - | 0 | - |
| - | - | 4.675E+04 | 982.5 | - | - | 0 | - |
| - | - | 2.745E+04 | 983.5 | - | - | 0 | - |
| - | - | 3.916E+04 | 1006 | - | - | 0 | - |
| - | - | 3.631E+05 | 1007 | - | - | 0 | - |
| - | - | 1.488E+05 | 1008 | - | - | 0 | - |
| - | - | 5.356E+04 | 1009 | - | - | 0 | - |
| 2 | w | 3.934E+05 | 1010 | 0.0003911 | 0.3874 | +1 | 11 |
| - | - | 1.885E+05 | 1011 | - | - | 0 | - |
| - | - | 1.009E+05 | 1012 | - | - | 0 | - |
| - | - | 3.751E+04 | 1014 | - | - | 0 | - |
| - | - | 2.825E+04 | 1015 | - | - | 0 | - |
| - | - | 4.225E+04 | 1019 | - | - | 0 | - |
| - | - | 2.136E+05 | 1021 | - | - | 0 | - |
| - | - | 4.77E+05 | 1022 | - | - | 0 | - |
| - | - | 3.208E+05 | 1023 | - | - | 0 | - |
| - | - | 1.592E+05 | 1024 | - | - | 0 | - |
| - | - | 3.911E+05 | 1031 | - | - | 0 | - |
| - | - | 3.256E+05 | 1032 | - | - | 0 | - |
| - | - | 1.636E+05 | 1033 | - | - | 0 | - |
| - | - | 3.42E+04 | 1034 | - | - | 0 | - |
| 11 | c | 5.158E+06 | 1049 | 0.0002782 | 0.2653 | +1 | 11 |
| - | - | 3.201E+06 | 1050 | - | - | 0 | - |
| - | - | 2.071E+06 | 1051 | - | - | 0 | - |
| - | - | 7.579E+05 | 1052 | - | - | 0 | - |
| 2 | z | 1.287E+06 | 1053 | 0.00366 | 3.477 | +1 | 11 |
| - | - | 7.943E+05 | 1054 | - | - | 0 | - |
| - | - | 2.494E+05 | 1055 | - | - | 0 | - |
| 11 | c | 1.365E+07 | 1066 | 0.0004625 | 0.434 | +1 | 11 |
| - | - | 8.48E+06 | 1067 | - | - | 0 | - |
| - | - | 2.968E+06 | 1068 | - | - | 0 | - |
| 2 | y | 4.522E+05 | 1069 | 0.02095 | 19.6 | +1 | 11 |
| - | - | 2.213E+05 | 1070 | - | - | 0 | - |
| - | - | 8.153E+04 | 1071 | - | - | 0 | - |
| - | - | 2.837E+04 | 1079 | - | - | 0 | - |
| - | - | 4.576E+04 | 1080 | - | - | 0 | - |
| - | - | 8.638E+05 | 1081 | - | - | 0 | - |
| - | - | 4.963E+05 | 1082 | - | - | 0 | - |
| - | - | 1.399E+05 | 1083 | - | - | 0 | - |
| - | - | 4.942E+04 | 1091 | - | - | 0 | - |
| - | - | 2.636E+06 | 1096 | - | - | 0 | - |
| - | - | 1.657E+06 | 1097 | - | - | 0 | - |
| - | - | 6.829E+05 | 1098 | - | - | 0 | - |
| - | - | 2.429E+05 | 1099 | - | - | 0 | - |
| - | - | 6.418E+04 | 1100 | - | - | 0 | - |
| - | - | 8.808E+04 | 1105 | - | - | 0 | - |
| - | - | 5.505E+04 | 1106 | - | - | 0 | - |
| - | - | 3.957E+04 | 1107 | - | - | 0 | - |
| - | - | 5E+04 | 1108 | - | - | 0 | - |
| - | - | 6.136E+04 | 1109 | - | - | 0 | - |
| - | - | 5.295E+04 | 1110 | - | - | 0 | - |
| - | - | 1.695E+05 | 1113 | - | - | 0 | - |
| - | - | 1.095E+05 | 1114 | - | - | 0 | - |
| - | - | 1.11E+05 | 1120 | - | - | 0 | - |
| - | - | 8.997E+04 | 1121 | - | - | 0 | - |
| - | - | 5.084E+04 | 1122 | - | - | 0 | - |
| - | - | 6.163E+05 | 1123 | - | - | 0 | - |
| - | - | 1.293E+06 | 1124 | - | - | 0 | - |
| - | - | 7.706E+05 | 1125 | - | - | 0 | - |
| - | - | 2.375E+05 | 1126 | - | - | 0 | - |
| - | - | 5.363E+04 | 1127 | - | - | 0 | - |
| - | - | 8.842E+04 | 1134 | - | - | 0 | - |
| - | - | 5.695E+04 | 1135 | - | - | 0 | - |
| - | - | 8.081E+04 | 1137 | - | - | 0 | - |
| - | - | 5.86E+04 | 1138 | - | - | 0 | - |
| - | - | 1.217E+06 | 1151 | - | - | 0 | - |
| - | - | 1.26E+07 | 1152 | - | - | 0 | - |
| - | - | 8.186E+06 | 1153 | - | - | 0 | - |
| - | - | 2.858E+06 | 1154 | - | - | 0 | - |
| - | - | 2.186E+05 | 1155 | - | - | 0 | - |
| - | - | 1.335E+07 | 1168 | - | - | 0 | - |
| - | - | 3.815E+07 | 1169 | - | - | 0 | - |
| - | - | 2.311E+07 | 1170 | - | - | 0 | - |
| - | - | 7.452E+06 | 1171 | - | - | 0 | - |
| - | - | 5.533E+05 | 1172 | - | - | 0 | - |
| - | - | 3.344E+04 | 1185 | - | - | 0 | - |
| - | - | 4.833E+04 | 1201 | - | - | 0 | - |
| - | - | 3.38E+04 | 3045 | - | - | 0 | - |

m/z Charge Intensity FragmentType MassShift Position
120.06582641601562 0 4847141.5 y 11
121.06914520263672 0 216086.56
157.09765625 0 36232.785
158.09263610839844 0 60697.19
159.11331176757812 0 23869.219
167.08177185058594 0 192793.89
167.62933349609375 0 18251.785
169.06109619140625 0 84497.3
169.09730529785156 0 22973.5
171.11317443847656 0 69881.54
173.45245361328125 0 56079.69
176.4202880859375 0 15603.08
184.10829162597656 0 38827.07
185.09226989746094 0 233937.36
185.1651611328125 0 5972305.5
186.08763122558594 0 670451.75
186.0959014892578 0 33469.516
186.16848754882812 0 705789.44
187.0912322998047 0 39909.527
187.1080322265625 0 102701.664
187.17185974121094 0 20766.545
188.1397247314453 0 61954.133
196.82284545898438 0 22673.385
201.12367248535156 0 307482.84 y Water loss 10
202.1187744140625 0 50075.32
202.12799072265625 0 28833.11
213.1600799560547 0 7503902.5
214.1634063720703 0 872596.9
215.16648864746094 0 27367.953
219.1342315673828 0 2734308.5 y 10
226.1552276611328 0 259052.53
227.15895080566406 0 29208.379
238.15631103515625 0 24126.553
245.40435791015625 0 20792.883
256.165771484375 0 178272.77
266.1501770019531 0 539676
267.15350341796875 0 70619.625
267.20611572265625 0 24534.502
270.18145751953125 0 870146.2
271.18475341796875 0 112639.664
284.1607971191406 0 628600.44
285.1566467285156 0 241246.44
286.1593322753906 0 39688.816
287.20819091796875 0 1582448 c 2
288.2112121582031 0 197639.67
297.193115234375 0 29453.191
299.1717834472656 0 909560.7
300.17510986328125 0 169957.73
306.1662902832031 0 116614.2 y 9
327.16693115234375 0 299383.47
327.2026062011719 0 22115.52
328.1691589355469 0 27754.037
337.187744140625 0 42901.223
341.2193298339844 0 26347.32
353.21820068359375 0 69127.7
355.1966857910156 0 36064.594
360.196533203125 0 50610.055
365.2185974121094 0 26178.566
368.2295227050781 0 734700.1
369.20318603515625 0 33928.652 z Ammonia loss 4
369.2326354980469 0 158890
370.2451171875 0 845573.56
371.2474670410156 0 131842.03
372.2509765625 0 25176.734
372.7087707519531 0 26839.33
375.7278747558594 0 45791.05
380.2306823730469 0 36448.13
381.21441650390625 0 69857.38
382.22442626953125 0 29470.947
382.7326354980469 0 29892.693
383.233642578125 0 130569.57 c Ammonia loss 7
383.7370910644531 0 60082.277
385.20849609375 0 260349.61 y Water loss 8
385.72418212890625 0 93364.195 y 4
394.2093505859375 0 25600.49
395.73162841796875 0 46869.207
396.2352600097656 0 25089.668
396.7450256347656 0 25200.92
398.2403869628906 0 4576195.5 c Ammonia loss 3
399.24322509765625 0 1081025.6
400.2453308105469 0 94813.24
401.216064453125 0 24074.48
403.21923828125 0 3979331.5 y 8
404.2224426269531 0 821946.8
404.7386169433594 0 115916.484
405.2264099121094 0 70456.69
405.7436218261719 0 22102.873
408.224609375 0 55713.664
409.7301940917969 0 218936.86
410.23175048828125 0 138529.66
410.7584533691406 0 31446.299
411.23394775390625 0 57622.516
412.25897216796875 0 28231.3
416.7379455566406 0 96064.836
417.2393798828125 0 28442.123
417.76434326171875 0 126587.664
418.2664794921875 0 56953.227
418.73529052734375 0 141127.8
418.7669372558594 0 32907.938
419.2348937988281 0 45607.117
419.7619934082031 0 27739.979
423.2441711425781 0 23871.4
423.27056884765625 0 27216.123
423.76849365234375 0 30548.4
425.7425231933594 0 47806.438
426.2366027832031 0 56739.285
430.75433349609375 0 38909.22
431.7613830566406 0 825213.5 c Ammonia loss 8
432.2646484375 0 474258.62
432.76861572265625 0 135431.1
439.7586669921875 0 44279.082
440.2552795410156 0 28553.475
440.7493896484375 0 174668.28 y Water loss 3
441.247314453125 0 90040.68 y Ammonia loss 3
444.76995849609375 0 74503.484
445.2711486816406 0 41652.48
448.751708984375 0 28732.564
449.7545166015625 0 60183.496 y 3
450.25299072265625 0 22776.354
450.755126953125 0 34743.777
451.7763977050781 0 39010.504
452.2749938964844 0 99935.055
452.7695007324219 0 184485.44
453.2689514160156 0 102447.39
453.77545166015625 0 95561.945
454.2733154296875 0 58998.13
457.263427734375 0 21944.428
457.7611999511719 0 90473.02
458.2612609863281 0 60637.504
458.76190185546875 0 39542.65
459.2645263671875 0 149912.22
459.764892578125 0 69002.64
460.29046630859375 0 290059.25
460.78973388671875 0 174797.4
461.28143310546875 0 380994.8
461.78216552734375 0 206644.58
462.2833251953125 0 61606.2
465.2830810546875 0 123354.83
466.2723388671875 0 3953313.8
466.77362060546875 0 2187830.8
467.27581787109375 0 732779.7
467.306640625 0 24378.021
467.77923583984375 0 92824.93
468.27056884765625 0 109426.25
468.7702331542969 0 70504.48
469.2598876953125 0 195160.6 y Water loss 2
469.2957763671875 0 319486.25
469.7596740722656 0 116886.53 y Ammonia loss 2
469.7974548339844 0 202010.62
470.29669189453125 0 52036.15
474.2564392089844 0 341242.28 y 7
474.785888671875 0 43956.72 c Water loss 9
475.277587890625 0 3047856.5 c Ammonia loss 9
475.7790222167969 0 1609638.9
476.28057861328125 0 462119.16
476.7845458984375 0 42853.754
478.26519775390625 0 187676.4 y 2
478.3026428222656 0 50786.117
478.7668151855469 0 114757.14
480.2877197265625 0 102828.54
480.784423828125 0 56711.906
482.27325439453125 0 122607.836
483.2721862792969 0 37859.535
489.2934265136719 0 130023.195
489.7958068847656 0 52843.55
490.2970275878906 0 28256.71
491.29754638671875 0 29744.744
493.2979431152344 0 71789.48
493.79998779296875 0 26595.854
494.29498291015625 0 26586.936
495.29278564453125 0 105865.38 c Ammonia loss 4
496.32110595703125 0 118737.375
497.3239440917969 0 40353.453
498.2873840332031 0 42515.72
501.80908203125 0 1374851.6
502.3096618652344 0 942929.5
502.8094787597656 0 225746.12
506.8007507324219 0 97108.07
507.2945251464844 0 257774.16
507.79742431640625 0 107932.52
508.2958679199219 0 56355.414
509.3102111816406 0 24950.51
509.8059997558594 0 39546.863
510.81427001953125 0 2689237.2
511.3156433105469 0 1661377.6
511.8169860839844 0 556936.56
512.322509765625 0 85808.57 c 4
513.3375854492188 0 30899.31
515.8065185546875 0 4693715
516.3074951171875 0 3306761
516.3539428710938 0 40991.79
516.8087158203125 0 1041953.2
517.306640625 0 75420.484
524.321533203125 0 41936.88 c Water loss 10
524.8116455078125 0 2910395 c Ammonia loss 10
525.313232421875 0 1896442.6
525.8116455078125 0 601534
526.303955078125 0 125421.96
526.8046264648438 0 48054.266 z 1
534.3045654296875 0 236261.6
534.80712890625 0 321583.1 y 1
535.307861328125 0 244908.27
535.810302734375 0 59761.445
536.3192749023438 0 41942.44
539.7989501953125 0 46069.23
540.3077392578125 0 74538.83
545.29345703125 0 78334.3 y 6
551.3406372070312 0 31627.068
552.3147583007812 0 725751.5
553.311767578125 0 783869.4
554.3167114257812 0 223670.19
556.3457641601562 0 924292.56
557.3489379882812 0 287085.78
558.3554077148438 0 31654.07
561.33740234375 0 44848.47
562.336669921875 0 33310.555
566.3306884765625 0 132146.36
566.8289794921875 0 194704.78
567.3302001953125 0 67946.445
568.3717041015625 0 43987.97
575.33544921875 0 3255737.8
575.8369140625 0 2173358.8
576.3383178710938 0 716694.06
576.8411865234375 0 74342.71
578.3888549804688 0 33942.273
579.3968505859375 0 72782.945
580.3433837890625 0 35668.31
580.3905639648438 0 41473.086
582.3231201171875 0 54260.824
583.341064453125 0 53922.05
584.3407592773438 0 3040067.2
584.8421020507812 0 2042044.2
585.1542358398438 0 22980.592
585.3433837890625 0 731361
585.8467407226562 0 49248.176
595.3546752929688 0 74977.19
596.3571166992188 0 22669.072
596.4013671875 0 183927.55
597.4063110351562 0 59218.336
605.3775024414062 0 92932.34
606.3829345703125 0 39992.94
608.3641357421875 0 74409.5
609.3677368164062 0 25593.102
623.3881225585938 0 653235.6 c Ammonia loss 5
624.3915405273438 0 219197.6
625.3953857421875 0 37074.758
633.3723754882812 0 781003.5
634.3755493164062 0 278682.84
635.3826904296875 0 27571.629
639.4068603515625 0 1658265.6
640.4143676757812 0 8686081 c 5
641.4174194335938 0 3071108.5
642.419921875 0 528709.1
650.3629150390625 0 131193.58
651.3826293945312 0 2120542
652.3851318359375 0 810779.06
653.387939453125 0 186903.73
657.36962890625 0 1698979.4 z 5
658.3760986328125 0 1956823.8
659.3799438476562 0 581402.3
660.3822631835938 0 86116.95
662.361083984375 0 25077.205
663.3826293945312 0 146010.56
664.3859252929688 0 64637.914
666.3953857421875 0 162197.72
667.438232421875 0 520341.2
668.4411010742188 0 209080
669.391845703125 0 31538.703
669.44873046875 0 42020.965
672.381103515625 0 201042.7
673.3883666992188 0 2089641.5 y 5
674.391357421875 0 752094.3
675.3933715820312 0 159374.64
676.4135131835938 0 31611.424
677.4246826171875 0 26402.633
680.3763427734375 0 27764.006
681.392333984375 0 74958.7
694.4249267578125 0 467775.97 c Ammonia loss 6
695.4283447265625 0 197630.98
704.3724975585938 0 32977.9
708.443603515625 0 38833.21
710.4434204101562 0 798871.3
711.4514770507812 0 5866639.5 c 6
712.4545288085938 0 2364505.8
713.45703125 0 416796.84
719.3840942382812 0 316866.56
720.3840942382812 0 109767.49
721.3851928710938 0 44576.043
722.4562377929688 0 34723.832
732.4432983398438 0 71411.41
733.4403686523438 0 51968.35
734.4185180664062 0 32302.238
736.43115234375 0 38676.383
737.394287109375 0 416881.56 z Ammonia loss 4
737.454345703125 0 88226.875
738.399169921875 0 187397.95
738.47509765625 0 2068361.2
739.4806518554688 0 1849609.6
740.4840087890625 0 484063.2
741.4871215820312 0 107562.27
748.4373779296875 0 38679.92
749.4361572265625 0 53933.605
750.4509887695312 0 157509.31
751.4505615234375 0 56215.58
752.431640625 0 517704.97 y Water loss 4
753.4337768554688 0 211364.98
754.4343872070312 0 52988.88 z 4
761.4288330078125 0 36820.547
762.4171752929688 0 67620.05
763.4292602539062 0 29028.98
764.45458984375 0 73965.516
765.4623413085938 0 5503219.5 c Ammonia loss 7
766.46533203125 0 2436018.2
767.468017578125 0 528596.75
768.4192504882812 0 31885.533
769.4329223632812 0 564712.3
770.4414672851562 0 12890696 y 4
771.4439697265625 0 5370771
772.4468994140625 0 1153917.6
773.4488525390625 0 60507.445
779.4414672851562 0 77895.195
780.4505615234375 0 47958.668
781.456298828125 0 175036.45
782.460205078125 0 60530.05
790.4595336914062 0 68839.89
791.45849609375 0 80570.78
793.4567260742188 0 53085.277
794.4625854492188 0 93808.94
795.4705200195312 0 39375.26
797.4312133789062 0 65471.684
798.435546875 0 417784.53
799.4393920898438 0 140247.98
800.4465942382812 0 38936.2
801.4270629882812 0 43731.957
802.4254760742188 0 24743.45
806.4619750976562 0 40972.812
807.493896484375 0 523703.8
808.4766235351562 0 461515.66
809.4771118164062 0 188858.8
810.4723510742188 0 28445.191
817.4496459960938 0 30319.383
818.45263671875 0 499294.4
819.4514770507812 0 247513.72
820.4452514648438 0 50488.668
820.5145874023438 0 95810.87
821.5126953125 0 65865.766
822.4711303710938 0 633390.5
823.474853515625 0 330522.94
824.4520874023438 0 3081140.2 w 3
825.4545288085938 0 1451987.6
826.4566650390625 0 393650.62
827.4606323242188 0 55927.613
832.4671020507812 0 139704.75
833.4692993164062 0 55539.336
834.476806640625 0 53188.156
835.5266723632812 0 472545.12
836.462646484375 0 1145282.5
836.5343627929688 0 178812.38
837.4658813476562 0 575385.3
837.5396118164062 0 57541.707
838.4716186523438 0 217307.11
839.475341796875 0 55521.062
840.4381103515625 0 34797.652
845.5173950195312 0 38969.516
850.4788208007812 0 168209.08
851.4816284179688 0 85591.8
852.4903564453125 0 48838.574
862.5138549804688 0 196709.39 c Ammonia loss 8
863.5227661132812 0 379247.75
864.529296875 0 381766.97
865.5309448242188 0 155168.88
866.534912109375 0 38086.42
868.4500122070312 0 30719.393
877.514404296875 0 198764.47
878.533447265625 0 9722805
879.539306640625 0 12262774 c 8
880.5428466796875 0 5138748.5
881.4716796875 0 524074.34 y Ammonia loss 3
881.5502319335938 0 1069080.4
882.4804077148438 0 5099625.5 z 3
883.4840087890625 0 2554278.5
884.4865112304688 0 620540
893.5111083984375 0 34345.375
896.496826171875 0 27951.494
897.4918212890625 0 937320.2
898.499267578125 0 6196692.5 y 3
899.502197265625 0 3048133.2
900.4039306640625 0 38746.11
900.5047607421875 0 770673.9
901.5048217773438 0 57059.87
902.5474243164062 0 49193.613
903.5480346679688 0 36065.41
906.5405883789062 0 60326.34
907.54833984375 0 99306.164
914.5040283203125 0 104722.695
915.5078125 0 58062.12
916.5242919921875 0 78776.01
917.5238037109375 0 103480.06
918.52001953125 0 46532.16
919.5164794921875 0 38071.93
920.4732666015625 0 36715.684
920.5559692382812 0 74826.28
921.477294921875 0 44171.332 z Water loss 2
921.5595703125 0 87093.2
931.536376953125 0 5656690
932.5384521484375 0 3047720
933.5441284179688 0 1124795.4
934.5510864257812 0 243096.64
935.5409545898438 0 104311.586
936.538818359375 0 39527.812
937.5106201171875 0 369660.7 y Water loss 2
938.5039672851562 0 278659.62 y Ammonia loss 2
939.50244140625 0 1428541.1 z 2
940.5052490234375 0 750080.06
941.5089111328125 0 231641.28
949.5472412109375 0 8919441 c Ammonia loss 9
950.5498657226562 0 4854426.5
951.553466796875 0 1409653.6
952.560546875 0 175843.11
955.5209350585938 0 7975503 y 2
956.5238037109375 0 4401978.5
957.5267944335938 0 1177510.9
958.5286865234375 0 55438.68
966.5731811523438 0 6977961 c 9
967.5761108398438 0 3929019.5
968.5784301757812 0 1162041
969.5809936523438 0 50223.727
982.51171875 0 46745.242
983.5138549804688 0 27447.688
1005.6143188476562 0 39158.113
1006.6177978515625 0 363082.88
1007.6190795898438 0 148848.56
1008.6198120117188 0 53555.86
1009.5316772460938 0 393371.06 w 1
1010.53564453125 0 188510.78
1011.5358276367188 0 100870.12
1013.5784301757812 0 37510.12
1014.57470703125 0 28251.812
1018.6049194335938 0 42254.6
1020.6192016601562 0 213595.36
1021.6267700195312 0 477027.8
1022.629150390625 0 320849.47
1023.639404296875 0 159204.86
1030.6041259765625 0 391084.84
1031.603515625 0 325629.72
1032.6060791015625 0 163640.17
1033.6295166015625 0 34203.035
1048.615234375 0 5157788.5 c Ammonia loss 10
1049.6180419921875 0 3200710.8
1050.6251220703125 0 2071179
1051.6324462890625 0 757877
1052.5897216796875 0 1286624.9 z 1
1053.58935546875 0 794320.06
1054.5919189453125 0 249381.58
1065.6419677734375 0 13647391 c 10
1066.644775390625 0 8479659
1067.6475830078125 0 2967690.8
1068.625732421875 0 452232.34 y 1
1069.6121826171875 0 221345.67
1070.610595703125 0 81533.766
1078.5885009765625 0 28373.371
1079.634033203125 0 45762.613
1080.614501953125 0 863787.94
1081.6180419921875 0 496311.66
1082.6168212890625 0 139855.44
1090.657958984375 0 49421.41
1095.59228515625 0 2635759.8
1096.5946044921875 0 1656684.5
1097.6099853515625 0 682926.44
1098.635009765625 0 242902.5
1099.6416015625 0 64180.867
1104.665771484375 0 88076.89
1105.6685791015625 0 55046.074
1106.6715087890625 0 39567.285
1107.65185546875 0 50004.3
1108.6195068359375 0 61362.89
1109.60595703125 0 52952.375
1112.6162109375 0 169502.3
1113.6219482421875 0 109546.43
1119.6546630859375 0 110962.2
1120.6505126953125 0 89974.305
1121.657470703125 0 50837.258
1122.6741943359375 0 616309.2
1123.66455078125 0 1293207.2
1124.6651611328125 0 770562.5
1125.6632080078125 0 237504.31
1126.6458740234375 0 53629.273
1133.6431884765625 0 88417.76
1134.6483154296875 0 56949.992
1136.632080078125 0 80813.73
1137.6397705078125 0 58596.566
1150.6697998046875 0 1216545.2
1151.6558837890625 0 12604719
1152.65771484375 0 8186145.5
1153.6600341796875 0 2858106.2
1154.6622314453125 0 218567.7
1167.673583984375 0 13346927
1168.6805419921875 0 38145990
1169.68359375 0 23109344
1170.6859130859375 0 7451690.5
1171.6893310546875 0 553315.75
1184.645751953125 0 33441.906
1200.6593017578125 0 48329.926
3044.831787109375 0 33797.96

Spectrum Details

|  |  |
| --- | --- |
| Matched peaks? Matched peaks The total absolute number of peaks matched. Additionally in brackets the total fraction of peaks matched and the total number of peaks is shown. | 57 (11.31% of 504) |
| FDR? FDR The false discovery rate estimated for this peptide. It is calculated by matching all theoretical fragments with a non-integer shift with the raw peaks for this spectrum. This is done with 40 different shifts. The resulting percentage is the average number of annotated peaks over the number of annotated peaks with the correct spectrum. | 0.42% |
| Satellite FDR? Satellite FDR See the FDR for details on its calculation. This satellite ion specific FDR only contains the satellite ions (d/w) for I/L/J positions. | 11.90% |
| PSM Score? PSM Score The PSM Score as given by Hecklib to this annotated spectrum. It is shown with three significant figures. | 570 |

#### Spectrum 4224? Spectrum 4224 The raw spectrum of this peptide as annotated by Hecklib. The fragments are coloured according to ion type (see legend). Any peaks with a star '\*' as text can be hovered over to see the full details, first the ion type second the mass shift type. By hovering over the amino acids in the peptide or ions in the legend the corresponding peaks are highlighted. By toggling the 'Unassigned' label you can turn the background (unassigned) peaks on or off in the plot. By updating the slider in the Ion legend you can update the spectrum to only show the top X% of the peaks with labels. The top X% means any peak that is within X% of the highest intensity. By dragging in the spectrum you can zoom in to a specific part of the spectrum and use 'Zoom Out' to get back to the original zoom level. The annotation of the spectrum is based on the given sequence in the peptides file and is done with different software so inconsistencies are likely. The peaks are annotated based on the given sequence, with 20 ppm tolerance.

Copy Data

##### Spectrum 4224 (TSV)

###### Preview

```
Loading example...
```

*Click on the button to copy the data to your clipboard.*

Mz MinMz MaxIntensity Max

WidthHeightPeptide font sizePeptide stroke widthSpectrum font sizeSpectrum stroke widthCompact peptide

Ion legend

wxyz

abcd

OtherUnassignedIon^Charge^_Position_Show for top:%

VLGQPKAAPSVT

01.34e+52.68e+54.01e+55.35e+5

Zoom Out

y^+1^_1_y^+1^_2_y^+1^_2_c^+1^_3_y^+1^_3_c^+2^_8_y^+1^_4_y^+2^_8_c^+1^_4_y^+1^_4_c^+2^_9_y^+2^_9_y^+2^_10_y^+2^_10_y^+1^_5_c^+2^_10_y^+2^_10_c^+2^_11_y^+2^_11_c^+1^_6_c^+1^_6_z^+1^_7_y^+1^_7_c^+1^_7_c^+1^_7_z^+1^_8_y^+1^_8_c^+1^_8_y^+1^_8_w^+1^_9_c^+1^_9_c^+1^_9_y^+1^_9_z^+1^_9_y^+1^_9_y^+1^_10_z^+1^_10_c^+1^_10_y^+1^_10_c^+1^_10_w^+1^_11_c^+1^_11_z^+1^_11_c^+1^_11_

0738147622142952

Fragment Matches Table

Show background peaks

| Position | Ion type | Intensity | mz Theoretical | mz Error (Th) | mz Error (ppm) | Charge | Series Number |
| --- | --- | --- | --- | --- | --- | --- | --- |
| 12 | y | 3.044E+04 | 120.1 | 0.0003602 | 3 | +1 | 1 |
| - | - | 943.6 | 121.1 | - | - | 0 | - |
| - | - | 379.9 | 122.9 | - | - | 0 | - |
| - | - | 437.2 | 126.4 | - | - | 0 | - |
| - | - | 446.7 | 129.1 | - | - | 0 | - |
| - | - | 1087 | 149 | - | - | 0 | - |
| - | - | 454 | 149.9 | - | - | 0 | - |
| - | - | 434.1 | 151.5 | - | - | 0 | - |
| - | - | 557.5 | 158.5 | - | - | 0 | - |
| - | - | 750.2 | 167.1 | - | - | 0 | - |
| - | - | 660.1 | 169.1 | - | - | 0 | - |
| - | - | 574.6 | 171.1 | - | - | 0 | - |
| - | - | 956.2 | 173.4 | - | - | 0 | - |
| - | - | 1852 | 185.1 | - | - | 0 | - |
| - | - | 3.816E+04 | 185.2 | - | - | 0 | - |
| - | - | 5160 | 186.1 | - | - | 0 | - |
| - | - | 3854 | 186.2 | - | - | 0 | - |
| - | - | 1177 | 187.1 | - | - | 0 | - |
| 11 | y | 2802 | 201.1 | 0.0003342 | 1.661 | +1 | 2 |
| - | - | 671.1 | 207.1 | - | - | 0 | - |
| - | - | 4.796E+04 | 213.2 | - | - | 0 | - |
| - | - | 5239 | 214.2 | - | - | 0 | - |
| - | - | 2247 | 215.1 | - | - | 0 | - |
| - | - | 799.6 | 217.1 | - | - | 0 | - |
| 11 | y | 1.676E+04 | 219.1 | 0.0004048 | 1.847 | +1 | 2 |
| - | - | 1861 | 226.2 | - | - | 0 | - |
| - | - | 755.6 | 256.2 | - | - | 0 | - |
| - | - | 3881 | 266.2 | - | - | 0 | - |
| - | - | 4666 | 270.2 | - | - | 0 | - |
| - | - | 3626 | 284.2 | - | - | 0 | - |
| - | - | 1824 | 285.2 | - | - | 0 | - |
| 3 | c | 9753 | 287.2 | 0.0006374 | 2.219 | +1 | 3 |
| - | - | 5379 | 299.2 | - | - | 0 | - |
| 10 | y | 927.8 | 306.2 | 0.0007556 | 2.468 | +1 | 3 |
| - | - | 2016 | 314.2 | - | - | 0 | - |
| - | - | 2164 | 327.2 | - | - | 0 | - |
| - | - | 6244 | 368.2 | - | - | 0 | - |
| - | - | 755.4 | 369.2 | - | - | 0 | - |
| - | - | 6000 | 370.2 | - | - | 0 | - |
| - | - | 7279 | 371.1 | - | - | 0 | - |
| - | - | 699.7 | 371.2 | - | - | 0 | - |
| 8 | c | 858.7 | 383.2 | 0.003404 | 8.881 | +2 | 8 |
| 9 | y | 1562 | 385.2 | 0.000335 | 0.8696 | +1 | 4 |
| 5 | y | 1106 | 385.7 | 0.0008338 | 2.162 | +2 | 8 |
| 4 | c | 2.944E+04 | 398.2 | 0.0007134 | 1.792 | +1 | 4 |
| - | - | 5333 | 399.2 | - | - | 0 | - |
| - | - | 715.1 | 400.2 | - | - | 0 | - |
| 9 | y | 2.519E+04 | 403.2 | 0.0006956 | 1.725 | +1 | 4 |
| - | - | 4628 | 404.2 | - | - | 0 | - |
| - | - | 1235 | 409.7 | - | - | 0 | - |
| - | - | 897.4 | 418.7 | - | - | 0 | - |
| - | - | 916.8 | 428.3 | - | - | 0 | - |
| 9 | c | 4862 | 431.8 | 0.0006405 | 1.483 | +2 | 9 |
| - | - | 3550 | 432.3 | - | - | 0 | - |
| - | - | 775.7 | 432.8 | - | - | 0 | - |
| 4 | y | 1245 | 440.7 | 0.0005177 | 1.175 | +2 | 9 |
| - | - | 601.4 | 451.8 | - | - | 0 | - |
| - | - | 845.9 | 452.3 | - | - | 0 | - |
| - | - | 1672 | 452.8 | - | - | 0 | - |
| - | - | 750.3 | 453.3 | - | - | 0 | - |
| - | - | 859.1 | 453.8 | - | - | 0 | - |
| - | - | 2533 | 460.3 | - | - | 0 | - |
| - | - | 1095 | 460.8 | - | - | 0 | - |
| - | - | 2606 | 461.3 | - | - | 0 | - |
| - | - | 1947 | 461.8 | - | - | 0 | - |
| - | - | 1098 | 465.3 | - | - | 0 | - |
| - | - | 2.702E+04 | 466.3 | - | - | 0 | - |
| - | - | 1.5E+04 | 466.8 | - | - | 0 | - |
| - | - | 3077 | 467.3 | - | - | 0 | - |
| - | - | 869.4 | 468.3 | - | - | 0 | - |
| 3 | y | 1310 | 469.3 | 0.003399 | 7.243 | +2 | 10 |
| - | - | 2994 | 469.3 | - | - | 0 | - |
| 3 | y | 1058 | 469.8 | 0.008919 | 18.99 | +2 | 10 |
| - | - | 733.9 | 469.8 | - | - | 0 | - |
| 8 | y | 2400 | 474.3 | 0.0004776 | 1.007 | +1 | 5 |
| 10 | c | 1.79E+04 | 475.3 | 0.0008311 | 1.749 | +2 | 10 |
| - | - | 1.032E+04 | 475.8 | - | - | 0 | - |
| - | - | 3599 | 476.3 | - | - | 0 | - |
| 3 | y | 1317 | 478.3 | 0.001779 | 3.719 | +2 | 10 |
| - | - | 909.9 | 480.3 | - | - | 0 | - |
| - | - | 1282 | 489.3 | - | - | 0 | - |
| - | - | 986.5 | 495.3 | - | - | 0 | - |
| - | - | 1614 | 496.3 | - | - | 0 | - |
| - | - | 8640 | 501.8 | - | - | 0 | - |
| - | - | 5578 | 502.3 | - | - | 0 | - |
| - | - | 1971 | 502.8 | - | - | 0 | - |
| - | - | 1643 | 507.3 | - | - | 0 | - |
| - | - | 1043 | 507.8 | - | - | 0 | - |
| - | - | 1.898E+04 | 510.8 | - | - | 0 | - |
| - | - | 1.248E+04 | 511.3 | - | - | 0 | - |
| - | - | 3800 | 511.8 | - | - | 0 | - |
| - | - | 787.6 | 515.3 | - | - | 0 | - |
| - | - | 2.757E+04 | 515.8 | - | - | 0 | - |
| - | - | 1.952E+04 | 516.3 | - | - | 0 | - |
| - | - | 5038 | 516.8 | - | - | 0 | - |
| 11 | c | 1.832E+04 | 524.8 | 0.0007733 | 1.474 | +2 | 11 |
| - | - | 1.018E+04 | 525.3 | - | - | 0 | - |
| - | - | 4019 | 525.8 | - | - | 0 | - |
| - | - | 1561 | 534.3 | - | - | 0 | - |
| 2 | y | 2314 | 534.8 | 0.0004876 | 0.9117 | +2 | 11 |
| - | - | 1749 | 535.3 | - | - | 0 | - |
| - | - | 610.3 | 549.8 | - | - | 0 | - |
| - | - | 5022 | 552.3 | - | - | 0 | - |
| - | - | 5271 | 553.3 | - | - | 0 | - |
| - | - | 1678 | 554.3 | - | - | 0 | - |
| - | - | 1.195E+04 | 556.3 | - | - | 0 | - |
| - | - | 1379 | 557.3 | - | - | 0 | - |
| - | - | 4188 | 557.3 | - | - | 0 | - |
| - | - | 728.3 | 566.3 | - | - | 0 | - |
| - | - | 1997 | 566.3 | - | - | 0 | - |
| - | - | 1225 | 566.8 | - | - | 0 | - |
| - | - | 2.085E+04 | 575.3 | - | - | 0 | - |
| - | - | 1.303E+04 | 575.8 | - | - | 0 | - |
| - | - | 5157 | 576.3 | - | - | 0 | - |
| - | - | 1473 | 579.4 | - | - | 0 | - |
| - | - | 1749 | 583.3 | - | - | 0 | - |
| - | - | 1.459E+04 | 584.3 | - | - | 0 | - |
| - | - | 9735 | 584.8 | - | - | 0 | - |
| - | - | 5962 | 585.2 | - | - | 0 | - |
| - | - | 4953 | 585.3 | - | - | 0 | - |
| - | - | 2198 | 585.3 | - | - | 0 | - |
| - | - | 2406 | 585.4 | - | - | 0 | - |
| - | - | 1007 | 595.4 | - | - | 0 | - |
| - | - | 2155 | 596.4 | - | - | 0 | - |
| - | - | 1330 | 597.4 | - | - | 0 | - |
| - | - | 1114 | 600.8 | - | - | 0 | - |
| - | - | 1007 | 608.4 | - | - | 0 | - |
| - | - | 614.9 | 618.1 | - | - | 0 | - |
| 6 | c | 4265 | 623.4 | 0.0006001 | 0.9626 | +1 | 6 |
| - | - | 1620 | 624.4 | - | - | 0 | - |
| - | - | 4908 | 633.4 | - | - | 0 | - |
| - | - | 1927 | 634.4 | - | - | 0 | - |
| - | - | 2.052E+04 | 639.4 | - | - | 0 | - |
| 6 | c | 1.115E+05 | 640.4 | 0.0004792 | 0.7483 | +1 | 6 |
| - | - | 3.951E+04 | 641.4 | - | - | 0 | - |
| - | - | 5739 | 642.4 | - | - | 0 | - |
| - | - | 1.405E+04 | 651.4 | - | - | 0 | - |
| - | - | 3288 | 651.9 | - | - | 0 | - |
| - | - | 5213 | 652.4 | - | - | 0 | - |
| - | - | 1121 | 653.4 | - | - | 0 | - |
| 6 | z | 2.364E+04 | 657.4 | 0.0005587 | 0.8498 | +1 | 7 |
| - | - | 2.411E+04 | 658.4 | - | - | 0 | - |
| - | - | 6630 | 659.4 | - | - | 0 | - |
| - | - | 1243 | 660.4 | - | - | 0 | - |
| - | - | 963.6 | 663.4 | - | - | 0 | - |
| - | - | 821.1 | 664.4 | - | - | 0 | - |
| - | - | 1450 | 666.4 | - | - | 0 | - |
| - | - | 7190 | 667.4 | - | - | 0 | - |
| - | - | 2973 | 668.4 | - | - | 0 | - |
| - | - | 2378 | 672.4 | - | - | 0 | - |
| 6 | y | 2.516E+04 | 673.4 | 0.0007555 | 1.122 | +1 | 7 |
| - | - | 9498 | 674.4 | - | - | 0 | - |
| - | - | 2003 | 675.4 | - | - | 0 | - |
| 7 | c | 2541 | 694.4 | 0.0003515 | 0.5062 | +1 | 7 |
| - | - | 1146 | 695.4 | - | - | 0 | - |
| - | - | 859.3 | 703.3 | - | - | 0 | - |
| - | - | 9417 | 710.4 | - | - | 0 | - |
| 7 | c | 7.654E+04 | 711.5 | 0.0003527 | 0.4958 | +1 | 7 |
| - | - | 3.059E+04 | 712.5 | - | - | 0 | - |
| - | - | 5475 | 713.5 | - | - | 0 | - |
| - | - | 2241 | 719.4 | - | - | 0 | - |
| - | - | 1185 | 720.4 | - | - | 0 | - |
| - | - | 797.3 | 732.4 | - | - | 0 | - |
| 5 | z | 2419 | 737.4 | 0.001242 | 1.684 | +1 | 8 |
| - | - | 1179 | 737.5 | - | - | 0 | - |
| - | - | 1193 | 738.4 | - | - | 0 | - |
| - | - | 2.918E+04 | 738.5 | - | - | 0 | - |
| - | - | 2.362E+04 | 739.5 | - | - | 0 | - |
| - | - | 7149 | 740.5 | - | - | 0 | - |
| - | - | 1211 | 741.5 | - | - | 0 | - |
| - | - | 1381 | 744.4 | - | - | 0 | - |
| - | - | 1902 | 745.4 | - | - | 0 | - |
| - | - | 743.4 | 746.4 | - | - | 0 | - |
| - | - | 1231 | 750.5 | - | - | 0 | - |
| 5 | y | 3764 | 752.4 | 0.0001839 | 0.2444 | +1 | 8 |
| - | - | 1471 | 753.4 | - | - | 0 | - |
| 8 | c | 3.849E+04 | 765.5 | 0.0005302 | 0.6927 | +1 | 8 |
| - | - | 1.618E+04 | 766.5 | - | - | 0 | - |
| - | - | 3349 | 767.5 | - | - | 0 | - |
| - | - | 8244 | 769.4 | - | - | 0 | - |
| 5 | y | 1.157E+05 | 770.4 | 0.000787 | 1.022 | +1 | 8 |
| - | - | 4.285E+04 | 771.4 | - | - | 0 | - |
| - | - | 1.144E+04 | 772.4 | - | - | 0 | - |
| - | - | 1810 | 781.5 | - | - | 0 | - |
| - | - | 3096 | 786.9 | - | - | 0 | - |
| - | - | 2026 | 787.4 | - | - | 0 | - |
| - | - | 1325 | 787.9 | - | - | 0 | - |
| - | - | 989.5 | 793.3 | - | - | 0 | - |
| - | - | 840.5 | 793.4 | - | - | 0 | - |
| - | - | 1081 | 794.5 | - | - | 0 | - |
| - | - | 1091 | 797.4 | - | - | 0 | - |
| - | - | 5563 | 798.4 | - | - | 0 | - |
| - | - | 2559 | 799.4 | - | - | 0 | - |
| - | - | 1022 | 806.5 | - | - | 0 | - |
| - | - | 8183 | 807.5 | - | - | 0 | - |
| - | - | 3558 | 808.5 | - | - | 0 | - |
| - | - | 1047 | 809.5 | - | - | 0 | - |
| - | - | 3399 | 810.5 | - | - | 0 | - |
| - | - | 1359 | 811.5 | - | - | 0 | - |
| - | - | 828.8 | 816.4 | - | - | 0 | - |
| - | - | 3978 | 818.5 | - | - | 0 | - |
| - | - | 1983 | 819.5 | - | - | 0 | - |
| - | - | 1442 | 820.5 | - | - | 0 | - |
| - | - | 8634 | 822.5 | - | - | 0 | - |
| - | - | 3093 | 823.5 | - | - | 0 | - |
| 4 | w | 4.104E+04 | 824.5 | 0.001148 | 1.392 | +1 | 9 |
| - | - | 1.908E+04 | 825.5 | - | - | 0 | - |
| - | - | 4403 | 826.5 | - | - | 0 | - |
| - | - | 1710 | 827.5 | - | - | 0 | - |
| - | - | 5539 | 835.5 | - | - | 0 | - |
| - | - | 6450 | 836.5 | - | - | 0 | - |
| - | - | 2667 | 836.5 | - | - | 0 | - |
| - | - | 3082 | 837.5 | - | - | 0 | - |
| - | - | 1145 | 837.5 | - | - | 0 | - |
| - | - | 2034 | 838.5 | - | - | 0 | - |
| - | - | 1588 | 848.5 | - | - | 0 | - |
| - | - | 1337 | 850.5 | - | - | 0 | - |
| - | - | 930.6 | 858.9 | - | - | 0 | - |
| 9 | c | 1653 | 862.5 | 0.003797 | 4.402 | +1 | 9 |
| - | - | 3954 | 863.5 | - | - | 0 | - |
| - | - | 1378 | 864.4 | - | - | 0 | - |
| - | - | 3988 | 864.5 | - | - | 0 | - |
| - | - | 1460 | 865.5 | - | - | 0 | - |
| - | - | 831.6 | 868.1 | - | - | 0 | - |
| - | - | 1048 | 868.5 | - | - | 0 | - |
| - | - | 1059 | 874.4 | - | - | 0 | - |
| - | - | 2239 | 877.4 | - | - | 0 | - |
| - | - | 2798 | 877.5 | - | - | 0 | - |
| - | - | 1.26E+05 | 878.5 | - | - | 0 | - |
| 9 | c | 1.576E+05 | 879.5 | 0.001573 | 1.789 | +1 | 9 |
| - | - | 6.134E+04 | 880.5 | - | - | 0 | - |
| 4 | y | 5827 | 881.5 | 0.002433 | 2.76 | +1 | 9 |
| - | - | 1.39E+04 | 881.5 | - | - | 0 | - |
| 4 | z | 6.698E+04 | 882.5 | 0.0001792 | 0.2031 | +1 | 9 |
| - | - | 3.117E+04 | 883.5 | - | - | 0 | - |
| - | - | 8871 | 884.5 | - | - | 0 | - |
| - | - | 1085 | 885.5 | - | - | 0 | - |
| - | - | 1.161E+04 | 897.5 | - | - | 0 | - |
| 4 | y | 7.868E+04 | 898.5 | 0.000254 | 0.2827 | +1 | 9 |
| - | - | 3.784E+04 | 899.5 | - | - | 0 | - |
| - | - | 9230 | 900.5 | - | - | 0 | - |
| - | - | 1180 | 901.5 | - | - | 0 | - |
| - | - | 1377 | 906.5 | - | - | 0 | - |
| - | - | 829 | 911.6 | - | - | 0 | - |
| - | - | 899 | 918.5 | - | - | 0 | - |
| - | - | 1339 | 922.4 | - | - | 0 | - |
| - | - | 1686 | 926.5 | - | - | 0 | - |
| - | - | 3.725E+04 | 931.5 | - | - | 0 | - |
| - | - | 2.031E+04 | 932.5 | - | - | 0 | - |
| - | - | 6424 | 933.5 | - | - | 0 | - |
| - | - | 1140 | 934.5 | - | - | 0 | - |
| - | - | 3200 | 936.4 | - | - | 0 | - |
| - | - | 1373 | 937.4 | - | - | 0 | - |
| 3 | y | 1905 | 937.5 | 0.003271 | 3.489 | +1 | 10 |
| - | - | 5357 | 938.5 | - | - | 0 | - |
| 3 | z | 1.713E+04 | 939.5 | 0.001848 | 1.967 | +1 | 10 |
| - | - | 9468 | 940.5 | - | - | 0 | - |
| - | - | 2682 | 941.5 | - | - | 0 | - |
| 10 | c | 5.911E+04 | 949.5 | 0.0006379 | 0.6718 | +1 | 10 |
| - | - | 2.995E+04 | 950.5 | - | - | 0 | - |
| - | - | 1.089E+04 | 951.6 | - | - | 0 | - |
| - | - | 2005 | 952.6 | - | - | 0 | - |
| 3 | y | 6.16E+04 | 955.5 | 3.048E-05 | 0.0319 | +1 | 10 |
| - | - | 2.787E+04 | 956.5 | - | - | 0 | - |
| - | - | 9550 | 957.5 | - | - | 0 | - |
| - | - | 1306 | 958.5 | - | - | 0 | - |
| - | - | 1318 | 958.8 | - | - | 0 | - |
| 10 | c | 9.552E+04 | 966.6 | 0.0003339 | 0.3454 | +1 | 10 |
| - | - | 5.246E+04 | 967.6 | - | - | 0 | - |
| - | - | 4312 | 967.8 | - | - | 0 | - |
| - | - | 3051 | 968.2 | - | - | 0 | - |
| - | - | 1687 | 968.5 | - | - | 0 | - |
| - | - | 1.428E+04 | 968.6 | - | - | 0 | - |
| - | - | 1810 | 969.6 | - | - | 0 | - |
| - | - | 2083 | 973.2 | - | - | 0 | - |
| - | - | 3577 | 973.5 | - | - | 0 | - |
| - | - | 4035 | 973.8 | - | - | 0 | - |
| - | - | 1626 | 974.2 | - | - | 0 | - |
| - | - | 821.4 | 974.5 | - | - | 0 | - |
| - | - | 1417 | 977.5 | - | - | 0 | - |
| - | - | 857.2 | 978.5 | - | - | 0 | - |
| - | - | 1413 | 982.5 | - | - | 0 | - |
| - | - | 1045 | 991 | - | - | 0 | - |
| - | - | 1748 | 991.5 | - | - | 0 | - |
| - | - | 1087 | 992 | - | - | 0 | - |
| - | - | 3842 | 1007 | - | - | 0 | - |
| - | - | 1813 | 1008 | - | - | 0 | - |
| 2 | w | 5294 | 1010 | 0.001185 | 1.173 | +1 | 11 |
| - | - | 3008 | 1011 | - | - | 0 | - |
| - | - | 934.5 | 1012 | - | - | 0 | - |
| - | - | 1548 | 1021 | - | - | 0 | - |
| - | - | 7207 | 1022 | - | - | 0 | - |
| - | - | 3425 | 1023 | - | - | 0 | - |
| - | - | 2021 | 1024 | - | - | 0 | - |
| - | - | 952.2 | 1025 | - | - | 0 | - |
| - | - | 2626 | 1031 | - | - | 0 | - |
| - | - | 2377 | 1032 | - | - | 0 | - |
| - | - | 1366 | 1038 | - | - | 0 | - |
| - | - | 1234 | 1039 | - | - | 0 | - |
| - | - | 883.7 | 1040 | - | - | 0 | - |
| 11 | c | 3.196E+04 | 1049 | 0.0002101 | 0.2004 | +1 | 11 |
| - | - | 1.868E+04 | 1050 | - | - | 0 | - |
| - | - | 2.181E+04 | 1051 | - | - | 0 | - |
| - | - | 8810 | 1052 | - | - | 0 | - |
| 2 | z | 1.817E+04 | 1053 | 0.002806 | 2.666 | +1 | 11 |
| - | - | 1.089E+04 | 1054 | - | - | 0 | - |
| - | - | 900.5 | 1054 | - | - | 0 | - |
| - | - | 3226 | 1055 | - | - | 0 | - |
| - | - | 1254 | 1056 | - | - | 0 | - |
| - | - | 1539 | 1056 | - | - | 0 | - |
| - | - | 913.9 | 1057 | - | - | 0 | - |
| 11 | c | 1.817E+05 | 1066 | 0.0005845 | 0.5485 | +1 | 11 |
| - | - | 1.114E+05 | 1067 | - | - | 0 | - |
| - | - | 4.413E+04 | 1068 | - | - | 0 | - |
| - | - | 7817 | 1069 | - | - | 0 | - |
| - | - | 1348 | 1079 | - | - | 0 | - |
| - | - | 1.01E+04 | 1081 | - | - | 0 | - |
| - | - | 7568 | 1082 | - | - | 0 | - |
| - | - | 2346 | 1083 | - | - | 0 | - |
| - | - | 3.334E+04 | 1096 | - | - | 0 | - |
| - | - | 2.152E+04 | 1097 | - | - | 0 | - |
| - | - | 9118 | 1098 | - | - | 0 | - |
| - | - | 3587 | 1099 | - | - | 0 | - |
| - | - | 1163 | 1100 | - | - | 0 | - |
| - | - | 710.7 | 1105 | - | - | 0 | - |
| - | - | 1066 | 1105 | - | - | 0 | - |
| - | - | 1002 | 1109 | - | - | 0 | - |
| - | - | 832.9 | 1110 | - | - | 0 | - |
| - | - | 771.8 | 1111 | - | - | 0 | - |
| - | - | 2337 | 1113 | - | - | 0 | - |
| - | - | 1286 | 1114 | - | - | 0 | - |
| - | - | 1601 | 1120 | - | - | 0 | - |
| - | - | 1168 | 1123 | - | - | 0 | - |
| - | - | 7850 | 1123 | - | - | 0 | - |
| - | - | 1.723E+04 | 1124 | - | - | 0 | - |
| - | - | 1.071E+04 | 1125 | - | - | 0 | - |
| - | - | 4000 | 1126 | - | - | 0 | - |
| - | - | 894 | 1134 | - | - | 0 | - |
| - | - | 938.2 | 1137 | - | - | 0 | - |
| - | - | 1656 | 1140 | - | - | 0 | - |
| - | - | 1005 | 1141 | - | - | 0 | - |
| - | - | 1.579E+04 | 1151 | - | - | 0 | - |
| - | - | 1.7E+05 | 1152 | - | - | 0 | - |
| - | - | 1.128E+05 | 1153 | - | - | 0 | - |
| - | - | 4.097E+04 | 1154 | - | - | 0 | - |
| - | - | 5864 | 1155 | - | - | 0 | - |
| - | - | 1.873E+05 | 1168 | - | - | 0 | - |
| - | - | 5.298E+05 | 1169 | - | - | 0 | - |
| - | - | 3.11E+05 | 1170 | - | - | 0 | - |
| - | - | 1.042E+05 | 1171 | - | - | 0 | - |
| - | - | 890.7 | 1172 | - | - | 0 | - |
| - | - | 1.366E+04 | 1172 | - | - | 0 | - |
| - | - | 1147 | 1180 | - | - | 0 | - |
| - | - | 2481 | 1201 | - | - | 0 | - |
| - | - | 1694 | 1202 | - | - | 0 | - |
| - | - | 1501 | 1223 | - | - | 0 | - |
| - | - | 1128 | 1224 | - | - | 0 | - |
| - | - | 961.1 | 1245 | - | - | 0 | - |
| - | - | 939.8 | 1246 | - | - | 0 | - |
| - | - | 791.7 | 1247 | - | - | 0 | - |
| - | - | 1042 | 1280 | - | - | 0 | - |
| - | - | 2128 | 1286 | - | - | 0 | - |
| - | - | 1155 | 1287 | - | - | 0 | - |
| - | - | 912.6 | 1293 | - | - | 0 | - |
| - | - | 2413 | 1301 | - | - | 0 | - |
| - | - | 5645 | 1302 | - | - | 0 | - |
| - | - | 1902 | 1302 | - | - | 0 | - |
| - | - | 5310 | 1303 | - | - | 0 | - |
| - | - | 2771 | 1304 | - | - | 0 | - |
| - | - | 2587 | 1358 | - | - | 0 | - |
| - | - | 2266 | 1359 | - | - | 0 | - |
| - | - | 1241 | 1359 | - | - | 0 | - |
| - | - | 1079 | 1360 | - | - | 0 | - |
| - | - | 1690 | 1360 | - | - | 0 | - |
| - | - | 989.2 | 1366 | - | - | 0 | - |
| - | - | 935.6 | 1367 | - | - | 0 | - |
| - | - | 897.5 | 1408 | - | - | 0 | - |
| - | - | 1108 | 1416 | - | - | 0 | - |
| - | - | 975.2 | 1416 | - | - | 0 | - |
| - | - | 1644 | 1429 | - | - | 0 | - |
| - | - | 2182 | 1429 | - | - | 0 | - |
| - | - | 1965 | 1430 | - | - | 0 | - |
| - | - | 2208 | 1430 | - | - | 0 | - |
| - | - | 2622 | 1431 | - | - | 0 | - |
| - | - | 1732 | 1431 | - | - | 0 | - |
| - | - | 979.1 | 1432 | - | - | 0 | - |
| - | - | 1108 | 1437 | - | - | 0 | - |
| - | - | 2865 | 1438 | - | - | 0 | - |
| - | - | 2748 | 1438 | - | - | 0 | - |
| - | - | 2584 | 1439 | - | - | 0 | - |
| - | - | 1114 | 1439 | - | - | 0 | - |
| - | - | 1679 | 1443 | - | - | 0 | - |
| - | - | 1985 | 1443 | - | - | 0 | - |
| - | - | 2749 | 1451 | - | - | 0 | - |
| - | - | 5521 | 1451 | - | - | 0 | - |
| - | - | 9310 | 1452 | - | - | 0 | - |
| - | - | 7177 | 1452 | - | - | 0 | - |
| - | - | 3517 | 1453 | - | - | 0 | - |
| - | - | 1624 | 1453 | - | - | 0 | - |
| - | - | 2161 | 1459 | - | - | 0 | - |
| - | - | 8379 | 1460 | - | - | 0 | - |
| - | - | 7851 | 1460 | - | - | 0 | - |
| - | - | 6510 | 1461 | - | - | 0 | - |
| - | - | 3384 | 1461 | - | - | 0 | - |
| - | - | 1634 | 1462 | - | - | 0 | - |
| - | - | 1047 | 1616 | - | - | 0 | - |
| - | - | 8055 | 1617 | - | - | 0 | - |
| - | - | 5412 | 1618 | - | - | 0 | - |
| - | - | 3399 | 1619 | - | - | 0 | - |
| - | - | 1143 | 1620 | - | - | 0 | - |
| - | - | 1234 | 1697 | - | - | 0 | - |
| - | - | 1098 | 1698 | - | - | 0 | - |
| - | - | 1429 | 1718 | - | - | 0 | - |
| - | - | 1282 | 1719 | - | - | 0 | - |
| - | - | 1077 | 1854 | - | - | 0 | - |
| - | - | 1051 | 1881 | - | - | 0 | - |
| - | - | 1340 | 1882 | - | - | 0 | - |
| - | - | 1038 | 1981 | - | - | 0 | - |
| - | - | 1692 | 1983 | - | - | 0 | - |
| - | - | 1410 | 1984 | - | - | 0 | - |
| - | - | 1249 | 2109 | - | - | 0 | - |
| - | - | 1054 | 2112 | - | - | 0 | - |
| - | - | 971.2 | 2113 | - | - | 0 | - |
| - | - | 903.8 | 2334 | - | - | 0 | - |
| - | - | 1736 | 2335 | - | - | 0 | - |
| - | - | 1434 | 2336 | - | - | 0 | - |
| - | - | 1320 | 2716 | - | - | 0 | - |
| - | - | 889.7 | 2874 | - | - | 0 | - |
| - | - | 2639 | 2902 | - | - | 0 | - |
| - | - | 2313 | 2903 | - | - | 0 | - |
| - | - | 1496 | 2904 | - | - | 0 | - |
| - | - | 1286 | 2918 | - | - | 0 | - |
| - | - | 2051 | 2919 | - | - | 0 | - |
| - | - | 3257 | 2920 | - | - | 0 | - |
| - | - | 2275 | 2921 | - | - | 0 | - |
| - | - | 1218 | 2923 | - | - | 0 | - |

m/z Charge Intensity FragmentType MassShift Position
120.06587982177734 0 30435.71 y 11
121.0694808959961 0 943.56396
122.86225128173828 0 379.87177
126.36900329589844 0 437.19147
129.10247802734375 0 446.7497
148.954345703125 0 1087.0524
149.9272003173828 0 454.00064
151.5273895263672 0 434.08505
158.5255889892578 0 557.45435
167.0818634033203 0 750.2369
169.0615692138672 0 660.0922
171.1132354736328 0 574.6274
173.4384765625 0 956.23444
185.09271240234375 0 1852.2211
185.16519165039062 0 38158.18
186.08773803710938 0 5159.6323
186.168701171875 0 3854.1028
187.10809326171875 0 1177.1119
201.1237030029297 0 2802.2922 y Water loss 10
207.0982208251953 0 671.06006
213.16017150878906 0 47959.54
214.16336059570312 0 5239.467
215.13938903808594 0 2246.825
217.11898803710938 0 799.5674
219.13433837890625 0 16763.719 y 10
226.15536499023438 0 1860.5035
256.16534423828125 0 755.6075
266.1502990722656 0 3880.6687
270.1815490722656 0 4666.38
284.16094970703125 0 3625.9338
285.156982421875 0 1824.4673
287.2084045410156 0 9752.871 c 2
299.1717224121094 0 5379.2065
306.1667175292969 0 927.7703 y 9
314.20794677734375 0 2016.2408
327.1674499511719 0 2164.0583
368.2296447753906 0 6243.5864
369.2325439453125 0 755.4404
370.2450256347656 0 6000.1025
371.0769958496094 0 7278.8125
371.24700927734375 0 699.6594
383.2311096191406 0 858.7471 c Ammonia loss 7
385.20849609375 0 1562.0657 y Water loss 8
385.72314453125 0 1106.4043 y 4
398.2405090332031 0 29440.158 c Ammonia loss 3
399.24383544921875 0 5333.184
400.2449951171875 0 715.13696
403.21942138671875 0 25193.182 y 8
404.2221984863281 0 4628.3804
409.73101806640625 0 1235.1855
418.7346496582031 0 897.403
428.26324462890625 0 916.7996
431.76153564453125 0 4862.348 c Ammonia loss 8
432.2653503417969 0 3550.423
432.7652587890625 0 775.68414
440.7474670410156 0 1244.9326 y Water loss 3
451.77447509765625 0 601.4431
452.2769775390625 0 845.9232
452.76885986328125 0 1672.1089
453.2708740234375 0 750.2556
453.7749328613281 0 859.06757
460.2900695800781 0 2532.5881
460.7876281738281 0 1094.8566
461.2815246582031 0 2606.1926
461.7813720703125 0 1947.0568
465.28179931640625 0 1098.4392
466.2725524902344 0 27015.404
466.7736511230469 0 15003.111
467.27679443359375 0 3076.65
468.2720031738281 0 869.3694
469.2621154785156 0 1310.3369 y Water loss 2
469.2965087890625 0 2994.134
469.7596435546875 0 1058.3579 y Ammonia loss 2
469.7969970703125 0 733.9136
474.2563171386719 0 2400.3396 y 7
475.2777404785156 0 17900.307 c Ammonia loss 9
475.779052734375 0 10318.136
476.2808532714844 0 3598.599
478.2657775878906 0 1317.4995 y 2
480.28857421875 0 909.87164
489.2920227050781 0 1281.61
495.31988525390625 0 986.50073
496.32403564453125 0 1613.796
501.8090515136719 0 8639.654
502.3097229003906 0 5578.1953
502.81317138671875 0 1971.3282
507.29736328125 0 1642.6138
507.7907409667969 0 1043.108
510.8144836425781 0 18975.312
511.3160095214844 0 12482.387
511.81707763671875 0 3799.9326
515.2822875976562 0 787.55237
515.8067626953125 0 27569.455
516.307861328125 0 19522.629
516.8090209960938 0 5037.796
524.8118896484375 0 18319.193 c Ammonia loss 10
525.3134155273438 0 10183.189
525.8125610351562 0 4018.6736
534.3047485351562 0 1561.385
534.8065185546875 0 2313.5037 y 1
535.30615234375 0 1748.8579
549.7825927734375 0 610.2873
552.3148803710938 0 5022.26
553.3124389648438 0 5270.558
554.3121337890625 0 1678.2815
556.3460693359375 0 11946.622
557.2904663085938 0 1378.6293
557.3485107421875 0 4188.4673
566.2767333984375 0 728.3253
566.3299560546875 0 1996.6086
566.8292236328125 0 1224.5789
575.3353271484375 0 20848.025
575.8372192382812 0 13028.5
576.338134765625 0 5157.1987
579.3970336914062 0 1472.8206
583.3228149414062 0 1748.9897
584.3438720703125 0 14592.511
584.8424682617188 0 9735.146
585.2091064453125 0 5961.5044
585.2830810546875 0 4953.031
585.3417358398438 0 2198.0728
585.3671875 0 2406.2888
595.3619384765625 0 1007.48
596.3995971679688 0 2155.4524
597.4026489257812 0 1329.5155
600.82958984375 0 1113.613
608.3633422851562 0 1007.47015
618.1279907226562 0 614.90204
623.3881225585938 0 4264.6895 c Ammonia loss 5
624.3916015625 0 1619.8069
633.3720703125 0 4908.37
634.3760375976562 0 1927.1091
639.406982421875 0 20521.328
640.41455078125 0 111544.7 c 5
641.4176635742188 0 39513.133
642.4207153320312 0 5738.724
651.3820190429688 0 14049.597
651.8538208007812 0 3287.5078
652.3840942382812 0 5213.3193
653.3867797851562 0 1120.8069
657.3697509765625 0 23643.145 z 5
658.3762817382812 0 24109.516
659.3798217773438 0 6630.483
660.3843994140625 0 1243.2185
663.3823852539062 0 963.5961
664.3865356445312 0 821.0848
666.3951416015625 0 1449.6083
667.438720703125 0 7189.7935
668.4410400390625 0 2973.433
672.3798217773438 0 2377.8535
673.388671875 0 25164.758 y 5
674.391357421875 0 9498.295
675.3916625976562 0 2002.6028
694.4249877929688 0 2540.5918 c Ammonia loss 6
695.4320068359375 0 1145.8191
703.3251342773438 0 859.26086
710.4439697265625 0 9416.786
711.4515380859375 0 76537.6 c 6
712.4548950195312 0 30588.396
713.4580078125 0 5474.832
719.3839721679688 0 2241.2385
720.3858642578125 0 1184.6416
732.4391479492188 0 797.3097
737.3941650390625 0 2419.0603 z Ammonia loss 4
737.4591674804688 0 1178.8418
738.4085693359375 0 1193.2192
738.4754638671875 0 29178.324
739.480712890625 0 23621.46
740.4841918945312 0 7148.8125
741.4863891601562 0 1211.0834
744.3907470703125 0 1381.4233
745.395751953125 0 1902.3867
746.3973999023438 0 743.42523
750.4505004882812 0 1231.4163
752.429931640625 0 3764.023 y Water loss 4
753.4339599609375 0 1470.9441
765.4622802734375 0 38488.81 c Ammonia loss 7
766.465576171875 0 16177.661
767.466796875 0 3349.4656
769.4337158203125 0 8243.824
770.4414672851562 0 115683.86 y 4
771.4445190429688 0 42845.504
772.4468994140625 0 11437.128
781.45703125 0 1809.9608
786.8699951171875 0 3096.082
787.3704833984375 0 2026.2734
787.87353515625 0 1325.3927
793.345703125 0 989.46545
793.44677734375 0 840.46515
794.4616088867188 0 1081.3612
797.4286499023438 0 1091.3585
798.4371948242188 0 5563.4834
799.4378051757812 0 2559.2344
806.4622192382812 0 1021.8653
807.4951782226562 0 8183.206
808.4863891601562 0 3557.7568
809.4774780273438 0 1047.1146
810.4575805664062 0 3398.9941
811.4581298828125 0 1359.0996
816.4037475585938 0 828.8297
818.4525756835938 0 3978.4675
819.4505615234375 0 1983.1981
820.5209350585938 0 1442.2131
822.4710083007812 0 8634.309
823.47314453125 0 3093.4023
824.452392578125 0 41042.6 w 3
825.4547119140625 0 19083.342
826.45654296875 0 4402.545
827.4580688476562 0 1710.3837
835.526611328125 0 5538.713
836.462158203125 0 6449.5864
836.535400390625 0 2666.5718
837.4651489257812 0 3082.2144
837.54443359375 0 1144.5934
838.4749755859375 0 2033.7014
848.4656372070312 0 1587.8835
850.4810180664062 0 1337.1909
858.9180908203125 0 930.6314
862.518310546875 0 1653.1378 c Ammonia loss 8
863.522216796875 0 3953.681
864.3836669921875 0 1378.4552
864.53076171875 0 3988.204
865.5349731445312 0 1459.9032
868.1246948242188 0 831.58997
868.4564819335938 0 1048.1255
874.4476318359375 0 1059.0194
877.409912109375 0 2238.9697
877.5169677734375 0 2798.4402
878.5335083007812 0 126002.66
879.5394897460938 0 157648.22 c 8
880.542724609375 0 61340.082
881.4702758789062 0 5827.478 y Ammonia loss 3
881.5497436523438 0 13898.538
882.480712890625 0 66979.33 z 3
883.4842529296875 0 31173.762
884.4866943359375 0 8871.402
885.4900512695312 0 1085.4614
897.4918212890625 0 11611.756
898.49951171875 0 78683.35 y 3
899.5026245117188 0 37840.684
900.5045776367188 0 9229.836
901.5089111328125 0 1180.1637
906.5424194335938 0 1377.0436
911.5591430664062 0 828.9942
918.5061645507812 0 898.95325
922.4060668945312 0 1339.1022
926.5093994140625 0 1685.7878
931.536376953125 0 37247.664
932.5386352539062 0 20314.97
933.5421752929688 0 6424.1597
934.5481567382812 0 1140.3093
936.4227294921875 0 3200.1558
937.422119140625 0 1372.8876
937.513427734375 0 1904.9062 y Water loss 2
938.513916015625 0 5357.197
939.5038452148438 0 17131.809 z 2
940.5071411132812 0 9467.805
941.5081176757812 0 2682.147
949.5471801757812 0 59105.33 c Ammonia loss 9
950.5499877929688 0 29950.379
951.5550537109375 0 10887.507
952.5650024414062 0 2004.7076
955.520751953125 0 61596.344 y 2
956.523681640625 0 27866.146
957.5264282226562 0 9550.466
958.5108642578125 0 1306.1608
958.827392578125 0 1317.6816
966.5734252929688 0 95516.7 c 9
967.575927734375 0 52463.992
967.8223266601562 0 4311.709
968.1614379882812 0 3050.704
968.4918823242188 0 1686.6172
968.5800170898438 0 14279.768
969.5762939453125 0 1810.4629
973.1644897460938 0 2082.6658
973.4971313476562 0 3576.77
973.8335571289062 0 4035.2368
974.1721801757812 0 1625.9121
974.5029907226562 0 821.3532
977.4693603515625 0 1417.2356
978.4736938476562 0 857.2413
982.5107421875 0 1412.5554
991.0069580078125 0 1045.3887
991.5352172851562 0 1747.8873
992.0463256835938 0 1087.1257
1006.6141357421875 0 3841.8008
1007.6148071289062 0 1813.2063
1009.532470703125 0 5293.807 w 1
1010.5382690429688 0 3007.51
1011.5382690429688 0 934.46045
1020.6244506835938 0 1547.5258
1021.6279907226562 0 7207.4683
1022.6307373046875 0 3424.8567
1023.6360473632812 0 2021.3923
1024.64892578125 0 952.20325
1030.604248046875 0 2626.166
1031.60302734375 0 2377.1008
1037.58349609375 0 1365.9366
1038.5843505859375 0 1233.96
1039.58544921875 0 883.6573
1048.61474609375 0 31956.602 c Ammonia loss 10
1049.6182861328125 0 18683.002
1050.6282958984375 0 21808.826
1051.6317138671875 0 8809.589
1052.5888671875 0 18169.379 z 1
1053.589599609375 0 10888.901
1054.0059814453125 0 900.5151
1054.592041015625 0 3225.689
1055.523681640625 0 1253.5054
1056.05517578125 0 1539.1545
1056.5572509765625 0 913.917
1065.64208984375 0 181650.81 c 10
1066.6448974609375 0 111374.6
1067.6474609375 0 44133.992
1068.646728515625 0 7816.5215
1078.5184326171875 0 1347.7418
1080.6165771484375 0 10104.998
1081.6209716796875 0 7568.367
1082.6270751953125 0 2346.341
1095.5921630859375 0 33340.434
1096.5950927734375 0 21518.553
1097.6092529296875 0 9118.099
1098.6268310546875 0 3587.3174
1099.6473388671875 0 1163.1526
1104.51171875 0 710.74774
1104.668212890625 0 1066.3496
1108.5982666015625 0 1001.757
1109.6090087890625 0 832.9363
1110.624267578125 0 771.76697
1112.6177978515625 0 2336.9268
1113.6204833984375 0 1286.0958
1119.6507568359375 0 1601.2253
1122.5274658203125 0 1168.0444
1122.6739501953125 0 7850.093
1123.666259765625 0 17226.213
1124.6656494140625 0 10714.031
1125.6632080078125 0 3999.788
1133.6322021484375 0 894.004
1136.630615234375 0 938.1554
1139.536865234375 0 1655.6232
1140.5528564453125 0 1004.71436
1150.6708984375 0 15785.86
1151.6556396484375 0 169993.78
1152.657958984375 0 112795.21
1153.660400390625 0 40973.957
1154.6644287109375 0 5864.3486
1167.673583984375 0 187320.64
1168.680419921875 0 529814.5
1169.68359375 0 310971.28
1170.685791015625 0 104244.26
1171.5435791015625 0 890.68317
1171.6898193359375 0 13655.515
1180.09619140625 0 1146.7992
1200.652099609375 0 2480.6697
1201.654541015625 0 1694.1609
1222.5711669921875 0 1500.5309
1223.5667724609375 0 1128.1204
1244.64208984375 0 961.10565
1246.127685546875 0 939.84094
1247.125 0 791.7006
1279.6654052734375 0 1041.8973
1285.6724853515625 0 2127.7043
1286.6763916015625 0 1155.3757
1293.1614990234375 0 912.56146
1301.168212890625 0 2413.3467
1301.6768798828125 0 5645.3823
1302.1669921875 0 1901.9353
1302.695556640625 0 5310.3784
1303.706787109375 0 2770.5718
1358.170654296875 0 2587.2156
1358.6707763671875 0 2265.6892
1359.16943359375 0 1240.9095
1359.68212890625 0 1079.254
1360.177001953125 0 1689.827
1366.2080078125 0 989.2379
1366.6973876953125 0 935.61926
1407.7115478515625 0 897.5355
1415.7313232421875 0 1108.2372
1416.22705078125 0 975.2229
1428.7265625 0 1643.7544
1429.22900390625 0 2182.2375
1429.7308349609375 0 1965.2173
1430.2254638671875 0 2208.1238
1430.7294921875 0 2621.9236
1431.2191162109375 0 1731.6378
1431.6973876953125 0 979.0979
1437.2427978515625 0 1107.5376
1437.7384033203125 0 2864.824
1438.2406005859375 0 2747.8567
1438.7386474609375 0 2583.8132
1439.22802734375 0 1113.8363
1442.720458984375 0 1679.0634
1443.2203369140625 0 1984.7572
1450.7353515625 0 2748.9263
1451.23291015625 0 5521.4546
1451.7332763671875 0 9310.381
1452.232666015625 0 7177.0522
1452.7381591796875 0 3517.2024
1453.244140625 0 1623.7874
1459.2431640625 0 2160.5583
1459.744384765625 0 8379.382
1460.24658203125 0 7850.957
1460.74609375 0 6509.7183
1461.248291015625 0 3384.2002
1461.7486572265625 0 1633.7489
1615.7659912109375 0 1046.7924
1616.791748046875 0 8055.24
1617.7919921875 0 5411.583
1618.793701171875 0 3399.269
1619.794189453125 0 1143.3452
1696.9322509765625 0 1234.3352
1697.933837890625 0 1098.1704
1717.8343505859375 0 1429.2421
1718.8299560546875 0 1282.0273
1854.036865234375 0 1076.7262
1880.8863525390625 0 1050.7717
1881.9091796875 0 1340.0806
1981.0015869140625 0 1038.248
1983.0765380859375 0 1692.3224
1984.0606689453125 0 1410.2838
2109.04541015625 0 1249.085
2112.09423828125 0 1053.513
2113.117431640625 0 971.2087
2334.126708984375 0 903.77704
2335.11474609375 0 1736.2267
2336.13525390625 0 1434.2063
2716.36669921875 0 1319.735
2874.460205078125 0 889.6578
2902.454833984375 0 2639.4932
2903.4677734375 0 2313.1091
2904.46484375 0 1496.1162
2918.472900390625 0 1286.464
2919.475830078125 0 2050.6436
2920.498291015625 0 3256.879
2921.49365234375 0 2275.386
2922.517578125 0 1217.5903

Spectrum Details

|  |  |
| --- | --- |
| Matched peaks? Matched peaks The total absolute number of peaks matched. Additionally in brackets the total fraction of peaks matched and the total number of peaks is shown. | 44 (10.09% of 436) |
| FDR? FDR The false discovery rate estimated for this peptide. It is calculated by matching all theoretical fragments with a non-integer shift with the raw peaks for this spectrum. This is done with 40 different shifts. The resulting percentage is the average number of annotated peaks over the number of annotated peaks with the correct spectrum. | 1.14% |
| Satellite FDR? Satellite FDR See the FDR for details on its calculation. This satellite ion specific FDR only contains the satellite ions (d/w) for I/L/J positions. | 11.90% |
| PSM Score? PSM Score The PSM Score as given by Hecklib to this annotated spectrum. It is shown with three significant figures. | 432 |

#### Spectrum 4298? Spectrum 4298 The raw spectrum of this peptide as annotated by Hecklib. The fragments are coloured according to ion type (see legend). Any peaks with a star '\*' as text can be hovered over to see the full details, first the ion type second the mass shift type. By hovering over the amino acids in the peptide or ions in the legend the corresponding peaks are highlighted. By toggling the 'Unassigned' label you can turn the background (unassigned) peaks on or off in the plot. By updating the slider in the Ion legend you can update the spectrum to only show the top X% of the peaks with labels. The top X% means any peak that is within X% of the highest intensity. By dragging in the spectrum you can zoom in to a specific part of the spectrum and use 'Zoom Out' to get back to the original zoom level. The annotation of the spectrum is based on the given sequence in the peptides file and is done with different software so inconsistencies are likely. The peaks are annotated based on the given sequence, with 20 ppm tolerance.

Copy Data

##### Spectrum 4298 (TSV)

###### Preview

```
Loading example...
```

*Click on the button to copy the data to your clipboard.*

Mz MinMz MaxIntensity Max

WidthHeightPeptide font sizePeptide stroke widthSpectrum font sizeSpectrum stroke widthCompact peptide

Ion legend

wxyz

abcd

OtherUnassignedIon^Charge^_Position_Show for top:%

VLGQPKAAPSVT

08.75e+41.75e+52.63e+53.50e+5

Zoom Out

y^+1^_1_y^+1^_2_y^+1^_2_c^+1^_3_c^+2^_8_y^+1^_4_y^+2^_8_c^+1^_4_y^+1^_4_c^+2^_9_y^+2^_9_y^+1^_5_c^+2^_10_c^+2^_11_y^+2^_11_c^+1^_6_c^+1^_6_z^+1^_7_y^+1^_7_c^+1^_7_c^+1^_7_z^+1^_8_z^+1^_8_y^+1^_8_c^+1^_8_y^+1^_8_w^+1^_9_c^+1^_9_c^+1^_9_y^+1^_9_z^+1^_9_y^+1^_9_z^+1^_10_y^+1^_10_z^+1^_10_c^+1^_10_y^+1^_10_c^+1^_10_w^+1^_11_c^+1^_11_z^+1^_11_c^+1^_11_

0738147522132951

Fragment Matches Table

Show background peaks

| Position | Ion type | Intensity | mz Theoretical | mz Error (Th) | mz Error (ppm) | Charge | Series Number |
| --- | --- | --- | --- | --- | --- | --- | --- |
| 12 | y | 1.883E+04 | 120.1 | 0.0002534 | 2.11 | +1 | 1 |
| - | - | 379.1 | 120.4 | - | - | 0 | - |
| - | - | 677.2 | 121.1 | - | - | 0 | - |
| - | - | 479.9 | 128.7 | - | - | 0 | - |
| - | - | 685.1 | 129.1 | - | - | 0 | - |
| - | - | 457.9 | 130.7 | - | - | 0 | - |
| - | - | 378.2 | 142.3 | - | - | 0 | - |
| - | - | 589.5 | 154.7 | - | - | 0 | - |
| - | - | 1191 | 167.1 | - | - | 0 | - |
| - | - | 1087 | 185.1 | - | - | 0 | - |
| - | - | 2.199E+04 | 185.2 | - | - | 0 | - |
| - | - | 2556 | 186.1 | - | - | 0 | - |
| - | - | 1626 | 186.2 | - | - | 0 | - |
| - | - | 517.7 | 187.1 | - | - | 0 | - |
| - | - | 456.7 | 190.4 | - | - | 0 | - |
| 11 | y | 859 | 201.1 | 0.0001663 | 0.8269 | +1 | 2 |
| - | - | 2.655E+04 | 213.2 | - | - | 0 | - |
| - | - | 3521 | 214.2 | - | - | 0 | - |
| - | - | 585.2 | 215.1 | - | - | 0 | - |
| - | - | 2148 | 217.1 | - | - | 0 | - |
| - | - | 507.1 | 217.6 | - | - | 0 | - |
| 11 | y | 9812 | 219.1 | 0.0002523 | 1.151 | +1 | 2 |
| - | - | 770 | 226.2 | - | - | 0 | - |
| - | - | 507.5 | 260.8 | - | - | 0 | - |
| - | - | 503.3 | 262.7 | - | - | 0 | - |
| - | - | 1933 | 266.1 | - | - | 0 | - |
| - | - | 3021 | 270.2 | - | - | 0 | - |
| - | - | 1376 | 284.2 | - | - | 0 | - |
| 3 | c | 7030 | 287.2 | 0.0002406 | 0.8379 | +1 | 3 |
| - | - | 3390 | 299.2 | - | - | 0 | - |
| - | - | 681.1 | 300.2 | - | - | 0 | - |
| - | - | 1106 | 314.2 | - | - | 0 | - |
| - | - | 755.1 | 327.2 | - | - | 0 | - |
| - | - | 3269 | 368.2 | - | - | 0 | - |
| - | - | 3306 | 370.2 | - | - | 0 | - |
| - | - | 2375 | 371.1 | - | - | 0 | - |
| 8 | c | 783.5 | 383.2 | 0.002397 | 6.253 | +2 | 8 |
| 9 | y | 1063 | 385.2 | 0.001495 | 3.88 | +1 | 4 |
| 5 | y | 641.6 | 385.7 | 0.0005897 | 1.529 | +2 | 8 |
| 4 | c | 1.745E+04 | 398.2 | 0.0004998 | 1.255 | +1 | 4 |
| - | - | 4096 | 399.2 | - | - | 0 | - |
| 9 | y | 1.455E+04 | 403.2 | 0.0003294 | 0.8169 | +1 | 4 |
| - | - | 3280 | 404.2 | - | - | 0 | - |
| - | - | 697.8 | 409.7 | - | - | 0 | - |
| - | - | 582.9 | 415.4 | - | - | 0 | - |
| - | - | 808.6 | 418.7 | - | - | 0 | - |
| - | - | 563.2 | 421.8 | - | - | 0 | - |
| - | - | 627.1 | 428.3 | - | - | 0 | - |
| 9 | c | 3644 | 431.8 | 0.0003658 | 0.8473 | +2 | 9 |
| - | - | 1681 | 432.3 | - | - | 0 | - |
| - | - | 814.8 | 432.8 | - | - | 0 | - |
| 4 | y | 567.4 | 441.2 | 0.008482 | 19.22 | +2 | 9 |
| - | - | 1139 | 460.3 | - | - | 0 | - |
| - | - | 1359 | 461.3 | - | - | 0 | - |
| - | - | 635.9 | 465.3 | - | - | 0 | - |
| - | - | 1.572E+04 | 466.3 | - | - | 0 | - |
| - | - | 8729 | 466.8 | - | - | 0 | - |
| - | - | 3421 | 467.3 | - | - | 0 | - |
| - | - | 1149 | 468.3 | - | - | 0 | - |
| - | - | 1078 | 469.3 | - | - | 0 | - |
| - | - | 675.2 | 469.8 | - | - | 0 | - |
| 8 | y | 957.3 | 474.3 | 0.0008957 | 1.889 | +1 | 5 |
| 10 | c | 1.16E+04 | 475.3 | 0.000709 | 1.492 | +2 | 10 |
| - | - | 6121 | 475.8 | - | - | 0 | - |
| - | - | 1756 | 476.3 | - | - | 0 | - |
| - | - | 757.7 | 478.8 | - | - | 0 | - |
| - | - | 720.8 | 482.3 | - | - | 0 | - |
| - | - | 722.6 | 489.3 | - | - | 0 | - |
| - | - | 799.1 | 496.3 | - | - | 0 | - |
| - | - | 5466 | 501.8 | - | - | 0 | - |
| - | - | 4025 | 502.3 | - | - | 0 | - |
| - | - | 665.8 | 502.8 | - | - | 0 | - |
| - | - | 674 | 507.3 | - | - | 0 | - |
| - | - | 740.5 | 507.8 | - | - | 0 | - |
| - | - | 8955 | 510.8 | - | - | 0 | - |
| - | - | 6281 | 511.3 | - | - | 0 | - |
| - | - | 2056 | 511.8 | - | - | 0 | - |
| - | - | 639.7 | 515.3 | - | - | 0 | - |
| - | - | 1.71E+04 | 515.8 | - | - | 0 | - |
| - | - | 1.097E+04 | 516.3 | - | - | 0 | - |
| - | - | 2513 | 516.8 | - | - | 0 | - |
| 11 | c | 1.057E+04 | 524.8 | 0.0004681 | 0.892 | +2 | 11 |
| - | - | 6855 | 525.3 | - | - | 0 | - |
| - | - | 2180 | 525.8 | - | - | 0 | - |
| - | - | 658.7 | 529.5 | - | - | 0 | - |
| - | - | 640 | 531.2 | - | - | 0 | - |
| 2 | y | 626.5 | 534.8 | 0.003051 | 5.705 | +2 | 11 |
| - | - | 2656 | 552.3 | - | - | 0 | - |
| - | - | 3466 | 553.3 | - | - | 0 | - |
| - | - | 873.7 | 554.3 | - | - | 0 | - |
| - | - | 8206 | 556.3 | - | - | 0 | - |
| - | - | 3071 | 557.3 | - | - | 0 | - |
| - | - | 891.4 | 566.3 | - | - | 0 | - |
| - | - | 1149 | 566.3 | - | - | 0 | - |
| - | - | 9244 | 575.3 | - | - | 0 | - |
| - | - | 8548 | 575.8 | - | - | 0 | - |
| - | - | 2209 | 576.3 | - | - | 0 | - |
| - | - | 885.2 | 576.8 | - | - | 0 | - |
| - | - | 1500 | 583.3 | - | - | 0 | - |
| - | - | 9480 | 584.4 | - | - | 0 | - |
| - | - | 4932 | 584.8 | - | - | 0 | - |
| - | - | 2087 | 585.2 | - | - | 0 | - |
| - | - | 896.2 | 585.3 | - | - | 0 | - |
| - | - | 1876 | 585.3 | - | - | 0 | - |
| - | - | 1936 | 585.4 | - | - | 0 | - |
| - | - | 1495 | 596.4 | - | - | 0 | - |
| - | - | 719 | 600.8 | - | - | 0 | - |
| - | - | 790.9 | 608.4 | - | - | 0 | - |
| 6 | c | 2271 | 623.4 | 0.0005391 | 0.8647 | +1 | 6 |
| - | - | 1388 | 624.4 | - | - | 0 | - |
| - | - | 2997 | 633.4 | - | - | 0 | - |
| - | - | 777 | 634.4 | - | - | 0 | - |
| - | - | 1.517E+04 | 639.4 | - | - | 0 | - |
| 6 | c | 7.181E+04 | 640.4 | 0.0002532 | 0.3954 | +1 | 6 |
| - | - | 2.622E+04 | 641.4 | - | - | 0 | - |
| - | - | 5065 | 642.4 | - | - | 0 | - |
| - | - | 7201 | 651.4 | - | - | 0 | - |
| - | - | 2307 | 651.9 | - | - | 0 | - |
| - | - | 1989 | 652.4 | - | - | 0 | - |
| 6 | z | 1.562E+04 | 657.4 | 9.343E-06 | 0.01421 | +1 | 7 |
| - | - | 1.67E+04 | 658.4 | - | - | 0 | - |
| - | - | 4549 | 659.4 | - | - | 0 | - |
| - | - | 586 | 661 | - | - | 0 | - |
| - | - | 1078 | 663.4 | - | - | 0 | - |
| - | - | 656.6 | 666.4 | - | - | 0 | - |
| - | - | 4291 | 667.4 | - | - | 0 | - |
| - | - | 1711 | 668.4 | - | - | 0 | - |
| - | - | 755.2 | 669.4 | - | - | 0 | - |
| - | - | 1337 | 672.4 | - | - | 0 | - |
| 6 | y | 1.624E+04 | 673.4 | 0.0001451 | 0.2155 | +1 | 7 |
| - | - | 5583 | 674.4 | - | - | 0 | - |
| - | - | 1259 | 675.4 | - | - | 0 | - |
| - | - | 636.3 | 681.4 | - | - | 0 | - |
| 7 | c | 2141 | 694.4 | 0.0003809 | 0.5485 | +1 | 7 |
| - | - | 803.9 | 704 | - | - | 0 | - |
| - | - | 1199 | 704.4 | - | - | 0 | - |
| - | - | 5984 | 710.4 | - | - | 0 | - |
| 7 | c | 5.087E+04 | 711.5 | 0.0001966 | 0.2763 | +1 | 7 |
| - | - | 1.881E+04 | 712.5 | - | - | 0 | - |
| - | - | 3828 | 713.5 | - | - | 0 | - |
| - | - | 1055 | 719.4 | - | - | 0 | - |
| - | - | 898.7 | 720.4 | - | - | 0 | - |
| 5 | z | 718 | 736.4 | 0.01378 | 18.71 | +1 | 8 |
| 5 | z | 1043 | 737.4 | 0.007407 | 10.04 | +1 | 8 |
| - | - | 997 | 738.4 | - | - | 0 | - |
| - | - | 1.692E+04 | 738.5 | - | - | 0 | - |
| - | - | 1.517E+04 | 739.5 | - | - | 0 | - |
| - | - | 4808 | 740.5 | - | - | 0 | - |
| - | - | 839.1 | 741.5 | - | - | 0 | - |
| - | - | 838.3 | 744.4 | - | - | 0 | - |
| - | - | 934.7 | 745.4 | - | - | 0 | - |
| 5 | y | 1819 | 752.4 | 0.00122 | 1.621 | +1 | 8 |
| 8 | c | 2.207E+04 | 765.5 | 4.196E-05 | 0.05482 | +1 | 8 |
| - | - | 8047 | 766.5 | - | - | 0 | - |
| - | - | 1805 | 767.5 | - | - | 0 | - |
| - | - | 3840 | 769.4 | - | - | 0 | - |
| 5 | y | 6.555E+04 | 770.4 | 0.0001767 | 0.2293 | +1 | 8 |
| - | - | 2.857E+04 | 771.4 | - | - | 0 | - |
| - | - | 6234 | 772.4 | - | - | 0 | - |
| - | - | 1089 | 781.5 | - | - | 0 | - |
| - | - | 913.4 | 782.5 | - | - | 0 | - |
| - | - | 2084 | 786.9 | - | - | 0 | - |
| - | - | 1481 | 787.4 | - | - | 0 | - |
| - | - | 878.7 | 787.9 | - | - | 0 | - |
| - | - | 726.3 | 788.4 | - | - | 0 | - |
| - | - | 816.4 | 793.3 | - | - | 0 | - |
| - | - | 842.6 | 793.5 | - | - | 0 | - |
| - | - | 2673 | 798.4 | - | - | 0 | - |
| - | - | 1466 | 799.4 | - | - | 0 | - |
| - | - | 5272 | 807.5 | - | - | 0 | - |
| - | - | 2545 | 808.5 | - | - | 0 | - |
| - | - | 1563 | 810.5 | - | - | 0 | - |
| - | - | 1142 | 811.5 | - | - | 0 | - |
| - | - | 796.1 | 812.5 | - | - | 0 | - |
| - | - | 1619 | 818.5 | - | - | 0 | - |
| - | - | 855.5 | 819.4 | - | - | 0 | - |
| - | - | 985.2 | 820.5 | - | - | 0 | - |
| - | - | 5401 | 822.5 | - | - | 0 | - |
| - | - | 644.7 | 823.4 | - | - | 0 | - |
| - | - | 2692 | 823.5 | - | - | 0 | - |
| 4 | w | 2.671E+04 | 824.5 | 0.0001101 | 0.1335 | +1 | 9 |
| - | - | 1.013E+04 | 825.5 | - | - | 0 | - |
| - | - | 734.8 | 826.4 | - | - | 0 | - |
| - | - | 3288 | 826.5 | - | - | 0 | - |
| - | - | 4038 | 835.5 | - | - | 0 | - |
| - | - | 4324 | 836.5 | - | - | 0 | - |
| - | - | 1605 | 836.5 | - | - | 0 | - |
| - | - | 2370 | 837.5 | - | - | 0 | - |
| - | - | 996.1 | 837.5 | - | - | 0 | - |
| - | - | 1542 | 838.5 | - | - | 0 | - |
| - | - | 1010 | 849 | - | - | 0 | - |
| - | - | 740.3 | 850.5 | - | - | 0 | - |
| 9 | c | 975.7 | 862.5 | 0.0007199 | 0.8347 | +1 | 9 |
| - | - | 2537 | 863.5 | - | - | 0 | - |
| - | - | 2551 | 864.5 | - | - | 0 | - |
| - | - | 1042 | 865.5 | - | - | 0 | - |
| - | - | 739 | 868.5 | - | - | 0 | - |
| - | - | 932.3 | 877.4 | - | - | 0 | - |
| - | - | 1344 | 877.5 | - | - | 0 | - |
| - | - | 693.8 | 878 | - | - | 0 | - |
| - | - | 8.375E+04 | 878.5 | - | - | 0 | - |
| 9 | c | 1.038E+05 | 879.5 | 0.002306 | 2.621 | +1 | 9 |
| - | - | 4.3E+04 | 880.5 | - | - | 0 | - |
| 4 | y | 2660 | 881.5 | 0.002982 | 3.383 | +1 | 9 |
| - | - | 1.04E+04 | 881.5 | - | - | 0 | - |
| 4 | z | 4.452E+04 | 882.5 | 0.0005532 | 0.6269 | +1 | 9 |
| - | - | 2.071E+04 | 883.5 | - | - | 0 | - |
| - | - | 5728 | 884.5 | - | - | 0 | - |
| - | - | 8525 | 897.5 | - | - | 0 | - |
| 4 | y | 4.845E+04 | 898.5 | 0.0007226 | 0.8042 | +1 | 9 |
| - | - | 2.468E+04 | 899.5 | - | - | 0 | - |
| - | - | 7823 | 900.5 | - | - | 0 | - |
| - | - | 1096 | 907.5 | - | - | 0 | - |
| 3 | z | 865.2 | 921.5 | 0.004006 | 4.347 | +1 | 10 |
| - | - | 1250 | 926.5 | - | - | 0 | - |
| - | - | 1656 | 927 | - | - | 0 | - |
| - | - | 1.975E+04 | 931.5 | - | - | 0 | - |
| - | - | 1.16E+04 | 932.5 | - | - | 0 | - |
| - | - | 4965 | 933.5 | - | - | 0 | - |
| - | - | 664.8 | 934.6 | - | - | 0 | - |
| - | - | 1772 | 936.4 | - | - | 0 | - |
| 3 | y | 1237 | 937.5 | 0.003759 | 4.01 | +1 | 10 |
| - | - | 803.4 | 938.4 | - | - | 0 | - |
| - | - | 2755 | 938.5 | - | - | 0 | - |
| 3 | z | 1.091E+04 | 939.5 | 0.000383 | 0.4076 | +1 | 10 |
| - | - | 5857 | 940.5 | - | - | 0 | - |
| - | - | 1322 | 941.5 | - | - | 0 | - |
| 10 | c | 3.219E+04 | 949.5 | 0.0002776 | 0.2924 | +1 | 10 |
| - | - | 1.724E+04 | 950.5 | - | - | 0 | - |
| - | - | 5616 | 951.6 | - | - | 0 | - |
| - | - | 1046 | 952.6 | - | - | 0 | - |
| 3 | y | 3.583E+04 | 955.5 | 0.0005188 | 0.543 | +1 | 10 |
| - | - | 1.698E+04 | 956.5 | - | - | 0 | - |
| - | - | 5857 | 957.5 | - | - | 0 | - |
| - | - | 733.3 | 958.5 | - | - | 0 | - |
| 10 | c | 6.236E+04 | 966.6 | 0.0007647 | 0.7912 | +1 | 10 |
| - | - | 3.174E+04 | 967.6 | - | - | 0 | - |
| - | - | 2404 | 967.8 | - | - | 0 | - |
| - | - | 1499 | 968.2 | - | - | 0 | - |
| - | - | 859 | 968.5 | - | - | 0 | - |
| - | - | 9585 | 968.6 | - | - | 0 | - |
| - | - | 1388 | 969.6 | - | - | 0 | - |
| - | - | 1854 | 973.2 | - | - | 0 | - |
| - | - | 1315 | 973.5 | - | - | 0 | - |
| - | - | 1131 | 973.8 | - | - | 0 | - |
| - | - | 1832 | 974.2 | - | - | 0 | - |
| - | - | 1188 | 991.5 | - | - | 0 | - |
| - | - | 3798 | 1007 | - | - | 0 | - |
| - | - | 1277 | 1008 | - | - | 0 | - |
| 2 | w | 3634 | 1010 | 0.0006963 | 0.6897 | +1 | 11 |
| - | - | 2032 | 1011 | - | - | 0 | - |
| - | - | 1030 | 1012 | - | - | 0 | - |
| - | - | 1092 | 1021 | - | - | 0 | - |
| - | - | 3215 | 1022 | - | - | 0 | - |
| - | - | 1686 | 1023 | - | - | 0 | - |
| - | - | 1078 | 1024 | - | - | 0 | - |
| - | - | 1771 | 1031 | - | - | 0 | - |
| - | - | 1732 | 1032 | - | - | 0 | - |
| - | - | 708 | 1033 | - | - | 0 | - |
| - | - | 721.1 | 1038 | - | - | 0 | - |
| - | - | 1112 | 1039 | - | - | 0 | - |
| 11 | c | 1.831E+04 | 1049 | 0.0006984 | 0.666 | +1 | 11 |
| - | - | 1.192E+04 | 1050 | - | - | 0 | - |
| - | - | 1.176E+04 | 1051 | - | - | 0 | - |
| - | - | 5865 | 1052 | - | - | 0 | - |
| 2 | z | 1.041E+04 | 1053 | 0.002318 | 2.202 | +1 | 11 |
| - | - | 7038 | 1054 | - | - | 0 | - |
| - | - | 2134 | 1055 | - | - | 0 | - |
| - | - | 753.2 | 1055 | - | - | 0 | - |
| - | - | 711.1 | 1056 | - | - | 0 | - |
| - | - | 1278 | 1056 | - | - | 0 | - |
| - | - | 1146 | 1057 | - | - | 0 | - |
| 11 | c | 1.214E+05 | 1066 | 0.0006362 | 0.597 | +1 | 11 |
| - | - | 7.503E+04 | 1067 | - | - | 0 | - |
| - | - | 2.375E+04 | 1068 | - | - | 0 | - |
| - | - | 5548 | 1069 | - | - | 0 | - |
| - | - | 661.7 | 1069 | - | - | 0 | - |
| - | - | 1999 | 1076 | - | - | 0 | - |
| - | - | 650.2 | 1077 | - | - | 0 | - |
| - | - | 1598 | 1080 | - | - | 0 | - |
| - | - | 6763 | 1081 | - | - | 0 | - |
| - | - | 3883 | 1082 | - | - | 0 | - |
| - | - | 2265 | 1083 | - | - | 0 | - |
| - | - | 2.378E+04 | 1096 | - | - | 0 | - |
| - | - | 1.426E+04 | 1097 | - | - | 0 | - |
| - | - | 6791 | 1098 | - | - | 0 | - |
| - | - | 1552 | 1099 | - | - | 0 | - |
| - | - | 1194 | 1105 | - | - | 0 | - |
| - | - | 1349 | 1113 | - | - | 0 | - |
| - | - | 954 | 1114 | - | - | 0 | - |
| - | - | 969.9 | 1120 | - | - | 0 | - |
| - | - | 4663 | 1123 | - | - | 0 | - |
| - | - | 676.9 | 1124 | - | - | 0 | - |
| - | - | 9100 | 1124 | - | - | 0 | - |
| - | - | 6053 | 1125 | - | - | 0 | - |
| - | - | 2203 | 1126 | - | - | 0 | - |
| - | - | 750.3 | 1133 | - | - | 0 | - |
| - | - | 786.2 | 1134 | - | - | 0 | - |
| - | - | 1012 | 1137 | - | - | 0 | - |
| - | - | 9113 | 1151 | - | - | 0 | - |
| - | - | 1.13E+05 | 1152 | - | - | 0 | - |
| - | - | 7.067E+04 | 1153 | - | - | 0 | - |
| - | - | 2.648E+04 | 1154 | - | - | 0 | - |
| - | - | 3510 | 1155 | - | - | 0 | - |
| - | - | 1201 | 1166 | - | - | 0 | - |
| - | - | 1.2E+05 | 1168 | - | - | 0 | - |
| - | - | 3.466E+05 | 1169 | - | - | 0 | - |
| - | - | 2.057E+05 | 1170 | - | - | 0 | - |
| - | - | 6.66E+04 | 1171 | - | - | 0 | - |
| - | - | 8785 | 1172 | - | - | 0 | - |
| - | - | 830.1 | 1186 | - | - | 0 | - |
| - | - | 1982 | 1201 | - | - | 0 | - |
| - | - | 1754 | 1202 | - | - | 0 | - |
| - | - | 864.7 | 1223 | - | - | 0 | - |
| - | - | 869.7 | 1224 | - | - | 0 | - |
| - | - | 805.9 | 1244 | - | - | 0 | - |
| - | - | 1334 | 1286 | - | - | 0 | - |
| - | - | 864.6 | 1287 | - | - | 0 | - |
| - | - | 1491 | 1301 | - | - | 0 | - |
| - | - | 4023 | 1302 | - | - | 0 | - |
| - | - | 1103 | 1302 | - | - | 0 | - |
| - | - | 3364 | 1303 | - | - | 0 | - |
| - | - | 1496 | 1304 | - | - | 0 | - |
| - | - | 1566 | 1359 | - | - | 0 | - |
| - | - | 1395 | 1359 | - | - | 0 | - |
| - | - | 1055 | 1360 | - | - | 0 | - |
| - | - | 881 | 1360 | - | - | 0 | - |
| - | - | 982.8 | 1416 | - | - | 0 | - |
| - | - | 811 | 1416 | - | - | 0 | - |
| - | - | 963.8 | 1429 | - | - | 0 | - |
| - | - | 955.5 | 1429 | - | - | 0 | - |
| - | - | 1481 | 1431 | - | - | 0 | - |
| - | - | 1598 | 1437 | - | - | 0 | - |
| - | - | 1865 | 1438 | - | - | 0 | - |
| - | - | 872.7 | 1438 | - | - | 0 | - |
| - | - | 1468 | 1443 | - | - | 0 | - |
| - | - | 1889 | 1443 | - | - | 0 | - |
| - | - | 1317 | 1444 | - | - | 0 | - |
| - | - | 1492 | 1451 | - | - | 0 | - |
| - | - | 5739 | 1451 | - | - | 0 | - |
| - | - | 7175 | 1452 | - | - | 0 | - |
| - | - | 3796 | 1452 | - | - | 0 | - |
| - | - | 3154 | 1453 | - | - | 0 | - |
| - | - | 940.8 | 1453 | - | - | 0 | - |
| - | - | 1058 | 1454 | - | - | 0 | - |
| - | - | 2614 | 1459 | - | - | 0 | - |
| - | - | 4502 | 1460 | - | - | 0 | - |
| - | - | 5974 | 1460 | - | - | 0 | - |
| - | - | 3891 | 1461 | - | - | 0 | - |
| - | - | 2595 | 1461 | - | - | 0 | - |
| - | - | 1444 | 1462 | - | - | 0 | - |
| - | - | 829.7 | 1609 | - | - | 0 | - |
| - | - | 5261 | 1617 | - | - | 0 | - |
| - | - | 4533 | 1618 | - | - | 0 | - |
| - | - | 2994 | 1619 | - | - | 0 | - |
| - | - | 803.8 | 1653 | - | - | 0 | - |
| - | - | 867.3 | 1700 | - | - | 0 | - |
| - | - | 796.4 | 1718 | - | - | 0 | - |
| - | - | 989.9 | 1719 | - | - | 0 | - |
| - | - | 1402 | 1750 | - | - | 0 | - |
| - | - | 1350 | 1751 | - | - | 0 | - |
| - | - | 991.2 | 1755 | - | - | 0 | - |
| - | - | 1527 | 1756 | - | - | 0 | - |
| - | - | 1409 | 1881 | - | - | 0 | - |
| - | - | 764.2 | 1882 | - | - | 0 | - |
| - | - | 1155 | 1983 | - | - | 0 | - |
| - | - | 911.4 | 2111 | - | - | 0 | - |
| - | - | 755.5 | 2112 | - | - | 0 | - |
| - | - | 962.4 | 2113 | - | - | 0 | - |
| - | - | 1078 | 2335 | - | - | 0 | - |
| - | - | 1365 | 2336 | - | - | 0 | - |
| - | - | 941.3 | 2337 | - | - | 0 | - |
| - | - | 777.7 | 2338 | - | - | 0 | - |
| - | - | 670.5 | 2887 | - | - | 0 | - |
| - | - | 780.3 | 2900 | - | - | 0 | - |
| - | - | 798.4 | 2901 | - | - | 0 | - |
| - | - | 1550 | 2902 | - | - | 0 | - |
| - | - | 2255 | 2903 | - | - | 0 | - |
| - | - | 1581 | 2904 | - | - | 0 | - |
| - | - | 1194 | 2918 | - | - | 0 | - |
| - | - | 2209 | 2919 | - | - | 0 | - |
| - | - | 2172 | 2920 | - | - | 0 | - |
| - | - | 2141 | 2921 | - | - | 0 | - |

m/z Charge Intensity FragmentType MassShift Position
120.0657730102539 0 18834.217 y 11
120.40908813476562 0 379.05292
121.06925201416016 0 677.1793
128.68333435058594 0 479.94122
129.10264587402344 0 685.1249
130.70169067382812 0 457.8504
142.31312561035156 0 378.16885
154.68235778808594 0 589.50385
167.08163452148438 0 1190.6072
185.0923614501953 0 1086.7517
185.16510009765625 0 21985.113
186.08763122558594 0 2555.7717
186.16815185546875 0 1626.1006
187.107177734375 0 517.65845
190.3984375 0 456.72073
201.12353515625 0 858.9516 y Water loss 10
213.16001892089844 0 26553.76
214.1633758544922 0 3521.171
215.13858032226562 0 585.24915
217.1184539794922 0 2148.177
217.57366943359375 0 507.11667
219.13418579101562 0 9812.012 y 10
226.15533447265625 0 770.02655
260.7999267578125 0 507.48505
262.7453308105469 0 503.29834
266.14996337890625 0 1932.8673
270.1813659667969 0 3021.3928
284.16064453125 0 1375.9902
287.2080078125 0 7030.2695 c 2
299.171630859375 0 3389.709
300.17413330078125 0 681.0845
314.209228515625 0 1106.3484
327.1653137207031 0 755.0638
368.2294616699219 0 3268.6736
370.2446594238281 0 3306.1045
371.07659912109375 0 2374.7676
383.23211669921875 0 783.4758 c Ammonia loss 7
385.20965576171875 0 1063.2157 y Water loss 8
385.723388671875 0 641.6371 y 4
398.24029541015625 0 17446.117 c Ammonia loss 3
399.24285888671875 0 4096.19
403.21905517578125 0 14554.624 y 8
404.2227783203125 0 3280.2114
409.7293395996094 0 697.839
415.40460205078125 0 582.8618
418.7373352050781 0 808.6443
421.8135681152344 0 563.1659
428.2615661621094 0 627.11444
431.7612609863281 0 3644.2131 c Ammonia loss 8
432.26519775390625 0 1680.9042
432.7676086425781 0 814.78
441.24847412109375 0 567.3732 y Ammonia loss 3
460.2896728515625 0 1138.5261
461.28021240234375 0 1359.0164
465.28253173828125 0 635.87036
466.2721862792969 0 15721.3
466.7733459472656 0 8728.763
467.2747802734375 0 3421.1326
468.2694396972656 0 1148.6884
469.29345703125 0 1078.2559
469.7603759765625 0 675.1695
474.25494384765625 0 957.29877 y 7
475.2776184082031 0 11604.957 c Ammonia loss 9
475.77886962890625 0 6120.5195
476.27899169921875 0 1755.5076
478.766845703125 0 757.68427
482.2740173339844 0 720.77704
489.29205322265625 0 722.64954
496.3268127441406 0 799.07666
501.80841064453125 0 5465.9116
502.3100891113281 0 4024.6514
502.810302734375 0 665.83545
507.29351806640625 0 673.9783
507.79443359375 0 740.4529
510.8138122558594 0 8955.461
511.3155517578125 0 6280.802
511.8171691894531 0 2055.5476
515.2818603515625 0 639.7344
515.8060913085938 0 17098.432
516.3070678710938 0 10966.969
516.8086547851562 0 2513.235
524.8115844726562 0 10567.769 c Ammonia loss 10
525.3132934570312 0 6854.8623
525.8116455078125 0 2180.4111
529.5469970703125 0 658.7469
531.1862182617188 0 640.0188
534.80908203125 0 626.5085 y 1
552.3154907226562 0 2656.0715
553.3120727539062 0 3466.089
554.3106689453125 0 873.73804
556.3468627929688 0 8206.11
557.3484497070312 0 3070.925
566.277587890625 0 891.36414
566.3292236328125 0 1148.5769
575.334228515625 0 9244.468
575.8363647460938 0 8547.573
576.3367919921875 0 2209.2966
576.8360595703125 0 885.1992
583.3182373046875 0 1499.8508
584.3500366210938 0 9479.586
584.8414916992188 0 4931.7563
585.2086181640625 0 2087.4587
585.2888793945312 0 896.2044
585.345947265625 0 1876.2722
585.3633422851562 0 1935.6067
596.3983764648438 0 1495.2048
600.8265991210938 0 718.98016
608.3654174804688 0 790.89966
623.3880615234375 0 2270.5251 c Ammonia loss 5
624.391357421875 0 1388.3943
633.371826171875 0 2996.8518
634.3761596679688 0 776.99255
639.40625 0 15167.283
640.413818359375 0 71810.695 c 5
641.4168090820312 0 26216.682
642.4197998046875 0 5064.774
651.3798828125 0 7201.025
651.8526000976562 0 2307.0305
652.3824462890625 0 1988.8772
657.3692016601562 0 15619.967 z 5
658.3753662109375 0 16699.416
659.3783569335938 0 4549.0977
661.0274047851562 0 586.0445
663.3860473632812 0 1077.9332
666.3871459960938 0 656.5831
667.4371337890625 0 4290.5513
668.440185546875 0 1710.5011
669.4411010742188 0 755.2025
672.380126953125 0 1337.1842
673.3880615234375 0 16236.181 y 5
674.3909912109375 0 5582.6226
675.3949584960938 0 1258.9656
681.3970947265625 0 636.25714
694.4242553710938 0 2141.189 c Ammonia loss 6
704.0402221679688 0 803.8687
704.3740844726562 0 1198.9476
710.4431762695312 0 5984.078
711.4509887695312 0 50870.773 c 6
712.453857421875 0 18806.426
713.4561767578125 0 3827.8086
719.3833618164062 0 1055.4878
720.3824462890625 0 898.7023
736.4251708984375 0 717.9732 z Water loss 4
737.3880004882812 0 1042.666 z Ammonia loss 4
738.40185546875 0 996.9788
738.4739990234375 0 16917.43
739.4801635742188 0 15171.17
740.484130859375 0 4808.0166
741.4827270507812 0 839.13806
744.385986328125 0 838.3108
745.3935546875 0 934.6504
752.4313354492188 0 1819.1206 y Water loss 4
765.4617919921875 0 22071.82 c Ammonia loss 7
766.46435546875 0 8047.373
767.4674682617188 0 1804.5894
769.43115234375 0 3840.4255
770.4408569335938 0 65554.664 y 4
771.4434814453125 0 28569.125
772.4459228515625 0 6233.9556
781.4613037109375 0 1089.135
782.4600219726562 0 913.40344
786.867919921875 0 2083.9019
787.371337890625 0 1481.0715
787.8713989257812 0 878.6878
788.3727416992188 0 726.2922
793.34033203125 0 816.44965
793.4584350585938 0 842.6318
798.4354858398438 0 2673.1597
799.434814453125 0 1466.3928
807.4937744140625 0 5272.288
808.4892578125 0 2545.4778
810.4591674804688 0 1562.8121
811.461181640625 0 1141.9252
812.4644165039062 0 796.08545
818.4503784179688 0 1618.8998
819.4451904296875 0 855.51044
820.5185546875 0 985.2114
822.4706420898438 0 5400.869
823.3931274414062 0 644.69403
823.4736328125 0 2692.02
824.4513549804688 0 26711.88 w 3
825.453369140625 0 10128.627
826.3787841796875 0 734.83453
826.4576416015625 0 3287.5972
835.5272216796875 0 4037.5334
836.4627685546875 0 4323.57
836.5342407226562 0 1604.5138
837.4649658203125 0 2369.983
837.5449829101562 0 996.0759
838.474365234375 0 1542.1562
848.9553833007812 0 1010.31006
850.4761962890625 0 740.3137
862.5137939453125 0 975.6811 c Ammonia loss 8
863.5235595703125 0 2536.9453
864.5291748046875 0 2551.413
865.5381469726562 0 1041.8983
868.457763671875 0 738.989
877.4105834960938 0 932.2911
877.5222778320312 0 1343.7811
877.9637451171875 0 693.8373
878.5326538085938 0 83748.375
879.5387573242188 0 103813.22 c 8
880.5419921875 0 42996.81
881.4697265625 0 2660.2554 y Ammonia loss 3
881.54833984375 0 10398.965
882.47998046875 0 44516.85 z 3
883.4830322265625 0 20705.53
884.484619140625 0 5728.2295
897.4923095703125 0 8524.517
898.49853515625 0 48446.79 y 3
899.50146484375 0 24679.668
900.5037231445312 0 7822.621
907.5472412109375 0 1095.8229
921.4874267578125 0 865.1533 z Water loss 2
926.5114135742188 0 1249.9276
927.012939453125 0 1656.3499
931.5350952148438 0 19745.166
932.5379638671875 0 11597.434
933.5428466796875 0 4964.625
934.5526123046875 0 664.7578
936.4193115234375 0 1772.4292
937.513916015625 0 1237.3679 y Water loss 2
938.4222412109375 0 803.36163
938.5142822265625 0 2754.6206
939.5023803710938 0 10905.039 z 2
940.5057983398438 0 5856.5596
941.5155029296875 0 1321.7487
949.5462646484375 0 32191.273 c Ammonia loss 9
950.5484619140625 0 17235.467
951.5536499023438 0 5615.6807
952.5570068359375 0 1045.6278
955.5202026367188 0 35830.465 y 2
956.5227661132812 0 16984.432
957.525634765625 0 5857.034
958.5230102539062 0 733.2845
966.5723266601562 0 62364.926 c 9
967.5751342773438 0 31736.527
967.8207397460938 0 2403.7122
968.15380859375 0 1498.6125
968.479736328125 0 859.02155
968.5783081054688 0 9585.458
969.579345703125 0 1388.4849
973.1637573242188 0 1853.9812
973.4998168945312 0 1314.8636
973.8338012695312 0 1131.0527
974.168701171875 0 1831.9753
991.52978515625 0 1187.7023
1006.6170043945312 0 3797.885
1007.6213989257812 0 1276.9609
1009.531982421875 0 3634.2407 w 1
1010.5338134765625 0 2032.1515
1011.5344848632812 0 1029.7189
1020.6234130859375 0 1091.8077
1021.62353515625 0 3215.4832
1022.6304321289062 0 1685.5308
1023.6466674804688 0 1078.3575
1030.60791015625 0 1771.0199
1031.6063232421875 0 1731.8385
1032.613037109375 0 708.0282
1037.5814208984375 0 721.11176
1038.585693359375 0 1112.0608
1048.6142578125 0 18310.967 c Ammonia loss 10
1049.6171875 0 11916.432
1050.627197265625 0 11758.86
1051.631591796875 0 5865.284
1052.58837890625 0 10411.299 z 1
1053.5885009765625 0 7037.9995
1054.59375 0 2134.122
1055.0146484375 0 753.2335
1055.537109375 0 711.05743
1056.0521240234375 0 1278.1772
1056.548095703125 0 1145.5983
1065.640869140625 0 121410.97 c 10
1066.6439208984375 0 75028.48
1067.647216796875 0 23746.709
1068.646240234375 0 5548.4204
1069.420654296875 0 661.726
1076.4022216796875 0 1999.2457
1077.404296875 0 650.20013
1080.39501953125 0 1598.2218
1080.6151123046875 0 6762.654
1081.6190185546875 0 3882.5977
1082.6173095703125 0 2265.1248
1095.591064453125 0 23783.441
1096.5943603515625 0 14260.881
1097.609375 0 6790.525
1098.6304931640625 0 1552.0128
1104.6610107421875 0 1194.1453
1112.6181640625 0 1348.6577
1113.615234375 0 953.97485
1119.6483154296875 0 969.87604
1122.6741943359375 0 4662.854
1123.5289306640625 0 676.8963
1123.6641845703125 0 9100.372
1124.664794921875 0 6053.052
1125.6656494140625 0 2202.8503
1133.3511962890625 0 750.2976
1134.343017578125 0 786.2076
1136.6219482421875 0 1012.2919
1150.6676025390625 0 9112.616
1151.654052734375 0 112976.41
1152.65673828125 0 70670.734
1153.6591796875 0 26478.469
1154.6612548828125 0 3510.1406
1166.49560546875 0 1200.7577
1167.672119140625 0 119987.37
1168.6788330078125 0 346581.3
1169.682373046875 0 205688.06
1170.6846923828125 0 66602.79
1171.688720703125 0 8784.866
1185.634033203125 0 830.1274
1200.650634765625 0 1982.3417
1201.656494140625 0 1754.1593
1222.5728759765625 0 864.7118
1223.5806884765625 0 869.74554
1243.66015625 0 805.92596
1285.66796875 0 1333.6147
1286.6767578125 0 864.5847
1301.159912109375 0 1491.0511
1301.6781005859375 0 4022.7832
1302.1671142578125 0 1103.4252
1302.689453125 0 3364.0383
1303.6956787109375 0 1496.2665
1358.6727294921875 0 1566.4985
1359.18017578125 0 1395.1704
1359.6729736328125 0 1055.0486
1360.168212890625 0 880.9732
1415.7113037109375 0 982.8489
1416.228271484375 0 810.9952
1428.7359619140625 0 963.8154
1429.2310791015625 0 955.5375
1430.732666015625 0 1480.8827
1437.2403564453125 0 1598.0897
1437.735595703125 0 1864.7499
1438.234130859375 0 872.67194
1442.7237548828125 0 1467.9987
1443.2220458984375 0 1889.0417
1443.722900390625 0 1316.754
1450.7335205078125 0 1491.6693
1451.2305908203125 0 5739.2866
1451.7315673828125 0 7175.495
1452.23828125 0 3795.8506
1452.7337646484375 0 3153.7969
1453.24267578125 0 940.829
1453.7242431640625 0 1058.3745
1459.240234375 0 2613.556
1459.74365234375 0 4502.04
1460.243408203125 0 5973.561
1460.74658203125 0 3891.4883
1461.247802734375 0 2594.5933
1461.7413330078125 0 1444.3557
1608.881591796875 0 829.695
1616.7860107421875 0 5261.3726
1617.7899169921875 0 4532.6187
1618.7930908203125 0 2994.249
1652.8876953125 0 803.7878
1699.9305419921875 0 867.32983
1717.8319091796875 0 796.36957
1718.8375244140625 0 989.94824
1749.76025390625 0 1401.7373
1750.7843017578125 0 1350.4976
1754.8857421875 0 991.239
1755.895751953125 0 1527.2125
1880.8988037109375 0 1408.8591
1881.892333984375 0 764.18994
1983.0615234375 0 1154.8091
2111.03564453125 0 911.44275
2112.10205078125 0 755.4641
2113.095458984375 0 962.436
2335.15673828125 0 1078.481
2336.1455078125 0 1364.9457
2337.138916015625 0 941.28296
2338.151123046875 0 777.65564
2887.476806640625 0 670.4682
2900.451904296875 0 780.3265
2901.4580078125 0 798.402
2902.472412109375 0 1550.1147
2903.4853515625 0 2254.7124
2904.457763671875 0 1581.0272
2918.485107421875 0 1193.8209
2919.483154296875 0 2209.1401
2920.495849609375 0 2172.422
2921.486328125 0 2140.5408

Spectrum Details

|  |  |
| --- | --- |
| Matched peaks? Matched peaks The total absolute number of peaks matched. Additionally in brackets the total fraction of peaks matched and the total number of peaks is shown. | 42 (10.97% of 383) |
| FDR? FDR The false discovery rate estimated for this peptide. It is calculated by matching all theoretical fragments with a non-integer shift with the raw peaks for this spectrum. This is done with 40 different shifts. The resulting percentage is the average number of annotated peaks over the number of annotated peaks with the correct spectrum. | 0.96% |
| Satellite FDR? Satellite FDR See the FDR for details on its calculation. This satellite ion specific FDR only contains the satellite ions (d/w) for I/L/J positions. | 9.52% |
| PSM Score? PSM Score The PSM Score as given by Hecklib to this annotated spectrum. It is shown with three significant figures. | 451 |

#### Spectrum 4546? Spectrum 4546 The raw spectrum of this peptide as annotated by Hecklib. The fragments are coloured according to ion type (see legend). Any peaks with a star '\*' as text can be hovered over to see the full details, first the ion type second the mass shift type. By hovering over the amino acids in the peptide or ions in the legend the corresponding peaks are highlighted. By toggling the 'Unassigned' label you can turn the background (unassigned) peaks on or off in the plot. By updating the slider in the Ion legend you can update the spectrum to only show the top X% of the peaks with labels. The top X% means any peak that is within X% of the highest intensity. By dragging in the spectrum you can zoom in to a specific part of the spectrum and use 'Zoom Out' to get back to the original zoom level. The annotation of the spectrum is based on the given sequence in the peptides file and is done with different software so inconsistencies are likely. The peaks are annotated based on the given sequence, with 20 ppm tolerance.

Copy Data

##### Spectrum 4546 (TSV)

###### Preview

```
Loading example...
```

*Click on the button to copy the data to your clipboard.*

Mz MinMz MaxIntensity Max

WidthHeightPeptide font sizePeptide stroke widthSpectrum font sizeSpectrum stroke widthCompact peptide

Ion legend

wxyz

abcd

OtherUnassignedIon^Charge^_Position_Show for top:%

VLGQPKAAPSVT

03.82e+47.64e+41.15e+51.53e+5

Zoom Out

y^+1^_1_y^+1^_2_y^+1^_2_c^+1^_3_c^+2^_8_c^+1^_4_y^+1^_4_c^+2^_9_y^+2^_9_y^+1^_5_c^+2^_10_c^+2^_11_y^+2^_11_c^+1^_6_z^+1^_7_y^+1^_7_c^+1^_7_c^+1^_7_y^+1^_8_c^+1^_8_y^+1^_8_w^+1^_9_z^+1^_9_c^+1^_9_y^+1^_9_z^+1^_9_y^+1^_9_z^+1^_10_c^+1^_10_y^+1^_10_c^+1^_10_w^+1^_11_c^+1^_11_z^+1^_11_c^+1^_11_

0591118217732364

Fragment Matches Table

Show background peaks

| Position | Ion type | Intensity | mz Theoretical | mz Error (Th) | mz Error (ppm) | Charge | Series Number |
| --- | --- | --- | --- | --- | --- | --- | --- |
| 12 | y | 7852 | 120.1 | 8.555E-05 | 0.7125 | +1 | 1 |
| - | - | 2055 | 120.1 | - | - | 0 | - |
| - | - | 444.6 | 121.1 | - | - | 0 | - |
| - | - | 2169 | 129.1 | - | - | 0 | - |
| - | - | 483.4 | 150.2 | - | - | 0 | - |
| - | - | 1105 | 173.5 | - | - | 0 | - |
| - | - | 558.6 | 175.9 | - | - | 0 | - |
| - | - | 462 | 179.1 | - | - | 0 | - |
| - | - | 464.7 | 179.2 | - | - | 0 | - |
| - | - | 709.5 | 185.1 | - | - | 0 | - |
| - | - | 8767 | 185.2 | - | - | 0 | - |
| - | - | 687.5 | 186.1 | - | - | 0 | - |
| - | - | 892.8 | 186.2 | - | - | 0 | - |
| 11 | y | 583.1 | 201.1 | 0.0004135 | 2.056 | +1 | 2 |
| - | - | 576.7 | 207.4 | - | - | 0 | - |
| - | - | 467 | 213 | - | - | 0 | - |
| - | - | 1.077E+04 | 213.2 | - | - | 0 | - |
| - | - | 970.2 | 214.2 | - | - | 0 | - |
| - | - | 796.2 | 215.1 | - | - | 0 | - |
| 11 | y | 3858 | 219.1 | 5.291E-05 | 0.2415 | +1 | 2 |
| - | - | 3301 | 219.1 | - | - | 0 | - |
| - | - | 978.5 | 234.1 | - | - | 0 | - |
| - | - | 1644 | 247.1 | - | - | 0 | - |
| - | - | 1239 | 251.2 | - | - | 0 | - |
| - | - | 958.3 | 270.2 | - | - | 0 | - |
| - | - | 1178 | 284.2 | - | - | 0 | - |
| 3 | c | 2143 | 287.2 | 0.0005153 | 1.794 | +1 | 3 |
| - | - | 945.8 | 299.2 | - | - | 0 | - |
| - | - | 579.6 | 330 | - | - | 0 | - |
| - | - | 1704 | 345.2 | - | - | 0 | - |
| - | - | 2442 | 346.2 | - | - | 0 | - |
| - | - | 860.8 | 350.2 | - | - | 0 | - |
| - | - | 912.6 | 355.2 | - | - | 0 | - |
| - | - | 1341 | 368.2 | - | - | 0 | - |
| - | - | 1341 | 370.2 | - | - | 0 | - |
| - | - | 1453 | 371.1 | - | - | 0 | - |
| - | - | 982.2 | 383.2 | - | - | 0 | - |
| 8 | c | 653.8 | 383.2 | 0.001693 | 4.417 | +2 | 8 |
| - | - | 634.5 | 392.1 | - | - | 0 | - |
| 4 | c | 6618 | 398.2 | 0.0003241 | 0.814 | +1 | 4 |
| - | - | 1171 | 399.2 | - | - | 0 | - |
| 9 | y | 5601 | 403.2 | 0.0001768 | 0.4385 | +1 | 4 |
| - | - | 825.8 | 404.2 | - | - | 0 | - |
| 9 | c | 824.4 | 431.8 | 0.001464 | 3.392 | +2 | 9 |
| - | - | 591 | 435.3 | - | - | 0 | - |
| - | - | 2558 | 440.3 | - | - | 0 | - |
| 4 | y | 1977 | 440.7 | 0.003144 | 7.134 | +2 | 9 |
| - | - | 4545 | 447.3 | - | - | 0 | - |
| - | - | 1595 | 448.3 | - | - | 0 | - |
| - | - | 988 | 460.8 | - | - | 0 | - |
| - | - | 771.3 | 466.2 | - | - | 0 | - |
| - | - | 6200 | 466.3 | - | - | 0 | - |
| - | - | 3230 | 466.8 | - | - | 0 | - |
| 8 | y | 839.8 | 474.3 | 0.001201 | 2.532 | +1 | 5 |
| 10 | c | 3500 | 475.3 | 0.0006337 | 1.333 | +2 | 10 |
| - | - | 1795 | 475.8 | - | - | 0 | - |
| - | - | 710 | 476.3 | - | - | 0 | - |
| - | - | 844.6 | 483.3 | - | - | 0 | - |
| - | - | 3679 | 484.2 | - | - | 0 | - |
| - | - | 1409 | 485.2 | - | - | 0 | - |
| - | - | 1178 | 501.3 | - | - | 0 | - |
| - | - | 2670 | 501.8 | - | - | 0 | - |
| - | - | 2434 | 504.3 | - | - | 0 | - |
| - | - | 5921 | 504.3 | - | - | 0 | - |
| - | - | 833.1 | 505.3 | - | - | 0 | - |
| - | - | 1896 | 505.3 | - | - | 0 | - |
| - | - | 3426 | 510.8 | - | - | 0 | - |
| - | - | 1900 | 511.3 | - | - | 0 | - |
| - | - | 1041 | 514.3 | - | - | 0 | - |
| - | - | 5463 | 515.8 | - | - | 0 | - |
| - | - | 3895 | 516.3 | - | - | 0 | - |
| - | - | 1147 | 516.8 | - | - | 0 | - |
| - | - | 800.1 | 518.3 | - | - | 0 | - |
| - | - | 707.2 | 519.3 | - | - | 0 | - |
| 11 | c | 3962 | 524.8 | 0.0009967 | 1.899 | +2 | 11 |
| - | - | 1753 | 525.3 | - | - | 0 | - |
| 2 | y | 661.3 | 525.8 | 0.009432 | 17.94 | +2 | 11 |
| - | - | 636.2 | 535.3 | - | - | 0 | - |
| - | - | 4463 | 536.3 | - | - | 0 | - |
| - | - | 1042 | 537.3 | - | - | 0 | - |
| - | - | 1474 | 552.3 | - | - | 0 | - |
| - | - | 1407 | 553.3 | - | - | 0 | - |
| - | - | 3485 | 556.3 | - | - | 0 | - |
| - | - | 740.9 | 557.3 | - | - | 0 | - |
| - | - | 2292 | 561.3 | - | - | 0 | - |
| - | - | 660.2 | 566.2 | - | - | 0 | - |
| - | - | 4034 | 575.3 | - | - | 0 | - |
| - | - | 3546 | 575.8 | - | - | 0 | - |
| - | - | 2628 | 576.3 | - | - | 0 | - |
| - | - | 815.6 | 583.3 | - | - | 0 | - |
| - | - | 1803 | 583.3 | - | - | 0 | - |
| - | - | 2302 | 584.3 | - | - | 0 | - |
| - | - | 1297 | 584.8 | - | - | 0 | - |
| - | - | 1697 | 585.2 | - | - | 0 | - |
| - | - | 668.6 | 585.3 | - | - | 0 | - |
| - | - | 1796 | 585.3 | - | - | 0 | - |
| - | - | 718.3 | 618.3 | - | - | 0 | - |
| - | - | 692.3 | 628.4 | - | - | 0 | - |
| - | - | 8135 | 633.4 | - | - | 0 | - |
| - | - | 2284 | 634.4 | - | - | 0 | - |
| - | - | 6648 | 639.4 | - | - | 0 | - |
| 6 | c | 3.21E+04 | 640.4 | 0.0009856 | 1.539 | +1 | 6 |
| - | - | 1.001E+04 | 641.4 | - | - | 0 | - |
| - | - | 2532 | 642.4 | - | - | 0 | - |
| - | - | 3269 | 651.4 | - | - | 0 | - |
| 6 | z | 8110 | 657.4 | 0.0004179 | 0.6357 | +1 | 7 |
| - | - | 6403 | 658.4 | - | - | 0 | - |
| - | - | 1923 | 659.4 | - | - | 0 | - |
| - | - | 656.1 | 661.4 | - | - | 0 | - |
| - | - | 6651 | 664.4 | - | - | 0 | - |
| - | - | 2607 | 665.4 | - | - | 0 | - |
| - | - | 992.8 | 666.4 | - | - | 0 | - |
| - | - | 2306 | 667.4 | - | - | 0 | - |
| - | - | 774.7 | 672.4 | - | - | 0 | - |
| 6 | y | 8369 | 673.4 | 0.0006483 | 0.9628 | +1 | 7 |
| - | - | 2334 | 674.4 | - | - | 0 | - |
| - | - | 628.6 | 683.4 | - | - | 0 | - |
| 7 | c | 848 | 694.4 | 0.0003809 | 0.5485 | +1 | 7 |
| - | - | 992.9 | 697.3 | - | - | 0 | - |
| - | - | 2878 | 700.4 | - | - | 0 | - |
| - | - | 1401 | 701.4 | - | - | 0 | - |
| - | - | 3530 | 710.4 | - | - | 0 | - |
| 7 | c | 2.243E+04 | 711.5 | 0.0006849 | 0.9626 | +1 | 7 |
| - | - | 8350 | 712.5 | - | - | 0 | - |
| - | - | 1931 | 713.5 | - | - | 0 | - |
| - | - | 8332 | 738.5 | - | - | 0 | - |
| - | - | 6506 | 739.5 | - | - | 0 | - |
| - | - | 1822 | 740.5 | - | - | 0 | - |
| - | - | 678.1 | 741.5 | - | - | 0 | - |
| 5 | y | 732.9 | 752.4 | 0.001586 | 2.108 | +1 | 8 |
| - | - | 849.3 | 753.4 | - | - | 0 | - |
| 8 | c | 7959 | 765.5 | 0.0008125 | 1.061 | +1 | 8 |
| - | - | 1071 | 765.9 | - | - | 0 | - |
| - | - | 593.7 | 766.4 | - | - | 0 | - |
| - | - | 3484 | 766.5 | - | - | 0 | - |
| - | - | 1478 | 769.4 | - | - | 0 | - |
| 5 | y | 2.707E+04 | 770.4 | 0.0005557 | 0.7213 | +1 | 8 |
| - | - | 1.077E+04 | 771.4 | - | - | 0 | - |
| - | - | 2595 | 772.4 | - | - | 0 | - |
| - | - | 8729 | 777.5 | - | - | 0 | - |
| - | - | 4050 | 778.5 | - | - | 0 | - |
| - | - | 1391 | 779.5 | - | - | 0 | - |
| - | - | 630.5 | 779.7 | - | - | 0 | - |
| - | - | 719.1 | 780.4 | - | - | 0 | - |
| - | - | 1873 | 798.4 | - | - | 0 | - |
| - | - | 2346 | 807.5 | - | - | 0 | - |
| - | - | 899.7 | 808.5 | - | - | 0 | - |
| - | - | 692.1 | 809.4 | - | - | 0 | - |
| - | - | 3303 | 810.4 | - | - | 0 | - |
| - | - | 849.5 | 811.4 | - | - | 0 | - |
| - | - | 1705 | 817.9 | - | - | 0 | - |
| - | - | 2524 | 822.5 | - | - | 0 | - |
| 4 | w | 1.287E+04 | 824.5 | 0.0003782 | 0.4588 | +1 | 9 |
| - | - | 6271 | 825.5 | - | - | 0 | - |
| - | - | 1236 | 826.5 | - | - | 0 | - |
| - | - | 1570 | 835.5 | - | - | 0 | - |
| - | - | 1189 | 836.5 | - | - | 0 | - |
| - | - | 826.4 | 836.5 | - | - | 0 | - |
| - | - | 1224 | 837.5 | - | - | 0 | - |
| - | - | 773.9 | 838.5 | - | - | 0 | - |
| - | - | 6058 | 863.5 | - | - | 0 | - |
| 4 | z | 3515 | 864.5 | 0.005007 | 5.792 | +1 | 9 |
| - | - | 828.5 | 865.5 | - | - | 0 | - |
| - | - | 805.4 | 867.4 | - | - | 0 | - |
| - | - | 1120 | 877.4 | - | - | 0 | - |
| - | - | 1269 | 877.9 | - | - | 0 | - |
| - | - | 883.4 | 878.4 | - | - | 0 | - |
| - | - | 3.702E+04 | 878.5 | - | - | 0 | - |
| 9 | c | 4.117E+04 | 879.5 | 0.005846 | 6.646 | +1 | 9 |
| - | - | 1.503E+04 | 880.5 | - | - | 0 | - |
| 4 | y | 2047 | 881.5 | 0.003976 | 4.511 | +1 | 9 |
| - | - | 2866 | 881.5 | - | - | 0 | - |
| 4 | z | 2.044E+04 | 882.5 | 0.0009805 | 1.111 | +1 | 9 |
| - | - | 8951 | 883.5 | - | - | 0 | - |
| - | - | 2692 | 884.5 | - | - | 0 | - |
| - | - | 606.5 | 888.5 | - | - | 0 | - |
| - | - | 3502 | 897.5 | - | - | 0 | - |
| 4 | y | 2.377E+04 | 898.5 | 0.001272 | 1.416 | +1 | 9 |
| - | - | 1.101E+04 | 899.5 | - | - | 0 | - |
| - | - | 3500 | 900.5 | - | - | 0 | - |
| - | - | 7506 | 931.5 | - | - | 0 | - |
| - | - | 4558 | 932.5 | - | - | 0 | - |
| - | - | 1640 | 933.5 | - | - | 0 | - |
| - | - | 683.3 | 934.6 | - | - | 0 | - |
| - | - | 961.4 | 938.4 | - | - | 0 | - |
| 3 | z | 4160 | 939.5 | 0.002181 | 2.321 | +1 | 10 |
| - | - | 2771 | 940.5 | - | - | 0 | - |
| 10 | c | 1.465E+04 | 949.5 | 0.0006439 | 0.6781 | +1 | 10 |
| - | - | 8804 | 950.5 | - | - | 0 | - |
| - | - | 2329 | 951.6 | - | - | 0 | - |
| 3 | y | 1.54E+04 | 955.5 | 0.001068 | 1.118 | +1 | 10 |
| - | - | 7470 | 956.5 | - | - | 0 | - |
| - | - | 1779 | 957.5 | - | - | 0 | - |
| 10 | c | 2.585E+04 | 966.6 | 0.001131 | 1.17 | +1 | 10 |
| - | - | 788.7 | 967.5 | - | - | 0 | - |
| - | - | 1.455E+04 | 967.6 | - | - | 0 | - |
| - | - | 1113 | 968.5 | - | - | 0 | - |
| - | - | 5436 | 968.6 | - | - | 0 | - |
| - | - | 1797 | 982.5 | - | - | 0 | - |
| - | - | 1005 | 983.5 | - | - | 0 | - |
| - | - | 992 | 1007 | - | - | 0 | - |
| 2 | w | 1543 | 1010 | 0.003881 | 3.845 | +1 | 11 |
| - | - | 1421 | 1011 | - | - | 0 | - |
| - | - | 763 | 1012 | - | - | 0 | - |
| - | - | 1847 | 1022 | - | - | 0 | - |
| - | - | 972.2 | 1024 | - | - | 0 | - |
| - | - | 906.4 | 1036 | - | - | 0 | - |
| 11 | c | 6878 | 1049 | 0.001553 | 1.481 | +1 | 11 |
| - | - | 5398 | 1050 | - | - | 0 | - |
| - | - | 5043 | 1051 | - | - | 0 | - |
| - | - | 1948 | 1052 | - | - | 0 | - |
| 2 | z | 4412 | 1053 | 0.0007306 | 0.6941 | +1 | 11 |
| - | - | 2698 | 1054 | - | - | 0 | - |
| - | - | 999.9 | 1055 | - | - | 0 | - |
| 11 | c | 5.352E+04 | 1066 | 0.001369 | 1.284 | +1 | 11 |
| - | - | 3.206E+04 | 1067 | - | - | 0 | - |
| - | - | 1.223E+04 | 1068 | - | - | 0 | - |
| - | - | 1770 | 1069 | - | - | 0 | - |
| - | - | 1014 | 1080 | - | - | 0 | - |
| - | - | 3236 | 1081 | - | - | 0 | - |
| - | - | 1492 | 1082 | - | - | 0 | - |
| - | - | 1128 | 1083 | - | - | 0 | - |
| - | - | 1.112E+04 | 1096 | - | - | 0 | - |
| - | - | 6186 | 1097 | - | - | 0 | - |
| - | - | 2719 | 1098 | - | - | 0 | - |
| - | - | 742.4 | 1099 | - | - | 0 | - |
| - | - | 1211 | 1105 | - | - | 0 | - |
| - | - | 689.5 | 1110 | - | - | 0 | - |
| - | - | 3193 | 1111 | - | - | 0 | - |
| - | - | 1732 | 1112 | - | - | 0 | - |
| - | - | 724.9 | 1123 | - | - | 0 | - |
| - | - | 2184 | 1123 | - | - | 0 | - |
| - | - | 4693 | 1124 | - | - | 0 | - |
| - | - | 2959 | 1125 | - | - | 0 | - |
| - | - | 1095 | 1126 | - | - | 0 | - |
| - | - | 1635 | 1131 | - | - | 0 | - |
| - | - | 2580 | 1132 | - | - | 0 | - |
| - | - | 1535 | 1132 | - | - | 0 | - |
| - | - | 1009 | 1133 | - | - | 0 | - |
| - | - | 871.3 | 1138 | - | - | 0 | - |
| - | - | 1033 | 1140 | - | - | 0 | - |
| - | - | 903.6 | 1140 | - | - | 0 | - |
| - | - | 721.7 | 1141 | - | - | 0 | - |
| - | - | 695.7 | 1147 | - | - | 0 | - |
| - | - | 1557 | 1147 | - | - | 0 | - |
| - | - | 1403 | 1150 | - | - | 0 | - |
| - | - | 5109 | 1151 | - | - | 0 | - |
| - | - | 4.998E+04 | 1152 | - | - | 0 | - |
| - | - | 3.088E+04 | 1153 | - | - | 0 | - |
| - | - | 1.041E+04 | 1154 | - | - | 0 | - |
| - | - | 1955 | 1155 | - | - | 0 | - |
| - | - | 779.5 | 1161 | - | - | 0 | - |
| - | - | 2219 | 1161 | - | - | 0 | - |
| - | - | 1543 | 1162 | - | - | 0 | - |
| - | - | 869.1 | 1162 | - | - | 0 | - |
| - | - | 1513 | 1168 | - | - | 0 | - |
| - | - | 5.012E+04 | 1168 | - | - | 0 | - |
| - | - | 1.512E+05 | 1169 | - | - | 0 | - |
| - | - | 9.1E+04 | 1170 | - | - | 0 | - |
| - | - | 1932 | 1170 | - | - | 0 | - |
| - | - | 2497 | 1171 | - | - | 0 | - |
| - | - | 3.113E+04 | 1171 | - | - | 0 | - |
| - | - | 1356 | 1171 | - | - | 0 | - |
| - | - | 4630 | 1172 | - | - | 0 | - |
| - | - | 908.1 | 1229 | - | - | 0 | - |
| - | - | 1036 | 1230 | - | - | 0 | - |
| - | - | 1563 | 1240 | - | - | 0 | - |
| - | - | 942.5 | 1241 | - | - | 0 | - |
| - | - | 785.4 | 1358 | - | - | 0 | - |
| - | - | 772.1 | 1359 | - | - | 0 | - |
| - | - | 1439 | 1400 | - | - | 0 | - |
| - | - | 1888 | 1403 | - | - | 0 | - |
| - | - | 1255 | 1404 | - | - | 0 | - |
| - | - | 1108 | 1414 | - | - | 0 | - |
| - | - | 984.5 | 1437 | - | - | 0 | - |
| - | - | 809.3 | 1438 | - | - | 0 | - |
| - | - | 1491 | 1452 | - | - | 0 | - |
| - | - | 1189 | 1452 | - | - | 0 | - |
| - | - | 1260 | 1460 | - | - | 0 | - |
| - | - | 1130 | 1461 | - | - | 0 | - |
| - | - | 1513 | 1529 | - | - | 0 | - |
| - | - | 1735 | 1530 | - | - | 0 | - |
| - | - | 1136 | 1531 | - | - | 0 | - |
| - | - | 1680 | 1532 | - | - | 0 | - |
| - | - | 1088 | 1617 | - | - | 0 | - |
| - | - | 1131 | 1618 | - | - | 0 | - |
| - | - | 2952 | 1635 | - | - | 0 | - |
| - | - | 1764 | 1636 | - | - | 0 | - |
| - | - | 790.3 | 1697 | - | - | 0 | - |
| - | - | 730.1 | 1708 | - | - | 0 | - |
| - | - | 699.3 | 1720 | - | - | 0 | - |
| - | - | 1163 | 1755 | - | - | 0 | - |
| - | - | 2577 | 1756 | - | - | 0 | - |
| - | - | 1670 | 1757 | - | - | 0 | - |
| - | - | 1158 | 2071 | - | - | 0 | - |
| - | - | 1516 | 2262 | - | - | 0 | - |
| - | - | 1946 | 2263 | - | - | 0 | - |
| - | - | 803.9 | 2265 | - | - | 0 | - |
| - | - | 878.5 | 2267 | - | - | 0 | - |
| - | - | 1124 | 2278 | - | - | 0 | - |
| - | - | 1354 | 2280 | - | - | 0 | - |
| - | - | 1218 | 2281 | - | - | 0 | - |
| - | - | 1415 | 2282 | - | - | 0 | - |
| - | - | 921.7 | 2294 | - | - | 0 | - |
| - | - | 791.2 | 2321 | - | - | 0 | - |
| - | - | 1196 | 2322 | - | - | 0 | - |
| - | - | 1369 | 2323 | - | - | 0 | - |
| - | - | 2357 | 2338 | - | - | 0 | - |
| - | - | 2255 | 2339 | - | - | 0 | - |
| - | - | 1457 | 2340 | - | - | 0 | - |

m/z Charge Intensity FragmentType MassShift Position
120.06560516357422 0 7852.054 y 11
120.08087921142578 0 2055.4485
121.06893920898438 0 444.61902
129.10220336914062 0 2168.7175
150.15570068359375 0 483.4093
173.45252990722656 0 1105.0558
175.89698791503906 0 558.5548
179.0678253173828 0 462.04703
179.19882202148438 0 464.73486
185.09210205078125 0 709.5207
185.1647491455078 0 8767.097
186.0870361328125 0 687.5221
186.16787719726562 0 892.84985
201.12295532226562 0 583.0755 y Water loss 10
207.4173583984375 0 576.6896
212.99876403808594 0 466.95776
213.1597137451172 0 10771.6875
214.16334533691406 0 970.17255
215.13938903808594 0 796.17755
219.13388061523438 0 3858.3896 y 10
219.1490936279297 0 3300.5942
234.1090545654297 0 978.51904
247.1447296142578 0 1643.7964
251.1500244140625 0 1239.0626
270.18060302734375 0 958.2552
284.1594543457031 0 1178.1267
287.2082824707031 0 2143.404 c 2
299.1705017089844 0 945.80566
330.03790283203125 0 579.6339
345.2256164550781 0 1704.2429
346.2329406738281 0 2441.9832
350.1512756347656 0 860.79144
355.1966857910156 0 912.62256
368.2275390625 0 1341.4348
370.24468994140625 0 1341.2532
371.0752868652344 0 1453.4468
383.19512939453125 0 982.15326
383.2362060546875 0 653.78534 c Ammonia loss 7
392.1387023925781 0 634.47095
398.2394714355469 0 6618.474 c Ammonia loss 3
399.24322509765625 0 1170.6003
403.2189025878906 0 5600.992 y 8
404.2198181152344 0 825.79407
431.7623596191406 0 824.3814 c Ammonia loss 8
435.2696838378906 0 591.03125
440.25103759765625 0 2557.528
440.7511291503906 0 1976.7177 y Water loss 3
447.2811279296875 0 4545.4
448.28387451171875 0 1595.215
460.78741455078125 0 987.9599
466.2347717285156 0 771.2964
466.2717590332031 0 6200.0054
466.77294921875 0 3230.19
474.254638671875 0 839.844 y 7
475.2762756347656 0 3499.6235 c Ammonia loss 9
475.7786560058594 0 1794.8135
476.2747802734375 0 709.9749
483.256591796875 0 844.584
484.2402648925781 0 3679.0017
485.24151611328125 0 1409.4808
501.26715087890625 0 1177.7035
501.8072509765625 0 2670.1172
504.2543640136719 0 2433.5605
504.3023986816406 0 5921.349
505.2578125 0 833.0668
505.3056335449219 0 1896.4329
510.8143310546875 0 3425.7678
511.31414794921875 0 1900.2766
514.2977294921875 0 1040.5734
515.8058471679688 0 5462.941
516.3070678710938 0 3895.449
516.8068237304688 0 1146.989
518.3035888671875 0 800.1024
519.3110961914062 0 707.1564
524.8101196289062 0 3961.8577 c Ammonia loss 10
525.31103515625 0 1752.8536
525.8101806640625 0 661.3196 y Water loss 1
535.271728515625 0 636.2122
536.2823486328125 0 4463.172
537.283935546875 0 1041.7081
552.3138427734375 0 1474.2175
553.3094482421875 0 1407.0161
556.3445434570312 0 3485.1494
557.3427734375 0 740.9123
561.3220825195312 0 2292.3928
566.2272338867188 0 660.1506
575.333251953125 0 4033.929
575.835693359375 0 3545.917
576.3360595703125 0 2628.1152
583.2657470703125 0 815.61316
583.3201904296875 0 1803.4099
584.34130859375 0 2301.6775
584.8436889648438 0 1297.1427
585.209228515625 0 1697.3507
585.2822875976562 0 668.6263
585.331787109375 0 1795.8732
618.3433227539062 0 718.2542
628.3867797851562 0 692.2832
633.3583984375 0 8134.9844
634.3615112304688 0 2283.608
639.4053344726562 0 6647.828
640.4130859375 0 32099.662 c 5
641.4156494140625 0 10010.488
642.4176025390625 0 2532.2175
651.3799438476562 0 3268.619
657.3687744140625 0 8109.553 z 5
658.3751831054688 0 6403.404
659.3799438476562 0 1923.3938
661.3656616210938 0 656.05225
664.3779907226562 0 6651.0273
665.3804931640625 0 2607.0208
666.3861083984375 0 992.78906
667.4365844726562 0 2306.4666
672.3795166015625 0 774.6748
673.3872680664062 0 8368.549 y 5
674.3900756835938 0 2334.379
683.376220703125 0 628.5847
694.4242553710938 0 847.9555 c Ammonia loss 6
697.3408203125 0 992.8788
700.3984375 0 2877.7654
701.3961791992188 0 1400.5095
710.4435424804688 0 3529.799
711.4505004882812 0 22433.727 c 6
712.45263671875 0 8350.385
713.456298828125 0 1931.0012
738.473876953125 0 8332.288
739.479248046875 0 6506.2676
740.4805908203125 0 1822.3469
741.4926147460938 0 678.084
752.4317016601562 0 732.9074 y Water loss 4
753.43359375 0 849.3133
765.4609375 0 7958.699 c Ammonia loss 7
765.8873901367188 0 1071.3591
766.4033203125 0 593.65845
766.4629516601562 0 3483.7305
769.4338989257812 0 1478.2723
770.4401245117188 0 27065.766 y 4
771.4432983398438 0 10770.844
772.447265625 0 2594.5752
777.4613037109375 0 8729.189
778.4639282226562 0 4050.1533
779.4693603515625 0 1391.3236
779.7125244140625 0 630.4919
780.4221801757812 0 719.0659
798.4359741210938 0 1872.8344
807.495361328125 0 2345.5
808.483154296875 0 899.7288
809.3750610351562 0 692.11786
810.4263916015625 0 3303.285
811.4309692382812 0 849.4964
817.8833618164062 0 1704.6506
822.4710083007812 0 2523.9429
824.4508666992188 0 12870.2295 w 3
825.4534912109375 0 6270.763
826.45361328125 0 1235.9008
835.5244750976562 0 1569.5573
836.4568481445312 0 1189.1139
836.5352783203125 0 826.39026
837.4586791992188 0 1223.695
838.4727172851562 0 773.9346
863.4747924804688 0 6057.617
864.4749755859375 0 3514.6514 z Water loss 3
865.4725952148438 0 828.5438
867.4186401367188 0 805.3998
877.3905639648438 0 1119.6241
877.8863525390625 0 1268.939
878.386474609375 0 883.44354
878.531982421875 0 37023.418
879.5352172851562 0 41172.816 c 8
880.5390625 0 15027.836
881.4766845703125 0 2046.6747 y Ammonia loss 3
881.5493774414062 0 2865.9739
882.4795532226562 0 20443.367 z 3
883.4818725585938 0 8951.02
884.4842529296875 0 2691.8982
888.4515380859375 0 606.48193
897.4886474609375 0 3502.2043
898.4979858398438 0 23771.633 y 3
899.5009155273438 0 11011.848
900.5030517578125 0 3500.2896
931.5350952148438 0 7506.327
932.5379638671875 0 4558.4106
933.5419311523438 0 1639.9828
934.5574951171875 0 683.2992
938.4274291992188 0 961.418
939.4998168945312 0 4160.2783 z 2
940.5050048828125 0 2770.8047
949.5458984375 0 14654.367 c Ammonia loss 9
950.548095703125 0 8804.259
951.5542602539062 0 2328.5981
955.5196533203125 0 15396.304 y 2
956.5230102539062 0 7470.3423
957.5223999023438 0 1779.104
966.5719604492188 0 25849.729 c 9
967.4622802734375 0 788.6555
967.5738525390625 0 14547.869
968.47265625 0 1113.022
968.5772094726562 0 5436.1826
982.476806640625 0 1796.5642
983.4801635742188 0 1004.93896
1006.6163330078125 0 992.0352
1009.5274047851562 0 1542.8268 w 1
1010.5302124023438 0 1420.9836
1011.5119018554688 0 762.98395
1021.6282958984375 0 1846.7638
1023.6414794921875 0 972.17993
1035.99365234375 0 906.413
1048.6134033203125 0 6877.9233 c Ammonia loss 10
1049.614990234375 0 5398.0894
1050.6241455078125 0 5043.2173
1051.630126953125 0 1948.2979
1052.5867919921875 0 4412.031 z 1
1053.5882568359375 0 2697.9148
1054.59521484375 0 999.8925
1065.64013671875 0 53516.285 c 10
1066.642578125 0 32060.822
1067.645263671875 0 12233.027
1068.642333984375 0 1769.6145
1079.6473388671875 0 1014.2104
1080.6168212890625 0 3235.6062
1081.6217041015625 0 1492.2222
1082.630126953125 0 1127.5381
1095.5904541015625 0 11121.586
1096.5946044921875 0 6186.3013
1097.6083984375 0 2719.4758
1098.648193359375 0 742.42236
1104.5240478515625 0 1210.9426
1109.58740234375 0 689.54004
1110.5670166015625 0 3193.1765
1111.5731201171875 0 1732.3275
1122.5447998046875 0 724.9472
1122.6761474609375 0 2183.7698
1123.661376953125 0 4692.659
1124.66259765625 0 2959.2913
1125.662109375 0 1094.9042
1131.0615234375 0 1635.2965
1131.5657958984375 0 2580.004
1132.0589599609375 0 1535.1582
1132.5653076171875 0 1008.69324
1138.066162109375 0 871.34576
1139.568603515625 0 1033.354
1140.0606689453125 0 903.55786
1140.580810546875 0 721.74774
1146.5572509765625 0 695.7309
1147.068359375 0 1557.1733
1149.5135498046875 0 1402.5612
1150.6663818359375 0 5108.718
1151.653564453125 0 49976.16
1152.6561279296875 0 30879.33
1153.6590576171875 0 10408.846
1154.6590576171875 0 1954.9031
1160.5784912109375 0 779.4707
1161.0662841796875 0 2219.1104
1161.5626220703125 0 1542.974
1162.06201171875 0 869.0514
1167.546875 0 1512.8113
1167.671630859375 0 50121.207
1168.6781005859375 0 151223.1
1169.6817626953125 0 90997.57
1170.075927734375 0 1932.104
1170.560546875 0 2496.889
1170.6845703125 0 31133.545
1171.0877685546875 0 1355.7328
1171.6866455078125 0 4630.3545
1228.57666015625 0 908.149
1229.5806884765625 0 1036.2946
1239.6121826171875 0 1562.7914
1240.6142578125 0 942.4721
1357.68408203125 0 785.38544
1358.686279296875 0 772.1011
1399.6822509765625 0 1439.0763
1402.6734619140625 0 1887.9023
1403.6708984375 0 1254.7927
1413.6961669921875 0 1107.5189
1437.224609375 0 984.5134
1437.728759765625 0 809.2947
1451.7294921875 0 1490.7533
1452.2352294921875 0 1188.6505
1460.2506103515625 0 1259.7125
1461.23828125 0 1130.0135
1528.7392578125 0 1512.9226
1529.72705078125 0 1735.234
1530.74951171875 0 1136.2784
1531.7559814453125 0 1680.4691
1616.7899169921875 0 1088.3694
1617.759765625 0 1131.3726
1634.7535400390625 0 2951.6213
1635.7608642578125 0 1764.4274
1696.8746337890625 0 790.3112
1707.78564453125 0 730.1361
1719.9180908203125 0 699.33356
1754.7781982421875 0 1163.4886
1755.789794921875 0 2576.6396
1756.7904052734375 0 1670.2589
2070.97900390625 0 1158.052
2262.120849609375 0 1515.5289
2263.13134765625 0 1946.0157
2265.121337890625 0 803.8522
2267.1328125 0 878.50806
2278.11767578125 0 1123.648
2280.136962890625 0 1353.8146
2281.126220703125 0 1217.561
2282.12890625 0 1414.9172
2294.13427734375 0 921.66864
2321.09912109375 0 791.2241
2322.110107421875 0 1196.419
2323.129638671875 0 1368.6526
2338.13818359375 0 2356.8406
2339.131103515625 0 2254.7488
2340.14453125 0 1456.724

Spectrum Details

|  |  |
| --- | --- |
| Matched peaks? Matched peaks The total absolute number of peaks matched. Additionally in brackets the total fraction of peaks matched and the total number of peaks is shown. | 35 (11.29% of 310) |
| FDR? FDR The false discovery rate estimated for this peptide. It is calculated by matching all theoretical fragments with a non-integer shift with the raw peaks for this spectrum. This is done with 40 different shifts. The resulting percentage is the average number of annotated peaks over the number of annotated peaks with the correct spectrum. | 0.82% |
| Satellite FDR? Satellite FDR See the FDR for details on its calculation. This satellite ion specific FDR only contains the satellite ions (d/w) for I/L/J positions. | 4.76% |
| PSM Score? PSM Score The PSM Score as given by Hecklib to this annotated spectrum. It is shown with three significant figures. | 341 |

#### Reverse Lookup? Reverse Lookup All places where this read could be placed.

| Group | Segment | Template | Template Part | Read Part | Score | Unique |
| --- | --- | --- | --- | --- | --- | --- |
| Homo sapiens Light Chain | IGLC | IGLC2 | [0..10] | [2..12] | 80 | False |
| Homo sapiens Light Chain | IGLC | IGLC6 | [0..10] | [2..12] | 80 | False |
| Homo sapiens Light Chain | IGLC | IGLC7 | [0..10] | [2..12] | 80 | False |

| Recombined | Template Part | Read Part | Score | Unique |
| --- | --- | --- | --- | --- |
| REC-0-1\_002 | [109..121] | [0..12] | 96 | True |

#### Meta Information from Multiple reads

##### Number of combined reads

4

##### Intensity

0.9781

##### TotalArea

9.979E+08

##### Changes to the peptide sequence

VLGQPKAAPSVT

J→LSupport for Leucine based on side chain ions (1 for L 0 for I) (Position: 2)

L→JEqual support for both Leucine and Isoleucine based on side chain ions (1 ions for both) (Position: 2)

#### Positional Score

Copy Data

##### Positional Score (TSV)

###### Preview

```
Loading example...
```

*Click on the button to copy the data to your clipboard.*

1001234567891011

Label Value
"0" 0.495
"1" 0.495
"2" 0.497
"3" 0.48
"4" 0.49
"5" 0.485
"6" 0.492
"7" 0.477
"8" 0.482
"9" 0.492
"10" 0.495
"11" 0.492

#### Meta Information from PEAKS

##### Scan Identifier

F1:4087

##### Original sequence

V

L

G

Q

P

K

A

A

P

S

V

T

##### Posttranslational Modifications

##### Source File

D:\separate\_stitch\_analyses\xle-disambiguation\raw\20210323\_F1\_UM1\_Peng0013\_SA\_F59\_ingel\_3ug\_ELA.raw

##### Fraction

1

##### Scan Feature

F1:7761

##### De Novo Score

99

##### ConfidenceScore

99

### m/z

584.3408

##### Mass

1166.6658

##### Charge

2

##### Retention Time

21.49

##### Predicted Retention Time

-

##### Area

9.979E+08

##### Parts Per Million

1.1

##### Fragmentation mode

ETHCD

##### Originating file

01 D:\separate\_stitch\_analyses\xle-disambiguation\20210325\_F59\_3ug\_DENOVO\_12.csv

#### Meta Information from PEAKS

##### Scan Identifier

F1:4224

##### Original sequence

V

L

G

Q

P

K

A

A

P

S

V

T

##### Posttranslational Modifications

##### Source File

D:\separate\_stitch\_analyses\xle-disambiguation\raw\20210323\_F1\_UM1\_Peng0013\_SA\_F59\_ingel\_3ug\_ELA.raw

##### Fraction

1

##### Scan Feature

-

##### De Novo Score

99

##### ConfidenceScore

99

### m/z

584.3412

##### Mass

1166.6658

##### Charge

2

##### Retention Time

22.26

##### Predicted Retention Time

-

##### Area

0

##### Parts Per Million

1.9

##### Fragmentation mode

ETHCD

##### Originating file

01 D:\separate\_stitch\_analyses\xle-disambiguation\20210325\_F59\_3ug\_DENOVO\_12.csv

#### Meta Information from PEAKS

##### Scan Identifier

F1:4298

##### Original sequence

V

L

G

Q

P

K

A

A

P

S

V

T

##### Posttranslational Modifications

##### Source File

D:\separate\_stitch\_analyses\xle-disambiguation\raw\20210323\_F1\_UM1\_Peng0013\_SA\_F59\_ingel\_3ug\_ELA.raw

##### Fraction

1

##### Scan Feature

-

##### De Novo Score

98

##### ConfidenceScore

98

### m/z

584.3409

##### Mass

1166.6658

##### Charge

2

##### Retention Time

22.69

##### Predicted Retention Time

-

##### Area

0

##### Parts Per Million

1.2

##### Fragmentation mode

ETHCD

##### Originating file

01 D:\separate\_stitch\_analyses\xle-disambiguation\20210325\_F59\_3ug\_DENOVO\_12.csv

#### Meta Information from PEAKS

##### Scan Identifier

F1:4546

##### Original sequence

V

L

G

Q

P

K

A

A

P

S

V

T

##### Posttranslational Modifications

##### Source File

D:\separate\_stitch\_analyses\xle-disambiguation\raw\20210323\_F1\_UM1\_Peng0013\_SA\_F59\_ingel\_3ug\_ELA.raw

##### Fraction

1

##### Scan Feature

-

##### De Novo Score

97

##### ConfidenceScore

97

### m/z

584.34

##### Mass

1166.6658

##### Charge

2

##### Retention Time

24.09

##### Predicted Retention Time

-

##### Area

0

##### Fragmentation mode

ETHCD

##### Originating file

01 D:\separate\_stitch\_analyses\xle-disambiguation\20210325\_F59\_3ug\_DENOVO\_12.csv
