## Supplementary material for "A handle on mass coincidence errors in *de novo* sequencing of antibodies by bottom-up proteomics": Combined_034.html

Details Combined\_034 | Stitch OverviewUndefined

### Read Combined\_034

#### Sequence (length=15)

VYTJPPSREEMTKNQ

#### Spectrum 3515? Spectrum 3515 The raw spectrum of this peptide as annotated by Hecklib. The fragments are coloured according to ion type (see legend). Any peaks with a star '\*' as text can be hovered over to see the full details, first the ion type second the mass shift type. By hovering over the amino acids in the peptide or ions in the legend the corresponding peaks are highlighted. By toggling the 'Unassigned' label you can turn the background (unassigned) peaks on or off in the plot. By updating the slider in the Ion legend you can update the spectrum to only show the top X% of the peaks with labels. The top X% means any peak that is within X% of the highest intensity. By dragging in the spectrum you can zoom in to a specific part of the spectrum and use 'Zoom Out' to get back to the original zoom level. The annotation of the spectrum is based on the given sequence in the peptides file and is done with different software so inconsistencies are likely. The peaks are annotated based on the given sequence, with 20 ppm tolerance.

Copy Data

##### Spectrum 3515 (TSV)

###### Preview

```
Loading example...
```

*Click on the button to copy the data to your clipboard.*

Mz MinMz MaxIntensity Max

WidthHeightPeptide font sizePeptide stroke widthSpectrum font sizeSpectrum stroke widthCompact peptide

Ion legend

wxyz

abcd

OtherUnassignedIon^Charge^_Position_Show for top:%

VYTJPPSREEMTKNQ

01.82e+43.64e+45.45e+47.27e+4

Zoom Out

y^+1^_1_y^+1^_1_y^+1^_2_y^+1^_2_w^+1^_3_z^+1^_3_y^+1^_3_y^+3^_11_w^+1^_4_w^+1^_4_w^+3^_12_y^+1^_4_z^+1^_4_y^+1^_4_y^+3^_13_y^+3^_13_y^+3^_14_y^+3^_14_z^+3^_14_z^+1^_5_y^+2^_10_z^+1^_5_y^+1^_5_y^+2^_11_y^+2^_11_z^+2^_11_y^+2^_11_c^+2^_11_c^+1^_6_w^+1^_6_w^+2^_12_z^+2^_12_y^+2^_12_z^+1^_6_y^+2^_13_y^+2^_13_z^+2^_13_y^+2^_13_c^+2^_13_c^+2^_13_c^+1^_7_c^+2^_13_w^+1^_7_c^+2^_14_y^+2^_14_y^+2^_14_z^+2^_14_y^+2^_14_z^+1^_7_y^+1^_7_c^+1^_8_z^+1^_8_y^+1^_8_c^+1^_9_z^+1^_9_y^+1^_9_c^+1^_10_y^+1^_10_y^+1^_11_c^+1^_11_z^+1^_12_c^+1^_12_y^+1^_12_z^+1^_13_y^+1^_13_c^+1^_13_c^+1^_13_c^+1^_13_c^+1^_14_z^+1^_14_

045891713751833

Fragment Matches Table

Show background peaks

| Position | Ion type | Intensity | mz Theoretical | mz Error (Th) | mz Error (ppm) | Charge | Series Number |
| --- | --- | --- | --- | --- | --- | --- | --- |
| - | - | 854.4 | 129.1 | - | - | 0 | - |
| 15 | y | 886.4 | 130 | 5.721E-05 | 0.4399 | +1 | 1 |
| - | - | 3528 | 136.1 | - | - | 0 | - |
| - | - | 426.3 | 144.2 | - | - | 0 | - |
| - | - | 479 | 145.7 | - | - | 0 | - |
| 15 | y | 6122 | 147.1 | 0.00014 | 0.9516 | +1 | 1 |
| - | - | 439.6 | 149 | - | - | 0 | - |
| - | - | 442.5 | 157 | - | - | 0 | - |
| - | - | 461 | 161.9 | - | - | 0 | - |
| - | - | 757.9 | 163.1 | - | - | 0 | - |
| - | - | 480 | 169.1 | - | - | 0 | - |
| - | - | 2253 | 173.4 | - | - | 0 | - |
| - | - | 621.5 | 183.1 | - | - | 0 | - |
| - | - | 561.7 | 184.1 | - | - | 0 | - |
| - | - | 1189 | 187.1 | - | - | 0 | - |
| - | - | 718.9 | 197.1 | - | - | 0 | - |
| - | - | 508 | 199.1 | - | - | 0 | - |
| - | - | 3040 | 212.1 | - | - | 0 | - |
| - | - | 1200 | 215.1 | - | - | 0 | - |
| - | - | 646.7 | 216.1 | - | - | 0 | - |
| - | - | 496.4 | 223.6 | - | - | 0 | - |
| - | - | 517.4 | 225.5 | - | - | 0 | - |
| - | - | 1.318E+04 | 235.1 | - | - | 0 | - |
| - | - | 2219 | 236.1 | - | - | 0 | - |
| - | - | 528.1 | 238 | - | - | 0 | - |
| 14 | y | 1030 | 244.1 | 0.0001761 | 0.7216 | +1 | 2 |
| - | - | 572.1 | 256.2 | - | - | 0 | - |
| 14 | y | 3818 | 261.1 | 0.0002665 | 1.021 | +1 | 2 |
| - | - | 1.051E+04 | 263.1 | - | - | 0 | - |
| - | - | 1230 | 264.1 | - | - | 0 | - |
| - | - | 612.4 | 298.1 | - | - | 0 | - |
| - | - | 2113 | 299.2 | - | - | 0 | - |
| 13 | w | 888.7 | 315.1 | 0.001005 | 3.188 | +1 | 3 |
| - | - | 1709 | 329.2 | - | - | 0 | - |
| - | - | 6263 | 346.2 | - | - | 0 | - |
| - | - | 924.4 | 347.2 | - | - | 0 | - |
| - | - | 979.8 | 360.2 | - | - | 0 | - |
| - | - | 8383 | 364.2 | - | - | 0 | - |
| - | - | 1600 | 365.2 | - | - | 0 | - |
| 13 | z | 2848 | 373.2 | 0.0003946 | 1.057 | +1 | 3 |
| - | - | 1048 | 374.2 | - | - | 0 | - |
| - | - | 1169 | 386.2 | - | - | 0 | - |
| 13 | y | 2541 | 389.2 | 0.0002905 | 0.7464 | +1 | 3 |
| - | - | 727.3 | 423.1 | - | - | 0 | - |
| - | - | 992.4 | 429.2 | - | - | 0 | - |
| - | - | 3635 | 430.2 | - | - | 0 | - |
| - | - | 1069 | 431.2 | - | - | 0 | - |
| - | - | 2986 | 431.3 | - | - | 0 | - |
| - | - | 642.9 | 432.3 | - | - | 0 | - |
| 5 | y | 4872 | 444.9 | 0.001642 | 3.69 | +3 | 11 |
| - | - | 3230 | 445.2 | - | - | 0 | - |
| - | - | 1238 | 445.5 | - | - | 0 | - |
| - | - | 609.3 | 446.2 | - | - | 0 | - |
| - | - | 1523 | 449.3 | - | - | 0 | - |
| - | - | 755.4 | 450.3 | - | - | 0 | - |
| 12 | w | 1412 | 457.2 | 0.000565 | 1.236 | +1 | 4 |
| 12 | w | 802.5 | 459.2 | 0.005248 | 11.43 | +1 | 4 |
| - | - | 6080 | 459.3 | - | - | 0 | - |
| - | - | 1887 | 460.3 | - | - | 0 | - |
| 4 | w | 703 | 472.2 | 0.003169 | 6.711 | +3 | 12 |
| 12 | y | 3762 | 473.2 | 0.00027 | 0.5706 | +1 | 4 |
| 12 | z | 1403 | 474.2 | 0.002693 | 5.679 | +1 | 4 |
| - | - | 1239 | 475.2 | - | - | 0 | - |
| - | - | 956.8 | 476.3 | - | - | 0 | - |
| - | - | 2154 | 477.3 | - | - | 0 | - |
| - | - | 966.1 | 478.3 | - | - | 0 | - |
| - | - | 595.1 | 481.7 | - | - | 0 | - |
| - | - | 667.7 | 488.2 | - | - | 0 | - |
| 12 | y | 1223 | 490.3 | 0.0002805 | 0.5721 | +1 | 4 |
| - | - | 620.3 | 491.3 | - | - | 0 | - |
| - | - | 847.7 | 498.2 | - | - | 0 | - |
| 3 | y | 1701 | 510.3 | 0.002916 | 5.714 | +3 | 13 |
| 3 | y | 1540 | 510.6 | 0.007522 | 14.73 | +3 | 13 |
| - | - | 686.4 | 514.3 | - | - | 0 | - |
| - | - | 1109 | 515.3 | - | - | 0 | - |
| - | - | 1797 | 531.3 | - | - | 0 | - |
| - | - | 650.7 | 532.3 | - | - | 0 | - |
| - | - | 1048 | 536.8 | - | - | 0 | - |
| - | - | 924.7 | 537.3 | - | - | 0 | - |
| - | - | 1496 | 541.3 | - | - | 0 | - |
| - | - | 582.3 | 552.6 | - | - | 0 | - |
| - | - | 642.1 | 552.9 | - | - | 0 | - |
| - | - | 1669 | 557.3 | - | - | 0 | - |
| - | - | 7262 | 558.3 | - | - | 0 | - |
| - | - | 1.083E+04 | 559.3 | - | - | 0 | - |
| - | - | 3208 | 560.3 | - | - | 0 | - |
| - | - | 816.2 | 560.8 | - | - | 0 | - |
| 2 | y | 1085 | 564.6 | 0.002172 | 3.846 | +3 | 14 |
| 2 | y | 1518 | 564.9 | 0.006198 | 10.97 | +3 | 14 |
| 2 | z | 1157 | 565.3 | 0.004057 | 7.177 | +3 | 14 |
| - | - | 546.1 | 565.6 | - | - | 0 | - |
| - | - | 8501 | 597.6 | - | - | 0 | - |
| - | - | 6941 | 598 | - | - | 0 | - |
| - | - | 5369 | 598.3 | - | - | 0 | - |
| - | - | 900.4 | 598.6 | - | - | 0 | - |
| - | - | 1404 | 602.8 | - | - | 0 | - |
| - | - | 685.9 | 603.3 | - | - | 0 | - |
| 11 | z | 1175 | 604.2 | 0.008172 | 13.52 | +1 | 5 |
| 6 | y | 1822 | 618.3 | 0.001751 | 2.833 | +2 | 10 |
| - | - | 824.1 | 618.8 | - | - | 0 | - |
| 11 | z | 3294 | 621.3 | 0.004328 | 6.966 | +1 | 5 |
| - | - | 1412 | 622.3 | - | - | 0 | - |
| - | - | 647.2 | 623.3 | - | - | 0 | - |
| - | - | 593.5 | 631.3 | - | - | 0 | - |
| - | - | 3292 | 634.8 | - | - | 0 | - |
| - | - | 1359 | 635.3 | - | - | 0 | - |
| - | - | 641.7 | 636.3 | - | - | 0 | - |
| 11 | y | 1956 | 637.3 | 0.006173 | 9.686 | +1 | 5 |
| - | - | 662.6 | 642.3 | - | - | 0 | - |
| - | - | 645.3 | 646.3 | - | - | 0 | - |
| 5 | y | 1218 | 657.8 | 0.002197 | 3.34 | +2 | 11 |
| 5 | y | 2463 | 658.3 | 0.003598 | 5.465 | +2 | 11 |
| 5 | z | 2030 | 658.8 | 0.0009253 | 1.404 | +2 | 11 |
| - | - | 923.9 | 659.3 | - | - | 0 | - |
| 5 | y | 5.154E+04 | 666.8 | 0.002286 | 3.428 | +2 | 11 |
| - | - | 3.68E+04 | 667.3 | - | - | 0 | - |
| - | - | 1.882E+04 | 667.8 | - | - | 0 | - |
| - | - | 5694 | 668.3 | - | - | 0 | - |
| 11 | c | 684 | 668.8 | 0.007502 | 11.22 | +2 | 11 |
| - | - | 754.2 | 679.3 | - | - | 0 | - |
| - | - | 824.2 | 686.3 | - | - | 0 | - |
| - | - | 3996 | 687.3 | - | - | 0 | - |
| - | - | 6014 | 687.4 | - | - | 0 | - |
| - | - | 764.7 | 687.8 | - | - | 0 | - |
| - | - | 1715 | 688.3 | - | - | 0 | - |
| 6 | c | 5815 | 688.4 | 0.001898 | 2.757 | +1 | 6 |
| - | - | 940.9 | 689.3 | - | - | 0 | - |
| - | - | 1992 | 689.4 | - | - | 0 | - |
| 10 | w | 2.167E+04 | 691.3 | 0.00458 | 6.626 | +1 | 6 |
| - | - | 8239 | 692.3 | - | - | 0 | - |
| - | - | 2071 | 693.3 | - | - | 0 | - |
| - | - | 1.508E+04 | 693.8 | - | - | 0 | - |
| - | - | 1.361E+04 | 694.3 | - | - | 0 | - |
| - | - | 6306 | 694.8 | - | - | 0 | - |
| - | - | 1591 | 695.3 | - | - | 0 | - |
| - | - | 637.6 | 695.8 | - | - | 0 | - |
| - | - | 1168 | 697.4 | - | - | 0 | - |
| 4 | w | 715.8 | 700.8 | 0.008222 | 11.73 | +2 | 12 |
| - | - | 1923 | 713.4 | - | - | 0 | - |
| - | - | 1227 | 714.4 | - | - | 0 | - |
| 4 | z | 1.765E+04 | 715.3 | 0.002209 | 3.088 | +2 | 12 |
| - | - | 1.255E+04 | 715.8 | - | - | 0 | - |
| - | - | 6466 | 716.3 | - | - | 0 | - |
| - | - | 1405 | 716.8 | - | - | 0 | - |
| 4 | y | 7990 | 723.4 | 0.001636 | 2.261 | +2 | 12 |
| - | - | 6938 | 723.9 | - | - | 0 | - |
| - | - | 4801 | 724.4 | - | - | 0 | - |
| - | - | 1392 | 724.9 | - | - | 0 | - |
| - | - | 657.4 | 729.9 | - | - | 0 | - |
| - | - | 2484 | 732.4 | - | - | 0 | - |
| - | - | 773.4 | 733.4 | - | - | 0 | - |
| - | - | 922.3 | 736.8 | - | - | 0 | - |
| - | - | 1063 | 737.3 | - | - | 0 | - |
| - | - | 5023 | 737.8 | - | - | 0 | - |
| - | - | 841.6 | 737.9 | - | - | 0 | - |
| - | - | 2833 | 738.3 | - | - | 0 | - |
| - | - | 2285 | 738.8 | - | - | 0 | - |
| - | - | 1757 | 741.9 | - | - | 0 | - |
| - | - | 1653 | 742.4 | - | - | 0 | - |
| - | - | 963.4 | 742.9 | - | - | 0 | - |
| 10 | z | 2.199E+04 | 750.3 | 0.004826 | 6.432 | +1 | 6 |
| - | - | 1.129E+04 | 751.3 | - | - | 0 | - |
| - | - | 3592 | 752.3 | - | - | 0 | - |
| - | - | 1033 | 753.3 | - | - | 0 | - |
| - | - | 605.5 | 754.9 | - | - | 0 | - |
| - | - | 1742 | 761.4 | - | - | 0 | - |
| - | - | 2426 | 761.9 | - | - | 0 | - |
| 3 | y | 1494 | 764.9 | 0.004897 | 6.403 | +2 | 13 |
| 3 | y | 4984 | 765.4 | 0.001325 | 1.731 | +2 | 13 |
| 3 | z | 9999 | 765.9 | 0.001502 | 1.961 | +2 | 13 |
| - | - | 9090 | 766.4 | - | - | 0 | - |
| - | - | 5410 | 766.9 | - | - | 0 | - |
| - | - | 1221 | 767.4 | - | - | 0 | - |
| - | - | 1483 | 773.4 | - | - | 0 | - |
| 3 | y | 3.471E+04 | 773.9 | 0.00215 | 2.778 | +2 | 13 |
| 13 | c | 2.895E+04 | 774.4 | 0.01397 | 18.04 | +2 | 13 |
| 13 | c | 1.304E+04 | 774.9 | 0.005917 | 7.636 | +2 | 13 |
| 7 | c | 1.211E+04 | 775.4 | 0.0005582 | 0.7199 | +1 | 7 |
| - | - | 2023 | 775.9 | - | - | 0 | - |
| - | - | 6509 | 776.4 | - | - | 0 | - |
| - | - | 1724 | 777.4 | - | - | 0 | - |
| - | - | 2606 | 777.9 | - | - | 0 | - |
| - | - | 1456 | 778.4 | - | - | 0 | - |
| - | - | 1104 | 778.9 | - | - | 0 | - |
| - | - | 765.8 | 779.4 | - | - | 0 | - |
| - | - | 5362 | 782.9 | - | - | 0 | - |
| 13 | c | 3.226E+04 | 783.4 | 0.002476 | 3.161 | +2 | 13 |
| - | - | 2.857E+04 | 783.9 | - | - | 0 | - |
| - | - | 1.323E+04 | 784.4 | - | - | 0 | - |
| - | - | 5306 | 784.9 | - | - | 0 | - |
| - | - | 863.5 | 785.4 | - | - | 0 | - |
| - | - | 2130 | 787.4 | - | - | 0 | - |
| - | - | 1748 | 787.9 | - | - | 0 | - |
| - | - | 916.8 | 788.4 | - | - | 0 | - |
| - | - | 774.4 | 817.9 | - | - | 0 | - |
| - | - | 1684 | 818.4 | - | - | 0 | - |
| - | - | 2642 | 818.9 | - | - | 0 | - |
| - | - | 886 | 819.9 | - | - | 0 | - |
| 9 | w | 5903 | 820.3 | 0.005566 | 6.785 | +1 | 7 |
| - | - | 1859 | 821.4 | - | - | 0 | - |
| - | - | 1557 | 832.9 | - | - | 0 | - |
| - | - | 1241 | 837.4 | - | - | 0 | - |
| - | - | 998.1 | 837.9 | - | - | 0 | - |
| - | - | 1063 | 839.4 | - | - | 0 | - |
| - | - | 1773 | 839.9 | - | - | 0 | - |
| 14 | c | 6.882E+04 | 840.4 | 0.00207 | 2.462 | +2 | 14 |
| - | - | 6.326E+04 | 840.9 | - | - | 0 | - |
| - | - | 3.769E+04 | 841.4 | - | - | 0 | - |
| - | - | 1.363E+04 | 841.9 | - | - | 0 | - |
| - | - | 2750 | 842.4 | - | - | 0 | - |
| 2 | y | 698.4 | 846.4 | 0.01025 | 12.11 | +2 | 14 |
| 2 | y | 997.3 | 846.9 | 0.01013 | 11.96 | +2 | 14 |
| 2 | z | 5232 | 847.4 | 0.001942 | 2.292 | +2 | 14 |
| - | - | 3829 | 847.9 | - | - | 0 | - |
| - | - | 2841 | 848.4 | - | - | 0 | - |
| - | - | 731.4 | 848.9 | - | - | 0 | - |
| - | - | 688.1 | 849.4 | - | - | 0 | - |
| - | - | 830.9 | 851.9 | - | - | 0 | - |
| - | - | 1778 | 853.4 | - | - | 0 | - |
| - | - | 2519 | 853.9 | - | - | 0 | - |
| - | - | 1609 | 854.4 | - | - | 0 | - |
| - | - | 1926 | 854.9 | - | - | 0 | - |
| 2 | y | 3415 | 855.4 | 0.002468 | 2.885 | +2 | 14 |
| - | - | 3703 | 855.9 | - | - | 0 | - |
| - | - | 1266 | 856.4 | - | - | 0 | - |
| - | - | 731.8 | 856.9 | - | - | 0 | - |
| - | - | 692.6 | 859.9 | - | - | 0 | - |
| - | - | 734.1 | 860.9 | - | - | 0 | - |
| - | - | 1668 | 861.4 | - | - | 0 | - |
| - | - | 1203 | 861.9 | - | - | 0 | - |
| - | - | 881.4 | 865.9 | - | - | 0 | - |
| - | - | 1251 | 867.9 | - | - | 0 | - |
| - | - | 2243 | 868.4 | - | - | 0 | - |
| - | - | 9566 | 868.9 | - | - | 0 | - |
| - | - | 1.004E+04 | 869.4 | - | - | 0 | - |
| - | - | 6485 | 869.9 | - | - | 0 | - |
| - | - | 2362 | 870.4 | - | - | 0 | - |
| - | - | 704.2 | 870.9 | - | - | 0 | - |
| - | - | 913.1 | 873.5 | - | - | 0 | - |
| - | - | 1563 | 873.9 | - | - | 0 | - |
| - | - | 1718 | 874.4 | - | - | 0 | - |
| - | - | 1378 | 874.9 | - | - | 0 | - |
| - | - | 978.7 | 875.4 | - | - | 0 | - |
| - | - | 6488 | 875.9 | - | - | 0 | - |
| - | - | 4067 | 876.4 | - | - | 0 | - |
| - | - | 4271 | 876.9 | - | - | 0 | - |
| - | - | 5072 | 877.4 | - | - | 0 | - |
| - | - | 3713 | 877.9 | - | - | 0 | - |
| - | - | 1085 | 878.4 | - | - | 0 | - |
| 9 | z | 1.005E+04 | 879.4 | 0.005079 | 5.776 | +1 | 7 |
| - | - | 5031 | 880.4 | - | - | 0 | - |
| - | - | 1818 | 881.4 | - | - | 0 | - |
| - | - | 1524 | 882.4 | - | - | 0 | - |
| - | - | 1.183E+04 | 882.9 | - | - | 0 | - |
| - | - | 1.034E+04 | 883.4 | - | - | 0 | - |
| - | - | 1.019E+04 | 883.9 | - | - | 0 | - |
| - | - | 4626 | 884.4 | - | - | 0 | - |
| - | - | 2356 | 884.9 | - | - | 0 | - |
| - | - | 818.3 | 885.4 | - | - | 0 | - |
| - | - | 1154 | 887.5 | - | - | 0 | - |
| - | - | 1631 | 887.9 | - | - | 0 | - |
| - | - | 2995 | 888.4 | - | - | 0 | - |
| - | - | 1099 | 888.5 | - | - | 0 | - |
| - | - | 1678 | 888.9 | - | - | 0 | - |
| - | - | 1706 | 889.4 | - | - | 0 | - |
| - | - | 1336 | 889.9 | - | - | 0 | - |
| - | - | 1030 | 890.4 | - | - | 0 | - |
| 9 | y | 1764 | 895.4 | 0.005215 | 5.825 | +1 | 7 |
| - | - | 1286 | 895.9 | - | - | 0 | - |
| - | - | 5187 | 896.4 | - | - | 0 | - |
| - | - | 4.874E+04 | 896.9 | - | - | 0 | - |
| - | - | 4.163E+04 | 897.4 | - | - | 0 | - |
| - | - | 2.894E+04 | 897.9 | - | - | 0 | - |
| - | - | 1.106E+04 | 898.4 | - | - | 0 | - |
| - | - | 5726 | 898.9 | - | - | 0 | - |
| - | - | 994.6 | 899.4 | - | - | 0 | - |
| - | - | 1101 | 904.4 | - | - | 0 | - |
| - | - | 3.817E+04 | 904.9 | - | - | 0 | - |
| - | - | 6.752E+04 | 905.4 | - | - | 0 | - |
| - | - | 5.119E+04 | 905.9 | - | - | 0 | - |
| - | - | 2.902E+04 | 906.5 | - | - | 0 | - |
| - | - | 1.056E+04 | 907 | - | - | 0 | - |
| - | - | 3782 | 907.5 | - | - | 0 | - |
| - | - | 1006 | 930.5 | - | - | 0 | - |
| 8 | c | 1.858E+04 | 931.5 | 0.000211 | 0.2265 | +1 | 8 |
| - | - | 1.066E+04 | 932.5 | - | - | 0 | - |
| - | - | 2160 | 933.5 | - | - | 0 | - |
| - | - | 885.8 | 957.5 | - | - | 0 | - |
| - | - | 1213 | 1007 | - | - | 0 | - |
| - | - | 1297 | 1009 | - | - | 0 | - |
| - | - | 898.8 | 1017 | - | - | 0 | - |
| - | - | 899.1 | 1018 | - | - | 0 | - |
| 8 | z | 1.375E+04 | 1035 | 0.004432 | 4.28 | +1 | 8 |
| - | - | 9429 | 1036 | - | - | 0 | - |
| - | - | 3742 | 1037 | - | - | 0 | - |
| - | - | 1512 | 1038 | - | - | 0 | - |
| - | - | 2694 | 1050 | - | - | 0 | - |
| 8 | y | 9746 | 1051 | 0.004141 | 3.938 | +1 | 8 |
| - | - | 5454 | 1052 | - | - | 0 | - |
| - | - | 2181 | 1053 | - | - | 0 | - |
| - | - | 3539 | 1060 | - | - | 0 | - |
| 9 | c | 4.883E+04 | 1061 | 0.0003236 | 0.3051 | +1 | 9 |
| - | - | 2.936E+04 | 1062 | - | - | 0 | - |
| - | - | 1.047E+04 | 1063 | - | - | 0 | - |
| - | - | 1596 | 1064 | - | - | 0 | - |
| - | - | 715.8 | 1087 | - | - | 0 | - |
| - | - | 808.8 | 1103 | - | - | 0 | - |
| - | - | 742.3 | 1104 | - | - | 0 | - |
| 7 | z | 4353 | 1122 | 0.005485 | 4.886 | +1 | 9 |
| - | - | 1.604E+04 | 1124 | - | - | 0 | - |
| - | - | 8283 | 1125 | - | - | 0 | - |
| - | - | 3140 | 1126 | - | - | 0 | - |
| 7 | y | 1800 | 1139 | 0.001409 | 1.238 | +1 | 9 |
| - | - | 1237 | 1146 | - | - | 0 | - |
| - | - | 2066 | 1189 | - | - | 0 | - |
| 10 | c | 3.581E+04 | 1190 | 0.0006804 | 0.5719 | +1 | 10 |
| - | - | 2.409E+04 | 1191 | - | - | 0 | - |
| - | - | 7490 | 1192 | - | - | 0 | - |
| - | - | 1463 | 1193 | - | - | 0 | - |
| - | - | 895 | 1205 | - | - | 0 | - |
| - | - | 2287 | 1206 | - | - | 0 | - |
| - | - | 1440 | 1207 | - | - | 0 | - |
| - | - | 1034 | 1208 | - | - | 0 | - |
| - | - | 809.1 | 1231 | - | - | 0 | - |
| 6 | y | 3013 | 1236 | 0.004187 | 3.389 | +1 | 10 |
| - | - | 1467 | 1237 | - | - | 0 | - |
| - | - | 3165 | 1293 | - | - | 0 | - |
| - | - | 2305 | 1294 | - | - | 0 | - |
| - | - | 1174 | 1295 | - | - | 0 | - |
| - | - | 2510 | 1301 | - | - | 0 | - |
| - | - | 1496 | 1302 | - | - | 0 | - |
| - | - | 854.6 | 1303 | - | - | 0 | - |
| - | - | 707.7 | 1323 | - | - | 0 | - |
| 5 | y | 2643 | 1333 | 0.006477 | 4.861 | +1 | 11 |
| - | - | 1105 | 1334 | - | - | 0 | - |
| - | - | 837.1 | 1335 | - | - | 0 | - |
| - | - | 4446 | 1336 | - | - | 0 | - |
| 11 | c | 1.636E+04 | 1337 | 0.002771 | 2.073 | +1 | 11 |
| - | - | 1.141E+04 | 1338 | - | - | 0 | - |
| - | - | 4950 | 1339 | - | - | 0 | - |
| - | - | 1681 | 1340 | - | - | 0 | - |
| - | - | 729.6 | 1341 | - | - | 0 | - |
| - | - | 1022 | 1402 | - | - | 0 | - |
| - | - | 1666 | 1403 | - | - | 0 | - |
| - | - | 838.3 | 1414 | - | - | 0 | - |
| - | - | 767.8 | 1415 | - | - | 0 | - |
| 4 | z | 5315 | 1430 | 0.003393 | 2.373 | +1 | 12 |
| - | - | 2.144E+04 | 1431 | - | - | 0 | - |
| - | - | 1.401E+04 | 1432 | - | - | 0 | - |
| - | - | 6218 | 1433 | - | - | 0 | - |
| - | - | 1793 | 1434 | - | - | 0 | - |
| - | - | 812.4 | 1437 | - | - | 0 | - |
| 12 | c | 2.615E+04 | 1438 | 0.002455 | 1.708 | +1 | 12 |
| - | - | 1.96E+04 | 1439 | - | - | 0 | - |
| - | - | 9543 | 1440 | - | - | 0 | - |
| - | - | 2933 | 1441 | - | - | 0 | - |
| 4 | y | 1078 | 1446 | 0.005299 | 3.665 | +1 | 12 |
| - | - | 869.1 | 1447 | - | - | 0 | - |
| - | - | 688.4 | 1500 | - | - | 0 | - |
| - | - | 777 | 1504 | - | - | 0 | - |
| - | - | 761.8 | 1509 | - | - | 0 | - |
| - | - | 1142 | 1511 | - | - | 0 | - |
| - | - | 1630 | 1523 | - | - | 0 | - |
| 3 | z | 1430 | 1531 | 0.004055 | 2.649 | +1 | 13 |
| - | - | 6129 | 1532 | - | - | 0 | - |
| - | - | 4781 | 1533 | - | - | 0 | - |
| - | - | 2072 | 1534 | - | - | 0 | - |
| 3 | y | 2968 | 1547 | 0.001608 | 1.04 | +1 | 13 |
| 13 | c | 3204 | 1548 | 0.02896 | 18.71 | +1 | 13 |
| 13 | c | 2314 | 1549 | 0.01127 | 7.277 | +1 | 13 |
| - | - | 1554 | 1550 | - | - | 0 | - |
| - | - | 1847 | 1565 | - | - | 0 | - |
| 13 | c | 1.612E+04 | 1566 | 0.002463 | 1.573 | +1 | 13 |
| - | - | 1.497E+04 | 1567 | - | - | 0 | - |
| - | - | 7669 | 1568 | - | - | 0 | - |
| - | - | 3600 | 1569 | - | - | 0 | - |
| - | - | 710.4 | 1570 | - | - | 0 | - |
| - | - | 663.3 | 1636 | - | - | 0 | - |
| - | - | 1695 | 1637 | - | - | 0 | - |
| - | - | 1508 | 1638 | - | - | 0 | - |
| - | - | 690.2 | 1639 | - | - | 0 | - |
| - | - | 2569 | 1664 | - | - | 0 | - |
| - | - | 2489 | 1665 | - | - | 0 | - |
| - | - | 1822 | 1666 | - | - | 0 | - |
| - | - | 827.8 | 1667 | - | - | 0 | - |
| 14 | c | 7064 | 1680 | 0.004824 | 2.872 | +1 | 14 |
| - | - | 1.536E+04 | 1681 | - | - | 0 | - |
| - | - | 1.296E+04 | 1682 | - | - | 0 | - |
| - | - | 7243 | 1683 | - | - | 0 | - |
| - | - | 1952 | 1684 | - | - | 0 | - |
| - | - | 839.1 | 1685 | - | - | 0 | - |
| 2 | z | 979.7 | 1694 | 0.0125 | 7.382 | +1 | 14 |
| - | - | 5343 | 1695 | - | - | 0 | - |
| - | - | 5370 | 1696 | - | - | 0 | - |
| - | - | 3032 | 1697 | - | - | 0 | - |
| - | - | 1524 | 1698 | - | - | 0 | - |
| - | - | 1065 | 1711 | - | - | 0 | - |
| - | - | 1129 | 1712 | - | - | 0 | - |
| - | - | 1574 | 1747 | - | - | 0 | - |
| - | - | 958.4 | 1748 | - | - | 0 | - |
| - | - | 3215 | 1749 | - | - | 0 | - |
| - | - | 2148 | 1750 | - | - | 0 | - |
| - | - | 2937 | 1751 | - | - | 0 | - |
| - | - | 5149 | 1752 | - | - | 0 | - |
| - | - | 5091 | 1753 | - | - | 0 | - |
| - | - | 2452 | 1754 | - | - | 0 | - |
| - | - | 813.3 | 1755 | - | - | 0 | - |
| - | - | 3955 | 1765 | - | - | 0 | - |
| - | - | 1.184E+04 | 1766 | - | - | 0 | - |
| - | - | 1.029E+04 | 1767 | - | - | 0 | - |
| - | - | 5393 | 1768 | - | - | 0 | - |
| - | - | 3335 | 1769 | - | - | 0 | - |
| - | - | 938.7 | 1770 | - | - | 0 | - |
| - | - | 1305 | 1777 | - | - | 0 | - |
| - | - | 1287 | 1778 | - | - | 0 | - |
| - | - | 1983 | 1782 | - | - | 0 | - |
| - | - | 6205 | 1783 | - | - | 0 | - |
| - | - | 5637 | 1784 | - | - | 0 | - |
| - | - | 3108 | 1785 | - | - | 0 | - |
| - | - | 979.7 | 1786 | - | - | 0 | - |
| - | - | 9669 | 1793 | - | - | 0 | - |
| - | - | 3.431E+04 | 1794 | - | - | 0 | - |
| - | - | 3.446E+04 | 1795 | - | - | 0 | - |
| - | - | 1.974E+04 | 1796 | - | - | 0 | - |
| - | - | 7520 | 1797 | - | - | 0 | - |
| - | - | 2061 | 1798 | - | - | 0 | - |
| - | - | 3764 | 1809 | - | - | 0 | - |
| - | - | 1.831E+04 | 1810 | - | - | 0 | - |
| - | - | 7.2E+04 | 1811 | - | - | 0 | - |
| - | - | 6.264E+04 | 1812 | - | - | 0 | - |
| - | - | 3.821E+04 | 1813 | - | - | 0 | - |
| - | - | 1.328E+04 | 1814 | - | - | 0 | - |
| - | - | 5191 | 1815 | - | - | 0 | - |

m/z Charge Intensity FragmentType MassShift Position
129.1021728515625 0 854.35065
130.0499267578125 0 886.44196 y Ammonia loss 14
136.07554626464844 0 3528.2031
144.16688537597656 0 426.28076
145.70657348632812 0 478.9856
147.07627868652344 0 6121.7 y 14
148.95562744140625 0 439.57086
156.98292541503906 0 442.5499
161.88966369628906 0 460.95517
163.07113647460938 0 757.8682
169.13369750976562 0 479.97064
173.43936157226562 0 2253.3857
183.11241149902344 0 621.5429
184.11338806152344 0 561.74994
187.14398193359375 0 1188.5011
197.1291046142578 0 718.8675
199.1319580078125 0 508.02505
212.13914489746094 0 3039.8381
215.13882446289062 0 1199.7769
216.09825134277344 0 646.686
223.62557983398438 0 496.38626
225.46670532226562 0 517.4184
235.1438446044922 0 13177.353
236.1470947265625 0 2218.5242
237.9993896484375 0 528.0807
244.09262084960938 0 1030.1577 y Ammonia loss 13
256.1659851074219 0 572.05536
261.11907958984375 0 3817.5303 y 13
263.1385803222656 0 10512.402
264.1424560546875 0 1230.0012
298.1025390625 0 612.3532
299.17132568359375 0 2112.8008
315.12890625 0 888.72906 w 12
329.18206787109375 0 1709.3549
346.17596435546875 0 6262.8047
347.1788024902344 0 924.401
360.1911315917969 0 979.7659
364.18621826171875 0 8382.901
365.18939208984375 0 1599.9902
373.1951904296875 0 2848.0122 z 12
374.1995849609375 0 1048.0083
386.2397155761719 0 1168.7892
389.214599609375 0 2541.083 y 12
423.0581970214844 0 727.2529
429.21875 0 992.44855
430.2298583984375 0 3634.753
431.2341613769531 0 1068.6991
431.265380859375 0 2985.6133
432.2709655761719 0 642.9024
444.87860107421875 0 4872.1885 y 4
445.212646484375 0 3229.7612
445.54541015625 0 1237.9222
446.1665954589844 0 609.2913
449.2760314941406 0 1523.1646
450.2814636230469 0 755.3528
457.2410888671875 0 1411.7357 w 11
459.22503662109375 0 802.4831 w 11
459.260009765625 0 6080.362
460.26416015625 0 1887.3057
472.2274169921875 0 703.0492 w 3
473.23516845703125 0 3762.3225 y Ammonia loss 11
474.2405700683594 0 1402.717 z 11
475.2492980957031 0 1238.859
476.2630310058594 0 956.81616
477.2701416015625 0 2153.854
478.2733154296875 0 966.1157
481.67926025390625 0 595.0982
488.220458984375 0 667.7275
490.26226806640625 0 1223.314 y 11
491.2682189941406 0 620.3063
498.23077392578125 0 847.73035
510.25360107421875 0 1701.2845 y Water loss 2
510.5862121582031 0 1540.2744 y Ammonia loss 2
514.2609252929688 0 686.39087
515.2785034179688 0 1108.9805
531.2638549804688 0 1797.4559
532.2669677734375 0 650.6668
536.7567749023438 0 1048.2512
537.259033203125 0 924.67065
541.2845458984375 0 1496.209
552.60498046875 0 582.2878
552.94384765625 0 642.1464
557.2789306640625 0 1669.3716
558.288330078125 0 7262.1055
559.2950439453125 0 10832.095
560.2979125976562 0 3208.4792
560.8043823242188 0 816.16284
564.6072998046875 0 1085.4609 y Water loss 1
564.9393310546875 0 1518.1166 y Ammonia loss 1
565.2731323242188 0 1157.2764 z 1
565.603515625 0 546.14264
597.6294555664062 0 8501.195
597.96337890625 0 6941.259
598.2982788085938 0 5369.333
598.6321411132812 0 900.3672
602.8279418945312 0 1403.9559
603.3228149414062 0 685.9276
604.25537109375 0 1175.1581 z Ammonia loss 10
618.2871704101562 0 1821.5242 y 5
618.7882690429688 0 824.0753
621.278076171875 0 3294.0647 z 10
622.2833862304688 0 1412.2357
623.28369140625 0 647.20667
631.3419799804688 0 593.45844
634.8145141601562 0 3292.4912
635.3162231445312 0 1358.6749
636.3218994140625 0 641.7383
637.2986450195312 0 1956.074 y 10
642.3390502929688 0 662.5675
646.2858276367188 0 645.2866
657.8087158203125 0 1218.4318 y Water loss 4
658.3021240234375 0 2463.0881 y Ammonia loss 4
658.801513671875 0 2030.0953 z 4
659.3031005859375 0 923.8724
666.8140869140625 0 51537.12 y 4
667.3153686523438 0 36801.25
667.8152465820312 0 18824.56
668.31591796875 0 5693.662
668.8219604492188 0 683.95764 c 10
679.3427734375 0 754.2449
686.3208618164062 0 824.1532
687.3268432617188 0 3996.0833
687.3950805664062 0 6013.6377
687.8150024414062 0 764.71814
688.334228515625 0 1714.7097
688.4009399414062 0 5815.2056 c 5
689.3439331054688 0 940.8956
689.4060668945312 0 1992.3099
691.3076171875 0 21671.074 w 9
692.3104858398438 0 8238.743
693.306396484375 0 2070.511
693.8200073242188 0 15077.573
694.3206176757812 0 13607.997
694.8217163085938 0 6305.6045
695.3226318359375 0 1591.0656
695.8261108398438 0 637.59283
697.437255859375 0 1168.374
700.8331298828125 0 715.76935 w 3
713.408447265625 0 1923.3585
714.4141845703125 0 1227.1256
715.3466796875 0 17651.025 z 3
715.84814453125 0 12546.541
716.3477783203125 0 6465.722
716.8488159179688 0 1404.743
723.35546875 0 7989.9043 y 3
723.8577880859375 0 6938.282
724.35888671875 0 4801.014
724.8564453125 0 1391.6407
729.8570556640625 0 657.3785
732.4283447265625 0 2484.2263
733.4303588867188 0 773.3787
736.847900390625 0 922.28253
737.3360595703125 0 1063.24
737.8387451171875 0 5023.016
737.9092407226562 0 841.5962
738.3405151367188 0 2832.8594
738.84326171875 0 2285.1704
741.87939453125 0 1757.3105
742.381103515625 0 1653.3319
742.8773193359375 0 963.415
750.3211669921875 0 21991.191 z 9
751.32568359375 0 11290.98
752.3246459960938 0 3592.1897
753.3319091796875 0 1032.527
754.8665771484375 0 605.45703
761.3972778320312 0 1741.6749
761.897216796875 0 2426.377
764.8674926757812 0 1494.3773 y Water loss 2
765.36572265625 0 4983.9756 y Ammonia loss 2
765.8698120117188 0 9998.924 z 2
766.3670043945312 0 9089.807
766.8733520507812 0 5410.3276
767.3648681640625 0 1221.2943
773.3753662109375 0 1483.3671
773.8798217773438 0 34711.227 y 2
774.3815307617188 0 28949.902 c Water loss 12
774.881591796875 0 13041.903 c Ammonia loss 12
775.4354248046875 0 12113.797 c 6
775.8853759765625 0 2023.4517
776.437744140625 0 6508.663
777.4378662109375 0 1723.586
777.8648681640625 0 2605.6545
778.365234375 0 1455.7328
778.865478515625 0 1104.4259
779.376708984375 0 765.77106
782.8993530273438 0 5361.816
783.4032592773438 0 32262.117 c 12
783.9039916992188 0 28567.447
784.4052734375 0 13227.938
784.9054565429688 0 5305.657
785.4043579101562 0 863.5071
787.3744506835938 0 2129.796
787.874267578125 0 1748.0951
788.3793334960938 0 916.78503
817.8994140625 0 774.3971
818.4138793945312 0 1684.3965
818.9161376953125 0 2642.2642
819.8681640625 0 885.99475
820.3511962890625 0 5903.4375 w 8
821.35546875 0 1858.5022
832.9178466796875 0 1556.5272
837.4408569335938 0 1240.888
837.9403076171875 0 998.11206
839.4158935546875 0 1063.0785
839.917724609375 0 1772.5974
840.42431640625 0 68815.33 c 13
840.9261474609375 0 63262.465
841.4266967773438 0 37686.49
841.9276733398438 0 13627.982
842.4298706054688 0 2749.983
846.414306640625 0 698.4031 y Water loss 1
846.9061889648438 0 997.29193 y Ammonia loss 1
847.4019165039062 0 5231.8647 z 1
847.9029541015625 0 3828.8208
848.4052734375 0 2840.818
848.9031982421875 0 731.4019
849.4144287109375 0 688.09436
851.9228515625 0 830.8514
853.4328002929688 0 1777.7219
853.929931640625 0 2519.2246
854.4306030273438 0 1609.4573
854.910888671875 0 1926.066
855.4118041992188 0 3415.1106 y 1
855.9115600585938 0 3702.5166
856.4122314453125 0 1266.4786
856.9122924804688 0 731.78314
859.9228515625 0 692.60876
860.92919921875 0 734.0828
861.4239501953125 0 1668.2732
861.91845703125 0 1202.8705
865.9359130859375 0 881.4325
867.9291381835938 0 1251.0479
868.4132690429688 0 2242.8892
868.9126586914062 0 9565.634
869.4169311523438 0 10035.458
869.9215698242188 0 6485.354
870.421630859375 0 2361.9468
870.9273071289062 0 704.20776
873.4532470703125 0 913.09863
873.9442138671875 0 1562.8584
874.4444580078125 0 1718.3353
874.9423828125 0 1377.6069
875.4429931640625 0 978.74097
875.9308471679688 0 6487.5967
876.4317626953125 0 4066.7969
876.9320068359375 0 4271.141
877.4219360351562 0 5071.788
877.9227905273438 0 3712.8757
878.4205932617188 0 1084.8519
879.364013671875 0 10047.9375 z 8
880.367431640625 0 5030.9404
881.367431640625 0 1817.5409
882.4379272460938 0 1524.4409
882.93896484375 0 11827.917
883.4407348632812 0 10344.792
883.9400634765625 0 10191.826
884.437255859375 0 4625.9653
884.936279296875 0 2356.0767
885.4386596679688 0 818.2945
887.5220336914062 0 1154.1865
887.9275512695312 0 1631.1078
888.4319458007812 0 2995.051
888.522705078125 0 1099.2677
888.9341430664062 0 1677.814
889.4297485351562 0 1706.4131
889.9290161132812 0 1336.4626
890.4268798828125 0 1030.3943
895.3828735351562 0 1764.0704 y 8
895.9344482421875 0 1285.7872
896.4412841796875 0 5186.7803
896.9390258789062 0 48741.242
897.4393920898438 0 41634.695
897.94091796875 0 28941.336
898.4418334960938 0 11056.316
898.9405517578125 0 5725.8325
899.4479370117188 0 994.57355
904.4390869140625 0 1100.5149
904.945556640625 0 38171.43
905.44873046875 0 67516.33
905.9495849609375 0 51187.31
906.450927734375 0 29020.584
906.951416015625 0 10562.239
907.4514770507812 0 3782.187
930.5205688476562 0 1005.55505
931.5357666015625 0 18582.607 c 7
932.5386352539062 0 10662.581
933.541748046875 0 2159.6367
957.5485229492188 0 885.8285
1006.852783203125 0 1212.6085
1009.4600219726562 0 1297.1741
1016.5665283203125 0 898.7893
1017.5736694335938 0 899.13403
1035.4644775390625 0 13745.191 z 7
1036.470458984375 0 9429.22
1037.4691162109375 0 3742.2122
1038.4676513671875 0 1511.9814
1050.4732666015625 0 2694.375
1051.48291015625 0 9746.112 y 7
1052.486572265625 0 5453.931
1053.4892578125 0 2180.6812
1059.5709228515625 0 3539.2703
1060.5782470703125 0 48831.312 c 8
1061.5810546875 0 29364.621
1062.584228515625 0 10473.278
1063.5860595703125 0 1596.49
1086.5987548828125 0 715.81476
1102.6087646484375 0 808.8253
1103.6231689453125 0 742.3131
1122.49755859375 0 4353.3594 z 6
1123.50390625 0 16035.449
1124.5057373046875 0 8282.528
1125.5079345703125 0 3139.6978
1138.51220703125 0 1800.1476 y 6
1145.61865234375 0 1236.8512
1188.619384765625 0 2065.9094
1189.6204833984375 0 35806.82 c 9
1190.6234130859375 0 24093.57
1191.6234130859375 0 7490.0933
1192.6312255859375 0 1463.1617
1204.6300048828125 0 894.9859
1205.645263671875 0 2287.069
1206.6513671875 0 1440.0072
1207.628173828125 0 1034.203
1230.6693115234375 0 809.11487
1235.5677490234375 0 3013.1995 y 5
1236.57373046875 0 1467.428
1292.6451416015625 0 3164.8257
1293.6510009765625 0 2304.7974
1294.64892578125 0 1173.6895
1300.6409912109375 0 2509.8357
1301.636962890625 0 1496.46
1302.6478271484375 0 854.60455
1322.6529541015625 0 707.7447
1332.622802734375 0 2642.5574 y 4
1333.6129150390625 0 1105.2428
1334.61669921875 0 837.09033
1335.6470947265625 0 4446.4463
1336.6544189453125 0 16362.034 c 10
1337.65771484375 0 11413.068
1338.6605224609375 0 4950.322
1339.6611328125 0 1680.6376
1340.6475830078125 0 729.63745
1401.7008056640625 0 1022.0345
1402.6964111328125 0 1665.5684
1413.6851806640625 0 838.29224
1414.690673828125 0 767.8217
1429.68505859375 0 5314.8604 z 3
1430.69189453125 0 21438.838
1431.693115234375 0 14007.286
1432.696044921875 0 6218.2144
1433.7008056640625 0 1792.7775
1436.6976318359375 0 812.43506
1437.7017822265625 0 26146.338 c 11
1438.7060546875 0 19602.45
1439.7071533203125 0 9543.067
1440.7061767578125 0 2932.8315
1445.7056884765625 0 1078.433 y 3
1446.70751953125 0 869.0983
1499.770751953125 0 688.429
1503.7584228515625 0 777.0438
1509.279541015625 0 761.82556
1510.759521484375 0 1142.2906
1522.7889404296875 0 1629.6724
1530.7333984375 0 1429.5339 z 2
1531.740234375 0 6128.606
1532.7452392578125 0 4781.134
1533.7366943359375 0 2071.8079
1546.7464599609375 0 2967.6006 y 2
1547.7547607421875 0 3203.8342 c Water loss 12
1548.7564697265625 0 2314.0378 c Ammonia loss 12
1549.76904296875 0 1553.9954
1564.7760009765625 0 1847.0686
1565.7967529296875 0 16117.725 c 12
1566.800537109375 0 14973.751
1567.8026123046875 0 7668.5967
1568.8033447265625 0 3600.4229
1569.7950439453125 0 710.4304
1635.8204345703125 0 663.25354
1636.8211669921875 0 1695.1815
1637.830810546875 0 1507.8976
1638.835205078125 0 690.18555
1663.829833984375 0 2569.0286
1664.826904296875 0 2488.9692
1665.8193359375 0 1822.4215
1666.8194580078125 0 827.8365
1679.842041015625 0 7063.9165 c 13
1680.847900390625 0 15355.125
1681.8502197265625 0 12957.687
1682.85009765625 0 7243.3364
1683.8548583984375 0 1952.0851
1684.8719482421875 0 839.1371
1693.80517578125 0 979.6953 z 1
1694.80224609375 0 5342.841
1695.804931640625 0 5370.189
1696.805908203125 0 3032.082
1697.80322265625 0 1523.912
1710.82373046875 0 1064.5867
1711.8212890625 0 1128.7009
1746.8978271484375 0 1574.0299
1747.8900146484375 0 958.373
1748.87890625 0 3215.1965
1749.8878173828125 0 2148.2085
1750.8531494140625 0 2936.793
1751.855712890625 0 5149.0938
1752.8612060546875 0 5091.309
1753.86083984375 0 2451.965
1754.856689453125 0 813.255
1764.884521484375 0 3955.3745
1765.8800048828125 0 11839.815
1766.8822021484375 0 10292.428
1767.8812255859375 0 5393.2964
1768.885009765625 0 3335.4187
1769.869384765625 0 938.66016
1776.8486328125 0 1304.8383
1777.8563232421875 0 1287.2893
1781.8870849609375 0 1983.4769
1782.901123046875 0 6204.9756
1783.9068603515625 0 5636.7886
1784.906494140625 0 3108.0789
1785.8946533203125 0 979.7453
1792.876953125 0 9669.064
1793.8741455078125 0 34308.2
1794.8770751953125 0 34457.88
1795.87841796875 0 19741.46
1796.88037109375 0 7519.639
1797.8914794921875 0 2061.1494
1808.8856201171875 0 3763.9019
1809.8917236328125 0 18307.207
1810.8983154296875 0 71995.54
1811.9022216796875 0 62637.066
1812.9027099609375 0 38207.18
1813.9034423828125 0 13277.759
1814.90478515625 0 5191.125

Spectrum Details

|  |  |
| --- | --- |
| Matched peaks? Matched peaks The total absolute number of peaks matched. Additionally in brackets the total fraction of peaks matched and the total number of peaks is shown. | 70 (16.17% of 433) |
| FDR? FDR The false discovery rate estimated for this peptide. It is calculated by matching all theoretical fragments with a non-integer shift with the raw peaks for this spectrum. This is done with 40 different shifts. The resulting percentage is the average number of annotated peaks over the number of annotated peaks with the correct spectrum. | 3.20% |
| Satellite FDR? Satellite FDR See the FDR for details on its calculation. This satellite ion specific FDR only contains the satellite ions (d/w) for I/L/J positions. | 0.00% |
| PSM Score? PSM Score The PSM Score as given by Hecklib to this annotated spectrum. It is shown with three significant figures. | 449 |

#### Spectrum 3810? Spectrum 3810 The raw spectrum of this peptide as annotated by Hecklib. The fragments are coloured according to ion type (see legend). Any peaks with a star '\*' as text can be hovered over to see the full details, first the ion type second the mass shift type. By hovering over the amino acids in the peptide or ions in the legend the corresponding peaks are highlighted. By toggling the 'Unassigned' label you can turn the background (unassigned) peaks on or off in the plot. By updating the slider in the Ion legend you can update the spectrum to only show the top X% of the peaks with labels. The top X% means any peak that is within X% of the highest intensity. By dragging in the spectrum you can zoom in to a specific part of the spectrum and use 'Zoom Out' to get back to the original zoom level. The annotation of the spectrum is based on the given sequence in the peptides file and is done with different software so inconsistencies are likely. The peaks are annotated based on the given sequence, with 20 ppm tolerance.

Copy Data

##### Spectrum 3810 (TSV)

###### Preview

```
Loading example...
```

*Click on the button to copy the data to your clipboard.*

Mz MinMz MaxIntensity Max

WidthHeightPeptide font sizePeptide stroke widthSpectrum font sizeSpectrum stroke widthCompact peptide

Ion legend

wxyz

abcd

OtherUnassignedIon^Charge^_Position_Show for top:%

VYTJPPSREEMTKNQ

01.07e+42.14e+43.21e+44.28e+4

Zoom Out

y^+1^_1_y^+1^_2_z^+1^_3_y^+1^_3_y^+3^_11_w^+1^_4_y^+1^_4_y^+1^_4_y^+3^_13_y^+3^_14_z^+3^_14_y^+2^_10_z^+1^_5_y^+1^_5_y^+2^_11_z^+2^_11_y^+2^_11_c^+1^_6_w^+1^_6_w^+2^_12_z^+2^_12_y^+2^_12_z^+1^_6_y^+2^_13_y^+2^_13_z^+2^_13_y^+2^_13_c^+2^_13_c^+1^_7_c^+2^_13_w^+1^_7_c^+2^_14_y^+2^_14_z^+2^_14_y^+2^_14_z^+1^_7_y^+1^_7_c^+1^_8_z^+1^_8_y^+1^_8_c^+1^_9_z^+1^_9_y^+1^_9_c^+1^_10_y^+1^_10_y^+1^_11_c^+1^_11_z^+1^_12_c^+1^_12_y^+1^_13_c^+1^_13_c^+1^_13_c^+1^_13_c^+1^_14_

0789157923683157

Fragment Matches Table

Show background peaks

| Position | Ion type | Intensity | mz Theoretical | mz Error (Th) | mz Error (ppm) | Charge | Series Number |
| --- | --- | --- | --- | --- | --- | --- | --- |
| - | - | 321.1 | 125 | - | - | 0 | - |
| - | - | 970.7 | 129.1 | - | - | 0 | - |
| - | - | 1885 | 136.1 | - | - | 0 | - |
| 15 | y | 3915 | 147.1 | 0.0003841 | 2.612 | +1 | 1 |
| - | - | 424.5 | 161.4 | - | - | 0 | - |
| - | - | 548.5 | 171.1 | - | - | 0 | - |
| - | - | 4577 | 173.4 | - | - | 0 | - |
| - | - | 789.3 | 183.1 | - | - | 0 | - |
| - | - | 436.1 | 184.5 | - | - | 0 | - |
| - | - | 662.2 | 187.1 | - | - | 0 | - |
| - | - | 3069 | 203.1 | - | - | 0 | - |
| - | - | 2181 | 212.1 | - | - | 0 | - |
| - | - | 646.7 | 215.1 | - | - | 0 | - |
| - | - | 7761 | 235.1 | - | - | 0 | - |
| - | - | 617.2 | 236.1 | - | - | 0 | - |
| - | - | 1400 | 251.2 | - | - | 0 | - |
| - | - | 914.3 | 260.1 | - | - | 0 | - |
| 14 | y | 2285 | 261.1 | 0.0007548 | 2.891 | +1 | 2 |
| - | - | 6019 | 263.1 | - | - | 0 | - |
| - | - | 655.8 | 264.1 | - | - | 0 | - |
| - | - | 599.1 | 281 | - | - | 0 | - |
| - | - | 1092 | 299.2 | - | - | 0 | - |
| - | - | 591.1 | 309.4 | - | - | 0 | - |
| - | - | 539 | 315.2 | - | - | 0 | - |
| - | - | 1934 | 317.1 | - | - | 0 | - |
| - | - | 569 | 320.2 | - | - | 0 | - |
| - | - | 942.9 | 329.2 | - | - | 0 | - |
| - | - | 3807 | 346.2 | - | - | 0 | - |
| - | - | 1014 | 347.2 | - | - | 0 | - |
| - | - | 4250 | 364.2 | - | - | 0 | - |
| - | - | 755.1 | 365.2 | - | - | 0 | - |
| 13 | z | 2246 | 373.2 | 0.0006082 | 1.63 | +1 | 3 |
| - | - | 741.3 | 374.2 | - | - | 0 | - |
| - | - | 1130 | 383.2 | - | - | 0 | - |
| - | - | 1045 | 386.2 | - | - | 0 | - |
| 13 | y | 1173 | 389.2 | 0.0004724 | 1.214 | +1 | 3 |
| - | - | 529 | 414.7 | - | - | 0 | - |
| - | - | 1850 | 430.2 | - | - | 0 | - |
| - | - | 1499 | 431.3 | - | - | 0 | - |
| 5 | y | 2523 | 444.9 | 0.001306 | 2.935 | +3 | 11 |
| - | - | 702.7 | 445.2 | - | - | 0 | - |
| 12 | w | 677.8 | 459.2 | 0.005798 | 12.62 | +1 | 4 |
| - | - | 2883 | 459.3 | - | - | 0 | - |
| - | - | 636.8 | 460.2 | - | - | 0 | - |
| - | - | 793.8 | 460.3 | - | - | 0 | - |
| 12 | y | 1774 | 473.2 | 0.0001267 | 0.2677 | +1 | 4 |
| - | - | 665.9 | 476.3 | - | - | 0 | - |
| - | - | 994.6 | 477.3 | - | - | 0 | - |
| 12 | y | 707.7 | 490.3 | 0.001379 | 2.813 | +1 | 4 |
| - | - | 704 | 498.2 | - | - | 0 | - |
| 3 | y | 689.7 | 510.3 | 0.001268 | 2.485 | +3 | 13 |
| - | - | 769.7 | 510.8 | - | - | 0 | - |
| - | - | 676.8 | 515.1 | - | - | 0 | - |
| - | - | 1508 | 515.3 | - | - | 0 | - |
| - | - | 823.3 | 531.3 | - | - | 0 | - |
| - | - | 773.6 | 536.8 | - | - | 0 | - |
| - | - | 726.6 | 541.3 | - | - | 0 | - |
| - | - | 676.2 | 544.3 | - | - | 0 | - |
| - | - | 796.2 | 552.9 | - | - | 0 | - |
| - | - | 984.7 | 557.3 | - | - | 0 | - |
| - | - | 3770 | 558.3 | - | - | 0 | - |
| - | - | 6198 | 559.3 | - | - | 0 | - |
| - | - | 1483 | 560.3 | - | - | 0 | - |
| 2 | y | 686.4 | 564.6 | 0.0008802 | 1.559 | +3 | 14 |
| 2 | z | 640.7 | 565.3 | 0.00363 | 6.422 | +3 | 14 |
| - | - | 629.1 | 567.3 | - | - | 0 | - |
| - | - | 7289 | 585.3 | - | - | 0 | - |
| - | - | 2731 | 586.3 | - | - | 0 | - |
| - | - | 4817 | 597.6 | - | - | 0 | - |
| - | - | 4544 | 598 | - | - | 0 | - |
| - | - | 2851 | 598.3 | - | - | 0 | - |
| - | - | 807.9 | 598.6 | - | - | 0 | - |
| - | - | 1029 | 602.3 | - | - | 0 | - |
| - | - | 4425 | 602.8 | - | - | 0 | - |
| - | - | 4663 | 603.3 | - | - | 0 | - |
| - | - | 730.5 | 604.3 | - | - | 0 | - |
| - | - | 1532 | 604.3 | - | - | 0 | - |
| - | - | 2545 | 605.3 | - | - | 0 | - |
| - | - | 889.6 | 609.3 | - | - | 0 | - |
| 6 | y | 1999 | 618.3 | 1.868E-05 | 0.03021 | +2 | 10 |
| - | - | 1047 | 618.8 | - | - | 0 | - |
| 11 | z | 1706 | 621.3 | 0.00506 | 8.145 | +1 | 5 |
| - | - | 839.5 | 622.3 | - | - | 0 | - |
| - | - | 1463 | 634.8 | - | - | 0 | - |
| - | - | 848.7 | 635.3 | - | - | 0 | - |
| 11 | y | 656.7 | 637.3 | 0.001945 | 3.052 | +1 | 5 |
| - | - | 831.1 | 638.3 | - | - | 0 | - |
| - | - | 608.3 | 644.3 | - | - | 0 | - |
| 5 | y | 954.9 | 658.3 | 0.006955 | 10.56 | +2 | 11 |
| 5 | z | 845.1 | 658.8 | 0.002329 | 3.535 | +2 | 11 |
| 5 | y | 3.075E+04 | 666.8 | 0.001126 | 1.689 | +2 | 11 |
| - | - | 2.056E+04 | 667.3 | - | - | 0 | - |
| - | - | 8707 | 667.8 | - | - | 0 | - |
| - | - | 3290 | 668.3 | - | - | 0 | - |
| - | - | 1667 | 687.3 | - | - | 0 | - |
| - | - | 2805 | 687.4 | - | - | 0 | - |
| - | - | 1538 | 688.3 | - | - | 0 | - |
| 6 | c | 3498 | 688.4 | 0.002387 | 3.467 | +1 | 6 |
| 10 | w | 1.245E+04 | 691.3 | 0.003543 | 5.125 | +1 | 6 |
| - | - | 3947 | 692.3 | - | - | 0 | - |
| - | - | 2048 | 693.3 | - | - | 0 | - |
| 4 | w | 9187 | 693.8 | 0.0009709 | 1.399 | +2 | 12 |
| - | - | 7403 | 694.3 | - | - | 0 | - |
| - | - | 2486 | 694.8 | - | - | 0 | - |
| - | - | 1324 | 695.3 | - | - | 0 | - |
| - | - | 1610 | 713.4 | - | - | 0 | - |
| 4 | z | 8455 | 715.3 | 0.001049 | 1.467 | +2 | 12 |
| - | - | 9311 | 715.8 | - | - | 0 | - |
| - | - | 3120 | 716.3 | - | - | 0 | - |
| - | - | 1008 | 716.8 | - | - | 0 | - |
| 4 | y | 4679 | 723.4 | 0.0009033 | 1.249 | +2 | 12 |
| - | - | 3822 | 723.9 | - | - | 0 | - |
| - | - | 1365 | 724.4 | - | - | 0 | - |
| - | - | 923.5 | 732.4 | - | - | 0 | - |
| - | - | 1753 | 737.8 | - | - | 0 | - |
| - | - | 691.4 | 738.3 | - | - | 0 | - |
| - | - | 727.7 | 738.8 | - | - | 0 | - |
| - | - | 654.6 | 739.3 | - | - | 0 | - |
| - | - | 943.4 | 741.9 | - | - | 0 | - |
| 10 | z | 1.156E+04 | 750.3 | 0.003544 | 4.723 | +1 | 6 |
| - | - | 5789 | 751.3 | - | - | 0 | - |
| - | - | 1925 | 752.3 | - | - | 0 | - |
| - | - | 991.4 | 761.4 | - | - | 0 | - |
| - | - | 1001 | 761.9 | - | - | 0 | - |
| 3 | y | 831.5 | 764.9 | 0.004226 | 5.525 | +2 | 13 |
| 3 | y | 3109 | 765.4 | 0.004011 | 5.24 | +2 | 13 |
| 3 | z | 5562 | 765.9 | 0.001502 | 1.961 | +2 | 13 |
| - | - | 4878 | 766.4 | - | - | 0 | - |
| - | - | 2760 | 766.9 | - | - | 0 | - |
| - | - | 1148 | 767.4 | - | - | 0 | - |
| 3 | y | 1.865E+04 | 773.9 | 0.001051 | 1.358 | +2 | 13 |
| - | - | 1.352E+04 | 774.4 | - | - | 0 | - |
| 13 | c | 9190 | 774.9 | 0.006893 | 8.896 | +2 | 13 |
| 7 | c | 7936 | 775.4 | 0.001151 | 1.484 | +1 | 7 |
| - | - | 1011 | 775.9 | - | - | 0 | - |
| - | - | 3617 | 776.4 | - | - | 0 | - |
| - | - | 1322 | 777.4 | - | - | 0 | - |
| - | - | 1701 | 777.9 | - | - | 0 | - |
| - | - | 1079 | 778.4 | - | - | 0 | - |
| - | - | 808 | 779.4 | - | - | 0 | - |
| - | - | 2117 | 782.9 | - | - | 0 | - |
| 13 | c | 1.578E+04 | 783.4 | 0.0004619 | 0.5897 | +2 | 13 |
| - | - | 1.447E+04 | 783.9 | - | - | 0 | - |
| - | - | 5822 | 784.4 | - | - | 0 | - |
| - | - | 2288 | 784.9 | - | - | 0 | - |
| - | - | 1027 | 785.4 | - | - | 0 | - |
| - | - | 1196 | 787.4 | - | - | 0 | - |
| - | - | 1006 | 795.4 | - | - | 0 | - |
| - | - | 1046 | 796.4 | - | - | 0 | - |
| - | - | 834.4 | 815.4 | - | - | 0 | - |
| - | - | 1036 | 818.9 | - | - | 0 | - |
| - | - | 788.5 | 819.4 | - | - | 0 | - |
| 9 | w | 3227 | 820.3 | 0.007764 | 9.464 | +1 | 7 |
| - | - | 657.2 | 820.9 | - | - | 0 | - |
| - | - | 1333 | 821.4 | - | - | 0 | - |
| - | - | 648.5 | 825.4 | - | - | 0 | - |
| - | - | 631.9 | 833.4 | - | - | 0 | - |
| - | - | 718.2 | 837.9 | - | - | 0 | - |
| - | - | 738.5 | 839.4 | - | - | 0 | - |
| 14 | c | 4.237E+04 | 840.4 | 0.0007878 | 0.9373 | +2 | 14 |
| - | - | 3.512E+04 | 840.9 | - | - | 0 | - |
| - | - | 1.549E+04 | 841.4 | - | - | 0 | - |
| - | - | 6148 | 841.9 | - | - | 0 | - |
| - | - | 1265 | 842.4 | - | - | 0 | - |
| - | - | 707.6 | 842.9 | - | - | 0 | - |
| 2 | y | 760.5 | 846.4 | 0.01123 | 13.27 | +2 | 14 |
| 2 | z | 3525 | 847.4 | 0.002552 | 3.012 | +2 | 14 |
| - | - | 1534 | 847.9 | - | - | 0 | - |
| - | - | 1639 | 848.4 | - | - | 0 | - |
| - | - | 933.5 | 848.9 | - | - | 0 | - |
| - | - | 1729 | 853.4 | - | - | 0 | - |
| - | - | 1231 | 853.9 | - | - | 0 | - |
| - | - | 1426 | 854.4 | - | - | 0 | - |
| 2 | y | 3105 | 855.4 | 0.0032 | 3.741 | +2 | 14 |
| - | - | 1781 | 855.9 | - | - | 0 | - |
| - | - | 1022 | 856.4 | - | - | 0 | - |
| - | - | 653.1 | 861.9 | - | - | 0 | - |
| - | - | 1853 | 868.4 | - | - | 0 | - |
| - | - | 5670 | 868.9 | - | - | 0 | - |
| - | - | 5191 | 869.4 | - | - | 0 | - |
| - | - | 4270 | 869.9 | - | - | 0 | - |
| - | - | 2659 | 870.4 | - | - | 0 | - |
| - | - | 881.2 | 870.9 | - | - | 0 | - |
| - | - | 1344 | 874.4 | - | - | 0 | - |
| - | - | 866.8 | 874.9 | - | - | 0 | - |
| - | - | 714.3 | 875.4 | - | - | 0 | - |
| - | - | 2483 | 875.9 | - | - | 0 | - |
| - | - | 3323 | 876.4 | - | - | 0 | - |
| - | - | 2182 | 876.9 | - | - | 0 | - |
| - | - | 1985 | 877.4 | - | - | 0 | - |
| - | - | 2072 | 877.9 | - | - | 0 | - |
| - | - | 1154 | 878.4 | - | - | 0 | - |
| 9 | z | 5393 | 879.4 | 0.003431 | 3.902 | +1 | 7 |
| - | - | 2145 | 880.4 | - | - | 0 | - |
| - | - | 9005 | 882.9 | - | - | 0 | - |
| - | - | 5555 | 883.4 | - | - | 0 | - |
| - | - | 4877 | 883.9 | - | - | 0 | - |
| - | - | 1959 | 884.4 | - | - | 0 | - |
| - | - | 1679 | 884.9 | - | - | 0 | - |
| - | - | 1515 | 887.9 | - | - | 0 | - |
| - | - | 1655 | 888.4 | - | - | 0 | - |
| - | - | 1643 | 888.9 | - | - | 0 | - |
| - | - | 1853 | 889.4 | - | - | 0 | - |
| - | - | 1044 | 890.4 | - | - | 0 | - |
| 9 | y | 799.4 | 895.4 | 0.003994 | 4.461 | +1 | 7 |
| - | - | 2474 | 896.4 | - | - | 0 | - |
| - | - | 2.586E+04 | 896.9 | - | - | 0 | - |
| - | - | 2.449E+04 | 897.4 | - | - | 0 | - |
| - | - | 1.205E+04 | 897.9 | - | - | 0 | - |
| - | - | 6601 | 898.4 | - | - | 0 | - |
| - | - | 2332 | 898.9 | - | - | 0 | - |
| - | - | 859.3 | 899.4 | - | - | 0 | - |
| - | - | 1354 | 904.4 | - | - | 0 | - |
| - | - | 1.813E+04 | 904.9 | - | - | 0 | - |
| - | - | 3.766E+04 | 905.4 | - | - | 0 | - |
| - | - | 2.863E+04 | 905.9 | - | - | 0 | - |
| - | - | 1.595E+04 | 906.4 | - | - | 0 | - |
| - | - | 7637 | 906.9 | - | - | 0 | - |
| - | - | 2086 | 907.4 | - | - | 0 | - |
| - | - | 1134 | 907.9 | - | - | 0 | - |
| 8 | c | 9128 | 931.5 | 0.001676 | 1.799 | +1 | 8 |
| - | - | 4199 | 932.5 | - | - | 0 | - |
| - | - | 945.4 | 933.5 | - | - | 0 | - |
| - | - | 979.3 | 957.5 | - | - | 0 | - |
| - | - | 1029 | 997.5 | - | - | 0 | - |
| - | - | 739.4 | 998.5 | - | - | 0 | - |
| - | - | 857.3 | 1017 | - | - | 0 | - |
| - | - | 774 | 1018 | - | - | 0 | - |
| 8 | z | 6895 | 1035 | 0.002967 | 2.866 | +1 | 8 |
| - | - | 4388 | 1036 | - | - | 0 | - |
| - | - | 2308 | 1037 | - | - | 0 | - |
| - | - | 830.6 | 1038 | - | - | 0 | - |
| 8 | y | 5914 | 1051 | 0.00231 | 2.197 | +1 | 8 |
| - | - | 3009 | 1052 | - | - | 0 | - |
| - | - | 1542 | 1053 | - | - | 0 | - |
| - | - | 1423 | 1060 | - | - | 0 | - |
| 9 | c | 2.508E+04 | 1061 | 0.001911 | 1.801 | +1 | 9 |
| - | - | 1.647E+04 | 1062 | - | - | 0 | - |
| - | - | 4976 | 1063 | - | - | 0 | - |
| 7 | z | 2551 | 1122 | 0.001701 | 1.515 | +1 | 9 |
| - | - | 7666 | 1124 | - | - | 0 | - |
| - | - | 4370 | 1125 | - | - | 0 | - |
| - | - | 1201 | 1126 | - | - | 0 | - |
| - | - | 742 | 1135 | - | - | 0 | - |
| 7 | y | 928.3 | 1139 | 0.004827 | 4.24 | +1 | 9 |
| - | - | 1641 | 1189 | - | - | 0 | - |
| 10 | c | 2.087E+04 | 1190 | 0.002878 | 2.419 | +1 | 10 |
| - | - | 1.336E+04 | 1191 | - | - | 0 | - |
| - | - | 4599 | 1192 | - | - | 0 | - |
| - | - | 1453 | 1193 | - | - | 0 | - |
| - | - | 698.4 | 1199 | - | - | 0 | - |
| - | - | 1946 | 1205 | - | - | 0 | - |
| - | - | 1760 | 1206 | - | - | 0 | - |
| - | - | 2162 | 1207 | - | - | 0 | - |
| 6 | y | 1066 | 1236 | 0.004309 | 3.488 | +1 | 10 |
| - | - | 787.6 | 1237 | - | - | 0 | - |
| - | - | 721.5 | 1278 | - | - | 0 | - |
| - | - | 1583 | 1293 | - | - | 0 | - |
| - | - | 899 | 1294 | - | - | 0 | - |
| - | - | 1135 | 1301 | - | - | 0 | - |
| - | - | 655.1 | 1303 | - | - | 0 | - |
| 5 | y | 1364 | 1333 | 0.004997 | 3.75 | +1 | 11 |
| - | - | 964.3 | 1334 | - | - | 0 | - |
| - | - | 2050 | 1336 | - | - | 0 | - |
| 11 | c | 8729 | 1337 | 0.002038 | 1.525 | +1 | 11 |
| - | - | 5492 | 1338 | - | - | 0 | - |
| - | - | 1864 | 1339 | - | - | 0 | - |
| - | - | 1294 | 1340 | - | - | 0 | - |
| - | - | 766.8 | 1403 | - | - | 0 | - |
| 4 | z | 2603 | 1430 | 2.482E-05 | 0.01736 | +1 | 12 |
| - | - | 1.204E+04 | 1431 | - | - | 0 | - |
| - | - | 7173 | 1432 | - | - | 0 | - |
| - | - | 4002 | 1433 | - | - | 0 | - |
| - | - | 1511 | 1434 | - | - | 0 | - |
| - | - | 740.9 | 1437 | - | - | 0 | - |
| 12 | c | 1.538E+04 | 1438 | 0.0009905 | 0.6889 | +1 | 12 |
| - | - | 9908 | 1439 | - | - | 0 | - |
| - | - | 4795 | 1440 | - | - | 0 | - |
| - | - | 1801 | 1441 | - | - | 0 | - |
| - | - | 849.7 | 1442 | - | - | 0 | - |
| - | - | 835.1 | 1447 | - | - | 0 | - |
| - | - | 977.1 | 1507 | - | - | 0 | - |
| - | - | 840.8 | 1523 | - | - | 0 | - |
| - | - | 2530 | 1532 | - | - | 0 | - |
| - | - | 2877 | 1533 | - | - | 0 | - |
| - | - | 1314 | 1534 | - | - | 0 | - |
| 3 | y | 1067 | 1547 | 0.001688 | 1.091 | +1 | 13 |
| 13 | c | 1897 | 1548 | 0.01895 | 12.25 | +1 | 13 |
| 13 | c | 819.1 | 1549 | 0.01054 | 6.805 | +1 | 13 |
| 13 | c | 9154 | 1566 | 0.002786 | 1.779 | +1 | 13 |
| - | - | 6914 | 1567 | - | - | 0 | - |
| - | - | 4855 | 1568 | - | - | 0 | - |
| - | - | 2219 | 1569 | - | - | 0 | - |
| - | - | 781.3 | 1638 | - | - | 0 | - |
| - | - | 1769 | 1664 | - | - | 0 | - |
| - | - | 1198 | 1665 | - | - | 0 | - |
| - | - | 754.8 | 1667 | - | - | 0 | - |
| 14 | c | 4212 | 1680 | 0.004824 | 2.872 | +1 | 14 |
| - | - | 9059 | 1681 | - | - | 0 | - |
| - | - | 7676 | 1682 | - | - | 0 | - |
| - | - | 3818 | 1683 | - | - | 0 | - |
| - | - | 1774 | 1684 | - | - | 0 | - |
| - | - | 3267 | 1695 | - | - | 0 | - |
| - | - | 2671 | 1696 | - | - | 0 | - |
| - | - | 1702 | 1697 | - | - | 0 | - |
| - | - | 1129 | 1749 | - | - | 0 | - |
| - | - | 3152 | 1752 | - | - | 0 | - |
| - | - | 1818 | 1753 | - | - | 0 | - |
| - | - | 984.4 | 1754 | - | - | 0 | - |
| - | - | 1867 | 1765 | - | - | 0 | - |
| - | - | 5556 | 1766 | - | - | 0 | - |
| - | - | 6703 | 1767 | - | - | 0 | - |
| - | - | 4351 | 1768 | - | - | 0 | - |
| - | - | 1424 | 1769 | - | - | 0 | - |
| - | - | 757.5 | 1770 | - | - | 0 | - |
| - | - | 956 | 1778 | - | - | 0 | - |
| - | - | 1038 | 1782 | - | - | 0 | - |
| - | - | 3086 | 1783 | - | - | 0 | - |
| - | - | 3025 | 1784 | - | - | 0 | - |
| - | - | 1698 | 1785 | - | - | 0 | - |
| - | - | 931.6 | 1786 | - | - | 0 | - |
| - | - | 5058 | 1793 | - | - | 0 | - |
| - | - | 1.98E+04 | 1794 | - | - | 0 | - |
| - | - | 1.56E+04 | 1795 | - | - | 0 | - |
| - | - | 1.063E+04 | 1796 | - | - | 0 | - |
| - | - | 5370 | 1797 | - | - | 0 | - |
| - | - | 1604 | 1798 | - | - | 0 | - |
| - | - | 842.7 | 1799 | - | - | 0 | - |
| - | - | 2246 | 1809 | - | - | 0 | - |
| - | - | 9783 | 1810 | - | - | 0 | - |
| - | - | 3.863E+04 | 1811 | - | - | 0 | - |
| - | - | 3.322E+04 | 1812 | - | - | 0 | - |
| - | - | 2.015E+04 | 1813 | - | - | 0 | - |
| - | - | 9895 | 1814 | - | - | 0 | - |
| - | - | 2975 | 1815 | - | - | 0 | - |
| - | - | 770.9 | 2773 | - | - | 0 | - |
| - | - | 700.2 | 3126 | - | - | 0 | - |

m/z Charge Intensity FragmentType MassShift Position
125.04859161376953 0 321.11786
129.10202026367188 0 970.6602
136.0753631591797 0 1885.0688
147.07603454589844 0 3914.928 y 14
161.4212646484375 0 424.5315
171.11216735839844 0 548.48016
173.4399871826172 0 4577.1953
183.1124725341797 0 789.3432
184.51876831054688 0 436.06873
187.1441192626953 0 662.22894
203.10215759277344 0 3069.0908
212.1384735107422 0 2181.428
215.13890075683594 0 646.7354
235.14349365234375 0 7760.5757
236.14781188964844 0 617.16345
251.1746063232422 0 1399.6257
260.1227111816406 0 914.31036
261.11859130859375 0 2284.5374 y 13
263.1383361816406 0 6018.944
264.14239501953125 0 655.7741
281.0367736816406 0 599.05023
299.171875 0 1092.269
309.36767578125 0 591.0699
315.1653137207031 0 539.0424
317.1446533203125 0 1934.3555
320.159423828125 0 569.03894
329.182373046875 0 942.9458
346.1756286621094 0 3806.6218
347.17889404296875 0 1013.59283
364.1858825683594 0 4250.049
365.18896484375 0 755.0614
373.1949768066406 0 2245.664 z 12
374.2012634277344 0 741.2678
383.22845458984375 0 1130.3647
386.24053955078125 0 1045.1418
389.2138366699219 0 1173.3601 y 12
414.7459716796875 0 529.0432
430.2293701171875 0 1850.0798
431.2651062011719 0 1498.9241
444.8782653808594 0 2522.671 y 4
445.2127685546875 0 702.6588
459.2255859375 0 677.8008 w 11
459.2592468261719 0 2882.6926
460.2231750488281 0 636.792
460.263671875 0 793.7681
473.2355651855469 0 1774.1879 y Ammonia loss 11
476.260986328125 0 665.9453
477.27032470703125 0 994.55963
490.26336669921875 0 707.72986 y 11
498.22930908203125 0 703.9787
510.251953125 0 689.6668 y Water loss 2
510.75103759765625 0 769.737
515.0819091796875 0 676.84863
515.2779541015625 0 1508.462
531.2626953125 0 823.2912
536.7568359375 0 773.60944
541.2855224609375 0 726.5683
544.26806640625 0 676.154
552.9403076171875 0 796.22943
557.2799072265625 0 984.72284
558.2860717773438 0 3770.4753
559.2937622070312 0 6198.3477
560.2974243164062 0 1483.1709
564.604248046875 0 686.40314 y Water loss 1
565.272705078125 0 640.6516 z 1
567.3112182617188 0 629.14124
585.3223876953125 0 7288.6406
586.325439453125 0 2730.6453
597.6283569335938 0 4816.856
597.9629516601562 0 4543.9106
598.29736328125 0 2851.2915
598.6290283203125 0 807.9109
602.320068359375 0 1028.716
602.8215942382812 0 4424.6143
603.32080078125 0 4663.237
604.2822265625 0 730.5443
604.329345703125 0 1531.5361
605.3140869140625 0 2544.7278
609.2970581054688 0 889.5539
618.285400390625 0 1998.7948 y 5
618.7899169921875 0 1046.9227
621.27880859375 0 1706.2898 z 10
622.28369140625 0 839.483
634.813232421875 0 1463.4302
635.3143310546875 0 848.65094
637.29052734375 0 656.7026 y 10
638.2996826171875 0 831.1025
644.3174438476562 0 608.2651
658.3054809570312 0 954.908 y Ammonia loss 4
658.8001098632812 0 845.0504 z 4
666.8129272460938 0 30752.805 y 4
667.3140869140625 0 20561.521
667.8145141601562 0 8706.869
668.31494140625 0 3290.48
687.3265380859375 0 1666.9829
687.3925170898438 0 2805.2825
688.336181640625 0 1538.1007
688.4004516601562 0 3498.475 c 5
691.3065795898438 0 12450.44 w 9
692.3090209960938 0 3947.0293
693.30908203125 0 2047.6296
693.8180541992188 0 9186.93 w 3
694.3192749023438 0 7403.3696
694.819091796875 0 2486.3062
695.3209838867188 0 1323.7546
713.4091796875 0 1610.1545
715.3455200195312 0 8454.584 z 3
715.84716796875 0 9310.859
716.3482666015625 0 3120.3403
716.8447265625 0 1007.6947
723.354736328125 0 4678.8184 y 3
723.8577880859375 0 3821.9478
724.35595703125 0 1365.4915
732.4266357421875 0 923.5211
737.8399047851562 0 1753.2095
738.3325805664062 0 691.447
738.8336791992188 0 727.7376
739.3408203125 0 654.56287
741.8778076171875 0 943.39294
750.3198852539062 0 11563.704 z 9
751.3238525390625 0 5789.476
752.322509765625 0 1925.4586
761.391845703125 0 991.4286
761.8985595703125 0 1001.06793
764.8681640625 0 831.5349 y Water loss 2
765.368408203125 0 3108.936 y Ammonia loss 2
765.8698120117188 0 5562.404 z 2
766.3674926757812 0 4877.547
766.8734741210938 0 2759.5715
767.3658447265625 0 1148.4968
773.8787231445312 0 18654.752 y 2
774.3795166015625 0 13521.314
774.880615234375 0 9189.929 c Ammonia loss 12
775.4337158203125 0 7935.5303 c 6
775.8836669921875 0 1010.64075
776.435791015625 0 3617.05
777.4423828125 0 1322.386
777.86181640625 0 1700.5767
778.363037109375 0 1079.4387
779.3697509765625 0 808.0282
782.8978881835938 0 2116.8464
783.4012451171875 0 15779.675 c 12
783.9028930664062 0 14472.55
784.4044799804688 0 5821.5234
784.905029296875 0 2287.7227
785.4016723632812 0 1027.18
787.371826171875 0 1195.5903
795.40673828125 0 1006.4933
796.4050903320312 0 1046.1398
815.4398193359375 0 834.35156
818.9138793945312 0 1036.0667
819.4210205078125 0 788.50714
820.3533935546875 0 3227.268 w 8
820.8714599609375 0 657.2021
821.351318359375 0 1333.1958
825.4111938476562 0 648.48376
833.4038696289062 0 631.923
837.9400024414062 0 718.16174
839.40625 0 738.482
840.4230346679688 0 42369.48 c 13
840.9246826171875 0 35115.92
841.42529296875 0 15486.45
841.9265747070312 0 6147.688
842.4254150390625 0 1264.9043
842.923583984375 0 707.5978
846.415283203125 0 760.5443 y Water loss 1
847.4025268554688 0 3525.2544 z 1
847.900390625 0 1534.4285
848.4013671875 0 1639.0706
848.8993530273438 0 933.52673
853.4287109375 0 1729.1936
853.9252319335938 0 1231.2793
854.4302978515625 0 1426.473
855.4125366210938 0 3104.7803 y 1
855.9100341796875 0 1781.2726
856.41015625 0 1021.77496
861.92041015625 0 653.0853
868.4149169921875 0 1853.2034
868.9111938476562 0 5670.4844
869.4196166992188 0 5191.1865
869.9216918945312 0 4269.979
870.41943359375 0 2658.6824
870.9285278320312 0 881.23517
874.4497680664062 0 1343.5641
874.939453125 0 866.78656
875.4296264648438 0 714.2574
875.9301147460938 0 2482.7375
876.4324340820312 0 3323.2932
876.9302368164062 0 2182.3728
877.4215698242188 0 1984.5072
877.9208374023438 0 2072.2808
878.4179077148438 0 1154.1257
879.3623657226562 0 5392.923 z 8
880.3660888671875 0 2145.2847
882.9380493164062 0 9004.682
883.4395141601562 0 5555.345
883.9371337890625 0 4877.413
884.4413452148438 0 1958.902
884.936767578125 0 1678.8047
887.9285278320312 0 1514.9178
888.4347534179688 0 1655.3925
888.9309692382812 0 1642.5656
889.4317016601562 0 1853.3695
890.4300537109375 0 1044.0917
895.3816528320312 0 799.4411 y 8
896.4368896484375 0 2473.8977
896.9375 0 25864.096
897.4382934570312 0 24485.914
897.9390869140625 0 12048.179
898.4384765625 0 6600.625
898.9412231445312 0 2332.4138
899.4393310546875 0 859.2574
904.4417114257812 0 1353.8329
904.9437866210938 0 18129.514
905.4464721679688 0 37660.35
905.9485473632812 0 28626.99
906.4493408203125 0 15948.164
906.9490966796875 0 7636.96
907.4468994140625 0 2085.963
907.945556640625 0 1133.6522
931.5343017578125 0 9128.155 c 7
932.5388793945312 0 4198.7007
933.5409545898438 0 945.448
957.5481567382812 0 979.25635
997.4546508789062 0 1028.9801
998.4699096679688 0 739.3617
1016.5584716796875 0 857.2855
1017.55908203125 0 773.9851
1035.4630126953125 0 6895.185 z 7
1036.468017578125 0 4388.128
1037.466796875 0 2308.248
1038.472900390625 0 830.572
1051.4810791015625 0 5914.0894 y 7
1052.48193359375 0 3008.8467
1053.4913330078125 0 1541.8352
1059.572998046875 0 1423.2344
1060.57666015625 0 25079.852 c 8
1061.5794677734375 0 16467.297
1062.5821533203125 0 4975.6655
1122.4937744140625 0 2551.4185 z 6
1123.5023193359375 0 7665.7026
1124.50537109375 0 4370.243
1125.5030517578125 0 1200.9126
1134.556396484375 0 741.96
1138.515625 0 928.28906 y 6
1188.6131591796875 0 1641.4886
1189.6182861328125 0 20869.654 c 9
1190.6209716796875 0 13363.757
1191.620361328125 0 4599.2607
1192.618896484375 0 1453.3417
1199.0926513671875 0 698.3517
1204.6368408203125 0 1945.5945
1205.6304931640625 0 1759.7935
1206.6336669921875 0 2161.9998
1235.56787109375 0 1065.8685 y 5
1236.5704345703125 0 787.6273
1277.622802734375 0 721.54913
1292.639892578125 0 1583.312
1293.6436767578125 0 899.0192
1300.6358642578125 0 1135.2838
1302.6500244140625 0 655.12726
1332.611328125 0 1364.2886 y 4
1333.6229248046875 0 964.33234
1335.6455078125 0 2050.4038
1336.6536865234375 0 8728.905 c 10
1337.65478515625 0 5492.1274
1338.658447265625 0 1864.0979
1339.6534423828125 0 1294.0247
1402.701416015625 0 766.8254
1429.681640625 0 2603.0776 z 3
1430.688720703125 0 12039.568
1431.69140625 0 7173.469
1432.6954345703125 0 4001.7751
1433.6895751953125 0 1511.1492
1436.7008056640625 0 740.8979
1437.7003173828125 0 15382.099 c 11
1438.702392578125 0 9908.376
1439.7066650390625 0 4795.2285
1440.7076416015625 0 1801.3394
1441.7125244140625 0 849.67017
1446.6865234375 0 835.0817
1507.262451171875 0 977.1192
1522.77490234375 0 840.8141
1531.7354736328125 0 2530.3289
1532.7374267578125 0 2877.2046
1533.7369384765625 0 1313.9409
1546.749755859375 0 1066.566 y 2
1547.7647705078125 0 1896.9734 c Water loss 12
1548.7572021484375 0 819.0743 c Ammonia loss 12
1565.79150390625 0 9153.561 c 12
1566.7987060546875 0 6914.4956
1567.7982177734375 0 4855.3735
1568.8017578125 0 2219.1729
1637.8201904296875 0 781.34467
1663.8165283203125 0 1768.564
1664.8140869140625 0 1198.162
1666.84228515625 0 754.7843
1679.842041015625 0 4211.6084 c 13
1680.8436279296875 0 9058.822
1681.8470458984375 0 7676.4106
1682.849853515625 0 3818.343
1683.8497314453125 0 1774.4562
1694.8016357421875 0 3266.901
1695.798095703125 0 2671.1138
1696.800048828125 0 1702.1716
1748.8804931640625 0 1129.062
1751.85302734375 0 3152.3228
1752.8525390625 0 1818.4247
1753.851806640625 0 984.4402
1764.88623046875 0 1866.8479
1765.8758544921875 0 5556.1367
1766.8779296875 0 6702.692
1767.8779296875 0 4350.505
1768.8707275390625 0 1423.5232
1769.86962890625 0 757.5008
1777.8865966796875 0 955.9692
1781.8975830078125 0 1038.3821
1782.9022216796875 0 3086.403
1783.9031982421875 0 3024.9272
1784.902587890625 0 1698.2162
1785.9080810546875 0 931.5988
1792.870361328125 0 5057.5063
1793.8717041015625 0 19800.658
1794.87353515625 0 15602.247
1795.8739013671875 0 10626.842
1796.8758544921875 0 5369.751
1797.88427734375 0 1604.2197
1798.86865234375 0 842.6608
1808.883544921875 0 2246.4436
1809.88671875 0 9782.982
1810.895751953125 0 38630.164
1811.897216796875 0 33219.445
1812.9000244140625 0 20147.791
1813.9017333984375 0 9894.698
1814.904296875 0 2975.2622
2772.798828125 0 770.9461
3126.201904296875 0 700.1712

Spectrum Details

|  |  |
| --- | --- |
| Matched peaks? Matched peaks The total absolute number of peaks matched. Additionally in brackets the total fraction of peaks matched and the total number of peaks is shown. | 54 (16.02% of 337) |
| FDR? FDR The false discovery rate estimated for this peptide. It is calculated by matching all theoretical fragments with a non-integer shift with the raw peaks for this spectrum. This is done with 40 different shifts. The resulting percentage is the average number of annotated peaks over the number of annotated peaks with the correct spectrum. | 2.82% |
| Satellite FDR? Satellite FDR See the FDR for details on its calculation. This satellite ion specific FDR only contains the satellite ions (d/w) for I/L/J positions. | 0.00% |
| PSM Score? PSM Score The PSM Score as given by Hecklib to this annotated spectrum. It is shown with three significant figures. | 334 |

#### Spectrum 3436? Spectrum 3436 The raw spectrum of this peptide as annotated by Hecklib. The fragments are coloured according to ion type (see legend). Any peaks with a star '\*' as text can be hovered over to see the full details, first the ion type second the mass shift type. By hovering over the amino acids in the peptide or ions in the legend the corresponding peaks are highlighted. By toggling the 'Unassigned' label you can turn the background (unassigned) peaks on or off in the plot. By updating the slider in the Ion legend you can update the spectrum to only show the top X% of the peaks with labels. The top X% means any peak that is within X% of the highest intensity. By dragging in the spectrum you can zoom in to a specific part of the spectrum and use 'Zoom Out' to get back to the original zoom level. The annotation of the spectrum is based on the given sequence in the peptides file and is done with different software so inconsistencies are likely. The peaks are annotated based on the given sequence, with 20 ppm tolerance.

Copy Data

##### Spectrum 3436 (TSV)

###### Preview

```
Loading example...
```

*Click on the button to copy the data to your clipboard.*

Mz MinMz MaxIntensity Max

WidthHeightPeptide font sizePeptide stroke widthSpectrum font sizeSpectrum stroke widthCompact peptide

Ion legend

wxyz

abcd

OtherUnassignedIon^Charge^_Position_Show for top:%

VYTJPPSREEMTKNQ

08.80e+31.76e+42.64e+43.52e+4

Zoom Out

y^+1^_1_y^+1^_2_y^+1^_2_z^+1^_3_y^+1^_3_y^+3^_11_w^+1^_4_w^+1^_4_y^+1^_4_z^+1^_4_y^+3^_13_y^+3^_13_y^+2^_10_z^+1^_5_y^+2^_11_z^+2^_11_y^+2^_11_c^+1^_6_w^+1^_6_w^+2^_12_z^+2^_12_y^+2^_12_z^+1^_6_y^+2^_13_z^+2^_13_y^+2^_13_c^+2^_13_c^+2^_13_c^+1^_7_c^+2^_13_w^+1^_7_c^+2^_14_y^+2^_14_z^+2^_14_y^+2^_14_z^+1^_7_c^+1^_8_z^+1^_8_y^+1^_8_c^+1^_9_z^+1^_9_c^+1^_10_y^+1^_10_c^+1^_11_z^+1^_12_c^+1^_12_z^+1^_13_y^+1^_13_c^+1^_13_c^+1^_13_c^+1^_13_c^+1^_14_z^+1^_14_

045991713761834

Fragment Matches Table

Show background peaks

| Position | Ion type | Intensity | mz Theoretical | mz Error (Th) | mz Error (ppm) | Charge | Series Number |
| --- | --- | --- | --- | --- | --- | --- | --- |
| - | - | 402 | 120.5 | - | - | 0 | - |
| - | - | 396.8 | 127.6 | - | - | 0 | - |
| - | - | 345.9 | 128.3 | - | - | 0 | - |
| - | - | 605.4 | 129.1 | - | - | 0 | - |
| - | - | 1105 | 136.1 | - | - | 0 | - |
| - | - | 417.1 | 144.2 | - | - | 0 | - |
| 15 | y | 2660 | 147.1 | 0.0002773 | 1.885 | +1 | 1 |
| - | - | 396.1 | 147.2 | - | - | 0 | - |
| - | - | 522.3 | 148.9 | - | - | 0 | - |
| - | - | 579.3 | 148.9 | - | - | 0 | - |
| - | - | 777.3 | 148.9 | - | - | 0 | - |
| - | - | 734.6 | 148.9 | - | - | 0 | - |
| - | - | 492.7 | 148.9 | - | - | 0 | - |
| - | - | 1278 | 148.9 | - | - | 0 | - |
| - | - | 1034 | 148.9 | - | - | 0 | - |
| - | - | 2119 | 148.9 | - | - | 0 | - |
| - | - | 3617 | 148.9 | - | - | 0 | - |
| - | - | 4415 | 149 | - | - | 0 | - |
| - | - | 2831 | 149 | - | - | 0 | - |
| - | - | 1549 | 149 | - | - | 0 | - |
| - | - | 1247 | 149 | - | - | 0 | - |
| - | - | 830.3 | 149 | - | - | 0 | - |
| - | - | 708.6 | 149 | - | - | 0 | - |
| - | - | 701 | 149 | - | - | 0 | - |
| - | - | 521.7 | 149 | - | - | 0 | - |
| - | - | 531.8 | 149 | - | - | 0 | - |
| - | - | 415.7 | 172.5 | - | - | 0 | - |
| - | - | 469.2 | 183.1 | - | - | 0 | - |
| - | - | 675.2 | 197.1 | - | - | 0 | - |
| - | - | 1801 | 212.1 | - | - | 0 | - |
| - | - | 1023 | 215.1 | - | - | 0 | - |
| - | - | 5883 | 235.1 | - | - | 0 | - |
| - | - | 845.9 | 236.1 | - | - | 0 | - |
| 14 | y | 650.7 | 244.1 | 0.0005424 | 2.222 | +1 | 2 |
| 14 | y | 1665 | 261.1 | 0.001029 | 3.942 | +1 | 2 |
| - | - | 4670 | 263.1 | - | - | 0 | - |
| - | - | 1226 | 299.2 | - | - | 0 | - |
| - | - | 553.2 | 305.4 | - | - | 0 | - |
| - | - | 542.5 | 324.4 | - | - | 0 | - |
| - | - | 690.2 | 329.2 | - | - | 0 | - |
| - | - | 2757 | 346.2 | - | - | 0 | - |
| - | - | 843.2 | 347.2 | - | - | 0 | - |
| - | - | 4141 | 364.2 | - | - | 0 | - |
| - | - | 1165 | 365.2 | - | - | 0 | - |
| - | - | 623.5 | 370.2 | - | - | 0 | - |
| 13 | z | 1974 | 373.2 | 0.0007913 | 2.12 | +1 | 3 |
| 13 | y | 1970 | 389.2 | 0.0007776 | 1.998 | +1 | 3 |
| - | - | 659.5 | 391.7 | - | - | 0 | - |
| - | - | 537.3 | 393.6 | - | - | 0 | - |
| - | - | 1429 | 430.2 | - | - | 0 | - |
| - | - | 1424 | 431.3 | - | - | 0 | - |
| - | - | 663.3 | 432.3 | - | - | 0 | - |
| 5 | y | 3024 | 444.9 | 0.001367 | 3.073 | +3 | 11 |
| - | - | 1407 | 445.2 | - | - | 0 | - |
| - | - | 625.9 | 445.5 | - | - | 0 | - |
| - | - | 826.8 | 449.3 | - | - | 0 | - |
| 12 | w | 952.1 | 457.2 | 0.0007778 | 1.701 | +1 | 4 |
| 12 | w | 528.3 | 459.2 | 0.003448 | 7.508 | +1 | 4 |
| - | - | 2026 | 459.3 | - | - | 0 | - |
| - | - | 532.8 | 460.3 | - | - | 0 | - |
| - | - | 552.4 | 472.2 | - | - | 0 | - |
| 12 | y | 1862 | 473.2 | 0.0009414 | 1.989 | +1 | 4 |
| 12 | z | 1118 | 474.2 | 0.003273 | 6.902 | +1 | 4 |
| - | - | 839.9 | 477.3 | - | - | 0 | - |
| - | - | 642.5 | 484.7 | - | - | 0 | - |
| 3 | y | 709.8 | 510.3 | 0.002702 | 5.296 | +3 | 13 |
| 3 | y | 754.4 | 510.6 | 0.008773 | 17.18 | +3 | 13 |
| - | - | 666.6 | 537.3 | - | - | 0 | - |
| - | - | 578.2 | 541.3 | - | - | 0 | - |
| - | - | 3423 | 558.3 | - | - | 0 | - |
| - | - | 4696 | 559.3 | - | - | 0 | - |
| - | - | 1140 | 560.3 | - | - | 0 | - |
| - | - | 1098 | 593.8 | - | - | 0 | - |
| - | - | 1882 | 594.3 | - | - | 0 | - |
| - | - | 5233 | 597.6 | - | - | 0 | - |
| - | - | 4479 | 598 | - | - | 0 | - |
| - | - | 1699 | 598.3 | - | - | 0 | - |
| - | - | 664.8 | 598.6 | - | - | 0 | - |
| 6 | y | 1018 | 618.3 | 0.0004696 | 0.7595 | +2 | 10 |
| 11 | z | 720.3 | 621.3 | 0.004694 | 7.556 | +1 | 5 |
| - | - | 2302 | 634.8 | - | - | 0 | - |
| - | - | 1122 | 635.3 | - | - | 0 | - |
| 5 | y | 1198 | 658.3 | 0.002072 | 3.147 | +2 | 11 |
| 5 | z | 1445 | 658.8 | 0.0006201 | 0.9413 | +2 | 11 |
| - | - | 1020 | 666.3 | - | - | 0 | - |
| 5 | y | 2.447E+04 | 666.8 | 0.001248 | 1.872 | +2 | 11 |
| - | - | 1.762E+04 | 667.3 | - | - | 0 | - |
| - | - | 9019 | 667.8 | - | - | 0 | - |
| - | - | 2991 | 668.3 | - | - | 0 | - |
| - | - | 1392 | 687.3 | - | - | 0 | - |
| - | - | 3638 | 687.4 | - | - | 0 | - |
| - | - | 997.5 | 688.3 | - | - | 0 | - |
| 6 | c | 2382 | 688.4 | 0.003058 | 4.442 | +1 | 6 |
| - | - | 814.1 | 689.4 | - | - | 0 | - |
| 10 | w | 1.114E+04 | 691.3 | 0.003726 | 5.39 | +1 | 6 |
| - | - | 2925 | 692.3 | - | - | 0 | - |
| - | - | 1575 | 693.3 | - | - | 0 | - |
| 4 | w | 9291 | 693.8 | 0.001459 | 2.103 | +2 | 12 |
| - | - | 6025 | 694.3 | - | - | 0 | - |
| - | - | 4189 | 694.8 | - | - | 0 | - |
| - | - | 885.7 | 695.3 | - | - | 0 | - |
| - | - | 1219 | 713.4 | - | - | 0 | - |
| 4 | z | 6821 | 715.3 | 0.001354 | 1.893 | +2 | 12 |
| - | - | 5980 | 715.8 | - | - | 0 | - |
| - | - | 1891 | 716.3 | - | - | 0 | - |
| - | - | 1480 | 716.8 | - | - | 0 | - |
| 4 | y | 4482 | 723.4 | 0.002612 | 3.611 | +2 | 12 |
| - | - | 3198 | 723.9 | - | - | 0 | - |
| - | - | 638 | 724.4 | - | - | 0 | - |
| - | - | 1371 | 732.4 | - | - | 0 | - |
| - | - | 607.1 | 733.4 | - | - | 0 | - |
| - | - | 1777 | 737.8 | - | - | 0 | - |
| - | - | 2060 | 738.3 | - | - | 0 | - |
| - | - | 840.1 | 738.8 | - | - | 0 | - |
| - | - | 838.3 | 741.9 | - | - | 0 | - |
| 10 | z | 1.151E+04 | 750.3 | 0.003422 | 4.561 | +1 | 6 |
| - | - | 5687 | 751.3 | - | - | 0 | - |
| - | - | 1389 | 752.3 | - | - | 0 | - |
| - | - | 757.7 | 761.4 | - | - | 0 | - |
| - | - | 931.3 | 761.9 | - | - | 0 | - |
| 3 | y | 2609 | 765.4 | 0.002302 | 3.007 | +2 | 13 |
| 3 | z | 4601 | 765.9 | 0.0006473 | 0.8452 | +2 | 13 |
| - | - | 3735 | 766.4 | - | - | 0 | - |
| - | - | 2352 | 766.9 | - | - | 0 | - |
| - | - | 941.2 | 767.4 | - | - | 0 | - |
| - | - | 688.9 | 772.4 | - | - | 0 | - |
| - | - | 674.8 | 773.4 | - | - | 0 | - |
| 3 | y | 1.81E+04 | 773.9 | 0.001051 | 1.358 | +2 | 13 |
| 13 | c | 1.463E+04 | 774.4 | 0.01507 | 19.46 | +2 | 13 |
| 13 | c | 7234 | 774.9 | 0.006405 | 8.266 | +2 | 13 |
| - | - | 1491 | 775.4 | - | - | 0 | - |
| 7 | c | 5191 | 775.4 | 0.001578 | 2.035 | +1 | 7 |
| - | - | 955.5 | 775.9 | - | - | 0 | - |
| - | - | 2908 | 776.4 | - | - | 0 | - |
| - | - | 995 | 777.4 | - | - | 0 | - |
| - | - | 1491 | 777.9 | - | - | 0 | - |
| - | - | 742.6 | 778.4 | - | - | 0 | - |
| - | - | 1087 | 778.9 | - | - | 0 | - |
| - | - | 2110 | 782.9 | - | - | 0 | - |
| 13 | c | 1.463E+04 | 783.4 | 0.001255 | 1.602 | +2 | 13 |
| - | - | 1.25E+04 | 783.9 | - | - | 0 | - |
| - | - | 7355 | 784.4 | - | - | 0 | - |
| - | - | 2548 | 784.9 | - | - | 0 | - |
| - | - | 793.6 | 785.4 | - | - | 0 | - |
| - | - | 1358 | 787.9 | - | - | 0 | - |
| - | - | 604.5 | 808.4 | - | - | 0 | - |
| - | - | 615.7 | 817.9 | - | - | 0 | - |
| - | - | 831.8 | 818.9 | - | - | 0 | - |
| 9 | w | 2104 | 820.3 | 0.003125 | 3.809 | +1 | 7 |
| - | - | 727.2 | 821.4 | - | - | 0 | - |
| - | - | 923.2 | 837.9 | - | - | 0 | - |
| 14 | c | 3.215E+04 | 840.4 | 0.001093 | 1.3 | +2 | 14 |
| - | - | 2.987E+04 | 840.9 | - | - | 0 | - |
| - | - | 694.9 | 841.1 | - | - | 0 | - |
| - | - | 1.688E+04 | 841.4 | - | - | 0 | - |
| - | - | 7569 | 841.9 | - | - | 0 | - |
| - | - | 1615 | 842.4 | - | - | 0 | - |
| - | - | 837.1 | 843.9 | - | - | 0 | - |
| 2 | y | 835 | 846.4 | 0.01282 | 15.14 | +2 | 14 |
| - | - | 861.3 | 846.9 | - | - | 0 | - |
| 2 | z | 2687 | 847.4 | 0.006459 | 7.622 | +2 | 14 |
| - | - | 3500 | 847.9 | - | - | 0 | - |
| - | - | 1973 | 848.4 | - | - | 0 | - |
| - | - | 835.9 | 848.9 | - | - | 0 | - |
| - | - | 1381 | 853.4 | - | - | 0 | - |
| - | - | 1011 | 853.9 | - | - | 0 | - |
| 2 | y | 2865 | 855.4 | 0.0002178 | 0.2547 | +2 | 14 |
| - | - | 2463 | 855.9 | - | - | 0 | - |
| - | - | 939.4 | 856.4 | - | - | 0 | - |
| - | - | 756.6 | 860.4 | - | - | 0 | - |
| - | - | 923.4 | 867.4 | - | - | 0 | - |
| - | - | 956.7 | 867.9 | - | - | 0 | - |
| - | - | 860.3 | 868.4 | - | - | 0 | - |
| - | - | 5068 | 868.9 | - | - | 0 | - |
| - | - | 5570 | 869.4 | - | - | 0 | - |
| - | - | 3340 | 869.9 | - | - | 0 | - |
| - | - | 1745 | 870.4 | - | - | 0 | - |
| - | - | 960.8 | 870.9 | - | - | 0 | - |
| - | - | 899 | 874.4 | - | - | 0 | - |
| - | - | 849.1 | 874.9 | - | - | 0 | - |
| - | - | 619.5 | 875.4 | - | - | 0 | - |
| - | - | 3691 | 875.9 | - | - | 0 | - |
| - | - | 3285 | 876.4 | - | - | 0 | - |
| - | - | 1402 | 876.9 | - | - | 0 | - |
| - | - | 2270 | 877.4 | - | - | 0 | - |
| - | - | 2222 | 877.9 | - | - | 0 | - |
| - | - | 742.5 | 878.9 | - | - | 0 | - |
| 9 | z | 3929 | 879.4 | 0.002211 | 2.514 | +1 | 7 |
| - | - | 1470 | 880.4 | - | - | 0 | - |
| - | - | 883.4 | 881.4 | - | - | 0 | - |
| - | - | 879.7 | 882.4 | - | - | 0 | - |
| - | - | 6329 | 882.9 | - | - | 0 | - |
| - | - | 5261 | 883.4 | - | - | 0 | - |
| - | - | 3659 | 883.9 | - | - | 0 | - |
| - | - | 1947 | 884.4 | - | - | 0 | - |
| - | - | 2102 | 884.9 | - | - | 0 | - |
| - | - | 869.3 | 888.4 | - | - | 0 | - |
| - | - | 671.8 | 888.9 | - | - | 0 | - |
| - | - | 1469 | 889.4 | - | - | 0 | - |
| - | - | 1360 | 889.9 | - | - | 0 | - |
| - | - | 2881 | 896.4 | - | - | 0 | - |
| - | - | 2.1E+04 | 896.9 | - | - | 0 | - |
| - | - | 2.447E+04 | 897.4 | - | - | 0 | - |
| - | - | 1.319E+04 | 897.9 | - | - | 0 | - |
| - | - | 6583 | 898.4 | - | - | 0 | - |
| - | - | 2679 | 898.9 | - | - | 0 | - |
| - | - | 995.7 | 899.4 | - | - | 0 | - |
| - | - | 1.478E+04 | 904.9 | - | - | 0 | - |
| - | - | 3.486E+04 | 905.4 | - | - | 0 | - |
| - | - | 2.553E+04 | 905.9 | - | - | 0 | - |
| - | - | 1.413E+04 | 906.4 | - | - | 0 | - |
| - | - | 5108 | 906.9 | - | - | 0 | - |
| - | - | 2319 | 907.4 | - | - | 0 | - |
| - | - | 790.6 | 930.5 | - | - | 0 | - |
| 8 | c | 8258 | 931.5 | 0.001432 | 1.537 | +1 | 8 |
| - | - | 3913 | 932.5 | - | - | 0 | - |
| - | - | 1584 | 933.5 | - | - | 0 | - |
| - | - | 815.4 | 1002 | - | - | 0 | - |
| - | - | 659.9 | 1017 | - | - | 0 | - |
| - | - | 737.1 | 1018 | - | - | 0 | - |
| 8 | z | 7422 | 1035 | 0.003211 | 3.102 | +1 | 8 |
| - | - | 4962 | 1036 | - | - | 0 | - |
| - | - | 1562 | 1037 | - | - | 0 | - |
| - | - | 962.4 | 1050 | - | - | 0 | - |
| 8 | y | 5241 | 1051 | 0.004263 | 4.054 | +1 | 8 |
| - | - | 2843 | 1052 | - | - | 0 | - |
| - | - | 1234 | 1053 | - | - | 0 | - |
| - | - | 1409 | 1060 | - | - | 0 | - |
| 9 | c | 2.473E+04 | 1061 | 0.001422 | 1.341 | +1 | 9 |
| - | - | 1.352E+04 | 1062 | - | - | 0 | - |
| - | - | 5153 | 1063 | - | - | 0 | - |
| - | - | 1038 | 1064 | - | - | 0 | - |
| - | - | 680.7 | 1088 | - | - | 0 | - |
| 7 | z | 2241 | 1122 | 0.002555 | 2.276 | +1 | 9 |
| - | - | 7401 | 1124 | - | - | 0 | - |
| - | - | 3887 | 1125 | - | - | 0 | - |
| - | - | 1533 | 1126 | - | - | 0 | - |
| - | - | 1098 | 1188 | - | - | 0 | - |
| - | - | 1100 | 1189 | - | - | 0 | - |
| 10 | c | 1.73E+04 | 1190 | 0.002023 | 1.701 | +1 | 10 |
| - | - | 1.089E+04 | 1191 | - | - | 0 | - |
| - | - | 4415 | 1192 | - | - | 0 | - |
| - | - | 804.5 | 1193 | - | - | 0 | - |
| - | - | 612.8 | 1198 | - | - | 0 | - |
| - | - | 734.3 | 1209 | - | - | 0 | - |
| - | - | 771.8 | 1231 | - | - | 0 | - |
| 6 | y | 1334 | 1236 | 0.005164 | 4.179 | +1 | 10 |
| - | - | 1135 | 1293 | - | - | 0 | - |
| - | - | 1130 | 1294 | - | - | 0 | - |
| - | - | 1119 | 1295 | - | - | 0 | - |
| - | - | 813.8 | 1301 | - | - | 0 | - |
| - | - | 799.5 | 1302 | - | - | 0 | - |
| - | - | 1068 | 1303 | - | - | 0 | - |
| - | - | 607.8 | 1322 | - | - | 0 | - |
| - | - | 811.9 | 1334 | - | - | 0 | - |
| - | - | 690.5 | 1335 | - | - | 0 | - |
| - | - | 1887 | 1336 | - | - | 0 | - |
| 11 | c | 8318 | 1337 | 0.002771 | 2.073 | +1 | 11 |
| - | - | 6372 | 1338 | - | - | 0 | - |
| - | - | 2258 | 1339 | - | - | 0 | - |
| - | - | 736.8 | 1395 | - | - | 0 | - |
| - | - | 619.5 | 1402 | - | - | 0 | - |
| - | - | 909.4 | 1403 | - | - | 0 | - |
| - | - | 636.4 | 1426 | - | - | 0 | - |
| 4 | z | 2506 | 1430 | 0.001562 | 1.093 | +1 | 12 |
| - | - | 1.1E+04 | 1431 | - | - | 0 | - |
| - | - | 6268 | 1432 | - | - | 0 | - |
| - | - | 3207 | 1433 | - | - | 0 | - |
| - | - | 1883 | 1434 | - | - | 0 | - |
| 12 | c | 1.294E+04 | 1438 | 0.001967 | 1.368 | +1 | 12 |
| - | - | 1.013E+04 | 1439 | - | - | 0 | - |
| - | - | 5104 | 1440 | - | - | 0 | - |
| - | - | 2033 | 1441 | - | - | 0 | - |
| - | - | 863.3 | 1447 | - | - | 0 | - |
| - | - | 889.5 | 1501 | - | - | 0 | - |
| - | - | 668.2 | 1502 | - | - | 0 | - |
| 3 | z | 997.2 | 1531 | 0.0008807 | 0.5753 | +1 | 13 |
| - | - | 2207 | 1532 | - | - | 0 | - |
| - | - | 2465 | 1533 | - | - | 0 | - |
| - | - | 1474 | 1534 | - | - | 0 | - |
| 3 | y | 1763 | 1547 | 0.005716 | 3.696 | +1 | 13 |
| 13 | c | 2133 | 1548 | 0.02603 | 16.82 | +1 | 13 |
| 13 | c | 1090 | 1549 | 0.009562 | 6.174 | +1 | 13 |
| - | - | 830.6 | 1565 | - | - | 0 | - |
| 13 | c | 8407 | 1566 | 0.000632 | 0.4036 | +1 | 13 |
| - | - | 7095 | 1567 | - | - | 0 | - |
| - | - | 3641 | 1568 | - | - | 0 | - |
| - | - | 1493 | 1569 | - | - | 0 | - |
| - | - | 715.4 | 1604 | - | - | 0 | - |
| - | - | 918.2 | 1637 | - | - | 0 | - |
| - | - | 856.8 | 1665 | - | - | 0 | - |
| 14 | c | 3038 | 1680 | 0.001284 | 0.7641 | +1 | 14 |
| - | - | 7710 | 1681 | - | - | 0 | - |
| - | - | 7037 | 1682 | - | - | 0 | - |
| - | - | 2596 | 1683 | - | - | 0 | - |
| - | - | 1034 | 1684 | - | - | 0 | - |
| 2 | z | 853 | 1694 | 0.0125 | 7.382 | +1 | 14 |
| - | - | 2744 | 1695 | - | - | 0 | - |
| - | - | 2813 | 1696 | - | - | 0 | - |
| - | - | 1029 | 1697 | - | - | 0 | - |
| - | - | 1019 | 1698 | - | - | 0 | - |
| - | - | 1054 | 1748 | - | - | 0 | - |
| - | - | 1411 | 1749 | - | - | 0 | - |
| - | - | 1305 | 1750 | - | - | 0 | - |
| - | - | 1960 | 1751 | - | - | 0 | - |
| - | - | 2523 | 1752 | - | - | 0 | - |
| - | - | 1821 | 1753 | - | - | 0 | - |
| - | - | 1069 | 1754 | - | - | 0 | - |
| - | - | 1934 | 1765 | - | - | 0 | - |
| - | - | 6159 | 1766 | - | - | 0 | - |
| - | - | 4248 | 1767 | - | - | 0 | - |
| - | - | 3310 | 1768 | - | - | 0 | - |
| - | - | 2321 | 1769 | - | - | 0 | - |
| - | - | 696.7 | 1777 | - | - | 0 | - |
| - | - | 754 | 1778 | - | - | 0 | - |
| - | - | 715.7 | 1782 | - | - | 0 | - |
| - | - | 2807 | 1783 | - | - | 0 | - |
| - | - | 2478 | 1784 | - | - | 0 | - |
| - | - | 1366 | 1785 | - | - | 0 | - |
| - | - | 4612 | 1793 | - | - | 0 | - |
| - | - | 1.694E+04 | 1794 | - | - | 0 | - |
| - | - | 1.344E+04 | 1795 | - | - | 0 | - |
| - | - | 8724 | 1796 | - | - | 0 | - |
| - | - | 4206 | 1797 | - | - | 0 | - |
| - | - | 1438 | 1798 | - | - | 0 | - |
| - | - | 1705 | 1809 | - | - | 0 | - |
| - | - | 1.026E+04 | 1810 | - | - | 0 | - |
| - | - | 3.401E+04 | 1811 | - | - | 0 | - |
| - | - | 2.9E+04 | 1812 | - | - | 0 | - |
| - | - | 1.81E+04 | 1813 | - | - | 0 | - |
| - | - | 7227 | 1814 | - | - | 0 | - |
| - | - | 2524 | 1815 | - | - | 0 | - |
| - | - | 955 | 1816 | - | - | 0 | - |

m/z Charge Intensity FragmentType MassShift Position
120.48392486572266 0 401.9664
127.6361083984375 0 396.81064
128.3043670654297 0 345.90112
129.10189819335938 0 605.37274
136.075439453125 0 1105.173
144.20785522460938 0 417.099
147.07614135742188 0 2659.7065 y 14
147.2150115966797 0 396.0761
148.86981201171875 0 522.2755
148.8838653564453 0 579.2821
148.89833068847656 0 777.26404
148.90541076660156 0 734.5643
148.91314697265625 0 492.71487
148.92039489746094 0 1277.936
148.9276580810547 0 1033.5681
148.9345245361328 0 2118.5344
148.94239807128906 0 3617.3926
148.95896911621094 0 4415.294
148.9668731689453 0 2831.3523
148.97402954101562 0 1549.4077
148.98130798339844 0 1246.5382
148.9881134033203 0 830.3487
148.9957733154297 0 708.64197
149.00265502929688 0 700.9556
149.01791381835938 0 521.6602
149.03201293945312 0 531.8363
172.53085327148438 0 415.71127
183.112548828125 0 469.15662
197.12808227539062 0 675.2361
212.13868713378906 0 1800.8011
215.13845825195312 0 1022.8861
235.14340209960938 0 5882.617
236.14752197265625 0 845.888
244.09225463867188 0 650.7471 y Ammonia loss 13
261.1183166503906 0 1665.2145 y 13
263.13848876953125 0 4670.3735
299.1711120605469 0 1225.6294
305.4007873535156 0 553.1678
324.38641357421875 0 542.5086
329.1801452636719 0 690.1894
346.1755676269531 0 2757.199
347.17962646484375 0 843.2124
364.1859130859375 0 4140.52
365.1883239746094 0 1165.3174
370.17010498046875 0 623.5316
373.1947937011719 0 1973.7748 z 12
389.2135314941406 0 1970.2937 y 12
391.7244873046875 0 659.5395
393.55291748046875 0 537.27795
430.2275695800781 0 1429.0253
431.2651672363281 0 1423.8484
432.2684020996094 0 663.3364
444.8783264160156 0 3024.2017 y 4
445.212646484375 0 1407.0897
445.5492248535156 0 625.8504
449.275390625 0 826.75464
457.23974609375 0 952.08765 w 11
459.2232360839844 0 528.27924 w 11
459.2593688964844 0 2026.3335
460.263916015625 0 532.8244
472.2288513183594 0 552.355
473.2344970703125 0 1861.9124 y Ammonia loss 11
474.239990234375 0 1117.7955 z 11
477.26727294921875 0 839.88293
484.6556701660156 0 642.4995
510.2533874511719 0 709.83057 y Water loss 2
510.58746337890625 0 754.4239 y Ammonia loss 2
537.26171875 0 666.598
541.2807006835938 0 578.232
558.2876586914062 0 3422.73
559.2941284179688 0 4696.3867
560.299560546875 0 1140.4321
593.7774658203125 0 1097.848
594.2832641601562 0 1882.1674
597.6286010742188 0 5233.346
597.9624633789062 0 4478.9155
598.2955322265625 0 1699.4988
598.6286010742188 0 664.76605
618.285888671875 0 1017.85223 y 5
621.2784423828125 0 720.29333 z 10
634.8136596679688 0 2302.4539
635.3132934570312 0 1122.118
658.3005981445312 0 1197.879 y Ammonia loss 4
658.8018188476562 0 1445.1306 z 4
666.3064575195312 0 1019.79126
666.8130493164062 0 24470.766 y 4
667.3147583007812 0 17617.334
667.8156127929688 0 9018.744
668.3145141601562 0 2990.8792
687.326904296875 0 1391.8156
687.3939819335938 0 3637.5718
688.3348388671875 0 997.4617
688.3997802734375 0 2381.9612 c 5
689.40478515625 0 814.1391
691.3067626953125 0 11139.439 w 9
692.3088989257812 0 2925.4849
693.3085327148438 0 1574.7305
693.8185424804688 0 9291.397 w 3
694.3192138671875 0 6025.403
694.8218383789062 0 4188.72
695.320068359375 0 885.73267
713.4102172851562 0 1219.2938
715.3458251953125 0 6820.875 z 3
715.84716796875 0 5980.275
716.3482055664062 0 1890.8818
716.8469848632812 0 1480.2002
723.3564453125 0 4482.461 y 3
723.8566284179688 0 3198.2224
724.35888671875 0 637.9682
732.4259033203125 0 1371.3878
733.4306640625 0 607.1387
737.8368530273438 0 1776.6382
738.3425903320312 0 2059.9639
738.8427734375 0 840.0988
741.8807983398438 0 838.3064
750.3197631835938 0 11513.031 z 9
751.3243408203125 0 5686.9785
752.3225708007812 0 1389.315
761.4019165039062 0 757.70966
761.8969116210938 0 931.2888
765.36669921875 0 2608.8342 y Ammonia loss 2
765.8689575195312 0 4600.981 z 2
766.3649291992188 0 3734.955
766.8727416992188 0 2352.0176
767.3583984375 0 941.17456
772.3888549804688 0 688.926
773.385986328125 0 674.81537
773.8787231445312 0 18104.367 y 2
774.3804321289062 0 14634.977 c Water loss 12
774.881103515625 0 7234.0557 c Ammonia loss 12
775.3693237304688 0 1490.5569
775.4332885742188 0 5190.792 c 6
775.8855590820312 0 955.49286
776.4367065429688 0 2908.0579
777.4397583007812 0 994.97363
777.8632202148438 0 1490.5718
778.361572265625 0 742.5722
778.8712768554688 0 1087.0215
782.9002075195312 0 2109.7068
783.4020385742188 0 14628.079 c 12
783.903564453125 0 12498.09
784.4036254882812 0 7355.493
784.9026489257812 0 2547.9033
785.4027709960938 0 793.582
787.868896484375 0 1358.1211
808.4307250976562 0 604.49225
817.9006958007812 0 615.65656
818.9147338867188 0 831.8212
820.3487548828125 0 2103.7607 w 8
821.35009765625 0 727.21625
837.9381103515625 0 923.2357
840.42333984375 0 32150.467 c 13
840.9251098632812 0 29868.188
841.062255859375 0 694.9034
841.4249877929688 0 16884.076
841.92724609375 0 7568.7886
842.427734375 0 1615.4202
843.9212036132812 0 837.0925
846.4168701171875 0 834.9673 y Water loss 1
846.9238891601562 0 861.2535
847.4064331054688 0 2687.3398 z 1
847.901611328125 0 3499.9622
848.4052124023438 0 1973.2695
848.9080810546875 0 835.89246
853.4318237304688 0 1380.8618
853.9295654296875 0 1010.6418
855.4091186523438 0 2864.5837 y 1
855.9088134765625 0 2462.8179
856.413330078125 0 939.41693
860.4221801757812 0 756.6331
867.4300537109375 0 923.4443
867.9218139648438 0 956.6884
868.4142456054688 0 860.2952
868.9130859375 0 5067.8115
869.4177856445312 0 5570.0596
869.9183959960938 0 3339.5479
870.416259765625 0 1745.1075
870.9182739257812 0 960.7872
874.4376220703125 0 898.9773
874.9498291015625 0 849.0863
875.4328002929688 0 619.4654
875.9307250976562 0 3691.0044
876.4295654296875 0 3284.8652
876.9326171875 0 1402.4022
877.4186401367188 0 2270.238
877.916259765625 0 2222.2551
878.9255981445312 0 742.4622
879.3611450195312 0 3928.876 z 8
880.3650512695312 0 1470.2677
881.3651733398438 0 883.3864
882.4434204101562 0 879.659
882.9381103515625 0 6328.9043
883.4397583007812 0 5260.9067
883.939453125 0 3659.1294
884.4386596679688 0 1946.7512
884.9332275390625 0 2101.7478
888.434326171875 0 869.33136
888.9382934570312 0 671.77325
889.4321899414062 0 1469.0574
889.93115234375 0 1359.9799
896.44189453125 0 2880.741
896.9379272460938 0 21003.268
897.4390258789062 0 24474.705
897.94091796875 0 13190.664
898.4395751953125 0 6583.0894
898.9390869140625 0 2678.603
899.4369506835938 0 995.66815
904.9442749023438 0 14784.182
905.4474487304688 0 34856.195
905.9484252929688 0 25526.998
906.4490356445312 0 14134.584
906.9495849609375 0 5107.569
907.4468994140625 0 2319.466
930.5301513671875 0 790.5648
931.5345458984375 0 8258.449 c 7
932.5374145507812 0 3912.606
933.5433959960938 0 1584.4259
1002.4976806640625 0 815.3737
1016.5548095703125 0 659.8565
1017.5538330078125 0 737.0916
1035.4632568359375 0 7422.1587 z 7
1036.4681396484375 0 4961.9634
1037.4700927734375 0 1561.7993
1050.4759521484375 0 962.3626
1051.4830322265625 0 5240.5664 y 7
1052.485107421875 0 2842.5857
1053.4884033203125 0 1233.6381
1059.567626953125 0 1408.6882
1060.5771484375 0 24730.531 c 8
1061.580322265625 0 13523.088
1062.5830078125 0 5153.175
1063.5792236328125 0 1037.579
1087.5941162109375 0 680.6745
1122.49462890625 0 2241.0103 z 6
1123.5028076171875 0 7400.975
1124.5064697265625 0 3886.9248
1125.509765625 0 1532.5798
1187.5546875 0 1097.8344
1188.602783203125 0 1099.5884
1189.619140625 0 17295.77 c 9
1190.6220703125 0 10891.5
1191.625 0 4414.673
1192.6341552734375 0 804.5271
1197.576904296875 0 612.7595
1208.5916748046875 0 734.3218
1230.6514892578125 0 771.8428
1235.5687255859375 0 1333.9026 y 5
1292.62939453125 0 1134.608
1293.6368408203125 0 1129.7816
1294.6524658203125 0 1118.9242
1300.639892578125 0 813.83875
1301.628662109375 0 799.50995
1302.6478271484375 0 1067.5061
1321.9447021484375 0 607.81134
1333.6317138671875 0 811.893
1334.6295166015625 0 690.5438
1335.6524658203125 0 1887.0938
1336.6544189453125 0 8318.189 c 10
1337.656005859375 0 6372.2
1338.6578369140625 0 2258.3228
1394.68505859375 0 736.842
1401.6839599609375 0 619.52374
1402.68115234375 0 909.44604
1426.0509033203125 0 636.39746
1429.6832275390625 0 2505.7083 z 3
1430.6910400390625 0 10999.705
1431.69189453125 0 6267.6367
1432.69873046875 0 3206.7166
1433.70166015625 0 1883.123
1437.7012939453125 0 12937.373 c 11
1438.7044677734375 0 10127.815
1439.706298828125 0 5104.2
1440.7095947265625 0 2032.8552
1446.714599609375 0 863.31854
1501.277099609375 0 889.5313
1501.761474609375 0 668.2379
1530.730224609375 0 997.2378 z 2
1531.7342529296875 0 2207.1987
1532.743408203125 0 2465.1738
1533.7445068359375 0 1474.4032
1546.7537841796875 0 1763.37 y 2
1547.7576904296875 0 2132.5312 c Water loss 12
1548.7581787109375 0 1089.8175 c Ammonia loss 12
1564.7757568359375 0 830.5542
1565.794921875 0 8406.795 c 12
1566.7982177734375 0 7095.1265
1567.799072265625 0 3640.504
1568.7923583984375 0 1493.0161
1603.548583984375 0 715.4392
1636.83251953125 0 918.2267
1664.8280029296875 0 856.7686
1679.8385009765625 0 3038.2163 c 13
1680.84521484375 0 7709.9937
1681.84912109375 0 7036.6543
1682.849853515625 0 2596.162
1683.85888671875 0 1033.6753
1693.80517578125 0 853.0091 z 1
1694.7933349609375 0 2743.691
1695.8031005859375 0 2812.585
1696.81982421875 0 1028.8491
1697.7967529296875 0 1018.79047
1747.8909912109375 0 1054.1063
1748.85302734375 0 1411.4287
1749.8905029296875 0 1304.7439
1750.86962890625 0 1959.538
1751.8516845703125 0 2522.7656
1752.8587646484375 0 1820.587
1753.8521728515625 0 1068.524
1764.882080078125 0 1934.4082
1765.87890625 0 6159.067
1766.8818359375 0 4247.988
1767.88671875 0 3309.937
1768.8887939453125 0 2321.0708
1776.8515625 0 696.7227
1777.8541259765625 0 754.0295
1781.8876953125 0 715.6835
1782.905517578125 0 2807.0347
1783.904052734375 0 2477.7866
1784.906494140625 0 1366.4944
1792.8731689453125 0 4611.7676
1793.8724365234375 0 16939.12
1794.8746337890625 0 13439.426
1795.8795166015625 0 8724.042
1796.875 0 4205.9067
1797.8748779296875 0 1438.34
1808.874267578125 0 1704.712
1809.8883056640625 0 10258.656
1810.8968505859375 0 34012.723
1811.8995361328125 0 29002.22
1812.90087890625 0 18101.148
1813.904052734375 0 7226.526
1814.9066162109375 0 2523.6675
1815.877197265625 0 954.99084

Spectrum Details

|  |  |
| --- | --- |
| Matched peaks? Matched peaks The total absolute number of peaks matched. Additionally in brackets the total fraction of peaks matched and the total number of peaks is shown. | 53 (15.92% of 333) |
| FDR? FDR The false discovery rate estimated for this peptide. It is calculated by matching all theoretical fragments with a non-integer shift with the raw peaks for this spectrum. This is done with 40 different shifts. The resulting percentage is the average number of annotated peaks over the number of annotated peaks with the correct spectrum. | 3.14% |
| Satellite FDR? Satellite FDR See the FDR for details on its calculation. This satellite ion specific FDR only contains the satellite ions (d/w) for I/L/J positions. | 0.00% |
| PSM Score? PSM Score The PSM Score as given by Hecklib to this annotated spectrum. It is shown with three significant figures. | 319 |

#### Spectrum 3646? Spectrum 3646 The raw spectrum of this peptide as annotated by Hecklib. The fragments are coloured according to ion type (see legend). Any peaks with a star '\*' as text can be hovered over to see the full details, first the ion type second the mass shift type. By hovering over the amino acids in the peptide or ions in the legend the corresponding peaks are highlighted. By toggling the 'Unassigned' label you can turn the background (unassigned) peaks on or off in the plot. By updating the slider in the Ion legend you can update the spectrum to only show the top X% of the peaks with labels. The top X% means any peak that is within X% of the highest intensity. By dragging in the spectrum you can zoom in to a specific part of the spectrum and use 'Zoom Out' to get back to the original zoom level. The annotation of the spectrum is based on the given sequence in the peptides file and is done with different software so inconsistencies are likely. The peaks are annotated based on the given sequence, with 20 ppm tolerance.

Copy Data

##### Spectrum 3646 (TSV)

###### Preview

```
Loading example...
```

*Click on the button to copy the data to your clipboard.*

Mz MinMz MaxIntensity Max

WidthHeightPeptide font sizePeptide stroke widthSpectrum font sizeSpectrum stroke widthCompact peptide

Ion legend

wxyz

abcd

OtherUnassignedIon^Charge^_Position_Show for top:%

VYTJPPSREEMTKNQ

01.36e+42.72e+44.09e+45.45e+4

Zoom Out

y^+1^_1_y^+1^_1_y^+1^_2_z^+1^_2_y^+1^_2_z^+1^_3_y^+1^_3_y^+3^_11_w^+1^_4_w^+1^_4_y^+1^_4_c^+3^_12_y^+1^_4_y^+3^_13_z^+3^_13_y^+3^_13_y^+3^_14_y^+3^_14_z^+3^_14_y^+2^_10_z^+1^_5_y^+1^_5_w^+2^_11_y^+2^_11_y^+2^_11_y^+2^_11_c^+1^_6_w^+1^_6_w^+2^_12_z^+2^_12_y^+2^_12_z^+1^_6_z^+2^_13_y^+2^_13_z^+2^_13_y^+2^_13_c^+2^_13_c^+2^_13_c^+1^_7_c^+2^_13_w^+1^_7_c^+2^_14_y^+2^_14_z^+2^_14_y^+2^_14_z^+1^_7_c^+1^_8_y^+1^_8_z^+1^_8_y^+1^_8_c^+1^_9_y^+1^_9_z^+1^_9_y^+1^_9_c^+1^_10_y^+1^_10_y^+1^_11_c^+1^_11_z^+1^_12_c^+1^_12_y^+1^_13_c^+1^_13_c^+1^_13_c^+1^_13_c^+1^_14_

045891713751833

Fragment Matches Table

Show background peaks

| Position | Ion type | Intensity | mz Theoretical | mz Error (Th) | mz Error (ppm) | Charge | Series Number |
| --- | --- | --- | --- | --- | --- | --- | --- |
| - | - | 448.7 | 125.1 | - | - | 0 | - |
| - | - | 842.4 | 126.1 | - | - | 0 | - |
| - | - | 480.5 | 128.1 | - | - | 0 | - |
| - | - | 615.8 | 129.1 | - | - | 0 | - |
| - | - | 375.3 | 129.7 | - | - | 0 | - |
| 15 | y | 616 | 130 | 0.0002174 | 1.672 | +1 | 1 |
| - | - | 432.3 | 130.4 | - | - | 0 | - |
| - | - | 3001 | 136.1 | - | - | 0 | - |
| - | - | 369.8 | 140.4 | - | - | 0 | - |
| 15 | y | 3393 | 147.1 | 0.0003231 | 2.197 | +1 | 1 |
| - | - | 373.1 | 147.4 | - | - | 0 | - |
| - | - | 775.1 | 168.1 | - | - | 0 | - |
| - | - | 589.1 | 170 | - | - | 0 | - |
| - | - | 1977 | 173.5 | - | - | 0 | - |
| - | - | 1992 | 175.1 | - | - | 0 | - |
| - | - | 523.4 | 183.1 | - | - | 0 | - |
| - | - | 1244 | 185.1 | - | - | 0 | - |
| - | - | 481.6 | 189.2 | - | - | 0 | - |
| - | - | 7326 | 199.2 | - | - | 0 | - |
| - | - | 874.1 | 200.2 | - | - | 0 | - |
| - | - | 2171 | 212.1 | - | - | 0 | - |
| - | - | 960.9 | 215.1 | - | - | 0 | - |
| - | - | 739.7 | 216.1 | - | - | 0 | - |
| - | - | 1727 | 217.1 | - | - | 0 | - |
| - | - | 4304 | 227.2 | - | - | 0 | - |
| - | - | 870.2 | 228.2 | - | - | 0 | - |
| - | - | 796.3 | 230.1 | - | - | 0 | - |
| - | - | 9541 | 235.1 | - | - | 0 | - |
| - | - | 1085 | 236.1 | - | - | 0 | - |
| - | - | 1508 | 243.2 | - | - | 0 | - |
| 14 | y | 782.9 | 244.1 | 0.0005118 | 2.097 | +1 | 2 |
| 14 | z | 657.3 | 245.1 | 0.0001582 | 0.6453 | +1 | 2 |
| - | - | 3707 | 245.1 | - | - | 0 | - |
| - | - | 597.1 | 255 | - | - | 0 | - |
| - | - | 674.6 | 259.2 | - | - | 0 | - |
| 14 | y | 3084 | 261.1 | 0.0005717 | 2.189 | +1 | 2 |
| - | - | 1437 | 262.2 | - | - | 0 | - |
| - | - | 687.5 | 263.1 | - | - | 0 | - |
| - | - | 7452 | 263.1 | - | - | 0 | - |
| - | - | 580.8 | 264.1 | - | - | 0 | - |
| - | - | 959 | 270.1 | - | - | 0 | - |
| - | - | 2051 | 299.2 | - | - | 0 | - |
| - | - | 1061 | 316.2 | - | - | 0 | - |
| - | - | 1389 | 329.2 | - | - | 0 | - |
| - | - | 642.7 | 332.2 | - | - | 0 | - |
| - | - | 1682 | 333.2 | - | - | 0 | - |
| - | - | 4739 | 346.2 | - | - | 0 | - |
| - | - | 663.5 | 347.2 | - | - | 0 | - |
| - | - | 933.8 | 349.2 | - | - | 0 | - |
| - | - | 1562 | 357.2 | - | - | 0 | - |
| - | - | 559.6 | 357.2 | - | - | 0 | - |
| - | - | 929.8 | 360.2 | - | - | 0 | - |
| - | - | 1780 | 361.2 | - | - | 0 | - |
| - | - | 7815 | 364.2 | - | - | 0 | - |
| - | - | 2258 | 365.2 | - | - | 0 | - |
| - | - | 628.9 | 366.2 | - | - | 0 | - |
| 13 | z | 3673 | 373.2 | 0.001371 | 3.674 | +1 | 3 |
| - | - | 669.1 | 374.2 | - | - | 0 | - |
| - | - | 859.4 | 386.2 | - | - | 0 | - |
| - | - | 1014 | 388.2 | - | - | 0 | - |
| 13 | y | 1562 | 389.2 | 0.0005945 | 1.527 | +1 | 3 |
| - | - | 624.3 | 391.7 | - | - | 0 | - |
| - | - | 971.1 | 415.2 | - | - | 0 | - |
| - | - | 1831 | 416.2 | - | - | 0 | - |
| - | - | 1108 | 417.2 | - | - | 0 | - |
| - | - | 867.2 | 422.7 | - | - | 0 | - |
| - | - | 1171 | 423.2 | - | - | 0 | - |
| - | - | 2380 | 430.2 | - | - | 0 | - |
| - | - | 970.9 | 431.2 | - | - | 0 | - |
| - | - | 1448 | 431.3 | - | - | 0 | - |
| - | - | 974.7 | 432.2 | - | - | 0 | - |
| 5 | y | 5547 | 444.9 | 0.001306 | 2.935 | +3 | 11 |
| - | - | 1873 | 445.2 | - | - | 0 | - |
| - | - | 969 | 445.5 | - | - | 0 | - |
| - | - | 642.5 | 445.7 | - | - | 0 | - |
| - | - | 653.3 | 446.2 | - | - | 0 | - |
| - | - | 951.4 | 448.3 | - | - | 0 | - |
| - | - | 703.7 | 450.7 | - | - | 0 | - |
| 12 | w | 1399 | 457.2 | 0.0006862 | 1.501 | +1 | 4 |
| 12 | w | 921.1 | 459.2 | 0.0004875 | 1.061 | +1 | 4 |
| - | - | 5785 | 459.3 | - | - | 0 | - |
| - | - | 1265 | 460.3 | - | - | 0 | - |
| - | - | 632 | 464.1 | - | - | 0 | - |
| - | - | 893 | 472.2 | - | - | 0 | - |
| 12 | y | 3633 | 473.2 | 0.000148 | 0.3127 | +1 | 4 |
| 12 | c | 1215 | 474.2 | 0.005692 | 12 | +3 | 12 |
| - | - | 824.5 | 475.3 | - | - | 0 | - |
| - | - | 2051 | 477.3 | - | - | 0 | - |
| - | - | 1814 | 489.3 | - | - | 0 | - |
| 12 | y | 2751 | 490.3 | 0.00149 | 3.038 | +1 | 4 |
| - | - | 898.9 | 498.2 | - | - | 0 | - |
| - | - | 597 | 504.3 | - | - | 0 | - |
| - | - | 4827 | 505.3 | - | - | 0 | - |
| - | - | 607.4 | 506.3 | - | - | 0 | - |
| - | - | 641.1 | 508.2 | - | - | 0 | - |
| 3 | y | 1918 | 510.3 | 0.0003522 | 0.6902 | +3 | 13 |
| - | - | 801.6 | 510.8 | - | - | 0 | - |
| 3 | z | 873.8 | 510.9 | 0.006663 | 13.04 | +3 | 13 |
| - | - | 631.6 | 511.3 | - | - | 0 | - |
| - | - | 599.7 | 513.8 | - | - | 0 | - |
| - | - | 708 | 515.1 | - | - | 0 | - |
| - | - | 792.5 | 515.3 | - | - | 0 | - |
| 3 | y | 1608 | 516.3 | 0.001216 | 2.356 | +3 | 13 |
| - | - | 1706 | 516.8 | - | - | 0 | - |
| - | - | 711.6 | 517.3 | - | - | 0 | - |
| - | - | 897.9 | 520.2 | - | - | 0 | - |
| - | - | 3163 | 524.3 | - | - | 0 | - |
| - | - | 763.3 | 524.8 | - | - | 0 | - |
| - | - | 1872 | 530.3 | - | - | 0 | - |
| - | - | 1160 | 530.6 | - | - | 0 | - |
| - | - | 1557 | 531.3 | - | - | 0 | - |
| - | - | 843.2 | 532.3 | - | - | 0 | - |
| - | - | 581.4 | 533.3 | - | - | 0 | - |
| - | - | 1130 | 536.8 | - | - | 0 | - |
| - | - | 1005 | 538.6 | - | - | 0 | - |
| - | - | 1204 | 541.3 | - | - | 0 | - |
| - | - | 698.4 | 542.3 | - | - | 0 | - |
| - | - | 831.6 | 543.3 | - | - | 0 | - |
| - | - | 811 | 550.8 | - | - | 0 | - |
| - | - | 754.3 | 551.3 | - | - | 0 | - |
| - | - | 5813 | 551.8 | - | - | 0 | - |
| - | - | 3459 | 552.3 | - | - | 0 | - |
| - | - | 921.5 | 552.8 | - | - | 0 | - |
| - | - | 1571 | 557.3 | - | - | 0 | - |
| - | - | 5890 | 558.3 | - | - | 0 | - |
| - | - | 3364 | 558.8 | - | - | 0 | - |
| - | - | 1.103E+04 | 559.3 | - | - | 0 | - |
| - | - | 674.2 | 559.3 | - | - | 0 | - |
| - | - | 1143 | 559.8 | - | - | 0 | - |
| - | - | 871.4 | 560.2 | - | - | 0 | - |
| - | - | 1957 | 560.3 | - | - | 0 | - |
| - | - | 799 | 561.3 | - | - | 0 | - |
| 2 | y | 748.1 | 564.6 | 0.002477 | 4.387 | +3 | 14 |
| 2 | y | 913.4 | 564.9 | 0.008273 | 14.64 | +3 | 14 |
| 2 | z | 583.8 | 565.3 | 0.00424 | 7.501 | +3 | 14 |
| - | - | 2918 | 565.8 | - | - | 0 | - |
| - | - | 2663 | 566.3 | - | - | 0 | - |
| - | - | 1395 | 566.8 | - | - | 0 | - |
| - | - | 705.4 | 571.3 | - | - | 0 | - |
| - | - | 1708 | 572.8 | - | - | 0 | - |
| - | - | 1338 | 573.3 | - | - | 0 | - |
| - | - | 1897 | 573.8 | - | - | 0 | - |
| - | - | 624.4 | 574.8 | - | - | 0 | - |
| - | - | 723.8 | 575.3 | - | - | 0 | - |
| - | - | 1675 | 576.3 | - | - | 0 | - |
| - | - | 1.351E+04 | 580.8 | - | - | 0 | - |
| - | - | 890.2 | 581.2 | - | - | 0 | - |
| - | - | 1.052E+04 | 581.3 | - | - | 0 | - |
| - | - | 7055 | 581.8 | - | - | 0 | - |
| - | - | 4206 | 582.3 | - | - | 0 | - |
| - | - | 766.3 | 584.3 | - | - | 0 | - |
| - | - | 766.2 | 584.8 | - | - | 0 | - |
| - | - | 1787 | 585.3 | - | - | 0 | - |
| - | - | 888.5 | 586.3 | - | - | 0 | - |
| - | - | 801.5 | 594.3 | - | - | 0 | - |
| - | - | 766.6 | 594.8 | - | - | 0 | - |
| - | - | 5468 | 597.6 | - | - | 0 | - |
| - | - | 6514 | 598 | - | - | 0 | - |
| - | - | 2654 | 598.3 | - | - | 0 | - |
| - | - | 1419 | 598.6 | - | - | 0 | - |
| - | - | 779.4 | 598.9 | - | - | 0 | - |
| - | - | 1108 | 600.3 | - | - | 0 | - |
| - | - | 1590 | 600.8 | - | - | 0 | - |
| - | - | 1187 | 601.3 | - | - | 0 | - |
| - | - | 1055 | 602.3 | - | - | 0 | - |
| - | - | 745.1 | 602.3 | - | - | 0 | - |
| - | - | 3173 | 602.8 | - | - | 0 | - |
| - | - | 917.6 | 602.8 | - | - | 0 | - |
| - | - | 4035 | 603.3 | - | - | 0 | - |
| - | - | 5578 | 603.3 | - | - | 0 | - |
| - | - | 1.681E+04 | 603.8 | - | - | 0 | - |
| - | - | 8885 | 604.3 | - | - | 0 | - |
| - | - | 1017 | 604.3 | - | - | 0 | - |
| - | - | 4124 | 604.8 | - | - | 0 | - |
| 6 | y | 1800 | 618.3 | 0.0007748 | 1.253 | +2 | 10 |
| - | - | 1218 | 618.8 | - | - | 0 | - |
| 11 | z | 1790 | 621.3 | 0.002558 | 4.117 | +1 | 5 |
| - | - | 1605 | 622.3 | - | - | 0 | - |
| - | - | 2722 | 630.3 | - | - | 0 | - |
| - | - | 2179 | 630.8 | - | - | 0 | - |
| - | - | 615.3 | 633.3 | - | - | 0 | - |
| - | - | 1813 | 634.8 | - | - | 0 | - |
| - | - | 1454 | 635.3 | - | - | 0 | - |
| - | - | 894.2 | 635.8 | - | - | 0 | - |
| 11 | y | 1239 | 637.3 | 0.00483 | 7.579 | +1 | 5 |
| - | - | 1124 | 637.8 | - | - | 0 | - |
| - | - | 2747 | 638.3 | - | - | 0 | - |
| - | - | 1939 | 638.8 | - | - | 0 | - |
| - | - | 1243 | 639.3 | - | - | 0 | - |
| - | - | 798.5 | 643.8 | - | - | 0 | - |
| - | - | 696.8 | 644.3 | - | - | 0 | - |
| 5 | w | 746.2 | 645.3 | 0.002878 | 4.46 | +2 | 11 |
| - | - | 2599 | 646.4 | - | - | 0 | - |
| - | - | 1423 | 647.4 | - | - | 0 | - |
| 5 | y | 823.3 | 657.8 | 0.002502 | 3.804 | +2 | 11 |
| 5 | y | 1968 | 658.3 | 0.007443 | 11.31 | +2 | 11 |
| - | - | 577 | 659.3 | - | - | 0 | - |
| - | - | 1465 | 665.8 | - | - | 0 | - |
| - | - | 1557 | 666.3 | - | - | 0 | - |
| 5 | y | 4.138E+04 | 666.8 | 0.001553 | 2.33 | +2 | 11 |
| - | - | 2.989E+04 | 667.3 | - | - | 0 | - |
| - | - | 445.9 | 667.4 | - | - | 0 | - |
| - | - | 1.624E+04 | 667.8 | - | - | 0 | - |
| - | - | 5210 | 668.3 | - | - | 0 | - |
| - | - | 1670 | 673.3 | - | - | 0 | - |
| - | - | 730.5 | 673.8 | - | - | 0 | - |
| - | - | 1650 | 674.3 | - | - | 0 | - |
| - | - | 1268 | 674.8 | - | - | 0 | - |
| - | - | 1012 | 675.3 | - | - | 0 | - |
| - | - | 1358 | 680.8 | - | - | 0 | - |
| - | - | 690.7 | 681.3 | - | - | 0 | - |
| - | - | 1.474E+04 | 681.8 | - | - | 0 | - |
| - | - | 1.076E+04 | 682.3 | - | - | 0 | - |
| - | - | 4338 | 682.8 | - | - | 0 | - |
| - | - | 2016 | 683.3 | - | - | 0 | - |
| - | - | 1148 | 683.8 | - | - | 0 | - |
| - | - | 870 | 686.3 | - | - | 0 | - |
| - | - | 4170 | 687.3 | - | - | 0 | - |
| - | - | 2557 | 687.4 | - | - | 0 | - |
| - | - | 1651 | 688.3 | - | - | 0 | - |
| 6 | c | 5170 | 688.4 | 0.002387 | 3.467 | +1 | 6 |
| - | - | 1702 | 689.4 | - | - | 0 | - |
| 10 | w | 1.771E+04 | 691.3 | 0.004336 | 6.272 | +1 | 6 |
| - | - | 5062 | 692.3 | - | - | 0 | - |
| - | - | 2162 | 693.3 | - | - | 0 | - |
| 4 | w | 1.082E+04 | 693.8 | 0.002069 | 2.983 | +2 | 12 |
| - | - | 1.094E+04 | 694.3 | - | - | 0 | - |
| - | - | 3453 | 694.8 | - | - | 0 | - |
| - | - | 2285 | 695.3 | - | - | 0 | - |
| - | - | 808.3 | 697.4 | - | - | 0 | - |
| - | - | 613.9 | 701.4 | - | - | 0 | - |
| - | - | 695.3 | 702.3 | - | - | 0 | - |
| - | - | 984.5 | 703.3 | - | - | 0 | - |
| - | - | 1780 | 713.4 | - | - | 0 | - |
| - | - | 755.8 | 714.4 | - | - | 0 | - |
| 4 | z | 1.16E+04 | 715.3 | 0.001049 | 1.467 | +2 | 12 |
| - | - | 773.5 | 715.4 | - | - | 0 | - |
| - | - | 1.033E+04 | 715.8 | - | - | 0 | - |
| - | - | 5626 | 716.3 | - | - | 0 | - |
| - | - | 2199 | 716.8 | - | - | 0 | - |
| - | - | 666.6 | 722.9 | - | - | 0 | - |
| 4 | y | 5629 | 723.4 | 0.0009033 | 1.249 | +2 | 12 |
| - | - | 5592 | 723.9 | - | - | 0 | - |
| - | - | 3104 | 724.4 | - | - | 0 | - |
| - | - | 991.4 | 729.9 | - | - | 0 | - |
| - | - | 828.6 | 730.3 | - | - | 0 | - |
| - | - | 1673 | 732.4 | - | - | 0 | - |
| - | - | 2357 | 737.8 | - | - | 0 | - |
| - | - | 1527 | 738.3 | - | - | 0 | - |
| - | - | 825.4 | 738.8 | - | - | 0 | - |
| - | - | 1244 | 741.9 | - | - | 0 | - |
| - | - | 1211 | 742.4 | - | - | 0 | - |
| - | - | 676.5 | 742.9 | - | - | 0 | - |
| - | - | 1018 | 743.4 | - | - | 0 | - |
| - | - | 803.8 | 744.4 | - | - | 0 | - |
| - | - | 681.7 | 744.8 | - | - | 0 | - |
| 10 | z | 1.633E+04 | 750.3 | 0.003971 | 5.293 | +1 | 6 |
| - | - | 7268 | 751.3 | - | - | 0 | - |
| - | - | 3018 | 752.3 | - | - | 0 | - |
| 3 | z | 753.6 | 757.4 | 0.007208 | 9.517 | +2 | 13 |
| - | - | 833.9 | 757.9 | - | - | 0 | - |
| - | - | 717 | 758.4 | - | - | 0 | - |
| - | - | 802.1 | 758.9 | - | - | 0 | - |
| - | - | 6589 | 761.4 | - | - | 0 | - |
| - | - | 631 | 761.9 | - | - | 0 | - |
| - | - | 2136 | 762.4 | - | - | 0 | - |
| 3 | y | 1.23E+04 | 765.4 | 0.0142 | 18.56 | +2 | 13 |
| 3 | z | 1.469E+04 | 765.9 | 0.00846 | 11.05 | +2 | 13 |
| - | - | 9430 | 766.4 | - | - | 0 | - |
| - | - | 6930 | 766.9 | - | - | 0 | - |
| - | - | 848.8 | 767.4 | - | - | 0 | - |
| - | - | 746.2 | 768.4 | - | - | 0 | - |
| - | - | 2123 | 773.4 | - | - | 0 | - |
| 3 | y | 2.583E+04 | 773.9 | 0.002943 | 3.803 | +2 | 13 |
| 13 | c | 2.029E+04 | 774.4 | 0.01336 | 17.25 | +2 | 13 |
| 13 | c | 9268 | 774.9 | 0.003903 | 5.036 | +2 | 13 |
| - | - | 915 | 775.4 | - | - | 0 | - |
| 7 | c | 8811 | 775.4 | 0.001822 | 2.35 | +1 | 7 |
| - | - | 881 | 775.9 | - | - | 0 | - |
| - | - | 4970 | 776.4 | - | - | 0 | - |
| - | - | 1798 | 777.4 | - | - | 0 | - |
| - | - | 2172 | 777.9 | - | - | 0 | - |
| - | - | 1124 | 778.4 | - | - | 0 | - |
| - | - | 1037 | 778.9 | - | - | 0 | - |
| - | - | 724.9 | 779.9 | - | - | 0 | - |
| - | - | 563.6 | 780.4 | - | - | 0 | - |
| - | - | 735.7 | 781.9 | - | - | 0 | - |
| - | - | 5486 | 782.9 | - | - | 0 | - |
| 13 | c | 2.539E+04 | 783.4 | 0.001744 | 2.226 | +2 | 13 |
| - | - | 1.851E+04 | 783.9 | - | - | 0 | - |
| - | - | 9853 | 784.4 | - | - | 0 | - |
| - | - | 3704 | 784.9 | - | - | 0 | - |
| - | - | 1931 | 785.4 | - | - | 0 | - |
| - | - | 984.2 | 786.4 | - | - | 0 | - |
| - | - | 9155 | 786.9 | - | - | 0 | - |
| - | - | 6689 | 787.4 | - | - | 0 | - |
| - | - | 4248 | 787.9 | - | - | 0 | - |
| - | - | 1762 | 788.4 | - | - | 0 | - |
| - | - | 638.6 | 788.9 | - | - | 0 | - |
| - | - | 678.5 | 793.9 | - | - | 0 | - |
| - | - | 1517 | 794.4 | - | - | 0 | - |
| - | - | 2.774E+04 | 794.9 | - | - | 0 | - |
| - | - | 2.961E+04 | 795.4 | - | - | 0 | - |
| - | - | 1.444E+04 | 795.9 | - | - | 0 | - |
| - | - | 7053 | 796.4 | - | - | 0 | - |
| - | - | 2305 | 796.9 | - | - | 0 | - |
| - | - | 871.6 | 797.4 | - | - | 0 | - |
| - | - | 2435 | 801.9 | - | - | 0 | - |
| - | - | 2584 | 802.4 | - | - | 0 | - |
| - | - | 1055 | 802.9 | - | - | 0 | - |
| - | - | 1670 | 803.4 | - | - | 0 | - |
| - | - | 3002 | 806.9 | - | - | 0 | - |
| - | - | 2060 | 807.4 | - | - | 0 | - |
| - | - | 794.6 | 808.9 | - | - | 0 | - |
| - | - | 1238 | 810.4 | - | - | 0 | - |
| - | - | 2748 | 810.9 | - | - | 0 | - |
| - | - | 4169 | 811.4 | - | - | 0 | - |
| - | - | 1583 | 811.9 | - | - | 0 | - |
| - | - | 712.5 | 812.4 | - | - | 0 | - |
| - | - | 898 | 814.4 | - | - | 0 | - |
| - | - | 883.3 | 818.4 | - | - | 0 | - |
| - | - | 1860 | 818.9 | - | - | 0 | - |
| - | - | 674.6 | 819.4 | - | - | 0 | - |
| - | - | 2961 | 819.9 | - | - | 0 | - |
| 9 | w | 5652 | 820.3 | 0.00874 | 10.65 | +1 | 7 |
| - | - | 794.1 | 820.9 | - | - | 0 | - |
| - | - | 1910 | 821.4 | - | - | 0 | - |
| - | - | 2267 | 821.9 | - | - | 0 | - |
| - | - | 1770 | 822.4 | - | - | 0 | - |
| - | - | 1934 | 822.9 | - | - | 0 | - |
| - | - | 899.1 | 824.9 | - | - | 0 | - |
| - | - | 849.4 | 825.4 | - | - | 0 | - |
| - | - | 752.6 | 826.4 | - | - | 0 | - |
| - | - | 1049 | 827.4 | - | - | 0 | - |
| - | - | 603.8 | 827.9 | - | - | 0 | - |
| - | - | 4511 | 829.4 | - | - | 0 | - |
| - | - | 2477 | 830.4 | - | - | 0 | - |
| - | - | 878.1 | 839.4 | - | - | 0 | - |
| - | - | 864.7 | 839.9 | - | - | 0 | - |
| 14 | c | 5.395E+04 | 840.4 | 0.001459 | 1.736 | +2 | 14 |
| - | - | 4.199E+04 | 840.9 | - | - | 0 | - |
| - | - | 2.629E+04 | 841.4 | - | - | 0 | - |
| - | - | 1.008E+04 | 841.9 | - | - | 0 | - |
| - | - | 3337 | 842.4 | - | - | 0 | - |
| - | - | 3245 | 843.4 | - | - | 0 | - |
| - | - | 1169 | 844.4 | - | - | 0 | - |
| - | - | 662.2 | 844.8 | - | - | 0 | - |
| - | - | 1755 | 845.4 | - | - | 0 | - |
| - | - | 606.6 | 846.5 | - | - | 0 | - |
| 2 | y | 710.2 | 846.9 | 0.008052 | 9.507 | +2 | 14 |
| 2 | z | 3498 | 847.4 | 0.002003 | 2.364 | +2 | 14 |
| - | - | 3970 | 847.9 | - | - | 0 | - |
| - | - | 2278 | 848.4 | - | - | 0 | - |
| - | - | 865.2 | 848.9 | - | - | 0 | - |
| - | - | 670.3 | 850.4 | - | - | 0 | - |
| - | - | 1206 | 853.4 | - | - | 0 | - |
| - | - | 1806 | 853.9 | - | - | 0 | - |
| - | - | 1189 | 854.4 | - | - | 0 | - |
| - | - | 1048 | 854.9 | - | - | 0 | - |
| 2 | y | 3415 | 855.4 | 0.003444 | 4.026 | +2 | 14 |
| - | - | 2002 | 855.9 | - | - | 0 | - |
| - | - | 850.3 | 856.4 | - | - | 0 | - |
| - | - | 814.4 | 859.4 | - | - | 0 | - |
| - | - | 778.9 | 860.4 | - | - | 0 | - |
| - | - | 1147 | 861.4 | - | - | 0 | - |
| - | - | 1313 | 867.4 | - | - | 0 | - |
| - | - | 772.1 | 867.9 | - | - | 0 | - |
| - | - | 1115 | 868.4 | - | - | 0 | - |
| - | - | 7864 | 868.9 | - | - | 0 | - |
| - | - | 7481 | 869.4 | - | - | 0 | - |
| - | - | 4907 | 869.9 | - | - | 0 | - |
| - | - | 1616 | 870.4 | - | - | 0 | - |
| - | - | 924 | 870.9 | - | - | 0 | - |
| - | - | 1461 | 873.9 | - | - | 0 | - |
| - | - | 1648 | 874.4 | - | - | 0 | - |
| - | - | 881.7 | 875.4 | - | - | 0 | - |
| - | - | 3405 | 875.9 | - | - | 0 | - |
| - | - | 4912 | 876.4 | - | - | 0 | - |
| - | - | 2517 | 876.9 | - | - | 0 | - |
| - | - | 2968 | 877.4 | - | - | 0 | - |
| - | - | 2012 | 877.9 | - | - | 0 | - |
| - | - | 2081 | 878.4 | - | - | 0 | - |
| - | - | 853.1 | 878.9 | - | - | 0 | - |
| 9 | z | 6325 | 879.4 | 0.006239 | 7.095 | +1 | 7 |
| - | - | 4446 | 880.4 | - | - | 0 | - |
| - | - | 1446 | 881.4 | - | - | 0 | - |
| - | - | 1881 | 882.4 | - | - | 0 | - |
| - | - | 1.047E+04 | 882.9 | - | - | 0 | - |
| - | - | 8733 | 883.4 | - | - | 0 | - |
| - | - | 6610 | 883.9 | - | - | 0 | - |
| - | - | 4555 | 884.4 | - | - | 0 | - |
| - | - | 1988 | 884.9 | - | - | 0 | - |
| - | - | 892.3 | 886.4 | - | - | 0 | - |
| - | - | 1578 | 887.9 | - | - | 0 | - |
| - | - | 2186 | 888.4 | - | - | 0 | - |
| - | - | 1194 | 888.9 | - | - | 0 | - |
| - | - | 2647 | 889.4 | - | - | 0 | - |
| - | - | 1516 | 889.9 | - | - | 0 | - |
| - | - | 3416 | 896.4 | - | - | 0 | - |
| - | - | 3.385E+04 | 896.9 | - | - | 0 | - |
| - | - | 3.344E+04 | 897.4 | - | - | 0 | - |
| - | - | 2.41E+04 | 897.9 | - | - | 0 | - |
| - | - | 1.141E+04 | 898.4 | - | - | 0 | - |
| - | - | 2995 | 898.9 | - | - | 0 | - |
| - | - | 987.7 | 900.5 | - | - | 0 | - |
| - | - | 933.8 | 901.5 | - | - | 0 | - |
| - | - | 895.7 | 902.5 | - | - | 0 | - |
| - | - | 1772 | 904.4 | - | - | 0 | - |
| - | - | 2.66E+04 | 904.9 | - | - | 0 | - |
| - | - | 5.15E+04 | 905.4 | - | - | 0 | - |
| - | - | 3.892E+04 | 905.9 | - | - | 0 | - |
| - | - | 2.271E+04 | 906.4 | - | - | 0 | - |
| - | - | 8334 | 907 | - | - | 0 | - |
| - | - | 1478 | 907.4 | - | - | 0 | - |
| 8 | c | 1.399E+04 | 931.5 | 0.0008214 | 0.8817 | +1 | 8 |
| - | - | 7441 | 932.5 | - | - | 0 | - |
| - | - | 2346 | 933.5 | - | - | 0 | - |
| - | - | 961.5 | 934.5 | - | - | 0 | - |
| - | - | 2334 | 944.5 | - | - | 0 | - |
| - | - | 2937 | 945.5 | - | - | 0 | - |
| - | - | 708 | 946.5 | - | - | 0 | - |
| - | - | 815 | 957.6 | - | - | 0 | - |
| - | - | 747.2 | 959.5 | - | - | 0 | - |
| - | - | 1526 | 960.5 | - | - | 0 | - |
| - | - | 1778 | 961.5 | - | - | 0 | - |
| - | - | 684.1 | 970.6 | - | - | 0 | - |
| - | - | 790.8 | 977.5 | - | - | 0 | - |
| - | - | 1474 | 989.5 | - | - | 0 | - |
| - | - | 963 | 990.5 | - | - | 0 | - |
| - | - | 903.3 | 1003 | - | - | 0 | - |
| - | - | 1242 | 1004 | - | - | 0 | - |
| - | - | 1229 | 1005 | - | - | 0 | - |
| - | - | 5136 | 1015 | - | - | 0 | - |
| - | - | 2790 | 1016 | - | - | 0 | - |
| - | - | 2333 | 1017 | - | - | 0 | - |
| - | - | 1442 | 1018 | - | - | 0 | - |
| - | - | 1513 | 1032 | - | - | 0 | - |
| - | - | 1648 | 1032 | - | - | 0 | - |
| 8 | y | 1027 | 1033 | 0.02056 | 19.9 | +1 | 8 |
| 8 | z | 1.026E+04 | 1035 | 0.00431 | 4.163 | +1 | 8 |
| - | - | 8618 | 1036 | - | - | 0 | - |
| - | - | 4486 | 1037 | - | - | 0 | - |
| - | - | 1985 | 1038 | - | - | 0 | - |
| - | - | 849.5 | 1045 | - | - | 0 | - |
| - | - | 1192 | 1046 | - | - | 0 | - |
| - | - | 1458 | 1048 | - | - | 0 | - |
| - | - | 793.8 | 1048 | - | - | 0 | - |
| - | - | 1208 | 1050 | - | - | 0 | - |
| 8 | y | 7223 | 1051 | 0.003652 | 3.474 | +1 | 8 |
| - | - | 3716 | 1052 | - | - | 0 | - |
| - | - | 1944 | 1053 | - | - | 0 | - |
| - | - | 795.7 | 1054 | - | - | 0 | - |
| - | - | 1770 | 1058 | - | - | 0 | - |
| - | - | 1043 | 1059 | - | - | 0 | - |
| - | - | 2311 | 1060 | - | - | 0 | - |
| 9 | c | 3.617E+04 | 1061 | 0.0006898 | 0.6504 | +1 | 9 |
| - | - | 2.293E+04 | 1062 | - | - | 0 | - |
| - | - | 8093 | 1063 | - | - | 0 | - |
| - | - | 950.1 | 1073 | - | - | 0 | - |
| - | - | 804.3 | 1075 | - | - | 0 | - |
| - | - | 1323 | 1076 | - | - | 0 | - |
| - | - | 1181 | 1092 | - | - | 0 | - |
| - | - | 630.6 | 1101 | - | - | 0 | - |
| - | - | 6281 | 1102 | - | - | 0 | - |
| - | - | 4328 | 1103 | - | - | 0 | - |
| - | - | 2329 | 1104 | - | - | 0 | - |
| - | - | 2227 | 1105 | - | - | 0 | - |
| - | - | 763.2 | 1106 | - | - | 0 | - |
| - | - | 919.2 | 1117 | - | - | 0 | - |
| - | - | 944.1 | 1118 | - | - | 0 | - |
| - | - | 809.3 | 1119 | - | - | 0 | - |
| 7 | y | 1485 | 1121 | 0.01991 | 17.77 | +1 | 9 |
| - | - | 683.2 | 1122 | - | - | 0 | - |
| 7 | z | 3149 | 1122 | 0.006461 | 5.756 | +1 | 9 |
| - | - | 1.058E+04 | 1124 | - | - | 0 | - |
| - | - | 4895 | 1125 | - | - | 0 | - |
| - | - | 2980 | 1126 | - | - | 0 | - |
| - | - | 763.1 | 1129 | - | - | 0 | - |
| - | - | 1835 | 1131 | - | - | 0 | - |
| - | - | 3847 | 1132 | - | - | 0 | - |
| - | - | 2060 | 1133 | - | - | 0 | - |
| - | - | 1578 | 1134 | - | - | 0 | - |
| 7 | y | 731.7 | 1139 | 0.004461 | 3.918 | +1 | 9 |
| - | - | 876.5 | 1140 | - | - | 0 | - |
| - | - | 1007 | 1144 | - | - | 0 | - |
| - | - | 3400 | 1145 | - | - | 0 | - |
| - | - | 5341 | 1146 | - | - | 0 | - |
| - | - | 1.499E+04 | 1147 | - | - | 0 | - |
| - | - | 8382 | 1148 | - | - | 0 | - |
| - | - | 2732 | 1149 | - | - | 0 | - |
| - | - | 2869 | 1159 | - | - | 0 | - |
| - | - | 1136 | 1160 | - | - | 0 | - |
| - | - | 1838 | 1161 | - | - | 0 | - |
| - | - | 6612 | 1162 | - | - | 0 | - |
| - | - | 5638 | 1163 | - | - | 0 | - |
| - | - | 4078 | 1164 | - | - | 0 | - |
| - | - | 2003 | 1165 | - | - | 0 | - |
| - | - | 1058 | 1188 | - | - | 0 | - |
| - | - | 2104 | 1189 | - | - | 0 | - |
| 10 | c | 2.751E+04 | 1190 | 0.002634 | 2.214 | +1 | 10 |
| - | - | 1.797E+04 | 1191 | - | - | 0 | - |
| - | - | 6633 | 1192 | - | - | 0 | - |
| - | - | 2123 | 1193 | - | - | 0 | - |
| - | - | 1153 | 1204 | - | - | 0 | - |
| - | - | 2694 | 1205 | - | - | 0 | - |
| - | - | 1013 | 1206 | - | - | 0 | - |
| - | - | 970.5 | 1206 | - | - | 0 | - |
| - | - | 3621 | 1207 | - | - | 0 | - |
| - | - | 2997 | 1208 | - | - | 0 | - |
| - | - | 1143 | 1209 | - | - | 0 | - |
| - | - | 762.9 | 1210 | - | - | 0 | - |
| - | - | 949.3 | 1214 | - | - | 0 | - |
| 6 | y | 2041 | 1236 | 0.004554 | 3.685 | +1 | 10 |
| - | - | 1640 | 1237 | - | - | 0 | - |
| - | - | 3561 | 1258 | - | - | 0 | - |
| - | - | 2424 | 1259 | - | - | 0 | - |
| - | - | 1591 | 1260 | - | - | 0 | - |
| - | - | 3128 | 1261 | - | - | 0 | - |
| - | - | 2571 | 1262 | - | - | 0 | - |
| - | - | 1123 | 1263 | - | - | 0 | - |
| - | - | 1426 | 1276 | - | - | 0 | - |
| - | - | 793.5 | 1278 | - | - | 0 | - |
| - | - | 2429 | 1293 | - | - | 0 | - |
| - | - | 2133 | 1294 | - | - | 0 | - |
| - | - | 737.8 | 1295 | - | - | 0 | - |
| - | - | 2131 | 1301 | - | - | 0 | - |
| - | - | 1862 | 1302 | - | - | 0 | - |
| - | - | 1171 | 1303 | - | - | 0 | - |
| 5 | y | 2407 | 1333 | 0.008064 | 6.051 | +1 | 11 |
| - | - | 1409 | 1334 | - | - | 0 | - |
| - | - | 774.1 | 1335 | - | - | 0 | - |
| - | - | 4081 | 1336 | - | - | 0 | - |
| 11 | c | 1.274E+04 | 1337 | 0.00155 | 1.159 | +1 | 11 |
| - | - | 8262 | 1338 | - | - | 0 | - |
| - | - | 3523 | 1339 | - | - | 0 | - |
| - | - | 6494 | 1345 | - | - | 0 | - |
| - | - | 4646 | 1346 | - | - | 0 | - |
| - | - | 2811 | 1347 | - | - | 0 | - |
| - | - | 2314 | 1348 | - | - | 0 | - |
| - | - | 3161 | 1349 | - | - | 0 | - |
| - | - | 729.4 | 1394 | - | - | 0 | - |
| - | - | 1392 | 1402 | - | - | 0 | - |
| - | - | 1683 | 1403 | - | - | 0 | - |
| 4 | z | 3870 | 1430 | 0.002661 | 1.861 | +1 | 12 |
| - | - | 1.492E+04 | 1431 | - | - | 0 | - |
| - | - | 1.359E+04 | 1432 | - | - | 0 | - |
| - | - | 9831 | 1433 | - | - | 0 | - |
| - | - | 4725 | 1434 | - | - | 0 | - |
| - | - | 1066 | 1435 | - | - | 0 | - |
| - | - | 1028 | 1437 | - | - | 0 | - |
| 12 | c | 1.889E+04 | 1438 | 0.003188 | 2.217 | +1 | 12 |
| - | - | 1.568E+04 | 1439 | - | - | 0 | - |
| - | - | 8341 | 1440 | - | - | 0 | - |
| - | - | 2128 | 1441 | - | - | 0 | - |
| - | - | 942.9 | 1456 | - | - | 0 | - |
| - | - | 784.1 | 1489 | - | - | 0 | - |
| - | - | 875.7 | 1490 | - | - | 0 | - |
| - | - | 935.9 | 1491 | - | - | 0 | - |
| - | - | 732.4 | 1492 | - | - | 0 | - |
| - | - | 1065 | 1523 | - | - | 0 | - |
| - | - | 2293 | 1531 | - | - | 0 | - |
| - | - | 5497 | 1532 | - | - | 0 | - |
| - | - | 4258 | 1533 | - | - | 0 | - |
| - | - | 2336 | 1534 | - | - | 0 | - |
| - | - | 1246 | 1535 | - | - | 0 | - |
| - | - | 900.9 | 1546 | - | - | 0 | - |
| 3 | y | 2745 | 1547 | 0.01048 | 6.774 | +1 | 13 |
| 13 | c | 2951 | 1548 | 0.01859 | 12.01 | +1 | 13 |
| 13 | c | 2299 | 1549 | 0.003703 | 2.391 | +1 | 13 |
| - | - | 853.5 | 1551 | - | - | 0 | - |
| - | - | 1002 | 1558 | - | - | 0 | - |
| - | - | 1054 | 1559 | - | - | 0 | - |
| - | - | 791.7 | 1560 | - | - | 0 | - |
| - | - | 1734 | 1563 | - | - | 0 | - |
| - | - | 728.6 | 1564 | - | - | 0 | - |
| - | - | 1538 | 1565 | - | - | 0 | - |
| 13 | c | 1.239E+04 | 1566 | 0.003318 | 2.119 | +1 | 13 |
| - | - | 8992 | 1567 | - | - | 0 | - |
| - | - | 5492 | 1568 | - | - | 0 | - |
| - | - | 2704 | 1569 | - | - | 0 | - |
| - | - | 715.7 | 1570 | - | - | 0 | - |
| - | - | 670.5 | 1572 | - | - | 0 | - |
| - | - | 3623 | 1573 | - | - | 0 | - |
| - | - | 1.291E+04 | 1574 | - | - | 0 | - |
| - | - | 1.011E+04 | 1575 | - | - | 0 | - |
| - | - | 4979 | 1576 | - | - | 0 | - |
| - | - | 2168 | 1577 | - | - | 0 | - |
| - | - | 1694 | 1578 | - | - | 0 | - |
| - | - | 1296 | 1588 | - | - | 0 | - |
| - | - | 3582 | 1589 | - | - | 0 | - |
| - | - | 1.22E+04 | 1590 | - | - | 0 | - |
| - | - | 1.796E+04 | 1591 | - | - | 0 | - |
| - | - | 1.163E+04 | 1592 | - | - | 0 | - |
| - | - | 5239 | 1593 | - | - | 0 | - |
| - | - | 1311 | 1594 | - | - | 0 | - |
| - | - | 943.6 | 1599 | - | - | 0 | - |
| - | - | 736.3 | 1604 | - | - | 0 | - |
| - | - | 1178 | 1605 | - | - | 0 | - |
| - | - | 1611 | 1606 | - | - | 0 | - |
| - | - | 892.7 | 1607 | - | - | 0 | - |
| - | - | 1415 | 1615 | - | - | 0 | - |
| - | - | 760.7 | 1617 | - | - | 0 | - |
| - | - | 1304 | 1621 | - | - | 0 | - |
| - | - | 1438 | 1622 | - | - | 0 | - |
| - | - | 1004 | 1623 | - | - | 0 | - |
| - | - | 930.2 | 1636 | - | - | 0 | - |
| - | - | 1554 | 1637 | - | - | 0 | - |
| - | - | 1473 | 1638 | - | - | 0 | - |
| - | - | 996.7 | 1639 | - | - | 0 | - |
| - | - | 810.8 | 1647 | - | - | 0 | - |
| - | - | 1002 | 1664 | - | - | 0 | - |
| - | - | 1739 | 1665 | - | - | 0 | - |
| - | - | 1187 | 1667 | - | - | 0 | - |
| 14 | c | 4514 | 1680 | 0.0007953 | 0.4735 | +1 | 14 |
| - | - | 1.165E+04 | 1681 | - | - | 0 | - |
| - | - | 1.01E+04 | 1682 | - | - | 0 | - |
| - | - | 4679 | 1683 | - | - | 0 | - |
| - | - | 1952 | 1684 | - | - | 0 | - |
| - | - | 4422 | 1695 | - | - | 0 | - |
| - | - | 4713 | 1696 | - | - | 0 | - |
| - | - | 1981 | 1697 | - | - | 0 | - |
| - | - | 1234 | 1698 | - | - | 0 | - |
| - | - | 782.1 | 1712 | - | - | 0 | - |
| - | - | 1006 | 1748 | - | - | 0 | - |
| - | - | 2218 | 1749 | - | - | 0 | - |
| - | - | 697.2 | 1750 | - | - | 0 | - |
| - | - | 2038 | 1751 | - | - | 0 | - |
| - | - | 3954 | 1752 | - | - | 0 | - |
| - | - | 3885 | 1753 | - | - | 0 | - |
| - | - | 2017 | 1754 | - | - | 0 | - |
| - | - | 960.5 | 1755 | - | - | 0 | - |
| - | - | 917.8 | 1764 | - | - | 0 | - |
| - | - | 2658 | 1765 | - | - | 0 | - |
| - | - | 8986 | 1766 | - | - | 0 | - |
| - | - | 8483 | 1767 | - | - | 0 | - |
| - | - | 6096 | 1768 | - | - | 0 | - |
| - | - | 1885 | 1769 | - | - | 0 | - |
| - | - | 995.9 | 1770 | - | - | 0 | - |
| - | - | 925.1 | 1777 | - | - | 0 | - |
| - | - | 1653 | 1778 | - | - | 0 | - |
| - | - | 1142 | 1779 | - | - | 0 | - |
| - | - | 4685 | 1783 | - | - | 0 | - |
| - | - | 3095 | 1784 | - | - | 0 | - |
| - | - | 1768 | 1785 | - | - | 0 | - |
| - | - | 1025 | 1786 | - | - | 0 | - |
| - | - | 1222 | 1792 | - | - | 0 | - |
| - | - | 6169 | 1793 | - | - | 0 | - |
| - | - | 2.799E+04 | 1794 | - | - | 0 | - |
| - | - | 2.371E+04 | 1795 | - | - | 0 | - |
| - | - | 1.416E+04 | 1796 | - | - | 0 | - |
| - | - | 6861 | 1797 | - | - | 0 | - |
| - | - | 2424 | 1798 | - | - | 0 | - |
| - | - | 3180 | 1809 | - | - | 0 | - |
| - | - | 1.44E+04 | 1810 | - | - | 0 | - |
| - | - | 5.366E+04 | 1811 | - | - | 0 | - |
| - | - | 4.754E+04 | 1812 | - | - | 0 | - |
| - | - | 2.542E+04 | 1813 | - | - | 0 | - |
| - | - | 1.23E+04 | 1814 | - | - | 0 | - |
| - | - | 4120 | 1815 | - | - | 0 | - |

m/z Charge Intensity FragmentType MassShift Position
125.10679626464844 0 448.66718
126.09136199951172 0 842.40656
128.08181762695312 0 480.50153
129.1023712158203 0 615.78455
129.7417449951172 0 375.27164
130.04965209960938 0 615.97797 y Ammonia loss 14
130.423828125 0 432.29166
136.07540893554688 0 3001.4316
140.36923217773438 0 369.82858
147.0760955810547 0 3392.8801 y 14
147.3645782470703 0 373.1096
168.11282348632812 0 775.1355
170.0467071533203 0 589.1325
173.45252990722656 0 1976.5721
175.11871337890625 0 1991.7743
183.1125946044922 0 523.35785
185.13951110839844 0 1243.566
189.219482421875 0 481.56625
199.1800537109375 0 7326.173
200.1839141845703 0 874.0904
212.13880920410156 0 2170.5105
215.1380615234375 0 960.9191
216.09768676757812 0 739.7168
217.08106994628906 0 1726.851
227.1748504638672 0 4304.185
228.17822265625 0 870.17944
230.14923095703125 0 796.2687
235.14361572265625 0 9541.058
236.1471710205078 0 1084.6084
243.16952514648438 0 1508.3525
244.09228515625 0 782.87274 y Ammonia loss 13
245.1004638671875 0 657.34247 z 13
245.12393188476562 0 3707.1016
255.04513549804688 0 597.1108
259.1656799316406 0 674.6237
261.1187744140625 0 3084.28 y 13
262.15008544921875 0 1436.9453
263.1256408691406 0 687.4745
263.1383972167969 0 7451.7026
264.1427307128906 0 580.79816
270.1205139160156 0 959.00714
299.17083740234375 0 2050.7463
316.1618957519531 0 1061.2125
329.18121337890625 0 1389.272
332.1573791503906 0 642.6921
333.1639709472656 0 1682.2157
346.1757507324219 0 4738.959
347.1781921386719 0 663.48285
349.1826171875 0 933.80774
357.1510314941406 0 1561.6355
357.1752624511719 0 559.6406
360.1921691894531 0 929.8208
361.1715087890625 0 1779.9731
364.18609619140625 0 7814.785
365.18914794921875 0 2258.2417
366.189697265625 0 628.8742
373.1942138671875 0 3673.3853 z 12
374.2003479003906 0 669.0689
386.2402648925781 0 859.3725
388.1814880371094 0 1013.7035
389.2137145996094 0 1561.6569 y 12
391.7252197265625 0 624.2979
415.2237243652344 0 971.1191
416.1769714355469 0 1831.3633
417.2088928222656 0 1107.8325
422.7166748046875 0 867.16504
423.2322998046875 0 1170.7063
430.2287902832031 0 2380.4587
431.2332458496094 0 970.8707
431.2647705078125 0 1448.0083
432.2326354980469 0 974.73883
444.8782653808594 0 5546.8267 y 4
445.2117919921875 0 1872.6151
445.5456848144531 0 969.01227
445.7349853515625 0 642.45483
446.2356872558594 0 653.28754
448.2523498535156 0 951.4162
450.74041748046875 0 703.6724
457.2398376464844 0 1398.9847 w 11
459.22027587890625 0 921.0528 w 11
459.2593078613281 0 5785.011
460.27154541015625 0 1265.465
464.1478576660156 0 632.00275
472.2266845703125 0 892.9777
473.23529052734375 0 3633.2817 y Ammonia loss 11
474.23480224609375 0 1215.3213 c Ammonia loss 11
475.2527770996094 0 824.4709
477.2701721191406 0 2051.3462
489.25360107421875 0 1814.4675
490.260498046875 0 2751.1013 y 11
498.2312927246094 0 898.9254
504.2650451660156 0 597.0278
505.2724914550781 0 4826.851
506.27496337890625 0 607.4465
508.2496337890625 0 641.0734
510.25103759765625 0 1918.2 y Water loss 2
510.7511291503906 0 801.60364
510.9212951660156 0 873.82196 z 2
511.2566223144531 0 631.565
513.765625 0 599.69934
515.0822143554688 0 708.02246
515.2778930664062 0 792.4862
516.2529907226562 0 1608.2 y 2
516.7542724609375 0 1706.1658
517.2551879882812 0 711.6153
520.2352294921875 0 897.93866
524.2620849609375 0 3162.7612
524.7630615234375 0 763.3349
530.280029296875 0 1872.1299
530.6148681640625 0 1160.4751
531.263671875 0 1557.0974
532.267578125 0 843.1851
533.270751953125 0 581.3595
536.7601928710938 0 1129.6428
538.5982055664062 0 1004.9567
541.2841186523438 0 1204.046
542.2753295898438 0 698.3821
543.2825317382812 0 831.5679
550.7538452148438 0 811.0082
551.2586669921875 0 754.2841
551.7647705078125 0 5812.7983
552.2658081054688 0 3459.3716
552.7682495117188 0 921.5161
557.2791137695312 0 1570.8579
558.2867431640625 0 5890.297
558.7890014648438 0 3364.285
559.2935791015625 0 11030.642
559.34423828125 0 674.2474
559.7957153320312 0 1143.0481
560.2320556640625 0 871.38116
560.295654296875 0 1956.9886
561.3053588867188 0 799.00305
564.6076049804688 0 748.0733 y Water loss 1
564.94140625 0 913.39026 y Ammonia loss 1
565.2733154296875 0 583.78656 z 1
565.7860107421875 0 2917.5354
566.28759765625 0 2663.355
566.791259765625 0 1394.8921
571.282470703125 0 705.41724
572.7885131835938 0 1707.8627
573.2847290039062 0 1338.2296
573.7957763671875 0 1897.0657
574.800048828125 0 624.42596
575.2785034179688 0 723.7583
576.2852783203125 0 1674.5468
580.7852783203125 0 13509.737
581.2384033203125 0 890.2289
581.2872924804688 0 10516.958
581.790771484375 0 7054.6724
582.2925415039062 0 4205.902
584.3086547851562 0 766.33215
584.8176879882812 0 766.20337
585.3202514648438 0 1787.3206
586.3253784179688 0 888.4667
594.2791137695312 0 801.5429
594.783203125 0 766.6237
597.6294555664062 0 5468.3457
597.9624633789062 0 6514.426
598.2976684570312 0 2653.5068
598.631103515625 0 1419.3086
598.9484252929688 0 779.4346
600.2718505859375 0 1107.6335
600.8023071289062 0 1590.101
601.3048706054688 0 1187.2882
602.2518920898438 0 1054.9044
602.3262329101562 0 745.14636
602.77978515625 0 3173.328
602.8299560546875 0 917.64355
603.2772216796875 0 4035.3496
603.3340454101562 0 5577.9634
603.7876586914062 0 16811.008
604.286865234375 0 8885.362
604.3384399414062 0 1016.8154
604.7924194335938 0 4124.0947
618.2861938476562 0 1799.7054 y 5
618.787109375 0 1218.4954
621.2763061523438 0 1789.7562 z 10
622.2841186523438 0 1604.6714
630.3069458007812 0 2721.9077
630.8106689453125 0 2178.5571
633.3013916015625 0 615.34625
634.81298828125 0 1812.7401
635.3163452148438 0 1453.517
635.8192138671875 0 894.15173
637.2973022460938 0 1239.2417 y 10
637.8111572265625 0 1123.7076
638.3155517578125 0 2746.8813
638.8157348632812 0 1939.1885
639.3140869140625 0 1243.1542
643.8193359375 0 798.4622
644.3223266601562 0 696.8162
645.2935791015625 0 746.1753 w 4
646.3760375976562 0 2598.8896
647.379638671875 0 1423.4076
657.8090209960938 0 823.3207 y Water loss 4
658.3059692382812 0 1968.1774 y Ammonia loss 4
659.3065185546875 0 576.9843
665.8108520507812 0 1464.7715
666.3126220703125 0 1557.2047
666.8133544921875 0 41378.656 y 4
667.314697265625 0 29890.273
667.3653564453125 0 445.949
667.815185546875 0 16238.435
668.31494140625 0 5209.6426
673.3208618164062 0 1670.1302
673.827392578125 0 730.46704
674.322021484375 0 1650.3651
674.8278198242188 0 1268.2799
675.3277587890625 0 1012.1397
680.8180541992188 0 1357.9607
681.3213500976562 0 690.68787
681.8330688476562 0 14739.344
682.3340454101562 0 10756.984
682.8358764648438 0 4338.2
683.3385009765625 0 2015.787
683.8434448242188 0 1147.5327
686.3145141601562 0 869.9514
687.3280639648438 0 4170.4995
687.3956298828125 0 2557.34
688.3333740234375 0 1650.5237
688.4004516601562 0 5170.1772 c 5
689.3529663085938 0 1702.2861
691.307373046875 0 17707.357 w 9
692.3084106445312 0 5062.306
693.3084716796875 0 2162.3645
693.8191528320312 0 10816.971 w 3
694.3197021484375 0 10935.204
694.8209228515625 0 3452.5212
695.3209228515625 0 2284.5632
697.4368896484375 0 808.32275
701.3508911132812 0 613.8606
702.33154296875 0 695.31586
703.331298828125 0 984.4899
713.41162109375 0 1779.9192
714.4130249023438 0 755.7602
715.3455200195312 0 11603.382 z 3
715.4149780273438 0 773.5386
715.8477783203125 0 10328.51
716.34814453125 0 5625.5786
716.848876953125 0 2198.5754
722.8509521484375 0 666.63696
723.354736328125 0 5628.779 y 3
723.8561401367188 0 5591.599
724.3557739257812 0 3103.7688
729.8618774414062 0 991.362
730.3356323242188 0 828.6051
732.4262084960938 0 1672.9468
737.8416137695312 0 2357.3757
738.3382568359375 0 1526.6835
738.8380737304688 0 825.39777
741.8819580078125 0 1244.4967
742.3812866210938 0 1210.9119
742.876708984375 0 676.5341
743.358154296875 0 1017.6272
744.3694458007812 0 803.79865
744.8447265625 0 681.6685
750.3203125 0 16329.181 z 9
751.3244018554688 0 7267.6777
752.3253173828125 0 3017.6208
757.3622436523438 0 753.6371 z Ammonia loss 2
757.8563842773438 0 833.891
758.3721923828125 0 716.9862
758.876220703125 0 802.1039
761.4013061523438 0 6588.6924
761.8958740234375 0 631.0151
762.4025268554688 0 2135.867
765.3786010742188 0 12296.835 y Ammonia loss 2
765.8767700195312 0 14690.146 z 2
766.3753051757812 0 9430.227
766.8764038085938 0 6930.185
767.3547973632812 0 848.81976
768.3533935546875 0 746.2315
773.3780517578125 0 2123.422
773.880615234375 0 25827.488 y 2
774.3821411132812 0 20294.486 c Water loss 12
774.8836059570312 0 9267.849 c Ammonia loss 12
775.362548828125 0 914.97253
775.4330444335938 0 8811.249 c 6
775.8839721679688 0 880.9903
776.436767578125 0 4969.823
777.4417724609375 0 1797.9854
777.8656616210938 0 2171.6099
778.361083984375 0 1124.0227
778.8661499023438 0 1036.6577
779.9078979492188 0 724.9256
780.4019775390625 0 563.60187
781.8738403320312 0 735.72864
782.8973388671875 0 5485.7705
783.4025268554688 0 25387.95 c 12
783.9036254882812 0 18505.293
784.4036865234375 0 9852.794
784.90576171875 0 3704.0737
785.4058227539062 0 1931.3898
786.410400390625 0 984.2293
786.9071044921875 0 9155.456
787.4052734375 0 6689.499
787.9075927734375 0 4248.099
788.4053955078125 0 1761.754
788.9193725585938 0 638.62524
793.9088745117188 0 678.47394
794.4027099609375 0 1517.293
794.9168701171875 0 27739.16
795.418701171875 0 29609.5
795.9193725585938 0 14441.472
796.4202270507812 0 7053.1294
796.9183959960938 0 2304.536
797.4218139648438 0 871.5552
801.904052734375 0 2435.3137
802.40478515625 0 2584.1191
802.9141845703125 0 1054.6569
803.4161987304688 0 1670.0619
806.8970336914062 0 3001.6587
807.3966674804688 0 2060.1304
808.8887329101562 0 794.5636
810.4063110351562 0 1238.094
810.9105224609375 0 2747.5706
811.4151611328125 0 4168.884
811.9030151367188 0 1583.0026
812.416748046875 0 712.47864
814.4304809570312 0 897.9996
818.3953857421875 0 883.3351
818.9031372070312 0 1860.3809
819.3798828125 0 674.6109
819.8765258789062 0 2961.0007
820.3543701171875 0 5651.9546 w 8
820.86865234375 0 794.1243
821.354736328125 0 1909.9896
821.8760986328125 0 2267.4927
822.3744506835938 0 1769.697
822.8828735351562 0 1933.7656
824.8973999023438 0 899.1393
825.4151000976562 0 849.4101
826.386474609375 0 752.6449
827.3804931640625 0 1048.9949
827.8750610351562 0 603.7654
829.43896484375 0 4511.2764
830.4375610351562 0 2477.1738
839.4196166992188 0 878.0536
839.926513671875 0 864.66583
840.4237060546875 0 53950.027 c 13
840.9254150390625 0 41992.363
841.4256591796875 0 26287.979
841.9267578125 0 10080.723
842.4282836914062 0 3337.3315
843.4188842773438 0 3245.009
844.4302368164062 0 1168.6473
844.8353271484375 0 662.1994
845.4341430664062 0 1754.7516
846.457275390625 0 606.6241
846.9041137695312 0 710.16833 y Ammonia loss 1
847.4019775390625 0 3497.6194 z 1
847.9026489257812 0 3969.6697
848.4041137695312 0 2278.4504
848.9035034179688 0 865.1928
850.4071044921875 0 670.3233
853.4381713867188 0 1205.8314
853.9277954101562 0 1806.3096
854.4303588867188 0 1188.5497
854.9196166992188 0 1048.4725
855.4127807617188 0 3414.68 y 1
855.9107666015625 0 2002.4005
856.41015625 0 850.29004
859.423828125 0 814.4193
860.42236328125 0 778.939
861.40966796875 0 1146.9059
867.4312133789062 0 1313.1329
867.9276733398438 0 772.1089
868.4176025390625 0 1114.671
868.9126586914062 0 7864.389
869.4194946289062 0 7481.0903
869.917724609375 0 4906.785
870.420654296875 0 1616.2947
870.9447021484375 0 923.96246
873.9456176757812 0 1461.1545
874.4390258789062 0 1648.2894
875.4310302734375 0 881.6549
875.9299926757812 0 3404.8518
876.4287109375 0 4911.7705
876.9324340820312 0 2516.7708
877.4248657226562 0 2967.5034
877.919921875 0 2011.939
878.422607421875 0 2081.2964
878.9197387695312 0 853.06714
879.3651733398438 0 6324.811 z 8
880.3655395507812 0 4445.559
881.3682250976562 0 1446.0277
882.4450073242188 0 1880.6339
882.9382934570312 0 10467.826
883.4395751953125 0 8732.828
883.93701171875 0 6610.2783
884.4373168945312 0 4555.491
884.9396362304688 0 1988.0361
886.3851928710938 0 892.3074
887.9340209960938 0 1577.5459
888.4315185546875 0 2185.659
888.93017578125 0 1194.0253
889.4276733398438 0 2647.3503
889.9259033203125 0 1516.4209
896.4380493164062 0 3416.1313
896.9381713867188 0 33851.35
897.4383544921875 0 33435.91
897.9400024414062 0 24099.709
898.439697265625 0 11409.087
898.9343872070312 0 2994.858
900.4795532226562 0 987.7165
901.47314453125 0 933.76434
902.4716796875 0 895.66705
904.4328002929688 0 1772.3617
904.9446411132812 0 26598.889
905.447509765625 0 51495.14
905.948974609375 0 38917.21
906.4492797851562 0 22708.72
906.951416015625 0 8334.407
907.4464721679688 0 1478.0656
931.53515625 0 13988.32 c 7
932.5386352539062 0 7441.2876
933.5391235351562 0 2346.369
934.5401611328125 0 961.5003
944.4652099609375 0 2334.2327
945.4597778320312 0 2937.0989
946.453857421875 0 707.9964
957.5595092773438 0 815.0163
959.4609985351562 0 747.19385
960.4678344726562 0 1525.5763
961.4678344726562 0 1778.2357
970.5503540039062 0 684.08527
977.4589233398438 0 790.81146
989.4945068359375 0 1473.8353
990.4981689453125 0 962.9764
1003.4873046875 0 903.3368
1004.4962768554688 0 1242.1871
1005.493408203125 0 1229.3737
1014.5570068359375 0 5135.541
1015.563232421875 0 2789.712
1016.5655517578125 0 2332.5085
1017.5673217773438 0 1441.7455
1031.5013427734375 0 1513.1877
1032.4949951171875 0 1648.4056
1033.48876953125 0 1027.0897 y Water loss 7
1035.46435546875 0 10256.493 z 7
1036.46826171875 0 8618.06
1037.46875 0 4485.913
1038.4715576171875 0 1985.3331
1045.4849853515625 0 849.5166
1046.4901123046875 0 1191.8394
1047.5048828125 0 1457.9244
1048.4937744140625 0 793.8446
1050.4754638671875 0 1207.761
1051.482421875 0 7222.7817 y 7
1052.48681640625 0 3716.14
1053.4832763671875 0 1944.1377
1054.4794921875 0 795.718
1057.5732421875 0 1769.5222
1058.5816650390625 0 1043.3175
1059.57275390625 0 2310.5332
1060.577880859375 0 36166.92 c 8
1061.580322265625 0 22934.102
1062.5828857421875 0 8092.6587
1072.503662109375 0 950.12494
1074.5185546875 0 804.28107
1075.530029296875 0 1322.5149
1091.5057373046875 0 1180.639
1101.467041015625 0 630.60034
1101.587646484375 0 6281.18
1102.5882568359375 0 4328.2275
1103.5313720703125 0 2329.43
1104.53759765625 0 2227.2256
1105.534423828125 0 763.1506
1117.4315185546875 0 919.1702
1117.5714111328125 0 944.11383
1118.56689453125 0 809.2577
1120.5201416015625 0 1485.25 y Water loss 6
1121.5108642578125 0 683.20087
1122.49853515625 0 3149.013 z 6
1123.5032958984375 0 10583.112
1124.506591796875 0 4895.012
1125.511474609375 0 2980.0132
1128.542724609375 0 763.0755
1130.558837890625 0 1834.8711
1131.5662841796875 0 3846.9917
1132.5684814453125 0 2060.3022
1133.572509765625 0 1578.4047
1138.5152587890625 0 731.6592 y 6
1139.51953125 0 876.5261
1143.5643310546875 0 1007.3373
1144.5582275390625 0 3399.6946
1145.5709228515625 0 5341.3496
1146.582275390625 0 14986.569
1147.5838623046875 0 8382.065
1148.5859375 0 2732.295
1158.6097412109375 0 2868.7012
1159.60693359375 0 1135.6731
1160.57177734375 0 1837.6505
1161.5723876953125 0 6612.2793
1162.5770263671875 0 5637.871
1163.5848388671875 0 4077.808
1164.57666015625 0 2003.4779
1187.569091796875 0 1058.2927
1188.5977783203125 0 2104.405
1189.6185302734375 0 27508.004 c 9
1190.6209716796875 0 17974.107
1191.6204833984375 0 6632.656
1192.6199951171875 0 2123.4727
1203.6090087890625 0 1153.0195
1204.598388671875 0 2693.628
1205.5523681640625 0 1012.76764
1205.6480712890625 0 970.511
1206.567626953125 0 3620.9207
1207.5703125 0 2996.9937
1208.5843505859375 0 1143.3159
1209.589599609375 0 762.93994
1213.6558837890625 0 949.2606
1235.568115234375 0 2041.1495 y 5
1236.5726318359375 0 1639.5974
1257.6812744140625 0 3560.653
1258.67578125 0 2424.0876
1259.6187744140625 0 1591.4103
1260.614501953125 0 3127.8809
1261.617919921875 0 2570.5547
1262.6162109375 0 1123.4392
1275.6217041015625 0 1426.204
1277.637451171875 0 793.5468
1292.6405029296875 0 2429.2246
1293.64404296875 0 2133.38
1294.64111328125 0 737.7956
1300.65087890625 0 2131.387
1301.6484375 0 1862.4988
1302.648681640625 0 1170.729
1332.6243896484375 0 2406.9617 y 4
1333.63037109375 0 1409.1511
1334.6285400390625 0 774.0765
1335.6458740234375 0 4080.6472
1336.6531982421875 0 12737.323 c 10
1337.65771484375 0 8261.505
1338.658935546875 0 3522.9304
1344.7099609375 0 6494.3433
1345.70849609375 0 4646.1562
1346.6590576171875 0 2811.207
1347.65087890625 0 2313.6855
1348.651123046875 0 3160.571
1393.6846923828125 0 729.39343
1401.6937255859375 0 1392.1307
1402.6903076171875 0 1683.0718
1429.684326171875 0 3870.3643 z 3
1430.6907958984375 0 14923.758
1431.7147216796875 0 13590.078
1432.724609375 0 9830.927
1433.734375 0 4724.728
1434.7513427734375 0 1065.5221
1436.7110595703125 0 1028.2964
1437.7025146484375 0 18885.213 c 11
1438.704345703125 0 15683.006
1439.7059326171875 0 8340.71
1440.7039794921875 0 2128.4124
1455.70556640625 0 942.8709
1488.7437744140625 0 784.1276
1489.744384765625 0 875.70624
1490.74853515625 0 935.9193
1491.7547607421875 0 732.36426
1522.7919921875 0 1064.8129
1530.77001953125 0 2292.657
1531.760498046875 0 5497.2676
1532.757080078125 0 4257.853
1533.75439453125 0 2335.742
1534.7781982421875 0 1246.0308
1545.8260498046875 0 900.9028
1546.758544921875 0 2744.7014 y 2
1547.76513671875 0 2951.1526 c Water loss 12
1548.7640380859375 0 2298.7944 c Ammonia loss 12
1550.7598876953125 0 853.46783
1557.789794921875 0 1001.96014
1558.8028564453125 0 1053.8724
1559.78515625 0 791.7462
1562.8404541015625 0 1733.8192
1563.8336181640625 0 728.6443
1564.79833984375 0 1537.6703
1565.797607421875 0 12387.43 c 12
1566.799072265625 0 8992.398
1567.8033447265625 0 5491.761
1568.7989501953125 0 2704.0698
1569.793701171875 0 715.6592
1571.7747802734375 0 670.4546
1572.813232421875 0 3623.1094
1573.81494140625 0 12911.002
1574.816162109375 0 10108.131
1575.8199462890625 0 4979.2153
1576.81396484375 0 2168.2512
1577.806640625 0 1693.809
1587.8128662109375 0 1296.0613
1588.8204345703125 0 3582.1445
1589.830810546875 0 12199.584
1590.8370361328125 0 17963.977
1591.8397216796875 0 11627.768
1592.8448486328125 0 5239.487
1593.8492431640625 0 1310.6588
1598.755615234375 0 943.58344
1603.79052734375 0 736.2522
1604.8018798828125 0 1177.6891
1605.8232421875 0 1611.0455
1606.8336181640625 0 892.7316
1614.7947998046875 0 1414.5186
1616.8140869140625 0 760.7035
1620.8233642578125 0 1304.3259
1621.8162841796875 0 1437.6683
1622.814697265625 0 1004.26324
1635.81884765625 0 930.2307
1636.8203125 0 1554.007
1637.8179931640625 0 1472.7339
1638.8182373046875 0 996.7185
1646.7481689453125 0 810.7863
1663.82568359375 0 1001.6818
1664.8255615234375 0 1739.2094
1666.8365478515625 0 1187.289
1679.8380126953125 0 4514.2446 c 13
1680.8468017578125 0 11649.309
1681.8485107421875 0 10100.423
1682.8548583984375 0 4678.846
1683.855224609375 0 1952.0295
1694.802001953125 0 4421.693
1695.80322265625 0 4712.7173
1696.8067626953125 0 1980.5735
1697.805419921875 0 1233.7253
1711.7698974609375 0 782.1261
1747.894775390625 0 1006.43695
1748.88671875 0 2217.5532
1749.8975830078125 0 697.2151
1750.8602294921875 0 2037.5596
1751.8525390625 0 3953.8528
1752.8642578125 0 3884.8389
1753.852783203125 0 2016.7913
1754.846435546875 0 960.4624
1763.880615234375 0 917.78516
1764.8812255859375 0 2658.007
1765.879150390625 0 8986.046
1766.8819580078125 0 8483.39
1767.882568359375 0 6096.0986
1768.879638671875 0 1884.7454
1769.87451171875 0 995.8617
1776.8663330078125 0 925.0747
1777.8560791015625 0 1653.3833
1778.843505859375 0 1142.498
1782.900146484375 0 4684.653
1783.9066162109375 0 3094.6858
1784.91357421875 0 1767.7834
1785.8974609375 0 1025.1348
1791.8758544921875 0 1222.4894
1792.874267578125 0 6168.6895
1793.873046875 0 27990.236
1794.875 0 23707.863
1795.8779296875 0 14159.263
1796.878173828125 0 6860.599
1797.880126953125 0 2424.4639
1808.8768310546875 0 3179.5356
1809.88720703125 0 14401.979
1810.8978271484375 0 53660.156
1811.900634765625 0 47537.71
1812.903076171875 0 25424.81
1813.8994140625 0 12303.475
1814.8887939453125 0 4119.533

Spectrum Details

|  |  |
| --- | --- |
| Matched peaks? Matched peaks The total absolute number of peaks matched. Additionally in brackets the total fraction of peaks matched and the total number of peaks is shown. | 65 (9.86% of 659) |
| FDR? FDR The false discovery rate estimated for this peptide. It is calculated by matching all theoretical fragments with a non-integer shift with the raw peaks for this spectrum. This is done with 40 different shifts. The resulting percentage is the average number of annotated peaks over the number of annotated peaks with the correct spectrum. | 4.95% |
| Satellite FDR? Satellite FDR See the FDR for details on its calculation. This satellite ion specific FDR only contains the satellite ions (d/w) for I/L/J positions. | 0.00% |
| PSM Score? PSM Score The PSM Score as given by Hecklib to this annotated spectrum. It is shown with three significant figures. | 415 |

#### Reverse Lookup? Reverse Lookup All places where this read could be placed.

| Group | Segment | Template | Template Part | Read Part | Score | Unique |
| --- | --- | --- | --- | --- | --- | --- |
| Homo sapiens Heavy Chain | IGHC | IGHG3 | [277..292] | [0..15] | 120 | False |
| Homo sapiens Heavy Chain | IGHC | IGHG2 | [226..241] | [0..15] | 120 | False |
| Homo sapiens Heavy Chain | IGHC | IGHG4 | [227..242] | [0..15] | 111 | False |

| Recombined | Template Part | Read Part | Score | Unique |
| --- | --- | --- | --- | --- |
| REC-0-1 | [355..370] | [0..15] | 106 | True |

#### Meta Information from Multiple reads

##### Number of combined reads

4

##### Intensity

0.6678

##### TotalArea

1.036E+08

##### Changes to the peptide sequence

VYTJPPSREEMTKNQ

L→JEqual support for both Leucine and Isoleucine based on side chain ions (1 ions for both) (Position: 4)

I→LSupport for Leucine based on side chain ions (1 for L 0 for I) (Position: 4)

L→ISupport for Isoleucine based on side chain ions (2 for I 1 for L) (Position: 4)

#### Positional Score

Copy Data

##### Positional Score (TSV)

###### Preview

```
Loading example...
```

*Click on the button to copy the data to your clipboard.*

1001234567891011121314

Label Value
"0" 0.5
"1" 0.5
"2" 0.497
"3" 0.475
"4" 0.45
"5" 0.485
"6" 0.497
"7" 0.497
"8" 0.497
"9" 0.495
"10" 0.487
"11" 0.495
"12" 0.497
"13" 0.495
"14" 0.495

#### Meta Information from PEAKS

##### Scan Identifier

F2:3515

##### Original sequence

V

Y

T

L

P

P

S

R

E

E

M

+15.99

T

K

N

Q

##### Posttranslational Modifications

Oxidation (M)

##### Source File

D:\separate\_stitch\_analyses\xle-disambiguation\raw\20210323\_F1\_UM1\_Peng0013\_SA\_F59\_ingel\_3ug\_TL.raw

##### Fraction

2

##### Scan Feature

F2:8363

##### De Novo Score

99

##### ConfidenceScore

99

### m/z

603.6332

##### Mass

1807.8774

##### Charge

3

##### Retention Time

19.29

##### Predicted Retention Time

-

##### Area

2.589E+07

##### Parts Per Million

0.1

##### Fragmentation mode

ETHCD

##### Originating file

01 D:\separate\_stitch\_analyses\xle-disambiguation\20210325\_F59\_3ug\_DENOVO\_12.csv

#### Meta Information from PEAKS

##### Scan Identifier

F2:3810

##### Original sequence

V

Y

T

L

P

P

S

R

E

E

M

+15.99

T

K

N

Q

##### Posttranslational Modifications

Oxidation (M)

##### Source File

D:\separate\_stitch\_analyses\xle-disambiguation\raw\20210323\_F1\_UM1\_Peng0013\_SA\_F59\_ingel\_3ug\_TL.raw

##### Fraction

2

##### Scan Feature

F2:8363

##### De Novo Score

99

##### ConfidenceScore

99

### m/z

603.6332

##### Mass

1807.8774

##### Charge

3

##### Retention Time

19.29

##### Predicted Retention Time

-

##### Area

2.589E+07

##### Parts Per Million

0.1

##### Fragmentation mode

ETHCD

##### Originating file

01 D:\separate\_stitch\_analyses\xle-disambiguation\20210325\_F59\_3ug\_DENOVO\_12.csv

#### Meta Information from PEAKS

##### Scan Identifier

F2:3436

##### Original sequence

V

Y

T

L

P

P

S

R

E

E

M

+15.99

T

K

N

Q

##### Posttranslational Modifications

Oxidation (M)

##### Source File

D:\separate\_stitch\_analyses\xle-disambiguation\raw\20210323\_F1\_UM1\_Peng0013\_SA\_F59\_ingel\_3ug\_TL.raw

##### Fraction

2

##### Scan Feature

F2:8363

##### De Novo Score

98

##### ConfidenceScore

98

### m/z

603.6332

##### Mass

1807.8774

##### Charge

3

##### Retention Time

19.29

##### Predicted Retention Time

-

##### Area

2.589E+07

##### Parts Per Million

0.1

##### Fragmentation mode

ETHCD

##### Originating file

01 D:\separate\_stitch\_analyses\xle-disambiguation\20210325\_F59\_3ug\_DENOVO\_12.csv

#### Meta Information from PEAKS

##### Scan Identifier

F2:3646

##### Original sequence

V

Y

T

L

P

P

S

R

E

E

M

+15.99

T

K

N

Q

##### Posttranslational Modifications

Oxidation (M)

##### Source File

D:\separate\_stitch\_analyses\xle-disambiguation\raw\20210323\_F1\_UM1\_Peng0013\_SA\_F59\_ingel\_3ug\_TL.raw

##### Fraction

2

##### Scan Feature

F2:8363

##### De Novo Score

97

##### ConfidenceScore

97

### m/z

603.6332

##### Mass

1807.8774

##### Charge

3

##### Retention Time

19.29

##### Predicted Retention Time

-

##### Area

2.589E+07

##### Parts Per Million

0.1

##### Fragmentation mode

ETHCD

##### Originating file

01 D:\separate\_stitch\_analyses\xle-disambiguation\20210325\_F59\_3ug\_DENOVO\_12.csv
