## Supplementary material for "A handle on mass coincidence errors in *de novo* sequencing of antibodies by bottom-up proteomics": Combined_035.html

Details Combined\_035 | Stitch OverviewUndefined

### Read Combined\_035

#### Sequence (length=10)

VLGQPKAAPS

#### Spectrum 2991? Spectrum 2991 The raw spectrum of this peptide as annotated by Hecklib. The fragments are coloured according to ion type (see legend). Any peaks with a star '\*' as text can be hovered over to see the full details, first the ion type second the mass shift type. By hovering over the amino acids in the peptide or ions in the legend the corresponding peaks are highlighted. By toggling the 'Unassigned' label you can turn the background (unassigned) peaks on or off in the plot. By updating the slider in the Ion legend you can update the spectrum to only show the top X% of the peaks with labels. The top X% means any peak that is within X% of the highest intensity. By dragging in the spectrum you can zoom in to a specific part of the spectrum and use 'Zoom Out' to get back to the original zoom level. The annotation of the spectrum is based on the given sequence in the peptides file and is done with different software so inconsistencies are likely. The peaks are annotated based on the given sequence, with 20 ppm tolerance.

Copy Data

##### Spectrum 2991 (TSV)

###### Preview

```
Loading example...
```

*Click on the button to copy the data to your clipboard.*

Mz MinMz MaxIntensity Max

WidthHeightPeptide font sizePeptide stroke widthSpectrum font sizeSpectrum stroke widthCompact peptide

Ion legend

wxyz

abcd

OtherUnassignedIon^Charge^_Position_Show for top:%

VLGQPKAAPS

01.70e+53.41e+55.11e+56.81e+5

Zoom Out

d^+1^_2_y^+1^_2_a^+1^_2_y^+1^_2_b^+1^_2_y^+2^_5_y^+1^_3_b^+1^_3_y^+1^_3_y^+2^_6_y^+1^_4_y^+2^_7_y^+1^_4_b^+2^_7_y^+2^_7_y^+2^_8_y^+2^_8_b^+1^_4_b^+2^_8_b^+1^_4_b^+2^_9_y^+2^_9_y^+2^_9_b^+2^_9_y^+2^_9_y^+1^_5_y^+1^_5_\*\*b^+1^_5_y^+1^_6_y^+1^_6_b^+1^_6_y^+1^_7_y^+1^_7_b^+1^_7_y^+1^_7_y^+1^_8_y^+1^_8_b^+1^_8_y^+1^_8_b^+1^_8_y^+1^_9_y^+1^_9_y^+1^_9_

0776155223283104

Fragment Matches Table

Show background peaks

| Position | Ion type | Intensity | mz Theoretical | mz Error (Th) | mz Error (ppm) | Charge | Series Number |
| --- | --- | --- | --- | --- | --- | --- | --- |
| - | - | 2004 | 124 | - | - | 0 | - |
| - | - | 496.2 | 125.1 | - | - | 0 | - |
| - | - | 1098 | 125.1 | - | - | 0 | - |
| - | - | 379 | 125.3 | - | - | 0 | - |
| - | - | 384.6 | 126.1 | - | - | 0 | - |
| - | - | 1411 | 127.1 | - | - | 0 | - |
| - | - | 435.5 | 127.1 | - | - | 0 | - |
| - | - | 455.8 | 128.1 | - | - | 0 | - |
| - | - | 362.6 | 128.1 | - | - | 0 | - |
| - | - | 1.442E+04 | 129.1 | - | - | 0 | - |
| - | - | 1.175E+05 | 129.1 | - | - | 0 | - |
| - | - | 1197 | 130.1 | - | - | 0 | - |
| - | - | 622 | 130.1 | - | - | 0 | - |
| - | - | 916.7 | 130.1 | - | - | 0 | - |
| - | - | 6680 | 130.1 | - | - | 0 | - |
| - | - | 407.2 | 132.5 | - | - | 0 | - |
| - | - | 357.7 | 132.7 | - | - | 0 | - |
| - | - | 364.6 | 134.6 | - | - | 0 | - |
| - | - | 1413 | 136.1 | - | - | 0 | - |
| - | - | 432.3 | 136.4 | - | - | 0 | - |
| - | - | 1007 | 137.1 | - | - | 0 | - |
| - | - | 3648 | 139.1 | - | - | 0 | - |
| - | - | 8523 | 141.1 | - | - | 0 | - |
| - | - | 8751 | 141.1 | - | - | 0 | - |
| - | - | 519.3 | 142.1 | - | - | 0 | - |
| - | - | 716.2 | 142.1 | - | - | 0 | - |
| - | - | 549.6 | 142.2 | - | - | 0 | - |
| - | - | 4849 | 143.1 | - | - | 0 | - |
| 2 | d | 5603 | 143.1 | 0.0004576 | 3.198 | +1 | 2 |
| - | - | 631.2 | 146.1 | - | - | 0 | - |
| - | - | 1017 | 147.1 | - | - | 0 | - |
| - | - | 979.3 | 149 | - | - | 0 | - |
| - | - | 7992 | 152 | - | - | 0 | - |
| - | - | 723.1 | 153 | - | - | 0 | - |
| - | - | 1734 | 153.1 | - | - | 0 | - |
| - | - | 2106 | 153.1 | - | - | 0 | - |
| - | - | 909.2 | 154.1 | - | - | 0 | - |
| - | - | 764.4 | 155.1 | - | - | 0 | - |
| - | - | 1.469E+04 | 155.1 | - | - | 0 | - |
| - | - | 950 | 156.1 | - | - | 0 | - |
| - | - | 3861 | 157.1 | - | - | 0 | - |
| - | - | 641.6 | 157.1 | - | - | 0 | - |
| - | - | 1342 | 157.2 | - | - | 0 | - |
| - | - | 6311 | 158.1 | - | - | 0 | - |
| - | - | 847 | 161.1 | - | - | 0 | - |
| - | - | 1175 | 163.1 | - | - | 0 | - |
| - | - | 2202 | 165.1 | - | - | 0 | - |
| - | - | 833 | 167.1 | - | - | 0 | - |
| - | - | 3.501E+04 | 169.1 | - | - | 0 | - |
| - | - | 1.266E+04 | 169.1 | - | - | 0 | - |
| - | - | 2781 | 169.1 | - | - | 0 | - |
| - | - | 2982 | 170.1 | - | - | 0 | - |
| - | - | 1211 | 170.1 | - | - | 0 | - |
| - | - | 2.082E+04 | 171.1 | - | - | 0 | - |
| - | - | 491.6 | 172.1 | - | - | 0 | - |
| - | - | 1833 | 172.1 | - | - | 0 | - |
| - | - | 1668 | 172.1 | - | - | 0 | - |
| - | - | 615.7 | 173.1 | - | - | 0 | - |
| - | - | 980.2 | 173.5 | - | - | 0 | - |
| - | - | 3786 | 180.1 | - | - | 0 | - |
| - | - | 9457 | 181.1 | - | - | 0 | - |
| - | - | 7666 | 181.1 | - | - | 0 | - |
| - | - | 1243 | 182.1 | - | - | 0 | - |
| - | - | 3.24E+04 | 182.1 | - | - | 0 | - |
| - | - | 3848 | 183.1 | - | - | 0 | - |
| - | - | 3603 | 183.1 | - | - | 0 | - |
| - | - | 2635 | 183.1 | - | - | 0 | - |
| - | - | 585.8 | 184.1 | - | - | 0 | - |
| - | - | 519.3 | 184.1 | - | - | 0 | - |
| 9 | y | 1.253E+04 | 185.1 | 0.0005369 | 2.901 | +1 | 2 |
| 2 | a | 6.747E+05 | 185.2 | 0.0005655 | 3.054 | +1 | 2 |
| - | - | 8.163E+04 | 186.1 | - | - | 0 | - |
| - | - | 666.9 | 186.1 | - | - | 0 | - |
| - | - | 6.893E+04 | 186.2 | - | - | 0 | - |
| - | - | 1056 | 187.1 | - | - | 0 | - |
| - | - | 6065 | 187.1 | - | - | 0 | - |
| - | - | 817.5 | 187.1 | - | - | 0 | - |
| - | - | 1022 | 187.1 | - | - | 0 | - |
| - | - | 2497 | 187.2 | - | - | 0 | - |
| - | - | 1.777E+04 | 188.1 | - | - | 0 | - |
| - | - | 1388 | 189.1 | - | - | 0 | - |
| - | - | 1964 | 191.1 | - | - | 0 | - |
| - | - | 1180 | 191.1 | - | - | 0 | - |
| - | - | 823.4 | 192.1 | - | - | 0 | - |
| - | - | 444.4 | 193.1 | - | - | 0 | - |
| - | - | 468.5 | 193.1 | - | - | 0 | - |
| - | - | 2210 | 193.1 | - | - | 0 | - |
| - | - | 2138 | 195.1 | - | - | 0 | - |
| - | - | 727.9 | 196.1 | - | - | 0 | - |
| - | - | 1673 | 197.1 | - | - | 0 | - |
| - | - | 3314 | 197.1 | - | - | 0 | - |
| - | - | 635.7 | 197.6 | - | - | 0 | - |
| - | - | 2021 | 198.1 | - | - | 0 | - |
| - | - | 1966 | 199.1 | - | - | 0 | - |
| - | - | 7476 | 200.1 | - | - | 0 | - |
| - | - | 1638 | 201.1 | - | - | 0 | - |
| - | - | 680 | 201.1 | - | - | 0 | - |
| 9 | y | 4.131E+05 | 203.1 | 0.0005465 | 2.691 | +1 | 2 |
| - | - | 3.601E+04 | 204.1 | - | - | 0 | - |
| - | - | 3981 | 205.1 | - | - | 0 | - |
| - | - | 1.033E+04 | 208.1 | - | - | 0 | - |
| - | - | 2252 | 208.1 | - | - | 0 | - |
| - | - | 8115 | 209.1 | - | - | 0 | - |
| - | - | 842.3 | 209.1 | - | - | 0 | - |
| - | - | 2382 | 209.1 | - | - | 0 | - |
| - | - | 4.163E+04 | 210.1 | - | - | 0 | - |
| - | - | 811.4 | 210.2 | - | - | 0 | - |
| - | - | 1175 | 211.1 | - | - | 0 | - |
| - | - | 4188 | 211.1 | - | - | 0 | - |
| - | - | 1652 | 211.1 | - | - | 0 | - |
| - | - | 2195 | 211.2 | - | - | 0 | - |
| - | - | 1391 | 212.1 | - | - | 0 | - |
| - | - | 764.2 | 212.1 | - | - | 0 | - |
| 2 | b | 3.44E+05 | 213.2 | 0.000585 | 2.744 | +1 | 2 |
| - | - | 1305 | 214.1 | - | - | 0 | - |
| - | - | 3.872E+04 | 214.2 | - | - | 0 | - |
| - | - | 1382 | 215.1 | - | - | 0 | - |
| - | - | 2768 | 215.2 | - | - | 0 | - |
| - | - | 538.2 | 217.2 | - | - | 0 | - |
| - | - | 2249 | 220.1 | - | - | 0 | - |
| - | - | 1140 | 221.1 | - | - | 0 | - |
| - | - | 1822 | 223.1 | - | - | 0 | - |
| - | - | 759.7 | 225.1 | - | - | 0 | - |
| - | - | 2.784E+05 | 226.2 | - | - | 0 | - |
| - | - | 3.19E+04 | 227.2 | - | - | 0 | - |
| 6 | y | 3.348E+04 | 228.1 | 0.0004977 | 2.182 | +2 | 5 |
| - | - | 1534 | 228.2 | - | - | 0 | - |
| - | - | 3838 | 229.1 | - | - | 0 | - |
| - | - | 794.8 | 235.1 | - | - | 0 | - |
| - | - | 2956 | 236.1 | - | - | 0 | - |
| - | - | 873.9 | 237.1 | - | - | 0 | - |
| - | - | 5037 | 238.1 | - | - | 0 | - |
| - | - | 1407 | 239.2 | - | - | 0 | - |
| - | - | 3918 | 240.1 | - | - | 0 | - |
| - | - | 1858 | 242.2 | - | - | 0 | - |
| - | - | 7976 | 243.2 | - | - | 0 | - |
| - | - | 888.1 | 244.2 | - | - | 0 | - |
| - | - | 746.3 | 248.1 | - | - | 0 | - |
| - | - | 3412 | 252.1 | - | - | 0 | - |
| - | - | 1.056E+04 | 252.2 | - | - | 0 | - |
| - | - | 558.7 | 253.1 | - | - | 0 | - |
| - | - | 3727 | 253.2 | - | - | 0 | - |
| - | - | 1271 | 254.1 | - | - | 0 | - |
| - | - | 1.091E+04 | 254.2 | - | - | 0 | - |
| - | - | 1095 | 255.2 | - | - | 0 | - |
| 8 | y | 5601 | 256.1 | 0.0006088 | 2.377 | +1 | 3 |
| - | - | 763.4 | 256.2 | - | - | 0 | - |
| - | - | 2863 | 257.1 | - | - | 0 | - |
| - | - | 1915 | 261.2 | - | - | 0 | - |
| - | - | 747.2 | 262.2 | - | - | 0 | - |
| - | - | 1787 | 264.1 | - | - | 0 | - |
| - | - | 2118 | 265.1 | - | - | 0 | - |
| - | - | 4154 | 266.1 | - | - | 0 | - |
| - | - | 799.6 | 266.1 | - | - | 0 | - |
| - | - | 1033 | 267.1 | - | - | 0 | - |
| - | - | 883.8 | 268.1 | - | - | 0 | - |
| - | - | 772.4 | 268.2 | - | - | 0 | - |
| - | - | 1126 | 269.2 | - | - | 0 | - |
| - | - | 1.398E+04 | 269.2 | - | - | 0 | - |
| - | - | 2009 | 270.1 | - | - | 0 | - |
| 3 | b | 4.103E+04 | 270.2 | 0.0005141 | 1.903 | +1 | 3 |
| - | - | 1693 | 270.2 | - | - | 0 | - |
| - | - | 1852 | 271.1 | - | - | 0 | - |
| - | - | 3E+04 | 271.2 | - | - | 0 | - |
| - | - | 2869 | 272.2 | - | - | 0 | - |
| 8 | y | 2.69E+04 | 274.1 | 0.0005116 | 1.866 | +1 | 3 |
| - | - | 966.7 | 274.2 | - | - | 0 | - |
| - | - | 757.3 | 274.7 | - | - | 0 | - |
| - | - | 2961 | 275.1 | - | - | 0 | - |
| - | - | 804 | 277.2 | - | - | 0 | - |
| - | - | 5.904E+04 | 279.2 | - | - | 0 | - |
| - | - | 816.6 | 280.2 | - | - | 0 | - |
| - | - | 6962 | 280.2 | - | - | 0 | - |
| - | - | 8356 | 281.2 | - | - | 0 | - |
| - | - | 768.1 | 281.2 | - | - | 0 | - |
| - | - | 1994 | 282.1 | - | - | 0 | - |
| - | - | 1091 | 282.2 | - | - | 0 | - |
| - | - | 6986 | 283.1 | - | - | 0 | - |
| - | - | 1091 | 283.2 | - | - | 0 | - |
| - | - | 1.05E+04 | 285.2 | - | - | 0 | - |
| 5 | y | 6644 | 285.7 | 0.0005718 | 2.001 | +2 | 6 |
| - | - | 1902 | 286.2 | - | - | 0 | - |
| - | - | 7.342E+04 | 287.2 | - | - | 0 | - |
| - | - | 988 | 288.2 | - | - | 0 | - |
| - | - | 1.138E+04 | 288.2 | - | - | 0 | - |
| - | - | 819.4 | 292.2 | - | - | 0 | - |
| - | - | 908.5 | 295.2 | - | - | 0 | - |
| - | - | 1314 | 295.2 | - | - | 0 | - |
| - | - | 812.4 | 296.2 | - | - | 0 | - |
| - | - | 1102 | 297.2 | - | - | 0 | - |
| - | - | 1443 | 297.2 | - | - | 0 | - |
| - | - | 1156 | 297.2 | - | - | 0 | - |
| - | - | 1177 | 297.2 | - | - | 0 | - |
| - | - | 1.133E+05 | 297.2 | - | - | 0 | - |
| - | - | 1.551E+04 | 298.2 | - | - | 0 | - |
| - | - | 2.093E+05 | 299.2 | - | - | 0 | - |
| - | - | 2.749E+04 | 300.2 | - | - | 0 | - |
| - | - | 3907 | 301.2 | - | - | 0 | - |
| - | - | 3939 | 309.2 | - | - | 0 | - |
| - | - | 989 | 309.2 | - | - | 0 | - |
| - | - | 1118 | 310.2 | - | - | 0 | - |
| - | - | 891.6 | 311.2 | - | - | 0 | - |
| - | - | 4918 | 314.2 | - | - | 0 | - |
| - | - | 1355 | 315.2 | - | - | 0 | - |
| - | - | 861.8 | 318.2 | - | - | 0 | - |
| - | - | 1344 | 319.2 | - | - | 0 | - |
| - | - | 655.9 | 319.7 | - | - | 0 | - |
| - | - | 1975 | 323.2 | - | - | 0 | - |
| - | - | 7623 | 323.2 | - | - | 0 | - |
| - | - | 899.4 | 324.2 | - | - | 0 | - |
| - | - | 5416 | 325.2 | - | - | 0 | - |
| - | - | 1515 | 325.7 | - | - | 0 | - |
| - | - | 912.1 | 326.2 | - | - | 0 | - |
| - | - | 1516 | 326.7 | - | - | 0 | - |
| 7 | y | 1.178E+04 | 327.2 | 0.0007569 | 2.314 | +1 | 4 |
| - | - | 696.1 | 327.2 | - | - | 0 | - |
| - | - | 6078 | 328.2 | - | - | 0 | - |
| - | - | 1223 | 328.2 | - | - | 0 | - |
| - | - | 685.8 | 329.2 | - | - | 0 | - |
| - | - | 673.7 | 331.7 | - | - | 0 | - |
| - | - | 4542 | 332.2 | - | - | 0 | - |
| - | - | 4348 | 336.2 | - | - | 0 | - |
| - | - | 8209 | 337.2 | - | - | 0 | - |
| - | - | 801.6 | 338.2 | - | - | 0 | - |
| - | - | 4.947E+04 | 340.2 | - | - | 0 | - |
| 4 | y | 1.611E+04 | 340.7 | 0.0007047 | 2.069 | +2 | 7 |
| - | - | 994.5 | 340.7 | - | - | 0 | - |
| - | - | 5020 | 341.2 | - | - | 0 | - |
| - | - | 7628 | 341.2 | - | - | 0 | - |
| - | - | 962 | 341.7 | - | - | 0 | - |
| - | - | 722.3 | 342.2 | - | - | 0 | - |
| 7 | y | 4796 | 345.2 | 0.0008734 | 2.53 | +1 | 4 |
| - | - | 955.1 | 345.2 | - | - | 0 | - |
| - | - | 839.6 | 346.2 | - | - | 0 | - |
| 7 | b | 1178 | 347.7 | 0.0001691 | 0.4864 | +2 | 7 |
| - | - | 890.2 | 349.2 | - | - | 0 | - |
| 4 | y | 2974 | 349.7 | 0.0007019 | 2.007 | +2 | 7 |
| - | - | 1027 | 350.2 | - | - | 0 | - |
| - | - | 3851 | 350.2 | - | - | 0 | - |
| - | - | 1497 | 351.2 | - | - | 0 | - |
| - | - | 3285 | 353.2 | - | - | 0 | - |
| - | - | 2771 | 354.2 | - | - | 0 | - |
| - | - | 1690 | 355.2 | - | - | 0 | - |
| - | - | 1136 | 359.7 | - | - | 0 | - |
| - | - | 973.4 | 360.7 | - | - | 0 | - |
| - | - | 2332 | 360.7 | - | - | 0 | - |
| - | - | 779.5 | 361.2 | - | - | 0 | - |
| - | - | 2558 | 363.2 | - | - | 0 | - |
| - | - | 3282 | 366.2 | - | - | 0 | - |
| - | - | 2.645E+05 | 368.2 | - | - | 0 | - |
| - | - | 1087 | 368.7 | - | - | 0 | - |
| - | - | 5.447E+04 | 369.2 | - | - | 0 | - |
| 3 | y | 1536 | 369.7 | 0.001139 | 3.081 | +2 | 8 |
| - | - | 2.208E+04 | 370.2 | - | - | 0 | - |
| - | - | 4050 | 371.2 | - | - | 0 | - |
| - | - | 799.7 | 372.2 | - | - | 0 | - |
| - | - | 812.9 | 372.2 | - | - | 0 | - |
| - | - | 1526 | 373.7 | - | - | 0 | - |
| - | - | 5806 | 376.2 | - | - | 0 | - |
| - | - | 2108 | 376.2 | - | - | 0 | - |
| - | - | 1217 | 378.2 | - | - | 0 | - |
| 3 | y | 4031 | 378.2 | 0.002391 | 6.321 | +2 | 8 |
| - | - | 889.1 | 378.7 | - | - | 0 | - |
| - | - | 5186 | 380.2 | - | - | 0 | - |
| 4 | b | 1082 | 381.2 | 0.002034 | 5.336 | +1 | 4 |
| - | - | 1161 | 381.2 | - | - | 0 | - |
| - | - | 5198 | 382.2 | - | - | 0 | - |
| - | - | 1680 | 382.7 | - | - | 0 | - |
| 8 | b | 4533 | 383.2 | 0.001296 | 3.382 | +2 | 8 |
| - | - | 1074 | 383.7 | - | - | 0 | - |
| - | - | 1367 | 384.3 | - | - | 0 | - |
| - | - | 890 | 385.3 | - | - | 0 | - |
| - | - | 1736 | 389.2 | - | - | 0 | - |
| - | - | 8490 | 390.2 | - | - | 0 | - |
| - | - | 901.6 | 391.2 | - | - | 0 | - |
| - | - | 1660 | 393.2 | - | - | 0 | - |
| - | - | 8980 | 394.2 | - | - | 0 | - |
| - | - | 4206 | 394.2 | - | - | 0 | - |
| - | - | 1981 | 395.2 | - | - | 0 | - |
| - | - | 1087 | 395.2 | - | - | 0 | - |
| - | - | 8465 | 396.2 | - | - | 0 | - |
| - | - | 832.4 | 396.7 | - | - | 0 | - |
| - | - | 1961 | 397.2 | - | - | 0 | - |
| - | - | 914.7 | 397.3 | - | - | 0 | - |
| 4 | b | 6.269E+04 | 398.2 | 0.0008966 | 2.251 | +1 | 4 |
| - | - | 1.245E+04 | 399.2 | - | - | 0 | - |
| - | - | 1951 | 400.2 | - | - | 0 | - |
| - | - | 3786 | 407.2 | - | - | 0 | - |
| - | - | 1.381E+04 | 408.2 | - | - | 0 | - |
| - | - | 2186 | 409.2 | - | - | 0 | - |
| - | - | 1.027E+04 | 411.2 | - | - | 0 | - |
| - | - | 1976 | 411.3 | - | - | 0 | - |
| - | - | 1858 | 412.2 | - | - | 0 | - |
| - | - | 1175 | 413.2 | - | - | 0 | - |
| - | - | 1079 | 413.3 | - | - | 0 | - |
| - | - | 1291 | 417.2 | - | - | 0 | - |
| - | - | 904.3 | 417.7 | - | - | 0 | - |
| - | - | 4921 | 417.8 | - | - | 0 | - |
| - | - | 1842 | 418.3 | - | - | 0 | - |
| - | - | 1590 | 420.2 | - | - | 0 | - |
| 9 | b | 1120 | 423.2 | 0.002013 | 4.756 | +2 | 9 |
| - | - | 1953 | 424.3 | - | - | 0 | - |
| - | - | 1107 | 425.2 | - | - | 0 | - |
| - | - | 2587 | 425.3 | - | - | 0 | - |
| 2 | y | 5400 | 425.7 | 0.0008584 | 2.016 | +2 | 9 |
| 2 | y | 4221 | 426.2 | 0.006562 | 15.39 | +2 | 9 |
| - | - | 2020 | 426.7 | - | - | 0 | - |
| - | - | 1187 | 429.2 | - | - | 0 | - |
| 9 | b | 9314 | 431.8 | 0.001068 | 2.473 | +2 | 9 |
| - | - | 4816 | 432.3 | - | - | 0 | - |
| - | - | 1321 | 432.8 | - | - | 0 | - |
| - | - | 1811 | 434.2 | - | - | 0 | - |
| 2 | y | 1.947E+04 | 434.7 | 0.0011 | 2.53 | +2 | 9 |
| - | - | 8413 | 435.3 | - | - | 0 | - |
| - | - | 2283 | 435.8 | - | - | 0 | - |
| - | - | 2573 | 437.3 | - | - | 0 | - |
| - | - | 5734 | 437.3 | - | - | 0 | - |
| - | - | 1052 | 438.3 | - | - | 0 | - |
| - | - | 2238 | 439.2 | - | - | 0 | - |
| - | - | 1860 | 439.3 | - | - | 0 | - |
| - | - | 5651 | 439.7 | - | - | 0 | - |
| - | - | 3192 | 440.2 | - | - | 0 | - |
| - | - | 1148 | 441.2 | - | - | 0 | - |
| - | - | 5169 | 442.3 | - | - | 0 | - |
| - | - | 1133 | 443.3 | - | - | 0 | - |
| - | - | 8559 | 447.2 | - | - | 0 | - |
| - | - | 1004 | 448.2 | - | - | 0 | - |
| - | - | 1194 | 449.2 | - | - | 0 | - |
| - | - | 3625 | 450.3 | - | - | 0 | - |
| - | - | 4548 | 451.3 | - | - | 0 | - |
| - | - | 954.9 | 452.3 | - | - | 0 | - |
| - | - | 1118 | 453.3 | - | - | 0 | - |
| - | - | 1.052E+04 | 454.3 | - | - | 0 | - |
| 6 | y | 817.8 | 455.3 | 0.002165 | 4.756 | +1 | 5 |
| - | - | 820.3 | 455.3 | - | - | 0 | - |
| - | - | 3246 | 456.3 | - | - | 0 | - |
| - | - | 989.8 | 457.8 | - | - | 0 | - |
| - | - | 697.1 | 463.3 | - | - | 0 | - |
| - | - | 3.006E+04 | 464.3 | - | - | 0 | - |
| - | - | 6347 | 465.2 | - | - | 0 | - |
| - | - | 2.228E+04 | 465.3 | - | - | 0 | - |
| - | - | 2234 | 466.2 | - | - | 0 | - |
| - | - | 5832 | 466.3 | - | - | 0 | - |
| - | - | 2423 | 466.8 | - | - | 0 | - |
| - | - | 1208 | 467.2 | - | - | 0 | - |
| - | - | 1067 | 467.3 | - | - | 0 | - |
| - | - | 1.131E+04 | 467.3 | - | - | 0 | - |
| - | - | 2261 | 468.3 | - | - | 0 | - |
| - | - | 1079 | 469.3 | - | - | 0 | - |
| 6 | y | 1.7E+04 | 473.3 | 0.001156 | 2.442 | +1 | 5 |
| - | - | 3966 | 474.3 | - | - | 0 | - |
| 0 | Precursor | 1.119E+04 | 475.3 | 0.001045 | 2.198 | +2 | -1 |
| - | - | 5972 | 475.8 | - | - | 0 | - |
| - | - | 1338 | 476.3 | - | - | 0 | - |
| - | - | 1115 | 477.3 | - | - | 0 | - |
| - | - | 7700 | 478.3 | - | - | 0 | - |
| - | - | 1.909E+04 | 479.3 | - | - | 0 | - |
| - | - | 1003 | 479.3 | - | - | 0 | - |
| - | - | 6626 | 480.3 | - | - | 0 | - |
| - | - | 1060 | 481.3 | - | - | 0 | - |
| - | - | 2526 | 481.3 | - | - | 0 | - |
| - | - | 7.613E+04 | 482.3 | - | - | 0 | - |
| - | - | 1.685E+04 | 483.3 | - | - | 0 | - |
| 0 | Precursor | 3.235E+04 | 484.3 | 0.001072 | 2.215 | +2 | -1 |
| - | - | 1.875E+04 | 484.8 | - | - | 0 | - |
| - | - | 955.8 | 485.3 | - | - | 0 | - |
| - | - | 5358 | 485.3 | - | - | 0 | - |
| - | - | 1131 | 485.4 | - | - | 0 | - |
| - | - | 1149 | 490.3 | - | - | 0 | - |
| - | - | 1883 | 491.3 | - | - | 0 | - |
| 5 | b | 4230 | 495.3 | 0.001386 | 2.798 | +1 | 5 |
| - | - | 6113 | 496.3 | - | - | 0 | - |
| - | - | 1083 | 497.3 | - | - | 0 | - |
| - | - | 1684 | 507.3 | - | - | 0 | - |
| - | - | 1.062E+04 | 508.3 | - | - | 0 | - |
| - | - | 3058 | 509.3 | - | - | 0 | - |
| - | - | 1618 | 509.3 | - | - | 0 | - |
| - | - | 4947 | 518.3 | - | - | 0 | - |
| - | - | 839.8 | 519.3 | - | - | 0 | - |
| - | - | 3417 | 524.3 | - | - | 0 | - |
| - | - | 1.96E+04 | 525.3 | - | - | 0 | - |
| - | - | 4037 | 526.3 | - | - | 0 | - |
| - | - | 1510 | 534.3 | - | - | 0 | - |
| - | - | 6111 | 535.3 | - | - | 0 | - |
| - | - | 2.459E+04 | 536.3 | - | - | 0 | - |
| - | - | 6398 | 537.3 | - | - | 0 | - |
| - | - | 1307 | 538.3 | - | - | 0 | - |
| - | - | 2387 | 540.3 | - | - | 0 | - |
| - | - | 1366 | 550.3 | - | - | 0 | - |
| - | - | 761.5 | 551.3 | - | - | 0 | - |
| - | - | 1078 | 551.3 | - | - | 0 | - |
| 5 | y | 1.848E+04 | 552.3 | 0.0009182 | 1.663 | +1 | 6 |
| - | - | 1296 | 552.4 | - | - | 0 | - |
| - | - | 2.22E+05 | 553.3 | - | - | 0 | - |
| - | - | 6.403E+04 | 554.3 | - | - | 0 | - |
| - | - | 1.22E+04 | 555.3 | - | - | 0 | - |
| - | - | 1260 | 555.4 | - | - | 0 | - |
| - | - | 805 | 556.3 | - | - | 0 | - |
| - | - | 1279 | 560.3 | - | - | 0 | - |
| - | - | 2005 | 564.4 | - | - | 0 | - |
| - | - | 937.2 | 565.4 | - | - | 0 | - |
| - | - | 3211 | 567.4 | - | - | 0 | - |
| - | - | 3133 | 568.3 | - | - | 0 | - |
| 5 | y | 4.711E+05 | 570.3 | 0.001218 | 2.135 | +1 | 6 |
| - | - | 1.428E+05 | 571.3 | - | - | 0 | - |
| - | - | 2.668E+04 | 572.3 | - | - | 0 | - |
| - | - | 2197 | 573.3 | - | - | 0 | - |
| - | - | 1687 | 576.3 | - | - | 0 | - |
| - | - | 759.2 | 577.3 | - | - | 0 | - |
| - | - | 7366 | 577.3 | - | - | 0 | - |
| - | - | 1941 | 578.3 | - | - | 0 | - |
| - | - | 2216 | 579.3 | - | - | 0 | - |
| - | - | 1038 | 580.3 | - | - | 0 | - |
| - | - | 1.05E+04 | 580.4 | - | - | 0 | - |
| - | - | 2191 | 581.4 | - | - | 0 | - |
| - | - | 811.9 | 583.4 | - | - | 0 | - |
| - | - | 1.332E+04 | 595.4 | - | - | 0 | - |
| - | - | 6275 | 596.4 | - | - | 0 | - |
| - | - | 1082 | 605.3 | - | - | 0 | - |
| - | - | 3839 | 621.4 | - | - | 0 | - |
| - | - | 3295 | 622.4 | - | - | 0 | - |
| 6 | b | 3675 | 623.4 | 0.002024 | 3.247 | +1 | 6 |
| - | - | 1583 | 624.4 | - | - | 0 | - |
| - | - | 835.6 | 631.4 | - | - | 0 | - |
| - | - | 2237 | 633.3 | - | - | 0 | - |
| - | - | 1150 | 637.4 | - | - | 0 | - |
| - | - | 6772 | 638.4 | - | - | 0 | - |
| - | - | 1550 | 639.4 | - | - | 0 | - |
| - | - | 686.6 | 640.4 | - | - | 0 | - |
| - | - | 1563 | 648.4 | - | - | 0 | - |
| - | - | 6094 | 649.4 | - | - | 0 | - |
| - | - | 1.796E+04 | 650.4 | - | - | 0 | - |
| - | - | 5629 | 651.4 | - | - | 0 | - |
| - | - | 3906 | 652.4 | - | - | 0 | - |
| - | - | 5219 | 654.4 | - | - | 0 | - |
| - | - | 1484 | 655.4 | - | - | 0 | - |
| - | - | 1125 | 659.4 | - | - | 0 | - |
| - | - | 981.3 | 662.4 | - | - | 0 | - |
| - | - | 5.315E+04 | 666.4 | - | - | 0 | - |
| - | - | 2.176E+04 | 667.4 | - | - | 0 | - |
| - | - | 4338 | 668.4 | - | - | 0 | - |
| - | - | 7695 | 676.4 | - | - | 0 | - |
| - | - | 3958 | 677.4 | - | - | 0 | - |
| - | - | 907.1 | 678.4 | - | - | 0 | - |
| 4 | y | 1.101E+04 | 680.4 | 0.001057 | 1.553 | +1 | 7 |
| 4 | y | 1.615E+04 | 681.4 | 0.004224 | 6.199 | +1 | 7 |
| - | - | 6692 | 682.4 | - | - | 0 | - |
| - | - | 1161 | 683.4 | - | - | 0 | - |
| 7 | b | 2.34E+04 | 694.4 | 0.0009009 | 1.297 | +1 | 7 |
| - | - | 1.033E+04 | 695.4 | - | - | 0 | - |
| - | - | 811.8 | 696.4 | - | - | 0 | - |
| 4 | y | 1.419E+04 | 698.4 | 0.001234 | 1.767 | +1 | 7 |
| - | - | 5285 | 699.4 | - | - | 0 | - |
| - | - | 904 | 700.4 | - | - | 0 | - |
| - | - | 4160 | 720.4 | - | - | 0 | - |
| - | - | 3415 | 720.4 | - | - | 0 | - |
| - | - | 1846 | 721.4 | - | - | 0 | - |
| - | - | 1140 | 721.4 | - | - | 0 | - |
| - | - | 2736 | 730.4 | - | - | 0 | - |
| - | - | 1367 | 731.4 | - | - | 0 | - |
| - | - | 1307 | 734.5 | - | - | 0 | - |
| - | - | 1043 | 735.5 | - | - | 0 | - |
| 3 | y | 1.457E+04 | 737.4 | 0.0004669 | 0.6331 | +1 | 8 |
| - | - | 1.637E+04 | 737.5 | - | - | 0 | - |
| 3 | y | 1.876E+04 | 738.4 | 0.004122 | 5.583 | +1 | 8 |
| - | - | 5624 | 738.5 | - | - | 0 | - |
| - | - | 6035 | 739.4 | - | - | 0 | - |
| - | - | 1594 | 739.5 | - | - | 0 | - |
| - | - | 951.3 | 740.4 | - | - | 0 | - |
| - | - | 846.7 | 746.4 | - | - | 0 | - |
| - | - | 3513 | 747.5 | - | - | 0 | - |
| 8 | b | 7580 | 748.4 | 0.003703 | 4.947 | +1 | 8 |
| - | - | 5479 | 749.4 | - | - | 0 | - |
| - | - | 1261 | 750.4 | - | - | 0 | - |
| - | - | 1376 | 753.4 | - | - | 0 | - |
| - | - | 1234 | 754.4 | - | - | 0 | - |
| 3 | y | 4.007E+05 | 755.4 | 0.001316 | 1.742 | +1 | 8 |
| - | - | 1.572E+05 | 756.4 | - | - | 0 | - |
| - | - | 3.581E+04 | 757.4 | - | - | 0 | - |
| - | - | 3782 | 758.4 | - | - | 0 | - |
| - | - | 6551 | 763.4 | - | - | 0 | - |
| - | - | 2456 | 764.5 | - | - | 0 | - |
| 8 | b | 2.128E+05 | 765.5 | 0.001019 | 1.331 | +1 | 8 |
| - | - | 9.011E+04 | 766.5 | - | - | 0 | - |
| - | - | 2.309E+04 | 767.5 | - | - | 0 | - |
| - | - | 1905 | 768.5 | - | - | 0 | - |
| - | - | 1767 | 791.5 | - | - | 0 | - |
| 2 | y | 1834 | 850.5 | 0.0004672 | 0.5494 | +1 | 9 |
| 2 | y | 3109 | 851.5 | 0.007827 | 9.192 | +1 | 9 |
| - | - | 1100 | 852.5 | - | - | 0 | - |
| - | - | 758.5 | 853.5 | - | - | 0 | - |
| 2 | y | 5.215E+04 | 868.5 | 0.000931 | 1.072 | +1 | 9 |
| - | - | 2.557E+04 | 869.5 | - | - | 0 | - |
| - | - | 7020 | 870.5 | - | - | 0 | - |
| - | - | 2158 | 878.5 | - | - | 0 | - |
| - | - | 1327 | 879.5 | - | - | 0 | - |
| - | - | 1051 | 880.5 | - | - | 0 | - |
| - | - | 1195 | 881.5 | - | - | 0 | - |
| - | - | 684.5 | 1785 | - | - | 0 | - |
| - | - | 791.3 | 2018 | - | - | 0 | - |
| - | - | 924 | 2993 | - | - | 0 | - |
| - | - | 891.4 | 3073 | - | - | 0 | - |

m/z Charge Intensity FragmentType MassShift Position
124.03974151611328 0 2004.4613
125.07144927978516 0 496.1844
125.10759735107422 0 1097.9637
125.32617950439453 0 379.01257
126.09187316894531 0 384.61084
127.08708953857422 0 1411.4152
127.1233139038086 0 435.54608
128.0713348388672 0 455.8358
128.1080322265625 0 362.59583
129.06629943847656 0 14421.9
129.10267639160156 0 117451.27
130.05043029785156 0 1196.8566
130.0698699951172 0 621.98444
130.1008758544922 0 916.6602
130.1060333251953 0 6679.749
132.4923095703125 0 407.20554
132.66871643066406 0 357.7294
134.64291381835938 0 364.56018
136.07608032226562 0 1413.2162
136.42433166503906 0 432.25372
137.10763549804688 0 1007.1392
139.08705139160156 0 3647.9968
141.06629943847656 0 8523.067
141.10264587402344 0 8750.823
142.10635375976562 0 519.32996
142.12315368652344 0 716.2495
142.1918487548828 0 549.56537
143.0818634033203 0 4848.7666
143.11834716796875 0 5602.716 d 1
146.09274291992188 0 631.1744
147.07723999023438 0 1016.5414
148.95384216308594 0 979.3023
152.03468322753906 0 7992.0303
153.0386505126953 0 723.14386
153.06614685058594 0 1733.886
153.1026611328125 0 2105.521
154.0867919921875 0 909.234
155.0819854736328 0 764.3899
155.11830139160156 0 14685.378
156.1217041015625 0 949.99365
157.0976104736328 0 3861.149
157.13404846191406 0 641.60175
157.1704559326172 0 1341.5625
158.0929412841797 0 6310.991
161.09237670898438 0 847.02563
163.07176208496094 0 1175.1013
165.1028289794922 0 2201.9897
167.11790466308594 0 833.01355
169.0612335205078 0 35006.92
169.09762573242188 0 12660.118
169.13397216796875 0 2780.7505
170.06455993652344 0 2982.22
170.10137939453125 0 1210.9753
171.11328125 0 20821.566
172.1082305908203 0 491.61157
172.1165771484375 0 1832.9358
172.14500427246094 0 1668.2185
173.12899780273438 0 615.70245
173.4522247314453 0 980.2331
180.11375427246094 0 3786.4841
181.09768676757812 0 9457.167
181.13401794433594 0 7665.7676
182.1016387939453 0 1242.5165
182.12930297851562 0 32404.688
183.11343383789062 0 3848.4148
183.1326904296875 0 3602.9817
183.14964294433594 0 2634.5742
184.10916137695312 0 585.8418
184.1173858642578 0 519.2828
185.0926055908203 0 12528.008 y Water loss 8
185.1654052734375 0 674656.06 a 1
186.08782958984375 0 81632.59
186.1239471435547 0 666.87256
186.16868591308594 0 68925.98
187.07188415527344 0 1055.7579
187.09112548828125 0 6064.738
187.1073760986328 0 817.5124
187.1447296142578 0 1022.1356
187.1715545654297 0 2497.3906
188.13980102539062 0 17766.168
189.1427459716797 0 1388.2223
191.08224487304688 0 1964.0771
191.1183319091797 0 1180.087
192.1136932373047 0 823.3567
193.0860595703125 0 444.4423
193.0881805419922 0 468.48257
193.0974578857422 0 2209.6833
195.11305236816406 0 2137.5837
196.1088104248047 0 727.9115
197.092529296875 0 1672.9963
197.12881469726562 0 3314.1843
197.62730407714844 0 635.6717
198.1243438720703 0 2020.8856
199.10842895507812 0 1966.0615
200.13973999023438 0 7475.8574
201.08738708496094 0 1638.2172
201.14414978027344 0 679.9802
203.10317993164062 0 413065.9 y 8
204.1064910888672 0 36006.42
205.1079559326172 0 3981.281
208.10855102539062 0 10329.81
208.14474487304688 0 2252.425
209.0926055908203 0 8115.2983
209.11178588867188 0 842.2719
209.1289825439453 0 2381.7654
210.12417602539062 0 41630.758
210.16055297851562 0 811.4087
211.1085205078125 0 1175.0714
211.12753295898438 0 4187.692
211.14456176757812 0 1651.6409
211.1810760498047 0 2195.01
212.1036834716797 0 1391.0582
212.13986206054688 0 764.1517
213.16033935546875 0 344045.25 b 1
214.11895751953125 0 1304.5038
214.16363525390625 0 38716.004
215.1389617919922 0 1381.8726
215.1656494140625 0 2768.2905
217.1678924560547 0 538.21954
220.10870361328125 0 2248.8708
221.0927734375 0 1139.8599
223.10836791992188 0 1822.1774
225.1352081298828 0 759.66327
226.1555938720703 0 278398.5
227.15895080566406 0 31895.207
228.134765625 0 33478.77 y Water loss 5
228.1619415283203 0 1534.124
229.1382293701172 0 3838.267
235.14405822753906 0 794.75977
236.1399383544922 0 2956.34
237.08871459960938 0 873.91766
238.1192169189453 0 5036.9077
239.17562866210938 0 1406.5398
240.135009765625 0 3917.5742
242.15025329589844 0 1858.3264
243.18211364746094 0 7975.926
244.1851043701172 0 888.1216
248.10299682617188 0 746.3232
252.13491821289062 0 3412.2368
252.1712646484375 0 10561.001
253.13763427734375 0 558.7073
253.1665496826172 0 3727.1301
254.11412048339844 0 1270.716
254.1505126953125 0 10914.806
255.15354919433594 0 1094.605
256.1297912597656 0 5601.2856 y Water loss 7
256.1663818359375 0 763.4088
257.12530517578125 0 2862.861
261.1718444824219 0 1914.7941
262.15533447265625 0 747.21857
264.1347961425781 0 1786.6489
265.13043212890625 0 2117.546
266.11407470703125 0 4153.719
266.1304931640625 0 799.61774
267.1457824707031 0 1033.0471
268.1283874511719 0 883.8059
268.165283203125 0 772.41547
269.16131591796875 0 1126.1343
269.1976318359375 0 13979.732
270.1462097167969 0 2009.0366
270.1817321777344 0 41031.863 b 2
270.2001647949219 0 1693.1606
271.1410827636719 0 1852.1619
271.1771545410156 0 30000.408
272.18035888671875 0 2868.6318
274.1402587890625 0 26900.156 y 7
274.16949462890625 0 966.65515
274.6732177734375 0 757.29565
275.1435241699219 0 2961.0508
277.1644287109375 0 804.02527
279.18212890625 0 59044.395
280.1677551269531 0 816.5664
280.1852722167969 0 6962
281.1614685058594 0 8355.784
281.1880187988281 0 768.06665
282.1452941894531 0 1994.01
282.1640930175781 0 1090.9722
283.14068603515625 0 6985.6895
283.17730712890625 0 1090.8572
285.1563415527344 0 10501.324
285.66650390625 0 6643.8433 y 4
286.1676330566406 0 1902.4674
287.20849609375 0 73422.62
288.17083740234375 0 988.0384
288.2117919921875 0 11375.619
292.1656799316406 0 819.3724
295.1768798828125 0 908.50635
295.2134704589844 0 1313.5933
296.1978759765625 0 812.43744
297.15533447265625 0 1102.2794
297.1603088378906 0 1442.7998
297.17236328125 0 1155.7863
297.17724609375 0 1176.6304
297.1928405761719 0 113342.33
298.19610595703125 0 15506.031
299.1721496582031 0 209266.39
300.1751403808594 0 27490.809
301.1773376464844 0 3906.9824
309.1561584472656 0 3939.1687
309.19171142578125 0 989.0006
310.2129821777344 0 1118.0515
311.1912536621094 0 891.6482
314.2196960449219 0 4917.6904
315.22161865234375 0 1354.7906
318.19281005859375 0 861.75757
319.17626953125 0 1344.3069
319.70361328125 0 655.8811
323.1729736328125 0 1974.5204
323.2085876464844 0 7623.1294
324.2123718261719 0 899.4476
325.1880187988281 0 5416.271
325.6860656738281 0 1515.1029
326.189697265625 0 912.1014
326.71112060546875 0 1516.4623
327.16705322265625 0 11775.81 y Water loss 6
327.2025451660156 0 696.11395
328.16412353515625 0 6078.4854
328.1977844238281 0 1223.2917
329.1675109863281 0 685.78644
331.6868591308594 0 673.6793
332.177734375 0 4541.846
336.2037048339844 0 4348.4355
337.1879577636719 0 8208.636
338.18994140625 0 801.59406
340.235107421875 0 49467.098
340.6906433105469 0 16112.064 y Water loss 3
340.712646484375 0 994.5123
341.19122314453125 0 5020.1206
341.23834228515625 0 7628.177
341.6872863769531 0 961.95776
342.243896484375 0 722.2894
345.177734375 0 4796.399 y 6
345.22674560546875 0 955.0805
346.1816101074219 0 839.6092
347.71612548828125 0 1177.7124 b 6
349.1864318847656 0 890.2142
349.6959228515625 0 2974.253 y 3
350.1974182128906 0 1026.9092
350.21929931640625 0 3851.3577
351.2030029296875 0 1497.1826
353.218994140625 0 3284.6584
354.2146911621094 0 2771.4805
355.2358703613281 0 1689.9264
359.7161865234375 0 1135.5594
360.6883850097656 0 973.4223
360.7239074707031 0 2331.6975
361.1883544921875 0 779.46387
363.2033996582031 0 2558.0288
366.2140808105469 0 3282.1763
368.22991943359375 0 264497.6
368.7331848144531 0 1086.953
369.233154296875 0 54470.887
369.6938171386719 0 1536.1765 y Ammonia loss 2
370.24468994140625 0 22084.684
371.2469177246094 0 4049.5908
372.2010192871094 0 799.7415
372.2459716796875 0 812.94635
373.714599609375 0 1526.1548
376.1988830566406 0 5806.2114
376.2444763183594 0 2107.5686
378.1802978515625 0 1216.667
378.2083435058594 0 4031.3933 y 2
378.7082824707031 0 889.1016
380.2301330566406 0 5185.8833
381.2112121582031 0 1081.5148 b Ammonia loss 3
381.23565673828125 0 1161.2677
382.2266845703125 0 5197.904
382.7284240722656 0 1680.455
383.2358093261719 0 4532.5137 b 7
383.7354431152344 0 1074.3885
384.2624816894531 0 1366.9619
385.2615966796875 0 890.01117
389.23065185546875 0 1735.5148
390.2142028808594 0 8490.224
391.2166748046875 0 901.62476
393.22637939453125 0 1659.9744
394.2094421386719 0 8980.37
394.24560546875 0 4205.636
395.2117919921875 0 1981.1561
395.24444580078125 0 1086.9589
396.2249755859375 0 8464.622
396.74444580078125 0 832.3757
397.22784423828125 0 1960.5732
397.2576599121094 0 914.7253
398.2406921386719 0 62689.395 b 3
399.2435302734375 0 12449.662
400.2473449707031 0 1950.686
407.2410583496094 0 3786.2014
408.2251892089844 0 13806.795
409.2275085449219 0 2186.393
411.2363586425781 0 10273.152
411.2721252441406 0 1975.7682
412.23858642578125 0 1857.7097
413.2152099609375 0 1175.1542
413.2503662109375 0 1079.2288
417.23040771484375 0 1290.6792
417.734375 0 904.25366
417.7645263671875 0 4920.6733
418.267333984375 0 1842.0212
420.223876953125 0 1590.1221
423.2496337890625 0 1120.0852 b Ammonia loss 8
424.25775146484375 0 1952.5021
425.2160949707031 0 1106.719
425.2518615722656 0 2587.1494
425.7435607910156 0 5399.596 y Water loss 1
426.24127197265625 0 4221.4062 y Ammonia loss 1
426.7422790527344 0 2019.8807
429.2273864746094 0 1187.0413
431.761962890625 0 9313.557 b 8
432.26336669921875 0 4815.629
432.76446533203125 0 1320.5319
434.24127197265625 0 1810.9601
434.74908447265625 0 19473.52 y 1
435.25006103515625 0 8413.326
435.7507629394531 0 2282.777
437.2519226074219 0 2572.9382
437.2884216308594 0 5734.1733
438.2900390625 0 1052.3949
439.2310485839844 0 2237.9844
439.3045654296875 0 1859.7654
439.74139404296875 0 5650.515
440.241943359375 0 3192.0378
441.24652099609375 0 1148.2479
442.2783203125 0 5169.2744
443.2804870605469 0 1133.4773
447.2360534667969 0 8558.703
448.24029541015625 0 1004.26953
449.2189636230469 0 1193.5642
450.2813720703125 0 3625.2698
451.26837158203125 0 4547.5127
452.2697448730469 0 954.8877
453.31805419921875 0 1117.9757
454.2784118652344 0 10517.624
455.25909423828125 0 817.8333 y Water loss 5
455.2850646972656 0 820.3096
456.25762939453125 0 3245.6655
457.7584533691406 0 989.7804
463.2674255371094 0 697.06256
464.2627258300781 0 30061.225
465.2475891113281 0 6347.444
465.2823181152344 0 22279.836
466.2456359863281 0 2234.494
466.2853088378906 0 5832.0835
466.76531982421875 0 2423.3804
467.226318359375 0 1208.0885
467.26544189453125 0 1066.6761
467.2982177734375 0 11310.318
468.3009338378906 0 2260.5576
469.2765197753906 0 1079.4741
473.2729797363281 0 17001.395 y 5
474.27532958984375 0 3966.3748
475.2779541015625 0 11191.804 Precursor Water loss
475.7792663574219 0 5972.148
476.27777099609375 0 1338.3153
477.28045654296875 0 1115.4905
478.2779541015625 0 7700.05
479.26263427734375 0 19092.508
479.296142578125 0 1002.94446
480.2653503417969 0 6625.672
481.268798828125 0 1060.1603
481.3138732910156 0 2526.0798
482.2734069824219 0 76133.1
483.2765197753906 0 16853.469
484.28326416015625 0 32354.445 Precursor
484.7848205566406 0 18749.53
485.2522888183594 0 955.8004
485.2868957519531 0 5358.008
485.35845947265625 0 1131.0558
490.27972412109375 0 1149.3552
491.2635498046875 0 1882.8058
495.2939453125 0 4230.0938 b 4
496.289794921875 0 6112.847
497.2940979003906 0 1082.6128
507.2960205078125 0 1683.6769
508.2891540527344 0 10621.365
509.2906799316406 0 3058.2888
509.3439025878906 0 1618.4484
518.2734375 0 4947.1885
519.2806396484375 0 839.77515
524.3203735351562 0 3417.3225
525.3155517578125 0 19599.387
526.3192138671875 0 4036.5085
534.3030395507812 0 1510.2301
535.2992553710938 0 6111.351
536.2841186523438 0 24591.959
537.287109375 0 6397.511
538.2923583984375 0 1307.4655
540.3162231445312 0 2387.4954
550.3359985351562 0 1365.5934
551.2924194335938 0 761.48926
551.3353271484375 0 1078.2452
552.31494140625 0 18480.39 y Water loss 4
552.3890991210938 0 1296.4685
553.310302734375 0 222006.47
554.3131713867188 0 64030.074
555.315185546875 0 12203.047
555.3633422851562 0 1260.196
556.3189697265625 0 805.02386
560.3211669921875 0 1278.5502
564.35009765625 0 2005.4276
565.3532104492188 0 937.18884
567.3616943359375 0 3211.3806
568.3098754882812 0 3132.7761
570.3258056640625 0 471058.22 y 4
571.3285522460938 0 142825.25
572.330810546875 0 26677.207
573.3345336914062 0 2197.4326
576.3150634765625 0 1687.2585
577.2655029296875 0 759.1902
577.3468627929688 0 7365.8823
578.3403930664062 0 1940.6255
579.3275146484375 0 2216.3909
580.331787109375 0 1037.9043
580.383056640625 0 10499.924
581.3858032226562 0 2190.9182
583.389892578125 0 811.8931
595.3571166992188 0 13315.922
596.3597412109375 0 6274.588
605.3456420898438 0 1081.6237
621.3721923828125 0 3839.0955
622.3711547851562 0 3294.6553
623.385498046875 0 3674.9143 b 5
624.3897705078125 0 1583.4384
631.3580932617188 0 835.63416
633.3375854492188 0 2236.6714
637.4058837890625 0 1149.5682
638.3998413085938 0 6771.792
639.4022827148438 0 1550.16
640.411865234375 0 686.5658
648.381103515625 0 1562.8004
649.3676147460938 0 6093.522
650.3635864257812 0 17955.871
651.3670043945312 0 5628.6465
652.3743286132812 0 3906.3296
654.4302978515625 0 5219.4844
655.4313354492188 0 1484.3477
659.3876342773438 0 1124.9156
662.3651733398438 0 981.3144
666.39453125 0 53149.062
667.3974609375 0 21759.947
668.400390625 0 4337.632
676.41552734375 0 7695.291
677.4171752929688 0 3957.751
678.4054565429688 0 907.08673
680.3736572265625 0 11014.825 y Water loss 3
681.36083984375 0 16152.848 y Ammonia loss 3
682.3618774414062 0 6692.485
683.3622436523438 0 1160.996
694.425537109375 0 23400.197 b 6
695.4287109375 0 10327.85
696.4212646484375 0 811.8197
698.3843994140625 0 14194.169 y 3
699.3871459960938 0 5285.3726
700.3936767578125 0 903.96423
720.3695678710938 0 4160.171
720.441650390625 0 3414.5137
721.3712768554688 0 1845.9827
721.4446411132812 0 1139.5568
730.42626953125 0 2735.5752
731.4271240234375 0 1366.6995
734.455810546875 0 1307.0215
735.4534912109375 0 1043.418
737.39453125 0 14572.435 y Water loss 2
737.4679565429688 0 16371.454
738.3822021484375 0 18761.523 y Ammonia loss 2
738.4711303710938 0 5623.9053
739.3854370117188 0 6034.722
739.4733276367188 0 1594.3257
740.387939453125 0 951.3466
746.4198608398438 0 846.6606
747.4530029296875 0 3513.419
748.4389038085938 0 7579.711 b Ammonia loss 7
749.4365234375 0 5479.2554
750.4385375976562 0 1260.7953
753.388916015625 0 1376.1725
754.3927001953125 0 1234.2388
755.4059448242188 0 400670.62 y 2
756.408447265625 0 157201.02
757.4111328125 0 35811.527
758.413818359375 0 3781.7004
763.4468994140625 0 6550.9624
764.4508056640625 0 2455.9421
765.4627685546875 0 212847.95 b 7
766.4656982421875 0 90112.3
767.4681396484375 0 23088.498
768.4707641601562 0 1904.7931
791.477783203125 0 1766.659
850.4776611328125 0 1833.6373 y Water loss 1
851.469970703125 0 3109.4543 y Ammonia loss 1
852.470947265625 0 1099.5619
853.4674682617188 0 758.48883
868.4896240234375 0 52150.617 y 1
869.4930419921875 0 25565.746
870.4963989257812 0 7019.9624
878.474853515625 0 2158.3594
879.4752197265625 0 1326.6405
880.525146484375 0 1050.8037
881.531005859375 0 1194.9897
1785.3184814453125 0 684.4972
2018.383056640625 0 791.34607
2992.51953125 0 923.98865
3073.169677734375 0 891.38116

Spectrum Details

|  |  |
| --- | --- |
| Matched peaks? Matched peaks The total absolute number of peaks matched. Additionally in brackets the total fraction of peaks matched and the total number of peaks is shown. | 45 (8.96% of 502) |
| FDR? FDR The false discovery rate estimated for this peptide. It is calculated by matching all theoretical fragments with a non-integer shift with the raw peaks for this spectrum. This is done with 40 different shifts. The resulting percentage is the average number of annotated peaks over the number of annotated peaks with the correct spectrum. | 0.05% |
| Satellite FDR? Satellite FDR See the FDR for details on its calculation. This satellite ion specific FDR only contains the satellite ions (d/w) for I/L/J positions. | 0.00% |
| PSM Score? PSM Score The PSM Score as given by Hecklib to this annotated spectrum. It is shown with three significant figures. | 476 |

#### Spectrum 2923? Spectrum 2923 The raw spectrum of this peptide as annotated by Hecklib. The fragments are coloured according to ion type (see legend). Any peaks with a star '\*' as text can be hovered over to see the full details, first the ion type second the mass shift type. By hovering over the amino acids in the peptide or ions in the legend the corresponding peaks are highlighted. By toggling the 'Unassigned' label you can turn the background (unassigned) peaks on or off in the plot. By updating the slider in the Ion legend you can update the spectrum to only show the top X% of the peaks with labels. The top X% means any peak that is within X% of the highest intensity. By dragging in the spectrum you can zoom in to a specific part of the spectrum and use 'Zoom Out' to get back to the original zoom level. The annotation of the spectrum is based on the given sequence in the peptides file and is done with different software so inconsistencies are likely. The peaks are annotated based on the given sequence, with 20 ppm tolerance.

Copy Data

##### Spectrum 2923 (TSV)

###### Preview

```
Loading example...
```

*Click on the button to copy the data to your clipboard.*

Mz MinMz MaxIntensity Max

WidthHeightPeptide font sizePeptide stroke widthSpectrum font sizeSpectrum stroke widthCompact peptide

Ion legend

wxyz

abcd

OtherUnassignedIon^Charge^_Position_Show for top:%

VLGQPKAAPS

03.88e+67.77e+61.17e+71.55e+7

Zoom Out

d^+1^_2_y^+1^_2_a^+1^_2_y^+1^_2_b^+1^_2_y^+2^_5_y^+1^_3_b^+1^_3_y^+1^_3_y^+2^_6_y^+2^_6_y^+1^_4_y^+2^_7_y^+1^_4_b^+2^_7_y^+2^_7_y^+2^_8_y^+2^_8_b^+2^_8_b^+1^_4_y^+2^_9_y^+2^_9_b^+2^_9_y^+2^_9_y^+1^_5_\*\*\*b^+1^_5_y^+1^_6_y^+1^_6_b^+1^_6_y^+1^_7_y^+1^_7_b^+1^_7_y^+1^_7_y^+1^_8_y^+1^_8_b^+1^_8_y^+1^_8_b^+1^_8_y^+1^_9_y^+1^_9_y^+1^_9_

0798159623933191

Fragment Matches Table

Show background peaks

| Position | Ion type | Intensity | mz Theoretical | mz Error (Th) | mz Error (ppm) | Charge | Series Number |
| --- | --- | --- | --- | --- | --- | --- | --- |
| - | - | 3.362E+04 | 124 | - | - | 0 | - |
| - | - | 1.975E+04 | 125.1 | - | - | 0 | - |
| - | - | 1.384E+04 | 126.7 | - | - | 0 | - |
| - | - | 2.812E+04 | 127.1 | - | - | 0 | - |
| - | - | 3.223E+05 | 129.1 | - | - | 0 | - |
| - | - | 2.687E+06 | 129.1 | - | - | 0 | - |
| - | - | 1.559E+05 | 130.1 | - | - | 0 | - |
| - | - | 2.665E+04 | 137.1 | - | - | 0 | - |
| - | - | 1.03E+05 | 139.1 | - | - | 0 | - |
| - | - | 1.716E+05 | 141.1 | - | - | 0 | - |
| - | - | 1.856E+05 | 141.1 | - | - | 0 | - |
| - | - | 2.819E+04 | 142.1 | - | - | 0 | - |
| - | - | 1.814E+04 | 142.1 | - | - | 0 | - |
| - | - | 8.713E+04 | 143.1 | - | - | 0 | - |
| 2 | d | 1.065E+05 | 143.1 | 0.0002593 | 1.812 | +1 | 2 |
| - | - | 1.293E+04 | 145.2 | - | - | 0 | - |
| - | - | 1.889E+05 | 152 | - | - | 0 | - |
| - | - | 1.358E+04 | 152.1 | - | - | 0 | - |
| - | - | 3.927E+04 | 153.1 | - | - | 0 | - |
| - | - | 5.54E+04 | 153.1 | - | - | 0 | - |
| - | - | 3.235E+05 | 155.1 | - | - | 0 | - |
| - | - | 1.79E+04 | 156.1 | - | - | 0 | - |
| - | - | 1.005E+05 | 157.1 | - | - | 0 | - |
| - | - | 4.233E+04 | 157.2 | - | - | 0 | - |
| - | - | 1.434E+04 | 157.3 | - | - | 0 | - |
| - | - | 1.339E+05 | 158.1 | - | - | 0 | - |
| - | - | 1.598E+04 | 161.1 | - | - | 0 | - |
| - | - | 1.562E+04 | 163.1 | - | - | 0 | - |
| - | - | 6.248E+04 | 165.1 | - | - | 0 | - |
| - | - | 2.645E+04 | 167.1 | - | - | 0 | - |
| - | - | 1.579E+04 | 168.5 | - | - | 0 | - |
| - | - | 8.12E+05 | 169.1 | - | - | 0 | - |
| - | - | 2.945E+05 | 169.1 | - | - | 0 | - |
| - | - | 4.921E+04 | 169.1 | - | - | 0 | - |
| - | - | 3.864E+04 | 170.1 | - | - | 0 | - |
| - | - | 1.748E+04 | 170.1 | - | - | 0 | - |
| - | - | 4.581E+05 | 171.1 | - | - | 0 | - |
| - | - | 6.154E+04 | 172.1 | - | - | 0 | - |
| - | - | 3.148E+04 | 172.1 | - | - | 0 | - |
| - | - | 1.643E+04 | 176.1 | - | - | 0 | - |
| - | - | 8.079E+04 | 180.1 | - | - | 0 | - |
| - | - | 2.398E+05 | 181.1 | - | - | 0 | - |
| - | - | 1.68E+05 | 181.1 | - | - | 0 | - |
| - | - | 2.102E+04 | 182.1 | - | - | 0 | - |
| - | - | 6.682E+05 | 182.1 | - | - | 0 | - |
| - | - | 1.125E+05 | 183.1 | - | - | 0 | - |
| - | - | 6.572E+04 | 183.1 | - | - | 0 | - |
| - | - | 3.892E+04 | 183.1 | - | - | 0 | - |
| 9 | y | 2.53E+05 | 185.1 | 0.0002012 | 1.087 | +1 | 2 |
| 2 | a | 1.539E+07 | 185.2 | 0.0003367 | 1.818 | +1 | 2 |
| - | - | 1.922E+06 | 186.1 | - | - | 0 | - |
| - | - | 1.54E+06 | 186.2 | - | - | 0 | - |
| - | - | 1.696E+05 | 187.1 | - | - | 0 | - |
| - | - | 2.524E+04 | 187.1 | - | - | 0 | - |
| - | - | 7.013E+04 | 187.2 | - | - | 0 | - |
| - | - | 4.301E+05 | 188.1 | - | - | 0 | - |
| - | - | 4.043E+04 | 191.1 | - | - | 0 | - |
| - | - | 2.077E+04 | 191.1 | - | - | 0 | - |
| - | - | 4.463E+04 | 193.1 | - | - | 0 | - |
| - | - | 2.774E+04 | 195.1 | - | - | 0 | - |
| - | - | 2.616E+04 | 196.1 | - | - | 0 | - |
| - | - | 2.744E+04 | 197.1 | - | - | 0 | - |
| - | - | 5.13E+04 | 197.1 | - | - | 0 | - |
| - | - | 5.138E+04 | 198.1 | - | - | 0 | - |
| - | - | 2.171E+04 | 198.2 | - | - | 0 | - |
| - | - | 1.847E+04 | 199.1 | - | - | 0 | - |
| - | - | 1.477E+05 | 200.1 | - | - | 0 | - |
| - | - | 4.086E+04 | 201.1 | - | - | 0 | - |
| - | - | 1.737E+04 | 201.1 | - | - | 0 | - |
| 9 | y | 9.596E+06 | 203.1 | 0.0002261 | 1.113 | +1 | 2 |
| - | - | 3.574E+04 | 204.1 | - | - | 0 | - |
| - | - | 8.312E+05 | 204.1 | - | - | 0 | - |
| - | - | 7.564E+04 | 205.1 | - | - | 0 | - |
| - | - | 2.298E+05 | 208.1 | - | - | 0 | - |
| - | - | 7.473E+04 | 208.1 | - | - | 0 | - |
| - | - | 1.76E+05 | 209.1 | - | - | 0 | - |
| - | - | 5.575E+04 | 209.1 | - | - | 0 | - |
| - | - | 8.317E+05 | 210.1 | - | - | 0 | - |
| - | - | 8.15E+04 | 211.1 | - | - | 0 | - |
| - | - | 3.416E+04 | 211.1 | - | - | 0 | - |
| - | - | 3.643E+04 | 211.2 | - | - | 0 | - |
| - | - | 6.695E+04 | 212.1 | - | - | 0 | - |
| 2 | b | 7.745E+06 | 213.2 | 0.0002951 | 1.384 | +1 | 2 |
| - | - | 2.984E+04 | 214.1 | - | - | 0 | - |
| - | - | 8.985E+05 | 214.2 | - | - | 0 | - |
| - | - | 3.972E+04 | 215.2 | - | - | 0 | - |
| - | - | 4.615E+04 | 220.1 | - | - | 0 | - |
| - | - | 4.602E+04 | 223.1 | - | - | 0 | - |
| - | - | 6.271E+06 | 226.2 | - | - | 0 | - |
| - | - | 6.883E+05 | 227.2 | - | - | 0 | - |
| 6 | y | 8.07E+05 | 228.1 | 0.0001926 | 0.8441 | +2 | 5 |
| - | - | 1.13E+04 | 228.1 | - | - | 0 | - |
| - | - | 5.303E+04 | 228.2 | - | - | 0 | - |
| - | - | 7.219E+04 | 229.1 | - | - | 0 | - |
| - | - | 5.38E+04 | 236.1 | - | - | 0 | - |
| - | - | 1.228E+05 | 238.1 | - | - | 0 | - |
| - | - | 2.3E+04 | 238.2 | - | - | 0 | - |
| - | - | 1.996E+04 | 239.2 | - | - | 0 | - |
| - | - | 5.541E+04 | 239.2 | - | - | 0 | - |
| - | - | 1.017E+05 | 240.1 | - | - | 0 | - |
| - | - | 5.84E+04 | 242.2 | - | - | 0 | - |
| - | - | 1.933E+05 | 243.2 | - | - | 0 | - |
| - | - | 8.226E+04 | 252.1 | - | - | 0 | - |
| - | - | 2.146E+05 | 252.2 | - | - | 0 | - |
| - | - | 6.571E+04 | 253.2 | - | - | 0 | - |
| - | - | 2.877E+05 | 254.2 | - | - | 0 | - |
| - | - | 2.103E+04 | 255.2 | - | - | 0 | - |
| 8 | y | 1.429E+05 | 256.1 | 0.0003036 | 1.185 | +1 | 3 |
| - | - | 5.039E+04 | 257.1 | - | - | 0 | - |
| - | - | 4.809E+04 | 261.2 | - | - | 0 | - |
| - | - | 1.879E+04 | 262.2 | - | - | 0 | - |
| - | - | 3.286E+04 | 264.1 | - | - | 0 | - |
| - | - | 4.43E+04 | 265.1 | - | - | 0 | - |
| - | - | 9.92E+04 | 266.1 | - | - | 0 | - |
| - | - | 2.109E+04 | 267.1 | - | - | 0 | - |
| - | - | 3.274E+04 | 268.1 | - | - | 0 | - |
| - | - | 2.564E+04 | 269.2 | - | - | 0 | - |
| - | - | 3.305E+05 | 269.2 | - | - | 0 | - |
| - | - | 3.381E+04 | 270.1 | - | - | 0 | - |
| 3 | b | 9.507E+05 | 270.2 | 0.0001174 | 0.4344 | +1 | 3 |
| - | - | 4.216E+04 | 270.2 | - | - | 0 | - |
| - | - | 4.957E+04 | 271.1 | - | - | 0 | - |
| - | - | 6.711E+05 | 271.2 | - | - | 0 | - |
| - | - | 9.739E+04 | 272.2 | - | - | 0 | - |
| 8 | y | 5.858E+05 | 274.1 | 0.0001454 | 0.5303 | +1 | 3 |
| - | - | 2.862E+04 | 274.7 | - | - | 0 | - |
| - | - | 7.502E+04 | 275.1 | - | - | 0 | - |
| 5 | y | 2.822E+04 | 276.7 | 6.631E-05 | 0.2397 | +2 | 6 |
| - | - | 1.201E+06 | 279.2 | - | - | 0 | - |
| - | - | 2.257E+04 | 280.2 | - | - | 0 | - |
| - | - | 1.665E+05 | 280.2 | - | - | 0 | - |
| - | - | 1.657E+05 | 281.2 | - | - | 0 | - |
| - | - | 3.021E+04 | 282.1 | - | - | 0 | - |
| - | - | 2.599E+04 | 282.2 | - | - | 0 | - |
| - | - | 1.739E+05 | 283.1 | - | - | 0 | - |
| - | - | 2.324E+04 | 284.1 | - | - | 0 | - |
| - | - | 2.737E+05 | 285.2 | - | - | 0 | - |
| 5 | y | 1.953E+05 | 285.7 | 2.244E-05 | 0.07855 | +2 | 6 |
| - | - | 5.507E+04 | 286.2 | - | - | 0 | - |
| - | - | 1.635E+06 | 287.2 | - | - | 0 | - |
| - | - | 2.42E+05 | 288.2 | - | - | 0 | - |
| - | - | 2.471E+04 | 295.2 | - | - | 0 | - |
| - | - | 2.104E+04 | 297.2 | - | - | 0 | - |
| - | - | 2.267E+04 | 297.2 | - | - | 0 | - |
| - | - | 2.28E+06 | 297.2 | - | - | 0 | - |
| - | - | 3.809E+05 | 298.2 | - | - | 0 | - |
| - | - | 4.638E+06 | 299.2 | - | - | 0 | - |
| - | - | 6.69E+05 | 300.2 | - | - | 0 | - |
| - | - | 5.475E+04 | 301.2 | - | - | 0 | - |
| - | - | 9.053E+04 | 309.2 | - | - | 0 | - |
| - | - | 2.054E+04 | 310.2 | - | - | 0 | - |
| - | - | 1.196E+05 | 314.2 | - | - | 0 | - |
| - | - | 2.453E+04 | 315.2 | - | - | 0 | - |
| - | - | 2.609E+04 | 319.2 | - | - | 0 | - |
| - | - | 6.427E+04 | 323.2 | - | - | 0 | - |
| - | - | 1.703E+05 | 323.2 | - | - | 0 | - |
| - | - | 2.49E+04 | 324.2 | - | - | 0 | - |
| - | - | 1.219E+05 | 325.2 | - | - | 0 | - |
| - | - | 6.258E+04 | 325.7 | - | - | 0 | - |
| - | - | 2.443E+04 | 326.7 | - | - | 0 | - |
| 7 | y | 3.3E+05 | 327.2 | 0.0001771 | 0.5413 | +1 | 4 |
| - | - | 8.025E+04 | 328.2 | - | - | 0 | - |
| - | - | 4.542E+04 | 331.7 | - | - | 0 | - |
| - | - | 1.253E+05 | 332.2 | - | - | 0 | - |
| - | - | 3.931E+04 | 332.7 | - | - | 0 | - |
| - | - | 3.057E+04 | 335.2 | - | - | 0 | - |
| - | - | 9.786E+04 | 336.2 | - | - | 0 | - |
| - | - | 1.852E+05 | 337.2 | - | - | 0 | - |
| - | - | 2.659E+04 | 338.2 | - | - | 0 | - |
| - | - | 1.168E+06 | 340.2 | - | - | 0 | - |
| 4 | y | 3.131E+05 | 340.7 | 0.0001554 | 0.4562 | +2 | 7 |
| - | - | 9.376E+04 | 341.2 | - | - | 0 | - |
| - | - | 2.942E+04 | 341.2 | - | - | 0 | - |
| - | - | 1.638E+05 | 341.2 | - | - | 0 | - |
| - | - | 2.71E+04 | 342.2 | - | - | 0 | - |
| 7 | y | 9.919E+04 | 345.2 | 0.000141 | 0.4084 | +1 | 4 |
| 7 | b | 3.725E+04 | 347.7 | 0.0008405 | 2.417 | +2 | 7 |
| - | - | 2.812E+04 | 349.2 | - | - | 0 | - |
| 4 | y | 9.582E+04 | 349.7 | 0.0006104 | 1.745 | +2 | 7 |
| - | - | 4.297E+04 | 350.2 | - | - | 0 | - |
| - | - | 9.605E+04 | 350.2 | - | - | 0 | - |
| - | - | 4.454E+04 | 351.2 | - | - | 0 | - |
| - | - | 1.885E+04 | 353.1 | - | - | 0 | - |
| - | - | 4.914E+04 | 353.2 | - | - | 0 | - |
| - | - | 7.521E+04 | 354.2 | - | - | 0 | - |
| - | - | 8.816E+04 | 360.7 | - | - | 0 | - |
| - | - | 2.139E+04 | 361.2 | - | - | 0 | - |
| - | - | 3.916E+04 | 361.2 | - | - | 0 | - |
| - | - | 6.813E+04 | 363.2 | - | - | 0 | - |
| - | - | 2.093E+04 | 365.2 | - | - | 0 | - |
| - | - | 4.624E+04 | 366.2 | - | - | 0 | - |
| - | - | 2.051E+04 | 366.2 | - | - | 0 | - |
| - | - | 6.074E+06 | 368.2 | - | - | 0 | - |
| - | - | 2.278E+04 | 368.7 | - | - | 0 | - |
| - | - | 1.167E+06 | 369.2 | - | - | 0 | - |
| 3 | y | 3.165E+04 | 369.7 | 4.029E-05 | 0.109 | +2 | 8 |
| - | - | 3.334E+04 | 369.7 | - | - | 0 | - |
| - | - | 4.935E+05 | 370.2 | - | - | 0 | - |
| - | - | 7.919E+04 | 371.2 | - | - | 0 | - |
| - | - | 9.269E+04 | 376.2 | - | - | 0 | - |
| - | - | 4.581E+04 | 376.2 | - | - | 0 | - |
| - | - | 3.355E+04 | 376.7 | - | - | 0 | - |
| - | - | 2.877E+04 | 377.2 | - | - | 0 | - |
| - | - | 2.409E+04 | 378.2 | - | - | 0 | - |
| 3 | y | 8.05E+04 | 378.2 | 0.001475 | 3.901 | +2 | 8 |
| - | - | 3.696E+04 | 378.7 | - | - | 0 | - |
| - | - | 2.02E+04 | 380.2 | - | - | 0 | - |
| - | - | 9.33E+04 | 380.2 | - | - | 0 | - |
| - | - | 1.321E+05 | 382.2 | - | - | 0 | - |
| - | - | 7.295E+04 | 382.7 | - | - | 0 | - |
| 8 | b | 1.04E+05 | 383.2 | 0.0004111 | 1.073 | +2 | 8 |
| - | - | 6.048E+04 | 383.7 | - | - | 0 | - |
| - | - | 3.961E+04 | 384.3 | - | - | 0 | - |
| - | - | 2.398E+04 | 386.2 | - | - | 0 | - |
| - | - | 3.714E+04 | 389.2 | - | - | 0 | - |
| - | - | 1.651E+05 | 390.2 | - | - | 0 | - |
| - | - | 2.839E+04 | 391.2 | - | - | 0 | - |
| - | - | 2.402E+04 | 393.2 | - | - | 0 | - |
| - | - | 2.091E+05 | 394.2 | - | - | 0 | - |
| - | - | 9.347E+04 | 394.2 | - | - | 0 | - |
| - | - | 2.967E+04 | 395.2 | - | - | 0 | - |
| - | - | 2.718E+04 | 395.2 | - | - | 0 | - |
| - | - | 3.539E+04 | 396.2 | - | - | 0 | - |
| - | - | 2.457E+05 | 396.2 | - | - | 0 | - |
| - | - | 3.72E+04 | 397.2 | - | - | 0 | - |
| 4 | b | 1.442E+06 | 398.2 | 0.0001031 | 0.2589 | +1 | 4 |
| - | - | 3.305E+05 | 399.2 | - | - | 0 | - |
| - | - | 5.405E+04 | 400.2 | - | - | 0 | - |
| - | - | 8.604E+04 | 407.2 | - | - | 0 | - |
| - | - | 2.4E+05 | 408.2 | - | - | 0 | - |
| - | - | 3.537E+04 | 409.2 | - | - | 0 | - |
| - | - | 2.135E+05 | 411.2 | - | - | 0 | - |
| - | - | 3.364E+04 | 411.3 | - | - | 0 | - |
| - | - | 3.202E+04 | 412.2 | - | - | 0 | - |
| - | - | 4.638E+04 | 413.2 | - | - | 0 | - |
| - | - | 1.041E+05 | 417.8 | - | - | 0 | - |
| - | - | 5.871E+04 | 418.3 | - | - | 0 | - |
| - | - | 2.933E+04 | 425.2 | - | - | 0 | - |
| - | - | 7.367E+04 | 425.3 | - | - | 0 | - |
| 2 | y | 1.562E+05 | 425.7 | 0.000126 | 0.2958 | +2 | 9 |
| 2 | y | 4.592E+04 | 426.2 | 0.0008235 | 1.932 | +2 | 9 |
| - | - | 2.673E+04 | 426.7 | - | - | 0 | - |
| - | - | 3.588E+04 | 429.2 | - | - | 0 | - |
| 9 | b | 1.993E+05 | 431.8 | 0.0001217 | 0.2818 | +2 | 9 |
| - | - | 1.512E+05 | 432.3 | - | - | 0 | - |
| - | - | 2.782E+04 | 432.8 | - | - | 0 | - |
| - | - | 3.674E+04 | 434.2 | - | - | 0 | - |
| 2 | y | 4.847E+05 | 434.7 | 0.0002757 | 0.6343 | +2 | 9 |
| - | - | 2.167E+05 | 435.2 | - | - | 0 | - |
| - | - | 6.858E+04 | 435.8 | - | - | 0 | - |
| - | - | 7.561E+04 | 437.3 | - | - | 0 | - |
| - | - | 9.241E+04 | 437.3 | - | - | 0 | - |
| - | - | 7.511E+04 | 439.2 | - | - | 0 | - |
| - | - | 5.187E+04 | 439.3 | - | - | 0 | - |
| - | - | 8.319E+04 | 439.7 | - | - | 0 | - |
| - | - | 7.609E+04 | 440.2 | - | - | 0 | - |
| - | - | 2.378E+04 | 440.3 | - | - | 0 | - |
| - | - | 3.578E+04 | 441.2 | - | - | 0 | - |
| - | - | 1.068E+05 | 442.3 | - | - | 0 | - |
| - | - | 1.791E+05 | 447.2 | - | - | 0 | - |
| - | - | 3.093E+04 | 449.2 | - | - | 0 | - |
| - | - | 4.946E+04 | 450.3 | - | - | 0 | - |
| - | - | 1.445E+05 | 451.3 | - | - | 0 | - |
| - | - | 2.72E+05 | 454.3 | - | - | 0 | - |
| - | - | 3.043E+04 | 455.3 | - | - | 0 | - |
| - | - | 7.341E+04 | 456.3 | - | - | 0 | - |
| - | - | 2.717E+04 | 457.8 | - | - | 0 | - |
| - | - | 6.275E+05 | 464.3 | - | - | 0 | - |
| - | - | 1.141E+05 | 465.2 | - | - | 0 | - |
| - | - | 5.083E+05 | 465.3 | - | - | 0 | - |
| - | - | 3.169E+04 | 466.2 | - | - | 0 | - |
| - | - | 1.131E+05 | 466.3 | - | - | 0 | - |
| - | - | 6.094E+04 | 466.8 | - | - | 0 | - |
| - | - | 2.823E+04 | 467.3 | - | - | 0 | - |
| - | - | 2.778E+05 | 467.3 | - | - | 0 | - |
| - | - | 8.717E+04 | 468.3 | - | - | 0 | - |
| - | - | 2.619E+04 | 469.3 | - | - | 0 | - |
| 6 | y | 3.898E+05 | 473.3 | 5.711E-05 | 0.1207 | +1 | 5 |
| - | - | 8.919E+04 | 474.3 | - | - | 0 | - |
| 0 | Precursor | 2.024E+05 | 475.3 | 0.0002208 | 0.4645 | +2 | -1 |
| 0 | Precursor | 1.366E+05 | 475.8 | 0.008549 | 17.97 | +2 | -1 |
| - | - | 5.324E+04 | 476.3 | - | - | 0 | - |
| - | - | 1.834E+05 | 478.3 | - | - | 0 | - |
| - | - | 4.148E+05 | 479.3 | - | - | 0 | - |
| - | - | 1.178E+05 | 480.3 | - | - | 0 | - |
| - | - | 5.638E+04 | 481.3 | - | - | 0 | - |
| - | - | 1.741E+06 | 482.3 | - | - | 0 | - |
| - | - | 4.134E+05 | 483.3 | - | - | 0 | - |
| - | - | 4.326E+04 | 484.2 | - | - | 0 | - |
| 0 | Precursor | 8.116E+05 | 484.3 | 5.671E-05 | 0.1171 | +2 | -1 |
| - | - | 4.191E+05 | 484.8 | - | - | 0 | - |
| - | - | 1.811E+05 | 485.3 | - | - | 0 | - |
| - | - | 2.945E+04 | 490.3 | - | - | 0 | - |
| - | - | 2.419E+04 | 492.3 | - | - | 0 | - |
| 5 | b | 6.705E+04 | 495.3 | 0.0009029 | 1.823 | +1 | 5 |
| - | - | 1.39E+05 | 496.3 | - | - | 0 | - |
| - | - | 3.103E+04 | 497.3 | - | - | 0 | - |
| - | - | 3.666E+04 | 507.3 | - | - | 0 | - |
| - | - | 1.846E+05 | 508.3 | - | - | 0 | - |
| - | - | 6.765E+04 | 509.3 | - | - | 0 | - |
| - | - | 1.028E+05 | 518.3 | - | - | 0 | - |
| - | - | 2.953E+04 | 519.3 | - | - | 0 | - |
| - | - | 1.093E+05 | 524.3 | - | - | 0 | - |
| - | - | 4.585E+05 | 525.3 | - | - | 0 | - |
| - | - | 1.133E+05 | 526.3 | - | - | 0 | - |
| - | - | 2.436E+04 | 527.3 | - | - | 0 | - |
| - | - | 3.22E+04 | 534.3 | - | - | 0 | - |
| - | - | 1.102E+05 | 535.3 | - | - | 0 | - |
| - | - | 5.912E+05 | 536.3 | - | - | 0 | - |
| - | - | 1.908E+05 | 537.3 | - | - | 0 | - |
| - | - | 4.243E+04 | 538.3 | - | - | 0 | - |
| - | - | 1.992E+04 | 539.3 | - | - | 0 | - |
| - | - | 2.816E+04 | 540.3 | - | - | 0 | - |
| - | - | 2.88E+04 | 551.3 | - | - | 0 | - |
| 5 | y | 3.914E+05 | 552.3 | 0.0003635 | 0.6581 | +1 | 6 |
| - | - | 3.698E+04 | 552.4 | - | - | 0 | - |
| - | - | 5.084E+06 | 553.3 | - | - | 0 | - |
| - | - | 1.49E+06 | 554.3 | - | - | 0 | - |
| - | - | 2.544E+05 | 555.3 | - | - | 0 | - |
| - | - | 4.134E+04 | 555.4 | - | - | 0 | - |
| - | - | 2.092E+04 | 556.3 | - | - | 0 | - |
| - | - | 3.378E+04 | 560.3 | - | - | 0 | - |
| - | - | 5.35E+04 | 564.4 | - | - | 0 | - |
| - | - | 2.167E+04 | 565.4 | - | - | 0 | - |
| - | - | 6.667E+04 | 567.4 | - | - | 0 | - |
| - | - | 6.913E+04 | 568.3 | - | - | 0 | - |
| 5 | y | 1.024E+07 | 570.3 | 6.392E-05 | 0.1121 | +1 | 6 |
| - | - | 3.199E+06 | 571.3 | - | - | 0 | - |
| - | - | 5.988E+05 | 572.3 | - | - | 0 | - |
| - | - | 4.254E+04 | 573.3 | - | - | 0 | - |
| - | - | 1.65E+05 | 577.3 | - | - | 0 | - |
| - | - | 7.025E+04 | 578.3 | - | - | 0 | - |
| - | - | 4.72E+04 | 579.3 | - | - | 0 | - |
| - | - | 2.446E+04 | 580.3 | - | - | 0 | - |
| - | - | 2.141E+05 | 580.4 | - | - | 0 | - |
| - | - | 6.067E+04 | 581.4 | - | - | 0 | - |
| - | - | 2.59E+05 | 595.4 | - | - | 0 | - |
| - | - | 1.013E+05 | 596.4 | - | - | 0 | - |
| - | - | 6.043E+04 | 621.4 | - | - | 0 | - |
| - | - | 5.526E+04 | 622.4 | - | - | 0 | - |
| 6 | b | 6.552E+04 | 623.4 | 0.0002949 | 0.4731 | +1 | 6 |
| - | - | 2.565E+04 | 631.4 | - | - | 0 | - |
| - | - | 2.933E+04 | 633.3 | - | - | 0 | - |
| - | - | 6.835E+04 | 637.4 | - | - | 0 | - |
| - | - | 1.351E+05 | 638.4 | - | - | 0 | - |
| - | - | 4.796E+04 | 639.4 | - | - | 0 | - |
| - | - | 2.284E+04 | 648.4 | - | - | 0 | - |
| - | - | 1.298E+05 | 649.4 | - | - | 0 | - |
| - | - | 4.017E+05 | 650.4 | - | - | 0 | - |
| - | - | 1.279E+05 | 651.4 | - | - | 0 | - |
| - | - | 6.223E+04 | 652.4 | - | - | 0 | - |
| - | - | 7.697E+04 | 654.4 | - | - | 0 | - |
| - | - | 3.522E+04 | 655.4 | - | - | 0 | - |
| - | - | 2.463E+04 | 660.4 | - | - | 0 | - |
| - | - | 1.286E+06 | 666.4 | - | - | 0 | - |
| - | - | 4.72E+05 | 667.4 | - | - | 0 | - |
| - | - | 8.184E+04 | 668.4 | - | - | 0 | - |
| - | - | 1.841E+05 | 676.4 | - | - | 0 | - |
| - | - | 8.257E+04 | 677.4 | - | - | 0 | - |
| 4 | y | 2.552E+05 | 680.4 | 4.208E-05 | 0.06185 | +1 | 7 |
| 4 | y | 3.723E+05 | 681.4 | 0.002026 | 2.974 | +1 | 7 |
| - | - | 1.153E+05 | 682.4 | - | - | 0 | - |
| 7 | b | 5.417E+05 | 694.4 | 0.000564 | 0.8122 | +1 | 7 |
| - | - | 1.825E+05 | 695.4 | - | - | 0 | - |
| - | - | 3.636E+04 | 696.4 | - | - | 0 | - |
| 4 | y | 2.608E+05 | 698.4 | 0.001513 | 2.166 | +1 | 7 |
| - | - | 1.344E+05 | 699.4 | - | - | 0 | - |
| - | - | 1.242E+05 | 720.4 | - | - | 0 | - |
| - | - | 6.415E+04 | 720.4 | - | - | 0 | - |
| - | - | 4.76E+04 | 721.4 | - | - | 0 | - |
| - | - | 2.664E+04 | 721.4 | - | - | 0 | - |
| - | - | 8.606E+04 | 730.4 | - | - | 0 | - |
| 3 | y | 3.204E+05 | 737.4 | 0.001852 | 2.512 | +1 | 8 |
| - | - | 3.637E+05 | 737.5 | - | - | 0 | - |
| 3 | y | 4.22E+05 | 738.4 | 0.003146 | 4.26 | +1 | 8 |
| - | - | 1.069E+05 | 738.5 | - | - | 0 | - |
| - | - | 1.57E+05 | 739.4 | - | - | 0 | - |
| - | - | 3.418E+04 | 739.5 | - | - | 0 | - |
| - | - | 3.051E+04 | 740.4 | - | - | 0 | - |
| - | - | 5.896E+04 | 747.4 | - | - | 0 | - |
| 8 | b | 1.724E+05 | 748.4 | 0.001078 | 1.441 | +1 | 8 |
| - | - | 1.063E+05 | 749.4 | - | - | 0 | - |
| - | - | 4.228E+04 | 750.4 | - | - | 0 | - |
| - | - | 3.467E+04 | 753.4 | - | - | 0 | - |
| - | - | 2.916E+04 | 754.4 | - | - | 0 | - |
| 3 | y | 8.858E+06 | 755.4 | 0.0009426 | 1.248 | +1 | 8 |
| - | - | 3.617E+06 | 756.4 | - | - | 0 | - |
| - | - | 8.593E+05 | 757.4 | - | - | 0 | - |
| - | - | 6.728E+04 | 758.4 | - | - | 0 | - |
| - | - | 1.369E+05 | 763.4 | - | - | 0 | - |
| - | - | 4.647E+04 | 764.4 | - | - | 0 | - |
| 8 | b | 5.026E+06 | 765.5 | 0.0007515 | 0.9818 | +1 | 8 |
| - | - | 2.099E+06 | 766.5 | - | - | 0 | - |
| - | - | 5.283E+05 | 767.5 | - | - | 0 | - |
| - | - | 4.283E+04 | 768.5 | - | - | 0 | - |
| - | - | 2.223E+04 | 791.5 | - | - | 0 | - |
| 2 | y | 3.909E+04 | 850.5 | 0.005716 | 6.721 | +1 | 9 |
| 2 | y | 8.669E+04 | 851.5 | 0.002089 | 2.454 | +1 | 9 |
| 2 | y | 1.208E+06 | 868.5 | 0.001388 | 1.599 | +1 | 9 |
| - | - | 5.732E+05 | 869.5 | - | - | 0 | - |
| - | - | 1.912E+05 | 870.5 | - | - | 0 | - |
| - | - | 2.474E+04 | 878.5 | - | - | 0 | - |
| - | - | 4.014E+04 | 880.5 | - | - | 0 | - |
| - | - | 2.016E+04 | 1269 | - | - | 0 | - |
| - | - | 2.191E+04 | 1298 | - | - | 0 | - |
| - | - | 2.268E+04 | 1451 | - | - | 0 | - |
| - | - | 2.277E+04 | 3159 | - | - | 0 | - |

m/z Charge Intensity FragmentType MassShift Position
124.03959655761719 0 33620.836
125.10774230957031 0 19754.688
126.71277618408203 0 13839.046
127.08695983886719 0 28118.783
129.0661163330078 0 322274.75
129.10250854492188 0 2686553.5
130.10589599609375 0 155886.42
137.10745239257812 0 26651.81
139.08694458007812 0 102997.46
141.066162109375 0 171619.95
141.10252380371094 0 185608.69
142.0693817138672 0 28191.156
142.08685302734375 0 18136.213
143.08168029785156 0 87129.18
143.11814880371094 0 106475.02 d 1
145.17193603515625 0 12927.268
152.03443908691406 0 188879.36
152.13128662109375 0 13575.505
153.06614685058594 0 39268.465
153.10235595703125 0 55397.34
155.1181182861328 0 323468.53
156.1213836669922 0 17899.021
157.09725952148438 0 100472.01
157.1702117919922 0 42331.637
157.2620849609375 0 14344.929
158.0926055908203 0 133879.52
161.09190368652344 0 15976.411
163.1231689453125 0 15623.743
165.10243225097656 0 62475.387
167.11842346191406 0 26448.344
168.5124053955078 0 15794.227
169.06101989746094 0 812025.4
169.0973663330078 0 294515.12
169.13412475585938 0 49211.555
170.0644989013672 0 38644.168
170.10107421875 0 17479.352
171.1129913330078 0 458065.4
172.11627197265625 0 61542.914
172.14483642578125 0 31481.342
176.11256408691406 0 16425.055
180.11328125 0 80788.445
181.0974884033203 0 239788.58
181.13377380371094 0 167976.11
182.1022186279297 0 21016.588
182.1290740966797 0 668206.94
183.11328125 0 112453.78
183.1323699951172 0 65720.445
183.14918518066406 0 38922.504
185.09226989746094 0 252988.61 y Water loss 8
185.16517639160156 0 15386081 a 1
186.08750915527344 0 1921999
186.16844177246094 0 1540489.1
187.09091186523438 0 169606
187.10800170898438 0 25238.518
187.17153930664062 0 70126.266
188.13955688476562 0 430085.66
191.08168029785156 0 40431.363
191.11766052246094 0 20771.629
193.0974884033203 0 44633.008
195.11354064941406 0 27735.73
196.10870361328125 0 26156.02
197.0920867919922 0 27439.23
197.12879943847656 0 51300.71
198.12399291992188 0 51383.86
198.15997314453125 0 21714.502
199.1074981689453 0 18467.037
200.13961791992188 0 147721.48
201.08706665039062 0 40856.195
201.1430206298828 0 17371.814
203.1028594970703 0 9595937 y 8
204.0964813232422 0 35739.24
204.10623168945312 0 831176.56
205.10755920410156 0 75643.71
208.10818481445312 0 229835.72
208.14459228515625 0 74727.11
209.09237670898438 0 175981.1
209.12860107421875 0 55750.09
210.12387084960938 0 831694.3
211.12722778320312 0 81504.4
211.14352416992188 0 34160.902
211.1801300048828 0 36428.06
212.10299682617188 0 66946.875
213.16004943847656 0 7744888.5 b 1
214.11875915527344 0 29839.521
214.16331481933594 0 898503.1
215.1658172607422 0 39720.117
220.10816955566406 0 46148.023
223.1079559326172 0 46022.91
226.15524291992188 0 6270895.5
227.1585235595703 0 688349
228.13446044921875 0 806963.25 y Water loss 5
228.14491271972656 0 11295.04
228.1612548828125 0 53026.08
229.137939453125 0 72193.734
236.13929748535156 0 53799.836
238.1189727783203 0 122834.21
238.15493774414062 0 22999.049
239.15060424804688 0 19963.527
239.1756591796875 0 55408.79
240.1343231201172 0 101712.06
242.1500701904297 0 58396.03
243.18177795410156 0 193275.58
252.134765625 0 82260.47
252.17086791992188 0 214569.47
253.166259765625 0 65712.875
254.1500244140625 0 287699.2
255.15078735351562 0 21028.754
256.1294860839844 0 142948.25 y Water loss 7
257.1247253417969 0 50389.664
261.17108154296875 0 48093.684
262.1563720703125 0 18794.1
264.13531494140625 0 32864.05
265.12945556640625 0 44301.844
266.1134948730469 0 99195.35
267.1470031738281 0 21090.613
268.129150390625 0 32739.703
269.1609802246094 0 25642.646
269.1972351074219 0 330514.25
270.1459655761719 0 33810.875
270.18133544921875 0 950692.1 b 2
270.1980285644531 0 42162.33
271.140380859375 0 49570.355
271.1767883300781 0 671128.44
272.1799621582031 0 97393.27
274.139892578125 0 585792.2 y 7
274.6726989746094 0 28622.04
275.1436462402344 0 75022.32
276.66058349609375 0 28217.848 y Water loss 4
279.1817321777344 0 1201005.8
280.1676025390625 0 22565.662
280.18463134765625 0 166517.7
281.16094970703125 0 165665.36
282.14617919921875 0 30206.416
282.16424560546875 0 25987.523
283.14019775390625 0 173940.77
284.1431884765625 0 23240.648
285.15594482421875 0 273725.78
285.66595458984375 0 195339.75 y 4
286.16754150390625 0 55070.984
287.2080383300781 0 1635033.6
288.21142578125 0 241992.47
295.213623046875 0 24710.498
297.1548767089844 0 21041.71
297.1595764160156 0 22672.092
297.1923828125 0 2279502.2
298.1955261230469 0 380927.84
299.1716613769531 0 4637800.5
300.1744689941406 0 668968.06
301.1763000488281 0 54750.1
309.15606689453125 0 90530.41
310.2115783691406 0 20539.168
314.2188720703125 0 119582.59
315.2236328125 0 24526.615
319.1747741699219 0 26092.738
323.17138671875 0 64274.367
323.2078552246094 0 170250.77
324.2125244140625 0 24895.26
325.187255859375 0 121913.484
325.68450927734375 0 62580.094
326.7111511230469 0 24432.416
327.1664733886719 0 330009.34 y Water loss 6
328.1634826660156 0 80252.09
331.6853332519531 0 45422.52
332.177490234375 0 125284.16
332.6777648925781 0 39306.535
335.20819091796875 0 30568.217
336.20294189453125 0 97861.88
337.187255859375 0 185217.92
338.1901550292969 0 26588.17
340.2345275878906 0 1168333.5
340.6900939941406 0 313087.5 y Water loss 3
341.1896667480469 0 93762.79
341.2176513671875 0 29416.033
341.23797607421875 0 163832.66
342.2424621582031 0 27101.725
345.177001953125 0 99189.9 y 6
347.716796875 0 37251.816 b 6
349.187744140625 0 28116.43
349.6958312988281 0 95821.07 y 3
350.19671630859375 0 42968.25
350.2187805175781 0 96045.59
351.20208740234375 0 44543.707
353.129638671875 0 18847.207
353.218017578125 0 49142.953
354.2143249511719 0 75212.45
360.72283935546875 0 88163.54
361.1814270019531 0 21385.182
361.22332763671875 0 39158.28
363.2027893066406 0 68131.58
365.2205810546875 0 20930.871
366.213134765625 0 46240.562
366.24969482421875 0 20507.309
368.22918701171875 0 6074268.5
368.734619140625 0 22775.49
369.2324523925781 0 1167039.5
369.6927185058594 0 31653.691 y Ammonia loss 2
369.7382507324219 0 33335.664
370.2438049316406 0 493514.47
371.2457580566406 0 79194.08
376.1980285644531 0 92685.18
376.2443542480469 0 45806.38
376.74566650390625 0 33546.434
377.20343017578125 0 28772.293
378.1792907714844 0 24090.17
378.2074279785156 0 80497.34 y 2
378.7069396972656 0 36958.062
380.1943359375 0 20201.645
380.2294616699219 0 93299.67
382.2269287109375 0 132074
382.72772216796875 0 72948.07
383.23492431640625 0 104030 b 7
383.7347717285156 0 60477.09
384.2622375488281 0 39607.723
386.2406921386719 0 23976.072
389.23004150390625 0 37142.434
390.2137756347656 0 165115.3
391.21759033203125 0 28392.354
393.22711181640625 0 24021.725
394.20892333984375 0 209100.48
394.2450256347656 0 93470.336
395.2132263183594 0 29665.783
395.2478942871094 0 27183.12
396.1919250488281 0 35391.562
396.2239074707031 0 245723.28
397.2274169921875 0 37202.164
398.2398986816406 0 1442311.1 b 3
399.2431335449219 0 330523.6
400.2441711425781 0 54047.406
407.2408752441406 0 86036.805
408.22467041015625 0 239983.25
409.22589111328125 0 35370.824
411.23504638671875 0 213477.94
411.26751708984375 0 33637.695
412.2358093261719 0 32019.984
413.2154541015625 0 46381.434
417.7640075683594 0 104072.305
418.2652587890625 0 58708.465
425.2165832519531 0 29325.268
425.25103759765625 0 73665.805
425.7428283691406 0 156243.47 y Water loss 1
426.23388671875 0 45918.39 y Ammonia loss 1
426.74176025390625 0 26727.887
429.22357177734375 0 35880.332
431.7610168457031 0 199253.75 b 8
432.2626037597656 0 151227.19
432.76470947265625 0 27823.146
434.2401123046875 0 36741.027
434.7482604980469 0 484704.94 y 1
435.24884033203125 0 216729.48
435.75091552734375 0 68577.44
437.2511901855469 0 75606.83
437.2874755859375 0 92412.47
439.23089599609375 0 75107.9
439.30267333984375 0 51868.09
439.7409362792969 0 83193.97
440.2395324707031 0 76094.07
440.3056335449219 0 23781.975
441.2442626953125 0 35782.914
442.27734375 0 106814.56
447.2354431152344 0 179106.8
449.2144775390625 0 30925.467
450.2772216796875 0 49460.72
451.2666015625 0 144495.36
454.27783203125 0 271977.72
455.2817687988281 0 30432.78
456.25714111328125 0 73414.82
457.7586364746094 0 27167.508
464.2617492675781 0 627477.8
465.2475280761719 0 114086.055
465.28143310546875 0 508268.88
466.24542236328125 0 31692.684
466.28411865234375 0 113131.01
466.76318359375 0 60939.727
467.2623596191406 0 28229.914
467.2974548339844 0 277845.84
468.2989501953125 0 87172.05
469.27728271484375 0 26194.588
473.2718811035156 0 389832.75 y 5
474.2748718261719 0 89190.41
475.2771301269531 0 202404.83 Precursor Water loss
475.7774658203125 0 136636.31 Precursor Ammonia loss
476.2776184082031 0 53237.09
478.2774658203125 0 183416.2
479.26190185546875 0 414802.72
480.26483154296875 0 117793.03
481.3121032714844 0 56384.055
482.2723083496094 0 1741092.5
483.2754211425781 0 413350.25
484.2442626953125 0 43257.06
484.2821350097656 0 811617.4 Precursor
484.78375244140625 0 419078.8
485.2853698730469 0 181142
490.2771911621094 0 29454.107
492.2635498046875 0 24192.33
495.2916564941406 0 67054.984 b 4
496.288330078125 0 139007.72
497.2931823730469 0 31026.967
507.2950744628906 0 36655.164
508.2884216308594 0 184569.88
509.2905578613281 0 67654.734
518.2714233398438 0 102758.97
519.2737426757812 0 29533.006
524.319091796875 0 109333.93
525.314453125 0 458464.88
526.3168334960938 0 113255.73
527.3215942382812 0 24357.809
534.3067016601562 0 32201.68
535.298095703125 0 110162.21
536.282958984375 0 591166.06
537.2860107421875 0 190827.86
538.2847900390625 0 42432.938
539.330810546875 0 19919.412
540.3154907226562 0 28162.33
551.332763671875 0 28798.756
552.3136596679688 0 391386.94 y Water loss 4
552.3882446289062 0 36975.68
553.3092041015625 0 5083961
554.3118896484375 0 1490345
555.314453125 0 254386.97
555.3599243164062 0 41337.87
556.3168334960938 0 20921.53
560.317626953125 0 33780.445
564.3509521484375 0 53502.332
565.3510131835938 0 21669.328
567.3594970703125 0 66670.625
568.3080444335938 0 69130.586
570.3245239257812 0 10237573 y 4
571.3272705078125 0 3199337
572.3297729492188 0 598815.75
573.330322265625 0 42541.168
577.3460083007812 0 165001.45
578.3432006835938 0 70251.78
579.326416015625 0 47200.176
580.3327026367188 0 24459.67
580.3812866210938 0 214066.58
581.3849487304688 0 60670.105
595.3558349609375 0 259026.53
596.359375 0 101281.086
621.3733520507812 0 60432.67
622.3679809570312 0 55260.754
623.3878173828125 0 65521.203 b 5
631.3587036132812 0 25645.51
633.3358154296875 0 29332.697
637.4016723632812 0 68349.33
638.3982543945312 0 135115.4
639.4017333984375 0 47955.703
648.3837890625 0 22839.63
649.364990234375 0 129801.73
650.3629760742188 0 401714.16
651.3655395507812 0 127921.305
652.3749389648438 0 62231.844
654.4288940429688 0 76966.6
655.4315795898438 0 35223.145
660.3931884765625 0 24630.512
666.3927001953125 0 1286384.6
667.395751953125 0 472005.03
668.398193359375 0 81837.09
676.4136962890625 0 184052.62
677.4149169921875 0 82570.875
680.37255859375 0 255211.4 y Water loss 3
681.358642578125 0 372305.28 y Ammonia loss 3
682.3594970703125 0 115278.83
694.424072265625 0 541695.1 b 6
695.4273071289062 0 182505.19
696.4296264648438 0 36361.79
698.3816528320312 0 260776.8 y 3
699.385498046875 0 134447.3
720.3668212890625 0 124193.18
720.4422607421875 0 64147.79
721.3687744140625 0 47595.215
721.441162109375 0 26643.885
730.42431640625 0 86057.5
737.3922119140625 0 320384.8 y Water loss 2
737.46630859375 0 363713.44
738.3812255859375 0 422029.47 y Ammonia loss 2
738.4695434570312 0 106939.71
739.3812255859375 0 156981.88
739.475341796875 0 34182.59
740.3847045898438 0 30510.717
747.449462890625 0 58960.3
748.436279296875 0 172359.58 b Ammonia loss 7
749.4353637695312 0 106317.875
750.4375 0 42279.215
753.38525390625 0 34671.71
754.3917236328125 0 29158.695
755.4036865234375 0 8857588 y 2
756.4066772460938 0 3616700.8
757.408935546875 0 859323.7
758.410400390625 0 67279.836
763.446044921875 0 136870.02
764.4465942382812 0 46474.36
765.4609985351562 0 5026282.5 b 7
766.463623046875 0 2099470.2
767.4666137695312 0 528316.7
768.4680786132812 0 42825.71
791.4734497070312 0 22225.863
850.472412109375 0 39088.746 y Water loss 1
851.4642333984375 0 86689.84 y Ammonia loss 1
868.4873046875 0 1208113.8 y 1
869.4899291992188 0 573211.8
870.4922485351562 0 191151.22
878.4721069335938 0 24741.365
880.5302124023438 0 40141.59
1268.6082763671875 0 20156.91
1298.07080078125 0 21912.977
1451.4495849609375 0 22679.053
3159.41259765625 0 22767.188

Spectrum Details

|  |  |
| --- | --- |
| Matched peaks? Matched peaks The total absolute number of peaks matched. Additionally in brackets the total fraction of peaks matched and the total number of peaks is shown. | 44 (10.81% of 407) |
| FDR? FDR The false discovery rate estimated for this peptide. It is calculated by matching all theoretical fragments with a non-integer shift with the raw peaks for this spectrum. This is done with 40 different shifts. The resulting percentage is the average number of annotated peaks over the number of annotated peaks with the correct spectrum. | 0.05% |
| Satellite FDR? Satellite FDR See the FDR for details on its calculation. This satellite ion specific FDR only contains the satellite ions (d/w) for I/L/J positions. | 0.00% |
| PSM Score? PSM Score The PSM Score as given by Hecklib to this annotated spectrum. It is shown with three significant figures. | 431 |

#### Spectrum 3209? Spectrum 3209 The raw spectrum of this peptide as annotated by Hecklib. The fragments are coloured according to ion type (see legend). Any peaks with a star '\*' as text can be hovered over to see the full details, first the ion type second the mass shift type. By hovering over the amino acids in the peptide or ions in the legend the corresponding peaks are highlighted. By toggling the 'Unassigned' label you can turn the background (unassigned) peaks on or off in the plot. By updating the slider in the Ion legend you can update the spectrum to only show the top X% of the peaks with labels. The top X% means any peak that is within X% of the highest intensity. By dragging in the spectrum you can zoom in to a specific part of the spectrum and use 'Zoom Out' to get back to the original zoom level. The annotation of the spectrum is based on the given sequence in the peptides file and is done with different software so inconsistencies are likely. The peaks are annotated based on the given sequence, with 20 ppm tolerance.

Copy Data

##### Spectrum 3209 (TSV)

###### Preview

```
Loading example...
```

*Click on the button to copy the data to your clipboard.*

Mz MinMz MaxIntensity Max

WidthHeightPeptide font sizePeptide stroke widthSpectrum font sizeSpectrum stroke widthCompact peptide

Ion legend

wxyz

abcd

OtherUnassignedIon^Charge^_Position_Show for top:%

VLGQPKAAPS

03.86e+47.71e+41.16e+51.54e+5

Zoom Out

y^+1^_2_y^+1^_3_y^+2^_6_c^+1^_3_z^+2^_7_y^+2^_7_y^+2^_8_c^+1^_4_c^+2^_9_y^+2^_9_z^+1^_5_y^+1^_5_c^+1^_5_y^+1^_6_z^+1^_6_y^+1^_6_w^+1^_7_c^+1^_6_y^+1^_7_z^+1^_7_c^+1^_7_y^+1^_7_c^+1^_7_y^+1^_8_z^+1^_8_y^+1^_8_c^+1^_8_w^+1^_9_y^+1^_9_z^+1^_9_y^+1^_9_c^+1^_9_

0758151522733031

Fragment Matches Table

Show background peaks

| Position | Ion type | Intensity | mz Theoretical | mz Error (Th) | mz Error (ppm) | Charge | Series Number |
| --- | --- | --- | --- | --- | --- | --- | --- |
| - | - | 399.9 | 121.3 | - | - | 0 | - |
| - | - | 398.2 | 122.1 | - | - | 0 | - |
| - | - | 432.8 | 123.7 | - | - | 0 | - |
| - | - | 518.7 | 144.2 | - | - | 0 | - |
| - | - | 541 | 148.9 | - | - | 0 | - |
| - | - | 723.2 | 148.9 | - | - | 0 | - |
| - | - | 553.5 | 148.9 | - | - | 0 | - |
| - | - | 747.1 | 148.9 | - | - | 0 | - |
| - | - | 1088 | 148.9 | - | - | 0 | - |
| - | - | 1191 | 148.9 | - | - | 0 | - |
| - | - | 1042 | 148.9 | - | - | 0 | - |
| - | - | 1820 | 148.9 | - | - | 0 | - |
| - | - | 3717 | 148.9 | - | - | 0 | - |
| - | - | 4764 | 149 | - | - | 0 | - |
| - | - | 2848 | 149 | - | - | 0 | - |
| - | - | 1483 | 149 | - | - | 0 | - |
| - | - | 1189 | 149 | - | - | 0 | - |
| - | - | 1183 | 149 | - | - | 0 | - |
| - | - | 902.2 | 149 | - | - | 0 | - |
| - | - | 696 | 149 | - | - | 0 | - |
| - | - | 568.7 | 149 | - | - | 0 | - |
| - | - | 796.4 | 149 | - | - | 0 | - |
| - | - | 496.3 | 149 | - | - | 0 | - |
| - | - | 463.4 | 149 | - | - | 0 | - |
| - | - | 502 | 149 | - | - | 0 | - |
| - | - | 507 | 169.1 | - | - | 0 | - |
| - | - | 985.1 | 173.4 | - | - | 0 | - |
| - | - | 428.5 | 173.7 | - | - | 0 | - |
| - | - | 3.177E+04 | 185.2 | - | - | 0 | - |
| - | - | 2728 | 186.1 | - | - | 0 | - |
| - | - | 3080 | 186.2 | - | - | 0 | - |
| - | - | 474.2 | 188.6 | - | - | 0 | - |
| - | - | 511.4 | 191.8 | - | - | 0 | - |
| 9 | y | 2.133E+04 | 203.1 | 2.773E-05 | 0.1365 | +1 | 2 |
| - | - | 1686 | 204.1 | - | - | 0 | - |
| - | - | 464.8 | 204.8 | - | - | 0 | - |
| - | - | 2.517E+04 | 213.2 | - | - | 0 | - |
| - | - | 2415 | 214.2 | - | - | 0 | - |
| - | - | 2422 | 226.2 | - | - | 0 | - |
| - | - | 553.6 | 244.4 | - | - | 0 | - |
| - | - | 539.9 | 245.9 | - | - | 0 | - |
| - | - | 2839 | 270.2 | - | - | 0 | - |
| - | - | 960.4 | 271.1 | - | - | 0 | - |
| 8 | y | 1095 | 274.1 | 7.199E-06 | 0.02626 | +1 | 3 |
| - | - | 568.2 | 283.1 | - | - | 0 | - |
| 5 | y | 623.8 | 285.7 | 0.0005412 | 1.895 | +2 | 6 |
| 3 | c | 3233 | 287.2 | 0.0003017 | 1.05 | +1 | 3 |
| - | - | 3347 | 299.2 | - | - | 0 | - |
| 4 | z | 574.7 | 332.7 | 0.00269 | 8.085 | +2 | 7 |
| 4 | y | 706.3 | 340.7 | 0.0007296 | 2.141 | +2 | 7 |
| - | - | 5853 | 368.2 | - | - | 0 | - |
| - | - | 1152 | 369.2 | - | - | 0 | - |
| - | - | 1323 | 370.2 | - | - | 0 | - |
| 3 | y | 582 | 378.2 | 0.0001422 | 0.376 | +2 | 8 |
| 4 | c | 4702 | 398.2 | 4.206E-05 | 0.1056 | +1 | 4 |
| - | - | 1084 | 399.2 | - | - | 0 | - |
| - | - | 580.4 | 401.6 | - | - | 0 | - |
| - | - | 679.5 | 418.3 | - | - | 0 | - |
| 9 | c | 621.8 | 431.8 | 0.0003353 | 0.7766 | +2 | 9 |
| 2 | y | 2137 | 434.7 | 0.0004872 | 1.121 | +2 | 9 |
| - | - | 933.5 | 435.2 | - | - | 0 | - |
| - | - | 624.5 | 452.7 | - | - | 0 | - |
| 6 | z | 3.253E+04 | 457.3 | 1.765E-05 | 0.0386 | +1 | 5 |
| - | - | 2.747E+04 | 458.3 | - | - | 0 | - |
| - | - | 6283 | 459.3 | - | - | 0 | - |
| - | - | 721.1 | 460.3 | - | - | 0 | - |
| - | - | 693.6 | 465.3 | - | - | 0 | - |
| - | - | 758.5 | 472.3 | - | - | 0 | - |
| 6 | y | 5283 | 473.3 | 0.000187 | 0.3952 | +1 | 5 |
| - | - | 1123 | 474.3 | - | - | 0 | - |
| - | - | 728.6 | 475.3 | - | - | 0 | - |
| - | - | 1932 | 482.3 | - | - | 0 | - |
| - | - | 1015 | 483.3 | - | - | 0 | - |
| - | - | 3465 | 484.3 | - | - | 0 | - |
| - | - | 1358 | 484.8 | - | - | 0 | - |
| - | - | 1409 | 485.3 | - | - | 0 | - |
| - | - | 1565 | 485.4 | - | - | 0 | - |
| - | - | 1151 | 489.3 | - | - | 0 | - |
| 5 | c | 1961 | 512.3 | 0.0001664 | 0.3247 | +1 | 5 |
| - | - | 850 | 513.3 | - | - | 0 | - |
| 5 | y | 1185 | 552.3 | 0.0009738 | 1.763 | +1 | 6 |
| - | - | 7179 | 553.3 | - | - | 0 | - |
| 5 | z | 2695 | 554.3 | 0.00258 | 4.655 | +1 | 6 |
| - | - | 1002 | 555.3 | - | - | 0 | - |
| - | - | 2029 | 568.3 | - | - | 0 | - |
| - | - | 1076 | 568.4 | - | - | 0 | - |
| - | - | 1.256E+04 | 569.3 | - | - | 0 | - |
| 5 | y | 6.197E+04 | 570.3 | 0.0005522 | 0.9682 | +1 | 6 |
| - | - | 1.608E+04 | 571.3 | - | - | 0 | - |
| - | - | 2470 | 572.3 | - | - | 0 | - |
| - | - | 673.9 | 597.4 | - | - | 0 | - |
| - | - | 555.8 | 600.5 | - | - | 0 | - |
| - | - | 1124 | 608.4 | - | - | 0 | - |
| - | - | 558.3 | 609.4 | - | - | 0 | - |
| 4 | w | 4.229E+04 | 624.3 | 0.0004357 | 0.6979 | +1 | 7 |
| - | - | 1.565E+04 | 625.3 | - | - | 0 | - |
| - | - | 3318 | 626.3 | - | - | 0 | - |
| - | - | 4419 | 639.4 | - | - | 0 | - |
| 6 | c | 6.252E+04 | 640.4 | 0.0008025 | 1.253 | +1 | 6 |
| - | - | 723 | 640.8 | - | - | 0 | - |
| - | - | 2.896E+04 | 641.4 | - | - | 0 | - |
| - | - | 6886 | 642.4 | - | - | 0 | - |
| - | - | 682.3 | 643.4 | - | - | 0 | - |
| - | - | 702.9 | 650.4 | - | - | 0 | - |
| - | - | 1522 | 666.4 | - | - | 0 | - |
| - | - | 637.3 | 667.4 | - | - | 0 | - |
| - | - | 1020 | 668.3 | - | - | 0 | - |
| - | - | 856.5 | 677.4 | - | - | 0 | - |
| 4 | y | 2333 | 681.4 | 0.0005004 | 0.7345 | +1 | 7 |
| - | - | 1112 | 682.3 | - | - | 0 | - |
| 4 | z | 2.088E+04 | 682.4 | 0.0006718 | 0.9845 | +1 | 7 |
| - | - | 1.011E+04 | 683.4 | - | - | 0 | - |
| - | - | 2858 | 684.4 | - | - | 0 | - |
| 7 | c | 1093 | 694.4 | 0.001968 | 2.834 | +1 | 7 |
| 4 | y | 9010 | 698.4 | 0.001146 | 1.641 | +1 | 7 |
| - | - | 4423 | 699.4 | - | - | 0 | - |
| - | - | 2114 | 700.4 | - | - | 0 | - |
| - | - | 652.5 | 701 | - | - | 0 | - |
| - | - | 2128 | 710.4 | - | - | 0 | - |
| 7 | c | 4.185E+04 | 711.5 | 0.0008069 | 1.134 | +1 | 7 |
| - | - | 2.263E+04 | 712.4 | - | - | 0 | - |
| - | - | 5609 | 713.4 | - | - | 0 | - |
| - | - | 847.7 | 721.4 | - | - | 0 | - |
| 3 | y | 1500 | 738.4 | 0.001864 | 2.524 | +1 | 8 |
| - | - | 1128 | 738.5 | - | - | 0 | - |
| 3 | z | 3340 | 739.4 | 0.001322 | 1.789 | +1 | 8 |
| - | - | 1624 | 740.4 | - | - | 0 | - |
| - | - | 862.5 | 741.4 | - | - | 0 | - |
| 3 | y | 4.035E+04 | 755.4 | 0.001004 | 1.329 | +1 | 8 |
| - | - | 2.375E+04 | 756.4 | - | - | 0 | - |
| - | - | 7853 | 757.4 | - | - | 0 | - |
| 8 | c | 1.52E+04 | 765.5 | 0.0007515 | 0.9818 | +1 | 8 |
| - | - | 6288 | 766.5 | - | - | 0 | - |
| - | - | 2120 | 767.5 | - | - | 0 | - |
| - | - | 750 | 781.4 | - | - | 0 | - |
| - | - | 619.2 | 793.4 | - | - | 0 | - |
| 2 | w | 9447 | 809.4 | 0.000704 | 0.8698 | +1 | 9 |
| - | - | 5393 | 810.4 | - | - | 0 | - |
| - | - | 1182 | 811.4 | - | - | 0 | - |
| - | - | 1034 | 820.5 | - | - | 0 | - |
| - | - | 736.6 | 821.5 | - | - | 0 | - |
| - | - | 2299 | 835.5 | - | - | 0 | - |
| - | - | 1165 | 836.5 | - | - | 0 | - |
| 2 | y | 873.1 | 851.5 | 0.01295 | 15.21 | +1 | 9 |
| 2 | z | 1.605E+04 | 852.5 | 0.0009749 | 1.144 | +1 | 9 |
| - | - | 9566 | 853.5 | - | - | 0 | - |
| - | - | 3331 | 854.5 | - | - | 0 | - |
| - | - | 615.4 | 855.5 | - | - | 0 | - |
| - | - | 930.4 | 864.5 | - | - | 0 | - |
| - | - | 712.5 | 865.5 | - | - | 0 | - |
| 2 | y | 3038 | 868.5 | 0.001877 | 2.161 | +1 | 9 |
| - | - | 2633 | 869.5 | - | - | 0 | - |
| - | - | 1055 | 870.5 | - | - | 0 | - |
| 9 | c | 5.366E+04 | 879.5 | 0.001512 | 1.719 | +1 | 9 |
| - | - | 3.356E+04 | 880.5 | - | - | 0 | - |
| - | - | 1.372E+04 | 881.5 | - | - | 0 | - |
| - | - | 2516 | 882.5 | - | - | 0 | - |
| - | - | 1241 | 889.5 | - | - | 0 | - |
| - | - | 7990 | 895.5 | - | - | 0 | - |
| - | - | 6701 | 896.5 | - | - | 0 | - |
| - | - | 1389 | 897.5 | - | - | 0 | - |
| - | - | 1606 | 897.5 | - | - | 0 | - |
| - | - | 1280 | 898.5 | - | - | 0 | - |
| - | - | 1529 | 906.6 | - | - | 0 | - |
| - | - | 1824 | 907.6 | - | - | 0 | - |
| - | - | 2981 | 912.5 | - | - | 0 | - |
| - | - | 1468 | 913.5 | - | - | 0 | - |
| - | - | 4011 | 922.6 | - | - | 0 | - |
| - | - | 8495 | 923.5 | - | - | 0 | - |
| - | - | 4874 | 924.5 | - | - | 0 | - |
| - | - | 1202 | 925.6 | - | - | 0 | - |
| - | - | 770.7 | 949.5 | - | - | 0 | - |
| - | - | 2649 | 950.5 | - | - | 0 | - |
| - | - | 5.272E+04 | 951.5 | - | - | 0 | - |
| - | - | 3.963E+04 | 952.5 | - | - | 0 | - |
| - | - | 1.407E+04 | 953.5 | - | - | 0 | - |
| - | - | 1689 | 954.5 | - | - | 0 | - |
| - | - | 683.7 | 959.9 | - | - | 0 | - |
| - | - | 624.7 | 965.4 | - | - | 0 | - |
| - | - | 588.4 | 966.4 | - | - | 0 | - |
| - | - | 3036 | 966.5 | - | - | 0 | - |
| - | - | 5.114E+04 | 967.6 | - | - | 0 | - |
| - | - | 1.527E+05 | 968.6 | - | - | 0 | - |
| - | - | 9.477E+04 | 969.6 | - | - | 0 | - |
| - | - | 3.081E+04 | 970.6 | - | - | 0 | - |
| - | - | 1421 | 971.4 | - | - | 0 | - |
| - | - | 2862 | 971.6 | - | - | 0 | - |
| - | - | 598.5 | 1013 | - | - | 0 | - |
| - | - | 686.4 | 1256 | - | - | 0 | - |
| - | - | 572.6 | 1270 | - | - | 0 | - |
| - | - | 648.8 | 1440 | - | - | 0 | - |
| - | - | 720.3 | 1451 | - | - | 0 | - |
| - | - | 725.4 | 1926 | - | - | 0 | - |
| - | - | 753.2 | 2321 | - | - | 0 | - |
| - | - | 649.1 | 2868 | - | - | 0 | - |
| - | - | 670 | 2966 | - | - | 0 | - |
| - | - | 666.8 | 3001 | - | - | 0 | - |

m/z Charge Intensity FragmentType MassShift Position
121.26897430419922 0 399.94296
122.05690002441406 0 398.21396
123.68619537353516 0 432.8424
144.22105407714844 0 518.6968
148.86915588378906 0 540.9978
148.88421630859375 0 723.1898
148.89083862304688 0 553.4592
148.90585327148438 0 747.1313
148.91275024414062 0 1088.3942
148.91970825195312 0 1191.4226
148.92782592773438 0 1041.5468
148.93418884277344 0 1819.8479
148.94200134277344 0 3717.199
148.95860290527344 0 4764.3247
148.96633911132812 0 2847.7136
148.9735107421875 0 1482.5248
148.9810028076172 0 1188.9152
148.98812866210938 0 1182.7748
148.99546813964844 0 902.17377
149.0027618408203 0 695.9736
149.01675415039062 0 568.6791
149.02398681640625 0 796.4259
149.03182983398438 0 496.29672
149.03903198242188 0 463.38724
149.0460662841797 0 502.0303
169.06051635742188 0 507.0384
173.44992065429688 0 985.10077
173.7084503173828 0 428.54535
185.1649169921875 0 31766.607
186.0873260498047 0 2727.5024
186.1683807373047 0 3079.9492
188.62820434570312 0 474.2298
191.760986328125 0 511.36826
203.1026611328125 0 21333.07 y 8
204.1060791015625 0 1686.0881
204.79876708984375 0 464.84503
213.15972900390625 0 25172.771
214.16314697265625 0 2415.0164
226.1553192138672 0 2421.9058
244.39085388183594 0 553.60547
245.94158935546875 0 539.9375
270.1812744140625 0 2838.7563
271.1400146484375 0 960.3866
274.1397399902344 0 1095.0277 y 7
283.14031982421875 0 568.15576
285.6664733886719 0 623.8161 y 4
287.20806884765625 0 3232.897 c 2
299.1714782714844 0 3346.91
332.6778869628906 0 574.68256 z Water loss 3
340.689208984375 0 706.2769 y Water loss 3
368.2289123535156 0 5852.9956
369.2328796386719 0 1152.4508
370.2439880371094 0 1322.9539
378.205810546875 0 581.96277 y 2
398.2398376464844 0 4702.2695 c Ammonia loss 3
399.2220764160156 0 1084.39
401.5960998535156 0 580.43176
418.2646789550781 0 679.54144
431.76123046875 0 621.78125 c Ammonia loss 8
434.74749755859375 0 2136.989 y 1
435.2428894042969 0 933.45276
452.6839599609375 0 624.5099
457.2530822753906 0 32529.293 z 5
458.259765625 0 27466.57
459.2628173828125 0 6282.591
460.2674865722656 0 721.0551
465.2834777832031 0 693.64703
472.26287841796875 0 758.4692
473.2716369628906 0 5282.7725 y 5
474.2737731933594 0 1122.6906
475.2784118652344 0 728.6115
482.2716064453125 0 1931.9781
483.2574462890625 0 1015.19415
484.28204345703125 0 3465.468
484.7825622558594 0 1358.1621
485.2785949707031 0 1408.5006
485.3578796386719 0 1565.3635
489.2787170410156 0 1151.2761
512.3192749023438 0 1960.6471 c 4
513.3184204101562 0 850.03595
552.3130493164062 0 1185.1483 y Water loss 4
553.309326171875 0 7178.713
554.3032836914062 0 2695.226 z 4
555.2942504882812 0 1002.11285
568.309326171875 0 2029.4865
568.3680419921875 0 1076.3746
569.3162841796875 0 12560.144
570.3240356445312 0 61973.938 y 4
571.32666015625 0 16078.374
572.3300170898438 0 2470.2385
597.3966064453125 0 673.8662
600.5344848632812 0 555.78326
608.3673706054688 0 1124.3665
609.3544311523438 0 558.3452
624.334716796875 0 42288.94 w 3
625.3372802734375 0 15650.648
626.3400268554688 0 3318.1584
639.4052734375 0 4418.522
640.4132690429688 0 62518.24 c 5
640.8295288085938 0 722.9935
641.408935546875 0 28959.76
642.4068603515625 0 6885.9097
643.4143676757812 0 682.27423
650.3582763671875 0 702.9038
666.3930053710938 0 1522.119
667.377685546875 0 637.33124
668.349853515625 0 1019.80817
677.3862915039062 0 856.50525
681.3571166992188 0 2332.964 y Ammonia loss 3
682.2975463867188 0 1111.5243
682.36376953125 0 20884.258 z 3
683.3594360351562 0 10105.874
684.3577270507812 0 2858.3958
694.4226684570312 0 1092.9921 c Ammonia loss 6
698.3820190429688 0 9009.526 y 3
699.3778076171875 0 4422.641
700.376708984375 0 2113.8083
700.952880859375 0 652.5094
710.4420776367188 0 2127.8083
711.4503784179688 0 41850.195 c 6
712.446044921875 0 22631.508
713.4474487304688 0 5609.2725
721.4439086914062 0 847.71234
738.3799438476562 0 1499.9978 y Ammonia loss 2
738.472900390625 0 1127.9822
739.3845825195312 0 3339.9812 z 2
740.3840942382812 0 1624.1561
741.388916015625 0 862.5359
755.4036254882812 0 40348.125 y 2
756.399658203125 0 23751.111
757.3973999023438 0 7852.8174
765.4609985351562 0 15200.873 c Ammonia loss 7
766.4601440429688 0 6288.198
767.4605712890625 0 2119.6064
781.4317016601562 0 749.9569
793.393310546875 0 619.21716
809.4144897460938 0 9447.477 w 1
810.4119262695312 0 5392.974
811.408447265625 0 1182.0669
820.5123291015625 0 1034.2557
821.5121459960938 0 736.5563
835.52490234375 0 2299.198
836.5216674804688 0 1165.0304
851.47509765625 0 873.1405 y Ammonia loss 1
852.468994140625 0 16049.611 z 1
853.466796875 0 9566.14
854.4669799804688 0 3330.9116
855.4664916992188 0 615.3697
864.52978515625 0 930.3909
865.5262451171875 0 712.4613
868.48681640625 0 3038.1338 y 1
869.4815673828125 0 2632.8196
870.4812622070312 0 1054.9933
879.53955078125 0 53661.33 c 8
880.5342407226562 0 33563.97
881.534423828125 0 13718.975
882.53466796875 0 2516.0134
889.539794921875 0 1240.9103
895.4740600585938 0 7990.3965
896.4702758789062 0 6701.087
897.4619750976562 0 1388.7196
897.5244750976562 0 1606.2697
898.5338134765625 0 1279.6619
906.5582275390625 0 1529.208
907.552978515625 0 1823.7477
912.5023803710938 0 2981.3167
913.4976806640625 0 1467.5486
922.56005859375 0 4011.4617
923.5445556640625 0 8494.708
924.549072265625 0 4873.9644
925.556396484375 0 1202.0925
949.5279541015625 0 770.72284
950.5478515625 0 2648.556
951.5371704101562 0 52721.047
952.5339965820312 0 39629.8
953.53515625 0 14071.53
954.5301513671875 0 1689.3527
959.939208984375 0 683.6701
965.4393310546875 0 624.6606
966.4385986328125 0 588.43304
966.5463256835938 0 3036.2603
967.5548095703125 0 51144.2
968.5610961914062 0 152708.98
969.5607299804688 0 94771.91
970.562255859375 0 30812.344
971.42333984375 0 1420.8489
971.563232421875 0 2861.7031
1013.1192626953125 0 598.4771
1255.8934326171875 0 686.3925
1269.863037109375 0 572.5687
1439.620361328125 0 648.75354
1450.733154296875 0 720.3024
1926.357421875 0 725.44495
2321.064697265625 0 753.23816
2867.5048828125 0 649.05536
2965.651611328125 0 669.98474
3000.662353515625 0 666.7645

Spectrum Details

|  |  |
| --- | --- |
| Matched peaks? Matched peaks The total absolute number of peaks matched. Additionally in brackets the total fraction of peaks matched and the total number of peaks is shown. | 32 (16.24% of 197) |
| FDR? FDR The false discovery rate estimated for this peptide. It is calculated by matching all theoretical fragments with a non-integer shift with the raw peaks for this spectrum. This is done with 40 different shifts. The resulting percentage is the average number of annotated peaks over the number of annotated peaks with the correct spectrum. | 0.89% |
| Satellite FDR? Satellite FDR See the FDR for details on its calculation. This satellite ion specific FDR only contains the satellite ions (d/w) for I/L/J positions. | 4.76% |
| PSM Score? PSM Score The PSM Score as given by Hecklib to this annotated spectrum. It is shown with three significant figures. | 316 |

#### Spectrum 3333? Spectrum 3333 The raw spectrum of this peptide as annotated by Hecklib. The fragments are coloured according to ion type (see legend). Any peaks with a star '\*' as text can be hovered over to see the full details, first the ion type second the mass shift type. By hovering over the amino acids in the peptide or ions in the legend the corresponding peaks are highlighted. By toggling the 'Unassigned' label you can turn the background (unassigned) peaks on or off in the plot. By updating the slider in the Ion legend you can update the spectrum to only show the top X% of the peaks with labels. The top X% means any peak that is within X% of the highest intensity. By dragging in the spectrum you can zoom in to a specific part of the spectrum and use 'Zoom Out' to get back to the original zoom level. The annotation of the spectrum is based on the given sequence in the peptides file and is done with different software so inconsistencies are likely. The peaks are annotated based on the given sequence, with 20 ppm tolerance.

Copy Data

##### Spectrum 3333 (TSV)

###### Preview

```
Loading example...
```

*Click on the button to copy the data to your clipboard.*

Mz MinMz MaxIntensity Max

WidthHeightPeptide font sizePeptide stroke widthSpectrum font sizeSpectrum stroke widthCompact peptide

Ion legend

wxyz

abcd

OtherUnassignedIon^Charge^_Position_Show for top:%

VLGQPKAAPS

02.18e+44.36e+46.54e+48.72e+4

Zoom Out

y^+2^_4_y^+1^_2_c^+1^_3_y^+2^_7_y^+2^_7_c^+1^_4_y^+2^_9_z^+1^_5_y^+1^_5_c^+1^_5_z^+1^_6_y^+1^_6_w^+1^_7_c^+1^_6_y^+1^_7_z^+1^_7_y^+1^_7_c^+1^_7_y^+1^_8_z^+1^_8_y^+1^_8_c^+1^_8_w^+1^_9_z^+1^_9_y^+1^_9_c^+1^_9_

0722144321652887

Fragment Matches Table

Show background peaks

| Position | Ion type | Intensity | mz Theoretical | mz Error (Th) | mz Error (ppm) | Charge | Series Number |
| --- | --- | --- | --- | --- | --- | --- | --- |
| - | - | 430 | 136.1 | - | - | 0 | - |
| - | - | 453.3 | 143.1 | - | - | 0 | - |
| - | - | 467.1 | 145.9 | - | - | 0 | - |
| - | - | 976.5 | 149 | - | - | 0 | - |
| - | - | 451.4 | 158.4 | - | - | 0 | - |
| - | - | 455.7 | 161.8 | - | - | 0 | - |
| - | - | 570.2 | 168.1 | - | - | 0 | - |
| 7 | y | 439.2 | 173.1 | 0.001224 | 7.069 | +2 | 4 |
| - | - | 1.453E+04 | 185.2 | - | - | 0 | - |
| - | - | 1228 | 186.1 | - | - | 0 | - |
| - | - | 502.8 | 186.1 | - | - | 0 | - |
| - | - | 1138 | 186.2 | - | - | 0 | - |
| - | - | 705.5 | 197.1 | - | - | 0 | - |
| 9 | y | 1.047E+04 | 203.1 | 0.0004453 | 2.192 | +1 | 2 |
| - | - | 609.9 | 204.1 | - | - | 0 | - |
| - | - | 671 | 213.1 | - | - | 0 | - |
| - | - | 1.079E+04 | 213.2 | - | - | 0 | - |
| - | - | 1100 | 214.2 | - | - | 0 | - |
| - | - | 1442 | 226.2 | - | - | 0 | - |
| - | - | 555.6 | 245 | - | - | 0 | - |
| - | - | 894.6 | 253.1 | - | - | 0 | - |
| - | - | 645.4 | 254.1 | - | - | 0 | - |
| - | - | 1295 | 270.2 | - | - | 0 | - |
| - | - | 1279 | 271.1 | - | - | 0 | - |
| 3 | c | 1244 | 287.2 | 0.0002171 | 0.756 | +1 | 3 |
| - | - | 1382 | 299.2 | - | - | 0 | - |
| - | - | 2728 | 325.2 | - | - | 0 | - |
| 4 | y | 694.5 | 340.7 | 0.001559 | 4.577 | +2 | 7 |
| - | - | 1015 | 343.2 | - | - | 0 | - |
| 4 | y | 686.3 | 349.7 | 0.0002746 | 0.7853 | +2 | 7 |
| - | - | 2318 | 368.2 | - | - | 0 | - |
| - | - | 673.3 | 369.2 | - | - | 0 | - |
| - | - | 659.9 | 370.2 | - | - | 0 | - |
| 4 | c | 2540 | 398.2 | 0.0001105 | 0.2775 | +1 | 4 |
| - | - | 794.1 | 399.2 | - | - | 0 | - |
| 2 | y | 778.7 | 434.7 | 0.001527 | 3.512 | +2 | 9 |
| - | - | 1131 | 435.2 | - | - | 0 | - |
| 6 | z | 1.661E+04 | 457.3 | 0.0002618 | 0.5725 | +1 | 5 |
| - | - | 1.381E+04 | 458.3 | - | - | 0 | - |
| - | - | 3555 | 459.3 | - | - | 0 | - |
| - | - | 682.1 | 464.1 | - | - | 0 | - |
| - | - | 522.2 | 465.2 | - | - | 0 | - |
| 6 | y | 1093 | 473.3 | 0.0007058 | 1.491 | +1 | 5 |
| - | - | 797.1 | 475.3 | - | - | 0 | - |
| - | - | 1429 | 482.1 | - | - | 0 | - |
| - | - | 1382 | 482.3 | - | - | 0 | - |
| - | - | 889.5 | 483.2 | - | - | 0 | - |
| - | - | 2316 | 484.3 | - | - | 0 | - |
| - | - | 781 | 484.8 | - | - | 0 | - |
| - | - | 1905 | 485.3 | - | - | 0 | - |
| - | - | 1594 | 485.4 | - | - | 0 | - |
| 5 | c | 1079 | 512.3 | 4.429E-05 | 0.08646 | +1 | 5 |
| - | - | 4323 | 553.3 | - | - | 0 | - |
| 5 | z | 1064 | 554.3 | 0.003584 | 6.467 | +1 | 6 |
| - | - | 5421 | 569.3 | - | - | 0 | - |
| 5 | y | 2.7E+04 | 570.3 | 0.0007353 | 1.289 | +1 | 6 |
| - | - | 7352 | 571.3 | - | - | 0 | - |
| - | - | 1925 | 572.3 | - | - | 0 | - |
| - | - | 772.1 | 596.4 | - | - | 0 | - |
| - | - | 1018 | 598.3 | - | - | 0 | - |
| - | - | 773.7 | 615.3 | - | - | 0 | - |
| - | - | 807.6 | 615.3 | - | - | 0 | - |
| 4 | w | 1.791E+04 | 624.3 | 0.0003747 | 0.6002 | +1 | 7 |
| - | - | 6614 | 625.3 | - | - | 0 | - |
| - | - | 1230 | 626.3 | - | - | 0 | - |
| - | - | 1978 | 639.4 | - | - | 0 | - |
| 6 | c | 3.481E+04 | 640.4 | 0.0006804 | 1.063 | +1 | 6 |
| - | - | 1.281E+04 | 641.4 | - | - | 0 | - |
| - | - | 2493 | 642.4 | - | - | 0 | - |
| - | - | 726 | 666.4 | - | - | 0 | - |
| - | - | 704.1 | 668.3 | - | - | 0 | - |
| - | - | 866.8 | 677.4 | - | - | 0 | - |
| 4 | y | 1221 | 681.4 | 0.0003173 | 0.4657 | +1 | 7 |
| 4 | z | 1.311E+04 | 682.4 | 0.0004887 | 0.7161 | +1 | 7 |
| - | - | 1382 | 683.3 | - | - | 0 | - |
| - | - | 3703 | 683.4 | - | - | 0 | - |
| - | - | 2159 | 683.8 | - | - | 0 | - |
| - | - | 649.9 | 684.3 | - | - | 0 | - |
| - | - | 1145 | 684.4 | - | - | 0 | - |
| - | - | 608.3 | 684.8 | - | - | 0 | - |
| - | - | 1254 | 696.4 | - | - | 0 | - |
| - | - | 1200 | 697.4 | - | - | 0 | - |
| 4 | y | 5733 | 698.4 | 0.0002918 | 0.4179 | +1 | 7 |
| - | - | 1640 | 699.4 | - | - | 0 | - |
| - | - | 655.1 | 701 | - | - | 0 | - |
| - | - | 1052 | 706.3 | - | - | 0 | - |
| - | - | 1079 | 710.4 | - | - | 0 | - |
| 7 | c | 2.527E+04 | 711.5 | 0.0006238 | 0.8768 | +1 | 7 |
| - | - | 8752 | 712.5 | - | - | 0 | - |
| - | - | 2456 | 713.5 | - | - | 0 | - |
| - | - | 1049 | 714.4 | - | - | 0 | - |
| - | - | 605.5 | 726.3 | - | - | 0 | - |
| 3 | y | 684.5 | 738.4 | 0.002348 | 3.179 | +1 | 8 |
| 3 | z | 1672 | 739.4 | 0.00413 | 5.586 | +1 | 8 |
| - | - | 954.6 | 740.4 | - | - | 0 | - |
| 3 | y | 2.416E+04 | 755.4 | 0.0006374 | 0.8438 | +1 | 8 |
| - | - | 9883 | 756.4 | - | - | 0 | - |
| - | - | 1959 | 757.4 | - | - | 0 | - |
| 8 | c | 8329 | 765.5 | 0.0005074 | 0.6628 | +1 | 8 |
| - | - | 3769 | 766.5 | - | - | 0 | - |
| - | - | 749.6 | 767.5 | - | - | 0 | - |
| - | - | 908.2 | 796.4 | - | - | 0 | - |
| 2 | w | 4266 | 809.4 | 0.000643 | 0.7944 | +1 | 9 |
| - | - | 2506 | 810.4 | - | - | 0 | - |
| - | - | 1696 | 812.4 | - | - | 0 | - |
| - | - | 1129 | 813.4 | - | - | 0 | - |
| - | - | 935.8 | 814.4 | - | - | 0 | - |
| - | - | 1008 | 820.5 | - | - | 0 | - |
| - | - | 706.1 | 821.5 | - | - | 0 | - |
| - | - | 1431 | 835.5 | - | - | 0 | - |
| - | - | 786.8 | 836.5 | - | - | 0 | - |
| 2 | z | 9004 | 852.5 | 0.0007917 | 0.9288 | +1 | 9 |
| - | - | 5121 | 853.5 | - | - | 0 | - |
| - | - | 739.7 | 854.5 | - | - | 0 | - |
| - | - | 2420 | 858.5 | - | - | 0 | - |
| - | - | 1166 | 864.5 | - | - | 0 | - |
| 2 | y | 1500 | 868.5 | 0.002365 | 2.723 | +1 | 9 |
| - | - | 1134 | 869.5 | - | - | 0 | - |
| 9 | c | 3.156E+04 | 879.5 | 0.0009629 | 1.095 | +1 | 9 |
| - | - | 1.68E+04 | 880.5 | - | - | 0 | - |
| - | - | 3725 | 881.5 | - | - | 0 | - |
| - | - | 961.3 | 889.5 | - | - | 0 | - |
| - | - | 4814 | 895.5 | - | - | 0 | - |
| - | - | 2150 | 896.5 | - | - | 0 | - |
| - | - | 1381 | 897.5 | - | - | 0 | - |
| - | - | 757.4 | 898.5 | - | - | 0 | - |
| - | - | 980.4 | 907.6 | - | - | 0 | - |
| - | - | 1790 | 912.5 | - | - | 0 | - |
| - | - | 1050 | 913.5 | - | - | 0 | - |
| - | - | 1975 | 922.6 | - | - | 0 | - |
| - | - | 4427 | 923.5 | - | - | 0 | - |
| - | - | 1517 | 924.6 | - | - | 0 | - |
| - | - | 1900 | 948.5 | - | - | 0 | - |
| - | - | 1735 | 950.5 | - | - | 0 | - |
| - | - | 3.158E+04 | 951.5 | - | - | 0 | - |
| - | - | 1.467E+04 | 952.5 | - | - | 0 | - |
| - | - | 4903 | 953.5 | - | - | 0 | - |
| - | - | 1211 | 966.6 | - | - | 0 | - |
| - | - | 2.663E+04 | 967.6 | - | - | 0 | - |
| - | - | 8.637E+04 | 968.6 | - | - | 0 | - |
| - | - | 3.925E+04 | 969.6 | - | - | 0 | - |
| - | - | 2061 | 970.4 | - | - | 0 | - |
| - | - | 1.145E+04 | 970.6 | - | - | 0 | - |
| - | - | 1477 | 971.5 | - | - | 0 | - |
| - | - | 949.1 | 971.6 | - | - | 0 | - |
| - | - | 729.8 | 1327 | - | - | 0 | - |
| - | - | 1409 | 1342 | - | - | 0 | - |
| - | - | 819 | 1343 | - | - | 0 | - |
| - | - | 1124 | 1349 | - | - | 0 | - |
| - | - | 1741 | 1350 | - | - | 0 | - |
| - | - | 848.2 | 1351 | - | - | 0 | - |
| - | - | 958 | 1366 | - | - | 0 | - |
| - | - | 2021 | 1367 | - | - | 0 | - |
| - | - | 1589 | 1368 | - | - | 0 | - |
| - | - | 811.2 | 1381 | - | - | 0 | - |
| - | - | 2746 | 1454 | - | - | 0 | - |
| - | - | 2086 | 1455 | - | - | 0 | - |
| - | - | 598.7 | 2738 | - | - | 0 | - |
| - | - | 740.1 | 2858 | - | - | 0 | - |

m/z Charge Intensity FragmentType MassShift Position
136.0751953125 0 430.02475
143.05747985839844 0 453.31335
145.86849975585938 0 467.11264
148.95458984375 0 976.5417
158.38876342773438 0 451.41492
161.8194580078125 0 455.66708
168.11090087890625 0 570.20044
173.09329223632812 0 439.2119 y 6
185.16455078125 0 14529.412
186.08714294433594 0 1227.9994
186.12197875976562 0 502.79437
186.16836547851562 0 1137.5443
197.12796020507812 0 705.4657
203.10218811035156 0 10465.025 y 8
204.10549926757812 0 609.94635
213.14947509765625 0 670.96906
213.15931701660156 0 10791.406
214.1628875732422 0 1099.6998
226.15493774414062 0 1441.975
245.0240478515625 0 555.57104
253.12911987304688 0 894.60077
254.1135711669922 0 645.3982
270.18096923828125 0 1295.0688
271.1396179199219 0 1279.1768
287.2075500488281 0 1244.3411 c 2
299.1708984375 0 1382.2468
325.1871643066406 0 2728.2996
340.6914978027344 0 694.4564 y Water loss 3
343.1974792480469 0 1014.53125
349.6949462890625 0 686.25726 y 3
368.2286071777344 0 2317.9092
369.2323913574219 0 673.3284
370.2448425292969 0 659.87683
398.23968505859375 0 2540.3975 c Ammonia loss 3
399.24163818359375 0 794.0994
434.74951171875 0 778.7234 y 1
435.2489013671875 0 1130.5321
457.2528381347656 0 16609.695 z 5
458.2594909667969 0 13805.186
459.2627258300781 0 3554.733
464.13238525390625 0 682.1152
465.24444580078125 0 522.219
473.2711181640625 0 1093.1766 y 5
475.2752380371094 0 797.0575
482.14520263671875 0 1429.2206
482.27166748046875 0 1382.2852
483.15521240234375 0 889.46344
484.2847900390625 0 2316.23
484.7817077636719 0 781.0374
485.2753601074219 0 1904.5963
485.35882568359375 0 1593.6313
512.3191528320312 0 1078.6025 c 4
553.3094482421875 0 4322.7866
554.3094482421875 0 1063.7174 z 4
569.3157348632812 0 5420.996
570.3238525390625 0 27001.857 y 4
571.3273315429688 0 7352.3315
572.328125 0 1925.1555
596.3995971679688 0 772.05566
598.3162841796875 0 1018.17725
615.2862548828125 0 773.73285
615.3453979492188 0 807.5533
624.3347778320312 0 17907.912 w 3
625.3378295898438 0 6614.387
626.3396606445312 0 1229.5787
639.4048461914062 0 1977.881
640.4133911132812 0 34807.83 c 5
641.4160766601562 0 12809.232
642.4183349609375 0 2492.83
666.392578125 0 726.0208
668.3482666015625 0 704.0952
677.38232421875 0 866.8214
681.35693359375 0 1220.7861 y Ammonia loss 3
682.3639526367188 0 13109.459 z 3
683.2916870117188 0 1381.8004
683.3660888671875 0 3702.886
683.7902221679688 0 2159.2104
684.2942504882812 0 649.9455
684.37158203125 0 1144.9185
684.7962036132812 0 608.3433
696.3843383789062 0 1254.0801
697.3853149414062 0 1200.3383
698.3828735351562 0 5732.8364 y 3
699.3876342773438 0 1639.9534
700.9638671875 0 655.13806
706.3092041015625 0 1052.2295
710.4423828125 0 1079.3237
711.4505615234375 0 25268.137 c 6
712.4530029296875 0 8751.722
713.4553833007812 0 2455.5408
714.4098510742188 0 1048.898
726.275634765625 0 605.53534
738.375732421875 0 684.48004 y Ammonia loss 2
739.3817749023438 0 1672.0522 z 2
740.3870849609375 0 954.56635
755.4039916992188 0 24163.951 y 2
756.4069213867188 0 9882.856
757.4094848632812 0 1958.8711
765.4612426757812 0 8329.466 c Ammonia loss 7
766.4644165039062 0 3768.588
767.4678955078125 0 749.563
796.3768920898438 0 908.20074
809.41455078125 0 4265.675 w 1
810.4175415039062 0 2505.7295
812.3943481445312 0 1696.0989
813.3958129882812 0 1129.1522
814.401611328125 0 935.84985
820.5093994140625 0 1008.32855
821.5174560546875 0 706.06604
835.52490234375 0 1430.9342
836.5247192382812 0 786.8479
852.4691772460938 0 9004.291 z 1
853.4708251953125 0 5121.302
854.467529296875 0 739.671
858.4652709960938 0 2420.4146
864.5315551757812 0 1166.3638
868.486328125 0 1500.0641 y 1
869.4927368164062 0 1133.7067
879.5401000976562 0 31557.035 c 8
880.5416259765625 0 16797.367
881.5432739257812 0 3724.8672
889.537841796875 0 961.33136
895.475830078125 0 4814.4165
896.4766845703125 0 2150.2905
897.52294921875 0 1380.9133
898.5321655273438 0 757.41766
907.5537109375 0 980.39185
912.5013427734375 0 1790.4706
913.5046997070312 0 1050.3274
922.559326171875 0 1975.0964
923.5479125976562 0 4427.496
924.5560302734375 0 1516.5835
948.4916381835938 0 1899.9691
950.5482788085938 0 1734.9677
951.5379638671875 0 31581.426
952.5405883789062 0 14670.525
953.5416259765625 0 4903.249
966.5513916015625 0 1210.8329
967.5567016601562 0 26627.873
968.5633544921875 0 86373.95
969.566162109375 0 39245.934
970.441650390625 0 2060.5708
970.5701904296875 0 11450.522
971.4505004882812 0 1476.7284
971.5715942382812 0 949.10065
1326.590576171875 0 729.8412
1341.6141357421875 0 1409.4985
1342.6251220703125 0 818.9682
1348.566162109375 0 1123.583
1349.5634765625 0 1741.3639
1350.5601806640625 0 848.2431
1365.583984375 0 958.03503
1366.5811767578125 0 2020.5519
1367.585693359375 0 1589.4263
1380.5557861328125 0 811.171
1453.552734375 0 2745.8228
1454.555419921875 0 2085.5996
2738.4619140625 0 598.74426
2858.38427734375 0 740.0574

Spectrum Details

|  |  |
| --- | --- |
| Matched peaks? Matched peaks The total absolute number of peaks matched. Additionally in brackets the total fraction of peaks matched and the total number of peaks is shown. | 26 (16.35% of 159) |
| FDR? FDR The false discovery rate estimated for this peptide. It is calculated by matching all theoretical fragments with a non-integer shift with the raw peaks for this spectrum. This is done with 40 different shifts. The resulting percentage is the average number of annotated peaks over the number of annotated peaks with the correct spectrum. | 0.92% |
| Satellite FDR? Satellite FDR See the FDR for details on its calculation. This satellite ion specific FDR only contains the satellite ions (d/w) for I/L/J positions. | 4.76% |
| PSM Score? PSM Score The PSM Score as given by Hecklib to this annotated spectrum. It is shown with three significant figures. | 245 |

#### Spectrum 3083? Spectrum 3083 The raw spectrum of this peptide as annotated by Hecklib. The fragments are coloured according to ion type (see legend). Any peaks with a star '\*' as text can be hovered over to see the full details, first the ion type second the mass shift type. By hovering over the amino acids in the peptide or ions in the legend the corresponding peaks are highlighted. By toggling the 'Unassigned' label you can turn the background (unassigned) peaks on or off in the plot. By updating the slider in the Ion legend you can update the spectrum to only show the top X% of the peaks with labels. The top X% means any peak that is within X% of the highest intensity. By dragging in the spectrum you can zoom in to a specific part of the spectrum and use 'Zoom Out' to get back to the original zoom level. The annotation of the spectrum is based on the given sequence in the peptides file and is done with different software so inconsistencies are likely. The peaks are annotated based on the given sequence, with 20 ppm tolerance.

Copy Data

##### Spectrum 3083 (TSV)

###### Preview

```
Loading example...
```

*Click on the button to copy the data to your clipboard.*

Mz MinMz MaxIntensity Max

WidthHeightPeptide font sizePeptide stroke widthSpectrum font sizeSpectrum stroke widthCompact peptide

Ion legend

wxyz

abcd

OtherUnassignedIon^Charge^_Position_Show for top:%

VLGQPKAAPS

07.91e+41.58e+52.37e+53.17e+5

Zoom Out

d^+1^_2_y^+1^_2_a^+1^_2_y^+1^_2_b^+1^_2_y^+2^_5_y^+1^_3_b^+1^_3_y^+1^_3_y^+2^_6_y^+1^_4_y^+2^_7_y^+1^_4_y^+2^_7_y^+2^_8_y^+2^_8_b^+2^_8_b^+1^_4_y^+2^_9_y^+2^_9_b^+2^_9_y^+2^_9_y^+1^_5_\*\*\*b^+1^_5_y^+1^_6_y^+1^_6_b^+1^_6_y^+1^_7_y^+1^_7_b^+1^_7_y^+1^_7_y^+1^_8_y^+1^_8_b^+1^_8_y^+1^_8_b^+1^_8_y^+1^_9_y^+1^_9_

0776155223273103

Fragment Matches Table

Show background peaks

| Position | Ion type | Intensity | mz Theoretical | mz Error (Th) | mz Error (ppm) | Charge | Series Number |
| --- | --- | --- | --- | --- | --- | --- | --- |
| - | - | 967.6 | 120.1 | - | - | 0 | - |
| - | - | 372.1 | 120.6 | - | - | 0 | - |
| - | - | 501.5 | 124 | - | - | 0 | - |
| - | - | 1776 | 125.1 | - | - | 0 | - |
| - | - | 799.7 | 127.1 | - | - | 0 | - |
| - | - | 1571 | 128.1 | - | - | 0 | - |
| - | - | 369 | 128.9 | - | - | 0 | - |
| - | - | 7266 | 129.1 | - | - | 0 | - |
| - | - | 6.14E+04 | 129.1 | - | - | 0 | - |
| - | - | 485.1 | 130.1 | - | - | 0 | - |
| - | - | 430.4 | 130.1 | - | - | 0 | - |
| - | - | 496.4 | 130.1 | - | - | 0 | - |
| - | - | 3833 | 130.1 | - | - | 0 | - |
| - | - | 401.4 | 132.9 | - | - | 0 | - |
| - | - | 5548 | 133.1 | - | - | 0 | - |
| - | - | 397.5 | 133.6 | - | - | 0 | - |
| - | - | 2347 | 136.1 | - | - | 0 | - |
| - | - | 449.6 | 137.8 | - | - | 0 | - |
| - | - | 470.6 | 138.1 | - | - | 0 | - |
| - | - | 1959 | 139.1 | - | - | 0 | - |
| - | - | 3742 | 141.1 | - | - | 0 | - |
| - | - | 4691 | 141.1 | - | - | 0 | - |
| - | - | 5071 | 142.1 | - | - | 0 | - |
| - | - | 2129 | 143.1 | - | - | 0 | - |
| 2 | d | 1796 | 143.1 | 0.0006713 | 4.69 | +1 | 2 |
| - | - | 814.3 | 143.1 | - | - | 0 | - |
| - | - | 612.6 | 147.1 | - | - | 0 | - |
| - | - | 529.6 | 148.9 | - | - | 0 | - |
| - | - | 514.9 | 151.1 | - | - | 0 | - |
| - | - | 3632 | 152 | - | - | 0 | - |
| - | - | 2726 | 152.1 | - | - | 0 | - |
| - | - | 942 | 153.1 | - | - | 0 | - |
| - | - | 1152 | 153.1 | - | - | 0 | - |
| - | - | 8321 | 155.1 | - | - | 0 | - |
| - | - | 584.8 | 156.1 | - | - | 0 | - |
| - | - | 2300 | 157.1 | - | - | 0 | - |
| - | - | 607 | 158.1 | - | - | 0 | - |
| - | - | 2914 | 158.1 | - | - | 0 | - |
| - | - | 484.9 | 161.1 | - | - | 0 | - |
| - | - | 535 | 163.1 | - | - | 0 | - |
| - | - | 1629 | 165.1 | - | - | 0 | - |
| - | - | 583.6 | 168.1 | - | - | 0 | - |
| - | - | 1.729E+04 | 169.1 | - | - | 0 | - |
| - | - | 6094 | 169.1 | - | - | 0 | - |
| - | - | 2430 | 169.1 | - | - | 0 | - |
| - | - | 1282 | 170.1 | - | - | 0 | - |
| - | - | 698.8 | 170.1 | - | - | 0 | - |
| - | - | 1.072E+04 | 171.1 | - | - | 0 | - |
| - | - | 1103 | 172.1 | - | - | 0 | - |
| - | - | 3735 | 173.1 | - | - | 0 | - |
| - | - | 1.031E+04 | 173.1 | - | - | 0 | - |
| - | - | 932.5 | 174.1 | - | - | 0 | - |
| - | - | 3091 | 175.1 | - | - | 0 | - |
| - | - | 1635 | 180.1 | - | - | 0 | - |
| - | - | 4593 | 181.1 | - | - | 0 | - |
| - | - | 3146 | 181.1 | - | - | 0 | - |
| - | - | 1.617E+04 | 182.1 | - | - | 0 | - |
| - | - | 1114 | 182.1 | - | - | 0 | - |
| - | - | 2184 | 183.1 | - | - | 0 | - |
| - | - | 1690 | 183.1 | - | - | 0 | - |
| - | - | 765.4 | 183.1 | - | - | 0 | - |
| 9 | y | 5829 | 185.1 | 0.0003385 | 1.829 | +1 | 2 |
| 2 | a | 3.134E+05 | 185.2 | 0.0004893 | 2.642 | +1 | 2 |
| - | - | 3.933E+04 | 186.1 | - | - | 0 | - |
| - | - | 3.218E+04 | 186.2 | - | - | 0 | - |
| - | - | 2802 | 187.1 | - | - | 0 | - |
| - | - | 954.4 | 187.1 | - | - | 0 | - |
| - | - | 8373 | 187.1 | - | - | 0 | - |
| - | - | 1159 | 187.2 | - | - | 0 | - |
| - | - | 8176 | 188.1 | - | - | 0 | - |
| - | - | 634.1 | 188.1 | - | - | 0 | - |
| - | - | 685 | 189.1 | - | - | 0 | - |
| - | - | 1511 | 190.1 | - | - | 0 | - |
| - | - | 748.6 | 191.1 | - | - | 0 | - |
| - | - | 1119 | 193.1 | - | - | 0 | - |
| - | - | 1257 | 195.1 | - | - | 0 | - |
| - | - | 853.3 | 197.1 | - | - | 0 | - |
| - | - | 2489 | 197.1 | - | - | 0 | - |
| - | - | 1580 | 198.1 | - | - | 0 | - |
| - | - | 1101 | 199.1 | - | - | 0 | - |
| - | - | 4119 | 200.1 | - | - | 0 | - |
| - | - | 1737 | 201.1 | - | - | 0 | - |
| 9 | y | 1.986E+05 | 203.1 | 0.0004092 | 2.015 | +1 | 2 |
| - | - | 1.632E+04 | 204.1 | - | - | 0 | - |
| - | - | 1775 | 205.1 | - | - | 0 | - |
| - | - | 4437 | 208.1 | - | - | 0 | - |
| - | - | 790.1 | 208.1 | - | - | 0 | - |
| - | - | 6014 | 209.1 | - | - | 0 | - |
| - | - | 650.8 | 209.1 | - | - | 0 | - |
| - | - | 570.5 | 210 | - | - | 0 | - |
| - | - | 1.767E+04 | 210.1 | - | - | 0 | - |
| - | - | 783.9 | 211.1 | - | - | 0 | - |
| - | - | 1503 | 211.1 | - | - | 0 | - |
| - | - | 1015 | 212.1 | - | - | 0 | - |
| 2 | b | 1.606E+05 | 213.2 | 0.0004629 | 2.172 | +1 | 2 |
| - | - | 728.3 | 214.1 | - | - | 0 | - |
| - | - | 1.853E+04 | 214.2 | - | - | 0 | - |
| - | - | 906.3 | 215.1 | - | - | 0 | - |
| - | - | 4040 | 215.1 | - | - | 0 | - |
| - | - | 1402 | 215.2 | - | - | 0 | - |
| - | - | 1485 | 220.1 | - | - | 0 | - |
| - | - | 884.1 | 223.1 | - | - | 0 | - |
| - | - | 999.1 | 224.1 | - | - | 0 | - |
| - | - | 3162 | 226.1 | - | - | 0 | - |
| - | - | 1.245E+05 | 226.2 | - | - | 0 | - |
| - | - | 920.7 | 227.1 | - | - | 0 | - |
| - | - | 1.472E+04 | 227.2 | - | - | 0 | - |
| 6 | y | 1.777E+04 | 228.1 | 0.0003757 | 1.647 | +2 | 5 |
| - | - | 1051 | 228.2 | - | - | 0 | - |
| - | - | 2133 | 229.1 | - | - | 0 | - |
| - | - | 1616 | 236.1 | - | - | 0 | - |
| - | - | 602.8 | 237.1 | - | - | 0 | - |
| - | - | 2762 | 238.1 | - | - | 0 | - |
| - | - | 937.2 | 239.2 | - | - | 0 | - |
| - | - | 2030 | 240.1 | - | - | 0 | - |
| - | - | 1118 | 242.2 | - | - | 0 | - |
| - | - | 4070 | 243.2 | - | - | 0 | - |
| - | - | 1843 | 244.1 | - | - | 0 | - |
| - | - | 851.3 | 244.1 | - | - | 0 | - |
| - | - | 521.2 | 248.1 | - | - | 0 | - |
| - | - | 1698 | 252.1 | - | - | 0 | - |
| - | - | 4774 | 252.2 | - | - | 0 | - |
| - | - | 796 | 253.2 | - | - | 0 | - |
| - | - | 1029 | 254.1 | - | - | 0 | - |
| - | - | 5983 | 254.2 | - | - | 0 | - |
| 8 | y | 2287 | 256.1 | 0.0004257 | 1.662 | +1 | 3 |
| - | - | 508.5 | 256.2 | - | - | 0 | - |
| - | - | 708.3 | 257.1 | - | - | 0 | - |
| - | - | 1117 | 261.2 | - | - | 0 | - |
| - | - | 557 | 263.4 | - | - | 0 | - |
| - | - | 797.7 | 264.1 | - | - | 0 | - |
| - | - | 607.1 | 265.1 | - | - | 0 | - |
| - | - | 2006 | 266.1 | - | - | 0 | - |
| - | - | 933.5 | 268.1 | - | - | 0 | - |
| - | - | 6129 | 269.2 | - | - | 0 | - |
| - | - | 1003 | 270.1 | - | - | 0 | - |
| 3 | b | 2.005E+04 | 270.2 | 0.0003615 | 1.338 | +1 | 3 |
| - | - | 664.6 | 270.2 | - | - | 0 | - |
| - | - | 1382 | 271.1 | - | - | 0 | - |
| - | - | 1.345E+04 | 271.2 | - | - | 0 | - |
| - | - | 2437 | 272.2 | - | - | 0 | - |
| 8 | y | 1.196E+04 | 274.1 | 0.0003285 | 1.198 | +1 | 3 |
| - | - | 758.7 | 274.2 | - | - | 0 | - |
| - | - | 1540 | 275.1 | - | - | 0 | - |
| - | - | 2.515E+04 | 279.2 | - | - | 0 | - |
| - | - | 653.7 | 280.2 | - | - | 0 | - |
| - | - | 3281 | 280.2 | - | - | 0 | - |
| - | - | 4566 | 281.2 | - | - | 0 | - |
| - | - | 611.1 | 282.1 | - | - | 0 | - |
| - | - | 1197 | 282.1 | - | - | 0 | - |
| - | - | 3711 | 283.1 | - | - | 0 | - |
| - | - | 659.1 | 284.1 | - | - | 0 | - |
| - | - | 5835 | 285.2 | - | - | 0 | - |
| 5 | y | 4383 | 285.7 | 0.0005107 | 1.788 | +2 | 6 |
| - | - | 3.342E+04 | 287.2 | - | - | 0 | - |
| - | - | 894.9 | 288.2 | - | - | 0 | - |
| - | - | 4728 | 288.2 | - | - | 0 | - |
| - | - | 640.1 | 295.2 | - | - | 0 | - |
| - | - | 1172 | 296.2 | - | - | 0 | - |
| - | - | 4.704E+04 | 297.2 | - | - | 0 | - |
| - | - | 8059 | 298.2 | - | - | 0 | - |
| - | - | 9.325E+04 | 299.2 | - | - | 0 | - |
| - | - | 925.7 | 300.1 | - | - | 0 | - |
| - | - | 1.292E+04 | 300.2 | - | - | 0 | - |
| - | - | 1515 | 301.2 | - | - | 0 | - |
| - | - | 1613 | 309.2 | - | - | 0 | - |
| - | - | 577.9 | 311.3 | - | - | 0 | - |
| - | - | 2514 | 314.2 | - | - | 0 | - |
| - | - | 662.9 | 323.2 | - | - | 0 | - |
| - | - | 3171 | 323.2 | - | - | 0 | - |
| - | - | 612.4 | 324.2 | - | - | 0 | - |
| - | - | 3109 | 325.2 | - | - | 0 | - |
| - | - | 591.7 | 325.2 | - | - | 0 | - |
| - | - | 706.4 | 325.7 | - | - | 0 | - |
| 7 | y | 5615 | 327.2 | 0.0006043 | 1.847 | +1 | 4 |
| - | - | 592.8 | 327.2 | - | - | 0 | - |
| - | - | 2134 | 328.2 | - | - | 0 | - |
| - | - | 673.2 | 331.7 | - | - | 0 | - |
| - | - | 2355 | 332.2 | - | - | 0 | - |
| - | - | 1099 | 332.7 | - | - | 0 | - |
| - | - | 1347 | 336.2 | - | - | 0 | - |
| - | - | 3581 | 337.2 | - | - | 0 | - |
| - | - | 846.2 | 338.2 | - | - | 0 | - |
| - | - | 2.459E+04 | 340.2 | - | - | 0 | - |
| 4 | y | 6974 | 340.7 | 0.0003996 | 1.173 | +2 | 7 |
| - | - | 2995 | 341.2 | - | - | 0 | - |
| - | - | 3951 | 341.2 | - | - | 0 | - |
| 7 | y | 1826 | 345.2 | 0.0005682 | 1.646 | +1 | 4 |
| 4 | y | 1999 | 349.7 | 3.048E-05 | 0.08715 | +2 | 7 |
| - | - | 1103 | 350.2 | - | - | 0 | - |
| - | - | 906.8 | 350.2 | - | - | 0 | - |
| - | - | 825.6 | 351.2 | - | - | 0 | - |
| - | - | 1665 | 353.2 | - | - | 0 | - |
| - | - | 1906 | 354.2 | - | - | 0 | - |
| - | - | 908.3 | 355.2 | - | - | 0 | - |
| - | - | 1165 | 360.7 | - | - | 0 | - |
| - | - | 1868 | 363.2 | - | - | 0 | - |
| - | - | 1030 | 366.2 | - | - | 0 | - |
| - | - | 588.5 | 366.2 | - | - | 0 | - |
| - | - | 1.232E+05 | 368.2 | - | - | 0 | - |
| - | - | 2.663E+04 | 369.2 | - | - | 0 | - |
| 3 | y | 1035 | 369.7 | 0.002543 | 6.878 | +2 | 8 |
| - | - | 914.5 | 369.7 | - | - | 0 | - |
| - | - | 837.9 | 370.2 | - | - | 0 | - |
| - | - | 1.023E+04 | 370.2 | - | - | 0 | - |
| - | - | 2042 | 371.2 | - | - | 0 | - |
| - | - | 2097 | 376.2 | - | - | 0 | - |
| - | - | 618.4 | 376.2 | - | - | 0 | - |
| 3 | y | 1683 | 378.2 | 0.001506 | 3.981 | +2 | 8 |
| - | - | 2630 | 380.2 | - | - | 0 | - |
| - | - | 2793 | 382.2 | - | - | 0 | - |
| - | - | 1615 | 382.7 | - | - | 0 | - |
| 8 | b | 1578 | 383.2 | 0.0002908 | 0.7589 | +2 | 8 |
| - | - | 762.6 | 383.7 | - | - | 0 | - |
| - | - | 767.3 | 384.3 | - | - | 0 | - |
| - | - | 4057 | 390.2 | - | - | 0 | - |
| - | - | 3488 | 394.2 | - | - | 0 | - |
| - | - | 2247 | 394.2 | - | - | 0 | - |
| - | - | 1011 | 395.2 | - | - | 0 | - |
| - | - | 3149 | 396.2 | - | - | 0 | - |
| - | - | 1189 | 397.2 | - | - | 0 | - |
| 4 | b | 3.104E+04 | 398.2 | 0.0006219 | 1.562 | +1 | 4 |
| - | - | 5426 | 399.2 | - | - | 0 | - |
| - | - | 666.3 | 400.2 | - | - | 0 | - |
| - | - | 561.8 | 406.2 | - | - | 0 | - |
| - | - | 1926 | 407.2 | - | - | 0 | - |
| - | - | 4740 | 408.2 | - | - | 0 | - |
| - | - | 1106 | 409.2 | - | - | 0 | - |
| - | - | 4446 | 411.2 | - | - | 0 | - |
| - | - | 1307 | 412.2 | - | - | 0 | - |
| - | - | 1927 | 417.8 | - | - | 0 | - |
| - | - | 1109 | 418.3 | - | - | 0 | - |
| - | - | 1837 | 425.3 | - | - | 0 | - |
| 2 | y | 3292 | 425.7 | 0.0005227 | 1.228 | +2 | 9 |
| 2 | y | 2098 | 426.2 | 0.007538 | 17.69 | +2 | 9 |
| - | - | 683.1 | 426.7 | - | - | 0 | - |
| 9 | b | 4483 | 431.8 | 0.0004879 | 1.13 | +2 | 9 |
| - | - | 2595 | 432.3 | - | - | 0 | - |
| - | - | 997.1 | 434.2 | - | - | 0 | - |
| 2 | y | 1.015E+04 | 434.7 | 0.0004894 | 1.126 | +2 | 9 |
| - | - | 4413 | 435.2 | - | - | 0 | - |
| - | - | 1574 | 435.8 | - | - | 0 | - |
| - | - | 1569 | 437.3 | - | - | 0 | - |
| - | - | 2011 | 437.3 | - | - | 0 | - |
| - | - | 1520 | 439.2 | - | - | 0 | - |
| - | - | 936.3 | 439.3 | - | - | 0 | - |
| - | - | 2737 | 439.7 | - | - | 0 | - |
| - | - | 1236 | 440.2 | - | - | 0 | - |
| - | - | 953.2 | 441.2 | - | - | 0 | - |
| - | - | 1964 | 442.3 | - | - | 0 | - |
| - | - | 3677 | 447.2 | - | - | 0 | - |
| - | - | 766 | 448.2 | - | - | 0 | - |
| - | - | 1094 | 449.2 | - | - | 0 | - |
| - | - | 2403 | 451.3 | - | - | 0 | - |
| - | - | 4188 | 454.3 | - | - | 0 | - |
| - | - | 810 | 455.3 | - | - | 0 | - |
| - | - | 1910 | 456.3 | - | - | 0 | - |
| - | - | 1033 | 458.7 | - | - | 0 | - |
| - | - | 1.428E+04 | 464.3 | - | - | 0 | - |
| - | - | 3295 | 465.2 | - | - | 0 | - |
| - | - | 9432 | 465.3 | - | - | 0 | - |
| - | - | 1199 | 466.2 | - | - | 0 | - |
| - | - | 2617 | 466.3 | - | - | 0 | - |
| - | - | 992.9 | 466.8 | - | - | 0 | - |
| - | - | 762.3 | 467.3 | - | - | 0 | - |
| - | - | 4300 | 467.3 | - | - | 0 | - |
| - | - | 1905 | 468.3 | - | - | 0 | - |
| 6 | y | 6852 | 473.3 | 0.0004538 | 0.9589 | +1 | 5 |
| - | - | 1727 | 474.3 | - | - | 0 | - |
| 0 | Precursor | 4400 | 475.3 | 0.000587 | 1.235 | +2 | -1 |
| 0 | Precursor | 786 | 475.8 | 0.007755 | 16.3 | +2 | -1 |
| - | - | 1175 | 476.3 | - | - | 0 | - |
| - | - | 842.8 | 477.6 | - | - | 0 | - |
| - | - | 3137 | 478.3 | - | - | 0 | - |
| - | - | 8590 | 479.3 | - | - | 0 | - |
| - | - | 1711 | 480.3 | - | - | 0 | - |
| - | - | 881.9 | 481.3 | - | - | 0 | - |
| - | - | 3.591E+04 | 482.3 | - | - | 0 | - |
| - | - | 7277 | 483.3 | - | - | 0 | - |
| - | - | 760.6 | 484.2 | - | - | 0 | - |
| 0 | Precursor | 1.545E+04 | 484.3 | 0.0003705 | 0.7651 | +2 | -1 |
| - | - | 8855 | 484.8 | - | - | 0 | - |
| - | - | 3735 | 485.3 | - | - | 0 | - |
| - | - | 1181 | 485.4 | - | - | 0 | - |
| - | - | 847.4 | 485.8 | - | - | 0 | - |
| - | - | 940.4 | 491.3 | - | - | 0 | - |
| 5 | b | 1374 | 495.3 | 0.0001705 | 0.3443 | +1 | 5 |
| - | - | 2897 | 496.3 | - | - | 0 | - |
| - | - | 951.7 | 502.2 | - | - | 0 | - |
| - | - | 4767 | 508.3 | - | - | 0 | - |
| - | - | 1694 | 509.3 | - | - | 0 | - |
| - | - | 1105 | 510.3 | - | - | 0 | - |
| - | - | 1013 | 510.8 | - | - | 0 | - |
| - | - | 1986 | 518.3 | - | - | 0 | - |
| - | - | 1865 | 522.8 | - | - | 0 | - |
| - | - | 1676 | 524.3 | - | - | 0 | - |
| - | - | 9183 | 525.3 | - | - | 0 | - |
| - | - | 2926 | 526.3 | - | - | 0 | - |
| - | - | 907.8 | 534.3 | - | - | 0 | - |
| - | - | 2566 | 535.3 | - | - | 0 | - |
| - | - | 1.246E+04 | 536.3 | - | - | 0 | - |
| - | - | 3489 | 537.3 | - | - | 0 | - |
| - | - | 972.9 | 540.3 | - | - | 0 | - |
| 5 | y | 7308 | 552.3 | 6.375E-05 | 0.1154 | +1 | 6 |
| - | - | 1.021E+05 | 553.3 | - | - | 0 | - |
| - | - | 3.067E+04 | 554.3 | - | - | 0 | - |
| - | - | 5382 | 555.3 | - | - | 0 | - |
| - | - | 991.7 | 555.4 | - | - | 0 | - |
| - | - | 1599 | 559.3 | - | - | 0 | - |
| - | - | 1048 | 567.4 | - | - | 0 | - |
| - | - | 1533 | 567.8 | - | - | 0 | - |
| - | - | 1028 | 568.3 | - | - | 0 | - |
| - | - | 1163 | 568.3 | - | - | 0 | - |
| 5 | y | 2.116E+05 | 570.3 | 0.0004244 | 0.7441 | +1 | 6 |
| - | - | 6.437E+04 | 571.3 | - | - | 0 | - |
| - | - | 1.227E+04 | 572.3 | - | - | 0 | - |
| - | - | 913.4 | 574.8 | - | - | 0 | - |
| - | - | 3996 | 577.3 | - | - | 0 | - |
| - | - | 952.5 | 578.3 | - | - | 0 | - |
| - | - | 1090 | 579.3 | - | - | 0 | - |
| - | - | 4043 | 580.4 | - | - | 0 | - |
| - | - | 1177 | 581.4 | - | - | 0 | - |
| - | - | 667.5 | 586.8 | - | - | 0 | - |
| - | - | 5223 | 595.4 | - | - | 0 | - |
| - | - | 1655 | 596.4 | - | - | 0 | - |
| - | - | 985 | 600.3 | - | - | 0 | - |
| - | - | 1091 | 600.8 | - | - | 0 | - |
| - | - | 4069 | 608.8 | - | - | 0 | - |
| - | - | 1305 | 609.3 | - | - | 0 | - |
| - | - | 8071 | 617.3 | - | - | 0 | - |
| - | - | 5438 | 617.8 | - | - | 0 | - |
| - | - | 2096 | 618.3 | - | - | 0 | - |
| - | - | 839.9 | 618.8 | - | - | 0 | - |
| - | - | 2042 | 621.4 | - | - | 0 | - |
| - | - | 1186 | 622.4 | - | - | 0 | - |
| 6 | b | 1420 | 623.4 | 0.001719 | 2.758 | +1 | 6 |
| - | - | 2117 | 638.4 | - | - | 0 | - |
| - | - | 961.6 | 639.4 | - | - | 0 | - |
| - | - | 635.6 | 640.2 | - | - | 0 | - |
| - | - | 3180 | 649.4 | - | - | 0 | - |
| - | - | 7190 | 650.4 | - | - | 0 | - |
| - | - | 818.1 | 651.3 | - | - | 0 | - |
| - | - | 3316 | 651.4 | - | - | 0 | - |
| - | - | 1585 | 652.4 | - | - | 0 | - |
| - | - | 1643 | 654.4 | - | - | 0 | - |
| - | - | 755.5 | 655.4 | - | - | 0 | - |
| - | - | 1038 | 658.3 | - | - | 0 | - |
| - | - | 7044 | 659.3 | - | - | 0 | - |
| - | - | 4053 | 659.8 | - | - | 0 | - |
| - | - | 1967 | 660.3 | - | - | 0 | - |
| - | - | 698.2 | 663.4 | - | - | 0 | - |
| - | - | 1134 | 664.3 | - | - | 0 | - |
| - | - | 819.7 | 664.8 | - | - | 0 | - |
| - | - | 2.355E+04 | 666.4 | - | - | 0 | - |
| - | - | 8559 | 667.4 | - | - | 0 | - |
| - | - | 8066 | 667.8 | - | - | 0 | - |
| - | - | 4795 | 668.3 | - | - | 0 | - |
| - | - | 1782 | 668.4 | - | - | 0 | - |
| - | - | 4302 | 668.8 | - | - | 0 | - |
| - | - | 668.4 | 669.3 | - | - | 0 | - |
| - | - | 1284 | 672.8 | - | - | 0 | - |
| - | - | 1704 | 673.3 | - | - | 0 | - |
| - | - | 3812 | 676.4 | - | - | 0 | - |
| - | - | 2118 | 677.4 | - | - | 0 | - |
| 4 | y | 4027 | 680.4 | 0.0005072 | 0.7455 | +1 | 7 |
| 4 | y | 7237 | 681.4 | 0.002759 | 4.049 | +1 | 7 |
| - | - | 2630 | 682.4 | - | - | 0 | - |
| 7 | b | 1.01E+04 | 694.4 | 1.466E-05 | 0.02111 | +1 | 7 |
| - | - | 4402 | 695.4 | - | - | 0 | - |
| - | - | 1516 | 696.4 | - | - | 0 | - |
| 4 | y | 6643 | 698.4 | 0.0001698 | 0.2431 | +1 | 7 |
| - | - | 2752 | 699.4 | - | - | 0 | - |
| - | - | 1626 | 720.4 | - | - | 0 | - |
| - | - | 900.3 | 720.4 | - | - | 0 | - |
| - | - | 688.9 | 721.4 | - | - | 0 | - |
| - | - | 960.3 | 730.4 | - | - | 0 | - |
| 3 | y | 5831 | 737.4 | 0.001364 | 1.85 | +1 | 8 |
| - | - | 6822 | 737.5 | - | - | 0 | - |
| 3 | y | 8269 | 738.4 | 0.002962 | 4.012 | +1 | 8 |
| - | - | 3318 | 738.5 | - | - | 0 | - |
| - | - | 2285 | 739.4 | - | - | 0 | - |
| - | - | 1394 | 747.5 | - | - | 0 | - |
| 8 | b | 3264 | 748.4 | 0.0012 | 1.604 | +1 | 8 |
| - | - | 2873 | 749.4 | - | - | 0 | - |
| - | - | 1034 | 750.4 | - | - | 0 | - |
| - | - | 1122 | 753.4 | - | - | 0 | - |
| 3 | y | 1.877E+05 | 755.4 | 0.0002712 | 0.359 | +1 | 8 |
| - | - | 7.429E+04 | 756.4 | - | - | 0 | - |
| - | - | 1.794E+04 | 757.4 | - | - | 0 | - |
| - | - | 1245 | 758.4 | - | - | 0 | - |
| - | - | 2325 | 763.4 | - | - | 0 | - |
| - | - | 1081 | 764.4 | - | - | 0 | - |
| 8 | b | 9.719E+04 | 765.5 | 0.0002022 | 0.2641 | +1 | 8 |
| - | - | 4.347E+04 | 766.5 | - | - | 0 | - |
| - | - | 9966 | 767.5 | - | - | 0 | - |
| - | - | 1099 | 768.5 | - | - | 0 | - |
| - | - | 1054 | 791.5 | - | - | 0 | - |
| - | - | 939.6 | 804.4 | - | - | 0 | - |
| - | - | 836.6 | 817.4 | - | - | 0 | - |
| - | - | 822 | 830.4 | - | - | 0 | - |
| 2 | y | 1067 | 851.5 | 0.000352 | 0.4134 | +1 | 9 |
| 2 | y | 2.235E+04 | 868.5 | 0.0004118 | 0.4742 | +1 | 9 |
| - | - | 1.17E+04 | 869.5 | - | - | 0 | - |
| - | - | 3553 | 870.5 | - | - | 0 | - |
| - | - | 1400 | 874.4 | - | - | 0 | - |
| - | - | 959.5 | 875.4 | - | - | 0 | - |
| - | - | 1054 | 878.5 | - | - | 0 | - |
| - | - | 2788 | 891.4 | - | - | 0 | - |
| - | - | 1209 | 892.4 | - | - | 0 | - |
| - | - | 1008 | 916.5 | - | - | 0 | - |
| - | - | 1041 | 917.5 | - | - | 0 | - |
| - | - | 765.5 | 945.5 | - | - | 0 | - |
| - | - | 1417 | 985.4 | - | - | 0 | - |
| - | - | 2051 | 1002 | - | - | 0 | - |
| - | - | 1035 | 1003 | - | - | 0 | - |
| - | - | 776.1 | 1046 | - | - | 0 | - |
| - | - | 898.8 | 1100 | - | - | 0 | - |
| - | - | 1962 | 1118 | - | - | 0 | - |
| - | - | 1480 | 1119 | - | - | 0 | - |
| - | - | 1032 | 1135 | - | - | 0 | - |
| - | - | 1499 | 1136 | - | - | 0 | - |
| - | - | 647.7 | 1156 | - | - | 0 | - |
| - | - | 668.6 | 2923 | - | - | 0 | - |
| - | - | 1018 | 3072 | - | - | 0 | - |

m/z Charge Intensity FragmentType MassShift Position
120.08126831054688 0 967.59033
120.62261962890625 0 372.07077
124.03963470458984 0 501.52423
125.10771179199219 0 1776.3557
127.08702850341797 0 799.7428
128.1073760986328 0 1571.2905
128.86654663085938 0 369.0227
129.06626892089844 0 7265.533
129.10263061523438 0 61401.91
130.065185546875 0 485.1126
130.07008361816406 0 430.38776
130.101318359375 0 496.4074
130.10597229003906 0 3833.317
132.91754150390625 0 401.4069
133.06117248535156 0 5548.238
133.61886596679688 0 397.5464
136.07606506347656 0 2347.1213
137.7515869140625 0 449.62878
138.06651306152344 0 470.57687
139.08705139160156 0 1958.5651
141.06631469726562 0 3741.7144
141.10263061523438 0 4690.6025
142.12306213378906 0 5071.0117
143.0818328857422 0 2128.7012
143.11856079101562 0 1796.1754 d 1
143.12619018554688 0 814.3272
147.07708740234375 0 612.6114
148.947021484375 0 529.60266
151.0870361328125 0 514.9024
152.03463745117188 0 3632.3555
152.10731506347656 0 2726.2705
153.06637573242188 0 942.0482
153.10272216796875 0 1152.3757
155.1182861328125 0 8320.704
156.0778045654297 0 584.81995
157.09751892089844 0 2299.9817
158.060791015625 0 606.98773
158.09286499023438 0 2914.2737
161.09317016601562 0 484.91592
163.07171630859375 0 534.9735
165.10269165039062 0 1629.104
168.07736206054688 0 583.60297
169.0611572265625 0 17294.738
169.09756469726562 0 6094.353
169.1339111328125 0 2429.6553
170.06472778320312 0 1281.8795
170.10118103027344 0 698.8316
171.1131591796875 0 10721.71
172.1168975830078 0 1102.8943
173.0561065673828 0 3734.9348
173.1288604736328 0 10309.724
174.13241577148438 0 932.5227
175.0870361328125 0 3091.1162
180.1132049560547 0 1634.8918
181.0975799560547 0 4593.3135
181.13375854492188 0 3145.9807
182.1292266845703 0 16173.089
182.13824462890625 0 1113.7225
183.1131591796875 0 2183.84
183.13259887695312 0 1690.4902
183.149658203125 0 765.3913
185.0924072265625 0 5828.803 y Water loss 8
185.1653289794922 0 313422.22 a 1
186.0877227783203 0 39331.27
186.16860961914062 0 32184.586
187.09103393554688 0 2802.1594
187.10794067382812 0 954.3641
187.14447021484375 0 8372.519
187.17196655273438 0 1159.4688
188.1397705078125 0 8176.1973
188.14840698242188 0 634.078
189.14334106445312 0 684.9774
190.08270263671875 0 1510.9637
191.0822296142578 0 748.61383
193.09754943847656 0 1118.5118
195.1133575439453 0 1256.8138
197.09239196777344 0 853.2969
197.12876892089844 0 2489.3643
198.1239776611328 0 1579.6793
199.1085968017578 0 1100.8895
200.13973999023438 0 4118.614
201.1237335205078 0 1737.3689
203.10304260253906 0 198623.25 y 8
204.10641479492188 0 16315.712
205.1080322265625 0 1775.0588
208.10838317871094 0 4436.706
208.14483642578125 0 790.14624
209.09254455566406 0 6014.0747
209.1285858154297 0 650.7833
210.02597045898438 0 570.47235
210.12408447265625 0 17667.83
211.10787963867188 0 783.89264
211.12734985351562 0 1503.2972
212.10321044921875 0 1014.9025
213.16021728515625 0 160598.83 b 1
214.1186981201172 0 728.2801
214.16351318359375 0 18527.145
215.10289001464844 0 906.3098
215.13929748535156 0 4040.3577
215.1665802001953 0 1402.0973
220.10821533203125 0 1484.699
223.10743713378906 0 884.07825
224.1400909423828 0 999.10126
226.11929321289062 0 3161.9773
226.1554412841797 0 124545.94
227.11410522460938 0 920.6787
227.15866088867188 0 14716.275
228.1346435546875 0 17766.596 y Water loss 5
228.16123962402344 0 1050.824
229.13829040527344 0 2132.9739
236.1396942138672 0 1616.2157
237.08724975585938 0 602.75085
238.11917114257812 0 2761.801
239.1754608154297 0 937.1867
240.13494873046875 0 2030.0013
242.150146484375 0 1118.4921
243.1817626953125 0 4070.2454
244.1294403076172 0 1842.9115
244.14210510253906 0 851.2817
248.10491943359375 0 521.1704
252.13453674316406 0 1697.7224
252.17112731933594 0 4773.832
253.1654510498047 0 795.97253
254.1138916015625 0 1029.1097
254.15028381347656 0 5982.699
256.1296081542969 0 2287.1794 y Water loss 7
256.1649475097656 0 508.47375
257.1248474121094 0 708.2594
261.1708984375 0 1116.698
263.4481506347656 0 557.0153
264.1355895996094 0 797.66064
265.1301574707031 0 607.1144
266.1138000488281 0 2006.4559
268.1293029785156 0 933.4968
269.197509765625 0 6129.0293
270.1452941894531 0 1002.7282
270.18157958984375 0 20053.303 b 2
270.1987609863281 0 664.6115
271.1407470703125 0 1382.0546
271.1770324707031 0 13449.8955
272.1802978515625 0 2436.6123
274.14007568359375 0 11955 y 7
274.1700439453125 0 758.7282
275.1432189941406 0 1540.3711
279.18194580078125 0 25149.934
280.1669616699219 0 653.7127
280.1851806640625 0 3281.427
281.1615905761719 0 4565.9175
282.12646484375 0 611.1495
282.1449890136719 0 1196.9825
283.1407470703125 0 3711.2915
284.0880432128906 0 659.0676
285.15631103515625 0 5835.471
285.66644287109375 0 4383.011 y 4
287.20831298828125 0 33420.668
288.16815185546875 0 894.8795
288.2115478515625 0 4727.9976
295.21356201171875 0 640.0997
296.1966857910156 0 1171.5869
297.1926574707031 0 47040.215
298.1959228515625 0 8058.8364
299.171875 0 93248.04
300.13134765625 0 925.6686
300.17474365234375 0 12916.64
301.1764831542969 0 1515.484
309.15582275390625 0 1613.4905
311.3487548828125 0 577.90814
314.2195739746094 0 2513.9482
323.17291259765625 0 662.8666
323.20819091796875 0 3171.0867
324.21258544921875 0 612.4117
325.1878662109375 0 3108.999
325.22222900390625 0 591.6725
325.6855773925781 0 706.3891
327.1669006347656 0 5615.385 y Water loss 6
327.2012634277344 0 592.78436
328.1636657714844 0 2133.576
331.6851501464844 0 673.17206
332.177490234375 0 2355.1577
332.6782531738281 0 1099.4702
336.2043762207031 0 1346.7319
337.1878662109375 0 3581.389
338.1882019042969 0 846.1825
340.23486328125 0 24591.125
340.6903381347656 0 6973.539 y Water loss 3
341.19061279296875 0 2994.5432
341.2385559082031 0 3950.8306
345.17742919921875 0 1826.0629 y 6
349.6951904296875 0 1998.8553 y 3
350.1972351074219 0 1103.4924
350.21759033203125 0 906.7978
351.201416015625 0 825.63794
353.21917724609375 0 1664.6838
354.2149658203125 0 1905.7737
355.2352294921875 0 908.3209
360.72454833984375 0 1164.514
363.2032470703125 0 1868.2676
366.2143249511719 0 1029.9135
366.2492370605469 0 588.54504
368.2295227050781 0 123213.12
369.23272705078125 0 26629.129
369.6952209472656 0 1035.2117 y Ammonia loss 2
369.74053955078125 0 914.4506
370.20928955078125 0 837.9121
370.2444763183594 0 10227.805
371.24652099609375 0 2042.3912
376.1983337402344 0 2096.8616
376.2406921386719 0 618.4113
378.20745849609375 0 1683.4495 y 2
380.2297668457031 0 2630.0374
382.2262878417969 0 2792.6023
382.72943115234375 0 1614.9857
383.2342224121094 0 1578.3008 b 7
383.7367248535156 0 762.5968
384.2624816894531 0 767.3342
390.2138671875 0 4057.2039
394.20867919921875 0 3487.5737
394.2447814941406 0 2247.2334
395.21148681640625 0 1010.6049
396.223876953125 0 3148.5173
397.2286071777344 0 1188.8376
398.24041748046875 0 31042.852 b 3
399.2433776855469 0 5425.8643
400.2440490722656 0 666.3235
406.20556640625 0 561.79913
407.2416687011719 0 1926.4033
408.22515869140625 0 4739.715
409.2270202636719 0 1105.7072
411.2350769042969 0 4446.065
412.2392272949219 0 1306.6198
417.76361083984375 0 1927.4907
418.2646789550781 0 1108.8707
425.251708984375 0 1836.8744
425.74322509765625 0 3292.3567 y Water loss 1
426.24224853515625 0 2097.6257 y Ammonia loss 1
426.74322509765625 0 683.0845
431.7613830566406 0 4482.572 b 8
432.2632751464844 0 2595.2114
434.2403869628906 0 997.13275
434.74847412109375 0 10148.332 y 1
435.2499084472656 0 4412.6465
435.7516174316406 0 1574.3224
437.2517395019531 0 1569.3182
437.28717041015625 0 2011.4679
439.2314147949219 0 1520.4535
439.3042907714844 0 936.2694
439.740966796875 0 2736.9915
440.2416076660156 0 1236.4103
441.2454833984375 0 953.1975
442.27801513671875 0 1964.0875
447.2358703613281 0 3676.828
448.2398681640625 0 765.97144
449.2153625488281 0 1093.9016
451.26715087890625 0 2403.3855
454.2777099609375 0 4187.8613
455.28387451171875 0 810.0312
456.25811767578125 0 1909.77
458.7362976074219 0 1033.0986
464.26220703125 0 14277.782
465.2471618652344 0 3294.5168
465.2820739746094 0 9431.57
466.2494812011719 0 1199.1615
466.2860107421875 0 2616.511
466.76458740234375 0 992.8681
467.2638854980469 0 762.33276
467.2976989746094 0 4299.775
468.2986145019531 0 1904.8539
473.27227783203125 0 6852.3105 y 5
474.27508544921875 0 1726.5251
475.2774963378906 0 4400.2363 Precursor Water loss
475.77667236328125 0 785.9971 Precursor Ammonia loss
476.28155517578125 0 1174.8058
477.57611083984375 0 842.8067
478.27783203125 0 3136.7024
479.2621765136719 0 8589.964
480.263916015625 0 1711.1772
481.3138427734375 0 881.94275
482.2727355957031 0 35906.703
483.27618408203125 0 7276.6665
484.2455749511719 0 760.5914
484.2825622558594 0 15446.211 Precursor
484.78448486328125 0 8854.782
485.28594970703125 0 3735.4563
485.35797119140625 0 1181.3593
485.7878723144531 0 847.3629
491.2645263671875 0 940.3873
495.2923889160156 0 1373.7742 b 4
496.2884216308594 0 2897.3545
502.2455139160156 0 951.6725
508.2887268066406 0 4766.8857
509.2915344238281 0 1693.9268
510.2546691894531 0 1104.9478
510.7552795410156 0 1013.1458
518.2723999023438 0 1986.3158
522.7662963867188 0 1865.4387
524.3177490234375 0 1675.888
525.3151245117188 0 9182.931
526.3176879882812 0 2926.0173
534.3037109375 0 907.809
535.299560546875 0 2566.4453
536.282958984375 0 12457.6
537.2852783203125 0 3489.2268
540.3160400390625 0 972.9458
552.3140869140625 0 7308.135 y Water loss 4
553.3096313476562 0 102139.15
554.3126220703125 0 30668.129
555.3147583007812 0 5382.2446
555.361572265625 0 991.6996
559.2530517578125 0 1598.9912
567.3627319335938 0 1047.6891
567.7679443359375 0 1532.574
568.2673950195312 0 1028.1887
568.3120727539062 0 1162.5709
570.3250122070312 0 211628.95 y 4
571.3278198242188 0 64365.355
572.329833984375 0 12266.822
574.8153686523438 0 913.419
577.344970703125 0 3996.4019
578.339111328125 0 952.5494
579.3233642578125 0 1089.6831
580.3818969726562 0 4042.553
581.3855590820312 0 1177.4882
586.778076171875 0 667.46344
595.3561401367188 0 5223.1855
596.3578491210938 0 1654.843
600.2821044921875 0 984.9535
600.7771606445312 0 1090.7579
608.7902221679688 0 4068.8235
609.2903442382812 0 1305.1487
617.3021240234375 0 8071.477
617.8038330078125 0 5437.9565
618.3037719726562 0 2095.8733
618.80517578125 0 839.9472
621.3712768554688 0 2042.3003
622.3677978515625 0 1186.3435
623.3858032226562 0 1419.5496 b 5
638.39990234375 0 2116.5042
639.4041748046875 0 961.6297
640.2451782226562 0 635.5577
649.369384765625 0 3180.0054
650.3622436523438 0 7190.159
651.3076782226562 0 818.1308
651.365966796875 0 3316.0066
652.3780517578125 0 1585.4834
654.4287109375 0 1643.4485
655.434326171875 0 755.54
658.3436889648438 0 1037.5364
659.3143920898438 0 7044.3257
659.8147583007812 0 4053.1707
660.3173828125 0 1966.5339
663.3519897460938 0 698.2345
664.3048706054688 0 1134.303
664.8018188476562 0 819.6675
666.3933715820312 0 23554.248
667.39697265625 0 8558.953
667.82568359375 0 8066.131
668.3267211914062 0 4795.227
668.3989868164062 0 1782.1396
668.8287963867188 0 4301.942
669.3262329101562 0 668.4412
672.8197631835938 0 1284.108
673.3175048828125 0 1703.6215
676.4149169921875 0 3812.2246
677.4173583984375 0 2117.7234
680.3731079101562 0 4026.7014 y Water loss 3
681.359375 0 7236.556 y Ammonia loss 3
682.3623657226562 0 2629.6885
694.4246215820312 0 10099.065 b 6
695.4281005859375 0 4401.958
696.4297485351562 0 1515.7722
698.3829956054688 0 6642.778 y 3
699.3847045898438 0 2751.7341
720.3661499023438 0 1625.9203
720.4409790039062 0 900.2702
721.3736572265625 0 688.90436
730.4222412109375 0 960.3423
737.3927001953125 0 5830.6694 y Water loss 2
737.4671630859375 0 6821.595
738.3810424804688 0 8269.462 y Ammonia loss 2
738.4695434570312 0 3317.6345
739.3831176757812 0 2285.4175
747.4520874023438 0 1394.2964
748.4364013671875 0 3263.7705 b Ammonia loss 7
749.4366455078125 0 2873.1438
750.4368286132812 0 1033.7456
753.3873291015625 0 1122.395
755.4043579101562 0 187660.14 y 2
756.4071044921875 0 74287.4
757.4100341796875 0 17941.283
758.4100952148438 0 1245.2396
763.4464111328125 0 2325.0642
764.4486083984375 0 1081.3573
765.4615478515625 0 97188.984 b 7
766.4644775390625 0 43466.938
767.4674072265625 0 9965.911
768.4743041992188 0 1098.9105
791.4759521484375 0 1053.9451
804.3781127929688 0 939.59784
817.3964233398438 0 836.6411
830.4214477539062 0 821.99585
851.4617919921875 0 1066.596 y Ammonia loss 1
868.48828125 0 22350.477 y 1
869.490478515625 0 11698.208
870.492431640625 0 3553.3247
874.3809814453125 0 1400.3815
875.3863525390625 0 959.456
878.4761962890625 0 1053.7773
891.40771484375 0 2788.0186
892.4134521484375 0 1208.6885
916.4577026367188 0 1007.70074
917.4647216796875 0 1040.8978
945.4617309570312 0 765.48694
985.4498901367188 0 1417.4547
1002.4736328125 0 2051.3877
1003.4686279296875 0 1035.2275
1045.528076171875 0 776.08636
1100.4700927734375 0 898.77734
1117.5030517578125 0 1961.9277
1118.50537109375 0 1480.1731
1134.532958984375 0 1032.1652
1135.5299072265625 0 1499.4653
1155.9202880859375 0 647.6556
2922.853515625 0 668.6286
3072.343505859375 0 1018.2688

Spectrum Details

|  |  |
| --- | --- |
| Matched peaks? Matched peaks The total absolute number of peaks matched. Additionally in brackets the total fraction of peaks matched and the total number of peaks is shown. | 41 (9.67% of 424) |
| FDR? FDR The false discovery rate estimated for this peptide. It is calculated by matching all theoretical fragments with a non-integer shift with the raw peaks for this spectrum. This is done with 40 different shifts. The resulting percentage is the average number of annotated peaks over the number of annotated peaks with the correct spectrum. | 0.46% |
| Satellite FDR? Satellite FDR See the FDR for details on its calculation. This satellite ion specific FDR only contains the satellite ions (d/w) for I/L/J positions. | 0.00% |
| PSM Score? PSM Score The PSM Score as given by Hecklib to this annotated spectrum. It is shown with three significant figures. | 408 |

#### Spectrum 3145? Spectrum 3145 The raw spectrum of this peptide as annotated by Hecklib. The fragments are coloured according to ion type (see legend). Any peaks with a star '\*' as text can be hovered over to see the full details, first the ion type second the mass shift type. By hovering over the amino acids in the peptide or ions in the legend the corresponding peaks are highlighted. By toggling the 'Unassigned' label you can turn the background (unassigned) peaks on or off in the plot. By updating the slider in the Ion legend you can update the spectrum to only show the top X% of the peaks with labels. The top X% means any peak that is within X% of the highest intensity. By dragging in the spectrum you can zoom in to a specific part of the spectrum and use 'Zoom Out' to get back to the original zoom level. The annotation of the spectrum is based on the given sequence in the peptides file and is done with different software so inconsistencies are likely. The peaks are annotated based on the given sequence, with 20 ppm tolerance.

Copy Data

##### Spectrum 3145 (TSV)

###### Preview

```
Loading example...
```

*Click on the button to copy the data to your clipboard.*

Mz MinMz MaxIntensity Max

WidthHeightPeptide font sizePeptide stroke widthSpectrum font sizeSpectrum stroke widthCompact peptide

Ion legend

wxyz

abcd

OtherUnassignedIon^Charge^_Position_Show for top:%

VLGQPKAAPS

05.60e+41.12e+51.68e+52.24e+5

Zoom Out

d^+1^_2_y^+1^_2_a^+1^_2_y^+1^_2_b^+1^_2_y^+2^_5_y^+1^_3_b^+1^_3_y^+1^_3_y^+2^_6_y^+1^_4_y^+2^_7_y^+1^_4_y^+2^_7_y^+2^_8_y^+2^_8_b^+2^_8_b^+1^_4_y^+2^_9_y^+2^_9_b^+2^_9_y^+2^_9_y^+1^_5_\*\*\*b^+1^_5_y^+1^_6_y^+1^_6_y^+1^_6_b^+1^_6_y^+1^_7_y^+1^_7_b^+1^_7_y^+1^_7_y^+1^_8_y^+1^_8_b^+1^_8_y^+1^_8_b^+1^_8_y^+1^_9_y^+1^_9_

0625124918742499

Fragment Matches Table

Show background peaks

| Position | Ion type | Intensity | mz Theoretical | mz Error (Th) | mz Error (ppm) | Charge | Series Number |
| --- | --- | --- | --- | --- | --- | --- | --- |
| - | - | 980.1 | 127.1 | - | - | 0 | - |
| - | - | 401.8 | 127.9 | - | - | 0 | - |
| - | - | 4951 | 129.1 | - | - | 0 | - |
| - | - | 4.048E+04 | 129.1 | - | - | 0 | - |
| - | - | 651.9 | 130 | - | - | 0 | - |
| - | - | 472.1 | 130.1 | - | - | 0 | - |
| - | - | 2968 | 130.1 | - | - | 0 | - |
| - | - | 373.7 | 133.1 | - | - | 0 | - |
| - | - | 398 | 135.1 | - | - | 0 | - |
| - | - | 1459 | 136.1 | - | - | 0 | - |
| - | - | 1401 | 139.1 | - | - | 0 | - |
| - | - | 3067 | 141.1 | - | - | 0 | - |
| - | - | 3242 | 141.1 | - | - | 0 | - |
| - | - | 1371 | 142.1 | - | - | 0 | - |
| - | - | 1715 | 143.1 | - | - | 0 | - |
| 2 | d | 1669 | 143.1 | 0.0001677 | 1.172 | +1 | 2 |
| - | - | 410.2 | 144.8 | - | - | 0 | - |
| - | - | 569.7 | 147.1 | - | - | 0 | - |
| - | - | 468.4 | 147.1 | - | - | 0 | - |
| - | - | 535.2 | 149 | - | - | 0 | - |
| - | - | 2856 | 152 | - | - | 0 | - |
| - | - | 455.6 | 152.1 | - | - | 0 | - |
| - | - | 722.8 | 153.1 | - | - | 0 | - |
| - | - | 591.6 | 153.1 | - | - | 0 | - |
| - | - | 5382 | 155.1 | - | - | 0 | - |
| - | - | 653.6 | 156.1 | - | - | 0 | - |
| - | - | 542.1 | 156.1 | - | - | 0 | - |
| - | - | 1712 | 157.1 | - | - | 0 | - |
| - | - | 949 | 157.2 | - | - | 0 | - |
| - | - | 2299 | 158.1 | - | - | 0 | - |
| - | - | 455.3 | 161.5 | - | - | 0 | - |
| - | - | 666.3 | 165.1 | - | - | 0 | - |
| - | - | 507.8 | 168.1 | - | - | 0 | - |
| - | - | 1.22E+04 | 169.1 | - | - | 0 | - |
| - | - | 3827 | 169.1 | - | - | 0 | - |
| - | - | 1333 | 169.1 | - | - | 0 | - |
| - | - | 766.3 | 170.1 | - | - | 0 | - |
| - | - | 473.9 | 170.3 | - | - | 0 | - |
| - | - | 6204 | 171.1 | - | - | 0 | - |
| - | - | 966.7 | 173.1 | - | - | 0 | - |
| - | - | 899.9 | 173.5 | - | - | 0 | - |
| - | - | 981.4 | 180.1 | - | - | 0 | - |
| - | - | 3524 | 181.1 | - | - | 0 | - |
| - | - | 2316 | 181.1 | - | - | 0 | - |
| - | - | 1.004E+04 | 182.1 | - | - | 0 | - |
| - | - | 928 | 182.1 | - | - | 0 | - |
| - | - | 2128 | 183.1 | - | - | 0 | - |
| - | - | 1341 | 183.1 | - | - | 0 | - |
| - | - | 1317 | 183.1 | - | - | 0 | - |
| 9 | y | 4292 | 185.1 | 0.0002012 | 1.087 | +1 | 2 |
| 2 | a | 2.217E+05 | 185.2 | 0.0002299 | 1.241 | +1 | 2 |
| - | - | 2.673E+04 | 186.1 | - | - | 0 | - |
| - | - | 667 | 186.1 | - | - | 0 | - |
| - | - | 2.288E+04 | 186.2 | - | - | 0 | - |
| - | - | 631.9 | 187.1 | - | - | 0 | - |
| - | - | 1651 | 187.1 | - | - | 0 | - |
| - | - | 1629 | 187.1 | - | - | 0 | - |
| - | - | 1225 | 187.2 | - | - | 0 | - |
| - | - | 5772 | 188.1 | - | - | 0 | - |
| - | - | 646.8 | 191.1 | - | - | 0 | - |
| - | - | 671.4 | 193.1 | - | - | 0 | - |
| - | - | 564.6 | 195.1 | - | - | 0 | - |
| - | - | 773.8 | 197.1 | - | - | 0 | - |
| - | - | 649.5 | 197.1 | - | - | 0 | - |
| - | - | 741.7 | 198.1 | - | - | 0 | - |
| - | - | 1006 | 199.1 | - | - | 0 | - |
| - | - | 3389 | 200.1 | - | - | 0 | - |
| - | - | 570.1 | 201.1 | - | - | 0 | - |
| - | - | 1453 | 202.1 | - | - | 0 | - |
| 9 | y | 1.426E+05 | 203.1 | 8.877E-05 | 0.4371 | +1 | 2 |
| - | - | 1.226E+04 | 204.1 | - | - | 0 | - |
| - | - | 969.1 | 205.1 | - | - | 0 | - |
| - | - | 2590 | 208.1 | - | - | 0 | - |
| - | - | 897.3 | 208.1 | - | - | 0 | - |
| - | - | 2889 | 209.1 | - | - | 0 | - |
| - | - | 772.8 | 209.1 | - | - | 0 | - |
| - | - | 1.232E+04 | 210.1 | - | - | 0 | - |
| - | - | 972.1 | 211.1 | - | - | 0 | - |
| - | - | 716.7 | 211.1 | - | - | 0 | - |
| - | - | 906.3 | 211.2 | - | - | 0 | - |
| - | - | 792.9 | 212.1 | - | - | 0 | - |
| 2 | b | 1.193E+05 | 213.2 | 0.000112 | 0.5253 | +1 | 2 |
| - | - | 1.371E+04 | 214.2 | - | - | 0 | - |
| - | - | 1165 | 215.1 | - | - | 0 | - |
| - | - | 910.4 | 215.2 | - | - | 0 | - |
| - | - | 684.7 | 223.1 | - | - | 0 | - |
| - | - | 1437 | 226.1 | - | - | 0 | - |
| - | - | 9.183E+04 | 226.2 | - | - | 0 | - |
| - | - | 1.034E+04 | 227.2 | - | - | 0 | - |
| 6 | y | 1.166E+04 | 228.1 | 0.000101 | 0.4428 | +2 | 5 |
| - | - | 1221 | 229.1 | - | - | 0 | - |
| - | - | 611.9 | 233.1 | - | - | 0 | - |
| - | - | 473 | 233.9 | - | - | 0 | - |
| - | - | 512.3 | 234.1 | - | - | 0 | - |
| - | - | 1212 | 236.1 | - | - | 0 | - |
| - | - | 1477 | 238.1 | - | - | 0 | - |
| - | - | 471.8 | 238.4 | - | - | 0 | - |
| - | - | 705.4 | 239.2 | - | - | 0 | - |
| - | - | 612.8 | 240.1 | - | - | 0 | - |
| - | - | 1967 | 240.1 | - | - | 0 | - |
| - | - | 507.2 | 241 | - | - | 0 | - |
| - | - | 1082 | 242.1 | - | - | 0 | - |
| - | - | 2567 | 243.2 | - | - | 0 | - |
| - | - | 1355 | 252.1 | - | - | 0 | - |
| - | - | 3683 | 252.2 | - | - | 0 | - |
| - | - | 765.3 | 253.2 | - | - | 0 | - |
| - | - | 989.4 | 254.1 | - | - | 0 | - |
| - | - | 3558 | 254.2 | - | - | 0 | - |
| 8 | y | 1932 | 256.1 | 0.0001542 | 0.602 | +1 | 3 |
| - | - | 1187 | 257.1 | - | - | 0 | - |
| - | - | 1406 | 266.1 | - | - | 0 | - |
| - | - | 4683 | 269.2 | - | - | 0 | - |
| 3 | b | 1.332E+04 | 270.2 | 6.573E-05 | 0.2433 | +1 | 3 |
| - | - | 1.015E+04 | 271.2 | - | - | 0 | - |
| - | - | 1577 | 272.2 | - | - | 0 | - |
| 8 | y | 8128 | 274.1 | 9.875E-05 | 0.3602 | +1 | 3 |
| - | - | 1127 | 275.1 | - | - | 0 | - |
| - | - | 598.8 | 277.2 | - | - | 0 | - |
| - | - | 1.828E+04 | 279.2 | - | - | 0 | - |
| - | - | 2436 | 280.2 | - | - | 0 | - |
| - | - | 2783 | 281.2 | - | - | 0 | - |
| - | - | 2674 | 283.1 | - | - | 0 | - |
| - | - | 1054 | 283.2 | - | - | 0 | - |
| - | - | 4816 | 285.2 | - | - | 0 | - |
| 5 | y | 1985 | 285.7 | 8.078E-06 | 0.02828 | +2 | 6 |
| - | - | 2.311E+04 | 287.2 | - | - | 0 | - |
| - | - | 654.6 | 288.2 | - | - | 0 | - |
| - | - | 3087 | 288.2 | - | - | 0 | - |
| - | - | 835.3 | 297.2 | - | - | 0 | - |
| - | - | 3.38E+04 | 297.2 | - | - | 0 | - |
| - | - | 5704 | 298.2 | - | - | 0 | - |
| - | - | 6.855E+04 | 299.2 | - | - | 0 | - |
| - | - | 1.005E+04 | 300.2 | - | - | 0 | - |
| - | - | 1095 | 309.2 | - | - | 0 | - |
| - | - | 1918 | 314.2 | - | - | 0 | - |
| - | - | 2022 | 323.2 | - | - | 0 | - |
| - | - | 2139 | 325.2 | - | - | 0 | - |
| 7 | y | 4265 | 327.2 | 0.0001586 | 0.4848 | +1 | 4 |
| - | - | 2419 | 328.2 | - | - | 0 | - |
| - | - | 1335 | 332.2 | - | - | 0 | - |
| - | - | 664.1 | 336.2 | - | - | 0 | - |
| - | - | 3193 | 337.2 | - | - | 0 | - |
| - | - | 1.722E+04 | 340.2 | - | - | 0 | - |
| 4 | y | 6894 | 340.7 | 3.336E-05 | 0.09792 | +2 | 7 |
| - | - | 2226 | 341.2 | - | - | 0 | - |
| - | - | 3632 | 341.2 | - | - | 0 | - |
| 7 | y | 885.6 | 345.2 | 0.0006293 | 1.823 | +1 | 4 |
| 4 | y | 848 | 349.7 | 0.0007934 | 2.269 | +2 | 7 |
| - | - | 1348 | 350.2 | - | - | 0 | - |
| - | - | 994.4 | 353.2 | - | - | 0 | - |
| - | - | 704.6 | 354.2 | - | - | 0 | - |
| - | - | 646 | 355.2 | - | - | 0 | - |
| - | - | 566.7 | 360.7 | - | - | 0 | - |
| - | - | 693.3 | 361.2 | - | - | 0 | - |
| - | - | 1470 | 363.2 | - | - | 0 | - |
| - | - | 9.09E+04 | 368.2 | - | - | 0 | - |
| - | - | 608.3 | 368.7 | - | - | 0 | - |
| - | - | 1.791E+04 | 369.2 | - | - | 0 | - |
| 3 | y | 632.9 | 369.7 | 0.002482 | 6.713 | +2 | 8 |
| - | - | 740 | 369.7 | - | - | 0 | - |
| - | - | 6568 | 370.2 | - | - | 0 | - |
| - | - | 1102 | 371.2 | - | - | 0 | - |
| - | - | 1746 | 376.2 | - | - | 0 | - |
| - | - | 619.1 | 376.2 | - | - | 0 | - |
| 3 | y | 1064 | 378.2 | 0.001353 | 3.578 | +2 | 8 |
| - | - | 601.7 | 378.7 | - | - | 0 | - |
| - | - | 1902 | 380.2 | - | - | 0 | - |
| 8 | b | 1075 | 383.2 | 4.669E-05 | 0.1218 | +2 | 8 |
| - | - | 798.6 | 383.7 | - | - | 0 | - |
| - | - | 1633 | 390.2 | - | - | 0 | - |
| - | - | 1865 | 394.2 | - | - | 0 | - |
| - | - | 993 | 394.2 | - | - | 0 | - |
| - | - | 3140 | 396.2 | - | - | 0 | - |
| 4 | b | 2.11E+04 | 398.2 | 0.0002021 | 0.5074 | +1 | 4 |
| - | - | 4257 | 399.2 | - | - | 0 | - |
| - | - | 1346 | 407.2 | - | - | 0 | - |
| - | - | 3365 | 408.2 | - | - | 0 | - |
| - | - | 706.4 | 409.2 | - | - | 0 | - |
| - | - | 2798 | 411.2 | - | - | 0 | - |
| - | - | 769.8 | 412.2 | - | - | 0 | - |
| - | - | 1747 | 417.8 | - | - | 0 | - |
| - | - | 617.7 | 418.3 | - | - | 0 | - |
| - | - | 1175 | 425.3 | - | - | 0 | - |
| 2 | y | 1917 | 425.7 | 0.0004844 | 1.138 | +2 | 9 |
| 2 | y | 947.1 | 426.2 | 0.004395 | 10.31 | +2 | 9 |
| 9 | b | 3076 | 431.8 | 0.0002445 | 0.5664 | +2 | 9 |
| - | - | 1896 | 432.3 | - | - | 0 | - |
| 2 | y | 6517 | 434.7 | 0.000121 | 0.2783 | +2 | 9 |
| - | - | 3433 | 435.2 | - | - | 0 | - |
| - | - | 1176 | 435.8 | - | - | 0 | - |
| - | - | 867.6 | 437.3 | - | - | 0 | - |
| - | - | 1533 | 437.3 | - | - | 0 | - |
| - | - | 635 | 439.2 | - | - | 0 | - |
| - | - | 567.3 | 439.3 | - | - | 0 | - |
| - | - | 1086 | 439.7 | - | - | 0 | - |
| - | - | 1136 | 440.2 | - | - | 0 | - |
| - | - | 1706 | 442.3 | - | - | 0 | - |
| - | - | 2765 | 447.2 | - | - | 0 | - |
| - | - | 965 | 448.2 | - | - | 0 | - |
| - | - | 690.7 | 449.2 | - | - | 0 | - |
| - | - | 1101 | 450.3 | - | - | 0 | - |
| - | - | 1338 | 451.3 | - | - | 0 | - |
| - | - | 3451 | 454.3 | - | - | 0 | - |
| - | - | 880.7 | 455.3 | - | - | 0 | - |
| - | - | 1178 | 456.3 | - | - | 0 | - |
| - | - | 1.021E+04 | 464.3 | - | - | 0 | - |
| - | - | 2067 | 465.2 | - | - | 0 | - |
| - | - | 7629 | 465.3 | - | - | 0 | - |
| - | - | 629.8 | 466.2 | - | - | 0 | - |
| - | - | 1270 | 466.3 | - | - | 0 | - |
| - | - | 888.8 | 467.2 | - | - | 0 | - |
| - | - | 3261 | 467.3 | - | - | 0 | - |
| - | - | 1170 | 468.3 | - | - | 0 | - |
| 6 | y | 5328 | 473.3 | 0.0002786 | 0.5886 | +1 | 5 |
| - | - | 890.2 | 474.3 | - | - | 0 | - |
| 0 | Precursor | 3825 | 475.3 | 0.000237 | 0.4987 | +2 | -1 |
| 0 | Precursor | 1686 | 475.8 | 0.00864 | 18.16 | +2 | -1 |
| - | - | 739.7 | 476.3 | - | - | 0 | - |
| - | - | 1875 | 478.3 | - | - | 0 | - |
| - | - | 7332 | 479.3 | - | - | 0 | - |
| - | - | 1478 | 480.3 | - | - | 0 | - |
| - | - | 1032 | 481.3 | - | - | 0 | - |
| - | - | 2.592E+04 | 482.3 | - | - | 0 | - |
| - | - | 6512 | 483.3 | - | - | 0 | - |
| - | - | 642.5 | 484.2 | - | - | 0 | - |
| 0 | Precursor | 9963 | 484.3 | 0.0002398 | 0.4952 | +2 | -1 |
| - | - | 6154 | 484.8 | - | - | 0 | - |
| - | - | 2951 | 485.3 | - | - | 0 | - |
| - | - | 779.2 | 490.3 | - | - | 0 | - |
| 5 | b | 1666 | 495.3 | 0.0005062 | 1.022 | +1 | 5 |
| - | - | 2136 | 496.3 | - | - | 0 | - |
| - | - | 3623 | 508.3 | - | - | 0 | - |
| - | - | 1872 | 518.3 | - | - | 0 | - |
| - | - | 1480 | 524.3 | - | - | 0 | - |
| - | - | 5555 | 525.3 | - | - | 0 | - |
| - | - | 1714 | 526.3 | - | - | 0 | - |
| - | - | 1418 | 535.3 | - | - | 0 | - |
| - | - | 8293 | 536.3 | - | - | 0 | - |
| - | - | 3046 | 537.3 | - | - | 0 | - |
| - | - | 853.1 | 540.3 | - | - | 0 | - |
| 5 | y | 6772 | 552.3 | 0.0004856 | 0.8791 | +1 | 6 |
| 5 | y | 7.378E+04 | 553.3 | 0.01086 | 19.63 | +1 | 6 |
| - | - | 2.18E+04 | 554.3 | - | - | 0 | - |
| - | - | 3728 | 555.3 | - | - | 0 | - |
| - | - | 1498 | 567.4 | - | - | 0 | - |
| - | - | 1459 | 568.3 | - | - | 0 | - |
| 5 | y | 1.442E+05 | 570.3 | 0.0004912 | 0.8612 | +1 | 6 |
| - | - | 4.197E+04 | 571.3 | - | - | 0 | - |
| - | - | 7818 | 572.3 | - | - | 0 | - |
| - | - | 3051 | 577.3 | - | - | 0 | - |
| - | - | 784.5 | 578.3 | - | - | 0 | - |
| - | - | 2989 | 580.4 | - | - | 0 | - |
| - | - | 1126 | 581.4 | - | - | 0 | - |
| - | - | 4056 | 595.4 | - | - | 0 | - |
| - | - | 1812 | 596.4 | - | - | 0 | - |
| - | - | 1587 | 621.4 | - | - | 0 | - |
| - | - | 881.4 | 622.4 | - | - | 0 | - |
| 6 | b | 610.5 | 623.4 | 0.004893 | 7.849 | +1 | 6 |
| - | - | 1813 | 638.4 | - | - | 0 | - |
| - | - | 2244 | 649.4 | - | - | 0 | - |
| - | - | 6337 | 650.4 | - | - | 0 | - |
| - | - | 2066 | 651.4 | - | - | 0 | - |
| - | - | 907.5 | 652.4 | - | - | 0 | - |
| - | - | 1452 | 654.4 | - | - | 0 | - |
| - | - | 1.845E+04 | 666.4 | - | - | 0 | - |
| - | - | 6107 | 667.4 | - | - | 0 | - |
| - | - | 1174 | 668.4 | - | - | 0 | - |
| - | - | 823.3 | 674 | - | - | 0 | - |
| - | - | 2831 | 676.4 | - | - | 0 | - |
| - | - | 840 | 677.4 | - | - | 0 | - |
| 4 | y | 3874 | 680.4 | 0.0003241 | 0.4764 | +1 | 7 |
| 4 | y | 5321 | 681.4 | 0.002209 | 3.243 | +1 | 7 |
| - | - | 2468 | 682.4 | - | - | 0 | - |
| - | - | 752.1 | 691.4 | - | - | 0 | - |
| 7 | b | 8082 | 694.4 | 0.0008692 | 1.252 | +1 | 7 |
| - | - | 3003 | 695.4 | - | - | 0 | - |
| 4 | y | 4179 | 698.4 | 0.002245 | 3.214 | +1 | 7 |
| - | - | 1637 | 699.4 | - | - | 0 | - |
| - | - | 1249 | 720.4 | - | - | 0 | - |
| - | - | 1068 | 720.4 | - | - | 0 | - |
| - | - | 1106 | 721.4 | - | - | 0 | - |
| - | - | 617.1 | 729.9 | - | - | 0 | - |
| - | - | 633.6 | 730.4 | - | - | 0 | - |
| 3 | y | 3752 | 737.4 | 0.001975 | 2.678 | +1 | 8 |
| - | - | 5781 | 737.5 | - | - | 0 | - |
| 3 | y | 5762 | 738.4 | 0.00162 | 2.194 | +1 | 8 |
| - | - | 1695 | 738.5 | - | - | 0 | - |
| - | - | 1732 | 739.4 | - | - | 0 | - |
| - | - | 1087 | 739.5 | - | - | 0 | - |
| - | - | 1145 | 747.5 | - | - | 0 | - |
| 8 | b | 2936 | 748.4 | 0.0008953 | 1.196 | +1 | 8 |
| - | - | 1217 | 749.4 | - | - | 0 | - |
| - | - | 700.4 | 750.4 | - | - | 0 | - |
| - | - | 785.6 | 754.4 | - | - | 0 | - |
| 3 | y | 1.251E+05 | 755.4 | 0.001553 | 2.056 | +1 | 8 |
| - | - | 5.394E+04 | 756.4 | - | - | 0 | - |
| - | - | 1.214E+04 | 757.4 | - | - | 0 | - |
| - | - | 1128 | 758.4 | - | - | 0 | - |
| - | - | 2115 | 763.4 | - | - | 0 | - |
| - | - | 1076 | 764.4 | - | - | 0 | - |
| 8 | b | 6.908E+04 | 765.5 | 0.001362 | 1.779 | +1 | 8 |
| - | - | 2.853E+04 | 766.5 | - | - | 0 | - |
| - | - | 6453 | 767.5 | - | - | 0 | - |
| 2 | y | 802.4 | 851.5 | 0.006911 | 8.117 | +1 | 9 |
| 2 | y | 1.672E+04 | 868.5 | 0.002365 | 2.723 | +1 | 9 |
| - | - | 653.9 | 868.7 | - | - | 0 | - |
| - | - | 6963 | 869.5 | - | - | 0 | - |
| - | - | 2012 | 870.5 | - | - | 0 | - |
| - | - | 857.3 | 878.5 | - | - | 0 | - |
| - | - | 664.9 | 1543 | - | - | 0 | - |
| - | - | 626.2 | 2128 | - | - | 0 | - |
| - | - | 701.6 | 2302 | - | - | 0 | - |
| - | - | 603.9 | 2404 | - | - | 0 | - |
| - | - | 611.4 | 2474 | - | - | 0 | - |

m/z Charge Intensity FragmentType MassShift Position
127.08671569824219 0 980.0743
127.89549255371094 0 401.78375
129.06605529785156 0 4951.2666
129.10244750976562 0 40483
130.04989624023438 0 651.87
130.1009979248047 0 472.12976
130.10580444335938 0 2968.0251
133.06077575683594 0 373.7472
135.09852600097656 0 398.04504
136.0757293701172 0 1458.7413
139.0868377685547 0 1401.1405
141.06600952148438 0 3067.3572
141.1024627685547 0 3241.5151
142.12281799316406 0 1370.7958
143.0816650390625 0 1715.469
143.11805725097656 0 1668.8182 d 1
144.82704162597656 0 410.15662
147.06549072265625 0 569.747
147.11270141601562 0 468.4127
149.02334594726562 0 535.2397
152.03433227539062 0 2856.3943
152.1072540283203 0 455.5554
153.06605529785156 0 722.7696
153.10218811035156 0 591.5618
155.11801147460938 0 5382.048
156.07786560058594 0 653.60187
156.12095642089844 0 542.11536
157.09742736816406 0 1712.4719
157.1699676513672 0 949.00397
158.09263610839844 0 2298.7627
161.51123046875 0 455.31073
165.102294921875 0 666.3322
168.07647705078125 0 507.8017
169.0609130859375 0 12203.512
169.0972442626953 0 3826.5635
169.133544921875 0 1333.0225
170.06396484375 0 766.3005
170.34129333496094 0 473.94083
171.11294555664062 0 6203.511
173.1289520263672 0 966.70514
173.451904296875 0 899.86487
180.1133575439453 0 981.3663
181.09713745117188 0 3524.0833
181.1335906982422 0 2316.4565
182.12892150878906 0 10042.603
182.1374053955078 0 927.9914
183.11312866210938 0 2128.1057
183.13232421875 0 1340.8433
183.14927673339844 0 1317.242
185.09226989746094 0 4291.7812 y Water loss 8
185.16506958007812 0 221709.98 a 1
186.08738708496094 0 26728.916
186.09442138671875 0 666.99005
186.16835021972656 0 22878.475
187.07162475585938 0 631.9333
187.09095764160156 0 1650.8743
187.14450073242188 0 1628.7181
187.1714630126953 0 1225.3788
188.13946533203125 0 5772.166
191.11753845214844 0 646.8232
193.0976104736328 0 671.4235
195.1131591796875 0 564.62744
197.0919647216797 0 773.7653
197.12802124023438 0 649.45966
198.12356567382812 0 741.71936
199.10784912109375 0 1005.81146
200.13934326171875 0 3388.7664
201.0870819091797 0 570.10803
202.1072540283203 0 1453.0166
203.10272216796875 0 142642.88 y 8
204.10618591308594 0 12257.7
205.1072998046875 0 969.1437
208.10812377929688 0 2590.371
208.1444091796875 0 897.2616
209.09205627441406 0 2888.6426
209.12803649902344 0 772.84125
210.123779296875 0 12318.835
211.12806701660156 0 972.0893
211.1443328857422 0 716.675
211.18040466308594 0 906.26624
212.10279846191406 0 792.85284
213.1598663330078 0 119281.21 b 1
214.16322326660156 0 13713.762
215.1390838623047 0 1164.775
215.16592407226562 0 910.38513
223.1080322265625 0 684.70776
226.11929321289062 0 1436.8905
226.1551055908203 0 91834.98
227.15843200683594 0 10338.058
228.13436889648438 0 11655.339 y Water loss 5
229.13775634765625 0 1220.7084
233.14405822753906 0 611.8719
233.9094696044922 0 472.98047
234.12315368652344 0 512.28253
236.1396484375 0 1211.9514
238.11849975585938 0 1477.2751
238.4471893310547 0 471.78714
239.17495727539062 0 705.41583
240.09788513183594 0 612.8311
240.13436889648438 0 1966.5659
240.95616149902344 0 507.20456
242.1497039794922 0 1082.4358
243.181396484375 0 2566.8237
252.1337127685547 0 1355.3057
252.1707763671875 0 3682.8206
253.1654815673828 0 765.3259
254.11355590820312 0 989.4147
254.1500244140625 0 3558.0027
256.1290283203125 0 1932.4161 y Water loss 7
257.12420654296875 0 1187.2842
266.1136474609375 0 1405.5033
269.197021484375 0 4682.634
270.18115234375 0 13320.189 b 2
271.17669677734375 0 10147.173
272.1793518066406 0 1577.0901
274.1396484375 0 8128.0127 y 7
275.1435241699219 0 1126.732
277.16412353515625 0 598.75073
279.18145751953125 0 18281.28
280.1849670410156 0 2435.9055
281.1607360839844 0 2783.4827
283.1399230957031 0 2673.6929
283.1763000488281 0 1053.8469
285.1556091308594 0 4816.4995
285.6659240722656 0 1984.6117 y 4
287.20782470703125 0 23106.352
288.16888427734375 0 654.6149
288.21112060546875 0 3087.1584
297.1722106933594 0 835.26306
297.1922302246094 0 33796.723
298.1952209472656 0 5704.0576
299.1714782714844 0 68545.695
300.17425537109375 0 10045.732
309.1560974121094 0 1095.4875
314.2185974121094 0 1917.6039
323.20831298828125 0 2021.7942
325.18695068359375 0 2139.3433
327.1661376953125 0 4264.7554 y Water loss 6
328.162353515625 0 2418.782
332.1780090332031 0 1334.6736
336.20404052734375 0 664.05505
337.1876525878906 0 3192.6057
340.2344055175781 0 17221.361
340.6899719238281 0 6893.564 y Water loss 3
341.1902770996094 0 2225.9697
341.2378234863281 0 3631.8037
345.177490234375 0 885.6139 y 6
349.6944274902344 0 848.01 y 3
350.21832275390625 0 1348.1339
353.2181396484375 0 994.43805
354.2137451171875 0 704.6224
355.23480224609375 0 646.0393
360.7248229980469 0 566.6738
361.22601318359375 0 693.2859
363.20233154296875 0 1470.0969
368.2289733886719 0 90895.78
368.7288513183594 0 608.27423
369.2325134277344 0 17908.428
369.6951599121094 0 632.9413 y Ammonia loss 2
369.7398986816406 0 739.9937
370.2436218261719 0 6568.3667
371.245849609375 0 1101.614
376.1971130371094 0 1746.4681
376.24066162109375 0 619.0908
378.2073059082031 0 1064.4473 y 2
378.7080383300781 0 601.71234
380.22930908203125 0 1902.3688
383.2344665527344 0 1075.0614 b 7
383.7362365722656 0 798.56067
390.2139587402344 0 1633.1196
394.20806884765625 0 1864.7941
394.2438049316406 0 992.97845
396.22393798828125 0 3139.5251
398.2395935058594 0 21097.467 b 3
399.2423095703125 0 4256.608
407.2400207519531 0 1345.7756
408.22418212890625 0 3364.9956
409.2246398925781 0 706.361
411.2349853515625 0 2798.3682
412.23638916015625 0 769.7615
417.7628479003906 0 1747.1573
418.2647399902344 0 617.718
425.2507019042969 0 1174.5557
425.7422180175781 0 1917.0713 y Water loss 1
426.2391052246094 0 947.14197 y Ammonia loss 1
431.7606506347656 0 3075.9727 b 8
432.26116943359375 0 1896.1852
434.74786376953125 0 6516.978 y 1
435.24969482421875 0 3432.8198
435.75042724609375 0 1176.0399
437.2540588378906 0 867.60913
437.2870178222656 0 1532.7979
439.2303161621094 0 634.9566
439.2996520996094 0 567.284
439.74151611328125 0 1085.508
440.2395935058594 0 1135.6091
442.27691650390625 0 1706.365
447.2348327636719 0 2764.6992
448.23907470703125 0 964.9961
449.2138977050781 0 690.74915
450.2806396484375 0 1100.8503
451.2663879394531 0 1337.7979
454.27764892578125 0 3451.2822
455.2786560058594 0 880.6716
456.2547607421875 0 1178.4917
464.2616271972656 0 10212.684
465.24609375 0 2067.44
465.2815246582031 0 7629.1787
466.24786376953125 0 629.81805
466.28460693359375 0 1269.7844
467.2239685058594 0 888.7601
467.297119140625 0 3260.6345
468.2976379394531 0 1170.2028
473.27154541015625 0 5328.3286 y 5
474.275390625 0 890.2454
475.27667236328125 0 3825.17 Precursor Water loss
475.7775573730469 0 1685.9086 Precursor Ammonia loss
476.27783203125 0 739.7298
478.278076171875 0 1875.36
479.26141357421875 0 7331.9766
480.26287841796875 0 1477.5629
481.3138427734375 0 1031.6144
482.2720031738281 0 25916.57
483.2756652832031 0 6511.6055
484.24530029296875 0 642.5317
484.2819519042969 0 9963.085 Precursor
484.7843322753906 0 6154.3623
485.28533935546875 0 2951.1487
490.27874755859375 0 779.167
495.29205322265625 0 1665.7289 b 4
496.2891540527344 0 2135.7158
508.2878723144531 0 3622.904
518.2720947265625 0 1871.5575
524.3179931640625 0 1480.3335
525.3142700195312 0 5554.7544
526.31640625 0 1713.6992
535.2987670898438 0 1418.2902
536.2825927734375 0 8293.301
537.28466796875 0 3045.5864
540.3136596679688 0 853.11237
552.3135375976562 0 6771.6514 y Water loss 4
553.3088989257812 0 73777.836 y Ammonia loss 4
554.3114013671875 0 21803.768
555.313720703125 0 3728.061
567.36181640625 0 1497.8839
568.3096313476562 0 1458.7709
570.3240966796875 0 144240.62 y 4
571.3270263671875 0 41970.832
572.3297729492188 0 7818.462
577.3441772460938 0 3051.0283
578.3323364257812 0 784.459
580.380126953125 0 2989.1377
581.3838500976562 0 1126.479
595.355224609375 0 4055.9082
596.3577880859375 0 1811.9332
621.3724365234375 0 1587.3955
622.369140625 0 881.40155
623.3826293945312 0 610.472 b 5
638.3975219726562 0 1813.0293
649.3646850585938 0 2243.871
650.3624267578125 0 6337.017
651.364013671875 0 2065.6318
652.3759155273438 0 907.54834
654.4300537109375 0 1452.3091
666.3922729492188 0 18453.203
667.3951416015625 0 6107.0923
668.4008178710938 0 1174.4354
673.9605102539062 0 823.3405
676.4119873046875 0 2830.566
677.416748046875 0 840.0139
680.3729248046875 0 3873.7217 y Water loss 3
681.3588256835938 0 5320.7007 y Ammonia loss 3
682.3583374023438 0 2467.8062
691.3826293945312 0 752.1379
694.4237670898438 0 8082.321 b 6
695.4266357421875 0 3003.369
698.3809204101562 0 4179.3003 y 3
699.3829345703125 0 1637.463
720.3668823242188 0 1249.3711
720.43701171875 0 1068.1396
721.3751831054688 0 1106.2253
729.9054565429688 0 617.09503
730.4243774414062 0 633.5986
737.39208984375 0 3752.2322 y Water loss 2
737.4658203125 0 5780.79
738.3796997070312 0 5762.3237 y Ammonia loss 2
738.4686889648438 0 1695.0084
739.381591796875 0 1731.6204
739.4673461914062 0 1086.7965
747.4546508789062 0 1144.9932
748.4360961914062 0 2936.3928 b Ammonia loss 7
749.4286499023438 0 1217.277
750.4400634765625 0 700.3936
754.3926391601562 0 785.6058
755.403076171875 0 125114.94 y 2
756.4060668945312 0 53938.613
757.40869140625 0 12137.741
758.4144287109375 0 1127.5056
763.4456176757812 0 2115.3054
764.4464721679688 0 1076.2157
765.4603881835938 0 69075.89 b 7
766.46337890625 0 28530.857
767.4654541015625 0 6453.4585
851.4690551757812 0 802.3557 y Ammonia loss 1
868.486328125 0 16724.443 y 1
868.6842651367188 0 653.94
869.4896850585938 0 6963.0664
870.4906005859375 0 2012.1578
878.46826171875 0 857.325
1542.707763671875 0 664.92224
2128.020263671875 0 626.1901
2302.48583984375 0 701.6377
2403.521728515625 0 603.9042
2474.044189453125 0 611.3718

Spectrum Details

|  |  |
| --- | --- |
| Matched peaks? Matched peaks The total absolute number of peaks matched. Additionally in brackets the total fraction of peaks matched and the total number of peaks is shown. | 42 (13.38% of 314) |
| FDR? FDR The false discovery rate estimated for this peptide. It is calculated by matching all theoretical fragments with a non-integer shift with the raw peaks for this spectrum. This is done with 40 different shifts. The resulting percentage is the average number of annotated peaks over the number of annotated peaks with the correct spectrum. | 0.11% |
| Satellite FDR? Satellite FDR See the FDR for details on its calculation. This satellite ion specific FDR only contains the satellite ions (d/w) for I/L/J positions. | 0.00% |
| PSM Score? PSM Score The PSM Score as given by Hecklib to this annotated spectrum. It is shown with three significant figures. | 431 |

#### Spectrum 3218? Spectrum 3218 The raw spectrum of this peptide as annotated by Hecklib. The fragments are coloured according to ion type (see legend). Any peaks with a star '\*' as text can be hovered over to see the full details, first the ion type second the mass shift type. By hovering over the amino acids in the peptide or ions in the legend the corresponding peaks are highlighted. By toggling the 'Unassigned' label you can turn the background (unassigned) peaks on or off in the plot. By updating the slider in the Ion legend you can update the spectrum to only show the top X% of the peaks with labels. The top X% means any peak that is within X% of the highest intensity. By dragging in the spectrum you can zoom in to a specific part of the spectrum and use 'Zoom Out' to get back to the original zoom level. The annotation of the spectrum is based on the given sequence in the peptides file and is done with different software so inconsistencies are likely. The peaks are annotated based on the given sequence, with 20 ppm tolerance.

Copy Data

##### Spectrum 3218 (TSV)

###### Preview

```
Loading example...
```

*Click on the button to copy the data to your clipboard.*

Mz MinMz MaxIntensity Max

WidthHeightPeptide font sizePeptide stroke widthSpectrum font sizeSpectrum stroke widthCompact peptide

Ion legend

wxyz

abcd

OtherUnassignedIon^Charge^_Position_Show for top:%

VLGQPKAAPS

01.31e+52.63e+53.94e+55.25e+5

Zoom Out

y^+1^_2_y^+1^_2_y^+1^_3_y^+2^_6_y^+2^_6_c^+1^_3_y^+1^_4_y^+2^_7_y^+2^_7_y^+2^_8_y^+2^_8_y^+2^_8_c^+2^_8_c^+1^_4_y^+2^_9_y^+2^_9_z^+2^_9_c^+2^_9_y^+2^_9_z^+1^_5_y^+1^_5_c^+1^_5_y^+1^_6_z^+1^_6_y^+1^_6_w^+1^_7_c^+1^_6_z^+1^_7_y^+1^_7_y^+1^_7_z^+1^_7_c^+1^_7_y^+1^_7_c^+1^_7_z^+1^_8_y^+1^_8_y^+1^_8_z^+1^_8_y^+1^_8_c^+1^_8_w^+1^_9_z^+1^_9_y^+1^_9_c^+1^_9_

049198114721962

Fragment Matches Table

Show background peaks

| Position | Ion type | Intensity | mz Theoretical | mz Error (Th) | mz Error (ppm) | Charge | Series Number |
| --- | --- | --- | --- | --- | --- | --- | --- |
| - | - | 352.6 | 121.7 | - | - | 0 | - |
| - | - | 434.5 | 125.4 | - | - | 0 | - |
| - | - | 436.1 | 128.3 | - | - | 0 | - |
| - | - | 353.4 | 128.6 | - | - | 0 | - |
| - | - | 550.4 | 129.1 | - | - | 0 | - |
| - | - | 933.1 | 136.1 | - | - | 0 | - |
| - | - | 535.5 | 138 | - | - | 0 | - |
| - | - | 386 | 141 | - | - | 0 | - |
| - | - | 643 | 148.9 | - | - | 0 | - |
| - | - | 579.6 | 148.9 | - | - | 0 | - |
| - | - | 653.8 | 148.9 | - | - | 0 | - |
| - | - | 1058 | 148.9 | - | - | 0 | - |
| - | - | 1300 | 148.9 | - | - | 0 | - |
| - | - | 1861 | 148.9 | - | - | 0 | - |
| - | - | 3454 | 148.9 | - | - | 0 | - |
| - | - | 4012 | 149 | - | - | 0 | - |
| - | - | 2540 | 149 | - | - | 0 | - |
| - | - | 1082 | 149 | - | - | 0 | - |
| - | - | 854.9 | 149 | - | - | 0 | - |
| - | - | 653.4 | 149 | - | - | 0 | - |
| - | - | 496.2 | 149 | - | - | 0 | - |
| - | - | 598.5 | 149 | - | - | 0 | - |
| - | - | 471.9 | 149 | - | - | 0 | - |
| - | - | 464.8 | 149 | - | - | 0 | - |
| - | - | 471 | 149.1 | - | - | 0 | - |
| - | - | 545.6 | 158.1 | - | - | 0 | - |
| - | - | 1647 | 169.1 | - | - | 0 | - |
| - | - | 974.7 | 171.1 | - | - | 0 | - |
| 9 | y | 1697 | 185.1 | 0.0004758 | 2.571 | +1 | 2 |
| - | - | 1.083E+05 | 185.2 | - | - | 0 | - |
| - | - | 8806 | 186.1 | - | - | 0 | - |
| - | - | 604.3 | 186.1 | - | - | 0 | - |
| - | - | 1.277E+04 | 186.2 | - | - | 0 | - |
| - | - | 664.3 | 187.1 | - | - | 0 | - |
| - | - | 466.3 | 187.2 | - | - | 0 | - |
| - | - | 1088 | 188.1 | - | - | 0 | - |
| - | - | 1302 | 190.1 | - | - | 0 | - |
| - | - | 543.7 | 191.6 | - | - | 0 | - |
| 9 | y | 7.626E+04 | 203.1 | 0.0005008 | 2.466 | +1 | 2 |
| - | - | 6422 | 204.1 | - | - | 0 | - |
| - | - | 543.2 | 205.1 | - | - | 0 | - |
| - | - | 8.156E+04 | 213.2 | - | - | 0 | - |
| - | - | 9045 | 214.2 | - | - | 0 | - |
| - | - | 699.7 | 215.2 | - | - | 0 | - |
| - | - | 7331 | 226.2 | - | - | 0 | - |
| - | - | 1055 | 227.2 | - | - | 0 | - |
| - | - | 502.3 | 229.2 | - | - | 0 | - |
| - | - | 565.6 | 243.2 | - | - | 0 | - |
| - | - | 1.196E+04 | 249.2 | - | - | 0 | - |
| - | - | 1609 | 250.2 | - | - | 0 | - |
| - | - | 8567 | 270.2 | - | - | 0 | - |
| - | - | 828.4 | 271.2 | - | - | 0 | - |
| 8 | y | 4720 | 274.1 | 0.0004506 | 1.644 | +1 | 3 |
| 5 | y | 1.977E+04 | 277.2 | 0.002585 | 9.328 | +2 | 6 |
| - | - | 2893 | 278.2 | - | - | 0 | - |
| - | - | 2498 | 285.2 | - | - | 0 | - |
| 5 | y | 932.3 | 285.7 | 0.001518 | 5.313 | +2 | 6 |
| 3 | c | 1.559E+04 | 287.2 | 0.0006984 | 2.432 | +1 | 3 |
| - | - | 2276 | 288.2 | - | - | 0 | - |
| - | - | 2213 | 297.2 | - | - | 0 | - |
| - | - | 1.301E+04 | 299.2 | - | - | 0 | - |
| - | - | 1831 | 300.2 | - | - | 0 | - |
| - | - | 1105 | 309.2 | - | - | 0 | - |
| - | - | 715.1 | 311.2 | - | - | 0 | - |
| 7 | y | 1339 | 327.2 | 0.001215 | 3.713 | +1 | 4 |
| - | - | 649 | 338.2 | - | - | 0 | - |
| - | - | 904.7 | 340.2 | - | - | 0 | - |
| 4 | y | 2518 | 340.7 | 0.0004301 | 1.262 | +2 | 7 |
| - | - | 729.6 | 341.2 | - | - | 0 | - |
| 4 | y | 894.3 | 349.7 | 6.099E-05 | 0.1744 | +2 | 7 |
| - | - | 695.3 | 350.2 | - | - | 0 | - |
| - | - | 602.9 | 351.1 | - | - | 0 | - |
| - | - | 568.9 | 361.2 | - | - | 0 | - |
| - | - | 2823 | 364.2 | - | - | 0 | - |
| - | - | 2.097E+04 | 368.2 | - | - | 0 | - |
| 3 | y | 1024 | 369.2 | 0.002149 | 5.822 | +2 | 8 |
| - | - | 3972 | 369.2 | - | - | 0 | - |
| 3 | y | 819.5 | 369.7 | 0.00297 | 8.034 | +2 | 8 |
| - | - | 5718 | 370.2 | - | - | 0 | - |
| - | - | 890.5 | 371.2 | - | - | 0 | - |
| - | - | 525.9 | 376.2 | - | - | 0 | - |
| 3 | y | 1878 | 378.2 | 0.0005292 | 1.399 | +2 | 8 |
| - | - | 911.7 | 382.2 | - | - | 0 | - |
| - | - | 603.6 | 382.7 | - | - | 0 | - |
| 8 | c | 929.6 | 383.2 | 0.001267 | 3.307 | +2 | 8 |
| - | - | 2611 | 396.2 | - | - | 0 | - |
| - | - | 908.3 | 396.2 | - | - | 0 | - |
| 4 | c | 2.295E+04 | 398.2 | 0.0008966 | 2.251 | +1 | 4 |
| - | - | 4616 | 399.2 | - | - | 0 | - |
| - | - | 521.3 | 407 | - | - | 0 | - |
| - | - | 1548 | 411.2 | - | - | 0 | - |
| - | - | 4532 | 414.2 | - | - | 0 | - |
| - | - | 1085 | 415.2 | - | - | 0 | - |
| - | - | 2222 | 417.8 | - | - | 0 | - |
| - | - | 569.5 | 418.3 | - | - | 0 | - |
| 2 | y | 1775 | 425.7 | 0.0006753 | 1.586 | +2 | 9 |
| 2 | y | 1410 | 426.2 | 0.006623 | 15.54 | +2 | 9 |
| 2 | z | 710.5 | 426.7 | 0.001856 | 4.349 | +2 | 9 |
| - | - | 1885 | 429.2 | - | - | 0 | - |
| 9 | c | 3430 | 431.8 | 0.0009762 | 2.261 | +2 | 9 |
| - | - | 1459 | 432.3 | - | - | 0 | - |
| 2 | y | 7503 | 434.7 | 0.0007335 | 1.687 | +2 | 9 |
| - | - | 4024 | 435.3 | - | - | 0 | - |
| - | - | 1143 | 435.8 | - | - | 0 | - |
| - | - | 1512 | 439.7 | - | - | 0 | - |
| - | - | 683.3 | 440.2 | - | - | 0 | - |
| 6 | z | 9.003E+04 | 457.3 | 0.001081 | 2.364 | +1 | 5 |
| - | - | 8.591E+04 | 458.3 | - | - | 0 | - |
| - | - | 1.831E+04 | 459.3 | - | - | 0 | - |
| - | - | 2397 | 460.3 | - | - | 0 | - |
| - | - | 1131 | 464.3 | - | - | 0 | - |
| - | - | 2566 | 465.3 | - | - | 0 | - |
| - | - | 1076 | 466.3 | - | - | 0 | - |
| - | - | 1069 | 466.8 | - | - | 0 | - |
| - | - | 1133 | 467.3 | - | - | 0 | - |
| 6 | y | 6096 | 473.3 | 0.0006064 | 1.281 | +1 | 5 |
| - | - | 1388 | 474.3 | - | - | 0 | - |
| - | - | 4328 | 475.3 | - | - | 0 | - |
| - | - | 2040 | 475.8 | - | - | 0 | - |
| - | - | 807.7 | 476.3 | - | - | 0 | - |
| - | - | 854.6 | 479.3 | - | - | 0 | - |
| - | - | 6759 | 482.3 | - | - | 0 | - |
| - | - | 1644 | 483.3 | - | - | 0 | - |
| - | - | 2.155E+04 | 484.3 | - | - | 0 | - |
| - | - | 1.093E+04 | 484.8 | - | - | 0 | - |
| - | - | 658.7 | 485.3 | - | - | 0 | - |
| - | - | 3352 | 485.3 | - | - | 0 | - |
| - | - | 2046 | 488.2 | - | - | 0 | - |
| - | - | 916.9 | 496.3 | - | - | 0 | - |
| - | - | 639.5 | 496.3 | - | - | 0 | - |
| - | - | 1038 | 499.3 | - | - | 0 | - |
| - | - | 971.7 | 511.3 | - | - | 0 | - |
| 5 | c | 6751 | 512.3 | 0.0007157 | 1.397 | +1 | 5 |
| - | - | 2549 | 513.3 | - | - | 0 | - |
| - | - | 900.2 | 525.3 | - | - | 0 | - |
| - | - | 706.9 | 530.3 | - | - | 0 | - |
| - | - | 1513 | 536.3 | - | - | 0 | - |
| - | - | 1191 | 537.3 | - | - | 0 | - |
| - | - | 746.5 | 538.3 | - | - | 0 | - |
| - | - | 643.6 | 547.9 | - | - | 0 | - |
| - | - | 1447 | 551.3 | - | - | 0 | - |
| 5 | y | 2512 | 552.3 | 0.0006131 | 1.11 | +1 | 6 |
| - | - | 3.333E+04 | 553.3 | - | - | 0 | - |
| 5 | z | 1.005E+04 | 554.3 | 0.007063 | 12.74 | +1 | 6 |
| - | - | 1875 | 555.3 | - | - | 0 | - |
| - | - | 602.9 | 558.4 | - | - | 0 | - |
| - | - | 3726 | 568.3 | - | - | 0 | - |
| - | - | 2290 | 568.4 | - | - | 0 | - |
| - | - | 590 | 568.4 | - | - | 0 | - |
| - | - | 3.956E+04 | 569.3 | - | - | 0 | - |
| 5 | y | 1.954E+05 | 570.3 | 0.0007295 | 1.279 | +1 | 6 |
| - | - | 830.7 | 570.9 | - | - | 0 | - |
| - | - | 5.54E+04 | 571.3 | - | - | 0 | - |
| - | - | 1.014E+04 | 572.3 | - | - | 0 | - |
| - | - | 799.7 | 580.4 | - | - | 0 | - |
| - | - | 2036 | 583.3 | - | - | 0 | - |
| - | - | 899.6 | 584.3 | - | - | 0 | - |
| - | - | 640.9 | 588.3 | - | - | 0 | - |
| - | - | 2173 | 595.3 | - | - | 0 | - |
| - | - | 2024 | 595.4 | - | - | 0 | - |
| - | - | 603.4 | 595.8 | - | - | 0 | - |
| - | - | 940.7 | 596.4 | - | - | 0 | - |
| - | - | 2702 | 596.4 | - | - | 0 | - |
| - | - | 1554 | 597.3 | - | - | 0 | - |
| - | - | 1701 | 597.4 | - | - | 0 | - |
| - | - | 2835 | 598.3 | - | - | 0 | - |
| - | - | 689.3 | 598.4 | - | - | 0 | - |
| - | - | 4936 | 608.4 | - | - | 0 | - |
| - | - | 1689 | 609.4 | - | - | 0 | - |
| - | - | 1576 | 617.3 | - | - | 0 | - |
| - | - | 640.7 | 620 | - | - | 0 | - |
| 4 | w | 1.039E+05 | 624.3 | 0.000846 | 1.355 | +1 | 7 |
| - | - | 3.548E+04 | 625.3 | - | - | 0 | - |
| - | - | 607.6 | 626 | - | - | 0 | - |
| - | - | 7908 | 626.3 | - | - | 0 | - |
| - | - | 374 | 639.4 | - | - | 0 | - |
| - | - | 1.321E+04 | 639.4 | - | - | 0 | - |
| 6 | c | 2.201E+05 | 640.4 | 0.001029 | 1.606 | +1 | 6 |
| - | - | 7.571E+04 | 641.4 | - | - | 0 | - |
| - | - | 866.7 | 642 | - | - | 0 | - |
| - | - | 1.475E+04 | 642.4 | - | - | 0 | - |
| - | - | 1059 | 643.4 | - | - | 0 | - |
| - | - | 615.2 | 646 | - | - | 0 | - |
| - | - | 1113 | 647 | - | - | 0 | - |
| - | - | 4645 | 647.3 | - | - | 0 | - |
| - | - | 4486 | 647.6 | - | - | 0 | - |
| - | - | 4192 | 650.4 | - | - | 0 | - |
| - | - | 1058 | 651.4 | - | - | 0 | - |
| - | - | 1446 | 652.4 | - | - | 0 | - |
| - | - | 763.9 | 653.4 | - | - | 0 | - |
| 4 | z | 665 | 664.4 | 0.003973 | 5.98 | +1 | 7 |
| - | - | 6923 | 666.4 | - | - | 0 | - |
| - | - | 2423 | 667.4 | - | - | 0 | - |
| - | - | 3240 | 668.3 | - | - | 0 | - |
| - | - | 1505 | 669.4 | - | - | 0 | - |
| - | - | 639.4 | 670.4 | - | - | 0 | - |
| - | - | 902.3 | 673.8 | - | - | 0 | - |
| - | - | 1545 | 676.8 | - | - | 0 | - |
| - | - | 1043 | 677.3 | - | - | 0 | - |
| - | - | 3655 | 677.4 | - | - | 0 | - |
| - | - | 1220 | 678.4 | - | - | 0 | - |
| 4 | y | 2194 | 680.4 | 0.0004462 | 0.6558 | +1 | 7 |
| 4 | y | 6585 | 681.4 | 0.001416 | 2.078 | +1 | 7 |
| 4 | z | 7.51E+04 | 682.4 | 0.000732 | 1.073 | +1 | 7 |
| - | - | 2.819E+04 | 683.4 | - | - | 0 | - |
| - | - | 6678 | 684.4 | - | - | 0 | - |
| - | - | 1375 | 688.3 | - | - | 0 | - |
| 7 | c | 4359 | 694.4 | 0.0005957 | 0.8578 | +1 | 7 |
| - | - | 558.4 | 695.3 | - | - | 0 | - |
| - | - | 1700 | 695.4 | - | - | 0 | - |
| - | - | 662.2 | 697.4 | - | - | 0 | - |
| 4 | y | 3.016E+04 | 698.4 | 0.0009289 | 1.33 | +1 | 7 |
| - | - | 9213 | 699.4 | - | - | 0 | - |
| - | - | 2251 | 700.4 | - | - | 0 | - |
| - | - | 7027 | 710.4 | - | - | 0 | - |
| 7 | c | 1.466E+05 | 711.5 | 0.000719 | 1.011 | +1 | 7 |
| - | - | 5.685E+04 | 712.5 | - | - | 0 | - |
| - | - | 1.31E+04 | 713.5 | - | - | 0 | - |
| - | - | 973.7 | 716.8 | - | - | 0 | - |
| - | - | 1300 | 720.4 | - | - | 0 | - |
| 3 | z | 1433 | 721.4 | 0.0008804 | 1.22 | +1 | 8 |
| - | - | 2838 | 721.4 | - | - | 0 | - |
| - | - | 1595 | 722.5 | - | - | 0 | - |
| - | - | 1466 | 724.8 | - | - | 0 | - |
| - | - | 1765 | 725.3 | - | - | 0 | - |
| - | - | 891.9 | 725.8 | - | - | 0 | - |
| - | - | 682.8 | 726.8 | - | - | 0 | - |
| - | - | 1896 | 727.3 | - | - | 0 | - |
| - | - | 826.8 | 727.8 | - | - | 0 | - |
| - | - | 770.2 | 730.4 | - | - | 0 | - |
| 3 | y | 3048 | 737.4 | 0.0001006 | 0.1365 | +1 | 8 |
| - | - | 2301 | 737.5 | - | - | 0 | - |
| 3 | y | 5139 | 738.4 | 0.003817 | 5.169 | +1 | 8 |
| - | - | 3706 | 738.5 | - | - | 0 | - |
| 3 | z | 1.269E+04 | 739.4 | 0.0003865 | 0.5227 | +1 | 8 |
| - | - | 850 | 739.5 | - | - | 0 | - |
| - | - | 6517 | 740.4 | - | - | 0 | - |
| - | - | 2073 | 741.4 | - | - | 0 | - |
| - | - | 1719 | 748.4 | - | - | 0 | - |
| - | - | 1346 | 749.4 | - | - | 0 | - |
| - | - | 3293 | 751.4 | - | - | 0 | - |
| - | - | 1363 | 752.4 | - | - | 0 | - |
| 3 | y | 1.711E+05 | 755.4 | 0.001011 | 1.338 | +1 | 8 |
| - | - | 6.725E+04 | 756.4 | - | - | 0 | - |
| - | - | 1.511E+04 | 757.4 | - | - | 0 | - |
| - | - | 968.4 | 758.4 | - | - | 0 | - |
| - | - | 846.9 | 763.5 | - | - | 0 | - |
| 8 | c | 6.971E+04 | 765.5 | 0.0008354 | 1.091 | +1 | 8 |
| - | - | 2.738E+04 | 766.5 | - | - | 0 | - |
| - | - | 928.6 | 767.4 | - | - | 0 | - |
| - | - | 6509 | 767.5 | - | - | 0 | - |
| - | - | 1807 | 781.4 | - | - | 0 | - |
| - | - | 1129 | 782.4 | - | - | 0 | - |
| - | - | 755.6 | 783.4 | - | - | 0 | - |
| - | - | 1245 | 790.5 | - | - | 0 | - |
| - | - | 2885 | 791.4 | - | - | 0 | - |
| - | - | 1114 | 791.5 | - | - | 0 | - |
| - | - | 1808 | 791.9 | - | - | 0 | - |
| - | - | 548.4 | 793.3 | - | - | 0 | - |
| - | - | 969.6 | 801.4 | - | - | 0 | - |
| - | - | 826.9 | 803.4 | - | - | 0 | - |
| - | - | 1138 | 803.9 | - | - | 0 | - |
| 2 | w | 2.891E+04 | 809.4 | 0.001005 | 1.242 | +1 | 9 |
| - | - | 1.406E+04 | 810.4 | - | - | 0 | - |
| - | - | 3129 | 811.4 | - | - | 0 | - |
| - | - | 3516 | 820.5 | - | - | 0 | - |
| - | - | 1606 | 821.5 | - | - | 0 | - |
| - | - | 1.049E+04 | 835.5 | - | - | 0 | - |
| - | - | 3708 | 836.5 | - | - | 0 | - |
| - | - | 1092 | 837.5 | - | - | 0 | - |
| 2 | z | 5.647E+04 | 852.5 | 0.0006731 | 0.7896 | +1 | 9 |
| - | - | 2.717E+04 | 853.5 | - | - | 0 | - |
| - | - | 6828 | 854.5 | - | - | 0 | - |
| - | - | 1723 | 855.4 | - | - | 0 | - |
| - | - | 2875 | 855.9 | - | - | 0 | - |
| - | - | 5080 | 864.5 | - | - | 0 | - |
| - | - | 2362 | 865.5 | - | - | 0 | - |
| 2 | y | 1.585E+04 | 868.5 | 0.0006258 | 0.7205 | +1 | 9 |
| - | - | 7452 | 869.5 | - | - | 0 | - |
| - | - | 1423 | 870.5 | - | - | 0 | - |
| 9 | c | 1.779E+05 | 879.5 | 0.0006851 | 0.7789 | +1 | 9 |
| - | - | 7.983E+04 | 880.5 | - | - | 0 | - |
| - | - | 2.588E+04 | 881.5 | - | - | 0 | - |
| - | - | 3454 | 882.6 | - | - | 0 | - |
| - | - | 880.6 | 888.4 | - | - | 0 | - |
| - | - | 821 | 888.6 | - | - | 0 | - |
| - | - | 4259 | 889.5 | - | - | 0 | - |
| - | - | 2160 | 890.5 | - | - | 0 | - |
| - | - | 2.834E+04 | 895.5 | - | - | 0 | - |
| - | - | 1.428E+04 | 896.5 | - | - | 0 | - |
| - | - | 6211 | 897.5 | - | - | 0 | - |
| - | - | 3274 | 898.5 | - | - | 0 | - |
| - | - | 1030 | 899.5 | - | - | 0 | - |
| - | - | 896.1 | 904.5 | - | - | 0 | - |
| - | - | 5280 | 906.6 | - | - | 0 | - |
| - | - | 4056 | 907.6 | - | - | 0 | - |
| - | - | 945.2 | 908.5 | - | - | 0 | - |
| - | - | 1534 | 908.6 | - | - | 0 | - |
| - | - | 9651 | 912.5 | - | - | 0 | - |
| - | - | 1105 | 913.4 | - | - | 0 | - |
| - | - | 3525 | 913.5 | - | - | 0 | - |
| - | - | 708.2 | 913.9 | - | - | 0 | - |
| - | - | 1514 | 914.5 | - | - | 0 | - |
| - | - | 1078 | 920.4 | - | - | 0 | - |
| - | - | 1.318E+04 | 922.6 | - | - | 0 | - |
| - | - | 2.712E+04 | 923.5 | - | - | 0 | - |
| - | - | 1.383E+04 | 924.6 | - | - | 0 | - |
| - | - | 3834 | 925.6 | - | - | 0 | - |
| - | - | 673.8 | 926.6 | - | - | 0 | - |
| - | - | 825.9 | 929.9 | - | - | 0 | - |
| - | - | 1498 | 933.5 | - | - | 0 | - |
| - | - | 1161 | 936.5 | - | - | 0 | - |
| - | - | 6313 | 938.5 | - | - | 0 | - |
| - | - | 3564 | 939.5 | - | - | 0 | - |
| - | - | 1087 | 940 | - | - | 0 | - |
| - | - | 793.3 | 940.5 | - | - | 0 | - |
| - | - | 1123 | 940.6 | - | - | 0 | - |
| - | - | 3788 | 948 | - | - | 0 | - |
| - | - | 2905 | 948.5 | - | - | 0 | - |
| - | - | 1060 | 949 | - | - | 0 | - |
| - | - | 556.7 | 950.4 | - | - | 0 | - |
| - | - | 9739 | 950.6 | - | - | 0 | - |
| - | - | 1.962E+05 | 951.5 | - | - | 0 | - |
| - | - | 1.011E+05 | 952.5 | - | - | 0 | - |
| - | - | 3.409E+04 | 953.5 | - | - | 0 | - |
| - | - | 830.8 | 953.7 | - | - | 0 | - |
| - | - | 802.9 | 954 | - | - | 0 | - |
| - | - | 3760 | 954.5 | - | - | 0 | - |
| - | - | 980.7 | 962 | - | - | 0 | - |
| - | - | 1.04E+04 | 962.5 | - | - | 0 | - |
| - | - | 7622 | 963 | - | - | 0 | - |
| - | - | 1059 | 963.5 | - | - | 0 | - |
| - | - | 1.583E+05 | 967.6 | - | - | 0 | - |
| - | - | 5.202E+05 | 968.6 | - | - | 0 | - |
| - | - | 2.598E+05 | 969.6 | - | - | 0 | - |
| - | - | 6194 | 970.5 | - | - | 0 | - |
| - | - | 7.696E+04 | 970.6 | - | - | 0 | - |
| - | - | 3.17E+04 | 971 | - | - | 0 | - |
| - | - | 2.573E+04 | 971.5 | - | - | 0 | - |
| - | - | 7465 | 971.6 | - | - | 0 | - |
| - | - | 968.7 | 983.5 | - | - | 0 | - |
| - | - | 872.8 | 984.5 | - | - | 0 | - |
| - | - | 623.2 | 995.5 | - | - | 0 | - |
| - | - | 996.8 | 1001 | - | - | 0 | - |
| - | - | 2249 | 1002 | - | - | 0 | - |
| - | - | 6795 | 1003 | - | - | 0 | - |
| - | - | 1814 | 1004 | - | - | 0 | - |
| - | - | 657.6 | 1053 | - | - | 0 | - |
| - | - | 659.7 | 1053 | - | - | 0 | - |
| - | - | 1063 | 1147 | - | - | 0 | - |
| - | - | 1364 | 1190 | - | - | 0 | - |
| - | - | 4464 | 1191 | - | - | 0 | - |
| - | - | 1418 | 1192 | - | - | 0 | - |
| - | - | 588.1 | 1234 | - | - | 0 | - |
| - | - | 780.7 | 1324 | - | - | 0 | - |
| - | - | 958 | 1325 | - | - | 0 | - |
| - | - | 670.2 | 1353 | - | - | 0 | - |
| - | - | 1467 | 1354 | - | - | 0 | - |
| - | - | 1168 | 1355 | - | - | 0 | - |
| - | - | 824.7 | 1392 | - | - | 0 | - |
| - | - | 660.1 | 1450 | - | - | 0 | - |
| - | - | 1605 | 1451 | - | - | 0 | - |
| - | - | 1250 | 1452 | - | - | 0 | - |
| - | - | 1463 | 1453 | - | - | 0 | - |
| - | - | 3199 | 1454 | - | - | 0 | - |
| - | - | 2102 | 1455 | - | - | 0 | - |
| - | - | 857.6 | 1566 | - | - | 0 | - |
| - | - | 1544 | 1582 | - | - | 0 | - |
| - | - | 2515 | 1583 | - | - | 0 | - |
| - | - | 899.5 | 1584 | - | - | 0 | - |
| - | - | 764.3 | 1592 | - | - | 0 | - |
| - | - | 1414 | 1711 | - | - | 0 | - |
| - | - | 1453 | 1712 | - | - | 0 | - |
| - | - | 938.9 | 1713 | - | - | 0 | - |
| - | - | 1464 | 1827 | - | - | 0 | - |
| - | - | 1344 | 1828 | - | - | 0 | - |
| - | - | 716.4 | 1879 | - | - | 0 | - |
| - | - | 1090 | 1895 | - | - | 0 | - |
| - | - | 2506 | 1896 | - | - | 0 | - |
| - | - | 2124 | 1897 | - | - | 0 | - |
| - | - | 780 | 1898 | - | - | 0 | - |
| - | - | 799.1 | 1909 | - | - | 0 | - |
| - | - | 2470 | 1913 | - | - | 0 | - |
| - | - | 2666 | 1914 | - | - | 0 | - |
| - | - | 2116 | 1924 | - | - | 0 | - |
| - | - | 4587 | 1925 | - | - | 0 | - |
| - | - | 3350 | 1926 | - | - | 0 | - |
| - | - | 802.1 | 1939 | - | - | 0 | - |
| - | - | 1136 | 1940 | - | - | 0 | - |
| - | - | 6672 | 1941 | - | - | 0 | - |
| - | - | 1.306E+04 | 1942 | - | - | 0 | - |
| - | - | 5580 | 1943 | - | - | 0 | - |

m/z Charge Intensity FragmentType MassShift Position
121.70883178710938 0 352.56442
125.43673706054688 0 434.49173
128.25965881347656 0 436.1094
128.581787109375 0 353.44742
129.1028594970703 0 550.377
136.07601928710938 0 933.08905
138.02232360839844 0 535.5267
140.96844482421875 0 385.97162
148.8994903564453 0 642.9769
148.9061737060547 0 579.6144
148.9137420654297 0 653.7703
148.92086791992188 0 1057.5118
148.92820739746094 0 1299.825
148.93504333496094 0 1861.0728
148.94288635253906 0 3453.5073
148.95944213867188 0 4011.944
148.9672393798828 0 2539.8462
148.9739227294922 0 1082.2914
148.98158264160156 0 854.88696
148.9891357421875 0 653.4176
149.0032958984375 0 496.22336
149.0101318359375 0 598.5096
149.02491760253906 0 471.9149
149.03240966796875 0 464.78424
149.05418395996094 0 471.0419
158.0927276611328 0 545.5701
169.06114196777344 0 1646.6832
171.113525390625 0 974.71216
185.09254455566406 0 1697.1075 y Water loss 8
185.1653289794922 0 108274.69
186.0878448486328 0 8806.249
186.09710693359375 0 604.311
186.16873168945312 0 12770.827
187.0912628173828 0 664.27747
187.17189025878906 0 466.2583
188.13983154296875 0 1087.9244
190.05364990234375 0 1302.1606
191.57740783691406 0 543.6764
203.10313415527344 0 76258.47 y 8
204.10650634765625 0 6421.874
205.10777282714844 0 543.1556
213.16024780273438 0 81563.914
214.16368103027344 0 9044.777
215.165771484375 0 699.6783
226.15553283691406 0 7330.9263
227.158935546875 0 1055.1852
229.17886352539062 0 502.3201
243.18161010742188 0 565.5975
249.16043090820312 0 11955.479
250.16360473632812 0 1608.6907
270.1816101074219 0 8566.778
271.186279296875 0 828.3905
274.14019775390625 0 4720.0557 y 7
277.1552429199219 0 19770.258 y Ammonia loss 4
278.15826416015625 0 2893.1067
285.1562805175781 0 2497.7173
285.6674499511719 0 932.3194 y 4
287.2084655761719 0 15590.147 c 2
288.2117919921875 0 2275.9617
297.1929016113281 0 2213.0706
299.172119140625 0 13014.85
300.1749267578125 0 1831.1602
309.2037048339844 0 1104.9863
311.2094421386719 0 715.0716
327.1675109863281 0 1338.5656 y Water loss 6
338.1988830566406 0 648.9867
340.2356262207031 0 904.73145
340.69036865234375 0 2517.7842 y Water loss 3
341.18988037109375 0 729.5862
349.6951599121094 0 894.313 y 3
350.19769287109375 0 695.3031
351.13983154296875 0 602.88245
361.2218322753906 0 568.938
364.1873779296875 0 2822.6353
368.22967529296875 0 20966.773
369.20281982421875 0 1023.68964 y Water loss 2
369.23358154296875 0 3971.6802
369.6956481933594 0 819.5174 y Ammonia loss 2
370.2452087402344 0 5718.0854
371.2478332519531 0 890.4801
376.2475891113281 0 525.8669
378.20648193359375 0 1877.6605 y 2
382.228515625 0 911.6977
382.7300109863281 0 603.6385
383.2332458496094 0 929.59686 c Ammonia loss 7
396.1590270996094 0 2610.7705
396.24237060546875 0 908.2871
398.2406921386719 0 22945.84 c Ammonia loss 3
399.24395751953125 0 4616.2104
406.9839172363281 0 521.27
411.23663330078125 0 1548.2946
414.1701354980469 0 4531.9614
415.1743469238281 0 1084.8477
417.76416015625 0 2221.902
418.2694091796875 0 569.47614
425.7433776855469 0 1774.9558 y Water loss 1
426.2413330078125 0 1410.4664 y Ammonia loss 1
426.740478515625 0 710.4827 z 1
429.2352294921875 0 1885.327
431.7618713378906 0 3430.0632 c Ammonia loss 8
432.2631530761719 0 1458.6628
434.74871826171875 0 7502.752 y 1
435.25091552734375 0 4024.303
435.75091552734375 0 1143.4211
439.7415771484375 0 1512.4482
440.24493408203125 0 683.3214
457.2541809082031 0 90025.71 z 5
458.2608947753906 0 85913.586
459.2641296386719 0 18308.783
460.26666259765625 0 2397.052
464.26214599609375 0 1130.5619
465.2831115722656 0 2566.1614
466.28900146484375 0 1076.4102
466.76409912109375 0 1069.1484
467.29833984375 0 1133.357
473.2724304199219 0 6095.843 y 5
474.2764892578125 0 1388.3973
475.27813720703125 0 4327.736
475.7796630859375 0 2039.6577
476.2784423828125 0 807.6835
479.2643737792969 0 854.59265
482.2734069824219 0 6758.627
483.2778625488281 0 1643.8562
484.2831726074219 0 21550.633
484.7846374511719 0 10933.02
485.251220703125 0 658.7149
485.28619384765625 0 3352.417
488.2306213378906 0 2045.6519
496.28948974609375 0 916.85895
496.32867431640625 0 639.54095
499.25341796875 0 1038.026
511.3114318847656 0 971.7215
512.31982421875 0 6751.0435 c 4
513.322998046875 0 2549.0698
525.3181762695312 0 900.22296
530.2811279296875 0 706.87415
536.2821655273438 0 1512.926
537.2906494140625 0 1191.1755
538.2955322265625 0 746.50354
547.9187622070312 0 643.6494
551.3438720703125 0 1447.1436
552.3146362304688 0 2511.8806 y Water loss 4
553.3102416992188 0 33330.633
554.3129272460938 0 10052.744 z 4
555.3153076171875 0 1875.1277
558.4146728515625 0 602.92834
568.31005859375 0 3725.8313
568.3694458007812 0 2289.978
568.4146118164062 0 590.0293
569.3175659179688 0 39562.9
570.3253173828125 0 195400.05 y 4
570.9447631835938 0 830.72107
571.3282470703125 0 55398.64
572.3310546875 0 10138.839
580.38427734375 0 799.6962
583.3447265625 0 2035.7285
584.3453369140625 0 899.60175
588.31103515625 0 640.9471
595.2813720703125 0 2173.4072
595.3556518554688 0 2023.7424
595.7814331054688 0 603.3717
596.3548583984375 0 940.6889
596.4022216796875 0 2701.5037
597.3123779296875 0 1554.499
597.408203125 0 1700.5784
598.3199462890625 0 2835.379
598.4102783203125 0 689.2918
608.3648071289062 0 4935.549
609.3681640625 0 1689.0201
617.272705078125 0 1575.5634
619.965087890625 0 640.71564
624.3359985351562 0 103925.78 w 3
625.3390502929688 0 35484.695
625.97412109375 0 607.62665
626.341064453125 0 7908.366
639.3615112304688 0 374.01004
639.407470703125 0 13209.603
640.4151000976562 0 220091.22 c 5
641.4178466796875 0 75714.32
641.9722900390625 0 866.67255
642.420654296875 0 14745.892
643.4229736328125 0 1059.3223
645.9816284179688 0 615.17847
646.97705078125 0 1112.9868
647.3125 0 4645.133
647.6473999023438 0 4485.7456
650.3638916015625 0 4191.6416
651.36376953125 0 1058.0034
652.3797607421875 0 1446.4445
653.375732421875 0 763.85394
664.3578491210938 0 664.9651 z Water loss 3
666.3947143554688 0 6923.1025
667.3959350585938 0 2423.2915
668.349365234375 0 3240.1963
669.3502197265625 0 1504.7881
670.3606567382812 0 639.446
673.8143310546875 0 902.33496
676.8150024414062 0 1545.1466
677.3172607421875 0 1042.6017
677.3883666992188 0 3655.3096
678.3903198242188 0 1219.7677
680.373046875 0 2193.7808 y Water loss 3
681.3580322265625 0 6585.093 y Ammonia loss 3
682.3651733398438 0 75104.586 z 3
683.368408203125 0 28190.902
684.3714599609375 0 6677.5303
688.3060913085938 0 1374.8441
694.4252319335938 0 4358.513 c Ammonia loss 6
695.3260498046875 0 558.3943
695.4287109375 0 1699.9338
697.3714599609375 0 662.16437
698.3840942382812 0 30159.5 y 3
699.3870239257812 0 9213.461
700.3881225585938 0 2251.33
710.4439086914062 0 7027.2446
711.451904296875 0 146584.02 c 6
712.4550170898438 0 56848.023
713.4576416015625 0 13095.065
716.8218994140625 0 973.708
720.3692626953125 0 1299.587
721.376220703125 0 1433.3151 z Water loss 2
721.4484252929688 0 2838.3564
722.4530639648438 0 1595.4352
724.8391723632812 0 1466.1395
725.3307495117188 0 1765.1378
725.8316040039062 0 891.85016
726.8455200195312 0 682.84863
727.3403930664062 0 1896.4081
727.8369750976562 0 826.82043
730.4224853515625 0 770.21136
737.3941650390625 0 3047.921 y Water loss 2
737.4661865234375 0 2301.1782
738.3818969726562 0 5138.847 y Ammonia loss 2
738.4735717773438 0 3706.0288
739.3862915039062 0 12693.902 z 2
739.4701538085938 0 849.9997
740.39111328125 0 6517.351
741.39404296875 0 2073.0752
748.4371948242188 0 1719.2612
749.439453125 0 1346.3956
751.37451171875 0 3293.0986
752.3778686523438 0 1362.9805
755.4056396484375 0 171116.95 y 2
756.408203125 0 67248.98
757.4108276367188 0 15111.452
758.411376953125 0 968.3664
763.4500732421875 0 846.85815
765.4625854492188 0 69714.81 c Ammonia loss 7
766.4657592773438 0 27382.262
767.3997802734375 0 928.6223
767.4684448242188 0 6509.021
781.4337158203125 0 1807.0883
782.3580322265625 0 1128.6848
783.4053955078125 0 755.62805
790.4706420898438 0 1245.0891
791.3662109375 0 2885.1995
791.4735717773438 0 1114.3264
791.8644409179688 0 1808.1273
793.317138671875 0 548.3791
801.3934936523438 0 969.6016
803.3881225585938 0 826.91595
803.8960571289062 0 1138.2432
809.4161987304688 0 28912.24 w 1
810.4189453125 0 14061.684
811.42236328125 0 3129.269
820.5170288085938 0 3516.1743
821.5201416015625 0 1605.5905
835.5284423828125 0 10486.581
836.5321655273438 0 3708.3518
837.53564453125 0 1092.4343
852.4706420898438 0 56468.98 z 1
853.473388671875 0 27166.65
854.476806640625 0 6827.532
855.4126586914062 0 1723.2585
855.9172973632812 0 2875.2617
864.530029296875 0 5079.8384
865.5311279296875 0 2362.2544
868.4893188476562 0 15853.359 y 1
869.4920043945312 0 7451.8013
870.4954223632812 0 1423.4386
879.541748046875 0 177866.75 c 8
880.54248046875 0 79826.78
881.5464477539062 0 25881.54
882.5506591796875 0 3453.6023
888.4338989257812 0 880.59906
888.555908203125 0 821.0326
889.5411376953125 0 4258.8525
890.5435791015625 0 2159.5132
895.4761962890625 0 28342.518
896.4796752929688 0 14279.006
897.5213012695312 0 6211.1206
898.5264892578125 0 3273.5562
899.5242309570312 0 1029.7072
904.5498046875 0 896.09534
906.5641479492188 0 5279.827
907.5589599609375 0 4055.7188
908.4652709960938 0 945.19275
908.5592041015625 0 1533.6913
912.5031127929688 0 9650.586
913.4199829101562 0 1105.2549
913.50732421875 0 3525.35
913.9315795898438 0 708.17535
914.5087890625 0 1513.6362
920.3993530273438 0 1077.6918
922.5594482421875 0 13179.351
923.5480346679688 0 27116.807
924.5557250976562 0 13834.839
925.5631103515625 0 3833.5872
926.5545043945312 0 673.81793
929.94140625 0 825.8719
933.5352783203125 0 1498.3882
936.5155639648438 0 1161.381
938.45166015625 0 6313.277
939.4594116210938 0 3564.1787
939.9603881835938 0 1087.1985
940.4677124023438 0 793.2934
940.5751953125 0 1123.326
947.9617309570312 0 3788.0583
948.4600219726562 0 2904.9502
948.9576416015625 0 1059.8948
950.4484252929688 0 556.6785
950.5549926757812 0 9738.832
951.539306640625 0 196165.02
952.5421142578125 0 101107.83
953.5443115234375 0 34092.22
953.6597290039062 0 830.76685
953.9571533203125 0 802.93665
954.5467529296875 0 3760.408
961.9605102539062 0 980.72076
962.4555053710938 0 10397.203
962.9574584960938 0 7622.1006
963.4603881835938 0 1059.3953
967.5578002929688 0 158323.53
968.5652465820312 0 520156.4
969.56787109375 0 259757.83
970.4673461914062 0 6193.6274
970.5709228515625 0 76957.57
970.9677124023438 0 31702.951
971.4691162109375 0 25728.21
971.5760498046875 0 7464.7617
983.5321655273438 0 968.6562
984.531494140625 0 872.8245
995.485107421875 0 623.2108
1000.5581665039062 0 996.7589
1002.4767456054688 0 2249.196
1003.4832763671875 0 6794.6157
1004.4848022460938 0 1813.8696
1052.501220703125 0 657.5973
1053.4951171875 0 659.7354
1146.5469970703125 0 1063.3224
1189.5543212890625 0 1364.0303
1190.5594482421875 0 4464.121
1191.5589599609375 0 1417.8688
1233.568359375 0 588.1391
1323.65673828125 0 780.6725
1324.6544189453125 0 957.9973
1352.6185302734375 0 670.2336
1353.618408203125 0 1467.1165
1354.63525390625 0 1167.6378
1391.6412353515625 0 824.6731
1449.68115234375 0 660.06195
1450.6656494140625 0 1605.358
1451.66552734375 0 1249.7809
1452.694091796875 0 1463.356
1453.68994140625 0 3198.9863
1454.6776123046875 0 2102.229
1565.686767578125 0 857.62646
1581.7130126953125 0 1544.0375
1582.724609375 0 2515.252
1583.730224609375 0 899.5055
1591.772705078125 0 764.3393
1710.8192138671875 0 1414.1885
1711.8311767578125 0 1453.3586
1712.814697265625 0 938.8924
1826.858642578125 0 1464.3362
1827.869873046875 0 1343.96
1878.9248046875 0 716.35645
1894.9202880859375 0 1089.9768
1895.9208984375 0 2505.9678
1896.92041015625 0 2124.1777
1897.9462890625 0 780.04987
1908.905029296875 0 799.14075
1912.9100341796875 0 2469.6147
1913.92333984375 0 2665.6636
1923.9085693359375 0 2115.534
1924.9091796875 0 4587.4517
1925.9056396484375 0 3349.683
1938.918701171875 0 802.14294
1939.9090576171875 0 1136.0221
1940.9237060546875 0 6672.292
1941.9248046875 0 13061.083
1942.9310302734375 0 5579.71

Spectrum Details

|  |  |
| --- | --- |
| Matched peaks? Matched peaks The total absolute number of peaks matched. Additionally in brackets the total fraction of peaks matched and the total number of peaks is shown. | 44 (11.22% of 392) |
| FDR? FDR The false discovery rate estimated for this peptide. It is calculated by matching all theoretical fragments with a non-integer shift with the raw peaks for this spectrum. This is done with 40 different shifts. The resulting percentage is the average number of annotated peaks over the number of annotated peaks with the correct spectrum. | 1.19% |
| Satellite FDR? Satellite FDR See the FDR for details on its calculation. This satellite ion specific FDR only contains the satellite ions (d/w) for I/L/J positions. | 7.14% |
| PSM Score? PSM Score The PSM Score as given by Hecklib to this annotated spectrum. It is shown with three significant figures. | 411 |

#### Spectrum 3487? Spectrum 3487 The raw spectrum of this peptide as annotated by Hecklib. The fragments are coloured according to ion type (see legend). Any peaks with a star '\*' as text can be hovered over to see the full details, first the ion type second the mass shift type. By hovering over the amino acids in the peptide or ions in the legend the corresponding peaks are highlighted. By toggling the 'Unassigned' label you can turn the background (unassigned) peaks on or off in the plot. By updating the slider in the Ion legend you can update the spectrum to only show the top X% of the peaks with labels. The top X% means any peak that is within X% of the highest intensity. By dragging in the spectrum you can zoom in to a specific part of the spectrum and use 'Zoom Out' to get back to the original zoom level. The annotation of the spectrum is based on the given sequence in the peptides file and is done with different software so inconsistencies are likely. The peaks are annotated based on the given sequence, with 20 ppm tolerance.

Copy Data

##### Spectrum 3487 (TSV)

###### Preview

```
Loading example...
```

*Click on the button to copy the data to your clipboard.*

Mz MinMz MaxIntensity Max

WidthHeightPeptide font sizePeptide stroke widthSpectrum font sizeSpectrum stroke widthCompact peptide

Ion legend

wxyz

abcd

OtherUnassignedIon^Charge^_Position_Show for top:%

VLGQPKAAPS

04.81e+49.61e+41.44e+51.92e+5

Zoom Out

y^+1^_2_y^+1^_3_y^+2^_6_c^+1^_3_y^+1^_4_y^+2^_7_y^+2^_8_c^+1^_4_y^+2^_9_c^+2^_9_y^+2^_9_z^+1^_5_y^+1^_5_c^+1^_5_z^+1^_6_y^+1^_6_c^+1^_6_w^+1^_7_c^+1^_6_y^+1^_7_y^+1^_7_z^+1^_7_c^+1^_7_y^+1^_7_c^+1^_7_z^+1^_8_y^+1^_8_y^+1^_8_z^+1^_8_y^+1^_8_c^+1^_8_w^+1^_9_z^+1^_9_y^+1^_9_c^+1^_9_

0767153323003067

Fragment Matches Table

Show background peaks

| Position | Ion type | Intensity | mz Theoretical | mz Error (Th) | mz Error (ppm) | Charge | Series Number |
| --- | --- | --- | --- | --- | --- | --- | --- |
| - | - | 373.6 | 122.5 | - | - | 0 | - |
| - | - | 494.3 | 136.1 | - | - | 0 | - |
| - | - | 575.7 | 158.1 | - | - | 0 | - |
| - | - | 435.6 | 160.3 | - | - | 0 | - |
| - | - | 1090 | 173.5 | - | - | 0 | - |
| - | - | 3.333E+04 | 185.2 | - | - | 0 | - |
| - | - | 3499 | 186.1 | - | - | 0 | - |
| - | - | 3226 | 186.2 | - | - | 0 | - |
| - | - | 2431 | 187.1 | - | - | 0 | - |
| - | - | 529.6 | 188.1 | - | - | 0 | - |
| - | - | 1074 | 190.1 | - | - | 0 | - |
| 9 | y | 2.509E+04 | 203.1 | 0.0002108 | 1.038 | +1 | 2 |
| - | - | 1193 | 204.1 | - | - | 0 | - |
| - | - | 2.667E+04 | 213.2 | - | - | 0 | - |
| - | - | 3197 | 214.2 | - | - | 0 | - |
| - | - | 4757 | 215.1 | - | - | 0 | - |
| - | - | 514.8 | 221.9 | - | - | 0 | - |
| - | - | 2477 | 226.2 | - | - | 0 | - |
| - | - | 746.8 | 229.2 | - | - | 0 | - |
| - | - | 6567 | 249.2 | - | - | 0 | - |
| - | - | 1541 | 250.2 | - | - | 0 | - |
| - | - | 610.7 | 255.1 | - | - | 0 | - |
| - | - | 3445 | 269.1 | - | - | 0 | - |
| - | - | 2175 | 270.2 | - | - | 0 | - |
| 8 | y | 1457 | 274.1 | 0.0003124 | 1.139 | +1 | 3 |
| 5 | y | 1.024E+04 | 277.2 | 0.002311 | 8.337 | +2 | 6 |
| - | - | 1668 | 278.2 | - | - | 0 | - |
| - | - | 594.1 | 283.1 | - | - | 0 | - |
| - | - | 901.1 | 285.2 | - | - | 0 | - |
| 3 | c | 4838 | 287.2 | 0.0003017 | 1.05 | +1 | 3 |
| - | - | 662.7 | 297.2 | - | - | 0 | - |
| - | - | 3875 | 299.2 | - | - | 0 | - |
| - | - | 758.3 | 300.2 | - | - | 0 | - |
| - | - | 1271 | 309.2 | - | - | 0 | - |
| - | - | 1643 | 314.2 | - | - | 0 | - |
| 7 | y | 690.1 | 327.2 | 0.0007569 | 2.314 | +1 | 4 |
| 4 | y | 907.7 | 340.7 | 0.0003691 | 1.083 | +2 | 7 |
| - | - | 551.1 | 354 | - | - | 0 | - |
| - | - | 906 | 364.2 | - | - | 0 | - |
| - | - | 6605 | 368.2 | - | - | 0 | - |
| - | - | 1179 | 369.2 | - | - | 0 | - |
| - | - | 2653 | 370.2 | - | - | 0 | - |
| 3 | y | 671.4 | 378.2 | 0.001384 | 3.658 | +2 | 8 |
| - | - | 922.1 | 396.2 | - | - | 0 | - |
| 4 | c | 6943 | 398.2 | 0.0003167 | 0.7953 | +1 | 4 |
| - | - | 1106 | 399.2 | - | - | 0 | - |
| - | - | 2021 | 414.2 | - | - | 0 | - |
| 2 | y | 1002 | 425.7 | 0.001347 | 3.163 | +2 | 9 |
| 9 | c | 625.9 | 431.8 | 0.004028 | 9.329 | +2 | 9 |
| - | - | 794.4 | 432.3 | - | - | 0 | - |
| 2 | y | 2856 | 434.7 | 0.0011 | 2.53 | +2 | 9 |
| - | - | 1577 | 435.3 | - | - | 0 | - |
| 6 | z | 3.428E+04 | 457.3 | 0.0004706 | 1.029 | +1 | 5 |
| - | - | 3.166E+04 | 458.3 | - | - | 0 | - |
| - | - | 6267 | 459.3 | - | - | 0 | - |
| - | - | 931 | 460.3 | - | - | 0 | - |
| - | - | 839.7 | 466.8 | - | - | 0 | - |
| 6 | y | 2712 | 473.3 | 0.0002481 | 0.5242 | +1 | 5 |
| - | - | 1779 | 475.3 | - | - | 0 | - |
| - | - | 1090 | 475.8 | - | - | 0 | - |
| - | - | 2637 | 482.3 | - | - | 0 | - |
| - | - | 1526 | 483.2 | - | - | 0 | - |
| - | - | 1340 | 484.2 | - | - | 0 | - |
| - | - | 756.6 | 484.3 | - | - | 0 | - |
| - | - | 5142 | 484.3 | - | - | 0 | - |
| - | - | 1826 | 484.8 | - | - | 0 | - |
| 5 | c | 3061 | 512.3 | 0.0005326 | 1.04 | +1 | 5 |
| - | - | 878.3 | 513.3 | - | - | 0 | - |
| - | - | 850.5 | 536.3 | - | - | 0 | - |
| - | - | 1.121E+04 | 553.3 | - | - | 0 | - |
| 5 | z | 2870 | 554.3 | 0.006148 | 11.09 | +1 | 6 |
| - | - | 1510 | 568.3 | - | - | 0 | - |
| - | - | 978.3 | 568.4 | - | - | 0 | - |
| - | - | 1.45E+04 | 569.3 | - | - | 0 | - |
| 5 | y | 6.564E+04 | 570.3 | 5.815E-05 | 0.102 | +1 | 6 |
| - | - | 1.941E+04 | 571.3 | - | - | 0 | - |
| - | - | 3099 | 572.3 | - | - | 0 | - |
| - | - | 708.8 | 583.3 | - | - | 0 | - |
| - | - | 562.9 | 593.5 | - | - | 0 | - |
| - | - | 1181 | 596.4 | - | - | 0 | - |
| - | - | 986.8 | 598.3 | - | - | 0 | - |
| - | - | 2088 | 608.4 | - | - | 0 | - |
| - | - | 1341 | 609.4 | - | - | 0 | - |
| - | - | 912.1 | 610.6 | - | - | 0 | - |
| 6 | c | 703.2 | 623.4 | 0.003957 | 6.348 | +1 | 6 |
| 4 | w | 3.961E+04 | 624.3 | 5.254E-05 | 0.08416 | +1 | 7 |
| - | - | 1.41E+04 | 625.3 | - | - | 0 | - |
| - | - | 2413 | 626.3 | - | - | 0 | - |
| - | - | 582 | 634.7 | - | - | 0 | - |
| - | - | 5518 | 639.4 | - | - | 0 | - |
| 6 | c | 8.495E+04 | 640.4 | 0.000113 | 0.1765 | +1 | 6 |
| - | - | 3.021E+04 | 641.4 | - | - | 0 | - |
| - | - | 5979 | 642.4 | - | - | 0 | - |
| - | - | 656 | 650.4 | - | - | 0 | - |
| - | - | 634.1 | 651.4 | - | - | 0 | - |
| - | - | 1920 | 666.4 | - | - | 0 | - |
| - | - | 1426 | 668.4 | - | - | 0 | - |
| - | - | 884 | 677.4 | - | - | 0 | - |
| - | - | 706 | 678.4 | - | - | 0 | - |
| 4 | y | 1642 | 680.4 | 0.0005914 | 0.8692 | +1 | 7 |
| 4 | y | 2166 | 681.4 | 0.002454 | 3.601 | +1 | 7 |
| 4 | z | 2.715E+04 | 682.4 | 6.141E-05 | 0.08999 | +1 | 7 |
| - | - | 1.004E+04 | 683.4 | - | - | 0 | - |
| - | - | 2147 | 684.4 | - | - | 0 | - |
| 7 | c | 1690 | 694.4 | 0.00148 | 2.131 | +1 | 7 |
| 4 | y | 1.173E+04 | 698.4 | 0.0001698 | 0.2431 | +1 | 7 |
| - | - | 3770 | 699.4 | - | - | 0 | - |
| - | - | 814.2 | 700.4 | - | - | 0 | - |
| - | - | 2731 | 710.4 | - | - | 0 | - |
| 7 | c | 5.724E+04 | 711.5 | 0.0001355 | 0.1905 | +1 | 7 |
| - | - | 2.316E+04 | 712.5 | - | - | 0 | - |
| - | - | 4936 | 713.5 | - | - | 0 | - |
| 3 | z | 720.7 | 721.4 | 0.0008286 | 1.149 | +1 | 8 |
| - | - | 1063 | 724.8 | - | - | 0 | - |
| - | - | 776.3 | 725.9 | - | - | 0 | - |
| 3 | y | 670.4 | 737.4 | 0.001975 | 2.678 | +1 | 8 |
| - | - | 615.7 | 737.5 | - | - | 0 | - |
| 3 | y | 699 | 738.4 | 0.001066 | 1.443 | +1 | 8 |
| - | - | 1116 | 738.5 | - | - | 0 | - |
| 3 | z | 4603 | 739.4 | 0.0008952 | 1.211 | +1 | 8 |
| - | - | 2923 | 740.4 | - | - | 0 | - |
| - | - | 642.9 | 741.4 | - | - | 0 | - |
| 3 | y | 5.41E+04 | 755.4 | 0.0002101 | 0.2782 | +1 | 8 |
| - | - | 2.101E+04 | 756.4 | - | - | 0 | - |
| - | - | 5457 | 757.4 | - | - | 0 | - |
| 8 | c | 2.187E+04 | 765.5 | 0.0001411 | 0.1844 | +1 | 8 |
| - | - | 8927 | 766.5 | - | - | 0 | - |
| - | - | 1646 | 767.5 | - | - | 0 | - |
| 2 | w | 1.247E+04 | 809.4 | 0.0002115 | 0.2613 | +1 | 9 |
| - | - | 4293 | 810.4 | - | - | 0 | - |
| - | - | 799.6 | 811.4 | - | - | 0 | - |
| - | - | 2115 | 820.5 | - | - | 0 | - |
| - | - | 706.6 | 821.5 | - | - | 0 | - |
| - | - | 3553 | 835.5 | - | - | 0 | - |
| - | - | 1405 | 836.5 | - | - | 0 | - |
| - | - | 852.9 | 851.4 | - | - | 0 | - |
| 2 | z | 2.115E+04 | 852.5 | 0.0004255 | 0.4992 | +1 | 9 |
| - | - | 7472 | 853.5 | - | - | 0 | - |
| - | - | 2894 | 854.5 | - | - | 0 | - |
| - | - | 1541 | 864.5 | - | - | 0 | - |
| - | - | 933 | 865.5 | - | - | 0 | - |
| 2 | y | 4708 | 868.5 | 0.0005037 | 0.58 | +1 | 9 |
| - | - | 1721 | 869.5 | - | - | 0 | - |
| - | - | 781.9 | 870.5 | - | - | 0 | - |
| 9 | c | 6.801E+04 | 879.5 | 0.0003525 | 0.4008 | +1 | 9 |
| - | - | 3.175E+04 | 880.5 | - | - | 0 | - |
| - | - | 8988 | 881.5 | - | - | 0 | - |
| - | - | 1548 | 882.5 | - | - | 0 | - |
| - | - | 1615 | 889.5 | - | - | 0 | - |
| - | - | 1156 | 890.5 | - | - | 0 | - |
| - | - | 1.163E+04 | 895.5 | - | - | 0 | - |
| - | - | 5619 | 896.5 | - | - | 0 | - |
| - | - | 2182 | 897.5 | - | - | 0 | - |
| - | - | 1316 | 898.5 | - | - | 0 | - |
| - | - | 1666 | 906.6 | - | - | 0 | - |
| - | - | 1623 | 907.6 | - | - | 0 | - |
| - | - | 769.8 | 908.6 | - | - | 0 | - |
| - | - | 4500 | 912.5 | - | - | 0 | - |
| - | - | 1504 | 913.5 | - | - | 0 | - |
| - | - | 2095 | 916 | - | - | 0 | - |
| - | - | 1796 | 916.5 | - | - | 0 | - |
| - | - | 992.4 | 917 | - | - | 0 | - |
| - | - | 912 | 920.4 | - | - | 0 | - |
| - | - | 4568 | 922.6 | - | - | 0 | - |
| - | - | 9973 | 923.5 | - | - | 0 | - |
| - | - | 5323 | 924.6 | - | - | 0 | - |
| - | - | 953.4 | 925.6 | - | - | 0 | - |
| - | - | 960.2 | 939.5 | - | - | 0 | - |
| - | - | 1983 | 947.5 | - | - | 0 | - |
| - | - | 661.6 | 950.4 | - | - | 0 | - |
| - | - | 3607 | 950.6 | - | - | 0 | - |
| - | - | 7.629E+04 | 951.5 | - | - | 0 | - |
| - | - | 3.806E+04 | 952.5 | - | - | 0 | - |
| - | - | 1.152E+04 | 953.5 | - | - | 0 | - |
| - | - | 1294 | 954.5 | - | - | 0 | - |
| - | - | 1125 | 965.5 | - | - | 0 | - |
| - | - | 6.202E+04 | 967.6 | - | - | 0 | - |
| - | - | 1.903E+05 | 968.6 | - | - | 0 | - |
| - | - | 9.405E+04 | 969.6 | - | - | 0 | - |
| - | - | 1324 | 970.5 | - | - | 0 | - |
| - | - | 2.824E+04 | 970.6 | - | - | 0 | - |
| - | - | 1395 | 971 | - | - | 0 | - |
| - | - | 1570 | 971.5 | - | - | 0 | - |
| - | - | 2916 | 971.6 | - | - | 0 | - |
| - | - | 611.5 | 1007 | - | - | 0 | - |
| - | - | 681.6 | 1392 | - | - | 0 | - |
| - | - | 1725 | 1434 | - | - | 0 | - |
| - | - | 866.2 | 1436 | - | - | 0 | - |
| - | - | 1392 | 1451 | - | - | 0 | - |
| - | - | 2086 | 1452 | - | - | 0 | - |
| - | - | 2049 | 1453 | - | - | 0 | - |
| - | - | 1236 | 1454 | - | - | 0 | - |
| - | - | 765.5 | 1458 | - | - | 0 | - |
| - | - | 1310 | 1832 | - | - | 0 | - |
| - | - | 827.9 | 1833 | - | - | 0 | - |
| - | - | 1218 | 1879 | - | - | 0 | - |
| - | - | 978.3 | 1880 | - | - | 0 | - |
| - | - | 758.1 | 1937 | - | - | 0 | - |
| - | - | 1381 | 1939 | - | - | 0 | - |
| - | - | 744.4 | 1940 | - | - | 0 | - |
| - | - | 1171 | 1941 | - | - | 0 | - |
| - | - | 1433 | 1942 | - | - | 0 | - |
| - | - | 732.7 | 2836 | - | - | 0 | - |
| - | - | 677.9 | 3036 | - | - | 0 | - |

m/z Charge Intensity FragmentType MassShift Position
122.48072814941406 0 373.55557
136.07591247558594 0 494.28088
158.09320068359375 0 575.6768
160.28651428222656 0 435.60007
173.45294189453125 0 1090.2505
185.16514587402344 0 33329.906
186.08775329589844 0 3499.208
186.16854858398438 0 3225.921
187.1444091796875 0 2430.9333
188.14755249023438 0 529.5761
190.052978515625 0 1073.7896
203.10284423828125 0 25092.898 y 8
204.10610961914062 0 1192.8475
213.16004943847656 0 26667.309
214.16336059570312 0 3197.4995
215.1393585205078 0 4757.4736
221.9279022216797 0 514.82776
226.155517578125 0 2476.8137
229.1787872314453 0 746.8446
249.16018676757812 0 6566.9106
250.16323852539062 0 1540.5436
255.0800018310547 0 610.6594
269.0804138183594 0 3445.3623
270.18133544921875 0 2175.2417
274.1394348144531 0 1457.049 y 7
277.15496826171875 0 10238.542 y Ammonia loss 4
278.15826416015625 0 1668.4377
283.142333984375 0 594.1414
285.15618896484375 0 901.0539
287.20806884765625 0 4837.6504 c 2
297.1919860839844 0 662.7276
299.1718444824219 0 3874.7031
300.17498779296875 0 758.29736
309.20367431640625 0 1270.6238
314.2082824707031 0 1643.1157
327.16705322265625 0 690.05334 y Water loss 6
340.6903076171875 0 907.7047 y Water loss 3
353.9639587402344 0 551.0686
364.18658447265625 0 905.9607
368.2292175292969 0 6605.266
369.23345947265625 0 1179.1808
370.2451171875 0 2652.8667
378.20733642578125 0 671.3842 y 2
396.1592102050781 0 922.1151
398.2401123046875 0 6942.9375 c Ammonia loss 3
399.2434997558594 0 1105.857
414.16961669921875 0 2021.0262
425.7440490722656 0 1002.10175 y Water loss 1
431.7649230957031 0 625.9466 c Ammonia loss 8
432.2660827636719 0 794.35895
434.74908447265625 0 2855.7012 y 1
435.2501220703125 0 1577.3845
457.2535705566406 0 34275.098 z 5
458.2602233886719 0 31656.691
459.26373291015625 0 6267.3896
460.26409912109375 0 931.0036
466.76104736328125 0 839.6866
473.2715759277344 0 2711.5476 y 5
475.27777099609375 0 1779.0841
475.78009033203125 0 1090.2881
482.2729797363281 0 2636.831
483.2130126953125 0 1526.2075
484.216064453125 0 1339.9822
484.2517395019531 0 756.644
484.2828369140625 0 5142.325
484.7837829589844 0 1826.403
512.3196411132812 0 3060.8455 c 4
513.3226928710938 0 878.2895
536.2822875976562 0 850.54083
553.3094482421875 0 11212.498
554.31201171875 0 2870.283 z 4
568.3109130859375 0 1510.1431
568.3713989257812 0 978.3107
569.3169555664062 0 14498.513
570.3246459960938 0 65639.69 y 4
571.32763671875 0 19414.504
572.331298828125 0 3098.842
583.342529296875 0 708.8286
593.480224609375 0 562.9051
596.4010009765625 0 1180.746
598.31787109375 0 986.8465
608.3649291992188 0 2087.5793
609.367919921875 0 1340.5012
610.640380859375 0 912.1196
623.3914794921875 0 703.1903 c Ammonia loss 5
624.335205078125 0 39614.05 w 3
625.3384399414062 0 14101.501
626.3407592773438 0 2412.539
634.7279052734375 0 581.97925
639.40625 0 5517.6045
640.4141845703125 0 84946.28 c 5
641.4170532226562 0 30214.527
642.4188842773438 0 5979.4697
650.3660278320312 0 655.95734
651.3651733398438 0 634.08374
666.3939208984375 0 1919.9692
668.3529663085938 0 1426.171
677.384765625 0 883.97485
678.392578125 0 705.9868
680.3720092773438 0 1641.8274 y Water loss 3
681.3590698242188 0 2166.2922 y Ammonia loss 3
682.3643798828125 0 27151.53 z 3
683.3677978515625 0 10041.827
684.3681640625 0 2147.0684
694.4231567382812 0 1690.0686 c Ammonia loss 6
698.3829956054688 0 11731.459 y 3
699.3867797851562 0 3770.0237
700.3895874023438 0 814.15704
710.4434204101562 0 2731.4373
711.4510498046875 0 57239.41 c 6
712.453857421875 0 23160.719
713.4553833007812 0 4935.64
721.37451171875 0 720.7176 z Water loss 2
724.8364868164062 0 1063.2482
725.86376953125 0 776.3171
737.39208984375 0 670.43146 y Water loss 2
737.46923828125 0 615.69714
738.3770141601562 0 698.9622 y Ammonia loss 2
738.47216796875 0 1115.9254
739.385009765625 0 4603.107 z 2
740.391845703125 0 2923.3315
741.3951416015625 0 642.85315
755.4044189453125 0 54099.29 y 2
756.4071044921875 0 21013.033
757.409423828125 0 5456.543
765.4616088867188 0 21868.32 c Ammonia loss 7
766.4641723632812 0 8926.52
767.4679565429688 0 1645.7828
809.4154052734375 0 12472.938 w 1
810.4178466796875 0 4292.8945
811.4228515625 0 799.6303
820.5175170898438 0 2114.782
821.5108642578125 0 706.55566
835.5250854492188 0 3552.9626
836.531005859375 0 1405.1049
851.3772583007812 0 852.8739
852.4695434570312 0 21154.691 z 1
853.472412109375 0 7471.968
854.4759521484375 0 2893.645
864.530029296875 0 1541.4158
865.5336303710938 0 933.02277
868.4891967773438 0 4708.1113 y 1
869.4931640625 0 1721.0782
870.483642578125 0 781.8683
879.5407104492188 0 68012.25 c 8
880.542236328125 0 31746.318
881.54296875 0 8987.903
882.5464477539062 0 1548.1874
889.5370483398438 0 1615.3398
890.5445556640625 0 1156.2793
895.4756469726562 0 11626.141
896.478515625 0 5619.2676
897.5175170898438 0 2181.554
898.5289916992188 0 1316.3335
906.5621948242188 0 1665.9191
907.556640625 0 1623.0203
908.561767578125 0 769.76807
912.4995727539062 0 4500.251
913.5062255859375 0 1503.7047
915.966064453125 0 2094.856
916.4679565429688 0 1795.9742
916.9715576171875 0 992.3648
920.3926391601562 0 911.9669
922.5586547851562 0 4568.3535
923.5474853515625 0 9972.503
924.553466796875 0 5323.2534
925.564453125 0 953.4457
939.4612426757812 0 960.2361
947.4716796875 0 1983.1764
950.4447631835938 0 661.5743
950.5532836914062 0 3606.7322
951.5380859375 0 76292.7
952.541015625 0 38059.984
953.5433959960938 0 11517.288
954.5491333007812 0 1294.3531
965.4725341796875 0 1125.3907
967.5565795898438 0 62022.273
968.5634765625 0 190344.1
969.5664672851562 0 94047.72
970.46240234375 0 1324.0558
970.5697021484375 0 28235.293
970.9699096679688 0 1395.2516
971.4555053710938 0 1569.9768
971.5735473632812 0 2915.8481
1006.6694946289062 0 611.5107
1391.6619873046875 0 681.6106
1433.64892578125 0 1725.3119
1435.70166015625 0 866.239
1450.6724853515625 0 1391.809
1451.6973876953125 0 2086.4697
1452.714599609375 0 2048.7808
1453.730712890625 0 1235.8041
1457.7381591796875 0 765.51306
1831.9296875 0 1310.3541
1832.940673828125 0 827.8697
1878.9093017578125 0 1218.479
1879.9088134765625 0 978.3069
1936.8955078125 0 758.0894
1938.932861328125 0 1380.7008
1939.9310302734375 0 744.42236
1940.901123046875 0 1171.3475
1941.92236328125 0 1432.6823
2835.701171875 0 732.70605
3036.319580078125 0 677.9018

Spectrum Details

|  |  |
| --- | --- |
| Matched peaks? Matched peaks The total absolute number of peaks matched. Additionally in brackets the total fraction of peaks matched and the total number of peaks is shown. | 35 (17.16% of 204) |
| FDR? FDR The false discovery rate estimated for this peptide. It is calculated by matching all theoretical fragments with a non-integer shift with the raw peaks for this spectrum. This is done with 40 different shifts. The resulting percentage is the average number of annotated peaks over the number of annotated peaks with the correct spectrum. | 0.61% |
| Satellite FDR? Satellite FDR See the FDR for details on its calculation. This satellite ion specific FDR only contains the satellite ions (d/w) for I/L/J positions. | 4.76% |
| PSM Score? PSM Score The PSM Score as given by Hecklib to this annotated spectrum. It is shown with three significant figures. | 372 |

#### Spectrum 2978? Spectrum 2978 The raw spectrum of this peptide as annotated by Hecklib. The fragments are coloured according to ion type (see legend). Any peaks with a star '\*' as text can be hovered over to see the full details, first the ion type second the mass shift type. By hovering over the amino acids in the peptide or ions in the legend the corresponding peaks are highlighted. By toggling the 'Unassigned' label you can turn the background (unassigned) peaks on or off in the plot. By updating the slider in the Ion legend you can update the spectrum to only show the top X% of the peaks with labels. The top X% means any peak that is within X% of the highest intensity. By dragging in the spectrum you can zoom in to a specific part of the spectrum and use 'Zoom Out' to get back to the original zoom level. The annotation of the spectrum is based on the given sequence in the peptides file and is done with different software so inconsistencies are likely. The peaks are annotated based on the given sequence, with 20 ppm tolerance.

Copy Data

##### Spectrum 2978 (TSV)

###### Preview

```
Loading example...
```

*Click on the button to copy the data to your clipboard.*

Mz MinMz MaxIntensity Max

WidthHeightPeptide font sizePeptide stroke widthSpectrum font sizeSpectrum stroke widthCompact peptide

Ion legend

wxyz

abcd

OtherUnassignedIon^Charge^_Position_Show for top:%

VLGQPKAAPS

01.38e+72.76e+74.15e+75.53e+7

Zoom Out

d^+1^_2_y^+1^_2_a^+1^_2_y^+1^_2_b^+1^_2_y^+2^_5_y^+1^_3_b^+1^_3_y^+1^_3_y^+2^_6_y^+1^_4_y^+2^_7_y^+1^_4_b^+2^_7_y^+2^_7_y^+2^_8_y^+2^_8_b^+1^_4_b^+2^_8_b^+1^_4_y^+2^_9_y^+2^_9_b^+2^_9_y^+2^_9_y^+1^_5_\*\*\*b^+1^_5_y^+1^_6_y^+1^_6_b^+1^_6_y^+1^_7_y^+1^_7_b^+1^_7_y^+1^_7_y^+1^_8_y^+1^_8_b^+1^_8_y^+1^_8_b^+1^_8_y^+1^_9_y^+1^_9_y^+1^_9_

0706141121172823

Fragment Matches Table

Show background peaks

| Position | Ion type | Intensity | mz Theoretical | mz Error (Th) | mz Error (ppm) | Charge | Series Number |
| --- | --- | --- | --- | --- | --- | --- | --- |
| - | - | 3.738E+04 | 120.1 | - | - | 0 | - |
| - | - | 4.298E+04 | 123.2 | - | - | 0 | - |
| - | - | 7.982E+04 | 124 | - | - | 0 | - |
| - | - | 4.1E+04 | 125.1 | - | - | 0 | - |
| - | - | 1.175E+06 | 129.1 | - | - | 0 | - |
| - | - | 9.378E+06 | 129.1 | - | - | 0 | - |
| - | - | 5.988E+04 | 130.1 | - | - | 0 | - |
| - | - | 8.711E+04 | 130.1 | - | - | 0 | - |
| - | - | 5.952E+05 | 130.1 | - | - | 0 | - |
| - | - | 6.308E+04 | 134 | - | - | 0 | - |
| - | - | 5.738E+05 | 136.1 | - | - | 0 | - |
| - | - | 6.122E+04 | 137.1 | - | - | 0 | - |
| - | - | 2.637E+05 | 139.1 | - | - | 0 | - |
| - | - | 5.307E+04 | 139.1 | - | - | 0 | - |
| - | - | 6.979E+05 | 141.1 | - | - | 0 | - |
| - | - | 7.789E+05 | 141.1 | - | - | 0 | - |
| - | - | 5.124E+04 | 142.1 | - | - | 0 | - |
| - | - | 3.816E+05 | 143.1 | - | - | 0 | - |
| 2 | d | 4.018E+05 | 143.1 | 0.0003203 | 2.238 | +1 | 2 |
| - | - | 4.824E+04 | 150 | - | - | 0 | - |
| - | - | 7.075E+05 | 152 | - | - | 0 | - |
| - | - | 1.431E+05 | 153.1 | - | - | 0 | - |
| - | - | 1.433E+05 | 153.1 | - | - | 0 | - |
| - | - | 1.211E+06 | 155.1 | - | - | 0 | - |
| - | - | 9.965E+04 | 156.1 | - | - | 0 | - |
| - | - | 2.916E+05 | 157.1 | - | - | 0 | - |
| - | - | 1.303E+05 | 157.2 | - | - | 0 | - |
| - | - | 4.08E+05 | 158.1 | - | - | 0 | - |
| - | - | 1.962E+05 | 165.1 | - | - | 0 | - |
| - | - | 7.655E+04 | 167.1 | - | - | 0 | - |
| - | - | 2.811E+06 | 169.1 | - | - | 0 | - |
| - | - | 1.026E+06 | 169.1 | - | - | 0 | - |
| - | - | 1.724E+05 | 169.1 | - | - | 0 | - |
| - | - | 1.996E+05 | 170.1 | - | - | 0 | - |
| - | - | 5.648E+04 | 170.1 | - | - | 0 | - |
| - | - | 1.737E+06 | 171.1 | - | - | 0 | - |
| - | - | 1.841E+05 | 172.1 | - | - | 0 | - |
| - | - | 1.179E+05 | 172.1 | - | - | 0 | - |
| - | - | 3.078E+05 | 180.1 | - | - | 0 | - |
| - | - | 8.571E+05 | 181.1 | - | - | 0 | - |
| - | - | 4.244E+05 | 181.1 | - | - | 0 | - |
| - | - | 5.724E+04 | 182.1 | - | - | 0 | - |
| - | - | 2.682E+06 | 182.1 | - | - | 0 | - |
| - | - | 2.707E+05 | 183.1 | - | - | 0 | - |
| - | - | 2.157E+05 | 183.1 | - | - | 0 | - |
| - | - | 4.306E+04 | 183.1 | - | - | 0 | - |
| 9 | y | 1.006E+06 | 185.1 | 0.0004301 | 2.324 | +1 | 2 |
| 2 | a | 5.472E+07 | 185.2 | 0.0004435 | 2.395 | +1 | 2 |
| - | - | 6.582E+06 | 186.1 | - | - | 0 | - |
| - | - | 5.619E+06 | 186.2 | - | - | 0 | - |
| - | - | 5.529E+04 | 187.1 | - | - | 0 | - |
| - | - | 4.42E+05 | 187.1 | - | - | 0 | - |
| - | - | 2.294E+05 | 187.2 | - | - | 0 | - |
| - | - | 1.519E+06 | 188.1 | - | - | 0 | - |
| - | - | 1.301E+05 | 189.1 | - | - | 0 | - |
| - | - | 1.308E+05 | 191.1 | - | - | 0 | - |
| - | - | 8.152E+04 | 191.1 | - | - | 0 | - |
| - | - | 1.689E+05 | 193.1 | - | - | 0 | - |
| - | - | 1.349E+05 | 195.1 | - | - | 0 | - |
| - | - | 8.693E+04 | 197.1 | - | - | 0 | - |
| - | - | 2.794E+05 | 197.1 | - | - | 0 | - |
| - | - | 1.883E+05 | 198.1 | - | - | 0 | - |
| - | - | 1.514E+05 | 199.1 | - | - | 0 | - |
| - | - | 5.467E+05 | 200.1 | - | - | 0 | - |
| - | - | 9.186E+04 | 201.1 | - | - | 0 | - |
| 9 | y | 3.424E+07 | 203.1 | 0.0003939 | 1.94 | +1 | 2 |
| - | - | 2.832E+06 | 204.1 | - | - | 0 | - |
| - | - | 2.793E+05 | 205.1 | - | - | 0 | - |
| - | - | 7.987E+05 | 208.1 | - | - | 0 | - |
| - | - | 2.229E+05 | 208.1 | - | - | 0 | - |
| - | - | 6.188E+05 | 209.1 | - | - | 0 | - |
| - | - | 8.254E+04 | 209.1 | - | - | 0 | - |
| - | - | 1.511E+05 | 209.1 | - | - | 0 | - |
| - | - | 8.558E+04 | 210.1 | - | - | 0 | - |
| - | - | 2.98E+06 | 210.1 | - | - | 0 | - |
| - | - | 7.508E+04 | 210.2 | - | - | 0 | - |
| - | - | 5.824E+04 | 211.1 | - | - | 0 | - |
| - | - | 2.622E+05 | 211.1 | - | - | 0 | - |
| - | - | 1.816E+05 | 211.2 | - | - | 0 | - |
| - | - | 1.822E+05 | 212.1 | - | - | 0 | - |
| 2 | b | 2.861E+07 | 213.2 | 0.0004324 | 2.029 | +1 | 2 |
| - | - | 3.417E+06 | 214.2 | - | - | 0 | - |
| - | - | 1.745E+05 | 215.2 | - | - | 0 | - |
| - | - | 6.825E+04 | 217.2 | - | - | 0 | - |
| - | - | 5.055E+04 | 219.1 | - | - | 0 | - |
| - | - | 2.052E+05 | 220.1 | - | - | 0 | - |
| - | - | 5.923E+04 | 221.1 | - | - | 0 | - |
| - | - | 1.402E+05 | 223.1 | - | - | 0 | - |
| - | - | 5.415E+04 | 225.1 | - | - | 0 | - |
| - | - | 6.519E+04 | 225.1 | - | - | 0 | - |
| - | - | 2.322E+07 | 226.2 | - | - | 0 | - |
| - | - | 2.626E+06 | 227.2 | - | - | 0 | - |
| 6 | y | 2.865E+06 | 228.1 | 0.0002994 | 1.312 | +2 | 5 |
| - | - | 1.797E+05 | 228.2 | - | - | 0 | - |
| - | - | 3.035E+05 | 229.1 | - | - | 0 | - |
| - | - | 3.247E+05 | 236.1 | - | - | 0 | - |
| - | - | 4.572E+05 | 238.1 | - | - | 0 | - |
| - | - | 1.397E+05 | 239.2 | - | - | 0 | - |
| - | - | 4.592E+05 | 240.1 | - | - | 0 | - |
| - | - | 7.705E+04 | 241.1 | - | - | 0 | - |
| - | - | 2.322E+05 | 242.2 | - | - | 0 | - |
| - | - | 7.829E+05 | 243.2 | - | - | 0 | - |
| - | - | 6.496E+04 | 248.1 | - | - | 0 | - |
| - | - | 4.022E+05 | 249.2 | - | - | 0 | - |
| - | - | 5.88E+04 | 250.2 | - | - | 0 | - |
| - | - | 1.649E+05 | 252.1 | - | - | 0 | - |
| - | - | 9.144E+05 | 252.2 | - | - | 0 | - |
| - | - | 2.232E+05 | 253.2 | - | - | 0 | - |
| - | - | 8.617E+05 | 254.2 | - | - | 0 | - |
| - | - | 9.483E+04 | 255.2 | - | - | 0 | - |
| 8 | y | 3.753E+05 | 256.1 | 8.996E-05 | 0.3512 | +1 | 3 |
| - | - | 1.9E+05 | 257.1 | - | - | 0 | - |
| - | - | 1.285E+05 | 261.2 | - | - | 0 | - |
| - | - | 1.113E+05 | 262.2 | - | - | 0 | - |
| - | - | 1.261E+05 | 264.1 | - | - | 0 | - |
| - | - | 1.499E+05 | 265.1 | - | - | 0 | - |
| - | - | 4.016E+05 | 266.1 | - | - | 0 | - |
| - | - | 1.071E+05 | 267.1 | - | - | 0 | - |
| - | - | 1.252E+05 | 268.1 | - | - | 0 | - |
| - | - | 7.167E+04 | 269.2 | - | - | 0 | - |
| - | - | 1.192E+06 | 269.2 | - | - | 0 | - |
| - | - | 1.042E+05 | 270.1 | - | - | 0 | - |
| 3 | b | 3.539E+06 | 270.2 | 0.00027 | 0.9992 | +1 | 3 |
| - | - | 1.488E+05 | 270.2 | - | - | 0 | - |
| - | - | 2.677E+06 | 271.2 | - | - | 0 | - |
| - | - | 3.053E+05 | 272.2 | - | - | 0 | - |
| 8 | y | 2.06E+06 | 274.1 | 0.0002675 | 0.9756 | +1 | 3 |
| - | - | 9.661E+04 | 274.2 | - | - | 0 | - |
| - | - | 2.354E+05 | 275.1 | - | - | 0 | - |
| - | - | 9.719E+04 | 276.1 | - | - | 0 | - |
| - | - | 4.628E+06 | 279.2 | - | - | 0 | - |
| - | - | 6.303E+04 | 280.2 | - | - | 0 | - |
| - | - | 6.238E+05 | 280.2 | - | - | 0 | - |
| - | - | 7.618E+05 | 281.2 | - | - | 0 | - |
| - | - | 1.859E+05 | 282.1 | - | - | 0 | - |
| - | - | 1.018E+05 | 282.2 | - | - | 0 | - |
| - | - | 7.137E+05 | 283.1 | - | - | 0 | - |
| - | - | 7.642E+04 | 284.2 | - | - | 0 | - |
| - | - | 9.417E+05 | 285.2 | - | - | 0 | - |
| 5 | y | 4.902E+05 | 285.7 | 0.0004497 | 1.574 | +2 | 6 |
| - | - | 2.265E+05 | 286.2 | - | - | 0 | - |
| - | - | 5.855E+06 | 287.2 | - | - | 0 | - |
| - | - | 1.332E+05 | 288.2 | - | - | 0 | - |
| - | - | 8.243E+05 | 288.2 | - | - | 0 | - |
| - | - | 6.452E+04 | 289.2 | - | - | 0 | - |
| - | - | 9.066E+04 | 295.2 | - | - | 0 | - |
| - | - | 7.875E+04 | 297.2 | - | - | 0 | - |
| - | - | 8.836E+06 | 297.2 | - | - | 0 | - |
| - | - | 1.195E+06 | 298.2 | - | - | 0 | - |
| - | - | 1.694E+07 | 299.2 | - | - | 0 | - |
| - | - | 2.321E+06 | 300.2 | - | - | 0 | - |
| - | - | 2.082E+05 | 301.2 | - | - | 0 | - |
| - | - | 7.163E+04 | 307.2 | - | - | 0 | - |
| - | - | 2.731E+05 | 309.2 | - | - | 0 | - |
| - | - | 5.795E+04 | 309.2 | - | - | 0 | - |
| - | - | 8.622E+04 | 311.2 | - | - | 0 | - |
| - | - | 7.068E+04 | 313.2 | - | - | 0 | - |
| - | - | 5.856E+04 | 313.2 | - | - | 0 | - |
| - | - | 4.808E+05 | 314.2 | - | - | 0 | - |
| - | - | 8.364E+04 | 315.2 | - | - | 0 | - |
| - | - | 1.311E+05 | 319.2 | - | - | 0 | - |
| - | - | 1.059E+05 | 323.2 | - | - | 0 | - |
| - | - | 5.943E+05 | 323.2 | - | - | 0 | - |
| - | - | 8.702E+04 | 324.2 | - | - | 0 | - |
| - | - | 1.049E+05 | 324.2 | - | - | 0 | - |
| - | - | 9.554E+04 | 325.2 | - | - | 0 | - |
| - | - | 4.429E+05 | 325.2 | - | - | 0 | - |
| - | - | 1.103E+05 | 325.7 | - | - | 0 | - |
| - | - | 1.048E+05 | 326.2 | - | - | 0 | - |
| 7 | y | 8.395E+05 | 327.2 | 0.0006043 | 1.847 | +1 | 4 |
| - | - | 4.968E+04 | 327.2 | - | - | 0 | - |
| - | - | 4.988E+05 | 328.2 | - | - | 0 | - |
| - | - | 5.737E+04 | 328.2 | - | - | 0 | - |
| - | - | 1.755E+05 | 331.7 | - | - | 0 | - |
| - | - | 3.374E+05 | 332.2 | - | - | 0 | - |
| - | - | 8.626E+04 | 332.7 | - | - | 0 | - |
| - | - | 5.145E+04 | 333.7 | - | - | 0 | - |
| - | - | 7.434E+04 | 335.2 | - | - | 0 | - |
| - | - | 3.633E+05 | 336.2 | - | - | 0 | - |
| - | - | 7.551E+05 | 337.2 | - | - | 0 | - |
| - | - | 7.205E+04 | 337.2 | - | - | 0 | - |
| - | - | 1.321E+05 | 338.2 | - | - | 0 | - |
| - | - | 5.135E+04 | 339.2 | - | - | 0 | - |
| - | - | 4.387E+06 | 340.2 | - | - | 0 | - |
| 4 | y | 1.475E+06 | 340.7 | 0.0005522 | 1.621 | +2 | 7 |
| - | - | 5.123E+05 | 341.2 | - | - | 0 | - |
| - | - | 7.289E+05 | 341.2 | - | - | 0 | - |
| - | - | 7.874E+04 | 341.7 | - | - | 0 | - |
| - | - | 4.752E+04 | 342.2 | - | - | 0 | - |
| 7 | y | 3.361E+05 | 345.2 | 4.942E-05 | 0.1432 | +1 | 4 |
| 7 | b | 9.967E+04 | 347.7 | 0.0008685 | 2.498 | +2 | 7 |
| - | - | 5.795E+04 | 349.2 | - | - | 0 | - |
| - | - | 8.519E+04 | 349.2 | - | - | 0 | - |
| 4 | y | 3.045E+05 | 349.7 | 0.0006409 | 1.833 | +2 | 7 |
| - | - | 2.752E+05 | 350.2 | - | - | 0 | - |
| - | - | 9.076E+04 | 351.2 | - | - | 0 | - |
| - | - | 2.703E+05 | 353.2 | - | - | 0 | - |
| - | - | 1.021E+05 | 354.2 | - | - | 0 | - |
| - | - | 3.137E+05 | 354.2 | - | - | 0 | - |
| - | - | 6.057E+04 | 356.2 | - | - | 0 | - |
| - | - | 1.043E+05 | 359.7 | - | - | 0 | - |
| - | - | 7.886E+04 | 360.7 | - | - | 0 | - |
| - | - | 1.668E+05 | 360.7 | - | - | 0 | - |
| - | - | 5.441E+04 | 361.2 | - | - | 0 | - |
| - | - | 1.498E+05 | 361.2 | - | - | 0 | - |
| - | - | 5.149E+04 | 361.7 | - | - | 0 | - |
| - | - | 2.518E+05 | 363.2 | - | - | 0 | - |
| - | - | 1.315E+05 | 366.2 | - | - | 0 | - |
| - | - | 2.203E+07 | 368.2 | - | - | 0 | - |
| - | - | 8.672E+04 | 368.7 | - | - | 0 | - |
| - | - | 4.471E+06 | 369.2 | - | - | 0 | - |
| 3 | y | 1.152E+05 | 369.7 | 0.0007422 | 2.008 | +2 | 8 |
| - | - | 1.749E+06 | 370.2 | - | - | 0 | - |
| - | - | 3.504E+05 | 371.2 | - | - | 0 | - |
| - | - | 5.337E+04 | 373.2 | - | - | 0 | - |
| - | - | 6.73E+04 | 375.2 | - | - | 0 | - |
| - | - | 5.264E+05 | 376.2 | - | - | 0 | - |
| - | - | 1.941E+05 | 376.2 | - | - | 0 | - |
| - | - | 1.244E+05 | 377.2 | - | - | 0 | - |
| - | - | 8.439E+04 | 378.2 | - | - | 0 | - |
| 3 | y | 2.451E+05 | 378.2 | 0.002757 | 7.29 | +2 | 8 |
| - | - | 1.693E+05 | 378.7 | - | - | 0 | - |
| - | - | 4.733E+05 | 380.2 | - | - | 0 | - |
| 4 | b | 1.42E+05 | 381.2 | 0.0004986 | 1.308 | +1 | 4 |
| - | - | 2.787E+05 | 382.2 | - | - | 0 | - |
| - | - | 1.574E+05 | 382.7 | - | - | 0 | - |
| 8 | b | 3.55E+05 | 383.2 | 0.0002603 | 0.6793 | +2 | 8 |
| - | - | 1.654E+05 | 383.7 | - | - | 0 | - |
| - | - | 1.008E+05 | 384.3 | - | - | 0 | - |
| - | - | 1.331E+05 | 389.2 | - | - | 0 | - |
| - | - | 5.704E+05 | 390.2 | - | - | 0 | - |
| - | - | 5.196E+04 | 390.2 | - | - | 0 | - |
| - | - | 9.451E+04 | 391.2 | - | - | 0 | - |
| - | - | 1.586E+05 | 393.2 | - | - | 0 | - |
| - | - | 6.055E+05 | 394.2 | - | - | 0 | - |
| - | - | 3.783E+05 | 394.2 | - | - | 0 | - |
| - | - | 1.214E+05 | 395.2 | - | - | 0 | - |
| - | - | 6.808E+04 | 395.2 | - | - | 0 | - |
| - | - | 7.954E+04 | 396.2 | - | - | 0 | - |
| - | - | 7.33E+05 | 396.2 | - | - | 0 | - |
| - | - | 9.679E+04 | 396.7 | - | - | 0 | - |
| - | - | 1.909E+05 | 397.2 | - | - | 0 | - |
| 4 | b | 5.171E+06 | 398.2 | 0.0005609 | 1.408 | +1 | 4 |
| - | - | 1.051E+06 | 399.2 | - | - | 0 | - |
| - | - | 1.884E+05 | 400.2 | - | - | 0 | - |
| - | - | 3.776E+05 | 407.2 | - | - | 0 | - |
| - | - | 9.871E+05 | 408.2 | - | - | 0 | - |
| - | - | 1.636E+05 | 409.2 | - | - | 0 | - |
| - | - | 6.655E+04 | 410.2 | - | - | 0 | - |
| - | - | 5.74E+04 | 410.3 | - | - | 0 | - |
| - | - | 7.63E+05 | 411.2 | - | - | 0 | - |
| - | - | 1.722E+05 | 412.2 | - | - | 0 | - |
| - | - | 1.052E+05 | 413.2 | - | - | 0 | - |
| - | - | 8.146E+04 | 417.2 | - | - | 0 | - |
| - | - | 4.278E+05 | 417.8 | - | - | 0 | - |
| - | - | 2.21E+05 | 418.3 | - | - | 0 | - |
| - | - | 8.44E+04 | 418.8 | - | - | 0 | - |
| - | - | 6.824E+04 | 419.2 | - | - | 0 | - |
| - | - | 1.822E+05 | 420.2 | - | - | 0 | - |
| - | - | 7.653E+04 | 421.2 | - | - | 0 | - |
| - | - | 7.978E+04 | 424.3 | - | - | 0 | - |
| - | - | 7.394E+04 | 425.2 | - | - | 0 | - |
| - | - | 3.301E+05 | 425.3 | - | - | 0 | - |
| 2 | y | 4.719E+05 | 425.7 | 0.0004311 | 1.013 | +2 | 9 |
| 2 | y | 2.46E+05 | 426.2 | 0.005097 | 11.96 | +2 | 9 |
| - | - | 1.074E+05 | 426.7 | - | - | 0 | - |
| - | - | 5.996E+04 | 427.2 | - | - | 0 | - |
| - | - | 8.494E+04 | 429.2 | - | - | 0 | - |
| 9 | b | 6.871E+05 | 431.8 | 0.0003963 | 0.918 | +2 | 9 |
| - | - | 3.022E+05 | 432.3 | - | - | 0 | - |
| - | - | 1.127E+05 | 432.8 | - | - | 0 | - |
| - | - | 1.446E+05 | 434.2 | - | - | 0 | - |
| 2 | y | 1.841E+06 | 434.7 | 0.0006725 | 1.547 | +2 | 9 |
| - | - | 7.579E+05 | 435.2 | - | - | 0 | - |
| - | - | 2.116E+05 | 435.8 | - | - | 0 | - |
| - | - | 2.238E+05 | 437.3 | - | - | 0 | - |
| - | - | 4.283E+05 | 437.3 | - | - | 0 | - |
| - | - | 1.329E+05 | 438.3 | - | - | 0 | - |
| - | - | 2.307E+05 | 439.2 | - | - | 0 | - |
| - | - | 2.497E+05 | 439.3 | - | - | 0 | - |
| - | - | 5.216E+05 | 439.7 | - | - | 0 | - |
| - | - | 2.476E+05 | 440.2 | - | - | 0 | - |
| - | - | 7.186E+04 | 440.3 | - | - | 0 | - |
| - | - | 1.504E+05 | 441.2 | - | - | 0 | - |
| - | - | 1.009E+05 | 441.3 | - | - | 0 | - |
| - | - | 8.241E+04 | 442.2 | - | - | 0 | - |
| - | - | 4.783E+05 | 442.3 | - | - | 0 | - |
| - | - | 1.225E+05 | 443.3 | - | - | 0 | - |
| - | - | 7.565E+05 | 447.2 | - | - | 0 | - |
| - | - | 1.308E+05 | 448.2 | - | - | 0 | - |
| - | - | 1.287E+05 | 449.2 | - | - | 0 | - |
| - | - | 2.927E+05 | 450.3 | - | - | 0 | - |
| - | - | 4.732E+05 | 451.3 | - | - | 0 | - |
| - | - | 6.213E+04 | 453.3 | - | - | 0 | - |
| - | - | 7.588E+04 | 453.3 | - | - | 0 | - |
| - | - | 8.57E+05 | 454.3 | - | - | 0 | - |
| - | - | 1.827E+05 | 455.3 | - | - | 0 | - |
| - | - | 2.96E+05 | 456.3 | - | - | 0 | - |
| - | - | 6.853E+04 | 457.3 | - | - | 0 | - |
| - | - | 5.616E+04 | 457.8 | - | - | 0 | - |
| - | - | 6.405E+04 | 463.3 | - | - | 0 | - |
| - | - | 2.636E+06 | 464.3 | - | - | 0 | - |
| - | - | 5.612E+05 | 465.2 | - | - | 0 | - |
| - | - | 1.815E+06 | 465.3 | - | - | 0 | - |
| - | - | 1.973E+05 | 466.2 | - | - | 0 | - |
| - | - | 5.053E+05 | 466.3 | - | - | 0 | - |
| - | - | 3.715E+05 | 466.8 | - | - | 0 | - |
| - | - | 1.064E+05 | 467.2 | - | - | 0 | - |
| - | - | 5.519E+04 | 467.3 | - | - | 0 | - |
| - | - | 8.594E+05 | 467.3 | - | - | 0 | - |
| - | - | 2.539E+05 | 468.3 | - | - | 0 | - |
| - | - | 9.754E+04 | 469.3 | - | - | 0 | - |
| - | - | 8.11E+04 | 470.3 | - | - | 0 | - |
| 6 | y | 1.376E+06 | 473.3 | 0.000759 | 1.604 | +1 | 5 |
| - | - | 2.504E+05 | 474.3 | - | - | 0 | - |
| 0 | Precursor | 6.959E+05 | 475.3 | 0.0006785 | 1.428 | +2 | -1 |
| 0 | Precursor | 3.691E+05 | 475.8 | 0.009312 | 19.57 | +2 | -1 |
| - | - | 9.573E+04 | 476.3 | - | - | 0 | - |
| - | - | 6.312E+04 | 477.3 | - | - | 0 | - |
| - | - | 4.845E+05 | 478.3 | - | - | 0 | - |
| - | - | 1.716E+06 | 479.3 | - | - | 0 | - |
| - | - | 3.742E+05 | 480.3 | - | - | 0 | - |
| - | - | 8.774E+04 | 480.3 | - | - | 0 | - |
| - | - | 8.469E+04 | 481.3 | - | - | 0 | - |
| - | - | 2.233E+05 | 481.3 | - | - | 0 | - |
| - | - | 6.176E+06 | 482.3 | - | - | 0 | - |
| - | - | 1.473E+06 | 483.3 | - | - | 0 | - |
| 0 | Precursor | 2.78E+06 | 484.3 | 0.0006452 | 1.332 | +2 | -1 |
| - | - | 1.23E+06 | 484.8 | - | - | 0 | - |
| - | - | 6.949E+04 | 485.3 | - | - | 0 | - |
| - | - | 5.515E+05 | 485.3 | - | - | 0 | - |
| - | - | 7.974E+04 | 490.3 | - | - | 0 | - |
| - | - | 1.385E+05 | 491.3 | - | - | 0 | - |
| - | - | 8.334E+04 | 492.3 | - | - | 0 | - |
| 5 | b | 2.964E+05 | 495.3 | 0.0005924 | 1.196 | +1 | 5 |
| - | - | 4.497E+05 | 496.3 | - | - | 0 | - |
| - | - | 6.181E+04 | 497.3 | - | - | 0 | - |
| - | - | 6.641E+04 | 505.3 | - | - | 0 | - |
| - | - | 1.553E+05 | 507.3 | - | - | 0 | - |
| - | - | 7.957E+05 | 508.3 | - | - | 0 | - |
| - | - | 2.983E+05 | 509.3 | - | - | 0 | - |
| - | - | 8.429E+04 | 509.3 | - | - | 0 | - |
| - | - | 5.159E+05 | 518.3 | - | - | 0 | - |
| - | - | 1.022E+05 | 519.3 | - | - | 0 | - |
| - | - | 2.776E+05 | 524.3 | - | - | 0 | - |
| - | - | 1.697E+06 | 525.3 | - | - | 0 | - |
| - | - | 4.874E+05 | 526.3 | - | - | 0 | - |
| - | - | 1.583E+05 | 534.3 | - | - | 0 | - |
| - | - | 5.234E+05 | 535.3 | - | - | 0 | - |
| - | - | 2.184E+06 | 536.3 | - | - | 0 | - |
| - | - | 6.174E+05 | 537.3 | - | - | 0 | - |
| - | - | 9.459E+04 | 538.3 | - | - | 0 | - |
| - | - | 8.216E+04 | 538.3 | - | - | 0 | - |
| - | - | 1.835E+05 | 540.3 | - | - | 0 | - |
| - | - | 7.132E+04 | 548.3 | - | - | 0 | - |
| - | - | 1.457E+05 | 550.3 | - | - | 0 | - |
| 5 | y | 1.43E+06 | 552.3 | 0.00043 | 0.7785 | +1 | 6 |
| - | - | 1.172E+05 | 552.4 | - | - | 0 | - |
| - | - | 1.909E+07 | 553.3 | - | - | 0 | - |
| - | - | 5.802E+06 | 554.3 | - | - | 0 | - |
| - | - | 7.653E+05 | 555.3 | - | - | 0 | - |
| - | - | 1.05E+05 | 555.4 | - | - | 0 | - |
| - | - | 6.983E+04 | 556.3 | - | - | 0 | - |
| - | - | 9.407E+04 | 560.3 | - | - | 0 | - |
| - | - | 1.686E+05 | 564.4 | - | - | 0 | - |
| - | - | 2.541E+05 | 567.4 | - | - | 0 | - |
| 5 | y | 3.892E+07 | 570.3 | 0.0006075 | 1.065 | +1 | 6 |
| - | - | 1.216E+07 | 571.3 | - | - | 0 | - |
| - | - | 1.997E+06 | 572.3 | - | - | 0 | - |
| - | - | 1.081E+05 | 573.3 | - | - | 0 | - |
| - | - | 6.967E+04 | 576.3 | - | - | 0 | - |
| - | - | 7.131E+05 | 577.3 | - | - | 0 | - |
| - | - | 2.236E+05 | 578.3 | - | - | 0 | - |
| - | - | 2.389E+05 | 579.3 | - | - | 0 | - |
| - | - | 1.055E+05 | 580.3 | - | - | 0 | - |
| - | - | 8.742E+05 | 580.4 | - | - | 0 | - |
| - | - | 2.292E+05 | 581.4 | - | - | 0 | - |
| - | - | 1.221E+05 | 586.8 | - | - | 0 | - |
| - | - | 1.346E+05 | 587.3 | - | - | 0 | - |
| - | - | 1.093E+06 | 595.4 | - | - | 0 | - |
| - | - | 3.823E+05 | 596.4 | - | - | 0 | - |
| - | - | 1.016E+05 | 605.3 | - | - | 0 | - |
| - | - | 2.466E+05 | 621.4 | - | - | 0 | - |
| - | - | 2.769E+05 | 622.4 | - | - | 0 | - |
| 6 | b | 3.005E+05 | 623.4 | 0.002391 | 3.835 | +1 | 6 |
| - | - | 1.211E+05 | 624.4 | - | - | 0 | - |
| - | - | 6.934E+04 | 631.4 | - | - | 0 | - |
| - | - | 1.07E+05 | 633.3 | - | - | 0 | - |
| - | - | 1.251E+05 | 637.4 | - | - | 0 | - |
| - | - | 4.519E+05 | 638.4 | - | - | 0 | - |
| - | - | 1.467E+05 | 639.4 | - | - | 0 | - |
| - | - | 6.678E+04 | 640.4 | - | - | 0 | - |
| - | - | 1.504E+05 | 648.4 | - | - | 0 | - |
| - | - | 6.274E+05 | 649.4 | - | - | 0 | - |
| - | - | 1.438E+06 | 650.4 | - | - | 0 | - |
| - | - | 5.944E+05 | 651.4 | - | - | 0 | - |
| - | - | 2.274E+05 | 652.4 | - | - | 0 | - |
| - | - | 6.415E+04 | 653.4 | - | - | 0 | - |
| - | - | 3.7E+05 | 654.4 | - | - | 0 | - |
| - | - | 1.092E+05 | 655.4 | - | - | 0 | - |
| - | - | 1.043E+05 | 659.4 | - | - | 0 | - |
| - | - | 5.207E+04 | 664.4 | - | - | 0 | - |
| - | - | 4.614E+06 | 666.4 | - | - | 0 | - |
| - | - | 1.769E+06 | 667.4 | - | - | 0 | - |
| - | - | 4.134E+05 | 668.4 | - | - | 0 | - |
| - | - | 6.915E+04 | 669.4 | - | - | 0 | - |
| - | - | 7.845E+05 | 676.4 | - | - | 0 | - |
| - | - | 2.847E+05 | 677.4 | - | - | 0 | - |
| - | - | 9.728E+04 | 678.4 | - | - | 0 | - |
| 4 | y | 8.602E+05 | 680.4 | 0.0005072 | 0.7455 | +1 | 7 |
| 4 | y | 1.223E+06 | 681.4 | 0.003796 | 5.572 | +1 | 7 |
| - | - | 4.399E+05 | 682.4 | - | - | 0 | - |
| - | - | 1.378E+05 | 683.4 | - | - | 0 | - |
| - | - | 1.001E+05 | 692.4 | - | - | 0 | - |
| 7 | b | 2.038E+06 | 694.4 | 0.0001684 | 0.2426 | +1 | 7 |
| - | - | 8.413E+05 | 695.4 | - | - | 0 | - |
| - | - | 1.953E+05 | 696.4 | - | - | 0 | - |
| 4 | y | 1.205E+06 | 698.4 | 0.0002308 | 0.3305 | +1 | 7 |
| - | - | 4.74E+05 | 699.4 | - | - | 0 | - |
| - | - | 9.351E+04 | 700.4 | - | - | 0 | - |
| - | - | 6.946E+04 | 719.4 | - | - | 0 | - |
| - | - | 2.676E+05 | 720.4 | - | - | 0 | - |
| - | - | 2.557E+05 | 720.4 | - | - | 0 | - |
| - | - | 1.255E+05 | 721.4 | - | - | 0 | - |
| - | - | 1.201E+05 | 721.4 | - | - | 0 | - |
| - | - | 1.932E+05 | 730.4 | - | - | 0 | - |
| - | - | 1.442E+05 | 731.4 | - | - | 0 | - |
| - | - | 1.022E+05 | 734.5 | - | - | 0 | - |
| - | - | 7.087E+04 | 735.5 | - | - | 0 | - |
| 3 | y | 1.136E+06 | 737.4 | 0.0001617 | 0.2193 | +1 | 8 |
| - | - | 1.192E+06 | 737.5 | - | - | 0 | - |
| 3 | y | 1.631E+06 | 738.4 | 0.004244 | 5.748 | +1 | 8 |
| - | - | 5.009E+05 | 738.5 | - | - | 0 | - |
| - | - | 5.153E+05 | 739.4 | - | - | 0 | - |
| - | - | 1.043E+05 | 739.5 | - | - | 0 | - |
| - | - | 1.411E+05 | 740.4 | - | - | 0 | - |
| - | - | 3.604E+05 | 747.4 | - | - | 0 | - |
| 8 | b | 6.256E+05 | 748.4 | 0.003276 | 4.377 | +1 | 8 |
| - | - | 5.04E+05 | 749.4 | - | - | 0 | - |
| - | - | 2.016E+05 | 750.4 | - | - | 0 | - |
| 3 | y | 3.317E+07 | 755.4 | 0.0003392 | 0.449 | +1 | 8 |
| - | - | 1.374E+07 | 756.4 | - | - | 0 | - |
| - | - | 3.102E+06 | 757.4 | - | - | 0 | - |
| - | - | 1.491E+05 | 758.4 | - | - | 0 | - |
| - | - | 4.578E+05 | 763.4 | - | - | 0 | - |
| - | - | 1.496E+05 | 764.4 | - | - | 0 | - |
| 8 | b | 1.786E+07 | 765.5 | 0.0002251 | 0.294 | +1 | 8 |
| - | - | 7.664E+06 | 766.5 | - | - | 0 | - |
| - | - | 1.848E+06 | 767.5 | - | - | 0 | - |
| - | - | 8.539E+04 | 768.5 | - | - | 0 | - |
| - | - | 1.456E+05 | 791.5 | - | - | 0 | - |
| - | - | 8.508E+04 | 792.5 | - | - | 0 | - |
| 2 | y | 1.601E+05 | 850.5 | 0.0002231 | 0.2623 | +1 | 9 |
| 2 | y | 2.62E+05 | 851.5 | 0.0009298 | 1.092 | +1 | 9 |
| - | - | 1.553E+05 | 852.5 | - | - | 0 | - |
| 2 | y | 4.625E+06 | 868.5 | 7.647E-05 | 0.08805 | +1 | 9 |
| - | - | 2.233E+06 | 869.5 | - | - | 0 | - |
| - | - | 6.446E+05 | 870.5 | - | - | 0 | - |
| - | - | 1.793E+05 | 878.5 | - | - | 0 | - |
| - | - | 1.244E+05 | 879.5 | - | - | 0 | - |
| - | - | 7.919E+04 | 897.4 | - | - | 0 | - |
| - | - | 5.018E+04 | 1205 | - | - | 0 | - |
| - | - | 5.536E+04 | 1509 | - | - | 0 | - |
| - | - | 6.434E+04 | 1572 | - | - | 0 | - |
| - | - | 7.644E+04 | 2736 | - | - | 0 | - |
| - | - | 5.774E+04 | 2795 | - | - | 0 | - |

m/z Charge Intensity FragmentType MassShift Position
120.06626892089844 0 37377.29
123.16212463378906 0 42984.02
124.03955078125 0 79816.97
125.10812377929688 0 41004.973
129.0662078857422 0 1174707
129.1026153564453 0 9377968
130.0697479248047 0 59883.973
130.10020446777344 0 87112.945
130.10595703125 0 595233.8
134.02752685546875 0 63082.34
136.07603454589844 0 573807.56
137.07965087890625 0 61217.168
139.08692932128906 0 263660.75
139.12362670898438 0 53067.082
141.0662078857422 0 697886.3
141.10260009765625 0 778851.25
142.08653259277344 0 51235.844
143.08181762695312 0 381561.1
143.1182098388672 0 401802.8 d 1
150.0166778564453 0 48244.848
152.03456115722656 0 707519.25
153.06626892089844 0 143098.11
153.10269165039062 0 143275.05
155.11819458007812 0 1211016.5
156.1216278076172 0 99649.27
157.09742736816406 0 291620.34
157.17031860351562 0 130315.73
158.09280395507812 0 408041.56
165.10260009765625 0 196222.95
167.11778259277344 0 76550.516
169.0611114501953 0 2810590.8
169.09750366210938 0 1026400.4
169.13385009765625 0 172368.02
170.0644073486328 0 199604.67
170.10098266601562 0 56481.297
171.11318969726562 0 1737076.9
172.11660766601562 0 184085.67
172.14479064941406 0 117855.31
180.11351013183594 0 307797.16
181.0976104736328 0 857147.94
181.1339111328125 0 424381.25
182.10121154785156 0 57236.953
182.1291961669922 0 2682204
183.113037109375 0 270747.84
183.132568359375 0 215710
183.14859008789062 0 43059.75
185.09249877929688 0 1006327.06 y Water loss 8
185.165283203125 0 54723596 a 1
186.08767700195312 0 6582167.5
186.1685791015625 0 5618954.5
187.07139587402344 0 55285.605
187.09104919433594 0 441969.47
187.17181396484375 0 229420.88
188.13970947265625 0 1519444.1
189.1430206298828 0 130075.695
191.08172607421875 0 130838.02
191.11746215820312 0 81524.19
193.09744262695312 0 168888.53
195.11325073242188 0 134902.94
197.0924530029297 0 86930.75
197.12860107421875 0 279363
198.12399291992188 0 188263.8
199.108154296875 0 151403.06
200.13978576660156 0 546676.25
201.1426239013672 0 91864.48
203.10302734375 0 34235960 y 8
204.1063690185547 0 2832131
205.10792541503906 0 279283.72
208.1083526611328 0 798679.9
208.1446990966797 0 222872.75
209.09251403808594 0 618840.6
209.1123046875 0 82540.94
209.12884521484375 0 151138.28
210.09664916992188 0 85584.02
210.1240234375 0 2979631
210.16030883789062 0 75082.48
211.10890197753906 0 58235.9
211.12742614746094 0 262211.72
211.180908203125 0 181571.89
212.1033172607422 0 182234.67
213.16018676757812 0 28605218 b 1
214.16346740722656 0 3416838.5
215.166015625 0 174463.78
217.16534423828125 0 68250.6
219.14784240722656 0 50552.613
220.10873413085938 0 205203.47
221.09292602539062 0 59233.848
223.10818481445312 0 140186.3
225.0994415283203 0 54146.34
225.1359100341797 0 65187.72
226.15542602539062 0 23223474
227.15875244140625 0 2626185.5
228.1345672607422 0 2864971.8 y Water loss 5
228.16123962402344 0 179746.78
229.13790893554688 0 303499.44
236.1398468017578 0 324693.75
238.1189727783203 0 457170.4
239.17552185058594 0 139685.28
240.13470458984375 0 459228.47
241.13807678222656 0 77046.125
242.15008544921875 0 232220.97
243.1819610595703 0 782851.8
248.10276794433594 0 64964.566
249.16001892089844 0 402194.38
250.1652374267578 0 58804.223
252.13461303710938 0 164908.73
252.17120361328125 0 914399
253.16622924804688 0 223206.45
254.1502685546875 0 861681.9
255.15386962890625 0 94829.82
256.1292724609375 0 375294.72 y Water loss 7
257.124755859375 0 189990.97
261.17108154296875 0 128543.695
262.1561584472656 0 111302.1
264.1349792480469 0 126071.555
265.1301574707031 0 149866.12
266.11395263671875 0 401583
267.1460266113281 0 107050.3
268.12945556640625 0 125179
269.1612243652344 0 71667.734
269.197509765625 0 1192283.8
270.146240234375 0 104171.57
270.1814880371094 0 3538993.8 b 2
270.19970703125 0 148754.38
271.1769714355469 0 2676604.8
272.1803894042969 0 305273.56
274.1400146484375 0 2059800.2 y 7
274.1694030761719 0 96614.64
275.14349365234375 0 235361
276.06439208984375 0 97194.914
279.1819152832031 0 4627789
280.16741943359375 0 63032.65
280.1852111816406 0 623763.6
281.1611022949219 0 761789.06
282.14508056640625 0 185868.34
282.16387939453125 0 101764.54
283.1405029296875 0 713674.1
284.1976623535156 0 76421.69
285.1561584472656 0 941714.94
285.6663818359375 0 490182.25 y 4
286.16790771484375 0 226515.61
287.208251953125 0 5854585.5
288.1692199707031 0 133246.72
288.21136474609375 0 824302.06
289.2126770019531 0 64519.348
295.21258544921875 0 90656.99
297.15362548828125 0 78745.05
297.192626953125 0 8835688
298.1957092285156 0 1194742.6
299.1719055175781 0 16940584
300.1746826171875 0 2320691.8
301.1757507324219 0 208196.67
307.1770324707031 0 71626.59
309.1560974121094 0 273052.94
309.1922912597656 0 57953.027
311.18963623046875 0 86215.01
313.18780517578125 0 70677.87
313.2242431640625 0 58558.07
314.2193298339844 0 480790.5
315.2217102050781 0 83644.33
319.1766662597656 0 131147.75
323.1720275878906 0 105853.52
323.20843505859375 0 594269.75
324.19287109375 0 87019.62
324.21173095703125 0 104907.38
325.1515808105469 0 95542.8
325.1877746582031 0 442900.25
325.68463134765625 0 110331.14
326.1899108886719 0 104795.7
327.1669006347656 0 839471.25 y Water loss 6
327.2020568847656 0 49679.34
328.16375732421875 0 498843.7
328.1959533691406 0 57373.08
331.68524169921875 0 175509.38
332.17822265625 0 337404.56
332.6795959472656 0 86257.39
333.69989013671875 0 51454.324
335.2081604003906 0 74342.125
336.2033996582031 0 363322.38
337.18768310546875 0 755098.4
337.20806884765625 0 72049.414
338.1899719238281 0 132139.52
339.16644287109375 0 51349.316
340.2348937988281 0 4386980.5
340.69049072265625 0 1475062.5 y Water loss 3
341.19091796875 0 512260.56
341.23809814453125 0 728946.25
341.6924133300781 0 78741.38
342.24078369140625 0 47524.895
345.1769104003906 0 336073.78 y 6
347.715087890625 0 99669 b 6
349.1894836425781 0 57947.348
349.2237548828125 0 85191.55
349.69586181640625 0 304547.78 y 3
350.21917724609375 0 275198.1
351.2026062011719 0 90762.55
353.2193298339844 0 270332.84
354.1787109375 0 102102.336
354.2146301269531 0 313671.5
356.1922912597656 0 60574.867
359.7160339355469 0 104309.27
360.68829345703125 0 78855.32
360.7239074707031 0 166803.2
361.18182373046875 0 54409.6
361.2247619628906 0 149755.12
361.7272644042969 0 51487.87
363.203857421875 0 251753.03
366.21319580078125 0 131523.25
368.2295837402344 0 22030324
368.72906494140625 0 86722.48
369.2327575683594 0 4470716.5
369.69342041015625 0 115161.81 y Ammonia loss 2
370.244384765625 0 1749087.2
371.2467041015625 0 350397.2
373.2245178222656 0 53371.016
375.21929931640625 0 67297.95
376.1982727050781 0 526435.94
376.2447204589844 0 194101.22
377.2026672363281 0 124398.23
378.1805419921875 0 84387.98
378.2087097167969 0 245114.64 y 2
378.7082214355469 0 169325.27
380.22955322265625 0 473262.44
381.2137451171875 0 142040.34 b Ammonia loss 3
382.2265319824219 0 278729.34
382.72894287109375 0 157420.97
383.2342529296875 0 354960.16 b 7
383.73590087890625 0 165429.53
384.26318359375 0 100825.82
389.22991943359375 0 133067.47
390.2141418457031 0 570404.25
390.2353515625 0 51957.79
391.2163391113281 0 94507.55
393.2250671386719 0 158564.11
394.208984375 0 605472.1
394.245361328125 0 378261.66
395.2118225097656 0 121357.37
395.2488098144531 0 68083.69
396.19146728515625 0 79543.44
396.2243347167969 0 733048.1
396.74334716796875 0 96792.48
397.227294921875 0 190914.22
398.2403564453125 0 5170885 b 3
399.2433166503906 0 1051104.5
400.2453308105469 0 188441.69
407.2411804199219 0 377606.25
408.22467041015625 0 987115.4
409.22711181640625 0 163609.83
410.2397766113281 0 66545.266
410.2776184082031 0 57399.43
411.23577880859375 0 763033.75
412.238037109375 0 172180.84
413.21502685546875 0 105223.35
417.2291259765625 0 81461.4
417.76416015625 0 427848.4
418.2654113769531 0 221005.6
418.76568603515625 0 84401.63
419.2414245605469 0 68244.17
420.22454833984375 0 182245.23
421.2249450683594 0 76528.8
424.25537109375 0 79776.445
425.2175598144531 0 73935.7
425.2514343261719 0 330064.28
425.7431335449219 0 471899.5 y Water loss 1
426.23980712890625 0 245971.56 y Ammonia loss 1
426.74005126953125 0 107409.086
427.2308654785156 0 59964.27
429.2239074707031 0 84944.055
431.76129150390625 0 687091.44 b 8
432.26318359375 0 302247.56
432.7637023925781 0 112681.78
434.2414245605469 0 144626.34
434.7486572265625 0 1840851.2 y 1
435.24945068359375 0 757913.5
435.7512512207031 0 211554.64
437.25140380859375 0 223781.3
437.28759765625 0 428346.84
438.2903137207031 0 132922
439.2314453125 0 230688.6
439.3036193847656 0 249718.6
439.74078369140625 0 521554.72
440.24017333984375 0 247617.94
440.3052062988281 0 71855.32
441.2464294433594 0 150355.11
441.2809143066406 0 100914.78
442.24658203125 0 82414.04
442.27801513671875 0 478280.03
443.27984619140625 0 122486.8
447.2359313964844 0 756485.56
448.2399597167969 0 130757.94
449.2158508300781 0 128666.734
450.28216552734375 0 292663.72
451.2669372558594 0 473209.72
453.2763671875 0 62130.77
453.32000732421875 0 75881.22
454.2779541015625 0 857006.06
455.2789611816406 0 182672.02
456.2576599121094 0 296042.28
457.26202392578125 0 68527.445
457.7578430175781 0 56155.418
463.266845703125 0 64049.01
464.2622985839844 0 2636148
465.246826171875 0 561227.3
465.2819519042969 0 1815174.2
466.24676513671875 0 197306.97
466.2849426269531 0 505336.84
466.7646789550781 0 371523.75
467.22564697265625 0 106403.87
467.2633056640625 0 55192.3
467.2977600097656 0 859382.44
468.3011779785156 0 253938.89
469.2758483886719 0 97537.56
470.2813415527344 0 81095.79
473.2725830078125 0 1376236.9 y 5
474.27587890625 0 250394.84
475.277587890625 0 695882.7 Precursor Water loss
475.7782287597656 0 369099.94 Precursor Ammonia loss
476.2799377441406 0 95729.516
477.28271484375 0 63117.47
478.2778015136719 0 484498.4
479.2622375488281 0 1716332.2
480.26458740234375 0 374242.2
480.2960510253906 0 87738.445
481.2712707519531 0 84686.47
481.3133544921875 0 223339.94
482.27294921875 0 6175889
483.27593994140625 0 1473386.5
484.2828369140625 0 2779540.5 Precursor
484.7845764160156 0 1229597.9
485.2514343261719 0 69492.87
485.28607177734375 0 551546.3
490.27679443359375 0 79743.95
491.2630615234375 0 138506.67
492.26171875 0 83336.21
495.29315185546875 0 296414.4 b 4
496.28857421875 0 449714.94
497.287841796875 0 61813.07
505.2789611816406 0 66407.05
507.2934875488281 0 155320.97
508.2885437011719 0 795695
509.2911071777344 0 298327.12
509.34576416015625 0 84285.83
518.272705078125 0 515856.2
519.2757568359375 0 102166.93
524.3196411132812 0 277587.66
525.3153686523438 0 1696623.4
526.318115234375 0 487352.1
534.304931640625 0 158283.69
535.298583984375 0 523394
536.283447265625 0 2183912.2
537.2861938476562 0 617438.75
538.2911987304688 0 94593.62
538.333740234375 0 82161.625
540.3151245117188 0 183509.97
548.3206787109375 0 71315.15
550.3360595703125 0 145664.06
552.314453125 0 1429504.5 y Water loss 4
552.3887939453125 0 117168.92
553.309814453125 0 19085506
554.312744140625 0 5801880
555.3151245117188 0 765271.4
555.3623046875 0 105023.79
556.3159790039062 0 69831.484
560.3189086914062 0 94074.4
564.3509521484375 0 168630.92
567.3618774414062 0 254066.42
570.3251953125 0 38918548 y 4
571.3280029296875 0 12162263
572.3302001953125 0 1997426
573.3322143554688 0 108100.336
576.3114013671875 0 69674.13
577.3458862304688 0 713059.4
578.3382568359375 0 223580.14
579.3272094726562 0 238919.36
580.3307495117188 0 105483.73
580.3818969726562 0 874220.94
581.3841552734375 0 229247.6
586.7523803710938 0 122112.61
587.2548828125 0 134627.34
595.3565063476562 0 1093281
596.3603515625 0 382340.8
605.3389892578125 0 101607.85
621.3709106445312 0 246551.1
622.3698120117188 0 276883.06
623.3851318359375 0 300501.9 b 5
624.38818359375 0 121114.305
631.35888671875 0 69336.34
633.3375244140625 0 107049.78
637.4047241210938 0 125144.76
638.4009399414062 0 451870.47
639.401123046875 0 146696.3
640.406005859375 0 66783.43
648.3832397460938 0 150401.33
649.3673095703125 0 627445.5
650.3634643554688 0 1438448.8
651.3658447265625 0 594398.2
652.3756103515625 0 227412.83
653.3875122070312 0 64152.332
654.4307250976562 0 370014.56
655.4342651367188 0 109154.97
659.3897094726562 0 104328.09
664.4149780273438 0 52073.2
666.393798828125 0 4613649
667.396728515625 0 1768665.2
668.4006958007812 0 413367.97
669.4071655273438 0 69147.27
676.4146728515625 0 784486.9
677.419189453125 0 284690.1
678.4125366210938 0 97279.44
680.3731079101562 0 860218.44 y Water loss 3
681.3604125976562 0 1223247.9 y Ammonia loss 3
682.3616333007812 0 439866.44
683.3573608398438 0 137795.73
692.411376953125 0 100098.63
694.4248046875 0 2037722.9 b 6
695.4277954101562 0 841297.44
696.4274291992188 0 195347.8
698.3829345703125 0 1205013.4 y 3
699.3858642578125 0 473987.16
700.394287109375 0 93505.29
719.3779296875 0 69464.01
720.3675537109375 0 267583.9
720.4408569335938 0 255698.52
721.3728637695312 0 125486.76
721.4451293945312 0 120077.38
730.423828125 0 193208.58
731.4293212890625 0 144241.47
734.4529418945312 0 102224.31
735.4573974609375 0 70873.6
737.3942260742188 0 1135696.6 y Water loss 2
737.4676513671875 0 1192005
738.38232421875 0 1630754.8 y Ammonia loss 2
738.469970703125 0 500912.1
739.3836669921875 0 515290.28
739.4725952148438 0 104288.59
740.3797607421875 0 141076.16
747.4496459960938 0 360357.88
748.4384765625 0 625639 b Ammonia loss 7
749.436767578125 0 504011.34
750.436279296875 0 201602.89
755.4049682617188 0 33168172 y 2
756.4076538085938 0 13740546
757.4100341796875 0 3101733.2
758.41357421875 0 149102.67
763.4467163085938 0 457784.4
764.4481201171875 0 149555.45
765.4619750976562 0 17863040 b 7
766.4647216796875 0 7663990.5
767.4672241210938 0 1847882.2
768.4671020507812 0 85393.875
791.4771728515625 0 145596.03
792.474365234375 0 85084.72
850.4779052734375 0 160064.73 y Water loss 1
851.4630737304688 0 262037.03 y Ammonia loss 1
852.4729614257812 0 155344.94
868.48876953125 0 4625412.5 y 1
869.4913940429688 0 2232531.8
870.4937744140625 0 644621.9
878.4755859375 0 179256.39
879.4741821289062 0 124375.266
897.4408569335938 0 79187.02
1204.86767578125 0 50176.293
1508.5477294921875 0 55360.883
1572.169921875 0 64342.895
2736.4716796875 0 76437.74
2795.043701171875 0 57743.207

Spectrum Details

|  |  |
| --- | --- |
| Matched peaks? Matched peaks The total absolute number of peaks matched. Additionally in brackets the total fraction of peaks matched and the total number of peaks is shown. | 44 (9.44% of 466) |
| FDR? FDR The false discovery rate estimated for this peptide. It is calculated by matching all theoretical fragments with a non-integer shift with the raw peaks for this spectrum. This is done with 40 different shifts. The resulting percentage is the average number of annotated peaks over the number of annotated peaks with the correct spectrum. | 0.11% |
| Satellite FDR? Satellite FDR See the FDR for details on its calculation. This satellite ion specific FDR only contains the satellite ions (d/w) for I/L/J positions. | 0.00% |
| PSM Score? PSM Score The PSM Score as given by Hecklib to this annotated spectrum. It is shown with three significant figures. | 453 |

#### Spectrum 3709? Spectrum 3709 The raw spectrum of this peptide as annotated by Hecklib. The fragments are coloured according to ion type (see legend). Any peaks with a star '\*' as text can be hovered over to see the full details, first the ion type second the mass shift type. By hovering over the amino acids in the peptide or ions in the legend the corresponding peaks are highlighted. By toggling the 'Unassigned' label you can turn the background (unassigned) peaks on or off in the plot. By updating the slider in the Ion legend you can update the spectrum to only show the top X% of the peaks with labels. The top X% means any peak that is within X% of the highest intensity. By dragging in the spectrum you can zoom in to a specific part of the spectrum and use 'Zoom Out' to get back to the original zoom level. The annotation of the spectrum is based on the given sequence in the peptides file and is done with different software so inconsistencies are likely. The peaks are annotated based on the given sequence, with 20 ppm tolerance.

Copy Data

##### Spectrum 3709 (TSV)

###### Preview

```
Loading example...
```

*Click on the button to copy the data to your clipboard.*

Mz MinMz MaxIntensity Max

WidthHeightPeptide font sizePeptide stroke widthSpectrum font sizeSpectrum stroke widthCompact peptide

Ion legend

wxyz

abcd

OtherUnassignedIon^Charge^_Position_Show for top:%

VLGQPKAAPS

03.13e+46.26e+49.40e+41.25e+5

Zoom Out

z^+1^_2_y^+1^_2_y^+1^_3_y^+2^_6_c^+1^_3_y^+2^_7_c^+1^_4_y^+2^_9_z^+1^_5_y^+1^_5_c^+1^_5_y^+1^_6_z^+1^_6_y^+1^_6_w^+1^_7_c^+1^_6_y^+1^_7_z^+1^_7_c^+1^_7_y^+1^_7_c^+1^_7_z^+1^_8_y^+1^_8_c^+1^_8_w^+1^_9_z^+1^_9_y^+1^_9_c^+1^_9_

0578115717352314

Fragment Matches Table

Show background peaks

| Position | Ion type | Intensity | mz Theoretical | mz Error (Th) | mz Error (ppm) | Charge | Series Number |
| --- | --- | --- | --- | --- | --- | --- | --- |
| - | - | 384.6 | 120.1 | - | - | 0 | - |
| - | - | 340.1 | 121.6 | - | - | 0 | - |
| - | - | 428.4 | 121.7 | - | - | 0 | - |
| - | - | 372.7 | 132.2 | - | - | 0 | - |
| - | - | 395.5 | 132.6 | - | - | 0 | - |
| - | - | 435.7 | 136.1 | - | - | 0 | - |
| - | - | 445.9 | 136.1 | - | - | 0 | - |
| - | - | 417.2 | 148.2 | - | - | 0 | - |
| - | - | 535 | 148.9 | - | - | 0 | - |
| - | - | 497.4 | 155.5 | - | - | 0 | - |
| - | - | 398.6 | 162.9 | - | - | 0 | - |
| - | - | 432 | 171.1 | - | - | 0 | - |
| - | - | 724.3 | 173.5 | - | - | 0 | - |
| - | - | 2.123E+04 | 185.2 | - | - | 0 | - |
| - | - | 2314 | 186.1 | - | - | 0 | - |
| - | - | 1682 | 186.2 | - | - | 0 | - |
| 9 | z | 471.3 | 187.1 | 0.001039 | 5.553 | +1 | 2 |
| - | - | 1625 | 187.1 | - | - | 0 | - |
| 9 | y | 1.596E+04 | 203.1 | 0.000104 | 0.5122 | +1 | 2 |
| - | - | 818.5 | 204.1 | - | - | 0 | - |
| - | - | 459.6 | 211.6 | - | - | 0 | - |
| - | - | 1.782E+04 | 213.2 | - | - | 0 | - |
| - | - | 1186 | 214.2 | - | - | 0 | - |
| - | - | 1790 | 215.1 | - | - | 0 | - |
| - | - | 1456 | 226.2 | - | - | 0 | - |
| - | - | 3856 | 249.2 | - | - | 0 | - |
| - | - | 986 | 269.1 | - | - | 0 | - |
| - | - | 2031 | 270.2 | - | - | 0 | - |
| 8 | y | 683.4 | 274.1 | 0.0001454 | 0.5303 | +1 | 3 |
| 5 | y | 6603 | 277.2 | 0.002128 | 7.676 | +2 | 6 |
| - | - | 1144 | 278.2 | - | - | 0 | - |
| - | - | 757.3 | 285.2 | - | - | 0 | - |
| 3 | c | 2744 | 287.2 | 0.0001491 | 0.5191 | +1 | 3 |
| - | - | 2465 | 299.2 | - | - | 0 | - |
| - | - | 1498 | 309.2 | - | - | 0 | - |
| 4 | y | 592.5 | 340.7 | 0.0007296 | 2.141 | +2 | 7 |
| - | - | 664.3 | 364.2 | - | - | 0 | - |
| - | - | 4586 | 368.2 | - | - | 0 | - |
| - | - | 727.9 | 369.2 | - | - | 0 | - |
| - | - | 1216 | 370.2 | - | - | 0 | - |
| - | - | 809.8 | 382.2 | - | - | 0 | - |
| 4 | c | 4245 | 398.2 | 0.0001641 | 0.4121 | +1 | 4 |
| - | - | 660.4 | 399.2 | - | - | 0 | - |
| - | - | 847.1 | 414.2 | - | - | 0 | - |
| - | - | 595.4 | 414.9 | - | - | 0 | - |
| 2 | y | 1382 | 434.7 | 0.001802 | 4.144 | +2 | 9 |
| - | - | 675.8 | 435.2 | - | - | 0 | - |
| 6 | z | 2.331E+04 | 457.3 | 0.000196 | 0.4286 | +1 | 5 |
| - | - | 2.074E+04 | 458.3 | - | - | 0 | - |
| - | - | 5286 | 459.3 | - | - | 0 | - |
| 6 | y | 1123 | 473.3 | 0.001072 | 2.265 | +1 | 5 |
| - | - | 611.9 | 475.3 | - | - | 0 | - |
| - | - | 1371 | 482.3 | - | - | 0 | - |
| - | - | 2456 | 484.2 | - | - | 0 | - |
| - | - | 2262 | 484.3 | - | - | 0 | - |
| - | - | 1812 | 484.8 | - | - | 0 | - |
| - | - | 660.9 | 485 | - | - | 0 | - |
| - | - | 1410 | 485.3 | - | - | 0 | - |
| - | - | 1196 | 500.2 | - | - | 0 | - |
| 5 | c | 1313 | 512.3 | 0.00136 | 2.654 | +1 | 5 |
| - | - | 1601 | 533.2 | - | - | 0 | - |
| - | - | 3423 | 534.2 | - | - | 0 | - |
| - | - | 873.9 | 535.2 | - | - | 0 | - |
| 5 | y | 874.2 | 552.3 | 0.001651 | 2.989 | +1 | 6 |
| - | - | 6148 | 553.3 | - | - | 0 | - |
| 5 | z | 1679 | 554.3 | 0.005904 | 10.65 | +1 | 6 |
| - | - | 1103 | 568.3 | - | - | 0 | - |
| - | - | 780.5 | 568.4 | - | - | 0 | - |
| - | - | 938.8 | 569.3 | - | - | 0 | - |
| - | - | 9275 | 569.3 | - | - | 0 | - |
| 5 | y | 4.077E+04 | 570.3 | 0.0005522 | 0.9682 | +1 | 6 |
| - | - | 1.2E+04 | 571.3 | - | - | 0 | - |
| - | - | 2381 | 572.3 | - | - | 0 | - |
| - | - | 790.4 | 598.3 | - | - | 0 | - |
| - | - | 1374 | 608.4 | - | - | 0 | - |
| - | - | 562.1 | 609.4 | - | - | 0 | - |
| 4 | w | 2.658E+04 | 624.3 | 0.0004357 | 0.6979 | +1 | 7 |
| - | - | 6480 | 625.3 | - | - | 0 | - |
| - | - | 2106 | 626.3 | - | - | 0 | - |
| - | - | 4311 | 639.4 | - | - | 0 | - |
| 6 | c | 4.857E+04 | 640.4 | 0.0006194 | 0.9672 | +1 | 6 |
| - | - | 1.694E+04 | 641.4 | - | - | 0 | - |
| - | - | 3999 | 642.4 | - | - | 0 | - |
| - | - | 956 | 650.4 | - | - | 0 | - |
| - | - | 714.4 | 651.3 | - | - | 0 | - |
| - | - | 786.5 | 652.4 | - | - | 0 | - |
| - | - | 933.5 | 666.4 | - | - | 0 | - |
| - | - | 730 | 667.4 | - | - | 0 | - |
| - | - | 898.5 | 668.3 | - | - | 0 | - |
| - | - | 945.9 | 677.4 | - | - | 0 | - |
| 4 | y | 1980 | 681.4 | 0.0004761 | 0.6988 | +1 | 7 |
| 4 | z | 1.755E+04 | 682.4 | 0.00116 | 1.7 | +1 | 7 |
| - | - | 6679 | 683.4 | - | - | 0 | - |
| - | - | 1705 | 684.4 | - | - | 0 | - |
| 7 | c | 973 | 694.4 | 0.0009009 | 1.297 | +1 | 7 |
| 4 | y | 6272 | 698.4 | 0.0007191 | 1.03 | +1 | 7 |
| - | - | 2547 | 699.4 | - | - | 0 | - |
| - | - | 1687 | 710.4 | - | - | 0 | - |
| 7 | c | 3.461E+04 | 711.5 | 0.0007459 | 1.048 | +1 | 7 |
| - | - | 1.479E+04 | 712.5 | - | - | 0 | - |
| - | - | 3314 | 713.5 | - | - | 0 | - |
| - | - | 699.9 | 721.4 | - | - | 0 | - |
| - | - | 797.2 | 722.4 | - | - | 0 | - |
| - | - | 573.8 | 724.3 | - | - | 0 | - |
| - | - | 684.7 | 738.4 | - | - | 0 | - |
| - | - | 1436 | 738.5 | - | - | 0 | - |
| 3 | z | 3911 | 739.4 | 0.001261 | 1.706 | +1 | 8 |
| - | - | 1316 | 740.4 | - | - | 0 | - |
| 3 | y | 3.46E+04 | 755.4 | 0.0008815 | 1.167 | +1 | 8 |
| - | - | 1.169E+04 | 756.4 | - | - | 0 | - |
| - | - | 3750 | 757.4 | - | - | 0 | - |
| 8 | c | 1.366E+04 | 765.5 | 0.0006294 | 0.8223 | +1 | 8 |
| - | - | 5781 | 766.5 | - | - | 0 | - |
| - | - | 651.4 | 781.4 | - | - | 0 | - |
| - | - | 917.9 | 790.5 | - | - | 0 | - |
| 2 | w | 8426 | 809.4 | 0.001131 | 1.398 | +1 | 9 |
| - | - | 3717 | 810.4 | - | - | 0 | - |
| - | - | 978.7 | 820.5 | - | - | 0 | - |
| - | - | 2924 | 835.5 | - | - | 0 | - |
| - | - | 1036 | 836.5 | - | - | 0 | - |
| 2 | z | 1.343E+04 | 852.5 | 0.00128 | 1.502 | +1 | 9 |
| - | - | 967 | 853.4 | - | - | 0 | - |
| - | - | 6033 | 853.5 | - | - | 0 | - |
| - | - | 1761 | 854.5 | - | - | 0 | - |
| - | - | 1026 | 864.5 | - | - | 0 | - |
| 2 | y | 2295 | 868.5 | 1.543E-05 | 0.01777 | +1 | 9 |
| - | - | 1156 | 869.5 | - | - | 0 | - |
| 9 | c | 4.245E+04 | 879.5 | 0.001512 | 1.719 | +1 | 9 |
| - | - | 2.09E+04 | 880.5 | - | - | 0 | - |
| - | - | 7084 | 881.5 | - | - | 0 | - |
| - | - | 1807 | 889.5 | - | - | 0 | - |
| - | - | 7703 | 895.5 | - | - | 0 | - |
| - | - | 3451 | 896.5 | - | - | 0 | - |
| - | - | 1509 | 897.5 | - | - | 0 | - |
| - | - | 1397 | 907.6 | - | - | 0 | - |
| - | - | 2815 | 912.5 | - | - | 0 | - |
| - | - | 1426 | 913.5 | - | - | 0 | - |
| - | - | 2596 | 922.6 | - | - | 0 | - |
| - | - | 7292 | 923.5 | - | - | 0 | - |
| - | - | 2682 | 924.6 | - | - | 0 | - |
| - | - | 1224 | 925.6 | - | - | 0 | - |
| - | - | 697.8 | 926.6 | - | - | 0 | - |
| - | - | 719.6 | 945.6 | - | - | 0 | - |
| - | - | 659.2 | 948.5 | - | - | 0 | - |
| - | - | 1869 | 950.4 | - | - | 0 | - |
| - | - | 2252 | 950.6 | - | - | 0 | - |
| - | - | 4.609E+04 | 951.5 | - | - | 0 | - |
| - | - | 2.262E+04 | 952.5 | - | - | 0 | - |
| - | - | 6869 | 953.5 | - | - | 0 | - |
| - | - | 939.2 | 954.6 | - | - | 0 | - |
| - | - | 826.5 | 966.4 | - | - | 0 | - |
| - | - | 1477 | 967.4 | - | - | 0 | - |
| - | - | 3.928E+04 | 967.6 | - | - | 0 | - |
| - | - | 1.241E+05 | 968.6 | - | - | 0 | - |
| - | - | 5.986E+04 | 969.6 | - | - | 0 | - |
| - | - | 734.8 | 970 | - | - | 0 | - |
| - | - | 1250 | 970.5 | - | - | 0 | - |
| - | - | 1.721E+04 | 970.6 | - | - | 0 | - |
| - | - | 748 | 971.5 | - | - | 0 | - |
| - | - | 1974 | 971.6 | - | - | 0 | - |
| - | - | 622.9 | 1283 | - | - | 0 | - |
| - | - | 857.5 | 1422 | - | - | 0 | - |
| - | - | 798.5 | 1434 | - | - | 0 | - |
| - | - | 697.9 | 1940 | - | - | 0 | - |
| - | - | 803.3 | 1941 | - | - | 0 | - |
| - | - | 690.9 | 2291 | - | - | 0 | - |

m/z Charge Intensity FragmentType MassShift Position
120.08114624023438 0 384.58218
121.61015319824219 0 340.05618
121.67631530761719 0 428.43954
132.16798400878906 0 372.74448
132.64341735839844 0 395.4709
136.07574462890625 0 435.67215
136.1212921142578 0 445.85834
148.1949005126953 0 417.19086
148.9463348388672 0 534.96655
155.52005004882812 0 497.36545
162.90072631835938 0 398.6232
171.112548828125 0 432.0137
173.452880859375 0 724.28156
185.16497802734375 0 21231.69
186.0875244140625 0 2313.9915
186.16839599609375 0 1681.8315
187.08287048339844 0 471.26996 z 8
187.144287109375 0 1625.3721
203.1027374267578 0 15957.724 y 8
204.1057891845703 0 818.4543
211.5884246826172 0 459.57132
213.15992736816406 0 17824.957
214.16366577148438 0 1185.6313
215.13888549804688 0 1789.8717
226.155517578125 0 1455.5563
249.1598358154297 0 3856.4802
269.0801696777344 0 985.9556
270.1811828613281 0 2031.1659
274.139892578125 0 683.35864 y 7
277.15478515625 0 6602.6455 y Ammonia loss 4
278.1582336425781 0 1143.968
285.1569519042969 0 757.3085
287.2079162597656 0 2743.9666 c 2
299.1715393066406 0 2464.5938
309.20379638671875 0 1497.5302
340.689208984375 0 592.538 y Water loss 3
364.1864929199219 0 664.25165
368.22906494140625 0 4586.141
369.2304992675781 0 727.8656
370.2441711425781 0 1216.0193
382.2273254394531 0 809.7726
398.2399597167969 0 4244.8306 c Ammonia loss 3
399.2428894042969 0 660.3973
414.1690979003906 0 847.1048
414.8851318359375 0 595.42566
434.7497863769531 0 1382.1855 y 1
435.2491760253906 0 675.7672
457.2532958984375 0 23305.969 z 5
458.26025390625 0 20738.12
459.26275634765625 0 5286.433
473.270751953125 0 1123.2936 y 5
475.2777099609375 0 611.8921
482.2726745605469 0 1371.0408
484.24896240234375 0 2456.2622
484.2831726074219 0 2261.6362
484.7837219238281 0 1812.3705
484.9815368652344 0 660.9211
485.2840881347656 0 1410.4855
500.2093811035156 0 1195.8634
512.3177490234375 0 1312.6212 c 4
533.1890869140625 0 1601.0659
534.1964721679688 0 3422.8325
535.1961669921875 0 873.94696
552.315673828125 0 874.213 y Water loss 4
553.3091430664062 0 6147.941
554.311767578125 0 1679.333 z 4
568.3099365234375 0 1102.7306
568.3682250976562 0 780.4567
569.2686157226562 0 938.7791
569.3167724609375 0 9274.6455
570.3240356445312 0 40772.03 y 4
571.3272094726562 0 11996.483
572.3290405273438 0 2381.025
598.3170166015625 0 790.39374
608.3634643554688 0 1373.7812
609.3699951171875 0 562.07104
624.334716796875 0 26577.996 w 3
625.33740234375 0 6480.1963
626.3422241210938 0 2106.3367
639.4059448242188 0 4310.553
640.4134521484375 0 48571.1 c 5
641.4168701171875 0 16935.602
642.4196166992188 0 3998.8162
650.3622436523438 0 956.0029
651.2850952148438 0 714.3949
652.3751831054688 0 786.5332
666.3959350585938 0 933.50775
667.3934936523438 0 729.97284
668.34912109375 0 898.4668
677.3810424804688 0 945.9143
681.3561401367188 0 1980.2013 y Ammonia loss 3
682.36328125 0 17549.393 z 3
683.366943359375 0 6679.2344
684.3684692382812 0 1705.4949
694.425537109375 0 972.95734 c Ammonia loss 6
698.3824462890625 0 6271.5425 y 3
699.3848266601562 0 2547.016
710.4420776367188 0 1686.5433
711.450439453125 0 34605.836 c 6
712.4532470703125 0 14793.448
713.456298828125 0 3313.921
721.4495239257812 0 699.8883
722.4468994140625 0 797.17816
724.3268432617188 0 573.8343
738.3938598632812 0 684.7263
738.4718017578125 0 1435.8029
739.3846435546875 0 3910.9382 z 2
740.392822265625 0 1316.4314
755.4037475585938 0 34601.824 y 2
756.4058837890625 0 11690.173
757.4088745117188 0 3749.6392
765.4611206054688 0 13662.767 c Ammonia loss 7
766.4634399414062 0 5780.822
781.4319458007812 0 651.3587
790.4714965820312 0 917.94324
809.4140625 0 8426.215 w 1
810.417236328125 0 3716.9265
820.5131225585938 0 978.6857
835.527587890625 0 2924.4778
836.529052734375 0 1036.4397
852.4686889648438 0 13427.254 z 1
853.3845825195312 0 966.9756
853.471435546875 0 6032.8955
854.4737548828125 0 1760.9989
864.5296630859375 0 1026.0034
868.4887084960938 0 2294.587 y 1
869.49462890625 0 1156.043
879.53955078125 0 42449.082 c 8
880.5407104492188 0 20900.54
881.5438842773438 0 7084.1367
889.5343627929688 0 1806.9197
895.4754028320312 0 7703.4116
896.4788818359375 0 3450.525
897.52392578125 0 1509.1361
907.5552368164062 0 1397.4984
912.5004272460938 0 2814.5518
913.5001220703125 0 1425.7242
922.5565795898438 0 2596.237
923.5453491210938 0 7291.9023
924.5562133789062 0 2682.13
925.5636596679688 0 1224.426
926.5589599609375 0 697.8164
945.6094360351562 0 719.5525
948.4506225585938 0 659.21106
950.38623046875 0 1869.2263
950.5534057617188 0 2252.48
951.5371704101562 0 46091.254
952.5401000976562 0 22622.377
953.5416259765625 0 6868.799
954.5521850585938 0 939.2279
966.4415893554688 0 826.524
967.4491577148438 0 1477.2378
967.555908203125 0 39279.465
968.562255859375 0 124054.38
969.5652465820312 0 59860.008
969.9766235351562 0 734.76306
970.4659423828125 0 1250.0911
970.5687866210938 0 17207.422
971.4633178710938 0 747.99
971.5709838867188 0 1974.1996
1282.6201171875 0 622.88965
1421.66162109375 0 857.54755
1433.650634765625 0 798.4778
1939.8787841796875 0 697.8961
1940.9267578125 0 803.3369
2290.730712890625 0 690.86487

Spectrum Details

|  |  |
| --- | --- |
| Matched peaks? Matched peaks The total absolute number of peaks matched. Additionally in brackets the total fraction of peaks matched and the total number of peaks is shown. | 28 (16.87% of 166) |
| FDR? FDR The false discovery rate estimated for this peptide. It is calculated by matching all theoretical fragments with a non-integer shift with the raw peaks for this spectrum. This is done with 40 different shifts. The resulting percentage is the average number of annotated peaks over the number of annotated peaks with the correct spectrum. | 0.94% |
| Satellite FDR? Satellite FDR See the FDR for details on its calculation. This satellite ion specific FDR only contains the satellite ions (d/w) for I/L/J positions. | 2.38% |
| PSM Score? PSM Score The PSM Score as given by Hecklib to this annotated spectrum. It is shown with three significant figures. | 298 |

#### Spectrum 3420? Spectrum 3420 The raw spectrum of this peptide as annotated by Hecklib. The fragments are coloured according to ion type (see legend). Any peaks with a star '\*' as text can be hovered over to see the full details, first the ion type second the mass shift type. By hovering over the amino acids in the peptide or ions in the legend the corresponding peaks are highlighted. By toggling the 'Unassigned' label you can turn the background (unassigned) peaks on or off in the plot. By updating the slider in the Ion legend you can update the spectrum to only show the top X% of the peaks with labels. The top X% means any peak that is within X% of the highest intensity. By dragging in the spectrum you can zoom in to a specific part of the spectrum and use 'Zoom Out' to get back to the original zoom level. The annotation of the spectrum is based on the given sequence in the peptides file and is done with different software so inconsistencies are likely. The peaks are annotated based on the given sequence, with 20 ppm tolerance.

Copy Data

##### Spectrum 3420 (TSV)

###### Preview

```
Loading example...
```

*Click on the button to copy the data to your clipboard.*

Mz MinMz MaxIntensity Max

WidthHeightPeptide font sizePeptide stroke widthSpectrum font sizeSpectrum stroke widthCompact peptide

Ion legend

wxyz

abcd

OtherUnassignedIon^Charge^_Position_Show for top:%

VLGQPKAAPS

06.31e+41.26e+51.89e+52.53e+5

Zoom Out

d^+1^_2_y^+2^_4_y^+1^_2_a^+1^_2_y^+1^_2_b^+1^_2_y^+2^_5_y^+1^_3_b^+1^_3_y^+1^_3_y^+2^_6_y^+2^_6_y^+1^_4_y^+2^_7_y^+1^_4_y^+2^_7_y^+2^_8_y^+2^_8_b^+1^_4_b^+2^_8_b^+1^_4_y^+2^_9_y^+2^_9_b^+2^_9_y^+2^_9_y^+1^_5_\*\*b^+1^_5_y^+1^_6_y^+1^_6_b^+1^_6_y^+1^_7_y^+1^_7_b^+1^_7_y^+1^_7_y^+1^_8_y^+1^_8_b^+1^_8_y^+1^_8_b^+1^_8_y^+1^_9_y^+1^_9_

0759151822783037

Fragment Matches Table

Show background peaks

| Position | Ion type | Intensity | mz Theoretical | mz Error (Th) | mz Error (ppm) | Charge | Series Number |
| --- | --- | --- | --- | --- | --- | --- | --- |
| - | - | 1493 | 120.1 | - | - | 0 | - |
| - | - | 1746 | 120.1 | - | - | 0 | - |
| - | - | 593.1 | 124 | - | - | 0 | - |
| - | - | 363.2 | 124.1 | - | - | 0 | - |
| - | - | 692.4 | 127.1 | - | - | 0 | - |
| - | - | 4562 | 129.1 | - | - | 0 | - |
| - | - | 5.389E+04 | 129.1 | - | - | 0 | - |
| - | - | 641.5 | 130.1 | - | - | 0 | - |
| - | - | 1113 | 130.1 | - | - | 0 | - |
| - | - | 3282 | 130.1 | - | - | 0 | - |
| - | - | 773.9 | 132.1 | - | - | 0 | - |
| - | - | 409.1 | 133.7 | - | - | 0 | - |
| - | - | 1061 | 134 | - | - | 0 | - |
| - | - | 2.308E+04 | 136.1 | - | - | 0 | - |
| - | - | 1764 | 137.1 | - | - | 0 | - |
| - | - | 548.5 | 137.1 | - | - | 0 | - |
| - | - | 644.1 | 138 | - | - | 0 | - |
| - | - | 420.1 | 138.1 | - | - | 0 | - |
| - | - | 1765 | 139.1 | - | - | 0 | - |
| - | - | 3451 | 141.1 | - | - | 0 | - |
| - | - | 4088 | 141.1 | - | - | 0 | - |
| - | - | 718.4 | 142.1 | - | - | 0 | - |
| - | - | 1249 | 143.1 | - | - | 0 | - |
| 2 | d | 2901 | 143.1 | 0.0004882 | 3.411 | +1 | 2 |
| - | - | 403.4 | 147 | - | - | 0 | - |
| - | - | 1356 | 147.1 | - | - | 0 | - |
| - | - | 486.1 | 148.9 | - | - | 0 | - |
| - | - | 635.6 | 149 | - | - | 0 | - |
| - | - | 3217 | 152 | - | - | 0 | - |
| - | - | 660.3 | 153.1 | - | - | 0 | - |
| - | - | 1161 | 153.1 | - | - | 0 | - |
| - | - | 5488 | 155.1 | - | - | 0 | - |
| - | - | 553.5 | 156.1 | - | - | 0 | - |
| - | - | 1874 | 157.1 | - | - | 0 | - |
| - | - | 1637 | 158.1 | - | - | 0 | - |
| - | - | 822.5 | 159.1 | - | - | 0 | - |
| - | - | 6074 | 159.1 | - | - | 0 | - |
| - | - | 1470 | 165.1 | - | - | 0 | - |
| - | - | 565.1 | 166.1 | - | - | 0 | - |
| - | - | 612.5 | 167.1 | - | - | 0 | - |
| - | - | 771.1 | 167.1 | - | - | 0 | - |
| - | - | 451 | 168 | - | - | 0 | - |
| - | - | 709.2 | 169.1 | - | - | 0 | - |
| - | - | 1.369E+04 | 169.1 | - | - | 0 | - |
| - | - | 3826 | 169.1 | - | - | 0 | - |
| - | - | 781.2 | 169.1 | - | - | 0 | - |
| - | - | 568.5 | 170.1 | - | - | 0 | - |
| - | - | 558.7 | 170.1 | - | - | 0 | - |
| - | - | 626.9 | 171.1 | - | - | 0 | - |
| - | - | 8586 | 171.1 | - | - | 0 | - |
| - | - | 861.8 | 172.1 | - | - | 0 | - |
| 7 | y | 2439 | 173.1 | 0.0005064 | 2.925 | +2 | 4 |
| - | - | 4938 | 173.1 | - | - | 0 | - |
| - | - | 1494 | 173.5 | - | - | 0 | - |
| - | - | 594.3 | 174.1 | - | - | 0 | - |
| - | - | 545.1 | 175.3 | - | - | 0 | - |
| - | - | 488.7 | 179 | - | - | 0 | - |
| - | - | 1807 | 180.1 | - | - | 0 | - |
| - | - | 4109 | 181.1 | - | - | 0 | - |
| - | - | 2517 | 181.1 | - | - | 0 | - |
| - | - | 1.266E+04 | 182.1 | - | - | 0 | - |
| - | - | 2757 | 183.1 | - | - | 0 | - |
| - | - | 1305 | 183.1 | - | - | 0 | - |
| - | - | 531.4 | 183.1 | - | - | 0 | - |
| - | - | 1386 | 185.1 | - | - | 0 | - |
| 9 | y | 5110 | 185.1 | 0.0004453 | 2.406 | +1 | 2 |
| 2 | a | 2.5E+05 | 185.2 | 0.0005198 | 2.807 | +1 | 2 |
| - | - | 3.086E+04 | 186.1 | - | - | 0 | - |
| - | - | 2.612E+04 | 186.2 | - | - | 0 | - |
| - | - | 1492 | 187.1 | - | - | 0 | - |
| - | - | 1745 | 187.1 | - | - | 0 | - |
| - | - | 1856 | 187.1 | - | - | 0 | - |
| - | - | 1.639E+04 | 187.1 | - | - | 0 | - |
| - | - | 986.9 | 187.2 | - | - | 0 | - |
| - | - | 6314 | 188.1 | - | - | 0 | - |
| - | - | 1237 | 188.1 | - | - | 0 | - |
| - | - | 834.5 | 191.1 | - | - | 0 | - |
| - | - | 1059 | 193.1 | - | - | 0 | - |
| - | - | 1004 | 195.1 | - | - | 0 | - |
| - | - | 1680 | 197.1 | - | - | 0 | - |
| - | - | 734.1 | 197.2 | - | - | 0 | - |
| - | - | 1174 | 199.1 | - | - | 0 | - |
| - | - | 3483 | 200.1 | - | - | 0 | - |
| - | - | 2246 | 201.1 | - | - | 0 | - |
| 9 | y | 1.472E+05 | 203.1 | 0.0004702 | 2.315 | +1 | 2 |
| - | - | 1.322E+04 | 204.1 | - | - | 0 | - |
| - | - | 675 | 204.1 | - | - | 0 | - |
| - | - | 678.8 | 205.1 | - | - | 0 | - |
| - | - | 642.7 | 207.1 | - | - | 0 | - |
| - | - | 859.7 | 207.1 | - | - | 0 | - |
| - | - | 3375 | 208.1 | - | - | 0 | - |
| - | - | 1161 | 208.1 | - | - | 0 | - |
| - | - | 3498 | 209.1 | - | - | 0 | - |
| - | - | 1032 | 209.1 | - | - | 0 | - |
| - | - | 581.7 | 210.1 | - | - | 0 | - |
| - | - | 1.4E+04 | 210.1 | - | - | 0 | - |
| - | - | 1762 | 211.1 | - | - | 0 | - |
| - | - | 621.6 | 211.2 | - | - | 0 | - |
| - | - | 1075 | 212.1 | - | - | 0 | - |
| - | - | 576 | 212.1 | - | - | 0 | - |
| 2 | b | 1.328E+05 | 213.2 | 0.0005087 | 2.387 | +1 | 2 |
| - | - | 1.523E+04 | 214.2 | - | - | 0 | - |
| - | - | 548.4 | 214.4 | - | - | 0 | - |
| - | - | 9788 | 215.1 | - | - | 0 | - |
| - | - | 874.2 | 215.2 | - | - | 0 | - |
| - | - | 1187 | 216.1 | - | - | 0 | - |
| - | - | 773.5 | 219.1 | - | - | 0 | - |
| - | - | 628.5 | 220.1 | - | - | 0 | - |
| - | - | 752.9 | 221.1 | - | - | 0 | - |
| - | - | 798.9 | 223.1 | - | - | 0 | - |
| - | - | 509.7 | 224.1 | - | - | 0 | - |
| - | - | 556.5 | 225.1 | - | - | 0 | - |
| - | - | 642.2 | 225.1 | - | - | 0 | - |
| - | - | 2040 | 226.1 | - | - | 0 | - |
| - | - | 9.912E+04 | 226.2 | - | - | 0 | - |
| - | - | 570 | 227.1 | - | - | 0 | - |
| - | - | 1.2E+04 | 227.2 | - | - | 0 | - |
| - | - | 774.2 | 228.1 | - | - | 0 | - |
| 6 | y | 1.324E+04 | 228.1 | 0.000452 | 1.981 | +2 | 5 |
| - | - | 761.4 | 229.1 | - | - | 0 | - |
| - | - | 1200 | 229.1 | - | - | 0 | - |
| - | - | 796.9 | 230.1 | - | - | 0 | - |
| - | - | 607.7 | 230.2 | - | - | 0 | - |
| - | - | 676.8 | 231 | - | - | 0 | - |
| - | - | 910 | 233.1 | - | - | 0 | - |
| - | - | 995.1 | 235.1 | - | - | 0 | - |
| - | - | 1188 | 236.1 | - | - | 0 | - |
| - | - | 1396 | 238.1 | - | - | 0 | - |
| - | - | 685.9 | 239.2 | - | - | 0 | - |
| - | - | 670.4 | 239.2 | - | - | 0 | - |
| - | - | 3287 | 240.1 | - | - | 0 | - |
| - | - | 861.8 | 242.1 | - | - | 0 | - |
| - | - | 3101 | 243.2 | - | - | 0 | - |
| - | - | 646.2 | 244.1 | - | - | 0 | - |
| - | - | 624.3 | 244.2 | - | - | 0 | - |
| - | - | 685.4 | 246.1 | - | - | 0 | - |
| - | - | 746.2 | 247.1 | - | - | 0 | - |
| - | - | 510.4 | 248.1 | - | - | 0 | - |
| - | - | 1.826E+04 | 249.2 | - | - | 0 | - |
| - | - | 2766 | 250.2 | - | - | 0 | - |
| - | - | 648.2 | 251.1 | - | - | 0 | - |
| - | - | 717.6 | 251.2 | - | - | 0 | - |
| - | - | 1545 | 252.1 | - | - | 0 | - |
| - | - | 1738 | 252.1 | - | - | 0 | - |
| - | - | 3271 | 252.2 | - | - | 0 | - |
| - | - | 1657 | 253.2 | - | - | 0 | - |
| - | - | 3951 | 254.2 | - | - | 0 | - |
| 8 | y | 1925 | 256.1 | 0.0006088 | 2.377 | +1 | 3 |
| - | - | 735.3 | 257.1 | - | - | 0 | - |
| - | - | 637.9 | 258.1 | - | - | 0 | - |
| - | - | 577.8 | 261.2 | - | - | 0 | - |
| - | - | 880 | 265.1 | - | - | 0 | - |
| - | - | 1660 | 266.1 | - | - | 0 | - |
| - | - | 984.9 | 267.1 | - | - | 0 | - |
| - | - | 580.7 | 268.1 | - | - | 0 | - |
| - | - | 870.2 | 269.1 | - | - | 0 | - |
| - | - | 6245 | 269.1 | - | - | 0 | - |
| - | - | 3412 | 269.2 | - | - | 0 | - |
| 3 | b | 1.401E+04 | 270.2 | 0.0004531 | 1.677 | +1 | 3 |
| - | - | 683.2 | 270.2 | - | - | 0 | - |
| - | - | 1.127E+04 | 271.2 | - | - | 0 | - |
| - | - | 1129 | 272.2 | - | - | 0 | - |
| - | - | 1235 | 273.1 | - | - | 0 | - |
| 8 | y | 9855 | 274.1 | 0.0004811 | 1.755 | +1 | 3 |
| - | - | 1236 | 275.1 | - | - | 0 | - |
| - | - | 847.8 | 276.1 | - | - | 0 | - |
| 5 | y | 8610 | 277.2 | 0.002555 | 9.218 | +2 | 6 |
| - | - | 1410 | 278.2 | - | - | 0 | - |
| - | - | 1.982E+04 | 279.2 | - | - | 0 | - |
| - | - | 2356 | 280.2 | - | - | 0 | - |
| - | - | 3367 | 281.2 | - | - | 0 | - |
| - | - | 3256 | 283.1 | - | - | 0 | - |
| - | - | 4133 | 285.2 | - | - | 0 | - |
| 5 | y | 2152 | 285.7 | 0.0005412 | 1.895 | +2 | 6 |
| - | - | 721 | 286.2 | - | - | 0 | - |
| - | - | 2.707E+04 | 287.2 | - | - | 0 | - |
| - | - | 3502 | 288.2 | - | - | 0 | - |
| - | - | 3.97E+04 | 297.2 | - | - | 0 | - |
| - | - | 5651 | 298.2 | - | - | 0 | - |
| - | - | 7.735E+04 | 299.2 | - | - | 0 | - |
| - | - | 9892 | 300.2 | - | - | 0 | - |
| - | - | 1886 | 301.1 | - | - | 0 | - |
| - | - | 601 | 301.2 | - | - | 0 | - |
| - | - | 1002 | 301.2 | - | - | 0 | - |
| - | - | 766.9 | 303.1 | - | - | 0 | - |
| - | - | 2184 | 309.2 | - | - | 0 | - |
| - | - | 1898 | 309.2 | - | - | 0 | - |
| - | - | 786.6 | 314.2 | - | - | 0 | - |
| - | - | 938.6 | 314.2 | - | - | 0 | - |
| - | - | 2970 | 323.2 | - | - | 0 | - |
| - | - | 649.8 | 323.2 | - | - | 0 | - |
| - | - | 2385 | 325.2 | - | - | 0 | - |
| - | - | 970.4 | 325.7 | - | - | 0 | - |
| 7 | y | 4444 | 327.2 | 0.001032 | 3.153 | +1 | 4 |
| - | - | 1646 | 328.2 | - | - | 0 | - |
| - | - | 614.4 | 331.7 | - | - | 0 | - |
| - | - | 1177 | 332.2 | - | - | 0 | - |
| - | - | 1720 | 336.2 | - | - | 0 | - |
| - | - | 2673 | 337.2 | - | - | 0 | - |
| - | - | 600.6 | 338.2 | - | - | 0 | - |
| - | - | 1.766E+04 | 340.2 | - | - | 0 | - |
| 4 | y | 5768 | 340.7 | 0.0006132 | 1.8 | +2 | 7 |
| - | - | 2028 | 341.2 | - | - | 0 | - |
| - | - | 2501 | 341.2 | - | - | 0 | - |
| 7 | y | 2118 | 345.2 | 0.0003241 | 0.9389 | +1 | 4 |
| - | - | 589 | 346.2 | - | - | 0 | - |
| - | - | 635.9 | 347.2 | - | - | 0 | - |
| - | - | 653.1 | 349.2 | - | - | 0 | - |
| 4 | y | 1352 | 349.7 | 0.0005799 | 1.658 | +2 | 7 |
| - | - | 1049 | 350.2 | - | - | 0 | - |
| - | - | 1742 | 350.2 | - | - | 0 | - |
| - | - | 1226 | 353.2 | - | - | 0 | - |
| - | - | 663.3 | 354.2 | - | - | 0 | - |
| - | - | 1437 | 354.2 | - | - | 0 | - |
| - | - | 719.1 | 359.2 | - | - | 0 | - |
| - | - | 655.4 | 360.7 | - | - | 0 | - |
| - | - | 1172 | 363.2 | - | - | 0 | - |
| - | - | 9.576E+04 | 368.2 | - | - | 0 | - |
| 3 | y | 1048 | 369.2 | 0.001478 | 4.003 | +2 | 8 |
| - | - | 1.993E+04 | 369.2 | - | - | 0 | - |
| - | - | 7893 | 370.2 | - | - | 0 | - |
| - | - | 1122 | 371.2 | - | - | 0 | - |
| - | - | 1857 | 376.2 | - | - | 0 | - |
| - | - | 787.9 | 376.2 | - | - | 0 | - |
| 3 | y | 1098 | 378.2 | 0.003978 | 10.52 | +2 | 8 |
| - | - | 729.3 | 380.2 | - | - | 0 | - |
| - | - | 2124 | 380.2 | - | - | 0 | - |
| 4 | b | 601.2 | 381.2 | 0.0002033 | 0.5332 | +1 | 4 |
| - | - | 968 | 382.2 | - | - | 0 | - |
| - | - | 2133 | 382.2 | - | - | 0 | - |
| - | - | 1274 | 382.7 | - | - | 0 | - |
| - | - | 552.4 | 383.2 | - | - | 0 | - |
| 8 | b | 1734 | 383.2 | 4.669E-05 | 0.1218 | +2 | 8 |
| - | - | 630 | 384.2 | - | - | 0 | - |
| - | - | 2400 | 390.2 | - | - | 0 | - |
| - | - | 913.4 | 393.2 | - | - | 0 | - |
| - | - | 3623 | 394.2 | - | - | 0 | - |
| - | - | 1670 | 394.2 | - | - | 0 | - |
| - | - | 694.8 | 395.2 | - | - | 0 | - |
| - | - | 587.2 | 395.2 | - | - | 0 | - |
| - | - | 807.2 | 396.2 | - | - | 0 | - |
| - | - | 2309 | 396.2 | - | - | 0 | - |
| 4 | b | 2.318E+04 | 398.2 | 0.000744 | 1.868 | +1 | 4 |
| - | - | 5204 | 399.2 | - | - | 0 | - |
| - | - | 1323 | 407.2 | - | - | 0 | - |
| - | - | 3449 | 408.2 | - | - | 0 | - |
| - | - | 612.1 | 409.2 | - | - | 0 | - |
| - | - | 3396 | 411.2 | - | - | 0 | - |
| - | - | 783.6 | 413.2 | - | - | 0 | - |
| - | - | 1446 | 414.2 | - | - | 0 | - |
| - | - | 652 | 417.2 | - | - | 0 | - |
| - | - | 1471 | 417.8 | - | - | 0 | - |
| - | - | 696.9 | 418.2 | - | - | 0 | - |
| - | - | 972.6 | 418.3 | - | - | 0 | - |
| - | - | 1131 | 420.2 | - | - | 0 | - |
| - | - | 1538 | 425.3 | - | - | 0 | - |
| 2 | y | 1794 | 425.7 | 0.001041 | 2.446 | +2 | 9 |
| 2 | y | 914.6 | 426.2 | 0.005585 | 13.1 | +2 | 9 |
| 9 | b | 3724 | 431.8 | 0.0004879 | 1.13 | +2 | 9 |
| - | - | 672.3 | 432.1 | - | - | 0 | - |
| - | - | 1905 | 432.3 | - | - | 0 | - |
| 2 | y | 7131 | 434.7 | 0.001039 | 2.389 | +2 | 9 |
| - | - | 3744 | 435.2 | - | - | 0 | - |
| - | - | 1058 | 435.8 | - | - | 0 | - |
| - | - | 1612 | 437.3 | - | - | 0 | - |
| - | - | 1710 | 437.3 | - | - | 0 | - |
| - | - | 762.2 | 439.2 | - | - | 0 | - |
| - | - | 2203 | 439.7 | - | - | 0 | - |
| - | - | 1104 | 440.2 | - | - | 0 | - |
| - | - | 1424 | 442.3 | - | - | 0 | - |
| - | - | 597.9 | 443.3 | - | - | 0 | - |
| - | - | 2354 | 447.2 | - | - | 0 | - |
| - | - | 920.4 | 450.3 | - | - | 0 | - |
| - | - | 1865 | 451.3 | - | - | 0 | - |
| - | - | 3351 | 454.3 | - | - | 0 | - |
| - | - | 630.3 | 456.2 | - | - | 0 | - |
| - | - | 640.6 | 456.2 | - | - | 0 | - |
| - | - | 1176 | 456.3 | - | - | 0 | - |
| - | - | 1.231E+04 | 464.3 | - | - | 0 | - |
| - | - | 1771 | 465.2 | - | - | 0 | - |
| - | - | 7998 | 465.3 | - | - | 0 | - |
| - | - | 951.1 | 466.2 | - | - | 0 | - |
| - | - | 1587 | 466.3 | - | - | 0 | - |
| - | - | 1546 | 466.8 | - | - | 0 | - |
| - | - | 696.6 | 467.2 | - | - | 0 | - |
| - | - | 3073 | 467.3 | - | - | 0 | - |
| - | - | 629.2 | 468.3 | - | - | 0 | - |
| 6 | y | 5548 | 473.3 | 0.001247 | 2.635 | +1 | 5 |
| - | - | 933.3 | 474.3 | - | - | 0 | - |
| 0 | Precursor | 4675 | 475.3 | 0.0007701 | 1.62 | +2 | -1 |
| - | - | 1993 | 475.8 | - | - | 0 | - |
| - | - | 770 | 476.3 | - | - | 0 | - |
| - | - | 2267 | 478.3 | - | - | 0 | - |
| - | - | 6142 | 479.3 | - | - | 0 | - |
| - | - | 1965 | 480.3 | - | - | 0 | - |
| - | - | 1169 | 481.3 | - | - | 0 | - |
| - | - | 2.499E+04 | 482.3 | - | - | 0 | - |
| - | - | 729.5 | 482.6 | - | - | 0 | - |
| - | - | 2140 | 482.9 | - | - | 0 | - |
| - | - | 867.4 | 483.2 | - | - | 0 | - |
| - | - | 7955 | 483.3 | - | - | 0 | - |
| - | - | 1215 | 484.2 | - | - | 0 | - |
| - | - | 1153 | 484.3 | - | - | 0 | - |
| 0 | Precursor | 1.079E+04 | 484.3 | 0.0007062 | 1.458 | +2 | -1 |
| - | - | 4756 | 484.8 | - | - | 0 | - |
| - | - | 851.5 | 484.9 | - | - | 0 | - |
| - | - | 2071 | 485.3 | - | - | 0 | - |
| - | - | 702 | 490.3 | - | - | 0 | - |
| 5 | b | 1132 | 495.3 | 0.000806 | 1.627 | +1 | 5 |
| - | - | 2236 | 496.3 | - | - | 0 | - |
| - | - | 4328 | 508.3 | - | - | 0 | - |
| - | - | 931 | 509.3 | - | - | 0 | - |
| - | - | 1940 | 518.3 | - | - | 0 | - |
| - | - | 1214 | 524.3 | - | - | 0 | - |
| - | - | 7332 | 525.3 | - | - | 0 | - |
| - | - | 1847 | 526.3 | - | - | 0 | - |
| - | - | 602.5 | 526.6 | - | - | 0 | - |
| - | - | 730.7 | 527.3 | - | - | 0 | - |
| - | - | 2119 | 535.3 | - | - | 0 | - |
| - | - | 9081 | 536.3 | - | - | 0 | - |
| - | - | 2446 | 537.3 | - | - | 0 | - |
| 5 | y | 6520 | 552.3 | 0.001101 | 1.994 | +1 | 6 |
| - | - | 8.087E+04 | 553.3 | - | - | 0 | - |
| - | - | 2.408E+04 | 554.3 | - | - | 0 | - |
| - | - | 3733 | 555.3 | - | - | 0 | - |
| - | - | 785.4 | 555.4 | - | - | 0 | - |
| - | - | 775.9 | 563.4 | - | - | 0 | - |
| 5 | y | 1.652E+05 | 570.3 | 0.0008516 | 1.493 | +1 | 6 |
| - | - | 5.036E+04 | 571.3 | - | - | 0 | - |
| - | - | 565.4 | 572.1 | - | - | 0 | - |
| - | - | 8942 | 572.3 | - | - | 0 | - |
| - | - | 815.3 | 573.3 | - | - | 0 | - |
| - | - | 2754 | 577.3 | - | - | 0 | - |
| - | - | 1604 | 577.8 | - | - | 0 | - |
| - | - | 1070 | 579.3 | - | - | 0 | - |
| - | - | 3417 | 580.4 | - | - | 0 | - |
| - | - | 969.8 | 581.6 | - | - | 0 | - |
| - | - | 937.8 | 585.3 | - | - | 0 | - |
| - | - | 2044 | 586.8 | - | - | 0 | - |
| - | - | 1234 | 588.8 | - | - | 0 | - |
| - | - | 5643 | 595.4 | - | - | 0 | - |
| - | - | 1234 | 596.4 | - | - | 0 | - |
| - | - | 827.4 | 603.3 | - | - | 0 | - |
| - | - | 840.4 | 621.4 | - | - | 0 | - |
| - | - | 1228 | 622.4 | - | - | 0 | - |
| 6 | b | 1259 | 623.4 | 0.003652 | 5.858 | +1 | 6 |
| - | - | 3702 | 628.3 | - | - | 0 | - |
| - | - | 2198 | 638.4 | - | - | 0 | - |
| - | - | 757.6 | 639.4 | - | - | 0 | - |
| - | - | 2125 | 649.4 | - | - | 0 | - |
| - | - | 5796 | 650.4 | - | - | 0 | - |
| - | - | 1567 | 651.4 | - | - | 0 | - |
| - | - | 1328 | 654.4 | - | - | 0 | - |
| - | - | 831.4 | 657.3 | - | - | 0 | - |
| - | - | 768.2 | 659.8 | - | - | 0 | - |
| - | - | 1.858E+04 | 666.4 | - | - | 0 | - |
| - | - | 6559 | 667.4 | - | - | 0 | - |
| - | - | 1163 | 668.4 | - | - | 0 | - |
| - | - | 2878 | 676.4 | - | - | 0 | - |
| - | - | 1327 | 677.4 | - | - | 0 | - |
| 4 | y | 3778 | 680.4 | 7.999E-05 | 0.1176 | +1 | 7 |
| 4 | y | 5150 | 681.4 | 0.004163 | 6.109 | +1 | 7 |
| - | - | 2239 | 682.4 | - | - | 0 | - |
| 7 | b | 9625 | 694.4 | 0.0009009 | 1.297 | +1 | 7 |
| - | - | 2998 | 695.4 | - | - | 0 | - |
| 4 | y | 5055 | 698.4 | 0.0004406 | 0.6309 | +1 | 7 |
| - | - | 1965 | 699.4 | - | - | 0 | - |
| - | - | 852.4 | 717.4 | - | - | 0 | - |
| - | - | 838.7 | 720.4 | - | - | 0 | - |
| - | - | 1212 | 720.4 | - | - | 0 | - |
| - | - | 785.7 | 721.4 | - | - | 0 | - |
| - | - | 722.5 | 722.4 | - | - | 0 | - |
| - | - | 1220 | 730.4 | - | - | 0 | - |
| 3 | y | 4372 | 737.4 | 0.0003266 | 0.4429 | +1 | 8 |
| - | - | 4144 | 737.5 | - | - | 0 | - |
| 3 | y | 5833 | 738.4 | 0.004427 | 5.996 | +1 | 8 |
| - | - | 1857 | 738.5 | - | - | 0 | - |
| - | - | 1850 | 739.4 | - | - | 0 | - |
| - | - | 1462 | 747.5 | - | - | 0 | - |
| 8 | b | 2651 | 748.4 | 0.002116 | 2.827 | +1 | 8 |
| - | - | 2140 | 749.4 | - | - | 0 | - |
| - | - | 856.8 | 750.4 | - | - | 0 | - |
| 3 | y | 1.413E+05 | 755.4 | 0.0006444 | 0.853 | +1 | 8 |
| - | - | 5.966E+04 | 756.4 | - | - | 0 | - |
| - | - | 1305 | 756.5 | - | - | 0 | - |
| - | - | 1.346E+04 | 757.4 | - | - | 0 | - |
| - | - | 1057 | 758.4 | - | - | 0 | - |
| - | - | 2208 | 763.4 | - | - | 0 | - |
| 8 | b | 7.395E+04 | 765.5 | 0.0005302 | 0.6927 | +1 | 8 |
| - | - | 3.308E+04 | 766.5 | - | - | 0 | - |
| - | - | 7094 | 767.5 | - | - | 0 | - |
| - | - | 749.3 | 800.4 | - | - | 0 | - |
| 2 | y | 964.1 | 851.5 | 0.003188 | 3.744 | +1 | 9 |
| 2 | y | 1.76E+04 | 868.5 | 0.0003816 | 0.4394 | +1 | 9 |
| - | - | 8695 | 869.5 | - | - | 0 | - |
| - | - | 3155 | 870.5 | - | - | 0 | - |
| - | - | 692.2 | 878.5 | - | - | 0 | - |
| - | - | 640.2 | 1269 | - | - | 0 | - |
| - | - | 815.4 | 1789 | - | - | 0 | - |
| - | - | 607.7 | 2616 | - | - | 0 | - |
| - | - | 763 | 3007 | - | - | 0 | - |

m/z Charge Intensity FragmentType MassShift Position
120.06590270996094 0 1492.7266
120.08112335205078 0 1746.2828
124.03980255126953 0 593.08594
124.07693481445312 0 363.17435
127.08708190917969 0 692.42957
129.0662384033203 0 4561.781
129.1026611328125 0 53886.77
130.0504608154297 0 641.466
130.08657836914062 0 1112.7197
130.10604858398438 0 3281.7007
132.10226440429688 0 773.9271
133.71316528320312 0 409.07822
134.02743530273438 0 1060.6729
136.0760955810547 0 23079.79
137.07937622070312 0 1764.1248
137.10787963867188 0 548.45374
138.02244567871094 0 644.10846
138.09210205078125 0 420.13522
139.0870819091797 0 1765.0889
141.0662384033203 0 3450.5872
141.10267639160156 0 4087.6606
142.12330627441406 0 718.36346
143.08213806152344 0 1249.3018
143.11837768554688 0 2901.4695 d 1
147.04444885253906 0 403.4117
147.11297607421875 0 1356.0491
148.90223693847656 0 486.09933
148.95404052734375 0 635.61115
152.0345458984375 0 3217.296
153.0666046142578 0 660.315
153.1026611328125 0 1160.8896
155.1182861328125 0 5487.598
156.1217498779297 0 553.4763
157.09767150878906 0 1873.918
158.09280395507812 0 1637.0862
159.09214782714844 0 822.48016
159.1132049560547 0 6073.5835
165.10284423828125 0 1469.7617
166.0617218017578 0 565.12915
167.08230590820312 0 612.48334
167.11810302734375 0 771.0819
168.03993225097656 0 451.0307
169.05294799804688 0 709.2246
169.06117248535156 0 13692.924
169.0975341796875 0 3826.1252
169.13381958007812 0 781.1725
170.0646514892578 0 568.53516
170.10162353515625 0 558.7418
171.07687377929688 0 626.9232
171.11322021484375 0 8586.112
172.1167449951172 0 861.841
173.0925750732422 0 2439.0642 y 6
173.12889099121094 0 4938.473
173.4514923095703 0 1493.7258
174.1313934326172 0 594.3212
175.2571258544922 0 545.1041
178.97166442871094 0 488.74005
180.11343383789062 0 1807.3693
181.09767150878906 0 4108.9985
181.1341094970703 0 2517.3582
182.1292266845703 0 12657.513
183.1134796142578 0 2757.1602
183.13253784179688 0 1305.4844
183.1495819091797 0 531.4333
185.07162475585938 0 1386.3762
185.09251403808594 0 5110.0054 y Water loss 8
185.1653594970703 0 250005.97 a 1
186.08778381347656 0 30860.588
186.16868591308594 0 26123.646
187.0717010498047 0 1492.0137
187.0910186767578 0 1744.8379
187.10806274414062 0 1855.9686
187.14454650878906 0 16388.348
187.1719512939453 0 986.9248
188.13980102539062 0 6314.135
188.148193359375 0 1237.0803
191.0821990966797 0 834.4575
193.09732055664062 0 1059.4514
195.11268615722656 0 1003.81354
197.12867736816406 0 1680.4878
197.16587829589844 0 734.0658
199.10816955566406 0 1173.6598
200.13980102539062 0 3482.8005
201.1238250732422 0 2246.1775
203.1031036376953 0 147245.25 y 8
204.1064453125 0 13215.1455
204.13499450683594 0 674.968
205.1080322265625 0 678.76556
207.0880126953125 0 642.7002
207.09869384765625 0 859.6974
208.1084442138672 0 3375.0156
208.14462280273438 0 1160.5541
209.09246826171875 0 3497.854
209.12905883789062 0 1031.6619
210.0959930419922 0 581.71155
210.12408447265625 0 13997.918
211.12750244140625 0 1761.7236
211.18057250976562 0 621.6499
212.10289001464844 0 1075.4874
212.13970947265625 0 575.98334
213.16026306152344 0 132817.28 b 1
214.16355895996094 0 15234.471
214.39450073242188 0 548.4445
215.139404296875 0 9788.304
215.166015625 0 874.1662
216.1437530517578 0 1187.0166
219.13442993164062 0 773.5212
220.10841369628906 0 628.5085
221.1040496826172 0 752.93243
223.1085205078125 0 798.911
224.1143798828125 0 509.73346
225.09967041015625 0 556.5107
225.12344360351562 0 642.16864
226.1190948486328 0 2039.9839
226.15548706054688 0 99116.3
227.1030731201172 0 570.0365
227.15887451171875 0 11997.194
228.1151123046875 0 774.245
228.1347198486328 0 13239.178 y Water loss 5
229.11924743652344 0 761.4371
229.13796997070312 0 1199.8564
230.11410522460938 0 796.89777
230.15460205078125 0 607.68756
231.0441131591797 0 676.76685
233.0929412841797 0 910.03394
235.0833740234375 0 995.103
236.1388702392578 0 1188.0858
238.1190948486328 0 1395.6572
239.15040588378906 0 685.9328
239.17633056640625 0 670.3891
240.13479614257812 0 3286.9668
242.1499786376953 0 861.8138
243.18212890625 0 3100.5488
244.12879943847656 0 646.21747
244.16720581054688 0 624.2586
246.12379455566406 0 685.411
247.10964965820312 0 746.2441
248.1044464111328 0 510.41235
249.16036987304688 0 18261.008
250.16358947753906 0 2765.8843
251.1060333251953 0 648.24225
251.1508331298828 0 717.57477
252.1095733642578 0 1544.6444
252.134765625 0 1737.6368
252.17105102539062 0 3271.4263
253.1669158935547 0 1657.2917
254.15029907226562 0 3951.1726
256.1297912597656 0 1924.7057 y Water loss 7
257.12481689453125 0 735.2936
258.14532470703125 0 637.87494
261.1739807128906 0 577.80255
265.1297607421875 0 880.04803
266.1136474609375 0 1660.0278
267.1087951660156 0 984.8884
268.1290588378906 0 580.72565
269.06378173828125 0 870.2167
269.08056640625 0 6245.34
269.19769287109375 0 3411.7954
270.1816711425781 0 14012.619 b 2
270.1980285644531 0 683.1689
271.1769104003906 0 11273.051
272.1802673339844 0 1128.9991
273.13555908203125 0 1234.8984
274.1402282714844 0 9854.95 y 7
275.1437072753906 0 1235.7645
276.0663146972656 0 847.76605
277.15521240234375 0 8609.96 y Ammonia loss 4
278.158447265625 0 1409.9307
279.18206787109375 0 19823.957
280.1854553222656 0 2356.214
281.16168212890625 0 3367.4565
283.1412353515625 0 3256.0151
285.15625 0 4132.959
285.6664733886719 0 2152.498 y 4
286.1690368652344 0 721.00354
287.2083435058594 0 27070.229
288.21136474609375 0 3502.0408
297.1927490234375 0 39697.953
298.1959533691406 0 5650.9795
299.1720275878906 0 77351.59
300.17486572265625 0 9892.347
301.130126953125 0 1885.8483
301.17413330078125 0 601.03754
301.1890869140625 0 1001.79114
303.0757751464844 0 766.9217
309.1563415527344 0 2183.6086
309.2042541503906 0 1897.6804
314.2055969238281 0 786.5903
314.2215576171875 0 938.6031
323.2080383300781 0 2969.9023
323.22723388671875 0 649.82855
325.18768310546875 0 2385.4224
325.6830749511719 0 970.35785
327.1673278808594 0 4443.961 y Water loss 6
328.1637268066406 0 1646.0131
331.6869201660156 0 614.3884
332.17767333984375 0 1176.8097
336.2035217285156 0 1720.0574
337.1876525878906 0 2672.5247
338.18927001953125 0 600.5532
340.2350769042969 0 17660.574
340.6905517578125 0 5768.379 y Water loss 3
341.1894226074219 0 2028.1012
341.2380065917969 0 2500.8694
345.17718505859375 0 2117.6755 y 6
346.17999267578125 0 589.01587
347.229736328125 0 635.9243
349.1885070800781 0 653.0784
349.69580078125 0 1351.9468 y 3
350.1973876953125 0 1049.4778
350.2189636230469 0 1741.8011
353.2181396484375 0 1225.5035
354.1778564453125 0 663.3344
354.2145690917969 0 1437.326
359.1925964355469 0 719.1236
360.7241516113281 0 655.41895
363.2017822265625 0 1172.2377
368.229736328125 0 95757.46
369.2021484375 0 1047.7782 y Water loss 2
369.2327575683594 0 19931.805
370.24420166015625 0 7892.6777
371.2474365234375 0 1122.3693
376.1982421875 0 1856.5728
376.24322509765625 0 787.891
378.2099304199219 0 1098.3302 y 2
380.2019348144531 0 729.253
380.22943115234375 0 2123.7363
381.2130432128906 0 601.16046 b Ammonia loss 3
382.16473388671875 0 968.0356
382.2272644042969 0 2133.2358
382.7279968261719 0 1274.3921
383.1988830566406 0 552.3935
383.2344665527344 0 1733.8527 b 7
384.2359924316406 0 630.03625
390.21405029296875 0 2400.1626
393.2251892089844 0 913.3856
394.2089538574219 0 3622.6829
394.2447509765625 0 1670.4607
395.21392822265625 0 694.8212
395.24737548828125 0 587.24146
396.158203125 0 807.1545
396.2246398925781 0 2308.924
398.24053955078125 0 23176.166 b 3
399.2437438964844 0 5203.761
407.24102783203125 0 1322.7834
408.2250671386719 0 3448.975
409.2283630371094 0 612.146
411.2359619140625 0 3396.2925
413.2087097167969 0 783.648
414.1697082519531 0 1446.1584
417.23052978515625 0 651.9688
417.7650146484375 0 1471.4338
418.1912536621094 0 696.9369
418.2649230957031 0 972.5959
420.2252197265625 0 1131.1475
425.25274658203125 0 1537.7764
425.7437438964844 0 1793.5763 y Water loss 1
426.24029541015625 0 914.641 y Ammonia loss 1
431.7613830566406 0 3723.6304 b 8
432.11834716796875 0 672.2619
432.2637939453125 0 1904.7064
434.7490234375 0 7131.1 y 1
435.2499084472656 0 3743.7175
435.7506408691406 0 1057.7876
437.2523498535156 0 1612.1794
437.2882080078125 0 1710.1404
439.2291564941406 0 762.2411
439.7401428222656 0 2203.2815
440.24200439453125 0 1103.9634
442.279052734375 0 1424.2041
443.2814025878906 0 597.8892
447.2355651855469 0 2353.8943
450.2843322753906 0 920.36444
451.268798828125 0 1864.7566
454.27783203125 0 3350.6218
456.166015625 0 630.3435
456.2217712402344 0 640.6083
456.2557373046875 0 1176.3982
464.2625427246094 0 12314.639
465.24774169921875 0 1770.7051
465.28216552734375 0 7997.9053
466.2410583496094 0 951.141
466.2837219238281 0 1587.2068
466.7635498046875 0 1545.6914
467.2278747558594 0 696.58936
467.29815673828125 0 3073.3818
468.29705810546875 0 629.2246
473.2730712890625 0 5548.199 y 5
474.2764892578125 0 933.3297
475.2776794433594 0 4675.0073 Precursor Water loss
475.7796630859375 0 1992.9707
476.2774963378906 0 769.9955
478.2779541015625 0 2266.6504
479.26263427734375 0 6142.4746
480.26458740234375 0 1965.0247
481.3132629394531 0 1168.9052
482.2730712890625 0 24985.145
482.5970458984375 0 729.5417
482.9324035644531 0 2139.5527
483.2132873535156 0 867.35144
483.2743225097656 0 7954.919
484.217041015625 0 1214.9447
484.252197265625 0 1153.1426
484.28289794921875 0 10794.062 Precursor
484.7855529785156 0 4756.258
484.91815185546875 0 851.5015
485.28656005859375 0 2071.3672
490.27764892578125 0 702.0438
495.2933654785156 0 1131.7152 b 4
496.28912353515625 0 2235.9346
508.288818359375 0 4328.0547
509.2924499511719 0 931.00024
518.2745971679688 0 1940.4401
524.3187255859375 0 1213.5957
525.3154296875 0 7331.7373
526.3171997070312 0 1847.4414
526.5963134765625 0 602.534
527.270263671875 0 730.7366
535.297607421875 0 2119.4355
536.28369140625 0 9080.577
537.2845458984375 0 2446.2168
552.3151245117188 0 6520.293 y Water loss 4
553.31005859375 0 80871.53
554.312744140625 0 24081.15
555.3151245117188 0 3733.1235
555.3595581054688 0 785.4243
563.39208984375 0 775.92883
570.325439453125 0 165217.81 y 4
571.3282470703125 0 50359.035
572.1226196289062 0 565.4146
572.3302612304688 0 8941.558
573.3331909179688 0 815.3325
577.3452758789062 0 2753.765
577.7897338867188 0 1604.421
579.3281860351562 0 1069.5481
580.3818359375 0 3417.2334
581.599609375 0 969.8444
585.273193359375 0 937.83624
586.7593383789062 0 2043.6101
588.7991333007812 0 1234.0413
595.3571166992188 0 5642.6494
596.3618774414062 0 1234.4811
603.3047485351562 0 827.44025
621.36962890625 0 840.4013
622.3695678710938 0 1228.1039
623.3911743164062 0 1259.1613 b 5
628.313720703125 0 3701.553
638.398681640625 0 2198.225
639.4050903320312 0 757.5661
649.3677978515625 0 2124.8428
650.3634033203125 0 5795.5312
651.3662719726562 0 1567.3553
654.4290771484375 0 1328.1246
657.2516479492188 0 831.4279
659.8489379882812 0 768.2117
666.3944091796875 0 18579.314
667.3970947265625 0 6558.957
668.3968505859375 0 1162.7362
676.4140014648438 0 2877.5159
677.4219970703125 0 1327.475
680.3726806640625 0 3778.2878 y Water loss 3
681.3607788085938 0 5149.73 y Ammonia loss 3
682.3629760742188 0 2238.9673
694.425537109375 0 9624.974 b 6
695.4275512695312 0 2998.0386
698.3836059570312 0 5054.618 y 3
699.3907470703125 0 1965.3412
717.3541870117188 0 852.44653
720.3716430664062 0 838.68964
720.440673828125 0 1212.2135
721.3751831054688 0 785.7018
722.3719482421875 0 722.47406
730.4276123046875 0 1219.9021
737.3937377929688 0 4371.574 y Water loss 2
737.468017578125 0 4144.486
738.3825073242188 0 5833.4536 y Ammonia loss 2
738.4710083007812 0 1856.7449
739.3832397460938 0 1849.6755
747.4556274414062 0 1462.4122
748.4373168945312 0 2650.8694 b Ammonia loss 7
749.4404907226562 0 2140.1504
750.4444580078125 0 856.7643
755.4052734375 0 141304.06 y 2
756.4076538085938 0 59661.54
756.489501953125 0 1304.797
757.4097290039062 0 13459.368
758.4090576171875 0 1057.0331
763.44677734375 0 2208.4617
765.4622802734375 0 73948.445 b 7
766.4653930664062 0 33078.547
767.4672241210938 0 7093.6445
800.4194946289062 0 749.2621
851.46533203125 0 964.14496 y Ammonia loss 1
868.4890747070312 0 17603.81 y 1
869.4918212890625 0 8694.6875
870.4966430664062 0 3154.6082
878.4678955078125 0 692.24567
1268.8427734375 0 640.2425
1788.76318359375 0 815.3587
2616.249267578125 0 607.7167
3006.836669921875 0 763.01636

Spectrum Details

|  |  |
| --- | --- |
| Matched peaks? Matched peaks The total absolute number of peaks matched. Additionally in brackets the total fraction of peaks matched and the total number of peaks is shown. | 43 (10.72% of 401) |
| FDR? FDR The false discovery rate estimated for this peptide. It is calculated by matching all theoretical fragments with a non-integer shift with the raw peaks for this spectrum. This is done with 40 different shifts. The resulting percentage is the average number of annotated peaks over the number of annotated peaks with the correct spectrum. | 0.61% |
| Satellite FDR? Satellite FDR See the FDR for details on its calculation. This satellite ion specific FDR only contains the satellite ions (d/w) for I/L/J positions. | 0.00% |
| PSM Score? PSM Score The PSM Score as given by Hecklib to this annotated spectrum. It is shown with three significant figures. | 431 |

#### Spectrum 3554? Spectrum 3554 The raw spectrum of this peptide as annotated by Hecklib. The fragments are coloured according to ion type (see legend). Any peaks with a star '\*' as text can be hovered over to see the full details, first the ion type second the mass shift type. By hovering over the amino acids in the peptide or ions in the legend the corresponding peaks are highlighted. By toggling the 'Unassigned' label you can turn the background (unassigned) peaks on or off in the plot. By updating the slider in the Ion legend you can update the spectrum to only show the top X% of the peaks with labels. The top X% means any peak that is within X% of the highest intensity. By dragging in the spectrum you can zoom in to a specific part of the spectrum and use 'Zoom Out' to get back to the original zoom level. The annotation of the spectrum is based on the given sequence in the peptides file and is done with different software so inconsistencies are likely. The peaks are annotated based on the given sequence, with 20 ppm tolerance.

Copy Data

##### Spectrum 3554 (TSV)

###### Preview

```
Loading example...
```

*Click on the button to copy the data to your clipboard.*

Mz MinMz MaxIntensity Max

WidthHeightPeptide font sizePeptide stroke widthSpectrum font sizeSpectrum stroke widthCompact peptide

Ion legend

wxyz

abcd

OtherUnassignedIon^Charge^_Position_Show for top:%

VLGQPKAAPS

04.48e+48.97e+41.34e+51.79e+5

Zoom Out

d^+1^_2_y^+2^_4_y^+1^_2_a^+1^_2_y^+1^_2_b^+1^_2_y^+2^_5_y^+1^_3_b^+1^_3_y^+1^_3_y^+2^_6_y^+2^_6_y^+1^_4_y^+2^_7_y^+2^_7_y^+1^_4_y^+2^_8_b^+1^_4_b^+2^_8_b^+1^_4_y^+2^_9_b^+2^_9_y^+2^_9_y^+1^_5_\*\*b^+1^_5_y^+1^_6_y^+1^_6_y^+1^_7_y^+1^_7_b^+1^_7_y^+1^_7_y^+1^_8_y^+1^_8_b^+1^_8_y^+1^_8_b^+1^_8_y^+1^_9_y^+1^_9_

0878175526333511

Fragment Matches Table

Show background peaks

| Position | Ion type | Intensity | mz Theoretical | mz Error (Th) | mz Error (ppm) | Charge | Series Number |
| --- | --- | --- | --- | --- | --- | --- | --- |
| - | - | 884.7 | 120 | - | - | 0 | - |
| - | - | 855.6 | 120.1 | - | - | 0 | - |
| - | - | 2113 | 120.1 | - | - | 0 | - |
| - | - | 319.1 | 120.8 | - | - | 0 | - |
| - | - | 366.9 | 121 | - | - | 0 | - |
| - | - | 337.2 | 125.9 | - | - | 0 | - |
| - | - | 459.1 | 127.1 | - | - | 0 | - |
| - | - | 3622 | 129.1 | - | - | 0 | - |
| - | - | 4.704E+04 | 129.1 | - | - | 0 | - |
| - | - | 688.6 | 130 | - | - | 0 | - |
| - | - | 557.5 | 130.1 | - | - | 0 | - |
| - | - | 2821 | 130.1 | - | - | 0 | - |
| - | - | 903.6 | 132.1 | - | - | 0 | - |
| - | - | 1204 | 134 | - | - | 0 | - |
| - | - | 1.753E+04 | 136.1 | - | - | 0 | - |
| - | - | 1734 | 137.1 | - | - | 0 | - |
| - | - | 2164 | 139.1 | - | - | 0 | - |
| - | - | 2482 | 141.1 | - | - | 0 | - |
| - | - | 3002 | 141.1 | - | - | 0 | - |
| - | - | 1167 | 143.1 | - | - | 0 | - |
| 2 | d | 1474 | 143.1 | 0.0003356 | 2.345 | +1 | 2 |
| - | - | 613.6 | 145.1 | - | - | 0 | - |
| - | - | 4627 | 146.1 | - | - | 0 | - |
| - | - | 453.7 | 147.1 | - | - | 0 | - |
| - | - | 466.1 | 148.8 | - | - | 0 | - |
| - | - | 516.8 | 148.8 | - | - | 0 | - |
| - | - | 433.7 | 148.9 | - | - | 0 | - |
| - | - | 554 | 148.9 | - | - | 0 | - |
| - | - | 534.3 | 148.9 | - | - | 0 | - |
| - | - | 580 | 148.9 | - | - | 0 | - |
| - | - | 704.7 | 148.9 | - | - | 0 | - |
| - | - | 973.6 | 148.9 | - | - | 0 | - |
| - | - | 1147 | 148.9 | - | - | 0 | - |
| - | - | 1637 | 148.9 | - | - | 0 | - |
| - | - | 3362 | 148.9 | - | - | 0 | - |
| - | - | 5269 | 149 | - | - | 0 | - |
| - | - | 3212 | 149 | - | - | 0 | - |
| - | - | 1261 | 149 | - | - | 0 | - |
| - | - | 1217 | 149 | - | - | 0 | - |
| - | - | 759.7 | 149 | - | - | 0 | - |
| - | - | 705.6 | 149 | - | - | 0 | - |
| - | - | 673.1 | 149 | - | - | 0 | - |
| - | - | 571.1 | 151.1 | - | - | 0 | - |
| - | - | 1896 | 152 | - | - | 0 | - |
| - | - | 638.4 | 153.1 | - | - | 0 | - |
| - | - | 803.1 | 153.1 | - | - | 0 | - |
| - | - | 4191 | 155.1 | - | - | 0 | - |
| - | - | 1558 | 157.1 | - | - | 0 | - |
| - | - | 507 | 158.1 | - | - | 0 | - |
| - | - | 1581 | 158.1 | - | - | 0 | - |
| - | - | 494.5 | 158.1 | - | - | 0 | - |
| - | - | 422.5 | 158.6 | - | - | 0 | - |
| - | - | 524.4 | 159.1 | - | - | 0 | - |
| - | - | 505.8 | 159.1 | - | - | 0 | - |
| - | - | 2571 | 159.1 | - | - | 0 | - |
| - | - | 881.8 | 165.1 | - | - | 0 | - |
| - | - | 979.1 | 167.1 | - | - | 0 | - |
| - | - | 9454 | 169.1 | - | - | 0 | - |
| - | - | 3223 | 169.1 | - | - | 0 | - |
| - | - | 891.1 | 169.1 | - | - | 0 | - |
| - | - | 702.2 | 171.1 | - | - | 0 | - |
| - | - | 6016 | 171.1 | - | - | 0 | - |
| - | - | 624.5 | 172.1 | - | - | 0 | - |
| - | - | 695 | 172.1 | - | - | 0 | - |
| 7 | y | 899 | 173.1 | 0.000369 | 2.132 | +2 | 4 |
| - | - | 2192 | 173.1 | - | - | 0 | - |
| - | - | 2643 | 174.1 | - | - | 0 | - |
| - | - | 916.5 | 175.1 | - | - | 0 | - |
| - | - | 540.3 | 176.1 | - | - | 0 | - |
| - | - | 1195 | 180.1 | - | - | 0 | - |
| - | - | 441.4 | 180.2 | - | - | 0 | - |
| - | - | 3005 | 181.1 | - | - | 0 | - |
| - | - | 1513 | 181.1 | - | - | 0 | - |
| - | - | 8696 | 182.1 | - | - | 0 | - |
| - | - | 1531 | 183.1 | - | - | 0 | - |
| - | - | 732.6 | 183.1 | - | - | 0 | - |
| - | - | 860.2 | 183.1 | - | - | 0 | - |
| - | - | 923.9 | 185.1 | - | - | 0 | - |
| 9 | y | 3485 | 185.1 | 0.0001707 | 0.9221 | +1 | 2 |
| - | - | 3299 | 185.1 | - | - | 0 | - |
| 2 | a | 1.775E+05 | 185.2 | 0.0003977 | 2.148 | +1 | 2 |
| - | - | 2.027E+04 | 186.1 | - | - | 0 | - |
| - | - | 803.2 | 186.1 | - | - | 0 | - |
| - | - | 966.7 | 186.1 | - | - | 0 | - |
| - | - | 2.041E+04 | 186.2 | - | - | 0 | - |
| - | - | 843.3 | 187.1 | - | - | 0 | - |
| - | - | 1467 | 187.1 | - | - | 0 | - |
| - | - | 4843 | 187.1 | - | - | 0 | - |
| - | - | 473.1 | 187.2 | - | - | 0 | - |
| - | - | 4228 | 188.1 | - | - | 0 | - |
| - | - | 637.2 | 190.1 | - | - | 0 | - |
| - | - | 977.6 | 191.1 | - | - | 0 | - |
| - | - | 626 | 191.1 | - | - | 0 | - |
| - | - | 1007 | 193.1 | - | - | 0 | - |
| - | - | 559.5 | 193.1 | - | - | 0 | - |
| - | - | 579.6 | 193.4 | - | - | 0 | - |
| - | - | 1034 | 195.1 | - | - | 0 | - |
| - | - | 489 | 197.1 | - | - | 0 | - |
| - | - | 1508 | 197.1 | - | - | 0 | - |
| - | - | 1125 | 198.1 | - | - | 0 | - |
| - | - | 2125 | 200.1 | - | - | 0 | - |
| - | - | 1193 | 201.1 | - | - | 0 | - |
| - | - | 528.2 | 202.1 | - | - | 0 | - |
| - | - | 664.1 | 202.1 | - | - | 0 | - |
| 9 | y | 1.118E+05 | 203.1 | 0.0003329 | 1.639 | +1 | 2 |
| - | - | 9459 | 204.1 | - | - | 0 | - |
| - | - | 696 | 205.1 | - | - | 0 | - |
| - | - | 2574 | 208.1 | - | - | 0 | - |
| - | - | 701.1 | 208.1 | - | - | 0 | - |
| - | - | 2075 | 209.1 | - | - | 0 | - |
| - | - | 1.019E+04 | 210.1 | - | - | 0 | - |
| - | - | 618.5 | 211.1 | - | - | 0 | - |
| - | - | 773.8 | 211.2 | - | - | 0 | - |
| 2 | b | 9.006E+04 | 213.2 | 0.0003714 | 1.742 | +1 | 2 |
| - | - | 9456 | 214.2 | - | - | 0 | - |
| - | - | 2941 | 215.1 | - | - | 0 | - |
| - | - | 843.6 | 215.2 | - | - | 0 | - |
| - | - | 662.8 | 219.1 | - | - | 0 | - |
| - | - | 531.9 | 221.1 | - | - | 0 | - |
| - | - | 635.5 | 221.1 | - | - | 0 | - |
| - | - | 575.2 | 223.1 | - | - | 0 | - |
| - | - | 7.351E+04 | 226.2 | - | - | 0 | - |
| - | - | 8015 | 227.2 | - | - | 0 | - |
| 6 | y | 9011 | 228.1 | 0.0003146 | 1.379 | +2 | 5 |
| - | - | 544.3 | 228.2 | - | - | 0 | - |
| - | - | 906.2 | 229.1 | - | - | 0 | - |
| - | - | 600.2 | 231 | - | - | 0 | - |
| - | - | 2136 | 233.1 | - | - | 0 | - |
| - | - | 814.1 | 235.1 | - | - | 0 | - |
| - | - | 881.6 | 236.1 | - | - | 0 | - |
| - | - | 1653 | 238.1 | - | - | 0 | - |
| - | - | 596.1 | 239.2 | - | - | 0 | - |
| - | - | 2441 | 240.1 | - | - | 0 | - |
| - | - | 881.6 | 241.1 | - | - | 0 | - |
| - | - | 763.4 | 242.1 | - | - | 0 | - |
| - | - | 754.5 | 242.2 | - | - | 0 | - |
| - | - | 2566 | 243.2 | - | - | 0 | - |
| - | - | 701.5 | 245.1 | - | - | 0 | - |
| - | - | 545.6 | 247.1 | - | - | 0 | - |
| - | - | 1.542E+04 | 249.2 | - | - | 0 | - |
| - | - | 2534 | 250.2 | - | - | 0 | - |
| - | - | 1046 | 251.2 | - | - | 0 | - |
| - | - | 2037 | 252.1 | - | - | 0 | - |
| - | - | 2315 | 252.2 | - | - | 0 | - |
| - | - | 1052 | 253.2 | - | - | 0 | - |
| - | - | 2694 | 254.2 | - | - | 0 | - |
| 8 | y | 1044 | 256.1 | 1.595E-06 | 0.006225 | +1 | 3 |
| - | - | 676.5 | 257.1 | - | - | 0 | - |
| - | - | 5246 | 261.1 | - | - | 0 | - |
| - | - | 612.2 | 261.2 | - | - | 0 | - |
| - | - | 866.7 | 262.1 | - | - | 0 | - |
| - | - | 1232 | 266.1 | - | - | 0 | - |
| - | - | 1678 | 269.1 | - | - | 0 | - |
| - | - | 3544 | 269.2 | - | - | 0 | - |
| - | - | 855.8 | 270.1 | - | - | 0 | - |
| 3 | b | 1.109E+04 | 270.2 | 0.0002089 | 0.7733 | +1 | 3 |
| - | - | 8630 | 271.2 | - | - | 0 | - |
| - | - | 1145 | 272.1 | - | - | 0 | - |
| - | - | 750.9 | 272.2 | - | - | 0 | - |
| 8 | y | 5915 | 274.1 | 0.0002369 | 0.8643 | +1 | 3 |
| - | - | 739.7 | 274.2 | - | - | 0 | - |
| - | - | 628.6 | 276.1 | - | - | 0 | - |
| 5 | y | 7964 | 277.2 | 0.00228 | 8.227 | +2 | 6 |
| - | - | 1091 | 278.2 | - | - | 0 | - |
| - | - | 1.392E+04 | 279.2 | - | - | 0 | - |
| - | - | 2359 | 280.2 | - | - | 0 | - |
| - | - | 2400 | 281.2 | - | - | 0 | - |
| - | - | 2169 | 283.1 | - | - | 0 | - |
| - | - | 2650 | 285.2 | - | - | 0 | - |
| 5 | y | 1621 | 285.7 | 0.0008159 | 2.856 | +2 | 6 |
| - | - | 1.772E+04 | 287.2 | - | - | 0 | - |
| - | - | 2119 | 288.2 | - | - | 0 | - |
| - | - | 1148 | 290.1 | - | - | 0 | - |
| - | - | 650.1 | 295.2 | - | - | 0 | - |
| - | - | 450.2 | 297.2 | - | - | 0 | - |
| - | - | 2.836E+04 | 297.2 | - | - | 0 | - |
| - | - | 4596 | 298.2 | - | - | 0 | - |
| - | - | 5.183E+04 | 299.2 | - | - | 0 | - |
| - | - | 544.8 | 299.4 | - | - | 0 | - |
| - | - | 6790 | 300.2 | - | - | 0 | - |
| - | - | 541.2 | 301.1 | - | - | 0 | - |
| - | - | 1764 | 309.2 | - | - | 0 | - |
| - | - | 891.3 | 314.2 | - | - | 0 | - |
| - | - | 2295 | 323.2 | - | - | 0 | - |
| - | - | 1554 | 325.2 | - | - | 0 | - |
| 7 | y | 2662 | 327.2 | 0.0004518 | 1.381 | +1 | 4 |
| - | - | 1602 | 328.2 | - | - | 0 | - |
| - | - | 1166 | 332.2 | - | - | 0 | - |
| - | - | 936.3 | 336.2 | - | - | 0 | - |
| - | - | 1397 | 337.2 | - | - | 0 | - |
| - | - | 638.8 | 339.2 | - | - | 0 | - |
| - | - | 1.378E+04 | 340.2 | - | - | 0 | - |
| 4 | y | 4841 | 340.7 | 0.000308 | 0.9041 | +2 | 7 |
| 4 | y | 1528 | 341.2 | 0.006805 | 19.94 | +2 | 7 |
| - | - | 2370 | 341.2 | - | - | 0 | - |
| 7 | y | 998.2 | 345.2 | 0.0003851 | 1.116 | +1 | 4 |
| - | - | 1367 | 347.7 | - | - | 0 | - |
| - | - | 878.2 | 349.7 | - | - | 0 | - |
| - | - | 613.9 | 350.2 | - | - | 0 | - |
| - | - | 885.1 | 354.2 | - | - | 0 | - |
| - | - | 702 | 360.7 | - | - | 0 | - |
| - | - | 6.919E+04 | 368.2 | - | - | 0 | - |
| - | - | 1.297E+04 | 369.2 | - | - | 0 | - |
| - | - | 4299 | 370.2 | - | - | 0 | - |
| - | - | 1018 | 371.2 | - | - | 0 | - |
| - | - | 1542 | 376.2 | - | - | 0 | - |
| - | - | 669.1 | 376.2 | - | - | 0 | - |
| 3 | y | 956.4 | 378.2 | 0.002635 | 6.967 | +2 | 8 |
| - | - | 1316 | 380.2 | - | - | 0 | - |
| - | - | 1128 | 381.2 | - | - | 0 | - |
| 4 | b | 689.2 | 381.2 | 0.002452 | 6.431 | +1 | 4 |
| - | - | 1146 | 382.2 | - | - | 0 | - |
| 8 | b | 951.7 | 383.2 | 0.0006876 | 1.794 | +2 | 8 |
| - | - | 622.1 | 383.7 | - | - | 0 | - |
| - | - | 693.4 | 389.2 | - | - | 0 | - |
| - | - | 753.4 | 389.2 | - | - | 0 | - |
| - | - | 1698 | 390.2 | - | - | 0 | - |
| - | - | 632.9 | 393.9 | - | - | 0 | - |
| - | - | 1962 | 394.2 | - | - | 0 | - |
| - | - | 643.7 | 394.2 | - | - | 0 | - |
| - | - | 853.1 | 395.2 | - | - | 0 | - |
| - | - | 661 | 396.2 | - | - | 0 | - |
| - | - | 2663 | 396.2 | - | - | 0 | - |
| - | - | 707.5 | 397.2 | - | - | 0 | - |
| 4 | b | 1.763E+04 | 398.2 | 0.0003472 | 0.8719 | +1 | 4 |
| - | - | 1008 | 399.2 | - | - | 0 | - |
| - | - | 4262 | 399.2 | - | - | 0 | - |
| - | - | 791.3 | 407.2 | - | - | 0 | - |
| - | - | 2496 | 408.2 | - | - | 0 | - |
| - | - | 1911 | 411.2 | - | - | 0 | - |
| - | - | 1360 | 414.2 | - | - | 0 | - |
| - | - | 1396 | 417.8 | - | - | 0 | - |
| - | - | 1192 | 425.3 | - | - | 0 | - |
| 2 | y | 2744 | 425.7 | 0.0005227 | 1.228 | +2 | 9 |
| - | - | 738.7 | 429.2 | - | - | 0 | - |
| 9 | b | 2734 | 431.8 | 0.0002751 | 0.637 | +2 | 9 |
| - | - | 875.4 | 432.3 | - | - | 0 | - |
| - | - | 592.9 | 434.2 | - | - | 0 | - |
| 2 | y | 5408 | 434.7 | 0.00113 | 2.6 | +2 | 9 |
| - | - | 1983 | 435.3 | - | - | 0 | - |
| - | - | 857.8 | 435.8 | - | - | 0 | - |
| - | - | 663.5 | 437.3 | - | - | 0 | - |
| - | - | 960.2 | 437.3 | - | - | 0 | - |
| - | - | 1563 | 439.7 | - | - | 0 | - |
| - | - | 748.8 | 442.3 | - | - | 0 | - |
| - | - | 1301 | 447.2 | - | - | 0 | - |
| - | - | 776.5 | 450.3 | - | - | 0 | - |
| - | - | 1025 | 451.3 | - | - | 0 | - |
| - | - | 3544 | 454.3 | - | - | 0 | - |
| - | - | 1293 | 456.3 | - | - | 0 | - |
| - | - | 7981 | 464.3 | - | - | 0 | - |
| - | - | 1277 | 465.2 | - | - | 0 | - |
| - | - | 4929 | 465.3 | - | - | 0 | - |
| - | - | 1148 | 466.2 | - | - | 0 | - |
| - | - | 2011 | 466.3 | - | - | 0 | - |
| - | - | 1076 | 466.8 | - | - | 0 | - |
| - | - | 2173 | 467.3 | - | - | 0 | - |
| 6 | y | 4151 | 473.3 | 5.711E-05 | 0.1207 | +1 | 5 |
| - | - | 775.9 | 474.3 | - | - | 0 | - |
| - | - | 700.2 | 475.2 | - | - | 0 | - |
| 0 | Precursor | 2865 | 475.3 | 0.0006175 | 1.299 | +2 | -1 |
| - | - | 980.6 | 475.8 | - | - | 0 | - |
| - | - | 1204 | 478.3 | - | - | 0 | - |
| - | - | 4294 | 479.3 | - | - | 0 | - |
| - | - | 1480 | 480.3 | - | - | 0 | - |
| - | - | 2.043E+04 | 482.3 | - | - | 0 | - |
| - | - | 4305 | 483.3 | - | - | 0 | - |
| - | - | 1006 | 484.2 | - | - | 0 | - |
| 0 | Precursor | 1.045E+04 | 484.3 | 2.619E-05 | 0.05408 | +2 | -1 |
| - | - | 4589 | 484.8 | - | - | 0 | - |
| - | - | 1205 | 485 | - | - | 0 | - |
| - | - | 1784 | 485.3 | - | - | 0 | - |
| 5 | b | 789.9 | 495.3 | 0.00188 | 3.795 | +1 | 5 |
| - | - | 1470 | 496.3 | - | - | 0 | - |
| - | - | 698.4 | 497.3 | - | - | 0 | - |
| - | - | 1055 | 504.2 | - | - | 0 | - |
| - | - | 700.4 | 506.8 | - | - | 0 | - |
| - | - | 753.6 | 507.3 | - | - | 0 | - |
| - | - | 2243 | 508.3 | - | - | 0 | - |
| - | - | 1052 | 518.3 | - | - | 0 | - |
| - | - | 3252 | 521.2 | - | - | 0 | - |
| - | - | 1185 | 522.3 | - | - | 0 | - |
| - | - | 1007 | 524.3 | - | - | 0 | - |
| - | - | 5007 | 525.3 | - | - | 0 | - |
| - | - | 1539 | 526.3 | - | - | 0 | - |
| - | - | 1559 | 535.3 | - | - | 0 | - |
| - | - | 6376 | 536.3 | - | - | 0 | - |
| - | - | 1910 | 537.3 | - | - | 0 | - |
| - | - | 801.1 | 547.3 | - | - | 0 | - |
| 5 | y | 4358 | 552.3 | 0.0003689 | 0.668 | +1 | 6 |
| - | - | 5.589E+04 | 553.3 | - | - | 0 | - |
| - | - | 1.702E+04 | 554.3 | - | - | 0 | - |
| - | - | 1179 | 555.3 | - | - | 0 | - |
| - | - | 2594 | 555.3 | - | - | 0 | - |
| - | - | 2197 | 563.8 | - | - | 0 | - |
| - | - | 1591 | 564.3 | - | - | 0 | - |
| - | - | 743.1 | 564.8 | - | - | 0 | - |
| - | - | 892 | 565.3 | - | - | 0 | - |
| 5 | y | 1.171E+05 | 570.3 | 0.0001802 | 0.316 | +1 | 6 |
| - | - | 3.699E+04 | 571.3 | - | - | 0 | - |
| - | - | 6867 | 572.3 | - | - | 0 | - |
| - | - | 999.3 | 573.3 | - | - | 0 | - |
| - | - | 2613 | 577.3 | - | - | 0 | - |
| - | - | 923.4 | 579.3 | - | - | 0 | - |
| - | - | 2226 | 580.4 | - | - | 0 | - |
| - | - | 1689 | 581.4 | - | - | 0 | - |
| - | - | 1385 | 586.8 | - | - | 0 | - |
| - | - | 709.8 | 587.3 | - | - | 0 | - |
| - | - | 3645 | 595.4 | - | - | 0 | - |
| - | - | 941.4 | 596.4 | - | - | 0 | - |
| - | - | 1228 | 601.3 | - | - | 0 | - |
| - | - | 1128 | 620.3 | - | - | 0 | - |
| - | - | 1060 | 622.4 | - | - | 0 | - |
| - | - | 1457 | 638.4 | - | - | 0 | - |
| - | - | 656.8 | 644.3 | - | - | 0 | - |
| - | - | 716.8 | 649.3 | - | - | 0 | - |
| - | - | 1498 | 649.4 | - | - | 0 | - |
| - | - | 4584 | 650.4 | - | - | 0 | - |
| - | - | 1547 | 651.4 | - | - | 0 | - |
| - | - | 1537 | 654.4 | - | - | 0 | - |
| - | - | 693 | 655.4 | - | - | 0 | - |
| - | - | 1.218E+04 | 666.4 | - | - | 0 | - |
| - | - | 4282 | 667.4 | - | - | 0 | - |
| - | - | 1490 | 668.4 | - | - | 0 | - |
| - | - | 2136 | 676.4 | - | - | 0 | - |
| - | - | 1172 | 677.4 | - | - | 0 | - |
| 4 | y | 2794 | 680.4 | 0.0007135 | 1.049 | +1 | 7 |
| 4 | y | 3715 | 681.4 | 0.003186 | 4.676 | +1 | 7 |
| - | - | 1188 | 682.4 | - | - | 0 | - |
| - | - | 691.8 | 694.3 | - | - | 0 | - |
| 7 | b | 4786 | 694.4 | 0.0001684 | 0.2426 | +1 | 7 |
| - | - | 2297 | 695.4 | - | - | 0 | - |
| 4 | y | 3598 | 698.4 | 0.001268 | 1.816 | +1 | 7 |
| - | - | 1067 | 699.4 | - | - | 0 | - |
| - | - | 1187 | 720.4 | - | - | 0 | - |
| - | - | 906 | 720.4 | - | - | 0 | - |
| 3 | y | 3192 | 737.4 | 0.0004487 | 0.6085 | +1 | 8 |
| - | - | 3842 | 737.5 | - | - | 0 | - |
| 3 | y | 4441 | 738.4 | 0.001925 | 2.607 | +1 | 8 |
| - | - | 844.7 | 738.5 | - | - | 0 | - |
| - | - | 1633 | 739.4 | - | - | 0 | - |
| - | - | 1359 | 747.5 | - | - | 0 | - |
| 8 | b | 1702 | 748.4 | 0.0008953 | 1.196 | +1 | 8 |
| - | - | 1296 | 749.4 | - | - | 0 | - |
| 3 | y | 1.01E+05 | 755.4 | 0.0007595 | 1.005 | +1 | 8 |
| - | - | 4.293E+04 | 756.4 | - | - | 0 | - |
| - | - | 9960 | 757.4 | - | - | 0 | - |
| - | - | 1210 | 763.4 | - | - | 0 | - |
| 8 | b | 5.232E+04 | 765.5 | 0.0005684 | 0.7425 | +1 | 8 |
| - | - | 2.438E+04 | 766.5 | - | - | 0 | - |
| - | - | 4992 | 767.5 | - | - | 0 | - |
| - | - | 624.7 | 798.4 | - | - | 0 | - |
| 2 | y | 747.6 | 851.5 | 0.002456 | 2.884 | +1 | 9 |
| 2 | y | 1.429E+04 | 868.5 | 0.0005949 | 0.685 | +1 | 9 |
| - | - | 6369 | 869.5 | - | - | 0 | - |
| - | - | 2364 | 870.5 | - | - | 0 | - |
| - | - | 720.4 | 2211 | - | - | 0 | - |
| - | - | 803.9 | 3039 | - | - | 0 | - |
| - | - | 726.2 | 3476 | - | - | 0 | - |

m/z Charge Intensity FragmentType MassShift Position
120.04475402832031 0 884.6894
120.06585693359375 0 855.6365
120.08110809326172 0 2112.8745
120.80268859863281 0 319.1084
121.03981018066406 0 366.91046
125.86412048339844 0 337.1606
127.0872573852539 0 459.14993
129.06614685058594 0 3621.7585
129.10260009765625 0 47038.613
130.0499267578125 0 688.6284
130.1002197265625 0 557.4827
130.1059112548828 0 2820.7488
132.10215759277344 0 903.58606
134.0275421142578 0 1204.1324
136.0760498046875 0 17531.607
137.07940673828125 0 1734.0431
139.0871124267578 0 2164.2727
141.0662078857422 0 2481.7913
141.10260009765625 0 3001.8987
143.08172607421875 0 1167.1936
143.11822509765625 0 1473.6691 d 1
145.0612335205078 0 613.6414
146.06036376953125 0 4626.6045
147.0769500732422 0 453.65387
148.83181762695312 0 466.05383
148.8387451171875 0 516.78424
148.87428283691406 0 433.69528
148.8822021484375 0 553.9674
148.89634704589844 0 534.3155
148.90394592285156 0 580.0272
148.9107666015625 0 704.6863
148.918212890625 0 973.5941
148.92588806152344 0 1147.2754
148.9325408935547 0 1637.0677
148.94024658203125 0 3361.703
148.9567413330078 0 5269.1333
148.9646759033203 0 3212.0146
148.97218322753906 0 1260.6787
148.979248046875 0 1217.0465
148.9863739013672 0 759.698
148.9939422607422 0 705.5954
149.00070190429688 0 673.0715
151.08641052246094 0 571.0627
152.0345001220703 0 1895.8978
153.0660858154297 0 638.406
153.10264587402344 0 803.0637
155.1182403564453 0 4191.394
157.09739685058594 0 1557.595
158.06007385253906 0 506.97916
158.09259033203125 0 1581.2504
158.0996856689453 0 494.45825
158.62063598632812 0 422.49512
159.07740783691406 0 524.40045
159.09170532226562 0 505.77737
159.1131134033203 0 2570.6235
165.1026153564453 0 881.7778
167.11819458007812 0 979.0903
169.06112670898438 0 9453.675
169.09750366210938 0 3223.2659
169.13414001464844 0 891.1441
171.07704162597656 0 702.21716
171.11318969726562 0 6016.0776
172.1160888671875 0 624.46106
172.1444549560547 0 695.0321
173.09243774414062 0 899.0147 y 6
173.1287841796875 0 2191.585
174.05526733398438 0 2643.1653
175.0870819091797 0 916.5043
176.08265686035156 0 540.3184
180.11373901367188 0 1195.066
180.1961669921875 0 441.40628
181.0976104736328 0 3004.6611
181.13397216796875 0 1512.8033
182.1291961669922 0 8695.86
183.11306762695312 0 1530.6537
183.1324462890625 0 732.5731
183.149169921875 0 860.16235
185.07138061523438 0 923.92114
185.0922393798828 0 3485.4543 y Water loss 8
185.12881469726562 0 3298.7354
185.1652374267578 0 177548.55 a 1
186.08767700195312 0 20271.488
186.0970916748047 0 803.2297
186.124267578125 0 966.6975
186.1685791015625 0 20406.77
187.09124755859375 0 843.2732
187.10791015625 0 1467.0847
187.14442443847656 0 4842.75
187.1722412109375 0 473.1069
188.13970947265625 0 4228.1655
190.05308532714844 0 637.19293
191.08212280273438 0 977.6491
191.1182098388672 0 625.9706
193.0975799560547 0 1007.35065
193.10739135742188 0 559.5046
193.39486694335938 0 579.57965
195.11329650878906 0 1033.7881
197.0930938720703 0 488.99533
197.12857055664062 0 1508.1823
198.1240234375 0 1124.9843
200.13975524902344 0 2124.8364
201.1240692138672 0 1193.2579
202.08206176757812 0 528.19824
202.1074676513672 0 664.0545
203.10296630859375 0 111804.516 y 8
204.10638427734375 0 9458.516
205.10842895507812 0 696.0221
208.10829162597656 0 2574.4214
208.14459228515625 0 701.1155
209.09266662597656 0 2074.7334
210.12399291992188 0 10194.258
211.12730407714844 0 618.542
211.18148803710938 0 773.77454
213.16012573242188 0 90061.32 b 1
214.16348266601562 0 9456.059
215.13914489746094 0 2940.8315
215.165771484375 0 843.5688
219.08004760742188 0 662.8216
221.09255981445312 0 531.88293
221.10330200195312 0 635.5499
223.10829162597656 0 575.24744
226.1553497314453 0 73514.84
227.15867614746094 0 8014.7495
228.13458251953125 0 9011.203 y Water loss 5
228.15989685058594 0 544.32184
229.13792419433594 0 906.2269
231.0442657470703 0 600.2116
233.14047241210938 0 2135.7795
235.08306884765625 0 814.0594
236.1393280029297 0 881.62177
238.1192169189453 0 1652.9497
239.1505584716797 0 596.109
240.13479614257812 0 2440.9958
241.13380432128906 0 881.55963
242.13552856445312 0 763.3676
242.15023803710938 0 754.5049
243.18174743652344 0 2566.0046
245.0953826904297 0 701.51154
247.1105499267578 0 545.56366
249.1601104736328 0 15419.257
250.16351318359375 0 2533.8552
251.1509246826172 0 1046.0529
252.10922241210938 0 2037.498
252.17111206054688 0 2314.8787
253.16615295410156 0 1051.6222
254.1501922607422 0 2693.633
256.1291809082031 0 1044.3467 y Water loss 7
257.12493896484375 0 676.5058
261.13458251953125 0 5245.638
261.1717224121094 0 612.17804
262.1387939453125 0 866.6768
266.11309814453125 0 1232.0156
269.0808410644531 0 1677.946
269.1972351074219 0 3544.2456
270.1456604003906 0 855.80035
270.1814270019531 0 11093.407 b 2
271.1770324707031 0 8630.142
272.1065673828125 0 1144.5977
272.1799621582031 0 750.86487
274.1399841308594 0 5914.6147 y 7
274.1727600097656 0 739.69055
276.06561279296875 0 628.57
277.1549377441406 0 7963.6055 y Ammonia loss 4
278.1598205566406 0 1091.4208
279.18182373046875 0 13915.586
280.1846923828125 0 2359.0613
281.160888671875 0 2400.3823
283.1412048339844 0 2168.5103
285.1561584472656 0 2649.8862
285.666748046875 0 1621.439 y 4
287.2082214355469 0 17724.307
288.211181640625 0 2119.044
290.11651611328125 0 1148.1184
295.2135925292969 0 650.1111
297.1759033203125 0 450.20795
297.1925048828125 0 28363.918
298.1959228515625 0 4595.954
299.1717834472656 0 51830.9
299.3655700683594 0 544.8469
300.17486572265625 0 6790.298
301.1461486816406 0 541.241
309.2040100097656 0 1763.7408
314.21844482421875 0 891.33276
323.2081298828125 0 2295.3984
325.1872253417969 0 1554.273
327.166748046875 0 2661.5068 y Water loss 6
328.1643981933594 0 1601.8511
332.177978515625 0 1165.7211
336.2046813964844 0 936.3191
337.1877136230469 0 1397.1481
339.16943359375 0 638.77405
340.2347717285156 0 13775.584
340.69024658203125 0 4840.5337 y Water loss 3
341.1887512207031 0 1528.3646 y Ammonia loss 3
341.2378234863281 0 2369.9475
345.17724609375 0 998.1627 y 6
347.6507873535156 0 1366.9452
349.67718505859375 0 878.1979
350.21905517578125 0 613.91766
354.2153625488281 0 885.10626
360.72271728515625 0 701.9757
368.2293395996094 0 69190.164
369.23248291015625 0 12965.486
370.24432373046875 0 4299.491
371.245361328125 0 1017.71173
376.1979675292969 0 1541.6255
376.2454833984375 0 669.1355
378.2085876464844 0 956.39764 y 2
380.2301025390625 0 1315.7942
381.15008544921875 0 1127.5101
381.2156982421875 0 689.24506 b Ammonia loss 3
382.2269287109375 0 1145.6372
383.23382568359375 0 951.66425 b 7
383.7338562011719 0 622.10095
389.1954345703125 0 693.3692
389.229248046875 0 753.44226
390.2130126953125 0 1698.3545
393.8704833984375 0 632.9474
394.2081604003906 0 1961.7845
394.2453308105469 0 643.69617
395.208740234375 0 853.0732
396.1594543457031 0 660.9761
396.2249450683594 0 2663.338
397.2265319824219 0 707.4828
398.2401428222656 0 17633.81 b 3
399.2120056152344 0 1008.1772
399.2431335449219 0 4261.5537
407.1994323730469 0 791.25366
408.2244567871094 0 2496.4792
411.23529052734375 0 1910.9833
414.1697692871094 0 1360.3645
417.7637939453125 0 1396.4633
425.2518310546875 0 1191.8019
425.74322509765625 0 2743.7285 y Water loss 1
429.2267150878906 0 738.70166
431.7606201171875 0 2733.9011 b 8
432.2616271972656 0 875.39575
434.239013671875 0 592.9486
434.7491149902344 0 5408.2095 y 1
435.25018310546875 0 1983.3953
435.75146484375 0 857.7784
437.25079345703125 0 663.46515
437.2865295410156 0 960.1637
439.7401123046875 0 1563.4266
442.2786865234375 0 748.77454
447.237060546875 0 1301.4938
450.2812805175781 0 776.5352
451.26605224609375 0 1024.8618
454.27789306640625 0 3543.9636
456.2562561035156 0 1292.9111
464.2619323730469 0 7981.314
465.2460632324219 0 1277.4724
465.2811584472656 0 4928.6064
466.2483825683594 0 1148.1221
466.2845764160156 0 2010.8644
466.7625427246094 0 1076.2476
467.29705810546875 0 2172.9587
473.2718811035156 0 4150.9663 y 5
474.27490234375 0 775.9134
475.2403259277344 0 700.1937
475.27752685546875 0 2864.6921 Precursor Water loss
475.7789306640625 0 980.6394
478.2795104980469 0 1203.7825
479.2624206542969 0 4294.488
480.26446533203125 0 1480.3699
482.27252197265625 0 20432.736
483.27520751953125 0 4304.7886
484.2493591308594 0 1006.1161
484.28216552734375 0 10454.198 Precursor
484.78399658203125 0 4589.3413
484.989990234375 0 1204.7058
485.28497314453125 0 1784.4728
495.2906799316406 0 789.8888 b 4
496.28924560546875 0 1469.6534
497.2931823730469 0 698.4387
504.2192077636719 0 1055.3564
506.76104736328125 0 700.3994
507.2602844238281 0 753.59216
508.2880859375 0 2242.7412
518.2732543945312 0 1052.2964
521.2465209960938 0 3251.7095
522.2501220703125 0 1184.942
524.3200073242188 0 1006.5561
525.3153686523438 0 5007.071
526.3171997070312 0 1538.9797
535.2969360351562 0 1559.4252
536.2830810546875 0 6376.3843
537.2864990234375 0 1909.9883
547.2628784179688 0 801.1362
552.3143920898438 0 4357.9214 y Water loss 4
553.3095092773438 0 55894.777
554.312255859375 0 17015
555.2701416015625 0 1178.5537
555.3157958984375 0 2593.8225
563.7818603515625 0 2197.4858
564.2821655273438 0 1591.3359
564.7816772460938 0 743.13043
565.2798461914062 0 891.95404
570.3247680664062 0 117130.82 y 4
571.3275756835938 0 36988.875
572.3296508789062 0 6867.0024
573.289306640625 0 999.3245
577.3460083007812 0 2613.438
579.3284912109375 0 923.3714
580.3822631835938 0 2225.9348
581.3856811523438 0 1689.2004
586.76171875 0 1384.6539
587.2560424804688 0 709.7638
595.3563232421875 0 3645.1526
596.3587646484375 0 941.3925
601.2817993164062 0 1227.783
620.3244018554688 0 1128.3335
622.3704223632812 0 1059.8416
638.3997192382812 0 1456.8732
644.2914428710938 0 656.75323
649.3038330078125 0 716.8407
649.3671264648438 0 1497.7084
650.3623046875 0 4584.2866
651.3665771484375 0 1547.1136
654.4284057617188 0 1537.1666
655.4334106445312 0 692.98157
666.3930053710938 0 12176.212
667.3958129882812 0 4281.663
668.3994140625 0 1489.9655
676.4138793945312 0 2136.373
677.4158935546875 0 1171.9554
680.3718872070312 0 2794.45 y Water loss 3
681.3598022460938 0 3714.8237 y Ammonia loss 3
682.3639526367188 0 1188.0217
694.2938842773438 0 691.833
694.4248046875 0 4786.163 b 6
695.4269409179688 0 2296.6624
698.3818969726562 0 3598.463 y 3
699.38525390625 0 1066.6613
720.3677978515625 0 1187.0686
720.4381713867188 0 905.9873
737.3936157226562 0 3191.8623 y Water loss 2
737.466796875 0 3842.4004
738.3800048828125 0 4441.1157 y Ammonia loss 2
738.462890625 0 844.71533
739.3850708007812 0 1632.7874
747.4505615234375 0 1359.3807
748.4360961914062 0 1702.168 b Ammonia loss 7
749.440673828125 0 1295.5305
755.4038696289062 0 100981.39 y 2
756.40673828125 0 42930.05
757.4091186523438 0 9959.702
763.44677734375 0 1210.4181
765.461181640625 0 52324.07 b 7
766.4639892578125 0 24384.467
767.465576171875 0 4991.5537
798.3820190429688 0 624.6804
851.464599609375 0 747.5528 y Ammonia loss 1
868.4880981445312 0 14285.8125 y 1
869.489990234375 0 6368.9814
870.4935913085938 0 2364.4377
2210.593994140625 0 720.37744
3039.089111328125 0 803.9205
3475.889892578125 0 726.2147

Spectrum Details

|  |  |
| --- | --- |
| Matched peaks? Matched peaks The total absolute number of peaks matched. Additionally in brackets the total fraction of peaks matched and the total number of peaks is shown. | 40 (11.14% of 359) |
| FDR? FDR The false discovery rate estimated for this peptide. It is calculated by matching all theoretical fragments with a non-integer shift with the raw peaks for this spectrum. This is done with 40 different shifts. The resulting percentage is the average number of annotated peaks over the number of annotated peaks with the correct spectrum. | 0.54% |
| Satellite FDR? Satellite FDR See the FDR for details on its calculation. This satellite ion specific FDR only contains the satellite ions (d/w) for I/L/J positions. | 0.00% |
| PSM Score? PSM Score The PSM Score as given by Hecklib to this annotated spectrum. It is shown with three significant figures. | 408 |

#### Spectrum 4216? Spectrum 4216 The raw spectrum of this peptide as annotated by Hecklib. The fragments are coloured according to ion type (see legend). Any peaks with a star '\*' as text can be hovered over to see the full details, first the ion type second the mass shift type. By hovering over the amino acids in the peptide or ions in the legend the corresponding peaks are highlighted. By toggling the 'Unassigned' label you can turn the background (unassigned) peaks on or off in the plot. By updating the slider in the Ion legend you can update the spectrum to only show the top X% of the peaks with labels. The top X% means any peak that is within X% of the highest intensity. By dragging in the spectrum you can zoom in to a specific part of the spectrum and use 'Zoom Out' to get back to the original zoom level. The annotation of the spectrum is based on the given sequence in the peptides file and is done with different software so inconsistencies are likely. The peaks are annotated based on the given sequence, with 20 ppm tolerance.

Copy Data

##### Spectrum 4216 (TSV)

###### Preview

```
Loading example...
```

*Click on the button to copy the data to your clipboard.*

Mz MinMz MaxIntensity Max

WidthHeightPeptide font sizePeptide stroke widthSpectrum font sizeSpectrum stroke widthCompact peptide

Ion legend

wxyz

abcd

OtherUnassignedIon^Charge^_Position_Show for top:%

VLGQPKAAPS

01.57e+43.14e+44.71e+46.28e+4

Zoom Out

y^+2^_4_y^+1^_2_y^+2^_6_c^+1^_3_c^+1^_4_z^+1^_5_y^+1^_5_c^+1^_5_y^+1^_6_z^+1^_6_y^+1^_6_w^+1^_7_c^+1^_6_z^+1^_7_y^+1^_7_c^+1^_7_z^+1^_8_y^+1^_8_c^+1^_8_w^+1^_9_z^+1^_9_y^+1^_9_c^+1^_9_

0636127119072542

Fragment Matches Table

Show background peaks

| Position | Ion type | Intensity | mz Theoretical | mz Error (Th) | mz Error (ppm) | Charge | Series Number |
| --- | --- | --- | --- | --- | --- | --- | --- |
| - | - | 635.8 | 129.1 | - | - | 0 | - |
| - | - | 562.7 | 136.1 | - | - | 0 | - |
| 7 | y | 543.4 | 173.1 | 2.819E-06 | 0.01628 | +2 | 4 |
| - | - | 2750 | 173.5 | - | - | 0 | - |
| - | - | 1.08E+04 | 185.2 | - | - | 0 | - |
| - | - | 529.3 | 185.3 | - | - | 0 | - |
| - | - | 1112 | 186.1 | - | - | 0 | - |
| - | - | 777.7 | 186.2 | - | - | 0 | - |
| - | - | 644.9 | 203.1 | - | - | 0 | - |
| 9 | y | 8593 | 203.1 | 0.000308 | 1.516 | +1 | 2 |
| - | - | 1021 | 204.1 | - | - | 0 | - |
| - | - | 482.5 | 211.7 | - | - | 0 | - |
| - | - | 8950 | 213.2 | - | - | 0 | - |
| - | - | 571.5 | 214.2 | - | - | 0 | - |
| - | - | 1145 | 215.1 | - | - | 0 | - |
| - | - | 1113 | 226.2 | - | - | 0 | - |
| - | - | 516.4 | 240.4 | - | - | 0 | - |
| - | - | 656.5 | 249.2 | - | - | 0 | - |
| - | - | 654.8 | 250.2 | - | - | 0 | - |
| - | - | 496.6 | 269.1 | - | - | 0 | - |
| - | - | 1062 | 270.2 | - | - | 0 | - |
| 5 | y | 1357 | 277.2 | 0.001883 | 6.796 | +2 | 6 |
| - | - | 491.4 | 278.4 | - | - | 0 | - |
| 3 | c | 1330 | 287.2 | 0.0003392 | 1.181 | +1 | 3 |
| - | - | 1188 | 299.2 | - | - | 0 | - |
| - | - | 1331 | 309.2 | - | - | 0 | - |
| - | - | 1882 | 368.2 | - | - | 0 | - |
| - | - | 609.2 | 369.2 | - | - | 0 | - |
| - | - | 909.7 | 370.2 | - | - | 0 | - |
| - | - | 1196 | 387.2 | - | - | 0 | - |
| 4 | c | 2242 | 398.2 | 0.000904 | 2.27 | +1 | 4 |
| - | - | 561 | 455.2 | - | - | 0 | - |
| 6 | z | 1.333E+04 | 457.3 | 0.0002618 | 0.5725 | +1 | 5 |
| - | - | 1.241E+04 | 458.3 | - | - | 0 | - |
| - | - | 2554 | 459.3 | - | - | 0 | - |
| - | - | 776.2 | 465.2 | - | - | 0 | - |
| 6 | y | 1166 | 473.3 | 0.0008584 | 1.814 | +1 | 5 |
| - | - | 3171 | 483.2 | - | - | 0 | - |
| - | - | 1970 | 484.2 | - | - | 0 | - |
| - | - | 1535 | 484.3 | - | - | 0 | - |
| - | - | 1357 | 484.8 | - | - | 0 | - |
| 5 | c | 716.4 | 512.3 | 0.0005936 | 1.159 | +1 | 5 |
| - | - | 628.8 | 519.3 | - | - | 0 | - |
| 5 | y | 3458 | 553.3 | 0.009762 | 17.64 | +1 | 6 |
| 5 | z | 597.3 | 554.3 | 0.003829 | 6.907 | +1 | 6 |
| - | - | 5589 | 569.3 | - | - | 0 | - |
| 5 | y | 2.117E+04 | 570.3 | 0.0007353 | 1.289 | +1 | 6 |
| - | - | 6361 | 571.3 | - | - | 0 | - |
| - | - | 1627 | 572.3 | - | - | 0 | - |
| 4 | w | 1.331E+04 | 624.3 | 0.0005578 | 0.8934 | +1 | 7 |
| - | - | 5700 | 625.3 | - | - | 0 | - |
| - | - | 718.8 | 626.3 | - | - | 0 | - |
| - | - | 1687 | 639.4 | - | - | 0 | - |
| 6 | c | 2.963E+04 | 640.4 | 0.0008635 | 1.348 | +1 | 6 |
| - | - | 9137 | 641.4 | - | - | 0 | - |
| - | - | 1454 | 642.4 | - | - | 0 | - |
| - | - | 607.6 | 651.4 | - | - | 0 | - |
| - | - | 1109 | 657.2 | - | - | 0 | - |
| - | - | 615.2 | 666.4 | - | - | 0 | - |
| 4 | z | 1E+04 | 682.4 | 0.00116 | 1.7 | +1 | 7 |
| - | - | 2847 | 683.4 | - | - | 0 | - |
| - | - | 749.9 | 684.4 | - | - | 0 | - |
| 4 | y | 4371 | 698.4 | 0.001635 | 2.341 | +1 | 7 |
| - | - | 849.9 | 699.4 | - | - | 0 | - |
| 7 | c | 1.966E+04 | 711.5 | 0.0006849 | 0.9626 | +1 | 7 |
| - | - | 7903 | 712.5 | - | - | 0 | - |
| - | - | 756.1 | 713.5 | - | - | 0 | - |
| - | - | 642.7 | 725.5 | - | - | 0 | - |
| 3 | z | 1334 | 739.4 | 0.002848 | 3.852 | +1 | 8 |
| - | - | 1090 | 740.4 | - | - | 0 | - |
| 3 | y | 1.697E+04 | 755.4 | 0.0005763 | 0.763 | +1 | 8 |
| - | - | 6691 | 756.4 | - | - | 0 | - |
| - | - | 1377 | 757.4 | - | - | 0 | - |
| 8 | c | 7024 | 765.5 | 0.0009956 | 1.301 | +1 | 8 |
| - | - | 2943 | 766.5 | - | - | 0 | - |
| - | - | 717.4 | 792.4 | - | - | 0 | - |
| 2 | w | 3514 | 809.4 | 0.0008871 | 1.096 | +1 | 9 |
| - | - | 2068 | 810.4 | - | - | 0 | - |
| - | - | 955.9 | 817.9 | - | - | 0 | - |
| - | - | 582.4 | 820.5 | - | - | 0 | - |
| - | - | 1123 | 835.5 | - | - | 0 | - |
| - | - | 648.6 | 836.5 | - | - | 0 | - |
| - | - | 1389 | 851.4 | - | - | 0 | - |
| 2 | z | 6199 | 852.5 | 0.001585 | 1.86 | +1 | 9 |
| - | - | 2995 | 853.5 | - | - | 0 | - |
| - | - | 1089 | 854.5 | - | - | 0 | - |
| - | - | 908.8 | 864.5 | - | - | 0 | - |
| 2 | y | 1565 | 868.5 | 0.00328 | 3.777 | +1 | 9 |
| - | - | 1146 | 878.4 | - | - | 0 | - |
| - | - | 929.4 | 879.4 | - | - | 0 | - |
| 9 | c | 2.317E+04 | 879.5 | 0.001329 | 1.511 | +1 | 9 |
| - | - | 9813 | 880.5 | - | - | 0 | - |
| - | - | 3525 | 881.5 | - | - | 0 | - |
| - | - | 4450 | 895.5 | - | - | 0 | - |
| - | - | 2169 | 896.5 | - | - | 0 | - |
| - | - | 942.5 | 897.5 | - | - | 0 | - |
| - | - | 559.5 | 898.5 | - | - | 0 | - |
| - | - | 885.2 | 904.4 | - | - | 0 | - |
| - | - | 1007 | 907.6 | - | - | 0 | - |
| - | - | 1174 | 912.5 | - | - | 0 | - |
| - | - | 1579 | 922.6 | - | - | 0 | - |
| - | - | 3056 | 923.5 | - | - | 0 | - |
| - | - | 1784 | 924.6 | - | - | 0 | - |
| - | - | 1599 | 950.6 | - | - | 0 | - |
| - | - | 1184 | 951.4 | - | - | 0 | - |
| - | - | 2.349E+04 | 951.5 | - | - | 0 | - |
| - | - | 912.5 | 952.4 | - | - | 0 | - |
| - | - | 1.256E+04 | 952.5 | - | - | 0 | - |
| - | - | 4116 | 953.5 | - | - | 0 | - |
| - | - | 1014 | 954.4 | - | - | 0 | - |
| - | - | 834 | 957 | - | - | 0 | - |
| - | - | 1606 | 965.5 | - | - | 0 | - |
| - | - | 1263 | 967.5 | - | - | 0 | - |
| - | - | 2.095E+04 | 967.6 | - | - | 0 | - |
| - | - | 596.3 | 968 | - | - | 0 | - |
| - | - | 6.217E+04 | 968.6 | - | - | 0 | - |
| - | - | 3.344E+04 | 969.6 | - | - | 0 | - |
| - | - | 1224 | 970.5 | - | - | 0 | - |
| - | - | 8636 | 970.6 | - | - | 0 | - |
| - | - | 972.5 | 971 | - | - | 0 | - |
| - | - | 1496 | 971.5 | - | - | 0 | - |
| - | - | 916.1 | 1237 | - | - | 0 | - |
| - | - | 840.7 | 1238 | - | - | 0 | - |
| - | - | 581.1 | 1240 | - | - | 0 | - |
| - | - | 865.1 | 1437 | - | - | 0 | - |
| - | - | 1219 | 1453 | - | - | 0 | - |
| - | - | 842.6 | 1454 | - | - | 0 | - |
| - | - | 862.6 | 1455 | - | - | 0 | - |
| - | - | 627.8 | 1550 | - | - | 0 | - |
| - | - | 737.2 | 1636 | - | - | 0 | - |
| - | - | 671.4 | 1729 | - | - | 0 | - |
| - | - | 602.2 | 1992 | - | - | 0 | - |
| - | - | 710.4 | 2517 | - | - | 0 | - |

m/z Charge Intensity FragmentType MassShift Position
129.10240173339844 0 635.8196
136.07554626464844 0 562.74634
173.09207153320312 0 543.3685 y 6
173.45098876953125 0 2750.1653
185.16464233398438 0 10797.147
185.3330535888672 0 529.2758
186.08738708496094 0 1111.5532
186.16830444335938 0 777.6673
203.09327697753906 0 644.8887
203.10232543945312 0 8593.44 y 8
204.106201171875 0 1021.13556
211.67843627929688 0 482.5418
213.1593780517578 0 8949.627
214.16336059570312 0 571.4666
215.13865661621094 0 1144.8044
226.1544189453125 0 1112.8877
240.4491729736328 0 516.36926
249.15884399414062 0 656.52563
250.1638641357422 0 654.7617
269.0811462402344 0 496.57773
270.18194580078125 0 1062.4552
277.154541015625 0 1357.48 y Ammonia loss 4
278.43853759765625 0 491.43814
287.2074279785156 0 1330.2817 c 2
299.172119140625 0 1188.3486
309.20343017578125 0 1330.9792
368.22857666015625 0 1882.4861
369.23077392578125 0 609.169
370.2464904785156 0 909.71387
387.187744140625 0 1196.0446
398.2388916015625 0 2241.646 c Ammonia loss 3
455.241943359375 0 561.0285
457.2528381347656 0 13332.274 z 5
458.25982666015625 0 12414.302
459.2626037597656 0 2554.387
465.2265319824219 0 776.15607
473.2709655761719 0 1165.8516 y 5
483.2366027832031 0 3171.3708
484.2477722167969 0 1970.0095
484.28277587890625 0 1534.6434
484.7832946777344 0 1356.9607
512.3197021484375 0 716.38074 c 4
519.29345703125 0 628.79517
553.3078002929688 0 3458.053 y Ammonia loss 4
554.3096923828125 0 597.25006 z 4
569.3163452148438 0 5588.6475
570.3238525390625 0 21173.906 y 4
571.3268432617188 0 6360.587
572.3263549804688 0 1627.3494
624.3345947265625 0 13311.478 w 3
625.3375244140625 0 5699.724
626.3411254882812 0 718.77734
639.4046020507812 0 1687.4712
640.4132080078125 0 29625.44 c 5
641.41650390625 0 9137.403
642.4204711914062 0 1453.6775
651.3642578125 0 607.5814
657.2491455078125 0 1109.255
666.3896484375 0 615.18756
682.36328125 0 10001.391 z 3
683.36572265625 0 2846.6282
684.371337890625 0 749.87604
698.3815307617188 0 4370.902 y 3
699.385009765625 0 849.9457
711.4505004882812 0 19661.42 c 6
712.4526977539062 0 7903.357
713.4541015625 0 756.14545
725.5031127929688 0 642.695
739.383056640625 0 1334.4159 z 2
740.388916015625 0 1090.3035
755.404052734375 0 16968.488 y 2
756.406494140625 0 6690.8354
757.4069213867188 0 1376.9238
765.4607543945312 0 7024.2524 c Ammonia loss 7
766.4628295898438 0 2943.2505
792.367431640625 0 717.3826
809.414306640625 0 3513.9663 w 1
810.4188232421875 0 2068.3826
817.8917846679688 0 955.8771
820.5188598632812 0 582.42615
835.5240478515625 0 1122.8737
836.5272216796875 0 648.61926
851.3790893554688 0 1388.7273
852.4683837890625 0 6198.679 z 1
853.4718017578125 0 2994.889
854.4773559570312 0 1088.5052
864.52783203125 0 908.8145
868.4854125976562 0 1565.1595 y 1
878.39794921875 0 1146.3964
879.394775390625 0 929.35077
879.5397338867188 0 23172.195 c 8
880.5405883789062 0 9812.738
881.5435180664062 0 3525.0627
895.4749755859375 0 4450.464
896.4794311523438 0 2169.2202
897.5185546875 0 942.49603
898.5005493164062 0 559.51526
904.3922729492188 0 885.2404
907.5585327148438 0 1006.8543
912.5018310546875 0 1173.8126
922.5557250976562 0 1579.2853
923.5443725585938 0 3056.462
924.554931640625 0 1783.7633
950.5519409179688 0 1598.7722
951.3650512695312 0 1183.8544
951.5372314453125 0 23494.826
952.370849609375 0 912.5094
952.540283203125 0 12563.981
953.5416870117188 0 4116.0825
954.4129638671875 0 1013.84985
956.9611206054688 0 833.9709
965.4610595703125 0 1606.2495
967.4556274414062 0 1263.407
967.5554809570312 0 20954.094
967.9735107421875 0 596.2732
968.5625 0 62170.508
969.56591796875 0 33435.83
970.471435546875 0 1223.7328
970.56982421875 0 8636.255
970.9676513671875 0 972.5269
971.4646606445312 0 1495.8157
1236.6094970703125 0 916.13184
1237.6011962890625 0 840.66675
1239.82763671875 0 581.112
1436.7174072265625 0 865.1133
1452.725341796875 0 1218.5535
1453.7587890625 0 842.6456
1454.7464599609375 0 862.5995
1549.782470703125 0 627.819
1635.77734375 0 737.19135
1728.5657958984375 0 671.4126
1992.128173828125 0 602.182
2517.287841796875 0 710.39246

Spectrum Details

|  |  |
| --- | --- |
| Matched peaks? Matched peaks The total absolute number of peaks matched. Additionally in brackets the total fraction of peaks matched and the total number of peaks is shown. | 23 (17.29% of 133) |
| FDR? FDR The false discovery rate estimated for this peptide. It is calculated by matching all theoretical fragments with a non-integer shift with the raw peaks for this spectrum. This is done with 40 different shifts. The resulting percentage is the average number of annotated peaks over the number of annotated peaks with the correct spectrum. | 0.62% |
| Satellite FDR? Satellite FDR See the FDR for details on its calculation. This satellite ion specific FDR only contains the satellite ions (d/w) for I/L/J positions. | 2.38% |
| PSM Score? PSM Score The PSM Score as given by Hecklib to this annotated spectrum. It is shown with three significant figures. | 229 |

#### Spectrum 4307? Spectrum 4307 The raw spectrum of this peptide as annotated by Hecklib. The fragments are coloured according to ion type (see legend). Any peaks with a star '\*' as text can be hovered over to see the full details, first the ion type second the mass shift type. By hovering over the amino acids in the peptide or ions in the legend the corresponding peaks are highlighted. By toggling the 'Unassigned' label you can turn the background (unassigned) peaks on or off in the plot. By updating the slider in the Ion legend you can update the spectrum to only show the top X% of the peaks with labels. The top X% means any peak that is within X% of the highest intensity. By dragging in the spectrum you can zoom in to a specific part of the spectrum and use 'Zoom Out' to get back to the original zoom level. The annotation of the spectrum is based on the given sequence in the peptides file and is done with different software so inconsistencies are likely. The peaks are annotated based on the given sequence, with 20 ppm tolerance.

Copy Data

##### Spectrum 4307 (TSV)

###### Preview

```
Loading example...
```

*Click on the button to copy the data to your clipboard.*

Mz MinMz MaxIntensity Max

WidthHeightPeptide font sizePeptide stroke widthSpectrum font sizeSpectrum stroke widthCompact peptide

Ion legend

wxyz

abcd

OtherUnassignedIon^Charge^_Position_Show for top:%

VLGQPKAAPS

01.59e+43.18e+44.77e+46.36e+4

Zoom Out

y^+1^_2_y^+2^_6_c^+1^_3_c^+1^_4_z^+1^_5_y^+1^_5_c^+1^_5_z^+1^_6_y^+1^_6_w^+1^_7_c^+1^_6_y^+1^_7_z^+1^_7_y^+1^_7_c^+1^_7_z^+1^_8_y^+1^_8_c^+1^_8_w^+1^_9_z^+1^_9_y^+1^_9_c^+1^_9_

0838167525133351

Fragment Matches Table

Show background peaks

| Position | Ion type | Intensity | mz Theoretical | mz Error (Th) | mz Error (ppm) | Charge | Series Number |
| --- | --- | --- | --- | --- | --- | --- | --- |
| - | - | 331.8 | 121.3 | - | - | 0 | - |
| - | - | 542.8 | 129.1 | - | - | 0 | - |
| - | - | 414.7 | 132.8 | - | - | 0 | - |
| - | - | 384.2 | 135 | - | - | 0 | - |
| - | - | 376.7 | 135.6 | - | - | 0 | - |
| - | - | 421.7 | 136.1 | - | - | 0 | - |
| - | - | 414.9 | 136.9 | - | - | 0 | - |
| - | - | 457.5 | 147.1 | - | - | 0 | - |
| - | - | 798.7 | 149 | - | - | 0 | - |
| - | - | 644.6 | 171.1 | - | - | 0 | - |
| - | - | 525.3 | 185.2 | - | - | 0 | - |
| - | - | 1.022E+04 | 185.2 | - | - | 0 | - |
| - | - | 698.1 | 186.1 | - | - | 0 | - |
| - | - | 992.2 | 186.2 | - | - | 0 | - |
| - | - | 679.1 | 199.1 | - | - | 0 | - |
| 9 | y | 7664 | 203.1 | 0.0002012 | 0.9904 | +1 | 2 |
| - | - | 474.8 | 207.1 | - | - | 0 | - |
| - | - | 8876 | 213.2 | - | - | 0 | - |
| - | - | 857.8 | 214.2 | - | - | 0 | - |
| - | - | 819.4 | 215.1 | - | - | 0 | - |
| - | - | 498.5 | 226.2 | - | - | 0 | - |
| - | - | 474.6 | 241.1 | - | - | 0 | - |
| - | - | 1311 | 270.2 | - | - | 0 | - |
| 5 | y | 1044 | 277.2 | 0.001853 | 6.685 | +2 | 6 |
| - | - | 661.2 | 283.1 | - | - | 0 | - |
| 3 | c | 918.7 | 287.2 | 2.701E-05 | 0.09406 | +1 | 3 |
| - | - | 988.2 | 299.2 | - | - | 0 | - |
| - | - | 2456 | 309.2 | - | - | 0 | - |
| - | - | 2597 | 368.2 | - | - | 0 | - |
| - | - | 699.3 | 369.2 | - | - | 0 | - |
| - | - | 589.1 | 370.2 | - | - | 0 | - |
| - | - | 903 | 387.2 | - | - | 0 | - |
| - | - | 671.4 | 390.6 | - | - | 0 | - |
| 4 | c | 2263 | 398.2 | 0.0001716 | 0.4308 | +1 | 4 |
| - | - | 652.2 | 399.8 | - | - | 0 | - |
| - | - | 531.3 | 412.3 | - | - | 0 | - |
| 6 | z | 1.173E+04 | 457.3 | 0.0002313 | 0.5058 | +1 | 5 |
| - | - | 9628 | 458.3 | - | - | 0 | - |
| - | - | 1939 | 459.3 | - | - | 0 | - |
| 6 | y | 1277 | 473.3 | 0.0002786 | 0.5886 | +1 | 5 |
| - | - | 1372 | 484.2 | - | - | 0 | - |
| - | - | 1384 | 484.3 | - | - | 0 | - |
| - | - | 664.2 | 484.3 | - | - | 0 | - |
| 5 | c | 938.5 | 512.3 | 0.003435 | 6.704 | +1 | 5 |
| - | - | 687 | 525.3 | - | - | 0 | - |
| - | - | 634.3 | 533.2 | - | - | 0 | - |
| - | - | 3195 | 553.3 | - | - | 0 | - |
| 5 | z | 752.2 | 554.3 | 0.004988 | 8.999 | +1 | 6 |
| - | - | 650.7 | 568.3 | - | - | 0 | - |
| - | - | 5265 | 569.3 | - | - | 0 | - |
| 5 | y | 1.854E+04 | 570.3 | 0.0007353 | 1.289 | +1 | 6 |
| - | - | 5498 | 571.3 | - | - | 0 | - |
| - | - | 847.7 | 572.3 | - | - | 0 | - |
| 4 | w | 1.257E+04 | 624.3 | 0.0001916 | 0.3069 | +1 | 7 |
| - | - | 4201 | 625.3 | - | - | 0 | - |
| - | - | 1017 | 626.3 | - | - | 0 | - |
| - | - | 1548 | 639.4 | - | - | 0 | - |
| 6 | c | 2.672E+04 | 640.4 | 0.0007415 | 1.158 | +1 | 6 |
| - | - | 8302 | 641.4 | - | - | 0 | - |
| - | - | 2620 | 642.4 | - | - | 0 | - |
| - | - | 1036 | 657.3 | - | - | 0 | - |
| - | - | 764.8 | 666.4 | - | - | 0 | - |
| 4 | y | 801.4 | 681.4 | 0.006116 | 8.976 | +1 | 7 |
| 4 | z | 7830 | 682.4 | 0.00116 | 1.7 | +1 | 7 |
| - | - | 2374 | 683.4 | - | - | 0 | - |
| 4 | y | 3772 | 698.4 | 0.000597 | 0.8548 | +1 | 7 |
| - | - | 853.5 | 699.4 | - | - | 0 | - |
| - | - | 732 | 709.3 | - | - | 0 | - |
| - | - | 625.9 | 710.4 | - | - | 0 | - |
| 7 | c | 1.868E+04 | 711.5 | 0.0006238 | 0.8768 | +1 | 7 |
| - | - | 7832 | 712.5 | - | - | 0 | - |
| - | - | 1543 | 713.5 | - | - | 0 | - |
| - | - | 609.2 | 721.4 | - | - | 0 | - |
| - | - | 541.6 | 722.5 | - | - | 0 | - |
| - | - | 1379 | 726.4 | - | - | 0 | - |
| - | - | 2105 | 726.9 | - | - | 0 | - |
| - | - | 1973 | 727.4 | - | - | 0 | - |
| - | - | 844.1 | 727.9 | - | - | 0 | - |
| 3 | z | 1685 | 739.4 | 0.0008342 | 1.128 | +1 | 8 |
| - | - | 1084 | 740.4 | - | - | 0 | - |
| - | - | 743.6 | 753.4 | - | - | 0 | - |
| - | - | 925 | 753.4 | - | - | 0 | - |
| 3 | y | 1.762E+04 | 755.4 | 0.0008815 | 1.167 | +1 | 8 |
| - | - | 6851 | 756.4 | - | - | 0 | - |
| - | - | 1619 | 757.4 | - | - | 0 | - |
| 8 | c | 6675 | 765.5 | 0.001301 | 1.699 | +1 | 8 |
| - | - | 2964 | 766.5 | - | - | 0 | - |
| - | - | 1008 | 771.4 | - | - | 0 | - |
| - | - | 903.4 | 781.3 | - | - | 0 | - |
| - | - | 694.1 | 781.4 | - | - | 0 | - |
| 2 | w | 3821 | 809.4 | 0.001497 | 1.85 | +1 | 9 |
| - | - | 1922 | 810.4 | - | - | 0 | - |
| - | - | 741.9 | 811.4 | - | - | 0 | - |
| - | - | 805 | 824.4 | - | - | 0 | - |
| - | - | 923.8 | 835.5 | - | - | 0 | - |
| - | - | 697.9 | 842.4 | - | - | 0 | - |
| - | - | 1784 | 851.4 | - | - | 0 | - |
| 2 | z | 6663 | 852.5 | 0.0008528 | 1 | +1 | 9 |
| - | - | 688 | 853.4 | - | - | 0 | - |
| - | - | 3677 | 853.5 | - | - | 0 | - |
| - | - | 954 | 854.5 | - | - | 0 | - |
| - | - | 929.7 | 864.4 | - | - | 0 | - |
| 2 | y | 1005 | 868.5 | 0.004989 | 5.745 | +1 | 9 |
| - | - | 1003 | 869.5 | - | - | 0 | - |
| 9 | c | 2.057E+04 | 879.5 | 0.001207 | 1.372 | +1 | 9 |
| - | - | 9350 | 880.5 | - | - | 0 | - |
| - | - | 2667 | 881.5 | - | - | 0 | - |
| - | - | 752.1 | 882.5 | - | - | 0 | - |
| - | - | 1018 | 883.5 | - | - | 0 | - |
| - | - | 3449 | 895.5 | - | - | 0 | - |
| - | - | 1800 | 896.5 | - | - | 0 | - |
| - | - | 3287 | 899.4 | - | - | 0 | - |
| - | - | 1361 | 900.4 | - | - | 0 | - |
| - | - | 891.1 | 909.4 | - | - | 0 | - |
| - | - | 623.3 | 910.4 | - | - | 0 | - |
| - | - | 1742 | 912.5 | - | - | 0 | - |
| - | - | 1664 | 922.6 | - | - | 0 | - |
| - | - | 3021 | 923.5 | - | - | 0 | - |
| - | - | 1086 | 924.4 | - | - | 0 | - |
| - | - | 2016 | 924.6 | - | - | 0 | - |
| - | - | 801.1 | 925.5 | - | - | 0 | - |
| - | - | 748.3 | 947.5 | - | - | 0 | - |
| - | - | 731.2 | 948.5 | - | - | 0 | - |
| - | - | 1516 | 951.4 | - | - | 0 | - |
| - | - | 2.506E+04 | 951.5 | - | - | 0 | - |
| - | - | 890.1 | 952.4 | - | - | 0 | - |
| - | - | 1.251E+04 | 952.5 | - | - | 0 | - |
| - | - | 891 | 953.4 | - | - | 0 | - |
| - | - | 4556 | 953.5 | - | - | 0 | - |
| - | - | 1137 | 954.4 | - | - | 0 | - |
| - | - | 778.9 | 961 | - | - | 0 | - |
| - | - | 758.9 | 961.5 | - | - | 0 | - |
| - | - | 1460 | 963 | - | - | 0 | - |
| - | - | 938.3 | 964.5 | - | - | 0 | - |
| - | - | 920.3 | 965.5 | - | - | 0 | - |
| - | - | 1025 | 967.4 | - | - | 0 | - |
| - | - | 2.006E+04 | 967.6 | - | - | 0 | - |
| - | - | 6.297E+04 | 968.6 | - | - | 0 | - |
| - | - | 1046 | 969 | - | - | 0 | - |
| - | - | 1782 | 969.5 | - | - | 0 | - |
| - | - | 3.036E+04 | 969.6 | - | - | 0 | - |
| - | - | 2729 | 970 | - | - | 0 | - |
| - | - | 4929 | 970.5 | - | - | 0 | - |
| - | - | 8374 | 970.6 | - | - | 0 | - |
| - | - | 1170 | 971 | - | - | 0 | - |
| - | - | 2559 | 971.5 | - | - | 0 | - |
| - | - | 1139 | 1095 | - | - | 0 | - |
| - | - | 674.5 | 1125 | - | - | 0 | - |
| - | - | 659.8 | 1224 | - | - | 0 | - |
| - | - | 941 | 1323 | - | - | 0 | - |
| - | - | 716.5 | 1395 | - | - | 0 | - |
| - | - | 970.6 | 1436 | - | - | 0 | - |
| - | - | 1805 | 1437 | - | - | 0 | - |
| - | - | 2391 | 1438 | - | - | 0 | - |
| - | - | 616.6 | 1452 | - | - | 0 | - |
| - | - | 2342 | 1453 | - | - | 0 | - |
| - | - | 3703 | 1454 | - | - | 0 | - |
| - | - | 1547 | 1455 | - | - | 0 | - |
| - | - | 972.4 | 1456 | - | - | 0 | - |
| - | - | 744.7 | 1810 | - | - | 0 | - |
| - | - | 762.2 | 1913 | - | - | 0 | - |
| - | - | 1053 | 1939 | - | - | 0 | - |
| - | - | 754.4 | 1940 | - | - | 0 | - |
| - | - | 1442 | 1941 | - | - | 0 | - |
| - | - | 935.6 | 1942 | - | - | 0 | - |
| - | - | 768.5 | 3043 | - | - | 0 | - |
| - | - | 656 | 3318 | - | - | 0 | - |

m/z Charge Intensity FragmentType MassShift Position
121.31905364990234 0 331.80447
129.10235595703125 0 542.79236
132.79330444335938 0 414.73932
135.02462768554688 0 384.22186
135.62376403808594 0 376.7093
136.07614135742188 0 421.72702
136.9151153564453 0 414.85062
147.11309814453125 0 457.45248
148.95472717285156 0 798.7039
171.11329650878906 0 644.60156
185.15597534179688 0 525.312
185.16470336914062 0 10224.675
186.0882568359375 0 698.07587
186.16851806640625 0 992.2468
199.14425659179688 0 679.06555
203.10243225097656 0 7664.0835 y 8
207.09796142578125 0 474.80408
213.1595001220703 0 8876.397
214.1632080078125 0 857.81445
215.13868713378906 0 819.4132
226.15496826171875 0 498.4928
241.06207275390625 0 474.56238
270.1806945800781 0 1310.8232
277.1545104980469 0 1044.2267 y Ammonia loss 4
283.14385986328125 0 661.15686
287.2077941894531 0 918.7273 c 2
299.1716613769531 0 988.1886
309.20343017578125 0 2455.6218
368.2284851074219 0 2597.0847
369.2310485839844 0 699.3017
370.208251953125 0 589.14014
387.18695068359375 0 902.9598
390.6120300292969 0 671.4111
398.2396240234375 0 2262.6892 c Ammonia loss 3
399.8357238769531 0 652.2227
412.2580261230469 0 531.2541
457.25286865234375 0 11731.971 z 5
458.2598876953125 0 9628.215
459.2632751464844 0 1938.7172
473.27154541015625 0 1276.727 y 5
484.24993896484375 0 1371.8964
484.28173828125 0 1384.4147
484.3177185058594 0 664.2457
512.315673828125 0 938.5028 c 4
525.3425903320312 0 686.9759
533.1919555664062 0 634.2521
553.3093872070312 0 3195.0898
554.3108520507812 0 752.2142 z 4
568.3098754882812 0 650.72943
569.31640625 0 5265.0547
570.3238525390625 0 18540.014 y 4
571.3272094726562 0 5497.522
572.3313598632812 0 847.662
624.3349609375 0 12568.544 w 3
625.337646484375 0 4201.194
626.340576171875 0 1017.29364
639.4074096679688 0 1548.2955
640.413330078125 0 26715.324 c 5
641.4163208007812 0 8301.86
642.4197998046875 0 2619.5527
657.25244140625 0 1035.5524
666.39501953125 0 764.7875
681.3627319335938 0 801.39886 y Ammonia loss 3
682.36328125 0 7829.7793 z 3
683.3646240234375 0 2374.1572
698.382568359375 0 3772.2283 y 3
699.384521484375 0 853.4723
709.3447265625 0 732.00525
710.4390869140625 0 625.8758
711.4505615234375 0 18684.932 c 6
712.452880859375 0 7831.985
713.454345703125 0 1542.9229
721.4493408203125 0 609.224
722.4548950195312 0 541.5581
726.3731079101562 0 1379.2793
726.874267578125 0 2104.6318
727.385009765625 0 1972.9537
727.881103515625 0 844.1263
739.3850708007812 0 1685.4698 z 2
740.3878784179688 0 1084.3265
753.3504028320312 0 743.62775
753.4223022460938 0 925.0445
755.4037475585938 0 17621.396 y 2
756.4063110351562 0 6850.7603
757.4056396484375 0 1619.1792
765.46044921875 0 6674.814 c Ammonia loss 7
766.464111328125 0 2964.3406
771.3541870117188 0 1008.4586
781.3491821289062 0 903.3716
781.4244384765625 0 694.1204
809.4136962890625 0 3820.9995 w 1
810.4179077148438 0 1922.4478
811.427490234375 0 741.912
824.4122924804688 0 805.047
835.52685546875 0 923.84314
842.4205322265625 0 697.94495
851.3764038085938 0 1784.045
852.4691162109375 0 6662.6826 z 1
853.3908081054688 0 688.04834
853.471435546875 0 3676.8145
854.4781494140625 0 953.9532
864.407470703125 0 929.67456
868.4837036132812 0 1005.4676 y 1
869.4942016601562 0 1003.39496
879.5398559570312 0 20569.52 c 8
880.5404052734375 0 9350.404
881.5408325195312 0 2666.885
882.467529296875 0 752.09985
883.47314453125 0 1018.01025
895.4730224609375 0 3448.8547
896.4743041992188 0 1800.0171
899.4453735351562 0 3286.6104
900.4412231445312 0 1360.8196
909.3748168945312 0 891.1317
910.3895263671875 0 623.3083
912.5025024414062 0 1741.6064
922.55810546875 0 1664.0703
923.5465698242188 0 3021.1108
924.4146118164062 0 1086.0607
924.55908203125 0 2015.7795
925.4500122070312 0 801.1076
947.451416015625 0 748.334
948.4578247070312 0 731.1783
951.3652954101562 0 1516.3862
951.5374755859375 0 25060.252
952.3709106445312 0 890.14886
952.5404052734375 0 12512.798
953.4396362304688 0 891.02795
953.5416259765625 0 4555.813
954.4171752929688 0 1136.8583
960.9744873046875 0 778.8728
961.4684448242188 0 758.8825
962.953125 0 1460.2236
964.4641723632812 0 938.30426
965.4734497070312 0 920.3405
967.3819580078125 0 1025.4618
967.55615234375 0 20061.688
968.5626220703125 0 62973.676
968.972900390625 0 1046.3354
969.4851684570312 0 1782.0198
969.5658569335938 0 30355.62
969.980224609375 0 2728.88
970.4622802734375 0 4928.869
970.5701904296875 0 8373.94
970.974365234375 0 1169.7776
971.4627075195312 0 2558.8179
1094.5303955078125 0 1139.0729
1124.7584228515625 0 674.5081
1223.601806640625 0 659.8062
1322.661865234375 0 941.0274
1394.722412109375 0 716.4533
1435.7181396484375 0 970.55225
1436.7176513671875 0 1804.6152
1437.7227783203125 0 2391.2568
1451.72998046875 0 616.6421
1452.749755859375 0 2342.1047
1453.7435302734375 0 3702.901
1454.7572021484375 0 1546.9854
1455.751708984375 0 972.3966
1810.4439697265625 0 744.687
1912.9102783203125 0 762.2229
1938.9228515625 0 1052.5247
1939.941162109375 0 754.4318
1940.9229736328125 0 1441.7832
1941.9124755859375 0 935.56775
3042.714111328125 0 768.5301
3317.55908203125 0 655.9731

Spectrum Details

|  |  |
| --- | --- |
| Matched peaks? Matched peaks The total absolute number of peaks matched. Additionally in brackets the total fraction of peaks matched and the total number of peaks is shown. | 22 (13.17% of 167) |
| FDR? FDR The false discovery rate estimated for this peptide. It is calculated by matching all theoretical fragments with a non-integer shift with the raw peaks for this spectrum. This is done with 40 different shifts. The resulting percentage is the average number of annotated peaks over the number of annotated peaks with the correct spectrum. | 1.19% |
| Satellite FDR? Satellite FDR See the FDR for details on its calculation. This satellite ion specific FDR only contains the satellite ions (d/w) for I/L/J positions. | 0.00% |
| PSM Score? PSM Score The PSM Score as given by Hecklib to this annotated spectrum. It is shown with three significant figures. | 229 |

#### Spectrum 4370? Spectrum 4370 The raw spectrum of this peptide as annotated by Hecklib. The fragments are coloured according to ion type (see legend). Any peaks with a star '\*' as text can be hovered over to see the full details, first the ion type second the mass shift type. By hovering over the amino acids in the peptide or ions in the legend the corresponding peaks are highlighted. By toggling the 'Unassigned' label you can turn the background (unassigned) peaks on or off in the plot. By updating the slider in the Ion legend you can update the spectrum to only show the top X% of the peaks with labels. The top X% means any peak that is within X% of the highest intensity. By dragging in the spectrum you can zoom in to a specific part of the spectrum and use 'Zoom Out' to get back to the original zoom level. The annotation of the spectrum is based on the given sequence in the peptides file and is done with different software so inconsistencies are likely. The peaks are annotated based on the given sequence, with 20 ppm tolerance.

Copy Data

##### Spectrum 4370 (TSV)

###### Preview

```
Loading example...
```

*Click on the button to copy the data to your clipboard.*

Mz MinMz MaxIntensity Max

WidthHeightPeptide font sizePeptide stroke widthSpectrum font sizeSpectrum stroke widthCompact peptide

Ion legend

wxyz

abcd

OtherUnassignedIon^Charge^_Position_Show for top:%

VLGQPKAAPS

01.42e+42.84e+44.26e+45.68e+4

Zoom Out

y^+1^_2_y^+2^_6_c^+1^_3_c^+1^_4_z^+1^_5_y^+1^_5_c^+1^_5_y^+1^_6_z^+1^_6_y^+1^_6_w^+1^_7_c^+1^_6_z^+1^_7_y^+1^_7_c^+1^_7_z^+1^_8_y^+1^_8_c^+1^_8_w^+1^_9_z^+1^_9_y^+1^_9_c^+1^_9_

0768153723053073

Fragment Matches Table

Show background peaks

| Position | Ion type | Intensity | mz Theoretical | mz Error (Th) | mz Error (ppm) | Charge | Series Number |
| --- | --- | --- | --- | --- | --- | --- | --- |
| - | - | 866.6 | 120.1 | - | - | 0 | - |
| - | - | 396.7 | 128.4 | - | - | 0 | - |
| - | - | 465.2 | 129.1 | - | - | 0 | - |
| - | - | 464.7 | 132.5 | - | - | 0 | - |
| - | - | 404.3 | 139.8 | - | - | 0 | - |
| - | - | 433.4 | 147.2 | - | - | 0 | - |
| - | - | 606.9 | 149 | - | - | 0 | - |
| - | - | 448.4 | 150.4 | - | - | 0 | - |
| - | - | 600.2 | 155.2 | - | - | 0 | - |
| - | - | 401 | 158.6 | - | - | 0 | - |
| - | - | 473.6 | 171.3 | - | - | 0 | - |
| - | - | 3702 | 173.4 | - | - | 0 | - |
| - | - | 1.051E+04 | 185.2 | - | - | 0 | - |
| - | - | 1141 | 186.1 | - | - | 0 | - |
| - | - | 909.5 | 186.2 | - | - | 0 | - |
| - | - | 465 | 188.5 | - | - | 0 | - |
| - | - | 779.9 | 199.1 | - | - | 0 | - |
| 9 | y | 7021 | 203.1 | 0.0001706 | 0.8401 | +1 | 2 |
| - | - | 859.4 | 204.1 | - | - | 0 | - |
| - | - | 7503 | 213.2 | - | - | 0 | - |
| - | - | 1027 | 214.2 | - | - | 0 | - |
| - | - | 1205 | 215.1 | - | - | 0 | - |
| - | - | 742.8 | 219.1 | - | - | 0 | - |
| - | - | 1173 | 226.2 | - | - | 0 | - |
| - | - | 784.4 | 249.2 | - | - | 0 | - |
| - | - | 683 | 270.2 | - | - | 0 | - |
| 5 | y | 864.2 | 277.2 | 0.001883 | 6.796 | +2 | 6 |
| 3 | c | 858 | 287.2 | 0.000858 | 2.987 | +1 | 3 |
| - | - | 862.7 | 299.2 | - | - | 0 | - |
| - | - | 2153 | 309.2 | - | - | 0 | - |
| - | - | 594.9 | 317.8 | - | - | 0 | - |
| - | - | 601.9 | 327 | - | - | 0 | - |
| - | - | 1849 | 368.2 | - | - | 0 | - |
| - | - | 590.9 | 370.2 | - | - | 0 | - |
| - | - | 973.6 | 372.2 | - | - | 0 | - |
| - | - | 780.1 | 373.2 | - | - | 0 | - |
| - | - | 982 | 387.2 | - | - | 0 | - |
| 4 | c | 1939 | 398.2 | 4.206E-05 | 0.1056 | +1 | 4 |
| - | - | 654.1 | 412.3 | - | - | 0 | - |
| 6 | z | 1.102E+04 | 457.3 | 0.0001397 | 0.3056 | +1 | 5 |
| - | - | 1.054E+04 | 458.3 | - | - | 0 | - |
| - | - | 2157 | 459.3 | - | - | 0 | - |
| 6 | y | 753.8 | 473.3 | 3.927E-06 | 0.008298 | +1 | 5 |
| - | - | 925.7 | 475.3 | - | - | 0 | - |
| - | - | 956.2 | 482.3 | - | - | 0 | - |
| - | - | 1122 | 484.2 | - | - | 0 | - |
| - | - | 1334 | 484.3 | - | - | 0 | - |
| - | - | 1342 | 484.8 | - | - | 0 | - |
| - | - | 685.8 | 508.3 | - | - | 0 | - |
| 5 | c | 802.5 | 512.3 | 0.0001998 | 0.3901 | +1 | 5 |
| - | - | 799.9 | 519.3 | - | - | 0 | - |
| 5 | y | 2323 | 553.3 | 0.01062 | 19.19 | +1 | 6 |
| 5 | z | 826.4 | 554.3 | 0.007247 | 13.07 | +1 | 6 |
| - | - | 3055 | 569.3 | - | - | 0 | - |
| 5 | y | 1.884E+04 | 570.3 | 0.0004301 | 0.7542 | +1 | 6 |
| - | - | 5718 | 571.3 | - | - | 0 | - |
| - | - | 757.1 | 572.3 | - | - | 0 | - |
| - | - | 1128 | 591.3 | - | - | 0 | - |
| - | - | 1063 | 591.3 | - | - | 0 | - |
| - | - | 637.7 | 592.3 | - | - | 0 | - |
| - | - | 564.8 | 600.3 | - | - | 0 | - |
| 4 | w | 1.187E+04 | 624.3 | 0.0003747 | 0.6002 | +1 | 7 |
| - | - | 4814 | 625.3 | - | - | 0 | - |
| - | - | 1142 | 626.3 | - | - | 0 | - |
| - | - | 1113 | 639.4 | - | - | 0 | - |
| 6 | c | 2.554E+04 | 640.4 | 0.0004363 | 0.6813 | +1 | 6 |
| - | - | 8378 | 641.4 | - | - | 0 | - |
| - | - | 1297 | 642.4 | - | - | 0 | - |
| - | - | 714.3 | 657.4 | - | - | 0 | - |
| - | - | 1454 | 664.4 | - | - | 0 | - |
| 4 | z | 6713 | 682.4 | 0.0006718 | 0.9845 | +1 | 7 |
| - | - | 3584 | 683.4 | - | - | 0 | - |
| - | - | 1302 | 684.4 | - | - | 0 | - |
| - | - | 862.7 | 686.4 | - | - | 0 | - |
| 4 | y | 3209 | 698.4 | 0.0009022 | 1.292 | +1 | 7 |
| - | - | 1475 | 699.4 | - | - | 0 | - |
| - | - | 1352 | 701.4 | - | - | 0 | - |
| - | - | 1586 | 702.4 | - | - | 0 | - |
| - | - | 1946 | 703.4 | - | - | 0 | - |
| - | - | 815.1 | 710.5 | - | - | 0 | - |
| 7 | c | 1.603E+04 | 711.5 | 0.0001355 | 0.1905 | +1 | 7 |
| - | - | 5799 | 712.5 | - | - | 0 | - |
| - | - | 1306 | 713.5 | - | - | 0 | - |
| - | - | 626.7 | 717.9 | - | - | 0 | - |
| - | - | 1185 | 720.4 | - | - | 0 | - |
| - | - | 806.6 | 723.3 | - | - | 0 | - |
| - | - | 870.8 | 726.4 | - | - | 0 | - |
| - | - | 1117 | 727.4 | - | - | 0 | - |
| 3 | z | 1439 | 739.4 | 0.000407 | 0.5504 | +1 | 8 |
| 3 | y | 1.819E+04 | 755.4 | 0.0004543 | 0.6014 | +1 | 8 |
| - | - | 6093 | 756.4 | - | - | 0 | - |
| - | - | 1516 | 757.4 | - | - | 0 | - |
| 8 | c | 6063 | 765.5 | 0.0002022 | 0.2641 | +1 | 8 |
| - | - | 2757 | 766.5 | - | - | 0 | - |
| - | - | 1043 | 767.5 | - | - | 0 | - |
| - | - | 608.4 | 771.4 | - | - | 0 | - |
| - | - | 597.7 | 773.2 | - | - | 0 | - |
| - | - | 788.4 | 778.5 | - | - | 0 | - |
| 2 | w | 3635 | 809.4 | 0.001615 | 1.996 | +1 | 9 |
| - | - | 1608 | 810.4 | - | - | 0 | - |
| - | - | 677.1 | 811.4 | - | - | 0 | - |
| - | - | 956.3 | 847.4 | - | - | 0 | - |
| 2 | z | 6006 | 852.5 | 0.0001238 | 0.1452 | +1 | 9 |
| - | - | 2968 | 853.5 | - | - | 0 | - |
| - | - | 829.6 | 854.5 | - | - | 0 | - |
| - | - | 2097 | 864.4 | - | - | 0 | - |
| - | - | 861.3 | 865.4 | - | - | 0 | - |
| 2 | y | 1140 | 868.5 | 0.002457 | 2.829 | +1 | 9 |
| - | - | 2124 | 878.5 | - | - | 0 | - |
| 9 | c | 2.08E+04 | 879.5 | 0.0007798 | 0.8866 | +1 | 9 |
| - | - | 8603 | 880.5 | - | - | 0 | - |
| - | - | 2408 | 881.5 | - | - | 0 | - |
| - | - | 3174 | 895.5 | - | - | 0 | - |
| - | - | 1804 | 896.5 | - | - | 0 | - |
| - | - | 979.6 | 897.5 | - | - | 0 | - |
| - | - | 2646 | 899.4 | - | - | 0 | - |
| - | - | 710 | 905.5 | - | - | 0 | - |
| - | - | 1889 | 906.5 | - | - | 0 | - |
| - | - | 583.8 | 907.4 | - | - | 0 | - |
| - | - | 1085 | 912.5 | - | - | 0 | - |
| - | - | 767.3 | 913.5 | - | - | 0 | - |
| - | - | 1682 | 922.6 | - | - | 0 | - |
| - | - | 2963 | 923.5 | - | - | 0 | - |
| - | - | 1480 | 924.6 | - | - | 0 | - |
| - | - | 975.1 | 938.4 | - | - | 0 | - |
| - | - | 4973 | 948.4 | - | - | 0 | - |
| - | - | 2413 | 949.4 | - | - | 0 | - |
| - | - | 744.6 | 950.5 | - | - | 0 | - |
| - | - | 1600 | 951.4 | - | - | 0 | - |
| - | - | 2.181E+04 | 951.5 | - | - | 0 | - |
| - | - | 1044 | 952.4 | - | - | 0 | - |
| - | - | 1.189E+04 | 952.5 | - | - | 0 | - |
| - | - | 1331 | 953.4 | - | - | 0 | - |
| - | - | 3589 | 953.5 | - | - | 0 | - |
| - | - | 1415 | 954.4 | - | - | 0 | - |
| - | - | 1690 | 964.5 | - | - | 0 | - |
| - | - | 2925 | 965.5 | - | - | 0 | - |
| - | - | 778.7 | 966.4 | - | - | 0 | - |
| - | - | 924.2 | 966.5 | - | - | 0 | - |
| - | - | 794.4 | 967.4 | - | - | 0 | - |
| - | - | 1.72E+04 | 967.6 | - | - | 0 | - |
| - | - | 5.627E+04 | 968.6 | - | - | 0 | - |
| - | - | 880.9 | 969.5 | - | - | 0 | - |
| - | - | 2.493E+04 | 969.6 | - | - | 0 | - |
| - | - | 2539 | 970.5 | - | - | 0 | - |
| - | - | 9976 | 970.6 | - | - | 0 | - |
| - | - | 1324 | 971.5 | - | - | 0 | - |
| - | - | 1083 | 971.6 | - | - | 0 | - |
| - | - | 729.4 | 1191 | - | - | 0 | - |
| - | - | 606.5 | 1288 | - | - | 0 | - |
| - | - | 700.1 | 1437 | - | - | 0 | - |
| - | - | 990.1 | 1454 | - | - | 0 | - |
| - | - | 780.1 | 1936 | - | - | 0 | - |
| - | - | 668.8 | 1940 | - | - | 0 | - |
| - | - | 680.3 | 3043 | - | - | 0 | - |

m/z Charge Intensity FragmentType MassShift Position
120.08059692382812 0 866.585
128.41888427734375 0 396.6885
129.1024627685547 0 465.16605
132.51597595214844 0 464.66455
139.77987670898438 0 404.3298
147.1928253173828 0 433.4109
148.9543914794922 0 606.92664
150.42498779296875 0 448.37695
155.24813842773438 0 600.2325
158.60122680664062 0 400.96545
171.26841735839844 0 473.6436
173.44053649902344 0 3702.4382
185.1646728515625 0 10506.529
186.08712768554688 0 1140.5421
186.16836547851562 0 909.51056
188.4987335205078 0 464.97305
199.1076202392578 0 779.8609
203.1024627685547 0 7021.148 y 8
204.10601806640625 0 859.4444
213.15951538085938 0 7503.1865
214.16342163085938 0 1026.5183
215.13900756835938 0 1205.205
219.1488494873047 0 742.8458
226.15480041503906 0 1173.0165
249.16024780273438 0 784.36597
270.1806335449219 0 683.03564
277.154541015625 0 864.1901 y Ammonia loss 4
287.2069091796875 0 857.96277 c 2
299.1709289550781 0 862.707
309.20379638671875 0 2153.0093
317.80352783203125 0 594.8532
327.0154113769531 0 601.9436
368.22930908203125 0 1849.1937
370.2443542480469 0 590.91364
372.2348327636719 0 973.6002
373.23919677734375 0 780.1103
387.1874694824219 0 982.02454
398.2398376464844 0 1938.9471 c Ammonia loss 3
412.25482177734375 0 654.1244
457.2529602050781 0 11016.108 z 5
458.26007080078125 0 10537.678
459.2623596191406 0 2156.6553
473.2718200683594 0 753.82965 y 5
475.27984619140625 0 925.70667
482.2739562988281 0 956.1513
484.2489929199219 0 1121.8302
484.2835388183594 0 1334.4902
484.78436279296875 0 1341.9346
508.3099670410156 0 685.8156
512.3189086914062 0 802.4892 c 4
519.2942504882812 0 799.94946
553.3086547851562 0 2322.5342 y Ammonia loss 4
554.3131103515625 0 826.3895 z 4
569.316650390625 0 3055.0327
570.3241577148438 0 18837.512 y 4
571.326904296875 0 5717.528
572.3284301757812 0 757.1048
591.285888671875 0 1127.9272
591.3355102539062 0 1063.3315
592.3377075195312 0 637.66595
600.3439331054688 0 564.7907
624.3347778320312 0 11868.555 w 3
625.3382568359375 0 4813.973
626.3395385742188 0 1142.182
639.4076538085938 0 1113.0726
640.4136352539062 0 25542.969 c 5
641.416259765625 0 8377.737
642.4207153320312 0 1297.3987
657.3884887695312 0 714.34174
664.3745727539062 0 1454.0543
682.36376953125 0 6712.9077 z 3
683.3681640625 0 3584.0183
684.3639526367188 0 1301.7778
686.3736572265625 0 862.7268
698.3822631835938 0 3209.291 y 3
699.3868408203125 0 1475.4579
701.3934326171875 0 1351.5392
702.3960571289062 0 1585.7667
703.3992309570312 0 1945.8407
710.4506225585938 0 815.0842
711.4510498046875 0 16031.785 c 6
712.453125 0 5798.527
713.4569702148438 0 1306.3738
717.8612060546875 0 626.691
720.3886108398438 0 1185.3536
723.2822265625 0 806.6021
726.3755493164062 0 870.7631
727.3953247070312 0 1117.096
739.385498046875 0 1438.9482 z 2
755.4041748046875 0 18188.152 y 2
756.406982421875 0 6093.3916
757.407958984375 0 1516.4172
765.4615478515625 0 6062.705 c Ammonia loss 7
766.4659423828125 0 2757.4792
767.4666748046875 0 1042.758
771.3511962890625 0 608.43744
773.2282104492188 0 597.6785
778.462158203125 0 788.3598
809.4168090820312 0 3635.1235 w 1
810.41796875 0 1608.2656
811.4229125976562 0 677.10443
847.3870849609375 0 956.2763
852.4700927734375 0 6006.261 z 1
853.4734497070312 0 2968.4055
854.4749755859375 0 829.6029
864.41162109375 0 2097.2651
865.4111938476562 0 861.27466
868.4911499023438 0 1139.5022 y 1
878.50732421875 0 2123.5732
879.540283203125 0 20798.123 c 8
880.5421142578125 0 8602.883
881.544677734375 0 2407.7405
895.4750366210938 0 3173.7874
896.4767456054688 0 1804.4678
897.5269165039062 0 979.64417
899.447998046875 0 2645.5708
905.4572143554688 0 709.9725
906.4556884765625 0 1889.3181
907.4488525390625 0 583.77045
912.5032958984375 0 1084.7738
913.5093994140625 0 767.33905
922.5596923828125 0 1681.7955
923.5482788085938 0 2962.5935
924.5523681640625 0 1479.9275
938.4346923828125 0 975.11346
948.447509765625 0 4973.2534
949.4471435546875 0 2413.2068
950.54736328125 0 744.60736
951.36474609375 0 1599.6005
951.5379028320312 0 21806.95
952.37841796875 0 1044.4675
952.5408935546875 0 11887.393
953.4329833984375 0 1330.9835
953.5440063476562 0 3589.111
954.4354248046875 0 1414.8595
964.469970703125 0 1689.7324
965.4767456054688 0 2925.3577
966.3607177734375 0 778.6986
966.506591796875 0 924.2101
967.3799438476562 0 794.4335
967.5560302734375 0 17195.387
968.5635375976562 0 56272.21
969.4629516601562 0 880.9136
969.5667724609375 0 24930.06
970.4631958007812 0 2539.3596
970.570068359375 0 9975.852
971.4612426757812 0 1324.4799
971.5702514648438 0 1082.7238
1190.5631103515625 0 729.38477
1288.4593505859375 0 606.5045
1436.7362060546875 0 700.0762
1453.736328125 0 990.1358
1935.933349609375 0 780.1017
1939.9185791015625 0 668.7524
3042.627685546875 0 680.26306

Spectrum Details

|  |  |
| --- | --- |
| Matched peaks? Matched peaks The total absolute number of peaks matched. Additionally in brackets the total fraction of peaks matched and the total number of peaks is shown. | 22 (14.19% of 155) |
| FDR? FDR The false discovery rate estimated for this peptide. It is calculated by matching all theoretical fragments with a non-integer shift with the raw peaks for this spectrum. This is done with 40 different shifts. The resulting percentage is the average number of annotated peaks over the number of annotated peaks with the correct spectrum. | 0.65% |
| Satellite FDR? Satellite FDR See the FDR for details on its calculation. This satellite ion specific FDR only contains the satellite ions (d/w) for I/L/J positions. | 0.00% |
| PSM Score? PSM Score The PSM Score as given by Hecklib to this annotated spectrum. It is shown with three significant figures. | 229 |

#### Reverse Lookup? Reverse Lookup All places where this read could be placed.

| Group | Segment | Template | Template Part | Read Part | Score | Unique |
| --- | --- | --- | --- | --- | --- | --- |
| Homo sapiens Light Chain | IGLC | IGLC2 | [0..8] | [2..10] | 64 | False |
| Homo sapiens Light Chain | IGLC | IGLC6 | [0..8] | [2..10] | 64 | False |
| Homo sapiens Light Chain | IGLC | IGLC7 | [0..8] | [2..10] | 64 | False |

| Recombined | Template Part | Read Part | Score | Unique |
| --- | --- | --- | --- | --- |
| REC-0-1\_002 | [109..119] | [0..10] | 80 | True |

#### Meta Information from Multiple reads

##### Number of combined reads

15

##### Intensity

0.9748

##### TotalArea

1.991E+09

##### Changes to the peptide sequence

VLGQPKAAPS

J→LSupport for Leucine based on side chain ions (1 for L 0 for I) (Position: 2)

L→JEqual support for both Leucine and Isoleucine based on side chain ions (1 ions for both) (Position: 2)

#### Positional Score

Copy Data

##### Positional Score (TSV)

###### Preview

```
Loading example...
```

*Click on the button to copy the data to your clipboard.*

100123456789

Label Value
"0" 0.863
"1" 0.859
"2" 0.857
"3" 0.825
"4" 0.843
"5" 0.843
"6" 0.854
"7" 0.859
"8" 0.849
"9" 0.852

#### Meta Information from PEAKS

##### Scan Identifier

F1:3218

##### Original sequence

V

L

G

Q

P

K

A

A

P

S

##### Posttranslational Modifications

##### Source File

D:\separate\_stitch\_analyses\xle-disambiguation\raw\20210323\_F1\_UM1\_Peng0013\_SA\_F59\_ingel\_3ug\_ELA.raw

##### Fraction

1

##### Scan Feature

-

##### De Novo Score

99

##### ConfidenceScore

99

### m/z

484.2834

##### Mass

966.5498

##### Charge

2

##### Retention Time

16.56

##### Predicted Retention Time

-

##### Area

0

##### Parts Per Million

2.4

##### Fragmentation mode

ETHCD

##### Originating file

01 D:\separate\_stitch\_analyses\xle-disambiguation\20210325\_F59\_3ug\_DENOVO\_12.csv

#### Meta Information from PEAKS

##### Scan Identifier

F1:3487

##### Original sequence

V

L

G

Q

P

K

A

A

P

S

##### Posttranslational Modifications

##### Source File

D:\separate\_stitch\_analyses\xle-disambiguation\raw\20210323\_F1\_UM1\_Peng0013\_SA\_F59\_ingel\_3ug\_ELA.raw

##### Fraction

1

##### Scan Feature

-

##### De Novo Score

98

##### ConfidenceScore

98

### m/z

484.2832

##### Mass

966.5498

##### Charge

2

##### Retention Time

18.06

##### Predicted Retention Time

-

##### Area

0

##### Parts Per Million

2.2

##### Fragmentation mode

ETHCD

##### Originating file

01 D:\separate\_stitch\_analyses\xle-disambiguation\20210325\_F59\_3ug\_DENOVO\_12.csv

#### Meta Information from PEAKS

##### Scan Identifier

F1:2978

##### Original sequence

V

L

G

Q

P

K

A

A

P

S

##### Posttranslational Modifications

##### Source File

D:\separate\_stitch\_analyses\xle-disambiguation\raw\20210323\_F1\_UM1\_Peng0013\_SA\_F59\_ingel\_3ug\_ELA.raw

##### Fraction

1

##### Scan Feature

F1:3457

##### De Novo Score

98

##### ConfidenceScore

98

### m/z

484.2831

##### Mass

966.5498

##### Charge

2

##### Retention Time

15.18

##### Predicted Retention Time

-

##### Area

1.235E+09

##### Parts Per Million

1.8

##### Fragmentation mode

HCD

##### Originating file

01 D:\separate\_stitch\_analyses\xle-disambiguation\20210325\_F59\_3ug\_DENOVO\_12.csv

#### Meta Information from PEAKS

##### Scan Identifier

F1:3709

##### Original sequence

V

L

G

Q

P

K

A

A

P

S

##### Posttranslational Modifications

##### Source File

D:\separate\_stitch\_analyses\xle-disambiguation\raw\20210323\_F1\_UM1\_Peng0013\_SA\_F59\_ingel\_3ug\_ELA.raw

##### Fraction

1

##### Scan Feature

-

##### De Novo Score

98

##### ConfidenceScore

98

### m/z

484.2828

##### Mass

966.5498

##### Charge

2

##### Retention Time

19.32

##### Predicted Retention Time

-

##### Area

0

##### Parts Per Million

1.4

##### Fragmentation mode

ETHCD

##### Originating file

01 D:\separate\_stitch\_analyses\xle-disambiguation\20210325\_F59\_3ug\_DENOVO\_12.csv

#### Meta Information from PEAKS

##### Scan Identifier

F2:2991

##### Original sequence

V

L

G

Q

P

K

A

A

P

S

##### Posttranslational Modifications

##### Source File

D:\separate\_stitch\_analyses\xle-disambiguation\raw\20210323\_F1\_UM1\_Peng0013\_SA\_F59\_ingel\_3ug\_TL.raw

##### Fraction

2

##### Scan Feature

F2:3463

##### De Novo Score

98

##### ConfidenceScore

98

### m/z

484.2833

##### Mass

966.5498

##### Charge

2

##### Retention Time

15.07

##### Predicted Retention Time

-

##### Area

3.782E+08

##### Parts Per Million

2.2

##### Fragmentation mode

HCD

##### Originating file

01 D:\separate\_stitch\_analyses\xle-disambiguation\20210325\_F59\_3ug\_DENOVO\_12.csv

#### Meta Information from PEAKS

##### Scan Identifier

F2:2923

##### Original sequence

V

L

G

Q

P

K

A

A

P

S

##### Posttranslational Modifications

##### Source File

D:\separate\_stitch\_analyses\xle-disambiguation\raw\20210323\_F1\_UM1\_Peng0013\_SA\_F59\_ingel\_3ug\_TL.raw

##### Fraction

2

##### Scan Feature

F2:3463

##### De Novo Score

98

##### ConfidenceScore

98

### m/z

484.2833

##### Mass

966.5498

##### Charge

2

##### Retention Time

15.07

##### Predicted Retention Time

-

##### Area

3.782E+08

##### Parts Per Million

2.2

##### Fragmentation mode

HCD

##### Originating file

01 D:\separate\_stitch\_analyses\xle-disambiguation\20210325\_F59\_3ug\_DENOVO\_12.csv

#### Meta Information from PEAKS

##### Scan Identifier

F2:3209

##### Original sequence

V

L

G

Q

P

K

A

A

P

S

##### Posttranslational Modifications

##### Source File

D:\separate\_stitch\_analyses\xle-disambiguation\raw\20210323\_F1\_UM1\_Peng0013\_SA\_F59\_ingel\_3ug\_TL.raw

##### Fraction

2

##### Scan Feature

-

##### De Novo Score

98

##### ConfidenceScore

98

### m/z

484.2823

##### Mass

966.5498

##### Charge

2

##### Retention Time

16.76

##### Predicted Retention Time

-

##### Area

0

##### Parts Per Million

0.2

##### Fragmentation mode

ETHCD

##### Originating file

01 D:\separate\_stitch\_analyses\xle-disambiguation\20210325\_F59\_3ug\_DENOVO\_12.csv

#### Meta Information from PEAKS

##### Scan Identifier

F1:3420

##### Original sequence

V

L

G

Q

P

K

A

A

P

S

##### Posttranslational Modifications

##### Source File

D:\separate\_stitch\_analyses\xle-disambiguation\raw\20210323\_F1\_UM1\_Peng0013\_SA\_F59\_ingel\_3ug\_ELA.raw

##### Fraction

1

##### Scan Feature

-

##### De Novo Score

98

##### ConfidenceScore

98

### m/z

484.283

##### Mass

966.5498

##### Charge

2

##### Retention Time

17.69

##### Predicted Retention Time

-

##### Area

0

##### Parts Per Million

1.6

##### Fragmentation mode

HCD

##### Originating file

01 D:\separate\_stitch\_analyses\xle-disambiguation\20210325\_F59\_3ug\_DENOVO\_12.csv

#### Meta Information from PEAKS

##### Scan Identifier

F1:3554

##### Original sequence

V

L

G

Q

P

K

A

A

P

S

##### Posttranslational Modifications

##### Source File

D:\separate\_stitch\_analyses\xle-disambiguation\raw\20210323\_F1\_UM1\_Peng0013\_SA\_F59\_ingel\_3ug\_ELA.raw

##### Fraction

1

##### Scan Feature

-

##### De Novo Score

98

##### ConfidenceScore

98

### m/z

484.2832

##### Mass

966.5498

##### Charge

2

##### Retention Time

18.43

##### Predicted Retention Time

-

##### Area

0

##### Parts Per Million

2.2

##### Fragmentation mode

HCD

##### Originating file

01 D:\separate\_stitch\_analyses\xle-disambiguation\20210325\_F59\_3ug\_DENOVO\_12.csv

#### Meta Information from PEAKS

##### Scan Identifier

F2:3333

##### Original sequence

V

L

G

Q

P

K

A

A

P

S

##### Posttranslational Modifications

##### Source File

D:\separate\_stitch\_analyses\xle-disambiguation\raw\20210323\_F1\_UM1\_Peng0013\_SA\_F59\_ingel\_3ug\_TL.raw

##### Fraction

2

##### Scan Feature

-

##### De Novo Score

98

##### ConfidenceScore

98

### m/z

484.2825

##### Mass

966.5498

##### Charge

2

##### Retention Time

17.49

##### Predicted Retention Time

-

##### Area

0

##### Parts Per Million

0.7

##### Fragmentation mode

ETHCD

##### Originating file

01 D:\separate\_stitch\_analyses\xle-disambiguation\20210325\_F59\_3ug\_DENOVO\_12.csv

#### Meta Information from PEAKS

##### Scan Identifier

F1:4216

##### Original sequence

V

L

G

Q

P

K

A

A

P

S

##### Posttranslational Modifications

##### Source File

D:\separate\_stitch\_analyses\xle-disambiguation\raw\20210323\_F1\_UM1\_Peng0013\_SA\_F59\_ingel\_3ug\_ELA.raw

##### Fraction

1

##### Scan Feature

-

##### De Novo Score

98

##### ConfidenceScore

98

### m/z

484.2828

##### Mass

966.5498

##### Charge

2

##### Retention Time

22.21

##### Predicted Retention Time

-

##### Area

0

##### Parts Per Million

1.2

##### Fragmentation mode

ETHCD

##### Originating file

01 D:\separate\_stitch\_analyses\xle-disambiguation\20210325\_F59\_3ug\_DENOVO\_12.csv

#### Meta Information from PEAKS

##### Scan Identifier

F2:3083

##### Original sequence

V

L

G

Q

P

K

A

A

P

S

##### Posttranslational Modifications

##### Source File

D:\separate\_stitch\_analyses\xle-disambiguation\raw\20210323\_F1\_UM1\_Peng0013\_SA\_F59\_ingel\_3ug\_TL.raw

##### Fraction

2

##### Scan Feature

-

##### De Novo Score

97

##### ConfidenceScore

97

### m/z

484.2831

##### Mass

966.5498

##### Charge

2

##### Retention Time

16.04

##### Predicted Retention Time

-

##### Area

0

##### Parts Per Million

2

##### Fragmentation mode

HCD

##### Originating file

01 D:\separate\_stitch\_analyses\xle-disambiguation\20210325\_F59\_3ug\_DENOVO\_12.csv

#### Meta Information from PEAKS

##### Scan Identifier

F1:4307

##### Original sequence

V

L

G

Q

P

K

A

A

P

S

##### Posttranslational Modifications

##### Source File

D:\separate\_stitch\_analyses\xle-disambiguation\raw\20210323\_F1\_UM1\_Peng0013\_SA\_F59\_ingel\_3ug\_ELA.raw

##### Fraction

1

##### Scan Feature

-

##### De Novo Score

97

##### ConfidenceScore

97

### m/z

484.2827

##### Mass

966.5498

##### Charge

2

##### Retention Time

22.74

##### Predicted Retention Time

-

##### Area

0

##### Parts Per Million

1.1

##### Fragmentation mode

ETHCD

##### Originating file

01 D:\separate\_stitch\_analyses\xle-disambiguation\20210325\_F59\_3ug\_DENOVO\_12.csv

#### Meta Information from PEAKS

##### Scan Identifier

F1:4370

##### Original sequence

V

L

G

Q

P

K

A

A

P

S

##### Posttranslational Modifications

##### Source File

D:\separate\_stitch\_analyses\xle-disambiguation\raw\20210323\_F1\_UM1\_Peng0013\_SA\_F59\_ingel\_3ug\_ELA.raw

##### Fraction

1

##### Scan Feature

-

##### De Novo Score

97

##### ConfidenceScore

97

### m/z

484.2833

##### Mass

966.5498

##### Charge

2

##### Retention Time

23.1

##### Predicted Retention Time

-

##### Area

0

##### Parts Per Million

2.2

##### Fragmentation mode

ETHCD

##### Originating file

01 D:\separate\_stitch\_analyses\xle-disambiguation\20210325\_F59\_3ug\_DENOVO\_12.csv

#### Meta Information from PEAKS

##### Scan Identifier

F2:3145

##### Original sequence

V

L

G

Q

P

K

A

A

P

S

##### Posttranslational Modifications

##### Source File

D:\separate\_stitch\_analyses\xle-disambiguation\raw\20210323\_F1\_UM1\_Peng0013\_SA\_F59\_ingel\_3ug\_TL.raw

##### Fraction

2

##### Scan Feature

-

##### De Novo Score

96

##### ConfidenceScore

96

### m/z

484.2825

##### Mass

966.5498

##### Charge

2

##### Retention Time

16.4

##### Predicted Retention Time

-

##### Area

0

##### Parts Per Million

0.7

##### Fragmentation mode

HCD

##### Originating file

01 D:\separate\_stitch\_analyses\xle-disambiguation\20210325\_F59\_3ug\_DENOVO\_12.csv
