## Supplementary material for "A handle on mass coincidence errors in *de novo* sequencing of antibodies by bottom-up proteomics": Combined_036.html

Details Combined\_036 | Stitch OverviewUndefined

### Read Combined\_036

#### Sequence (length=6)

VJHQDW

#### Spectrum 4662? Spectrum 4662 The raw spectrum of this peptide as annotated by Hecklib. The fragments are coloured according to ion type (see legend). Any peaks with a star '\*' as text can be hovered over to see the full details, first the ion type second the mass shift type. By hovering over the amino acids in the peptide or ions in the legend the corresponding peaks are highlighted. By toggling the 'Unassigned' label you can turn the background (unassigned) peaks on or off in the plot. By updating the slider in the Ion legend you can update the spectrum to only show the top X% of the peaks with labels. The top X% means any peak that is within X% of the highest intensity. By dragging in the spectrum you can zoom in to a specific part of the spectrum and use 'Zoom Out' to get back to the original zoom level. The annotation of the spectrum is based on the given sequence in the peptides file and is done with different software so inconsistencies are likely. The peaks are annotated based on the given sequence, with 20 ppm tolerance.

Copy Data

##### Spectrum 4662 (TSV)

###### Preview

```
Loading example...
```

*Click on the button to copy the data to your clipboard.*

Mz MinMz MaxIntensity Max

WidthHeightPeptide font sizePeptide stroke widthSpectrum font sizeSpectrum stroke widthCompact peptide

Ion legend

wxyz

abcd

OtherUnassignedIon^Charge^_Position_Show for top:%

VJHQDW

01.43e+52.87e+54.30e+55.74e+5

Zoom Out

a^+1^_2_y^+1^_1_b^+1^_2_b^+2^_4_y^+2^_4_b^+2^_5_y^+2^_4_b^+2^_5_y^+1^_2_y^+1^_2_a^+1^_3_y^+2^_5_y^+2^_5_y^+2^_5_b^+1^_3_\*\*\*y^+1^_3_y^+1^_3_y^+1^_3_b^+1^_4_b^+1^_4_y^+1^_4_y^+1^_4_b^+1^_5_b^+1^_5_y^+1^_4_b^+1^_5_y^+1^_5_y^+1^_5_y^+1^_5_

0791158223743165

Fragment Matches Table

Show background peaks

| Position | Ion type | Intensity | mz Theoretical | mz Error (Th) | mz Error (ppm) | Charge | Series Number |
| --- | --- | --- | --- | --- | --- | --- | --- |
| - | - | 1195 | 120.1 | - | - | 0 | - |
| - | - | 1695 | 120.1 | - | - | 0 | - |
| - | - | 343.2 | 120.7 | - | - | 0 | - |
| - | - | 2646 | 121 | - | - | 0 | - |
| - | - | 6443 | 122.1 | - | - | 0 | - |
| - | - | 4717 | 123.1 | - | - | 0 | - |
| - | - | 848 | 123.1 | - | - | 0 | - |
| - | - | 601.4 | 124.1 | - | - | 0 | - |
| - | - | 494.4 | 125.1 | - | - | 0 | - |
| - | - | 1760 | 125.1 | - | - | 0 | - |
| - | - | 795.2 | 127.1 | - | - | 0 | - |
| - | - | 1384 | 128.1 | - | - | 0 | - |
| - | - | 1589 | 129.1 | - | - | 0 | - |
| - | - | 4214 | 129.1 | - | - | 0 | - |
| - | - | 2341 | 129.1 | - | - | 0 | - |
| - | - | 1.972E+05 | 130.1 | - | - | 0 | - |
| - | - | 1.847E+04 | 131.1 | - | - | 0 | - |
| - | - | 507.2 | 132 | - | - | 0 | - |
| - | - | 5.873E+04 | 132.1 | - | - | 0 | - |
| - | - | 3410 | 133.1 | - | - | 0 | - |
| - | - | 4795 | 133.1 | - | - | 0 | - |
| - | - | 522.3 | 134 | - | - | 0 | - |
| - | - | 585.5 | 136 | - | - | 0 | - |
| - | - | 2991 | 136.1 | - | - | 0 | - |
| - | - | 1.746E+04 | 138.1 | - | - | 0 | - |
| - | - | 521.8 | 138.1 | - | - | 0 | - |
| - | - | 2149 | 139.1 | - | - | 0 | - |
| - | - | 1066 | 139.1 | - | - | 0 | - |
| - | - | 481.2 | 139.1 | - | - | 0 | - |
| - | - | 699.4 | 140.1 | - | - | 0 | - |
| - | - | 451.3 | 140.1 | - | - | 0 | - |
| - | - | 805.4 | 140.1 | - | - | 0 | - |
| - | - | 1967 | 141.1 | - | - | 0 | - |
| - | - | 9516 | 142.1 | - | - | 0 | - |
| - | - | 6223 | 143 | - | - | 0 | - |
| - | - | 629 | 143.1 | - | - | 0 | - |
| - | - | 2171 | 143.1 | - | - | 0 | - |
| - | - | 606.2 | 144 | - | - | 0 | - |
| - | - | 2.08E+04 | 144.1 | - | - | 0 | - |
| - | - | 490.9 | 145 | - | - | 0 | - |
| - | - | 2088 | 145.1 | - | - | 0 | - |
| - | - | 1.064E+05 | 146.1 | - | - | 0 | - |
| - | - | 9369 | 147.1 | - | - | 0 | - |
| - | - | 705.7 | 148.1 | - | - | 0 | - |
| - | - | 565 | 152.1 | - | - | 0 | - |
| - | - | 1052 | 153.1 | - | - | 0 | - |
| - | - | 1827 | 154.1 | - | - | 0 | - |
| - | - | 4960 | 154.1 | - | - | 0 | - |
| - | - | 4288 | 155.1 | - | - | 0 | - |
| - | - | 8.779E+04 | 155.1 | - | - | 0 | - |
| - | - | 3822 | 156.1 | - | - | 0 | - |
| - | - | 1158 | 156.1 | - | - | 0 | - |
| - | - | 4835 | 156.1 | - | - | 0 | - |
| - | - | 541.6 | 156.1 | - | - | 0 | - |
| - | - | 683.2 | 158.1 | - | - | 0 | - |
| - | - | 8103 | 158.1 | - | - | 0 | - |
| - | - | 4.414E+05 | 159.1 | - | - | 0 | - |
| - | - | 7345 | 160.1 | - | - | 0 | - |
| - | - | 3041 | 160.1 | - | - | 0 | - |
| - | - | 4.616E+04 | 160.1 | - | - | 0 | - |
| - | - | 468.7 | 161.1 | - | - | 0 | - |
| - | - | 1721 | 161.1 | - | - | 0 | - |
| - | - | 568.2 | 164.1 | - | - | 0 | - |
| - | - | 2867 | 164.1 | - | - | 0 | - |
| - | - | 3536 | 165.1 | - | - | 0 | - |
| - | - | 7.694E+04 | 166.1 | - | - | 0 | - |
| - | - | 760.4 | 167.1 | - | - | 0 | - |
| - | - | 5657 | 167.1 | - | - | 0 | - |
| - | - | 888.6 | 167.1 | - | - | 0 | - |
| - | - | 9358 | 169.1 | - | - | 0 | - |
| - | - | 3.603E+04 | 170.1 | - | - | 0 | - |
| - | - | 3949 | 171.1 | - | - | 0 | - |
| - | - | 738.4 | 171.1 | - | - | 0 | - |
| - | - | 730 | 171.1 | - | - | 0 | - |
| - | - | 1630 | 172.1 | - | - | 0 | - |
| - | - | 449.4 | 173.3 | - | - | 0 | - |
| - | - | 1243 | 175 | - | - | 0 | - |
| - | - | 4287 | 175.1 | - | - | 0 | - |
| - | - | 4399 | 176.1 | - | - | 0 | - |
| - | - | 3.928E+04 | 178.1 | - | - | 0 | - |
| - | - | 3477 | 179.1 | - | - | 0 | - |
| - | - | 928 | 180.1 | - | - | 0 | - |
| - | - | 1.527E+04 | 181.1 | - | - | 0 | - |
| - | - | 1213 | 182 | - | - | 0 | - |
| - | - | 1709 | 183.1 | - | - | 0 | - |
| - | - | 748.3 | 183.1 | - | - | 0 | - |
| - | - | 599.2 | 183.2 | - | - | 0 | - |
| - | - | 957.5 | 184.1 | - | - | 0 | - |
| 2 | a | 1.346E+05 | 185.2 | 0.0005503 | 2.972 | +1 | 2 |
| - | - | 1.492E+04 | 186.2 | - | - | 0 | - |
| - | - | 2.388E+04 | 187.1 | - | - | 0 | - |
| - | - | 1136 | 187.1 | - | - | 0 | - |
| - | - | 857.8 | 187.2 | - | - | 0 | - |
| - | - | 5.682E+05 | 188.1 | - | - | 0 | - |
| - | - | 6.577E+04 | 189.1 | - | - | 0 | - |
| - | - | 4324 | 190.1 | - | - | 0 | - |
| - | - | 8770 | 190.1 | - | - | 0 | - |
| - | - | 1050 | 191.1 | - | - | 0 | - |
| - | - | 959.9 | 194.1 | - | - | 0 | - |
| - | - | 1273 | 195.1 | - | - | 0 | - |
| - | - | 1412 | 195.1 | - | - | 0 | - |
| - | - | 770.8 | 196.1 | - | - | 0 | - |
| - | - | 663.6 | 196.1 | - | - | 0 | - |
| - | - | 786.8 | 197.1 | - | - | 0 | - |
| - | - | 605.9 | 197.1 | - | - | 0 | - |
| - | - | 562.2 | 197.1 | - | - | 0 | - |
| - | - | 1.702E+04 | 198.1 | - | - | 0 | - |
| - | - | 7489 | 199.1 | - | - | 0 | - |
| - | - | 792.3 | 199.1 | - | - | 0 | - |
| - | - | 734.7 | 200.1 | - | - | 0 | - |
| - | - | 856.1 | 200.1 | - | - | 0 | - |
| - | - | 789.9 | 202.1 | - | - | 0 | - |
| - | - | 1921 | 203.1 | - | - | 0 | - |
| - | - | 1448 | 204.1 | - | - | 0 | - |
| - | - | 966.9 | 204.1 | - | - | 0 | - |
| 6 | y | 1.642E+05 | 205.1 | 0.0005479 | 2.672 | +1 | 1 |
| - | - | 4354 | 205.1 | - | - | 0 | - |
| - | - | 1.759E+04 | 206.1 | - | - | 0 | - |
| - | - | 4297 | 206.1 | - | - | 0 | - |
| - | - | 1066 | 207.1 | - | - | 0 | - |
| - | - | 723.7 | 207.1 | - | - | 0 | - |
| - | - | 5043 | 207.2 | - | - | 0 | - |
| - | - | 2302 | 209.1 | - | - | 0 | - |
| - | - | 828.5 | 211.1 | - | - | 0 | - |
| - | - | 2126 | 212.2 | - | - | 0 | - |
| - | - | 4745 | 213.1 | - | - | 0 | - |
| 2 | b | 2.622E+04 | 213.2 | 0.000524 | 2.458 | +1 | 2 |
| - | - | 535.4 | 214.1 | - | - | 0 | - |
| - | - | 620.3 | 214.1 | - | - | 0 | - |
| - | - | 3405 | 214.2 | - | - | 0 | - |
| - | - | 1771 | 215.1 | - | - | 0 | - |
| - | - | 8752 | 216.1 | - | - | 0 | - |
| - | - | 653.6 | 217.1 | - | - | 0 | - |
| - | - | 1190 | 217.1 | - | - | 0 | - |
| - | - | 5831 | 220.1 | - | - | 0 | - |
| - | - | 3339 | 221.1 | - | - | 0 | - |
| - | - | 1.728E+04 | 221.1 | - | - | 0 | - |
| - | - | 1140 | 221.1 | - | - | 0 | - |
| - | - | 811.1 | 222.1 | - | - | 0 | - |
| - | - | 1168 | 223.1 | - | - | 0 | - |
| - | - | 5.547E+04 | 223.2 | - | - | 0 | - |
| - | - | 587.7 | 224.1 | - | - | 0 | - |
| - | - | 5474 | 224.2 | - | - | 0 | - |
| - | - | 1620 | 225.6 | - | - | 0 | - |
| - | - | 2.707E+04 | 226.1 | - | - | 0 | - |
| - | - | 614 | 226.2 | - | - | 0 | - |
| - | - | 2.131E+04 | 227.1 | - | - | 0 | - |
| - | - | 2588 | 227.1 | - | - | 0 | - |
| - | - | 1023 | 227.2 | - | - | 0 | - |
| - | - | 1649 | 228.1 | - | - | 0 | - |
| - | - | 697 | 229.1 | - | - | 0 | - |
| - | - | 616.8 | 229.6 | - | - | 0 | - |
| - | - | 1210 | 230.1 | - | - | 0 | - |
| - | - | 2916 | 231.1 | - | - | 0 | - |
| - | - | 7560 | 232.1 | - | - | 0 | - |
| - | - | 2409 | 232.6 | - | - | 0 | - |
| - | - | 1254 | 233.1 | - | - | 0 | - |
| - | - | 2.456E+04 | 233.1 | - | - | 0 | - |
| - | - | 8.51E+04 | 234.1 | - | - | 0 | - |
| - | - | 2415 | 234.1 | - | - | 0 | - |
| - | - | 9820 | 235.1 | - | - | 0 | - |
| - | - | 2249 | 235.2 | - | - | 0 | - |
| - | - | 649.7 | 238.1 | - | - | 0 | - |
| - | - | 5074 | 238.1 | - | - | 0 | - |
| - | - | 1987 | 238.6 | - | - | 0 | - |
| - | - | 5532 | 239.1 | - | - | 0 | - |
| - | - | 742 | 239.1 | - | - | 0 | - |
| 4 | b | 2194 | 239.6 | 0.0004401 | 1.837 | +2 | 4 |
| - | - | 1073 | 240.1 | - | - | 0 | - |
| - | - | 1302 | 241.1 | - | - | 0 | - |
| - | - | 990.1 | 241.1 | - | - | 0 | - |
| - | - | 828.2 | 242.2 | - | - | 0 | - |
| - | - | 5415 | 243.1 | - | - | 0 | - |
| - | - | 4832 | 243.1 | - | - | 0 | - |
| - | - | 986.6 | 243.6 | - | - | 0 | - |
| - | - | 4.54E+04 | 244.1 | - | - | 0 | - |
| - | - | 2420 | 245.1 | - | - | 0 | - |
| - | - | 3091 | 245.1 | - | - | 0 | - |
| - | - | 4843 | 246.1 | - | - | 0 | - |
| - | - | 712.4 | 247.1 | - | - | 0 | - |
| - | - | 3.266E+04 | 248.1 | - | - | 0 | - |
| - | - | 5.962E+04 | 249.1 | - | - | 0 | - |
| - | - | 2715 | 249.1 | - | - | 0 | - |
| - | - | 1392 | 249.1 | - | - | 0 | - |
| - | - | 2497 | 249.2 | - | - | 0 | - |
| - | - | 8325 | 250.1 | - | - | 0 | - |
| - | - | 1607 | 250.2 | - | - | 0 | - |
| - | - | 2.752E+05 | 251.2 | - | - | 0 | - |
| - | - | 3.629E+04 | 252.2 | - | - | 0 | - |
| - | - | 604.7 | 252.2 | - | - | 0 | - |
| - | - | 1.117E+04 | 252.6 | - | - | 0 | - |
| - | - | 1639 | 252.7 | - | - | 0 | - |
| - | - | 3606 | 253.1 | - | - | 0 | - |
| - | - | 2142 | 253.2 | - | - | 0 | - |
| - | - | 1178 | 254.1 | - | - | 0 | - |
| - | - | 4014 | 254.7 | - | - | 0 | - |
| - | - | 986.1 | 255.2 | - | - | 0 | - |
| - | - | 616.1 | 255.2 | - | - | 0 | - |
| - | - | 5418 | 256.1 | - | - | 0 | - |
| - | - | 1519 | 257.1 | - | - | 0 | - |
| - | - | 1041 | 257.1 | - | - | 0 | - |
| - | - | 921 | 258.1 | - | - | 0 | - |
| - | - | 2261 | 258.1 | - | - | 0 | - |
| - | - | 5597 | 260.1 | - | - | 0 | - |
| - | - | 8044 | 261.1 | - | - | 0 | - |
| - | - | 7192 | 261.1 | - | - | 0 | - |
| - | - | 6066 | 261.2 | - | - | 0 | - |
| - | - | 7355 | 261.6 | - | - | 0 | - |
| - | - | 854.4 | 261.7 | - | - | 0 | - |
| - | - | 1394 | 262.1 | - | - | 0 | - |
| - | - | 2004 | 262.1 | - | - | 0 | - |
| - | - | 1136 | 262.1 | - | - | 0 | - |
| - | - | 4673 | 265.1 | - | - | 0 | - |
| - | - | 600 | 265.6 | - | - | 0 | - |
| - | - | 1.501E+05 | 266.1 | - | - | 0 | - |
| - | - | 1677 | 267.1 | - | - | 0 | - |
| - | - | 2532 | 267.1 | - | - | 0 | - |
| - | - | 1.589E+04 | 267.1 | - | - | 0 | - |
| - | - | 1351 | 268.1 | - | - | 0 | - |
| - | - | 4102 | 268.2 | - | - | 0 | - |
| - | - | 8.03E+04 | 270.1 | - | - | 0 | - |
| - | - | 2.214E+04 | 270.6 | - | - | 0 | - |
| - | - | 3573 | 271.1 | - | - | 0 | - |
| - | - | 9729 | 274.1 | - | - | 0 | - |
| - | - | 2.653E+04 | 274.1 | - | - | 0 | - |
| - | - | 3130 | 274.6 | - | - | 0 | - |
| - | - | 1279 | 275.1 | - | - | 0 | - |
| - | - | 3564 | 275.1 | - | - | 0 | - |
| - | - | 743.1 | 275.1 | - | - | 0 | - |
| - | - | 598.1 | 275.6 | - | - | 0 | - |
| - | - | 5271 | 276.1 | - | - | 0 | - |
| - | - | 2240 | 277.1 | - | - | 0 | - |
| - | - | 588.1 | 277.1 | - | - | 0 | - |
| - | - | 1939 | 277.2 | - | - | 0 | - |
| - | - | 1430 | 277.2 | - | - | 0 | - |
| - | - | 1537 | 278.2 | - | - | 0 | - |
| - | - | 1.026E+04 | 279.1 | - | - | 0 | - |
| - | - | 737.4 | 279.2 | - | - | 0 | - |
| - | - | 1351 | 280.1 | - | - | 0 | - |
| - | - | 1948 | 283.2 | - | - | 0 | - |
| - | - | 658.4 | 283.7 | - | - | 0 | - |
| - | - | 1252 | 284.1 | - | - | 0 | - |
| - | - | 8317 | 284.1 | - | - | 0 | - |
| 3 | y | 5410 | 284.1 | 0.0006629 | 2.333 | +2 | 4 |
| - | - | 1565 | 284.1 | - | - | 0 | - |
| - | - | 2484 | 284.6 | - | - | 0 | - |
| - | - | 9449 | 285.1 | - | - | 0 | - |
| - | - | 1450 | 285.1 | - | - | 0 | - |
| - | - | 2341 | 286.1 | - | - | 0 | - |
| - | - | 2645 | 287.2 | - | - | 0 | - |
| 5 | b | 3405 | 288.6 | 0.0006399 | 2.217 | +2 | 5 |
| - | - | 705.8 | 289.1 | - | - | 0 | - |
| - | - | 1337 | 289.2 | - | - | 0 | - |
| - | - | 1412 | 292.1 | - | - | 0 | - |
| 3 | y | 2134 | 293.1 | 0.0008432 | 2.877 | +2 | 4 |
| - | - | 686.4 | 293.6 | - | - | 0 | - |
| - | - | 5524 | 294.1 | - | - | 0 | - |
| - | - | 761 | 295.1 | - | - | 0 | - |
| - | - | 1760 | 296.2 | - | - | 0 | - |
| - | - | 689.2 | 296.7 | - | - | 0 | - |
| 5 | b | 2.681E+04 | 297.2 | 0.000671 | 2.258 | +2 | 5 |
| - | - | 7563 | 297.7 | - | - | 0 | - |
| - | - | 1119 | 298.1 | - | - | 0 | - |
| - | - | 1964 | 298.2 | - | - | 0 | - |
| - | - | 951.6 | 299.1 | - | - | 0 | - |
| - | - | 2728 | 300.1 | - | - | 0 | - |
| - | - | 6062 | 301.1 | - | - | 0 | - |
| - | - | 3969 | 301.1 | - | - | 0 | - |
| 5 | y | 1.123E+04 | 302.1 | 0.0007254 | 2.401 | +1 | 2 |
| - | - | 4505 | 302.1 | - | - | 0 | - |
| - | - | 790 | 302.1 | - | - | 0 | - |
| - | - | 600.5 | 302.2 | - | - | 0 | - |
| - | - | 1808 | 303.1 | - | - | 0 | - |
| - | - | 788.8 | 303.1 | - | - | 0 | - |
| - | - | 963.2 | 304.1 | - | - | 0 | - |
| - | - | 3958 | 304.2 | - | - | 0 | - |
| - | - | 1393 | 305.2 | - | - | 0 | - |
| - | - | 753.7 | 306.2 | - | - | 0 | - |
| - | - | 1.335E+04 | 306.2 | - | - | 0 | - |
| - | - | 2150 | 307.2 | - | - | 0 | - |
| - | - | 862.8 | 309.1 | - | - | 0 | - |
| - | - | 2193 | 309.2 | - | - | 0 | - |
| - | - | 5411 | 310.2 | - | - | 0 | - |
| - | - | 1630 | 310.7 | - | - | 0 | - |
| - | - | 1319 | 310.7 | - | - | 0 | - |
| - | - | 784.5 | 315.2 | - | - | 0 | - |
| - | - | 1681 | 316.2 | - | - | 0 | - |
| - | - | 1038 | 317.1 | - | - | 0 | - |
| - | - | 879.4 | 317.2 | - | - | 0 | - |
| - | - | 642.7 | 317.2 | - | - | 0 | - |
| - | - | 1105 | 317.7 | - | - | 0 | - |
| - | - | 4312 | 318.1 | - | - | 0 | - |
| - | - | 4170 | 318.2 | - | - | 0 | - |
| - | - | 1776 | 318.7 | - | - | 0 | - |
| - | - | 1113 | 319.1 | - | - | 0 | - |
| - | - | 3093 | 319.1 | - | - | 0 | - |
| - | - | 6183 | 319.2 | - | - | 0 | - |
| 5 | y | 4.01E+04 | 320.1 | 0.0007808 | 2.439 | +1 | 2 |
| - | - | 6276 | 321.1 | - | - | 0 | - |
| - | - | 896.8 | 322.1 | - | - | 0 | - |
| 3 | a | 2771 | 322.2 | 0.0003695 | 1.147 | +1 | 3 |
| - | - | 675.7 | 323.1 | - | - | 0 | - |
| - | - | 1835 | 323.2 | - | - | 0 | - |
| - | - | 3928 | 326.1 | - | - | 0 | - |
| - | - | 702 | 326.2 | - | - | 0 | - |
| - | - | 2355 | 326.2 | - | - | 0 | - |
| - | - | 2.581E+04 | 326.7 | - | - | 0 | - |
| - | - | 9963 | 327.2 | - | - | 0 | - |
| - | - | 815.7 | 327.2 | - | - | 0 | - |
| - | - | 2220 | 327.7 | - | - | 0 | - |
| - | - | 1783 | 328.1 | - | - | 0 | - |
| - | - | 1529 | 331.7 | - | - | 0 | - |
| - | - | 1185 | 332.1 | - | - | 0 | - |
| - | - | 3.763E+04 | 332.2 | - | - | 0 | - |
| - | - | 814.5 | 333.2 | - | - | 0 | - |
| - | - | 7327 | 333.2 | - | - | 0 | - |
| - | - | 816.3 | 335.1 | - | - | 0 | - |
| - | - | 2747 | 336.1 | - | - | 0 | - |
| - | - | 1454 | 337.2 | - | - | 0 | - |
| 2 | y | 2534 | 340.7 | 0.001722 | 5.054 | +2 | 5 |
| 2 | y | 1124 | 341.2 | 0.003977 | 11.66 | +2 | 5 |
| - | - | 1090 | 343.2 | - | - | 0 | - |
| - | - | 6587 | 344.2 | - | - | 0 | - |
| - | - | 896.3 | 344.7 | - | - | 0 | - |
| - | - | 1.816E+04 | 345.1 | - | - | 0 | - |
| - | - | 980.8 | 345.2 | - | - | 0 | - |
| - | - | 636.3 | 345.2 | - | - | 0 | - |
| - | - | 6.777E+04 | 346.1 | - | - | 0 | - |
| - | - | 811.2 | 347.1 | - | - | 0 | - |
| - | - | 1.066E+04 | 347.1 | - | - | 0 | - |
| - | - | 1796 | 348.1 | - | - | 0 | - |
| - | - | 902.2 | 348.2 | - | - | 0 | - |
| - | - | 1430 | 349.1 | - | - | 0 | - |
| 2 | y | 1553 | 349.7 | 0.001414 | 4.043 | +2 | 5 |
| 3 | b | 1.599E+05 | 350.2 | 0.0006941 | 1.982 | +1 | 3 |
| - | - | 757.2 | 351.1 | - | - | 0 | - |
| - | - | 3.033E+04 | 351.2 | - | - | 0 | - |
| - | - | 3732 | 352.2 | - | - | 0 | - |
| - | - | 2092 | 353.7 | - | - | 0 | - |
| - | - | 1003 | 354.2 | - | - | 0 | - |
| - | - | 2855 | 354.2 | - | - | 0 | - |
| - | - | 887.9 | 355.2 | - | - | 0 | - |
| - | - | 7599 | 358.7 | - | - | 0 | - |
| - | - | 3668 | 359.2 | - | - | 0 | - |
| - | - | 1865 | 359.7 | - | - | 0 | - |
| - | - | 1213 | 360.2 | - | - | 0 | - |
| - | - | 7032 | 361.2 | - | - | 0 | - |
| - | - | 8511 | 362.2 | - | - | 0 | - |
| - | - | 724.5 | 362.2 | - | - | 0 | - |
| - | - | 7.029E+04 | 363.1 | - | - | 0 | - |
| - | - | 1996 | 363.2 | - | - | 0 | - |
| - | - | 3.389E+04 | 364.1 | - | - | 0 | - |
| - | - | 7886 | 364.1 | - | - | 0 | - |
| - | - | 6344 | 365.1 | - | - | 0 | - |
| - | - | 669 | 366.1 | - | - | 0 | - |
| - | - | 1699 | 366.2 | - | - | 0 | - |
| - | - | 7210 | 367.1 | - | - | 0 | - |
| - | - | 4905 | 367.2 | - | - | 0 | - |
| - | - | 2.292E+04 | 367.2 | - | - | 0 | - |
| - | - | 4.12E+04 | 367.7 | - | - | 0 | - |
| - | - | 1137 | 368.1 | - | - | 0 | - |
| - | - | 1.782E+04 | 368.2 | - | - | 0 | - |
| - | - | 4233 | 368.2 | - | - | 0 | - |
| - | - | 5131 | 368.7 | - | - | 0 | - |
| - | - | 638.7 | 369.2 | - | - | 0 | - |
| - | - | 1751 | 372.7 | - | - | 0 | - |
| - | - | 685.9 | 374.1 | - | - | 0 | - |
| - | - | 9.609E+04 | 376.2 | - | - | 0 | - |
| - | - | 4.535E+04 | 376.7 | - | - | 0 | - |
| - | - | 1.18E+04 | 377.2 | - | - | 0 | - |
| - | - | 925.7 | 377.7 | - | - | 0 | - |
| - | - | 7.905E+04 | 379.2 | - | - | 0 | - |
| - | - | 910.1 | 380.2 | - | - | 0 | - |
| - | - | 739.1 | 380.2 | - | - | 0 | - |
| - | - | 1.578E+04 | 380.2 | - | - | 0 | - |
| - | - | 2.025E+05 | 381.2 | - | - | 0 | - |
| - | - | 3907 | 381.7 | - | - | 0 | - |
| - | - | 3.446E+04 | 382.2 | - | - | 0 | - |
| - | - | 1017 | 382.2 | - | - | 0 | - |
| - | - | 5303 | 383.2 | - | - | 0 | - |
| - | - | 4152 | 384.2 | - | - | 0 | - |
| - | - | 9790 | 385.2 | - | - | 0 | - |
| - | - | 1794 | 386.2 | - | - | 0 | - |
| - | - | 749.7 | 386.2 | - | - | 0 | - |
| - | - | 869.9 | 388.2 | - | - | 0 | - |
| - | - | 3035 | 389.2 | - | - | 0 | - |
| - | - | 762 | 390.1 | - | - | 0 | - |
| 0 | Precursor | 1.411E+04 | 390.2 | 0.0006568 | 1.683 | +2 | -1 |
| 0 | Precursor | 1.365E+04 | 390.7 | 0.002423 | 6.203 | +2 | -1 |
| - | - | 4450 | 391.1 | - | - | 0 | - |
| - | - | 4037 | 391.2 | - | - | 0 | - |
| - | - | 1271 | 391.7 | - | - | 0 | - |
| - | - | 2488 | 392.1 | - | - | 0 | - |
| - | - | 888.9 | 393.2 | - | - | 0 | - |
| - | - | 1386 | 394.2 | - | - | 0 | - |
| - | - | 6610 | 395.1 | - | - | 0 | - |
| - | - | 1883 | 396.1 | - | - | 0 | - |
| - | - | 8578 | 398.2 | - | - | 0 | - |
| - | - | 5540 | 398.2 | - | - | 0 | - |
| - | - | 8943 | 399.2 | - | - | 0 | - |
| 0 | Precursor | 3.979E+04 | 399.2 | 0.0008677 | 2.173 | +2 | -1 |
| - | - | 1.913E+04 | 399.7 | - | - | 0 | - |
| - | - | 1221 | 400.2 | - | - | 0 | - |
| - | - | 6056 | 400.2 | - | - | 0 | - |
| - | - | 9608 | 402.2 | - | - | 0 | - |
| - | - | 1222 | 403.1 | - | - | 0 | - |
| - | - | 1564 | 403.2 | - | - | 0 | - |
| - | - | 2196 | 405.2 | - | - | 0 | - |
| - | - | 9711 | 407.2 | - | - | 0 | - |
| - | - | 1870 | 408.2 | - | - | 0 | - |
| - | - | 8106 | 409.1 | - | - | 0 | - |
| - | - | 1455 | 410.1 | - | - | 0 | - |
| - | - | 5828 | 412.2 | - | - | 0 | - |
| - | - | 6061 | 413.1 | - | - | 0 | - |
| - | - | 1245 | 414.1 | - | - | 0 | - |
| - | - | 3575 | 416.2 | - | - | 0 | - |
| - | - | 1482 | 417.2 | - | - | 0 | - |
| - | - | 7755 | 421.1 | - | - | 0 | - |
| - | - | 1884 | 422.2 | - | - | 0 | - |
| - | - | 691.5 | 422.3 | - | - | 0 | - |
| 4 | y | 5.474E+04 | 430.2 | 0.0008332 | 1.937 | +1 | 3 |
| 4 | y | 7544 | 431.2 | 0.0008216 | 1.906 | +1 | 3 |
| - | - | 7392 | 431.2 | - | - | 0 | - |
| - | - | 1459 | 432.2 | - | - | 0 | - |
| - | - | 994.4 | 432.2 | - | - | 0 | - |
| - | - | 2975 | 433.3 | - | - | 0 | - |
| - | - | 1754 | 435.2 | - | - | 0 | - |
| - | - | 2328 | 438.2 | - | - | 0 | - |
| - | - | 2384 | 439.2 | - | - | 0 | - |
| - | - | 683.6 | 441.2 | - | - | 0 | - |
| - | - | 1119 | 443.2 | - | - | 0 | - |
| 4 | y | 3.697E+04 | 448.2 | 0.0008276 | 1.846 | +1 | 3 |
| - | - | 9551 | 449.2 | - | - | 0 | - |
| - | - | 1389 | 450.2 | - | - | 0 | - |
| - | - | 5567 | 450.3 | - | - | 0 | - |
| - | - | 1135 | 451.3 | - | - | 0 | - |
| - | - | 1.204E+04 | 456.2 | - | - | 0 | - |
| - | - | 2634 | 457.2 | - | - | 0 | - |
| - | - | 4555 | 459.2 | - | - | 0 | - |
| - | - | 1192 | 460.3 | - | - | 0 | - |
| 4 | b | 8182 | 461.3 | 0.0008923 | 1.935 | +1 | 4 |
| - | - | 1964 | 462.3 | - | - | 0 | - |
| - | - | 2153 | 463.2 | - | - | 0 | - |
| - | - | 836.9 | 465.2 | - | - | 0 | - |
| - | - | 767.8 | 468.2 | - | - | 0 | - |
| - | - | 3285 | 476.2 | - | - | 0 | - |
| - | - | 1.478E+04 | 477.2 | - | - | 0 | - |
| - | - | 3472 | 478.2 | - | - | 0 | - |
| 4 | b | 3.634E+04 | 478.3 | 0.000985 | 2.06 | +1 | 4 |
| - | - | 752.6 | 479.2 | - | - | 0 | - |
| - | - | 8504 | 479.3 | - | - | 0 | - |
| - | - | 1190 | 480.2 | - | - | 0 | - |
| - | - | 1489 | 480.3 | - | - | 0 | - |
| - | - | 765.5 | 484.2 | - | - | 0 | - |
| - | - | 1619 | 486.2 | - | - | 0 | - |
| - | - | 1157 | 487.2 | - | - | 0 | - |
| - | - | 5400 | 491.3 | - | - | 0 | - |
| - | - | 764.5 | 492.3 | - | - | 0 | - |
| - | - | 9.726E+04 | 494.2 | - | - | 0 | - |
| - | - | 2.274E+04 | 495.2 | - | - | 0 | - |
| - | - | 3994 | 496.2 | - | - | 0 | - |
| - | - | 959.4 | 496.3 | - | - | 0 | - |
| - | - | 6696 | 504.2 | - | - | 0 | - |
| - | - | 1862 | 504.3 | - | - | 0 | - |
| - | - | 3075 | 505.2 | - | - | 0 | - |
| - | - | 805.2 | 505.2 | - | - | 0 | - |
| - | - | 1150 | 506.2 | - | - | 0 | - |
| - | - | 1406 | 511.3 | - | - | 0 | - |
| - | - | 3773 | 512.2 | - | - | 0 | - |
| - | - | 1864 | 514.2 | - | - | 0 | - |
| - | - | 3553 | 521.2 | - | - | 0 | - |
| - | - | 2257 | 521.3 | - | - | 0 | - |
| - | - | 8421 | 522.2 | - | - | 0 | - |
| - | - | 925.3 | 522.3 | - | - | 0 | - |
| - | - | 5302 | 523.2 | - | - | 0 | - |
| - | - | 2563 | 524.2 | - | - | 0 | - |
| - | - | 919 | 530.2 | - | - | 0 | - |
| - | - | 4896 | 532.2 | - | - | 0 | - |
| - | - | 1406 | 533.2 | - | - | 0 | - |
| - | - | 3.176E+04 | 539.2 | - | - | 0 | - |
| - | - | 9342 | 540.2 | - | - | 0 | - |
| - | - | 1753 | 541.2 | - | - | 0 | - |
| - | - | 1817 | 547.3 | - | - | 0 | - |
| - | - | 4482 | 548.3 | - | - | 0 | - |
| - | - | 6010 | 549.2 | - | - | 0 | - |
| - | - | 1154 | 549.3 | - | - | 0 | - |
| - | - | 3.668E+04 | 550.2 | - | - | 0 | - |
| - | - | 1.11E+04 | 551.2 | - | - | 0 | - |
| - | - | 707.7 | 551.3 | - | - | 0 | - |
| - | - | 1320 | 552.2 | - | - | 0 | - |
| - | - | 726.4 | 552.2 | - | - | 0 | - |
| - | - | 6788 | 558.3 | - | - | 0 | - |
| - | - | 2112 | 559.3 | - | - | 0 | - |
| - | - | 974.2 | 562.3 | - | - | 0 | - |
| - | - | 651.9 | 563.3 | - | - | 0 | - |
| - | - | 3595 | 565.3 | - | - | 0 | - |
| - | - | 672.2 | 566.3 | - | - | 0 | - |
| 3 | y | 1.432E+04 | 567.2 | 0.0005456 | 0.9618 | +1 | 4 |
| 3 | y | 1.387E+04 | 568.2 | 0.003102 | 5.46 | +1 | 4 |
| - | - | 4455 | 569.2 | - | - | 0 | - |
| - | - | 1054 | 570.2 | - | - | 0 | - |
| 5 | b | 6257 | 575.3 | 0.0004453 | 0.7741 | +1 | 5 |
| 5 | b | 4.303E+04 | 576.3 | 0.0008658 | 1.502 | +1 | 5 |
| - | - | 1.277E+04 | 577.3 | - | - | 0 | - |
| - | - | 699.5 | 578.2 | - | - | 0 | - |
| - | - | 3425 | 578.3 | - | - | 0 | - |
| - | - | 680 | 583.2 | - | - | 0 | - |
| 3 | y | 1.088E+05 | 585.2 | 0.0007231 | 1.236 | +1 | 4 |
| - | - | 3.165E+04 | 586.2 | - | - | 0 | - |
| - | - | 5981 | 587.2 | - | - | 0 | - |
| - | - | 1302 | 592.3 | - | - | 0 | - |
| 5 | b | 2.044E+05 | 593.3 | 0.0007449 | 1.255 | +1 | 5 |
| - | - | 6.542E+04 | 594.3 | - | - | 0 | - |
| - | - | 2838 | 595.2 | - | - | 0 | - |
| - | - | 1.36E+04 | 595.3 | - | - | 0 | - |
| - | - | 909.9 | 596.2 | - | - | 0 | - |
| - | - | 651.7 | 603.3 | - | - | 0 | - |
| - | - | 7873 | 610.3 | - | - | 0 | - |
| - | - | 1.195E+04 | 611.3 | - | - | 0 | - |
| - | - | 4338 | 612.3 | - | - | 0 | - |
| - | - | 1161 | 617.3 | - | - | 0 | - |
| - | - | 955 | 620.3 | - | - | 0 | - |
| - | - | 702.5 | 622.3 | - | - | 0 | - |
| - | - | 2257 | 633.3 | - | - | 0 | - |
| - | - | 1782 | 634.3 | - | - | 0 | - |
| - | - | 3271 | 635.3 | - | - | 0 | - |
| - | - | 2214 | 636.3 | - | - | 0 | - |
| - | - | 1585 | 637.3 | - | - | 0 | - |
| - | - | 2270 | 645.3 | - | - | 0 | - |
| - | - | 1028 | 646.3 | - | - | 0 | - |
| - | - | 4071 | 650.3 | - | - | 0 | - |
| - | - | 4203 | 651.3 | - | - | 0 | - |
| - | - | 1.187E+04 | 652.3 | - | - | 0 | - |
| - | - | 4209 | 653.3 | - | - | 0 | - |
| - | - | 716.6 | 654.3 | - | - | 0 | - |
| - | - | 6908 | 662.3 | - | - | 0 | - |
| - | - | 1.138E+04 | 663.3 | - | - | 0 | - |
| - | - | 4715 | 664.3 | - | - | 0 | - |
| - | - | 858.7 | 665.3 | - | - | 0 | - |
| - | - | 1.066E+04 | 668.3 | - | - | 0 | - |
| - | - | 4090 | 669.3 | - | - | 0 | - |
| - | - | 1244 | 670.3 | - | - | 0 | - |
| 2 | y | 2.039E+04 | 680.3 | 0.000588 | 0.8644 | +1 | 5 |
| 2 | y | 4.091E+04 | 681.3 | 0.002656 | 3.899 | +1 | 5 |
| - | - | 1.59E+04 | 682.3 | - | - | 0 | - |
| - | - | 3784 | 683.3 | - | - | 0 | - |
| 2 | y | 1.275E+05 | 698.3 | 0.0005824 | 0.834 | +1 | 5 |
| - | - | 4.666E+04 | 699.3 | - | - | 0 | - |
| - | - | 1.269E+04 | 700.3 | - | - | 0 | - |
| - | - | 1036 | 701.3 | - | - | 0 | - |
| - | - | 851.8 | 708.3 | - | - | 0 | - |
| - | - | 1854 | 709.3 | - | - | 0 | - |
| - | - | 703.2 | 2316 | - | - | 0 | - |
| - | - | 742.7 | 3133 | - | - | 0 | - |

m/z Charge Intensity FragmentType MassShift Position
120.05602264404297 0 1195.167
120.0811767578125 0 1695.063
120.6639175415039 0 343.21985
121.04004669189453 0 2645.79
122.07173156738281 0 6443.049
123.05573272705078 0 4716.882
123.11721801757812 0 847.96906
124.0874252319336 0 601.39575
125.07110595703125 0 494.40747
125.10783386230469 0 1759.9045
127.08682250976562 0 795.1518
128.0824432373047 0 1384.338
129.0551300048828 0 1589.1324
129.06625366210938 0 4213.6577
129.10272216796875 0 2340.9072
130.0656280517578 0 197177.25
131.06895446777344 0 18469.838
132.04429626464844 0 507.2117
132.0812530517578 0 58732.727
133.06134033203125 0 3409.989
133.0847625732422 0 4794.5356
134.0450897216797 0 522.25104
136.0397186279297 0 585.53516
136.07614135742188 0 2991.2183
138.06668090820312 0 17456.238
138.0918731689453 0 521.7571
139.0506134033203 0 2148.6196
139.07025146484375 0 1066.4917
139.0984344482422 0 481.1886
140.0823516845703 0 699.35913
140.10707092285156 0 451.29034
140.14414978027344 0 805.3904
141.10267639160156 0 1967.1072
142.0656280517578 0 9515.991
143.0455780029297 0 6222.6006
143.0682373046875 0 629.0419
143.07354736328125 0 2170.6724
144.04434204101562 0 606.24506
144.0812530517578 0 20800.021
145.0484161376953 0 490.93005
145.0845184326172 0 2087.7588
146.060546875 0 106426.53
147.0640411376953 0 9369.12
148.05079650878906 0 705.65076
152.08287048339844 0 565.0438
153.10267639160156 0 1052.1517
154.0613250732422 0 1827.0024
154.09799194335938 0 4959.874
155.08175659179688 0 4287.525
155.09323120117188 0 87788.7
156.0771942138672 0 3822.4177
156.09039306640625 0 1157.6238
156.09664916992188 0 4834.581
156.11361694335938 0 541.5967
158.06024169921875 0 683.22327
158.08436584472656 0 8102.505
159.09222412109375 0 441436.47
160.07614135742188 0 7345.211
160.08909606933594 0 3040.5063
160.09555053710938 0 46164.926
161.080078125 0 468.67917
161.0985870361328 0 1720.7242
164.08213806152344 0 568.15283
164.11868286132812 0 2867.4473
165.07763671875 0 3536.178
166.0615997314453 0 76938.57
167.05776977539062 0 760.4223
167.06495666503906 0 5656.858
167.093017578125 0 888.6005
169.07647705078125 0 9358.388
170.06051635742188 0 36031.574
171.0638885498047 0 3948.5186
171.0769805908203 0 738.36334
171.09207153320312 0 730.037
172.10861206054688 0 1630.0493
173.3477325439453 0 449.36307
175.01614379882812 0 1243.1565
175.0983428955078 0 4286.984
176.0823974609375 0 4399.282
178.1343994140625 0 39278.215
179.1376953125 0 3476.5923
180.07745361328125 0 927.9557
181.06129455566406 0 15266.011
182.04547119140625 0 1212.7845
183.0883026123047 0 1708.6675
183.11325073242188 0 748.3455
183.1500701904297 0 599.23193
184.12673950195312 0 957.4561
185.16539001464844 0 134640.53 a 1
186.16879272460938 0 14924.536
187.08714294433594 0 23880.645
187.10816955566406 0 1135.578
187.17132568359375 0 857.7596
188.0712127685547 0 568211
189.07447814941406 0 65768.06
190.07705688476562 0 4323.879
190.13433837890625 0 8769.9375
191.13754272460938 0 1049.9332
194.0926513671875 0 959.85565
195.09193420410156 0 1272.9619
195.14976501464844 0 1412.3868
196.11280822753906 0 770.79626
196.13412475585938 0 663.61304
197.07156372070312 0 786.8366
197.10723876953125 0 605.8701
197.13619995117188 0 562.21246
198.0877227783203 0 17019.504
199.07188415527344 0 7489.251
199.09031677246094 0 792.3203
200.10289001464844 0 734.721
200.1433563232422 0 856.09045
202.10887145996094 0 789.94183
203.0929718017578 0 1921.2561
204.07725524902344 0 1447.8015
204.13482666015625 0 966.8742
205.0977020263672 0 164178.84 y 5
205.14527893066406 0 4354.242
206.10104370117188 0 17586.117
206.129150390625 0 4297.0396
207.1039581298828 0 1065.6519
207.13259887695312 0 723.6697
207.16085815429688 0 5043.473
209.05580139160156 0 2302.0095
211.0871124267578 0 828.5004
212.17636108398438 0 2126.4126
213.13926696777344 0 4744.596
213.1602783203125 0 26220.484 b 1
214.09808349609375 0 535.42346
214.14329528808594 0 620.31586
214.16371154785156 0 3405.3772
215.08212280273438 0 1771.4397
216.09841918945312 0 8752.404
217.10098266601562 0 653.60095
217.1329803466797 0 1190.1259
220.11972045898438 0 5830.604
221.0711669921875 0 3339.307
221.1038360595703 0 17279.129
221.1230010986328 0 1139.6482
222.10755920410156 0 811.0636
223.0876922607422 0 1167.794
223.15589904785156 0 55465.04
224.11862182617188 0 587.7489
224.15927124023438 0 5473.999
225.64524841308594 0 1619.5549
226.08277893066406 0 27065.21
226.15675354003906 0 613.97015
227.06678771972656 0 21310.357
227.0864715576172 0 2587.508
227.1547088623047 0 1023.0349
228.07052612304688 0 1648.9283
229.0970458984375 0 696.9534
229.60032653808594 0 616.7942
230.1036376953125 0 1210.4939
231.08811950683594 0 2916.135
232.14503479003906 0 7559.807
232.64674377441406 0 2408.6794
233.1048583984375 0 1254.0261
233.14027404785156 0 24560.066
234.1243133544922 0 85103.01
234.14500427246094 0 2415.027
235.1276397705078 0 9820.198
235.15611267089844 0 2248.7832
238.09817504882812 0 649.7114
238.13038635253906 0 5074.102
238.606201171875 0 1986.953
239.08187866210938 0 5532.334
239.1143341064453 0 742.0048
239.6427001953125 0 2194.0776 b 3
240.0856475830078 0 1072.7327
241.0975799560547 0 1301.5658
241.1339569091797 0 990.14105
242.15069580078125 0 828.1586
243.07701110839844 0 5415.198
243.1096954345703 0 4832.003
243.5982208251953 0 986.634
244.0934600830078 0 45395.984
245.0770721435547 0 2420.051
245.0966796875 0 3090.609
246.135986328125 0 4842.951
247.11830139160156 0 712.41486
248.1148681640625 0 32661.389
249.0988311767578 0 59624.062
249.11656188964844 0 2714.6047
249.13539123535156 0 1391.6981
249.15980529785156 0 2496.9795
250.1021270751953 0 8325.323
250.16616821289062 0 1606.8828
251.1509552001953 0 275172.44
252.15411376953125 0 36288.773
252.1707763671875 0 604.74384
252.60369873046875 0 11170.253
252.6505889892578 0 1638.7957
253.10523986816406 0 3606.176
253.15573120117188 0 2141.5713
254.0780029296875 0 1177.928
254.65562438964844 0 4014.3257
255.15774536132812 0 986.09686
255.19473266601562 0 616.09467
256.1085510253906 0 5418.0967
257.0923156738281 0 1519.0266
257.11138916015625 0 1040.6249
258.0993347167969 0 920.9856
258.1239929199219 0 2260.924
260.1035461425781 0 5596.802
261.1181640625 0 8044.3604
261.1351318359375 0 7192.2886
261.1639099121094 0 6065.6265
261.6089782714844 0 7354.9424
261.66448974609375 0 854.4101
262.050537109375 0 1394.0579
262.1047058105469 0 2003.8909
262.1386413574219 0 1136.2505
265.1412048339844 0 4672.7114
265.6407470703125 0 599.9559
266.12530517578125 0 150119.52
267.0764465332031 0 1677.2947
267.10906982421875 0 2532.389
267.1282653808594 0 15886.864
268.1306457519531 0 1350.8776
268.17724609375 0 4101.829
270.1221923828125 0 80296.89
270.62359619140625 0 22139.23
271.12451171875 0 3573.269
274.1180725097656 0 9728.516
274.1304931640625 0 26534.773
274.6455383300781 0 3130.2239
275.1192321777344 0 1279.4514
275.1337890625 0 3564.234
275.1481628417969 0 743.0708
275.60601806640625 0 598.0959
276.109375 0 5270.53
277.0937194824219 0 2239.9204
277.10943603515625 0 588.082
277.155517578125 0 1939.2494
277.2023620605469 0 1430.0345
278.1615295410156 0 1536.9606
279.1457824707031 0 10262.386
279.16302490234375 0 737.4185
280.1490173339844 0 1350.5012
283.154296875 0 1948.3328
283.6602783203125 0 658.40436
284.08697509765625 0 1252.0897
284.103515625 0 8316.958
284.11981201171875 0 5410.426 y Water loss 2
284.13604736328125 0 1564.6224
284.6207580566406 0 2483.8018
285.08758544921875 0 9449.07
285.10589599609375 0 1449.5756
286.09228515625 0 2340.941
287.1871032714844 0 2645.1575
288.6430969238281 0 3405.2988 b Ammonia loss 4
289.1440734863281 0 705.7912
289.2022399902344 0 1337.314
292.14111328125 0 1411.8958
293.1252746582031 0 2134.0444 y 2
293.6248779296875 0 686.39716
294.12030029296875 0 5524.322
295.1228332519531 0 760.975
296.17431640625 0 1759.6239
296.67291259765625 0 689.153
297.1564025878906 0 26809.582 b 4
297.6576232910156 0 7563.109
298.14141845703125 0 1118.966
298.1593322753906 0 1964.2468
299.1251220703125 0 951.58685
300.1099548339844 0 2727.6348
301.1148376464844 0 6062.23
301.14105224609375 0 3969.0178
302.1142578125 0 11228.459 y Water loss 4
302.1278076171875 0 4505.1143
302.1478271484375 0 790.02106
302.16436767578125 0 600.53906
303.11785888671875 0 1807.6254
303.1301574707031 0 788.77026
304.10577392578125 0 963.21625
304.21368408203125 0 3957.6467
305.1973876953125 0 1392.6174
306.1618957519531 0 753.72894
306.2294921875 0 13348.145
307.2329406738281 0 2150.3193
309.1454772949219 0 862.80005
309.1674499511719 0 2193.1008
310.1721496582031 0 5411.377
310.66168212890625 0 1629.9265
310.67529296875 0 1319.2246
315.1834716796875 0 784.4869
316.1778564453125 0 1681.0693
317.1369934082031 0 1038.2457
317.1612243652344 0 879.39557
317.18060302734375 0 642.7055
317.65985107421875 0 1104.7938
318.1199951171875 0 4312.2803
318.15179443359375 0 4169.6157
318.6530456542969 0 1775.9055
319.1051940917969 0 1113.4758
319.1248779296875 0 3092.809
319.1520690917969 0 6183.1045
320.1248779296875 0 40096.793 y 4
321.12835693359375 0 6275.6177
322.132080078125 0 896.8087
322.22412109375 0 2770.7085 a 2
323.1433410644531 0 675.72943
323.1800537109375 0 1834.8877
326.13519287109375 0 3928.2012
326.1573791503906 0 701.99457
326.2229309082031 0 2354.8787
326.66448974609375 0 25809.258
327.1656188964844 0 9963.22
327.2255859375 0 815.7299
327.66748046875 0 2219.6853
328.1041259765625 0 1783.2583
331.65643310546875 0 1528.8643
332.1492919921875 0 1185.3848
332.2088317871094 0 37629.793
333.1925964355469 0 814.52936
333.21185302734375 0 7326.8037
335.1479797363281 0 816.27356
336.1307373046875 0 2747.3604
337.16162109375 0 1453.7468
340.66290283203125 0 2534.2888 y Water loss 1
341.15716552734375 0 1124.4081 y Ammonia loss 1
343.16217041015625 0 1089.6805
344.17205810546875 0 6587.0244
344.68280029296875 0 896.2546
345.13128662109375 0 18160.05
345.1549377441406 0 980.7759
345.1788024902344 0 636.3244
346.115234375 0 67766.15
347.0960998535156 0 811.1603
347.1182861328125 0 10660.275
348.120849609375 0 1796.1943
348.1673278808594 0 902.1908
349.1296691894531 0 1430.247
349.6678771972656 0 1553.2158 y 1
350.2193603515625 0 159894.53 b 2
351.14691162109375 0 757.2156
351.2223205566406 0 30333.654
352.224853515625 0 3731.9458
353.6874694824219 0 2092.4526
354.1920471191406 0 1003.1615
354.2178649902344 0 2854.8315
355.2193298339844 0 887.86725
358.6800537109375 0 7598.681
359.1756896972656 0 3668.354
359.6757507324219 0 1865.133
360.2047119140625 0 1213.2297
361.1990661621094 0 7032.2344
362.1828918457031 0 8510.588
362.2021484375 0 724.5029
363.1418151855469 0 70285.875
363.18524169921875 0 1996.309
364.1255187988281 0 33891.887
364.14593505859375 0 7885.592
365.1288146972656 0 6343.9727
366.1321105957031 0 668.9738
366.15618896484375 0 1698.5723
367.1404724121094 0 7209.7827
367.1933288574219 0 4904.727
367.2458190917969 0 22915.842
367.6854553222656 0 41199.344
368.14691162109375 0 1137.2097
368.1866760253906 0 17815.121
368.2484130859375 0 4232.842
368.6875915527344 0 5130.7544
369.15081787109375 0 638.6693
372.6776123046875 0 1751.1743
374.10882568359375 0 685.87946
376.19854736328125 0 96094.29
376.6991882324219 0 45348.266
377.1994323730469 0 11798.372
377.6949157714844 0 925.6559
379.2095642089844 0 79048.64
380.16748046875 0 910.12744
380.189697265625 0 739.05493
380.2126159667969 0 15782.128
381.1523742675781 0 202540.34
381.68682861328125 0 3906.6704
382.1551208496094 0 34461.617
382.1836242675781 0 1017.17474
383.15753173828125 0 5302.655
384.1669006347656 0 4152.4233
385.1512145996094 0 9790.395
386.15447998046875 0 1793.6511
386.181396484375 0 749.65436
388.2347717285156 0 869.8786
389.19329833984375 0 3035.1174
390.14862060546875 0 762.0312
390.196044921875 0 14113.089 Precursor Water loss
390.6898193359375 0 13653.954 Precursor Ammonia loss
391.1367492675781 0 4449.912
391.19146728515625 0 4037.4175
391.69287109375 0 1270.6093
392.1212158203125 0 2487.9846
393.153076171875 0 888.8722
394.1520080566406 0 1386.0574
395.1356506347656 0 6609.62
396.13885498046875 0 1882.5535
398.178955078125 0 8578.394
398.2393798828125 0 5539.5557
399.1635437011719 0 8942.55
399.2015380859375 0 39794.645 Precursor
399.70306396484375 0 19127.08
400.1662292480469 0 1221.2677
400.2051086425781 0 6055.642
402.1778259277344 0 9608.35
403.1368713378906 0 1221.9817
403.1803283691406 0 1564.4807
405.2259216308594 0 2196.293
407.20452880859375 0 9711.181
408.2076416015625 0 1870.4214
409.1474304199219 0 8106.1187
410.1487731933594 0 1455.3325
412.16253662109375 0 5828.0405
413.1463623046875 0 6061.19
414.1488952636719 0 1245.3608
416.2298889160156 0 3575.049
417.2333679199219 0 1481.7087
421.14739990234375 0 7755.1997
422.1504821777344 0 1883.8098
422.25201416015625 0 691.50116
430.1729431152344 0 54739.258 y Water loss 3
431.1553039550781 0 7543.549 y Ammonia loss 3
431.17803955078125 0 7391.717
432.15667724609375 0 1458.8745
432.18365478515625 0 994.39056
433.2563171386719 0 2975.21
435.1997375488281 0 1753.5422
438.17437744140625 0 2327.9385
439.15869140625 0 2383.6387
441.1889953613281 0 683.61035
443.2410888671875 0 1118.605
448.1835021972656 0 36970.54 y 3
449.1865539550781 0 9550.889
450.1883544921875 0 1388.7576
450.2832946777344 0 5567.192
451.28765869140625 0 1134.8762
456.18450927734375 0 12044.661
457.18743896484375 0 2634.2522
459.1993713378906 0 4555.008
460.2673645019531 0 1192.2468
461.2515869140625 0 8182.296 b Ammonia loss 3
462.25396728515625 0 1964.1267
463.1942138671875 0 2153.317
465.2473449707031 0 836.865
468.18707275390625 0 767.80286
476.2255554199219 0 3284.5852
477.210205078125 0 14782.035
478.2138366699219 0 3471.7935
478.2782287597656 0 36335.977 b 3
479.2208251953125 0 752.58704
479.2807312011719 0 8504.074
480.2200012207031 0 1189.9731
480.28424072265625 0 1489.2213
484.2203369140625 0 765.46533
486.1896667480469 0 1618.5105
487.1934814453125 0 1156.9581
491.27703857421875 0 5399.788
492.2815246582031 0 764.5083
494.2365417480469 0 97256.83
495.2391662597656 0 22739.75
496.2413635253906 0 3993.761
496.28436279296875 0 959.40314
504.20025634765625 0 6695.706
504.293212890625 0 1862.3474
505.2038269042969 0 3075.447
505.23748779296875 0 805.1559
506.21148681640625 0 1150.2556
511.2641296386719 0 1405.5564
512.2474975585938 0 3773.4387
514.1829223632812 0 1863.5598
521.2273559570312 0 3553.426
521.3190307617188 0 2257.237
522.2303466796875 0 8420.517
522.3240356445312 0 925.3348
523.2396240234375 0 5301.941
524.22705078125 0 2562.943
530.2258911132812 0 918.95245
532.195556640625 0 4896.3843
533.1987915039062 0 1405.8354
539.2366333007812 0 31764.854
540.2385864257812 0 9341.887
541.2415771484375 0 1752.5793
547.2994995117188 0 1816.7124
548.2832641601562 0 4482.044
549.22119140625 0 6010.092
549.3193969726562 0 1154.1561
550.20556640625 0 36675.137
551.2086181640625 0 11104.1455
551.2536010742188 0 707.7241
552.2055053710938 0 1319.8613
552.2484130859375 0 726.4317
558.2676391601562 0 6788.2544
559.2703247070312 0 2112.3179
562.2646484375 0 974.1713
563.2600708007812 0 651.9324
565.31005859375 0 3595.379
566.3092041015625 0 672.1757
567.2315673828125 0 14321.342 y Water loss 2
568.2181396484375 0 13874.346 y Ammonia loss 2
569.2199096679688 0 4454.801
570.2219848632812 0 1054.1571
575.2940673828125 0 6257.4355 b Water loss 4
576.2785034179688 0 43028.207 b Ammonia loss 4
577.2814331054688 0 12767.766
578.2032470703125 0 699.5323
578.2841796875 0 3425.0776
583.2279663085938 0 680.017
585.2423095703125 0 108754.59 y 2
586.2449340820312 0 31653.854
587.2481689453125 0 5981.2935
592.3233642578125 0 1301.618
593.304931640625 0 204357.7 b 4
594.3069458007812 0 65417.63
595.2262573242188 0 2837.762
595.3088989257812 0 13602.501
596.2290649414062 0 909.9097
603.2791137695312 0 651.65155
610.3314208984375 0 7872.728
611.3169555664062 0 11950.901
612.3191528320312 0 4338.2983
617.282958984375 0 1161.3384
620.3159790039062 0 954.9522
622.331787109375 0 702.4684
633.2998657226562 0 2257.1125
634.3057250976562 0 1782.4158
635.2954711914062 0 3270.6155
636.3236083984375 0 2213.9023
637.314453125 0 1584.5225
645.2796630859375 0 2270.4558
646.2763671875 0 1027.5453
650.3267211914062 0 4070.6597
651.3157348632812 0 4203.3403
652.3196411132812 0 11872.585
653.3212890625 0 4209.1997
654.330322265625 0 716.60175
662.30517578125 0 6907.7456
663.291259765625 0 11380.537
664.2938842773438 0 4715.2905
665.2954711914062 0 858.7427
668.3368530273438 0 10656.495
669.339599609375 0 4090.0876
670.3395385742188 0 1243.818
680.315673828125 0 20392.572 y Water loss 1
681.3017578125 0 40905.438 y Ammonia loss 1
682.3040771484375 0 15901.703
683.3057861328125 0 3783.8906
698.3262329101562 0 127529.07 y 1
699.3289794921875 0 46657.387
700.3319091796875 0 12687.746
701.3345336914062 0 1036.2024
708.3082885742188 0 851.83514
709.2986450195312 0 1854.4811
2316.32470703125 0 703.2344
3133.494140625 0 742.6552

Spectrum Details

|  |  |
| --- | --- |
| Matched peaks? Matched peaks The total absolute number of peaks matched. Additionally in brackets the total fraction of peaks matched and the total number of peaks is shown. | 32 (5.78% of 554) |
| FDR? FDR The false discovery rate estimated for this peptide. It is calculated by matching all theoretical fragments with a non-integer shift with the raw peaks for this spectrum. This is done with 40 different shifts. The resulting percentage is the average number of annotated peaks over the number of annotated peaks with the correct spectrum. | 1.26% |
| Satellite FDR? Satellite FDR See the FDR for details on its calculation. This satellite ion specific FDR only contains the satellite ions (d/w) for I/L/J positions. | - |
| PSM Score? PSM Score The PSM Score as given by Hecklib to this annotated spectrum. It is shown with three significant figures. | 387 |

#### Spectrum 4739? Spectrum 4739 The raw spectrum of this peptide as annotated by Hecklib. The fragments are coloured according to ion type (see legend). Any peaks with a star '\*' as text can be hovered over to see the full details, first the ion type second the mass shift type. By hovering over the amino acids in the peptide or ions in the legend the corresponding peaks are highlighted. By toggling the 'Unassigned' label you can turn the background (unassigned) peaks on or off in the plot. By updating the slider in the Ion legend you can update the spectrum to only show the top X% of the peaks with labels. The top X% means any peak that is within X% of the highest intensity. By dragging in the spectrum you can zoom in to a specific part of the spectrum and use 'Zoom Out' to get back to the original zoom level. The annotation of the spectrum is based on the given sequence in the peptides file and is done with different software so inconsistencies are likely. The peaks are annotated based on the given sequence, with 20 ppm tolerance.

Copy Data

##### Spectrum 4739 (TSV)

###### Preview

```
Loading example...
```

*Click on the button to copy the data to your clipboard.*

Mz MinMz MaxIntensity Max

WidthHeightPeptide font sizePeptide stroke widthSpectrum font sizeSpectrum stroke widthCompact peptide

Ion legend

wxyz

abcd

OtherUnassignedIon^Charge^_Position_Show for top:%

VJHQDW

01.54e+43.08e+44.62e+46.16e+4

Zoom Out

d^+1^_2_a^+1^_2_y^+1^_1_b^+1^_2_y^+2^_4_b^+2^_5_y^+1^_2_y^+1^_2_b^+1^_3_\*\*\*y^+1^_3_y^+1^_3_y^+1^_3_b^+1^_4_b^+1^_4_y^+1^_4_y^+1^_4_b^+1^_5_b^+1^_5_y^+1^_4_b^+1^_5_y^+1^_5_y^+1^_5_y^+1^_5_

0844168725313374

Fragment Matches Table

Show background peaks

| Position | Ion type | Intensity | mz Theoretical | mz Error (Th) | mz Error (ppm) | Charge | Series Number |
| --- | --- | --- | --- | --- | --- | --- | --- |
| - | - | 369.9 | 120.1 | - | - | 0 | - |
| - | - | 426.5 | 120.1 | - | - | 0 | - |
| - | - | 2845 | 120.1 | - | - | 0 | - |
| - | - | 372.6 | 121.5 | - | - | 0 | - |
| - | - | 1068 | 122.1 | - | - | 0 | - |
| - | - | 344.5 | 123.1 | - | - | 0 | - |
| - | - | 712.7 | 123.1 | - | - | 0 | - |
| - | - | 554.1 | 125.1 | - | - | 0 | - |
| - | - | 616.3 | 126.1 | - | - | 0 | - |
| - | - | 477.6 | 127.1 | - | - | 0 | - |
| - | - | 499.1 | 128.1 | - | - | 0 | - |
| - | - | 652.1 | 128.1 | - | - | 0 | - |
| - | - | 3776 | 129.1 | - | - | 0 | - |
| - | - | 611.2 | 129.1 | - | - | 0 | - |
| - | - | 3669 | 129.1 | - | - | 0 | - |
| - | - | 2.222E+04 | 130.1 | - | - | 0 | - |
| - | - | 549.4 | 130.1 | - | - | 0 | - |
| - | - | 2077 | 131.1 | - | - | 0 | - |
| - | - | 6664 | 132.1 | - | - | 0 | - |
| - | - | 373.8 | 132.9 | - | - | 0 | - |
| - | - | 1308 | 133.1 | - | - | 0 | - |
| - | - | 1.359E+04 | 133.1 | - | - | 0 | - |
| - | - | 485.5 | 134.1 | - | - | 0 | - |
| - | - | 416.9 | 135.9 | - | - | 0 | - |
| - | - | 640.9 | 136 | - | - | 0 | - |
| - | - | 1431 | 136.1 | - | - | 0 | - |
| - | - | 421.1 | 137.1 | - | - | 0 | - |
| - | - | 434.4 | 138 | - | - | 0 | - |
| - | - | 2922 | 138.1 | - | - | 0 | - |
| - | - | 883.3 | 139.1 | - | - | 0 | - |
| - | - | 440.7 | 139.1 | - | - | 0 | - |
| - | - | 1162 | 140.1 | - | - | 0 | - |
| - | - | 1630 | 141.1 | - | - | 0 | - |
| - | - | 1194 | 142.1 | - | - | 0 | - |
| - | - | 525 | 142.1 | - | - | 0 | - |
| - | - | 471.4 | 142.1 | - | - | 0 | - |
| - | - | 807.2 | 143 | - | - | 0 | - |
| - | - | 480.9 | 143.1 | - | - | 0 | - |
| 2 | d | 499.2 | 143.1 | 0.0001222 | 0.8538 | +1 | 2 |
| - | - | 1792 | 144.1 | - | - | 0 | - |
| - | - | 1235 | 144.1 | - | - | 0 | - |
| - | - | 393.7 | 144.4 | - | - | 0 | - |
| - | - | 528.1 | 145.1 | - | - | 0 | - |
| - | - | 456.4 | 145.1 | - | - | 0 | - |
| - | - | 471.7 | 145.4 | - | - | 0 | - |
| - | - | 1.044E+04 | 146.1 | - | - | 0 | - |
| - | - | 694.1 | 146.1 | - | - | 0 | - |
| - | - | 3541 | 147.1 | - | - | 0 | - |
| - | - | 656.3 | 148.9 | - | - | 0 | - |
| - | - | 574.4 | 153.1 | - | - | 0 | - |
| - | - | 693.9 | 154.1 | - | - | 0 | - |
| - | - | 857.5 | 155.1 | - | - | 0 | - |
| - | - | 9716 | 155.1 | - | - | 0 | - |
| - | - | 812.1 | 155.1 | - | - | 0 | - |
| - | - | 1051 | 156.1 | - | - | 0 | - |
| - | - | 802.1 | 156.1 | - | - | 0 | - |
| - | - | 456 | 157.1 | - | - | 0 | - |
| - | - | 559 | 157.1 | - | - | 0 | - |
| - | - | 616.6 | 157.1 | - | - | 0 | - |
| - | - | 1442 | 158.1 | - | - | 0 | - |
| - | - | 4.873E+04 | 159.1 | - | - | 0 | - |
| - | - | 1284 | 160.1 | - | - | 0 | - |
| - | - | 5169 | 160.1 | - | - | 0 | - |
| - | - | 651.2 | 161.1 | - | - | 0 | - |
| - | - | 442.1 | 162.2 | - | - | 0 | - |
| - | - | 644.2 | 163 | - | - | 0 | - |
| - | - | 393.4 | 163.8 | - | - | 0 | - |
| - | - | 449.1 | 165.1 | - | - | 0 | - |
| - | - | 8123 | 166.1 | - | - | 0 | - |
| - | - | 620.6 | 167.1 | - | - | 0 | - |
| - | - | 775.5 | 169.1 | - | - | 0 | - |
| - | - | 1286 | 169.1 | - | - | 0 | - |
| - | - | 3959 | 170.1 | - | - | 0 | - |
| - | - | 614.1 | 171.1 | - | - | 0 | - |
| - | - | 1689 | 173.5 | - | - | 0 | - |
| - | - | 802 | 175 | - | - | 0 | - |
| - | - | 2001 | 175 | - | - | 0 | - |
| - | - | 612.5 | 175 | - | - | 0 | - |
| - | - | 1291 | 175.1 | - | - | 0 | - |
| - | - | 1131 | 176.1 | - | - | 0 | - |
| - | - | 4078 | 177.1 | - | - | 0 | - |
| - | - | 4586 | 178.1 | - | - | 0 | - |
| - | - | 596.6 | 179.1 | - | - | 0 | - |
| - | - | 1694 | 181.1 | - | - | 0 | - |
| - | - | 549.6 | 183.1 | - | - | 0 | - |
| - | - | 781.2 | 183.1 | - | - | 0 | - |
| 2 | a | 1.57E+04 | 185.2 | 0.0003061 | 1.653 | +1 | 2 |
| - | - | 1710 | 186.2 | - | - | 0 | - |
| - | - | 2211 | 187.1 | - | - | 0 | - |
| - | - | 922.4 | 187.1 | - | - | 0 | - |
| - | - | 502.6 | 188 | - | - | 0 | - |
| - | - | 6.094E+04 | 188.1 | - | - | 0 | - |
| - | - | 7351 | 189.1 | - | - | 0 | - |
| - | - | 497.7 | 190.1 | - | - | 0 | - |
| - | - | 1634 | 198.1 | - | - | 0 | - |
| - | - | 553.7 | 198.9 | - | - | 0 | - |
| - | - | 2760 | 199.1 | - | - | 0 | - |
| - | - | 1412 | 202.1 | - | - | 0 | - |
| - | - | 1083 | 204.1 | - | - | 0 | - |
| - | - | 861.1 | 205.1 | - | - | 0 | - |
| 6 | y | 1.74E+04 | 205.1 | 0.0002885 | 1.407 | +1 | 1 |
| - | - | 1116 | 205.1 | - | - | 0 | - |
| - | - | 586.4 | 205.1 | - | - | 0 | - |
| - | - | 1538 | 206.1 | - | - | 0 | - |
| - | - | 759 | 207.2 | - | - | 0 | - |
| - | - | 784.7 | 213.1 | - | - | 0 | - |
| 2 | b | 2994 | 213.2 | 0.000173 | 0.8117 | +1 | 2 |
| - | - | 664.6 | 214.1 | - | - | 0 | - |
| - | - | 540.3 | 215.1 | - | - | 0 | - |
| - | - | 828 | 216.1 | - | - | 0 | - |
| - | - | 670.2 | 218 | - | - | 0 | - |
| - | - | 702.2 | 220.2 | - | - | 0 | - |
| - | - | 615.7 | 221.1 | - | - | 0 | - |
| - | - | 1138 | 221.1 | - | - | 0 | - |
| - | - | 5731 | 223.2 | - | - | 0 | - |
| - | - | 2693 | 226.1 | - | - | 0 | - |
| - | - | 2163 | 227.1 | - | - | 0 | - |
| - | - | 702.9 | 228.2 | - | - | 0 | - |
| - | - | 713.3 | 230 | - | - | 0 | - |
| - | - | 793.8 | 231.1 | - | - | 0 | - |
| - | - | 1459 | 232.1 | - | - | 0 | - |
| - | - | 3017 | 233.1 | - | - | 0 | - |
| - | - | 8878 | 234.1 | - | - | 0 | - |
| - | - | 1067 | 235.1 | - | - | 0 | - |
| - | - | 752.4 | 238.1 | - | - | 0 | - |
| - | - | 1071 | 239.1 | - | - | 0 | - |
| - | - | 510.1 | 240.1 | - | - | 0 | - |
| - | - | 577 | 243.1 | - | - | 0 | - |
| - | - | 4534 | 244.1 | - | - | 0 | - |
| - | - | 2986 | 248.1 | - | - | 0 | - |
| - | - | 5302 | 249.1 | - | - | 0 | - |
| - | - | 2.7E+04 | 251.2 | - | - | 0 | - |
| - | - | 3507 | 252.2 | - | - | 0 | - |
| - | - | 624.6 | 254.7 | - | - | 0 | - |
| - | - | 1225 | 255.2 | - | - | 0 | - |
| - | - | 693 | 256.1 | - | - | 0 | - |
| - | - | 705 | 261.1 | - | - | 0 | - |
| - | - | 1003 | 261.1 | - | - | 0 | - |
| - | - | 701.6 | 261.2 | - | - | 0 | - |
| - | - | 2797 | 262.1 | - | - | 0 | - |
| - | - | 1.522E+04 | 266.1 | - | - | 0 | - |
| - | - | 2299 | 267.1 | - | - | 0 | - |
| - | - | 9060 | 270.1 | - | - | 0 | - |
| - | - | 2244 | 270.6 | - | - | 0 | - |
| - | - | 735.4 | 271.1 | - | - | 0 | - |
| - | - | 961.6 | 274.1 | - | - | 0 | - |
| - | - | 3387 | 274.1 | - | - | 0 | - |
| - | - | 727.3 | 276.1 | - | - | 0 | - |
| - | - | 1533 | 279.1 | - | - | 0 | - |
| - | - | 1107 | 283.2 | - | - | 0 | - |
| - | - | 666 | 284.1 | - | - | 0 | - |
| 3 | y | 815.5 | 284.1 | 0.0002221 | 0.7818 | +2 | 4 |
| - | - | 758.2 | 285.1 | - | - | 0 | - |
| - | - | 531.5 | 291.1 | - | - | 0 | - |
| 5 | b | 2082 | 297.2 | 0.000153 | 0.5149 | +2 | 5 |
| - | - | 947.2 | 297.7 | - | - | 0 | - |
| - | - | 1148 | 299.2 | - | - | 0 | - |
| 5 | y | 904.4 | 302.1 | 0.001716 | 5.68 | +1 | 2 |
| - | - | 1093 | 306.2 | - | - | 0 | - |
| - | - | 891.2 | 318.2 | - | - | 0 | - |
| 5 | y | 3954 | 320.1 | 0.0002315 | 0.7231 | +1 | 2 |
| - | - | 550.7 | 321.1 | - | - | 0 | - |
| - | - | 2483 | 326.7 | - | - | 0 | - |
| - | - | 3984 | 327.2 | - | - | 0 | - |
| - | - | 4014 | 332.2 | - | - | 0 | - |
| - | - | 1092 | 333.2 | - | - | 0 | - |
| - | - | 2944 | 345.1 | - | - | 0 | - |
| - | - | 557.3 | 345.2 | - | - | 0 | - |
| - | - | 7196 | 346.1 | - | - | 0 | - |
| - | - | 869.3 | 347.1 | - | - | 0 | - |
| 3 | b | 1.582E+04 | 350.2 | 8.379E-05 | 0.2392 | +1 | 3 |
| - | - | 3614 | 351.2 | - | - | 0 | - |
| - | - | 990.5 | 355.2 | - | - | 0 | - |
| - | - | 658.1 | 361.2 | - | - | 0 | - |
| - | - | 973.2 | 362.2 | - | - | 0 | - |
| - | - | 7080 | 363.1 | - | - | 0 | - |
| - | - | 3785 | 364.1 | - | - | 0 | - |
| - | - | 704.4 | 367.1 | - | - | 0 | - |
| - | - | 668.3 | 367.2 | - | - | 0 | - |
| - | - | 2564 | 367.2 | - | - | 0 | - |
| - | - | 5086 | 367.7 | - | - | 0 | - |
| - | - | 2819 | 368.2 | - | - | 0 | - |
| - | - | 1147 | 371.2 | - | - | 0 | - |
| - | - | 9081 | 376.2 | - | - | 0 | - |
| - | - | 3967 | 376.7 | - | - | 0 | - |
| - | - | 833 | 377.2 | - | - | 0 | - |
| - | - | 8249 | 379.2 | - | - | 0 | - |
| - | - | 649.2 | 380.2 | - | - | 0 | - |
| - | - | 1.94E+04 | 381.2 | - | - | 0 | - |
| - | - | 3041 | 382.2 | - | - | 0 | - |
| - | - | 836.2 | 385.1 | - | - | 0 | - |
| 0 | Precursor | 1159 | 390.2 | 0.0001686 | 0.432 | +2 | -1 |
| 0 | Precursor | 859.8 | 390.7 | 0.005323 | 13.62 | +2 | -1 |
| - | - | 713.3 | 398.2 | - | - | 0 | - |
| - | - | 6631 | 398.2 | - | - | 0 | - |
| - | - | 1296 | 399.2 | - | - | 0 | - |
| 0 | Precursor | 3812 | 399.2 | 1.317E-05 | 0.03299 | +2 | -1 |
| - | - | 1097 | 399.2 | - | - | 0 | - |
| - | - | 1468 | 399.3 | - | - | 0 | - |
| - | - | 2099 | 399.7 | - | - | 0 | - |
| - | - | 1036 | 400.2 | - | - | 0 | - |
| - | - | 1022 | 400.3 | - | - | 0 | - |
| - | - | 1300 | 402.2 | - | - | 0 | - |
| - | - | 1187 | 407.2 | - | - | 0 | - |
| - | - | 736 | 413.1 | - | - | 0 | - |
| - | - | 1015 | 421.1 | - | - | 0 | - |
| 4 | y | 5578 | 430.2 | 0.0001129 | 0.2624 | +1 | 3 |
| 4 | y | 800.1 | 431.2 | 0.004087 | 9.479 | +1 | 3 |
| - | - | 792.3 | 431.2 | - | - | 0 | - |
| 4 | y | 4441 | 448.2 | 0.0009496 | 2.119 | +1 | 3 |
| - | - | 893.8 | 449.2 | - | - | 0 | - |
| - | - | 1383 | 456.2 | - | - | 0 | - |
| - | - | 952.3 | 459.2 | - | - | 0 | - |
| 4 | b | 955.6 | 461.3 | 6.832E-05 | 0.1481 | +1 | 4 |
| - | - | 1592 | 477.2 | - | - | 0 | - |
| 4 | b | 2618 | 478.3 | 0.0007714 | 1.613 | +1 | 4 |
| - | - | 728.4 | 479.3 | - | - | 0 | - |
| - | - | 983 | 491.3 | - | - | 0 | - |
| - | - | 1.019E+04 | 494.2 | - | - | 0 | - |
| - | - | 2356 | 495.2 | - | - | 0 | - |
| - | - | 1081 | 522.2 | - | - | 0 | - |
| - | - | 3139 | 539.2 | - | - | 0 | - |
| - | - | 1144 | 540.2 | - | - | 0 | - |
| - | - | 3667 | 550.2 | - | - | 0 | - |
| - | - | 1322 | 551.2 | - | - | 0 | - |
| - | - | 846.7 | 558.3 | - | - | 0 | - |
| 3 | y | 869.3 | 567.2 | 0.00275 | 4.849 | +1 | 4 |
| 3 | y | 1560 | 568.2 | 0.001821 | 3.204 | +1 | 4 |
| 5 | b | 842 | 575.3 | 0.003461 | 6.016 | +1 | 5 |
| 5 | b | 3228 | 576.3 | 0.0002554 | 0.4432 | +1 | 5 |
| - | - | 805.8 | 577.3 | - | - | 0 | - |
| 3 | y | 1.001E+04 | 585.2 | 0.0004976 | 0.8503 | +1 | 4 |
| - | - | 2891 | 586.2 | - | - | 0 | - |
| 5 | b | 1.885E+04 | 593.3 | 0.0002927 | 0.4934 | +1 | 5 |
| - | - | 4755 | 594.3 | - | - | 0 | - |
| - | - | 1337 | 595.3 | - | - | 0 | - |
| - | - | 941.1 | 610.3 | - | - | 0 | - |
| - | - | 1049 | 611.3 | - | - | 0 | - |
| - | - | 694.8 | 636.3 | - | - | 0 | - |
| - | - | 831.6 | 652.3 | - | - | 0 | - |
| - | - | 951.2 | 663.3 | - | - | 0 | - |
| - | - | 743.1 | 668.3 | - | - | 0 | - |
| 2 | y | 2116 | 680.3 | 0.001304 | 1.917 | +1 | 5 |
| 2 | y | 3768 | 681.3 | 3.192E-05 | 0.04685 | +1 | 5 |
| - | - | 1344 | 682.3 | - | - | 0 | - |
| 2 | y | 1.205E+04 | 698.3 | 0.0006993 | 1.001 | +1 | 5 |
| - | - | 4810 | 699.3 | - | - | 0 | - |
| - | - | 755.9 | 700.3 | - | - | 0 | - |
| - | - | 746.2 | 1313 | - | - | 0 | - |
| - | - | 632.3 | 1827 | - | - | 0 | - |
| - | - | 705.2 | 3019 | - | - | 0 | - |
| - | - | 673.3 | 3341 | - | - | 0 | - |

m/z Charge Intensity FragmentType MassShift Position
120.05565643310547 0 369.85767
120.0657958984375 0 426.53787
120.08109283447266 0 2844.7668
121.50505828857422 0 372.58185
122.07156372070312 0 1068.0482
123.05572509765625 0 344.4737
123.11714935302734 0 712.6883
125.07132720947266 0 554.13226
126.1024398803711 0 616.33276
127.08687591552734 0 477.64194
128.08206176757812 0 499.1461
128.10740661621094 0 652.11957
129.05496215820312 0 3776.4954
129.06619262695312 0 611.17487
129.1025848388672 0 3669.3562
130.06544494628906 0 22220.7
130.0863494873047 0 549.4144
131.06906127929688 0 2077.313
132.0811004638672 0 6664.4165
132.94447326660156 0 373.75916
133.06118774414062 0 1307.9622
133.0862274169922 0 13586.877
134.08958435058594 0 485.52563
135.85374450683594 0 416.8662
136.01866149902344 0 640.86505
136.0758514404297 0 1430.9784
137.0830078125 0 421.0925
138.0345916748047 0 434.38077
138.06658935546875 0 2921.6226
139.05047607421875 0 883.313
139.08712768554688 0 440.65567
140.08218383789062 0 1161.9417
141.1024932861328 0 1630.3982
142.06564331054688 0 1193.5333
142.10586547851562 0 524.95245
142.12257385253906 0 471.3676
143.04537963867188 0 807.1591
143.0819854736328 0 480.87164
143.11776733398438 0 499.18195 d 1
144.0812225341797 0 1792.0548
144.10226440429688 0 1234.8646
144.4281768798828 0 393.7122
145.0501251220703 0 528.0873
145.0862274169922 0 456.3647
145.39300537109375 0 471.67154
146.06036376953125 0 10444.194
146.0669403076172 0 694.0582
147.0653076171875 0 3541.424
148.947265625 0 656.27374
153.05386352539062 0 574.44324
154.09799194335938 0 693.87317
155.08212280273438 0 857.53406
155.09307861328125 0 9715.853
155.11830139160156 0 812.1382
156.0771026611328 0 1050.5825
156.0963134765625 0 802.1279
157.08734130859375 0 455.9892
157.0976104736328 0 559.02155
157.13388061523438 0 616.5555
158.08413696289062 0 1441.5815
159.09197998046875 0 48727.215
160.0758514404297 0 1283.8927
160.09552001953125 0 5169.187
161.08084106445312 0 651.17413
162.16761779785156 0 442.0818
163.02919006347656 0 644.16705
163.78176879882812 0 393.37274
165.10191345214844 0 449.13074
166.0613555908203 0 8123.0547
167.064697265625 0 620.6457
169.07655334472656 0 775.5368
169.0975341796875 0 1285.6665
170.0604248046875 0 3958.8726
171.07693481445312 0 614.0737
173.45127868652344 0 1689.2881
175.00686645507812 0 802.0297
175.015869140625 0 2001.0452
175.0292205810547 0 612.48627
175.09715270996094 0 1291.4862
176.08241271972656 0 1130.7462
177.11244201660156 0 4078.2634
178.13421630859375 0 4586.3296
179.09271240234375 0 596.61725
181.06100463867188 0 1693.9164
183.08853149414062 0 549.5505
183.1498260498047 0 781.1625
185.16514587402344 0 15703.527 a 1
186.1685333251953 0 1710.3442
187.08705139160156 0 2210.8914
187.10791015625 0 922.35425
187.9763946533203 0 502.64603
188.07089233398438 0 60942.1
189.0742950439453 0 7350.8213
190.13365173339844 0 497.65903
198.08767700195312 0 1633.8768
198.86712646484375 0 553.7318
199.07257080078125 0 2760.4004
202.10752868652344 0 1411.5709
204.1234588623047 0 1083.1299
205.08628845214844 0 861.055
205.09744262695312 0 17404.824 y 5
205.10693359375 0 1115.9923
205.14500427246094 0 586.41656
206.1007843017578 0 1537.9651
207.16094970703125 0 759.00806
213.13925170898438 0 784.742
213.15992736816406 0 2993.7576 b 1
214.11880493164062 0 664.56696
215.103759765625 0 540.2819
216.09828186035156 0 827.9585
218.03494262695312 0 670.2245
220.15476989746094 0 702.19946
221.0716094970703 0 615.74835
221.103515625 0 1138.0786
223.15557861328125 0 5731.002
226.08253479003906 0 2692.8674
227.06643676757812 0 2162.666
228.16009521484375 0 702.89954
230.0364227294922 0 713.27893
231.08822631835938 0 793.79736
232.14450073242188 0 1458.751
233.13990783691406 0 3017.2166
234.1239471435547 0 8878.059
235.12767028808594 0 1066.906
238.13023376464844 0 752.4384
239.1497039794922 0 1070.5891
240.13461303710938 0 510.10226
243.1091766357422 0 576.9724
244.09310913085938 0 4533.53
248.11453247070312 0 2986.0747
249.0984344482422 0 5301.529
251.15054321289062 0 27004.453
252.1535186767578 0 3507.0864
254.6553192138672 0 624.63916
255.1962432861328 0 1225.0808
256.1070251464844 0 693.0017
261.1173400878906 0 705.0258
261.1343078613281 0 1002.59863
261.1626892089844 0 701.61163
262.0500183105469 0 2796.838
266.1249694824219 0 15221.965
267.12811279296875 0 2299.4478
270.12176513671875 0 9059.86
270.623291015625 0 2244.136
271.1240539550781 0 735.4462
274.1154479980469 0 961.6453
274.1300964355469 0 3387.4966
276.1083679199219 0 727.29895
279.1451416015625 0 1533.2905
283.1751708984375 0 1107.0153
284.1029968261719 0 665.95
284.1189270019531 0 815.54956 y Water loss 2
285.0869445800781 0 758.2012
291.1188659667969 0 531.46655
297.15557861328125 0 2081.9353 b 4
297.65716552734375 0 947.1813
299.16168212890625 0 1147.758
302.11181640625 0 904.37445 y Water loss 4
306.2292175292969 0 1092.682
318.1513671875 0 891.162
320.12432861328125 0 3954.2715 y 4
321.126708984375 0 550.74286
326.66461181640625 0 2482.8857
327.2016296386719 0 3983.5984
332.20849609375 0 4014.0645
333.2129821777344 0 1091.6864
345.1305236816406 0 2944.0947
345.15240478515625 0 557.2842
346.11492919921875 0 7195.6816
347.1171569824219 0 869.2599
350.21875 0 15818.601 b 2
351.2212829589844 0 3614.308
355.1968078613281 0 990.495
361.1987609863281 0 658.1096
362.180908203125 0 973.2185
363.1413879394531 0 7079.8784
364.12518310546875 0 3784.6719
367.14056396484375 0 704.4363
367.1922912597656 0 668.3127
367.2446594238281 0 2563.862
367.6850891113281 0 5086.108
368.184814453125 0 2818.8752
371.22601318359375 0 1147.3298
376.1980895996094 0 9081.215
376.69915771484375 0 3967.057
377.2007751464844 0 833.0042
379.20880126953125 0 8249.286
380.2120666503906 0 649.19446
381.15179443359375 0 19401.479
382.1544494628906 0 3041.3884
385.14984130859375 0 836.2276
390.195556640625 0 1159.0856 Precursor Water loss
390.6927185058594 0 859.7598 Precursor Ammonia loss
398.1767272949219 0 713.28107
398.23858642578125 0 6630.7485
399.16241455078125 0 1295.9583
399.20068359375 0 3812.2686 Precursor
399.2439880371094 0 1097.4958
399.29638671875 0 1468.032
399.70233154296875 0 2098.7295
400.218017578125 0 1036.0023
400.2510070800781 0 1022.1218
402.17755126953125 0 1299.6758
407.20404052734375 0 1187.1713
413.1457214355469 0 735.9654
421.1480407714844 0 1015.4003
430.1719970703125 0 5578.4463 y Water loss 3
431.15203857421875 0 800.13947 y Ammonia loss 3
431.1802978515625 0 792.3353
448.1836242675781 0 4441.126 y 3
449.1849365234375 0 893.79083
456.1833190917969 0 1383.286
459.1998291015625 0 952.3337
461.2507629394531 0 955.59937 b Ammonia loss 3
477.2098388671875 0 1591.6582
478.27801513671875 0 2617.5388 b 3
479.2801208496094 0 728.4267
491.2785339355469 0 982.9952
494.23577880859375 0 10189.419
495.23760986328125 0 2355.709
522.2315063476562 0 1080.8356
539.2354736328125 0 3139.2114
540.2380981445312 0 1144.3054
550.2037963867188 0 3666.586
551.2069091796875 0 1321.8362
558.2713012695312 0 846.7397
567.228271484375 0 869.3264 y Water loss 2
568.2168579101562 0 1560.3444 y Ammonia loss 2
575.2901611328125 0 841.988 b Water loss 4
576.2778930664062 0 3228.2896 b Ammonia loss 4
577.2793579101562 0 805.79034
585.2410888671875 0 10009.472 y 2
586.245361328125 0 2891.117
593.3038940429688 0 18851.361 b 4
594.3053588867188 0 4754.9517
595.308349609375 0 1337.4324
610.3327026367188 0 941.07104
611.316650390625 0 1048.7712
636.327880859375 0 694.7888
652.319091796875 0 831.6346
663.2900390625 0 951.2116
668.3330688476562 0 743.05896
680.3137817382812 0 2116.0085 y Water loss 1
681.2991333007812 0 3768.0425 y Ammonia loss 1
682.302734375 0 1344.0635
698.324951171875 0 12054.487 y 1
699.3282470703125 0 4810.257
700.3309326171875 0 755.8669
1313.334716796875 0 746.19946
1827.022705078125 0 632.2542
3018.55419921875 0 705.19434
3340.970458984375 0 673.2638

Spectrum Details

|  |  |
| --- | --- |
| Matched peaks? Matched peaks The total absolute number of peaks matched. Additionally in brackets the total fraction of peaks matched and the total number of peaks is shown. | 26 (10.32% of 252) |
| FDR? FDR The false discovery rate estimated for this peptide. It is calculated by matching all theoretical fragments with a non-integer shift with the raw peaks for this spectrum. This is done with 40 different shifts. The resulting percentage is the average number of annotated peaks over the number of annotated peaks with the correct spectrum. | 0.46% |
| Satellite FDR? Satellite FDR See the FDR for details on its calculation. This satellite ion specific FDR only contains the satellite ions (d/w) for I/L/J positions. | 0.00% |
| PSM Score? PSM Score The PSM Score as given by Hecklib to this annotated spectrum. It is shown with three significant figures. | 360 |

#### Spectrum 4673? Spectrum 4673 The raw spectrum of this peptide as annotated by Hecklib. The fragments are coloured according to ion type (see legend). Any peaks with a star '\*' as text can be hovered over to see the full details, first the ion type second the mass shift type. By hovering over the amino acids in the peptide or ions in the legend the corresponding peaks are highlighted. By toggling the 'Unassigned' label you can turn the background (unassigned) peaks on or off in the plot. By updating the slider in the Ion legend you can update the spectrum to only show the top X% of the peaks with labels. The top X% means any peak that is within X% of the highest intensity. By dragging in the spectrum you can zoom in to a specific part of the spectrum and use 'Zoom Out' to get back to the original zoom level. The annotation of the spectrum is based on the given sequence in the peptides file and is done with different software so inconsistencies are likely. The peaks are annotated based on the given sequence, with 20 ppm tolerance.

Copy Data

##### Spectrum 4673 (TSV)

###### Preview

```
Loading example...
```

*Click on the button to copy the data to your clipboard.*

Mz MinMz MaxIntensity Max

WidthHeightPeptide font sizePeptide stroke widthSpectrum font sizeSpectrum stroke widthCompact peptide

Ion legend

wxyz

abcd

OtherUnassignedIon^Charge^_Position_Show for top:%

VJHQDW

02.01e+44.02e+46.03e+48.04e+4

Zoom Out

a^+2^_3_a^+1^_2_b^+1^_2_b^+2^_5_y^+2^_4_y^+1^_2_a^+1^_3_y^+1^_2_b^+1^_3_\*\*y^+1^_3_y^+1^_3_b^+1^_4_y^+1^_3_b^+1^_4_b^+1^_5_b^+1^_5_y^+1^_4_y^+1^_4_b^+1^_5_y^+1^_4_y^+1^_5_y^+1^_5_y^+1^_5_

0763152522883051

Fragment Matches Table

Show background peaks

| Position | Ion type | Intensity | mz Theoretical | mz Error (Th) | mz Error (ppm) | Charge | Series Number |
| --- | --- | --- | --- | --- | --- | --- | --- |
| - | - | 787.3 | 120.1 | - | - | 0 | - |
| - | - | 2708 | 120.1 | - | - | 0 | - |
| - | - | 797.9 | 121 | - | - | 0 | - |
| - | - | 1646 | 122.1 | - | - | 0 | - |
| - | - | 886.7 | 123.1 | - | - | 0 | - |
| - | - | 1140 | 123.1 | - | - | 0 | - |
| - | - | 554.4 | 126.1 | - | - | 0 | - |
| - | - | 544.6 | 127.1 | - | - | 0 | - |
| - | - | 724.6 | 128 | - | - | 0 | - |
| - | - | 603.4 | 128.1 | - | - | 0 | - |
| - | - | 1127 | 129.1 | - | - | 0 | - |
| - | - | 3009 | 129.1 | - | - | 0 | - |
| - | - | 429.1 | 130 | - | - | 0 | - |
| - | - | 2.349E+04 | 130.1 | - | - | 0 | - |
| - | - | 523.7 | 130.1 | - | - | 0 | - |
| - | - | 2116 | 131.1 | - | - | 0 | - |
| - | - | 4599 | 132 | - | - | 0 | - |
| - | - | 2087 | 132.1 | - | - | 0 | - |
| - | - | 602 | 133 | - | - | 0 | - |
| - | - | 2959 | 133.1 | - | - | 0 | - |
| - | - | 811.8 | 133.1 | - | - | 0 | - |
| - | - | 1135 | 134 | - | - | 0 | - |
| - | - | 1061 | 134.1 | - | - | 0 | - |
| - | - | 939.7 | 136.1 | - | - | 0 | - |
| - | - | 5184 | 136.1 | - | - | 0 | - |
| - | - | 521.4 | 137.1 | - | - | 0 | - |
| - | - | 3981 | 138.1 | - | - | 0 | - |
| - | - | 881.8 | 139.1 | - | - | 0 | - |
| - | - | 831.5 | 140.1 | - | - | 0 | - |
| - | - | 932.5 | 143 | - | - | 0 | - |
| - | - | 7.137E+04 | 146.1 | - | - | 0 | - |
| - | - | 5696 | 147.1 | - | - | 0 | - |
| - | - | 5344 | 148 | - | - | 0 | - |
| - | - | 4970 | 148.1 | - | - | 0 | - |
| - | - | 455.1 | 148.8 | - | - | 0 | - |
| - | - | 544.8 | 148.9 | - | - | 0 | - |
| - | - | 689.3 | 148.9 | - | - | 0 | - |
| - | - | 761.7 | 148.9 | - | - | 0 | - |
| - | - | 985 | 148.9 | - | - | 0 | - |
| - | - | 1305 | 148.9 | - | - | 0 | - |
| - | - | 1492 | 148.9 | - | - | 0 | - |
| - | - | 2903 | 148.9 | - | - | 0 | - |
| - | - | 4236 | 148.9 | - | - | 0 | - |
| - | - | 4166 | 149 | - | - | 0 | - |
| - | - | 2318 | 149 | - | - | 0 | - |
| - | - | 1468 | 149 | - | - | 0 | - |
| - | - | 1147 | 149 | - | - | 0 | - |
| - | - | 653.7 | 149 | - | - | 0 | - |
| - | - | 806.1 | 149 | - | - | 0 | - |
| - | - | 731.6 | 149 | - | - | 0 | - |
| - | - | 687.3 | 149 | - | - | 0 | - |
| - | - | 1447 | 149 | - | - | 0 | - |
| - | - | 633.1 | 149 | - | - | 0 | - |
| - | - | 456.9 | 149.1 | - | - | 0 | - |
| - | - | 852.6 | 151.1 | - | - | 0 | - |
| - | - | 650.1 | 152.1 | - | - | 0 | - |
| - | - | 696.7 | 154.1 | - | - | 0 | - |
| - | - | 1196 | 154.1 | - | - | 0 | - |
| - | - | 1850 | 155.1 | - | - | 0 | - |
| - | - | 1.656E+04 | 155.1 | - | - | 0 | - |
| - | - | 1338 | 156.1 | - | - | 0 | - |
| - | - | 1228 | 156.1 | - | - | 0 | - |
| - | - | 5444 | 157.1 | - | - | 0 | - |
| - | - | 3.93E+04 | 158.1 | - | - | 0 | - |
| - | - | 586.1 | 159.1 | - | - | 0 | - |
| - | - | 3018 | 159.1 | - | - | 0 | - |
| - | - | 1.224E+04 | 159.1 | - | - | 0 | - |
| - | - | 510.1 | 160.1 | - | - | 0 | - |
| - | - | 1593 | 160.1 | - | - | 0 | - |
| - | - | 1586 | 160.1 | - | - | 0 | - |
| - | - | 3957 | 161.1 | - | - | 0 | - |
| 3 | a | 516.1 | 161.6 | 0.0002849 | 1.763 | +2 | 3 |
| - | - | 5633 | 162.1 | - | - | 0 | - |
| - | - | 615.7 | 162.1 | - | - | 0 | - |
| - | - | 537.1 | 163 | - | - | 0 | - |
| - | - | 489.9 | 163.1 | - | - | 0 | - |
| - | - | 861.2 | 164.1 | - | - | 0 | - |
| - | - | 1003 | 165.1 | - | - | 0 | - |
| - | - | 1.333E+04 | 166.1 | - | - | 0 | - |
| - | - | 743.6 | 167 | - | - | 0 | - |
| - | - | 1176 | 167.1 | - | - | 0 | - |
| - | - | 497.8 | 169.7 | - | - | 0 | - |
| - | - | 1199 | 170.1 | - | - | 0 | - |
| - | - | 7376 | 173.1 | - | - | 0 | - |
| - | - | 5306 | 174.1 | - | - | 0 | - |
| - | - | 896.6 | 174.1 | - | - | 0 | - |
| - | - | 714.6 | 174.1 | - | - | 0 | - |
| - | - | 7.959E+04 | 175.1 | - | - | 0 | - |
| - | - | 1443 | 176.1 | - | - | 0 | - |
| - | - | 8306 | 176.1 | - | - | 0 | - |
| - | - | 528.7 | 177.1 | - | - | 0 | - |
| - | - | 676.7 | 178.1 | - | - | 0 | - |
| - | - | 6644 | 178.1 | - | - | 0 | - |
| - | - | 545.6 | 179.1 | - | - | 0 | - |
| - | - | 544.1 | 179.1 | - | - | 0 | - |
| - | - | 678.3 | 181.1 | - | - | 0 | - |
| - | - | 3738 | 181.1 | - | - | 0 | - |
| - | - | 567.2 | 183.1 | - | - | 0 | - |
| - | - | 739.7 | 183.1 | - | - | 0 | - |
| - | - | 758.7 | 185 | - | - | 0 | - |
| - | - | 1924 | 185.1 | - | - | 0 | - |
| 2 | a | 2.718E+04 | 185.2 | 0.0002604 | 1.406 | +1 | 2 |
| - | - | 4045 | 186.1 | - | - | 0 | - |
| - | - | 2338 | 186.2 | - | - | 0 | - |
| - | - | 691.1 | 187.1 | - | - | 0 | - |
| - | - | 887.8 | 187.1 | - | - | 0 | - |
| - | - | 542.9 | 187.1 | - | - | 0 | - |
| - | - | 615.5 | 187.1 | - | - | 0 | - |
| - | - | 1.09E+04 | 188.1 | - | - | 0 | - |
| - | - | 1836 | 189.1 | - | - | 0 | - |
| - | - | 1983 | 190.1 | - | - | 0 | - |
| - | - | 8786 | 191.1 | - | - | 0 | - |
| - | - | 566.3 | 197.1 | - | - | 0 | - |
| - | - | 2950 | 197.1 | - | - | 0 | - |
| - | - | 1374 | 197.6 | - | - | 0 | - |
| - | - | 3808 | 198.1 | - | - | 0 | - |
| - | - | 1558 | 199.1 | - | - | 0 | - |
| - | - | 466.1 | 199.1 | - | - | 0 | - |
| - | - | 1798 | 201.1 | - | - | 0 | - |
| - | - | 941 | 201.1 | - | - | 0 | - |
| - | - | 2391 | 202.1 | - | - | 0 | - |
| - | - | 1842 | 203.1 | - | - | 0 | - |
| - | - | 1.035E+04 | 203.1 | - | - | 0 | - |
| - | - | 1130 | 203.1 | - | - | 0 | - |
| - | - | 3.445E+04 | 204.1 | - | - | 0 | - |
| - | - | 1135 | 204.1 | - | - | 0 | - |
| - | - | 766.1 | 204.1 | - | - | 0 | - |
| - | - | 3185 | 205.1 | - | - | 0 | - |
| - | - | 657.5 | 205.1 | - | - | 0 | - |
| - | - | 3229 | 205.1 | - | - | 0 | - |
| - | - | 596.5 | 206.1 | - | - | 0 | - |
| - | - | 865 | 206.1 | - | - | 0 | - |
| - | - | 572.3 | 207.2 | - | - | 0 | - |
| - | - | 1026 | 211.1 | - | - | 0 | - |
| - | - | 551.2 | 211.2 | - | - | 0 | - |
| 2 | b | 6052 | 213.2 | 0.0002188 | 1.026 | +1 | 2 |
| - | - | 942.7 | 214.2 | - | - | 0 | - |
| - | - | 592.8 | 215.1 | - | - | 0 | - |
| - | - | 532 | 215.1 | - | - | 0 | - |
| - | - | 2507 | 216.1 | - | - | 0 | - |
| - | - | 574.2 | 216.1 | - | - | 0 | - |
| - | - | 952.5 | 218.1 | - | - | 0 | - |
| - | - | 6518 | 219.1 | - | - | 0 | - |
| - | - | 3676 | 219.1 | - | - | 0 | - |
| - | - | 1480 | 220.1 | - | - | 0 | - |
| - | - | 1074 | 220.1 | - | - | 0 | - |
| - | - | 3.063E+04 | 221.1 | - | - | 0 | - |
| - | - | 2714 | 221.1 | - | - | 0 | - |
| - | - | 3178 | 222.1 | - | - | 0 | - |
| - | - | 1.133E+04 | 223.2 | - | - | 0 | - |
| - | - | 1232 | 224.2 | - | - | 0 | - |
| - | - | 702.4 | 225.6 | - | - | 0 | - |
| - | - | 482.2 | 226 | - | - | 0 | - |
| - | - | 4591 | 226.1 | - | - | 0 | - |
| - | - | 715 | 226.1 | - | - | 0 | - |
| - | - | 4542 | 227.1 | - | - | 0 | - |
| - | - | 822.5 | 227.1 | - | - | 0 | - |
| - | - | 581.2 | 227.1 | - | - | 0 | - |
| - | - | 846.9 | 229.1 | - | - | 0 | - |
| - | - | 759.9 | 232.1 | - | - | 0 | - |
| - | - | 5371 | 233.1 | - | - | 0 | - |
| - | - | 1.536E+04 | 234.1 | - | - | 0 | - |
| - | - | 630.7 | 235.1 | - | - | 0 | - |
| - | - | 808.6 | 235.1 | - | - | 0 | - |
| - | - | 2084 | 235.1 | - | - | 0 | - |
| - | - | 917.2 | 237.1 | - | - | 0 | - |
| - | - | 1194 | 238.1 | - | - | 0 | - |
| - | - | 1696 | 239.2 | - | - | 0 | - |
| - | - | 612.1 | 240.1 | - | - | 0 | - |
| - | - | 525.4 | 241.5 | - | - | 0 | - |
| - | - | 1370 | 243.1 | - | - | 0 | - |
| - | - | 8098 | 244.1 | - | - | 0 | - |
| - | - | 1371 | 244.1 | - | - | 0 | - |
| - | - | 672.3 | 244.7 | - | - | 0 | - |
| - | - | 695 | 245.1 | - | - | 0 | - |
| - | - | 865.6 | 246.1 | - | - | 0 | - |
| - | - | 2539 | 247.1 | - | - | 0 | - |
| - | - | 5855 | 248.1 | - | - | 0 | - |
| - | - | 1.075E+04 | 249.1 | - | - | 0 | - |
| - | - | 1606 | 249.1 | - | - | 0 | - |
| - | - | 1384 | 250.1 | - | - | 0 | - |
| - | - | 1820 | 251 | - | - | 0 | - |
| - | - | 4.355E+04 | 251.2 | - | - | 0 | - |
| - | - | 6976 | 252.2 | - | - | 0 | - |
| - | - | 1058 | 252.7 | - | - | 0 | - |
| - | - | 733.5 | 254.6 | - | - | 0 | - |
| - | - | 1488 | 255.1 | - | - | 0 | - |
| - | - | 721.8 | 256.2 | - | - | 0 | - |
| - | - | 1019 | 260.6 | - | - | 0 | - |
| - | - | 1541 | 261.1 | - | - | 0 | - |
| - | - | 1560 | 261.1 | - | - | 0 | - |
| - | - | 1348 | 261.2 | - | - | 0 | - |
| - | - | 1133 | 261.7 | - | - | 0 | - |
| - | - | 706.3 | 262.1 | - | - | 0 | - |
| - | - | 686.7 | 265.1 | - | - | 0 | - |
| - | - | 2.926E+04 | 266.1 | - | - | 0 | - |
| - | - | 3286 | 267.1 | - | - | 0 | - |
| - | - | 992.2 | 267.1 | - | - | 0 | - |
| - | - | 715.8 | 268.2 | - | - | 0 | - |
| - | - | 1145 | 269.1 | - | - | 0 | - |
| - | - | 971.9 | 269.6 | - | - | 0 | - |
| - | - | 4162 | 270.1 | - | - | 0 | - |
| - | - | 1031 | 270.6 | - | - | 0 | - |
| - | - | 1224 | 272.1 | - | - | 0 | - |
| - | - | 3125 | 273.1 | - | - | 0 | - |
| - | - | 5086 | 274.1 | - | - | 0 | - |
| - | - | 842 | 274.7 | - | - | 0 | - |
| - | - | 653.9 | 275.1 | - | - | 0 | - |
| - | - | 864.3 | 275.2 | - | - | 0 | - |
| - | - | 9293 | 278.1 | - | - | 0 | - |
| - | - | 3209 | 278.6 | - | - | 0 | - |
| - | - | 2913 | 279.1 | - | - | 0 | - |
| - | - | 945.7 | 282.1 | - | - | 0 | - |
| - | - | 609.4 | 283.1 | - | - | 0 | - |
| - | - | 9022 | 283.2 | - | - | 0 | - |
| - | - | 4304 | 283.7 | - | - | 0 | - |
| - | - | 635.1 | 284.1 | - | - | 0 | - |
| - | - | 746.5 | 284.2 | - | - | 0 | - |
| - | - | 1809 | 290.1 | - | - | 0 | - |
| - | - | 1876 | 290.2 | - | - | 0 | - |
| - | - | 1009 | 291.2 | - | - | 0 | - |
| - | - | 1243 | 294.1 | - | - | 0 | - |
| - | - | 1075 | 294.2 | - | - | 0 | - |
| 5 | b | 2573 | 297.2 | 0.0001217 | 0.4094 | +2 | 5 |
| - | - | 1523 | 297.7 | - | - | 0 | - |
| - | - | 593 | 300.1 | - | - | 0 | - |
| - | - | 633.5 | 301.1 | - | - | 0 | - |
| 3 | y | 1589 | 301.1 | 0.0007486 | 2.486 | +2 | 4 |
| - | - | 1038 | 301.1 | - | - | 0 | - |
| - | - | 2184 | 302.1 | - | - | 0 | - |
| - | - | 941.2 | 304.2 | - | - | 0 | - |
| - | - | 2846 | 306.2 | - | - | 0 | - |
| - | - | 1356 | 307.6 | - | - | 0 | - |
| - | - | 756.2 | 309.2 | - | - | 0 | - |
| - | - | 553.4 | 317.1 | - | - | 0 | - |
| 5 | y | 1956 | 318.1 | 0.005843 | 18.37 | +1 | 2 |
| - | - | 939.6 | 318.1 | - | - | 0 | - |
| - | - | 797.9 | 318.2 | - | - | 0 | - |
| - | - | 639.5 | 318.6 | - | - | 0 | - |
| - | - | 855 | 319.1 | - | - | 0 | - |
| - | - | 1416 | 319.2 | - | - | 0 | - |
| - | - | 810.9 | 320.1 | - | - | 0 | - |
| 3 | a | 1677 | 322.2 | 6.433E-05 | 0.1996 | +1 | 3 |
| - | - | 776 | 325.2 | - | - | 0 | - |
| - | - | 890.1 | 327.2 | - | - | 0 | - |
| - | - | 642.8 | 329.5 | - | - | 0 | - |
| - | - | 707.7 | 331.1 | - | - | 0 | - |
| - | - | 7201 | 332.2 | - | - | 0 | - |
| - | - | 1517 | 333.2 | - | - | 0 | - |
| - | - | 1342 | 334.2 | - | - | 0 | - |
| - | - | 3468 | 334.7 | - | - | 0 | - |
| - | - | 3403 | 335.7 | - | - | 0 | - |
| 5 | y | 3738 | 336.1 | 0.005166 | 15.37 | +1 | 2 |
| - | - | 1741 | 336.2 | - | - | 0 | - |
| - | - | 1183 | 344.2 | - | - | 0 | - |
| - | - | 3022 | 345.1 | - | - | 0 | - |
| - | - | 1.231E+04 | 346.1 | - | - | 0 | - |
| - | - | 1793 | 347.1 | - | - | 0 | - |
| 3 | b | 3.105E+04 | 350.2 | 2.275E-05 | 0.06497 | +1 | 3 |
| - | - | 7108 | 351.2 | - | - | 0 | - |
| - | - | 958.2 | 352.2 | - | - | 0 | - |
| - | - | 4802 | 353.2 | - | - | 0 | - |
| - | - | 573.6 | 357.4 | - | - | 0 | - |
| - | - | 858.7 | 361.2 | - | - | 0 | - |
| - | - | 1050 | 362.2 | - | - | 0 | - |
| - | - | 1.476E+04 | 363.1 | - | - | 0 | - |
| - | - | 4530 | 364.1 | - | - | 0 | - |
| - | - | 1708 | 364.1 | - | - | 0 | - |
| - | - | 1220 | 365.1 | - | - | 0 | - |
| - | - | 914.3 | 366.7 | - | - | 0 | - |
| - | - | 720 | 367.2 | - | - | 0 | - |
| - | - | 4790 | 367.2 | - | - | 0 | - |
| - | - | 1042 | 374.7 | - | - | 0 | - |
| - | - | 847.9 | 375.2 | - | - | 0 | - |
| - | - | 5215 | 375.7 | - | - | 0 | - |
| - | - | 803.2 | 376.2 | - | - | 0 | - |
| - | - | 1.222E+04 | 379.2 | - | - | 0 | - |
| - | - | 2326 | 380.2 | - | - | 0 | - |
| - | - | 3.697E+04 | 381.2 | - | - | 0 | - |
| - | - | 6386 | 382.2 | - | - | 0 | - |
| - | - | 985.3 | 383.1 | - | - | 0 | - |
| - | - | 1129 | 383.2 | - | - | 0 | - |
| - | - | 1914 | 383.2 | - | - | 0 | - |
| - | - | 1506 | 383.7 | - | - | 0 | - |
| - | - | 9868 | 384.2 | - | - | 0 | - |
| - | - | 3940 | 384.7 | - | - | 0 | - |
| - | - | 1162 | 385.2 | - | - | 0 | - |
| - | - | 672.3 | 389.2 | - | - | 0 | - |
| - | - | 793.2 | 391.1 | - | - | 0 | - |
| - | - | 2163 | 392.2 | - | - | 0 | - |
| - | - | 1391 | 392.7 | - | - | 0 | - |
| - | - | 1959 | 393.3 | - | - | 0 | - |
| - | - | 1801 | 397.1 | - | - | 0 | - |
| - | - | 768.7 | 397.2 | - | - | 0 | - |
| - | - | 843.5 | 397.7 | - | - | 0 | - |
| - | - | 4016 | 398.2 | - | - | 0 | - |
| 0 | Precursor | 2018 | 398.7 | 0.005165 | 12.96 | +2 | -1 |
| - | - | 1135 | 401.1 | - | - | 0 | - |
| - | - | 872 | 405.2 | - | - | 0 | - |
| - | - | 1246 | 406.2 | - | - | 0 | - |
| - | - | 1651 | 406.7 | - | - | 0 | - |
| 0 | Precursor | 6304 | 407.2 | 0.003731 | 9.164 | +2 | -1 |
| - | - | 2296 | 407.7 | - | - | 0 | - |
| - | - | 981.3 | 409.1 | - | - | 0 | - |
| - | - | 1175 | 411.1 | - | - | 0 | - |
| - | - | 802.5 | 416.2 | - | - | 0 | - |
| - | - | 2016 | 418.2 | - | - | 0 | - |
| - | - | 1308 | 428.2 | - | - | 0 | - |
| - | - | 1981 | 429.1 | - | - | 0 | - |
| 4 | y | 7877 | 446.2 | 0.004791 | 10.74 | +1 | 3 |
| 4 | y | 1048 | 447.1 | 0.001671 | 3.737 | +1 | 3 |
| - | - | 612.9 | 448.2 | - | - | 0 | - |
| - | - | 949 | 450.3 | - | - | 0 | - |
| - | - | 897.4 | 455.2 | - | - | 0 | - |
| - | - | 1396 | 456.2 | - | - | 0 | - |
| - | - | 821.6 | 459.2 | - | - | 0 | - |
| 4 | b | 787.1 | 461.3 | 0.0007702 | 1.67 | +1 | 4 |
| - | - | 989.2 | 463.2 | - | - | 0 | - |
| 4 | y | 3380 | 464.2 | 0.004541 | 9.783 | +1 | 3 |
| - | - | 1169 | 465.2 | - | - | 0 | - |
| - | - | 3271 | 466.2 | - | - | 0 | - |
| - | - | 1176 | 467.2 | - | - | 0 | - |
| - | - | 589.8 | 471.8 | - | - | 0 | - |
| - | - | 1117 | 476.2 | - | - | 0 | - |
| - | - | 1790 | 477.2 | - | - | 0 | - |
| - | - | 600.9 | 478.2 | - | - | 0 | - |
| 4 | b | 6898 | 478.3 | 0.0003272 | 0.6842 | +1 | 4 |
| - | - | 2242 | 479.3 | - | - | 0 | - |
| - | - | 1235 | 480.3 | - | - | 0 | - |
| - | - | 1360 | 488.3 | - | - | 0 | - |
| - | - | 860.1 | 490.2 | - | - | 0 | - |
| - | - | 1.465E+04 | 494.2 | - | - | 0 | - |
| - | - | 545.6 | 494.3 | - | - | 0 | - |
| - | - | 3783 | 495.2 | - | - | 0 | - |
| - | - | 728 | 504.2 | - | - | 0 | - |
| - | - | 824.4 | 510.2 | - | - | 0 | - |
| - | - | 764.8 | 511.3 | - | - | 0 | - |
| - | - | 972 | 515.2 | - | - | 0 | - |
| - | - | 1797 | 516.2 | - | - | 0 | - |
| - | - | 1936 | 516.3 | - | - | 0 | - |
| - | - | 739.3 | 517.3 | - | - | 0 | - |
| - | - | 1460 | 520.2 | - | - | 0 | - |
| - | - | 1557 | 522.2 | - | - | 0 | - |
| - | - | 689.9 | 537.2 | - | - | 0 | - |
| - | - | 1196 | 538.2 | - | - | 0 | - |
| - | - | 2036 | 539.2 | - | - | 0 | - |
| - | - | 610.4 | 547.3 | - | - | 0 | - |
| - | - | 1097 | 548.2 | - | - | 0 | - |
| - | - | 594.2 | 550.5 | - | - | 0 | - |
| - | - | 4390 | 555.2 | - | - | 0 | - |
| - | - | 793.1 | 556.2 | - | - | 0 | - |
| - | - | 5460 | 565.3 | - | - | 0 | - |
| - | - | 2988 | 566.2 | - | - | 0 | - |
| - | - | 1498 | 566.3 | - | - | 0 | - |
| - | - | 1472 | 568.3 | - | - | 0 | - |
| - | - | 786.2 | 569.3 | - | - | 0 | - |
| 5 | b | 909.2 | 575.3 | 0.0004453 | 0.7741 | +1 | 5 |
| 5 | b | 5531 | 576.3 | 0.0006601 | 1.145 | +1 | 5 |
| - | - | 1143 | 577.3 | - | - | 0 | - |
| - | - | 587.4 | 578.8 | - | - | 0 | - |
| 3 | y | 3275 | 583.2 | 0.005236 | 8.977 | +1 | 4 |
| 3 | y | 1757 | 584.2 | 0.009684 | 16.58 | +1 | 4 |
| - | - | 1135 | 591.3 | - | - | 0 | - |
| 5 | b | 2.952E+04 | 593.3 | 0.0005369 | 0.9049 | +1 | 5 |
| - | - | 1250 | 593.8 | - | - | 0 | - |
| - | - | 9516 | 594.3 | - | - | 0 | - |
| - | - | 1594 | 595.3 | - | - | 0 | - |
| - | - | 891.3 | 599.2 | - | - | 0 | - |
| 3 | y | 1.179E+04 | 601.2 | 0.004681 | 7.785 | +1 | 4 |
| - | - | 4122 | 602.2 | - | - | 0 | - |
| - | - | 1541 | 609.3 | - | - | 0 | - |
| - | - | 4691 | 610.3 | - | - | 0 | - |
| - | - | 1628 | 611.3 | - | - | 0 | - |
| - | - | 771.6 | 650.3 | - | - | 0 | - |
| - | - | 668.8 | 651.3 | - | - | 0 | - |
| - | - | 1216 | 667.3 | - | - | 0 | - |
| - | - | 2714 | 668.3 | - | - | 0 | - |
| - | - | 1477 | 679.3 | - | - | 0 | - |
| - | - | 2778 | 694.3 | - | - | 0 | - |
| - | - | 1436 | 695.3 | - | - | 0 | - |
| 2 | y | 3694 | 696.3 | 0.003935 | 5.652 | +1 | 5 |
| 2 | y | 4156 | 697.3 | 0.007407 | 10.62 | +1 | 5 |
| - | - | 1208 | 698.3 | - | - | 0 | - |
| - | - | 1306 | 712.3 | - | - | 0 | - |
| - | - | 1216 | 713.3 | - | - | 0 | - |
| 2 | y | 1.343E+04 | 714.3 | 0.003746 | 5.245 | +1 | 5 |
| - | - | 699.9 | 714.7 | - | - | 0 | - |
| - | - | 5204 | 715.3 | - | - | 0 | - |
| - | - | 1395 | 716.3 | - | - | 0 | - |
| - | - | 703.6 | 725.3 | - | - | 0 | - |
| - | - | 580.5 | 827.2 | - | - | 0 | - |
| - | - | 601.7 | 861.6 | - | - | 0 | - |
| - | - | 633 | 866.4 | - | - | 0 | - |
| - | - | 588.6 | 1229 | - | - | 0 | - |
| - | - | 593.2 | 1792 | - | - | 0 | - |
| - | - | 631.3 | 2284 | - | - | 0 | - |
| - | - | 763.4 | 3020 | - | - | 0 | - |

m/z Charge Intensity FragmentType MassShift Position
120.0660629272461 0 787.3473
120.08106231689453 0 2707.7883
121.03990936279297 0 797.90643
122.07160186767578 0 1646.4391
123.05558776855469 0 886.68994
123.11710357666016 0 1139.5032
126.06649017333984 0 554.44025
127.08697509765625 0 544.6127
128.0498504638672 0 724.57446
128.08241271972656 0 603.3965
129.0662384033203 0 1126.8597
129.1025390625 0 3009.2915
130.0349884033203 0 429.13318
130.06541442871094 0 23489.055
130.08682250976562 0 523.733
131.06883239746094 0 2115.9788
132.04469299316406 0 4598.8267
132.08106994628906 0 2086.5908
133.0478515625 0 602.03357
133.05262756347656 0 2958.7625
133.06106567382812 0 811.7708
134.0448455810547 0 1134.5316
134.06048583984375 0 1061.0376
136.0508575439453 0 939.7326
136.07595825195312 0 5184.3223
137.05914306640625 0 521.42065
138.06646728515625 0 3981.1523
139.05035400390625 0 881.8334
140.08180236816406 0 831.48663
143.0454864501953 0 932.5337
146.0603485107422 0 71367.92
147.06370544433594 0 5695.5327
148.03964233398438 0 5344.301
148.07594299316406 0 4969.648
148.79222106933594 0 455.1173
148.89285278320312 0 544.7827
148.9003143310547 0 689.2592
148.90731811523438 0 761.6792
148.9143524169922 0 985.0143
148.921630859375 0 1305.1938
148.92877197265625 0 1491.9644
148.93609619140625 0 2902.9666
148.9439239501953 0 4236.3936
148.96046447753906 0 4166.092
148.96832275390625 0 2318.121
148.97543334960938 0 1468.2352
148.98268127441406 0 1146.6305
148.9898681640625 0 653.72595
148.99710083007812 0 806.1049
149.0042266845703 0 731.6101
149.01185607910156 0 687.33276
149.0240020751953 0 1446.6199
149.0331573486328 0 633.14777
149.0550537109375 0 456.92102
151.07553100585938 0 852.618
152.10736083984375 0 650.0851
154.06126403808594 0 696.6894
154.097412109375 0 1195.9641
155.0816192626953 0 1849.7255
155.09298706054688 0 16557.729
156.07691955566406 0 1338.1326
156.09652709960938 0 1228.304
157.07623291015625 0 5443.6245
158.06028747558594 0 39295.773
159.05728149414062 0 586.10614
159.06370544433594 0 3017.9773
159.09190368652344 0 12236.782
160.06455993652344 0 510.13962
160.0759735107422 0 1592.8987
160.0952606201172 0 1586.0194
161.07118225097656 0 3956.7607
161.6157989501953 0 516.1041 a 2
162.0551300048828 0 5633.381
162.09121704101562 0 615.7285
163.02981567382812 0 537.1149
163.0592498779297 0 489.86722
164.08229064941406 0 861.2341
165.07766723632812 0 1003.14026
166.06137084960938 0 13328.693
167.0341796875 0 743.617
167.06442260742188 0 1175.5461
169.72938537597656 0 497.82492
170.0600128173828 0 1198.9436
173.07113647460938 0 7375.621
174.05528259277344 0 5306.2773
174.07855224609375 0 896.5808
174.1026611328125 0 714.5679
175.08688354492188 0 79585.16
176.0826416015625 0 1442.8052
176.09031677246094 0 8306.169
177.09268188476562 0 528.6576
178.0500030517578 0 676.7014
178.13406372070312 0 6644.289
179.0927734375 0 545.6119
179.13803100585938 0 544.102
181.05303955078125 0 678.3441
181.0610809326172 0 3737.6506
183.11288452148438 0 567.1824
183.14947509765625 0 739.7103
185.0470428466797 0 758.663
185.0711669921875 0 1924.3152
185.16510009765625 0 27176.934 a 1
186.05520629882812 0 4044.6514
186.16848754882812 0 2337.529
187.06338500976562 0 691.1125
187.0872802734375 0 887.83966
187.1075897216797 0 542.942
187.14535522460938 0 615.5058
188.07080078125 0 10900.205
189.07431030273438 0 1835.737
190.13421630859375 0 1983.1929
191.08175659179688 0 8785.908
197.103515625 0 566.2899
197.13409423828125 0 2950.1365
197.63592529296875 0 1373.6925
198.08750915527344 0 3807.7073
199.071044921875 0 1557.9452
199.08985900878906 0 466.0532
201.06640625 0 1797.7697
201.1243896484375 0 941.0277
202.0501708984375 0 2390.8376
203.05775451660156 0 1841.6196
203.08164978027344 0 10348.781
203.0924530029297 0 1129.6772
204.06573486328125 0 34450.08
204.08372497558594 0 1134.7861
204.11317443847656 0 766.111
205.06918334960938 0 3184.89
205.0874481201172 0 657.4561
205.0972442626953 0 3228.8176
206.0605926513672 0 596.47876
206.1291961669922 0 864.9552
207.16053771972656 0 572.3197
211.14437866210938 0 1026.31
211.18020629882812 0 551.23114
213.15997314453125 0 6051.715 b 1
214.16383361816406 0 942.7466
215.11822509765625 0 592.78107
215.13909912109375 0 532.02716
216.0981903076172 0 2507.0051
216.12306213378906 0 574.2273
218.06919860839844 0 952.5379
219.07669067382812 0 6518.2334
219.1242218017578 0 3675.8647
220.08056640625 0 1479.5338
220.11936950683594 0 1074.443
221.09228515625 0 30630.83
221.10330200195312 0 2713.9211
222.09576416015625 0 3178.2068
223.1555633544922 0 11326.085
224.15872192382812 0 1231.8881
225.64505004882812 0 702.3546
226.04150390625 0 482.17224
226.08251953125 0 4590.783
226.14675903320312 0 715.02167
227.06655883789062 0 4542.4526
227.0857391357422 0 822.47064
227.11817932128906 0 581.18665
229.134033203125 0 846.92377
232.10809326171875 0 759.8841
233.13999938964844 0 5371.1475
234.12393188476562 0 15360.333
235.0759735107422 0 630.73755
235.09617614746094 0 808.60535
235.12744140625 0 2083.556
237.09083557128906 0 917.20245
238.13038635253906 0 1194.2743
239.15020751953125 0 1695.78
240.1422119140625 0 612.1333
241.51878356933594 0 525.4392
243.10906982421875 0 1369.7388
244.0929718017578 0 8097.956
244.144775390625 0 1370.9961
244.6524658203125 0 672.2921
245.1283416748047 0 694.9855
246.13450622558594 0 865.6047
247.1193084716797 0 2539.2554
248.1144256591797 0 5854.707
249.09844970703125 0 10745.201
249.1353302001953 0 1606.3875
250.1015167236328 0 1383.8663
251.04690551757812 0 1820.3627
251.15048217773438 0 43547.402
252.1538848876953 0 6975.803
252.65020751953125 0 1057.9565
254.6469268798828 0 733.5258
255.07586669921875 0 1488.1608
256.17755126953125 0 721.78937
260.6000671386719 0 1018.518
261.1200866699219 0 1540.8568
261.1346740722656 0 1559.8595
261.1634521484375 0 1348.2487
261.6646423339844 0 1132.5721
262.1036376953125 0 706.3177
265.1417541503906 0 686.66034
266.1248474121094 0 29261.861
267.1282043457031 0 3286.4917
267.1438903808594 0 992.2289
268.1767272949219 0 715.7707
269.1137390136719 0 1145.3351
269.6058654785156 0 971.86365
270.1219482421875 0 4162.229
270.6221923828125 0 1031.2466
272.1033630371094 0 1224.115
273.1233825683594 0 3124.7375
274.129638671875 0 5085.607
274.658447265625 0 841.99493
275.1329345703125 0 653.89606
275.1750793457031 0 864.2896
278.11920166015625 0 9293.157
278.6206970214844 0 3208.619
279.14532470703125 0 2912.8213
282.12115478515625 0 945.7136
283.11199951171875 0 609.391
283.1584167480469 0 9022.071
283.6601867675781 0 4304.063
284.13568115234375 0 635.1163
284.16229248046875 0 746.52936
290.113525390625 0 1808.6982
290.1504211425781 0 1875.6047
291.154052734375 0 1008.51
294.1195983886719 0 1242.6791
294.19268798828125 0 1075.169
297.1558532714844 0 2573.3298 b 4
297.6573486328125 0 1522.7052
300.0970153808594 0 593.00775
301.082275390625 0 633.5496
301.1186828613281 0 1589.3112 y 2
301.1411437988281 0 1038.3473
302.12469482421875 0 2184.1821
304.2135009765625 0 941.15204
306.2292175292969 0 2845.881
307.6200866699219 0 1355.8447
309.16668701171875 0 756.1952
317.13568115234375 0 553.38715
318.109375 0 1955.6549 y Water loss 4
318.13433837890625 0 939.60297
318.1699523925781 0 797.93146
318.6328125 0 639.53735
319.125732421875 0 854.9893
319.1514892578125 0 1416.08
320.1227111816406 0 810.9051
322.22381591796875 0 1676.6586 a 2
325.1630554199219 0 775.9907
327.1661682128906 0 890.1362
329.47344970703125 0 642.7568
331.13763427734375 0 707.74005
332.20831298828125 0 7200.5225
333.2120361328125 0 1517.2076
334.1679992675781 0 1342.3125
334.6623229980469 0 3468.139
335.65093994140625 0 3402.629
336.1192626953125 0 3737.8357 y 4
336.1529846191406 0 1740.9253
344.1709289550781 0 1182.9958
345.1311950683594 0 3022.2134
346.11492919921875 0 12308.281
347.11767578125 0 1792.7134
350.21868896484375 0 31045.275 b 2
351.2215881347656 0 7108.0796
352.223388671875 0 958.2401
353.1572265625 0 4802.034
357.3723449707031 0 573.58215
361.19805908203125 0 858.70294
362.18182373046875 0 1050.2079
363.1410217285156 0 14764.53
364.125 0 4529.806
364.1451110839844 0 1707.8484
365.1263732910156 0 1220.3352
366.67706298828125 0 914.27234
367.17938232421875 0 719.956
367.2452697753906 0 4790.222
374.6752624511719 0 1042.0392
375.1797790527344 0 847.8889
375.6826477050781 0 5215.0005
376.2015686035156 0 803.2134
379.2086486816406 0 12224.599
380.2115478515625 0 2325.9324
381.1517639160156 0 36974.637
382.1547546386719 0 6386.245
383.1333312988281 0 985.25793
383.15875244140625 0 1128.9342
383.1874084472656 0 1914.2252
383.6824951171875 0 1506.0295
384.1949462890625 0 9868.203
384.6953125 0 3939.5227
385.19342041015625 0 1162.1326
389.2257385253906 0 672.2891
391.1373596191406 0 793.15234
392.1934814453125 0 2162.873
392.6947326660156 0 1391.1666
393.2613220214844 0 1959.3599
397.147216796875 0 1801.3834
397.18658447265625 0 768.717
397.6790771484375 0 843.4592
398.17950439453125 0 4015.8115
398.68756103515625 0 2017.5632 Precursor Ammonia loss
401.1453857421875 0 1134.518
405.17724609375 0 871.9537
406.1897888183594 0 1245.984
406.69146728515625 0 1651.1917
407.19940185546875 0 6304.042 Precursor
407.69989013671875 0 2295.8577
409.1470642089844 0 981.3411
411.1306457519531 0 1174.866
416.2292785644531 0 802.5389
418.17193603515625 0 2015.6602
428.15679931640625 0 1307.6984
429.1408386230469 0 1980.5021
446.1669006347656 0 7877.142 y Water loss 3
447.1477966308594 0 1048.3263 y Ammonia loss 3
448.22705078125 0 612.8572
450.2825927734375 0 949.0349
455.17535400390625 0 897.3917
456.1855163574219 0 1396.3326
459.1981506347656 0 821.61676
461.25146484375 0 787.11487 b Ammonia loss 3
463.1960144042969 0 989.2256
464.1772155761719 0 3380.2913 y 3
465.1802978515625 0 1169.2512
466.2410583496094 0 3271.055
467.2447204589844 0 1175.9818
471.8440246582031 0 589.83887
476.2244873046875 0 1117.4033
477.2100830078125 0 1789.8689
478.2118225097656 0 600.884
478.27691650390625 0 6897.7085 b 3
479.2803649902344 0 2241.5925
480.2571716308594 0 1234.8633
488.2979736328125 0 1359.5757
490.2041015625 0 860.05414
494.2356872558594 0 14652.163
494.2722473144531 0 545.6256
495.2392578125 0 3782.6875
504.21942138671875 0 727.98114
510.23028564453125 0 824.4121
511.2653503417969 0 764.7812
515.2232666015625 0 971.9888
516.2110595703125 0 1797.0583
516.2928466796875 0 1936.2566
517.297119140625 0 739.2913
520.1942749023438 0 1460.1482
522.2301635742188 0 1557.3357
537.2169189453125 0 689.89075
538.2032470703125 0 1196.3091
539.2356567382812 0 2035.5049
547.2985229492188 0 610.3528
548.190185546875 0 1097.4113
550.5447998046875 0 594.2341
555.2313842773438 0 4389.8223
556.2359008789062 0 793.13684
565.3094482421875 0 5459.8906
566.2003173828125 0 2987.5308
566.3121948242188 0 1497.5383
568.259521484375 0 1472.1527
569.2659301757812 0 786.18384
575.2940673828125 0 909.211 b Water loss 4
576.2769775390625 0 5531.44 b Ammonia loss 4
577.2815551757812 0 1142.5416
578.8295288085938 0 587.3531
583.2262573242188 0 3275.4255 y Water loss 2
584.2147216796875 0 1757.0325 y Ammonia loss 2
591.2894897460938 0 1134.8513
593.3036499023438 0 29522.744 b 4
593.7853393554688 0 1249.6992
594.3054809570312 0 9516.38
595.3072509765625 0 1593.7456
599.220703125 0 891.32135
601.2362670898438 0 11788.33 y 2
602.2378540039062 0 4122.4316
609.300537109375 0 1541.0127
610.3298950195312 0 4691.308
611.3261108398438 0 1627.6099
650.3060913085938 0 771.6009
651.30029296875 0 668.7917
667.3294067382812 0 1215.8046
668.3212280273438 0 2713.5825
679.2833251953125 0 1476.923
694.292724609375 0 2777.5015
695.2926635742188 0 1435.6299
696.3090209960938 0 3693.5012 y Water loss 1
697.2965087890625 0 4156.323 y Ammonia loss 1
698.3029174804688 0 1207.8662
712.3026733398438 0 1306.3417
713.3056640625 0 1215.8896
714.3193969726562 0 13432.948 y 1
714.7156982421875 0 699.93823
715.3224487304688 0 5204.172
716.32421875 0 1394.9637
725.2942504882812 0 703.5507
827.2239379882812 0 580.5435
861.5789794921875 0 601.6907
866.4373779296875 0 633.00397
1228.8580322265625 0 588.61896
1791.9805908203125 0 593.2453
2284.4931640625 0 631.2626
3020.478759765625 0 763.35333

Spectrum Details

|  |  |
| --- | --- |
| Matched peaks? Matched peaks The total absolute number of peaks matched. Additionally in brackets the total fraction of peaks matched and the total number of peaks is shown. | 25 (6.30% of 397) |
| FDR? FDR The false discovery rate estimated for this peptide. It is calculated by matching all theoretical fragments with a non-integer shift with the raw peaks for this spectrum. This is done with 40 different shifts. The resulting percentage is the average number of annotated peaks over the number of annotated peaks with the correct spectrum. | 3.05% |
| Satellite FDR? Satellite FDR See the FDR for details on its calculation. This satellite ion specific FDR only contains the satellite ions (d/w) for I/L/J positions. | - |
| PSM Score? PSM Score The PSM Score as given by Hecklib to this annotated spectrum. It is shown with three significant figures. | 360 |

#### Spectrum 4808? Spectrum 4808 The raw spectrum of this peptide as annotated by Hecklib. The fragments are coloured according to ion type (see legend). Any peaks with a star '\*' as text can be hovered over to see the full details, first the ion type second the mass shift type. By hovering over the amino acids in the peptide or ions in the legend the corresponding peaks are highlighted. By toggling the 'Unassigned' label you can turn the background (unassigned) peaks on or off in the plot. By updating the slider in the Ion legend you can update the spectrum to only show the top X% of the peaks with labels. The top X% means any peak that is within X% of the highest intensity. By dragging in the spectrum you can zoom in to a specific part of the spectrum and use 'Zoom Out' to get back to the original zoom level. The annotation of the spectrum is based on the given sequence in the peptides file and is done with different software so inconsistencies are likely. The peaks are annotated based on the given sequence, with 20 ppm tolerance.

Copy Data

##### Spectrum 4808 (TSV)

###### Preview

```
Loading example...
```

*Click on the button to copy the data to your clipboard.*

Mz MinMz MaxIntensity Max

WidthHeightPeptide font sizePeptide stroke widthSpectrum font sizeSpectrum stroke widthCompact peptide

Ion legend

wxyz

abcd

OtherUnassignedIon^Charge^_Position_Show for top:%

VJHQDW

06.27e+31.25e+41.88e+42.51e+4

Zoom Out

d^+1^_2_a^+1^_2_y^+1^_1_b^+1^_2_b^+2^_5_y^+1^_2_b^+1^_3_\*y^+1^_3_y^+1^_3_b^+1^_4_y^+1^_4_b^+1^_5_y^+1^_4_b^+1^_5_y^+1^_5_y^+1^_5_y^+1^_5_

0577115417312308

Fragment Matches Table

Show background peaks

| Position | Ion type | Intensity | mz Theoretical | mz Error (Th) | mz Error (ppm) | Charge | Series Number |
| --- | --- | --- | --- | --- | --- | --- | --- |
| - | - | 1.057E+04 | 120.1 | - | - | 0 | - |
| - | - | 727.9 | 121.1 | - | - | 0 | - |
| - | - | 416.3 | 123 | - | - | 0 | - |
| - | - | 1068 | 123.1 | - | - | 0 | - |
| - | - | 412.7 | 124.1 | - | - | 0 | - |
| - | - | 516.5 | 126.1 | - | - | 0 | - |
| - | - | 3206 | 129.1 | - | - | 0 | - |
| - | - | 936.6 | 129.1 | - | - | 0 | - |
| - | - | 7109 | 129.1 | - | - | 0 | - |
| - | - | 599.4 | 130.1 | - | - | 0 | - |
| - | - | 9228 | 130.1 | - | - | 0 | - |
| - | - | 780.7 | 131.1 | - | - | 0 | - |
| - | - | 2848 | 132.1 | - | - | 0 | - |
| - | - | 448.7 | 132.1 | - | - | 0 | - |
| - | - | 903.6 | 133.1 | - | - | 0 | - |
| - | - | 3500 | 133.1 | - | - | 0 | - |
| - | - | 9051 | 136.1 | - | - | 0 | - |
| - | - | 386.3 | 136.3 | - | - | 0 | - |
| - | - | 1069 | 138.1 | - | - | 0 | - |
| - | - | 680.6 | 140.1 | - | - | 0 | - |
| - | - | 434.6 | 143.1 | - | - | 0 | - |
| 2 | d | 597.6 | 143.1 | 1.539E-05 | 0.1075 | +1 | 2 |
| - | - | 580.2 | 144.1 | - | - | 0 | - |
| - | - | 509 | 144.1 | - | - | 0 | - |
| - | - | 416.6 | 145 | - | - | 0 | - |
| - | - | 4021 | 146.1 | - | - | 0 | - |
| - | - | 3005 | 147.1 | - | - | 0 | - |
| - | - | 525.3 | 148 | - | - | 0 | - |
| - | - | 855.6 | 148.9 | - | - | 0 | - |
| - | - | 443.2 | 149 | - | - | 0 | - |
| - | - | 518.8 | 151.1 | - | - | 0 | - |
| - | - | 424.8 | 152.1 | - | - | 0 | - |
| - | - | 436.7 | 154.1 | - | - | 0 | - |
| - | - | 3530 | 155.1 | - | - | 0 | - |
| - | - | 427.5 | 155.6 | - | - | 0 | - |
| - | - | 811.7 | 156.1 | - | - | 0 | - |
| - | - | 490 | 158.4 | - | - | 0 | - |
| - | - | 2.068E+04 | 159.1 | - | - | 0 | - |
| - | - | 1631 | 160.1 | - | - | 0 | - |
| - | - | 462.4 | 161.1 | - | - | 0 | - |
| - | - | 3947 | 166.1 | - | - | 0 | - |
| - | - | 442 | 166.1 | - | - | 0 | - |
| - | - | 1457 | 170.1 | - | - | 0 | - |
| - | - | 515.6 | 172.1 | - | - | 0 | - |
| - | - | 590.2 | 173.1 | - | - | 0 | - |
| - | - | 1482 | 173.4 | - | - | 0 | - |
| - | - | 536.9 | 174.5 | - | - | 0 | - |
| - | - | 1518 | 175 | - | - | 0 | - |
| - | - | 1062 | 175 | - | - | 0 | - |
| - | - | 1202 | 177.1 | - | - | 0 | - |
| - | - | 1751 | 178.1 | - | - | 0 | - |
| - | - | 938.1 | 181.1 | - | - | 0 | - |
| 2 | a | 7128 | 185.2 | 0.0003519 | 1.901 | +1 | 2 |
| - | - | 716.3 | 187 | - | - | 0 | - |
| - | - | 1275 | 187.1 | - | - | 0 | - |
| - | - | 2.483E+04 | 188.1 | - | - | 0 | - |
| - | - | 560.6 | 189.1 | - | - | 0 | - |
| - | - | 3228 | 189.1 | - | - | 0 | - |
| - | - | 695.8 | 191 | - | - | 0 | - |
| - | - | 2559 | 199.1 | - | - | 0 | - |
| - | - | 928.7 | 200.1 | - | - | 0 | - |
| - | - | 614.2 | 202 | - | - | 0 | - |
| - | - | 1602 | 202.1 | - | - | 0 | - |
| 6 | y | 7527 | 205.1 | 0.0002428 | 1.184 | +1 | 1 |
| - | - | 782.2 | 205.1 | - | - | 0 | - |
| 2 | b | 1959 | 213.2 | 0.0002341 | 1.098 | +1 | 2 |
| - | - | 554.6 | 214 | - | - | 0 | - |
| - | - | 907.9 | 215.1 | - | - | 0 | - |
| - | - | 552.3 | 220.1 | - | - | 0 | - |
| - | - | 684.8 | 221.1 | - | - | 0 | - |
| - | - | 3159 | 223.2 | - | - | 0 | - |
| - | - | 641.7 | 226.1 | - | - | 0 | - |
| - | - | 994.3 | 226.2 | - | - | 0 | - |
| - | - | 759.4 | 227.1 | - | - | 0 | - |
| - | - | 623.4 | 233.1 | - | - | 0 | - |
| - | - | 934.1 | 233.2 | - | - | 0 | - |
| - | - | 4005 | 234.1 | - | - | 0 | - |
| - | - | 548.6 | 237.1 | - | - | 0 | - |
| - | - | 649.3 | 239.2 | - | - | 0 | - |
| - | - | 1615 | 244.1 | - | - | 0 | - |
| - | - | 1150 | 248.1 | - | - | 0 | - |
| - | - | 2316 | 249.1 | - | - | 0 | - |
| - | - | 750.2 | 249.1 | - | - | 0 | - |
| - | - | 1.056E+04 | 251.2 | - | - | 0 | - |
| - | - | 1240 | 252.2 | - | - | 0 | - |
| - | - | 580.9 | 252.6 | - | - | 0 | - |
| - | - | 732.1 | 255.2 | - | - | 0 | - |
| - | - | 665.5 | 261.1 | - | - | 0 | - |
| - | - | 610.5 | 261.1 | - | - | 0 | - |
| - | - | 2777 | 262.1 | - | - | 0 | - |
| - | - | 723.4 | 262.1 | - | - | 0 | - |
| - | - | 6325 | 266.1 | - | - | 0 | - |
| - | - | 746.3 | 267.1 | - | - | 0 | - |
| - | - | 4065 | 270.1 | - | - | 0 | - |
| - | - | 1295 | 274.1 | - | - | 0 | - |
| - | - | 3082 | 283.1 | - | - | 0 | - |
| - | - | 642.7 | 285.1 | - | - | 0 | - |
| 5 | b | 1535 | 297.2 | 0.0007625 | 2.566 | +2 | 5 |
| - | - | 581.2 | 299.1 | - | - | 0 | - |
| - | - | 582.2 | 306.2 | - | - | 0 | - |
| 5 | y | 891.9 | 320.1 | 0.0004756 | 1.486 | +1 | 2 |
| - | - | 1007 | 326.7 | - | - | 0 | - |
| - | - | 1350 | 327.2 | - | - | 0 | - |
| - | - | 1501 | 332.2 | - | - | 0 | - |
| - | - | 1337 | 345.1 | - | - | 0 | - |
| - | - | 3138 | 346.1 | - | - | 0 | - |
| 3 | b | 6839 | 350.2 | 0.000389 | 1.111 | +1 | 3 |
| - | - | 959.2 | 351.2 | - | - | 0 | - |
| - | - | 3913 | 363.1 | - | - | 0 | - |
| - | - | 1381 | 364.1 | - | - | 0 | - |
| - | - | 1053 | 367.2 | - | - | 0 | - |
| - | - | 1455 | 367.7 | - | - | 0 | - |
| - | - | 605.5 | 368.2 | - | - | 0 | - |
| - | - | 2523 | 376.2 | - | - | 0 | - |
| - | - | 1494 | 376.7 | - | - | 0 | - |
| - | - | 3568 | 379.2 | - | - | 0 | - |
| - | - | 6815 | 381.2 | - | - | 0 | - |
| - | - | 1544 | 382.2 | - | - | 0 | - |
| - | - | 633.6 | 382.2 | - | - | 0 | - |
| - | - | 2788 | 398.2 | - | - | 0 | - |
| 0 | Precursor | 1093 | 399.2 | 0.00136 | 3.407 | +2 | -1 |
| 4 | y | 2277 | 430.2 | 0.0007416 | 1.724 | +1 | 3 |
| 4 | y | 1373 | 448.2 | 0.001224 | 2.732 | +1 | 3 |
| 4 | b | 1673 | 478.3 | 0.00129 | 2.698 | +1 | 4 |
| - | - | 536.1 | 479.3 | - | - | 0 | - |
| - | - | 3573 | 494.2 | - | - | 0 | - |
| - | - | 1116 | 539.2 | - | - | 0 | - |
| - | - | 528.5 | 547.9 | - | - | 0 | - |
| - | - | 654.6 | 549.3 | - | - | 0 | - |
| - | - | 1451 | 550.2 | - | - | 0 | - |
| 3 | y | 670.8 | 568.2 | 0.002797 | 4.923 | +1 | 4 |
| 5 | b | 1647 | 576.3 | 0.00111 | 1.926 | +1 | 5 |
| 3 | y | 3308 | 585.2 | 0.001211 | 2.07 | +1 | 4 |
| - | - | 949.8 | 586.2 | - | - | 0 | - |
| 5 | b | 8427 | 593.3 | 0.0003787 | 0.6382 | +1 | 5 |
| - | - | 2656 | 594.3 | - | - | 0 | - |
| - | - | 752.6 | 611.3 | - | - | 0 | - |
| - | - | 569.5 | 652.3 | - | - | 0 | - |
| 2 | y | 766.5 | 680.3 | 0.0003275 | 0.4814 | +1 | 5 |
| 2 | y | 985.6 | 681.3 | 0.001558 | 2.287 | +1 | 5 |
| - | - | 600.3 | 685.5 | - | - | 0 | - |
| 2 | y | 4808 | 698.3 | 0.0001552 | 0.2222 | +1 | 5 |
| - | - | 1343 | 699.3 | - | - | 0 | - |
| - | - | 591.6 | 718.5 | - | - | 0 | - |
| - | - | 2124 | 775.3 | - | - | 0 | - |
| - | - | 756.6 | 938.3 | - | - | 0 | - |
| - | - | 734.1 | 1232 | - | - | 0 | - |
| - | - | 678.5 | 1614 | - | - | 0 | - |
| - | - | 644.3 | 1711 | - | - | 0 | - |
| - | - | 633.4 | 1848 | - | - | 0 | - |
| - | - | 714.1 | 2038 | - | - | 0 | - |
| - | - | 673.5 | 2285 | - | - | 0 | - |

m/z Charge Intensity FragmentType MassShift Position
120.08100891113281 0 10566.206
121.0845718383789 0 727.8927
123.04410552978516 0 416.26395
123.1170883178711 0 1067.6941
124.08734893798828 0 412.7352
126.10321807861328 0 516.4581
129.0548858642578 0 3205.5464
129.0663299560547 0 936.6121
129.1024932861328 0 7109.1445
130.06080627441406 0 599.4358
130.06536865234375 0 9227.92
131.06919860839844 0 780.73285
132.08106994628906 0 2848.3967
132.1020965576172 0 448.68167
133.0608367919922 0 903.6266
133.08609008789062 0 3499.9773
136.075927734375 0 9051.193
136.27139282226562 0 386.34256
138.06639099121094 0 1069.1149
140.0818634033203 0 680.5841
143.09483337402344 0 434.58167
143.1178741455078 0 597.6446 d 1
144.08164978027344 0 580.2311
144.1022491455078 0 509.04517
145.04994201660156 0 416.6353
146.0602264404297 0 4021.192
147.06515502929688 0 3005.3545
148.01873779296875 0 525.26184
148.9470977783203 0 855.56976
149.0243377685547 0 443.16272
151.07614135742188 0 518.84515
152.1071319580078 0 424.81976
154.0612335205078 0 436.66492
155.0929412841797 0 3529.6416
155.63375854492188 0 427.45676
156.0769805908203 0 811.6729
158.41778564453125 0 489.99353
159.09190368652344 0 20675.062
160.09535217285156 0 1631.468
161.08128356933594 0 462.35077
166.06134033203125 0 3947.057
166.0867919921875 0 442.02405
170.06024169921875 0 1456.6833
172.0723114013672 0 515.59625
173.12879943847656 0 590.16327
173.43980407714844 0 1482.3798
174.47137451171875 0 536.8781
175.01576232910156 0 1517.5562
175.0297088623047 0 1062.2004
177.1124725341797 0 1202.3872
178.1339569091797 0 1750.6842
181.06112670898438 0 938.13806
185.16519165039062 0 7127.9336 a 1
187.02964782714844 0 716.3291
187.08670043945312 0 1274.8358
188.0708465576172 0 24828.24
189.0662078857422 0 560.6361
189.0744171142578 0 3228.221
191.02474975585938 0 695.7949
199.0731658935547 0 2559.3792
200.13983154296875 0 928.7311
202.04055786132812 0 614.1969
202.10780334472656 0 1601.871
205.09739685058594 0 7526.7583 y 5
205.10733032226562 0 782.2084
213.1599884033203 0 1959.4474 b 1
214.04092407226562 0 554.56726
215.13909912109375 0 907.93494
220.05093383789062 0 552.339
221.1036376953125 0 684.80084
223.15576171875 0 3158.5417
226.08253479003906 0 641.6639
226.1552276611328 0 994.3144
227.06594848632812 0 759.39197
233.13995361328125 0 623.4188
233.16554260253906 0 934.09216
234.12400817871094 0 4004.979
237.0896759033203 0 548.64746
239.15005493164062 0 649.29584
244.0930938720703 0 1614.7897
248.1148681640625 0 1150.2833
249.09901428222656 0 2315.5105
249.13479614257812 0 750.19507
251.15061950683594 0 10556.079
252.1537628173828 0 1239.8695
252.6028594970703 0 580.8959
255.19485473632812 0 732.1198
261.1151123046875 0 665.45325
261.1333923339844 0 610.5485
262.0505676269531 0 2777.0562
262.1393127441406 0 723.4299
266.1249694824219 0 6324.8027
267.12786865234375 0 746.3123
270.1219482421875 0 4065.311
274.1308898925781 0 1294.6133
283.1444396972656 0 3081.912
285.07373046875 0 642.70776
297.156494140625 0 1535.3173 b 4
299.1393737792969 0 581.2009
306.2291259765625 0 582.1561
320.12457275390625 0 891.9165 y 4
326.6639709472656 0 1006.9116
327.2024230957031 0 1350.493
332.20892333984375 0 1501.4225
345.1308288574219 0 1336.9011
346.1150817871094 0 3138.0674
350.21905517578125 0 6839.046 b 2
351.2225341796875 0 959.1997
363.1418151855469 0 3913.2495
364.125 0 1381.4342
367.2457580566406 0 1053.1366
367.68499755859375 0 1454.9343
368.1864929199219 0 605.4753
376.1981506347656 0 2522.7349
376.6986083984375 0 1494.2766
379.2091064453125 0 3568.0444
381.15203857421875 0 6815.005
382.15753173828125 0 1544.1351
382.24530029296875 0 633.586
398.23876953125 0 2788.2988
399.1993103027344 0 1092.8425 Precursor
430.1728515625 0 2277.3901 y Water loss 3
448.18389892578125 0 1372.9977 y 3
478.2785339355469 0 1673.0101 b 3
479.2811584472656 0 536.05896
494.23651123046875 0 3573.2527
539.236328125 0 1116.3187
547.87109375 0 528.4749
549.3392944335938 0 654.58167
550.2055053710938 0 1450.9874
568.2178344726562 0 670.79224 y Ammonia loss 2
576.2787475585938 0 1647.1451 b Ammonia loss 4
585.2427978515625 0 3307.6309 y 2
586.2450561523438 0 949.7979
593.3045654296875 0 8426.797 b 4
594.3038940429688 0 2655.9329
611.319580078125 0 752.5886
652.3096313476562 0 569.45844
680.3147583007812 0 766.46155 y Water loss 1
681.3006591796875 0 985.63367 y Ammonia loss 1
685.4815673828125 0 600.2728
698.3258056640625 0 4807.749 y 1
699.3279418945312 0 1342.8887
718.501953125 0 591.6092
775.2736206054688 0 2123.5208
938.3427734375 0 756.5624
1232.212158203125 0 734.05273
1613.57080078125 0 678.53937
1711.256591796875 0 644.28455
1847.962646484375 0 633.383
2037.6663818359375 0 714.06525
2284.794189453125 0 673.4676

Spectrum Details

|  |  |
| --- | --- |
| Matched peaks? Matched peaks The total absolute number of peaks matched. Additionally in brackets the total fraction of peaks matched and the total number of peaks is shown. | 18 (11.84% of 152) |
| FDR? FDR The false discovery rate estimated for this peptide. It is calculated by matching all theoretical fragments with a non-integer shift with the raw peaks for this spectrum. This is done with 40 different shifts. The resulting percentage is the average number of annotated peaks over the number of annotated peaks with the correct spectrum. | 0.66% |
| Satellite FDR? Satellite FDR See the FDR for details on its calculation. This satellite ion specific FDR only contains the satellite ions (d/w) for I/L/J positions. | 0.00% |
| PSM Score? PSM Score The PSM Score as given by Hecklib to this annotated spectrum. It is shown with three significant figures. | 239 |

#### Spectrum 4874? Spectrum 4874 The raw spectrum of this peptide as annotated by Hecklib. The fragments are coloured according to ion type (see legend). Any peaks with a star '\*' as text can be hovered over to see the full details, first the ion type second the mass shift type. By hovering over the amino acids in the peptide or ions in the legend the corresponding peaks are highlighted. By toggling the 'Unassigned' label you can turn the background (unassigned) peaks on or off in the plot. By updating the slider in the Ion legend you can update the spectrum to only show the top X% of the peaks with labels. The top X% means any peak that is within X% of the highest intensity. By dragging in the spectrum you can zoom in to a specific part of the spectrum and use 'Zoom Out' to get back to the original zoom level. The annotation of the spectrum is based on the given sequence in the peptides file and is done with different software so inconsistencies are likely. The peaks are annotated based on the given sequence, with 20 ppm tolerance.

Copy Data

##### Spectrum 4874 (TSV)

###### Preview

```
Loading example...
```

*Click on the button to copy the data to your clipboard.*

Mz MinMz MaxIntensity Max

WidthHeightPeptide font sizePeptide stroke widthSpectrum font sizeSpectrum stroke widthCompact peptide

Ion legend

wxyz

abcd

OtherUnassignedIon^Charge^_Position_Show for top:%

VJHQDW

04.22e+38.45e+31.27e+41.69e+4

Zoom Out

a^+1^_2_y^+1^_1_b^+1^_2_y^+1^_2_b^+1^_3_\*\*y^+1^_3_y^+1^_3_b^+1^_4_b^+1^_5_y^+1^_4_b^+1^_5_y^+1^_5_y^+1^_5_y^+1^_5_

0536107116072143

Fragment Matches Table

Show background peaks

| Position | Ion type | Intensity | mz Theoretical | mz Error (Th) | mz Error (ppm) | Charge | Series Number |
| --- | --- | --- | --- | --- | --- | --- | --- |
| - | - | 4249 | 120.1 | - | - | 0 | - |
| - | - | 325.4 | 120.7 | - | - | 0 | - |
| - | - | 569.2 | 122.1 | - | - | 0 | - |
| - | - | 1015 | 123.1 | - | - | 0 | - |
| - | - | 598 | 125.1 | - | - | 0 | - |
| - | - | 725 | 126.1 | - | - | 0 | - |
| - | - | 442.3 | 127.4 | - | - | 0 | - |
| - | - | 468 | 127.7 | - | - | 0 | - |
| - | - | 388.9 | 128.1 | - | - | 0 | - |
| - | - | 430.9 | 128.1 | - | - | 0 | - |
| - | - | 3710 | 129.1 | - | - | 0 | - |
| - | - | 428.5 | 129.1 | - | - | 0 | - |
| - | - | 3903 | 129.1 | - | - | 0 | - |
| - | - | 5550 | 130.1 | - | - | 0 | - |
| - | - | 468.2 | 131 | - | - | 0 | - |
| - | - | 576.1 | 131.1 | - | - | 0 | - |
| - | - | 1743 | 132.1 | - | - | 0 | - |
| - | - | 855.5 | 133.1 | - | - | 0 | - |
| - | - | 3851 | 133.1 | - | - | 0 | - |
| - | - | 400.9 | 135.6 | - | - | 0 | - |
| - | - | 1018 | 136 | - | - | 0 | - |
| - | - | 2789 | 136.1 | - | - | 0 | - |
| - | - | 1005 | 138.1 | - | - | 0 | - |
| - | - | 483.2 | 139.1 | - | - | 0 | - |
| - | - | 1247 | 140.1 | - | - | 0 | - |
| - | - | 631 | 141.1 | - | - | 0 | - |
| - | - | 528.4 | 145 | - | - | 0 | - |
| - | - | 7846 | 145 | - | - | 0 | - |
| - | - | 3594 | 146.1 | - | - | 0 | - |
| - | - | 2978 | 147.1 | - | - | 0 | - |
| - | - | 413.7 | 148.5 | - | - | 0 | - |
| - | - | 561.4 | 148.9 | - | - | 0 | - |
| - | - | 392.1 | 149.5 | - | - | 0 | - |
| - | - | 462.7 | 155.1 | - | - | 0 | - |
| - | - | 578 | 155.1 | - | - | 0 | - |
| - | - | 2544 | 155.1 | - | - | 0 | - |
| - | - | 903.9 | 156.1 | - | - | 0 | - |
| - | - | 637.3 | 159 | - | - | 0 | - |
| - | - | 1.406E+04 | 159.1 | - | - | 0 | - |
| - | - | 950.5 | 160.1 | - | - | 0 | - |
| - | - | 630.2 | 161.1 | - | - | 0 | - |
| - | - | 735.1 | 162.1 | - | - | 0 | - |
| - | - | 882.6 | 163 | - | - | 0 | - |
| - | - | 2686 | 166.1 | - | - | 0 | - |
| - | - | 450.2 | 174.1 | - | - | 0 | - |
| - | - | 738.1 | 175 | - | - | 0 | - |
| - | - | 2422 | 175 | - | - | 0 | - |
| - | - | 1437 | 175 | - | - | 0 | - |
| - | - | 1738 | 178.1 | - | - | 0 | - |
| - | - | 801.3 | 181.1 | - | - | 0 | - |
| - | - | 539.1 | 182.1 | - | - | 0 | - |
| 2 | a | 4633 | 185.2 | 0.0003672 | 1.983 | +1 | 2 |
| - | - | 806.9 | 186.2 | - | - | 0 | - |
| - | - | 720.2 | 187.1 | - | - | 0 | - |
| - | - | 1.673E+04 | 188.1 | - | - | 0 | - |
| - | - | 2362 | 189.1 | - | - | 0 | - |
| - | - | 925 | 191.1 | - | - | 0 | - |
| - | - | 768.8 | 195.1 | - | - | 0 | - |
| - | - | 603 | 196.1 | - | - | 0 | - |
| - | - | 590.2 | 198.1 | - | - | 0 | - |
| - | - | 2614 | 199.1 | - | - | 0 | - |
| - | - | 1262 | 202.1 | - | - | 0 | - |
| - | - | 596.8 | 203 | - | - | 0 | - |
| 6 | y | 4798 | 205.1 | 7.491E-05 | 0.3652 | +1 | 1 |
| - | - | 903.8 | 205.1 | - | - | 0 | - |
| - | - | 747.9 | 206.1 | - | - | 0 | - |
| - | - | 499.4 | 212 | - | - | 0 | - |
| 2 | b | 1004 | 213.2 | 0.0004324 | 2.029 | +1 | 2 |
| - | - | 842.3 | 214 | - | - | 0 | - |
| - | - | 1838 | 223.2 | - | - | 0 | - |
| - | - | 877.6 | 226.1 | - | - | 0 | - |
| - | - | 1211 | 226.2 | - | - | 0 | - |
| - | - | 614.6 | 227.1 | - | - | 0 | - |
| - | - | 712.1 | 230 | - | - | 0 | - |
| - | - | 1396 | 233.1 | - | - | 0 | - |
| - | - | 911.8 | 233.1 | - | - | 0 | - |
| - | - | 737 | 233.2 | - | - | 0 | - |
| - | - | 2144 | 234.1 | - | - | 0 | - |
| - | - | 1315 | 239.1 | - | - | 0 | - |
| - | - | 729.5 | 244.1 | - | - | 0 | - |
| - | - | 583.2 | 248 | - | - | 0 | - |
| - | - | 779.4 | 248.1 | - | - | 0 | - |
| - | - | 1447 | 249.1 | - | - | 0 | - |
| - | - | 709.8 | 249.1 | - | - | 0 | - |
| - | - | 7398 | 251.2 | - | - | 0 | - |
| - | - | 1052 | 252.2 | - | - | 0 | - |
| - | - | 694.4 | 252.6 | - | - | 0 | - |
| - | - | 1041 | 255.2 | - | - | 0 | - |
| - | - | 1364 | 255.2 | - | - | 0 | - |
| - | - | 2616 | 262.1 | - | - | 0 | - |
| - | - | 4391 | 266.1 | - | - | 0 | - |
| - | - | 583.2 | 267.1 | - | - | 0 | - |
| - | - | 2209 | 270.1 | - | - | 0 | - |
| - | - | 1169 | 274.1 | - | - | 0 | - |
| - | - | 2014 | 277.1 | - | - | 0 | - |
| - | - | 1827 | 283.2 | - | - | 0 | - |
| - | - | 613.6 | 284.2 | - | - | 0 | - |
| - | - | 781.3 | 299.2 | - | - | 0 | - |
| - | - | 541.3 | 319.2 | - | - | 0 | - |
| 5 | y | 728.5 | 320.1 | 0.0007451 | 2.327 | +1 | 2 |
| - | - | 934.1 | 327.2 | - | - | 0 | - |
| - | - | 593.7 | 327.4 | - | - | 0 | - |
| - | - | 641.4 | 332.2 | - | - | 0 | - |
| - | - | 1250 | 345 | - | - | 0 | - |
| - | - | 2162 | 346.1 | - | - | 0 | - |
| - | - | 710.1 | 350.2 | - | - | 0 | - |
| 3 | b | 3357 | 350.2 | 0.0008162 | 2.331 | +1 | 3 |
| - | - | 989.2 | 351.2 | - | - | 0 | - |
| - | - | 2004 | 363.1 | - | - | 0 | - |
| - | - | 769 | 363.2 | - | - | 0 | - |
| - | - | 2077 | 365.2 | - | - | 0 | - |
| - | - | 764.2 | 367.2 | - | - | 0 | - |
| - | - | 1451 | 367.7 | - | - | 0 | - |
| - | - | 1948 | 376.2 | - | - | 0 | - |
| - | - | 669.5 | 376.7 | - | - | 0 | - |
| - | - | 594.7 | 377.8 | - | - | 0 | - |
| - | - | 1393 | 379.2 | - | - | 0 | - |
| - | - | 5526 | 381.2 | - | - | 0 | - |
| - | - | 1057 | 383.2 | - | - | 0 | - |
| 0 | Precursor | 609.1 | 390.2 | 0.003312 | 8.488 | +2 | -1 |
| - | - | 990.7 | 398.2 | - | - | 0 | - |
| - | - | 1112 | 398.2 | - | - | 0 | - |
| 0 | Precursor | 1447 | 399.2 | 0.002459 | 6.159 | +2 | -1 |
| 4 | y | 1548 | 430.2 | 0.001749 | 4.065 | +1 | 3 |
| 4 | y | 1008 | 448.2 | 0.001072 | 2.391 | +1 | 3 |
| 4 | b | 584.4 | 478.3 | 0.002555 | 5.342 | +1 | 4 |
| - | - | 625.4 | 479.8 | - | - | 0 | - |
| - | - | 2199 | 494.2 | - | - | 0 | - |
| - | - | 687.7 | 539.2 | - | - | 0 | - |
| - | - | 874.6 | 550.2 | - | - | 0 | - |
| 5 | b | 746.2 | 576.3 | 0.001537 | 2.667 | +1 | 5 |
| - | - | 608.1 | 581.8 | - | - | 0 | - |
| 3 | y | 2204 | 585.2 | 0.0001925 | 0.3288 | +1 | 4 |
| - | - | 910 | 586.2 | - | - | 0 | - |
| 5 | b | 4548 | 593.3 | 0.002088 | 3.519 | +1 | 5 |
| - | - | 1431 | 594.3 | - | - | 0 | - |
| 2 | y | 781.9 | 680.3 | 0.002053 | 3.018 | +1 | 5 |
| 2 | y | 978.4 | 681.3 | 0.001067 | 1.566 | +1 | 5 |
| 2 | y | 3430 | 698.3 | 0.001681 | 2.407 | +1 | 5 |
| - | - | 2049 | 699.3 | - | - | 0 | - |
| - | - | 745.4 | 1478 | - | - | 0 | - |
| - | - | 716.4 | 2122 | - | - | 0 | - |

m/z Charge Intensity FragmentType MassShift Position
120.08094787597656 0 4249.431
120.73297882080078 0 325.4479
122.07154846191406 0 569.24963
123.1170425415039 0 1015.32043
125.07112121582031 0 598.0217
126.10267639160156 0 725.0349
127.40895080566406 0 442.31012
127.71942901611328 0 468.03174
128.0706329345703 0 388.94968
128.08206176757812 0 430.9158
129.05474853515625 0 3710.4053
129.0660400390625 0 428.54657
129.1024169921875 0 3903.0903
130.0653076171875 0 5549.5933
131.0448760986328 0 468.1629
131.0691375732422 0 576.0975
132.0809326171875 0 1743.0099
133.06092834472656 0 855.5035
133.0860137939453 0 3851.0913
135.5779571533203 0 400.9057
136.01856994628906 0 1017.84705
136.07583618164062 0 2788.9146
138.0664520263672 0 1004.556
139.07496643066406 0 483.22208
140.08201599121094 0 1247.234
141.06619262695312 0 630.96747
145.04364013671875 0 528.3525
145.04969787597656 0 7846.143
146.0601806640625 0 3593.8828
147.06532287597656 0 2978.0667
148.5200653076172 0 413.7385
148.947021484375 0 561.3963
149.47874450683594 0 392.05017
155.05874633789062 0 462.7248
155.08177185058594 0 578.03656
155.09317016601562 0 2544.2625
156.07705688476562 0 903.88226
159.0348358154297 0 637.3326
159.0918426513672 0 14057.062
160.09535217285156 0 950.4715
161.095458984375 0 630.1951
162.0548095703125 0 735.14667
163.02914428710938 0 882.6411
166.06109619140625 0 2685.5264
174.0555877685547 0 450.23425
175.006103515625 0 738.1166
175.01577758789062 0 2422.2195
175.0297393798828 0 1437.0046
178.13397216796875 0 1737.768
181.06076049804688 0 801.34784
182.08114624023438 0 539.14374
185.1652069091797 0 4632.717 a 1
186.1685028076172 0 806.9363
187.08642578125 0 720.199
188.07078552246094 0 16729.736
189.0747528076172 0 2362.474
191.11795043945312 0 925.03015
195.123046875 0 768.8495
196.1453399658203 0 603.04816
198.0878143310547 0 590.22833
199.07308959960938 0 2613.574
202.10739135742188 0 1262.1746
203.02444458007812 0 596.775
205.09722900390625 0 4797.9365 y 5
205.107177734375 0 903.7719
206.10121154785156 0 747.90985
212.0340118408203 0 499.40668
213.16018676757812 0 1003.9654 b 1
214.0404052734375 0 842.3311
223.15574645996094 0 1837.5723
226.0823516845703 0 877.57947
226.15567016601562 0 1210.5062
227.103759765625 0 614.64636
230.03494262695312 0 712.05066
233.102294921875 0 1396.0255
233.1395721435547 0 911.76337
233.16481018066406 0 737.041
234.12403869628906 0 2143.6538
239.14987182617188 0 1314.9056
244.09286499023438 0 729.4572
248.0466766357422 0 583.2452
248.1147003173828 0 779.39233
249.09823608398438 0 1446.8113
249.13536071777344 0 709.77454
251.1506805419922 0 7398.0684
252.15354919433594 0 1051.7885
252.6030731201172 0 694.38403
255.17059326171875 0 1041.4802
255.19601440429688 0 1364.4852
262.0501403808594 0 2616.425
266.1248474121094 0 4390.7256
267.12530517578125 0 583.1686
270.1220703125 0 2208.9773
274.1306457519531 0 1168.8136
277.1284484863281 0 2014.1993
283.1755676269531 0 1827.3245
284.1850280761719 0 613.59546
299.1617431640625 0 781.2987
319.1531982421875 0 541.2686
320.12335205078125 0 728.5137 y 4
327.2012939453125 0 934.13635
327.40875244140625 0 593.6598
332.2088623046875 0 641.3611
344.9755859375 0 1250.1897
346.1146240234375 0 2161.6453
350.1966552734375 0 710.094
350.219482421875 0 3357.213 b 2
351.2216796875 0 989.2183
363.14178466796875 0 2004.2053
363.1938171386719 0 769.0313
365.18084716796875 0 2076.653
367.2467041015625 0 764.21704
367.68536376953125 0 1451.3076
376.19793701171875 0 1947.9941
376.6969299316406 0 669.48956
377.8024597167969 0 594.7155
379.20916748046875 0 1393.0063
381.1523132324219 0 5526.2397
383.19232177734375 0 1056.5052
390.1986999511719 0 609.1479 Precursor Water loss
398.192626953125 0 990.73303
398.2388610839844 0 1111.6033
399.1982116699219 0 1446.5814 Precursor
430.1738586425781 0 1547.7186 y Water loss 3
448.1837463378906 0 1007.81085 y 3
478.2746887207031 0 584.3642 b 3
479.7606201171875 0 625.413
494.2361145019531 0 2198.918
539.23974609375 0 687.6674
550.2052001953125 0 874.57544
576.2791748046875 0 746.18536 b Ammonia loss 4
581.7853393554688 0 608.09186
585.2413940429688 0 2203.684 y 2
586.2449951171875 0 910.0054
593.3062744140625 0 4547.8296 b 4
594.3062133789062 0 1430.9651
680.317138671875 0 781.86884 y Water loss 1
681.2980346679688 0 978.39404 y Ammonia loss 1
698.3273315429688 0 3429.5889 y 1
699.3290405273438 0 2049.4731
1478.0513916015625 0 745.4416
2121.716552734375 0 716.3866

Spectrum Details

|  |  |
| --- | --- |
| Matched peaks? Matched peaks The total absolute number of peaks matched. Additionally in brackets the total fraction of peaks matched and the total number of peaks is shown. | 16 (11.27% of 142) |
| FDR? FDR The false discovery rate estimated for this peptide. It is calculated by matching all theoretical fragments with a non-integer shift with the raw peaks for this spectrum. This is done with 40 different shifts. The resulting percentage is the average number of annotated peaks over the number of annotated peaks with the correct spectrum. | 0.15% |
| Satellite FDR? Satellite FDR See the FDR for details on its calculation. This satellite ion specific FDR only contains the satellite ions (d/w) for I/L/J positions. | - |
| PSM Score? PSM Score The PSM Score as given by Hecklib to this annotated spectrum. It is shown with three significant figures. | 217 |

#### Spectrum 4545? Spectrum 4545 The raw spectrum of this peptide as annotated by Hecklib. The fragments are coloured according to ion type (see legend). Any peaks with a star '\*' as text can be hovered over to see the full details, first the ion type second the mass shift type. By hovering over the amino acids in the peptide or ions in the legend the corresponding peaks are highlighted. By toggling the 'Unassigned' label you can turn the background (unassigned) peaks on or off in the plot. By updating the slider in the Ion legend you can update the spectrum to only show the top X% of the peaks with labels. The top X% means any peak that is within X% of the highest intensity. By dragging in the spectrum you can zoom in to a specific part of the spectrum and use 'Zoom Out' to get back to the original zoom level. The annotation of the spectrum is based on the given sequence in the peptides file and is done with different software so inconsistencies are likely. The peaks are annotated based on the given sequence, with 20 ppm tolerance.

Copy Data

##### Spectrum 4545 (TSV)

###### Preview

```
Loading example...
```

*Click on the button to copy the data to your clipboard.*

Mz MinMz MaxIntensity Max

WidthHeightPeptide font sizePeptide stroke widthSpectrum font sizeSpectrum stroke widthCompact peptide

Ion legend

wxyz

abcd

OtherUnassignedIon^Charge^_Position_Show for top:%

VJHQDW

04.01e+48.03e+41.20e+51.61e+5

Zoom Out

a^+1^_2_y^+1^_1_b^+1^_2_b^+2^_4_y^+2^_4_b^+2^_5_b^+2^_5_y^+1^_2_y^+1^_2_a^+1^_3_y^+2^_5_y^+2^_5_b^+1^_3_\*\*\*y^+1^_3_y^+1^_3_y^+1^_3_b^+1^_4_b^+1^_4_y^+1^_4_y^+1^_4_b^+1^_5_b^+1^_5_y^+1^_4_b^+1^_5_y^+1^_5_y^+1^_5_y^+1^_5_

0874174726213495

Fragment Matches Table

Show background peaks

| Position | Ion type | Intensity | mz Theoretical | mz Error (Th) | mz Error (ppm) | Charge | Series Number |
| --- | --- | --- | --- | --- | --- | --- | --- |
| - | - | 3374 | 120.1 | - | - | 0 | - |
| - | - | 770.5 | 121 | - | - | 0 | - |
| - | - | 585.2 | 121.1 | - | - | 0 | - |
| - | - | 339.1 | 121.2 | - | - | 0 | - |
| - | - | 1949 | 122.1 | - | - | 0 | - |
| - | - | 1586 | 123 | - | - | 0 | - |
| - | - | 1524 | 123.1 | - | - | 0 | - |
| - | - | 435.1 | 124 | - | - | 0 | - |
| - | - | 393.8 | 124.2 | - | - | 0 | - |
| - | - | 848.3 | 125.1 | - | - | 0 | - |
| - | - | 720.9 | 126 | - | - | 0 | - |
| - | - | 640.1 | 128.1 | - | - | 0 | - |
| - | - | 3142 | 129.1 | - | - | 0 | - |
| - | - | 1350 | 129.1 | - | - | 0 | - |
| - | - | 8352 | 129.1 | - | - | 0 | - |
| - | - | 5.423E+04 | 130.1 | - | - | 0 | - |
| - | - | 772 | 130.1 | - | - | 0 | - |
| - | - | 5218 | 131.1 | - | - | 0 | - |
| - | - | 1.808E+04 | 132.1 | - | - | 0 | - |
| - | - | 469.1 | 133.1 | - | - | 0 | - |
| - | - | 1722 | 133.1 | - | - | 0 | - |
| - | - | 382.5 | 134 | - | - | 0 | - |
| - | - | 2147 | 134 | - | - | 0 | - |
| - | - | 481.9 | 134 | - | - | 0 | - |
| - | - | 1.276E+04 | 136.1 | - | - | 0 | - |
| - | - | 1348 | 137.1 | - | - | 0 | - |
| - | - | 5269 | 138.1 | - | - | 0 | - |
| - | - | 894.9 | 139.1 | - | - | 0 | - |
| - | - | 619.1 | 139.1 | - | - | 0 | - |
| - | - | 1144 | 141.1 | - | - | 0 | - |
| - | - | 469.6 | 141.8 | - | - | 0 | - |
| - | - | 2392 | 142.1 | - | - | 0 | - |
| - | - | 2045 | 143 | - | - | 0 | - |
| - | - | 753.2 | 143.1 | - | - | 0 | - |
| - | - | 5360 | 144.1 | - | - | 0 | - |
| - | - | 2.777E+04 | 146.1 | - | - | 0 | - |
| - | - | 1842 | 147 | - | - | 0 | - |
| - | - | 4102 | 147.1 | - | - | 0 | - |
| - | - | 534.6 | 147.1 | - | - | 0 | - |
| - | - | 519.9 | 148.1 | - | - | 0 | - |
| - | - | 910.5 | 148.9 | - | - | 0 | - |
| - | - | 455.7 | 149.7 | - | - | 0 | - |
| - | - | 499.6 | 153.1 | - | - | 0 | - |
| - | - | 982.9 | 154.1 | - | - | 0 | - |
| - | - | 1718 | 155.1 | - | - | 0 | - |
| - | - | 2.36E+04 | 155.1 | - | - | 0 | - |
| - | - | 1142 | 155.1 | - | - | 0 | - |
| - | - | 1385 | 156.1 | - | - | 0 | - |
| - | - | 1475 | 156.1 | - | - | 0 | - |
| - | - | 1495 | 157.1 | - | - | 0 | - |
| - | - | 643.9 | 158 | - | - | 0 | - |
| - | - | 2325 | 158.1 | - | - | 0 | - |
| - | - | 1.191E+05 | 159.1 | - | - | 0 | - |
| - | - | 2089 | 159.1 | - | - | 0 | - |
| - | - | 2761 | 160.1 | - | - | 0 | - |
| - | - | 419.2 | 160.1 | - | - | 0 | - |
| - | - | 1.158E+04 | 160.1 | - | - | 0 | - |
| - | - | 752.9 | 164.1 | - | - | 0 | - |
| - | - | 7240 | 165.1 | - | - | 0 | - |
| - | - | 693.5 | 165.1 | - | - | 0 | - |
| - | - | 2.268E+04 | 166.1 | - | - | 0 | - |
| - | - | 661.1 | 167.1 | - | - | 0 | - |
| - | - | 2364 | 169.1 | - | - | 0 | - |
| - | - | 472 | 169.1 | - | - | 0 | - |
| - | - | 1.004E+04 | 170.1 | - | - | 0 | - |
| - | - | 516.5 | 171.1 | - | - | 0 | - |
| - | - | 3081 | 173.5 | - | - | 0 | - |
| - | - | 841.7 | 174 | - | - | 0 | - |
| - | - | 2322 | 175 | - | - | 0 | - |
| - | - | 715.3 | 175.1 | - | - | 0 | - |
| - | - | 572 | 175.1 | - | - | 0 | - |
| - | - | 778.3 | 176 | - | - | 0 | - |
| - | - | 1429 | 176.1 | - | - | 0 | - |
| - | - | 558.8 | 177.1 | - | - | 0 | - |
| - | - | 1.004E+04 | 178.1 | - | - | 0 | - |
| - | - | 979.2 | 179.1 | - | - | 0 | - |
| - | - | 4510 | 181.1 | - | - | 0 | - |
| - | - | 724.1 | 181.1 | - | - | 0 | - |
| - | - | 8875 | 182.1 | - | - | 0 | - |
| - | - | 831 | 183.1 | - | - | 0 | - |
| - | - | 860.5 | 183.1 | - | - | 0 | - |
| - | - | 657 | 183.1 | - | - | 0 | - |
| - | - | 2358 | 183.1 | - | - | 0 | - |
| - | - | 708 | 185.1 | - | - | 0 | - |
| 2 | a | 3.48E+04 | 185.2 | 0.0003519 | 1.901 | +1 | 2 |
| - | - | 438.2 | 186 | - | - | 0 | - |
| - | - | 3900 | 186.2 | - | - | 0 | - |
| - | - | 6311 | 187.1 | - | - | 0 | - |
| - | - | 1.589E+05 | 188.1 | - | - | 0 | - |
| - | - | 1.678E+04 | 189.1 | - | - | 0 | - |
| - | - | 445.8 | 190.1 | - | - | 0 | - |
| - | - | 780.7 | 190.1 | - | - | 0 | - |
| - | - | 2325 | 190.1 | - | - | 0 | - |
| - | - | 485.4 | 190.8 | - | - | 0 | - |
| - | - | 4763 | 198.1 | - | - | 0 | - |
| - | - | 2613 | 199.1 | - | - | 0 | - |
| - | - | 719.2 | 200.1 | - | - | 0 | - |
| - | - | 625.3 | 201.1 | - | - | 0 | - |
| - | - | 600.6 | 202.1 | - | - | 0 | - |
| - | - | 616.9 | 203.1 | - | - | 0 | - |
| 6 | y | 4.275E+04 | 205.1 | 0.0002885 | 1.407 | +1 | 1 |
| - | - | 1111 | 205.1 | - | - | 0 | - |
| - | - | 428 | 206.1 | - | - | 0 | - |
| - | - | 4514 | 206.1 | - | - | 0 | - |
| - | - | 1115 | 206.1 | - | - | 0 | - |
| - | - | 1448 | 207.2 | - | - | 0 | - |
| - | - | 842.7 | 209.1 | - | - | 0 | - |
| - | - | 474 | 211.5 | - | - | 0 | - |
| - | - | 827.5 | 212.2 | - | - | 0 | - |
| - | - | 1480 | 213.1 | - | - | 0 | - |
| 2 | b | 7940 | 213.2 | 0.0002951 | 1.384 | +1 | 2 |
| - | - | 906.5 | 214.2 | - | - | 0 | - |
| - | - | 1764 | 216.1 | - | - | 0 | - |
| - | - | 634.8 | 216.1 | - | - | 0 | - |
| - | - | 1267 | 220.1 | - | - | 0 | - |
| - | - | 1818 | 221.1 | - | - | 0 | - |
| - | - | 775.2 | 221.1 | - | - | 0 | - |
| - | - | 4557 | 221.1 | - | - | 0 | - |
| - | - | 500.5 | 222.1 | - | - | 0 | - |
| - | - | 580.3 | 222.1 | - | - | 0 | - |
| - | - | 510.9 | 223.1 | - | - | 0 | - |
| - | - | 528.9 | 223.1 | - | - | 0 | - |
| - | - | 1.53E+04 | 223.2 | - | - | 0 | - |
| - | - | 1712 | 224.2 | - | - | 0 | - |
| - | - | 685.1 | 225.6 | - | - | 0 | - |
| - | - | 7541 | 226.1 | - | - | 0 | - |
| - | - | 861.9 | 226.1 | - | - | 0 | - |
| - | - | 4446 | 227.1 | - | - | 0 | - |
| - | - | 753.7 | 227.1 | - | - | 0 | - |
| - | - | 884.5 | 227.2 | - | - | 0 | - |
| - | - | 1560 | 230.1 | - | - | 0 | - |
| - | - | 601.9 | 230.1 | - | - | 0 | - |
| - | - | 555.3 | 231 | - | - | 0 | - |
| - | - | 1132 | 231.1 | - | - | 0 | - |
| - | - | 2343 | 232.1 | - | - | 0 | - |
| - | - | 806.7 | 233.1 | - | - | 0 | - |
| - | - | 6020 | 233.1 | - | - | 0 | - |
| - | - | 2.126E+04 | 234.1 | - | - | 0 | - |
| - | - | 1072 | 234.1 | - | - | 0 | - |
| - | - | 1095 | 235.1 | - | - | 0 | - |
| - | - | 2434 | 235.1 | - | - | 0 | - |
| - | - | 800.7 | 235.2 | - | - | 0 | - |
| - | - | 1619 | 238.1 | - | - | 0 | - |
| - | - | 1517 | 239.1 | - | - | 0 | - |
| 4 | b | 1046 | 239.6 | 0.0001855 | 0.7741 | +2 | 4 |
| - | - | 1136 | 240.1 | - | - | 0 | - |
| - | - | 1081 | 243.1 | - | - | 0 | - |
| - | - | 1054 | 243.1 | - | - | 0 | - |
| - | - | 1.265E+04 | 244.1 | - | - | 0 | - |
| - | - | 765.7 | 245.1 | - | - | 0 | - |
| - | - | 1888 | 246.1 | - | - | 0 | - |
| - | - | 594.6 | 247.1 | - | - | 0 | - |
| - | - | 7501 | 248.1 | - | - | 0 | - |
| - | - | 1.753E+04 | 249.1 | - | - | 0 | - |
| - | - | 1998 | 250.1 | - | - | 0 | - |
| - | - | 7.339E+04 | 251.2 | - | - | 0 | - |
| - | - | 1202 | 252.1 | - | - | 0 | - |
| - | - | 8791 | 252.2 | - | - | 0 | - |
| - | - | 3204 | 252.6 | - | - | 0 | - |
| - | - | 863.2 | 252.7 | - | - | 0 | - |
| - | - | 871.2 | 253.1 | - | - | 0 | - |
| - | - | 902 | 253.1 | - | - | 0 | - |
| - | - | 681.4 | 254.1 | - | - | 0 | - |
| - | - | 1807 | 255.2 | - | - | 0 | - |
| - | - | 1716 | 256.1 | - | - | 0 | - |
| - | - | 1370 | 260.1 | - | - | 0 | - |
| - | - | 1531 | 261.1 | - | - | 0 | - |
| - | - | 1908 | 261.1 | - | - | 0 | - |
| - | - | 1359 | 261.2 | - | - | 0 | - |
| - | - | 1907 | 261.6 | - | - | 0 | - |
| - | - | 763.4 | 261.7 | - | - | 0 | - |
| - | - | 3544 | 262.1 | - | - | 0 | - |
| - | - | 842.1 | 262.1 | - | - | 0 | - |
| - | - | 929.6 | 265.1 | - | - | 0 | - |
| - | - | 3.744E+04 | 266.1 | - | - | 0 | - |
| - | - | 1197 | 267.1 | - | - | 0 | - |
| - | - | 3955 | 267.1 | - | - | 0 | - |
| - | - | 940.9 | 268.2 | - | - | 0 | - |
| - | - | 772.1 | 269.1 | - | - | 0 | - |
| - | - | 667 | 269.2 | - | - | 0 | - |
| - | - | 2.039E+04 | 270.1 | - | - | 0 | - |
| - | - | 7280 | 270.6 | - | - | 0 | - |
| - | - | 1054 | 271.1 | - | - | 0 | - |
| - | - | 2314 | 274.1 | - | - | 0 | - |
| - | - | 6566 | 274.1 | - | - | 0 | - |
| - | - | 964.9 | 275.1 | - | - | 0 | - |
| - | - | 1142 | 276.1 | - | - | 0 | - |
| - | - | 3294 | 279.1 | - | - | 0 | - |
| - | - | 735.1 | 280.1 | - | - | 0 | - |
| - | - | 632.7 | 280.1 | - | - | 0 | - |
| - | - | 524.8 | 283.2 | - | - | 0 | - |
| - | - | 2071 | 284.1 | - | - | 0 | - |
| 3 | y | 1409 | 284.1 | 0.0002051 | 0.7219 | +2 | 4 |
| - | - | 2594 | 285.1 | - | - | 0 | - |
| - | - | 759.3 | 286.1 | - | - | 0 | - |
| - | - | 811.7 | 287.2 | - | - | 0 | - |
| 5 | b | 706.7 | 288.6 | 0.0003652 | 1.265 | +2 | 5 |
| - | - | 1361 | 294.1 | - | - | 0 | - |
| - | - | 969.4 | 296.2 | - | - | 0 | - |
| - | - | 813.5 | 296.2 | - | - | 0 | - |
| 5 | b | 5981 | 297.2 | 0.0003963 | 1.334 | +2 | 5 |
| - | - | 2153 | 297.7 | - | - | 0 | - |
| - | - | 753.4 | 298.1 | - | - | 0 | - |
| - | - | 794.4 | 300.1 | - | - | 0 | - |
| - | - | 1163 | 301.1 | - | - | 0 | - |
| - | - | 916.1 | 301.1 | - | - | 0 | - |
| 5 | y | 2656 | 302.1 | 0.0003286 | 1.088 | +1 | 2 |
| - | - | 871.6 | 302.1 | - | - | 0 | - |
| - | - | 678.1 | 302.1 | - | - | 0 | - |
| - | - | 1151 | 304.2 | - | - | 0 | - |
| - | - | 4188 | 306.2 | - | - | 0 | - |
| - | - | 767.4 | 309.2 | - | - | 0 | - |
| - | - | 1513 | 310.2 | - | - | 0 | - |
| - | - | 2216 | 313.2 | - | - | 0 | - |
| - | - | 749.6 | 316.1 | - | - | 0 | - |
| - | - | 776.1 | 318.1 | - | - | 0 | - |
| - | - | 981.3 | 318.2 | - | - | 0 | - |
| 5 | y | 1.049E+04 | 320.1 | 0.0004451 | 1.39 | +1 | 2 |
| - | - | 897.1 | 320.1 | - | - | 0 | - |
| - | - | 1728 | 321.1 | - | - | 0 | - |
| 3 | a | 745.3 | 322.2 | 0.0001188 | 0.3686 | +1 | 3 |
| - | - | 716.7 | 326.1 | - | - | 0 | - |
| - | - | 692.6 | 326.2 | - | - | 0 | - |
| - | - | 7906 | 326.7 | - | - | 0 | - |
| - | - | 1005 | 327.1 | - | - | 0 | - |
| - | - | 2228 | 327.2 | - | - | 0 | - |
| - | - | 624.9 | 327.7 | - | - | 0 | - |
| - | - | 1.071E+04 | 332.2 | - | - | 0 | - |
| - | - | 1950 | 333.2 | - | - | 0 | - |
| - | - | 889.3 | 336.1 | - | - | 0 | - |
| 2 | y | 765.8 | 340.7 | 0.001874 | 5.502 | +2 | 5 |
| - | - | 836.4 | 343.2 | - | - | 0 | - |
| - | - | 2057 | 344.2 | - | - | 0 | - |
| - | - | 4271 | 345.1 | - | - | 0 | - |
| - | - | 1.71E+04 | 346.1 | - | - | 0 | - |
| - | - | 899.1 | 346.1 | - | - | 0 | - |
| - | - | 1706 | 347.1 | - | - | 0 | - |
| 2 | y | 846.6 | 349.7 | 0.0008644 | 2.472 | +2 | 5 |
| 3 | b | 4.191E+04 | 350.2 | 0.0002669 | 0.7621 | +1 | 3 |
| - | - | 7393 | 351.2 | - | - | 0 | - |
| - | - | 852.9 | 352.2 | - | - | 0 | - |
| - | - | 1094 | 353.7 | - | - | 0 | - |
| - | - | 658.4 | 354.2 | - | - | 0 | - |
| - | - | 787 | 354.2 | - | - | 0 | - |
| - | - | 1691 | 358.7 | - | - | 0 | - |
| - | - | 728.9 | 359.2 | - | - | 0 | - |
| - | - | 668.4 | 360.2 | - | - | 0 | - |
| - | - | 2506 | 361.2 | - | - | 0 | - |
| - | - | 2222 | 362.2 | - | - | 0 | - |
| - | - | 1.919E+04 | 363.1 | - | - | 0 | - |
| - | - | 914.5 | 363.2 | - | - | 0 | - |
| - | - | 9211 | 364.1 | - | - | 0 | - |
| - | - | 1902 | 364.1 | - | - | 0 | - |
| - | - | 1402 | 365.1 | - | - | 0 | - |
| - | - | 1809 | 367.1 | - | - | 0 | - |
| - | - | 1420 | 367.2 | - | - | 0 | - |
| - | - | 6404 | 367.2 | - | - | 0 | - |
| - | - | 9259 | 367.7 | - | - | 0 | - |
| - | - | 5162 | 368.2 | - | - | 0 | - |
| - | - | 824.2 | 368.2 | - | - | 0 | - |
| - | - | 661.7 | 368.7 | - | - | 0 | - |
| - | - | 2.615E+04 | 376.2 | - | - | 0 | - |
| - | - | 640.6 | 376.6 | - | - | 0 | - |
| - | - | 9655 | 376.7 | - | - | 0 | - |
| - | - | 2726 | 377.2 | - | - | 0 | - |
| - | - | 751.3 | 379.2 | - | - | 0 | - |
| - | - | 2.053E+04 | 379.2 | - | - | 0 | - |
| - | - | 4347 | 380.2 | - | - | 0 | - |
| - | - | 5.225E+04 | 381.2 | - | - | 0 | - |
| - | - | 1212 | 381.7 | - | - | 0 | - |
| - | - | 9378 | 382.2 | - | - | 0 | - |
| - | - | 1081 | 383.2 | - | - | 0 | - |
| - | - | 875 | 384.2 | - | - | 0 | - |
| - | - | 2236 | 385.2 | - | - | 0 | - |
| - | - | 2574 | 389.2 | - | - | 0 | - |
| 0 | Precursor | 4508 | 390.2 | 0.0004127 | 1.058 | +2 | -1 |
| 0 | Precursor | 1784 | 390.7 | 0.004865 | 12.45 | +2 | -1 |
| - | - | 1644 | 391.1 | - | - | 0 | - |
| - | - | 1177 | 391.2 | - | - | 0 | - |
| - | - | 617.9 | 394.2 | - | - | 0 | - |
| - | - | 1632 | 395.1 | - | - | 0 | - |
| - | - | 2718 | 398.2 | - | - | 0 | - |
| - | - | 2619 | 399.2 | - | - | 0 | - |
| 0 | Precursor | 1.109E+04 | 399.2 | 0.0002268 | 0.5681 | +2 | -1 |
| - | - | 5017 | 399.7 | - | - | 0 | - |
| - | - | 607.5 | 400.2 | - | - | 0 | - |
| - | - | 1434 | 400.2 | - | - | 0 | - |
| - | - | 2947 | 402.2 | - | - | 0 | - |
| - | - | 840.2 | 403.1 | - | - | 0 | - |
| - | - | 656.1 | 406.7 | - | - | 0 | - |
| - | - | 2148 | 407.2 | - | - | 0 | - |
| - | - | 2005 | 409.1 | - | - | 0 | - |
| - | - | 1201 | 410.3 | - | - | 0 | - |
| - | - | 1398 | 412.2 | - | - | 0 | - |
| - | - | 1784 | 413.1 | - | - | 0 | - |
| - | - | 1185 | 416.2 | - | - | 0 | - |
| - | - | 1430 | 421.1 | - | - | 0 | - |
| 4 | y | 1.572E+04 | 430.2 | 0.0003754 | 0.8727 | +1 | 3 |
| 4 | y | 2236 | 431.2 | 0.002378 | 5.515 | +1 | 3 |
| - | - | 2644 | 431.2 | - | - | 0 | - |
| - | - | 552.8 | 431.4 | - | - | 0 | - |
| - | - | 1616 | 432.1 | - | - | 0 | - |
| - | - | 960.3 | 433.3 | - | - | 0 | - |
| 4 | y | 9570 | 448.2 | 0.0005529 | 1.234 | +1 | 3 |
| - | - | 2178 | 449.2 | - | - | 0 | - |
| - | - | 1329 | 450.3 | - | - | 0 | - |
| - | - | 3542 | 456.2 | - | - | 0 | - |
| - | - | 653.5 | 457.2 | - | - | 0 | - |
| - | - | 1140 | 459.2 | - | - | 0 | - |
| 4 | b | 1802 | 461.3 | 0.0006787 | 1.471 | +1 | 4 |
| - | - | 726.3 | 476.2 | - | - | 0 | - |
| - | - | 3544 | 477.2 | - | - | 0 | - |
| - | - | 1198 | 478.2 | - | - | 0 | - |
| 4 | b | 9263 | 478.3 | 0.0003136 | 0.6558 | +1 | 4 |
| - | - | 1791 | 479.3 | - | - | 0 | - |
| - | - | 617.2 | 480.2 | - | - | 0 | - |
| - | - | 764 | 480.3 | - | - | 0 | - |
| - | - | 1088 | 481.3 | - | - | 0 | - |
| - | - | 1746 | 491.3 | - | - | 0 | - |
| - | - | 2.384E+04 | 494.2 | - | - | 0 | - |
| - | - | 4924 | 495.2 | - | - | 0 | - |
| - | - | 1073 | 496.2 | - | - | 0 | - |
| - | - | 1059 | 500.2 | - | - | 0 | - |
| - | - | 1715 | 504.2 | - | - | 0 | - |
| - | - | 959.9 | 505.2 | - | - | 0 | - |
| - | - | 765.7 | 509.3 | - | - | 0 | - |
| - | - | 1254 | 512.2 | - | - | 0 | - |
| - | - | 798 | 514.2 | - | - | 0 | - |
| - | - | 629.8 | 517.2 | - | - | 0 | - |
| - | - | 2004 | 522.2 | - | - | 0 | - |
| - | - | 1889 | 523.2 | - | - | 0 | - |
| - | - | 1358 | 528.2 | - | - | 0 | - |
| - | - | 1074 | 532.2 | - | - | 0 | - |
| - | - | 8504 | 539.2 | - | - | 0 | - |
| - | - | 1986 | 540.2 | - | - | 0 | - |
| - | - | 1344 | 545.2 | - | - | 0 | - |
| - | - | 655.7 | 547.3 | - | - | 0 | - |
| - | - | 998.6 | 548.3 | - | - | 0 | - |
| - | - | 747.2 | 549.2 | - | - | 0 | - |
| - | - | 1.001E+04 | 550.2 | - | - | 0 | - |
| - | - | 2349 | 551.2 | - | - | 0 | - |
| - | - | 744.6 | 557.8 | - | - | 0 | - |
| - | - | 1128 | 558.3 | - | - | 0 | - |
| - | - | 669.5 | 558.3 | - | - | 0 | - |
| - | - | 1219 | 565.3 | - | - | 0 | - |
| 3 | y | 3510 | 567.2 | 0.0001183 | 0.2086 | +1 | 4 |
| 3 | y | 3438 | 568.2 | 0.002309 | 4.063 | +1 | 4 |
| - | - | 1621 | 569.2 | - | - | 0 | - |
| - | - | 846.8 | 571.2 | - | - | 0 | - |
| 5 | b | 1559 | 575.3 | 0.00163 | 2.833 | +1 | 5 |
| 5 | b | 9982 | 576.3 | 0.0001108 | 0.1923 | +1 | 5 |
| - | - | 3997 | 577.3 | - | - | 0 | - |
| 3 | y | 2.635E+04 | 585.2 | 9.351E-06 | 0.01598 | +1 | 4 |
| - | - | 8346 | 586.2 | - | - | 0 | - |
| - | - | 1169 | 587.2 | - | - | 0 | - |
| 5 | b | 4.718E+04 | 593.3 | 0.0001096 | 0.1847 | +1 | 5 |
| - | - | 1.565E+04 | 594.3 | - | - | 0 | - |
| - | - | 895.4 | 595.2 | - | - | 0 | - |
| - | - | 3203 | 595.3 | - | - | 0 | - |
| - | - | 1827 | 610.3 | - | - | 0 | - |
| - | - | 2841 | 611.3 | - | - | 0 | - |
| - | - | 664.1 | 615.6 | - | - | 0 | - |
| - | - | 780.9 | 633.3 | - | - | 0 | - |
| - | - | 783.6 | 634.3 | - | - | 0 | - |
| - | - | 637.2 | 645.3 | - | - | 0 | - |
| - | - | 624.7 | 647.4 | - | - | 0 | - |
| - | - | 994.3 | 650.3 | - | - | 0 | - |
| - | - | 1030 | 651.3 | - | - | 0 | - |
| - | - | 2149 | 652.3 | - | - | 0 | - |
| - | - | 1122 | 653.3 | - | - | 0 | - |
| - | - | 1457 | 662.3 | - | - | 0 | - |
| - | - | 1867 | 663.3 | - | - | 0 | - |
| - | - | 1066 | 664.3 | - | - | 0 | - |
| - | - | 2045 | 668.3 | - | - | 0 | - |
| - | - | 1140 | 669.3 | - | - | 0 | - |
| 2 | y | 4312 | 680.3 | 0.0003439 | 0.5055 | +1 | 5 |
| 2 | y | 9990 | 681.3 | 0.001131 | 1.659 | +1 | 5 |
| - | - | 3535 | 682.3 | - | - | 0 | - |
| - | - | 1343 | 683.3 | - | - | 0 | - |
| 2 | y | 3.107E+04 | 698.3 | 0.0004552 | 0.6518 | +1 | 5 |
| - | - | 1.176E+04 | 699.3 | - | - | 0 | - |
| - | - | 2528 | 700.3 | - | - | 0 | - |
| - | - | 602.4 | 887.4 | - | - | 0 | - |
| - | - | 585.4 | 1065 | - | - | 0 | - |
| - | - | 666.2 | 1711 | - | - | 0 | - |
| - | - | 642 | 2897 | - | - | 0 | - |
| - | - | 719.8 | 3191 | - | - | 0 | - |
| - | - | 656.9 | 3460 | - | - | 0 | - |

m/z Charge Intensity FragmentType MassShift Position
120.08109283447266 0 3373.8582
121.0401611328125 0 770.5353
121.08489227294922 0 585.2176
121.21875762939453 0 339.0621
122.07160186767578 0 1949.4805
123.04447174072266 0 1585.8158
123.0555419921875 0 1524.3452
124.0473403930664 0 435.08322
124.2462387084961 0 393.83115
125.10757446289062 0 848.27875
126.03767395019531 0 720.9029
128.0531005859375 0 640.12933
129.05499267578125 0 3142.198
129.066162109375 0 1349.8909
129.10255432128906 0 8352.063
130.06549072265625 0 54225.395
130.1060791015625 0 771.9661
131.06883239746094 0 5217.8374
132.08111572265625 0 18081.88
133.061279296875 0 469.11072
133.08453369140625 0 1722.0919
133.9642791748047 0 382.50903
134.0274200439453 0 2146.849
134.04510498046875 0 481.85
136.0760498046875 0 12759.898
137.0791778564453 0 1348.3434
138.0665283203125 0 5268.9434
139.0504913330078 0 894.9431
139.08657836914062 0 619.08984
141.1024932861328 0 1143.6564
141.8125457763672 0 469.59067
142.06541442871094 0 2392.252
143.04544067382812 0 2045.345
143.0733642578125 0 753.15814
144.08116149902344 0 5359.9697
146.06039428710938 0 27773.42
147.04444885253906 0 1842.1909
147.06454467773438 0 4102.189
147.1134033203125 0 534.58575
148.05087280273438 0 519.91846
148.94715881347656 0 910.46075
149.7315673828125 0 455.66852
153.1028594970703 0 499.61935
154.0980682373047 0 982.8778
155.08155822753906 0 1717.5087
155.09307861328125 0 23598.809
155.11827087402344 0 1142.2675
156.07737731933594 0 1385.4788
156.09628295898438 0 1475.3267
157.06130981445312 0 1494.8635
158.0273895263672 0 643.9323
158.0843048095703 0 2325.2295
159.09205627441406 0 119142.63
159.11288452148438 0 2089.4785
160.0760040283203 0 2761.0188
160.0887908935547 0 419.2351
160.0953826904297 0 11581.725
164.11868286132812 0 752.9068
165.054931640625 0 7240.254
165.0771942138672 0 693.45337
166.06143188476562 0 22675.045
167.06515502929688 0 661.14
169.0762939453125 0 2364.002
169.09771728515625 0 472.01755
170.06036376953125 0 10036.033
171.06344604492188 0 516.48303
173.45066833496094 0 3080.503
174.0372314453125 0 841.685
175.0157928466797 0 2322.3796
175.07151794433594 0 715.30725
175.09793090820312 0 572.0026
176.0379180908203 0 778.28455
176.0823974609375 0 1428.9413
177.1025848388672 0 558.8158
178.1342010498047 0 10039.505
179.13768005371094 0 979.20184
181.06106567382812 0 4509.514
181.1339569091797 0 724.07776
182.0814971923828 0 8875.353
183.07640075683594 0 830.99384
183.08529663085938 0 860.4995
183.1131134033203 0 657.0316
183.1496124267578 0 2358.4453
185.0564727783203 0 708.0268
185.16519165039062 0 34796.87 a 1
186.02276611328125 0 438.22824
186.1685333251953 0 3900.1414
187.0869140625 0 6311.3965
188.0709686279297 0 158922.81
189.07432556152344 0 16784.193
190.06051635742188 0 445.7581
190.07723999023438 0 780.7474
190.13418579101562 0 2324.6455
190.7676544189453 0 485.3784
198.08770751953125 0 4763.14
199.07217407226562 0 2612.8408
200.14041137695312 0 719.22327
201.0875701904297 0 625.33057
202.107421875 0 600.6206
203.09349060058594 0 616.89856
205.09744262695312 0 42754.36 y 5
205.1447296142578 0 1110.5394
206.09268188476562 0 428.04425
206.10092163085938 0 4514.2695
206.12911987304688 0 1114.9314
207.1605224609375 0 1447.6486
209.05612182617188 0 842.69196
211.45343017578125 0 474.03262
212.17649841308594 0 827.4807
213.13870239257812 0 1480.4391
213.16004943847656 0 7940.145 b 1
214.16311645507812 0 906.53033
216.09837341308594 0 1763.6754
216.13401794433594 0 634.8197
220.119384765625 0 1266.9471
221.0595703125 0 1817.5323
221.0712432861328 0 775.19226
221.10374450683594 0 4557.432
222.08607482910156 0 500.48065
222.10699462890625 0 580.2961
223.05592346191406 0 510.87488
223.144287109375 0 528.9437
223.1556854248047 0 15299.812
224.15940856933594 0 1712.2882
225.64584350585938 0 685.14734
226.0825958251953 0 7541.3857
226.09474182128906 0 861.8801
227.06678771972656 0 4446.407
227.08615112304688 0 753.72876
227.1553955078125 0 884.50464
230.07762145996094 0 1560.4506
230.10336303710938 0 601.8762
231.04534912109375 0 555.2835
231.0877227783203 0 1132.4675
232.14450073242188 0 2343.154
233.1039581298828 0 806.67017
233.13995361328125 0 6019.842
234.12408447265625 0 21255.768
234.14389038085938 0 1072.2461
235.0820770263672 0 1095.4344
235.12693786621094 0 2433.543
235.15631103515625 0 800.6767
238.13014221191406 0 1618.779
239.0823211669922 0 1516.8242
239.64207458496094 0 1046.1162 b 3
240.13436889648438 0 1136.2963
243.07708740234375 0 1081.1527
243.10906982421875 0 1054.477
244.09315490722656 0 12649.634
245.09669494628906 0 765.6907
246.13558959960938 0 1888.136
247.11166381835938 0 594.62524
248.1145477294922 0 7500.6533
249.0985565185547 0 17530.504
250.10125732421875 0 1997.94
251.15065002441406 0 73386.42
252.10916137695312 0 1202.0868
252.1538848876953 0 8791.054
252.60328674316406 0 3203.604
252.6506805419922 0 863.1727
253.1039581298828 0 871.1569
253.11827087402344 0 902.03876
254.078125 0 681.35535
255.19589233398438 0 1807.2089
256.1083679199219 0 1715.8351
260.10333251953125 0 1369.6251
261.1179504394531 0 1530.9426
261.1344299316406 0 1908.0452
261.1636047363281 0 1359.3065
261.60858154296875 0 1906.6052
261.6647644042969 0 763.38104
262.05047607421875 0 3543.5889
262.10406494140625 0 842.0987
265.1401062011719 0 929.6436
266.1249694824219 0 37439.742
267.10894775390625 0 1197.2806
267.12786865234375 0 3954.9497
268.1767883300781 0 940.9327
269.1058044433594 0 772.1419
269.16131591796875 0 666.9712
270.1218566894531 0 20386.467
270.6232604980469 0 7279.951
271.1255187988281 0 1054.2214
274.1177978515625 0 2314.1067
274.13018798828125 0 6565.603
275.1343078613281 0 964.88434
276.109375 0 1142.2744
279.1455993652344 0 3293.6968
280.1291809082031 0 735.05194
280.1475830078125 0 632.7437
283.1524353027344 0 524.7583
284.1035461425781 0 2070.698
284.1193542480469 0 1408.9095 y Water loss 2
285.0876770019531 0 2594.4136
286.0904846191406 0 759.3171
287.1866455078125 0 811.6557
288.642822265625 0 706.71436 b Ammonia loss 4
294.12054443359375 0 1360.648
296.17333984375 0 969.43
296.1981201171875 0 813.52124
297.1561279296875 0 5981.1675 b 4
297.6574401855469 0 2152.905
298.1393737792969 0 753.4353
300.1104431152344 0 794.38654
301.11334228515625 0 1163.1095
301.1412353515625 0 916.0941
302.1138610839844 0 2655.818 y Water loss 4
302.12908935546875 0 871.6498
302.1439514160156 0 678.10114
304.2128601074219 0 1151.1049
306.2292785644531 0 4187.529
309.16741943359375 0 767.38306
310.1722717285156 0 1513.2449
313.223876953125 0 2215.729
316.1146240234375 0 749.56146
318.1202697753906 0 776.09717
318.1507873535156 0 981.3196
320.1245422363281 0 10489.768 y 4
320.1451110839844 0 897.05914
321.12744140625 0 1727.552
322.2236328125 0 745.29987 a 2
326.1354064941406 0 716.7113
326.2228088378906 0 692.5705
326.6640625 0 7905.5415
327.0943603515625 0 1005.1195
327.16552734375 0 2227.9536
327.6667175292969 0 624.90955
332.2084655761719 0 10707.761
333.2128601074219 0 1949.6312
336.12969970703125 0 889.2716
340.6630554199219 0 765.8326 y Water loss 1
343.1614685058594 0 836.4392
344.17181396484375 0 2057.192
345.1309814453125 0 4271.3174
346.1148681640625 0 17104.562
346.1376037597656 0 899.05304
347.1177673339844 0 1706.2145
349.6673278808594 0 846.6261 y 1
350.21893310546875 0 41909.55 b 2
351.2218017578125 0 7393.493
352.2247619628906 0 852.8544
353.6885986328125 0 1093.6072
354.1922607421875 0 658.38
354.2175598144531 0 787.0095
358.67987060546875 0 1691.0569
359.1775817871094 0 728.9214
360.203125 0 668.36914
361.1982727050781 0 2506.291
362.18212890625 0 2222.3325
363.14111328125 0 19188.045
363.16668701171875 0 914.5488
364.1252136230469 0 9211.432
364.14593505859375 0 1902.1257
365.12823486328125 0 1402.2885
367.13983154296875 0 1808.7914
367.192138671875 0 1419.7329
367.2454528808594 0 6403.6523
367.6852111816406 0 9258.59
368.18609619140625 0 5162.2544
368.2486267089844 0 824.1643
368.6846008300781 0 661.6678
376.197998046875 0 26154.744
376.5605773925781 0 640.59
376.6986999511719 0 9654.81
377.1982727050781 0 2726.0066
379.1805419921875 0 751.3115
379.20892333984375 0 20527.168
380.2120056152344 0 4346.7144
381.1518859863281 0 52249.457
381.68963623046875 0 1212.2386
382.15484619140625 0 9378.488
383.1573791503906 0 1080.9926
384.1648864746094 0 875.018
385.1508483886719 0 2236.3782
389.1913757324219 0 2573.8818
390.19580078125 0 4508.3975 Precursor Water loss
390.6922607421875 0 1784.0536 Precursor Ammonia loss
391.1368103027344 0 1644.1147
391.18841552734375 0 1177.3081
394.151123046875 0 617.8768
395.1353759765625 0 1632.3763
398.1784973144531 0 2718.1418
399.1623840332031 0 2619.2349
399.2008972167969 0 11091.711 Precursor
399.702880859375 0 5016.527
400.1654052734375 0 607.5494
400.20269775390625 0 1434.1473
402.1775207519531 0 2946.5105
403.1357727050781 0 840.16486
406.7015686035156 0 656.05347
407.20379638671875 0 2148.0593
409.1470947265625 0 2005.4572
410.2773132324219 0 1201.1251
412.1607360839844 0 1398.1274
413.14581298828125 0 1784.2493
416.23004150390625 0 1185.2611
421.1458435058594 0 1429.6667
430.1724853515625 0 15724.095 y Water loss 3
431.15374755859375 0 2236.3533 y Ammonia loss 3
431.1778869628906 0 2644.2979
431.371826171875 0 552.8438
432.1355895996094 0 1616.2104
433.2547912597656 0 960.276
448.1832275390625 0 9569.762 y 3
449.1861877441406 0 2178.0618
450.2823486328125 0 1328.7554
456.1836853027344 0 3541.805
457.1875915527344 0 653.50934
459.19952392578125 0 1139.6301
461.2513732910156 0 1802.0354 b Ammonia loss 3
476.2239074707031 0 726.27936
477.2095642089844 0 3544.18
478.212158203125 0 1198.1174
478.2775573730469 0 9263.425 b 3
479.27862548828125 0 1791.4792
480.2190856933594 0 617.2169
480.28228759765625 0 764.03925
481.3127746582031 0 1088.0442
491.2777404785156 0 1745.5862
494.2359313964844 0 23841.965
495.23834228515625 0 4923.724
496.2405700683594 0 1073.1078
500.2005920410156 0 1058.5088
504.2005920410156 0 1714.8517
505.1992492675781 0 959.937
509.31768798828125 0 765.6942
512.2477416992188 0 1253.6052
514.1812133789062 0 798.0017
517.2225341796875 0 629.759
522.2279663085938 0 2003.8356
523.2355346679688 0 1889.4518
528.19482421875 0 1357.6366
532.196044921875 0 1074.1296
539.236572265625 0 8503.572
540.2369995117188 0 1986.3478
545.222900390625 0 1344.43
547.30029296875 0 655.6641
548.2837524414062 0 998.5634
549.220703125 0 747.1698
550.20458984375 0 10007.781
551.2062377929688 0 2349.4426
557.843505859375 0 744.635
558.265625 0 1128.123
558.3475341796875 0 669.5326
565.3085327148438 0 1219.0696
567.2311401367188 0 3509.8638 y Water loss 2
568.2173461914062 0 3437.7683 y Ammonia loss 2
569.2177734375 0 1621.3436
571.2382202148438 0 846.8269
575.2919921875 0 1559.355 b Water loss 4
576.2775268554688 0 9982.132 b Ammonia loss 4
577.2799072265625 0 3997.3845
585.2415771484375 0 26346.076 y 2
586.244873046875 0 8345.611
587.244384765625 0 1168.5061
593.3040771484375 0 47179.445 b 4
594.3060913085938 0 15648.307
595.2279663085938 0 895.3954
595.307861328125 0 3203.1204
610.3301391601562 0 1826.6168
611.315185546875 0 2840.6467
615.599853515625 0 664.06366
633.2988891601562 0 780.8787
634.3045654296875 0 783.6463
645.2745971679688 0 637.204
647.4092407226562 0 624.69226
650.326171875 0 994.30554
651.3191528320312 0 1029.5818
652.3184204101562 0 2148.7358
653.3209228515625 0 1121.5154
662.3025512695312 0 1456.9866
663.2943725585938 0 1867.0898
664.2940063476562 0 1066.3641
668.3339233398438 0 2044.845
669.3384399414062 0 1139.9547
680.3154296875 0 4311.6123 y Water loss 1
681.3002319335938 0 9989.747 y Ammonia loss 1
682.3041381835938 0 3534.5276
683.3047485351562 0 1343.3396
698.3251953125 0 31068.56 y 1
699.3286743164062 0 11755.318
700.332275390625 0 2527.7449
887.444091796875 0 602.38904
1064.7264404296875 0 585.4301
1710.762451171875 0 666.15814
2897.47998046875 0 642.00836
3190.5908203125 0 719.7771
3460.318603515625 0 656.91754

Spectrum Details

|  |  |
| --- | --- |
| Matched peaks? Matched peaks The total absolute number of peaks matched. Additionally in brackets the total fraction of peaks matched and the total number of peaks is shown. | 30 (7.73% of 388) |
| FDR? FDR The false discovery rate estimated for this peptide. It is calculated by matching all theoretical fragments with a non-integer shift with the raw peaks for this spectrum. This is done with 40 different shifts. The resulting percentage is the average number of annotated peaks over the number of annotated peaks with the correct spectrum. | 1.03% |
| Satellite FDR? Satellite FDR See the FDR for details on its calculation. This satellite ion specific FDR only contains the satellite ions (d/w) for I/L/J positions. | - |
| PSM Score? PSM Score The PSM Score as given by Hecklib to this annotated spectrum. It is shown with three significant figures. | 387 |

#### Reverse Lookup? Reverse Lookup All places where this read could be placed.

| Group | Segment | Template | Template Part | Read Part | Score | Unique |
| --- | --- | --- | --- | --- | --- | --- |
| Homo sapiens Heavy Chain | IGHC | IGHG1 | [190..196] | [0..6] | 48 | False |
| Homo sapiens Heavy Chain | IGHC | IGHG3 | [237..243] | [0..6] | 48 | False |
| Homo sapiens Heavy Chain | IGHC | IGHG4 | [187..193] | [0..6] | 48 | False |

| Recombined | Template Part | Read Part | Score | Unique |
| --- | --- | --- | --- | --- |
| REC-0-1 | [315..321] | [0..6] | 48 | True |

#### Meta Information from Multiple reads

##### Number of combined reads

6

##### Intensity

0.7284

##### TotalArea

5.567E+06

##### Changes to the peptide sequence

VJHQDW

L→JNo support for either Leucine or Isoleucine based on side chain ions (Position: 2)

J→LSupport for Leucine based on side chain ions (1 for L 0 for I) (Position: 2)

L→JNo support for either Leucine or Isoleucine based on side chain ions (Position: 2)

#### Positional Score

Copy Data

##### Positional Score (TSV)

###### Preview

```
Loading example...
```

*Click on the button to copy the data to your clipboard.*

10012345

Label Value
"0" 0.655
"1" 0.658
"2" 0.65
"3" 0.625
"4" 0.655
"5" 0.66

#### Meta Information from PEAKS

##### Scan Identifier

F2:4662

##### Original sequence

V

L

H

Q

D

W

##### Posttranslational Modifications

##### Source File

D:\separate\_stitch\_analyses\xle-disambiguation\raw\20210323\_F1\_UM1\_Peng0013\_SA\_F59\_ingel\_3ug\_TL.raw

##### Fraction

2

##### Scan Feature

F2:790

##### De Novo Score

99

##### ConfidenceScore

99

### m/z

399.2009

##### Mass

796.3868

##### Charge

2

##### Retention Time

25.05

##### Predicted Retention Time

-

##### Area

4.066E+06

##### Parts Per Million

0.6

##### Fragmentation mode

HCD

##### Originating file

01 D:\separate\_stitch\_analyses\xle-disambiguation\20210325\_F59\_3ug\_DENOVO\_12.csv

#### Meta Information from PEAKS

##### Scan Identifier

F2:4739

##### Original sequence

V

L

H

Q

D

W

##### Posttranslational Modifications

##### Source File

D:\separate\_stitch\_analyses\xle-disambiguation\raw\20210323\_F1\_UM1\_Peng0013\_SA\_F59\_ingel\_3ug\_TL.raw

##### Fraction

2

##### Scan Feature

-

##### De Novo Score

98

##### ConfidenceScore

98

### m/z

399.2013

##### Mass

796.3868

##### Charge

2

##### Retention Time

25.42

##### Predicted Retention Time

-

##### Area

0

##### Parts Per Million

1.7

##### Fragmentation mode

HCD

##### Originating file

01 D:\separate\_stitch\_analyses\xle-disambiguation\20210325\_F59\_3ug\_DENOVO\_12.csv

#### Meta Information from PEAKS

##### Scan Identifier

F3:4545

##### Original sequence

V

L

H

Q

D

W

##### Posttranslational Modifications

##### Source File

D:\separate\_stitch\_analyses\xle-disambiguation\raw\20210323\_F1\_UM1\_Peng0013\_SA\_F59\_ingel\_3ug\_chymo.raw

##### Fraction

3

##### Scan Feature

F3:699

##### De Novo Score

98

##### ConfidenceScore

98

### m/z

399.2011

##### Mass

796.3868

##### Charge

2

##### Retention Time

25.09

##### Predicted Retention Time

-

##### Area

1.23E+06

##### Parts Per Million

1.2

##### Fragmentation mode

HCD

##### Originating file

01 D:\separate\_stitch\_analyses\xle-disambiguation\20210325\_F59\_3ug\_DENOVO\_12.csv

#### Meta Information from PEAKS

##### Scan Identifier

F2:4673

##### Original sequence

V

L

H

Q

D

W

+15.99

##### Posttranslational Modifications

Oxidation (HW)

##### Source File

D:\separate\_stitch\_analyses\xle-disambiguation\raw\20210323\_F1\_UM1\_Peng0013\_SA\_F59\_ingel\_3ug\_TL.raw

##### Fraction

2

##### Scan Feature

F2:943

##### De Novo Score

97

##### ConfidenceScore

97

### m/z

407.1978

##### Mass

812.3817

##### Charge

2

##### Retention Time

25.05

##### Predicted Retention Time

-

##### Area

2.718E+05

##### Fragmentation mode

HCD

##### Originating file

01 D:\separate\_stitch\_analyses\xle-disambiguation\20210325\_F59\_3ug\_DENOVO\_12.csv

#### Meta Information from PEAKS

##### Scan Identifier

F2:4808

##### Original sequence

V

L

H

Q

D

W

##### Posttranslational Modifications

##### Source File

D:\separate\_stitch\_analyses\xle-disambiguation\raw\20210323\_F1\_UM1\_Peng0013\_SA\_F59\_ingel\_3ug\_TL.raw

##### Fraction

2

##### Scan Feature

-

##### De Novo Score

97

##### ConfidenceScore

97

### m/z

399.2016

##### Mass

796.3868

##### Charge

2

##### Retention Time

25.81

##### Predicted Retention Time

-

##### Area

0

##### Parts Per Million

2.4

##### Fragmentation mode

HCD

##### Originating file

01 D:\separate\_stitch\_analyses\xle-disambiguation\20210325\_F59\_3ug\_DENOVO\_12.csv

#### Meta Information from PEAKS

##### Scan Identifier

F2:4874

##### Original sequence

V

L

H

Q

D

W

##### Posttranslational Modifications

##### Source File

D:\separate\_stitch\_analyses\xle-disambiguation\raw\20210323\_F1\_UM1\_Peng0013\_SA\_F59\_ingel\_3ug\_TL.raw

##### Fraction

2

##### Scan Feature

-

##### De Novo Score

96

##### ConfidenceScore

96

### m/z

399.2011

##### Mass

796.3868

##### Charge

2

##### Retention Time

26.2

##### Predicted Retention Time

-

##### Area

0

##### Parts Per Million

1

##### Fragmentation mode

HCD

##### Originating file

01 D:\separate\_stitch\_analyses\xle-disambiguation\20210325\_F59\_3ug\_DENOVO\_12.csv
