## Supplementary material for "A handle on mass coincidence errors in *de novo* sequencing of antibodies by bottom-up proteomics": Combined_037.html

Details Combined\_037 | Stitch OverviewUndefined

### Read Combined\_037

#### Sequence (length=15)

JFPPSSEEJQANKAT

#### Spectrum 6210? Spectrum 6210 The raw spectrum of this peptide as annotated by Hecklib. The fragments are coloured according to ion type (see legend). Any peaks with a star '\*' as text can be hovered over to see the full details, first the ion type second the mass shift type. By hovering over the amino acids in the peptide or ions in the legend the corresponding peaks are highlighted. By toggling the 'Unassigned' label you can turn the background (unassigned) peaks on or off in the plot. By updating the slider in the Ion legend you can update the spectrum to only show the top X% of the peaks with labels. The top X% means any peak that is within X% of the highest intensity. By dragging in the spectrum you can zoom in to a specific part of the spectrum and use 'Zoom Out' to get back to the original zoom level. The annotation of the spectrum is based on the given sequence in the peptides file and is done with different software so inconsistencies are likely. The peaks are annotated based on the given sequence, with 20 ppm tolerance.

Copy Data

##### Spectrum 6210 (TSV)

###### Preview

```
Loading example...
```

*Click on the button to copy the data to your clipboard.*

Mz MinMz MaxIntensity Max

WidthHeightPeptide font sizePeptide stroke widthSpectrum font sizeSpectrum stroke widthCompact peptide

Ion legend

wxyz

abcd

OtherUnassignedIon^Charge^_Position_Show for top:%

JFPPSSEEJQANKAT

02.04e+54.08e+56.12e+58.16e+5

Zoom Out

y^+1^_2_y^+1^_3_y^+1^_4_y^+1^_5_y^+1^_6_z^+1^_6_y^+1^_6_y^+2^_12_y^+2^_13_y^+2^_13_z^+2^_13_y^+2^_13_z^+1^_7_y^+1^_7_y^+2^_14_z^+2^_14_c^+2^_14_c^+1^_7_y^+2^_14_z^+1^_8_y^+1^_8_c^+1^_8_w^+1^_9_z^+1^_9_y^+1^_9_c^+1^_9_y^+1^_10_y^+1^_10_z^+1^_10_y^+1^_10_c^+1^_10_c^+1^_10_y^+1^_11_z^+1^_11_y^+1^_11_c^+1^_11_c^+1^_11_y^+1^_12_c^+1^_12_c^+1^_12_y^+1^_13_y^+1^_13_c^+1^_13_z^+1^_14_c^+1^_14_

0776155123273103

Fragment Matches Table

Show background peaks

| Position | Ion type | Intensity | mz Theoretical | mz Error (Th) | mz Error (ppm) | Charge | Series Number |
| --- | --- | --- | --- | --- | --- | --- | --- |
| - | - | 3560 | 120.1 | - | - | 0 | - |
| - | - | 5316 | 129.1 | - | - | 0 | - |
| - | - | 597.9 | 144.7 | - | - | 0 | - |
| - | - | 1219 | 149 | - | - | 0 | - |
| - | - | 578.9 | 159.1 | - | - | 0 | - |
| - | - | 3221 | 173.4 | - | - | 0 | - |
| 14 | y | 1964 | 191.1 | 0.0003329 | 1.742 | +1 | 2 |
| - | - | 732.7 | 215.5 | - | - | 0 | - |
| - | - | 738 | 220.2 | - | - | 0 | - |
| - | - | 1015 | 221.1 | - | - | 0 | - |
| - | - | 1382 | 226.1 | - | - | 0 | - |
| - | - | 789.6 | 232.5 | - | - | 0 | - |
| - | - | 3.427E+04 | 233.2 | - | - | 0 | - |
| - | - | 5502 | 234.2 | - | - | 0 | - |
| - | - | 2445 | 243.1 | - | - | 0 | - |
| - | - | 4.134E+04 | 261.2 | - | - | 0 | - |
| - | - | 5079 | 262.2 | - | - | 0 | - |
| - | - | 798.3 | 276.7 | - | - | 0 | - |
| - | - | 2349 | 282.1 | - | - | 0 | - |
| - | - | 2249 | 297.2 | - | - | 0 | - |
| - | - | 1306 | 299.1 | - | - | 0 | - |
| - | - | 2289 | 314.2 | - | - | 0 | - |
| 13 | y | 4283 | 319.2 | 4.899E-06 | 0.01535 | +1 | 3 |
| - | - | 5216 | 355.1 | - | - | 0 | - |
| - | - | 1965 | 356.1 | - | - | 0 | - |
| - | - | 7179 | 358.2 | - | - | 0 | - |
| - | - | 1107 | 359.2 | - | - | 0 | - |
| - | - | 984.4 | 385.2 | - | - | 0 | - |
| - | - | 913.5 | 414.7 | - | - | 0 | - |
| - | - | 1122 | 425.2 | - | - | 0 | - |
| 12 | y | 4589 | 433.2 | 0.0001683 | 0.3884 | +1 | 4 |
| - | - | 1589 | 455.3 | - | - | 0 | - |
| - | - | 1029 | 470.7 | - | - | 0 | - |
| - | - | 984.9 | 479.8 | - | - | 0 | - |
| - | - | 1051 | 498.2 | - | - | 0 | - |
| 11 | y | 5038 | 504.3 | 0.0009878 | 1.959 | +1 | 5 |
| - | - | 1621 | 505.3 | - | - | 0 | - |
| - | - | 1002 | 542.3 | - | - | 0 | - |
| - | - | 1160 | 582.3 | - | - | 0 | - |
| - | - | 5373 | 582.8 | - | - | 0 | - |
| - | - | 2220 | 583.3 | - | - | 0 | - |
| - | - | 4714 | 591.3 | - | - | 0 | - |
| - | - | 2901 | 591.8 | - | - | 0 | - |
| - | - | 1714 | 612.8 | - | - | 0 | - |
| - | - | 1895 | 613.3 | - | - | 0 | - |
| 10 | y | 1436 | 615.3 | 0.002537 | 4.123 | +1 | 6 |
| 10 | z | 5173 | 616.3 | 0.0004745 | 0.7699 | +1 | 6 |
| - | - | 8403 | 617.3 | - | - | 0 | - |
| - | - | 7389 | 617.8 | - | - | 0 | - |
| - | - | 6974 | 618.3 | - | - | 0 | - |
| - | - | 3291 | 618.8 | - | - | 0 | - |
| - | - | 4.311E+04 | 626.8 | - | - | 0 | - |
| - | - | 1543 | 627.3 | - | - | 0 | - |
| - | - | 2.793E+04 | 627.3 | - | - | 0 | - |
| - | - | 7141 | 627.8 | - | - | 0 | - |
| - | - | 1663 | 629.3 | - | - | 0 | - |
| 10 | y | 5451 | 632.3 | 0.0003997 | 0.6322 | +1 | 6 |
| - | - | 1482 | 633.3 | - | - | 0 | - |
| 4 | y | 2709 | 637.8 | 0.0009385 | 1.471 | +2 | 12 |
| - | - | 2722 | 638.3 | - | - | 0 | - |
| - | - | 3249 | 655.3 | - | - | 0 | - |
| - | - | 2629 | 655.8 | - | - | 0 | - |
| - | - | 3964 | 663.3 | - | - | 0 | - |
| - | - | 1939 | 663.8 | - | - | 0 | - |
| - | - | 2409 | 664.3 | - | - | 0 | - |
| - | - | 1167 | 664.8 | - | - | 0 | - |
| - | - | 1378 | 668.8 | - | - | 0 | - |
| 3 | y | 2.004E+04 | 677.3 | 0.0004926 | 0.7273 | +2 | 13 |
| 3 | y | 2.215E+04 | 677.8 | 0.006218 | 9.173 | +2 | 13 |
| 3 | z | 1.111E+04 | 678.3 | 0.001024 | 1.509 | +2 | 13 |
| - | - | 5346 | 678.8 | - | - | 0 | - |
| - | - | 1384 | 679.3 | - | - | 0 | - |
| - | - | 4130 | 685.3 | - | - | 0 | - |
| - | - | 3666 | 685.8 | - | - | 0 | - |
| 3 | y | 7.23E+05 | 686.3 | 0.0001454 | 0.2119 | +2 | 13 |
| - | - | 5.42E+05 | 686.8 | - | - | 0 | - |
| - | - | 1.894E+05 | 687.3 | - | - | 0 | - |
| - | - | 2.02E+04 | 687.8 | - | - | 0 | - |
| - | - | 1211 | 700.8 | - | - | 0 | - |
| 9 | z | 2.162E+04 | 729.4 | 0.00031 | 0.425 | +1 | 7 |
| - | - | 1.809E+04 | 730.4 | - | - | 0 | - |
| - | - | 5802 | 731.4 | - | - | 0 | - |
| - | - | 2694 | 740.3 | - | - | 0 | - |
| 9 | y | 5282 | 745.4 | 0.00123 | 1.65 | +1 | 7 |
| 2 | y | 1189 | 751.4 | 0.005214 | 6.939 | +2 | 14 |
| 2 | z | 924.7 | 751.9 | 0.01326 | 17.64 | +2 | 14 |
| 14 | c | 5062 | 756.9 | 0.001501 | 1.983 | +2 | 14 |
| 7 | c | 3769 | 757.4 | 0.01003 | 13.25 | +1 | 7 |
| - | - | 1539 | 758.4 | - | - | 0 | - |
| 2 | y | 8653 | 759.9 | 0.0009727 | 1.28 | +2 | 14 |
| - | - | 8318 | 760.4 | - | - | 0 | - |
| - | - | 3615 | 760.9 | - | - | 0 | - |
| - | - | 2100 | 802.4 | - | - | 0 | - |
| - | - | 3562 | 807.4 | - | - | 0 | - |
| - | - | 4499 | 807.9 | - | - | 0 | - |
| - | - | 2357 | 808.4 | - | - | 0 | - |
| - | - | 2196 | 815.4 | - | - | 0 | - |
| - | - | 4271 | 815.9 | - | - | 0 | - |
| - | - | 4.984E+04 | 816.4 | - | - | 0 | - |
| - | - | 4.882E+04 | 816.9 | - | - | 0 | - |
| - | - | 2.688E+04 | 817.4 | - | - | 0 | - |
| - | - | 3995 | 817.9 | - | - | 0 | - |
| 8 | z | 5.123E+04 | 858.4 | 0.0001784 | 0.2079 | +1 | 8 |
| - | - | 5.714E+04 | 859.4 | - | - | 0 | - |
| - | - | 1.889E+04 | 860.5 | - | - | 0 | - |
| - | - | 3545 | 861.5 | - | - | 0 | - |
| - | - | 5185 | 868.4 | - | - | 0 | - |
| - | - | 2866 | 869.4 | - | - | 0 | - |
| - | - | 2340 | 873.5 | - | - | 0 | - |
| 8 | y | 5227 | 874.5 | 0.001996 | 2.282 | +1 | 8 |
| - | - | 2208 | 875.5 | - | - | 0 | - |
| - | - | 1899 | 886.5 | - | - | 0 | - |
| - | - | 1143 | 887.4 | - | - | 0 | - |
| - | - | 2948 | 903.4 | - | - | 0 | - |
| 8 | c | 7321 | 904.4 | 0.002537 | 2.805 | +1 | 8 |
| - | - | 2297 | 905.4 | - | - | 0 | - |
| 7 | w | 1073 | 928.5 | 0.009046 | 9.743 | +1 | 9 |
| - | - | 1296 | 929.5 | - | - | 0 | - |
| - | - | 5159 | 939.4 | - | - | 0 | - |
| - | - | 1818 | 940.4 | - | - | 0 | - |
| - | - | 1997 | 960.5 | - | - | 0 | - |
| - | - | 1048 | 961.5 | - | - | 0 | - |
| - | - | 1508 | 971.5 | - | - | 0 | - |
| - | - | 3300 | 973.5 | - | - | 0 | - |
| - | - | 1696 | 974.5 | - | - | 0 | - |
| 7 | z | 5.929E+04 | 987.5 | 0.0002911 | 0.2948 | +1 | 9 |
| - | - | 6.494E+04 | 988.5 | - | - | 0 | - |
| - | - | 2.528E+04 | 989.5 | - | - | 0 | - |
| - | - | 4668 | 990.5 | - | - | 0 | - |
| - | - | 2905 | 1000 | - | - | 0 | - |
| - | - | 1570 | 1002 | - | - | 0 | - |
| - | - | 5598 | 1002 | - | - | 0 | - |
| 7 | y | 5935 | 1004 | 0.003756 | 3.743 | +1 | 9 |
| - | - | 2427 | 1005 | - | - | 0 | - |
| - | - | 1.52E+04 | 1017 | - | - | 0 | - |
| 9 | c | 2.137E+04 | 1018 | 0.001945 | 1.911 | +1 | 9 |
| - | - | 1.044E+04 | 1019 | - | - | 0 | - |
| - | - | 1403 | 1020 | - | - | 0 | - |
| - | - | 5308 | 1053 | - | - | 0 | - |
| - | - | 3657 | 1054 | - | - | 0 | - |
| - | - | 1204 | 1061 | - | - | 0 | - |
| - | - | 1122 | 1070 | - | - | 0 | - |
| 6 | y | 2253 | 1073 | 0.01021 | 9.515 | +1 | 10 |
| 6 | y | 2968 | 1074 | 0.002605 | 2.427 | +1 | 10 |
| 6 | z | 6505 | 1075 | 0.000215 | 0.2001 | +1 | 10 |
| - | - | 8.353E+04 | 1076 | - | - | 0 | - |
| - | - | 4.202E+04 | 1077 | - | - | 0 | - |
| - | - | 1.098E+04 | 1078 | - | - | 0 | - |
| - | - | 1268 | 1085 | - | - | 0 | - |
| - | - | 9514 | 1090 | - | - | 0 | - |
| 6 | y | 1.87E+04 | 1091 | 0.002093 | 1.92 | +1 | 10 |
| - | - | 9046 | 1092 | - | - | 0 | - |
| - | - | 1894 | 1093 | - | - | 0 | - |
| - | - | 3247 | 1102 | - | - | 0 | - |
| - | - | 2790 | 1103 | - | - | 0 | - |
| - | - | 2969 | 1118 | - | - | 0 | - |
| - | - | 2367 | 1119 | - | - | 0 | - |
| 10 | c | 3062 | 1129 | 0.002967 | 2.629 | +1 | 10 |
| - | - | 2062 | 1130 | - | - | 0 | - |
| - | - | 1425 | 1131 | - | - | 0 | - |
| - | - | 2.664E+04 | 1145 | - | - | 0 | - |
| 10 | c | 1.295E+05 | 1146 | 0.001318 | 1.151 | +1 | 10 |
| - | - | 7.616E+04 | 1147 | - | - | 0 | - |
| - | - | 2.125E+04 | 1148 | - | - | 0 | - |
| - | - | 2362 | 1149 | - | - | 0 | - |
| 5 | y | 1790 | 1160 | 0.01159 | 9.999 | +1 | 11 |
| 5 | z | 7680 | 1162 | 0.0009934 | 0.8552 | +1 | 11 |
| - | - | 4.055E+04 | 1163 | - | - | 0 | - |
| - | - | 2.246E+04 | 1164 | - | - | 0 | - |
| - | - | 6064 | 1165 | - | - | 0 | - |
| - | - | 3489 | 1173 | - | - | 0 | - |
| - | - | 4037 | 1174 | - | - | 0 | - |
| - | - | 2831 | 1175 | - | - | 0 | - |
| - | - | 1622 | 1177 | - | - | 0 | - |
| 5 | y | 8840 | 1178 | 0.001651 | 1.402 | +1 | 11 |
| - | - | 4176 | 1179 | - | - | 0 | - |
| - | - | 3059 | 1180 | - | - | 0 | - |
| - | - | 2649 | 1182 | - | - | 0 | - |
| - | - | 1700 | 1183 | - | - | 0 | - |
| 11 | c | 2229 | 1200 | 0.0008124 | 0.6772 | +1 | 11 |
| - | - | 5.664E+04 | 1216 | - | - | 0 | - |
| 11 | c | 7.484E+04 | 1217 | 0.00352 | 2.893 | +1 | 11 |
| - | - | 4.052E+04 | 1218 | - | - | 0 | - |
| - | - | 1.001E+04 | 1219 | - | - | 0 | - |
| - | - | 1120 | 1247 | - | - | 0 | - |
| - | - | 1657 | 1253 | - | - | 0 | - |
| - | - | 3867 | 1274 | - | - | 0 | - |
| 4 | y | 3.552E+04 | 1275 | 0.001681 | 1.318 | +1 | 12 |
| - | - | 2.252E+04 | 1276 | - | - | 0 | - |
| - | - | 7032 | 1277 | - | - | 0 | - |
| - | - | 4212 | 1287 | - | - | 0 | - |
| - | - | 5018 | 1288 | - | - | 0 | - |
| - | - | 2705 | 1289 | - | - | 0 | - |
| 12 | c | 2457 | 1314 | 0.003907 | 2.974 | +1 | 12 |
| - | - | 1539 | 1315 | - | - | 0 | - |
| - | - | 6452 | 1330 | - | - | 0 | - |
| 12 | c | 1.188E+05 | 1331 | 0.00238 | 1.789 | +1 | 12 |
| - | - | 8.643E+04 | 1332 | - | - | 0 | - |
| - | - | 1461 | 1332 | - | - | 0 | - |
| - | - | 2.986E+04 | 1333 | - | - | 0 | - |
| - | - | 3714 | 1334 | - | - | 0 | - |
| 3 | y | 1480 | 1355 | 0.0005471 | 0.4039 | +1 | 13 |
| - | - | 1214 | 1359 | - | - | 0 | - |
| - | - | 7045 | 1371 | - | - | 0 | - |
| 3 | y | 8.083E+04 | 1372 | 0.002687 | 1.959 | +1 | 13 |
| - | - | 5.529E+04 | 1373 | - | - | 0 | - |
| - | - | 1.989E+04 | 1374 | - | - | 0 | - |
| - | - | 2504 | 1375 | - | - | 0 | - |
| - | - | 3624 | 1400 | - | - | 0 | - |
| - | - | 2218 | 1401 | - | - | 0 | - |
| - | - | 5025 | 1402 | - | - | 0 | - |
| - | - | 3033 | 1403 | - | - | 0 | - |
| - | - | 2.229E+04 | 1415 | - | - | 0 | - |
| - | - | 1.53E+04 | 1416 | - | - | 0 | - |
| - | - | 5794 | 1417 | - | - | 0 | - |
| - | - | 3452 | 1431 | - | - | 0 | - |
| - | - | 3884 | 1432 | - | - | 0 | - |
| - | - | 1729 | 1433 | - | - | 0 | - |
| 13 | c | 7.026E+04 | 1459 | 0.003349 | 2.296 | +1 | 13 |
| - | - | 5.878E+04 | 1460 | - | - | 0 | - |
| - | - | 2.287E+04 | 1461 | - | - | 0 | - |
| - | - | 2660 | 1462 | - | - | 0 | - |
| - | - | 1316 | 1472 | - | - | 0 | - |
| - | - | 8330 | 1486 | - | - | 0 | - |
| - | - | 1.177E+04 | 1487 | - | - | 0 | - |
| - | - | 9097 | 1488 | - | - | 0 | - |
| - | - | 3881 | 1489 | - | - | 0 | - |
| - | - | 1686 | 1490 | - | - | 0 | - |
| 2 | z | 4.115E+04 | 1503 | 0.002938 | 1.955 | +1 | 14 |
| - | - | 3.303E+04 | 1504 | - | - | 0 | - |
| - | - | 1.831E+04 | 1505 | - | - | 0 | - |
| - | - | 2724 | 1506 | - | - | 0 | - |
| - | - | 8278 | 1515 | - | - | 0 | - |
| - | - | 7578 | 1516 | - | - | 0 | - |
| - | - | 3664 | 1517 | - | - | 0 | - |
| - | - | 1748 | 1529 | - | - | 0 | - |
| 14 | c | 9.914E+04 | 1530 | 0.002865 | 1.873 | +1 | 14 |
| - | - | 9.021E+04 | 1531 | - | - | 0 | - |
| - | - | 6.133E+04 | 1532 | - | - | 0 | - |
| - | - | 2.288E+04 | 1533 | - | - | 0 | - |
| - | - | 6966 | 1534 | - | - | 0 | - |
| - | - | 1.703E+04 | 1544 | - | - | 0 | - |
| - | - | 1.654E+04 | 1545 | - | - | 0 | - |
| - | - | 9742 | 1546 | - | - | 0 | - |
| - | - | 1623 | 1547 | - | - | 0 | - |
| - | - | 8551 | 1560 | - | - | 0 | - |
| - | - | 1.637E+04 | 1561 | - | - | 0 | - |
| - | - | 1.277E+04 | 1562 | - | - | 0 | - |
| - | - | 7543 | 1563 | - | - | 0 | - |
| - | - | 2544 | 1564 | - | - | 0 | - |
| - | - | 5075 | 1569 | - | - | 0 | - |
| - | - | 5318 | 1570 | - | - | 0 | - |
| - | - | 1.218E+04 | 1571 | - | - | 0 | - |
| - | - | 8789 | 1572 | - | - | 0 | - |
| - | - | 5150 | 1573 | - | - | 0 | - |
| - | - | 7498 | 1574 | - | - | 0 | - |
| - | - | 7839 | 1575 | - | - | 0 | - |
| - | - | 4535 | 1576 | - | - | 0 | - |
| - | - | 1.287E+04 | 1577 | - | - | 0 | - |
| - | - | 1.107E+04 | 1578 | - | - | 0 | - |
| - | - | 5798 | 1579 | - | - | 0 | - |
| - | - | 1156 | 1580 | - | - | 0 | - |
| - | - | 1462 | 1586 | - | - | 0 | - |
| - | - | 1.653E+04 | 1587 | - | - | 0 | - |
| - | - | 1.069E+05 | 1588 | - | - | 0 | - |
| - | - | 9.583E+04 | 1589 | - | - | 0 | - |
| - | - | 4.372E+04 | 1590 | - | - | 0 | - |
| - | - | 7789 | 1591 | - | - | 0 | - |
| - | - | 2497 | 1597 | - | - | 0 | - |
| - | - | 3227 | 1598 | - | - | 0 | - |
| - | - | 2254 | 1599 | - | - | 0 | - |
| - | - | 6280 | 1605 | - | - | 0 | - |
| - | - | 6694 | 1606 | - | - | 0 | - |
| - | - | 4430 | 1607 | - | - | 0 | - |
| - | - | 2018 | 1614 | - | - | 0 | - |
| - | - | 3.905E+04 | 1615 | - | - | 0 | - |
| - | - | 1.355E+05 | 1616 | - | - | 0 | - |
| - | - | 1.177E+05 | 1617 | - | - | 0 | - |
| - | - | 4.994E+04 | 1618 | - | - | 0 | - |
| - | - | 8934 | 1619 | - | - | 0 | - |
| - | - | 3176 | 1630 | - | - | 0 | - |
| - | - | 1.585E+04 | 1631 | - | - | 0 | - |
| - | - | 2.874E+05 | 1632 | - | - | 0 | - |
| - | - | 8.08E+05 | 1633 | - | - | 0 | - |
| - | - | 6.334E+05 | 1634 | - | - | 0 | - |
| - | - | 2.559E+05 | 1635 | - | - | 0 | - |
| - | - | 3.242E+04 | 1636 | - | - | 0 | - |
| - | - | 1914 | 1665 | - | - | 0 | - |
| - | - | 1324 | 1666 | - | - | 0 | - |
| - | - | 1287 | 3072 | - | - | 0 | - |

m/z Charge Intensity FragmentType MassShift Position
120.08092498779297 0 3560.3572
129.1023712158203 0 5315.704
144.67819213867188 0 597.8718
148.95399475097656 0 1219.0833
159.13232421875 0 578.934
173.43902587890625 0 3220.7668
191.10296630859375 0 1963.8988 y 13
215.4649658203125 0 732.7497
220.17019653320312 0 737.997
221.0845947265625 0 1014.692
226.11892700195312 0 1382.3739
232.5020751953125 0 789.6
233.16485595703125 0 34267.992
234.16822814941406 0 5502.273
243.14483642578125 0 2444.725
261.15966796875 0 41343.387
262.1630859375 0 5078.891
276.6896057128906 0 798.28015
282.1448974609375 0 2349.422
297.1553039550781 0 2248.8704
299.06207275390625 0 1305.6812
314.181884765625 0 2288.8452
319.1976013183594 0 4282.808 y 12
355.0698547363281 0 5216.172
356.0686340332031 0 1964.5889
358.2125549316406 0 7179.317
359.21588134765625 0 1106.9569
385.21875 0 984.3877
414.7284240722656 0 913.5329
425.212158203125 0 1121.5238
433.2406921386719 0 4588.885 y 11
455.2651672363281 0 1589.1852
470.72369384765625 0 1028.884
479.7829284667969 0 984.86017
498.2174072265625 0 1051.0751
504.27862548828125 0 5037.9766 y 10
505.2801208496094 0 1621.192
542.301513671875 0 1001.688
582.2844848632812 0 1160.1871
582.7815551757812 0 5372.9585
583.2799682617188 0 2220.1711
591.2920532226562 0 4714.4517
591.795654296875 0 2901.2458
612.8146362304688 0 1714.4479
613.313232421875 0 1894.8552
615.30712890625 0 1435.6481 y Ammonia loss 9
616.3170166015625 0 5173.207 z 9
617.3253173828125 0 8402.915
617.8059692382812 0 7388.787
618.3076782226562 0 6973.698
618.806640625 0 3291.3115
626.8119506835938 0 43109.234
627.26318359375 0 1543.4492
627.3134155273438 0 27932.367
627.8151245117188 0 7141.0044
629.33154296875 0 1662.5967
632.3358154296875 0 5451.0933 y 9
633.3388671875 0 1482.2268
637.8138427734375 0 2708.6362 y 3
638.3153686523438 0 2721.7278
655.3406372070312 0 3249.3274
655.84130859375 0 2628.7502
663.3389282226562 0 3963.6401
663.8424682617188 0 1939.3567
664.342041015625 0 2409.2537
664.8413696289062 0 1167.3411
668.8223266601562 0 1378.0865
677.3353881835938 0 20044.375 y Water loss 2
677.8341064453125 0 22153.66 y Ammonia loss 2
678.3328247070312 0 11110.012 z 2
678.8304443359375 0 5346.208
679.3319702148438 0 1384.3177
685.3336181640625 0 4130.199
685.835205078125 0 3665.603
686.34130859375 0 722954.56 y 2
686.8427734375 0 542035.7
687.3436279296875 0 189426.36
687.8446044921875 0 20200.672
700.849365234375 0 1210.7955
729.4012451171875 0 21623.729 z 8
730.40673828125 0 18086.88
731.4093017578125 0 5802.131
740.3439331054688 0 2694.434
745.4215087890625 0 5281.6196 y 8
751.3673095703125 0 1189.4109 y Ammonia loss 1
751.8792724609375 0 924.67755 z 1
756.8867797851562 0 5061.983 c Ammonia loss 13
757.39794921875 0 3769.1428 c Water loss 6
758.3650512695312 0 1538.7506
759.8763427734375 0 8653.0205 y 1
760.376708984375 0 8318.32
760.8764038085938 0 3614.5276
802.3787231445312 0 2100.401
807.409912109375 0 3561.9507
807.9114379882812 0 4499.46
808.4095458984375 0 2356.9226
815.423583984375 0 2195.7913
815.9224853515625 0 4271.0444
816.41796875 0 49839.598
816.9190673828125 0 48821.07
817.420166015625 0 26878.062
817.9236450195312 0 3995.1948
858.4439697265625 0 51234.902 z 7
859.4495239257812 0 57138.32
860.4529418945312 0 18887.008
861.4568481445312 0 3545.2798
868.4046020507812 0 5184.5576
869.4097900390625 0 2866.1675
873.455322265625 0 2340.119
874.4608764648438 0 5227.4985 y 7
875.4662475585938 0 2207.8398
886.459228515625 0 1898.6956
887.4218139648438 0 1143.3004
903.4337768554688 0 2947.7397
904.4385375976562 0 7321.32 c 7
905.4426879882812 0 2296.833
928.4824829101562 0 1073.0717 w 6
929.4710693359375 0 1295.9299
939.442626953125 0 5159.382
940.446044921875 0 1818.2567
960.4568481445312 0 1997.3164
961.4686279296875 0 1048.244
971.4852294921875 0 1507.9336
973.5006103515625 0 3300.3408
974.4937744140625 0 1696.138
987.4864501953125 0 59292.406 z 6
988.4917602539062 0 64942.867
989.4954223632812 0 25283.11
990.4976196289062 0 4668.32
1000.49755859375 0 2905.1333
1001.50048828125 0 1570.1393
1002.4993896484375 0 5597.5093
1003.501708984375 0 5935.1074 y 6
1004.5054931640625 0 2427.4487
1016.51708984375 0 15195.429
1017.523193359375 0 21367.035 c 8
1018.5253295898438 0 10435.494
1019.5326538085938 0 1403.2542
1053.4820556640625 0 5307.8164
1054.4859619140625 0 3656.6223
1060.5399169921875 0 1203.855
1070.4986572265625 0 1122.438
1072.5167236328125 0 2252.8235 y Water loss 5
1073.5135498046875 0 2967.7834 y Ammonia loss 5
1074.5185546875 0 6504.5547 z 5
1075.5262451171875 0 83529.49
1076.528564453125 0 42016.79
1077.5311279296875 0 10980.454
1084.528076171875 0 1267.6631
1089.5277099609375 0 9513.749
1090.535400390625 0 18695.406 y 5
1091.537353515625 0 9046.211
1092.543701171875 0 1893.6481
1101.5745849609375 0 3247.0217
1102.5697021484375 0 2790.053
1117.531005859375 0 2969.0767
1118.5357666015625 0 2367.4768
1128.55419921875 0 3062.4458 c Ammonia loss 9
1129.5537109375 0 2062.0896
1130.5535888671875 0 1424.5841
1144.5755615234375 0 26640.604
1145.5823974609375 0 129500.86 c 9
1146.5860595703125 0 76160.64
1147.588623046875 0 21250.656
1148.592041015625 0 2361.995
1159.54736328125 0 1789.9745 y Water loss 4
1161.5498046875 0 7679.7905 z 4
1162.5567626953125 0 40545.176
1163.5589599609375 0 22456.344
1164.5648193359375 0 6064.182
1172.610107421875 0 3488.7603
1173.6102294921875 0 4036.6055
1174.6209716796875 0 2831.225
1176.5657958984375 0 1621.899
1177.56787109375 0 8840.319 y 4
1178.5736083984375 0 4176.1426
1179.57373046875 0 3058.786
1181.5811767578125 0 2649.484
1182.5831298828125 0 1699.6475
1199.5950927734375 0 2229.3699 c Ammonia loss 10
1215.611572265625 0 56635.082
1216.6173095703125 0 74835.49 c 10
1217.620361328125 0 40524.98
1218.623779296875 0 10005.256
1247.150146484375 0 1120.0664
1252.624755859375 0 1657.085
1273.61181640625 0 3866.7402
1274.62060546875 0 35516.33 y 3
1275.623291015625 0 22515.875
1276.62353515625 0 7031.545
1286.6488037109375 0 4211.884
1287.6505126953125 0 5017.6597
1288.6494140625 0 2704.8828
1313.63330078125 0 2457.2385 c Ammonia loss 11
1314.628173828125 0 1538.683
1329.6510009765625 0 6451.705
1330.661376953125 0 118753.17 c 11
1331.66455078125 0 86433.74
1331.856689453125 0 1460.9125
1332.6671142578125 0 29857.516
1333.6708984375 0 3714.413
1354.6490478515625 0 1480.4615 y Ammonia loss 2
1358.688720703125 0 1214.254
1370.6658935546875 0 7045.151
1371.67236328125 0 80827.83 y 2
1372.6761474609375 0 55287.2
1373.678955078125 0 19886.387
1374.678466796875 0 2503.903
1399.6656494140625 0 3623.6196
1400.6719970703125 0 2217.9116
1401.69091796875 0 5024.8423
1402.7005615234375 0 3033.3687
1414.74169921875 0 22290.771
1415.744140625 0 15296.566
1416.7501220703125 0 5794.391
1430.7076416015625 0 3452.157
1431.7052001953125 0 3884.0654
1432.7060546875 0 1728.7897
1458.75537109375 0 70260.73 c 12
1459.7574462890625 0 58780.6
1460.759765625 0 22865.451
1461.7645263671875 0 2660.014
1471.785400390625 0 1316.3195
1485.77978515625 0 8329.776
1486.783447265625 0 11771.764
1487.7891845703125 0 9096.84
1488.7958984375 0 3881.0793
1489.8035888671875 0 1686.334
1502.7218017578125 0 41149.51 z 1
1503.724365234375 0 33034.707
1504.72705078125 0 18310.53
1505.7330322265625 0 2724.2478
1514.78271484375 0 8277.998
1515.781982421875 0 7578.2188
1516.7835693359375 0 3664.252
1528.8153076171875 0 1747.7068
1529.79296875 0 99138.54 c 13
1530.7957763671875 0 90205.08
1531.7989501953125 0 61333.926
1532.7987060546875 0 22883.37
1533.8074951171875 0 6966.259
1543.785888671875 0 17025.723
1544.784912109375 0 16544.562
1545.784423828125 0 9741.54
1546.789306640625 0 1622.9409
1559.7623291015625 0 8551.053
1560.79150390625 0 16373.834
1561.794189453125 0 12769.245
1562.7982177734375 0 7542.975
1563.7921142578125 0 2543.9224
1568.8197021484375 0 5074.753
1569.815673828125 0 5318.2554
1570.8272705078125 0 12181.699
1571.8280029296875 0 8789.235
1572.80859375 0 5149.7285
1573.787353515625 0 7497.6064
1574.788330078125 0 7839.463
1575.7958984375 0 4535.2354
1576.7698974609375 0 12871.889
1577.7730712890625 0 11070.712
1578.7762451171875 0 5798.217
1579.7576904296875 0 1155.7727
1585.8348388671875 0 1461.8479
1586.82373046875 0 16531.477
1587.8125 0 106904.07
1588.8150634765625 0 95828.42
1589.8138427734375 0 43723.85
1590.8116455078125 0 7789.0107
1596.7943115234375 0 2497.091
1597.8094482421875 0 3226.7825
1598.7978515625 0 2254.3687
1604.835693359375 0 6280.016
1605.8370361328125 0 6693.853
1606.838623046875 0 4430.347
1613.77783203125 0 2017.5543
1614.820556640625 0 39047.5
1615.8099365234375 0 135521.05
1616.8104248046875 0 117728.05
1617.8104248046875 0 49937.992
1618.8128662109375 0 8934.11
1629.801025390625 0 3176.104
1630.8140869140625 0 15848.38
1631.82373046875 0 287379.03
1632.8310546875 0 808016.2
1633.8341064453125 0 633441.1
1634.83642578125 0 255938.58
1635.839599609375 0 32417.857
1664.82275390625 0 1913.9661
1665.8101806640625 0 1323.8489
3071.947265625 0 1287.0513

Spectrum Details

|  |  |
| --- | --- |
| Matched peaks? Matched peaks The total absolute number of peaks matched. Additionally in brackets the total fraction of peaks matched and the total number of peaks is shown. | 45 (15.52% of 290) |
| FDR? FDR The false discovery rate estimated for this peptide. It is calculated by matching all theoretical fragments with a non-integer shift with the raw peaks for this spectrum. This is done with 40 different shifts. The resulting percentage is the average number of annotated peaks over the number of annotated peaks with the correct spectrum. | 0.11% |
| Satellite FDR? Satellite FDR See the FDR for details on its calculation. This satellite ion specific FDR only contains the satellite ions (d/w) for I/L/J positions. | - |
| PSM Score? PSM Score The PSM Score as given by Hecklib to this annotated spectrum. It is shown with three significant figures. | 419 |

#### Spectrum 6152? Spectrum 6152 The raw spectrum of this peptide as annotated by Hecklib. The fragments are coloured according to ion type (see legend). Any peaks with a star '\*' as text can be hovered over to see the full details, first the ion type second the mass shift type. By hovering over the amino acids in the peptide or ions in the legend the corresponding peaks are highlighted. By toggling the 'Unassigned' label you can turn the background (unassigned) peaks on or off in the plot. By updating the slider in the Ion legend you can update the spectrum to only show the top X% of the peaks with labels. The top X% means any peak that is within X% of the highest intensity. By dragging in the spectrum you can zoom in to a specific part of the spectrum and use 'Zoom Out' to get back to the original zoom level. The annotation of the spectrum is based on the given sequence in the peptides file and is done with different software so inconsistencies are likely. The peaks are annotated based on the given sequence, with 20 ppm tolerance.

Copy Data

##### Spectrum 6152 (TSV)

###### Preview

```
Loading example...
```

*Click on the button to copy the data to your clipboard.*

Mz MinMz MaxIntensity Max

WidthHeightPeptide font sizePeptide stroke widthSpectrum font sizeSpectrum stroke widthCompact peptide

Ion legend

wxyz

abcd

OtherUnassignedIon^Charge^_Position_Show for top:%

JFPPSSEEJQANKAT

02.14e+54.28e+56.42e+58.56e+5

Zoom Out

y^+1^_2_y^+1^_3_y^+1^_4_y^+1^_5_y^+1^_6_z^+1^_6_y^+1^_6_y^+2^_12_z^+2^_13_y^+2^_13_y^+2^_13_z^+2^_13_y^+2^_13_c^+2^_13_z^+1^_7_y^+1^_7_y^+2^_14_c^+2^_14_y^+2^_14_w^+1^_8_z^+1^_8_y^+1^_8_c^+1^_8_w^+1^_9_z^+1^_9_y^+1^_9_c^+1^_9_y^+1^_10_y^+1^_10_z^+1^_10_y^+1^_10_c^+1^_10_c^+1^_10_y^+1^_11_y^+1^_11_z^+1^_11_y^+1^_11_c^+1^_11_c^+1^_11_z^+1^_12_y^+1^_12_c^+1^_12_w^+1^_13_c^+1^_12_y^+1^_13_y^+1^_13_y^+1^_13_c^+1^_13_z^+1^_14_y^+1^_14_c^+1^_14_

0799159823973196

Fragment Matches Table

Show background peaks

| Position | Ion type | Intensity | mz Theoretical | mz Error (Th) | mz Error (ppm) | Charge | Series Number |
| --- | --- | --- | --- | --- | --- | --- | --- |
| - | - | 3805 | 120.1 | - | - | 0 | - |
| - | - | 536.3 | 129.1 | - | - | 0 | - |
| - | - | 6569 | 129.1 | - | - | 0 | - |
| - | - | 746 | 148.9 | - | - | 0 | - |
| - | - | 506 | 148.9 | - | - | 0 | - |
| - | - | 755.9 | 148.9 | - | - | 0 | - |
| - | - | 888.8 | 148.9 | - | - | 0 | - |
| - | - | 1123 | 148.9 | - | - | 0 | - |
| - | - | 1732 | 148.9 | - | - | 0 | - |
| - | - | 1692 | 148.9 | - | - | 0 | - |
| - | - | 4137 | 148.9 | - | - | 0 | - |
| - | - | 7522 | 148.9 | - | - | 0 | - |
| - | - | 4560 | 149 | - | - | 0 | - |
| - | - | 2055 | 149 | - | - | 0 | - |
| - | - | 1801 | 149 | - | - | 0 | - |
| - | - | 1530 | 149 | - | - | 0 | - |
| - | - | 869.2 | 149 | - | - | 0 | - |
| - | - | 799.9 | 149 | - | - | 0 | - |
| - | - | 789.3 | 149 | - | - | 0 | - |
| - | - | 541.2 | 149 | - | - | 0 | - |
| - | - | 651.6 | 149.1 | - | - | 0 | - |
| - | - | 669.4 | 155.1 | - | - | 0 | - |
| - | - | 570.6 | 168.7 | - | - | 0 | - |
| - | - | 779.8 | 185.1 | - | - | 0 | - |
| 14 | y | 1475 | 191.1 | 1.804E-05 | 0.09442 | +1 | 2 |
| - | - | 611.8 | 200.1 | - | - | 0 | - |
| - | - | 631.2 | 209.7 | - | - | 0 | - |
| - | - | 646.6 | 217.1 | - | - | 0 | - |
| - | - | 678.4 | 220.1 | - | - | 0 | - |
| - | - | 663.6 | 221.1 | - | - | 0 | - |
| - | - | 1706 | 226.1 | - | - | 0 | - |
| - | - | 724.9 | 229 | - | - | 0 | - |
| - | - | 576.5 | 232 | - | - | 0 | - |
| - | - | 3.929E+04 | 233.2 | - | - | 0 | - |
| - | - | 4386 | 234.2 | - | - | 0 | - |
| - | - | 1577 | 243.1 | - | - | 0 | - |
| - | - | 735 | 258.4 | - | - | 0 | - |
| - | - | 4.46E+04 | 261.2 | - | - | 0 | - |
| - | - | 7220 | 262.2 | - | - | 0 | - |
| - | - | 1645 | 282.1 | - | - | 0 | - |
| - | - | 2204 | 297.2 | - | - | 0 | - |
| - | - | 2249 | 299.1 | - | - | 0 | - |
| - | - | 1883 | 313.2 | - | - | 0 | - |
| - | - | 2274 | 314.2 | - | - | 0 | - |
| 13 | y | 4662 | 319.2 | 0.0003101 | 0.9714 | +1 | 3 |
| - | - | 969.4 | 320.2 | - | - | 0 | - |
| - | - | 798 | 341 | - | - | 0 | - |
| - | - | 3975 | 355.1 | - | - | 0 | - |
| - | - | 1992 | 356.1 | - | - | 0 | - |
| - | - | 1134 | 357.1 | - | - | 0 | - |
| - | - | 7344 | 358.2 | - | - | 0 | - |
| - | - | 1159 | 359.2 | - | - | 0 | - |
| - | - | 783.3 | 361.2 | - | - | 0 | - |
| 12 | y | 3775 | 433.2 | 0.0004429 | 1.022 | +1 | 4 |
| - | - | 1807 | 455.3 | - | - | 0 | - |
| - | - | 1677 | 498.2 | - | - | 0 | - |
| 11 | y | 6845 | 504.3 | 0.000538 | 1.067 | +1 | 5 |
| - | - | 959.8 | 505.3 | - | - | 0 | - |
| - | - | 1011 | 582.3 | - | - | 0 | - |
| - | - | 5971 | 582.8 | - | - | 0 | - |
| - | - | 3481 | 583.3 | - | - | 0 | - |
| - | - | 1154 | 583.8 | - | - | 0 | - |
| - | - | 5279 | 591.3 | - | - | 0 | - |
| - | - | 3462 | 591.8 | - | - | 0 | - |
| - | - | 1035 | 612.8 | - | - | 0 | - |
| - | - | 1505 | 613.3 | - | - | 0 | - |
| 10 | y | 920.8 | 615.3 | 0.001133 | 1.842 | +1 | 6 |
| 10 | z | 5751 | 616.3 | 0.0003524 | 0.5718 | +1 | 6 |
| - | - | 7367 | 617.3 | - | - | 0 | - |
| - | - | 7277 | 617.8 | - | - | 0 | - |
| - | - | 7809 | 618.3 | - | - | 0 | - |
| - | - | 2138 | 618.8 | - | - | 0 | - |
| - | - | 4.71E+04 | 626.8 | - | - | 0 | - |
| - | - | 2008 | 627.3 | - | - | 0 | - |
| - | - | 2.788E+04 | 627.3 | - | - | 0 | - |
| - | - | 8286 | 627.8 | - | - | 0 | - |
| - | - | 1214 | 628.3 | - | - | 0 | - |
| - | - | 1149 | 629.3 | - | - | 0 | - |
| 10 | y | 5693 | 632.3 | 9.456E-05 | 0.1495 | +1 | 6 |
| - | - | 1090 | 633.3 | - | - | 0 | - |
| - | - | 1078 | 635.8 | - | - | 0 | - |
| 4 | y | 3296 | 637.8 | 0.0006485 | 1.017 | +2 | 12 |
| - | - | 3031 | 655.3 | - | - | 0 | - |
| - | - | 3290 | 655.8 | - | - | 0 | - |
| - | - | 1416 | 656.3 | - | - | 0 | - |
| - | - | 2815 | 663.3 | - | - | 0 | - |
| - | - | 3974 | 663.8 | - | - | 0 | - |
| - | - | 2855 | 664.3 | - | - | 0 | - |
| - | - | 2164 | 668.8 | - | - | 0 | - |
| 3 | z | 1360 | 669.3 | 0.002666 | 3.983 | +2 | 13 |
| 3 | y | 1.748E+04 | 677.3 | 0.0002398 | 0.354 | +2 | 13 |
| 3 | y | 2.534E+04 | 677.8 | 0.005607 | 8.273 | +2 | 13 |
| 3 | z | 1.032E+04 | 678.3 | 0.001939 | 2.859 | +2 | 13 |
| - | - | 3842 | 678.8 | - | - | 0 | - |
| - | - | 3705 | 685.3 | - | - | 0 | - |
| - | - | 5038 | 685.8 | - | - | 0 | - |
| 3 | y | 7.915E+05 | 686.3 | 0.0003896 | 0.5676 | +2 | 13 |
| - | - | 5.774E+05 | 686.8 | - | - | 0 | - |
| - | - | 2.142E+05 | 687.3 | - | - | 0 | - |
| - | - | 2.141E+04 | 687.8 | - | - | 0 | - |
| - | - | 1258 | 700.3 | - | - | 0 | - |
| - | - | 1377 | 700.8 | - | - | 0 | - |
| 13 | c | 939.7 | 721.4 | 0.002774 | 3.845 | +2 | 13 |
| 9 | z | 2.413E+04 | 729.4 | 6.582E-05 | 0.09024 | +1 | 7 |
| - | - | 1.814E+04 | 730.4 | - | - | 0 | - |
| - | - | 6643 | 731.4 | - | - | 0 | - |
| - | - | 3270 | 740.3 | - | - | 0 | - |
| 9 | y | 4806 | 745.4 | 5.21E-05 | 0.06989 | +1 | 7 |
| - | - | 1148 | 746.4 | - | - | 0 | - |
| - | - | 1451 | 748.4 | - | - | 0 | - |
| - | - | 966.4 | 748.9 | - | - | 0 | - |
| 2 | y | 1291 | 751.4 | 0.00204 | 2.715 | +2 | 14 |
| 14 | c | 5764 | 756.9 | 0.0007073 | 0.9345 | +2 | 14 |
| - | - | 4240 | 757.4 | - | - | 0 | - |
| - | - | 2033 | 757.9 | - | - | 0 | - |
| - | - | 1120 | 758.4 | - | - | 0 | - |
| - | - | 1174 | 759.4 | - | - | 0 | - |
| 2 | y | 1.156E+04 | 759.9 | 0.0005454 | 0.7178 | +2 | 14 |
| - | - | 1.033E+04 | 760.4 | - | - | 0 | - |
| - | - | 5128 | 760.9 | - | - | 0 | - |
| 8 | w | 1274 | 799.4 | 0.006772 | 8.47 | +1 | 8 |
| - | - | 2514 | 802.4 | - | - | 0 | - |
| - | - | 3308 | 807.4 | - | - | 0 | - |
| - | - | 4215 | 807.9 | - | - | 0 | - |
| - | - | 2944 | 808.4 | - | - | 0 | - |
| - | - | 979.5 | 808.9 | - | - | 0 | - |
| - | - | 1197 | 815.4 | - | - | 0 | - |
| - | - | 4752 | 815.9 | - | - | 0 | - |
| - | - | 5.854E+04 | 816.4 | - | - | 0 | - |
| - | - | 6.547E+04 | 816.9 | - | - | 0 | - |
| - | - | 2.571E+04 | 817.4 | - | - | 0 | - |
| - | - | 3151 | 817.9 | - | - | 0 | - |
| 8 | z | 5.412E+04 | 858.4 | 5.638E-05 | 0.06567 | +1 | 8 |
| - | - | 5.524E+04 | 859.4 | - | - | 0 | - |
| - | - | 1.766E+04 | 860.5 | - | - | 0 | - |
| - | - | 2624 | 861.5 | - | - | 0 | - |
| - | - | 6436 | 868.4 | - | - | 0 | - |
| - | - | 2412 | 869.4 | - | - | 0 | - |
| - | - | 2285 | 873.5 | - | - | 0 | - |
| 8 | y | 6232 | 874.5 | 0.001141 | 1.305 | +1 | 8 |
| - | - | 2609 | 875.5 | - | - | 0 | - |
| - | - | 1924 | 886.5 | - | - | 0 | - |
| - | - | 2468 | 903.4 | - | - | 0 | - |
| 8 | c | 6613 | 904.4 | 0.0009497 | 1.05 | +1 | 8 |
| - | - | 2886 | 905.4 | - | - | 0 | - |
| - | - | 1022 | 906.5 | - | - | 0 | - |
| 7 | w | 1849 | 928.5 | 0.004835 | 5.207 | +1 | 9 |
| - | - | 1137 | 929.5 | - | - | 0 | - |
| - | - | 5754 | 939.4 | - | - | 0 | - |
| - | - | 3088 | 940.4 | - | - | 0 | - |
| - | - | 1624 | 956.5 | - | - | 0 | - |
| - | - | 1928 | 960.5 | - | - | 0 | - |
| - | - | 3100 | 973.5 | - | - | 0 | - |
| - | - | 2421 | 974.5 | - | - | 0 | - |
| - | - | 959.5 | 975.5 | - | - | 0 | - |
| 7 | z | 6.212E+04 | 987.5 | 0.0004131 | 0.4184 | +1 | 9 |
| - | - | 6.815E+04 | 988.5 | - | - | 0 | - |
| - | - | 2.775E+04 | 989.5 | - | - | 0 | - |
| - | - | 4724 | 990.5 | - | - | 0 | - |
| - | - | 3498 | 1000 | - | - | 0 | - |
| - | - | 1847 | 1001 | - | - | 0 | - |
| - | - | 4940 | 1002 | - | - | 0 | - |
| 7 | y | 5962 | 1004 | 0.004306 | 4.291 | +1 | 9 |
| - | - | 2073 | 1005 | - | - | 0 | - |
| - | - | 1267 | 1006 | - | - | 0 | - |
| - | - | 1.838E+04 | 1017 | - | - | 0 | - |
| 9 | c | 2.212E+04 | 1018 | 0.001762 | 1.731 | +1 | 9 |
| - | - | 1.084E+04 | 1019 | - | - | 0 | - |
| - | - | 1415 | 1020 | - | - | 0 | - |
| - | - | 1216 | 1031 | - | - | 0 | - |
| - | - | 6617 | 1053 | - | - | 0 | - |
| - | - | 3723 | 1054 | - | - | 0 | - |
| - | - | 1510 | 1071 | - | - | 0 | - |
| 6 | y | 1952 | 1073 | 0.02144 | 19.99 | +1 | 10 |
| 6 | y | 2885 | 1074 | 8.039E-05 | 0.07489 | +1 | 10 |
| 6 | z | 9356 | 1075 | 0.0009474 | 0.8817 | +1 | 10 |
| - | - | 8.653E+04 | 1076 | - | - | 0 | - |
| - | - | 4.605E+04 | 1077 | - | - | 0 | - |
| - | - | 1.067E+04 | 1078 | - | - | 0 | - |
| - | - | 1.091E+04 | 1090 | - | - | 0 | - |
| 6 | y | 1.881E+04 | 1091 | 0.001727 | 1.584 | +1 | 10 |
| - | - | 9435 | 1092 | - | - | 0 | - |
| - | - | 2800 | 1093 | - | - | 0 | - |
| - | - | 2429 | 1102 | - | - | 0 | - |
| - | - | 3195 | 1103 | - | - | 0 | - |
| - | - | 1368 | 1117 | - | - | 0 | - |
| - | - | 1466 | 1118 | - | - | 0 | - |
| - | - | 2162 | 1119 | - | - | 0 | - |
| - | - | 1062 | 1120 | - | - | 0 | - |
| 10 | c | 5554 | 1129 | 0.002357 | 2.088 | +1 | 10 |
| - | - | 3374 | 1130 | - | - | 0 | - |
| - | - | 3.02E+04 | 1145 | - | - | 0 | - |
| 10 | c | 1.29E+05 | 1146 | 0.001074 | 0.9376 | +1 | 10 |
| - | - | 7.495E+04 | 1147 | - | - | 0 | - |
| - | - | 2.168E+04 | 1148 | - | - | 0 | - |
| - | - | 2150 | 1149 | - | - | 0 | - |
| - | - | 1363 | 1156 | - | - | 0 | - |
| 5 | y | 1404 | 1160 | 0.01172 | 10.1 | +1 | 11 |
| 5 | y | 2105 | 1161 | 0.002437 | 2.1 | +1 | 11 |
| 5 | z | 7216 | 1162 | 0.0005051 | 0.4349 | +1 | 11 |
| - | - | 4.488E+04 | 1163 | - | - | 0 | - |
| - | - | 2.561E+04 | 1164 | - | - | 0 | - |
| - | - | 7037 | 1165 | - | - | 0 | - |
| - | - | 3210 | 1173 | - | - | 0 | - |
| - | - | 4163 | 1174 | - | - | 0 | - |
| - | - | 2731 | 1175 | - | - | 0 | - |
| - | - | 1521 | 1177 | - | - | 0 | - |
| 5 | y | 1.105E+04 | 1178 | 0.001163 | 0.9874 | +1 | 11 |
| - | - | 6340 | 1179 | - | - | 0 | - |
| - | - | 1751 | 1180 | - | - | 0 | - |
| - | - | 2796 | 1182 | - | - | 0 | - |
| - | - | 1690 | 1183 | - | - | 0 | - |
| 11 | c | 2733 | 1200 | 0.0002862 | 0.2386 | +1 | 11 |
| - | - | 2008 | 1201 | - | - | 0 | - |
| - | - | 6.226E+04 | 1216 | - | - | 0 | - |
| 11 | c | 7.966E+04 | 1217 | 0.00291 | 2.391 | +1 | 11 |
| - | - | 4.148E+04 | 1218 | - | - | 0 | - |
| - | - | 1.11E+04 | 1219 | - | - | 0 | - |
| - | - | 1546 | 1253 | - | - | 0 | - |
| 4 | z | 1131 | 1259 | 0.001267 | 1.007 | +1 | 12 |
| - | - | 3772 | 1274 | - | - | 0 | - |
| 4 | y | 3.897E+04 | 1275 | 0.001558 | 1.223 | +1 | 12 |
| - | - | 2.903E+04 | 1276 | - | - | 0 | - |
| - | - | 7975 | 1277 | - | - | 0 | - |
| - | - | 4346 | 1287 | - | - | 0 | - |
| - | - | 6857 | 1288 | - | - | 0 | - |
| - | - | 2936 | 1289 | - | - | 0 | - |
| 12 | c | 2240 | 1314 | 0.001954 | 1.487 | +1 | 12 |
| - | - | 2271 | 1315 | - | - | 0 | - |
| - | - | 1488 | 1320 | - | - | 0 | - |
| - | - | 1261 | 1321 | - | - | 0 | - |
| 3 | w | 1160 | 1329 | 0.01363 | 10.26 | +1 | 13 |
| - | - | 6588 | 1330 | - | - | 0 | - |
| 12 | c | 1.217E+05 | 1331 | 0.001892 | 1.422 | +1 | 12 |
| - | - | 9.187E+04 | 1332 | - | - | 0 | - |
| - | - | 3.372E+04 | 1333 | - | - | 0 | - |
| - | - | 3601 | 1334 | - | - | 0 | - |
| 3 | y | 1394 | 1354 | 0.004948 | 3.656 | +1 | 13 |
| 3 | y | 1352 | 1355 | 0.002138 | 1.579 | +1 | 13 |
| - | - | 1206 | 1358 | - | - | 0 | - |
| - | - | 6016 | 1371 | - | - | 0 | - |
| 3 | y | 8.495E+04 | 1372 | 0.002442 | 1.781 | +1 | 13 |
| - | - | 6.418E+04 | 1373 | - | - | 0 | - |
| - | - | 2.386E+04 | 1374 | - | - | 0 | - |
| - | - | 2317 | 1375 | - | - | 0 | - |
| - | - | 2650 | 1400 | - | - | 0 | - |
| - | - | 2785 | 1401 | - | - | 0 | - |
| - | - | 5398 | 1402 | - | - | 0 | - |
| - | - | 3127 | 1403 | - | - | 0 | - |
| - | - | 2.205E+04 | 1415 | - | - | 0 | - |
| - | - | 1.652E+04 | 1416 | - | - | 0 | - |
| - | - | 7609 | 1417 | - | - | 0 | - |
| - | - | 3546 | 1431 | - | - | 0 | - |
| - | - | 4569 | 1432 | - | - | 0 | - |
| - | - | 1927 | 1433 | - | - | 0 | - |
| - | - | 1233 | 1444 | - | - | 0 | - |
| - | - | 1361 | 1457 | - | - | 0 | - |
| 13 | c | 7.125E+04 | 1459 | 0.002861 | 1.961 | +1 | 13 |
| - | - | 6.155E+04 | 1460 | - | - | 0 | - |
| - | - | 2.561E+04 | 1461 | - | - | 0 | - |
| - | - | 3788 | 1462 | - | - | 0 | - |
| - | - | 1.051E+04 | 1486 | - | - | 0 | - |
| - | - | 9620 | 1487 | - | - | 0 | - |
| - | - | 9314 | 1488 | - | - | 0 | - |
| - | - | 6868 | 1489 | - | - | 0 | - |
| - | - | 2369 | 1490 | - | - | 0 | - |
| 2 | z | 4.154E+04 | 1503 | 0.003182 | 2.118 | +1 | 14 |
| - | - | 3.926E+04 | 1504 | - | - | 0 | - |
| - | - | 1.508E+04 | 1505 | - | - | 0 | - |
| - | - | 1852 | 1506 | - | - | 0 | - |
| - | - | 8640 | 1515 | - | - | 0 | - |
| - | - | 7087 | 1516 | - | - | 0 | - |
| - | - | 4391 | 1517 | - | - | 0 | - |
| 2 | y | 1087 | 1519 | 0.0102 | 6.715 | +1 | 14 |
| - | - | 1022 | 1520 | - | - | 0 | - |
| - | - | 1405 | 1529 | - | - | 0 | - |
| 14 | c | 1.05E+05 | 1530 | 0.002377 | 1.554 | +1 | 14 |
| - | - | 9.619E+04 | 1531 | - | - | 0 | - |
| - | - | 6.252E+04 | 1532 | - | - | 0 | - |
| - | - | 2.396E+04 | 1533 | - | - | 0 | - |
| - | - | 6911 | 1534 | - | - | 0 | - |
| - | - | 1277 | 1535 | - | - | 0 | - |
| - | - | 1.831E+04 | 1544 | - | - | 0 | - |
| - | - | 1.71E+04 | 1545 | - | - | 0 | - |
| - | - | 9782 | 1546 | - | - | 0 | - |
| - | - | 1699 | 1547 | - | - | 0 | - |
| - | - | 9080 | 1560 | - | - | 0 | - |
| - | - | 1.42E+04 | 1561 | - | - | 0 | - |
| - | - | 1.606E+04 | 1562 | - | - | 0 | - |
| - | - | 8438 | 1563 | - | - | 0 | - |
| - | - | 3284 | 1564 | - | - | 0 | - |
| - | - | 5826 | 1569 | - | - | 0 | - |
| - | - | 7834 | 1570 | - | - | 0 | - |
| - | - | 1.502E+04 | 1571 | - | - | 0 | - |
| - | - | 9437 | 1572 | - | - | 0 | - |
| - | - | 5879 | 1573 | - | - | 0 | - |
| - | - | 8576 | 1574 | - | - | 0 | - |
| - | - | 7945 | 1575 | - | - | 0 | - |
| - | - | 4097 | 1576 | - | - | 0 | - |
| - | - | 1.445E+04 | 1577 | - | - | 0 | - |
| - | - | 1.061E+04 | 1578 | - | - | 0 | - |
| - | - | 5644 | 1579 | - | - | 0 | - |
| - | - | 2216 | 1586 | - | - | 0 | - |
| - | - | 1.922E+04 | 1587 | - | - | 0 | - |
| - | - | 1.147E+05 | 1588 | - | - | 0 | - |
| - | - | 9.977E+04 | 1589 | - | - | 0 | - |
| - | - | 4.986E+04 | 1590 | - | - | 0 | - |
| - | - | 8675 | 1591 | - | - | 0 | - |
| - | - | 1466 | 1592 | - | - | 0 | - |
| - | - | 1872 | 1597 | - | - | 0 | - |
| - | - | 3406 | 1598 | - | - | 0 | - |
| - | - | 2638 | 1599 | - | - | 0 | - |
| - | - | 8873 | 1605 | - | - | 0 | - |
| - | - | 7346 | 1606 | - | - | 0 | - |
| - | - | 2303 | 1607 | - | - | 0 | - |
| - | - | 3042 | 1614 | - | - | 0 | - |
| - | - | 4.268E+04 | 1615 | - | - | 0 | - |
| - | - | 1.414E+05 | 1616 | - | - | 0 | - |
| - | - | 1.251E+05 | 1617 | - | - | 0 | - |
| - | - | 5.457E+04 | 1618 | - | - | 0 | - |
| - | - | 8093 | 1619 | - | - | 0 | - |
| - | - | 3884 | 1630 | - | - | 0 | - |
| - | - | 1.374E+04 | 1631 | - | - | 0 | - |
| - | - | 3.053E+05 | 1632 | - | - | 0 | - |
| - | - | 8.475E+05 | 1633 | - | - | 0 | - |
| - | - | 6.612E+05 | 1634 | - | - | 0 | - |
| - | - | 2.672E+05 | 1635 | - | - | 0 | - |
| - | - | 3.841E+04 | 1636 | - | - | 0 | - |
| - | - | 2125 | 1666 | - | - | 0 | - |
| - | - | 815 | 3165 | - | - | 0 | - |

m/z Charge Intensity FragmentType MassShift Position
120.08100891113281 0 3804.8862
129.09854125976562 0 536.3497
129.10243225097656 0 6568.8965
148.8592529296875 0 746.0131
148.8802947998047 0 505.9828
148.9020233154297 0 755.93176
148.9092559814453 0 888.7833
148.9166717529297 0 1123.0438
148.92388916015625 0 1731.8778
148.93116760253906 0 1692.4438
148.9385986328125 0 4137.2725
148.94627380371094 0 7521.8735
148.96278381347656 0 4559.909
148.97047424316406 0 2055.3838
148.97756958007812 0 1800.865
148.9849090576172 0 1529.8077
148.99179077148438 0 869.1559
148.99964904785156 0 799.8916
149.0061492919922 0 789.2876
149.01400756835938 0 541.1548
149.06427001953125 0 651.5889
155.11767578125 0 669.4399
168.74513244628906 0 570.5898
185.1286163330078 0 779.77094
191.1026153564453 0 1475.2557 y 13
200.14080810546875 0 611.7702
209.68812561035156 0 631.20355
217.13377380371094 0 646.57635
220.0535125732422 0 678.40106
221.083251953125 0 663.62006
226.1188201904297 0 1705.5009
228.98890686035156 0 724.93567
231.9686737060547 0 576.4611
233.1649169921875 0 39286.086
234.16822814941406 0 4385.7085
243.1456756591797 0 1576.6938
258.369140625 0 734.98126
261.1596984863281 0 44604.98
262.1629333496094 0 7219.533
282.14556884765625 0 1645.1486
297.15576171875 0 2203.5042
299.0619812011719 0 2248.8137
313.1872253417969 0 1883.2853
314.1827087402344 0 2274.2192
319.1979064941406 0 4662.442 y 12
320.2034606933594 0 969.4185
341.0160217285156 0 797.95856
355.0699462890625 0 3974.6016
356.0714416503906 0 1992.3647
357.06927490234375 0 1133.6051
358.2126770019531 0 7344.317
359.2149658203125 0 1158.6891
361.17645263671875 0 783.26294
433.240966796875 0 3774.56 y 11
455.26507568359375 0 1807.4255
498.2189636230469 0 1677.0016
504.277099609375 0 6844.5156 y 10
505.28302001953125 0 959.8406
582.2874755859375 0 1011.3485
582.7821044921875 0 5971.1646
583.2825927734375 0 3480.5332
583.784912109375 0 1154.3756
591.2920532226562 0 5279.2
591.7957153320312 0 3461.8975
612.8157348632812 0 1035.2765
613.3145141601562 0 1505.1136
615.3085327148438 0 920.77765 y Ammonia loss 9
616.317138671875 0 5751.341 z 9
617.3255004882812 0 7367.3237
617.8071899414062 0 7277.283
618.3093872070312 0 7808.9814
618.8067626953125 0 2138.3772
626.8121948242188 0 47103.508
627.261962890625 0 2007.9912
627.3136596679688 0 27881.93
627.81396484375 0 8286.349
628.3181762695312 0 1213.6641
629.3282470703125 0 1148.8048
632.3361206054688 0 5692.591 y 9
633.34228515625 0 1089.6187
635.8228149414062 0 1078.3729
637.8154296875 0 3295.9307 y 3
655.3402099609375 0 3031.0337
655.84228515625 0 3289.565
656.344482421875 0 1415.6964
663.338134765625 0 2815.3113
663.8378295898438 0 3974.349
664.33740234375 0 2855.028
668.8239135742188 0 2163.5188
669.3238525390625 0 1359.7185 z Water loss 2
677.3361206054688 0 17482.701 y Water loss 2
677.83349609375 0 25337.234 y Ammonia loss 2
678.333740234375 0 10321.998 z 2
678.8311157226562 0 3842.065
685.3327026367188 0 3704.9568
685.834716796875 0 5037.8467
686.341552734375 0 791483.6 y 2
686.8428955078125 0 577403.1
687.3438110351562 0 214226.77
687.8445434570312 0 21409.33
700.3401489257812 0 1257.8824
700.8494873046875 0 1377.2449
721.3724975585938 0 939.70184 c Ammonia loss 12
729.4014892578125 0 24132.98 z 8
730.4067993164062 0 18140.016
731.4102783203125 0 6642.624
740.3451538085938 0 3270.4036
745.4202270507812 0 4805.691 y 8
746.42578125 0 1147.586
748.3726196289062 0 1450.5938
748.8803100585938 0 966.35986
751.3641357421875 0 1291.3796 y Ammonia loss 1
756.8875732421875 0 5763.818 c Ammonia loss 13
757.416259765625 0 4240.1245
757.8914794921875 0 2033.1321
758.3598022460938 0 1120.0039
759.3772583007812 0 1173.6948
759.8759155273438 0 11559.474 y 1
760.377197265625 0 10325.732
760.8772583007812 0 5127.7466
799.424072265625 0 1273.709 w 7
802.3824462890625 0 2514.0222
807.4120483398438 0 3307.9192
807.9124145507812 0 4215.352
808.4135131835938 0 2944.3179
808.9138793945312 0 979.54193
815.4180297851562 0 1197.4908
815.9227294921875 0 4751.5327
816.4178466796875 0 58536.527
816.9191284179688 0 65465.324
817.4201049804688 0 25708.654
817.9234619140625 0 3150.6448
858.444091796875 0 54116.227 z 7
859.44970703125 0 55243.21
860.4535522460938 0 17655.062
861.4534301757812 0 2624.3245
868.4039306640625 0 6436.073
869.4083251953125 0 2411.5623
873.4573974609375 0 2284.7776
874.4617309570312 0 6231.69 y 7
875.4613647460938 0 2608.6836
886.4609375 0 1923.6748
903.4315795898438 0 2468.0833
904.4401245117188 0 6612.5874 c 7
905.4449462890625 0 2886.3179
906.4561157226562 0 1021.8689
928.478271484375 0 1849.2317 w 6
929.465087890625 0 1136.9642
939.4424438476562 0 5753.5103
940.444580078125 0 3087.7693
956.4639892578125 0 1623.952
960.4591674804688 0 1928.3281
973.497314453125 0 3099.6062
974.4982299804688 0 2420.8298
975.5147705078125 0 959.5106
987.486328125 0 62116.508 z 6
988.4917602539062 0 68146.88
989.49462890625 0 27746.578
990.4971923828125 0 4723.8022
1000.4925537109375 0 3497.5618
1001.4957275390625 0 1847.2162
1002.4999389648438 0 4940.3535
1003.5011596679688 0 5962.481 y 6
1004.5050048828125 0 2073.101
1005.516357421875 0 1266.5692
1016.5166015625 0 18381.104
1017.5233764648438 0 22115.543 c 8
1018.5256958007812 0 10843.656
1019.5300903320312 0 1415.4805
1031.4971923828125 0 1215.6278
1053.4832763671875 0 6617.2686
1054.4884033203125 0 3722.9944
1070.5072021484375 0 1510.2731
1072.5054931640625 0 1951.878 y Water loss 5
1073.5108642578125 0 2884.5906 y Ammonia loss 5
1074.517822265625 0 9355.69 z 5
1075.526123046875 0 86532.57
1076.529052734375 0 46046.008
1077.5306396484375 0 10667.38
1089.529296875 0 10905.3
1090.5357666015625 0 18814.174 y 5
1091.5401611328125 0 9434.812
1092.543701171875 0 2800.1357
1101.5673828125 0 2429.3875
1102.568603515625 0 3194.7004
1116.525146484375 0 1368.1687
1117.535400390625 0 1466.1316
1118.5311279296875 0 2161.7224
1119.5433349609375 0 1062.333
1128.5548095703125 0 5553.8 c Ammonia loss 9
1129.560302734375 0 3373.8926
1144.5755615234375 0 30198.91
1145.5826416015625 0 128975.86 c 9
1146.5860595703125 0 74947.02
1147.589111328125 0 21683.182
1148.59716796875 0 2149.7224
1155.56298828125 0 1362.5565
1159.5472412109375 0 1404.08 y Water loss 4
1160.54541015625 0 2105.1277 y Ammonia loss 4
1161.55029296875 0 7216.008 z 4
1162.5567626953125 0 44875.535
1163.560302734375 0 25610.531
1164.5640869140625 0 7037.331
1172.6103515625 0 3209.6843
1173.610595703125 0 4163.106
1174.61572265625 0 2731.0066
1176.5643310546875 0 1521.4404
1177.568359375 0 11047.759 y 4
1178.569580078125 0 6339.7534
1179.5712890625 0 1751.4333
1181.5775146484375 0 2796.266
1182.5872802734375 0 1690.0758
1199.593994140625 0 2733.1575 c Ammonia loss 10
1200.5921630859375 0 2008.1179
1215.61181640625 0 62257.523
1216.617919921875 0 79663.805 c 10
1217.6209716796875 0 41484.977
1218.6234130859375 0 11100.165
1252.614990234375 0 1546.0524
1258.602294921875 0 1130.9874 z 3
1273.6097412109375 0 3772.1062
1274.6207275390625 0 38972.27 y 3
1275.6229248046875 0 29028.512
1276.6229248046875 0 7975.058
1286.6461181640625 0 4346.1973
1287.6529541015625 0 6856.8813
1288.656005859375 0 2936.1309
1313.63525390625 0 2239.757 c Ammonia loss 11
1314.634521484375 0 2270.7852
1319.5823974609375 0 1488.409
1320.5989990234375 0 1260.5798
1328.646484375 0 1160.2705 w 2
1329.6500244140625 0 6587.9297
1330.661865234375 0 121669.664 c 11
1331.6650390625 0 91870.28
1332.6676025390625 0 33718.098
1333.6728515625 0 3600.829
1353.66943359375 0 1394.3522 y Water loss 2
1354.6463623046875 0 1352.0957 y Ammonia loss 2
1357.6650390625 0 1206.2974
1370.6630859375 0 6016.34
1371.672607421875 0 84950.6 y 2
1372.676025390625 0 64183.945
1373.6790771484375 0 23856.385
1374.6776123046875 0 2317.3972
1399.667236328125 0 2649.9302
1400.6634521484375 0 2784.9065
1401.6915283203125 0 5397.8794
1402.7025146484375 0 3126.7742
1414.7420654296875 0 22051.299
1415.7462158203125 0 16524.988
1416.7464599609375 0 7608.943
1430.69921875 0 3546.236
1431.690673828125 0 4568.74
1432.69873046875 0 1926.5573
1443.7276611328125 0 1232.8312
1456.7401123046875 0 1360.9047
1458.755859375 0 71249.5 c 12
1459.75830078125 0 61548.742
1460.760009765625 0 25607.842
1461.7611083984375 0 3787.8962
1485.7769775390625 0 10507.745
1486.7822265625 0 9620.226
1487.7874755859375 0 9314.144
1488.7926025390625 0 6868.2515
1489.80126953125 0 2368.7468
1502.7215576171875 0 41543.26 z 1
1503.7249755859375 0 39259.85
1504.7279052734375 0 15083.925
1505.726806640625 0 1852.1095
1514.7833251953125 0 8640.396
1515.7841796875 0 7086.9897
1516.7880859375 0 4391.1406
1518.753662109375 0 1087.0645 y 1
1519.7379150390625 0 1021.55286
1528.7974853515625 0 1404.8127
1529.79345703125 0 104984.38 c 13
1530.7955322265625 0 96189.24
1531.7984619140625 0 62520.707
1532.803466796875 0 23956.088
1533.8052978515625 0 6910.976
1534.7852783203125 0 1277.2917
1543.784912109375 0 18305.434
1544.7833251953125 0 17097.613
1545.78564453125 0 9782.173
1546.7828369140625 0 1699.2487
1559.7596435546875 0 9080.021
1560.7923583984375 0 14204.482
1561.795654296875 0 16060.313
1562.799560546875 0 8438.315
1563.802490234375 0 3284.3567
1568.8187255859375 0 5826.1934
1569.8172607421875 0 7834.3584
1570.8255615234375 0 15017.3125
1571.82666015625 0 9437.244
1572.80517578125 0 5878.9434
1573.7861328125 0 8575.936
1574.7872314453125 0 7944.6655
1575.78564453125 0 4096.817
1576.7733154296875 0 14451.822
1577.771728515625 0 10606.061
1578.777099609375 0 5643.5474
1585.815185546875 0 2216.0662
1586.8226318359375 0 19220.473
1587.8125 0 114746.34
1588.8148193359375 0 99772.305
1589.814697265625 0 49855.09
1590.809814453125 0 8675.3545
1591.7938232421875 0 1466.3446
1596.8084716796875 0 1871.5944
1597.8046875 0 3405.8127
1598.8072509765625 0 2637.5566
1604.831298828125 0 8872.65
1605.83349609375 0 7345.5454
1606.8341064453125 0 2303.138
1613.7933349609375 0 3042.05
1614.820068359375 0 42682.832
1615.810546875 0 141382.56
1616.8104248046875 0 125077.29
1617.81005859375 0 54566.77
1618.805908203125 0 8093.087
1629.8035888671875 0 3883.6184
1630.8165283203125 0 13740.591
1631.823974609375 0 305300.1
1632.8310546875 0 847530.8
1633.834228515625 0 661195.06
1634.8355712890625 0 267246.88
1635.83740234375 0 38410.62
1665.8135986328125 0 2125.3225
3164.677734375 0 814.98706

Spectrum Details

|  |  |
| --- | --- |
| Matched peaks? Matched peaks The total absolute number of peaks matched. Additionally in brackets the total fraction of peaks matched and the total number of peaks is shown. | 51 (15.45% of 330) |
| FDR? FDR The false discovery rate estimated for this peptide. It is calculated by matching all theoretical fragments with a non-integer shift with the raw peaks for this spectrum. This is done with 40 different shifts. The resulting percentage is the average number of annotated peaks over the number of annotated peaks with the correct spectrum. | 0.23% |
| Satellite FDR? Satellite FDR See the FDR for details on its calculation. This satellite ion specific FDR only contains the satellite ions (d/w) for I/L/J positions. | - |
| PSM Score? PSM Score The PSM Score as given by Hecklib to this annotated spectrum. It is shown with three significant figures. | 470 |

#### Spectrum 6090? Spectrum 6090 The raw spectrum of this peptide as annotated by Hecklib. The fragments are coloured according to ion type (see legend). Any peaks with a star '\*' as text can be hovered over to see the full details, first the ion type second the mass shift type. By hovering over the amino acids in the peptide or ions in the legend the corresponding peaks are highlighted. By toggling the 'Unassigned' label you can turn the background (unassigned) peaks on or off in the plot. By updating the slider in the Ion legend you can update the spectrum to only show the top X% of the peaks with labels. The top X% means any peak that is within X% of the highest intensity. By dragging in the spectrum you can zoom in to a specific part of the spectrum and use 'Zoom Out' to get back to the original zoom level. The annotation of the spectrum is based on the given sequence in the peptides file and is done with different software so inconsistencies are likely. The peaks are annotated based on the given sequence, with 20 ppm tolerance.

Copy Data

##### Spectrum 6090 (TSV)

###### Preview

```
Loading example...
```

*Click on the button to copy the data to your clipboard.*

Mz MinMz MaxIntensity Max

WidthHeightPeptide font sizePeptide stroke widthSpectrum font sizeSpectrum stroke widthCompact peptide

Ion legend

wxyz

abcd

OtherUnassignedIon^Charge^_Position_Show for top:%

JFPPSSEEJQANKAT

01.82e+53.63e+55.45e+57.26e+5

Zoom Out

y^+1^_2_y^+1^_3_y^+1^_4_y^+1^_5_y^+1^_6_z^+1^_6_y^+2^_12_y^+1^_6_y^+2^_12_c^+2^_12_z^+2^_13_y^+2^_13_y^+2^_13_z^+2^_13_y^+2^_13_c^+2^_13_z^+1^_7_y^+1^_7_y^+2^_14_c^+2^_14_c^+1^_7_y^+2^_14_z^+1^_8_y^+1^_8_c^+1^_8_w^+1^_9_z^+1^_9_y^+1^_9_c^+1^_9_y^+1^_10_y^+1^_10_z^+1^_10_y^+1^_10_c^+1^_10_c^+1^_10_y^+1^_11_y^+1^_11_z^+1^_11_y^+1^_11_c^+1^_11_c^+1^_11_y^+1^_12_c^+1^_12_c^+1^_12_y^+1^_13_c^+1^_13_z^+1^_14_c^+1^_14_

042184112621682

Fragment Matches Table

Show background peaks

| Position | Ion type | Intensity | mz Theoretical | mz Error (Th) | mz Error (ppm) | Charge | Series Number |
| --- | --- | --- | --- | --- | --- | --- | --- |
| - | - | 2662 | 120.1 | - | - | 0 | - |
| - | - | 418.2 | 120.4 | - | - | 0 | - |
| - | - | 511.6 | 121.1 | - | - | 0 | - |
| - | - | 427.7 | 126 | - | - | 0 | - |
| - | - | 5468 | 129.1 | - | - | 0 | - |
| - | - | 448 | 129.6 | - | - | 0 | - |
| - | - | 376.8 | 133.1 | - | - | 0 | - |
| - | - | 4326 | 133.1 | - | - | 0 | - |
| - | - | 4146 | 136.1 | - | - | 0 | - |
| - | - | 498.3 | 155.1 | - | - | 0 | - |
| - | - | 511.7 | 188.8 | - | - | 0 | - |
| 14 | y | 1259 | 191.1 | 1.804E-05 | 0.09442 | +1 | 2 |
| - | - | 581.2 | 223.8 | - | - | 0 | - |
| - | - | 2248 | 226.1 | - | - | 0 | - |
| - | - | 521.7 | 226.4 | - | - | 0 | - |
| - | - | 3.602E+04 | 233.2 | - | - | 0 | - |
| - | - | 5189 | 234.2 | - | - | 0 | - |
| - | - | 1573 | 239.1 | - | - | 0 | - |
| - | - | 1671 | 243.1 | - | - | 0 | - |
| - | - | 922.8 | 245.1 | - | - | 0 | - |
| - | - | 3.924E+04 | 261.2 | - | - | 0 | - |
| - | - | 5369 | 262.2 | - | - | 0 | - |
| - | - | 1059 | 278.1 | - | - | 0 | - |
| - | - | 1817 | 282.1 | - | - | 0 | - |
| - | - | 3156 | 296.1 | - | - | 0 | - |
| - | - | 2462 | 297.2 | - | - | 0 | - |
| - | - | 1790 | 299.1 | - | - | 0 | - |
| - | - | 1300 | 310.2 | - | - | 0 | - |
| - | - | 1899 | 314.2 | - | - | 0 | - |
| 13 | y | 4972 | 319.2 | 0.0003406 | 1.067 | +1 | 3 |
| - | - | 967 | 320.2 | - | - | 0 | - |
| - | - | 740.1 | 353.1 | - | - | 0 | - |
| - | - | 5224 | 355.1 | - | - | 0 | - |
| - | - | 1537 | 356.1 | - | - | 0 | - |
| - | - | 7671 | 358.2 | - | - | 0 | - |
| - | - | 1826 | 359.2 | - | - | 0 | - |
| - | - | 2599 | 410.2 | - | - | 0 | - |
| - | - | 1005 | 425.2 | - | - | 0 | - |
| - | - | 683.2 | 427.3 | - | - | 0 | - |
| 12 | y | 4103 | 433.2 | 0.0002903 | 0.6702 | +1 | 4 |
| - | - | 747.1 | 434.2 | - | - | 0 | - |
| - | - | 1287 | 455.3 | - | - | 0 | - |
| - | - | 743.5 | 470.2 | - | - | 0 | - |
| - | - | 1160 | 498.2 | - | - | 0 | - |
| 11 | y | 5891 | 504.3 | 0.0002249 | 0.446 | +1 | 5 |
| - | - | 888.2 | 505.3 | - | - | 0 | - |
| - | - | 754.3 | 509.2 | - | - | 0 | - |
| - | - | 1177 | 582.3 | - | - | 0 | - |
| - | - | 4994 | 582.8 | - | - | 0 | - |
| - | - | 2944 | 583.3 | - | - | 0 | - |
| - | - | 1459 | 583.8 | - | - | 0 | - |
| - | - | 5841 | 591.3 | - | - | 0 | - |
| - | - | 4052 | 591.8 | - | - | 0 | - |
| - | - | 1288 | 592.3 | - | - | 0 | - |
| - | - | 1047 | 609.8 | - | - | 0 | - |
| - | - | 1707 | 612.8 | - | - | 0 | - |
| - | - | 891.9 | 613.3 | - | - | 0 | - |
| 10 | y | 664.8 | 615.3 | 0.0015 | 2.437 | +1 | 6 |
| 10 | z | 4785 | 616.3 | 0.0002914 | 0.4728 | +1 | 6 |
| - | - | 7506 | 617.3 | - | - | 0 | - |
| - | - | 5896 | 617.8 | - | - | 0 | - |
| - | - | 6370 | 618.3 | - | - | 0 | - |
| - | - | 2492 | 618.8 | - | - | 0 | - |
| - | - | 4.283E+04 | 626.8 | - | - | 0 | - |
| - | - | 1933 | 627.3 | - | - | 0 | - |
| - | - | 2.824E+04 | 627.3 | - | - | 0 | - |
| - | - | 7180 | 627.8 | - | - | 0 | - |
| 4 | y | 1083 | 629.3 | 0.007148 | 11.36 | +2 | 12 |
| 10 | y | 5656 | 632.3 | 0.0003997 | 0.6322 | +1 | 6 |
| - | - | 1064 | 633.3 | - | - | 0 | - |
| - | - | 1823 | 635.8 | - | - | 0 | - |
| 4 | y | 2201 | 637.8 | 0.001503 | 2.356 | +2 | 12 |
| - | - | 1275 | 638.3 | - | - | 0 | - |
| - | - | 4386 | 655.3 | - | - | 0 | - |
| - | - | 2901 | 655.8 | - | - | 0 | - |
| 12 | c | 981.5 | 656.8 | 0.01017 | 15.48 | +2 | 12 |
| - | - | 2704 | 663.3 | - | - | 0 | - |
| - | - | 2688 | 663.8 | - | - | 0 | - |
| - | - | 2039 | 664.3 | - | - | 0 | - |
| - | - | 1225 | 664.8 | - | - | 0 | - |
| - | - | 786.6 | 668.3 | - | - | 0 | - |
| - | - | 1639 | 668.8 | - | - | 0 | - |
| 3 | z | 1223 | 669.3 | 0.003765 | 5.625 | +2 | 13 |
| 3 | y | 1.809E+04 | 677.3 | 0.000667 | 0.9848 | +2 | 13 |
| 3 | y | 2.347E+04 | 677.8 | 0.006889 | 10.16 | +2 | 13 |
| 3 | z | 1.125E+04 | 678.3 | 0.001878 | 2.769 | +2 | 13 |
| - | - | 4110 | 678.8 | - | - | 0 | - |
| - | - | 1516 | 679.3 | - | - | 0 | - |
| - | - | 4506 | 685.3 | - | - | 0 | - |
| - | - | 4276 | 685.8 | - | - | 0 | - |
| 3 | y | 7.192E+05 | 686.3 | 0.0006337 | 0.9233 | +2 | 13 |
| - | - | 5.228E+05 | 686.8 | - | - | 0 | - |
| - | - | 1.862E+05 | 687.3 | - | - | 0 | - |
| - | - | 2.066E+04 | 687.8 | - | - | 0 | - |
| - | - | 1062 | 711.3 | - | - | 0 | - |
| - | - | 798.2 | 712.9 | - | - | 0 | - |
| 13 | c | 895.7 | 721.4 | 0.004544 | 6.299 | +2 | 13 |
| 9 | z | 1.907E+04 | 729.4 | 0.0001879 | 0.2576 | +1 | 7 |
| - | - | 1.697E+04 | 730.4 | - | - | 0 | - |
| - | - | 804 | 730.5 | - | - | 0 | - |
| - | - | 4282 | 731.4 | - | - | 0 | - |
| - | - | 976.3 | 732.4 | - | - | 0 | - |
| - | - | 3545 | 740.3 | - | - | 0 | - |
| - | - | 817.6 | 741.3 | - | - | 0 | - |
| 9 | y | 4255 | 745.4 | 0.001108 | 1.486 | +1 | 7 |
| - | - | 1342 | 746.4 | - | - | 0 | - |
| - | - | 1109 | 748.4 | - | - | 0 | - |
| 2 | y | 3384 | 751.4 | 0.003139 | 4.177 | +2 | 14 |
| - | - | 925.9 | 752.4 | - | - | 0 | - |
| 14 | c | 5010 | 756.9 | 0.0007683 | 1.015 | +2 | 14 |
| 7 | c | 3446 | 757.4 | 0.01138 | 15.02 | +1 | 7 |
| - | - | 1124 | 757.9 | - | - | 0 | - |
| - | - | 1231 | 758.4 | - | - | 0 | - |
| 2 | y | 1.049E+04 | 759.9 | 0.0004844 | 0.6374 | +2 | 14 |
| - | - | 1.001E+04 | 760.4 | - | - | 0 | - |
| - | - | 3533 | 760.9 | - | - | 0 | - |
| - | - | 938.9 | 798.3 | - | - | 0 | - |
| - | - | 1487 | 802.4 | - | - | 0 | - |
| - | - | 841.5 | 803.4 | - | - | 0 | - |
| - | - | 3421 | 807.4 | - | - | 0 | - |
| - | - | 3918 | 807.9 | - | - | 0 | - |
| - | - | 2064 | 808.4 | - | - | 0 | - |
| - | - | 841.2 | 808.9 | - | - | 0 | - |
| - | - | 765.1 | 814.9 | - | - | 0 | - |
| - | - | 2344 | 815.4 | - | - | 0 | - |
| - | - | 1.279E+04 | 815.9 | - | - | 0 | - |
| - | - | 6.651E+04 | 816.4 | - | - | 0 | - |
| - | - | 5.367E+04 | 816.9 | - | - | 0 | - |
| - | - | 2.359E+04 | 817.4 | - | - | 0 | - |
| - | - | 3282 | 817.9 | - | - | 0 | - |
| - | - | 949.6 | 851.4 | - | - | 0 | - |
| - | - | 2423 | 852.4 | - | - | 0 | - |
| 8 | z | 4.404E+04 | 858.4 | 6.569E-05 | 0.07653 | +1 | 8 |
| - | - | 5.014E+04 | 859.5 | - | - | 0 | - |
| - | - | 1.68E+04 | 860.5 | - | - | 0 | - |
| - | - | 3324 | 861.5 | - | - | 0 | - |
| - | - | 6849 | 868.4 | - | - | 0 | - |
| - | - | 2483 | 869.4 | - | - | 0 | - |
| - | - | 2366 | 873.5 | - | - | 0 | - |
| 8 | y | 5401 | 874.5 | 0.00163 | 1.864 | +1 | 8 |
| - | - | 2959 | 875.5 | - | - | 0 | - |
| - | - | 1030 | 886.5 | - | - | 0 | - |
| - | - | 1266 | 887.4 | - | - | 0 | - |
| - | - | 901.3 | 888.4 | - | - | 0 | - |
| - | - | 1308 | 899.4 | - | - | 0 | - |
| - | - | 2612 | 903.4 | - | - | 0 | - |
| 8 | c | 6386 | 904.4 | 0.003025 | 3.344 | +1 | 8 |
| - | - | 2919 | 905.4 | - | - | 0 | - |
| 7 | w | 816.1 | 928.5 | 0.01185 | 12.77 | +1 | 9 |
| - | - | 1220 | 934.5 | - | - | 0 | - |
| - | - | 1492 | 938.4 | - | - | 0 | - |
| - | - | 8939 | 939.4 | - | - | 0 | - |
| - | - | 4398 | 940.4 | - | - | 0 | - |
| - | - | 964.2 | 941.4 | - | - | 0 | - |
| - | - | 910.3 | 944.5 | - | - | 0 | - |
| - | - | 708.4 | 945.5 | - | - | 0 | - |
| - | - | 872 | 955.4 | - | - | 0 | - |
| - | - | 1490 | 956.4 | - | - | 0 | - |
| - | - | 1057 | 957.5 | - | - | 0 | - |
| - | - | 897.5 | 960.5 | - | - | 0 | - |
| - | - | 816.7 | 961.5 | - | - | 0 | - |
| - | - | 2871 | 973.5 | - | - | 0 | - |
| - | - | 1442 | 974.5 | - | - | 0 | - |
| 7 | z | 5.32E+04 | 987.5 | 0.0002582 | 0.2615 | +1 | 9 |
| - | - | 5.598E+04 | 988.5 | - | - | 0 | - |
| - | - | 2.125E+04 | 989.5 | - | - | 0 | - |
| - | - | 4327 | 990.5 | - | - | 0 | - |
| - | - | 3509 | 1000 | - | - | 0 | - |
| - | - | 1376 | 1002 | - | - | 0 | - |
| - | - | 4864 | 1002 | - | - | 0 | - |
| 7 | y | 4112 | 1004 | 0.001559 | 1.554 | +1 | 9 |
| - | - | 2339 | 1005 | - | - | 0 | - |
| - | - | 1.266E+04 | 1017 | - | - | 0 | - |
| 9 | c | 2.109E+04 | 1018 | 0.001273 | 1.251 | +1 | 9 |
| - | - | 8821 | 1019 | - | - | 0 | - |
| - | - | 729.2 | 1020 | - | - | 0 | - |
| - | - | 1762 | 1025 | - | - | 0 | - |
| - | - | 4533 | 1026 | - | - | 0 | - |
| - | - | 1577 | 1027 | - | - | 0 | - |
| - | - | 1185 | 1031 | - | - | 0 | - |
| - | - | 1070 | 1032 | - | - | 0 | - |
| - | - | 6306 | 1053 | - | - | 0 | - |
| - | - | 2957 | 1054 | - | - | 0 | - |
| - | - | 1105 | 1057 | - | - | 0 | - |
| - | - | 854.5 | 1071 | - | - | 0 | - |
| 6 | y | 1576 | 1073 | 0.005567 | 5.19 | +1 | 10 |
| 6 | y | 2675 | 1074 | 0.0008962 | 0.8348 | +1 | 10 |
| 6 | z | 8216 | 1075 | 0.001006 | 0.936 | +1 | 10 |
| - | - | 7.147E+04 | 1076 | - | - | 0 | - |
| - | - | 3.477E+04 | 1077 | - | - | 0 | - |
| - | - | 1134 | 1077 | - | - | 0 | - |
| - | - | 9908 | 1078 | - | - | 0 | - |
| - | - | 757 | 1079 | - | - | 0 | - |
| - | - | 1.103E+04 | 1090 | - | - | 0 | - |
| 6 | y | 1.863E+04 | 1091 | 0.001849 | 1.696 | +1 | 10 |
| - | - | 7590 | 1092 | - | - | 0 | - |
| - | - | 1549 | 1093 | - | - | 0 | - |
| - | - | 995.6 | 1097 | - | - | 0 | - |
| - | - | 1893 | 1102 | - | - | 0 | - |
| - | - | 3348 | 1103 | - | - | 0 | - |
| - | - | 2043 | 1111 | - | - | 0 | - |
| - | - | 1089 | 1112 | - | - | 0 | - |
| - | - | 2118 | 1118 | - | - | 0 | - |
| - | - | 2144 | 1119 | - | - | 0 | - |
| - | - | 1390 | 1120 | - | - | 0 | - |
| - | - | 1401 | 1124 | - | - | 0 | - |
| 10 | c | 3033 | 1129 | 0.002479 | 2.197 | +1 | 10 |
| - | - | 2501 | 1130 | - | - | 0 | - |
| - | - | 2000 | 1131 | - | - | 0 | - |
| - | - | 1051 | 1132 | - | - | 0 | - |
| - | - | 1927 | 1140 | - | - | 0 | - |
| - | - | 3074 | 1142 | - | - | 0 | - |
| - | - | 1626 | 1143 | - | - | 0 | - |
| - | - | 862.6 | 1144 | - | - | 0 | - |
| - | - | 2.444E+04 | 1145 | - | - | 0 | - |
| 10 | c | 1.126E+05 | 1146 | 0.0008299 | 0.7245 | +1 | 10 |
| - | - | 6.712E+04 | 1147 | - | - | 0 | - |
| - | - | 1.985E+04 | 1148 | - | - | 0 | - |
| - | - | 1616 | 1149 | - | - | 0 | - |
| - | - | 1218 | 1156 | - | - | 0 | - |
| 5 | y | 1609 | 1160 | 0.01306 | 11.26 | +1 | 11 |
| 5 | y | 1898 | 1161 | 0.01037 | 8.937 | +1 | 11 |
| 5 | z | 5646 | 1162 | 0.001204 | 1.036 | +1 | 11 |
| - | - | 3.673E+04 | 1163 | - | - | 0 | - |
| - | - | 2.061E+04 | 1164 | - | - | 0 | - |
| - | - | 5493 | 1165 | - | - | 0 | - |
| - | - | 1217 | 1166 | - | - | 0 | - |
| - | - | 2705 | 1173 | - | - | 0 | - |
| - | - | 3648 | 1174 | - | - | 0 | - |
| - | - | 1809 | 1175 | - | - | 0 | - |
| - | - | 1235 | 1177 | - | - | 0 | - |
| 5 | y | 1.004E+04 | 1178 | 0.001895 | 1.609 | +1 | 11 |
| - | - | 4781 | 1179 | - | - | 0 | - |
| - | - | 1563 | 1180 | - | - | 0 | - |
| - | - | 1911 | 1182 | - | - | 0 | - |
| - | - | 1847 | 1183 | - | - | 0 | - |
| - | - | 766.9 | 1197 | - | - | 0 | - |
| - | - | 981.3 | 1198 | - | - | 0 | - |
| 11 | c | 2095 | 1200 | 0.003704 | 3.088 | +1 | 11 |
| - | - | 1629 | 1201 | - | - | 0 | - |
| - | - | 1101 | 1206 | - | - | 0 | - |
| - | - | 4.927E+04 | 1216 | - | - | 0 | - |
| 11 | c | 6.54E+04 | 1217 | 0.002543 | 2.09 | +1 | 11 |
| - | - | 3.1E+04 | 1218 | - | - | 0 | - |
| - | - | 7559 | 1219 | - | - | 0 | - |
| - | - | 1163 | 1224 | - | - | 0 | - |
| - | - | 1655 | 1240 | - | - | 0 | - |
| - | - | 1454 | 1241 | - | - | 0 | - |
| - | - | 1166 | 1253 | - | - | 0 | - |
| - | - | 2041 | 1260 | - | - | 0 | - |
| - | - | 1531 | 1261 | - | - | 0 | - |
| - | - | 3204 | 1274 | - | - | 0 | - |
| 4 | y | 3.791E+04 | 1275 | 0.0009481 | 0.7438 | +1 | 12 |
| - | - | 2.385E+04 | 1276 | - | - | 0 | - |
| - | - | 8492 | 1277 | - | - | 0 | - |
| - | - | 984 | 1278 | - | - | 0 | - |
| - | - | 4377 | 1287 | - | - | 0 | - |
| - | - | 4149 | 1288 | - | - | 0 | - |
| - | - | 2092 | 1289 | - | - | 0 | - |
| - | - | 2588 | 1297 | - | - | 0 | - |
| - | - | 2579 | 1298 | - | - | 0 | - |
| - | - | 1311 | 1299 | - | - | 0 | - |
| - | - | 1687 | 1304 | - | - | 0 | - |
| - | - | 2263 | 1305 | - | - | 0 | - |
| - | - | 960.6 | 1306 | - | - | 0 | - |
| - | - | 1672 | 1312 | - | - | 0 | - |
| - | - | 1319 | 1313 | - | - | 0 | - |
| 12 | c | 2535 | 1314 | 0.003419 | 2.602 | +1 | 12 |
| - | - | 1527 | 1315 | - | - | 0 | - |
| - | - | 1039 | 1329 | - | - | 0 | - |
| - | - | 6466 | 1330 | - | - | 0 | - |
| 12 | c | 1.064E+05 | 1331 | 0.001159 | 0.8712 | +1 | 12 |
| - | - | 7.876E+04 | 1332 | - | - | 0 | - |
| - | - | 2.797E+04 | 1333 | - | - | 0 | - |
| - | - | 3158 | 1334 | - | - | 0 | - |
| - | - | 2122 | 1338 | - | - | 0 | - |
| - | - | 1270 | 1339 | - | - | 0 | - |
| - | - | 1119 | 1347 | - | - | 0 | - |
| - | - | 7215 | 1355 | - | - | 0 | - |
| - | - | 3937 | 1356 | - | - | 0 | - |
| - | - | 1623 | 1357 | - | - | 0 | - |
| - | - | 1171 | 1358 | - | - | 0 | - |
| - | - | 5028 | 1371 | - | - | 0 | - |
| 3 | y | 7.248E+04 | 1372 | 0.001832 | 1.336 | +1 | 13 |
| - | - | 5.47E+04 | 1373 | - | - | 0 | - |
| - | - | 1.701E+04 | 1374 | - | - | 0 | - |
| - | - | 1921 | 1375 | - | - | 0 | - |
| - | - | 1078 | 1386 | - | - | 0 | - |
| - | - | 1246 | 1387 | - | - | 0 | - |
| - | - | 2926 | 1400 | - | - | 0 | - |
| - | - | 2611 | 1401 | - | - | 0 | - |
| - | - | 5508 | 1402 | - | - | 0 | - |
| - | - | 4379 | 1403 | - | - | 0 | - |
| - | - | 1497 | 1404 | - | - | 0 | - |
| - | - | 1.847E+04 | 1415 | - | - | 0 | - |
| - | - | 1.489E+04 | 1416 | - | - | 0 | - |
| - | - | 6250 | 1417 | - | - | 0 | - |
| - | - | 3695 | 1431 | - | - | 0 | - |
| - | - | 3917 | 1432 | - | - | 0 | - |
| - | - | 2137 | 1433 | - | - | 0 | - |
| 13 | c | 6.32E+04 | 1459 | 0.002372 | 1.626 | +1 | 13 |
| - | - | 5.318E+04 | 1460 | - | - | 0 | - |
| - | - | 2.235E+04 | 1461 | - | - | 0 | - |
| - | - | 2607 | 1462 | - | - | 0 | - |
| - | - | 1711 | 1474 | - | - | 0 | - |
| - | - | 1262 | 1475 | - | - | 0 | - |
| - | - | 8293 | 1486 | - | - | 0 | - |
| - | - | 9750 | 1487 | - | - | 0 | - |
| - | - | 7908 | 1488 | - | - | 0 | - |
| - | - | 4759 | 1489 | - | - | 0 | - |
| - | - | 1218 | 1490 | - | - | 0 | - |
| 2 | z | 3.713E+04 | 1503 | 0.002328 | 1.549 | +1 | 14 |
| - | - | 3.331E+04 | 1504 | - | - | 0 | - |
| - | - | 1.244E+04 | 1505 | - | - | 0 | - |
| - | - | 6831 | 1515 | - | - | 0 | - |
| - | - | 6920 | 1516 | - | - | 0 | - |
| - | - | 3370 | 1517 | - | - | 0 | - |
| - | - | 1.576E+04 | 1518 | - | - | 0 | - |
| - | - | 1.345E+04 | 1519 | - | - | 0 | - |
| - | - | 3747 | 1520 | - | - | 0 | - |
| - | - | 1091 | 1529 | - | - | 0 | - |
| 14 | c | 9.072E+04 | 1530 | 0.001278 | 0.8354 | +1 | 14 |
| - | - | 8.376E+04 | 1531 | - | - | 0 | - |
| - | - | 5.152E+04 | 1532 | - | - | 0 | - |
| - | - | 1.923E+04 | 1533 | - | - | 0 | - |
| - | - | 5387 | 1534 | - | - | 0 | - |
| - | - | 4574 | 1543 | - | - | 0 | - |
| - | - | 1.647E+04 | 1544 | - | - | 0 | - |
| - | - | 1.515E+04 | 1545 | - | - | 0 | - |
| - | - | 7815 | 1546 | - | - | 0 | - |
| - | - | 2111 | 1547 | - | - | 0 | - |
| - | - | 8565 | 1560 | - | - | 0 | - |
| - | - | 1.096E+04 | 1561 | - | - | 0 | - |
| - | - | 1.355E+04 | 1562 | - | - | 0 | - |
| - | - | 7536 | 1563 | - | - | 0 | - |
| - | - | 2303 | 1564 | - | - | 0 | - |
| - | - | 5583 | 1569 | - | - | 0 | - |
| - | - | 6239 | 1570 | - | - | 0 | - |
| - | - | 9922 | 1571 | - | - | 0 | - |
| - | - | 7683 | 1572 | - | - | 0 | - |
| - | - | 6006 | 1573 | - | - | 0 | - |
| - | - | 7446 | 1574 | - | - | 0 | - |
| - | - | 7372 | 1575 | - | - | 0 | - |
| - | - | 6688 | 1576 | - | - | 0 | - |
| - | - | 1.115E+04 | 1577 | - | - | 0 | - |
| - | - | 9753 | 1578 | - | - | 0 | - |
| - | - | 3982 | 1579 | - | - | 0 | - |
| - | - | 911.7 | 1585 | - | - | 0 | - |
| - | - | 1948 | 1586 | - | - | 0 | - |
| - | - | 1.425E+04 | 1587 | - | - | 0 | - |
| - | - | 9.688E+04 | 1588 | - | - | 0 | - |
| - | - | 8.455E+04 | 1589 | - | - | 0 | - |
| - | - | 4.015E+04 | 1590 | - | - | 0 | - |
| - | - | 6631 | 1591 | - | - | 0 | - |
| - | - | 1276 | 1592 | - | - | 0 | - |
| - | - | 1175 | 1597 | - | - | 0 | - |
| - | - | 2611 | 1598 | - | - | 0 | - |
| - | - | 2027 | 1599 | - | - | 0 | - |
| - | - | 6697 | 1605 | - | - | 0 | - |
| - | - | 4833 | 1606 | - | - | 0 | - |
| - | - | 2648 | 1607 | - | - | 0 | - |
| - | - | 1093 | 1613 | - | - | 0 | - |
| - | - | 3213 | 1614 | - | - | 0 | - |
| - | - | 3.398E+04 | 1615 | - | - | 0 | - |
| - | - | 1.245E+05 | 1616 | - | - | 0 | - |
| - | - | 1.016E+05 | 1617 | - | - | 0 | - |
| - | - | 4.282E+04 | 1618 | - | - | 0 | - |
| - | - | 6364 | 1619 | - | - | 0 | - |
| - | - | 1151 | 1620 | - | - | 0 | - |
| - | - | 2973 | 1630 | - | - | 0 | - |
| - | - | 1.366E+04 | 1631 | - | - | 0 | - |
| - | - | 2.601E+05 | 1632 | - | - | 0 | - |
| - | - | 7.153E+05 | 1633 | - | - | 0 | - |
| - | - | 5.331E+05 | 1634 | - | - | 0 | - |
| - | - | 2.208E+05 | 1635 | - | - | 0 | - |
| - | - | 2.946E+04 | 1636 | - | - | 0 | - |
| - | - | 1455 | 1665 | - | - | 0 | - |
| - | - | 1192 | 1666 | - | - | 0 | - |

m/z Charge Intensity FragmentType MassShift Position
120.08090209960938 0 2661.704
120.35990142822266 0 418.1652
121.08427429199219 0 511.64563
126.03141784667969 0 427.68637
129.10240173339844 0 5467.5312
129.558349609375 0 447.98965
133.05691528320312 0 376.755
133.06088256835938 0 4325.7354
136.0758514404297 0 4146.241
155.1178436279297 0 498.33545
188.76084899902344 0 511.6843
191.1026153564453 0 1259.4319 y 13
223.77708435058594 0 581.1697
226.11856079101562 0 2248.2756
226.44581604003906 0 521.6801
233.16494750976562 0 36021.312
234.1683807373047 0 5189.2793
239.09500122070312 0 1573.0035
243.14524841308594 0 1671.0421
245.1288299560547 0 922.8478
261.1597595214844 0 39237.715
262.16326904296875 0 5368.915
278.1133728027344 0 1059.2783
282.1448669433594 0 1816.8516
296.12445068359375 0 3155.8135
297.1556396484375 0 2461.512
299.0626525878906 0 1789.6205
310.2122497558594 0 1300.4592
314.1826477050781 0 1898.5203
319.19793701171875 0 4971.8037 y 12
320.2019348144531 0 966.9819
353.1455078125 0 740.0891
355.0699768066406 0 5223.5034
356.07110595703125 0 1537.2638
358.2125549316406 0 7671.236
359.21502685546875 0 1826.2891
410.16705322265625 0 2599.2722
425.2127685546875 0 1004.9474
427.2694091796875 0 683.1869
433.2408142089844 0 4102.9634 y 11
434.2424011230469 0 747.1461
455.2646789550781 0 1287.0349
470.2222595214844 0 743.4753
498.218505859375 0 1160.3999
504.2778625488281 0 5891.352 y 10
505.2835388183594 0 888.2467
509.2370300292969 0 754.25366
582.2874755859375 0 1176.6982
582.7822265625 0 4994.433
583.2817993164062 0 2944.2407
583.7838745117188 0 1458.7418
591.2935791015625 0 5841.4697
591.794677734375 0 4052.03
592.2978515625 0 1287.6976
609.794921875 0 1047.2327
612.8165283203125 0 1706.932
613.3102416992188 0 891.91144
615.3081665039062 0 664.7532 y Ammonia loss 9
616.3171997070312 0 4784.614 z 9
617.3248291015625 0 7506.096
617.8068237304688 0 5895.5
618.3082885742188 0 6370.481
618.8040161132812 0 2492.255
626.8123779296875 0 42828.363
627.2632446289062 0 1932.9215
627.3135375976562 0 28235.365
627.8150634765625 0 7179.966
629.3086547851562 0 1082.8842 y Ammonia loss 3
632.3358154296875 0 5656.4814 y 9
633.3286743164062 0 1063.8772
635.8179931640625 0 1823.1685
637.8162841796875 0 2200.7444 y 3
638.316162109375 0 1274.6127
655.3421020507812 0 4386.4766
655.8424682617188 0 2900.898
656.820068359375 0 981.46423 c Water loss 11
663.3399047851562 0 2704.1353
663.84033203125 0 2687.8682
664.3381958007812 0 2039.1881
664.8403930664062 0 1224.6462
668.3280639648438 0 786.5796
668.82568359375 0 1638.6761
669.32275390625 0 1222.8846 z Water loss 2
677.3365478515625 0 18088.209 y Water loss 2
677.8347778320312 0 23471.451 y Ammonia loss 2
678.3336791992188 0 11245.183 z 2
678.8307495117188 0 4110.281
679.3340454101562 0 1516.1498
685.33544921875 0 4506.2275
685.8346557617188 0 4275.996
686.341796875 0 719224.06 y 2
686.8432006835938 0 522761.97
687.3441162109375 0 186181.25
687.8453979492188 0 20656.363
711.2909545898438 0 1061.7712
712.8521728515625 0 798.23376
721.374267578125 0 895.7267 c Ammonia loss 12
729.4013671875 0 19069.299 z 8
730.4078369140625 0 16967.348
730.483154296875 0 804.0075
731.412353515625 0 4282.2603
732.4070434570312 0 976.25995
740.3453369140625 0 3544.6602
741.346923828125 0 817.59906
745.42138671875 0 4255.263 y 8
746.4295654296875 0 1342.3943
748.3790283203125 0 1108.7191
751.365234375 0 3383.5603 y Ammonia loss 1
752.374267578125 0 925.91064
756.8875122070312 0 5010.498 c Ammonia loss 13
757.3992919921875 0 3446.2832 c Water loss 6
757.9000854492188 0 1124.465
758.4281616210938 0 1230.8435
759.8758544921875 0 10488.785 y 1
760.3770751953125 0 10009.729
760.8781127929688 0 3533.018
798.3223876953125 0 938.928
802.38720703125 0 1487.0065
803.37255859375 0 841.50665
807.4094848632812 0 3420.6714
807.912353515625 0 3918.2527
808.4108276367188 0 2064.144
808.9132080078125 0 841.1712
814.91943359375 0 765.07544
815.4174194335938 0 2343.9675
815.923583984375 0 12790.222
816.4192504882812 0 66514.805
816.919921875 0 53668.598
817.4208374023438 0 23591.941
817.9227294921875 0 3282.3308
851.3927001953125 0 949.6349
852.4108276367188 0 2423.3694
858.4442138671875 0 44044.957 z 7
859.4501953125 0 50143.95
860.4534912109375 0 16795.568
861.4544067382812 0 3324.303
868.4035034179688 0 6849.214
869.408935546875 0 2483.2632
873.455810546875 0 2366.3716
874.4612426757812 0 5401.3457 y 7
875.466552734375 0 2959.3684
886.4674072265625 0 1029.8568
887.4169921875 0 1265.5953
888.4130859375 0 901.2641
899.3685913085938 0 1307.7281
903.4317626953125 0 2611.716
904.4380493164062 0 6386.3384 c 7
905.4401245117188 0 2918.805
928.4852905273438 0 816.1345 w 6
934.4805297851562 0 1219.646
938.4403686523438 0 1492.1665
939.4434204101562 0 8938.513
940.4466552734375 0 4398.286
941.4464111328125 0 964.23883
944.4855346679688 0 910.29504
945.4817504882812 0 708.43713
955.380126953125 0 872.0453
956.3873901367188 0 1490.0914
957.4730224609375 0 1056.6788
960.4562377929688 0 897.4782
961.4791870117188 0 816.74585
973.4996337890625 0 2870.558
974.5032348632812 0 1442.2754
987.4869995117188 0 53202.586 z 6
988.4923095703125 0 55983
989.4955444335938 0 21248.15
990.497802734375 0 4326.703
1000.4970703125 0 3509.1892
1001.504638671875 0 1375.567
1002.498779296875 0 4864.087
1003.50390625 0 4111.5645 y 6
1004.503173828125 0 2338.829
1016.5164184570312 0 12664.447
1017.5238647460938 0 21089.951 c 8
1018.5260009765625 0 8821.168
1019.543212890625 0 729.1909
1025.46875 0 1762.4276
1026.4766845703125 0 4533.1367
1027.4759521484375 0 1577.0039
1031.4898681640625 0 1185.3306
1032.4913330078125 0 1070.3676
1053.4827880859375 0 6306.042
1054.4840087890625 0 2957.0957
1057.4381103515625 0 1105.3475
1070.5042724609375 0 854.4649
1072.5213623046875 0 1576.1544 y Water loss 5
1073.5118408203125 0 2674.7046 y Ammonia loss 5
1074.519775390625 0 8215.934 z 5
1075.5269775390625 0 71474.18
1076.529541015625 0 34772.25
1076.66357421875 0 1134.4424
1077.5306396484375 0 9908.073
1078.5250244140625 0 756.9575
1089.5303955078125 0 11025.378
1090.53564453125 0 18631.24 y 5
1091.5390625 0 7590.241
1092.54345703125 0 1548.5896
1097.478271484375 0 995.58344
1101.569091796875 0 1892.9172
1102.5709228515625 0 3347.887
1111.4556884765625 0 2042.5852
1112.4404296875 0 1089.3671
1117.5299072265625 0 2117.6096
1118.5338134765625 0 2144.1787
1119.5421142578125 0 1390.4612
1124.4844970703125 0 1400.667
1128.5546875 0 3033.4807 c Ammonia loss 9
1129.5594482421875 0 2500.7312
1130.5728759765625 0 2000.3679
1131.5672607421875 0 1050.5859
1140.494873046875 0 1927.1952
1141.5042724609375 0 3074.3962
1142.5045166015625 0 1625.6346
1144.42919921875 0 862.5763
1144.576171875 0 24441.342
1145.5828857421875 0 112591.69 c 9
1146.586181640625 0 67121.67
1147.589599609375 0 19854.717
1148.599365234375 0 1616.2203
1155.564208984375 0 1218.091
1159.5458984375 0 1608.5328 y Water loss 4
1160.5533447265625 0 1897.6193 y Ammonia loss 4
1161.552001953125 0 5645.698 z 4
1162.5577392578125 0 36726.54
1163.5606689453125 0 20613.49
1164.5655517578125 0 5493.2793
1165.57470703125 0 1217.4004
1172.6103515625 0 2704.5732
1173.610595703125 0 3648.0857
1174.6181640625 0 1809.0477
1176.5565185546875 0 1235.4135
1177.567626953125 0 10039.705 y 4
1178.572021484375 0 4780.6904
1179.57080078125 0 1562.7163
1181.5679931640625 0 1910.9652
1182.5867919921875 0 1846.691
1196.55615234375 0 766.9271
1197.5662841796875 0 981.32166
1199.590576171875 0 2094.5823 c Ammonia loss 10
1200.6005859375 0 1628.7357
1205.54833984375 0 1101.3534
1215.6124267578125 0 49266.477
1216.6182861328125 0 65403.707 c 10
1217.6214599609375 0 31002.531
1218.6236572265625 0 7558.593
1223.553466796875 0 1163.1002
1239.5654296875 0 1654.7216
1240.5623779296875 0 1453.6995
1252.6221923828125 0 1165.5032
1259.6102294921875 0 2041.4623
1260.61083984375 0 1530.9276
1273.611328125 0 3204.125
1274.621337890625 0 37909.79 y 3
1275.6241455078125 0 23847.879
1276.62646484375 0 8491.674
1277.639892578125 0 983.9663
1286.650146484375 0 4376.511
1287.651611328125 0 4148.784
1288.6627197265625 0 2091.8423
1296.5953369140625 0 2587.5066
1297.596923828125 0 2578.5098
1298.60302734375 0 1311.0084
1304.479248046875 0 1687.0883
1305.4873046875 0 2262.6333
1306.49755859375 0 960.5928
1311.6070556640625 0 1671.5332
1312.60595703125 0 1319.3402
1313.6337890625 0 2534.875 c Ammonia loss 11
1314.642578125 0 1526.8615
1328.6649169921875 0 1038.7466
1329.6527099609375 0 6466.464
1330.66259765625 0 106421.64 c 11
1331.66552734375 0 78761.52
1332.6673583984375 0 27974.408
1333.672607421875 0 3158.2615
1337.589111328125 0 2121.6016
1338.5946044921875 0 1270.293
1346.64501953125 0 1118.6108
1354.6171875 0 7215.3433
1355.6170654296875 0 3936.6956
1356.649169921875 0 1623.4138
1357.67724609375 0 1170.9037
1370.6622314453125 0 5027.625
1371.6732177734375 0 72479.7 y 2
1372.6767578125 0 54695.418
1373.68017578125 0 17008.822
1374.677490234375 0 1920.7274
1385.699462890625 0 1077.9475
1386.709716796875 0 1246.1475
1399.6651611328125 0 2925.547
1400.6737060546875 0 2610.957
1401.6927490234375 0 5507.9473
1402.701416015625 0 4379.3564
1403.6988525390625 0 1497.175
1414.7427978515625 0 18466.342
1415.7457275390625 0 14887.589
1416.7503662109375 0 6249.5605
1430.7010498046875 0 3694.9902
1431.69970703125 0 3916.61
1432.698974609375 0 2136.9646
1458.75634765625 0 63196.293 c 12
1459.758544921875 0 53180.457
1460.7606201171875 0 22348.852
1461.765380859375 0 2607.142
1473.74365234375 0 1711.4976
1474.7469482421875 0 1262.0913
1485.78076171875 0 8292.691
1486.782958984375 0 9750.276
1487.7890625 0 7907.7944
1488.7984619140625 0 4758.782
1489.80126953125 0 1218.3011
1502.722412109375 0 37125.15 z 1
1503.7257080078125 0 33312.113
1504.7298583984375 0 12443.725
1514.783935546875 0 6830.5176
1515.7852783203125 0 6920.216
1516.78173828125 0 3369.6877
1517.6783447265625 0 15760.439
1518.68212890625 0 13447.162
1519.686279296875 0 3747.0164
1528.79248046875 0 1090.5587
1529.7945556640625 0 90717.36 c 13
1530.79638671875 0 83760.44
1531.7998046875 0 51518.734
1532.803955078125 0 19225.748
1533.8045654296875 0 5387.3384
1542.7310791015625 0 4574.3765
1543.7750244140625 0 16469.867
1544.7816162109375 0 15145.309
1545.787841796875 0 7815.3877
1546.77587890625 0 2111.427
1559.7630615234375 0 8565.395
1560.7913818359375 0 10957.11
1561.7952880859375 0 13552.468
1562.80078125 0 7535.98
1563.8057861328125 0 2302.5007
1568.81884765625 0 5583.3813
1569.81689453125 0 6239.086
1570.824462890625 0 9922.416
1571.8271484375 0 7683.477
1572.8001708984375 0 6005.632
1573.793212890625 0 7446.39
1574.7054443359375 0 7371.6436
1575.70556640625 0 6687.5586
1576.7647705078125 0 11150.084
1577.7718505859375 0 9752.781
1578.7767333984375 0 3981.62
1584.7603759765625 0 911.7247
1585.8109130859375 0 1948.3945
1586.82373046875 0 14250.361
1587.8143310546875 0 96882.625
1588.815673828125 0 84553.33
1589.8153076171875 0 40147.043
1590.80859375 0 6631.1045
1591.7952880859375 0 1276.2628
1596.7965087890625 0 1174.9818
1597.8028564453125 0 2611.25
1598.7977294921875 0 2026.8706
1604.8385009765625 0 6696.7295
1605.8363037109375 0 4832.633
1606.8367919921875 0 2648.078
1612.77001953125 0 1093.4589
1613.801025390625 0 3212.763
1614.821044921875 0 33975.75
1615.8106689453125 0 124467.39
1616.810791015625 0 101637.516
1617.8115234375 0 42819.13
1618.8092041015625 0 6364.44
1619.7882080078125 0 1150.5317
1629.8123779296875 0 2972.756
1630.8214111328125 0 13658.311
1631.8253173828125 0 260091.1
1632.83203125 0 715332.25
1633.8350830078125 0 533117.25
1634.8370361328125 0 220813.45
1635.8394775390625 0 29458.637
1664.817138671875 0 1454.6975
1665.81884765625 0 1191.6123

Spectrum Details

|  |  |
| --- | --- |
| Matched peaks? Matched peaks The total absolute number of peaks matched. Additionally in brackets the total fraction of peaks matched and the total number of peaks is shown. | 48 (12.70% of 378) |
| FDR? FDR The false discovery rate estimated for this peptide. It is calculated by matching all theoretical fragments with a non-integer shift with the raw peaks for this spectrum. This is done with 40 different shifts. The resulting percentage is the average number of annotated peaks over the number of annotated peaks with the correct spectrum. | 0.99% |
| Satellite FDR? Satellite FDR See the FDR for details on its calculation. This satellite ion specific FDR only contains the satellite ions (d/w) for I/L/J positions. | - |
| PSM Score? PSM Score The PSM Score as given by Hecklib to this annotated spectrum. It is shown with three significant figures. | 419 |

#### Spectrum 6239? Spectrum 6239 The raw spectrum of this peptide as annotated by Hecklib. The fragments are coloured according to ion type (see legend). Any peaks with a star '\*' as text can be hovered over to see the full details, first the ion type second the mass shift type. By hovering over the amino acids in the peptide or ions in the legend the corresponding peaks are highlighted. By toggling the 'Unassigned' label you can turn the background (unassigned) peaks on or off in the plot. By updating the slider in the Ion legend you can update the spectrum to only show the top X% of the peaks with labels. The top X% means any peak that is within X% of the highest intensity. By dragging in the spectrum you can zoom in to a specific part of the spectrum and use 'Zoom Out' to get back to the original zoom level. The annotation of the spectrum is based on the given sequence in the peptides file and is done with different software so inconsistencies are likely. The peaks are annotated based on the given sequence, with 20 ppm tolerance.

Copy Data

##### Spectrum 6239 (TSV)

###### Preview

```
Loading example...
```

*Click on the button to copy the data to your clipboard.*

Mz MinMz MaxIntensity Max

WidthHeightPeptide font sizePeptide stroke widthSpectrum font sizeSpectrum stroke widthCompact peptide

Ion legend

wxyz

abcd

OtherUnassignedIon^Charge^_Position_Show for top:%

JFPPSSEEJQANKAT

05.89e+31.18e+41.77e+42.36e+4

Zoom Out

y^+1^_1_w^+2^_4_y^+1^_2_y^+3^_7_z^+1^_3_y^+1^_3_y^+2^_7_y^+1^_4_y^+3^_13_y^+3^_13_c^+1^_4_z^+1^_5_y^+1^_5_w^+1^_6_c^+1^_5_c^+2^_11_c^+2^_11_z^+1^_6_c^+1^_6_c^+2^_12_y^+2^_13_y^+2^_13_z^+2^_13_y^+2^_13_z^+1^_7_c^+2^_13_z^+2^_14_z^+2^_14_y^+1^_7_z^+2^_14_c^+2^_14_c^+2^_14_c^+1^_7_w^+1^_8_z^+1^_8_c^+1^_8_w^+1^_9_z^+1^_9_c^+1^_9_z^+1^_10_y^+1^_10_c^+1^_10_z^+1^_11_y^+1^_11_c^+1^_11_y^+1^_12_c^+1^_12_y^+1^_13_c^+1^_13_z^+1^_14_c^+1^_14_

0798159623943192

Fragment Matches Table

Show background peaks

| Position | Ion type | Intensity | mz Theoretical | mz Error (Th) | mz Error (ppm) | Charge | Series Number |
| --- | --- | --- | --- | --- | --- | --- | --- |
| 15 | y | 2441 | 120.1 | 0.0001281 | 1.067 | +1 | 1 |
| - | - | 1912 | 120.1 | - | - | 0 | - |
| - | - | 347.4 | 124.7 | - | - | 0 | - |
| - | - | 734.1 | 128.1 | - | - | 0 | - |
| - | - | 1835 | 129.1 | - | - | 0 | - |
| - | - | 577.8 | 131.1 | - | - | 0 | - |
| - | - | 497.8 | 142.1 | - | - | 0 | - |
| - | - | 491.6 | 158.1 | - | - | 0 | - |
| - | - | 463.4 | 170.7 | - | - | 0 | - |
| - | - | 467.6 | 173.1 | - | - | 0 | - |
| - | - | 1065 | 173.5 | - | - | 0 | - |
| - | - | 488.1 | 175.1 | - | - | 0 | - |
| - | - | 513.5 | 180.7 | - | - | 0 | - |
| - | - | 780.7 | 183.1 | - | - | 0 | - |
| - | - | 1013 | 185.1 | - | - | 0 | - |
| 12 | w | 1523 | 187.1 | 2.225E-05 | 0.1189 | +2 | 4 |
| - | - | 503.4 | 191.1 | - | - | 0 | - |
| 14 | y | 1226 | 191.1 | 9.434E-05 | 0.4937 | +1 | 2 |
| - | - | 493.9 | 191.8 | - | - | 0 | - |
| - | - | 1921 | 193.1 | - | - | 0 | - |
| - | - | 523.2 | 201.1 | - | - | 0 | - |
| - | - | 646.8 | 201.1 | - | - | 0 | - |
| - | - | 2947 | 212.1 | - | - | 0 | - |
| - | - | 762.1 | 221.1 | - | - | 0 | - |
| - | - | 901.5 | 227.1 | - | - | 0 | - |
| - | - | 574.7 | 228.1 | - | - | 0 | - |
| - | - | 5243 | 233.2 | - | - | 0 | - |
| - | - | 641.5 | 234.2 | - | - | 0 | - |
| 9 | y | 547.3 | 243.1 | 0.004436 | 18.25 | +3 | 7 |
| - | - | 664 | 245.1 | - | - | 0 | - |
| - | - | 942.8 | 254.1 | - | - | 0 | - |
| - | - | 862.7 | 256.2 | - | - | 0 | - |
| - | - | 3515 | 261.2 | - | - | 0 | - |
| - | - | 839.2 | 263.1 | - | - | 0 | - |
| - | - | 1545 | 282.1 | - | - | 0 | - |
| - | - | 1.296E+04 | 299.2 | - | - | 0 | - |
| - | - | 480.2 | 299.6 | - | - | 0 | - |
| - | - | 1632 | 300.2 | - | - | 0 | - |
| 13 | z | 2705 | 303.2 | 0.0004971 | 1.64 | +1 | 3 |
| - | - | 979.5 | 304.2 | - | - | 0 | - |
| - | - | 672.1 | 309.6 | - | - | 0 | - |
| - | - | 699.2 | 313.7 | - | - | 0 | - |
| - | - | 848.1 | 314.2 | - | - | 0 | - |
| 13 | y | 1068 | 319.2 | 0.001287 | 4.031 | +1 | 3 |
| - | - | 695.7 | 325.2 | - | - | 0 | - |
| - | - | 508.4 | 334.7 | - | - | 0 | - |
| - | - | 1394 | 341.1 | - | - | 0 | - |
| - | - | 582.7 | 341.2 | - | - | 0 | - |
| - | - | 560.2 | 343.2 | - | - | 0 | - |
| - | - | 976.9 | 358.2 | - | - | 0 | - |
| - | - | 1335 | 359.2 | - | - | 0 | - |
| - | - | 1554 | 369.2 | - | - | 0 | - |
| 9 | y | 3310 | 373.2 | 0.005587 | 14.97 | +2 | 7 |
| - | - | 581.8 | 374.8 | - | - | 0 | - |
| - | - | 7630 | 377.2 | - | - | 0 | - |
| - | - | 977.9 | 378.2 | - | - | 0 | - |
| - | - | 8035 | 386.2 | - | - | 0 | - |
| - | - | 1173 | 387.2 | - | - | 0 | - |
| - | - | 766.5 | 387.2 | - | - | 0 | - |
| - | - | 1.162E+04 | 405.2 | - | - | 0 | - |
| - | - | 1512 | 406.2 | - | - | 0 | - |
| - | - | 1019 | 418.2 | - | - | 0 | - |
| - | - | 590.4 | 423.7 | - | - | 0 | - |
| - | - | 647.4 | 431.2 | - | - | 0 | - |
| 12 | y | 839.4 | 433.2 | 0.0006566 | 1.515 | +1 | 4 |
| - | - | 4011 | 444.2 | - | - | 0 | - |
| - | - | 903.8 | 445.2 | - | - | 0 | - |
| 3 | y | 1430 | 451.9 | 0.0001147 | 0.2539 | +3 | 13 |
| 3 | y | 648.5 | 452.2 | 0.003561 | 7.874 | +3 | 13 |
| - | - | 2024 | 468.3 | - | - | 0 | - |
| - | - | 865.2 | 471.3 | - | - | 0 | - |
| 4 | c | 883 | 472.3 | 0.00167 | 3.536 | +1 | 4 |
| - | - | 1618 | 476.2 | - | - | 0 | - |
| - | - | 1519 | 486.3 | - | - | 0 | - |
| 11 | z | 3072 | 488.3 | 0.0003943 | 0.8075 | +1 | 5 |
| - | - | 1130 | 489.3 | - | - | 0 | - |
| - | - | 946.4 | 497.3 | - | - | 0 | - |
| - | - | 1089 | 503.3 | - | - | 0 | - |
| - | - | 2929 | 504.2 | - | - | 0 | - |
| 11 | y | 1870 | 504.3 | 0.0001639 | 0.3249 | +1 | 5 |
| - | - | 844.4 | 505.2 | - | - | 0 | - |
| - | - | 752.8 | 505.3 | - | - | 0 | - |
| - | - | 562.3 | 508.6 | - | - | 0 | - |
| - | - | 1159 | 515.2 | - | - | 0 | - |
| - | - | 635 | 515.3 | - | - | 0 | - |
| - | - | 923 | 528.3 | - | - | 0 | - |
| - | - | 762.6 | 543.3 | - | - | 0 | - |
| - | - | 901.8 | 544.3 | - | - | 0 | - |
| - | - | 591.1 | 545.3 | - | - | 0 | - |
| - | - | 646.5 | 545.8 | - | - | 0 | - |
| - | - | 946.7 | 546.3 | - | - | 0 | - |
| - | - | 863.7 | 546.3 | - | - | 0 | - |
| - | - | 1835 | 551.3 | - | - | 0 | - |
| - | - | 871.4 | 555.3 | - | - | 0 | - |
| 10 | w | 4343 | 558.3 | 0.0004216 | 0.7551 | +1 | 6 |
| - | - | 1073 | 559.3 | - | - | 0 | - |
| 5 | c | 1547 | 559.3 | 0.001031 | 1.843 | +1 | 5 |
| - | - | 682.7 | 560.3 | - | - | 0 | - |
| - | - | 2536 | 569.3 | - | - | 0 | - |
| - | - | 795 | 570.3 | - | - | 0 | - |
| - | - | 1675 | 587.3 | - | - | 0 | - |
| - | - | 758.5 | 597.3 | - | - | 0 | - |
| 11 | c | 4608 | 599.8 | 0.002313 | 3.856 | +2 | 11 |
| 11 | c | 2513 | 600.3 | 0.007632 | 12.71 | +2 | 11 |
| - | - | 1207 | 600.8 | - | - | 0 | - |
| - | - | 655.4 | 603.4 | - | - | 0 | - |
| - | - | 4152 | 604.4 | - | - | 0 | - |
| - | - | 1505 | 605.3 | - | - | 0 | - |
| - | - | 1316 | 605.4 | - | - | 0 | - |
| 10 | z | 3755 | 616.3 | 0.0005966 | 0.968 | +1 | 6 |
| - | - | 981.2 | 617.3 | - | - | 0 | - |
| - | - | 808 | 617.8 | - | - | 0 | - |
| - | - | 1323 | 618.3 | - | - | 0 | - |
| - | - | 3001 | 626.8 | - | - | 0 | - |
| - | - | 2752 | 627.3 | - | - | 0 | - |
| - | - | 1301 | 627.3 | - | - | 0 | - |
| - | - | 893.4 | 627.8 | - | - | 0 | - |
| - | - | 1537 | 632.4 | - | - | 0 | - |
| - | - | 1305 | 635.3 | - | - | 0 | - |
| - | - | 1384 | 635.8 | - | - | 0 | - |
| 6 | c | 7172 | 646.4 | 0.0001911 | 0.2957 | +1 | 6 |
| - | - | 3792 | 647.4 | - | - | 0 | - |
| - | - | 1477 | 650.4 | - | - | 0 | - |
| - | - | 839.8 | 651.4 | - | - | 0 | - |
| - | - | 1004 | 655.3 | - | - | 0 | - |
| - | - | 825.5 | 655.8 | - | - | 0 | - |
| 12 | c | 1061 | 656.8 | 0.0008204 | 1.249 | +2 | 12 |
| - | - | 833.1 | 659.4 | - | - | 0 | - |
| - | - | 688.7 | 664.3 | - | - | 0 | - |
| - | - | 1030 | 664.8 | - | - | 0 | - |
| - | - | 1396 | 665.3 | - | - | 0 | - |
| - | - | 3351 | 668.4 | - | - | 0 | - |
| - | - | 655.9 | 669.4 | - | - | 0 | - |
| - | - | 1088 | 670.4 | - | - | 0 | - |
| 3 | y | 1891 | 677.3 | 0.0009112 | 1.345 | +2 | 13 |
| 3 | y | 1194 | 677.8 | 0.00695 | 10.25 | +2 | 13 |
| 3 | z | 1101 | 678.3 | 0.001357 | 2 | +2 | 13 |
| - | - | 1518 | 681.8 | - | - | 0 | - |
| - | - | 1224 | 682.3 | - | - | 0 | - |
| - | - | 1536 | 685.8 | - | - | 0 | - |
| 3 | y | 2.232E+04 | 686.3 | 0.001183 | 1.724 | +2 | 13 |
| - | - | 2648 | 686.4 | - | - | 0 | - |
| - | - | 1.185E+04 | 686.8 | - | - | 0 | - |
| - | - | 9078 | 687.3 | - | - | 0 | - |
| - | - | 1064 | 687.4 | - | - | 0 | - |
| - | - | 1432 | 687.8 | - | - | 0 | - |
| - | - | 1025 | 692.3 | - | - | 0 | - |
| - | - | 784.7 | 692.8 | - | - | 0 | - |
| - | - | 2338 | 696.4 | - | - | 0 | - |
| - | - | 737.8 | 697.4 | - | - | 0 | - |
| - | - | 786.6 | 698.4 | - | - | 0 | - |
| - | - | 1203 | 704.4 | - | - | 0 | - |
| - | - | 2988 | 714.4 | - | - | 0 | - |
| - | - | 840.8 | 715.4 | - | - | 0 | - |
| 9 | z | 4992 | 729.4 | 0.0002489 | 0.3413 | +1 | 7 |
| 13 | c | 9522 | 729.9 | 0.000674 | 0.9234 | +2 | 13 |
| - | - | 9350 | 730.4 | - | - | 0 | - |
| - | - | 3668 | 730.9 | - | - | 0 | - |
| - | - | 1135 | 731.4 | - | - | 0 | - |
| 2 | z | 1502 | 742.9 | 0.002632 | 3.544 | +2 | 14 |
| 2 | z | 1113 | 743.4 | 0.007313 | 9.838 | +2 | 14 |
| - | - | 1086 | 743.9 | - | - | 0 | - |
| - | - | 770.6 | 744.4 | - | - | 0 | - |
| 9 | y | 1125 | 745.4 | 0.001291 | 1.731 | +1 | 7 |
| - | - | 711.8 | 749.4 | - | - | 0 | - |
| 2 | z | 2366 | 751.9 | 0.0008958 | 1.191 | +2 | 14 |
| - | - | 1018 | 752.4 | - | - | 0 | - |
| - | - | 719.5 | 752.9 | - | - | 0 | - |
| 14 | c | 1096 | 756.9 | 0.005701 | 7.533 | +2 | 14 |
| - | - | 953.5 | 757.4 | - | - | 0 | - |
| - | - | 666.1 | 757.9 | - | - | 0 | - |
| - | - | 900.1 | 758.4 | - | - | 0 | - |
| - | - | 1037 | 758.9 | - | - | 0 | - |
| - | - | 1302 | 764.9 | - | - | 0 | - |
| 14 | c | 1.291E+04 | 765.4 | 0.0002489 | 0.3252 | +2 | 14 |
| - | - | 1.055E+04 | 765.9 | - | - | 0 | - |
| - | - | 7797 | 766.4 | - | - | 0 | - |
| - | - | 1615 | 766.9 | - | - | 0 | - |
| - | - | 960.8 | 767.4 | - | - | 0 | - |
| - | - | 861.5 | 772.4 | - | - | 0 | - |
| 7 | c | 1.553E+04 | 775.4 | 0.0001656 | 0.2136 | +1 | 7 |
| - | - | 6481 | 776.4 | - | - | 0 | - |
| - | - | 2442 | 777.4 | - | - | 0 | - |
| - | - | 2553 | 779.4 | - | - | 0 | - |
| - | - | 2318 | 779.9 | - | - | 0 | - |
| - | - | 873.7 | 780.4 | - | - | 0 | - |
| - | - | 2202 | 780.9 | - | - | 0 | - |
| - | - | 2461 | 781.4 | - | - | 0 | - |
| - | - | 910.7 | 781.9 | - | - | 0 | - |
| - | - | 905.5 | 785.4 | - | - | 0 | - |
| - | - | 1013 | 786.4 | - | - | 0 | - |
| - | - | 707.2 | 786.9 | - | - | 0 | - |
| - | - | 1861 | 787.4 | - | - | 0 | - |
| - | - | 1047 | 788.4 | - | - | 0 | - |
| - | - | 832.2 | 789.4 | - | - | 0 | - |
| - | - | 3655 | 794.4 | - | - | 0 | - |
| - | - | 4841 | 794.9 | - | - | 0 | - |
| - | - | 2172 | 795.4 | - | - | 0 | - |
| - | - | 1154 | 795.9 | - | - | 0 | - |
| 8 | w | 2970 | 799.4 | 0.005429 | 6.791 | +1 | 8 |
| - | - | 1701 | 801.4 | - | - | 0 | - |
| - | - | 963.9 | 802.4 | - | - | 0 | - |
| - | - | 2378 | 807.4 | - | - | 0 | - |
| - | - | 3472 | 807.9 | - | - | 0 | - |
| - | - | 2283 | 808.4 | - | - | 0 | - |
| - | - | 1658 | 808.9 | - | - | 0 | - |
| - | - | 908.6 | 809.9 | - | - | 0 | - |
| - | - | 1496 | 815.9 | - | - | 0 | - |
| - | - | 1.356E+04 | 816.4 | - | - | 0 | - |
| - | - | 2.334E+04 | 816.9 | - | - | 0 | - |
| - | - | 1.659E+04 | 817.4 | - | - | 0 | - |
| - | - | 8395 | 817.9 | - | - | 0 | - |
| - | - | 3001 | 818.4 | - | - | 0 | - |
| - | - | 1748 | 828.5 | - | - | 0 | - |
| - | - | 710.6 | 829.4 | - | - | 0 | - |
| - | - | 723.4 | 856.4 | - | - | 0 | - |
| 8 | z | 7691 | 858.4 | 0.0007888 | 0.9189 | +1 | 8 |
| - | - | 3317 | 859.4 | - | - | 0 | - |
| - | - | 791 | 860.4 | - | - | 0 | - |
| - | - | 890.9 | 874.4 | - | - | 0 | - |
| - | - | 1231 | 875.4 | - | - | 0 | - |
| - | - | 1045 | 886.5 | - | - | 0 | - |
| - | - | 827.3 | 887.4 | - | - | 0 | - |
| - | - | 1364 | 887.5 | - | - | 0 | - |
| - | - | 810.8 | 890.4 | - | - | 0 | - |
| 8 | c | 1.522E+04 | 904.4 | 3.413E-05 | 0.03774 | +1 | 8 |
| - | - | 8437 | 905.4 | - | - | 0 | - |
| - | - | 2309 | 906.4 | - | - | 0 | - |
| - | - | 668.4 | 907.4 | - | - | 0 | - |
| - | - | 5909 | 915.5 | - | - | 0 | - |
| - | - | 1948 | 916.5 | - | - | 0 | - |
| 7 | w | 1865 | 928.5 | 0.002184 | 2.353 | +1 | 9 |
| - | - | 1683 | 929.5 | - | - | 0 | - |
| - | - | 1320 | 943.4 | - | - | 0 | - |
| - | - | 944.1 | 944.5 | - | - | 0 | - |
| - | - | 589.7 | 952.4 | - | - | 0 | - |
| - | - | 807 | 961.5 | - | - | 0 | - |
| - | - | 5326 | 973.5 | - | - | 0 | - |
| - | - | 1614 | 974.5 | - | - | 0 | - |
| - | - | 1755 | 974.6 | - | - | 0 | - |
| 7 | z | 1.008E+04 | 987.5 | 0.0004413 | 0.4469 | +1 | 9 |
| - | - | 6056 | 988.5 | - | - | 0 | - |
| - | - | 1857 | 989.5 | - | - | 0 | - |
| - | - | 780.4 | 990.5 | - | - | 0 | - |
| - | - | 7147 | 1003 | - | - | 0 | - |
| - | - | 2717 | 1004 | - | - | 0 | - |
| - | - | 1596 | 1005 | - | - | 0 | - |
| - | - | 779.1 | 1006 | - | - | 0 | - |
| 9 | c | 8869 | 1018 | 0.0001304 | 0.1282 | +1 | 9 |
| - | - | 6207 | 1019 | - | - | 0 | - |
| - | - | 1469 | 1020 | - | - | 0 | - |
| - | - | 3348 | 1046 | - | - | 0 | - |
| - | - | 1756 | 1047 | - | - | 0 | - |
| - | - | 886.8 | 1057 | - | - | 0 | - |
| - | - | 612.6 | 1073 | - | - | 0 | - |
| - | - | 1581 | 1074 | - | - | 0 | - |
| 6 | z | 7662 | 1075 | 0.007598 | 7.071 | +1 | 10 |
| - | - | 6739 | 1076 | - | - | 0 | - |
| - | - | 2471 | 1077 | - | - | 0 | - |
| - | - | 1067 | 1078 | - | - | 0 | - |
| - | - | 1078 | 1089 | - | - | 0 | - |
| - | - | 2326 | 1090 | - | - | 0 | - |
| - | - | 695.4 | 1090 | - | - | 0 | - |
| 6 | y | 7543 | 1091 | 0.007672 | 7.035 | +1 | 10 |
| - | - | 1.751E+04 | 1092 | - | - | 0 | - |
| - | - | 8589 | 1093 | - | - | 0 | - |
| - | - | 800.4 | 1102 | - | - | 0 | - |
| - | - | 1453 | 1145 | - | - | 0 | - |
| 10 | c | 1.606E+04 | 1146 | 0.0002196 | 0.1917 | +1 | 10 |
| - | - | 9346 | 1147 | - | - | 0 | - |
| - | - | 4063 | 1148 | - | - | 0 | - |
| - | - | 1056 | 1149 | - | - | 0 | - |
| 5 | z | 2036 | 1162 | 0.006975 | 6.005 | +1 | 11 |
| - | - | 2638 | 1163 | - | - | 0 | - |
| - | - | 1373 | 1164 | - | - | 0 | - |
| 5 | y | 943 | 1178 | 0.002988 | 2.537 | +1 | 11 |
| - | - | 899.1 | 1216 | - | - | 0 | - |
| 11 | c | 9493 | 1217 | 0.0003461 | 0.2844 | +1 | 11 |
| - | - | 7691 | 1218 | - | - | 0 | - |
| - | - | 3786 | 1219 | - | - | 0 | - |
| - | - | 884.2 | 1220 | - | - | 0 | - |
| 4 | y | 1391 | 1275 | 0.001859 | 1.459 | +1 | 12 |
| - | - | 1527 | 1276 | - | - | 0 | - |
| - | - | 1603 | 1330 | - | - | 0 | - |
| 12 | c | 1.257E+04 | 1331 | 0.001892 | 1.422 | +1 | 12 |
| - | - | 8958 | 1332 | - | - | 0 | - |
| - | - | 3298 | 1333 | - | - | 0 | - |
| 3 | y | 820.2 | 1372 | 0.0008555 | 0.6237 | +1 | 13 |
| - | - | 963.9 | 1373 | - | - | 0 | - |
| - | - | 809.1 | 1387 | - | - | 0 | - |
| - | - | 1207 | 1415 | - | - | 0 | - |
| - | - | 1337 | 1416 | - | - | 0 | - |
| 13 | c | 1946 | 1459 | 0.01653 | 11.33 | +1 | 13 |
| - | - | 4522 | 1460 | - | - | 0 | - |
| - | - | 2972 | 1461 | - | - | 0 | - |
| - | - | 1419 | 1462 | - | - | 0 | - |
| - | - | 1675 | 1474 | - | - | 0 | - |
| - | - | 1142 | 1475 | - | - | 0 | - |
| - | - | 1234 | 1486 | - | - | 0 | - |
| - | - | 1037 | 1487 | - | - | 0 | - |
| - | - | 1113 | 1488 | - | - | 0 | - |
| 2 | z | 1642 | 1503 | 0.001351 | 0.8991 | +1 | 14 |
| - | - | 5395 | 1504 | - | - | 0 | - |
| - | - | 4259 | 1505 | - | - | 0 | - |
| - | - | 1688 | 1506 | - | - | 0 | - |
| - | - | 992.5 | 1514 | - | - | 0 | - |
| 14 | c | 762.6 | 1530 | 0.01043 | 6.82 | +1 | 14 |
| - | - | 4290 | 1531 | - | - | 0 | - |
| - | - | 3375 | 1532 | - | - | 0 | - |
| - | - | 1979 | 1533 | - | - | 0 | - |
| - | - | 1143 | 1571 | - | - | 0 | - |
| - | - | 1699 | 1572 | - | - | 0 | - |
| - | - | 816.4 | 1573 | - | - | 0 | - |
| - | - | 1165 | 1588 | - | - | 0 | - |
| - | - | 6590 | 1589 | - | - | 0 | - |
| - | - | 6021 | 1590 | - | - | 0 | - |
| - | - | 2220 | 1591 | - | - | 0 | - |
| - | - | 937.6 | 1592 | - | - | 0 | - |
| - | - | 1018 | 1606 | - | - | 0 | - |
| - | - | 864.8 | 1607 | - | - | 0 | - |
| - | - | 1173 | 1614 | - | - | 0 | - |
| - | - | 2980 | 1615 | - | - | 0 | - |
| - | - | 5379 | 1616 | - | - | 0 | - |
| - | - | 1.211E+04 | 1617 | - | - | 0 | - |
| - | - | 9861 | 1618 | - | - | 0 | - |
| - | - | 4944 | 1619 | - | - | 0 | - |
| - | - | 1636 | 1620 | - | - | 0 | - |
| - | - | 738.6 | 1621 | - | - | 0 | - |
| - | - | 1572 | 1632 | - | - | 0 | - |
| - | - | 6771 | 1633 | - | - | 0 | - |
| - | - | 1.943E+04 | 1634 | - | - | 0 | - |
| - | - | 1.347E+04 | 1635 | - | - | 0 | - |
| - | - | 7712 | 1636 | - | - | 0 | - |
| - | - | 2953 | 1637 | - | - | 0 | - |
| - | - | 722.9 | 2505 | - | - | 0 | - |
| - | - | 698.3 | 2791 | - | - | 0 | - |
| - | - | 779.7 | 3161 | - | - | 0 | - |

m/z Charge Intensity FragmentType MassShift Position
120.06539154052734 0 2440.646 y 14
120.08076477050781 0 1912.4932
124.66665649414062 0 347.41397
128.10696411132812 0 734.1074
129.10220336914062 0 1834.9635
131.11773681640625 0 577.7673
142.0978546142578 0 497.80228
158.09295654296875 0 491.58878
170.72450256347656 0 463.36234
173.1287078857422 0 467.5505
173.45188903808594 0 1064.958
175.07130432128906 0 488.10892
180.7196807861328 0 513.51044
183.1129913330078 0 780.65576
185.0919189453125 0 1013.4069
187.10769653320312 0 1523.322 w 11
191.09327697753906 0 503.4357
191.1025390625 0 1225.8329 y 13
191.776123046875 0 493.91852
193.08180236816406 0 1921.166
201.0967559814453 0 523.18445
201.12356567382812 0 646.8487
212.13914489746094 0 2946.6743
221.0765838623047 0 762.1154
227.10247802734375 0 901.4579
228.13485717773438 0 574.7484
233.16468811035156 0 5243.3174
234.1677703857422 0 641.52014
243.14585876464844 0 547.3083 y Water loss 8
245.12490844726562 0 664.03723
254.1136932373047 0 942.77216
256.1656799316406 0 862.6682
261.1595458984375 0 3515.2341
263.13958740234375 0 839.1844
282.1445617675781 0 1545.4241
299.1712341308594 0 12955.1455
299.5751037597656 0 480.2013
300.1739196777344 0 1632.086
303.1783752441406 0 2704.9282 z 12
304.18359375 0 979.4756
309.5939636230469 0 672.11615
313.6622314453125 0 699.1546
314.1812744140625 0 848.10876
319.1988830566406 0 1068.2474 y 12
325.1862487792969 0 695.6909
334.7031555175781 0 508.43024
341.1455993652344 0 1393.5425
341.21722412109375 0 582.67737
343.1978454589844 0 560.22046
358.21173095703125 0 976.9033
359.1562805175781 0 1335.2239
369.2116394042969 0 1554.2551
373.20819091796875 0 3309.5557 y 8
374.83819580078125 0 581.75964
377.1661682128906 0 7629.9746
378.1675109863281 0 977.91705
386.20330810546875 0 8034.6733
387.2052001953125 0 1173.1611
387.2310485839844 0 766.53265
405.1614685058594 0 11621.153
406.1632385253906 0 1511.9454
418.2292175292969 0 1019.4299
423.7115783691406 0 590.3507
431.2279968261719 0 647.4409
433.2411804199219 0 839.3964 y 11
444.24542236328125 0 4011.0056
445.2468566894531 0 903.77374
451.89312744140625 0 1430.3838 y Water loss 2
452.2245788574219 0 648.5027 y Ammonia loss 2
468.2814636230469 0 2024.4478
471.28125 0 865.24207
472.2901611328125 0 882.9507 c 3
476.2353515625 0 1617.6833
486.29193115234375 0 1519.2291
488.2593078613281 0 3071.742 z 10
489.26141357421875 0 1130.3251
497.2982482910156 0 946.3929
503.3119201660156 0 1088.9901
504.2300109863281 0 2928.9197
504.2778015136719 0 1870.0671 y 10
505.2313537597656 0 844.4164
505.2789306640625 0 752.7951
508.5723571777344 0 562.3368
515.24560546875 0 1158.6025
515.2926635742188 0 635.03326
528.27294921875 0 923.0324
543.3148193359375 0 762.61664
544.3172607421875 0 901.81805
545.2974853515625 0 591.0813
545.7922973632812 0 646.4613
546.2954711914062 0 946.719
546.3468627929688 0 863.6618
551.317626953125 0 1835.179
555.3138427734375 0 871.3678
558.2877807617188 0 4342.73 w 9
559.284423828125 0 1072.8488
559.3248901367188 0 1547.2897 c 4
560.3268432617188 0 682.68097
569.3285522460938 0 2535.793
570.330078125 0 794.98914
587.34130859375 0 1674.7625
597.3250732421875 0 758.5379
599.8064575195312 0 4608.2407 c Water loss 10
600.3084106445312 0 2513.2485 c Ammonia loss 10
600.8084716796875 0 1206.6187
603.35595703125 0 655.3696
604.3660888671875 0 4151.792
605.2772216796875 0 1505.0695
605.3699951171875 0 1315.7505
616.31689453125 0 3754.522 z 9
617.3194580078125 0 981.1721
617.8062133789062 0 807.9857
618.306396484375 0 1323.1587
626.8124389648438 0 3000.5095
627.2621459960938 0 2752.1333
627.3152465820312 0 1300.5632
627.8174438476562 0 893.36456
632.3580322265625 0 1536.8301
635.3245239257812 0 1305.0844
635.826416015625 0 1383.9237
646.3560791015625 0 7172.4253 c 5
647.3584594726562 0 3791.879
650.3875732421875 0 1477.1846
651.388427734375 0 839.7511
655.3399658203125 0 1003.6376
655.837158203125 0 825.504
656.8310546875 0 1061.0767 c Water loss 11
659.4199829101562 0 833.1398
664.32763671875 0 688.70123
664.836669921875 0 1029.5951
665.3416137695312 0 1396.0187
668.3969116210938 0 3351.1902
669.4020385742188 0 655.8636
670.3943481445312 0 1087.7454
677.3367919921875 0 1890.9684 y Water loss 2
677.8348388671875 0 1193.9417 y Ammonia loss 2
678.3304443359375 0 1100.7292 z 2
681.8316040039062 0 1518.0929
682.3319702148438 0 1224.0435
685.8383178710938 0 1535.7596
686.3423461914062 0 22315.676 y 2
686.4071044921875 0 2648.0334
686.84228515625 0 11847.778
687.3453979492188 0 9077.603
687.4116821289062 0 1064.0831
687.8458251953125 0 1432.3422
692.3372802734375 0 1025.1237
692.8367309570312 0 784.6914
696.393798828125 0 2338.3025
697.3984375 0 737.7565
698.3870849609375 0 786.5948
704.3504638671875 0 1202.8265
714.4029541015625 0 2988.4363
715.4043579101562 0 840.8141
729.4013061523438 0 4992.38 z 8
729.88232421875 0 9521.68 c 12
730.3856201171875 0 9349.657
730.8836059570312 0 3667.738
731.3876342773438 0 1134.802
742.8580932617188 0 1502.4406 z Water loss 1
743.3600463867188 0 1112.5801 z Ammonia loss 1
743.9003295898438 0 1086.1193
744.4019165039062 0 770.5917
745.4215698242188 0 1124.614 y 8
749.3801879882812 0 711.7845
751.8651123046875 0 2366.436 z 1
752.370361328125 0 1017.91223
752.8684692382812 0 719.48956
756.8939819335938 0 1096.2959 c Ammonia loss 13
757.4140014648438 0 953.5033
757.8944091796875 0 666.1065
758.3916625976562 0 900.1361
758.8949584960938 0 1036.5581
764.9038696289062 0 1301.6569
765.4013061523438 0 12907.373 c 13
765.9033813476562 0 10547.173
766.4044189453125 0 7796.769
766.9029541015625 0 1614.7758
767.410888671875 0 960.7586
772.401611328125 0 861.54376
775.3983154296875 0 15531.743 c 6
776.4024658203125 0 6480.6274
777.4033203125 0 2441.6606
779.3921508789062 0 2553.0774
779.8939819335938 0 2318.362
780.3922729492188 0 873.7371
780.9059448242188 0 2201.6016
781.412109375 0 2460.9106
781.9058227539062 0 910.70764
785.4140625 0 905.47095
786.4058227539062 0 1012.5909
786.8782958984375 0 707.1611
787.4022827148438 0 1861.4122
788.4033203125 0 1047.3915
789.3883056640625 0 832.17316
794.4115600585938 0 3654.5671
794.9134521484375 0 4841.379
795.4139404296875 0 2172.345
795.9133911132812 0 1154.4395
799.4254150390625 0 2969.9087 w 7
801.414306640625 0 1701.39
802.3799438476562 0 963.87054
807.4222412109375 0 2378.2803
807.9180908203125 0 3471.712
808.412353515625 0 2282.659
808.9152221679688 0 1658.1012
809.9071655273438 0 908.582
815.9197387695312 0 1496.2849
816.4178466796875 0 13560.453
816.9197998046875 0 23340.47
817.4215087890625 0 16589.375
817.9210815429688 0 8395.35
818.4244995117188 0 3000.9578
828.4876098632812 0 1747.7518
829.4246826171875 0 710.58026
856.4137573242188 0 723.4177
858.443359375 0 7691.141 z 7
859.4465942382812 0 3317.0996
860.4465942382812 0 791.0253
874.4307861328125 0 890.93823
875.4418334960938 0 1230.824
886.4613037109375 0 1045.3064
887.4105224609375 0 827.2796
887.51904296875 0 1364.1597
890.449951171875 0 810.81793
904.4410400390625 0 15222.993 c 7
905.4437255859375 0 8436.972
906.4464111328125 0 2308.9363
907.44580078125 0 668.3778
915.5142822265625 0 5909.145
916.5140380859375 0 1947.6982
928.4712524414062 0 1864.7172 w 6
929.4735717773438 0 1683.2303
943.4483032226562 0 1320.4275
944.4581298828125 0 944.06586
952.385498046875 0 589.737
961.4616088867188 0 806.98883
973.496337890625 0 5325.5312
974.4873657226562 0 1614.2312
974.55908203125 0 1755.4314
987.4871826171875 0 10084.235 z 6
988.4895629882812 0 6055.885
989.4908447265625 0 1856.9843
990.4959716796875 0 780.4418
1002.544677734375 0 7146.934
1003.5375366210938 0 2717.1665
1004.5042724609375 0 1595.727
1006.4412231445312 0 779.1208
1017.5252685546875 0 8869.483 c 8
1018.52880859375 0 6207.463
1019.5277709960938 0 1468.8933
1045.56396484375 0 3348.1985
1046.5703125 0 1756.1886
1056.538818359375 0 886.7902
1072.502197265625 0 612.64386
1073.556396484375 0 1580.7457
1074.5263671875 0 7661.687 z 5
1075.526123046875 0 6739.3228
1076.525634765625 0 2471.3062
1077.5198974609375 0 1067.1117
1088.5479736328125 0 1077.9486
1089.5322265625 0 2326.4546
1090.011962890625 0 695.3506
1090.545166015625 0 7542.7656 y 5
1091.565185546875 0 17513.982
1092.5677490234375 0 8588.855
1101.5643310546875 0 800.4491
1144.5885009765625 0 1453.0021
1145.58349609375 0 16064.155 c 9
1146.5867919921875 0 9346.437
1147.5889892578125 0 4062.986
1148.59521484375 0 1055.6747
1161.5438232421875 0 2036.0896 z 4
1162.5523681640625 0 2638.2793
1163.5555419921875 0 1373.022
1177.572509765625 0 942.9803 y 4
1215.6011962890625 0 899.0961
1216.6204833984375 0 9493.197 c 10
1217.622314453125 0 7691.2896
1218.6253662109375 0 3785.803
1219.6297607421875 0 884.23755
1274.6241455078125 0 1390.937 y 3
1275.62353515625 0 1526.8359
1329.642578125 0 1602.9763
1330.661865234375 0 12569.81 c 11
1331.6629638671875 0 8957.894
1332.67138671875 0 3297.8499
1371.6741943359375 0 820.2286 y 2
1372.6834716796875 0 963.93427
1386.7144775390625 0 809.0765
1414.7325439453125 0 1206.7639
1415.7376708984375 0 1336.6918
1458.7421875 0 1945.5273 c 12
1459.7626953125 0 4522.235
1460.7607421875 0 2972.287
1461.7738037109375 0 1418.8086
1473.7459716796875 0 1675.1073
1474.7564697265625 0 1142.3492
1485.76904296875 0 1234.1531
1486.7796630859375 0 1037.4167
1487.7955322265625 0 1112.9453
1502.723388671875 0 1641.56 z 1
1503.7313232421875 0 5395.342
1504.735107421875 0 4258.857
1505.730712890625 0 1687.7117
1513.7529296875 0 992.5404
1529.785400390625 0 762.62463 c 13
1530.798095703125 0 4289.8296
1531.80322265625 0 3374.5334
1532.8056640625 0 1979.0812
1570.8050537109375 0 1143.1615
1571.802490234375 0 1698.8964
1572.802001953125 0 816.4139
1587.8292236328125 0 1164.7877
1588.8193359375 0 6589.747
1589.8240966796875 0 6020.8335
1590.8228759765625 0 2220.3677
1591.830078125 0 937.5875
1605.8466796875 0 1017.791
1606.84228515625 0 864.75745
1613.8326416015625 0 1173.4824
1614.841552734375 0 2980.2004
1615.83154296875 0 5379.0537
1616.817626953125 0 12106.9
1617.8155517578125 0 9860.93
1618.8226318359375 0 4943.549
1619.8148193359375 0 1636.1437
1620.800048828125 0 738.5758
1631.8173828125 0 1571.6871
1632.8330078125 0 6770.9062
1633.8389892578125 0 19428.268
1634.8431396484375 0 13465.39
1635.84716796875 0 7712.1196
1636.8486328125 0 2953.186
2504.835693359375 0 722.86487
2791.415283203125 0 698.3109
3160.68408203125 0 779.6919

Spectrum Details

|  |  |
| --- | --- |
| Matched peaks? Matched peaks The total absolute number of peaks matched. Additionally in brackets the total fraction of peaks matched and the total number of peaks is shown. | 51 (15.13% of 337) |
| FDR? FDR The false discovery rate estimated for this peptide. It is calculated by matching all theoretical fragments with a non-integer shift with the raw peaks for this spectrum. This is done with 40 different shifts. The resulting percentage is the average number of annotated peaks over the number of annotated peaks with the correct spectrum. | 0.23% |
| Satellite FDR? Satellite FDR See the FDR for details on its calculation. This satellite ion specific FDR only contains the satellite ions (d/w) for I/L/J positions. | - |
| PSM Score? PSM Score The PSM Score as given by Hecklib to this annotated spectrum. It is shown with three significant figures. | 337 |

#### Reverse Lookup? Reverse Lookup All places where this read could be placed.

| Group | Segment | Template | Template Part | Read Part | Score | Unique |
| --- | --- | --- | --- | --- | --- | --- |
| Homo sapiens Light Chain | IGLC | IGLC2 | [10..25] | [0..15] | 120 | False |
| Homo sapiens Light Chain | IGLC | IGLC3 | [8..23] | [0..15] | 120 | False |
| Homo sapiens Light Chain | IGLC | IGLC6 | [10..25] | [0..15] | 120 | False |
| Homo sapiens Light Chain | IGLC | IGLC7 | [10..25] | [0..15] | 120 | False |

| Recombined | Template Part | Read Part | Score | Unique |
| --- | --- | --- | --- | --- |
| REC-0-1\_002 | [121..136] | [0..15] | 120 | True |

#### Meta Information from Multiple reads

##### Number of combined reads

4

##### Intensity

0.7103

##### TotalArea

2.893E+08

##### Changes to the peptide sequence

JFPPSSEEJQANKAT

L→JNo support for either Leucine or Isoleucine based on side chain ions (Position: 9)

L→JNo support for either Leucine or Isoleucine based on side chain ions (Position: 1)

#### Positional Score

Copy Data

##### Positional Score (TSV)

###### Preview

```
Loading example...
```

*Click on the button to copy the data to your clipboard.*

1001234567891011121314

Label Value
"0" 0.49
"1" 0.48
"2" 0.492
"3" 0.495
"4" 0.495
"5" 0.49
"6" 0.495
"7" 0.497
"8" 0.497
"9" 0.487
"10" 0.495
"11" 0.492
"12" 0.497
"13" 0.495
"14" 0.49

#### Meta Information from PEAKS

##### Scan Identifier

F2:6210

##### Original sequence

L

F

P

P

S

S

E

E

L

Q

A

N

K

A

T

##### Posttranslational Modifications

##### Source File

D:\separate\_stitch\_analyses\xle-disambiguation\raw\20210323\_F1\_UM1\_Peng0013\_SA\_F59\_ingel\_3ug\_TL.raw

##### Fraction

2

##### Scan Feature

F2:15589

##### De Novo Score

99

##### ConfidenceScore

99

### m/z

816.4174

##### Mass

1630.8203

##### Charge

2

##### Retention Time

33.88

##### Predicted Retention Time

-

##### Area

9.418E+07

##### Fragmentation mode

ETHCD

##### Originating file

01 D:\separate\_stitch\_analyses\xle-disambiguation\20210325\_F59\_3ug\_DENOVO\_12.csv

#### Meta Information from PEAKS

##### Scan Identifier

F2:6152

##### Original sequence

L

F

P

P

S

S

E

E

L

Q

A

N

K

A

T

##### Posttranslational Modifications

##### Source File

D:\separate\_stitch\_analyses\xle-disambiguation\raw\20210323\_F1\_UM1\_Peng0013\_SA\_F59\_ingel\_3ug\_TL.raw

##### Fraction

2

##### Scan Feature

F2:15589

##### De Novo Score

98

##### ConfidenceScore

98

### m/z

816.4174

##### Mass

1630.8203

##### Charge

2

##### Retention Time

33.88

##### Predicted Retention Time

-

##### Area

9.418E+07

##### Fragmentation mode

ETHCD

##### Originating file

01 D:\separate\_stitch\_analyses\xle-disambiguation\20210325\_F59\_3ug\_DENOVO\_12.csv

#### Meta Information from PEAKS

##### Scan Identifier

F2:6090

##### Original sequence

L

F

P

P

S

S

E

E

L

Q

A

N

K

A

T

##### Posttranslational Modifications

##### Source File

D:\separate\_stitch\_analyses\xle-disambiguation\raw\20210323\_F1\_UM1\_Peng0013\_SA\_F59\_ingel\_3ug\_TL.raw

##### Fraction

2

##### Scan Feature

F2:15589

##### De Novo Score

98

##### ConfidenceScore

98

### m/z

816.4174

##### Mass

1630.8203

##### Charge

2

##### Retention Time

33.88

##### Predicted Retention Time

-

##### Area

9.418E+07

##### Fragmentation mode

ETHCD

##### Originating file

01 D:\separate\_stitch\_analyses\xle-disambiguation\20210325\_F59\_3ug\_DENOVO\_12.csv

#### Meta Information from PEAKS

##### Scan Identifier

F2:6239

##### Original sequence

L

F

P

P

S

S

E

E

L

Q

A

N

K

A

T

##### Posttranslational Modifications

##### Source File

D:\separate\_stitch\_analyses\xle-disambiguation\raw\20210323\_F1\_UM1\_Peng0013\_SA\_F59\_ingel\_3ug\_TL.raw

##### Fraction

2

##### Scan Feature

F2:5997

##### De Novo Score

98

##### ConfidenceScore

98

### m/z

544.6152

##### Mass

1630.8203

##### Charge

3

##### Retention Time

33.72

##### Predicted Retention Time

-

##### Area

6.806E+06

##### Parts Per Million

2.2

##### Fragmentation mode

ETHCD

##### Originating file

01 D:\separate\_stitch\_analyses\xle-disambiguation\20210325\_F59\_3ug\_DENOVO\_12.csv
