## Supplementary material for "A handle on mass coincidence errors in *de novo* sequencing of antibodies by bottom-up proteomics": Combined_038.html

Details Combined\_038 | Stitch OverviewUndefined

### Read Combined\_038

#### Sequence (length=12)

TISRDDAKNSJY

#### Spectrum 3647? Spectrum 3647 The raw spectrum of this peptide as annotated by Hecklib. The fragments are coloured according to ion type (see legend). Any peaks with a star '\*' as text can be hovered over to see the full details, first the ion type second the mass shift type. By hovering over the amino acids in the peptide or ions in the legend the corresponding peaks are highlighted. By toggling the 'Unassigned' label you can turn the background (unassigned) peaks on or off in the plot. By updating the slider in the Ion legend you can update the spectrum to only show the top X% of the peaks with labels. The top X% means any peak that is within X% of the highest intensity. By dragging in the spectrum you can zoom in to a specific part of the spectrum and use 'Zoom Out' to get back to the original zoom level. The annotation of the spectrum is based on the given sequence in the peptides file and is done with different software so inconsistencies are likely. The peaks are annotated based on the given sequence, with 20 ppm tolerance.

Copy Data

##### Spectrum 3647 (TSV)

###### Preview

```
Loading example...
```

*Click on the button to copy the data to your clipboard.*

Mz MinMz MaxIntensity Max

WidthHeightPeptide font sizePeptide stroke widthSpectrum font sizeSpectrum stroke widthCompact peptide

Ion legend

wxyz

abcd

OtherUnassignedIon^Charge^_Position_Show for top:%

TISRDDAKNSJY

04.61e+59.22e+51.38e+61.84e+6

Zoom Out

y^+1^_1_c^+1^_2_c^+2^_5_y^+1^_2_c^+1^_3_c^+2^_6_z^+1^_3_y^+1^_3_c^+3^_11_c^+2^_8_c^+2^_8_c^+2^_8_c^+1^_4_z^+1^_4_y^+2^_8_c^+1^_4_z^+1^_4_y^+1^_4_c^+2^_9_c^+2^_9_c^+2^_9_z^+2^_9_y^+2^_9_c^+2^_10_c^+2^_10_c^+2^_10_z^+2^_10_c^+1^_5_y^+2^_10_y^+2^_10_z^+2^_10_y^+2^_10_c^+1^_5_c^+2^_11_c^+2^_11_z^+1^_5_c^+2^_11_w^+2^_11_y^+1^_5_w^+2^_11_y^+2^_11_z^+2^_11_y^+2^_11_z^+1^_6_z^+1^_6_c^+1^_6_y^+1^_6_c^+1^_6_w^+1^_7_c^+1^_7_c^+1^_7_z^+1^_7_y^+1^_7_c^+1^_8_c^+1^_8_c^+1^_8_z^+1^_8_y^+1^_8_c^+1^_9_c^+1^_9_c^+1^_9_z^+1^_9_y^+1^_9_c^+1^_10_c^+1^_10_z^+1^_10_y^+1^_10_z^+1^_10_y^+1^_10_c^+1^_11_c^+1^_11_z^+1^_11_y^+1^_11_

0588117517632350

Fragment Matches Table

Show background peaks

| Position | Ion type | Intensity | mz Theoretical | mz Error (Th) | mz Error (ppm) | Charge | Series Number |
| --- | --- | --- | --- | --- | --- | --- | --- |
| - | - | 2697 | 123.4 | - | - | 0 | - |
| - | - | 3814 | 129.1 | - | - | 0 | - |
| - | - | 4481 | 131.1 | - | - | 0 | - |
| - | - | 6.25E+04 | 136.1 | - | - | 0 | - |
| - | - | 5958 | 137.1 | - | - | 0 | - |
| - | - | 3478 | 141.6 | - | - | 0 | - |
| - | - | 1.452E+04 | 142.1 | - | - | 0 | - |
| - | - | 8897 | 143.1 | - | - | 0 | - |
| - | - | 4729 | 158.1 | - | - | 0 | - |
| - | - | 6.171E+04 | 165.1 | - | - | 0 | - |
| - | - | 9295 | 166.1 | - | - | 0 | - |
| - | - | 9363 | 169.1 | - | - | 0 | - |
| - | - | 6827 | 173.1 | - | - | 0 | - |
| 12 | y | 2.367E+05 | 182.1 | 0.0004038 | 2.218 | +1 | 1 |
| - | - | 2.744E+04 | 183.1 | - | - | 0 | - |
| - | - | 4404 | 186.1 | - | - | 0 | - |
| - | - | 7.416E+05 | 187.1 | - | - | 0 | - |
| - | - | 6.826E+04 | 188.1 | - | - | 0 | - |
| - | - | 3930 | 189.1 | - | - | 0 | - |
| - | - | 3779 | 199.5 | - | - | 0 | - |
| - | - | 7118 | 200.1 | - | - | 0 | - |
| - | - | 2.8E+04 | 201.1 | - | - | 0 | - |
| - | - | 6197 | 201.1 | - | - | 0 | - |
| - | - | 7675 | 202.1 | - | - | 0 | - |
| - | - | 4572 | 202.1 | - | - | 0 | - |
| - | - | 1.973E+05 | 215.1 | - | - | 0 | - |
| - | - | 1.522E+04 | 216.1 | - | - | 0 | - |
| - | - | 9524 | 229.1 | - | - | 0 | - |
| 2 | c | 1.7E+04 | 232.2 | 0.0006002 | 2.585 | +1 | 2 |
| - | - | 3918 | 235.1 | - | - | 0 | - |
| - | - | 6796 | 244.1 | - | - | 0 | - |
| - | - | 5922 | 272.2 | - | - | 0 | - |
| - | - | 4041 | 274.6 | - | - | 0 | - |
| - | - | 1.163E+04 | 281.1 | - | - | 0 | - |
| - | - | 5.435E+04 | 282.1 | - | - | 0 | - |
| - | - | 8900 | 283.1 | - | - | 0 | - |
| 5 | c | 6565 | 287.2 | 0.001138 | 3.965 | +2 | 5 |
| - | - | 8282 | 287.2 | - | - | 0 | - |
| 11 | y | 3.077E+04 | 295.2 | 0.0006294 | 2.132 | +1 | 2 |
| - | - | 4984 | 296.2 | - | - | 0 | - |
| - | - | 9483 | 297.2 | - | - | 0 | - |
| - | - | 1.729E+04 | 299.1 | - | - | 0 | - |
| - | - | 1.883E+04 | 299.2 | - | - | 0 | - |
| - | - | 1.239E+05 | 300.2 | - | - | 0 | - |
| - | - | 2.046E+04 | 301.2 | - | - | 0 | - |
| - | - | 4427 | 310.1 | - | - | 0 | - |
| - | - | 2.221E+04 | 315.2 | - | - | 0 | - |
| - | - | 8.862E+04 | 317.2 | - | - | 0 | - |
| - | - | 1.184E+04 | 318.2 | - | - | 0 | - |
| 3 | c | 5.678E+04 | 319.2 | 0.0007373 | 2.31 | +1 | 3 |
| - | - | 7286 | 320.2 | - | - | 0 | - |
| - | - | 4479 | 328.8 | - | - | 0 | - |
| - | - | 9068 | 335.7 | - | - | 0 | - |
| - | - | 3999 | 336.2 | - | - | 0 | - |
| - | - | 9673 | 341 | - | - | 0 | - |
| 6 | c | 4563 | 344.7 | 8.762E-05 | 0.2542 | +2 | 6 |
| - | - | 1.167E+04 | 358.2 | - | - | 0 | - |
| - | - | 3.705E+04 | 359 | - | - | 0 | - |
| - | - | 1.181E+04 | 359.2 | - | - | 0 | - |
| 10 | z | 5447 | 366.2 | 0.001882 | 5.139 | +1 | 3 |
| - | - | 4460 | 367.2 | - | - | 0 | - |
| - | - | 1.606E+04 | 373.2 | - | - | 0 | - |
| - | - | 1.593E+04 | 374.2 | - | - | 0 | - |
| - | - | 6288 | 375.2 | - | - | 0 | - |
| - | - | 7085 | 377.2 | - | - | 0 | - |
| 10 | y | 2.941E+04 | 382.2 | 0.0001867 | 0.4884 | +1 | 3 |
| - | - | 5955 | 383.2 | - | - | 0 | - |
| - | - | 6715 | 385.2 | - | - | 0 | - |
| - | - | 2.374E+04 | 385.9 | - | - | 0 | - |
| - | - | 5.251E+04 | 386.2 | - | - | 0 | - |
| - | - | 5985 | 386.5 | - | - | 0 | - |
| - | - | 1.159E+04 | 387.2 | - | - | 0 | - |
| - | - | 6528 | 391.6 | - | - | 0 | - |
| - | - | 4.266E+04 | 391.9 | - | - | 0 | - |
| - | - | 2.74E+04 | 392.2 | - | - | 0 | - |
| - | - | 1.217E+04 | 392.5 | - | - | 0 | - |
| - | - | 5901 | 394.2 | - | - | 0 | - |
| - | - | 1.33E+04 | 395.2 | - | - | 0 | - |
| - | - | 7888 | 400.3 | - | - | 0 | - |
| 11 | c | 7097 | 401.2 | 0.002784 | 6.938 | +3 | 11 |
| - | - | 1.819E+04 | 415.3 | - | - | 0 | - |
| - | - | 7209 | 416.3 | - | - | 0 | - |
| - | - | 7522 | 423.7 | - | - | 0 | - |
| - | - | 4334 | 424.2 | - | - | 0 | - |
| - | - | 6567 | 426.2 | - | - | 0 | - |
| - | - | 9.203E+04 | 429.1 | - | - | 0 | - |
| - | - | 4722 | 430.1 | - | - | 0 | - |
| - | - | 2.674E+04 | 430.2 | - | - | 0 | - |
| - | - | 1.315E+04 | 430.3 | - | - | 0 | - |
| - | - | 6768 | 430.8 | - | - | 0 | - |
| - | - | 6512 | 431.2 | - | - | 0 | - |
| - | - | 1.288E+04 | 431.3 | - | - | 0 | - |
| - | - | 1.312E+04 | 437.7 | - | - | 0 | - |
| - | - | 3.223E+04 | 443.2 | - | - | 0 | - |
| - | - | 1.003E+04 | 443.3 | - | - | 0 | - |
| 8 | c | 3.417E+04 | 443.7 | 0.001677 | 3.78 | +2 | 8 |
| 8 | c | 1.169E+04 | 444.2 | 0.00369 | 8.307 | +2 | 8 |
| - | - | 8168 | 444.7 | - | - | 0 | - |
| - | - | 5.071E+04 | 445.1 | - | - | 0 | - |
| - | - | 5485 | 445.2 | - | - | 0 | - |
| - | - | 5321 | 446.1 | - | - | 0 | - |
| - | - | 3.174E+04 | 446.2 | - | - | 0 | - |
| - | - | 5302 | 447.2 | - | - | 0 | - |
| - | - | 1.57E+04 | 451.8 | - | - | 0 | - |
| - | - | 1.003E+05 | 452.2 | - | - | 0 | - |
| 8 | c | 4.25E+05 | 452.7 | 0.0008528 | 1.884 | +2 | 8 |
| - | - | 1.816E+05 | 453.2 | - | - | 0 | - |
| - | - | 5.524E+04 | 453.7 | - | - | 0 | - |
| - | - | 1.181E+04 | 454.3 | - | - | 0 | - |
| - | - | 1.154E+04 | 455.6 | - | - | 0 | - |
| - | - | 1.186E+04 | 455.9 | - | - | 0 | - |
| 4 | c | 1.886E+04 | 458.3 | 0.00015 | 0.3273 | +1 | 4 |
| - | - | 9445 | 458.8 | - | - | 0 | - |
| - | - | 2.028E+04 | 461.6 | - | - | 0 | - |
| - | - | 9081 | 461.9 | - | - | 0 | - |
| 9 | z | 7467 | 462.2 | 0.001153 | 2.495 | +1 | 4 |
| 5 | y | 5091 | 463.2 | 0.003622 | 7.82 | +2 | 8 |
| - | - | 5943 | 466.3 | - | - | 0 | - |
| - | - | 6626 | 467.2 | - | - | 0 | - |
| - | - | 1.179E+04 | 471.2 | - | - | 0 | - |
| - | - | 1.436E+04 | 471.7 | - | - | 0 | - |
| - | - | 7309 | 472.2 | - | - | 0 | - |
| - | - | 8596 | 472.7 | - | - | 0 | - |
| - | - | 5.768E+04 | 474.2 | - | - | 0 | - |
| - | - | 7813 | 474.3 | - | - | 0 | - |
| 4 | c | 1.479E+06 | 475.3 | 0.001006 | 2.116 | +1 | 4 |
| - | - | 3.15E+05 | 476.3 | - | - | 0 | - |
| - | - | 5986 | 476.4 | - | - | 0 | - |
| - | - | 7454 | 476.7 | - | - | 0 | - |
| - | - | 4.87E+04 | 477.3 | - | - | 0 | - |
| - | - | 5477 | 478.3 | - | - | 0 | - |
| - | - | 2.611E+04 | 479.8 | - | - | 0 | - |
| 9 | z | 2.411E+04 | 480.2 | 0.001342 | 2.795 | +1 | 4 |
| - | - | 1.1E+05 | 480.3 | - | - | 0 | - |
| - | - | 6.524E+04 | 480.8 | - | - | 0 | - |
| - | - | 1.342E+05 | 481.2 | - | - | 0 | - |
| - | - | 2.289E+04 | 482.2 | - | - | 0 | - |
| - | - | 4551 | 482.3 | - | - | 0 | - |
| - | - | 2.287E+04 | 485.2 | - | - | 0 | - |
| - | - | 2.321E+04 | 485.7 | - | - | 0 | - |
| - | - | 1.143E+04 | 486.2 | - | - | 0 | - |
| - | - | 8828 | 486.3 | - | - | 0 | - |
| - | - | 1.411E+04 | 487.3 | - | - | 0 | - |
| - | - | 3.574E+04 | 487.8 | - | - | 0 | - |
| - | - | 2.291E+04 | 488.3 | - | - | 0 | - |
| - | - | 5386 | 488.8 | - | - | 0 | - |
| - | - | 7794 | 492.3 | - | - | 0 | - |
| - | - | 6.051E+04 | 494.2 | - | - | 0 | - |
| - | - | 2.301E+04 | 494.7 | - | - | 0 | - |
| - | - | 6171 | 495.2 | - | - | 0 | - |
| - | - | 5932 | 495.7 | - | - | 0 | - |
| 9 | y | 7.227E+04 | 496.2 | 0.0004416 | 0.8899 | +1 | 4 |
| - | - | 2.477E+04 | 497.2 | - | - | 0 | - |
| - | - | 1.934E+04 | 499.8 | - | - | 0 | - |
| - | - | 1.233E+05 | 500.3 | - | - | 0 | - |
| 9 | c | 7.405E+04 | 500.8 | 0.001535 | 3.065 | +2 | 9 |
| 9 | c | 5.018E+04 | 501.3 | 0.003864 | 7.708 | +2 | 9 |
| - | - | 2.642E+04 | 501.8 | - | - | 0 | - |
| - | - | 1.937E+05 | 502.3 | - | - | 0 | - |
| - | - | 1.429E+04 | 502.8 | - | - | 0 | - |
| - | - | 3.632E+04 | 503.3 | - | - | 0 | - |
| - | - | 4973 | 504.3 | - | - | 0 | - |
| - | - | 7157 | 508.3 | - | - | 0 | - |
| - | - | 3.796E+04 | 508.8 | - | - | 0 | - |
| - | - | 1.99E+05 | 509.3 | - | - | 0 | - |
| 9 | c | 6.069E+05 | 509.8 | 0.0007208 | 1.414 | +2 | 9 |
| - | - | 2.852E+05 | 510.3 | - | - | 0 | - |
| - | - | 7.652E+04 | 510.8 | - | - | 0 | - |
| - | - | 2.19E+04 | 511.3 | - | - | 0 | - |
| - | - | 6423 | 514.3 | - | - | 0 | - |
| - | - | 5794 | 516.3 | - | - | 0 | - |
| - | - | 8415 | 521.3 | - | - | 0 | - |
| - | - | 2.153E+04 | 522.3 | - | - | 0 | - |
| - | - | 1.438E+04 | 522.8 | - | - | 0 | - |
| - | - | 1.413E+04 | 523.3 | - | - | 0 | - |
| - | - | 7711 | 523.8 | - | - | 0 | - |
| - | - | 2.981E+04 | 526.7 | - | - | 0 | - |
| - | - | 1.918E+04 | 527.3 | - | - | 0 | - |
| - | - | 8365 | 527.7 | - | - | 0 | - |
| - | - | 1.627E+04 | 530.3 | - | - | 0 | - |
| - | - | 1.578E+04 | 530.8 | - | - | 0 | - |
| - | - | 7284 | 531.8 | - | - | 0 | - |
| - | - | 9202 | 532.8 | - | - | 0 | - |
| 4 | z | 5.987E+04 | 533.3 | 0.001186 | 2.225 | +2 | 9 |
| - | - | 3.922E+04 | 533.8 | - | - | 0 | - |
| - | - | 1.638E+04 | 534.3 | - | - | 0 | - |
| - | - | 1.849E+04 | 535.8 | - | - | 0 | - |
| - | - | 1.606E+04 | 536.3 | - | - | 0 | - |
| - | - | 5262 | 540.8 | - | - | 0 | - |
| 4 | y | 1.385E+04 | 541.3 | 0.002017 | 3.727 | +2 | 9 |
| - | - | 5269 | 541.8 | - | - | 0 | - |
| - | - | 6053 | 542.3 | - | - | 0 | - |
| 10 | c | 1.095E+04 | 544.3 | 0.00168 | 3.086 | +2 | 10 |
| - | - | 5244 | 544.3 | - | - | 0 | - |
| 10 | c | 5.607E+04 | 544.8 | 0.002772 | 5.089 | +2 | 10 |
| - | - | 6769 | 545.2 | - | - | 0 | - |
| - | - | 2.531E+04 | 545.3 | - | - | 0 | - |
| - | - | 2.678E+04 | 545.3 | - | - | 0 | - |
| - | - | 7074 | 545.8 | - | - | 0 | - |
| - | - | 7986 | 546.3 | - | - | 0 | - |
| - | - | 2.861E+04 | 546.3 | - | - | 0 | - |
| - | - | 7863 | 547.3 | - | - | 0 | - |
| - | - | 1.189E+04 | 548.3 | - | - | 0 | - |
| - | - | 8.11E+04 | 548.7 | - | - | 0 | - |
| - | - | 5.177E+04 | 549.2 | - | - | 0 | - |
| - | - | 1.808E+04 | 549.7 | - | - | 0 | - |
| - | - | 1.699E+04 | 550.8 | - | - | 0 | - |
| - | - | 6440 | 552.3 | - | - | 0 | - |
| - | - | 3.847E+04 | 552.8 | - | - | 0 | - |
| 10 | c | 8.209E+05 | 553.3 | 0.0009725 | 1.758 | +2 | 10 |
| - | - | 4.375E+05 | 553.8 | - | - | 0 | - |
| - | - | 1.557E+05 | 554.3 | - | - | 0 | - |
| - | - | 3.166E+04 | 554.8 | - | - | 0 | - |
| - | - | 1.475E+04 | 555.3 | - | - | 0 | - |
| - | - | 1.449E+04 | 562.3 | - | - | 0 | - |
| - | - | 5815 | 563.3 | - | - | 0 | - |
| - | - | 6.146E+04 | 564.3 | - | - | 0 | - |
| - | - | 8172 | 565.3 | - | - | 0 | - |
| - | - | 2.122E+04 | 565.3 | - | - | 0 | - |
| - | - | 3.69E+04 | 566.3 | - | - | 0 | - |
| - | - | 1.371E+04 | 566.8 | - | - | 0 | - |
| - | - | 6335 | 567.3 | - | - | 0 | - |
| 3 | z | 1.089E+04 | 568.3 | 0.00687 | 12.09 | +2 | 10 |
| - | - | 9032 | 568.8 | - | - | 0 | - |
| - | - | 9678 | 569.3 | - | - | 0 | - |
| 5 | c | 2.638E+04 | 573.3 | 0.003694 | 6.443 | +1 | 5 |
| - | - | 9249 | 574.3 | - | - | 0 | - |
| 3 | y | 2.421E+04 | 575.8 | 0.0002918 | 0.5069 | +2 | 10 |
| 3 | y | 4.111E+04 | 576.3 | 0.005747 | 9.973 | +2 | 10 |
| 3 | z | 1.31E+05 | 576.8 | 0.001102 | 1.911 | +2 | 10 |
| - | - | 8.145E+04 | 577.3 | - | - | 0 | - |
| - | - | 2.983E+04 | 577.8 | - | - | 0 | - |
| - | - | 2.28E+04 | 578.3 | - | - | 0 | - |
| - | - | 2.47E+04 | 578.8 | - | - | 0 | - |
| - | - | 1.288E+04 | 579.3 | - | - | 0 | - |
| - | - | 2.063E+04 | 579.8 | - | - | 0 | - |
| - | - | 1.841E+04 | 580.3 | - | - | 0 | - |
| - | - | 7070 | 581.3 | - | - | 0 | - |
| - | - | 1.621E+04 | 583.8 | - | - | 0 | - |
| - | - | 6.332E+04 | 584.3 | - | - | 0 | - |
| 3 | y | 8.337E+05 | 584.8 | 0.0009566 | 1.636 | +2 | 10 |
| - | - | 4.792E+05 | 585.3 | - | - | 0 | - |
| - | - | 1.701E+05 | 585.8 | - | - | 0 | - |
| - | - | 5.173E+04 | 586.3 | - | - | 0 | - |
| - | - | 7691 | 586.8 | - | - | 0 | - |
| - | - | 4.735E+04 | 587.3 | - | - | 0 | - |
| - | - | 5.283E+04 | 587.8 | - | - | 0 | - |
| - | - | 8135 | 588.3 | - | - | 0 | - |
| - | - | 2.164E+04 | 588.3 | - | - | 0 | - |
| - | - | 3.727E+04 | 588.8 | - | - | 0 | - |
| - | - | 7916 | 588.8 | - | - | 0 | - |
| - | - | 1.51E+04 | 589.3 | - | - | 0 | - |
| - | - | 8560 | 589.3 | - | - | 0 | - |
| - | - | 7099 | 589.8 | - | - | 0 | - |
| 5 | c | 1.825E+06 | 590.3 | 0.0008266 | 1.4 | +1 | 5 |
| - | - | 5.179E+05 | 591.3 | - | - | 0 | - |
| - | - | 6641 | 591.4 | - | - | 0 | - |
| - | - | 9.093E+04 | 592.3 | - | - | 0 | - |
| - | - | 3.776E+04 | 592.8 | - | - | 0 | - |
| - | - | 1.994E+04 | 593.3 | - | - | 0 | - |
| - | - | 5686 | 593.8 | - | - | 0 | - |
| - | - | 9659 | 598.3 | - | - | 0 | - |
| - | - | 6649 | 598.8 | - | - | 0 | - |
| 11 | c | 1.758E+04 | 600.8 | 0.001231 | 2.049 | +2 | 11 |
| 11 | c | 8.424E+04 | 601.3 | 0.002427 | 4.037 | +2 | 11 |
| - | - | 5.518E+04 | 601.8 | - | - | 0 | - |
| - | - | 3.578E+04 | 602.3 | - | - | 0 | - |
| - | - | 7883 | 602.8 | - | - | 0 | - |
| - | - | 6216 | 602.8 | - | - | 0 | - |
| - | - | 7172 | 605.3 | - | - | 0 | - |
| - | - | 1.931E+04 | 607.8 | - | - | 0 | - |
| 8 | z | 1.702E+05 | 608.3 | 0.001137 | 1.87 | +1 | 5 |
| - | - | 5811 | 608.8 | - | - | 0 | - |
| - | - | 1.658E+05 | 609.3 | - | - | 0 | - |
| 11 | c | 1.04E+06 | 609.8 | 0.001299 | 2.13 | +2 | 11 |
| - | - | 5.72E+05 | 610.3 | - | - | 0 | - |
| - | - | 2.168E+05 | 610.8 | - | - | 0 | - |
| - | - | 5.698E+04 | 611.3 | - | - | 0 | - |
| - | - | 1.883E+04 | 611.8 | - | - | 0 | - |
| - | - | 5707 | 613.3 | - | - | 0 | - |
| - | - | 5258 | 616.3 | - | - | 0 | - |
| 2 | w | 9.613E+04 | 618.8 | 0.001155 | 1.866 | +2 | 11 |
| - | - | 6.122E+04 | 619.3 | - | - | 0 | - |
| - | - | 1.956E+04 | 619.8 | - | - | 0 | - |
| - | - | 6015 | 621.8 | - | - | 0 | - |
| - | - | 6742 | 622.3 | - | - | 0 | - |
| 8 | y | 1.271E+05 | 624.3 | 0.0005408 | 0.8662 | +1 | 5 |
| - | - | 8415 | 624.8 | - | - | 0 | - |
| - | - | 3.873E+04 | 625.3 | - | - | 0 | - |
| 2 | w | 9874 | 625.8 | 0.0004446 | 0.7104 | +2 | 11 |
| - | - | 1.146E+04 | 625.9 | - | - | 0 | - |
| - | - | 7536 | 626.3 | - | - | 0 | - |
| - | - | 5944 | 626.4 | - | - | 0 | - |
| - | - | 5466 | 626.9 | - | - | 0 | - |
| - | - | 1.565E+04 | 629.3 | - | - | 0 | - |
| - | - | 8523 | 630.3 | - | - | 0 | - |
| - | - | 4.328E+04 | 630.8 | - | - | 0 | - |
| - | - | 2.297E+04 | 631.3 | - | - | 0 | - |
| - | - | 1.946E+04 | 631.8 | - | - | 0 | - |
| - | - | 8114 | 632.3 | - | - | 0 | - |
| 2 | y | 1.426E+04 | 632.8 | 0.003693 | 5.836 | +2 | 11 |
| 2 | z | 4.184E+04 | 633.3 | 0.0003413 | 0.5389 | +2 | 11 |
| - | - | 5.235E+04 | 633.8 | - | - | 0 | - |
| - | - | 2.249E+04 | 634.3 | - | - | 0 | - |
| - | - | 1.432E+04 | 634.8 | - | - | 0 | - |
| - | - | 6337 | 635.4 | - | - | 0 | - |
| - | - | 2.222E+04 | 636.3 | - | - | 0 | - |
| - | - | 6028 | 637.4 | - | - | 0 | - |
| - | - | 3.462E+04 | 638.3 | - | - | 0 | - |
| - | - | 3.044E+04 | 638.8 | - | - | 0 | - |
| - | - | 1.226E+04 | 639.3 | - | - | 0 | - |
| - | - | 2.336E+04 | 639.8 | - | - | 0 | - |
| - | - | 4.807E+04 | 640.3 | - | - | 0 | - |
| - | - | 3.408E+04 | 640.8 | - | - | 0 | - |
| 2 | y | 8.879E+04 | 641.3 | 0.0007337 | 1.144 | +2 | 11 |
| - | - | 6.308E+04 | 641.8 | - | - | 0 | - |
| - | - | 2.075E+04 | 642.3 | - | - | 0 | - |
| - | - | 7614 | 642.8 | - | - | 0 | - |
| - | - | 7646 | 644.3 | - | - | 0 | - |
| - | - | 1.108E+04 | 645.3 | - | - | 0 | - |
| - | - | 1.844E+05 | 647.3 | - | - | 0 | - |
| - | - | 4.918E+04 | 647.8 | - | - | 0 | - |
| - | - | 6.152E+04 | 648.3 | - | - | 0 | - |
| - | - | 1.104E+04 | 648.9 | - | - | 0 | - |
| - | - | 1.542E+04 | 649.3 | - | - | 0 | - |
| - | - | 1.135E+04 | 652.8 | - | - | 0 | - |
| - | - | 4.737E+04 | 653.3 | - | - | 0 | - |
| - | - | 2.829E+04 | 653.8 | - | - | 0 | - |
| - | - | 2.413E+04 | 654.3 | - | - | 0 | - |
| - | - | 1.146E+04 | 654.8 | - | - | 0 | - |
| - | - | 2.464E+04 | 655.3 | - | - | 0 | - |
| - | - | 1.078E+05 | 655.8 | - | - | 0 | - |
| - | - | 6.659E+04 | 656.3 | - | - | 0 | - |
| - | - | 2.223E+04 | 656.8 | - | - | 0 | - |
| - | - | 1.485E+04 | 657.3 | - | - | 0 | - |
| - | - | 1.771E+04 | 659.3 | - | - | 0 | - |
| - | - | 4.151E+04 | 660.3 | - | - | 0 | - |
| - | - | 2.974E+04 | 660.8 | - | - | 0 | - |
| 7 | z | 9.501E+04 | 661.3 | 0.003702 | 5.598 | +1 | 6 |
| - | - | 2.07E+05 | 661.8 | - | - | 0 | - |
| - | - | 2.227E+05 | 662.3 | - | - | 0 | - |
| - | - | 1.637E+05 | 662.8 | - | - | 0 | - |
| - | - | 8.004E+04 | 663.3 | - | - | 0 | - |
| - | - | 2.356E+04 | 663.8 | - | - | 0 | - |
| - | - | 7769 | 664.3 | - | - | 0 | - |
| - | - | 5.517E+04 | 669.3 | - | - | 0 | - |
| - | - | 3.537E+05 | 669.8 | - | - | 0 | - |
| - | - | 2.473E+05 | 670.3 | - | - | 0 | - |
| - | - | 3.767E+05 | 670.8 | - | - | 0 | - |
| - | - | 1.893E+05 | 671.3 | - | - | 0 | - |
| - | - | 6.619E+04 | 671.8 | - | - | 0 | - |
| - | - | 3.55E+04 | 672.3 | - | - | 0 | - |
| - | - | 2.262E+04 | 673.3 | - | - | 0 | - |
| - | - | 3.552E+04 | 674.3 | - | - | 0 | - |
| - | - | 7.236E+04 | 674.8 | - | - | 0 | - |
| - | - | 1.706E+05 | 675.3 | - | - | 0 | - |
| - | - | 9.359E+04 | 675.8 | - | - | 0 | - |
| - | - | 8.674E+04 | 676.3 | - | - | 0 | - |
| - | - | 3.628E+04 | 676.8 | - | - | 0 | - |
| - | - | 2.281E+04 | 677.3 | - | - | 0 | - |
| - | - | 7913 | 677.8 | - | - | 0 | - |
| 7 | z | 5.221E+05 | 679.4 | 0.0008889 | 1.308 | +1 | 6 |
| - | - | 1.798E+05 | 680.4 | - | - | 0 | - |
| - | - | 4.301E+04 | 681.4 | - | - | 0 | - |
| - | - | 8807 | 682.4 | - | - | 0 | - |
| - | - | 5297 | 682.9 | - | - | 0 | - |
| - | - | 9.118E+04 | 683.3 | - | - | 0 | - |
| - | - | 9.314E+05 | 683.8 | - | - | 0 | - |
| - | - | 5.708E+05 | 684.3 | - | - | 0 | - |
| - | - | 2.664E+05 | 684.8 | - | - | 0 | - |
| - | - | 8.003E+04 | 685.3 | - | - | 0 | - |
| - | - | 2.056E+04 | 685.8 | - | - | 0 | - |
| 6 | c | 2.567E+04 | 688.3 | 0.003362 | 4.885 | +1 | 6 |
| - | - | 7456 | 689.3 | - | - | 0 | - |
| - | - | 8050 | 690.3 | - | - | 0 | - |
| - | - | 6.348E+04 | 691.4 | - | - | 0 | - |
| - | - | 9.797E+05 | 691.9 | - | - | 0 | - |
| - | - | 6.249E+05 | 692.4 | - | - | 0 | - |
| - | - | 2.613E+05 | 692.9 | - | - | 0 | - |
| - | - | 6.624E+04 | 693.4 | - | - | 0 | - |
| - | - | 1.353E+04 | 693.9 | - | - | 0 | - |
| 7 | y | 7.281E+04 | 695.4 | 0.0008416 | 1.21 | +1 | 6 |
| - | - | 2.857E+04 | 696.4 | - | - | 0 | - |
| - | - | 6935 | 701.4 | - | - | 0 | - |
| - | - | 5867 | 702.4 | - | - | 0 | - |
| - | - | 6.927E+04 | 704.4 | - | - | 0 | - |
| 6 | c | 5.06E+05 | 705.4 | 0.00141 | 2 | +1 | 6 |
| - | - | 1.608E+05 | 706.4 | - | - | 0 | - |
| - | - | 4.316E+04 | 707.4 | - | - | 0 | - |
| - | - | 7856 | 708.4 | - | - | 0 | - |
| - | - | 8733 | 711.3 | - | - | 0 | - |
| - | - | 5433 | 712.3 | - | - | 0 | - |
| - | - | 5930 | 722.4 | - | - | 0 | - |
| - | - | 7469 | 731.4 | - | - | 0 | - |
| - | - | 8.121E+04 | 732.4 | - | - | 0 | - |
| - | - | 3.046E+04 | 733.4 | - | - | 0 | - |
| - | - | 7490 | 734.4 | - | - | 0 | - |
| - | - | 2.409E+05 | 737.4 | - | - | 0 | - |
| - | - | 4714 | 738.3 | - | - | 0 | - |
| - | - | 8.775E+04 | 738.4 | - | - | 0 | - |
| - | - | 1.927E+04 | 739.4 | - | - | 0 | - |
| 6 | w | 3.359E+04 | 749.4 | 0.001263 | 1.686 | +1 | 7 |
| - | - | 4.503E+04 | 750.4 | - | - | 0 | - |
| - | - | 1.72E+04 | 751.4 | - | - | 0 | - |
| 7 | c | 3.17E+04 | 759.4 | 0.001832 | 2.413 | +1 | 7 |
| - | - | 1.169E+04 | 760.4 | - | - | 0 | - |
| - | - | 8349 | 761.4 | - | - | 0 | - |
| - | - | 6056 | 764.4 | - | - | 0 | - |
| - | - | 5835 | 774.4 | - | - | 0 | - |
| - | - | 7.808E+04 | 775.4 | - | - | 0 | - |
| 7 | c | 6.707E+05 | 776.4 | 0.0009177 | 1.182 | +1 | 7 |
| - | - | 2.244E+05 | 777.4 | - | - | 0 | - |
| - | - | 5.381E+04 | 778.4 | - | - | 0 | - |
| - | - | 1.116E+04 | 779.4 | - | - | 0 | - |
| - | - | 1.187E+04 | 787.4 | - | - | 0 | - |
| - | - | 4493 | 788.3 | - | - | 0 | - |
| - | - | 2.518E+04 | 788.4 | - | - | 0 | - |
| - | - | 2.492E+04 | 789.4 | - | - | 0 | - |
| - | - | 9288 | 790.4 | - | - | 0 | - |
| - | - | 1.667E+05 | 793.4 | - | - | 0 | - |
| 6 | z | 1.397E+06 | 794.4 | 0.001534 | 1.931 | +1 | 7 |
| - | - | 5.189E+05 | 795.4 | - | - | 0 | - |
| - | - | 1.508E+05 | 796.4 | - | - | 0 | - |
| - | - | 2.064E+04 | 797.4 | - | - | 0 | - |
| - | - | 6348 | 803.4 | - | - | 0 | - |
| 6 | y | 5.111E+04 | 810.4 | 0.0008151 | 1.006 | +1 | 7 |
| - | - | 2.714E+04 | 811.4 | - | - | 0 | - |
| - | - | 8184 | 812.4 | - | - | 0 | - |
| - | - | 9714 | 816.5 | - | - | 0 | - |
| - | - | 7.035E+04 | 821.4 | - | - | 0 | - |
| - | - | 2.56E+04 | 822.4 | - | - | 0 | - |
| - | - | 1.053E+04 | 823.4 | - | - | 0 | - |
| - | - | 4901 | 826.4 | - | - | 0 | - |
| - | - | 5146 | 837.4 | - | - | 0 | - |
| - | - | 7991 | 842.5 | - | - | 0 | - |
| - | - | 7053 | 844.5 | - | - | 0 | - |
| - | - | 2.321E+04 | 859.5 | - | - | 0 | - |
| - | - | 1.028E+05 | 860.5 | - | - | 0 | - |
| - | - | 8.45E+04 | 861.5 | - | - | 0 | - |
| - | - | 2.583E+04 | 862.5 | - | - | 0 | - |
| - | - | 5625 | 863.5 | - | - | 0 | - |
| - | - | 4.942E+04 | 864.4 | - | - | 0 | - |
| - | - | 4.18E+05 | 865.4 | - | - | 0 | - |
| - | - | 1.871E+05 | 866.4 | - | - | 0 | - |
| - | - | 5.69E+04 | 867.4 | - | - | 0 | - |
| - | - | 1.62E+04 | 868.4 | - | - | 0 | - |
| - | - | 8064 | 869.4 | - | - | 0 | - |
| - | - | 1.175E+04 | 874.4 | - | - | 0 | - |
| - | - | 1.82E+04 | 875.4 | - | - | 0 | - |
| - | - | 6640 | 876.4 | - | - | 0 | - |
| - | - | 4.626E+04 | 885.5 | - | - | 0 | - |
| 8 | c | 5.358E+04 | 886.5 | 0.003341 | 3.769 | +1 | 8 |
| 8 | c | 5.683E+04 | 887.5 | 0.004769 | 5.374 | +1 | 8 |
| - | - | 1.878E+04 | 888.5 | - | - | 0 | - |
| - | - | 1.638E+04 | 889.5 | - | - | 0 | - |
| - | - | 1.448E+04 | 891.4 | - | - | 0 | - |
| - | - | 6976 | 900.5 | - | - | 0 | - |
| - | - | 3.128E+04 | 901.5 | - | - | 0 | - |
| - | - | 3.904E+04 | 902.5 | - | - | 0 | - |
| - | - | 2.472E+05 | 903.5 | - | - | 0 | - |
| 8 | c | 1.01E+06 | 904.5 | 0.0007423 | 0.8207 | +1 | 8 |
| - | - | 4.227E+05 | 905.5 | - | - | 0 | - |
| - | - | 1.17E+05 | 906.5 | - | - | 0 | - |
| - | - | 3.146E+04 | 907.5 | - | - | 0 | - |
| - | - | 4.298E+04 | 908.4 | - | - | 0 | - |
| 5 | z | 6.96E+05 | 909.4 | 0.0007138 | 0.7849 | +1 | 8 |
| - | - | 3.124E+05 | 910.4 | - | - | 0 | - |
| - | - | 1.041E+05 | 911.4 | - | - | 0 | - |
| - | - | 2.206E+04 | 912.4 | - | - | 0 | - |
| 5 | y | 1.129E+05 | 925.4 | 0.0006054 | 0.6542 | +1 | 8 |
| - | - | 4.915E+04 | 926.4 | - | - | 0 | - |
| - | - | 1.007E+04 | 927.4 | - | - | 0 | - |
| - | - | 6022 | 928.4 | - | - | 0 | - |
| - | - | 5949 | 952.4 | - | - | 0 | - |
| - | - | 6103 | 958.5 | - | - | 0 | - |
| - | - | 9156 | 959.5 | - | - | 0 | - |
| - | - | 1.601E+04 | 969.5 | - | - | 0 | - |
| - | - | 6227 | 970.5 | - | - | 0 | - |
| - | - | 1.771E+04 | 974.5 | - | - | 0 | - |
| - | - | 2.878E+04 | 975.5 | - | - | 0 | - |
| - | - | 1.596E+04 | 976.5 | - | - | 0 | - |
| - | - | 5549 | 977.5 | - | - | 0 | - |
| - | - | 9152 | 986.5 | - | - | 0 | - |
| - | - | 4.257E+04 | 987.5 | - | - | 0 | - |
| - | - | 4.853E+04 | 988.5 | - | - | 0 | - |
| - | - | 2.79E+04 | 989.5 | - | - | 0 | - |
| - | - | 7878 | 990.5 | - | - | 0 | - |
| - | - | 7582 | 999.5 | - | - | 0 | - |
| 9 | c | 2.562E+04 | 1001 | 0.005062 | 5.059 | +1 | 9 |
| 9 | c | 3.44E+04 | 1002 | 0.009266 | 9.252 | +1 | 9 |
| - | - | 1.842E+04 | 1003 | - | - | 0 | - |
| - | - | 2.615E+04 | 1004 | - | - | 0 | - |
| - | - | 1.697E+04 | 1005 | - | - | 0 | - |
| - | - | 7922 | 1006 | - | - | 0 | - |
| - | - | 7.633E+04 | 1018 | - | - | 0 | - |
| 9 | c | 4.34E+05 | 1019 | 0.001089 | 1.069 | +1 | 9 |
| - | - | 2.104E+05 | 1020 | - | - | 0 | - |
| - | - | 7.759E+04 | 1021 | - | - | 0 | - |
| - | - | 1.769E+04 | 1022 | - | - | 0 | - |
| - | - | 1.316E+04 | 1023 | - | - | 0 | - |
| - | - | 9072 | 1023 | - | - | 0 | - |
| - | - | 7740 | 1045 | - | - | 0 | - |
| - | - | 7257 | 1046 | - | - | 0 | - |
| - | - | 1.008E+04 | 1047 | - | - | 0 | - |
| - | - | 1.029E+04 | 1048 | - | - | 0 | - |
| - | - | 1.517E+04 | 1049 | - | - | 0 | - |
| - | - | 9080 | 1051 | - | - | 0 | - |
| - | - | 1.64E+04 | 1062 | - | - | 0 | - |
| - | - | 2.179E+04 | 1063 | - | - | 0 | - |
| - | - | 2.585E+04 | 1064 | - | - | 0 | - |
| - | - | 3.478E+04 | 1065 | - | - | 0 | - |
| 4 | z | 3.595E+05 | 1066 | 0.001226 | 1.151 | +1 | 9 |
| - | - | 2.801E+05 | 1067 | - | - | 0 | - |
| - | - | 1.102E+05 | 1068 | - | - | 0 | - |
| - | - | 3.746E+04 | 1069 | - | - | 0 | - |
| - | - | 7173 | 1070 | - | - | 0 | - |
| - | - | 1.674E+04 | 1081 | - | - | 0 | - |
| 4 | y | 3.789E+04 | 1082 | 0.0007742 | 0.7158 | +1 | 9 |
| - | - | 1.696E+04 | 1083 | - | - | 0 | - |
| - | - | 6182 | 1084 | - | - | 0 | - |
| 10 | c | 1.288E+04 | 1089 | 0.007084 | 6.508 | +1 | 10 |
| - | - | 6.746E+04 | 1090 | - | - | 0 | - |
| - | - | 3.17E+04 | 1091 | - | - | 0 | - |
| - | - | 1.066E+04 | 1092 | - | - | 0 | - |
| - | - | 2.041E+04 | 1096 | - | - | 0 | - |
| - | - | 2.559E+04 | 1097 | - | - | 0 | - |
| - | - | 7430 | 1098 | - | - | 0 | - |
| - | - | 6901 | 1102 | - | - | 0 | - |
| - | - | 1.521E+04 | 1105 | - | - | 0 | - |
| 10 | c | 2.063E+05 | 1106 | 0.0005545 | 0.5015 | +1 | 10 |
| - | - | 1.4E+05 | 1107 | - | - | 0 | - |
| - | - | 6.479E+04 | 1108 | - | - | 0 | - |
| - | - | 2.095E+04 | 1109 | - | - | 0 | - |
| - | - | 1.152E+04 | 1111 | - | - | 0 | - |
| - | - | 8466 | 1127 | - | - | 0 | - |
| 3 | z | 9138 | 1136 | 0.01992 | 17.54 | +1 | 10 |
| - | - | 1.605E+04 | 1137 | - | - | 0 | - |
| - | - | 9260 | 1138 | - | - | 0 | - |
| 3 | y | 1.767E+04 | 1152 | 0.01889 | 16.41 | +1 | 10 |
| 3 | z | 1.919E+05 | 1153 | 0.002157 | 1.871 | +1 | 10 |
| - | - | 2.362E+05 | 1154 | - | - | 0 | - |
| - | - | 1.084E+05 | 1155 | - | - | 0 | - |
| - | - | 3.398E+04 | 1156 | - | - | 0 | - |
| - | - | 5733 | 1157 | - | - | 0 | - |
| - | - | 1.09E+04 | 1159 | - | - | 0 | - |
| - | - | 7434 | 1160 | - | - | 0 | - |
| - | - | 1.138E+04 | 1161 | - | - | 0 | - |
| 3 | y | 4.118E+04 | 1169 | 0.0008201 | 0.7018 | +1 | 10 |
| - | - | 3.379E+04 | 1170 | - | - | 0 | - |
| - | - | 1.91E+04 | 1171 | - | - | 0 | - |
| - | - | 3.288E+04 | 1175 | - | - | 0 | - |
| - | - | 6.215E+04 | 1176 | - | - | 0 | - |
| - | - | 4.495E+04 | 1177 | - | - | 0 | - |
| - | - | 2.125E+04 | 1178 | - | - | 0 | - |
| - | - | 8743 | 1186 | - | - | 0 | - |
| - | - | 6332 | 1187 | - | - | 0 | - |
| 11 | c | 1.398E+04 | 1202 | 0.01909 | 15.89 | +1 | 11 |
| - | - | 8.627E+04 | 1203 | - | - | 0 | - |
| - | - | 6.256E+04 | 1204 | - | - | 0 | - |
| - | - | 2.085E+04 | 1205 | - | - | 0 | - |
| - | - | 6647 | 1206 | - | - | 0 | - |
| - | - | 6779 | 1218 | - | - | 0 | - |
| 11 | c | 1.127E+05 | 1219 | 0.0002575 | 0.2113 | +1 | 11 |
| - | - | 1.494E+05 | 1220 | - | - | 0 | - |
| - | - | 7.298E+04 | 1221 | - | - | 0 | - |
| - | - | 2.608E+04 | 1222 | - | - | 0 | - |
| - | - | 6956 | 1223 | - | - | 0 | - |
| - | - | 6327 | 1252 | - | - | 0 | - |
| 2 | z | 1.455E+04 | 1266 | 0.005129 | 4.053 | +1 | 11 |
| - | - | 7.142E+04 | 1267 | - | - | 0 | - |
| - | - | 5.657E+04 | 1268 | - | - | 0 | - |
| - | - | 2.348E+04 | 1269 | - | - | 0 | - |
| - | - | 8361 | 1281 | - | - | 0 | - |
| 2 | y | 1.431E+04 | 1282 | 0.007401 | 5.775 | +1 | 11 |
| - | - | 8077 | 1283 | - | - | 0 | - |
| - | - | 8214 | 1296 | - | - | 0 | - |
| - | - | 1.643E+04 | 1306 | - | - | 0 | - |
| - | - | 7.2E+04 | 1307 | - | - | 0 | - |
| - | - | 7.872E+04 | 1308 | - | - | 0 | - |
| - | - | 4.686E+04 | 1309 | - | - | 0 | - |
| - | - | 2.252E+04 | 1310 | - | - | 0 | - |
| - | - | 1.079E+04 | 1311 | - | - | 0 | - |
| - | - | 2.677E+04 | 1322 | - | - | 0 | - |
| - | - | 5.284E+04 | 1323 | - | - | 0 | - |
| - | - | 5.732E+04 | 1324 | - | - | 0 | - |
| - | - | 2.615E+05 | 1325 | - | - | 0 | - |
| - | - | 2.435E+05 | 1326 | - | - | 0 | - |
| - | - | 1.184E+05 | 1327 | - | - | 0 | - |
| - | - | 3.685E+04 | 1328 | - | - | 0 | - |
| - | - | 1.087E+04 | 1329 | - | - | 0 | - |
| - | - | 8772 | 1338 | - | - | 0 | - |
| - | - | 6.482E+04 | 1339 | - | - | 0 | - |
| - | - | 2.59E+05 | 1340 | - | - | 0 | - |
| - | - | 1.724E+05 | 1341 | - | - | 0 | - |
| - | - | 1.154E+05 | 1342 | - | - | 0 | - |
| - | - | 4.192E+04 | 1343 | - | - | 0 | - |
| - | - | 1.232E+04 | 1344 | - | - | 0 | - |
| - | - | 6.887E+04 | 1350 | - | - | 0 | - |
| - | - | 1.18E+05 | 1351 | - | - | 0 | - |
| - | - | 6.782E+04 | 1352 | - | - | 0 | - |
| - | - | 2.575E+04 | 1353 | - | - | 0 | - |
| - | - | 1.31E+04 | 1356 | - | - | 0 | - |
| - | - | 3.697E+04 | 1357 | - | - | 0 | - |
| - | - | 1.666E+04 | 1358 | - | - | 0 | - |
| - | - | 2.431E+04 | 1366 | - | - | 0 | - |
| - | - | 2.165E+05 | 1367 | - | - | 0 | - |
| - | - | 9.644E+05 | 1368 | - | - | 0 | - |
| - | - | 5.997E+05 | 1369 | - | - | 0 | - |
| - | - | 2.541E+05 | 1370 | - | - | 0 | - |
| - | - | 7.72E+04 | 1371 | - | - | 0 | - |
| - | - | 8909 | 1372 | - | - | 0 | - |
| - | - | 5.618E+04 | 1383 | - | - | 0 | - |
| - | - | 1.701E+05 | 1384 | - | - | 0 | - |
| - | - | 2.617E+05 | 1385 | - | - | 0 | - |
| - | - | 1.409E+05 | 1386 | - | - | 0 | - |
| - | - | 5.757E+04 | 1387 | - | - | 0 | - |
| - | - | 1.373E+04 | 1388 | - | - | 0 | - |
| - | - | 6127 | 2253 | - | - | 0 | - |
| - | - | 5466 | 2327 | - | - | 0 | - |

m/z Charge Intensity FragmentType MassShift Position
123.38087463378906 0 2696.6694
129.10232543945312 0 3814.4187
131.11827087402344 0 4481.443
136.07603454589844 0 62502.145
137.07943725585938 0 5957.847
141.60659790039062 0 3477.9
142.1230010986328 0 14524.388
143.11819458007812 0 8897.202
158.11769104003906 0 4729.261
165.05494689941406 0 61706.938
166.05838012695312 0 9295.298
169.13369750976562 0 9363.259
173.1284637451172 0 6827.043
182.08157348632812 0 236655.4 y 11
183.0850067138672 0 27443.064
186.1125946044922 0 4404.429
187.14451599121094 0 741600.1
188.14785766601562 0 68262.9
189.14956665039062 0 3930.147
199.54624938964844 0 3779.463
200.13986206054688 0 7118.4404
201.12371826171875 0 27995.898
201.1472625732422 0 6197.0054
202.0825958251953 0 7674.8286
202.14212036132812 0 4572.4854
215.139404296875 0 197309.72
216.14259338378906 0 15217.43
229.12916564941406 0 9524.36
232.16616821289062 0 16998.01 c 1
235.1436767578125 0 3917.5886
244.14112854003906 0 6796.4487
272.1605529785156 0 5922.198
274.5708923339844 0 4041.4438
281.1371765136719 0 11630.131
282.1453552246094 0 54347.1
283.14801025390625 0 8899.818
287.1543273925781 0 6565.48 c Ammonia loss 4
287.17095947265625 0 8282.118
295.1658630371094 0 30768.105 y 10
296.1697998046875 0 4983.8955
297.1553039550781 0 9482.873
299.1483459472656 0 17291.44
299.1718444824219 0 18831.674
300.15582275390625 0 123893.08
301.1596374511719 0 20455.482
310.11553955078125 0 4426.612
315.1665344238281 0 22209.818
317.18267822265625 0 88616.32
318.1854553222656 0 11842.835
319.1983337402344 0 56782.637 c 2
320.2032470703125 0 7285.6406
328.8470764160156 0 4479.325
335.66082763671875 0 9067.8545
336.1611022949219 0 3998.6611
341.0179443359375 0 9673.209
344.666748046875 0 4563.2773 c Ammonia loss 5
358.1968688964844 0 11674.35
359.02880859375 0 37045.383
359.1678161621094 0 11812.099
366.180419921875 0 5447.148 z 9
367.1879577636719 0 4459.978
373.1893005371094 0 16058.27
374.1802062988281 0 15928.836
375.1816101074219 0 6288.4263
377.1793212890625 0 7085.472
382.19744873046875 0 29408.404 y 9
383.20166015625 0 5954.968
385.2197265625 0 6714.8784
385.8765563964844 0 23743.463
386.2050476074219 0 52508.008
386.5425109863281 0 5984.611
387.2098693847656 0 11587.039
391.5500183105469 0 6527.596
391.88092041015625 0 42664.78
392.2134704589844 0 27401.285
392.54901123046875 0 12171.972
394.1893005371094 0 5901.07
395.20257568359375 0 13295.319
400.2558898925781 0 7887.985
401.2133483886719 0 7097.4185 c Ammonia loss 10
415.2668151855469 0 18190.857
416.2718811035156 0 7209.365
423.7060852050781 0 7522.3447
424.16162109375 0 4334.164
426.2327880859375 0 6567.3833
429.089599609375 0 92033.875
430.08740234375 0 4721.621
430.2462463378906 0 26738.363
430.2794189453125 0 13152.043
430.75048828125 0 6767.577
431.2234802246094 0 6512.214
431.28131103515625 0 12876.704
437.70587158203125 0 13118.934
443.2373352050781 0 32234.328
443.26629638671875 0 10025.519
443.739013671875 0 34172.594 c Water loss 7
444.23638916015625 0 11685.609 c Ammonia loss 7
444.7411804199219 0 8167.916
445.1209411621094 0 50706.844
445.23504638671875 0 5485.2275
446.12176513671875 0 5320.764
446.2127380371094 0 31740.953
447.2159729003906 0 5302.0625
451.75146484375 0 15700.758
452.2452697753906 0 100274.195
452.746826171875 0 424966.5 c 7
453.24810791015625 0 181640.53
453.7492980957031 0 55235.54
454.25140380859375 0 11809.155
455.56658935546875 0 11540.239
455.89874267578125 0 11862.276
458.2723083496094 0 18857.127 c Ammonia loss 3
458.7587585449219 0 9445.177
461.56982421875 0 20283.168
461.904052734375 0 9080.941
462.2097473144531 0 7467.3267 z Water loss 8
463.2203369140625 0 5090.6523 y 4
466.2608642578125 0 5943.424
467.223876953125 0 6626.4067
471.2442932128906 0 11794.312
471.7389831542969 0 14360.262
472.2449645996094 0 7308.5386
472.74444580078125 0 8596.393
474.1951904296875 0 57676.496
474.2937316894531 0 7813.3545
475.2997131347656 0 1478998.9 c 3
476.3025207519531 0 315019.8
476.36077880859375 0 5985.6514
476.7310791015625 0 7454.1777
477.3045959472656 0 48695.4
478.264404296875 0 5477.2456
479.7559814453125 0 26109.48
480.2201232910156 0 24105.045 z 8
480.2520446777344 0 110021.055
480.7523498535156 0 65240.844
481.2297668457031 0 134176.94
482.2322082519531 0 22891.19
482.26873779296875 0 4551.2695
485.2426452636719 0 22868.361
485.73809814453125 0 23214.889
486.23876953125 0 11431.502
486.30157470703125 0 8827.88
487.2658996582031 0 14110.059
487.76153564453125 0 35737.844
488.2630920410156 0 22910.139
488.7684020996094 0 5385.7773
492.25177001953125 0 7794.292
494.2474670410156 0 60507.79
494.7484436035156 0 23014.73
495.2496643066406 0 6170.8486
495.7205810546875 0 5931.7085
496.2406311035156 0 72269.48 y 8
497.2441101074219 0 24769.89
499.766845703125 0 19343.125
500.2594299316406 0 123327.484
500.7606201171875 0 74052.94 c Water loss 8
501.2580261230469 0 50180.83 c Ammonia loss 8
501.75946044921875 0 26424.416
502.27410888671875 0 193682.95
502.76263427734375 0 14286.65
503.2763366699219 0 36323.758
504.2835998535156 0 4973.05
508.2642822265625 0 7156.586
508.77130126953125 0 37955.91
509.26678466796875 0 199023.62
509.7681579589844 0 606872.2 c 8
510.2695617675781 0 285186.78
510.77056884765625 0 76523.23
511.2672424316406 0 21904.193
514.2998046875 0 6423.001
516.3170776367188 0 5793.6016
521.271484375 0 8414.894
522.2716674804688 0 21525.256
522.770751953125 0 14376.572
523.2689819335938 0 14125.497
523.77099609375 0 7711.119
526.7490234375 0 29809.223
527.2510375976562 0 19184.209
527.7495727539062 0 8365.241
530.2941284179688 0 16269.501
530.7723999023438 0 15776.089
531.78076171875 0 7283.584
532.766357421875 0 9202.102
533.2590942382812 0 59866.477 z 3
533.75927734375 0 39222.38
534.2605590820312 0 16377.274
535.765869140625 0 18488.969
536.265869140625 0 16061.599
540.7683715820312 0 5261.5146
541.269287109375 0 13852.833 y 3
541.77490234375 0 5268.738
542.2772827148438 0 6052.5054
544.2764892578125 0 10946.0625 c Water loss 9
544.3203125 0 5244.2417
544.77294921875 0 56069.785 c Ammonia loss 9
545.2288208007812 0 6768.557
545.2721557617188 0 25307.877
545.3056640625 0 26778.988
545.7744750976562 0 7074.093
546.254150390625 0 7985.5693
546.3114013671875 0 28614.377
547.315673828125 0 7862.6895
548.2531127929688 0 11891.682
548.7437744140625 0 81103.734
549.2447509765625 0 51766.45
549.74560546875 0 18075.508
550.7879028320312 0 16988.234
552.253662109375 0 6439.9
552.7896728515625 0 38466.656
553.284423828125 0 820922.2 c 9
553.78564453125 0 437518.6
554.2868041992188 0 155713.47
554.7830200195312 0 31657.148
555.2796020507812 0 14745.756
562.2836303710938 0 14486.904
563.3209838867188 0 5814.8438
564.304443359375 0 61459.332
565.2625122070312 0 8171.791
565.3088989257812 0 21219.16
566.2933959960938 0 36901.363
566.794677734375 0 13711.285
567.27099609375 0 6334.5977
568.2675170898438 0 10887.965 z Ammonia loss 2
568.7686767578125 0 9031.527
569.269775390625 0 9677.975
573.3027954101562 0 26379.926 c Ammonia loss 4
574.301513671875 0 9248.8
575.7777099609375 0 24208.137 y Water loss 2
576.2757568359375 0 41108.53 y Ammonia loss 2
576.7750244140625 0 130957.98 z 2
577.276611328125 0 81449.81
577.7776489257812 0 29834.457
578.3101806640625 0 22803.104
578.8099975585938 0 24700.037
579.3082885742188 0 12884.656
579.81005859375 0 20625.48
580.310546875 0 18407.36
581.2613525390625 0 7069.7075
583.794189453125 0 16205.736
584.2901000976562 0 63318.33
584.7842407226562 0 833694.4 y 2
585.2854614257812 0 479183.88
585.7869262695312 0 170066.98
586.2881469726562 0 51730.992
586.78125 0 7690.6943
587.3163452148438 0 47351.684
587.8189086914062 0 52826.508
588.270263671875 0 8135.136
588.3204345703125 0 21642.049
588.768798828125 0 37269.617
588.8195190429688 0 7916.2334
589.2697143554688 0 15103.076
589.3165893554688 0 8559.827
589.768798828125 0 7098.693
590.3264770507812 0 1825179.6 c 4
591.3290405273438 0 517871.1
591.3873901367188 0 6640.6733
592.3295288085938 0 90926.3
592.8065185546875 0 37755.094
593.3086547851562 0 19938.037
593.8030395507812 0 5685.9805
598.2793579101562 0 9659.024
598.77734375 0 6649.127
600.8189697265625 0 17577.988 c Water loss 10
601.3146362304688 0 84235.25 c Ammonia loss 10
601.8152465820312 0 55180.742
602.3179931640625 0 35776.793
602.7838134765625 0 7882.881
602.8326416015625 0 6216.0767
605.2930297851562 0 7172.1807
607.8275756835938 0 19314.955
608.3175659179688 0 170161.86 z 7
608.8275146484375 0 5811.1343
609.3309326171875 0 165792.58
609.8267822265625 0 1040325 c 10
610.327880859375 0 571990.06
610.8285522460938 0 216799.75
611.3289794921875 0 56983.19
611.8302001953125 0 18831.281
613.3110961914062 0 5706.842
616.339599609375 0 5257.9
618.7975463867188 0 96128.87 w 1
619.2990112304688 0 61220.8
619.80322265625 0 19558.764
621.814208984375 0 6015.1777
622.30908203125 0 6741.5825
624.335693359375 0 127119.016 y 7
624.8152465820312 0 8415.191
625.3379516601562 0 38728.703
625.8037719726562 0 9874.032 w 1
625.8541259765625 0 11460.949
626.2978515625 0 7535.801
626.3543090820312 0 5943.7617
626.8576049804688 0 5466.244
629.3173828125 0 15653.701
630.3255004882812 0 8523.071
630.8202514648438 0 43276.89
631.3234252929688 0 22966.219
631.8392333984375 0 19462.594
632.3468017578125 0 8114.46
632.8157348632812 0 14259.864 y Ammonia loss 1
633.3156127929688 0 41839.125 z 1
633.814208984375 0 52351.336
634.3167724609375 0 22492.92
634.8175659179688 0 14323.62
635.3673706054688 0 6337.1807
636.3487548828125 0 22220.807
637.3514404296875 0 6027.823
638.3258666992188 0 34624.984
638.828369140625 0 30441.854
639.3301391601562 0 12263.796
639.8472900390625 0 23358.104
640.346923828125 0 48074.13
640.844482421875 0 34078.16
641.3260498046875 0 88792.65 y 1
641.827392578125 0 63081.996
642.3256225585938 0 20747.34
642.8340454101562 0 7613.9297
644.3439331054688 0 7645.8433
645.3236083984375 0 11082.5205
647.323974609375 0 184389.84
647.8483276367188 0 49178.984
648.3313598632812 0 61520.74
648.850341796875 0 11043.081
649.332763671875 0 15419.24
652.8358154296875 0 11352.327
653.3379516601562 0 47372.598
653.8357543945312 0 28294.332
654.3328247070312 0 24127.094
654.8342895507812 0 11461.497
655.318115234375 0 24639.81
655.8101196289062 0 107805.66
656.3114624023438 0 66587.46
656.8134155273438 0 22231.957
657.3168334960938 0 14848.149
659.348388671875 0 17710.95
660.340087890625 0 41506.695
660.8428955078125 0 29738.947
661.3466796875 0 95014.47 z Water loss 6
661.8457641601562 0 207032.44
662.3430786132812 0 222659.9
662.8406982421875 0 163723.8
663.3422241210938 0 80036.97
663.840576171875 0 23563.25
664.3380126953125 0 7769.332
669.3267822265625 0 55165.836
669.842529296875 0 353682.03
670.3438110351562 0 247343.23
670.8411865234375 0 376742.75
671.3408813476562 0 189289.27
671.8424072265625 0 66191.17
672.3428344726562 0 35502.363
673.3272705078125 0 22621.234
674.34130859375 0 35517.95
674.838134765625 0 72358.68
675.3377685546875 0 170583.4
675.8395385742188 0 93587.695
676.3335571289062 0 86743.48
676.83203125 0 36282.543
677.3351440429688 0 22812.695
677.835205078125 0 7913.279
679.3544311523438 0 522114.16 z 6
680.356689453125 0 179830.84
681.359130859375 0 43011.027
682.3567504882812 0 8807.326
682.85986328125 0 5296.681
683.34716796875 0 91176.875
683.8410034179688 0 931369.2
684.3424072265625 0 570754.56
684.8434448242188 0 266355.03
685.3446044921875 0 80026.46
685.8453369140625 0 20562.012
688.3294067382812 0 25671.271 c Ammonia loss 5
689.3306274414062 0 7456.2275
690.3392333984375 0 8050.384
691.357421875 0 63481.92
691.8501586914062 0 979710.4
692.3517456054688 0 624930.5
692.8524780273438 0 261328.23
693.353759765625 0 66238.79
693.8553466796875 0 13527.529
695.3731079101562 0 72813.625 y 6
696.3742065429688 0 28574.703
701.353515625 0 6934.94
702.3592529296875 0 5867.0176
704.3688354492188 0 69270.195
705.35400390625 0 506046.72 c 5
706.3561401367188 0 160772.12
707.3583374023438 0 43157.266
708.3622436523438 0 7856.3315
711.340087890625 0 8733.201
712.3466186523438 0 5433.4775
722.363037109375 0 5929.5337
731.3897094726562 0 7469.4497
732.3765869140625 0 81208.1
733.3797607421875 0 30464.85
734.382080078125 0 7490.04
737.3836059570312 0 240938.75
738.3201293945312 0 4713.793
738.3867797851562 0 87745.11
739.3883666992188 0 19265.572
749.3840942382812 0 33589.66 w 5
750.3878173828125 0 45028.492
751.3836059570312 0 17196.14
759.364990234375 0 31698.475 c Ammonia loss 6
760.366943359375 0 11690.88
761.3723754882812 0 8349.475
764.399169921875 0 6056.09
774.380126953125 0 5834.8955
775.4049682617188 0 78081.42
776.390625 0 670670.94 c 6
777.3933715820312 0 224380.14
778.3950805664062 0 53809.86
779.4005737304688 0 11164.673
787.3717651367188 0 11865.127
788.2900390625 0 4493.419
788.3768920898438 0 25182.4
789.3839721679688 0 24920.43
790.3826293945312 0 9288.1045
793.3970947265625 0 166730.62
794.3820190429688 0 1396572.2 z 5
795.3843994140625 0 518885.25
796.3865966796875 0 150822.31
797.38623046875 0 20636.045
803.3681640625 0 6348.366
810.4000244140625 0 51107.125 y 5
811.4006958007812 0 27138.35
812.400146484375 0 8183.9683
816.4877319335938 0 9713.551
821.427490234375 0 70352.945
822.4307250976562 0 25598.117
823.43408203125 0 10532.794
826.3701782226562 0 4901.33
837.3914794921875 0 5145.671
842.4598999023438 0 7990.9043
844.453369140625 0 7053.4316
859.48828125 0 23210.29
860.473876953125 0 102831.83
861.4783935546875 0 84498.414
862.4782104492188 0 25834.256
863.4874267578125 0 5625.2495
864.4337158203125 0 49422.11
865.4189453125 0 417966.47
866.4208984375 0 187134.69
867.4246215820312 0 56898.055
868.4278564453125 0 16202.673
869.4484252929688 0 8064.164
874.4061279296875 0 11751.853
875.410400390625 0 18199.074
876.410888671875 0 6639.941
885.4660034179688 0 46262.754
886.4707641601562 0 53579.73 c Water loss 7
887.462890625 0 56829.633 c Ammonia loss 7
888.4657592773438 0 18784.637
889.4729614257812 0 16379.566
891.4292602539062 0 14477.704
900.4668579101562 0 6975.5337
901.4636840820312 0 31278.52
902.4823608398438 0 39035.734
903.4832763671875 0 247219.2
904.4854125976562 0 1009521.3 c 7
905.48779296875 0 422690.38
906.489990234375 0 117008.555
907.494384765625 0 31461.715
908.423583984375 0 42980.098
909.4081420898438 0 695959.56 z 4
910.4109497070312 0 312438.94
911.4132690429688 0 104133.42
912.4148559570312 0 22056.94
925.4267578125 0 112941.01 y 4
926.4297485351562 0 49146.97
927.4301147460938 0 10074.971
928.4335327148438 0 6022.2744
952.44287109375 0 5949.4375
958.492431640625 0 6102.9585
959.5020751953125 0 9155.739
969.4771728515625 0 16006.632
970.463134765625 0 6226.928
974.5169677734375 0 17714.941
975.5194702148438 0 28784.328
976.5214233398438 0 15964.592
977.519287109375 0 5549.3105
986.5001220703125 0 9151.875
987.489990234375 0 42570.574
988.493408203125 0 48530.19
989.495849609375 0 27897.344
990.4973754882812 0 7878.3623
999.5184326171875 0 7581.776
1000.5220947265625 0 25622.426 c Water loss 8
1001.5103149414062 0 34399.36 c Ammonia loss 8
1002.5126342773438 0 18416.986
1003.5162963867188 0 26154.32
1004.51513671875 0 16969.598
1005.517333984375 0 7922.2324
1017.5320434570312 0 76332.28
1018.5286865234375 0 434032.1 c 8
1019.531494140625 0 210439.03
1020.5331420898438 0 77587.664
1021.530029296875 0 17687.68
1022.51708984375 0 13155.104
1023.4927368164062 0 9072.108
1044.5516357421875 0 7739.6646
1045.5260009765625 0 7256.607
1046.546142578125 0 10078.948
1047.534912109375 0 10290.793
1049.494384765625 0 15172.468
1050.5010986328125 0 9079.844
1061.5472412109375 0 16398.049
1062.55517578125 0 21789.2
1063.546142578125 0 25853.756
1064.5225830078125 0 34781.027
1065.509765625 0 359518.7 z 3
1066.5142822265625 0 280129.75
1067.516845703125 0 110214.99
1068.5203857421875 0 37463.434
1069.52880859375 0 7173.0176
1080.5247802734375 0 16737.697
1081.5264892578125 0 37891.31 y 3
1082.5284423828125 0 16959.73
1083.531982421875 0 6181.744
1088.5401611328125 0 12877.851 c Ammonia loss 9
1089.5430908203125 0 67455.85
1090.5439453125 0 31703.295
1091.546142578125 0 10659.728
1096.4769287109375 0 20413.174
1097.4837646484375 0 25592.37
1098.4910888671875 0 7429.6465
1101.5750732421875 0 6901.117
1104.5753173828125 0 15211.249
1105.5601806640625 0 206280.73 c 9
1106.564697265625 0 139976.8
1107.56640625 0 64792.008
1108.564208984375 0 20945.967
1110.525390625 0 11519.14
1126.5537109375 0 8466.205
1135.533935546875 0 9138.017 z Ammonia loss 2
1136.5299072265625 0 16054.436
1137.5355224609375 0 9259.722
1151.5516357421875 0 17671.043 y Ammonia loss 2
1152.542724609375 0 191947.67 z 2
1153.5477294921875 0 236233.81
1154.549560546875 0 108371.37
1155.5543212890625 0 33978.71
1156.5455322265625 0 5732.55
1158.619384765625 0 10901.748
1159.6187744140625 0 7434.373
1160.6177978515625 0 11375.872
1168.5584716796875 0 41183.883 y 2
1169.564208984375 0 33793.547
1170.568603515625 0 19103.484
1174.6326904296875 0 32876.152
1175.6414794921875 0 62154.184
1176.6373291015625 0 44946.312
1177.638427734375 0 21254.018
1185.6080322265625 0 8742.63
1186.6116943359375 0 6331.9653
1201.63623046875 0 13981.418 c Ammonia loss 10
1202.625732421875 0 86272.58
1203.628662109375 0 62562.133
1204.632568359375 0 20853.586
1205.628173828125 0 6646.894
1217.6693115234375 0 6778.974
1218.6434326171875 0 112670.12 c 10
1219.648681640625 0 149444.25
1220.65185546875 0 72979.4
1221.654541015625 0 26081.658
1222.6572265625 0 6956.2593
1251.5877685546875 0 6326.993
1265.6297607421875 0 14548.679 z 1
1266.630615234375 0 71424.97
1267.6341552734375 0 56573.996
1268.630126953125 0 23475.223
1280.64404296875 0 8361.233
1281.6507568359375 0 14306.504 y 1
1282.6483154296875 0 8076.97
1295.6251220703125 0 8214.351
1305.6748046875 0 16432.908
1306.642578125 0 71998.76
1307.6435546875 0 78722.2
1308.6478271484375 0 46858.633
1309.6419677734375 0 22517.977
1310.6478271484375 0 10793.206
1321.6839599609375 0 26770.055
1322.6817626953125 0 52843.863
1323.6778564453125 0 57315.617
1324.66552734375 0 261543.62
1325.666015625 0 243535.98
1326.6676025390625 0 118387.4
1327.6680908203125 0 36850.484
1328.6732177734375 0 10865.029
1337.691650390625 0 8771.713
1338.6929931640625 0 64818.754
1339.6861572265625 0 259005.17
1340.6871337890625 0 172379.36
1341.6845703125 0 115360.836
1342.687255859375 0 41921.234
1343.685791015625 0 12320.5
1349.6697998046875 0 68866.41
1350.6588134765625 0 117952.13
1351.65966796875 0 67818.2
1352.6575927734375 0 25748.232
1355.7071533203125 0 13099.844
1356.7098388671875 0 36969.06
1357.718505859375 0 16664.062
1365.686767578125 0 24313.953
1366.681640625 0 216510.36
1367.68017578125 0 964439.9
1368.681640625 0 599734.5
1369.6842041015625 0 254142.95
1370.6861572265625 0 77201.37
1371.693603515625 0 8909.366
1382.6932373046875 0 56178.273
1383.698974609375 0 170111
1384.7041015625 0 261694.3
1385.7059326171875 0 140898.62
1386.7081298828125 0 57570.38
1387.71044921875 0 13727.7
2253.1826171875 0 6126.8086
2326.7724609375 0 5465.9175

Spectrum Details

|  |  |
| --- | --- |
| Matched peaks? Matched peaks The total absolute number of peaks matched. Additionally in brackets the total fraction of peaks matched and the total number of peaks is shown. | 73 (11.79% of 619) |
| FDR? FDR The false discovery rate estimated for this peptide. It is calculated by matching all theoretical fragments with a non-integer shift with the raw peaks for this spectrum. This is done with 40 different shifts. The resulting percentage is the average number of annotated peaks over the number of annotated peaks with the correct spectrum. | 1.70% |
| Satellite FDR? Satellite FDR See the FDR for details on its calculation. This satellite ion specific FDR only contains the satellite ions (d/w) for I/L/J positions. | 0.00% |
| PSM Score? PSM Score The PSM Score as given by Hecklib to this annotated spectrum. It is shown with three significant figures. | 640 |

#### Spectrum 3545? Spectrum 3545 The raw spectrum of this peptide as annotated by Hecklib. The fragments are coloured according to ion type (see legend). Any peaks with a star '\*' as text can be hovered over to see the full details, first the ion type second the mass shift type. By hovering over the amino acids in the peptide or ions in the legend the corresponding peaks are highlighted. By toggling the 'Unassigned' label you can turn the background (unassigned) peaks on or off in the plot. By updating the slider in the Ion legend you can update the spectrum to only show the top X% of the peaks with labels. The top X% means any peak that is within X% of the highest intensity. By dragging in the spectrum you can zoom in to a specific part of the spectrum and use 'Zoom Out' to get back to the original zoom level. The annotation of the spectrum is based on the given sequence in the peptides file and is done with different software so inconsistencies are likely. The peaks are annotated based on the given sequence, with 20 ppm tolerance.

Copy Data

##### Spectrum 3545 (TSV)

###### Preview

```
Loading example...
```

*Click on the button to copy the data to your clipboard.*

Mz MinMz MaxIntensity Max

WidthHeightPeptide font sizePeptide stroke widthSpectrum font sizeSpectrum stroke widthCompact peptide

Ion legend

wxyz

abcd

OtherUnassignedIon^Charge^_Position_Show for top:%

TISRDDAKNSJY

03.19e+56.37e+59.56e+51.27e+6

Zoom Out

c^+3^_4_y^+1^_1_w^+3^_5_c^+2^_4_c^+1^_2_z^+1^_2_c^+2^_5_y^+1^_2_c^+1^_3_c^+3^_8_c^+1^_3_c^+2^_6_c^+2^_6_c^+3^_10_c^+3^_10_y^+1^_3_c^+3^_11_c^+3^_11_y^+3^_11_c^+2^_8_c^+2^_8_c^+2^_8_y^+2^_8_c^+1^_4_z^+1^_4_y^+2^_8_c^+1^_4_y^+1^_4_c^+2^_9_c^+2^_9_c^+2^_9_z^+2^_9_y^+2^_9_c^+2^_10_c^+2^_10_w^+1^_5_c^+2^_10_z^+2^_10_w^+2^_10_c^+1^_5_c^+1^_5_y^+2^_10_y^+2^_10_z^+2^_10_y^+2^_10_c^+1^_5_c^+2^_11_c^+2^_11_z^+1^_5_c^+2^_11_w^+2^_11_y^+1^_5_w^+2^_11_y^+2^_11_y^+2^_11_z^+2^_11_y^+2^_11_z^+1^_6_y^+1^_6_z^+1^_6_c^+1^_6_c^+1^_6_y^+1^_6_c^+1^_6_w^+1^_7_c^+1^_7_c^+1^_7_c^+1^_7_z^+1^_7_y^+1^_7_c^+1^_8_c^+1^_8_c^+1^_8_z^+1^_8_y^+1^_8_w^+1^_9_c^+1^_9_c^+1^_9_z^+1^_9_y^+1^_9_c^+1^_10_c^+1^_10_z^+1^_10_y^+1^_10_c^+1^_11_c^+1^_11_y^+1^_11_

035070010501401

Fragment Matches Table

Show background peaks

| Position | Ion type | Intensity | mz Theoretical | mz Error (Th) | mz Error (ppm) | Charge | Series Number |
| --- | --- | --- | --- | --- | --- | --- | --- |
| - | - | 5688 | 123 | - | - | 0 | - |
| - | - | 1488 | 123.1 | - | - | 0 | - |
| - | - | 860.3 | 124.1 | - | - | 0 | - |
| - | - | 1030 | 126.1 | - | - | 0 | - |
| - | - | 1377 | 126.1 | - | - | 0 | - |
| - | - | 1.235E+04 | 129.1 | - | - | 0 | - |
| - | - | 939.2 | 130.1 | - | - | 0 | - |
| - | - | 802.9 | 130.9 | - | - | 0 | - |
| - | - | 2016 | 131.1 | - | - | 0 | - |
| - | - | 1485 | 132.1 | - | - | 0 | - |
| - | - | 1297 | 133.1 | - | - | 0 | - |
| - | - | 2.078E+05 | 136.1 | - | - | 0 | - |
| - | - | 947.1 | 136.8 | - | - | 0 | - |
| - | - | 1989 | 137.1 | - | - | 0 | - |
| - | - | 1.885E+04 | 137.1 | - | - | 0 | - |
| - | - | 1.639E+04 | 142.1 | - | - | 0 | - |
| - | - | 1.81E+04 | 143.1 | - | - | 0 | - |
| - | - | 1986 | 143.1 | - | - | 0 | - |
| - | - | 5102 | 147 | - | - | 0 | - |
| - | - | 1330 | 149 | - | - | 0 | - |
| - | - | 910.1 | 149.1 | - | - | 0 | - |
| 4 | c | 1070 | 153.1 | 0.002236 | 14.6 | +3 | 4 |
| - | - | 1256 | 155.1 | - | - | 0 | - |
| - | - | 8184 | 155.1 | - | - | 0 | - |
| - | - | 1078 | 156.1 | - | - | 0 | - |
| - | - | 4097 | 157.1 | - | - | 0 | - |
| - | - | 1421 | 157.1 | - | - | 0 | - |
| - | - | 1544 | 158.1 | - | - | 0 | - |
| - | - | 1.856E+05 | 165.1 | - | - | 0 | - |
| - | - | 1.746E+04 | 166.1 | - | - | 0 | - |
| - | - | 1156 | 168.1 | - | - | 0 | - |
| - | - | 1.325E+04 | 169.1 | - | - | 0 | - |
| - | - | 3115 | 171.1 | - | - | 0 | - |
| - | - | 1910 | 171.1 | - | - | 0 | - |
| - | - | 2.051E+04 | 173.1 | - | - | 0 | - |
| - | - | 5001 | 173.5 | - | - | 0 | - |
| - | - | 4922 | 174.1 | - | - | 0 | - |
| - | - | 1337 | 174.1 | - | - | 0 | - |
| - | - | 1838 | 175.1 | - | - | 0 | - |
| - | - | 1037 | 181.1 | - | - | 0 | - |
| 12 | y | 6.584E+05 | 182.1 | 0.0006785 | 3.726 | +1 | 1 |
| - | - | 5.956E+04 | 183.1 | - | - | 0 | - |
| - | - | 3027 | 183.1 | - | - | 0 | - |
| - | - | 3780 | 184.1 | - | - | 0 | - |
| 8 | w | 4727 | 184.1 | 0.0005451 | 2.961 | +3 | 5 |
| - | - | 1939 | 184.1 | - | - | 0 | - |
| - | - | 1761 | 185.1 | - | - | 0 | - |
| - | - | 3241 | 186.1 | - | - | 0 | - |
| - | - | 1.197E+06 | 187.1 | - | - | 0 | - |
| - | - | 1.119E+05 | 188.1 | - | - | 0 | - |
| - | - | 6606 | 189.2 | - | - | 0 | - |
| - | - | 6919 | 197.1 | - | - | 0 | - |
| - | - | 4465 | 197.2 | - | - | 0 | - |
| - | - | 5095 | 200.1 | - | - | 0 | - |
| - | - | 2.537E+04 | 201.1 | - | - | 0 | - |
| - | - | 1699 | 201.1 | - | - | 0 | - |
| - | - | 1494 | 201.1 | - | - | 0 | - |
| - | - | 2.235E+04 | 202.1 | - | - | 0 | - |
| - | - | 1417 | 202.1 | - | - | 0 | - |
| - | - | 1121 | 203.1 | - | - | 0 | - |
| - | - | 1160 | 203.1 | - | - | 0 | - |
| - | - | 2672 | 203.1 | - | - | 0 | - |
| - | - | 1473 | 210.1 | - | - | 0 | - |
| - | - | 1353 | 213.1 | - | - | 0 | - |
| - | - | 2.91E+05 | 215.1 | - | - | 0 | - |
| - | - | 3.082E+04 | 216.1 | - | - | 0 | - |
| - | - | 2641 | 217.1 | - | - | 0 | - |
| - | - | 1162 | 221.1 | - | - | 0 | - |
| - | - | 1218 | 223.1 | - | - | 0 | - |
| - | - | 1164 | 225 | - | - | 0 | - |
| - | - | 4144 | 227.1 | - | - | 0 | - |
| - | - | 1.047E+04 | 229.1 | - | - | 0 | - |
| 4 | c | 2696 | 229.6 | 0.001854 | 8.072 | +2 | 4 |
| - | - | 1681 | 230.1 | - | - | 0 | - |
| 2 | c | 7164 | 232.2 | 0.0009359 | 4.031 | +1 | 2 |
| - | - | 1693 | 243.1 | - | - | 0 | - |
| - | - | 1822 | 244.1 | - | - | 0 | - |
| - | - | 6267 | 244.1 | - | - | 0 | - |
| - | - | 2109 | 245.2 | - | - | 0 | - |
| - | - | 1919 | 249.2 | - | - | 0 | - |
| - | - | 1277 | 253.1 | - | - | 0 | - |
| - | - | 1320 | 261.2 | - | - | 0 | - |
| - | - | 2515 | 262.2 | - | - | 0 | - |
| - | - | 6253 | 266.2 | - | - | 0 | - |
| - | - | 1890 | 267.2 | - | - | 0 | - |
| - | - | 2597 | 268.1 | - | - | 0 | - |
| - | - | 3796 | 269.2 | - | - | 0 | - |
| - | - | 6027 | 270.1 | - | - | 0 | - |
| - | - | 1538 | 274.2 | - | - | 0 | - |
| 11 | z | 2155 | 279.1 | 0.0005851 | 2.096 | +1 | 2 |
| - | - | 2048 | 280.1 | - | - | 0 | - |
| - | - | 3242 | 281.1 | - | - | 0 | - |
| - | - | 1.893E+04 | 282.1 | - | - | 0 | - |
| - | - | 1541 | 283.1 | - | - | 0 | - |
| - | - | 3318 | 283.1 | - | - | 0 | - |
| - | - | 1419 | 285.2 | - | - | 0 | - |
| 5 | c | 2205 | 287.2 | 0.002664 | 9.279 | +2 | 5 |
| - | - | 6804 | 287.2 | - | - | 0 | - |
| 11 | y | 3.637E+04 | 295.2 | 0.0009346 | 3.166 | +1 | 2 |
| - | - | 5197 | 296.2 | - | - | 0 | - |
| - | - | 1.456E+04 | 297.2 | - | - | 0 | - |
| - | - | 8400 | 298.1 | - | - | 0 | - |
| - | - | 2967 | 298.2 | - | - | 0 | - |
| - | - | 7662 | 299.1 | - | - | 0 | - |
| - | - | 6601 | 299.2 | - | - | 0 | - |
| - | - | 5.152E+04 | 300.2 | - | - | 0 | - |
| - | - | 2915 | 301.1 | - | - | 0 | - |
| - | - | 8520 | 301.2 | - | - | 0 | - |
| 3 | c | 1335 | 301.2 | 0.0007124 | 2.365 | +1 | 3 |
| 8 | c | 7712 | 302.2 | 0.005559 | 18.4 | +3 | 8 |
| - | - | 1310 | 303.2 | - | - | 0 | - |
| - | - | 3975 | 310.1 | - | - | 0 | - |
| - | - | 1471 | 313.1 | - | - | 0 | - |
| - | - | 3.666E+04 | 315.2 | - | - | 0 | - |
| - | - | 2076 | 315.2 | - | - | 0 | - |
| - | - | 2193 | 316.2 | - | - | 0 | - |
| - | - | 4881 | 316.2 | - | - | 0 | - |
| - | - | 2.953E+04 | 317.2 | - | - | 0 | - |
| - | - | 3711 | 318.2 | - | - | 0 | - |
| 3 | c | 3.946E+04 | 319.2 | 0.001134 | 3.553 | +1 | 3 |
| - | - | 5272 | 320.2 | - | - | 0 | - |
| - | - | 1138 | 329.5 | - | - | 0 | - |
| - | - | 5064 | 330.2 | - | - | 0 | - |
| - | - | 3129 | 335.7 | - | - | 0 | - |
| - | - | 1257 | 335.7 | - | - | 0 | - |
| - | - | 3154 | 337.2 | - | - | 0 | - |
| - | - | 7366 | 341 | - | - | 0 | - |
| - | - | 8603 | 342.2 | - | - | 0 | - |
| - | - | 6168 | 343.2 | - | - | 0 | - |
| 6 | c | 1489 | 344.2 | 0.002166 | 6.294 | +2 | 6 |
| 6 | c | 4504 | 344.7 | 0.001125 | 3.265 | +2 | 6 |
| - | - | 2157 | 345.2 | - | - | 0 | - |
| - | - | 1476 | 351.5 | - | - | 0 | - |
| - | - | 1157 | 352.2 | - | - | 0 | - |
| - | - | 1413 | 352.5 | - | - | 0 | - |
| - | - | 1401 | 354.2 | - | - | 0 | - |
| - | - | 1153 | 356.2 | - | - | 0 | - |
| - | - | 2504 | 357.2 | - | - | 0 | - |
| - | - | 1138 | 357.2 | - | - | 0 | - |
| - | - | 4326 | 357.5 | - | - | 0 | - |
| - | - | 3268 | 357.8 | - | - | 0 | - |
| - | - | 1871 | 357.9 | - | - | 0 | - |
| - | - | 3155 | 358.2 | - | - | 0 | - |
| - | - | 3154 | 358.5 | - | - | 0 | - |
| - | - | 1679 | 358.9 | - | - | 0 | - |
| - | - | 1.59E+04 | 359 | - | - | 0 | - |
| - | - | 1077 | 359.1 | - | - | 0 | - |
| - | - | 7155 | 359.2 | - | - | 0 | - |
| - | - | 1489 | 360.2 | - | - | 0 | - |
| 10 | c | 1994 | 363.2 | 0.001215 | 3.347 | +3 | 10 |
| 10 | c | 6551 | 363.5 | 0.00277 | 7.619 | +3 | 10 |
| - | - | 4554 | 363.9 | - | - | 0 | - |
| - | - | 4510 | 366.2 | - | - | 0 | - |
| - | - | 5689 | 367.2 | - | - | 0 | - |
| - | - | 1813 | 370.1 | - | - | 0 | - |
| - | - | 5419 | 373.2 | - | - | 0 | - |
| - | - | 1541 | 373.7 | - | - | 0 | - |
| - | - | 6312 | 374.2 | - | - | 0 | - |
| - | - | 1509 | 375.9 | - | - | 0 | - |
| - | - | 1455 | 377.2 | - | - | 0 | - |
| - | - | 1566 | 379.5 | - | - | 0 | - |
| - | - | 7053 | 379.9 | - | - | 0 | - |
| - | - | 5710 | 380.2 | - | - | 0 | - |
| - | - | 1417 | 380.2 | - | - | 0 | - |
| - | - | 2154 | 380.5 | - | - | 0 | - |
| - | - | 1329 | 380.7 | - | - | 0 | - |
| - | - | 1427 | 380.9 | - | - | 0 | - |
| - | - | 1775 | 381.2 | - | - | 0 | - |
| 10 | y | 1.362E+04 | 382.2 | 0.001194 | 3.123 | +1 | 3 |
| - | - | 3203 | 383.2 | - | - | 0 | - |
| - | - | 3837 | 385.2 | - | - | 0 | - |
| - | - | 4189 | 385.2 | - | - | 0 | - |
| - | - | 1.093E+04 | 385.5 | - | - | 0 | - |
| - | - | 3635 | 385.7 | - | - | 0 | - |
| - | - | 6.216E+04 | 385.9 | - | - | 0 | - |
| - | - | 3.868E+04 | 386.2 | - | - | 0 | - |
| - | - | 1.56E+04 | 386.5 | - | - | 0 | - |
| - | - | 5893 | 386.9 | - | - | 0 | - |
| - | - | 3653 | 387.2 | - | - | 0 | - |
| - | - | 8324 | 387.2 | - | - | 0 | - |
| - | - | 1520 | 387.5 | - | - | 0 | - |
| - | - | 1623 | 388.2 | - | - | 0 | - |
| - | - | 1298 | 389.2 | - | - | 0 | - |
| - | - | 1751 | 389.5 | - | - | 0 | - |
| - | - | 1631 | 389.9 | - | - | 0 | - |
| - | - | 3.51E+04 | 391.6 | - | - | 0 | - |
| - | - | 1.439E+05 | 391.9 | - | - | 0 | - |
| - | - | 8.346E+04 | 392.2 | - | - | 0 | - |
| - | - | 1684 | 392.2 | - | - | 0 | - |
| - | - | 1155 | 392.3 | - | - | 0 | - |
| - | - | 3.276E+04 | 392.5 | - | - | 0 | - |
| - | - | 6978 | 392.9 | - | - | 0 | - |
| - | - | 3137 | 393.2 | - | - | 0 | - |
| - | - | 3900 | 393.7 | - | - | 0 | - |
| - | - | 1.131E+04 | 394.2 | - | - | 0 | - |
| - | - | 1145 | 394.2 | - | - | 0 | - |
| - | - | 2122 | 394.7 | - | - | 0 | - |
| - | - | 8255 | 395.2 | - | - | 0 | - |
| - | - | 3877 | 395.5 | - | - | 0 | - |
| - | - | 1919 | 395.9 | - | - | 0 | - |
| - | - | 4479 | 398.2 | - | - | 0 | - |
| - | - | 2690 | 400.3 | - | - | 0 | - |
| 11 | c | 1834 | 400.9 | 0.001169 | 2.915 | +3 | 11 |
| 11 | c | 6169 | 401.2 | 0.003028 | 7.547 | +3 | 11 |
| - | - | 2735 | 401.5 | - | - | 0 | - |
| - | - | 1652 | 401.9 | - | - | 0 | - |
| - | - | 1483 | 406.7 | - | - | 0 | - |
| - | - | 9938 | 408.2 | - | - | 0 | - |
| - | - | 1.055E+04 | 415.3 | - | - | 0 | - |
| - | - | 1892 | 415.7 | - | - | 0 | - |
| - | - | 2143 | 416.2 | - | - | 0 | - |
| - | - | 1943 | 416.3 | - | - | 0 | - |
| - | - | 1508 | 419.2 | - | - | 0 | - |
| - | - | 1842 | 419.7 | - | - | 0 | - |
| - | - | 1287 | 420.2 | - | - | 0 | - |
| - | - | 2446 | 423.7 | - | - | 0 | - |
| - | - | 7556 | 424.7 | - | - | 0 | - |
| - | - | 1.074E+04 | 425.3 | - | - | 0 | - |
| - | - | 1.399E+04 | 426.2 | - | - | 0 | - |
| - | - | 2899 | 427.2 | - | - | 0 | - |
| 2 | y | 2237 | 427.9 | 0.00136 | 3.177 | +3 | 11 |
| - | - | 1710 | 428.2 | - | - | 0 | - |
| - | - | 6210 | 428.7 | - | - | 0 | - |
| - | - | 5.623E+04 | 429.1 | - | - | 0 | - |
| - | - | 7545 | 429.2 | - | - | 0 | - |
| - | - | 3811 | 429.2 | - | - | 0 | - |
| - | - | 2471 | 429.7 | - | - | 0 | - |
| - | - | 7081 | 429.7 | - | - | 0 | - |
| - | - | 2801 | 429.8 | - | - | 0 | - |
| - | - | 1841 | 430.2 | - | - | 0 | - |
| - | - | 1494 | 430.2 | - | - | 0 | - |
| - | - | 6229 | 430.2 | - | - | 0 | - |
| - | - | 9308 | 430.3 | - | - | 0 | - |
| - | - | 2723 | 430.7 | - | - | 0 | - |
| - | - | 1668 | 431.2 | - | - | 0 | - |
| - | - | 4922 | 431.2 | - | - | 0 | - |
| - | - | 4370 | 431.3 | - | - | 0 | - |
| - | - | 3836 | 435.2 | - | - | 0 | - |
| - | - | 2040 | 436.2 | - | - | 0 | - |
| - | - | 1869 | 436.7 | - | - | 0 | - |
| - | - | 7142 | 437.2 | - | - | 0 | - |
| - | - | 2.191E+04 | 437.7 | - | - | 0 | - |
| - | - | 6031 | 438.2 | - | - | 0 | - |
| - | - | 1.64E+04 | 438.7 | - | - | 0 | - |
| - | - | 1.032E+04 | 439.2 | - | - | 0 | - |
| - | - | 1645 | 439.7 | - | - | 0 | - |
| - | - | 2417 | 440.6 | - | - | 0 | - |
| - | - | 1803 | 440.9 | - | - | 0 | - |
| - | - | 1.036E+04 | 442.7 | - | - | 0 | - |
| - | - | 8769 | 443.2 | - | - | 0 | - |
| - | - | 2.897E+04 | 443.3 | - | - | 0 | - |
| 8 | c | 1.685E+04 | 443.7 | 0.0008556 | 1.928 | +2 | 8 |
| 8 | c | 1.27E+04 | 444.2 | 0.007017 | 15.8 | +2 | 8 |
| - | - | 5455 | 444.7 | - | - | 0 | - |
| - | - | 2.995E+04 | 445.1 | - | - | 0 | - |
| - | - | 1822 | 445.2 | - | - | 0 | - |
| - | - | 1936 | 446.1 | - | - | 0 | - |
| - | - | 9431 | 446.2 | - | - | 0 | - |
| - | - | 1931 | 446.6 | - | - | 0 | - |
| - | - | 2178 | 446.9 | - | - | 0 | - |
| - | - | 1654 | 449.6 | - | - | 0 | - |
| - | - | 1314 | 450.2 | - | - | 0 | - |
| - | - | 1295 | 450.6 | - | - | 0 | - |
| - | - | 4428 | 450.7 | - | - | 0 | - |
| - | - | 1351 | 451.2 | - | - | 0 | - |
| - | - | 2.909E+04 | 451.8 | - | - | 0 | - |
| - | - | 8.692E+04 | 452.3 | - | - | 0 | - |
| 8 | c | 1.517E+05 | 452.7 | 0.00302 | 6.669 | +2 | 8 |
| - | - | 5.342E+04 | 453.2 | - | - | 0 | - |
| - | - | 1.358E+04 | 453.8 | - | - | 0 | - |
| 5 | y | 1516 | 454.2 | 0.007715 | 16.98 | +2 | 8 |
| - | - | 2641 | 454.3 | - | - | 0 | - |
| - | - | 4355 | 455.2 | - | - | 0 | - |
| - | - | 2.711E+04 | 455.6 | - | - | 0 | - |
| - | - | 2.241E+04 | 455.9 | - | - | 0 | - |
| - | - | 8295 | 456.2 | - | - | 0 | - |
| - | - | 2230 | 456.6 | - | - | 0 | - |
| - | - | 2073 | 457.2 | - | - | 0 | - |
| 4 | c | 1.086E+04 | 458.3 | 0.001035 | 2.259 | +1 | 4 |
| - | - | 2011 | 458.8 | - | - | 0 | - |
| - | - | 2991 | 459.3 | - | - | 0 | - |
| - | - | 7593 | 460.7 | - | - | 0 | - |
| - | - | 5687 | 461.2 | - | - | 0 | - |
| - | - | 4.025E+04 | 461.6 | - | - | 0 | - |
| - | - | 2.891E+04 | 461.9 | - | - | 0 | - |
| 9 | z | 1528 | 462.2 | 0.004785 | 10.35 | +1 | 4 |
| - | - | 1.585E+04 | 462.2 | - | - | 0 | - |
| - | - | 3546 | 462.6 | - | - | 0 | - |
| - | - | 7789 | 462.7 | - | - | 0 | - |
| 5 | y | 5026 | 463.2 | 0.007315 | 15.79 | +2 | 8 |
| - | - | 1318 | 465.7 | - | - | 0 | - |
| - | - | 3152 | 467.2 | - | - | 0 | - |
| - | - | 5644 | 467.7 | - | - | 0 | - |
| - | - | 1759 | 468.2 | - | - | 0 | - |
| - | - | 3108 | 469.7 | - | - | 0 | - |
| - | - | 2440 | 470.2 | - | - | 0 | - |
| - | - | 4931 | 470.8 | - | - | 0 | - |
| - | - | 2.956E+04 | 471.2 | - | - | 0 | - |
| - | - | 2.555E+04 | 471.7 | - | - | 0 | - |
| - | - | 9403 | 472.2 | - | - | 0 | - |
| - | - | 6289 | 472.7 | - | - | 0 | - |
| - | - | 3022 | 473.2 | - | - | 0 | - |
| - | - | 4195 | 473.2 | - | - | 0 | - |
| - | - | 2.672E+04 | 474.2 | - | - | 0 | - |
| - | - | 1578 | 474.3 | - | - | 0 | - |
| 4 | c | 5.335E+05 | 475.3 | 0.001677 | 3.528 | +1 | 4 |
| - | - | 3277 | 476.2 | - | - | 0 | - |
| - | - | 1.098E+05 | 476.3 | - | - | 0 | - |
| - | - | 1.728E+04 | 476.7 | - | - | 0 | - |
| - | - | 6376 | 477.2 | - | - | 0 | - |
| - | - | 1.91E+04 | 477.3 | - | - | 0 | - |
| - | - | 3698 | 477.7 | - | - | 0 | - |
| - | - | 1991 | 478.3 | - | - | 0 | - |
| - | - | 5801 | 478.7 | - | - | 0 | - |
| - | - | 2527 | 479.3 | - | - | 0 | - |
| - | - | 5.946E+04 | 479.8 | - | - | 0 | - |
| - | - | 2.258E+05 | 480.3 | - | - | 0 | - |
| - | - | 1.089E+05 | 480.8 | - | - | 0 | - |
| - | - | 4.069E+04 | 481.2 | - | - | 0 | - |
| - | - | 1.927E+04 | 481.3 | - | - | 0 | - |
| - | - | 7664 | 481.8 | - | - | 0 | - |
| - | - | 1.467E+04 | 482.2 | - | - | 0 | - |
| - | - | 4473 | 483.2 | - | - | 0 | - |
| - | - | 1452 | 484.2 | - | - | 0 | - |
| - | - | 6636 | 484.8 | - | - | 0 | - |
| - | - | 4.272E+04 | 485.2 | - | - | 0 | - |
| - | - | 3.856E+04 | 485.7 | - | - | 0 | - |
| - | - | 1.562E+04 | 486.2 | - | - | 0 | - |
| - | - | 4542 | 486.3 | - | - | 0 | - |
| - | - | 5506 | 486.7 | - | - | 0 | - |
| - | - | 3614 | 486.8 | - | - | 0 | - |
| - | - | 2.096E+04 | 487.3 | - | - | 0 | - |
| - | - | 1.825E+04 | 487.8 | - | - | 0 | - |
| - | - | 1.191E+04 | 488.3 | - | - | 0 | - |
| - | - | 2914 | 488.8 | - | - | 0 | - |
| - | - | 2471 | 489.3 | - | - | 0 | - |
| - | - | 2434 | 491.3 | - | - | 0 | - |
| - | - | 4998 | 491.8 | - | - | 0 | - |
| - | - | 2.062E+04 | 492.3 | - | - | 0 | - |
| - | - | 1.102E+04 | 492.7 | - | - | 0 | - |
| - | - | 3247 | 493.3 | - | - | 0 | - |
| - | - | 1.545E+04 | 493.8 | - | - | 0 | - |
| - | - | 8.35E+04 | 494.2 | - | - | 0 | - |
| - | - | 4.075E+04 | 494.7 | - | - | 0 | - |
| - | - | 1.032E+04 | 495.2 | - | - | 0 | - |
| - | - | 3762 | 495.7 | - | - | 0 | - |
| - | - | 2464 | 495.8 | - | - | 0 | - |
| 9 | y | 3.382E+04 | 496.2 | 0.001143 | 2.304 | +1 | 4 |
| - | - | 2128 | 496.3 | - | - | 0 | - |
| - | - | 7170 | 497.2 | - | - | 0 | - |
| - | - | 1423 | 498.3 | - | - | 0 | - |
| - | - | 2.156E+04 | 499.8 | - | - | 0 | - |
| - | - | 6.511E+04 | 500.3 | - | - | 0 | - |
| 9 | c | 4.156E+04 | 500.8 | 0.0009983 | 1.994 | +2 | 9 |
| 9 | c | 5.972E+04 | 501.3 | 0.003314 | 6.612 | +2 | 9 |
| - | - | 3.332E+04 | 501.8 | - | - | 0 | - |
| - | - | 5.161E+04 | 502.3 | - | - | 0 | - |
| - | - | 9865 | 502.8 | - | - | 0 | - |
| - | - | 1.086E+04 | 503.3 | - | - | 0 | - |
| - | - | 8679 | 503.3 | - | - | 0 | - |
| - | - | 5602 | 503.8 | - | - | 0 | - |
| - | - | 1250 | 504.3 | - | - | 0 | - |
| - | - | 1139 | 504.3 | - | - | 0 | - |
| - | - | 1486 | 504.8 | - | - | 0 | - |
| - | - | 3771 | 508.3 | - | - | 0 | - |
| - | - | 4.344E+04 | 508.8 | - | - | 0 | - |
| - | - | 1.72E+05 | 509.3 | - | - | 0 | - |
| 9 | c | 3.702E+05 | 509.8 | 0.001667 | 3.27 | +2 | 9 |
| - | - | 1.594E+05 | 510.3 | - | - | 0 | - |
| - | - | 5.386E+04 | 510.8 | - | - | 0 | - |
| - | - | 1.699E+04 | 511.3 | - | - | 0 | - |
| - | - | 3102 | 511.8 | - | - | 0 | - |
| - | - | 2114 | 512.3 | - | - | 0 | - |
| - | - | 3116 | 513.3 | - | - | 0 | - |
| - | - | 1333 | 513.8 | - | - | 0 | - |
| - | - | 9579 | 514.3 | - | - | 0 | - |
| - | - | 1684 | 515.3 | - | - | 0 | - |
| - | - | 2008 | 517.8 | - | - | 0 | - |
| - | - | 3222 | 518.2 | - | - | 0 | - |
| - | - | 2145 | 521.8 | - | - | 0 | - |
| - | - | 6789 | 522.3 | - | - | 0 | - |
| - | - | 4429 | 522.8 | - | - | 0 | - |
| - | - | 1E+04 | 523.3 | - | - | 0 | - |
| - | - | 3382 | 523.8 | - | - | 0 | - |
| - | - | 1761 | 524.3 | - | - | 0 | - |
| - | - | 1365 | 525.7 | - | - | 0 | - |
| - | - | 7786 | 526.3 | - | - | 0 | - |
| - | - | 1.227E+04 | 526.8 | - | - | 0 | - |
| - | - | 1.515E+04 | 527.3 | - | - | 0 | - |
| - | - | 3479 | 527.8 | - | - | 0 | - |
| - | - | 1985 | 527.8 | - | - | 0 | - |
| - | - | 2334 | 528.3 | - | - | 0 | - |
| - | - | 2979 | 528.3 | - | - | 0 | - |
| - | - | 2494 | 529.3 | - | - | 0 | - |
| - | - | 1.477E+04 | 530.3 | - | - | 0 | - |
| - | - | 1.43E+04 | 530.8 | - | - | 0 | - |
| - | - | 7256 | 531.3 | - | - | 0 | - |
| - | - | 5273 | 531.8 | - | - | 0 | - |
| - | - | 7366 | 532.8 | - | - | 0 | - |
| 4 | z | 3.252E+04 | 533.3 | 0.0032 | 6.002 | +2 | 9 |
| - | - | 1.47E+04 | 533.8 | - | - | 0 | - |
| - | - | 7326 | 534.3 | - | - | 0 | - |
| - | - | 3214 | 534.3 | - | - | 0 | - |
| - | - | 3240 | 534.8 | - | - | 0 | - |
| - | - | 1.277E+04 | 535.3 | - | - | 0 | - |
| - | - | 4.778E+04 | 535.8 | - | - | 0 | - |
| - | - | 3.172E+04 | 536.3 | - | - | 0 | - |
| - | - | 1.943E+04 | 536.8 | - | - | 0 | - |
| - | - | 1.385E+04 | 537.3 | - | - | 0 | - |
| - | - | 4534 | 537.8 | - | - | 0 | - |
| - | - | 2592 | 540.3 | - | - | 0 | - |
| - | - | 6625 | 540.8 | - | - | 0 | - |
| 4 | y | 1.272E+04 | 541.3 | 0.002383 | 4.403 | +2 | 9 |
| - | - | 1.13E+04 | 541.8 | - | - | 0 | - |
| - | - | 1.048E+04 | 542.3 | - | - | 0 | - |
| - | - | 3797 | 542.8 | - | - | 0 | - |
| - | - | 2433 | 543.3 | - | - | 0 | - |
| - | - | 1758 | 543.8 | - | - | 0 | - |
| 10 | c | 3.208E+04 | 544.3 | 0.002593 | 4.764 | +2 | 10 |
| 10 | c | 1.146E+05 | 544.8 | 0.00265 | 4.865 | +2 | 10 |
| - | - | 1.091E+04 | 545.2 | - | - | 0 | - |
| - | - | 4.386E+04 | 545.3 | - | - | 0 | - |
| - | - | 1.948E+04 | 545.3 | - | - | 0 | - |
| - | - | 2.088E+04 | 545.8 | - | - | 0 | - |
| - | - | 3735 | 546.2 | - | - | 0 | - |
| - | - | 2493 | 546.3 | - | - | 0 | - |
| - | - | 1.652E+04 | 546.3 | - | - | 0 | - |
| - | - | 2515 | 546.8 | - | - | 0 | - |
| - | - | 3103 | 547.3 | - | - | 0 | - |
| - | - | 1860 | 547.8 | - | - | 0 | - |
| - | - | 8041 | 548.3 | - | - | 0 | - |
| - | - | 3.482E+04 | 548.7 | - | - | 0 | - |
| - | - | 2574 | 548.8 | - | - | 0 | - |
| - | - | 1.758E+04 | 549.2 | - | - | 0 | - |
| - | - | 2680 | 549.3 | - | - | 0 | - |
| - | - | 6923 | 549.7 | - | - | 0 | - |
| 8 | w | 2260 | 550.3 | 0.001992 | 3.621 | +1 | 5 |
| - | - | 2536 | 550.3 | - | - | 0 | - |
| - | - | 1.941E+04 | 550.8 | - | - | 0 | - |
| - | - | 8829 | 551.3 | - | - | 0 | - |
| - | - | 3411 | 551.8 | - | - | 0 | - |
| - | - | 2005 | 552.3 | - | - | 0 | - |
| - | - | 1985 | 552.3 | - | - | 0 | - |
| - | - | 5.279E+04 | 552.8 | - | - | 0 | - |
| 10 | c | 3.065E+05 | 553.3 | 0.002132 | 3.854 | +2 | 10 |
| - | - | 1.604E+05 | 553.8 | - | - | 0 | - |
| - | - | 5.972E+04 | 554.3 | - | - | 0 | - |
| - | - | 2.281E+04 | 554.8 | - | - | 0 | - |
| - | - | 7975 | 555.3 | - | - | 0 | - |
| - | - | 2061 | 555.8 | - | - | 0 | - |
| - | - | 2344 | 556.3 | - | - | 0 | - |
| - | - | 1382 | 556.8 | - | - | 0 | - |
| - | - | 3960 | 557.3 | - | - | 0 | - |
| - | - | 6021 | 557.8 | - | - | 0 | - |
| - | - | 4078 | 558.3 | - | - | 0 | - |
| - | - | 1846 | 559.8 | - | - | 0 | - |
| - | - | 2023 | 560.8 | - | - | 0 | - |
| - | - | 9465 | 561.3 | - | - | 0 | - |
| - | - | 1.377E+04 | 561.8 | - | - | 0 | - |
| - | - | 8973 | 562.3 | - | - | 0 | - |
| - | - | 4534 | 562.8 | - | - | 0 | - |
| - | - | 2354 | 563.3 | - | - | 0 | - |
| - | - | 1769 | 563.3 | - | - | 0 | - |
| - | - | 1.769E+04 | 564.3 | - | - | 0 | - |
| - | - | 2108 | 565.3 | - | - | 0 | - |
| - | - | 1.012E+04 | 565.3 | - | - | 0 | - |
| - | - | 3731 | 565.8 | - | - | 0 | - |
| - | - | 2.372E+04 | 566.3 | - | - | 0 | - |
| - | - | 1.209E+04 | 566.8 | - | - | 0 | - |
| - | - | 8919 | 567.3 | - | - | 0 | - |
| 3 | z | 5600 | 567.8 | 0.004432 | 7.805 | +2 | 10 |
| 3 | w | 7538 | 568.3 | 0.003632 | 6.391 | +2 | 10 |
| - | - | 1.215E+04 | 568.8 | - | - | 0 | - |
| - | - | 6352 | 569.3 | - | - | 0 | - |
| - | - | 5730 | 569.3 | - | - | 0 | - |
| - | - | 2.063E+04 | 569.8 | - | - | 0 | - |
| - | - | 1.543E+04 | 570.3 | - | - | 0 | - |
| - | - | 6451 | 570.8 | - | - | 0 | - |
| - | - | 4027 | 571.3 | - | - | 0 | - |
| 5 | c | 2022 | 572.3 | 0.00309 | 5.4 | +1 | 5 |
| - | - | 2317 | 572.8 | - | - | 0 | - |
| 5 | c | 1.885E+04 | 573.3 | 0.003755 | 6.55 | +1 | 5 |
| - | - | 7139 | 574.3 | - | - | 0 | - |
| - | - | 1.172E+04 | 574.8 | - | - | 0 | - |
| - | - | 1.333E+04 | 575.3 | - | - | 0 | - |
| 3 | y | 4.027E+04 | 575.8 | 0.002516 | 4.369 | +2 | 10 |
| 3 | y | 6.411E+04 | 576.3 | 0.00825 | 14.32 | +2 | 10 |
| 3 | z | 1.037E+05 | 576.8 | 0.002811 | 4.874 | +2 | 10 |
| - | - | 5.676E+04 | 577.3 | - | - | 0 | - |
| - | - | 2.317E+04 | 577.8 | - | - | 0 | - |
| - | - | 3898 | 577.8 | - | - | 0 | - |
| - | - | 5.273E+04 | 578.3 | - | - | 0 | - |
| - | - | 5.444E+04 | 578.8 | - | - | 0 | - |
| - | - | 2.755E+04 | 579.3 | - | - | 0 | - |
| - | - | 1.496E+04 | 579.8 | - | - | 0 | - |
| - | - | 9364 | 580.3 | - | - | 0 | - |
| - | - | 2106 | 581.3 | - | - | 0 | - |
| - | - | 4587 | 582.8 | - | - | 0 | - |
| - | - | 2.528E+04 | 583.3 | - | - | 0 | - |
| - | - | 4.544E+04 | 583.8 | - | - | 0 | - |
| - | - | 2.464E+05 | 584.3 | - | - | 0 | - |
| 3 | y | 1.261E+06 | 584.8 | 0.00291 | 4.976 | +2 | 10 |
| - | - | 7.16E+05 | 585.3 | - | - | 0 | - |
| - | - | 2.685E+05 | 585.8 | - | - | 0 | - |
| - | - | 7.089E+04 | 586.3 | - | - | 0 | - |
| - | - | 7173 | 586.8 | - | - | 0 | - |
| - | - | 2.558E+04 | 586.8 | - | - | 0 | - |
| - | - | 1.438E+05 | 587.3 | - | - | 0 | - |
| - | - | 9.875E+04 | 587.8 | - | - | 0 | - |
| - | - | 4.274E+04 | 588.3 | - | - | 0 | - |
| - | - | 1.663E+04 | 588.8 | - | - | 0 | - |
| - | - | 9281 | 588.8 | - | - | 0 | - |
| - | - | 1.229E+04 | 589.3 | - | - | 0 | - |
| - | - | 4196 | 589.3 | - | - | 0 | - |
| - | - | 4319 | 589.8 | - | - | 0 | - |
| 5 | c | 9.486E+05 | 590.3 | 0.00162 | 2.744 | +1 | 5 |
| - | - | 2.761E+05 | 591.3 | - | - | 0 | - |
| - | - | 1.51E+04 | 591.8 | - | - | 0 | - |
| - | - | 9.483E+04 | 592.3 | - | - | 0 | - |
| - | - | 1.121E+05 | 592.8 | - | - | 0 | - |
| - | - | 4.433E+04 | 593.3 | - | - | 0 | - |
| - | - | 1.833E+04 | 593.8 | - | - | 0 | - |
| - | - | 4171 | 594.3 | - | - | 0 | - |
| - | - | 5969 | 598.3 | - | - | 0 | - |
| - | - | 3358 | 598.8 | - | - | 0 | - |
| 11 | c | 4.43E+04 | 600.8 | 0.001393 | 2.319 | +2 | 11 |
| 11 | c | 1.754E+05 | 601.3 | 0.003038 | 5.052 | +2 | 11 |
| - | - | 1.033E+05 | 601.8 | - | - | 0 | - |
| - | - | 4.529E+04 | 602.3 | - | - | 0 | - |
| - | - | 1.197E+04 | 602.8 | - | - | 0 | - |
| - | - | 3714 | 603.3 | - | - | 0 | - |
| - | - | 4407 | 603.3 | - | - | 0 | - |
| - | - | 3012 | 603.8 | - | - | 0 | - |
| - | - | 1797 | 604.3 | - | - | 0 | - |
| - | - | 2786 | 605.3 | - | - | 0 | - |
| - | - | 2864 | 607.3 | - | - | 0 | - |
| - | - | 6971 | 607.8 | - | - | 0 | - |
| 8 | z | 5.479E+04 | 608.3 | 0.002419 | 3.977 | +1 | 5 |
| - | - | 3071 | 608.8 | - | - | 0 | - |
| - | - | 1.313E+05 | 609.3 | - | - | 0 | - |
| 11 | c | 7.327E+05 | 609.8 | 0.002703 | 4.432 | +2 | 11 |
| - | - | 4.159E+05 | 610.3 | - | - | 0 | - |
| - | - | 1.669E+05 | 610.8 | - | - | 0 | - |
| - | - | 4.607E+04 | 611.3 | - | - | 0 | - |
| - | - | 9343 | 611.8 | - | - | 0 | - |
| - | - | 7348 | 616.3 | - | - | 0 | - |
| - | - | 5615 | 617.3 | - | - | 0 | - |
| - | - | 9810 | 618.3 | - | - | 0 | - |
| 2 | w | 3.416E+04 | 618.8 | 0.003047 | 4.924 | +2 | 11 |
| - | - | 2.162E+04 | 619.3 | - | - | 0 | - |
| - | - | 9789 | 619.8 | - | - | 0 | - |
| - | - | 2257 | 621.3 | - | - | 0 | - |
| - | - | 3989 | 621.8 | - | - | 0 | - |
| - | - | 2483 | 622.3 | - | - | 0 | - |
| - | - | 1731 | 622.8 | - | - | 0 | - |
| - | - | 1608 | 623.3 | - | - | 0 | - |
| 8 | y | 9.692E+04 | 624.3 | 0.0017 | 2.724 | +1 | 5 |
| - | - | 3047 | 624.8 | - | - | 0 | - |
| - | - | 3.682E+04 | 625.3 | - | - | 0 | - |
| 2 | w | 4251 | 625.8 | 0.001936 | 3.093 | +2 | 11 |
| - | - | 5624 | 625.9 | - | - | 0 | - |
| - | - | 2402 | 626.3 | - | - | 0 | - |
| - | - | 8355 | 626.3 | - | - | 0 | - |
| - | - | 3243 | 626.8 | - | - | 0 | - |
| - | - | 2523 | 629.3 | - | - | 0 | - |
| - | - | 6236 | 630.3 | - | - | 0 | - |
| - | - | 2.606E+04 | 630.8 | - | - | 0 | - |
| - | - | 1.659E+04 | 631.3 | - | - | 0 | - |
| - | - | 8478 | 631.8 | - | - | 0 | - |
| 2 | y | 8262 | 632.3 | 0.003453 | 5.46 | +2 | 11 |
| 2 | y | 9226 | 632.8 | 0.005646 | 8.923 | +2 | 11 |
| 2 | z | 2.373E+04 | 633.3 | 0.003016 | 4.762 | +2 | 11 |
| - | - | 2.972E+04 | 633.8 | - | - | 0 | - |
| - | - | 1.559E+04 | 634.3 | - | - | 0 | - |
| - | - | 4196 | 634.8 | - | - | 0 | - |
| - | - | 2306 | 635.4 | - | - | 0 | - |
| - | - | 1.618E+04 | 636.4 | - | - | 0 | - |
| - | - | 6343 | 637.4 | - | - | 0 | - |
| - | - | 1861 | 637.8 | - | - | 0 | - |
| - | - | 1.816E+04 | 638.3 | - | - | 0 | - |
| - | - | 1.266E+04 | 638.8 | - | - | 0 | - |
| - | - | 7549 | 639.3 | - | - | 0 | - |
| - | - | 8815 | 639.8 | - | - | 0 | - |
| - | - | 1.218E+04 | 640.3 | - | - | 0 | - |
| - | - | 2.653E+04 | 640.8 | - | - | 0 | - |
| 2 | y | 1.616E+05 | 641.3 | 0.002321 | 3.618 | +2 | 11 |
| - | - | 9.776E+04 | 641.8 | - | - | 0 | - |
| - | - | 4.607E+04 | 642.3 | - | - | 0 | - |
| - | - | 8396 | 642.8 | - | - | 0 | - |
| - | - | 4949 | 643.3 | - | - | 0 | - |
| - | - | 6852 | 644.3 | - | - | 0 | - |
| - | - | 6700 | 645.3 | - | - | 0 | - |
| - | - | 6465 | 646.3 | - | - | 0 | - |
| - | - | 4846 | 646.8 | - | - | 0 | - |
| - | - | 9390 | 647.3 | - | - | 0 | - |
| - | - | 1.012E+05 | 647.8 | - | - | 0 | - |
| - | - | 6.107E+04 | 648.4 | - | - | 0 | - |
| - | - | 2.505E+04 | 648.9 | - | - | 0 | - |
| - | - | 9355 | 649.3 | - | - | 0 | - |
| - | - | 2624 | 649.9 | - | - | 0 | - |
| - | - | 5922 | 650.3 | - | - | 0 | - |
| - | - | 8788 | 652.8 | - | - | 0 | - |
| - | - | 2.426E+04 | 653.3 | - | - | 0 | - |
| - | - | 2.253E+04 | 653.8 | - | - | 0 | - |
| - | - | 1.282E+04 | 654.3 | - | - | 0 | - |
| - | - | 5618 | 654.8 | - | - | 0 | - |
| - | - | 1.815E+04 | 655.3 | - | - | 0 | - |
| - | - | 5.594E+04 | 655.8 | - | - | 0 | - |
| - | - | 3.106E+04 | 656.3 | - | - | 0 | - |
| - | - | 1.26E+04 | 656.8 | - | - | 0 | - |
| - | - | 3384 | 657.3 | - | - | 0 | - |
| - | - | 2007 | 658.3 | - | - | 0 | - |
| - | - | 1.336E+04 | 659.3 | - | - | 0 | - |
| - | - | 5.508E+04 | 660.3 | - | - | 0 | - |
| - | - | 3.759E+04 | 660.8 | - | - | 0 | - |
| 7 | z | 8.21E+04 | 661.3 | 0.00535 | 8.09 | +1 | 6 |
| - | - | 1.291E+05 | 661.8 | - | - | 0 | - |
| - | - | 1.974E+05 | 662.3 | - | - | 0 | - |
| - | - | 1.289E+05 | 662.8 | - | - | 0 | - |
| - | - | 5.662E+04 | 663.3 | - | - | 0 | - |
| - | - | 1.941E+04 | 663.8 | - | - | 0 | - |
| - | - | 4483 | 664.3 | - | - | 0 | - |
| - | - | 9773 | 668.8 | - | - | 0 | - |
| - | - | 2.914E+04 | 669.3 | - | - | 0 | - |
| - | - | 2.753E+04 | 669.3 | - | - | 0 | - |
| - | - | 7.502E+04 | 669.8 | - | - | 0 | - |
| - | - | 5.849E+04 | 670.3 | - | - | 0 | - |
| - | - | 1.112E+05 | 670.8 | - | - | 0 | - |
| - | - | 6.837E+04 | 671.3 | - | - | 0 | - |
| - | - | 2.835E+04 | 671.8 | - | - | 0 | - |
| - | - | 2.128E+04 | 672.3 | - | - | 0 | - |
| - | - | 2084 | 672.8 | - | - | 0 | - |
| - | - | 3.255E+04 | 673.3 | - | - | 0 | - |
| - | - | 2886 | 673.9 | - | - | 0 | - |
| - | - | 2.18E+04 | 674.3 | - | - | 0 | - |
| - | - | 3.852E+04 | 674.8 | - | - | 0 | - |
| - | - | 8.472E+04 | 675.3 | - | - | 0 | - |
| - | - | 5.722E+04 | 675.8 | - | - | 0 | - |
| - | - | 6.377E+04 | 676.3 | - | - | 0 | - |
| - | - | 3.395E+04 | 676.8 | - | - | 0 | - |
| - | - | 1.291E+04 | 677.3 | - | - | 0 | - |
| - | - | 4234 | 677.8 | - | - | 0 | - |
| 7 | y | 2122 | 678.3 | 0.003188 | 4.7 | +1 | 6 |
| 7 | z | 2.235E+05 | 679.4 | 0.001988 | 2.926 | +1 | 6 |
| - | - | 8.373E+04 | 680.4 | - | - | 0 | - |
| - | - | 2.172E+04 | 681.4 | - | - | 0 | - |
| - | - | 2525 | 681.9 | - | - | 0 | - |
| - | - | 4206 | 682.4 | - | - | 0 | - |
| - | - | 2100 | 682.9 | - | - | 0 | - |
| - | - | 1.37E+05 | 683.3 | - | - | 0 | - |
| - | - | 4.457E+05 | 683.8 | - | - | 0 | - |
| - | - | 2.733E+05 | 684.3 | - | - | 0 | - |
| - | - | 1.222E+05 | 684.8 | - | - | 0 | - |
| - | - | 4.041E+04 | 685.3 | - | - | 0 | - |
| - | - | 6268 | 685.8 | - | - | 0 | - |
| 6 | c | 7379 | 687.3 | 0.005078 | 7.388 | +1 | 6 |
| 6 | c | 3.932E+04 | 688.3 | 0.003179 | 4.619 | +1 | 6 |
| - | - | 4413 | 688.4 | - | - | 0 | - |
| - | - | 1.295E+04 | 689.3 | - | - | 0 | - |
| - | - | 1520 | 689.4 | - | - | 0 | - |
| - | - | 3815 | 690.3 | - | - | 0 | - |
| - | - | 7.653E+04 | 691.4 | - | - | 0 | - |
| - | - | 4.303E+05 | 691.9 | - | - | 0 | - |
| - | - | 3.072E+05 | 692.4 | - | - | 0 | - |
| - | - | 1.407E+05 | 692.9 | - | - | 0 | - |
| - | - | 4.487E+04 | 693.4 | - | - | 0 | - |
| - | - | 1E+04 | 693.9 | - | - | 0 | - |
| - | - | 1.241E+04 | 694.3 | - | - | 0 | - |
| 7 | y | 5.15E+04 | 695.4 | 0.002001 | 2.878 | +1 | 6 |
| - | - | 2.155E+04 | 696.4 | - | - | 0 | - |
| - | - | 5309 | 697.4 | - | - | 0 | - |
| - | - | 7.007E+04 | 704.4 | - | - | 0 | - |
| 6 | c | 1.988E+05 | 705.4 | 0.003119 | 4.422 | +1 | 6 |
| - | - | 5.948E+04 | 706.4 | - | - | 0 | - |
| - | - | 1.464E+04 | 707.4 | - | - | 0 | - |
| - | - | 1913 | 708.4 | - | - | 0 | - |
| - | - | 1623 | 709.3 | - | - | 0 | - |
| - | - | 3506 | 711.3 | - | - | 0 | - |
| - | - | 1990 | 714.4 | - | - | 0 | - |
| - | - | 2138 | 723.3 | - | - | 0 | - |
| - | - | 3166 | 730.4 | - | - | 0 | - |
| - | - | 1.141E+04 | 731.4 | - | - | 0 | - |
| - | - | 3.544E+04 | 732.4 | - | - | 0 | - |
| - | - | 1.194E+04 | 733.4 | - | - | 0 | - |
| - | - | 3652 | 734.4 | - | - | 0 | - |
| - | - | 2699 | 737.3 | - | - | 0 | - |
| - | - | 4444 | 738.3 | - | - | 0 | - |
| - | - | 3621 | 740.4 | - | - | 0 | - |
| - | - | 8118 | 741.4 | - | - | 0 | - |
| - | - | 3338 | 742.4 | - | - | 0 | - |
| - | - | 5386 | 744.4 | - | - | 0 | - |
| - | - | 1775 | 745.4 | - | - | 0 | - |
| 6 | w | 2.809E+04 | 749.4 | 0.002606 | 3.478 | +1 | 7 |
| - | - | 4.527E+05 | 750.4 | - | - | 0 | - |
| - | - | 1.718E+05 | 751.4 | - | - | 0 | - |
| - | - | 4.384E+04 | 752.4 | - | - | 0 | - |
| - | - | 8229 | 753.4 | - | - | 0 | - |
| - | - | 1782 | 757.4 | - | - | 0 | - |
| 7 | c | 3.253E+04 | 758.4 | 0.002571 | 3.39 | +1 | 7 |
| 7 | c | 6.869E+04 | 759.4 | 0.004884 | 6.431 | +1 | 7 |
| - | - | 2.735E+04 | 760.4 | - | - | 0 | - |
| - | - | 7044 | 761.4 | - | - | 0 | - |
| - | - | 1877 | 762.4 | - | - | 0 | - |
| - | - | 3643 | 769.4 | - | - | 0 | - |
| - | - | 4684 | 770.4 | - | - | 0 | - |
| - | - | 7.225E+04 | 775.4 | - | - | 0 | - |
| 7 | c | 2.172E+05 | 776.4 | 0.003786 | 4.877 | +1 | 7 |
| - | - | 7.437E+04 | 777.4 | - | - | 0 | - |
| - | - | 2.114E+04 | 778.4 | - | - | 0 | - |
| - | - | 3766 | 779.4 | - | - | 0 | - |
| - | - | 5029 | 782.4 | - | - | 0 | - |
| - | - | 1970 | 783.4 | - | - | 0 | - |
| - | - | 7934 | 786.4 | - | - | 0 | - |
| - | - | 1.436E+04 | 787.4 | - | - | 0 | - |
| - | - | 9012 | 788.4 | - | - | 0 | - |
| - | - | 3878 | 789.4 | - | - | 0 | - |
| - | - | 1.245E+05 | 793.4 | - | - | 0 | - |
| 6 | z | 1.413E+05 | 794.4 | 0.007393 | 9.307 | +1 | 7 |
| - | - | 5.736E+04 | 795.4 | - | - | 0 | - |
| - | - | 1.662E+04 | 796.4 | - | - | 0 | - |
| - | - | 3402 | 797.4 | - | - | 0 | - |
| - | - | 2358 | 804.4 | - | - | 0 | - |
| - | - | 2989 | 809.4 | - | - | 0 | - |
| 6 | y | 3.094E+04 | 810.4 | 0.002402 | 2.964 | +1 | 7 |
| - | - | 1.358E+04 | 811.4 | - | - | 0 | - |
| - | - | 3134 | 812.4 | - | - | 0 | - |
| - | - | 2314 | 815.5 | - | - | 0 | - |
| - | - | 3413 | 816.5 | - | - | 0 | - |
| - | - | 6605 | 821.4 | - | - | 0 | - |
| - | - | 3979 | 822.4 | - | - | 0 | - |
| - | - | 1978 | 823.4 | - | - | 0 | - |
| - | - | 6130 | 832.4 | - | - | 0 | - |
| - | - | 3776 | 833.5 | - | - | 0 | - |
| - | - | 2163 | 838.4 | - | - | 0 | - |
| - | - | 2470 | 842.5 | - | - | 0 | - |
| - | - | 1972 | 843.4 | - | - | 0 | - |
| - | - | 6017 | 853.3 | - | - | 0 | - |
| - | - | 1723 | 854.3 | - | - | 0 | - |
| - | - | 4793 | 856.4 | - | - | 0 | - |
| - | - | 6421 | 857.4 | - | - | 0 | - |
| - | - | 2042 | 858.4 | - | - | 0 | - |
| - | - | 2025 | 858.5 | - | - | 0 | - |
| - | - | 2.382E+04 | 859.5 | - | - | 0 | - |
| - | - | 2.984E+04 | 860.5 | - | - | 0 | - |
| - | - | 1.374E+04 | 861.5 | - | - | 0 | - |
| - | - | 3214 | 862.5 | - | - | 0 | - |
| - | - | 3.64E+04 | 864.4 | - | - | 0 | - |
| - | - | 1.497E+05 | 865.4 | - | - | 0 | - |
| - | - | 6.944E+04 | 866.4 | - | - | 0 | - |
| - | - | 2.197E+04 | 867.4 | - | - | 0 | - |
| - | - | 3336 | 868.4 | - | - | 0 | - |
| - | - | 6417 | 869.5 | - | - | 0 | - |
| - | - | 2653 | 870.5 | - | - | 0 | - |
| - | - | 4345 | 873.4 | - | - | 0 | - |
| - | - | 1.502E+04 | 874.4 | - | - | 0 | - |
| - | - | 9078 | 875.4 | - | - | 0 | - |
| - | - | 2303 | 876.4 | - | - | 0 | - |
| - | - | 2696 | 883.4 | - | - | 0 | - |
| - | - | 3457 | 884.5 | - | - | 0 | - |
| - | - | 1.957E+04 | 885.5 | - | - | 0 | - |
| 8 | c | 2.494E+04 | 886.5 | 0.001083 | 1.222 | +1 | 8 |
| 8 | c | 5.548E+04 | 887.5 | 0.004647 | 5.237 | +1 | 8 |
| - | - | 2.357E+04 | 888.5 | - | - | 0 | - |
| - | - | 7508 | 889.5 | - | - | 0 | - |
| - | - | 2455 | 897.4 | - | - | 0 | - |
| - | - | 7071 | 900.5 | - | - | 0 | - |
| - | - | 7096 | 901.5 | - | - | 0 | - |
| - | - | 1.968E+04 | 902.5 | - | - | 0 | - |
| - | - | 7.317E+04 | 903.5 | - | - | 0 | - |
| 8 | c | 1.695E+05 | 904.5 | 0.003306 | 3.655 | +1 | 8 |
| - | - | 7.117E+04 | 905.5 | - | - | 0 | - |
| - | - | 2.092E+04 | 906.5 | - | - | 0 | - |
| - | - | 4143 | 907.5 | - | - | 0 | - |
| - | - | 3.349E+04 | 908.4 | - | - | 0 | - |
| 5 | z | 1.789E+05 | 909.4 | 0.003155 | 3.469 | +1 | 8 |
| - | - | 7.799E+04 | 910.4 | - | - | 0 | - |
| - | - | 2.576E+04 | 911.4 | - | - | 0 | - |
| - | - | 5976 | 912.4 | - | - | 0 | - |
| - | - | 1925 | 920.5 | - | - | 0 | - |
| - | - | 3960 | 921.5 | - | - | 0 | - |
| - | - | 4380 | 924.4 | - | - | 0 | - |
| 5 | y | 1.489E+04 | 925.4 | 0.003352 | 3.622 | +1 | 8 |
| - | - | 6490 | 926.4 | - | - | 0 | - |
| - | - | 3439 | 927.4 | - | - | 0 | - |
| - | - | 1952 | 951.5 | - | - | 0 | - |
| - | - | 1.013E+04 | 952.5 | - | - | 0 | - |
| - | - | 4612 | 953.5 | - | - | 0 | - |
| - | - | 2496 | 956.5 | - | - | 0 | - |
| - | - | 1862 | 958.5 | - | - | 0 | - |
| - | - | 3634 | 959.5 | - | - | 0 | - |
| - | - | 5450 | 968.5 | - | - | 0 | - |
| - | - | 2.461E+04 | 969.5 | - | - | 0 | - |
| - | - | 2.582E+04 | 970.5 | - | - | 0 | - |
| - | - | 9173 | 971.5 | - | - | 0 | - |
| - | - | 4572 | 972.5 | - | - | 0 | - |
| - | - | 2348 | 974.5 | - | - | 0 | - |
| 4 | w | 3285 | 979.4 | 0.003468 | 3.541 | +1 | 9 |
| - | - | 2577 | 980.4 | - | - | 0 | - |
| - | - | 1730 | 983.5 | - | - | 0 | - |
| - | - | 1.548E+04 | 986.5 | - | - | 0 | - |
| - | - | 8.543E+04 | 987.5 | - | - | 0 | - |
| - | - | 4.572E+04 | 988.5 | - | - | 0 | - |
| - | - | 1.646E+04 | 989.5 | - | - | 0 | - |
| - | - | 3952 | 990.5 | - | - | 0 | - |
| - | - | 1.083E+04 | 1001 | - | - | 0 | - |
| 9 | c | 1.536E+04 | 1002 | 0.009449 | 9.435 | +1 | 9 |
| - | - | 9024 | 1003 | - | - | 0 | - |
| - | - | 4266 | 1004 | - | - | 0 | - |
| - | - | 3070 | 1005 | - | - | 0 | - |
| - | - | 2388 | 1006 | - | - | 0 | - |
| - | - | 2.088E+04 | 1018 | - | - | 0 | - |
| 9 | c | 7.457E+04 | 1019 | 0.002981 | 2.927 | +1 | 9 |
| - | - | 3.243E+04 | 1020 | - | - | 0 | - |
| - | - | 1.21E+04 | 1021 | - | - | 0 | - |
| - | - | 1.266E+04 | 1022 | - | - | 0 | - |
| - | - | 6141 | 1023 | - | - | 0 | - |
| - | - | 2068 | 1053 | - | - | 0 | - |
| - | - | 1701 | 1064 | - | - | 0 | - |
| - | - | 1.311E+04 | 1065 | - | - | 0 | - |
| 4 | z | 1.296E+05 | 1066 | 0.003424 | 3.213 | +1 | 9 |
| - | - | 8.672E+04 | 1067 | - | - | 0 | - |
| - | - | 3.351E+04 | 1068 | - | - | 0 | - |
| - | - | 9783 | 1069 | - | - | 0 | - |
| - | - | 3686 | 1070 | - | - | 0 | - |
| - | - | 2091 | 1071 | - | - | 0 | - |
| - | - | 3237 | 1081 | - | - | 0 | - |
| 4 | y | 6151 | 1082 | 0.006145 | 5.682 | +1 | 9 |
| - | - | 3739 | 1083 | - | - | 0 | - |
| - | - | 2387 | 1084 | - | - | 0 | - |
| 10 | c | 2074 | 1089 | 0.01209 | 11.11 | +1 | 10 |
| - | - | 3054 | 1090 | - | - | 0 | - |
| - | - | 3248 | 1091 | - | - | 0 | - |
| - | - | 2461 | 1096 | - | - | 0 | - |
| - | - | 3777 | 1097 | - | - | 0 | - |
| - | - | 1747 | 1105 | - | - | 0 | - |
| 10 | c | 1.808E+04 | 1106 | 0.003728 | 3.372 | +1 | 10 |
| - | - | 1.137E+04 | 1107 | - | - | 0 | - |
| - | - | 5056 | 1108 | - | - | 0 | - |
| - | - | 2098 | 1109 | - | - | 0 | - |
| - | - | 6977 | 1152 | - | - | 0 | - |
| 3 | z | 5.815E+04 | 1153 | 0.004476 | 3.884 | +1 | 10 |
| - | - | 5.014E+04 | 1154 | - | - | 0 | - |
| - | - | 2.459E+04 | 1155 | - | - | 0 | - |
| - | - | 6856 | 1156 | - | - | 0 | - |
| - | - | 2930 | 1159 | - | - | 0 | - |
| - | - | 3141 | 1160 | - | - | 0 | - |
| - | - | 2536 | 1161 | - | - | 0 | - |
| 3 | y | 1730 | 1169 | 0.01505 | 12.88 | +1 | 10 |
| - | - | 5010 | 1175 | - | - | 0 | - |
| - | - | 6990 | 1176 | - | - | 0 | - |
| - | - | 4420 | 1177 | - | - | 0 | - |
| - | - | 2152 | 1178 | - | - | 0 | - |
| 11 | c | 3255 | 1202 | 0.01262 | 10.5 | +1 | 11 |
| - | - | 1.098E+04 | 1203 | - | - | 0 | - |
| - | - | 5806 | 1204 | - | - | 0 | - |
| - | - | 2830 | 1205 | - | - | 0 | - |
| 11 | c | 9878 | 1219 | 0.005968 | 4.897 | +1 | 11 |
| - | - | 1.656E+04 | 1220 | - | - | 0 | - |
| - | - | 7765 | 1221 | - | - | 0 | - |
| - | - | 3397 | 1222 | - | - | 0 | - |
| 2 | y | 1719 | 1264 | 0.02114 | 16.73 | +1 | 11 |
| - | - | 3828 | 1267 | - | - | 0 | - |
| - | - | 4121 | 1268 | - | - | 0 | - |
| - | - | 4495 | 1281 | - | - | 0 | - |
| - | - | 1656 | 1283 | - | - | 0 | - |
| - | - | 3402 | 1296 | - | - | 0 | - |
| - | - | 3489 | 1297 | - | - | 0 | - |
| - | - | 2328 | 1306 | - | - | 0 | - |
| - | - | 2.194E+04 | 1307 | - | - | 0 | - |
| - | - | 1.709E+04 | 1308 | - | - | 0 | - |
| - | - | 8417 | 1309 | - | - | 0 | - |
| - | - | 2641 | 1310 | - | - | 0 | - |
| - | - | 5140 | 1323 | - | - | 0 | - |
| - | - | 6980 | 1324 | - | - | 0 | - |
| - | - | 3.071E+04 | 1325 | - | - | 0 | - |
| - | - | 2.367E+04 | 1326 | - | - | 0 | - |
| - | - | 1.165E+04 | 1327 | - | - | 0 | - |
| - | - | 3860 | 1328 | - | - | 0 | - |
| - | - | 6437 | 1339 | - | - | 0 | - |
| - | - | 2.039E+04 | 1340 | - | - | 0 | - |
| - | - | 1.377E+04 | 1341 | - | - | 0 | - |
| - | - | 6754 | 1342 | - | - | 0 | - |
| - | - | 2270 | 1343 | - | - | 0 | - |
| - | - | 4017 | 1350 | - | - | 0 | - |
| - | - | 7545 | 1351 | - | - | 0 | - |
| - | - | 4970 | 1352 | - | - | 0 | - |
| - | - | 2945 | 1353 | - | - | 0 | - |
| - | - | 2064 | 1357 | - | - | 0 | - |
| - | - | 2582 | 1366 | - | - | 0 | - |
| - | - | 2.314E+04 | 1367 | - | - | 0 | - |
| - | - | 7.506E+04 | 1368 | - | - | 0 | - |
| - | - | 4.611E+04 | 1369 | - | - | 0 | - |
| - | - | 2.153E+04 | 1370 | - | - | 0 | - |
| - | - | 5987 | 1371 | - | - | 0 | - |
| - | - | 5056 | 1383 | - | - | 0 | - |
| - | - | 1.242E+04 | 1384 | - | - | 0 | - |
| - | - | 1.529E+04 | 1385 | - | - | 0 | - |
| - | - | 1.165E+04 | 1386 | - | - | 0 | - |
| - | - | 3358 | 1387 | - | - | 0 | - |

m/z Charge Intensity FragmentType MassShift Position
123.04463958740234 0 5688.333
123.10481262207031 0 1487.634
124.11229705810547 0 860.28125
126.09197998046875 0 1030.2057
126.12842559814453 0 1376.971
129.10279846191406 0 12346.256
130.0503692626953 0 939.24347
130.85960388183594 0 802.8925
131.1183624267578 0 2016.1449
132.07737731933594 0 1485.0049
133.0614471435547 0 1297.4526
136.07623291015625 0 207790.9
136.824951171875 0 947.05524
137.07371520996094 0 1989.2871
137.07958984375 0 18851.215
142.12319946289062 0 16394.21
143.1184844970703 0 18096.912
143.1254119873047 0 1986.371
147.0447540283203 0 5102.0796
149.0453643798828 0 1330.418
149.07156372070312 0 910.05475
153.10313415527344 0 1070.0906 c Water loss 3
155.1121063232422 0 1256.4261
155.11837768554688 0 8183.541
156.1220703125 0 1077.506
157.0614013671875 0 4096.809
157.13417053222656 0 1421.4192
158.11801147460938 0 1544.24
165.0552215576172 0 185609.88
166.05859375 0 17462.621
168.1266632080078 0 1156.2385
169.13412475585938 0 13254.409
171.1132049560547 0 3114.8667
171.12567138671875 0 1910.061
173.12904357910156 0 20508.254
173.45040893554688 0 5000.601
174.087890625 0 4922.19
174.13267517089844 0 1336.797
175.07205200195312 0 1838.3586
181.09799194335938 0 1037.4498
182.08184814453125 0 658374.9 y 11
183.08514404296875 0 59562.508
183.11375427246094 0 3026.647
184.07225036621094 0 3780.1938
184.087890625 0 4726.787 w 7
184.0970916748047 0 1939.3632
185.12890625 0 1760.7672
186.1130828857422 0 3240.9438
187.144775390625 0 1196529.5
188.1481170654297 0 111850.36
189.15029907226562 0 6605.66
197.1291046142578 0 6918.763
197.16531372070312 0 4464.965
200.1399688720703 0 5095.494
201.1239471435547 0 25367.027
201.1349334716797 0 1699.3296
201.14755249023438 0 1494.2601
202.08282470703125 0 22351.074
202.12754821777344 0 1417.2251
203.06683349609375 0 1120.6222
203.08642578125 0 1160.2206
203.1399383544922 0 2671.8206
210.12454223632812 0 1473.3073
213.14797973632812 0 1352.7225
215.1396942138672 0 291015.72
216.14315795898438 0 30818.078
217.14569091796875 0 2641.047
221.08534240722656 0 1162.4572
223.0822296142578 0 1217.5878
225.04373168945312 0 1163.7458
227.11448669433594 0 4143.515
229.13021850585938 0 10468.804
229.64157104492188 0 2695.7903 c Ammonia loss 3
230.14285278320312 0 1680.809
232.16650390625 0 7163.614 c 1
243.14613342285156 0 1693.124
244.12905883789062 0 1821.6261
244.14157104492188 0 6266.962
245.16207885742188 0 2108.7976
249.1608428955078 0 1919.0299
253.1189422607422 0 1277.077
261.1591796875 0 1320.1376
262.1512756347656 0 2514.8884
266.18701171875 0 6252.9575
267.1903991699219 0 1890.2017
268.1298828125 0 2596.9124
269.161376953125 0 3795.743
270.1456298828125 0 6026.9614
274.1761474609375 0 1538.1665
279.1470947265625 0 2155.406 z 10
280.13067626953125 0 2047.5872
281.1383361816406 0 3242.1301
282.1457214355469 0 18932.848
283.12939453125 0 1540.7896
283.1490173339844 0 3317.9636
285.1563415527344 0 1418.7279
287.1558532714844 0 2205.3018 c Ammonia loss 4
287.1722717285156 0 6803.9443
295.1661682128906 0 36373.617 y 10
296.1698913574219 0 5196.653
297.15673828125 0 14555.053
298.140869140625 0 8400.396
298.15972900390625 0 2967.3853
299.148193359375 0 7661.6943
299.1723937988281 0 6600.682
300.15631103515625 0 51522.977
301.1402282714844 0 2914.8862
301.15985107421875 0 8519.669
301.187744140625 0 1335.3903 c Water loss 2
302.1719665527344 0 7711.894 c 7
303.17559814453125 0 1309.6862
310.11724853515625 0 3975.0437
313.1492919921875 0 1470.5619
315.16741943359375 0 36660.4
315.18817138671875 0 2076.2705
316.15118408203125 0 2192.6167
316.1703796386719 0 4881.464
317.1830139160156 0 29531.697
318.18646240234375 0 3710.7856
319.19873046875 0 39463.707 c 2
320.2020568847656 0 5271.929
329.5185852050781 0 1137.7983
330.1778564453125 0 5064.303
335.6632385253906 0 3129.049
335.73736572265625 0 1257.1179
337.1680908203125 0 3153.9106
341.01861572265625 0 7365.568
342.214599609375 0 8602.645
343.2215576171875 0 6168.2407
344.17681884765625 0 1488.6599 c Water loss 5
344.66778564453125 0 4503.952 c Ammonia loss 5
345.1690673828125 0 2157.1045
351.51177978515625 0 1476.212
352.1730651855469 0 1157.2808
352.52886962890625 0 1413.0096
354.1870422363281 0 1400.9575
356.1705627441406 0 1153.249
357.1847839355469 0 2503.9648
357.22552490234375 0 1138.1647
357.51519775390625 0 4325.529
357.84375 0 3267.829
357.8697814941406 0 1870.6703
358.1976013183594 0 3154.9653
358.5311584472656 0 3154.3564
358.8663330078125 0 1678.7673
359.02947998046875 0 15896.223
359.1454772949219 0 1077.4783
359.16845703125 0 7154.5977
360.17138671875 0 1488.6681
363.1890869140625 0 1993.6697 c Water loss 9
363.5186462402344 0 6551.225 c Ammonia loss 9
363.8507995605469 0 4553.8076
366.18621826171875 0 4510.143
367.18707275390625 0 5688.9062
370.1234130859375 0 1812.675
373.19012451171875 0 5419.006
373.6901550292969 0 1541.2144
374.1806335449219 0 6311.9297
375.8630676269531 0 1509.3002
377.1803894042969 0 1455.1769
379.5446472167969 0 1565.7711
379.87353515625 0 7052.841
380.2032775878906 0 5710.406
380.2294921875 0 1417.0763
380.5397033691406 0 2153.9177
380.6881408691406 0 1329.0344
380.8727111816406 0 1426.5237
381.21368408203125 0 1775.0709
382.1984558105469 0 13621.221 y 9
383.2029724121094 0 3202.8188
385.1846008300781 0 3837.2488
385.2200012207031 0 4188.62
385.5484924316406 0 10926.926
385.68206787109375 0 3634.7136
385.8772277832031 0 62158.18
386.20928955078125 0 38684.72
386.5432434082031 0 15597.241
386.877197265625 0 5892.9097
387.2112731933594 0 3652.9094
387.23614501953125 0 8324.104
387.54156494140625 0 1519.6044
388.2391052246094 0 1623.3799
389.2066955566406 0 1297.5804
389.5335388183594 0 1751.3903
389.8708801269531 0 1631.367
391.5520935058594 0 35095.54
391.8808288574219 0 143902.38
392.2146301269531 0 83464.4
392.24908447265625 0 1683.783
392.2549743652344 0 1155.46
392.5487365722656 0 32764.828
392.88330078125 0 6978.057
393.2179260253906 0 3137.103
393.7091979980469 0 3900.3872
394.1893615722656 0 11305.093
394.21209716796875 0 1144.5516
394.6887512207031 0 2122.1682
395.2062072753906 0 8254.657
395.5423889160156 0 3876.7578
395.87579345703125 0 1918.6029
398.2411804199219 0 4478.7227
400.2571716308594 0 2689.9636
400.88372802734375 0 1833.8572 c Water loss 10
401.2135925292969 0 6168.7617 c Ammonia loss 10
401.5454406738281 0 2735.2373
401.8835754394531 0 1652.3008
406.6875915527344 0 1482.9158
408.2253112792969 0 9937.501
415.2681579589844 0 10552.215
415.7107849121094 0 1891.5023
416.2022705078125 0 2143.25
416.2728271484375 0 1942.5575
419.2150573730469 0 1508.3174
419.7085266113281 0 1841.9082
420.18548583984375 0 1286.623
423.7083740234375 0 2446.2087
424.7367858886719 0 7556.395
425.2509765625 0 10740.006
426.2362365722656 0 13992.799
427.2384948730469 0 2899.3438
427.8873291015625 0 2237.1504 y 1
428.2162170410156 0 1709.8674
428.70074462890625 0 6209.7983
429.0902099609375 0 56231.14
429.1958312988281 0 7544.684
429.24835205078125 0 3811.1826
429.69720458984375 0 2471.084
429.7285461425781 0 7081.162
429.75701904296875 0 2801.3728
430.201904296875 0 1840.7603
430.22650146484375 0 1493.9893
430.247802734375 0 6228.9565
430.27880859375 0 9308.136
430.7450256347656 0 2722.8396
431.1950378417969 0 1668.3937
431.22674560546875 0 4922.015
431.2807312011719 0 4369.9385
435.23175048828125 0 3836.333
436.2347717285156 0 2040.2773
436.73492431640625 0 1869.2418
437.2157897949219 0 7141.898
437.7066955566406 0 21909.043
438.2082824707031 0 6030.7515
438.7336730957031 0 16399.277
439.2351379394531 0 10317.816
439.7385559082031 0 1645.373
440.5625305175781 0 2416.8242
440.8922119140625 0 1802.5057
442.74627685546875 0 10357.224
443.23712158203125 0 8769.095
443.2625427246094 0 28965.541
443.7415466308594 0 16848.459 c Water loss 7
444.2397155761719 0 12704.311 c Ammonia loss 7
444.7374572753906 0 5455.462
445.12158203125 0 29947.164
445.2406311035156 0 1822.2645
446.1204833984375 0 1936.4982
446.2137145996094 0 9430.848
446.5662536621094 0 1930.5776
446.9031677246094 0 2177.5217
449.5633239746094 0 1654.0098
450.2313232421875 0 1314.0867
450.5598449707031 0 1295.4606
450.73248291015625 0 4427.798
451.2329406738281 0 1351.1487
451.7516784667969 0 29092.623
452.2532653808594 0 86921.266
452.7489929199219 0 151669.55 c 7
453.24951171875 0 53424.9
453.7507629394531 0 13584.012
454.2191467285156 0 1515.565 y Water loss 4
454.2543640136719 0 2640.682
455.23779296875 0 4354.523
455.5670471191406 0 27114.41
455.9007873535156 0 22407.893
456.23223876953125 0 8294.978
456.5670471191406 0 2230.4597
457.24176025390625 0 2073.395
458.273193359375 0 10858.298 c Ammonia loss 3
458.760986328125 0 2010.516
459.27447509765625 0 2991.4937
460.7293701171875 0 7592.6587
461.2384338378906 0 5687.3115
461.57049560546875 0 40247.703
461.90447998046875 0 28906.516
462.20611572265625 0 1528.0298 z Water loss 8
462.2387390136719 0 15852.726
462.5706787109375 0 3546.0786
462.7335205078125 0 7788.67
463.2240295410156 0 5025.9053 y 4
465.7472839355469 0 1318.4489
467.22479248046875 0 3151.7075
467.72369384765625 0 5644.3823
468.2191162109375 0 1758.9635
469.7406005859375 0 3107.9917
470.2405700683594 0 2439.7917
470.752685546875 0 4930.8403
471.24609375 0 29563.713
471.741455078125 0 25554.441
472.2428283691406 0 9402.781
472.7438049316406 0 6289.4062
473.21160888671875 0 3022.1057
473.2440490722656 0 4194.6963
474.19586181640625 0 26723.805
474.255859375 0 1577.986
475.3003845214844 0 533455.44 c 3
476.2388916015625 0 3276.9163
476.3030090332031 0 109751.9
476.730224609375 0 17280.9
477.2311706542969 0 6376.151
477.3053283691406 0 19101.375
477.7334289550781 0 3698.4236
478.31201171875 0 1990.5366
478.74969482421875 0 5801.2036
479.25787353515625 0 2526.6167
479.7582092285156 0 59456.152
480.25146484375 0 225782.08
480.75244140625 0 108895.73
481.2303161621094 0 40690.34
481.2579040527344 0 19273.457
481.7529296875 0 7663.9653
482.2330017089844 0 14667.46
483.2388916015625 0 4473.4155
484.2361755371094 0 1452.3992
484.75091552734375 0 6635.962
485.24334716796875 0 42724.555
485.73876953125 0 38562.777
486.2386169433594 0 15624.441
486.3039245605469 0 4542.2197
486.7371826171875 0 5506.1846
486.77532958984375 0 3614.2617
487.2672119140625 0 20964.787
487.76397705078125 0 18254.377
488.2641906738281 0 11905.989
488.76739501953125 0 2913.6624
489.2653503417969 0 2470.537
491.27984619140625 0 2434.2065
491.76739501953125 0 4998.176
492.2513732910156 0 20623.459
492.7493896484375 0 11015.055
493.2523193359375 0 3247.3506
493.75604248046875 0 15447.64
494.24859619140625 0 83495.31
494.7492980957031 0 40754.492
495.24835205078125 0 10320.403
495.7219543457031 0 3762.0679
495.7572326660156 0 2463.8582
496.2413330078125 0 33820.45 y 8
496.2858581542969 0 2127.6755
497.2409973144531 0 7170.3604
498.2821960449219 0 1422.8013
499.76788330078125 0 21562.668
500.26104736328125 0 65109.316
500.7631530761719 0 41562.867 c Water loss 8
501.2574768066406 0 59718.766 c Ammonia loss 8
501.75836181640625 0 33322.598
502.2738037109375 0 51608.336
502.76348876953125 0 9864.902
503.2566223144531 0 10859.744
503.27850341796875 0 8678.503
503.757080078125 0 5602.0957
504.2623291015625 0 1249.6931
504.2822570800781 0 1139.0465
504.7519836425781 0 1485.6786
508.2637634277344 0 3771.16
508.7730407714844 0 43437.098
509.26800537109375 0 171958.3
509.76910400390625 0 370209.38 c 8
510.2699890136719 0 159417.72
510.77093505859375 0 53858.938
511.2690734863281 0 16987.59
511.7672424316406 0 3101.6472
512.2659912109375 0 2113.9795
513.2568359375 0 3116.2542
513.75146484375 0 1332.6016
514.2998046875 0 9579.382
515.2947998046875 0 1684.2946
517.7628173828125 0 2007.643
518.24853515625 0 3222.3508
521.774658203125 0 2144.9202
522.2664794921875 0 6789.4805
522.7691040039062 0 4428.8633
523.2684936523438 0 10000.919
523.7720947265625 0 3381.993
524.2640991210938 0 1761.4279
525.7494506835938 0 1364.8792
526.2505493164062 0 7785.867
526.7557373046875 0 12272.971
527.2538452148438 0 15150.955
527.7526245117188 0 3478.5247
527.7925415039062 0 1985.3279
528.2559814453125 0 2333.5857
528.2999267578125 0 2979.329
529.31005859375 0 2493.9395
530.29345703125 0 14767.552
530.7744750976562 0 14295.622
531.2759399414062 0 7255.9673
531.7760620117188 0 5273.2285
532.7667846679688 0 7365.608
533.2611083984375 0 32523.34 z 3
533.7623291015625 0 14700.823
534.2621459960938 0 7326.427
534.3069458007812 0 3214.2693
534.7950439453125 0 3240.029
535.283203125 0 12772.538
535.7672729492188 0 47784.203
536.2653198242188 0 31720.354
536.7923583984375 0 19432.896
537.2952270507812 0 13846.342
537.7935791015625 0 4534.4795
540.2763061523438 0 2592.0654
540.7739868164062 0 6625.182
541.2696533203125 0 12716.817 y 3
541.7756958007812 0 11301.133
542.2802734375 0 10482.398
542.7838745117188 0 3796.54
543.2835693359375 0 2433.11
543.7759399414062 0 1758.4468
544.28076171875 0 32077.775 c Water loss 9
544.7728271484375 0 114568.51 c Ammonia loss 9
545.2313232421875 0 10909.179
545.272216796875 0 43860.44
545.3075561523438 0 19483.604
545.7747802734375 0 20882.098
546.2398071289062 0 3734.857
546.2703857421875 0 2492.5232
546.3121948242188 0 16515.336
546.779052734375 0 2515.152
547.314697265625 0 3103.4558
547.7701416015625 0 1859.616
548.2540283203125 0 8040.857
548.7449340820312 0 34821.81
548.7928466796875 0 2574.1326
549.2459716796875 0 17582.193
549.289794921875 0 2679.9326
549.7473754882812 0 6923.127
550.2527465820312 0 2259.6194 w 7
550.298828125 0 2536.2556
550.790283203125 0 19413.475
551.2896118164062 0 8829.166
551.7900390625 0 3411.206
552.2525634765625 0 2005.3983
552.294189453125 0 1984.8683
552.7921142578125 0 52790.98
553.2855834960938 0 306476 c 9
553.7859497070312 0 160405.4
554.2863159179688 0 59721.09
554.7823486328125 0 22806.248
555.280517578125 0 7974.683
555.784912109375 0 2061.3105
556.2974243164062 0 2344.4978
556.7900390625 0 1382.2212
557.3035888671875 0 3959.6814
557.7987060546875 0 6021.208
558.2996215820312 0 4078.092
559.7972412109375 0 1846.3312
560.7948608398438 0 2023.0802
561.2835083007812 0 9465.47
561.78369140625 0 13771.481
562.283203125 0 8972.717
562.7845458984375 0 4533.702
563.2614135742188 0 2354.191
563.319091796875 0 1768.7609
564.3050537109375 0 17693.277
565.26220703125 0 2107.747
565.3089599609375 0 10119.162
565.8007202148438 0 3730.9248
566.29296875 0 23715.537
566.78662109375 0 12091.357
567.2715454101562 0 8919.203
567.7730712890625 0 5600.3096 z Water loss 2
568.2761840820312 0 7537.518 w 2
568.772216796875 0 12150.104
569.2723388671875 0 6351.9688
569.3095703125 0 5729.9663
569.7999877929688 0 20631.68
570.302734375 0 15428.782
570.8017578125 0 6451.4116
571.30224609375 0 4027.1401
572.3181762695312 0 2022.0195 c Water loss 4
572.8043212890625 0 2317.1304
573.3028564453125 0 18845.662 c Ammonia loss 4
574.30078125 0 7139.237
574.7911987304688 0 11722.668
575.2889404296875 0 13325.151
575.780517578125 0 40268.09 y Water loss 2
576.2782592773438 0 64108.902 y Ammonia loss 2
576.7767333984375 0 103725.59 z 2
577.2774658203125 0 56764.016
577.7782592773438 0 23169.838
577.818359375 0 3897.7197
578.3119506835938 0 52734.87
578.8082275390625 0 54437.523
579.3082885742188 0 27554.527
579.81005859375 0 14957.19
580.311279296875 0 9364.201
581.3126220703125 0 2105.9236
582.8114013671875 0 4586.7134
583.304443359375 0 25276.271
583.797607421875 0 45435.723
584.2936401367188 0 246380.78
584.7861938476562 0 1261471.9 y 2
585.286376953125 0 715995.3
585.7872314453125 0 268540.78
586.2885131835938 0 70892.03
586.7853393554688 0 7173.0986
586.82470703125 0 25578.998
587.3173828125 0 143792.06
587.8181762695312 0 98751.336
588.3195190429688 0 42736.414
588.7699584960938 0 16633.014
588.821044921875 0 9280.624
589.2706298828125 0 12289.813
589.3204956054688 0 4196.136
589.7724609375 0 4318.6255
590.3272705078125 0 948641.7 c 4
591.3300170898438 0 276094.97
591.8157958984375 0 15096.54
592.3121948242188 0 94826.91
592.8059692382812 0 112088.42
593.306396484375 0 44332.39
593.8071899414062 0 18331.531
594.3087768554688 0 4171.162
598.2792358398438 0 5968.666
598.7764282226562 0 3357.936
600.8215942382812 0 44299.094 c Water loss 10
601.3152465820312 0 175360.14 c Ammonia loss 10
601.81591796875 0 103270.75
602.3167724609375 0 45293.336
602.8173828125 0 11968.114
603.296875 0 3714.4646
603.316650390625 0 4406.8657
603.7930297851562 0 3011.6367
604.3011474609375 0 1796.965
605.2950439453125 0 2785.61
607.32373046875 0 2864.2979
607.8280029296875 0 6970.5713
608.31884765625 0 54790.125 z 7
608.8265991210938 0 3071.3425
609.3341064453125 0 131344.7
609.8281860351562 0 732736.2 c 10
610.3284301757812 0 415918.1
610.8294067382812 0 166909.77
611.3291015625 0 46072.316
611.8294067382812 0 9343.281
616.3421630859375 0 7348.4917
617.3463745117188 0 5615.2783
618.3049926757812 0 9810.319
618.7994384765625 0 34157.504 w 1
619.2996215820312 0 21624.96
619.8018798828125 0 9789.242
621.3197021484375 0 2256.7092
621.8173217773438 0 3988.928
622.3118286132812 0 2482.8215
622.81298828125 0 1731.1724
623.2974243164062 0 1608.0801
624.3368530273438 0 96917.28 y 7
624.8154296875 0 3047.4668
625.3365478515625 0 36824.316
625.80615234375 0 4251.2764 w 1
625.8550415039062 0 5623.99
626.3012084960938 0 2401.6072
626.3483276367188 0 8354.886
626.80419921875 0 3242.7285
629.3302612304688 0 2523.4014
630.3272705078125 0 6236.1772
630.8214111328125 0 26057.938
631.3228149414062 0 16593.318
631.8285522460938 0 8478.102
632.323486328125 0 8261.817 y Water loss 1
632.8176879882812 0 9225.861 y Ammonia loss 1
633.3189697265625 0 23725.398 z 1
633.8170166015625 0 29717.162
634.3168334960938 0 15594.966
634.8176879882812 0 4196.3496
635.3582153320312 0 2305.8157
636.3501586914062 0 16175.169
637.3522338867188 0 6342.789
637.8316040039062 0 1860.77
638.3275146484375 0 18156.377
638.8348388671875 0 12662.98
639.3406372070312 0 7548.5986
639.8435668945312 0 8814.653
640.3464965820312 0 12178.81
640.8363647460938 0 26531.959
641.32763671875 0 161574.52 y 1
641.8286743164062 0 97762.22
642.3297119140625 0 46070.137
642.8299560546875 0 8396.474
643.3289794921875 0 4948.889
644.3463745117188 0 6851.741
645.3245239257812 0 6699.768
646.3218383789062 0 6465.1943
646.814697265625 0 4845.886
647.3472900390625 0 9390.374
647.8494262695312 0 101152.45
648.3502807617188 0 61066.543
648.8515625 0 25050.416
649.3447875976562 0 9355.155
649.8572998046875 0 2623.6223
650.3374633789062 0 5921.9634
652.8445434570312 0 8787.6875
653.3390502929688 0 24261.17
653.8373413085938 0 22530.848
654.334716796875 0 12816.332
654.8311157226562 0 5618.3955
655.320556640625 0 18154.451
655.8120727539062 0 55944.547
656.312255859375 0 31063.08
656.8137817382812 0 12600.653
657.3242797851562 0 3383.9668
658.3142700195312 0 2006.5176
659.3488159179688 0 13355.09
660.336669921875 0 55083.78
660.8438110351562 0 37586.363
661.3483276367188 0 82095.16 z Water loss 6
661.8475952148438 0 129071.04
662.3453979492188 0 197441.17
662.8442993164062 0 128916.78
663.3442993164062 0 56618.977
663.844970703125 0 19407.9
664.3426513671875 0 4482.796
668.833740234375 0 9772.815
669.3297119140625 0 29139.014
669.34130859375 0 27532.271
669.8433837890625 0 75021.86
670.3455200195312 0 58486.227
670.8422241210938 0 111248.914
671.3409423828125 0 68366.24
671.8429565429688 0 28348.707
672.34375 0 21275.465
672.841796875 0 2083.586
673.3289794921875 0 32548.19
673.850341796875 0 2885.7437
674.33935546875 0 21799.203
674.8405151367188 0 38515.844
675.3402709960938 0 84724.82
675.8410034179688 0 57218.93
676.3347778320312 0 63771.344
676.832275390625 0 33952.508
677.3331909179688 0 12911.889
677.8356323242188 0 4233.886
678.342529296875 0 2121.5146 y Ammonia loss 6
679.3555297851562 0 223452.14 z 6
680.3550415039062 0 83725.11
681.3563232421875 0 21719.912
681.879150390625 0 2525.087
682.3591918945312 0 4205.8325
682.855224609375 0 2099.8604
683.3497924804688 0 136999.83
683.8436889648438 0 445699.72
684.3438720703125 0 273317.47
684.8441772460938 0 122179.695
685.344970703125 0 40411.863
685.8438110351562 0 6267.6616
687.3471069335938 0 7378.5537 c Water loss 5
688.3292236328125 0 39324.04 c Ammonia loss 5
688.3890991210938 0 4413.2036
689.3311767578125 0 12953.326
689.393310546875 0 1520.4836
690.3375854492188 0 3814.8567
691.3587036132812 0 76529.46
691.8526000976562 0 430292.7
692.3527221679688 0 307201.8
692.853759765625 0 140697.62
693.3541870117188 0 44866.41
693.8561401367188 0 10003.769
694.3307495117188 0 12412.051
695.374267578125 0 51499.027 y 6
696.3729858398438 0 21547.686
697.3709106445312 0 5308.8594
704.3703002929688 0 70068.14
705.355712890625 0 198807.23 c 5
706.358642578125 0 59479.75
707.3650512695312 0 14643.515
708.3989868164062 0 1913.327
709.3282470703125 0 1622.7024
711.3440551757812 0 3506.3862
714.3682861328125 0 1990.2538
723.3389282226562 0 2137.6665
730.3864135742188 0 3166.0522
731.3912353515625 0 11410.027
732.3795166015625 0 35442.52
733.38232421875 0 11937.253
734.3825073242188 0 3651.5217
737.3353271484375 0 2699.1414
738.3211669921875 0 4444.4165
740.3707275390625 0 3620.7559
741.3573608398438 0 8117.529
742.3585815429688 0 3338.3103
744.3563232421875 0 5385.6475
745.3623657226562 0 1774.601
749.3854370117188 0 28085.646 w 5
750.392333984375 0 452691.3
751.3950805664062 0 171821.19
752.3977661132812 0 43843.78
753.4000854492188 0 8229.274
757.402099609375 0 1782.0809
758.3817138671875 0 32525.154 c Water loss 6
759.3680419921875 0 68690.59 c Ammonia loss 6
760.3703002929688 0 27345.586
761.3721923828125 0 7044.4995
762.3731689453125 0 1877.3324
769.362060546875 0 3643.2795
770.3512573242188 0 4684.3213
775.4076538085938 0 72251.35
776.3934936523438 0 217193.12 c 6
777.3950805664062 0 74374.3
778.39697265625 0 21144.291
779.398681640625 0 3765.6707
782.3824462890625 0 5029.14
783.387939453125 0 1969.9385
786.4075927734375 0 7934.095
787.371826171875 0 14364.291
788.37451171875 0 9012.271
789.3792724609375 0 3878.2983
793.3985595703125 0 124526.55
794.3878784179688 0 141345.42 z 5
795.383544921875 0 57362.25
796.3837890625 0 16622.537
797.381591796875 0 3401.941
804.3987426757812 0 2357.7156
809.3543701171875 0 2989.1143
810.401611328125 0 30935.076 y 5
811.4007568359375 0 13582.506
812.406494140625 0 3133.766
815.4934692382812 0 2313.854
816.4774169921875 0 3413.3298
821.4230346679688 0 6605.265
822.4258422851562 0 3978.822
823.4288940429688 0 1978.2231
832.4464111328125 0 6130.0195
833.4501953125 0 3776.0522
838.4026489257812 0 2163.2021
842.4678955078125 0 2470.2441
843.4485473632812 0 1972.0463
853.3486328125 0 6016.8374
854.3442993164062 0 1723.1658
856.3980712890625 0 4792.639
857.3806762695312 0 6421.1577
858.378173828125 0 2041.9497
858.4876098632812 0 2025.1277
859.4869384765625 0 23819.125
860.4780883789062 0 29844.361
861.4801025390625 0 13738.161
862.4830322265625 0 3214.3618
864.4353637695312 0 36396.906
865.4209594726562 0 149707.53
866.4220581054688 0 69438.5
867.4241943359375 0 21971.066
868.42529296875 0 3336.4133
869.4517822265625 0 6416.8125
870.4524536132812 0 2652.7
873.4197387695312 0 4344.7485
874.4055786132812 0 15018.158
875.4082641601562 0 9078.498
876.408935546875 0 2302.8557
883.4302368164062 0 2696.167
884.4798583984375 0 3456.5012
885.4697265625 0 19570.465
886.4730224609375 0 24940.188 c Water loss 7
887.4627685546875 0 55479.11 c Ammonia loss 7
888.4644775390625 0 23565.787
889.4669189453125 0 7508.184
897.40576171875 0 2455.322
900.4608764648438 0 7070.6816
901.460205078125 0 7095.9014
902.4912109375 0 19677.754
903.4923095703125 0 73171.99
904.4879760742188 0 169469.62 c 7
905.4902954101562 0 71167.586
906.4910888671875 0 20915.576
907.493896484375 0 4142.9277
908.425048828125 0 33489.633
909.4105834960938 0 178918.97 z 4
910.412353515625 0 77987.78
911.4143676757812 0 25759.832
912.4165649414062 0 5976.2324
920.45263671875 0 1924.6241
921.4577026367188 0 3960.3518
924.4444580078125 0 4380.462
925.4295043945312 0 14892.42 y 4
926.4290161132812 0 6490.1704
927.4320068359375 0 3438.7612
951.4616088867188 0 1951.6991
952.4509887695312 0 10127.174
953.453369140625 0 4612.251
956.4906616210938 0 2495.814
958.4887084960938 0 1861.9644
959.4934692382812 0 3633.908
968.4911499023438 0 5449.748
969.4786376953125 0 24610.922
970.468505859375 0 25817.307
971.4702758789062 0 9173.425
972.4690551757812 0 4572.1094
974.5126342773438 0 2348.4592
979.440185546875 0 3285.129 w 3
980.4481811523438 0 2577.208
983.5 0 1730.0492
986.5038452148438 0 15478.774
987.4892578125 0 85428.16
988.4904174804688 0 45721.07
989.49169921875 0 16463.783
990.4918823242188 0 3952.367
1000.5387573242188 0 10827.154
1001.510498046875 0 15359.542 c Ammonia loss 8
1002.507080078125 0 9023.713
1003.51416015625 0 4265.7725
1004.508544921875 0 3069.624
1005.511962890625 0 2388.2512
1017.5380859375 0 20883.365
1018.5305786132812 0 74571.84 c 8
1019.531494140625 0 32432.852
1020.5335693359375 0 12096.977
1021.52490234375 0 12655.153
1022.517822265625 0 6140.54
1052.532470703125 0 2067.994
1063.5479736328125 0 1701.0526
1064.52783203125 0 13108.04
1065.511962890625 0 129564.98 z 3
1066.513427734375 0 86720.94
1067.5152587890625 0 33508.285
1068.5208740234375 0 9782.895
1069.561767578125 0 3686.1865
1070.5537109375 0 2090.7178
1080.52294921875 0 3236.673
1081.5211181640625 0 6150.6763 y 3
1082.5244140625 0 3739.182
1083.5367431640625 0 2386.5383
1088.545166015625 0 2073.5542 c Ammonia loss 9
1089.55029296875 0 3053.7268
1090.5391845703125 0 3247.7107
1096.4866943359375 0 2460.6836
1097.488037109375 0 3777.232
1104.59033203125 0 1747.3907
1105.5633544921875 0 18080.361 c 9
1106.5640869140625 0 11368.782
1107.5662841796875 0 5055.5835
1108.5587158203125 0 2098.4502
1151.5570068359375 0 6977.2505
1152.5450439453125 0 58147.812 z 2
1153.546875 0 50137.78
1154.55078125 0 24589.639
1155.5543212890625 0 6856.443
1158.6160888671875 0 2929.936
1159.630126953125 0 3141.435
1160.62060546875 0 2536.3608
1168.5743408203125 0 1730.4321 y 2
1174.6363525390625 0 5010.406
1175.644775390625 0 6989.511
1176.6353759765625 0 4420.131
1177.6444091796875 0 2152.478
1201.6297607421875 0 3254.916 c Ammonia loss 10
1202.6268310546875 0 10981.377
1203.6268310546875 0 5805.7754
1204.6419677734375 0 2830.2734
1218.649658203125 0 9878.135 c 10
1219.6519775390625 0 16561.756
1220.65380859375 0 7764.9795
1221.6546630859375 0 3396.7976
1263.6539306640625 0 1718.93 y Water loss 1
1266.63427734375 0 3828.336
1267.620849609375 0 4121.253
1280.618896484375 0 4494.9395
1282.602294921875 0 1655.5939
1295.620849609375 0 3401.903
1296.6119384765625 0 3489.2122
1305.6630859375 0 2327.6223
1306.642822265625 0 21942.55
1307.645751953125 0 17087.34
1308.6500244140625 0 8417.055
1309.6461181640625 0 2641.4417
1322.681640625 0 5139.937
1323.67822265625 0 6979.8584
1324.6673583984375 0 30714.588
1325.66796875 0 23671.072
1326.6644287109375 0 11647.175
1327.665771484375 0 3859.512
1338.696044921875 0 6437.4175
1339.690673828125 0 20386.8
1340.689453125 0 13772.37
1341.692626953125 0 6753.9404
1342.683837890625 0 2269.6177
1349.661376953125 0 4017.309
1350.6661376953125 0 7545.22
1351.6632080078125 0 4970.013
1352.658935546875 0 2944.5415
1356.71826171875 0 2063.739
1365.6917724609375 0 2582.3562
1366.69091796875 0 23144.611
1367.68310546875 0 75064.55
1368.6849365234375 0 46105.92
1369.6842041015625 0 21532.117
1370.68310546875 0 5987.3794
1382.6944580078125 0 5055.677
1383.7021484375 0 12422.199
1384.70751953125 0 15285.19
1385.7064208984375 0 11654.716
1386.7083740234375 0 3358.4246

Spectrum Details

|  |  |
| --- | --- |
| Matched peaks? Matched peaks The total absolute number of peaks matched. Additionally in brackets the total fraction of peaks matched and the total number of peaks is shown. | 87 (9.68% of 899) |
| FDR? FDR The false discovery rate estimated for this peptide. It is calculated by matching all theoretical fragments with a non-integer shift with the raw peaks for this spectrum. This is done with 40 different shifts. The resulting percentage is the average number of annotated peaks over the number of annotated peaks with the correct spectrum. | 1.48% |
| Satellite FDR? Satellite FDR See the FDR for details on its calculation. This satellite ion specific FDR only contains the satellite ions (d/w) for I/L/J positions. | 0.00% |
| PSM Score? PSM Score The PSM Score as given by Hecklib to this annotated spectrum. It is shown with three significant figures. | 640 |

#### Reverse Lookup? Reverse Lookup All places where this read could be placed.

| Group | Segment | Template | Template Part | Read Part | Score | Unique |
| --- | --- | --- | --- | --- | --- | --- |
| Homo sapiens Heavy Chain | IGHV | IGHV3-9 | [68..80] | [0..12] | 91 | False |
| Homo sapiens Heavy Chain | IGHV | IGHV3-48 | [68..80] | [0..12] | 91 | False |
| Homo sapiens Heavy Chain | IGHV | IGHV3-21 | [68..80] | [0..12] | 91 | False |
| Homo sapiens Heavy Chain | IGHV | IGHV3-7 | [68..80] | [0..12] | 91 | False |
| Homo sapiens Heavy Chain | IGHV | IGHV3-43 | [68..80] | [0..12] | 82 | False |
| Homo sapiens Heavy Chain | IGHV | IGHV3-13 | [67..79] | [0..12] | 82 | False |
| Homo sapiens Heavy Chain | IGHV | IGHV3-11 | [68..80] | [0..12] | 91 | False |
| Homo sapiens Heavy Chain | IGHV | IGHV3-74 | [68..80] | [0..12] | 82 | False |
| Homo sapiens Heavy Chain | IGHV | IGHV3-20 | [68..80] | [0..12] | 91 | False |
| Homo sapiens Heavy Chain | IGHV | IGHV3-72 | [70..82] | [0..12] | 87 | False |

| Recombined | Template Part | Read Part | Score | Unique |
| --- | --- | --- | --- | --- |
| REC-0-1 | [68..80] | [0..12] | 91 | True |

#### Meta Information from Multiple reads

##### Number of combined reads

2

##### Intensity

0.7249

##### TotalArea

1.226E+08

##### Changes to the peptide sequence

TISRDDAKNSJY

L→JNo support for either Leucine or Isoleucine based on side chain ions (Position: 11)

L→ISupport for Isoleucine based on side chain ions (2 for I 0 for L) (Position: 2)

#### Positional Score

Copy Data

##### Positional Score (TSV)

###### Preview

```
Loading example...
```

*Click on the button to copy the data to your clipboard.*

0001234567891011

Label Value
"0" 0
"1" 0
"2" 0
"3" 0
"4" 0
"5" 0
"6" 0
"7" 0
"8" 0
"9" 0
"10" 0
"11" 0

#### Meta Information from PEAKS

##### Scan Identifier

F3:3647

##### Original sequence

T

L

S

R

D

D

A

K

N

S

L

Y

##### Posttranslational Modifications

##### Source File

D:\separate\_stitch\_analyses\xle-disambiguation\raw\20210323\_F1\_UM1\_Peng0013\_SA\_F59\_ingel\_3ug\_chymo.raw

##### Fraction

3

##### Scan Feature

F3:2691

##### De Novo Score

99

##### ConfidenceScore

99

### m/z

461.5691

##### Mass

1381.6838

##### Charge

3

##### Retention Time

19.96

##### Predicted Retention Time

-

##### Area

6.915E+07

##### Parts Per Million

1.2

##### Fragmentation mode

ETHCD

##### Originating file

01 D:\separate\_stitch\_analyses\xle-disambiguation\20210325\_F59\_3ug\_DENOVO\_12.csv

#### Meta Information from PEAKS

##### Scan Identifier

F3:3545

##### Original sequence

T

L

S

R

D

D

A

K

N

S

L

Y

##### Posttranslational Modifications

##### Source File

D:\separate\_stitch\_analyses\xle-disambiguation\raw\20210323\_F1\_UM1\_Peng0013\_SA\_F59\_ingel\_3ug\_chymo.raw

##### Fraction

3

##### Scan Feature

F3:2692

##### De Novo Score

98

##### ConfidenceScore

98

### m/z

461.5702

##### Mass

1381.6838

##### Charge

3

##### Retention Time

19.23

##### Predicted Retention Time

-

##### Area

5.34E+07

##### Parts Per Million

3.6

##### Fragmentation mode

ETHCD

##### Originating file

01 D:\separate\_stitch\_analyses\xle-disambiguation\20210325\_F59\_3ug\_DENOVO\_12.csv
