## Supplementary material for "A handle on mass coincidence errors in *de novo* sequencing of antibodies by bottom-up proteomics": Combined_039.html

Details Combined\_039 | Stitch OverviewUndefined

### Read Combined\_039

#### Sequence (length=14)

TJSGJQAEDESMYF

#### Spectrum 8780? Spectrum 8780 The raw spectrum of this peptide as annotated by Hecklib. The fragments are coloured according to ion type (see legend). Any peaks with a star '\*' as text can be hovered over to see the full details, first the ion type second the mass shift type. By hovering over the amino acids in the peptide or ions in the legend the corresponding peaks are highlighted. By toggling the 'Unassigned' label you can turn the background (unassigned) peaks on or off in the plot. By updating the slider in the Ion legend you can update the spectrum to only show the top X% of the peaks with labels. The top X% means any peak that is within X% of the highest intensity. By dragging in the spectrum you can zoom in to a specific part of the spectrum and use 'Zoom Out' to get back to the original zoom level. The annotation of the spectrum is based on the given sequence in the peptides file and is done with different software so inconsistencies are likely. The peaks are annotated based on the given sequence, with 20 ppm tolerance.

Copy Data

##### Spectrum 8780 (TSV)

###### Preview

```
Loading example...
```

*Click on the button to copy the data to your clipboard.*

Mz MinMz MaxIntensity Max

WidthHeightPeptide font sizePeptide stroke widthSpectrum font sizeSpectrum stroke widthCompact peptide

Ion legend

wxyz

abcd

OtherUnassignedIon^Charge^_Position_Show for top:%

TJSGJQAEDESMYF

05.49e+41.10e+51.65e+52.20e+5

Zoom Out

y^+1^_1_y^+1^_2_c^+2^_8_y^+1^_3_c^+1^_5_c^+2^_10_y^+1^_4_c^+2^_11_c^+1^_6_c^+1^_6_c^+2^_12_c^+1^_7_c^+1^_7_y^+1^_5_y^+1^_6_z^+1^_6_c^+1^_8_y^+1^_6_c^+1^_8_c^+1^_9_c^+1^_9_y^+1^_7_y^+1^_8_z^+1^_8_y^+1^_8_c^+1^_10_c^+1^_10_c^+1^_10_y^+1^_9_z^+1^_9_c^+1^_11_c^+1^_11_y^+1^_9_c^+1^_11_z^+1^_10_y^+1^_10_z^+1^_11_c^+1^_12_z^+1^_11_c^+1^_12_y^+1^_11_z^+1^_12_z^+1^_12_y^+1^_12_y^+1^_12_z^+1^_12_y^+1^_12_c^+1^_13_c^+1^_13_z^+1^_13_z^+1^_13_z^+1^_13_

0609121818272435

Fragment Matches Table

Show background peaks

| Position | Ion type | Intensity | mz Theoretical | mz Error (Th) | mz Error (ppm) | Charge | Series Number |
| --- | --- | --- | --- | --- | --- | --- | --- |
| - | - | 5160 | 120.1 | - | - | 0 | - |
| - | - | 494.4 | 121.1 | - | - | 0 | - |
| - | - | 357 | 124.3 | - | - | 0 | - |
| - | - | 452.5 | 124.3 | - | - | 0 | - |
| - | - | 337.1 | 125.6 | - | - | 0 | - |
| - | - | 408.9 | 125.8 | - | - | 0 | - |
| - | - | 5.61E+04 | 136.1 | - | - | 0 | - |
| - | - | 4057 | 137.1 | - | - | 0 | - |
| - | - | 454.6 | 148.2 | - | - | 0 | - |
| - | - | 480.3 | 148.9 | - | - | 0 | - |
| - | - | 527.5 | 148.9 | - | - | 0 | - |
| - | - | 676.6 | 148.9 | - | - | 0 | - |
| - | - | 767.8 | 148.9 | - | - | 0 | - |
| - | - | 722.4 | 148.9 | - | - | 0 | - |
| - | - | 1325 | 148.9 | - | - | 0 | - |
| - | - | 1147 | 148.9 | - | - | 0 | - |
| - | - | 3104 | 148.9 | - | - | 0 | - |
| - | - | 5871 | 148.9 | - | - | 0 | - |
| - | - | 3992 | 149 | - | - | 0 | - |
| - | - | 1698 | 149 | - | - | 0 | - |
| - | - | 1219 | 149 | - | - | 0 | - |
| - | - | 1108 | 149 | - | - | 0 | - |
| - | - | 782.1 | 149 | - | - | 0 | - |
| - | - | 590.8 | 149 | - | - | 0 | - |
| - | - | 708.9 | 149 | - | - | 0 | - |
| - | - | 643.7 | 149 | - | - | 0 | - |
| - | - | 585.5 | 149 | - | - | 0 | - |
| - | - | 697.6 | 166.1 | - | - | 0 | - |
| 14 | y | 1.022E+04 | 166.1 | 5.816E-05 | 0.3502 | +1 | 1 |
| - | - | 749.8 | 173.1 | - | - | 0 | - |
| - | - | 1.016E+04 | 187.1 | - | - | 0 | - |
| - | - | 505.2 | 187.7 | - | - | 0 | - |
| - | - | 754.1 | 188.1 | - | - | 0 | - |
| - | - | 591.3 | 201.1 | - | - | 0 | - |
| - | - | 490.8 | 203.3 | - | - | 0 | - |
| - | - | 548.5 | 212.6 | - | - | 0 | - |
| - | - | 711.6 | 215.1 | - | - | 0 | - |
| - | - | 9728 | 215.1 | - | - | 0 | - |
| - | - | 4728 | 217.1 | - | - | 0 | - |
| - | - | 4992 | 235.1 | - | - | 0 | - |
| - | - | 593.4 | 236.1 | - | - | 0 | - |
| - | - | 587.3 | 242.1 | - | - | 0 | - |
| - | - | 646.2 | 245.1 | - | - | 0 | - |
| - | - | 1956 | 258.1 | - | - | 0 | - |
| - | - | 2743 | 283.1 | - | - | 0 | - |
| - | - | 580.1 | 283.1 | - | - | 0 | - |
| - | - | 1002 | 299.2 | - | - | 0 | - |
| - | - | 3772 | 302.2 | - | - | 0 | - |
| - | - | 662.2 | 303.2 | - | - | 0 | - |
| - | - | 2442 | 311.1 | - | - | 0 | - |
| - | - | 1540 | 312.1 | - | - | 0 | - |
| - | - | 902.4 | 314.1 | - | - | 0 | - |
| - | - | 2288 | 318.1 | - | - | 0 | - |
| - | - | 642.3 | 327.1 | - | - | 0 | - |
| 13 | y | 5.36E+04 | 329.1 | 0.000139 | 0.4223 | +1 | 2 |
| - | - | 9957 | 330.2 | - | - | 0 | - |
| - | - | 518.4 | 331.2 | - | - | 0 | - |
| - | - | 3285 | 334.1 | - | - | 0 | - |
| - | - | 637.1 | 335.1 | - | - | 0 | - |
| - | - | 5541 | 341.2 | - | - | 0 | - |
| - | - | 774.6 | 342.2 | - | - | 0 | - |
| - | - | 1573 | 346.1 | - | - | 0 | - |
| - | - | 4298 | 359.2 | - | - | 0 | - |
| - | - | 937 | 364.1 | - | - | 0 | - |
| - | - | 1178 | 370.1 | - | - | 0 | - |
| - | - | 617.7 | 371.2 | - | - | 0 | - |
| - | - | 4396 | 386.2 | - | - | 0 | - |
| - | - | 662.6 | 387.2 | - | - | 0 | - |
| - | - | 724.8 | 397.2 | - | - | 0 | - |
| - | - | 3931 | 398.1 | - | - | 0 | - |
| 8 | c | 556.2 | 400.2 | 0.002477 | 6.189 | +2 | 8 |
| - | - | 831.1 | 426.3 | - | - | 0 | - |
| - | - | 576.9 | 431.2 | - | - | 0 | - |
| - | - | 565.6 | 432.2 | - | - | 0 | - |
| - | - | 4500 | 444.3 | - | - | 0 | - |
| - | - | 588.4 | 445.3 | - | - | 0 | - |
| - | - | 7849 | 454.3 | - | - | 0 | - |
| - | - | 2022 | 455.3 | - | - | 0 | - |
| - | - | 1852 | 457.2 | - | - | 0 | - |
| - | - | 1282 | 460.2 | - | - | 0 | - |
| - | - | 5387 | 461.1 | - | - | 0 | - |
| - | - | 576.8 | 462.1 | - | - | 0 | - |
| - | - | 898 | 469.2 | - | - | 0 | - |
| - | - | 7355 | 472.3 | - | - | 0 | - |
| - | - | 2073 | 473.3 | - | - | 0 | - |
| 12 | y | 8057 | 476.2 | 0.004716 | 9.903 | +1 | 3 |
| - | - | 1637 | 477.2 | - | - | 0 | - |
| - | - | 3465 | 479.1 | - | - | 0 | - |
| - | - | 1078 | 487.3 | - | - | 0 | - |
| 5 | c | 1314 | 489.3 | 0.001244 | 2.543 | +1 | 5 |
| - | - | 504.9 | 500.6 | - | - | 0 | - |
| - | - | 590 | 509.2 | - | - | 0 | - |
| - | - | 1138 | 514.2 | - | - | 0 | - |
| 10 | c | 1477 | 522.7 | 0.001146 | 2.192 | +2 | 10 |
| - | - | 1017 | 523.2 | - | - | 0 | - |
| - | - | 826.3 | 524.2 | - | - | 0 | - |
| - | - | 992.4 | 536.2 | - | - | 0 | - |
| - | - | 1336 | 540.3 | - | - | 0 | - |
| - | - | 561.2 | 548.2 | - | - | 0 | - |
| - | - | 5091 | 557.3 | - | - | 0 | - |
| - | - | 2352 | 557.8 | - | - | 0 | - |
| 11 | y | 7368 | 563.2 | 0.004273 | 7.587 | +1 | 4 |
| - | - | 1950 | 564.2 | - | - | 0 | - |
| 11 | c | 2408 | 566.3 | 0.0003756 | 0.6633 | +2 | 11 |
| - | - | 902.7 | 566.8 | - | - | 0 | - |
| - | - | 567 | 571.2 | - | - | 0 | - |
| - | - | 1214 | 572.2 | - | - | 0 | - |
| - | - | 1642 | 573.2 | - | - | 0 | - |
| - | - | 871.9 | 574.4 | - | - | 0 | - |
| - | - | 6150 | 582.3 | - | - | 0 | - |
| - | - | 1989 | 583.3 | - | - | 0 | - |
| - | - | 2431 | 586.3 | - | - | 0 | - |
| - | - | 3577 | 590.2 | - | - | 0 | - |
| - | - | 602.3 | 591.2 | - | - | 0 | - |
| - | - | 1214 | 598.8 | - | - | 0 | - |
| - | - | 769.9 | 599.3 | - | - | 0 | - |
| 6 | c | 1.251E+04 | 600.3 | 0.0003137 | 0.5225 | +1 | 6 |
| - | - | 3380 | 601.3 | - | - | 0 | - |
| - | - | 691.4 | 602.3 | - | - | 0 | - |
| - | - | 1474 | 608.2 | - | - | 0 | - |
| - | - | 890.7 | 614.2 | - | - | 0 | - |
| - | - | 802.5 | 615.2 | - | - | 0 | - |
| - | - | 2431 | 616.4 | - | - | 0 | - |
| 6 | c | 2.105E+04 | 617.4 | 0.0004345 | 0.7039 | +1 | 6 |
| - | - | 5769 | 618.4 | - | - | 0 | - |
| - | - | 860.2 | 619.4 | - | - | 0 | - |
| - | - | 2215 | 621.8 | - | - | 0 | - |
| - | - | 2741 | 622.3 | - | - | 0 | - |
| - | - | 911.3 | 622.8 | - | - | 0 | - |
| - | - | 1.493E+04 | 630.8 | - | - | 0 | - |
| - | - | 8858 | 631.3 | - | - | 0 | - |
| - | - | 2996 | 631.8 | - | - | 0 | - |
| 12 | c | 2505 | 639.8 | 0.0009837 | 1.538 | +2 | 12 |
| - | - | 953.4 | 640.3 | - | - | 0 | - |
| - | - | 4427 | 642.2 | - | - | 0 | - |
| - | - | 1117 | 643.2 | - | - | 0 | - |
| - | - | 1067 | 643.4 | - | - | 0 | - |
| - | - | 1465 | 644.4 | - | - | 0 | - |
| - | - | 3132 | 645.4 | - | - | 0 | - |
| - | - | 6732 | 653.4 | - | - | 0 | - |
| - | - | 2758 | 654.4 | - | - | 0 | - |
| - | - | 623.9 | 655.4 | - | - | 0 | - |
| - | - | 4367 | 661.2 | - | - | 0 | - |
| - | - | 1232 | 662.2 | - | - | 0 | - |
| - | - | 1078 | 664.8 | - | - | 0 | - |
| - | - | 1055 | 665.3 | - | - | 0 | - |
| - | - | 700.5 | 666.3 | - | - | 0 | - |
| - | - | 1455 | 669.3 | - | - | 0 | - |
| 7 | c | 8305 | 671.4 | 0.000257 | 0.3829 | +1 | 7 |
| - | - | 2709 | 672.4 | - | - | 0 | - |
| - | - | 857.9 | 673.4 | - | - | 0 | - |
| - | - | 1034 | 675.3 | - | - | 0 | - |
| - | - | 1080 | 675.8 | - | - | 0 | - |
| - | - | 2946 | 679.2 | - | - | 0 | - |
| - | - | 604.1 | 680.2 | - | - | 0 | - |
| - | - | 1104 | 687.3 | - | - | 0 | - |
| - | - | 8323 | 687.4 | - | - | 0 | - |
| 7 | c | 6.035E+04 | 688.4 | 0.0006221 | 0.9036 | +1 | 7 |
| - | - | 1.818E+04 | 689.4 | - | - | 0 | - |
| - | - | 3031 | 690.4 | - | - | 0 | - |
| 10 | y | 888.3 | 692.3 | 0.007517 | 10.86 | +1 | 5 |
| - | - | 2155 | 701.3 | - | - | 0 | - |
| - | - | 835.4 | 712.3 | - | - | 0 | - |
| - | - | 740.9 | 712.8 | - | - | 0 | - |
| - | - | 859.3 | 713.4 | - | - | 0 | - |
| - | - | 2046 | 714.3 | - | - | 0 | - |
| - | - | 1228 | 714.4 | - | - | 0 | - |
| - | - | 1019 | 715.3 | - | - | 0 | - |
| - | - | 723.5 | 715.4 | - | - | 0 | - |
| - | - | 1096 | 743.3 | - | - | 0 | - |
| - | - | 1452 | 753.2 | - | - | 0 | - |
| - | - | 694.3 | 760.4 | - | - | 0 | - |
| - | - | 652.1 | 763.3 | - | - | 0 | - |
| - | - | 628.9 | 763.8 | - | - | 0 | - |
| - | - | 2833 | 771.3 | - | - | 0 | - |
| - | - | 1755 | 771.9 | - | - | 0 | - |
| - | - | 1492 | 772.3 | - | - | 0 | - |
| - | - | 899 | 772.3 | - | - | 0 | - |
| - | - | 5088 | 773.4 | - | - | 0 | - |
| - | - | 5274 | 774.4 | - | - | 0 | - |
| - | - | 644.7 | 775.3 | - | - | 0 | - |
| - | - | 1840 | 775.4 | - | - | 0 | - |
| - | - | 1251 | 776.3 | - | - | 0 | - |
| - | - | 9882 | 777.3 | - | - | 0 | - |
| - | - | 3217 | 778.3 | - | - | 0 | - |
| - | - | 844.2 | 779.3 | - | - | 0 | - |
| - | - | 6630 | 782.4 | - | - | 0 | - |
| - | - | 2621 | 783.4 | - | - | 0 | - |
| - | - | 1247 | 784.4 | - | - | 0 | - |
| 9 | y | 2249 | 789.3 | 0.00111 | 1.406 | +1 | 6 |
| - | - | 1404 | 790.3 | - | - | 0 | - |
| 9 | z | 830.8 | 791.3 | 0.001068 | 1.35 | +1 | 6 |
| - | - | 3514 | 793.3 | - | - | 0 | - |
| - | - | 2367 | 794.3 | - | - | 0 | - |
| 8 | c | 1.1E+04 | 800.4 | 0.0006138 | 0.7669 | +1 | 8 |
| - | - | 3672 | 801.4 | - | - | 0 | - |
| - | - | 2700 | 802.4 | - | - | 0 | - |
| - | - | 3344 | 802.9 | - | - | 0 | - |
| - | - | 3647 | 803.4 | - | - | 0 | - |
| 9 | y | 5755 | 807.3 | 0.004012 | 4.97 | +1 | 6 |
| - | - | 1730 | 808.3 | - | - | 0 | - |
| - | - | 754.1 | 809.3 | - | - | 0 | - |
| - | - | 800.5 | 812.3 | - | - | 0 | - |
| - | - | 1357 | 812.4 | - | - | 0 | - |
| - | - | 2067 | 813.4 | - | - | 0 | - |
| - | - | 1113 | 814.3 | - | - | 0 | - |
| - | - | 1394 | 814.4 | - | - | 0 | - |
| - | - | 2.129E+04 | 816.4 | - | - | 0 | - |
| 8 | c | 3.506E+04 | 817.4 | 0.001772 | 2.168 | +1 | 8 |
| - | - | 1.07E+04 | 818.4 | - | - | 0 | - |
| - | - | 1966 | 819.4 | - | - | 0 | - |
| - | - | 1149 | 824.3 | - | - | 0 | - |
| - | - | 3295 | 830.4 | - | - | 0 | - |
| - | - | 1296 | 831.4 | - | - | 0 | - |
| - | - | 4475 | 842.3 | - | - | 0 | - |
| - | - | 1695 | 843.3 | - | - | 0 | - |
| - | - | 1098 | 872.3 | - | - | 0 | - |
| - | - | 1071 | 887.4 | - | - | 0 | - |
| - | - | 7483 | 888.5 | - | - | 0 | - |
| - | - | 5458 | 889.5 | - | - | 0 | - |
| - | - | 2029 | 890.5 | - | - | 0 | - |
| - | - | 695.7 | 893.3 | - | - | 0 | - |
| - | - | 729.9 | 897.4 | - | - | 0 | - |
| - | - | 835.2 | 898.4 | - | - | 0 | - |
| 9 | c | 5675 | 915.4 | 0.0007624 | 0.8328 | +1 | 9 |
| - | - | 2029 | 916.4 | - | - | 0 | - |
| - | - | 685.7 | 925.3 | - | - | 0 | - |
| - | - | 9187 | 931.5 | - | - | 0 | - |
| 9 | c | 5.319E+04 | 932.5 | 0.001616 | 1.733 | +1 | 9 |
| - | - | 2.218E+04 | 933.5 | - | - | 0 | - |
| - | - | 4726 | 934.5 | - | - | 0 | - |
| - | - | 2358 | 935.3 | - | - | 0 | - |
| 8 | y | 7587 | 936.3 | 0.002862 | 3.056 | +1 | 7 |
| - | - | 3503 | 937.3 | - | - | 0 | - |
| - | - | 1097 | 938.3 | - | - | 0 | - |
| - | - | 685.6 | 942.4 | - | - | 0 | - |
| - | - | 1903 | 943.4 | - | - | 0 | - |
| - | - | 1294 | 958.5 | - | - | 0 | - |
| - | - | 944.3 | 964.5 | - | - | 0 | - |
| - | - | 1023 | 970.3 | - | - | 0 | - |
| - | - | 1415 | 971.4 | - | - | 0 | - |
| - | - | 949.8 | 979.4 | - | - | 0 | - |
| - | - | 1845 | 984.5 | - | - | 0 | - |
| - | - | 841.2 | 988.5 | - | - | 0 | - |
| 7 | y | 1168 | 989.4 | 0.001093 | 1.105 | +1 | 8 |
| 7 | z | 1417 | 991.3 | 0.001379 | 1.391 | +1 | 8 |
| - | - | 966.9 | 992.4 | - | - | 0 | - |
| - | - | 890.9 | 999.5 | - | - | 0 | - |
| - | - | 606.4 | 1000 | - | - | 0 | - |
| - | - | 631.3 | 1001 | - | - | 0 | - |
| - | - | 2062 | 1006 | - | - | 0 | - |
| 7 | y | 1.078E+04 | 1007 | 0.003162 | 3.139 | +1 | 8 |
| - | - | 4895 | 1008 | - | - | 0 | - |
| - | - | 2071 | 1009 | - | - | 0 | - |
| - | - | 1030 | 1011 | - | - | 0 | - |
| - | - | 7325 | 1017 | - | - | 0 | - |
| - | - | 3852 | 1019 | - | - | 0 | - |
| - | - | 1377 | 1020 | - | - | 0 | - |
| - | - | 936.5 | 1023 | - | - | 0 | - |
| - | - | 5006 | 1026 | - | - | 0 | - |
| - | - | 3561 | 1027 | - | - | 0 | - |
| - | - | 1130 | 1028 | - | - | 0 | - |
| - | - | 2475 | 1029 | - | - | 0 | - |
| - | - | 1275 | 1030 | - | - | 0 | - |
| - | - | 2232 | 1042 | - | - | 0 | - |
| 10 | c | 1749 | 1044 | 0.0121 | 11.59 | +1 | 10 |
| 10 | c | 1.603E+04 | 1044 | 0.001119 | 1.072 | +1 | 10 |
| - | - | 7646 | 1045 | - | - | 0 | - |
| - | - | 1967 | 1046 | - | - | 0 | - |
| - | - | 1518 | 1047 | - | - | 0 | - |
| - | - | 2554 | 1047 | - | - | 0 | - |
| - | - | 2312 | 1048 | - | - | 0 | - |
| - | - | 1033 | 1049 | - | - | 0 | - |
| - | - | 757.1 | 1054 | - | - | 0 | - |
| - | - | 5.713E+04 | 1061 | - | - | 0 | - |
| 10 | c | 4.405E+04 | 1062 | 0.004719 | 4.446 | +1 | 10 |
| - | - | 1.547E+04 | 1063 | - | - | 0 | - |
| - | - | 3198 | 1064 | - | - | 0 | - |
| - | - | 2934 | 1064 | - | - | 0 | - |
| - | - | 2139 | 1065 | - | - | 0 | - |
| - | - | 3532 | 1066 | - | - | 0 | - |
| - | - | 1058 | 1067 | - | - | 0 | - |
| - | - | 631.4 | 1068 | - | - | 0 | - |
| - | - | 1240 | 1071 | - | - | 0 | - |
| - | - | 800.4 | 1087 | - | - | 0 | - |
| - | - | 2604 | 1088 | - | - | 0 | - |
| - | - | 2039 | 1089 | - | - | 0 | - |
| - | - | 1151 | 1091 | - | - | 0 | - |
| - | - | 637.9 | 1092 | - | - | 0 | - |
| - | - | 2590 | 1095 | - | - | 0 | - |
| - | - | 1362 | 1096 | - | - | 0 | - |
| - | - | 1748 | 1106 | - | - | 0 | - |
| - | - | 1.315E+04 | 1114 | - | - | 0 | - |
| - | - | 8257 | 1115 | - | - | 0 | - |
| - | - | 1831 | 1116 | - | - | 0 | - |
| 6 | y | 681 | 1118 | 0.01618 | 14.47 | +1 | 9 |
| 6 | z | 2925 | 1119 | 0.004447 | 3.972 | +1 | 9 |
| - | - | 2198 | 1120 | - | - | 0 | - |
| - | - | 683.8 | 1121 | - | - | 0 | - |
| - | - | 1205 | 1130 | - | - | 0 | - |
| 11 | c | 1197 | 1131 | 0.01349 | 11.93 | +1 | 11 |
| 11 | c | 1.154E+04 | 1132 | 0.000799 | 0.7061 | +1 | 11 |
| - | - | 6008 | 1133 | - | - | 0 | - |
| - | - | 1821 | 1134 | - | - | 0 | - |
| - | - | 7951 | 1134 | - | - | 0 | - |
| 6 | y | 1.068E+04 | 1135 | 0.001714 | 1.509 | +1 | 9 |
| - | - | 6144 | 1136 | - | - | 0 | - |
| - | - | 2332 | 1137 | - | - | 0 | - |
| - | - | 3.314E+04 | 1148 | - | - | 0 | - |
| 11 | c | 8.23E+04 | 1149 | 0.002202 | 1.917 | +1 | 11 |
| - | - | 4.008E+04 | 1150 | - | - | 0 | - |
| - | - | 8845 | 1151 | - | - | 0 | - |
| - | - | 975.4 | 1162 | - | - | 0 | - |
| - | - | 919.9 | 1165 | - | - | 0 | - |
| - | - | 2709 | 1180 | - | - | 0 | - |
| - | - | 1902 | 1181 | - | - | 0 | - |
| - | - | 1043 | 1183 | - | - | 0 | - |
| - | - | 761.9 | 1184 | - | - | 0 | - |
| - | - | 995.5 | 1196 | - | - | 0 | - |
| - | - | 1364 | 1197 | - | - | 0 | - |
| - | - | 1110 | 1198 | - | - | 0 | - |
| - | - | 831.8 | 1200 | - | - | 0 | - |
| - | - | 1574 | 1204 | - | - | 0 | - |
| - | - | 1536 | 1205 | - | - | 0 | - |
| - | - | 1454 | 1211 | - | - | 0 | - |
| - | - | 1226 | 1212 | - | - | 0 | - |
| - | - | 748.5 | 1215 | - | - | 0 | - |
| - | - | 1895 | 1227 | - | - | 0 | - |
| - | - | 819.6 | 1228 | - | - | 0 | - |
| - | - | 728.2 | 1229 | - | - | 0 | - |
| 5 | z | 1.248E+04 | 1232 | 0.004123 | 3.345 | +1 | 10 |
| - | - | 1.002E+04 | 1233 | - | - | 0 | - |
| - | - | 5320 | 1234 | - | - | 0 | - |
| - | - | 5133 | 1236 | - | - | 0 | - |
| - | - | 3287 | 1237 | - | - | 0 | - |
| - | - | 1705 | 1238 | - | - | 0 | - |
| - | - | 4025 | 1243 | - | - | 0 | - |
| - | - | 3853 | 1244 | - | - | 0 | - |
| - | - | 2168 | 1245 | - | - | 0 | - |
| - | - | 1092 | 1246 | - | - | 0 | - |
| - | - | 1129 | 1247 | - | - | 0 | - |
| - | - | 1.934E+04 | 1248 | - | - | 0 | - |
| 5 | y | 1.612E+04 | 1249 | 0.001418 | 1.135 | +1 | 10 |
| - | - | 6316 | 1250 | - | - | 0 | - |
| - | - | 1842 | 1251 | - | - | 0 | - |
| - | - | 6305 | 1252 | - | - | 0 | - |
| - | - | 4146 | 1253 | - | - | 0 | - |
| - | - | 2072 | 1254 | - | - | 0 | - |
| - | - | 1.987E+04 | 1261 | - | - | 0 | - |
| - | - | 1.56E+04 | 1262 | - | - | 0 | - |
| - | - | 1.527E+04 | 1263 | - | - | 0 | - |
| - | - | 1248 | 1263 | - | - | 0 | - |
| - | - | 7749 | 1264 | - | - | 0 | - |
| - | - | 1008 | 1264 | - | - | 0 | - |
| - | - | 2794 | 1265 | - | - | 0 | - |
| 4 | z | 740.3 | 1271 | 0.001647 | 1.295 | +1 | 11 |
| - | - | 702.3 | 1273 | - | - | 0 | - |
| 12 | c | 2.825E+04 | 1279 | 0.001797 | 1.406 | +1 | 12 |
| - | - | 1.823E+04 | 1280 | - | - | 0 | - |
| - | - | 9918 | 1281 | - | - | 0 | - |
| - | - | 3631 | 1282 | - | - | 0 | - |
| - | - | 1062 | 1287 | - | - | 0 | - |
| 4 | z | 7308 | 1290 | 0.003533 | 2.74 | +1 | 11 |
| - | - | 1.307E+04 | 1291 | - | - | 0 | - |
| - | - | 9007 | 1292 | - | - | 0 | - |
| - | - | 3403 | 1293 | - | - | 0 | - |
| 12 | c | 6.931E+04 | 1296 | 0.001982 | 1.53 | +1 | 12 |
| - | - | 4.338E+04 | 1297 | - | - | 0 | - |
| - | - | 859.8 | 1297 | - | - | 0 | - |
| - | - | 1.581E+04 | 1298 | - | - | 0 | - |
| - | - | 1623 | 1299 | - | - | 0 | - |
| - | - | 1.41E+04 | 1305 | - | - | 0 | - |
| 4 | y | 1.73E+04 | 1306 | 0.0009087 | 0.696 | +1 | 11 |
| - | - | 9188 | 1307 | - | - | 0 | - |
| - | - | 2360 | 1308 | - | - | 0 | - |
| - | - | 1054 | 1311 | - | - | 0 | - |
| - | - | 844.8 | 1312 | - | - | 0 | - |
| - | - | 933.1 | 1325 | - | - | 0 | - |
| - | - | 1047 | 1326 | - | - | 0 | - |
| - | - | 1721 | 1329 | - | - | 0 | - |
| - | - | 1524 | 1330 | - | - | 0 | - |
| - | - | 828.8 | 1344 | - | - | 0 | - |
| - | - | 985.2 | 1352 | - | - | 0 | - |
| 3 | z | 4680 | 1359 | 0.002455 | 1.807 | +1 | 12 |
| 3 | z | 4107 | 1360 | 0.02198 | 16.17 | +1 | 12 |
| - | - | 2228 | 1361 | - | - | 0 | - |
| - | - | 5000 | 1362 | - | - | 0 | - |
| - | - | 4429 | 1363 | - | - | 0 | - |
| - | - | 2501 | 1364 | - | - | 0 | - |
| 3 | y | 2254 | 1375 | 0.01358 | 9.882 | +1 | 12 |
| 3 | y | 1820 | 1376 | 0.00826 | 6.005 | +1 | 12 |
| 3 | z | 3.499E+04 | 1377 | 0.001778 | 1.292 | +1 | 12 |
| - | - | 2.684E+04 | 1378 | - | - | 0 | - |
| - | - | 1.026E+04 | 1379 | - | - | 0 | - |
| - | - | 1769 | 1380 | - | - | 0 | - |
| - | - | 1334 | 1392 | - | - | 0 | - |
| 3 | y | 2.74E+04 | 1393 | 0.001609 | 1.155 | +1 | 12 |
| - | - | 2.208E+04 | 1394 | - | - | 0 | - |
| - | - | 8971 | 1395 | - | - | 0 | - |
| - | - | 1470 | 1396 | - | - | 0 | - |
| - | - | 914.9 | 1414 | - | - | 0 | - |
| - | - | 5317 | 1415 | - | - | 0 | - |
| - | - | 4990 | 1416 | - | - | 0 | - |
| - | - | 2014 | 1417 | - | - | 0 | - |
| - | - | 2072 | 1419 | - | - | 0 | - |
| - | - | 1569 | 1420 | - | - | 0 | - |
| - | - | 1549 | 1424 | - | - | 0 | - |
| - | - | 1515 | 1425 | - | - | 0 | - |
| - | - | 7047 | 1426 | - | - | 0 | - |
| - | - | 1.345E+04 | 1427 | - | - | 0 | - |
| - | - | 2.815E+04 | 1428 | - | - | 0 | - |
| - | - | 1.998E+04 | 1429 | - | - | 0 | - |
| - | - | 7096 | 1430 | - | - | 0 | - |
| - | - | 1574 | 1440 | - | - | 0 | - |
| - | - | 2061 | 1441 | - | - | 0 | - |
| 13 | c | 3120 | 1442 | 0.001091 | 0.7568 | +1 | 13 |
| - | - | 7125 | 1443 | - | - | 0 | - |
| - | - | 1.414E+04 | 1444 | - | - | 0 | - |
| - | - | 8091 | 1445 | - | - | 0 | - |
| - | - | 4223 | 1446 | - | - | 0 | - |
| - | - | 1008 | 1447 | - | - | 0 | - |
| 13 | c | 8.663E+04 | 1459 | 0.001153 | 0.7906 | +1 | 13 |
| - | - | 6.61E+04 | 1460 | - | - | 0 | - |
| - | - | 2.899E+04 | 1461 | - | - | 0 | - |
| - | - | 3782 | 1462 | - | - | 0 | - |
| 2 | z | 2448 | 1472 | 0.007503 | 5.098 | +1 | 13 |
| 2 | z | 2162 | 1473 | 0.02373 | 16.12 | +1 | 13 |
| - | - | 1667 | 1474 | - | - | 0 | - |
| 2 | z | 3.017E+04 | 1490 | 0.0009661 | 0.6486 | +1 | 13 |
| - | - | 2.684E+04 | 1491 | - | - | 0 | - |
| - | - | 1.205E+04 | 1492 | - | - | 0 | - |
| - | - | 2019 | 1493 | - | - | 0 | - |
| - | - | 6224 | 1507 | - | - | 0 | - |
| - | - | 5577 | 1508 | - | - | 0 | - |
| - | - | 2109 | 1509 | - | - | 0 | - |
| - | - | 5609 | 1516 | - | - | 0 | - |
| - | - | 5177 | 1517 | - | - | 0 | - |
| - | - | 2515 | 1518 | - | - | 0 | - |
| - | - | 1024 | 1519 | - | - | 0 | - |
| - | - | 1049 | 1520 | - | - | 0 | - |
| - | - | 1617 | 1521 | - | - | 0 | - |
| - | - | 1925 | 1527 | - | - | 0 | - |
| - | - | 2860 | 1528 | - | - | 0 | - |
| - | - | 2025 | 1529 | - | - | 0 | - |
| - | - | 959.1 | 1530 | - | - | 0 | - |
| - | - | 2845 | 1535 | - | - | 0 | - |
| - | - | 4337 | 1536 | - | - | 0 | - |
| - | - | 4451 | 1537 | - | - | 0 | - |
| - | - | 2908 | 1538 | - | - | 0 | - |
| - | - | 1401 | 1539 | - | - | 0 | - |
| - | - | 6485 | 1544 | - | - | 0 | - |
| - | - | 9177 | 1545 | - | - | 0 | - |
| - | - | 1.354E+04 | 1546 | - | - | 0 | - |
| - | - | 9881 | 1547 | - | - | 0 | - |
| - | - | 9121 | 1548 | - | - | 0 | - |
| - | - | 7049 | 1549 | - | - | 0 | - |
| - | - | 5354 | 1550 | - | - | 0 | - |
| - | - | 1628 | 1551 | - | - | 0 | - |
| - | - | 4351 | 1552 | - | - | 0 | - |
| - | - | 2713 | 1553 | - | - | 0 | - |
| - | - | 1889 | 1554 | - | - | 0 | - |
| - | - | 853.1 | 1555 | - | - | 0 | - |
| - | - | 5923 | 1562 | - | - | 0 | - |
| - | - | 3.314E+04 | 1563 | - | - | 0 | - |
| - | - | 3.011E+04 | 1564 | - | - | 0 | - |
| - | - | 1.677E+04 | 1565 | - | - | 0 | - |
| - | - | 4458 | 1566 | - | - | 0 | - |
| - | - | 985.3 | 1567 | - | - | 0 | - |
| - | - | 2288 | 1572 | - | - | 0 | - |
| - | - | 1.366E+04 | 1573 | - | - | 0 | - |
| - | - | 1.282E+04 | 1574 | - | - | 0 | - |
| - | - | 7368 | 1575 | - | - | 0 | - |
| - | - | 977.8 | 1576 | - | - | 0 | - |
| - | - | 872 | 1577 | - | - | 0 | - |
| - | - | 1.348E+04 | 1580 | - | - | 0 | - |
| - | - | 1.06E+04 | 1581 | - | - | 0 | - |
| - | - | 5215 | 1582 | - | - | 0 | - |
| - | - | 2432 | 1588 | - | - | 0 | - |
| - | - | 5764 | 1589 | - | - | 0 | - |
| - | - | 4.853E+04 | 1590 | - | - | 0 | - |
| - | - | 1.851E+05 | 1591 | - | - | 0 | - |
| - | - | 1.502E+05 | 1592 | - | - | 0 | - |
| - | - | 7.183E+04 | 1593 | - | - | 0 | - |
| - | - | 1.332E+04 | 1594 | - | - | 0 | - |
| - | - | 2123 | 1595 | - | - | 0 | - |
| - | - | 1624 | 1604 | - | - | 0 | - |
| - | - | 5195 | 1605 | - | - | 0 | - |
| - | - | 5698 | 1606 | - | - | 0 | - |
| - | - | 6.112E+04 | 1607 | - | - | 0 | - |
| - | - | 2.174E+05 | 1608 | - | - | 0 | - |
| - | - | 1.698E+05 | 1609 | - | - | 0 | - |
| - | - | 7.905E+04 | 1610 | - | - | 0 | - |
| - | - | 9450 | 1611 | - | - | 0 | - |
| - | - | 796.8 | 1622 | - | - | 0 | - |
| - | - | 953.3 | 1642 | - | - | 0 | - |
| - | - | 794.9 | 2046 | - | - | 0 | - |
| - | - | 1121 | 2047 | - | - | 0 | - |
| - | - | 890.9 | 2377 | - | - | 0 | - |
| - | - | 767.6 | 2391 | - | - | 0 | - |
| - | - | 712.8 | 2392 | - | - | 0 | - |
| - | - | 1592 | 2396 | - | - | 0 | - |
| - | - | 1008 | 2397 | - | - | 0 | - |
| - | - | 1197 | 2407 | - | - | 0 | - |
| - | - | 3436 | 2408 | - | - | 0 | - |
| - | - | 3712 | 2409 | - | - | 0 | - |
| - | - | 1627 | 2410 | - | - | 0 | - |
| - | - | 1414 | 2411 | - | - | 0 | - |

m/z Charge Intensity FragmentType MassShift Position
120.08076477050781 0 5160.044
121.08429718017578 0 494.4195
124.26469421386719 0 357.03516
124.27033996582031 0 452.45636
125.60387420654297 0 337.05667
125.77693176269531 0 408.94217
136.07569885253906 0 56104.363
137.07901000976562 0 4057.425
148.24473571777344 0 454.6431
148.8871612548828 0 480.2692
148.89453125 0 527.535
148.90199279785156 0 676.5651
148.90896606445312 0 767.8132
148.91600036621094 0 722.37695
148.92355346679688 0 1324.5182
148.93075561523438 0 1146.9901
148.93821716308594 0 3104.457
148.9460906982422 0 5871.481
148.9626007080078 0 3991.6934
148.97036743164062 0 1698.1556
148.9775390625 0 1218.9049
148.98484802246094 0 1108.4994
148.99209594726562 0 782.1338
148.9991912841797 0 590.7813
149.00674438476562 0 708.9457
149.0137481689453 0 643.6942
149.02134704589844 0 585.497
166.0789337158203 0 697.6038
166.08619689941406 0 10218.138 y 13
173.12841796875 0 749.7629
187.1439666748047 0 10164.191
187.69822692871094 0 505.17828
188.1477813720703 0 754.0652
201.1227569580078 0 591.29553
203.3450927734375 0 490.77972
212.55889892578125 0 548.528
215.12779235839844 0 711.6251
215.13880920410156 0 9728.493
217.06398010253906 0 4728.4253
235.0745391845703 0 4991.779
236.0772247314453 0 593.42285
242.14889526367188 0 587.3407
245.07717895507812 0 646.2022
258.1447448730469 0 1955.9402
283.1109924316406 0 2742.7092
283.1432189941406 0 580.0941
299.1709899902344 0 1001.694
302.17083740234375 0 3771.7764
303.17462158203125 0 662.1974
311.1057434082031 0 2441.806
312.1223449707031 0 1539.8488
314.0976867675781 0 902.4152
318.14483642578125 0 2288.3154
327.0982971191406 0 642.34546
329.1494445800781 0 53598.22 y 12
330.15252685546875 0 9956.603
331.1552429199219 0 518.3703
334.1395568847656 0 3284.9473
335.14276123046875 0 637.0541
341.1817932128906 0 5541.3384
342.1850891113281 0 774.5558
346.10650634765625 0 1573.1729
359.1922607421875 0 4298.1523
364.1173400878906 0 937.0139
370.142578125 0 1178.4314
371.2274475097656 0 617.69226
386.20306396484375 0 4395.6147
387.2042236328125 0 662.60144
397.2071228027344 0 724.78516
398.1377868652344 0 3930.771
400.2165832519531 0 556.2457 c Water loss 7
426.2727355957031 0 831.08563
431.19561767578125 0 576.87744
432.1855773925781 0 565.616
444.2820129394531 0 4500.396
445.28619384765625 0 588.4364
454.26580810546875 0 7849.2065
455.2677917480469 0 2021.5956
457.2409973144531 0 1851.9436
460.18292236328125 0 1282.0024
461.13360595703125 0 5386.597
462.1338806152344 0 576.8499
469.24163818359375 0 897.9714
472.27630615234375 0 7355.461
473.279541015625 0 2072.9995
476.1847839355469 0 8057.1035 y 11
477.18646240234375 0 1636.5833
479.14361572265625 0 3465.0222
487.25048828125 0 1078.2139
489.3018798828125 0 1313.7577 c 4
500.6255798339844 0 504.94858
509.1685485839844 0 590.0034
514.176513671875 0 1138.0737
522.7446899414062 0 1477.3469 c Ammonia loss 9
523.2484741210938 0 1017.4152
524.2134399414062 0 826.27386
536.2159423828125 0 992.3855
540.277099609375 0 1335.628
548.248779296875 0 561.2291
557.2562866210938 0 5090.9307
557.75732421875 0 2351.5247
563.2163696289062 0 7367.6274 y 10
564.2203369140625 0 1949.6115
566.261474609375 0 2407.7158 c Ammonia loss 10
566.762939453125 0 902.7355
571.2479858398438 0 567.0028
572.1647338867188 0 1214.1741
573.2179565429688 0 1642.1086
574.356201171875 0 871.90515
582.3236694335938 0 6149.9863
583.32177734375 0 1988.8278
586.28173828125 0 2430.5837
590.17626953125 0 3576.9055
591.1783447265625 0 602.2602
598.7743530273438 0 1213.6978
599.2734375 0 769.948
600.3348388671875 0 12513.471 c Ammonia loss 5
601.3364868164062 0 3379.6467
602.343017578125 0 691.4246
608.1871948242188 0 1473.9316
614.2115478515625 0 890.6526
615.216064453125 0 802.5454
616.3538818359375 0 2431.0266
617.3612670898438 0 21047.55 c 5
618.3641967773438 0 5768.9507
619.363037109375 0 860.1572
621.7705688476562 0 2214.7715
622.2662963867188 0 2740.9019
622.7694702148438 0 911.2885
630.7737426757812 0 14926.583
631.2747192382812 0 8857.826
631.7747802734375 0 2996.32
639.778076171875 0 2504.5251 c Ammonia loss 11
640.2810668945312 0 953.4055
642.2078857421875 0 4426.8574
643.20751953125 0 1116.8894
643.378662109375 0 1067.1357
644.3832397460938 0 1465.4352
645.3909301757812 0 3132.1104
653.361083984375 0 6732.2354
654.3612060546875 0 2758.1926
655.3599853515625 0 623.9351
661.21337890625 0 4366.6777
662.215087890625 0 1231.9745
664.7811279296875 0 1077.5913
665.284912109375 0 1055.4097
666.3030395507812 0 700.5114
669.3214111328125 0 1454.8716
671.3720092773438 0 8304.792 c Ammonia loss 6
672.3743896484375 0 2708.8904
673.3845825195312 0 857.8999
675.3147583007812 0 1034.1089
675.8136596679688 0 1080.2905
679.2229614257812 0 2946.2986
680.229736328125 0 604.0628
687.3294677734375 0 1104.332
687.3910522460938 0 8322.99
688.398193359375 0 60351.44 c 6
689.400634765625 0 18182.34
690.4036865234375 0 3031.164
692.26220703125 0 888.2887 y 9
701.3093872070312 0 2155.0186
712.3020629882812 0 835.40125
712.7966918945312 0 740.9211
713.3505249023438 0 859.2598
714.33154296875 0 2045.5865
714.4095458984375 0 1227.8138
715.3388671875 0 1019.06915
715.4188232421875 0 723.4587
743.2549438476562 0 1095.5382
753.2372436523438 0 1451.9624
760.37548828125 0 694.3034
763.3432006835938 0 652.06964
763.84130859375 0 628.94403
771.2532348632812 0 2833.3105
771.8504028320312 0 1754.6761
772.251708984375 0 1492.4062
772.3473510742188 0 898.9692
773.4274291992188 0 5088.352
774.4321899414062 0 5273.8677
775.30810546875 0 644.7294
775.4371948242188 0 1839.6246
776.3095092773438 0 1250.9636
777.319091796875 0 9881.662
778.3211669921875 0 3216.9172
779.3230590820312 0 844.1992
782.4038696289062 0 6629.968
783.4024047851562 0 2620.7385
784.3977661132812 0 1246.6554
789.2699584960938 0 2248.9856 y Water loss 8
790.2744750976562 0 1404.143
791.2618408203125 0 830.8217 z 8
793.3154907226562 0 3513.8567
794.32080078125 0 2366.6775
800.4142456054688 0 10996.157 c Ammonia loss 7
801.4175415039062 0 3672.1467
802.3934936523438 0 2700.3853
802.895263671875 0 3344.061
803.3956298828125 0 3647.0278
807.28564453125 0 5755.0625 y 8
808.2881469726562 0 1730.3082
809.2982788085938 0 754.1071
812.3368530273438 0 800.47504
812.4147338867188 0 1357.4843
813.4030151367188 0 2066.544
814.29150390625 0 1112.9103
814.4010009765625 0 1393.5171
816.4327392578125 0 21294.395
817.4396362304688 0 35061.938 c 7
818.4429321289062 0 10704.407
819.4423828125 0 1965.7502
824.27880859375 0 1149.4524
830.35107421875 0 3294.6624
831.35302734375 0 1295.932
842.2857666015625 0 4474.8765
843.2881469726562 0 1694.9694
872.3306274414062 0 1097.8423
887.436767578125 0 1070.6418
888.4530639648438 0 7483.189
889.4601440429688 0 5458.4336
890.4618530273438 0 2029.2957
893.343505859375 0 695.72955
897.437255859375 0 729.88574
898.41162109375 0 835.22705
915.4410400390625 0 5675.1704 c Ammonia loss 8
916.4417114257812 0 2028.7642
925.346923828125 0 685.65753
931.4583740234375 0 9187.066
932.4667358398438 0 53191.223 c 8
933.4698486328125 0 22175.045
934.4718627929688 0 4725.519
935.3238525390625 0 2358.0144
936.3270874023438 0 7587.2417 y 7
937.3319091796875 0 3503.1868
938.335205078125 0 1097.1678
942.3572387695312 0 685.59155
943.3631591796875 0 1903.1409
958.4832153320312 0 1294.4166
964.4974365234375 0 944.32684
970.3436889648438 0 1022.8619
971.352294921875 0 1414.9087
979.4042358398438 0 949.80273
984.4799194335938 0 1844.611
988.490234375 0 841.2407
989.3518676757812 0 1167.941 y Water loss 6
991.343994140625 0 1417.3451 z 6
992.35205078125 0 966.8731
999.4668579101562 0 890.9191
1000.455322265625 0 606.4393
1001.4744873046875 0 631.2857
1006.3560180664062 0 2061.6338
1007.364501953125 0 10784.594 y 6
1008.3671264648438 0 4895.126
1009.3673095703125 0 2071.4702
1010.5447998046875 0 1030.0923
1017.4954833984375 0 7325.143
1018.5010375976562 0 3852.3247
1019.5013427734375 0 1376.5566
1023.049560546875 0 936.5254
1026.4735107421875 0 5006.2896
1027.466796875 0 3561.2114
1028.46484375 0 1130.4989
1029.4703369140625 0 2474.9397
1030.474853515625 0 1275.4448
1042.4912109375 0 2231.842
1043.48828125 0 1749.2771 c Water loss 9
1044.4832763671875 0 16026.675 c Ammonia loss 9
1045.4862060546875 0 7646.068
1046.4083251953125 0 1966.8723
1046.5032958984375 0 1518.3232
1047.40869140625 0 2553.7039
1048.416748046875 0 2311.8333
1049.428955078125 0 1033.1409
1054.401123046875 0 757.0733
1060.501953125 0 57126.04
1061.5062255859375 0 44050.734 c 9
1062.5103759765625 0 15465.69
1063.5166015625 0 3198.1838
1064.4193115234375 0 2933.8335
1065.4256591796875 0 2139.0613
1066.4326171875 0 3532.4663
1067.43359375 0 1058.0005
1068.4263916015625 0 631.41156
1071.424560546875 0 1239.9546
1086.521240234375 0 800.3843
1087.526123046875 0 2603.9587
1088.52880859375 0 2039.3062
1091.469482421875 0 1151.0588
1092.470458984375 0 637.9141
1095.4912109375 0 2590.0754
1096.494384765625 0 1362.2104
1105.5316162109375 0 1747.9099
1113.5045166015625 0 13153.929
1114.5032958984375 0 8257.267
1115.5 0 1830.6572
1118.4095458984375 0 680.9782 y Ammonia loss 5
1119.4056396484375 0 2925.2007 z 5
1120.4058837890625 0 2198.0332
1121.415283203125 0 683.80444
1129.5335693359375 0 1205.4875
1130.5189208984375 0 1197.0894 c Water loss 10
1131.515625 0 11537.649 c Ammonia loss 10
1132.51708984375 0 6008.3
1133.5206298828125 0 1821.0183
1134.41552734375 0 7950.5825
1135.421630859375 0 10675.831 y 5
1136.423828125 0 6143.608
1137.4312744140625 0 2332.0518
1147.5330810546875 0 33142.98
1148.540771484375 0 82297.6 c 10
1149.5440673828125 0 40080.12
1150.54443359375 0 8845.028
1161.507568359375 0 975.3724
1165.462890625 0 919.93774
1179.5216064453125 0 2709.0776
1180.52392578125 0 1902.2191
1182.5877685546875 0 1042.8097
1183.57958984375 0 761.9458
1196.122314453125 0 995.5315
1196.538330078125 0 1363.9976
1198.1153564453125 0 1109.6296
1199.5009765625 0 831.8483
1204.13525390625 0 1573.5377
1204.6334228515625 0 1535.9111
1211.4588623046875 0 1453.7133
1212.4676513671875 0 1226.2185
1214.543701171875 0 748.46857
1227.4793701171875 0 1895.2589
1228.0643310546875 0 819.55853
1229.48828125 0 728.2056
1232.4893798828125 0 12482.459 z 4
1233.4921875 0 10022.107
1234.499267578125 0 5319.774
1235.537841796875 0 5132.785
1236.5465087890625 0 3286.794
1237.551025390625 0 1705.2365
1242.5294189453125 0 4025.1309
1243.5269775390625 0 3853.3047
1244.52392578125 0 2168.24
1245.5203857421875 0 1092.2095
1246.521484375 0 1128.5853
1247.5003662109375 0 19343.37
1248.5025634765625 0 16121.065 y 4
1249.5047607421875 0 6316.2036
1250.5301513671875 0 1841.522
1251.561279296875 0 6305.2124
1252.5654296875 0 4146.2007
1253.570556640625 0 2071.8677
1260.5391845703125 0 19866.877
1261.5399169921875 0 15601.266
1262.5467529296875 0 15265.963
1262.68310546875 0 1248.3997
1263.5518798828125 0 7749.077
1263.690673828125 0 1008.3113
1264.55517578125 0 2794.155
1271.497802734375 0 740.3289 z Water loss 3
1272.5233154296875 0 702.2523
1278.5487060546875 0 28250.998 c Ammonia loss 11
1279.55126953125 0 18234.223
1280.558349609375 0 9917.613
1281.5687255859375 0 3630.5825
1286.5079345703125 0 1062.4293
1289.51025390625 0 7307.9277 z 3
1290.5159912109375 0 13068.561
1291.5185546875 0 9006.691
1292.52099609375 0 3403.1765
1295.575439453125 0 69305.6 c 11
1296.5782470703125 0 43375.57
1296.758056640625 0 859.8358
1297.5787353515625 0 15812.448
1298.583740234375 0 1622.8811
1304.5196533203125 0 14103.049
1305.5245361328125 0 17299.139 y 3
1306.527099609375 0 9187.935
1307.533203125 0 2359.6147
1310.5482177734375 0 1053.5692
1311.69287109375 0 844.8212
1324.560302734375 0 933.0587
1325.536865234375 0 1047.036
1328.5640869140625 0 1720.6641
1329.5543212890625 0 1524.0508
1343.5885009765625 0 828.7644
1351.632568359375 0 985.2295
1358.5306396484375 0 4680.045 z Water loss 2
1359.5341796875 0 4106.7334 z Ammonia loss 2
1360.54052734375 0 2227.52
1361.5888671875 0 5000.3486
1362.595458984375 0 4428.6587
1363.6053466796875 0 2500.7878
1374.5333251953125 0 2253.7175 y Water loss 2
1375.5391845703125 0 1819.6947 y Ammonia loss 2
1376.54052734375 0 34985.47 z 2
1377.544189453125 0 26839.08
1378.546142578125 0 10258.948
1379.5546875 0 1768.9854
1391.5489501953125 0 1333.5927
1392.55908203125 0 27397.885 y 2
1393.5614013671875 0 22079.352
1394.564208984375 0 8970.892
1395.5625 0 1470.4825
1413.622314453125 0 914.8756
1414.6226806640625 0 5316.9033
1415.626953125 0 4990.292
1416.6273193359375 0 2014.0367
1418.590576171875 0 2072.031
1419.58935546875 0 1569.0457
1423.6090087890625 0 1548.8971
1424.600341796875 0 1514.5403
1425.596435546875 0 7047.126
1426.6571044921875 0 13453.955
1427.656494140625 0 28153.916
1428.6591796875 0 19984.602
1429.6632080078125 0 7095.741
1439.702880859375 0 1574.4612
1440.724365234375 0 2061.0535
1441.611328125 0 3120.1294 c Ammonia loss 12
1442.656982421875 0 7125.4077
1443.6502685546875 0 14138.775
1444.6544189453125 0 8090.7124
1445.655517578125 0 4223.297
1446.660888671875 0 1007.94464
1458.637939453125 0 86626.43 c 12
1459.6412353515625 0 66104.54
1460.6417236328125 0 28990.84
1461.6436767578125 0 3781.706
1471.6197509765625 0 2447.8916 z Water loss 1
1472.6199951171875 0 2161.7122 z Ammonia loss 1
1473.6337890625 0 1667.205
1489.623779296875 0 30174.734 z 1
1490.6268310546875 0 26839.533
1491.6285400390625 0 12051.625
1492.640380859375 0 2019.4691
1506.6170654296875 0 6223.867
1507.6212158203125 0 5576.7036
1508.61376953125 0 2109.4675
1515.666748046875 0 5608.5024
1516.6707763671875 0 5177.3584
1517.67236328125 0 2515.2717
1518.6700439453125 0 1024.0577
1519.6475830078125 0 1048.9877
1520.67236328125 0 1617.3008
1526.6781005859375 0 1925.1315
1527.6771240234375 0 2859.5994
1528.6849365234375 0 2025.1835
1529.6842041015625 0 959.1302
1534.638671875 0 2844.5457
1535.6639404296875 0 4337.2075
1536.668212890625 0 4450.8057
1537.6715087890625 0 2907.5908
1538.66552734375 0 1401.0741
1543.7010498046875 0 6485.183
1544.6947021484375 0 9177.139
1545.67578125 0 13540.75
1546.6778564453125 0 9880.538
1547.67724609375 0 9121.471
1548.680419921875 0 7049.339
1549.6815185546875 0 5354.476
1550.6812744140625 0 1627.8992
1551.6358642578125 0 4350.887
1552.6431884765625 0 2713.1504
1553.6431884765625 0 1889.3925
1554.6507568359375 0 853.1496
1561.6947021484375 0 5922.7266
1562.680419921875 0 33135.457
1563.68310546875 0 30111.674
1564.6856689453125 0 16769.42
1565.6790771484375 0 4457.7524
1566.6688232421875 0 985.31396
1571.6846923828125 0 2287.985
1572.6669921875 0 13663.721
1573.671630859375 0 12820.275
1574.6749267578125 0 7367.7417
1575.6669921875 0 977.8168
1576.7696533203125 0 872.0497
1579.703369140625 0 13475.684
1580.708251953125 0 10600.214
1581.70556640625 0 5214.57
1587.7501220703125 0 2432.2312
1588.723876953125 0 5764.132
1589.690673828125 0 48532.324
1590.689697265625 0 185129.58
1591.69140625 0 150220.28
1592.6915283203125 0 71830.59
1593.6884765625 0 13315.702
1594.67919921875 0 2123.4924
1603.77734375 0 1624.2551
1604.7747802734375 0 5194.7207
1605.7728271484375 0 5698.0527
1606.6912841796875 0 61118.01
1607.6966552734375 0 217397.1
1608.7000732421875 0 169762.75
1609.7027587890625 0 79052.65
1610.703857421875 0 9449.638
1621.8094482421875 0 796.81903
1641.7078857421875 0 953.34344
2046.096435546875 0 794.9357
2047.0963134765625 0 1121.3691
2377.2001953125 0 890.92596
2391.228271484375 0 767.59863
2392.223388671875 0 712.8391
2396.210693359375 0 1592.0297
2397.22265625 0 1008.3764
2407.2333984375 0 1196.9081
2408.256103515625 0 3436.157
2409.251220703125 0 3711.8132
2410.260009765625 0 1626.9526
2411.268798828125 0 1414.0549

Spectrum Details

|  |  |
| --- | --- |
| Matched peaks? Matched peaks The total absolute number of peaks matched. Additionally in brackets the total fraction of peaks matched and the total number of peaks is shown. | 52 (10.26% of 507) |
| FDR? FDR The false discovery rate estimated for this peptide. It is calculated by matching all theoretical fragments with a non-integer shift with the raw peaks for this spectrum. This is done with 40 different shifts. The resulting percentage is the average number of annotated peaks over the number of annotated peaks with the correct spectrum. | 5.54% |
| Satellite FDR? Satellite FDR See the FDR for details on its calculation. This satellite ion specific FDR only contains the satellite ions (d/w) for I/L/J positions. | ∞ |
| PSM Score? PSM Score The PSM Score as given by Hecklib to this annotated spectrum. It is shown with three significant figures. | 709 |

#### Spectrum 8663? Spectrum 8663 The raw spectrum of this peptide as annotated by Hecklib. The fragments are coloured according to ion type (see legend). Any peaks with a star '\*' as text can be hovered over to see the full details, first the ion type second the mass shift type. By hovering over the amino acids in the peptide or ions in the legend the corresponding peaks are highlighted. By toggling the 'Unassigned' label you can turn the background (unassigned) peaks on or off in the plot. By updating the slider in the Ion legend you can update the spectrum to only show the top X% of the peaks with labels. The top X% means any peak that is within X% of the highest intensity. By dragging in the spectrum you can zoom in to a specific part of the spectrum and use 'Zoom Out' to get back to the original zoom level. The annotation of the spectrum is based on the given sequence in the peptides file and is done with different software so inconsistencies are likely. The peaks are annotated based on the given sequence, with 20 ppm tolerance.

Copy Data

##### Spectrum 8663 (TSV)

###### Preview

```
Loading example...
```

*Click on the button to copy the data to your clipboard.*

Mz MinMz MaxIntensity Max

WidthHeightPeptide font sizePeptide stroke widthSpectrum font sizeSpectrum stroke widthCompact peptide

Ion legend

wxyz

abcd

OtherUnassignedIon^Charge^_Position_Show for top:%

TJSGJQAEDESMYF

09.69e+51.94e+62.91e+63.87e+6

Zoom Out

y^+1^_1_y^+1^_2_w^+1^_3_y^+1^_3_c^+1^_5_y^+1^_4_c^+2^_11_c^+1^_6_c^+1^_6_c^+2^_12_c^+1^_7_c^+1^_7_y^+1^_5_z^+1^_6_y^+1^_6_c^+1^_8_y^+1^_6_c^+1^_8_c^+1^_9_c^+1^_9_y^+1^_7_z^+1^_8_y^+1^_8_c^+1^_10_c^+1^_10_c^+1^_10_y^+1^_9_z^+1^_9_c^+1^_11_c^+1^_11_y^+1^_9_c^+1^_11_z^+1^_10_y^+1^_10_z^+1^_11_c^+1^_12_y^+1^_11_z^+1^_11_c^+1^_12_y^+1^_11_z^+1^_12_z^+1^_12_y^+1^_12_y^+1^_12_z^+1^_12_y^+1^_12_c^+1^_13_c^+1^_13_z^+1^_13_z^+1^_13_

046893614041872

Fragment Matches Table

Show background peaks

| Position | Ion type | Intensity | mz Theoretical | mz Error (Th) | mz Error (ppm) | Charge | Series Number |
| --- | --- | --- | --- | --- | --- | --- | --- |
| - | - | 8.042E+04 | 120.1 | - | - | 0 | - |
| - | - | 9.955E+05 | 136.1 | - | - | 0 | - |
| - | - | 7.017E+04 | 137.1 | - | - | 0 | - |
| - | - | 9488 | 138.2 | - | - | 0 | - |
| - | - | 1.105E+04 | 146.7 | - | - | 0 | - |
| - | - | 1.935E+04 | 148.9 | - | - | 0 | - |
| - | - | 1.086E+04 | 158.7 | - | - | 0 | - |
| - | - | 1.097E+04 | 158.7 | - | - | 0 | - |
| - | - | 1.189E+04 | 166.1 | - | - | 0 | - |
| 14 | y | 1.913E+05 | 166.1 | 5.816E-05 | 0.3502 | +1 | 1 |
| - | - | 1.535E+04 | 167.1 | - | - | 0 | - |
| - | - | 1.402E+04 | 168.1 | - | - | 0 | - |
| - | - | 1.265E+04 | 173.4 | - | - | 0 | - |
| - | - | 3.271E+04 | 173.4 | - | - | 0 | - |
| - | - | 1.82E+04 | 173.5 | - | - | 0 | - |
| - | - | 1.105E+04 | 178.6 | - | - | 0 | - |
| - | - | 1.157E+04 | 182.6 | - | - | 0 | - |
| - | - | 1.982E+05 | 187.1 | - | - | 0 | - |
| - | - | 1.316E+04 | 195.3 | - | - | 0 | - |
| - | - | 1.294E+04 | 201 | - | - | 0 | - |
| - | - | 1.21E+04 | 208.2 | - | - | 0 | - |
| - | - | 1.851E+05 | 215.1 | - | - | 0 | - |
| - | - | 7.533E+04 | 217.1 | - | - | 0 | - |
| - | - | 1.334E+04 | 226.4 | - | - | 0 | - |
| - | - | 1.12E+05 | 235.1 | - | - | 0 | - |
| - | - | 3.068E+04 | 258.1 | - | - | 0 | - |
| - | - | 1.349E+04 | 262.1 | - | - | 0 | - |
| - | - | 4.646E+04 | 283.1 | - | - | 0 | - |
| - | - | 1.6E+04 | 284.1 | - | - | 0 | - |
| - | - | 5.378E+04 | 302.2 | - | - | 0 | - |
| - | - | 5.082E+04 | 311.1 | - | - | 0 | - |
| - | - | 2.143E+04 | 312.1 | - | - | 0 | - |
| 13 | y | 1.016E+06 | 329.1 | 4.409E-05 | 0.134 | +1 | 2 |
| - | - | 1.787E+05 | 330.2 | - | - | 0 | - |
| - | - | 2.838E+04 | 334.1 | - | - | 0 | - |
| - | - | 9.021E+04 | 341.2 | - | - | 0 | - |
| - | - | 3.067E+04 | 346.1 | - | - | 0 | - |
| - | - | 1.42E+04 | 358.1 | - | - | 0 | - |
| - | - | 9.108E+04 | 359.2 | - | - | 0 | - |
| - | - | 2.144E+04 | 360.2 | - | - | 0 | - |
| - | - | 1.584E+04 | 370.1 | - | - | 0 | - |
| - | - | 6.339E+04 | 386.2 | - | - | 0 | - |
| 12 | w | 1.529E+04 | 399.2 | 0.007784 | 19.5 | +1 | 3 |
| - | - | 1.708E+04 | 412.2 | - | - | 0 | - |
| - | - | 1.494E+04 | 424.7 | - | - | 0 | - |
| - | - | 1.538E+04 | 426.3 | - | - | 0 | - |
| - | - | 1.398E+04 | 434.1 | - | - | 0 | - |
| - | - | 1.466E+04 | 443.1 | - | - | 0 | - |
| - | - | 1.667E+04 | 444.2 | - | - | 0 | - |
| - | - | 8.895E+04 | 444.3 | - | - | 0 | - |
| - | - | 1.483E+05 | 454.3 | - | - | 0 | - |
| - | - | 2.095E+04 | 455.3 | - | - | 0 | - |
| - | - | 3.645E+04 | 457.2 | - | - | 0 | - |
| - | - | 8.229E+04 | 461.1 | - | - | 0 | - |
| - | - | 2.431E+04 | 462.1 | - | - | 0 | - |
| - | - | 2.245E+04 | 469.2 | - | - | 0 | - |
| - | - | 1.605E+04 | 469.4 | - | - | 0 | - |
| - | - | 1.171E+05 | 472.3 | - | - | 0 | - |
| - | - | 3.485E+04 | 473.3 | - | - | 0 | - |
| 12 | y | 1.337E+05 | 476.2 | 0.004929 | 10.35 | +1 | 3 |
| - | - | 3.02E+04 | 477.2 | - | - | 0 | - |
| - | - | 5.171E+04 | 479.1 | - | - | 0 | - |
| 5 | c | 1.588E+04 | 489.3 | 0.002357 | 4.817 | +1 | 5 |
| - | - | 1.938E+04 | 509.2 | - | - | 0 | - |
| - | - | 1.78E+04 | 523.2 | - | - | 0 | - |
| - | - | 2.389E+04 | 523.7 | - | - | 0 | - |
| - | - | 1.677E+04 | 556.3 | - | - | 0 | - |
| - | - | 1.139E+05 | 557.3 | - | - | 0 | - |
| - | - | 6.445E+04 | 557.8 | - | - | 0 | - |
| 11 | y | 1.169E+05 | 563.2 | 0.004456 | 7.912 | +1 | 4 |
| - | - | 3.965E+04 | 564.2 | - | - | 0 | - |
| 11 | c | 2.837E+04 | 566.3 | 0.001047 | 1.849 | +2 | 11 |
| - | - | 1.959E+04 | 572.2 | - | - | 0 | - |
| - | - | 2.454E+04 | 572.3 | - | - | 0 | - |
| - | - | 2.497E+04 | 573.2 | - | - | 0 | - |
| - | - | 1.837E+04 | 574.4 | - | - | 0 | - |
| - | - | 1.112E+05 | 582.3 | - | - | 0 | - |
| - | - | 3.265E+04 | 583.3 | - | - | 0 | - |
| - | - | 1.667E+04 | 584.3 | - | - | 0 | - |
| - | - | 5.23E+04 | 586.3 | - | - | 0 | - |
| - | - | 4.646E+04 | 590.2 | - | - | 0 | - |
| 6 | c | 1.999E+05 | 600.3 | 0.0001746 | 0.2909 | +1 | 6 |
| - | - | 4.634E+04 | 601.3 | - | - | 0 | - |
| - | - | 2.785E+04 | 608.2 | - | - | 0 | - |
| - | - | 3.73E+04 | 614.2 | - | - | 0 | - |
| - | - | 5.113E+04 | 616.4 | - | - | 0 | - |
| 6 | c | 4.077E+05 | 617.4 | 0.0001148 | 0.1859 | +1 | 6 |
| - | - | 1.292E+05 | 618.4 | - | - | 0 | - |
| - | - | 2.176E+04 | 619.4 | - | - | 0 | - |
| - | - | 4.512E+04 | 621.8 | - | - | 0 | - |
| - | - | 4.235E+04 | 622.3 | - | - | 0 | - |
| - | - | 1.925E+04 | 622.8 | - | - | 0 | - |
| - | - | 1.73E+04 | 624.2 | - | - | 0 | - |
| - | - | 1.604E+04 | 628.9 | - | - | 0 | - |
| - | - | 1.853E+05 | 630.8 | - | - | 0 | - |
| - | - | 1.332E+05 | 631.3 | - | - | 0 | - |
| - | - | 7.462E+04 | 631.8 | - | - | 0 | - |
| - | - | 2.704E+04 | 635.4 | - | - | 0 | - |
| 12 | c | 4.507E+04 | 639.8 | 0.002326 | 3.636 | +2 | 12 |
| - | - | 1.76E+04 | 640.3 | - | - | 0 | - |
| - | - | 6.924E+04 | 642.2 | - | - | 0 | - |
| - | - | 2.157E+04 | 643.2 | - | - | 0 | - |
| - | - | 4.748E+04 | 644.4 | - | - | 0 | - |
| - | - | 8.313E+04 | 645.4 | - | - | 0 | - |
| - | - | 2.735E+04 | 646.4 | - | - | 0 | - |
| - | - | 1.187E+05 | 653.4 | - | - | 0 | - |
| - | - | 3.652E+04 | 654.4 | - | - | 0 | - |
| - | - | 1.641E+04 | 655.4 | - | - | 0 | - |
| - | - | 6.124E+04 | 661.2 | - | - | 0 | - |
| - | - | 1.702E+04 | 662.2 | - | - | 0 | - |
| - | - | 1.768E+04 | 669.3 | - | - | 0 | - |
| 7 | c | 1.331E+05 | 671.4 | 0.0009637 | 1.435 | +1 | 7 |
| - | - | 4.458E+04 | 672.4 | - | - | 0 | - |
| - | - | 5.611E+04 | 679.2 | - | - | 0 | - |
| - | - | 2.333E+04 | 687.3 | - | - | 0 | - |
| - | - | 1.602E+05 | 687.4 | - | - | 0 | - |
| 7 | c | 1.201E+06 | 688.4 | 4.932E-05 | 0.07165 | +1 | 7 |
| - | - | 3.58E+05 | 689.4 | - | - | 0 | - |
| - | - | 3.908E+04 | 690.4 | - | - | 0 | - |
| 10 | y | 2.769E+04 | 692.3 | 0.003733 | 5.393 | +1 | 5 |
| - | - | 3.03E+04 | 701.3 | - | - | 0 | - |
| - | - | 1.559E+04 | 714.4 | - | - | 0 | - |
| - | - | 1.83E+04 | 753.2 | - | - | 0 | - |
| - | - | 3.818E+04 | 760.4 | - | - | 0 | - |
| - | - | 2.146E+04 | 763.3 | - | - | 0 | - |
| - | - | 4.148E+04 | 771.3 | - | - | 0 | - |
| - | - | 3.922E+04 | 771.9 | - | - | 0 | - |
| - | - | 2.71E+04 | 772.3 | - | - | 0 | - |
| - | - | 2.916E+04 | 772.4 | - | - | 0 | - |
| 9 | z | 1.742E+04 | 773.3 | 0.006372 | 8.241 | +1 | 6 |
| - | - | 1.061E+05 | 773.4 | - | - | 0 | - |
| - | - | 8.577E+04 | 774.4 | - | - | 0 | - |
| - | - | 1.02E+05 | 782.4 | - | - | 0 | - |
| - | - | 3.441E+04 | 783.4 | - | - | 0 | - |
| - | - | 2.804E+04 | 784.4 | - | - | 0 | - |
| 9 | y | 3.49E+04 | 789.3 | 0.001576 | 1.997 | +1 | 6 |
| - | - | 2.328E+04 | 790.3 | - | - | 0 | - |
| - | - | 4.796E+04 | 793.3 | - | - | 0 | - |
| - | - | 3.033E+04 | 794.3 | - | - | 0 | - |
| 8 | c | 1.824E+05 | 800.4 | 3.462E-06 | 0.004325 | +1 | 8 |
| - | - | 7.644E+04 | 801.4 | - | - | 0 | - |
| 9 | y | 1.119E+05 | 807.3 | 0.003218 | 3.987 | +1 | 6 |
| - | - | 3.437E+04 | 808.3 | - | - | 0 | - |
| - | - | 2.301E+04 | 809.3 | - | - | 0 | - |
| - | - | 1.896E+04 | 813.3 | - | - | 0 | - |
| - | - | 2.004E+04 | 814.3 | - | - | 0 | - |
| - | - | 3.579E+05 | 816.4 | - | - | 0 | - |
| 8 | c | 6.607E+05 | 817.4 | 0.001101 | 1.347 | +1 | 8 |
| - | - | 2.557E+05 | 818.4 | - | - | 0 | - |
| - | - | 4.239E+04 | 819.4 | - | - | 0 | - |
| - | - | 1.596E+04 | 825.3 | - | - | 0 | - |
| - | - | 7.905E+04 | 830.4 | - | - | 0 | - |
| - | - | 1.728E+04 | 831.4 | - | - | 0 | - |
| - | - | 7.799E+04 | 842.3 | - | - | 0 | - |
| - | - | 2.872E+04 | 843.3 | - | - | 0 | - |
| - | - | 1.092E+05 | 888.5 | - | - | 0 | - |
| - | - | 7.957E+04 | 889.5 | - | - | 0 | - |
| - | - | 2.54E+04 | 890.5 | - | - | 0 | - |
| - | - | 1.782E+04 | 899.4 | - | - | 0 | - |
| 9 | c | 1.204E+05 | 915.4 | 0.0005793 | 0.6328 | +1 | 9 |
| - | - | 2.79E+04 | 916.4 | - | - | 0 | - |
| - | - | 1.82E+05 | 931.5 | - | - | 0 | - |
| 9 | c | 1.041E+06 | 932.5 | 0.0002119 | 0.2273 | +1 | 9 |
| - | - | 4.741E+05 | 933.5 | - | - | 0 | - |
| - | - | 1.119E+05 | 934.5 | - | - | 0 | - |
| - | - | 2.934E+04 | 935.3 | - | - | 0 | - |
| 8 | y | 1.562E+05 | 936.3 | 0.004204 | 4.49 | +1 | 7 |
| - | - | 7.56E+04 | 937.3 | - | - | 0 | - |
| - | - | 2.099E+04 | 938.3 | - | - | 0 | - |
| - | - | 4.041E+04 | 943.4 | - | - | 0 | - |
| - | - | 2.513E+04 | 944.4 | - | - | 0 | - |
| - | - | 3.663E+04 | 979.4 | - | - | 0 | - |
| 7 | z | 1.984E+04 | 991.3 | 0.009069 | 9.148 | +1 | 8 |
| - | - | 1.701E+04 | 999.5 | - | - | 0 | - |
| - | - | 1.942E+04 | 1004 | - | - | 0 | - |
| - | - | 4.073E+04 | 1006 | - | - | 0 | - |
| 7 | y | 2.112E+05 | 1007 | 0.004566 | 4.533 | +1 | 8 |
| - | - | 1.344E+05 | 1008 | - | - | 0 | - |
| - | - | 2.652E+04 | 1009 | - | - | 0 | - |
| - | - | 1.455E+05 | 1017 | - | - | 0 | - |
| - | - | 8.488E+04 | 1019 | - | - | 0 | - |
| - | - | 3.537E+04 | 1020 | - | - | 0 | - |
| - | - | 9.463E+04 | 1026 | - | - | 0 | - |
| - | - | 6.034E+04 | 1027 | - | - | 0 | - |
| - | - | 3.726E+04 | 1028 | - | - | 0 | - |
| - | - | 5.535E+04 | 1029 | - | - | 0 | - |
| - | - | 2.88E+04 | 1030 | - | - | 0 | - |
| - | - | 5.347E+04 | 1042 | - | - | 0 | - |
| 10 | c | 3.908E+04 | 1044 | 0.009901 | 9.489 | +1 | 10 |
| 10 | c | 2.961E+05 | 1044 | 0.0003457 | 0.3309 | +1 | 10 |
| - | - | 1.368E+05 | 1045 | - | - | 0 | - |
| - | - | 2.887E+04 | 1046 | - | - | 0 | - |
| - | - | 2.14E+04 | 1047 | - | - | 0 | - |
| - | - | 2.515E+04 | 1047 | - | - | 0 | - |
| - | - | 3.366E+04 | 1048 | - | - | 0 | - |
| - | - | 1.089E+06 | 1061 | - | - | 0 | - |
| 10 | c | 9.262E+05 | 1062 | 0.002522 | 2.376 | +1 | 10 |
| - | - | 3.098E+05 | 1063 | - | - | 0 | - |
| - | - | 4.484E+04 | 1064 | - | - | 0 | - |
| - | - | 7.771E+04 | 1064 | - | - | 0 | - |
| - | - | 6.453E+04 | 1066 | - | - | 0 | - |
| - | - | 2.966E+04 | 1067 | - | - | 0 | - |
| - | - | 2.501E+04 | 1068 | - | - | 0 | - |
| - | - | 2.025E+04 | 1071 | - | - | 0 | - |
| - | - | 1.873E+04 | 1072 | - | - | 0 | - |
| - | - | 4.888E+04 | 1088 | - | - | 0 | - |
| - | - | 2.389E+04 | 1089 | - | - | 0 | - |
| - | - | 2.933E+04 | 1095 | - | - | 0 | - |
| - | - | 4.231E+04 | 1106 | - | - | 0 | - |
| - | - | 2.329E+05 | 1114 | - | - | 0 | - |
| - | - | 1.217E+05 | 1115 | - | - | 0 | - |
| - | - | 3.699E+04 | 1116 | - | - | 0 | - |
| 6 | y | 1.715E+04 | 1118 | 0.01838 | 16.43 | +1 | 9 |
| 6 | z | 5.12E+04 | 1119 | 0.007132 | 6.371 | +1 | 9 |
| - | - | 4.778E+04 | 1120 | - | - | 0 | - |
| - | - | 1.968E+04 | 1125 | - | - | 0 | - |
| 11 | c | 2.112E+04 | 1131 | 0.01739 | 15.39 | +1 | 11 |
| 11 | c | 1.929E+05 | 1132 | 0.0004327 | 0.3824 | +1 | 11 |
| - | - | 1.109E+05 | 1133 | - | - | 0 | - |
| - | - | 3.799E+04 | 1134 | - | - | 0 | - |
| - | - | 1.501E+05 | 1134 | - | - | 0 | - |
| 6 | y | 1.583E+05 | 1135 | 0.003056 | 2.692 | +1 | 9 |
| - | - | 9.948E+04 | 1136 | - | - | 0 | - |
| - | - | 3.339E+04 | 1137 | - | - | 0 | - |
| - | - | 5.757E+05 | 1148 | - | - | 0 | - |
| 11 | c | 1.563E+06 | 1149 | 0.0007367 | 0.6414 | +1 | 11 |
| - | - | 7.59E+05 | 1150 | - | - | 0 | - |
| - | - | 1.922E+05 | 1151 | - | - | 0 | - |
| - | - | 3.406E+04 | 1180 | - | - | 0 | - |
| - | - | 3.481E+04 | 1181 | - | - | 0 | - |
| - | - | 1.669E+04 | 1190 | - | - | 0 | - |
| - | - | 2.084E+04 | 1197 | - | - | 0 | - |
| - | - | 2.428E+04 | 1198 | - | - | 0 | - |
| - | - | 1.869E+04 | 1200 | - | - | 0 | - |
| - | - | 2.42E+04 | 1212 | - | - | 0 | - |
| - | - | 2.841E+04 | 1225 | - | - | 0 | - |
| - | - | 1.82E+04 | 1229 | - | - | 0 | - |
| 5 | z | 2.627E+05 | 1232 | 0.004977 | 4.038 | +1 | 10 |
| - | - | 2.309E+05 | 1233 | - | - | 0 | - |
| - | - | 1.041E+05 | 1235 | - | - | 0 | - |
| - | - | 9.712E+04 | 1236 | - | - | 0 | - |
| - | - | 6.286E+04 | 1237 | - | - | 0 | - |
| - | - | 8.568E+04 | 1243 | - | - | 0 | - |
| - | - | 8.27E+04 | 1244 | - | - | 0 | - |
| - | - | 3.961E+04 | 1245 | - | - | 0 | - |
| - | - | 2.731E+04 | 1246 | - | - | 0 | - |
| - | - | 3.486E+05 | 1248 | - | - | 0 | - |
| 5 | y | 2.861E+05 | 1249 | 7.487E-05 | 0.05997 | +1 | 10 |
| - | - | 1.309E+05 | 1250 | - | - | 0 | - |
| - | - | 1.327E+05 | 1252 | - | - | 0 | - |
| - | - | 1.283E+05 | 1253 | - | - | 0 | - |
| - | - | 3.516E+04 | 1254 | - | - | 0 | - |
| - | - | 3.39E+05 | 1261 | - | - | 0 | - |
| - | - | 2.287E+05 | 1262 | - | - | 0 | - |
| - | - | 3E+05 | 1263 | - | - | 0 | - |
| - | - | 1.368E+05 | 1264 | - | - | 0 | - |
| - | - | 7.39E+04 | 1265 | - | - | 0 | - |
| 4 | z | 2.321E+04 | 1271 | 0.001039 | 0.8171 | +1 | 11 |
| 12 | c | 4.396E+05 | 1279 | 0.004605 | 3.602 | +1 | 12 |
| - | - | 3.059E+05 | 1280 | - | - | 0 | - |
| - | - | 2.145E+05 | 1281 | - | - | 0 | - |
| - | - | 6.614E+04 | 1282 | - | - | 0 | - |
| - | - | 2.243E+04 | 1283 | - | - | 0 | - |
| - | - | 2.36E+04 | 1287 | - | - | 0 | - |
| 4 | y | 2.511E+04 | 1288 | 0.001208 | 0.9385 | +1 | 11 |
| 4 | z | 1.129E+05 | 1290 | 0.006707 | 5.201 | +1 | 11 |
| - | - | 2.467E+05 | 1291 | - | - | 0 | - |
| - | - | 1.621E+05 | 1292 | - | - | 0 | - |
| - | - | 4.175E+04 | 1293 | - | - | 0 | - |
| - | - | 1.934E+04 | 1295 | - | - | 0 | - |
| 12 | c | 1.278E+06 | 1296 | 0.003813 | 2.943 | +1 | 12 |
| - | - | 8.184E+05 | 1297 | - | - | 0 | - |
| - | - | 3.084E+05 | 1298 | - | - | 0 | - |
| - | - | 2.023E+04 | 1299 | - | - | 0 | - |
| - | - | 3.017E+05 | 1305 | - | - | 0 | - |
| 4 | y | 2.674E+05 | 1306 | 0.0005561 | 0.426 | +1 | 11 |
| - | - | 1.452E+05 | 1307 | - | - | 0 | - |
| - | - | 4.176E+04 | 1308 | - | - | 0 | - |
| - | - | 2.669E+04 | 1325 | - | - | 0 | - |
| - | - | 2.063E+04 | 1326 | - | - | 0 | - |
| - | - | 2.816E+04 | 1329 | - | - | 0 | - |
| - | - | 3.538E+04 | 1330 | - | - | 0 | - |
| - | - | 1.662E+04 | 1352 | - | - | 0 | - |
| 3 | z | 1.215E+05 | 1359 | 0.006239 | 4.593 | +1 | 12 |
| 3 | z | 7.188E+04 | 1360 | 0.02125 | 15.63 | +1 | 12 |
| - | - | 2.49E+04 | 1361 | - | - | 0 | - |
| - | - | 8.258E+04 | 1362 | - | - | 0 | - |
| - | - | 8.189E+04 | 1363 | - | - | 0 | - |
| - | - | 3.067E+04 | 1364 | - | - | 0 | - |
| 3 | y | 3.418E+04 | 1375 | 0.02079 | 15.12 | +1 | 12 |
| 3 | y | 3.671E+04 | 1376 | 0.01424 | 10.35 | +1 | 12 |
| 3 | z | 6.855E+05 | 1377 | 0.003731 | 2.711 | +1 | 12 |
| - | - | 4.468E+05 | 1378 | - | - | 0 | - |
| - | - | 1.881E+05 | 1379 | - | - | 0 | - |
| - | - | 4.604E+04 | 1380 | - | - | 0 | - |
| - | - | 3.138E+04 | 1392 | - | - | 0 | - |
| 3 | y | 4.977E+05 | 1393 | 0.002952 | 2.12 | +1 | 12 |
| - | - | 3.771E+05 | 1394 | - | - | 0 | - |
| - | - | 1.545E+05 | 1395 | - | - | 0 | - |
| - | - | 2.809E+04 | 1396 | - | - | 0 | - |
| - | - | 1.288E+05 | 1415 | - | - | 0 | - |
| - | - | 1.823E+04 | 1415 | - | - | 0 | - |
| - | - | 1.038E+05 | 1416 | - | - | 0 | - |
| - | - | 2.702E+04 | 1417 | - | - | 0 | - |
| - | - | 2.031E+04 | 1418 | - | - | 0 | - |
| - | - | 3.703E+04 | 1419 | - | - | 0 | - |
| - | - | 2.889E+04 | 1420 | - | - | 0 | - |
| - | - | 1.387E+05 | 1426 | - | - | 0 | - |
| - | - | 9.029E+04 | 1427 | - | - | 0 | - |
| - | - | 5.807E+04 | 1428 | - | - | 0 | - |
| - | - | 2.076E+04 | 1429 | - | - | 0 | - |
| 13 | c | 4.155E+04 | 1442 | 0.002678 | 1.858 | +1 | 13 |
| - | - | 2.819E+05 | 1443 | - | - | 0 | - |
| - | - | 2.257E+05 | 1444 | - | - | 0 | - |
| - | - | 1.029E+05 | 1445 | - | - | 0 | - |
| - | - | 5.159E+04 | 1446 | - | - | 0 | - |
| 13 | c | 1.523E+06 | 1459 | 0.003839 | 2.632 | +1 | 13 |
| - | - | 1.167E+06 | 1460 | - | - | 0 | - |
| - | - | 3.145E+04 | 1460 | - | - | 0 | - |
| - | - | 5.093E+05 | 1461 | - | - | 0 | - |
| - | - | 7.98E+04 | 1462 | - | - | 0 | - |
| 2 | z | 5.003E+04 | 1472 | 0.008601 | 5.845 | +1 | 13 |
| - | - | 4.883E+04 | 1473 | - | - | 0 | - |
| 2 | z | 5.422E+05 | 1490 | 0.00353 | 2.369 | +1 | 13 |
| - | - | 4.444E+05 | 1491 | - | - | 0 | - |
| - | - | 2.267E+05 | 1492 | - | - | 0 | - |
| - | - | 2.637E+04 | 1493 | - | - | 0 | - |
| - | - | 1.318E+05 | 1507 | - | - | 0 | - |
| - | - | 9.308E+04 | 1508 | - | - | 0 | - |
| - | - | 3.898E+04 | 1509 | - | - | 0 | - |
| - | - | 9.811E+04 | 1516 | - | - | 0 | - |
| - | - | 7.066E+04 | 1517 | - | - | 0 | - |
| - | - | 5.845E+04 | 1518 | - | - | 0 | - |
| - | - | 2.091E+04 | 1519 | - | - | 0 | - |
| - | - | 2.132E+04 | 1520 | - | - | 0 | - |
| - | - | 2.451E+04 | 1522 | - | - | 0 | - |
| - | - | 5.775E+04 | 1528 | - | - | 0 | - |
| - | - | 3.359E+04 | 1529 | - | - | 0 | - |
| - | - | 2.01E+04 | 1534 | - | - | 0 | - |
| - | - | 4.67E+04 | 1535 | - | - | 0 | - |
| - | - | 7.385E+04 | 1536 | - | - | 0 | - |
| - | - | 8.577E+04 | 1537 | - | - | 0 | - |
| - | - | 4.52E+04 | 1538 | - | - | 0 | - |
| - | - | 1.295E+05 | 1544 | - | - | 0 | - |
| - | - | 2.051E+05 | 1545 | - | - | 0 | - |
| - | - | 2.129E+05 | 1546 | - | - | 0 | - |
| - | - | 1.681E+05 | 1547 | - | - | 0 | - |
| - | - | 1.603E+05 | 1548 | - | - | 0 | - |
| - | - | 1.33E+05 | 1549 | - | - | 0 | - |
| - | - | 9.662E+04 | 1550 | - | - | 0 | - |
| - | - | 4.232E+04 | 1551 | - | - | 0 | - |
| - | - | 8.352E+04 | 1552 | - | - | 0 | - |
| - | - | 4.886E+04 | 1553 | - | - | 0 | - |
| - | - | 2.219E+04 | 1554 | - | - | 0 | - |
| - | - | 2.254E+04 | 1555 | - | - | 0 | - |
| - | - | 9.178E+04 | 1562 | - | - | 0 | - |
| - | - | 6.257E+05 | 1563 | - | - | 0 | - |
| - | - | 5.593E+05 | 1564 | - | - | 0 | - |
| - | - | 2.513E+05 | 1565 | - | - | 0 | - |
| - | - | 6.642E+04 | 1566 | - | - | 0 | - |
| - | - | 2.734E+04 | 1572 | - | - | 0 | - |
| - | - | 2.539E+05 | 1573 | - | - | 0 | - |
| - | - | 2.341E+05 | 1574 | - | - | 0 | - |
| - | - | 1.117E+05 | 1575 | - | - | 0 | - |
| - | - | 1.911E+05 | 1580 | - | - | 0 | - |
| - | - | 1.974E+05 | 1581 | - | - | 0 | - |
| - | - | 8.805E+04 | 1582 | - | - | 0 | - |
| - | - | 2.509E+04 | 1583 | - | - | 0 | - |
| - | - | 7.361E+05 | 1590 | - | - | 0 | - |
| - | - | 3.313E+06 | 1591 | - | - | 0 | - |
| - | - | 2.653E+06 | 1592 | - | - | 0 | - |
| - | - | 1.282E+06 | 1593 | - | - | 0 | - |
| - | - | 2.624E+05 | 1594 | - | - | 0 | - |
| - | - | 2.28E+04 | 1595 | - | - | 0 | - |
| - | - | 1.784E+04 | 1606 | - | - | 0 | - |
| - | - | 1.179E+06 | 1607 | - | - | 0 | - |
| - | - | 3.837E+06 | 1608 | - | - | 0 | - |
| - | - | 3.125E+06 | 1609 | - | - | 0 | - |
| - | - | 1.303E+06 | 1610 | - | - | 0 | - |
| - | - | 1.773E+05 | 1611 | - | - | 0 | - |
| - | - | 1.729E+04 | 1853 | - | - | 0 | - |

m/z Charge Intensity FragmentType MassShift Position
120.08071899414062 0 80419.414
136.07571411132812 0 995520.06
137.0791015625 0 70172.51
138.20120239257812 0 9487.889
146.68736267089844 0 11049
148.9458770751953 0 19352.072
158.69459533691406 0 10860.744
158.74205017089844 0 10971.931
166.053466796875 0 11887.791
166.08619689941406 0 191277.44 y 13
167.089599609375 0 15350.835
168.10528564453125 0 14024.266
173.42652893066406 0 12648.412
173.43621826171875 0 32714.393
173.45870971679688 0 18195.543
178.58126831054688 0 11052.745
182.56863403320312 0 11571.387
187.14410400390625 0 198169.72
195.28610229492188 0 13162.997
201.0308074951172 0 12939.768
208.2481689453125 0 12102.113
215.1388397216797 0 185050.2
217.06410217285156 0 75334.87
226.3524169921875 0 13336.549
235.07455444335938 0 111995.73
258.14501953125 0 30684.006
262.1293640136719 0 13487.093
283.1106872558594 0 46456.477
284.114990234375 0 15997.158
302.1709289550781 0 53781.188
311.10601806640625 0 50820.332
312.12237548828125 0 21429.96
329.1496276855469 0 1016019.56 y 12
330.1529541015625 0 178667.06
334.1402587890625 0 28383.258
341.1820373535156 0 90211.56
346.10723876953125 0 30666.688
358.0752868652344 0 14198.858
359.1919250488281 0 91084.805
360.1955871582031 0 21438.49
370.1402893066406 0 15839.375
386.2030944824219 0 63385.02
399.1423645019531 0 15291.979 w 11
412.1858825683594 0 17079.998
424.66082763671875 0 14943.955
426.271240234375 0 15384.046
434.0911865234375 0 13977.985
443.121337890625 0 14660.889
444.17120361328125 0 16669.756
444.2823486328125 0 88952.336
454.2658996582031 0 148278.22
455.2674255371094 0 20949.754
457.2408142089844 0 36454.668
461.13421630859375 0 82293.445
462.1346740722656 0 24312.504
469.2393493652344 0 22451.562
469.3835754394531 0 16052.516
472.2764892578125 0 117055.586
473.27923583984375 0 34849.082
476.18499755859375 0 133681.42 y 11
477.1880798339844 0 30202.375
479.1444091796875 0 51709.96
489.30548095703125 0 15877.547 c 4
509.17279052734375 0 19381.592
523.248291015625 0 17797.129
523.7069702148438 0 23886.918
556.3042602539062 0 16769.553
557.2564086914062 0 113907.24
557.7576293945312 0 64452.23
563.216552734375 0 116869.21 y 10
564.2211303710938 0 39650.785
566.2608032226562 0 28373.857 c Ammonia loss 10
572.1663208007812 0 19591.875
572.3416137695312 0 24535.078
573.2161254882812 0 24972.057
574.3560791015625 0 18372.854
582.32373046875 0 111213.14
583.3251953125 0 32652.883
584.3357543945312 0 16668.03
586.2838134765625 0 52303.09
590.1762084960938 0 46456.137
600.3353271484375 0 199889.27 c Ammonia loss 5
601.337646484375 0 46337.36
608.1879272460938 0 27846.25
614.213134765625 0 37300.67
616.3530883789062 0 51130.266
617.36181640625 0 407734.84 c 5
618.3646240234375 0 129228.28
619.3697509765625 0 21764.775
621.7697143554688 0 45121.85
622.2669677734375 0 42350.293
622.7622680664062 0 19252.3
624.1962890625 0 17304.432
628.9099731445312 0 16044.674
630.7742309570312 0 185339.83
631.27587890625 0 133234
631.774169921875 0 74616.42
635.353271484375 0 27040.611
639.7794189453125 0 45072.086 c Ammonia loss 11
640.2788696289062 0 17601.844
642.2086181640625 0 69244.36
643.2091674804688 0 21567.797
644.3851318359375 0 47479.41
645.3916015625 0 83132.73
646.3956298828125 0 27350.746
653.3624267578125 0 118719.1
654.3655395507812 0 36515.63
655.3619995117188 0 16410.719
661.2142333984375 0 61238.074
662.2138671875 0 17022.035
669.3184814453125 0 17681.602
671.3732299804688 0 133140.83 c Ammonia loss 6
672.3746948242188 0 44576.54
679.2257080078125 0 56109.547
687.3280639648438 0 23331.02
687.3909301757812 0 160168.25
688.3988647460938 0 1201182 c 6
689.4016723632812 0 358019.84
690.4024658203125 0 39079.766
692.2584228515625 0 27694.562 y 9
701.3099975585938 0 30301.96
714.4113159179688 0 15592.364
753.2386474609375 0 18301.59
760.3693237304688 0 38179.19
763.3405151367188 0 21463.555
771.2506103515625 0 41475.195
771.8530883789062 0 39219.793
772.2536010742188 0 27098.998
772.4241333007812 0 29157.785
773.2459716796875 0 17415.785 z Water loss 8
773.42724609375 0 106131.734
774.4332275390625 0 85774.58
782.40478515625 0 102008.09
783.4035034179688 0 34414.625
784.4002685546875 0 28038.953
789.2726440429688 0 34904.438 y Water loss 8
790.2743530273438 0 23282.049
793.31591796875 0 47955.74
794.31884765625 0 30330.873
800.4148559570312 0 182445.5 c Ammonia loss 7
801.4193115234375 0 76439.7
807.2848510742188 0 111855.89 y 8
808.2868041992188 0 34368.555
809.2877197265625 0 23007.332
813.3303833007812 0 18961.81
814.2904663085938 0 20038.594
816.4332885742188 0 357927.1
817.4403076171875 0 660684.1 c 7
818.4439697265625 0 255737.03
819.4463500976562 0 42385.55
825.2865600585938 0 15956.217
830.3517456054688 0 79053.77
831.3673095703125 0 17278.748
842.2888793945312 0 77988.29
843.2871704101562 0 28715.467
888.4547119140625 0 109208.836
889.4607543945312 0 79569.96
890.464111328125 0 25397.555
899.3739624023438 0 17820.5
915.4412231445312 0 120438.59 c Ammonia loss 8
916.442138671875 0 27904.266
931.4599609375 0 182022.53
932.4681396484375 0 1040777.5 c 8
933.4708251953125 0 474057.5
934.4729614257812 0 111882.99
935.3255615234375 0 29336.633
936.3284301757812 0 156155.11 y 7
937.3318481445312 0 75597.48
938.3306884765625 0 20992.117
943.366455078125 0 40410.375
944.3602294921875 0 25126.34
979.4030151367188 0 36628.99
991.3516845703125 0 19837.46 z 6
999.46044921875 0 17006.545
1004.4280395507812 0 19416.338
1006.3619995117188 0 40726.047
1007.3659057617188 0 211198.17 y 6
1008.3695068359375 0 134379.19
1009.3680419921875 0 26515.44
1017.4970703125 0 145462.67
1018.5018310546875 0 84878.68
1019.5094604492188 0 35374.324
1026.4754638671875 0 94631.45
1027.471435546875 0 60335.81
1028.46630859375 0 37264.242
1029.4752197265625 0 55349.633
1030.47265625 0 28803.533
1042.4910888671875 0 53471.43
1043.490478515625 0 39084.4 c Water loss 9
1044.4847412109375 0 296100.75 c Ammonia loss 9
1045.4881591796875 0 136833.33
1046.3955078125 0 28870.342
1046.5093994140625 0 21400.465
1047.408447265625 0 25153.068
1048.419921875 0 33659.043
1060.5032958984375 0 1089279.4
1061.5084228515625 0 926244.44 c 9
1062.5123291015625 0 309756.28
1063.5164794921875 0 44841.74
1064.4208984375 0 77706.46
1066.4302978515625 0 64531.062
1067.42919921875 0 29663.584
1068.4217529296875 0 25009.734
1071.427490234375 0 20254.035
1072.430419921875 0 18726.215
1087.5228271484375 0 48877.42
1088.5328369140625 0 23887.074
1095.498046875 0 29333.625
1105.5352783203125 0 42307.734
1113.505615234375 0 232866.89
1114.5048828125 0 121650.57
1115.5123291015625 0 36987.67
1118.4117431640625 0 17147.312 y Ammonia loss 5
1119.4083251953125 0 51200.902 z 5
1120.41748046875 0 47781.49
1125.4405517578125 0 19678.666
1130.5150146484375 0 21122.73 c Water loss 10
1131.5159912109375 0 192910.48 c Ammonia loss 10
1132.518310546875 0 110909.37
1133.5177001953125 0 37992.51
1134.4185791015625 0 150077.11
1135.4229736328125 0 158303.22 y 5
1136.423828125 0 99484.32
1137.4307861328125 0 33393.53
1147.5347900390625 0 575688.9
1148.542236328125 0 1562509.4 c 10
1149.5452880859375 0 758963.56
1150.5489501953125 0 192232.8
1179.51318359375 0 34055.355
1180.5205078125 0 34812.68
1189.59033203125 0 16685.258
1196.548583984375 0 20841.426
1197.541015625 0 24281.889
1199.548828125 0 18692.945
1212.4737548828125 0 24201.504
1224.5123291015625 0 28405.236
1229.4847412109375 0 18202.994
1232.490234375 0 262658.4 z 4
1233.4940185546875 0 230927.42
1234.5035400390625 0 104065.016
1235.5411376953125 0 97122.65
1236.5523681640625 0 62860.098
1242.529296875 0 85680.39
1243.528564453125 0 82696.25
1244.5245361328125 0 39609.215
1245.5269775390625 0 27306.734
1247.500732421875 0 348591.6
1248.50390625 0 286081.2 y 4
1249.507568359375 0 130865.63
1251.5634765625 0 132712.62
1252.5689697265625 0 128329.32
1253.5682373046875 0 35157.035
1260.5411376953125 0 338958.22
1261.5418701171875 0 228728.03
1262.5506591796875 0 300008.44
1263.55615234375 0 136752.48
1264.55859375 0 73895.03
1271.4951171875 0 23209.816 z Water loss 3
1278.551513671875 0 439605.97 c Ammonia loss 11
1279.5548095703125 0 305941.06
1280.56005859375 0 214542.08
1281.567138671875 0 66143.68
1282.5758056640625 0 22434.059
1286.517822265625 0 23598.945
1287.513671875 0 25106.62 y Water loss 3
1289.513427734375 0 112945.586 z 3
1290.51806640625 0 246683.55
1291.5208740234375 0 162073.77
1292.51904296875 0 41754.273
1294.57177734375 0 19338.432
1295.5772705078125 0 1278481.6 c 11
1296.5797119140625 0 818413.25
1297.581298828125 0 308400.8
1298.58447265625 0 20228.234
1304.5213623046875 0 301666.22
1305.5260009765625 0 267360.12 y 3
1306.5289306640625 0 145174.19
1307.526123046875 0 41756.613
1324.536376953125 0 26692.955
1325.5394287109375 0 20626.611
1328.5506591796875 0 28158.516
1329.5670166015625 0 35376.35
1351.6231689453125 0 16621.953
1358.534423828125 0 121544.39 z Water loss 2
1359.533447265625 0 71884.57 z Ammonia loss 2
1360.5335693359375 0 24897.965
1361.5970458984375 0 82584.25
1362.5975341796875 0 81894.86
1363.6251220703125 0 30673.959
1374.526123046875 0 34176.465 y Water loss 2
1375.545166015625 0 36707.242 y Ammonia loss 2
1376.54248046875 0 685521.7 z 2
1377.5458984375 0 446821.4
1378.5479736328125 0 188143.9
1379.5538330078125 0 46039.777
1391.5509033203125 0 31375.057
1392.5604248046875 0 497653.78 y 2
1393.5645751953125 0 377112.34
1394.56298828125 0 154489.48
1395.5545654296875 0 28090.295
1414.6273193359375 0 128782.055
1415.4234619140625 0 18233.807
1415.627685546875 0 103830.93
1416.6395263671875 0 27015.146
1417.627197265625 0 20308.438
1418.5850830078125 0 37030.35
1419.5987548828125 0 28890.1
1425.5966796875 0 138741.84
1426.6019287109375 0 90288.95
1427.6224365234375 0 58072.54
1428.6441650390625 0 20758.02
1441.6129150390625 0 41548.32 c Ammonia loss 12
1442.6585693359375 0 281939.78
1443.6624755859375 0 225656.81
1444.6695556640625 0 102858.516
1445.659912109375 0 51591.324
1458.640625 0 1523400.8 c 12
1459.643798828125 0 1167269.1
1459.855224609375 0 31446.412
1460.64453125 0 509290.5
1461.640869140625 0 79800.4
1471.620849609375 0 50031.86 z Water loss 1
1472.6258544921875 0 48832.85
1489.6263427734375 0 542185.8 z 1
1490.6285400390625 0 444379.7
1491.6318359375 0 226659.81
1492.6376953125 0 26366.264
1506.616455078125 0 131785.95
1507.6214599609375 0 93081.9
1508.63330078125 0 38979.633
1515.66943359375 0 98108.26
1516.6741943359375 0 70660.87
1517.67822265625 0 58449.324
1518.63525390625 0 20906.045
1519.652099609375 0 21315.824
1521.6798095703125 0 24511.92
1527.680908203125 0 57753.28
1528.6859130859375 0 33593.863
1533.6439208984375 0 20102.24
1534.643310546875 0 46696.914
1535.65478515625 0 73849.65
1536.6689453125 0 85774.48
1537.66845703125 0 45203.945
1543.701904296875 0 129547.266
1544.6895751953125 0 205060.72
1545.6795654296875 0 212876.25
1546.682861328125 0 168067.67
1547.6826171875 0 160310.78
1548.6844482421875 0 132960.2
1549.68505859375 0 96620.24
1550.6748046875 0 42322.79
1551.636962890625 0 83522.43
1552.6436767578125 0 48858.047
1553.6431884765625 0 22192.734
1554.6431884765625 0 22537.822
1561.697998046875 0 91784.28
1562.6820068359375 0 625705.06
1563.6865234375 0 559254.3
1564.6885986328125 0 251251.48
1565.6837158203125 0 66422.58
1571.681640625 0 27335.83
1572.6671142578125 0 253924.4
1573.6685791015625 0 234052.06
1574.67041015625 0 111735.05
1579.7049560546875 0 191102.03
1580.70849609375 0 197445.14
1581.7122802734375 0 88049.195
1582.720703125 0 25092.898
1589.6903076171875 0 736065.94
1590.6910400390625 0 3312669
1591.6932373046875 0 2653294
1592.69287109375 0 1281966.8
1593.6942138671875 0 262388.66
1594.6695556640625 0 22796.078
1605.76220703125 0 17841.594
1606.6929931640625 0 1178904.6
1607.698974609375 0 3836504.2
1608.7022705078125 0 3125204.5
1609.7049560546875 0 1303446.1
1610.7069091796875 0 177279.9
1853.2088623046875 0 17292.076

Spectrum Details

|  |  |
| --- | --- |
| Matched peaks? Matched peaks The total absolute number of peaks matched. Additionally in brackets the total fraction of peaks matched and the total number of peaks is shown. | 50 (13.12% of 381) |
| FDR? FDR The false discovery rate estimated for this peptide. It is calculated by matching all theoretical fragments with a non-integer shift with the raw peaks for this spectrum. This is done with 40 different shifts. The resulting percentage is the average number of annotated peaks over the number of annotated peaks with the correct spectrum. | 5.14% |
| Satellite FDR? Satellite FDR See the FDR for details on its calculation. This satellite ion specific FDR only contains the satellite ions (d/w) for I/L/J positions. | ∞ |
| PSM Score? PSM Score The PSM Score as given by Hecklib to this annotated spectrum. It is shown with three significant figures. | 688 |

#### Spectrum 9004? Spectrum 9004 The raw spectrum of this peptide as annotated by Hecklib. The fragments are coloured according to ion type (see legend). Any peaks with a star '\*' as text can be hovered over to see the full details, first the ion type second the mass shift type. By hovering over the amino acids in the peptide or ions in the legend the corresponding peaks are highlighted. By toggling the 'Unassigned' label you can turn the background (unassigned) peaks on or off in the plot. By updating the slider in the Ion legend you can update the spectrum to only show the top X% of the peaks with labels. The top X% means any peak that is within X% of the highest intensity. By dragging in the spectrum you can zoom in to a specific part of the spectrum and use 'Zoom Out' to get back to the original zoom level. The annotation of the spectrum is based on the given sequence in the peptides file and is done with different software so inconsistencies are likely. The peaks are annotated based on the given sequence, with 20 ppm tolerance.

Copy Data

##### Spectrum 9004 (TSV)

###### Preview

```
Loading example...
```

*Click on the button to copy the data to your clipboard.*

Mz MinMz MaxIntensity Max

WidthHeightPeptide font sizePeptide stroke widthSpectrum font sizeSpectrum stroke widthCompact peptide

Ion legend

wxyz

abcd

OtherUnassignedIon^Charge^_Position_Show for top:%

TJSGJQAEDESMYF

01.50e+43.00e+44.51e+46.01e+4

Zoom Out

y^+1^_1_y^+1^_2_y^+1^_3_y^+1^_4_c^+2^_11_c^+1^_6_c^+1^_6_c^+1^_7_c^+1^_7_c^+1^_8_y^+1^_6_c^+1^_8_c^+1^_9_c^+1^_9_y^+1^_7_y^+1^_8_c^+1^_10_c^+1^_10_z^+1^_9_c^+1^_11_c^+1^_11_y^+1^_9_c^+1^_11_z^+1^_10_y^+1^_10_c^+1^_12_z^+1^_11_c^+1^_12_y^+1^_11_z^+1^_12_z^+1^_12_y^+1^_12_c^+1^_13_c^+1^_13_z^+1^_13_

0688137720652753

Fragment Matches Table

Show background peaks

| Position | Ion type | Intensity | mz Theoretical | mz Error (Th) | mz Error (ppm) | Charge | Series Number |
| --- | --- | --- | --- | --- | --- | --- | --- |
| - | - | 1519 | 120.1 | - | - | 0 | - |
| - | - | 395.5 | 124.6 | - | - | 0 | - |
| - | - | 372.6 | 126.2 | - | - | 0 | - |
| - | - | 408 | 133.5 | - | - | 0 | - |
| - | - | 385.6 | 135.8 | - | - | 0 | - |
| - | - | 1.355E+04 | 136.1 | - | - | 0 | - |
| - | - | 743 | 137.1 | - | - | 0 | - |
| - | - | 393.4 | 143.2 | - | - | 0 | - |
| - | - | 867.1 | 148.9 | - | - | 0 | - |
| - | - | 569.5 | 150.5 | - | - | 0 | - |
| 14 | y | 2425 | 166.1 | 0.0002775 | 1.671 | +1 | 1 |
| - | - | 499.5 | 171 | - | - | 0 | - |
| - | - | 467.3 | 172.4 | - | - | 0 | - |
| - | - | 732.2 | 173.1 | - | - | 0 | - |
| - | - | 1761 | 173.5 | - | - | 0 | - |
| - | - | 521 | 174.5 | - | - | 0 | - |
| - | - | 2819 | 187.1 | - | - | 0 | - |
| - | - | 521.9 | 188.1 | - | - | 0 | - |
| - | - | 1304 | 201.1 | - | - | 0 | - |
| - | - | 2101 | 215.1 | - | - | 0 | - |
| - | - | 749.5 | 217.1 | - | - | 0 | - |
| - | - | 1518 | 235.1 | - | - | 0 | - |
| - | - | 732.9 | 283.1 | - | - | 0 | - |
| - | - | 561.5 | 288.3 | - | - | 0 | - |
| - | - | 649.6 | 299.2 | - | - | 0 | - |
| - | - | 965.8 | 302.2 | - | - | 0 | - |
| - | - | 983.1 | 318.1 | - | - | 0 | - |
| 13 | y | 1.183E+04 | 329.1 | 0.0001356 | 0.4121 | +1 | 2 |
| - | - | 2200 | 330.2 | - | - | 0 | - |
| - | - | 556.3 | 332.9 | - | - | 0 | - |
| - | - | 1363 | 334.1 | - | - | 0 | - |
| - | - | 1342 | 341.2 | - | - | 0 | - |
| - | - | 601.5 | 346.1 | - | - | 0 | - |
| - | - | 782.6 | 359.2 | - | - | 0 | - |
| - | - | 743.3 | 376.2 | - | - | 0 | - |
| - | - | 1025 | 386.2 | - | - | 0 | - |
| - | - | 586.7 | 414.5 | - | - | 0 | - |
| - | - | 1148 | 444.3 | - | - | 0 | - |
| - | - | 589.1 | 452.5 | - | - | 0 | - |
| - | - | 1576 | 454.3 | - | - | 0 | - |
| - | - | 1330 | 461.1 | - | - | 0 | - |
| - | - | 1686 | 472.3 | - | - | 0 | - |
| 12 | y | 1507 | 476.2 | 0.005692 | 11.95 | +1 | 3 |
| - | - | 605.9 | 477.2 | - | - | 0 | - |
| - | - | 764.5 | 479.1 | - | - | 0 | - |
| - | - | 553.2 | 479.8 | - | - | 0 | - |
| - | - | 1554 | 557.3 | - | - | 0 | - |
| 11 | y | 1755 | 563.2 | 0.006226 | 11.05 | +1 | 4 |
| 11 | c | 786.4 | 566.3 | 0.001944 | 3.433 | +2 | 11 |
| - | - | 729.2 | 573.2 | - | - | 0 | - |
| - | - | 1731 | 582.3 | - | - | 0 | - |
| 6 | c | 2199 | 600.3 | 8.491E-06 | 0.01414 | +1 | 6 |
| - | - | 1020 | 601.2 | - | - | 0 | - |
| - | - | 810.3 | 601.3 | - | - | 0 | - |
| - | - | 661.3 | 614.2 | - | - | 0 | - |
| 6 | c | 5541 | 617.4 | 5.374E-05 | 0.08704 | +1 | 6 |
| - | - | 1990 | 618.4 | - | - | 0 | - |
| - | - | 840.1 | 622.3 | - | - | 0 | - |
| - | - | 2787 | 630.8 | - | - | 0 | - |
| - | - | 1583 | 631.3 | - | - | 0 | - |
| - | - | 1348 | 631.8 | - | - | 0 | - |
| - | - | 1208 | 642.2 | - | - | 0 | - |
| - | - | 812.4 | 644.4 | - | - | 0 | - |
| - | - | 918.2 | 645.4 | - | - | 0 | - |
| - | - | 1237 | 653.4 | - | - | 0 | - |
| - | - | 893 | 661.2 | - | - | 0 | - |
| - | - | 572.7 | 664.8 | - | - | 0 | - |
| 7 | c | 1492 | 671.4 | 0.000257 | 0.3829 | +1 | 7 |
| - | - | 797.9 | 679.2 | - | - | 0 | - |
| - | - | 2441 | 687.4 | - | - | 0 | - |
| 7 | c | 1.528E+04 | 688.4 | 0.0003169 | 0.4603 | +1 | 7 |
| - | - | 4587 | 689.4 | - | - | 0 | - |
| - | - | 912.7 | 690.4 | - | - | 0 | - |
| - | - | 589.2 | 696 | - | - | 0 | - |
| - | - | 730.7 | 714.3 | - | - | 0 | - |
| - | - | 619.2 | 772.4 | - | - | 0 | - |
| - | - | 1529 | 773.4 | - | - | 0 | - |
| - | - | 780.8 | 774.4 | - | - | 0 | - |
| - | - | 867.6 | 776.3 | - | - | 0 | - |
| - | - | 3563 | 777.3 | - | - | 0 | - |
| - | - | 1236 | 778.3 | - | - | 0 | - |
| - | - | 1616 | 782.4 | - | - | 0 | - |
| - | - | 716.3 | 783.4 | - | - | 0 | - |
| - | - | 697.7 | 785.4 | - | - | 0 | - |
| - | - | 1532 | 793.3 | - | - | 0 | - |
| 8 | c | 2287 | 800.4 | 0.002262 | 2.826 | +1 | 8 |
| - | - | 662.1 | 801.4 | - | - | 0 | - |
| - | - | 2816 | 802.4 | - | - | 0 | - |
| - | - | 3374 | 802.9 | - | - | 0 | - |
| - | - | 2012 | 803.4 | - | - | 0 | - |
| - | - | 1209 | 803.9 | - | - | 0 | - |
| - | - | 624.1 | 804.4 | - | - | 0 | - |
| 9 | y | 1883 | 807.3 | 0.003951 | 4.894 | +1 | 6 |
| - | - | 740.5 | 812.4 | - | - | 0 | - |
| - | - | 1684 | 813.4 | - | - | 0 | - |
| - | - | 736.7 | 814.4 | - | - | 0 | - |
| - | - | 5765 | 816.4 | - | - | 0 | - |
| 8 | c | 9448 | 817.4 | 0.0005516 | 0.6748 | +1 | 8 |
| - | - | 3750 | 818.4 | - | - | 0 | - |
| - | - | 700 | 819.4 | - | - | 0 | - |
| - | - | 1073 | 842.3 | - | - | 0 | - |
| - | - | 1937 | 888.5 | - | - | 0 | - |
| - | - | 1746 | 889.5 | - | - | 0 | - |
| 9 | c | 1403 | 915.4 | 0.001008 | 1.101 | +1 | 9 |
| - | - | 2216 | 931.5 | - | - | 0 | - |
| 9 | c | 1.51E+04 | 932.5 | 0.001127 | 1.209 | +1 | 9 |
| - | - | 6701 | 933.5 | - | - | 0 | - |
| - | - | 1709 | 934.5 | - | - | 0 | - |
| 8 | y | 1577 | 936.3 | 0.005303 | 5.664 | +1 | 7 |
| - | - | 764 | 937.3 | - | - | 0 | - |
| - | - | 577.3 | 970.5 | - | - | 0 | - |
| 7 | y | 3575 | 1007 | 0.003712 | 3.685 | +1 | 8 |
| - | - | 1522 | 1008 | - | - | 0 | - |
| - | - | 794.5 | 1012 | - | - | 0 | - |
| - | - | 1818 | 1017 | - | - | 0 | - |
| - | - | 1007 | 1019 | - | - | 0 | - |
| - | - | 1129 | 1026 | - | - | 0 | - |
| - | - | 846.7 | 1029 | - | - | 0 | - |
| 10 | c | 3908 | 1044 | 0.000875 | 0.8378 | +1 | 10 |
| - | - | 1693 | 1045 | - | - | 0 | - |
| - | - | 1.505E+04 | 1061 | - | - | 0 | - |
| 10 | c | 1.261E+04 | 1062 | 0.003254 | 3.066 | +1 | 10 |
| - | - | 4955 | 1063 | - | - | 0 | - |
| - | - | 823 | 1064 | - | - | 0 | - |
| - | - | 681.5 | 1064 | - | - | 0 | - |
| - | - | 691.7 | 1066 | - | - | 0 | - |
| - | - | 963.4 | 1072 | - | - | 0 | - |
| - | - | 989.2 | 1088 | - | - | 0 | - |
| - | - | 2836 | 1114 | - | - | 0 | - |
| - | - | 2122 | 1115 | - | - | 0 | - |
| 6 | z | 700.6 | 1119 | 0.002249 | 2.009 | +1 | 9 |
| - | - | 1115 | 1120 | - | - | 0 | - |
| 11 | c | 957.8 | 1131 | 0.007628 | 6.747 | +1 | 11 |
| 11 | c | 2703 | 1132 | 0.002264 | 2.001 | +1 | 11 |
| - | - | 1753 | 1133 | - | - | 0 | - |
| - | - | 2206 | 1134 | - | - | 0 | - |
| 6 | y | 2974 | 1135 | 0.00501 | 4.412 | +1 | 9 |
| - | - | 748.3 | 1136 | - | - | 0 | - |
| - | - | 1784 | 1137 | - | - | 0 | - |
| - | - | 1464 | 1138 | - | - | 0 | - |
| - | - | 8411 | 1148 | - | - | 0 | - |
| 11 | c | 2.236E+04 | 1149 | 0.002324 | 2.023 | +1 | 11 |
| - | - | 1.134E+04 | 1150 | - | - | 0 | - |
| - | - | 3309 | 1151 | - | - | 0 | - |
| - | - | 1173 | 1180 | - | - | 0 | - |
| - | - | 1309 | 1183 | - | - | 0 | - |
| - | - | 659.7 | 1187 | - | - | 0 | - |
| - | - | 1136 | 1188 | - | - | 0 | - |
| - | - | 928.5 | 1196 | - | - | 0 | - |
| - | - | 991.6 | 1204 | - | - | 0 | - |
| - | - | 1101 | 1205 | - | - | 0 | - |
| 5 | z | 3457 | 1232 | 0.003268 | 2.652 | +1 | 10 |
| - | - | 3242 | 1233 | - | - | 0 | - |
| - | - | 982 | 1235 | - | - | 0 | - |
| - | - | 1797 | 1236 | - | - | 0 | - |
| - | - | 1020 | 1244 | - | - | 0 | - |
| - | - | 882.6 | 1247 | - | - | 0 | - |
| - | - | 5071 | 1248 | - | - | 0 | - |
| 5 | y | 4340 | 1249 | 0.0005355 | 0.4289 | +1 | 10 |
| - | - | 1659 | 1250 | - | - | 0 | - |
| - | - | 1734 | 1252 | - | - | 0 | - |
| - | - | 1185 | 1253 | - | - | 0 | - |
| - | - | 1051 | 1254 | - | - | 0 | - |
| - | - | 4581 | 1261 | - | - | 0 | - |
| - | - | 4093 | 1262 | - | - | 0 | - |
| - | - | 3721 | 1263 | - | - | 0 | - |
| - | - | 2100 | 1264 | - | - | 0 | - |
| - | - | 736 | 1276 | - | - | 0 | - |
| 12 | c | 5305 | 1279 | 0.001065 | 0.833 | +1 | 12 |
| - | - | 3982 | 1280 | - | - | 0 | - |
| - | - | 2344 | 1281 | - | - | 0 | - |
| - | - | 1379 | 1282 | - | - | 0 | - |
| 4 | z | 1890 | 1290 | 0.0002372 | 0.184 | +1 | 11 |
| - | - | 2876 | 1291 | - | - | 0 | - |
| - | - | 2325 | 1292 | - | - | 0 | - |
| - | - | 729.4 | 1295 | - | - | 0 | - |
| 12 | c | 1.745E+04 | 1296 | 0.002958 | 2.283 | +1 | 12 |
| - | - | 1.135E+04 | 1297 | - | - | 0 | - |
| - | - | 4346 | 1298 | - | - | 0 | - |
| - | - | 3773 | 1305 | - | - | 0 | - |
| 4 | y | 3828 | 1306 | 0.001166 | 0.8935 | +1 | 11 |
| - | - | 2013 | 1307 | - | - | 0 | - |
| - | - | 1048 | 1311 | - | - | 0 | - |
| - | - | 788.4 | 1316 | - | - | 0 | - |
| 3 | z | 1668 | 1359 | 0.002428 | 1.787 | +1 | 12 |
| - | - | 1075 | 1362 | - | - | 0 | - |
| - | - | 1291 | 1363 | - | - | 0 | - |
| 3 | z | 9077 | 1377 | 0.0019 | 1.38 | +1 | 12 |
| - | - | 6949 | 1378 | - | - | 0 | - |
| - | - | 3188 | 1379 | - | - | 0 | - |
| 3 | y | 6720 | 1393 | 0.001975 | 1.418 | +1 | 12 |
| - | - | 4863 | 1394 | - | - | 0 | - |
| - | - | 3054 | 1395 | - | - | 0 | - |
| - | - | 926.6 | 1396 | - | - | 0 | - |
| - | - | 1797 | 1415 | - | - | 0 | - |
| - | - | 1417 | 1416 | - | - | 0 | - |
| - | - | 1789 | 1426 | - | - | 0 | - |
| - | - | 2951 | 1427 | - | - | 0 | - |
| - | - | 1.244E+04 | 1428 | - | - | 0 | - |
| - | - | 8530 | 1429 | - | - | 0 | - |
| - | - | 2688 | 1430 | - | - | 0 | - |
| - | - | 1256 | 1440 | - | - | 0 | - |
| - | - | 1633 | 1441 | - | - | 0 | - |
| 13 | c | 885 | 1442 | 0.002312 | 1.604 | +1 | 13 |
| - | - | 3397 | 1443 | - | - | 0 | - |
| - | - | 8108 | 1444 | - | - | 0 | - |
| - | - | 8149 | 1445 | - | - | 0 | - |
| - | - | 3887 | 1446 | - | - | 0 | - |
| - | - | 1397 | 1447 | - | - | 0 | - |
| 13 | c | 2.493E+04 | 1459 | 0.001031 | 0.7069 | +1 | 13 |
| - | - | 1.669E+04 | 1460 | - | - | 0 | - |
| - | - | 7249 | 1461 | - | - | 0 | - |
| - | - | 1356 | 1462 | - | - | 0 | - |
| 2 | z | 8020 | 1490 | 0.002065 | 1.386 | +1 | 13 |
| - | - | 6958 | 1491 | - | - | 0 | - |
| - | - | 3425 | 1492 | - | - | 0 | - |
| - | - | 2189 | 1507 | - | - | 0 | - |
| - | - | 1199 | 1508 | - | - | 0 | - |
| - | - | 920.2 | 1509 | - | - | 0 | - |
| - | - | 1182 | 1516 | - | - | 0 | - |
| - | - | 849.2 | 1518 | - | - | 0 | - |
| - | - | 783.9 | 1529 | - | - | 0 | - |
| - | - | 743 | 1536 | - | - | 0 | - |
| - | - | 1840 | 1537 | - | - | 0 | - |
| - | - | 2325 | 1544 | - | - | 0 | - |
| - | - | 2542 | 1545 | - | - | 0 | - |
| - | - | 3956 | 1546 | - | - | 0 | - |
| - | - | 2624 | 1547 | - | - | 0 | - |
| - | - | 2112 | 1548 | - | - | 0 | - |
| - | - | 1437 | 1549 | - | - | 0 | - |
| - | - | 1491 | 1550 | - | - | 0 | - |
| - | - | 1412 | 1552 | - | - | 0 | - |
| - | - | 1415 | 1553 | - | - | 0 | - |
| - | - | 787.2 | 1560 | - | - | 0 | - |
| - | - | 1478 | 1561 | - | - | 0 | - |
| - | - | 1895 | 1562 | - | - | 0 | - |
| - | - | 8784 | 1563 | - | - | 0 | - |
| - | - | 7962 | 1564 | - | - | 0 | - |
| - | - | 3595 | 1565 | - | - | 0 | - |
| - | - | 1122 | 1566 | - | - | 0 | - |
| - | - | 1114 | 1571 | - | - | 0 | - |
| - | - | 1417 | 1572 | - | - | 0 | - |
| - | - | 5318 | 1573 | - | - | 0 | - |
| - | - | 4548 | 1574 | - | - | 0 | - |
| - | - | 2482 | 1575 | - | - | 0 | - |
| - | - | 672.2 | 1578 | - | - | 0 | - |
| - | - | 3575 | 1580 | - | - | 0 | - |
| - | - | 3032 | 1581 | - | - | 0 | - |
| - | - | 1407 | 1582 | - | - | 0 | - |
| - | - | 2634 | 1588 | - | - | 0 | - |
| - | - | 2818 | 1589 | - | - | 0 | - |
| - | - | 1.589E+04 | 1590 | - | - | 0 | - |
| - | - | 5.124E+04 | 1591 | - | - | 0 | - |
| - | - | 3.91E+04 | 1592 | - | - | 0 | - |
| - | - | 2.288E+04 | 1593 | - | - | 0 | - |
| - | - | 4423 | 1594 | - | - | 0 | - |
| - | - | 2159 | 1604 | - | - | 0 | - |
| - | - | 6122 | 1605 | - | - | 0 | - |
| - | - | 6861 | 1606 | - | - | 0 | - |
| - | - | 1.636E+04 | 1607 | - | - | 0 | - |
| - | - | 5.949E+04 | 1608 | - | - | 0 | - |
| - | - | 4.552E+04 | 1609 | - | - | 0 | - |
| - | - | 2.42E+04 | 1610 | - | - | 0 | - |
| - | - | 2902 | 1611 | - | - | 0 | - |
| - | - | 757.7 | 1676 | - | - | 0 | - |
| - | - | 753.7 | 1798 | - | - | 0 | - |
| - | - | 751 | 2046 | - | - | 0 | - |
| - | - | 677.3 | 2407 | - | - | 0 | - |
| - | - | 2575 | 2408 | - | - | 0 | - |
| - | - | 3103 | 2409 | - | - | 0 | - |
| - | - | 3044 | 2410 | - | - | 0 | - |
| - | - | 1250 | 2411 | - | - | 0 | - |
| - | - | 831.1 | 2413 | - | - | 0 | - |
| - | - | 740.3 | 2726 | - | - | 0 | - |

m/z Charge Intensity FragmentType MassShift Position
120.08097839355469 0 1518.505
124.59553527832031 0 395.50186
126.20233154296875 0 372.5629
133.54322814941406 0 408.0214
135.84185791015625 0 385.5987
136.0758514404297 0 13548.397
137.07919311523438 0 743.0172
143.1610870361328 0 393.3708
148.94717407226562 0 867.14124
150.51876831054688 0 569.5066
166.08653259277344 0 2424.9119 y 13
170.96099853515625 0 499.48337
172.41180419921875 0 467.2964
173.12889099121094 0 732.1928
173.45240783691406 0 1761.1538
174.5266571044922 0 521.03735
187.14422607421875 0 2818.527
188.14698791503906 0 521.8686
201.1234588623047 0 1303.7485
215.13902282714844 0 2100.76
217.0641632080078 0 749.4715
235.07460021972656 0 1518.2258
283.1126403808594 0 732.89233
288.30413818359375 0 561.5151
299.1712646484375 0 649.5757
302.1709289550781 0 965.8447
318.14501953125 0 983.12537
329.14971923828125 0 11832.605 y 12
330.1530456542969 0 2199.9873
332.8993835449219 0 556.29297
334.13983154296875 0 1363.3219
341.18267822265625 0 1341.7286
346.13970947265625 0 601.5062
359.1930236816406 0 782.64044
376.1861572265625 0 743.25977
386.20361328125 0 1024.6833
414.51446533203125 0 586.67255
444.2825622558594 0 1147.5731
452.4518127441406 0 589.0592
454.265625 0 1575.807
461.1328430175781 0 1329.6509
472.2765808105469 0 1686.0659
476.1857604980469 0 1506.5482 y 11
477.1872863769531 0 605.8654
479.14324951171875 0 764.5022
479.765625 0 553.2143
557.2571411132812 0 1553.862
563.2183227539062 0 1754.7559 y 10
566.2637939453125 0 786.36847 c Ammonia loss 10
573.2164306640625 0 729.1592
582.3237915039062 0 1730.8779
600.3351440429688 0 2198.7458 c Ammonia loss 5
601.247314453125 0 1020.24835
601.3397216796875 0 810.26825
614.2132568359375 0 661.2838
617.3617553710938 0 5541.136 c 5
618.3622436523438 0 1989.92
622.2656860351562 0 840.1044
630.7742919921875 0 2787.1245
631.2769165039062 0 1583.273
631.7750244140625 0 1347.9174
642.205322265625 0 1208.4396
644.3850708007812 0 812.4034
645.38818359375 0 918.15857
653.3631591796875 0 1237.41
661.2130126953125 0 893.02
664.7821655273438 0 572.67584
671.3720092773438 0 1492.3815 c Ammonia loss 6
679.2205810546875 0 797.91943
687.3905029296875 0 2440.7236
688.3984985351562 0 15283.49 c 6
689.4010620117188 0 4586.9326
690.4066162109375 0 912.71497
696.0443725585938 0 589.1883
714.3299560546875 0 730.69275
772.3541870117188 0 619.24866
773.4315185546875 0 1528.8854
774.4326782226562 0 780.75366
776.3045043945312 0 867.57184
777.3194580078125 0 3563.339
778.3231811523438 0 1235.6478
782.4057006835938 0 1615.5184
783.4066772460938 0 716.3009
785.3993530273438 0 697.73303
793.3179931640625 0 1531.6343
800.41259765625 0 2286.8794 c Ammonia loss 7
801.41845703125 0 662.0944
802.3932495117188 0 2816.037
802.8947143554688 0 3374.1062
803.3942260742188 0 2012.1221
803.8939208984375 0 1208.755
804.3858642578125 0 624.1044
807.2855834960938 0 1882.8213 y 8
812.4147338867188 0 740.4595
813.3997802734375 0 1684.0581
814.3935546875 0 736.6738
816.4337768554688 0 5765.463
817.4408569335938 0 9448.049 c 7
818.4439697265625 0 3749.674
819.4432373046875 0 700.01514
842.2848510742188 0 1072.8804
888.4526977539062 0 1936.617
889.4619750976562 0 1746.406
915.4428100585938 0 1403.4893 c Ammonia loss 8
931.4595947265625 0 2215.5483
932.4672241210938 0 15102.193 c 8
933.469482421875 0 6701.1733
934.4740600585938 0 1708.8478
936.3295288085938 0 1577.03 y 7
937.3294067382812 0 763.9839
970.4723510742188 0 577.32227
1007.3650512695312 0 3575.1675 y 6
1008.3722534179688 0 1522.3508
1011.5465087890625 0 794.5234
1017.499755859375 0 1818.225
1018.5049438476562 0 1006.97577
1026.4696044921875 0 1129.2634
1029.4742431640625 0 846.69836
1044.4835205078125 0 3907.6194 c Ammonia loss 9
1045.486572265625 0 1693.2458
1060.5023193359375 0 15045.72
1061.5076904296875 0 12608.674 c 9
1062.510986328125 0 4955.283
1063.51025390625 0 822.97595
1064.4285888671875 0 681.50977
1066.4429931640625 0 691.6733
1071.535888671875 0 963.3806
1087.5128173828125 0 989.1779
1113.5035400390625 0 2835.6018
1114.5054931640625 0 2122.339
1119.4034423828125 0 700.6038 z 5
1120.412353515625 0 1115.0549
1130.5247802734375 0 957.75385 c Water loss 10
1131.51416015625 0 2703.279 c Ammonia loss 10
1132.515869140625 0 1752.8573
1134.4156494140625 0 2206.4902
1135.4249267578125 0 2974.3826 y 5
1136.3978271484375 0 748.3091
1136.5166015625 0 1783.6542
1137.5064697265625 0 1464.45
1147.5345458984375 0 8410.526
1148.5406494140625 0 22360.885 c 10
1149.544189453125 0 11338.313
1150.5469970703125 0 3309.3237
1179.5162353515625 0 1173.3536
1182.600341796875 0 1308.7455
1186.55810546875 0 659.72656
1187.5604248046875 0 1135.7195
1195.6148681640625 0 928.509
1204.1309814453125 0 991.6158
1204.634521484375 0 1100.5679
1232.488525390625 0 3457.1208 z 4
1233.4923095703125 0 3241.8875
1234.5003662109375 0 982.03357
1235.5401611328125 0 1797.132
1243.5220947265625 0 1019.65906
1246.516845703125 0 882.6256
1247.5003662109375 0 5070.6343
1248.5045166015625 0 4340.194 y 4
1249.5057373046875 0 1659.4907
1251.5614013671875 0 1733.7957
1252.568115234375 0 1184.7394
1253.56689453125 0 1050.6274
1260.5386962890625 0 4581.028
1261.537841796875 0 4092.8428
1262.5482177734375 0 3720.799
1263.55126953125 0 2099.675
1275.5501708984375 0 735.9502
1278.5479736328125 0 5305.3516 c Ammonia loss 11
1279.549560546875 0 3981.573
1280.5623779296875 0 2343.7566
1281.5653076171875 0 1379.4199
1289.5069580078125 0 1890.4403 z 3
1290.515869140625 0 2875.6104
1291.518798828125 0 2324.6992
1294.5604248046875 0 729.4025
1295.576416015625 0 17454.713 c 11
1296.5787353515625 0 11349.903
1297.579833984375 0 4345.8374
1304.5220947265625 0 3772.5068
1305.526611328125 0 3827.9163 y 3
1306.523193359375 0 2013.038
1310.6763916015625 0 1048.146
1315.6573486328125 0 788.3944
1358.5257568359375 0 1667.8169 z Water loss 2
1361.58837890625 0 1075.3438
1362.5909423828125 0 1291.1705
1376.5406494140625 0 9077.456 z 2
1377.5447998046875 0 6948.9375
1378.5467529296875 0 3188.4106
1392.5594482421875 0 6720.4595 y 2
1393.565673828125 0 4862.9883
1394.562744140625 0 3054.2979
1395.5667724609375 0 926.6156
1414.62548828125 0 1796.5817
1415.6358642578125 0 1416.8636
1425.5977783203125 0 1788.5387
1426.653076171875 0 2951.2117
1427.656005859375 0 12437.234
1428.6552734375 0 8529.88
1429.6597900390625 0 2687.807
1439.722900390625 0 1256.3116
1440.729248046875 0 1632.7758
1441.612548828125 0 884.97833 c Ammonia loss 12
1442.662841796875 0 3397.4355
1443.650390625 0 8108.4365
1444.6627197265625 0 8149.042
1445.6590576171875 0 3887.1646
1446.65283203125 0 1396.7399
1458.6378173828125 0 24926.12 c 12
1459.6409912109375 0 16685.473
1460.64404296875 0 7248.6763
1461.6419677734375 0 1355.9935
1489.6248779296875 0 8019.599 z 1
1490.6282958984375 0 6957.9077
1491.632080078125 0 3424.633
1506.6077880859375 0 2189.2766
1507.6285400390625 0 1198.7715
1508.6478271484375 0 920.2095
1515.66552734375 0 1181.748
1517.69482421875 0 849.23303
1528.6854248046875 0 783.92816
1535.656494140625 0 743.01636
1536.6661376953125 0 1839.8416
1543.70361328125 0 2325.1147
1544.6988525390625 0 2542.051
1545.6878662109375 0 3955.5605
1546.683349609375 0 2624.1758
1547.6767578125 0 2112.473
1548.6943359375 0 1437.3964
1549.700927734375 0 1490.7803
1551.6429443359375 0 1411.6807
1552.6385498046875 0 1415.4907
1559.7537841796875 0 787.1956
1560.7657470703125 0 1477.7645
1561.71240234375 0 1895.4504
1562.682373046875 0 8783.543
1563.68505859375 0 7962.194
1564.698486328125 0 3595.1362
1565.6796875 0 1122.4059
1570.7164306640625 0 1113.6748
1571.6953125 0 1416.5417
1572.68115234375 0 5317.6143
1573.6800537109375 0 4548.0977
1574.688720703125 0 2482.4653
1577.8070068359375 0 672.2029
1579.7020263671875 0 3575.4663
1580.707763671875 0 3031.8264
1581.7122802734375 0 1407.4436
1587.7373046875 0 2633.6091
1588.72998046875 0 2818.0251
1589.6947021484375 0 15893.502
1590.6900634765625 0 51238.88
1591.692138671875 0 39101.93
1592.6966552734375 0 22880.188
1593.6939697265625 0 4423.135
1603.775634765625 0 2158.671
1604.77783203125 0 6121.776
1605.7784423828125 0 6860.553
1606.6961669921875 0 16362.013
1607.697265625 0 59490.445
1608.703857421875 0 45523.543
1609.7132568359375 0 24201.258
1610.724365234375 0 2902.3926
1676.17138671875 0 757.73
1798.2376708984375 0 753.7382
2046.0787353515625 0 750.9772
2407.19482421875 0 677.263
2408.2607421875 0 2575.1296
2409.25830078125 0 3102.8557
2410.259521484375 0 3044.2456
2411.26318359375 0 1250.3359
2413.222412109375 0 831.12695
2725.857421875 0 740.2943

Spectrum Details

|  |  |
| --- | --- |
| Matched peaks? Matched peaks The total absolute number of peaks matched. Additionally in brackets the total fraction of peaks matched and the total number of peaks is shown. | 35 (12.77% of 274) |
| FDR? FDR The false discovery rate estimated for this peptide. It is calculated by matching all theoretical fragments with a non-integer shift with the raw peaks for this spectrum. This is done with 40 different shifts. The resulting percentage is the average number of annotated peaks over the number of annotated peaks with the correct spectrum. | 4.90% |
| Satellite FDR? Satellite FDR See the FDR for details on its calculation. This satellite ion specific FDR only contains the satellite ions (d/w) for I/L/J positions. | ∞ |
| PSM Score? PSM Score The PSM Score as given by Hecklib to this annotated spectrum. It is shown with three significant figures. | 438 |

#### Reverse Lookup? Reverse Lookup All places where this read could be placed.

| Group | Segment | Template | Template Part | Read Part | Score | Unique |
| --- | --- | --- | --- | --- | --- | --- |
| Homo sapiens Light Chain | IGLV | IGLV2-14 | [75..89] | [0..14] | 85 | False |
| Homo sapiens Light Chain | IGLV | IGLV2-23 | [75..89] | [0..14] | 85 | False |
| Homo sapiens Light Chain | IGLV | IGLV2-18 | [75..89] | [0..14] | 85 | False |
| Homo sapiens Light Chain | IGLV | IGLV2-11 | [75..89] | [0..14] | 85 | False |

| Recombined | Template Part | Read Part | Score | Unique |
| --- | --- | --- | --- | --- |
| REC-0-1\_002 | [75..89] | [0..14] | 112 | True |

#### Meta Information from Multiple reads

##### Number of combined reads

3

##### Intensity

0.9022

##### TotalArea

8.273E+08

##### Changes to the peptide sequence

TJSGJQAEDESMYF

L→JNo support for either Leucine or Isoleucine based on side chain ions (Position: 5)

L→JNo support for either Leucine or Isoleucine based on side chain ions (Position: 2)

#### Positional Score

Copy Data

##### Positional Score (TSV)

###### Preview

```
Loading example...
```

*Click on the button to copy the data to your clipboard.*

10012345678910111213

Label Value
"0" 0.32
"1" 0.32
"2" 0.323
"3" 0.32
"4" 0.32
"5" 0.307
"6" 0.333
"7" 0.333
"8" 0.333
"9" 0.333
"10" 0.333
"11" 0.333
"12" 0.333
"13" 0.333

#### Meta Information from PEAKS

##### Scan Identifier

F3:8780

##### Original sequence

T

L

S

G

L

Q

A

E

D

E

S

M

+15.99

Y

F

##### Posttranslational Modifications

Oxidation (M)

##### Source File

D:\separate\_stitch\_analyses\xle-disambiguation\raw\20210323\_F1\_UM1\_Peng0013\_SA\_F59\_ingel\_3ug\_chymo.raw

##### Fraction

3

##### Scan Feature

F3:14509

##### De Novo Score

99

##### ConfidenceScore

99

### m/z

803.8518

##### Mass

1605.6868

##### Charge

2

##### Retention Time

48.36

##### Predicted Retention Time

-

##### Area

4.137E+08

##### Parts Per Million

1.4

##### Fragmentation mode

ETHCD

##### Originating file

01 D:\separate\_stitch\_analyses\xle-disambiguation\20210325\_F59\_3ug\_DENOVO\_12.csv

#### Meta Information from PEAKS

##### Scan Identifier

F3:8663

##### Original sequence

T

L

S

G

L

Q

A

E

D

E

S

M

+15.99

Y

F

##### Posttranslational Modifications

Oxidation (M)

##### Source File

D:\separate\_stitch\_analyses\xle-disambiguation\raw\20210323\_F1\_UM1\_Peng0013\_SA\_F59\_ingel\_3ug\_chymo.raw

##### Fraction

3

##### Scan Feature

F3:14509

##### De Novo Score

98

##### ConfidenceScore

98

### m/z

803.8518

##### Mass

1605.6868

##### Charge

2

##### Retention Time

48.36

##### Predicted Retention Time

-

##### Area

4.137E+08

##### Parts Per Million

1.4

##### Fragmentation mode

ETHCD

##### Originating file

01 D:\separate\_stitch\_analyses\xle-disambiguation\20210325\_F59\_3ug\_DENOVO\_12.csv

#### Meta Information from PEAKS

##### Scan Identifier

F3:9004

##### Original sequence

T

L

S

G

L

Q

A

E

D

E

S

M

+15.99

Y

F

##### Posttranslational Modifications

Oxidation (M)

##### Source File

D:\separate\_stitch\_analyses\xle-disambiguation\raw\20210323\_F1\_UM1\_Peng0013\_SA\_F59\_ingel\_3ug\_chymo.raw

##### Fraction

3

##### Scan Feature

-

##### De Novo Score

98

##### ConfidenceScore

98

### m/z

803.8524

##### Mass

1605.6868

##### Charge

2

##### Retention Time

50.42

##### Predicted Retention Time

-

##### Area

0

##### Parts Per Million

2.2

##### Fragmentation mode

ETHCD

##### Originating file

01 D:\separate\_stitch\_analyses\xle-disambiguation\20210325\_F59\_3ug\_DENOVO\_12.csv
