## Supplementary material for "A handle on mass coincidence errors in *de novo* sequencing of antibodies by bottom-up proteomics": Combined_040.html

Details Combined\_040 | Stitch OverviewUndefined

### Read Combined\_040

#### Sequence (length=15)

TLPPSREEMTKNQKJ

#### Spectrum 4527? Spectrum 4527 The raw spectrum of this peptide as annotated by Hecklib. The fragments are coloured according to ion type (see legend). Any peaks with a star '\*' as text can be hovered over to see the full details, first the ion type second the mass shift type. By hovering over the amino acids in the peptide or ions in the legend the corresponding peaks are highlighted. By toggling the 'Unassigned' label you can turn the background (unassigned) peaks on or off in the plot. By updating the slider in the Ion legend you can update the spectrum to only show the top X% of the peaks with labels. The top X% means any peak that is within X% of the highest intensity. By dragging in the spectrum you can zoom in to a specific part of the spectrum and use 'Zoom Out' to get back to the original zoom level. The annotation of the spectrum is based on the given sequence in the peptides file and is done with different software so inconsistencies are likely. The peaks are annotated based on the given sequence, with 20 ppm tolerance.

Copy Data

##### Spectrum 4527 (TSV)

###### Preview

```
Loading example...
```

*Click on the button to copy the data to your clipboard.*

Mz MinMz MaxIntensity Max

WidthHeightPeptide font sizePeptide stroke widthSpectrum font sizeSpectrum stroke widthCompact peptide

Ion legend

wxyz

abcd

OtherUnassignedIon^Charge^_Position_Show for top:%

TLPPSREEMTKNQKJ

05.52e+41.10e+51.66e+52.21e+5

Zoom Out

y^+1^_1_y^+1^_2_c^+1^_4_c^+1^_4_z^+1^_3_y^+1^_3_z^+3^_12_z^+3^_12_c^+1^_5_y^+3^_13_y^+3^_13_z^+3^_13_z^+1^_4_y^+3^_13_y^+1^_4_y^+2^_9_c^+2^_11_c^+2^_11_c^+2^_11_z^+2^_10_z^+2^_10_c^+1^_6_z^+1^_5_y^+2^_10_y^+1^_5_c^+2^_12_c^+2^_12_z^+2^_11_c^+2^_12_y^+2^_11_z^+2^_11_y^+2^_11_w^+2^_12_z^+2^_12_z^+2^_12_w^+1^_6_w^+1^_6_z^+2^_12_c^+2^_13_y^+2^_12_y^+1^_6_c^+2^_13_y^+1^_6_w^+2^_13_c^+1^_7_y^+2^_13_y^+2^_13_z^+2^_13_y^+2^_13_w^+2^_14_c^+2^_14_y^+2^_14_z^+2^_14_c^+2^_14_y^+2^_14_z^+1^_7_z^+1^_7_c^+1^_8_y^+1^_7_w^+1^_8_z^+1^_8_y^+1^_8_c^+1^_9_w^+1^_9_c^+1^_10_y^+1^_9_z^+1^_9_y^+1^_9_c^+1^_11_c^+1^_11_y^+1^_10_z^+1^_10_y^+1^_10_z^+1^_11_c^+1^_12_y^+1^_11_z^+1^_11_y^+1^_11_c^+1^_13_y^+1^_12_c^+1^_13_y^+1^_13_z^+1^_13_y^+1^_13_z^+1^_14_c^+1^_14_

0779155823373116

Fragment Matches Table

Show background peaks

| Position | Ion type | Intensity | mz Theoretical | mz Error (Th) | mz Error (ppm) | Charge | Series Number |
| --- | --- | --- | --- | --- | --- | --- | --- |
| 15 | y | 1781 | 132.1 | 0.000283 | 2.142 | +1 | 1 |
| - | - | 736.1 | 143.1 | - | - | 0 | - |
| - | - | 537.4 | 143.7 | - | - | 0 | - |
| - | - | 983.8 | 149 | - | - | 0 | - |
| - | - | 530.7 | 159.4 | - | - | 0 | - |
| - | - | 660.9 | 162.7 | - | - | 0 | - |
| - | - | 663.5 | 173.1 | - | - | 0 | - |
| - | - | 2629 | 173.5 | - | - | 0 | - |
| - | - | 6.047E+04 | 187.1 | - | - | 0 | - |
| - | - | 5641 | 188.1 | - | - | 0 | - |
| - | - | 1180 | 201.1 | - | - | 0 | - |
| - | - | 4080 | 212.1 | - | - | 0 | - |
| - | - | 3.811E+04 | 215.1 | - | - | 0 | - |
| - | - | 3449 | 216.1 | - | - | 0 | - |
| - | - | 4972 | 219.1 | - | - | 0 | - |
| - | - | 700.1 | 220.1 | - | - | 0 | - |
| - | - | 861.9 | 230.2 | - | - | 0 | - |
| - | - | 1452 | 256.2 | - | - | 0 | - |
| - | - | 726.4 | 292.7 | - | - | 0 | - |
| - | - | 8690 | 299.2 | - | - | 0 | - |
| - | - | 859.7 | 300.2 | - | - | 0 | - |
| - | - | 1509 | 312.2 | - | - | 0 | - |
| - | - | 754.8 | 313.2 | - | - | 0 | - |
| 14 | y | 1195 | 318.2 | 0.003072 | 9.653 | +1 | 2 |
| - | - | 1995 | 326.2 | - | - | 0 | - |
| - | - | 2143 | 327.2 | - | - | 0 | - |
| - | - | 1544 | 344.2 | - | - | 0 | - |
| - | - | 655.2 | 369.1 | - | - | 0 | - |
| - | - | 1570 | 382.3 | - | - | 0 | - |
| - | - | 848.5 | 395.2 | - | - | 0 | - |
| - | - | 1755 | 396.2 | - | - | 0 | - |
| - | - | 2307 | 407.3 | - | - | 0 | - |
| 4 | c | 925 | 408.3 | 0.004092 | 10.02 | +1 | 4 |
| - | - | 1163 | 412.2 | - | - | 0 | - |
| - | - | 1595 | 414.2 | - | - | 0 | - |
| - | - | 794.4 | 425.2 | - | - | 0 | - |
| - | - | 3502 | 425.3 | - | - | 0 | - |
| 4 | c | 2427 | 426.3 | 0.0002361 | 0.5538 | +1 | 4 |
| - | - | 1789 | 427.3 | - | - | 0 | - |
| - | - | 1283 | 428.3 | - | - | 0 | - |
| - | - | 764.4 | 430.1 | - | - | 0 | - |
| 13 | z | 1569 | 430.2 | 0.004229 | 9.829 | +1 | 3 |
| - | - | 998.3 | 431.3 | - | - | 0 | - |
| 13 | y | 4214 | 446.3 | 0.003788 | 8.488 | +1 | 3 |
| - | - | 1018 | 447.3 | - | - | 0 | - |
| - | - | 1387 | 462.9 | - | - | 0 | - |
| - | - | 1082 | 466.2 | - | - | 0 | - |
| - | - | 1695 | 466.6 | - | - | 0 | - |
| - | - | 4585 | 470.3 | - | - | 0 | - |
| - | - | 1211 | 471.3 | - | - | 0 | - |
| - | - | 2779 | 471.9 | - | - | 0 | - |
| - | - | 1344 | 472.2 | - | - | 0 | - |
| - | - | 832.2 | 472.3 | - | - | 0 | - |
| - | - | 2120 | 472.6 | - | - | 0 | - |
| - | - | 1593 | 491.6 | - | - | 0 | - |
| - | - | 725 | 491.9 | - | - | 0 | - |
| - | - | 2753 | 494.3 | - | - | 0 | - |
| - | - | 1107 | 494.9 | - | - | 0 | - |
| - | - | 2819 | 495.2 | - | - | 0 | - |
| - | - | 858.4 | 495.6 | - | - | 0 | - |
| 4 | z | 3013 | 500.9 | 0.003499 | 6.985 | +3 | 12 |
| 4 | z | 3136 | 501.2 | 0.002633 | 5.253 | +3 | 12 |
| - | - | 1256 | 501.6 | - | - | 0 | - |
| - | - | 1349 | 512.3 | - | - | 0 | - |
| 5 | c | 3.614E+04 | 513.3 | 0.0007699 | 1.5 | +1 | 5 |
| - | - | 9497 | 514.3 | - | - | 0 | - |
| - | - | 2695 | 515.3 | - | - | 0 | - |
| - | - | 4198 | 523.3 | - | - | 0 | - |
| - | - | 3480 | 523.6 | - | - | 0 | - |
| - | - | 2034 | 523.9 | - | - | 0 | - |
| - | - | 1435 | 524.3 | - | - | 0 | - |
| - | - | 942.6 | 528.3 | - | - | 0 | - |
| 3 | y | 2337 | 538.6 | 0.002111 | 3.919 | +3 | 13 |
| 3 | y | 3357 | 538.9 | 0.005466 | 10.14 | +3 | 13 |
| 3 | z | 1194 | 539.3 | 0.002959 | 5.487 | +3 | 13 |
| - | - | 2138 | 539.6 | - | - | 0 | - |
| - | - | 1616 | 539.9 | - | - | 0 | - |
| - | - | 1112 | 540.3 | - | - | 0 | - |
| - | - | 884.4 | 541.6 | - | - | 0 | - |
| - | - | 1092 | 541.9 | - | - | 0 | - |
| 12 | z | 2935 | 544.3 | 0.003089 | 5.675 | +1 | 4 |
| 3 | y | 2.186E+05 | 544.6 | 0.001092 | 2.005 | +3 | 13 |
| - | - | 1.922E+05 | 544.9 | - | - | 0 | - |
| - | - | 1.132E+05 | 545.3 | - | - | 0 | - |
| - | - | 4.35E+04 | 545.6 | - | - | 0 | - |
| - | - | 1.182E+04 | 545.9 | - | - | 0 | - |
| - | - | 1999 | 546.3 | - | - | 0 | - |
| - | - | 893.1 | 546.8 | - | - | 0 | - |
| - | - | 834.6 | 547.3 | - | - | 0 | - |
| - | - | 1429 | 552.8 | - | - | 0 | - |
| - | - | 1175 | 557.9 | - | - | 0 | - |
| 12 | y | 8894 | 560.3 | 0.003747 | 6.687 | +1 | 4 |
| - | - | 3280 | 561.3 | - | - | 0 | - |
| - | - | 2067 | 561.8 | - | - | 0 | - |
| - | - | 3182 | 562.3 | - | - | 0 | - |
| - | - | 1576 | 562.8 | - | - | 0 | - |
| - | - | 1191 | 571.2 | - | - | 0 | - |
| - | - | 942.8 | 588.3 | - | - | 0 | - |
| - | - | 921.6 | 590.3 | - | - | 0 | - |
| - | - | 1309 | 590.8 | - | - | 0 | - |
| 7 | y | 926.1 | 597.8 | 0.00849 | 14.2 | +2 | 9 |
| - | - | 886.8 | 598.2 | - | - | 0 | - |
| - | - | 1252 | 598.3 | - | - | 0 | - |
| - | - | 1644 | 599.2 | - | - | 0 | - |
| - | - | 1172 | 607.4 | - | - | 0 | - |
| - | - | 1125 | 610 | - | - | 0 | - |
| - | - | 1365 | 614.4 | - | - | 0 | - |
| - | - | 2203 | 615.3 | - | - | 0 | - |
| - | - | 811.8 | 615.9 | - | - | 0 | - |
| - | - | 2874 | 616.3 | - | - | 0 | - |
| - | - | 1443 | 625.4 | - | - | 0 | - |
| - | - | 2745 | 626.4 | - | - | 0 | - |
| - | - | 815.4 | 627.4 | - | - | 0 | - |
| - | - | 1.29E+04 | 628.4 | - | - | 0 | - |
| - | - | 5726 | 629.3 | - | - | 0 | - |
| - | - | 4372 | 629.4 | - | - | 0 | - |
| - | - | 1540 | 630.3 | - | - | 0 | - |
| - | - | 1030 | 631.3 | - | - | 0 | - |
| - | - | 1359 | 637.3 | - | - | 0 | - |
| - | - | 2075 | 642.8 | - | - | 0 | - |
| 11 | c | 1411 | 643.3 | 0.00303 | 4.711 | +2 | 11 |
| 11 | c | 2055 | 643.8 | 0.005816 | 9.034 | +2 | 11 |
| - | - | 1857 | 647.3 | - | - | 0 | - |
| - | - | 1788 | 647.8 | - | - | 0 | - |
| - | - | 1164 | 648.3 | - | - | 0 | - |
| - | - | 731.5 | 648.8 | - | - | 0 | - |
| - | - | 4935 | 651.8 | - | - | 0 | - |
| 11 | c | 3.105E+04 | 652.3 | 0.003223 | 4.94 | +2 | 11 |
| - | - | 2.006E+04 | 652.8 | - | - | 0 | - |
| - | - | 8918 | 653.3 | - | - | 0 | - |
| - | - | 1239 | 653.8 | - | - | 0 | - |
| - | - | 1421 | 657.8 | - | - | 0 | - |
| 6 | z | 2074 | 658.8 | 0.007157 | 10.86 | +2 | 10 |
| 6 | z | 2644 | 667.8 | 0.002636 | 3.947 | +2 | 10 |
| - | - | 3030 | 668.3 | - | - | 0 | - |
| - | - | 1194 | 668.8 | - | - | 0 | - |
| 6 | c | 7.846E+04 | 669.4 | 0.00116 | 1.733 | +1 | 6 |
| - | - | 2.639E+04 | 670.4 | - | - | 0 | - |
| - | - | 7229 | 671.4 | - | - | 0 | - |
| 11 | z | 2.982E+04 | 672.4 | 0.003081 | 4.583 | +1 | 5 |
| - | - | 1.1E+04 | 673.4 | - | - | 0 | - |
| - | - | 3010 | 674.4 | - | - | 0 | - |
| - | - | 3283 | 675.3 | - | - | 0 | - |
| 6 | y | 4545 | 675.8 | 0.0003542 | 0.5241 | +2 | 10 |
| - | - | 4719 | 676.3 | - | - | 0 | - |
| - | - | 1354 | 676.8 | - | - | 0 | - |
| - | - | 2265 | 680.8 | - | - | 0 | - |
| - | - | 2294 | 681.3 | - | - | 0 | - |
| - | - | 1109 | 681.8 | - | - | 0 | - |
| - | - | 1257 | 686.9 | - | - | 0 | - |
| - | - | 3050 | 687.3 | - | - | 0 | - |
| - | - | 2299 | 687.9 | - | - | 0 | - |
| - | - | 797.8 | 688.3 | - | - | 0 | - |
| 11 | y | 8204 | 688.4 | 0.003678 | 5.343 | +1 | 5 |
| - | - | 1939 | 689.4 | - | - | 0 | - |
| - | - | 1911 | 695.4 | - | - | 0 | - |
| - | - | 3442 | 699.9 | - | - | 0 | - |
| 12 | c | 2510 | 700.4 | 0.00478 | 6.825 | +2 | 12 |
| 12 | c | 1478 | 700.8 | 0.007241 | 10.33 | +2 | 12 |
| 5 | z | 1996 | 702.3 | 0.01394 | 19.85 | +2 | 11 |
| - | - | 1.089E+04 | 707.3 | - | - | 0 | - |
| - | - | 6533 | 707.8 | - | - | 0 | - |
| - | - | 4102 | 708.3 | - | - | 0 | - |
| - | - | 1.223E+04 | 708.9 | - | - | 0 | - |
| 12 | c | 9.491E+04 | 709.4 | 0.002938 | 4.142 | +2 | 12 |
| - | - | 7.149E+04 | 709.9 | - | - | 0 | - |
| 5 | y | 3.819E+04 | 710.3 | 0.01269 | 17.87 | +2 | 11 |
| - | - | 1.192E+04 | 710.9 | - | - | 0 | - |
| 5 | z | 3650 | 711.3 | 0.01122 | 15.77 | +2 | 11 |
| - | - | 6974 | 711.8 | - | - | 0 | - |
| - | - | 5131 | 712.4 | - | - | 0 | - |
| - | - | 3502 | 712.9 | - | - | 0 | - |
| - | - | 1161 | 713.4 | - | - | 0 | - |
| - | - | 1121 | 717.4 | - | - | 0 | - |
| 5 | y | 6666 | 719.4 | 0.001552 | 2.157 | +2 | 11 |
| - | - | 2586 | 719.9 | - | - | 0 | - |
| - | - | 2492 | 720.4 | - | - | 0 | - |
| - | - | 917.9 | 723.4 | - | - | 0 | - |
| - | - | 2513 | 729.4 | - | - | 0 | - |
| - | - | 3350 | 737.9 | - | - | 0 | - |
| - | - | 3076 | 738.4 | - | - | 0 | - |
| - | - | 2994 | 738.9 | - | - | 0 | - |
| - | - | 970.4 | 739.4 | - | - | 0 | - |
| 4 | w | 1129 | 746.4 | 0.002068 | 2.771 | +2 | 12 |
| - | - | 1184 | 746.9 | - | - | 0 | - |
| - | - | 1529 | 747.4 | - | - | 0 | - |
| - | - | 864.8 | 748.9 | - | - | 0 | - |
| - | - | 934.2 | 749.9 | - | - | 0 | - |
| 4 | z | 5325 | 750.9 | 0.005059 | 6.737 | +2 | 12 |
| 4 | z | 3808 | 751.4 | 0.004215 | 5.61 | +2 | 12 |
| - | - | 3910 | 751.9 | - | - | 0 | - |
| - | - | 1021 | 752.4 | - | - | 0 | - |
| - | - | 1378 | 754.4 | - | - | 0 | - |
| - | - | 1608 | 755.4 | - | - | 0 | - |
| - | - | 2490 | 755.9 | - | - | 0 | - |
| 10 | w | 5973 | 756.4 | 0.004502 | 5.952 | +1 | 6 |
| - | - | 1316 | 756.9 | - | - | 0 | - |
| - | - | 2266 | 757.4 | - | - | 0 | - |
| 10 | w | 9541 | 758.4 | 0.003054 | 4.026 | +1 | 6 |
| - | - | 4389 | 759.4 | - | - | 0 | - |
| 4 | z | 988 | 759.9 | 0.002345 | 3.087 | +2 | 12 |
| - | - | 805.8 | 760.4 | - | - | 0 | - |
| - | - | 1229 | 760.9 | - | - | 0 | - |
| 13 | c | 1272 | 764.4 | 0.007802 | 10.21 | +2 | 13 |
| - | - | 946.8 | 765.4 | - | - | 0 | - |
| - | - | 1519 | 765.9 | - | - | 0 | - |
| - | - | 1057 | 766.4 | - | - | 0 | - |
| - | - | 871.5 | 766.9 | - | - | 0 | - |
| - | - | 1225 | 767.4 | - | - | 0 | - |
| 4 | y | 3.599E+04 | 767.9 | 0.001537 | 2.002 | +2 | 12 |
| - | - | 2.663E+04 | 768.4 | - | - | 0 | - |
| - | - | 1.681E+04 | 768.9 | - | - | 0 | - |
| - | - | 5593 | 769.4 | - | - | 0 | - |
| - | - | 1001 | 769.9 | - | - | 0 | - |
| - | - | 3340 | 771.4 | - | - | 0 | - |
| 10 | y | 7910 | 772.4 | 0.003994 | 5.171 | +1 | 6 |
| - | - | 3590 | 772.9 | - | - | 0 | - |
| 13 | c | 4.326E+04 | 773.4 | 0.004716 | 6.098 | +2 | 13 |
| - | - | 4.558E+04 | 773.9 | - | - | 0 | - |
| - | - | 1.99E+04 | 774.4 | - | - | 0 | - |
| - | - | 1.103E+04 | 774.4 | - | - | 0 | - |
| - | - | 9927 | 774.9 | - | - | 0 | - |
| - | - | 5600 | 775.4 | - | - | 0 | - |
| - | - | 1498 | 776.4 | - | - | 0 | - |
| - | - | 1332 | 779.4 | - | - | 0 | - |
| - | - | 952.2 | 779.9 | - | - | 0 | - |
| - | - | 1309 | 782.4 | - | - | 0 | - |
| - | - | 4483 | 784.4 | - | - | 0 | - |
| - | - | 4242 | 784.9 | - | - | 0 | - |
| - | - | 2199 | 785.4 | - | - | 0 | - |
| - | - | 894 | 785.9 | - | - | 0 | - |
| - | - | 1192 | 787.4 | - | - | 0 | - |
| - | - | 962.3 | 787.9 | - | - | 0 | - |
| - | - | 1677 | 788.4 | - | - | 0 | - |
| 10 | y | 9567 | 789.5 | 0.004664 | 5.909 | +1 | 6 |
| - | - | 1218 | 789.9 | - | - | 0 | - |
| - | - | 4174 | 790.4 | - | - | 0 | - |
| - | - | 1026 | 790.9 | - | - | 0 | - |
| - | - | 1168 | 791.9 | - | - | 0 | - |
| - | - | 1278 | 792.4 | - | - | 0 | - |
| - | - | 2170 | 792.9 | - | - | 0 | - |
| - | - | 3634 | 793.4 | - | - | 0 | - |
| - | - | 2297 | 793.9 | - | - | 0 | - |
| - | - | 1949 | 794.4 | - | - | 0 | - |
| 3 | w | 2947 | 794.9 | 0.004434 | 5.578 | +2 | 13 |
| - | - | 5534 | 795.4 | - | - | 0 | - |
| - | - | 2790 | 795.9 | - | - | 0 | - |
| - | - | 1940 | 797.4 | - | - | 0 | - |
| 7 | c | 1.219E+05 | 798.4 | 0.001353 | 1.694 | +1 | 7 |
| - | - | 4.673E+04 | 799.5 | - | - | 0 | - |
| - | - | 1.127E+04 | 800.5 | - | - | 0 | - |
| - | - | 3468 | 800.9 | - | - | 0 | - |
| - | - | 2776 | 801.4 | - | - | 0 | - |
| - | - | 3654 | 801.9 | - | - | 0 | - |
| - | - | 2140 | 802.4 | - | - | 0 | - |
| - | - | 957.9 | 803.4 | - | - | 0 | - |
| - | - | 3712 | 803.9 | - | - | 0 | - |
| - | - | 2906 | 804.4 | - | - | 0 | - |
| - | - | 1164 | 804.9 | - | - | 0 | - |
| - | - | 3028 | 805.4 | - | - | 0 | - |
| - | - | 1024 | 805.9 | - | - | 0 | - |
| 3 | y | 3400 | 807.4 | 0.00241 | 2.985 | +2 | 13 |
| 3 | y | 3403 | 807.9 | 0.006313 | 7.814 | +2 | 13 |
| 3 | z | 3564 | 808.4 | 0.002034 | 2.517 | +2 | 13 |
| - | - | 1160 | 809.4 | - | - | 0 | - |
| - | - | 7156 | 811.9 | - | - | 0 | - |
| - | - | 8114 | 812.4 | - | - | 0 | - |
| - | - | 5537 | 812.9 | - | - | 0 | - |
| - | - | 6071 | 813.4 | - | - | 0 | - |
| - | - | 5031 | 813.9 | - | - | 0 | - |
| - | - | 4655 | 814.4 | - | - | 0 | - |
| - | - | 3475 | 814.9 | - | - | 0 | - |
| - | - | 2736 | 815.4 | - | - | 0 | - |
| - | - | 2389 | 815.9 | - | - | 0 | - |
| 3 | y | 1.287E+05 | 816.4 | 0.001461 | 1.79 | +2 | 13 |
| - | - | 1.154E+05 | 816.9 | - | - | 0 | - |
| - | - | 6.384E+04 | 817.4 | - | - | 0 | - |
| - | - | 2.392E+04 | 817.9 | - | - | 0 | - |
| - | - | 6034 | 818.4 | - | - | 0 | - |
| - | - | 1.176E+04 | 822.4 | - | - | 0 | - |
| - | - | 5.816E+04 | 822.9 | - | - | 0 | - |
| - | - | 4.91E+04 | 823.4 | - | - | 0 | - |
| - | - | 2.816E+04 | 823.9 | - | - | 0 | - |
| - | - | 7855 | 824.5 | - | - | 0 | - |
| - | - | 3334 | 824.9 | - | - | 0 | - |
| - | - | 3950 | 825.5 | - | - | 0 | - |
| - | - | 1016 | 826.5 | - | - | 0 | - |
| - | - | 2071 | 828.9 | - | - | 0 | - |
| - | - | 1414 | 829.4 | - | - | 0 | - |
| - | - | 1143 | 829.9 | - | - | 0 | - |
| - | - | 2158 | 830.4 | - | - | 0 | - |
| - | - | 3328 | 830.9 | - | - | 0 | - |
| - | - | 4946 | 835.4 | - | - | 0 | - |
| - | - | 5203 | 835.9 | - | - | 0 | - |
| - | - | 5143 | 836.4 | - | - | 0 | - |
| - | - | 4487 | 836.9 | - | - | 0 | - |
| - | - | 2947 | 837.4 | - | - | 0 | - |
| - | - | 2091 | 837.9 | - | - | 0 | - |
| - | - | 910.2 | 838.4 | - | - | 0 | - |
| - | - | 1589 | 840.5 | - | - | 0 | - |
| 2 | w | 4.98E+04 | 843.4 | 0.001367 | 1.621 | +2 | 14 |
| - | - | 4.513E+04 | 843.9 | - | - | 0 | - |
| - | - | 2.705E+04 | 844.4 | - | - | 0 | - |
| - | - | 9153 | 844.9 | - | - | 0 | - |
| - | - | 3803 | 845.4 | - | - | 0 | - |
| - | - | 1839 | 849.4 | - | - | 0 | - |
| - | - | 1115 | 850.4 | - | - | 0 | - |
| - | - | 1575 | 851.4 | - | - | 0 | - |
| - | - | 2427 | 856 | - | - | 0 | - |
| - | - | 4435 | 856.5 | - | - | 0 | - |
| - | - | 2194 | 857 | - | - | 0 | - |
| - | - | 1.489E+04 | 857.5 | - | - | 0 | - |
| 14 | c | 1935 | 857.9 | 0.009577 | 11.16 | +2 | 14 |
| - | - | 4.439E+04 | 858.5 | - | - | 0 | - |
| - | - | 1972 | 858.9 | - | - | 0 | - |
| - | - | 1.987E+04 | 859.5 | - | - | 0 | - |
| - | - | 1200 | 859.9 | - | - | 0 | - |
| - | - | 5127 | 860.5 | - | - | 0 | - |
| - | - | 1117 | 863.5 | - | - | 0 | - |
| 2 | y | 1191 | 863.9 | 0.00591 | 6.841 | +2 | 14 |
| 2 | z | 1.955E+04 | 864.9 | 0.001179 | 1.363 | +2 | 14 |
| - | - | 1.881E+04 | 865.4 | - | - | 0 | - |
| - | - | 1.327E+04 | 865.9 | - | - | 0 | - |
| 14 | c | 1.261E+05 | 866.4 | 0.001551 | 1.79 | +2 | 14 |
| - | - | 1.225E+05 | 866.9 | - | - | 0 | - |
| - | - | 7.532E+04 | 867.4 | - | - | 0 | - |
| - | - | 3.024E+04 | 867.9 | - | - | 0 | - |
| - | - | 7133 | 868.4 | - | - | 0 | - |
| - | - | 2102 | 870 | - | - | 0 | - |
| - | - | 2660 | 870.5 | - | - | 0 | - |
| - | - | 1147 | 871.5 | - | - | 0 | - |
| - | - | 3834 | 872 | - | - | 0 | - |
| - | - | 4225 | 872.5 | - | - | 0 | - |
| 2 | y | 4307 | 872.9 | 0.00661 | 7.572 | +2 | 14 |
| - | - | 2503 | 873.4 | - | - | 0 | - |
| - | - | 2972 | 873.9 | - | - | 0 | - |
| - | - | 2830 | 874.4 | - | - | 0 | - |
| - | - | 3251 | 878 | - | - | 0 | - |
| - | - | 3476 | 878.5 | - | - | 0 | - |
| - | - | 3472 | 878.9 | - | - | 0 | - |
| - | - | 5820 | 879.4 | - | - | 0 | - |
| - | - | 4421 | 879.9 | - | - | 0 | - |
| - | - | 3422 | 880.4 | - | - | 0 | - |
| - | - | 1260 | 880.9 | - | - | 0 | - |
| - | - | 4388 | 883.5 | - | - | 0 | - |
| - | - | 6135 | 884.5 | - | - | 0 | - |
| - | - | 2878 | 885 | - | - | 0 | - |
| - | - | 2973 | 885.5 | - | - | 0 | - |
| - | - | 3845 | 886 | - | - | 0 | - |
| - | - | 4674 | 886.5 | - | - | 0 | - |
| - | - | 4635 | 886.9 | - | - | 0 | - |
| - | - | 3.502E+04 | 887.4 | - | - | 0 | - |
| - | - | 3.796E+04 | 887.9 | - | - | 0 | - |
| - | - | 2.492E+04 | 888.4 | - | - | 0 | - |
| - | - | 1.09E+04 | 888.9 | - | - | 0 | - |
| - | - | 4761 | 889.5 | - | - | 0 | - |
| - | - | 3324 | 892 | - | - | 0 | - |
| - | - | 1.011E+04 | 892.5 | - | - | 0 | - |
| - | - | 1.196E+04 | 893 | - | - | 0 | - |
| - | - | 1.131E+04 | 893.5 | - | - | 0 | - |
| - | - | 9792 | 894 | - | - | 0 | - |
| - | - | 2.157E+04 | 894.5 | - | - | 0 | - |
| - | - | 1.815E+04 | 895 | - | - | 0 | - |
| - | - | 9130 | 895.5 | - | - | 0 | - |
| - | - | 1.312E+04 | 896 | - | - | 0 | - |
| - | - | 1.242E+04 | 896.4 | - | - | 0 | - |
| - | - | 6796 | 896.9 | - | - | 0 | - |
| - | - | 3566 | 897.4 | - | - | 0 | - |
| - | - | 1018 | 900.5 | - | - | 0 | - |
| - | - | 1555 | 901 | - | - | 0 | - |
| - | - | 2.56E+04 | 901.5 | - | - | 0 | - |
| - | - | 2.477E+04 | 902 | - | - | 0 | - |
| 9 | z | 2.476E+04 | 902.5 | 0.01257 | 13.93 | +1 | 7 |
| - | - | 1.532E+04 | 903 | - | - | 0 | - |
| - | - | 9264 | 903.5 | - | - | 0 | - |
| - | - | 8695 | 906 | - | - | 0 | - |
| - | - | 2.398E+04 | 906.5 | - | - | 0 | - |
| - | - | 1.985E+04 | 907 | - | - | 0 | - |
| - | - | 1.418E+04 | 907.5 | - | - | 0 | - |
| - | - | 6875 | 908 | - | - | 0 | - |
| - | - | 3334 | 908.5 | - | - | 0 | - |
| - | - | 3.103E+04 | 915 | - | - | 0 | - |
| - | - | 1.252E+05 | 915.5 | - | - | 0 | - |
| - | - | 1.17E+05 | 916 | - | - | 0 | - |
| - | - | 7.804E+04 | 916.5 | - | - | 0 | - |
| - | - | 3.37E+04 | 917 | - | - | 0 | - |
| - | - | 1.218E+04 | 917.5 | - | - | 0 | - |
| 9 | z | 1.289E+04 | 920.5 | 0.001702 | 1.85 | +1 | 7 |
| - | - | 9228 | 921.5 | - | - | 0 | - |
| - | - | 3746 | 922.5 | - | - | 0 | - |
| - | - | 8.404E+04 | 923.5 | - | - | 0 | - |
| - | - | 1.632E+05 | 924 | - | - | 0 | - |
| - | - | 1.338E+05 | 924.5 | - | - | 0 | - |
| - | - | 7.693E+04 | 925 | - | - | 0 | - |
| - | - | 3.191E+04 | 925.5 | - | - | 0 | - |
| - | - | 7928 | 926 | - | - | 0 | - |
| - | - | 1634 | 926.5 | - | - | 0 | - |
| 8 | c | 9.509E+04 | 927.5 | 0.0009961 | 1.074 | +1 | 8 |
| - | - | 4.423E+04 | 928.5 | - | - | 0 | - |
| - | - | 1.365E+04 | 929.5 | - | - | 0 | - |
| - | - | 2588 | 930.5 | - | - | 0 | - |
| 9 | y | 1.389E+04 | 936.5 | 0.0006785 | 0.7246 | +1 | 7 |
| - | - | 7732 | 937.5 | - | - | 0 | - |
| - | - | 2487 | 938.5 | - | - | 0 | - |
| - | - | 1474 | 953.5 | - | - | 0 | - |
| - | - | 2810 | 968.5 | - | - | 0 | - |
| - | - | 2088 | 969.5 | - | - | 0 | - |
| - | - | 2164 | 978.5 | - | - | 0 | - |
| - | - | 1042 | 979.5 | - | - | 0 | - |
| - | - | 3516 | 986.5 | - | - | 0 | - |
| - | - | 3236 | 987.5 | - | - | 0 | - |
| - | - | 3445 | 988.5 | - | - | 0 | - |
| - | - | 1884 | 989.5 | - | - | 0 | - |
| 8 | w | 1.959E+04 | 990.5 | 0.002687 | 2.713 | +1 | 8 |
| - | - | 1.013E+04 | 991.5 | - | - | 0 | - |
| - | - | 5286 | 992.5 | - | - | 0 | - |
| - | - | 1491 | 993.5 | - | - | 0 | - |
| - | - | 1240 | 994.4 | - | - | 0 | - |
| - | - | 1421 | 1013 | - | - | 0 | - |
| - | - | 8940 | 1031 | - | - | 0 | - |
| - | - | 6453 | 1032 | - | - | 0 | - |
| - | - | 3920 | 1033 | - | - | 0 | - |
| - | - | 1181 | 1034 | - | - | 0 | - |
| - | - | 1446 | 1046 | - | - | 0 | - |
| 8 | z | 5.836E+04 | 1050 | 0.001712 | 1.631 | +1 | 8 |
| - | - | 3.949E+04 | 1051 | - | - | 0 | - |
| - | - | 1.711E+04 | 1052 | - | - | 0 | - |
| - | - | 4743 | 1053 | - | - | 0 | - |
| - | - | 1751 | 1054 | - | - | 0 | - |
| 8 | y | 3559 | 1066 | 0.001054 | 0.9894 | +1 | 8 |
| - | - | 2275 | 1067 | - | - | 0 | - |
| - | - | 1876 | 1068 | - | - | 0 | - |
| - | - | 1568 | 1074 | - | - | 0 | - |
| 9 | c | 7.645E+04 | 1075 | 0.006095 | 5.672 | +1 | 9 |
| - | - | 4.312E+04 | 1076 | - | - | 0 | - |
| - | - | 1.979E+04 | 1077 | - | - | 0 | - |
| - | - | 6035 | 1078 | - | - | 0 | - |
| - | - | 1033 | 1108 | - | - | 0 | - |
| - | - | 2343 | 1114 | - | - | 0 | - |
| - | - | 1503 | 1115 | - | - | 0 | - |
| 7 | w | 8508 | 1120 | 0.0004993 | 0.446 | +1 | 9 |
| - | - | 4767 | 1121 | - | - | 0 | - |
| - | - | 2316 | 1122 | - | - | 0 | - |
| - | - | 2129 | 1122 | - | - | 0 | - |
| - | - | 2277 | 1132 | - | - | 0 | - |
| - | - | 1701 | 1133 | - | - | 0 | - |
| - | - | 2151 | 1134 | - | - | 0 | - |
| - | - | 2134 | 1135 | - | - | 0 | - |
| - | - | 1622 | 1145 | - | - | 0 | - |
| 10 | c | 7.317E+04 | 1176 | 0.00578 | 4.917 | +1 | 10 |
| 7 | y | 4.399E+04 | 1177 | 0.01981 | 16.83 | +1 | 9 |
| - | - | 1.886E+04 | 1178 | - | - | 0 | - |
| 7 | z | 5.106E+04 | 1179 | 0.003552 | 3.014 | +1 | 9 |
| - | - | 3.047E+04 | 1180 | - | - | 0 | - |
| - | - | 1.434E+04 | 1181 | - | - | 0 | - |
| - | - | 4510 | 1182 | - | - | 0 | - |
| - | - | 1416 | 1194 | - | - | 0 | - |
| 7 | y | 6061 | 1195 | 8.707E-05 | 0.07289 | +1 | 9 |
| - | - | 4356 | 1196 | - | - | 0 | - |
| - | - | 1668 | 1197 | - | - | 0 | - |
| - | - | 1968 | 1221 | - | - | 0 | - |
| - | - | 1845 | 1222 | - | - | 0 | - |
| - | - | 1303 | 1223 | - | - | 0 | - |
| - | - | 1138 | 1224 | - | - | 0 | - |
| - | - | 1373 | 1232 | - | - | 0 | - |
| - | - | 1376 | 1232 | - | - | 0 | - |
| - | - | 1061 | 1233 | - | - | 0 | - |
| - | - | 1397 | 1233 | - | - | 0 | - |
| - | - | 4865 | 1260 | - | - | 0 | - |
| - | - | 3566 | 1261 | - | - | 0 | - |
| - | - | 2474 | 1262 | - | - | 0 | - |
| - | - | 1213 | 1276 | - | - | 0 | - |
| - | - | 1333 | 1278 | - | - | 0 | - |
| - | - | 1976 | 1279 | - | - | 0 | - |
| - | - | 2644 | 1285 | - | - | 0 | - |
| 11 | c | 1491 | 1287 | 0.01415 | 11 | +1 | 11 |
| - | - | 8102 | 1303 | - | - | 0 | - |
| 11 | c | 1.98E+04 | 1304 | 0.003346 | 2.567 | +1 | 11 |
| - | - | 1.311E+04 | 1305 | - | - | 0 | - |
| - | - | 6118 | 1306 | - | - | 0 | - |
| - | - | 1707 | 1319 | - | - | 0 | - |
| - | - | 2038 | 1320 | - | - | 0 | - |
| - | - | 1988 | 1321 | - | - | 0 | - |
| 6 | y | 1794 | 1334 | 0.007312 | 5.483 | +1 | 10 |
| 6 | z | 3.125E+04 | 1335 | 0.0005859 | 0.439 | +1 | 10 |
| - | - | 4.422E+04 | 1336 | - | - | 0 | - |
| - | - | 2.712E+04 | 1337 | - | - | 0 | - |
| - | - | 1.123E+04 | 1338 | - | - | 0 | - |
| - | - | 3746 | 1339 | - | - | 0 | - |
| - | - | 5361 | 1350 | - | - | 0 | - |
| 6 | y | 1.335E+04 | 1351 | 0.001048 | 0.7762 | +1 | 10 |
| - | - | 8740 | 1352 | - | - | 0 | - |
| - | - | 3601 | 1353 | - | - | 0 | - |
| - | - | 1430 | 1354 | - | - | 0 | - |
| - | - | 1310 | 1374 | - | - | 0 | - |
| - | - | 2287 | 1375 | - | - | 0 | - |
| - | - | 1817 | 1376 | - | - | 0 | - |
| - | - | 1036 | 1377 | - | - | 0 | - |
| - | - | 1187 | 1402 | - | - | 0 | - |
| 5 | z | 1360 | 1404 | 0.01367 | 9.737 | +1 | 11 |
| - | - | 3012 | 1417 | - | - | 0 | - |
| 12 | c | 3.823E+04 | 1418 | 0.004364 | 3.078 | +1 | 12 |
| - | - | 2.951E+04 | 1419 | - | - | 0 | - |
| 5 | y | 1.511E+04 | 1420 | 0.02326 | 16.39 | +1 | 11 |
| - | - | 5635 | 1421 | - | - | 0 | - |
| 5 | z | 3906 | 1422 | 0.008719 | 6.133 | +1 | 11 |
| - | - | 1.734E+04 | 1423 | - | - | 0 | - |
| - | - | 1.327E+04 | 1424 | - | - | 0 | - |
| - | - | 7699 | 1425 | - | - | 0 | - |
| - | - | 3043 | 1426 | - | - | 0 | - |
| 5 | y | 3545 | 1438 | 0.002446 | 1.701 | +1 | 11 |
| - | - | 2444 | 1439 | - | - | 0 | - |
| - | - | 2302 | 1440 | - | - | 0 | - |
| - | - | 1259 | 1476 | - | - | 0 | - |
| - | - | 1283 | 1477 | - | - | 0 | - |
| - | - | 1501 | 1503 | - | - | 0 | - |
| - | - | 1219 | 1504 | - | - | 0 | - |
| 13 | c | 1278 | 1528 | 0.01873 | 12.26 | +1 | 13 |
| - | - | 1338 | 1530 | - | - | 0 | - |
| 4 | y | 1.05E+04 | 1535 | 9.687E-05 | 0.06312 | +1 | 12 |
| - | - | 6847 | 1536 | - | - | 0 | - |
| - | - | 5564 | 1537 | - | - | 0 | - |
| - | - | 1852 | 1538 | - | - | 0 | - |
| 13 | c | 1.963E+04 | 1546 | 0.004014 | 2.597 | +1 | 13 |
| - | - | 1.982E+04 | 1547 | - | - | 0 | - |
| - | - | 1.093E+04 | 1548 | - | - | 0 | - |
| - | - | 6087 | 1549 | - | - | 0 | - |
| - | - | 1778 | 1550 | - | - | 0 | - |
| - | - | 1494 | 1566 | - | - | 0 | - |
| - | - | 2532 | 1602 | - | - | 0 | - |
| - | - | 1717 | 1603 | - | - | 0 | - |
| - | - | 3045 | 1608 | - | - | 0 | - |
| - | - | 1852 | 1609 | - | - | 0 | - |
| - | - | 1655 | 1610 | - | - | 0 | - |
| 3 | y | 4070 | 1615 | 0.02442 | 15.12 | +1 | 13 |
| 3 | z | 4251 | 1616 | 0.01684 | 10.42 | +1 | 13 |
| - | - | 2291 | 1617 | - | - | 0 | - |
| - | - | 1543 | 1624 | - | - | 0 | - |
| - | - | 5078 | 1625 | - | - | 0 | - |
| - | - | 3814 | 1626 | - | - | 0 | - |
| - | - | 4478 | 1627 | - | - | 0 | - |
| - | - | 1949 | 1628 | - | - | 0 | - |
| - | - | 2486 | 1629 | - | - | 0 | - |
| - | - | 2263 | 1630 | - | - | 0 | - |
| - | - | 1611 | 1631 | - | - | 0 | - |
| 3 | y | 2085 | 1632 | 0.007392 | 4.53 | +1 | 13 |
| - | - | 2450 | 1633 | - | - | 0 | - |
| - | - | 1343 | 1634 | - | - | 0 | - |
| - | - | 1.712E+04 | 1645 | - | - | 0 | - |
| - | - | 2.278E+04 | 1646 | - | - | 0 | - |
| - | - | 1.676E+04 | 1647 | - | - | 0 | - |
| - | - | 8844 | 1648 | - | - | 0 | - |
| - | - | 3919 | 1649 | - | - | 0 | - |
| - | - | 2869 | 1671 | - | - | 0 | - |
| - | - | 1968 | 1672 | - | - | 0 | - |
| - | - | 1587 | 1674 | - | - | 0 | - |
| - | - | 1292 | 1675 | - | - | 0 | - |
| - | - | 2045 | 1685 | - | - | 0 | - |
| - | - | 2228 | 1686 | - | - | 0 | - |
| - | - | 2635 | 1688 | - | - | 0 | - |
| - | - | 7114 | 1689 | - | - | 0 | - |
| - | - | 5132 | 1690 | - | - | 0 | - |
| - | - | 2593 | 1691 | - | - | 0 | - |
| - | - | 3759 | 1716 | - | - | 0 | - |
| - | - | 3401 | 1717 | - | - | 0 | - |
| - | - | 2382 | 1718 | - | - | 0 | - |
| 2 | z | 6385 | 1729 | 0.001064 | 0.6152 | +1 | 14 |
| - | - | 3.61E+04 | 1730 | - | - | 0 | - |
| - | - | 3.209E+04 | 1731 | - | - | 0 | - |
| 14 | c | 2.951E+04 | 1732 | 0.00513 | 2.962 | +1 | 14 |
| - | - | 3.913E+04 | 1733 | - | - | 0 | - |
| - | - | 2.828E+04 | 1734 | - | - | 0 | - |
| - | - | 1.62E+04 | 1735 | - | - | 0 | - |
| - | - | 5527 | 1736 | - | - | 0 | - |
| - | - | 1361 | 1748 | - | - | 0 | - |
| - | - | 1311 | 1749 | - | - | 0 | - |
| - | - | 1841 | 1770 | - | - | 0 | - |
| - | - | 3202 | 1771 | - | - | 0 | - |
| - | - | 2238 | 1772 | - | - | 0 | - |
| - | - | 1508 | 1773 | - | - | 0 | - |
| - | - | 2772 | 1784 | - | - | 0 | - |
| - | - | 3696 | 1785 | - | - | 0 | - |
| - | - | 7388 | 1786 | - | - | 0 | - |
| - | - | 6517 | 1787 | - | - | 0 | - |
| - | - | 7520 | 1788 | - | - | 0 | - |
| - | - | 1.227E+04 | 1789 | - | - | 0 | - |
| - | - | 1.167E+04 | 1790 | - | - | 0 | - |
| - | - | 6602 | 1791 | - | - | 0 | - |
| - | - | 2825 | 1792 | - | - | 0 | - |
| - | - | 1026 | 1800 | - | - | 0 | - |
| - | - | 6562 | 1802 | - | - | 0 | - |
| - | - | 2.455E+04 | 1803 | - | - | 0 | - |
| - | - | 2.333E+04 | 1804 | - | - | 0 | - |
| - | - | 1.545E+04 | 1805 | - | - | 0 | - |
| - | - | 6039 | 1806 | - | - | 0 | - |
| - | - | 1950 | 1807 | - | - | 0 | - |
| - | - | 1428 | 1812 | - | - | 0 | - |
| - | - | 2942 | 1813 | - | - | 0 | - |
| - | - | 5898 | 1814 | - | - | 0 | - |
| - | - | 5966 | 1815 | - | - | 0 | - |
| - | - | 2402 | 1816 | - | - | 0 | - |
| - | - | 1726 | 1817 | - | - | 0 | - |
| - | - | 1174 | 1818 | - | - | 0 | - |
| - | - | 2633 | 1819 | - | - | 0 | - |
| - | - | 1.577E+04 | 1820 | - | - | 0 | - |
| - | - | 1.478E+04 | 1821 | - | - | 0 | - |
| - | - | 7205 | 1822 | - | - | 0 | - |
| - | - | 3746 | 1823 | - | - | 0 | - |
| - | - | 2114 | 1829 | - | - | 0 | - |
| - | - | 2.138E+04 | 1830 | - | - | 0 | - |
| - | - | 7.431E+04 | 1831 | - | - | 0 | - |
| - | - | 6.662E+04 | 1832 | - | - | 0 | - |
| - | - | 4.15E+04 | 1833 | - | - | 0 | - |
| - | - | 1.697E+04 | 1834 | - | - | 0 | - |
| - | - | 5436 | 1835 | - | - | 0 | - |
| - | - | 7314 | 1846 | - | - | 0 | - |
| - | - | 4.437E+04 | 1847 | - | - | 0 | - |
| - | - | 1.917E+05 | 1848 | - | - | 0 | - |
| - | - | 1.896E+05 | 1849 | - | - | 0 | - |
| - | - | 1.123E+05 | 1850 | - | - | 0 | - |
| - | - | 4.768E+04 | 1851 | - | - | 0 | - |
| - | - | 1.472E+04 | 1852 | - | - | 0 | - |
| - | - | 1141 | 3085 | - | - | 0 | - |

m/z Charge Intensity FragmentType MassShift Position
132.10218811035156 0 1781.4822 y 14
143.1177520751953 0 736.0863
143.6822967529297 0 537.4266
148.95401000976562 0 983.7532
159.38150024414062 0 530.71796
162.653564453125 0 660.8836
173.1287078857422 0 663.47675
173.45162963867188 0 2628.7366
187.14450073242188 0 60469.785
188.14791870117188 0 5640.5107
201.12303161621094 0 1180.2795
212.13983154296875 0 4080.3135
215.13941955566406 0 38112.414
216.1427459716797 0 3449.4573
219.1344451904297 0 4971.575
220.13795471191406 0 700.11957
230.15045166015625 0 861.85236
256.1663513183594 0 1452.1493
292.6524658203125 0 726.4487
299.171875 0 8689.505
300.17401123046875 0 859.73114
312.1929931640625 0 1509.3821
313.1976318359375 0 754.84845
318.20379638671875 0 1194.6602 y 13
326.1584167480469 0 1995.1587
327.16595458984375 0 2143.303
344.1936340332031 0 1544.0238
369.0833740234375 0 655.1532
382.2578125 0 1569.886
395.1816711425781 0 848.47076
396.1886291503906 0 1755.4772
407.2530517578125 0 2306.9678
408.2564392089844 0 925.03815 c Water loss 3
412.2333068847656 0 1163.3218
414.1988525390625 0 1594.5215
425.23199462890625 0 794.3753
425.2637939453125 0 3502.209
426.2713317871094 0 2427.051 c 3
427.2685852050781 0 1788.9749
428.2730712890625 0 1283.4548
430.09100341796875 0 764.37756
430.24249267578125 0 1568.6768 z 12
431.25091552734375 0 998.28455
446.26165771484375 0 4213.6157 y 12
447.2650146484375 0 1018.11066
462.903076171875 0 1387.4865
466.22528076171875 0 1081.9106
466.5586242675781 0 1695.18
470.29754638671875 0 4585.235
471.3021240234375 0 1211.4911
471.89990234375 0 2779.1072
472.2318115234375 0 1343.9811
472.2915344238281 0 832.19275
472.5663757324219 0 2120.052
491.5782470703125 0 1593.2777
491.9105224609375 0 724.99384
494.28643798828125 0 2753.4746
494.9050598144531 0 1106.5886
495.23712158203125 0 2819.3525
495.57275390625 0 858.3784
500.90911865234375 0 3013.4756 z Water loss 3
501.2432556152344 0 3136.2026 z Ammonia loss 3
501.57427978515625 0 1255.986
512.2959594726562 0 1349.2383
513.3038940429688 0 36137.01 c 4
514.30615234375 0 9497.466
515.3086547851562 0 2694.6282
523.2741088867188 0 4198.022
523.6090087890625 0 3479.8608
523.9396362304688 0 2034.1918
524.2737426757812 0 1435.3495
528.3153686523438 0 942.58344
538.605224609375 0 2336.9192 y Water loss 2
538.9365844726562 0 3357.2966 y Ammonia loss 2
539.27001953125 0 1193.5798 z 2
539.6015014648438 0 2137.728
539.9385375976562 0 1616.3258
540.2669067382812 0 1111.8071
541.610595703125 0 884.3651
541.9435424804688 0 1091.6086
544.2865600585938 0 2935.2864 z 11
544.6077270507812 0 218591.28 y 2
544.9415893554688 0 192213.61
545.2756958007812 0 113228.39
545.6089477539062 0 43497.766
545.9424438476562 0 11820.642
546.2958984375 0 1998.9888
546.7721557617188 0 893.124
547.2775268554688 0 834.6401
552.7769775390625 0 1428.6963
557.9176635742188 0 1174.5088
560.3046264648438 0 8893.874 y 11
561.3060302734375 0 3279.8691
561.783935546875 0 2067.3813
562.28564453125 0 3182.2126
562.78759765625 0 1576.0184
571.2379150390625 0 1191.0636
588.2632446289062 0 942.80914
590.2913208007812 0 921.62415
590.7908325195312 0 1308.588
597.7954711914062 0 926.1004 y 6
598.2481689453125 0 886.7696
598.2999267578125 0 1252.4932
599.23193359375 0 1643.7318
607.382080078125 0 1172.4564
609.9783325195312 0 1125.3169
614.3508911132812 0 1364.8744
615.34326171875 0 2202.6301
615.8567504882812 0 811.8198
616.2584838867188 0 2874.0847
625.3909301757812 0 1442.9434
626.3985595703125 0 2745.4583
627.4031372070312 0 815.427
628.3670043945312 0 12904.686
629.3237915039062 0 5726.479
629.3717651367188 0 4371.901
630.325927734375 0 1539.893
631.3363037109375 0 1029.7368
637.3306884765625 0 1358.9484
642.8291625976562 0 2074.7913
643.3265991210938 0 1410.7959 c Water loss 10
643.8274536132812 0 2055.152 c Ammonia loss 10
647.33251953125 0 1857.4465
647.8323974609375 0 1788.2722
648.3237915039062 0 1164.0756
648.8389282226562 0 731.4878
651.8347778320312 0 4934.781
652.338134765625 0 31050.256 c 10
652.83935546875 0 20062.059
653.3397827148438 0 8917.666
653.842041015625 0 1238.5028
657.8104858398438 0 1421.1665
658.8157348632812 0 2073.7546 z Water loss 5
667.830810546875 0 2643.9753 z 5
668.330810546875 0 3029.7014
668.83056640625 0 1193.8954
669.4053955078125 0 78458.94 c 5
670.4081420898438 0 26389.877
671.4109497070312 0 7229.272
672.3815307617188 0 29819.648 z 10
673.3818969726562 0 10998.885
674.3856811523438 0 3010.375
675.3389892578125 0 3283.439
675.837890625 0 4545.0894 y 5
676.33935546875 0 4719.467
676.8389282226562 0 1353.6963
680.8331298828125 0 2264.5737
681.3363037109375 0 2294.3633
681.8369750976562 0 1108.6307
686.8524169921875 0 1256.9681
687.3488159179688 0 3049.6494
687.8548583984375 0 2299.4834
688.3453979492188 0 797.8075
688.399658203125 0 8203.7705 y 10
689.399658203125 0 1938.7432
695.42236328125 0 1910.6602
699.851806640625 0 3441.8271
700.3463134765625 0 2510.4592 c Water loss 11
700.850341796875 0 1478.2218 c Ammonia loss 11
702.3528442382812 0 1996.22 z Water loss 4
707.3436889648438 0 10887.616
707.8446044921875 0 6532.6133
708.3450927734375 0 4102.0737
708.854736328125 0 12227.087
709.3593139648438 0 94910.64 c 11
709.8606567382812 0 71488.99
710.3609619140625 0 38189.613 y Water loss 4
710.861083984375 0 11917.755
711.3554077148438 0 3649.686 z 4
711.8489379882812 0 6973.6504
712.3512573242188 0 5130.6626
712.8509521484375 0 3502.2979
713.35546875 0 1161.0881
717.3717651367188 0 1121.0643
719.3551025390625 0 6665.793 y 4
719.85498046875 0 2585.5894
720.357666015625 0 2492.0686
723.3607788085938 0 917.9475
729.4159545898438 0 2512.5088
737.8826904296875 0 3349.9727
738.3870239257812 0 3075.606
738.886474609375 0 2993.6587
739.3865356445312 0 970.4471
746.3609008789062 0 1128.8906 w 3
746.8637084960938 0 1183.9832
747.358642578125 0 1528.6056
748.8677368164062 0 864.75256
749.8541259765625 0 934.19403
750.8602294921875 0 5324.9424 z Water loss 3
751.3615112304688 0 3807.636 z Ammonia loss 3
751.8668212890625 0 3909.604
752.3751831054688 0 1020.9031
754.4337158203125 0 1377.8702
755.3655395507812 0 1608.0104
755.8709106445312 0 2490.032
756.425048828125 0 5972.544 w 9
756.8745727539062 0 1315.5668
757.4293823242188 0 2265.6724
758.40576171875 0 9540.772 w 9
759.4083251953125 0 4388.788
759.8682250976562 0 987.9747 z 3
760.3827514648438 0 805.815
760.8687744140625 0 1229.4175
764.38818359375 0 1271.7097 c Water loss 12
765.3980102539062 0 946.81036
765.891357421875 0 1518.9022
766.3936157226562 0 1057.2319
766.873046875 0 871.4757
767.379150390625 0 1225.1442
767.8814697265625 0 35993.695 y 3
768.3822631835938 0 26625.318
768.883056640625 0 16810.771
769.3836059570312 0 5593.046
769.8862915039062 0 1000.51434
771.413330078125 0 3339.8154
772.4204711914062 0 7909.6484 y Ammonia loss 9
772.88427734375 0 3589.6794
773.390380859375 0 43259.695 c 12
773.8904418945312 0 45582.637
774.3901977539062 0 19899.416
774.4293212890625 0 11030.795
774.8906860351562 0 9926.549
775.4306640625 0 5599.884
776.4317626953125 0 1498.1425
779.4257202148438 0 1331.5944
779.8795166015625 0 952.17883
782.4066162109375 0 1308.8878
784.4078979492188 0 4483.3296
784.9086303710938 0 4242.066
785.4091796875 0 2199.3506
785.9078979492188 0 893.9519
787.440673828125 0 1191.7983
787.8914794921875 0 962.3257
788.4468383789062 0 1676.9406
789.4463500976562 0 9567.404 y 9
789.8805541992188 0 1218.1302
790.4481201171875 0 4173.5474
790.895263671875 0 1025.7529
791.9110717773438 0 1168.4254
792.4053344726562 0 1278.2407
792.9136352539062 0 2170.1353
793.40966796875 0 3634.352
793.908935546875 0 2297.0388
794.39990234375 0 1948.8179
794.8896484375 0 2946.6157 w 2
795.387451171875 0 5534.235
795.8909301757812 0 2790.0796
797.4324340820312 0 1939.8473
798.4481811523438 0 121925.04 c 6
799.45068359375 0 46734.85
800.4522705078125 0 11266.632
800.9172973632812 0 3468.0933
801.4202270507812 0 2776.1765
801.9172973632812 0 3654.176
802.419189453125 0 2139.7104
803.41552734375 0 957.8767
803.9036254882812 0 3711.8909
804.4038696289062 0 2905.8494
804.9041137695312 0 1163.9137
805.4019165039062 0 3028.3872
805.9154052734375 0 1023.9374
807.4034423828125 0 3399.7378 y Water loss 2
807.8993530273438 0 3402.67 y Ammonia loss 2
808.3989868164062 0 3564.234 z 2
809.3983764648438 0 1159.8228
811.9116821289062 0 7156.4556
812.4132690429688 0 8113.9346
812.9134521484375 0 5536.82
813.419189453125 0 6071.2437
813.9122924804688 0 5031.317
814.4158325195312 0 4654.592
814.9157104492188 0 3474.817
815.4078369140625 0 2735.628
815.9116821289062 0 2389.196
816.4077758789062 0 128685.375 y 2
816.9083862304688 0 115356.87
817.408935546875 0 63837.125
817.9089965820312 0 23920.957
818.4113159179688 0 6033.5674
822.4193115234375 0 11756.562
822.9227905273438 0 58161.492
823.4236450195312 0 49095.27
823.9241333007812 0 28161.988
824.4534912109375 0 7855.0317
824.9267578125 0 3334.4587
825.4661865234375 0 3949.8572
826.4649658203125 0 1015.522
828.9290771484375 0 2070.8892
829.4347534179688 0 1414.4114
829.9215698242188 0 1142.8423
830.4447021484375 0 2157.5032
830.9070434570312 0 3328.1084
835.4284057617188 0 4946.3755
835.9285278320312 0 5203.137
836.4275512695312 0 5142.9673
836.9146118164062 0 4486.8276
837.4121704101562 0 2946.6462
837.9110107421875 0 2091.2876
838.4378662109375 0 910.1686
840.4717407226562 0 1589.2987
843.4129638671875 0 49803.418 w 1
843.9133911132812 0 45133.22
844.4149169921875 0 27054.459
844.9153442382812 0 9152.561
845.4203491210938 0 3802.525
849.4197998046875 0 1839.1427
850.4212036132812 0 1115.014
851.4176025390625 0 1575.2386
855.9672241210938 0 2426.606
856.465087890625 0 4434.639
856.960205078125 0 2194.1865
857.4730834960938 0 14885.377
857.9344482421875 0 1935.3599 c Ammonia loss 13
858.4805908203125 0 44391.734
858.927001953125 0 1971.8186
859.4810180664062 0 19874.07
859.9353637695312 0 1200.13
860.4824829101562 0 5127.3916
863.4598388671875 0 1117.3485
863.948974609375 0 1191.4277 y Water loss 1
864.9378051757812 0 19546.95 z 1
865.441650390625 0 18805.611
865.944580078125 0 13269.303
866.439697265625 0 126132.7 c 13
866.9401245117188 0 122478.68
867.440673828125 0 75316.766
867.9415893554688 0 30244.068
868.4403076171875 0 7132.705
869.972412109375 0 2102.0696
870.4717407226562 0 2660.223
871.4646606445312 0 1147.3693
871.9576416015625 0 3833.6387
872.4532470703125 0 4224.963
872.9549560546875 0 4306.5835 y 1
873.4432983398438 0 2503.4465
873.9362182617188 0 2972.1228
874.438232421875 0 2829.5354
877.9609375 0 3250.713
878.4593505859375 0 3476.0447
878.9348754882812 0 3471.997
879.4484252929688 0 5819.757
879.9459228515625 0 4420.998
880.4478759765625 0 3422.3616
880.945556640625 0 1260.1611
883.475830078125 0 4388.227
884.4758911132812 0 6134.713
884.9688110351562 0 2877.767
885.4672241210938 0 2973.3418
885.954345703125 0 3844.9783
886.4555053710938 0 4674.379
886.9473266601562 0 4634.6924
887.4349975585938 0 35018.33
887.937744140625 0 37958.793
888.4403076171875 0 24922.676
888.9446411132812 0 10903.964
889.45263671875 0 4761.145
891.973388671875 0 3324.249
892.4683227539062 0 10110.406
892.9647827148438 0 11958.755
893.4613037109375 0 11314.1
893.9550170898438 0 9792.053
894.4576416015625 0 21568.8
894.9581298828125 0 18145.16
895.4589233398438 0 9129.966
895.9505615234375 0 13115.301
896.447998046875 0 12415.269
896.94873046875 0 6796.262
897.445556640625 0 3566.4072
900.4608154296875 0 1018.2305
900.97119140625 0 1555.1311
901.4684448242188 0 25595.113
901.9678344726562 0 24771.525
902.4647827148438 0 24760.717 z Water loss 8
902.9638061523438 0 15317.908
903.4631958007812 0 9263.581
905.96728515625 0 8694.559
906.4625854492188 0 23981.049
906.9603881835938 0 19845.844
907.4595336914062 0 14176.983
907.958984375 0 6875.0635
908.4571533203125 0 3334.2358
914.9719848632812 0 31027.99
915.4683227539062 0 125189.836
915.9689331054688 0 117042.53
916.4683227539062 0 78041.86
916.9686279296875 0 33703.285
917.4699096679688 0 12179.546
920.4644775390625 0 12892.726 z 8
921.4668579101562 0 9228.399
922.46923828125 0 3745.8735
923.473388671875 0 84038
923.9757690429688 0 163215.55
924.4764404296875 0 133821.73
924.977783203125 0 76927.46
925.4786376953125 0 31905.133
925.9805297851562 0 7928.1035
926.4827270507812 0 1633.9448
927.4904174804688 0 95094.56 c 7
928.4932861328125 0 44234.035
929.4965209960938 0 13649.58
930.4959106445312 0 2587.603
936.482177734375 0 13893.231 y 8
937.4856567382812 0 7731.657
938.486328125 0 2486.5962
953.5123291015625 0 1474.4434
968.5272827148438 0 2809.7654
969.5280151367188 0 2088.2346
978.4653930664062 0 2164.1042
979.4603881835938 0 1041.6217
986.5147705078125 0 3516.3938
987.5153198242188 0 3235.578
988.510009765625 0 3445.4553
989.5107421875 0 1884.3478
990.4947509765625 0 19585.725 w 7
991.494140625 0 10132.111
992.4952392578125 0 5286.466
993.4535522460938 0 1491.0732
994.43505859375 0 1239.8685
1012.5053100585938 0 1421.479
1030.5123291015625 0 8940.0205
1031.5155029296875 0 6452.827
1032.52197265625 0 3919.9084
1033.51611328125 0 1181.3788
1045.5291748046875 0 1445.7765
1049.507080078125 0 58359.1 z 7
1050.5084228515625 0 39486.082
1051.509521484375 0 17109.291
1052.5096435546875 0 4742.7427
1053.5115966796875 0 1750.5842
1065.525146484375 0 3558.5725 y 7
1066.5208740234375 0 2274.6929
1067.52783203125 0 1876.0802
1073.5250244140625 0 1567.532
1074.5260009765625 0 76451.27 c 8
1075.5286865234375 0 43119.125
1076.529541015625 0 19786.69
1077.5306396484375 0 6034.694
1107.53076171875 0 1033.2179
1113.547119140625 0 2342.5107
1114.552490234375 0 1502.8987
1119.53515625 0 8508.324 w 6
1120.5357666015625 0 4767.033
1121.53662109375 0 2316.1643
1122.4901123046875 0 2129.4702
1131.55859375 0 2276.9753
1132.5618896484375 0 1700.5045
1133.5604248046875 0 2151.0044
1134.5556640625 0 2134.2502
1144.5872802734375 0 1621.7584
1175.5733642578125 0 73167.37 c 9
1176.575927734375 0 43993.8 y Water loss 6
1177.5767822265625 0 18856.662
1178.551513671875 0 51059.773 z 6
1179.550537109375 0 30471.148
1180.55029296875 0 14338.122
1181.5537109375 0 4510.19
1193.5592041015625 0 1416.4995
1194.5667724609375 0 6061.47 y 6
1195.5684814453125 0 4355.771
1196.5614013671875 0 1667.6514
1220.580078125 0 1968.0168
1221.5810546875 0 1844.7961
1222.5828857421875 0 1303.1736
1223.57958984375 0 1138.1962
1231.556396484375 0 1373.0399
1232.43408203125 0 1375.689
1232.5972900390625 0 1060.8658
1233.427001953125 0 1397.3394
1259.652099609375 0 4864.602
1260.6558837890625 0 3566.0627
1261.66748046875 0 2473.684
1275.6424560546875 0 1212.9291
1277.6259765625 0 1333.0808
1278.6207275390625 0 1975.8994
1284.6492919921875 0 2643.6938
1286.650146484375 0 1491.2925 c Ammonia loss 10
1302.6597900390625 0 8101.8364
1303.6658935546875 0 19796.887 c 10
1304.670166015625 0 13112.693
1305.6693115234375 0 6117.6685
1318.6473388671875 0 1707.0388
1319.6484375 0 2037.5607
1320.6568603515625 0 1987.5222
1333.6485595703125 0 1793.5476 y Ammonia loss 5
1334.649658203125 0 31254.977 z 5
1335.6546630859375 0 44216.652
1336.655517578125 0 27118.05
1337.6580810546875 0 11229.151
1338.662353515625 0 3746.4421
1349.6590576171875 0 5361.1313
1350.666748046875 0 13350.82 y 5
1351.6669921875 0 8739.79
1352.6697998046875 0 3600.8577
1353.6702880859375 0 1429.9015
1373.69287109375 0 1310.4365
1374.7030029296875 0 2287.1587
1375.703369140625 0 1817.0848
1376.6995849609375 0 1035.6456
1401.700439453125 0 1187.4207
1403.6842041015625 0 1359.7911 z Water loss 4
1416.7037353515625 0 3012.3423
1417.7098388671875 0 38234.63 c 11
1418.7105712890625 0 29512.936
1419.7125244140625 0 15108.057 y Water loss 4
1420.706787109375 0 5635.0527
1421.6898193359375 0 3906.1047 z 4
1422.6881103515625 0 17339.979
1423.6905517578125 0 13267.32
1424.6932373046875 0 7698.7056
1425.6929931640625 0 3043.1729
1437.7022705078125 0 3545.276 y 4
1438.69580078125 0 2443.907
1439.7047119140625 0 2302.098
1475.7576904296875 0 1259.0636
1476.7550048828125 0 1282.7292
1502.7510986328125 0 1501.118
1503.7652587890625 0 1219.4603
1527.772216796875 0 1277.6976 c Water loss 12
1529.7833251953125 0 1338.0948
1534.752685546875 0 10502.035 y 3
1535.757080078125 0 6847.2915
1536.761474609375 0 5563.6733
1537.760986328125 0 1852.2069
1545.76806640625 0 19626.893 c 12
1546.7701416015625 0 19823.96
1547.771484375 0 10926.81
1548.7733154296875 0 6087.4106
1549.771484375 0 1777.6318
1565.7977294921875 0 1494.401
1601.828857421875 0 2532.0098
1602.816162109375 0 1716.9594
1607.7962646484375 0 3044.758
1608.8033447265625 0 1851.7783
1609.8109130859375 0 1654.8591
1614.80322265625 0 4069.7322 y Ammonia loss 2
1615.803466796875 0 4251.2305 z 2
1616.8121337890625 0 2290.6401
1623.82177734375 0 1542.8907
1624.8282470703125 0 5077.518
1625.826416015625 0 3813.637
1626.8375244140625 0 4477.9756
1627.8258056640625 0 1949.4586
1628.8341064453125 0 2486.2002
1629.8094482421875 0 2263.077
1630.83251953125 0 1610.6959
1631.812744140625 0 2084.8984 y 2
1632.811767578125 0 2449.6245
1633.8037109375 0 1342.7711
1644.8372802734375 0 17116.598
1645.841552734375 0 22783
1646.84326171875 0 16758.248
1647.8424072265625 0 8843.767
1648.837890625 0 3918.6812
1670.8424072265625 0 2868.7585
1671.8453369140625 0 1968.2937
1673.8646240234375 0 1586.5028
1674.8477783203125 0 1291.6698
1684.861328125 0 2044.963
1685.86083984375 0 2227.705
1687.8587646484375 0 2635.1294
1688.86279296875 0 7114.3037
1689.8648681640625 0 5132.2334
1690.8671875 0 2593.328
1715.861328125 0 3759.3267
1716.849365234375 0 3401.4246
1717.8670654296875 0 2381.5051
1728.86962890625 0 6385.0356 z 1
1729.877685546875 0 36100.996
1730.878662109375 0 32091.166
1731.8741455078125 0 29509.809 c 13
1732.8751220703125 0 39132.8
1733.8780517578125 0 28282.502
1734.878173828125 0 16198.468
1735.88037109375 0 5527.476
1747.8480224609375 0 1360.7502
1748.864990234375 0 1311.036
1769.903564453125 0 1840.8997
1770.88623046875 0 3202.1433
1771.8824462890625 0 2237.9697
1772.8809814453125 0 1508.0283
1783.9525146484375 0 2771.791
1784.9512939453125 0 3695.6594
1785.927978515625 0 7387.65
1786.9144287109375 0 6516.578
1787.9105224609375 0 7520.3506
1788.913330078125 0 12265.129
1789.9154052734375 0 11667.556
1790.9136962890625 0 6601.912
1791.9058837890625 0 2824.5813
1799.9052734375 0 1026.3174
1801.9365234375 0 6561.772
1802.93212890625 0 24548.62
1803.933349609375 0 23328.055
1804.9324951171875 0 15445.582
1805.931640625 0 6039.061
1806.92236328125 0 1950.4761
1811.90673828125 0 1428.3817
1812.9195556640625 0 2942.2227
1813.9119873046875 0 5898.1963
1814.9078369140625 0 5965.827
1815.9224853515625 0 2401.562
1816.9061279296875 0 1726.2323
1817.91064453125 0 1174.2681
1818.94970703125 0 2633.148
1819.958984375 0 15768.686
1820.9571533203125 0 14776.216
1821.9581298828125 0 7204.5093
1822.9617919921875 0 3746.262
1828.9149169921875 0 2114.2192
1829.9295654296875 0 21384.771
1830.929443359375 0 74308.73
1831.9300537109375 0 66617.67
1832.931396484375 0 41497.42
1833.930908203125 0 16972.076
1834.9332275390625 0 5435.9126
1845.9339599609375 0 7313.8174
1846.9429931640625 0 44367.195
1847.950927734375 0 191696
1848.9532470703125 0 189633.3
1849.954833984375 0 112325.6
1850.95556640625 0 47680.406
1851.958740234375 0 14720.464
3084.695556640625 0 1141.0854

Spectrum Details

|  |  |
| --- | --- |
| Matched peaks? Matched peaks The total absolute number of peaks matched. Additionally in brackets the total fraction of peaks matched and the total number of peaks is shown. | 86 (13.80% of 623) |
| FDR? FDR The false discovery rate estimated for this peptide. It is calculated by matching all theoretical fragments with a non-integer shift with the raw peaks for this spectrum. This is done with 40 different shifts. The resulting percentage is the average number of annotated peaks over the number of annotated peaks with the correct spectrum. | 0.17% |
| Satellite FDR? Satellite FDR See the FDR for details on its calculation. This satellite ion specific FDR only contains the satellite ions (d/w) for I/L/J positions. | 0.00% |
| PSM Score? PSM Score The PSM Score as given by Hecklib to this annotated spectrum. It is shown with three significant figures. | 584 |

#### Spectrum 4335? Spectrum 4335 The raw spectrum of this peptide as annotated by Hecklib. The fragments are coloured according to ion type (see legend). Any peaks with a star '\*' as text can be hovered over to see the full details, first the ion type second the mass shift type. By hovering over the amino acids in the peptide or ions in the legend the corresponding peaks are highlighted. By toggling the 'Unassigned' label you can turn the background (unassigned) peaks on or off in the plot. By updating the slider in the Ion legend you can update the spectrum to only show the top X% of the peaks with labels. The top X% means any peak that is within X% of the highest intensity. By dragging in the spectrum you can zoom in to a specific part of the spectrum and use 'Zoom Out' to get back to the original zoom level. The annotation of the spectrum is based on the given sequence in the peptides file and is done with different software so inconsistencies are likely. The peaks are annotated based on the given sequence, with 20 ppm tolerance.

Copy Data

##### Spectrum 4335 (TSV)

###### Preview

```
Loading example...
```

*Click on the button to copy the data to your clipboard.*

Mz MinMz MaxIntensity Max

WidthHeightPeptide font sizePeptide stroke widthSpectrum font sizeSpectrum stroke widthCompact peptide

Ion legend

wxyz

abcd

OtherUnassignedIon^Charge^_Position_Show for top:%

TLPPSREEMTKNQKJ

01.59e+53.18e+54.78e+56.37e+5

Zoom Out

y^+1^_1_y^+1^_2_c^+1^_4_c^+1^_4_z^+1^_3_y^+1^_3_c^+1^_5_z^+3^_12_z^+3^_12_c^+1^_5_w^+3^_13_y^+3^_13_y^+3^_13_z^+3^_13_z^+1^_4_y^+3^_13_y^+1^_4_y^+3^_14_c^+2^_11_c^+2^_11_c^+2^_11_z^+2^_10_z^+2^_10_c^+1^_6_z^+1^_5_y^+2^_10_y^+1^_5_c^+2^_12_c^+2^_12_z^+2^_11_c^+2^_12_y^+2^_11_z^+2^_11_y^+2^_11_w^+2^_12_z^+2^_12_z^+2^_12_z^+1^_6_w^+1^_6_w^+1^_6_z^+2^_12_c^+2^_13_y^+2^_12_y^+1^_6_c^+2^_13_y^+1^_6_w^+2^_13_c^+1^_7_y^+2^_13_y^+2^_13_z^+2^_13_y^+2^_13_w^+2^_14_c^+2^_14_y^+2^_14_y^+2^_14_z^+2^_14_c^+2^_14_y^+2^_14_z^+1^_7_z^+1^_7_c^+1^_8_y^+1^_7_w^+1^_8_z^+1^_8_z^+1^_8_y^+1^_8_c^+1^_9_w^+1^_9_c^+1^_10_y^+1^_9_z^+1^_9_y^+1^_9_c^+1^_11_c^+1^_11_c^+1^_11_y^+1^_10_z^+1^_10_y^+1^_10_w^+1^_11_c^+1^_12_y^+1^_11_z^+1^_11_y^+1^_11_c^+1^_13_y^+1^_12_c^+1^_13_y^+1^_13_z^+1^_13_y^+1^_13_c^+1^_14_z^+1^_14_c^+1^_14_

0599119717962394

Fragment Matches Table

Show background peaks

| Position | Ion type | Intensity | mz Theoretical | mz Error (Th) | mz Error (ppm) | Charge | Series Number |
| --- | --- | --- | --- | --- | --- | --- | --- |
| - | - | 885.7 | 127.7 | - | - | 0 | - |
| 15 | y | 4574 | 132.1 | 0.0002982 | 2.258 | +1 | 1 |
| - | - | 1638 | 143.1 | - | - | 0 | - |
| - | - | 1014 | 143.5 | - | - | 0 | - |
| - | - | 945.7 | 145.5 | - | - | 0 | - |
| - | - | 1229 | 149 | - | - | 0 | - |
| - | - | 1125 | 152 | - | - | 0 | - |
| - | - | 1070 | 155.1 | - | - | 0 | - |
| - | - | 1428 | 169.1 | - | - | 0 | - |
| - | - | 1177 | 172.2 | - | - | 0 | - |
| - | - | 5017 | 173.5 | - | - | 0 | - |
| - | - | 1036 | 175.6 | - | - | 0 | - |
| - | - | 1434 | 185.1 | - | - | 0 | - |
| - | - | 1.611E+05 | 187.1 | - | - | 0 | - |
| - | - | 1.554E+04 | 188.1 | - | - | 0 | - |
| - | - | 1792 | 200.1 | - | - | 0 | - |
| - | - | 1.043E+04 | 212.1 | - | - | 0 | - |
| - | - | 1692 | 213.1 | - | - | 0 | - |
| - | - | 1.007E+05 | 215.1 | - | - | 0 | - |
| - | - | 1.106E+04 | 216.1 | - | - | 0 | - |
| - | - | 1.573E+04 | 219.1 | - | - | 0 | - |
| - | - | 1652 | 220.1 | - | - | 0 | - |
| - | - | 1469 | 230.2 | - | - | 0 | - |
| - | - | 5020 | 256.2 | - | - | 0 | - |
| - | - | 1205 | 297.7 | - | - | 0 | - |
| - | - | 2.358E+04 | 299.2 | - | - | 0 | - |
| - | - | 4904 | 300.2 | - | - | 0 | - |
| - | - | 4866 | 312.2 | - | - | 0 | - |
| 14 | y | 2906 | 318.2 | 0.004262 | 13.39 | +1 | 2 |
| - | - | 8200 | 326.2 | - | - | 0 | - |
| - | - | 5796 | 327.2 | - | - | 0 | - |
| - | - | 1476 | 328.2 | - | - | 0 | - |
| - | - | 1687 | 341.2 | - | - | 0 | - |
| - | - | 3083 | 344.2 | - | - | 0 | - |
| - | - | 3340 | 382.3 | - | - | 0 | - |
| - | - | 1485 | 386.2 | - | - | 0 | - |
| - | - | 1808 | 396.2 | - | - | 0 | - |
| - | - | 1545 | 397.2 | - | - | 0 | - |
| - | - | 6562 | 407.3 | - | - | 0 | - |
| 4 | c | 3021 | 408.3 | 0.003634 | 8.901 | +1 | 4 |
| - | - | 1966 | 413.2 | - | - | 0 | - |
| - | - | 2634 | 414.2 | - | - | 0 | - |
| - | - | 1409 | 415.2 | - | - | 0 | - |
| - | - | 1794 | 415.2 | - | - | 0 | - |
| - | - | 8992 | 425.3 | - | - | 0 | - |
| 4 | c | 6539 | 426.3 | 0.0002971 | 0.697 | +1 | 4 |
| - | - | 3580 | 427.3 | - | - | 0 | - |
| - | - | 2384 | 428.3 | - | - | 0 | - |
| 13 | z | 4628 | 430.2 | 0.003802 | 8.836 | +1 | 3 |
| - | - | 2323 | 431.2 | - | - | 0 | - |
| 13 | y | 1.053E+04 | 446.3 | 0.003544 | 7.941 | +1 | 3 |
| - | - | 2655 | 447.3 | - | - | 0 | - |
| - | - | 1752 | 462.9 | - | - | 0 | - |
| - | - | 2884 | 466.2 | - | - | 0 | - |
| - | - | 2780 | 466.6 | - | - | 0 | - |
| - | - | 1.084E+04 | 470.3 | - | - | 0 | - |
| - | - | 2417 | 471.3 | - | - | 0 | - |
| - | - | 1.27E+04 | 471.9 | - | - | 0 | - |
| - | - | 1.108E+04 | 472.2 | - | - | 0 | - |
| - | - | 2425 | 472.3 | - | - | 0 | - |
| - | - | 3409 | 472.6 | - | - | 0 | - |
| - | - | 1675 | 485.6 | - | - | 0 | - |
| - | - | 2942 | 491.6 | - | - | 0 | - |
| - | - | 2578 | 491.9 | - | - | 0 | - |
| - | - | 5861 | 494.3 | - | - | 0 | - |
| - | - | 3123 | 495.2 | - | - | 0 | - |
| 5 | c | 2119 | 495.3 | 0.004596 | 9.278 | +1 | 5 |
| 4 | z | 7541 | 500.9 | 0.003316 | 6.62 | +3 | 12 |
| 4 | z | 7337 | 501.2 | 0.002572 | 5.131 | +3 | 12 |
| - | - | 4268 | 501.6 | - | - | 0 | - |
| - | - | 2593 | 512.3 | - | - | 0 | - |
| - | - | 1729 | 512.6 | - | - | 0 | - |
| 5 | c | 1.025E+05 | 513.3 | 0.000953 | 1.857 | +1 | 5 |
| - | - | 1360 | 514.3 | - | - | 0 | - |
| - | - | 2.547E+04 | 514.3 | - | - | 0 | - |
| - | - | 4762 | 515.3 | - | - | 0 | - |
| - | - | 2959 | 520.2 | - | - | 0 | - |
| - | - | 9180 | 523.3 | - | - | 0 | - |
| - | - | 6152 | 523.6 | - | - | 0 | - |
| - | - | 3470 | 523.9 | - | - | 0 | - |
| - | - | 2400 | 526.3 | - | - | 0 | - |
| - | - | 1807 | 529.3 | - | - | 0 | - |
| - | - | 2017 | 529.6 | - | - | 0 | - |
| - | - | 1726 | 529.9 | - | - | 0 | - |
| 3 | w | 1705 | 530.3 | 0.008831 | 16.65 | +3 | 13 |
| - | - | 2232 | 532.9 | - | - | 0 | - |
| - | - | 2147 | 536.8 | - | - | 0 | - |
| 3 | y | 7926 | 538.6 | 0.001928 | 3.579 | +3 | 13 |
| 3 | y | 7264 | 538.9 | 0.006625 | 12.29 | +3 | 13 |
| 3 | z | 2608 | 539.3 | 0.0005177 | 0.9599 | +3 | 13 |
| - | - | 4056 | 539.6 | - | - | 0 | - |
| - | - | 4285 | 539.9 | - | - | 0 | - |
| 12 | z | 8556 | 544.3 | 0.005713 | 10.5 | +1 | 4 |
| 3 | y | 6.306E+05 | 544.6 | 0.001275 | 2.341 | +3 | 13 |
| - | - | 5.338E+05 | 544.9 | - | - | 0 | - |
| - | - | 3.043E+05 | 545.3 | - | - | 0 | - |
| - | - | 1.152E+05 | 545.6 | - | - | 0 | - |
| - | - | 3.001E+04 | 545.9 | - | - | 0 | - |
| - | - | 7874 | 546.3 | - | - | 0 | - |
| 12 | y | 2.125E+04 | 560.3 | 0.00338 | 6.033 | +1 | 4 |
| - | - | 6404 | 561.3 | - | - | 0 | - |
| - | - | 1623 | 565.8 | - | - | 0 | - |
| - | - | 2164 | 571.3 | - | - | 0 | - |
| - | - | 2912 | 573.3 | - | - | 0 | - |
| 2 | y | 1905 | 582.3 | 0.0004957 | 0.8513 | +3 | 14 |
| - | - | 1473 | 607.3 | - | - | 0 | - |
| - | - | 2057 | 607.4 | - | - | 0 | - |
| - | - | 1463 | 613.3 | - | - | 0 | - |
| - | - | 5721 | 614.4 | - | - | 0 | - |
| - | - | 7856 | 615.3 | - | - | 0 | - |
| - | - | 2441 | 616.3 | - | - | 0 | - |
| - | - | 4939 | 625.4 | - | - | 0 | - |
| - | - | 7105 | 626.4 | - | - | 0 | - |
| - | - | 2193 | 627.4 | - | - | 0 | - |
| - | - | 3.272E+04 | 628.4 | - | - | 0 | - |
| - | - | 1.1E+04 | 629.3 | - | - | 0 | - |
| - | - | 9445 | 629.4 | - | - | 0 | - |
| - | - | 6721 | 630.3 | - | - | 0 | - |
| - | - | 2112 | 630.4 | - | - | 0 | - |
| - | - | 4159 | 642.8 | - | - | 0 | - |
| 11 | c | 4826 | 643.3 | 0.0009553 | 1.485 | +2 | 11 |
| 11 | c | 2757 | 643.8 | 0.006549 | 10.17 | +2 | 11 |
| - | - | 1.155E+04 | 651.8 | - | - | 0 | - |
| 11 | c | 6.442E+04 | 652.3 | 0.003162 | 4.847 | +2 | 11 |
| - | - | 4.387E+04 | 652.8 | - | - | 0 | - |
| - | - | 2.162E+04 | 653.3 | - | - | 0 | - |
| - | - | 6881 | 653.8 | - | - | 0 | - |
| - | - | 2790 | 657.8 | - | - | 0 | - |
| - | - | 3551 | 658.3 | - | - | 0 | - |
| 6 | z | 2121 | 659.3 | 0.001873 | 2.84 | +2 | 10 |
| 6 | z | 4851 | 667.8 | 0.0004766 | 0.7137 | +2 | 10 |
| - | - | 6901 | 668.3 | - | - | 0 | - |
| - | - | 3487 | 668.8 | - | - | 0 | - |
| 6 | c | 1.859E+05 | 669.4 | 0.001343 | 2.007 | +1 | 6 |
| - | - | 7.229E+04 | 670.4 | - | - | 0 | - |
| - | - | 1.529E+04 | 671.4 | - | - | 0 | - |
| 11 | z | 7.26E+04 | 672.4 | 0.003386 | 5.036 | +1 | 5 |
| - | - | 2.321E+04 | 673.4 | - | - | 0 | - |
| - | - | 5963 | 674.4 | - | - | 0 | - |
| - | - | 5477 | 675.3 | - | - | 0 | - |
| 6 | y | 1.124E+04 | 675.8 | 0.001087 | 1.608 | +2 | 10 |
| - | - | 5494 | 676.3 | - | - | 0 | - |
| - | - | 4888 | 676.8 | - | - | 0 | - |
| - | - | 4461 | 680.8 | - | - | 0 | - |
| - | - | 1604 | 681.3 | - | - | 0 | - |
| - | - | 4257 | 686.8 | - | - | 0 | - |
| - | - | 4549 | 687.4 | - | - | 0 | - |
| - | - | 6300 | 687.9 | - | - | 0 | - |
| 11 | y | 1.986E+04 | 688.4 | 0.003129 | 4.545 | +1 | 5 |
| - | - | 6349 | 689.4 | - | - | 0 | - |
| - | - | 1818 | 690.4 | - | - | 0 | - |
| - | - | 5156 | 695.4 | - | - | 0 | - |
| - | - | 8575 | 699.8 | - | - | 0 | - |
| 12 | c | 5362 | 700.4 | 0.0006523 | 0.9314 | +2 | 12 |
| 12 | c | 3732 | 700.8 | 0.01157 | 16.51 | +2 | 12 |
| - | - | 2938 | 701.8 | - | - | 0 | - |
| 5 | z | 2822 | 702.3 | 0.01223 | 17.41 | +2 | 11 |
| - | - | 2009 | 703.4 | - | - | 0 | - |
| - | - | 2.522E+04 | 707.3 | - | - | 0 | - |
| - | - | 1.762E+04 | 707.8 | - | - | 0 | - |
| - | - | 1.112E+04 | 708.3 | - | - | 0 | - |
| - | - | 2.683E+04 | 708.9 | - | - | 0 | - |
| 12 | c | 2.398E+05 | 709.4 | 0.003182 | 4.486 | +2 | 12 |
| - | - | 1.827E+05 | 709.9 | - | - | 0 | - |
| 5 | y | 8.439E+04 | 710.3 | 0.01324 | 18.64 | +2 | 11 |
| - | - | 2.829E+04 | 710.9 | - | - | 0 | - |
| 5 | z | 9244 | 711.3 | 0.005482 | 7.706 | +2 | 11 |
| - | - | 1.854E+04 | 711.8 | - | - | 0 | - |
| - | - | 6631 | 712.3 | - | - | 0 | - |
| - | - | 3883 | 712.9 | - | - | 0 | - |
| - | - | 2399 | 713.4 | - | - | 0 | - |
| - | - | 1753 | 716.4 | - | - | 0 | - |
| - | - | 3467 | 717.4 | - | - | 0 | - |
| - | - | 3547 | 718.9 | - | - | 0 | - |
| 5 | y | 9812 | 719.4 | 0.001918 | 2.666 | +2 | 11 |
| - | - | 8019 | 719.9 | - | - | 0 | - |
| - | - | 3997 | 720.4 | - | - | 0 | - |
| - | - | 2064 | 722.4 | - | - | 0 | - |
| - | - | 2272 | 722.9 | - | - | 0 | - |
| - | - | 6589 | 729.4 | - | - | 0 | - |
| - | - | 2309 | 730.4 | - | - | 0 | - |
| - | - | 2005 | 743.4 | - | - | 0 | - |
| 4 | w | 2686 | 746.4 | 0.0008008 | 1.073 | +2 | 12 |
| - | - | 2345 | 747.4 | - | - | 0 | - |
| 4 | z | 1.344E+04 | 750.9 | 0.005547 | 7.387 | +2 | 12 |
| 4 | z | 1.239E+04 | 751.4 | 0.00806 | 10.73 | +2 | 12 |
| - | - | 6129 | 751.9 | - | - | 0 | - |
| - | - | 1625 | 752.4 | - | - | 0 | - |
| - | - | 4948 | 754.4 | - | - | 0 | - |
| 10 | z | 2169 | 755.4 | 0.01309 | 17.33 | +1 | 6 |
| 10 | w | 1.363E+04 | 756.4 | 0.002061 | 2.724 | +1 | 6 |
| - | - | 5185 | 757.4 | - | - | 0 | - |
| 10 | w | 2.451E+04 | 758.4 | 0.002749 | 3.624 | +1 | 6 |
| - | - | 8154 | 759.4 | - | - | 0 | - |
| 4 | z | 4296 | 759.9 | 0.0032 | 4.211 | +2 | 12 |
| - | - | 1601 | 760.9 | - | - | 0 | - |
| - | - | 1768 | 763.9 | - | - | 0 | - |
| 13 | c | 2900 | 764.9 | 0.009324 | 12.19 | +2 | 13 |
| - | - | 5058 | 765.4 | - | - | 0 | - |
| - | - | 2236 | 766.4 | - | - | 0 | - |
| - | - | 1988 | 766.9 | - | - | 0 | - |
| - | - | 4137 | 767.4 | - | - | 0 | - |
| 4 | y | 8.067E+04 | 767.9 | 0.001415 | 1.843 | +2 | 12 |
| - | - | 7.9E+04 | 768.4 | - | - | 0 | - |
| - | - | 3.263E+04 | 768.9 | - | - | 0 | - |
| - | - | 1735 | 769.3 | - | - | 0 | - |
| - | - | 1.298E+04 | 769.4 | - | - | 0 | - |
| - | - | 2377 | 769.9 | - | - | 0 | - |
| - | - | 8611 | 771.4 | - | - | 0 | - |
| 10 | y | 1.76E+04 | 772.4 | 0.004116 | 5.329 | +1 | 6 |
| - | - | 1.003E+04 | 772.9 | - | - | 0 | - |
| 13 | c | 1.043E+05 | 773.4 | 0.004533 | 5.862 | +2 | 13 |
| - | - | 1.279E+05 | 773.9 | - | - | 0 | - |
| - | - | 3.923E+04 | 774.4 | - | - | 0 | - |
| - | - | 2.702E+04 | 774.4 | - | - | 0 | - |
| - | - | 2.182E+04 | 774.9 | - | - | 0 | - |
| - | - | 1.379E+04 | 775.4 | - | - | 0 | - |
| - | - | 2802 | 779.4 | - | - | 0 | - |
| - | - | 2611 | 781.9 | - | - | 0 | - |
| - | - | 2877 | 782.4 | - | - | 0 | - |
| - | - | 1.583E+04 | 784.4 | - | - | 0 | - |
| - | - | 1.025E+04 | 784.9 | - | - | 0 | - |
| - | - | 5007 | 785.4 | - | - | 0 | - |
| - | - | 4226 | 787.4 | - | - | 0 | - |
| - | - | 2831 | 788.4 | - | - | 0 | - |
| 10 | y | 2.557E+04 | 789.5 | 0.003261 | 4.13 | +1 | 6 |
| - | - | 8856 | 790.4 | - | - | 0 | - |
| - | - | 3538 | 791.4 | - | - | 0 | - |
| - | - | 4697 | 791.9 | - | - | 0 | - |
| - | - | 2277 | 792.4 | - | - | 0 | - |
| - | - | 5924 | 792.9 | - | - | 0 | - |
| - | - | 8949 | 793.4 | - | - | 0 | - |
| - | - | 5838 | 793.9 | - | - | 0 | - |
| - | - | 2516 | 794.4 | - | - | 0 | - |
| 3 | w | 1.128E+04 | 794.9 | 0.004189 | 5.27 | +2 | 13 |
| - | - | 7907 | 795.4 | - | - | 0 | - |
| - | - | 4272 | 795.9 | - | - | 0 | - |
| - | - | 4452 | 797.4 | - | - | 0 | - |
| 7 | c | 2.757E+05 | 798.4 | 0.001475 | 1.847 | +1 | 7 |
| - | - | 1.134E+05 | 799.5 | - | - | 0 | - |
| - | - | 2.858E+04 | 800.5 | - | - | 0 | - |
| - | - | 1.011E+04 | 800.9 | - | - | 0 | - |
| - | - | 7462 | 801.4 | - | - | 0 | - |
| - | - | 9303 | 801.9 | - | - | 0 | - |
| - | - | 6327 | 802.4 | - | - | 0 | - |
| - | - | 3606 | 805.4 | - | - | 0 | - |
| - | - | 1806 | 806.4 | - | - | 0 | - |
| 3 | y | 4763 | 807.4 | 0.001008 | 1.248 | +2 | 13 |
| 3 | y | 1.146E+04 | 807.9 | 0.008693 | 10.76 | +2 | 13 |
| 3 | z | 7073 | 808.4 | 0.001201 | 1.485 | +2 | 13 |
| - | - | 5016 | 808.9 | - | - | 0 | - |
| - | - | 3473 | 809.4 | - | - | 0 | - |
| - | - | 7325 | 813.4 | - | - | 0 | - |
| - | - | 1.062E+04 | 813.9 | - | - | 0 | - |
| - | - | 1.143E+04 | 814.4 | - | - | 0 | - |
| - | - | 7313 | 814.9 | - | - | 0 | - |
| - | - | 7338 | 815.4 | - | - | 0 | - |
| - | - | 5036 | 815.9 | - | - | 0 | - |
| 3 | y | 3.549E+05 | 816.4 | 0.001583 | 1.94 | +2 | 13 |
| - | - | 3.417E+05 | 816.9 | - | - | 0 | - |
| - | - | 1.741E+05 | 817.4 | - | - | 0 | - |
| - | - | 6.856E+04 | 817.9 | - | - | 0 | - |
| - | - | 1.55E+04 | 818.4 | - | - | 0 | - |
| - | - | 2.908E+04 | 822.4 | - | - | 0 | - |
| - | - | 1.488E+05 | 822.9 | - | - | 0 | - |
| - | - | 1.182E+05 | 823.4 | - | - | 0 | - |
| - | - | 7.525E+04 | 823.9 | - | - | 0 | - |
| - | - | 2.151E+04 | 824.4 | - | - | 0 | - |
| - | - | 7440 | 824.9 | - | - | 0 | - |
| - | - | 1.038E+04 | 825.5 | - | - | 0 | - |
| - | - | 2343 | 826.5 | - | - | 0 | - |
| - | - | 2261 | 828.9 | - | - | 0 | - |
| - | - | 5119 | 829.4 | - | - | 0 | - |
| - | - | 1968 | 829.9 | - | - | 0 | - |
| - | - | 4524 | 830.4 | - | - | 0 | - |
| - | - | 7851 | 830.9 | - | - | 0 | - |
| - | - | 5228 | 831.4 | - | - | 0 | - |
| - | - | 2798 | 831.9 | - | - | 0 | - |
| - | - | 1.161E+04 | 835.4 | - | - | 0 | - |
| - | - | 9899 | 835.9 | - | - | 0 | - |
| - | - | 7636 | 836.4 | - | - | 0 | - |
| - | - | 9142 | 836.9 | - | - | 0 | - |
| - | - | 8841 | 837.4 | - | - | 0 | - |
| - | - | 4656 | 837.9 | - | - | 0 | - |
| - | - | 1730 | 838.4 | - | - | 0 | - |
| - | - | 2728 | 840.5 | - | - | 0 | - |
| - | - | 2510 | 841.5 | - | - | 0 | - |
| 2 | w | 1.122E+05 | 843.4 | 0.001489 | 1.766 | +2 | 14 |
| - | - | 1.042E+05 | 843.9 | - | - | 0 | - |
| - | - | 6.955E+04 | 844.4 | - | - | 0 | - |
| - | - | 2.952E+04 | 844.9 | - | - | 0 | - |
| - | - | 8173 | 845.4 | - | - | 0 | - |
| - | - | 1687 | 845.9 | - | - | 0 | - |
| - | - | 2883 | 849.4 | - | - | 0 | - |
| - | - | 2310 | 850.4 | - | - | 0 | - |
| - | - | 2795 | 851.4 | - | - | 0 | - |
| - | - | 2016 | 852.4 | - | - | 0 | - |
| - | - | 3908 | 856 | - | - | 0 | - |
| - | - | 1.333E+04 | 856.5 | - | - | 0 | - |
| - | - | 4151 | 857 | - | - | 0 | - |
| - | - | 3.496E+04 | 857.5 | - | - | 0 | - |
| 14 | c | 3907 | 857.9 | 0.003717 | 4.333 | +2 | 14 |
| - | - | 1.059E+05 | 858.5 | - | - | 0 | - |
| - | - | 5733 | 858.9 | - | - | 0 | - |
| - | - | 4.581E+04 | 859.5 | - | - | 0 | - |
| - | - | 2283 | 859.9 | - | - | 0 | - |
| - | - | 1.275E+04 | 860.5 | - | - | 0 | - |
| - | - | 4815 | 863.4 | - | - | 0 | - |
| 2 | y | 2797 | 863.9 | 0.007742 | 8.961 | +2 | 14 |
| 2 | y | 3995 | 864.4 | 0.0001354 | 0.1566 | +2 | 14 |
| 2 | z | 5.085E+04 | 864.9 | 0.001118 | 1.293 | +2 | 14 |
| - | - | 5.2E+04 | 865.4 | - | - | 0 | - |
| - | - | 2.323E+04 | 865.9 | - | - | 0 | - |
| 14 | c | 3.342E+05 | 866.4 | 0.00149 | 1.72 | +2 | 14 |
| - | - | 2.912E+05 | 866.9 | - | - | 0 | - |
| - | - | 1.663E+05 | 867.4 | - | - | 0 | - |
| - | - | 6.709E+04 | 867.9 | - | - | 0 | - |
| - | - | 1.781E+04 | 868.4 | - | - | 0 | - |
| - | - | 5710 | 870 | - | - | 0 | - |
| - | - | 6882 | 870.5 | - | - | 0 | - |
| - | - | 4005 | 871 | - | - | 0 | - |
| - | - | 2468 | 871.4 | - | - | 0 | - |
| - | - | 7117 | 872 | - | - | 0 | - |
| - | - | 7216 | 872.5 | - | - | 0 | - |
| 2 | y | 8365 | 872.9 | 0.001849 | 2.118 | +2 | 14 |
| - | - | 7994 | 873.4 | - | - | 0 | - |
| - | - | 6316 | 873.9 | - | - | 0 | - |
| - | - | 2907 | 874.4 | - | - | 0 | - |
| - | - | 1.163E+04 | 878 | - | - | 0 | - |
| - | - | 7663 | 878.5 | - | - | 0 | - |
| - | - | 9172 | 878.9 | - | - | 0 | - |
| - | - | 1.195E+04 | 879.4 | - | - | 0 | - |
| - | - | 1.396E+04 | 879.9 | - | - | 0 | - |
| - | - | 8258 | 880.4 | - | - | 0 | - |
| - | - | 4098 | 881 | - | - | 0 | - |
| - | - | 2546 | 881.4 | - | - | 0 | - |
| - | - | 1.288E+04 | 883.5 | - | - | 0 | - |
| - | - | 3638 | 884 | - | - | 0 | - |
| - | - | 9394 | 884.5 | - | - | 0 | - |
| - | - | 6468 | 885 | - | - | 0 | - |
| - | - | 6410 | 885.5 | - | - | 0 | - |
| - | - | 8713 | 886 | - | - | 0 | - |
| - | - | 1.104E+04 | 886.5 | - | - | 0 | - |
| - | - | 8271 | 887 | - | - | 0 | - |
| - | - | 8.775E+04 | 887.4 | - | - | 0 | - |
| - | - | 8.406E+04 | 887.9 | - | - | 0 | - |
| - | - | 6.192E+04 | 888.4 | - | - | 0 | - |
| - | - | 2.318E+04 | 888.9 | - | - | 0 | - |
| - | - | 7306 | 889.5 | - | - | 0 | - |
| - | - | 1.028E+04 | 892 | - | - | 0 | - |
| - | - | 2.517E+04 | 892.5 | - | - | 0 | - |
| - | - | 2.677E+04 | 893 | - | - | 0 | - |
| - | - | 2.597E+04 | 893.5 | - | - | 0 | - |
| - | - | 2.348E+04 | 894 | - | - | 0 | - |
| - | - | 5.04E+04 | 894.5 | - | - | 0 | - |
| - | - | 4.732E+04 | 895 | - | - | 0 | - |
| - | - | 2.474E+04 | 895.5 | - | - | 0 | - |
| - | - | 4.422E+04 | 895.9 | - | - | 0 | - |
| - | - | 3.492E+04 | 896.4 | - | - | 0 | - |
| - | - | 1.565E+04 | 896.9 | - | - | 0 | - |
| - | - | 4575 | 897.5 | - | - | 0 | - |
| - | - | 4962 | 901 | - | - | 0 | - |
| - | - | 6.508E+04 | 901.5 | - | - | 0 | - |
| - | - | 6.719E+04 | 902 | - | - | 0 | - |
| 9 | z | 6.101E+04 | 902.5 | 0.01245 | 13.8 | +1 | 7 |
| - | - | 3.39E+04 | 903 | - | - | 0 | - |
| - | - | 1.653E+04 | 903.5 | - | - | 0 | - |
| - | - | 6060 | 904 | - | - | 0 | - |
| - | - | 2738 | 904.5 | - | - | 0 | - |
| - | - | 1.754E+04 | 906 | - | - | 0 | - |
| - | - | 5.592E+04 | 906.5 | - | - | 0 | - |
| - | - | 5.128E+04 | 907 | - | - | 0 | - |
| - | - | 3.346E+04 | 907.5 | - | - | 0 | - |
| - | - | 1.014E+04 | 908 | - | - | 0 | - |
| - | - | 7553 | 908.5 | - | - | 0 | - |
| - | - | 2635 | 909 | - | - | 0 | - |
| - | - | 8.02E+04 | 915 | - | - | 0 | - |
| - | - | 3.208E+05 | 915.5 | - | - | 0 | - |
| - | - | 2.894E+05 | 916 | - | - | 0 | - |
| - | - | 1.814E+05 | 916.5 | - | - | 0 | - |
| - | - | 7.725E+04 | 917 | - | - | 0 | - |
| - | - | 2.848E+04 | 917.5 | - | - | 0 | - |
| - | - | 2254 | 918 | - | - | 0 | - |
| 9 | z | 3.279E+04 | 920.5 | 0.002008 | 2.181 | +1 | 7 |
| - | - | 2.152E+04 | 921.5 | - | - | 0 | - |
| - | - | 1.157E+04 | 922.5 | - | - | 0 | - |
| - | - | 2.125E+05 | 923.5 | - | - | 0 | - |
| - | - | 4.174E+05 | 924 | - | - | 0 | - |
| - | - | 3.192E+05 | 924.5 | - | - | 0 | - |
| - | - | 1.857E+05 | 925 | - | - | 0 | - |
| - | - | 6.485E+04 | 925.5 | - | - | 0 | - |
| - | - | 1.839E+04 | 926 | - | - | 0 | - |
| - | - | 3868 | 926.5 | - | - | 0 | - |
| 8 | c | 2.276E+05 | 927.5 | 0.001362 | 1.469 | +1 | 8 |
| - | - | 1.128E+05 | 928.5 | - | - | 0 | - |
| - | - | 3.574E+04 | 929.5 | - | - | 0 | - |
| - | - | 5198 | 930.5 | - | - | 0 | - |
| 9 | y | 4.328E+04 | 936.5 | 0.001289 | 1.376 | +1 | 7 |
| - | - | 1.899E+04 | 937.5 | - | - | 0 | - |
| - | - | 6647 | 938.5 | - | - | 0 | - |
| - | - | 2457 | 955.5 | - | - | 0 | - |
| - | - | 2471 | 967.5 | - | - | 0 | - |
| - | - | 8298 | 968.5 | - | - | 0 | - |
| - | - | 5023 | 969.5 | - | - | 0 | - |
| - | - | 1910 | 970.5 | - | - | 0 | - |
| - | - | 3958 | 978.5 | - | - | 0 | - |
| - | - | 2121 | 979.5 | - | - | 0 | - |
| - | - | 9807 | 986.5 | - | - | 0 | - |
| - | - | 7206 | 987.5 | - | - | 0 | - |
| 8 | w | 4.199E+04 | 990.5 | 0.001772 | 1.789 | +1 | 8 |
| - | - | 2.31E+04 | 991.5 | - | - | 0 | - |
| - | - | 9718 | 992.5 | - | - | 0 | - |
| - | - | 5730 | 993.5 | - | - | 0 | - |
| - | - | 4158 | 994.4 | - | - | 0 | - |
| - | - | 3742 | 1013 | - | - | 0 | - |
| - | - | 1848 | 1013 | - | - | 0 | - |
| - | - | 2083 | 1015 | - | - | 0 | - |
| - | - | 2.047E+04 | 1031 | - | - | 0 | - |
| 8 | z | 1.141E+04 | 1031 | 0.02033 | 19.71 | +1 | 8 |
| - | - | 7994 | 1033 | - | - | 0 | - |
| - | - | 3459 | 1034 | - | - | 0 | - |
| - | - | 2469 | 1034 | - | - | 0 | - |
| 8 | z | 1.47E+05 | 1050 | 0.001834 | 1.747 | +1 | 8 |
| - | - | 8.991E+04 | 1051 | - | - | 0 | - |
| - | - | 3.578E+04 | 1052 | - | - | 0 | - |
| - | - | 1.021E+04 | 1053 | - | - | 0 | - |
| - | - | 2179 | 1054 | - | - | 0 | - |
| - | - | 1879 | 1058 | - | - | 0 | - |
| - | - | 2866 | 1060 | - | - | 0 | - |
| 8 | y | 8808 | 1066 | 0.000688 | 0.6457 | +1 | 8 |
| - | - | 6123 | 1067 | - | - | 0 | - |
| - | - | 2067 | 1068 | - | - | 0 | - |
| - | - | 3763 | 1074 | - | - | 0 | - |
| 9 | c | 1.811E+05 | 1075 | 0.006217 | 5.786 | +1 | 9 |
| - | - | 9.771E+04 | 1076 | - | - | 0 | - |
| - | - | 3.902E+04 | 1077 | - | - | 0 | - |
| - | - | 1.198E+04 | 1078 | - | - | 0 | - |
| - | - | 1995 | 1079 | - | - | 0 | - |
| - | - | 2375 | 1107 | - | - | 0 | - |
| - | - | 2813 | 1108 | - | - | 0 | - |
| - | - | 5974 | 1114 | - | - | 0 | - |
| - | - | 4305 | 1115 | - | - | 0 | - |
| - | - | 3979 | 1116 | - | - | 0 | - |
| 7 | w | 1.894E+04 | 1120 | 0.003307 | 2.954 | +1 | 9 |
| - | - | 1.234E+04 | 1121 | - | - | 0 | - |
| - | - | 6374 | 1122 | - | - | 0 | - |
| - | - | 2410 | 1122 | - | - | 0 | - |
| - | - | 3208 | 1123 | - | - | 0 | - |
| - | - | 6357 | 1132 | - | - | 0 | - |
| - | - | 5147 | 1133 | - | - | 0 | - |
| - | - | 4191 | 1134 | - | - | 0 | - |
| - | - | 5458 | 1135 | - | - | 0 | - |
| - | - | 4676 | 1136 | - | - | 0 | - |
| 10 | c | 1.531E+05 | 1176 | 0.006024 | 5.124 | +1 | 10 |
| 7 | y | 9.97E+04 | 1177 | 0.02005 | 17.04 | +1 | 9 |
| - | - | 4.646E+04 | 1178 | - | - | 0 | - |
| 7 | z | 1.191E+05 | 1179 | 0.00343 | 2.911 | +1 | 9 |
| - | - | 7.708E+04 | 1180 | - | - | 0 | - |
| - | - | 3.399E+04 | 1181 | - | - | 0 | - |
| - | - | 9386 | 1182 | - | - | 0 | - |
| - | - | 3156 | 1194 | - | - | 0 | - |
| 7 | y | 1.454E+04 | 1195 | 0.003383 | 2.832 | +1 | 9 |
| - | - | 9889 | 1196 | - | - | 0 | - |
| - | - | 4051 | 1197 | - | - | 0 | - |
| - | - | 2157 | 1213 | - | - | 0 | - |
| - | - | 2119 | 1216 | - | - | 0 | - |
| - | - | 3860 | 1222 | - | - | 0 | - |
| - | - | 2155 | 1223 | - | - | 0 | - |
| - | - | 5490 | 1230 | - | - | 0 | - |
| - | - | 3533 | 1231 | - | - | 0 | - |
| - | - | 2988 | 1233 | - | - | 0 | - |
| - | - | 9959 | 1260 | - | - | 0 | - |
| - | - | 7567 | 1261 | - | - | 0 | - |
| - | - | 4317 | 1262 | - | - | 0 | - |
| - | - | 4050 | 1263 | - | - | 0 | - |
| - | - | 3329 | 1264 | - | - | 0 | - |
| - | - | 3649 | 1276 | - | - | 0 | - |
| - | - | 2127 | 1277 | - | - | 0 | - |
| - | - | 4127 | 1285 | - | - | 0 | - |
| 11 | c | 4443 | 1286 | 0.007329 | 5.701 | +1 | 11 |
| 11 | c | 2766 | 1287 | 0.01171 | 9.099 | +1 | 11 |
| - | - | 2462 | 1292 | - | - | 0 | - |
| - | - | 2488 | 1302 | - | - | 0 | - |
| - | - | 1.774E+04 | 1303 | - | - | 0 | - |
| 11 | c | 4.184E+04 | 1304 | 0.004201 | 3.222 | +1 | 11 |
| - | - | 2.802E+04 | 1305 | - | - | 0 | - |
| - | - | 1.435E+04 | 1306 | - | - | 0 | - |
| - | - | 3428 | 1307 | - | - | 0 | - |
| - | - | 2107 | 1309 | - | - | 0 | - |
| - | - | 2345 | 1320 | - | - | 0 | - |
| 6 | y | 3407 | 1334 | 0.001233 | 0.9243 | +1 | 10 |
| 6 | z | 7.423E+04 | 1335 | 0.0007079 | 0.5304 | +1 | 10 |
| - | - | 1.058E+05 | 1336 | - | - | 0 | - |
| - | - | 6.215E+04 | 1337 | - | - | 0 | - |
| - | - | 2.497E+04 | 1338 | - | - | 0 | - |
| - | - | 6002 | 1339 | - | - | 0 | - |
| - | - | 1.462E+04 | 1350 | - | - | 0 | - |
| 6 | y | 3.029E+04 | 1351 | 0.000438 | 0.3243 | +1 | 10 |
| - | - | 2.321E+04 | 1352 | - | - | 0 | - |
| - | - | 8911 | 1353 | - | - | 0 | - |
| - | - | 4510 | 1354 | - | - | 0 | - |
| - | - | 6469 | 1374 | - | - | 0 | - |
| - | - | 6813 | 1375 | - | - | 0 | - |
| - | - | 3913 | 1376 | - | - | 0 | - |
| - | - | 2095 | 1403 | - | - | 0 | - |
| - | - | 2752 | 1404 | - | - | 0 | - |
| 5 | w | 2024 | 1405 | 0.008284 | 5.898 | +1 | 11 |
| - | - | 4177 | 1417 | - | - | 0 | - |
| 12 | c | 8.165E+04 | 1418 | 0.00412 | 2.906 | +1 | 12 |
| - | - | 5.962E+04 | 1419 | - | - | 0 | - |
| 5 | y | 3.022E+04 | 1420 | 0.02412 | 16.99 | +1 | 11 |
| - | - | 8880 | 1421 | - | - | 0 | - |
| 5 | z | 8213 | 1422 | 0.007986 | 5.617 | +1 | 11 |
| - | - | 4.063E+04 | 1423 | - | - | 0 | - |
| - | - | 2.561E+04 | 1424 | - | - | 0 | - |
| - | - | 1.517E+04 | 1425 | - | - | 0 | - |
| - | - | 4200 | 1426 | - | - | 0 | - |
| 5 | y | 7222 | 1438 | 0.0009723 | 0.6763 | +1 | 11 |
| - | - | 6619 | 1439 | - | - | 0 | - |
| - | - | 3429 | 1440 | - | - | 0 | - |
| - | - | 2847 | 1503 | - | - | 0 | - |
| 13 | c | 3544 | 1528 | 0.01299 | 8.504 | +1 | 13 |
| - | - | 3836 | 1529 | - | - | 0 | - |
| - | - | 3217 | 1530 | - | - | 0 | - |
| - | - | 1827 | 1531 | - | - | 0 | - |
| 4 | y | 2.154E+04 | 1535 | 0.002538 | 1.654 | +1 | 12 |
| - | - | 1.599E+04 | 1536 | - | - | 0 | - |
| - | - | 9043 | 1537 | - | - | 0 | - |
| - | - | 6200 | 1538 | - | - | 0 | - |
| - | - | 1728 | 1539 | - | - | 0 | - |
| 13 | c | 4.667E+04 | 1546 | 0.003526 | 2.281 | +1 | 13 |
| - | - | 4.135E+04 | 1547 | - | - | 0 | - |
| - | - | 2.632E+04 | 1548 | - | - | 0 | - |
| - | - | 1.264E+04 | 1549 | - | - | 0 | - |
| - | - | 4773 | 1550 | - | - | 0 | - |
| - | - | 3368 | 1601 | - | - | 0 | - |
| - | - | 4038 | 1602 | - | - | 0 | - |
| - | - | 4255 | 1603 | - | - | 0 | - |
| - | - | 2449 | 1604 | - | - | 0 | - |
| 3 | y | 8387 | 1615 | 0.0243 | 15.05 | +1 | 13 |
| 3 | z | 9997 | 1616 | 0.01525 | 9.439 | +1 | 13 |
| - | - | 5737 | 1617 | - | - | 0 | - |
| - | - | 1955 | 1618 | - | - | 0 | - |
| - | - | 1972 | 1619 | - | - | 0 | - |
| - | - | 1675 | 1627 | - | - | 0 | - |
| - | - | 2559 | 1628 | - | - | 0 | - |
| - | - | 5563 | 1629 | - | - | 0 | - |
| - | - | 6074 | 1630 | - | - | 0 | - |
| - | - | 2523 | 1631 | - | - | 0 | - |
| 3 | y | 4908 | 1632 | 0.01569 | 9.617 | +1 | 13 |
| - | - | 4999 | 1633 | - | - | 0 | - |
| - | - | 4065 | 1634 | - | - | 0 | - |
| - | - | 3178 | 1644 | - | - | 0 | - |
| - | - | 3.503E+04 | 1645 | - | - | 0 | - |
| - | - | 4.466E+04 | 1646 | - | - | 0 | - |
| - | - | 3.31E+04 | 1647 | - | - | 0 | - |
| - | - | 1.647E+04 | 1648 | - | - | 0 | - |
| - | - | 4437 | 1649 | - | - | 0 | - |
| - | - | 6760 | 1671 | - | - | 0 | - |
| - | - | 7215 | 1672 | - | - | 0 | - |
| - | - | 2758 | 1673 | - | - | 0 | - |
| - | - | 2403 | 1674 | - | - | 0 | - |
| - | - | 3008 | 1685 | - | - | 0 | - |
| - | - | 3582 | 1686 | - | - | 0 | - |
| - | - | 2332 | 1687 | - | - | 0 | - |
| - | - | 7677 | 1688 | - | - | 0 | - |
| - | - | 1.593E+04 | 1689 | - | - | 0 | - |
| - | - | 1.192E+04 | 1690 | - | - | 0 | - |
| - | - | 6393 | 1691 | - | - | 0 | - |
| - | - | 2165 | 1702 | - | - | 0 | - |
| - | - | 3250 | 1713 | - | - | 0 | - |
| 14 | c | 2585 | 1715 | 0.01459 | 8.508 | +1 | 14 |
| - | - | 7130 | 1716 | - | - | 0 | - |
| - | - | 9270 | 1717 | - | - | 0 | - |
| - | - | 5499 | 1718 | - | - | 0 | - |
| - | - | 2718 | 1719 | - | - | 0 | - |
| 2 | z | 8164 | 1729 | 0.0005234 | 0.3027 | +1 | 14 |
| - | - | 7.311E+04 | 1730 | - | - | 0 | - |
| - | - | 6.163E+04 | 1731 | - | - | 0 | - |
| 14 | c | 6.237E+04 | 1732 | 0.005618 | 3.244 | +1 | 14 |
| - | - | 8.135E+04 | 1733 | - | - | 0 | - |
| - | - | 5.883E+04 | 1734 | - | - | 0 | - |
| - | - | 3.225E+04 | 1735 | - | - | 0 | - |
| - | - | 1.174E+04 | 1736 | - | - | 0 | - |
| - | - | 2480 | 1749 | - | - | 0 | - |
| - | - | 5914 | 1770 | - | - | 0 | - |
| - | - | 4345 | 1771 | - | - | 0 | - |
| - | - | 5464 | 1772 | - | - | 0 | - |
| - | - | 4086 | 1773 | - | - | 0 | - |
| - | - | 3506 | 1776 | - | - | 0 | - |
| - | - | 5211 | 1784 | - | - | 0 | - |
| - | - | 6522 | 1785 | - | - | 0 | - |
| - | - | 1.191E+04 | 1786 | - | - | 0 | - |
| - | - | 1.147E+04 | 1787 | - | - | 0 | - |
| - | - | 1.427E+04 | 1788 | - | - | 0 | - |
| - | - | 2.692E+04 | 1789 | - | - | 0 | - |
| - | - | 2.247E+04 | 1790 | - | - | 0 | - |
| - | - | 1.129E+04 | 1791 | - | - | 0 | - |
| - | - | 5824 | 1792 | - | - | 0 | - |
| - | - | 3060 | 1793 | - | - | 0 | - |
| - | - | 1.236E+04 | 1802 | - | - | 0 | - |
| - | - | 4.503E+04 | 1803 | - | - | 0 | - |
| - | - | 4.461E+04 | 1804 | - | - | 0 | - |
| - | - | 2.716E+04 | 1805 | - | - | 0 | - |
| - | - | 1.016E+04 | 1806 | - | - | 0 | - |
| - | - | 3830 | 1807 | - | - | 0 | - |
| - | - | 6611 | 1813 | - | - | 0 | - |
| - | - | 1.547E+04 | 1814 | - | - | 0 | - |
| - | - | 9129 | 1815 | - | - | 0 | - |
| - | - | 5488 | 1816 | - | - | 0 | - |
| - | - | 2148 | 1817 | - | - | 0 | - |
| - | - | 7488 | 1819 | - | - | 0 | - |
| - | - | 3.283E+04 | 1820 | - | - | 0 | - |
| - | - | 3.326E+04 | 1821 | - | - | 0 | - |
| - | - | 1.96E+04 | 1822 | - | - | 0 | - |
| - | - | 8350 | 1823 | - | - | 0 | - |
| - | - | 2227 | 1829 | - | - | 0 | - |
| - | - | 4.554E+04 | 1830 | - | - | 0 | - |
| - | - | 1.415E+05 | 1831 | - | - | 0 | - |
| - | - | 1.364E+05 | 1832 | - | - | 0 | - |
| - | - | 8.369E+04 | 1833 | - | - | 0 | - |
| - | - | 3.151E+04 | 1834 | - | - | 0 | - |
| - | - | 1.33E+04 | 1835 | - | - | 0 | - |
| - | - | 1.598E+04 | 1846 | - | - | 0 | - |
| - | - | 9.198E+04 | 1847 | - | - | 0 | - |
| - | - | 4.213E+05 | 1848 | - | - | 0 | - |
| - | - | 3.793E+05 | 1849 | - | - | 0 | - |
| - | - | 2.209E+05 | 1850 | - | - | 0 | - |
| - | - | 9.167E+04 | 1851 | - | - | 0 | - |
| - | - | 2.778E+04 | 1852 | - | - | 0 | - |
| - | - | 2480 | 2051 | - | - | 0 | - |
| - | - | 1691 | 2371 | - | - | 0 | - |

m/z Charge Intensity FragmentType MassShift Position
127.7265625 0 885.69965
132.10220336914062 0 4573.738 y 14
143.11825561523438 0 1638.1431
143.5453643798828 0 1014.4709
145.4530487060547 0 945.7113
148.9545440673828 0 1228.5065
152.000244140625 0 1125.136
155.1202392578125 0 1069.8375
169.1339111328125 0 1428.1115
172.19448852539062 0 1177.4983
173.45143127441406 0 5017.223
175.59312438964844 0 1035.9548
185.12908935546875 0 1434.0367
187.14454650878906 0 161137.48
188.14784240722656 0 15542.747
200.14013671875 0 1791.9231
212.139892578125 0 10434.928
213.1239471435547 0 1691.5936
215.1394500732422 0 100714.7
216.14273071289062 0 11058.194
219.13433837890625 0 15726.778
220.13796997070312 0 1651.6001
230.15061950683594 0 1468.9355
256.1661682128906 0 5019.7104
297.69146728515625 0 1205.1559
299.17205810546875 0 23581.56
300.1753845214844 0 4904.1426
312.1925354003906 0 4866.496
318.2026062011719 0 2906.4062 y 13
326.1590576171875 0 8199.638
327.1664123535156 0 5796.4907
328.1976318359375 0 1475.5422
341.1812438964844 0 1686.6272
344.1929016113281 0 3082.5078
382.25860595703125 0 3340.3137
386.2420654296875 0 1485.3801
396.18890380859375 0 1807.5269
397.195068359375 0 1544.6687
407.2535705566406 0 6561.762
408.25689697265625 0 3020.9373 c Water loss 3
413.238037109375 0 1966.3992
414.1974182128906 0 2633.9214
415.1655578613281 0 1409.466
415.2007141113281 0 1794.0007
425.263427734375 0 8991.566
426.2713928222656 0 6538.578 c 3
427.2684020996094 0 3579.5505
428.2776184082031 0 2383.6565
430.242919921875 0 4627.937 z 12
431.2486572265625 0 2322.631
446.26190185546875 0 10530.659 y 12
447.2645263671875 0 2654.6099
462.9010314941406 0 1751.5802
466.2218017578125 0 2884.3374
466.5567932128906 0 2779.9775
470.2993469238281 0 10844.931
471.30328369140625 0 2417.2332
471.8994140625 0 12699.869
472.23291015625 0 11075.653
472.28564453125 0 2424.8235
472.5667419433594 0 3408.614
485.5751037597656 0 1674.5227
491.57733154296875 0 2941.63
491.9123840332031 0 2577.668
494.2854919433594 0 5860.8438
495.23907470703125 0 3122.535
495.2879638671875 0 2119.299 c Water loss 4
500.9093017578125 0 7541.279 z Water loss 3
501.2431945800781 0 7337.109 z Ammonia loss 3
501.5775451660156 0 4268.182
512.2978515625 0 2592.961
512.5885620117188 0 1729.4792
513.3040771484375 0 102504.086 c 4
514.26708984375 0 1359.5878
514.3072509765625 0 25471.78
515.3097534179688 0 4761.918
520.2310791015625 0 2959.1467
523.27490234375 0 9180.109
523.608642578125 0 6152.3057
523.9408569335938 0 3469.6619
526.2720947265625 0 2400.008
529.318603515625 0 1807.2496
529.6107177734375 0 2017.422
529.9329833984375 0 1725.5348
530.26806640625 0 1704.7478 w 2
532.9299926757812 0 2231.5034
536.7569580078125 0 2146.7886
538.6050415039062 0 7925.8936 y Water loss 2
538.937744140625 0 7264.3584 y Ammonia loss 2
539.267578125 0 2607.9575 z 2
539.6016845703125 0 4055.9983
539.9352416992188 0 4284.8013
544.283935546875 0 8555.921 z 11
544.60791015625 0 630558.25 y 2
544.9419555664062 0 533815.3
545.2761840820312 0 304308.6
545.609619140625 0 115211.58
545.9434204101562 0 30011.273
546.2967529296875 0 7873.522
560.3049926757812 0 21248.639 y 11
561.3067016601562 0 6404.453
565.763427734375 0 1622.7645
571.3194580078125 0 2163.7195
573.3363647460938 0 2911.5261
582.3018188476562 0 1904.8242 y 1
607.33056640625 0 1473.0311
607.38037109375 0 2056.5518
613.3103637695312 0 1462.8859
614.3507080078125 0 5721.341
615.348388671875 0 7855.899
616.317626953125 0 2440.5269
625.3900756835938 0 4939.1304
626.397216796875 0 7105.07
627.4052734375 0 2192.8618
628.3679809570312 0 32719.547
629.3250122070312 0 10996.266
629.372314453125 0 9445.496
630.3275146484375 0 6721.4204
630.3780517578125 0 2112.1707
642.8299560546875 0 4159.0127
643.3286743164062 0 4825.7837 c Water loss 10
643.8281860351562 0 2756.8833 c Ammonia loss 10
651.8348388671875 0 11551.76
652.3380737304688 0 64416.137 c 10
652.8395385742188 0 43866.14
653.3399047851562 0 21615.55
653.8408813476562 0 6881.3813
657.8115844726562 0 2790.3274
658.3141479492188 0 3550.7085
659.3167724609375 0 2121.0027 z Ammonia loss 5
667.8276977539062 0 4851.102 z 5
668.3322143554688 0 6901.0024
668.8333129882812 0 3487.0693
669.4055786132812 0 185936.38 c 5
670.408203125 0 72293.25
671.4107055664062 0 15294.913
672.3812255859375 0 72597.96 z 10
673.3831787109375 0 23208.545
674.3844604492188 0 5962.9546
675.3380126953125 0 5476.584
675.838623046875 0 11236.749 y 5
676.3391723632812 0 5494.443
676.8388671875 0 4888.122
680.832763671875 0 4460.739
681.336181640625 0 1604.1339
686.8494873046875 0 4257.02
687.3525390625 0 4548.9873
687.8550415039062 0 6300.1807
688.4002075195312 0 19864.016 y 10
689.40234375 0 6348.905
690.4042358398438 0 1817.7767
695.4210205078125 0 5155.7695
699.8499145507812 0 8574.659
700.3517456054688 0 5361.6865 c Water loss 11
700.8546752929688 0 3732.142 c Ammonia loss 11
701.8453369140625 0 2937.6714
702.3511352539062 0 2822.3572 z Water loss 4
703.3501586914062 0 2008.9597
707.34423828125 0 25217.809
707.8452758789062 0 17623.73
708.3456420898438 0 11119.599
708.8553466796875 0 26833.13
709.3595581054688 0 239781.03 c 11
709.8609619140625 0 182680.39
710.3615112304688 0 84394.62 y Water loss 4
710.8616333007812 0 28287.254
711.3496704101562 0 9244.265 z 4
711.8485717773438 0 18536.324
712.3489990234375 0 6631.318
712.8516235351562 0 3883.1448
713.3530883789062 0 2399.1897
716.3869018554688 0 1752.9938
717.3678588867188 0 3467.0493
718.8533935546875 0 3546.8467
719.35546875 0 9811.678 y 4
719.8568115234375 0 8018.596
720.364990234375 0 3997.3374
722.3665771484375 0 2064.0564
722.8685302734375 0 2271.5818
729.416259765625 0 6588.9434
730.4266357421875 0 2308.589
743.3823852539062 0 2004.9536
746.3580322265625 0 2686.2502 w 3
747.3643188476562 0 2345.3198
750.8597412109375 0 13441.833 z Water loss 3
751.3653564453125 0 12391.164 z Ammonia loss 3
751.8696899414062 0 6129.363
752.3751831054688 0 1624.8499
754.4337158203125 0 4948.267
755.434814453125 0 2168.5342 z Water loss 9
756.427490234375 0 13627.107 w 9
757.430419921875 0 5185.0923
758.4060668945312 0 24509.379 w 9
759.4071655273438 0 8153.6055
759.8673706054688 0 4296.1265 z 3
760.8577880859375 0 1601.3065
763.8889770507812 0 1767.8395
764.8817138671875 0 2899.7273 c Ammonia loss 12
765.3887939453125 0 5057.5996
766.4009399414062 0 2236.085
766.874755859375 0 1988.1316
767.3756713867188 0 4136.629
767.88134765625 0 80671.24 y 3
768.3824462890625 0 79003.65
768.8839111328125 0 32627.074
769.2647705078125 0 1734.8224
769.384765625 0 12984.92
769.8848266601562 0 2376.7166
771.4129028320312 0 8611.006
772.4203491210938 0 17602.06 y Ammonia loss 9
772.884521484375 0 10028.397
773.3901977539062 0 104279.05 c 12
773.890625 0 127918.87
774.3892822265625 0 39230.535
774.4354858398438 0 27021.564
774.8925170898438 0 21821.916
775.4328002929688 0 13791.255
779.4271850585938 0 2801.5388
781.8772583007812 0 2611.1135
782.3811645507812 0 2877.21
784.408203125 0 15833.729
784.9095458984375 0 10249.21
785.4130249023438 0 5007.385
787.4415283203125 0 4225.5664
788.4442749023438 0 2830.959
789.44775390625 0 25574.232 y 9
790.4497680664062 0 8856.441
791.4461669921875 0 3537.7004
791.9135131835938 0 4697.143
792.4094848632812 0 2277.2795
792.9107666015625 0 5924.4214
793.4104614257812 0 8949.297
793.9080810546875 0 5837.7725
794.408935546875 0 2516.0388
794.889404296875 0 11278.849 w 2
795.3892211914062 0 7906.7373
795.8896484375 0 4271.7964
797.4376831054688 0 4451.829
798.4483032226562 0 275741.88 c 6
799.4507446289062 0 113398.25
800.4527587890625 0 28583.879
800.9129028320312 0 10114.893
801.4190063476562 0 7461.5728
801.9199829101562 0 9302.596
802.419921875 0 6327.2373
805.4072265625 0 3606.421
806.3936767578125 0 1806.055
807.4000244140625 0 4763.266 y Water loss 2
807.9017333984375 0 11456.996 y Ammonia loss 2
808.395751953125 0 7072.517 z 2
808.8974609375 0 5016.1626
809.3991088867188 0 3472.7803
813.4188232421875 0 7324.7256
813.9164428710938 0 10619.96
814.41552734375 0 11433.175
814.9111328125 0 7312.506
815.4121704101562 0 7337.7046
815.9118041992188 0 5035.6533
816.4078979492188 0 354851.03 y 2
816.908935546875 0 341663.12
817.4096069335938 0 174050.95
817.9107666015625 0 68558.04
818.4129028320312 0 15495.797
822.4193725585938 0 29079.682
822.9229736328125 0 148771.61
823.4243774414062 0 118217.96
823.9248046875 0 75248.62
824.4395141601562 0 21508.652
824.9288940429688 0 7440.308
825.4672241210938 0 10376.644
826.4723510742188 0 2343.4453
828.9281616210938 0 2260.8657
829.4341430664062 0 5118.97
829.9195556640625 0 1968.0934
830.4451904296875 0 4524.4136
830.9094848632812 0 7850.796
831.412353515625 0 5228.092
831.911376953125 0 2798.479
835.4288330078125 0 11606.012
835.927490234375 0 9899.075
836.4301147460938 0 7636.0303
836.9139404296875 0 9142.237
837.4152221679688 0 8841.06
837.9138793945312 0 4656.204
838.4060668945312 0 1730.2506
840.4711303710938 0 2728.1802
841.4752807617188 0 2510.3816
843.4130859375 0 112191.97 w 1
843.9143676757812 0 104179.78
844.4158325195312 0 69552.65
844.9195556640625 0 29523.686
845.4239501953125 0 8172.571
845.943359375 0 1686.8148
849.4248657226562 0 2883.3845
850.4248657226562 0 2309.655
851.4180297851562 0 2794.7363
852.4214477539062 0 2015.5782
855.9696655273438 0 3908.4949
856.464599609375 0 13331.343
856.962646484375 0 4151.1587
857.4727783203125 0 34962.336
857.9285888671875 0 3907.216 c Ammonia loss 13
858.4807739257812 0 105872.95
858.928955078125 0 5733.0254
859.4827880859375 0 45808.98
859.9381103515625 0 2283.0176
860.4862060546875 0 12754.713
863.447021484375 0 4814.6216
863.9508056640625 0 2796.9333 y Water loss 1
864.4349365234375 0 3995.4385 y Ammonia loss 1
864.9378662109375 0 50851.934 z 1
865.4417114257812 0 51999.88
865.9427490234375 0 23230.48
866.4396362304688 0 334221.53 c 13
866.9405517578125 0 291156.6
867.4412841796875 0 166338
867.9417724609375 0 67086.84
868.4429931640625 0 17813.352
869.968994140625 0 5709.6455
870.4728393554688 0 6881.7812
870.9623413085938 0 4004.914
871.4478759765625 0 2468.0835
871.9584350585938 0 7117.1294
872.4556884765625 0 7215.6104
872.9501953125 0 8365.198 y 1
873.439453125 0 7993.62
873.939453125 0 6316.3975
874.43603515625 0 2907.2441
877.9598388671875 0 11626.177
878.4575805664062 0 7663.4624
878.9423217773438 0 9172.329
879.444580078125 0 11949.928
879.9481201171875 0 13959.816
880.4400634765625 0 8258.388
880.9542236328125 0 4098.42
881.4488525390625 0 2546.0571
883.474853515625 0 12879.11
883.96630859375 0 3638.252
884.4754028320312 0 9394.323
884.9664916992188 0 6468.2627
885.4647827148438 0 6410.4272
885.958740234375 0 8712.716
886.4555053710938 0 11042.473
886.9506225585938 0 8270.916
887.4346313476562 0 87748.37
887.9385375976562 0 84056.28
888.4411010742188 0 61921.1
888.9475708007812 0 23175.27
889.4595947265625 0 7306.294
891.972900390625 0 10283.879
892.4698486328125 0 25174.867
892.9632568359375 0 26770.342
893.4620361328125 0 25965.207
893.953857421875 0 23475.223
894.4573364257812 0 50401.664
894.9589233398438 0 47316.727
895.4609985351562 0 24736.277
895.947998046875 0 44221.27
896.448486328125 0 34924.082
896.9481201171875 0 15651.289
897.4515991210938 0 4575.48
900.9705200195312 0 4962.0786
901.46728515625 0 65080.156
901.96826171875 0 67192.19
902.4646606445312 0 61014.16 z Water loss 8
902.964599609375 0 33901.32
903.4622802734375 0 16528.748
903.9608764648438 0 6060.351
904.462158203125 0 2737.573
905.96630859375 0 17538.234
906.4624633789062 0 55921.406
906.9603881835938 0 51282.008
907.4598999023438 0 33460.23
907.9603271484375 0 10142.685
908.4620361328125 0 7553.4473
908.967041015625 0 2635.3318
914.9725341796875 0 80199.3
915.468994140625 0 320810.53
915.9696044921875 0 289372.44
916.4689331054688 0 181368.14
916.9701538085938 0 77253.4
917.469970703125 0 28480.494
917.9603271484375 0 2253.6038
920.4647827148438 0 32793.43 z 8
921.4674682617188 0 21524.736
922.46923828125 0 11573.529
923.4736328125 0 212549.98
923.9761352539062 0 417401.88
924.4773559570312 0 319220.7
924.978759765625 0 185693.08
925.4791259765625 0 64845.375
925.9803466796875 0 18387.123
926.4804077148438 0 3868.2297
927.4907836914062 0 227616.52 c 7
928.4935302734375 0 112757.32
929.496337890625 0 35737.527
930.49658203125 0 5197.788
936.4827880859375 0 43275.02 y 8
937.4848022460938 0 18987.291
938.4852905273438 0 6646.6543
955.5147705078125 0 2457.318
967.5253295898438 0 2471.2764
968.5302734375 0 8298.019
969.5326538085938 0 5022.9224
970.5385131835938 0 1909.6395
978.4702758789062 0 3957.9534
979.4668579101562 0 2120.9856
986.516845703125 0 9806.979
987.518798828125 0 7205.8154
990.4938354492188 0 41993.37 w 7
991.4949340820312 0 23098.592
992.4957885742188 0 9717.808
993.4508666992188 0 5729.609
994.445068359375 0 4158.4893
1012.50146484375 0 3741.6704
1013.4940795898438 0 1847.826
1014.5108032226562 0 2082.7004
1030.5130615234375 0 20472.55
1031.51513671875 0 11413.837 z Water loss 7
1032.5213623046875 0 7994.1704
1033.5189208984375 0 3458.6501
1034.49560546875 0 2468.9946
1049.5072021484375 0 146995.17 z 7
1050.5098876953125 0 89914.04
1051.5106201171875 0 35777.703
1052.5140380859375 0 10214.248
1053.50927734375 0 2178.7454
1058.4949951171875 0 1878.6102
1059.5101318359375 0 2866.3608
1065.5247802734375 0 8807.62 y 7
1066.52978515625 0 6122.7603
1067.5303955078125 0 2066.994
1073.52001953125 0 3762.5056
1074.526123046875 0 181112.67 c 8
1075.5289306640625 0 97714.54
1076.5308837890625 0 39023.004
1077.53076171875 0 11976.461
1078.5438232421875 0 1994.9746
1106.5303955078125 0 2375.031
1107.5255126953125 0 2812.867
1113.546630859375 0 5974.3906
1114.5479736328125 0 4304.5845
1115.5496826171875 0 3978.5938
1119.5379638671875 0 18936.346 w 6
1120.536376953125 0 12344.563
1121.5406494140625 0 6374.3306
1122.4923095703125 0 2410.2566
1123.492919921875 0 3207.5757
1131.5618896484375 0 6357.025
1132.568603515625 0 5146.87
1133.560546875 0 4191.2617
1134.557861328125 0 5458.1562
1135.556884765625 0 4676.4487
1175.5736083984375 0 153111.23 c 9
1176.576171875 0 99701.75 y Water loss 6
1177.577392578125 0 46456.88
1178.5513916015625 0 119079.14 z 6
1179.552734375 0 77076.11
1180.5533447265625 0 33989.223
1181.5521240234375 0 9385.894
1193.569580078125 0 3155.704
1194.570068359375 0 14541.805 y 6
1195.5684814453125 0 9889.4
1196.5748291015625 0 4051.452
1213.4866943359375 0 2156.7476
1215.6541748046875 0 2119.1987
1221.580810546875 0 3859.5432
1222.5928955078125 0 2154.8735
1229.507568359375 0 5490.382
1230.5118408203125 0 3532.568
1232.6712646484375 0 2988.178
1259.654296875 0 9959.004
1260.6568603515625 0 7566.744
1261.6611328125 0 4317.2925
1262.644775390625 0 4049.9312
1263.6240234375 0 3328.8528
1275.637939453125 0 3649.2256
1276.642578125 0 2127.106
1284.648681640625 0 4127.2944
1285.6446533203125 0 4442.838 c Water loss 10
1286.647705078125 0 2766.1077 c Ammonia loss 10
1291.6302490234375 0 2462.4487
1301.635986328125 0 2488.2317
1302.6575927734375 0 17739.303
1303.666748046875 0 41840.742 c 10
1304.66943359375 0 28023.926
1305.669677734375 0 14346.86
1306.6768798828125 0 3428.3447
1308.6229248046875 0 2107.4558
1319.6407470703125 0 2344.933
1333.6400146484375 0 3406.6626 y Ammonia loss 5
1334.6497802734375 0 74229.414 z 5
1335.655029296875 0 105793.33
1336.6566162109375 0 62146.03
1337.6583251953125 0 24965.865
1338.658447265625 0 6001.8784
1349.6602783203125 0 14619.661
1350.6673583984375 0 30290.12 y 5
1351.6693115234375 0 23208.057
1352.6739501953125 0 8910.955
1353.674560546875 0 4509.8354
1373.696044921875 0 6468.8877
1374.7061767578125 0 6812.717
1375.6995849609375 0 3913.4402
1402.6761474609375 0 2095.4539
1403.698974609375 0 2751.6648
1404.6866455078125 0 2024.3066 w 4
1416.6995849609375 0 4176.544
1417.7095947265625 0 81650.83 c 11
1418.7117919921875 0 59623.543
1419.71337890625 0 30222.555 y Water loss 4
1420.708251953125 0 8879.94
1421.6890869140625 0 8212.888 z 4
1422.688720703125 0 40627.355
1423.6900634765625 0 25614.959
1424.692626953125 0 15165.511
1425.6932373046875 0 4199.6064
1437.6988525390625 0 7221.87 y 4
1438.6993408203125 0 6619.4634
1439.7042236328125 0 3429.4893
1502.7568359375 0 2847.1768
1527.7664794921875 0 3543.7188 c Water loss 12
1528.774169921875 0 3835.5103
1529.7652587890625 0 3216.7126
1530.76953125 0 1826.5992
1534.755126953125 0 21536.703 y 3
1535.7557373046875 0 15992.154
1536.7626953125 0 9043.339
1537.7640380859375 0 6199.6543
1538.746826171875 0 1727.6572
1545.767578125 0 46668.11 c 12
1546.7725830078125 0 41353.504
1547.7716064453125 0 26317.777
1548.7772216796875 0 12639.379
1549.78466796875 0 4773.175
1600.8284912109375 0 3367.6401
1601.83447265625 0 4038.1304
1602.829833984375 0 4255.3657
1603.8287353515625 0 2448.8818
1614.8031005859375 0 8387.158 y Ammonia loss 2
1615.8018798828125 0 9996.782 z 2
1616.8082275390625 0 5737.1113
1617.808837890625 0 1955.3845
1618.826416015625 0 1972.4556
1626.800537109375 0 1674.5886
1627.8258056640625 0 2559.4084
1628.8248291015625 0 5562.815
1629.8193359375 0 6074.03
1630.7955322265625 0 2522.6035
1631.821044921875 0 4908.105 y 2
1632.808349609375 0 4999.4106
1633.8135986328125 0 4064.6333
1643.8436279296875 0 3178.2695
1644.8370361328125 0 35028.062
1645.8419189453125 0 44656.87
1646.843017578125 0 33100.02
1647.8477783203125 0 16473.15
1648.8458251953125 0 4436.531
1670.8553466796875 0 6760.2656
1671.844970703125 0 7214.737
1672.8460693359375 0 2758.1252
1673.857421875 0 2402.5574
1684.8448486328125 0 3007.7222
1685.867919921875 0 3582.3762
1686.8572998046875 0 2332.3875
1687.851806640625 0 7676.6924
1688.862060546875 0 15926.756
1689.8680419921875 0 11916.034
1690.86767578125 0 6393.3477
1701.8934326171875 0 2165.1304
1712.86865234375 0 3249.6494
1714.8570556640625 0 2585.153 c Ammonia loss 13
1715.8656005859375 0 7129.6997
1716.8524169921875 0 9269.682
1717.870361328125 0 5498.782
1718.8616943359375 0 2717.575
1728.8712158203125 0 8164.4185 z 1
1729.8773193359375 0 73112.055
1730.8807373046875 0 61631.094
1731.8746337890625 0 62368.86 c 13
1732.876220703125 0 81353.734
1733.8792724609375 0 58827.31
1734.877685546875 0 32246.451
1735.882568359375 0 11740.466
1748.857421875 0 2480.085
1769.89453125 0 5914.433
1770.8876953125 0 4344.665
1771.885498046875 0 5464.372
1772.90283203125 0 4086.4521
1775.9066162109375 0 3506.3684
1783.9580078125 0 5211.093
1784.947021484375 0 6521.736
1785.92626953125 0 11907.958
1786.9185791015625 0 11470.731
1787.9158935546875 0 14270.42
1788.911376953125 0 26921.393
1789.914794921875 0 22470.68
1790.9122314453125 0 11293.626
1791.923828125 0 5823.689
1792.9239501953125 0 3059.58
1801.9283447265625 0 12363.65
1802.9322509765625 0 45033.168
1803.935302734375 0 44612.9
1804.9345703125 0 27157.25
1805.932373046875 0 10155.0205
1806.9293212890625 0 3829.5781
1812.92333984375 0 6611.063
1813.9093017578125 0 15466.105
1814.917236328125 0 9128.573
1815.9095458984375 0 5488.4336
1816.922119140625 0 2148.4915
1818.943359375 0 7488.2524
1819.9564208984375 0 32834.094
1820.961181640625 0 33261.54
1821.963623046875 0 19598.855
1822.95947265625 0 8350.074
1828.9161376953125 0 2227.2288
1829.931396484375 0 45543.742
1830.92919921875 0 141499.84
1831.931396484375 0 136360.58
1832.9329833984375 0 83692.71
1833.9342041015625 0 31505.96
1834.9320068359375 0 13300.035
1845.940185546875 0 15976.809
1846.9434814453125 0 91979.164
1847.9510498046875 0 421289.12
1848.953857421875 0 379297.53
1849.9560546875 0 220890.45
1850.9561767578125 0 91668.5
1851.95947265625 0 27778.77
2051.005615234375 0 2480.0747
2370.64306640625 0 1690.6371

Spectrum Details

|  |  |
| --- | --- |
| Matched peaks? Matched peaks The total absolute number of peaks matched. Additionally in brackets the total fraction of peaks matched and the total number of peaks is shown. | 93 (14.72% of 632) |
| FDR? FDR The false discovery rate estimated for this peptide. It is calculated by matching all theoretical fragments with a non-integer shift with the raw peaks for this spectrum. This is done with 40 different shifts. The resulting percentage is the average number of annotated peaks over the number of annotated peaks with the correct spectrum. | 0.23% |
| Satellite FDR? Satellite FDR See the FDR for details on its calculation. This satellite ion specific FDR only contains the satellite ions (d/w) for I/L/J positions. | 2.38% |
| PSM Score? PSM Score The PSM Score as given by Hecklib to this annotated spectrum. It is shown with three significant figures. | 659 |

#### Spectrum 4114? Spectrum 4114 The raw spectrum of this peptide as annotated by Hecklib. The fragments are coloured according to ion type (see legend). Any peaks with a star '\*' as text can be hovered over to see the full details, first the ion type second the mass shift type. By hovering over the amino acids in the peptide or ions in the legend the corresponding peaks are highlighted. By toggling the 'Unassigned' label you can turn the background (unassigned) peaks on or off in the plot. By updating the slider in the Ion legend you can update the spectrum to only show the top X% of the peaks with labels. The top X% means any peak that is within X% of the highest intensity. By dragging in the spectrum you can zoom in to a specific part of the spectrum and use 'Zoom Out' to get back to the original zoom level. The annotation of the spectrum is based on the given sequence in the peptides file and is done with different software so inconsistencies are likely. The peaks are annotated based on the given sequence, with 20 ppm tolerance.

Copy Data

##### Spectrum 4114 (TSV)

###### Preview

```
Loading example...
```

*Click on the button to copy the data to your clipboard.*

Mz MinMz MaxIntensity Max

WidthHeightPeptide font sizePeptide stroke widthSpectrum font sizeSpectrum stroke widthCompact peptide

Ion legend

wxyz

abcd

OtherUnassignedIon^Charge^_Position_Show for top:%

TLPPSREEMTKNQKJ

01.35e+42.69e+44.04e+45.39e+4

Zoom Out

y^+1^_3_z^+3^_12_z^+3^_12_c^+1^_5_y^+3^_13_z^+1^_4_y^+3^_13_y^+1^_4_c^+2^_11_c^+1^_6_z^+1^_5_y^+2^_10_y^+1^_5_c^+2^_12_c^+2^_12_y^+2^_11_y^+2^_11_w^+1^_6_w^+1^_6_y^+2^_12_y^+1^_6_c^+2^_13_y^+1^_6_c^+1^_7_y^+2^_13_z^+2^_13_y^+2^_13_w^+2^_14_z^+2^_14_c^+2^_14_z^+1^_7_z^+1^_7_c^+1^_8_y^+1^_7_w^+1^_8_z^+1^_8_z^+1^_8_y^+1^_8_c^+1^_9_w^+1^_9_c^+1^_10_y^+1^_9_z^+1^_9_y^+1^_9_c^+1^_11_z^+1^_10_y^+1^_10_c^+1^_12_y^+1^_11_z^+1^_11_y^+1^_12_c^+1^_13_y^+1^_13_z^+1^_13_y^+1^_13_z^+1^_14_c^+1^_14_

046893514031870

Fragment Matches Table

Show background peaks

| Position | Ion type | Intensity | mz Theoretical | mz Error (Th) | mz Error (ppm) | Charge | Series Number |
| --- | --- | --- | --- | --- | --- | --- | --- |
| - | - | 422.2 | 124.4 | - | - | 0 | - |
| - | - | 373.4 | 128.2 | - | - | 0 | - |
| - | - | 436.2 | 162.9 | - | - | 0 | - |
| - | - | 569.3 | 173.1 | - | - | 0 | - |
| - | - | 1341 | 173.4 | - | - | 0 | - |
| - | - | 828 | 187.1 | - | - | 0 | - |
| - | - | 1.088E+04 | 187.1 | - | - | 0 | - |
| - | - | 930.1 | 188.1 | - | - | 0 | - |
| - | - | 459.4 | 198.9 | - | - | 0 | - |
| - | - | 1014 | 212.1 | - | - | 0 | - |
| - | - | 6586 | 215.1 | - | - | 0 | - |
| - | - | 805.1 | 219.1 | - | - | 0 | - |
| - | - | 542.5 | 224 | - | - | 0 | - |
| - | - | 612.9 | 243.6 | - | - | 0 | - |
| - | - | 591.9 | 254.1 | - | - | 0 | - |
| - | - | 802.7 | 256.2 | - | - | 0 | - |
| - | - | 557.2 | 270.9 | - | - | 0 | - |
| - | - | 2016 | 299.2 | - | - | 0 | - |
| - | - | 834.9 | 326.2 | - | - | 0 | - |
| - | - | 823.5 | 327.2 | - | - | 0 | - |
| - | - | 611.1 | 364.8 | - | - | 0 | - |
| - | - | 709.9 | 382.2 | - | - | 0 | - |
| - | - | 692.6 | 395.2 | - | - | 0 | - |
| - | - | 563.7 | 404.5 | - | - | 0 | - |
| - | - | 869.5 | 425.3 | - | - | 0 | - |
| 13 | y | 1351 | 446.3 | 0.005039 | 11.29 | +1 | 3 |
| - | - | 811.1 | 470.3 | - | - | 0 | - |
| - | - | 1402 | 471.9 | - | - | 0 | - |
| 4 | z | 692 | 500.9 | 0.002126 | 4.244 | +3 | 12 |
| 4 | z | 887.1 | 501.2 | 0.0001914 | 0.3819 | +3 | 12 |
| 5 | c | 7979 | 513.3 | 0.0002677 | 0.5215 | +1 | 5 |
| - | - | 2137 | 514.3 | - | - | 0 | - |
| - | - | 844.4 | 515.3 | - | - | 0 | - |
| - | - | 769.8 | 517.2 | - | - | 0 | - |
| - | - | 930.8 | 523.3 | - | - | 0 | - |
| - | - | 585.5 | 529.6 | - | - | 0 | - |
| 3 | y | 1241 | 538.6 | 0.002416 | 4.486 | +3 | 13 |
| 12 | z | 964.9 | 544.3 | 0.003516 | 6.46 | +1 | 4 |
| 3 | y | 4.533E+04 | 544.6 | 6.776E-05 | 0.1244 | +3 | 13 |
| - | - | 4.022E+04 | 544.9 | - | - | 0 | - |
| - | - | 2.227E+04 | 545.3 | - | - | 0 | - |
| - | - | 7876 | 545.6 | - | - | 0 | - |
| - | - | 1455 | 545.9 | - | - | 0 | - |
| - | - | 604.9 | 553.3 | - | - | 0 | - |
| 12 | y | 1581 | 560.3 | 0.005883 | 10.5 | +1 | 4 |
| - | - | 907.9 | 598.8 | - | - | 0 | - |
| - | - | 1161 | 614.3 | - | - | 0 | - |
| - | - | 1489 | 615.4 | - | - | 0 | - |
| - | - | 658.1 | 626.4 | - | - | 0 | - |
| - | - | 3127 | 628.4 | - | - | 0 | - |
| - | - | 757.1 | 629.3 | - | - | 0 | - |
| - | - | 916.5 | 629.4 | - | - | 0 | - |
| - | - | 580.9 | 642.8 | - | - | 0 | - |
| - | - | 1413 | 651.8 | - | - | 0 | - |
| 11 | c | 6689 | 652.3 | 0.001331 | 2.04 | +2 | 11 |
| - | - | 4170 | 652.8 | - | - | 0 | - |
| - | - | 2037 | 653.3 | - | - | 0 | - |
| - | - | 730.2 | 653.8 | - | - | 0 | - |
| 6 | c | 1.964E+04 | 669.4 | 0.0006707 | 1.002 | +1 | 6 |
| - | - | 6169 | 670.4 | - | - | 0 | - |
| - | - | 1689 | 671.4 | - | - | 0 | - |
| 11 | z | 6816 | 672.4 | 0.005462 | 8.123 | +1 | 5 |
| - | - | 2938 | 673.4 | - | - | 0 | - |
| - | - | 1827 | 675.3 | - | - | 0 | - |
| 6 | y | 959.3 | 675.8 | 0.0007444 | 1.102 | +2 | 10 |
| - | - | 1051 | 687.4 | - | - | 0 | - |
| 11 | y | 1968 | 688.4 | 0.0038 | 5.52 | +1 | 5 |
| - | - | 681.6 | 689.4 | - | - | 0 | - |
| - | - | 689 | 695.4 | - | - | 0 | - |
| 12 | c | 1201 | 700.4 | 0.005146 | 7.348 | +2 | 12 |
| - | - | 597.8 | 701.9 | - | - | 0 | - |
| - | - | 1875 | 707.3 | - | - | 0 | - |
| - | - | 1834 | 707.8 | - | - | 0 | - |
| - | - | 1003 | 708.9 | - | - | 0 | - |
| 12 | c | 2.37E+04 | 709.4 | 0.001596 | 2.249 | +2 | 12 |
| - | - | 1.566E+04 | 709.9 | - | - | 0 | - |
| 5 | y | 7267 | 710.3 | 0.01092 | 15.38 | +2 | 11 |
| - | - | 1998 | 710.9 | - | - | 0 | - |
| - | - | 1352 | 711.4 | - | - | 0 | - |
| 5 | y | 1289 | 719.4 | 0.001927 | 2.679 | +2 | 11 |
| - | - | 1157 | 719.9 | - | - | 0 | - |
| 10 | w | 1415 | 756.4 | 0.004807 | 6.355 | +1 | 6 |
| 10 | w | 1895 | 758.4 | 0.00458 | 6.038 | +1 | 6 |
| - | - | 953.3 | 759.4 | - | - | 0 | - |
| 4 | y | 7858 | 767.9 | 0.0006601 | 0.8596 | +2 | 12 |
| - | - | 7039 | 768.4 | - | - | 0 | - |
| - | - | 2557 | 768.9 | - | - | 0 | - |
| - | - | 792.2 | 769.4 | - | - | 0 | - |
| 10 | y | 1447 | 772.4 | 0.007534 | 9.754 | +1 | 6 |
| - | - | 798.7 | 772.9 | - | - | 0 | - |
| 13 | c | 8939 | 773.4 | 0.002153 | 2.784 | +2 | 13 |
| - | - | 1.074E+04 | 773.9 | - | - | 0 | - |
| - | - | 4734 | 774.4 | - | - | 0 | - |
| - | - | 2040 | 774.4 | - | - | 0 | - |
| - | - | 2288 | 774.9 | - | - | 0 | - |
| - | - | 1670 | 775.4 | - | - | 0 | - |
| - | - | 741.2 | 784.4 | - | - | 0 | - |
| - | - | 701.8 | 784.9 | - | - | 0 | - |
| - | - | 602.7 | 785.4 | - | - | 0 | - |
| 10 | y | 2398 | 789.5 | 0.006923 | 8.769 | +1 | 6 |
| - | - | 1049 | 790.4 | - | - | 0 | - |
| - | - | 579.7 | 795.9 | - | - | 0 | - |
| 7 | c | 2.545E+04 | 798.4 | 0.0008444 | 1.058 | +1 | 7 |
| - | - | 736.5 | 799.4 | - | - | 0 | - |
| - | - | 1.073E+04 | 799.4 | - | - | 0 | - |
| - | - | 3077 | 800.5 | - | - | 0 | - |
| - | - | 1173 | 800.9 | - | - | 0 | - |
| - | - | 833.2 | 801.4 | - | - | 0 | - |
| - | - | 952.2 | 801.9 | - | - | 0 | - |
| - | - | 827.4 | 802.4 | - | - | 0 | - |
| - | - | 654.5 | 805.4 | - | - | 0 | - |
| 3 | y | 895.7 | 807.9 | 0.004665 | 5.774 | +2 | 13 |
| 3 | z | 879.2 | 808.4 | 0.003133 | 3.876 | +2 | 13 |
| - | - | 1253 | 813.4 | - | - | 0 | - |
| - | - | 764.9 | 813.9 | - | - | 0 | - |
| - | - | 694 | 814.4 | - | - | 0 | - |
| - | - | 804.9 | 815.4 | - | - | 0 | - |
| 3 | y | 2.485E+04 | 816.4 | 0.0002476 | 0.3033 | +2 | 13 |
| - | - | 2.047E+04 | 816.9 | - | - | 0 | - |
| - | - | 1.284E+04 | 817.4 | - | - | 0 | - |
| - | - | 3574 | 817.9 | - | - | 0 | - |
| - | - | 1117 | 818.4 | - | - | 0 | - |
| - | - | 1904 | 822.4 | - | - | 0 | - |
| - | - | 1.283E+04 | 822.9 | - | - | 0 | - |
| - | - | 9415 | 823.4 | - | - | 0 | - |
| - | - | 5996 | 823.9 | - | - | 0 | - |
| - | - | 3221 | 824.4 | - | - | 0 | - |
| - | - | 696.1 | 824.9 | - | - | 0 | - |
| - | - | 831.9 | 825.5 | - | - | 0 | - |
| - | - | 589.4 | 830.4 | - | - | 0 | - |
| - | - | 1115 | 830.9 | - | - | 0 | - |
| - | - | 1408 | 835.4 | - | - | 0 | - |
| - | - | 742.8 | 835.9 | - | - | 0 | - |
| - | - | 932.8 | 836.4 | - | - | 0 | - |
| - | - | 955.2 | 836.9 | - | - | 0 | - |
| 2 | w | 1.217E+04 | 843.4 | 9.781E-05 | 0.116 | +2 | 14 |
| - | - | 1.133E+04 | 843.9 | - | - | 0 | - |
| - | - | 7188 | 844.4 | - | - | 0 | - |
| - | - | 1819 | 844.9 | - | - | 0 | - |
| - | - | 1107 | 845.4 | - | - | 0 | - |
| - | - | 742 | 849.4 | - | - | 0 | - |
| - | - | 774.9 | 856.5 | - | - | 0 | - |
| - | - | 3763 | 857.5 | - | - | 0 | - |
| - | - | 8359 | 858.5 | - | - | 0 | - |
| - | - | 798.5 | 858.9 | - | - | 0 | - |
| - | - | 5127 | 859.5 | - | - | 0 | - |
| - | - | 1382 | 860.5 | - | - | 0 | - |
| 2 | z | 3137 | 864.9 | 0.0002027 | 0.2344 | +2 | 14 |
| - | - | 4523 | 865.4 | - | - | 0 | - |
| - | - | 3013 | 865.9 | - | - | 0 | - |
| 14 | c | 2.61E+04 | 866.4 | 0.0002797 | 0.3228 | +2 | 14 |
| - | - | 2.631E+04 | 866.9 | - | - | 0 | - |
| - | - | 1.433E+04 | 867.4 | - | - | 0 | - |
| - | - | 5727 | 867.9 | - | - | 0 | - |
| - | - | 1867 | 868.4 | - | - | 0 | - |
| - | - | 687.5 | 872 | - | - | 0 | - |
| - | - | 1379 | 873.4 | - | - | 0 | - |
| - | - | 937.3 | 878 | - | - | 0 | - |
| - | - | 1325 | 878.5 | - | - | 0 | - |
| - | - | 978.5 | 879.4 | - | - | 0 | - |
| - | - | 575.8 | 879.9 | - | - | 0 | - |
| - | - | 1176 | 880.4 | - | - | 0 | - |
| - | - | 921.8 | 883.5 | - | - | 0 | - |
| - | - | 1391 | 884.5 | - | - | 0 | - |
| - | - | 705.5 | 886.5 | - | - | 0 | - |
| - | - | 1191 | 886.9 | - | - | 0 | - |
| - | - | 9570 | 887.4 | - | - | 0 | - |
| - | - | 7641 | 887.9 | - | - | 0 | - |
| - | - | 4590 | 888.4 | - | - | 0 | - |
| - | - | 2136 | 889 | - | - | 0 | - |
| - | - | 808 | 889.5 | - | - | 0 | - |
| - | - | 1302 | 892 | - | - | 0 | - |
| - | - | 2231 | 892.5 | - | - | 0 | - |
| - | - | 1840 | 893 | - | - | 0 | - |
| - | - | 2415 | 893.5 | - | - | 0 | - |
| - | - | 2906 | 894 | - | - | 0 | - |
| - | - | 3458 | 894.5 | - | - | 0 | - |
| - | - | 2382 | 895 | - | - | 0 | - |
| - | - | 2213 | 895.5 | - | - | 0 | - |
| - | - | 3750 | 895.9 | - | - | 0 | - |
| - | - | 2217 | 896.4 | - | - | 0 | - |
| - | - | 2119 | 896.9 | - | - | 0 | - |
| - | - | 6543 | 901.5 | - | - | 0 | - |
| - | - | 6043 | 902 | - | - | 0 | - |
| 9 | z | 5834 | 902.5 | 0.01111 | 12.31 | +1 | 7 |
| - | - | 3329 | 903 | - | - | 0 | - |
| - | - | 2059 | 903.5 | - | - | 0 | - |
| - | - | 1634 | 906 | - | - | 0 | - |
| - | - | 3798 | 906.5 | - | - | 0 | - |
| - | - | 4377 | 907 | - | - | 0 | - |
| - | - | 2279 | 907.5 | - | - | 0 | - |
| - | - | 1253 | 908 | - | - | 0 | - |
| - | - | 619.2 | 908.5 | - | - | 0 | - |
| - | - | 6965 | 915 | - | - | 0 | - |
| - | - | 2.738E+04 | 915.5 | - | - | 0 | - |
| - | - | 3.055E+04 | 916 | - | - | 0 | - |
| - | - | 1.646E+04 | 916.5 | - | - | 0 | - |
| - | - | 6324 | 917 | - | - | 0 | - |
| - | - | 1832 | 917.5 | - | - | 0 | - |
| 9 | z | 2885 | 920.5 | 0.001105 | 1.201 | +1 | 7 |
| - | - | 1985 | 921.5 | - | - | 0 | - |
| - | - | 952.5 | 922.5 | - | - | 0 | - |
| - | - | 2.033E+04 | 923.5 | - | - | 0 | - |
| - | - | 3.584E+04 | 924 | - | - | 0 | - |
| - | - | 2.966E+04 | 924.5 | - | - | 0 | - |
| - | - | 1.328E+04 | 925 | - | - | 0 | - |
| - | - | 6161 | 925.5 | - | - | 0 | - |
| - | - | 1758 | 926 | - | - | 0 | - |
| 8 | c | 2.179E+04 | 927.5 | 0.001323 | 1.427 | +1 | 8 |
| - | - | 1.021E+04 | 928.5 | - | - | 0 | - |
| - | - | 2828 | 929.5 | - | - | 0 | - |
| 9 | y | 2819 | 936.5 | 0.0006642 | 0.7093 | +1 | 7 |
| - | - | 962 | 937.5 | - | - | 0 | - |
| - | - | 727.6 | 969.4 | - | - | 0 | - |
| 8 | w | 4796 | 990.5 | 0.0006088 | 0.6146 | +1 | 8 |
| - | - | 2161 | 991.5 | - | - | 0 | - |
| - | - | 2026 | 1031 | - | - | 0 | - |
| 8 | z | 1269 | 1031 | 0.01655 | 16.04 | +1 | 8 |
| 8 | z | 1.383E+04 | 1050 | 0.0001192 | 0.1136 | +1 | 8 |
| - | - | 7670 | 1051 | - | - | 0 | - |
| - | - | 3312 | 1052 | - | - | 0 | - |
| - | - | 1133 | 1053 | - | - | 0 | - |
| 8 | y | 992.3 | 1066 | 0.001265 | 1.187 | +1 | 8 |
| 9 | c | 1.852E+04 | 1075 | 0.004264 | 3.968 | +1 | 9 |
| - | - | 1.116E+04 | 1076 | - | - | 0 | - |
| - | - | 3594 | 1077 | - | - | 0 | - |
| - | - | 872.6 | 1078 | - | - | 0 | - |
| 7 | w | 1646 | 1120 | 0.002552 | 2.28 | +1 | 9 |
| - | - | 1182 | 1121 | - | - | 0 | - |
| - | - | 915.7 | 1132 | - | - | 0 | - |
| - | - | 901.6 | 1135 | - | - | 0 | - |
| 10 | c | 1.433E+04 | 1176 | 0.003827 | 3.255 | +1 | 10 |
| 7 | y | 1.026E+04 | 1177 | 0.01785 | 15.17 | +1 | 9 |
| - | - | 3757 | 1178 | - | - | 0 | - |
| 7 | z | 1.125E+04 | 1179 | 0.0008668 | 0.7355 | +1 | 9 |
| - | - | 7046 | 1180 | - | - | 0 | - |
| - | - | 2954 | 1181 | - | - | 0 | - |
| - | - | 931.7 | 1182 | - | - | 0 | - |
| - | - | 790.1 | 1194 | - | - | 0 | - |
| 7 | y | 1661 | 1195 | 0.006016 | 5.037 | +1 | 9 |
| - | - | 840.2 | 1196 | - | - | 0 | - |
| - | - | 2267 | 1214 | - | - | 0 | - |
| - | - | 1274 | 1215 | - | - | 0 | - |
| - | - | 912.8 | 1216 | - | - | 0 | - |
| - | - | 1126 | 1217 | - | - | 0 | - |
| - | - | 671.3 | 1221 | - | - | 0 | - |
| - | - | 665.2 | 1222 | - | - | 0 | - |
| - | - | 1432 | 1303 | - | - | 0 | - |
| 11 | c | 4359 | 1304 | 0.002858 | 2.192 | +1 | 11 |
| - | - | 2847 | 1305 | - | - | 0 | - |
| 6 | z | 5963 | 1335 | 0.002466 | 1.848 | +1 | 10 |
| - | - | 1.04E+04 | 1336 | - | - | 0 | - |
| - | - | 6648 | 1337 | - | - | 0 | - |
| - | - | 2532 | 1338 | - | - | 0 | - |
| - | - | 943.5 | 1339 | - | - | 0 | - |
| - | - | 1389 | 1350 | - | - | 0 | - |
| 6 | y | 3898 | 1351 | 0.002757 | 2.041 | +1 | 10 |
| - | - | 2346 | 1352 | - | - | 0 | - |
| - | - | 830.5 | 1417 | - | - | 0 | - |
| 12 | c | 9836 | 1418 | 0.002655 | 1.873 | +1 | 12 |
| - | - | 6509 | 1419 | - | - | 0 | - |
| 5 | y | 3486 | 1420 | 0.02265 | 15.96 | +1 | 11 |
| - | - | 1173 | 1421 | - | - | 0 | - |
| 5 | z | 950 | 1422 | 0.02202 | 15.49 | +1 | 11 |
| - | - | 3637 | 1423 | - | - | 0 | - |
| - | - | 2666 | 1424 | - | - | 0 | - |
| - | - | 1082 | 1425 | - | - | 0 | - |
| - | - | 720 | 1520 | - | - | 0 | - |
| 4 | y | 2301 | 1535 | 0.0002189 | 0.1427 | +1 | 12 |
| - | - | 1870 | 1536 | - | - | 0 | - |
| - | - | 1365 | 1537 | - | - | 0 | - |
| 13 | c | 6273 | 1546 | 0.001357 | 0.8779 | +1 | 13 |
| - | - | 6345 | 1547 | - | - | 0 | - |
| - | - | 2587 | 1548 | - | - | 0 | - |
| - | - | 902.6 | 1549 | - | - | 0 | - |
| - | - | 882.4 | 1602 | - | - | 0 | - |
| 3 | y | 1065 | 1615 | 0.02039 | 12.63 | +1 | 13 |
| 3 | z | 1076 | 1616 | 0.01476 | 9.137 | +1 | 13 |
| - | - | 1049 | 1629 | - | - | 0 | - |
| - | - | 706.1 | 1630 | - | - | 0 | - |
| 3 | y | 723.2 | 1632 | 0.008612 | 5.278 | +1 | 13 |
| - | - | 807.5 | 1633 | - | - | 0 | - |
| - | - | 4285 | 1645 | - | - | 0 | - |
| - | - | 4739 | 1646 | - | - | 0 | - |
| - | - | 4362 | 1647 | - | - | 0 | - |
| - | - | 1722 | 1648 | - | - | 0 | - |
| - | - | 843.2 | 1671 | - | - | 0 | - |
| - | - | 1004 | 1672 | - | - | 0 | - |
| - | - | 1176 | 1673 | - | - | 0 | - |
| - | - | 1175 | 1688 | - | - | 0 | - |
| - | - | 1596 | 1689 | - | - | 0 | - |
| - | - | 1131 | 1690 | - | - | 0 | - |
| - | - | 1196 | 1716 | - | - | 0 | - |
| - | - | 740 | 1718 | - | - | 0 | - |
| 2 | z | 1485 | 1729 | 0.001674 | 0.9682 | +1 | 14 |
| - | - | 7302 | 1730 | - | - | 0 | - |
| - | - | 8131 | 1731 | - | - | 0 | - |
| 14 | c | 7041 | 1732 | 0.004276 | 2.469 | +1 | 14 |
| - | - | 9763 | 1733 | - | - | 0 | - |
| - | - | 7098 | 1734 | - | - | 0 | - |
| - | - | 3329 | 1735 | - | - | 0 | - |
| - | - | 1493 | 1736 | - | - | 0 | - |
| - | - | 807.6 | 1784 | - | - | 0 | - |
| - | - | 1903 | 1786 | - | - | 0 | - |
| - | - | 1055 | 1787 | - | - | 0 | - |
| - | - | 1745 | 1788 | - | - | 0 | - |
| - | - | 3441 | 1789 | - | - | 0 | - |
| - | - | 2242 | 1790 | - | - | 0 | - |
| - | - | 2058 | 1791 | - | - | 0 | - |
| - | - | 1363 | 1802 | - | - | 0 | - |
| - | - | 5363 | 1803 | - | - | 0 | - |
| - | - | 6102 | 1804 | - | - | 0 | - |
| - | - | 2728 | 1805 | - | - | 0 | - |
| - | - | 1092 | 1806 | - | - | 0 | - |
| - | - | 1567 | 1814 | - | - | 0 | - |
| - | - | 1589 | 1815 | - | - | 0 | - |
| - | - | 1061 | 1816 | - | - | 0 | - |
| - | - | 4267 | 1820 | - | - | 0 | - |
| - | - | 3893 | 1821 | - | - | 0 | - |
| - | - | 2302 | 1822 | - | - | 0 | - |
| - | - | 4420 | 1830 | - | - | 0 | - |
| - | - | 1.783E+04 | 1831 | - | - | 0 | - |
| - | - | 1.796E+04 | 1832 | - | - | 0 | - |
| - | - | 9260 | 1833 | - | - | 0 | - |
| - | - | 3955 | 1834 | - | - | 0 | - |
| - | - | 807 | 1835 | - | - | 0 | - |
| - | - | 2241 | 1846 | - | - | 0 | - |
| - | - | 1.102E+04 | 1847 | - | - | 0 | - |
| - | - | 5.336E+04 | 1848 | - | - | 0 | - |
| - | - | 4.858E+04 | 1849 | - | - | 0 | - |
| - | - | 2.64E+04 | 1850 | - | - | 0 | - |
| - | - | 1.187E+04 | 1851 | - | - | 0 | - |
| - | - | 2673 | 1852 | - | - | 0 | - |

m/z Charge Intensity FragmentType MassShift Position
124.4217758178711 0 422.19672
128.1709442138672 0 373.35443
162.87222290039062 0 436.17444
173.0546112060547 0 569.3303
173.43896484375 0 1341.0754
187.13552856445312 0 827.9665
187.14390563964844 0 10877.812
188.1474151611328 0 930.0573
198.9484100341797 0 459.3628
212.1393280029297 0 1013.53406
215.13877868652344 0 6586.4097
219.13369750976562 0 805.143
223.98277282714844 0 542.5279
243.63925170898438 0 612.9141
254.1490936279297 0 591.92413
256.1651306152344 0 802.66284
270.89434814453125 0 557.2197
299.1716613769531 0 2016.4805
326.1582336425781 0 834.89307
327.1656188964844 0 823.4612
364.7673645019531 0 611.07513
382.2075500488281 0 709.897
395.1811828613281 0 692.6002
404.5128173828125 0 563.6818
425.2624206542969 0 869.52
446.2604064941406 0 1350.7446 y 12
470.29705810546875 0 811.0645
471.89654541015625 0 1401.6714
500.9104919433594 0 692.04095 z Water loss 3
501.2408142089844 0 887.087 z Ammonia loss 3
513.3028564453125 0 7978.5205 c 4
514.306640625 0 2136.6963
515.3074340820312 0 844.3956
517.169677734375 0 769.8032
523.2719116210938 0 930.82916
529.6067504882812 0 585.4702
538.6055297851562 0 1240.8108 y Water loss 2
544.2861328125 0 964.9405 z 11
544.6065673828125 0 45329.734 y 2
544.9407958984375 0 40223.285
545.275390625 0 22265.252
545.608154296875 0 7875.842
545.9428100585938 0 1455.0952
553.2569580078125 0 604.8568
560.302490234375 0 1581.0315 y 11
598.7772216796875 0 907.9143
614.34912109375 0 1161.2833
615.3502197265625 0 1489.4994
626.3999633789062 0 658.05756
628.3665161132812 0 3126.9814
629.322998046875 0 757.1239
629.3724975585938 0 916.46497
642.8282470703125 0 580.94165
651.8306884765625 0 1413.063
652.3362426757812 0 6688.586 c 10
652.8383178710938 0 4169.9507
653.33935546875 0 2036.659
653.840087890625 0 730.163
669.403564453125 0 19637.297 c 5
670.4064331054688 0 6168.8555
671.4077758789062 0 1688.8168
672.379150390625 0 6815.8413 z 10
673.3833618164062 0 2937.6277
675.33544921875 0 1826.8838
675.8367919921875 0 959.27496 y 5
687.351806640625 0 1051.4313
688.3995361328125 0 1968.2935 y 10
689.4032592773438 0 681.5786
695.4194946289062 0 689.0205
700.345947265625 0 1201.3138 c Water loss 11
701.85107421875 0 597.84705
707.342041015625 0 1874.8478
707.8440551757812 0 1833.8949
708.8565673828125 0 1002.89795
709.3579711914062 0 23701.81 c 11
709.8598022460938 0 15658.008
710.3591918945312 0 7267.4136 y Water loss 4
710.8604125976562 0 1997.8574
711.3585815429688 0 1352.0133
719.3516235351562 0 1289.1943 y 4
719.8564453125 0 1157.2092
756.4247436523438 0 1414.6157 w 9
758.4042358398438 0 1895.0477 w 9
759.4107055664062 0 953.31946
767.8792724609375 0 7858.2188 y 3
768.3806762695312 0 7038.9316
768.881591796875 0 2557.0188
769.3804321289062 0 792.2247
772.4169311523438 0 1447.2246 y Ammonia loss 9
772.8914794921875 0 798.7304
773.3878173828125 0 8939.033 c 12
773.8887939453125 0 10735.206
774.3889770507812 0 4734.2134
774.4356689453125 0 2040.024
774.8909912109375 0 2287.928
775.4346313476562 0 1669.926
784.4027099609375 0 741.1999
784.9107055664062 0 701.8158
785.41015625 0 602.73474
789.444091796875 0 2398.299 y 9
790.4476928710938 0 1048.792
795.8798217773438 0 579.6671
798.4459838867188 0 25448.26 c 6
799.3665161132812 0 736.53925
799.4492797851562 0 10732.508
800.4506225585938 0 3076.8386
800.9118041992188 0 1172.6733
801.4197387695312 0 833.2145
801.918212890625 0 952.2241
802.4210205078125 0 827.41797
805.3932495117188 0 654.5142
807.897705078125 0 895.7302 y Ammonia loss 2
808.4000854492188 0 879.168 z 2
813.4121704101562 0 1252.9149
813.9141845703125 0 764.9378
814.4072875976562 0 693.9585
815.406982421875 0 804.8875
816.4060668945312 0 24845.574 y 2
816.9074096679688 0 20466.85
817.4081420898438 0 12839.705
817.9092407226562 0 3573.6448
818.4098510742188 0 1116.9854
822.4178466796875 0 1904.2864
822.921142578125 0 12832.69
823.4229736328125 0 9415.086
823.9252319335938 0 5995.5664
824.4367065429688 0 3220.6028
824.923583984375 0 696.1025
825.4637451171875 0 831.8564
830.44580078125 0 589.42303
830.9067993164062 0 1114.6785
835.4232177734375 0 1408.2864
835.9222412109375 0 742.81586
836.427490234375 0 932.8213
836.9149780273438 0 955.178
843.4114990234375 0 12167.091 w 1
843.9124145507812 0 11325.86
844.41357421875 0 7188.497
844.9176635742188 0 1818.9319
845.4218139648438 0 1106.7178
849.4149169921875 0 742.0004
856.46240234375 0 774.89984
857.4708251953125 0 3762.5764
858.478759765625 0 8359.318
858.9321899414062 0 798.544
859.4832763671875 0 5126.5054
860.4827880859375 0 1381.6213
864.9387817382812 0 3137.1497 z 1
865.4391479492188 0 4523.096
865.9426879882812 0 3013.0686
866.4378662109375 0 26095.484 c 13
866.9392700195312 0 26306.91
867.4395751953125 0 14325.165
867.9415893554688 0 5726.6675
868.4391479492188 0 1867.0172
871.966064453125 0 687.52277
873.435302734375 0 1379.2997
877.9603271484375 0 937.3333
878.460205078125 0 1324.5994
879.43017578125 0 978.4904
879.9266967773438 0 575.79596
880.43798828125 0 1175.8368
883.4671630859375 0 921.83826
884.4743041992188 0 1391.199
886.4542236328125 0 705.5052
886.9467163085938 0 1191.4343
887.4336547851562 0 9569.632
887.936279296875 0 7640.5996
888.4411010742188 0 4589.7427
888.950927734375 0 2135.523
889.4594116210938 0 807.9847
891.969482421875 0 1301.603
892.460693359375 0 2231.4233
892.9639282226562 0 1840.1318
893.4588623046875 0 2415.4568
893.95361328125 0 2905.7673
894.4533081054688 0 3458.0754
894.9606323242188 0 2382.006
895.460693359375 0 2212.7153
895.94921875 0 3749.9915
896.4451293945312 0 2217.2964
896.9463500976562 0 2119.27
901.4650268554688 0 6543.3535
901.9662475585938 0 6042.5425
902.4633178710938 0 5833.895 z Water loss 8
902.96044921875 0 3329.313
903.460205078125 0 2059.4036
905.9668579101562 0 1634.1078
906.460693359375 0 3797.5122
906.9612426757812 0 4376.513
907.456298828125 0 2279.019
907.96142578125 0 1252.7324
908.4628295898438 0 619.2022
914.9694213867188 0 6965.2153
915.4669799804688 0 27379.5
915.9672241210938 0 30545.979
916.4678955078125 0 16461.422
916.9674682617188 0 6324.259
917.4673461914062 0 1831.8832
920.461669921875 0 2885.3008 z 8
921.4649658203125 0 1985.2239
922.4747924804688 0 952.47675
923.4718017578125 0 20329.434
923.9740600585938 0 35840.074
924.4752807617188 0 29662.154
924.9773559570312 0 13280.454
925.476806640625 0 6161.1016
925.9835815429688 0 1757.6721
927.4880981445312 0 21794.203 c 7
928.4906616210938 0 10210.265
929.4910278320312 0 2828.2812
936.4808349609375 0 2818.909 y 8
937.4852294921875 0 962.0351
969.4468994140625 0 727.6485
990.491455078125 0 4796.0674 w 7
991.494140625 0 2161.4893
1030.508056640625 0 2026.2911
1031.5113525390625 0 1269.3424 z Water loss 7
1049.5052490234375 0 13830.004 z 7
1050.5084228515625 0 7669.511
1051.5101318359375 0 3311.8916
1052.5078125 0 1132.808
1065.5228271484375 0 992.28894 y 7
1074.524169921875 0 18524.057 c 8
1075.5267333984375 0 11159.598
1076.5284423828125 0 3593.9658
1077.53271484375 0 872.5544
1119.5321044921875 0 1646.111 w 6
1120.54345703125 0 1181.8083
1131.56103515625 0 915.7356
1134.5513916015625 0 901.58026
1175.5714111328125 0 14332.991 c 9
1176.573974609375 0 10260.197 y Water loss 6
1177.575927734375 0 3757.0269
1178.548828125 0 11246.39 z 6
1179.5516357421875 0 7046.499
1180.550048828125 0 2953.8245
1181.5623779296875 0 931.7137
1193.5570068359375 0 790.1332
1194.5606689453125 0 1661.2012 y 6
1195.5574951171875 0 840.2497
1213.5438232421875 0 2267.3738
1214.545654296875 0 1274.1735
1215.540283203125 0 912.8174
1216.578125 0 1125.7932
1220.577880859375 0 671.291
1221.5902099609375 0 665.15173
1302.6632080078125 0 1431.8077
1303.6654052734375 0 4359.453 c 10
1304.6636962890625 0 2846.9521
1334.6466064453125 0 5962.7236 z 5
1335.6531982421875 0 10404.56
1336.6534423828125 0 6647.778
1337.6629638671875 0 2532.2568
1338.6588134765625 0 943.4757
1349.646728515625 0 1389.2745
1350.6650390625 0 3898.2598 y 5
1351.666259765625 0 2345.596
1416.703125 0 830.54315
1417.7081298828125 0 9835.537 c 11
1418.7103271484375 0 6509.319
1419.7119140625 0 3485.8008 y Water loss 4
1420.714599609375 0 1172.7513
1421.703125 0 949.9564 z 4
1422.6827392578125 0 3636.5083
1423.68701171875 0 2666.4888
1424.696533203125 0 1082.2734
1520.2586669921875 0 719.9776
1534.7528076171875 0 2300.6313 y 3
1535.7569580078125 0 1869.6859
1536.75830078125 0 1364.892
1545.7626953125 0 6272.89 c 12
1546.7708740234375 0 6345.077
1547.7696533203125 0 2586.652
1548.7864990234375 0 902.6252
1601.8321533203125 0 882.36316
1614.7991943359375 0 1065.4426 y Ammonia loss 2
1615.8013916015625 0 1075.8314 z 2
1628.829833984375 0 1049.0223
1629.8310546875 0 706.11127
1631.81396484375 0 723.2237 y 2
1632.801513671875 0 807.5244
1644.8336181640625 0 4285.442
1645.8392333984375 0 4738.9727
1646.8431396484375 0 4361.9736
1647.8463134765625 0 1722.4708
1670.845947265625 0 843.159
1671.8316650390625 0 1003.87555
1672.850830078125 0 1176.0011
1687.8563232421875 0 1174.5815
1688.8658447265625 0 1596.0537
1689.85595703125 0 1130.5347
1715.84716796875 0 1196.2844
1717.8818359375 0 740.01416
1728.8690185546875 0 1484.994 z 1
1729.8760986328125 0 7301.904
1730.880126953125 0 8130.5264
1731.873291015625 0 7040.803 c 13
1732.873046875 0 9763.143
1733.8797607421875 0 7098.375
1734.882568359375 0 3328.7878
1735.8787841796875 0 1493.2308
1783.9381103515625 0 807.559
1785.92333984375 0 1903.374
1786.9281005859375 0 1054.6423
1787.9141845703125 0 1745.3395
1788.91015625 0 3441.36
1789.9151611328125 0 2242.29
1790.9034423828125 0 2057.6445
1801.9371337890625 0 1362.7361
1802.931640625 0 5362.812
1803.9320068359375 0 6101.8213
1804.9285888671875 0 2727.6465
1805.949951171875 0 1092.1035
1813.91015625 0 1567.3857
1814.9171142578125 0 1588.7076
1815.91064453125 0 1061.0322
1819.9527587890625 0 4267.312
1820.9625244140625 0 3893.4177
1821.9532470703125 0 2302.2722
1829.928955078125 0 4420.2344
1830.925537109375 0 17833.645
1831.929443359375 0 17964.34
1832.9310302734375 0 9259.851
1833.933349609375 0 3955.4697
1834.933837890625 0 807.0004
1845.939453125 0 2240.898
1846.9412841796875 0 11018.01
1847.9488525390625 0 53360.625
1848.951171875 0 48584.58
1849.9537353515625 0 26397.264
1850.9527587890625 0 11865.503
1851.9537353515625 0 2672.565

Spectrum Details

|  |  |
| --- | --- |
| Matched peaks? Matched peaks The total absolute number of peaks matched. Additionally in brackets the total fraction of peaks matched and the total number of peaks is shown. | 57 (17.12% of 333) |
| FDR? FDR The false discovery rate estimated for this peptide. It is calculated by matching all theoretical fragments with a non-integer shift with the raw peaks for this spectrum. This is done with 40 different shifts. The resulting percentage is the average number of annotated peaks over the number of annotated peaks with the correct spectrum. | 0.17% |
| Satellite FDR? Satellite FDR See the FDR for details on its calculation. This satellite ion specific FDR only contains the satellite ions (d/w) for I/L/J positions. | 0.00% |
| PSM Score? PSM Score The PSM Score as given by Hecklib to this annotated spectrum. It is shown with three significant figures. | 424 |

#### Spectrum 4656? Spectrum 4656 The raw spectrum of this peptide as annotated by Hecklib. The fragments are coloured according to ion type (see legend). Any peaks with a star '\*' as text can be hovered over to see the full details, first the ion type second the mass shift type. By hovering over the amino acids in the peptide or ions in the legend the corresponding peaks are highlighted. By toggling the 'Unassigned' label you can turn the background (unassigned) peaks on or off in the plot. By updating the slider in the Ion legend you can update the spectrum to only show the top X% of the peaks with labels. The top X% means any peak that is within X% of the highest intensity. By dragging in the spectrum you can zoom in to a specific part of the spectrum and use 'Zoom Out' to get back to the original zoom level. The annotation of the spectrum is based on the given sequence in the peptides file and is done with different software so inconsistencies are likely. The peaks are annotated based on the given sequence, with 20 ppm tolerance.

Copy Data

##### Spectrum 4656 (TSV)

###### Preview

```
Loading example...
```

*Click on the button to copy the data to your clipboard.*

Mz MinMz MaxIntensity Max

WidthHeightPeptide font sizePeptide stroke widthSpectrum font sizeSpectrum stroke widthCompact peptide

Ion legend

wxyz

abcd

OtherUnassignedIon^Charge^_Position_Show for top:%

TLPPSREEMTKNQKJ

03.19e+46.37e+49.56e+41.27e+5

Zoom Out

y^+1^_1_c^+1^_4_z^+1^_3_y^+1^_3_z^+3^_12_z^+3^_12_c^+1^_5_y^+3^_13_y^+3^_13_z^+3^_13_z^+1^_4_y^+3^_13_y^+1^_4_c^+2^_11_c^+2^_11_z^+2^_10_c^+1^_6_z^+1^_5_y^+2^_10_y^+1^_5_c^+2^_12_c^+2^_12_z^+2^_11_c^+2^_12_y^+2^_11_z^+2^_11_y^+2^_11_z^+2^_12_z^+2^_12_w^+1^_6_w^+1^_6_y^+2^_12_y^+1^_6_c^+2^_13_y^+1^_6_w^+2^_13_c^+1^_7_y^+2^_13_y^+2^_13_z^+2^_13_y^+2^_13_w^+2^_14_y^+2^_14_z^+2^_14_c^+2^_14_y^+2^_14_z^+1^_7_z^+1^_7_c^+1^_8_y^+1^_7_w^+1^_8_z^+1^_8_z^+1^_8_y^+1^_8_c^+1^_9_w^+1^_9_c^+1^_10_y^+1^_9_z^+1^_9_y^+1^_9_c^+1^_11_c^+1^_11_c^+1^_11_y^+1^_10_z^+1^_10_y^+1^_10_z^+1^_11_c^+1^_12_y^+1^_11_z^+1^_11_y^+1^_11_c^+1^_13_c^+1^_13_y^+1^_12_c^+1^_13_y^+1^_13_z^+1^_13_y^+1^_13_y^+1^_14_z^+1^_14_c^+1^_14_

046893614041871

Fragment Matches Table

Show background peaks

| Position | Ion type | Intensity | mz Theoretical | mz Error (Th) | mz Error (ppm) | Charge | Series Number |
| --- | --- | --- | --- | --- | --- | --- | --- |
| 15 | y | 1077 | 132.1 | 0.0002067 | 1.565 | +1 | 1 |
| - | - | 416.5 | 149.4 | - | - | 0 | - |
| - | - | 457.6 | 166.1 | - | - | 0 | - |
| - | - | 807.2 | 173.4 | - | - | 0 | - |
| - | - | 455 | 177.6 | - | - | 0 | - |
| - | - | 476.8 | 186.1 | - | - | 0 | - |
| - | - | 3.713E+04 | 187.1 | - | - | 0 | - |
| - | - | 3168 | 188.1 | - | - | 0 | - |
| - | - | 626.6 | 201.1 | - | - | 0 | - |
| - | - | 2674 | 212.1 | - | - | 0 | - |
| - | - | 550.7 | 212.1 | - | - | 0 | - |
| - | - | 2.281E+04 | 215.1 | - | - | 0 | - |
| - | - | 2588 | 216.1 | - | - | 0 | - |
| - | - | 3277 | 219.1 | - | - | 0 | - |
| - | - | 5201 | 221.1 | - | - | 0 | - |
| - | - | 561.8 | 227.4 | - | - | 0 | - |
| - | - | 712.8 | 238.2 | - | - | 0 | - |
| - | - | 817.9 | 247.1 | - | - | 0 | - |
| - | - | 2538 | 249.1 | - | - | 0 | - |
| - | - | 1015 | 256.2 | - | - | 0 | - |
| - | - | 481.6 | 268.6 | - | - | 0 | - |
| - | - | 518.3 | 278.5 | - | - | 0 | - |
| - | - | 6559 | 299.2 | - | - | 0 | - |
| - | - | 904.7 | 312.2 | - | - | 0 | - |
| - | - | 1958 | 326.2 | - | - | 0 | - |
| - | - | 967.9 | 327.2 | - | - | 0 | - |
| - | - | 651.3 | 344.2 | - | - | 0 | - |
| - | - | 2123 | 371.2 | - | - | 0 | - |
| - | - | 640.3 | 374.1 | - | - | 0 | - |
| - | - | 664.5 | 382.3 | - | - | 0 | - |
| - | - | 602.8 | 386.2 | - | - | 0 | - |
| - | - | 583.2 | 395.2 | - | - | 0 | - |
| - | - | 996.8 | 407.3 | - | - | 0 | - |
| - | - | 689.1 | 414.2 | - | - | 0 | - |
| - | - | 658.5 | 415.5 | - | - | 0 | - |
| - | - | 2023 | 425.3 | - | - | 0 | - |
| 4 | c | 1621 | 426.3 | 0.00071 | 1.666 | +1 | 4 |
| 13 | z | 907.2 | 430.2 | 0.002947 | 6.85 | +1 | 3 |
| - | - | 623.8 | 431.2 | - | - | 0 | - |
| - | - | 961.2 | 435.2 | - | - | 0 | - |
| 13 | y | 3235 | 446.3 | 0.003696 | 8.283 | +1 | 3 |
| - | - | 861.6 | 451.3 | - | - | 0 | - |
| - | - | 653.9 | 466.6 | - | - | 0 | - |
| - | - | 2952 | 470.3 | - | - | 0 | - |
| - | - | 609 | 471.3 | - | - | 0 | - |
| - | - | 2137 | 471.9 | - | - | 0 | - |
| - | - | 1423 | 472.2 | - | - | 0 | - |
| - | - | 1083 | 472.3 | - | - | 0 | - |
| - | - | 637.9 | 491.6 | - | - | 0 | - |
| - | - | 1189 | 494.3 | - | - | 0 | - |
| - | - | 667 | 495.6 | - | - | 0 | - |
| - | - | 1003 | 500.2 | - | - | 0 | - |
| 4 | z | 2046 | 500.9 | 0.002767 | 5.523 | +3 | 12 |
| 4 | z | 1617 | 501.2 | 0.004128 | 8.236 | +3 | 12 |
| - | - | 1313 | 501.6 | - | - | 0 | - |
| 5 | c | 2.518E+04 | 513.3 | 0.000892 | 1.738 | +1 | 5 |
| - | - | 6291 | 514.3 | - | - | 0 | - |
| - | - | 1334 | 515.3 | - | - | 0 | - |
| - | - | 2575 | 523.3 | - | - | 0 | - |
| - | - | 2277 | 523.6 | - | - | 0 | - |
| - | - | 1609 | 523.9 | - | - | 0 | - |
| 3 | y | 1392 | 538.6 | 0.001501 | 2.786 | +3 | 13 |
| 3 | y | 3447 | 538.9 | 0.00687 | 12.75 | +3 | 13 |
| 3 | z | 1363 | 539.3 | 0.004668 | 8.656 | +3 | 13 |
| - | - | 1306 | 539.6 | - | - | 0 | - |
| - | - | 901.6 | 539.9 | - | - | 0 | - |
| - | - | 608.6 | 540.3 | - | - | 0 | - |
| 12 | z | 2758 | 544.3 | 0.004859 | 8.927 | +1 | 4 |
| 3 | y | 1.233E+05 | 544.6 | 0.0009698 | 1.781 | +3 | 13 |
| - | - | 1.146E+05 | 544.9 | - | - | 0 | - |
| - | - | 6.395E+04 | 545.3 | - | - | 0 | - |
| - | - | 2.617E+04 | 545.6 | - | - | 0 | - |
| - | - | 9304 | 545.9 | - | - | 0 | - |
| - | - | 1913 | 546.3 | - | - | 0 | - |
| 12 | y | 5326 | 560.3 | 0.003441 | 6.142 | +1 | 4 |
| - | - | 1005 | 561.3 | - | - | 0 | - |
| - | - | 565.7 | 576.2 | - | - | 0 | - |
| - | - | 864.3 | 592.2 | - | - | 0 | - |
| - | - | 572.1 | 595.3 | - | - | 0 | - |
| - | - | 781.6 | 607.4 | - | - | 0 | - |
| - | - | 772.4 | 608.2 | - | - | 0 | - |
| - | - | 539.3 | 608.4 | - | - | 0 | - |
| - | - | 1184 | 614.4 | - | - | 0 | - |
| - | - | 926 | 615.4 | - | - | 0 | - |
| - | - | 1609 | 616.3 | - | - | 0 | - |
| - | - | 990.5 | 616.8 | - | - | 0 | - |
| - | - | 1917 | 617.2 | - | - | 0 | - |
| - | - | 6528 | 617.3 | - | - | 0 | - |
| - | - | 532.7 | 617.4 | - | - | 0 | - |
| - | - | 1092 | 617.7 | - | - | 0 | - |
| - | - | 566.4 | 625.1 | - | - | 0 | - |
| - | - | 1523 | 625.4 | - | - | 0 | - |
| - | - | 1441 | 626.4 | - | - | 0 | - |
| - | - | 7964 | 628.4 | - | - | 0 | - |
| - | - | 2809 | 629.3 | - | - | 0 | - |
| - | - | 3432 | 629.4 | - | - | 0 | - |
| - | - | 1264 | 630.3 | - | - | 0 | - |
| - | - | 854.7 | 631.3 | - | - | 0 | - |
| - | - | 1095 | 642.8 | - | - | 0 | - |
| 11 | c | 1204 | 643.3 | 0.002481 | 3.857 | +2 | 11 |
| - | - | 3633 | 651.8 | - | - | 0 | - |
| 11 | c | 2.131E+04 | 652.3 | 0.003101 | 4.753 | +2 | 11 |
| - | - | 1.343E+04 | 652.8 | - | - | 0 | - |
| - | - | 5377 | 653.3 | - | - | 0 | - |
| - | - | 2026 | 653.8 | - | - | 0 | - |
| - | - | 710.4 | 658.3 | - | - | 0 | - |
| 6 | z | 1192 | 667.8 | 0.00111 | 1.663 | +2 | 10 |
| - | - | 1606 | 668.3 | - | - | 0 | - |
| - | - | 779.4 | 668.8 | - | - | 0 | - |
| 6 | c | 5.091E+04 | 669.4 | 0.001099 | 1.642 | +1 | 6 |
| - | - | 1.901E+04 | 670.4 | - | - | 0 | - |
| - | - | 3578 | 671.4 | - | - | 0 | - |
| 11 | z | 1.614E+04 | 672.4 | 0.003325 | 4.946 | +1 | 5 |
| - | - | 6961 | 673.4 | - | - | 0 | - |
| - | - | 2135 | 674.4 | - | - | 0 | - |
| - | - | 1464 | 675.3 | - | - | 0 | - |
| 6 | y | 2640 | 675.8 | 0.0007444 | 1.102 | +2 | 10 |
| - | - | 1058 | 676.3 | - | - | 0 | - |
| - | - | 729.8 | 676.8 | - | - | 0 | - |
| - | - | 2105 | 680.8 | - | - | 0 | - |
| - | - | 845.7 | 681.3 | - | - | 0 | - |
| - | - | 1636 | 687.4 | - | - | 0 | - |
| - | - | 1509 | 687.9 | - | - | 0 | - |
| - | - | 605.7 | 688.3 | - | - | 0 | - |
| 11 | y | 5421 | 688.4 | 0.003678 | 5.343 | +1 | 5 |
| - | - | 1166 | 689.4 | - | - | 0 | - |
| - | - | 1223 | 695.4 | - | - | 0 | - |
| - | - | 740.8 | 696.4 | - | - | 0 | - |
| - | - | 791.7 | 698.8 | - | - | 0 | - |
| - | - | 1867 | 699.9 | - | - | 0 | - |
| 12 | c | 978.4 | 700.4 | 0.003742 | 5.343 | +2 | 12 |
| 12 | c | 842.9 | 700.8 | 0.002236 | 3.19 | +2 | 12 |
| - | - | 741.8 | 701.8 | - | - | 0 | - |
| 5 | z | 1376 | 702.3 | 0.008262 | 11.76 | +2 | 11 |
| - | - | 6797 | 707.3 | - | - | 0 | - |
| - | - | 4265 | 707.8 | - | - | 0 | - |
| - | - | 3389 | 708.3 | - | - | 0 | - |
| - | - | 5272 | 708.9 | - | - | 0 | - |
| 12 | c | 5.65E+04 | 709.4 | 0.002877 | 4.056 | +2 | 12 |
| - | - | 4.353E+04 | 709.9 | - | - | 0 | - |
| 5 | y | 2.108E+04 | 710.3 | 0.01257 | 17.7 | +2 | 11 |
| - | - | 7673 | 710.9 | - | - | 0 | - |
| 5 | z | 4114 | 711.3 | 0.006458 | 9.079 | +2 | 11 |
| - | - | 3366 | 711.8 | - | - | 0 | - |
| - | - | 2333 | 712.3 | - | - | 0 | - |
| - | - | 890.7 | 712.8 | - | - | 0 | - |
| 5 | y | 3467 | 719.4 | 0.002101 | 2.921 | +2 | 11 |
| - | - | 2008 | 719.9 | - | - | 0 | - |
| - | - | 1648 | 720.4 | - | - | 0 | - |
| - | - | 2402 | 729.4 | - | - | 0 | - |
| - | - | 1109 | 739.3 | - | - | 0 | - |
| - | - | 836.5 | 747.4 | - | - | 0 | - |
| 4 | z | 1189 | 750.9 | 0.002251 | 2.998 | +2 | 12 |
| 4 | z | 1927 | 751.4 | 0.00629 | 8.372 | +2 | 12 |
| - | - | 1120 | 751.9 | - | - | 0 | - |
| - | - | 1125 | 754.4 | - | - | 0 | - |
| - | - | 706.9 | 755.3 | - | - | 0 | - |
| - | - | 783.2 | 755.4 | - | - | 0 | - |
| 10 | w | 5052 | 756.4 | 0.002 | 2.643 | +1 | 6 |
| - | - | 1412 | 757.4 | - | - | 0 | - |
| 10 | w | 5581 | 758.4 | 0.002993 | 3.946 | +1 | 6 |
| - | - | 2198 | 759.4 | - | - | 0 | - |
| - | - | 653.4 | 760.4 | - | - | 0 | - |
| - | - | 1252 | 765.4 | - | - | 0 | - |
| - | - | 712 | 765.9 | - | - | 0 | - |
| 4 | y | 1.777E+04 | 767.9 | 0.0009268 | 1.207 | +2 | 12 |
| - | - | 1.788E+04 | 768.4 | - | - | 0 | - |
| - | - | 1.014E+04 | 768.9 | - | - | 0 | - |
| - | - | 3126 | 769.4 | - | - | 0 | - |
| - | - | 1651 | 769.9 | - | - | 0 | - |
| - | - | 1385 | 771.4 | - | - | 0 | - |
| 10 | y | 4628 | 772.4 | 0.003384 | 4.381 | +1 | 6 |
| - | - | 2317 | 772.9 | - | - | 0 | - |
| 13 | c | 2.659E+04 | 773.4 | 0.004533 | 5.862 | +2 | 13 |
| - | - | 3.367E+04 | 773.9 | - | - | 0 | - |
| - | - | 1.368E+04 | 774.4 | - | - | 0 | - |
| - | - | 6774 | 774.9 | - | - | 0 | - |
| - | - | 3281 | 775.4 | - | - | 0 | - |
| - | - | 597.6 | 779.4 | - | - | 0 | - |
| - | - | 875.2 | 781.9 | - | - | 0 | - |
| - | - | 3345 | 784.4 | - | - | 0 | - |
| - | - | 2647 | 784.9 | - | - | 0 | - |
| - | - | 1178 | 785.4 | - | - | 0 | - |
| - | - | 1043 | 785.9 | - | - | 0 | - |
| - | - | 1229 | 788.4 | - | - | 0 | - |
| 10 | y | 7288 | 789.5 | 0.00381 | 4.826 | +1 | 6 |
| - | - | 2720 | 790.4 | - | - | 0 | - |
| - | - | 734.1 | 791.4 | - | - | 0 | - |
| - | - | 1371 | 792.4 | - | - | 0 | - |
| - | - | 1195 | 792.9 | - | - | 0 | - |
| - | - | 951.8 | 793.4 | - | - | 0 | - |
| - | - | 2244 | 793.9 | - | - | 0 | - |
| - | - | 1864 | 794.4 | - | - | 0 | - |
| 3 | w | 2634 | 794.9 | 0.004434 | 5.578 | +2 | 13 |
| - | - | 3279 | 795.4 | - | - | 0 | - |
| - | - | 1908 | 795.9 | - | - | 0 | - |
| - | - | 1212 | 797.4 | - | - | 0 | - |
| 7 | c | 7.074E+04 | 798.4 | 0.001109 | 1.389 | +1 | 7 |
| - | - | 2.984E+04 | 799.5 | - | - | 0 | - |
| - | - | 7781 | 800.5 | - | - | 0 | - |
| - | - | 1691 | 800.9 | - | - | 0 | - |
| - | - | 3212 | 801.4 | - | - | 0 | - |
| - | - | 2095 | 801.9 | - | - | 0 | - |
| - | - | 957.8 | 802.4 | - | - | 0 | - |
| - | - | 655.5 | 805.4 | - | - | 0 | - |
| 3 | y | 1840 | 807.4 | 0.001191 | 1.475 | +2 | 13 |
| 3 | y | 1765 | 807.9 | 0.004177 | 5.17 | +2 | 13 |
| 3 | z | 2577 | 808.4 | 0.003499 | 4.329 | +2 | 13 |
| - | - | 1359 | 808.9 | - | - | 0 | - |
| - | - | 2208 | 813.4 | - | - | 0 | - |
| - | - | 2758 | 813.9 | - | - | 0 | - |
| - | - | 1613 | 814.4 | - | - | 0 | - |
| - | - | 2212 | 814.9 | - | - | 0 | - |
| - | - | 1737 | 815.4 | - | - | 0 | - |
| - | - | 1032 | 815.9 | - | - | 0 | - |
| 3 | y | 6.754E+04 | 816.4 | 0.0014 | 1.715 | +2 | 13 |
| - | - | 6.788E+04 | 816.9 | - | - | 0 | - |
| - | - | 3.905E+04 | 817.4 | - | - | 0 | - |
| - | - | 1.534E+04 | 817.9 | - | - | 0 | - |
| - | - | 5471 | 818.4 | - | - | 0 | - |
| - | - | 1055 | 818.9 | - | - | 0 | - |
| - | - | 7583 | 822.4 | - | - | 0 | - |
| - | - | 3.794E+04 | 822.9 | - | - | 0 | - |
| - | - | 3.526E+04 | 823.4 | - | - | 0 | - |
| - | - | 1.918E+04 | 823.9 | - | - | 0 | - |
| - | - | 5682 | 824.4 | - | - | 0 | - |
| - | - | 1926 | 824.9 | - | - | 0 | - |
| - | - | 3215 | 825.5 | - | - | 0 | - |
| - | - | 1180 | 826.5 | - | - | 0 | - |
| - | - | 1358 | 828.9 | - | - | 0 | - |
| - | - | 1388 | 829.4 | - | - | 0 | - |
| - | - | 796.2 | 830.5 | - | - | 0 | - |
| - | - | 2095 | 830.9 | - | - | 0 | - |
| - | - | 1933 | 831.4 | - | - | 0 | - |
| - | - | 3879 | 835.4 | - | - | 0 | - |
| - | - | 3081 | 835.9 | - | - | 0 | - |
| - | - | 2399 | 836.4 | - | - | 0 | - |
| - | - | 2669 | 836.9 | - | - | 0 | - |
| - | - | 998.7 | 837.4 | - | - | 0 | - |
| - | - | 878.5 | 840.5 | - | - | 0 | - |
| 2 | w | 2.914E+04 | 843.4 | 0.001062 | 1.259 | +2 | 14 |
| - | - | 2.726E+04 | 843.9 | - | - | 0 | - |
| - | - | 1.592E+04 | 844.4 | - | - | 0 | - |
| - | - | 6914 | 844.9 | - | - | 0 | - |
| - | - | 3221 | 845.4 | - | - | 0 | - |
| - | - | 989.1 | 845.9 | - | - | 0 | - |
| - | - | 968 | 849.4 | - | - | 0 | - |
| - | - | 770.4 | 850.4 | - | - | 0 | - |
| - | - | 1043 | 854.3 | - | - | 0 | - |
| - | - | 1772 | 855.3 | - | - | 0 | - |
| - | - | 710.8 | 856 | - | - | 0 | - |
| - | - | 3311 | 856.5 | - | - | 0 | - |
| - | - | 1708 | 857 | - | - | 0 | - |
| - | - | 8336 | 857.5 | - | - | 0 | - |
| - | - | 2.572E+04 | 858.5 | - | - | 0 | - |
| - | - | 2987 | 858.9 | - | - | 0 | - |
| - | - | 1.336E+04 | 859.5 | - | - | 0 | - |
| - | - | 906.1 | 859.9 | - | - | 0 | - |
| - | - | 3774 | 860.5 | - | - | 0 | - |
| - | - | 1420 | 861.5 | - | - | 0 | - |
| - | - | 814.9 | 863.5 | - | - | 0 | - |
| 2 | y | 889.6 | 863.9 | 0.003225 | 3.733 | +2 | 14 |
| 2 | z | 1.189E+04 | 864.9 | 0.0004687 | 0.5419 | +2 | 14 |
| - | - | 1.257E+04 | 865.4 | - | - | 0 | - |
| - | - | 7352 | 865.9 | - | - | 0 | - |
| 14 | c | 7.49E+04 | 866.4 | 0.0008799 | 1.016 | +2 | 14 |
| - | - | 6.831E+04 | 866.9 | - | - | 0 | - |
| - | - | 4.416E+04 | 867.4 | - | - | 0 | - |
| - | - | 2.125E+04 | 867.9 | - | - | 0 | - |
| - | - | 5954 | 868.4 | - | - | 0 | - |
| - | - | 2039 | 868.9 | - | - | 0 | - |
| - | - | 1113 | 870.3 | - | - | 0 | - |
| - | - | 2032 | 870.5 | - | - | 0 | - |
| - | - | 788.9 | 871 | - | - | 0 | - |
| - | - | 673.7 | 871.5 | - | - | 0 | - |
| - | - | 1965 | 872 | - | - | 0 | - |
| - | - | 2863 | 872.5 | - | - | 0 | - |
| 2 | y | 1929 | 872.9 | 0.004656 | 5.334 | +2 | 14 |
| - | - | 1493 | 873.4 | - | - | 0 | - |
| - | - | 1947 | 873.9 | - | - | 0 | - |
| - | - | 1026 | 874.4 | - | - | 0 | - |
| - | - | 2184 | 878 | - | - | 0 | - |
| - | - | 2234 | 878.5 | - | - | 0 | - |
| - | - | 1642 | 878.9 | - | - | 0 | - |
| - | - | 3540 | 879.4 | - | - | 0 | - |
| - | - | 3010 | 879.9 | - | - | 0 | - |
| - | - | 3329 | 880.4 | - | - | 0 | - |
| - | - | 1103 | 880.9 | - | - | 0 | - |
| - | - | 2923 | 883.5 | - | - | 0 | - |
| - | - | 2732 | 884.5 | - | - | 0 | - |
| - | - | 941.3 | 885 | - | - | 0 | - |
| - | - | 2399 | 885.5 | - | - | 0 | - |
| - | - | 2239 | 886 | - | - | 0 | - |
| - | - | 2257 | 886.5 | - | - | 0 | - |
| - | - | 2624 | 886.9 | - | - | 0 | - |
| - | - | 2.192E+04 | 887.4 | - | - | 0 | - |
| - | - | 2.464E+04 | 887.9 | - | - | 0 | - |
| - | - | 1.453E+04 | 888.4 | - | - | 0 | - |
| - | - | 5967 | 888.9 | - | - | 0 | - |
| - | - | 3102 | 889.4 | - | - | 0 | - |
| - | - | 1121 | 889.9 | - | - | 0 | - |
| - | - | 4080 | 892 | - | - | 0 | - |
| - | - | 4312 | 892.5 | - | - | 0 | - |
| - | - | 7322 | 893 | - | - | 0 | - |
| - | - | 7003 | 893.5 | - | - | 0 | - |
| - | - | 5952 | 894 | - | - | 0 | - |
| - | - | 1.27E+04 | 894.5 | - | - | 0 | - |
| - | - | 1.264E+04 | 895 | - | - | 0 | - |
| - | - | 7679 | 895.5 | - | - | 0 | - |
| - | - | 1.02E+04 | 895.9 | - | - | 0 | - |
| - | - | 7442 | 896.4 | - | - | 0 | - |
| - | - | 5438 | 896.9 | - | - | 0 | - |
| - | - | 1509 | 897.5 | - | - | 0 | - |
| - | - | 729.2 | 898 | - | - | 0 | - |
| - | - | 1466 | 901 | - | - | 0 | - |
| - | - | 1.566E+04 | 901.5 | - | - | 0 | - |
| - | - | 1.596E+04 | 902 | - | - | 0 | - |
| 9 | z | 1.225E+04 | 902.5 | 0.01312 | 14.54 | +1 | 7 |
| - | - | 9884 | 903 | - | - | 0 | - |
| - | - | 6644 | 903.5 | - | - | 0 | - |
| - | - | 1965 | 904 | - | - | 0 | - |
| - | - | 4731 | 906 | - | - | 0 | - |
| - | - | 1.383E+04 | 906.5 | - | - | 0 | - |
| - | - | 1.136E+04 | 907 | - | - | 0 | - |
| - | - | 8261 | 907.5 | - | - | 0 | - |
| - | - | 4968 | 908 | - | - | 0 | - |
| - | - | 2314 | 908.5 | - | - | 0 | - |
| - | - | 942.7 | 908.9 | - | - | 0 | - |
| - | - | 1.918E+04 | 915 | - | - | 0 | - |
| - | - | 7.559E+04 | 915.5 | - | - | 0 | - |
| - | - | 7.552E+04 | 916 | - | - | 0 | - |
| - | - | 4.69E+04 | 916.5 | - | - | 0 | - |
| - | - | 2.413E+04 | 917 | - | - | 0 | - |
| - | - | 7806 | 917.5 | - | - | 0 | - |
| - | - | 2640 | 918 | - | - | 0 | - |
| 9 | z | 8598 | 920.5 | 0.002008 | 2.181 | +1 | 7 |
| - | - | 5943 | 921.5 | - | - | 0 | - |
| - | - | 2754 | 922.5 | - | - | 0 | - |
| - | - | 809.6 | 923 | - | - | 0 | - |
| - | - | 4.907E+04 | 923.5 | - | - | 0 | - |
| - | - | 1.014E+05 | 924 | - | - | 0 | - |
| - | - | 7.813E+04 | 924.5 | - | - | 0 | - |
| - | - | 4.678E+04 | 925 | - | - | 0 | - |
| - | - | 2.092E+04 | 925.5 | - | - | 0 | - |
| - | - | 5925 | 926 | - | - | 0 | - |
| - | - | 2369 | 926.5 | - | - | 0 | - |
| 8 | c | 6.277E+04 | 927.5 | 0.001179 | 1.271 | +1 | 8 |
| - | - | 2.998E+04 | 928.5 | - | - | 0 | - |
| - | - | 917.9 | 928.6 | - | - | 0 | - |
| - | - | 8052 | 929.5 | - | - | 0 | - |
| - | - | 1826 | 930.5 | - | - | 0 | - |
| 9 | y | 9116 | 936.5 | 0.0001903 | 0.2032 | +1 | 7 |
| - | - | 5128 | 937.5 | - | - | 0 | - |
| - | - | 1811 | 938.5 | - | - | 0 | - |
| - | - | 1388 | 942.5 | - | - | 0 | - |
| - | - | 2382 | 968.5 | - | - | 0 | - |
| - | - | 1289 | 969.5 | - | - | 0 | - |
| - | - | 1747 | 978.5 | - | - | 0 | - |
| - | - | 868.3 | 979.5 | - | - | 0 | - |
| - | - | 6457 | 985.3 | - | - | 0 | - |
| - | - | 1738 | 986.3 | - | - | 0 | - |
| - | - | 1992 | 986.5 | - | - | 0 | - |
| - | - | 1162 | 987.5 | - | - | 0 | - |
| 8 | w | 1.096E+04 | 990.5 | 0.001222 | 1.234 | +1 | 8 |
| - | - | 6279 | 991.5 | - | - | 0 | - |
| - | - | 2722 | 992.5 | - | - | 0 | - |
| - | - | 1022 | 993.5 | - | - | 0 | - |
| - | - | 1743 | 994.5 | - | - | 0 | - |
| - | - | 1006 | 1014 | - | - | 0 | - |
| - | - | 7018 | 1031 | - | - | 0 | - |
| 8 | z | 5286 | 1031 | 0.02033 | 19.71 | +1 | 8 |
| - | - | 2256 | 1033 | - | - | 0 | - |
| - | - | 1327 | 1040 | - | - | 0 | - |
| 8 | z | 3.58E+04 | 1050 | 0.00159 | 1.515 | +1 | 8 |
| - | - | 2.322E+04 | 1051 | - | - | 0 | - |
| - | - | 1.148E+04 | 1052 | - | - | 0 | - |
| - | - | 3367 | 1053 | - | - | 0 | - |
| - | - | 767.1 | 1054 | - | - | 0 | - |
| 8 | y | 2260 | 1066 | 0.004594 | 4.312 | +1 | 8 |
| - | - | 1765 | 1067 | - | - | 0 | - |
| - | - | 1515 | 1074 | - | - | 0 | - |
| 9 | c | 4.616E+04 | 1075 | 0.005973 | 5.559 | +1 | 9 |
| - | - | 2.547E+04 | 1076 | - | - | 0 | - |
| - | - | 1.201E+04 | 1077 | - | - | 0 | - |
| - | - | 2662 | 1078 | - | - | 0 | - |
| - | - | 1394 | 1114 | - | - | 0 | - |
| - | - | 1069 | 1115 | - | - | 0 | - |
| 7 | w | 3978 | 1120 | 0.003429 | 3.063 | +1 | 9 |
| - | - | 2898 | 1121 | - | - | 0 | - |
| - | - | 1480 | 1122 | - | - | 0 | - |
| - | - | 728.6 | 1123 | - | - | 0 | - |
| - | - | 743.4 | 1123 | - | - | 0 | - |
| - | - | 1619 | 1132 | - | - | 0 | - |
| - | - | 1289 | 1133 | - | - | 0 | - |
| - | - | 1220 | 1134 | - | - | 0 | - |
| - | - | 831.2 | 1135 | - | - | 0 | - |
| - | - | 1289 | 1174 | - | - | 0 | - |
| 10 | c | 4.215E+04 | 1176 | 0.00578 | 4.917 | +1 | 10 |
| 7 | y | 2.716E+04 | 1177 | 0.01956 | 16.63 | +1 | 9 |
| - | - | 1.342E+04 | 1178 | - | - | 0 | - |
| 7 | z | 3.348E+04 | 1179 | 0.004407 | 3.739 | +1 | 9 |
| - | - | 2.015E+04 | 1180 | - | - | 0 | - |
| - | - | 9659 | 1181 | - | - | 0 | - |
| - | - | 2615 | 1182 | - | - | 0 | - |
| 7 | y | 2804 | 1195 | 0.0001571 | 0.1315 | +1 | 9 |
| - | - | 2304 | 1196 | - | - | 0 | - |
| - | - | 869.6 | 1197 | - | - | 0 | - |
| - | - | 1560 | 1216 | - | - | 0 | - |
| - | - | 800.8 | 1217 | - | - | 0 | - |
| - | - | 4203 | 1217 | - | - | 0 | - |
| - | - | 2155 | 1218 | - | - | 0 | - |
| - | - | 1485 | 1222 | - | - | 0 | - |
| - | - | 814.1 | 1223 | - | - | 0 | - |
| - | - | 905 | 1232 | - | - | 0 | - |
| - | - | 944.2 | 1233 | - | - | 0 | - |
| - | - | 1832 | 1233 | - | - | 0 | - |
| - | - | 7412 | 1234 | - | - | 0 | - |
| - | - | 1032 | 1235 | - | - | 0 | - |
| - | - | 3865 | 1235 | - | - | 0 | - |
| - | - | 2278 | 1260 | - | - | 0 | - |
| - | - | 1832 | 1261 | - | - | 0 | - |
| - | - | 932.4 | 1264 | - | - | 0 | - |
| 11 | c | 1268 | 1286 | 0.01917 | 14.91 | +1 | 11 |
| 11 | c | 1124 | 1287 | 0.01403 | 10.9 | +1 | 11 |
| - | - | 978 | 1288 | - | - | 0 | - |
| - | - | 4982 | 1303 | - | - | 0 | - |
| 11 | c | 1.239E+04 | 1304 | 0.004445 | 3.409 | +1 | 11 |
| - | - | 8167 | 1305 | - | - | 0 | - |
| - | - | 2929 | 1306 | - | - | 0 | - |
| - | - | 1828 | 1307 | - | - | 0 | - |
| 6 | y | 1081 | 1334 | 0.005627 | 4.219 | +1 | 10 |
| 6 | z | 2.031E+04 | 1335 | 0.0004638 | 0.3475 | +1 | 10 |
| - | - | 968.7 | 1335 | - | - | 0 | - |
| - | - | 2.951E+04 | 1336 | - | - | 0 | - |
| - | - | 1.636E+04 | 1337 | - | - | 0 | - |
| - | - | 8186 | 1338 | - | - | 0 | - |
| - | - | 2436 | 1339 | - | - | 0 | - |
| - | - | 1045 | 1340 | - | - | 0 | - |
| - | - | 3353 | 1350 | - | - | 0 | - |
| 6 | y | 9100 | 1351 | 0.001781 | 1.318 | +1 | 10 |
| - | - | 5304 | 1352 | - | - | 0 | - |
| - | - | 2738 | 1353 | - | - | 0 | - |
| - | - | 2185 | 1374 | - | - | 0 | - |
| - | - | 1438 | 1375 | - | - | 0 | - |
| - | - | 1634 | 1376 | - | - | 0 | - |
| 5 | z | 1100 | 1404 | 0.009884 | 7.041 | +1 | 11 |
| - | - | 1397 | 1417 | - | - | 0 | - |
| 12 | c | 2.579E+04 | 1418 | 0.004608 | 3.25 | +1 | 12 |
| - | - | 1.805E+04 | 1419 | - | - | 0 | - |
| 5 | y | 9857 | 1420 | 0.02314 | 16.3 | +1 | 11 |
| - | - | 4742 | 1421 | - | - | 0 | - |
| 5 | z | 3264 | 1422 | 0.018 | 12.66 | +1 | 11 |
| - | - | 1.077E+04 | 1423 | - | - | 0 | - |
| - | - | 7400 | 1424 | - | - | 0 | - |
| - | - | 4135 | 1425 | - | - | 0 | - |
| - | - | 1240 | 1426 | - | - | 0 | - |
| - | - | 743.6 | 1437 | - | - | 0 | - |
| 5 | y | 3090 | 1438 | 0.000484 | 0.3367 | +1 | 11 |
| - | - | 2347 | 1439 | - | - | 0 | - |
| 13 | c | 1076 | 1528 | 0.02337 | 15.3 | +1 | 13 |
| 13 | c | 957.3 | 1529 | 0.02849 | 18.63 | +1 | 13 |
| - | - | 1488 | 1530 | - | - | 0 | - |
| - | - | 748.4 | 1531 | - | - | 0 | - |
| 4 | y | 6129 | 1535 | 0.0008293 | 0.5403 | +1 | 12 |
| - | - | 5015 | 1536 | - | - | 0 | - |
| - | - | 3249 | 1537 | - | - | 0 | - |
| - | - | 764.3 | 1539 | - | - | 0 | - |
| 13 | c | 1.306E+04 | 1546 | 0.004136 | 2.676 | +1 | 13 |
| - | - | 1.275E+04 | 1547 | - | - | 0 | - |
| - | - | 9383 | 1548 | - | - | 0 | - |
| - | - | 3427 | 1549 | - | - | 0 | - |
| - | - | 1195 | 1550 | - | - | 0 | - |
| - | - | 1068 | 1601 | - | - | 0 | - |
| - | - | 1909 | 1602 | - | - | 0 | - |
| - | - | 1587 | 1603 | - | - | 0 | - |
| 3 | y | 3186 | 1615 | 0.02064 | 12.78 | +1 | 13 |
| 3 | z | 2476 | 1616 | 0.01464 | 9.061 | +1 | 13 |
| - | - | 1969 | 1617 | - | - | 0 | - |
| - | - | 1164 | 1618 | - | - | 0 | - |
| - | - | 782.2 | 1628 | - | - | 0 | - |
| - | - | 2084 | 1629 | - | - | 0 | - |
| - | - | 1681 | 1630 | - | - | 0 | - |
| - | - | 1262 | 1631 | - | - | 0 | - |
| 3 | y | 1011 | 1632 | 0.00849 | 5.203 | +1 | 13 |
| - | - | 1201 | 1633 | - | - | 0 | - |
| - | - | 1424 | 1634 | - | - | 0 | - |
| - | - | 1022 | 1644 | - | - | 0 | - |
| - | - | 1.247E+04 | 1645 | - | - | 0 | - |
| - | - | 1.573E+04 | 1646 | - | - | 0 | - |
| - | - | 1.137E+04 | 1647 | - | - | 0 | - |
| - | - | 5685 | 1648 | - | - | 0 | - |
| - | - | 2397 | 1649 | - | - | 0 | - |
| - | - | 1432 | 1671 | - | - | 0 | - |
| - | - | 1916 | 1672 | - | - | 0 | - |
| - | - | 1528 | 1673 | - | - | 0 | - |
| - | - | 1105 | 1674 | - | - | 0 | - |
| - | - | 968.4 | 1685 | - | - | 0 | - |
| - | - | 1010 | 1686 | - | - | 0 | - |
| - | - | 906.6 | 1687 | - | - | 0 | - |
| - | - | 1470 | 1688 | - | - | 0 | - |
| - | - | 5111 | 1689 | - | - | 0 | - |
| - | - | 3776 | 1690 | - | - | 0 | - |
| - | - | 1886 | 1691 | - | - | 0 | - |
| - | - | 816 | 1692 | - | - | 0 | - |
| - | - | 732.4 | 1705 | - | - | 0 | - |
| - | - | 2934 | 1716 | - | - | 0 | - |
| - | - | 1834 | 1717 | - | - | 0 | - |
| - | - | 1539 | 1718 | - | - | 0 | - |
| - | - | 1075 | 1719 | - | - | 0 | - |
| 2 | y | 1053 | 1728 | 0.0103 | 5.962 | +1 | 14 |
| 2 | z | 4171 | 1729 | 0.0003311 | 0.1915 | +1 | 14 |
| - | - | 2.291E+04 | 1730 | - | - | 0 | - |
| - | - | 1.793E+04 | 1731 | - | - | 0 | - |
| 14 | c | 2.132E+04 | 1732 | 0.004642 | 2.68 | +1 | 14 |
| - | - | 2.552E+04 | 1733 | - | - | 0 | - |
| - | - | 1.991E+04 | 1734 | - | - | 0 | - |
| - | - | 1.121E+04 | 1735 | - | - | 0 | - |
| - | - | 3386 | 1736 | - | - | 0 | - |
| - | - | 1283 | 1737 | - | - | 0 | - |
| - | - | 1088 | 1748 | - | - | 0 | - |
| - | - | 1193 | 1770 | - | - | 0 | - |
| - | - | 1862 | 1772 | - | - | 0 | - |
| - | - | 896.5 | 1775 | - | - | 0 | - |
| - | - | 834.3 | 1776 | - | - | 0 | - |
| - | - | 1939 | 1784 | - | - | 0 | - |
| - | - | 2638 | 1785 | - | - | 0 | - |
| - | - | 4389 | 1786 | - | - | 0 | - |
| - | - | 4299 | 1787 | - | - | 0 | - |
| - | - | 3979 | 1788 | - | - | 0 | - |
| - | - | 8071 | 1789 | - | - | 0 | - |
| - | - | 7146 | 1790 | - | - | 0 | - |
| - | - | 4998 | 1791 | - | - | 0 | - |
| - | - | 2241 | 1792 | - | - | 0 | - |
| - | - | 1016 | 1793 | - | - | 0 | - |
| - | - | 4226 | 1802 | - | - | 0 | - |
| - | - | 1.533E+04 | 1803 | - | - | 0 | - |
| - | - | 1.386E+04 | 1804 | - | - | 0 | - |
| - | - | 9342 | 1805 | - | - | 0 | - |
| - | - | 4134 | 1806 | - | - | 0 | - |
| - | - | 1359 | 1807 | - | - | 0 | - |
| - | - | 1679 | 1813 | - | - | 0 | - |
| - | - | 4693 | 1814 | - | - | 0 | - |
| - | - | 3248 | 1815 | - | - | 0 | - |
| - | - | 2414 | 1816 | - | - | 0 | - |
| - | - | 2477 | 1817 | - | - | 0 | - |
| - | - | 1670 | 1819 | - | - | 0 | - |
| - | - | 9853 | 1820 | - | - | 0 | - |
| - | - | 1.086E+04 | 1821 | - | - | 0 | - |
| - | - | 5536 | 1822 | - | - | 0 | - |
| - | - | 3669 | 1823 | - | - | 0 | - |
| - | - | 850.9 | 1824 | - | - | 0 | - |
| - | - | 1520 | 1829 | - | - | 0 | - |
| - | - | 1.212E+04 | 1830 | - | - | 0 | - |
| - | - | 4.659E+04 | 1831 | - | - | 0 | - |
| - | - | 4.821E+04 | 1832 | - | - | 0 | - |
| - | - | 3.152E+04 | 1833 | - | - | 0 | - |
| - | - | 1.473E+04 | 1834 | - | - | 0 | - |
| - | - | 4841 | 1835 | - | - | 0 | - |
| - | - | 1409 | 1836 | - | - | 0 | - |
| - | - | 5054 | 1846 | - | - | 0 | - |
| - | - | 2.874E+04 | 1847 | - | - | 0 | - |
| - | - | 1.236E+05 | 1848 | - | - | 0 | - |
| - | - | 1.261E+05 | 1849 | - | - | 0 | - |
| - | - | 7.478E+04 | 1850 | - | - | 0 | - |
| - | - | 3.399E+04 | 1851 | - | - | 0 | - |
| - | - | 1.184E+04 | 1852 | - | - | 0 | - |
| - | - | 3110 | 1853 | - | - | 0 | - |

m/z Charge Intensity FragmentType MassShift Position
132.10211181640625 0 1076.8431 y 14
149.36911010742188 0 416.51498
166.0865478515625 0 457.60898
173.4385223388672 0 807.16974
177.6192169189453 0 454.97803
186.12391662597656 0 476.8071
187.1444091796875 0 37126.29
188.1478271484375 0 3167.7083
201.1233673095703 0 626.5853
212.13963317871094 0 2674.0134
212.14999389648438 0 550.6589
215.13929748535156 0 22811.957
216.142578125 0 2587.6633
219.13449096679688 0 3277.3218
221.1287384033203 0 5201.3677
227.4397430419922 0 561.8149
238.1553192138672 0 712.76733
247.129150390625 0 817.9479
249.1234588623047 0 2538.436
256.165283203125 0 1014.56036
268.5540771484375 0 481.565
278.45977783203125 0 518.325
299.171875 0 6559.4478
312.1925354003906 0 904.6692
326.1595153808594 0 1957.7622
327.1656188964844 0 967.9069
344.19219970703125 0 651.3255
371.1927490234375 0 2123.4038
374.1437072753906 0 640.29065
382.2589111328125 0 664.45276
386.2418518066406 0 602.81165
395.1802062988281 0 583.1975
407.2523498535156 0 996.7572
414.1988525390625 0 689.1071
415.537353515625 0 658.529
425.2640380859375 0 2022.5442
426.2703857421875 0 1620.5779 c 3
430.2437744140625 0 907.20447 z 12
431.2489929199219 0 623.8388
435.15093994140625 0 961.2249
446.2617492675781 0 3235.1008 y 12
451.2782897949219 0 861.57947
466.5572814941406 0 653.8628
470.2981872558594 0 2952.2046
471.3034973144531 0 608.9917
471.89776611328125 0 2137.4163
472.2320251464844 0 1423.4653
472.2901611328125 0 1083.1256
491.5772705078125 0 637.91144
494.2843322753906 0 1188.9932
495.5674743652344 0 667.032
500.2375793457031 0 1003.14453
500.90985107421875 0 2046.2153 z Water loss 3
501.2447509765625 0 1617.1543 z Ammonia loss 3
501.5762634277344 0 1312.8868
513.3040161132812 0 25178.48 c 4
514.3070068359375 0 6290.916
515.3075561523438 0 1334.4867
523.2742309570312 0 2575.2988
523.609130859375 0 2277.0513
523.9420776367188 0 1608.9563
538.6046142578125 0 1391.8191 y Water loss 2
538.93798828125 0 3446.54 y Ammonia loss 2
539.271728515625 0 1362.9597 z 2
539.6014404296875 0 1306.1007
539.9266357421875 0 901.59784
540.2691040039062 0 608.59485
544.2847900390625 0 2757.84 z 11
544.6076049804688 0 123279.44 y 2
544.9413452148438 0 114621.805
545.2754516601562 0 63947.13
545.6083984375 0 26165.771
545.9423217773438 0 9304.458
546.2839965820312 0 1913.2664
560.304931640625 0 5325.9272 y 11
561.3078002929688 0 1005.1671
576.1981811523438 0 565.70667
592.2220458984375 0 864.2626
595.3016357421875 0 572.0916
607.3843383789062 0 781.6346
608.2244873046875 0 772.3826
608.3693237304688 0 539.27014
614.3530883789062 0 1183.549
615.3530883789062 0 925.95776
616.3163452148438 0 1609.264
616.7919311523438 0 990.5111
617.2294311523438 0 1916.6898
617.3147583007812 0 6527.7964
617.35693359375 0 532.6796
617.7322387695312 0 1092.2029
625.1087646484375 0 566.3662
625.3901977539062 0 1523.1532
626.3977661132812 0 1440.8306
628.3673706054688 0 7963.764
629.32470703125 0 2809.424
629.3726806640625 0 3432.239
630.3295288085938 0 1264.496
631.3339233398438 0 854.6703
642.8301391601562 0 1094.5565
643.3271484375 0 1203.6473 c Water loss 10
651.8353271484375 0 3633.1746
652.3380126953125 0 21309.96 c 10
652.8394165039062 0 13431.042
653.339599609375 0 5376.9287
653.841552734375 0 2025.9299
658.3148803710938 0 710.37427
667.8292846679688 0 1192.4352 z 5
668.3322143554688 0 1605.6389
668.82763671875 0 779.4244
669.4053344726562 0 50912.87 c 5
670.4075317382812 0 19006.148
671.4112548828125 0 3577.618
672.3812866210938 0 16141.554 z 10
673.3818969726562 0 6960.5527
674.3849487304688 0 2134.5022
675.3373413085938 0 1464.126
675.8367919921875 0 2639.9092 y 5
676.341064453125 0 1058.476
676.8397216796875 0 729.7942
680.8343505859375 0 2105.4763
681.3353271484375 0 845.6646
687.3519287109375 0 1635.8824
687.852783203125 0 1509.1239
688.3436279296875 0 605.73114
688.399658203125 0 5421.315 y 10
689.4012451171875 0 1165.6665
695.4219360351562 0 1223.4828
696.4228515625 0 740.80945
698.8336791992188 0 791.6819
699.8501586914062 0 1867.4307
700.3473510742188 0 978.4053 c Water loss 11
700.8453369140625 0 842.92365 c Ammonia loss 11
701.8491821289062 0 741.781
702.34716796875 0 1375.8551 z Water loss 4
707.3442993164062 0 6797.375
707.8450317382812 0 4265.332
708.3427124023438 0 3388.7542
708.8538818359375 0 5272.1104
709.3592529296875 0 56497.258 c 11
709.860107421875 0 43533.098
710.36083984375 0 21083.205 y Water loss 4
710.8599853515625 0 7672.753
711.3506469726562 0 4113.914 z 4
711.8482055664062 0 3365.989
712.3473510742188 0 2333.019
712.849853515625 0 890.69824
719.3556518554688 0 3466.7778 y 4
719.8547973632812 0 2008.4509
720.3550415039062 0 1648.3292
729.4136962890625 0 2402.193
739.2640991210938 0 1109.3407
747.3624267578125 0 836.5093
750.863037109375 0 1188.6566 z Water loss 3
751.3635864257812 0 1927.2661 z Ammonia loss 3
751.874267578125 0 1120.1012
754.4285888671875 0 1125.1444
755.284912109375 0 706.8831
755.4368896484375 0 783.24786
756.4275512695312 0 5051.804 w 9
757.42578125 0 1411.8713
758.4058227539062 0 5580.565 w 9
759.404052734375 0 2198.4395
760.4154052734375 0 653.36426
765.395263671875 0 1252.2212
765.8892822265625 0 711.9791
767.880859375 0 17765.129 y 3
768.3815307617188 0 17880.553
768.8829956054688 0 10138.959
769.3822631835938 0 3125.8833
769.8823852539062 0 1650.5181
771.4151611328125 0 1384.644
772.4210815429688 0 4628.2954 y Ammonia loss 9
772.8871459960938 0 2317.2803
773.3901977539062 0 26586.916 c 12
773.8899536132812 0 33669.664
774.3914184570312 0 13678.364
774.8916015625 0 6773.8047
775.43212890625 0 3280.5354
779.4208984375 0 597.6446
781.8810424804688 0 875.2
784.4077758789062 0 3344.5317
784.9075927734375 0 2646.6494
785.4065551757812 0 1178.1367
785.9066162109375 0 1042.6539
788.4380493164062 0 1229.0383
789.4472045898438 0 7288.2505 y 9
790.44921875 0 2719.9504
791.4466552734375 0 734.14215
792.4056396484375 0 1371.2705
792.9070434570312 0 1195.2402
793.4058227539062 0 951.821
793.9122314453125 0 2244.3523
794.4046020507812 0 1863.7373
794.8896484375 0 2633.9404 w 2
795.3907470703125 0 3278.596
795.8887329101562 0 1908.4023
797.436279296875 0 1211.8708
798.4479370117188 0 70737.15 c 6
799.4505615234375 0 29843.79
800.4528198242188 0 7781.2275
800.9127197265625 0 1690.7988
801.4183959960938 0 3211.6533
801.9180297851562 0 2094.6216
802.4219360351562 0 957.7696
805.4111328125 0 655.5278
807.3998413085938 0 1839.8159 y Water loss 2
807.897216796875 0 1765.4392 y Ammonia loss 2
808.4004516601562 0 2576.5466 z 2
808.89501953125 0 1359.3953
813.4185180664062 0 2207.6716
813.911376953125 0 2757.8025
814.4078979492188 0 1613.1882
814.911376953125 0 2212.33
815.4050903320312 0 1737.0364
815.9130249023438 0 1032.3516
816.40771484375 0 67538.81 y 2
816.9082641601562 0 67881.04
817.4083862304688 0 39049.35
817.9092407226562 0 15340.745
818.40869140625 0 5471.2573
818.9099731445312 0 1054.9563
822.4193115234375 0 7583.312
822.9226684570312 0 37944.14
823.4231567382812 0 35259.297
823.9237670898438 0 19178.105
824.4320678710938 0 5681.6064
824.9268798828125 0 1926.4526
825.4639892578125 0 3215.3308
826.47021484375 0 1180.3818
828.9299926757812 0 1357.8384
829.4281616210938 0 1388.1599
830.452880859375 0 796.2394
830.9097900390625 0 2095.1807
831.4074096679688 0 1933.029
835.4271240234375 0 3879.478
835.9265747070312 0 3081.3977
836.4302368164062 0 2399.2976
836.9163208007812 0 2668.878
837.4198608398438 0 998.65546
840.4705810546875 0 878.46783
843.4126586914062 0 29137.662 w 1
843.9135131835938 0 27256.734
844.4146118164062 0 15916.485
844.9180297851562 0 6914.361
845.4222412109375 0 3221.0115
845.9227294921875 0 989.1405
849.4261474609375 0 968.0177
850.40673828125 0 770.38226
854.2901611328125 0 1043.3535
855.2988891601562 0 1772.3911
855.9601440429688 0 710.80206
856.4614868164062 0 3310.59
856.9556274414062 0 1708.343
857.4723510742188 0 8335.994
858.4800415039062 0 25720.326
858.9339599609375 0 2986.9768
859.479248046875 0 13363.574
859.9266967773438 0 906.1304
860.48388671875 0 3773.8013
861.4898071289062 0 1419.9331
863.4567260742188 0 814.8688
863.9462890625 0 889.6232 y Water loss 1
864.939453125 0 11887.969 z 1
865.4417114257812 0 12569.272
865.942138671875 0 7351.6445
866.4390258789062 0 74901.44 c 13
866.9397583007812 0 68305.914
867.4403076171875 0 44164.63
867.940673828125 0 21250.734
868.4419555664062 0 5953.8154
868.9430541992188 0 2039.3353
870.3145141601562 0 1112.8184
870.4667358398438 0 2031.7462
870.967041015625 0 788.9013
871.471435546875 0 673.68884
871.9561767578125 0 1965.1552
872.460205078125 0 2863.071
872.9530029296875 0 1929.2017 y 1
873.4435424804688 0 1493.3766
873.9385375976562 0 1947.2285
874.4387817382812 0 1025.6093
877.9598388671875 0 2183.6982
878.4560546875 0 2233.8193
878.9281005859375 0 1641.7144
879.4475708007812 0 3540.3857
879.9498291015625 0 3010.039
880.449951171875 0 3328.937
880.9482421875 0 1102.5669
883.4743041992188 0 2922.6294
884.471923828125 0 2731.7568
884.9669799804688 0 941.3373
885.4649047851562 0 2399.3027
885.9546508789062 0 2238.9321
886.4527587890625 0 2257.4456
886.94677734375 0 2623.5042
887.435546875 0 21918.158
887.9364624023438 0 24640.898
888.4408569335938 0 14533.228
888.9407348632812 0 5966.754
889.4449462890625 0 3102.4634
889.9436645507812 0 1120.7766
891.9713745117188 0 4080.2898
892.4688110351562 0 4312.433
892.9630126953125 0 7322.2505
893.4606323242188 0 7002.6797
893.9544677734375 0 5951.7295
894.4580078125 0 12702.107
894.959228515625 0 12643.269
895.4598388671875 0 7679.167
895.949951171875 0 10202.367
896.448486328125 0 7442.0767
896.9473876953125 0 5437.8267
897.4515991210938 0 1508.6113
897.9545288085938 0 729.1955
900.9737548828125 0 1466.3341
901.4674072265625 0 15659.226
901.9678344726562 0 15961.461
902.46533203125 0 12245.275 z Water loss 8
902.9636840820312 0 9883.77
903.46484375 0 6643.6094
903.9589233398438 0 1965.307
905.967529296875 0 4730.653
906.4613037109375 0 13832.769
906.9596557617188 0 11364.162
907.4602661132812 0 8261.493
907.9580688476562 0 4967.746
908.45556640625 0 2314.032
908.9495849609375 0 942.67444
914.9730224609375 0 19177.354
915.46826171875 0 75586.81
915.9688110351562 0 75517.02
916.4680786132812 0 46898.19
916.9688110351562 0 24128.83
917.4669799804688 0 7806.1494
917.9679565429688 0 2639.654
920.4647827148438 0 8597.617 z 8
921.466552734375 0 5943.4673
922.4668579101562 0 2754.4136
922.967041015625 0 809.57635
923.4733276367188 0 49071.03
923.9752807617188 0 101407.35
924.4759521484375 0 78130.02
924.9776000976562 0 46777.406
925.4776000976562 0 20920.043
925.9786987304688 0 5924.84
926.4800415039062 0 2368.8845
927.4906005859375 0 62770.203 c 7
928.4933471679688 0 29979.816
928.5993041992188 0 917.9438
929.495361328125 0 8052.3403
930.5018310546875 0 1825.8457
936.481689453125 0 9116.413 y 8
937.483154296875 0 5127.616
938.4859008789062 0 1810.5706
942.4667358398438 0 1388.3666
968.530029296875 0 2382.3955
969.53369140625 0 1288.7487
978.4678955078125 0 1747.4475
979.4712524414062 0 868.3182
985.3370971679688 0 6456.87
986.3379516601562 0 1738.0154
986.5173950195312 0 1991.8278
987.5152587890625 0 1161.6968
990.4932861328125 0 10958.686 w 7
991.4947509765625 0 6279.0073
992.4943237304688 0 2722.0107
993.4551391601562 0 1021.7231
994.451416015625 0 1743.251
1014.4019165039062 0 1005.7925
1030.5137939453125 0 7017.5225
1031.51513671875 0 5286.34 z Water loss 7
1032.5208740234375 0 2256.016
1040.418212890625 0 1326.5171
1049.5069580078125 0 35800.53 z 7
1050.5076904296875 0 23218.152
1051.5069580078125 0 11484.826
1052.5067138671875 0 3366.8027
1053.5128173828125 0 767.1096
1065.5286865234375 0 2260.015 y 7
1066.525634765625 0 1764.6925
1073.516357421875 0 1514.631
1074.52587890625 0 46160.977 c 8
1075.5286865234375 0 25469.814
1076.5289306640625 0 12005.744
1077.53076171875 0 2661.8657
1113.548583984375 0 1393.8362
1114.5400390625 0 1069.3884
1119.5380859375 0 3977.968 w 6
1120.5316162109375 0 2898.1992
1121.537353515625 0 1479.6863
1122.519775390625 0 728.5631
1123.4866943359375 0 743.4254
1131.5609130859375 0 1618.772
1132.56201171875 0 1288.9457
1133.5654296875 0 1219.6035
1134.5556640625 0 831.23627
1174.439453125 0 1288.8052
1175.5733642578125 0 42150.484 c 9
1176.57568359375 0 27159.645 y Water loss 6
1177.5772705078125 0 13415.719
1178.5523681640625 0 33479.605 z 6
1179.54931640625 0 20150.424
1180.5506591796875 0 9658.958
1181.5511474609375 0 2615.1382
1194.5665283203125 0 2804.0918 y 6
1195.572998046875 0 2303.9714
1196.5643310546875 0 869.60095
1216.45166015625 0 1560.4373
1216.5810546875 0 800.79614
1217.44091796875 0 4203.226
1218.441162109375 0 2154.8433
1221.583984375 0 1484.8064
1222.5728759765625 0 814.1196
1231.56298828125 0 904.9879
1232.5653076171875 0 944.20667
1233.4473876953125 0 1832.478
1234.458251953125 0 7412.3604
1234.5914306640625 0 1031.9978
1235.462646484375 0 3865.389
1259.653076171875 0 2278.4133
1260.6597900390625 0 1832.2542
1263.6339111328125 0 932.4017
1285.6328125 0 1267.528 c Water loss 10
1286.6500244140625 0 1124.425 c Ammonia loss 10
1287.6353759765625 0 977.9506
1302.65576171875 0 4982.46
1303.6669921875 0 12391.819 c 10
1304.67041015625 0 8167.484
1305.6748046875 0 2929.0117
1306.67138671875 0 1828.0063
1333.6356201171875 0 1081.2043 y Ammonia loss 5
1334.6495361328125 0 20305.977 z 5
1334.832763671875 0 968.71967
1335.6544189453125 0 29514.01
1336.655029296875 0 16358.307
1337.657470703125 0 8185.8315
1338.6600341796875 0 2436.3665
1339.65625 0 1045.1787
1349.659423828125 0 3352.569
1350.666015625 0 9099.786 y 5
1351.668701171875 0 5303.7827
1352.6695556640625 0 2737.8928
1373.69140625 0 2185.137
1374.708740234375 0 1437.503
1375.69921875 0 1633.537
1403.680419921875 0 1100.0743 z Water loss 4
1416.6890869140625 0 1396.6897
1417.7100830078125 0 25794.01 c 11
1418.710205078125 0 18054.965
1419.71240234375 0 9857.212 y Water loss 4
1420.7095947265625 0 4741.771
1421.6990966796875 0 3264.3584 z 4
1422.68798828125 0 10772.172
1423.6929931640625 0 7399.577
1424.687744140625 0 4134.9404
1425.6883544921875 0 1239.5323
1436.697265625 0 743.6183
1437.6993408203125 0 3090.0864 y 4
1438.7069091796875 0 2346.8047
1527.77685546875 0 1076.3843 c Water loss 12
1528.7659912109375 0 957.3086 c Ammonia loss 12
1529.7567138671875 0 1488.3452
1530.76171875 0 748.3957
1534.75341796875 0 6129.274 y 3
1535.757080078125 0 5014.6353
1536.76171875 0 3249.1912
1538.7716064453125 0 764.28424
1545.7681884765625 0 13062.754 c 12
1546.7706298828125 0 12746.034
1547.7711181640625 0 9383.14
1548.770751953125 0 3427.4666
1549.7862548828125 0 1194.6093
1600.8177490234375 0 1067.7611
1601.828125 0 1908.6787
1602.829345703125 0 1586.9882
1614.7994384765625 0 3186.3684 y Ammonia loss 2
1615.80126953125 0 2475.9675 z 2
1616.81103515625 0 1969.0118
1617.8272705078125 0 1164.1854
1627.827392578125 0 782.1816
1628.8289794921875 0 2083.5947
1629.8184814453125 0 1680.659
1630.8089599609375 0 1261.6353
1631.8138427734375 0 1010.6033 y 2
1632.80810546875 0 1200.9447
1633.814453125 0 1423.6533
1643.84423828125 0 1021.83057
1644.837646484375 0 12473.693
1645.8394775390625 0 15729.418
1646.84375 0 11371.8545
1647.843994140625 0 5684.512
1648.8526611328125 0 2397.0566
1670.8463134765625 0 1431.8445
1671.845703125 0 1916.3992
1672.855712890625 0 1528.393
1673.8414306640625 0 1104.9802
1684.8409423828125 0 968.4318
1685.87353515625 0 1010.3193
1686.86181640625 0 906.6105
1687.8548583984375 0 1469.9712
1688.8648681640625 0 5110.9893
1689.8614501953125 0 3776.4358
1690.85595703125 0 1885.8779
1691.8507080078125 0 815.95636
1704.88037109375 0 732.434
1715.8623046875 0 2934.411
1716.8624267578125 0 1833.9983
1717.8564453125 0 1539.2588
1718.8658447265625 0 1074.8627
1727.8731689453125 0 1052.911 y Ammonia loss 1
1728.870361328125 0 4170.871 z 1
1729.878173828125 0 22907.86
1730.878662109375 0 17929.969
1731.8736572265625 0 21318.316 c 13
1732.876708984375 0 25518.764
1733.8765869140625 0 19912.715
1734.876708984375 0 11206.439
1735.87451171875 0 3385.9548
1736.873046875 0 1282.5208
1747.8863525390625 0 1087.5012
1769.9080810546875 0 1193.1912
1771.892822265625 0 1861.617
1774.8970947265625 0 896.5212
1775.8880615234375 0 834.30444
1783.955322265625 0 1939.4553
1784.9417724609375 0 2638.261
1785.9228515625 0 4389.3037
1786.923828125 0 4298.592
1787.91259765625 0 3978.7375
1788.9130859375 0 8070.9077
1789.9156494140625 0 7145.731
1790.9122314453125 0 4997.6963
1791.9156494140625 0 2241.3345
1792.90771484375 0 1016.1771
1801.9315185546875 0 4225.6123
1802.93115234375 0 15327.291
1803.93359375 0 13863.688
1804.933349609375 0 9341.842
1805.93017578125 0 4133.842
1806.9442138671875 0 1358.9636
1812.920654296875 0 1679.2659
1813.9130859375 0 4692.535
1814.9061279296875 0 3248.284
1815.902099609375 0 2413.8245
1816.9129638671875 0 2476.6792
1818.9476318359375 0 1670.4375
1819.9552001953125 0 9853.159
1820.9580078125 0 10855.279
1821.9617919921875 0 5536.2964
1822.9603271484375 0 3668.743
1823.953125 0 850.85004
1828.9127197265625 0 1519.8901
1829.928955078125 0 12119.265
1830.9293212890625 0 46590.25
1831.929931640625 0 48209.25
1832.9310302734375 0 31516.82
1833.93115234375 0 14731.395
1834.9315185546875 0 4840.8037
1835.9307861328125 0 1409.4141
1845.9404296875 0 5053.6274
1846.943603515625 0 28741.104
1847.9510498046875 0 123575.9
1848.9525146484375 0 126141.34
1849.953857421875 0 74780.25
1850.9544677734375 0 33985.86
1851.95654296875 0 11844.657
1852.95263671875 0 3109.5312

Spectrum Details

|  |  |
| --- | --- |
| Matched peaks? Matched peaks The total absolute number of peaks matched. Additionally in brackets the total fraction of peaks matched and the total number of peaks is shown. | 81 (14.29% of 567) |
| FDR? FDR The false discovery rate estimated for this peptide. It is calculated by matching all theoretical fragments with a non-integer shift with the raw peaks for this spectrum. This is done with 40 different shifts. The resulting percentage is the average number of annotated peaks over the number of annotated peaks with the correct spectrum. | 0.73% |
| Satellite FDR? Satellite FDR See the FDR for details on its calculation. This satellite ion specific FDR only contains the satellite ions (d/w) for I/L/J positions. | 0.00% |
| PSM Score? PSM Score The PSM Score as given by Hecklib to this annotated spectrum. It is shown with three significant figures. | 621 |

#### Spectrum 4786? Spectrum 4786 The raw spectrum of this peptide as annotated by Hecklib. The fragments are coloured according to ion type (see legend). Any peaks with a star '\*' as text can be hovered over to see the full details, first the ion type second the mass shift type. By hovering over the amino acids in the peptide or ions in the legend the corresponding peaks are highlighted. By toggling the 'Unassigned' label you can turn the background (unassigned) peaks on or off in the plot. By updating the slider in the Ion legend you can update the spectrum to only show the top X% of the peaks with labels. The top X% means any peak that is within X% of the highest intensity. By dragging in the spectrum you can zoom in to a specific part of the spectrum and use 'Zoom Out' to get back to the original zoom level. The annotation of the spectrum is based on the given sequence in the peptides file and is done with different software so inconsistencies are likely. The peaks are annotated based on the given sequence, with 20 ppm tolerance.

Copy Data

##### Spectrum 4786 (TSV)

###### Preview

```
Loading example...
```

*Click on the button to copy the data to your clipboard.*

Mz MinMz MaxIntensity Max

WidthHeightPeptide font sizePeptide stroke widthSpectrum font sizeSpectrum stroke widthCompact peptide

Ion legend

wxyz

abcd

OtherUnassignedIon^Charge^_Position_Show for top:%

TLPPSREEMTKNQKJ

02.01e+44.02e+46.03e+48.05e+4

Zoom Out

y^+1^_3_z^+3^_12_z^+3^_12_c^+1^_5_y^+3^_13_z^+3^_13_z^+1^_4_y^+3^_13_y^+1^_4_c^+2^_11_c^+2^_11_c^+1^_6_z^+1^_5_y^+2^_10_y^+1^_5_c^+2^_12_c^+2^_12_c^+2^_12_y^+2^_11_z^+2^_11_y^+2^_11_z^+2^_12_z^+2^_12_w^+1^_6_w^+1^_6_y^+2^_12_y^+1^_6_c^+2^_13_y^+1^_6_w^+2^_13_c^+1^_7_z^+2^_13_y^+2^_13_w^+2^_14_z^+2^_14_c^+2^_14_y^+2^_14_z^+1^_7_z^+1^_7_c^+1^_8_y^+1^_7_w^+1^_8_z^+1^_8_z^+1^_8_y^+1^_8_c^+1^_9_w^+1^_9_c^+1^_10_y^+1^_9_z^+1^_9_y^+1^_9_c^+1^_11_c^+1^_11_z^+1^_10_y^+1^_10_c^+1^_12_y^+1^_11_z^+1^_11_y^+1^_11_y^+1^_12_c^+1^_13_y^+1^_13_z^+1^_13_y^+1^_13_z^+1^_14_c^+1^_14_

046893614041871

Fragment Matches Table

Show background peaks

| Position | Ion type | Intensity | mz Theoretical | mz Error (Th) | mz Error (ppm) | Charge | Series Number |
| --- | --- | --- | --- | --- | --- | --- | --- |
| - | - | 396.7 | 132.9 | - | - | 0 | - |
| - | - | 1376 | 173.4 | - | - | 0 | - |
| - | - | 500.2 | 178.1 | - | - | 0 | - |
| - | - | 1.864E+04 | 187.1 | - | - | 0 | - |
| - | - | 1484 | 188.1 | - | - | 0 | - |
| - | - | 1801 | 212.1 | - | - | 0 | - |
| - | - | 1.061E+04 | 215.1 | - | - | 0 | - |
| - | - | 1112 | 216.1 | - | - | 0 | - |
| - | - | 2176 | 219.1 | - | - | 0 | - |
| - | - | 2143 | 221.1 | - | - | 0 | - |
| - | - | 642.1 | 249.1 | - | - | 0 | - |
| - | - | 592.7 | 256.2 | - | - | 0 | - |
| - | - | 2789 | 299.2 | - | - | 0 | - |
| - | - | 669.6 | 313.2 | - | - | 0 | - |
| - | - | 1366 | 326.2 | - | - | 0 | - |
| - | - | 1513 | 327.2 | - | - | 0 | - |
| - | - | 653.4 | 407.3 | - | - | 0 | - |
| - | - | 1587 | 425.3 | - | - | 0 | - |
| 13 | y | 1368 | 446.3 | 0.004124 | 9.24 | +1 | 3 |
| - | - | 673.3 | 456.2 | - | - | 0 | - |
| - | - | 606.7 | 464.2 | - | - | 0 | - |
| - | - | 674.9 | 465.2 | - | - | 0 | - |
| - | - | 829.5 | 466.2 | - | - | 0 | - |
| - | - | 1697 | 470.3 | - | - | 0 | - |
| - | - | 640 | 471.3 | - | - | 0 | - |
| - | - | 1646 | 471.9 | - | - | 0 | - |
| - | - | 2150 | 472.2 | - | - | 0 | - |
| - | - | 786.1 | 483.2 | - | - | 0 | - |
| - | - | 1725 | 484.2 | - | - | 0 | - |
| - | - | 619.4 | 491.6 | - | - | 0 | - |
| - | - | 673.9 | 494.3 | - | - | 0 | - |
| 4 | z | 1705 | 500.9 | 0.004506 | 8.996 | +3 | 12 |
| 4 | z | 662.3 | 501.2 | 0.002236 | 4.461 | +3 | 12 |
| 5 | c | 1.36E+04 | 513.3 | 0.0003427 | 0.6676 | +1 | 5 |
| - | - | 4119 | 514.3 | - | - | 0 | - |
| - | - | 1072 | 515.3 | - | - | 0 | - |
| - | - | 1193 | 523.3 | - | - | 0 | - |
| - | - | 607.6 | 529.3 | - | - | 0 | - |
| 3 | y | 717.5 | 538.6 | 0.001246 | 2.313 | +3 | 13 |
| 3 | z | 802.2 | 539.3 | 0.002044 | 3.789 | +3 | 13 |
| 12 | z | 965.8 | 544.3 | 0.006263 | 11.51 | +1 | 4 |
| 3 | y | 6.377E+04 | 544.6 | 0.0002984 | 0.548 | +3 | 13 |
| - | - | 5.798E+04 | 544.9 | - | - | 0 | - |
| - | - | 3.311E+04 | 545.3 | - | - | 0 | - |
| - | - | 1.683E+04 | 545.6 | - | - | 0 | - |
| - | - | 4043 | 545.9 | - | - | 0 | - |
| 12 | y | 3383 | 560.3 | 0.00454 | 8.103 | +1 | 4 |
| - | - | 745.8 | 561.8 | - | - | 0 | - |
| - | - | 5375 | 615.3 | - | - | 0 | - |
| - | - | 771.3 | 616.3 | - | - | 0 | - |
| - | - | 1089 | 617.3 | - | - | 0 | - |
| - | - | 4888 | 628.4 | - | - | 0 | - |
| - | - | 1580 | 629.3 | - | - | 0 | - |
| - | - | 1845 | 629.4 | - | - | 0 | - |
| - | - | 792.2 | 630.4 | - | - | 0 | - |
| 11 | c | 715.6 | 643.8 | 0.001142 | 1.773 | +2 | 11 |
| - | - | 675 | 644.8 | - | - | 0 | - |
| - | - | 1689 | 651.8 | - | - | 0 | - |
| 11 | c | 1.165E+04 | 652.3 | 0.002002 | 3.069 | +2 | 11 |
| - | - | 8752 | 652.8 | - | - | 0 | - |
| - | - | 3104 | 653.3 | - | - | 0 | - |
| - | - | 986.2 | 653.8 | - | - | 0 | - |
| - | - | 727.1 | 668.3 | - | - | 0 | - |
| - | - | 650.4 | 668.8 | - | - | 0 | - |
| 6 | c | 2.943E+04 | 669.4 | 0.0001824 | 0.2725 | +1 | 6 |
| - | - | 1.071E+04 | 670.4 | - | - | 0 | - |
| - | - | 2649 | 671.4 | - | - | 0 | - |
| 11 | z | 8775 | 672.4 | 0.005034 | 7.487 | +1 | 5 |
| - | - | 4062 | 673.4 | - | - | 0 | - |
| - | - | 942.9 | 674.4 | - | - | 0 | - |
| - | - | 678.2 | 675.3 | - | - | 0 | - |
| 6 | y | 1552 | 675.8 | 0.0001341 | 0.1984 | +2 | 10 |
| - | - | 849.1 | 676.3 | - | - | 0 | - |
| - | - | 1153 | 681.3 | - | - | 0 | - |
| - | - | 845.6 | 687.4 | - | - | 0 | - |
| 11 | y | 2888 | 688.4 | 0.004716 | 6.85 | +1 | 5 |
| - | - | 1362 | 689.4 | - | - | 0 | - |
| - | - | 790 | 699.9 | - | - | 0 | - |
| 12 | c | 1334 | 700.4 | 0.001545 | 2.206 | +2 | 12 |
| 12 | c | 1067 | 700.8 | 0.009377 | 13.38 | +2 | 12 |
| - | - | 2485 | 707.3 | - | - | 0 | - |
| - | - | 1751 | 707.8 | - | - | 0 | - |
| - | - | 1583 | 708.3 | - | - | 0 | - |
| - | - | 2645 | 708.9 | - | - | 0 | - |
| 12 | c | 3.549E+04 | 709.4 | 0.002023 | 2.852 | +2 | 12 |
| - | - | 2.604E+04 | 709.9 | - | - | 0 | - |
| 5 | y | 1.23E+04 | 710.3 | 0.01123 | 15.81 | +2 | 11 |
| - | - | 4623 | 710.9 | - | - | 0 | - |
| 5 | z | 1520 | 711.3 | 0.005421 | 7.62 | +2 | 11 |
| - | - | 2190 | 711.8 | - | - | 0 | - |
| - | - | 1193 | 712.3 | - | - | 0 | - |
| - | - | 605 | 716.3 | - | - | 0 | - |
| 5 | y | 1373 | 719.4 | 0.001735 | 2.412 | +2 | 11 |
| - | - | 768.9 | 719.9 | - | - | 0 | - |
| - | - | 1040 | 729.4 | - | - | 0 | - |
| - | - | 737.6 | 746.9 | - | - | 0 | - |
| 4 | z | 1184 | 750.9 | 0.007378 | 9.826 | +2 | 12 |
| 4 | z | 888.1 | 751.4 | 0.005253 | 6.991 | +2 | 12 |
| - | - | 865.1 | 751.9 | - | - | 0 | - |
| 10 | w | 1876 | 756.4 | 0.002976 | 3.934 | +1 | 6 |
| - | - | 1220 | 757.4 | - | - | 0 | - |
| 10 | w | 4011 | 758.4 | 0.00342 | 4.509 | +1 | 6 |
| - | - | 1394 | 759.4 | - | - | 0 | - |
| - | - | 940.2 | 765.4 | - | - | 0 | - |
| - | - | 614.9 | 766.4 | - | - | 0 | - |
| - | - | 825 | 767.4 | - | - | 0 | - |
| 4 | y | 1.136E+04 | 767.9 | 0.0001108 | 0.1443 | +2 | 12 |
| - | - | 9605 | 768.4 | - | - | 0 | - |
| - | - | 4671 | 768.9 | - | - | 0 | - |
| - | - | 829.4 | 769.4 | - | - | 0 | - |
| - | - | 800.7 | 769.9 | - | - | 0 | - |
| 10 | y | 3786 | 772.4 | 0.0101 | 13.07 | +1 | 6 |
| - | - | 1694 | 772.9 | - | - | 0 | - |
| 13 | c | 1.36E+04 | 773.4 | 0.003191 | 4.125 | +2 | 13 |
| - | - | 1.5E+04 | 773.9 | - | - | 0 | - |
| - | - | 7176 | 774.4 | - | - | 0 | - |
| - | - | 5100 | 774.4 | - | - | 0 | - |
| - | - | 3151 | 774.9 | - | - | 0 | - |
| - | - | 1876 | 775.4 | - | - | 0 | - |
| - | - | 1415 | 784.4 | - | - | 0 | - |
| - | - | 1596 | 784.9 | - | - | 0 | - |
| - | - | 878.6 | 785.4 | - | - | 0 | - |
| - | - | 734 | 785.9 | - | - | 0 | - |
| 10 | y | 3634 | 789.5 | 0.005519 | 6.991 | +1 | 6 |
| - | - | 1951 | 790.4 | - | - | 0 | - |
| - | - | 695.8 | 791.4 | - | - | 0 | - |
| - | - | 816.8 | 792.9 | - | - | 0 | - |
| - | - | 1445 | 793.4 | - | - | 0 | - |
| - | - | 865.7 | 793.9 | - | - | 0 | - |
| 3 | w | 1693 | 794.9 | 0.00248 | 3.12 | +2 | 13 |
| - | - | 1210 | 795.4 | - | - | 0 | - |
| - | - | 990.5 | 795.9 | - | - | 0 | - |
| 7 | c | 4.435E+04 | 798.4 | 0.000173 | 0.2166 | +1 | 7 |
| - | - | 1.815E+04 | 799.4 | - | - | 0 | - |
| - | - | 4820 | 800.5 | - | - | 0 | - |
| - | - | 1137 | 800.9 | - | - | 0 | - |
| - | - | 1759 | 801.4 | - | - | 0 | - |
| - | - | 1092 | 801.9 | - | - | 0 | - |
| - | - | 1465 | 802.4 | - | - | 0 | - |
| 3 | z | 917.1 | 808.4 | 0.005635 | 6.971 | +2 | 13 |
| - | - | 748.5 | 808.9 | - | - | 0 | - |
| - | - | 1203 | 813.4 | - | - | 0 | - |
| - | - | 1943 | 813.9 | - | - | 0 | - |
| - | - | 688.6 | 814.4 | - | - | 0 | - |
| - | - | 1287 | 814.9 | - | - | 0 | - |
| - | - | 1140 | 815.9 | - | - | 0 | - |
| 3 | y | 3.887E+04 | 816.4 | 0.0002407 | 0.2948 | +2 | 13 |
| - | - | 3.627E+04 | 816.9 | - | - | 0 | - |
| - | - | 1.662E+04 | 817.4 | - | - | 0 | - |
| - | - | 7594 | 817.9 | - | - | 0 | - |
| - | - | 2498 | 818.4 | - | - | 0 | - |
| - | - | 925.9 | 818.9 | - | - | 0 | - |
| - | - | 3364 | 822.4 | - | - | 0 | - |
| - | - | 1.993E+04 | 822.9 | - | - | 0 | - |
| - | - | 1.69E+04 | 823.4 | - | - | 0 | - |
| - | - | 1.069E+04 | 823.9 | - | - | 0 | - |
| - | - | 3473 | 824.4 | - | - | 0 | - |
| - | - | 1273 | 824.9 | - | - | 0 | - |
| - | - | 1739 | 825.5 | - | - | 0 | - |
| - | - | 693.8 | 828.9 | - | - | 0 | - |
| - | - | 1567 | 835.4 | - | - | 0 | - |
| - | - | 2004 | 835.9 | - | - | 0 | - |
| - | - | 1142 | 836.4 | - | - | 0 | - |
| - | - | 1555 | 836.9 | - | - | 0 | - |
| - | - | 1512 | 837.4 | - | - | 0 | - |
| 2 | w | 1.639E+04 | 843.4 | 0.0001463 | 0.1735 | +2 | 14 |
| - | - | 1.726E+04 | 843.9 | - | - | 0 | - |
| - | - | 1.129E+04 | 844.4 | - | - | 0 | - |
| - | - | 5138 | 844.9 | - | - | 0 | - |
| - | - | 1305 | 845.4 | - | - | 0 | - |
| - | - | 982.9 | 854.3 | - | - | 0 | - |
| - | - | 2024 | 856.5 | - | - | 0 | - |
| - | - | 992.4 | 857 | - | - | 0 | - |
| - | - | 3968 | 857.5 | - | - | 0 | - |
| - | - | 1.418E+04 | 858.5 | - | - | 0 | - |
| - | - | 860.7 | 858.9 | - | - | 0 | - |
| - | - | 6583 | 859.5 | - | - | 0 | - |
| - | - | 2186 | 860.5 | - | - | 0 | - |
| 2 | z | 6079 | 864.9 | 0.001912 | 2.21 | +2 | 14 |
| - | - | 7328 | 865.4 | - | - | 0 | - |
| - | - | 3645 | 865.9 | - | - | 0 | - |
| 14 | c | 4.089E+04 | 866.4 | 3.559E-05 | 0.04107 | +2 | 14 |
| - | - | 4.083E+04 | 866.9 | - | - | 0 | - |
| - | - | 2.375E+04 | 867.4 | - | - | 0 | - |
| - | - | 1.09E+04 | 867.9 | - | - | 0 | - |
| - | - | 3639 | 868.4 | - | - | 0 | - |
| - | - | 1052 | 872 | - | - | 0 | - |
| - | - | 1401 | 872.4 | - | - | 0 | - |
| 2 | y | 1460 | 872.9 | 0.005999 | 6.872 | +2 | 14 |
| - | - | 731.9 | 873.4 | - | - | 0 | - |
| - | - | 1356 | 873.9 | - | - | 0 | - |
| - | - | 976.7 | 878 | - | - | 0 | - |
| - | - | 1134 | 878.5 | - | - | 0 | - |
| - | - | 1949 | 879.5 | - | - | 0 | - |
| - | - | 1687 | 879.9 | - | - | 0 | - |
| - | - | 2150 | 880.4 | - | - | 0 | - |
| - | - | 935.9 | 880.9 | - | - | 0 | - |
| - | - | 1912 | 883.5 | - | - | 0 | - |
| - | - | 1789 | 884.5 | - | - | 0 | - |
| - | - | 1416 | 885.5 | - | - | 0 | - |
| - | - | 1133 | 886.5 | - | - | 0 | - |
| - | - | 1693 | 887 | - | - | 0 | - |
| - | - | 1.427E+04 | 887.4 | - | - | 0 | - |
| - | - | 1.108E+04 | 887.9 | - | - | 0 | - |
| - | - | 9285 | 888.4 | - | - | 0 | - |
| - | - | 3438 | 888.9 | - | - | 0 | - |
| - | - | 1244 | 889.4 | - | - | 0 | - |
| - | - | 862 | 889.9 | - | - | 0 | - |
| - | - | 1891 | 892 | - | - | 0 | - |
| - | - | 2633 | 892.5 | - | - | 0 | - |
| - | - | 4158 | 893 | - | - | 0 | - |
| - | - | 2892 | 893.5 | - | - | 0 | - |
| - | - | 3673 | 894 | - | - | 0 | - |
| - | - | 7629 | 894.5 | - | - | 0 | - |
| - | - | 6826 | 895 | - | - | 0 | - |
| - | - | 3701 | 895.5 | - | - | 0 | - |
| - | - | 3956 | 896 | - | - | 0 | - |
| - | - | 4319 | 896.4 | - | - | 0 | - |
| - | - | 3004 | 896.9 | - | - | 0 | - |
| - | - | 1847 | 897.5 | - | - | 0 | - |
| - | - | 1005 | 901 | - | - | 0 | - |
| - | - | 8538 | 901.5 | - | - | 0 | - |
| - | - | 9616 | 902 | - | - | 0 | - |
| 9 | z | 8883 | 902.5 | 0.01215 | 13.46 | +1 | 7 |
| - | - | 4463 | 903 | - | - | 0 | - |
| - | - | 3151 | 903.5 | - | - | 0 | - |
| - | - | 2051 | 906 | - | - | 0 | - |
| - | - | 7110 | 906.5 | - | - | 0 | - |
| - | - | 7691 | 907 | - | - | 0 | - |
| - | - | 4816 | 907.5 | - | - | 0 | - |
| - | - | 2011 | 908 | - | - | 0 | - |
| - | - | 1332 | 908.5 | - | - | 0 | - |
| - | - | 9462 | 915 | - | - | 0 | - |
| - | - | 4.197E+04 | 915.5 | - | - | 0 | - |
| - | - | 4.259E+04 | 916 | - | - | 0 | - |
| - | - | 2.712E+04 | 916.5 | - | - | 0 | - |
| - | - | 1.145E+04 | 917 | - | - | 0 | - |
| - | - | 5350 | 917.5 | - | - | 0 | - |
| - | - | 1736 | 918 | - | - | 0 | - |
| 9 | z | 3966 | 920.5 | 0.001397 | 1.518 | +1 | 7 |
| - | - | 3578 | 921.5 | - | - | 0 | - |
| - | - | 1446 | 922.5 | - | - | 0 | - |
| - | - | 2.807E+04 | 923.5 | - | - | 0 | - |
| - | - | 5.547E+04 | 924 | - | - | 0 | - |
| - | - | 4.108E+04 | 924.5 | - | - | 0 | - |
| - | - | 2.3E+04 | 925 | - | - | 0 | - |
| - | - | 1.068E+04 | 925.5 | - | - | 0 | - |
| - | - | 2763 | 926 | - | - | 0 | - |
| - | - | 2270 | 926.5 | - | - | 0 | - |
| 8 | c | 3.242E+04 | 927.5 | 0.0005908 | 0.637 | +1 | 8 |
| - | - | 1.733E+04 | 928.5 | - | - | 0 | - |
| - | - | 5266 | 929.5 | - | - | 0 | - |
| - | - | 811.1 | 930.5 | - | - | 0 | - |
| 9 | y | 6438 | 936.5 | 0.0001149 | 0.1227 | +1 | 7 |
| - | - | 3244 | 937.5 | - | - | 0 | - |
| - | - | 851 | 938.5 | - | - | 0 | - |
| - | - | 928.3 | 968.5 | - | - | 0 | - |
| - | - | 809.2 | 969.5 | - | - | 0 | - |
| - | - | 1094 | 978.5 | - | - | 0 | - |
| - | - | 1373 | 985.3 | - | - | 0 | - |
| - | - | 1010 | 986.5 | - | - | 0 | - |
| 8 | w | 5868 | 990.5 | 6.258E-05 | 0.06318 | +1 | 8 |
| - | - | 3447 | 991.5 | - | - | 0 | - |
| - | - | 1122 | 992.5 | - | - | 0 | - |
| - | - | 3726 | 1031 | - | - | 0 | - |
| 8 | z | 2807 | 1031 | 0.01569 | 15.22 | +1 | 8 |
| - | - | 907.3 | 1033 | - | - | 0 | - |
| 8 | z | 2.189E+04 | 1050 | 2.87E-06 | 0.002735 | +1 | 8 |
| - | - | 1.428E+04 | 1051 | - | - | 0 | - |
| - | - | 5958 | 1052 | - | - | 0 | - |
| - | - | 1955 | 1053 | - | - | 0 | - |
| 8 | y | 1048 | 1066 | 0.001998 | 1.875 | +1 | 8 |
| - | - | 1057 | 1067 | - | - | 0 | - |
| 9 | c | 2.605E+04 | 1075 | 0.004508 | 4.195 | +1 | 9 |
| - | - | 1.535E+04 | 1076 | - | - | 0 | - |
| - | - | 6571 | 1077 | - | - | 0 | - |
| - | - | 2075 | 1078 | - | - | 0 | - |
| - | - | 766.5 | 1114 | - | - | 0 | - |
| - | - | 819.1 | 1115 | - | - | 0 | - |
| 7 | w | 2484 | 1120 | 0.003551 | 3.172 | +1 | 9 |
| - | - | 1779 | 1121 | - | - | 0 | - |
| - | - | 879.6 | 1133 | - | - | 0 | - |
| - | - | 800.7 | 1135 | - | - | 0 | - |
| - | - | 723.6 | 1146 | - | - | 0 | - |
| - | - | 751.4 | 1174 | - | - | 0 | - |
| 10 | c | 2.274E+04 | 1176 | 0.003949 | 3.359 | +1 | 10 |
| 7 | y | 1.445E+04 | 1177 | 0.01798 | 15.28 | +1 | 9 |
| - | - | 6615 | 1178 | - | - | 0 | - |
| 7 | z | 1.691E+04 | 1179 | 0.002576 | 2.186 | +1 | 9 |
| - | - | 1.177E+04 | 1180 | - | - | 0 | - |
| - | - | 5792 | 1181 | - | - | 0 | - |
| - | - | 2178 | 1182 | - | - | 0 | - |
| 7 | y | 1548 | 1195 | 0.005284 | 4.423 | +1 | 9 |
| - | - | 1614 | 1196 | - | - | 0 | - |
| - | - | 1406 | 1216 | - | - | 0 | - |
| - | - | 877.5 | 1217 | - | - | 0 | - |
| - | - | 926.7 | 1224 | - | - | 0 | - |
| - | - | 780.4 | 1232 | - | - | 0 | - |
| - | - | 3290 | 1234 | - | - | 0 | - |
| - | - | 1367 | 1235 | - | - | 0 | - |
| - | - | 804.1 | 1260 | - | - | 0 | - |
| - | - | 986.1 | 1261 | - | - | 0 | - |
| - | - | 1225 | 1262 | - | - | 0 | - |
| 11 | c | 1085 | 1286 | 0.008916 | 6.935 | +1 | 11 |
| - | - | 3120 | 1303 | - | - | 0 | - |
| 11 | c | 7113 | 1304 | 0.001881 | 1.443 | +1 | 11 |
| - | - | 4384 | 1305 | - | - | 0 | - |
| - | - | 2456 | 1306 | - | - | 0 | - |
| 6 | z | 1.025E+04 | 1335 | 0.001611 | 1.207 | +1 | 10 |
| - | - | 1.633E+04 | 1336 | - | - | 0 | - |
| - | - | 1.031E+04 | 1337 | - | - | 0 | - |
| - | - | 4143 | 1338 | - | - | 0 | - |
| - | - | 2220 | 1350 | - | - | 0 | - |
| 6 | y | 5840 | 1351 | 0.003124 | 2.313 | +1 | 10 |
| - | - | 3482 | 1352 | - | - | 0 | - |
| - | - | 1335 | 1353 | - | - | 0 | - |
| - | - | 1081 | 1354 | - | - | 0 | - |
| - | - | 981.8 | 1374 | - | - | 0 | - |
| - | - | 1384 | 1375 | - | - | 0 | - |
| 12 | c | 1.343E+04 | 1418 | 0.001678 | 1.184 | +1 | 12 |
| - | - | 9976 | 1419 | - | - | 0 | - |
| 5 | y | 6436 | 1420 | 0.02058 | 14.5 | +1 | 11 |
| - | - | 2859 | 1421 | - | - | 0 | - |
| 5 | z | 1927 | 1422 | 0.01067 | 7.506 | +1 | 11 |
| - | - | 5162 | 1423 | - | - | 0 | - |
| - | - | 5458 | 1424 | - | - | 0 | - |
| - | - | 3243 | 1425 | - | - | 0 | - |
| - | - | 897.9 | 1426 | - | - | 0 | - |
| 5 | y | 1161 | 1438 | 0.01209 | 8.409 | +1 | 11 |
| - | - | 1062 | 1439 | - | - | 0 | - |
| 4 | y | 4002 | 1535 | 0.002833 | 1.846 | +1 | 12 |
| - | - | 2967 | 1536 | - | - | 0 | - |
| - | - | 1688 | 1537 | - | - | 0 | - |
| - | - | 1214 | 1538 | - | - | 0 | - |
| 13 | c | 9318 | 1546 | 0.003892 | 2.518 | +1 | 13 |
| - | - | 7483 | 1547 | - | - | 0 | - |
| - | - | 4755 | 1548 | - | - | 0 | - |
| - | - | 2077 | 1549 | - | - | 0 | - |
| - | - | 1272 | 1550 | - | - | 0 | - |
| - | - | 873.6 | 1603 | - | - | 0 | - |
| 3 | y | 1888 | 1615 | 0.01246 | 7.714 | +1 | 13 |
| 3 | z | 1331 | 1616 | 0.01305 | 8.079 | +1 | 13 |
| - | - | 1734 | 1617 | - | - | 0 | - |
| - | - | 754.8 | 1618 | - | - | 0 | - |
| - | - | 1517 | 1629 | - | - | 0 | - |
| - | - | 975.4 | 1630 | - | - | 0 | - |
| 3 | y | 832.5 | 1632 | 0.004815 | 2.951 | +1 | 13 |
| - | - | 1108 | 1633 | - | - | 0 | - |
| - | - | 7247 | 1645 | - | - | 0 | - |
| - | - | 7847 | 1646 | - | - | 0 | - |
| - | - | 6999 | 1647 | - | - | 0 | - |
| - | - | 2906 | 1648 | - | - | 0 | - |
| - | - | 1431 | 1649 | - | - | 0 | - |
| - | - | 1164 | 1671 | - | - | 0 | - |
| - | - | 1280 | 1672 | - | - | 0 | - |
| - | - | 854.3 | 1686 | - | - | 0 | - |
| - | - | 719 | 1687 | - | - | 0 | - |
| - | - | 859.2 | 1688 | - | - | 0 | - |
| - | - | 2273 | 1689 | - | - | 0 | - |
| - | - | 2063 | 1690 | - | - | 0 | - |
| - | - | 1218 | 1691 | - | - | 0 | - |
| - | - | 1585 | 1716 | - | - | 0 | - |
| - | - | 1088 | 1717 | - | - | 0 | - |
| - | - | 1188 | 1718 | - | - | 0 | - |
| 2 | z | 2074 | 1729 | 0.00443 | 2.562 | +1 | 14 |
| - | - | 1.165E+04 | 1730 | - | - | 0 | - |
| - | - | 1.346E+04 | 1731 | - | - | 0 | - |
| 14 | c | 1.182E+04 | 1732 | 0.002933 | 1.693 | +1 | 14 |
| - | - | 1.633E+04 | 1733 | - | - | 0 | - |
| - | - | 1.251E+04 | 1734 | - | - | 0 | - |
| - | - | 5871 | 1735 | - | - | 0 | - |
| - | - | 1399 | 1736 | - | - | 0 | - |
| - | - | 951.9 | 1771 | - | - | 0 | - |
| - | - | 1065 | 1772 | - | - | 0 | - |
| - | - | 715.6 | 1774 | - | - | 0 | - |
| - | - | 1411 | 1784 | - | - | 0 | - |
| - | - | 1754 | 1785 | - | - | 0 | - |
| - | - | 1985 | 1786 | - | - | 0 | - |
| - | - | 2283 | 1787 | - | - | 0 | - |
| - | - | 2261 | 1788 | - | - | 0 | - |
| - | - | 5638 | 1789 | - | - | 0 | - |
| - | - | 3669 | 1790 | - | - | 0 | - |
| - | - | 2214 | 1791 | - | - | 0 | - |
| - | - | 1134 | 1792 | - | - | 0 | - |
| - | - | 862.2 | 1793 | - | - | 0 | - |
| - | - | 2493 | 1802 | - | - | 0 | - |
| - | - | 8471 | 1803 | - | - | 0 | - |
| - | - | 6375 | 1804 | - | - | 0 | - |
| - | - | 6454 | 1805 | - | - | 0 | - |
| - | - | 2187 | 1806 | - | - | 0 | - |
| - | - | 716 | 1807 | - | - | 0 | - |
| - | - | 843.3 | 1808 | - | - | 0 | - |
| - | - | 1732 | 1813 | - | - | 0 | - |
| - | - | 2645 | 1814 | - | - | 0 | - |
| - | - | 2345 | 1815 | - | - | 0 | - |
| - | - | 1779 | 1816 | - | - | 0 | - |
| - | - | 821.4 | 1817 | - | - | 0 | - |
| - | - | 919.6 | 1819 | - | - | 0 | - |
| - | - | 6995 | 1820 | - | - | 0 | - |
| - | - | 5496 | 1821 | - | - | 0 | - |
| - | - | 3129 | 1822 | - | - | 0 | - |
| - | - | 1827 | 1823 | - | - | 0 | - |
| - | - | 7452 | 1830 | - | - | 0 | - |
| - | - | 2.809E+04 | 1831 | - | - | 0 | - |
| - | - | 2.691E+04 | 1832 | - | - | 0 | - |
| - | - | 1.937E+04 | 1833 | - | - | 0 | - |
| - | - | 7292 | 1834 | - | - | 0 | - |
| - | - | 2255 | 1835 | - | - | 0 | - |
| - | - | 1395 | 1836 | - | - | 0 | - |
| - | - | 3359 | 1846 | - | - | 0 | - |
| - | - | 1.807E+04 | 1847 | - | - | 0 | - |
| - | - | 7.966E+04 | 1848 | - | - | 0 | - |
| - | - | 7.356E+04 | 1849 | - | - | 0 | - |
| - | - | 4.272E+04 | 1850 | - | - | 0 | - |
| - | - | 1.987E+04 | 1851 | - | - | 0 | - |
| - | - | 6391 | 1852 | - | - | 0 | - |
| - | - | 1730 | 1853 | - | - | 0 | - |

m/z Charge Intensity FragmentType MassShift Position
132.8741455078125 0 396.66086
173.4386749267578 0 1376.0005
178.1350555419922 0 500.2045
187.14427185058594 0 18637.576
188.14768981933594 0 1483.7285
212.13929748535156 0 1801.2335
215.1391143798828 0 10613.598
216.1426544189453 0 1111.8672
219.1341552734375 0 2176.0671
221.12832641601562 0 2142.8506
249.1233367919922 0 642.067
256.1639709472656 0 592.72906
299.17181396484375 0 2788.875
313.1971130371094 0 669.6021
326.1583557128906 0 1365.579
327.1667175292969 0 1513.1328
407.2522888183594 0 653.3683
425.26373291015625 0 1587.221
446.2613220214844 0 1368.4281 y 12
456.2279357910156 0 673.30774
464.2059020996094 0 606.6783
465.2062683105469 0 674.9477
466.2100524902344 0 829.52747
470.2981872558594 0 1697.2501
471.2995300292969 0 639.95087
471.8979797363281 0 1646.0176
472.2317199707031 0 2150.1716
483.2172546386719 0 786.1065
484.2203369140625 0 1724.5878
491.5782470703125 0 619.37494
494.2841491699219 0 673.9405
500.9081115722656 0 1705.021 z Water loss 3
501.24285888671875 0 662.29926 z Ammonia loss 3
513.303466796875 0 13597.353 c 4
514.3052978515625 0 4119.0835
515.3090209960938 0 1071.933
523.2725830078125 0 1192.5795
529.2678833007812 0 607.61957
538.6018676757812 0 717.54095 y Water loss 2
539.2691040039062 0 802.2002 z 2
544.2833862304688 0 965.8499 z 11
544.60693359375 0 63765.332 y 2
544.9406127929688 0 57978.72
545.274658203125 0 33113.566
545.6077270507812 0 16825.541
545.9415893554688 0 4042.8303
560.3038330078125 0 3383.4336 y 11
561.7817993164062 0 745.7737
615.3150024414062 0 5375.418
616.3203735351562 0 771.27966
617.3173217773438 0 1088.7102
628.366455078125 0 4887.638
629.3252563476562 0 1579.6849
629.3724975585938 0 1845.149
630.3759765625 0 792.22375
643.8204956054688 0 715.6318 c Ammonia loss 10
644.8335571289062 0 674.9535
651.8336791992188 0 1688.5795
652.3369140625 0 11646.646 c 10
652.8385009765625 0 8751.726
653.3391723632812 0 3103.62
653.8377685546875 0 986.24646
668.330810546875 0 727.0812
668.8309936523438 0 650.35645
669.404052734375 0 29430.004 c 5
670.4074096679688 0 10709.133
671.4099731445312 0 2649.2246
672.3795776367188 0 8775.402 z 10
673.380126953125 0 4061.6777
674.3841552734375 0 942.9321
675.3333740234375 0 678.24054
675.83740234375 0 1552.0815 y 5
676.3403930664062 0 849.12305
681.336669921875 0 1153.2847
687.352294921875 0 845.62506
688.3986206054688 0 2888.0728 y 10
689.40185546875 0 1361.8063
699.8505859375 0 789.9791
700.3495483398438 0 1334.2838 c Water loss 11
700.8524780273438 0 1066.8062 c Ammonia loss 11
707.3434448242188 0 2485.1663
707.8451538085938 0 1750.7025
708.3447875976562 0 1583.1636
708.8514404296875 0 2644.8977
709.3583984375 0 35494.24 c 11
709.8594360351562 0 26038.004
710.3594970703125 0 12297.038 y Water loss 4
710.85791015625 0 4622.5864
711.349609375 0 1519.8286 z 4
711.848876953125 0 2189.676
712.3473510742188 0 1193.2296
716.3323974609375 0 604.9507
719.3552856445312 0 1373.246 y 4
719.858154296875 0 768.8778
729.4143676757812 0 1040.3652
746.8548583984375 0 737.63556
750.85791015625 0 1183.5247 z Water loss 3
751.362548828125 0 888.06323 z Ammonia loss 3
751.8682250976562 0 865.1132
756.4265747070312 0 1876.3107 w 9
757.4221801757812 0 1219.7236
758.4053955078125 0 4010.6018 w 9
759.4058837890625 0 1393.614
765.3939819335938 0 940.2146
766.3956909179688 0 614.9221
767.3739624023438 0 825.00635
767.8798217773438 0 11358.848 y 3
768.3802490234375 0 9604.655
768.8815307617188 0 4670.9907
769.3776245117188 0 829.4275
769.8851318359375 0 800.6834
772.4143676757812 0 3785.8938 y Ammonia loss 9
772.8870239257812 0 1694.1411
773.3888549804688 0 13604.281 c 12
773.888671875 0 15001.917
774.3883666992188 0 7175.953
774.4318237304688 0 5099.9473
774.8871459960938 0 3150.6462
775.4342041015625 0 1875.9523
784.4083251953125 0 1415.3442
784.9097900390625 0 1596.3962
785.404052734375 0 878.57367
785.910888671875 0 733.96564
789.4454956054688 0 3633.6995 y 9
790.4469604492188 0 1951.3268
791.44677734375 0 695.7696
792.9113159179688 0 816.7625
793.404541015625 0 1444.5254
793.9109497070312 0 865.6937
794.8876953125 0 1692.9772 w 2
795.3856201171875 0 1210.2825
795.8804321289062 0 990.5249
798.4466552734375 0 44347.035 c 6
799.4496459960938 0 18146.03
800.451171875 0 4819.703
800.909912109375 0 1136.513
801.412109375 0 1759.4004
801.9122314453125 0 1092.2141
802.4140625 0 1464.5104
808.402587890625 0 917.12354 z 2
808.8930053710938 0 748.47974
813.4122924804688 0 1203.0626
813.9124145507812 0 1942.9808
814.4081420898438 0 688.6338
814.9077758789062 0 1286.6595
815.9124145507812 0 1139.6735
816.4065551757812 0 38874.656 y 2
816.9071044921875 0 36266.805
817.4077758789062 0 16618.455
817.90869140625 0 7593.6465
818.40869140625 0 2497.641
818.9053955078125 0 925.9457
822.4172973632812 0 3364.306
822.9215698242188 0 19934.342
823.4219360351562 0 16898.062
823.9231567382812 0 10688.321
824.4368896484375 0 3472.6255
824.9287719726562 0 1273.4093
825.467041015625 0 1738.6691
828.927978515625 0 693.79504
835.4301147460938 0 1566.5532
835.9290771484375 0 2003.7811
836.4247436523438 0 1142.3333
836.922119140625 0 1554.5607
837.4137573242188 0 1511.6443
843.4117431640625 0 16385.164 w 1
843.9117431640625 0 17260.846
844.4130249023438 0 11287.54
844.9142456054688 0 5137.896
845.4125366210938 0 1304.9727
854.2954711914062 0 982.877
856.46240234375 0 2024.3641
856.9639892578125 0 992.4148
857.47119140625 0 3967.7366
858.4793090820312 0 14180.399
858.9240112304688 0 860.6708
859.4776611328125 0 6582.598
860.47900390625 0 2186.1448
864.9370727539062 0 6078.504 z 1
865.4407348632812 0 7327.8936
865.941650390625 0 3644.9705
866.4381103515625 0 40890.773 c 13
866.9387817382812 0 40834.785
867.4390869140625 0 23751.715
867.9396362304688 0 10898.763
868.4379272460938 0 3638.7017
871.9548950195312 0 1051.9515
872.448486328125 0 1401.1658
872.954345703125 0 1460.3477 y 1
873.4371337890625 0 731.8791
873.9334716796875 0 1356.0602
877.952392578125 0 976.69324
878.4512329101562 0 1133.5168
879.4503173828125 0 1949.3196
879.9371337890625 0 1686.8839
880.4481201171875 0 2150.384
880.9396362304688 0 935.8539
883.4747314453125 0 1911.6852
884.473876953125 0 1788.9502
885.4661865234375 0 1416.194
886.4575805664062 0 1133.3855
886.9517822265625 0 1692.7477
887.4332275390625 0 14271.868
887.9346313476562 0 11081.934
888.4390258789062 0 9284.769
888.9453125 0 3438.483
889.4429321289062 0 1243.6837
889.9407958984375 0 861.97614
891.971923828125 0 1891.1028
892.4659423828125 0 2632.687
892.9618530273438 0 4157.536
893.45849609375 0 2891.7495
893.9536743164062 0 3673.1465
894.4559936523438 0 7628.587
894.958251953125 0 6825.911
895.4584350585938 0 3700.849
895.95166015625 0 3956.1885
896.4442138671875 0 4318.89
896.947265625 0 3003.9468
897.4522094726562 0 1847.113
900.9725341796875 0 1005.017
901.4654541015625 0 8537.697
901.9661254882812 0 9615.89
902.46435546875 0 8883.344 z Water loss 8
902.9619140625 0 4462.5913
903.4589233398438 0 3151.0293
905.964599609375 0 2051.3054
906.4608154296875 0 7110.073
906.958740234375 0 7690.8496
907.458251953125 0 4816.199
907.9580688476562 0 2011.2297
908.4586181640625 0 1332.3933
914.9712524414062 0 9462.3545
915.466796875 0 41968.805
915.967041015625 0 42585.57
916.466552734375 0 27117.953
916.9664306640625 0 11450.275
917.4666137695312 0 5350.175
917.9725952148438 0 1735.645
920.4641723632812 0 3965.9675 z 8
921.46484375 0 3578.4937
922.4696655273438 0 1445.9141
923.47216796875 0 28065.86
923.973876953125 0 55472.58
924.4744262695312 0 41078.56
924.974853515625 0 23003.799
925.47607421875 0 10678.318
925.9783935546875 0 2762.6628
926.4804077148438 0 2269.5571
927.4888305664062 0 32423.89 c 7
928.4923706054688 0 17325.635
929.4955444335938 0 5265.6455
930.493896484375 0 811.05444
936.4813842773438 0 6437.9756 y 8
937.4808959960938 0 3244.4622
938.4810791015625 0 850.9623
968.5289916992188 0 928.2761
969.5319213867188 0 809.1756
978.4611206054688 0 1093.5704
985.3358764648438 0 1372.541
986.5133056640625 0 1009.7985
990.4921264648438 0 5867.773 w 7
991.4871215820312 0 3446.6377
992.49267578125 0 1122.0258
1030.511962890625 0 3725.8416
1031.510498046875 0 2807.3289 z Water loss 7
1032.5205078125 0 907.29877
1049.50537109375 0 21887.066 z 7
1050.5054931640625 0 14278.593
1051.50390625 0 5958.2393
1052.5076904296875 0 1955.2448
1065.5220947265625 0 1048.287 y 7
1066.51708984375 0 1056.831
1074.5244140625 0 26046.648 c 8
1075.5272216796875 0 15353.094
1076.52685546875 0 6570.661
1077.5355224609375 0 2075.4104
1113.5543212890625 0 766.483
1114.5589599609375 0 819.0956
1119.5382080078125 0 2483.787 w 6
1120.5240478515625 0 1778.9358
1132.5625 0 879.6391
1134.54541015625 0 800.73663
1146.1224365234375 0 723.62573
1174.4295654296875 0 751.3718
1175.571533203125 0 22740.5 c 9
1176.5740966796875 0 14447.592 y Water loss 6
1177.5760498046875 0 6615.1987
1178.550537109375 0 16906.48 z 6
1179.548583984375 0 11767.039
1180.5484619140625 0 5791.8535
1181.54541015625 0 2177.935
1194.5614013671875 0 1548.4807 y 6
1195.5679931640625 0 1614.2596
1216.443359375 0 1405.5903
1217.43896484375 0 877.49915
1223.595703125 0 926.6666
1232.4302978515625 0 780.3962
1234.4576416015625 0 3290.1418
1235.4541015625 0 1366.7921
1259.6494140625 0 804.143
1260.6541748046875 0 986.14044
1261.66015625 0 1224.748
1285.64306640625 0 1085.2848 c Water loss 10
1302.65673828125 0 3120.3828
1303.6644287109375 0 7112.924 c 10
1304.667724609375 0 4384.3022
1305.6729736328125 0 2456.04
1334.6474609375 0 10254.056 z 5
1335.651611328125 0 16334.266
1336.6552734375 0 10310.346
1337.6539306640625 0 4142.7666
1349.6575927734375 0 2220.306
1350.6646728515625 0 5839.5127 y 5
1351.66552734375 0 3482.1912
1352.6702880859375 0 1335.1614
1353.6768798828125 0 1081.479
1373.691162109375 0 981.78656
1374.706298828125 0 1383.6276
1417.7071533203125 0 13429.345 c 11
1418.70947265625 0 9975.644
1419.7098388671875 0 6435.8594 y Water loss 4
1420.7047119140625 0 2858.695
1421.6917724609375 0 1927.2996 z 4
1422.6868896484375 0 5162.472
1423.68603515625 0 5458.2334
1424.6912841796875 0 3242.7793
1425.688720703125 0 897.8787
1437.7119140625 0 1161.4839 y 4
1438.68310546875 0 1062.0736
1534.749755859375 0 4002.0066 y 3
1535.755859375 0 2967.2695
1536.7598876953125 0 1688.0164
1537.7591552734375 0 1214.4904
1545.7679443359375 0 9318.003 c 12
1546.7646484375 0 7483.0933
1547.76708984375 0 4755.403
1548.767333984375 0 2076.892
1549.76611328125 0 1271.521
1602.826904296875 0 873.60657
1614.791259765625 0 1888.3726 y Ammonia loss 2
1615.7996826171875 0 1330.7029 z 2
1616.802734375 0 1733.6174
1617.826416015625 0 754.83905
1628.8209228515625 0 1516.6382
1629.8096923828125 0 975.43976
1631.800537109375 0 832.53455 y 2
1632.8089599609375 0 1107.6665
1644.8328857421875 0 7247.306
1645.8397216796875 0 7847.486
1646.84033203125 0 6999.41
1647.8431396484375 0 2906.22
1648.84130859375 0 1431.4246
1670.8564453125 0 1164.189
1671.8499755859375 0 1280.1882
1685.86181640625 0 854.3384
1686.8927001953125 0 719.02905
1687.8553466796875 0 859.1552
1688.8486328125 0 2273.2327
1689.8565673828125 0 2062.883
1690.85400390625 0 1217.5028
1715.8575439453125 0 1585.2356
1716.8448486328125 0 1088.2417
1717.858154296875 0 1188.0107
1728.8751220703125 0 2073.9302 z 1
1729.8759765625 0 11649.883
1730.8736572265625 0 13460.843
1731.8719482421875 0 11819.259 c 13
1732.87109375 0 16325.777
1733.8741455078125 0 12506.25
1734.8743896484375 0 5871.13
1735.873046875 0 1398.6409
1770.8792724609375 0 951.8724
1771.890625 0 1064.6248
1773.8834228515625 0 715.62726
1783.9498291015625 0 1411.4209
1784.9310302734375 0 1753.8921
1785.9202880859375 0 1985.4388
1786.907958984375 0 2283.2039
1787.913330078125 0 2260.9019
1788.9073486328125 0 5637.925
1789.9072265625 0 3669.092
1790.90380859375 0 2214.3826
1791.92724609375 0 1134.2004
1792.90087890625 0 862.17944
1801.930908203125 0 2492.7385
1802.9305419921875 0 8471.252
1803.9306640625 0 6374.8706
1804.930419921875 0 6453.7627
1805.92236328125 0 2187.2708
1806.937744140625 0 716.0206
1807.94677734375 0 843.27783
1812.9190673828125 0 1732.1476
1813.9010009765625 0 2645.103
1814.9066162109375 0 2344.7212
1815.90576171875 0 1778.9166
1816.9288330078125 0 821.42676
1818.955810546875 0 919.5815
1819.951904296875 0 6995.336
1820.955322265625 0 5495.555
1821.94921875 0 3128.5547
1822.961181640625 0 1827.3241
1829.9219970703125 0 7452.305
1830.9248046875 0 28092.463
1831.9266357421875 0 26914.082
1832.925048828125 0 19371.387
1833.9254150390625 0 7291.642
1834.9222412109375 0 2255.219
1835.926513671875 0 1394.7047
1845.9306640625 0 3358.9746
1846.9395751953125 0 18065.137
1847.947265625 0 79655.555
1848.9483642578125 0 73563.195
1849.9512939453125 0 42723.285
1850.9517822265625 0 19867.115
1851.955322265625 0 6390.887
1852.9573974609375 0 1730.4554

Spectrum Details

|  |  |
| --- | --- |
| Matched peaks? Matched peaks The total absolute number of peaks matched. Additionally in brackets the total fraction of peaks matched and the total number of peaks is shown. | 66 (15.83% of 417) |
| FDR? FDR The false discovery rate estimated for this peptide. It is calculated by matching all theoretical fragments with a non-integer shift with the raw peaks for this spectrum. This is done with 40 different shifts. The resulting percentage is the average number of annotated peaks over the number of annotated peaks with the correct spectrum. | 0.18% |
| Satellite FDR? Satellite FDR See the FDR for details on its calculation. This satellite ion specific FDR only contains the satellite ions (d/w) for I/L/J positions. | 0.00% |
| PSM Score? PSM Score The PSM Score as given by Hecklib to this annotated spectrum. It is shown with three significant figures. | 458 |

#### Spectrum 4405? Spectrum 4405 The raw spectrum of this peptide as annotated by Hecklib. The fragments are coloured according to ion type (see legend). Any peaks with a star '\*' as text can be hovered over to see the full details, first the ion type second the mass shift type. By hovering over the amino acids in the peptide or ions in the legend the corresponding peaks are highlighted. By toggling the 'Unassigned' label you can turn the background (unassigned) peaks on or off in the plot. By updating the slider in the Ion legend you can update the spectrum to only show the top X% of the peaks with labels. The top X% means any peak that is within X% of the highest intensity. By dragging in the spectrum you can zoom in to a specific part of the spectrum and use 'Zoom Out' to get back to the original zoom level. The annotation of the spectrum is based on the given sequence in the peptides file and is done with different software so inconsistencies are likely. The peaks are annotated based on the given sequence, with 20 ppm tolerance.

Copy Data

##### Spectrum 4405 (TSV)

###### Preview

```
Loading example...
```

*Click on the button to copy the data to your clipboard.*

Mz MinMz MaxIntensity Max

WidthHeightPeptide font sizePeptide stroke widthSpectrum font sizeSpectrum stroke widthCompact peptide

Ion legend

wxyz

abcd

OtherUnassignedIon^Charge^_Position_Show for top:%

TLPPSREEMTKNQKJ

01.03e+52.06e+53.09e+54.12e+5

Zoom Out

y^+1^_1_c^+1^_4_z^+1^_3_y^+1^_3_z^+3^_12_z^+3^_12_y^+3^_12_c^+1^_5_y^+3^_13_y^+3^_13_z^+3^_13_z^+1^_4_y^+3^_13_y^+1^_4_c^+2^_11_c^+2^_11_z^+2^_10_z^+2^_10_c^+1^_6_z^+1^_5_y^+2^_10_y^+1^_5_c^+2^_12_c^+2^_12_z^+2^_11_c^+2^_12_y^+2^_11_z^+2^_11_y^+2^_11_z^+2^_12_z^+2^_12_z^+1^_6_w^+1^_6_w^+1^_6_y^+2^_12_z^+2^_12_c^+2^_13_c^+2^_13_y^+2^_12_y^+1^_6_c^+2^_13_y^+1^_6_w^+2^_13_c^+1^_7_y^+2^_13_y^+2^_13_z^+2^_13_y^+2^_13_w^+2^_14_c^+2^_14_y^+2^_14_z^+2^_14_c^+2^_14_y^+2^_14_z^+1^_7_y^+1^_7_z^+1^_7_c^+1^_8_y^+1^_7_w^+1^_8_z^+1^_8_y^+1^_8_c^+1^_9_w^+1^_9_c^+1^_10_y^+1^_9_z^+1^_9_y^+1^_9_c^+1^_11_c^+1^_11_c^+1^_11_z^+1^_10_y^+1^_10_c^+1^_12_y^+1^_11_z^+1^_11_y^+1^_11_c^+1^_13_y^+1^_12_c^+1^_13_y^+1^_13_z^+1^_13_y^+1^_13_z^+1^_14_c^+1^_14_

0779155723363115

Fragment Matches Table

Show background peaks

| Position | Ion type | Intensity | mz Theoretical | mz Error (Th) | mz Error (ppm) | Charge | Series Number |
| --- | --- | --- | --- | --- | --- | --- | --- |
| 15 | y | 4370 | 132.1 | 0.0003593 | 2.72 | +1 | 1 |
| - | - | 1465 | 143.1 | - | - | 0 | - |
| - | - | 1018 | 148.9 | - | - | 0 | - |
| - | - | 803 | 149.4 | - | - | 0 | - |
| - | - | 971.2 | 157.2 | - | - | 0 | - |
| - | - | 904 | 161.2 | - | - | 0 | - |
| - | - | 5637 | 173.4 | - | - | 0 | - |
| - | - | 1465 | 185.1 | - | - | 0 | - |
| - | - | 1.151E+05 | 187.1 | - | - | 0 | - |
| - | - | 1.068E+04 | 188.1 | - | - | 0 | - |
| - | - | 865.3 | 195.4 | - | - | 0 | - |
| - | - | 1259 | 212.1 | - | - | 0 | - |
| - | - | 6611 | 212.1 | - | - | 0 | - |
| - | - | 7.67E+04 | 215.1 | - | - | 0 | - |
| - | - | 7256 | 216.1 | - | - | 0 | - |
| - | - | 1.138E+04 | 219.1 | - | - | 0 | - |
| - | - | 1539 | 238.2 | - | - | 0 | - |
| - | - | 2354 | 256.2 | - | - | 0 | - |
| - | - | 1034 | 291.8 | - | - | 0 | - |
| - | - | 1.764E+04 | 299.2 | - | - | 0 | - |
| - | - | 2386 | 300.2 | - | - | 0 | - |
| - | - | 3124 | 312.2 | - | - | 0 | - |
| - | - | 1221 | 313.2 | - | - | 0 | - |
| - | - | 4808 | 326.2 | - | - | 0 | - |
| - | - | 5137 | 327.2 | - | - | 0 | - |
| - | - | 2482 | 344.2 | - | - | 0 | - |
| - | - | 1691 | 382.3 | - | - | 0 | - |
| - | - | 1070 | 383.3 | - | - | 0 | - |
| - | - | 2074 | 396.2 | - | - | 0 | - |
| - | - | 1384 | 402.7 | - | - | 0 | - |
| - | - | 5272 | 407.3 | - | - | 0 | - |
| - | - | 1795 | 412.2 | - | - | 0 | - |
| - | - | 1841 | 414.2 | - | - | 0 | - |
| - | - | 1680 | 425.2 | - | - | 0 | - |
| - | - | 5805 | 425.3 | - | - | 0 | - |
| 4 | c | 4228 | 426.3 | 0.0009236 | 2.167 | +1 | 4 |
| - | - | 3091 | 427.3 | - | - | 0 | - |
| - | - | 1715 | 428.3 | - | - | 0 | - |
| 13 | z | 3150 | 430.2 | 0.003466 | 8.056 | +1 | 3 |
| - | - | 1351 | 431.2 | - | - | 0 | - |
| - | - | 1004 | 444.2 | - | - | 0 | - |
| 13 | y | 8539 | 446.3 | 0.003605 | 8.078 | +1 | 3 |
| - | - | 2506 | 447.3 | - | - | 0 | - |
| - | - | 960.7 | 454.2 | - | - | 0 | - |
| - | - | 3023 | 462.9 | - | - | 0 | - |
| - | - | 1191 | 463.6 | - | - | 0 | - |
| - | - | 3146 | 466.2 | - | - | 0 | - |
| - | - | 9639 | 470.3 | - | - | 0 | - |
| - | - | 2322 | 471.3 | - | - | 0 | - |
| - | - | 8543 | 471.9 | - | - | 0 | - |
| - | - | 7482 | 472.2 | - | - | 0 | - |
| - | - | 1648 | 472.3 | - | - | 0 | - |
| - | - | 2326 | 472.6 | - | - | 0 | - |
| - | - | 2187 | 491.6 | - | - | 0 | - |
| - | - | 1786 | 491.9 | - | - | 0 | - |
| - | - | 3316 | 494.3 | - | - | 0 | - |
| - | - | 1541 | 494.9 | - | - | 0 | - |
| - | - | 1788 | 495.6 | - | - | 0 | - |
| - | - | 1328 | 495.9 | - | - | 0 | - |
| 4 | z | 5920 | 500.9 | 0.002919 | 5.828 | +3 | 12 |
| 4 | z | 3471 | 501.2 | 0.003854 | 7.688 | +3 | 12 |
| - | - | 2688 | 501.6 | - | - | 0 | - |
| - | - | 2270 | 501.9 | - | - | 0 | - |
| - | - | 1212 | 504.6 | - | - | 0 | - |
| - | - | 1626 | 507.2 | - | - | 0 | - |
| 4 | y | 1718 | 512.3 | 0.00161 | 3.144 | +3 | 12 |
| - | - | 3101 | 512.3 | - | - | 0 | - |
| 5 | c | 6.63E+04 | 513.3 | 0.000831 | 1.619 | +1 | 5 |
| - | - | 1.976E+04 | 514.3 | - | - | 0 | - |
| - | - | 4287 | 515.3 | - | - | 0 | - |
| - | - | 1296 | 520.2 | - | - | 0 | - |
| - | - | 7173 | 523.3 | - | - | 0 | - |
| - | - | 8673 | 523.6 | - | - | 0 | - |
| - | - | 2510 | 523.9 | - | - | 0 | - |
| - | - | 1654 | 528.3 | - | - | 0 | - |
| - | - | 2019 | 529.3 | - | - | 0 | - |
| - | - | 2243 | 529.9 | - | - | 0 | - |
| - | - | 2154 | 536.8 | - | - | 0 | - |
| 3 | y | 6892 | 538.6 | 0.001379 | 2.56 | +3 | 13 |
| 3 | y | 5539 | 538.9 | 0.006564 | 12.18 | +3 | 13 |
| 3 | z | 5317 | 539.3 | 0.003753 | 6.959 | +3 | 13 |
| - | - | 2706 | 539.6 | - | - | 0 | - |
| - | - | 2073 | 539.9 | - | - | 0 | - |
| 12 | z | 7893 | 544.3 | 0.004554 | 8.367 | +1 | 4 |
| 3 | y | 4.078E+05 | 544.6 | 0.001092 | 2.005 | +3 | 13 |
| - | - | 3.721E+05 | 544.9 | - | - | 0 | - |
| - | - | 1.985E+05 | 545.3 | - | - | 0 | - |
| - | - | 8.495E+04 | 545.6 | - | - | 0 | - |
| - | - | 2.128E+04 | 545.9 | - | - | 0 | - |
| - | - | 5227 | 546.3 | - | - | 0 | - |
| 12 | y | 1.603E+04 | 560.3 | 0.00393 | 7.014 | +1 | 4 |
| - | - | 3048 | 561.3 | - | - | 0 | - |
| - | - | 1248 | 564.2 | - | - | 0 | - |
| - | - | 2055 | 582.6 | - | - | 0 | - |
| - | - | 1991 | 596.3 | - | - | 0 | - |
| - | - | 1440 | 607.4 | - | - | 0 | - |
| - | - | 2455 | 614.4 | - | - | 0 | - |
| - | - | 4588 | 615.3 | - | - | 0 | - |
| - | - | 2196 | 616.3 | - | - | 0 | - |
| - | - | 1178 | 617.3 | - | - | 0 | - |
| - | - | 2363 | 625.4 | - | - | 0 | - |
| - | - | 4272 | 626.4 | - | - | 0 | - |
| - | - | 1917 | 627.4 | - | - | 0 | - |
| - | - | 2.368E+04 | 628.4 | - | - | 0 | - |
| - | - | 7514 | 629.3 | - | - | 0 | - |
| - | - | 7718 | 629.4 | - | - | 0 | - |
| - | - | 4556 | 630.3 | - | - | 0 | - |
| - | - | 4770 | 642.8 | - | - | 0 | - |
| 11 | c | 3844 | 643.3 | 0.002542 | 3.952 | +2 | 11 |
| - | - | 1384 | 643.8 | - | - | 0 | - |
| - | - | 1211 | 644.3 | - | - | 0 | - |
| - | - | 1.217E+04 | 651.8 | - | - | 0 | - |
| 11 | c | 5.142E+04 | 652.3 | 0.003223 | 4.94 | +2 | 11 |
| - | - | 3.738E+04 | 652.8 | - | - | 0 | - |
| - | - | 1.313E+04 | 653.3 | - | - | 0 | - |
| - | - | 4932 | 653.8 | - | - | 0 | - |
| - | - | 2111 | 657.8 | - | - | 0 | - |
| - | - | 2191 | 658.3 | - | - | 0 | - |
| 6 | z | 3361 | 658.8 | 0.008134 | 12.35 | +2 | 10 |
| 6 | z | 2705 | 667.8 | 0.0002935 | 0.4395 | +2 | 10 |
| - | - | 4870 | 668.3 | - | - | 0 | - |
| - | - | 1942 | 668.8 | - | - | 0 | - |
| 6 | c | 1.358E+05 | 669.4 | 0.001099 | 1.642 | +1 | 6 |
| - | - | 1404 | 670.3 | - | - | 0 | - |
| - | - | 4.903E+04 | 670.4 | - | - | 0 | - |
| - | - | 1.17E+04 | 671.4 | - | - | 0 | - |
| 11 | z | 4.954E+04 | 672.4 | 0.003203 | 4.764 | +1 | 5 |
| - | - | 2.118E+04 | 673.4 | - | - | 0 | - |
| - | - | 4612 | 674.4 | - | - | 0 | - |
| - | - | 6232 | 675.3 | - | - | 0 | - |
| 6 | y | 7490 | 675.8 | 0.002246 | 3.324 | +2 | 10 |
| - | - | 3935 | 676.3 | - | - | 0 | - |
| - | - | 1962 | 676.8 | - | - | 0 | - |
| - | - | 3587 | 680.8 | - | - | 0 | - |
| - | - | 1779 | 681.3 | - | - | 0 | - |
| - | - | 1455 | 686.8 | - | - | 0 | - |
| - | - | 3539 | 687.4 | - | - | 0 | - |
| - | - | 3202 | 687.9 | - | - | 0 | - |
| 11 | y | 1.551E+04 | 688.4 | 0.003983 | 5.786 | +1 | 5 |
| - | - | 5272 | 689.4 | - | - | 0 | - |
| - | - | 1465 | 693.3 | - | - | 0 | - |
| - | - | 3577 | 695.4 | - | - | 0 | - |
| - | - | 1218 | 696.3 | - | - | 0 | - |
| - | - | 4229 | 699.9 | - | - | 0 | - |
| 12 | c | 6234 | 700.4 | 0.0002633 | 0.3759 | +2 | 12 |
| 12 | c | 3017 | 700.8 | 0.002724 | 3.887 | +2 | 12 |
| - | - | 1779 | 701.4 | - | - | 0 | - |
| - | - | 2240 | 701.8 | - | - | 0 | - |
| 5 | z | 2916 | 702.3 | 0.01357 | 19.32 | +2 | 11 |
| - | - | 1908 | 703.3 | - | - | 0 | - |
| - | - | 2.412E+04 | 707.3 | - | - | 0 | - |
| - | - | 1.463E+04 | 707.8 | - | - | 0 | - |
| - | - | 5000 | 708.3 | - | - | 0 | - |
| - | - | 1.993E+04 | 708.9 | - | - | 0 | - |
| 12 | c | 1.808E+05 | 709.4 | 0.002877 | 4.056 | +2 | 12 |
| - | - | 1.274E+05 | 709.9 | - | - | 0 | - |
| 5 | y | 5.659E+04 | 710.3 | 0.01294 | 18.21 | +2 | 11 |
| - | - | 2.333E+04 | 710.9 | - | - | 0 | - |
| 5 | z | 8204 | 711.3 | 0.009083 | 12.77 | +2 | 11 |
| - | - | 9810 | 711.8 | - | - | 0 | - |
| - | - | 8228 | 712.3 | - | - | 0 | - |
| - | - | 4556 | 712.8 | - | - | 0 | - |
| - | - | 1816 | 715.4 | - | - | 0 | - |
| - | - | 1726 | 717.4 | - | - | 0 | - |
| - | - | 1612 | 718.9 | - | - | 0 | - |
| 5 | y | 1.02E+04 | 719.4 | 0.0005753 | 0.7998 | +2 | 11 |
| - | - | 7565 | 719.9 | - | - | 0 | - |
| - | - | 2840 | 720.4 | - | - | 0 | - |
| - | - | 3431 | 722.4 | - | - | 0 | - |
| - | - | 3049 | 722.9 | - | - | 0 | - |
| - | - | 1328 | 723.4 | - | - | 0 | - |
| - | - | 4381 | 729.4 | - | - | 0 | - |
| - | - | 1478 | 730.4 | - | - | 0 | - |
| - | - | 1831 | 735.9 | - | - | 0 | - |
| - | - | 1803 | 742.4 | - | - | 0 | - |
| - | - | 1469 | 743.9 | - | - | 0 | - |
| - | - | 2801 | 746.9 | - | - | 0 | - |
| 4 | z | 6839 | 750.9 | 0.004692 | 6.249 | +2 | 12 |
| 4 | z | 6327 | 751.4 | 0.006046 | 8.047 | +2 | 12 |
| - | - | 5284 | 751.9 | - | - | 0 | - |
| - | - | 1657 | 752.4 | - | - | 0 | - |
| - | - | 3770 | 754.4 | - | - | 0 | - |
| 10 | z | 2857 | 755.4 | 0.01461 | 19.35 | +1 | 6 |
| 10 | w | 1.147E+04 | 756.4 | 0.002549 | 3.37 | +1 | 6 |
| - | - | 4785 | 757.4 | - | - | 0 | - |
| 10 | w | 1.746E+04 | 758.4 | 0.00281 | 3.705 | +1 | 6 |
| 4 | y | 1405 | 758.9 | 0.00374 | 4.928 | +2 | 12 |
| - | - | 6806 | 759.4 | - | - | 0 | - |
| 4 | z | 2500 | 759.9 | 0.005336 | 7.022 | +2 | 12 |
| - | - | 1574 | 760.4 | - | - | 0 | - |
| 13 | c | 2367 | 764.4 | 0.003468 | 4.537 | +2 | 13 |
| 13 | c | 3791 | 764.9 | 0.004807 | 6.285 | +2 | 13 |
| - | - | 1777 | 765.4 | - | - | 0 | - |
| - | - | 3026 | 765.9 | - | - | 0 | - |
| - | - | 1528 | 766.4 | - | - | 0 | - |
| - | - | 1695 | 767.4 | - | - | 0 | - |
| 4 | y | 7.058E+04 | 767.9 | 0.00111 | 1.445 | +2 | 12 |
| - | - | 5.557E+04 | 768.4 | - | - | 0 | - |
| - | - | 2.252E+04 | 768.9 | - | - | 0 | - |
| - | - | 8332 | 769.4 | - | - | 0 | - |
| - | - | 3332 | 769.9 | - | - | 0 | - |
| - | - | 5829 | 771.4 | - | - | 0 | - |
| 10 | y | 1.647E+04 | 772.4 | 0.004483 | 5.803 | +1 | 6 |
| - | - | 4894 | 772.9 | - | - | 0 | - |
| 13 | c | 8.227E+04 | 773.4 | 0.004961 | 6.414 | +2 | 13 |
| - | - | 8.309E+04 | 773.9 | - | - | 0 | - |
| - | - | 2.951E+04 | 774.4 | - | - | 0 | - |
| - | - | 2.022E+04 | 774.4 | - | - | 0 | - |
| - | - | 1.828E+04 | 774.9 | - | - | 0 | - |
| - | - | 4787 | 775.4 | - | - | 0 | - |
| - | - | 7810 | 775.4 | - | - | 0 | - |
| - | - | 3374 | 779.4 | - | - | 0 | - |
| - | - | 2289 | 782.4 | - | - | 0 | - |
| - | - | 1594 | 783.4 | - | - | 0 | - |
| - | - | 1.191E+04 | 784.4 | - | - | 0 | - |
| - | - | 8719 | 784.9 | - | - | 0 | - |
| - | - | 3585 | 785.4 | - | - | 0 | - |
| - | - | 3708 | 787.4 | - | - | 0 | - |
| - | - | 2331 | 788.4 | - | - | 0 | - |
| 10 | y | 1.97E+04 | 789.5 | 0.0032 | 4.053 | +1 | 6 |
| - | - | 8250 | 790.4 | - | - | 0 | - |
| - | - | 2266 | 791.4 | - | - | 0 | - |
| - | - | 1456 | 791.9 | - | - | 0 | - |
| - | - | 2390 | 792.4 | - | - | 0 | - |
| - | - | 5378 | 792.9 | - | - | 0 | - |
| - | - | 6151 | 793.4 | - | - | 0 | - |
| - | - | 3664 | 793.9 | - | - | 0 | - |
| - | - | 2148 | 794.4 | - | - | 0 | - |
| 3 | w | 6994 | 794.9 | 0.004983 | 6.269 | +2 | 13 |
| - | - | 5618 | 795.4 | - | - | 0 | - |
| - | - | 3016 | 795.9 | - | - | 0 | - |
| - | - | 4588 | 797.4 | - | - | 0 | - |
| 7 | c | 1.993E+05 | 798.4 | 0.00117 | 1.465 | +1 | 7 |
| - | - | 8.893E+04 | 799.5 | - | - | 0 | - |
| - | - | 2.076E+04 | 800.5 | - | - | 0 | - |
| - | - | 7967 | 800.9 | - | - | 0 | - |
| - | - | 3552 | 801.4 | - | - | 0 | - |
| - | - | 5630 | 801.9 | - | - | 0 | - |
| - | - | 2965 | 802.4 | - | - | 0 | - |
| - | - | 1428 | 802.9 | - | - | 0 | - |
| - | - | 2275 | 805.4 | - | - | 0 | - |
| 3 | y | 3362 | 807.4 | 9.09E-05 | 0.1126 | +2 | 13 |
| 3 | y | 7183 | 807.9 | 0.006557 | 8.116 | +2 | 13 |
| 3 | z | 3643 | 808.4 | 0.003682 | 4.555 | +2 | 13 |
| - | - | 2771 | 808.9 | - | - | 0 | - |
| - | - | 2164 | 809.4 | - | - | 0 | - |
| - | - | 1546 | 811.5 | - | - | 0 | - |
| - | - | 8110 | 813.4 | - | - | 0 | - |
| - | - | 8129 | 813.9 | - | - | 0 | - |
| - | - | 6668 | 814.4 | - | - | 0 | - |
| - | - | 4536 | 814.9 | - | - | 0 | - |
| - | - | 3155 | 815.4 | - | - | 0 | - |
| - | - | 1846 | 815.5 | - | - | 0 | - |
| - | - | 4511 | 815.9 | - | - | 0 | - |
| 3 | y | 2.554E+05 | 816.4 | 0.001339 | 1.641 | +2 | 13 |
| - | - | 2.261E+05 | 816.9 | - | - | 0 | - |
| - | - | 1.159E+05 | 817.4 | - | - | 0 | - |
| - | - | 5.706E+04 | 817.9 | - | - | 0 | - |
| - | - | 1.592E+04 | 818.4 | - | - | 0 | - |
| - | - | 2375 | 818.9 | - | - | 0 | - |
| - | - | 2.267E+04 | 822.4 | - | - | 0 | - |
| - | - | 9.823E+04 | 822.9 | - | - | 0 | - |
| - | - | 8.479E+04 | 823.4 | - | - | 0 | - |
| - | - | 1744 | 823.5 | - | - | 0 | - |
| - | - | 5.386E+04 | 823.9 | - | - | 0 | - |
| - | - | 2.434E+04 | 824.4 | - | - | 0 | - |
| - | - | 7886 | 824.9 | - | - | 0 | - |
| - | - | 1.025E+04 | 825.5 | - | - | 0 | - |
| - | - | 3194 | 826.5 | - | - | 0 | - |
| - | - | 3303 | 828.9 | - | - | 0 | - |
| - | - | 1468 | 829.4 | - | - | 0 | - |
| - | - | 1251 | 829.9 | - | - | 0 | - |
| - | - | 2211 | 830.5 | - | - | 0 | - |
| - | - | 4765 | 830.9 | - | - | 0 | - |
| - | - | 3186 | 831.4 | - | - | 0 | - |
| - | - | 2281 | 831.9 | - | - | 0 | - |
| - | - | 9036 | 835.4 | - | - | 0 | - |
| - | - | 7800 | 835.9 | - | - | 0 | - |
| - | - | 5353 | 836.4 | - | - | 0 | - |
| - | - | 5756 | 836.9 | - | - | 0 | - |
| - | - | 6022 | 837.4 | - | - | 0 | - |
| - | - | 2994 | 837.9 | - | - | 0 | - |
| - | - | 3294 | 840.5 | - | - | 0 | - |
| 2 | w | 7.712E+04 | 843.4 | 0.001428 | 1.693 | +2 | 14 |
| - | - | 7.206E+04 | 843.9 | - | - | 0 | - |
| - | - | 4.441E+04 | 844.4 | - | - | 0 | - |
| - | - | 2.167E+04 | 844.9 | - | - | 0 | - |
| - | - | 7182 | 845.4 | - | - | 0 | - |
| - | - | 1357 | 845.9 | - | - | 0 | - |
| - | - | 1728 | 849.4 | - | - | 0 | - |
| - | - | 3290 | 850.4 | - | - | 0 | - |
| - | - | 1685 | 856 | - | - | 0 | - |
| - | - | 8913 | 856.5 | - | - | 0 | - |
| - | - | 4113 | 857 | - | - | 0 | - |
| - | - | 2.799E+04 | 857.5 | - | - | 0 | - |
| 14 | c | 1656 | 857.9 | 0.002558 | 2.981 | +2 | 14 |
| - | - | 7.346E+04 | 858.5 | - | - | 0 | - |
| - | - | 5440 | 858.9 | - | - | 0 | - |
| - | - | 3.542E+04 | 859.5 | - | - | 0 | - |
| - | - | 1.064E+04 | 860.5 | - | - | 0 | - |
| - | - | 3539 | 861.5 | - | - | 0 | - |
| 2 | y | 1547 | 863.9 | 0.007558 | 8.749 | +2 | 14 |
| 2 | z | 3.475E+04 | 864.9 | 0.0008131 | 0.94 | +2 | 14 |
| - | - | 3.847E+04 | 865.4 | - | - | 0 | - |
| - | - | 2.033E+04 | 865.9 | - | - | 0 | - |
| 14 | c | 2.202E+05 | 866.4 | 0.001063 | 1.227 | +2 | 14 |
| - | - | 2.001E+05 | 866.9 | - | - | 0 | - |
| - | - | 1.25E+05 | 867.4 | - | - | 0 | - |
| - | - | 5.104E+04 | 867.9 | - | - | 0 | - |
| - | - | 1.564E+04 | 868.4 | - | - | 0 | - |
| - | - | 3889 | 868.9 | - | - | 0 | - |
| - | - | 4058 | 870 | - | - | 0 | - |
| - | - | 3274 | 870.5 | - | - | 0 | - |
| - | - | 2596 | 871.5 | - | - | 0 | - |
| - | - | 7303 | 872 | - | - | 0 | - |
| - | - | 5671 | 872.5 | - | - | 0 | - |
| 2 | y | 5517 | 872.9 | 0.004046 | 4.635 | +2 | 14 |
| - | - | 6168 | 873.4 | - | - | 0 | - |
| - | - | 4916 | 873.9 | - | - | 0 | - |
| - | - | 3812 | 874.4 | - | - | 0 | - |
| - | - | 5555 | 878 | - | - | 0 | - |
| - | - | 6103 | 878.5 | - | - | 0 | - |
| - | - | 5496 | 878.9 | - | - | 0 | - |
| - | - | 1.142E+04 | 879.4 | - | - | 0 | - |
| - | - | 9279 | 879.9 | - | - | 0 | - |
| - | - | 7165 | 880.4 | - | - | 0 | - |
| - | - | 4007 | 880.9 | - | - | 0 | - |
| - | - | 2043 | 882.5 | - | - | 0 | - |
| - | - | 8957 | 883.5 | - | - | 0 | - |
| - | - | 8260 | 884.5 | - | - | 0 | - |
| - | - | 2617 | 885 | - | - | 0 | - |
| - | - | 3794 | 885.5 | - | - | 0 | - |
| - | - | 8152 | 886 | - | - | 0 | - |
| - | - | 9109 | 886.5 | - | - | 0 | - |
| - | - | 8785 | 886.9 | - | - | 0 | - |
| - | - | 7.264E+04 | 887.4 | - | - | 0 | - |
| - | - | 6.898E+04 | 887.9 | - | - | 0 | - |
| - | - | 4.168E+04 | 888.4 | - | - | 0 | - |
| - | - | 2.003E+04 | 888.9 | - | - | 0 | - |
| - | - | 7313 | 889.4 | - | - | 0 | - |
| - | - | 2343 | 890 | - | - | 0 | - |
| - | - | 8555 | 892 | - | - | 0 | - |
| - | - | 1.677E+04 | 892.5 | - | - | 0 | - |
| - | - | 1.791E+04 | 893 | - | - | 0 | - |
| - | - | 2.06E+04 | 893.5 | - | - | 0 | - |
| - | - | 1.852E+04 | 894 | - | - | 0 | - |
| - | - | 4.073E+04 | 894.5 | - | - | 0 | - |
| - | - | 3.413E+04 | 895 | - | - | 0 | - |
| - | - | 2.098E+04 | 895.5 | - | - | 0 | - |
| - | - | 2.556E+04 | 895.9 | - | - | 0 | - |
| - | - | 2.532E+04 | 896.4 | - | - | 0 | - |
| - | - | 1.632E+04 | 896.9 | - | - | 0 | - |
| - | - | 5252 | 897.4 | - | - | 0 | - |
| - | - | 4549 | 901 | - | - | 0 | - |
| - | - | 4.775E+04 | 901.5 | - | - | 0 | - |
| - | - | 4.557E+04 | 902 | - | - | 0 | - |
| 9 | z | 4.358E+04 | 902.5 | 0.01343 | 14.88 | +1 | 7 |
| - | - | 2.921E+04 | 903 | - | - | 0 | - |
| - | - | 1.439E+04 | 903.5 | - | - | 0 | - |
| - | - | 7005 | 904 | - | - | 0 | - |
| - | - | 1637 | 904.5 | - | - | 0 | - |
| - | - | 1.356E+04 | 906 | - | - | 0 | - |
| - | - | 3.608E+04 | 906.5 | - | - | 0 | - |
| - | - | 3.688E+04 | 907 | - | - | 0 | - |
| - | - | 2.558E+04 | 907.5 | - | - | 0 | - |
| - | - | 1.076E+04 | 908 | - | - | 0 | - |
| - | - | 5204 | 908.5 | - | - | 0 | - |
| - | - | 2877 | 909 | - | - | 0 | - |
| - | - | 1563 | 909.5 | - | - | 0 | - |
| - | - | 5.787E+04 | 915 | - | - | 0 | - |
| - | - | 2.291E+05 | 915.5 | - | - | 0 | - |
| - | - | 2.109E+05 | 916 | - | - | 0 | - |
| - | - | 1.342E+05 | 916.5 | - | - | 0 | - |
| - | - | 6.146E+04 | 917 | - | - | 0 | - |
| - | - | 2.162E+04 | 917.5 | - | - | 0 | - |
| - | - | 6859 | 918 | - | - | 0 | - |
| 9 | y | 1760 | 918.5 | 0.01152 | 12.55 | +1 | 7 |
| 9 | z | 2.692E+04 | 920.5 | 0.002069 | 2.247 | +1 | 7 |
| - | - | 1.813E+04 | 921.5 | - | - | 0 | - |
| - | - | 7924 | 922.5 | - | - | 0 | - |
| - | - | 1.5E+05 | 923.5 | - | - | 0 | - |
| - | - | 2.953E+05 | 924 | - | - | 0 | - |
| - | - | 2.42E+05 | 924.5 | - | - | 0 | - |
| - | - | 1.262E+05 | 925 | - | - | 0 | - |
| - | - | 5.607E+04 | 925.5 | - | - | 0 | - |
| - | - | 1.626E+04 | 926 | - | - | 0 | - |
| - | - | 6297 | 926.5 | - | - | 0 | - |
| 8 | c | 1.622E+05 | 927.5 | 0.001118 | 1.206 | +1 | 8 |
| - | - | 8.504E+04 | 928.5 | - | - | 0 | - |
| - | - | 2.898E+04 | 929.5 | - | - | 0 | - |
| - | - | 7598 | 930.5 | - | - | 0 | - |
| 9 | y | 2.872E+04 | 936.5 | 0.001228 | 1.311 | +1 | 7 |
| - | - | 1.184E+04 | 937.5 | - | - | 0 | - |
| - | - | 4992 | 938.5 | - | - | 0 | - |
| - | - | 1724 | 953.5 | - | - | 0 | - |
| - | - | 6793 | 968.5 | - | - | 0 | - |
| - | - | 3491 | 969.5 | - | - | 0 | - |
| - | - | 1581 | 975.5 | - | - | 0 | - |
| - | - | 3981 | 978.5 | - | - | 0 | - |
| - | - | 5560 | 986.5 | - | - | 0 | - |
| - | - | 4677 | 987.5 | - | - | 0 | - |
| 8 | w | 3.168E+04 | 990.5 | 0.001772 | 1.789 | +1 | 8 |
| - | - | 1.912E+04 | 991.5 | - | - | 0 | - |
| - | - | 7784 | 992.5 | - | - | 0 | - |
| - | - | 3132 | 993.5 | - | - | 0 | - |
| - | - | 2254 | 994.4 | - | - | 0 | - |
| - | - | 1536 | 1013 | - | - | 0 | - |
| - | - | 1.673E+04 | 1031 | - | - | 0 | - |
| - | - | 1.175E+04 | 1032 | - | - | 0 | - |
| - | - | 5710 | 1033 | - | - | 0 | - |
| - | - | 1601 | 1034 | - | - | 0 | - |
| 8 | z | 1.031E+05 | 1050 | 0.001834 | 1.747 | +1 | 8 |
| - | - | 6.705E+04 | 1051 | - | - | 0 | - |
| - | - | 3.103E+04 | 1052 | - | - | 0 | - |
| - | - | 1.109E+04 | 1053 | - | - | 0 | - |
| 8 | y | 6708 | 1066 | 0.0009321 | 0.8748 | +1 | 8 |
| - | - | 4826 | 1067 | - | - | 0 | - |
| - | - | 1738 | 1074 | - | - | 0 | - |
| 9 | c | 1.349E+05 | 1075 | 0.005973 | 5.559 | +1 | 9 |
| - | - | 7.711E+04 | 1076 | - | - | 0 | - |
| - | - | 3.481E+04 | 1077 | - | - | 0 | - |
| - | - | 1E+04 | 1078 | - | - | 0 | - |
| - | - | 2376 | 1079 | - | - | 0 | - |
| - | - | 2436 | 1107 | - | - | 0 | - |
| - | - | 2241 | 1108 | - | - | 0 | - |
| - | - | 2994 | 1114 | - | - | 0 | - |
| - | - | 2494 | 1115 | - | - | 0 | - |
| - | - | 1542 | 1116 | - | - | 0 | - |
| 7 | w | 1.486E+04 | 1120 | 0.002574 | 2.3 | +1 | 9 |
| - | - | 7809 | 1121 | - | - | 0 | - |
| - | - | 3672 | 1122 | - | - | 0 | - |
| - | - | 3100 | 1122 | - | - | 0 | - |
| - | - | 5236 | 1132 | - | - | 0 | - |
| - | - | 4928 | 1133 | - | - | 0 | - |
| - | - | 4084 | 1134 | - | - | 0 | - |
| - | - | 4068 | 1135 | - | - | 0 | - |
| - | - | 1797 | 1137 | - | - | 0 | - |
| 10 | c | 1.202E+05 | 1176 | 0.005536 | 4.709 | +1 | 10 |
| 7 | y | 8.211E+04 | 1177 | 0.01993 | 16.94 | +1 | 9 |
| - | - | 3.258E+04 | 1178 | - | - | 0 | - |
| 7 | z | 8.705E+04 | 1179 | 0.003064 | 2.6 | +1 | 9 |
| - | - | 5.494E+04 | 1180 | - | - | 0 | - |
| - | - | 2.197E+04 | 1181 | - | - | 0 | - |
| - | - | 7499 | 1182 | - | - | 0 | - |
| - | - | 2423 | 1183 | - | - | 0 | - |
| - | - | 2271 | 1194 | - | - | 0 | - |
| 7 | y | 9068 | 1195 | 0.0008895 | 0.7446 | +1 | 9 |
| - | - | 4553 | 1196 | - | - | 0 | - |
| - | - | 2232 | 1197 | - | - | 0 | - |
| - | - | 3418 | 1221 | - | - | 0 | - |
| - | - | 4528 | 1222 | - | - | 0 | - |
| - | - | 1884 | 1223 | - | - | 0 | - |
| - | - | 1707 | 1234 | - | - | 0 | - |
| - | - | 2262 | 1235 | - | - | 0 | - |
| - | - | 1617 | 1243 | - | - | 0 | - |
| - | - | 7313 | 1260 | - | - | 0 | - |
| - | - | 6066 | 1261 | - | - | 0 | - |
| - | - | 1946 | 1263 | - | - | 0 | - |
| - | - | 2440 | 1276 | - | - | 0 | - |
| - | - | 1961 | 1277 | - | - | 0 | - |
| - | - | 2575 | 1285 | - | - | 0 | - |
| 11 | c | 2983 | 1286 | 0.007207 | 5.606 | +1 | 11 |
| 11 | c | 2989 | 1287 | 0.009265 | 7.201 | +1 | 11 |
| - | - | 1726 | 1290 | - | - | 0 | - |
| - | - | 1.346E+04 | 1303 | - | - | 0 | - |
| 11 | c | 2.988E+04 | 1304 | 0.002247 | 1.724 | +1 | 11 |
| - | - | 2.016E+04 | 1305 | - | - | 0 | - |
| - | - | 9008 | 1306 | - | - | 0 | - |
| - | - | 2203 | 1307 | - | - | 0 | - |
| - | - | 1912 | 1308 | - | - | 0 | - |
| - | - | 1941 | 1309 | - | - | 0 | - |
| - | - | 1757 | 1322 | - | - | 0 | - |
| 6 | z | 5.22E+04 | 1335 | 0.0007079 | 0.5304 | +1 | 10 |
| - | - | 7.371E+04 | 1336 | - | - | 0 | - |
| - | - | 2301 | 1336 | - | - | 0 | - |
| - | - | 4.612E+04 | 1337 | - | - | 0 | - |
| - | - | 1.955E+04 | 1338 | - | - | 0 | - |
| - | - | 6244 | 1339 | - | - | 0 | - |
| - | - | 1768 | 1340 | - | - | 0 | - |
| - | - | 1.213E+04 | 1350 | - | - | 0 | - |
| 6 | y | 2.287E+04 | 1351 | 0.002635 | 1.951 | +1 | 10 |
| - | - | 1.348E+04 | 1352 | - | - | 0 | - |
| - | - | 6383 | 1353 | - | - | 0 | - |
| - | - | 3336 | 1354 | - | - | 0 | - |
| - | - | 1638 | 1361 | - | - | 0 | - |
| - | - | 3831 | 1374 | - | - | 0 | - |
| - | - | 5873 | 1375 | - | - | 0 | - |
| - | - | 3666 | 1376 | - | - | 0 | - |
| - | - | 3970 | 1377 | - | - | 0 | - |
| - | - | 1792 | 1402 | - | - | 0 | - |
| - | - | 3905 | 1417 | - | - | 0 | - |
| 12 | c | 6.082E+04 | 1418 | 0.003632 | 2.562 | +1 | 12 |
| - | - | 4.659E+04 | 1419 | - | - | 0 | - |
| 5 | y | 2.24E+04 | 1420 | 0.024 | 16.9 | +1 | 11 |
| - | - | 1.052E+04 | 1421 | - | - | 0 | - |
| 5 | z | 5537 | 1422 | 0.01116 | 7.85 | +1 | 11 |
| - | - | 2.766E+04 | 1423 | - | - | 0 | - |
| - | - | 2.314E+04 | 1424 | - | - | 0 | - |
| - | - | 1.055E+04 | 1425 | - | - | 0 | - |
| - | - | 4571 | 1426 | - | - | 0 | - |
| 5 | y | 6717 | 1438 | 0.00317 | 2.205 | +1 | 11 |
| - | - | 4603 | 1439 | - | - | 0 | - |
| - | - | 2157 | 1440 | - | - | 0 | - |
| - | - | 3257 | 1503 | - | - | 0 | - |
| - | - | 1962 | 1504 | - | - | 0 | - |
| - | - | 1492 | 1528 | - | - | 0 | - |
| 13 | c | 2102 | 1529 | 0.02812 | 18.4 | +1 | 13 |
| - | - | 2297 | 1530 | - | - | 0 | - |
| - | - | 1748 | 1531 | - | - | 0 | - |
| 4 | y | 1.661E+04 | 1535 | 0.0001473 | 0.09596 | +1 | 12 |
| - | - | 1.276E+04 | 1536 | - | - | 0 | - |
| - | - | 7541 | 1537 | - | - | 0 | - |
| - | - | 3699 | 1538 | - | - | 0 | - |
| 13 | c | 3.547E+04 | 1546 | 0.002915 | 1.886 | +1 | 13 |
| - | - | 3.275E+04 | 1547 | - | - | 0 | - |
| - | - | 1.763E+04 | 1548 | - | - | 0 | - |
| - | - | 8745 | 1549 | - | - | 0 | - |
| - | - | 2495 | 1550 | - | - | 0 | - |
| - | - | 1666 | 1584 | - | - | 0 | - |
| - | - | 1439 | 1601 | - | - | 0 | - |
| - | - | 3570 | 1602 | - | - | 0 | - |
| - | - | 4767 | 1603 | - | - | 0 | - |
| - | - | 1436 | 1604 | - | - | 0 | - |
| 3 | y | 6827 | 1615 | 0.02125 | 13.16 | +1 | 13 |
| 3 | z | 6126 | 1616 | 0.0194 | 12.01 | +1 | 13 |
| - | - | 3674 | 1617 | - | - | 0 | - |
| - | - | 1916 | 1618 | - | - | 0 | - |
| - | - | 1850 | 1619 | - | - | 0 | - |
| - | - | 1717 | 1628 | - | - | 0 | - |
| - | - | 4165 | 1629 | - | - | 0 | - |
| - | - | 3606 | 1630 | - | - | 0 | - |
| - | - | 1899 | 1631 | - | - | 0 | - |
| 3 | y | 2631 | 1632 | 0.01508 | 9.243 | +1 | 13 |
| - | - | 4196 | 1633 | - | - | 0 | - |
| - | - | 3274 | 1634 | - | - | 0 | - |
| - | - | 1641 | 1635 | - | - | 0 | - |
| - | - | 3175 | 1644 | - | - | 0 | - |
| - | - | 2.189E+04 | 1645 | - | - | 0 | - |
| - | - | 3.626E+04 | 1646 | - | - | 0 | - |
| - | - | 2.697E+04 | 1647 | - | - | 0 | - |
| - | - | 1.088E+04 | 1648 | - | - | 0 | - |
| - | - | 4558 | 1649 | - | - | 0 | - |
| - | - | 4821 | 1671 | - | - | 0 | - |
| - | - | 5600 | 1672 | - | - | 0 | - |
| - | - | 3270 | 1673 | - | - | 0 | - |
| - | - | 2341 | 1674 | - | - | 0 | - |
| - | - | 1806 | 1685 | - | - | 0 | - |
| - | - | 3740 | 1686 | - | - | 0 | - |
| - | - | 1529 | 1687 | - | - | 0 | - |
| - | - | 6448 | 1688 | - | - | 0 | - |
| - | - | 1.161E+04 | 1689 | - | - | 0 | - |
| - | - | 7372 | 1690 | - | - | 0 | - |
| - | - | 4190 | 1691 | - | - | 0 | - |
| - | - | 1947 | 1692 | - | - | 0 | - |
| - | - | 1483 | 1693 | - | - | 0 | - |
| - | - | 1490 | 1702 | - | - | 0 | - |
| - | - | 6363 | 1716 | - | - | 0 | - |
| - | - | 7762 | 1717 | - | - | 0 | - |
| - | - | 4427 | 1718 | - | - | 0 | - |
| - | - | 2789 | 1719 | - | - | 0 | - |
| - | - | 1690 | 1720 | - | - | 0 | - |
| 2 | z | 8155 | 1729 | 0.00143 | 0.827 | +1 | 14 |
| - | - | 5.133E+04 | 1730 | - | - | 0 | - |
| - | - | 5.314E+04 | 1731 | - | - | 0 | - |
| 14 | c | 4.937E+04 | 1732 | 0.005374 | 3.103 | +1 | 14 |
| - | - | 6.689E+04 | 1733 | - | - | 0 | - |
| - | - | 4.877E+04 | 1734 | - | - | 0 | - |
| - | - | 2.312E+04 | 1735 | - | - | 0 | - |
| - | - | 8072 | 1736 | - | - | 0 | - |
| - | - | 2731 | 1737 | - | - | 0 | - |
| - | - | 2921 | 1770 | - | - | 0 | - |
| - | - | 4866 | 1771 | - | - | 0 | - |
| - | - | 4161 | 1772 | - | - | 0 | - |
| - | - | 2787 | 1773 | - | - | 0 | - |
| - | - | 1944 | 1774 | - | - | 0 | - |
| - | - | 3080 | 1775 | - | - | 0 | - |
| - | - | 1952 | 1776 | - | - | 0 | - |
| - | - | 4785 | 1784 | - | - | 0 | - |
| - | - | 7002 | 1785 | - | - | 0 | - |
| - | - | 9831 | 1786 | - | - | 0 | - |
| - | - | 8758 | 1787 | - | - | 0 | - |
| - | - | 1.055E+04 | 1788 | - | - | 0 | - |
| - | - | 2.16E+04 | 1789 | - | - | 0 | - |
| - | - | 1.518E+04 | 1790 | - | - | 0 | - |
| - | - | 8750 | 1791 | - | - | 0 | - |
| - | - | 2961 | 1792 | - | - | 0 | - |
| - | - | 2411 | 1793 | - | - | 0 | - |
| - | - | 9651 | 1802 | - | - | 0 | - |
| - | - | 3.629E+04 | 1803 | - | - | 0 | - |
| - | - | 3.494E+04 | 1804 | - | - | 0 | - |
| - | - | 2.095E+04 | 1805 | - | - | 0 | - |
| - | - | 8782 | 1806 | - | - | 0 | - |
| - | - | 3622 | 1807 | - | - | 0 | - |
| - | - | 1696 | 1812 | - | - | 0 | - |
| - | - | 5301 | 1813 | - | - | 0 | - |
| - | - | 8396 | 1814 | - | - | 0 | - |
| - | - | 9749 | 1815 | - | - | 0 | - |
| - | - | 6634 | 1816 | - | - | 0 | - |
| - | - | 3387 | 1817 | - | - | 0 | - |
| - | - | 5296 | 1819 | - | - | 0 | - |
| - | - | 2.483E+04 | 1820 | - | - | 0 | - |
| - | - | 2.63E+04 | 1821 | - | - | 0 | - |
| - | - | 1.271E+04 | 1822 | - | - | 0 | - |
| - | - | 4401 | 1823 | - | - | 0 | - |
| - | - | 2828 | 1829 | - | - | 0 | - |
| - | - | 3.888E+04 | 1830 | - | - | 0 | - |
| - | - | 1.143E+05 | 1831 | - | - | 0 | - |
| - | - | 1.095E+05 | 1832 | - | - | 0 | - |
| - | - | 6.6E+04 | 1833 | - | - | 0 | - |
| - | - | 2.738E+04 | 1834 | - | - | 0 | - |
| - | - | 9376 | 1835 | - | - | 0 | - |
| - | - | 3139 | 1836 | - | - | 0 | - |
| - | - | 1.175E+04 | 1846 | - | - | 0 | - |
| - | - | 6.804E+04 | 1847 | - | - | 0 | - |
| - | - | 3.194E+05 | 1848 | - | - | 0 | - |
| - | - | 3.006E+05 | 1849 | - | - | 0 | - |
| - | - | 1.759E+05 | 1850 | - | - | 0 | - |
| - | - | 8.082E+04 | 1851 | - | - | 0 | - |
| - | - | 2.173E+04 | 1852 | - | - | 0 | - |
| - | - | 5150 | 1853 | - | - | 0 | - |
| - | - | 1422 | 3084 | - | - | 0 | - |

m/z Charge Intensity FragmentType MassShift Position
132.10226440429688 0 4369.8325 y 14
143.11827087402344 0 1465.0345
148.94638061523438 0 1017.7535
149.38064575195312 0 802.95483
157.2446746826172 0 971.17194
161.23483276367188 0 904.03094
173.440185546875 0 5636.8223
185.12908935546875 0 1464.8668
187.1444854736328 0 115095.23
188.14793395996094 0 10682.358
195.35203552246094 0 865.3015
212.11509704589844 0 1259.2618
212.13973999023438 0 6610.7554
215.13937377929688 0 76700.65
216.14244079589844 0 7256.11
219.13438415527344 0 11384.63
238.15528869628906 0 1539.3298
256.1661682128906 0 2353.8547
291.7546081542969 0 1034.2533
299.1719970703125 0 17639.37
300.1756286621094 0 2385.5496
312.19256591796875 0 3123.7512
313.20013427734375 0 1220.7051
326.1593933105469 0 4808.4077
327.1664733886719 0 5136.977
344.19342041015625 0 2481.6758
382.2579345703125 0 1691.1437
383.2626953125 0 1069.6239
396.18768310546875 0 2074.4963
402.6925964355469 0 1383.739
407.2536315917969 0 5272.1865
412.23388671875 0 1795.36
414.198974609375 0 1840.5754
425.2320861816406 0 1679.8508
425.26446533203125 0 5804.567
426.2701721191406 0 4228.4663 c 3
427.2675476074219 0 3091.3784
428.2755126953125 0 1715.3004
430.2432556152344 0 3150.2866 z 12
431.2477111816406 0 1351.064
444.1805725097656 0 1003.91943
446.2618408203125 0 8539.317 y 12
447.2648620605469 0 2505.965
454.1549987792969 0 960.7168
462.9015197753906 0 3023.0603
463.563720703125 0 1190.7454
466.2236633300781 0 3145.6917
470.2978210449219 0 9638.615
471.30059814453125 0 2322.378
471.8988342285156 0 8543.248
472.2333984375 0 7482.2905
472.2914123535156 0 1647.8475
472.5659484863281 0 2326.1519
491.57537841796875 0 2187.3079
491.91180419921875 0 1785.8378
494.28619384765625 0 3315.6309
494.9047546386719 0 1541.0121
495.5693359375 0 1788.1072
495.906005859375 0 1327.8229
500.9096984863281 0 5920.134 z Water loss 3
501.2444763183594 0 3471.4565 z Ammonia loss 3
501.57861328125 0 2688.0898
501.9116516113281 0 2269.7737
504.6270751953125 0 1211.8723
507.244140625 0 1626.3314
512.25732421875 0 1718.2355 y 3
512.2973022460938 0 3100.836
513.303955078125 0 66299.69 c 4
514.306640625 0 19758.016
515.30908203125 0 4286.9814
520.2332763671875 0 1295.8278
523.27490234375 0 7172.794
523.6090087890625 0 8673.411
523.94140625 0 2510.4377
528.3147583007812 0 1653.9259
529.3207397460938 0 2019.2365
529.9329223632812 0 2243.198
536.7604370117188 0 2154.4124
538.6044921875 0 6891.9546 y Water loss 2
538.9376831054688 0 5539.346 y Ammonia loss 2
539.2708129882812 0 5316.686 z 2
539.6006469726562 0 2706.4285
539.9331665039062 0 2073.1938
544.2850952148438 0 7892.617 z 11
544.6077270507812 0 407830.62 y 2
544.9417724609375 0 372121.8
545.2761840820312 0 198514.48
545.6091918945312 0 84953.97
545.9425659179688 0 21278.709
546.2777709960938 0 5227.262
560.304443359375 0 16026.734 y 11
561.3070068359375 0 3047.876
564.2418823242188 0 1247.5055
582.6340942382812 0 2055.33
596.3148803710938 0 1990.9078
607.3826904296875 0 1440.3207
614.3529663085938 0 2455.4377
615.3477783203125 0 4588.457
616.3263549804688 0 2195.8215
617.3125 0 1177.5742
625.389404296875 0 2362.6306
626.3995971679688 0 4271.9346
627.40234375 0 1916.9777
628.3673706054688 0 23681.807
629.3242797851562 0 7513.831
629.3729248046875 0 7717.7275
630.3280029296875 0 4556.416
642.8295288085938 0 4769.62
643.3270874023438 0 3843.5889 c Water loss 10
643.8356323242188 0 1383.6143
644.3257446289062 0 1210.6718
651.8350219726562 0 12165.1455
652.338134765625 0 51418.363 c 10
652.839599609375 0 37378.613
653.3401489257812 0 13133.018
653.8399658203125 0 4931.6235
657.8107299804688 0 2110.8708
658.3076171875 0 2191.329
658.8147583007812 0 3360.7812 z Water loss 5
667.827880859375 0 2705.0642 z 5
668.33251953125 0 4870.006
668.8326416015625 0 1942.0221
669.4053344726562 0 135809.42 c 5
670.3477172851562 0 1403.6079
670.4080200195312 0 49030.6
671.4114990234375 0 11699.511
672.3814086914062 0 49536.09 z 10
673.3828735351562 0 21183.662
674.3853149414062 0 4612.1924
675.3355102539062 0 6231.5015
675.8397827148438 0 7490.2095 y 5
676.3408203125 0 3935.129
676.838134765625 0 1962.1407
680.8365478515625 0 3586.5317
681.3404541015625 0 1779.3579
686.849609375 0 1454.5347
687.3526611328125 0 3539.2954
687.8511352539062 0 3202.32
688.3993530273438 0 15514.386 y 10
689.401123046875 0 5271.8975
693.3452758789062 0 1464.648
695.4208984375 0 3577.3218
696.32958984375 0 1218.3774
699.8511962890625 0 4229.3384
700.350830078125 0 6234.1235 c Water loss 11
700.8458251953125 0 3017.3877 c Ammonia loss 11
701.3507080078125 0 1779.1221
701.8413696289062 0 2239.596
702.3524780273438 0 2915.5193 z Water loss 4
703.34326171875 0 1907.5275
707.3440551757812 0 24118.416
707.8453979492188 0 14629.679
708.3482055664062 0 4999.758
708.8553466796875 0 19934.533
709.3592529296875 0 180832.86 c 11
709.8607788085938 0 127404.47
710.3612060546875 0 56593.56 y Water loss 4
710.8612670898438 0 23325.809
711.353271484375 0 8203.555 z 4
711.8496704101562 0 9809.626
712.3486938476562 0 8227.618
712.8497924804688 0 4555.866
715.40234375 0 1815.5645
717.3629760742188 0 1725.6857
718.8555908203125 0 1611.6664
719.3541259765625 0 10198.792 y 4
719.854248046875 0 7565.1377
720.3609619140625 0 2839.6309
722.3653564453125 0 3431.4268
722.868896484375 0 3048.5337
723.364501953125 0 1327.6667
729.4158325195312 0 4381.459
730.4263305664062 0 1478.3734
735.8884887695312 0 1831.481
742.3615112304688 0 1802.5621
743.8861083984375 0 1469.3226
746.8634033203125 0 2800.7925
750.860595703125 0 6839.0396 z Water loss 3
751.3633422851562 0 6326.849 z Ammonia loss 3
751.8682250976562 0 5284.4253
752.3704223632812 0 1656.516
754.4315795898438 0 3770.3289
755.4363403320312 0 2857.0935 z Water loss 9
756.427001953125 0 11473.3955 w 9
757.430419921875 0 4784.9497
758.406005859375 0 17461.488 w 9
758.8709106445312 0 1405.316 y Water loss 3
759.4085083007812 0 6806.4087
759.865234375 0 2500.079 z 3
760.3822631835938 0 1573.6094
764.3838500976562 0 2366.6135 c Water loss 12
764.877197265625 0 3790.983 c Ammonia loss 12
765.3920288085938 0 1776.5535
765.8972778320312 0 3025.5103
766.3998413085938 0 1527.7909
767.3792114257812 0 1695.0276
767.8810424804688 0 70579.484 y 3
768.382080078125 0 55565.04
768.8826293945312 0 22518.398
769.38330078125 0 8332.022
769.8880004882812 0 3331.934
771.413330078125 0 5828.821
772.4199829101562 0 16469.93 y Ammonia loss 9
772.8871459960938 0 4894.4707
773.390625 0 82265.58 c 12
773.8903198242188 0 83085.18
774.3898315429688 0 29505.234
774.43115234375 0 20216.736
774.8912963867188 0 18279.447
775.3975830078125 0 4787.358
775.4366455078125 0 7809.9775
779.4263916015625 0 3373.9192
782.378173828125 0 2288.973
783.3826293945312 0 1593.9554
784.4091186523438 0 11914.629
784.9099731445312 0 8718.585
785.4140014648438 0 3584.8928
787.4453125 0 3708.4785
788.446044921875 0 2331.1953
789.4478149414062 0 19704.035 y 9
790.4483032226562 0 8250.173
791.447265625 0 2266.1194
791.9195556640625 0 1456.2334
792.4058837890625 0 2389.9624
792.9111328125 0 5377.6714
793.4095458984375 0 6150.5073
793.9099731445312 0 3664.2122
794.407958984375 0 2148.3948
794.8901977539062 0 6994.392 w 2
795.3895874023438 0 5617.9575
795.89013671875 0 3016.3118
797.431640625 0 4587.532
798.447998046875 0 199326.38 c 6
799.4505615234375 0 88933.61
800.4531860351562 0 20764.24
800.9130859375 0 7967.382
801.419677734375 0 3551.8013
801.9190063476562 0 5629.9604
802.4141845703125 0 2965.1792
802.92626953125 0 1428.3904
805.3993530273438 0 2275.0007
807.401123046875 0 3361.9438 y Water loss 2
807.8995971679688 0 7182.8193 y Ammonia loss 2
808.400634765625 0 3643.0562 z 2
808.9019165039062 0 2771.4685
809.4025268554688 0 2164.1953
811.4569091796875 0 1545.5216
813.4195556640625 0 8110.0977
813.9124145507812 0 8128.5103
814.4144287109375 0 6667.694
814.9104614257812 0 4535.928
815.4078979492188 0 3154.9214
815.480224609375 0 1846.2806
815.9107055664062 0 4511.103
816.4076538085938 0 255368.98 y 2
816.90869140625 0 226138.88
817.4096069335938 0 115930.375
817.9097290039062 0 57064.18
818.4111328125 0 15916.776
818.9102783203125 0 2374.8691
822.4188842773438 0 22670.992
822.9227294921875 0 98230.47
823.4236450195312 0 84793.17
823.5162353515625 0 1743.6661
823.925048828125 0 53858.008
824.4385375976562 0 24342.963
824.9278564453125 0 7886.1553
825.4618530273438 0 10254.856
826.468505859375 0 3194.3281
828.9284057617188 0 3303.0352
829.43701171875 0 1467.8018
829.9345703125 0 1251.4346
830.4592895507812 0 2211.3408
830.91015625 0 4764.641
831.4097900390625 0 3186.0127
831.9135131835938 0 2280.745
835.42919921875 0 9036.5
835.9297485351562 0 7800.006
836.423828125 0 5353.1455
836.9171142578125 0 5756.36
837.411376953125 0 6021.759
837.9161987304688 0 2994.3064
840.4698486328125 0 3293.8945
843.4130249023438 0 77123.15 w 1
843.9136352539062 0 72061.08
844.4154052734375 0 44409.9
844.917724609375 0 21670.504
845.4234008789062 0 7182.4634
845.9175415039062 0 1356.8398
849.4208374023438 0 1727.9917
850.4229736328125 0 3289.7563
855.96435546875 0 1684.8971
856.461669921875 0 8912.796
856.9608764648438 0 4112.749
857.4729614257812 0 27991.06
857.9274291992188 0 1656.2573 c Ammonia loss 13
858.4804077148438 0 73457.79
858.9337768554688 0 5439.559
859.4817504882812 0 35417.73
860.4859619140625 0 10640.992
861.4883422851562 0 3538.7832
863.9506225585938 0 1547.0852 y Water loss 1
864.9381713867188 0 34754.91 z 1
865.4415283203125 0 38467.016
865.9423217773438 0 20333.082
866.439208984375 0 220157.86 c 13
866.9403686523438 0 200099.83
867.4407348632812 0 124977.08
867.9415283203125 0 51038.7
868.4425048828125 0 15636.835
868.9480590820312 0 3888.7642
869.9688110351562 0 4058.3262
870.4724731445312 0 3274.4138
871.4617919921875 0 2595.5552
871.9561767578125 0 7303.4854
872.4579467773438 0 5671.362
872.952392578125 0 5516.9 y 1
873.4415893554688 0 6168.38
873.9402465820312 0 4916.415
874.4371948242188 0 3811.937
877.9580688476562 0 5555.497
878.4533081054688 0 6102.509
878.9299926757812 0 5496.336
879.4448852539062 0 11421.541
879.9488525390625 0 9279.012
880.4453125 0 7164.701
880.9485473632812 0 4007.2498
882.4652099609375 0 2043.2567
883.4757080078125 0 8957.178
884.4747924804688 0 8260.307
884.9671020507812 0 2616.8323
885.4668579101562 0 3793.6626
885.9583129882812 0 8152.4463
886.454345703125 0 9108.815
886.9471435546875 0 8784.647
887.4346313476562 0 72639.51
887.9386596679688 0 68976.85
888.4415283203125 0 41681.137
888.9454345703125 0 20030.586
889.448974609375 0 7313.231
889.9524536132812 0 2342.904
891.9746704101562 0 8554.797
892.4700927734375 0 16765.656
892.9653930664062 0 17906.662
893.4619750976562 0 20597.484
893.9544067382812 0 18523.059
894.4566650390625 0 40731.97
894.959228515625 0 34131.36
895.46044921875 0 20975.395
895.9494018554688 0 25557.623
896.448486328125 0 25317.328
896.9482421875 0 16316.019
897.4474487304688 0 5252.1885
900.9749755859375 0 4548.5674
901.4673461914062 0 47750.406
901.968017578125 0 45568.56
902.4656372070312 0 43581.523 z Water loss 8
902.9653930664062 0 29212.746
903.4625244140625 0 14391.053
903.9638061523438 0 7004.544
904.454833984375 0 1637.0135
905.9667358398438 0 13561.702
906.4609375 0 36076.19
906.9609375 0 36875.844
907.4591674804688 0 25581.373
907.9598388671875 0 10762.15
908.4617919921875 0 5203.9297
908.9658813476562 0 2877.3445
909.45751953125 0 1562.824
914.9715576171875 0 57869.45
915.4681396484375 0 229086.9
915.9690551757812 0 210897.34
916.4689331054688 0 134230.06
916.9696655273438 0 61459.543
917.4700317382812 0 21616.96
917.9705810546875 0 6858.803
918.4594116210938 0 1759.806 y Water loss 8
920.46484375 0 26920.375 z 8
921.4671020507812 0 18129.625
922.4691162109375 0 7924.114
923.473388671875 0 150029.64
923.9758911132812 0 295254.22
924.4768676757812 0 242039.56
924.9779663085938 0 126249.74
925.4786376953125 0 56071.56
925.9797973632812 0 16260.709
926.4796752929688 0 6297.0654
927.4905395507812 0 162161.3 c 7
928.4933471679688 0 85038.51
929.4959106445312 0 28975.54
930.4969482421875 0 7597.6914
936.4827270507812 0 28723.25 y 8
937.4848022460938 0 11836.62
938.4868774414062 0 4992.0205
953.5101928710938 0 1723.579
968.529541015625 0 6793.2603
969.53369140625 0 3490.799
975.452880859375 0 1580.8049
978.4702758789062 0 3981.4404
986.5136108398438 0 5560.036
987.5189819335938 0 4676.8774
990.4938354492188 0 31684.293 w 7
991.494873046875 0 19115.602
992.4954223632812 0 7783.6685
993.4725952148438 0 3132.373
994.446044921875 0 2254.0364
1012.5076904296875 0 1536.1079
1030.511962890625 0 16733.318
1031.5162353515625 0 11750.768
1032.516845703125 0 5710.281
1033.5167236328125 0 1600.8375
1049.5072021484375 0 103100.195 z 7
1050.5086669921875 0 67053.82
1051.5101318359375 0 31027.857
1052.510498046875 0 11094.901
1065.5250244140625 0 6708.17 y 7
1066.5272216796875 0 4826.0557
1073.5152587890625 0 1737.9337
1074.52587890625 0 134872.78 c 8
1075.528564453125 0 77112.15
1076.5296630859375 0 34811.17
1077.5302734375 0 10000.797
1078.537353515625 0 2375.964
1106.5301513671875 0 2436.2014
1107.514404296875 0 2240.916
1113.5540771484375 0 2994.2993
1114.5560302734375 0 2493.5916
1115.544677734375 0 1541.9303
1119.5372314453125 0 14862.695 w 6
1120.53857421875 0 7809.1484
1121.5382080078125 0 3672.037
1122.4864501953125 0 3099.7437
1131.5587158203125 0 5236.133
1132.5621337890625 0 4927.9634
1133.5584716796875 0 4084.1892
1134.552978515625 0 4067.7307
1136.5565185546875 0 1797.3851
1175.5731201171875 0 120215.99 c 9
1176.5760498046875 0 82109.31 y Water loss 6
1177.5771484375 0 32575.977
1178.551025390625 0 87046.35 z 6
1179.55126953125 0 54941.93
1180.5509033203125 0 21973.637
1181.5518798828125 0 7499.1816
1182.54638671875 0 2422.5144
1193.5565185546875 0 2271.1794
1194.5657958984375 0 9068.277 y 6
1195.572265625 0 4552.841
1196.56689453125 0 2232.3284
1220.580810546875 0 3418.4805
1221.5802001953125 0 4527.64
1222.5855712890625 0 1883.6321
1233.574951171875 0 1706.8638
1234.593017578125 0 2261.7717
1242.6444091796875 0 1617.3502
1259.658447265625 0 7313.192
1260.656982421875 0 6066.103
1262.6358642578125 0 1945.8009
1275.62841796875 0 2440.4294
1276.6375732421875 0 1961.1519
1284.6500244140625 0 2574.7043
1285.644775390625 0 2983.0352 c Water loss 10
1286.645263671875 0 2988.8354 c Ammonia loss 10
1289.6336669921875 0 1726.1737
1302.65869140625 0 13464.361
1303.664794921875 0 29880.787 c 10
1304.6685791015625 0 20160.258
1305.6708984375 0 9008.09
1306.6585693359375 0 2202.757
1307.674560546875 0 1912.1869
1308.625 0 1940.675
1321.65576171875 0 1757.4514
1334.6497802734375 0 52198.844 z 5
1335.6551513671875 0 73712.25
1336.4730224609375 0 2300.6006
1336.656494140625 0 46124.164
1337.6573486328125 0 19549.928
1338.65966796875 0 6244.2935
1339.6605224609375 0 1768.2194
1349.6588134765625 0 12128.802
1350.6651611328125 0 22871.357 y 5
1351.6683349609375 0 13481.152
1352.66943359375 0 6383.3994
1353.6632080078125 0 3336.097
1360.668212890625 0 1637.5468
1373.6937255859375 0 3830.9934
1374.7020263671875 0 5873.4414
1375.705322265625 0 3665.9966
1376.705322265625 0 3970.1458
1401.7021484375 0 1791.8751
1416.7015380859375 0 3905.477
1417.7091064453125 0 60816.3 c 11
1418.711181640625 0 46586.766
1419.7132568359375 0 22395.355 y Water loss 4
1420.7119140625 0 10519.937
1421.6922607421875 0 5537.374 z 4
1422.6878662109375 0 27661.604
1423.6905517578125 0 23138.135
1424.6920166015625 0 10554.02
1425.6961669921875 0 4570.958
1437.6966552734375 0 6716.579 y 4
1438.69970703125 0 4602.584
1439.695556640625 0 2156.735
1502.7652587890625 0 3256.6716
1503.7276611328125 0 1962.0632
1527.7880859375 0 1492.0042
1528.765625 0 2102.473 c Ammonia loss 12
1529.75927734375 0 2297.3818
1530.7554931640625 0 1748.4913
1534.75244140625 0 16609.11 y 3
1535.7525634765625 0 12758.149
1536.758056640625 0 7540.8516
1537.7611083984375 0 3698.9346
1545.7669677734375 0 35465.758 c 12
1546.7706298828125 0 32751.867
1547.771728515625 0 17627.65
1548.776611328125 0 8745.438
1549.7877197265625 0 2494.6946
1583.8529052734375 0 1666.4995
1600.8074951171875 0 1439.499
1601.8260498046875 0 3570.065
1602.83056640625 0 4766.8145
1603.8280029296875 0 1436.2551
1614.800048828125 0 6826.866 y Ammonia loss 2
1615.8060302734375 0 6126.254 z 2
1616.81982421875 0 3674.2476
1617.8114013671875 0 1915.8306
1618.8172607421875 0 1849.854
1627.8101806640625 0 1716.8378
1628.8165283203125 0 4164.7686
1629.8194580078125 0 3605.9165
1630.8162841796875 0 1898.5403
1631.8204345703125 0 2630.86 y 2
1632.812744140625 0 4196.388
1633.8209228515625 0 3274.4465
1634.80908203125 0 1641.1913
1643.83154296875 0 3175.4338
1644.83642578125 0 21886.152
1645.8408203125 0 36260.355
1646.8447265625 0 26970.074
1647.843994140625 0 10877.364
1648.8431396484375 0 4558.4004
1670.849365234375 0 4820.842
1671.8502197265625 0 5599.747
1672.84521484375 0 3269.703
1673.8375244140625 0 2341.4138
1684.8621826171875 0 1806.191
1685.855224609375 0 3740.2593
1686.8485107421875 0 1528.9353
1687.8592529296875 0 6447.735
1688.862548828125 0 11611.262
1689.8641357421875 0 7372.182
1690.86962890625 0 4189.6616
1691.8634033203125 0 1947.4136
1692.900146484375 0 1482.6637
1701.900390625 0 1489.6282
1715.8597412109375 0 6362.9736
1716.8612060546875 0 7761.6025
1717.856201171875 0 4426.778
1718.880615234375 0 2788.6628
1719.8712158203125 0 1689.9243
1728.8692626953125 0 8155.106 z 1
1729.8773193359375 0 51325.156
1730.880126953125 0 53143.25
1731.8743896484375 0 49372.86 c 13
1732.8760986328125 0 66893.984
1733.8775634765625 0 48769.39
1734.877685546875 0 23124.582
1735.88037109375 0 8071.7686
1736.87451171875 0 2731.2393
1769.8900146484375 0 2920.7317
1770.8863525390625 0 4866.02
1771.8956298828125 0 4160.8413
1772.8814697265625 0 2787.3623
1773.872802734375 0 1944.4426
1774.8929443359375 0 3079.9155
1775.9144287109375 0 1951.9032
1783.9466552734375 0 4784.5435
1784.9385986328125 0 7002.0566
1785.9183349609375 0 9830.823
1786.9193115234375 0 8757.792
1787.9075927734375 0 10546.974
1788.913818359375 0 21604.846
1789.91552734375 0 15175.852
1790.919189453125 0 8750.366
1791.9163818359375 0 2961.164
1792.91943359375 0 2411.3794
1801.9315185546875 0 9651.435
1802.9334716796875 0 36290.67
1803.933349609375 0 34943.44
1804.9364013671875 0 20950.453
1805.9365234375 0 8782.267
1806.9346923828125 0 3622.4802
1811.960205078125 0 1695.7858
1812.922119140625 0 5301.4375
1813.9114990234375 0 8395.679
1814.91064453125 0 9748.529
1815.9095458984375 0 6634.3022
1816.91162109375 0 3387.2026
1818.95068359375 0 5296.129
1819.9544677734375 0 24831.086
1820.9595947265625 0 26301.11
1821.9625244140625 0 12712.472
1822.95849609375 0 4400.967
1828.93408203125 0 2828.44
1829.9312744140625 0 38879.586
1830.9296875 0 114267.1
1831.9305419921875 0 109524.164
1832.9327392578125 0 66002.57
1833.9315185546875 0 27382.883
1834.93359375 0 9375.537
1835.938232421875 0 3139.3792
1845.9365234375 0 11751.241
1846.9432373046875 0 68037.21
1847.9510498046875 0 319371.28
1848.9532470703125 0 300592.5
1849.9552001953125 0 175895.94
1850.9552001953125 0 80822.67
1851.957275390625 0 21726.514
1852.9578857421875 0 5149.5107
3083.882080078125 0 1422.3395

Spectrum Details

|  |  |
| --- | --- |
| Matched peaks? Matched peaks The total absolute number of peaks matched. Additionally in brackets the total fraction of peaks matched and the total number of peaks is shown. | 85 (13.69% of 621) |
| FDR? FDR The false discovery rate estimated for this peptide. It is calculated by matching all theoretical fragments with a non-integer shift with the raw peaks for this spectrum. This is done with 40 different shifts. The resulting percentage is the average number of annotated peaks over the number of annotated peaks with the correct spectrum. | 0.39% |
| Satellite FDR? Satellite FDR See the FDR for details on its calculation. This satellite ion specific FDR only contains the satellite ions (d/w) for I/L/J positions. | 4.76% |
| PSM Score? PSM Score The PSM Score as given by Hecklib to this annotated spectrum. It is shown with three significant figures. | 565 |

#### Spectrum 5470? Spectrum 5470 The raw spectrum of this peptide as annotated by Hecklib. The fragments are coloured according to ion type (see legend). Any peaks with a star '\*' as text can be hovered over to see the full details, first the ion type second the mass shift type. By hovering over the amino acids in the peptide or ions in the legend the corresponding peaks are highlighted. By toggling the 'Unassigned' label you can turn the background (unassigned) peaks on or off in the plot. By updating the slider in the Ion legend you can update the spectrum to only show the top X% of the peaks with labels. The top X% means any peak that is within X% of the highest intensity. By dragging in the spectrum you can zoom in to a specific part of the spectrum and use 'Zoom Out' to get back to the original zoom level. The annotation of the spectrum is based on the given sequence in the peptides file and is done with different software so inconsistencies are likely. The peaks are annotated based on the given sequence, with 20 ppm tolerance.

Copy Data

##### Spectrum 5470 (TSV)

###### Preview

```
Loading example...
```

*Click on the button to copy the data to your clipboard.*

Mz MinMz MaxIntensity Max

WidthHeightPeptide font sizePeptide stroke widthSpectrum font sizeSpectrum stroke widthCompact peptide

Ion legend

wxyz

abcd

OtherUnassignedIon^Charge^_Position_Show for top:%

TLPPSREEMTKNQKJ

01.15e+42.31e+43.46e+44.62e+4

Zoom Out

y^+1^_3_z^+3^_11_z^+3^_12_c^+1^_5_y^+3^_13_y^+1^_4_c^+2^_11_c^+1^_6_z^+1^_5_y^+1^_5_c^+2^_12_y^+2^_11_z^+2^_11_y^+2^_11_z^+2^_12_w^+1^_6_w^+1^_6_y^+2^_12_c^+2^_13_y^+1^_6_z^+1^_6_w^+2^_13_y^+1^_6_c^+1^_7_y^+2^_13_y^+2^_13_w^+2^_14_w^+1^_7_z^+2^_14_c^+2^_14_z^+1^_7_z^+1^_7_z^+1^_7_c^+1^_8_y^+1^_7_c^+1^_8_w^+1^_8_z^+1^_8_z^+1^_8_c^+1^_9_w^+1^_9_c^+1^_10_y^+1^_9_z^+1^_9_y^+1^_9_c^+1^_11_z^+1^_10_y^+1^_10_c^+1^_12_y^+1^_11_y^+1^_11_c^+1^_13_y^+1^_12_c^+1^_13_y^+1^_13_z^+1^_13_z^+1^_14_c^+1^_14_

0619123818572476

Fragment Matches Table

Show background peaks

| Position | Ion type | Intensity | mz Theoretical | mz Error (Th) | mz Error (ppm) | Charge | Series Number |
| --- | --- | --- | --- | --- | --- | --- | --- |
| - | - | 468.2 | 120.1 | - | - | 0 | - |
| - | - | 400.2 | 123.9 | - | - | 0 | - |
| - | - | 987.9 | 133.1 | - | - | 0 | - |
| - | - | 958.2 | 148.9 | - | - | 0 | - |
| - | - | 864 | 149 | - | - | 0 | - |
| - | - | 511.1 | 159.3 | - | - | 0 | - |
| - | - | 2263 | 173.5 | - | - | 0 | - |
| - | - | 758.9 | 187.1 | - | - | 0 | - |
| - | - | 1.068E+04 | 187.1 | - | - | 0 | - |
| - | - | 896.4 | 188.1 | - | - | 0 | - |
| - | - | 3812 | 203.1 | - | - | 0 | - |
| - | - | 734.3 | 204.1 | - | - | 0 | - |
| - | - | 874.5 | 204.1 | - | - | 0 | - |
| - | - | 6784 | 215.1 | - | - | 0 | - |
| - | - | 1191 | 219.1 | - | - | 0 | - |
| - | - | 3364 | 221.1 | - | - | 0 | - |
| - | - | 3288 | 221.1 | - | - | 0 | - |
| - | - | 721.5 | 222.1 | - | - | 0 | - |
| - | - | 645.5 | 222.1 | - | - | 0 | - |
| - | - | 954.7 | 223.1 | - | - | 0 | - |
| - | - | 748.6 | 225 | - | - | 0 | - |
| - | - | 611.7 | 230.1 | - | - | 0 | - |
| - | - | 6171 | 239.1 | - | - | 0 | - |
| - | - | 994.1 | 240.1 | - | - | 0 | - |
| - | - | 1128 | 241.1 | - | - | 0 | - |
| - | - | 1098 | 249.1 | - | - | 0 | - |
| - | - | 849.2 | 256.2 | - | - | 0 | - |
| - | - | 1453 | 281.1 | - | - | 0 | - |
| - | - | 504.2 | 282.1 | - | - | 0 | - |
| - | - | 813.5 | 295.1 | - | - | 0 | - |
| - | - | 2722 | 295.2 | - | - | 0 | - |
| - | - | 547.4 | 296.1 | - | - | 0 | - |
| - | - | 796 | 297.1 | - | - | 0 | - |
| - | - | 604.8 | 298.1 | - | - | 0 | - |
| - | - | 4747 | 299.1 | - | - | 0 | - |
| - | - | 2719 | 299.1 | - | - | 0 | - |
| - | - | 2926 | 299.2 | - | - | 0 | - |
| - | - | 1076 | 300.1 | - | - | 0 | - |
| - | - | 812 | 301.1 | - | - | 0 | - |
| - | - | 1883 | 317.1 | - | - | 0 | - |
| - | - | 619.8 | 327.2 | - | - | 0 | - |
| - | - | 575.8 | 331.9 | - | - | 0 | - |
| - | - | 521.4 | 335 | - | - | 0 | - |
| - | - | 3.797E+04 | 355.1 | - | - | 0 | - |
| - | - | 1.174E+04 | 356.1 | - | - | 0 | - |
| - | - | 5144 | 357.1 | - | - | 0 | - |
| - | - | 2625 | 371.1 | - | - | 0 | - |
| - | - | 670.1 | 375.2 | - | - | 0 | - |
| - | - | 1313 | 382.2 | - | - | 0 | - |
| - | - | 1138 | 391.2 | - | - | 0 | - |
| - | - | 598.8 | 409.2 | - | - | 0 | - |
| - | - | 2053 | 415 | - | - | 0 | - |
| - | - | 713.8 | 419 | - | - | 0 | - |
| - | - | 785.5 | 431.2 | - | - | 0 | - |
| - | - | 1549 | 434.2 | - | - | 0 | - |
| - | - | 2345 | 444.2 | - | - | 0 | - |
| 13 | y | 1240 | 446.3 | 0.004887 | 10.95 | +1 | 3 |
| - | - | 724.1 | 447.3 | - | - | 0 | - |
| - | - | 2778 | 452.2 | - | - | 0 | - |
| - | - | 619.1 | 461.3 | - | - | 0 | - |
| - | - | 7732 | 462.2 | - | - | 0 | - |
| - | - | 1120 | 463.2 | - | - | 0 | - |
| - | - | 978.9 | 466.6 | - | - | 0 | - |
| 5 | z | 1979 | 469.2 | 0.006153 | 13.11 | +3 | 11 |
| - | - | 1044 | 470.3 | - | - | 0 | - |
| - | - | 7294 | 480.2 | - | - | 0 | - |
| - | - | 1238 | 481.2 | - | - | 0 | - |
| - | - | 810.5 | 489.1 | - | - | 0 | - |
| - | - | 718.1 | 490.1 | - | - | 0 | - |
| - | - | 764 | 494.3 | - | - | 0 | - |
| 4 | z | 1897 | 495.9 | 0.00167 | 3.367 | +3 | 12 |
| - | - | 700.2 | 496.2 | - | - | 0 | - |
| 5 | c | 6263 | 513.3 | 3.75E-05 | 0.07305 | +1 | 5 |
| - | - | 1404 | 514.3 | - | - | 0 | - |
| 3 | y | 3.188E+04 | 539.3 | 0.001855 | 3.44 | +3 | 13 |
| - | - | 2.791E+04 | 539.6 | - | - | 0 | - |
| - | - | 1.629E+04 | 539.9 | - | - | 0 | - |
| - | - | 7871 | 540.3 | - | - | 0 | - |
| - | - | 1979 | 540.6 | - | - | 0 | - |
| - | - | 2939 | 545.3 | - | - | 0 | - |
| - | - | 710 | 546.3 | - | - | 0 | - |
| - | - | 1256 | 554.2 | - | - | 0 | - |
| - | - | 623.6 | 555.7 | - | - | 0 | - |
| 12 | y | 3762 | 560.3 | 0.00454 | 8.103 | +1 | 4 |
| - | - | 840.5 | 561.3 | - | - | 0 | - |
| - | - | 816.6 | 562.7 | - | - | 0 | - |
| - | - | 1185 | 575.3 | - | - | 0 | - |
| - | - | 1397 | 576.2 | - | - | 0 | - |
| - | - | 1096 | 576.7 | - | - | 0 | - |
| - | - | 1488 | 577.1 | - | - | 0 | - |
| - | - | 1136 | 578.1 | - | - | 0 | - |
| - | - | 1046 | 584.7 | - | - | 0 | - |
| - | - | 1011 | 585.8 | - | - | 0 | - |
| - | - | 1491 | 593.3 | - | - | 0 | - |
| - | - | 799.3 | 602.3 | - | - | 0 | - |
| - | - | 9798 | 610.2 | - | - | 0 | - |
| - | - | 666.7 | 610.3 | - | - | 0 | - |
| - | - | 6854 | 611.2 | - | - | 0 | - |
| - | - | 1.279E+04 | 611.3 | - | - | 0 | - |
| - | - | 706.3 | 611.7 | - | - | 0 | - |
| - | - | 2236 | 611.8 | - | - | 0 | - |
| - | - | 3276 | 612.2 | - | - | 0 | - |
| - | - | 2083 | 612.3 | - | - | 0 | - |
| - | - | 1060 | 626.4 | - | - | 0 | - |
| - | - | 2908 | 628.4 | - | - | 0 | - |
| - | - | 855 | 629.4 | - | - | 0 | - |
| - | - | 908 | 630.3 | - | - | 0 | - |
| 11 | c | 4453 | 644.3 | 0.0003734 | 0.5795 | +2 | 11 |
| - | - | 2508 | 644.8 | - | - | 0 | - |
| - | - | 1023 | 645.3 | - | - | 0 | - |
| - | - | 1279 | 645.8 | - | - | 0 | - |
| - | - | 867.3 | 668.3 | - | - | 0 | - |
| 6 | c | 1.323E+04 | 669.4 | 0.0009759 | 1.458 | +1 | 6 |
| - | - | 4574 | 670.4 | - | - | 0 | - |
| - | - | 1172 | 671.4 | - | - | 0 | - |
| 11 | z | 4034 | 672.4 | 0.004851 | 7.215 | +1 | 5 |
| - | - | 1123 | 673.4 | - | - | 0 | - |
| 11 | y | 2324 | 688.4 | 0.005814 | 8.446 | +1 | 5 |
| - | - | 691.5 | 689.4 | - | - | 0 | - |
| - | - | 2057 | 699.3 | - | - | 0 | - |
| - | - | 1365 | 699.8 | - | - | 0 | - |
| - | - | 1474 | 700.9 | - | - | 0 | - |
| 12 | c | 1.863E+04 | 701.4 | 0.001024 | 1.46 | +2 | 12 |
| - | - | 1.368E+04 | 701.9 | - | - | 0 | - |
| 5 | y | 6016 | 702.4 | 0.008792 | 12.52 | +2 | 11 |
| - | - | 1986 | 702.9 | - | - | 0 | - |
| 5 | z | 905.5 | 703.3 | 0.007501 | 10.66 | +2 | 11 |
| - | - | 820.4 | 704.3 | - | - | 0 | - |
| 5 | y | 1006 | 711.4 | 0.003753 | 5.276 | +2 | 11 |
| - | - | 1370 | 714.4 | - | - | 0 | - |
| - | - | 911.5 | 729.4 | - | - | 0 | - |
| - | - | 663 | 734.4 | - | - | 0 | - |
| 4 | z | 736.9 | 742.9 | 0.0136 | 18.31 | +2 | 12 |
| - | - | 985.5 | 743.9 | - | - | 0 | - |
| - | - | 618.1 | 748.4 | - | - | 0 | - |
| 10 | w | 1106 | 756.4 | 0.004014 | 5.306 | +1 | 6 |
| - | - | 762.1 | 757.4 | - | - | 0 | - |
| 10 | w | 2463 | 758.4 | 0.003847 | 5.073 | +1 | 6 |
| 4 | y | 6517 | 759.9 | 0.003036 | 3.995 | +2 | 12 |
| - | - | 4266 | 760.4 | - | - | 0 | - |
| - | - | 3697 | 760.9 | - | - | 0 | - |
| - | - | 1262 | 761.4 | - | - | 0 | - |
| 13 | c | 1.386E+04 | 765.4 | 0.001443 | 1.886 | +2 | 13 |
| - | - | 1.114E+04 | 765.9 | - | - | 0 | - |
| - | - | 5442 | 766.4 | - | - | 0 | - |
| - | - | 1431 | 766.9 | - | - | 0 | - |
| 10 | y | 745.6 | 772.4 | 0.01345 | 17.42 | +1 | 6 |
| 10 | z | 7787 | 773.4 | 0.005289 | 6.838 | +1 | 6 |
| - | - | 709.4 | 774.3 | - | - | 0 | - |
| - | - | 4913 | 774.4 | - | - | 0 | - |
| - | - | 1377 | 775.4 | - | - | 0 | - |
| - | - | 960.7 | 777.9 | - | - | 0 | - |
| 3 | w | 617 | 786.9 | 0.0005932 | 0.7538 | +2 | 13 |
| - | - | 912.9 | 787.4 | - | - | 0 | - |
| 10 | y | 2480 | 789.5 | 0.004909 | 6.218 | +1 | 6 |
| - | - | 1126 | 790.4 | - | - | 0 | - |
| - | - | 700.6 | 792.4 | - | - | 0 | - |
| - | - | 768.2 | 792.9 | - | - | 0 | - |
| - | - | 720.4 | 793.4 | - | - | 0 | - |
| 7 | c | 2.851E+04 | 798.4 | 0.0007833 | 0.9811 | +1 | 7 |
| - | - | 1.339E+04 | 799.4 | - | - | 0 | - |
| 3 | y | 646.6 | 799.9 | 0.01218 | 15.22 | +2 | 13 |
| - | - | 3308 | 800.5 | - | - | 0 | - |
| - | - | 1831 | 805.4 | - | - | 0 | - |
| - | - | 1064 | 805.9 | - | - | 0 | - |
| - | - | 753.5 | 807.4 | - | - | 0 | - |
| - | - | 774.7 | 807.9 | - | - | 0 | - |
| 3 | y | 2.542E+04 | 808.4 | 0.002989 | 3.698 | +2 | 13 |
| - | - | 2.004E+04 | 808.9 | - | - | 0 | - |
| - | - | 1.307E+04 | 809.4 | - | - | 0 | - |
| - | - | 4581 | 809.9 | - | - | 0 | - |
| - | - | 1150 | 810.4 | - | - | 0 | - |
| - | - | 798.7 | 810.9 | - | - | 0 | - |
| - | - | 804.3 | 814.4 | - | - | 0 | - |
| - | - | 1.074E+04 | 814.9 | - | - | 0 | - |
| - | - | 1.291E+04 | 815.4 | - | - | 0 | - |
| - | - | 6210 | 815.9 | - | - | 0 | - |
| - | - | 3573 | 816.4 | - | - | 0 | - |
| - | - | 1907 | 824.5 | - | - | 0 | - |
| - | - | 1002 | 825.5 | - | - | 0 | - |
| - | - | 940.5 | 827.4 | - | - | 0 | - |
| - | - | 865.5 | 827.9 | - | - | 0 | - |
| 2 | w | 9730 | 835.4 | 0.002473 | 2.961 | +2 | 14 |
| - | - | 8997 | 835.9 | - | - | 0 | - |
| - | - | 6054 | 836.4 | - | - | 0 | - |
| - | - | 2374 | 836.9 | - | - | 0 | - |
| 9 | w | 5540 | 843.5 | 0.004731 | 5.609 | +1 | 7 |
| - | - | 2472 | 844.5 | - | - | 0 | - |
| 2 | z | 3135 | 856.9 | 0.003372 | 3.935 | +2 | 14 |
| - | - | 4083 | 857.4 | - | - | 0 | - |
| - | - | 2402 | 857.9 | - | - | 0 | - |
| 14 | c | 2.627E+04 | 858.4 | 0.002716 | 3.164 | +2 | 14 |
| - | - | 2.239E+04 | 858.9 | - | - | 0 | - |
| - | - | 1.252E+04 | 859.4 | - | - | 0 | - |
| - | - | 5914 | 859.9 | - | - | 0 | - |
| - | - | 1769 | 860.4 | - | - | 0 | - |
| - | - | 1206 | 863 | - | - | 0 | - |
| - | - | 2005 | 863.5 | - | - | 0 | - |
| - | - | 738.3 | 864 | - | - | 0 | - |
| - | - | 700.2 | 864.5 | - | - | 0 | - |
| - | - | 1160 | 865.4 | - | - | 0 | - |
| - | - | 1109 | 870.4 | - | - | 0 | - |
| - | - | 652.6 | 870.9 | - | - | 0 | - |
| - | - | 1059 | 871.4 | - | - | 0 | - |
| - | - | 1927 | 872 | - | - | 0 | - |
| - | - | 1054 | 872.4 | - | - | 0 | - |
| - | - | 1794 | 879 | - | - | 0 | - |
| - | - | 1.044E+04 | 879.4 | - | - | 0 | - |
| - | - | 1.075E+04 | 879.9 | - | - | 0 | - |
| - | - | 7967 | 880.4 | - | - | 0 | - |
| - | - | 3346 | 881 | - | - | 0 | - |
| - | - | 1220 | 881.5 | - | - | 0 | - |
| - | - | 981.7 | 883.5 | - | - | 0 | - |
| - | - | 932.6 | 884.5 | - | - | 0 | - |
| - | - | 1674 | 885 | - | - | 0 | - |
| - | - | 2725 | 885.5 | - | - | 0 | - |
| - | - | 2400 | 886 | - | - | 0 | - |
| 9 | z | 6787 | 886.5 | 0.00225 | 2.538 | +1 | 7 |
| - | - | 4288 | 887 | - | - | 0 | - |
| 9 | z | 1901 | 887.4 | 0.007692 | 8.668 | +1 | 7 |
| - | - | 1953 | 887.9 | - | - | 0 | - |
| - | - | 1624 | 888.4 | - | - | 0 | - |
| - | - | 1721 | 888.9 | - | - | 0 | - |
| - | - | 3954 | 893.5 | - | - | 0 | - |
| - | - | 3941 | 894 | - | - | 0 | - |
| - | - | 3102 | 894.5 | - | - | 0 | - |
| - | - | 2498 | 895 | - | - | 0 | - |
| - | - | 793.1 | 895.5 | - | - | 0 | - |
| - | - | 1799 | 898 | - | - | 0 | - |
| - | - | 4536 | 898.5 | - | - | 0 | - |
| - | - | 3835 | 899 | - | - | 0 | - |
| - | - | 4181 | 899.5 | - | - | 0 | - |
| - | - | 1733 | 900 | - | - | 0 | - |
| 9 | z | 7053 | 904.5 | 0.00665 | 7.352 | +1 | 7 |
| - | - | 3967 | 905.5 | - | - | 0 | - |
| - | - | 1459 | 906.5 | - | - | 0 | - |
| - | - | 3840 | 907 | - | - | 0 | - |
| - | - | 2.397E+04 | 907.5 | - | - | 0 | - |
| - | - | 1.966E+04 | 908 | - | - | 0 | - |
| - | - | 1.511E+04 | 908.5 | - | - | 0 | - |
| - | - | 6094 | 909 | - | - | 0 | - |
| 8 | c | 2039 | 909.5 | 0.01298 | 14.27 | +1 | 8 |
| - | - | 1.762E+04 | 915.5 | - | - | 0 | - |
| - | - | 3.285E+04 | 916 | - | - | 0 | - |
| - | - | 2.665E+04 | 916.5 | - | - | 0 | - |
| - | - | 1.255E+04 | 917 | - | - | 0 | - |
| - | - | 6359 | 917.5 | - | - | 0 | - |
| - | - | 1762 | 918 | - | - | 0 | - |
| 9 | y | 3647 | 920.5 | 0.004866 | 5.286 | +1 | 7 |
| - | - | 1665 | 921.5 | - | - | 0 | - |
| 8 | c | 1.582E+04 | 927.5 | 0.001079 | 1.163 | +1 | 8 |
| - | - | 8459 | 928.5 | - | - | 0 | - |
| - | - | 2336 | 929.5 | - | - | 0 | - |
| - | - | 747.4 | 960.5 | - | - | 0 | - |
| - | - | 713.9 | 969.3 | - | - | 0 | - |
| - | - | 1479 | 970.3 | - | - | 0 | - |
| - | - | 882.9 | 971.3 | - | - | 0 | - |
| - | - | 1278 | 972.5 | - | - | 0 | - |
| - | - | 1437 | 973.5 | - | - | 0 | - |
| 8 | w | 3172 | 974.5 | 0.004383 | 4.498 | +1 | 8 |
| - | - | 2301 | 975.5 | - | - | 0 | - |
| - | - | 895.6 | 976.5 | - | - | 0 | - |
| - | - | 1168 | 986.5 | - | - | 0 | - |
| - | - | 1209 | 997.4 | - | - | 0 | - |
| - | - | 1695 | 998.4 | - | - | 0 | - |
| - | - | 861.4 | 1014 | - | - | 0 | - |
| - | - | 2232 | 1015 | - | - | 0 | - |
| - | - | 826.6 | 1015 | - | - | 0 | - |
| 8 | z | 843.2 | 1016 | 0.006183 | 6.088 | +1 | 8 |
| - | - | 696.9 | 1017 | - | - | 0 | - |
| 8 | z | 1.547E+04 | 1034 | 0.006213 | 6.011 | +1 | 8 |
| - | - | 1.042E+04 | 1035 | - | - | 0 | - |
| - | - | 3296 | 1036 | - | - | 0 | - |
| - | - | 871.9 | 1037 | - | - | 0 | - |
| - | - | 707.1 | 1038 | - | - | 0 | - |
| - | - | 1556 | 1039 | - | - | 0 | - |
| 9 | c | 1.384E+04 | 1059 | 0.001341 | 1.267 | +1 | 9 |
| - | - | 7987 | 1060 | - | - | 0 | - |
| - | - | 4785 | 1061 | - | - | 0 | - |
| - | - | 713.4 | 1062 | - | - | 0 | - |
| 7 | w | 1522 | 1104 | 0.009135 | 8.277 | +1 | 9 |
| - | - | 1083 | 1105 | - | - | 0 | - |
| - | - | 662.9 | 1106 | - | - | 0 | - |
| - | - | 831.6 | 1121 | - | - | 0 | - |
| - | - | 1278 | 1157 | - | - | 0 | - |
| 10 | c | 1.127E+04 | 1160 | 0.001046 | 0.9023 | +1 | 10 |
| 7 | y | 8505 | 1161 | 0.01176 | 10.13 | +1 | 9 |
| - | - | 3398 | 1162 | - | - | 0 | - |
| 7 | z | 8994 | 1163 | 0.00364 | 3.131 | +1 | 9 |
| - | - | 5702 | 1164 | - | - | 0 | - |
| - | - | 1635 | 1165 | - | - | 0 | - |
| - | - | 985.1 | 1166 | - | - | 0 | - |
| - | - | 3234 | 1168 | - | - | 0 | - |
| - | - | 1473 | 1169 | - | - | 0 | - |
| - | - | 1020 | 1170 | - | - | 0 | - |
| - | - | 1207 | 1172 | - | - | 0 | - |
| - | - | 1588 | 1174 | - | - | 0 | - |
| - | - | 827.3 | 1175 | - | - | 0 | - |
| - | - | 799.9 | 1177 | - | - | 0 | - |
| 7 | y | 1923 | 1179 | 0.002955 | 2.507 | +1 | 9 |
| - | - | 1905 | 1180 | - | - | 0 | - |
| - | - | 2626 | 1185 | - | - | 0 | - |
| - | - | 2553 | 1186 | - | - | 0 | - |
| - | - | 5358 | 1187 | - | - | 0 | - |
| - | - | 3214 | 1189 | - | - | 0 | - |
| - | - | 1238 | 1190 | - | - | 0 | - |
| - | - | 1301 | 1201 | - | - | 0 | - |
| - | - | 949.4 | 1202 | - | - | 0 | - |
| - | - | 1018 | 1203 | - | - | 0 | - |
| - | - | 819.8 | 1204 | - | - | 0 | - |
| - | - | 825.7 | 1206 | - | - | 0 | - |
| - | - | 780 | 1207 | - | - | 0 | - |
| - | - | 2169 | 1218 | - | - | 0 | - |
| - | - | 2755 | 1219 | - | - | 0 | - |
| - | - | 1927 | 1220 | - | - | 0 | - |
| - | - | 2537 | 1220 | - | - | 0 | - |
| - | - | 1692 | 1221 | - | - | 0 | - |
| - | - | 2279 | 1221 | - | - | 0 | - |
| - | - | 832.4 | 1222 | - | - | 0 | - |
| - | - | 1001 | 1223 | - | - | 0 | - |
| - | - | 1399 | 1224 | - | - | 0 | - |
| - | - | 3275 | 1224 | - | - | 0 | - |
| - | - | 1902 | 1225 | - | - | 0 | - |
| - | - | 1857 | 1225 | - | - | 0 | - |
| - | - | 1052 | 1244 | - | - | 0 | - |
| - | - | 1860 | 1245 | - | - | 0 | - |
| - | - | 1540 | 1287 | - | - | 0 | - |
| 11 | c | 4214 | 1288 | 0.001771 | 1.375 | +1 | 11 |
| - | - | 2435 | 1289 | - | - | 0 | - |
| - | - | 1019 | 1290 | - | - | 0 | - |
| 6 | z | 5432 | 1319 | 0.006851 | 5.195 | +1 | 10 |
| - | - | 8778 | 1320 | - | - | 0 | - |
| - | - | 6141 | 1321 | - | - | 0 | - |
| - | - | 3280 | 1322 | - | - | 0 | - |
| - | - | 952.6 | 1334 | - | - | 0 | - |
| 6 | y | 4670 | 1335 | 0.008607 | 6.449 | +1 | 10 |
| - | - | 2581 | 1336 | - | - | 0 | - |
| - | - | 1590 | 1337 | - | - | 0 | - |
| - | - | 755.2 | 1358 | - | - | 0 | - |
| - | - | 933.8 | 1359 | - | - | 0 | - |
| 12 | c | 7422 | 1402 | 0.002706 | 1.931 | +1 | 12 |
| - | - | 7136 | 1403 | - | - | 0 | - |
| 5 | y | 3243 | 1404 | 0.01693 | 12.06 | +1 | 11 |
| 5 | y | 1205 | 1405 | 0.02473 | 17.61 | +1 | 11 |
| - | - | 2268 | 1407 | - | - | 0 | - |
| - | - | 1818 | 1408 | - | - | 0 | - |
| - | - | 759.1 | 1486 | - | - | 0 | - |
| 13 | c | 781.8 | 1512 | 0.005921 | 3.917 | +1 | 13 |
| 4 | y | 2477 | 1519 | 0.006851 | 4.511 | +1 | 12 |
| - | - | 1997 | 1520 | - | - | 0 | - |
| - | - | 866.3 | 1521 | - | - | 0 | - |
| - | - | 729.4 | 1529 | - | - | 0 | - |
| 13 | c | 4810 | 1530 | 0.003423 | 2.237 | +1 | 13 |
| - | - | 5150 | 1531 | - | - | 0 | - |
| - | - | 2663 | 1532 | - | - | 0 | - |
| - | - | 1782 | 1533 | - | - | 0 | - |
| 3 | y | 832.4 | 1599 | 0.0006253 | 0.3911 | +1 | 13 |
| 3 | z | 1192 | 1600 | 0.002078 | 1.299 | +1 | 13 |
| - | - | 4448 | 1629 | - | - | 0 | - |
| - | - | 5831 | 1630 | - | - | 0 | - |
| - | - | 3166 | 1631 | - | - | 0 | - |
| - | - | 2328 | 1632 | - | - | 0 | - |
| - | - | 2209 | 1673 | - | - | 0 | - |
| - | - | 1122 | 1674 | - | - | 0 | - |
| - | - | 1035 | 1675 | - | - | 0 | - |
| - | - | 896.5 | 1701 | - | - | 0 | - |
| 2 | z | 1068 | 1713 | 0.008988 | 5.248 | +1 | 14 |
| - | - | 9123 | 1714 | - | - | 0 | - |
| - | - | 9946 | 1715 | - | - | 0 | - |
| 14 | c | 6485 | 1716 | 0.001696 | 0.9885 | +1 | 14 |
| - | - | 9962 | 1717 | - | - | 0 | - |
| - | - | 6755 | 1718 | - | - | 0 | - |
| - | - | 3559 | 1719 | - | - | 0 | - |
| - | - | 844.4 | 1720 | - | - | 0 | - |
| - | - | 762.2 | 1755 | - | - | 0 | - |
| - | - | 920.9 | 1756 | - | - | 0 | - |
| - | - | 1722 | 1770 | - | - | 0 | - |
| - | - | 1251 | 1771 | - | - | 0 | - |
| - | - | 2066 | 1772 | - | - | 0 | - |
| - | - | 3258 | 1773 | - | - | 0 | - |
| - | - | 2559 | 1774 | - | - | 0 | - |
| - | - | 1497 | 1775 | - | - | 0 | - |
| - | - | 1131 | 1786 | - | - | 0 | - |
| - | - | 5091 | 1787 | - | - | 0 | - |
| - | - | 3864 | 1788 | - | - | 0 | - |
| - | - | 3173 | 1789 | - | - | 0 | - |
| - | - | 813.5 | 1790 | - | - | 0 | - |
| - | - | 944.6 | 1797 | - | - | 0 | - |
| - | - | 1158 | 1798 | - | - | 0 | - |
| - | - | 1091 | 1799 | - | - | 0 | - |
| - | - | 4658 | 1804 | - | - | 0 | - |
| - | - | 3955 | 1805 | - | - | 0 | - |
| - | - | 2645 | 1806 | - | - | 0 | - |
| - | - | 4150 | 1814 | - | - | 0 | - |
| - | - | 1.965E+04 | 1815 | - | - | 0 | - |
| - | - | 1.804E+04 | 1816 | - | - | 0 | - |
| - | - | 1.02E+04 | 1817 | - | - | 0 | - |
| - | - | 4113 | 1818 | - | - | 0 | - |
| - | - | 1803 | 1819 | - | - | 0 | - |
| - | - | 2002 | 1830 | - | - | 0 | - |
| - | - | 1.189E+04 | 1831 | - | - | 0 | - |
| - | - | 4.571E+04 | 1832 | - | - | 0 | - |
| - | - | 4.186E+04 | 1833 | - | - | 0 | - |
| - | - | 2.457E+04 | 1834 | - | - | 0 | - |
| - | - | 1.085E+04 | 1835 | - | - | 0 | - |
| - | - | 3907 | 1836 | - | - | 0 | - |
| - | - | 882.3 | 1837 | - | - | 0 | - |
| - | - | 752.1 | 2451 | - | - | 0 | - |

m/z Charge Intensity FragmentType MassShift Position
120.08076477050781 0 468.2171
123.94280242919922 0 400.1626
133.06082153320312 0 987.9295
148.9470672607422 0 958.2109
149.0448455810547 0 864.04034
159.2568359375 0 511.07138
173.45144653320312 0 2263.1543
187.13551330566406 0 758.8526
187.14401245117188 0 10675.203
188.14752197265625 0 896.44684
203.08143615722656 0 3812.1567
204.08543395996094 0 734.33057
204.09754943847656 0 874.4533
215.1388702392578 0 6783.8687
219.13389587402344 0 1190.7122
221.0842742919922 0 3363.6028
221.12826538085938 0 3287.8186
222.08441162109375 0 721.49634
222.1326904296875 0 645.48444
223.08116149902344 0 954.6881
225.04263305664062 0 748.5634
230.1134033203125 0 611.7097
239.09486389160156 0 6170.719
240.0967254638672 0 994.1072
241.09164428710938 0 1128.0359
249.1229705810547 0 1097.5627
256.1653747558594 0 849.20197
281.0516662597656 0 1453.1957
282.05340576171875 0 504.21747
295.1021728515625 0 813.4982
295.16510009765625 0 2721.9644
296.10186767578125 0 547.3702
297.0997314453125 0 795.9969
298.101806640625 0 604.8215
299.0616149902344 0 4746.888
299.13543701171875 0 2718.5767
299.17120361328125 0 2925.8442
300.0613098144531 0 1076.3054
301.05914306640625 0 811.9845
317.1460266113281 0 1882.7584
327.1660461425781 0 619.8012
331.9129943847656 0 575.7777
334.9867248535156 0 521.38855
355.0696105957031 0 37965.496
356.07025146484375 0 11735.093
357.0675354003906 0 5143.901
371.1007995605469 0 2624.625
375.1639709472656 0 670.113
382.19622802734375 0 1312.5326
391.1595153808594 0 1138.3818
409.17010498046875 0 598.80615
415.0364685058594 0 2052.981
418.99285888671875 0 713.8311
431.19110107421875 0 785.49414
434.20343017578125 0 1548.9132
444.1871032714844 0 2345.3643
446.26055908203125 0 1240.266 y 12
447.26275634765625 0 724.1326
452.2137145996094 0 2777.5352
461.2709045410156 0 619.1381
462.197998046875 0 7732.235
463.20172119140625 0 1120.3958
466.5666198730469 0 978.91473
469.22906494140625 0 1979.4375 z 4
470.2958068847656 0 1043.5037
480.20892333984375 0 7294.272
481.20989990234375 0 1237.9812
489.0556945800781 0 810.50397
490.0593566894531 0 718.09717
494.2838134765625 0 764.0467
495.91229248046875 0 1897.0735 z Ammonia loss 3
496.2449035644531 0 700.16656
513.3031616210938 0 6263.4893 c 4
514.30517578125 0 1403.6724
539.2747802734375 0 31876.268 y 2
539.6087646484375 0 27912.367
539.9425659179688 0 16291.49
540.2763061523438 0 7870.6245
540.6082153320312 0 1979.0385
545.2926025390625 0 2939.2783
546.2938842773438 0 710.0499
554.2195434570312 0 1256.0153
555.7311401367188 0 623.646
560.3038330078125 0 3762.127 y 11
561.3085327148438 0 840.5339
562.7351684570312 0 816.64325
575.2823486328125 0 1184.6217
576.2280883789062 0 1397.1766
576.7326049804688 0 1096.1072
577.1259155273438 0 1488.1436
578.130126953125 0 1136.4537
584.7445678710938 0 1046.1263
585.7515869140625 0 1010.88214
593.2907104492188 0 1491.3318
602.3107299804688 0 799.31506
610.1840209960938 0 9798.25
610.3276977539062 0 666.71027
611.1843872070312 0 6853.8633
611.3031616210938 0 12790.935
611.7325439453125 0 706.2625
611.8374633789062 0 2235.6104
612.1827392578125 0 3276.4001
612.30029296875 0 2083.4504
626.397216796875 0 1059.5033
628.3665771484375 0 2907.6265
629.3699951171875 0 855.00024
630.3253784179688 0 907.97217
644.3395385742188 0 4452.766 c 10
644.8418579101562 0 2507.6672
645.3402709960938 0 1022.54175
645.8424072265625 0 1278.5895
668.33544921875 0 867.3359
669.4032592773438 0 13232.457 c 5
670.406982421875 0 4573.5225
671.4114379882812 0 1172.4172
672.3797607421875 0 4033.5447 z 10
673.3804321289062 0 1123.1069
688.3975219726562 0 2323.7935 y 10
689.4060668945312 0 691.4669
699.3448486328125 0 2056.5017
699.8468627929688 0 1365.462
700.8544921875 0 1473.617
701.3603515625 0 18625.418 c 11
701.8615112304688 0 13675.255
702.362060546875 0 6016.109 y Water loss 4
702.8612060546875 0 1986.4108
703.356689453125 0 905.53064 z 4
704.3477172851562 0 820.42395
711.3547973632812 0 1006.15546 y 4
714.368896484375 0 1369.5394
729.4151611328125 0 911.4624
734.368408203125 0 662.9684
742.856689453125 0 736.87244 z Water loss 3
743.87939453125 0 985.46295
748.3613891601562 0 618.09393
756.425537109375 0 1105.9535 w 9
757.4268798828125 0 762.11414
758.4049682617188 0 2462.603 w 9
759.8818969726562 0 6517.4805 y 3
760.3851318359375 0 4265.9863
760.8842163085938 0 3696.796
761.3837890625 0 1261.6798
765.3892211914062 0 13862.128 c 12
765.8909912109375 0 11143.53
766.3915405273438 0 5442.084
766.8942260742188 0 1430.9127
772.4110107421875 0 745.63696 y Ammonia loss 9
773.427001953125 0 7786.546 z 9
774.3465576171875 0 709.4313
774.4305419921875 0 4912.748
775.437255859375 0 1377.2546
777.9051513671875 0 960.7152
786.8908081054688 0 617.0425 w 2
787.3851928710938 0 912.8829
789.4461059570312 0 2480.3171 y 9
790.4449462890625 0 1125.855
792.4114379882812 0 700.58417
792.9154663085938 0 768.208
793.415771484375 0 720.4086
798.446044921875 0 28514.428 c 6
799.4489135742188 0 13388.353
799.9102172851562 0 646.5552 y Ammonia loss 2
800.451416015625 0 3308.0334
805.416748046875 0 1831.1589
805.9197998046875 0 1063.9911
807.4141845703125 0 753.53516
807.9100952148438 0 774.65326
808.4083251953125 0 25416.867 y 2
808.9091186523438 0 20036.781
809.409912109375 0 13073.489
809.91015625 0 4581.2793
810.4157104492188 0 1150.2041
810.906982421875 0 798.70337
814.4219970703125 0 804.3043
814.9237670898438 0 10740.2295
815.4246826171875 0 12910.38
815.9264526367188 0 6209.9014
816.4266967773438 0 3572.574
824.462158203125 0 1906.902
825.4690551757812 0 1002.39825
827.427734375 0 940.4866
827.9236450195312 0 865.49194
835.4141235351562 0 9729.744 w 1
835.9154052734375 0 8997.003
836.416259765625 0 6053.792
836.9217529296875 0 2374.084
843.4568481445312 0 5540.493 w 8
844.4596557617188 0 2471.7903
856.9406127929688 0 3135.3823 z 1
857.443115234375 0 4083.351
857.944580078125 0 2402.1292
858.4404296875 0 26271.24 c 13
858.9406127929688 0 22389.596
859.4423828125 0 12520.027
859.941650390625 0 5914.19
860.44873046875 0 1769.1544
862.9595336914062 0 1206.0521
863.4627685546875 0 2005.0392
863.9678955078125 0 738.2985
864.4523315429688 0 700.17316
865.4400634765625 0 1160.3069
870.4421997070312 0 1108.7886
870.949951171875 0 652.5759
871.447998046875 0 1059.2537
871.9517822265625 0 1926.8944
872.4461669921875 0 1054.4918
878.9578857421875 0 1793.6384
879.4353637695312 0 10437.27
879.9373779296875 0 10752.967
880.4385986328125 0 7966.8994
880.9502563476562 0 3345.7305
881.450439453125 0 1219.949
883.4750366210938 0 981.69775
884.4703369140625 0 932.6129
884.967041015625 0 1674.3356
885.466796875 0 2725.382
885.961669921875 0 2399.5938
886.4599609375 0 6786.5625 z Water loss 8
886.9588012695312 0 4287.786
887.4539184570312 0 1900.8662 z Ammonia loss 8
887.94970703125 0 1952.6573
888.4478759765625 0 1624.3983
888.9479370117188 0 1721.3396
893.467041015625 0 3954.1533
893.9686279296875 0 3940.8296
894.462890625 0 3102.2422
894.9627685546875 0 2497.945
895.4620971679688 0 793.14435
897.9677124023438 0 1798.5645
898.4628295898438 0 4535.978
898.9618530273438 0 3835.2236
899.4595336914062 0 4180.513
899.9580078125 0 1733.0498
904.4661254882812 0 7053.2188 z 8
905.4671020507812 0 3966.9846
906.4678955078125 0 1459.0707
906.9730834960938 0 3840.344
907.4656982421875 0 23967.58
907.966796875 0 19655.256
908.4674682617188 0 15110.69
908.9686889648438 0 6094.216
909.4658813476562 0 2038.6173 c Water loss 7
915.473388671875 0 17620.03
915.9761352539062 0 32852.863
916.4776611328125 0 26653.113
916.9785766601562 0 12551.227
917.47998046875 0 6358.883
917.9856567382812 0 1761.5565
920.4866333007812 0 3646.7656 y 8
921.486572265625 0 1664.8021
927.4883422851562 0 15821.839 c 7
928.490966796875 0 8458.635
929.4922485351562 0 2336.1455
960.4959716796875 0 747.4154
969.3493041992188 0 713.9026
970.3462524414062 0 1478.719
971.3428344726562 0 882.9183
972.4928588867188 0 1278.2927
973.49365234375 0 1437.2065
974.4976806640625 0 3171.9958 w 7
975.4996337890625 0 2301.189
976.4934692382812 0 895.58435
986.51220703125 0 1168.0608
997.3735961914062 0 1209.0002
998.3712158203125 0 1694.699
1014.4056396484375 0 861.4403
1014.5160522460938 0 2232.0073
1015.3937377929688 0 826.6438
1015.510986328125 0 843.2252 z Water loss 7
1016.5136108398438 0 696.867
1033.5091552734375 0 15472.271 z 7
1034.511474609375 0 10417.456
1035.513427734375 0 3296.0974
1036.5101318359375 0 871.9057
1037.5216064453125 0 707.05035
1039.445068359375 0 1556.1691
1058.528564453125 0 13835.216 c 8
1059.5328369140625 0 7986.998
1060.53173828125 0 4785.073
1061.522705078125 0 713.3701
1103.5355224609375 0 1521.5654 w 6
1104.53271484375 0 1082.6559
1105.5379638671875 0 662.9196
1120.5546875 0 831.642
1157.4652099609375 0 1278.231
1159.5765380859375 0 11269.826 c 9
1160.577880859375 0 8504.958 y Water loss 6
1161.5787353515625 0 3397.6116
1162.5543212890625 0 8994.5 z 6
1163.5535888671875 0 5701.6797
1164.559814453125 0 1634.8925
1165.54931640625 0 985.0812
1168.4808349609375 0 3234.3943
1169.4903564453125 0 1473.2688
1170.4986572265625 0 1020.30853
1172.4998779296875 0 1207.4237
1173.5023193359375 0 1587.639
1174.5103759765625 0 827.2978
1176.5050048828125 0 799.882
1178.57373046875 0 1922.733 y 6
1179.5745849609375 0 1905.4182
1185.4833984375 0 2625.5967
1186.488525390625 0 2553.1018
1187.497802734375 0 5357.8633
1188.501220703125 0 3213.6284
1189.5008544921875 0 1238.4299
1201.437744140625 0 1301.4094
1202.4312744140625 0 949.414
1203.469970703125 0 1017.7499
1204.4735107421875 0 819.75616
1205.5997314453125 0 825.67334
1206.5357666015625 0 780.0382
1218.4580078125 0 2168.735
1219.46044921875 0 2755.0852
1219.58203125 0 1927.1376
1220.462158203125 0 2537.1921
1220.591064453125 0 1691.6184
1221.2012939453125 0 2278.7412
1222.2022705078125 0 832.4343
1222.63037109375 0 1001.49866
1223.538818359375 0 1398.5739
1223.675048828125 0 3274.9678
1224.5499267578125 0 1902.1707
1224.67431640625 0 1856.9595
1243.6466064453125 0 1052.4955
1244.652099609375 0 1859.6753
1286.6578369140625 0 1540.1296
1287.6707763671875 0 4213.5044 c 10
1288.6729736328125 0 2434.9868
1289.676025390625 0 1019.3151
1318.6522216796875 0 5432.4595 z 5
1319.657958984375 0 8777.689
1320.6575927734375 0 6140.537
1321.6629638671875 0 3280.4473
1333.6717529296875 0 952.577
1334.669189453125 0 4669.6353 y 5
1335.665283203125 0 2581.461
1336.678955078125 0 1589.8245
1357.7015380859375 0 755.1591
1358.6986083984375 0 933.7736
1401.7127685546875 0 7421.951 c 11
1402.713134765625 0 7135.7217
1403.7161865234375 0 3242.8667 y Water loss 4
1404.7080078125 0 1204.9653 y Ammonia loss 4
1406.6912841796875 0 2268.1323
1407.6846923828125 0 1818.2015
1485.7491455078125 0 759.11914
1511.7694091796875 0 781.79565 c Water loss 12
1518.7557373046875 0 2476.8127 y 3
1519.76171875 0 1997.0287
1520.7388916015625 0 866.2934
1528.7474365234375 0 729.39215
1529.7706298828125 0 4810.055 c 12
1530.775146484375 0 5150.2397
1531.7740478515625 0 2663.1902
1532.7845458984375 0 1782.0098
1598.7894287109375 0 832.393 y Ammonia loss 2
1599.7987060546875 0 1192.4812 z 2
1628.835693359375 0 4448.0806
1629.8438720703125 0 5831.2896
1630.842041015625 0 3166.163
1631.84130859375 0 2327.676
1672.8677978515625 0 2209.2874
1673.87451171875 0 1121.731
1674.8602294921875 0 1034.7173
1700.856689453125 0 896.5317
1712.8717041015625 0 1067.7573 z 1
1713.8782958984375 0 9123.293
1714.881103515625 0 9946.011
1715.8773193359375 0 6485.348 c 13
1716.8765869140625 0 9962.413
1717.8763427734375 0 6755.203
1718.8802490234375 0 3559.1365
1719.886474609375 0 844.44824
1754.8900146484375 0 762.2256
1755.8880615234375 0 920.9353
1769.91796875 0 1722.4657
1770.9010009765625 0 1251.4889
1771.9176025390625 0 2065.889
1772.912109375 0 3257.626
1773.921142578125 0 2558.7136
1774.9031982421875 0 1497.36
1785.943115234375 0 1130.8661
1786.9317626953125 0 5091.428
1787.930419921875 0 3864.3645
1788.9268798828125 0 3173.2153
1789.920166015625 0 813.5136
1796.9222412109375 0 944.5589
1797.911865234375 0 1157.8018
1798.89697265625 0 1091.2855
1803.9605712890625 0 4657.8174
1804.956787109375 0 3955.0894
1805.9569091796875 0 2645.1948
1813.92529296875 0 4149.507
1814.9288330078125 0 19650.688
1815.9281005859375 0 18042.975
1816.93359375 0 10200.825
1817.931640625 0 4113.387
1818.9324951171875 0 1802.9149
1829.9365234375 0 2002.3967
1830.9442138671875 0 11887.704
1831.951904296875 0 45709.74
1832.95458984375 0 41857.273
1833.956787109375 0 24573.283
1834.959716796875 0 10849.258
1835.9644775390625 0 3906.9714
1836.9642333984375 0 882.2889
2451.15576171875 0 752.1199

Spectrum Details

|  |  |
| --- | --- |
| Matched peaks? Matched peaks The total absolute number of peaks matched. Additionally in brackets the total fraction of peaks matched and the total number of peaks is shown. | 58 (14.22% of 408) |
| FDR? FDR The false discovery rate estimated for this peptide. It is calculated by matching all theoretical fragments with a non-integer shift with the raw peaks for this spectrum. This is done with 40 different shifts. The resulting percentage is the average number of annotated peaks over the number of annotated peaks with the correct spectrum. | 1.68% |
| Satellite FDR? Satellite FDR See the FDR for details on its calculation. This satellite ion specific FDR only contains the satellite ions (d/w) for I/L/J positions. | 0.00% |
| PSM Score? PSM Score The PSM Score as given by Hecklib to this annotated spectrum. It is shown with three significant figures. | 441 |

#### Spectrum 4991? Spectrum 4991 The raw spectrum of this peptide as annotated by Hecklib. The fragments are coloured according to ion type (see legend). Any peaks with a star '\*' as text can be hovered over to see the full details, first the ion type second the mass shift type. By hovering over the amino acids in the peptide or ions in the legend the corresponding peaks are highlighted. By toggling the 'Unassigned' label you can turn the background (unassigned) peaks on or off in the plot. By updating the slider in the Ion legend you can update the spectrum to only show the top X% of the peaks with labels. The top X% means any peak that is within X% of the highest intensity. By dragging in the spectrum you can zoom in to a specific part of the spectrum and use 'Zoom Out' to get back to the original zoom level. The annotation of the spectrum is based on the given sequence in the peptides file and is done with different software so inconsistencies are likely. The peaks are annotated based on the given sequence, with 20 ppm tolerance.

Copy Data

##### Spectrum 4991 (TSV)

###### Preview

```
Loading example...
```

*Click on the button to copy the data to your clipboard.*

Mz MinMz MaxIntensity Max

WidthHeightPeptide font sizePeptide stroke widthSpectrum font sizeSpectrum stroke widthCompact peptide

Ion legend

wxyz

abcd

OtherUnassignedIon^Charge^_Position_Show for top:%

TLPPSREEMTKNQKJ

07.94e+31.59e+42.38e+43.17e+4

Zoom Out

y^+1^_3_c^+1^_5_y^+3^_13_c^+2^_11_c^+1^_6_z^+1^_5_y^+1^_5_c^+2^_12_y^+2^_11_z^+2^_11_y^+2^_11_w^+1^_6_w^+1^_6_y^+2^_12_y^+1^_6_c^+2^_13_y^+1^_6_c^+1^_7_y^+2^_13_w^+2^_14_y^+2^_14_z^+2^_14_c^+2^_14_z^+1^_7_z^+1^_7_c^+1^_8_y^+1^_7_w^+1^_8_z^+1^_8_c^+1^_9_w^+1^_9_c^+1^_10_y^+1^_9_z^+1^_9_c^+1^_11_c^+1^_11_y^+1^_10_z^+1^_10_y^+1^_10_c^+1^_12_y^+1^_11_y^+1^_11_y^+1^_12_c^+1^_13_z^+1^_14_c^+1^_14_

0779155923383117

Fragment Matches Table

Show background peaks

| Position | Ion type | Intensity | mz Theoretical | mz Error (Th) | mz Error (ppm) | Charge | Series Number |
| --- | --- | --- | --- | --- | --- | --- | --- |
| - | - | 356.8 | 120.2 | - | - | 0 | - |
| - | - | 398.2 | 127.8 | - | - | 0 | - |
| - | - | 1486 | 129.1 | - | - | 0 | - |
| - | - | 374.4 | 138.3 | - | - | 0 | - |
| - | - | 417.9 | 143.9 | - | - | 0 | - |
| - | - | 442.8 | 150.9 | - | - | 0 | - |
| - | - | 464.8 | 156.3 | - | - | 0 | - |
| - | - | 458.6 | 176.2 | - | - | 0 | - |
| - | - | 448.3 | 182.2 | - | - | 0 | - |
| - | - | 593.6 | 185.2 | - | - | 0 | - |
| - | - | 6249 | 187.1 | - | - | 0 | - |
| - | - | 735.5 | 188.1 | - | - | 0 | - |
| - | - | 675.6 | 212.1 | - | - | 0 | - |
| - | - | 523.5 | 215.1 | - | - | 0 | - |
| - | - | 4466 | 215.1 | - | - | 0 | - |
| - | - | 522 | 216.1 | - | - | 0 | - |
| - | - | 6929 | 221.1 | - | - | 0 | - |
| - | - | 2794 | 249.1 | - | - | 0 | - |
| - | - | 850.6 | 299.2 | - | - | 0 | - |
| - | - | 613.7 | 363.1 | - | - | 0 | - |
| - | - | 1202 | 364.2 | - | - | 0 | - |
| - | - | 600.2 | 407.3 | - | - | 0 | - |
| - | - | 574.5 | 412.6 | - | - | 0 | - |
| - | - | 828.4 | 431.2 | - | - | 0 | - |
| 13 | y | 864.8 | 446.3 | 0.004337 | 9.719 | +1 | 3 |
| - | - | 619.2 | 472.2 | - | - | 0 | - |
| - | - | 1698 | 488.3 | - | - | 0 | - |
| - | - | 983.6 | 499.2 | - | - | 0 | - |
| 5 | c | 5433 | 513.3 | 3.75E-05 | 0.07305 | +1 | 5 |
| - | - | 1482 | 514.3 | - | - | 0 | - |
| 3 | y | 2.421E+04 | 544.6 | 6.728E-06 | 0.01235 | +3 | 13 |
| - | - | 2.285E+04 | 544.9 | - | - | 0 | - |
| - | - | 999.2 | 545 | - | - | 0 | - |
| - | - | 1.629E+04 | 545.3 | - | - | 0 | - |
| - | - | 4461 | 545.6 | - | - | 0 | - |
| - | - | 1568 | 545.9 | - | - | 0 | - |
| - | - | 778.1 | 549.3 | - | - | 0 | - |
| - | - | 1197 | 576.2 | - | - | 0 | - |
| - | - | 1122 | 592.2 | - | - | 0 | - |
| - | - | 960.4 | 592.9 | - | - | 0 | - |
| - | - | 665.2 | 593.4 | - | - | 0 | - |
| - | - | 1796 | 608.2 | - | - | 0 | - |
| - | - | 992.5 | 614.4 | - | - | 0 | - |
| - | - | 809 | 616.3 | - | - | 0 | - |
| - | - | 2947 | 617.2 | - | - | 0 | - |
| - | - | 796.5 | 617.4 | - | - | 0 | - |
| - | - | 1568 | 628.4 | - | - | 0 | - |
| - | - | 1078 | 629.4 | - | - | 0 | - |
| - | - | 642.7 | 644.4 | - | - | 0 | - |
| - | - | 949.1 | 651.8 | - | - | 0 | - |
| 11 | c | 4648 | 652.3 | 0.001331 | 2.04 | +2 | 11 |
| - | - | 2597 | 652.8 | - | - | 0 | - |
| - | - | 1251 | 653.3 | - | - | 0 | - |
| 6 | c | 1.134E+04 | 669.4 | 6.871E-07 | 0.001026 | +1 | 6 |
| - | - | 4058 | 670.4 | - | - | 0 | - |
| - | - | 1071 | 671.4 | - | - | 0 | - |
| 11 | z | 4164 | 672.4 | 0.004058 | 6.035 | +1 | 5 |
| - | - | 2775 | 673.4 | - | - | 0 | - |
| - | - | 1144 | 673.9 | - | - | 0 | - |
| 11 | y | 1640 | 688.4 | 0.004838 | 7.027 | +1 | 5 |
| - | - | 683.6 | 689.4 | - | - | 0 | - |
| - | - | 859 | 707.3 | - | - | 0 | - |
| - | - | 782.6 | 708.3 | - | - | 0 | - |
| - | - | 1224 | 708.9 | - | - | 0 | - |
| 12 | c | 1.261E+04 | 709.4 | 0.002328 | 3.282 | +2 | 12 |
| - | - | 9559 | 709.9 | - | - | 0 | - |
| 5 | y | 5986 | 710.3 | 0.01257 | 17.7 | +2 | 11 |
| - | - | 1859 | 710.9 | - | - | 0 | - |
| 5 | z | 673.3 | 711.3 | 0.01189 | 16.72 | +2 | 11 |
| 5 | y | 714.7 | 719.4 | 0.0005843 | 0.8123 | +2 | 11 |
| - | - | 959.6 | 729.4 | - | - | 0 | - |
| - | - | 1456 | 739.3 | - | - | 0 | - |
| - | - | 3597 | 743.5 | - | - | 0 | - |
| - | - | 770.2 | 744.5 | - | - | 0 | - |
| - | - | 1390 | 755.3 | - | - | 0 | - |
| 10 | w | 1156 | 756.4 | 0.008469 | 11.2 | +1 | 6 |
| 10 | w | 1990 | 758.4 | 0.0009175 | 1.21 | +1 | 6 |
| - | - | 600.2 | 759.4 | - | - | 0 | - |
| - | - | 787.4 | 764.3 | - | - | 0 | - |
| 4 | y | 4300 | 767.9 | 0.0001334 | 0.1737 | +2 | 12 |
| - | - | 3468 | 768.4 | - | - | 0 | - |
| - | - | 1999 | 768.9 | - | - | 0 | - |
| - | - | 816.9 | 769.4 | - | - | 0 | - |
| 10 | y | 868.6 | 772.4 | 0.004605 | 5.961 | +1 | 6 |
| 13 | c | 6541 | 773.4 | 0.004961 | 6.414 | +2 | 13 |
| - | - | 8226 | 773.9 | - | - | 0 | - |
| - | - | 4237 | 774.4 | - | - | 0 | - |
| - | - | 1611 | 774.9 | - | - | 0 | - |
| - | - | 924.3 | 775.4 | - | - | 0 | - |
| - | - | 825.6 | 779.8 | - | - | 0 | - |
| - | - | 1524 | 780.3 | - | - | 0 | - |
| - | - | 588.3 | 782.4 | - | - | 0 | - |
| 10 | y | 1307 | 789.5 | 0.0032 | 4.053 | +1 | 6 |
| - | - | 1206 | 793.4 | - | - | 0 | - |
| 7 | c | 1.571E+04 | 798.4 | 0.0004782 | 0.5989 | +1 | 7 |
| - | - | 7065 | 799.4 | - | - | 0 | - |
| - | - | 1664 | 800.5 | - | - | 0 | - |
| - | - | 1111 | 801.9 | - | - | 0 | - |
| - | - | 886.9 | 810.3 | - | - | 0 | - |
| 3 | y | 1.492E+04 | 816.4 | 0.0006069 | 0.7434 | +2 | 13 |
| - | - | 1.105E+04 | 816.9 | - | - | 0 | - |
| - | - | 7482 | 817.4 | - | - | 0 | - |
| - | - | 2563 | 817.9 | - | - | 0 | - |
| - | - | 1774 | 822.4 | - | - | 0 | - |
| - | - | 7668 | 822.9 | - | - | 0 | - |
| - | - | 6791 | 823.4 | - | - | 0 | - |
| - | - | 2592 | 823.9 | - | - | 0 | - |
| - | - | 1054 | 824.4 | - | - | 0 | - |
| - | - | 726.5 | 824.9 | - | - | 0 | - |
| - | - | 863.5 | 825.5 | - | - | 0 | - |
| - | - | 1262 | 835.9 | - | - | 0 | - |
| - | - | 743.3 | 836.9 | - | - | 0 | - |
| 2 | w | 6828 | 843.4 | 0.0001588 | 0.1883 | +2 | 14 |
| - | - | 6795 | 843.9 | - | - | 0 | - |
| - | - | 4060 | 844.4 | - | - | 0 | - |
| - | - | 1153 | 844.9 | - | - | 0 | - |
| - | - | 923.6 | 845.4 | - | - | 0 | - |
| - | - | 1187 | 848.5 | - | - | 0 | - |
| - | - | 1616 | 854.3 | - | - | 0 | - |
| - | - | 1735 | 855.3 | - | - | 0 | - |
| - | - | 1706 | 857.5 | - | - | 0 | - |
| - | - | 5643 | 858.5 | - | - | 0 | - |
| - | - | 2648 | 859.5 | - | - | 0 | - |
| - | - | 836.8 | 860.5 | - | - | 0 | - |
| 2 | y | 804 | 863.9 | 0.008962 | 10.37 | +2 | 14 |
| 2 | z | 2452 | 864.9 | 0.003377 | 3.904 | +2 | 14 |
| - | - | 3399 | 865.4 | - | - | 0 | - |
| - | - | 1550 | 865.9 | - | - | 0 | - |
| 14 | c | 1.982E+04 | 866.4 | 0.0002187 | 0.2524 | +2 | 14 |
| - | - | 1.923E+04 | 866.9 | - | - | 0 | - |
| - | - | 9953 | 867.4 | - | - | 0 | - |
| - | - | 3490 | 867.9 | - | - | 0 | - |
| - | - | 883.3 | 868.4 | - | - | 0 | - |
| - | - | 1534 | 870.3 | - | - | 0 | - |
| - | - | 842.8 | 872.5 | - | - | 0 | - |
| - | - | 1020 | 873.4 | - | - | 0 | - |
| - | - | 751.2 | 878.9 | - | - | 0 | - |
| - | - | 1391 | 885.5 | - | - | 0 | - |
| - | - | 1010 | 886.4 | - | - | 0 | - |
| - | - | 4037 | 887.4 | - | - | 0 | - |
| - | - | 4718 | 887.9 | - | - | 0 | - |
| - | - | 3946 | 888.4 | - | - | 0 | - |
| - | - | 1199 | 888.9 | - | - | 0 | - |
| - | - | 1060 | 892.5 | - | - | 0 | - |
| - | - | 1526 | 893 | - | - | 0 | - |
| - | - | 1945 | 893.5 | - | - | 0 | - |
| - | - | 1434 | 894 | - | - | 0 | - |
| - | - | 3070 | 894.5 | - | - | 0 | - |
| - | - | 3017 | 895 | - | - | 0 | - |
| - | - | 849.8 | 895.5 | - | - | 0 | - |
| - | - | 1702 | 895.9 | - | - | 0 | - |
| - | - | 2134 | 896.4 | - | - | 0 | - |
| - | - | 1507 | 896.9 | - | - | 0 | - |
| - | - | 724.1 | 897.4 | - | - | 0 | - |
| - | - | 3576 | 901.5 | - | - | 0 | - |
| - | - | 4237 | 902 | - | - | 0 | - |
| 9 | z | 3851 | 902.5 | 0.01227 | 13.59 | +1 | 7 |
| - | - | 2462 | 903 | - | - | 0 | - |
| - | - | 1786 | 903.5 | - | - | 0 | - |
| - | - | 890.6 | 904 | - | - | 0 | - |
| - | - | 844.5 | 904.4 | - | - | 0 | - |
| - | - | 849.3 | 906 | - | - | 0 | - |
| - | - | 2474 | 906.5 | - | - | 0 | - |
| - | - | 2926 | 907 | - | - | 0 | - |
| - | - | 2091 | 907.5 | - | - | 0 | - |
| - | - | 768 | 908 | - | - | 0 | - |
| - | - | 5749 | 915 | - | - | 0 | - |
| - | - | 1.841E+04 | 915.5 | - | - | 0 | - |
| - | - | 1.843E+04 | 916 | - | - | 0 | - |
| - | - | 1.138E+04 | 916.5 | - | - | 0 | - |
| - | - | 3895 | 917 | - | - | 0 | - |
| - | - | 1219 | 917.5 | - | - | 0 | - |
| 9 | z | 2167 | 920.5 | 0.0001155 | 0.1255 | +1 | 7 |
| - | - | 1645 | 921.5 | - | - | 0 | - |
| - | - | 1009 | 922.5 | - | - | 0 | - |
| - | - | 1.137E+04 | 923.5 | - | - | 0 | - |
| - | - | 2.103E+04 | 924 | - | - | 0 | - |
| - | - | 1.923E+04 | 924.5 | - | - | 0 | - |
| - | - | 8892 | 925 | - | - | 0 | - |
| - | - | 1291 | 925.3 | - | - | 0 | - |
| - | - | 3631 | 925.5 | - | - | 0 | - |
| - | - | 1163 | 926 | - | - | 0 | - |
| 8 | c | 1.227E+04 | 927.5 | 0.0007739 | 0.8344 | +1 | 8 |
| - | - | 6507 | 928.5 | - | - | 0 | - |
| - | - | 1804 | 929.5 | - | - | 0 | - |
| 9 | y | 1888 | 936.5 | 0.001045 | 1.116 | +1 | 7 |
| - | - | 1159 | 937.5 | - | - | 0 | - |
| - | - | 921.6 | 967.3 | - | - | 0 | - |
| - | - | 9814 | 985.3 | - | - | 0 | - |
| - | - | 996.5 | 989.5 | - | - | 0 | - |
| 8 | w | 2771 | 990.5 | 0.001039 | 1.049 | +1 | 8 |
| - | - | 1523 | 991.5 | - | - | 0 | - |
| - | - | 828.9 | 1001 | - | - | 0 | - |
| - | - | 1091 | 1002 | - | - | 0 | - |
| - | - | 825.9 | 1002 | - | - | 0 | - |
| - | - | 1364 | 1003 | - | - | 0 | - |
| - | - | 2044 | 1008 | - | - | 0 | - |
| - | - | 2843 | 1009 | - | - | 0 | - |
| - | - | 1197 | 1009 | - | - | 0 | - |
| - | - | 1685 | 1014 | - | - | 0 | - |
| - | - | 2245 | 1023 | - | - | 0 | - |
| - | - | 4429 | 1023 | - | - | 0 | - |
| - | - | 860.2 | 1023 | - | - | 0 | - |
| - | - | 938.6 | 1028 | - | - | 0 | - |
| - | - | 2966 | 1028 | - | - | 0 | - |
| - | - | 3558 | 1029 | - | - | 0 | - |
| - | - | 1011 | 1029 | - | - | 0 | - |
| - | - | 1315 | 1031 | - | - | 0 | - |
| - | - | 830.5 | 1032 | - | - | 0 | - |
| - | - | 735.3 | 1033 | - | - | 0 | - |
| 8 | z | 7564 | 1050 | 0.001224 | 1.166 | +1 | 8 |
| - | - | 4983 | 1051 | - | - | 0 | - |
| - | - | 2717 | 1052 | - | - | 0 | - |
| - | - | 1043 | 1053 | - | - | 0 | - |
| 9 | c | 1.11E+04 | 1075 | 0.005118 | 4.763 | +1 | 9 |
| - | - | 6527 | 1076 | - | - | 0 | - |
| - | - | 2948 | 1077 | - | - | 0 | - |
| 7 | w | 1478 | 1120 | 0.001332 | 1.19 | +1 | 9 |
| - | - | 861.8 | 1121 | - | - | 0 | - |
| - | - | 1360 | 1171 | - | - | 0 | - |
| - | - | 1540 | 1172 | - | - | 0 | - |
| - | - | 1758 | 1174 | - | - | 0 | - |
| 10 | c | 8799 | 1176 | 0.003705 | 3.151 | +1 | 10 |
| 7 | y | 6398 | 1177 | 0.01907 | 16.21 | +1 | 9 |
| - | - | 3069 | 1178 | - | - | 0 | - |
| 7 | z | 5834 | 1179 | 0.003919 | 3.325 | +1 | 9 |
| - | - | 4675 | 1180 | - | - | 0 | - |
| - | - | 2097 | 1181 | - | - | 0 | - |
| - | - | 909.1 | 1182 | - | - | 0 | - |
| - | - | 729.7 | 1215 | - | - | 0 | - |
| - | - | 4869 | 1216 | - | - | 0 | - |
| - | - | 8163 | 1217 | - | - | 0 | - |
| - | - | 831 | 1218 | - | - | 0 | - |
| - | - | 804 | 1221 | - | - | 0 | - |
| - | - | 825.5 | 1230 | - | - | 0 | - |
| - | - | 739.8 | 1233 | - | - | 0 | - |
| - | - | 3306 | 1233 | - | - | 0 | - |
| - | - | 716 | 1234 | - | - | 0 | - |
| - | - | 1.28E+04 | 1234 | - | - | 0 | - |
| - | - | 886.9 | 1235 | - | - | 0 | - |
| 11 | c | 920.6 | 1286 | 0.005376 | 4.182 | +1 | 11 |
| - | - | 1381 | 1299 | - | - | 0 | - |
| - | - | 860.9 | 1303 | - | - | 0 | - |
| 11 | c | 3513 | 1304 | 0.002858 | 2.192 | +1 | 11 |
| - | - | 1834 | 1305 | - | - | 0 | - |
| - | - | 1153 | 1333 | - | - | 0 | - |
| 6 | y | 1254 | 1334 | 0.002209 | 1.657 | +1 | 10 |
| - | - | 881 | 1334 | - | - | 0 | - |
| 6 | z | 4384 | 1335 | 0.002832 | 2.122 | +1 | 10 |
| - | - | 6492 | 1336 | - | - | 0 | - |
| - | - | 3023 | 1337 | - | - | 0 | - |
| - | - | 1413 | 1338 | - | - | 0 | - |
| 6 | y | 2193 | 1351 | 0.001027 | 0.7602 | +1 | 10 |
| - | - | 1660 | 1352 | - | - | 0 | - |
| - | - | 3073 | 1363 | - | - | 0 | - |
| - | - | 3254 | 1363 | - | - | 0 | - |
| - | - | 1113 | 1364 | - | - | 0 | - |
| - | - | 945.6 | 1399 | - | - | 0 | - |
| 12 | c | 6174 | 1418 | 0.003509 | 2.475 | +1 | 12 |
| - | - | 5004 | 1419 | - | - | 0 | - |
| 5 | y | 2476 | 1420 | 0.02522 | 17.76 | +1 | 11 |
| - | - | 1777 | 1420 | - | - | 0 | - |
| 5 | y | 2051 | 1421 | 0.0003021 | 0.2126 | +1 | 11 |
| - | - | 2488 | 1423 | - | - | 0 | - |
| - | - | 1726 | 1424 | - | - | 0 | - |
| - | - | 1142 | 1475 | - | - | 0 | - |
| - | - | 900.3 | 1484 | - | - | 0 | - |
| - | - | 854.9 | 1489 | - | - | 0 | - |
| - | - | 1411 | 1489 | - | - | 0 | - |
| - | - | 1316 | 1496 | - | - | 0 | - |
| - | - | 1051 | 1504 | - | - | 0 | - |
| - | - | 2611 | 1505 | - | - | 0 | - |
| - | - | 2123 | 1505 | - | - | 0 | - |
| - | - | 1052 | 1512 | - | - | 0 | - |
| - | - | 2797 | 1513 | - | - | 0 | - |
| - | - | 3792 | 1513 | - | - | 0 | - |
| - | - | 2802 | 1514 | - | - | 0 | - |
| - | - | 1226 | 1514 | - | - | 0 | - |
| - | - | 886.7 | 1520 | - | - | 0 | - |
| - | - | 841.3 | 1525 | - | - | 0 | - |
| - | - | 1007 | 1534 | - | - | 0 | - |
| - | - | 3186 | 1534 | - | - | 0 | - |
| 4 | y | 4251 | 1535 | 0.01345 | 8.766 | +1 | 12 |
| - | - | 960.2 | 1535 | - | - | 0 | - |
| - | - | 1294 | 1536 | - | - | 0 | - |
| - | - | 966.1 | 1538 | - | - | 0 | - |
| - | - | 801.6 | 1538 | - | - | 0 | - |
| - | - | 1416 | 1542 | - | - | 0 | - |
| - | - | 3418 | 1543 | - | - | 0 | - |
| - | - | 2751 | 1543 | - | - | 0 | - |
| 13 | c | 2468 | 1546 | 0.004624 | 2.992 | +1 | 13 |
| - | - | 3613 | 1547 | - | - | 0 | - |
| - | - | 1989 | 1548 | - | - | 0 | - |
| - | - | 1063 | 1549 | - | - | 0 | - |
| - | - | 710.3 | 1560 | - | - | 0 | - |
| - | - | 2432 | 1645 | - | - | 0 | - |
| - | - | 3580 | 1646 | - | - | 0 | - |
| - | - | 2789 | 1647 | - | - | 0 | - |
| - | - | 738.2 | 1648 | - | - | 0 | - |
| - | - | 925.5 | 1689 | - | - | 0 | - |
| - | - | 1178 | 1690 | - | - | 0 | - |
| - | - | 861.3 | 1702 | - | - | 0 | - |
| 2 | z | 1341 | 1729 | 0.006557 | 3.792 | +1 | 14 |
| - | - | 4773 | 1730 | - | - | 0 | - |
| - | - | 4400 | 1731 | - | - | 0 | - |
| 14 | c | 5010 | 1732 | 0.002811 | 1.623 | +1 | 14 |
| - | - | 6585 | 1733 | - | - | 0 | - |
| - | - | 5366 | 1734 | - | - | 0 | - |
| - | - | 2575 | 1735 | - | - | 0 | - |
| - | - | 1690 | 1736 | - | - | 0 | - |
| - | - | 1549 | 1739 | - | - | 0 | - |
| - | - | 678.6 | 1775 | - | - | 0 | - |
| - | - | 1198 | 1786 | - | - | 0 | - |
| - | - | 1232 | 1787 | - | - | 0 | - |
| - | - | 1085 | 1788 | - | - | 0 | - |
| - | - | 1494 | 1789 | - | - | 0 | - |
| - | - | 1579 | 1790 | - | - | 0 | - |
| - | - | 1374 | 1802 | - | - | 0 | - |
| - | - | 3073 | 1803 | - | - | 0 | - |
| - | - | 2855 | 1804 | - | - | 0 | - |
| - | - | 2005 | 1805 | - | - | 0 | - |
| - | - | 1030 | 1806 | - | - | 0 | - |
| - | - | 853.5 | 1813 | - | - | 0 | - |
| - | - | 858.7 | 1814 | - | - | 0 | - |
| - | - | 966.3 | 1815 | - | - | 0 | - |
| - | - | 878.8 | 1816 | - | - | 0 | - |
| - | - | 2131 | 1820 | - | - | 0 | - |
| - | - | 2107 | 1821 | - | - | 0 | - |
| - | - | 1552 | 1822 | - | - | 0 | - |
| - | - | 886 | 1829 | - | - | 0 | - |
| - | - | 3790 | 1830 | - | - | 0 | - |
| - | - | 1.167E+04 | 1831 | - | - | 0 | - |
| - | - | 1.096E+04 | 1832 | - | - | 0 | - |
| - | - | 6653 | 1833 | - | - | 0 | - |
| - | - | 2437 | 1834 | - | - | 0 | - |
| - | - | 2085 | 1846 | - | - | 0 | - |
| - | - | 7357 | 1847 | - | - | 0 | - |
| - | - | 3.143E+04 | 1848 | - | - | 0 | - |
| - | - | 2.955E+04 | 1849 | - | - | 0 | - |
| - | - | 1.952E+04 | 1850 | - | - | 0 | - |
| - | - | 7572 | 1851 | - | - | 0 | - |
| - | - | 1753 | 1852 | - | - | 0 | - |
| - | - | 842.9 | 1877 | - | - | 0 | - |
| - | - | 712.4 | 3086 | - | - | 0 | - |

m/z Charge Intensity FragmentType MassShift Position
120.20887756347656 0 356.7529
127.79176330566406 0 398.15152
129.1021728515625 0 1485.6655
138.3363037109375 0 374.408
143.85897827148438 0 417.86588
150.88975524902344 0 442.79117
156.318603515625 0 464.751
176.18553161621094 0 458.58154
182.23130798339844 0 448.34305
185.2146759033203 0 593.64923
187.14405822753906 0 6248.886
188.14698791503906 0 735.5104
212.13998413085938 0 675.6135
215.1017303466797 0 523.48535
215.1388397216797 0 4465.6763
216.14224243164062 0 521.9591
221.12828063964844 0 6929.033
249.12322998046875 0 2793.7688
299.1712646484375 0 850.5815
363.1045227050781 0 613.6923
364.1507873535156 0 1202.4669
407.2521667480469 0 600.24817
412.6099548339844 0 574.46576
431.19183349609375 0 828.41187
446.2611083984375 0 864.7543 y 12
472.233154296875 0 619.2486
488.3191833496094 0 1698.1879
499.1745300292969 0 983.56854
513.3031616210938 0 5432.757 c 4
514.304931640625 0 1481.563
544.6066284179688 0 24209.95 y 2
544.9407348632812 0 22854.715
544.9888916015625 0 999.23175
545.2750854492188 0 16291.428
545.6068725585938 0 4461.1274
545.939453125 0 1568.3928
549.343017578125 0 778.1003
576.2001342773438 0 1196.5992
592.21826171875 0 1122.1947
592.853515625 0 960.39526
593.3641357421875 0 665.1843
608.22412109375 0 1796.4121
614.3890380859375 0 992.5334
616.2744750976562 0 809.0329
617.2275390625 0 2946.5994
617.3518676757812 0 796.53564
628.3663940429688 0 1568.4371
629.3722534179688 0 1078.0823
644.4217529296875 0 642.74945
651.8341064453125 0 949.13855
652.3362426757812 0 4648.383 c 10
652.839111328125 0 2596.7651
653.3392333984375 0 1250.526
669.4042358398438 0 11343.967 c 5
670.406494140625 0 4058.499
671.4102783203125 0 1071.0555
672.3805541992188 0 4164.3833 z 10
673.3710327148438 0 2774.8218
673.86669921875 0 1144.401
688.3984985351562 0 1639.6832 y 10
689.399169921875 0 683.6258
707.3435668945312 0 858.98956
708.3422241210938 0 782.58466
708.8560791015625 0 1223.72
709.3587036132812 0 12605.959 c 11
709.8592529296875 0 9559.276
710.36083984375 0 5985.967 y Water loss 4
710.8623657226562 0 1858.5214
711.3560791015625 0 673.28906 z 4
719.3529663085938 0 714.6921 y 4
729.4144897460938 0 959.5548
739.263916015625 0 1455.6123
743.4880981445312 0 3596.8594
744.4934692382812 0 770.1865
755.2799072265625 0 1390.3802
756.4210815429688 0 1155.8496 w 9
758.4078979492188 0 1989.6715 w 9
759.3951416015625 0 600.15027
764.2869262695312 0 787.44495
767.8800659179688 0 4300.015 y 3
768.3832397460938 0 3468.2656
768.879638671875 0 1998.6583
769.3795776367188 0 816.86255
772.4198608398438 0 868.60144 y Ammonia loss 9
773.390625 0 6541.442 c 12
773.8892211914062 0 8225.968
774.3903198242188 0 4237.301
774.8914794921875 0 1610.9828
775.4308471679688 0 924.26025
779.8134765625 0 825.6311
780.3177490234375 0 1523.5745
782.3589477539062 0 588.3232
789.4478149414062 0 1306.8463 y 9
793.4085693359375 0 1205.5751
798.4463500976562 0 15707.133 c 6
799.449951171875 0 7064.9565
800.451904296875 0 1663.8716
801.9208374023438 0 1110.7874
810.302978515625 0 886.8909
816.4069213867188 0 14921.483 y 2
816.90673828125 0 11053.752
817.4070434570312 0 7481.555
817.9071655273438 0 2563.4895
822.4176635742188 0 1773.6918
822.9219970703125 0 7668.389
823.4215698242188 0 6790.8433
823.9224243164062 0 2591.5247
824.4253540039062 0 1053.719
824.9249267578125 0 726.5306
825.469970703125 0 863.4696
835.9263305664062 0 1261.6222
836.9351806640625 0 743.2939
843.4114379882812 0 6827.761 w 1
843.912109375 0 6794.8853
844.4127807617188 0 4059.5908
844.9154052734375 0 1153.2626
845.4447631835938 0 923.6427
848.4647216796875 0 1187.4318
854.2905883789062 0 1616.1211
855.296875 0 1734.8673
857.4691772460938 0 1706.2872
858.4791870117188 0 5642.687
859.4801635742188 0 2647.988
860.4743041992188 0 836.776
863.9520263671875 0 804.00214 y Water loss 1
864.9356079101562 0 2452.1045 z 1
865.4397583007812 0 3399.2788
865.9363403320312 0 1549.5804
866.4379272460938 0 19821.082 c 13
866.9391479492188 0 19229.736
867.4386596679688 0 9953.486
867.9400634765625 0 3490.2366
868.4426879882812 0 883.253
870.3084106445312 0 1534.2999
872.4522705078125 0 842.8158
873.444091796875 0 1019.6126
878.9498291015625 0 751.23755
885.4638671875 0 1390.9557
886.443115234375 0 1010.2431
887.4328002929688 0 4036.8699
887.9376220703125 0 4718.0605
888.443115234375 0 3945.9138
888.9442138671875 0 1199.437
892.466796875 0 1059.9785
892.9640502929688 0 1525.7244
893.4555053710938 0 1945.0762
893.9691772460938 0 1434.024
894.4571533203125 0 3070.0044
894.9581298828125 0 3016.6682
895.46142578125 0 849.7896
895.9464111328125 0 1702.0582
896.4454956054688 0 2134.0513
896.9427490234375 0 1507.3325
897.4483642578125 0 724.1407
901.4662475585938 0 3576.4973
901.9672241210938 0 4237.4854
902.4644775390625 0 3850.8113 z Water loss 8
902.9629516601562 0 2462.4697
903.4600830078125 0 1786.3848
903.9501342773438 0 890.5566
904.4075317382812 0 844.4601
905.9635009765625 0 849.28
906.4592895507812 0 2474.2996
906.9600830078125 0 2925.7656
907.4566650390625 0 2090.554
907.965087890625 0 767.9785
914.9705810546875 0 5749.3994
915.467529296875 0 18407.982
915.966552734375 0 18431.633
916.4676513671875 0 11378.923
916.9663696289062 0 3894.506
917.4632568359375 0 1219.1205
920.462890625 0 2167.3552 z 8
921.4661865234375 0 1645.3676
922.4699096679688 0 1009.23486
923.471923828125 0 11365.732
923.974609375 0 21028.451
924.4749145507812 0 19233.611
924.9756469726562 0 8892.1875
925.3255004882812 0 1291.486
925.4783935546875 0 3630.9712
925.9757690429688 0 1163.2158
927.4886474609375 0 12271.764 c 7
928.4927978515625 0 6507.2075
929.495849609375 0 1804.1372
936.4825439453125 0 1888.3601 y 8
937.47998046875 0 1159.1068
967.32470703125 0 921.62366
985.3359985351562 0 9813.714
989.492919921875 0 996.5341
990.4931030273438 0 2771.3967 w 7
991.4916381835938 0 1523.1204
1001.4178466796875 0 828.93933
1001.9153442382812 0 1091.4033
1002.4937133789062 0 825.9186
1003.1558227539062 0 1364.2522
1008.4952392578125 0 2043.9331
1008.8264770507812 0 2843.0325
1009.1560668945312 0 1197.2667
1014.3970336914062 0 1684.7158
1022.8245239257812 0 2244.8179
1023.1578369140625 0 4428.811
1023.4898681640625 0 860.2154
1028.162841796875 0 938.57697
1028.498779296875 0 2966.3054
1028.837158203125 0 3557.762
1029.162353515625 0 1011.1799
1030.5057373046875 0 1314.5271
1031.5166015625 0 830.51227
1032.5006103515625 0 735.2508
1049.506591796875 0 7563.8027 z 7
1050.5076904296875 0 4982.9805
1051.5068359375 0 2716.6624
1052.50830078125 0 1043.0021
1074.5250244140625 0 11101.058 c 8
1075.5272216796875 0 6527.201
1076.5274658203125 0 2948.1365
1119.5333251953125 0 1478.1716 w 6
1120.5250244140625 0 861.7758
1171.0068359375 0 1359.6324
1171.51416015625 0 1539.5956
1174.43701171875 0 1758.3013
1175.5712890625 0 8798.71 c 9
1176.5751953125 0 6398.3413 y Water loss 6
1177.5787353515625 0 3069.367
1178.5518798828125 0 5833.519 z 6
1179.55078125 0 4675.478
1180.5498046875 0 2097.1172
1181.5546875 0 909.14905
1214.5330810546875 0 729.6755
1216.446044921875 0 4868.969
1217.4361572265625 0 8162.631
1217.579345703125 0 831.0137
1220.5623779296875 0 803.95917
1229.638671875 0 825.4762
1232.6243896484375 0 739.79816
1233.449951171875 0 3305.6494
1233.6044921875 0 715.9691
1234.458984375 0 12797.1
1234.611328125 0 886.93884
1285.6466064453125 0 920.63 c Water loss 10
1299.0931396484375 0 1380.8855
1302.6510009765625 0 860.87897
1303.6654052734375 0 3513.252 c 10
1304.669921875 0 1834.4584
1333.1319580078125 0 1153.2042
1333.6390380859375 0 1253.7664 y Ammonia loss 5
1334.1337890625 0 880.9638
1334.646240234375 0 4384.274 z 5
1335.65380859375 0 6492.3213
1336.6488037109375 0 3022.968
1337.6531982421875 0 1412.8351
1350.6688232421875 0 2193.1807 y 5
1351.6600341796875 0 1660.3323
1362.6334228515625 0 3072.634
1363.141845703125 0 3253.787
1363.6468505859375 0 1112.9392
1399.1566162109375 0 945.64233
1417.708984375 0 6173.914 c 11
1418.7115478515625 0 5004.2397
1419.7144775390625 0 2475.901 y Water loss 4
1420.154541015625 0 1776.5212
1420.6729736328125 0 2050.5347 y Ammonia loss 4
1422.6890869140625 0 2488.264
1423.692138671875 0 1726.184
1475.22509765625 0 1142.2899
1483.73291015625 0 900.32654
1488.7464599609375 0 854.8536
1489.2474365234375 0 1411.1986
1496.2279052734375 0 1316.2711
1504.2310791015625 0 1051.4442
1504.7353515625 0 2611.065
1505.2222900390625 0 2122.891
1512.2298583984375 0 1052.2942
1512.7330322265625 0 2796.5635
1513.234130859375 0 3792.2222
1513.7415771484375 0 2802.207
1514.2442626953125 0 1226.1835
1520.248779296875 0 886.6923
1525.2598876953125 0 841.3384
1533.7325439453125 0 1006.51495
1534.23291015625 0 3186.3694
1534.7391357421875 0 4251.3276 y 3
1535.2608642578125 0 960.2397
1535.7705078125 0 1293.7941
1537.7564697265625 0 966.1163
1538.2354736328125 0 801.5801
1542.249267578125 0 1416.0791
1542.7474365234375 0 3417.7065
1543.2498779296875 0 2750.636
1545.7686767578125 0 2467.7024 c 12
1546.7711181640625 0 3612.5698
1547.772705078125 0 1988.542
1548.7705078125 0 1063.1367
1559.620849609375 0 710.29944
1644.831787109375 0 2432.1177
1645.8380126953125 0 3580.4417
1646.8438720703125 0 2789.3242
1647.85009765625 0 738.23895
1688.8673095703125 0 925.463
1689.8739013671875 0 1178.2662
1701.87744140625 0 861.3065
1728.8641357421875 0 1341.3422 z 1
1729.8731689453125 0 4773.034
1730.8792724609375 0 4399.589
1731.871826171875 0 5009.9253 c 13
1732.874755859375 0 6585.3066
1733.872802734375 0 5365.9688
1734.872802734375 0 2574.5532
1735.878662109375 0 1689.8702
1738.7637939453125 0 1549.3342
1774.8846435546875 0 678.5742
1785.92626953125 0 1197.951
1786.9437255859375 0 1232.1514
1787.900390625 0 1084.9991
1788.9202880859375 0 1493.761
1789.9254150390625 0 1579.0162
1801.926513671875 0 1374.1971
1802.931396484375 0 3072.8176
1803.92822265625 0 2854.9111
1804.939208984375 0 2004.9825
1805.9193115234375 0 1029.5311
1812.9232177734375 0 853.4526
1813.9180908203125 0 858.695
1814.9234619140625 0 966.2551
1815.8974609375 0 878.7807
1819.955810546875 0 2130.6572
1820.95361328125 0 2106.7705
1821.949462890625 0 1551.8467
1828.923095703125 0 886.0452
1829.931640625 0 3790.2148
1830.9273681640625 0 11670.757
1831.92724609375 0 10963.767
1832.9266357421875 0 6653.4346
1833.928955078125 0 2437.1292
1845.939697265625 0 2084.9414
1846.943115234375 0 7356.96
1847.9501953125 0 31430.496
1848.951416015625 0 29548.479
1849.9537353515625 0 19520.072
1850.955322265625 0 7572.119
1851.9549560546875 0 1752.8944
1876.719482421875 0 842.86615
3086.466552734375 0 712.3916

Spectrum Details

|  |  |
| --- | --- |
| Matched peaks? Matched peaks The total absolute number of peaks matched. Additionally in brackets the total fraction of peaks matched and the total number of peaks is shown. | 46 (13.37% of 344) |
| FDR? FDR The false discovery rate estimated for this peptide. It is calculated by matching all theoretical fragments with a non-integer shift with the raw peaks for this spectrum. This is done with 40 different shifts. The resulting percentage is the average number of annotated peaks over the number of annotated peaks with the correct spectrum. | 1.14% |
| Satellite FDR? Satellite FDR See the FDR for details on its calculation. This satellite ion specific FDR only contains the satellite ions (d/w) for I/L/J positions. | 0.00% |
| PSM Score? PSM Score The PSM Score as given by Hecklib to this annotated spectrum. It is shown with three significant figures. | 325 |

#### Reverse Lookup? Reverse Lookup All places where this read could be placed.

| Group | Segment | Template | Template Part | Read Part | Score | Unique |
| --- | --- | --- | --- | --- | --- | --- |
| Homo sapiens Heavy Chain | IGHC | IGHG3 | [279..294] | [0..15] | 102 | False |
| Homo sapiens Heavy Chain | IGHC | IGHG2 | [228..243] | [0..15] | 102 | False |
| Homo sapiens Heavy Chain | IGHC | IGHG4 | [229..244] | [0..15] | 93 | False |

| Recombined | Template Part | Read Part | Score | Unique |
| --- | --- | --- | --- | --- |
| REC-0-1 | [357..372] | [0..15] | 88 | True |

#### Meta Information from Multiple reads

##### Number of combined reads

8

##### Intensity

0.813

##### TotalArea

2.21E+09

##### Changes to the peptide sequence

TLPPSREEMTKNQKJ

J→LSupport for Leucine based on side chain ions (1 for L 0 for I) (Position: 2)

L→JEqual support for both Leucine and Isoleucine based on side chain ions (1 ions for both) (Position: 2)

J→LSupport for Leucine based on side chain ions (1 for L 0 for I) (Position: 2)

L→JEqual support for both Leucine and Isoleucine based on side chain ions (1 ions for both) (Position: 2)

J→LSupport for Leucine based on side chain ions (1 for L 0 for I) (Position: 2)

L→JNo support for either Leucine or Isoleucine based on side chain ions (Position: 15)

L→JEqual support for both Leucine and Isoleucine based on side chain ions (1 ions for both) (Position: 2)

#### Positional Score

Copy Data

##### Positional Score (TSV)

###### Preview

```
Loading example...
```

*Click on the button to copy the data to your clipboard.*

1001234567891011121314

Label Value
"0" 0.715
"1" 0.699
"2" 0.711
"3" 0.711
"4" 0.721
"5" 0.743
"6" 0.749
"7" 0.749
"8" 0.746
"9" 0.749
"10" 0.745
"11" 0.725
"12" 0.685
"13" 0.729
"14" 0.75

#### Meta Information from PEAKS

##### Scan Identifier

F3:4527

##### Original sequence

T

L

P

P

S

R

E

E

M

+15.99

T

K

N

Q

K

+58.01

L

##### Posttranslational Modifications

Oxidation (M); Carboxymethyl (KW X@N-term)

##### Source File

D:\separate\_stitch\_analyses\xle-disambiguation\raw\20210323\_F1\_UM1\_Peng0013\_SA\_F59\_ingel\_3ug\_chymo.raw

##### Fraction

3

##### Scan Feature

F3:8858

##### De Novo Score

98

##### ConfidenceScore

98

### m/z

615.9847

##### Mass

1844.9302

##### Charge

3

##### Retention Time

23.56

##### Predicted Retention Time

-

##### Area

3.122E+08

##### Parts Per Million

1.2

##### Fragmentation mode

ETHCD

##### Originating file

01 D:\separate\_stitch\_analyses\xle-disambiguation\20210325\_F59\_3ug\_DENOVO\_12.csv

#### Meta Information from PEAKS

##### Scan Identifier

F3:4335

##### Original sequence

T

L

P

P

S

R

E

E

M

+15.99

T

K

N

Q

K

+58.01

L

##### Posttranslational Modifications

Oxidation (M); Carboxymethyl (KW X@N-term)

##### Source File

D:\separate\_stitch\_analyses\xle-disambiguation\raw\20210323\_F1\_UM1\_Peng0013\_SA\_F59\_ingel\_3ug\_chymo.raw

##### Fraction

3

##### Scan Feature

F3:8858

##### De Novo Score

98

##### ConfidenceScore

98

### m/z

615.9847

##### Mass

1844.9302

##### Charge

3

##### Retention Time

23.56

##### Predicted Retention Time

-

##### Area

3.122E+08

##### Parts Per Million

1.2

##### Fragmentation mode

ETHCD

##### Originating file

01 D:\separate\_stitch\_analyses\xle-disambiguation\20210325\_F59\_3ug\_DENOVO\_12.csv

#### Meta Information from PEAKS

##### Scan Identifier

F3:4114

##### Original sequence

T

L

P

P

S

R

E

E

M

+15.99

T

K

N

Q

K

+58.01

L

##### Posttranslational Modifications

Oxidation (M); Carboxymethyl (KW X@N-term)

##### Source File

D:\separate\_stitch\_analyses\xle-disambiguation\raw\20210323\_F1\_UM1\_Peng0013\_SA\_F59\_ingel\_3ug\_chymo.raw

##### Fraction

3

##### Scan Feature

F3:8858

##### De Novo Score

98

##### ConfidenceScore

98

### m/z

615.9847

##### Mass

1844.9302

##### Charge

3

##### Retention Time

23.56

##### Predicted Retention Time

-

##### Area

3.122E+08

##### Parts Per Million

1.2

##### Fragmentation mode

ETHCD

##### Originating file

01 D:\separate\_stitch\_analyses\xle-disambiguation\20210325\_F59\_3ug\_DENOVO\_12.csv

#### Meta Information from PEAKS

##### Scan Identifier

F3:4656

##### Original sequence

T

L

P

P

S

R

E

E

M

+15.99

T

K

N

Q

K

+58.01

L

##### Posttranslational Modifications

Oxidation (M); Carboxymethyl (KW X@N-term)

##### Source File

D:\separate\_stitch\_analyses\xle-disambiguation\raw\20210323\_F1\_UM1\_Peng0013\_SA\_F59\_ingel\_3ug\_chymo.raw

##### Fraction

3

##### Scan Feature

F3:8858

##### De Novo Score

97

##### ConfidenceScore

97

### m/z

615.9847

##### Mass

1844.9302

##### Charge

3

##### Retention Time

23.56

##### Predicted Retention Time

-

##### Area

3.122E+08

##### Parts Per Million

1.2

##### Fragmentation mode

ETHCD

##### Originating file

01 D:\separate\_stitch\_analyses\xle-disambiguation\20210325\_F59\_3ug\_DENOVO\_12.csv

#### Meta Information from PEAKS

##### Scan Identifier

F3:4786

##### Original sequence

T

L

P

P

S

R

E

E

M

+15.99

T

K

N

Q

K

+58.01

L

##### Posttranslational Modifications

Oxidation (M); Carboxymethyl (KW X@N-term)

##### Source File

D:\separate\_stitch\_analyses\xle-disambiguation\raw\20210323\_F1\_UM1\_Peng0013\_SA\_F59\_ingel\_3ug\_chymo.raw

##### Fraction

3

##### Scan Feature

F3:8858

##### De Novo Score

97

##### ConfidenceScore

97

### m/z

615.9847

##### Mass

1844.9302

##### Charge

3

##### Retention Time

23.56

##### Predicted Retention Time

-

##### Area

3.122E+08

##### Parts Per Million

1.2

##### Fragmentation mode

ETHCD

##### Originating file

01 D:\separate\_stitch\_analyses\xle-disambiguation\20210325\_F59\_3ug\_DENOVO\_12.csv

#### Meta Information from PEAKS

##### Scan Identifier

F3:4405

##### Original sequence

T

L

P

P

S

R

E

E

M

+15.99

T

K

N

Q

K

+58.01

L

##### Posttranslational Modifications

Oxidation (M); Carboxymethyl (KW X@N-term)

##### Source File

D:\separate\_stitch\_analyses\xle-disambiguation\raw\20210323\_F1\_UM1\_Peng0013\_SA\_F59\_ingel\_3ug\_chymo.raw

##### Fraction

3

##### Scan Feature

F3:8858

##### De Novo Score

97

##### ConfidenceScore

97

### m/z

615.9847

##### Mass

1844.9302

##### Charge

3

##### Retention Time

23.56

##### Predicted Retention Time

-

##### Area

3.122E+08

##### Parts Per Million

1.2

##### Fragmentation mode

ETHCD

##### Originating file

01 D:\separate\_stitch\_analyses\xle-disambiguation\20210325\_F59\_3ug\_DENOVO\_12.csv

#### Meta Information from PEAKS

##### Scan Identifier

F3:5470

##### Original sequence

T

L

P

P

S

R

E

E

M

T

K

N

Q

K

+58.01

L

##### Posttranslational Modifications

Carboxymethyl (KW X@N-term)

##### Source File

D:\separate\_stitch\_analyses\xle-disambiguation\raw\20210323\_F1\_UM1\_Peng0013\_SA\_F59\_ingel\_3ug\_chymo.raw

##### Fraction

3

##### Scan Feature

F3:8647

##### De Novo Score

95

##### ConfidenceScore

95

### m/z

610.6514

##### Mass

1828.9353

##### Charge

3

##### Retention Time

29.73

##### Predicted Retention Time

-

##### Area

2.399E+07

##### Fragmentation mode

ETHCD

##### Originating file

01 D:\separate\_stitch\_analyses\xle-disambiguation\20210325\_F59\_3ug\_DENOVO\_12.csv

#### Meta Information from PEAKS

##### Scan Identifier

F3:4991

##### Original sequence

T

L

P

P

S

R

E

E

M

+15.99

T

K

N

Q

K

+58.01

L

##### Posttranslational Modifications

Oxidation (M); Carboxymethyl (KW X@N-term)

##### Source File

D:\separate\_stitch\_analyses\xle-disambiguation\raw\20210323\_F1\_UM1\_Peng0013\_SA\_F59\_ingel\_3ug\_chymo.raw

##### Fraction

3

##### Scan Feature

F3:8858

##### De Novo Score

95

##### ConfidenceScore

95

### m/z

615.9847

##### Mass

1844.9302

##### Charge

3

##### Retention Time

23.56

##### Predicted Retention Time

-

##### Area

3.122E+08

##### Parts Per Million

1.2

##### Fragmentation mode

ETHCD

##### Originating file

01 D:\separate\_stitch\_analyses\xle-disambiguation\20210325\_F59\_3ug\_DENOVO\_12.csv
