## Supplementary material for "A handle on mass coincidence errors in *de novo* sequencing of antibodies by bottom-up proteomics": Combined_041.html

Details Combined\_041 | Stitch OverviewUndefined

### Read Combined\_041

#### Sequence (length=12)

SYLSJTPEQWKS

#### Spectrum 8765? Spectrum 8765 The raw spectrum of this peptide as annotated by Hecklib. The fragments are coloured according to ion type (see legend). Any peaks with a star '\*' as text can be hovered over to see the full details, first the ion type second the mass shift type. By hovering over the amino acids in the peptide or ions in the legend the corresponding peaks are highlighted. By toggling the 'Unassigned' label you can turn the background (unassigned) peaks on or off in the plot. By updating the slider in the Ion legend you can update the spectrum to only show the top X% of the peaks with labels. The top X% means any peak that is within X% of the highest intensity. By dragging in the spectrum you can zoom in to a specific part of the spectrum and use 'Zoom Out' to get back to the original zoom level. The annotation of the spectrum is based on the given sequence in the peptides file and is done with different software so inconsistencies are likely. The peaks are annotated based on the given sequence, with 20 ppm tolerance.

Copy Data

##### Spectrum 8765 (TSV)

###### Preview

```
Loading example...
```

*Click on the button to copy the data to your clipboard.*

Mz MinMz MaxIntensity Max

WidthHeightPeptide font sizePeptide stroke widthSpectrum font sizeSpectrum stroke widthCompact peptide

Ion legend

wxyz

abcd

OtherUnassignedIon^Charge^_Position_Show for top:%

SYLSJTPEQWKS

08.81e+51.76e+62.64e+63.52e+6

Zoom Out

y^+1^_2_y^+1^_2_y^+2^_6_y^+1^_3_y^+2^_7_y^+2^_8_y^+2^_9_y^+1^_4_y^+1^_4_z^+1^_4_y^+2^_9_y^+1^_4_y^+2^_10_y^+2^_10_z^+2^_10_y^+2^_10_w^+1^_5_z^+1^_5_c^+2^_11_y^+2^_11_y^+1^_5_y^+1^_6_y^+1^_6_y^+1^_6_w^+1^_7_y^+1^_7_z^+1^_7_y^+1^_7_c^+1^_8_w^+1^_8_y^+1^_8_y^+1^_8_z^+1^_8_y^+1^_8_c^+1^_9_c^+1^_9_c^+1^_9_y^+1^_9_y^+1^_9_z^+1^_9_y^+1^_9_w^+1^_10_z^+1^_10_z^+1^_10_y^+1^_10_y^+1^_10_z^+1^_10_y^+1^_10_c^+1^_10_c^+1^_10_c^+1^_10_z^+1^_11_c^+1^_11_y^+1^_11_

0560112116812242

Fragment Matches Table

Show background peaks

| Position | Ion type | Intensity | mz Theoretical | mz Error (Th) | mz Error (ppm) | Charge | Series Number |
| --- | --- | --- | --- | --- | --- | --- | --- |
| - | - | 2378 | 120.5 | - | - | 0 | - |
| - | - | 3053 | 122 | - | - | 0 | - |
| - | - | 2555 | 122.6 | - | - | 0 | - |
| - | - | 2682 | 127.5 | - | - | 0 | - |
| - | - | 3053 | 128.2 | - | - | 0 | - |
| - | - | 5.206E+04 | 129.1 | - | - | 0 | - |
| - | - | 4057 | 130.1 | - | - | 0 | - |
| - | - | 2765 | 131.1 | - | - | 0 | - |
| - | - | 3519 | 132.5 | - | - | 0 | - |
| - | - | 2617 | 135 | - | - | 0 | - |
| - | - | 3344 | 159.1 | - | - | 0 | - |
| - | - | 1.095E+04 | 173.1 | - | - | 0 | - |
| - | - | 1.295E+04 | 173.4 | - | - | 0 | - |
| - | - | 3810 | 194.8 | - | - | 0 | - |
| - | - | 3433 | 195.7 | - | - | 0 | - |
| - | - | 4.733E+04 | 201.1 | - | - | 0 | - |
| - | - | 7226 | 215.1 | - | - | 0 | - |
| 11 | y | 4857 | 217.1 | 0.001239 | 5.705 | +1 | 2 |
| - | - | 3.262E+05 | 223.1 | - | - | 0 | - |
| - | - | 4.322E+04 | 224.1 | - | - | 0 | - |
| - | - | 4265 | 232.9 | - | - | 0 | - |
| 11 | y | 1.795E+05 | 234.1 | 0.0003243 | 1.385 | +1 | 2 |
| - | - | 2.118E+04 | 235.1 | - | - | 0 | - |
| - | - | 7374 | 249.2 | - | - | 0 | - |
| - | - | 6.415E+05 | 251.1 | - | - | 0 | - |
| - | - | 7.448E+04 | 252.1 | - | - | 0 | - |
| - | - | 5461 | 253.1 | - | - | 0 | - |
| - | - | 3647 | 258.9 | - | - | 0 | - |
| - | - | 3973 | 266.6 | - | - | 0 | - |
| - | - | 4836 | 270.1 | - | - | 0 | - |
| - | - | 1.044E+04 | 277.2 | - | - | 0 | - |
| - | - | 8209 | 284.2 | - | - | 0 | - |
| - | - | 5384 | 288.2 | - | - | 0 | - |
| - | - | 2.781E+04 | 296.2 | - | - | 0 | - |
| - | - | 9689 | 298.2 | - | - | 0 | - |
| - | - | 2.147E+04 | 302.2 | - | - | 0 | - |
| - | - | 3680 | 304.8 | - | - | 0 | - |
| - | - | 9934 | 314.2 | - | - | 0 | - |
| - | - | 1.911E+04 | 315.2 | - | - | 0 | - |
| - | - | 4792 | 316.2 | - | - | 0 | - |
| - | - | 3.297E+04 | 319.2 | - | - | 0 | - |
| - | - | 4958 | 320.1 | - | - | 0 | - |
| - | - | 6460 | 320.2 | - | - | 0 | - |
| - | - | 2.154E+04 | 346.2 | - | - | 0 | - |
| - | - | 5449 | 353.2 | - | - | 0 | - |
| - | - | 3.116E+04 | 355.2 | - | - | 0 | - |
| - | - | 2.616E+05 | 364.2 | - | - | 0 | - |
| - | - | 4.477E+04 | 365.2 | - | - | 0 | - |
| - | - | 6165 | 365.2 | - | - | 0 | - |
| - | - | 6295 | 370.2 | - | - | 0 | - |
| - | - | 4754 | 371.2 | - | - | 0 | - |
| - | - | 7150 | 372.2 | - | - | 0 | - |
| - | - | 6365 | 387.3 | - | - | 0 | - |
| 7 | y | 1.032E+04 | 387.7 | 0.0003149 | 0.8124 | +2 | 6 |
| - | - | 7494 | 388.2 | - | - | 0 | - |
| - | - | 7375 | 392.2 | - | - | 0 | - |
| - | - | 1.74E+04 | 397.2 | - | - | 0 | - |
| - | - | 4665 | 398.2 | - | - | 0 | - |
| - | - | 1.223E+04 | 405.2 | - | - | 0 | - |
| - | - | 1.278E+04 | 406.2 | - | - | 0 | - |
| - | - | 5.381E+04 | 415.2 | - | - | 0 | - |
| - | - | 4.435E+04 | 415.3 | - | - | 0 | - |
| - | - | 1.101E+04 | 416.2 | - | - | 0 | - |
| - | - | 1.002E+04 | 416.3 | - | - | 0 | - |
| 10 | y | 8.756E+04 | 420.2 | 0.0003418 | 0.8133 | +1 | 3 |
| - | - | 1.475E+04 | 421.2 | - | - | 0 | - |
| - | - | 1.077E+04 | 423.2 | - | - | 0 | - |
| - | - | 1.534E+04 | 432.3 | - | - | 0 | - |
| - | - | 1.978E+05 | 433.2 | - | - | 0 | - |
| - | - | 4.763E+04 | 434.2 | - | - | 0 | - |
| - | - | 6176 | 437.2 | - | - | 0 | - |
| 6 | y | 7141 | 438.2 | 0.001256 | 2.866 | +2 | 7 |
| - | - | 4798 | 443.2 | - | - | 0 | - |
| - | - | 5627 | 447.2 | - | - | 0 | - |
| - | - | 1.342E+05 | 451.2 | - | - | 0 | - |
| - | - | 2.932E+04 | 452.2 | - | - | 0 | - |
| - | - | 1.399E+04 | 456.2 | - | - | 0 | - |
| - | - | 5209 | 465.2 | - | - | 0 | - |
| 5 | y | 9883 | 494.8 | 0.0005844 | 1.181 | +2 | 8 |
| - | - | 1.081E+04 | 495.3 | - | - | 0 | - |
| - | - | 5010 | 500.3 | - | - | 0 | - |
| - | - | 8070 | 501.3 | - | - | 0 | - |
| - | - | 7203 | 513.2 | - | - | 0 | - |
| - | - | 5.199E+04 | 516.2 | - | - | 0 | - |
| - | - | 1.774E+04 | 517.2 | - | - | 0 | - |
| - | - | 3.167E+04 | 518.3 | - | - | 0 | - |
| - | - | 4588 | 519.3 | - | - | 0 | - |
| - | - | 4.23E+04 | 528.3 | - | - | 0 | - |
| 4 | y | 1.954E+04 | 529.3 | 0.006292 | 11.89 | +2 | 9 |
| - | - | 4389 | 529.8 | - | - | 0 | - |
| 9 | y | 1.441E+04 | 530.3 | 0.002744 | 5.175 | +1 | 4 |
| 9 | y | 8126 | 531.3 | 0.002127 | 4.003 | +1 | 4 |
| 9 | z | 1.343E+05 | 532.3 | 0.0005884 | 1.106 | +1 | 4 |
| - | - | 2.252E+05 | 533.3 | - | - | 0 | - |
| - | - | 4.891E+04 | 534.3 | - | - | 0 | - |
| - | - | 1.548E+04 | 535.3 | - | - | 0 | - |
| - | - | 3.926E+04 | 536.3 | - | - | 0 | - |
| - | - | 6321 | 537.3 | - | - | 0 | - |
| 4 | y | 7656 | 538.3 | 0.001193 | 2.217 | +2 | 9 |
| - | - | 3.102E+04 | 541.2 | - | - | 0 | - |
| - | - | 1.323E+04 | 542.2 | - | - | 0 | - |
| - | - | 6188 | 542.3 | - | - | 0 | - |
| - | - | 1.384E+05 | 546.3 | - | - | 0 | - |
| - | - | 5.144E+04 | 547.3 | - | - | 0 | - |
| 9 | y | 4.646E+04 | 548.3 | 0.001823 | 3.325 | +1 | 4 |
| - | - | 1.356E+04 | 549.3 | - | - | 0 | - |
| - | - | 2.146E+04 | 552.3 | - | - | 0 | - |
| - | - | 6623 | 553.3 | - | - | 0 | - |
| - | - | 5.75E+04 | 564.3 | - | - | 0 | - |
| - | - | 1.944E+04 | 565.3 | - | - | 0 | - |
| 3 | y | 4.136E+04 | 585.8 | 0.0008205 | 1.401 | +2 | 10 |
| 3 | y | 3.077E+04 | 586.3 | 0.009362 | 15.97 | +2 | 10 |
| 3 | z | 1.217E+04 | 586.8 | 0.00667 | 11.37 | +2 | 10 |
| 3 | y | 9.762E+04 | 594.8 | 0.0007261 | 1.221 | +2 | 10 |
| - | - | 6.313E+04 | 595.3 | - | - | 0 | - |
| - | - | 1.98E+04 | 595.8 | - | - | 0 | - |
| - | - | 4300 | 596.3 | - | - | 0 | - |
| - | - | 4629 | 601.3 | - | - | 0 | - |
| 8 | w | 1.232E+04 | 602.3 | 0.002672 | 4.436 | +1 | 5 |
| - | - | 5026 | 603.3 | - | - | 0 | - |
| - | - | 6814 | 611.3 | - | - | 0 | - |
| - | - | 7808 | 619.3 | - | - | 0 | - |
| - | - | 1.319E+04 | 620.3 | - | - | 0 | - |
| - | - | 4.596E+04 | 629.3 | - | - | 0 | - |
| - | - | 1.643E+04 | 630.3 | - | - | 0 | - |
| - | - | 2.267E+04 | 637.4 | - | - | 0 | - |
| - | - | 7981 | 638.4 | - | - | 0 | - |
| - | - | 2.412E+04 | 642.3 | - | - | 0 | - |
| - | - | 1.109E+05 | 647.3 | - | - | 0 | - |
| - | - | 4.119E+04 | 648.3 | - | - | 0 | - |
| - | - | 8689 | 649.3 | - | - | 0 | - |
| - | - | 5806 | 656.3 | - | - | 0 | - |
| - | - | 4497 | 657.3 | - | - | 0 | - |
| - | - | 1.179E+04 | 658.3 | - | - | 0 | - |
| - | - | 3485 | 661.3 | - | - | 0 | - |
| 8 | z | 1.722E+05 | 661.3 | 0.000781 | 1.181 | +1 | 5 |
| - | - | 2.362E+05 | 662.3 | - | - | 0 | - |
| - | - | 7.996E+04 | 663.3 | - | - | 0 | - |
| - | - | 1.153E+04 | 664.3 | - | - | 0 | - |
| - | - | 1.153E+05 | 665.4 | - | - | 0 | - |
| - | - | 3.702E+04 | 666.4 | - | - | 0 | - |
| 11 | c | 1.49E+04 | 667.3 | 0.007364 | 11.04 | +2 | 11 |
| - | - | 9.372E+04 | 669.3 | - | - | 0 | - |
| - | - | 2.872E+04 | 670.3 | - | - | 0 | - |
| - | - | 6276 | 671.3 | - | - | 0 | - |
| 2 | y | 7210 | 676.3 | 0.0007868 | 1.163 | +2 | 11 |
| 8 | y | 3.124E+04 | 677.3 | 0.0002454 | 0.3623 | +1 | 5 |
| - | - | 6920 | 678.3 | - | - | 0 | - |
| - | - | 8534 | 701.9 | - | - | 0 | - |
| - | - | 1.162E+04 | 702.4 | - | - | 0 | - |
| - | - | 1.099E+05 | 710.9 | - | - | 0 | - |
| - | - | 1.135E+05 | 711.4 | - | - | 0 | - |
| - | - | 4.538E+04 | 711.9 | - | - | 0 | - |
| - | - | 6097 | 712.4 | - | - | 0 | - |
| - | - | 2.585E+04 | 719.9 | - | - | 0 | - |
| - | - | 3.397E+04 | 720.4 | - | - | 0 | - |
| - | - | 1.067E+04 | 720.9 | - | - | 0 | - |
| - | - | 7742 | 755.4 | - | - | 0 | - |
| 7 | y | 1.166E+04 | 756.4 | 0.001686 | 2.23 | +1 | 6 |
| 7 | y | 8178 | 757.4 | 0.004365 | 5.764 | +1 | 6 |
| - | - | 5.1E+04 | 770.4 | - | - | 0 | - |
| - | - | 1.726E+04 | 771.4 | - | - | 0 | - |
| - | - | 5211 | 772.4 | - | - | 0 | - |
| - | - | 4.432E+04 | 773.4 | - | - | 0 | - |
| 7 | y | 8.899E+05 | 774.4 | 0.000277 | 0.3576 | +1 | 6 |
| - | - | 3.472E+05 | 775.4 | - | - | 0 | - |
| - | - | 8.002E+04 | 776.4 | - | - | 0 | - |
| - | - | 1.073E+04 | 801.4 | - | - | 0 | - |
| - | - | 3.368E+04 | 802.4 | - | - | 0 | - |
| - | - | 1.174E+04 | 803.4 | - | - | 0 | - |
| - | - | 6241 | 804.4 | - | - | 0 | - |
| - | - | 4796 | 815.4 | - | - | 0 | - |
| - | - | 9334 | 824.4 | - | - | 0 | - |
| - | - | 1.004E+04 | 831.4 | - | - | 0 | - |
| - | - | 6533 | 832.4 | - | - | 0 | - |
| 6 | w | 1.183E+04 | 842.4 | 0.003237 | 3.843 | +1 | 7 |
| - | - | 6183 | 843.4 | - | - | 0 | - |
| - | - | 6.294E+04 | 857.4 | - | - | 0 | - |
| 6 | y | 4.105E+05 | 858.4 | 0.0002657 | 0.3096 | +1 | 7 |
| 6 | z | 2.545E+05 | 859.4 | 0.003531 | 4.109 | +1 | 7 |
| - | - | 6.649E+05 | 860.4 | - | - | 0 | - |
| - | - | 2.778E+05 | 861.4 | - | - | 0 | - |
| - | - | 6.626E+04 | 862.4 | - | - | 0 | - |
| - | - | 5137 | 863.4 | - | - | 0 | - |
| - | - | 1.275E+04 | 874.4 | - | - | 0 | - |
| 6 | y | 6.84E+05 | 875.4 | 0.0006942 | 0.793 | +1 | 7 |
| - | - | 3.214E+05 | 876.4 | - | - | 0 | - |
| - | - | 7.517E+04 | 877.4 | - | - | 0 | - |
| - | - | 9675 | 878.4 | - | - | 0 | - |
| - | - | 4264 | 883.5 | - | - | 0 | - |
| - | - | 6624 | 891.4 | - | - | 0 | - |
| - | - | 2.558E+04 | 907.5 | - | - | 0 | - |
| 8 | c | 6.59E+04 | 908.5 | 0.0002674 | 0.2943 | +1 | 8 |
| - | - | 2.789E+04 | 909.5 | - | - | 0 | - |
| - | - | 6060 | 910.5 | - | - | 0 | - |
| - | - | 4567 | 928.5 | - | - | 0 | - |
| 5 | w | 2.671E+04 | 929.4 | 0.002214 | 2.383 | +1 | 8 |
| - | - | 1.527E+04 | 930.4 | - | - | 0 | - |
| - | - | 1.041E+04 | 934.5 | - | - | 0 | - |
| - | - | 1.003E+04 | 955.5 | - | - | 0 | - |
| 5 | y | 2.671E+04 | 970.5 | 0.001028 | 1.059 | +1 | 8 |
| 5 | y | 1.449E+04 | 971.5 | 0.0166 | 17.09 | +1 | 8 |
| 5 | z | 7.623E+05 | 972.5 | 0.0002346 | 0.2413 | +1 | 8 |
| - | - | 5.482E+05 | 973.5 | - | - | 0 | - |
| - | - | 1.928E+05 | 974.5 | - | - | 0 | - |
| - | - | 3.242E+04 | 975.5 | - | - | 0 | - |
| - | - | 5.266E+04 | 987.5 | - | - | 0 | - |
| 5 | y | 2.89E+05 | 988.5 | 0.003353 | 3.392 | +1 | 8 |
| - | - | 1.503E+05 | 989.5 | - | - | 0 | - |
| - | - | 3.969E+04 | 990.5 | - | - | 0 | - |
| - | - | 9.821E+04 | 992.5 | - | - | 0 | - |
| - | - | 9.085E+04 | 993.5 | - | - | 0 | - |
| - | - | 3.188E+04 | 994.5 | - | - | 0 | - |
| - | - | 7773 | 995.5 | - | - | 0 | - |
| - | - | 1.785E+04 | 1001 | - | - | 0 | - |
| - | - | 1.153E+04 | 1002 | - | - | 0 | - |
| - | - | 2.928E+04 | 1003 | - | - | 0 | - |
| - | - | 1.344E+04 | 1004 | - | - | 0 | - |
| - | - | 1.158E+04 | 1005 | - | - | 0 | - |
| - | - | 9526 | 1016 | - | - | 0 | - |
| - | - | 1.224E+04 | 1018 | - | - | 0 | - |
| 9 | c | 1.671E+04 | 1019 | 0.01447 | 14.2 | +1 | 9 |
| 9 | c | 2.437E+04 | 1020 | 0.004081 | 4.003 | +1 | 9 |
| - | - | 1.156E+04 | 1021 | - | - | 0 | - |
| - | - | 3.553E+04 | 1036 | - | - | 0 | - |
| 9 | c | 1.345E+05 | 1037 | 0.0002982 | 0.2877 | +1 | 9 |
| - | - | 7.131E+04 | 1038 | - | - | 0 | - |
| - | - | 1.906E+04 | 1039 | - | - | 0 | - |
| 4 | y | 4.856E+04 | 1058 | 0.00614 | 5.806 | +1 | 9 |
| 4 | y | 4.749E+04 | 1059 | 0.009967 | 9.416 | +1 | 9 |
| 4 | z | 2.945E+05 | 1060 | 0.0006769 | 0.6389 | +1 | 9 |
| - | - | 7.548E+05 | 1061 | - | - | 0 | - |
| - | - | 3.957E+05 | 1062 | - | - | 0 | - |
| - | - | 1.172E+05 | 1063 | - | - | 0 | - |
| - | - | 1.933E+04 | 1064 | - | - | 0 | - |
| - | - | 1.976E+04 | 1070 | - | - | 0 | - |
| - | - | 1.11E+04 | 1071 | - | - | 0 | - |
| - | - | 6307 | 1074 | - | - | 0 | - |
| - | - | 5.467E+04 | 1075 | - | - | 0 | - |
| 4 | y | 7.789E+05 | 1076 | 0.0001413 | 0.1314 | +1 | 9 |
| - | - | 4.935E+05 | 1077 | - | - | 0 | - |
| - | - | 1.461E+05 | 1078 | - | - | 0 | - |
| - | - | 5717 | 1078 | - | - | 0 | - |
| - | - | 1.627E+04 | 1079 | - | - | 0 | - |
| - | - | 6521 | 1084 | - | - | 0 | - |
| - | - | 8205 | 1101 | - | - | 0 | - |
| - | - | 1.642E+04 | 1102 | - | - | 0 | - |
| - | - | 1.06E+04 | 1103 | - | - | 0 | - |
| - | - | 7126 | 1108 | - | - | 0 | - |
| - | - | 7193 | 1111 | - | - | 0 | - |
| - | - | 1.059E+04 | 1114 | - | - | 0 | - |
| - | - | 5580 | 1115 | - | - | 0 | - |
| 3 | w | 1.1E+05 | 1130 | 0.0008071 | 0.7146 | +1 | 10 |
| - | - | 7.208E+04 | 1131 | - | - | 0 | - |
| - | - | 2.109E+04 | 1132 | - | - | 0 | - |
| 3 | z | 6841 | 1155 | 0.01006 | 8.716 | +1 | 10 |
| 3 | z | 1.191E+04 | 1156 | 0.02043 | 17.68 | +1 | 10 |
| - | - | 3.11E+04 | 1161 | - | - | 0 | - |
| - | - | 2.986E+04 | 1162 | - | - | 0 | - |
| - | - | 1.885E+04 | 1163 | - | - | 0 | - |
| - | - | 5539 | 1170 | - | - | 0 | - |
| 3 | y | 2.119E+04 | 1171 | 0.004754 | 4.061 | +1 | 10 |
| 3 | y | 1.263E+04 | 1172 | 0.01672 | 14.27 | +1 | 10 |
| 3 | z | 2.627E+05 | 1173 | 0.0002311 | 0.1971 | +1 | 10 |
| - | - | 1.64E+05 | 1174 | - | - | 0 | - |
| - | - | 6.17E+04 | 1175 | - | - | 0 | - |
| - | - | 1.038E+04 | 1176 | - | - | 0 | - |
| - | - | 6569 | 1178 | - | - | 0 | - |
| - | - | 2.28E+05 | 1179 | - | - | 0 | - |
| - | - | 1.952E+05 | 1180 | - | - | 0 | - |
| - | - | 7.282E+04 | 1181 | - | - | 0 | - |
| - | - | 7328 | 1182 | - | - | 0 | - |
| - | - | 4.214E+04 | 1188 | - | - | 0 | - |
| 3 | y | 2.75E+05 | 1189 | 0.002013 | 1.694 | +1 | 10 |
| - | - | 1.836E+05 | 1190 | - | - | 0 | - |
| - | - | 5.911E+04 | 1191 | - | - | 0 | - |
| - | - | 9723 | 1192 | - | - | 0 | - |
| - | - | 1.447E+04 | 1204 | - | - | 0 | - |
| 10 | c | 2.583E+04 | 1205 | 0.01376 | 11.43 | +1 | 10 |
| 10 | c | 6.994E+04 | 1206 | 0.0002687 | 0.2229 | +1 | 10 |
| - | - | 4.442E+04 | 1207 | - | - | 0 | - |
| - | - | 1.235E+04 | 1208 | - | - | 0 | - |
| - | - | 1.068E+05 | 1222 | - | - | 0 | - |
| 10 | c | 8.896E+05 | 1223 | 0.0007677 | 0.6279 | +1 | 10 |
| - | - | 6.1E+05 | 1224 | - | - | 0 | - |
| - | - | 2.103E+05 | 1225 | - | - | 0 | - |
| - | - | 3.239E+04 | 1226 | - | - | 0 | - |
| - | - | 9054 | 1280 | - | - | 0 | - |
| - | - | 5499 | 1281 | - | - | 0 | - |
| - | - | 6571 | 1293 | - | - | 0 | - |
| - | - | 2.2E+04 | 1307 | - | - | 0 | - |
| - | - | 1.878E+04 | 1308 | - | - | 0 | - |
| - | - | 8490 | 1309 | - | - | 0 | - |
| - | - | 8044 | 1323 | - | - | 0 | - |
| 2 | z | 1.17E+05 | 1336 | 0.004407 | 3.3 | +1 | 11 |
| - | - | 9.291E+04 | 1337 | - | - | 0 | - |
| - | - | 4.062E+04 | 1338 | - | - | 0 | - |
| - | - | 1.055E+04 | 1339 | - | - | 0 | - |
| - | - | 9663 | 1349 | - | - | 0 | - |
| - | - | 7924 | 1350 | - | - | 0 | - |
| 11 | c | 7.357E+05 | 1351 | 0.001248 | 0.9242 | +1 | 11 |
| 2 | y | 6.111E+05 | 1352 | 0.01645 | 12.17 | +1 | 11 |
| - | - | 2.738E+05 | 1353 | - | - | 0 | - |
| - | - | 5.07E+04 | 1354 | - | - | 0 | - |
| - | - | 1.277E+04 | 1355 | - | - | 0 | - |
| - | - | 5638 | 1361 | - | - | 0 | - |
| - | - | 4.856E+04 | 1367 | - | - | 0 | - |
| - | - | 5.216E+04 | 1368 | - | - | 0 | - |
| - | - | 4.839E+04 | 1369 | - | - | 0 | - |
| - | - | 2.261E+04 | 1370 | - | - | 0 | - |
| - | - | 8262 | 1371 | - | - | 0 | - |
| - | - | 1.6E+04 | 1379 | - | - | 0 | - |
| - | - | 2.169E+04 | 1380 | - | - | 0 | - |
| - | - | 3.04E+04 | 1381 | - | - | 0 | - |
| - | - | 2.919E+04 | 1382 | - | - | 0 | - |
| - | - | 1.553E+04 | 1383 | - | - | 0 | - |
| - | - | 4.83E+04 | 1384 | - | - | 0 | - |
| - | - | 4.086E+04 | 1385 | - | - | 0 | - |
| - | - | 1.408E+04 | 1386 | - | - | 0 | - |
| - | - | 8565 | 1394 | - | - | 0 | - |
| - | - | 4.695E+04 | 1395 | - | - | 0 | - |
| - | - | 4.089E+04 | 1396 | - | - | 0 | - |
| - | - | 2.562E+04 | 1397 | - | - | 0 | - |
| - | - | 9688 | 1398 | - | - | 0 | - |
| - | - | 5606 | 1407 | - | - | 0 | - |
| - | - | 6746 | 1411 | - | - | 0 | - |
| - | - | 9.788E+04 | 1412 | - | - | 0 | - |
| - | - | 8.222E+04 | 1413 | - | - | 0 | - |
| - | - | 3.246E+04 | 1414 | - | - | 0 | - |
| - | - | 6933 | 1415 | - | - | 0 | - |
| - | - | 1.502E+04 | 1421 | - | - | 0 | - |
| - | - | 9.273E+04 | 1422 | - | - | 0 | - |
| - | - | 1.273E+06 | 1423 | - | - | 0 | - |
| - | - | 1.042E+06 | 1424 | - | - | 0 | - |
| - | - | 4.453E+05 | 1425 | - | - | 0 | - |
| - | - | 7.099E+04 | 1426 | - | - | 0 | - |
| - | - | 4.211E+04 | 1438 | - | - | 0 | - |
| - | - | 1.183E+06 | 1439 | - | - | 0 | - |
| - | - | 3.489E+06 | 1440 | - | - | 0 | - |
| - | - | 2.475E+06 | 1441 | - | - | 0 | - |
| - | - | 9.713E+05 | 1442 | - | - | 0 | - |
| - | - | 1.445E+05 | 1443 | - | - | 0 | - |
| - | - | 7435 | 1455 | - | - | 0 | - |
| - | - | 8814 | 1472 | - | - | 0 | - |
| - | - | 4992 | 2167 | - | - | 0 | - |
| - | - | 4656 | 2220 | - | - | 0 | - |

m/z Charge Intensity FragmentType MassShift Position
120.52883911132812 0 2377.8816
121.99918365478516 0 3052.8147
122.5577392578125 0 2555
127.48995208740234 0 2682.3352
128.244384765625 0 3053.3435
129.10247802734375 0 52055.13
130.10598754882812 0 4057.2397
131.1025390625 0 2764.9456
132.45458984375 0 3519.1038
134.97653198242188 0 2617.2212
159.0917205810547 0 3344.2744
173.1284637451172 0 10946.288
173.43898010253906 0 12952.119
194.77151489257812 0 3810.1243
195.70477294921875 0 3433.08
201.12355041503906 0 47327.14
215.13890075683594 0 7225.638
217.11952209472656 0 4857.1426 y Ammonia loss 10
223.1079864501953 0 326159
224.11129760742188 0 43223.707
232.936279296875 0 4264.77
234.14515686035156 0 179457.62 y 10
235.14840698242188 0 21180.238
249.16099548339844 0 7373.9077
251.1029510498047 0 641499.2
252.106201171875 0 74484.914
253.1085662841797 0 5460.5127
258.9385681152344 0 3646.8044
266.612060546875 0 3973.4722
270.1449279785156 0 4836.3535
277.1552734375 0 10439.977
284.1605224609375 0 8208.939
288.1547546386719 0 5384.3667
296.1971435546875 0 27805.434
298.1558837890625 0 9689.1045
302.1713562011719 0 21465.707
304.80364990234375 0 3679.9006
314.2080993652344 0 9934.409
315.1822204589844 0 19112.936
316.1849060058594 0 4792.0195
319.16552734375 0 32965.637
320.12451171875 0 4958.4644
320.1686706542969 0 6459.786
346.1766052246094 0 21543.31
353.1815490722656 0 5448.634
355.1614685058594 0 31163.852
364.1869201660156 0 261569.7
365.1900634765625 0 44774.184
365.216064453125 0 6164.573
370.1775207519531 0 6295.326
371.19219970703125 0 4754.1514
372.1885681152344 0 7149.857
387.2608947753906 0 6365.151
387.6929931640625 0 10317.318 y 6
388.19305419921875 0 7493.6807
392.1817932128906 0 7374.649
397.2455749511719 0 17403.014
398.2494812011719 0 4665.2314
405.2131042480469 0 12228.204
406.1975402832031 0 12780.667
415.19805908203125 0 53809.434
415.25567626953125 0 44347.62
416.1994323730469 0 11009.068
416.26025390625 0 10024.903
420.2244873046875 0 87555.01 y 9
421.22698974609375 0 14752.543
423.2248229980469 0 10773.558
432.2507019042969 0 15342.98
433.2088928222656 0 197799.12
434.2120666503906 0 47631.88
437.24224853515625 0 6175.683
438.2177734375 0 7140.9575 y 5
443.24114990234375 0 4798.3726
447.22491455078125 0 5627.106
451.2192687988281 0 134170.06
452.22235107421875 0 29320.62
456.2104797363281 0 13992.977
465.2356262207031 0 5208.888
494.7579650878906 0 9883.35 y 4
495.260009765625 0 10806.503
500.28643798828125 0 5010.336
501.27154541015625 0 8070.286
513.2460327148438 0 7203.2295
516.2454833984375 0 51988.363
517.2477416992188 0 17737.121
518.2971801757812 0 31672.717
519.29931640625 0 4588.3945
528.2822875976562 0 42303.066
529.2755737304688 0 19542.867 y Water loss 3
529.7720947265625 0 4389.4976
530.27490234375 0 14414.962 y Water loss 8
531.25830078125 0 8126.3945 y Ammonia loss 8
532.2645874023438 0 134262.81 z 8
533.2716674804688 0 225174.94
534.2666625976562 0 48914.816
535.2628784179688 0 15484.059
536.308349609375 0 39257.85
537.3145141601562 0 6320.8594
538.2757568359375 0 7656.366 y 3
541.240966796875 0 31018.459
542.2446899414062 0 13231.93
542.2936401367188 0 6187.86
546.2926025390625 0 138438.3
547.2959594726562 0 51438.973
548.2845458984375 0 46459.19 y 8
549.2867431640625 0 13563.674
552.2666625976562 0 21460.166
553.2706909179688 0 6622.745
564.3035888671875 0 57503.902
565.3074340820312 0 19444.697
585.8121337890625 0 41362.824 y Water loss 2
586.3126831054688 0 30773.22 y Ammonia loss 2
586.8139038085938 0 12167.001 z 2
594.8173217773438 0 97621.94 y 2
595.3187255859375 0 63129.08
595.8201293945312 0 19798.818
596.3226928710938 0 4300.432
601.333984375 0 4628.882
602.2959594726562 0 12317.381 w 7
603.3206176757812 0 5026.024
611.3231201171875 0 6814.4683
619.3456420898438 0 7808.071
620.33203125 0 13186.737
629.3294677734375 0 45963.652
630.3322143554688 0 16427.346
637.3566284179688 0 22668.742
638.35986328125 0 7980.851
642.288330078125 0 24123.082
647.3403930664062 0 110946.34
648.3436889648438 0 41185.96
649.3428344726562 0 8688.91
656.3299560546875 0 5806.1636
657.3316040039062 0 4496.558
658.3385620117188 0 11793.562
661.2588500976562 0 3484.6494
661.307373046875 0 172207.52 z 7
662.3136596679688 0 236167.6
663.3171997070312 0 79955.03
664.32080078125 0 11525.29
665.3509521484375 0 115334.66
666.3538818359375 0 37015.223
667.350341796875 0 14895.087 c Ammonia loss 10
669.3363647460938 0 93723.94
670.3389892578125 0 28717.65
671.3423461914062 0 6276.3745
676.3474731445312 0 7210.372 y 1
677.3255615234375 0 31238.463 y 7
678.3263549804688 0 6919.775
701.85498046875 0 8533.894
702.3556518554688 0 11616.876
710.8591918945312 0 109919.78
711.3607177734375 0 113507.125
711.861572265625 0 45375.4
712.363037109375 0 6096.5156
719.865478515625 0 25854.227
720.3671264648438 0 33970.465
720.8682861328125 0 10674.506
755.37109375 0 7742.002
756.3692016601562 0 11655.737 y Water loss 6
757.3558959960938 0 8178.1753 y Ammonia loss 6
770.384033203125 0 50998.805
771.38818359375 0 17257.06
772.3805541992188 0 5211.089
773.3715209960938 0 44324.035
774.3783569335938 0 889932.2 y 6
775.3814086914062 0 347189.75
776.3842163085938 0 80019.16
801.3663330078125 0 10731.923
802.3724365234375 0 33677.85
803.3742065429688 0 11739.878
804.3834838867188 0 6240.5645
815.4057006835938 0 4796.3594
824.3958740234375 0 9333.51
831.3981323242188 0 10040.714
832.4014892578125 0 6532.8726
842.4075317382812 0 11827.369 w 5
843.4110107421875 0 6182.587
857.396240234375 0 62939.227
858.3994750976562 0 410492.7 y Ammonia loss 5
859.4035034179688 0 254500.34 z 5
860.41455078125 0 664943.75
861.4178466796875 0 277762.56
862.4215087890625 0 66263.11
863.4266357421875 0 5137.1787
874.4197387695312 0 12751.034
875.4264526367188 0 684036.9 y 5
876.42919921875 0 321429.4
877.4317016601562 0 75168.055
878.4342651367188 0 9675.285
883.4671020507812 0 4263.711
891.4439086914062 0 6624.034
907.4630126953125 0 25578.398
908.4721069335938 0 65897.76 c 7
909.4743041992188 0 27891.578
910.4829711914062 0 6059.5786
928.4913330078125 0 4566.538
929.4385375976562 0 26710.447 w 4
930.445068359375 0 15268.806
934.487548828125 0 10411.948
955.492919921875 0 10032.308
970.4982299804688 0 26711.96 y Water loss 4
971.4998779296875 0 14487.708 y Ammonia loss 4
972.4913330078125 0 762253.7 z 4
973.4957885742188 0 548159.44
974.49951171875 0 192750.92
975.50146484375 0 32420.367
987.5021362304688 0 52660.23
988.5064697265625 0 288999.28 y 4
989.509033203125 0 150265.16
990.5115966796875 0 39692.867
992.5178833007812 0 98212.53
993.52294921875 0 90849.93
994.525390625 0 31882.994
995.531005859375 0 7772.67
1001.4956665039062 0 17851.889
1002.4929809570312 0 11531.077
1003.4623413085938 0 29284.426
1004.461181640625 0 13435.412
1005.4680786132812 0 11578.375
1016.4785766601562 0 9525.845
1017.5128173828125 0 12236.367
1018.5059204101562 0 16708.459 c Water loss 8
1019.5084838867188 0 24369.768 c Ammonia loss 8
1020.5086669921875 0 11562.571
1035.5234375 0 35532.418
1036.53125 0 134522.53 c 8
1037.533935546875 0 71312.67
1038.537353515625 0 19061.371
1057.525146484375 0 48563.32 y Water loss 3
1058.5252685546875 0 47491.633 y Ammonia loss 3
1059.5238037109375 0 294455.8 z 3
1060.5301513671875 0 754810.56
1061.53369140625 0 395734.97
1062.5382080078125 0 117194.945
1063.541748046875 0 19328.97
1069.5784912109375 0 19760.135
1070.5831298828125 0 11100.402
1073.51806640625 0 6307.323
1074.53369140625 0 54671.54
1075.5419921875 0 778925.25 y 3
1076.544921875 0 493548.16
1077.547119140625 0 146129.38
1077.6854248046875 0 5716.5806
1078.552490234375 0 16274.913
1083.5831298828125 0 6521.341
1100.582763671875 0 8205.474
1101.5736083984375 0 16415.127
1102.578857421875 0 10600.473
1107.56396484375 0 7125.959
1110.6053466796875 0 7193.3916
1113.593505859375 0 10588.909
1114.598388671875 0 5579.846
1129.55322265625 0 110016.71 w 2
1130.5552978515625 0 72084.63
1131.55615234375 0 21092.148
1154.606689453125 0 6840.6655 z Water loss 2
1155.60107421875 0 11906.563 z Ammonia loss 2
1160.5853271484375 0 31098.291
1161.5770263671875 0 29862.133
1162.5750732421875 0 18846.963
1169.564208984375 0 5539.0864
1170.610595703125 0 21190.658 y Water loss 2
1171.6160888671875 0 12633.598 y Ammonia loss 2
1172.607421875 0 262741.47 z 2
1173.610107421875 0 164033.62
1174.6153564453125 0 61698.727
1175.6173095703125 0 10384.928
1177.5875244140625 0 6569.294
1178.596435546875 0 227954.39
1179.600341796875 0 195188.89
1180.6033935546875 0 72815.07
1181.60009765625 0 7328.287
1187.5753173828125 0 42141.945
1188.6239013671875 0 275044 y 2
1189.6282958984375 0 183649.2
1190.631103515625 0 59113.137
1191.629638671875 0 9723.451
1203.594482421875 0 14466.168
1204.5859375 0 25826.584 c Water loss 9
1205.583984375 0 69944.81 c Ammonia loss 9
1206.587890625 0 44424.957
1207.5887451171875 0 12349.171
1221.6011962890625 0 106822.53
1222.6094970703125 0 889595.75 c 9
1223.6123046875 0 610004.8
1224.61474609375 0 210320.83
1225.61572265625 0 32385.844
1279.6025390625 0 9054.255
1280.60986328125 0 5499.2324
1292.67138671875 0 6571.2603
1306.691650390625 0 22000.371
1307.696044921875 0 18780.527
1308.6900634765625 0 8489.808
1322.6822509765625 0 8043.8623
1335.6749267578125 0 116956.21 z 1
1336.677490234375 0 92905.016
1337.677734375 0 40618.67
1338.6737060546875 0 10554.047
1348.6806640625 0 9662.798
1349.691162109375 0 7923.7295
1350.7039794921875 0 735715.8 c 10
1351.7056884765625 0 611097.3 y 1
1352.7078857421875 0 273795.4
1353.7027587890625 0 50704.832
1354.7113037109375 0 12768.671
1360.6954345703125 0 5638.0566
1366.6385498046875 0 48563.785
1367.6666259765625 0 52157.535
1368.6829833984375 0 48387.555
1369.6884765625 0 22605.928
1370.6973876953125 0 8262.135
1378.6900634765625 0 15998.907
1379.6907958984375 0 21690.275
1380.7027587890625 0 30399.742
1381.7080078125 0 29192.771
1382.70751953125 0 15532.961
1383.669189453125 0 48302.484
1384.6707763671875 0 40862.46
1385.6712646484375 0 14079.357
1393.721435546875 0 8565.124
1394.710205078125 0 46949.05
1395.7105712890625 0 40894.71
1396.7135009765625 0 25620.086
1397.7208251953125 0 9688.149
1406.7119140625 0 5606.224
1410.7196044921875 0 6746.0195
1411.7322998046875 0 97880.12
1412.735107421875 0 82224.195
1413.7357177734375 0 32462.244
1414.7431640625 0 6932.6694
1420.682373046875 0 15024.572
1421.712890625 0 92734.266
1422.701904296875 0 1273070.1
1423.7047119140625 0 1042472.7
1424.70654296875 0 445292.8
1425.7099609375 0 70985.63
1437.708251953125 0 42109.13
1438.7197265625 0 1183161.5
1439.7264404296875 0 3488555.8
1440.7301025390625 0 2475101.5
1441.732666015625 0 971264
1442.73388671875 0 144548.78
1454.6953125 0 7435.4106
1471.7242431640625 0 8814.475
2167.393798828125 0 4992.0776
2219.796142578125 0 4656.1924

Spectrum Details

|  |  |
| --- | --- |
| Matched peaks? Matched peaks The total absolute number of peaks matched. Additionally in brackets the total fraction of peaks matched and the total number of peaks is shown. | 54 (15.61% of 346) |
| FDR? FDR The false discovery rate estimated for this peptide. It is calculated by matching all theoretical fragments with a non-integer shift with the raw peaks for this spectrum. This is done with 40 different shifts. The resulting percentage is the average number of annotated peaks over the number of annotated peaks with the correct spectrum. | 0.00% |
| Satellite FDR? Satellite FDR See the FDR for details on its calculation. This satellite ion specific FDR only contains the satellite ions (d/w) for I/L/J positions. | 0.00% |
| PSM Score? PSM Score The PSM Score as given by Hecklib to this annotated spectrum. It is shown with three significant figures. | 533 |

#### Spectrum 8833? Spectrum 8833 The raw spectrum of this peptide as annotated by Hecklib. The fragments are coloured according to ion type (see legend). Any peaks with a star '\*' as text can be hovered over to see the full details, first the ion type second the mass shift type. By hovering over the amino acids in the peptide or ions in the legend the corresponding peaks are highlighted. By toggling the 'Unassigned' label you can turn the background (unassigned) peaks on or off in the plot. By updating the slider in the Ion legend you can update the spectrum to only show the top X% of the peaks with labels. The top X% means any peak that is within X% of the highest intensity. By dragging in the spectrum you can zoom in to a specific part of the spectrum and use 'Zoom Out' to get back to the original zoom level. The annotation of the spectrum is based on the given sequence in the peptides file and is done with different software so inconsistencies are likely. The peaks are annotated based on the given sequence, with 20 ppm tolerance.

Copy Data

##### Spectrum 8833 (TSV)

###### Preview

```
Loading example...
```

*Click on the button to copy the data to your clipboard.*

Mz MinMz MaxIntensity Max

WidthHeightPeptide font sizePeptide stroke widthSpectrum font sizeSpectrum stroke widthCompact peptide

Ion legend

wxyz

abcd

OtherUnassignedIon^Charge^_Position_Show for top:%

SYLSJTPEQWKS

04.51e+49.02e+41.35e+51.80e+5

Zoom Out

y^+1^_2_y^+1^_3_w^+1^_4_y^+2^_8_y^+2^_9_z^+1^_4_y^+1^_4_y^+2^_10_y^+2^_10_z^+2^_10_y^+2^_10_z^+1^_5_y^+1^_5_y^+1^_6_y^+1^_6_y^+1^_7_z^+1^_7_y^+1^_7_c^+1^_8_w^+1^_8_y^+1^_8_y^+1^_8_z^+1^_8_y^+1^_8_c^+1^_9_c^+1^_9_c^+1^_9_y^+1^_9_y^+1^_9_z^+1^_9_y^+1^_9_w^+1^_10_z^+1^_10_y^+1^_10_z^+1^_10_y^+1^_10_c^+1^_10_c^+1^_10_c^+1^_10_z^+1^_11_c^+1^_11_y^+1^_11_

0700140021002800

Fragment Matches Table

Show background peaks

| Position | Ion type | Intensity | mz Theoretical | mz Error (Th) | mz Error (ppm) | Charge | Series Number |
| --- | --- | --- | --- | --- | --- | --- | --- |
| - | - | 2678 | 129.1 | - | - | 0 | - |
| - | - | 1313 | 136.1 | - | - | 0 | - |
| - | - | 495.4 | 140.4 | - | - | 0 | - |
| - | - | 381.3 | 144.9 | - | - | 0 | - |
| - | - | 449.3 | 145 | - | - | 0 | - |
| - | - | 382.1 | 146.4 | - | - | 0 | - |
| - | - | 408.5 | 149 | - | - | 0 | - |
| - | - | 443 | 162.3 | - | - | 0 | - |
| - | - | 441 | 164 | - | - | 0 | - |
| - | - | 1917 | 173.1 | - | - | 0 | - |
| - | - | 705.5 | 173.1 | - | - | 0 | - |
| - | - | 667.7 | 187.1 | - | - | 0 | - |
| - | - | 417 | 189.6 | - | - | 0 | - |
| - | - | 524.3 | 190.7 | - | - | 0 | - |
| - | - | 2967 | 201.1 | - | - | 0 | - |
| - | - | 896.6 | 215.1 | - | - | 0 | - |
| - | - | 552.8 | 218 | - | - | 0 | - |
| - | - | 1.404E+04 | 223.1 | - | - | 0 | - |
| - | - | 886.6 | 224.1 | - | - | 0 | - |
| - | - | 1361 | 227.1 | - | - | 0 | - |
| 11 | y | 5727 | 234.1 | 0.0002175 | 0.9288 | +1 | 2 |
| - | - | 622.9 | 245.2 | - | - | 0 | - |
| - | - | 2.428E+04 | 251.1 | - | - | 0 | - |
| - | - | 2954 | 252.1 | - | - | 0 | - |
| - | - | 1471 | 296.2 | - | - | 0 | - |
| - | - | 721.1 | 302.2 | - | - | 0 | - |
| - | - | 1211 | 315.2 | - | - | 0 | - |
| - | - | 1433 | 319.2 | - | - | 0 | - |
| - | - | 646.1 | 340.2 | - | - | 0 | - |
| - | - | 1325 | 355.2 | - | - | 0 | - |
| - | - | 9790 | 364.2 | - | - | 0 | - |
| - | - | 2163 | 365.2 | - | - | 0 | - |
| - | - | 904.6 | 397.2 | - | - | 0 | - |
| - | - | 498.6 | 405.4 | - | - | 0 | - |
| - | - | 525.4 | 415.2 | - | - | 0 | - |
| - | - | 1471 | 415.2 | - | - | 0 | - |
| - | - | 1535 | 415.3 | - | - | 0 | - |
| 10 | y | 3435 | 420.2 | 0.0002686 | 0.6391 | +1 | 3 |
| - | - | 619.2 | 421.2 | - | - | 0 | - |
| - | - | 620.9 | 423.2 | - | - | 0 | - |
| - | - | 649.3 | 424.1 | - | - | 0 | - |
| - | - | 8134 | 433.2 | - | - | 0 | - |
| - | - | 1878 | 434.2 | - | - | 0 | - |
| - | - | 6189 | 451.2 | - | - | 0 | - |
| - | - | 1450 | 452.2 | - | - | 0 | - |
| - | - | 1812 | 471.3 | - | - | 0 | - |
| - | - | 1183 | 472.3 | - | - | 0 | - |
| 9 | w | 589.1 | 474.2 | 0.000854 | 1.801 | +1 | 4 |
| - | - | 853.5 | 477.3 | - | - | 0 | - |
| 5 | y | 618 | 494.8 | 0.0003007 | 0.6077 | +2 | 8 |
| - | - | 673.7 | 498.2 | - | - | 0 | - |
| - | - | 1772 | 516.2 | - | - | 0 | - |
| - | - | 1441 | 518.3 | - | - | 0 | - |
| - | - | 1418 | 528.3 | - | - | 0 | - |
| 4 | y | 645.1 | 529.3 | 0.003485 | 6.584 | +2 | 9 |
| 9 | z | 7179 | 532.3 | 0.0002833 | 0.5322 | +1 | 4 |
| - | - | 1.268E+04 | 533.3 | - | - | 0 | - |
| - | - | 3457 | 534.3 | - | - | 0 | - |
| - | - | 1241 | 536.3 | - | - | 0 | - |
| - | - | 1721 | 541.2 | - | - | 0 | - |
| - | - | 5084 | 546.3 | - | - | 0 | - |
| - | - | 2232 | 547.3 | - | - | 0 | - |
| 9 | y | 1594 | 548.3 | 0.001945 | 3.547 | +1 | 4 |
| - | - | 638 | 552.3 | - | - | 0 | - |
| - | - | 2613 | 564.3 | - | - | 0 | - |
| 3 | y | 2384 | 585.8 | 8.807E-05 | 0.1503 | +2 | 10 |
| 3 | y | 1650 | 586.3 | 0.008874 | 15.14 | +2 | 10 |
| 3 | z | 602.6 | 586.8 | 0.005022 | 8.559 | +2 | 10 |
| 3 | y | 3903 | 594.8 | 0.0002379 | 0.3999 | +2 | 10 |
| - | - | 2086 | 595.3 | - | - | 0 | - |
| - | - | 926.1 | 595.8 | - | - | 0 | - |
| - | - | 833.9 | 603.3 | - | - | 0 | - |
| - | - | 2184 | 629.3 | - | - | 0 | - |
| - | - | 1352 | 630.3 | - | - | 0 | - |
| - | - | 866.6 | 637.4 | - | - | 0 | - |
| - | - | 1046 | 639.4 | - | - | 0 | - |
| - | - | 4290 | 647.3 | - | - | 0 | - |
| - | - | 1476 | 648.3 | - | - | 0 | - |
| - | - | 1398 | 653.3 | - | - | 0 | - |
| 8 | z | 7794 | 661.3 | 0.0004758 | 0.7195 | +1 | 5 |
| - | - | 1.085E+04 | 662.3 | - | - | 0 | - |
| - | - | 3398 | 663.3 | - | - | 0 | - |
| - | - | 4408 | 665.4 | - | - | 0 | - |
| - | - | 1172 | 666.4 | - | - | 0 | - |
| - | - | 4224 | 669.3 | - | - | 0 | - |
| - | - | 1348 | 670.3 | - | - | 0 | - |
| - | - | 664.3 | 671.3 | - | - | 0 | - |
| 8 | y | 1525 | 677.3 | 0.001222 | 1.804 | +1 | 5 |
| - | - | 688.2 | 702.3 | - | - | 0 | - |
| - | - | 4164 | 710.9 | - | - | 0 | - |
| - | - | 5990 | 711.4 | - | - | 0 | - |
| - | - | 2222 | 711.9 | - | - | 0 | - |
| - | - | 763.5 | 714.4 | - | - | 0 | - |
| - | - | 885.4 | 719.4 | - | - | 0 | - |
| 7 | y | 670.5 | 756.4 | 0.002403 | 3.177 | +1 | 6 |
| - | - | 1842 | 770.4 | - | - | 0 | - |
| - | - | 1985 | 773.4 | - | - | 0 | - |
| 7 | y | 3.798E+04 | 774.4 | 0.0003944 | 0.5094 | +1 | 6 |
| - | - | 1.702E+04 | 775.4 | - | - | 0 | - |
| - | - | 3947 | 776.4 | - | - | 0 | - |
| - | - | 1065 | 800.4 | - | - | 0 | - |
| - | - | 1438 | 802.4 | - | - | 0 | - |
| - | - | 553.1 | 804.4 | - | - | 0 | - |
| - | - | 629.5 | 830.5 | - | - | 0 | - |
| - | - | 766.6 | 843.4 | - | - | 0 | - |
| - | - | 3427 | 857.4 | - | - | 0 | - |
| 6 | y | 2.245E+04 | 858.4 | 0.0001615 | 0.1881 | +1 | 7 |
| 6 | z | 1.31E+04 | 859.4 | 0.003592 | 4.18 | +1 | 7 |
| - | - | 3.396E+04 | 860.4 | - | - | 0 | - |
| - | - | 1.325E+04 | 861.4 | - | - | 0 | - |
| - | - | 3116 | 862.4 | - | - | 0 | - |
| 6 | y | 2.882E+04 | 875.4 | 0.0004045 | 0.462 | +1 | 7 |
| - | - | 1.319E+04 | 876.4 | - | - | 0 | - |
| - | - | 3447 | 877.4 | - | - | 0 | - |
| - | - | 708.5 | 878.4 | - | - | 0 | - |
| - | - | 715.8 | 889.4 | - | - | 0 | - |
| - | - | 931.8 | 902.4 | - | - | 0 | - |
| - | - | 1146 | 907.5 | - | - | 0 | - |
| 8 | c | 2303 | 908.5 | 0.0007556 | 0.8318 | +1 | 8 |
| - | - | 1280 | 909.5 | - | - | 0 | - |
| - | - | 673.7 | 911.3 | - | - | 0 | - |
| - | - | 787.6 | 912.3 | - | - | 0 | - |
| 5 | w | 1476 | 929.4 | 0.0001393 | 0.1498 | +1 | 8 |
| - | - | 830.3 | 930.4 | - | - | 0 | - |
| - | - | 1206 | 947.3 | - | - | 0 | - |
| - | - | 1754 | 948.3 | - | - | 0 | - |
| - | - | 890.8 | 949.3 | - | - | 0 | - |
| 5 | y | 953.1 | 970.5 | 0.002371 | 2.443 | +1 | 8 |
| 5 | y | 1853 | 971.5 | 0.0069 | 7.103 | +1 | 8 |
| 5 | z | 4.04E+04 | 972.5 | 0.000864 | 0.8884 | +1 | 8 |
| - | - | 2.828E+04 | 973.5 | - | - | 0 | - |
| - | - | 8796 | 974.5 | - | - | 0 | - |
| - | - | 1494 | 975.5 | - | - | 0 | - |
| - | - | 2650 | 987.5 | - | - | 0 | - |
| 5 | y | 1.369E+04 | 988.5 | 0.005001 | 5.059 | +1 | 8 |
| - | - | 8289 | 989.5 | - | - | 0 | - |
| - | - | 1383 | 990.5 | - | - | 0 | - |
| - | - | 6014 | 992.5 | - | - | 0 | - |
| - | - | 4419 | 993.5 | - | - | 0 | - |
| - | - | 1758 | 994.5 | - | - | 0 | - |
| - | - | 849.7 | 995.5 | - | - | 0 | - |
| - | - | 660.1 | 1001 | - | - | 0 | - |
| - | - | 872.3 | 1003 | - | - | 0 | - |
| - | - | 960.4 | 1004 | - | - | 0 | - |
| - | - | 931.2 | 1018 | - | - | 0 | - |
| 9 | c | 2061 | 1019 | 0.002504 | 2.458 | +1 | 9 |
| 9 | c | 1419 | 1020 | 0.01055 | 10.35 | +1 | 9 |
| - | - | 1408 | 1036 | - | - | 0 | - |
| 9 | c | 7594 | 1037 | 0.001655 | 1.597 | +1 | 9 |
| - | - | 3353 | 1038 | - | - | 0 | - |
| - | - | 1048 | 1039 | - | - | 0 | - |
| 4 | y | 2238 | 1058 | 0.00614 | 5.806 | +1 | 9 |
| 4 | y | 2224 | 1059 | 0.01021 | 9.646 | +1 | 9 |
| 4 | z | 1.445E+04 | 1060 | 0.0005549 | 0.5237 | +1 | 9 |
| - | - | 3.992E+04 | 1061 | - | - | 0 | - |
| - | - | 883.3 | 1061 | - | - | 0 | - |
| - | - | 2.085E+04 | 1062 | - | - | 0 | - |
| - | - | 6418 | 1063 | - | - | 0 | - |
| - | - | 2048 | 1064 | - | - | 0 | - |
| - | - | 3151 | 1075 | - | - | 0 | - |
| 4 | y | 3.477E+04 | 1076 | 0.001079 | 1.004 | +1 | 9 |
| - | - | 2.009E+04 | 1077 | - | - | 0 | - |
| - | - | 6400 | 1078 | - | - | 0 | - |
| - | - | 1138 | 1079 | - | - | 0 | - |
| - | - | 626.8 | 1080 | - | - | 0 | - |
| - | - | 959.6 | 1081 | - | - | 0 | - |
| - | - | 2037 | 1082 | - | - | 0 | - |
| - | - | 769.2 | 1087 | - | - | 0 | - |
| - | - | 1102 | 1088 | - | - | 0 | - |
| - | - | 964.8 | 1103 | - | - | 0 | - |
| - | - | 870.6 | 1114 | - | - | 0 | - |
| - | - | 785.1 | 1117 | - | - | 0 | - |
| - | - | 1009 | 1120 | - | - | 0 | - |
| 3 | w | 7006 | 1130 | 0.0009019 | 0.7984 | +1 | 10 |
| - | - | 3394 | 1131 | - | - | 0 | - |
| - | - | 1094 | 1132 | - | - | 0 | - |
| 3 | z | 823.8 | 1156 | 0.008226 | 7.118 | +1 | 10 |
| - | - | 713.8 | 1158 | - | - | 0 | - |
| - | - | 1634 | 1162 | - | - | 0 | - |
| - | - | 968.8 | 1163 | - | - | 0 | - |
| 3 | y | 819.3 | 1171 | 0.01672 | 14.28 | +1 | 10 |
| 3 | z | 1.267E+04 | 1173 | 1.302E-05 | 0.01111 | +1 | 10 |
| - | - | 742 | 1173 | - | - | 0 | - |
| - | - | 8967 | 1174 | - | - | 0 | - |
| - | - | 3691 | 1175 | - | - | 0 | - |
| - | - | 1.067E+04 | 1179 | - | - | 0 | - |
| - | - | 8414 | 1180 | - | - | 0 | - |
| - | - | 3636 | 1181 | - | - | 0 | - |
| - | - | 1207 | 1188 | - | - | 0 | - |
| 3 | y | 1.306E+04 | 1189 | 0.003356 | 2.824 | +1 | 10 |
| - | - | 7801 | 1190 | - | - | 0 | - |
| - | - | 2678 | 1191 | - | - | 0 | - |
| 10 | c | 1142 | 1205 | 0.02145 | 17.81 | +1 | 10 |
| 10 | c | 3181 | 1206 | 0.001123 | 0.9317 | +1 | 10 |
| - | - | 1359 | 1207 | - | - | 0 | - |
| - | - | 1161 | 1209 | - | - | 0 | - |
| - | - | 4663 | 1222 | - | - | 0 | - |
| 10 | c | 4.561E+04 | 1223 | 0.001744 | 1.427 | +1 | 10 |
| - | - | 2.92E+04 | 1224 | - | - | 0 | - |
| - | - | 1.023E+04 | 1225 | - | - | 0 | - |
| - | - | 1428 | 1226 | - | - | 0 | - |
| - | - | 730.7 | 1240 | - | - | 0 | - |
| - | - | 758 | 1252 | - | - | 0 | - |
| - | - | 761.9 | 1269 | - | - | 0 | - |
| - | - | 819.3 | 1280 | - | - | 0 | - |
| - | - | 1190 | 1307 | - | - | 0 | - |
| - | - | 835.8 | 1308 | - | - | 0 | - |
| - | - | 1132 | 1323 | - | - | 0 | - |
| - | - | 1289 | 1328 | - | - | 0 | - |
| - | - | 942.8 | 1329 | - | - | 0 | - |
| 2 | z | 6513 | 1336 | 0.003187 | 2.386 | +1 | 11 |
| - | - | 4004 | 1337 | - | - | 0 | - |
| - | - | 1981 | 1338 | - | - | 0 | - |
| - | - | 4034 | 1345 | - | - | 0 | - |
| - | - | 4171 | 1346 | - | - | 0 | - |
| - | - | 2446 | 1347 | - | - | 0 | - |
| - | - | 767.5 | 1348 | - | - | 0 | - |
| - | - | 847.9 | 1350 | - | - | 0 | - |
| 11 | c | 3.827E+04 | 1351 | 0.002225 | 1.647 | +1 | 11 |
| 2 | y | 2.892E+04 | 1352 | 0.01486 | 10.99 | +1 | 11 |
| - | - | 1.346E+04 | 1353 | - | - | 0 | - |
| - | - | 3309 | 1354 | - | - | 0 | - |
| - | - | 1192 | 1363 | - | - | 0 | - |
| - | - | 1120 | 1364 | - | - | 0 | - |
| - | - | 1417 | 1365 | - | - | 0 | - |
| - | - | 912.8 | 1366 | - | - | 0 | - |
| - | - | 4224 | 1367 | - | - | 0 | - |
| - | - | 2862 | 1368 | - | - | 0 | - |
| - | - | 2237 | 1369 | - | - | 0 | - |
| - | - | 1164 | 1370 | - | - | 0 | - |
| - | - | 2461 | 1378 | - | - | 0 | - |
| - | - | 4165 | 1379 | - | - | 0 | - |
| - | - | 2705 | 1380 | - | - | 0 | - |
| - | - | 786.8 | 1381 | - | - | 0 | - |
| - | - | 809.7 | 1381 | - | - | 0 | - |
| - | - | 1002 | 1382 | - | - | 0 | - |
| - | - | 3067 | 1384 | - | - | 0 | - |
| - | - | 2539 | 1385 | - | - | 0 | - |
| - | - | 1427 | 1386 | - | - | 0 | - |
| - | - | 2166 | 1395 | - | - | 0 | - |
| - | - | 1858 | 1396 | - | - | 0 | - |
| - | - | 1970 | 1397 | - | - | 0 | - |
| - | - | 777.8 | 1406 | - | - | 0 | - |
| - | - | 5556 | 1412 | - | - | 0 | - |
| - | - | 3266 | 1413 | - | - | 0 | - |
| - | - | 2570 | 1414 | - | - | 0 | - |
| - | - | 1144 | 1415 | - | - | 0 | - |
| - | - | 1258 | 1420 | - | - | 0 | - |
| - | - | 2757 | 1421 | - | - | 0 | - |
| - | - | 5649 | 1422 | - | - | 0 | - |
| - | - | 6.839E+04 | 1423 | - | - | 0 | - |
| - | - | 5.428E+04 | 1424 | - | - | 0 | - |
| - | - | 2.455E+04 | 1425 | - | - | 0 | - |
| - | - | 5413 | 1426 | - | - | 0 | - |
| - | - | 3330 | 1437 | - | - | 0 | - |
| - | - | 4098 | 1437 | - | - | 0 | - |
| - | - | 6953 | 1438 | - | - | 0 | - |
| - | - | 6.4E+04 | 1439 | - | - | 0 | - |
| - | - | 1.785E+05 | 1440 | - | - | 0 | - |
| - | - | 1.331E+05 | 1441 | - | - | 0 | - |
| - | - | 1132 | 1441 | - | - | 0 | - |
| - | - | 5.447E+04 | 1442 | - | - | 0 | - |
| - | - | 8093 | 1443 | - | - | 0 | - |
| - | - | 910.8 | 2082 | - | - | 0 | - |
| - | - | 1093 | 2083 | - | - | 0 | - |
| - | - | 697.1 | 2138 | - | - | 0 | - |
| - | - | 699 | 2140 | - | - | 0 | - |
| - | - | 1161 | 2141 | - | - | 0 | - |
| - | - | 906.2 | 2156 | - | - | 0 | - |
| - | - | 901.3 | 2158 | - | - | 0 | - |
| - | - | 958.4 | 2160 | - | - | 0 | - |
| - | - | 1001 | 2161 | - | - | 0 | - |
| - | - | 2922 | 2162 | - | - | 0 | - |
| - | - | 1549 | 2163 | - | - | 0 | - |
| - | - | 732.4 | 2753 | - | - | 0 | - |
| - | - | 762.6 | 2773 | - | - | 0 | - |

m/z Charge Intensity FragmentType MassShift Position
129.1024627685547 0 2677.979
136.0758514404297 0 1313.2668
140.42457580566406 0 495.37985
144.91195678710938 0 381.29037
144.95071411132812 0 449.32666
146.3795166015625 0 382.1387
148.95513916015625 0 408.45676
162.26541137695312 0 443.01193
163.9969024658203 0 441.04462
173.09213256835938 0 1916.9365
173.12815856933594 0 705.5377
187.1083526611328 0 667.65717
189.56056213378906 0 417.02103
190.65142822265625 0 524.3409
201.12339782714844 0 2967.1443
215.13929748535156 0 896.58527
218.04129028320312 0 552.79694
223.1078338623047 0 14044.46
224.11134338378906 0 886.5673
227.10279846191406 0 1360.9121
234.14505004882812 0 5727.2964 y 10
245.1857452392578 0 622.8611
251.1027374267578 0 24284.973
252.10618591308594 0 2954.2693
296.1974182128906 0 1471.0239
302.1715393066406 0 721.09467
315.18182373046875 0 1210.8787
319.164794921875 0 1433.4961
340.18597412109375 0 646.14886
355.16156005859375 0 1324.59
364.1866760253906 0 9789.82
365.19000244140625 0 2162.9922
397.2422180175781 0 904.64435
405.37554931640625 0 498.63522
415.1663513183594 0 525.404
415.1991882324219 0 1470.738
415.2552795410156 0 1535.0765
420.223876953125 0 3434.7036 y 9
421.2265625 0 619.15894
423.2251281738281 0 620.9433
424.1302490234375 0 649.3193
433.20867919921875 0 8134.2227
434.21185302734375 0 1878.2922
451.21917724609375 0 6189.0103
452.2217712402344 0 1450.2468
471.28070068359375 0 1811.6569
472.2845458984375 0 1183.3083
474.2338562011719 0 589.0525 w 8
477.2695007324219 0 853.4713
494.75885009765625 0 617.95483 y 4
498.23223876953125 0 673.70233
516.2456665039062 0 1772.3933
518.2982788085938 0 1440.7444
528.2799682617188 0 1418.2216
529.2727661132812 0 645.0809 y Water loss 3
532.2642822265625 0 7178.851 z 8
533.2716064453125 0 12682.397
534.2722778320312 0 3456.6108
536.3090209960938 0 1241.3195
541.24169921875 0 1721.3167
546.2924194335938 0 5084.2144
547.294677734375 0 2231.6472
548.28466796875 0 1593.6996 y 8
552.2681274414062 0 638.0386
564.3026123046875 0 2612.909
585.8114013671875 0 2384.3525 y Water loss 2
586.3121948242188 0 1649.574 y Ammonia loss 2
586.812255859375 0 602.6098 z 2
594.8168334960938 0 3903.3267 y 2
595.3179931640625 0 2085.9382
595.8202514648438 0 926.075
603.310546875 0 833.8894
629.3292236328125 0 2183.5847
630.33251953125 0 1352.0825
637.3552856445312 0 866.6243
639.3716430664062 0 1046.2999
647.34033203125 0 4290.268
648.3419799804688 0 1476.2457
653.269287109375 0 1398.4979
661.3070678710938 0 7793.5776 z 7
662.3134155273438 0 10854.618
663.3179321289062 0 3398.3787
665.35107421875 0 4407.9023
666.3526000976562 0 1172.3644
669.3357543945312 0 4223.5977
670.3427124023438 0 1348.0358
671.3436889648438 0 664.2851
677.3265380859375 0 1525.2246 y 7
702.3467407226562 0 688.20166
710.8585205078125 0 4163.6997
711.3596801757812 0 5989.798
711.8597412109375 0 2222.1401
714.3650512695312 0 763.45074
719.3599243164062 0 885.4347
756.3651123046875 0 670.53424 y Water loss 6
770.3834228515625 0 1841.9849
773.3710327148438 0 1985.0376
774.377685546875 0 37976.145 y 6
775.380859375 0 17024.943
776.3841552734375 0 3947.3406
800.390625 0 1065.4612
802.37548828125 0 1437.6001
804.385986328125 0 553.09344
830.4771728515625 0 629.5014
843.4193115234375 0 766.5733
857.397216796875 0 3426.6973
858.3990478515625 0 22447.84 y Ammonia loss 5
859.4034423828125 0 13104.202 z 5
860.4138793945312 0 33958.105
861.4171142578125 0 13245.934
862.419677734375 0 3115.5955
875.4253540039062 0 28823.617 y 5
876.4286499023438 0 13192.374
877.43115234375 0 3447.2805
878.443115234375 0 708.53827
889.4486694335938 0 715.8141
902.4329223632812 0 931.752
907.4619140625 0 1146.3622
908.4716186523438 0 2302.5078 c 7
909.4785766601562 0 1280.2448
911.3319091796875 0 673.70996
912.3416137695312 0 787.6289
929.4364624023438 0 1476.4292 w 4
930.4412231445312 0 830.33887
947.337158203125 0 1205.7012
948.340576171875 0 1754.2396
949.3458251953125 0 890.8161
970.4968872070312 0 953.1003 y Water loss 4
971.4901733398438 0 1852.5712 y Ammonia loss 4
972.490234375 0 40396.406 z 4
973.4949951171875 0 28275.893
974.498779296875 0 8796.167
975.508544921875 0 1493.6129
987.5014038085938 0 2649.6143
988.5048217773438 0 13685.075 y 4
989.5092163085938 0 8288.648
990.515625 0 1382.5446
992.5166015625 0 6014.2144
993.52197265625 0 4419.425
994.5242309570312 0 1758.2611
995.4893188476562 0 849.71606
1001.4909057617188 0 660.09937
1003.4617309570312 0 872.3288
1004.4637451171875 0 960.44586
1017.5096435546875 0 931.18866
1018.5178833007812 0 2060.87 c Water loss 8
1019.5149536132812 0 1419.3108 c Ammonia loss 8
1035.5235595703125 0 1407.614
1036.529296875 0 7593.647 c 8
1037.53125 0 3353.0283
1038.5408935546875 0 1047.8655
1057.525146484375 0 2237.9377 y Water loss 3
1058.5255126953125 0 2223.932 y Ammonia loss 3
1059.523681640625 0 14454.908 z 3
1060.529296875 0 39915.06
1061.402099609375 0 883.25726
1061.5330810546875 0 20848.719
1062.537109375 0 6417.6587
1063.5421142578125 0 2048.2722
1074.532958984375 0 3151.476
1075.540771484375 0 34772.586 y 3
1076.5435791015625 0 20092.158
1077.5462646484375 0 6399.773
1078.5467529296875 0 1137.5863
1079.5244140625 0 626.76056
1081.053955078125 0 959.5713
1081.5540771484375 0 2037.3428
1086.53125 0 769.217
1087.5347900390625 0 1102.1023
1102.549560546875 0 964.8159
1113.59521484375 0 870.59955
1116.54296875 0 785.1057
1119.5775146484375 0 1009.1175
1129.551513671875 0 7006.1265 w 2
1130.556640625 0 3393.568
1131.559326171875 0 1094.3002
1155.5888671875 0 823.80963 z Ammonia loss 2
1157.5821533203125 0 713.7975
1161.5765380859375 0 1634.3859
1162.5765380859375 0 968.82214
1170.5986328125 0 819.2988 y Water loss 2
1172.607177734375 0 12674.553 z 2
1173.4503173828125 0 742.01086
1173.60986328125 0 8966.664
1174.613037109375 0 3691.3196
1178.594970703125 0 10671.477
1179.5986328125 0 8413.688
1180.6005859375 0 3635.7566
1187.5726318359375 0 1207.2526
1188.62255859375 0 13058.636 y 2
1189.6265869140625 0 7801.1323
1190.6297607421875 0 2677.7307
1204.5782470703125 0 1142.199 c Water loss 9
1205.5848388671875 0 3181.4602 c Ammonia loss 9
1206.592041015625 0 1358.7971
1209.4805908203125 0 1161.446
1221.6029052734375 0 4662.6025
1222.6085205078125 0 45609.16 c 9
1223.6107177734375 0 29195.81
1224.6141357421875 0 10232.906
1225.6192626953125 0 1428.267
1239.517822265625 0 730.67676
1251.6063232421875 0 757.97186
1268.609375 0 761.92084
1279.5972900390625 0 819.31226
1306.686279296875 0 1189.894
1307.6949462890625 0 835.7626
1322.54541015625 0 1131.9146
1327.6905517578125 0 1289.3499
1328.6802978515625 0 942.8139
1335.6737060546875 0 6513.0493 z 1
1336.6793212890625 0 4003.536
1337.67919921875 0 1980.7343
1344.6939697265625 0 4034.0657
1345.696044921875 0 4171.32
1346.6923828125 0 2445.8806
1347.6961669921875 0 767.5445
1349.61572265625 0 847.9179
1350.7030029296875 0 38270.95 c 10
1351.7041015625 0 28922.674 y 1
1352.70556640625 0 13457.736
1353.7098388671875 0 3308.7239
1362.6844482421875 0 1191.5973
1363.6883544921875 0 1120.3938
1364.6737060546875 0 1416.7906
1365.678466796875 0 912.7835
1366.640380859375 0 4224.2256
1367.6805419921875 0 2861.7139
1368.6868896484375 0 2236.5122
1369.683349609375 0 1164.4529
1377.6082763671875 0 2461.3564
1378.6156005859375 0 4164.922
1379.6214599609375 0 2704.9177
1380.605224609375 0 786.7855
1380.7158203125 0 809.7018
1381.71533203125 0 1001.5109
1383.6651611328125 0 3067.4375
1384.659912109375 0 2539.2974
1385.6671142578125 0 1426.6232
1394.704833984375 0 2165.7078
1395.704833984375 0 1858.0073
1396.690673828125 0 1969.5236
1405.7008056640625 0 777.7644
1411.7310791015625 0 5556.377
1412.7359619140625 0 3266.2356
1413.731689453125 0 2569.7573
1414.73876953125 0 1143.9365
1419.6873779296875 0 1258.0964
1420.6839599609375 0 2757.302
1421.704833984375 0 5648.687
1422.701171875 0 68394.68
1423.7022705078125 0 54279.94
1424.703857421875 0 24545.812
1425.7047119140625 0 5412.7085
1436.6383056640625 0 3330.1155
1436.719482421875 0 4098.0415
1437.6766357421875 0 6953.24
1438.71728515625 0 63998.89
1439.72412109375 0 178534.4
1440.7283935546875 0 133064.75
1441.151123046875 0 1132.1793
1441.7301025390625 0 54474.01
1442.7301025390625 0 8092.8003
2082.049560546875 0 910.8066
2083.07421875 0 1093.4674
2138.074462890625 0 697.1266
2140.08056640625 0 699.0252
2141.096923828125 0 1161.1165
2156.06982421875 0 906.191
2158.09814453125 0 901.28876
2160.0986328125 0 958.39746
2161.11376953125 0 1001.0058
2162.10009765625 0 2922.425
2163.09765625 0 1549.1862
2753.00390625 0 732.40485
2772.650146484375 0 762.6444

Spectrum Details

|  |  |
| --- | --- |
| Matched peaks? Matched peaks The total absolute number of peaks matched. Additionally in brackets the total fraction of peaks matched and the total number of peaks is shown. | 42 (15.22% of 276) |
| FDR? FDR The false discovery rate estimated for this peptide. It is calculated by matching all theoretical fragments with a non-integer shift with the raw peaks for this spectrum. This is done with 40 different shifts. The resulting percentage is the average number of annotated peaks over the number of annotated peaks with the correct spectrum. | 0.45% |
| Satellite FDR? Satellite FDR See the FDR for details on its calculation. This satellite ion specific FDR only contains the satellite ions (d/w) for I/L/J positions. | 2.38% |
| PSM Score? PSM Score The PSM Score as given by Hecklib to this annotated spectrum. It is shown with three significant figures. | 422 |

#### Spectrum 8777? Spectrum 8777 The raw spectrum of this peptide as annotated by Hecklib. The fragments are coloured according to ion type (see legend). Any peaks with a star '\*' as text can be hovered over to see the full details, first the ion type second the mass shift type. By hovering over the amino acids in the peptide or ions in the legend the corresponding peaks are highlighted. By toggling the 'Unassigned' label you can turn the background (unassigned) peaks on or off in the plot. By updating the slider in the Ion legend you can update the spectrum to only show the top X% of the peaks with labels. The top X% means any peak that is within X% of the highest intensity. By dragging in the spectrum you can zoom in to a specific part of the spectrum and use 'Zoom Out' to get back to the original zoom level. The annotation of the spectrum is based on the given sequence in the peptides file and is done with different software so inconsistencies are likely. The peaks are annotated based on the given sequence, with 20 ppm tolerance.

Copy Data

##### Spectrum 8777 (TSV)

###### Preview

```
Loading example...
```

*Click on the button to copy the data to your clipboard.*

Mz MinMz MaxIntensity Max

WidthHeightPeptide font sizePeptide stroke widthSpectrum font sizeSpectrum stroke widthCompact peptide

Ion legend

wxyz

abcd

OtherUnassignedIon^Charge^_Position_Show for top:%

SYLSJTPEQWKS

01.41e+32.81e+34.22e+35.62e+3

Zoom Out

w^+1^_2_z^+3^_5_y^+2^_6_y^+1^_3_y^+1^_3_w^+1^_4_w^+2^_8_w^+1^_5_c^+2^_11_c^+2^_11_y^+2^_11_y^+1^_6_c^+1^_7_y^+1^_7_z^+1^_7_y^+1^_7_c^+1^_8_z^+1^_8_y^+1^_8_c^+1^_9_z^+1^_9_y^+1^_9_z^+1^_10_c^+1^_10_y^+1^_11_z^+1^_11_c^+1^_11_y^+1^_11_

0867173326003466

Fragment Matches Table

Show background peaks

| Position | Ion type | Intensity | mz Theoretical | mz Error (Th) | mz Error (ppm) | Charge | Series Number |
| --- | --- | --- | --- | --- | --- | --- | --- |
| - | - | 478.4 | 120.1 | - | - | 0 | - |
| - | - | 340.9 | 120.1 | - | - | 0 | - |
| - | - | 354 | 126.1 | - | - | 0 | - |
| - | - | 437.6 | 127.1 | - | - | 0 | - |
| - | - | 985.8 | 129.1 | - | - | 0 | - |
| - | - | 672.7 | 130.1 | - | - | 0 | - |
| - | - | 923.7 | 130.1 | - | - | 0 | - |
| - | - | 397.7 | 130.9 | - | - | 0 | - |
| - | - | 398.7 | 132.1 | - | - | 0 | - |
| - | - | 921.1 | 133.1 | - | - | 0 | - |
| - | - | 474.8 | 135.1 | - | - | 0 | - |
| - | - | 1221 | 136.1 | - | - | 0 | - |
| - | - | 495.2 | 138.1 | - | - | 0 | - |
| - | - | 1463 | 140.1 | - | - | 0 | - |
| - | - | 557.3 | 142.1 | - | - | 0 | - |
| - | - | 500.6 | 146.1 | - | - | 0 | - |
| - | - | 670.7 | 149.1 | - | - | 0 | - |
| - | - | 748.4 | 151.1 | - | - | 0 | - |
| - | - | 520.4 | 153.1 | - | - | 0 | - |
| - | - | 1214 | 156.1 | - | - | 0 | - |
| - | - | 740.9 | 158.1 | - | - | 0 | - |
| 11 | w | 848 | 160.1 | 0.0001126 | 0.7037 | +1 | 2 |
| - | - | 573.3 | 173.4 | - | - | 0 | - |
| - | - | 489.5 | 174.1 | - | - | 0 | - |
| - | - | 622.3 | 178.3 | - | - | 0 | - |
| - | - | 483.9 | 184.1 | - | - | 0 | - |
| - | - | 662.3 | 201.1 | - | - | 0 | - |
| - | - | 467.1 | 213.9 | - | - | 0 | - |
| - | - | 582.5 | 219.1 | - | - | 0 | - |
| - | - | 557.8 | 219.5 | - | - | 0 | - |
| 8 | z | 564 | 221.1 | 0.003319 | 15.01 | +3 | 5 |
| - | - | 998 | 223.1 | - | - | 0 | - |
| - | - | 3351 | 223.1 | - | - | 0 | - |
| - | - | 562.9 | 231.1 | - | - | 0 | - |
| - | - | 534.8 | 236.7 | - | - | 0 | - |
| - | - | 2454 | 251.1 | - | - | 0 | - |
| - | - | 503.6 | 282.3 | - | - | 0 | - |
| - | - | 1770 | 288.2 | - | - | 0 | - |
| - | - | 707.1 | 297.1 | - | - | 0 | - |
| - | - | 1010 | 299.1 | - | - | 0 | - |
| - | - | 662.8 | 309.2 | - | - | 0 | - |
| - | - | 530.7 | 317.3 | - | - | 0 | - |
| - | - | 653.6 | 360 | - | - | 0 | - |
| - | - | 1725 | 364.2 | - | - | 0 | - |
| - | - | 579.3 | 369.7 | - | - | 0 | - |
| - | - | 668.4 | 372.2 | - | - | 0 | - |
| - | - | 596.7 | 387.2 | - | - | 0 | - |
| 7 | y | 1696 | 387.7 | 0.0005395 | 1.392 | +2 | 6 |
| - | - | 780.2 | 388.2 | - | - | 0 | - |
| - | - | 611 | 388.7 | - | - | 0 | - |
| - | - | 592.5 | 395.7 | - | - | 0 | - |
| - | - | 761.5 | 402.2 | - | - | 0 | - |
| 10 | y | 862.7 | 403.2 | 0.001744 | 4.326 | +1 | 3 |
| 10 | y | 780.7 | 420.2 | 0.001654 | 3.936 | +1 | 3 |
| - | - | 1286 | 430.1 | - | - | 0 | - |
| - | - | 1136 | 431.1 | - | - | 0 | - |
| - | - | 683.4 | 432.1 | - | - | 0 | - |
| - | - | 700 | 433.2 | - | - | 0 | - |
| - | - | 705.1 | 437.9 | - | - | 0 | - |
| - | - | 919.4 | 446.1 | - | - | 0 | - |
| - | - | 782.2 | 451.2 | - | - | 0 | - |
| 9 | w | 3636 | 474.2 | 0.000916 | 1.932 | +1 | 4 |
| - | - | 651.1 | 475.2 | - | - | 0 | - |
| 5 | w | 876.7 | 479.2 | 0.005195 | 10.84 | +2 | 8 |
| - | - | 803.9 | 481.2 | - | - | 0 | - |
| - | - | 542.7 | 492.2 | - | - | 0 | - |
| - | - | 1176 | 546.3 | - | - | 0 | - |
| - | - | 880.7 | 564.3 | - | - | 0 | - |
| - | - | 616.3 | 585.4 | - | - | 0 | - |
| 8 | w | 2579 | 602.3 | 1.379E-05 | 0.0229 | +1 | 5 |
| - | - | 1012 | 654.8 | - | - | 0 | - |
| - | - | 1647 | 665.4 | - | - | 0 | - |
| 11 | c | 835.4 | 667.3 | 0.004679 | 7.011 | +2 | 11 |
| 11 | c | 1708 | 675.9 | 0.001597 | 2.363 | +2 | 11 |
| 2 | y | 1493 | 676.3 | 0.01227 | 18.15 | +2 | 11 |
| - | - | 580 | 676.9 | - | - | 0 | - |
| - | - | 661.9 | 710.9 | - | - | 0 | - |
| - | - | 1143 | 711.3 | - | - | 0 | - |
| - | - | 888 | 711.9 | - | - | 0 | - |
| - | - | 884 | 719.4 | - | - | 0 | - |
| - | - | 2232 | 719.9 | - | - | 0 | - |
| - | - | 3934 | 720.4 | - | - | 0 | - |
| - | - | 1726 | 720.9 | - | - | 0 | - |
| - | - | 1145 | 721.4 | - | - | 0 | - |
| - | - | 1187 | 721.9 | - | - | 0 | - |
| - | - | 581.8 | 735.9 | - | - | 0 | - |
| 7 | y | 2029 | 774.4 | 0.0006386 | 0.8246 | +1 | 6 |
| 7 | c | 2125 | 779.4 | 0.001742 | 2.235 | +1 | 7 |
| - | - | 819.2 | 780.4 | - | - | 0 | - |
| - | - | 941.7 | 787.4 | - | - | 0 | - |
| - | - | 1285 | 810.4 | - | - | 0 | - |
| - | - | 651.5 | 841.5 | - | - | 0 | - |
| 6 | y | 748 | 858.4 | 0.003745 | 4.362 | +1 | 7 |
| 6 | z | 2031 | 859.4 | 0.0003753 | 0.4367 | +1 | 7 |
| - | - | 1147 | 860.4 | - | - | 0 | - |
| - | - | 636.9 | 870.3 | - | - | 0 | - |
| 6 | y | 981.3 | 875.4 | 0.00132 | 1.508 | +1 | 7 |
| 8 | c | 3233 | 908.5 | 0.0001599 | 0.176 | +1 | 8 |
| - | - | 1663 | 909.5 | - | - | 0 | - |
| - | - | 705.4 | 910.5 | - | - | 0 | - |
| - | - | 934 | 932 | - | - | 0 | - |
| - | - | 809.5 | 958.5 | - | - | 0 | - |
| - | - | 850.3 | 960.3 | - | - | 0 | - |
| - | - | 682.2 | 961 | - | - | 0 | - |
| - | - | 679.7 | 961.5 | - | - | 0 | - |
| - | - | 916.1 | 962 | - | - | 0 | - |
| 5 | z | 2450 | 972.5 | 0.000784 | 0.8061 | +1 | 8 |
| - | - | 1924 | 973.5 | - | - | 0 | - |
| - | - | 758.8 | 974.5 | - | - | 0 | - |
| - | - | 719.1 | 982.5 | - | - | 0 | - |
| 5 | y | 1040 | 988.5 | 0.01538 | 15.56 | +1 | 8 |
| - | - | 778.3 | 992.5 | - | - | 0 | - |
| 9 | c | 3905 | 1037 | 0.0004203 | 0.4054 | +1 | 9 |
| - | - | 1919 | 1038 | - | - | 0 | - |
| 4 | z | 754.8 | 1060 | 0.00445 | 4.2 | +1 | 9 |
| - | - | 958.3 | 1061 | - | - | 0 | - |
| 4 | y | 763 | 1076 | 0.004619 | 4.295 | +1 | 9 |
| 3 | z | 1612 | 1173 | 0.001696 | 1.446 | +1 | 10 |
| - | - | 1070 | 1174 | - | - | 0 | - |
| - | - | 816.1 | 1175 | - | - | 0 | - |
| 10 | c | 5568 | 1223 | 0.001012 | 0.8276 | +1 | 10 |
| - | - | 4075 | 1224 | - | - | 0 | - |
| - | - | 1579 | 1225 | - | - | 0 | - |
| 2 | y | 798.4 | 1335 | 0.009913 | 7.427 | +1 | 11 |
| 2 | z | 800.9 | 1336 | 0.006116 | 4.579 | +1 | 11 |
| - | - | 1055 | 1337 | - | - | 0 | - |
| 11 | c | 777 | 1351 | 0.002536 | 1.877 | +1 | 11 |
| 2 | y | 821.4 | 1352 | 0.02523 | 18.67 | +1 | 11 |
| - | - | 880.8 | 1354 | - | - | 0 | - |
| - | - | 666.7 | 1406 | - | - | 0 | - |
| - | - | 1411 | 1423 | - | - | 0 | - |
| - | - | 4619 | 1424 | - | - | 0 | - |
| - | - | 3481 | 1425 | - | - | 0 | - |
| - | - | 1695 | 1426 | - | - | 0 | - |
| - | - | 1509 | 1439 | - | - | 0 | - |
| - | - | 2125 | 1440 | - | - | 0 | - |
| - | - | 4147 | 1441 | - | - | 0 | - |
| - | - | 1929 | 1442 | - | - | 0 | - |
| - | - | 1793 | 1443 | - | - | 0 | - |
| - | - | 1071 | 3037 | - | - | 0 | - |
| - | - | 714 | 3432 | - | - | 0 | - |

m/z Charge Intensity FragmentType MassShift Position
120.08094024658203 0 478.3711
120.08586120605469 0 340.85754
126.10227966308594 0 354.03412
127.07586669921875 0 437.6351
129.10243225097656 0 985.82996
130.06076049804688 0 672.7014
130.06549072265625 0 923.7243
130.86029052734375 0 397.69214
132.10186767578125 0 398.69473
133.06072998046875 0 921.0522
135.07611083984375 0 474.80026
136.0758819580078 0 1220.5121
138.0659637451172 0 495.18747
140.0820770263672 0 1462.7927
142.09783935546875 0 557.261
146.0563201904297 0 500.5805
149.055908203125 0 670.7455
151.0758056640625 0 748.39935
153.07762145996094 0 520.39343
156.0768280029297 0 1213.9673
158.09262084960938 0 740.8758
160.060546875 0 848.0258 w 10
173.43797302246094 0 573.3219
174.08740234375 0 489.55
178.32504272460938 0 622.296
184.0715789794922 0 483.89685
201.1238250732422 0 662.25275
213.87562561035156 0 467.11548
219.13475036621094 0 582.4609
219.51747131347656 0 557.8228
221.10372924804688 0 563.951 z 7
223.06390380859375 0 998.0114
223.1078643798828 0 3350.9727
231.11094665527344 0 562.8548
236.66900634765625 0 534.8106
251.1029510498047 0 2454.0708
282.2513732910156 0 503.57462
288.1550598144531 0 1770.0967
297.0812072753906 0 707.14417
299.0616455078125 0 1009.9031
309.2034912109375 0 662.75507
317.2511291503906 0 530.71045
360.0284729003906 0 653.6263
364.18695068359375 0 1724.8715
369.7227783203125 0 579.2976
372.1901550292969 0 668.37616
387.19268798828125 0 596.7461
387.692138671875 0 1695.9163 y 6
388.19537353515625 0 780.2492
388.6944885253906 0 611.04156
395.6551513671875 0 592.5154
402.1891784667969 0 761.4765
403.1993408203125 0 862.67255 y Ammonia loss 9
420.2257995605469 0 780.67896 y 9
430.0890197753906 0 1286.426
431.08880615234375 0 1135.8843
432.0890197753906 0 683.3518
433.2075500488281 0 700.0442
437.91607666015625 0 705.1247
446.1205749511719 0 919.3691
451.2193908691406 0 782.2251
474.2356262207031 0 3636.3037 w 8
475.2403259277344 0 651.05096
479.2426452636719 0 876.722 w 4
481.2347106933594 0 803.8631
492.1941223144531 0 542.6852
546.2920532226562 0 1175.6727
564.302001953125 0 880.7236
585.3716430664062 0 616.27893
602.2932739257812 0 2579.418 w 7
654.8416748046875 0 1011.51886
665.3511962890625 0 1647.3495
667.34765625 0 835.42694 c Ammonia loss 10
675.8578491210938 0 1708.0812 c 10
676.3605346679688 0 1493.4421 y 1
676.8636474609375 0 580.03735
710.8513793945312 0 661.8508
711.3489990234375 0 1143.4305
711.8522338867188 0 888.0021
719.35693359375 0 883.96936
719.86669921875 0 2232.451
720.3731689453125 0 3933.653
720.8765869140625 0 1725.5422
721.3704223632812 0 1145.4763
721.863037109375 0 1187.1833
735.868896484375 0 581.7677
774.37744140625 0 2028.7626 y 6
779.4280395507812 0 2124.6587 c 6
780.4308471679688 0 819.2246
787.4039916992188 0 941.6773
810.4234008789062 0 1284.8762
841.4946899414062 0 651.5046
858.4029541015625 0 748.0054 y Ammonia loss 5
859.4074096679688 0 2031.3239 z 5
860.4105834960938 0 1146.7955
870.28271484375 0 636.91583
875.4244384765625 0 981.2553 y 5
908.4725341796875 0 3232.6982 c 7
909.4737548828125 0 1663.0951
910.4729614257812 0 705.4063
931.9752807617188 0 934.0212
958.544189453125 0 809.5113
960.3429565429688 0 850.27234
960.9916381835938 0 682.1504
961.4898681640625 0 679.6659
961.9758911132812 0 916.14545
972.4918823242188 0 2450.1406 z 4
973.4917602539062 0 1924.3589
974.4948120117188 0 758.7611
982.4768676757812 0 719.14496
988.4944458007812 0 1040.1694 y 4
992.514404296875 0 778.2781
1036.5313720703125 0 3905.0076 c 8
1037.5301513671875 0 1918.7389
1059.5186767578125 0 754.7749 z 3
1060.525390625 0 958.3102
1075.5372314453125 0 763.01184 y 3
1172.60888671875 0 1612.1248 z 2
1173.6243896484375 0 1069.8804
1174.6151123046875 0 816.1199
1222.6092529296875 0 5567.601 c 9
1223.610595703125 0 4074.5427
1224.6187744140625 0 1579.0872
1334.672607421875 0 798.4215 y Ammonia loss 1
1335.6766357421875 0 800.89606 z 1
1336.686767578125 0 1054.839
1350.707763671875 0 777.0492 c 10
1351.7144775390625 0 821.36084 y 1
1353.7144775390625 0 880.7623
1405.699462890625 0 666.7127
1422.7010498046875 0 1411.1351
1423.7100830078125 0 4618.6865
1424.7152099609375 0 3481.1382
1425.71826171875 0 1694.6655
1438.728759765625 0 1509.4747
1439.73095703125 0 2125.3035
1440.7457275390625 0 4146.7915
1441.7535400390625 0 1929.1412
1442.7476806640625 0 1792.549
3036.781005859375 0 1070.9703
3431.826171875 0 713.9683

Spectrum Details

|  |  |
| --- | --- |
| Matched peaks? Matched peaks The total absolute number of peaks matched. Additionally in brackets the total fraction of peaks matched and the total number of peaks is shown. | 28 (19.86% of 141) |
| FDR? FDR The false discovery rate estimated for this peptide. It is calculated by matching all theoretical fragments with a non-integer shift with the raw peaks for this spectrum. This is done with 40 different shifts. The resulting percentage is the average number of annotated peaks over the number of annotated peaks with the correct spectrum. | 0.43% |
| Satellite FDR? Satellite FDR See the FDR for details on its calculation. This satellite ion specific FDR only contains the satellite ions (d/w) for I/L/J positions. | 2.38% |
| PSM Score? PSM Score The PSM Score as given by Hecklib to this annotated spectrum. It is shown with three significant figures. | 194 |

#### Spectrum 8905? Spectrum 8905 The raw spectrum of this peptide as annotated by Hecklib. The fragments are coloured according to ion type (see legend). Any peaks with a star '\*' as text can be hovered over to see the full details, first the ion type second the mass shift type. By hovering over the amino acids in the peptide or ions in the legend the corresponding peaks are highlighted. By toggling the 'Unassigned' label you can turn the background (unassigned) peaks on or off in the plot. By updating the slider in the Ion legend you can update the spectrum to only show the top X% of the peaks with labels. The top X% means any peak that is within X% of the highest intensity. By dragging in the spectrum you can zoom in to a specific part of the spectrum and use 'Zoom Out' to get back to the original zoom level. The annotation of the spectrum is based on the given sequence in the peptides file and is done with different software so inconsistencies are likely. The peaks are annotated based on the given sequence, with 20 ppm tolerance.

Copy Data

##### Spectrum 8905 (TSV)

###### Preview

```
Loading example...
```

*Click on the button to copy the data to your clipboard.*

Mz MinMz MaxIntensity Max

WidthHeightPeptide font sizePeptide stroke widthSpectrum font sizeSpectrum stroke widthCompact peptide

Ion legend

wxyz

abcd

OtherUnassignedIon^Charge^_Position_Show for top:%

SYLSJTPEQWKS

01.69e+43.39e+45.08e+46.77e+4

Zoom Out

y^+1^_2_y^+1^_3_z^+1^_4_z^+1^_5_y^+1^_6_y^+1^_7_z^+1^_7_y^+1^_7_c^+1^_8_z^+1^_8_y^+1^_8_c^+1^_9_c^+1^_9_y^+1^_9_z^+1^_9_y^+1^_9_w^+1^_10_z^+1^_10_y^+1^_10_c^+1^_10_z^+1^_11_c^+1^_11_y^+1^_11_

0546109216392185

Fragment Matches Table

Show background peaks

| Position | Ion type | Intensity | mz Theoretical | mz Error (Th) | mz Error (ppm) | Charge | Series Number |
| --- | --- | --- | --- | --- | --- | --- | --- |
| - | - | 428 | 127.6 | - | - | 0 | - |
| - | - | 940.5 | 129.1 | - | - | 0 | - |
| - | - | 499.5 | 148.9 | - | - | 0 | - |
| - | - | 643.7 | 148.9 | - | - | 0 | - |
| - | - | 979.1 | 148.9 | - | - | 0 | - |
| - | - | 1125 | 148.9 | - | - | 0 | - |
| - | - | 1127 | 148.9 | - | - | 0 | - |
| - | - | 1573 | 148.9 | - | - | 0 | - |
| - | - | 3396 | 148.9 | - | - | 0 | - |
| - | - | 4388 | 149 | - | - | 0 | - |
| - | - | 2954 | 149 | - | - | 0 | - |
| - | - | 1116 | 149 | - | - | 0 | - |
| - | - | 959.7 | 149 | - | - | 0 | - |
| - | - | 909.7 | 149 | - | - | 0 | - |
| - | - | 674.5 | 149 | - | - | 0 | - |
| - | - | 531.6 | 149 | - | - | 0 | - |
| - | - | 472.2 | 149 | - | - | 0 | - |
| - | - | 529.2 | 149 | - | - | 0 | - |
| - | - | 503.7 | 157.9 | - | - | 0 | - |
| - | - | 435 | 168.5 | - | - | 0 | - |
| - | - | 2084 | 173.1 | - | - | 0 | - |
| - | - | 533.2 | 173.5 | - | - | 0 | - |
| - | - | 452.5 | 195.9 | - | - | 0 | - |
| - | - | 742.9 | 201.1 | - | - | 0 | - |
| - | - | 743.9 | 215.1 | - | - | 0 | - |
| - | - | 5373 | 223.1 | - | - | 0 | - |
| - | - | 644 | 224.1 | - | - | 0 | - |
| - | - | 597.3 | 227.1 | - | - | 0 | - |
| 11 | y | 2550 | 234.1 | 0.0003396 | 1.45 | +1 | 2 |
| - | - | 8556 | 251.1 | - | - | 0 | - |
| - | - | 1207 | 252.1 | - | - | 0 | - |
| - | - | 799.2 | 296.2 | - | - | 0 | - |
| - | - | 602 | 343.4 | - | - | 0 | - |
| - | - | 819.7 | 346.2 | - | - | 0 | - |
| - | - | 607.1 | 355.2 | - | - | 0 | - |
| - | - | 4984 | 364.2 | - | - | 0 | - |
| - | - | 557.5 | 409.5 | - | - | 0 | - |
| - | - | 954 | 415.2 | - | - | 0 | - |
| - | - | 938.3 | 415.3 | - | - | 0 | - |
| 10 | y | 1071 | 420.2 | 0.001288 | 3.065 | +1 | 3 |
| - | - | 3319 | 433.2 | - | - | 0 | - |
| - | - | 1197 | 434.2 | - | - | 0 | - |
| - | - | 1769 | 451.2 | - | - | 0 | - |
| - | - | 1162 | 471.3 | - | - | 0 | - |
| - | - | 778 | 528.3 | - | - | 0 | - |
| 9 | z | 2451 | 532.3 | 0.000205 | 0.3852 | +1 | 4 |
| - | - | 4241 | 533.3 | - | - | 0 | - |
| - | - | 936.3 | 534.3 | - | - | 0 | - |
| - | - | 675.9 | 536.3 | - | - | 0 | - |
| - | - | 884.5 | 541.2 | - | - | 0 | - |
| - | - | 1870 | 546.3 | - | - | 0 | - |
| - | - | 813.6 | 595.3 | - | - | 0 | - |
| - | - | 1128 | 629.3 | - | - | 0 | - |
| - | - | 2294 | 647.3 | - | - | 0 | - |
| 8 | z | 2903 | 661.3 | 7.351E-05 | 0.1112 | +1 | 5 |
| - | - | 4235 | 662.3 | - | - | 0 | - |
| - | - | 1536 | 663.3 | - | - | 0 | - |
| - | - | 1617 | 665.3 | - | - | 0 | - |
| - | - | 787.8 | 666.4 | - | - | 0 | - |
| - | - | 1638 | 669.3 | - | - | 0 | - |
| - | - | 1741 | 710.9 | - | - | 0 | - |
| - | - | 941.1 | 711.4 | - | - | 0 | - |
| - | - | 633.9 | 714.4 | - | - | 0 | - |
| - | - | 641.5 | 720.4 | - | - | 0 | - |
| - | - | 670.1 | 773.4 | - | - | 0 | - |
| 7 | y | 1.432E+04 | 774.4 | 0.0002113 | 0.2729 | +1 | 6 |
| - | - | 5979 | 775.4 | - | - | 0 | - |
| - | - | 819 | 776.4 | - | - | 0 | - |
| - | - | 936.2 | 857.4 | - | - | 0 | - |
| 6 | y | 7620 | 858.4 | 0.0002047 | 0.2385 | +1 | 7 |
| 6 | z | 5330 | 859.4 | 0.002921 | 3.398 | +1 | 7 |
| - | - | 1.338E+04 | 860.4 | - | - | 0 | - |
| - | - | 4517 | 861.4 | - | - | 0 | - |
| - | - | 2255 | 862.4 | - | - | 0 | - |
| 6 | y | 1.094E+04 | 875.4 | 0.00045 | 0.5141 | +1 | 7 |
| - | - | 4194 | 876.4 | - | - | 0 | - |
| - | - | 1388 | 877.4 | - | - | 0 | - |
| 8 | c | 1368 | 908.5 | 0.001061 | 1.168 | +1 | 8 |
| - | - | 750.8 | 947.3 | - | - | 0 | - |
| 5 | z | 1.491E+04 | 972.5 | 0.001413 | 1.453 | +1 | 8 |
| - | - | 1.066E+04 | 973.5 | - | - | 0 | - |
| - | - | 2757 | 974.5 | - | - | 0 | - |
| - | - | 692.4 | 975.5 | - | - | 0 | - |
| - | - | 1313 | 987.5 | - | - | 0 | - |
| 5 | y | 5467 | 988.5 | 0.006038 | 6.108 | +1 | 8 |
| - | - | 2298 | 989.5 | - | - | 0 | - |
| - | - | 863.3 | 990.5 | - | - | 0 | - |
| - | - | 2057 | 992.5 | - | - | 0 | - |
| - | - | 2326 | 993.5 | - | - | 0 | - |
| - | - | 958.9 | 1003 | - | - | 0 | - |
| 9 | c | 786.6 | 1020 | 0.005363 | 5.26 | +1 | 9 |
| 9 | c | 2724 | 1037 | 0.0005563 | 0.5367 | +1 | 9 |
| - | - | 1672 | 1038 | - | - | 0 | - |
| 4 | y | 1449 | 1058 | 0.0004024 | 0.3805 | +1 | 9 |
| 4 | z | 5600 | 1060 | 0.0005549 | 0.5237 | +1 | 9 |
| - | - | 1.392E+04 | 1061 | - | - | 0 | - |
| - | - | 7453 | 1062 | - | - | 0 | - |
| - | - | 1906 | 1063 | - | - | 0 | - |
| - | - | 855 | 1069 | - | - | 0 | - |
| - | - | 945.9 | 1075 | - | - | 0 | - |
| 4 | y | 1.191E+04 | 1076 | 0.0007132 | 0.6631 | +1 | 9 |
| - | - | 7081 | 1077 | - | - | 0 | - |
| - | - | 2463 | 1078 | - | - | 0 | - |
| - | - | 649.5 | 1081 | - | - | 0 | - |
| - | - | 735 | 1082 | - | - | 0 | - |
| - | - | 797.7 | 1086 | - | - | 0 | - |
| 3 | w | 2497 | 1130 | 0.0004409 | 0.3903 | +1 | 10 |
| - | - | 1676 | 1131 | - | - | 0 | - |
| - | - | 1184 | 1157 | - | - | 0 | - |
| - | - | 774 | 1162 | - | - | 0 | - |
| 3 | z | 4865 | 1173 | 0.0002572 | 0.2193 | +1 | 10 |
| - | - | 2513 | 1174 | - | - | 0 | - |
| - | - | 1510 | 1175 | - | - | 0 | - |
| - | - | 4402 | 1179 | - | - | 0 | - |
| - | - | 3000 | 1180 | - | - | 0 | - |
| - | - | 1094 | 1181 | - | - | 0 | - |
| 3 | y | 4947 | 1189 | 0.004943 | 4.159 | +1 | 10 |
| - | - | 2603 | 1190 | - | - | 0 | - |
| - | - | 1350 | 1191 | - | - | 0 | - |
| - | - | 1923 | 1222 | - | - | 0 | - |
| 10 | c | 1.731E+04 | 1223 | 0.002355 | 1.926 | +1 | 10 |
| - | - | 1.278E+04 | 1224 | - | - | 0 | - |
| - | - | 4124 | 1225 | - | - | 0 | - |
| 2 | z | 3366 | 1336 | 0.004285 | 3.208 | +1 | 11 |
| - | - | 1889 | 1337 | - | - | 0 | - |
| - | - | 823.4 | 1338 | - | - | 0 | - |
| 11 | c | 1.312E+04 | 1351 | 0.002957 | 2.189 | +1 | 11 |
| 2 | y | 1.176E+04 | 1352 | 0.01547 | 11.44 | +1 | 11 |
| - | - | 4461 | 1353 | - | - | 0 | - |
| - | - | 1063 | 1354 | - | - | 0 | - |
| - | - | 1438 | 1367 | - | - | 0 | - |
| - | - | 819.2 | 1368 | - | - | 0 | - |
| - | - | 790.3 | 1378 | - | - | 0 | - |
| - | - | 1165 | 1379 | - | - | 0 | - |
| - | - | 1172 | 1380 | - | - | 0 | - |
| - | - | 1715 | 1384 | - | - | 0 | - |
| - | - | 1007 | 1396 | - | - | 0 | - |
| - | - | 719.4 | 1397 | - | - | 0 | - |
| - | - | 736.8 | 1406 | - | - | 0 | - |
| - | - | 1680 | 1412 | - | - | 0 | - |
| - | - | 1832 | 1413 | - | - | 0 | - |
| - | - | 1126 | 1414 | - | - | 0 | - |
| - | - | 649.9 | 1421 | - | - | 0 | - |
| - | - | 2905 | 1422 | - | - | 0 | - |
| - | - | 2.411E+04 | 1423 | - | - | 0 | - |
| - | - | 2.082E+04 | 1424 | - | - | 0 | - |
| - | - | 8858 | 1425 | - | - | 0 | - |
| - | - | 1070 | 1426 | - | - | 0 | - |
| - | - | 1736 | 1437 | - | - | 0 | - |
| - | - | 3548 | 1438 | - | - | 0 | - |
| - | - | 2.677E+04 | 1439 | - | - | 0 | - |
| - | - | 6.703E+04 | 1440 | - | - | 0 | - |
| - | - | 5.123E+04 | 1441 | - | - | 0 | - |
| - | - | 4978 | 1442 | - | - | 0 | - |
| - | - | 1.743E+04 | 1442 | - | - | 0 | - |
| - | - | 2481 | 1443 | - | - | 0 | - |
| - | - | 2091 | 1443 | - | - | 0 | - |
| - | - | 746.3 | 1549 | - | - | 0 | - |
| - | - | 1337 | 2156 | - | - | 0 | - |
| - | - | 1032 | 2157 | - | - | 0 | - |
| - | - | 715.5 | 2158 | - | - | 0 | - |
| - | - | 844.1 | 2159 | - | - | 0 | - |
| - | - | 732.9 | 2161 | - | - | 0 | - |
| - | - | 1052 | 2162 | - | - | 0 | - |
| - | - | 1376 | 2163 | - | - | 0 | - |

m/z Charge Intensity FragmentType MassShift Position
127.62824249267578 0 428.0412
129.10235595703125 0 940.53345
148.8976287841797 0 499.54376
148.90524291992188 0 643.66614
148.91209411621094 0 979.1333
148.91925048828125 0 1124.858
148.9264373779297 0 1127.2004
148.93353271484375 0 1573.4893
148.94131469726562 0 3396.2446
148.95791625976562 0 4388.072
148.96571350097656 0 2954.3218
148.97267150878906 0 1115.7479
148.98011779785156 0 959.71265
148.98744201660156 0 909.70184
148.99452209472656 0 674.52057
149.00169372558594 0 531.5969
149.00868225097656 0 472.22726
149.03822326660156 0 529.2219
157.9376678466797 0 503.7418
168.4743194580078 0 434.9712
173.09230041503906 0 2083.902
173.45069885253906 0 533.15186
195.9469451904297 0 452.49976
201.1235809326172 0 742.91223
215.13858032226562 0 743.9213
223.10787963867188 0 5373.4624
224.1112823486328 0 643.9971
227.10256958007812 0 597.3443
234.14517211914062 0 2550.1414 y 10
251.10275268554688 0 8555.568
252.10646057128906 0 1207.0444
296.19842529296875 0 799.1784
343.40399169921875 0 602.00226
346.17724609375 0 819.74963
355.1621398925781 0 607.118
364.18670654296875 0 4984.0464
409.4647216796875 0 557.4703
415.1973876953125 0 953.97687
415.25555419921875 0 938.3208
420.2254333496094 0 1071.0458 y 9
433.2084655761719 0 3318.6777
434.2101745605469 0 1196.8306
451.21844482421875 0 1769.0209
471.28192138671875 0 1162.0419
528.2809448242188 0 778.02185
532.2637939453125 0 2450.7732 z 8
533.2711791992188 0 4241.425
534.2626953125 0 936.25824
536.3079223632812 0 675.9179
541.2380981445312 0 884.54724
546.2925415039062 0 1870.4188
595.319091796875 0 813.5876
629.3291015625 0 1127.9277
647.3406982421875 0 2294.086
661.3065185546875 0 2902.504 z 7
662.3136596679688 0 4235.126
663.3148193359375 0 1535.7998
665.3482666015625 0 1616.7637
666.354736328125 0 787.8227
669.3380126953125 0 1638.1848
710.8583984375 0 1741.1268
711.3619995117188 0 941.1265
714.3590698242188 0 633.8667
720.3890991210938 0 641.49884
773.3712768554688 0 670.1086
774.3778686523438 0 14317.437 y 6
775.3817749023438 0 5978.9536
776.3826293945312 0 819.0376
857.3937377929688 0 936.2364
858.3994140625 0 7619.5884 y Ammonia loss 5
859.4041137695312 0 5330.4365 z 5
860.4140625 0 13377.252
861.4182739257812 0 4517.3003
862.4200439453125 0 2254.5532
875.4262084960938 0 10940.795 y 5
876.4279174804688 0 4193.7
877.4331665039062 0 1387.7063
908.4713134765625 0 1367.5895 c 7
947.3412475585938 0 750.81433
972.4896850585938 0 14905.415 z 4
973.4939575195312 0 10658.562
974.5008544921875 0 2756.7815
975.49951171875 0 692.43475
987.5052490234375 0 1313.0314
988.5037841796875 0 5467.383 y 4
989.508056640625 0 2297.798
990.5107421875 0 863.2787
992.5162963867188 0 2056.6223
993.5233154296875 0 2325.6733
1003.4562377929688 0 958.8753
1019.509765625 0 786.6273 c Ammonia loss 8
1036.5303955078125 0 2724.0325 c 8
1037.5377197265625 0 1671.5105
1057.5308837890625 0 1448.7185 y Water loss 3
1059.523681640625 0 5599.5176 z 3
1060.5296630859375 0 13919.126
1061.5321044921875 0 7452.999
1062.5406494140625 0 1906.4435
1068.54052734375 0 855.03375
1074.53466796875 0 945.9235
1075.5411376953125 0 11912.165 y 3
1076.54296875 0 7081.034
1077.542724609375 0 2463.1404
1081.426513671875 0 649.5216
1081.5582275390625 0 735.0168
1085.5531005859375 0 797.72437
1129.5528564453125 0 2496.5032 w 2
1130.55810546875 0 1676.0016
1156.5849609375 0 1184.5
1161.5767822265625 0 773.9955
1172.60693359375 0 4864.793 z 2
1173.612060546875 0 2513.1362
1174.608642578125 0 1510.003
1178.5955810546875 0 4402.477
1179.598876953125 0 3000.0007
1180.61181640625 0 1093.6956
1188.6209716796875 0 4947.3667 y 2
1189.627197265625 0 2602.5232
1190.6234130859375 0 1349.8945
1221.599853515625 0 1922.6359
1222.60791015625 0 17306.643 c 9
1223.6104736328125 0 12782.289
1224.6141357421875 0 4123.579
1335.6748046875 0 3366.0037 z 1
1336.6788330078125 0 1889.1865
1337.6776123046875 0 823.35175
1350.7022705078125 0 13120.41 c 10
1351.7047119140625 0 11759.42 y 1
1352.7073974609375 0 4461.457
1353.7044677734375 0 1063.0336
1366.6466064453125 0 1437.897
1367.6654052734375 0 819.19763
1377.60595703125 0 790.25543
1378.602783203125 0 1164.9264
1379.6212158203125 0 1172.2102
1383.6666259765625 0 1715.0204
1395.686279296875 0 1006.8544
1396.7115478515625 0 719.4097
1405.6988525390625 0 736.8481
1411.7318115234375 0 1679.6935
1412.725830078125 0 1831.7698
1413.735595703125 0 1126.2977
1420.690185546875 0 649.88574
1421.7003173828125 0 2905.0808
1422.700439453125 0 24109.898
1423.7020263671875 0 20823.648
1424.702880859375 0 8857.721
1425.716064453125 0 1070.1754
1436.713623046875 0 1735.9095
1437.658447265625 0 3547.646
1438.7177734375 0 26770.611
1439.72314453125 0 67032.98
1440.7266845703125 0 51225.508
1441.6007080078125 0 4977.6724
1441.728759765625 0 17431.566
1442.6124267578125 0 2481.116
1442.7476806640625 0 2090.7036
1549.29443359375 0 746.2878
2156.087890625 0 1336.5907
2157.061767578125 0 1031.607
2158.121826171875 0 715.5431
2159.11865234375 0 844.07886
2161.1416015625 0 732.88306
2162.111083984375 0 1051.8414
2163.091064453125 0 1375.537

Spectrum Details

|  |  |
| --- | --- |
| Matched peaks? Matched peaks The total absolute number of peaks matched. Additionally in brackets the total fraction of peaks matched and the total number of peaks is shown. | 23 (13.94% of 165) |
| FDR? FDR The false discovery rate estimated for this peptide. It is calculated by matching all theoretical fragments with a non-integer shift with the raw peaks for this spectrum. This is done with 40 different shifts. The resulting percentage is the average number of annotated peaks over the number of annotated peaks with the correct spectrum. | 0.10% |
| Satellite FDR? Satellite FDR See the FDR for details on its calculation. This satellite ion specific FDR only contains the satellite ions (d/w) for I/L/J positions. | 0.00% |
| PSM Score? PSM Score The PSM Score as given by Hecklib to this annotated spectrum. It is shown with three significant figures. | 239 |

#### Reverse Lookup? Reverse Lookup All places where this read could be placed.

| Group | Segment | Template | Template Part | Read Part | Score | Unique |
| --- | --- | --- | --- | --- | --- | --- |
| Homo sapiens Light Chain | IGLC | IGLC2 | [69..81] | [0..12] | 96 | False |
| Homo sapiens Light Chain | IGLC | IGLC3 | [67..79] | [0..12] | 96 | False |
| Homo sapiens Light Chain | IGLC | IGLC6 | [69..81] | [0..12] | 96 | False |
| Homo sapiens Light Chain | IGLC | IGLC7 | [69..81] | [0..12] | 96 | False |

| Recombined | Template Part | Read Part | Score | Unique |
| --- | --- | --- | --- | --- |
| REC-0-1\_002 | [180..192] | [0..12] | 96 | True |

#### Meta Information from Multiple reads

##### Number of combined reads

4

##### Intensity

0.6387

##### TotalArea

1.94E+08

##### Changes to the peptide sequence

SYLSJTPEQWKS

I→JNo support for either Leucine or Isoleucine based on side chain ions (Position: 5)

J→LSupport for Leucine based on side chain ions (1 for L 0 for I) (Position: 3)

L→ISupport for Isoleucine based on side chain ions (1 for I 0 for L) (Position: 5)

L→JEqual support for both Leucine and Isoleucine based on side chain ions (1 ions for both) (Position: 3)

#### Positional Score

Copy Data

##### Positional Score (TSV)

###### Preview

```
Loading example...
```

*Click on the button to copy the data to your clipboard.*

1001234567891011

Label Value
"0" 0.5
"1" 0.497
"2" 0.497
"3" 0.497
"4" 0.497
"5" 0.482
"6" 0.445
"7" 0.448
"8" 0.45
"9" 0.48
"10" 0.497
"11" 0.497

#### Meta Information from PEAKS

##### Scan Identifier

F1:8765

##### Original sequence

S

Y

L

S

L

T

P

E

Q

W

K

S

##### Posttranslational Modifications

##### Source File

D:\separate\_stitch\_analyses\xle-disambiguation\raw\20210323\_F1\_UM1\_Peng0013\_SA\_F59\_ingel\_3ug\_ELA.raw

##### Fraction

1

##### Scan Feature

F1:13758

##### De Novo Score

99

##### ConfidenceScore

99

### m/z

719.8654

##### Mass

1437.7139

##### Charge

2

##### Retention Time

48.13

##### Predicted Retention Time

-

##### Area

6.456E+07

##### Parts Per Million

1.6

##### Fragmentation mode

ETHCD

##### Originating file

01 D:\separate\_stitch\_analyses\xle-disambiguation\20210325\_F59\_3ug\_DENOVO\_12.csv

#### Meta Information from PEAKS

##### Scan Identifier

F1:8833

##### Original sequence

S

Y

L

S

L

T

P

E

Q

W

K

S

##### Posttranslational Modifications

##### Source File

D:\separate\_stitch\_analyses\xle-disambiguation\raw\20210323\_F1\_UM1\_Peng0013\_SA\_F59\_ingel\_3ug\_ELA.raw

##### Fraction

1

##### Scan Feature

F1:13758

##### De Novo Score

98

##### ConfidenceScore

98

### m/z

719.8654

##### Mass

1437.7139

##### Charge

2

##### Retention Time

48.13

##### Predicted Retention Time

-

##### Area

6.456E+07

##### Parts Per Million

1.6

##### Fragmentation mode

ETHCD

##### Originating file

01 D:\separate\_stitch\_analyses\xle-disambiguation\20210325\_F59\_3ug\_DENOVO\_12.csv

#### Meta Information from PEAKS

##### Scan Identifier

F1:8777

##### Original sequence

S

Y

L

S

L

T

P

E

Q

W

K

S

##### Posttranslational Modifications

##### Source File

D:\separate\_stitch\_analyses\xle-disambiguation\raw\20210323\_F1\_UM1\_Peng0013\_SA\_F59\_ingel\_3ug\_ELA.raw

##### Fraction

1

##### Scan Feature

F1:3290

##### De Novo Score

96

##### ConfidenceScore

96

### m/z

480.2455

##### Mass

1437.7139

##### Charge

3

##### Retention Time

48.13

##### Predicted Retention Time

-

##### Area

2.926E+05

##### Parts Per Million

0.7

##### Fragmentation mode

ETHCD

##### Originating file

01 D:\separate\_stitch\_analyses\xle-disambiguation\20210325\_F59\_3ug\_DENOVO\_12.csv

#### Meta Information from PEAKS

##### Scan Identifier

F1:8905

##### Original sequence

S

Y

L

S

L

T

P

E

Q

W

K

S

##### Posttranslational Modifications

##### Source File

D:\separate\_stitch\_analyses\xle-disambiguation\raw\20210323\_F1\_UM1\_Peng0013\_SA\_F59\_ingel\_3ug\_ELA.raw

##### Fraction

1

##### Scan Feature

F1:13758

##### De Novo Score

96

##### ConfidenceScore

96

### m/z

719.8654

##### Mass

1437.7139

##### Charge

2

##### Retention Time

48.13

##### Predicted Retention Time

-

##### Area

6.456E+07

##### Parts Per Million

1.6

##### Fragmentation mode

ETHCD

##### Originating file

01 D:\separate\_stitch\_analyses\xle-disambiguation\20210325\_F59\_3ug\_DENOVO\_12.csv
