## Supplementary material for "A handle on mass coincidence errors in *de novo* sequencing of antibodies by bottom-up proteomics": Combined_042.html

Details Combined\_042 | Stitch OverviewUndefined

### Read Combined\_042

#### Sequence (length=10)

EVSJQDKTGF

#### Spectrum 7326? Spectrum 7326 The raw spectrum of this peptide as annotated by Hecklib. The fragments are coloured according to ion type (see legend). Any peaks with a star '\*' as text can be hovered over to see the full details, first the ion type second the mass shift type. By hovering over the amino acids in the peptide or ions in the legend the corresponding peaks are highlighted. By toggling the 'Unassigned' label you can turn the background (unassigned) peaks on or off in the plot. By updating the slider in the Ion legend you can update the spectrum to only show the top X% of the peaks with labels. The top X% means any peak that is within X% of the highest intensity. By dragging in the spectrum you can zoom in to a specific part of the spectrum and use 'Zoom Out' to get back to the original zoom level. The annotation of the spectrum is based on the given sequence in the peptides file and is done with different software so inconsistencies are likely. The peaks are annotated based on the given sequence, with 20 ppm tolerance.

Copy Data

##### Spectrum 7326 (TSV)

###### Preview

```
Loading example...
```

*Click on the button to copy the data to your clipboard.*

Mz MinMz MaxIntensity Max

WidthHeightPeptide font sizePeptide stroke widthSpectrum font sizeSpectrum stroke widthCompact peptide

Ion legend

wxyz

abcd

OtherUnassignedIon^Charge^_Position_Show for top:%

EVSJQDKTGF

01.55e+53.10e+54.65e+56.21e+5

Zoom Out

y^+1^_1_d^+1^_2_a^+1^_2_b^+1^_2_y^+1^_2_a^+1^_3_a^+1^_3_b^+1^_3_y^+2^_5_b^+1^_3_y^+1^_3_b^+2^_6_y^+1^_3_y^+2^_6_y^+2^_6_b^+2^_7_b^+2^_7_b^+1^_4_y^+2^_7_y^+2^_7_y^+2^_7_b^+1^_4_b^+2^_8_y^+1^_4_y^+1^_4_y^+2^_8_y^+2^_8_b^+2^_8_y^+2^_8_y^+1^_4_b^+2^_9_b^+2^_9_y^+2^_9_y^+2^_9_b^+1^_5_b^+1^_5_b^+1^_5_\*\*y^+1^_5_\*y^+1^_5_b^+1^_6_b^+1^_6_b^+1^_6_y^+1^_6_y^+1^_6_y^+1^_6_b^+1^_7_b^+1^_7_b^+1^_7_y^+1^_7_y^+1^_7_y^+1^_7_b^+1^_8_b^+1^_8_y^+1^_8_y^+1^_8_b^+1^_8_y^+1^_8_b^+1^_9_b^+1^_9_b^+1^_9_

0796159223893185

Fragment Matches Table

Show background peaks

| Position | Ion type | Intensity | mz Theoretical | mz Error (Th) | mz Error (ppm) | Charge | Series Number |
| --- | --- | --- | --- | --- | --- | --- | --- |
| - | - | 3.536E+05 | 120.1 | - | - | 0 | - |
| - | - | 2.749E+04 | 121.1 | - | - | 0 | - |
| - | - | 674.3 | 122.1 | - | - | 0 | - |
| - | - | 378 | 122.6 | - | - | 0 | - |
| - | - | 1344 | 123.1 | - | - | 0 | - |
| - | - | 980.7 | 124.1 | - | - | 0 | - |
| - | - | 421.1 | 124.1 | - | - | 0 | - |
| - | - | 934.4 | 125 | - | - | 0 | - |
| - | - | 3916 | 126.1 | - | - | 0 | - |
| - | - | 2135 | 127.1 | - | - | 0 | - |
| - | - | 2.222E+04 | 127.1 | - | - | 0 | - |
| - | - | 1633 | 128.1 | - | - | 0 | - |
| - | - | 1.449E+04 | 128.1 | - | - | 0 | - |
| - | - | 1761 | 128.1 | - | - | 0 | - |
| - | - | 5733 | 129.1 | - | - | 0 | - |
| - | - | 3.664E+05 | 129.1 | - | - | 0 | - |
| - | - | 1264 | 130.1 | - | - | 0 | - |
| - | - | 2749 | 130.1 | - | - | 0 | - |
| - | - | 2541 | 130.1 | - | - | 0 | - |
| - | - | 3285 | 130.1 | - | - | 0 | - |
| - | - | 2.151E+04 | 130.1 | - | - | 0 | - |
| - | - | 6287 | 131 | - | - | 0 | - |
| - | - | 1.445E+04 | 131.1 | - | - | 0 | - |
| - | - | 2.633E+04 | 131.1 | - | - | 0 | - |
| - | - | 624.7 | 132.1 | - | - | 0 | - |
| - | - | 638.4 | 132.1 | - | - | 0 | - |
| - | - | 1701 | 132.1 | - | - | 0 | - |
| - | - | 1940 | 132.1 | - | - | 0 | - |
| - | - | 5288 | 133.1 | - | - | 0 | - |
| - | - | 5681 | 136.1 | - | - | 0 | - |
| - | - | 886.2 | 137.1 | - | - | 0 | - |
| - | - | 1.447E+04 | 138.1 | - | - | 0 | - |
| - | - | 2065 | 139.1 | - | - | 0 | - |
| - | - | 8302 | 139.1 | - | - | 0 | - |
| - | - | 998.3 | 139.1 | - | - | 0 | - |
| - | - | 1047 | 139.1 | - | - | 0 | - |
| - | - | 920.6 | 140.1 | - | - | 0 | - |
| - | - | 2713 | 140.1 | - | - | 0 | - |
| - | - | 5827 | 141.1 | - | - | 0 | - |
| - | - | 1.305E+04 | 141.1 | - | - | 0 | - |
| - | - | 2463 | 142.1 | - | - | 0 | - |
| - | - | 630.4 | 142.1 | - | - | 0 | - |
| - | - | 612.1 | 142.1 | - | - | 0 | - |
| - | - | 605.9 | 143 | - | - | 0 | - |
| - | - | 536.5 | 143.1 | - | - | 0 | - |
| - | - | 1499 | 143.1 | - | - | 0 | - |
| - | - | 863.2 | 147.1 | - | - | 0 | - |
| - | - | 2129 | 147.1 | - | - | 0 | - |
| - | - | 517.4 | 148.1 | - | - | 0 | - |
| - | - | 992.8 | 149 | - | - | 0 | - |
| - | - | 524.4 | 149 | - | - | 0 | - |
| - | - | 466.4 | 149.1 | - | - | 0 | - |
| - | - | 6354 | 149.1 | - | - | 0 | - |
| - | - | 2064 | 149.1 | - | - | 0 | - |
| - | - | 646.3 | 150.1 | - | - | 0 | - |
| - | - | 524.7 | 150.1 | - | - | 0 | - |
| - | - | 1.736E+04 | 151.1 | - | - | 0 | - |
| - | - | 1716 | 152.1 | - | - | 0 | - |
| - | - | 1138 | 152.1 | - | - | 0 | - |
| - | - | 1095 | 153.1 | - | - | 0 | - |
| - | - | 1075 | 153.1 | - | - | 0 | - |
| - | - | 8742 | 154.1 | - | - | 0 | - |
| - | - | 5494 | 155.1 | - | - | 0 | - |
| - | - | 6.098E+04 | 155.1 | - | - | 0 | - |
| - | - | 595.3 | 156.1 | - | - | 0 | - |
| - | - | 550.1 | 156.1 | - | - | 0 | - |
| - | - | 1849 | 156.1 | - | - | 0 | - |
| - | - | 649 | 156.1 | - | - | 0 | - |
| - | - | 5153 | 156.1 | - | - | 0 | - |
| - | - | 1039 | 157.1 | - | - | 0 | - |
| - | - | 1883 | 157.1 | - | - | 0 | - |
| - | - | 5.405E+04 | 159.1 | - | - | 0 | - |
| - | - | 7102 | 159.1 | - | - | 0 | - |
| - | - | 1503 | 159.1 | - | - | 0 | - |
| - | - | 3000 | 160.1 | - | - | 0 | - |
| - | - | 1066 | 163.1 | - | - | 0 | - |
| - | - | 2764 | 165.1 | - | - | 0 | - |
| 10 | y | 4.35E+05 | 166.1 | 0.000598 | 3.6 | +1 | 1 |
| - | - | 754.7 | 167.1 | - | - | 0 | - |
| - | - | 2619 | 167.1 | - | - | 0 | - |
| - | - | 3.997E+04 | 167.1 | - | - | 0 | - |
| - | - | 1.747E+04 | 167.1 | - | - | 0 | - |
| - | - | 913.2 | 167.1 | - | - | 0 | - |
| - | - | 1582 | 168.1 | - | - | 0 | - |
| - | - | 1.464E+04 | 168.1 | - | - | 0 | - |
| - | - | 1467 | 168.1 | - | - | 0 | - |
| - | - | 977.8 | 168.2 | - | - | 0 | - |
| - | - | 1178 | 169.1 | - | - | 0 | - |
| 2 | d | 1964 | 169.1 | 0.0001388 | 0.821 | +1 | 2 |
| - | - | 1416 | 169.1 | - | - | 0 | - |
| - | - | 1581 | 169.1 | - | - | 0 | - |
| - | - | 1960 | 171.1 | - | - | 0 | - |
| - | - | 684.7 | 171.1 | - | - | 0 | - |
| - | - | 736.6 | 172.1 | - | - | 0 | - |
| - | - | 5.337E+05 | 173.1 | - | - | 0 | - |
| - | - | 2242 | 173.5 | - | - | 0 | - |
| - | - | 4.378E+04 | 174.1 | - | - | 0 | - |
| - | - | 1549 | 175.1 | - | - | 0 | - |
| - | - | 9.007E+04 | 177.1 | - | - | 0 | - |
| - | - | 8753 | 178.1 | - | - | 0 | - |
| - | - | 723.7 | 179.1 | - | - | 0 | - |
| - | - | 1219 | 179.1 | - | - | 0 | - |
| - | - | 2306 | 180.1 | - | - | 0 | - |
| - | - | 1891 | 180.1 | - | - | 0 | - |
| - | - | 1.821E+04 | 181.1 | - | - | 0 | - |
| - | - | 1919 | 181.1 | - | - | 0 | - |
| - | - | 433.3 | 181.1 | - | - | 0 | - |
| - | - | 917.2 | 182 | - | - | 0 | - |
| - | - | 1013 | 182.1 | - | - | 0 | - |
| - | - | 971.3 | 182.1 | - | - | 0 | - |
| - | - | 2010 | 182.1 | - | - | 0 | - |
| 2 | a | 2.683E+05 | 183.1 | 9.551E-05 | 0.5216 | +1 | 2 |
| - | - | 1.455E+04 | 184.1 | - | - | 0 | - |
| - | - | 2.239E+04 | 184.1 | - | - | 0 | - |
| - | - | 2663 | 184.1 | - | - | 0 | - |
| - | - | 923.1 | 185.1 | - | - | 0 | - |
| - | - | 938.9 | 185.1 | - | - | 0 | - |
| - | - | 2.32E+04 | 185.1 | - | - | 0 | - |
| - | - | 4627 | 186.1 | - | - | 0 | - |
| - | - | 980.3 | 186.2 | - | - | 0 | - |
| - | - | 693.4 | 187.1 | - | - | 0 | - |
| - | - | 1.191E+04 | 187.1 | - | - | 0 | - |
| - | - | 1618 | 188.1 | - | - | 0 | - |
| - | - | 1364 | 188.1 | - | - | 0 | - |
| - | - | 1659 | 191.1 | - | - | 0 | - |
| - | - | 3318 | 193.1 | - | - | 0 | - |
| - | - | 367.2 | 194.1 | - | - | 0 | - |
| - | - | 2.001E+04 | 194.1 | - | - | 0 | - |
| - | - | 4108 | 195.1 | - | - | 0 | - |
| - | - | 2251 | 195.1 | - | - | 0 | - |
| - | - | 4167 | 196.1 | - | - | 0 | - |
| - | - | 5123 | 197.1 | - | - | 0 | - |
| - | - | 3082 | 197.1 | - | - | 0 | - |
| - | - | 3.844E+04 | 198.1 | - | - | 0 | - |
| - | - | 1.078E+04 | 198.1 | - | - | 0 | - |
| - | - | 1.06E+04 | 199.1 | - | - | 0 | - |
| - | - | 3395 | 199.1 | - | - | 0 | - |
| - | - | 3274 | 199.1 | - | - | 0 | - |
| - | - | 909.3 | 199.1 | - | - | 0 | - |
| - | - | 504.3 | 199.2 | - | - | 0 | - |
| - | - | 1380 | 200.1 | - | - | 0 | - |
| - | - | 1.604E+05 | 201.1 | - | - | 0 | - |
| - | - | 1.629E+04 | 202.1 | - | - | 0 | - |
| - | - | 4543 | 202.2 | - | - | 0 | - |
| - | - | 857.3 | 203.1 | - | - | 0 | - |
| - | - | 696 | 203.1 | - | - | 0 | - |
| - | - | 1020 | 203.1 | - | - | 0 | - |
| - | - | 588.3 | 203.2 | - | - | 0 | - |
| - | - | 1316 | 204.1 | - | - | 0 | - |
| - | - | 1007 | 204.1 | - | - | 0 | - |
| - | - | 1.127E+04 | 205.1 | - | - | 0 | - |
| - | - | 1002 | 206.1 | - | - | 0 | - |
| - | - | 618.9 | 207.1 | - | - | 0 | - |
| - | - | 720.3 | 208.1 | - | - | 0 | - |
| - | - | 6420 | 208.1 | - | - | 0 | - |
| - | - | 6300 | 208.1 | - | - | 0 | - |
| - | - | 5826 | 209.1 | - | - | 0 | - |
| - | - | 5.355E+04 | 209.1 | - | - | 0 | - |
| - | - | 616.4 | 210.1 | - | - | 0 | - |
| - | - | 555.1 | 210.1 | - | - | 0 | - |
| - | - | 5008 | 210.1 | - | - | 0 | - |
| - | - | 1915 | 210.1 | - | - | 0 | - |
| 2 | b | 9.553E+04 | 211.1 | 8.445E-05 | 0.4001 | +1 | 2 |
| - | - | 1.034E+04 | 212.1 | - | - | 0 | - |
| - | - | 1.281E+04 | 212.1 | - | - | 0 | - |
| - | - | 1101 | 213.1 | - | - | 0 | - |
| - | - | 5576 | 213.1 | - | - | 0 | - |
| - | - | 971.5 | 213.1 | - | - | 0 | - |
| - | - | 4124 | 214.1 | - | - | 0 | - |
| - | - | 2748 | 214.2 | - | - | 0 | - |
| - | - | 4106 | 215.1 | - | - | 0 | - |
| - | - | 1.305E+04 | 216.1 | - | - | 0 | - |
| - | - | 884.8 | 217.1 | - | - | 0 | - |
| - | - | 507 | 217.7 | - | - | 0 | - |
| - | - | 9519 | 218.2 | - | - | 0 | - |
| - | - | 983.6 | 219.2 | - | - | 0 | - |
| - | - | 3546 | 222.1 | - | - | 0 | - |
| 9 | y | 2.527E+05 | 223.1 | 0.0006949 | 3.115 | +1 | 2 |
| - | - | 3.062E+04 | 224.1 | - | - | 0 | - |
| - | - | 3957 | 224.1 | - | - | 0 | - |
| - | - | 1408 | 225 | - | - | 0 | - |
| - | - | 3759 | 225.1 | - | - | 0 | - |
| - | - | 1689 | 225.1 | - | - | 0 | - |
| - | - | 1.333E+04 | 225.1 | - | - | 0 | - |
| - | - | 586.3 | 225.1 | - | - | 0 | - |
| - | - | 7.623E+04 | 226.1 | - | - | 0 | - |
| - | - | 1.609E+05 | 226.1 | - | - | 0 | - |
| - | - | 3.44E+04 | 227.1 | - | - | 0 | - |
| - | - | 7320 | 227.1 | - | - | 0 | - |
| - | - | 1.101E+04 | 227.1 | - | - | 0 | - |
| - | - | 1.503E+04 | 227.1 | - | - | 0 | - |
| - | - | 3110 | 228.1 | - | - | 0 | - |
| - | - | 634.2 | 228.1 | - | - | 0 | - |
| - | - | 846.4 | 228.1 | - | - | 0 | - |
| - | - | 1036 | 228.1 | - | - | 0 | - |
| - | - | 1806 | 228.1 | - | - | 0 | - |
| - | - | 489.3 | 229.1 | - | - | 0 | - |
| - | - | 3927 | 229.1 | - | - | 0 | - |
| - | - | 705.7 | 229.2 | - | - | 0 | - |
| - | - | 814.3 | 230.1 | - | - | 0 | - |
| - | - | 1.917E+04 | 230.2 | - | - | 0 | - |
| - | - | 812 | 231.1 | - | - | 0 | - |
| - | - | 3425 | 231.2 | - | - | 0 | - |
| - | - | 1041 | 233.1 | - | - | 0 | - |
| - | - | 2484 | 234.1 | - | - | 0 | - |
| - | - | 2349 | 236.1 | - | - | 0 | - |
| - | - | 727.3 | 237.1 | - | - | 0 | - |
| - | - | 1804 | 238.1 | - | - | 0 | - |
| - | - | 2071 | 239.1 | - | - | 0 | - |
| - | - | 720.7 | 239.1 | - | - | 0 | - |
| - | - | 1173 | 239.2 | - | - | 0 | - |
| - | - | 1061 | 240.1 | - | - | 0 | - |
| - | - | 6110 | 240.1 | - | - | 0 | - |
| - | - | 1731 | 241.1 | - | - | 0 | - |
| - | - | 808.1 | 241.1 | - | - | 0 | - |
| - | - | 8.374E+04 | 242.2 | - | - | 0 | - |
| - | - | 1.071E+04 | 243.1 | - | - | 0 | - |
| - | - | 9197 | 243.2 | - | - | 0 | - |
| - | - | 7.361E+04 | 244.1 | - | - | 0 | - |
| - | - | 1.435E+05 | 244.1 | - | - | 0 | - |
| - | - | 1034 | 245.1 | - | - | 0 | - |
| - | - | 6011 | 245.1 | - | - | 0 | - |
| - | - | 1.498E+04 | 245.1 | - | - | 0 | - |
| - | - | 890.6 | 246.1 | - | - | 0 | - |
| - | - | 680.5 | 246.1 | - | - | 0 | - |
| - | - | 2779 | 248.1 | - | - | 0 | - |
| - | - | 808.2 | 248.1 | - | - | 0 | - |
| - | - | 1529 | 248.1 | - | - | 0 | - |
| - | - | 2825 | 248.2 | - | - | 0 | - |
| - | - | 2318 | 249.1 | - | - | 0 | - |
| - | - | 1259 | 250.2 | - | - | 0 | - |
| - | - | 1332 | 251.2 | - | - | 0 | - |
| - | - | 879.8 | 251.2 | - | - | 0 | - |
| 3 | a | 6647 | 252.1 | 0.0001619 | 0.6423 | +1 | 3 |
| - | - | 4135 | 253.1 | - | - | 0 | - |
| - | - | 772.5 | 254.1 | - | - | 0 | - |
| - | - | 3947 | 254.1 | - | - | 0 | - |
| - | - | 3070 | 254.2 | - | - | 0 | - |
| - | - | 933.3 | 255.1 | - | - | 0 | - |
| - | - | 1150 | 256.1 | - | - | 0 | - |
| - | - | 1.005E+04 | 258.1 | - | - | 0 | - |
| - | - | 1375 | 259.1 | - | - | 0 | - |
| - | - | 1281 | 259.2 | - | - | 0 | - |
| - | - | 602.7 | 260.1 | - | - | 0 | - |
| - | - | 5479 | 261.1 | - | - | 0 | - |
| - | - | 669.9 | 261.2 | - | - | 0 | - |
| - | - | 510.8 | 262.1 | - | - | 0 | - |
| - | - | 1739 | 262.1 | - | - | 0 | - |
| - | - | 2128 | 262.1 | - | - | 0 | - |
| - | - | 705.1 | 263.1 | - | - | 0 | - |
| - | - | 1852 | 264.1 | - | - | 0 | - |
| - | - | 5771 | 265.1 | - | - | 0 | - |
| - | - | 1241 | 265.2 | - | - | 0 | - |
| - | - | 1264 | 265.6 | - | - | 0 | - |
| - | - | 6839 | 266.1 | - | - | 0 | - |
| - | - | 1217 | 266.1 | - | - | 0 | - |
| - | - | 3402 | 266.2 | - | - | 0 | - |
| - | - | 1116 | 267.1 | - | - | 0 | - |
| - | - | 2380 | 268.1 | - | - | 0 | - |
| - | - | 5154 | 269.2 | - | - | 0 | - |
| 3 | a | 2.658E+04 | 270.1 | 0.0001106 | 0.4093 | +1 | 3 |
| - | - | 2935 | 271.1 | - | - | 0 | - |
| - | - | 992.4 | 273.2 | - | - | 0 | - |
| - | - | 4386 | 274.1 | - | - | 0 | - |
| - | - | 1652 | 276.1 | - | - | 0 | - |
| - | - | 582.1 | 277.1 | - | - | 0 | - |
| - | - | 5000 | 278.2 | - | - | 0 | - |
| - | - | 6360 | 279.1 | - | - | 0 | - |
| 3 | b | 7.95E+04 | 280.1 | 0.0002577 | 0.9199 | +1 | 3 |
| - | - | 1.07E+04 | 281.1 | - | - | 0 | - |
| - | - | 1201 | 281.2 | - | - | 0 | - |
| - | - | 548.8 | 282.1 | - | - | 0 | - |
| - | - | 3001 | 282.1 | - | - | 0 | - |
| - | - | 2325 | 282.2 | - | - | 0 | - |
| - | - | 4.794E+04 | 283.1 | - | - | 0 | - |
| - | - | 2066 | 284.1 | - | - | 0 | - |
| 6 | y | 6008 | 284.1 | 0.001967 | 6.922 | +2 | 5 |
| - | - | 943.1 | 284.2 | - | - | 0 | - |
| - | - | 545 | 285 | - | - | 0 | - |
| - | - | 1673 | 286.1 | - | - | 0 | - |
| - | - | 9795 | 287.2 | - | - | 0 | - |
| - | - | 1100 | 288.1 | - | - | 0 | - |
| - | - | 860.7 | 288.2 | - | - | 0 | - |
| - | - | 1404 | 288.2 | - | - | 0 | - |
| - | - | 3887 | 290.1 | - | - | 0 | - |
| - | - | 931 | 291.1 | - | - | 0 | - |
| - | - | 6061 | 291.1 | - | - | 0 | - |
| - | - | 1.016E+04 | 292.1 | - | - | 0 | - |
| - | - | 703.1 | 292.1 | - | - | 0 | - |
| - | - | 583.3 | 293.1 | - | - | 0 | - |
| - | - | 2309 | 293.2 | - | - | 0 | - |
| - | - | 5266 | 294.1 | - | - | 0 | - |
| - | - | 590.4 | 295.1 | - | - | 0 | - |
| - | - | 616.7 | 296.1 | - | - | 0 | - |
| - | - | 707.2 | 297.1 | - | - | 0 | - |
| - | - | 1.761E+04 | 297.2 | - | - | 0 | - |
| 3 | b | 1.955E+05 | 298.1 | 0.0004047 | 1.357 | +1 | 3 |
| - | - | 1139 | 299.1 | - | - | 0 | - |
| - | - | 2.79E+04 | 299.1 | - | - | 0 | - |
| - | - | 2445 | 299.2 | - | - | 0 | - |
| - | - | 2629 | 300.1 | - | - | 0 | - |
| - | - | 773.6 | 300.2 | - | - | 0 | - |
| - | - | 9540 | 301.2 | - | - | 0 | - |
| - | - | 1535 | 301.2 | - | - | 0 | - |
| - | - | 1836 | 302.2 | - | - | 0 | - |
| - | - | 916.5 | 304.1 | - | - | 0 | - |
| - | - | 622.2 | 305.1 | - | - | 0 | - |
| - | - | 753.1 | 305.1 | - | - | 0 | - |
| - | - | 8425 | 305.2 | - | - | 0 | - |
| 8 | y | 1.665E+04 | 306.1 | 0.0009194 | 3.003 | +1 | 3 |
| - | - | 1014 | 306.2 | - | - | 0 | - |
| - | - | 2341 | 307.1 | - | - | 0 | - |
| - | - | 2.22E+04 | 309.2 | - | - | 0 | - |
| - | - | 6896 | 310.1 | - | - | 0 | - |
| - | - | 3315 | 310.2 | - | - | 0 | - |
| - | - | 3615 | 311.1 | - | - | 0 | - |
| - | - | 5644 | 311.2 | - | - | 0 | - |
| - | - | 1300 | 312.2 | - | - | 0 | - |
| - | - | 928.9 | 312.2 | - | - | 0 | - |
| - | - | 8090 | 315.2 | - | - | 0 | - |
| - | - | 1096 | 316.1 | - | - | 0 | - |
| - | - | 2789 | 316.2 | - | - | 0 | - |
| - | - | 1804 | 316.7 | - | - | 0 | - |
| - | - | 5755 | 317.2 | - | - | 0 | - |
| - | - | 1135 | 318.2 | - | - | 0 | - |
| - | - | 928.4 | 318.2 | - | - | 0 | - |
| 6 | b | 1.28E+04 | 319.1 | 0.004382 | 13.73 | +2 | 6 |
| - | - | 1014 | 320.1 | - | - | 0 | - |
| - | - | 2281 | 320.1 | - | - | 0 | - |
| - | - | 1853 | 321.2 | - | - | 0 | - |
| - | - | 2351 | 322.1 | - | - | 0 | - |
| - | - | 1519 | 323.2 | - | - | 0 | - |
| - | - | 679.9 | 323.2 | - | - | 0 | - |
| - | - | 743 | 323.7 | - | - | 0 | - |
| 8 | y | 7.86E+04 | 324.2 | 0.001036 | 3.196 | +1 | 3 |
| - | - | 1.197E+04 | 325.2 | - | - | 0 | - |
| - | - | 1505 | 326.2 | - | - | 0 | - |
| - | - | 3.2E+04 | 327.2 | - | - | 0 | - |
| - | - | 5134 | 328.2 | - | - | 0 | - |
| - | - | 6359 | 328.2 | - | - | 0 | - |
| - | - | 821.1 | 328.7 | - | - | 0 | - |
| - | - | 2499 | 329.1 | - | - | 0 | - |
| - | - | 6.072E+04 | 329.2 | - | - | 0 | - |
| - | - | 1.022E+04 | 330.2 | - | - | 0 | - |
| - | - | 718.8 | 330.7 | - | - | 0 | - |
| - | - | 744.3 | 331.2 | - | - | 0 | - |
| - | - | 1504 | 334.1 | - | - | 0 | - |
| - | - | 1.697E+04 | 336.2 | - | - | 0 | - |
| - | - | 6.077E+04 | 337.2 | - | - | 0 | - |
| - | - | 2481 | 338.1 | - | - | 0 | - |
| - | - | 1.164E+04 | 338.2 | - | - | 0 | - |
| 5 | y | 5.348E+04 | 339.2 | 0.0008485 | 2.502 | +2 | 6 |
| - | - | 1.524E+04 | 339.7 | - | - | 0 | - |
| - | - | 8919 | 340.2 | - | - | 0 | - |
| - | - | 2113 | 340.2 | - | - | 0 | - |
| - | - | 1260 | 340.7 | - | - | 0 | - |
| - | - | 2190 | 341.2 | - | - | 0 | - |
| - | - | 777.5 | 341.2 | - | - | 0 | - |
| - | - | 665.2 | 343.2 | - | - | 0 | - |
| - | - | 820.6 | 343.7 | - | - | 0 | - |
| - | - | 6.612E+04 | 345.2 | - | - | 0 | - |
| - | - | 1.039E+04 | 346.2 | - | - | 0 | - |
| - | - | 1645 | 347.2 | - | - | 0 | - |
| - | - | 2549 | 347.2 | - | - | 0 | - |
| 5 | y | 6.35E+04 | 348.2 | 0.0009067 | 2.604 | +2 | 6 |
| - | - | 2.656E+04 | 348.7 | - | - | 0 | - |
| - | - | 3728 | 349.2 | - | - | 0 | - |
| - | - | 4426 | 349.2 | - | - | 0 | - |
| - | - | 3640 | 351.1 | - | - | 0 | - |
| - | - | 9.076E+04 | 354.2 | - | - | 0 | - |
| - | - | 1612 | 355.1 | - | - | 0 | - |
| - | - | 4.371E+04 | 355.2 | - | - | 0 | - |
| - | - | 1.138E+04 | 355.2 | - | - | 0 | - |
| - | - | 6917 | 356.2 | - | - | 0 | - |
| - | - | 2012 | 356.2 | - | - | 0 | - |
| - | - | 1407 | 356.7 | - | - | 0 | - |
| - | - | 2.667E+04 | 357.2 | - | - | 0 | - |
| - | - | 5228 | 358.2 | - | - | 0 | - |
| - | - | 626.9 | 361.2 | - | - | 0 | - |
| - | - | 1.674E+05 | 365.2 | - | - | 0 | - |
| - | - | 3369 | 366.2 | - | - | 0 | - |
| - | - | 3.39E+04 | 366.2 | - | - | 0 | - |
| - | - | 724.2 | 367.1 | - | - | 0 | - |
| - | - | 5583 | 367.2 | - | - | 0 | - |
| - | - | 4123 | 367.2 | - | - | 0 | - |
| - | - | 3728 | 368.2 | - | - | 0 | - |
| - | - | 733 | 371.2 | - | - | 0 | - |
| - | - | 5.431E+04 | 372.2 | - | - | 0 | - |
| - | - | 5933 | 373.2 | - | - | 0 | - |
| - | - | 8691 | 373.2 | - | - | 0 | - |
| - | - | 845.2 | 373.7 | - | - | 0 | - |
| - | - | 1150 | 374.2 | - | - | 0 | - |
| - | - | 2576 | 374.2 | - | - | 0 | - |
| - | - | 2798 | 375.2 | - | - | 0 | - |
| - | - | 1487 | 375.2 | - | - | 0 | - |
| - | - | 7302 | 376.2 | - | - | 0 | - |
| - | - | 1053 | 377.2 | - | - | 0 | - |
| - | - | 981.9 | 381.2 | - | - | 0 | - |
| - | - | 1213 | 381.7 | - | - | 0 | - |
| - | - | 1809 | 382.2 | - | - | 0 | - |
| 7 | b | 2428 | 382.7 | 0.0004473 | 1.169 | +2 | 7 |
| - | - | 1.932E+05 | 383.2 | - | - | 0 | - |
| - | - | 897 | 384.2 | - | - | 0 | - |
| - | - | 1.42E+04 | 384.2 | - | - | 0 | - |
| - | - | 3.918E+04 | 384.2 | - | - | 0 | - |
| - | - | 3262 | 385.2 | - | - | 0 | - |
| - | - | 2387 | 385.2 | - | - | 0 | - |
| - | - | 4093 | 385.2 | - | - | 0 | - |
| - | - | 743.6 | 386.2 | - | - | 0 | - |
| - | - | 3391 | 387.2 | - | - | 0 | - |
| - | - | 3450 | 387.7 | - | - | 0 | - |
| - | - | 1059 | 388.2 | - | - | 0 | - |
| - | - | 1655 | 389.2 | - | - | 0 | - |
| - | - | 1101 | 390.2 | - | - | 0 | - |
| 7 | b | 1761 | 391.7 | 0.0001088 | 0.2778 | +2 | 7 |
| - | - | 1550 | 392.2 | - | - | 0 | - |
| - | - | 1.367E+04 | 393.2 | - | - | 0 | - |
| 4 | b | 1.699E+05 | 393.2 | 0.0005443 | 1.384 | +1 | 4 |
| - | - | 5.733E+04 | 394.2 | - | - | 0 | - |
| - | - | 3.888E+04 | 394.2 | - | - | 0 | - |
| - | - | 1.095E+04 | 395.2 | - | - | 0 | - |
| - | - | 3745 | 395.2 | - | - | 0 | - |
| 4 | y | 2866 | 395.7 | 0.0009308 | 2.352 | +2 | 7 |
| 4 | y | 2.793E+04 | 396.2 | 0.001355 | 3.419 | +2 | 7 |
| - | - | 1.215E+04 | 396.7 | - | - | 0 | - |
| - | - | 1337 | 397.2 | - | - | 0 | - |
| - | - | 1124 | 399.2 | - | - | 0 | - |
| - | - | 2617 | 401.2 | - | - | 0 | - |
| - | - | 2.887E+04 | 402.2 | - | - | 0 | - |
| - | - | 5229 | 403.2 | - | - | 0 | - |
| 4 | y | 8.735E+04 | 404.7 | 0.001081 | 2.67 | +2 | 7 |
| - | - | 4.025E+04 | 405.2 | - | - | 0 | - |
| - | - | 9779 | 405.7 | - | - | 0 | - |
| - | - | 1100 | 406.2 | - | - | 0 | - |
| - | - | 2402 | 406.2 | - | - | 0 | - |
| - | - | 1132 | 408.2 | - | - | 0 | - |
| - | - | 2309 | 408.2 | - | - | 0 | - |
| - | - | 1404 | 409.2 | - | - | 0 | - |
| - | - | 1893 | 409.2 | - | - | 0 | - |
| - | - | 1084 | 409.7 | - | - | 0 | - |
| - | - | 5891 | 410.2 | - | - | 0 | - |
| - | - | 1652 | 410.2 | - | - | 0 | - |
| - | - | 1.435E+04 | 411.2 | - | - | 0 | - |
| 4 | b | 1.484E+05 | 411.2 | 0.0008134 | 1.978 | +1 | 4 |
| - | - | 1.714E+04 | 412.2 | - | - | 0 | - |
| - | - | 3.297E+04 | 412.2 | - | - | 0 | - |
| - | - | 3481 | 413.2 | - | - | 0 | - |
| - | - | 4745 | 413.2 | - | - | 0 | - |
| - | - | 643.1 | 414.2 | - | - | 0 | - |
| - | - | 604.1 | 414.2 | - | - | 0 | - |
| - | - | 786.3 | 416.2 | - | - | 0 | - |
| - | - | 1984 | 416.7 | - | - | 0 | - |
| - | - | 4808 | 417.2 | - | - | 0 | - |
| - | - | 888.9 | 418.2 | - | - | 0 | - |
| - | - | 1.363E+04 | 419.2 | - | - | 0 | - |
| - | - | 2335 | 419.2 | - | - | 0 | - |
| - | - | 2640 | 419.7 | - | - | 0 | - |
| - | - | 2.994E+04 | 420.2 | - | - | 0 | - |
| - | - | 5298 | 420.2 | - | - | 0 | - |
| - | - | 1517 | 420.7 | - | - | 0 | - |
| - | - | 5931 | 421.2 | - | - | 0 | - |
| - | - | 961.1 | 421.7 | - | - | 0 | - |
| - | - | 973.3 | 422.2 | - | - | 0 | - |
| - | - | 913.5 | 423.2 | - | - | 0 | - |
| - | - | 604.5 | 424.2 | - | - | 0 | - |
| - | - | 3144 | 424.2 | - | - | 0 | - |
| - | - | 2840 | 424.7 | - | - | 0 | - |
| - | - | 3308 | 425.2 | - | - | 0 | - |
| - | - | 824.5 | 425.7 | - | - | 0 | - |
| - | - | 2042 | 426.2 | - | - | 0 | - |
| - | - | 1.162E+04 | 427.2 | - | - | 0 | - |
| - | - | 4771 | 427.2 | - | - | 0 | - |
| - | - | 1580 | 428.2 | - | - | 0 | - |
| - | - | 1.314E+04 | 428.2 | - | - | 0 | - |
| - | - | 3.324E+04 | 428.3 | - | - | 0 | - |
| - | - | 749.6 | 428.7 | - | - | 0 | - |
| - | - | 6931 | 429.2 | - | - | 0 | - |
| - | - | 9120 | 429.3 | - | - | 0 | - |
| - | - | 2339 | 430.2 | - | - | 0 | - |
| - | - | 2090 | 430.2 | - | - | 0 | - |
| - | - | 4601 | 430.7 | - | - | 0 | - |
| - | - | 1936 | 431.2 | - | - | 0 | - |
| - | - | 752.4 | 431.7 | - | - | 0 | - |
| - | - | 1497 | 432.2 | - | - | 0 | - |
| 8 | b | 2.273E+04 | 433.2 | 0.000839 | 1.937 | +2 | 8 |
| - | - | 8560 | 433.7 | - | - | 0 | - |
| 7 | y | 1.49E+04 | 434.2 | 0.0004083 | 0.9402 | +1 | 4 |
| 7 | y | 2714 | 435.2 | 0.002202 | 5.059 | +1 | 4 |
| - | - | 2259 | 435.2 | - | - | 0 | - |
| - | - | 1009 | 436.2 | - | - | 0 | - |
| - | - | 1.576E+04 | 437.2 | - | - | 0 | - |
| - | - | 4.05E+04 | 438.2 | - | - | 0 | - |
| - | - | 7513 | 439.2 | - | - | 0 | - |
| 3 | y | 7894 | 439.2 | 0.001793 | 4.082 | +2 | 8 |
| 3 | y | 4.034E+04 | 439.7 | 0.001972 | 4.486 | +2 | 8 |
| - | - | 882.9 | 440.2 | - | - | 0 | - |
| - | - | 1.875E+04 | 440.2 | - | - | 0 | - |
| - | - | 5720 | 440.7 | - | - | 0 | - |
| 8 | b | 2949 | 442.2 | 0.0008362 | 1.891 | +2 | 8 |
| - | - | 1509 | 442.7 | - | - | 0 | - |
| - | - | 2.565E+04 | 444.2 | - | - | 0 | - |
| - | - | 4982 | 445.2 | - | - | 0 | - |
| - | - | 4345 | 445.2 | - | - | 0 | - |
| - | - | 1346 | 446.2 | - | - | 0 | - |
| - | - | 1215 | 447.7 | - | - | 0 | - |
| 3 | y | 5.773E+04 | 448.2 | 0.001363 | 3.04 | +2 | 8 |
| - | - | 2.866E+04 | 448.7 | - | - | 0 | - |
| - | - | 9782 | 449.2 | - | - | 0 | - |
| - | - | 1238 | 449.7 | - | - | 0 | - |
| - | - | 1652 | 450.2 | - | - | 0 | - |
| - | - | 6206 | 450.2 | - | - | 0 | - |
| - | - | 2129 | 451.2 | - | - | 0 | - |
| 7 | y | 1.822E+05 | 452.3 | 0.001349 | 2.982 | +1 | 4 |
| - | - | 1583 | 452.7 | - | - | 0 | - |
| - | - | 4.686E+04 | 453.3 | - | - | 0 | - |
| - | - | 995.9 | 453.7 | - | - | 0 | - |
| - | - | 7217 | 454.3 | - | - | 0 | - |
| - | - | 954.3 | 455.2 | - | - | 0 | - |
| - | - | 3.436E+04 | 455.2 | - | - | 0 | - |
| - | - | 8.857E+04 | 456.2 | - | - | 0 | - |
| - | - | 2.189E+04 | 457.2 | - | - | 0 | - |
| - | - | 3177 | 458.2 | - | - | 0 | - |
| - | - | 752.5 | 461.2 | - | - | 0 | - |
| - | - | 835.5 | 461.2 | - | - | 0 | - |
| 9 | b | 1.213E+04 | 461.7 | 0.001002 | 2.17 | +2 | 9 |
| - | - | 1.108E+04 | 462.2 | - | - | 0 | - |
| - | - | 1850 | 462.7 | - | - | 0 | - |
| - | - | 677.1 | 465.2 | - | - | 0 | - |
| - | - | 679.1 | 466.2 | - | - | 0 | - |
| - | - | 2970 | 467.2 | - | - | 0 | - |
| - | - | 1534 | 467.3 | - | - | 0 | - |
| - | - | 3085 | 468.2 | - | - | 0 | - |
| - | - | 2669 | 468.2 | - | - | 0 | - |
| - | - | 5022 | 469.2 | - | - | 0 | - |
| - | - | 872.1 | 470.2 | - | - | 0 | - |
| - | - | 1526 | 470.2 | - | - | 0 | - |
| 9 | b | 3538 | 470.7 | 0.0002362 | 0.5018 | +2 | 9 |
| - | - | 1524 | 471.2 | - | - | 0 | - |
| - | - | 1061 | 471.7 | - | - | 0 | - |
| - | - | 4.933E+04 | 473.2 | - | - | 0 | - |
| - | - | 1.215E+04 | 474.2 | - | - | 0 | - |
| - | - | 1650 | 475.2 | - | - | 0 | - |
| - | - | 3318 | 477.2 | - | - | 0 | - |
| - | - | 823.4 | 478.2 | - | - | 0 | - |
| - | - | 811.3 | 483.2 | - | - | 0 | - |
| - | - | 1512 | 484.3 | - | - | 0 | - |
| - | - | 4912 | 485.2 | - | - | 0 | - |
| - | - | 9131 | 485.3 | - | - | 0 | - |
| - | - | 4217 | 486.2 | - | - | 0 | - |
| - | - | 2021 | 486.3 | - | - | 0 | - |
| 2 | y | 1723 | 489.3 | 0.0009381 | 1.917 | +2 | 9 |
| - | - | 839.1 | 492.2 | - | - | 0 | - |
| - | - | 4674 | 494.2 | - | - | 0 | - |
| - | - | 2.13E+04 | 495.2 | - | - | 0 | - |
| - | - | 5198 | 496.2 | - | - | 0 | - |
| 2 | y | 1630 | 497.8 | 0.0003589 | 0.721 | +2 | 9 |
| - | - | 832 | 498.3 | - | - | 0 | - |
| - | - | 838.6 | 498.8 | - | - | 0 | - |
| - | - | 1017 | 502.2 | - | - | 0 | - |
| - | - | 2178 | 503.3 | - | - | 0 | - |
| - | - | 4308 | 505.2 | - | - | 0 | - |
| - | - | 692.3 | 506.2 | - | - | 0 | - |
| - | - | 1425 | 511.3 | - | - | 0 | - |
| - | - | 2.406E+04 | 512.2 | - | - | 0 | - |
| - | - | 789.6 | 512.8 | - | - | 0 | - |
| - | - | 4.867E+04 | 513.2 | - | - | 0 | - |
| - | - | 1558 | 513.8 | - | - | 0 | - |
| - | - | 1.471E+04 | 514.2 | - | - | 0 | - |
| - | - | 1922 | 515.2 | - | - | 0 | - |
| 5 | b | 1.401E+04 | 521.3 | 1.122E-05 | 0.02153 | +1 | 5 |
| 5 | b | 1.622E+04 | 522.3 | 0.008122 | 15.55 | +1 | 5 |
| - | - | 8666 | 522.8 | - | - | 0 | - |
| - | - | 4265 | 523.3 | - | - | 0 | - |
| - | - | 2270 | 524.3 | - | - | 0 | - |
| - | - | 4.083E+04 | 530.3 | - | - | 0 | - |
| - | - | 1.484E+04 | 531.3 | - | - | 0 | - |
| - | - | 8173 | 532.2 | - | - | 0 | - |
| - | - | 1988 | 533.3 | - | - | 0 | - |
| - | - | 3747 | 534.3 | - | - | 0 | - |
| - | - | 5115 | 535.3 | - | - | 0 | - |
| - | - | 956 | 535.3 | - | - | 0 | - |
| - | - | 5262 | 535.8 | - | - | 0 | - |
| - | - | 3002 | 536.3 | - | - | 0 | - |
| - | - | 898.5 | 536.8 | - | - | 0 | - |
| - | - | 1569 | 537.3 | - | - | 0 | - |
| - | - | 932.1 | 538.3 | - | - | 0 | - |
| 5 | b | 4.487E+04 | 539.3 | 0.000616 | 1.142 | +1 | 5 |
| - | - | 1267 | 540.2 | - | - | 0 | - |
| - | - | 1.243E+04 | 540.3 | - | - | 0 | - |
| - | - | 1019 | 541.2 | - | - | 0 | - |
| - | - | 2384 | 541.3 | - | - | 0 | - |
| - | - | 1413 | 542.3 | - | - | 0 | - |
| 0 | Precursor | 1.371E+04 | 544.3 | 0.0003005 | 0.5521 | +2 | -1 |
| 0 | Precursor | 7901 | 544.8 | 0.009635 | 17.69 | +2 | -1 |
| - | - | 2845 | 545.3 | - | - | 0 | - |
| - | - | 996.1 | 545.8 | - | - | 0 | - |
| - | - | 3114 | 548.3 | - | - | 0 | - |
| 6 | y | 4.678E+04 | 549.3 | 0.001267 | 2.306 | +1 | 5 |
| - | - | 1.204E+04 | 550.3 | - | - | 0 | - |
| - | - | 3765 | 551.3 | - | - | 0 | - |
| - | - | 2483 | 552.3 | - | - | 0 | - |
| - | - | 5033 | 552.8 | - | - | 0 | - |
| 0 | Precursor | 5051 | 553.3 | 0.0006944 | 1.255 | +2 | -1 |
| - | - | 1078 | 553.6 | - | - | 0 | - |
| - | - | 1609 | 553.8 | - | - | 0 | - |
| - | - | 974.1 | 554 | - | - | 0 | - |
| - | - | 2566 | 554.3 | - | - | 0 | - |
| - | - | 2621 | 555.3 | - | - | 0 | - |
| - | - | 956.2 | 556.3 | - | - | 0 | - |
| - | - | 1157 | 558.3 | - | - | 0 | - |
| - | - | 1050 | 558.3 | - | - | 0 | - |
| - | - | 1604 | 559.3 | - | - | 0 | - |
| - | - | 791.1 | 562.3 | - | - | 0 | - |
| 6 | y | 6.145E+05 | 567.3 | 0.001139 | 2.008 | +1 | 5 |
| - | - | 1.892E+05 | 568.3 | - | - | 0 | - |
| - | - | 3.497E+04 | 569.3 | - | - | 0 | - |
| - | - | 2466 | 570.3 | - | - | 0 | - |
| - | - | 1656 | 571.3 | - | - | 0 | - |
| - | - | 1037 | 572.3 | - | - | 0 | - |
| - | - | 9181 | 572.3 | - | - | 0 | - |
| - | - | 2633 | 573.3 | - | - | 0 | - |
| - | - | 1789 | 577.3 | - | - | 0 | - |
| - | - | 948.9 | 578.3 | - | - | 0 | - |
| - | - | 1.223E+04 | 586.3 | - | - | 0 | - |
| - | - | 3350 | 587.3 | - | - | 0 | - |
| - | - | 770.7 | 588.3 | - | - | 0 | - |
| - | - | 1229 | 593.3 | - | - | 0 | - |
| - | - | 838.2 | 594.3 | - | - | 0 | - |
| - | - | 2801 | 608.3 | - | - | 0 | - |
| - | - | 1095 | 609.3 | - | - | 0 | - |
| - | - | 1022 | 611.3 | - | - | 0 | - |
| - | - | 2825 | 611.3 | - | - | 0 | - |
| - | - | 1175 | 612.3 | - | - | 0 | - |
| - | - | 1718 | 614.3 | - | - | 0 | - |
| - | - | 2265 | 615.3 | - | - | 0 | - |
| - | - | 2237 | 616.3 | - | - | 0 | - |
| - | - | 1408 | 616.3 | - | - | 0 | - |
| - | - | 980.9 | 617.3 | - | - | 0 | - |
| - | - | 760.7 | 618.3 | - | - | 0 | - |
| - | - | 944.9 | 619.3 | - | - | 0 | - |
| - | - | 2574 | 625.3 | - | - | 0 | - |
| - | - | 1758 | 626.3 | - | - | 0 | - |
| - | - | 1528 | 629.3 | - | - | 0 | - |
| - | - | 2300 | 631.3 | - | - | 0 | - |
| - | - | 8817 | 632.3 | - | - | 0 | - |
| - | - | 7811 | 633.3 | - | - | 0 | - |
| - | - | 4532 | 634.3 | - | - | 0 | - |
| - | - | 772.1 | 635.3 | - | - | 0 | - |
| 6 | b | 6756 | 636.3 | 0.000534 | 0.8392 | +1 | 6 |
| 6 | b | 2609 | 637.3 | 0.00126 | 1.977 | +1 | 6 |
| - | - | 4389 | 637.3 | - | - | 0 | - |
| - | - | 2496 | 638.3 | - | - | 0 | - |
| - | - | 2252 | 641.3 | - | - | 0 | - |
| - | - | 6380 | 642.3 | - | - | 0 | - |
| - | - | 1917 | 643.3 | - | - | 0 | - |
| - | - | 7572 | 643.3 | - | - | 0 | - |
| - | - | 2732 | 644.3 | - | - | 0 | - |
| - | - | 2705 | 645.4 | - | - | 0 | - |
| - | - | 982.4 | 646.4 | - | - | 0 | - |
| - | - | 9694 | 649.3 | - | - | 0 | - |
| - | - | 3086 | 650.3 | - | - | 0 | - |
| - | - | 3422 | 651.3 | - | - | 0 | - |
| - | - | 1030 | 652.3 | - | - | 0 | - |
| - | - | 1199 | 653.3 | - | - | 0 | - |
| 6 | b | 3.254E+04 | 654.3 | 0.0007725 | 1.181 | +1 | 6 |
| - | - | 8831 | 655.3 | - | - | 0 | - |
| - | - | 4266 | 656.3 | - | - | 0 | - |
| - | - | 1524 | 657.3 | - | - | 0 | - |
| - | - | 1.66E+04 | 659.3 | - | - | 0 | - |
| - | - | 3.029E+04 | 660.3 | - | - | 0 | - |
| - | - | 1.197E+04 | 661.3 | - | - | 0 | - |
| - | - | 2476 | 662.3 | - | - | 0 | - |
| - | - | 1675 | 671.3 | - | - | 0 | - |
| - | - | 1.306E+04 | 673.4 | - | - | 0 | - |
| - | - | 4325 | 674.4 | - | - | 0 | - |
| - | - | 1048 | 675.4 | - | - | 0 | - |
| 5 | y | 1.852E+05 | 677.3 | 0.001283 | 1.894 | +1 | 6 |
| 5 | y | 4.195E+05 | 678.3 | 0.003351 | 4.941 | +1 | 6 |
| - | - | 1.499E+05 | 679.3 | - | - | 0 | - |
| - | - | 3.296E+04 | 680.3 | - | - | 0 | - |
| - | - | 2011 | 681.3 | - | - | 0 | - |
| - | - | 1463 | 686.3 | - | - | 0 | - |
| - | - | 742.8 | 694.4 | - | - | 0 | - |
| 5 | y | 5.395E+05 | 695.3 | 0.00146 | 2.1 | +1 | 6 |
| - | - | 2.076E+05 | 696.3 | - | - | 0 | - |
| - | - | 1396 | 696.5 | - | - | 0 | - |
| - | - | 4.294E+04 | 697.3 | - | - | 0 | - |
| - | - | 3339 | 698.3 | - | - | 0 | - |
| - | - | 3861 | 705.3 | - | - | 0 | - |
| - | - | 2372 | 706.3 | - | - | 0 | - |
| - | - | 6139 | 712.4 | - | - | 0 | - |
| - | - | 4740 | 713.4 | - | - | 0 | - |
| - | - | 1756 | 714.4 | - | - | 0 | - |
| - | - | 1.232E+04 | 730.4 | - | - | 0 | - |
| - | - | 5400 | 731.4 | - | - | 0 | - |
| - | - | 1467 | 732.4 | - | - | 0 | - |
| - | - | 1248 | 746.4 | - | - | 0 | - |
| - | - | 1647 | 747.4 | - | - | 0 | - |
| - | - | 1394 | 748.4 | - | - | 0 | - |
| - | - | 1177 | 749.4 | - | - | 0 | - |
| - | - | 1362 | 755.4 | - | - | 0 | - |
| - | - | 889 | 756.4 | - | - | 0 | - |
| 7 | b | 5651 | 764.4 | 0.0007858 | 1.028 | +1 | 7 |
| 7 | b | 3278 | 765.4 | 0.01323 | 17.29 | +1 | 7 |
| - | - | 2631 | 766.4 | - | - | 0 | - |
| - | - | 1.508E+04 | 773.4 | - | - | 0 | - |
| - | - | 5545 | 774.4 | - | - | 0 | - |
| - | - | 1190 | 775.4 | - | - | 0 | - |
| 7 | b | 1.237E+04 | 782.4 | 0.0005361 | 0.6852 | +1 | 7 |
| - | - | 5818 | 783.4 | - | - | 0 | - |
| - | - | 1360 | 784.4 | - | - | 0 | - |
| 4 | y | 4407 | 790.4 | 0.0002268 | 0.287 | +1 | 7 |
| 4 | y | 3644 | 791.4 | 0.01285 | 16.24 | +1 | 7 |
| - | - | 1139 | 794.4 | - | - | 0 | - |
| - | - | 919.8 | 798.5 | - | - | 0 | - |
| - | - | 830 | 800.4 | - | - | 0 | - |
| - | - | 1978 | 803.4 | - | - | 0 | - |
| - | - | 2798 | 804.4 | - | - | 0 | - |
| - | - | 1347 | 805.4 | - | - | 0 | - |
| 4 | y | 5.248E+04 | 808.4 | 0.0008316 | 1.029 | +1 | 7 |
| - | - | 2.678E+04 | 809.4 | - | - | 0 | - |
| - | - | 5557 | 810.4 | - | - | 0 | - |
| - | - | 830.3 | 811.4 | - | - | 0 | - |
| - | - | 680.4 | 813.4 | - | - | 0 | - |
| - | - | 735 | 818.4 | - | - | 0 | - |
| - | - | 652.3 | 819.4 | - | - | 0 | - |
| - | - | 3.293E+04 | 821.4 | - | - | 0 | - |
| - | - | 1.354E+04 | 822.4 | - | - | 0 | - |
| - | - | 4058 | 823.4 | - | - | 0 | - |
| - | - | 1900 | 829.4 | - | - | 0 | - |
| - | - | 2171 | 830.4 | - | - | 0 | - |
| - | - | 997.7 | 832.4 | - | - | 0 | - |
| - | - | 738.9 | 833.4 | - | - | 0 | - |
| - | - | 683.1 | 836.3 | - | - | 0 | - |
| - | - | 1681 | 837.4 | - | - | 0 | - |
| - | - | 2024 | 838.4 | - | - | 0 | - |
| - | - | 7298 | 839.4 | - | - | 0 | - |
| - | - | 1805 | 840.4 | - | - | 0 | - |
| - | - | 1.755E+04 | 847.4 | - | - | 0 | - |
| - | - | 1.072E+04 | 848.4 | - | - | 0 | - |
| - | - | 3686 | 849.4 | - | - | 0 | - |
| - | - | 677 | 850.4 | - | - | 0 | - |
| - | - | 8649 | 855.5 | - | - | 0 | - |
| - | - | 4781 | 856.5 | - | - | 0 | - |
| - | - | 1262 | 857.5 | - | - | 0 | - |
| - | - | 1856 | 859.4 | - | - | 0 | - |
| - | - | 1508 | 860.4 | - | - | 0 | - |
| 8 | b | 3.889E+04 | 865.4 | 0.0001397 | 0.1614 | +1 | 8 |
| 8 | b | 2.373E+04 | 866.4 | 0.01487 | 17.16 | +1 | 8 |
| - | - | 7835 | 867.4 | - | - | 0 | - |
| - | - | 1154 | 868.4 | - | - | 0 | - |
| 3 | y | 5694 | 877.4 | 0.0002464 | 0.2808 | +1 | 8 |
| 3 | y | 8180 | 878.4 | 0.006217 | 7.077 | +1 | 8 |
| - | - | 5114 | 879.4 | - | - | 0 | - |
| - | - | 1448 | 880.4 | - | - | 0 | - |
| 8 | b | 1.132E+05 | 883.5 | 9.883E-05 | 0.1119 | +1 | 8 |
| - | - | 5.714E+04 | 884.5 | - | - | 0 | - |
| - | - | 1.618E+04 | 885.5 | - | - | 0 | - |
| - | - | 1128 | 886.5 | - | - | 0 | - |
| - | - | 837.3 | 887.4 | - | - | 0 | - |
| - | - | 739.8 | 894.5 | - | - | 0 | - |
| 3 | y | 3.88E+04 | 895.5 | 0.0006635 | 0.741 | +1 | 8 |
| - | - | 2.512E+04 | 896.5 | - | - | 0 | - |
| - | - | 7648 | 897.5 | - | - | 0 | - |
| - | - | 1619 | 898.5 | - | - | 0 | - |
| - | - | 1936 | 904.5 | - | - | 0 | - |
| - | - | 4668 | 905.4 | - | - | 0 | - |
| - | - | 2882 | 906.4 | - | - | 0 | - |
| - | - | 1133 | 907.4 | - | - | 0 | - |
| - | - | 3351 | 912.5 | - | - | 0 | - |
| - | - | 1130 | 913.5 | - | - | 0 | - |
| 9 | b | 3.714E+04 | 922.5 | 0.0002472 | 0.2679 | +1 | 9 |
| 9 | b | 2.636E+04 | 923.4 | 0.01287 | 13.94 | +1 | 9 |
| - | - | 9465 | 924.5 | - | - | 0 | - |
| - | - | 1572 | 925.5 | - | - | 0 | - |
| 9 | b | 1.517E+05 | 940.5 | 6.363E-05 | 0.06766 | +1 | 9 |
| - | - | 8.137E+04 | 941.5 | - | - | 0 | - |
| - | - | 2.497E+04 | 942.5 | - | - | 0 | - |
| - | - | 3529 | 943.5 | - | - | 0 | - |
| - | - | 3941 | 958.5 | - | - | 0 | - |
| - | - | 1711 | 959.5 | - | - | 0 | - |
| - | - | 718.5 | 2121 | - | - | 0 | - |
| - | - | 740.6 | 2555 | - | - | 0 | - |
| - | - | 955.3 | 3082 | - | - | 0 | - |
| - | - | 700.4 | 3153 | - | - | 0 | - |

m/z Charge Intensity FragmentType MassShift Position
120.08131408691406 0 353629.38
121.08464050292969 0 27487.703
122.08795166015625 0 674.3431
122.5987548828125 0 378.0374
123.09230041503906 0 1343.5732
124.07633209228516 0 980.7039
124.1127700805664 0 421.0689
125.03520965576172 0 934.3852
126.09184265136719 0 3916.3564
127.08727264404297 0 2135.2566
127.12350463867188 0 22220.014
128.08250427246094 0 1633.0051
128.10751342773438 0 14492.297
128.12689208984375 0 1760.7278
129.0663604736328 0 5732.8193
129.10279846191406 0 366393.06
130.05050659179688 0 1263.514
130.06568908691406 0 2748.876
130.08680725097656 0 2540.6387
130.0998992919922 0 3285.2515
130.10609436035156 0 21509.895
131.04965209960938 0 6287.1816
131.08201599121094 0 14448.647
131.11839294433594 0 26325.559
132.05294799804688 0 624.74384
132.08583068847656 0 638.39667
132.10240173339844 0 1701.3325
132.12181091308594 0 1939.5073
133.06129455566406 0 5287.861
136.07620239257812 0 5680.5625
137.10797119140625 0 886.22314
138.09190368652344 0 14474.801
139.05064392089844 0 2064.7124
139.08717346191406 0 8302.366
139.0945587158203 0 998.3297
139.12342834472656 0 1046.5206
140.09046936035156 0 920.5948
140.10745239257812 0 2712.5781
141.06643676757812 0 5826.947
141.10276794433594 0 13054.014
142.08685302734375 0 2462.5159
142.1060028076172 0 630.38995
142.12286376953125 0 612.0502
143.0455322265625 0 605.8611
143.0819854736328 0 536.50543
143.118408203125 0 1498.7136
147.0771942138672 0 863.2177
147.1133575439453 0 2129.1104
148.07676696777344 0 517.3615
148.9546356201172 0 992.7877
149.04551696777344 0 524.369
149.05520629882812 0 466.3869
149.06021118164062 0 6353.845
149.1079864501953 0 2064.3584
150.06398010253906 0 646.3318
150.0918731689453 0 524.65753
151.08717346191406 0 17356.5
152.09060668945312 0 1715.9982
152.10772705078125 0 1137.8328
153.06642150878906 0 1094.5719
153.10256958007812 0 1074.7404
154.0980987548828 0 8742.023
155.08206176757812 0 5493.6924
155.11842346191406 0 60976.348
156.0780487060547 0 595.26
156.08538818359375 0 550.1438
156.10255432128906 0 1848.7173
156.11492919921875 0 648.98267
156.12179565429688 0 5152.95
157.06146240234375 0 1039.1047
157.0977325439453 0 1883.1868
159.07696533203125 0 54046.242
159.0922393798828 0 7101.6084
159.11331176757812 0 1503.4489
160.08035278320312 0 3000.2085
163.0870819091797 0 1065.6854
165.10287475585938 0 2763.5864
166.08685302734375 0 434961.06 y 9
167.05636596679688 0 754.7132
167.08285522460938 0 2618.6218
167.0902099609375 0 39973
167.118408203125 0 17468.637
167.12661743164062 0 913.1745
168.0924072265625 0 1582.0808
168.11367797851562 0 14637.382
168.121826171875 0 1466.7905
168.15045166015625 0 977.7515
169.06185913085938 0 1177.7709
169.0975799560547 0 1963.5048 d 1
169.1171875 0 1415.7079
169.1340789794922 0 1581.0734
171.07711791992188 0 1959.8204
171.14950561523438 0 684.66486
172.108642578125 0 736.57574
173.12905883789062 0 533729.5
173.4521942138672 0 2241.5386
174.13246154785156 0 43780.637
175.13409423828125 0 1548.5485
177.10284423828125 0 90071.06
178.10623168945312 0 8752.663
179.08177185058594 0 723.68634
179.11859130859375 0 1218.5022
180.07749938964844 0 2306.1885
180.1139373779297 0 1891.0039
181.06134033203125 0 18212.357
181.09780883789062 0 1919.3923
181.10623168945312 0 433.34692
182.0454559326172 0 917.17346
182.06539916992188 0 1012.75525
182.0818634033203 0 971.3303
182.12937927246094 0 2010.2911
183.11346435546875 0 268344.78 a 1
184.10867309570312 0 14553.465
184.11691284179688 0 22391.79
184.1450653076172 0 2663.3203
185.09263610839844 0 923.11694
185.11190795898438 0 938.91504
185.12899780273438 0 23200.13
186.12474060058594 0 4627.266
186.16064453125 0 980.301
187.0869140625 0 693.36163
187.1083221435547 0 11911.812
188.0713653564453 0 1618.3015
188.1116943359375 0 1363.895
191.08233642578125 0 1658.529
193.09774780273438 0 3317.6287
194.12155151367188 0 367.23688
194.12942504882812 0 20012.793
195.11346435546875 0 4108.149
195.13287353515625 0 2251.1956
196.10865783691406 0 4167.285
197.0926055908203 0 5123.359
197.1290283203125 0 3082.0955
198.087890625 0 38435.176
198.124267578125 0 10781.204
199.0718536376953 0 10597.594
199.09130859375 0 3394.9844
199.10826110839844 0 3274.154
199.12704467773438 0 909.34467
199.1794891357422 0 504.33023
200.14007568359375 0 1380.3433
201.1239471435547 0 160446.17
202.12733459472656 0 16292.867
202.15562438964844 0 4543.1694
203.0820770263672 0 857.3005
203.11729431152344 0 695.9561
203.12844848632812 0 1019.6735
203.15821838378906 0 588.2625
204.11370849609375 0 1316.306
204.1353759765625 0 1006.5565
205.0977325439453 0 11274.762
206.1017303466797 0 1001.5883
207.11343383789062 0 618.94055
208.07293701171875 0 720.3219
208.0972900390625 0 6419.6294
208.10862731933594 0 6299.6167
209.05625915527344 0 5825.9897
209.09274291992188 0 53548.8
210.078125 0 616.3943
210.08668518066406 0 555.13116
210.09616088867188 0 5007.657
210.1243896484375 0 1914.706
211.10836791992188 0 95525.75 b 1
212.1117401123047 0 10344.15
212.1399688720703 0 12805.63
213.11399841308594 0 1101.2025
213.12411499023438 0 5576.22
213.14381408691406 0 971.5311
214.11915588378906 0 4124.2437
214.15562438964844 0 2747.7058
215.10316467285156 0 4106.47
216.09860229492188 0 13054.645
217.10145568847656 0 884.81213
217.71360778808594 0 506.97614
218.15061950683594 0 9519.247
219.15380859375 0 983.59
222.1241912841797 0 3546.0574
223.10841369628906 0 252667.92 y 8
224.11184692382812 0 30624.219
224.1400909423828 0 3956.9922
225.0440216064453 0 1407.7755
225.09889221191406 0 3758.9978
225.11376953125 0 1688.5022
225.1241455078125 0 13326.261
225.13555908203125 0 586.31714
226.08299255371094 0 76225.984
226.1193084716797 0 160944.95
227.0669403076172 0 34403.613
227.08645629882812 0 7320.2627
227.10317993164062 0 11014.727
227.12261962890625 0 15028.132
228.07032775878906 0 3109.6887
228.08795166015625 0 634.1591
228.10777282714844 0 846.36285
228.12396240234375 0 1036.0383
228.1349334716797 0 1805.5913
229.0730438232422 0 489.34946
229.11903381347656 0 3926.6257
229.1551971435547 0 705.6707
230.12991333007812 0 814.2767
230.15057373046875 0 19172.395
231.13693237304688 0 811.99316
231.15164184570312 0 3424.5369
233.1288604736328 0 1041.3667
234.12445068359375 0 2483.5254
236.10391235351562 0 2348.873
237.12368774414062 0 727.30566
238.1192626953125 0 1804.0743
239.09539794921875 0 2071.0894
239.1394500732422 0 720.7295
239.1514129638672 0 1173.4451
240.0962677001953 0 1060.6067
240.13504028320312 0 6109.502
241.09292602539062 0 1731.0151
241.1386260986328 0 808.1488
242.1506805419922 0 83739.4
243.10971069335938 0 10709.695
243.15408325195312 0 9197.102
244.09361267089844 0 73611.91
244.12989807128906 0 143505.94
245.07754516601562 0 1033.9807
245.0965576171875 0 6011.048
245.13320922851562 0 14978.952
246.12330627441406 0 890.564
246.13584899902344 0 680.4619
248.10348510742188 0 2778.8972
248.1160125732422 0 808.2132
248.1405487060547 0 1529.3505
248.16116333007812 0 2824.6646
249.12411499023438 0 2318.221
250.1560516357422 0 1259.3253
251.15184020996094 0 1331.7351
251.17608642578125 0 879.77264
252.13499450683594 0 6646.775 a Water loss 2
253.11898803710938 0 4134.9043
254.0774383544922 0 772.50696
254.1140899658203 0 3946.8613
254.1870574951172 0 3069.7065
255.14492797851562 0 933.26587
256.1302795410156 0 1149.9749
258.1455078125 0 10053.11
259.1486511230469 0 1374.8027
259.17755126953125 0 1281.3574
260.1394958496094 0 602.6609
261.1203918457031 0 5479.239
261.1560974121094 0 669.8794
262.10406494140625 0 510.77914
262.12005615234375 0 1738.5996
262.140380859375 0 2127.6104
263.1036682128906 0 705.0507
264.1354064941406 0 1852.2347
265.1304016113281 0 5770.5483
265.16632080078125 0 1241.3739
265.6336364746094 0 1264.0443
266.1141357421875 0 6839.427
266.1335144042969 0 1216.5244
266.1506652832031 0 3402.1313
267.11859130859375 0 1115.9646
268.130126953125 0 2380.4817
269.16162109375 0 5153.6567
270.1455078125 0 26579.879 a 2
271.148681640625 0 2935.1677
273.1564025878906 0 992.3641
274.11944580078125 0 4386.275
276.135009765625 0 1652.3607
277.1352844238281 0 582.0951
278.1504821777344 0 5000.4146
279.1346740722656 0 6359.86
280.1300048828125 0 79500.81 b Water loss 2
281.1331787109375 0 10702.462
281.160888671875 0 1200.7827
282.110107421875 0 548.8386
282.1452331542969 0 3000.5222
282.1824035644531 0 2324.955
283.140869140625 0 47943.863
284.1251220703125 0 2066.0618
284.1442565917969 0 6007.8945 y 5
284.1611022949219 0 943.1327
285.00836181640625 0 544.9781
286.1408386230469 0 1672.5583
287.1721496582031 0 9795.341
288.1347351074219 0 1100.3086
288.1563415527344 0 860.6623
288.17510986328125 0 1403.9331
290.1144104003906 0 3887.191
291.1187438964844 0 931.01306
291.1457214355469 0 6061.418
292.1299743652344 0 10160.8
292.1474609375 0 703.1304
293.1317138671875 0 583.2923
293.161865234375 0 2309.271
294.1460266113281 0 5265.8574
295.1495056152344 0 590.43665
296.1063232421875 0 616.68286
297.1019592285156 0 707.23254
297.15667724609375 0 17607.557
298.1407165527344 0 195461.05 b 2
299.06353759765625 0 1138.572
299.1437683105469 0 27900.484
299.1728210449219 0 2445.4023
300.1465148925781 0 2628.5435
300.1748962402344 0 773.5777
301.1513366699219 0 9539.669
301.18743896484375 0 1535.0093
302.154296875 0 1835.8427
304.1295471191406 0 916.46497
305.1123962402344 0 622.1509
305.1302490234375 0 753.0746
305.18292236328125 0 8425.002
306.145751953125 0 16648.787 y Water loss 7
306.18603515625 0 1014.19165
307.1485900878906 0 2341.021
309.15673828125 0 22200.867
310.14056396484375 0 6895.9287
310.15972900390625 0 3315.2642
311.13702392578125 0 3615.4468
311.1722717285156 0 5644.433
312.1566467285156 0 1300.4072
312.17578125 0 928.9488
315.16748046875 0 8090.082
316.13079833984375 0 1096.3575
316.16644287109375 0 2789.4395
316.6578063964844 0 1803.884
317.18280029296875 0 5755.491
318.1582336425781 0 1134.7423
318.1847229003906 0 928.36847
319.14093017578125 0 12799.931 b Ammonia loss 5
320.1246337890625 0 1014.4938
320.1441650390625 0 2281.155
321.1556701660156 0 1853.2408
322.14093017578125 0 2350.9775
323.1732177734375 0 1519.4645
323.2109680175781 0 679.8516
323.685302734375 0 743.0321
324.15643310546875 0 78601.37 y 7
325.1595764160156 0 11968.099
326.1618347167969 0 1505.2462
327.16729736328125 0 31998.133
328.1510009765625 0 5134.3926
328.1722106933594 0 6359.459
328.6737060546875 0 821.1464
329.1490173339844 0 2498.6892
329.1829528808594 0 60722.445
330.1860046386719 0 10220.526
330.655029296875 0 718.8341
331.18634033203125 0 744.32745
334.14031982421875 0 1503.8071
336.1675720214844 0 16968.809
337.15167236328125 0 60765.8
338.13494873046875 0 2481.426
338.1548767089844 0 11635.783
339.1671447753906 0 53480.63 y Water loss 4
339.6651611328125 0 15236.467
340.15228271484375 0 8919.227
340.1708068847656 0 2113.0425
340.6634521484375 0 1260.1399
341.15435791015625 0 2189.8464
341.1822204589844 0 777.5248
343.160400390625 0 665.20374
343.6798400878906 0 820.59796
345.1777648925781 0 66117.77
346.180419921875 0 10385.192
347.1829528808594 0 1645.1421
347.2092590332031 0 2549.0469
348.1724853515625 0 63502.566 y 4
348.673828125 0 26562.885
349.1512451171875 0 3728.1855
349.1753845214844 0 4425.754
351.13482666015625 0 3639.7402
354.1781311035156 0 90756.97
355.0705261230469 0 1611.7427
355.1618957519531 0 43713.383
355.181884765625 0 11382.522
356.1646728515625 0 6916.9775
356.1907653808594 0 2011.5112
356.6858215332031 0 1407.3434
357.1778564453125 0 26666.697
358.1800231933594 0 5227.624
361.1921691894531 0 626.89075
365.2192077636719 0 167354.17
366.1780090332031 0 3369.4985
366.2225341796875 0 33899.25
367.12493896484375 0 724.219
367.1620178222656 0 5582.974
367.2252197265625 0 4122.9873
368.16180419921875 0 3728.125
371.2049560546875 0 732.9801
372.188720703125 0 54311.25
373.1719970703125 0 5932.627
373.1916198730469 0 8690.604
373.6966247558594 0 845.216
374.1741027832031 0 1149.6793
374.2024230957031 0 2575.666
375.1753845214844 0 2798.0317
375.2059631347656 0 1486.6082
376.1622009277344 0 7302.266
377.16644287109375 0 1052.997
381.1786193847656 0 981.9064
381.71173095703125 0 1212.8582
382.1734313964844 0 1808.8649
382.70123291015625 0 2427.9092 b Water loss 6
383.22991943359375 0 193150.61
384.1596374511719 0 896.9793
384.1885681152344 0 14204.05
384.2328796386719 0 39175.51
385.1725158691406 0 3261.8613
385.1922912597656 0 2387.038
385.2354736328125 0 4092.8274
386.1719970703125 0 743.55237
387.19793701171875 0 3391.2168
387.69342041015625 0 3450.0671
388.1911315917969 0 1059.2661
389.2197570800781 0 1655.2844
390.2181701660156 0 1101.2131
391.7061767578125 0 1761.4904 b 6
392.1953430175781 0 1549.8065
393.18890380859375 0 13668.244
393.21435546875 0 169943.67 b Water loss 3
394.173095703125 0 57327.883
394.2176208496094 0 38881.41
395.1762390136719 0 10950.457
395.2206726074219 0 3744.6777
395.7092590332031 0 2865.9192 y Water loss 3
396.2016906738281 0 27934.209 y Ammonia loss 3
396.7030029296875 0 12154.502
397.2039794921875 0 1336.9838
399.2025146484375 0 1123.5734
401.1932373046875 0 2616.59
402.1995544433594 0 28874.535
403.2021484375 0 5229.2983
404.7146911621094 0 87347.97 y 3
405.21624755859375 0 40248.16
405.71734619140625 0 9779.218
406.21722412109375 0 1099.8026
406.2459716796875 0 2402.3457
408.189208984375 0 1132.3885
408.22479248046875 0 2308.5176
409.1736755371094 0 1403.7032
409.2228088378906 0 1892.5316
409.7080078125 0 1083.827
410.20477294921875 0 5891.417
410.24005126953125 0 1652.4535
411.20001220703125 0 14346.203
411.2251892089844 0 148438.81 b 3
412.183837890625 0 17139.604
412.2279052734375 0 32973.797
413.18646240234375 0 3481.0806
413.2296142578125 0 4745.35
414.1885681152344 0 643.1484
414.2298278808594 0 604.13715
416.2322692871094 0 786.2954
416.7151184082031 0 1983.563
417.2135009765625 0 4808.398
418.2082214355469 0 888.89606
419.2044677734375 0 13632.684
419.2291259765625 0 2335.215
419.7271728515625 0 2640.2837
420.1888732910156 0 29942.127
420.2113037109375 0 5298.4004
420.720458984375 0 1516.8942
421.1913757324219 0 5930.653
421.7064514160156 0 961.0911
422.2043151855469 0 973.33673
423.22735595703125 0 913.52966
424.1850280761719 0 604.54047
424.2204284667969 0 3143.5771
424.71734619140625 0 2840.224
425.2267150878906 0 3307.546
425.7303771972656 0 824.46204
426.19915771484375 0 2041.7795
427.1835632324219 0 11620.592
427.23077392578125 0 4770.685
428.1844482421875 0 1579.9514
428.2162170410156 0 13143.879
428.251708984375 0 33240.35
428.7342834472656 0 749.5736
429.2142639160156 0 6930.67
429.25439453125 0 9119.917
430.1941223144531 0 2339.3706
430.2199401855469 0 2089.9797
430.7132873535156 0 4601.0825
431.21380615234375 0 1935.8654
431.7154846191406 0 752.436
432.22509765625 0 1497.229
433.2254638671875 0 22728.404 b Water loss 7
433.7269287109375 0 8559.795
434.2402038574219 0 14895.257 y Water loss 6
435.22601318359375 0 2714.1936 y Ammonia loss 6
435.24462890625 0 2259.1118
436.228515625 0 1008.91205
437.2156677246094 0 15761.46
438.19970703125 0 40496.938
439.2004699707031 0 7513.325
439.22613525390625 0 7894.4395 y Water loss 2
439.71832275390625 0 40339.48 y Ammonia loss 2
440.18359375 0 882.9335
440.21917724609375 0 18749.002
440.72039794921875 0 5719.8315
442.2307434082031 0 2949.1611 b 7
442.7317199707031 0 1508.8558
444.21063232421875 0 25645.234
445.21270751953125 0 4982.1914
445.24212646484375 0 4345.0957
446.2442932128906 0 1346.2354
447.7384338378906 0 1214.6002
448.2309875488281 0 57734.055 y 2
448.73223876953125 0 28662.068
449.2336730957031 0 9781.985
449.7341613769531 0 1237.7467
450.2005920410156 0 1651.5262
450.23626708984375 0 6205.6196
451.2351379394531 0 2129.0254
452.251708984375 0 182212.39 y 6
452.73114013671875 0 1583.0511
453.2547302246094 0 46864.14
453.72491455078125 0 995.89264
454.2574768066406 0 7217.117
455.1967468261719 0 954.2812
455.2261962890625 0 34355.45
456.21026611328125 0 88574
457.21319580078125 0 21888.21
458.21490478515625 0 3176.579
461.15399169921875 0 752.5204
461.2368469238281 0 835.53265
461.7363586425781 0 12129.764 b Water loss 8
462.23675537109375 0 11079.879
462.7378234863281 0 1850.4337
465.2115478515625 0 677.1002
466.19482421875 0 679.05707
467.22601318359375 0 2970.032
467.26190185546875 0 1533.8323
468.2165832519531 0 3084.8508
468.2477722167969 0 2669.0825
469.2061462402344 0 5021.568
470.2067565917969 0 872.119
470.24212646484375 0 1525.6891
470.7408752441406 0 3538.2175 b 8
471.2435607910156 0 1523.9083
471.7442932128906 0 1061.1283
473.2367858886719 0 49332.984
474.2392272949219 0 12153.48
475.24176025390625 0 1649.5002
477.21014404296875 0 3317.9006
478.1925048828125 0 823.3777
483.2219543457031 0 811.28296
484.25262451171875 0 1511.8678
485.2364501953125 0 4912.419
485.2734069824219 0 9130.8955
486.2345275878906 0 4216.805
486.2765197753906 0 2021.1298
489.2514953613281 0 1723.3374 y Ammonia loss 1
492.2490539550781 0 839.10376
494.2373962402344 0 4673.571
495.22137451171875 0 21296.709
496.22320556640625 0 5198.4575
497.7641906738281 0 1630.0732 y 1
498.2676086425781 0 832.0167
498.76922607421875 0 838.6496
502.2301330566406 0 1017.1387
503.2633361816406 0 2177.9973
505.24224853515625 0 4308.178
506.2468566894531 0 692.30835
511.2889099121094 0 1424.7688
512.2476196289062 0 24059.744
512.763671875 0 789.647
513.2322998046875 0 48674.14
513.7504272460938 0 1558.2598
514.2344360351562 0 14708.537
515.234130859375 0 1921.9596
521.2723999023438 0 14008.352 b Water loss 4
522.2645263671875 0 16223.481 b Ammonia loss 4
522.7637329101562 0 8665.9375
523.2579956054688 0 4265.4976
524.2830810546875 0 2270.0105
530.2581787109375 0 40825.95
531.2540893554688 0 14841.366
532.24560546875 0 8172.9507
533.270751953125 0 1987.8473
534.304931640625 0 3746.5015
535.27001953125 0 5114.8423
535.309326171875 0 956.0166
535.7662353515625 0 5262.4355
536.2662963867188 0 3002.1477
536.7644653320312 0 898.4702
537.2698364257812 0 1568.5677
538.2694702148438 0 932.10675
539.2835693359375 0 44865.805 b 4
540.2431030273438 0 1266.9171
540.28662109375 0 12430.853
541.2425537109375 0 1019.2964
541.2901611328125 0 2384.1255
542.2930908203125 0 1412.9706
544.275146484375 0 13707.279 Precursor Water loss
544.7764892578125 0 7901.224 Precursor Ammonia loss
545.2771606445312 0 2845.288
545.77490234375 0 996.1122
548.2693481445312 0 3114.024
549.2680053710938 0 46780.195 y Water loss 5
550.2681274414062 0 12038.488
551.2825317382812 0 3765.1526
552.2828369140625 0 2483.0579
552.7780151367188 0 5032.794
553.2808227539062 0 5050.9097 Precursor
553.6392822265625 0 1077.8123
553.7845458984375 0 1608.6132
553.9732055664062 0 974.05585
554.2965698242188 0 2565.6448
555.2783203125 0 2620.9434
556.280517578125 0 956.16693
558.2608032226562 0 1156.9202
558.326171875 0 1049.9784
559.2542114257812 0 1604.4503
562.3017578125 0 791.1492
567.2784423828125 0 614503.1 y 5
568.2814331054688 0 189193.2
569.2839965820312 0 34967.62
570.2872924804688 0 2465.815
571.25341796875 0 1655.5337
572.2559204101562 0 1036.9784
572.3049926757812 0 9180.61
573.309326171875 0 2632.8
577.26171875 0 1789.4083
578.2667236328125 0 948.922
586.3203125 0 12227.555
587.32373046875 0 3349.8591
588.3268432617188 0 770.6584
593.307861328125 0 1229.3174
594.2928466796875 0 838.2133
608.3051147460938 0 2800.5757
609.3076171875 0 1095.313
611.267578125 0 1022.0775
611.3165893554688 0 2824.645
612.3158569335938 0 1174.8744
614.294677734375 0 1717.6414
615.2882690429688 0 2264.6418
616.2722778320312 0 2236.7593
616.3170166015625 0 1408.04
617.2744750976562 0 980.9276
618.286865234375 0 760.6688
619.3204956054688 0 944.8821
625.3312377929688 0 2573.985
626.3201293945312 0 1757.9863
629.33056640625 0 1527.9854
631.322265625 0 2299.6785
632.3062133789062 0 8816.941
633.3024291992188 0 7810.68
634.2900390625 0 4532.0205
635.2880859375 0 772.08734
636.2998657226562 0 6756.35 b Water loss 5
637.2846069335938 0 2609.3853 b Ammonia loss 5
637.3306884765625 0 4388.8413
638.3253173828125 0 2495.7073
641.3068237304688 0 2251.5496
642.2915649414062 0 6379.9165
643.2858276367188 0 1916.9358
643.3425903320312 0 7571.8833
644.3433837890625 0 2731.7112
645.3545532226562 0 2704.7014
646.3633422851562 0 982.3824
649.33203125 0 9694.474
650.33447265625 0 3086.2295
651.3125610351562 0 3421.613
652.3135375976562 0 1030.0985
653.3231811523438 0 1199.1193
654.3106689453125 0 32536.535 b 5
655.3150024414062 0 8831.058
656.3311157226562 0 4265.862
657.3374633789062 0 1523.9867
659.3157348632812 0 16599.666
660.3026123046875 0 30288.486
661.3028564453125 0 11967.599
662.302734375 0 2476.191
671.339111328125 0 1674.9862
673.3528442382812 0 13061.464
674.3555908203125 0 4325.0923
675.3597412109375 0 1047.8519
677.3265991210938 0 185201.4 y Water loss 4
678.3126831054688 0 419508.3 y Ammonia loss 4
679.3148803710938 0 149899.62
680.3165893554688 0 32963.06
681.3193969726562 0 2010.6158
686.3475341796875 0 1463.21
694.3528442382812 0 742.78723
695.3373413085938 0 539533.3 y 4
696.340087890625 0 207645.03
696.4808349609375 0 1395.687
697.3421630859375 0 42938.125
698.3444213867188 0 3339.4617
705.3212890625 0 3861.2305
706.31396484375 0 2371.7058
712.3634033203125 0 6139.1353
713.3556518554688 0 4740.2393
714.3535766601562 0 1755.8536
730.3740844726562 0 12321.099
731.3761596679688 0 5400.0005
732.3797607421875 0 1467.3215
746.3845825195312 0 1247.6357
747.3739013671875 0 1646.536
748.3877563476562 0 1393.6005
749.3896484375 0 1177.1375
755.3745727539062 0 1361.576
756.3789672851562 0 888.9992
764.3950805664062 0 5651.237 b Water loss 6
765.3915405273438 0 3277.855 b Ammonia loss 6
766.404541015625 0 2631.3875
773.3840942382812 0 15075.809
774.3864135742188 0 5545.499
775.3912353515625 0 1190.1777
782.4053955078125 0 12371.058 b 6
783.4083862304688 0 5818.458
784.408935546875 0 1360.2799
790.4096069335938 0 4406.759 y Water loss 3
791.40625 0 3644.2168 y Ammonia loss 3
794.4083251953125 0 1138.572
798.4710693359375 0 919.7828
800.4112548828125 0 830.0049
803.4042358398438 0 1978.2377
804.396484375 0 2797.8364
805.396240234375 0 1347.3115
808.4207763671875 0 52475.867 y 3
809.4238891601562 0 26776.951
810.4266357421875 0 5556.6685
811.428466796875 0 830.32214
813.3991088867188 0 680.3808
818.4043579101562 0 734.99915
819.4114990234375 0 652.275
821.4161987304688 0 32929.24
822.41845703125 0 13537.267
823.4208984375 0 4057.7615
829.4197387695312 0 1900.2853
830.4125366210938 0 2170.6992
832.4219970703125 0 997.72864
833.4163818359375 0 738.90594
836.26513671875 0 683.1321
837.4476928710938 0 1681.4446
838.4403076171875 0 2024.3973
839.4304809570312 0 7297.765
840.4302978515625 0 1804.6495
847.4319458007812 0 17550.604
848.4290771484375 0 10715.877
849.4293212890625 0 3685.8074
850.4408569335938 0 677.01666
855.4573974609375 0 8648.973
856.4601440429688 0 4780.8496
857.464111328125 0 1261.9415
859.4304809570312 0 1855.6024
860.4193725585938 0 1507.922
865.4418334960938 0 38887.05 b Water loss 7
866.4408569335938 0 23732.426 b Ammonia loss 7
867.4417114257812 0 7834.6416
868.4479370117188 0 1153.6204
877.441162109375 0 5694.4355 y Water loss 2
878.431640625 0 8180.0596 y Ammonia loss 2
879.4331665039062 0 5113.936
880.4345092773438 0 1447.5493
883.45263671875 0 113241.19 b 7
884.4554443359375 0 57141.883
885.4581298828125 0 16175.458
886.4581909179688 0 1127.6826
887.4295654296875 0 837.2725
894.4655151367188 0 739.8093
895.45263671875 0 38796.062 y 2
896.453857421875 0 25116.29
897.4547119140625 0 7648.0273
898.4530639648438 0 1619.1866
904.4549560546875 0 1936.0677
905.4409790039062 0 4668.0093
906.441650390625 0 2882.4998
907.4400024414062 0 1133.2278
912.4765625 0 3351.1763
913.4837036132812 0 1129.5393
922.4636840820312 0 37143.688 b Water loss 8
923.4603271484375 0 26360.133 b Ammonia loss 8
924.460693359375 0 9465.166
925.4571533203125 0 1571.6951
940.4739379882812 0 151668.81 b 8
941.4772338867188 0 81372.51
942.4796142578125 0 24970.998
943.4820556640625 0 3529.2295
958.4844360351562 0 3940.502
959.48779296875 0 1710.7133
2120.70068359375 0 718.49347
2555.42529296875 0 740.5633
3081.777587890625 0 955.2641
3153.3310546875 0 700.37555

Spectrum Details

|  |  |
| --- | --- |
| Matched peaks? Matched peaks The total absolute number of peaks matched. Additionally in brackets the total fraction of peaks matched and the total number of peaks is shown. | 63 (8.03% of 785) |
| FDR? FDR The false discovery rate estimated for this peptide. It is calculated by matching all theoretical fragments with a non-integer shift with the raw peaks for this spectrum. This is done with 40 different shifts. The resulting percentage is the average number of annotated peaks over the number of annotated peaks with the correct spectrum. | 0.00% |
| Satellite FDR? Satellite FDR See the FDR for details on its calculation. This satellite ion specific FDR only contains the satellite ions (d/w) for I/L/J positions. | - |
| PSM Score? PSM Score The PSM Score as given by Hecklib to this annotated spectrum. It is shown with three significant figures. | 742 |

#### Spectrum 5620? Spectrum 5620 The raw spectrum of this peptide as annotated by Hecklib. The fragments are coloured according to ion type (see legend). Any peaks with a star '\*' as text can be hovered over to see the full details, first the ion type second the mass shift type. By hovering over the amino acids in the peptide or ions in the legend the corresponding peaks are highlighted. By toggling the 'Unassigned' label you can turn the background (unassigned) peaks on or off in the plot. By updating the slider in the Ion legend you can update the spectrum to only show the top X% of the peaks with labels. The top X% means any peak that is within X% of the highest intensity. By dragging in the spectrum you can zoom in to a specific part of the spectrum and use 'Zoom Out' to get back to the original zoom level. The annotation of the spectrum is based on the given sequence in the peptides file and is done with different software so inconsistencies are likely. The peaks are annotated based on the given sequence, with 20 ppm tolerance.

Copy Data

##### Spectrum 5620 (TSV)

###### Preview

```
Loading example...
```

*Click on the button to copy the data to your clipboard.*

Mz MinMz MaxIntensity Max

WidthHeightPeptide font sizePeptide stroke widthSpectrum font sizeSpectrum stroke widthCompact peptide

Ion legend

wxyz

abcd

OtherUnassignedIon^Charge^_Position_Show for top:%

EVSJQDKTGF

08.98e+41.80e+52.70e+53.59e+5

Zoom Out

y^+1^_1_a^+1^_2_d^+1^_2_a^+1^_2_b^+1^_2_y^+1^_2_b^+1^_2_a^+1^_3_b^+2^_5_y^+2^_5_a^+1^_3_b^+1^_3_y^+1^_3_b^+1^_3_y^+1^_3_b^+2^_6_y^+2^_6_y^+2^_6_y^+2^_6_b^+2^_7_y^+2^_7_y^+2^_7_y^+2^_7_b^+1^_4_b^+1^_4_y^+1^_4_y^+2^_8_y^+2^_8_b^+2^_8_y^+2^_8_b^+2^_8_y^+1^_4_b^+2^_9_y^+2^_9_b^+1^_5_y^+1^_5_\*b^+1^_5_\*y^+1^_5_b^+1^_6_b^+1^_6_y^+1^_6_y^+1^_6_y^+1^_6_b^+1^_7_y^+1^_7_b^+1^_7_y^+1^_7_y^+1^_8_y^+1^_8_b^+1^_8_b^+1^_8_y^+1^_8_b^+1^_8_b^+1^_9_b^+1^_9_b^+1^_9_y^+1^_9_y^+1^_9_

049899614941992

Fragment Matches Table

Show background peaks

| Position | Ion type | Intensity | mz Theoretical | mz Error (Th) | mz Error (ppm) | Charge | Series Number |
| --- | --- | --- | --- | --- | --- | --- | --- |
| - | - | 8.507E+04 | 120.1 | - | - | 0 | - |
| - | - | 7089 | 121.1 | - | - | 0 | - |
| - | - | 513.4 | 123.1 | - | - | 0 | - |
| - | - | 491.4 | 124.1 | - | - | 0 | - |
| - | - | 400.6 | 125.1 | - | - | 0 | - |
| - | - | 1398 | 126.1 | - | - | 0 | - |
| - | - | 458.4 | 127.1 | - | - | 0 | - |
| - | - | 503.6 | 127.1 | - | - | 0 | - |
| - | - | 3060 | 127.1 | - | - | 0 | - |
| - | - | 1190 | 128.1 | - | - | 0 | - |
| - | - | 4138 | 128.1 | - | - | 0 | - |
| - | - | 418.3 | 129 | - | - | 0 | - |
| - | - | 558.4 | 129.1 | - | - | 0 | - |
| - | - | 1.041E+04 | 129.1 | - | - | 0 | - |
| - | - | 1.45E+05 | 129.1 | - | - | 0 | - |
| - | - | 1420 | 130.1 | - | - | 0 | - |
| - | - | 462.9 | 130.1 | - | - | 0 | - |
| - | - | 702.7 | 130.1 | - | - | 0 | - |
| - | - | 9483 | 130.1 | - | - | 0 | - |
| - | - | 1055 | 131 | - | - | 0 | - |
| - | - | 3961 | 131.1 | - | - | 0 | - |
| - | - | 7705 | 131.1 | - | - | 0 | - |
| - | - | 564.9 | 132.1 | - | - | 0 | - |
| - | - | 1653 | 133.1 | - | - | 0 | - |
| - | - | 1462 | 133.1 | - | - | 0 | - |
| - | - | 2454 | 136.1 | - | - | 0 | - |
| - | - | 407.6 | 137.1 | - | - | 0 | - |
| - | - | 3.402E+04 | 138.1 | - | - | 0 | - |
| - | - | 700 | 139.1 | - | - | 0 | - |
| - | - | 3639 | 139.1 | - | - | 0 | - |
| - | - | 2496 | 139.1 | - | - | 0 | - |
| - | - | 606.4 | 139.1 | - | - | 0 | - |
| - | - | 1418 | 140.1 | - | - | 0 | - |
| - | - | 2681 | 141.1 | - | - | 0 | - |
| - | - | 6246 | 141.1 | - | - | 0 | - |
| - | - | 721.6 | 143.1 | - | - | 0 | - |
| - | - | 511.7 | 146.1 | - | - | 0 | - |
| - | - | 923 | 147.1 | - | - | 0 | - |
| - | - | 1475 | 147.1 | - | - | 0 | - |
| - | - | 558.2 | 148.9 | - | - | 0 | - |
| - | - | 556.8 | 148.9 | - | - | 0 | - |
| - | - | 541.7 | 148.9 | - | - | 0 | - |
| - | - | 765.2 | 148.9 | - | - | 0 | - |
| - | - | 791.5 | 148.9 | - | - | 0 | - |
| - | - | 1204 | 148.9 | - | - | 0 | - |
| - | - | 1417 | 148.9 | - | - | 0 | - |
| - | - | 3064 | 148.9 | - | - | 0 | - |
| - | - | 4838 | 148.9 | - | - | 0 | - |
| - | - | 3767 | 149 | - | - | 0 | - |
| - | - | 2155 | 149 | - | - | 0 | - |
| - | - | 1323 | 149 | - | - | 0 | - |
| - | - | 1109 | 149 | - | - | 0 | - |
| - | - | 672.6 | 149 | - | - | 0 | - |
| - | - | 752 | 149 | - | - | 0 | - |
| - | - | 524.2 | 149 | - | - | 0 | - |
| - | - | 432.9 | 149 | - | - | 0 | - |
| - | - | 638.5 | 149 | - | - | 0 | - |
| - | - | 1109 | 149.1 | - | - | 0 | - |
| - | - | 1042 | 149.1 | - | - | 0 | - |
| - | - | 6226 | 151.1 | - | - | 0 | - |
| - | - | 540.3 | 152.1 | - | - | 0 | - |
| - | - | 773.6 | 152.1 | - | - | 0 | - |
| - | - | 764.2 | 153.1 | - | - | 0 | - |
| - | - | 3149 | 154.1 | - | - | 0 | - |
| - | - | 755.5 | 155.1 | - | - | 0 | - |
| - | - | 2591 | 155.1 | - | - | 0 | - |
| - | - | 3.492E+04 | 155.1 | - | - | 0 | - |
| - | - | 2.584E+04 | 156.1 | - | - | 0 | - |
| - | - | 3378 | 156.1 | - | - | 0 | - |
| - | - | 727.4 | 157.1 | - | - | 0 | - |
| - | - | 1063 | 157.1 | - | - | 0 | - |
| - | - | 1991 | 157.1 | - | - | 0 | - |
| - | - | 1333 | 157.1 | - | - | 0 | - |
| - | - | 1.886E+04 | 159.1 | - | - | 0 | - |
| - | - | 1465 | 159.1 | - | - | 0 | - |
| - | - | 3261 | 159.1 | - | - | 0 | - |
| - | - | 483.7 | 159.1 | - | - | 0 | - |
| - | - | 1561 | 160.1 | - | - | 0 | - |
| - | - | 1093 | 165.1 | - | - | 0 | - |
| - | - | 478.2 | 165.1 | - | - | 0 | - |
| - | - | 1086 | 165.1 | - | - | 0 | - |
| 10 | y | 1.164E+05 | 166.1 | 0.0004301 | 2.59 | +1 | 1 |
| - | - | 629.9 | 166.1 | - | - | 0 | - |
| - | - | 916.4 | 167.1 | - | - | 0 | - |
| - | - | 1.16E+04 | 167.1 | - | - | 0 | - |
| - | - | 512.3 | 167.1 | - | - | 0 | - |
| - | - | 6949 | 167.1 | - | - | 0 | - |
| - | - | 5798 | 168.1 | - | - | 0 | - |
| - | - | 854.8 | 168.1 | - | - | 0 | - |
| - | - | 3155 | 169.1 | - | - | 0 | - |
| - | - | 754 | 169.1 | - | - | 0 | - |
| - | - | 922.3 | 171.1 | - | - | 0 | - |
| - | - | 1001 | 172.1 | - | - | 0 | - |
| - | - | 1.724E+05 | 173.1 | - | - | 0 | - |
| - | - | 541.9 | 173.4 | - | - | 0 | - |
| - | - | 1.45E+04 | 174.1 | - | - | 0 | - |
| - | - | 1296 | 177.1 | - | - | 0 | - |
| - | - | 2.46E+04 | 177.1 | - | - | 0 | - |
| - | - | 1286 | 177.1 | - | - | 0 | - |
| - | - | 2379 | 178.1 | - | - | 0 | - |
| - | - | 791.9 | 179.1 | - | - | 0 | - |
| - | - | 7188 | 181.1 | - | - | 0 | - |
| - | - | 1068 | 181.1 | - | - | 0 | - |
| - | - | 559.5 | 182.1 | - | - | 0 | - |
| - | - | 1122 | 182.1 | - | - | 0 | - |
| - | - | 7.284E+04 | 182.1 | - | - | 0 | - |
| 2 | a | 2.311E+05 | 183.1 | 0.0004618 | 2.522 | +1 | 2 |
| - | - | 8365 | 183.1 | - | - | 0 | - |
| - | - | 2921 | 184.1 | - | - | 0 | - |
| - | - | 4472 | 184.1 | - | - | 0 | - |
| - | - | 2.084E+04 | 184.1 | - | - | 0 | - |
| - | - | 605.8 | 184.1 | - | - | 0 | - |
| - | - | 1157 | 184.1 | - | - | 0 | - |
| - | - | 1146 | 185.1 | - | - | 0 | - |
| - | - | 9514 | 185.1 | - | - | 0 | - |
| - | - | 737.9 | 186.1 | - | - | 0 | - |
| - | - | 574.5 | 186.2 | - | - | 0 | - |
| 2 | d | 8527 | 187.1 | 0.0003592 | 1.92 | +1 | 2 |
| - | - | 1069 | 187.1 | - | - | 0 | - |
| - | - | 1374 | 188.1 | - | - | 0 | - |
| - | - | 1121 | 188.1 | - | - | 0 | - |
| - | - | 766.2 | 191.1 | - | - | 0 | - |
| - | - | 968.5 | 193.1 | - | - | 0 | - |
| - | - | 8959 | 194.1 | - | - | 0 | - |
| - | - | 1569 | 195.1 | - | - | 0 | - |
| - | - | 966.6 | 195.1 | - | - | 0 | - |
| - | - | 1024 | 196.1 | - | - | 0 | - |
| - | - | 3676 | 197.1 | - | - | 0 | - |
| - | - | 2005 | 197.1 | - | - | 0 | - |
| - | - | 1.17E+04 | 198.1 | - | - | 0 | - |
| - | - | 4393 | 198.1 | - | - | 0 | - |
| - | - | 4722 | 199.1 | - | - | 0 | - |
| - | - | 1419 | 199.1 | - | - | 0 | - |
| - | - | 1.13E+05 | 200.1 | - | - | 0 | - |
| 2 | a | 3.558E+05 | 201.1 | 0.0004105 | 2.041 | +1 | 2 |
| - | - | 1.214E+04 | 201.1 | - | - | 0 | - |
| - | - | 3.232E+04 | 202.1 | - | - | 0 | - |
| - | - | 632.2 | 202.1 | - | - | 0 | - |
| - | - | 1533 | 202.2 | - | - | 0 | - |
| - | - | 2298 | 203.1 | - | - | 0 | - |
| - | - | 3317 | 205.1 | - | - | 0 | - |
| - | - | 1571 | 208.1 | - | - | 0 | - |
| - | - | 1805 | 208.1 | - | - | 0 | - |
| - | - | 1500 | 209.1 | - | - | 0 | - |
| - | - | 1.609E+04 | 209.1 | - | - | 0 | - |
| - | - | 1910 | 210.1 | - | - | 0 | - |
| - | - | 3.184E+04 | 210.1 | - | - | 0 | - |
| 2 | b | 7.084E+04 | 211.1 | 0.000405 | 1.918 | +1 | 2 |
| - | - | 3153 | 211.1 | - | - | 0 | - |
| - | - | 730 | 212.1 | - | - | 0 | - |
| - | - | 7418 | 212.1 | - | - | 0 | - |
| - | - | 5380 | 212.1 | - | - | 0 | - |
| - | - | 567.6 | 213.1 | - | - | 0 | - |
| - | - | 1931 | 213.1 | - | - | 0 | - |
| - | - | 726.6 | 213.1 | - | - | 0 | - |
| - | - | 1475 | 214.1 | - | - | 0 | - |
| - | - | 634.8 | 214.2 | - | - | 0 | - |
| - | - | 1561 | 215.1 | - | - | 0 | - |
| - | - | 1036 | 215.1 | - | - | 0 | - |
| - | - | 6129 | 216.1 | - | - | 0 | - |
| - | - | 5831 | 218.2 | - | - | 0 | - |
| - | - | 1008 | 219.1 | - | - | 0 | - |
| - | - | 1391 | 222.1 | - | - | 0 | - |
| 9 | y | 7.227E+04 | 223.1 | 0.0004355 | 1.952 | +1 | 2 |
| - | - | 9264 | 224.1 | - | - | 0 | - |
| - | - | 912.1 | 225.1 | - | - | 0 | - |
| - | - | 1070 | 225.1 | - | - | 0 | - |
| - | - | 4700 | 225.1 | - | - | 0 | - |
| - | - | 485 | 225.5 | - | - | 0 | - |
| - | - | 2.911E+04 | 226.1 | - | - | 0 | - |
| - | - | 5.786E+04 | 226.1 | - | - | 0 | - |
| - | - | 1.906E+04 | 227.1 | - | - | 0 | - |
| - | - | 2988 | 227.1 | - | - | 0 | - |
| - | - | 2920 | 227.1 | - | - | 0 | - |
| - | - | 4737 | 227.1 | - | - | 0 | - |
| - | - | 2127 | 228.1 | - | - | 0 | - |
| - | - | 4.189E+04 | 228.1 | - | - | 0 | - |
| 2 | b | 7.147E+04 | 229.1 | 0.0004452 | 1.943 | +1 | 2 |
| - | - | 3703 | 229.1 | - | - | 0 | - |
| - | - | 6414 | 230.1 | - | - | 0 | - |
| - | - | 7668 | 230.2 | - | - | 0 | - |
| - | - | 824.4 | 231.1 | - | - | 0 | - |
| - | - | 853.8 | 231.2 | - | - | 0 | - |
| - | - | 1082 | 234.1 | - | - | 0 | - |
| - | - | 780.4 | 236.1 | - | - | 0 | - |
| - | - | 779.1 | 238.1 | - | - | 0 | - |
| - | - | 658 | 238.2 | - | - | 0 | - |
| - | - | 1499 | 239.2 | - | - | 0 | - |
| - | - | 2989 | 240.1 | - | - | 0 | - |
| - | - | 2.526E+04 | 242.2 | - | - | 0 | - |
| - | - | 4688 | 243.1 | - | - | 0 | - |
| - | - | 2374 | 243.2 | - | - | 0 | - |
| - | - | 1208 | 244.1 | - | - | 0 | - |
| - | - | 4.596E+04 | 244.1 | - | - | 0 | - |
| - | - | 5.269E+04 | 244.1 | - | - | 0 | - |
| - | - | 1180 | 245.1 | - | - | 0 | - |
| - | - | 3875 | 245.1 | - | - | 0 | - |
| - | - | 6516 | 245.1 | - | - | 0 | - |
| - | - | 796.9 | 246.1 | - | - | 0 | - |
| - | - | 1078 | 248.1 | - | - | 0 | - |
| - | - | 785.1 | 248.1 | - | - | 0 | - |
| - | - | 837.1 | 248.2 | - | - | 0 | - |
| - | - | 589.6 | 249.1 | - | - | 0 | - |
| - | - | 525.4 | 250.2 | - | - | 0 | - |
| - | - | 1423 | 251.2 | - | - | 0 | - |
| - | - | 3129 | 252.1 | - | - | 0 | - |
| - | - | 2831 | 253.1 | - | - | 0 | - |
| - | - | 1307 | 254.1 | - | - | 0 | - |
| - | - | 1363 | 254.2 | - | - | 0 | - |
| - | - | 582.7 | 258.1 | - | - | 0 | - |
| - | - | 3414 | 258.1 | - | - | 0 | - |
| - | - | 3361 | 261.1 | - | - | 0 | - |
| - | - | 833.6 | 261.1 | - | - | 0 | - |
| - | - | 952.7 | 262.1 | - | - | 0 | - |
| - | - | 942.5 | 262.1 | - | - | 0 | - |
| - | - | 1019 | 264.1 | - | - | 0 | - |
| - | - | 1644 | 265.1 | - | - | 0 | - |
| - | - | 2692 | 266.1 | - | - | 0 | - |
| - | - | 1880 | 266.2 | - | - | 0 | - |
| - | - | 1720 | 267.1 | - | - | 0 | - |
| - | - | 1164 | 268.1 | - | - | 0 | - |
| - | - | 597.7 | 268.1 | - | - | 0 | - |
| - | - | 1864 | 269.2 | - | - | 0 | - |
| 3 | a | 9959 | 270.1 | 0.0004006 | 1.483 | +1 | 3 |
| - | - | 669.1 | 271.1 | - | - | 0 | - |
| - | - | 974.5 | 274.1 | - | - | 0 | - |
| - | - | 952.7 | 276.1 | - | - | 0 | - |
| - | - | 2086 | 278.2 | - | - | 0 | - |
| - | - | 1309 | 279.1 | - | - | 0 | - |
| 5 | b | 4412 | 279.2 | 0.00418 | 14.97 | +2 | 5 |
| - | - | 2.832E+04 | 280.1 | - | - | 0 | - |
| - | - | 3241 | 281.1 | - | - | 0 | - |
| - | - | 670.3 | 281.2 | - | - | 0 | - |
| - | - | 1650 | 282.1 | - | - | 0 | - |
| - | - | 4431 | 282.2 | - | - | 0 | - |
| - | - | 1.482E+04 | 283.1 | - | - | 0 | - |
| - | - | 569.1 | 284.1 | - | - | 0 | - |
| 6 | y | 2171 | 284.1 | 0.001448 | 5.096 | +2 | 5 |
| - | - | 786.6 | 284.2 | - | - | 0 | - |
| - | - | 611.2 | 285.2 | - | - | 0 | - |
| - | - | 3086 | 287.2 | - | - | 0 | - |
| 3 | a | 766.7 | 288.2 | 0.000368 | 1.277 | +1 | 3 |
| - | - | 1124 | 290.1 | - | - | 0 | - |
| - | - | 1162 | 291.1 | - | - | 0 | - |
| - | - | 2561 | 292.1 | - | - | 0 | - |
| - | - | 698.7 | 292.1 | - | - | 0 | - |
| - | - | 2480 | 294.1 | - | - | 0 | - |
| - | - | 2.53E+04 | 297.2 | - | - | 0 | - |
| - | - | 511.3 | 297.9 | - | - | 0 | - |
| 3 | b | 8.034E+04 | 298.1 | 0.0006032 | 2.023 | +1 | 3 |
| - | - | 4195 | 298.2 | - | - | 0 | - |
| - | - | 1.105E+04 | 299.1 | - | - | 0 | - |
| - | - | 1029 | 299.2 | - | - | 0 | - |
| - | - | 749.8 | 300.2 | - | - | 0 | - |
| - | - | 2325 | 301.2 | - | - | 0 | - |
| - | - | 992.1 | 301.2 | - | - | 0 | - |
| - | - | 745.4 | 302.2 | - | - | 0 | - |
| - | - | 3106 | 305.2 | - | - | 0 | - |
| 8 | y | 5609 | 306.1 | 0.0005227 | 1.707 | +1 | 3 |
| - | - | 8511 | 309.2 | - | - | 0 | - |
| - | - | 3660 | 310.1 | - | - | 0 | - |
| - | - | 954.5 | 310.2 | - | - | 0 | - |
| - | - | 2026 | 310.2 | - | - | 0 | - |
| - | - | 1173 | 311.1 | - | - | 0 | - |
| - | - | 3724 | 311.2 | - | - | 0 | - |
| - | - | 954.3 | 312.2 | - | - | 0 | - |
| - | - | 1.091E+04 | 315.2 | - | - | 0 | - |
| - | - | 1216 | 316.1 | - | - | 0 | - |
| 3 | b | 2.547E+04 | 316.2 | 0.0005365 | 1.697 | +1 | 3 |
| - | - | 1960 | 316.2 | - | - | 0 | - |
| - | - | 3740 | 317.2 | - | - | 0 | - |
| - | - | 2056 | 317.2 | - | - | 0 | - |
| - | - | 732.4 | 317.2 | - | - | 0 | - |
| - | - | 4919 | 319.1 | - | - | 0 | - |
| - | - | 1167 | 320.1 | - | - | 0 | - |
| - | - | 1452 | 322.1 | - | - | 0 | - |
| 8 | y | 2.713E+04 | 324.2 | 0.0006696 | 2.066 | +1 | 3 |
| - | - | 4752 | 325.2 | - | - | 0 | - |
| - | - | 9810 | 327.2 | - | - | 0 | - |
| 6 | b | 1381 | 328.2 | 0.0004755 | 1.449 | +2 | 6 |
| - | - | 1852 | 328.2 | - | - | 0 | - |
| - | - | 2.935E+04 | 329.2 | - | - | 0 | - |
| - | - | 1047 | 330.2 | - | - | 0 | - |
| - | - | 5522 | 330.2 | - | - | 0 | - |
| - | - | 6941 | 336.2 | - | - | 0 | - |
| - | - | 2.7E+04 | 337.2 | - | - | 0 | - |
| - | - | 1385 | 337.2 | - | - | 0 | - |
| - | - | 915.7 | 338.1 | - | - | 0 | - |
| - | - | 4741 | 338.2 | - | - | 0 | - |
| 5 | y | 1.474E+04 | 339.2 | 0.0004823 | 1.422 | +2 | 6 |
| 5 | y | 4522 | 339.7 | 0.006765 | 19.92 | +2 | 6 |
| - | - | 4717 | 340.2 | - | - | 0 | - |
| - | - | 927.1 | 341.2 | - | - | 0 | - |
| - | - | 569.4 | 343.7 | - | - | 0 | - |
| - | - | 1.892E+04 | 345.2 | - | - | 0 | - |
| - | - | 3235 | 346.2 | - | - | 0 | - |
| 5 | y | 1.745E+04 | 348.2 | 0.0005405 | 1.552 | +2 | 6 |
| - | - | 5709 | 348.7 | - | - | 0 | - |
| - | - | 1115 | 349.2 | - | - | 0 | - |
| - | - | 3.984E+04 | 354.2 | - | - | 0 | - |
| - | - | 1.86E+04 | 355.2 | - | - | 0 | - |
| - | - | 5362 | 355.2 | - | - | 0 | - |
| - | - | 2370 | 356.2 | - | - | 0 | - |
| - | - | 926.6 | 356.2 | - | - | 0 | - |
| - | - | 936.7 | 357.2 | - | - | 0 | - |
| - | - | 1.297E+04 | 357.2 | - | - | 0 | - |
| - | - | 2039 | 358.2 | - | - | 0 | - |
| - | - | 606.3 | 360.2 | - | - | 0 | - |
| - | - | 588 | 362 | - | - | 0 | - |
| - | - | 2193 | 364.2 | - | - | 0 | - |
| - | - | 3.7E+04 | 365.2 | - | - | 0 | - |
| - | - | 1067 | 366.2 | - | - | 0 | - |
| - | - | 6364 | 366.2 | - | - | 0 | - |
| - | - | 2480 | 367.2 | - | - | 0 | - |
| - | - | 690.2 | 371.2 | - | - | 0 | - |
| - | - | 2.781E+04 | 372.2 | - | - | 0 | - |
| - | - | 1164 | 373.2 | - | - | 0 | - |
| - | - | 4421 | 373.2 | - | - | 0 | - |
| - | - | 1104 | 374.2 | - | - | 0 | - |
| - | - | 848.3 | 375.2 | - | - | 0 | - |
| - | - | 2080 | 376.2 | - | - | 0 | - |
| - | - | 705.3 | 377.2 | - | - | 0 | - |
| - | - | 778.8 | 381.2 | - | - | 0 | - |
| - | - | 754.3 | 382.2 | - | - | 0 | - |
| - | - | 853.7 | 382.2 | - | - | 0 | - |
| - | - | 2735 | 382.2 | - | - | 0 | - |
| - | - | 776.8 | 382.7 | - | - | 0 | - |
| - | - | 4.185E+04 | 383.2 | - | - | 0 | - |
| - | - | 4427 | 384.2 | - | - | 0 | - |
| - | - | 8484 | 384.2 | - | - | 0 | - |
| - | - | 2028 | 385.2 | - | - | 0 | - |
| - | - | 927.7 | 385.2 | - | - | 0 | - |
| - | - | 1181 | 387.2 | - | - | 0 | - |
| 7 | b | 897.8 | 391.7 | 0.0006658 | 1.7 | +2 | 7 |
| - | - | 654.1 | 392.2 | - | - | 0 | - |
| - | - | 4372 | 392.2 | - | - | 0 | - |
| - | - | 4557 | 393.2 | - | - | 0 | - |
| - | - | 3.039E+04 | 393.2 | - | - | 0 | - |
| - | - | 1.71E+04 | 394.2 | - | - | 0 | - |
| - | - | 6053 | 394.2 | - | - | 0 | - |
| - | - | 2844 | 395.2 | - | - | 0 | - |
| - | - | 1315 | 395.2 | - | - | 0 | - |
| 4 | y | 1053 | 395.7 | 0.001114 | 2.815 | +2 | 7 |
| 4 | y | 5328 | 396.2 | 0.0005611 | 1.416 | +2 | 7 |
| - | - | 2767 | 396.7 | - | - | 0 | - |
| - | - | 722.4 | 399.2 | - | - | 0 | - |
| - | - | 1017 | 401.2 | - | - | 0 | - |
| - | - | 3746 | 401.2 | - | - | 0 | - |
| - | - | 8318 | 402.2 | - | - | 0 | - |
| - | - | 1195 | 403.2 | - | - | 0 | - |
| 4 | y | 2.04E+04 | 404.7 | 0.0006533 | 1.614 | +2 | 7 |
| - | - | 9657 | 405.2 | - | - | 0 | - |
| - | - | 1893 | 405.7 | - | - | 0 | - |
| - | - | 1107 | 408.2 | - | - | 0 | - |
| - | - | 841.9 | 409.2 | - | - | 0 | - |
| - | - | 729 | 410.2 | - | - | 0 | - |
| - | - | 2096 | 410.2 | - | - | 0 | - |
| - | - | 1.065E+04 | 410.2 | - | - | 0 | - |
| - | - | 7424 | 410.7 | - | - | 0 | - |
| - | - | 4712 | 411.2 | - | - | 0 | - |
| 4 | b | 4.046E+04 | 411.2 | 0.0004015 | 0.9763 | +1 | 4 |
| - | - | 1109 | 411.7 | - | - | 0 | - |
| - | - | 4401 | 412.2 | - | - | 0 | - |
| - | - | 8432 | 412.2 | - | - | 0 | - |
| - | - | 993.6 | 413.2 | - | - | 0 | - |
| - | - | 1385 | 413.2 | - | - | 0 | - |
| - | - | 855.5 | 415.2 | - | - | 0 | - |
| - | - | 710.3 | 416.2 | - | - | 0 | - |
| - | - | 3455 | 419.2 | - | - | 0 | - |
| - | - | 1224 | 419.2 | - | - | 0 | - |
| - | - | 1.045E+04 | 420.2 | - | - | 0 | - |
| - | - | 1743 | 421.2 | - | - | 0 | - |
| - | - | 973.3 | 422.2 | - | - | 0 | - |
| - | - | 1818 | 423.7 | - | - | 0 | - |
| - | - | 923.4 | 424.2 | - | - | 0 | - |
| - | - | 1116 | 424.2 | - | - | 0 | - |
| - | - | 867.2 | 425.2 | - | - | 0 | - |
| - | - | 2627 | 426.2 | - | - | 0 | - |
| - | - | 3916 | 427.2 | - | - | 0 | - |
| - | - | 1741 | 427.2 | - | - | 0 | - |
| - | - | 1928 | 427.3 | - | - | 0 | - |
| - | - | 1443 | 427.7 | - | - | 0 | - |
| - | - | 4864 | 428.2 | - | - | 0 | - |
| - | - | 1.048E+04 | 428.3 | - | - | 0 | - |
| - | - | 1361 | 429.2 | - | - | 0 | - |
| 4 | b | 2506 | 429.2 | 0.00122 | 2.842 | +1 | 4 |
| - | - | 894.4 | 430.2 | - | - | 0 | - |
| - | - | 1168 | 430.7 | - | - | 0 | - |
| - | - | 966.5 | 431.2 | - | - | 0 | - |
| - | - | 4370 | 432.7 | - | - | 0 | - |
| - | - | 6307 | 433.2 | - | - | 0 | - |
| - | - | 2263 | 433.7 | - | - | 0 | - |
| 7 | y | 6592 | 434.2 | 0.0005914 | 1.362 | +1 | 4 |
| - | - | 936.1 | 435.2 | - | - | 0 | - |
| - | - | 6871 | 437.2 | - | - | 0 | - |
| - | - | 9221 | 438.2 | - | - | 0 | - |
| - | - | 2081 | 439.2 | - | - | 0 | - |
| 3 | y | 3947 | 439.2 | 0.001152 | 2.623 | +2 | 8 |
| 3 | y | 1.156E+04 | 439.7 | 0.001911 | 4.347 | +2 | 8 |
| - | - | 3860 | 440.2 | - | - | 0 | - |
| - | - | 1524 | 440.7 | - | - | 0 | - |
| - | - | 7803 | 441.7 | - | - | 0 | - |
| 8 | b | 4189 | 442.2 | 0.008443 | 19.09 | +2 | 8 |
| - | - | 2117 | 442.7 | - | - | 0 | - |
| - | - | 2.504E+04 | 444.2 | - | - | 0 | - |
| - | - | 4926 | 445.2 | - | - | 0 | - |
| - | - | 1010 | 445.2 | - | - | 0 | - |
| - | - | 691.6 | 445.3 | - | - | 0 | - |
| - | - | 702.2 | 446.2 | - | - | 0 | - |
| - | - | 905.7 | 446.3 | - | - | 0 | - |
| - | - | 892.1 | 447.7 | - | - | 0 | - |
| 3 | y | 2.248E+04 | 448.2 | 0.0005998 | 1.338 | +2 | 8 |
| - | - | 1168 | 448.3 | - | - | 0 | - |
| - | - | 1.265E+04 | 448.7 | - | - | 0 | - |
| - | - | 2732 | 449.2 | - | - | 0 | - |
| - | - | 951.4 | 450.2 | - | - | 0 | - |
| - | - | 2342 | 450.2 | - | - | 0 | - |
| - | - | 1150 | 450.7 | - | - | 0 | - |
| 8 | b | 653.2 | 451.2 | 0.00255 | 5.651 | +2 | 8 |
| 7 | y | 6.743E+04 | 452.3 | 0.0006468 | 1.43 | +1 | 4 |
| - | - | 1184 | 452.7 | - | - | 0 | - |
| - | - | 1.743E+04 | 453.3 | - | - | 0 | - |
| - | - | 1994 | 454.3 | - | - | 0 | - |
| - | - | 9728 | 455.2 | - | - | 0 | - |
| - | - | 2.784E+04 | 456.2 | - | - | 0 | - |
| - | - | 6090 | 457.2 | - | - | 0 | - |
| - | - | 1861 | 458.2 | - | - | 0 | - |
| - | - | 790.3 | 459.2 | - | - | 0 | - |
| - | - | 3080 | 461.2 | - | - | 0 | - |
| - | - | 4067 | 461.7 | - | - | 0 | - |
| - | - | 3148 | 462.2 | - | - | 0 | - |
| - | - | 848.2 | 467.2 | - | - | 0 | - |
| - | - | 893.2 | 467.3 | - | - | 0 | - |
| - | - | 1237 | 468.2 | - | - | 0 | - |
| - | - | 2508 | 468.2 | - | - | 0 | - |
| - | - | 799.3 | 469.2 | - | - | 0 | - |
| - | - | 6665 | 470.2 | - | - | 0 | - |
| 9 | b | 4348 | 470.7 | 0.006958 | 14.78 | +2 | 9 |
| - | - | 1669 | 471.3 | - | - | 0 | - |
| - | - | 1.524E+04 | 473.2 | - | - | 0 | - |
| - | - | 3845 | 474.2 | - | - | 0 | - |
| - | - | 963.5 | 477.2 | - | - | 0 | - |
| - | - | 712.3 | 478.2 | - | - | 0 | - |
| - | - | 4686 | 479.3 | - | - | 0 | - |
| - | - | 2502 | 479.8 | - | - | 0 | - |
| - | - | 652.2 | 480.7 | - | - | 0 | - |
| - | - | 714 | 483.2 | - | - | 0 | - |
| - | - | 1521 | 485.2 | - | - | 0 | - |
| - | - | 6227 | 485.3 | - | - | 0 | - |
| - | - | 987 | 486.2 | - | - | 0 | - |
| - | - | 696.3 | 486.3 | - | - | 0 | - |
| - | - | 725.7 | 488.3 | - | - | 0 | - |
| - | - | 1370 | 494.2 | - | - | 0 | - |
| - | - | 6147 | 495.2 | - | - | 0 | - |
| - | - | 1573 | 496.2 | - | - | 0 | - |
| 2 | y | 5373 | 497.8 | 0.0009387 | 1.886 | +2 | 9 |
| - | - | 2497 | 498.3 | - | - | 0 | - |
| - | - | 1178 | 498.8 | - | - | 0 | - |
| - | - | 2246 | 502.8 | - | - | 0 | - |
| - | - | 1514 | 503.3 | - | - | 0 | - |
| - | - | 765.4 | 505.2 | - | - | 0 | - |
| - | - | 2145 | 510.3 | - | - | 0 | - |
| - | - | 5567 | 512.2 | - | - | 0 | - |
| - | - | 1.154E+04 | 513.2 | - | - | 0 | - |
| - | - | 4114 | 514.2 | - | - | 0 | - |
| - | - | 877.1 | 520.3 | - | - | 0 | - |
| - | - | 4472 | 521.3 | - | - | 0 | - |
| - | - | 735.2 | 521.8 | - | - | 0 | - |
| - | - | 3328 | 522.3 | - | - | 0 | - |
| - | - | 817.1 | 522.8 | - | - | 0 | - |
| - | - | 1127 | 523.3 | - | - | 0 | - |
| - | - | 1498 | 524.3 | - | - | 0 | - |
| - | - | 722.6 | 526.8 | - | - | 0 | - |
| - | - | 1031 | 529.3 | - | - | 0 | - |
| - | - | 933.2 | 529.8 | - | - | 0 | - |
| - | - | 1.278E+04 | 530.3 | - | - | 0 | - |
| - | - | 5734 | 531.3 | - | - | 0 | - |
| - | - | 2410 | 532.2 | - | - | 0 | - |
| - | - | 1847 | 534.3 | - | - | 0 | - |
| - | - | 2235 | 535.3 | - | - | 0 | - |
| - | - | 1524 | 535.8 | - | - | 0 | - |
| - | - | 1501 | 536.3 | - | - | 0 | - |
| - | - | 1049 | 536.8 | - | - | 0 | - |
| - | - | 952.7 | 537.3 | - | - | 0 | - |
| - | - | 8982 | 538.3 | - | - | 0 | - |
| 5 | b | 1.149E+04 | 539.3 | 0.001852 | 3.434 | +1 | 5 |
| - | - | 897.2 | 539.8 | - | - | 0 | - |
| - | - | 3072 | 540.3 | - | - | 0 | - |
| - | - | 811.1 | 541.3 | - | - | 0 | - |
| - | - | 1005 | 542.3 | - | - | 0 | - |
| - | - | 977.7 | 543.3 | - | - | 0 | - |
| - | - | 8028 | 543.8 | - | - | 0 | - |
| - | - | 1.024E+04 | 544.3 | - | - | 0 | - |
| - | - | 7578 | 544.8 | - | - | 0 | - |
| - | - | 1874 | 545.3 | - | - | 0 | - |
| - | - | 884.5 | 548.3 | - | - | 0 | - |
| 6 | y | 1.314E+04 | 549.3 | 0.0006564 | 1.195 | +1 | 5 |
| - | - | 2290 | 550.3 | - | - | 0 | - |
| - | - | 1881 | 550.3 | - | - | 0 | - |
| - | - | 859.6 | 551.3 | - | - | 0 | - |
| - | - | 922.7 | 551.7 | - | - | 0 | - |
| - | - | 2364 | 552.2 | - | - | 0 | - |
| - | - | 1.664E+05 | 552.8 | - | - | 0 | - |
| 0 | Precursor | 1.119E+05 | 553.3 | 0.009155 | 16.55 | +2 | -1 |
| - | - | 4.283E+04 | 553.8 | - | - | 0 | - |
| - | - | 1.424E+04 | 554.3 | - | - | 0 | - |
| - | - | 1050 | 554.8 | - | - | 0 | - |
| - | - | 2913 | 555.3 | - | - | 0 | - |
| - | - | 1606 | 556.3 | - | - | 0 | - |
| 5 | b | 2911 | 557.3 | 0.0003206 | 0.5752 | +1 | 5 |
| - | - | 969.5 | 558.3 | - | - | 0 | - |
| - | - | 2.637E+04 | 561.8 | - | - | 0 | - |
| 0 | Precursor | 2.419E+04 | 562.3 | 0.005399 | 9.602 | +2 | -1 |
| - | - | 1.328E+04 | 562.8 | - | - | 0 | - |
| - | - | 774.2 | 562.9 | - | - | 0 | - |
| - | - | 3931 | 563.3 | - | - | 0 | - |
| 6 | y | 1.714E+05 | 567.3 | 0.0002846 | 0.5017 | +1 | 5 |
| - | - | 4.865E+04 | 568.3 | - | - | 0 | - |
| - | - | 8316 | 569.3 | - | - | 0 | - |
| - | - | 787.7 | 570.3 | - | - | 0 | - |
| - | - | 2.276E+04 | 572.3 | - | - | 0 | - |
| - | - | 5993 | 573.3 | - | - | 0 | - |
| - | - | 1019 | 574.3 | - | - | 0 | - |
| - | - | 716.5 | 577.3 | - | - | 0 | - |
| - | - | 8540 | 586.3 | - | - | 0 | - |
| - | - | 2682 | 587.3 | - | - | 0 | - |
| - | - | 1745 | 593.3 | - | - | 0 | - |
| - | - | 984.9 | 608.3 | - | - | 0 | - |
| - | - | 746.1 | 610.3 | - | - | 0 | - |
| - | - | 1.012E+04 | 611.3 | - | - | 0 | - |
| - | - | 2988 | 612.3 | - | - | 0 | - |
| - | - | 614.8 | 614.3 | - | - | 0 | - |
| - | - | 1919 | 620.3 | - | - | 0 | - |
| - | - | 1962 | 625.3 | - | - | 0 | - |
| - | - | 645.4 | 626.3 | - | - | 0 | - |
| - | - | 972.3 | 628.3 | - | - | 0 | - |
| - | - | 2964 | 629.3 | - | - | 0 | - |
| - | - | 2168 | 632.3 | - | - | 0 | - |
| - | - | 2373 | 633.3 | - | - | 0 | - |
| - | - | 821.1 | 634.3 | - | - | 0 | - |
| - | - | 791.4 | 635.3 | - | - | 0 | - |
| - | - | 2795 | 636.3 | - | - | 0 | - |
| - | - | 1.173E+04 | 637.3 | - | - | 0 | - |
| - | - | 4451 | 638.3 | - | - | 0 | - |
| - | - | 753.4 | 639.3 | - | - | 0 | - |
| - | - | 882.3 | 641.3 | - | - | 0 | - |
| - | - | 5056 | 641.9 | - | - | 0 | - |
| - | - | 1497 | 642.3 | - | - | 0 | - |
| - | - | 3979 | 642.4 | - | - | 0 | - |
| - | - | 964.7 | 643.3 | - | - | 0 | - |
| - | - | 4741 | 643.3 | - | - | 0 | - |
| - | - | 1023 | 644.3 | - | - | 0 | - |
| - | - | 3506 | 645.4 | - | - | 0 | - |
| - | - | 4067 | 649.3 | - | - | 0 | - |
| - | - | 1916 | 650.3 | - | - | 0 | - |
| - | - | 748.4 | 651.3 | - | - | 0 | - |
| - | - | 1.486E+04 | 653.3 | - | - | 0 | - |
| 6 | b | 8375 | 654.3 | 0.007319 | 11.19 | +1 | 6 |
| - | - | 867.9 | 654.4 | - | - | 0 | - |
| - | - | 9436 | 655.3 | - | - | 0 | - |
| - | - | 4762 | 656.3 | - | - | 0 | - |
| - | - | 1136 | 657.3 | - | - | 0 | - |
| - | - | 5630 | 659.3 | - | - | 0 | - |
| - | - | 8296 | 660.3 | - | - | 0 | - |
| - | - | 2612 | 661.3 | - | - | 0 | - |
| - | - | 1126 | 662.3 | - | - | 0 | - |
| - | - | 1088 | 668.3 | - | - | 0 | - |
| - | - | 3062 | 670.4 | - | - | 0 | - |
| - | - | 1738 | 671.4 | - | - | 0 | - |
| 6 | b | 1472 | 672.3 | 0.001293 | 1.923 | +1 | 6 |
| - | - | 4.032E+04 | 673.4 | - | - | 0 | - |
| - | - | 1.543E+04 | 674.4 | - | - | 0 | - |
| - | - | 2360 | 675.4 | - | - | 0 | - |
| 5 | y | 5.431E+04 | 677.3 | 0.0002454 | 0.3623 | +1 | 6 |
| 5 | y | 1.161E+05 | 678.3 | 0.002192 | 3.231 | +1 | 6 |
| - | - | 3.828E+04 | 679.3 | - | - | 0 | - |
| - | - | 9013 | 680.3 | - | - | 0 | - |
| - | - | 1083 | 681.3 | - | - | 0 | - |
| - | - | 1170 | 686.3 | - | - | 0 | - |
| 5 | y | 1.74E+05 | 695.3 | 0.0001788 | 0.2571 | +1 | 6 |
| - | - | 6.287E+04 | 696.3 | - | - | 0 | - |
| - | - | 1.384E+04 | 697.3 | - | - | 0 | - |
| - | - | 1685 | 698.3 | - | - | 0 | - |
| - | - | 810.9 | 705.3 | - | - | 0 | - |
| - | - | 1138 | 706.3 | - | - | 0 | - |
| - | - | 802.1 | 711.9 | - | - | 0 | - |
| - | - | 7135 | 712.4 | - | - | 0 | - |
| - | - | 2398 | 713.4 | - | - | 0 | - |
| - | - | 1.776E+04 | 730.4 | - | - | 0 | - |
| - | - | 6939 | 731.4 | - | - | 0 | - |
| - | - | 1180 | 732.4 | - | - | 0 | - |
| - | - | 959.2 | 746.4 | - | - | 0 | - |
| - | - | 2991 | 748.4 | - | - | 0 | - |
| - | - | 1104 | 749.4 | - | - | 0 | - |
| - | - | 704.1 | 755.9 | - | - | 0 | - |
| - | - | 1668 | 764.4 | - | - | 0 | - |
| - | - | 1130 | 765.4 | - | - | 0 | - |
| - | - | 1547 | 772.4 | - | - | 0 | - |
| - | - | 4624 | 773.4 | - | - | 0 | - |
| - | - | 2067 | 774.4 | - | - | 0 | - |
| - | - | 9262 | 781.4 | - | - | 0 | - |
| 7 | b | 6570 | 782.4 | 0.01001 | 12.8 | +1 | 7 |
| - | - | 2081 | 783.4 | - | - | 0 | - |
| 4 | y | 2684 | 790.4 | 0.00136 | 1.721 | +1 | 7 |
| - | - | 1608 | 791.4 | - | - | 0 | - |
| - | - | 1432 | 799.4 | - | - | 0 | - |
| 7 | b | 2805 | 800.4 | 0.0009731 | 1.216 | +1 | 7 |
| - | - | 864.6 | 801.4 | - | - | 0 | - |
| 4 | y | 5.502E+04 | 808.4 | 0.0003281 | 0.4059 | +1 | 7 |
| - | - | 2.293E+04 | 809.4 | - | - | 0 | - |
| - | - | 5361 | 810.4 | - | - | 0 | - |
| - | - | 850 | 811.4 | - | - | 0 | - |
| - | - | 936.6 | 818.4 | - | - | 0 | - |
| - | - | 4278 | 821.4 | - | - | 0 | - |
| - | - | 1980 | 822.4 | - | - | 0 | - |
| - | - | 968.5 | 823.4 | - | - | 0 | - |
| - | - | 1539 | 829.4 | - | - | 0 | - |
| - | - | 1793 | 833.4 | - | - | 0 | - |
| - | - | 791.6 | 838.4 | - | - | 0 | - |
| - | - | 2163 | 839.4 | - | - | 0 | - |
| - | - | 845.7 | 840.4 | - | - | 0 | - |
| - | - | 799.5 | 842.4 | - | - | 0 | - |
| - | - | 1536 | 846.4 | - | - | 0 | - |
| - | - | 3570 | 847.4 | - | - | 0 | - |
| - | - | 1479 | 848.4 | - | - | 0 | - |
| - | - | 1589 | 849.4 | - | - | 0 | - |
| - | - | 1210 | 855.5 | - | - | 0 | - |
| - | - | 2533 | 859.4 | - | - | 0 | - |
| - | - | 2672 | 860.4 | - | - | 0 | - |
| - | - | 8993 | 865.4 | - | - | 0 | - |
| - | - | 4165 | 866.4 | - | - | 0 | - |
| - | - | 1531 | 867.4 | - | - | 0 | - |
| - | - | 1038 | 877.3 | - | - | 0 | - |
| 3 | y | 1.477E+04 | 877.4 | 0.0004295 | 0.4895 | +1 | 8 |
| 3 | y | 1.128E+04 | 878.4 | 0.009268 | 10.55 | +1 | 8 |
| - | - | 3841 | 879.4 | - | - | 0 | - |
| - | - | 1377 | 880.4 | - | - | 0 | - |
| 8 | b | 2.517E+04 | 883.5 | 0.0002363 | 0.2674 | +1 | 8 |
| 8 | b | 1.259E+04 | 884.4 | 0.01756 | 19.86 | +1 | 8 |
| - | - | 2846 | 885.5 | - | - | 0 | - |
| 3 | y | 2.428E+05 | 895.5 | 0.0006182 | 0.6904 | +1 | 8 |
| - | - | 1.211E+05 | 896.5 | - | - | 0 | - |
| - | - | 3.475E+04 | 897.5 | - | - | 0 | - |
| - | - | 1261 | 897.6 | - | - | 0 | - |
| - | - | 4216 | 898.5 | - | - | 0 | - |
| - | - | 2020 | 900.5 | - | - | 0 | - |
| 8 | b | 8885 | 901.5 | 0.001696 | 1.881 | +1 | 8 |
| - | - | 4020 | 902.5 | - | - | 0 | - |
| - | - | 1676 | 903.5 | - | - | 0 | - |
| - | - | 2245 | 905.4 | - | - | 0 | - |
| - | - | 847.7 | 906.4 | - | - | 0 | - |
| - | - | 5084 | 922.5 | - | - | 0 | - |
| - | - | 4617 | 923.5 | - | - | 0 | - |
| - | - | 2338 | 924.5 | - | - | 0 | - |
| 9 | b | 3.256E+04 | 940.5 | 0.0005976 | 0.6354 | +1 | 9 |
| 9 | b | 1.616E+04 | 941.5 | 0.01819 | 19.33 | +1 | 9 |
| - | - | 4289 | 942.5 | - | - | 0 | - |
| - | - | 2873 | 957.5 | - | - | 0 | - |
| 9 | b | 1.336E+04 | 958.5 | 0.0009837 | 1.026 | +1 | 9 |
| - | - | 6778 | 959.5 | - | - | 0 | - |
| - | - | 1994 | 960.5 | - | - | 0 | - |
| 2 | y | 1180 | 976.5 | 0.009456 | 9.684 | +1 | 9 |
| 2 | y | 6524 | 994.5 | 0.0006117 | 0.6151 | +1 | 9 |
| - | - | 2641 | 995.5 | - | - | 0 | - |
| - | - | 1678 | 996.5 | - | - | 0 | - |
| - | - | 1063 | 1005 | - | - | 0 | - |
| - | - | 695.3 | 1401 | - | - | 0 | - |
| - | - | 657.6 | 1659 | - | - | 0 | - |
| - | - | 624.7 | 1720 | - | - | 0 | - |
| - | - | 643.7 | 1972 | - | - | 0 | - |

m/z Charge Intensity FragmentType MassShift Position
120.08119201660156 0 85070.4
121.08452606201172 0 7089.4683
123.09194946289062 0 513.4157
124.07605743408203 0 491.40692
125.07051849365234 0 400.64758
126.09173583984375 0 1397.5781
127.05069732666016 0 458.35806
127.08706665039062 0 503.60324
127.12340545654297 0 3060.1855
128.0823516845703 0 1189.892
128.10733032226562 0 4138.035
129.018798828125 0 418.26956
129.0608367919922 0 558.35596
129.06625366210938 0 10410.242
129.1026611328125 0 145041.62
130.05030822753906 0 1420.3185
130.08648681640625 0 462.93555
130.10020446777344 0 702.70856
130.10601806640625 0 9483.365
131.04949951171875 0 1054.8263
131.08187866210938 0 3961.1821
131.1182403564453 0 7704.729
132.12188720703125 0 564.93774
133.06117248535156 0 1652.8987
133.08616638183594 0 1462.479
136.07611083984375 0 2453.5366
137.10797119140625 0 407.62216
138.0917510986328 0 34017.375
139.050537109375 0 700.00183
139.08702087402344 0 3638.9072
139.09506225585938 0 2495.6697
139.1237030029297 0 606.36456
140.1073760986328 0 1417.6055
141.06625366210938 0 2680.802
141.10260009765625 0 6245.835
143.11830139160156 0 721.62213
146.06056213378906 0 511.70886
147.07676696777344 0 922.954
147.11305236816406 0 1475.3795
148.8858642578125 0 558.1879
148.89305114746094 0 556.81274
148.90045166015625 0 541.68726
148.90773010253906 0 765.2023
148.91519165039062 0 791.5454
148.9223175048828 0 1203.6688
148.92953491210938 0 1416.857
148.93685913085938 0 3064.137
148.94476318359375 0 4838.2524
148.96145629882812 0 3766.5757
148.96929931640625 0 2155.2766
148.97637939453125 0 1323.3351
148.98336791992188 0 1108.7013
148.99090576171875 0 672.60144
148.99856567382812 0 752.0176
149.00564575195312 0 524.2051
149.01303100585938 0 432.9466
149.03466796875 0 638.4678
149.06008911132812 0 1108.5842
149.1075439453125 0 1042.3931
151.0869598388672 0 6225.5825
152.09010314941406 0 540.31024
152.10745239257812 0 773.62335
153.10269165039062 0 764.1821
154.0978546142578 0 3148.985
155.0707244873047 0 755.48047
155.08187866210938 0 2591.0571
155.11827087402344 0 34923.117
156.10231018066406 0 25838.818
156.1216583251953 0 3378.2773
157.06124877929688 0 727.3585
157.09823608398438 0 1063.0643
157.10585021972656 0 1990.9036
157.13375854492188 0 1332.5264
159.07681274414062 0 18857.553
159.09217834472656 0 1464.5486
159.11317443847656 0 3260.7773
159.1238250732422 0 483.68875
160.0800323486328 0 1560.5154
165.054931640625 0 1092.6636
165.06625366210938 0 478.1982
165.10256958007812 0 1085.6583
166.08668518066406 0 116387.58 y 9
166.1341094970703 0 629.86255
167.0825958251953 0 916.40424
167.09002685546875 0 11596.809
167.1028594970703 0 512.2965
167.1182861328125 0 6948.734
168.11355590820312 0 5797.906
168.12124633789062 0 854.8353
169.0976104736328 0 3155.43
169.13389587402344 0 753.95905
171.07701110839844 0 922.30414
172.1085968017578 0 1001.3066
173.12890625 0 172376
173.4407501220703 0 541.93805
174.13226318359375 0 14503.135
177.08798217773438 0 1295.5012
177.10264587402344 0 24596.941
177.1116180419922 0 1286.4478
178.10623168945312 0 2378.7139
179.08226013183594 0 791.8751
181.06130981445312 0 7188.4575
181.09739685058594 0 1068.1846
182.0648193359375 0 559.48987
182.08180236816406 0 1122.3711
182.12925720214844 0 72844.49
183.11326599121094 0 231052.36 a Water loss 1
183.1325225830078 0 8364.882
184.0972900390625 0 2921.3435
184.10862731933594 0 4472.4395
184.11669921875 0 20843.066
184.13525390625 0 605.8321
184.14453125 0 1157.1941
185.11953735351562 0 1146.1526
185.12889099121094 0 9514.261
186.12672424316406 0 737.9431
186.16123962402344 0 574.4844
187.1080780029297 0 8527.165 d 1
187.14479064941406 0 1069.4122
188.07101440429688 0 1374.3313
188.1118927001953 0 1121.229
191.08218383789062 0 766.1501
193.09759521484375 0 968.45294
194.12918090820312 0 8959.089
195.1129608154297 0 1568.7253
195.1326446533203 0 966.55634
196.1083984375 0 1024.1915
197.09230041503906 0 3675.5366
197.1287078857422 0 2005.013
198.08767700195312 0 11703.537
198.1241455078125 0 4392.559
199.07183837890625 0 4721.5605
199.10809326171875 0 1418.8467
200.13973999023438 0 113040.53
201.123779296875 0 355805.4 a 1
201.1424102783203 0 12140.659
202.1271514892578 0 32318.764
202.14358520507812 0 632.2451
202.1557159423828 0 1533.276
203.12872314453125 0 2298.0098
205.09756469726562 0 3317.1836
208.09719848632812 0 1570.9889
208.10855102539062 0 1805.175
209.0560302734375 0 1500.0864
209.0924835205078 0 16089.605
210.0956573486328 0 1910.0143
210.12411499023438 0 31836.834
211.10812377929688 0 70839.6 b Water loss 1
211.1271514892578 0 3153.1633
212.10244750976562 0 730.0092
212.11146545410156 0 7417.8955
212.13973999023438 0 5380.028
213.113525390625 0 567.55914
213.12376403808594 0 1930.5677
213.1435089111328 0 726.6067
214.1189422607422 0 1475.0093
214.1562042236328 0 634.7794
215.1031036376953 0 1560.9382
215.13954162597656 0 1036.495
216.0981903076172 0 6128.5767
218.15023803710938 0 5830.5786
219.1349334716797 0 1008.4789
222.12380981445312 0 1390.7683
223.108154296875 0 72270.03 y 8
224.1113739013672 0 9263.947
225.09837341308594 0 912.0604
225.11288452148438 0 1070.37
225.12391662597656 0 4700.1426
225.4588623046875 0 484.95187
226.0826873779297 0 29106.238
226.11904907226562 0 57863.54
227.06671142578125 0 19061.787
227.08668518066406 0 2987.7014
227.10287475585938 0 2920.2878
227.122314453125 0 4736.5933
228.07005310058594 0 2127.427
228.13470458984375 0 41890.723
229.1187286376953 0 71471.555 b 1
229.1387481689453 0 3703.442
230.12216186523438 0 6414.3813
230.15040588378906 0 7667.9014
231.12387084960938 0 824.3808
231.152587890625 0 853.8383
234.12374877929688 0 1082.3458
236.10337829589844 0 780.3748
238.1188507080078 0 779.10376
238.1558380126953 0 658.0461
239.1504364013672 0 1499.467
240.13461303710938 0 2989.0215
242.150390625 0 25262.967
243.10928344726562 0 4688.1294
243.15374755859375 0 2374.1294
244.07870483398438 0 1208.0392
244.0932159423828 0 45959.05
244.12965393066406 0 52686.207
245.0774688720703 0 1180.0621
245.09654235839844 0 3874.8215
245.1329345703125 0 6515.5986
246.1349639892578 0 796.88257
248.10317993164062 0 1077.5934
248.13990783691406 0 785.06683
248.16082763671875 0 837.1476
249.12322998046875 0 589.5852
250.15431213378906 0 525.4444
251.1509552001953 0 1422.7245
252.13458251953125 0 3128.5022
253.11891174316406 0 2830.573
254.1133270263672 0 1307.1589
254.1868438720703 0 1363.2402
258.1086120605469 0 582.733
258.1451110839844 0 3414.3777
261.1199645996094 0 3361.4314
261.13427734375 0 833.6012
262.1191101074219 0 952.7277
262.1402587890625 0 942.4908
264.134765625 0 1018.6868
265.1297607421875 0 1644.4141
266.1138916015625 0 2691.965
266.1500549316406 0 1880.3651
267.1453552246094 0 1719.8643
268.1291198730469 0 1164.2916
268.1492614746094 0 597.74
269.1612243652344 0 1864.0154
270.1452331542969 0 9959.447 a Water loss 2
271.1488342285156 0 669.1484
274.1189880371094 0 974.4519
276.1346435546875 0 952.69244
278.1507568359375 0 2085.7234
279.1317443847656 0 1308.8885
279.14593505859375 0 4412.417 b 4
280.12969970703125 0 28324.945
281.1326904296875 0 3240.712
281.16192626953125 0 670.2967
282.14471435546875 0 1650.0452
282.181884765625 0 4430.737
283.1405029296875 0 14823.641
284.1246643066406 0 569.0906
284.14373779296875 0 2171.2507 y 5
284.16094970703125 0 786.5635
285.1927490234375 0 611.2468
287.1719055175781 0 3086.3174
288.155029296875 0 766.662 a 2
290.1144104003906 0 1124.2887
291.14642333984375 0 1162.0957
292.12939453125 0 2561.4875
292.1471252441406 0 698.6719
294.1448974609375 0 2480.3267
297.15631103515625 0 25297.014
297.8763122558594 0 511.27597
298.1403503417969 0 80338.46 b Water loss 2
298.1578063964844 0 4194.861
299.1433410644531 0 11050.885
299.1728210449219 0 1029.1665
300.1576232910156 0 749.7852
301.15130615234375 0 2324.872
301.1876220703125 0 992.0718
302.1545715332031 0 745.3556
305.1825256347656 0 3106.168
306.1453552246094 0 5608.648 y Water loss 7
309.15625 0 8510.902
310.14031982421875 0 3659.8804
310.15936279296875 0 954.4606
310.1768798828125 0 2025.5062
311.13629150390625 0 1173.1681
311.1719665527344 0 3724.021
312.1568298339844 0 954.27954
315.1668701171875 0 10913.115
316.1299743652344 0 1216.2004
316.1508483886719 0 25472.215 b 2
316.1692810058594 0 1960.3131
317.1539001464844 0 3739.7163
317.182861328125 0 2055.518
317.21856689453125 0 732.35297
319.1405029296875 0 4918.654
320.1437683105469 0 1166.7084
322.1401062011719 0 1452.1179
324.15606689453125 0 27128.094 y 7
325.1590576171875 0 4752.408
327.1668395996094 0 9810.384
328.1507873535156 0 1381.4893 b Ammonia loss 5
328.1707458496094 0 1851.7249
329.18255615234375 0 29353.615
330.1656799316406 0 1047.2198
330.1858825683594 0 5521.9927
336.167236328125 0 6940.7266
337.1512145996094 0 27003.58
337.1692810058594 0 1384.7825
338.1341857910156 0 915.7469
338.1544494628906 0 4741.2666
339.1667785644531 0 14741.341 y Water loss 4
339.6650695800781 0 4522.0464 y Ammonia loss 4
340.152099609375 0 4717.1787
341.1822814941406 0 927.05676
343.6765441894531 0 569.4147
345.1773376464844 0 18916.1
346.1800231933594 0 3235.112
348.172119140625 0 17450.36 y 4
348.67333984375 0 5709.437
349.1502685546875 0 1115.0134
354.1775817871094 0 39840.133
355.1614685058594 0 18598.623
355.181884765625 0 5361.9263
356.16375732421875 0 2369.7166
356.18951416015625 0 926.64246
357.15435791015625 0 936.69824
357.17718505859375 0 12973.993
358.1789245605469 0 2038.7034
360.1888427734375 0 606.29236
362.0348815917969 0 588.00464
364.234619140625 0 2193.341
365.21875 0 36996.824
366.1778869628906 0 1066.7856
366.2218933105469 0 6363.8203
367.1612854003906 0 2479.675
371.20611572265625 0 690.15814
372.1881103515625 0 27807.846
373.1686096191406 0 1164.2712
373.19171142578125 0 4420.7373
374.2001037597656 0 1104.2605
375.176025390625 0 848.34125
376.161865234375 0 2079.847
377.21844482421875 0 705.3182
381.17681884765625 0 778.7935
382.17388916015625 0 754.346
382.2091369628906 0 853.7245
382.2456359863281 0 2734.5308
382.7005920410156 0 776.7775
383.2294921875 0 41853.35
384.18841552734375 0 4426.797
384.232421875 0 8483.985
385.1722717285156 0 2028.3041
385.2341613769531 0 927.6988
387.19818115234375 0 1180.9711
391.7064514160156 0 897.7752 b Water loss 6
392.2058410644531 0 654.09546
392.2301330566406 0 4372.089
393.18695068359375 0 4556.5957
393.21392822265625 0 30391.426
394.1727294921875 0 17099.92
394.21722412109375 0 6053.4126
395.17572021484375 0 2844.1904
395.2191467285156 0 1315.0886
395.70721435546875 0 1053.0593 y Water loss 3
396.2008972167969 0 5328.371 y Ammonia loss 3
396.702880859375 0 2766.8179
399.2036437988281 0 722.3506
401.1921081542969 0 1017.34247
401.2402648925781 0 3746.2031
402.19927978515625 0 8317.577
403.20306396484375 0 1194.7561
404.7142639160156 0 20403.105 y 3
405.2156677246094 0 9656.534
405.7163391113281 0 1892.5546
408.2255859375 0 1106.9222
409.17242431640625 0 841.928
410.1770935058594 0 728.9506
410.20465087890625 0 2096.2104
410.2401428222656 0 10649.77
410.72003173828125 0 7423.5835
411.1984558105469 0 4712.2676
411.2242126464844 0 40455.4 b Water loss 3
411.7223815917969 0 1109.2667
412.1828918457031 0 4401.131
412.22735595703125 0 8432.012
413.1869201660156 0 993.6367
413.22930908203125 0 1384.5742
415.22357177734375 0 855.5475
416.2312927246094 0 710.34106
419.2037048339844 0 3455.334
419.2306213378906 0 1224.1655
420.1883544921875 0 10446.339
421.1890869140625 0 1743.3235
422.1958923339844 0 973.3447
423.7283020019531 0 1817.8202
424.1835632324219 0 923.3775
424.22235107421875 0 1115.9827
425.22149658203125 0 867.1882
426.1986083984375 0 2627.2153
427.1827087402344 0 3915.9102
427.2304382324219 0 1741.3751
427.2678527832031 0 1927.9933
427.74017333984375 0 1442.9364
428.215576171875 0 4864.464
428.2509460449219 0 10475.853
429.2094421386719 0 1361.2025
429.235595703125 0 2506.3123 b 3
430.2381286621094 0 894.3573
430.71075439453125 0 1167.8986
431.2127990722656 0 966.5172
432.7330322265625 0 4369.6064
433.2272033691406 0 6306.769
433.72808837890625 0 2263.4072
434.2403869628906 0 6592.448 y Water loss 6
435.2413330078125 0 936.10895
437.2146911621094 0 6871.1973
438.1993408203125 0 9220.746
439.1977844238281 0 2081.1729
439.2254943847656 0 3946.968 y Water loss 2
439.71826171875 0 11558.549 y Ammonia loss 2
440.2186584472656 0 3860.205
440.7214050292969 0 1524.341
441.7384338378906 0 7802.5596
442.2380676269531 0 4189.4097 b Water loss 7
442.7386779785156 0 2117.2056
444.2097473144531 0 25035.76
445.2127685546875 0 4925.678
445.24249267578125 0 1009.9931
445.2745666503906 0 691.6383
446.21630859375 0 702.17737
446.2612609863281 0 905.6513
447.7427978515625 0 892.05304
448.230224609375 0 22482.83 y 2
448.2666320800781 0 1168.1696
448.7314758300781 0 12645.34
449.23223876953125 0 2732.4666
450.20166015625 0 951.4043
450.2355651855469 0 2342.0354
450.7439270019531 0 1150.1855
451.2374572753906 0 653.2029 b 7
452.2510070800781 0 67426.37 y 6
452.7312316894531 0 1184.361
453.2539978027344 0 17434.398
454.2569580078125 0 1994.4764
455.2257080078125 0 9727.925
456.20965576171875 0 27840.168
457.2115478515625 0 6089.937
458.22265625 0 1860.8506
459.2262878417969 0 790.29376
461.2400817871094 0 3079.9443
461.7370910644531 0 4067.3054
462.2357177734375 0 3147.6333
467.22747802734375 0 848.1839
467.26220703125 0 893.1789
468.2120361328125 0 1237.0222
468.2464294433594 0 2508.4443
469.2065124511719 0 799.2965
470.2489929199219 0 6664.923
470.747314453125 0 4348.141 b Water loss 8
471.2690734863281 0 1668.8422
473.236083984375 0 15243.486
474.23883056640625 0 3844.5898
477.21112060546875 0 963.5415
478.1899719238281 0 712.3197
479.2544250488281 0 4686.153
479.75543212890625 0 2501.8645
480.7371826171875 0 652.1993
483.22216796875 0 713.9953
485.2364501953125 0 1520.9188
485.2726745605469 0 6227.012
486.23638916015625 0 986.9696
486.27520751953125 0 696.29254
488.2590026855469 0 725.74243
494.2335510253906 0 1370.1216
495.2208251953125 0 6146.6064
496.22479248046875 0 1572.744
497.7647705078125 0 5372.782 y 1
498.2655944824219 0 2496.805
498.7677307128906 0 1178.4368
502.75714111328125 0 2245.5598
503.25982666015625 0 1513.9254
505.242919921875 0 765.42487
510.3042297363281 0 2144.6643
512.246826171875 0 5566.7397
513.2315063476562 0 11536.232
514.233642578125 0 4113.585
520.2882080078125 0 877.08215
521.27197265625 0 4471.6226
521.769775390625 0 735.1773
522.263427734375 0 3327.6658
522.7634887695312 0 817.14813
523.2532958984375 0 1127.2344
524.2838745117188 0 1497.9998
526.7574462890625 0 722.61237
529.2994995117188 0 1030.5537
529.784912109375 0 933.19025
530.2576293945312 0 12779.061
531.2568359375 0 5734.209
532.2433471679688 0 2409.9175
534.302490234375 0 1846.9435
535.2694702148438 0 2234.9116
535.7654418945312 0 1523.6746
536.2640380859375 0 1501.4268
536.7642822265625 0 1048.8138
537.262451171875 0 952.6521
538.298828125 0 8981.558
539.2842407226562 0 11493.224 b Water loss 4
539.8138427734375 0 897.1506
540.2860717773438 0 3071.6199
541.2861328125 0 811.14246
542.2945556640625 0 1005.26056
543.2777709960938 0 977.6873
543.7831420898438 0 8028.1084
544.2791748046875 0 10236.312
544.7763061523438 0 7578.325
545.2771606445312 0 1873.521
548.2677001953125 0 884.475
549.2673950195312 0 13143.391 y Water loss 5
550.2640991210938 0 2289.7273
550.3030395507812 0 1880.5154
551.286376953125 0 859.6325
551.6914672851562 0 922.7464
552.1898193359375 0 2364.1067
552.7883911132812 0 166444.25
553.2890014648438 0 111852.664 Precursor Water loss
553.7897338867188 0 42834.625
554.2921142578125 0 14244.257
554.78759765625 0 1050.2664
555.2789916992188 0 2913.4854
556.308837890625 0 1606.2373
557.2932739257812 0 2910.6082 b 4
558.2948608398438 0 969.4855
561.793701171875 0 26369.678
562.29052734375 0 24192.668 Precursor
562.7891845703125 0 13276.446
562.9349975585938 0 774.18866
563.2923583984375 0 3931.1917
567.277587890625 0 171422.12 y 5
568.280517578125 0 48647.535
569.2840576171875 0 8315.57
570.2906494140625 0 787.69403
572.3042602539062 0 22763.23
573.3072509765625 0 5992.5283
574.3086547851562 0 1019.32544
577.26025390625 0 716.491
586.3198852539062 0 8539.583
587.322021484375 0 2681.5425
593.3025512695312 0 1745.0071
608.3078002929688 0 984.9147
610.3238525390625 0 746.101
611.3150634765625 0 10120.392
612.3156127929688 0 2988.0413
614.306884765625 0 614.7698
620.30615234375 0 1918.9053
625.3331298828125 0 1962.3453
626.3253173828125 0 645.4456
628.3287353515625 0 972.3173
629.3247680664062 0 2963.5693
632.3057250976562 0 2168.1907
633.3019409179688 0 2372.8213
634.2833251953125 0 821.0697
635.31787109375 0 791.3757
636.2996826171875 0 2794.6772
637.3306884765625 0 11732.304
638.3265380859375 0 4450.8467
639.32763671875 0 753.38855
641.3006591796875 0 882.25916
641.8546752929688 0 5056.1367
642.2908325195312 0 1496.8007
642.3565673828125 0 3978.5415
643.2898559570312 0 964.66254
643.3419799804688 0 4741.4473
644.3455200195312 0 1022.9395
645.3575439453125 0 3505.841
649.3309936523438 0 4067.2366
650.3330688476562 0 1916.429
651.3180541992188 0 748.3833
653.325439453125 0 14859.831
654.316650390625 0 8375.39 b Water loss 5
654.3836669921875 0 867.89154
655.3388671875 0 9436.204
656.334716796875 0 4762.1606
657.3314208984375 0 1135.5026
659.3145751953125 0 5630.0015
660.3016967773438 0 8296.052
661.300537109375 0 2612.2495
662.3018188476562 0 1126.0383
668.3389282226562 0 1087.7416
670.352294921875 0 3062.0295
671.3511352539062 0 1738.0886
672.318603515625 0 1471.8757 b 5
673.3519897460938 0 40322.66
674.3543701171875 0 15433.037
675.3571166992188 0 2359.6575
677.3255615234375 0 54311.855 y Water loss 4
678.3115234375 0 116118.23 y Ammonia loss 4
679.3136596679688 0 38282.453
680.3158569335938 0 9012.983
681.3169555664062 0 1082.7391
686.3482055664062 0 1169.9679
695.3360595703125 0 174018.28 y 4
696.3390502929688 0 62872.742
697.3416748046875 0 13837.321
698.3428955078125 0 1684.6135
705.317626953125 0 810.93494
706.3185424804688 0 1138.1012
711.8501586914062 0 802.0533
712.3616943359375 0 7134.5737
713.3582763671875 0 2398.2666
730.3729858398438 0 17755.297
731.3760375976562 0 6938.848
732.3739624023438 0 1179.8196
746.3832397460938 0 959.19855
748.3822021484375 0 2991.044
749.3875122070312 0 1104.0677
755.8648071289062 0 704.10376
764.3959350585938 0 1668.0946
765.3910522460938 0 1130.3065
772.418212890625 0 1546.9528
773.38330078125 0 4624.406
774.3862915039062 0 2066.835
781.4210205078125 0 9262.496
782.414306640625 0 6570.0864 b Water loss 6
783.4140014648438 0 2081.4053
790.4080200195312 0 2683.836 y Water loss 3
791.4110107421875 0 1607.7692
799.4282836914062 0 1431.7604
800.4158325195312 0 2805.3984 b 6
801.4159545898438 0 864.60895
808.4196166992188 0 55023.344 y 3
809.4225463867188 0 22925.834
810.4261474609375 0 5360.9766
811.4266967773438 0 850.0426
818.4039916992188 0 936.59045
821.4166259765625 0 4278.1494
822.4178466796875 0 1979.8097
823.4205932617188 0 968.5413
829.4332885742188 0 1538.9053
833.4172973632812 0 1792.9904
838.439453125 0 791.60956
839.43017578125 0 2163.2603
840.4323120117188 0 845.7373
842.4006958007812 0 799.5052
846.4459228515625 0 1535.6993
847.436279296875 0 3569.773
848.4303588867188 0 1478.8219
849.4392700195312 0 1589.0924
855.4533081054688 0 1209.8265
859.4308471679688 0 2532.5215
860.422119140625 0 2672.3386
865.441162109375 0 8993.046
866.4395751953125 0 4164.6533
867.446044921875 0 1530.9768
877.343017578125 0 1037.8479
877.4409790039062 0 14766.155 y Water loss 2
878.4346923828125 0 11275.332 y Ammonia loss 2
879.4343872070312 0 3841.0068
880.4358520507812 0 1377.3192
883.4522094726562 0 25174.455 b Water loss 7
884.4535522460938 0 12588.61 b Ammonia loss 7
885.4572143554688 0 2846.4478
895.4513549804688 0 242835.28 y 2
896.4544677734375 0 121106.81
897.4569702148438 0 34751.797
897.5609130859375 0 1260.5836
898.4607543945312 0 4216.0576
900.4776611328125 0 2020.4874
901.4642333984375 0 8885.482 b 7
902.4669189453125 0 4020.0085
903.4654541015625 0 1676.2327
905.4362182617188 0 2245.3442
906.4424438476562 0 847.7385
922.4623413085938 0 5083.8774
923.4561157226562 0 4616.59
924.4564208984375 0 2338.3079
940.4728393554688 0 32558.23 b Water loss 8
941.4756469726562 0 16161.896 b Ammonia loss 8
942.4783325195312 0 4289.3223
957.4996337890625 0 2873.1099
958.4849853515625 0 13357.101 b 8
959.485595703125 0 6777.8125
960.4866943359375 0 1993.7926
976.5003662109375 0 1180.4551 y Water loss 1
994.519775390625 0 6523.7646 y 1
995.5237426757812 0 2640.6868
996.5272827148438 0 1677.8564
1004.5051879882812 0 1063.3611
1401.0146484375 0 695.33966
1658.7186279296875 0 657.5535
1720.025634765625 0 624.66595
1971.9923095703125 0 643.7003

Spectrum Details

|  |  |
| --- | --- |
| Matched peaks? Matched peaks The total absolute number of peaks matched. Additionally in brackets the total fraction of peaks matched and the total number of peaks is shown. | 60 (8.96% of 670) |
| FDR? FDR The false discovery rate estimated for this peptide. It is calculated by matching all theoretical fragments with a non-integer shift with the raw peaks for this spectrum. This is done with 40 different shifts. The resulting percentage is the average number of annotated peaks over the number of annotated peaks with the correct spectrum. | 0.36% |
| Satellite FDR? Satellite FDR See the FDR for details on its calculation. This satellite ion specific FDR only contains the satellite ions (d/w) for I/L/J positions. | - |
| PSM Score? PSM Score The PSM Score as given by Hecklib to this annotated spectrum. It is shown with three significant figures. | 700 |

#### Spectrum 5655? Spectrum 5655 The raw spectrum of this peptide as annotated by Hecklib. The fragments are coloured according to ion type (see legend). Any peaks with a star '\*' as text can be hovered over to see the full details, first the ion type second the mass shift type. By hovering over the amino acids in the peptide or ions in the legend the corresponding peaks are highlighted. By toggling the 'Unassigned' label you can turn the background (unassigned) peaks on or off in the plot. By updating the slider in the Ion legend you can update the spectrum to only show the top X% of the peaks with labels. The top X% means any peak that is within X% of the highest intensity. By dragging in the spectrum you can zoom in to a specific part of the spectrum and use 'Zoom Out' to get back to the original zoom level. The annotation of the spectrum is based on the given sequence in the peptides file and is done with different software so inconsistencies are likely. The peaks are annotated based on the given sequence, with 20 ppm tolerance.

Copy Data

##### Spectrum 5655 (TSV)

###### Preview

```
Loading example...
```

*Click on the button to copy the data to your clipboard.*

Mz MinMz MaxIntensity Max

WidthHeightPeptide font sizePeptide stroke widthSpectrum font sizeSpectrum stroke widthCompact peptide

Ion legend

wxyz

abcd

OtherUnassignedIon^Charge^_Position_Show for top:%

EVSJQDKTGF

04.59e+49.18e+41.38e+51.84e+5

Zoom Out

y^+1^_1_a^+1^_2_d^+1^_2_a^+1^_2_b^+1^_2_y^+1^_2_b^+1^_2_a^+1^_3_b^+2^_5_y^+2^_5_b^+1^_3_y^+1^_3_b^+1^_3_y^+1^_3_b^+2^_6_y^+2^_6_y^+2^_6_y^+2^_7_y^+2^_7_b^+1^_4_y^+1^_4_y^+2^_8_y^+2^_8_b^+2^_8_y^+2^_8_y^+1^_4_b^+2^_9_b^+2^_9_y^+2^_9_b^+1^_5_y^+1^_5_\*\*y^+1^_5_b^+1^_6_y^+1^_6_y^+1^_6_y^+1^_6_b^+1^_7_y^+1^_7_y^+1^_7_y^+1^_8_y^+1^_8_b^+1^_8_b^+1^_8_y^+1^_8_b^+1^_8_b^+1^_9_b^+1^_9_b^+1^_9_y^+1^_9_

0779155923383118

Fragment Matches Table

Show background peaks

| Position | Ion type | Intensity | mz Theoretical | mz Error (Th) | mz Error (ppm) | Charge | Series Number |
| --- | --- | --- | --- | --- | --- | --- | --- |
| - | - | 4.104E+04 | 120.1 | - | - | 0 | - |
| - | - | 396.5 | 120.2 | - | - | 0 | - |
| - | - | 2546 | 121.1 | - | - | 0 | - |
| - | - | 336.6 | 122.4 | - | - | 0 | - |
| - | - | 464.9 | 125.1 | - | - | 0 | - |
| - | - | 690.5 | 126.1 | - | - | 0 | - |
| - | - | 386 | 126.7 | - | - | 0 | - |
| - | - | 487.5 | 127.1 | - | - | 0 | - |
| - | - | 1633 | 127.1 | - | - | 0 | - |
| - | - | 420.3 | 128.1 | - | - | 0 | - |
| - | - | 2982 | 128.1 | - | - | 0 | - |
| - | - | 8227 | 129.1 | - | - | 0 | - |
| - | - | 7.047E+04 | 129.1 | - | - | 0 | - |
| - | - | 451.4 | 130 | - | - | 0 | - |
| - | - | 755.9 | 130.1 | - | - | 0 | - |
| - | - | 4431 | 130.1 | - | - | 0 | - |
| - | - | 1057 | 131 | - | - | 0 | - |
| - | - | 2931 | 131.1 | - | - | 0 | - |
| - | - | 3947 | 131.1 | - | - | 0 | - |
| - | - | 886.1 | 132.1 | - | - | 0 | - |
| - | - | 1049 | 133.1 | - | - | 0 | - |
| - | - | 1211 | 133.1 | - | - | 0 | - |
| - | - | 3890 | 136.1 | - | - | 0 | - |
| - | - | 406.5 | 137.1 | - | - | 0 | - |
| - | - | 2.102E+04 | 138.1 | - | - | 0 | - |
| - | - | 388.9 | 139.1 | - | - | 0 | - |
| - | - | 1604 | 139.1 | - | - | 0 | - |
| - | - | 1656 | 139.1 | - | - | 0 | - |
| - | - | 465.5 | 140.1 | - | - | 0 | - |
| - | - | 1320 | 141.1 | - | - | 0 | - |
| - | - | 3647 | 141.1 | - | - | 0 | - |
| - | - | 589.4 | 148.9 | - | - | 0 | - |
| - | - | 689.3 | 149.1 | - | - | 0 | - |
| - | - | 471.3 | 150.1 | - | - | 0 | - |
| - | - | 3089 | 151.1 | - | - | 0 | - |
| - | - | 697.4 | 153.1 | - | - | 0 | - |
| - | - | 1240 | 154.1 | - | - | 0 | - |
| - | - | 1300 | 155.1 | - | - | 0 | - |
| - | - | 2.566E+04 | 155.1 | - | - | 0 | - |
| - | - | 496.9 | 156.1 | - | - | 0 | - |
| - | - | 1783 | 156.1 | - | - | 0 | - |
| - | - | 1756 | 156.1 | - | - | 0 | - |
| - | - | 507.3 | 157.1 | - | - | 0 | - |
| - | - | 628.2 | 157.1 | - | - | 0 | - |
| - | - | 522.8 | 157.1 | - | - | 0 | - |
| - | - | 1.081E+04 | 159.1 | - | - | 0 | - |
| - | - | 502 | 159.1 | - | - | 0 | - |
| - | - | 1190 | 159.1 | - | - | 0 | - |
| - | - | 939.1 | 165.1 | - | - | 0 | - |
| - | - | 1254 | 165.1 | - | - | 0 | - |
| 10 | y | 5.294E+04 | 166.1 | 0.0001402 | 0.8442 | +1 | 1 |
| - | - | 991.3 | 167.1 | - | - | 0 | - |
| - | - | 5522 | 167.1 | - | - | 0 | - |
| - | - | 4852 | 167.1 | - | - | 0 | - |
| - | - | 3474 | 168.1 | - | - | 0 | - |
| - | - | 1718 | 169.1 | - | - | 0 | - |
| - | - | 722.1 | 169.1 | - | - | 0 | - |
| - | - | 581 | 171.1 | - | - | 0 | - |
| - | - | 7.195E+04 | 173.1 | - | - | 0 | - |
| - | - | 1540 | 173.5 | - | - | 0 | - |
| - | - | 781.7 | 174.1 | - | - | 0 | - |
| - | - | 5643 | 174.1 | - | - | 0 | - |
| - | - | 1226 | 175.1 | - | - | 0 | - |
| - | - | 442.3 | 177.1 | - | - | 0 | - |
| - | - | 1.061E+04 | 177.1 | - | - | 0 | - |
| - | - | 380.6 | 177.1 | - | - | 0 | - |
| - | - | 1058 | 178.1 | - | - | 0 | - |
| - | - | 3076 | 181.1 | - | - | 0 | - |
| - | - | 1578 | 182.1 | - | - | 0 | - |
| - | - | 8.129E+04 | 182.1 | - | - | 0 | - |
| 2 | a | 1.199E+05 | 183.1 | 0.0001261 | 0.6889 | +1 | 2 |
| - | - | 8123 | 183.1 | - | - | 0 | - |
| - | - | 587.3 | 184.1 | - | - | 0 | - |
| - | - | 2056 | 184.1 | - | - | 0 | - |
| - | - | 1.093E+04 | 184.1 | - | - | 0 | - |
| - | - | 607.8 | 184.1 | - | - | 0 | - |
| - | - | 3762 | 185.1 | - | - | 0 | - |
| - | - | 790.3 | 186.1 | - | - | 0 | - |
| 2 | d | 4974 | 187.1 | 0.0001914 | 1.023 | +1 | 2 |
| - | - | 1530 | 187.1 | - | - | 0 | - |
| - | - | 1246 | 188.1 | - | - | 0 | - |
| - | - | 4912 | 194.1 | - | - | 0 | - |
| - | - | 1283 | 195.1 | - | - | 0 | - |
| - | - | 553.8 | 196.1 | - | - | 0 | - |
| - | - | 757.1 | 197.1 | - | - | 0 | - |
| - | - | 1335 | 197.1 | - | - | 0 | - |
| - | - | 4930 | 198.1 | - | - | 0 | - |
| - | - | 1702 | 198.1 | - | - | 0 | - |
| - | - | 2659 | 199.1 | - | - | 0 | - |
| - | - | 1050 | 199.1 | - | - | 0 | - |
| - | - | 1.269E+05 | 200.1 | - | - | 0 | - |
| 2 | a | 4.622E+04 | 201.1 | 5.95E-05 | 0.2958 | +1 | 2 |
| - | - | 1.184E+04 | 201.1 | - | - | 0 | - |
| - | - | 3252 | 202.1 | - | - | 0 | - |
| - | - | 2099 | 205.1 | - | - | 0 | - |
| - | - | 830 | 208.1 | - | - | 0 | - |
| - | - | 830.8 | 208.1 | - | - | 0 | - |
| - | - | 982.6 | 209.1 | - | - | 0 | - |
| - | - | 8630 | 209.1 | - | - | 0 | - |
| - | - | 3.363E+04 | 210.1 | - | - | 0 | - |
| 2 | b | 2.253E+04 | 211.1 | 2.353E-05 | 0.1115 | +1 | 2 |
| - | - | 4122 | 211.1 | - | - | 0 | - |
| - | - | 560.6 | 211.1 | - | - | 0 | - |
| - | - | 927.8 | 211.1 | - | - | 0 | - |
| - | - | 1537 | 212.1 | - | - | 0 | - |
| - | - | 3196 | 212.1 | - | - | 0 | - |
| - | - | 851.3 | 213.1 | - | - | 0 | - |
| - | - | 625.7 | 213.1 | - | - | 0 | - |
| - | - | 853.7 | 215.1 | - | - | 0 | - |
| - | - | 2425 | 216.1 | - | - | 0 | - |
| - | - | 1995 | 218.1 | - | - | 0 | - |
| - | - | 1397 | 222.1 | - | - | 0 | - |
| 9 | y | 3.62E+04 | 223.1 | 8.457E-05 | 0.379 | +1 | 2 |
| - | - | 3506 | 224.1 | - | - | 0 | - |
| - | - | 3258 | 225.1 | - | - | 0 | - |
| - | - | 1.146E+04 | 226.1 | - | - | 0 | - |
| - | - | 2.798E+04 | 226.1 | - | - | 0 | - |
| - | - | 7928 | 227.1 | - | - | 0 | - |
| - | - | 1203 | 227.1 | - | - | 0 | - |
| - | - | 1807 | 227.1 | - | - | 0 | - |
| - | - | 2989 | 227.1 | - | - | 0 | - |
| - | - | 4.448E+04 | 228.1 | - | - | 0 | - |
| 2 | b | 5955 | 229.1 | 7.896E-05 | 0.3446 | +1 | 2 |
| - | - | 4720 | 229.1 | - | - | 0 | - |
| - | - | 3119 | 230.1 | - | - | 0 | - |
| - | - | 1196 | 234.1 | - | - | 0 | - |
| - | - | 695.6 | 238.1 | - | - | 0 | - |
| - | - | 1350 | 239.2 | - | - | 0 | - |
| - | - | 3000 | 240.1 | - | - | 0 | - |
| - | - | 1.082E+04 | 242.1 | - | - | 0 | - |
| - | - | 404.3 | 242.2 | - | - | 0 | - |
| - | - | 1502 | 243.1 | - | - | 0 | - |
| - | - | 810.7 | 243.2 | - | - | 0 | - |
| - | - | 863.9 | 244.1 | - | - | 0 | - |
| - | - | 2.153E+04 | 244.1 | - | - | 0 | - |
| - | - | 2.502E+04 | 244.1 | - | - | 0 | - |
| - | - | 1922 | 245.1 | - | - | 0 | - |
| - | - | 2125 | 245.1 | - | - | 0 | - |
| - | - | 761.8 | 249.1 | - | - | 0 | - |
| - | - | 587.9 | 250.2 | - | - | 0 | - |
| - | - | 961.7 | 251.2 | - | - | 0 | - |
| - | - | 1669 | 252.1 | - | - | 0 | - |
| - | - | 925.7 | 253.1 | - | - | 0 | - |
| - | - | 1861 | 258.1 | - | - | 0 | - |
| - | - | 1157 | 261.1 | - | - | 0 | - |
| - | - | 646.8 | 262.1 | - | - | 0 | - |
| - | - | 905.8 | 265.1 | - | - | 0 | - |
| - | - | 1377 | 266.1 | - | - | 0 | - |
| - | - | 1526 | 267.1 | - | - | 0 | - |
| - | - | 694.6 | 268.1 | - | - | 0 | - |
| - | - | 1127 | 269.2 | - | - | 0 | - |
| 3 | a | 4346 | 270.1 | 0.0003013 | 1.115 | +1 | 3 |
| - | - | 1831 | 279.1 | - | - | 0 | - |
| 5 | b | 4316 | 279.2 | 0.004699 | 16.83 | +2 | 5 |
| - | - | 7105 | 280.1 | - | - | 0 | - |
| - | - | 952.8 | 280.1 | - | - | 0 | - |
| - | - | 728.1 | 281.1 | - | - | 0 | - |
| - | - | 2307 | 282.2 | - | - | 0 | - |
| - | - | 6682 | 283.1 | - | - | 0 | - |
| 6 | y | 1237 | 284.1 | 0.001478 | 5.203 | +2 | 5 |
| - | - | 2315 | 287.2 | - | - | 0 | - |
| - | - | 809.5 | 290.1 | - | - | 0 | - |
| - | - | 1053 | 291.1 | - | - | 0 | - |
| - | - | 1437 | 292.1 | - | - | 0 | - |
| - | - | 714.1 | 294.1 | - | - | 0 | - |
| - | - | 2.537E+04 | 297.2 | - | - | 0 | - |
| 3 | b | 2.337E+04 | 298.1 | 6.823E-05 | 0.2289 | +1 | 3 |
| - | - | 2992 | 298.2 | - | - | 0 | - |
| - | - | 3755 | 299.1 | - | - | 0 | - |
| - | - | 687.8 | 300.2 | - | - | 0 | - |
| - | - | 996.5 | 305.2 | - | - | 0 | - |
| 8 | y | 2815 | 306.1 | 0.0001792 | 0.5855 | +1 | 3 |
| - | - | 3937 | 309.2 | - | - | 0 | - |
| - | - | 1611 | 310.1 | - | - | 0 | - |
| - | - | 751 | 310.2 | - | - | 0 | - |
| - | - | 1661 | 311.2 | - | - | 0 | - |
| - | - | 1.083E+04 | 315.2 | - | - | 0 | - |
| 3 | b | 1749 | 316.2 | 0.0005621 | 1.778 | +1 | 3 |
| - | - | 1790 | 316.2 | - | - | 0 | - |
| - | - | 1319 | 317.2 | - | - | 0 | - |
| - | - | 2539 | 319.1 | - | - | 0 | - |
| 8 | y | 1.318E+04 | 324.2 | 5.929E-05 | 0.1829 | +1 | 3 |
| - | - | 1803 | 325.2 | - | - | 0 | - |
| - | - | 4052 | 327.2 | - | - | 0 | - |
| 6 | b | 897.4 | 328.2 | 7.875E-05 | 0.24 | +2 | 6 |
| - | - | 771.1 | 328.2 | - | - | 0 | - |
| - | - | 649.1 | 329.1 | - | - | 0 | - |
| - | - | 1.229E+04 | 329.2 | - | - | 0 | - |
| - | - | 1558 | 330.2 | - | - | 0 | - |
| - | - | 2653 | 336.2 | - | - | 0 | - |
| - | - | 1.271E+04 | 337.2 | - | - | 0 | - |
| - | - | 728.3 | 337.2 | - | - | 0 | - |
| - | - | 2425 | 338.2 | - | - | 0 | - |
| 5 | y | 4830 | 339.2 | 5.503E-05 | 0.1622 | +2 | 6 |
| - | - | 1030 | 339.7 | - | - | 0 | - |
| - | - | 1674 | 340.2 | - | - | 0 | - |
| - | - | 9956 | 345.2 | - | - | 0 | - |
| - | - | 1934 | 346.2 | - | - | 0 | - |
| 5 | y | 7203 | 348.2 | 0.0005581 | 1.603 | +2 | 6 |
| - | - | 2633 | 348.7 | - | - | 0 | - |
| - | - | 1.737E+04 | 354.2 | - | - | 0 | - |
| - | - | 6939 | 355.2 | - | - | 0 | - |
| - | - | 1788 | 355.2 | - | - | 0 | - |
| - | - | 978.8 | 356.2 | - | - | 0 | - |
| - | - | 6362 | 357.2 | - | - | 0 | - |
| - | - | 2782 | 364.2 | - | - | 0 | - |
| - | - | 1.397E+04 | 365.2 | - | - | 0 | - |
| - | - | 782.1 | 366.2 | - | - | 0 | - |
| - | - | 2919 | 366.2 | - | - | 0 | - |
| - | - | 1426 | 367.2 | - | - | 0 | - |
| - | - | 987.9 | 371.2 | - | - | 0 | - |
| - | - | 1.43E+04 | 372.2 | - | - | 0 | - |
| - | - | 2411 | 373.2 | - | - | 0 | - |
| - | - | 741.3 | 376.2 | - | - | 0 | - |
| - | - | 965.9 | 380.2 | - | - | 0 | - |
| - | - | 602.3 | 382.2 | - | - | 0 | - |
| - | - | 3460 | 382.2 | - | - | 0 | - |
| - | - | 1.682E+04 | 383.2 | - | - | 0 | - |
| - | - | 1486 | 384.2 | - | - | 0 | - |
| - | - | 2418 | 384.2 | - | - | 0 | - |
| - | - | 6593 | 392.2 | - | - | 0 | - |
| - | - | 1619 | 393.2 | - | - | 0 | - |
| - | - | 1.364E+04 | 393.2 | - | - | 0 | - |
| - | - | 6968 | 394.2 | - | - | 0 | - |
| - | - | 2548 | 394.2 | - | - | 0 | - |
| - | - | 1258 | 395.2 | - | - | 0 | - |
| 4 | y | 1825 | 396.2 | 0.0005306 | 1.339 | +2 | 7 |
| - | - | 1344 | 396.7 | - | - | 0 | - |
| - | - | 967.4 | 400.3 | - | - | 0 | - |
| - | - | 3413 | 402.2 | - | - | 0 | - |
| 4 | y | 8715 | 404.7 | 0.0001707 | 0.4217 | +2 | 7 |
| - | - | 5706 | 405.2 | - | - | 0 | - |
| - | - | 1010 | 405.7 | - | - | 0 | - |
| - | - | 1130 | 410.2 | - | - | 0 | - |
| - | - | 1.078E+04 | 410.2 | - | - | 0 | - |
| - | - | 7865 | 410.7 | - | - | 0 | - |
| - | - | 1315 | 411.2 | - | - | 0 | - |
| 4 | b | 1.834E+04 | 411.2 | 0.0007277 | 1.77 | +1 | 4 |
| - | - | 1100 | 411.7 | - | - | 0 | - |
| - | - | 737.8 | 412.2 | - | - | 0 | - |
| - | - | 2650 | 412.2 | - | - | 0 | - |
| - | - | 2948 | 412.2 | - | - | 0 | - |
| - | - | 620.2 | 418.2 | - | - | 0 | - |
| - | - | 1213 | 419.2 | - | - | 0 | - |
| - | - | 594 | 419.7 | - | - | 0 | - |
| - | - | 2702 | 420.2 | - | - | 0 | - |
| - | - | 1656 | 423.7 | - | - | 0 | - |
| - | - | 1658 | 424.2 | - | - | 0 | - |
| - | - | 1075 | 426.2 | - | - | 0 | - |
| - | - | 1446 | 427.2 | - | - | 0 | - |
| - | - | 1066 | 427.2 | - | - | 0 | - |
| - | - | 2170 | 427.3 | - | - | 0 | - |
| - | - | 1772 | 427.7 | - | - | 0 | - |
| - | - | 753.9 | 428.2 | - | - | 0 | - |
| - | - | 1515 | 428.2 | - | - | 0 | - |
| - | - | 5550 | 428.2 | - | - | 0 | - |
| - | - | 721 | 429.3 | - | - | 0 | - |
| - | - | 871.1 | 430.7 | - | - | 0 | - |
| - | - | 6253 | 432.7 | - | - | 0 | - |
| - | - | 2471 | 433.2 | - | - | 0 | - |
| - | - | 786.8 | 433.7 | - | - | 0 | - |
| 7 | y | 2957 | 434.2 | 1.897E-05 | 0.04369 | +1 | 4 |
| - | - | 1618 | 435.2 | - | - | 0 | - |
| - | - | 1239 | 435.7 | - | - | 0 | - |
| - | - | 2836 | 437.2 | - | - | 0 | - |
| - | - | 4346 | 438.2 | - | - | 0 | - |
| - | - | 1201 | 439.2 | - | - | 0 | - |
| 3 | y | 1110 | 439.2 | 0.002983 | 6.791 | +2 | 8 |
| 3 | y | 4704 | 439.7 | 0.00185 | 4.208 | +2 | 8 |
| - | - | 1759 | 440.2 | - | - | 0 | - |
| - | - | 935.9 | 440.7 | - | - | 0 | - |
| - | - | 7027 | 441.7 | - | - | 0 | - |
| 8 | b | 4201 | 442.2 | 0.008504 | 19.23 | +2 | 8 |
| - | - | 874.6 | 442.7 | - | - | 0 | - |
| - | - | 1.32E+04 | 444.2 | - | - | 0 | - |
| - | - | 2246 | 445.2 | - | - | 0 | - |
| 3 | y | 8692 | 448.2 | 5.046E-05 | 0.1126 | +2 | 8 |
| - | - | 1173 | 448.3 | - | - | 0 | - |
| - | - | 4549 | 448.7 | - | - | 0 | - |
| - | - | 1028 | 450.2 | - | - | 0 | - |
| 7 | y | 3.126E+04 | 452.3 | 0.0002382 | 0.5267 | +1 | 4 |
| - | - | 5960 | 453.3 | - | - | 0 | - |
| - | - | 950.9 | 454.3 | - | - | 0 | - |
| - | - | 4332 | 455.2 | - | - | 0 | - |
| - | - | 8507 | 456.2 | - | - | 0 | - |
| - | - | 723.3 | 457.2 | - | - | 0 | - |
| - | - | 2846 | 457.2 | - | - | 0 | - |
| - | - | 2387 | 461.2 | - | - | 0 | - |
| - | - | 3516 | 461.7 | - | - | 0 | - |
| - | - | 1907 | 462.2 | - | - | 0 | - |
| - | - | 770.4 | 468.2 | - | - | 0 | - |
| - | - | 5310 | 470.2 | - | - | 0 | - |
| 9 | b | 2859 | 470.7 | 0.007965 | 16.92 | +2 | 9 |
| - | - | 1829 | 471.2 | - | - | 0 | - |
| - | - | 5580 | 473.2 | - | - | 0 | - |
| - | - | 1563 | 474.2 | - | - | 0 | - |
| - | - | 6067 | 479.3 | - | - | 0 | - |
| 9 | b | 2484 | 479.7 | 0.009 | 18.76 | +2 | 9 |
| - | - | 1589 | 480.3 | - | - | 0 | - |
| - | - | 1109 | 485.2 | - | - | 0 | - |
| - | - | 3212 | 485.3 | - | - | 0 | - |
| - | - | 690.4 | 486.2 | - | - | 0 | - |
| - | - | 1692 | 495.2 | - | - | 0 | - |
| 2 | y | 2064 | 497.8 | 0.001136 | 2.283 | +2 | 9 |
| - | - | 1364 | 498.3 | - | - | 0 | - |
| - | - | 1915 | 510.3 | - | - | 0 | - |
| - | - | 2221 | 512.2 | - | - | 0 | - |
| - | - | 4487 | 513.2 | - | - | 0 | - |
| - | - | 1659 | 514.2 | - | - | 0 | - |
| - | - | 983.6 | 521.3 | - | - | 0 | - |
| - | - | 1395 | 522.3 | - | - | 0 | - |
| - | - | 1050 | 522.7 | - | - | 0 | - |
| - | - | 873.6 | 522.8 | - | - | 0 | - |
| - | - | 1241 | 524.3 | - | - | 0 | - |
| - | - | 1093 | 529.8 | - | - | 0 | - |
| - | - | 4322 | 530.3 | - | - | 0 | - |
| - | - | 742.1 | 530.8 | - | - | 0 | - |
| - | - | 1641 | 531.3 | - | - | 0 | - |
| - | - | 1115 | 532.2 | - | - | 0 | - |
| - | - | 3895 | 534.3 | - | - | 0 | - |
| - | - | 1093 | 534.8 | - | - | 0 | - |
| - | - | 1138 | 535.3 | - | - | 0 | - |
| - | - | 941.1 | 535.3 | - | - | 0 | - |
| - | - | 1313 | 535.8 | - | - | 0 | - |
| - | - | 1.022E+04 | 538.3 | - | - | 0 | - |
| 5 | b | 5171 | 539.3 | 0.003317 | 6.151 | +1 | 5 |
| - | - | 802.1 | 542.7 | - | - | 0 | - |
| - | - | 2680 | 543.3 | - | - | 0 | - |
| - | - | 1.194E+04 | 543.8 | - | - | 0 | - |
| - | - | 9841 | 544.3 | - | - | 0 | - |
| - | - | 3893 | 544.8 | - | - | 0 | - |
| - | - | 1726 | 545.3 | - | - | 0 | - |
| - | - | 650.5 | 545.8 | - | - | 0 | - |
| 6 | y | 5594 | 549.3 | 4.605E-05 | 0.08384 | +1 | 5 |
| - | - | 1182 | 550.3 | - | - | 0 | - |
| - | - | 5973 | 551.7 | - | - | 0 | - |
| - | - | 3168 | 552.2 | - | - | 0 | - |
| - | - | 1.818E+05 | 552.8 | - | - | 0 | - |
| 0 | Precursor | 1.104E+05 | 553.3 | 0.008423 | 15.22 | +2 | -1 |
| - | - | 4.116E+04 | 553.8 | - | - | 0 | - |
| - | - | 7517 | 554.3 | - | - | 0 | - |
| - | - | 1384 | 555.3 | - | - | 0 | - |
| - | - | 1809 | 556.3 | - | - | 0 | - |
| - | - | 653.7 | 558.3 | - | - | 0 | - |
| - | - | 2.671E+04 | 561.8 | - | - | 0 | - |
| 0 | Precursor | 1.7E+04 | 562.3 | 0.008451 | 15.03 | +2 | -1 |
| - | - | 687.4 | 562.7 | - | - | 0 | - |
| - | - | 5932 | 562.8 | - | - | 0 | - |
| - | - | 1098 | 563.3 | - | - | 0 | - |
| 6 | y | 7.441E+04 | 567.3 | 0.0005699 | 1.005 | +1 | 5 |
| - | - | 1.955E+04 | 568.3 | - | - | 0 | - |
| - | - | 3662 | 569.3 | - | - | 0 | - |
| - | - | 1.054E+04 | 572.3 | - | - | 0 | - |
| - | - | 2234 | 573.3 | - | - | 0 | - |
| - | - | 790.9 | 574.3 | - | - | 0 | - |
| - | - | 873.5 | 583.3 | - | - | 0 | - |
| - | - | 4819 | 586.3 | - | - | 0 | - |
| - | - | 1025 | 587.3 | - | - | 0 | - |
| - | - | 788.4 | 597.9 | - | - | 0 | - |
| - | - | 639.9 | 599.6 | - | - | 0 | - |
| - | - | 609.7 | 601.8 | - | - | 0 | - |
| - | - | 640.6 | 606.3 | - | - | 0 | - |
| - | - | 959.5 | 608.3 | - | - | 0 | - |
| - | - | 5617 | 611.3 | - | - | 0 | - |
| - | - | 1724 | 612.3 | - | - | 0 | - |
| - | - | 977 | 625.3 | - | - | 0 | - |
| - | - | 623.4 | 626.3 | - | - | 0 | - |
| - | - | 1242 | 629.3 | - | - | 0 | - |
| - | - | 647.4 | 632.6 | - | - | 0 | - |
| - | - | 912.2 | 632.9 | - | - | 0 | - |
| - | - | 2167 | 636.3 | - | - | 0 | - |
| - | - | 5201 | 637.3 | - | - | 0 | - |
| - | - | 1724 | 638.3 | - | - | 0 | - |
| - | - | 2043 | 641.9 | - | - | 0 | - |
| - | - | 899.5 | 642.3 | - | - | 0 | - |
| - | - | 1981 | 642.4 | - | - | 0 | - |
| - | - | 1685 | 643.3 | - | - | 0 | - |
| - | - | 1700 | 645.4 | - | - | 0 | - |
| - | - | 6678 | 649.3 | - | - | 0 | - |
| - | - | 2437 | 650.3 | - | - | 0 | - |
| - | - | 1.443E+04 | 653.3 | - | - | 0 | - |
| 6 | b | 5587 | 654.3 | 0.01068 | 16.32 | +1 | 6 |
| - | - | 5192 | 655.3 | - | - | 0 | - |
| - | - | 2244 | 656.3 | - | - | 0 | - |
| - | - | 3083 | 659.3 | - | - | 0 | - |
| - | - | 3840 | 660.3 | - | - | 0 | - |
| - | - | 1728 | 661.3 | - | - | 0 | - |
| - | - | 4604 | 670.4 | - | - | 0 | - |
| - | - | 1826 | 671.3 | - | - | 0 | - |
| - | - | 1.971E+04 | 673.4 | - | - | 0 | - |
| - | - | 7843 | 674.4 | - | - | 0 | - |
| - | - | 1022 | 675.4 | - | - | 0 | - |
| 5 | y | 1.928E+04 | 677.3 | 0.001036 | 1.53 | +1 | 6 |
| 5 | y | 4.55E+04 | 678.3 | 0.0005438 | 0.8017 | +1 | 6 |
| - | - | 1.616E+04 | 679.3 | - | - | 0 | - |
| - | - | 2247 | 680.3 | - | - | 0 | - |
| - | - | 940.6 | 686.3 | - | - | 0 | - |
| 5 | y | 6.21E+04 | 695.3 | 0.0009199 | 1.323 | +1 | 6 |
| - | - | 2.59E+04 | 696.3 | - | - | 0 | - |
| - | - | 4744 | 697.3 | - | - | 0 | - |
| - | - | 1497 | 711.9 | - | - | 0 | - |
| - | - | 3214 | 712.4 | - | - | 0 | - |
| - | - | 1215 | 713.4 | - | - | 0 | - |
| - | - | 680 | 713.5 | - | - | 0 | - |
| - | - | 8230 | 730.4 | - | - | 0 | - |
| - | - | 2633 | 731.4 | - | - | 0 | - |
| - | - | 1662 | 748.4 | - | - | 0 | - |
| - | - | 1024 | 755.4 | - | - | 0 | - |
| - | - | 674.7 | 763.4 | - | - | 0 | - |
| - | - | 891.2 | 764.4 | - | - | 0 | - |
| - | - | 684.9 | 765.4 | - | - | 0 | - |
| - | - | 964.5 | 772.4 | - | - | 0 | - |
| - | - | 1807 | 773.4 | - | - | 0 | - |
| - | - | 898.4 | 774.4 | - | - | 0 | - |
| - | - | 1.024E+04 | 781.4 | - | - | 0 | - |
| 7 | b | 5040 | 782.4 | 0.01392 | 17.79 | +1 | 7 |
| - | - | 1067 | 783.4 | - | - | 0 | - |
| 4 | y | 1171 | 790.4 | 0.005572 | 7.049 | +1 | 7 |
| - | - | 1455 | 799.4 | - | - | 0 | - |
| 4 | y | 1.979E+04 | 808.4 | 0.001366 | 1.689 | +1 | 7 |
| - | - | 8573 | 809.4 | - | - | 0 | - |
| - | - | 2836 | 810.4 | - | - | 0 | - |
| - | - | 2545 | 821.4 | - | - | 0 | - |
| - | - | 1216 | 822.4 | - | - | 0 | - |
| - | - | 1034 | 839.4 | - | - | 0 | - |
| - | - | 1592 | 846.4 | - | - | 0 | - |
| - | - | 1716 | 847.4 | - | - | 0 | - |
| - | - | 784.9 | 848.4 | - | - | 0 | - |
| - | - | 1053 | 849.4 | - | - | 0 | - |
| - | - | 786.9 | 855.5 | - | - | 0 | - |
| - | - | 1544 | 859.4 | - | - | 0 | - |
| - | - | 908.8 | 860.4 | - | - | 0 | - |
| - | - | 663.4 | 864.5 | - | - | 0 | - |
| - | - | 4030 | 865.4 | - | - | 0 | - |
| - | - | 1471 | 866.4 | - | - | 0 | - |
| 3 | y | 6013 | 877.4 | 0.002627 | 2.994 | +1 | 8 |
| 3 | y | 4767 | 878.4 | 0.008048 | 9.161 | +1 | 8 |
| - | - | 1627 | 879.4 | - | - | 0 | - |
| 8 | b | 1.114E+04 | 883.5 | 0.0006182 | 0.6998 | +1 | 8 |
| 8 | b | 4808 | 884.4 | 0.01634 | 18.48 | +1 | 8 |
| 3 | y | 1.131E+05 | 895.5 | 0.002266 | 2.531 | +1 | 8 |
| - | - | 5.309E+04 | 896.5 | - | - | 0 | - |
| - | - | 1.379E+04 | 897.5 | - | - | 0 | - |
| - | - | 772.3 | 898.5 | - | - | 0 | - |
| - | - | 2919 | 900.5 | - | - | 0 | - |
| 8 | b | 1761 | 901.5 | 0.01061 | 11.77 | +1 | 8 |
| - | - | 718.6 | 902.5 | - | - | 0 | - |
| - | - | 2722 | 922.5 | - | - | 0 | - |
| - | - | 2897 | 923.5 | - | - | 0 | - |
| - | - | 937.1 | 924.5 | - | - | 0 | - |
| - | - | 1832 | 939.5 | - | - | 0 | - |
| 9 | b | 1.223E+04 | 940.5 | 0.0009028 | 0.9599 | +1 | 9 |
| 9 | b | 7245 | 941.5 | 0.01716 | 18.22 | +1 | 9 |
| - | - | 1914 | 942.5 | - | - | 0 | - |
| - | - | 3474 | 957.5 | - | - | 0 | - |
| 9 | b | 2177 | 958.5 | 0.005683 | 5.93 | +1 | 9 |
| - | - | 987.1 | 959.5 | - | - | 0 | - |
| 2 | y | 6405 | 994.5 | 0.002199 | 2.211 | +1 | 9 |
| - | - | 2569 | 995.5 | - | - | 0 | - |
| - | - | 647 | 1712 | - | - | 0 | - |
| - | - | 828.6 | 2886 | - | - | 0 | - |
| - | - | 830.6 | 3087 | - | - | 0 | - |

m/z Charge Intensity FragmentType MassShift Position
120.08097839355469 0 41038.445
120.2103500366211 0 396.5366
121.08429718017578 0 2545.8594
122.44593811035156 0 336.64847
125.10733795166016 0 464.9029
126.09142303466797 0 690.4737
126.6981430053711 0 386.02765
127.08714294433594 0 487.4773
127.12320709228516 0 1632.7842
128.07066345214844 0 420.29025
128.107177734375 0 2981.941
129.06602478027344 0 8227.337
129.10243225097656 0 70467.89
130.04995727539062 0 451.3657
130.10009765625 0 755.8686
130.10572814941406 0 4431.445
131.04920959472656 0 1056.7667
131.0816650390625 0 2931.277
131.11807250976562 0 3946.649
132.10227966308594 0 886.0639
133.0610809326172 0 1049.0758
133.0861053466797 0 1210.8982
136.07582092285156 0 3890.4546
137.07919311523438 0 406.45377
138.0915069580078 0 21021.377
139.05010986328125 0 388.90808
139.0867462158203 0 1604.0583
139.09487915039062 0 1655.6877
140.1072998046875 0 465.4892
141.06600952148438 0 1320.294
141.10238647460938 0 3646.8752
148.94747924804688 0 589.4145
149.0599822998047 0 689.31494
150.0911865234375 0 471.32346
151.08689880371094 0 3089.4138
153.10208129882812 0 697.3817
154.09774780273438 0 1239.6897
155.08172607421875 0 1300.1377
155.11802673339844 0 25658.852
156.07643127441406 0 496.8988
156.10206604003906 0 1782.9536
156.12124633789062 0 1756.183
157.06044006347656 0 507.3002
157.0972137451172 0 628.15985
157.13352966308594 0 522.84753
159.0765380859375 0 10808.762
159.09144592285156 0 501.9635
159.11282348632812 0 1190.2192
165.05474853515625 0 939.0559
165.10256958007812 0 1254.0507
166.08639526367188 0 52943.55 y 9
167.08262634277344 0 991.3103
167.08984375 0 5521.7534
167.11793518066406 0 4852.2515
168.1133575439453 0 3473.6777
169.09722900390625 0 1717.9204
169.13368225097656 0 722.1104
171.11305236816406 0 581.0282
173.12857055664062 0 71953.25
173.4518280029297 0 1539.8889
174.0552978515625 0 781.65533
174.1319580078125 0 5643.146
175.08645629882812 0 1225.7274
177.08798217773438 0 442.33978
177.1023406982422 0 10614.82
177.11093139648438 0 380.5701
178.10577392578125 0 1057.578
181.06094360351562 0 3076.15
182.08155822753906 0 1577.8917
182.12892150878906 0 81292.13
183.11293029785156 0 119906.61 a Water loss 1
183.13233947753906 0 8122.538
184.09767150878906 0 587.2814
184.1081085205078 0 2055.766
184.11631774902344 0 10932.106
184.1442108154297 0 607.771
185.12860107421875 0 3762.356
186.12376403808594 0 790.3274
187.10791015625 0 4974.438 d 1
187.14447021484375 0 1530.0243
188.07054138183594 0 1246.3453
194.12887573242188 0 4911.7285
195.11265563964844 0 1283.3336
196.10787963867188 0 553.83105
197.09152221679688 0 757.1113
197.1282501220703 0 1334.7208
198.0873260498047 0 4929.6274
198.1236572265625 0 1702.0803
199.07131958007812 0 2658.6716
199.10772705078125 0 1049.8236
200.1394500732422 0 126861.54
201.12342834472656 0 46221.113 a 1
201.14271545410156 0 11842.241
202.1268768310547 0 3252.023
205.09707641601562 0 2098.8723
208.09654235839844 0 829.95667
208.10763549804688 0 830.7584
209.055419921875 0 982.5759
209.09214782714844 0 8630.13
210.123779296875 0 33627.914
211.1077423095703 0 22529.572 b Water loss 1
211.12693786621094 0 4121.6553
211.1375732421875 0 560.6197
211.14390563964844 0 927.80255
212.11099243164062 0 1537.4303
212.13929748535156 0 3195.5688
213.12376403808594 0 851.336
213.14283752441406 0 625.7055
215.13943481445312 0 853.71436
216.09796142578125 0 2424.922
218.14996337890625 0 1994.585
222.12376403808594 0 1397.0941
223.10780334472656 0 36204.92 y 8
224.1109161376953 0 3506.2437
225.12353515625 0 3258.3018
226.0823211669922 0 11455.077
226.11865234375 0 27976.467
227.06631469726562 0 7928.3613
227.0862579345703 0 1202.9406
227.10240173339844 0 1806.512
227.12222290039062 0 2988.6953
228.1343536376953 0 44484.465
229.1183624267578 0 5955.429 b 1
229.13772583007812 0 4719.906
230.14991760253906 0 3118.8203
234.12326049804688 0 1196.4703
238.1185302734375 0 695.58
239.1503143310547 0 1349.6937
240.13421630859375 0 3000.2852
242.14993286132812 0 10816.553
242.1627197265625 0 404.32056
243.1089630126953 0 1501.7147
243.15406799316406 0 810.68854
244.0786895751953 0 863.9011
244.0928497314453 0 21530.34
244.1291961669922 0 25015.398
245.09642028808594 0 1922.3788
245.13233947753906 0 2125.0686
249.12271118164062 0 761.82587
250.15505981445312 0 587.9262
251.15016174316406 0 961.74207
252.13436889648438 0 1669.4584
253.117431640625 0 925.70636
258.1446533203125 0 1860.5697
261.1197204589844 0 1157.3787
262.1193542480469 0 646.75867
265.1296081542969 0 905.83167
266.113525390625 0 1377.0131
267.1455993652344 0 1526.2252
268.1298828125 0 694.56177
269.1602783203125 0 1127.2013
270.14453125 0 4346.149 a Water loss 2
279.1323547363281 0 1831.3037
279.1454162597656 0 4315.6143 b 4
280.1292724609375 0 7104.7407
280.1464538574219 0 952.8187
281.1328125 0 728.1461
282.181396484375 0 2306.9463
283.1401062011719 0 6682.4175
284.1437683105469 0 1236.5116 y 5
287.17138671875 0 2315.4119
290.1144104003906 0 809.5412
291.14422607421875 0 1053.4417
292.1297912597656 0 1437.3903
294.1440124511719 0 714.1205
297.1557312011719 0 25371.709
298.1396789550781 0 23366.285 b Water loss 2
298.15850830078125 0 2992.4077
299.1424865722656 0 3754.7898
300.154541015625 0 687.7924
305.1812438964844 0 996.5039
306.1446533203125 0 2815.0593 y Water loss 7
309.15570068359375 0 3937.3186
310.13983154296875 0 1611.0183
310.1593017578125 0 750.95215
311.17181396484375 0 1661.4764
315.16632080078125 0 10831.883
316.1497497558594 0 1748.7477 b 2
316.1701965332031 0 1789.6431
317.182861328125 0 1319.1719
319.1396484375 0 2538.6482
324.15545654296875 0 13182.594 y 7
325.1580505371094 0 1802.9891
327.1661682128906 0 4051.6052
328.150390625 0 897.3728 b Ammonia loss 5
328.1704406738281 0 771.09216
329.1488037109375 0 649.05225
329.1819763183594 0 12287.9
330.18414306640625 0 1558.4523
336.1665344238281 0 2653.3438
337.15057373046875 0 12712.938
337.17041015625 0 728.2784
338.15325927734375 0 2425.1736
339.1663513183594 0 4830.005 y Water loss 4
339.6659851074219 0 1030.3231
340.1505126953125 0 1673.8475
345.1768493652344 0 9955.713
346.18011474609375 0 1934.419
348.1710205078125 0 7202.919 y 4
348.6725769042969 0 2632.5715
354.17694091796875 0 17369.629
355.1608581542969 0 6938.9917
355.18084716796875 0 1788.1997
356.1636657714844 0 978.79205
357.1764831542969 0 6361.6387
364.23382568359375 0 2781.602
365.21807861328125 0 13965.454
366.1992492675781 0 782.1233
366.2218933105469 0 2919.3003
367.1613464355469 0 1426.357
371.2031555175781 0 987.8982
372.18743896484375 0 14300.174
373.1896057128906 0 2410.5557
376.16021728515625 0 741.273
380.2305603027344 0 965.9452
382.2179870605469 0 602.3377
382.2444763183594 0 3459.888
383.2284240722656 0 16818.748
384.1870422363281 0 1485.9097
384.2320861816406 0 2418.335
392.2288513183594 0 6593.1367
393.1859436035156 0 1618.9453
393.21319580078125 0 13641.6045
394.1718444824219 0 6967.7812
394.21539306640625 0 2548.1782
395.17431640625 0 1258.3843
396.20086669921875 0 1824.9933 y Ammonia loss 3
396.70263671875 0 1343.5789
400.2568359375 0 967.4422
402.1995544433594 0 3412.9585
404.71343994140625 0 8714.714 y 3
405.2147216796875 0 5705.783
405.7156066894531 0 1010.01465
410.2086181640625 0 1129.9714
410.23980712890625 0 10784.21
410.7190856933594 0 7865.0176
411.2019348144531 0 1315.4532
411.22308349609375 0 18338.975 b Water loss 3
411.72314453125 0 1100.0676
412.1548767089844 0 737.7971
412.1830749511719 0 2650.3179
412.2254943847656 0 2947.9307
418.2087707519531 0 620.2171
419.2038269042969 0 1212.5603
419.72369384765625 0 593.9559
420.1867980957031 0 2701.8337
423.7284851074219 0 1656.446
424.2253112792969 0 1658.4396
426.19677734375 0 1075.3597
427.1832580566406 0 1445.8088
427.23248291015625 0 1066.1451
427.267333984375 0 2170.0942
427.74072265625 0 1771.9629
428.1597595214844 0 753.8719
428.2145080566406 0 1515.4805
428.2493591308594 0 5549.6553
429.2550354003906 0 721.0443
430.7116394042969 0 871.0982
432.732177734375 0 6252.8223
433.2279052734375 0 2470.984
433.7276916503906 0 786.7516
434.2397766113281 0 2957.295 y Water loss 6
435.23406982421875 0 1618.1213
435.7265930175781 0 1238.6516
437.2144775390625 0 2835.5244
438.1982116699219 0 4346.239
439.1971130371094 0 1201.2499
439.2273254394531 0 1109.6499 y Water loss 2
439.71820068359375 0 4704.097 y Ammonia loss 2
440.2191467285156 0 1759.0669
440.7184143066406 0 935.93427
441.73779296875 0 7027.174
442.2381286621094 0 4200.656 b Water loss 7
442.73699951171875 0 874.56854
444.2088928222656 0 13196.139
445.2115783691406 0 2246.0684
448.22967529296875 0 8692.436 y 2
448.26275634765625 0 1173.3237
448.7314147949219 0 4548.717
450.234375 0 1028.0715
452.2501220703125 0 31255.883 y 6
453.25341796875 0 5959.5073
454.2578430175781 0 950.90906
455.22418212890625 0 4331.6406
456.20904541015625 0 8506.9
457.17578125 0 723.27435
457.2107849121094 0 2846.4155
461.24139404296875 0 2387.2964
461.73870849609375 0 3515.5913
462.23736572265625 0 1906.5619
468.2466735839844 0 770.43176
470.247802734375 0 5309.8926
470.7483215332031 0 2859.0166 b Water loss 8
471.24517822265625 0 1828.7953
473.23529052734375 0 5580.131
474.23712158203125 0 1563.0062
479.25341796875 0 6067.3984
479.754638671875 0 2483.793 b 8
480.25421142578125 0 1589.07
485.2367858886719 0 1109.2106
485.2726745605469 0 3212.351
486.2349548339844 0 690.4086
495.2188720703125 0 1691.8153
497.7626953125 0 2063.809 y 1
498.2630920410156 0 1364.4346
510.3026428222656 0 1914.864
512.2469482421875 0 2221.2979
513.2303466796875 0 4486.9634
514.232666015625 0 1658.6049
521.2692260742188 0 983.59875
522.2684326171875 0 1395.0417
522.685791015625 0 1050.2794
522.76123046875 0 873.5921
524.2820434570312 0 1241.4414
529.7791748046875 0 1093.2136
530.256591796875 0 4322.2217
530.7755126953125 0 742.1039
531.2578735351562 0 1641.293
532.2430419921875 0 1115.1705
534.3027954101562 0 3894.5703
534.7785034179688 0 1092.8721
535.268798828125 0 1138.4233
535.310791015625 0 941.09534
535.7735595703125 0 1312.8723
538.2977905273438 0 10224.109
539.2857055664062 0 5170.852 b Water loss 4
542.6848754882812 0 802.1213
543.301025390625 0 2680.4148
543.7821044921875 0 11936.649
544.2793579101562 0 9841.461
544.7776489257812 0 3893.2344
545.274169921875 0 1725.5457
545.763916015625 0 650.4544
549.2667846679688 0 5593.522 y Water loss 5
550.2625732421875 0 1181.7993
551.6881713867188 0 5972.7124
552.18994140625 0 3168.1484
552.7872314453125 0 181823.92
553.2882690429688 0 110375.19 Precursor Water loss
553.7896118164062 0 41158.77
554.2911376953125 0 7517.067
555.27978515625 0 1383.7473
556.3074951171875 0 1809.1404
558.2523193359375 0 653.66956
561.7922973632812 0 26708.635
562.2935791015625 0 16999.871 Precursor
562.6538696289062 0 687.37494
562.7953491210938 0 5931.7935
563.291259765625 0 1097.8713
567.2767333984375 0 74414.664 y 5
568.2797241210938 0 19552.523
569.281982421875 0 3661.7593
572.3033447265625 0 10535.349
573.3079833984375 0 2233.9275
574.3104858398438 0 790.9325
583.2706909179688 0 873.48425
586.3189697265625 0 4818.724
587.3223266601562 0 1024.902
597.9470825195312 0 788.44244
599.5799560546875 0 639.92487
601.78564453125 0 609.6987
606.2868041992188 0 640.6317
608.30322265625 0 959.5062
611.314208984375 0 5617.227
612.3182983398438 0 1723.8367
625.3302001953125 0 976.9676
626.3251342773438 0 623.4038
629.325439453125 0 1241.7179
632.6143798828125 0 647.42126
632.8663940429688 0 912.1833
636.3001098632812 0 2167.2134
637.3289794921875 0 5201.285
638.3262939453125 0 1723.5425
641.8541870117188 0 2042.5955
642.2920532226562 0 899.5413
642.357177734375 0 1980.6064
643.3419189453125 0 1685.189
645.3558959960938 0 1700.1968
649.3297119140625 0 6678.37
650.3311767578125 0 2437.3857
653.32421875 0 14425.559
654.3200073242188 0 5587.4243 b Water loss 5
655.338134765625 0 5191.5347
656.3346557617188 0 2244.235
659.3135375976562 0 3083.4282
660.3005981445312 0 3839.6284
661.3001708984375 0 1728.4329
670.3515014648438 0 4603.775
671.3435668945312 0 1825.7509
673.3505859375 0 19708.37
674.3535766601562 0 7842.8726
675.357177734375 0 1022.2504
677.3242797851562 0 19278.318 y Water loss 4
678.3098754882812 0 45498.367 y Ammonia loss 4
679.31201171875 0 16159.849
680.3118286132812 0 2246.8218
686.3427124023438 0 940.63184
695.3349609375 0 62097.043 y 4
696.337890625 0 25898.602
697.3392333984375 0 4744.2227
711.8506469726562 0 1497.1343
712.3591918945312 0 3213.9104
713.3576049804688 0 1214.9889
713.4730224609375 0 679.97363
730.3715209960938 0 8229.915
731.3758544921875 0 2633.3708
748.3770141601562 0 1661.7717
755.364501953125 0 1023.7949
763.412353515625 0 674.7345
764.3942260742188 0 891.2326
765.3888549804688 0 684.8922
772.414794921875 0 964.4954
773.3798217773438 0 1806.8992
774.3851318359375 0 898.4257
781.4192504882812 0 10238.523
782.418212890625 0 5039.8384 b Water loss 6
783.4216918945312 0 1067.3136
790.40380859375 0 1170.6033 y Water loss 3
799.428955078125 0 1455.1321
808.4185791015625 0 19791.602 y 3
809.4212646484375 0 8573.446
810.423583984375 0 2835.7085
821.4136352539062 0 2545.0464
822.4166259765625 0 1215.5386
839.4327392578125 0 1033.8188
846.4453735351562 0 1592.1633
847.4363403320312 0 1715.7866
848.4242553710938 0 784.8692
849.44580078125 0 1053.142
855.4554443359375 0 786.94336
859.4270629882812 0 1543.6753
860.4227905273438 0 908.7671
864.4523315429688 0 663.39496
865.4408569335938 0 4030.0613
866.4392700195312 0 1471.382
877.4387817382812 0 6013.189 y Water loss 2
878.4334716796875 0 4766.9077 y Ammonia loss 2
879.4329223632812 0 1627.4672
883.4513549804688 0 11135.846 b Water loss 7
884.4523315429688 0 4808.356 b Ammonia loss 7
895.44970703125 0 113142.36 y 2
896.4525146484375 0 53093.6
897.4553833007812 0 13793.896
898.458984375 0 772.3009
900.4762573242188 0 2918.853
901.47314453125 0 1761.4036 b 7
902.4700317382812 0 718.59875
922.4625244140625 0 2721.7998
923.4553833007812 0 2897.2053
924.4518432617188 0 937.1225
939.4832153320312 0 1832.1586
940.4725341796875 0 12226.186 b Water loss 8
941.474609375 0 7245.441 b Ammonia loss 8
942.4803466796875 0 1913.9521
957.4954223632812 0 3474.07
958.4896850585938 0 2177.2788 b 8
959.5023193359375 0 987.10864
994.5181884765625 0 6404.813 y 1
995.52001953125 0 2569.4287
1711.5938720703125 0 647.03876
2885.630859375 0 828.56195
3086.826904296875 0 830.62335

Spectrum Details

|  |  |
| --- | --- |
| Matched peaks? Matched peaks The total absolute number of peaks matched. Additionally in brackets the total fraction of peaks matched and the total number of peaks is shown. | 51 (11.04% of 462) |
| FDR? FDR The false discovery rate estimated for this peptide. It is calculated by matching all theoretical fragments with a non-integer shift with the raw peaks for this spectrum. This is done with 40 different shifts. The resulting percentage is the average number of annotated peaks over the number of annotated peaks with the correct spectrum. | 0.09% |
| Satellite FDR? Satellite FDR See the FDR for details on its calculation. This satellite ion specific FDR only contains the satellite ions (d/w) for I/L/J positions. | - |
| PSM Score? PSM Score The PSM Score as given by Hecklib to this annotated spectrum. It is shown with three significant figures. | 554 |

#### Reverse Lookup? Reverse Lookup All places where this read could be placed.

| Group | Segment | Template | Template Part | Read Part | Score | Unique |
| --- | --- | --- | --- | --- | --- | --- |
| Homo sapiens Heavy Chain | IGHC | IGHG4 | [72..82] | [0..10] | 34 | False |
| Homo sapiens Heavy Chain | IGHC | IGHD | [179..189] | [0..10] | 34 | False |
| Decoy | Decoy | TRYP | [22..32] | [0..10] | 34 | False |

| Recombined | Template Part | Read Part | Score | Unique |
| --- | --- | --- | --- | --- |
| TRYP | [22..32] | [0..10] | 34 | True |

#### Meta Information from Multiple reads

##### Number of combined reads

3

##### Intensity

0.5901

##### TotalArea

1.143E+08

##### Changes to the peptide sequence

EVSJQDKTGF

L→JNo support for either Leucine or Isoleucine based on side chain ions (Position: 4)

#### Positional Score

Copy Data

##### Positional Score (TSV)

###### Preview

```
Loading example...
```

*Click on the button to copy the data to your clipboard.*

100123456789

Label Value
"0" 0.32
"1" 0.317
"2" 0.33
"3" 0.33
"4" 0.327
"5" 0.33
"6" 0.327
"7" 0.327
"8" 0.33
"9" 0.333

#### Meta Information from PEAKS

##### Scan Identifier

F3:7326

##### Original sequence

E

-18.01

V

S

L

Q

D

K

T

G

F

##### Posttranslational Modifications

Pyro-glu from E

##### Source File

D:\separate\_stitch\_analyses\xle-disambiguation\raw\20210323\_F1\_UM1\_Peng0013\_SA\_F59\_ingel\_3ug\_chymo.raw

##### Fraction

3

##### Scan Feature

F3:6472

##### De Novo Score

98

##### ConfidenceScore

98

### m/z

553.2811

##### Mass

1104.5452

##### Charge

2

##### Retention Time

40.88

##### Predicted Retention Time

-

##### Area

1.095E+08

##### Parts Per Million

2.3

##### Fragmentation mode

HCD

##### Originating file

01 D:\separate\_stitch\_analyses\xle-disambiguation\20210325\_F59\_3ug\_DENOVO\_12.csv

#### Meta Information from PEAKS

##### Scan Identifier

F3:5620

##### Original sequence

E

V

S

L

Q

D

K

T

G

F

##### Posttranslational Modifications

##### Source File

D:\separate\_stitch\_analyses\xle-disambiguation\raw\20210323\_F1\_UM1\_Peng0013\_SA\_F59\_ingel\_3ug\_chymo.raw

##### Fraction

3

##### Scan Feature

F3:6817

##### De Novo Score

98

##### ConfidenceScore

98

### m/z

562.2872

##### Mass

1122.5557

##### Charge

2

##### Retention Time

31.11

##### Predicted Retention Time

-

##### Area

3.299E+06

##### Parts Per Million

3.8

##### Fragmentation mode

HCD

##### Originating file

01 D:\separate\_stitch\_analyses\xle-disambiguation\20210325\_F59\_3ug\_DENOVO\_12.csv

#### Meta Information from PEAKS

##### Scan Identifier

F3:5655

##### Original sequence

E

V

S

L

Q

D

K

T

G

F

##### Posttranslational Modifications

##### Source File

D:\separate\_stitch\_analyses\xle-disambiguation\raw\20210323\_F1\_UM1\_Peng0013\_SA\_F59\_ingel\_3ug\_chymo.raw

##### Fraction

3

##### Scan Feature

F3:6823

##### De Novo Score

98

##### ConfidenceScore

98

### m/z

562.2943

##### Mass

1122.5557

##### Charge

2

##### Retention Time

31.41

##### Predicted Retention Time

-

##### Area

1.479E+06

##### Parts Per Million

16.4

##### Fragmentation mode

HCD

##### Originating file

01 D:\separate\_stitch\_analyses\xle-disambiguation\20210325\_F59\_3ug\_DENOVO\_12.csv
