## Supplementary material for "A handle on mass coincidence errors in *de novo* sequencing of antibodies by bottom-up proteomics": Combined_043.html

Details Combined\_043 | Stitch OverviewUndefined

### Read Combined\_043

#### Sequence (length=11)

VLQSSGJYSJS

#### Spectrum 7916? Spectrum 7916 The raw spectrum of this peptide as annotated by Hecklib. The fragments are coloured according to ion type (see legend). Any peaks with a star '\*' as text can be hovered over to see the full details, first the ion type second the mass shift type. By hovering over the amino acids in the peptide or ions in the legend the corresponding peaks are highlighted. By toggling the 'Unassigned' label you can turn the background (unassigned) peaks on or off in the plot. By updating the slider in the Ion legend you can update the spectrum to only show the top X% of the peaks with labels. The top X% means any peak that is within X% of the highest intensity. By dragging in the spectrum you can zoom in to a specific part of the spectrum and use 'Zoom Out' to get back to the original zoom level. The annotation of the spectrum is based on the given sequence in the peptides file and is done with different software so inconsistencies are likely. The peaks are annotated based on the given sequence, with 20 ppm tolerance.

Copy Data

##### Spectrum 7916 (TSV)

###### Preview

```
Loading example...
```

*Click on the button to copy the data to your clipboard.*

Mz MinMz MaxIntensity Max

WidthHeightPeptide font sizePeptide stroke widthSpectrum font sizeSpectrum stroke widthCompact peptide

Ion legend

wxyz

abcd

OtherUnassignedIon^Charge^_Position_Show for top:%

VLQSSGJYSJS

08.03e+41.61e+52.41e+53.21e+5

Zoom Out

y^+1^_2_y^+1^_2_y^+1^_3_y^+1^_3_c^+1^_3_c^+2^_7_c^+1^_3_c^+2^_8_c^+1^_4_c^+1^_4_y^+1^_4_y^+2^_9_c^+2^_9_y^+1^_4_c^+1^_5_c^+1^_5_c^+1^_5_c^+1^_6_c^+1^_6_y^+1^_5_c^+1^_6_z^+1^_6_y^+1^_6_y^+1^_6_c^+1^_7_c^+1^_7_z^+1^_7_c^+1^_7_y^+1^_7_z^+1^_7_y^+1^_7_z^+1^_8_w^+1^_8_y^+1^_8_z^+1^_8_y^+1^_8_c^+1^_8_c^+1^_8_c^+1^_8_w^+1^_9_z^+1^_9_y^+1^_9_z^+1^_9_c^+1^_9_c^+1^_9_y^+1^_9_c^+1^_9_w^+1^_10_z^+1^_10_z^+1^_10_y^+1^_10_z^+1^_10_c^+1^_10_c^+1^_10_y^+1^_10_c^+1^_10_

0698139520932791

Fragment Matches Table

Show background peaks

| Position | Ion type | Intensity | mz Theoretical | mz Error (Th) | mz Error (ppm) | Charge | Series Number |
| --- | --- | --- | --- | --- | --- | --- | --- |
| - | - | 638.3 | 123.1 | - | - | 0 | - |
| - | - | 692.7 | 126.7 | - | - | 0 | - |
| - | - | 671.7 | 127.5 | - | - | 0 | - |
| - | - | 1.678E+04 | 136.1 | - | - | 0 | - |
| - | - | 1291 | 137.1 | - | - | 0 | - |
| - | - | 623.5 | 142.4 | - | - | 0 | - |
| - | - | 655 | 144.1 | - | - | 0 | - |
| - | - | 605.1 | 144.2 | - | - | 0 | - |
| - | - | 675.8 | 148.9 | - | - | 0 | - |
| - | - | 660.9 | 153.9 | - | - | 0 | - |
| - | - | 988.8 | 155.1 | - | - | 0 | - |
| - | - | 782.1 | 166.6 | - | - | 0 | - |
| - | - | 1156 | 173.1 | - | - | 0 | - |
| - | - | 1.117E+04 | 173.1 | - | - | 0 | - |
| - | - | 5555 | 173.4 | - | - | 0 | - |
| - | - | 926.3 | 183.1 | - | - | 0 | - |
| - | - | 1.814E+04 | 185.2 | - | - | 0 | - |
| - | - | 1532 | 186.2 | - | - | 0 | - |
| 10 | y | 2.072E+04 | 201.1 | 6.257E-05 | 0.3111 | +1 | 2 |
| - | - | 2467 | 202.1 | - | - | 0 | - |
| - | - | 782.7 | 209.2 | - | - | 0 | - |
| - | - | 1005 | 210.1 | - | - | 0 | - |
| - | - | 1329 | 213.1 | - | - | 0 | - |
| - | - | 2.244E+04 | 213.2 | - | - | 0 | - |
| - | - | 1368 | 213.2 | - | - | 0 | - |
| - | - | 2333 | 214.2 | - | - | 0 | - |
| - | - | 2028 | 216.1 | - | - | 0 | - |
| 10 | y | 2.678E+04 | 219.1 | 6.817E-05 | 0.3111 | +1 | 2 |
| - | - | 1998 | 220.1 | - | - | 0 | - |
| - | - | 883.3 | 221.1 | - | - | 0 | - |
| - | - | 788.9 | 223.4 | - | - | 0 | - |
| - | - | 9296 | 233.1 | - | - | 0 | - |
| - | - | 4257 | 242.1 | - | - | 0 | - |
| - | - | 2226 | 251.1 | - | - | 0 | - |
| - | - | 998.8 | 256.2 | - | - | 0 | - |
| - | - | 806.1 | 260.2 | - | - | 0 | - |
| - | - | 1042 | 273.2 | - | - | 0 | - |
| - | - | 1089 | 282.8 | - | - | 0 | - |
| 9 | y | 5582 | 288.2 | 0.0001814 | 0.6294 | +1 | 3 |
| 9 | y | 7493 | 306.2 | 9.89E-05 | 0.323 | +1 | 3 |
| - | - | 949.8 | 307.2 | - | - | 0 | - |
| - | - | 1643 | 314.2 | - | - | 0 | - |
| 3 | c | 1.673E+04 | 341.2 | 0.000174 | 0.5099 | +1 | 3 |
| - | - | 3448 | 342.2 | - | - | 0 | - |
| 7 | c | 1177 | 343.2 | 0.0002796 | 0.8146 | +2 | 7 |
| - | - | 4365 | 345.2 | - | - | 0 | - |
| - | - | 5148 | 346.2 | - | - | 0 | - |
| - | - | 3531 | 347.2 | - | - | 0 | - |
| - | - | 1375 | 353.2 | - | - | 0 | - |
| - | - | 3491 | 355.1 | - | - | 0 | - |
| - | - | 1760 | 356.1 | - | - | 0 | - |
| - | - | 1633 | 357.2 | - | - | 0 | - |
| 3 | c | 1.537E+04 | 358.2 | 0.0001605 | 0.448 | +1 | 3 |
| - | - | 2224 | 359.2 | - | - | 0 | - |
| - | - | 1698 | 364.2 | - | - | 0 | - |
| - | - | 991.4 | 386.6 | - | - | 0 | - |
| - | - | 1194 | 387.2 | - | - | 0 | - |
| - | - | 981.7 | 395.7 | - | - | 0 | - |
| - | - | 1396 | 403.2 | - | - | 0 | - |
| - | - | 3550 | 409.7 | - | - | 0 | - |
| - | - | 1647 | 410.2 | - | - | 0 | - |
| - | - | 1387 | 410.2 | - | - | 0 | - |
| - | - | 1106 | 410.7 | - | - | 0 | - |
| - | - | 1523 | 411.2 | - | - | 0 | - |
| - | - | 1282 | 414.2 | - | - | 0 | - |
| - | - | 1133 | 415.2 | - | - | 0 | - |
| - | - | 1910 | 421.2 | - | - | 0 | - |
| 8 | c | 4549 | 424.7 | 0.0004146 | 0.9762 | +2 | 8 |
| - | - | 1683 | 425.2 | - | - | 0 | - |
| - | - | 2015 | 426.2 | - | - | 0 | - |
| - | - | 1579 | 426.3 | - | - | 0 | - |
| 4 | c | 5922 | 428.3 | 9.749E-05 | 0.2277 | +1 | 4 |
| - | - | 993.8 | 429.3 | - | - | 0 | - |
| - | - | 3174 | 430.3 | - | - | 0 | - |
| - | - | 1.352E+04 | 444.3 | - | - | 0 | - |
| - | - | 3349 | 445.2 | - | - | 0 | - |
| 4 | c | 6.064E+04 | 445.3 | 0.0001597 | 0.3587 | +1 | 4 |
| - | - | 2734 | 445.7 | - | - | 0 | - |
| - | - | 1.389E+04 | 446.3 | - | - | 0 | - |
| - | - | 1249 | 447.3 | - | - | 0 | - |
| - | - | 3278 | 450.2 | - | - | 0 | - |
| - | - | 3705 | 450.7 | - | - | 0 | - |
| 8 | y | 3532 | 451.2 | 0.00448 | 9.928 | +1 | 4 |
| - | - | 1925 | 453.2 | - | - | 0 | - |
| - | - | 2390 | 454.2 | - | - | 0 | - |
| - | - | 1782 | 456.2 | - | - | 0 | - |
| - | - | 1233 | 457.2 | - | - | 0 | - |
| - | - | 1112 | 457.2 | - | - | 0 | - |
| - | - | 2.967E+04 | 459.2 | - | - | 0 | - |
| - | - | 1.512E+04 | 459.7 | - | - | 0 | - |
| - | - | 6253 | 460.2 | - | - | 0 | - |
| 3 | y | 5767 | 462.2 | 0.001006 | 2.177 | +2 | 9 |
| 9 | c | 8220 | 468.2 | 0.0003916 | 0.8363 | +2 | 9 |
| - | - | 6148 | 468.7 | - | - | 0 | - |
| 8 | y | 8185 | 469.2 | 0.000629 | 1.34 | +1 | 4 |
| - | - | 1757 | 470.2 | - | - | 0 | - |
| - | - | 2582 | 471.3 | - | - | 0 | - |
| - | - | 1053 | 471.7 | - | - | 0 | - |
| - | - | 1437 | 473.2 | - | - | 0 | - |
| - | - | 4940 | 475.3 | - | - | 0 | - |
| - | - | 1069 | 476.3 | - | - | 0 | - |
| - | - | 1261 | 479.3 | - | - | 0 | - |
| - | - | 1425 | 480.2 | - | - | 0 | - |
| - | - | 3448 | 489.3 | - | - | 0 | - |
| - | - | 2427 | 490.2 | - | - | 0 | - |
| - | - | 4428 | 497.3 | - | - | 0 | - |
| - | - | 1259 | 498.3 | - | - | 0 | - |
| - | - | 1698 | 501.8 | - | - | 0 | - |
| - | - | 904.6 | 502.3 | - | - | 0 | - |
| - | - | 5626 | 508.2 | - | - | 0 | - |
| - | - | 4104 | 513.3 | - | - | 0 | - |
| 5 | c | 1156 | 514.3 | 0.0008877 | 1.726 | +1 | 5 |
| 5 | c | 4394 | 515.3 | 0.0006924 | 1.344 | +1 | 5 |
| - | - | 4356 | 515.8 | - | - | 0 | - |
| - | - | 5222 | 516.3 | - | - | 0 | - |
| - | - | 1934 | 527.3 | - | - | 0 | - |
| - | - | 1.842E+04 | 531.3 | - | - | 0 | - |
| 5 | c | 5.147E+04 | 532.3 | 9.989E-05 | 0.1877 | +1 | 5 |
| - | - | 1.472E+04 | 533.3 | - | - | 0 | - |
| - | - | 849.6 | 534.3 | - | - | 0 | - |
| - | - | 857.9 | 534.3 | - | - | 0 | - |
| - | - | 1214 | 539.3 | - | - | 0 | - |
| - | - | 1743 | 545.3 | - | - | 0 | - |
| - | - | 1.733E+04 | 546.3 | - | - | 0 | - |
| - | - | 4596 | 547.3 | - | - | 0 | - |
| - | - | 3503 | 554.3 | - | - | 0 | - |
| - | - | 1620 | 555.3 | - | - | 0 | - |
| - | - | 1906 | 557.3 | - | - | 0 | - |
| - | - | 1163 | 558.3 | - | - | 0 | - |
| - | - | 1135 | 559.3 | - | - | 0 | - |
| - | - | 1469 | 559.3 | - | - | 0 | - |
| - | - | 1496 | 570.3 | - | - | 0 | - |
| 6 | c | 4130 | 571.3 | 0.01008 | 17.65 | +1 | 6 |
| 6 | c | 1.082E+04 | 572.3 | 0.0008961 | 1.566 | +1 | 6 |
| - | - | 3308 | 573.3 | - | - | 0 | - |
| - | - | 1164 | 574.4 | - | - | 0 | - |
| - | - | 3126 | 577.3 | - | - | 0 | - |
| - | - | 1074 | 578.3 | - | - | 0 | - |
| - | - | 3802 | 578.3 | - | - | 0 | - |
| 7 | y | 1155 | 582.3 | 0.001038 | 1.782 | +1 | 5 |
| - | - | 1827 | 586.3 | - | - | 0 | - |
| - | - | 3.298E+04 | 588.3 | - | - | 0 | - |
| 6 | c | 1.438E+05 | 589.3 | 0.000409 | 0.6941 | +1 | 6 |
| - | - | 4.327E+04 | 590.3 | - | - | 0 | - |
| - | - | 6465 | 591.3 | - | - | 0 | - |
| - | - | 2149 | 595.3 | - | - | 0 | - |
| - | - | 1247 | 603.3 | - | - | 0 | - |
| 6 | z | 2266 | 605.3 | 0.001844 | 3.046 | +1 | 6 |
| - | - | 1849 | 607.3 | - | - | 0 | - |
| - | - | 1177 | 618.3 | - | - | 0 | - |
| - | - | 3468 | 619.3 | - | - | 0 | - |
| - | - | 1253 | 620.3 | - | - | 0 | - |
| 6 | y | 3657 | 621.3 | 0.0003315 | 0.5335 | +1 | 6 |
| - | - | 2292 | 636.3 | - | - | 0 | - |
| - | - | 1782 | 637.3 | - | - | 0 | - |
| - | - | 6816 | 638.3 | - | - | 0 | - |
| 6 | y | 3540 | 639.3 | 0.003397 | 5.314 | +1 | 6 |
| - | - | 1674 | 641.3 | - | - | 0 | - |
| - | - | 2379 | 649.4 | - | - | 0 | - |
| - | - | 1313 | 650.4 | - | - | 0 | - |
| - | - | 2037 | 655.3 | - | - | 0 | - |
| - | - | 4755 | 657.4 | - | - | 0 | - |
| - | - | 2.56E+04 | 658.4 | - | - | 0 | - |
| - | - | 2.765E+04 | 659.4 | - | - | 0 | - |
| - | - | 6665 | 660.4 | - | - | 0 | - |
| - | - | 1656 | 661.4 | - | - | 0 | - |
| - | - | 7072 | 667.4 | - | - | 0 | - |
| - | - | 5049 | 668.4 | - | - | 0 | - |
| - | - | 4414 | 669.4 | - | - | 0 | - |
| - | - | 2777 | 670.4 | - | - | 0 | - |
| 7 | c | 2500 | 684.4 | 0.004914 | 7.18 | +1 | 7 |
| 7 | c | 5.587E+04 | 685.4 | 0.0003893 | 0.568 | +1 | 7 |
| - | - | 1.952E+04 | 686.4 | - | - | 0 | - |
| - | - | 8278 | 687.4 | - | - | 0 | - |
| - | - | 1348 | 688.3 | - | - | 0 | - |
| - | - | 1239 | 688.4 | - | - | 0 | - |
| - | - | 2637 | 690.3 | - | - | 0 | - |
| 5 | z | 3708 | 692.3 | 0.002164 | 3.125 | +1 | 7 |
| - | - | 2027 | 694.3 | - | - | 0 | - |
| - | - | 3.402E+04 | 701.4 | - | - | 0 | - |
| 7 | c | 1.196E+05 | 702.4 | 0.0001463 | 0.2083 | +1 | 7 |
| - | - | 4.276E+04 | 703.4 | - | - | 0 | - |
| - | - | 7868 | 704.4 | - | - | 0 | - |
| - | - | 7121 | 705.3 | - | - | 0 | - |
| - | - | 3860 | 706.3 | - | - | 0 | - |
| 5 | y | 3814 | 708.4 | 0.01284 | 18.12 | +1 | 7 |
| - | - | 1990 | 709.3 | - | - | 0 | - |
| 5 | z | 9455 | 710.3 | 0.001365 | 1.921 | +1 | 7 |
| - | - | 5584 | 711.4 | - | - | 0 | - |
| - | - | 1147 | 717.3 | - | - | 0 | - |
| - | - | 1637 | 720.4 | - | - | 0 | - |
| - | - | 3972 | 723.3 | - | - | 0 | - |
| - | - | 1956 | 724.3 | - | - | 0 | - |
| - | - | 1.949E+04 | 725.4 | - | - | 0 | - |
| 5 | y | 1.78E+04 | 726.4 | 0.002223 | 3.06 | +1 | 7 |
| - | - | 4616 | 727.4 | - | - | 0 | - |
| - | - | 2508 | 733.4 | - | - | 0 | - |
| - | - | 3378 | 734.4 | - | - | 0 | - |
| - | - | 1199 | 735.4 | - | - | 0 | - |
| - | - | 1824 | 737.4 | - | - | 0 | - |
| - | - | 1521 | 738.4 | - | - | 0 | - |
| - | - | 2145 | 749.4 | - | - | 0 | - |
| - | - | 1388 | 750.4 | - | - | 0 | - |
| - | - | 3607 | 751.4 | - | - | 0 | - |
| - | - | 1219 | 752.4 | - | - | 0 | - |
| - | - | 8740 | 754.4 | - | - | 0 | - |
| - | - | 4977 | 755.4 | - | - | 0 | - |
| - | - | 2242 | 756.4 | - | - | 0 | - |
| - | - | 1325 | 772.4 | - | - | 0 | - |
| 4 | z | 6651 | 779.4 | 0.000592 | 0.7596 | +1 | 8 |
| 4 | w | 2499 | 780.4 | 0.00351 | 4.498 | +1 | 8 |
| - | - | 1052 | 781.4 | - | - | 0 | - |
| - | - | 1751 | 789.4 | - | - | 0 | - |
| - | - | 1825 | 790.4 | - | - | 0 | - |
| - | - | 1294 | 791.4 | - | - | 0 | - |
| - | - | 1823 | 794.4 | - | - | 0 | - |
| 4 | y | 5496 | 795.4 | 0.01075 | 13.51 | +1 | 8 |
| - | - | 2769 | 796.4 | - | - | 0 | - |
| 4 | z | 9852 | 797.4 | 0.0004033 | 0.5057 | +1 | 8 |
| - | - | 4291 | 798.4 | - | - | 0 | - |
| - | - | 4359 | 802.4 | - | - | 0 | - |
| - | - | 1445 | 803.4 | - | - | 0 | - |
| - | - | 1241 | 805.4 | - | - | 0 | - |
| - | - | 1700 | 807.4 | - | - | 0 | - |
| - | - | 1523 | 808.4 | - | - | 0 | - |
| - | - | 2373 | 809.4 | - | - | 0 | - |
| - | - | 1.655E+04 | 812.4 | - | - | 0 | - |
| 4 | y | 2.139E+04 | 813.4 | 0.00117 | 1.438 | +1 | 8 |
| - | - | 8445 | 814.4 | - | - | 0 | - |
| - | - | 1313 | 815.4 | - | - | 0 | - |
| - | - | 3737 | 818.4 | - | - | 0 | - |
| - | - | 2051 | 819.4 | - | - | 0 | - |
| - | - | 1.045E+04 | 820.4 | - | - | 0 | - |
| - | - | 5138 | 821.4 | - | - | 0 | - |
| - | - | 4531 | 821.5 | - | - | 0 | - |
| - | - | 2653 | 822.4 | - | - | 0 | - |
| - | - | 2338 | 822.5 | - | - | 0 | - |
| - | - | 1940 | 823.5 | - | - | 0 | - |
| - | - | 1046 | 826.5 | - | - | 0 | - |
| - | - | 6027 | 830.4 | - | - | 0 | - |
| - | - | 5891 | 831.4 | - | - | 0 | - |
| - | - | 6579 | 832.4 | - | - | 0 | - |
| - | - | 3978 | 833.4 | - | - | 0 | - |
| - | - | 1500 | 834.4 | - | - | 0 | - |
| - | - | 1370 | 836.4 | - | - | 0 | - |
| - | - | 2073 | 838.4 | - | - | 0 | - |
| - | - | 1890 | 839.4 | - | - | 0 | - |
| - | - | 4349 | 846.5 | - | - | 0 | - |
| 8 | c | 3888 | 847.5 | 0.01484 | 17.51 | +1 | 8 |
| 8 | c | 4.529E+04 | 848.5 | 0.001148 | 1.353 | +1 | 8 |
| - | - | 2.08E+04 | 849.5 | - | - | 0 | - |
| - | - | 7626 | 850.5 | - | - | 0 | - |
| - | - | 1899 | 851.5 | - | - | 0 | - |
| - | - | 1.231E+04 | 864.5 | - | - | 0 | - |
| 8 | c | 1.214E+05 | 865.5 | 0.0004775 | 0.5517 | +1 | 8 |
| - | - | 1383 | 865.9 | - | - | 0 | - |
| - | - | 6.025E+04 | 866.5 | - | - | 0 | - |
| 3 | w | 1188 | 867.4 | 0.000321 | 0.3701 | +1 | 9 |
| - | - | 2.076E+04 | 867.5 | - | - | 0 | - |
| - | - | 5820 | 868.5 | - | - | 0 | - |
| - | - | 3424 | 869.4 | - | - | 0 | - |
| - | - | 1826 | 870.4 | - | - | 0 | - |
| - | - | 1567 | 889.5 | - | - | 0 | - |
| - | - | 3354 | 890.4 | - | - | 0 | - |
| - | - | 2242 | 891.4 | - | - | 0 | - |
| - | - | 5903 | 899.5 | - | - | 0 | - |
| - | - | 3982 | 900.5 | - | - | 0 | - |
| - | - | 1307 | 901.5 | - | - | 0 | - |
| - | - | 2353 | 906.4 | - | - | 0 | - |
| 3 | z | 2.124E+04 | 907.4 | 0.000181 | 0.1994 | +1 | 9 |
| - | - | 7040 | 908.4 | - | - | 0 | - |
| - | - | 1937 | 909.4 | - | - | 0 | - |
| - | - | 1608 | 909.5 | - | - | 0 | - |
| - | - | 1552 | 910.5 | - | - | 0 | - |
| - | - | 1.203E+04 | 915.5 | - | - | 0 | - |
| - | - | 9273 | 916.5 | - | - | 0 | - |
| - | - | 5.208E+04 | 917.5 | - | - | 0 | - |
| - | - | 2.734E+04 | 918.5 | - | - | 0 | - |
| - | - | 1.032E+04 | 919.5 | - | - | 0 | - |
| - | - | 3672 | 920.5 | - | - | 0 | - |
| 3 | y | 3805 | 923.4 | 0.001697 | 1.838 | +1 | 9 |
| - | - | 2943 | 924.5 | - | - | 0 | - |
| 3 | z | 3.753E+04 | 925.4 | 0.0008467 | 0.915 | +1 | 9 |
| - | - | 1.925E+04 | 926.4 | - | - | 0 | - |
| - | - | 5134 | 927.4 | - | - | 0 | - |
| - | - | 3.398E+04 | 933.5 | - | - | 0 | - |
| 9 | c | 1.957E+04 | 934.5 | 0.0147 | 15.73 | +1 | 9 |
| 9 | c | 4.047E+04 | 935.5 | 0.001041 | 1.112 | +1 | 9 |
| - | - | 2.117E+04 | 936.5 | - | - | 0 | - |
| - | - | 4899 | 937.5 | - | - | 0 | - |
| 3 | y | 1.008E+04 | 941.5 | 0.0004943 | 0.525 | +1 | 9 |
| - | - | 4508 | 942.5 | - | - | 0 | - |
| - | - | 4302 | 951.5 | - | - | 0 | - |
| 9 | c | 2.121E+05 | 952.5 | 0.0006756 | 0.7093 | +1 | 9 |
| - | - | 1.12E+05 | 953.5 | - | - | 0 | - |
| - | - | 3.191E+04 | 954.5 | - | - | 0 | - |
| - | - | 4055 | 955.5 | - | - | 0 | - |
| - | - | 2580 | 976.5 | - | - | 0 | - |
| - | - | 1969 | 977.5 | - | - | 0 | - |
| - | - | 5209 | 982.5 | - | - | 0 | - |
| - | - | 1401 | 983.5 | - | - | 0 | - |
| 2 | w | 1.414E+04 | 995.5 | 0.00116 | 1.165 | +1 | 10 |
| - | - | 5720 | 996.5 | - | - | 0 | - |
| - | - | 2918 | 997.5 | - | - | 0 | - |
| - | - | 3451 | 1004 | - | - | 0 | - |
| - | - | 2529 | 1005 | - | - | 0 | - |
| - | - | 1408 | 1006 | - | - | 0 | - |
| - | - | 2.709E+04 | 1015 | - | - | 0 | - |
| - | - | 1.574E+04 | 1016 | - | - | 0 | - |
| - | - | 5413 | 1017 | - | - | 0 | - |
| 2 | z | 1.582E+04 | 1021 | 0.001322 | 1.295 | +1 | 10 |
| 2 | z | 3690 | 1021 | 0.004794 | 4.693 | +1 | 10 |
| - | - | 1.152E+04 | 1022 | - | - | 0 | - |
| - | - | 7258 | 1023 | - | - | 0 | - |
| - | - | 2287 | 1024 | - | - | 0 | - |
| - | - | 7829 | 1031 | - | - | 0 | - |
| - | - | 5684 | 1032 | - | - | 0 | - |
| - | - | 3.456E+04 | 1033 | - | - | 0 | - |
| - | - | 1.807E+04 | 1034 | - | - | 0 | - |
| - | - | 5596 | 1035 | - | - | 0 | - |
| 2 | y | 1454 | 1037 | 0.01545 | 14.9 | +1 | 10 |
| 2 | z | 1.255E+05 | 1039 | 0.001133 | 1.091 | +1 | 10 |
| - | - | 7.193E+04 | 1040 | - | - | 0 | - |
| - | - | 2.676E+04 | 1041 | - | - | 0 | - |
| - | - | 2714 | 1042 | - | - | 0 | - |
| 10 | c | 2033 | 1048 | 0.01362 | 13 | +1 | 10 |
| 10 | c | 2.209E+04 | 1049 | 0.001877 | 1.79 | +1 | 10 |
| - | - | 1.362E+04 | 1050 | - | - | 0 | - |
| - | - | 5160 | 1051 | - | - | 0 | - |
| - | - | 7497 | 1054 | - | - | 0 | - |
| 2 | y | 4987 | 1055 | 0.00514 | 4.874 | +1 | 10 |
| - | - | 1267 | 1056 | - | - | 0 | - |
| - | - | 2286 | 1064 | - | - | 0 | - |
| - | - | 1809 | 1065 | - | - | 0 | - |
| 10 | c | 2.719E+05 | 1066 | 0.001206 | 1.132 | +1 | 10 |
| - | - | 1.638E+05 | 1067 | - | - | 0 | - |
| - | - | 5.566E+04 | 1068 | - | - | 0 | - |
| - | - | 8632 | 1069 | - | - | 0 | - |
| - | - | 1570 | 1071 | - | - | 0 | - |
| - | - | 2.007E+04 | 1082 | - | - | 0 | - |
| - | - | 1.13E+04 | 1083 | - | - | 0 | - |
| - | - | 4479 | 1084 | - | - | 0 | - |
| - | - | 1409 | 1092 | - | - | 0 | - |
| - | - | 2825 | 1093 | - | - | 0 | - |
| - | - | 9502 | 1099 | - | - | 0 | - |
| - | - | 6483 | 1100 | - | - | 0 | - |
| - | - | 4540 | 1109 | - | - | 0 | - |
| - | - | 1.874E+04 | 1110 | - | - | 0 | - |
| - | - | 1.184E+04 | 1111 | - | - | 0 | - |
| - | - | 4320 | 1112 | - | - | 0 | - |
| - | - | 1207 | 1113 | - | - | 0 | - |
| - | - | 6023 | 1120 | - | - | 0 | - |
| - | - | 5259 | 1121 | - | - | 0 | - |
| - | - | 1.028E+04 | 1127 | - | - | 0 | - |
| - | - | 5504 | 1128 | - | - | 0 | - |
| - | - | 3154 | 1129 | - | - | 0 | - |
| - | - | 7875 | 1137 | - | - | 0 | - |
| - | - | 3.701E+04 | 1138 | - | - | 0 | - |
| - | - | 2.266E+04 | 1139 | - | - | 0 | - |
| - | - | 1.054E+04 | 1140 | - | - | 0 | - |
| - | - | 1721 | 1141 | - | - | 0 | - |
| - | - | 6345 | 1153 | - | - | 0 | - |
| - | - | 9.724E+04 | 1154 | - | - | 0 | - |
| - | - | 3.18E+05 | 1155 | - | - | 0 | - |
| - | - | 1.816E+05 | 1156 | - | - | 0 | - |
| - | - | 6.257E+04 | 1157 | - | - | 0 | - |
| - | - | 8796 | 1158 | - | - | 0 | - |
| - | - | 1117 | 1430 | - | - | 0 | - |
| - | - | 1372 | 2763 | - | - | 0 | - |

m/z Charge Intensity FragmentType MassShift Position
123.08396911621094 0 638.2524
126.67093658447266 0 692.69336
127.5208740234375 0 671.6571
136.07569885253906 0 16784.094
137.07907104492188 0 1291.3887
142.4107208251953 0 623.4874
144.0757598876953 0 655.03674
144.19461059570312 0 605.1317
148.9473419189453 0 675.8121
153.8539581298828 0 660.86707
155.1179656982422 0 988.76544
166.60458374023438 0 782.1402
173.12124633789062 0 1155.7776
173.12855529785156 0 11170.388
173.4403839111328 0 5555.253
183.11288452148438 0 926.3448
185.1647491455078 0 18137.094
186.16830444335938 0 1531.9009
201.12330627441406 0 20717.697 y Water loss 9
202.12672424316406 0 2466.539
209.23284912109375 0 782.6914
210.14297485351562 0 1004.786
213.1491241455078 0 1329.1874
213.1597137451172 0 22441.281
213.17161560058594 0 1367.9149
214.1632080078125 0 2333.3342
216.09793090820312 0 2028.3989
219.1338653564453 0 26775.021 y 9
220.13783264160156 0 1997.9288
221.1393280029297 0 883.3386
223.42100524902344 0 788.93915
233.09197998046875 0 9295.5205
242.14999389648438 0 4257.221
251.10340881347656 0 2225.763
256.20159912109375 0 998.7704
260.2278137207031 0 806.1284
273.1990966796875 0 1042.1031
282.7813415527344 0 1089.0685
288.15557861328125 0 5582.2803 y Water loss 8
306.1658630371094 0 7492.6133 y 8
307.1703186035156 0 949.76636
314.2313537597656 0 1642.9733
341.218505859375 0 16734.049 c Ammonia loss 2
342.2216491699219 0 3448.4412
343.1978759765625 0 1176.621 c Ammonia loss 6
345.1769104003906 0 4364.8604
346.1766662597656 0 5148.355
347.1925964355469 0 3530.956
353.1654968261719 0 1374.6172
355.0699768066406 0 3491.404
356.0697937011719 0 1760.0098
357.23663330078125 0 1633.3701
358.2447204589844 0 15374.154 c 2
359.24786376953125 0 2224.1858
364.18634033203125 0 1697.8438
386.5823669433594 0 991.40564
387.2228698730469 0 1193.7422
395.7102355957031 0 981.7372
403.1982421875 0 1395.5715
409.70587158203125 0 3550.109
410.20880126953125 0 1647.3396
410.23785400390625 0 1387.2762
410.7331237792969 0 1106.3558
411.2301330566406 0 1522.7252
414.197021484375 0 1281.5962
415.2051086425781 0 1132.9354
421.2082824707031 0 1909.5708
424.72967529296875 0 4548.952 c Ammonia loss 7
425.2305908203125 0 1682.6616
426.2325439453125 0 2014.8104
426.2621765136719 0 1579.4412
428.2504577636719 0 5922.0747 c Ammonia loss 3
429.2532043457031 0 993.84656
430.26556396484375 0 3174.468
444.2695617675781 0 13515.489
445.244873046875 0 3349.1995
445.2770690917969 0 60637.59 c 3
445.7437438964844 0 2733.7236
446.2796325683594 0 13892.638
447.2798156738281 0 1249.1704
450.2352600097656 0 3277.9365
450.72930908203125 0 3705.4863
451.22320556640625 0 3531.9275 y Water loss 7
453.22161865234375 0 1925.156
454.2490539550781 0 2390.3193
456.2092590332031 0 1781.8386
457.2093811035156 0 1233.011
457.2416076660156 0 1111.983
459.240234375 0 29668.174
459.7420349121094 0 15119.201
460.2424011230469 0 6253.1094
462.22808837890625 0 5766.873 y Water loss 2
468.24566650390625 0 8219.823 c Ammonia loss 8
468.7474365234375 0 6147.959
469.22991943359375 0 8185.1753 y 7
470.23150634765625 0 1756.6987
471.2939147949219 0 2582.0408
471.676513671875 0 1053.1952
473.2358093261719 0 1437.2485
475.25128173828125 0 4940.4624
476.2549133300781 0 1068.8026
479.2621154785156 0 1260.8606
480.2444152832031 0 1424.9506
489.3030090332031 0 3447.965
490.2303771972656 0 2427.0806
497.2724304199219 0 4428.349
498.27471923828125 0 1259.1206
501.7844543457031 0 1697.9227
502.2745666503906 0 904.61
508.2406921386719 0 5625.93
513.2900390625 0 4103.725
514.2974853515625 0 1155.6906 c Water loss 4
515.2830810546875 0 4393.526 c Ammonia loss 4
515.7820434570312 0 4356.149
516.283447265625 0 5221.619
527.2828979492188 0 1933.8363
531.30126953125 0 18423.646
532.308837890625 0 51472.793 c 4
533.3118286132812 0 14724.397
534.2994384765625 0 849.5551
534.3110961914062 0 857.8669
539.3195190429688 0 1214.3213
545.3176879882812 0 1742.7422
546.324951171875 0 17331.604
547.326904296875 0 4596.226
554.294189453125 0 3503.4194
555.27880859375 0 1619.8418
557.3306274414062 0 1906.1353
558.3275756835938 0 1163.1837
559.2509155273438 0 1134.6613
559.2984619140625 0 1468.688
570.3126831054688 0 1495.5762
571.3097534179688 0 4129.79 c Water loss 5
572.3047485351562 0 10824.113 c Ammonia loss 5
573.3082885742188 0 3307.5474
574.35546875 0 1164.0223
577.26220703125 0 3126.391
578.2677001953125 0 1073.9984
578.3284912109375 0 3801.802
582.3123168945312 0 1155.008 y 6
586.3204345703125 0 1826.5876
588.3263549804688 0 32975.22
589.330810546875 0 143752.19 c 5
590.3336181640625 0 43271.945
591.336181640625 0 6465.1587
595.2708740234375 0 2149.261
603.3114624023438 0 1246.6003
605.307373046875 0 2265.8247 z Water loss 5
607.3089599609375 0 1849.2332
618.286865234375 0 1176.6462
619.2733154296875 0 3467.5298
620.2719116210938 0 1253.4635
621.3245849609375 0 3657.3835 y Water loss 5
636.3018188476562 0 2292.4858
637.2987670898438 0 1781.516
638.32275390625 0 6815.8477
639.3314208984375 0 3540.1616 y 5
641.326416015625 0 1674.2914
649.3663940429688 0 2378.8801
650.3635864257812 0 1312.599
655.3402099609375 0 2036.9913
657.393310546875 0 4755.111
658.40087890625 0 25599.766
659.4075317382812 0 27652.91
660.4116821289062 0 6665.1494
661.4126586914062 0 1656.0221
667.37841796875 0 7072.1323
668.3660888671875 0 5049.127
669.3692016601562 0 4414.031
670.3770141601562 0 2777.4814
684.3989868164062 0 2500.0208 c Water loss 6
685.3883056640625 0 55868.93 c Ammonia loss 6
686.39111328125 0 19515.295
687.3984375 0 8278.002
688.30224609375 0 1347.6791
688.4109497070312 0 1239.2275
690.34130859375 0 2637.1843
692.3397216796875 0 3708.287 z Water loss 4
694.3436889648438 0 2027.41
701.4071655273438 0 34017.324
702.4146118164062 0 119591.95 c 6
703.417236328125 0 42757.215
704.4208984375 0 7868.4927
705.3213500976562 0 7120.553
706.3203735351562 0 3859.9253
708.3434448242188 0 3814.2312 y Water loss 4
709.3487548828125 0 1990.0348
710.3494873046875 0 9455.361 z 4
711.3553466796875 0 5583.5093
717.3446655273438 0 1147.3981
720.3919067382812 0 1636.8862
723.3325805664062 0 3972.368
724.33251953125 0 1956.0966
725.359619140625 0 19492.963
726.3646240234375 0 17800.463 y 4
727.3718872070312 0 4615.867
733.3651123046875 0 2507.6902
734.3699951171875 0 3377.7278
735.3763427734375 0 1199.2504
737.3881225585938 0 1823.5048
738.39111328125 0 1520.8712
749.3843994140625 0 2145.2798
750.3892822265625 0 1388.3668
751.3991088867188 0 3607.1338
752.4044189453125 0 1218.9875
754.4095458984375 0 8739.902
755.4118041992188 0 4976.993
756.4148559570312 0 2242.3003
772.4223022460938 0 1324.9825
779.3701782226562 0 6650.966 z Water loss 3
780.3739013671875 0 2498.5513 w 3
781.3729248046875 0 1052.3058
789.413818359375 0 1750.6295
790.4132080078125 0 1825.3456
791.4114990234375 0 1294.0474
794.386962890625 0 1823.3326
795.3775634765625 0 5496.3896 y Water loss 3
796.3815307617188 0 2768.8738
797.3805541992188 0 9852.259 z 3
798.3867797851562 0 4290.9824
802.3845825195312 0 4359.3833
803.3876342773438 0 1444.9623
805.3983154296875 0 1240.6663
807.4313354492188 0 1699.9459
808.4124755859375 0 1523.2666
809.4246215820312 0 2372.9949
812.392578125 0 16546.885
813.397705078125 0 21392.822 y 3
814.40185546875 0 8444.724
815.4067993164062 0 1313.0867
818.4058227539062 0 3737.4148
819.4056396484375 0 2050.835
820.39599609375 0 10445.446
821.3998413085938 0 5137.78
821.4655151367188 0 4530.599
822.4014892578125 0 2653.2202
822.4713745117188 0 2338.2405
823.4783935546875 0 1940.3978
826.4550170898438 0 1046.2987
830.4427490234375 0 6026.9873
831.437744140625 0 5890.7188
832.4341430664062 0 6579.0107
833.4393310546875 0 3978.437
834.444091796875 0 1500.2295
836.4183349609375 0 1369.7863
838.4371337890625 0 2073.3982
839.432373046875 0 1890.4429
846.4583740234375 0 4349.3384
847.452392578125 0 3888.477 c Water loss 7
848.452392578125 0 45292.89 c Ammonia loss 7
849.454833984375 0 20800.504
850.460693359375 0 7625.911
851.4691162109375 0 1898.6866
864.469970703125 0 12309.644
865.478271484375 0 121403.805 c 7
865.9105224609375 0 1383.432
866.4810180664062 0 60252.258
867.4091186523438 0 1187.7125 w 2
867.48779296875 0 20759.219
868.4923706054688 0 5820.1475
869.3772583007812 0 3424.3123
870.3807373046875 0 1825.5503
889.47998046875 0 1566.7595
890.4232788085938 0 3354.0781
891.415283203125 0 2241.858
899.4611206054688 0 5903.437
900.458251953125 0 3982.008
901.4613647460938 0 1307.2615
906.4207153320312 0 2353.2742
907.4283447265625 0 21243.295 z Water loss 2
908.4310302734375 0 7040.4272
909.4265747070312 0 1936.9631
909.5142822265625 0 1607.7566
910.5120849609375 0 1551.705
915.468994140625 0 12027.912
916.4741821289062 0 9272.8
917.4737548828125 0 52076.445
918.4749145507812 0 27337.033
919.4747314453125 0 10321.028
920.4718627929688 0 3672.1233
923.4451904296875 0 3805.0247 y Water loss 2
924.4506225585938 0 2943.4197
925.4395751953125 0 37533.664 z 2
926.4425048828125 0 19246.176
927.4451293945312 0 5133.6655
933.4808959960938 0 33980.76
934.4845581054688 0 19574.63 c Water loss 8
935.4843139648438 0 40473.75 c Ammonia loss 8
936.4873657226562 0 21166.062
937.4880981445312 0 4899.2183
941.4579467773438 0 10081.205 y 2
942.4618530273438 0 4508.043
951.5074462890625 0 4302.172
952.510498046875 0 212124.64 c 8
953.5133666992188 0 112049.31
954.51611328125 0 31912.453
955.5198364257812 0 4054.6035
976.4880981445312 0 2579.5623
977.4744873046875 0 1969.0292
982.4612426757812 0 5208.6084
983.466064453125 0 1401.1405
995.4691772460938 0 14141.058 w 1
996.4727172851562 0 5719.9575
997.4732666015625 0 2917.6929
1003.572509765625 0 3451.041
1004.5687255859375 0 2528.7502
1005.5673217773438 0 1408.1262
1014.5386962890625 0 27091.7
1015.5419921875 0 15738.091
1016.5454711914062 0 5413.335
1020.5135498046875 0 15815.657 z Water loss 1
1021.5010375976562 0 3690.0383 z Ammonia loss 1
1021.5800170898438 0 11521.369
1022.5852661132812 0 7258.1323
1023.590087890625 0 2286.7466
1030.557861328125 0 7829.0947
1031.55908203125 0 5683.903
1032.55029296875 0 34558.707
1033.552734375 0 18065.125
1034.553955078125 0 5596.087
1036.5155029296875 0 1453.9698 y Water loss 1
1038.52392578125 0 125507.19 z 1
1039.5267333984375 0 71927.71
1040.5296630859375 0 26759.193
1041.5316162109375 0 2713.7715
1047.5697021484375 0 2032.6989 c Water loss 9
1048.5692138671875 0 22089.256 c Ammonia loss 9
1049.5731201171875 0 13621.052
1050.5780029296875 0 5160.3657
1053.5345458984375 0 7496.843
1054.536376953125 0 4987.495 y 1
1055.5474853515625 0 1266.7272
1063.5765380859375 0 2286.27
1064.583740234375 0 1809.1565
1065.5950927734375 0 271853 c 9
1066.5975341796875 0 163845.05
1067.6007080078125 0 55660.53
1068.602294921875 0 8631.539
1070.5054931640625 0 1570.2915
1081.53466796875 0 20069.412
1082.5369873046875 0 11303.951
1083.548828125 0 4479.2544
1091.5865478515625 0 1408.6293
1092.5987548828125 0 2824.5867
1098.554931640625 0 9502.343
1099.55810546875 0 6482.7173
1108.60791015625 0 4540.472
1109.599853515625 0 18737.732
1110.6044921875 0 11844.898
1111.595703125 0 4319.6685
1112.576416015625 0 1207.404
1119.580810546875 0 6023.1255
1120.579833984375 0 5258.786
1126.62353515625 0 10280.699
1127.6268310546875 0 5504.2964
1128.632080078125 0 3154.3904
1136.6026611328125 0 7875.3564
1137.59326171875 0 37009.555
1138.5943603515625 0 22655.941
1139.595947265625 0 10538.591
1140.5980224609375 0 1721.1466
1152.5966796875 0 6344.796
1153.6099853515625 0 97241.17
1154.6171875 0 318019.22
1155.620361328125 0 181601.45
1156.623291015625 0 62574.215
1157.62158203125 0 8795.76
1429.6817626953125 0 1117.2146
2763.035888671875 0 1371.7291

Spectrum Details

|  |  |
| --- | --- |
| Matched peaks? Matched peaks The total absolute number of peaks matched. Additionally in brackets the total fraction of peaks matched and the total number of peaks is shown. | 56 (15.18% of 369) |
| FDR? FDR The false discovery rate estimated for this peptide. It is calculated by matching all theoretical fragments with a non-integer shift with the raw peaks for this spectrum. This is done with 40 different shifts. The resulting percentage is the average number of annotated peaks over the number of annotated peaks with the correct spectrum. | 0.26% |
| Satellite FDR? Satellite FDR See the FDR for details on its calculation. This satellite ion specific FDR only contains the satellite ions (d/w) for I/L/J positions. | 9.52% |
| PSM Score? PSM Score The PSM Score as given by Hecklib to this annotated spectrum. It is shown with three significant figures. | 816 |

#### Spectrum 8039? Spectrum 8039 The raw spectrum of this peptide as annotated by Hecklib. The fragments are coloured according to ion type (see legend). Any peaks with a star '\*' as text can be hovered over to see the full details, first the ion type second the mass shift type. By hovering over the amino acids in the peptide or ions in the legend the corresponding peaks are highlighted. By toggling the 'Unassigned' label you can turn the background (unassigned) peaks on or off in the plot. By updating the slider in the Ion legend you can update the spectrum to only show the top X% of the peaks with labels. The top X% means any peak that is within X% of the highest intensity. By dragging in the spectrum you can zoom in to a specific part of the spectrum and use 'Zoom Out' to get back to the original zoom level. The annotation of the spectrum is based on the given sequence in the peptides file and is done with different software so inconsistencies are likely. The peaks are annotated based on the given sequence, with 20 ppm tolerance.

Copy Data

##### Spectrum 8039 (TSV)

###### Preview

```
Loading example...
```

*Click on the button to copy the data to your clipboard.*

Mz MinMz MaxIntensity Max

WidthHeightPeptide font sizePeptide stroke widthSpectrum font sizeSpectrum stroke widthCompact peptide

Ion legend

wxyz

abcd

OtherUnassignedIon^Charge^_Position_Show for top:%

VLQSSGJYSJS

08.90e+31.78e+42.67e+43.56e+4

Zoom Out

y^+1^_2_y^+1^_2_y^+1^_3_y^+1^_3_c^+1^_3_c^+1^_3_c^+1^_4_y^+1^_4_c^+2^_9_y^+1^_4_c^+1^_5_c^+1^_6_c^+1^_6_y^+1^_6_c^+1^_7_c^+1^_7_z^+1^_7_y^+1^_7_z^+1^_8_z^+1^_8_y^+1^_8_c^+1^_8_c^+1^_8_z^+1^_9_z^+1^_9_z^+1^_9_c^+1^_9_c^+1^_9_y^+1^_9_c^+1^_9_w^+1^_10_z^+1^_10_z^+1^_10_c^+1^_10_y^+1^_10_c^+1^_10_

049599014851980

Fragment Matches Table

Show background peaks

| Position | Ion type | Intensity | mz Theoretical | mz Error (Th) | mz Error (ppm) | Charge | Series Number |
| --- | --- | --- | --- | --- | --- | --- | --- |
| - | - | 350.6 | 120.7 | - | - | 0 | - |
| - | - | 339.7 | 129.1 | - | - | 0 | - |
| - | - | 548.9 | 133.1 | - | - | 0 | - |
| - | - | 2747 | 136.1 | - | - | 0 | - |
| - | - | 494.8 | 172.5 | - | - | 0 | - |
| - | - | 1116 | 173.1 | - | - | 0 | - |
| - | - | 568.2 | 175.1 | - | - | 0 | - |
| - | - | 1605 | 185.2 | - | - | 0 | - |
| - | - | 475.4 | 198.9 | - | - | 0 | - |
| - | - | 474.1 | 201.1 | - | - | 0 | - |
| 10 | y | 2393 | 201.1 | 0.0001511 | 0.7511 | +1 | 2 |
| - | - | 1617 | 203.1 | - | - | 0 | - |
| - | - | 3071 | 213.2 | - | - | 0 | - |
| - | - | 906.1 | 215.1 | - | - | 0 | - |
| - | - | 553.4 | 218.2 | - | - | 0 | - |
| 10 | y | 3527 | 219.1 | 9.967E-05 | 0.4549 | +1 | 2 |
| - | - | 786.5 | 221.1 | - | - | 0 | - |
| - | - | 814 | 233.1 | - | - | 0 | - |
| - | - | 744 | 239.1 | - | - | 0 | - |
| - | - | 553.2 | 245.4 | - | - | 0 | - |
| - | - | 935.6 | 249.1 | - | - | 0 | - |
| 9 | y | 1149 | 288.2 | 0.0002729 | 0.9471 | +1 | 3 |
| - | - | 1299 | 299.1 | - | - | 0 | - |
| 9 | y | 597.6 | 306.2 | 0.0007093 | 2.317 | +1 | 3 |
| - | - | 572.9 | 334.2 | - | - | 0 | - |
| - | - | 1825 | 336.2 | - | - | 0 | - |
| 3 | c | 1489 | 341.2 | 0.0002961 | 0.8677 | +1 | 3 |
| - | - | 549.2 | 345.2 | - | - | 0 | - |
| - | - | 903 | 346.2 | - | - | 0 | - |
| - | - | 5293 | 355.1 | - | - | 0 | - |
| - | - | 1168 | 356.1 | - | - | 0 | - |
| 3 | c | 1900 | 358.2 | 0.00013 | 0.3628 | +1 | 3 |
| - | - | 720.2 | 402.3 | - | - | 0 | - |
| - | - | 573.5 | 415.7 | - | - | 0 | - |
| - | - | 2419 | 444.3 | - | - | 0 | - |
| 4 | c | 6791 | 445.3 | 0.0001149 | 0.2581 | +1 | 4 |
| - | - | 1240 | 446.3 | - | - | 0 | - |
| - | - | 517.2 | 450.7 | - | - | 0 | - |
| - | - | 591.8 | 451.1 | - | - | 0 | - |
| 8 | y | 851.1 | 451.2 | 0.001397 | 3.097 | +1 | 4 |
| - | - | 671.5 | 453.7 | - | - | 0 | - |
| - | - | 1210 | 457.2 | - | - | 0 | - |
| - | - | 2800 | 459.2 | - | - | 0 | - |
| - | - | 1599 | 459.7 | - | - | 0 | - |
| 9 | c | 1634 | 468.2 | 9.667E-05 | 0.2065 | +2 | 9 |
| - | - | 968.9 | 468.7 | - | - | 0 | - |
| 8 | y | 640.8 | 469.2 | 0.000751 | 1.601 | +1 | 4 |
| - | - | 596.1 | 502.5 | - | - | 0 | - |
| - | - | 854.6 | 508.2 | - | - | 0 | - |
| - | - | 559 | 513.3 | - | - | 0 | - |
| - | - | 547 | 515.8 | - | - | 0 | - |
| - | - | 911.3 | 516.3 | - | - | 0 | - |
| - | - | 2432 | 531.3 | - | - | 0 | - |
| 5 | c | 5456 | 532.3 | 0.001015 | 1.908 | +1 | 5 |
| - | - | 2077 | 533.3 | - | - | 0 | - |
| - | - | 1276 | 541.3 | - | - | 0 | - |
| - | - | 3255 | 546.3 | - | - | 0 | - |
| - | - | 1620 | 559.3 | - | - | 0 | - |
| - | - | 622.2 | 560.3 | - | - | 0 | - |
| 6 | c | 1094 | 572.3 | 0.0005077 | 0.8871 | +1 | 6 |
| - | - | 707.7 | 576.9 | - | - | 0 | - |
| - | - | 891 | 577.3 | - | - | 0 | - |
| - | - | 1060 | 579.3 | - | - | 0 | - |
| - | - | 1883 | 585.3 | - | - | 0 | - |
| - | - | 3406 | 588.3 | - | - | 0 | - |
| 6 | c | 1.56E+04 | 589.3 | 1.821E-05 | 0.03089 | +1 | 6 |
| - | - | 5442 | 590.3 | - | - | 0 | - |
| - | - | 690.8 | 608.6 | - | - | 0 | - |
| - | - | 627.7 | 621.8 | - | - | 0 | - |
| - | - | 898 | 638.3 | - | - | 0 | - |
| 6 | y | 1092 | 639.3 | 0.0001428 | 0.2233 | +1 | 6 |
| - | - | 795.9 | 657.4 | - | - | 0 | - |
| - | - | 3038 | 658.4 | - | - | 0 | - |
| - | - | 2943 | 659.4 | - | - | 0 | - |
| - | - | 697.5 | 660.4 | - | - | 0 | - |
| - | - | 594 | 669.4 | - | - | 0 | - |
| 7 | c | 6239 | 685.4 | 0.0002672 | 0.3899 | +1 | 7 |
| - | - | 2266 | 686.4 | - | - | 0 | - |
| - | - | 935.7 | 687.4 | - | - | 0 | - |
| - | - | 874.5 | 698.4 | - | - | 0 | - |
| - | - | 4255 | 701.4 | - | - | 0 | - |
| 7 | c | 1.173E+04 | 702.4 | 0.001196 | 1.703 | +1 | 7 |
| - | - | 5378 | 703.4 | - | - | 0 | - |
| - | - | 797.3 | 704.4 | - | - | 0 | - |
| - | - | 924.1 | 705.3 | - | - | 0 | - |
| 5 | z | 953 | 710.3 | 0.001138 | 1.602 | +1 | 7 |
| - | - | 649.7 | 716.3 | - | - | 0 | - |
| - | - | 2479 | 725.4 | - | - | 0 | - |
| 5 | y | 2578 | 726.4 | 0.002223 | 3.06 | +1 | 7 |
| - | - | 665.7 | 727.4 | - | - | 0 | - |
| - | - | 606.8 | 732.4 | - | - | 0 | - |
| - | - | 1105 | 733.4 | - | - | 0 | - |
| - | - | 865.9 | 751.4 | - | - | 0 | - |
| - | - | 943.8 | 754.4 | - | - | 0 | - |
| - | - | 851.8 | 759.4 | - | - | 0 | - |
| - | - | 823 | 778.9 | - | - | 0 | - |
| 4 | z | 1610 | 779.4 | 0.001727 | 2.216 | +1 | 8 |
| - | - | 625.6 | 792.4 | - | - | 0 | - |
| - | - | 794.2 | 793.9 | - | - | 0 | - |
| 4 | z | 1331 | 797.4 | 0.0004512 | 0.5659 | +1 | 8 |
| - | - | 915 | 798.4 | - | - | 0 | - |
| - | - | 716.1 | 801.3 | - | - | 0 | - |
| - | - | 641.7 | 808.4 | - | - | 0 | - |
| - | - | 2180 | 812.4 | - | - | 0 | - |
| 4 | y | 2520 | 813.4 | 0.001536 | 1.889 | +1 | 8 |
| - | - | 665.9 | 814.4 | - | - | 0 | - |
| - | - | 1803 | 820.4 | - | - | 0 | - |
| - | - | 796.9 | 822.5 | - | - | 0 | - |
| - | - | 1097 | 826.5 | - | - | 0 | - |
| - | - | 747.7 | 829.9 | - | - | 0 | - |
| - | - | 1038 | 830.4 | - | - | 0 | - |
| - | - | 1196 | 830.9 | - | - | 0 | - |
| - | - | 1586 | 846.5 | - | - | 0 | - |
| 8 | c | 5512 | 848.5 | 0.0009035 | 1.065 | +1 | 8 |
| - | - | 3144 | 849.5 | - | - | 0 | - |
| - | - | 1770 | 864.5 | - | - | 0 | - |
| 8 | c | 1.245E+04 | 865.5 | 0.001109 | 1.282 | +1 | 8 |
| - | - | 915.7 | 865.9 | - | - | 0 | - |
| - | - | 7095 | 866.5 | - | - | 0 | - |
| - | - | 978.3 | 866.9 | - | - | 0 | - |
| - | - | 2282 | 867.5 | - | - | 0 | - |
| - | - | 847.5 | 868.5 | - | - | 0 | - |
| - | - | 681 | 869.4 | - | - | 0 | - |
| - | - | 1129 | 886.4 | - | - | 0 | - |
| - | - | 657.8 | 889.5 | - | - | 0 | - |
| - | - | 1034 | 890.5 | - | - | 0 | - |
| - | - | 851.1 | 902.5 | - | - | 0 | - |
| 3 | z | 2005 | 907.4 | 0.001955 | 2.155 | +1 | 9 |
| 3 | z | 922.7 | 908.4 | 0.01653 | 18.2 | +1 | 9 |
| - | - | 1697 | 915.5 | - | - | 0 | - |
| - | - | 1092 | 916.5 | - | - | 0 | - |
| - | - | 6404 | 917.5 | - | - | 0 | - |
| - | - | 3719 | 918.5 | - | - | 0 | - |
| - | - | 3145 | 919.5 | - | - | 0 | - |
| - | - | 951.2 | 920.5 | - | - | 0 | - |
| 3 | z | 5013 | 925.4 | 0.0003129 | 0.3381 | +1 | 9 |
| - | - | 2714 | 926.4 | - | - | 0 | - |
| - | - | 4004 | 933.5 | - | - | 0 | - |
| 9 | c | 2216 | 934.5 | 0.01336 | 14.29 | +1 | 9 |
| 9 | c | 5897 | 935.5 | 0.001401 | 1.497 | +1 | 9 |
| - | - | 2504 | 936.5 | - | - | 0 | - |
| 3 | y | 926.7 | 941.5 | 0.001593 | 1.692 | +1 | 9 |
| - | - | 779.7 | 942.5 | - | - | 0 | - |
| 9 | c | 2.374E+04 | 952.5 | 0.0005451 | 0.5723 | +1 | 9 |
| - | - | 1.321E+04 | 953.5 | - | - | 0 | - |
| - | - | 3968 | 954.5 | - | - | 0 | - |
| - | - | 671.5 | 955.5 | - | - | 0 | - |
| - | - | 1177 | 960.5 | - | - | 0 | - |
| - | - | 902.6 | 961.5 | - | - | 0 | - |
| - | - | 844.3 | 973.5 | - | - | 0 | - |
| - | - | 2164 | 975.5 | - | - | 0 | - |
| - | - | 1088 | 976.5 | - | - | 0 | - |
| - | - | 832.8 | 977.5 | - | - | 0 | - |
| 2 | w | 2484 | 995.5 | 0.0006107 | 0.6135 | +1 | 10 |
| - | - | 2641 | 1015 | - | - | 0 | - |
| - | - | 1853 | 1016 | - | - | 0 | - |
| 2 | z | 1501 | 1021 | 0.003439 | 3.37 | +1 | 10 |
| - | - | 951.2 | 1022 | - | - | 0 | - |
| - | - | 857.7 | 1024 | - | - | 0 | - |
| - | - | 1047 | 1031 | - | - | 0 | - |
| - | - | 3781 | 1033 | - | - | 0 | - |
| - | - | 2341 | 1034 | - | - | 0 | - |
| 2 | z | 1.584E+04 | 1039 | 0.0004536 | 0.4367 | +1 | 10 |
| - | - | 9128 | 1040 | - | - | 0 | - |
| - | - | 2428 | 1041 | - | - | 0 | - |
| 10 | c | 2331 | 1049 | 0.008744 | 8.339 | +1 | 10 |
| - | - | 1362 | 1050 | - | - | 0 | - |
| - | - | 929.8 | 1051 | - | - | 0 | - |
| 2 | y | 696.9 | 1055 | 0.002307 | 2.187 | +1 | 10 |
| 10 | c | 3.524E+04 | 1066 | 0.001479 | 1.388 | +1 | 10 |
| - | - | 1.809E+04 | 1067 | - | - | 0 | - |
| - | - | 7674 | 1068 | - | - | 0 | - |
| - | - | 1794 | 1069 | - | - | 0 | - |
| - | - | 1656 | 1082 | - | - | 0 | - |
| - | - | 920.8 | 1083 | - | - | 0 | - |
| - | - | 1370 | 1099 | - | - | 0 | - |
| - | - | 706.6 | 1101 | - | - | 0 | - |
| - | - | 1423 | 1105 | - | - | 0 | - |
| - | - | 890.8 | 1106 | - | - | 0 | - |
| - | - | 730.4 | 1107 | - | - | 0 | - |
| - | - | 2323 | 1110 | - | - | 0 | - |
| - | - | 1600 | 1111 | - | - | 0 | - |
| - | - | 881.3 | 1120 | - | - | 0 | - |
| - | - | 1211 | 1128 | - | - | 0 | - |
| - | - | 1781 | 1135 | - | - | 0 | - |
| - | - | 991.9 | 1136 | - | - | 0 | - |
| - | - | 758.7 | 1137 | - | - | 0 | - |
| - | - | 3205 | 1138 | - | - | 0 | - |
| - | - | 3147 | 1139 | - | - | 0 | - |
| - | - | 1956 | 1140 | - | - | 0 | - |
| - | - | 1841 | 1151 | - | - | 0 | - |
| - | - | 3775 | 1152 | - | - | 0 | - |
| - | - | 3055 | 1153 | - | - | 0 | - |
| - | - | 1.427E+04 | 1154 | - | - | 0 | - |
| - | - | 3.451E+04 | 1155 | - | - | 0 | - |
| - | - | 2.704E+04 | 1156 | - | - | 0 | - |
| - | - | 1.44E+04 | 1157 | - | - | 0 | - |
| - | - | 2520 | 1158 | - | - | 0 | - |
| - | - | 1396 | 1218 | - | - | 0 | - |
| - | - | 800.8 | 1219 | - | - | 0 | - |
| - | - | 753.6 | 1347 | - | - | 0 | - |
| - | - | 1584 | 1367 | - | - | 0 | - |
| - | - | 1829 | 1368 | - | - | 0 | - |
| - | - | 1207 | 1369 | - | - | 0 | - |
| - | - | 1805 | 1402 | - | - | 0 | - |
| - | - | 2180 | 1403 | - | - | 0 | - |
| - | - | 1450 | 1404 | - | - | 0 | - |
| - | - | 1343 | 1417 | - | - | 0 | - |
| - | - | 1887 | 1531 | - | - | 0 | - |
| - | - | 1858 | 1532 | - | - | 0 | - |
| - | - | 1217 | 1533 | - | - | 0 | - |
| - | - | 1839 | 1559 | - | - | 0 | - |
| - | - | 746.1 | 1560 | - | - | 0 | - |
| - | - | 902.1 | 1633 | - | - | 0 | - |
| - | - | 946.1 | 1634 | - | - | 0 | - |
| - | - | 894.6 | 1660 | - | - | 0 | - |
| - | - | 776.5 | 1688 | - | - | 0 | - |
| - | - | 719.6 | 1699 | - | - | 0 | - |
| - | - | 960.6 | 1711 | - | - | 0 | - |
| - | - | 969.4 | 1712 | - | - | 0 | - |
| - | - | 1402 | 1713 | - | - | 0 | - |
| - | - | 920 | 1714 | - | - | 0 | - |
| - | - | 1571 | 1716 | - | - | 0 | - |
| - | - | 1591 | 1717 | - | - | 0 | - |
| - | - | 849 | 1718 | - | - | 0 | - |
| - | - | 1226 | 1730 | - | - | 0 | - |
| - | - | 945.3 | 1731 | - | - | 0 | - |
| - | - | 952.7 | 1732 | - | - | 0 | - |
| - | - | 2788 | 1733 | - | - | 0 | - |
| - | - | 1737 | 1734 | - | - | 0 | - |
| - | - | 2126 | 1735 | - | - | 0 | - |
| - | - | 845.7 | 1736 | - | - | 0 | - |
| - | - | 802.7 | 1748 | - | - | 0 | - |
| - | - | 722.2 | 1960 | - | - | 0 | - |

m/z Charge Intensity FragmentType MassShift Position
120.68168640136719 0 350.5567
129.05490112304688 0 339.6928
133.08612060546875 0 548.9466
136.07583618164062 0 2747.2615
172.46243286132812 0 494.76468
173.12840270996094 0 1115.9519
175.08656311035156 0 568.1838
185.1651153564453 0 1604.7693
198.89085388183594 0 475.37125
201.1138458251953 0 474.09348
201.12351989746094 0 2392.724 y Water loss 9
203.10272216796875 0 1616.6305
213.15988159179688 0 3070.6802
215.13912963867188 0 906.0621
218.15057373046875 0 553.41394
219.134033203125 0 3527.2236 y 9
221.0845184326172 0 786.4574
233.09207153320312 0 813.9603
239.09535217285156 0 743.959
245.35455322265625 0 553.176
249.12245178222656 0 935.5688
288.1556701660156 0 1149.312 y Water loss 8
299.06158447265625 0 1299.0518
306.1652526855469 0 597.58203 y 8
334.178466796875 0 572.93915
336.15496826171875 0 1824.6548
341.2186279296875 0 1489.0676 c Ammonia loss 2
345.1758728027344 0 549.2009
346.1774597167969 0 902.9896
355.06982421875 0 5292.5137
356.07086181640625 0 1168.0311
358.2447509765625 0 1899.6152 c 2
402.27008056640625 0 720.17267
415.7211608886719 0 573.4544
444.2697448730469 0 2419.011
445.27679443359375 0 6790.7046 c 3
446.2795715332031 0 1239.8567
450.7281494140625 0 517.2
451.0502624511719 0 591.8409
451.2201232910156 0 851.0537 y Water loss 7
453.7261047363281 0 671.51337
457.2410888671875 0 1209.5939
459.240234375 0 2799.5774
459.74163818359375 0 1599.2694
468.24517822265625 0 1634.2808 c Ammonia loss 8
468.7473449707031 0 968.8694
469.23004150390625 0 640.82355 y 7
502.5058288574219 0 596.0994
508.2405090332031 0 854.5728
513.28955078125 0 558.9521
515.7833251953125 0 547.0081
516.2813110351562 0 911.3324
531.2999267578125 0 2431.604
532.3079223632812 0 5456.0713 c 4
533.3116455078125 0 2076.6707
541.2872314453125 0 1276.2826
546.3238525390625 0 3255.019
559.2965698242188 0 1619.8119
560.3051147460938 0 622.1531
572.3033447265625 0 1094.4482 c Ammonia loss 5
576.8734741210938 0 707.7064
577.297607421875 0 891.04297
579.306396484375 0 1060.2521
585.2984619140625 0 1882.6814
588.3257446289062 0 3405.964
589.3303833007812 0 15601.164 c 5
590.33349609375 0 5441.6733
608.5692749023438 0 690.84344
621.8132934570312 0 627.68665
638.3153686523438 0 898.0211
639.3349609375 0 1091.789 y 5
657.3912353515625 0 795.9147
658.4003295898438 0 3037.981
659.4075317382812 0 2943.0295
660.408447265625 0 697.47705
669.3726806640625 0 594.04706
685.38818359375 0 6238.8994 c Ammonia loss 6
686.390869140625 0 2266.1243
687.4036865234375 0 935.7061
698.3801879882812 0 874.479
701.40576171875 0 4255.173
702.4132690429688 0 11732.47 c 6
703.4154052734375 0 5378.4053
704.4160766601562 0 797.3309
705.3220825195312 0 924.09
710.3469848632812 0 953.0333 z 4
716.346435546875 0 649.73376
725.3582763671875 0 2478.5063
726.3646240234375 0 2577.6301 y 4
727.3781127929688 0 665.6781
732.376953125 0 606.8415
733.3759765625 0 1105.312
751.3946533203125 0 865.8726
754.4115600585938 0 943.8498
759.4373779296875 0 851.8453
778.8806762695312 0 822.9538
779.3678588867188 0 1610.1012 z Water loss 3
792.3618774414062 0 625.6316
793.8600463867188 0 794.1885
797.3796997070312 0 1330.6648 z 3
798.3812866210938 0 914.9677
801.3494873046875 0 716.0717
808.416015625 0 641.74695
812.3916625976562 0 2180.0234
813.3973388671875 0 2519.8928 y 3
814.3988647460938 0 665.8703
820.3972778320312 0 1802.6737
822.4713745117188 0 796.89575
826.4580078125 0 1097.0349
829.9376831054688 0 747.67883
830.4349975585938 0 1037.8104
830.9434204101562 0 1195.634
846.466064453125 0 1586.4161
848.4521484375 0 5512.2285 c Ammonia loss 7
849.4527587890625 0 3143.7695
864.4685668945312 0 1769.8927
865.4766845703125 0 12450.621 c 7
865.944580078125 0 915.74097
866.4762573242188 0 7094.8164
866.9487915039062 0 978.30615
867.4841918945312 0 2281.7766
868.4857177734375 0 847.5187
869.3728637695312 0 681.0208
886.4375 0 1128.9264
889.4633178710938 0 657.83026
890.464111328125 0 1034.251
902.4542236328125 0 851.06616
907.4262084960938 0 2005.4885 z Water loss 2
908.4287109375 0 922.699 z Ammonia loss 2
915.469970703125 0 1697.2269
916.4769897460938 0 1092.0524
917.4722900390625 0 6404.433
918.47265625 0 3719.4773
919.4813232421875 0 3144.764
920.4891357421875 0 951.2046
925.4384155273438 0 5013.401 z 2
926.439697265625 0 2714.196
933.4790649414062 0 4003.6824
934.4859008789062 0 2216.4414 c Water loss 8
935.4818725585938 0 5896.7017 c Ammonia loss 8
936.48779296875 0 2504.3105
941.4590454101562 0 926.6907 y 2
942.4520874023438 0 779.72394
952.50927734375 0 23737.803 c 8
953.5121459960938 0 13213.567
954.5148315429688 0 3968.048
955.5155639648438 0 671.46594
960.495361328125 0 1176.5569
961.4852905273438 0 902.6249
973.46533203125 0 844.3137
975.513671875 0 2163.548
976.5062866210938 0 1088.1699
977.515380859375 0 832.81854
995.4686279296875 0 2483.7778 w 1
1014.5377197265625 0 2640.9192
1015.540283203125 0 1852.5249
1020.5087890625 0 1500.7716 z Water loss 1
1021.5850219726562 0 951.16815
1023.557373046875 0 857.6756
1030.555419921875 0 1046.6855
1032.548583984375 0 3780.8357
1033.5517578125 0 2341.2078
1038.5223388671875 0 15838.671 z 1
1039.5255126953125 0 9128.025
1040.5277099609375 0 2428.2415
1048.55859375 0 2330.6877 c Ammonia loss 9
1049.5760498046875 0 1361.9918
1050.5718994140625 0 929.8419
1054.5438232421875 0 696.86 y 1
1065.5924072265625 0 35235.426 c 9
1066.59619140625 0 18089.873
1067.5968017578125 0 7674.3604
1068.5987548828125 0 1794.2017
1081.53173828125 0 1656.341
1082.55126953125 0 920.75916
1098.5499267578125 0 1370.1017
1100.5616455078125 0 706.58417
1104.5550537109375 0 1423.3452
1105.552001953125 0 890.8021
1106.5548095703125 0 730.37274
1109.5992431640625 0 2322.9822
1110.59716796875 0 1600.1991
1119.5806884765625 0 881.3392
1127.6285400390625 0 1210.6694
1134.5540771484375 0 1780.8142
1135.5611572265625 0 991.90424
1136.5699462890625 0 758.73254
1137.5888671875 0 3204.976
1138.5914306640625 0 3146.8057
1139.5833740234375 0 1956.0024
1150.5701904296875 0 1840.719
1151.5738525390625 0 3775.068
1152.583984375 0 3054.5708
1153.6063232421875 0 14274.261
1154.6146240234375 0 34506.605
1155.6180419921875 0 27038.812
1156.6207275390625 0 14399.142
1157.6123046875 0 2520.2393
1217.6424560546875 0 1396.4348
1218.6400146484375 0 800.79736
1346.700927734375 0 753.57324
1366.669677734375 0 1584.029
1367.6732177734375 0 1828.8116
1368.67333984375 0 1207.4496
1401.6717529296875 0 1805.1902
1402.677490234375 0 2179.7952
1403.680419921875 0 1450.3604
1416.7403564453125 0 1343.3142
1530.777099609375 0 1886.6018
1531.7742919921875 0 1858.3547
1532.7724609375 0 1216.8591
1558.75439453125 0 1839.3993
1559.788818359375 0 746.1235
1632.6417236328125 0 902.0751
1633.650146484375 0 946.11334
1659.863525390625 0 894.6015
1687.8865966796875 0 776.4715
1698.77294921875 0 719.597
1710.748046875 0 960.5525
1711.7596435546875 0 969.39386
1712.7396240234375 0 1402.1693
1713.7257080078125 0 920.0271
1715.87060546875 0 1570.6018
1716.8980712890625 0 1590.7637
1717.8748779296875 0 849.02844
1729.8033447265625 0 1226.1865
1730.78955078125 0 945.32916
1731.8507080078125 0 952.74384
1732.8946533203125 0 2788.362
1733.90087890625 0 1736.6963
1734.9068603515625 0 2126.4434
1735.9169921875 0 845.6769
1748.46142578125 0 802.6549
1960.3363037109375 0 722.2289

Spectrum Details

|  |  |
| --- | --- |
| Matched peaks? Matched peaks The total absolute number of peaks matched. Additionally in brackets the total fraction of peaks matched and the total number of peaks is shown. | 36 (15.38% of 234) |
| FDR? FDR The false discovery rate estimated for this peptide. It is calculated by matching all theoretical fragments with a non-integer shift with the raw peaks for this spectrum. This is done with 40 different shifts. The resulting percentage is the average number of annotated peaks over the number of annotated peaks with the correct spectrum. | 0.46% |
| Satellite FDR? Satellite FDR See the FDR for details on its calculation. This satellite ion specific FDR only contains the satellite ions (d/w) for I/L/J positions. | 2.38% |
| PSM Score? PSM Score The PSM Score as given by Hecklib to this annotated spectrum. It is shown with three significant figures. | 487 |

#### Reverse Lookup? Reverse Lookup All places where this read could be placed.

| Group | Segment | Template | Template Part | Read Part | Score | Unique |
| --- | --- | --- | --- | --- | --- | --- |
| Homo sapiens Heavy Chain | IGHC | IGHG1 | [55..66] | [0..11] | 88 | False |
| Homo sapiens Heavy Chain | IGHC | IGHG3 | [55..66] | [0..11] | 88 | False |
| Homo sapiens Heavy Chain | IGHC | IGHG2 | [55..66] | [0..11] | 88 | False |
| Homo sapiens Heavy Chain | IGHC | IGHG4 | [55..66] | [0..11] | 88 | False |

| Recombined | Template Part | Read Part | Score | Unique |
| --- | --- | --- | --- | --- |
| REC-0-1 | [177..188] | [0..11] | 88 | True |

#### Meta Information from Multiple reads

##### Number of combined reads

2

##### Intensity

0.7514

##### TotalArea

1.803E+08

##### Changes to the peptide sequence

VLQSSGJYSJS

L→JNo support for either Leucine or Isoleucine based on side chain ions (Position: 10)

L→JNo support for either Leucine or Isoleucine based on side chain ions (Position: 7)

#### Positional Score

Copy Data

##### Positional Score (TSV)

###### Preview

```
Loading example...
```

*Click on the button to copy the data to your clipboard.*

00012345678910

Label Value
"0" 0
"1" 0
"2" 0
"3" 0
"4" 0
"5" 0
"6" 0
"7" 0
"8" 0
"9" 0
"10" 0

#### Meta Information from PEAKS

##### Scan Identifier

F1:7916

##### Original sequence

V

L

Q

S

S

G

L

Y

S

L

S

##### Posttranslational Modifications

##### Source File

D:\separate\_stitch\_analyses\xle-disambiguation\raw\20210323\_F1\_UM1\_Peng0013\_SA\_F59\_ingel\_3ug\_ELA.raw

##### Fraction

1

##### Scan Feature

F1:7458

##### De Novo Score

99

##### ConfidenceScore

99

### m/z

577.3096

##### Mass

1152.6025

##### Charge

2

##### Retention Time

42.88

##### Predicted Retention Time

-

##### Area

9.016E+07

##### Parts Per Million

1.8

##### Fragmentation mode

ETHCD

##### Originating file

01 D:\separate\_stitch\_analyses\xle-disambiguation\20210325\_F59\_3ug\_DENOVO\_12.csv

#### Meta Information from PEAKS

##### Scan Identifier

F1:8039

##### Original sequence

V

L

Q

S

S

G

L

Y

S

L

S

##### Posttranslational Modifications

##### Source File

D:\separate\_stitch\_analyses\xle-disambiguation\raw\20210323\_F1\_UM1\_Peng0013\_SA\_F59\_ingel\_3ug\_ELA.raw

##### Fraction

1

##### Scan Feature

F1:7458

##### De Novo Score

98

##### ConfidenceScore

98

### m/z

577.3096

##### Mass

1152.6025

##### Charge

2

##### Retention Time

42.88

##### Predicted Retention Time

-

##### Area

9.016E+07

##### Parts Per Million

1.8

##### Fragmentation mode

ETHCD

##### Originating file

01 D:\separate\_stitch\_analyses\xle-disambiguation\20210325\_F59\_3ug\_DENOVO\_12.csv
