## Supplementary material for "A handle on mass coincidence errors in *de novo* sequencing of antibodies by bottom-up proteomics": Combined_044.html

Details Combined\_044 | Stitch OverviewUndefined

### Read Combined\_044

#### Sequence (length=11)

SFVVFGGGTKJ

#### Spectrum 9569? Spectrum 9569 The raw spectrum of this peptide as annotated by Hecklib. The fragments are coloured according to ion type (see legend). Any peaks with a star '\*' as text can be hovered over to see the full details, first the ion type second the mass shift type. By hovering over the amino acids in the peptide or ions in the legend the corresponding peaks are highlighted. By toggling the 'Unassigned' label you can turn the background (unassigned) peaks on or off in the plot. By updating the slider in the Ion legend you can update the spectrum to only show the top X% of the peaks with labels. The top X% means any peak that is within X% of the highest intensity. By dragging in the spectrum you can zoom in to a specific part of the spectrum and use 'Zoom Out' to get back to the original zoom level. The annotation of the spectrum is based on the given sequence in the peptides file and is done with different software so inconsistencies are likely. The peaks are annotated based on the given sequence, with 20 ppm tolerance.

Copy Data

##### Spectrum 9569 (TSV)

###### Preview

```
Loading example...
```

*Click on the button to copy the data to your clipboard.*

Mz MinMz MaxIntensity Max

WidthHeightPeptide font sizePeptide stroke widthSpectrum font sizeSpectrum stroke widthCompact peptide

Ion legend

wxyz

abcd

OtherUnassignedIon^Charge^_Position_Show for top:%

SFVVFGGGTKJ

01.28e+52.57e+53.85e+55.13e+5

Zoom Out

y^+2^_2_y^+1^_1_a^+1^_2_a^+1^_2_b^+2^_4_b^+1^_2_b^+1^_2_y^+1^_2_y^+1^_2_a^+1^_3_a^+1^_3_b^+1^_3_b^+1^_3_y^+2^_7_y^+1^_3_y^+1^_3_y^+1^_4_b^+1^_4_y^+1^_4_b^+1^_4_y^+2^_9_y^+1^_5_y^+1^_5_b^+2^_10_b^+2^_10_b^+2^_10_y^+2^_10_y^+1^_6_y^+1^_6_\*\*\*b^+1^_5_b^+1^_5_b^+1^_6_b^+1^_6_y^+1^_7_b^+1^_7_y^+1^_7_b^+1^_7_b^+1^_8_y^+1^_8_y^+1^_8_b^+1^_9_b^+1^_9_y^+1^_9_y^+1^_9_b^+1^_10_b^+1^_10_b^+1^_10_y^+1^_10_

0837167525123349

Fragment Matches Table

Show background peaks

| Position | Ion type | Intensity | mz Theoretical | mz Error (Th) | mz Error (ppm) | Charge | Series Number |
| --- | --- | --- | --- | --- | --- | --- | --- |
| - | - | 1.053E+05 | 120.1 | - | - | 0 | - |
| - | - | 854.9 | 121.1 | - | - | 0 | - |
| - | - | 8117 | 121.1 | - | - | 0 | - |
| 10 | y | 505.7 | 122.1 | 0.0008003 | 6.555 | +2 | 2 |
| - | - | 402.7 | 124 | - | - | 0 | - |
| - | - | 562.5 | 125.1 | - | - | 0 | - |
| - | - | 1547 | 127.1 | - | - | 0 | - |
| - | - | 432.9 | 127.1 | - | - | 0 | - |
| - | - | 1968 | 128.1 | - | - | 0 | - |
| - | - | 723.3 | 129.1 | - | - | 0 | - |
| - | - | 1.441E+05 | 129.1 | - | - | 0 | - |
| - | - | 576.9 | 130.1 | - | - | 0 | - |
| - | - | 1263 | 130.1 | - | - | 0 | - |
| - | - | 1066 | 130.1 | - | - | 0 | - |
| - | - | 8666 | 130.1 | - | - | 0 | - |
| - | - | 674.6 | 131 | - | - | 0 | - |
| - | - | 5153 | 131.1 | - | - | 0 | - |
| - | - | 559.8 | 132.1 | - | - | 0 | - |
| - | - | 1791 | 132.1 | - | - | 0 | - |
| 11 | y | 1.409E+04 | 132.1 | 0.0003745 | 2.835 | +1 | 1 |
| - | - | 1202 | 133.1 | - | - | 0 | - |
| - | - | 1228 | 134.1 | - | - | 0 | - |
| - | - | 2232 | 136.1 | - | - | 0 | - |
| - | - | 864 | 139.1 | - | - | 0 | - |
| - | - | 1124 | 140.1 | - | - | 0 | - |
| - | - | 2658 | 141.1 | - | - | 0 | - |
| - | - | 831.4 | 141.1 | - | - | 0 | - |
| - | - | 365 | 141.9 | - | - | 0 | - |
| - | - | 436.9 | 142.1 | - | - | 0 | - |
| - | - | 438.6 | 143.1 | - | - | 0 | - |
| - | - | 1.087E+04 | 144.1 | - | - | 0 | - |
| - | - | 937.2 | 145.1 | - | - | 0 | - |
| - | - | 507.1 | 146.1 | - | - | 0 | - |
| - | - | 6565 | 147.1 | - | - | 0 | - |
| - | - | 557.4 | 148.1 | - | - | 0 | - |
| - | - | 405.5 | 152.5 | - | - | 0 | - |
| - | - | 751.1 | 153.1 | - | - | 0 | - |
| - | - | 433.8 | 153.1 | - | - | 0 | - |
| - | - | 845.9 | 154.1 | - | - | 0 | - |
| - | - | 1678 | 155.1 | - | - | 0 | - |
| - | - | 2805 | 155.1 | - | - | 0 | - |
| - | - | 642.6 | 156.1 | - | - | 0 | - |
| - | - | 477.2 | 157.1 | - | - | 0 | - |
| - | - | 437.1 | 157.5 | - | - | 0 | - |
| - | - | 986.7 | 158.1 | - | - | 0 | - |
| - | - | 4357 | 159.1 | - | - | 0 | - |
| - | - | 951.6 | 160.1 | - | - | 0 | - |
| - | - | 3.115E+04 | 162.1 | - | - | 0 | - |
| - | - | 2947 | 163.1 | - | - | 0 | - |
| - | - | 629.5 | 166.1 | - | - | 0 | - |
| - | - | 717.4 | 167.1 | - | - | 0 | - |
| - | - | 644.2 | 167.1 | - | - | 0 | - |
| - | - | 480.9 | 167.8 | - | - | 0 | - |
| - | - | 689.3 | 169.1 | - | - | 0 | - |
| - | - | 1082 | 171.1 | - | - | 0 | - |
| - | - | 4.672E+04 | 171.1 | - | - | 0 | - |
| - | - | 5458 | 172.1 | - | - | 0 | - |
| - | - | 4447 | 172.2 | - | - | 0 | - |
| - | - | 464 | 172.4 | - | - | 0 | - |
| - | - | 2859 | 173.4 | - | - | 0 | - |
| - | - | 745.1 | 174.1 | - | - | 0 | - |
| - | - | 485 | 174.5 | - | - | 0 | - |
| - | - | 5095 | 176.1 | - | - | 0 | - |
| - | - | 6289 | 177.1 | - | - | 0 | - |
| - | - | 833.9 | 177.1 | - | - | 0 | - |
| - | - | 493.4 | 181.1 | - | - | 0 | - |
| - | - | 467.7 | 183.1 | - | - | 0 | - |
| - | - | 486.1 | 184.1 | - | - | 0 | - |
| - | - | 1684 | 185.1 | - | - | 0 | - |
| - | - | 2253 | 186.1 | - | - | 0 | - |
| - | - | 1179 | 187.1 | - | - | 0 | - |
| - | - | 1323 | 188.1 | - | - | 0 | - |
| 2 | a | 1503 | 189.1 | 0.0003718 | 1.966 | +1 | 2 |
| - | - | 798.7 | 190.1 | - | - | 0 | - |
| - | - | 1902 | 195.1 | - | - | 0 | - |
| - | - | 687.3 | 196.1 | - | - | 0 | - |
| - | - | 465.9 | 197.1 | - | - | 0 | - |
| - | - | 1383 | 197.2 | - | - | 0 | - |
| - | - | 4217 | 198.1 | - | - | 0 | - |
| - | - | 2.761E+04 | 199.1 | - | - | 0 | - |
| - | - | 3836 | 200.1 | - | - | 0 | - |
| - | - | 1020 | 203.1 | - | - | 0 | - |
| - | - | 1.076E+04 | 205.1 | - | - | 0 | - |
| - | - | 1248 | 206.1 | - | - | 0 | - |
| 2 | a | 5.082E+05 | 207.1 | 0.0005076 | 2.451 | +1 | 2 |
| 4 | b | 5.852E+04 | 208.1 | 0.004083 | 19.62 | +2 | 4 |
| - | - | 1063 | 209.1 | - | - | 0 | - |
| - | - | 3781 | 209.1 | - | - | 0 | - |
| - | - | 1562 | 210.1 | - | - | 0 | - |
| - | - | 1201 | 211.1 | - | - | 0 | - |
| - | - | 1.572E+04 | 212.1 | - | - | 0 | - |
| - | - | 984.3 | 213.1 | - | - | 0 | - |
| - | - | 1313 | 213.1 | - | - | 0 | - |
| - | - | 587.2 | 214.2 | - | - | 0 | - |
| - | - | 544.7 | 215.1 | - | - | 0 | - |
| - | - | 899.3 | 215.1 | - | - | 0 | - |
| - | - | 1421 | 215.1 | - | - | 0 | - |
| - | - | 4174 | 216.1 | - | - | 0 | - |
| 2 | b | 3424 | 217.1 | 0.0005327 | 2.454 | +1 | 2 |
| - | - | 2.663E+04 | 219.1 | - | - | 0 | - |
| - | - | 2439 | 220.2 | - | - | 0 | - |
| - | - | 802.2 | 221.1 | - | - | 0 | - |
| - | - | 530.1 | 224.1 | - | - | 0 | - |
| - | - | 672.9 | 224.2 | - | - | 0 | - |
| - | - | 2018 | 225.1 | - | - | 0 | - |
| - | - | 666.9 | 225.2 | - | - | 0 | - |
| - | - | 1217 | 226.1 | - | - | 0 | - |
| - | - | 530.5 | 227.1 | - | - | 0 | - |
| - | - | 584.2 | 227.1 | - | - | 0 | - |
| - | - | 5888 | 227.1 | - | - | 0 | - |
| - | - | 2692 | 228.1 | - | - | 0 | - |
| - | - | 1210 | 229.1 | - | - | 0 | - |
| - | - | 604.1 | 229.1 | - | - | 0 | - |
| - | - | 2.733E+04 | 230.2 | - | - | 0 | - |
| - | - | 2219 | 231.1 | - | - | 0 | - |
| - | - | 2728 | 231.2 | - | - | 0 | - |
| - | - | 618.9 | 233.1 | - | - | 0 | - |
| - | - | 831.7 | 234.1 | - | - | 0 | - |
| 2 | b | 3.382E+05 | 235.1 | 0.0005423 | 2.307 | +1 | 2 |
| - | - | 4.411E+04 | 236.1 | - | - | 0 | - |
| - | - | 1400 | 237.1 | - | - | 0 | - |
| - | - | 3791 | 237.1 | - | - | 0 | - |
| - | - | 868.2 | 241.2 | - | - | 0 | - |
| - | - | 1635 | 242.2 | - | - | 0 | - |
| - | - | 2803 | 243.1 | - | - | 0 | - |
| - | - | 2194 | 243.1 | - | - | 0 | - |
| 10 | y | 1.13E+04 | 243.2 | 0.0003963 | 1.63 | +1 | 2 |
| - | - | 1855 | 244.1 | - | - | 0 | - |
| - | - | 1048 | 244.2 | - | - | 0 | - |
| - | - | 4668 | 245.1 | - | - | 0 | - |
| - | - | 1785 | 246.1 | - | - | 0 | - |
| - | - | 1.655E+04 | 247.1 | - | - | 0 | - |
| - | - | 1683 | 248.1 | - | - | 0 | - |
| - | - | 1262 | 248.2 | - | - | 0 | - |
| - | - | 1192 | 251.2 | - | - | 0 | - |
| - | - | 1.587E+04 | 255.1 | - | - | 0 | - |
| - | - | 1752 | 256.1 | - | - | 0 | - |
| - | - | 1760 | 259.1 | - | - | 0 | - |
| 10 | y | 4.751E+04 | 260.2 | 0.0003365 | 1.293 | +1 | 2 |
| - | - | 2753 | 261.1 | - | - | 0 | - |
| - | - | 2848 | 261.2 | - | - | 0 | - |
| - | - | 6326 | 261.2 | - | - | 0 | - |
| - | - | 6306 | 262.1 | - | - | 0 | - |
| - | - | 882.3 | 263.1 | - | - | 0 | - |
| - | - | 1.852E+04 | 269.2 | - | - | 0 | - |
| - | - | 2267 | 270.2 | - | - | 0 | - |
| - | - | 1116 | 270.2 | - | - | 0 | - |
| - | - | 1240 | 271.1 | - | - | 0 | - |
| - | - | 971.6 | 272.1 | - | - | 0 | - |
| - | - | 1.785E+04 | 273.1 | - | - | 0 | - |
| - | - | 1549 | 273.2 | - | - | 0 | - |
| - | - | 5766 | 274.1 | - | - | 0 | - |
| - | - | 885 | 275.1 | - | - | 0 | - |
| - | - | 2060 | 275.2 | - | - | 0 | - |
| - | - | 745.7 | 279.1 | - | - | 0 | - |
| - | - | 1377 | 282.2 | - | - | 0 | - |
| - | - | 3376 | 287.2 | - | - | 0 | - |
| - | - | 566 | 288.1 | - | - | 0 | - |
| 3 | a | 2197 | 288.2 | 0.001801 | 6.251 | +1 | 3 |
| - | - | 1.967E+04 | 289.2 | - | - | 0 | - |
| - | - | 3125 | 290.2 | - | - | 0 | - |
| - | - | 4095 | 291.1 | - | - | 0 | - |
| - | - | 824.7 | 292.1 | - | - | 0 | - |
| - | - | 607.1 | 295.1 | - | - | 0 | - |
| - | - | 853 | 297.2 | - | - | 0 | - |
| - | - | 1397 | 300.1 | - | - | 0 | - |
| - | - | 874.3 | 300.2 | - | - | 0 | - |
| - | - | 2534 | 301.2 | - | - | 0 | - |
| - | - | 761.2 | 303.2 | - | - | 0 | - |
| - | - | 2450 | 304.2 | - | - | 0 | - |
| - | - | 1128 | 305.2 | - | - | 0 | - |
| 3 | a | 1251 | 306.2 | 0.001063 | 3.473 | +1 | 3 |
| - | - | 2171 | 309.2 | - | - | 0 | - |
| - | - | 902.1 | 313.2 | - | - | 0 | - |
| 3 | b | 8813 | 316.2 | 0.0006002 | 1.898 | +1 | 3 |
| - | - | 779.2 | 317.2 | - | - | 0 | - |
| - | - | 1480 | 318.1 | - | - | 0 | - |
| - | - | 8442 | 319.1 | - | - | 0 | - |
| - | - | 925.1 | 320.1 | - | - | 0 | - |
| - | - | 1.353E+04 | 326.2 | - | - | 0 | - |
| - | - | 641.9 | 327.2 | - | - | 0 | - |
| - | - | 1839 | 327.2 | - | - | 0 | - |
| - | - | 898.7 | 328.1 | - | - | 0 | - |
| - | - | 756.9 | 330.2 | - | - | 0 | - |
| - | - | 664.2 | 332.2 | - | - | 0 | - |
| - | - | 1501 | 333.2 | - | - | 0 | - |
| 3 | b | 1.72E+05 | 334.2 | 0.0007472 | 2.236 | +1 | 3 |
| - | - | 3.201E+04 | 335.2 | - | - | 0 | - |
| - | - | 2895 | 336.2 | - | - | 0 | - |
| - | - | 3405 | 337.2 | - | - | 0 | - |
| - | - | 1596 | 339.2 | - | - | 0 | - |
| 5 | y | 4202 | 340.2 | 0.0003434 | 1.009 | +2 | 7 |
| - | - | 1816 | 340.7 | - | - | 0 | - |
| 9 | y | 2116 | 343.2 | 0.0004846 | 1.412 | +1 | 3 |
| - | - | 3220 | 344.2 | - | - | 0 | - |
| - | - | 754.4 | 344.2 | - | - | 0 | - |
| - | - | 683.2 | 345.2 | - | - | 0 | - |
| - | - | 1145 | 345.2 | - | - | 0 | - |
| - | - | 659.8 | 346.2 | - | - | 0 | - |
| - | - | 4423 | 346.2 | - | - | 0 | - |
| - | - | 844.2 | 347.2 | - | - | 0 | - |
| - | - | 3325 | 348.2 | - | - | 0 | - |
| - | - | 2791 | 354.2 | - | - | 0 | - |
| - | - | 813.8 | 356.2 | - | - | 0 | - |
| - | - | 5167 | 357.2 | - | - | 0 | - |
| - | - | 754.3 | 358.2 | - | - | 0 | - |
| - | - | 968.2 | 360.2 | - | - | 0 | - |
| - | - | 1420 | 361.2 | - | - | 0 | - |
| 9 | y | 1.081E+04 | 361.2 | 0.000418 | 1.157 | +1 | 3 |
| - | - | 1219 | 362.2 | - | - | 0 | - |
| - | - | 2136 | 362.2 | - | - | 0 | - |
| - | - | 666.4 | 363.2 | - | - | 0 | - |
| - | - | 3091 | 364.2 | - | - | 0 | - |
| - | - | 647.7 | 364.6 | - | - | 0 | - |
| - | - | 874.2 | 365.2 | - | - | 0 | - |
| - | - | 2593 | 365.2 | - | - | 0 | - |
| - | - | 800.4 | 365.2 | - | - | 0 | - |
| - | - | 793.1 | 368.2 | - | - | 0 | - |
| - | - | 1934 | 372.2 | - | - | 0 | - |
| - | - | 2667 | 373.2 | - | - | 0 | - |
| - | - | 935.2 | 373.7 | - | - | 0 | - |
| - | - | 5661 | 374.2 | - | - | 0 | - |
| - | - | 952.8 | 374.2 | - | - | 0 | - |
| - | - | 3355 | 375.2 | - | - | 0 | - |
| - | - | 1208 | 376.2 | - | - | 0 | - |
| - | - | 2882 | 382.2 | - | - | 0 | - |
| - | - | 821.4 | 382.2 | - | - | 0 | - |
| - | - | 822.3 | 383.2 | - | - | 0 | - |
| - | - | 1.608E+04 | 383.2 | - | - | 0 | - |
| - | - | 733.6 | 384.2 | - | - | 0 | - |
| - | - | 2901 | 384.2 | - | - | 0 | - |
| - | - | 1457 | 385.2 | - | - | 0 | - |
| - | - | 1338 | 387.2 | - | - | 0 | - |
| - | - | 2228 | 388.2 | - | - | 0 | - |
| - | - | 3878 | 390.2 | - | - | 0 | - |
| - | - | 7349 | 392.2 | - | - | 0 | - |
| - | - | 1474 | 393.2 | - | - | 0 | - |
| 8 | y | 2405 | 400.3 | 0.0007189 | 1.796 | +1 | 4 |
| - | - | 1.343E+04 | 401.2 | - | - | 0 | - |
| - | - | 2.149E+04 | 402.2 | - | - | 0 | - |
| - | - | 1518 | 402.2 | - | - | 0 | - |
| - | - | 4887 | 403.2 | - | - | 0 | - |
| - | - | 1178 | 403.2 | - | - | 0 | - |
| - | - | 1.11E+04 | 405.3 | - | - | 0 | - |
| - | - | 3033 | 406.3 | - | - | 0 | - |
| - | - | 601.8 | 413.3 | - | - | 0 | - |
| 4 | b | 4623 | 415.2 | 0.0005762 | 1.388 | +1 | 4 |
| - | - | 1312 | 416.2 | - | - | 0 | - |
| - | - | 3000 | 418.2 | - | - | 0 | - |
| 8 | y | 9769 | 418.3 | 0.0009574 | 2.289 | +1 | 4 |
| - | - | 5242 | 419.2 | - | - | 0 | - |
| - | - | 2636 | 419.3 | - | - | 0 | - |
| - | - | 1.216E+04 | 420.2 | - | - | 0 | - |
| - | - | 894.8 | 420.2 | - | - | 0 | - |
| - | - | 2757 | 421.2 | - | - | 0 | - |
| 4 | b | 3.427E+04 | 433.2 | 0.0007842 | 1.81 | +1 | 4 |
| - | - | 8603 | 434.2 | - | - | 0 | - |
| - | - | 1199 | 435.2 | - | - | 0 | - |
| - | - | 4040 | 436.2 | - | - | 0 | - |
| - | - | 516.2 | 437.2 | - | - | 0 | - |
| 3 | y | 7909 | 439.3 | 0.001204 | 2.742 | +2 | 9 |
| - | - | 3118 | 439.8 | - | - | 0 | - |
| - | - | 1390 | 440.3 | - | - | 0 | - |
| - | - | 515.3 | 440.4 | - | - | 0 | - |
| - | - | 1088 | 445.2 | - | - | 0 | - |
| - | - | 610.5 | 446.2 | - | - | 0 | - |
| - | - | 622.9 | 451.2 | - | - | 0 | - |
| - | - | 2631 | 453.2 | - | - | 0 | - |
| - | - | 1504 | 454.3 | - | - | 0 | - |
| - | - | 1762 | 456.2 | - | - | 0 | - |
| 7 | y | 2899 | 457.3 | 0.001441 | 3.152 | +1 | 5 |
| - | - | 1178 | 458.3 | - | - | 0 | - |
| - | - | 2682 | 463.2 | - | - | 0 | - |
| - | - | 960.5 | 464.2 | - | - | 0 | - |
| - | - | 672.1 | 470.3 | - | - | 0 | - |
| - | - | 709.5 | 471.2 | - | - | 0 | - |
| - | - | 1081 | 472.3 | - | - | 0 | - |
| - | - | 2756 | 473.3 | - | - | 0 | - |
| - | - | 876 | 474.2 | - | - | 0 | - |
| 7 | y | 2.431E+04 | 475.3 | 0.0008255 | 1.737 | +1 | 5 |
| - | - | 5821 | 476.3 | - | - | 0 | - |
| - | - | 724.8 | 477.3 | - | - | 0 | - |
| - | - | 6603 | 481.2 | - | - | 0 | - |
| 10 | b | 886.5 | 481.8 | 0.0006502 | 1.35 | +2 | 10 |
| 10 | b | 2958 | 482.3 | 0.0005858 | 1.215 | +2 | 10 |
| - | - | 695.2 | 483.2 | - | - | 0 | - |
| - | - | 940.2 | 486.2 | - | - | 0 | - |
| - | - | 1120 | 486.3 | - | - | 0 | - |
| - | - | 1393 | 489.3 | - | - | 0 | - |
| - | - | 921.3 | 490.3 | - | - | 0 | - |
| 10 | b | 840 | 490.8 | 0.003913 | 7.973 | +2 | 10 |
| - | - | 3806 | 491.3 | - | - | 0 | - |
| - | - | 1583 | 492.3 | - | - | 0 | - |
| - | - | 754.9 | 495.2 | - | - | 0 | - |
| - | - | 2796 | 496.3 | - | - | 0 | - |
| - | - | 1578 | 497.3 | - | - | 0 | - |
| - | - | 841 | 499.3 | - | - | 0 | - |
| - | - | 9908 | 501.2 | - | - | 0 | - |
| - | - | 2706 | 502.2 | - | - | 0 | - |
| - | - | 650.8 | 503.2 | - | - | 0 | - |
| - | - | 939.2 | 510.3 | - | - | 0 | - |
| - | - | 520.3 | 510.7 | - | - | 0 | - |
| - | - | 907.5 | 511.3 | - | - | 0 | - |
| - | - | 2466 | 512.3 | - | - | 0 | - |
| 2 | y | 1044 | 512.8 | 0.002001 | 3.902 | +2 | 10 |
| 6 | y | 6732 | 514.3 | 0.0002719 | 0.5287 | +1 | 6 |
| - | - | 2385 | 515.3 | - | - | 0 | - |
| - | - | 2476 | 517.3 | - | - | 0 | - |
| - | - | 5788 | 519.3 | - | - | 0 | - |
| - | - | 2916 | 520.3 | - | - | 0 | - |
| - | - | 1.24E+04 | 530.3 | - | - | 0 | - |
| - | - | 2865 | 531.3 | - | - | 0 | - |
| 6 | y | 1.105E+05 | 532.3 | 0.0005715 | 1.074 | +1 | 6 |
| - | - | 2.802E+04 | 533.3 | - | - | 0 | - |
| - | - | 5711 | 534.3 | - | - | 0 | - |
| - | - | 2272 | 535.3 | - | - | 0 | - |
| - | - | 1103 | 536.3 | - | - | 0 | - |
| - | - | 1341 | 538.3 | - | - | 0 | - |
| 0 | Precursor | 756.3 | 547.3 | 0.004519 | 8.257 | +2 | -1 |
| 0 | Precursor | 913.9 | 547.8 | 0.009759 | 17.82 | +2 | -1 |
| - | - | 2.068E+04 | 548.3 | - | - | 0 | - |
| - | - | 6960 | 549.3 | - | - | 0 | - |
| - | - | 894.2 | 550.3 | - | - | 0 | - |
| - | - | 4312 | 552.3 | - | - | 0 | - |
| - | - | 1357 | 553.3 | - | - | 0 | - |
| - | - | 605.1 | 554.6 | - | - | 0 | - |
| - | - | 2059 | 555 | - | - | 0 | - |
| - | - | 1423 | 555.3 | - | - | 0 | - |
| - | - | 1915 | 556.2 | - | - | 0 | - |
| 0 | Precursor | 2115 | 556.3 | 0.0005132 | 0.9226 | +2 | -1 |
| 5 | b | 3425 | 562.3 | 0.001254 | 2.23 | +1 | 5 |
| - | - | 675.7 | 563.3 | - | - | 0 | - |
| - | - | 5366 | 566.3 | - | - | 0 | - |
| - | - | 1951 | 567.3 | - | - | 0 | - |
| - | - | 890 | 568.3 | - | - | 0 | - |
| - | - | 1437 | 573.3 | - | - | 0 | - |
| 5 | b | 8504 | 580.3 | 0.0003329 | 0.5737 | +1 | 5 |
| - | - | 2115 | 581.3 | - | - | 0 | - |
| - | - | 1919 | 590.3 | - | - | 0 | - |
| - | - | 771.3 | 592.3 | - | - | 0 | - |
| - | - | 618.9 | 595.3 | - | - | 0 | - |
| - | - | 3002 | 600.3 | - | - | 0 | - |
| - | - | 1181 | 601.3 | - | - | 0 | - |
| - | - | 953.3 | 609.3 | - | - | 0 | - |
| - | - | 1168 | 611.3 | - | - | 0 | - |
| - | - | 972.1 | 617.3 | - | - | 0 | - |
| - | - | 3414 | 618.3 | - | - | 0 | - |
| 6 | b | 2598 | 619.3 | 0.003045 | 4.916 | +1 | 6 |
| - | - | 987.7 | 620.3 | - | - | 0 | - |
| - | - | 768.8 | 625.3 | - | - | 0 | - |
| - | - | 4976 | 629.3 | - | - | 0 | - |
| - | - | 1789 | 630.3 | - | - | 0 | - |
| - | - | 1111 | 633.4 | - | - | 0 | - |
| 6 | b | 2490 | 637.3 | 0.0003178 | 0.4987 | +1 | 6 |
| - | - | 582.4 | 638.3 | - | - | 0 | - |
| - | - | 3962 | 643.4 | - | - | 0 | - |
| - | - | 867.4 | 644.3 | - | - | 0 | - |
| - | - | 1.022E+04 | 647.4 | - | - | 0 | - |
| - | - | 4080 | 648.4 | - | - | 0 | - |
| - | - | 983.8 | 649.4 | - | - | 0 | - |
| 5 | y | 1.04E+04 | 661.4 | 0.0004005 | 0.6055 | +1 | 7 |
| - | - | 4102 | 662.4 | - | - | 0 | - |
| - | - | 1164 | 663.4 | - | - | 0 | - |
| - | - | 4107 | 665.4 | - | - | 0 | - |
| - | - | 2092 | 666.4 | - | - | 0 | - |
| 7 | b | 732.9 | 676.3 | 0.001417 | 2.096 | +1 | 7 |
| - | - | 809.8 | 677.4 | - | - | 0 | - |
| 5 | y | 2.178E+05 | 679.4 | 0.0004559 | 0.6711 | +1 | 7 |
| - | - | 8.102E+04 | 680.4 | - | - | 0 | - |
| - | - | 1.724E+04 | 681.4 | - | - | 0 | - |
| - | - | 1433 | 682.4 | - | - | 0 | - |
| - | - | 655 | 683.4 | - | - | 0 | - |
| 7 | b | 1415 | 694.4 | 0.0006634 | 0.9554 | +1 | 7 |
| - | - | 674.1 | 711.4 | - | - | 0 | - |
| - | - | 914.7 | 716.4 | - | - | 0 | - |
| - | - | 3775 | 728.4 | - | - | 0 | - |
| - | - | 1705 | 729.4 | - | - | 0 | - |
| - | - | 856.1 | 732.4 | - | - | 0 | - |
| - | - | 2809 | 742.4 | - | - | 0 | - |
| - | - | 1429 | 743.4 | - | - | 0 | - |
| - | - | 687.1 | 744.4 | - | - | 0 | - |
| - | - | 5962 | 746.4 | - | - | 0 | - |
| - | - | 2640 | 747.4 | - | - | 0 | - |
| 8 | b | 1933 | 751.4 | 0.002043 | 2.719 | +1 | 8 |
| - | - | 783.7 | 752.4 | - | - | 0 | - |
| 4 | y | 8298 | 760.4 | 0.0007732 | 1.017 | +1 | 8 |
| - | - | 2892 | 761.4 | - | - | 0 | - |
| - | - | 778.9 | 762.4 | - | - | 0 | - |
| - | - | 1778 | 764.4 | - | - | 0 | - |
| - | - | 1021 | 765.4 | - | - | 0 | - |
| 4 | y | 2.023E+05 | 778.4 | 0.0006455 | 0.8292 | +1 | 8 |
| - | - | 8.661E+04 | 779.4 | - | - | 0 | - |
| - | - | 2.022E+04 | 780.5 | - | - | 0 | - |
| - | - | 2970 | 781.5 | - | - | 0 | - |
| - | - | 609.2 | 807.4 | - | - | 0 | - |
| - | - | 851.9 | 824.4 | - | - | 0 | - |
| 9 | b | 1919 | 834.4 | 0.001489 | 1.785 | +1 | 9 |
| - | - | 1546 | 835.4 | - | - | 0 | - |
| - | - | 1498 | 841.5 | - | - | 0 | - |
| - | - | 669.2 | 842.5 | - | - | 0 | - |
| 9 | b | 4090 | 852.4 | 0.0001034 | 0.1213 | +1 | 9 |
| - | - | 2913 | 853.4 | - | - | 0 | - |
| 3 | y | 5751 | 859.5 | 0.0005021 | 0.5841 | +1 | 9 |
| - | - | 3378 | 860.5 | - | - | 0 | - |
| - | - | 707.4 | 876.5 | - | - | 0 | - |
| 3 | y | 1.618E+05 | 877.5 | 0.0001637 | 0.1866 | +1 | 9 |
| - | - | 8.112E+04 | 878.5 | - | - | 0 | - |
| - | - | 2.061E+04 | 879.5 | - | - | 0 | - |
| - | - | 3862 | 880.5 | - | - | 0 | - |
| - | - | 635.5 | 893.5 | - | - | 0 | - |
| 10 | b | 1327 | 962.5 | 0.001009 | 1.048 | +1 | 10 |
| 10 | b | 940.8 | 963.5 | 0.0137 | 14.22 | +1 | 10 |
| 10 | b | 3274 | 980.5 | 0.001561 | 1.592 | +1 | 10 |
| - | - | 1858 | 981.5 | - | - | 0 | - |
| 2 | y | 1167 | 1025 | 0.003832 | 3.74 | +1 | 10 |
| - | - | 727.3 | 1026 | - | - | 0 | - |
| - | - | 615.4 | 1163 | - | - | 0 | - |
| - | - | 672.5 | 1599 | - | - | 0 | - |
| - | - | 701.6 | 1805 | - | - | 0 | - |
| - | - | 662.9 | 1829 | - | - | 0 | - |
| - | - | 569.9 | 1848 | - | - | 0 | - |
| - | - | 708.5 | 2023 | - | - | 0 | - |
| - | - | 671.2 | 2046 | - | - | 0 | - |
| - | - | 593.3 | 2146 | - | - | 0 | - |
| - | - | 626.1 | 3316 | - | - | 0 | - |

m/z Charge Intensity FragmentType MassShift Position
120.08118438720703 0 105320.71
121.07968139648438 0 854.8749
121.08451080322266 0 8117.0713
122.08799743652344 0 505.66205 y Ammonia loss 9
124.0398178100586 0 402.66202
125.07124328613281 0 562.5068
127.05069732666016 0 1547.0654
127.08665466308594 0 432.9343
128.1073760986328 0 1967.8152
129.06610107421875 0 723.25433
129.10267639160156 0 144050.53
130.0612030029297 0 576.9291
130.08663940429688 0 1263.4539
130.1001739501953 0 1066.1477
130.10598754882812 0 8666.172
131.0452117919922 0 674.60675
131.08189392089844 0 5153.173
132.07640075683594 0 559.7969
132.0812225341797 0 1790.956
132.10227966308594 0 14089.878 y 10
133.1056365966797 0 1202.0829
134.09664916992188 0 1227.5356
136.07608032226562 0 2232.1924
139.0869903564453 0 864.0328
140.08245849609375 0 1123.7308
141.06625366210938 0 2657.853
141.1024932861328 0 831.3538
141.94659423828125 0 364.9836
142.0609588623047 0 436.90152
143.0816192626953 0 438.63968
144.08116149902344 0 10873.494
145.08462524414062 0 937.23804
146.09640502929688 0 507.06824
147.11322021484375 0 6564.8545
148.07635498046875 0 557.43506
152.45999145507812 0 405.50842
153.06661987304688 0 751.0954
153.10206604003906 0 433.78073
154.06167602539062 0 845.88135
155.08181762695312 0 1678.1199
155.11825561523438 0 2804.9614
156.0769500732422 0 642.6449
157.096923828125 0 477.2418
157.46932983398438 0 437.13358
158.092529296875 0 986.749
159.07679748535156 0 4357.051
160.07568359375 0 951.63513
162.09170532226562 0 31152.533
163.09523010253906 0 2946.8071
166.0987091064453 0 629.53534
167.09317016601562 0 717.429
167.118408203125 0 644.18317
167.8037567138672 0 480.90845
169.09774780273438 0 689.257
171.07691955566406 0 1082.0575
171.1495819091797 0 46722.43
172.0720672607422 0 5458.1187
172.15306091308594 0 4447.4873
172.3776397705078 0 463.9935
173.43983459472656 0 2858.6123
174.127685546875 0 745.06714
174.52102661132812 0 484.99103
176.1073760986328 0 5094.5835
177.1026611328125 0 6289.2446
177.11062622070312 0 833.93964
181.09776306152344 0 493.42587
183.14984130859375 0 467.72214
184.1450958251953 0 486.11002
185.12892150878906 0 1683.9977
186.12425231933594 0 2253.0986
187.1077117919922 0 1178.7667
188.103271484375 0 1323.1006
189.10186767578125 0 1503.3223 a Water loss 1
190.08631896972656 0 798.65393
195.11337280273438 0 1902.0562
196.10797119140625 0 687.317
197.10366821289062 0 465.9175
197.16519165039062 0 1382.5864
198.08778381347656 0 4217.485
199.1444854736328 0 27607.576
200.1479949951172 0 3836.1328
203.11851501464844 0 1020.1923
205.09756469726562 0 10762.19
206.10130310058594 0 1247.8909
207.11331176757812 0 508239.8 a 1
208.11654663085938 0 58523.35 b Water loss 3
209.1041259765625 0 1062.959
209.11895751953125 0 3781.4126
210.08799743652344 0 1561.7693
211.14450073242188 0 1201.249
212.1397247314453 0 15716.756
213.12359619140625 0 984.3259
213.1431121826172 0 1312.8943
214.1904754638672 0 587.15796
215.10324096679688 0 544.6833
215.1146240234375 0 899.2736
215.1402130126953 0 1420.9498
216.09828186035156 0 4173.7964
217.09768676757812 0 3424.4956 b Water loss 1
219.1495819091797 0 26625.096
220.15313720703125 0 2438.9536
221.12828063964844 0 802.2066
224.14097595214844 0 530.13617
224.17684936523438 0 672.9455
225.13490295410156 0 2017.888
225.1608123779297 0 666.875
226.1188507080078 0 1217.299
227.08062744140625 0 530.53876
227.1028594970703 0 584.1837
227.11448669433594 0 5888.4126
228.0982666015625 0 2691.5464
229.093505859375 0 1210.3662
229.11972045898438 0 604.1357
230.15036010742188 0 27327.945
231.11318969726562 0 2218.568
231.15304565429688 0 2727.7056
233.1286163330078 0 618.86597
234.12290954589844 0 831.7285
235.10826110839844 0 338169.06 b 1
236.1115264892578 0 44112.156
237.09854125976562 0 1399.5293
237.11370849609375 0 3791.4644
241.15513610839844 0 868.2185
242.1869659423828 0 1634.6511
243.11318969726562 0 2803.3015
243.14564514160156 0 2194.2542
243.17071533203125 0 11295.033 y Ammonia loss 9
244.10862731933594 0 1854.5863
244.1740264892578 0 1048.2139
245.12513732910156 0 4667.717
246.1239013671875 0 1785.4926
247.1445770263672 0 16546.01
248.1475067138672 0 1682.5286
248.1611328125 0 1261.8427
251.1504669189453 0 1192.2549
255.10926818847656 0 15874.156
256.1126708984375 0 1752.2292
259.1445007324219 0 1759.8162
260.19720458984375 0 47506.84 y 9
261.12359619140625 0 2752.8267
261.1601257324219 0 2847.9756
261.20068359375 0 6325.7534
262.11907958984375 0 6306.4985
263.1224365234375 0 882.34015
269.1612548828125 0 18518.373
270.1644287109375 0 2266.5647
270.18157958984375 0 1116.2219
271.1398620605469 0 1239.726
272.1377258300781 0 971.5618
273.1197204589844 0 17848.732
273.196533203125 0 1549.4669
274.11993408203125 0 5766.272
275.12298583984375 0 885.01764
275.1759948730469 0 2060.161
279.14520263671875 0 745.7146
282.1561584472656 0 1377.1809
287.1719665527344 0 3376.3625
288.1337890625 0 566.02844
288.1724548339844 0 2196.8105 a Water loss 2
289.1551818847656 0 19668.414
290.15826416015625 0 3125.1238
291.1453857421875 0 4094.765
292.129150390625 0 824.73413
295.0776062011719 0 607.0991
297.1558532714844 0 852.9892
300.1344909667969 0 1396.8745
300.1670227050781 0 874.33374
301.1917724609375 0 2534.2156
303.17132568359375 0 761.17816
304.166748046875 0 2449.63
305.1835021972656 0 1128.3574
306.1822814941406 0 1250.9769 a 2
309.15960693359375 0 2171.4404
313.1883850097656 0 902.05176
316.1661682128906 0 8812.95 b Water loss 2
317.1678161621094 0 779.2192
318.14520263671875 0 1480.4391
319.14080810546875 0 8441.936
320.1441345214844 0 925.13043
326.1830139160156 0 13531.923
327.1662902832031 0 641.9167
327.18585205078125 0 1838.9432
328.129638671875 0 898.6992
330.1561279296875 0 756.9378
332.16192626953125 0 664.1859
333.19317626953125 0 1501.4001
334.1768798828125 0 171959.14 b 2
335.179931640625 0 32007.047
336.182373046875 0 2895.2224
337.15472412109375 0 3404.5996
339.17816162109375 0 1595.6377
340.1926574707031 0 4202.1313 y 4
340.6950378417969 0 1816.2748
343.2344665527344 0 2116.2783 y Water loss 8
344.1933898925781 0 3220.2185
344.2384338378906 0 754.4171
345.15472412109375 0 683.1643
345.1955261230469 0 1145.3086
346.1771545410156 0 659.77954
346.2131042480469 0 4423.198
347.21539306640625 0 844.2341
348.1671447753906 0 3324.5276
354.177734375 0 2790.9517
356.1729431152344 0 813.81177
357.15631103515625 0 5166.7065
358.1580505371094 0 754.31946
360.189697265625 0 968.186
361.1876220703125 0 1420.1538
361.2449645996094 0 10808.774 y 8
362.2034606933594 0 1218.5883
362.2486877441406 0 2136.289
363.2044677734375 0 666.40155
364.16595458984375 0 3091.0593
364.6256408691406 0 647.7258
365.16973876953125 0 874.21344
365.19317626953125 0 2593.4316
365.2173767089844 0 800.3598
368.1911926269531 0 793.05566
372.18865966796875 0 1933.844
373.1877136230469 0 2667.1907
373.7127380371094 0 935.20197
374.18316650390625 0 5661.159
374.2086486816406 0 952.79236
375.1668395996094 0 3354.544
376.1639099121094 0 1207.5042
382.176513671875 0 2882.2932
382.2440185546875 0 821.41724
383.17584228515625 0 822.2518
383.2042541503906 0 16077.936
384.1645202636719 0 733.61255
384.2076416015625 0 2900.6328
385.15093994140625 0 1456.7942
387.23944091796875 0 1337.5543
388.2236633300781 0 2228.3843
390.2145690917969 0 3877.9724
392.1936950683594 0 7348.9756
393.1939697265625 0 1473.9862
400.25616455078125 0 2404.9458 y Water loss 7
401.2149658203125 0 13431.079
402.17779541015625 0 21485.396
402.21527099609375 0 1518.419
403.18109130859375 0 4887.327
403.2348327636719 0 1177.7935
405.250244140625 0 11096.445
406.2531433105469 0 3032.9944
413.2525329589844 0 601.7961
415.23455810546875 0 4622.676 b Water loss 3
416.2381286621094 0 1312.0245
418.2101135253906 0 2999.5225
418.2669677734375 0 9768.866 y 7
419.224853515625 0 5242.454
419.2701110839844 0 2636.009
420.1885070800781 0 12159.133
420.2226257324219 0 894.7502
421.1910095214844 0 2757.4329
433.2453308105469 0 34268.062 b 3
434.24853515625 0 8602.974
435.24835205078125 0 1199.0486
436.2235107421875 0 4039.9363
437.2270812988281 0 516.15015
439.2619323730469 0 7908.8125 y 2
439.7627868652344 0 3118.2952
440.2663879394531 0 1390.1316
440.4487609863281 0 515.30884
445.2479553222656 0 1087.7606
446.2442321777344 0 610.48627
451.23382568359375 0 622.937
453.2496643066406 0 2630.8728
454.2538146972656 0 1503.8682
456.2254943847656 0 1761.969
457.2783508300781 0 2898.794 y Water loss 6
458.2802734375 0 1178.0302
463.2347412109375 0 2681.917
464.2372741699219 0 960.51245
470.27325439453125 0 672.1362
471.19842529296875 0 709.5106
472.2566223144531 0 1080.7375
473.2516174316406 0 2756.4675
474.2329406738281 0 876.03986
475.2882995605469 0 24313.002 y 6
476.29132080078125 0 5820.7744
477.2916564941406 0 724.8001
481.2453918457031 0 6603.048
481.7590026855469 0 886.51495 b Water loss 9
482.2509460449219 0 2958.1177 b Ammonia loss 9
483.2359924316406 0 695.1623
486.2477111816406 0 940.1837
486.3040466308594 0 1120.3855
489.282470703125 0 1393.102
490.26617431640625 0 921.33636
490.7675476074219 0 839.98645 b 9
491.2625732421875 0 3806.3984
492.26287841796875 0 1583.3364
495.2365417480469 0 754.942
496.2883605957031 0 2795.8901
497.274169921875 0 1577.8552
499.2664489746094 0 840.9773
501.24615478515625 0 9908.251
502.2485656738281 0 2706.0195
503.24969482421875 0 650.7526
510.27239990234375 0 939.1988
510.6837158203125 0 520.2828
511.2735290527344 0 907.529
512.2625732421875 0 2466.27
512.7969360351562 0 1044.0013 y 1
514.2986450195312 0 6731.783 y Water loss 5
515.301513671875 0 2384.758
517.2776489257812 0 2476.0244
519.2567749023438 0 5787.7793
520.259521484375 0 2915.783
530.273193359375 0 12401.204
531.276123046875 0 2864.6267
532.3095092773438 0 110548.28 y 5
533.3123168945312 0 28015.654
534.3134155273438 0 5711.38
535.29443359375 0 2272.4329
536.2940673828125 0 1103.1663
538.2652587890625 0 1340.7134
547.3011474609375 0 756.28455 Precursor Water loss
547.8074340820312 0 913.87274 Precursor Ammonia loss
548.283447265625 0 20680.953
549.2864990234375 0 6959.5195
550.2872924804688 0 894.21356
552.3184204101562 0 4311.903
553.3211059570312 0 1356.9833
554.643310546875 0 605.121
554.9682006835938 0 2059.4265
555.3008422851562 0 1422.7335
556.1905517578125 0 1914.8265
556.3114624023438 0 2115.4722 Precursor
562.3036499023438 0 3425.4778 b Water loss 4
563.3050537109375 0 675.6748
566.294677734375 0 5365.805
567.2965698242188 0 1950.7417
568.2998657226562 0 889.98895
573.3046875 0 1436.6256
580.3132934570312 0 8503.923 b 4
581.3173217773438 0 2115.4688
590.3297119140625 0 1918.9723
592.3130493164062 0 771.2609
595.29931640625 0 618.9186
600.314697265625 0 3001.6978
601.3167114257812 0 1181.1943
609.342529296875 0 953.27124
611.3343505859375 0 1168.017
617.3419799804688 0 972.09143
618.3263549804688 0 3413.9438
619.326904296875 0 2598.1343 b Water loss 5
620.3277587890625 0 987.67285
625.3464965820312 0 768.82495
629.3409423828125 0 4975.936
630.3419799804688 0 1788.545
633.373779296875 0 1110.6465
637.3341064453125 0 2490.2458 b 5
638.3442993164062 0 582.36993
643.3570556640625 0 3962.0647
644.347900390625 0 867.3538
647.3511352539062 0 10217.342
648.3543701171875 0 4080.341
649.3614501953125 0 983.7734
661.3671875 0 10395.574 y Water loss 4
662.36962890625 0 4102.0225
663.3688354492188 0 1164.2448
665.3623046875 0 4106.692
666.3656616210938 0 2092.2175
676.3467407226562 0 732.8909 b Water loss 6
677.3585815429688 0 809.82086
679.3778076171875 0 217822.67 y 4
680.3809204101562 0 81022.375
681.383056640625 0 17237.19
682.3869018554688 0 1432.5731
683.357666015625 0 655.0255
694.355224609375 0 1415.1093 b 6
711.4082641601562 0 674.1013
716.4153442382812 0 914.7198
728.40966796875 0 3775.498
729.41064453125 0 1705.0321
732.4386596679688 0 856.13116
742.426025390625 0 2809.3718
743.4152221679688 0 1428.6389
744.4168090820312 0 687.05145
746.4197998046875 0 5962.0576
747.4234619140625 0 2640.3877
751.37939453125 0 1932.5757 b 7
752.38623046875 0 783.6708
760.4359741210938 0 8298.358 y Water loss 3
761.4364624023438 0 2892.3809
762.434326171875 0 778.94617
764.4315795898438 0 1778.2632
765.4329833984375 0 1021.3574
778.4464111328125 0 202291.39 y 3
779.4491577148438 0 86613
780.4515380859375 0 20222.188
781.4547119140625 0 2969.716
807.4058227539062 0 609.2193
824.4273681640625 0 851.90894
834.4159545898438 0 1919.2491 b Water loss 8
835.4159545898438 0 1546.2067
841.4925537109375 0 1498.1309
842.4927978515625 0 669.22034
852.4249267578125 0 4090.0188 b 8
853.4269409179688 0 2912.664
859.5031127929688 0 5750.784 y Water loss 2
860.5073852539062 0 3378.428
876.5017700195312 0 707.3628
877.5143432617188 0 161811.73 y 2
878.51708984375 0 81116.336
879.5198364257812 0 20606.494
880.5222778320312 0 3862.3293
893.4801025390625 0 635.497
962.5104370117188 0 1326.6433 b Water loss 9
963.5071411132812 0 940.7959 b Ammonia loss 9
980.5184326171875 0 3273.7253 b 9
981.5226440429688 0 1858.4258
1024.58642578125 0 1167.1202 y 1
1025.594970703125 0 727.3272
1162.73779296875 0 615.3953
1599.24951171875 0 672.50885
1805.3421630859375 0 701.6443
1829.3453369140625 0 662.8683
1848.0787353515625 0 569.9236
2023.11669921875 0 708.5366
2046.300048828125 0 671.2254
2146.477783203125 0 593.26416
3315.84375 0 626.1413

Spectrum Details

|  |  |
| --- | --- |
| Matched peaks? Matched peaks The total absolute number of peaks matched. Additionally in brackets the total fraction of peaks matched and the total number of peaks is shown. | 51 (12.00% of 425) |
| FDR? FDR The false discovery rate estimated for this peptide. It is calculated by matching all theoretical fragments with a non-integer shift with the raw peaks for this spectrum. This is done with 40 different shifts. The resulting percentage is the average number of annotated peaks over the number of annotated peaks with the correct spectrum. | 0.51% |
| Satellite FDR? Satellite FDR See the FDR for details on its calculation. This satellite ion specific FDR only contains the satellite ions (d/w) for I/L/J positions. | - |
| PSM Score? PSM Score The PSM Score as given by Hecklib to this annotated spectrum. It is shown with three significant figures. | 681 |

#### Spectrum 9687? Spectrum 9687 The raw spectrum of this peptide as annotated by Hecklib. The fragments are coloured according to ion type (see legend). Any peaks with a star '\*' as text can be hovered over to see the full details, first the ion type second the mass shift type. By hovering over the amino acids in the peptide or ions in the legend the corresponding peaks are highlighted. By toggling the 'Unassigned' label you can turn the background (unassigned) peaks on or off in the plot. By updating the slider in the Ion legend you can update the spectrum to only show the top X% of the peaks with labels. The top X% means any peak that is within X% of the highest intensity. By dragging in the spectrum you can zoom in to a specific part of the spectrum and use 'Zoom Out' to get back to the original zoom level. The annotation of the spectrum is based on the given sequence in the peptides file and is done with different software so inconsistencies are likely. The peaks are annotated based on the given sequence, with 20 ppm tolerance.

Copy Data

##### Spectrum 9687 (TSV)

###### Preview

```
Loading example...
```

*Click on the button to copy the data to your clipboard.*

Mz MinMz MaxIntensity Max

WidthHeightPeptide font sizePeptide stroke widthSpectrum font sizeSpectrum stroke widthCompact peptide

Ion legend

wxyz

abcd

OtherUnassignedIon^Charge^_Position_Show for top:%

SFVVFGGGTKJ

03.16e+46.32e+49.48e+41.26e+5

Zoom Out

y^+1^_1_y^+2^_4_a^+1^_2_b^+2^_4_b^+1^_2_b^+1^_2_y^+1^_2_y^+1^_2_b^+1^_3_b^+1^_3_y^+2^_7_y^+1^_3_y^+1^_3_b^+1^_4_y^+1^_4_b^+1^_4_y^+2^_9_y^+1^_5_y^+1^_5_y^+1^_6_y^+1^_6_b^+1^_5_b^+1^_5_y^+1^_7_y^+1^_7_y^+1^_8_y^+1^_8_b^+1^_9_y^+1^_9_y^+1^_9_y^+1^_10_

039278411761568

Fragment Matches Table

Show background peaks

| Position | Ion type | Intensity | mz Theoretical | mz Error (Th) | mz Error (ppm) | Charge | Series Number |
| --- | --- | --- | --- | --- | --- | --- | --- |
| - | - | 3.042E+04 | 120.1 | - | - | 0 | - |
| - | - | 2385 | 121.1 | - | - | 0 | - |
| - | - | 340.9 | 121.4 | - | - | 0 | - |
| - | - | 441.7 | 125.1 | - | - | 0 | - |
| - | - | 400.7 | 125.7 | - | - | 0 | - |
| - | - | 464.1 | 126.1 | - | - | 0 | - |
| - | - | 517.4 | 126.1 | - | - | 0 | - |
| - | - | 636.2 | 127.1 | - | - | 0 | - |
| - | - | 565 | 127.1 | - | - | 0 | - |
| - | - | 586.3 | 128 | - | - | 0 | - |
| - | - | 1015 | 128.1 | - | - | 0 | - |
| - | - | 863.2 | 129.1 | - | - | 0 | - |
| - | - | 3.809E+04 | 129.1 | - | - | 0 | - |
| - | - | 426.1 | 130.1 | - | - | 0 | - |
| - | - | 607.6 | 130.1 | - | - | 0 | - |
| - | - | 536.5 | 130.1 | - | - | 0 | - |
| - | - | 2005 | 130.1 | - | - | 0 | - |
| - | - | 468.1 | 131 | - | - | 0 | - |
| - | - | 638.5 | 131.1 | - | - | 0 | - |
| - | - | 1160 | 131.1 | - | - | 0 | - |
| - | - | 1298 | 132.1 | - | - | 0 | - |
| 11 | y | 3765 | 132.1 | 0.000344 | 2.604 | +1 | 1 |
| - | - | 461.2 | 132.2 | - | - | 0 | - |
| - | - | 524.2 | 133.1 | - | - | 0 | - |
| - | - | 6811 | 133.1 | - | - | 0 | - |
| - | - | 550.2 | 133.1 | - | - | 0 | - |
| - | - | 439.6 | 134.5 | - | - | 0 | - |
| - | - | 4613 | 136.1 | - | - | 0 | - |
| - | - | 875.3 | 138.1 | - | - | 0 | - |
| - | - | 744.1 | 139.1 | - | - | 0 | - |
| - | - | 890.9 | 140.1 | - | - | 0 | - |
| - | - | 789.6 | 141.1 | - | - | 0 | - |
| - | - | 430.7 | 141.1 | - | - | 0 | - |
| - | - | 3059 | 144.1 | - | - | 0 | - |
| - | - | 640.3 | 145.1 | - | - | 0 | - |
| - | - | 522.3 | 145.1 | - | - | 0 | - |
| - | - | 463.5 | 146.1 | - | - | 0 | - |
| - | - | 2482 | 147.1 | - | - | 0 | - |
| - | - | 488.2 | 152.6 | - | - | 0 | - |
| - | - | 1448 | 155.1 | - | - | 0 | - |
| - | - | 960.9 | 155.1 | - | - | 0 | - |
| - | - | 1054 | 156.1 | - | - | 0 | - |
| - | - | 699.8 | 158.1 | - | - | 0 | - |
| - | - | 1082 | 159.1 | - | - | 0 | - |
| - | - | 617.3 | 159.1 | - | - | 0 | - |
| - | - | 504.9 | 159.2 | - | - | 0 | - |
| - | - | 7845 | 162.1 | - | - | 0 | - |
| - | - | 470.9 | 165.8 | - | - | 0 | - |
| - | - | 438.3 | 166.8 | - | - | 0 | - |
| - | - | 693.2 | 169.1 | - | - | 0 | - |
| - | - | 1.327E+04 | 171.1 | - | - | 0 | - |
| - | - | 1795 | 172.1 | - | - | 0 | - |
| - | - | 702.1 | 172.1 | - | - | 0 | - |
| - | - | 1288 | 172.2 | - | - | 0 | - |
| - | - | 1477 | 173.1 | - | - | 0 | - |
| - | - | 595.1 | 175.1 | - | - | 0 | - |
| - | - | 1497 | 176.1 | - | - | 0 | - |
| - | - | 1340 | 177.1 | - | - | 0 | - |
| - | - | 2381 | 177.1 | - | - | 0 | - |
| - | - | 663.1 | 181.1 | - | - | 0 | - |
| - | - | 595.4 | 183.1 | - | - | 0 | - |
| - | - | 563.5 | 183.1 | - | - | 0 | - |
| - | - | 551.2 | 184.1 | - | - | 0 | - |
| - | - | 758.1 | 185.1 | - | - | 0 | - |
| - | - | 597.1 | 185.2 | - | - | 0 | - |
| - | - | 690.6 | 186.1 | - | - | 0 | - |
| - | - | 532.7 | 187.1 | - | - | 0 | - |
| - | - | 600.3 | 187.1 | - | - | 0 | - |
| - | - | 718.5 | 187.1 | - | - | 0 | - |
| - | - | 865.4 | 187.1 | - | - | 0 | - |
| - | - | 565.2 | 189.1 | - | - | 0 | - |
| - | - | 699.7 | 195.1 | - | - | 0 | - |
| - | - | 604.6 | 197.2 | - | - | 0 | - |
| - | - | 816.1 | 198.1 | - | - | 0 | - |
| - | - | 7586 | 199.1 | - | - | 0 | - |
| 8 | y | 1361 | 201.1 | 0.0004715 | 2.344 | +2 | 4 |
| - | - | 2845 | 203.1 | - | - | 0 | - |
| - | - | 3372 | 205.1 | - | - | 0 | - |
| 2 | a | 1.251E+05 | 207.1 | 0.0003703 | 1.788 | +1 | 2 |
| 4 | b | 1.231E+04 | 208.1 | 0.004098 | 19.69 | +2 | 4 |
| - | - | 637.3 | 209.1 | - | - | 0 | - |
| - | - | 4258 | 212.1 | - | - | 0 | - |
| - | - | 679.2 | 215.1 | - | - | 0 | - |
| - | - | 1291 | 215.1 | - | - | 0 | - |
| - | - | 883.4 | 216.1 | - | - | 0 | - |
| 2 | b | 1404 | 217.1 | 0.0002275 | 1.048 | +1 | 2 |
| - | - | 6348 | 219.1 | - | - | 0 | - |
| - | - | 1133 | 220.2 | - | - | 0 | - |
| - | - | 769.4 | 221.1 | - | - | 0 | - |
| - | - | 616 | 222.1 | - | - | 0 | - |
| - | - | 2424 | 226.2 | - | - | 0 | - |
| - | - | 741.5 | 227.1 | - | - | 0 | - |
| - | - | 1558 | 227.1 | - | - | 0 | - |
| - | - | 576.5 | 228.1 | - | - | 0 | - |
| - | - | 680.7 | 229.1 | - | - | 0 | - |
| - | - | 7087 | 230.2 | - | - | 0 | - |
| - | - | 924.6 | 231.2 | - | - | 0 | - |
| 2 | b | 8.204E+04 | 235.1 | 0.0003745 | 1.593 | +1 | 2 |
| - | - | 1.1E+04 | 236.1 | - | - | 0 | - |
| - | - | 866 | 237.1 | - | - | 0 | - |
| - | - | 578.5 | 239.1 | - | - | 0 | - |
| - | - | 471.8 | 243.1 | - | - | 0 | - |
| - | - | 1065 | 243.1 | - | - | 0 | - |
| 10 | y | 2697 | 243.2 | 0.0002742 | 1.128 | +1 | 2 |
| - | - | 581.4 | 244.2 | - | - | 0 | - |
| - | - | 758.2 | 245.1 | - | - | 0 | - |
| - | - | 1961 | 246.1 | - | - | 0 | - |
| - | - | 948.5 | 247.1 | - | - | 0 | - |
| - | - | 4423 | 247.1 | - | - | 0 | - |
| - | - | 3357 | 255.1 | - | - | 0 | - |
| - | - | 606.3 | 258.1 | - | - | 0 | - |
| 10 | y | 1.186E+04 | 260.2 | 0.0002449 | 0.9412 | +1 | 2 |
| - | - | 1105 | 261.1 | - | - | 0 | - |
| - | - | 721.6 | 261.2 | - | - | 0 | - |
| - | - | 1913 | 261.2 | - | - | 0 | - |
| - | - | 1627 | 262.1 | - | - | 0 | - |
| - | - | 3691 | 269.2 | - | - | 0 | - |
| - | - | 4672 | 273.1 | - | - | 0 | - |
| - | - | 638.2 | 273.2 | - | - | 0 | - |
| - | - | 2426 | 274.1 | - | - | 0 | - |
| - | - | 533.6 | 282.2 | - | - | 0 | - |
| - | - | 805.2 | 283.2 | - | - | 0 | - |
| - | - | 1439 | 287.2 | - | - | 0 | - |
| - | - | 3962 | 289.2 | - | - | 0 | - |
| - | - | 580.1 | 289.3 | - | - | 0 | - |
| - | - | 931.8 | 290.2 | - | - | 0 | - |
| - | - | 817.6 | 291.1 | - | - | 0 | - |
| - | - | 578.2 | 297.2 | - | - | 0 | - |
| - | - | 785.8 | 301.2 | - | - | 0 | - |
| - | - | 776.2 | 304.2 | - | - | 0 | - |
| 3 | b | 2260 | 316.2 | 0.0005697 | 1.802 | +1 | 3 |
| - | - | 1193 | 319.1 | - | - | 0 | - |
| - | - | 3185 | 326.2 | - | - | 0 | - |
| - | - | 664 | 327.2 | - | - | 0 | - |
| 3 | b | 3.972E+04 | 334.2 | 0.0005336 | 1.597 | +1 | 3 |
| - | - | 6069 | 335.2 | - | - | 0 | - |
| 5 | y | 960.3 | 340.2 | 0.0005416 | 1.592 | +2 | 7 |
| 9 | y | 758.5 | 343.2 | 0.001125 | 3.279 | +1 | 3 |
| - | - | 1099 | 346.2 | - | - | 0 | - |
| - | - | 650.4 | 348.2 | - | - | 0 | - |
| - | - | 1327 | 357.2 | - | - | 0 | - |
| 9 | y | 1924 | 361.2 | 7.029E-05 | 0.1946 | +1 | 3 |
| - | - | 1075 | 364.2 | - | - | 0 | - |
| - | - | 699 | 366.2 | - | - | 0 | - |
| - | - | 975.8 | 368.2 | - | - | 0 | - |
| - | - | 1758 | 374.2 | - | - | 0 | - |
| - | - | 860.6 | 375.2 | - | - | 0 | - |
| - | - | 584.9 | 382.2 | - | - | 0 | - |
| - | - | 3443 | 383.2 | - | - | 0 | - |
| - | - | 728.5 | 384.2 | - | - | 0 | - |
| - | - | 763.8 | 388.2 | - | - | 0 | - |
| - | - | 1782 | 392.2 | - | - | 0 | - |
| - | - | 3379 | 401.2 | - | - | 0 | - |
| - | - | 3644 | 402.2 | - | - | 0 | - |
| - | - | 621.1 | 402.2 | - | - | 0 | - |
| - | - | 2023 | 405.2 | - | - | 0 | - |
| - | - | 969.9 | 406.3 | - | - | 0 | - |
| 4 | b | 1416 | 415.2 | 0.0004541 | 1.094 | +1 | 4 |
| - | - | 1196 | 418.2 | - | - | 0 | - |
| 8 | y | 3664 | 418.3 | 4.191E-05 | 0.1002 | +1 | 4 |
| - | - | 1225 | 419.2 | - | - | 0 | - |
| - | - | 2277 | 420.2 | - | - | 0 | - |
| 4 | b | 8067 | 433.2 | 0.0005401 | 1.247 | +1 | 4 |
| - | - | 2362 | 434.2 | - | - | 0 | - |
| - | - | 737.7 | 436.2 | - | - | 0 | - |
| 3 | y | 1579 | 439.3 | 0.002547 | 5.799 | +2 | 9 |
| - | - | 1098 | 453.2 | - | - | 0 | - |
| - | - | 749.4 | 456.2 | - | - | 0 | - |
| 7 | y | 724.3 | 457.3 | 0.001061 | 2.32 | +1 | 5 |
| - | - | 854.2 | 463.2 | - | - | 0 | - |
| 7 | y | 5402 | 475.3 | 0.0007645 | 1.608 | +1 | 5 |
| - | - | 1048 | 476.3 | - | - | 0 | - |
| - | - | 1377 | 481.2 | - | - | 0 | - |
| - | - | 1909 | 501.2 | - | - | 0 | - |
| 6 | y | 1758 | 514.3 | 2.778E-05 | 0.05402 | +1 | 6 |
| - | - | 1727 | 519.3 | - | - | 0 | - |
| - | - | 2822 | 530.3 | - | - | 0 | - |
| - | - | 1022 | 531.3 | - | - | 0 | - |
| 6 | y | 2.286E+04 | 532.3 | 0.0001442 | 0.271 | +1 | 6 |
| - | - | 6586 | 533.3 | - | - | 0 | - |
| - | - | 1079 | 534.3 | - | - | 0 | - |
| - | - | 752.6 | 535.3 | - | - | 0 | - |
| - | - | 5307 | 548.3 | - | - | 0 | - |
| - | - | 1878 | 549.3 | - | - | 0 | - |
| - | - | 1214 | 552.3 | - | - | 0 | - |
| - | - | 837.5 | 554.6 | - | - | 0 | - |
| - | - | 2081 | 555 | - | - | 0 | - |
| - | - | 2313 | 555.3 | - | - | 0 | - |
| - | - | 2506 | 556.4 | - | - | 0 | - |
| - | - | 914 | 557.3 | - | - | 0 | - |
| 5 | b | 1352 | 562.3 | 0.002225 | 3.957 | +1 | 5 |
| - | - | 1360 | 566.3 | - | - | 0 | - |
| 5 | b | 1835 | 580.3 | 0.0003329 | 0.5737 | +1 | 5 |
| - | - | 658.1 | 581.3 | - | - | 0 | - |
| - | - | 639.3 | 599.6 | - | - | 0 | - |
| - | - | 861.8 | 600.3 | - | - | 0 | - |
| - | - | 807.3 | 610.8 | - | - | 0 | - |
| - | - | 672.8 | 613.1 | - | - | 0 | - |
| - | - | 575.1 | 616.6 | - | - | 0 | - |
| - | - | 673.8 | 617.3 | - | - | 0 | - |
| - | - | 932.7 | 618.3 | - | - | 0 | - |
| - | - | 643 | 629.3 | - | - | 0 | - |
| - | - | 2160 | 629.3 | - | - | 0 | - |
| - | - | 1066 | 630.3 | - | - | 0 | - |
| - | - | 780.8 | 643.4 | - | - | 0 | - |
| - | - | 603.6 | 644.4 | - | - | 0 | - |
| - | - | 1890 | 647.3 | - | - | 0 | - |
| 5 | y | 2250 | 661.4 | 0.0003319 | 0.5019 | +1 | 7 |
| - | - | 972.8 | 662.4 | - | - | 0 | - |
| - | - | 663.5 | 666.4 | - | - | 0 | - |
| - | - | 618 | 677.4 | - | - | 0 | - |
| 5 | y | 4.487E+04 | 679.4 | 3.236E-05 | 0.04764 | +1 | 7 |
| - | - | 1.858E+04 | 680.4 | - | - | 0 | - |
| - | - | 4115 | 681.4 | - | - | 0 | - |
| - | - | 611.5 | 694.9 | - | - | 0 | - |
| - | - | 614.2 | 695.4 | - | - | 0 | - |
| - | - | 935.1 | 728.4 | - | - | 0 | - |
| - | - | 637.2 | 742.4 | - | - | 0 | - |
| - | - | 1764 | 746.4 | - | - | 0 | - |
| - | - | 725.6 | 747.4 | - | - | 0 | - |
| 4 | y | 2085 | 760.4 | 0.002584 | 3.398 | +1 | 8 |
| 4 | y | 4.638E+04 | 778.4 | 0.000209 | 0.2685 | +1 | 8 |
| - | - | 1.923E+04 | 779.4 | - | - | 0 | - |
| - | - | 4463 | 780.5 | - | - | 0 | - |
| - | - | 655.3 | 781.5 | - | - | 0 | - |
| 9 | b | 685.6 | 834.4 | 0.003259 | 3.906 | +1 | 9 |
| - | - | 906 | 853.4 | - | - | 0 | - |
| 3 | y | 958.1 | 859.5 | 0.001573 | 1.83 | +1 | 9 |
| - | - | 622.3 | 860.5 | - | - | 0 | - |
| 3 | y | 3.544E+04 | 877.5 | 0.0009959 | 1.135 | +1 | 9 |
| - | - | 1.812E+04 | 878.5 | - | - | 0 | - |
| - | - | 4834 | 879.5 | - | - | 0 | - |
| - | - | 629.9 | 971.4 | - | - | 0 | - |
| 2 | y | 677.9 | 1025 | 0.00105 | 1.025 | +1 | 10 |
| - | - | 561.3 | 1552 | - | - | 0 | - |

m/z Charge Intensity FragmentType MassShift Position
120.08111572265625 0 30416.6
121.08448028564453 0 2385.0764
121.3980712890625 0 340.8632
125.0710220336914 0 441.6922
125.70158386230469 0 400.74066
126.06633758544922 0 464.13586
126.1030044555664 0 517.3743
127.05075073242188 0 636.16205
127.0870590209961 0 564.9815
128.0459442138672 0 586.3386
128.1073455810547 0 1014.79114
129.06631469726562 0 863.17017
129.10256958007812 0 38087.312
130.0612335205078 0 426.11078
130.06602478027344 0 607.5579
130.08670043945312 0 536.53754
130.10586547851562 0 2004.7515
131.0455322265625 0 468.14148
131.07044982910156 0 638.4993
131.08154296875 0 1160.0911
132.0811004638672 0 1298.1416
132.1022491455078 0 3764.505 y 10
132.23167419433594 0 461.21564
133.06065368652344 0 524.1512
133.08621215820312 0 6810.524
133.10507202148438 0 550.24585
134.4835968017578 0 439.56943
136.07598876953125 0 4612.887
138.0663604736328 0 875.278
139.08688354492188 0 744.1004
140.0821075439453 0 890.90265
141.0660400390625 0 789.55597
141.14273071289062 0 430.70694
144.08114624023438 0 3058.8562
145.06100463867188 0 640.2675
145.0852813720703 0 522.3019
146.0558319091797 0 463.48926
147.1131134033203 0 2481.99
152.6229705810547 0 488.17874
155.08192443847656 0 1447.6031
155.1183319091797 0 960.86615
156.07704162597656 0 1054.2172
158.09197998046875 0 699.7701
159.07650756835938 0 1082.4424
159.09194946289062 0 617.2673
159.17169189453125 0 504.91327
162.0916748046875 0 7845.437
165.75393676757812 0 470.8829
166.78184509277344 0 438.28677
169.09768676757812 0 693.2011
171.1495361328125 0 13274.872
172.07203674316406 0 1794.5502
172.10853576660156 0 702.11896
172.15277099609375 0 1287.712
173.1287384033203 0 1477.3274
175.1190185546875 0 595.11395
176.10733032226562 0 1497.3318
177.1027069091797 0 1340.3402
177.11251831054688 0 2381.2002
181.0611114501953 0 663.1226
183.11289978027344 0 595.39777
183.14990234375 0 563.4556
184.0729217529297 0 551.1507
185.12850952148438 0 758.05853
185.16539001464844 0 597.07776
186.12432861328125 0 690.58417
187.0724334716797 0 532.7266
187.08724975585938 0 600.2844
187.0969696044922 0 718.5099
187.1078643798828 0 865.3968
189.1121826171875 0 565.20026
195.11367797851562 0 699.67413
197.1651611328125 0 604.5661
198.08810424804688 0 816.09894
199.14447021484375 0 7586.327
201.12384033203125 0 1360.977 y Ammonia loss 7
203.10301208496094 0 2845.4827
205.09751892089844 0 3371.6543
207.11317443847656 0 125148.83 a 1
208.1165313720703 0 12307.264 b Water loss 3
209.11956787109375 0 637.2804
212.13963317871094 0 4257.753
215.11415100097656 0 679.1836
215.1394500732422 0 1291.3413
216.09744262695312 0 883.4317
217.09738159179688 0 1403.556 b Water loss 1
219.1494903564453 0 6348.3486
220.15335083007812 0 1132.6521
221.1038055419922 0 769.4177
222.12387084960938 0 616.00476
226.15570068359375 0 2423.6125
227.10281372070312 0 741.51447
227.11422729492188 0 1557.6149
228.0982666015625 0 576.5067
229.11904907226562 0 680.7113
230.15023803710938 0 7086.847
231.1520538330078 0 924.5604
235.10809326171875 0 82036.04 b 1
236.1114501953125 0 10998.639
237.11277770996094 0 866.01013
239.14923095703125 0 578.53033
243.11231994628906 0 471.77902
243.1455841064453 0 1065.1033
243.17059326171875 0 2696.7302 y Ammonia loss 9
244.17591857910156 0 581.38043
245.12600708007812 0 758.1998
246.12371826171875 0 1960.5753
247.12994384765625 0 948.54407
247.14431762695312 0 4423.3345
255.10928344726562 0 3357.3728
258.0982666015625 0 606.3251
260.1971130371094 0 11859.949 y 9
261.1229248046875 0 1105.392
261.1601867675781 0 721.5756
261.200439453125 0 1913.2441
262.1188659667969 0 1627.3854
269.1612243652344 0 3690.8923
273.1195373535156 0 4672.3066
273.1958312988281 0 638.2426
274.119140625 0 2425.6116
282.15673828125 0 533.5813
283.1759948730469 0 805.20917
287.171142578125 0 1439.4551
289.1549377441406 0 3961.8452
289.3199462890625 0 580.14417
290.1575622558594 0 931.79333
291.1459655761719 0 817.6497
297.1925964355469 0 578.2219
301.1921691894531 0 785.80536
304.16546630859375 0 776.22626
316.1661376953125 0 2260.107 b Water loss 2
319.1398010253906 0 1193.3859
326.1826171875 0 3184.8003
327.1658630371094 0 664.0173
334.1766662597656 0 39717.062 b 2
335.1795654296875 0 6068.7466
340.1917724609375 0 960.29956 y 4
343.235107421875 0 758.52625 y Water loss 8
346.21282958984375 0 1099.3923
348.16693115234375 0 650.3993
357.1557312011719 0 1326.5585
361.2444763183594 0 1924.017 y 8
364.16534423828125 0 1075.1997
366.17608642578125 0 699.028
368.2293701171875 0 975.77246
374.18182373046875 0 1757.7825
375.1662902832031 0 860.56555
382.17718505859375 0 584.89575
383.2035827636719 0 3443.1404
384.2053527832031 0 728.48987
388.22381591796875 0 763.80646
392.1936950683594 0 1782.4015
401.21490478515625 0 3379.4873
402.1771240234375 0 3644.4507
402.2165832519531 0 621.07007
405.24969482421875 0 2022.7556
406.25396728515625 0 969.8919
415.23443603515625 0 1415.5342 b Water loss 3
418.2132263183594 0 1196.4824
418.26605224609375 0 3663.6936 y 7
419.2238464355469 0 1224.5098
420.18890380859375 0 2277.3948
433.2450866699219 0 8067.3745 b 3
434.24822998046875 0 2362.449
436.2248229980469 0 737.6785
439.2632751464844 0 1579.1771 y 2
453.24951171875 0 1098.4579
456.22283935546875 0 749.3621
457.2758483886719 0 724.2578 y Water loss 6
463.235107421875 0 854.1995
475.2882385253906 0 5401.8364 y 6
476.291259765625 0 1048.3916
481.2440490722656 0 1376.661
501.2455749511719 0 1908.8145
514.2984008789062 0 1757.6487 y Water loss 5
519.2551879882812 0 1727.3008
530.27294921875 0 2821.5498
531.2772827148438 0 1021.7124
532.30908203125 0 22862.775 y 5
533.3119506835938 0 6585.9795
534.3115234375 0 1078.6497
535.2896118164062 0 752.55005
548.282958984375 0 5306.762
549.28662109375 0 1878.2716
552.3168334960938 0 1213.8966
554.6312866210938 0 837.5023
554.9700317382812 0 2081.016
555.3035278320312 0 2312.9976
556.3814086914062 0 2505.8533
557.3108520507812 0 914.0442
562.3001708984375 0 1352.2213 b Water loss 4
566.2938842773438 0 1360.469
580.3132934570312 0 1834.7789 b 4
581.3145751953125 0 658.0684
599.6431274414062 0 639.3106
600.3119506835938 0 861.8034
610.7868041992188 0 807.3024
613.1491088867188 0 672.77686
616.5615844726562 0 575.11707
617.344970703125 0 673.7885
618.3212890625 0 932.6594
629.2843017578125 0 643.03625
629.3423461914062 0 2159.912
630.3433837890625 0 1066.4381
643.3562622070312 0 780.8263
644.3506469726562 0 603.63513
647.3490600585938 0 1889.9286
661.366455078125 0 2249.566 y Water loss 4
662.369140625 0 972.7935
666.3645629882812 0 663.46967
677.3573608398438 0 618.04926
679.3773193359375 0 44871.582 y 4
680.3804931640625 0 18582.916
681.382568359375 0 4114.9883
694.8950805664062 0 611.4725
695.3976440429688 0 614.2243
728.4074096679688 0 935.1049
742.4262084960938 0 637.1529
746.4161987304688 0 1764.0629
747.4169921875 0 725.64557
760.4326171875 0 2085.0256 y Water loss 3
778.445556640625 0 46384.445 y 3
779.44873046875 0 19232.482
780.4505004882812 0 4463.4854
781.455078125 0 655.27325
834.417724609375 0 685.579 b Water loss 8
853.4315795898438 0 906.0274
859.5051879882812 0 958.1491 y Water loss 2
860.51220703125 0 622.31616
877.51318359375 0 35436.156 y 2
878.5162353515625 0 18122.717
879.5203247070312 0 4833.9717
971.3637084960938 0 629.8817
1024.58154296875 0 677.8815 y 1
1552.4715576171875 0 561.2611

Spectrum Details

|  |  |
| --- | --- |
| Matched peaks? Matched peaks The total absolute number of peaks matched. Additionally in brackets the total fraction of peaks matched and the total number of peaks is shown. | 31 (13.19% of 235) |
| FDR? FDR The false discovery rate estimated for this peptide. It is calculated by matching all theoretical fragments with a non-integer shift with the raw peaks for this spectrum. This is done with 40 different shifts. The resulting percentage is the average number of annotated peaks over the number of annotated peaks with the correct spectrum. | 0.46% |
| Satellite FDR? Satellite FDR See the FDR for details on its calculation. This satellite ion specific FDR only contains the satellite ions (d/w) for I/L/J positions. | - |
| PSM Score? PSM Score The PSM Score as given by Hecklib to this annotated spectrum. It is shown with three significant figures. | 391 |

#### Reverse Lookup? Reverse Lookup All places where this read could be placed.

| Group | Segment | Template | Template Part | Read Part | Score | Unique |
| --- | --- | --- | --- | --- | --- | --- |
| Homo sapiens Light Chain | IGLJ | IGLJ2 | [0..9] | [2..11] | 72 | True |

| Recombined | Template Part | Read Part | Score | Unique |
| --- | --- | --- | --- | --- |
| REC-0-1\_002 | [98..108] | [0..11] | 79 | True |

#### Meta Information from Multiple reads

##### Number of combined reads

2

##### Intensity

0.6912

##### TotalArea

7.338E+07

##### Changes to the peptide sequence

SFVVFGGGTKJ

L→JNo support for either Leucine or Isoleucine based on side chain ions (Position: 11)

#### Positional Score

Copy Data

##### Positional Score (TSV)

###### Preview

```
Loading example...
```

*Click on the button to copy the data to your clipboard.*

00012345678910

Label Value
"0" 0
"1" 0
"2" 0
"3" 0
"4" 0
"5" 0
"6" 0
"7" 0
"8" 0
"9" 0
"10" 0

#### Meta Information from PEAKS

##### Scan Identifier

F1:9569

##### Original sequence

S

F

V

V

F

G

G

G

T

K

L

##### Posttranslational Modifications

##### Source File

D:\separate\_stitch\_analyses\xle-disambiguation\raw\20210323\_F1\_UM1\_Peng0013\_SA\_F59\_ingel\_3ug\_ELA.raw

##### Fraction

1

##### Scan Feature

F1:6501

##### De Novo Score

99

##### ConfidenceScore

99

### m/z

556.3126

##### Mass

1110.6074

##### Charge

2

##### Retention Time

52.71

##### Predicted Retention Time

-

##### Area

3.669E+07

##### Parts Per Million

2.8

##### Fragmentation mode

HCD

##### Originating file

01 D:\separate\_stitch\_analyses\xle-disambiguation\20210325\_F59\_3ug\_DENOVO\_12.csv

#### Meta Information from PEAKS

##### Scan Identifier

F1:9687

##### Original sequence

S

F

V

V

F

G

G

G

T

K

L

##### Posttranslational Modifications

##### Source File

D:\separate\_stitch\_analyses\xle-disambiguation\raw\20210323\_F1\_UM1\_Peng0013\_SA\_F59\_ingel\_3ug\_ELA.raw

##### Fraction

1

##### Scan Feature

F1:6501

##### De Novo Score

98

##### ConfidenceScore

98

### m/z

556.3126

##### Mass

1110.6074

##### Charge

2

##### Retention Time

52.71

##### Predicted Retention Time

-

##### Area

3.669E+07

##### Parts Per Million

2.8

##### Fragmentation mode

HCD

##### Originating file

01 D:\separate\_stitch\_analyses\xle-disambiguation\20210325\_F59\_3ug\_DENOVO\_12.csv
