## Supplementary material for "A handle on mass coincidence errors in *de novo* sequencing of antibodies by bottom-up proteomics": Combined_045.html

Details Combined\_045 | Stitch OverviewUndefined

### Read Combined\_045

#### Sequence (length=12)

TFPAVJQSSGJY

#### Spectrum 11139? Spectrum 11139 The raw spectrum of this peptide as annotated by Hecklib. The fragments are coloured according to ion type (see legend). Any peaks with a star '\*' as text can be hovered over to see the full details, first the ion type second the mass shift type. By hovering over the amino acids in the peptide or ions in the legend the corresponding peaks are highlighted. By toggling the 'Unassigned' label you can turn the background (unassigned) peaks on or off in the plot. By updating the slider in the Ion legend you can update the spectrum to only show the top X% of the peaks with labels. The top X% means any peak that is within X% of the highest intensity. By dragging in the spectrum you can zoom in to a specific part of the spectrum and use 'Zoom Out' to get back to the original zoom level. The annotation of the spectrum is based on the given sequence in the peptides file and is done with different software so inconsistencies are likely. The peaks are annotated based on the given sequence, with 20 ppm tolerance.

Copy Data

##### Spectrum 11139 (TSV)

###### Preview

```
Loading example...
```

*Click on the button to copy the data to your clipboard.*

Mz MinMz MaxIntensity Max

WidthHeightPeptide font sizePeptide stroke widthSpectrum font sizeSpectrum stroke widthCompact peptide

Ion legend

wxyz

abcd

OtherUnassignedIon^Charge^_Position_Show for top:%

TFPAVJQSSGJY

02.65e+45.30e+47.95e+41.06e+5

Zoom Out

y^+1^_1_y^+1^_2_y^+1^_3_z^+2^_8_w^+2^_8_c^+2^_8_y^+2^_8_y^+1^_4_y^+1^_5_y^+1^_5_c^+1^_5_y^+1^_6_c^+1^_6_y^+1^_6_y^+1^_7_z^+1^_7_c^+1^_7_c^+1^_7_y^+1^_7_c^+1^_7_c^+1^_8_c^+1^_8_z^+1^_8_c^+1^_8_y^+1^_8_z^+1^_9_z^+1^_9_z^+1^_9_c^+1^_9_c^+1^_9_y^+1^_9_c^+1^_9_c^+1^_10_c^+1^_10_y^+1^_10_z^+1^_10_y^+1^_10_c^+1^_11_c^+1^_11_z^+1^_11_z^+1^_11_z^+1^_11_

0794158723813174

Fragment Matches Table

Show background peaks

| Position | Ion type | Intensity | mz Theoretical | mz Error (Th) | mz Error (ppm) | Charge | Series Number |
| --- | --- | --- | --- | --- | --- | --- | --- |
| - | - | 632.1 | 120.1 | - | - | 0 | - |
| - | - | 322.7 | 121.5 | - | - | 0 | - |
| - | - | 411.3 | 131.9 | - | - | 0 | - |
| - | - | 503.2 | 134.7 | - | - | 0 | - |
| - | - | 5193 | 136.1 | - | - | 0 | - |
| - | - | 505.2 | 137.1 | - | - | 0 | - |
| - | - | 462.4 | 142 | - | - | 0 | - |
| - | - | 615.7 | 143.1 | - | - | 0 | - |
| - | - | 535.3 | 146.2 | - | - | 0 | - |
| - | - | 953.2 | 149 | - | - | 0 | - |
| - | - | 4286 | 165.1 | - | - | 0 | - |
| - | - | 525 | 168.1 | - | - | 0 | - |
| - | - | 2332 | 169.1 | - | - | 0 | - |
| - | - | 1364 | 171.1 | - | - | 0 | - |
| - | - | 419.4 | 171.6 | - | - | 0 | - |
| - | - | 2599 | 173.4 | - | - | 0 | - |
| 12 | y | 3.153E+04 | 182.1 | 0.0001139 | 0.6255 | +1 | 1 |
| - | - | 2373 | 183.1 | - | - | 0 | - |
| - | - | 484.5 | 186.1 | - | - | 0 | - |
| - | - | 562.2 | 214.1 | - | - | 0 | - |
| - | - | 1.081E+04 | 221.1 | - | - | 0 | - |
| - | - | 940.9 | 222.1 | - | - | 0 | - |
| - | - | 510.7 | 230.9 | - | - | 0 | - |
| - | - | 647.3 | 232.1 | - | - | 0 | - |
| - | - | 432.8 | 234.9 | - | - | 0 | - |
| - | - | 967.1 | 240.2 | - | - | 0 | - |
| - | - | 7728 | 249.1 | - | - | 0 | - |
| - | - | 1679 | 258.1 | - | - | 0 | - |
| - | - | 510.7 | 259.1 | - | - | 0 | - |
| - | - | 541.8 | 267.9 | - | - | 0 | - |
| - | - | 5199 | 268.2 | - | - | 0 | - |
| - | - | 782 | 269.2 | - | - | 0 | - |
| 11 | y | 4150 | 295.2 | 4.201E-05 | 0.1423 | +1 | 2 |
| - | - | 594.1 | 308.1 | - | - | 0 | - |
| - | - | 3431 | 317.2 | - | - | 0 | - |
| - | - | 549.6 | 318.2 | - | - | 0 | - |
| - | - | 495.9 | 343.4 | - | - | 0 | - |
| - | - | 4372 | 345.2 | - | - | 0 | - |
| - | - | 1341 | 346.2 | - | - | 0 | - |
| - | - | 729.1 | 351.2 | - | - | 0 | - |
| 10 | y | 1722 | 352.2 | 5.187E-05 | 0.1473 | +1 | 3 |
| - | - | 2428 | 361.7 | - | - | 0 | - |
| - | - | 750.4 | 362.2 | - | - | 0 | - |
| - | - | 2308 | 381.2 | - | - | 0 | - |
| - | - | 585.3 | 398.3 | - | - | 0 | - |
| - | - | 1504 | 399.2 | - | - | 0 | - |
| - | - | 1636 | 414.3 | - | - | 0 | - |
| 5 | z | 3066 | 417.2 | 0.001411 | 3.383 | +2 | 8 |
| 5 | w | 578.9 | 418.2 | 0.005551 | 13.27 | +2 | 8 |
| 8 | c | 599.2 | 422.7 | 0.001579 | 3.735 | +2 | 8 |
| 5 | y | 574.2 | 425.2 | 0.006608 | 15.54 | +2 | 8 |
| - | - | 1023 | 428.2 | - | - | 0 | - |
| 9 | y | 1154 | 439.2 | 0.0001158 | 0.2636 | +1 | 4 |
| - | - | 1065 | 455.2 | - | - | 0 | - |
| - | - | 1844 | 456.2 | - | - | 0 | - |
| - | - | 750.1 | 457.2 | - | - | 0 | - |
| - | - | 1870 | 473.2 | - | - | 0 | - |
| - | - | 644.6 | 482.3 | - | - | 0 | - |
| - | - | 2871 | 485.8 | - | - | 0 | - |
| - | - | 1371 | 486.3 | - | - | 0 | - |
| - | - | 1178 | 488.3 | - | - | 0 | - |
| - | - | 4609 | 498.3 | - | - | 0 | - |
| - | - | 724.6 | 499.3 | - | - | 0 | - |
| 8 | y | 1154 | 508.2 | 0.00154 | 3.03 | +1 | 5 |
| - | - | 4962 | 509.3 | - | - | 0 | - |
| - | - | 1653 | 510.3 | - | - | 0 | - |
| - | - | 7073 | 516.3 | - | - | 0 | - |
| - | - | 2611 | 517.3 | - | - | 0 | - |
| 8 | y | 5039 | 526.3 | 0.000388 | 0.7373 | +1 | 5 |
| - | - | 1655 | 526.3 | - | - | 0 | - |
| - | - | 1401 | 527.3 | - | - | 0 | - |
| - | - | 5482 | 532.3 | - | - | 0 | - |
| 5 | c | 1.813E+04 | 533.3 | 0.0003482 | 0.6529 | +1 | 5 |
| - | - | 5428 | 534.3 | - | - | 0 | - |
| - | - | 753.5 | 535.3 | - | - | 0 | - |
| - | - | 629.5 | 568.3 | - | - | 0 | - |
| - | - | 627.3 | 569.3 | - | - | 0 | - |
| - | - | 532.9 | 584.3 | - | - | 0 | - |
| - | - | 2289 | 586.3 | - | - | 0 | - |
| - | - | 1285 | 596.3 | - | - | 0 | - |
| - | - | 712.5 | 601.4 | - | - | 0 | - |
| - | - | 1734 | 602.4 | - | - | 0 | - |
| - | - | 9265 | 603.4 | - | - | 0 | - |
| - | - | 3245 | 604.4 | - | - | 0 | - |
| - | - | 546.2 | 609.3 | - | - | 0 | - |
| - | - | 4840 | 611.4 | - | - | 0 | - |
| - | - | 2065 | 612.3 | - | - | 0 | - |
| - | - | 687.1 | 613.4 | - | - | 0 | - |
| - | - | 6782 | 629.4 | - | - | 0 | - |
| - | - | 2802 | 630.4 | - | - | 0 | - |
| - | - | 981.2 | 631.4 | - | - | 0 | - |
| - | - | 651.1 | 635.3 | - | - | 0 | - |
| 7 | y | 1957 | 636.3 | 0.002624 | 4.125 | +1 | 6 |
| - | - | 1981 | 641.9 | - | - | 0 | - |
| - | - | 3810 | 642.4 | - | - | 0 | - |
| - | - | 954.9 | 642.9 | - | - | 0 | - |
| - | - | 1.182E+04 | 645.4 | - | - | 0 | - |
| 6 | c | 3.832E+04 | 646.4 | 0.0006719 | 1.039 | +1 | 6 |
| - | - | 1.332E+04 | 647.4 | - | - | 0 | - |
| - | - | 2163 | 648.4 | - | - | 0 | - |
| - | - | 5317 | 653.3 | - | - | 0 | - |
| 7 | y | 1.727E+04 | 654.3 | 0.001104 | 1.688 | +1 | 6 |
| - | - | 6229 | 655.3 | - | - | 0 | - |
| - | - | 1280 | 656.3 | - | - | 0 | - |
| - | - | 2015 | 672.4 | - | - | 0 | - |
| - | - | 805.6 | 685.4 | - | - | 0 | - |
| - | - | 867.7 | 700.4 | - | - | 0 | - |
| - | - | 1095 | 702.4 | - | - | 0 | - |
| - | - | 767.1 | 712.4 | - | - | 0 | - |
| - | - | 2857 | 722.4 | - | - | 0 | - |
| - | - | 1455 | 723.4 | - | - | 0 | - |
| - | - | 946.2 | 727.4 | - | - | 0 | - |
| - | - | 6164 | 730.4 | - | - | 0 | - |
| - | - | 1.44E+04 | 731.4 | - | - | 0 | - |
| - | - | 5086 | 732.4 | - | - | 0 | - |
| - | - | 869.9 | 733.4 | - | - | 0 | - |
| - | - | 3073 | 739.4 | - | - | 0 | - |
| - | - | 6898 | 740.4 | - | - | 0 | - |
| - | - | 4052 | 741.4 | - | - | 0 | - |
| - | - | 1889 | 742.4 | - | - | 0 | - |
| 6 | y | 669.7 | 750.4 | 0.002355 | 3.139 | +1 | 7 |
| 6 | z | 3508 | 751.4 | 0.0005261 | 0.7002 | +1 | 7 |
| - | - | 1761 | 752.4 | - | - | 0 | - |
| - | - | 800.7 | 753.4 | - | - | 0 | - |
| - | - | 2049 | 755.4 | - | - | 0 | - |
| 7 | c | 868.5 | 756.4 | 0.01438 | 19.01 | +1 | 7 |
| 7 | c | 8618 | 757.4 | 0.0002296 | 0.3032 | +1 | 7 |
| - | - | 5301 | 758.4 | - | - | 0 | - |
| - | - | 3287 | 759.4 | - | - | 0 | - |
| - | - | 1794 | 766.4 | - | - | 0 | - |
| 6 | y | 1.012E+04 | 767.4 | 0.001489 | 1.94 | +1 | 7 |
| - | - | 4109 | 768.4 | - | - | 0 | - |
| - | - | 1.854E+04 | 773.4 | - | - | 0 | - |
| 7 | c | 2.711E+04 | 774.5 | 0.002853 | 3.684 | +1 | 7 |
| - | - | 1.095E+04 | 775.5 | - | - | 0 | - |
| - | - | 2627 | 776.5 | - | - | 0 | - |
| - | - | 939 | 799.5 | - | - | 0 | - |
| - | - | 1485 | 800.5 | - | - | 0 | - |
| - | - | 683.7 | 809.4 | - | - | 0 | - |
| - | - | 1097 | 817.5 | - | - | 0 | - |
| - | - | 858.8 | 819.5 | - | - | 0 | - |
| - | - | 1646 | 825.5 | - | - | 0 | - |
| - | - | 905.7 | 826.4 | - | - | 0 | - |
| - | - | 1166 | 827.4 | - | - | 0 | - |
| - | - | 1005 | 835.5 | - | - | 0 | - |
| - | - | 2242 | 842.5 | - | - | 0 | - |
| 8 | c | 1099 | 843.5 | 0.009302 | 11.03 | +1 | 8 |
| 8 | c | 4413 | 844.5 | 0.0001535 | 0.1818 | +1 | 8 |
| - | - | 1936 | 845.5 | - | - | 0 | - |
| - | - | 905.5 | 846.5 | - | - | 0 | - |
| 5 | z | 3291 | 850.4 | 0.0002755 | 0.3239 | +1 | 8 |
| - | - | 2317 | 851.4 | - | - | 0 | - |
| - | - | 7665 | 853.5 | - | - | 0 | - |
| - | - | 3235 | 854.5 | - | - | 0 | - |
| - | - | 1.345E+04 | 860.5 | - | - | 0 | - |
| 8 | c | 1.551E+04 | 861.5 | 0.004303 | 4.995 | +1 | 8 |
| - | - | 720.1 | 861.6 | - | - | 0 | - |
| - | - | 5599 | 862.5 | - | - | 0 | - |
| - | - | 1875 | 863.5 | - | - | 0 | - |
| - | - | 722.6 | 865.4 | - | - | 0 | - |
| 5 | y | 4666 | 866.5 | 0.0009942 | 1.147 | +1 | 8 |
| - | - | 1856 | 867.5 | - | - | 0 | - |
| 4 | z | 2422 | 903.5 | 0.000275 | 0.3044 | +1 | 9 |
| 4 | z | 2436 | 904.5 | 0.01498 | 16.56 | +1 | 9 |
| - | - | 1433 | 913.5 | - | - | 0 | - |
| - | - | 769.9 | 914.5 | - | - | 0 | - |
| - | - | 733.1 | 919.4 | - | - | 0 | - |
| 4 | z | 2.049E+04 | 921.5 | 0.001562 | 1.695 | +1 | 9 |
| - | - | 1.202E+04 | 922.5 | - | - | 0 | - |
| - | - | 3663 | 923.5 | - | - | 0 | - |
| - | - | 788.3 | 924.5 | - | - | 0 | - |
| - | - | 1331 | 929.5 | - | - | 0 | - |
| 9 | c | 1798 | 930.5 | 0.009104 | 9.784 | +1 | 9 |
| 9 | c | 3396 | 931.5 | 0.0005657 | 0.6074 | +1 | 9 |
| - | - | 1830 | 932.5 | - | - | 0 | - |
| 4 | y | 1625 | 937.5 | 0.002908 | 3.102 | +1 | 9 |
| - | - | 1.476E+04 | 947.5 | - | - | 0 | - |
| 9 | c | 3.325E+04 | 948.5 | 0.002213 | 2.333 | +1 | 9 |
| - | - | 1.523E+04 | 949.5 | - | - | 0 | - |
| - | - | 3682 | 950.5 | - | - | 0 | - |
| - | - | 929.5 | 952.5 | - | - | 0 | - |
| - | - | 1608 | 953.5 | - | - | 0 | - |
| - | - | 841.4 | 955.5 | - | - | 0 | - |
| - | - | 5950 | 970.5 | - | - | 0 | - |
| - | - | 3786 | 971.5 | - | - | 0 | - |
| - | - | 1196 | 972.5 | - | - | 0 | - |
| - | - | 8387 | 984.5 | - | - | 0 | - |
| - | - | 5374 | 985.5 | - | - | 0 | - |
| - | - | 1460 | 986.5 | - | - | 0 | - |
| 10 | c | 7512 | 988.5 | 0.0006672 | 0.6749 | +1 | 10 |
| - | - | 4061 | 989.5 | - | - | 0 | - |
| - | - | 918.8 | 990.5 | - | - | 0 | - |
| 10 | c | 9.966E+04 | 1006 | 0.001215 | 1.209 | +1 | 10 |
| - | - | 5.667E+04 | 1007 | - | - | 0 | - |
| - | - | 1.746E+04 | 1008 | - | - | 0 | - |
| - | - | 1698 | 1009 | - | - | 0 | - |
| - | - | 749 | 1013 | - | - | 0 | - |
| - | - | 1074 | 1016 | - | - | 0 | - |
| 3 | y | 1241 | 1017 | 0.01201 | 11.81 | +1 | 10 |
| 3 | z | 751.3 | 1019 | 0.003292 | 3.232 | +1 | 10 |
| - | - | 817.7 | 1034 | - | - | 0 | - |
| 3 | y | 6290 | 1035 | 0.002005 | 1.938 | +1 | 10 |
| - | - | 3790 | 1036 | - | - | 0 | - |
| - | - | 689.2 | 1058 | - | - | 0 | - |
| - | - | 5161 | 1069 | - | - | 0 | - |
| - | - | 3394 | 1070 | - | - | 0 | - |
| - | - | 1022 | 1071 | - | - | 0 | - |
| - | - | 1729 | 1074 | - | - | 0 | - |
| - | - | 5087 | 1075 | - | - | 0 | - |
| - | - | 2883 | 1076 | - | - | 0 | - |
| - | - | 1109 | 1077 | - | - | 0 | - |
| - | - | 2820 | 1084 | - | - | 0 | - |
| - | - | 1732 | 1085 | - | - | 0 | - |
| - | - | 1.304E+04 | 1086 | - | - | 0 | - |
| - | - | 8405 | 1087 | - | - | 0 | - |
| - | - | 2911 | 1088 | - | - | 0 | - |
| - | - | 834.1 | 1095 | - | - | 0 | - |
| - | - | 1455 | 1096 | - | - | 0 | - |
| 11 | c | 8136 | 1102 | 0.0007468 | 0.6779 | +1 | 11 |
| - | - | 3085 | 1103 | - | - | 0 | - |
| - | - | 3321 | 1104 | - | - | 0 | - |
| - | - | 1042 | 1105 | - | - | 0 | - |
| 11 | c | 8.559E+04 | 1119 | 0.002271 | 2.031 | +1 | 11 |
| - | - | 5.681E+04 | 1120 | - | - | 0 | - |
| - | - | 1.978E+04 | 1121 | - | - | 0 | - |
| - | - | 1698 | 1122 | - | - | 0 | - |
| - | - | 1215 | 1133 | - | - | 0 | - |
| - | - | 805.2 | 1134 | - | - | 0 | - |
| 2 | z | 2148 | 1148 | 0.001335 | 1.163 | +1 | 11 |
| 2 | z | 1029 | 1149 | 0.01795 | 15.62 | +1 | 11 |
| 2 | z | 1.845E+04 | 1166 | 0.002134 | 1.831 | +1 | 11 |
| - | - | 1.451E+04 | 1167 | - | - | 0 | - |
| - | - | 4958 | 1168 | - | - | 0 | - |
| - | - | 1620 | 1196 | - | - | 0 | - |
| - | - | 1114 | 1205 | - | - | 0 | - |
| - | - | 1398 | 1206 | - | - | 0 | - |
| - | - | 2557 | 1211 | - | - | 0 | - |
| - | - | 1601 | 1212 | - | - | 0 | - |
| - | - | 2405 | 1213 | - | - | 0 | - |
| - | - | 991 | 1214 | - | - | 0 | - |
| - | - | 3763 | 1222 | - | - | 0 | - |
| - | - | 2571 | 1223 | - | - | 0 | - |
| - | - | 973.1 | 1224 | - | - | 0 | - |
| - | - | 1576 | 1228 | - | - | 0 | - |
| - | - | 1165 | 1229 | - | - | 0 | - |
| - | - | 1.278E+04 | 1239 | - | - | 0 | - |
| - | - | 9406 | 1240 | - | - | 0 | - |
| - | - | 5540 | 1241 | - | - | 0 | - |
| - | - | 2199 | 1242 | - | - | 0 | - |
| - | - | 8307 | 1249 | - | - | 0 | - |
| - | - | 6877 | 1250 | - | - | 0 | - |
| - | - | 2019 | 1251 | - | - | 0 | - |
| - | - | 4684 | 1256 | - | - | 0 | - |
| - | - | 4238 | 1257 | - | - | 0 | - |
| - | - | 1374 | 1258 | - | - | 0 | - |
| - | - | 2081 | 1266 | - | - | 0 | - |
| - | - | 1.44E+04 | 1267 | - | - | 0 | - |
| - | - | 1.127E+04 | 1268 | - | - | 0 | - |
| - | - | 3904 | 1269 | - | - | 0 | - |
| - | - | 763.8 | 1281 | - | - | 0 | - |
| - | - | 739.7 | 1282 | - | - | 0 | - |
| - | - | 2.521E+04 | 1283 | - | - | 0 | - |
| - | - | 1.049E+05 | 1284 | - | - | 0 | - |
| - | - | 7.142E+04 | 1285 | - | - | 0 | - |
| - | - | 2.893E+04 | 1286 | - | - | 0 | - |
| - | - | 3280 | 1287 | - | - | 0 | - |
| - | - | 743.2 | 1911 | - | - | 0 | - |
| - | - | 655.2 | 2190 | - | - | 0 | - |
| - | - | 737.8 | 3080 | - | - | 0 | - |
| - | - | 798.8 | 3081 | - | - | 0 | - |
| - | - | 650.7 | 3143 | - | - | 0 | - |

m/z Charge Intensity FragmentType MassShift Position
120.08087921142578 0 632.1291
121.47071838378906 0 322.71402
131.93402099609375 0 411.28604
134.74502563476562 0 503.15765
136.07579040527344 0 5193.121
137.07920837402344 0 505.16754
141.9900360107422 0 462.36667
143.11807250976562 0 615.6765
146.21578979492188 0 535.26117
148.95460510253906 0 953.1686
165.05477905273438 0 4286.4673
168.12098693847656 0 524.96716
169.0972137451172 0 2332.0085
171.11300659179688 0 1363.7734
171.55931091308594 0 419.4396
173.4394073486328 0 2598.9292
182.08128356933594 0 31525.857 y 11
183.08456420898438 0 2372.6885
186.12257385253906 0 484.52145
214.0830078125 0 562.1711
221.12850952148438 0 10805.618
222.1315155029297 0 940.9317
230.89662170410156 0 510.6546
232.0924835205078 0 647.3363
234.9329071044922 0 432.78824
240.1703643798828 0 967.0959
249.12350463867188 0 7727.852
258.1448974609375 0 1679.4059
259.1496276855469 0 510.67776
267.9453125 0 541.8423
268.1656494140625 0 5198.963
269.1686096191406 0 782.0062
295.1651916503906 0 4149.657 y 10
308.12548828125 0 594.0562
317.1820983886719 0 3430.662
318.18414306640625 0 549.6078
343.37255859375 0 495.86942
345.1770324707031 0 4371.9067
346.1778259277344 0 1341.3832
351.20330810546875 0 729.0545
352.1866455078125 0 1721.7473 y 9
361.6949157714844 0 2427.7092
362.197509765625 0 750.3927
381.24951171875 0 2307.854
398.27642822265625 0 585.32263
399.203369140625 0 1504.407
414.2713317871094 0 1636.234
417.21331787109375 0 3066.109 z Ammonia loss 4
418.218994140625 0 578.93207 w 4
422.730224609375 0 599.1985 c Ammonia loss 7
425.21466064453125 0 574.22455 y Ammonia loss 4
428.21246337890625 0 1023.35394
439.2188415527344 0 1154.1969 y 8
455.22528076171875 0 1064.61
456.2093505859375 0 1844.0914
457.2449645996094 0 750.05646
473.23651123046875 0 1869.7925
482.297119140625 0 644.64856
485.7524108886719 0 2870.9565
486.2543640136719 0 1370.9513
488.28460693359375 0 1178.1084
498.27093505859375 0 4609.162
499.271484375 0 724.62933
508.2417297363281 0 1154.0618 y Water loss 7
509.308349609375 0 4962.0913
510.3094787597656 0 1652.588
516.2817993164062 0 7073.0645
517.2847900390625 0 2610.5706
526.2503662109375 0 5039.289 y 7
526.333740234375 0 1655.014
527.2540893554688 0 1400.593
532.300537109375 0 5481.816
533.307861328125 0 18129.67 c 4
534.310791015625 0 5428.461
535.313232421875 0 753.4793
568.3087768554688 0 629.5078
569.2911987304688 0 627.2995
584.3482666015625 0 532.938
586.3176879882812 0 2288.5017
596.3402709960938 0 1285.4597
601.3685302734375 0 712.5315
602.3773193359375 0 1734.3922
603.3855590820312 0 9264.64
604.3887939453125 0 3245.1667
609.3038330078125 0 546.2362
611.3545532226562 0 4839.642
612.3455200195312 0 2065.3052
613.3604125976562 0 687.08374
629.36474609375 0 6782.336
630.3679809570312 0 2801.8164
631.3743896484375 0 981.1945
635.2842407226562 0 651.1091
636.296142578125 0 1957.4125 y Water loss 6
641.8541870117188 0 1981.1643
642.35498046875 0 3810.0571
642.8572998046875 0 954.93915
645.3838500976562 0 11824.368
646.3916015625 0 38318.9 c 5
647.3944702148438 0 13323.563
648.3971557617188 0 2163.42
653.3019409179688 0 5316.557
654.3082275390625 0 17274.78 y 6
655.3115844726562 0 6228.552
656.312744140625 0 1280.3661
672.40869140625 0 2015.4204
685.3848876953125 0 805.55493
700.3970947265625 0 867.7018
702.4036254882812 0 1094.739
712.3909301757812 0 767.13025
722.3820190429688 0 2856.5742
723.383056640625 0 1454.6108
727.3872680664062 0 946.23303
730.4364013671875 0 6163.756
731.4432373046875 0 14395.65
732.4464111328125 0 5085.505
733.4481201171875 0 869.9251
739.4110107421875 0 3072.6099
740.3992309570312 0 6897.99
741.4009399414062 0 4051.9216
742.4075927734375 0 1888.6594
750.3692016601562 0 669.6557 y Ammonia loss 5
751.3741455078125 0 3508.3237 z 5
752.3798217773438 0 1760.8201
753.3789672851562 0 800.71735
755.42919921875 0 2048.9382
756.4259033203125 0 868.5188 c Water loss 6
757.424072265625 0 8618.276 c Ammonia loss 6
758.4288940429688 0 5301.2876
759.4367065429688 0 3287.0442
766.3861083984375 0 1794.293
767.3919067382812 0 10123.975 y 5
768.3944702148438 0 4108.954
773.4421997070312 0 18539.229
774.447998046875 0 27108.896 c 6
775.4517211914062 0 10954.893
776.4563598632812 0 2627.2314
799.4550170898438 0 938.9519
800.461669921875 0 1485.3525
809.4319458007812 0 683.6955
817.4639892578125 0 1096.6505
819.4775390625 0 858.804
825.4835815429688 0 1646.3439
826.444091796875 0 905.65186
827.4384155273438 0 1165.7883
835.464111328125 0 1004.9108
842.4616088867188 0 2242.1162
843.4630126953125 0 1099.3303 c Water loss 7
844.4561767578125 0 4412.7153 c Ammonia loss 7
845.4607543945312 0 1935.931
846.477294921875 0 905.48016
850.4428100585938 0 3291.4456 z 4
851.4457397460938 0 2316.6257
853.4776611328125 0 7664.9575
854.4771118164062 0 3235.0183
860.4735107421875 0 13445.153
861.4785766601562 0 15508.274 c 7
861.6229248046875 0 720.07336
862.4830932617188 0 5599.294
863.4905395507812 0 1875.1923
865.44921875 0 722.6498
866.4608154296875 0 4665.9194 y 4
867.4639282226562 0 1856.3225
903.4699096679688 0 2421.5051 z Water loss 3
904.4686279296875 0 2435.994 z Ammonia loss 3
913.4798583984375 0 1432.7589
914.4639892578125 0 769.9392
919.40576171875 0 733.10034
921.4786376953125 0 20486.35 z 3
922.4813842773438 0 12020.884
923.4841918945312 0 3662.6838
924.491943359375 0 788.34924
929.4928588867188 0 1330.7657
930.4952392578125 0 1798.2716 c Water loss 8
931.48779296875 0 3396.4756 c Ammonia loss 8
932.4878540039062 0 1830.3881
937.5018310546875 0 1624.7682 y 3
947.5057983398438 0 14758.801
948.5126953125 0 33247.062 c 8
949.5149536132812 0 15226.999
950.5182495117188 0 3681.878
952.48193359375 0 929.4659
953.47412109375 0 1608.2445
955.4901123046875 0 841.3921
970.497802734375 0 5949.8525
971.498046875 0 3785.5095
972.504150390625 0 1196.0616
984.5272827148438 0 8387.105
985.5303955078125 0 5373.6206
986.5353393554688 0 1460.1516
988.5091552734375 0 7511.976 c Ammonia loss 9
989.5105590820312 0 4060.9026
990.5162963867188 0 918.76196
1005.53515625 0 99661.664 c 9
1006.5372314453125 0 56671.598
1007.5397338867188 0 17459.998
1008.5402221679688 0 1697.5527
1012.5140991210938 0 748.9633
1015.534423828125 0 1073.6969
1016.5291137695312 0 1241.2014 y Water loss 2
1018.5362548828125 0 751.3221 z 2
1033.550048828125 0 817.68866
1034.5496826171875 0 6289.558 y 2
1035.551513671875 0 3789.5442
1057.58154296875 0 689.2425
1068.570556640625 0 5161.0786
1069.574951171875 0 3393.5732
1070.5828857421875 0 1021.8443
1073.5933837890625 0 1729.405
1074.60400390625 0 5087.3735
1075.6064453125 0 2882.8164
1076.6138916015625 0 1109.3208
1083.5809326171875 0 2820.2695
1084.5804443359375 0 1732.4891
1085.58837890625 0 13042.546
1086.5921630859375 0 8405.197
1087.58984375 0 2911.038
1094.5584716796875 0 834.14575
1095.6402587890625 0 1454.6757
1101.5931396484375 0 8135.9814 c Ammonia loss 10
1102.59375 0 3085.0684
1103.6024169921875 0 3321.321
1104.6070556640625 0 1041.9965
1118.6181640625 0 85590.52 c 10
1119.620849609375 0 56805.402
1120.623291015625 0 19776.352
1121.6220703125 0 1698.1624
1132.631591796875 0 1214.7109
1133.6422119140625 0 805.19336
1147.5894775390625 0 2147.9841 z Water loss 1
1148.5927734375 0 1028.5966 z Ammonia loss 1
1165.5992431640625 0 18445.361 z 1
1166.6014404296875 0 14511.553
1167.6051025390625 0 4957.916
1195.6085205078125 0 1619.8019
1204.6373291015625 0 1114.3143
1205.6492919921875 0 1397.8822
1210.5919189453125 0 2556.7761
1211.59228515625 0 1600.6389
1212.6258544921875 0 2405.102
1213.639404296875 0 991.04236
1221.6695556640625 0 3763.0085
1222.6717529296875 0 2571.0605
1223.6778564453125 0 973.0861
1227.6070556640625 0 1576.1173
1228.61376953125 0 1165.4557
1238.6507568359375 0 12779.215
1239.6556396484375 0 9405.655
1240.6431884765625 0 5539.8
1241.6304931640625 0 2198.888
1248.6356201171875 0 8307.391
1249.6383056640625 0 6877.1807
1250.642822265625 0 2019.339
1255.6768798828125 0 4684.081
1256.6778564453125 0 4237.513
1257.689208984375 0 1374.4174
1265.657958984375 0 2081.1665
1266.6485595703125 0 14403.311
1267.6505126953125 0 11266.061
1268.64892578125 0 3904.4028
1280.7220458984375 0 763.75934
1281.7587890625 0 739.7401
1282.665283203125 0 25207.879
1283.6715087890625 0 104898.83
1284.6744384765625 0 71415.09
1285.6783447265625 0 28925.555
1286.6793212890625 0 3279.5356
1910.918212890625 0 743.1831
2189.730224609375 0 655.19836
3080.22607421875 0 737.7763
3080.944091796875 0 798.8204
3142.61865234375 0 650.7437

Spectrum Details

|  |  |
| --- | --- |
| Matched peaks? Matched peaks The total absolute number of peaks matched. Additionally in brackets the total fraction of peaks matched and the total number of peaks is shown. | 42 (15.50% of 271) |
| FDR? FDR The false discovery rate estimated for this peptide. It is calculated by matching all theoretical fragments with a non-integer shift with the raw peaks for this spectrum. This is done with 40 different shifts. The resulting percentage is the average number of annotated peaks over the number of annotated peaks with the correct spectrum. | 1.42% |
| Satellite FDR? Satellite FDR See the FDR for details on its calculation. This satellite ion specific FDR only contains the satellite ions (d/w) for I/L/J positions. | ∞ |
| PSM Score? PSM Score The PSM Score as given by Hecklib to this annotated spectrum. It is shown with three significant figures. | 546 |

#### Spectrum 10666? Spectrum 10666 The raw spectrum of this peptide as annotated by Hecklib. The fragments are coloured according to ion type (see legend). Any peaks with a star '\*' as text can be hovered over to see the full details, first the ion type second the mass shift type. By hovering over the amino acids in the peptide or ions in the legend the corresponding peaks are highlighted. By toggling the 'Unassigned' label you can turn the background (unassigned) peaks on or off in the plot. By updating the slider in the Ion legend you can update the spectrum to only show the top X% of the peaks with labels. The top X% means any peak that is within X% of the highest intensity. By dragging in the spectrum you can zoom in to a specific part of the spectrum and use 'Zoom Out' to get back to the original zoom level. The annotation of the spectrum is based on the given sequence in the peptides file and is done with different software so inconsistencies are likely. The peaks are annotated based on the given sequence, with 20 ppm tolerance.

Copy Data

##### Spectrum 10666 (TSV)

###### Preview

```
Loading example...
```

*Click on the button to copy the data to your clipboard.*

Mz MinMz MaxIntensity Max

WidthHeightPeptide font sizePeptide stroke widthSpectrum font sizeSpectrum stroke widthCompact peptide

Ion legend

wxyz

abcd

OtherUnassignedIon^Charge^_Position_Show for top:%

TFPAVJQSSGJY

01.44e+62.89e+64.33e+65.77e+6

Zoom Out

y^+1^_1_y^+1^_2_y^+1^_3_c^+2^_7_z^+2^_8_w^+2^_8_y^+1^_4_c^+2^_8_c^+1^_4_y^+1^_4_c^+2^_10_y^+1^_5_y^+2^_10_y^+1^_5_c^+1^_5_c^+2^_11_y^+1^_6_z^+1^_6_c^+1^_6_y^+1^_6_z^+1^_7_y^+1^_7_y^+1^_7_z^+1^_7_c^+1^_7_c^+1^_7_y^+1^_7_c^+1^_7_z^+1^_8_c^+1^_8_c^+1^_8_z^+1^_8_c^+1^_8_y^+1^_8_z^+1^_9_z^+1^_9_z^+1^_9_c^+1^_9_c^+1^_9_y^+1^_9_c^+1^_9_c^+1^_10_c^+1^_10_w^+1^_10_c^+1^_10_y^+1^_10_y^+1^_10_z^+1^_10_y^+1^_10_c^+1^_11_c^+1^_11_z^+1^_11_z^+1^_11_z^+1^_11_y^+1^_11_

038276511471530

Fragment Matches Table

Show background peaks

| Position | Ion type | Intensity | mz Theoretical | mz Error (Th) | mz Error (ppm) | Charge | Series Number |
| --- | --- | --- | --- | --- | --- | --- | --- |
| - | - | 1.469E+04 | 120.1 | - | - | 0 | - |
| - | - | 5656 | 120.9 | - | - | 0 | - |
| - | - | 5064 | 121.8 | - | - | 0 | - |
| - | - | 1.122E+04 | 123 | - | - | 0 | - |
| - | - | 6683 | 125.7 | - | - | 0 | - |
| - | - | 5942 | 128.9 | - | - | 0 | - |
| - | - | 6406 | 130.6 | - | - | 0 | - |
| - | - | 6356 | 135.4 | - | - | 0 | - |
| - | - | 3.725E+05 | 136.1 | - | - | 0 | - |
| - | - | 2.989E+04 | 137.1 | - | - | 0 | - |
| - | - | 1.191E+04 | 141.1 | - | - | 0 | - |
| - | - | 6.211E+04 | 143.1 | - | - | 0 | - |
| - | - | 3.355E+05 | 165.1 | - | - | 0 | - |
| - | - | 3.707E+04 | 166.1 | - | - | 0 | - |
| - | - | 1.525E+05 | 169.1 | - | - | 0 | - |
| - | - | 8.71E+04 | 171.1 | - | - | 0 | - |
| - | - | 1.418E+04 | 175.1 | - | - | 0 | - |
| - | - | 7470 | 177.8 | - | - | 0 | - |
| 12 | y | 2.188E+06 | 182.1 | 0.0004343 | 2.385 | +1 | 1 |
| - | - | 2.041E+05 | 183.1 | - | - | 0 | - |
| - | - | 2.456E+04 | 186.1 | - | - | 0 | - |
| - | - | 7370 | 189.5 | - | - | 0 | - |
| - | - | 8236 | 196 | - | - | 0 | - |
| - | - | 7674 | 200 | - | - | 0 | - |
| - | - | 9953 | 213.2 | - | - | 0 | - |
| - | - | 4.924E+04 | 214.1 | - | - | 0 | - |
| - | - | 2.156E+04 | 216.1 | - | - | 0 | - |
| - | - | 8667 | 218 | - | - | 0 | - |
| - | - | 8.072E+05 | 221.1 | - | - | 0 | - |
| - | - | 9.765E+04 | 222.1 | - | - | 0 | - |
| - | - | 7.949E+04 | 232.1 | - | - | 0 | - |
| - | - | 7.34E+04 | 240.2 | - | - | 0 | - |
| - | - | 4.061E+04 | 242.2 | - | - | 0 | - |
| - | - | 5.919E+05 | 249.1 | - | - | 0 | - |
| - | - | 6.883E+04 | 250.1 | - | - | 0 | - |
| - | - | 1.042E+05 | 258.1 | - | - | 0 | - |
| - | - | 1.178E+04 | 259.1 | - | - | 0 | - |
| - | - | 3.68E+05 | 268.2 | - | - | 0 | - |
| - | - | 6.398E+04 | 269.2 | - | - | 0 | - |
| - | - | 1.7E+04 | 285.2 | - | - | 0 | - |
| - | - | 1.096E+04 | 286.2 | - | - | 0 | - |
| 11 | y | 3.058E+05 | 295.2 | 0.0005683 | 1.926 | +1 | 2 |
| - | - | 4.57E+04 | 296.2 | - | - | 0 | - |
| - | - | 1.01E+04 | 299.2 | - | - | 0 | - |
| - | - | 1.009E+04 | 300.2 | - | - | 0 | - |
| - | - | 2.152E+04 | 316.2 | - | - | 0 | - |
| - | - | 2.356E+05 | 317.2 | - | - | 0 | - |
| - | - | 2.235E+04 | 318.2 | - | - | 0 | - |
| - | - | 8314 | 318.5 | - | - | 0 | - |
| - | - | 1.327E+04 | 324.2 | - | - | 0 | - |
| - | - | 3.061E+04 | 327.2 | - | - | 0 | - |
| - | - | 1.825E+04 | 328.2 | - | - | 0 | - |
| - | - | 1.078E+04 | 329.2 | - | - | 0 | - |
| - | - | 2.042E+04 | 333.2 | - | - | 0 | - |
| - | - | 2.017E+04 | 341.2 | - | - | 0 | - |
| - | - | 1.996E+04 | 342.1 | - | - | 0 | - |
| - | - | 4.727E+04 | 343.1 | - | - | 0 | - |
| - | - | 8571 | 344.2 | - | - | 0 | - |
| - | - | 4.689E+05 | 345.2 | - | - | 0 | - |
| - | - | 1.211E+05 | 346.2 | - | - | 0 | - |
| - | - | 1.172E+04 | 347.2 | - | - | 0 | - |
| - | - | 4.589E+04 | 351.2 | - | - | 0 | - |
| 10 | y | 1.087E+05 | 352.2 | 0.0003143 | 0.8925 | +1 | 3 |
| - | - | 1.457E+04 | 352.2 | - | - | 0 | - |
| - | - | 1.154E+04 | 352.7 | - | - | 0 | - |
| - | - | 2.447E+04 | 353.2 | - | - | 0 | - |
| - | - | 3.708E+04 | 353.3 | - | - | 0 | - |
| - | - | 1.406E+04 | 360.2 | - | - | 0 | - |
| - | - | 1.454E+05 | 361.7 | - | - | 0 | - |
| - | - | 6.525E+04 | 362.2 | - | - | 0 | - |
| - | - | 9411 | 362.7 | - | - | 0 | - |
| - | - | 4.346E+04 | 369.2 | - | - | 0 | - |
| - | - | 2.707E+04 | 370.7 | - | - | 0 | - |
| - | - | 2.893E+04 | 371.2 | - | - | 0 | - |
| - | - | 1.136E+04 | 377.2 | - | - | 0 | - |
| 7 | c | 1.849E+04 | 379.2 | 0.00171 | 4.508 | +2 | 7 |
| - | - | 1.568E+05 | 381.3 | - | - | 0 | - |
| - | - | 3.236E+04 | 382.3 | - | - | 0 | - |
| - | - | 1.154E+04 | 387.2 | - | - | 0 | - |
| - | - | 2.281E+04 | 389.2 | - | - | 0 | - |
| - | - | 8861 | 395.2 | - | - | 0 | - |
| - | - | 2.189E+04 | 398.2 | - | - | 0 | - |
| - | - | 2.832E+04 | 398.3 | - | - | 0 | - |
| - | - | 8.032E+04 | 399.2 | - | - | 0 | - |
| - | - | 1.006E+04 | 400.2 | - | - | 0 | - |
| - | - | 1.104E+04 | 400.2 | - | - | 0 | - |
| - | - | 1.061E+04 | 410.2 | - | - | 0 | - |
| - | - | 1.157E+04 | 412.3 | - | - | 0 | - |
| - | - | 1.8E+04 | 413.2 | - | - | 0 | - |
| - | - | 7.017E+04 | 414.3 | - | - | 0 | - |
| - | - | 3.608E+04 | 415.2 | - | - | 0 | - |
| - | - | 1.771E+04 | 415.3 | - | - | 0 | - |
| - | - | 1.469E+04 | 416.2 | - | - | 0 | - |
| 5 | z | 2.283E+05 | 417.2 | 0.002113 | 5.065 | +2 | 8 |
| 5 | w | 5.355E+04 | 418.2 | 0.00375 | 8.967 | +2 | 8 |
| - | - | 9214 | 420.9 | - | - | 0 | - |
| 9 | y | 1.053E+04 | 421.2 | 0.001525 | 3.621 | +1 | 4 |
| 8 | c | 3.845E+04 | 422.7 | 0.0008016 | 1.896 | +2 | 8 |
| - | - | 1.381E+04 | 423.2 | - | - | 0 | - |
| - | - | 2.78E+04 | 427.2 | - | - | 0 | - |
| - | - | 2.518E+04 | 427.7 | - | - | 0 | - |
| - | - | 6.685E+04 | 428.2 | - | - | 0 | - |
| - | - | 1.738E+04 | 428.2 | - | - | 0 | - |
| - | - | 1.861E+04 | 429.2 | - | - | 0 | - |
| - | - | 1.648E+04 | 433.2 | - | - | 0 | - |
| 4 | c | 2.389E+04 | 434.2 | 0.0003167 | 0.7294 | +1 | 4 |
| - | - | 1.198E+04 | 438.2 | - | - | 0 | - |
| 9 | y | 9.702E+04 | 439.2 | 0.0009703 | 2.209 | +1 | 4 |
| - | - | 2.49E+04 | 440.2 | - | - | 0 | - |
| - | - | 1.224E+04 | 440.3 | - | - | 0 | - |
| - | - | 4.336E+04 | 445.2 | - | - | 0 | - |
| - | - | 4.941E+04 | 448.2 | - | - | 0 | - |
| - | - | 5.761E+04 | 455.2 | - | - | 0 | - |
| - | - | 1.182E+05 | 456.2 | - | - | 0 | - |
| - | - | 2.74E+04 | 457.2 | - | - | 0 | - |
| - | - | 4.27E+04 | 457.2 | - | - | 0 | - |
| - | - | 1.236E+04 | 457.7 | - | - | 0 | - |
| - | - | 1.648E+04 | 458.3 | - | - | 0 | - |
| - | - | 4.221E+04 | 464.3 | - | - | 0 | - |
| - | - | 1.841E+04 | 470.3 | - | - | 0 | - |
| - | - | 1.419E+04 | 471.3 | - | - | 0 | - |
| - | - | 1.082E+04 | 472.2 | - | - | 0 | - |
| - | - | 3.535E+04 | 472.3 | - | - | 0 | - |
| - | - | 1.506E+05 | 473.2 | - | - | 0 | - |
| - | - | 3.543E+04 | 474.2 | - | - | 0 | - |
| - | - | 1.042E+04 | 475.3 | - | - | 0 | - |
| - | - | 2.532E+04 | 476.7 | - | - | 0 | - |
| - | - | 2.252E+04 | 477.2 | - | - | 0 | - |
| - | - | 1.344E+04 | 481.3 | - | - | 0 | - |
| - | - | 3.311E+04 | 482.3 | - | - | 0 | - |
| - | - | 1.529E+04 | 483.3 | - | - | 0 | - |
| - | - | 1.698E+05 | 485.8 | - | - | 0 | - |
| - | - | 6.922E+04 | 486.3 | - | - | 0 | - |
| - | - | 3.24E+04 | 486.8 | - | - | 0 | - |
| - | - | 1.725E+05 | 488.3 | - | - | 0 | - |
| - | - | 4.18E+04 | 489.3 | - | - | 0 | - |
| - | - | 1.724E+04 | 490.2 | - | - | 0 | - |
| - | - | 2.752E+04 | 490.3 | - | - | 0 | - |
| - | - | 9589 | 491.3 | - | - | 0 | - |
| - | - | 9226 | 492.3 | - | - | 0 | - |
| 10 | c | 1.513E+04 | 494.8 | 9.608E-05 | 0.1942 | +2 | 10 |
| - | - | 3.116E+05 | 498.3 | - | - | 0 | - |
| - | - | 6.341E+04 | 499.3 | - | - | 0 | - |
| - | - | 2.597E+04 | 499.8 | - | - | 0 | - |
| - | - | 3.46E+04 | 500.3 | - | - | 0 | - |
| - | - | 2.285E+04 | 501.3 | - | - | 0 | - |
| - | - | 1.027E+04 | 502.3 | - | - | 0 | - |
| 8 | y | 4.655E+04 | 508.2 | 0.0001059 | 0.2084 | +1 | 5 |
| - | - | 1.33E+04 | 508.3 | - | - | 0 | - |
| 3 | y | 2.421E+04 | 508.8 | 6.01E-05 | 0.1181 | +2 | 10 |
| - | - | 4.03E+05 | 509.3 | - | - | 0 | - |
| - | - | 1.18E+05 | 510.3 | - | - | 0 | - |
| - | - | 1.305E+04 | 511.3 | - | - | 0 | - |
| - | - | 2.865E+04 | 515.3 | - | - | 0 | - |
| - | - | 5.242E+05 | 516.3 | - | - | 0 | - |
| - | - | 1.414E+05 | 517.3 | - | - | 0 | - |
| - | - | 2.384E+04 | 518.3 | - | - | 0 | - |
| - | - | 1.348E+04 | 525.3 | - | - | 0 | - |
| 8 | y | 3.807E+05 | 526.3 | 0.0007106 | 1.35 | +1 | 5 |
| - | - | 9.886E+04 | 526.3 | - | - | 0 | - |
| - | - | 9.516E+04 | 527.3 | - | - | 0 | - |
| - | - | 1.557E+04 | 527.3 | - | - | 0 | - |
| - | - | 1.214E+04 | 528.3 | - | - | 0 | - |
| - | - | 1.54E+04 | 530.3 | - | - | 0 | - |
| - | - | 2.822E+05 | 532.3 | - | - | 0 | - |
| 5 | c | 9.544E+05 | 533.3 | 0.0003842 | 0.7205 | +1 | 5 |
| - | - | 2.825E+05 | 534.3 | - | - | 0 | - |
| - | - | 4.3E+04 | 535.3 | - | - | 0 | - |
| - | - | 1.049E+04 | 541.3 | - | - | 0 | - |
| - | - | 1.089E+04 | 550.3 | - | - | 0 | - |
| 11 | c | 9115 | 551.3 | 0.0092 | 16.69 | +2 | 11 |
| - | - | 1.31E+04 | 552.3 | - | - | 0 | - |
| - | - | 9217 | 557.3 | - | - | 0 | - |
| - | - | 4.317E+04 | 558.3 | - | - | 0 | - |
| - | - | 1.594E+04 | 559.3 | - | - | 0 | - |
| - | - | 4.649E+04 | 568.3 | - | - | 0 | - |
| - | - | 2.495E+04 | 569.3 | - | - | 0 | - |
| - | - | 1.264E+04 | 570.3 | - | - | 0 | - |
| - | - | 1.005E+04 | 573.2 | - | - | 0 | - |
| - | - | 1.173E+04 | 577.3 | - | - | 0 | - |
| - | - | 2.447E+04 | 578.3 | - | - | 0 | - |
| - | - | 2.336E+04 | 583.4 | - | - | 0 | - |
| - | - | 3.406E+04 | 584.4 | - | - | 0 | - |
| - | - | 2.7E+04 | 585.3 | - | - | 0 | - |
| - | - | 1.796E+05 | 586.3 | - | - | 0 | - |
| - | - | 5.048E+04 | 587.3 | - | - | 0 | - |
| - | - | 2.097E+04 | 588.3 | - | - | 0 | - |
| - | - | 2.696E+04 | 589.3 | - | - | 0 | - |
| - | - | 1.021E+04 | 590.3 | - | - | 0 | - |
| - | - | 1.634E+04 | 592.3 | - | - | 0 | - |
| - | - | 1.21E+04 | 593.3 | - | - | 0 | - |
| - | - | 7E+04 | 596.3 | - | - | 0 | - |
| - | - | 2.662E+04 | 597.3 | - | - | 0 | - |
| - | - | 6.002E+04 | 601.4 | - | - | 0 | - |
| - | - | 1.022E+05 | 602.4 | - | - | 0 | - |
| - | - | 4.627E+05 | 603.4 | - | - | 0 | - |
| - | - | 1.763E+05 | 604.4 | - | - | 0 | - |
| - | - | 2.2E+04 | 605.4 | - | - | 0 | - |
| - | - | 1.408E+04 | 610.4 | - | - | 0 | - |
| - | - | 3.094E+05 | 611.4 | - | - | 0 | - |
| - | - | 1.199E+05 | 612.4 | - | - | 0 | - |
| - | - | 7.208E+04 | 613.4 | - | - | 0 | - |
| - | - | 1.433E+04 | 618.3 | - | - | 0 | - |
| - | - | 1.514E+04 | 627.3 | - | - | 0 | - |
| - | - | 2.539E+04 | 628.4 | - | - | 0 | - |
| - | - | 4.651E+05 | 629.4 | - | - | 0 | - |
| - | - | 1.637E+05 | 630.4 | - | - | 0 | - |
| - | - | 2.746E+04 | 631.4 | - | - | 0 | - |
| - | - | 4.063E+04 | 635.3 | - | - | 0 | - |
| 7 | y | 1.392E+05 | 636.3 | 0.0002442 | 0.3838 | +1 | 6 |
| - | - | 3.944E+04 | 637.3 | - | - | 0 | - |
| 7 | z | 1.983E+04 | 638.3 | 0.001568 | 2.456 | +1 | 6 |
| - | - | 1.017E+04 | 639.3 | - | - | 0 | - |
| - | - | 3.181E+04 | 640.4 | - | - | 0 | - |
| - | - | 2.7E+04 | 641.4 | - | - | 0 | - |
| - | - | 1.588E+04 | 642.3 | - | - | 0 | - |
| - | - | 1.342E+04 | 643.3 | - | - | 0 | - |
| - | - | 6.094E+05 | 645.4 | - | - | 0 | - |
| 6 | c | 1.928E+06 | 646.4 | 0.0001826 | 0.2824 | +1 | 6 |
| - | - | 1.599E+04 | 647.3 | - | - | 0 | - |
| - | - | 6.743E+05 | 647.4 | - | - | 0 | - |
| - | - | 1.071E+05 | 648.4 | - | - | 0 | - |
| - | - | 3.057E+05 | 653.3 | - | - | 0 | - |
| 7 | y | 1.094E+06 | 654.3 | 0.0002386 | 0.3647 | +1 | 6 |
| - | - | 3.709E+05 | 655.3 | - | - | 0 | - |
| - | - | 1.11E+04 | 655.4 | - | - | 0 | - |
| - | - | 7.771E+04 | 656.3 | - | - | 0 | - |
| - | - | 3.192E+04 | 656.4 | - | - | 0 | - |
| - | - | 1.083E+04 | 657.4 | - | - | 0 | - |
| - | - | 1.582E+04 | 659.4 | - | - | 0 | - |
| - | - | 3.501E+04 | 665.4 | - | - | 0 | - |
| - | - | 2.014E+04 | 666.4 | - | - | 0 | - |
| - | - | 1.567E+04 | 667.4 | - | - | 0 | - |
| - | - | 1.122E+04 | 668.4 | - | - | 0 | - |
| - | - | 1.399E+04 | 669.4 | - | - | 0 | - |
| - | - | 1.018E+05 | 672.4 | - | - | 0 | - |
| - | - | 4.009E+04 | 673.4 | - | - | 0 | - |
| - | - | 2.781E+04 | 674.4 | - | - | 0 | - |
| - | - | 1.516E+04 | 675.4 | - | - | 0 | - |
| - | - | 4.578E+04 | 683.4 | - | - | 0 | - |
| - | - | 2.055E+04 | 684.4 | - | - | 0 | - |
| - | - | 5.55E+04 | 685.4 | - | - | 0 | - |
| - | - | 3.79E+04 | 686.4 | - | - | 0 | - |
| - | - | 2.351E+04 | 687.4 | - | - | 0 | - |
| - | - | 9967 | 697.4 | - | - | 0 | - |
| - | - | 2.243E+04 | 699.4 | - | - | 0 | - |
| - | - | 5.215E+04 | 700.4 | - | - | 0 | - |
| - | - | 1.474E+04 | 701.4 | - | - | 0 | - |
| - | - | 2.704E+04 | 702.4 | - | - | 0 | - |
| - | - | 2.84E+04 | 703.3 | - | - | 0 | - |
| - | - | 1.31E+04 | 703.4 | - | - | 0 | - |
| - | - | 2.993E+04 | 704.4 | - | - | 0 | - |
| - | - | 1.451E+04 | 705.4 | - | - | 0 | - |
| - | - | 9171 | 706.4 | - | - | 0 | - |
| - | - | 1.947E+04 | 710.4 | - | - | 0 | - |
| - | - | 4.091E+04 | 712.4 | - | - | 0 | - |
| - | - | 2.939E+04 | 713.4 | - | - | 0 | - |
| - | - | 1.677E+04 | 714.4 | - | - | 0 | - |
| - | - | 1.301E+04 | 715.4 | - | - | 0 | - |
| - | - | 1.5E+04 | 721.3 | - | - | 0 | - |
| - | - | 1.713E+05 | 722.4 | - | - | 0 | - |
| - | - | 7.308E+04 | 723.4 | - | - | 0 | - |
| - | - | 1.841E+04 | 724.4 | - | - | 0 | - |
| - | - | 3.758E+04 | 727.4 | - | - | 0 | - |
| - | - | 2.954E+04 | 729.4 | - | - | 0 | - |
| - | - | 1.64E+04 | 730.4 | - | - | 0 | - |
| - | - | 3.357E+05 | 730.4 | - | - | 0 | - |
| - | - | 7.456E+05 | 731.4 | - | - | 0 | - |
| - | - | 3.062E+05 | 732.4 | - | - | 0 | - |
| 6 | z | 1.403E+04 | 733.4 | 0.00955 | 13.02 | +1 | 7 |
| - | - | 4.942E+04 | 733.5 | - | - | 0 | - |
| - | - | 1.339E+05 | 739.4 | - | - | 0 | - |
| - | - | 4.884E+05 | 740.4 | - | - | 0 | - |
| - | - | 2.136E+05 | 741.4 | - | - | 0 | - |
| - | - | 1.229E+05 | 742.4 | - | - | 0 | - |
| - | - | 4.009E+04 | 743.4 | - | - | 0 | - |
| - | - | 1E+04 | 747.4 | - | - | 0 | - |
| 6 | y | 4.411E+04 | 749.4 | 0.0002016 | 0.269 | +1 | 7 |
| 6 | y | 3.463E+04 | 750.4 | 0.009008 | 12 | +1 | 7 |
| 6 | z | 1.971E+05 | 751.4 | 0.0005115 | 0.6807 | +1 | 7 |
| - | - | 1.122E+05 | 752.4 | - | - | 0 | - |
| - | - | 2.918E+04 | 753.4 | - | - | 0 | - |
| - | - | 1.158E+05 | 755.4 | - | - | 0 | - |
| 7 | c | 9.221E+04 | 756.4 | 0.01097 | 14.5 | +1 | 7 |
| 7 | c | 6.784E+05 | 757.4 | 0.0007469 | 0.9862 | +1 | 7 |
| - | - | 3.237E+05 | 758.4 | - | - | 0 | - |
| - | - | 1.807E+05 | 759.4 | - | - | 0 | - |
| - | - | 4.978E+04 | 760.4 | - | - | 0 | - |
| - | - | 1.056E+05 | 766.4 | - | - | 0 | - |
| 6 | y | 7.647E+05 | 767.4 | 0.00022 | 0.2867 | +1 | 7 |
| - | - | 2.937E+05 | 768.4 | - | - | 0 | - |
| - | - | 6.502E+04 | 769.4 | - | - | 0 | - |
| - | - | 1.12E+06 | 773.4 | - | - | 0 | - |
| 7 | c | 1.454E+06 | 774.5 | 0.001205 | 1.556 | +1 | 7 |
| - | - | 5.801E+05 | 775.5 | - | - | 0 | - |
| - | - | 1.301E+05 | 776.5 | - | - | 0 | - |
| - | - | 2.468E+04 | 777.5 | - | - | 0 | - |
| - | - | 1.039E+04 | 781.5 | - | - | 0 | - |
| - | - | 1.313E+04 | 789.4 | - | - | 0 | - |
| - | - | 3.185E+04 | 794.4 | - | - | 0 | - |
| - | - | 4.407E+04 | 799.5 | - | - | 0 | - |
| - | - | 8.699E+04 | 800.5 | - | - | 0 | - |
| - | - | 4.626E+04 | 801.5 | - | - | 0 | - |
| - | - | 1.684E+04 | 802.5 | - | - | 0 | - |
| - | - | 1.531E+04 | 808.4 | - | - | 0 | - |
| - | - | 1.128E+04 | 809.4 | - | - | 0 | - |
| - | - | 1.265E+04 | 812.4 | - | - | 0 | - |
| - | - | 2.164E+04 | 814.4 | - | - | 0 | - |
| - | - | 1.989E+04 | 815.4 | - | - | 0 | - |
| - | - | 1.725E+04 | 816.4 | - | - | 0 | - |
| - | - | 4.737E+04 | 817.5 | - | - | 0 | - |
| - | - | 3.968E+04 | 818.5 | - | - | 0 | - |
| - | - | 2.541E+04 | 819.5 | - | - | 0 | - |
| - | - | 1.04E+04 | 823.4 | - | - | 0 | - |
| - | - | 1.001E+05 | 825.5 | - | - | 0 | - |
| - | - | 5.067E+04 | 826.4 | - | - | 0 | - |
| - | - | 7.606E+04 | 827.4 | - | - | 0 | - |
| - | - | 2.655E+04 | 828.4 | - | - | 0 | - |
| - | - | 2.499E+04 | 829.4 | - | - | 0 | - |
| - | - | 1.84E+04 | 830.4 | - | - | 0 | - |
| 5 | z | 1.186E+04 | 832.4 | 0.004796 | 5.761 | +1 | 8 |
| - | - | 1.748E+04 | 834.4 | - | - | 0 | - |
| - | - | 8.397E+04 | 835.5 | - | - | 0 | - |
| - | - | 7.418E+04 | 836.5 | - | - | 0 | - |
| - | - | 1.69E+04 | 837.5 | - | - | 0 | - |
| - | - | 1.25E+05 | 842.5 | - | - | 0 | - |
| 8 | c | 8.444E+04 | 843.5 | 0.007898 | 9.364 | +1 | 8 |
| 8 | c | 2.735E+05 | 844.5 | 0.001311 | 1.553 | +1 | 8 |
| - | - | 1.172E+05 | 845.5 | - | - | 0 | - |
| - | - | 6E+04 | 846.5 | - | - | 0 | - |
| - | - | 2.286E+04 | 847.5 | - | - | 0 | - |
| 5 | z | 2.015E+05 | 850.4 | 0.000518 | 0.6091 | +1 | 8 |
| - | - | 9.691E+04 | 851.4 | - | - | 0 | - |
| - | - | 1.69E+04 | 852.5 | - | - | 0 | - |
| - | - | 5.036E+05 | 853.5 | - | - | 0 | - |
| - | - | 2.344E+05 | 854.5 | - | - | 0 | - |
| - | - | 5.935E+04 | 855.5 | - | - | 0 | - |
| - | - | 7.814E+05 | 860.5 | - | - | 0 | - |
| 8 | c | 8.591E+05 | 861.5 | 0.001678 | 1.948 | +1 | 8 |
| - | - | 3.446E+05 | 862.5 | - | - | 0 | - |
| - | - | 1.162E+05 | 863.5 | - | - | 0 | - |
| - | - | 2.735E+04 | 864.5 | - | - | 0 | - |
| - | - | 4.542E+04 | 865.5 | - | - | 0 | - |
| 5 | y | 2.119E+05 | 866.5 | 0.0001397 | 0.1612 | +1 | 8 |
| - | - | 7.889E+04 | 867.5 | - | - | 0 | - |
| - | - | 2.419E+04 | 868.5 | - | - | 0 | - |
| - | - | 1.607E+04 | 871.4 | - | - | 0 | - |
| - | - | 2.053E+04 | 872.4 | - | - | 0 | - |
| - | - | 2.552E+04 | 887.5 | - | - | 0 | - |
| - | - | 2.608E+04 | 889.5 | - | - | 0 | - |
| - | - | 2.859E+04 | 891.4 | - | - | 0 | - |
| - | - | 2.156E+04 | 892.5 | - | - | 0 | - |
| - | - | 1.325E+04 | 893.5 | - | - | 0 | - |
| - | - | 4.758E+04 | 895.5 | - | - | 0 | - |
| - | - | 5.76E+04 | 896.5 | - | - | 0 | - |
| - | - | 2.804E+04 | 897.5 | - | - | 0 | - |
| - | - | 2.109E+04 | 898.5 | - | - | 0 | - |
| - | - | 1.207E+04 | 899.5 | - | - | 0 | - |
| 4 | z | 1.542E+05 | 903.5 | 0.001252 | 1.385 | +1 | 9 |
| 4 | z | 7.944E+04 | 904.5 | 0.01779 | 19.66 | +1 | 9 |
| - | - | 2.701E+04 | 905.5 | - | - | 0 | - |
| - | - | 5.893E+04 | 913.5 | - | - | 0 | - |
| - | - | 6.795E+04 | 914.5 | - | - | 0 | - |
| - | - | 2.752E+04 | 915.5 | - | - | 0 | - |
| 4 | z | 1.227E+06 | 921.5 | 0.0001473 | 0.1599 | +1 | 9 |
| - | - | 7.806E+05 | 922.5 | - | - | 0 | - |
| - | - | 2.233E+05 | 923.5 | - | - | 0 | - |
| - | - | 3.273E+04 | 924.5 | - | - | 0 | - |
| - | - | 1.05E+05 | 929.5 | - | - | 0 | - |
| 9 | c | 6.788E+04 | 930.5 | 0.008982 | 9.653 | +1 | 9 |
| 9 | c | 2.287E+05 | 931.5 | 0.000716 | 0.7687 | +1 | 9 |
| - | - | 1.058E+05 | 932.5 | - | - | 0 | - |
| - | - | 3.169E+04 | 933.5 | - | - | 0 | - |
| - | - | 1.101E+04 | 934.5 | - | - | 0 | - |
| - | - | 1.385E+04 | 935.5 | - | - | 0 | - |
| - | - | 5.276E+04 | 936.5 | - | - | 0 | - |
| 4 | y | 9.962E+04 | 937.5 | 0.001382 | 1.474 | +1 | 9 |
| - | - | 4.195E+04 | 938.5 | - | - | 0 | - |
| - | - | 1.265E+04 | 939.5 | - | - | 0 | - |
| - | - | 1.525E+04 | 942.5 | - | - | 0 | - |
| - | - | 1.326E+04 | 945.5 | - | - | 0 | - |
| - | - | 7.783E+05 | 947.5 | - | - | 0 | - |
| 9 | c | 1.889E+06 | 948.5 | 0.0006256 | 0.6595 | +1 | 9 |
| - | - | 9.355E+05 | 949.5 | - | - | 0 | - |
| - | - | 2.228E+05 | 950.5 | - | - | 0 | - |
| - | - | 1.346E+04 | 951.5 | - | - | 0 | - |
| - | - | 9.626E+04 | 952.5 | - | - | 0 | - |
| - | - | 9.022E+04 | 953.5 | - | - | 0 | - |
| - | - | 4.848E+04 | 954.5 | - | - | 0 | - |
| - | - | 2.205E+04 | 955.5 | - | - | 0 | - |
| - | - | 2.372E+04 | 960.5 | - | - | 0 | - |
| - | - | 5.302E+04 | 961.5 | - | - | 0 | - |
| - | - | 4.488E+04 | 962.5 | - | - | 0 | - |
| - | - | 1.576E+04 | 963.5 | - | - | 0 | - |
| - | - | 4.052E+04 | 967.5 | - | - | 0 | - |
| - | - | 2.043E+04 | 968.5 | - | - | 0 | - |
| - | - | 4.915E+05 | 970.5 | - | - | 0 | - |
| - | - | 2.976E+05 | 971.5 | - | - | 0 | - |
| - | - | 1.047E+05 | 972.5 | - | - | 0 | - |
| - | - | 1.993E+04 | 973.5 | - | - | 0 | - |
| - | - | 2.453E+04 | 977.5 | - | - | 0 | - |
| - | - | 4.6E+05 | 984.5 | - | - | 0 | - |
| - | - | 3.047E+05 | 985.5 | - | - | 0 | - |
| - | - | 1.055E+05 | 986.5 | - | - | 0 | - |
| 10 | c | 3.362E+04 | 987.5 | 0.005992 | 6.068 | +1 | 10 |
| 10 | c | 5.734E+05 | 988.5 | 0.0004315 | 0.4365 | +1 | 10 |
| - | - | 3.076E+05 | 989.5 | - | - | 0 | - |
| - | - | 9.445E+04 | 990.5 | - | - | 0 | - |
| 3 | w | 1.412E+04 | 991.5 | 0.01627 | 16.41 | +1 | 10 |
| - | - | 2.432E+04 | 998.5 | - | - | 0 | - |
| - | - | 1.342E+04 | 1001 | - | - | 0 | - |
| - | - | 1.232E+04 | 1002 | - | - | 0 | - |
| - | - | 3.185E+04 | 1003 | - | - | 0 | - |
| - | - | 2.343E+04 | 1004 | - | - | 0 | - |
| 10 | c | 5.469E+06 | 1006 | 0.0007378 | 0.7338 | +1 | 10 |
| - | - | 3.117E+06 | 1007 | - | - | 0 | - |
| - | - | 8.432E+05 | 1008 | - | - | 0 | - |
| - | - | 8.12E+04 | 1009 | - | - | 0 | - |
| - | - | 5.882E+04 | 1016 | - | - | 0 | - |
| 3 | y | 8.064E+04 | 1017 | 0.002731 | 2.687 | +1 | 10 |
| 3 | y | 5.053E+04 | 1018 | 0.01643 | 16.14 | +1 | 10 |
| 3 | z | 2.087E+04 | 1019 | 0.007625 | 7.486 | +1 | 10 |
| - | - | 1.284E+04 | 1030 | - | - | 0 | - |
| - | - | 3.16E+04 | 1032 | - | - | 0 | - |
| - | - | 1.607E+04 | 1033 | - | - | 0 | - |
| - | - | 4.342E+04 | 1034 | - | - | 0 | - |
| 3 | y | 5.15E+05 | 1035 | 0.0002957 | 0.2858 | +1 | 10 |
| - | - | 2.785E+05 | 1036 | - | - | 0 | - |
| - | - | 6.291E+04 | 1037 | - | - | 0 | - |
| - | - | 1.574E+04 | 1039 | - | - | 0 | - |
| - | - | 2.169E+04 | 1042 | - | - | 0 | - |
| - | - | 2.296E+04 | 1057 | - | - | 0 | - |
| - | - | 5.687E+04 | 1058 | - | - | 0 | - |
| - | - | 3.693E+04 | 1059 | - | - | 0 | - |
| - | - | 2.142E+04 | 1063 | - | - | 0 | - |
| - | - | 1.905E+04 | 1066 | - | - | 0 | - |
| - | - | 3.366E+04 | 1067 | - | - | 0 | - |
| - | - | 2.224E+04 | 1068 | - | - | 0 | - |
| - | - | 2.532E+05 | 1069 | - | - | 0 | - |
| - | - | 1.62E+05 | 1070 | - | - | 0 | - |
| - | - | 5.904E+04 | 1071 | - | - | 0 | - |
| - | - | 1.063E+05 | 1074 | - | - | 0 | - |
| - | - | 2.637E+05 | 1075 | - | - | 0 | - |
| - | - | 1.74E+05 | 1076 | - | - | 0 | - |
| - | - | 4.584E+04 | 1077 | - | - | 0 | - |
| - | - | 2.422E+05 | 1084 | - | - | 0 | - |
| - | - | 1.758E+05 | 1085 | - | - | 0 | - |
| - | - | 7.136E+05 | 1086 | - | - | 0 | - |
| - | - | 5.234E+05 | 1087 | - | - | 0 | - |
| - | - | 1.56E+05 | 1088 | - | - | 0 | - |
| - | - | 2.456E+04 | 1089 | - | - | 0 | - |
| - | - | 3.493E+04 | 1095 | - | - | 0 | - |
| - | - | 2.947E+04 | 1096 | - | - | 0 | - |
| 11 | c | 4.493E+05 | 1102 | 0.000596 | 0.541 | +1 | 11 |
| - | - | 2.949E+05 | 1103 | - | - | 0 | - |
| - | - | 1.906E+05 | 1104 | - | - | 0 | - |
| - | - | 7.668E+04 | 1105 | - | - | 0 | - |
| - | - | 3.168E+04 | 1106 | - | - | 0 | - |
| - | - | 1.561E+04 | 1110 | - | - | 0 | - |
| - | - | 2.434E+04 | 1115 | - | - | 0 | - |
| - | - | 2.434E+04 | 1116 | - | - | 0 | - |
| 11 | c | 4.851E+06 | 1119 | 4.788E-05 | 0.0428 | +1 | 11 |
| - | - | 3.109E+06 | 1120 | - | - | 0 | - |
| - | - | 1.009E+06 | 1121 | - | - | 0 | - |
| - | - | 1.08E+05 | 1122 | - | - | 0 | - |
| - | - | 7.945E+04 | 1133 | - | - | 0 | - |
| - | - | 4.992E+04 | 1134 | - | - | 0 | - |
| - | - | 1.909E+04 | 1135 | - | - | 0 | - |
| - | - | 4.882E+04 | 1143 | - | - | 0 | - |
| - | - | 2.746E+04 | 1144 | - | - | 0 | - |
| 2 | z | 1.112E+05 | 1148 | 7.9E-06 | 0.006884 | +1 | 11 |
| 2 | z | 7.004E+04 | 1149 | 0.01721 | 14.99 | +1 | 11 |
| - | - | 2.441E+04 | 1150 | - | - | 0 | - |
| 2 | z | 1.085E+06 | 1166 | 5.874E-05 | 0.05039 | +1 | 11 |
| - | - | 7.888E+05 | 1167 | - | - | 0 | - |
| - | - | 2.764E+05 | 1168 | - | - | 0 | - |
| - | - | 3.036E+04 | 1169 | - | - | 0 | - |
| - | - | 1.55E+04 | 1176 | - | - | 0 | - |
| - | - | 1.734E+04 | 1177 | - | - | 0 | - |
| - | - | 2.088E+04 | 1178 | - | - | 0 | - |
| 2 | y | 1.592E+04 | 1182 | 0.01085 | 9.181 | +1 | 11 |
| - | - | 2.343E+04 | 1183 | - | - | 0 | - |
| - | - | 1.75E+04 | 1184 | - | - | 0 | - |
| - | - | 4.645E+04 | 1196 | - | - | 0 | - |
| - | - | 2.902E+04 | 1197 | - | - | 0 | - |
| - | - | 2.641E+04 | 1198 | - | - | 0 | - |
| - | - | 1.662E+04 | 1199 | - | - | 0 | - |
| - | - | 7.296E+04 | 1205 | - | - | 0 | - |
| - | - | 5.937E+04 | 1206 | - | - | 0 | - |
| - | - | 3.199E+04 | 1207 | - | - | 0 | - |
| - | - | 1.445E+05 | 1211 | - | - | 0 | - |
| - | - | 1.063E+05 | 1212 | - | - | 0 | - |
| - | - | 1.099E+05 | 1213 | - | - | 0 | - |
| - | - | 7.219E+04 | 1214 | - | - | 0 | - |
| - | - | 3.26E+04 | 1215 | - | - | 0 | - |
| - | - | 2.219E+04 | 1221 | - | - | 0 | - |
| - | - | 2.119E+05 | 1222 | - | - | 0 | - |
| - | - | 1.724E+05 | 1223 | - | - | 0 | - |
| - | - | 5.755E+04 | 1224 | - | - | 0 | - |
| - | - | 9.07E+04 | 1228 | - | - | 0 | - |
| - | - | 6.651E+04 | 1229 | - | - | 0 | - |
| - | - | 2.328E+04 | 1230 | - | - | 0 | - |
| - | - | 1.202E+04 | 1231 | - | - | 0 | - |
| - | - | 3.005E+04 | 1238 | - | - | 0 | - |
| - | - | 6.89E+05 | 1239 | - | - | 0 | - |
| - | - | 5.451E+05 | 1240 | - | - | 0 | - |
| - | - | 2.664E+05 | 1241 | - | - | 0 | - |
| - | - | 9.831E+04 | 1242 | - | - | 0 | - |
| - | - | 3.554E+04 | 1243 | - | - | 0 | - |
| - | - | 5.068E+05 | 1249 | - | - | 0 | - |
| - | - | 3.973E+05 | 1250 | - | - | 0 | - |
| - | - | 1.124E+05 | 1251 | - | - | 0 | - |
| - | - | 3.561E+04 | 1252 | - | - | 0 | - |
| - | - | 2.203E+04 | 1253 | - | - | 0 | - |
| - | - | 2.853E+05 | 1256 | - | - | 0 | - |
| - | - | 2.394E+05 | 1257 | - | - | 0 | - |
| - | - | 8.254E+04 | 1258 | - | - | 0 | - |
| - | - | 1.38E+04 | 1259 | - | - | 0 | - |
| - | - | 1.004E+05 | 1266 | - | - | 0 | - |
| - | - | 7.239E+05 | 1267 | - | - | 0 | - |
| - | - | 5.373E+05 | 1268 | - | - | 0 | - |
| - | - | 1.884E+05 | 1269 | - | - | 0 | - |
| - | - | 2.41E+04 | 1270 | - | - | 0 | - |
| - | - | 7.891E+04 | 1282 | - | - | 0 | - |
| - | - | 1.334E+06 | 1283 | - | - | 0 | - |
| - | - | 5.717E+06 | 1284 | - | - | 0 | - |
| - | - | 3.971E+06 | 1285 | - | - | 0 | - |
| - | - | 1.444E+06 | 1286 | - | - | 0 | - |
| - | - | 1.557E+05 | 1287 | - | - | 0 | - |
| - | - | 1.709E+04 | 1316 | - | - | 0 | - |
| - | - | 2.372E+04 | 1317 | - | - | 0 | - |
| - | - | 1.332E+04 | 1318 | - | - | 0 | - |
| - | - | 1.2E+04 | 1515 | - | - | 0 | - |

m/z Charge Intensity FragmentType MassShift Position
120.081298828125 0 14685.79
120.94535064697266 0 5656.1255
121.77444458007812 0 5064.1885
123.04434204101562 0 11218.942
125.7257080078125 0 6683.122
128.92152404785156 0 5942.37
130.5825653076172 0 6406.4146
135.36268615722656 0 6356.179
136.07603454589844 0 372524.16
137.0794677734375 0 29890.201
141.10252380371094 0 11909.876
143.1181640625 0 62105.824
165.05494689941406 0 335494.6
166.0582733154297 0 37065.4
169.0975341796875 0 152459.69
171.11314392089844 0 87099.805
175.07196044921875 0 14176.31
177.7549591064453 0 7469.8413
182.08160400390625 0 2187959.8 y 11
183.0849151611328 0 204093.27
186.1238555908203 0 24560.69
189.5147705078125 0 7369.645
195.97044372558594 0 8236.008
199.99818420410156 0 7673.765
213.1599884033203 0 9952.664
214.0824432373047 0 49241.836
216.09820556640625 0 21561.83
218.00706481933594 0 8666.624
221.1288604736328 0 807182.8
222.13221740722656 0 97646.84
232.0933837890625 0 79493.36
240.17103576660156 0 73402.04
242.15042114257812 0 40609.11
249.12379455566406 0 591937.2
250.12701416015625 0 68830.234
258.1451110839844 0 104177.22
259.1477966308594 0 11776.855
268.1658630371094 0 367972.4
269.1692199707031 0 63979.766
285.1927795410156 0 16998.496
286.1767272949219 0 10964.009
295.1658020019531 0 305804.47 y 10
296.169189453125 0 45701.484
299.1730041503906 0 10095.528
300.1562805175781 0 10089.309
316.1657409667969 0 21519.473
317.1827087402344 0 235629.4
318.1850891113281 0 22349.516
318.53564453125 0 8314.187
324.1792297363281 0 13265.935
327.1671447753906 0 30609.43
328.1670227050781 0 18252.184
329.1834716796875 0 10777.348
333.18560791015625 0 20420.29
341.2187805175781 0 20165.555
342.14166259765625 0 19963.334
343.1252136230469 0 47274.223
344.1972351074219 0 8571.287
345.1773986816406 0 468890.8
346.178466796875 0 121052.95
347.1803283691406 0 11724.935
351.20294189453125 0 45890.8
352.18701171875 0 108737.36 y 9
352.2091064453125 0 14574.714
352.69122314453125 0 11541.093
353.18951416015625 0 24469.953
353.2554016113281 0 37079.176
360.1518249511719 0 14055.208
361.6954650878906 0 145424.98
362.1977233886719 0 65249.434
362.6972961425781 0 9410.52
369.21405029296875 0 43464.39
370.7001647949219 0 27074.967
371.20172119140625 0 28932.746
377.18035888671875 0 11357.04
379.2174987792969 0 18486.404 c Ammonia loss 6
381.2500915527344 0 156815.67
382.25360107421875 0 32357.557
387.23883056640625 0 11542.081
389.2192687988281 0 22811.291
395.2370300292969 0 8861.028
398.2044372558594 0 21888.422
398.2764587402344 0 28321.557
399.20330810546875 0 80320.69
400.18267822265625 0 10059.215
400.2082214355469 0 11037.837
410.24066162109375 0 10613.779
412.2596435546875 0 11565.402
413.2463684082031 0 17999.668
414.2720642089844 0 70166.6
415.2344970703125 0 36083.844
415.27459716796875 0 17712.6
416.2132263183594 0 14686.711
417.2140197753906 0 228313.89 z Ammonia loss 4
418.2171936035156 0 53553.832 w 4
420.8575134277344 0 9214.488
421.2096862792969 0 10532.63 y Water loss 8
422.73260498046875 0 38449.43 c Ammonia loss 7
423.2312316894531 0 13809.367
427.24237060546875 0 27804.67
427.74359130859375 0 25179.479
428.2147521972656 0 66852.516
428.2478332519531 0 17382.645
429.21636962890625 0 18614.244
433.241455078125 0 16475.107
434.2401123046875 0 23887.049 c 3
438.19842529296875 0 11980.646
439.2196960449219 0 97021.32 y 8
440.2218933105469 0 24898.717
440.2527770996094 0 12240.618
445.2419128417969 0 43361.18
448.2376403808594 0 49412.945
455.225341796875 0 57614.91
456.2099914550781 0 118186.67
457.2114562988281 0 27403.62
457.2434997558594 0 42696.605
457.7438049316406 0 12358.04
458.26080322265625 0 16482.426
464.287841796875 0 42207.02
470.2759704589844 0 18412.52
471.2586364746094 0 14186.34
472.2225036621094 0 10820.76
472.2574462890625 0 35346.055
473.2364807128906 0 150627.84
474.2389831542969 0 35432.027
475.28680419921875 0 10415.311
476.7474670410156 0 25317.26
477.2465515136719 0 22523.98
481.31439208984375 0 13443.551
482.29974365234375 0 33108.54
483.2987976074219 0 15294.189
485.7537841796875 0 169810.92
486.2546691894531 0 69216.695
486.7566833496094 0 32399.223
488.28759765625 0 172461.64
489.2909851074219 0 41797.195
490.22784423828125 0 17241.814
490.26397705078125 0 27522.521
491.2684020996094 0 9589.212
492.2808532714844 0 9226.036
494.7584533691406 0 15133.483 c Ammonia loss 9
498.2716064453125 0 311563.8
499.2742919921875 0 63407.504
499.7688293457031 0 25968.463
500.2670593261719 0 34603.5
501.3045959472656 0 22852.121
502.3076477050781 0 10272.18
508.24029541015625 0 46547.465 y Water loss 7
508.3030090332031 0 13302.878
508.7741394042969 0 24206.191 y Water loss 2
509.3088684082031 0 403043.5
510.3118591308594 0 117954.234
511.3163146972656 0 13047.614
515.2762451171875 0 28647.521
516.2822875976562 0 524181.7
517.2849731445312 0 141403.47
518.2865600585938 0 23841.023
525.3292236328125 0 13478.79
526.25146484375 0 380653.97 y 7
526.3356323242188 0 98859.93
527.2544555664062 0 95157.52
527.3366088867188 0 15573.578
528.318359375 0 12140.355
530.297607421875 0 15399.278
532.3016357421875 0 282248.47
533.30859375 0 954444.44 c 4
534.3116455078125 0 282474.6
535.3135986328125 0 42995.887
541.2988891601562 0 10491.504
550.296142578125 0 10888.583
551.2913818359375 0 9115.208 c Ammonia loss 10
552.347900390625 0 13098.829
557.3325805664062 0 9217.319
558.3253173828125 0 43170.28
559.3292236328125 0 15942.734
568.3094482421875 0 46490.28
569.3006591796875 0 24953.494
570.2996826171875 0 12639.329
573.2266235351562 0 10045.616
577.3001708984375 0 11725.514
578.3296508789062 0 24472.578
583.36083984375 0 23359.594
584.3623046875 0 34057.836
585.3452758789062 0 27004.203
586.3201904296875 0 179562.77
587.3226318359375 0 50477.34
588.330810546875 0 20972.684
589.326416015625 0 26958.621
590.3323364257812 0 10205.672
592.3438110351562 0 16337.741
593.3479614257812 0 12103.296
596.3410034179688 0 70002.29
597.3427734375 0 26623.668
601.371826171875 0 60019.223
602.378173828125 0 102190.12
603.386962890625 0 462681.9
604.3901977539062 0 176271.75
605.3912963867188 0 22003.162
610.3541259765625 0 14078.591
611.3550415039062 0 309443.28
612.3527221679688 0 119865.71
613.3604736328125 0 72084.23
618.2921752929688 0 14330.619
627.3142700195312 0 15144.385
628.36083984375 0 25387.584
629.3661499023438 0 465071.12
630.369873046875 0 163691.16
631.3749389648438 0 27461.678
635.2914428710938 0 40634.406
636.2990112304688 0 139153.78 y Water loss 6
637.2958374023438 0 39442.633
638.2921752929688 0 19828.426 z 6
639.296875 0 10168.374
640.3598022460938 0 31812.229
641.3621215820312 0 26999.08
642.3197631835938 0 15879.574
643.3308715820312 0 13418.709
645.385009765625 0 609364.06
646.3924560546875 0 1928408.9 c 5
647.3323364257812 0 15989.721
647.3955078125 0 674251.2
648.3990478515625 0 107107.266
653.302001953125 0 305692.78
654.3095703125 0 1093807.4 y 6
655.3126831054688 0 370884.9
655.3702392578125 0 11097.432
656.3141479492188 0 77713.516
656.3787231445312 0 31915.664
657.386962890625 0 10829.402
659.3991088867188 0 15817.411
665.3634033203125 0 35006.207
666.3653564453125 0 20139.799
667.375732421875 0 15673.3125
668.3701782226562 0 11220.978
669.3667602539062 0 13990.901
672.408935546875 0 101761.945
673.4100952148438 0 40085.566
674.3739624023438 0 27809.043
675.3801879882812 0 15160.188
683.373779296875 0 45777.08
684.3781127929688 0 20552.064
685.3881225585938 0 55498.89
686.3912353515625 0 37898.94
687.394775390625 0 23506.04
697.3787841796875 0 9967.202
699.3916625976562 0 22427.586
700.397705078125 0 52149.043
701.40185546875 0 14735.502
702.40771484375 0 27039.914
703.34130859375 0 28402.842
703.4075317382812 0 13096.72
704.3720092773438 0 29927.234
705.3764038085938 0 14512.089
706.370849609375 0 9171.018
710.3609008789062 0 19469.797
712.3935546875 0 40910.6
713.4034423828125 0 29386.023
714.4158325195312 0 16769.441
715.4212036132812 0 13005.224
721.3499145507812 0 14996.959
722.38427734375 0 171286.5
723.3812866210938 0 73079.39
724.3802490234375 0 18409.912
727.3897094726562 0 37582.484
729.4288330078125 0 29537.826
730.3638305664062 0 16398.512
730.4375 0 335747.34
731.4445190429688 0 745571.44
732.4486694335938 0 306165.75
733.3736572265625 0 14033.743 z Water loss 5
733.4512329101562 0 49424.047
739.4129028320312 0 133923.16
740.3987426757812 0 488423.53
741.403564453125 0 213553
742.4115600585938 0 122943.5
743.414794921875 0 40091.53
747.3628540039062 0 10003.512
749.3826293945312 0 44109.18 y Water loss 5
750.3758544921875 0 34627.367 y Ammonia loss 5
751.3751831054688 0 197134.81 z 5
752.3809204101562 0 112191.12
753.3826904296875 0 29178.373
755.4317016601562 0 115814.695
756.4293212890625 0 92207.95 c Water loss 6
757.425048828125 0 678396.94 c Ammonia loss 6
758.4308471679688 0 323696.78
759.4381103515625 0 180698.81
760.4430541992188 0 49781.53
766.3869018554688 0 105630.16
767.3936157226562 0 764652.56 y 5
768.3966674804688 0 293728.3
769.3995971679688 0 65017.867
773.4432983398438 0 1119666
774.4496459960938 0 1454230.6 c 6
775.45361328125 0 580147.3
776.4594116210938 0 130074.24
777.4650268554688 0 24679.76
781.457763671875 0 10394.647
789.4340209960938 0 13125.77
794.382080078125 0 31846.152
799.4551391601562 0 44069.207
800.464599609375 0 86994.17
801.464111328125 0 46255.543
802.4677124023438 0 16841.77
808.4402465820312 0 15310.521
809.4395141601562 0 11276.874
812.4367065429688 0 12654.928
814.424560546875 0 21635.27
815.4306030273438 0 19887.834
816.428466796875 0 17251.338
817.4640502929688 0 47366.72
818.4725341796875 0 39678.13
819.478515625 0 25405.918
823.4342041015625 0 10404.452
825.4832763671875 0 100107.04
826.4461669921875 0 50671.637
827.4376831054688 0 76064.336
828.4386596679688 0 26549.992
829.4429931640625 0 24989.324
830.4445190429688 0 18401.52
832.4373168945312 0 11859.451 z Water loss 4
834.4368896484375 0 17479.908
835.464111328125 0 83970.93
836.4609985351562 0 74178.336
837.4571533203125 0 16898.182
842.4625244140625 0 125041.95
843.4644165039062 0 84442.86 c Water loss 7
844.4576416015625 0 273521.44 c Ammonia loss 7
845.4598999023438 0 117208.3
846.4699096679688 0 59995.906
847.4722900390625 0 22859.23
850.443603515625 0 201462.9 z 4
851.447021484375 0 96907.234
852.4535522460938 0 16897.87
853.478271484375 0 503597.2
854.4813842773438 0 234366.17
855.4837646484375 0 59352.906
860.4749145507812 0 781373.4
861.481201171875 0 859058.8 c 7
862.4844970703125 0 344644.84
863.4932250976562 0 116217.12
864.5003051757812 0 27353.453
865.45068359375 0 45424.965
866.461669921875 0 211893.66 y 4
867.4638671875 0 78892.03
868.465576171875 0 24190.008
871.4306640625 0 16074.4375
872.4443359375 0 20531.889
887.4930419921875 0 25515.635
889.4780883789062 0 26079.537
891.4431762695312 0 28591.969
892.4545288085938 0 21559.217
893.4613647460938 0 13248.734
895.4671630859375 0 47580.805
896.4544067382812 0 57603.55
897.4534912109375 0 28042.129
898.4661865234375 0 21091.713
899.4830932617188 0 12070.033
903.4708862304688 0 154177.69 z Water loss 3
904.471435546875 0 79435.85 z Ammonia loss 3
905.4747924804688 0 27007.023
913.4779663085938 0 58926.465
914.471923828125 0 67945.75
915.4796142578125 0 27521.363
921.4803466796875 0 1226660.8 z 3
922.4842529296875 0 780593.5
923.4883422851562 0 223307.45
924.4909057617188 0 32726.15
929.4966430664062 0 105036.03
930.495361328125 0 67876 c Water loss 8
931.4890747070312 0 228672.12 c Ammonia loss 8
932.4920654296875 0 105784.945
933.4971313476562 0 31694.562
934.4832763671875 0 11009.977
935.4688110351562 0 13849.056
936.5094604492188 0 52756.227
937.5003051757812 0 99622.61 y 3
938.5008544921875 0 41952.996
939.4991455078125 0 12647.122
942.4998779296875 0 15248.113
945.492919921875 0 13256.44
947.5077514648438 0 778320.1
948.5142822265625 0 1889101.2 c 8
949.517578125 0 935475.2
950.5206909179688 0 222847.55
951.5264282226562 0 13459.266
952.4904174804688 0 96264.95
953.4838256835938 0 90224.44
954.495849609375 0 48476.285
955.499755859375 0 22052.814
960.5170288085938 0 23716.482
961.5233154296875 0 53020.98
962.5267944335938 0 44884.066
963.5349731445312 0 15759.551
967.5254516601562 0 40516.766
968.5292358398438 0 20427.744
970.4998168945312 0 491463.25
971.5010375976562 0 297561.4
972.502685546875 0 104702.83
973.510009765625 0 19930.91
977.4837036132812 0 24532.041
984.5296630859375 0 460046.53
985.5328979492188 0 304716.25
986.5347900390625 0 105498.664
987.5317993164062 0 33624.027 c Water loss 9
988.51025390625 0 573448.06 c Ammonia loss 9
989.5130004882812 0 307609.5
990.5164184570312 0 94453.42
991.5257568359375 0 14117.585 w 2
998.5265502929688 0 24319.266
1000.5467529296875 0 13416.655
1001.5264892578125 0 12321.98
1002.5534057617188 0 31847.639
1003.5607299804688 0 23433.098
1005.537109375 0 5468797 c 9
1006.5394897460938 0 3117353.2
1007.5426025390625 0 843162.25
1008.54638671875 0 81203.02
1015.536376953125 0 58815.797
1016.5383911132812 0 80635.4 y Water loss 2
1017.5415649414062 0 50528.727 y Ammonia loss 2
1018.5405883789062 0 20866.512 z 2
1029.5166015625 0 12835.581
1031.552978515625 0 31595.396
1032.55419921875 0 16074.911
1033.549560546875 0 43420.11
1034.5513916015625 0 515032.78 y 2
1035.5556640625 0 278497.78
1036.5584716796875 0 62912.875
1038.571044921875 0 15742.992
1041.5604248046875 0 21689.51
1056.590576171875 0 22960.02
1057.58251953125 0 56873.445
1058.581298828125 0 36928.79
1062.5439453125 0 21422.81
1065.5654296875 0 19054.531
1066.5645751953125 0 33662.42
1067.565673828125 0 22238.416
1068.5732421875 0 253237.11
1069.5760498046875 0 161999.55
1070.575927734375 0 59036.477
1073.6004638671875 0 106327.21
1074.606201171875 0 263672.72
1075.6092529296875 0 173975.72
1076.612060546875 0 45843.47
1083.5833740234375 0 242181.69
1084.585205078125 0 175835.14
1085.5906982421875 0 713594.5
1086.59326171875 0 523393.84
1087.5960693359375 0 156016.89
1088.5994873046875 0 24556.271
1094.563232421875 0 34933.72
1095.5712890625 0 29472.29
1101.594482421875 0 449282.66 c Ammonia loss 10
1102.5966796875 0 294941.16
1103.6033935546875 0 190641.9
1104.6114501953125 0 76677.82
1105.6175537109375 0 31683.068
1109.5316162109375 0 15611.873
1114.625 0 24339.865
1115.6160888671875 0 24342.404
1118.6204833984375 0 4851336 c 10
1119.62353515625 0 3108983.5
1120.6258544921875 0 1008986
1121.6278076171875 0 107954.195
1132.6341552734375 0 79445.34
1133.6407470703125 0 49918.64
1134.6424560546875 0 19091.285
1142.5987548828125 0 48818.082
1143.597412109375 0 27463.637
1147.5908203125 0 111183.016 z Water loss 1
1148.592041015625 0 70035.375 z Ammonia loss 1
1149.5960693359375 0 24412.59
1165.601318359375 0 1084525.2 z 1
1166.6044921875 0 788796.5
1167.6070556640625 0 276383.62
1168.6063232421875 0 30356.98
1175.616943359375 0 15498.545
1176.622314453125 0 17343.402
1177.6259765625 0 20884.264
1181.6092529296875 0 15916.023 y 1
1182.59375 0 23428.287
1183.594970703125 0 17501.357
1195.6097412109375 0 46448.91
1196.6143798828125 0 29021.094
1197.5977783203125 0 26411.22
1198.5958251953125 0 16621.348
1204.6483154296875 0 72962.61
1205.6513671875 0 59370.176
1206.65478515625 0 31990.111
1210.5921630859375 0 144519.9
1211.5926513671875 0 106250.31
1212.62890625 0 109859.27
1213.640869140625 0 72188.18
1214.6422119140625 0 32602.143
1220.645751953125 0 22188.365
1221.672119140625 0 211910.81
1222.6749267578125 0 172416.73
1223.6783447265625 0 57549.875
1227.61181640625 0 90699.31
1228.615478515625 0 66509.125
1229.617431640625 0 23276.01
1230.6376953125 0 12022.412
1237.6650390625 0 30053.67
1238.653564453125 0 688990.7
1239.656982421875 0 545087.94
1240.64892578125 0 266441.66
1241.6298828125 0 98311.2
1242.6209716796875 0 35542.22
1248.637451171875 0 506829.12
1249.6397705078125 0 397258.1
1250.6435546875 0 112398.766
1251.63427734375 0 35608.36
1252.63720703125 0 22032.836
1255.6795654296875 0 285286.3
1256.6826171875 0 239447.33
1257.685302734375 0 82541.22
1258.687255859375 0 13804.233
1265.6650390625 0 100422
1266.64990234375 0 723902.9
1267.65234375 0 537292.4
1268.6534423828125 0 188409.86
1269.65771484375 0 24097.354
1281.7618408203125 0 78906.625
1282.6669921875 0 1333850.6
1283.6741943359375 0 5717163
1284.6767578125 0 3971463
1285.6802978515625 0 1444481.6
1286.6826171875 0 155654.89
1315.6658935546875 0 17090.357
1316.669677734375 0 23724.346
1317.67529296875 0 13318.308
1514.7232666015625 0 12002.948

Spectrum Details

|  |  |
| --- | --- |
| Matched peaks? Matched peaks The total absolute number of peaks matched. Additionally in brackets the total fraction of peaks matched and the total number of peaks is shown. | 55 (10.32% of 533) |
| FDR? FDR The false discovery rate estimated for this peptide. It is calculated by matching all theoretical fragments with a non-integer shift with the raw peaks for this spectrum. This is done with 40 different shifts. The resulting percentage is the average number of annotated peaks over the number of annotated peaks with the correct spectrum. | 2.03% |
| Satellite FDR? Satellite FDR See the FDR for details on its calculation. This satellite ion specific FDR only contains the satellite ions (d/w) for I/L/J positions. | ∞ |
| PSM Score? PSM Score The PSM Score as given by Hecklib to this annotated spectrum. It is shown with three significant figures. | 736 |

#### Spectrum 11052? Spectrum 11052 The raw spectrum of this peptide as annotated by Hecklib. The fragments are coloured according to ion type (see legend). Any peaks with a star '\*' as text can be hovered over to see the full details, first the ion type second the mass shift type. By hovering over the amino acids in the peptide or ions in the legend the corresponding peaks are highlighted. By toggling the 'Unassigned' label you can turn the background (unassigned) peaks on or off in the plot. By updating the slider in the Ion legend you can update the spectrum to only show the top X% of the peaks with labels. The top X% means any peak that is within X% of the highest intensity. By dragging in the spectrum you can zoom in to a specific part of the spectrum and use 'Zoom Out' to get back to the original zoom level. The annotation of the spectrum is based on the given sequence in the peptides file and is done with different software so inconsistencies are likely. The peaks are annotated based on the given sequence, with 20 ppm tolerance.

Copy Data

##### Spectrum 11052 (TSV)

###### Preview

```
Loading example...
```

*Click on the button to copy the data to your clipboard.*

Mz MinMz MaxIntensity Max

WidthHeightPeptide font sizePeptide stroke widthSpectrum font sizeSpectrum stroke widthCompact peptide

Ion legend

wxyz

abcd

OtherUnassignedIon^Charge^_Position_Show for top:%

TFPAVJQSSGJY

03.06e+46.13e+49.19e+41.23e+5

Zoom Out

y^+1^_1_y^+1^_2_y^+1^_3_c^+2^_7_z^+2^_8_w^+2^_8_y^+1^_4_y^+1^_5_y^+1^_5_c^+1^_5_y^+1^_6_c^+1^_6_y^+1^_6_y^+1^_7_z^+1^_7_c^+1^_7_c^+1^_7_y^+1^_7_c^+1^_7_c^+1^_8_c^+1^_8_z^+1^_8_c^+1^_8_y^+1^_8_z^+1^_9_z^+1^_9_z^+1^_9_c^+1^_9_c^+1^_9_y^+1^_9_c^+1^_9_c^+1^_10_c^+1^_10_c^+1^_10_y^+1^_10_y^+1^_10_c^+1^_11_c^+1^_11_z^+1^_11_z^+1^_11_z^+1^_11_

0610122018302440

Fragment Matches Table

Show background peaks

| Position | Ion type | Intensity | mz Theoretical | mz Error (Th) | mz Error (ppm) | Charge | Series Number |
| --- | --- | --- | --- | --- | --- | --- | --- |
| - | - | 758.8 | 120.1 | - | - | 0 | - |
| - | - | 726.1 | 123 | - | - | 0 | - |
| - | - | 388.1 | 127.5 | - | - | 0 | - |
| - | - | 387.8 | 131.7 | - | - | 0 | - |
| - | - | 8158 | 136.1 | - | - | 0 | - |
| - | - | 686.5 | 137.1 | - | - | 0 | - |
| - | - | 1159 | 143.1 | - | - | 0 | - |
| - | - | 928.5 | 148.9 | - | - | 0 | - |
| - | - | 6109 | 165.1 | - | - | 0 | - |
| - | - | 985.9 | 166.1 | - | - | 0 | - |
| - | - | 3558 | 169.1 | - | - | 0 | - |
| - | - | 1640 | 171.1 | - | - | 0 | - |
| - | - | 500 | 172.1 | - | - | 0 | - |
| - | - | 538.3 | 173.1 | - | - | 0 | - |
| - | - | 484.2 | 173.5 | - | - | 0 | - |
| 12 | y | 4.226E+04 | 182.1 | 0.0001749 | 0.9607 | +1 | 1 |
| - | - | 493.7 | 182.3 | - | - | 0 | - |
| - | - | 4709 | 183.1 | - | - | 0 | - |
| - | - | 524.9 | 184.1 | - | - | 0 | - |
| - | - | 688 | 186.1 | - | - | 0 | - |
| - | - | 554.6 | 213.6 | - | - | 0 | - |
| - | - | 946.3 | 214.1 | - | - | 0 | - |
| - | - | 611.6 | 216.1 | - | - | 0 | - |
| - | - | 551.5 | 216.8 | - | - | 0 | - |
| - | - | 1.705E+04 | 221.1 | - | - | 0 | - |
| - | - | 522.2 | 222 | - | - | 0 | - |
| - | - | 2090 | 222.1 | - | - | 0 | - |
| - | - | 1935 | 232.1 | - | - | 0 | - |
| - | - | 1558 | 240.2 | - | - | 0 | - |
| - | - | 458.8 | 240.5 | - | - | 0 | - |
| - | - | 595.7 | 242.1 | - | - | 0 | - |
| - | - | 1.223E+04 | 249.1 | - | - | 0 | - |
| - | - | 1552 | 250.1 | - | - | 0 | - |
| - | - | 549.4 | 253.5 | - | - | 0 | - |
| - | - | 2285 | 258.1 | - | - | 0 | - |
| - | - | 8923 | 268.2 | - | - | 0 | - |
| - | - | 1257 | 269.2 | - | - | 0 | - |
| - | - | 518.6 | 281 | - | - | 0 | - |
| - | - | 497.4 | 285.2 | - | - | 0 | - |
| 11 | y | 5928 | 295.2 | 0.0003242 | 1.098 | +1 | 2 |
| - | - | 632 | 296.2 | - | - | 0 | - |
| - | - | 506.9 | 307.9 | - | - | 0 | - |
| - | - | 4839 | 317.2 | - | - | 0 | - |
| - | - | 881.1 | 327.2 | - | - | 0 | - |
| - | - | 918.2 | 343.1 | - | - | 0 | - |
| - | - | 8180 | 345.2 | - | - | 0 | - |
| - | - | 2053 | 346.2 | - | - | 0 | - |
| - | - | 946.3 | 351.2 | - | - | 0 | - |
| 10 | y | 1752 | 352.2 | 0.0004181 | 1.187 | +1 | 3 |
| - | - | 643.7 | 353.2 | - | - | 0 | - |
| - | - | 1099 | 353.3 | - | - | 0 | - |
| - | - | 580.3 | 360.2 | - | - | 0 | - |
| - | - | 2385 | 361.7 | - | - | 0 | - |
| - | - | 595.7 | 362.2 | - | - | 0 | - |
| - | - | 743.7 | 369.2 | - | - | 0 | - |
| 7 | c | 710.8 | 379.2 | 0.00183 | 4.827 | +2 | 7 |
| - | - | 3021 | 381.2 | - | - | 0 | - |
| - | - | 664.5 | 382.3 | - | - | 0 | - |
| - | - | 890.5 | 389.2 | - | - | 0 | - |
| - | - | 1218 | 398.3 | - | - | 0 | - |
| - | - | 2021 | 399.2 | - | - | 0 | - |
| - | - | 608.5 | 412.6 | - | - | 0 | - |
| - | - | 669.5 | 413.3 | - | - | 0 | - |
| - | - | 1685 | 414.3 | - | - | 0 | - |
| 5 | z | 5472 | 417.2 | 0.001595 | 3.822 | +2 | 8 |
| 5 | w | 1320 | 418.2 | 0.00314 | 7.508 | +2 | 8 |
| - | - | 1385 | 428.2 | - | - | 0 | - |
| 9 | y | 2344 | 439.2 | 0.0003904 | 0.8889 | +1 | 4 |
| - | - | 862.2 | 445.2 | - | - | 0 | - |
| - | - | 687.6 | 448.2 | - | - | 0 | - |
| - | - | 849.1 | 455.2 | - | - | 0 | - |
| - | - | 3230 | 456.2 | - | - | 0 | - |
| - | - | 697.6 | 457.2 | - | - | 0 | - |
| - | - | 616.9 | 458.3 | - | - | 0 | - |
| - | - | 700.6 | 472.3 | - | - | 0 | - |
| - | - | 2950 | 473.2 | - | - | 0 | - |
| - | - | 2691 | 485.8 | - | - | 0 | - |
| - | - | 1508 | 486.3 | - | - | 0 | - |
| - | - | 765.3 | 486.8 | - | - | 0 | - |
| - | - | 3088 | 488.3 | - | - | 0 | - |
| - | - | 932.1 | 489.3 | - | - | 0 | - |
| - | - | 728.1 | 490.3 | - | - | 0 | - |
| - | - | 5823 | 498.3 | - | - | 0 | - |
| - | - | 1043 | 499.3 | - | - | 0 | - |
| - | - | 838 | 499.8 | - | - | 0 | - |
| - | - | 700.1 | 500.3 | - | - | 0 | - |
| - | - | 824.3 | 501.3 | - | - | 0 | - |
| 8 | y | 664.9 | 508.2 | 0.001695 | 3.334 | +1 | 5 |
| - | - | 7622 | 509.3 | - | - | 0 | - |
| - | - | 1773 | 510.3 | - | - | 0 | - |
| - | - | 1215 | 515.3 | - | - | 0 | - |
| - | - | 9600 | 516.3 | - | - | 0 | - |
| - | - | 3631 | 517.3 | - | - | 0 | - |
| - | - | 702.3 | 518.3 | - | - | 0 | - |
| 8 | y | 6565 | 526.3 | 0.0001613 | 0.3066 | +1 | 5 |
| - | - | 2597 | 526.3 | - | - | 0 | - |
| - | - | 2121 | 527.3 | - | - | 0 | - |
| - | - | 622.9 | 527.3 | - | - | 0 | - |
| - | - | 6733 | 532.3 | - | - | 0 | - |
| 5 | c | 2.486E+04 | 533.3 | 0.0004092 | 0.7673 | +1 | 5 |
| - | - | 8154 | 534.3 | - | - | 0 | - |
| - | - | 1905 | 535.3 | - | - | 0 | - |
| - | - | 872 | 558.3 | - | - | 0 | - |
| - | - | 1023 | 568.3 | - | - | 0 | - |
| - | - | 1037 | 569.3 | - | - | 0 | - |
| - | - | 928.8 | 584.4 | - | - | 0 | - |
| - | - | 767.3 | 585.3 | - | - | 0 | - |
| - | - | 4112 | 586.3 | - | - | 0 | - |
| - | - | 984.4 | 587.3 | - | - | 0 | - |
| - | - | 1646 | 596.3 | - | - | 0 | - |
| - | - | 1399 | 601.4 | - | - | 0 | - |
| - | - | 2416 | 602.4 | - | - | 0 | - |
| - | - | 1.073E+04 | 603.4 | - | - | 0 | - |
| - | - | 4159 | 604.4 | - | - | 0 | - |
| - | - | 5387 | 611.4 | - | - | 0 | - |
| - | - | 3515 | 612.3 | - | - | 0 | - |
| - | - | 1043 | 613.4 | - | - | 0 | - |
| - | - | 603.1 | 614.4 | - | - | 0 | - |
| - | - | 9352 | 629.4 | - | - | 0 | - |
| - | - | 2861 | 630.4 | - | - | 0 | - |
| - | - | 1168 | 635.3 | - | - | 0 | - |
| 7 | y | 2988 | 636.3 | 0.0001221 | 0.1919 | +1 | 6 |
| - | - | 1065 | 637.3 | - | - | 0 | - |
| - | - | 997.8 | 640.4 | - | - | 0 | - |
| - | - | 3319 | 641.9 | - | - | 0 | - |
| - | - | 2165 | 642.4 | - | - | 0 | - |
| - | - | 649.2 | 643.3 | - | - | 0 | - |
| - | - | 1.505E+04 | 645.4 | - | - | 0 | - |
| 6 | c | 4.65E+04 | 646.4 | 0.0006109 | 0.9451 | +1 | 6 |
| - | - | 2.082E+04 | 647.4 | - | - | 0 | - |
| - | - | 4538 | 648.4 | - | - | 0 | - |
| - | - | 6614 | 653.3 | - | - | 0 | - |
| 7 | y | 2.291E+04 | 654.3 | 0.0006769 | 1.035 | +1 | 6 |
| - | - | 7787 | 655.3 | - | - | 0 | - |
| - | - | 2102 | 656.3 | - | - | 0 | - |
| - | - | 821.5 | 656.4 | - | - | 0 | - |
| - | - | 644.6 | 669.4 | - | - | 0 | - |
| - | - | 3330 | 672.4 | - | - | 0 | - |
| - | - | 1422 | 673.4 | - | - | 0 | - |
| - | - | 891.2 | 674.4 | - | - | 0 | - |
| - | - | 1014 | 683.4 | - | - | 0 | - |
| - | - | 1142 | 685.4 | - | - | 0 | - |
| - | - | 1513 | 700.4 | - | - | 0 | - |
| - | - | 788.9 | 701.4 | - | - | 0 | - |
| - | - | 967.5 | 712.4 | - | - | 0 | - |
| - | - | 2770 | 722.4 | - | - | 0 | - |
| - | - | 1489 | 723.4 | - | - | 0 | - |
| - | - | 773 | 727.4 | - | - | 0 | - |
| - | - | 614 | 728.4 | - | - | 0 | - |
| - | - | 7320 | 730.4 | - | - | 0 | - |
| - | - | 1.72E+04 | 731.4 | - | - | 0 | - |
| - | - | 7786 | 732.4 | - | - | 0 | - |
| - | - | 2793 | 733.4 | - | - | 0 | - |
| - | - | 2522 | 739.4 | - | - | 0 | - |
| - | - | 1.054E+04 | 740.4 | - | - | 0 | - |
| - | - | 4351 | 741.4 | - | - | 0 | - |
| - | - | 2192 | 742.4 | - | - | 0 | - |
| - | - | 866 | 743.4 | - | - | 0 | - |
| 6 | y | 1421 | 749.4 | 0.003498 | 4.667 | +1 | 7 |
| 6 | z | 3176 | 751.4 | 0.0009534 | 1.269 | +1 | 7 |
| - | - | 4004 | 752.4 | - | - | 0 | - |
| - | - | 833.1 | 753.4 | - | - | 0 | - |
| - | - | 2141 | 755.4 | - | - | 0 | - |
| 7 | c | 2035 | 756.4 | 0.01219 | 16.11 | +1 | 7 |
| 7 | c | 1.365E+04 | 757.4 | 4.652E-05 | 0.06142 | +1 | 7 |
| - | - | 7579 | 758.4 | - | - | 0 | - |
| - | - | 3403 | 759.4 | - | - | 0 | - |
| - | - | 1835 | 760.4 | - | - | 0 | - |
| - | - | 2548 | 766.4 | - | - | 0 | - |
| 6 | y | 1.617E+04 | 767.4 | 0.001611 | 2.099 | +1 | 7 |
| - | - | 5841 | 768.4 | - | - | 0 | - |
| - | - | 1644 | 769.4 | - | - | 0 | - |
| - | - | 2.441E+04 | 773.4 | - | - | 0 | - |
| 7 | c | 3.582E+04 | 774.5 | 0.003341 | 4.314 | +1 | 7 |
| - | - | 1.674E+04 | 775.4 | - | - | 0 | - |
| - | - | 4224 | 776.5 | - | - | 0 | - |
| - | - | 791.5 | 777.5 | - | - | 0 | - |
| - | - | 1348 | 799.5 | - | - | 0 | - |
| - | - | 1878 | 800.5 | - | - | 0 | - |
| - | - | 1871 | 801.5 | - | - | 0 | - |
| - | - | 755.8 | 802.5 | - | - | 0 | - |
| - | - | 889.2 | 817.5 | - | - | 0 | - |
| - | - | 1011 | 818.5 | - | - | 0 | - |
| - | - | 658 | 819.5 | - | - | 0 | - |
| - | - | 2328 | 825.5 | - | - | 0 | - |
| - | - | 1078 | 826.4 | - | - | 0 | - |
| - | - | 1603 | 827.4 | - | - | 0 | - |
| - | - | 1463 | 828.4 | - | - | 0 | - |
| - | - | 758.7 | 829.4 | - | - | 0 | - |
| - | - | 666 | 829.6 | - | - | 0 | - |
| - | - | 1594 | 835.5 | - | - | 0 | - |
| - | - | 1504 | 836.5 | - | - | 0 | - |
| - | - | 2988 | 842.5 | - | - | 0 | - |
| 8 | c | 2094 | 843.5 | 0.01144 | 13.56 | +1 | 8 |
| 8 | c | 5431 | 844.5 | 0.0001516 | 0.1796 | +1 | 8 |
| - | - | 2877 | 845.5 | - | - | 0 | - |
| - | - | 1126 | 846.5 | - | - | 0 | - |
| - | - | 731.9 | 847.5 | - | - | 0 | - |
| 5 | z | 5061 | 850.4 | 0.001128 | 1.327 | +1 | 8 |
| - | - | 2660 | 851.4 | - | - | 0 | - |
| - | - | 8854 | 853.5 | - | - | 0 | - |
| - | - | 5092 | 854.5 | - | - | 0 | - |
| - | - | 1832 | 855.5 | - | - | 0 | - |
| - | - | 1.607E+04 | 860.5 | - | - | 0 | - |
| 8 | c | 1.951E+04 | 861.5 | 0.003875 | 4.499 | +1 | 8 |
| - | - | 1.036E+04 | 862.5 | - | - | 0 | - |
| - | - | 4159 | 863.5 | - | - | 0 | - |
| - | - | 783.6 | 864.5 | - | - | 0 | - |
| - | - | 712 | 865.4 | - | - | 0 | - |
| 5 | y | 5286 | 866.5 | 0.000689 | 0.7952 | +1 | 8 |
| - | - | 3205 | 867.5 | - | - | 0 | - |
| - | - | 803.3 | 868.5 | - | - | 0 | - |
| - | - | 1159 | 896.5 | - | - | 0 | - |
| - | - | 1410 | 897.4 | - | - | 0 | - |
| - | - | 863 | 898.5 | - | - | 0 | - |
| 4 | z | 3098 | 903.5 | 0.00113 | 1.25 | +1 | 9 |
| 4 | z | 2453 | 904.5 | 0.01272 | 14.06 | +1 | 9 |
| - | - | 1543 | 913.5 | - | - | 0 | - |
| - | - | 1455 | 914.5 | - | - | 0 | - |
| - | - | 731.8 | 915.5 | - | - | 0 | - |
| 4 | z | 2.826E+04 | 921.5 | 0.001012 | 1.099 | +1 | 9 |
| - | - | 1.86E+04 | 922.5 | - | - | 0 | - |
| - | - | 6764 | 923.5 | - | - | 0 | - |
| - | - | 2084 | 924.5 | - | - | 0 | - |
| - | - | 2259 | 929.5 | - | - | 0 | - |
| 9 | c | 1973 | 930.5 | 0.01051 | 11.29 | +1 | 9 |
| 9 | c | 5091 | 931.5 | 0.001786 | 1.918 | +1 | 9 |
| - | - | 2796 | 932.5 | - | - | 0 | - |
| - | - | 1062 | 933.5 | - | - | 0 | - |
| 4 | y | 2058 | 937.5 | 0.0007545 | 0.8048 | +1 | 9 |
| - | - | 1669 | 938.5 | - | - | 0 | - |
| - | - | 696.5 | 946.5 | - | - | 0 | - |
| - | - | 1.816E+04 | 947.5 | - | - | 0 | - |
| 9 | c | 4.387E+04 | 948.5 | 0.002762 | 2.912 | +1 | 9 |
| - | - | 2.453E+04 | 949.5 | - | - | 0 | - |
| - | - | 7901 | 950.5 | - | - | 0 | - |
| - | - | 1663 | 951.5 | - | - | 0 | - |
| - | - | 1614 | 952.5 | - | - | 0 | - |
| - | - | 1885 | 953.5 | - | - | 0 | - |
| - | - | 766 | 954.5 | - | - | 0 | - |
| - | - | 1161 | 955.5 | - | - | 0 | - |
| - | - | 1494 | 961.5 | - | - | 0 | - |
| - | - | 1033 | 962.5 | - | - | 0 | - |
| - | - | 770.5 | 963.5 | - | - | 0 | - |
| - | - | 774.1 | 967.5 | - | - | 0 | - |
| - | - | 8415 | 970.5 | - | - | 0 | - |
| - | - | 6144 | 971.5 | - | - | 0 | - |
| - | - | 1788 | 972.5 | - | - | 0 | - |
| - | - | 934.8 | 973.5 | - | - | 0 | - |
| - | - | 9186 | 984.5 | - | - | 0 | - |
| - | - | 6234 | 985.5 | - | - | 0 | - |
| - | - | 2892 | 986.5 | - | - | 0 | - |
| 10 | c | 1150 | 987.5 | 0.0007434 | 0.7528 | +1 | 10 |
| 10 | c | 8833 | 988.5 | 0.0004841 | 0.4897 | +1 | 10 |
| - | - | 6244 | 989.5 | - | - | 0 | - |
| - | - | 1873 | 990.5 | - | - | 0 | - |
| - | - | 844.6 | 1004 | - | - | 0 | - |
| 10 | c | 1.207E+05 | 1006 | 0.0008491 | 0.8444 | +1 | 10 |
| - | - | 8.097E+04 | 1007 | - | - | 0 | - |
| - | - | 3.051E+04 | 1008 | - | - | 0 | - |
| - | - | 7912 | 1009 | - | - | 0 | - |
| - | - | 793.6 | 1010 | - | - | 0 | - |
| - | - | 1768 | 1016 | - | - | 0 | - |
| 3 | y | 2039 | 1017 | 0.0009004 | 0.8857 | +1 | 10 |
| - | - | 1941 | 1034 | - | - | 0 | - |
| 3 | y | 8592 | 1035 | 0.001394 | 1.348 | +1 | 10 |
| - | - | 6069 | 1036 | - | - | 0 | - |
| - | - | 1774 | 1037 | - | - | 0 | - |
| - | - | 1052 | 1038 | - | - | 0 | - |
| - | - | 1580 | 1058 | - | - | 0 | - |
| - | - | 1800 | 1059 | - | - | 0 | - |
| - | - | 757.4 | 1067 | - | - | 0 | - |
| - | - | 5180 | 1069 | - | - | 0 | - |
| - | - | 4137 | 1070 | - | - | 0 | - |
| - | - | 1806 | 1071 | - | - | 0 | - |
| - | - | 641.8 | 1072 | - | - | 0 | - |
| - | - | 2274 | 1074 | - | - | 0 | - |
| - | - | 6507 | 1075 | - | - | 0 | - |
| - | - | 3737 | 1076 | - | - | 0 | - |
| - | - | 1556 | 1077 | - | - | 0 | - |
| - | - | 3461 | 1084 | - | - | 0 | - |
| - | - | 4349 | 1085 | - | - | 0 | - |
| - | - | 1.59E+04 | 1086 | - | - | 0 | - |
| - | - | 1.102E+04 | 1087 | - | - | 0 | - |
| - | - | 4864 | 1088 | - | - | 0 | - |
| - | - | 1721 | 1089 | - | - | 0 | - |
| - | - | 785.1 | 1095 | - | - | 0 | - |
| 11 | c | 1.122E+04 | 1102 | 0.001723 | 1.564 | +1 | 11 |
| - | - | 7828 | 1103 | - | - | 0 | - |
| - | - | 4402 | 1104 | - | - | 0 | - |
| - | - | 2167 | 1105 | - | - | 0 | - |
| 11 | c | 1.119E+05 | 1119 | 0.001783 | 1.594 | +1 | 11 |
| - | - | 7.714E+04 | 1120 | - | - | 0 | - |
| - | - | 3.466E+04 | 1121 | - | - | 0 | - |
| - | - | 9532 | 1122 | - | - | 0 | - |
| - | - | 1689 | 1123 | - | - | 0 | - |
| - | - | 1534 | 1133 | - | - | 0 | - |
| - | - | 1531 | 1134 | - | - | 0 | - |
| - | - | 1455 | 1143 | - | - | 0 | - |
| 2 | z | 2517 | 1148 | 0.002189 | 1.908 | +1 | 11 |
| 2 | z | 1487 | 1149 | 0.0166 | 14.46 | +1 | 11 |
| - | - | 836.4 | 1150 | - | - | 0 | - |
| 2 | z | 2.117E+04 | 1166 | 0.001646 | 1.412 | +1 | 11 |
| - | - | 1.847E+04 | 1167 | - | - | 0 | - |
| - | - | 8175 | 1168 | - | - | 0 | - |
| - | - | 2275 | 1169 | - | - | 0 | - |
| - | - | 834.8 | 1176 | - | - | 0 | - |
| - | - | 996.2 | 1183 | - | - | 0 | - |
| - | - | 845.6 | 1196 | - | - | 0 | - |
| - | - | 1780 | 1205 | - | - | 0 | - |
| - | - | 1554 | 1206 | - | - | 0 | - |
| - | - | 3806 | 1211 | - | - | 0 | - |
| - | - | 2534 | 1212 | - | - | 0 | - |
| - | - | 2908 | 1213 | - | - | 0 | - |
| - | - | 935.3 | 1214 | - | - | 0 | - |
| - | - | 1113 | 1221 | - | - | 0 | - |
| - | - | 3379 | 1222 | - | - | 0 | - |
| - | - | 3836 | 1223 | - | - | 0 | - |
| - | - | 1141 | 1224 | - | - | 0 | - |
| - | - | 962.8 | 1225 | - | - | 0 | - |
| - | - | 1009 | 1226 | - | - | 0 | - |
| - | - | 742.3 | 1227 | - | - | 0 | - |
| - | - | 2915 | 1228 | - | - | 0 | - |
| - | - | 1830 | 1229 | - | - | 0 | - |
| - | - | 1.401E+04 | 1239 | - | - | 0 | - |
| - | - | 1.126E+04 | 1240 | - | - | 0 | - |
| - | - | 7039 | 1241 | - | - | 0 | - |
| - | - | 3245 | 1242 | - | - | 0 | - |
| - | - | 1059 | 1243 | - | - | 0 | - |
| - | - | 1.088E+04 | 1249 | - | - | 0 | - |
| - | - | 8270 | 1250 | - | - | 0 | - |
| - | - | 3458 | 1251 | - | - | 0 | - |
| - | - | 1403 | 1252 | - | - | 0 | - |
| - | - | 7097 | 1256 | - | - | 0 | - |
| - | - | 5168 | 1257 | - | - | 0 | - |
| - | - | 3201 | 1258 | - | - | 0 | - |
| - | - | 2323 | 1266 | - | - | 0 | - |
| - | - | 1.674E+04 | 1267 | - | - | 0 | - |
| - | - | 1.324E+04 | 1268 | - | - | 0 | - |
| - | - | 6231 | 1269 | - | - | 0 | - |
| - | - | 1834 | 1270 | - | - | 0 | - |
| - | - | 2.871E+04 | 1283 | - | - | 0 | - |
| - | - | 1.213E+05 | 1284 | - | - | 0 | - |
| - | - | 1.017E+05 | 1285 | - | - | 0 | - |
| - | - | 4.625E+04 | 1286 | - | - | 0 | - |
| - | - | 1.434E+04 | 1287 | - | - | 0 | - |
| - | - | 3626 | 1288 | - | - | 0 | - |
| - | - | 1048 | 1316 | - | - | 0 | - |
| - | - | 679 | 1459 | - | - | 0 | - |
| - | - | 728.7 | 1547 | - | - | 0 | - |
| - | - | 661.9 | 1896 | - | - | 0 | - |
| - | - | 1993 | 1931 | - | - | 0 | - |
| - | - | 1363 | 1932 | - | - | 0 | - |
| - | - | 876.4 | 2214 | - | - | 0 | - |
| - | - | 690.7 | 2415 | - | - | 0 | - |

m/z Charge Intensity FragmentType MassShift Position
120.08092498779297 0 758.8091
123.04409790039062 0 726.0871
127.50553894042969 0 388.13855
131.7090606689453 0 387.83246
136.07589721679688 0 8157.7393
137.0791778564453 0 686.4628
143.11830139160156 0 1158.9738
148.9473419189453 0 928.46716
165.0547637939453 0 6109.435
166.05856323242188 0 985.8809
169.09725952148438 0 3557.9775
171.11314392089844 0 1640.2434
172.11715698242188 0 499.991
173.14251708984375 0 538.3382
173.45388793945312 0 484.20346
182.0813446044922 0 42258.383 y 11
182.33665466308594 0 493.6881
183.08468627929688 0 4709.489
184.08660888671875 0 524.89856
186.12362670898438 0 688.0123
213.6310577392578 0 554.6048
214.08206176757812 0 946.3187
216.0980682373047 0 611.55853
216.84202575683594 0 551.4898
221.1285858154297 0 17048.895
221.9576416015625 0 522.2463
222.13230895996094 0 2089.607
232.09307861328125 0 1935.3293
240.17105102539062 0 1558.0626
240.53451538085938 0 458.7831
242.1499481201172 0 595.6685
249.12356567382812 0 12230.887
250.126708984375 0 1551.8845
253.5421600341797 0 549.3586
258.1448974609375 0 2285.067
268.16552734375 0 8923.444
269.1685791015625 0 1257.1083
281.0037841796875 0 518.6274
285.19207763671875 0 497.3812
295.1655578613281 0 5927.9395 y 10
296.1680908203125 0 631.95905
307.8754577636719 0 506.93674
317.1822204589844 0 4838.575
327.1673889160156 0 881.0915
343.1247253417969 0 918.2402
345.177001953125 0 8180.3223
346.17742919921875 0 2053.4814
351.2024230957031 0 946.31384
352.186279296875 0 1752.134 y 9
353.1861572265625 0 643.72363
353.2547302246094 0 1099.0105
360.15179443359375 0 580.323
361.6952819824219 0 2385.2134
362.1959533691406 0 595.6866
369.2132263183594 0 743.6805
379.2139587402344 0 710.7785 c Ammonia loss 6
381.2496643066406 0 3021.363
382.25543212890625 0 664.49634
389.2176818847656 0 890.50806
398.27679443359375 0 1218.1833
399.2033386230469 0 2021.2997
412.64202880859375 0 608.48126
413.2667541503906 0 669.51373
414.2712097167969 0 1685.4617
417.2135009765625 0 5471.763 z Ammonia loss 4
418.2165832519531 0 1319.8113 w 4
428.21392822265625 0 1385.1135
439.2191162109375 0 2343.6277 y 8
445.24127197265625 0 862.17163
448.2364196777344 0 687.59174
455.2242126464844 0 849.1299
456.2097473144531 0 3230.178
457.2093811035156 0 697.60345
458.2615661621094 0 616.8506
472.255126953125 0 700.6309
473.2360534667969 0 2949.8137
485.75396728515625 0 2691.0461
486.2532653808594 0 1508.3821
486.7516784667969 0 765.3253
488.28607177734375 0 3087.5266
489.2877197265625 0 932.12964
490.2632141113281 0 728.05853
498.2708435058594 0 5823.483
499.2759094238281 0 1042.6782
499.77069091796875 0 838.0128
500.261962890625 0 700.0766
501.306396484375 0 824.2957
508.2384948730469 0 664.917 y Water loss 7
509.3083801269531 0 7621.9497
510.3096008300781 0 1772.9998
515.2806396484375 0 1214.5266
516.281494140625 0 9599.546
517.2847290039062 0 3630.824
518.2852172851562 0 702.2739
526.2509155273438 0 6564.705 y 7
526.3335571289062 0 2596.5364
527.253173828125 0 2121.1638
527.34228515625 0 622.85315
532.3007202148438 0 6733.425
533.3078002929688 0 24860.348 c 4
534.3107299804688 0 8154.304
535.3148193359375 0 1904.7372
558.324951171875 0 872.0229
568.3084716796875 0 1022.8001
569.2965087890625 0 1036.599
584.362548828125 0 928.7652
585.3441772460938 0 767.30164
586.3194580078125 0 4112.1294
587.324462890625 0 984.388
596.3414916992188 0 1646.4293
601.3711547851562 0 1399.2566
602.3781127929688 0 2415.585
603.3864135742188 0 10733.45
604.3900756835938 0 4159.1465
611.355224609375 0 5386.852
612.3468627929688 0 3515.357
613.3643798828125 0 1042.5352
614.3729248046875 0 603.1245
629.3653564453125 0 9352.3
630.3680419921875 0 2861.088
635.29052734375 0 1168.1985
636.2988891601562 0 2988.3389 y Water loss 6
637.2987670898438 0 1064.5642
640.3519287109375 0 997.8368
641.8535766601562 0 3319.1504
642.357421875 0 2165.355
643.3446655273438 0 649.1651
645.3839721679688 0 15051.649
646.3916625976562 0 46501.49 c 5
647.3944702148438 0 20822.998
648.3969116210938 0 4538.1445
653.3009033203125 0 6613.501
654.3086547851562 0 22909.984 y 6
655.3106689453125 0 7786.8945
656.3118896484375 0 2102.1655
656.3717041015625 0 821.5188
669.37158203125 0 644.5902
672.4080810546875 0 3330.1199
673.4096069335938 0 1422.3252
674.3734741210938 0 891.1721
683.3728637695312 0 1013.5854
685.3912353515625 0 1141.5876
700.3982543945312 0 1513.2446
701.4034423828125 0 788.89813
712.3955078125 0 967.50995
722.3823852539062 0 2770.2559
723.37158203125 0 1488.7657
727.386474609375 0 772.998
728.3846435546875 0 614.0186
730.4360961914062 0 7319.7646
731.44287109375 0 17202.04
732.4462890625 0 7786.188
733.448974609375 0 2792.7651
739.4113159179688 0 2522.3196
740.3988647460938 0 10536.867
741.4026489257812 0 4350.5776
742.4052734375 0 2191.5068
743.4068603515625 0 865.9909
749.3793334960938 0 1421.0063 y Water loss 5
751.3737182617188 0 3176.4624 z 5
752.3787841796875 0 4003.664
753.3827514648438 0 833.0952
755.4306640625 0 2140.6877
756.4281005859375 0 2034.9907 c Water loss 6
757.4242553710938 0 13646.716 c Ammonia loss 6
758.4268798828125 0 7579.4116
759.4346313476562 0 3402.8596
760.435791015625 0 1834.5912
766.3867797851562 0 2548.3386
767.3917846679688 0 16167.171 y 5
768.39306640625 0 5841.0967
769.398193359375 0 1643.9741
773.4425048828125 0 24413.09
774.447509765625 0 35815.777 c 6
775.4495239257812 0 16740.883
776.4541625976562 0 4224.1885
777.4593505859375 0 791.49896
799.4620361328125 0 1347.5974
800.4620971679688 0 1878.2642
801.4645385742188 0 1870.7384
802.4552001953125 0 755.7941
817.45947265625 0 889.2131
818.4735107421875 0 1011.2442
819.4780883789062 0 657.9512
825.4827270507812 0 2327.7551
826.4428100585938 0 1078.014
827.43408203125 0 1602.6035
828.4328002929688 0 1463.1138
829.4409790039062 0 758.65814
829.5822143554688 0 665.9617
835.4662475585938 0 1593.916
836.4575805664062 0 1504.4312
842.462890625 0 2988.0564
843.4608764648438 0 2093.7632 c Water loss 7
844.4564819335938 0 5430.607 c Ammonia loss 7
845.454345703125 0 2877.157
846.4647216796875 0 1126.1968
847.4663696289062 0 731.8552
850.4442138671875 0 5060.846 z 4
851.4446411132812 0 2660.2456
853.4776000976562 0 8853.58
854.4776000976562 0 5092.4805
855.4778442382812 0 1832.3268
860.4738159179688 0 16067.776
861.47900390625 0 19512.943 c 7
862.4811401367188 0 10360.3125
863.4869995117188 0 4159.053
864.4945068359375 0 783.61975
865.4454345703125 0 711.98553
866.4611206054688 0 5285.87 y 4
867.4603881835938 0 3205.135
868.4653930664062 0 803.3005
896.4551391601562 0 1159.3768
897.4490356445312 0 1410.4684
898.45751953125 0 863.0311
903.4707641601562 0 3097.5005 z Water loss 3
904.4663696289062 0 2453.3489 z Ammonia loss 3
913.474853515625 0 1542.6904
914.4713134765625 0 1455.225
915.4634399414062 0 731.8338
921.4791870117188 0 28260.486 z 3
922.4812622070312 0 18604.078
923.4813842773438 0 6764.3936
924.4854736328125 0 2083.9302
929.496337890625 0 2259.361
930.4938354492188 0 1973.376 c Water loss 8
931.486572265625 0 5091.448 c Ammonia loss 8
932.4866333007812 0 2795.5034
933.49267578125 0 1061.6844
937.4981689453125 0 2057.7078 y 3
938.49658203125 0 1668.6962
946.47998046875 0 696.4563
947.505615234375 0 18158.611
948.5121459960938 0 43870.06 c 8
949.5133666992188 0 24527.154
950.5154418945312 0 7900.9497
951.5225830078125 0 1663.4456
952.4881591796875 0 1614.2418
953.479736328125 0 1884.8735
954.49267578125 0 765.98145
955.4927368164062 0 1160.812
961.5200805664062 0 1494.1952
962.5142211914062 0 1033.0154
963.531982421875 0 770.5161
967.5179443359375 0 774.0982
970.498291015625 0 8414.753
971.495849609375 0 6144.1387
972.4947509765625 0 1788.0287
973.5010375976562 0 934.7626
984.5283203125 0 9185.759
985.5294799804688 0 6233.98
986.531982421875 0 2891.8186
987.5265502929688 0 1149.6 c Water loss 9
988.5093383789062 0 8833.001 c Ammonia loss 9
989.5096435546875 0 6244.4165
990.5099487304688 0 1873.0635
1003.5661010742188 0 844.55145
1005.5355224609375 0 120703.625 c 9
1006.5360717773438 0 80965.28
1007.5368041992188 0 30513.264
1008.537353515625 0 7912.2764
1009.5397338867188 0 793.6001
1015.5296020507812 0 1767.9133
1016.5402221679688 0 2038.7668 y Water loss 2
1033.5452880859375 0 1940.7242
1034.55029296875 0 8591.989 y 2
1035.551513671875 0 6069.4727
1036.551513671875 0 1773.6622
1037.555908203125 0 1051.5651
1057.58154296875 0 1579.5597
1058.5755615234375 0 1800.1196
1066.5599365234375 0 757.39984
1068.5731201171875 0 5180.337
1069.5723876953125 0 4136.563
1070.5728759765625 0 1805.612
1071.587646484375 0 641.8416
1073.59814453125 0 2273.7366
1074.604248046875 0 6507.046
1075.6029052734375 0 3737.0781
1076.6046142578125 0 1556.4271
1083.5831298828125 0 3461.0208
1084.579833984375 0 4348.875
1085.588623046875 0 15895.77
1086.5867919921875 0 11017.683
1087.586669921875 0 4864.301
1088.5799560546875 0 1721.1157
1094.5634765625 0 785.14166
1101.5921630859375 0 11224.73 c Ammonia loss 10
1102.5936279296875 0 7827.823
1103.5997314453125 0 4402.3667
1104.6016845703125 0 2167.121
1118.61865234375 0 111864.26 c 10
1119.61962890625 0 77136.69
1120.6219482421875 0 34659.97
1121.6236572265625 0 9532.288
1122.623779296875 0 1689.3972
1132.6312255859375 0 1533.575
1133.63623046875 0 1531.099
1142.599853515625 0 1455.0463
1147.588623046875 0 2516.965 z Water loss 1
1148.5914306640625 0 1487.0372 z Ammonia loss 1
1149.5921630859375 0 836.3959
1165.5997314453125 0 21167.521 z 1
1166.6005859375 0 18466.86
1167.6043701171875 0 8174.7437
1168.6051025390625 0 2274.7512
1175.6204833984375 0 834.82806
1182.606201171875 0 996.15894
1195.6134033203125 0 845.5503
1204.6494140625 0 1780.1129
1205.650634765625 0 1554.4335
1210.5869140625 0 3806.0854
1211.59130859375 0 2533.7322
1212.626220703125 0 2907.7102
1213.628173828125 0 935.30005
1220.63671875 0 1113.3132
1221.6649169921875 0 3379.3572
1222.674072265625 0 3835.604
1223.669677734375 0 1140.8761
1224.6619873046875 0 962.82184
1225.6512451171875 0 1009.199
1226.6483154296875 0 742.33105
1227.6109619140625 0 2914.8875
1228.60693359375 0 1829.827
1238.652099609375 0 14009.954
1239.6546630859375 0 11258.645
1240.648193359375 0 7038.5283
1241.63818359375 0 3244.7212
1242.637939453125 0 1058.9116
1248.63623046875 0 10883.623
1249.63623046875 0 8270.251
1250.6412353515625 0 3457.5562
1251.6351318359375 0 1403.4377
1255.6768798828125 0 7097.132
1256.67822265625 0 5167.8477
1257.6798095703125 0 3201.4036
1265.6583251953125 0 2322.6438
1266.648681640625 0 16741.006
1267.6495361328125 0 13237.368
1268.648681640625 0 6231.485
1269.64697265625 0 1834.1943
1282.6650390625 0 28714.754
1283.6717529296875 0 121299.914
1284.6732177734375 0 101721.195
1285.675537109375 0 46252.516
1286.677490234375 0 14335.918
1287.683837890625 0 3625.653
1315.6533203125 0 1047.6423
1459.267578125 0 678.9553
1547.414794921875 0 728.7321
1896.0091552734375 0 661.8782
1930.9658203125 0 1993.3121
1931.9849853515625 0 1362.5457
2214.275390625 0 876.3797
2415.450927734375 0 690.70056

Spectrum Details

|  |  |
| --- | --- |
| Matched peaks? Matched peaks The total absolute number of peaks matched. Additionally in brackets the total fraction of peaks matched and the total number of peaks is shown. | 41 (11.55% of 355) |
| FDR? FDR The false discovery rate estimated for this peptide. It is calculated by matching all theoretical fragments with a non-integer shift with the raw peaks for this spectrum. This is done with 40 different shifts. The resulting percentage is the average number of annotated peaks over the number of annotated peaks with the correct spectrum. | 1.45% |
| Satellite FDR? Satellite FDR See the FDR for details on its calculation. This satellite ion specific FDR only contains the satellite ions (d/w) for I/L/J positions. | ∞ |
| PSM Score? PSM Score The PSM Score as given by Hecklib to this annotated spectrum. It is shown with three significant figures. | 546 |

#### Spectrum 11433? Spectrum 11433 The raw spectrum of this peptide as annotated by Hecklib. The fragments are coloured according to ion type (see legend). Any peaks with a star '\*' as text can be hovered over to see the full details, first the ion type second the mass shift type. By hovering over the amino acids in the peptide or ions in the legend the corresponding peaks are highlighted. By toggling the 'Unassigned' label you can turn the background (unassigned) peaks on or off in the plot. By updating the slider in the Ion legend you can update the spectrum to only show the top X% of the peaks with labels. The top X% means any peak that is within X% of the highest intensity. By dragging in the spectrum you can zoom in to a specific part of the spectrum and use 'Zoom Out' to get back to the original zoom level. The annotation of the spectrum is based on the given sequence in the peptides file and is done with different software so inconsistencies are likely. The peaks are annotated based on the given sequence, with 20 ppm tolerance.

Copy Data

##### Spectrum 11433 (TSV)

###### Preview

```
Loading example...
```

*Click on the button to copy the data to your clipboard.*

Mz MinMz MaxIntensity Max

WidthHeightPeptide font sizePeptide stroke widthSpectrum font sizeSpectrum stroke widthCompact peptide

Ion legend

wxyz

abcd

OtherUnassignedIon^Charge^_Position_Show for top:%

TFPAVJQSSGJY

01.22e+42.44e+43.66e+44.88e+4

Zoom Out

y^+1^_1_y^+1^_2_y^+1^_3_z^+2^_8_y^+1^_5_c^+1^_5_y^+1^_6_c^+1^_6_y^+1^_6_z^+1^_7_c^+1^_7_c^+1^_7_y^+1^_7_c^+1^_7_c^+1^_8_c^+1^_8_z^+1^_8_c^+1^_8_y^+1^_8_z^+1^_9_z^+1^_9_z^+1^_9_c^+1^_9_y^+1^_9_c^+1^_9_c^+1^_10_c^+1^_10_c^+1^_10_y^+1^_10_c^+1^_11_c^+1^_11_z^+1^_11_

03256509751300

Fragment Matches Table

Show background peaks

| Position | Ion type | Intensity | mz Theoretical | mz Error (Th) | mz Error (ppm) | Charge | Series Number |
| --- | --- | --- | --- | --- | --- | --- | --- |
| - | - | 389.7 | 135.7 | - | - | 0 | - |
| - | - | 2662 | 136.1 | - | - | 0 | - |
| - | - | 496.4 | 141.1 | - | - | 0 | - |
| - | - | 747.5 | 143.1 | - | - | 0 | - |
| - | - | 485.3 | 149 | - | - | 0 | - |
| - | - | 435.4 | 156.1 | - | - | 0 | - |
| - | - | 484 | 162.8 | - | - | 0 | - |
| - | - | 500.7 | 163.5 | - | - | 0 | - |
| - | - | 2118 | 165.1 | - | - | 0 | - |
| - | - | 1585 | 169.1 | - | - | 0 | - |
| - | - | 624.3 | 171.1 | - | - | 0 | - |
| - | - | 2700 | 173.4 | - | - | 0 | - |
| - | - | 560.9 | 174.1 | - | - | 0 | - |
| 12 | y | 1.336E+04 | 182.1 | 5.286E-05 | 0.2903 | +1 | 1 |
| - | - | 1135 | 183.1 | - | - | 0 | - |
| - | - | 579.2 | 211.2 | - | - | 0 | - |
| - | - | 546.4 | 216.2 | - | - | 0 | - |
| - | - | 4969 | 221.1 | - | - | 0 | - |
| - | - | 574.8 | 222.1 | - | - | 0 | - |
| - | - | 601.4 | 232.1 | - | - | 0 | - |
| - | - | 701.5 | 240.2 | - | - | 0 | - |
| - | - | 3829 | 249.1 | - | - | 0 | - |
| - | - | 718.8 | 258.1 | - | - | 0 | - |
| - | - | 475.2 | 258.2 | - | - | 0 | - |
| - | - | 2488 | 268.2 | - | - | 0 | - |
| 11 | y | 2222 | 295.2 | 8.006E-05 | 0.2712 | +1 | 2 |
| - | - | 717.4 | 308.1 | - | - | 0 | - |
| - | - | 1323 | 317.2 | - | - | 0 | - |
| - | - | 511.8 | 328.9 | - | - | 0 | - |
| - | - | 3214 | 345.2 | - | - | 0 | - |
| - | - | 862.3 | 346.2 | - | - | 0 | - |
| 10 | y | 740.1 | 352.2 | 8.239E-05 | 0.2339 | +1 | 3 |
| - | - | 1333 | 361.7 | - | - | 0 | - |
| - | - | 780.8 | 381.2 | - | - | 0 | - |
| - | - | 707.6 | 399.2 | - | - | 0 | - |
| - | - | 678.6 | 405.2 | - | - | 0 | - |
| - | - | 553.5 | 413.3 | - | - | 0 | - |
| - | - | 860.9 | 414.3 | - | - | 0 | - |
| 5 | z | 1136 | 417.2 | 0.001595 | 3.822 | +2 | 8 |
| - | - | 645.3 | 425.2 | - | - | 0 | - |
| - | - | 758.7 | 456.2 | - | - | 0 | - |
| - | - | 1171 | 473.2 | - | - | 0 | - |
| - | - | 791.9 | 485.8 | - | - | 0 | - |
| - | - | 583.3 | 486.3 | - | - | 0 | - |
| - | - | 2106 | 498.3 | - | - | 0 | - |
| - | - | 2455 | 509.3 | - | - | 0 | - |
| - | - | 787.3 | 510.3 | - | - | 0 | - |
| - | - | 2428 | 516.3 | - | - | 0 | - |
| - | - | 1488 | 517.3 | - | - | 0 | - |
| 8 | y | 2826 | 526.3 | 0.0001613 | 0.3066 | +1 | 5 |
| - | - | 730.4 | 526.3 | - | - | 0 | - |
| - | - | 3009 | 532.3 | - | - | 0 | - |
| 5 | c | 8386 | 533.3 | 0.0004092 | 0.7673 | +1 | 5 |
| - | - | 1667 | 534.3 | - | - | 0 | - |
| - | - | 861.5 | 558.3 | - | - | 0 | - |
| - | - | 1110 | 586.3 | - | - | 0 | - |
| - | - | 560.2 | 587.3 | - | - | 0 | - |
| - | - | 641.9 | 596.3 | - | - | 0 | - |
| - | - | 1523 | 602.4 | - | - | 0 | - |
| - | - | 3851 | 603.4 | - | - | 0 | - |
| - | - | 1475 | 604.4 | - | - | 0 | - |
| - | - | 1451 | 611.4 | - | - | 0 | - |
| - | - | 909.2 | 612.4 | - | - | 0 | - |
| - | - | 940.6 | 613.4 | - | - | 0 | - |
| - | - | 3225 | 629.4 | - | - | 0 | - |
| - | - | 1229 | 630.4 | - | - | 0 | - |
| 7 | y | 1253 | 636.3 | 0.004028 | 6.331 | +1 | 6 |
| - | - | 569.7 | 637.3 | - | - | 0 | - |
| - | - | 2707 | 641.9 | - | - | 0 | - |
| - | - | 2009 | 642.4 | - | - | 0 | - |
| - | - | 1447 | 642.9 | - | - | 0 | - |
| - | - | 5612 | 645.4 | - | - | 0 | - |
| 6 | c | 1.692E+04 | 646.4 | 0.000733 | 1.134 | +1 | 6 |
| - | - | 6424 | 647.4 | - | - | 0 | - |
| - | - | 2457 | 653.3 | - | - | 0 | - |
| 7 | y | 8666 | 654.3 | 0.0003718 | 0.5682 | +1 | 6 |
| - | - | 3666 | 655.3 | - | - | 0 | - |
| - | - | 908.6 | 672.4 | - | - | 0 | - |
| - | - | 791.3 | 683.4 | - | - | 0 | - |
| - | - | 641.1 | 700.4 | - | - | 0 | - |
| - | - | 635.4 | 722.4 | - | - | 0 | - |
| - | - | 2436 | 730.4 | - | - | 0 | - |
| - | - | 6366 | 731.4 | - | - | 0 | - |
| - | - | 2938 | 732.4 | - | - | 0 | - |
| - | - | 2897 | 740.4 | - | - | 0 | - |
| - | - | 1469 | 741.4 | - | - | 0 | - |
| - | - | 935.8 | 742.4 | - | - | 0 | - |
| 6 | z | 2017 | 751.4 | 0.001869 | 2.487 | +1 | 7 |
| - | - | 1552 | 752.4 | - | - | 0 | - |
| 7 | c | 864.3 | 756.4 | 0.01286 | 17 | +1 | 7 |
| 7 | c | 4407 | 757.4 | 0.0004418 | 0.5832 | +1 | 7 |
| - | - | 1840 | 758.4 | - | - | 0 | - |
| - | - | 1595 | 759.4 | - | - | 0 | - |
| - | - | 1388 | 766.4 | - | - | 0 | - |
| 6 | y | 4402 | 767.4 | 0.00155 | 2.02 | +1 | 7 |
| - | - | 809 | 768.4 | - | - | 0 | - |
| - | - | 8613 | 773.4 | - | - | 0 | - |
| 7 | c | 1.148E+04 | 774.5 | 0.001693 | 2.186 | +1 | 7 |
| - | - | 6040 | 775.5 | - | - | 0 | - |
| - | - | 1393 | 776.5 | - | - | 0 | - |
| - | - | 921.8 | 799.5 | - | - | 0 | - |
| - | - | 1065 | 800.5 | - | - | 0 | - |
| - | - | 777.9 | 801.5 | - | - | 0 | - |
| - | - | 796.5 | 818.5 | - | - | 0 | - |
| - | - | 584.6 | 837.7 | - | - | 0 | - |
| - | - | 1445 | 842.5 | - | - | 0 | - |
| 8 | c | 587.7 | 843.5 | 0.0104 | 12.33 | +1 | 8 |
| 8 | c | 2155 | 844.5 | 0.0008249 | 0.9769 | +1 | 8 |
| - | - | 955.1 | 845.5 | - | - | 0 | - |
| - | - | 673.1 | 846.5 | - | - | 0 | - |
| 5 | z | 1392 | 850.4 | 0.0006417 | 0.7545 | +1 | 8 |
| - | - | 816.6 | 851.4 | - | - | 0 | - |
| - | - | 3107 | 853.5 | - | - | 0 | - |
| - | - | 907.1 | 854.5 | - | - | 0 | - |
| - | - | 7182 | 860.5 | - | - | 0 | - |
| 8 | c | 7867 | 861.5 | 0.001556 | 1.806 | +1 | 8 |
| - | - | 2637 | 862.5 | - | - | 0 | - |
| - | - | 764.2 | 863.5 | - | - | 0 | - |
| - | - | 764.6 | 865.4 | - | - | 0 | - |
| 5 | y | 1574 | 866.5 | 0.002668 | 3.079 | +1 | 8 |
| 4 | z | 1083 | 903.5 | 0.004181 | 4.628 | +1 | 9 |
| 4 | z | 973 | 904.5 | 0.01498 | 16.56 | +1 | 9 |
| 4 | z | 1.083E+04 | 921.5 | 0.001195 | 1.297 | +1 | 9 |
| - | - | 5867 | 922.5 | - | - | 0 | - |
| - | - | 2800 | 923.5 | - | - | 0 | - |
| - | - | 955.7 | 929.5 | - | - | 0 | - |
| 9 | c | 1260 | 931.5 | 0.000716 | 0.7687 | +1 | 9 |
| - | - | 1408 | 932.5 | - | - | 0 | - |
| 4 | y | 847.9 | 937.5 | 0.003579 | 3.818 | +1 | 9 |
| - | - | 6932 | 947.5 | - | - | 0 | - |
| 9 | c | 1.686E+04 | 948.5 | 0.001724 | 1.818 | +1 | 9 |
| - | - | 7931 | 949.5 | - | - | 0 | - |
| - | - | 1816 | 950.5 | - | - | 0 | - |
| - | - | 1083 | 953.5 | - | - | 0 | - |
| - | - | 3034 | 970.5 | - | - | 0 | - |
| - | - | 1649 | 971.5 | - | - | 0 | - |
| - | - | 3875 | 984.5 | - | - | 0 | - |
| - | - | 2522 | 985.5 | - | - | 0 | - |
| - | - | 1414 | 986.5 | - | - | 0 | - |
| 10 | c | 724.9 | 987.5 | 0.006664 | 6.748 | +1 | 10 |
| 10 | c | 3695 | 988.5 | 0.001042 | 1.054 | +1 | 10 |
| - | - | 1809 | 989.5 | - | - | 0 | - |
| 10 | c | 4.735E+04 | 1006 | 0.0005439 | 0.5409 | +1 | 10 |
| - | - | 2.847E+04 | 1007 | - | - | 0 | - |
| - | - | 8321 | 1008 | - | - | 0 | - |
| 3 | y | 2896 | 1035 | 0.002371 | 2.292 | +1 | 10 |
| - | - | 1862 | 1036 | - | - | 0 | - |
| - | - | 2479 | 1069 | - | - | 0 | - |
| - | - | 1055 | 1070 | - | - | 0 | - |
| - | - | 899.6 | 1074 | - | - | 0 | - |
| - | - | 2233 | 1075 | - | - | 0 | - |
| - | - | 1042 | 1076 | - | - | 0 | - |
| - | - | 1227 | 1084 | - | - | 0 | - |
| - | - | 1012 | 1085 | - | - | 0 | - |
| - | - | 6729 | 1086 | - | - | 0 | - |
| - | - | 4226 | 1087 | - | - | 0 | - |
| - | - | 884.2 | 1088 | - | - | 0 | - |
| - | - | 932.8 | 1096 | - | - | 0 | - |
| 11 | c | 3200 | 1102 | 0.001113 | 1.01 | +1 | 11 |
| - | - | 1993 | 1103 | - | - | 0 | - |
| - | - | 918.8 | 1104 | - | - | 0 | - |
| - | - | 718.5 | 1105 | - | - | 0 | - |
| 11 | c | 4.552E+04 | 1119 | 0.001295 | 1.158 | +1 | 11 |
| - | - | 2.808E+04 | 1120 | - | - | 0 | - |
| - | - | 7639 | 1121 | - | - | 0 | - |
| - | - | 903 | 1122 | - | - | 0 | - |
| - | - | 828.7 | 1133 | - | - | 0 | - |
| 2 | z | 8695 | 1166 | 0.002134 | 1.831 | +1 | 11 |
| - | - | 6478 | 1167 | - | - | 0 | - |
| - | - | 2473 | 1168 | - | - | 0 | - |
| - | - | 5413 | 1208 | - | - | 0 | - |
| - | - | 4099 | 1209 | - | - | 0 | - |
| - | - | 1188 | 1210 | - | - | 0 | - |
| - | - | 908.7 | 1211 | - | - | 0 | - |
| - | - | 1347 | 1213 | - | - | 0 | - |
| - | - | 1538 | 1222 | - | - | 0 | - |
| - | - | 1179 | 1223 | - | - | 0 | - |
| - | - | 1091 | 1226 | - | - | 0 | - |
| - | - | 7332 | 1239 | - | - | 0 | - |
| - | - | 5081 | 1240 | - | - | 0 | - |
| - | - | 2374 | 1241 | - | - | 0 | - |
| - | - | 733.5 | 1242 | - | - | 0 | - |
| - | - | 671.5 | 1243 | - | - | 0 | - |
| - | - | 4125 | 1249 | - | - | 0 | - |
| - | - | 3601 | 1250 | - | - | 0 | - |
| - | - | 965.8 | 1251 | - | - | 0 | - |
| - | - | 2186 | 1256 | - | - | 0 | - |
| - | - | 2472 | 1257 | - | - | 0 | - |
| - | - | 1053 | 1258 | - | - | 0 | - |
| - | - | 1916 | 1266 | - | - | 0 | - |
| - | - | 7334 | 1267 | - | - | 0 | - |
| - | - | 6170 | 1268 | - | - | 0 | - |
| - | - | 1896 | 1269 | - | - | 0 | - |
| - | - | 1321 | 1282 | - | - | 0 | - |
| - | - | 6007 | 1283 | - | - | 0 | - |
| - | - | 1.162E+04 | 1283 | - | - | 0 | - |
| - | - | 4.835E+04 | 1284 | - | - | 0 | - |
| - | - | 3.386E+04 | 1285 | - | - | 0 | - |
| - | - | 1.271E+04 | 1286 | - | - | 0 | - |
| - | - | 1667 | 1287 | - | - | 0 | - |

m/z Charge Intensity FragmentType MassShift Position
135.73480224609375 0 389.69476
136.07574462890625 0 2661.979
141.10227966308594 0 496.43695
143.1180877685547 0 747.5428
148.9545135498047 0 485.25262
156.13970947265625 0 435.3812
162.84864807128906 0 483.99054
163.50135803222656 0 500.66315
165.05477905273438 0 2118.4202
169.09742736816406 0 1585.098
171.1124267578125 0 624.33594
173.43917846679688 0 2699.9463
174.0907745361328 0 560.8584
182.0812225341797 0 13356.4795 y 11
183.08448791503906 0 1134.6334
211.17828369140625 0 579.21246
216.19375610351562 0 546.43634
221.12843322753906 0 4968.702
222.13218688964844 0 574.79913
232.09243774414062 0 601.4176
240.17010498046875 0 701.5373
249.12351989746094 0 3828.566
258.1444396972656 0 718.7999
258.23822021484375 0 475.24088
268.1654052734375 0 2488.2898
295.1653137207031 0 2222.423 y 10
308.1236877441406 0 717.3795
317.1819763183594 0 1322.8542
328.9121398925781 0 511.83328
345.17694091796875 0 3213.8743
346.1777648925781 0 862.3254
352.1866149902344 0 740.08405 y 9
361.69281005859375 0 1332.5797
381.2489318847656 0 780.79553
399.2025146484375 0 707.60156
405.24560546875 0 678.58167
413.2705993652344 0 553.4976
414.26788330078125 0 860.89935
417.2135009765625 0 1136.1665 z Ammonia loss 4
425.2121887207031 0 645.2977
456.2086486816406 0 758.6838
473.2372741699219 0 1170.8607
485.7531433105469 0 791.93823
486.2505798339844 0 583.3262
498.27069091796875 0 2105.6836
509.30902099609375 0 2454.6091
510.3115539550781 0 787.28455
516.2821044921875 0 2427.6052
517.28466796875 0 1488.248
526.2509155273438 0 2825.8665 y 7
526.3372192382812 0 730.4061
532.2988891601562 0 3009.2932
533.3078002929688 0 8386.079 c 4
534.310546875 0 1666.8485
558.3240966796875 0 861.51483
586.3176879882812 0 1110.2704
587.3248291015625 0 560.20935
596.3397827148438 0 641.8886
602.3778686523438 0 1523.1729
603.3860473632812 0 3850.6802
604.3909301757812 0 1475.4099
611.3544311523438 0 1451.4434
612.3500366210938 0 909.1756
613.3672485351562 0 940.6231
629.3658447265625 0 3224.544
630.3683471679688 0 1229.1409
636.2947387695312 0 1253.4163 y Water loss 6
637.298095703125 0 569.732
641.8540649414062 0 2706.7942
642.3546142578125 0 2008.882
642.8593139648438 0 1446.5114
645.3843994140625 0 5612.233
646.3915405273438 0 16924.38 c 5
647.3946533203125 0 6424.3965
653.30029296875 0 2457.1821
654.3089599609375 0 8665.698 y 6
655.311767578125 0 3665.9905
672.4070434570312 0 908.6436
683.37109375 0 791.2999
700.3917236328125 0 641.09625
722.3834838867188 0 635.38116
730.4369506835938 0 2435.738
731.4441528320312 0 6365.553
732.44775390625 0 2938.098
740.4009399414062 0 2897.053
741.4017944335938 0 1469.0593
742.401123046875 0 935.76044
751.372802734375 0 2017.4332 z 5
752.3790283203125 0 1552.0183
756.4274291992188 0 864.2563 c Water loss 6
757.4247436523438 0 4407.0635 c Ammonia loss 6
758.4312744140625 0 1839.8921
759.4356689453125 0 1594.7429
766.3843994140625 0 1388.3821
767.391845703125 0 4402.313 y 5
768.3956298828125 0 808.9964
773.4426879882812 0 8612.774
774.4491577148438 0 11479.477 c 6
775.4529418945312 0 6039.7793
776.4594116210938 0 1393.405
799.456787109375 0 921.77
800.45751953125 0 1065.0438
801.4557495117188 0 777.9181
818.4688720703125 0 796.54706
837.6954345703125 0 584.6097
842.4646606445312 0 1444.6965
843.4619140625 0 587.667 c Water loss 7
844.4555053710938 0 2154.878 c Ammonia loss 7
845.4591064453125 0 955.1032
846.46533203125 0 673.10846
850.4424438476562 0 1392.1864 z 4
851.4454956054688 0 816.58514
853.4796752929688 0 3106.7258
854.483642578125 0 907.0622
860.4738159179688 0 7182.2817
861.4813232421875 0 7866.8564 c 7
862.48388671875 0 2637.0493
863.4939575195312 0 764.1697
865.4491577148438 0 764.62067
866.4644775390625 0 1574.1968 y 4
903.4738159179688 0 1082.7744 z Water loss 3
904.4686279296875 0 973.0264 z Ammonia loss 3
921.47900390625 0 10828.446 z 3
922.4835205078125 0 5867.242
923.4846801757812 0 2800.229
929.4976196289062 0 955.7349
931.4890747070312 0 1259.5186 c Ammonia loss 8
932.4928588867188 0 1408.3914
937.5025024414062 0 847.8847 y 3
947.5070190429688 0 6932.398
948.51318359375 0 16863.277 c 8
949.5164184570312 0 7931.005
950.5226440429688 0 1815.535
953.4895629882812 0 1082.8846
970.4977416992188 0 3033.8384
971.4998168945312 0 1649.1344
984.526611328125 0 3875.12
985.5291137695312 0 2522.2808
986.5338134765625 0 1413.6838
987.532470703125 0 724.9005 c Water loss 9
988.5108642578125 0 3695.026 c Ammonia loss 9
989.513916015625 0 1808.6107
1005.5358276367188 0 47351.906 c 9
1006.5383911132812 0 28471.812
1007.5413208007812 0 8321.103
1034.54931640625 0 2895.6445 y 2
1035.552001953125 0 1861.8894
1068.5716552734375 0 2478.5479
1069.57666015625 0 1055.0525
1073.599853515625 0 899.62305
1074.60400390625 0 2232.627
1075.603515625 0 1042.1744
1083.572265625 0 1226.6871
1084.5804443359375 0 1012.4832
1085.5892333984375 0 6729.4526
1086.5927734375 0 4225.8765
1087.59716796875 0 884.1715
1095.6376953125 0 932.83203
1101.5927734375 0 3200.272 c Ammonia loss 10
1102.594482421875 0 1992.8622
1103.6077880859375 0 918.7584
1104.6104736328125 0 718.50824
1118.619140625 0 45523.93 c 10
1119.6220703125 0 28084.662
1120.625244140625 0 7638.507
1121.623291015625 0 902.98816
1132.6339111328125 0 828.65186
1165.5992431640625 0 8694.848 z 1
1166.6038818359375 0 6477.952
1167.606201171875 0 2472.9658
1207.7257080078125 0 5412.8613
1208.7264404296875 0 4098.8955
1209.7303466796875 0 1187.8009
1210.58740234375 0 908.7119
1212.6295166015625 0 1347.4565
1221.6719970703125 0 1537.733
1222.67626953125 0 1178.9863
1225.7537841796875 0 1090.8428
1238.6529541015625 0 7331.8936
1239.6561279296875 0 5080.662
1240.650390625 0 2373.843
1241.637939453125 0 733.4668
1242.642333984375 0 671.5204
1248.6373291015625 0 4125.3257
1249.640380859375 0 3601.3628
1250.6409912109375 0 965.78015
1255.679443359375 0 2186.144
1256.6912841796875 0 2472.102
1257.6910400390625 0 1052.9358
1265.7178955078125 0 1915.6687
1266.654296875 0 7334.2817
1267.6552734375 0 6170.2324
1268.6578369140625 0 1895.9857
1281.764404296875 0 1320.9989
1282.6522216796875 0 6007.2275
1282.756103515625 0 11617.43
1283.6759033203125 0 48349.953
1284.677734375 0 33862.457
1285.680908203125 0 12705.581
1286.674072265625 0 1666.9032

Spectrum Details

|  |  |
| --- | --- |
| Matched peaks? Matched peaks The total absolute number of peaks matched. Additionally in brackets the total fraction of peaks matched and the total number of peaks is shown. | 32 (16.00% of 200) |
| FDR? FDR The false discovery rate estimated for this peptide. It is calculated by matching all theoretical fragments with a non-integer shift with the raw peaks for this spectrum. This is done with 40 different shifts. The resulting percentage is the average number of annotated peaks over the number of annotated peaks with the correct spectrum. | 1.26% |
| Satellite FDR? Satellite FDR See the FDR for details on its calculation. This satellite ion specific FDR only contains the satellite ions (d/w) for I/L/J positions. | ∞ |
| PSM Score? PSM Score The PSM Score as given by Hecklib to this annotated spectrum. It is shown with three significant figures. | 411 |

#### Spectrum 11765? Spectrum 11765 The raw spectrum of this peptide as annotated by Hecklib. The fragments are coloured according to ion type (see legend). Any peaks with a star '\*' as text can be hovered over to see the full details, first the ion type second the mass shift type. By hovering over the amino acids in the peptide or ions in the legend the corresponding peaks are highlighted. By toggling the 'Unassigned' label you can turn the background (unassigned) peaks on or off in the plot. By updating the slider in the Ion legend you can update the spectrum to only show the top X% of the peaks with labels. The top X% means any peak that is within X% of the highest intensity. By dragging in the spectrum you can zoom in to a specific part of the spectrum and use 'Zoom Out' to get back to the original zoom level. The annotation of the spectrum is based on the given sequence in the peptides file and is done with different software so inconsistencies are likely. The peaks are annotated based on the given sequence, with 20 ppm tolerance.

Copy Data

##### Spectrum 11765 (TSV)

###### Preview

```
Loading example...
```

*Click on the button to copy the data to your clipboard.*

Mz MinMz MaxIntensity Max

WidthHeightPeptide font sizePeptide stroke widthSpectrum font sizeSpectrum stroke widthCompact peptide

Ion legend

wxyz

abcd

OtherUnassignedIon^Charge^_Position_Show for top:%

TFPAVJQSSGJY

07.60e+31.52e+42.28e+43.04e+4

Zoom Out

y^+1^_1_y^+1^_2_z^+2^_8_y^+2^_8_y^+1^_5_c^+1^_5_c^+1^_6_y^+1^_6_z^+1^_7_c^+1^_7_y^+1^_7_c^+1^_7_c^+1^_8_z^+1^_8_c^+1^_8_y^+1^_8_z^+1^_9_c^+1^_9_c^+1^_9_c^+1^_9_c^+1^_10_c^+1^_10_y^+1^_10_c^+1^_11_c^+1^_11_z^+1^_11_

0777155423313108

Fragment Matches Table

Show background peaks

| Position | Ion type | Intensity | mz Theoretical | mz Error (Th) | mz Error (ppm) | Charge | Series Number |
| --- | --- | --- | --- | --- | --- | --- | --- |
| - | - | 366.8 | 127.5 | - | - | 0 | - |
| - | - | 406.3 | 132.9 | - | - | 0 | - |
| - | - | 1460 | 136.1 | - | - | 0 | - |
| - | - | 397.2 | 145.3 | - | - | 0 | - |
| - | - | 555.1 | 148.9 | - | - | 0 | - |
| - | - | 579.7 | 148.9 | - | - | 0 | - |
| - | - | 640.9 | 148.9 | - | - | 0 | - |
| - | - | 788 | 148.9 | - | - | 0 | - |
| - | - | 671 | 148.9 | - | - | 0 | - |
| - | - | 746.3 | 148.9 | - | - | 0 | - |
| - | - | 2333 | 148.9 | - | - | 0 | - |
| - | - | 1940 | 148.9 | - | - | 0 | - |
| - | - | 4048 | 149 | - | - | 0 | - |
| - | - | 2423 | 149 | - | - | 0 | - |
| - | - | 1416 | 149 | - | - | 0 | - |
| - | - | 662.6 | 149 | - | - | 0 | - |
| - | - | 927.7 | 149 | - | - | 0 | - |
| - | - | 473.5 | 149 | - | - | 0 | - |
| - | - | 368.9 | 159.2 | - | - | 0 | - |
| - | - | 418.1 | 161.2 | - | - | 0 | - |
| - | - | 515.1 | 162.7 | - | - | 0 | - |
| - | - | 1217 | 165.1 | - | - | 0 | - |
| - | - | 539.6 | 169.1 | - | - | 0 | - |
| - | - | 468.6 | 173.2 | - | - | 0 | - |
| 12 | y | 8774 | 182.1 | 6.812E-05 | 0.3741 | +1 | 1 |
| - | - | 654.3 | 183.1 | - | - | 0 | - |
| - | - | 460 | 185.1 | - | - | 0 | - |
| - | - | 636.6 | 187.2 | - | - | 0 | - |
| - | - | 441.8 | 192.5 | - | - | 0 | - |
| - | - | 692.5 | 205.1 | - | - | 0 | - |
| - | - | 640 | 221.1 | - | - | 0 | - |
| - | - | 2977 | 221.1 | - | - | 0 | - |
| - | - | 506.3 | 222.1 | - | - | 0 | - |
| - | - | 2676 | 249.1 | - | - | 0 | - |
| - | - | 1031 | 252.2 | - | - | 0 | - |
| - | - | 580.6 | 258.1 | - | - | 0 | - |
| - | - | 1285 | 268.2 | - | - | 0 | - |
| 11 | y | 1196 | 295.2 | 0.0001411 | 0.478 | +1 | 2 |
| - | - | 1046 | 317.2 | - | - | 0 | - |
| - | - | 509.1 | 320.5 | - | - | 0 | - |
| - | - | 504.8 | 336.1 | - | - | 0 | - |
| - | - | 1909 | 345.2 | - | - | 0 | - |
| - | - | 540.2 | 345.8 | - | - | 0 | - |
| - | - | 723.6 | 361.7 | - | - | 0 | - |
| - | - | 888 | 381.2 | - | - | 0 | - |
| 5 | z | 961.3 | 417.2 | 0.0004654 | 1.115 | +2 | 8 |
| - | - | 729.7 | 420.1 | - | - | 0 | - |
| 5 | y | 610.7 | 425.2 | 0.007768 | 18.27 | +2 | 8 |
| - | - | 612.9 | 433.4 | - | - | 0 | - |
| - | - | 652.7 | 470.1 | - | - | 0 | - |
| - | - | 722.3 | 485.8 | - | - | 0 | - |
| - | - | 583.6 | 486.2 | - | - | 0 | - |
| - | - | 882.6 | 498.3 | - | - | 0 | - |
| - | - | 1339 | 509.3 | - | - | 0 | - |
| - | - | 2483 | 516.3 | - | - | 0 | - |
| - | - | 864.5 | 517.3 | - | - | 0 | - |
| 8 | y | 1875 | 526.3 | 0.001077 | 2.046 | +1 | 5 |
| - | - | 1646 | 532.3 | - | - | 0 | - |
| 5 | c | 5415 | 533.3 | 4.3E-05 | 0.08062 | +1 | 5 |
| - | - | 1991 | 534.3 | - | - | 0 | - |
| - | - | 3249 | 603.4 | - | - | 0 | - |
| - | - | 911.8 | 611.4 | - | - | 0 | - |
| - | - | 1995 | 629.4 | - | - | 0 | - |
| - | - | 1214 | 630.4 | - | - | 0 | - |
| - | - | 657.1 | 640.3 | - | - | 0 | - |
| - | - | 961.8 | 641.3 | - | - | 0 | - |
| - | - | 3213 | 641.9 | - | - | 0 | - |
| - | - | 1781 | 642.4 | - | - | 0 | - |
| - | - | 763.7 | 642.9 | - | - | 0 | - |
| - | - | 3683 | 645.4 | - | - | 0 | - |
| 6 | c | 1.078E+04 | 646.4 | 0.0005498 | 0.8506 | +1 | 6 |
| - | - | 4160 | 647.4 | - | - | 0 | - |
| - | - | 1233 | 653.3 | - | - | 0 | - |
| 7 | y | 5353 | 654.3 | 0.0009211 | 1.408 | +1 | 6 |
| - | - | 1467 | 655.3 | - | - | 0 | - |
| - | - | 989.5 | 672.4 | - | - | 0 | - |
| - | - | 1304 | 730.4 | - | - | 0 | - |
| - | - | 4171 | 731.4 | - | - | 0 | - |
| - | - | 2239 | 732.4 | - | - | 0 | - |
| - | - | 674.8 | 733.5 | - | - | 0 | - |
| - | - | 1202 | 740.4 | - | - | 0 | - |
| 6 | z | 657.1 | 751.4 | 0.0001599 | 0.2128 | +1 | 7 |
| - | - | 965.7 | 752.4 | - | - | 0 | - |
| 7 | c | 3112 | 757.4 | 0.00145 | 1.915 | +1 | 7 |
| - | - | 769.5 | 758.4 | - | - | 0 | - |
| - | - | 793.2 | 759.4 | - | - | 0 | - |
| 6 | y | 2887 | 767.4 | 0.0009396 | 1.224 | +1 | 7 |
| - | - | 1556 | 768.4 | - | - | 0 | - |
| - | - | 5987 | 773.4 | - | - | 0 | - |
| 7 | c | 6651 | 774.5 | 0.001754 | 2.265 | +1 | 7 |
| - | - | 2121 | 775.5 | - | - | 0 | - |
| 8 | c | 1091 | 844.5 | 0.003264 | 3.866 | +1 | 8 |
| - | - | 617.4 | 845.5 | - | - | 0 | - |
| 5 | z | 1343 | 850.4 | 0.003142 | 3.695 | +1 | 8 |
| - | - | 880.2 | 851.4 | - | - | 0 | - |
| - | - | 1261 | 853.5 | - | - | 0 | - |
| - | - | 771.2 | 854.5 | - | - | 0 | - |
| - | - | 4293 | 860.5 | - | - | 0 | - |
| 8 | c | 5152 | 861.5 | 0.003021 | 3.507 | +1 | 8 |
| - | - | 2064 | 862.5 | - | - | 0 | - |
| - | - | 864.7 | 863.5 | - | - | 0 | - |
| 5 | y | 978.9 | 866.5 | 0.005572 | 6.431 | +1 | 8 |
| - | - | 767.8 | 874.4 | - | - | 0 | - |
| 4 | z | 7690 | 921.5 | 0.0008902 | 0.9661 | +1 | 9 |
| - | - | 4337 | 922.5 | - | - | 0 | - |
| - | - | 636.7 | 923.5 | - | - | 0 | - |
| 9 | c | 680.8 | 930.5 | 0.006418 | 6.898 | +1 | 9 |
| 9 | c | 1271 | 931.5 | 0.0008381 | 0.8997 | +1 | 9 |
| - | - | 3671 | 947.5 | - | - | 0 | - |
| 9 | c | 9128 | 948.5 | 0.001785 | 1.882 | +1 | 9 |
| - | - | 5513 | 949.5 | - | - | 0 | - |
| - | - | 1285 | 950.5 | - | - | 0 | - |
| - | - | 784.4 | 961.5 | - | - | 0 | - |
| - | - | 783.8 | 966.6 | - | - | 0 | - |
| - | - | 1496 | 970.5 | - | - | 0 | - |
| - | - | 1046 | 971.5 | - | - | 0 | - |
| - | - | 2401 | 984.5 | - | - | 0 | - |
| - | - | 1047 | 985.5 | - | - | 0 | - |
| - | - | 763.6 | 986.5 | - | - | 0 | - |
| 10 | c | 1542 | 988.5 | 0.001522 | 1.539 | +1 | 10 |
| - | - | 866.5 | 989.5 | - | - | 0 | - |
| - | - | 743.8 | 1003 | - | - | 0 | - |
| 10 | c | 3.009E+04 | 1006 | 0.0004218 | 0.4195 | +1 | 10 |
| - | - | 1.703E+04 | 1007 | - | - | 0 | - |
| - | - | 4700 | 1008 | - | - | 0 | - |
| - | - | 687.6 | 1013 | - | - | 0 | - |
| 3 | y | 1837 | 1035 | 0.000906 | 0.8757 | +1 | 10 |
| - | - | 1082 | 1036 | - | - | 0 | - |
| - | - | 1026 | 1069 | - | - | 0 | - |
| - | - | 724 | 1070 | - | - | 0 | - |
| - | - | 937.7 | 1075 | - | - | 0 | - |
| - | - | 1092 | 1084 | - | - | 0 | - |
| - | - | 3426 | 1086 | - | - | 0 | - |
| - | - | 2127 | 1087 | - | - | 0 | - |
| - | - | 1118 | 1088 | - | - | 0 | - |
| - | - | 921.8 | 1096 | - | - | 0 | - |
| 11 | c | 1927 | 1102 | 0.006728 | 6.108 | +1 | 11 |
| - | - | 795.4 | 1103 | - | - | 0 | - |
| - | - | 692.4 | 1104 | - | - | 0 | - |
| 11 | c | 2.776E+04 | 1119 | 0.001417 | 1.267 | +1 | 11 |
| - | - | 1.542E+04 | 1120 | - | - | 0 | - |
| - | - | 6171 | 1121 | - | - | 0 | - |
| - | - | 719.9 | 1122 | - | - | 0 | - |
| - | - | 1527 | 1150 | - | - | 0 | - |
| - | - | 978.7 | 1151 | - | - | 0 | - |
| 2 | z | 4868 | 1166 | 0.00189 | 1.621 | +1 | 11 |
| - | - | 3385 | 1167 | - | - | 0 | - |
| - | - | 2105 | 1168 | - | - | 0 | - |
| - | - | 1183 | 1211 | - | - | 0 | - |
| - | - | 823.4 | 1212 | - | - | 0 | - |
| - | - | 636.2 | 1213 | - | - | 0 | - |
| - | - | 1388 | 1222 | - | - | 0 | - |
| - | - | 1266 | 1223 | - | - | 0 | - |
| - | - | 1387 | 1228 | - | - | 0 | - |
| - | - | 3238 | 1239 | - | - | 0 | - |
| - | - | 2921 | 1240 | - | - | 0 | - |
| - | - | 917.5 | 1241 | - | - | 0 | - |
| - | - | 701.7 | 1242 | - | - | 0 | - |
| - | - | 2627 | 1249 | - | - | 0 | - |
| - | - | 1733 | 1250 | - | - | 0 | - |
| - | - | 897.6 | 1256 | - | - | 0 | - |
| - | - | 896 | 1257 | - | - | 0 | - |
| - | - | 4004 | 1267 | - | - | 0 | - |
| - | - | 3037 | 1268 | - | - | 0 | - |
| - | - | 1017 | 1269 | - | - | 0 | - |
| - | - | 1582 | 1281 | - | - | 0 | - |
| - | - | 792.7 | 1282 | - | - | 0 | - |
| - | - | 6650 | 1283 | - | - | 0 | - |
| - | - | 2.981E+04 | 1284 | - | - | 0 | - |
| - | - | 2.107E+04 | 1285 | - | - | 0 | - |
| - | - | 9051 | 1286 | - | - | 0 | - |
| - | - | 1848 | 1287 | - | - | 0 | - |
| - | - | 717.1 | 1940 | - | - | 0 | - |
| - | - | 606.5 | 2146 | - | - | 0 | - |
| - | - | 654.3 | 2791 | - | - | 0 | - |
| - | - | 717.5 | 3012 | - | - | 0 | - |
| - | - | 587.9 | 3078 | - | - | 0 | - |

m/z Charge Intensity FragmentType MassShift Position
127.47740936279297 0 366.83676
132.87123107910156 0 406.2864
136.0759735107422 0 1460.068
145.32708740234375 0 397.1983
148.88426208496094 0 555.14734
148.89552307128906 0 579.7052
148.9111785888672 0 640.85645
148.92198181152344 0 788.0043
148.92762756347656 0 670.99664
148.93289184570312 0 746.2752
148.93832397460938 0 2333.1003
148.94390869140625 0 1940.0925
148.95640563964844 0 4047.9646
148.9617919921875 0 2422.8872
148.96778869628906 0 1415.9749
148.97312927246094 0 662.6351
148.9835968017578 0 927.7068
149.0051727294922 0 473.48706
159.191650390625 0 368.87314
161.21189880371094 0 418.12637
162.70843505859375 0 515.1311
165.05450439453125 0 1217.0668
169.09754943847656 0 539.64465
173.20106506347656 0 468.61667
182.08123779296875 0 8774.439 y 11
183.08436584472656 0 654.26074
185.10513305664062 0 459.98526
187.15740966796875 0 636.60974
192.53207397460938 0 441.77374
205.0972137451172 0 692.4772
221.09132385253906 0 639.99536
221.12843322753906 0 2976.8853
222.13209533691406 0 506.3216
249.1233367919922 0 2675.9353
252.1564483642578 0 1031.2458
258.1448669433594 0 580.58673
268.1653137207031 0 1284.9888
295.1653747558594 0 1195.7009 y 10
317.1833190917969 0 1045.7802
320.5458068847656 0 509.07614
336.1405944824219 0 504.78815
345.1761474609375 0 1908.9855
345.8378601074219 0 540.1521
361.6944885253906 0 723.648
381.249755859375 0 887.964
417.2123718261719 0 961.34814 z Ammonia loss 4
420.1432800292969 0 729.6666
425.2135009765625 0 610.7436 y Ammonia loss 4
433.3656921386719 0 612.9107
470.0723571777344 0 652.6629
485.75341796875 0 722.3281
486.1805114746094 0 583.5788
498.2684631347656 0 882.63184
509.3089294433594 0 1339.2327
516.2809448242188 0 2482.7363
517.2847900390625 0 864.52814
526.2518310546875 0 1874.9038 y 7
532.3019409179688 0 1645.9957
533.3081665039062 0 5415.2754 c 4
534.3095703125 0 1990.9963
603.3865356445312 0 3248.9404
611.3546142578125 0 911.75726
629.364013671875 0 1994.6377
630.36669921875 0 1213.6958
640.345947265625 0 657.1099
641.2565307617188 0 961.7745
641.854248046875 0 3212.9143
642.3558959960938 0 1780.6715
642.8611450195312 0 763.6571
645.3845825195312 0 3682.6501
646.3917236328125 0 10782.282 c 5
647.3944702148438 0 4160.411
653.3007202148438 0 1232.9099
654.3084106445312 0 5352.857 y 6
655.3140869140625 0 1466.926
672.4067993164062 0 989.5024
730.4371948242188 0 1304.4055
731.443359375 0 4170.857
732.4478149414062 0 2239.4912
733.452880859375 0 674.7794
740.3975830078125 0 1201.9095
751.37451171875 0 657.08466 z 5
752.3826293945312 0 965.6937
757.4228515625 0 3112.1963 c Ammonia loss 6
758.4281616210938 0 769.455
759.4339599609375 0 793.2427
767.3924560546875 0 2886.999 y 5
768.3960571289062 0 1555.5724
773.4424438476562 0 5987.2144
774.4490966796875 0 6651.378 c 6
775.45263671875 0 2120.6768
844.4595947265625 0 1090.9888 c Ammonia loss 7
845.4571533203125 0 617.3858
850.4462280273438 0 1343.4495 z 4
851.4451904296875 0 880.19055
853.4801635742188 0 1261.4762
854.4841918945312 0 771.2066
860.4727172851562 0 4293.4277
861.4798583984375 0 5152.4873 c 7
862.48291015625 0 2063.6472
863.4954223632812 0 864.6731
866.4562377929688 0 978.91907 y 4
874.4317626953125 0 767.75024
921.4793090820312 0 7690.492 z 3
922.4830932617188 0 4336.917
923.493896484375 0 636.7257
930.4979248046875 0 680.83154 c Water loss 8
931.4891967773438 0 1270.7036 c Ammonia loss 8
947.507568359375 0 3671.1875
948.5131225585938 0 9127.759 c 8
949.51611328125 0 5512.6665
950.5206909179688 0 1285.4905
961.5169677734375 0 784.40283
966.5897216796875 0 783.7544
970.5003051757812 0 1495.6443
971.4981079101562 0 1046.4852
984.525634765625 0 2401.217
985.5343017578125 0 1047.1324
986.5352172851562 0 763.6398
988.50830078125 0 1542.2617 c Ammonia loss 9
989.5030517578125 0 866.4544
1002.5325317382812 0 743.8023
1005.5359497070312 0 30085.244 c 9
1006.5375366210938 0 17030.943
1007.541015625 0 4700.092
1012.51171875 0 687.5707
1034.55078125 0 1837.0217 y 2
1035.552978515625 0 1081.5684
1068.5704345703125 0 1026.1123
1069.57470703125 0 723.9531
1074.6041259765625 0 937.6836
1083.5843505859375 0 1092.0242
1085.5888671875 0 3425.7168
1086.5908203125 0 2126.9495
1087.5904541015625 0 1117.6742
1095.6436767578125 0 921.7651
1101.587158203125 0 1927.0621 c Ammonia loss 10
1102.59375 0 795.40393
1103.60400390625 0 692.3802
1118.6190185546875 0 27757.785 c 10
1119.6224365234375 0 15424.285
1120.6248779296875 0 6170.796
1121.6317138671875 0 719.9272
1150.4189453125 0 1526.6923
1151.4281005859375 0 978.72266
1165.5994873046875 0 4868.3833 z 1
1166.6007080078125 0 3384.754
1167.6036376953125 0 2104.561
1210.5928955078125 0 1182.628
1211.5908203125 0 823.4095
1212.60595703125 0 636.19244
1221.67529296875 0 1388.0789
1222.6748046875 0 1265.6494
1227.611083984375 0 1387.1469
1238.6492919921875 0 3237.5115
1239.6614990234375 0 2920.8313
1240.646240234375 0 917.45636
1241.6419677734375 0 701.6878
1248.6351318359375 0 2627.115
1249.638671875 0 1733.3391
1255.6768798828125 0 897.6055
1256.6834716796875 0 896.0225
1266.6544189453125 0 4003.9553
1267.6650390625 0 3036.594
1268.658935546875 0 1016.59357
1280.690673828125 0 1582.2228
1281.647216796875 0 792.73663
1282.67236328125 0 6649.509
1283.674072265625 0 29814.232
1284.6767578125 0 21071.21
1285.6817626953125 0 9051.443
1286.6873779296875 0 1847.722
1939.5191650390625 0 717.0863
2146.165771484375 0 606.5446
2790.76611328125 0 654.274
3011.6748046875 0 717.5226
3077.6259765625 0 587.87634

Spectrum Details

|  |  |
| --- | --- |
| Matched peaks? Matched peaks The total absolute number of peaks matched. Additionally in brackets the total fraction of peaks matched and the total number of peaks is shown. | 26 (14.69% of 177) |
| FDR? FDR The false discovery rate estimated for this peptide. It is calculated by matching all theoretical fragments with a non-integer shift with the raw peaks for this spectrum. This is done with 40 different shifts. The resulting percentage is the average number of annotated peaks over the number of annotated peaks with the correct spectrum. | 1.65% |
| Satellite FDR? Satellite FDR See the FDR for details on its calculation. This satellite ion specific FDR only contains the satellite ions (d/w) for I/L/J positions. | ∞ |
| PSM Score? PSM Score The PSM Score as given by Hecklib to this annotated spectrum. It is shown with three significant figures. | 288 |

#### Spectrum 10897? Spectrum 10897 The raw spectrum of this peptide as annotated by Hecklib. The fragments are coloured according to ion type (see legend). Any peaks with a star '\*' as text can be hovered over to see the full details, first the ion type second the mass shift type. By hovering over the amino acids in the peptide or ions in the legend the corresponding peaks are highlighted. By toggling the 'Unassigned' label you can turn the background (unassigned) peaks on or off in the plot. By updating the slider in the Ion legend you can update the spectrum to only show the top X% of the peaks with labels. The top X% means any peak that is within X% of the highest intensity. By dragging in the spectrum you can zoom in to a specific part of the spectrum and use 'Zoom Out' to get back to the original zoom level. The annotation of the spectrum is based on the given sequence in the peptides file and is done with different software so inconsistencies are likely. The peaks are annotated based on the given sequence, with 20 ppm tolerance.

Copy Data

##### Spectrum 10897 (TSV)

###### Preview

```
Loading example...
```

*Click on the button to copy the data to your clipboard.*

Mz MinMz MaxIntensity Max

WidthHeightPeptide font sizePeptide stroke widthSpectrum font sizeSpectrum stroke widthCompact peptide

Ion legend

wxyz

abcd

OtherUnassignedIon^Charge^_Position_Show for top:%

TFPAVJQSSGJY

01.01e+52.01e+53.02e+54.03e+5

Zoom Out

y^+1^_1_a^+1^_2_y^+2^_4_a^+1^_2_b^+1^_2_b^+1^_2_d^+1^_3_y^+1^_2_a^+1^_3_b^+2^_6_b^+2^_6_a^+1^_3_b^+1^_3_b^+1^_3_y^+1^_3_b^+2^_7_b^+2^_7_b^+2^_7_b^+1^_4_b^+1^_4_y^+1^_4_b^+2^_8_y^+1^_4_b^+2^_9_b^+2^_9_b^+2^_10_b^+1^_5_y^+1^_5_y^+2^_10_b^+1^_5_y^+1^_5_b^+2^_11_b^+1^_6_b^+1^_6_y^+1^_6_y^+1^_6_b^+1^_7_b^+1^_7_y^+1^_7_y^+1^_7_b^+1^_7_y^+1^_7_b^+1^_8_b^+1^_8_y^+1^_8_y^+1^_8_b^+1^_9_b^+1^_9_b^+1^_9_y^+1^_9_b^+1^_10_b^+1^_10_b^+1^_10_y^+1^_10_y^+1^_10_y^+1^_10_b^+1^_11_b^+1^_11_b^+1^_11_

0793158523783170

Fragment Matches Table

Show background peaks

| Position | Ion type | Intensity | mz Theoretical | mz Error (Th) | mz Error (ppm) | Charge | Series Number |
| --- | --- | --- | --- | --- | --- | --- | --- |
| - | - | 1195 | 120.1 | - | - | 0 | - |
| - | - | 4.751E+04 | 120.1 | - | - | 0 | - |
| - | - | 468.2 | 120.8 | - | - | 0 | - |
| - | - | 709.5 | 121.1 | - | - | 0 | - |
| - | - | 4209 | 121.1 | - | - | 0 | - |
| - | - | 458.2 | 122.9 | - | - | 0 | - |
| - | - | 2.541E+04 | 123 | - | - | 0 | - |
| - | - | 1554 | 124 | - | - | 0 | - |
| - | - | 516.3 | 125.1 | - | - | 0 | - |
| - | - | 1612 | 126.1 | - | - | 0 | - |
| - | - | 2247 | 127.1 | - | - | 0 | - |
| - | - | 1123 | 127.1 | - | - | 0 | - |
| - | - | 4575 | 129.1 | - | - | 0 | - |
| - | - | 2838 | 129.1 | - | - | 0 | - |
| - | - | 616.4 | 130 | - | - | 0 | - |
| - | - | 1679 | 130.1 | - | - | 0 | - |
| - | - | 481.8 | 130.1 | - | - | 0 | - |
| - | - | 9032 | 131.1 | - | - | 0 | - |
| - | - | 3015 | 132.1 | - | - | 0 | - |
| - | - | 607.1 | 132.1 | - | - | 0 | - |
| - | - | 573.5 | 133.1 | - | - | 0 | - |
| - | - | 511.5 | 135.7 | - | - | 0 | - |
| - | - | 2.14E+05 | 136.1 | - | - | 0 | - |
| - | - | 559.8 | 137.1 | - | - | 0 | - |
| - | - | 878.3 | 137.1 | - | - | 0 | - |
| - | - | 1.594E+04 | 137.1 | - | - | 0 | - |
| - | - | 619.1 | 138.1 | - | - | 0 | - |
| - | - | 1308 | 139.1 | - | - | 0 | - |
| - | - | 1578 | 140.1 | - | - | 0 | - |
| - | - | 6.023E+04 | 141.1 | - | - | 0 | - |
| - | - | 612.3 | 142.1 | - | - | 0 | - |
| - | - | 3777 | 142.1 | - | - | 0 | - |
| - | - | 5.447E+04 | 143.1 | - | - | 0 | - |
| - | - | 703.7 | 144.1 | - | - | 0 | - |
| - | - | 3042 | 144.1 | - | - | 0 | - |
| - | - | 448 | 144.2 | - | - | 0 | - |
| - | - | 4298 | 145.1 | - | - | 0 | - |
| - | - | 602.4 | 146.1 | - | - | 0 | - |
| - | - | 1.941E+04 | 147 | - | - | 0 | - |
| - | - | 2798 | 147.1 | - | - | 0 | - |
| - | - | 1676 | 148 | - | - | 0 | - |
| - | - | 488.4 | 149 | - | - | 0 | - |
| - | - | 3025 | 153.1 | - | - | 0 | - |
| - | - | 1090 | 154.2 | - | - | 0 | - |
| - | - | 1503 | 155.1 | - | - | 0 | - |
| - | - | 953.3 | 155.1 | - | - | 0 | - |
| - | - | 1098 | 156.1 | - | - | 0 | - |
| - | - | 1.54E+04 | 157.1 | - | - | 0 | - |
| - | - | 816.2 | 157.1 | - | - | 0 | - |
| - | - | 1.019E+04 | 158.1 | - | - | 0 | - |
| - | - | 1152 | 159.1 | - | - | 0 | - |
| - | - | 1042 | 160.1 | - | - | 0 | - |
| - | - | 577.7 | 160.1 | - | - | 0 | - |
| - | - | 3744 | 160.1 | - | - | 0 | - |
| - | - | 906.6 | 162.1 | - | - | 0 | - |
| - | - | 2.069E+05 | 165.1 | - | - | 0 | - |
| - | - | 1.893E+04 | 166.1 | - | - | 0 | - |
| - | - | 1042 | 166.1 | - | - | 0 | - |
| - | - | 783.5 | 167.1 | - | - | 0 | - |
| - | - | 692.2 | 167.1 | - | - | 0 | - |
| - | - | 859 | 168.1 | - | - | 0 | - |
| - | - | 3386 | 169.1 | - | - | 0 | - |
| - | - | 2.415E+05 | 169.1 | - | - | 0 | - |
| - | - | 4961 | 170.1 | - | - | 0 | - |
| - | - | 1.979E+04 | 170.1 | - | - | 0 | - |
| - | - | 3452 | 171.1 | - | - | 0 | - |
| - | - | 2.509E+04 | 171.1 | - | - | 0 | - |
| - | - | 1302 | 172.1 | - | - | 0 | - |
| - | - | 597 | 173.1 | - | - | 0 | - |
| - | - | 2965 | 173.5 | - | - | 0 | - |
| - | - | 5937 | 175.1 | - | - | 0 | - |
| - | - | 534.5 | 176 | - | - | 0 | - |
| - | - | 1.958E+04 | 176.1 | - | - | 0 | - |
| - | - | 1006 | 177.1 | - | - | 0 | - |
| - | - | 1.282E+04 | 177.1 | - | - | 0 | - |
| - | - | 2505 | 177.1 | - | - | 0 | - |
| - | - | 1106 | 177.1 | - | - | 0 | - |
| - | - | 1768 | 178.1 | - | - | 0 | - |
| - | - | 2099 | 180.1 | - | - | 0 | - |
| - | - | 7101 | 181.1 | - | - | 0 | - |
| - | - | 1799 | 181.1 | - | - | 0 | - |
| 12 | y | 3.987E+05 | 182.1 | 0.000358 | 1.966 | +1 | 1 |
| - | - | 3.344E+04 | 183.1 | - | - | 0 | - |
| - | - | 1313 | 183.1 | - | - | 0 | - |
| - | - | 837.6 | 183.1 | - | - | 0 | - |
| - | - | 2640 | 184.1 | - | - | 0 | - |
| - | - | 1162 | 185.1 | - | - | 0 | - |
| - | - | 1.181E+04 | 185.2 | - | - | 0 | - |
| - | - | 758.1 | 186.1 | - | - | 0 | - |
| - | - | 1.234E+04 | 186.1 | - | - | 0 | - |
| - | - | 874.5 | 186.2 | - | - | 0 | - |
| - | - | 1639 | 187.1 | - | - | 0 | - |
| - | - | 709.3 | 187.1 | - | - | 0 | - |
| - | - | 3590 | 188.1 | - | - | 0 | - |
| - | - | 1637 | 189.1 | - | - | 0 | - |
| - | - | 906.5 | 191.1 | - | - | 0 | - |
| - | - | 1218 | 193.1 | - | - | 0 | - |
| - | - | 556.1 | 194.1 | - | - | 0 | - |
| - | - | 835.5 | 195.1 | - | - | 0 | - |
| - | - | 2415 | 196.1 | - | - | 0 | - |
| - | - | 955.8 | 197.1 | - | - | 0 | - |
| - | - | 3057 | 197.1 | - | - | 0 | - |
| - | - | 1.164E+04 | 198.1 | - | - | 0 | - |
| - | - | 639.1 | 198.1 | - | - | 0 | - |
| - | - | 5441 | 199.1 | - | - | 0 | - |
| - | - | 1074 | 199.1 | - | - | 0 | - |
| - | - | 1101 | 199.1 | - | - | 0 | - |
| - | - | 1734 | 199.2 | - | - | 0 | - |
| - | - | 1961 | 201.1 | - | - | 0 | - |
| 2 | a | 6209 | 203.1 | 0.000244 | 1.201 | +1 | 2 |
| - | - | 771.9 | 204.1 | - | - | 0 | - |
| - | - | 1278 | 205.1 | - | - | 0 | - |
| - | - | 664.3 | 206.1 | - | - | 0 | - |
| - | - | 911.4 | 207.1 | - | - | 0 | - |
| - | - | 2662 | 208.1 | - | - | 0 | - |
| - | - | 1009 | 208.1 | - | - | 0 | - |
| - | - | 949.1 | 209.2 | - | - | 0 | - |
| 9 | y | 1494 | 211.1 | 0.0003134 | 1.485 | +2 | 4 |
| - | - | 1804 | 211.1 | - | - | 0 | - |
| - | - | 880.9 | 211.2 | - | - | 0 | - |
| - | - | 1800 | 212.1 | - | - | 0 | - |
| - | - | 666.1 | 213.1 | - | - | 0 | - |
| - | - | 5469 | 213.1 | - | - | 0 | - |
| - | - | 1.14E+04 | 213.2 | - | - | 0 | - |
| - | - | 4.81E+04 | 214.1 | - | - | 0 | - |
| - | - | 1421 | 214.2 | - | - | 0 | - |
| - | - | 834.8 | 214.2 | - | - | 0 | - |
| - | - | 3334 | 215.1 | - | - | 0 | - |
| - | - | 1494 | 215.1 | - | - | 0 | - |
| - | - | 1087 | 216.1 | - | - | 0 | - |
| - | - | 1.411E+04 | 216.1 | - | - | 0 | - |
| - | - | 1065 | 217.1 | - | - | 0 | - |
| - | - | 5537 | 217.1 | - | - | 0 | - |
| - | - | 821.3 | 219.1 | - | - | 0 | - |
| 2 | a | 3.335E+05 | 221.1 | 0.0003452 | 1.561 | +1 | 2 |
| - | - | 4.02E+04 | 222.1 | - | - | 0 | - |
| - | - | 2667 | 223.1 | - | - | 0 | - |
| - | - | 2846 | 223.1 | - | - | 0 | - |
| - | - | 3636 | 223.1 | - | - | 0 | - |
| - | - | 3707 | 224.1 | - | - | 0 | - |
| - | - | 8350 | 225.1 | - | - | 0 | - |
| - | - | 4472 | 226.1 | - | - | 0 | - |
| - | - | 1252 | 228.1 | - | - | 0 | - |
| - | - | 832.4 | 230.1 | - | - | 0 | - |
| - | - | 3551 | 230.2 | - | - | 0 | - |
| - | - | 1455 | 230.2 | - | - | 0 | - |
| 2 | b | 3852 | 231.1 | 0.0003398 | 1.47 | +1 | 2 |
| - | - | 3.884E+04 | 232.1 | - | - | 0 | - |
| - | - | 2674 | 233.1 | - | - | 0 | - |
| - | - | 976.8 | 233.1 | - | - | 0 | - |
| - | - | 982.6 | 233.2 | - | - | 0 | - |
| - | - | 777.6 | 234.1 | - | - | 0 | - |
| - | - | 905.6 | 235.1 | - | - | 0 | - |
| - | - | 819 | 237.2 | - | - | 0 | - |
| - | - | 1814 | 238.1 | - | - | 0 | - |
| - | - | 897 | 239.2 | - | - | 0 | - |
| - | - | 1718 | 239.2 | - | - | 0 | - |
| - | - | 2738 | 240.1 | - | - | 0 | - |
| - | - | 2778 | 240.1 | - | - | 0 | - |
| - | - | 3.771E+04 | 240.2 | - | - | 0 | - |
| - | - | 1618 | 241.2 | - | - | 0 | - |
| - | - | 4465 | 241.2 | - | - | 0 | - |
| - | - | 1259 | 242.1 | - | - | 0 | - |
| - | - | 4.006E+04 | 242.2 | - | - | 0 | - |
| - | - | 1425 | 243.1 | - | - | 0 | - |
| - | - | 4100 | 243.2 | - | - | 0 | - |
| - | - | 568.8 | 244.1 | - | - | 0 | - |
| - | - | 780.1 | 244.2 | - | - | 0 | - |
| - | - | 1.079E+04 | 245.1 | - | - | 0 | - |
| - | - | 700.3 | 246.1 | - | - | 0 | - |
| - | - | 1770 | 246.1 | - | - | 0 | - |
| 2 | b | 1.117E+05 | 249.1 | 0.0003189 | 1.28 | +1 | 2 |
| - | - | 3025 | 249.2 | - | - | 0 | - |
| - | - | 9661 | 250.1 | - | - | 0 | - |
| - | - | 1.403E+04 | 250.1 | - | - | 0 | - |
| - | - | 1438 | 251.1 | - | - | 0 | - |
| - | - | 896.2 | 251.2 | - | - | 0 | - |
| - | - | 4810 | 252.1 | - | - | 0 | - |
| - | - | 3271 | 254.2 | - | - | 0 | - |
| - | - | 6177 | 255.2 | - | - | 0 | - |
| - | - | 1815 | 255.7 | - | - | 0 | - |
| - | - | 4548 | 256.1 | - | - | 0 | - |
| - | - | 613.4 | 256.2 | - | - | 0 | - |
| - | - | 1158 | 258.1 | - | - | 0 | - |
| - | - | 2.445E+04 | 258.1 | - | - | 0 | - |
| - | - | 3478 | 259.1 | - | - | 0 | - |
| - | - | 693.8 | 261.1 | - | - | 0 | - |
| - | - | 892.8 | 262.1 | - | - | 0 | - |
| - | - | 777.6 | 262.2 | - | - | 0 | - |
| - | - | 882.9 | 263.1 | - | - | 0 | - |
| - | - | 981.8 | 263.1 | - | - | 0 | - |
| - | - | 2002 | 266.1 | - | - | 0 | - |
| - | - | 6067 | 267.1 | - | - | 0 | - |
| - | - | 6082 | 268.1 | - | - | 0 | - |
| - | - | 2.271E+05 | 268.2 | - | - | 0 | - |
| - | - | 2.994E+04 | 269.2 | - | - | 0 | - |
| - | - | 4282 | 270.1 | - | - | 0 | - |
| - | - | 2145 | 270.2 | - | - | 0 | - |
| - | - | 1996 | 271.1 | - | - | 0 | - |
| - | - | 831.2 | 272.2 | - | - | 0 | - |
| - | - | 1204 | 273.2 | - | - | 0 | - |
| - | - | 689 | 274.1 | - | - | 0 | - |
| - | - | 704.1 | 274.5 | - | - | 0 | - |
| - | - | 2060 | 275.1 | - | - | 0 | - |
| - | - | 1953 | 275.2 | - | - | 0 | - |
| - | - | 983.2 | 276.1 | - | - | 0 | - |
| - | - | 1223 | 277.1 | - | - | 0 | - |
| - | - | 1588 | 279.1 | - | - | 0 | - |
| - | - | 978.2 | 279.1 | - | - | 0 | - |
| - | - | 1830 | 280.1 | - | - | 0 | - |
| - | - | 2397 | 280.2 | - | - | 0 | - |
| - | - | 1788 | 281.2 | - | - | 0 | - |
| - | - | 3043 | 282.1 | - | - | 0 | - |
| - | - | 5751 | 282.2 | - | - | 0 | - |
| - | - | 1104 | 282.2 | - | - | 0 | - |
| - | - | 742.5 | 284.1 | - | - | 0 | - |
| - | - | 2488 | 284.2 | - | - | 0 | - |
| - | - | 3793 | 284.2 | - | - | 0 | - |
| - | - | 5589 | 285.1 | - | - | 0 | - |
| - | - | 2074 | 285.2 | - | - | 0 | - |
| - | - | 4664 | 285.2 | - | - | 0 | - |
| - | - | 2936 | 286.1 | - | - | 0 | - |
| - | - | 850.9 | 286.1 | - | - | 0 | - |
| - | - | 3190 | 286.2 | - | - | 0 | - |
| - | - | 525.6 | 286.2 | - | - | 0 | - |
| - | - | 635 | 287.1 | - | - | 0 | - |
| - | - | 789.7 | 288.1 | - | - | 0 | - |
| - | - | 2842 | 289.2 | - | - | 0 | - |
| - | - | 2172 | 289.7 | - | - | 0 | - |
| - | - | 1019 | 290.2 | - | - | 0 | - |
| - | - | 612.8 | 290.2 | - | - | 0 | - |
| - | - | 664.7 | 291.1 | - | - | 0 | - |
| 3 | d | 1177 | 292.2 | 4.067E-05 | 0.1392 | +1 | 3 |
| - | - | 1452 | 294.1 | - | - | 0 | - |
| - | - | 3027 | 294.1 | - | - | 0 | - |
| - | - | 1500 | 294.2 | - | - | 0 | - |
| - | - | 1.015E+04 | 295.1 | - | - | 0 | - |
| 11 | y | 5.624E+04 | 295.2 | 0.0004158 | 1.409 | +1 | 2 |
| - | - | 615.2 | 296.1 | - | - | 0 | - |
| - | - | 1208 | 296.1 | - | - | 0 | - |
| - | - | 8818 | 296.2 | - | - | 0 | - |
| - | - | 5432 | 296.2 | - | - | 0 | - |
| - | - | 3854 | 297.1 | - | - | 0 | - |
| - | - | 2259 | 297.2 | - | - | 0 | - |
| - | - | 777.6 | 297.2 | - | - | 0 | - |
| - | - | 2241 | 297.2 | - | - | 0 | - |
| - | - | 1916 | 298.2 | - | - | 0 | - |
| - | - | 3089 | 298.7 | - | - | 0 | - |
| - | - | 8567 | 299.2 | - | - | 0 | - |
| - | - | 3894 | 300.2 | - | - | 0 | - |
| 3 | a | 1438 | 300.2 | 0.003358 | 11.19 | +1 | 3 |
| - | - | 685.4 | 301.2 | - | - | 0 | - |
| - | - | 990.5 | 301.2 | - | - | 0 | - |
| - | - | 862 | 302.1 | - | - | 0 | - |
| - | - | 959.3 | 302.2 | - | - | 0 | - |
| - | - | 3157 | 303.1 | - | - | 0 | - |
| - | - | 2156 | 303.2 | - | - | 0 | - |
| - | - | 760.4 | 304.1 | - | - | 0 | - |
| 6 | b | 1831 | 306.2 | 0.00027 | 0.8817 | +2 | 6 |
| - | - | 1689 | 307.1 | - | - | 0 | - |
| - | - | 871.7 | 307.1 | - | - | 0 | - |
| - | - | 950 | 308.1 | - | - | 0 | - |
| - | - | 2747 | 309.2 | - | - | 0 | - |
| - | - | 602 | 309.2 | - | - | 0 | - |
| - | - | 1588 | 310.1 | - | - | 0 | - |
| - | - | 946.2 | 310.2 | - | - | 0 | - |
| - | - | 1638 | 310.2 | - | - | 0 | - |
| - | - | 3917 | 310.2 | - | - | 0 | - |
| - | - | 955.8 | 311.1 | - | - | 0 | - |
| - | - | 4817 | 311.2 | - | - | 0 | - |
| - | - | 1189 | 312.1 | - | - | 0 | - |
| - | - | 3723 | 312.2 | - | - | 0 | - |
| - | - | 934.1 | 312.2 | - | - | 0 | - |
| - | - | 782.9 | 312.2 | - | - | 0 | - |
| - | - | 775.2 | 313.1 | - | - | 0 | - |
| - | - | 3598 | 313.2 | - | - | 0 | - |
| - | - | 978.9 | 314.2 | - | - | 0 | - |
| - | - | 574.7 | 314.2 | - | - | 0 | - |
| - | - | 2589 | 315.1 | - | - | 0 | - |
| 6 | b | 1551 | 315.2 | 0.0007554 | 2.397 | +2 | 6 |
| - | - | 3193 | 315.7 | - | - | 0 | - |
| - | - | 937.1 | 315.7 | - | - | 0 | - |
| - | - | 1.566E+04 | 316.2 | - | - | 0 | - |
| - | - | 677.1 | 316.2 | - | - | 0 | - |
| - | - | 5.994E+04 | 317.2 | - | - | 0 | - |
| - | - | 1545 | 318.1 | - | - | 0 | - |
| 3 | a | 8584 | 318.2 | 0.004603 | 14.47 | +1 | 3 |
| - | - | 1006 | 319.2 | - | - | 0 | - |
| - | - | 884.3 | 320.2 | - | - | 0 | - |
| - | - | 752.8 | 320.2 | - | - | 0 | - |
| - | - | 631.8 | 323.2 | - | - | 0 | - |
| - | - | 1099 | 323.2 | - | - | 0 | - |
| - | - | 4943 | 324.1 | - | - | 0 | - |
| - | - | 1.41E+04 | 324.2 | - | - | 0 | - |
| - | - | 4933 | 324.7 | - | - | 0 | - |
| - | - | 1.43E+04 | 325.1 | - | - | 0 | - |
| - | - | 1693 | 325.2 | - | - | 0 | - |
| - | - | 7081 | 325.2 | - | - | 0 | - |
| - | - | 2138 | 326.1 | - | - | 0 | - |
| - | - | 864.3 | 326.2 | - | - | 0 | - |
| - | - | 9303 | 327.2 | - | - | 0 | - |
| - | - | 1644 | 327.2 | - | - | 0 | - |
| 3 | b | 1.134E+04 | 328.2 | 0.0009359 | 2.852 | +1 | 3 |
| - | - | 1.06E+04 | 329.2 | - | - | 0 | - |
| - | - | 843 | 330.1 | - | - | 0 | - |
| - | - | 3682 | 330.2 | - | - | 0 | - |
| - | - | 658.6 | 331.2 | - | - | 0 | - |
| - | - | 799.2 | 333.2 | - | - | 0 | - |
| - | - | 3913 | 333.2 | - | - | 0 | - |
| - | - | 879 | 333.7 | - | - | 0 | - |
| - | - | 1532 | 334.2 | - | - | 0 | - |
| - | - | 1364 | 336.2 | - | - | 0 | - |
| - | - | 1735 | 337.2 | - | - | 0 | - |
| - | - | 1781 | 337.2 | - | - | 0 | - |
| - | - | 994.9 | 339.2 | - | - | 0 | - |
| - | - | 2384 | 339.2 | - | - | 0 | - |
| - | - | 1687 | 341.2 | - | - | 0 | - |
| - | - | 1.144E+04 | 341.2 | - | - | 0 | - |
| - | - | 1.227E+04 | 342.1 | - | - | 0 | - |
| - | - | 2953 | 342.2 | - | - | 0 | - |
| - | - | 2.432E+04 | 343.1 | - | - | 0 | - |
| - | - | 1996 | 343.1 | - | - | 0 | - |
| - | - | 1574 | 343.2 | - | - | 0 | - |
| - | - | 3188 | 344.1 | - | - | 0 | - |
| - | - | 4206 | 344.2 | - | - | 0 | - |
| - | - | 6.614E+04 | 345.2 | - | - | 0 | - |
| 3 | b | 2.641E+04 | 346.2 | 0.00151 | 4.362 | +1 | 3 |
| - | - | 717.4 | 347.1 | - | - | 0 | - |
| - | - | 5218 | 347.2 | - | - | 0 | - |
| - | - | 2300 | 348.2 | - | - | 0 | - |
| - | - | 745.1 | 349.2 | - | - | 0 | - |
| - | - | 1063 | 350.2 | - | - | 0 | - |
| - | - | 1226 | 351.2 | - | - | 0 | - |
| - | - | 2.046E+04 | 351.2 | - | - | 0 | - |
| - | - | 593.1 | 352.1 | - | - | 0 | - |
| 10 | y | 1.438E+04 | 352.2 | 0.0003143 | 0.8925 | +1 | 3 |
| - | - | 2812 | 352.2 | - | - | 0 | - |
| - | - | 3329 | 352.7 | - | - | 0 | - |
| - | - | 5224 | 353.2 | - | - | 0 | - |
| - | - | 3.216E+04 | 353.3 | - | - | 0 | - |
| - | - | 1148 | 353.7 | - | - | 0 | - |
| - | - | 5486 | 354.3 | - | - | 0 | - |
| - | - | 1149 | 355.2 | - | - | 0 | - |
| - | - | 3266 | 355.2 | - | - | 0 | - |
| - | - | 828.9 | 355.3 | - | - | 0 | - |
| - | - | 1402 | 356.2 | - | - | 0 | - |
| - | - | 1091 | 357.2 | - | - | 0 | - |
| - | - | 1189 | 358.2 | - | - | 0 | - |
| - | - | 672 | 358.2 | - | - | 0 | - |
| - | - | 2524 | 359.2 | - | - | 0 | - |
| - | - | 7348 | 360.2 | - | - | 0 | - |
| - | - | 953.6 | 360.2 | - | - | 0 | - |
| - | - | 1453 | 361.2 | - | - | 0 | - |
| - | - | 2.6E+04 | 361.7 | - | - | 0 | - |
| - | - | 9997 | 362.2 | - | - | 0 | - |
| - | - | 1423 | 362.7 | - | - | 0 | - |
| - | - | 1276 | 363.2 | - | - | 0 | - |
| - | - | 1600 | 364.2 | - | - | 0 | - |
| - | - | 2344 | 365.1 | - | - | 0 | - |
| - | - | 696.8 | 365.2 | - | - | 0 | - |
| - | - | 2929 | 365.2 | - | - | 0 | - |
| - | - | 1337 | 366.2 | - | - | 0 | - |
| - | - | 1942 | 367.2 | - | - | 0 | - |
| - | - | 2721 | 367.2 | - | - | 0 | - |
| - | - | 1216 | 368.2 | - | - | 0 | - |
| - | - | 756.4 | 369.2 | - | - | 0 | - |
| - | - | 845.9 | 369.2 | - | - | 0 | - |
| - | - | 1.105E+04 | 369.2 | - | - | 0 | - |
| - | - | 1654 | 370.1 | - | - | 0 | - |
| 7 | b | 6876 | 370.2 | 0.00333 | 8.994 | +2 | 7 |
| 7 | b | 3782 | 370.7 | 0.00177 | 4.775 | +2 | 7 |
| - | - | 1287 | 371.2 | - | - | 0 | - |
| - | - | 2430 | 372.2 | - | - | 0 | - |
| - | - | 1985 | 373.2 | - | - | 0 | - |
| - | - | 1242 | 373.2 | - | - | 0 | - |
| - | - | 1808 | 375.2 | - | - | 0 | - |
| - | - | 1940 | 377.2 | - | - | 0 | - |
| - | - | 1944 | 378.2 | - | - | 0 | - |
| - | - | 4572 | 379.2 | - | - | 0 | - |
| 7 | b | 3093 | 379.2 | 0.00174 | 4.589 | +2 | 7 |
| - | - | 1413 | 379.7 | - | - | 0 | - |
| - | - | 3987 | 380.2 | - | - | 0 | - |
| - | - | 6.399E+04 | 381.2 | - | - | 0 | - |
| - | - | 3131 | 382.2 | - | - | 0 | - |
| - | - | 1.208E+04 | 382.3 | - | - | 0 | - |
| - | - | 916 | 383.2 | - | - | 0 | - |
| - | - | 2982 | 383.2 | - | - | 0 | - |
| - | - | 1181 | 383.3 | - | - | 0 | - |
| - | - | 987.3 | 384.2 | - | - | 0 | - |
| - | - | 1449 | 384.2 | - | - | 0 | - |
| - | - | 1683 | 385.1 | - | - | 0 | - |
| - | - | 1171 | 386.2 | - | - | 0 | - |
| - | - | 1181 | 387.2 | - | - | 0 | - |
| - | - | 9985 | 387.2 | - | - | 0 | - |
| - | - | 1661 | 388.2 | - | - | 0 | - |
| - | - | 7018 | 389.2 | - | - | 0 | - |
| - | - | 839.2 | 390.2 | - | - | 0 | - |
| - | - | 1604 | 392.2 | - | - | 0 | - |
| - | - | 1888 | 393.2 | - | - | 0 | - |
| - | - | 731.4 | 393.3 | - | - | 0 | - |
| - | - | 1.163E+04 | 394.2 | - | - | 0 | - |
| - | - | 1027 | 395.2 | - | - | 0 | - |
| - | - | 1338 | 395.2 | - | - | 0 | - |
| - | - | 2276 | 396.2 | - | - | 0 | - |
| - | - | 1150 | 397.2 | - | - | 0 | - |
| - | - | 5462 | 398.2 | - | - | 0 | - |
| - | - | 763.6 | 398.2 | - | - | 0 | - |
| - | - | 1.841E+04 | 398.3 | - | - | 0 | - |
| 4 | b | 2.703E+04 | 399.2 | 0.0005042 | 1.263 | +1 | 4 |
| - | - | 1161 | 399.3 | - | - | 0 | - |
| - | - | 2613 | 399.3 | - | - | 0 | - |
| - | - | 5249 | 400.2 | - | - | 0 | - |
| - | - | 1028 | 401.2 | - | - | 0 | - |
| - | - | 1204 | 404.2 | - | - | 0 | - |
| - | - | 1037 | 405.2 | - | - | 0 | - |
| - | - | 963.1 | 406.2 | - | - | 0 | - |
| - | - | 732.9 | 407.2 | - | - | 0 | - |
| - | - | 960.3 | 408.2 | - | - | 0 | - |
| - | - | 1183 | 408.2 | - | - | 0 | - |
| - | - | 960.3 | 409.2 | - | - | 0 | - |
| - | - | 4401 | 410.2 | - | - | 0 | - |
| - | - | 4891 | 410.2 | - | - | 0 | - |
| - | - | 2084 | 410.3 | - | - | 0 | - |
| - | - | 809.2 | 411.2 | - | - | 0 | - |
| - | - | 1254 | 412.2 | - | - | 0 | - |
| - | - | 4473 | 412.2 | - | - | 0 | - |
| - | - | 2592 | 412.3 | - | - | 0 | - |
| - | - | 2247 | 413.2 | - | - | 0 | - |
| - | - | 935.8 | 413.7 | - | - | 0 | - |
| - | - | 852.3 | 414.2 | - | - | 0 | - |
| - | - | 1.23E+04 | 415.2 | - | - | 0 | - |
| - | - | 1036 | 415.3 | - | - | 0 | - |
| - | - | 2533 | 416.2 | - | - | 0 | - |
| - | - | 1191 | 416.2 | - | - | 0 | - |
| 4 | b | 4.329E+04 | 417.2 | 0.0004986 | 1.195 | +1 | 4 |
| - | - | 8715 | 418.2 | - | - | 0 | - |
| - | - | 1896 | 419.2 | - | - | 0 | - |
| - | - | 1712 | 420.2 | - | - | 0 | - |
| - | - | 1050 | 420.2 | - | - | 0 | - |
| 9 | y | 2121 | 421.2 | 0.0004571 | 1.085 | +1 | 4 |
| - | - | 1313 | 422.2 | - | - | 0 | - |
| 8 | b | 4868 | 422.7 | 0.0002523 | 0.5968 | +2 | 8 |
| - | - | 2006 | 423.2 | - | - | 0 | - |
| - | - | 2642 | 424.2 | - | - | 0 | - |
| - | - | 1028 | 424.3 | - | - | 0 | - |
| - | - | 1226 | 426.2 | - | - | 0 | - |
| - | - | 2007 | 426.2 | - | - | 0 | - |
| - | - | 1058 | 427.2 | - | - | 0 | - |
| - | - | 1253 | 427.2 | - | - | 0 | - |
| - | - | 7121 | 427.2 | - | - | 0 | - |
| - | - | 1743 | 427.7 | - | - | 0 | - |
| - | - | 1.562E+04 | 428.2 | - | - | 0 | - |
| - | - | 3793 | 428.2 | - | - | 0 | - |
| - | - | 5472 | 429.2 | - | - | 0 | - |
| - | - | 1606 | 429.3 | - | - | 0 | - |
| - | - | 1772 | 430.3 | - | - | 0 | - |
| - | - | 2277 | 433.2 | - | - | 0 | - |
| - | - | 934 | 434.2 | - | - | 0 | - |
| - | - | 1446 | 435.2 | - | - | 0 | - |
| - | - | 1660 | 436.2 | - | - | 0 | - |
| - | - | 3580 | 437.2 | - | - | 0 | - |
| - | - | 6429 | 438.2 | - | - | 0 | - |
| - | - | 1760 | 438.2 | - | - | 0 | - |
| - | - | 3310 | 438.3 | - | - | 0 | - |
| 9 | y | 1.233E+04 | 439.2 | 0.0004209 | 0.9584 | +1 | 4 |
| - | - | 942.5 | 440.2 | - | - | 0 | - |
| - | - | 1923 | 440.2 | - | - | 0 | - |
| - | - | 3809 | 440.3 | - | - | 0 | - |
| - | - | 1700 | 441.2 | - | - | 0 | - |
| - | - | 970.6 | 441.3 | - | - | 0 | - |
| - | - | 1234 | 442.2 | - | - | 0 | - |
| - | - | 1383 | 443.2 | - | - | 0 | - |
| - | - | 773.7 | 443.2 | - | - | 0 | - |
| - | - | 2319 | 443.3 | - | - | 0 | - |
| - | - | 984 | 444.2 | - | - | 0 | - |
| - | - | 2771 | 444.3 | - | - | 0 | - |
| - | - | 1.19E+04 | 445.2 | - | - | 0 | - |
| - | - | 1565 | 446.2 | - | - | 0 | - |
| - | - | 2123 | 446.2 | - | - | 0 | - |
| - | - | 5135 | 448.2 | - | - | 0 | - |
| - | - | 2495 | 448.7 | - | - | 0 | - |
| - | - | 1136 | 449.2 | - | - | 0 | - |
| - | - | 1313 | 450.3 | - | - | 0 | - |
| - | - | 1192 | 452.3 | - | - | 0 | - |
| - | - | 3008 | 453.2 | - | - | 0 | - |
| - | - | 1080 | 454.3 | - | - | 0 | - |
| - | - | 965.4 | 455.2 | - | - | 0 | - |
| - | - | 1.671E+04 | 455.2 | - | - | 0 | - |
| - | - | 956.2 | 455.3 | - | - | 0 | - |
| - | - | 2.153E+04 | 456.2 | - | - | 0 | - |
| - | - | 5156 | 457.2 | - | - | 0 | - |
| 9 | b | 1373 | 457.2 | 0.0006529 | 1.428 | +2 | 9 |
| 9 | b | 1149 | 457.7 | 0.008987 | 19.63 | +2 | 9 |
| - | - | 5810 | 458.3 | - | - | 0 | - |
| - | - | 1176 | 459.3 | - | - | 0 | - |
| - | - | 860.1 | 460.2 | - | - | 0 | - |
| - | - | 1146 | 461.3 | - | - | 0 | - |
| - | - | 2031 | 462.2 | - | - | 0 | - |
| - | - | 1088 | 462.2 | - | - | 0 | - |
| - | - | 3135 | 463.2 | - | - | 0 | - |
| - | - | 938.1 | 463.3 | - | - | 0 | - |
| - | - | 2449 | 464.2 | - | - | 0 | - |
| - | - | 1.311E+04 | 464.3 | - | - | 0 | - |
| - | - | 1151 | 465.2 | - | - | 0 | - |
| - | - | 1404 | 465.3 | - | - | 0 | - |
| - | - | 3473 | 465.3 | - | - | 0 | - |
| - | - | 1741 | 466.2 | - | - | 0 | - |
| - | - | 1807 | 468.3 | - | - | 0 | - |
| - | - | 887.6 | 469.3 | - | - | 0 | - |
| - | - | 6688 | 470.3 | - | - | 0 | - |
| - | - | 4289 | 471.3 | - | - | 0 | - |
| - | - | 989.9 | 472.2 | - | - | 0 | - |
| - | - | 7576 | 472.3 | - | - | 0 | - |
| - | - | 2.743E+04 | 473.2 | - | - | 0 | - |
| - | - | 6248 | 474.2 | - | - | 0 | - |
| - | - | 2028 | 475.3 | - | - | 0 | - |
| - | - | 758.8 | 476.3 | - | - | 0 | - |
| - | - | 2405 | 476.7 | - | - | 0 | - |
| - | - | 2584 | 477.2 | - | - | 0 | - |
| - | - | 1426 | 479.3 | - | - | 0 | - |
| - | - | 2537 | 480.2 | - | - | 0 | - |
| - | - | 4040 | 481.2 | - | - | 0 | - |
| - | - | 5524 | 481.3 | - | - | 0 | - |
| - | - | 1667 | 482.2 | - | - | 0 | - |
| - | - | 6524 | 482.3 | - | - | 0 | - |
| - | - | 1326 | 483.2 | - | - | 0 | - |
| - | - | 1059 | 483.3 | - | - | 0 | - |
| - | - | 4557 | 483.3 | - | - | 0 | - |
| - | - | 1023 | 484.3 | - | - | 0 | - |
| - | - | 3033 | 485.3 | - | - | 0 | - |
| 10 | b | 1.306E+04 | 485.8 | 0.0005781 | 1.19 | +2 | 10 |
| - | - | 6955 | 486.3 | - | - | 0 | - |
| - | - | 1467 | 486.8 | - | - | 0 | - |
| - | - | 867.4 | 488.3 | - | - | 0 | - |
| - | - | 2.483E+04 | 488.3 | - | - | 0 | - |
| - | - | 6493 | 489.3 | - | - | 0 | - |
| - | - | 2607 | 490.2 | - | - | 0 | - |
| - | - | 1697 | 490.3 | - | - | 0 | - |
| - | - | 750.2 | 491.3 | - | - | 0 | - |
| - | - | 3817 | 491.3 | - | - | 0 | - |
| - | - | 2848 | 492.3 | - | - | 0 | - |
| - | - | 1893 | 493.2 | - | - | 0 | - |
| - | - | 1459 | 493.3 | - | - | 0 | - |
| - | - | 859.6 | 494.2 | - | - | 0 | - |
| - | - | 1609 | 495.3 | - | - | 0 | - |
| - | - | 2244 | 497.3 | - | - | 0 | - |
| - | - | 1438 | 497.3 | - | - | 0 | - |
| 5 | b | 5.693E+04 | 498.3 | 0.0005107 | 1.025 | +1 | 5 |
| - | - | 1.36E+04 | 499.3 | - | - | 0 | - |
| - | - | 1700 | 499.8 | - | - | 0 | - |
| - | - | 3361 | 500.3 | - | - | 0 | - |
| - | - | 3068 | 500.3 | - | - | 0 | - |
| - | - | 968.5 | 500.8 | - | - | 0 | - |
| - | - | 968.9 | 501.3 | - | - | 0 | - |
| - | - | 4940 | 507.3 | - | - | 0 | - |
| 8 | y | 7102 | 508.2 | 0.0006247 | 1.229 | +1 | 5 |
| - | - | 1306 | 508.3 | - | - | 0 | - |
| 3 | y | 2718 | 508.8 | 0.00113 | 2.221 | +2 | 10 |
| - | - | 1917 | 509.2 | - | - | 0 | - |
| - | - | 7.359E+04 | 509.3 | - | - | 0 | - |
| - | - | 1031 | 510.2 | - | - | 0 | - |
| - | - | 2.025E+04 | 510.3 | - | - | 0 | - |
| - | - | 2942 | 511.2 | - | - | 0 | - |
| - | - | 3474 | 511.3 | - | - | 0 | - |
| - | - | 950.1 | 512.2 | - | - | 0 | - |
| - | - | 1494 | 512.3 | - | - | 0 | - |
| - | - | 1643 | 513.3 | - | - | 0 | - |
| - | - | 1398 | 515.3 | - | - | 0 | - |
| 5 | b | 6.553E+04 | 516.3 | 0.0003831 | 0.742 | +1 | 5 |
| - | - | 2.099E+04 | 517.3 | - | - | 0 | - |
| - | - | 3337 | 518.3 | - | - | 0 | - |
| - | - | 1654 | 519.3 | - | - | 0 | - |
| - | - | 1663 | 523.3 | - | - | 0 | - |
| - | - | 854.7 | 524.3 | - | - | 0 | - |
| - | - | 3369 | 525.3 | - | - | 0 | - |
| 8 | y | 4.504E+04 | 526.3 | 0.0005275 | 1.002 | +1 | 5 |
| - | - | 1.06E+04 | 527.3 | - | - | 0 | - |
| - | - | 3453 | 528.3 | - | - | 0 | - |
| - | - | 4776 | 528.3 | - | - | 0 | - |
| - | - | 1650 | 529.3 | - | - | 0 | - |
| - | - | 885.9 | 530.3 | - | - | 0 | - |
| - | - | 2201 | 530.3 | - | - | 0 | - |
| - | - | 916.2 | 536.3 | - | - | 0 | - |
| - | - | 1403 | 537.3 | - | - | 0 | - |
| - | - | 1159 | 538.3 | - | - | 0 | - |
| - | - | 1804 | 539.2 | - | - | 0 | - |
| - | - | 2142 | 540.3 | - | - | 0 | - |
| - | - | 2718 | 541.3 | - | - | 0 | - |
| 11 | b | 1467 | 542.3 | 0.001027 | 1.893 | +2 | 11 |
| - | - | 2102 | 543.3 | - | - | 0 | - |
| - | - | 702.8 | 544.3 | - | - | 0 | - |
| - | - | 808.8 | 544.3 | - | - | 0 | - |
| - | - | 1344 | 548.8 | - | - | 0 | - |
| - | - | 1387 | 549.3 | - | - | 0 | - |
| - | - | 2783 | 550.3 | - | - | 0 | - |
| - | - | 2341 | 551.3 | - | - | 0 | - |
| - | - | 1354 | 551.3 | - | - | 0 | - |
| - | - | 2468 | 554.3 | - | - | 0 | - |
| - | - | 1138 | 555.3 | - | - | 0 | - |
| - | - | 1658 | 556.3 | - | - | 0 | - |
| - | - | 1071 | 557.3 | - | - | 0 | - |
| - | - | 868.6 | 557.3 | - | - | 0 | - |
| - | - | 1.048E+04 | 558.3 | - | - | 0 | - |
| - | - | 3025 | 559.3 | - | - | 0 | - |
| - | - | 1157 | 560.3 | - | - | 0 | - |
| - | - | 3525 | 566.3 | - | - | 0 | - |
| - | - | 983.4 | 567.3 | - | - | 0 | - |
| - | - | 8007 | 568.3 | - | - | 0 | - |
| - | - | 4210 | 569.3 | - | - | 0 | - |
| - | - | 2167 | 570.3 | - | - | 0 | - |
| - | - | 1040 | 571.3 | - | - | 0 | - |
| - | - | 1444 | 572.2 | - | - | 0 | - |
| - | - | 2087 | 572.3 | - | - | 0 | - |
| - | - | 922.9 | 573.2 | - | - | 0 | - |
| - | - | 2180 | 573.3 | - | - | 0 | - |
| - | - | 817.7 | 574.3 | - | - | 0 | - |
| - | - | 1356 | 575.3 | - | - | 0 | - |
| - | - | 2804 | 576.3 | - | - | 0 | - |
| - | - | 2671 | 577.3 | - | - | 0 | - |
| - | - | 844.3 | 577.4 | - | - | 0 | - |
| - | - | 7207 | 578.3 | - | - | 0 | - |
| - | - | 3357 | 579.3 | - | - | 0 | - |
| - | - | 933.8 | 580.3 | - | - | 0 | - |
| - | - | 1073 | 582.3 | - | - | 0 | - |
| - | - | 727.5 | 583.3 | - | - | 0 | - |
| - | - | 4449 | 583.4 | - | - | 0 | - |
| - | - | 8327 | 584.3 | - | - | 0 | - |
| - | - | 885.2 | 585.3 | - | - | 0 | - |
| - | - | 3842 | 585.3 | - | - | 0 | - |
| - | - | 2.447E+04 | 586.3 | - | - | 0 | - |
| - | - | 7018 | 587.3 | - | - | 0 | - |
| - | - | 1986 | 590.3 | - | - | 0 | - |
| - | - | 5959 | 592.3 | - | - | 0 | - |
| - | - | 1175 | 593.3 | - | - | 0 | - |
| - | - | 2440 | 593.3 | - | - | 0 | - |
| - | - | 3550 | 594.3 | - | - | 0 | - |
| - | - | 1727 | 595.3 | - | - | 0 | - |
| - | - | 2.021E+04 | 596.3 | - | - | 0 | - |
| - | - | 6115 | 597.3 | - | - | 0 | - |
| - | - | 1279 | 598.3 | - | - | 0 | - |
| - | - | 8198 | 601.4 | - | - | 0 | - |
| - | - | 3098 | 602.4 | - | - | 0 | - |
| - | - | 1770 | 603.3 | - | - | 0 | - |
| - | - | 2083 | 609.3 | - | - | 0 | - |
| - | - | 3778 | 610.3 | - | - | 0 | - |
| - | - | 2451 | 610.4 | - | - | 0 | - |
| 6 | b | 4.782E+04 | 611.4 | 0.0003091 | 0.5055 | +1 | 6 |
| - | - | 1.483E+04 | 612.4 | - | - | 0 | - |
| - | - | 2600 | 613.4 | - | - | 0 | - |
| - | - | 2346 | 618.3 | - | - | 0 | - |
| - | - | 1944 | 619.3 | - | - | 0 | - |
| - | - | 1198 | 623.3 | - | - | 0 | - |
| - | - | 1087 | 624.3 | - | - | 0 | - |
| - | - | 950.9 | 625.3 | - | - | 0 | - |
| - | - | 2860 | 627.3 | - | - | 0 | - |
| - | - | 1217 | 628.3 | - | - | 0 | - |
| - | - | 1017 | 628.4 | - | - | 0 | - |
| 6 | b | 4.253E+04 | 629.4 | 5.931E-05 | 0.09424 | +1 | 6 |
| - | - | 1.586E+04 | 630.4 | - | - | 0 | - |
| - | - | 856.3 | 630.6 | - | - | 0 | - |
| - | - | 2493 | 631.4 | - | - | 0 | - |
| - | - | 1080 | 635.4 | - | - | 0 | - |
| 7 | y | 2.184E+04 | 636.3 | 0.0004273 | 0.6716 | +1 | 6 |
| - | - | 1068 | 636.4 | - | - | 0 | - |
| - | - | 7533 | 637.3 | - | - | 0 | - |
| - | - | 1490 | 638.3 | - | - | 0 | - |
| - | - | 1151 | 640.4 | - | - | 0 | - |
| - | - | 2027 | 641.3 | - | - | 0 | - |
| - | - | 1.354E+04 | 641.9 | - | - | 0 | - |
| - | - | 9034 | 642.4 | - | - | 0 | - |
| - | - | 4225 | 642.9 | - | - | 0 | - |
| - | - | 2202 | 643.3 | - | - | 0 | - |
| - | - | 1872 | 644.3 | - | - | 0 | - |
| - | - | 985 | 646.3 | - | - | 0 | - |
| - | - | 2616 | 646.4 | - | - | 0 | - |
| - | - | 4534 | 647.4 | - | - | 0 | - |
| - | - | 2221 | 648.3 | - | - | 0 | - |
| - | - | 1320 | 649.4 | - | - | 0 | - |
| - | - | 810.6 | 651.3 | - | - | 0 | - |
| 7 | y | 4.47E+04 | 654.3 | 0.0002996 | 0.4579 | +1 | 6 |
| - | - | 1.377E+04 | 655.3 | - | - | 0 | - |
| - | - | 3188 | 656.3 | - | - | 0 | - |
| - | - | 3611 | 656.4 | - | - | 0 | - |
| - | - | 2517 | 657.4 | - | - | 0 | - |
| - | - | 1073 | 658.4 | - | - | 0 | - |
| - | - | 993.4 | 659.4 | - | - | 0 | - |
| - | - | 1449 | 660.3 | - | - | 0 | - |
| - | - | 6894 | 665.4 | - | - | 0 | - |
| - | - | 2896 | 666.4 | - | - | 0 | - |
| - | - | 2391 | 667.4 | - | - | 0 | - |
| - | - | 2472 | 668.4 | - | - | 0 | - |
| - | - | 1982 | 669.4 | - | - | 0 | - |
| - | - | 890.7 | 672.3 | - | - | 0 | - |
| - | - | 887 | 677.4 | - | - | 0 | - |
| - | - | 1071 | 678.4 | - | - | 0 | - |
| - | - | 769.3 | 682.6 | - | - | 0 | - |
| - | - | 8111 | 683.4 | - | - | 0 | - |
| - | - | 2572 | 684.4 | - | - | 0 | - |
| - | - | 1653 | 685.3 | - | - | 0 | - |
| - | - | 7815 | 685.4 | - | - | 0 | - |
| - | - | 1311 | 686.3 | - | - | 0 | - |
| - | - | 2663 | 686.4 | - | - | 0 | - |
| - | - | 918.8 | 687.4 | - | - | 0 | - |
| - | - | 1643 | 694.4 | - | - | 0 | - |
| - | - | 2690 | 695.4 | - | - | 0 | - |
| - | - | 1363 | 696.4 | - | - | 0 | - |
| - | - | 1271 | 697.4 | - | - | 0 | - |
| - | - | 1109 | 698.4 | - | - | 0 | - |
| - | - | 4533 | 703.3 | - | - | 0 | - |
| - | - | 9179 | 704.4 | - | - | 0 | - |
| - | - | 5653 | 705.4 | - | - | 0 | - |
| - | - | 1883 | 706.4 | - | - | 0 | - |
| - | - | 1436 | 711.4 | - | - | 0 | - |
| - | - | 5439 | 712.4 | - | - | 0 | - |
| - | - | 2507 | 713.4 | - | - | 0 | - |
| - | - | 975.9 | 719.4 | - | - | 0 | - |
| - | - | 2323 | 721.4 | - | - | 0 | - |
| - | - | 3.846E+04 | 722.4 | - | - | 0 | - |
| - | - | 1.84E+04 | 723.4 | - | - | 0 | - |
| - | - | 4426 | 724.4 | - | - | 0 | - |
| - | - | 1788 | 725.4 | - | - | 0 | - |
| - | - | 829.4 | 726.4 | - | - | 0 | - |
| - | - | 2763 | 729.4 | - | - | 0 | - |
| - | - | 4707 | 729.4 | - | - | 0 | - |
| - | - | 925.3 | 730.4 | - | - | 0 | - |
| - | - | 1481 | 730.4 | - | - | 0 | - |
| - | - | 2402 | 731.4 | - | - | 0 | - |
| - | - | 1669 | 732.4 | - | - | 0 | - |
| - | - | 915.6 | 737.4 | - | - | 0 | - |
| 7 | b | 1.719E+04 | 739.4 | 0.0002032 | 0.2748 | +1 | 7 |
| 7 | b | 6.55E+04 | 740.4 | 0.002428 | 3.279 | +1 | 7 |
| - | - | 2.455E+04 | 741.4 | - | - | 0 | - |
| - | - | 4716 | 742.4 | - | - | 0 | - |
| - | - | 1001 | 743.4 | - | - | 0 | - |
| - | - | 1227 | 747.4 | - | - | 0 | - |
| 6 | y | 6398 | 749.4 | 0.0005308 | 0.7083 | +1 | 7 |
| 6 | y | 3860 | 750.4 | 0.01072 | 14.28 | +1 | 7 |
| - | - | 1221 | 751.4 | - | - | 0 | - |
| - | - | 1044 | 754.4 | - | - | 0 | - |
| - | - | 3881 | 755.4 | - | - | 0 | - |
| - | - | 2511 | 756.4 | - | - | 0 | - |
| 7 | b | 3.236E+04 | 757.4 | 0.0002296 | 0.3032 | +1 | 7 |
| - | - | 918.5 | 758.4 | - | - | 0 | - |
| - | - | 1.185E+04 | 758.4 | - | - | 0 | - |
| - | - | 1742 | 759.4 | - | - | 0 | - |
| - | - | 1193 | 766.4 | - | - | 0 | - |
| 6 | y | 5.183E+04 | 767.4 | 0.00022 | 0.2867 | +1 | 7 |
| - | - | 2.169E+04 | 768.4 | - | - | 0 | - |
| - | - | 4942 | 769.4 | - | - | 0 | - |
| - | - | 860.8 | 770.4 | - | - | 0 | - |
| - | - | 885.6 | 773.4 | - | - | 0 | - |
| - | - | 808.4 | 784.4 | - | - | 0 | - |
| - | - | 1644 | 785.4 | - | - | 0 | - |
| - | - | 912.8 | 786.4 | - | - | 0 | - |
| - | - | 941.3 | 796.4 | - | - | 0 | - |
| - | - | 825.7 | 802.4 | - | - | 0 | - |
| - | - | 1166 | 805.4 | - | - | 0 | - |
| - | - | 961.3 | 807.5 | - | - | 0 | - |
| - | - | 4542 | 808.4 | - | - | 0 | - |
| - | - | 2163 | 809.4 | - | - | 0 | - |
| - | - | 865.3 | 813.4 | - | - | 0 | - |
| - | - | 2351 | 816.4 | - | - | 0 | - |
| - | - | 3422 | 817.5 | - | - | 0 | - |
| - | - | 2139 | 818.4 | - | - | 0 | - |
| - | - | 1396 | 819.5 | - | - | 0 | - |
| - | - | 4642 | 823.4 | - | - | 0 | - |
| - | - | 3290 | 824.4 | - | - | 0 | - |
| - | - | 1190 | 825.4 | - | - | 0 | - |
| - | - | 2.078E+04 | 825.5 | - | - | 0 | - |
| 8 | b | 6454 | 826.4 | 0.002355 | 2.849 | +1 | 8 |
| - | - | 5616 | 826.5 | - | - | 0 | - |
| - | - | 3686 | 827.5 | - | - | 0 | - |
| - | - | 906.5 | 828.4 | - | - | 0 | - |
| - | - | 3154 | 834.4 | - | - | 0 | - |
| - | - | 1.234E+04 | 835.5 | - | - | 0 | - |
| - | - | 8435 | 836.5 | - | - | 0 | - |
| - | - | 3913 | 837.5 | - | - | 0 | - |
| - | - | 1042 | 838.5 | - | - | 0 | - |
| - | - | 1987 | 841.4 | - | - | 0 | - |
| - | - | 3860 | 842.4 | - | - | 0 | - |
| - | - | 887.4 | 843.4 | - | - | 0 | - |
| 8 | b | 1.253E+04 | 844.5 | 0.0007029 | 0.8323 | +1 | 8 |
| - | - | 4981 | 845.5 | - | - | 0 | - |
| - | - | 1453 | 846.5 | - | - | 0 | - |
| 5 | y | 1112 | 848.5 | 0.001355 | 1.597 | +1 | 8 |
| - | - | 5.865E+04 | 853.5 | - | - | 0 | - |
| - | - | 2.634E+04 | 854.5 | - | - | 0 | - |
| - | - | 7185 | 855.5 | - | - | 0 | - |
| - | - | 2076 | 860.5 | - | - | 0 | - |
| - | - | 1805 | 861.5 | - | - | 0 | - |
| 5 | y | 5850 | 866.5 | 0.0004449 | 0.5134 | +1 | 8 |
| - | - | 3454 | 867.5 | - | - | 0 | - |
| - | - | 1763 | 868.5 | - | - | 0 | - |
| - | - | 1341 | 869.5 | - | - | 0 | - |
| - | - | 1238 | 877.5 | - | - | 0 | - |
| - | - | 1227 | 887.5 | - | - | 0 | - |
| - | - | 960.6 | 888.5 | - | - | 0 | - |
| - | - | 910.2 | 889.4 | - | - | 0 | - |
| - | - | 4397 | 895.5 | - | - | 0 | - |
| - | - | 1887 | 896.5 | - | - | 0 | - |
| - | - | 989.7 | 903.5 | - | - | 0 | - |
| 9 | b | 4944 | 913.5 | 0.000255 | 0.2791 | +1 | 9 |
| 9 | b | 4089 | 914.5 | 0.0178 | 19.47 | +1 | 9 |
| - | - | 2410 | 915.5 | - | - | 0 | - |
| - | - | 1016 | 925.5 | - | - | 0 | - |
| - | - | 1464 | 926.5 | - | - | 0 | - |
| 9 | b | 5592 | 931.5 | 0.0008099 | 0.8695 | +1 | 9 |
| - | - | 1766 | 932.5 | - | - | 0 | - |
| - | - | 1164 | 933.5 | - | - | 0 | - |
| - | - | 1155 | 934.5 | - | - | 0 | - |
| - | - | 1417 | 935.5 | - | - | 0 | - |
| - | - | 5216 | 936.5 | - | - | 0 | - |
| 4 | y | 3519 | 937.5 | 0.01524 | 16.25 | +1 | 9 |
| - | - | 1841 | 938.5 | - | - | 0 | - |
| - | - | 3320 | 942.5 | - | - | 0 | - |
| - | - | 1806 | 943.5 | - | - | 0 | - |
| - | - | 1952 | 944.5 | - | - | 0 | - |
| - | - | 930.3 | 945.5 | - | - | 0 | - |
| - | - | 1.376E+04 | 952.5 | - | - | 0 | - |
| - | - | 9310 | 953.5 | - | - | 0 | - |
| - | - | 4508 | 954.5 | - | - | 0 | - |
| - | - | 1707 | 955.5 | - | - | 0 | - |
| - | - | 1200 | 956.5 | - | - | 0 | - |
| - | - | 3642 | 960.5 | - | - | 0 | - |
| - | - | 2273 | 961.5 | - | - | 0 | - |
| 10 | b | 6.414E+04 | 970.5 | 0.0004174 | 0.4301 | +1 | 10 |
| 10 | b | 3.961E+04 | 971.5 | 0.01691 | 17.41 | +1 | 10 |
| - | - | 9929 | 972.5 | - | - | 0 | - |
| - | - | 2057 | 973.5 | - | - | 0 | - |
| - | - | 1185 | 980.5 | - | - | 0 | - |
| - | - | 993.3 | 981.5 | - | - | 0 | - |
| - | - | 1055 | 982.5 | - | - | 0 | - |
| 10 | b | 5.781E+04 | 988.5 | 0.0004841 | 0.4897 | +1 | 10 |
| - | - | 3.066E+04 | 989.5 | - | - | 0 | - |
| - | - | 9101 | 990.5 | - | - | 0 | - |
| - | - | 2408 | 998.5 | - | - | 0 | - |
| - | - | 1417 | 999.5 | - | - | 0 | - |
| - | - | 2008 | 1001 | - | - | 0 | - |
| - | - | 1127 | 1002 | - | - | 0 | - |
| - | - | 888.2 | 1003 | - | - | 0 | - |
| - | - | 976.5 | 1004 | - | - | 0 | - |
| 3 | y | 6308 | 1017 | 0.0005342 | 0.5255 | +1 | 10 |
| 3 | y | 5001 | 1018 | 0.01502 | 14.76 | +1 | 10 |
| - | - | 2036 | 1019 | - | - | 0 | - |
| 3 | y | 4.299E+04 | 1035 | 0.0005398 | 0.5218 | +1 | 10 |
| - | - | 2.642E+04 | 1036 | - | - | 0 | - |
| - | - | 8393 | 1037 | - | - | 0 | - |
| - | - | 877.1 | 1038 | - | - | 0 | - |
| - | - | 957.3 | 1039 | - | - | 0 | - |
| - | - | 2004 | 1056 | - | - | 0 | - |
| - | - | 2058 | 1057 | - | - | 0 | - |
| - | - | 1319 | 1058 | - | - | 0 | - |
| - | - | 4074 | 1066 | - | - | 0 | - |
| - | - | 4381 | 1067 | - | - | 0 | - |
| - | - | 2022 | 1068 | - | - | 0 | - |
| - | - | 8194 | 1074 | - | - | 0 | - |
| - | - | 5520 | 1075 | - | - | 0 | - |
| - | - | 1717 | 1076 | - | - | 0 | - |
| 11 | b | 2.818E+04 | 1084 | 0.0006801 | 0.6277 | +1 | 11 |
| 11 | b | 2.25E+04 | 1085 | 0.01567 | 14.45 | +1 | 11 |
| - | - | 7810 | 1086 | - | - | 0 | - |
| - | - | 1051 | 1087 | - | - | 0 | - |
| 11 | b | 3.064E+04 | 1102 | 0.0005026 | 0.4563 | +1 | 11 |
| - | - | 1450 | 1102 | - | - | 0 | - |
| - | - | 2.126E+04 | 1103 | - | - | 0 | - |
| - | - | 5946 | 1104 | - | - | 0 | - |
| - | - | 863.3 | 1821 | - | - | 0 | - |
| - | - | 733.2 | 2233 | - | - | 0 | - |
| - | - | 849.9 | 2341 | - | - | 0 | - |
| - | - | 749.8 | 3139 | - | - | 0 | - |

m/z Charge Intensity FragmentType MassShift Position
120.05299377441406 0 1195.0677
120.08106231689453 0 47512.883
120.76138305664062 0 468.19653
121.06510925292969 0 709.46954
121.08436584472656 0 4208.819
122.90186309814453 0 458.2497
123.04434204101562 0 25410.729
124.0477294921875 0 1553.516
125.10774993896484 0 516.26355
126.05514526367188 0 1612.213
127.05052947998047 0 2247.4153
127.08692169189453 0 1123.4746
129.06607055664062 0 4575.209
129.10250854492188 0 2838.1333
130.0497283935547 0 616.39606
130.06521606445312 0 1678.5962
130.08689880371094 0 481.76926
131.11817932128906 0 9032.119
132.0811309814453 0 3015.1987
132.1023712158203 0 607.08075
133.0854949951172 0 573.5054
135.7295379638672 0 511.48108
136.07601928710938 0 213959.78
137.05984497070312 0 559.84235
137.07347106933594 0 878.3122
137.079345703125 0 15939.784
138.1282501220703 0 619.1209
139.05038452148438 0 1308.3066
140.1437225341797 0 1578.4271
141.10252380371094 0 60234.98
142.1005096435547 0 612.33185
142.10592651367188 0 3777.3286
143.1181640625 0 54469.777
144.11587524414062 0 703.6949
144.1215362548828 0 3041.577
144.18992614746094 0 447.96948
145.06109619140625 0 4298.4736
146.09742736816406 0 602.4109
147.04429626464844 0 19409.29
147.07672119140625 0 2798.1948
148.04769897460938 0 1675.7054
148.95526123046875 0 488.36624
153.06622314453125 0 3024.5225
154.15951538085938 0 1089.8666
155.0816192626953 0 1503.495
155.1180877685547 0 953.3183
156.07717895507812 0 1098.2876
157.06103515625 0 15400.682
157.09710693359375 0 816.1809
158.09664916992188 0 10185.7295
159.0998077392578 0 1152.4761
160.075927734375 0 1041.7888
160.10598754882812 0 577.73785
160.11241149902344 0 3744.2764
162.0876922607422 0 906.5908
165.05491638183594 0 206915.62
166.0582733154297 0 18925.16
166.0865020751953 0 1042.2657
167.06069946289062 0 783.4713
167.08209228515625 0 692.15015
168.0769500732422 0 859.0255
169.06103515625 0 3386.3325
169.09747314453125 0 241534.97
170.09292602539062 0 4960.9834
170.100830078125 0 19786.41
171.0767822265625 0 3451.6287
171.11305236816406 0 25089.115
172.11660766601562 0 1301.5203
173.12864685058594 0 597.00684
173.4513397216797 0 2965.281
175.07151794433594 0 5936.501
176.0387725830078 0 534.4655
176.1072998046875 0 19576.379
177.0944061279297 0 1006.0283
177.10247802734375 0 12824.878
177.1111297607422 0 2505.4558
177.13119506835938 0 1105.8827
178.10595703125 0 1768.1272
180.07711791992188 0 2099.3894
181.06112670898438 0 7100.6807
181.097412109375 0 1799.2738
182.08152770996094 0 398706.16 y 11
183.08480834960938 0 33440.668
183.11317443847656 0 1312.8707
183.1497344970703 0 837.5786
184.0866241455078 0 2639.77
185.12814331054688 0 1162.4419
185.1651153564453 0 11806.298
186.0911407470703 0 758.1127
186.1239776611328 0 12339.76
186.16864013671875 0 874.4866
187.10781860351562 0 1638.9243
187.128173828125 0 709.2684
188.10337829589844 0 3589.5183
189.123779296875 0 1637.3984
191.11770629882812 0 906.49066
193.09722900390625 0 1218.1501
194.09190368652344 0 556.11334
195.11297607421875 0 835.4571
196.0721435546875 0 2415.188
197.05581665039062 0 955.7511
197.12863159179688 0 3057.378
198.08755493164062 0 11639.224
198.12367248535156 0 639.12744
199.0715789794922 0 5440.961
199.09146118164062 0 1073.9406
199.10781860351562 0 1101.301
199.1808624267578 0 1733.8794
201.12384033203125 0 1961.456
203.11813354492188 0 6209.11 a Water loss 1
204.1215057373047 0 771.914
205.0973663330078 0 1277.7681
206.09262084960938 0 664.31903
207.1131134033203 0 911.393
208.09707641601562 0 2661.59
208.10714721679688 0 1009.1078
209.16494750976562 0 949.0687
211.1080322265625 0 1493.7605 y Water loss 8
211.14425659179688 0 1804.1804
211.180419921875 0 880.9474
212.1031036376953 0 1799.5912
213.08700561523438 0 666.0732
213.1236114501953 0 5469.231
213.16009521484375 0 11398.919
214.08250427246094 0 48097.656
214.15451049804688 0 1421.0607
214.1647491455078 0 834.8234
215.08560180664062 0 3334.2544
215.13919067382812 0 1494.0168
216.08653259277344 0 1086.5101
216.09812927246094 0 14111.388
217.10189819335938 0 1065.1122
217.1337890625 0 5536.7373
219.13478088378906 0 821.3421
221.12879943847656 0 333462.6 a 1
222.13204956054688 0 40197.246
223.10800170898438 0 2666.9785
223.13421630859375 0 2846.465
223.14466857910156 0 3636.127
224.10324096679688 0 3706.7495
225.1236114501953 0 8350.297
226.11895751953125 0 4472.2246
228.0983428955078 0 1251.7347
230.11351013183594 0 832.3955
230.15061950683594 0 3550.7659
230.1868438720703 0 1454.9994
231.11314392089844 0 3852.2444 b Water loss 1
232.09315490722656 0 38836.94
233.0968780517578 0 2674.1665
233.12872314453125 0 976.8334
233.16567993164062 0 982.584
234.12393188476562 0 777.57745
235.10833740234375 0 905.6391
237.16015625 0 818.9725
238.11891174316406 0 1814.1
239.15097045898438 0 897.0481
239.1754150390625 0 1718.2075
240.0983123779297 0 2738.004
240.13441467285156 0 2777.5002
240.17095947265625 0 37713.105
241.16038513183594 0 1618.0317
241.17449951171875 0 4465.0166
242.11390686035156 0 1258.6823
242.15025329589844 0 40056.688
243.13430786132812 0 1425.066
243.1530303955078 0 4100.1733
244.12942504882812 0 568.7979
244.15335083007812 0 780.0851
245.12879943847656 0 10792.893
246.10931396484375 0 700.3374
246.1320343017578 0 1769.8202
249.12368774414062 0 111703.67 b 1
249.1603546142578 0 3025.0544
250.08255004882812 0 9660.982
250.1269989013672 0 14033.312
251.08648681640625 0 1438.2577
251.17608642578125 0 896.17017
252.1343994140625 0 4809.8643
254.15013122558594 0 3271.2886
255.15780639648438 0 6177.3926
255.65985107421875 0 1814.9692
256.1294250488281 0 4547.5693
256.1630859375 0 613.3725
258.1099548339844 0 1158.2505
258.14501953125 0 24450.34
259.14776611328125 0 3477.7068
261.1234130859375 0 693.83545
262.0830078125 0 892.754
262.15570068359375 0 777.60315
263.1022033691406 0 882.947
263.1388244628906 0 981.799
266.1495361328125 0 2001.7386
267.10906982421875 0 6066.656
268.09320068359375 0 6081.7544
268.1658020019531 0 227143.34
269.1689147949219 0 29937.617
270.1448059082031 0 4282.3223
270.1717224121094 0 2145.2017
271.1443176269531 0 1995.5046
272.16162109375 0 831.2324
273.1589660644531 0 1203.6705
274.1187744140625 0 688.9848
274.4502258300781 0 704.1036
275.1387634277344 0 2059.7778
275.1755065917969 0 1953.3115
276.1355895996094 0 983.20013
277.1181640625 0 1223.2375
279.1094665527344 0 1587.8157
279.1454772949219 0 978.20123
280.0931091308594 0 1830.0844
280.16607666015625 0 2397.3557
281.1616516113281 0 1788.3499
282.1454162597656 0 3043.1475
282.1814880371094 0 5750.675
282.2177734375 0 1103.7721
284.13916015625 0 742.48425
284.1610412597656 0 2488.0537
284.19720458984375 0 3792.9124
285.1200256347656 0 5589.0537
285.1585388183594 0 2073.8486
285.1926574707031 0 4664.3857
286.1040344238281 0 2935.9055
286.12139892578125 0 850.86487
286.17681884765625 0 3190.4417
286.1953125 0 525.6406
287.10748291015625 0 634.9988
288.1344909667969 0 789.6735
289.1914367675781 0 2841.9673
289.6690368652344 0 2172.3281
290.17047119140625 0 1019.4912
290.1934509277344 0 612.7684
291.132080078125 0 664.7124
292.16552734375 0 1177.4104 d 2
294.10931396484375 0 1451.8806
294.145263671875 0 3027.254
294.1817321777344 0 1500.0466
295.1407470703125 0 10148.054
295.1656494140625 0 56240.66 y 10
296.1231994628906 0 615.24884
296.1428527832031 0 1207.8778
296.1689147949219 0 8817.757
296.1976013183594 0 5432.3306
297.1197509765625 0 3854.2334
297.15618896484375 0 2259.1252
297.1740417480469 0 777.5671
297.1941223144531 0 2240.6155
298.1551818847656 0 1915.5193
298.6743469238281 0 3089.4297
299.17205810546875 0 8566.579
300.15576171875 0 3894.1406
300.17401123046875 0 1438.4501 a Water loss 2
301.1878356933594 0 685.38
301.2219543457031 0 990.50854
302.1498107910156 0 862.034
302.18646240234375 0 959.3311
303.13043212890625 0 3156.8103
303.1708068847656 0 2155.98
304.1321105957031 0 760.36017
306.1814880371094 0 1831.2798 b Water loss 5
307.1042785644531 0 1689.2404
307.140380859375 0 871.65594
308.12420654296875 0 949.9596
309.1559753417969 0 2746.894
309.2362365722656 0 601.98267
310.1390380859375 0 1588.1803
310.1588134765625 0 946.16986
310.17755126953125 0 1638.3762
310.2129211425781 0 3916.7969
311.1364440917969 0 955.77826
311.17236328125 0 4817.06
312.1344299316406 0 1188.8986
312.1559753417969 0 3723.3435
312.1739807128906 0 934.097
312.1919860839844 0 782.8877
313.1157531738281 0 775.2201
313.15142822265625 0 3598.1118
314.15081787109375 0 978.88153
314.20611572265625 0 574.7402
315.13043212890625 0 2589.1462
315.187255859375 0 1551.4541 b 5
315.6661071777344 0 3192.8518
315.686279296875 0 937.08875
316.1661682128906 0 15663.923
316.19873046875 0 677.05084
317.1826477050781 0 59935.934
318.1447448730469 0 1545.2263
318.1858215332031 0 8583.746 a 2
319.18743896484375 0 1005.6855
320.1592712402344 0 884.2908
320.19439697265625 0 752.8138
323.1714172363281 0 631.837
323.2082824707031 0 1099.4773
324.1307678222656 0 4942.9443
324.17962646484375 0 14100.39
324.68048095703125 0 4932.647
325.11468505859375 0 14298.251
325.1512451171875 0 1692.8976
325.18707275390625 0 7080.649
326.1178894042969 0 2138.3928
326.1529235839844 0 864.27765
327.1667785644531 0 9303.077
327.2030029296875 0 1643.7474
328.16650390625 0 11344.227 b Water loss 2
329.1824035644531 0 10602.95
330.1454162597656 0 842.9993
330.1835021972656 0 3682.0818
331.1612854003906 0 658.60486
333.16265869140625 0 799.235
333.1865539550781 0 3912.5652
333.6852722167969 0 878.9621
334.17669677734375 0 1532.1641
336.1554260253906 0 1364.4233
337.1522521972656 0 1735.1528
337.1875915527344 0 1780.7318
339.1672058105469 0 994.8896
339.202880859375 0 2384.0156
341.18243408203125 0 1686.6521
341.21875 0 11437.607
342.1414794921875 0 12270.282
342.2207946777344 0 2952.6475
343.1251525878906 0 24323.805
343.14520263671875 0 1995.8894
343.19744873046875 0 1574.0266
344.128173828125 0 3188.2952
344.1973571777344 0 4205.8975
345.1772155761719 0 66142.555
346.1776428222656 0 26412.506 b 2
347.1346435546875 0 717.4217
347.18023681640625 0 5217.9937
348.19195556640625 0 2299.727
349.1934814453125 0 745.0676
350.21795654296875 0 1063.2056
351.1669921875 0 1226.3215
351.2029724121094 0 20461.527
352.1270446777344 0 593.09033
352.18701171875 0 14379.983 y 9
352.2064514160156 0 2811.8213
352.6901550292969 0 3328.7905
353.18719482421875 0 5223.955
353.25506591796875 0 32164.223
353.6828308105469 0 1148
354.2577819824219 0 5486.426
355.16082763671875 0 1149.3107
355.1981506347656 0 3266.4011
355.2607727050781 0 828.87854
356.19976806640625 0 1401.7732
357.1770935058594 0 1090.9403
358.17626953125 0 1188.8407
358.2098083496094 0 672.0293
359.1706848144531 0 2524.3345
360.1517639160156 0 7348.3306
360.17529296875 0 953.5834
361.1545715332031 0 1453.0236
361.6954650878906 0 25996.326
362.19830322265625 0 9996.737
362.6960144042969 0 1422.99
363.1662292480469 0 1275.8942
364.1617431640625 0 1599.9534
365.1455078125 0 2343.6184
365.19549560546875 0 696.7977
365.2189636230469 0 2929.3806
366.2138977050781 0 1337.2213
367.16107177734375 0 1941.5352
367.2345886230469 0 2721.213
368.1931457519531 0 1215.693
369.17779541015625 0 756.3953
369.1845397949219 0 845.9094
369.21331787109375 0 11046.223
370.13714599609375 0 1654.3905
370.2138366699219 0 6876.39 b Water loss 6
370.70074462890625 0 3782.2676 b Ammonia loss 6
371.2069091796875 0 1287.0919
372.1915283203125 0 2429.8389
373.1880187988281 0 1984.7295
373.2224426269531 0 1241.9895
375.1669616699219 0 1807.9305
377.179931640625 0 1940.1714
378.2140197753906 0 1943.9468
379.1970520019531 0 4571.562
379.217529296875 0 3092.7788 b 6
379.71746826171875 0 1413.3363
380.1949462890625 0 3987.1345
381.2499084472656 0 63992.48
382.1733093261719 0 3130.7275
382.2528381347656 0 12080.65
383.1989440917969 0 915.96576
383.22930908203125 0 2981.5574
383.2555847167969 0 1180.7454
384.1888122558594 0 987.3304
384.2282409667969 0 1448.6415
385.1287841796875 0 1682.6184
386.2039489746094 0 1171.3774
387.16802978515625 0 1180.5774
387.2393798828125 0 9985.271
388.24249267578125 0 1661.4169
389.2188720703125 0 7018.2266
390.2213134765625 0 839.20734
392.19207763671875 0 1603.7777
393.1763000488281 0 1888.4491
393.2500915527344 0 731.3625
394.2091369628906 0 11629.277
395.2104187011719 0 1026.755
395.23553466796875 0 1337.7965
396.2244567871094 0 2276.0496
397.246826171875 0 1150.2955
398.20391845703125 0 5461.619
398.2469177246094 0 763.55005
398.2767028808594 0 18407.387
399.20318603515625 0 27029.77 b Water loss 3
399.2558288574219 0 1161.2706
399.28094482421875 0 2613.2864
400.2064208984375 0 5248.7236
401.2184143066406 0 1027.6171
404.2400817871094 0 1203.8213
405.1776428222656 0 1036.9791
406.2124328613281 0 963.0672
407.1917419433594 0 732.94867
408.1521911621094 0 960.3002
408.22406005859375 0 1183.4882
409.2218322753906 0 960.25977
410.20416259765625 0 4401.388
410.240234375 0 4890.986
410.2762756347656 0 2083.5298
411.24383544921875 0 809.1993
412.1856384277344 0 1253.8083
412.21978759765625 0 4472.5654
412.2555847167969 0 2592.1716
413.2449645996094 0 2247.3271
413.7486877441406 0 935.8055
414.19830322265625 0 852.33905
415.2345886230469 0 12303.824
415.2669982910156 0 1036.2931
416.2131652832031 0 2533.176
416.2378234863281 0 1190.5961
417.2137451171875 0 43293.47 b 3
418.2165222167969 0 8714.671
419.2267761230469 0 1896.491
420.1893005371094 0 1711.9655
420.224853515625 0 1049.8774
421.2086181640625 0 2121.0693 y Water loss 8
422.24072265625 0 1312.9409
422.7320556640625 0 4867.774 b 7
423.2295227050781 0 2005.5176
424.2204895019531 0 2641.613
424.2549133300781 0 1027.6125
426.1629638671875 0 1225.6565
426.23468017578125 0 2007.3383
427.1644287109375 0 1058.0685
427.1987609863281 0 1253.0267
427.2337951660156 0 7120.705
427.74432373046875 0 1742.76
428.21453857421875 0 15622.241
428.2491760253906 0 3792.6157
429.2162170410156 0 5471.545
429.2516174316406 0 1605.7074
430.2644348144531 0 1772.1156
433.24554443359375 0 2277.1045
434.2441711425781 0 933.9945
435.2357482910156 0 1445.6841
436.2201843261719 0 1659.7877
437.21600341796875 0 3579.5112
438.1991882324219 0 6428.519
438.23541259765625 0 1760.209
438.2718811035156 0 3310.2832
439.2191467285156 0 12327.557 y 8
440.17877197265625 0 942.4726
440.2214050292969 0 1923.1511
440.2516784667969 0 3808.6335
441.2169494628906 0 1700.3987
441.2533874511719 0 970.6214
442.22955322265625 0 1233.7606
443.1923828125 0 1383.2081
443.2335205078125 0 773.74585
443.26654052734375 0 2319.3027
444.191162109375 0 984.0254
444.26055908203125 0 2771.1262
445.24151611328125 0 11902.601
446.20404052734375 0 1565.1061
446.243896484375 0 2123.3235
448.2374267578125 0 5134.729
448.7396545410156 0 2494.5544
449.2410583496094 0 1135.6702
450.2743225097656 0 1313.4462
452.2524108886719 0 1192.0376
453.24847412109375 0 3008.2722
454.2679443359375 0 1080.165
455.1922607421875 0 965.4084
455.22576904296875 0 16711.318
455.2622375488281 0 956.2093
456.2099609375 0 21527.584
457.2116394042969 0 5155.875
457.24188232421875 0 1372.7003 b Water loss 8
457.7435302734375 0 1149.4519 b Ammonia loss 8
458.2608947753906 0 5809.639
459.26397705078125 0 1176.1517
460.2193908691406 0 860.1362
461.2742614746094 0 1145.9323
462.198486328125 0 2030.5938
462.23406982421875 0 1087.9805
463.2309265136719 0 3135.1216
463.3036193847656 0 938.08124
464.2153015136719 0 2448.6477
464.28753662109375 0 13106.125
465.2130432128906 0 1150.9476
465.25054931640625 0 1403.7544
465.29119873046875 0 3473.1545
466.22991943359375 0 1741.2195
468.2816162109375 0 1806.9104
469.2845764160156 0 887.60803
470.2764892578125 0 6687.636
471.260009765625 0 4288.938
472.2190246582031 0 989.85657
472.2569580078125 0 7575.6333
473.23626708984375 0 27427.896
474.2388000488281 0 6247.532
475.2876892089844 0 2027.7268
476.2928161621094 0 758.80865
476.74981689453125 0 2405.4683
477.2455139160156 0 2583.8572
479.2622375488281 0 1425.5242
480.2471923828125 0 2537.4062
481.2429504394531 0 4040.2124
481.3141784667969 0 5524.359
482.2425842285156 0 1666.7891
482.2987976074219 0 6523.9307
483.22100830078125 0 1326.1322
483.2589111328125 0 1058.5562
483.2984619140625 0 4556.9272
484.3040466308594 0 1023.48596
485.2757873535156 0 3032.844
485.75384521484375 0 13057.952 b Water loss 9
486.2555847167969 0 6954.6807
486.756591796875 0 1467.2899
488.2522888183594 0 867.4141
488.2874755859375 0 24831.281
489.2903137207031 0 6492.669
490.2296142578125 0 2607.3625
490.2640380859375 0 1696.7256
491.2649230957031 0 750.1946
491.2982482910156 0 3817.0303
492.2827453613281 0 2848.414
493.2042541503906 0 1892.8374
493.280517578125 0 1459.3374
494.2099609375 0 859.61035
495.25860595703125 0 1609.1787
497.2704162597656 0 2243.8357
497.308837890625 0 1438.4454
498.2716064453125 0 56933.605 b Water loss 4
499.2751159667969 0 13596.358
499.7694091796875 0 1700.4105
500.275634765625 0 3360.8877
500.3246154785156 0 3068.421
500.77349853515625 0 968.4982
501.3272399902344 0 968.8554
507.2934875488281 0 4940.284
508.2408142089844 0 7101.9736 y Water loss 7
508.2892150878906 0 1306.3812
508.77532958984375 0 2717.949 y Water loss 2
509.239501953125 0 1916.5952
509.3088073730469 0 73592.4
510.2379455566406 0 1030.5159
510.3115234375 0 20253.002
511.2153015136719 0 2942.0256
511.31439208984375 0 3474.3955
512.2183227539062 0 950.1069
512.2887573242188 0 1493.5673
513.2676391601562 0 1642.5027
515.2827758789062 0 1397.9218
516.2820434570312 0 65529.58 b 4
517.2852172851562 0 20988.467
518.287109375 0 3336.5886
519.2910766601562 0 1653.8655
523.287353515625 0 1663.4958
524.2879638671875 0 854.6932
525.3035888671875 0 3369.1
526.2512817382812 0 45044.637 y 7
527.2539672851562 0 10600.742
528.2501220703125 0 3452.9448
528.3193359375 0 4775.563
529.3230590820312 0 1649.5851
530.2562255859375 0 885.89386
530.2969360351562 0 2201.3982
536.280029296875 0 916.17957
537.3058471679688 0 1402.6385
538.2633666992188 0 1158.6163
539.246826171875 0 1803.9708
540.3154296875 0 2141.958
541.2947998046875 0 2718.382
542.2963256835938 0 1466.6234 b Water loss 10
543.2576904296875 0 2102.1938
544.2531127929688 0 702.78345
544.3101196289062 0 808.81696
548.8145751953125 0 1343.6814
549.3069458007812 0 1386.7139
550.2982177734375 0 2783.1375
551.2828979492188 0 2341.3198
551.3214111328125 0 1354.4889
554.2931518554688 0 2467.9856
555.2825317382812 0 1137.5695
556.27685546875 0 1658.026
557.2728271484375 0 1071.0477
557.3395385742188 0 868.62994
558.324462890625 0 10481.688
559.3285522460938 0 3025.171
560.3199462890625 0 1157.3187
566.33349609375 0 3524.7441
567.3292236328125 0 983.4406
568.3092041015625 0 8006.5186
569.300048828125 0 4210.0986
570.2930908203125 0 2166.6604
571.3272705078125 0 1040.2911
572.2491455078125 0 1443.5726
572.3040161132812 0 2087.1765
573.2290649414062 0 922.9007
573.30615234375 0 2180.3826
574.2807006835938 0 817.7485
575.2811279296875 0 1356.152
576.3158569335938 0 2804.0957
577.3013916015625 0 2670.647
577.3723754882812 0 844.2805
578.329833984375 0 7207.0195
579.318115234375 0 3356.824
580.3163452148438 0 933.7722
582.2887573242188 0 1073.4333
583.2890625 0 727.5219
583.360107421875 0 4449.4375
584.3449096679688 0 8326.959
585.2982788085938 0 885.24023
585.3440551757812 0 3841.6326
586.3201293945312 0 24474.453
587.3226928710938 0 7017.8706
590.2571411132812 0 1986.1221
592.3463745117188 0 5959.2095
593.29248046875 0 1174.8402
593.3477783203125 0 2439.8882
594.3242797851562 0 3550.1304
595.32861328125 0 1727.2854
596.3406372070312 0 20212.59
597.3435668945312 0 6114.7607
598.3439331054688 0 1279.4292
601.3715209960938 0 8197.599
602.3734130859375 0 3098.0474
603.314697265625 0 1770.0746
609.2994384765625 0 2082.7202
610.2859497070312 0 3777.99
610.3564453125 0 2451.104
611.35546875 0 47821.598 b Water loss 5
612.3565063476562 0 14828.885
613.359619140625 0 2600.247
618.2896728515625 0 2345.6206
619.2753295898438 0 1944.3271
623.3101806640625 0 1198.1777
624.3037109375 0 1086.7904
625.3038940429688 0 950.8831
627.3157348632812 0 2860.1477
628.31640625 0 1216.6166
628.3794555664062 0 1016.79266
629.3657836914062 0 42528.38 b 5
630.3693237304688 0 15863.905
630.637939453125 0 856.3143
631.371337890625 0 2493.3628
635.3534545898438 0 1080.0881
636.2991943359375 0 21844.76 y Water loss 6
636.3555908203125 0 1068.418
637.2996215820312 0 7533.4204
638.2978515625 0 1489.8466
640.3656005859375 0 1151.3287
641.3269653320312 0 2027.1981
641.8545532226562 0 13540.921
642.3548583984375 0 9033.58
642.8572387695312 0 4225.464
643.333984375 0 2202.0762
644.3394165039062 0 1871.6566
646.3154907226562 0 985.04517
646.38916015625 0 2615.532
647.3507080078125 0 4534.042
648.3370361328125 0 2221.2446
649.36767578125 0 1320.0784
651.3460693359375 0 810.6415
654.3096313476562 0 44699.01 y 6
655.312744140625 0 13765.923
656.3143310546875 0 3188.1963
656.37744140625 0 3611.2234
657.38623046875 0 2517.2737
658.3511352539062 0 1073.0074
659.3602905273438 0 993.40265
660.3348999023438 0 1448.9554
665.362548828125 0 6894.415
666.3617553710938 0 2896.4143
667.3748168945312 0 2391.3794
668.3718872070312 0 2472.0703
669.3594360351562 0 1981.9806
672.33251953125 0 890.6553
677.3662719726562 0 887.03217
678.3505249023438 0 1070.5563
682.6473388671875 0 769.2559
683.37255859375 0 8111.2793
684.3767700195312 0 2572.3848
685.3284301757812 0 1652.6866
685.388671875 0 7814.595
686.3175659179688 0 1311.2308
686.3897705078125 0 2662.7769
687.36376953125 0 918.7887
694.3896484375 0 1642.6265
695.3753662109375 0 2690.391
696.3763427734375 0 1362.9719
697.4301147460938 0 1271.4042
698.3849487304688 0 1108.9153
703.3414306640625 0 4532.5044
704.372314453125 0 9179.324
705.3640747070312 0 5653.321
706.3701171875 0 1883.2521
711.4166259765625 0 1435.5881
712.4009399414062 0 5438.5483
713.3973999023438 0 2507.1428
719.371337890625 0 975.87024
721.353271484375 0 2323.234
722.3834838867188 0 38457.203
723.381591796875 0 18399.186
724.3810424804688 0 4425.98
725.4198608398438 0 1788.3546
726.4221801757812 0 829.3976
729.3560791015625 0 2763.2832
729.4302368164062 0 4707.0356
730.365966796875 0 925.3086
730.4326782226562 0 1481.2614
731.37353515625 0 2402.0303
732.361083984375 0 1668.6798
737.3929443359375 0 915.62366
739.4139404296875 0 17187.889 b Water loss 6
740.3953247070312 0 65498.625 b Ammonia loss 6
741.3975830078125 0 24551.506
742.3988037109375 0 4715.5015
743.40478515625 0 1000.67865
747.3659057617188 0 1227.2543
749.3833618164062 0 6398.278 y Water loss 5
750.3775634765625 0 3860.1816 y Ammonia loss 5
751.3779907226562 0 1220.5046
754.4097900390625 0 1043.8175
755.4113159179688 0 3881.0737
756.4149780273438 0 2510.5164
757.424072265625 0 32356.756 b 6
758.3541870117188 0 918.4805
758.4273681640625 0 11849.867
759.4291381835938 0 1742.3971
766.411376953125 0 1193.3733
767.3936157226562 0 51826.84 y 5
768.396728515625 0 21688.014
769.3991088867188 0 4942.0947
770.4095458984375 0 860.80634
773.4133911132812 0 885.5633
784.4157104492188 0 808.40814
785.4210205078125 0 1643.6104
786.4281005859375 0 912.7512
796.4239501953125 0 941.2746
802.4158935546875 0 825.7468
805.4227905273438 0 1165.8037
807.470458984375 0 961.25104
808.442626953125 0 4542.4746
809.4474487304688 0 2163.232
813.4415283203125 0 865.3081
816.431396484375 0 2350.8513
817.4534301757812 0 3422.0945
818.449951171875 0 2139.1064
819.4519653320312 0 1395.685
823.4318237304688 0 4641.865
824.4326171875 0 3289.7588
825.403564453125 0 1190.4204
825.4833984375 0 20777.223
826.4481201171875 0 6453.514 b Water loss 7
826.477783203125 0 5616.3975
827.4502563476562 0 3685.9001
828.4479370117188 0 906.54004
834.435546875 0 3154.0208
835.4652709960938 0 12339.407
836.4627685546875 0 8434.995
837.4613037109375 0 3913.2505
838.45947265625 0 1042.2931
841.4437866210938 0 1987.3934
842.44140625 0 3860.2656
843.4465942382812 0 887.41437
844.4556274414062 0 12530.132 b 7
845.4591674804688 0 4981.061
846.45947265625 0 1452.8303
848.4498901367188 0 1112.2847 y Water loss 4
853.4779052734375 0 58651.977
854.4808349609375 0 26344.467
855.4838256835938 0 7185.258
860.4532470703125 0 2076.0093
861.4551391601562 0 1805.3911
866.4613647460938 0 5850.0244 y 4
867.4657592773438 0 3454.1067
868.470703125 0 1762.549
869.4556884765625 0 1340.8584
877.4707641601562 0 1238.1014
887.4630737304688 0 1226.8573
888.4723510742188 0 960.6057
889.43701171875 0 910.17505
895.4682006835938 0 4396.923
896.4649658203125 0 1887.4493
903.4921875 0 989.6546
913.4775390625 0 4943.923 b Water loss 8
914.4796142578125 0 4089.0532 b Ammonia loss 8
915.4846801757812 0 2409.7063
925.4880981445312 0 1016.30646
926.4713745117188 0 1463.7941
931.487548828125 0 5591.9946 b 8
932.4896850585938 0 1766.187
933.4961547851562 0 1164.4862
934.4828491210938 0 1155.4836
935.4700317382812 0 1417.1919
936.5135498046875 0 5215.939
937.51416015625 0 3518.6323 y 3
938.5106201171875 0 1840.88
942.5032348632812 0 3319.7725
943.4953002929688 0 1806.3734
944.4918212890625 0 1952.3505
945.474365234375 0 930.2587
952.4883422851562 0 13756.079
953.4855346679688 0 9309.608
954.4967651367188 0 4508.3833
955.524169921875 0 1706.9421
956.5291748046875 0 1199.9966
960.51611328125 0 3641.6143
961.5145874023438 0 2273.1882
970.4988403320312 0 64137.965 b Water loss 9
971.5001831054688 0 39611.85 b Ammonia loss 9
972.5015258789062 0 9928.812
973.515625 0 2056.8394
980.516845703125 0 1184.8899
981.5319213867188 0 993.33563
982.5360717773438 0 1054.5929
988.5093383789062 0 57808.34 b 9
989.5125122070312 0 30656.12
990.5150146484375 0 9100.561
998.529052734375 0 2407.814
999.5364379882812 0 1417.3567
1000.5396728515625 0 2007.588
1001.5401611328125 0 1127.1075
1002.5162963867188 0 888.198
1003.5286254882812 0 976.5403
1016.5405883789062 0 6307.6284 y Water loss 2
1017.5401611328125 0 5000.988 y Ammonia loss 2
1018.5464477539062 0 2035.6986
1034.5511474609375 0 42992.715 y 2
1035.5540771484375 0 26424.86
1036.5562744140625 0 8392.901
1037.56591796875 0 877.1226
1038.554931640625 0 957.263
1055.5863037109375 0 2004.0748
1056.583251953125 0 2058.0188
1057.576171875 0 1318.9585
1065.5728759765625 0 4073.5613
1066.5667724609375 0 4381.1265
1067.5714111328125 0 2021.9261
1073.5970458984375 0 8194.122
1074.6015625 0 5520.3154
1075.600341796875 0 1716.9353
1083.5826416015625 0 28176.326 b Water loss 10
1084.5830078125 0 22496.67 b Ammonia loss 10
1085.584716796875 0 7810.01
1086.5931396484375 0 1051.4493
1101.5933837890625 0 30636.566 b 10
1101.7318115234375 0 1450.056
1102.5965576171875 0 21260.418
1103.598388671875 0 5945.505
1820.7415771484375 0 863.2662
2232.537353515625 0 733.2228
2340.9404296875 0 849.8642
3138.666259765625 0 749.81433

Spectrum Details

|  |  |
| --- | --- |
| Matched peaks? Matched peaks The total absolute number of peaks matched. Additionally in brackets the total fraction of peaks matched and the total number of peaks is shown. | 59 (6.78% of 870) |
| FDR? FDR The false discovery rate estimated for this peptide. It is calculated by matching all theoretical fragments with a non-integer shift with the raw peaks for this spectrum. This is done with 40 different shifts. The resulting percentage is the average number of annotated peaks over the number of annotated peaks with the correct spectrum. | 0.16% |
| Satellite FDR? Satellite FDR See the FDR for details on its calculation. This satellite ion specific FDR only contains the satellite ions (d/w) for I/L/J positions. | - |
| PSM Score? PSM Score The PSM Score as given by Hecklib to this annotated spectrum. It is shown with three significant figures. | 808 |

#### Spectrum 11657? Spectrum 11657 The raw spectrum of this peptide as annotated by Hecklib. The fragments are coloured according to ion type (see legend). Any peaks with a star '\*' as text can be hovered over to see the full details, first the ion type second the mass shift type. By hovering over the amino acids in the peptide or ions in the legend the corresponding peaks are highlighted. By toggling the 'Unassigned' label you can turn the background (unassigned) peaks on or off in the plot. By updating the slider in the Ion legend you can update the spectrum to only show the top X% of the peaks with labels. The top X% means any peak that is within X% of the highest intensity. By dragging in the spectrum you can zoom in to a specific part of the spectrum and use 'Zoom Out' to get back to the original zoom level. The annotation of the spectrum is based on the given sequence in the peptides file and is done with different software so inconsistencies are likely. The peaks are annotated based on the given sequence, with 20 ppm tolerance.

Copy Data

##### Spectrum 11657 (TSV)

###### Preview

```
Loading example...
```

*Click on the button to copy the data to your clipboard.*

Mz MinMz MaxIntensity Max

WidthHeightPeptide font sizePeptide stroke widthSpectrum font sizeSpectrum stroke widthCompact peptide

Ion legend

wxyz

abcd

OtherUnassignedIon^Charge^_Position_Show for top:%

TFPAVJQSSGJY

08.88e+31.78e+42.66e+43.55e+4

Zoom Out

y^+1^_1_y^+1^_2_z^+2^_8_y^+1^_4_y^+1^_5_c^+1^_5_c^+1^_6_y^+1^_6_z^+1^_7_c^+1^_7_y^+1^_7_c^+1^_7_c^+1^_8_z^+1^_8_c^+1^_8_y^+1^_8_z^+1^_9_z^+1^_9_z^+1^_9_c^+1^_9_c^+1^_9_c^+1^_10_c^+1^_10_y^+1^_10_c^+1^_11_c^+1^_11_z^+1^_11_z^+1^_11_

0778155523333110

Fragment Matches Table

Show background peaks

| Position | Ion type | Intensity | mz Theoretical | mz Error (Th) | mz Error (ppm) | Charge | Series Number |
| --- | --- | --- | --- | --- | --- | --- | --- |
| - | - | 1658 | 136.1 | - | - | 0 | - |
| - | - | 599.2 | 143.1 | - | - | 0 | - |
| - | - | 467.4 | 145.5 | - | - | 0 | - |
| - | - | 569 | 148.9 | - | - | 0 | - |
| - | - | 1578 | 165.1 | - | - | 0 | - |
| - | - | 459 | 167.7 | - | - | 0 | - |
| - | - | 552.8 | 169.1 | - | - | 0 | - |
| - | - | 631.5 | 171.1 | - | - | 0 | - |
| - | - | 1863 | 173.5 | - | - | 0 | - |
| 12 | y | 9654 | 182.1 | 5.286E-05 | 0.2903 | +1 | 1 |
| - | - | 1175 | 183.1 | - | - | 0 | - |
| - | - | 424.6 | 183.1 | - | - | 0 | - |
| - | - | 3794 | 221.1 | - | - | 0 | - |
| - | - | 493.7 | 239.1 | - | - | 0 | - |
| - | - | 538 | 242.1 | - | - | 0 | - |
| - | - | 2938 | 249.1 | - | - | 0 | - |
| - | - | 1457 | 252.2 | - | - | 0 | - |
| - | - | 548 | 266.3 | - | - | 0 | - |
| - | - | 1985 | 268.2 | - | - | 0 | - |
| 11 | y | 1636 | 295.2 | 1.903E-05 | 0.06446 | +1 | 2 |
| - | - | 616.7 | 308.1 | - | - | 0 | - |
| - | - | 1161 | 317.2 | - | - | 0 | - |
| - | - | 2328 | 345.2 | - | - | 0 | - |
| - | - | 555.2 | 348.5 | - | - | 0 | - |
| - | - | 693 | 381.2 | - | - | 0 | - |
| - | - | 654 | 413.3 | - | - | 0 | - |
| 5 | z | 800.2 | 417.2 | 0.00193 | 4.626 | +2 | 8 |
| 9 | y | 671.7 | 439.2 | 0.001184 | 2.695 | +1 | 4 |
| - | - | 1244 | 498.3 | - | - | 0 | - |
| - | - | 2083 | 509.3 | - | - | 0 | - |
| - | - | 1571 | 516.3 | - | - | 0 | - |
| 8 | y | 1230 | 526.3 | 0.002725 | 5.178 | +1 | 5 |
| - | - | 713.7 | 526.3 | - | - | 0 | - |
| - | - | 1915 | 532.3 | - | - | 0 | - |
| 5 | c | 6456 | 533.3 | 0.001142 | 2.141 | +1 | 5 |
| - | - | 1835 | 534.3 | - | - | 0 | - |
| - | - | 717.9 | 570.4 | - | - | 0 | - |
| - | - | 577.5 | 596.3 | - | - | 0 | - |
| - | - | 2656 | 603.4 | - | - | 0 | - |
| - | - | 755.6 | 604.4 | - | - | 0 | - |
| - | - | 1364 | 611.4 | - | - | 0 | - |
| - | - | 652.2 | 612.4 | - | - | 0 | - |
| - | - | 720.7 | 628.4 | - | - | 0 | - |
| - | - | 2134 | 629.4 | - | - | 0 | - |
| - | - | 785.4 | 641.3 | - | - | 0 | - |
| - | - | 2799 | 641.9 | - | - | 0 | - |
| - | - | 1703 | 642.4 | - | - | 0 | - |
| - | - | 878 | 642.9 | - | - | 0 | - |
| - | - | 3840 | 645.4 | - | - | 0 | - |
| 6 | c | 1.211E+04 | 646.4 | 0.0004278 | 0.6618 | +1 | 6 |
| - | - | 4404 | 647.4 | - | - | 0 | - |
| - | - | 1879 | 653.3 | - | - | 0 | - |
| 7 | y | 6347 | 654.3 | 0.0003718 | 0.5682 | +1 | 6 |
| - | - | 1572 | 655.3 | - | - | 0 | - |
| - | - | 850.9 | 673.4 | - | - | 0 | - |
| - | - | 756.5 | 693.9 | - | - | 0 | - |
| - | - | 2338 | 730.4 | - | - | 0 | - |
| - | - | 4814 | 731.4 | - | - | 0 | - |
| - | - | 2080 | 732.4 | - | - | 0 | - |
| - | - | 1983 | 740.4 | - | - | 0 | - |
| - | - | 1018 | 741.4 | - | - | 0 | - |
| - | - | 870.3 | 742.4 | - | - | 0 | - |
| 6 | z | 1163 | 751.4 | 0.0007556 | 1.006 | +1 | 7 |
| - | - | 686.1 | 752.4 | - | - | 0 | - |
| - | - | 782 | 756.4 | - | - | 0 | - |
| 7 | c | 2935 | 757.4 | 0.001328 | 1.754 | +1 | 7 |
| - | - | 1757 | 758.4 | - | - | 0 | - |
| - | - | 1485 | 759.4 | - | - | 0 | - |
| - | - | 853.1 | 766.4 | - | - | 0 | - |
| 6 | y | 3289 | 767.4 | 0.002527 | 3.292 | +1 | 7 |
| - | - | 1465 | 768.4 | - | - | 0 | - |
| - | - | 6854 | 773.4 | - | - | 0 | - |
| 7 | c | 9572 | 774.5 | 0.002304 | 2.975 | +1 | 7 |
| - | - | 3082 | 775.5 | - | - | 0 | - |
| - | - | 596.4 | 795.3 | - | - | 0 | - |
| - | - | 764.8 | 825.5 | - | - | 0 | - |
| - | - | 970.2 | 842.5 | - | - | 0 | - |
| 8 | c | 1714 | 844.5 | 0.001372 | 1.625 | +1 | 8 |
| 5 | z | 974.4 | 850.4 | 0.003266 | 3.841 | +1 | 8 |
| - | - | 2055 | 853.5 | - | - | 0 | - |
| - | - | 1522 | 854.5 | - | - | 0 | - |
| - | - | 4385 | 860.5 | - | - | 0 | - |
| 8 | c | 4941 | 861.5 | 0.004059 | 4.711 | +1 | 8 |
| - | - | 1761 | 862.5 | - | - | 0 | - |
| 5 | y | 1235 | 866.5 | 0.003436 | 3.965 | +1 | 8 |
| - | - | 745.8 | 867.5 | - | - | 0 | - |
| 4 | z | 661.3 | 903.5 | 0.0004581 | 0.5071 | +1 | 9 |
| 4 | z | 648 | 904.5 | 0.007531 | 8.327 | +1 | 9 |
| 4 | z | 8291 | 921.5 | 0.001134 | 1.231 | +1 | 9 |
| - | - | 4223 | 922.5 | - | - | 0 | - |
| - | - | 2148 | 923.5 | - | - | 0 | - |
| 9 | c | 829.8 | 931.5 | 0.006669 | 7.16 | +1 | 9 |
| - | - | 734.3 | 932.5 | - | - | 0 | - |
| - | - | 4861 | 947.5 | - | - | 0 | - |
| 9 | c | 1.208E+04 | 948.5 | 0.001602 | 1.689 | +1 | 9 |
| - | - | 4527 | 949.5 | - | - | 0 | - |
| - | - | 1018 | 950.5 | - | - | 0 | - |
| - | - | 932.7 | 966.6 | - | - | 0 | - |
| - | - | 2206 | 970.5 | - | - | 0 | - |
| - | - | 992.7 | 971.5 | - | - | 0 | - |
| - | - | 2915 | 984.5 | - | - | 0 | - |
| - | - | 2016 | 985.5 | - | - | 0 | - |
| 10 | c | 3061 | 988.5 | 0.001522 | 1.539 | +1 | 10 |
| - | - | 1649 | 989.5 | - | - | 0 | - |
| 10 | c | 3.48E+04 | 1006 | 0.0003608 | 0.3588 | +1 | 10 |
| - | - | 1.993E+04 | 1007 | - | - | 0 | - |
| - | - | 5346 | 1008 | - | - | 0 | - |
| - | - | 965.3 | 1009 | - | - | 0 | - |
| 3 | y | 1954 | 1035 | 0.0001926 | 0.1862 | +1 | 10 |
| - | - | 1186 | 1036 | - | - | 0 | - |
| - | - | 2323 | 1069 | - | - | 0 | - |
| - | - | 1372 | 1070 | - | - | 0 | - |
| - | - | 1602 | 1075 | - | - | 0 | - |
| - | - | 1374 | 1076 | - | - | 0 | - |
| - | - | 1085 | 1084 | - | - | 0 | - |
| - | - | 957 | 1085 | - | - | 0 | - |
| - | - | 5141 | 1086 | - | - | 0 | - |
| - | - | 4069 | 1087 | - | - | 0 | - |
| - | - | 810 | 1096 | - | - | 0 | - |
| - | - | 659.7 | 1097 | - | - | 0 | - |
| 11 | c | 2383 | 1102 | 0.0006247 | 0.5671 | +1 | 11 |
| - | - | 2029 | 1103 | - | - | 0 | - |
| - | - | 1062 | 1104 | - | - | 0 | - |
| 11 | c | 3.001E+04 | 1119 | 0.001173 | 1.048 | +1 | 11 |
| - | - | 2.108E+04 | 1120 | - | - | 0 | - |
| - | - | 7076 | 1121 | - | - | 0 | - |
| - | - | 1007 | 1122 | - | - | 0 | - |
| - | - | 747.1 | 1133 | - | - | 0 | - |
| 2 | z | 683 | 1148 | 0.002922 | 2.546 | +1 | 11 |
| 2 | z | 7854 | 1166 | 0.001646 | 1.412 | +1 | 11 |
| - | - | 3841 | 1167 | - | - | 0 | - |
| - | - | 877.7 | 1168 | - | - | 0 | - |
| - | - | 646.8 | 1213 | - | - | 0 | - |
| - | - | 865.9 | 1222 | - | - | 0 | - |
| - | - | 887.4 | 1223 | - | - | 0 | - |
| - | - | 4611 | 1239 | - | - | 0 | - |
| - | - | 3144 | 1240 | - | - | 0 | - |
| - | - | 2014 | 1241 | - | - | 0 | - |
| - | - | 3138 | 1249 | - | - | 0 | - |
| - | - | 1982 | 1250 | - | - | 0 | - |
| - | - | 1055 | 1251 | - | - | 0 | - |
| - | - | 1556 | 1256 | - | - | 0 | - |
| - | - | 1348 | 1257 | - | - | 0 | - |
| - | - | 1164 | 1258 | - | - | 0 | - |
| - | - | 900.2 | 1266 | - | - | 0 | - |
| - | - | 3943 | 1267 | - | - | 0 | - |
| - | - | 3705 | 1268 | - | - | 0 | - |
| - | - | 1511 | 1269 | - | - | 0 | - |
| - | - | 930.8 | 1281 | - | - | 0 | - |
| - | - | 5988 | 1283 | - | - | 0 | - |
| - | - | 3945 | 1283 | - | - | 0 | - |
| - | - | 3.516E+04 | 1284 | - | - | 0 | - |
| - | - | 2.358E+04 | 1285 | - | - | 0 | - |
| - | - | 9988 | 1286 | - | - | 0 | - |
| - | - | 1435 | 1287 | - | - | 0 | - |
| - | - | 733.3 | 1925 | - | - | 0 | - |
| - | - | 676.2 | 3080 | - | - | 0 | - |

m/z Charge Intensity FragmentType MassShift Position
136.07582092285156 0 1658.1606
143.117919921875 0 599.2373
145.49256896972656 0 467.35785
148.94757080078125 0 569.0077
165.05462646484375 0 1577.8611
167.73068237304688 0 459.00342
169.09646606445312 0 552.81964
171.11329650878906 0 631.53296
173.45164489746094 0 1863.1184
182.0812225341797 0 9653.769 y 11
183.08468627929688 0 1174.7798
183.09390258789062 0 424.56808
221.12844848632812 0 3793.9087
239.13052368164062 0 493.7447
242.1494140625 0 538.04645
249.1236572265625 0 2937.8672
252.1571044921875 0 1457.3387
266.2837219238281 0 547.98206
268.16510009765625 0 1984.907
295.1652526855469 0 1636.0228 y 10
308.1249084472656 0 616.68994
317.1814880371094 0 1160.7635
345.1764831542969 0 2327.694
348.4671325683594 0 555.2136
381.24951171875 0 693.0019
413.2664794921875 0 653.96265
417.2138366699219 0 800.22156 z Ammonia loss 4
439.21990966796875 0 671.7106 y 8
498.269775390625 0 1244.0789
509.3084411621094 0 2083.4934
516.2828369140625 0 1570.6283
526.2534790039062 0 1229.6614 y 7
526.335205078125 0 713.6958
532.2996826171875 0 1915.0347
533.3070678710938 0 6456.262 c 4
534.3106079101562 0 1834.5024
570.35302734375 0 717.8562
596.3367919921875 0 577.5345
603.3872680664062 0 2656.244
604.3920288085938 0 755.55383
611.3547973632812 0 1364.2435
612.35400390625 0 652.23035
628.3616333007812 0 720.7209
629.3653564453125 0 2133.7961
641.2523193359375 0 785.4189
641.8544921875 0 2799.1836
642.3555908203125 0 1702.9087
642.8551025390625 0 877.9959
645.3834838867188 0 3839.8225
646.391845703125 0 12108.853 c 5
647.3942260742188 0 4403.621
653.2994995117188 0 1878.8254
654.3089599609375 0 6347.406 y 6
655.3125 0 1572.1882
673.4117431640625 0 850.94696
693.9342651367188 0 756.5457
730.4364013671875 0 2337.786
731.4440307617188 0 4814.4976
732.446044921875 0 2079.8738
740.395751953125 0 1982.8151
741.4027099609375 0 1018.2688
742.4103393554688 0 870.30975
751.3754272460938 0 1162.6404 z 5
752.3856201171875 0 686.1143
756.4193115234375 0 781.974
757.4229736328125 0 2935.055 c Ammonia loss 6
758.4286499023438 0 1757.1135
759.4317016601562 0 1484.5599
766.3827514648438 0 853.0582
767.390869140625 0 3289.1865 y 5
768.3960571289062 0 1465.3555
773.4431762695312 0 6854.403
774.4485473632812 0 9571.752 c 6
775.451904296875 0 3082.0159
795.3279418945312 0 596.42194
825.4832763671875 0 764.78876
842.4663696289062 0 970.1755
844.4577026367188 0 1713.6521 c Ammonia loss 7
850.4398193359375 0 974.40857 z 4
853.4818725585938 0 2054.742
854.4810791015625 0 1521.6626
860.4740600585938 0 4385.2285
861.4788208007812 0 4941.2217 c 7
862.4832763671875 0 1760.9764
866.4583740234375 0 1234.6515 y 4
867.4694213867188 0 745.8484
903.4700927734375 0 661.283 z Water loss 3
904.461181640625 0 648.0162 z Ammonia loss 3
921.4790649414062 0 8290.949 z 3
922.4820556640625 0 4223.3105
923.48388671875 0 2148.0576
931.481689453125 0 829.84955 c Ammonia loss 8
932.488525390625 0 734.3017
947.506103515625 0 4860.5356
948.5133056640625 0 12081.266 c 8
949.5148315429688 0 4526.772
950.5177001953125 0 1018.26575
966.590576171875 0 932.67487
970.4983520507812 0 2205.8945
971.4972534179688 0 992.7293
984.5281372070312 0 2914.8398
985.5330810546875 0 2015.7625
988.50830078125 0 3061.0024 c Ammonia loss 9
989.5093994140625 0 1648.9143
1005.5360107421875 0 34800.34 c 9
1006.5383911132812 0 19930.02
1007.541015625 0 5345.919
1008.5404052734375 0 965.2783
1034.5518798828125 0 1954.26 y 2
1035.551513671875 0 1186.4268
1068.571044921875 0 2322.5679
1069.578369140625 0 1372.1165
1074.6048583984375 0 1601.8643
1075.611083984375 0 1373.6991
1083.5775146484375 0 1084.8308
1084.580078125 0 956.9532
1085.5875244140625 0 5140.9634
1086.594482421875 0 4068.933
1095.6429443359375 0 809.96875
1096.633056640625 0 659.74976
1101.59326171875 0 2383.1685 c Ammonia loss 10
1102.5985107421875 0 2029.0662
1103.604736328125 0 1062.3317
1118.6192626953125 0 30010.436 c 10
1119.6219482421875 0 21081.479
1120.6246337890625 0 7075.544
1122.4271240234375 0 1007.29724
1132.624755859375 0 747.1091
1147.587890625 0 683.00653 z Water loss 1
1165.5997314453125 0 7854.348 z 1
1166.6048583984375 0 3841.0662
1167.5946044921875 0 877.7069
1212.6097412109375 0 646.7564
1221.6871337890625 0 865.8836
1222.6717529296875 0 887.37555
1238.6544189453125 0 4610.9795
1239.65576171875 0 3143.7725
1240.648193359375 0 2014.3438
1248.6361083984375 0 3138.1636
1249.6390380859375 0 1982.4995
1250.6444091796875 0 1054.5527
1255.6871337890625 0 1556.1161
1256.6768798828125 0 1347.7888
1257.6719970703125 0 1163.9977
1265.715087890625 0 900.23486
1266.65771484375 0 3942.5618
1267.655029296875 0 3704.6733
1268.6580810546875 0 1510.583
1280.6953125 0 930.75226
1282.665283203125 0 5988.1704
1282.7618408203125 0 3945.2002
1283.67529296875 0 35163.71
1284.67724609375 0 23580.084
1285.680419921875 0 9987.769
1286.6832275390625 0 1434.5278
1924.9017333984375 0 733.2726
3079.55908203125 0 676.16187

Spectrum Details

|  |  |
| --- | --- |
| Matched peaks? Matched peaks The total absolute number of peaks matched. Additionally in brackets the total fraction of peaks matched and the total number of peaks is shown. | 28 (17.83% of 157) |
| FDR? FDR The false discovery rate estimated for this peptide. It is calculated by matching all theoretical fragments with a non-integer shift with the raw peaks for this spectrum. This is done with 40 different shifts. The resulting percentage is the average number of annotated peaks over the number of annotated peaks with the correct spectrum. | 1.36% |
| Satellite FDR? Satellite FDR See the FDR for details on its calculation. This satellite ion specific FDR only contains the satellite ions (d/w) for I/L/J positions. | ∞ |
| PSM Score? PSM Score The PSM Score as given by Hecklib to this annotated spectrum. It is shown with three significant figures. | 339 |

#### Spectrum 11194? Spectrum 11194 The raw spectrum of this peptide as annotated by Hecklib. The fragments are coloured according to ion type (see legend). Any peaks with a star '\*' as text can be hovered over to see the full details, first the ion type second the mass shift type. By hovering over the amino acids in the peptide or ions in the legend the corresponding peaks are highlighted. By toggling the 'Unassigned' label you can turn the background (unassigned) peaks on or off in the plot. By updating the slider in the Ion legend you can update the spectrum to only show the top X% of the peaks with labels. The top X% means any peak that is within X% of the highest intensity. By dragging in the spectrum you can zoom in to a specific part of the spectrum and use 'Zoom Out' to get back to the original zoom level. The annotation of the spectrum is based on the given sequence in the peptides file and is done with different software so inconsistencies are likely. The peaks are annotated based on the given sequence, with 20 ppm tolerance.

Copy Data

##### Spectrum 11194 (TSV)

###### Preview

```
Loading example...
```

*Click on the button to copy the data to your clipboard.*

Mz MinMz MaxIntensity Max

WidthHeightPeptide font sizePeptide stroke widthSpectrum font sizeSpectrum stroke widthCompact peptide

Ion legend

wxyz

abcd

OtherUnassignedIon^Charge^_Position_Show for top:%

TFPAVJQSSGJY

03.37e+46.75e+41.01e+51.35e+5

Zoom Out

y^+1^_1_a^+1^_2_a^+1^_2_b^+1^_2_b^+1^_2_d^+1^_3_y^+1^_2_b^+2^_6_a^+1^_3_b^+1^_3_b^+1^_3_y^+1^_3_b^+2^_7_b^+2^_7_b^+1^_4_b^+1^_4_b^+2^_8_y^+2^_8_y^+1^_4_b^+2^_9_b^+2^_10_b^+2^_10_b^+1^_5_y^+1^_5_y^+2^_10_b^+1^_5_y^+1^_5_b^+1^_6_b^+1^_6_y^+1^_6_y^+1^_6_b^+1^_7_b^+1^_7_y^+1^_7_y^+1^_7_b^+1^_7_y^+1^_7_b^+1^_8_b^+1^_8_y^+1^_8_b^+1^_9_b^+1^_9_b^+1^_9_y^+1^_9_b^+1^_10_b^+1^_10_b^+1^_10_y^+1^_10_y^+1^_10_y^+1^_10_b^+1^_11_b^+1^_11_b^+1^_11_

0778155723353114

Fragment Matches Table

Show background peaks

| Position | Ion type | Intensity | mz Theoretical | mz Error (Th) | mz Error (ppm) | Charge | Series Number |
| --- | --- | --- | --- | --- | --- | --- | --- |
| - | - | 406.2 | 120.1 | - | - | 0 | - |
| - | - | 1.68E+04 | 120.1 | - | - | 0 | - |
| - | - | 1531 | 121.1 | - | - | 0 | - |
| - | - | 734.8 | 123 | - | - | 0 | - |
| - | - | 7485 | 123 | - | - | 0 | - |
| - | - | 737.4 | 124 | - | - | 0 | - |
| - | - | 382.4 | 126.1 | - | - | 0 | - |
| - | - | 644.3 | 127.1 | - | - | 0 | - |
| - | - | 407.9 | 128.1 | - | - | 0 | - |
| - | - | 2166 | 129.1 | - | - | 0 | - |
| - | - | 2441 | 129.1 | - | - | 0 | - |
| - | - | 890.5 | 130.1 | - | - | 0 | - |
| - | - | 3365 | 131.1 | - | - | 0 | - |
| - | - | 556.3 | 132.1 | - | - | 0 | - |
| - | - | 658.7 | 132.1 | - | - | 0 | - |
| - | - | 506.2 | 133.1 | - | - | 0 | - |
| - | - | 442.6 | 134.4 | - | - | 0 | - |
| - | - | 434.2 | 134.5 | - | - | 0 | - |
| - | - | 7.207E+04 | 136.1 | - | - | 0 | - |
| - | - | 5793 | 137.1 | - | - | 0 | - |
| - | - | 389.3 | 138.5 | - | - | 0 | - |
| - | - | 608.1 | 139.1 | - | - | 0 | - |
| - | - | 600.8 | 140.1 | - | - | 0 | - |
| - | - | 1.949E+04 | 141.1 | - | - | 0 | - |
| - | - | 949.6 | 142.1 | - | - | 0 | - |
| - | - | 1.982E+04 | 143.1 | - | - | 0 | - |
| - | - | 1242 | 144.1 | - | - | 0 | - |
| - | - | 1866 | 145.1 | - | - | 0 | - |
| - | - | 1049 | 146.1 | - | - | 0 | - |
| - | - | 799.7 | 147 | - | - | 0 | - |
| - | - | 7199 | 147 | - | - | 0 | - |
| - | - | 1165 | 147.1 | - | - | 0 | - |
| - | - | 462.6 | 148 | - | - | 0 | - |
| - | - | 707.3 | 153.1 | - | - | 0 | - |
| - | - | 659.5 | 155.1 | - | - | 0 | - |
| - | - | 5446 | 157.1 | - | - | 0 | - |
| - | - | 3734 | 158.1 | - | - | 0 | - |
| - | - | 564.3 | 159.1 | - | - | 0 | - |
| - | - | 552.5 | 159.1 | - | - | 0 | - |
| - | - | 508.6 | 159.1 | - | - | 0 | - |
| - | - | 1625 | 160.1 | - | - | 0 | - |
| - | - | 6.693E+04 | 165.1 | - | - | 0 | - |
| - | - | 6244 | 166.1 | - | - | 0 | - |
| - | - | 839.4 | 166.1 | - | - | 0 | - |
| - | - | 1173 | 169.1 | - | - | 0 | - |
| - | - | 8.163E+04 | 169.1 | - | - | 0 | - |
| - | - | 1180 | 169.1 | - | - | 0 | - |
| - | - | 1607 | 170.1 | - | - | 0 | - |
| - | - | 7222 | 170.1 | - | - | 0 | - |
| - | - | 1148 | 171.1 | - | - | 0 | - |
| - | - | 8775 | 171.1 | - | - | 0 | - |
| - | - | 599.2 | 172.1 | - | - | 0 | - |
| - | - | 1663 | 173.5 | - | - | 0 | - |
| - | - | 1778 | 175.1 | - | - | 0 | - |
| - | - | 739.5 | 175.1 | - | - | 0 | - |
| - | - | 7009 | 176.1 | - | - | 0 | - |
| - | - | 4414 | 177.1 | - | - | 0 | - |
| - | - | 946.5 | 177.1 | - | - | 0 | - |
| - | - | 656.5 | 178.1 | - | - | 0 | - |
| - | - | 747.7 | 180.1 | - | - | 0 | - |
| - | - | 2125 | 181.1 | - | - | 0 | - |
| - | - | 751.9 | 181.1 | - | - | 0 | - |
| 12 | y | 1.336E+05 | 182.1 | 0.000358 | 1.966 | +1 | 1 |
| - | - | 591 | 183.1 | - | - | 0 | - |
| - | - | 1.067E+04 | 183.1 | - | - | 0 | - |
| - | - | 1035 | 184.1 | - | - | 0 | - |
| - | - | 579.1 | 184.1 | - | - | 0 | - |
| - | - | 3096 | 185.2 | - | - | 0 | - |
| - | - | 3925 | 186.1 | - | - | 0 | - |
| - | - | 1144 | 188.1 | - | - | 0 | - |
| - | - | 462.1 | 193.1 | - | - | 0 | - |
| - | - | 666.1 | 196.1 | - | - | 0 | - |
| - | - | 741.6 | 197.1 | - | - | 0 | - |
| - | - | 5033 | 198.1 | - | - | 0 | - |
| - | - | 1945 | 199.1 | - | - | 0 | - |
| - | - | 959.8 | 201.1 | - | - | 0 | - |
| 2 | a | 2714 | 203.1 | 0.0002288 | 1.126 | +1 | 2 |
| - | - | 969.8 | 205.1 | - | - | 0 | - |
| - | - | 1170 | 208.1 | - | - | 0 | - |
| - | - | 495.4 | 209.1 | - | - | 0 | - |
| - | - | 517.7 | 209.8 | - | - | 0 | - |
| - | - | 873.9 | 211.1 | - | - | 0 | - |
| - | - | 778.6 | 212.1 | - | - | 0 | - |
| - | - | 481.8 | 213.1 | - | - | 0 | - |
| - | - | 1283 | 213.1 | - | - | 0 | - |
| - | - | 3864 | 213.2 | - | - | 0 | - |
| - | - | 1.702E+04 | 214.1 | - | - | 0 | - |
| - | - | 666.9 | 214.2 | - | - | 0 | - |
| - | - | 709.9 | 214.2 | - | - | 0 | - |
| - | - | 1191 | 215.1 | - | - | 0 | - |
| - | - | 632 | 215.1 | - | - | 0 | - |
| - | - | 5069 | 216.1 | - | - | 0 | - |
| - | - | 1496 | 217.1 | - | - | 0 | - |
| - | - | 1934 | 221.1 | - | - | 0 | - |
| 2 | a | 1.118E+05 | 221.1 | 0.0003452 | 1.561 | +1 | 2 |
| - | - | 1.311E+04 | 222.1 | - | - | 0 | - |
| - | - | 911.2 | 223.1 | - | - | 0 | - |
| - | - | 921.6 | 223.1 | - | - | 0 | - |
| - | - | 1471 | 224.1 | - | - | 0 | - |
| - | - | 2980 | 225.1 | - | - | 0 | - |
| - | - | 2112 | 226.1 | - | - | 0 | - |
| - | - | 840.9 | 226.2 | - | - | 0 | - |
| - | - | 1362 | 230.2 | - | - | 0 | - |
| 2 | b | 1718 | 231.1 | 4.071E-06 | 0.01761 | +1 | 2 |
| - | - | 1.293E+04 | 232.1 | - | - | 0 | - |
| - | - | 643.4 | 233.2 | - | - | 0 | - |
| - | - | 868.9 | 238.1 | - | - | 0 | - |
| - | - | 783.1 | 239.2 | - | - | 0 | - |
| - | - | 958.7 | 240.1 | - | - | 0 | - |
| - | - | 1109 | 240.1 | - | - | 0 | - |
| - | - | 1.237E+04 | 240.2 | - | - | 0 | - |
| - | - | 789.4 | 241.2 | - | - | 0 | - |
| - | - | 1396 | 241.2 | - | - | 0 | - |
| - | - | 1.475E+04 | 242.2 | - | - | 0 | - |
| - | - | 600.2 | 243.1 | - | - | 0 | - |
| - | - | 1026 | 243.2 | - | - | 0 | - |
| - | - | 3239 | 245.1 | - | - | 0 | - |
| 2 | b | 3.645E+04 | 249.1 | 0.0003342 | 1.341 | +1 | 2 |
| - | - | 1289 | 249.2 | - | - | 0 | - |
| - | - | 2950 | 250.1 | - | - | 0 | - |
| - | - | 4467 | 250.1 | - | - | 0 | - |
| - | - | 657.4 | 251.1 | - | - | 0 | - |
| - | - | 1057 | 252.1 | - | - | 0 | - |
| - | - | 933.6 | 254.2 | - | - | 0 | - |
| - | - | 2291 | 255.2 | - | - | 0 | - |
| - | - | 1035 | 256.1 | - | - | 0 | - |
| - | - | 6776 | 258.1 | - | - | 0 | - |
| - | - | 805.1 | 259.1 | - | - | 0 | - |
| - | - | 672.5 | 259.2 | - | - | 0 | - |
| - | - | 579.1 | 262.1 | - | - | 0 | - |
| - | - | 642.1 | 262.2 | - | - | 0 | - |
| - | - | 1598 | 267.1 | - | - | 0 | - |
| - | - | 2535 | 268.1 | - | - | 0 | - |
| - | - | 7.361E+04 | 268.2 | - | - | 0 | - |
| - | - | 9542 | 269.2 | - | - | 0 | - |
| - | - | 2021 | 270.1 | - | - | 0 | - |
| - | - | 615.1 | 270.2 | - | - | 0 | - |
| - | - | 1030 | 271.1 | - | - | 0 | - |
| - | - | 658.5 | 273.2 | - | - | 0 | - |
| - | - | 872.5 | 275.1 | - | - | 0 | - |
| - | - | 959.3 | 275.2 | - | - | 0 | - |
| - | - | 665.1 | 275.7 | - | - | 0 | - |
| - | - | 920.9 | 280.1 | - | - | 0 | - |
| - | - | 975.7 | 280.2 | - | - | 0 | - |
| - | - | 1041 | 282.1 | - | - | 0 | - |
| - | - | 2276 | 282.2 | - | - | 0 | - |
| - | - | 533.5 | 284.2 | - | - | 0 | - |
| - | - | 1381 | 284.2 | - | - | 0 | - |
| - | - | 606.5 | 284.7 | - | - | 0 | - |
| - | - | 1427 | 285.1 | - | - | 0 | - |
| - | - | 1734 | 285.2 | - | - | 0 | - |
| - | - | 695.5 | 286.1 | - | - | 0 | - |
| - | - | 1004 | 286.2 | - | - | 0 | - |
| - | - | 875.1 | 287.2 | - | - | 0 | - |
| - | - | 509.9 | 288.2 | - | - | 0 | - |
| - | - | 794.3 | 289.2 | - | - | 0 | - |
| - | - | 769.6 | 289.7 | - | - | 0 | - |
| 3 | d | 585.8 | 292.2 | 0.0004171 | 1.428 | +1 | 3 |
| - | - | 548.4 | 294.1 | - | - | 0 | - |
| - | - | 3010 | 295.1 | - | - | 0 | - |
| 11 | y | 1.975E+04 | 295.2 | 0.0005073 | 1.719 | +1 | 2 |
| - | - | 2315 | 296.2 | - | - | 0 | - |
| - | - | 1764 | 296.2 | - | - | 0 | - |
| - | - | 1464 | 297.1 | - | - | 0 | - |
| - | - | 755.9 | 297.2 | - | - | 0 | - |
| - | - | 727.2 | 297.2 | - | - | 0 | - |
| - | - | 2713 | 299.2 | - | - | 0 | - |
| - | - | 1637 | 300.2 | - | - | 0 | - |
| - | - | 1216 | 303.1 | - | - | 0 | - |
| 6 | b | 738.6 | 306.2 | 0.000392 | 1.28 | +2 | 6 |
| - | - | 964.4 | 307.1 | - | - | 0 | - |
| - | - | 767.3 | 309.2 | - | - | 0 | - |
| - | - | 884.1 | 310.1 | - | - | 0 | - |
| - | - | 853.7 | 310.2 | - | - | 0 | - |
| - | - | 1825 | 310.2 | - | - | 0 | - |
| - | - | 1707 | 311.2 | - | - | 0 | - |
| - | - | 1347 | 312.2 | - | - | 0 | - |
| - | - | 1125 | 313.2 | - | - | 0 | - |
| - | - | 758.6 | 315.1 | - | - | 0 | - |
| - | - | 1349 | 315.7 | - | - | 0 | - |
| - | - | 4945 | 316.2 | - | - | 0 | - |
| - | - | 1.72E+04 | 317.2 | - | - | 0 | - |
| 3 | a | 3189 | 318.2 | 0.004848 | 15.24 | +1 | 3 |
| - | - | 1728 | 324.1 | - | - | 0 | - |
| - | - | 3856 | 324.2 | - | - | 0 | - |
| - | - | 1232 | 324.7 | - | - | 0 | - |
| - | - | 5627 | 325.1 | - | - | 0 | - |
| - | - | 714 | 325.2 | - | - | 0 | - |
| - | - | 2286 | 325.2 | - | - | 0 | - |
| - | - | 721.5 | 326.1 | - | - | 0 | - |
| - | - | 3808 | 327.2 | - | - | 0 | - |
| - | - | 842.3 | 327.2 | - | - | 0 | - |
| 3 | b | 2941 | 328.2 | 0.0008443 | 2.573 | +1 | 3 |
| - | - | 3894 | 329.2 | - | - | 0 | - |
| - | - | 704.3 | 330.2 | - | - | 0 | - |
| - | - | 1691 | 333.2 | - | - | 0 | - |
| - | - | 1018 | 339.2 | - | - | 0 | - |
| - | - | 663.9 | 341.2 | - | - | 0 | - |
| - | - | 5522 | 341.2 | - | - | 0 | - |
| - | - | 3905 | 342.1 | - | - | 0 | - |
| - | - | 7056 | 343.1 | - | - | 0 | - |
| - | - | 769.8 | 343.2 | - | - | 0 | - |
| - | - | 1403 | 344.2 | - | - | 0 | - |
| - | - | 2.192E+04 | 345.2 | - | - | 0 | - |
| 3 | b | 9597 | 346.2 | 0.001693 | 4.891 | +1 | 3 |
| - | - | 994.4 | 347.2 | - | - | 0 | - |
| - | - | 6807 | 351.2 | - | - | 0 | - |
| 10 | y | 3969 | 352.2 | 0.0004059 | 1.152 | +1 | 3 |
| - | - | 653 | 352.2 | - | - | 0 | - |
| - | - | 594.7 | 352.7 | - | - | 0 | - |
| - | - | 1757 | 353.2 | - | - | 0 | - |
| - | - | 1.025E+04 | 353.3 | - | - | 0 | - |
| - | - | 1534 | 354.3 | - | - | 0 | - |
| - | - | 807.9 | 355.2 | - | - | 0 | - |
| - | - | 1688 | 360.2 | - | - | 0 | - |
| - | - | 7484 | 361.7 | - | - | 0 | - |
| - | - | 3418 | 362.2 | - | - | 0 | - |
| - | - | 754.5 | 362.7 | - | - | 0 | - |
| - | - | 1026 | 363.2 | - | - | 0 | - |
| - | - | 1043 | 365.1 | - | - | 0 | - |
| - | - | 992.2 | 365.2 | - | - | 0 | - |
| - | - | 3214 | 369.2 | - | - | 0 | - |
| - | - | 704.3 | 370.1 | - | - | 0 | - |
| 7 | b | 2154 | 370.2 | 0.003421 | 9.242 | +2 | 7 |
| 7 | b | 993.1 | 370.7 | 0.002502 | 6.75 | +2 | 7 |
| - | - | 691.2 | 372.2 | - | - | 0 | - |
| - | - | 838.3 | 373.2 | - | - | 0 | - |
| - | - | 635 | 378.2 | - | - | 0 | - |
| - | - | 1500 | 379.2 | - | - | 0 | - |
| - | - | 1110 | 380.2 | - | - | 0 | - |
| - | - | 2.111E+04 | 381.2 | - | - | 0 | - |
| - | - | 1117 | 382.2 | - | - | 0 | - |
| - | - | 4047 | 382.3 | - | - | 0 | - |
| - | - | 775.5 | 383.2 | - | - | 0 | - |
| - | - | 1030 | 384.2 | - | - | 0 | - |
| - | - | 737.2 | 384.2 | - | - | 0 | - |
| - | - | 3179 | 387.2 | - | - | 0 | - |
| - | - | 2104 | 389.2 | - | - | 0 | - |
| - | - | 1243 | 393.2 | - | - | 0 | - |
| - | - | 3522 | 394.2 | - | - | 0 | - |
| - | - | 712 | 396.2 | - | - | 0 | - |
| - | - | 1272 | 398.2 | - | - | 0 | - |
| - | - | 907.2 | 398.2 | - | - | 0 | - |
| - | - | 5757 | 398.3 | - | - | 0 | - |
| 4 | b | 6814 | 399.2 | 0.0003822 | 0.9573 | +1 | 4 |
| - | - | 1003 | 399.3 | - | - | 0 | - |
| - | - | 1341 | 400.2 | - | - | 0 | - |
| - | - | 857.8 | 401.2 | - | - | 0 | - |
| - | - | 560 | 407.2 | - | - | 0 | - |
| - | - | 934.4 | 408.2 | - | - | 0 | - |
| - | - | 1513 | 410.2 | - | - | 0 | - |
| - | - | 1545 | 410.2 | - | - | 0 | - |
| - | - | 962.1 | 412.2 | - | - | 0 | - |
| - | - | 759.7 | 412.3 | - | - | 0 | - |
| - | - | 2622 | 415.2 | - | - | 0 | - |
| - | - | 1088 | 416.2 | - | - | 0 | - |
| 4 | b | 1.343E+04 | 417.2 | 0.0005597 | 1.341 | +1 | 4 |
| - | - | 2251 | 418.2 | - | - | 0 | - |
| - | - | 1225 | 420.2 | - | - | 0 | - |
| 8 | b | 1971 | 422.7 | 0.0008932 | 2.113 | +2 | 8 |
| - | - | 1031 | 424.2 | - | - | 0 | - |
| 5 | y | 1253 | 425.2 | 0.003831 | 9.009 | +2 | 8 |
| - | - | 1058 | 426.2 | - | - | 0 | - |
| - | - | 1835 | 427.2 | - | - | 0 | - |
| - | - | 5205 | 428.2 | - | - | 0 | - |
| - | - | 810.5 | 428.2 | - | - | 0 | - |
| - | - | 1308 | 429.2 | - | - | 0 | - |
| - | - | 990 | 433.2 | - | - | 0 | - |
| - | - | 1334 | 437.2 | - | - | 0 | - |
| - | - | 1676 | 438.2 | - | - | 0 | - |
| - | - | 975.3 | 438.3 | - | - | 0 | - |
| 9 | y | 3586 | 439.2 | 0.0001768 | 0.4025 | +1 | 4 |
| - | - | 1053 | 440.2 | - | - | 0 | - |
| - | - | 1070 | 444.3 | - | - | 0 | - |
| - | - | 3803 | 445.2 | - | - | 0 | - |
| - | - | 652.1 | 446.2 | - | - | 0 | - |
| - | - | 1589 | 448.2 | - | - | 0 | - |
| - | - | 1165 | 448.7 | - | - | 0 | - |
| - | - | 877.7 | 453.2 | - | - | 0 | - |
| - | - | 6387 | 455.2 | - | - | 0 | - |
| - | - | 4889 | 456.2 | - | - | 0 | - |
| - | - | 1650 | 457.2 | - | - | 0 | - |
| 9 | b | 887.1 | 457.2 | 0.001575 | 3.444 | +2 | 9 |
| - | - | 1790 | 458.3 | - | - | 0 | - |
| - | - | 683 | 462.2 | - | - | 0 | - |
| - | - | 1094 | 464.2 | - | - | 0 | - |
| - | - | 3909 | 464.3 | - | - | 0 | - |
| - | - | 1136 | 465.3 | - | - | 0 | - |
| - | - | 730.3 | 466.2 | - | - | 0 | - |
| - | - | 2050 | 470.3 | - | - | 0 | - |
| - | - | 1191 | 471.3 | - | - | 0 | - |
| - | - | 2004 | 472.3 | - | - | 0 | - |
| - | - | 7387 | 473.2 | - | - | 0 | - |
| - | - | 2932 | 474.2 | - | - | 0 | - |
| - | - | 1128 | 480.2 | - | - | 0 | - |
| - | - | 1169 | 481.2 | - | - | 0 | - |
| - | - | 1159 | 481.3 | - | - | 0 | - |
| - | - | 1959 | 482.3 | - | - | 0 | - |
| - | - | 1382 | 483.3 | - | - | 0 | - |
| 10 | b | 3924 | 485.8 | 0.0006697 | 1.379 | +2 | 10 |
| 10 | b | 1668 | 486.2 | 0.007624 | 15.68 | +2 | 10 |
| - | - | 7327 | 488.3 | - | - | 0 | - |
| - | - | 1864 | 489.3 | - | - | 0 | - |
| - | - | 833.7 | 490.2 | - | - | 0 | - |
| - | - | 734.6 | 491.3 | - | - | 0 | - |
| - | - | 1206 | 492.3 | - | - | 0 | - |
| - | - | 851 | 497.3 | - | - | 0 | - |
| 5 | b | 1.779E+04 | 498.3 | 0.0003887 | 0.78 | +1 | 5 |
| - | - | 4229 | 499.3 | - | - | 0 | - |
| - | - | 902.1 | 499.8 | - | - | 0 | - |
| - | - | 1015 | 500.3 | - | - | 0 | - |
| - | - | 2196 | 507.3 | - | - | 0 | - |
| 8 | y | 2166 | 508.2 | 0.0002585 | 0.5086 | +1 | 5 |
| 3 | y | 925.4 | 508.8 | 0.0001535 | 0.3017 | +2 | 10 |
| - | - | 850.6 | 509.2 | - | - | 0 | - |
| - | - | 2.298E+04 | 509.3 | - | - | 0 | - |
| - | - | 6475 | 510.3 | - | - | 0 | - |
| - | - | 1030 | 511.3 | - | - | 0 | - |
| - | - | 864.5 | 512.2 | - | - | 0 | - |
| - | - | 1174 | 512.3 | - | - | 0 | - |
| 5 | b | 1.984E+04 | 516.3 | 0.0005051 | 0.9784 | +1 | 5 |
| - | - | 7064 | 517.3 | - | - | 0 | - |
| - | - | 959.2 | 518.3 | - | - | 0 | - |
| - | - | 961.2 | 525.3 | - | - | 0 | - |
| 8 | y | 1.337E+04 | 526.3 | 0.0005886 | 1.118 | +1 | 5 |
| - | - | 2385 | 527.3 | - | - | 0 | - |
| - | - | 1428 | 528.3 | - | - | 0 | - |
| - | - | 802.1 | 530.3 | - | - | 0 | - |
| - | - | 714.8 | 531.3 | - | - | 0 | - |
| - | - | 928 | 538.3 | - | - | 0 | - |
| - | - | 779.1 | 539.8 | - | - | 0 | - |
| - | - | 1285 | 540.3 | - | - | 0 | - |
| - | - | 687.2 | 541.3 | - | - | 0 | - |
| - | - | 770.7 | 543.3 | - | - | 0 | - |
| - | - | 615.7 | 549.8 | - | - | 0 | - |
| - | - | 908.7 | 551.3 | - | - | 0 | - |
| - | - | 859.8 | 554.3 | - | - | 0 | - |
| - | - | 611.8 | 557.3 | - | - | 0 | - |
| - | - | 3949 | 558.3 | - | - | 0 | - |
| - | - | 1199 | 566.3 | - | - | 0 | - |
| - | - | 2468 | 568.3 | - | - | 0 | - |
| - | - | 600.6 | 569.3 | - | - | 0 | - |
| - | - | 951.3 | 572.3 | - | - | 0 | - |
| - | - | 639.4 | 573.2 | - | - | 0 | - |
| - | - | 704 | 575.3 | - | - | 0 | - |
| - | - | 940.4 | 577.3 | - | - | 0 | - |
| - | - | 3133 | 578.3 | - | - | 0 | - |
| - | - | 1220 | 578.8 | - | - | 0 | - |
| - | - | 1060 | 579.3 | - | - | 0 | - |
| - | - | 1101 | 583.4 | - | - | 0 | - |
| - | - | 2252 | 584.3 | - | - | 0 | - |
| - | - | 1424 | 585.3 | - | - | 0 | - |
| - | - | 7434 | 586.3 | - | - | 0 | - |
| - | - | 2802 | 587.3 | - | - | 0 | - |
| - | - | 1568 | 592.3 | - | - | 0 | - |
| - | - | 857.4 | 593.3 | - | - | 0 | - |
| - | - | 1209 | 594.3 | - | - | 0 | - |
| - | - | 6847 | 596.3 | - | - | 0 | - |
| - | - | 1771 | 597.3 | - | - | 0 | - |
| - | - | 2743 | 601.4 | - | - | 0 | - |
| - | - | 1462 | 602.4 | - | - | 0 | - |
| - | - | 783.5 | 609.3 | - | - | 0 | - |
| - | - | 908.5 | 610.3 | - | - | 0 | - |
| 6 | b | 1.319E+04 | 611.4 | 0.0006142 | 1.005 | +1 | 6 |
| - | - | 4470 | 612.4 | - | - | 0 | - |
| - | - | 857.8 | 613.4 | - | - | 0 | - |
| - | - | 785.6 | 618.3 | - | - | 0 | - |
| - | - | 800.3 | 624.3 | - | - | 0 | - |
| 6 | b | 1.162E+04 | 629.4 | 5.931E-05 | 0.09424 | +1 | 6 |
| - | - | 4394 | 630.4 | - | - | 0 | - |
| - | - | 821.4 | 631.4 | - | - | 0 | - |
| 7 | y | 7344 | 636.3 | 0.0004883 | 0.7675 | +1 | 6 |
| - | - | 1049 | 637.3 | - | - | 0 | - |
| - | - | 1.374E+04 | 641.9 | - | - | 0 | - |
| - | - | 8869 | 642.4 | - | - | 0 | - |
| - | - | 4348 | 642.9 | - | - | 0 | - |
| - | - | 1126 | 643.4 | - | - | 0 | - |
| - | - | 617.7 | 644.3 | - | - | 0 | - |
| - | - | 1161 | 646.4 | - | - | 0 | - |
| - | - | 1499 | 647.3 | - | - | 0 | - |
| - | - | 1075 | 648.3 | - | - | 0 | - |
| 7 | y | 1.509E+04 | 654.3 | 0.0007269 | 1.111 | +1 | 6 |
| - | - | 4637 | 655.3 | - | - | 0 | - |
| - | - | 881.4 | 656.3 | - | - | 0 | - |
| - | - | 1054 | 656.4 | - | - | 0 | - |
| - | - | 1215 | 657.4 | - | - | 0 | - |
| - | - | 2023 | 665.4 | - | - | 0 | - |
| - | - | 962.6 | 667.4 | - | - | 0 | - |
| - | - | 719.7 | 668.4 | - | - | 0 | - |
| - | - | 925.6 | 672.3 | - | - | 0 | - |
| - | - | 2429 | 683.4 | - | - | 0 | - |
| - | - | 737.1 | 684.4 | - | - | 0 | - |
| - | - | 821.7 | 685.3 | - | - | 0 | - |
| - | - | 1835 | 685.4 | - | - | 0 | - |
| - | - | 865 | 686.4 | - | - | 0 | - |
| - | - | 1281 | 695.4 | - | - | 0 | - |
| - | - | 1286 | 703.3 | - | - | 0 | - |
| - | - | 2128 | 704.4 | - | - | 0 | - |
| - | - | 1639 | 705.4 | - | - | 0 | - |
| - | - | 681.4 | 706.4 | - | - | 0 | - |
| - | - | 1643 | 712.4 | - | - | 0 | - |
| - | - | 1396 | 713.4 | - | - | 0 | - |
| - | - | 1.274E+04 | 722.4 | - | - | 0 | - |
| - | - | 5918 | 723.4 | - | - | 0 | - |
| - | - | 1467 | 724.4 | - | - | 0 | - |
| - | - | 889.2 | 726.4 | - | - | 0 | - |
| - | - | 1114 | 729.4 | - | - | 0 | - |
| 7 | b | 4381 | 739.4 | 0.0006513 | 0.8808 | +1 | 7 |
| 7 | b | 1.903E+04 | 740.4 | 0.002184 | 2.95 | +1 | 7 |
| - | - | 8652 | 741.4 | - | - | 0 | - |
| - | - | 754 | 742.4 | - | - | 0 | - |
| 6 | y | 1573 | 749.4 | 0.000836 | 1.116 | +1 | 7 |
| 6 | y | 1682 | 750.4 | 0.007238 | 9.646 | +1 | 7 |
| - | - | 952 | 755.4 | - | - | 0 | - |
| - | - | 785.1 | 756.4 | - | - | 0 | - |
| 7 | b | 9255 | 757.4 | 0.0004738 | 0.6255 | +1 | 7 |
| - | - | 3974 | 758.4 | - | - | 0 | - |
| - | - | 778.2 | 765 | - | - | 0 | - |
| - | - | 894.2 | 766.4 | - | - | 0 | - |
| 6 | y | 1.491E+04 | 767.4 | 0.000159 | 0.2072 | +1 | 7 |
| - | - | 6273 | 768.4 | - | - | 0 | - |
| - | - | 1422 | 769.4 | - | - | 0 | - |
| - | - | 1231 | 808.4 | - | - | 0 | - |
| - | - | 823.4 | 809.4 | - | - | 0 | - |
| - | - | 692.9 | 816.4 | - | - | 0 | - |
| - | - | 1113 | 817.5 | - | - | 0 | - |
| - | - | 1039 | 824.4 | - | - | 0 | - |
| - | - | 6263 | 825.5 | - | - | 0 | - |
| 8 | b | 1992 | 826.4 | 0.007726 | 9.348 | +1 | 8 |
| - | - | 782.1 | 830.4 | - | - | 0 | - |
| - | - | 3519 | 835.5 | - | - | 0 | - |
| - | - | 3052 | 836.5 | - | - | 0 | - |
| - | - | 955 | 837.5 | - | - | 0 | - |
| - | - | 820.4 | 841.4 | - | - | 0 | - |
| - | - | 1514 | 842.4 | - | - | 0 | - |
| 8 | b | 3639 | 844.5 | 0.0005198 | 0.6155 | +1 | 8 |
| - | - | 1928 | 845.5 | - | - | 0 | - |
| - | - | 2.032E+04 | 853.5 | - | - | 0 | - |
| - | - | 9044 | 854.5 | - | - | 0 | - |
| - | - | 2534 | 855.5 | - | - | 0 | - |
| 5 | y | 1668 | 866.5 | 0.001177 | 1.359 | +1 | 8 |
| - | - | 754.8 | 895.5 | - | - | 0 | - |
| - | - | 645.5 | 896.5 | - | - | 0 | - |
| 9 | b | 1166 | 913.5 | 0.01063 | 11.64 | +1 | 9 |
| 9 | b | 1171 | 914.5 | 0.01256 | 13.73 | +1 | 9 |
| 9 | b | 2171 | 931.5 | 0.001387 | 1.489 | +1 | 9 |
| - | - | 1188 | 932.5 | - | - | 0 | - |
| - | - | 764.2 | 935.5 | - | - | 0 | - |
| - | - | 1657 | 936.5 | - | - | 0 | - |
| 4 | y | 1047 | 937.5 | 0.01188 | 12.67 | +1 | 9 |
| - | - | 933.8 | 942.5 | - | - | 0 | - |
| - | - | 745.3 | 944.5 | - | - | 0 | - |
| - | - | 3784 | 952.5 | - | - | 0 | - |
| - | - | 3220 | 953.5 | - | - | 0 | - |
| - | - | 1093 | 954.5 | - | - | 0 | - |
| - | - | 1739 | 960.5 | - | - | 0 | - |
| - | - | 808.4 | 961.5 | - | - | 0 | - |
| 10 | b | 1.916E+04 | 970.5 | 0.0001733 | 0.1785 | +1 | 10 |
| 10 | b | 1.2E+04 | 971.5 | 0.01593 | 16.4 | +1 | 10 |
| - | - | 3979 | 972.5 | - | - | 0 | - |
| - | - | 906.1 | 973.5 | - | - | 0 | - |
| 10 | b | 1.79E+04 | 988.5 | 0.0001789 | 0.181 | +1 | 10 |
| - | - | 9190 | 989.5 | - | - | 0 | - |
| - | - | 2314 | 990.5 | - | - | 0 | - |
| 3 | y | 1802 | 1017 | 0.003433 | 3.377 | +1 | 10 |
| 3 | y | 1530 | 1018 | 0.008004 | 7.866 | +1 | 10 |
| 3 | y | 1.258E+04 | 1035 | 0.001028 | 0.9937 | +1 | 10 |
| - | - | 7545 | 1036 | - | - | 0 | - |
| - | - | 2440 | 1037 | - | - | 0 | - |
| - | - | 865.6 | 1057 | - | - | 0 | - |
| - | - | 1190 | 1066 | - | - | 0 | - |
| - | - | 1939 | 1067 | - | - | 0 | - |
| - | - | 2267 | 1074 | - | - | 0 | - |
| - | - | 1333 | 1075 | - | - | 0 | - |
| 11 | b | 8429 | 1084 | 0.0009243 | 0.853 | +1 | 11 |
| 11 | b | 4916 | 1085 | 0.01457 | 13.44 | +1 | 11 |
| - | - | 2806 | 1086 | - | - | 0 | - |
| 11 | b | 8284 | 1102 | 0.0002298 | 0.2086 | +1 | 11 |
| - | - | 5728 | 1103 | - | - | 0 | - |
| - | - | 1744 | 1104 | - | - | 0 | - |
| - | - | 728.3 | 1212 | - | - | 0 | - |
| - | - | 607.7 | 2353 | - | - | 0 | - |
| - | - | 641.4 | 2511 | - | - | 0 | - |
| - | - | 736.4 | 3083 | - | - | 0 | - |

m/z Charge Intensity FragmentType MassShift Position
120.05274963378906 0 406.19394
120.08109283447266 0 16801.633
121.0843276977539 0 1531.0687
123.0395736694336 0 734.76373
123.04438018798828 0 7485.4297
124.04774475097656 0 737.4206
126.05525207519531 0 382.4083
127.05039978027344 0 644.3441
128.08758544921875 0 407.91327
129.0662841796875 0 2165.9202
129.1025848388672 0 2441.3757
130.06533813476562 0 890.53345
131.11814880371094 0 3364.7744
132.08152770996094 0 556.28015
132.1021728515625 0 658.7022
133.08570861816406 0 506.2071
134.3588104248047 0 442.59595
134.522216796875 0 434.19098
136.07601928710938 0 72072.34
137.07933044433594 0 5792.8613
138.46229553222656 0 389.27615
139.05078125 0 608.0995
140.14346313476562 0 600.8002
141.1025390625 0 19488.926
142.1057586669922 0 949.5717
143.11817932128906 0 19823.129
144.1216278076172 0 1242.1638
145.06112670898438 0 1866.0027
146.09689331054688 0 1048.5127
147.03858947753906 0 799.72925
147.0443572998047 0 7198.8135
147.07652282714844 0 1165.048
148.04771423339844 0 462.6158
153.06607055664062 0 707.3231
155.08187866210938 0 659.47614
157.06105041503906 0 5445.507
158.0965118408203 0 3734.4595
159.0923614501953 0 564.315
159.0996856689453 0 552.4883
159.13748168945312 0 508.6154
160.1123809814453 0 1624.5972
165.054931640625 0 66926.68
166.0582733154297 0 6243.926
166.08697509765625 0 839.42377
169.06101989746094 0 1173.3209
169.0974578857422 0 81633.05
169.13375854492188 0 1180.1077
170.09278869628906 0 1607.3778
170.10089111328125 0 7222.054
171.07640075683594 0 1148.3655
171.1131134033203 0 8774.672
172.11642456054688 0 599.2238
173.4522247314453 0 1663.1327
175.0717315673828 0 1778.0892
175.08700561523438 0 739.47565
176.10728454589844 0 7009.238
177.10263061523438 0 4414.1846
177.11033630371094 0 946.48926
178.10606384277344 0 656.516
180.0778350830078 0 747.6725
181.06109619140625 0 2125.1418
181.09716796875 0 751.92126
182.08152770996094 0 133570.34 y 11
183.07595825195312 0 591.0111
183.08482360839844 0 10670.948
184.0733184814453 0 1034.6676
184.08677673339844 0 579.07306
185.16514587402344 0 3096.3962
186.12400817871094 0 3924.649
188.10321044921875 0 1144.2365
193.1345977783203 0 462.1435
196.0714569091797 0 666.0786
197.1284637451172 0 741.5597
198.08758544921875 0 5032.7344
199.07147216796875 0 1945.239
201.12339782714844 0 959.77905
203.1181182861328 0 2713.8086 a Water loss 1
205.0972137451172 0 969.7785
208.0977325439453 0 1170.3522
209.0803985595703 0 495.35687
209.831787109375 0 517.7201
211.14385986328125 0 873.9443
212.1039276123047 0 778.5954
213.08753967285156 0 481.81198
213.1234130859375 0 1282.6229
213.1602325439453 0 3864.2673
214.08248901367188 0 17020.963
214.1543731689453 0 666.92096
214.16456604003906 0 709.9347
215.08612060546875 0 1191.406
215.1387176513672 0 632.0462
216.0982666015625 0 5068.5435
217.13343811035156 0 1495.6492
221.0926971435547 0 1933.5686
221.12879943847656 0 111832.05 a 1
222.13211059570312 0 13109.065
223.13377380371094 0 911.2475
223.14505004882812 0 921.5797
224.10311889648438 0 1470.9957
225.12356567382812 0 2979.817
226.11868286132812 0 2111.859
226.15541076660156 0 840.87006
230.15110778808594 0 1361.898
231.11280822753906 0 1717.9866 b Water loss 1
232.0931854248047 0 12932.659
233.16619873046875 0 643.3836
238.11866760253906 0 868.9112
239.1516876220703 0 783.14905
240.09848022460938 0 958.703
240.13426208496094 0 1109.2977
240.17098999023438 0 12368.17
241.16099548339844 0 789.3988
241.17384338378906 0 1396.4655
242.1503143310547 0 14746.778
243.13458251953125 0 600.20776
243.15243530273438 0 1025.7733
245.1288604736328 0 3238.6416
249.1237030029297 0 36448.062 b 1
249.15992736816406 0 1288.8718
250.0828094482422 0 2950.0183
250.126953125 0 4467.375
251.12794494628906 0 657.438
252.13461303710938 0 1056.9426
254.1501922607422 0 933.5764
255.1579132080078 0 2291.1003
256.1294250488281 0 1035.3138
258.14520263671875 0 6776.1445
259.1475524902344 0 805.10364
259.180908203125 0 672.48895
262.0830078125 0 579.1409
262.15618896484375 0 642.06323
267.1086730957031 0 1598.1056
268.0928039550781 0 2535.3645
268.16583251953125 0 73614.81
269.1688537597656 0 9542.205
270.1450500488281 0 2020.746
270.17242431640625 0 615.08765
271.1445007324219 0 1030.2069
273.1608581542969 0 658.53094
275.1392822265625 0 872.48975
275.175048828125 0 959.34534
275.6720886230469 0 665.06964
280.1290283203125 0 920.8859
280.1649169921875 0 975.6611
282.144287109375 0 1041.4109
282.1813049316406 0 2276.423
284.1607971191406 0 533.48846
284.19696044921875 0 1381.0388
284.6777038574219 0 606.45465
285.11993408203125 0 1426.5789
285.1932067871094 0 1733.6106
286.1035461425781 0 695.5327
286.1763000488281 0 1004.1804
287.17578125 0 875.1016
288.17291259765625 0 509.9148
289.1917724609375 0 794.30133
289.6700439453125 0 769.5748
292.1659851074219 0 585.7516 d 2
294.14459228515625 0 548.3682
295.1409606933594 0 3009.8516
295.1657409667969 0 19749.154 y 10
296.1683654785156 0 2315.3232
296.197021484375 0 1763.9982
297.11944580078125 0 1463.6188
297.1556396484375 0 755.8653
297.194580078125 0 727.15533
299.1720275878906 0 2713.2056
300.156005859375 0 1636.999
303.1299743652344 0 1216.4576
306.1816101074219 0 738.64795 b Water loss 5
307.1042785644531 0 964.4468
309.1553649902344 0 767.3261
310.14013671875 0 884.1024
310.1758117675781 0 853.6718
310.2125244140625 0 1824.9237
311.1721496582031 0 1707.4728
312.1559753417969 0 1346.6268
313.1514587402344 0 1125.3667
315.130126953125 0 758.5648
315.6671142578125 0 1348.5935
316.1660461425781 0 4945.1084
317.1827697753906 0 17196.219
318.1860656738281 0 3188.9807 a 2
324.1312561035156 0 1728.466
324.1797180175781 0 3856.0356
324.68145751953125 0 1231.8306
325.114501953125 0 5627.1846
325.15179443359375 0 713.9534
325.187744140625 0 2286.1375
326.11688232421875 0 721.5197
327.1671447753906 0 3807.9668
327.2021484375 0 842.3182
328.1664123535156 0 2940.7395 b Water loss 2
329.18267822265625 0 3893.843
330.18231201171875 0 704.3418
333.1866149902344 0 1691.1068
339.2027587890625 0 1018.41675
341.18243408203125 0 663.9375
341.218994140625 0 5521.6055
342.14129638671875 0 3905.271
343.1252136230469 0 7055.8594
343.1988220214844 0 769.80304
344.1967468261719 0 1402.7836
345.1773376464844 0 21922.824
346.1778259277344 0 9596.566 b 2
347.1789245605469 0 994.3635
351.203369140625 0 6806.855
352.1871032714844 0 3969.418 y 9
352.20867919921875 0 652.9686
352.69140625 0 594.6772
353.1834716796875 0 1757.0023
353.25506591796875 0 10246.785
354.25750732421875 0 1534.2728
355.1974182128906 0 807.9003
360.15234375 0 1687.8732
361.6954650878906 0 7483.8384
362.1985168457031 0 3417.5264
362.6983337402344 0 754.4977
363.1665344238281 0 1026.1152
365.1475830078125 0 1042.7579
365.21868896484375 0 992.2196
369.2137451171875 0 3213.9224
370.1373596191406 0 704.2644
370.21392822265625 0 2154.0613 b Water loss 6
370.70001220703125 0 993.1338 b Ammonia loss 6
372.19134521484375 0 691.2275
373.1893310546875 0 838.34827
378.2138671875 0 635.04193
379.1972961425781 0 1500.2261
380.1967468261719 0 1110.0525
381.2498779296875 0 21110.893
382.1723327636719 0 1116.6588
382.2530517578125 0 4047.1057
383.22967529296875 0 775.54083
384.18817138671875 0 1030.1626
384.2304992675781 0 737.1817
387.2393798828125 0 3179.1746
389.218994140625 0 2103.714
393.177734375 0 1243.3037
394.2085266113281 0 3522.0908
396.22576904296875 0 712.03357
398.2054748535156 0 1271.6802
398.24725341796875 0 907.1933
398.2768249511719 0 5756.7686
399.20306396484375 0 6813.653 b Water loss 3
399.2817077636719 0 1003.1356
400.2080383300781 0 1341.3185
401.2162170410156 0 857.81335
407.1954650878906 0 560.0118
408.2237854003906 0 934.3596
410.2029113769531 0 1513.229
410.2401428222656 0 1544.7213
412.2205505371094 0 962.1486
412.25543212890625 0 759.7069
415.2346496582031 0 2621.6716
416.2403259277344 0 1088.1208
417.21380615234375 0 13430.73 b 3
418.2164306640625 0 2251.179
420.18975830078125 0 1225.2716
422.7326965332031 0 1970.5813 b 7
424.21990966796875 0 1031.1294
425.2174377441406 0 1252.5939 y Ammonia loss 4
426.23638916015625 0 1058.2997
427.23516845703125 0 1834.508
428.21466064453125 0 5204.5894
428.24542236328125 0 810.4528
429.2163391113281 0 1308.144
433.2444152832031 0 989.97906
437.2150573730469 0 1334.3577
438.1986083984375 0 1676.2184
438.2715759277344 0 975.3493
439.2189025878906 0 3586.252 y 8
440.22247314453125 0 1052.8342
444.260986328125 0 1069.6049
445.2423400878906 0 3803.4976
446.2399597167969 0 652.1267
448.2373046875 0 1588.8573
448.7381591796875 0 1165.3729
453.24920654296875 0 877.73517
455.2257385253906 0 6387.299
456.2098083496094 0 4889.0747
457.2113037109375 0 1649.88
457.2441101074219 0 887.13367 b Water loss 8
458.26202392578125 0 1790.1486
462.19921875 0 683.013
464.2142333984375 0 1094.4177
464.2880859375 0 3908.5457
465.291259765625 0 1135.6407
466.2320556640625 0 730.3332
470.2765808105469 0 2049.7808
471.2559509277344 0 1190.987
472.25701904296875 0 2003.8962
473.23614501953125 0 7387.1113
474.23919677734375 0 2932.1494
480.2458190917969 0 1128.4253
481.24114990234375 0 1169.1565
481.3135070800781 0 1158.5787
482.2988586425781 0 1959.0835
483.2984313964844 0 1381.6649
485.7539367675781 0 3924.2388 b Water loss 9
486.2528991699219 0 1668.3982 b Ammonia loss 9
488.28741455078125 0 7326.7427
489.2890319824219 0 1863.8252
490.2279052734375 0 833.68933
491.2982177734375 0 734.61633
492.28155517578125 0 1205.944
497.2727355957031 0 850.9733
498.271484375 0 17789.32 b Water loss 4
499.275146484375 0 4228.9517
499.7704772949219 0 902.14557
500.3225402832031 0 1015.0497
507.2935791015625 0 2196.2488
508.2404479980469 0 2165.8455 y Water loss 7
508.77435302734375 0 925.37885 y Water loss 2
509.2419738769531 0 850.60815
509.3088684082031 0 22984.387
510.3116760253906 0 6474.9517
511.31494140625 0 1030.0406
512.2146606445312 0 864.53815
512.28857421875 0 1173.9403
516.2821655273438 0 19843.465 b 4
517.2850341796875 0 7064.4097
518.2882080078125 0 959.17163
525.3032836914062 0 961.1882
526.2513427734375 0 13370.923 y 7
527.2547607421875 0 2385.0908
528.319580078125 0 1428.1383
530.2992553710938 0 802.05304
531.2984619140625 0 714.8161
538.2608642578125 0 927.96063
539.8102416992188 0 779.093
540.3156127929688 0 1284.7301
541.3012084960938 0 687.15234
543.2559814453125 0 770.6705
549.8176879882812 0 615.7372
551.2815551757812 0 908.73157
554.2965087890625 0 859.7676
557.339599609375 0 611.83154
558.3251953125 0 3949.2004
566.3309326171875 0 1199.2979
568.3098754882812 0 2468.008
569.309326171875 0 600.6445
572.302978515625 0 951.3463
573.2278442382812 0 639.4334
575.2811279296875 0 704.00195
577.3024291992188 0 940.3942
578.3302001953125 0 3133.2249
578.8102416992188 0 1220.2826
579.3258056640625 0 1060.2771
583.3623046875 0 1100.6439
584.3462524414062 0 2251.5244
585.3482055664062 0 1424.2142
586.3197631835938 0 7433.6216
587.3233642578125 0 2801.8447
592.34375 0 1567.8982
593.3469848632812 0 857.4303
594.3245849609375 0 1208.8679
596.3406372070312 0 6846.6895
597.3430786132812 0 1770.8124
601.3713989257812 0 2743.3877
602.3731689453125 0 1462.094
609.3001098632812 0 783.4887
610.28759765625 0 908.4552
611.3557739257812 0 13186.244 b Water loss 5
612.3556518554688 0 4470.106
613.3656616210938 0 857.7663
618.2877197265625 0 785.59424
624.3119506835938 0 800.3322
629.3657836914062 0 11618.244 b 5
630.3690795898438 0 4394.2847
631.37353515625 0 821.4376
636.2992553710938 0 7344.06 y Water loss 6
637.2985229492188 0 1048.8499
641.8544921875 0 13743.981
642.3558349609375 0 8868.674
642.8578491210938 0 4347.823
643.3506469726562 0 1126.2501
644.3353881835938 0 617.65204
646.3890991210938 0 1160.8733
647.349609375 0 1498.7695
648.34326171875 0 1075.0387
654.31005859375 0 15087.51 y 6
655.3124389648438 0 4637.192
656.3157958984375 0 881.44684
656.3763427734375 0 1054.2372
657.3882446289062 0 1215.0728
665.3619995117188 0 2023.1549
667.3805541992188 0 962.61597
668.3685913085938 0 719.68066
672.3344116210938 0 925.63696
683.3709106445312 0 2428.853
684.3741455078125 0 737.0744
685.32666015625 0 821.6515
685.3888549804688 0 1835.4188
686.3888549804688 0 864.9503
695.3759155273438 0 1280.7607
703.33642578125 0 1286.4199
704.3715209960938 0 2128.2837
705.3695068359375 0 1639.021
706.375 0 681.38086
712.4013061523438 0 1643.1622
713.403076171875 0 1396.2668
722.3834838867188 0 12737.546
723.381103515625 0 5918.04
724.3764038085938 0 1466.8534
726.4197387695312 0 889.15985
729.4279174804688 0 1114.2332
739.4130859375 0 4381.4746 b Water loss 6
740.3955688476562 0 19030.885 b Ammonia loss 6
741.3975830078125 0 8651.813
742.3938598632812 0 753.9978
749.3836669921875 0 1572.8715 y Water loss 5
750.3740844726562 0 1681.7292 y Ammonia loss 5
755.4126586914062 0 952.0251
756.4220581054688 0 785.0647
757.423828125 0 9255.182 b 6
758.42724609375 0 3973.6318
764.9608764648438 0 778.2194
766.4140625 0 894.1662
767.3935546875 0 14911.288 y 5
768.3971557617188 0 6272.757
769.40185546875 0 1422.2662
808.4474487304688 0 1231.4271
809.442138671875 0 823.3616
816.4322509765625 0 692.9373
817.451171875 0 1112.8965
824.4234619140625 0 1038.681
825.4832153320312 0 6263.4727
826.4534912109375 0 1992.3262 b Water loss 7
830.4385375976562 0 782.13556
835.4673461914062 0 3519.4878
836.4607543945312 0 3052.3896
837.4544677734375 0 955.05
841.447509765625 0 820.38336
842.4445190429688 0 1514.3411
844.455810546875 0 3639.0598 b 7
845.459716796875 0 1927.8325
853.4779052734375 0 20322.527
854.480224609375 0 9043.655
855.4808349609375 0 2533.6316
866.4606323242188 0 1667.9747 y 4
895.465576171875 0 754.7573
896.4677124023438 0 645.48975
913.4671630859375 0 1165.7076 b Water loss 8
914.474365234375 0 1170.5328 b Ammonia loss 8
931.48974609375 0 2170.7678 b 8
932.4891357421875 0 1187.5317
935.4617919921875 0 764.1581
936.513916015625 0 1657.003
937.5108032226562 0 1046.9474 y 3
942.5045776367188 0 933.8308
944.4879150390625 0 745.3274
952.4862670898438 0 3784.3145
953.4861450195312 0 3219.8088
954.4946899414062 0 1092.6757
960.515625 0 1738.998
961.5106811523438 0 808.4393
970.4990844726562 0 19160.36 b Water loss 9
971.4992065429688 0 11995.358 b Ammonia loss 9
972.4994506835938 0 3978.6511
973.5198974609375 0 906.13324
988.5096435546875 0 17903.75 b 9
989.5123291015625 0 9190.189
990.5138549804688 0 2313.5474
1016.5445556640625 0 1801.739 y Water loss 2
1017.5331420898438 0 1530.4739 y Ammonia loss 2
1034.5506591796875 0 12578.9 y 2
1035.5538330078125 0 7545.3423
1036.5595703125 0 2440.015
1056.58203125 0 865.58276
1065.569091796875 0 1189.887
1066.5701904296875 0 1939.4452
1073.598388671875 0 2267.3174
1074.59716796875 0 1332.7272
1083.5823974609375 0 8429.002 b Water loss 10
1084.5819091796875 0 4915.673 b Ammonia loss 10
1085.5849609375 0 2806.1675
1101.5941162109375 0 8283.63 b 10
1102.5948486328125 0 5727.6724
1103.5987548828125 0 1744.1918
1211.806884765625 0 728.29395
2352.51708984375 0 607.7312
2511.380615234375 0 641.35315
3083.14111328125 0 736.35583

Spectrum Details

|  |  |
| --- | --- |
| Matched peaks? Matched peaks The total absolute number of peaks matched. Additionally in brackets the total fraction of peaks matched and the total number of peaks is shown. | 53 (10.95% of 484) |
| FDR? FDR The false discovery rate estimated for this peptide. It is calculated by matching all theoretical fragments with a non-integer shift with the raw peaks for this spectrum. This is done with 40 different shifts. The resulting percentage is the average number of annotated peaks over the number of annotated peaks with the correct spectrum. | 0.13% |
| Satellite FDR? Satellite FDR See the FDR for details on its calculation. This satellite ion specific FDR only contains the satellite ions (d/w) for I/L/J positions. | - |
| PSM Score? PSM Score The PSM Score as given by Hecklib to this annotated spectrum. It is shown with three significant figures. | 732 |

#### Reverse Lookup? Reverse Lookup All places where this read could be placed.

| Group | Segment | Template | Template Part | Read Part | Score | Unique |
| --- | --- | --- | --- | --- | --- | --- |
| Homo sapiens Heavy Chain | IGHC | IGHG1 | [51..63] | [0..12] | 96 | False |
| Homo sapiens Heavy Chain | IGHC | IGHG3 | [51..63] | [0..12] | 96 | False |
| Homo sapiens Heavy Chain | IGHC | IGHG2 | [51..63] | [0..12] | 96 | False |
| Homo sapiens Heavy Chain | IGHC | IGHG4 | [51..63] | [0..12] | 96 | False |

| Recombined | Template Part | Read Part | Score | Unique |
| --- | --- | --- | --- | --- |
| REC-0-1 | [173..185] | [0..12] | 96 | True |

#### Meta Information from Multiple reads

##### Number of combined reads

8

##### Intensity

0.977

##### TotalArea

1.871E+09

##### Changes to the peptide sequence

TFPAVJQSSGJY

L→JNo support for either Leucine or Isoleucine based on side chain ions (Position: 11)

L→JNo support for either Leucine or Isoleucine based on side chain ions (Position: 6)

#### Positional Score

Copy Data

##### Positional Score (TSV)

###### Preview

```
Loading example...
```

*Click on the button to copy the data to your clipboard.*

1001234567891011

Label Value
"0" 0.722
"1" 0.675
"2" 0.711
"3" 0.716
"4" 0.741
"5" 0.749
"6" 0.736
"7" 0.746
"8" 0.748
"9" 0.746
"10" 0.75
"11" 0.75

#### Meta Information from PEAKS

##### Scan Identifier

F3:11139

##### Original sequence

T

F

P

A

V

L

Q

S

S

G

L

Y

##### Posttranslational Modifications

##### Source File

D:\separate\_stitch\_analyses\xle-disambiguation\raw\20210323\_F1\_UM1\_Peng0013\_SA\_F59\_ingel\_3ug\_chymo.raw

##### Fraction

3

##### Scan Feature

-

##### De Novo Score

99

##### ConfidenceScore

99

### m/z

641.8378

##### Mass

1281.6604

##### Charge

2

##### Retention Time

63.02

##### Predicted Retention Time

-

##### Area

0

##### Parts Per Million

0.5

##### Fragmentation mode

ETHCD

##### Originating file

01 D:\separate\_stitch\_analyses\xle-disambiguation\20210325\_F59\_3ug\_DENOVO\_12.csv

#### Meta Information from PEAKS

##### Scan Identifier

F3:10666

##### Original sequence

T

F

P

A

V

L

Q

S

S

G

L

Y

##### Posttranslational Modifications

##### Source File

D:\separate\_stitch\_analyses\xle-disambiguation\raw\20210323\_F1\_UM1\_Peng0013\_SA\_F59\_ingel\_3ug\_chymo.raw

##### Fraction

3

##### Scan Feature

F3:9878

##### De Novo Score

98

##### ConfidenceScore

98

### m/z

641.8386

##### Mass

1281.6604

##### Charge

2

##### Retention Time

60.4

##### Predicted Retention Time

-

##### Area

9.355E+08

##### Parts Per Million

1.8

##### Fragmentation mode

ETHCD

##### Originating file

01 D:\separate\_stitch\_analyses\xle-disambiguation\20210325\_F59\_3ug\_DENOVO\_12.csv

#### Meta Information from PEAKS

##### Scan Identifier

F3:11052

##### Original sequence

T

F

P

A

V

L

Q

S

S

G

L

Y

##### Posttranslational Modifications

##### Source File

D:\separate\_stitch\_analyses\xle-disambiguation\raw\20210323\_F1\_UM1\_Peng0013\_SA\_F59\_ingel\_3ug\_chymo.raw

##### Fraction

3

##### Scan Feature

-

##### De Novo Score

98

##### ConfidenceScore

98

### m/z

641.838

##### Mass

1281.6604

##### Charge

2

##### Retention Time

62.5

##### Predicted Retention Time

-

##### Area

0

##### Parts Per Million

0.8

##### Fragmentation mode

ETHCD

##### Originating file

01 D:\separate\_stitch\_analyses\xle-disambiguation\20210325\_F59\_3ug\_DENOVO\_12.csv

#### Meta Information from PEAKS

##### Scan Identifier

F3:11433

##### Original sequence

T

F

P

A

V

L

Q

S

S

G

L

Y

##### Posttranslational Modifications

##### Source File

D:\separate\_stitch\_analyses\xle-disambiguation\raw\20210323\_F1\_UM1\_Peng0013\_SA\_F59\_ingel\_3ug\_chymo.raw

##### Fraction

3

##### Scan Feature

-

##### De Novo Score

98

##### ConfidenceScore

98

### m/z

641.8381

##### Mass

1281.6604

##### Charge

2

##### Retention Time

64.75

##### Predicted Retention Time

-

##### Area

0

##### Parts Per Million

0.9

##### Fragmentation mode

ETHCD

##### Originating file

01 D:\separate\_stitch\_analyses\xle-disambiguation\20210325\_F59\_3ug\_DENOVO\_12.csv

#### Meta Information from PEAKS

##### Scan Identifier

F3:11765

##### Original sequence

T

F

P

A

V

L

Q

S

S

G

L

Y

##### Posttranslational Modifications

##### Source File

D:\separate\_stitch\_analyses\xle-disambiguation\raw\20210323\_F1\_UM1\_Peng0013\_SA\_F59\_ingel\_3ug\_chymo.raw

##### Fraction

3

##### Scan Feature

-

##### De Novo Score

97

##### ConfidenceScore

97

### m/z

641.8381

##### Mass

1281.6604

##### Charge

2

##### Retention Time

66.63

##### Predicted Retention Time

-

##### Area

0

##### Parts Per Million

1

##### Fragmentation mode

ETHCD

##### Originating file

01 D:\separate\_stitch\_analyses\xle-disambiguation\20210325\_F59\_3ug\_DENOVO\_12.csv

#### Meta Information from PEAKS

##### Scan Identifier

F3:10897

##### Original sequence

T

F

P

A

V

L

Q

S

S

G

L

Y

##### Posttranslational Modifications

##### Source File

D:\separate\_stitch\_analyses\xle-disambiguation\raw\20210323\_F1\_UM1\_Peng0013\_SA\_F59\_ingel\_3ug\_chymo.raw

##### Fraction

3

##### Scan Feature

F3:9878

##### De Novo Score

97

##### ConfidenceScore

97

### m/z

641.8386

##### Mass

1281.6604

##### Charge

2

##### Retention Time

60.4

##### Predicted Retention Time

-

##### Area

9.355E+08

##### Parts Per Million

1.8

##### Fragmentation mode

HCD

##### Originating file

01 D:\separate\_stitch\_analyses\xle-disambiguation\20210325\_F59\_3ug\_DENOVO\_12.csv

#### Meta Information from PEAKS

##### Scan Identifier

F3:11657

##### Original sequence

T

F

P

A

V

L

Q

S

S

G

L

Y

##### Posttranslational Modifications

##### Source File

D:\separate\_stitch\_analyses\xle-disambiguation\raw\20210323\_F1\_UM1\_Peng0013\_SA\_F59\_ingel\_3ug\_chymo.raw

##### Fraction

3

##### Scan Feature

-

##### De Novo Score

97

##### ConfidenceScore

97

### m/z

641.8383

##### Mass

1281.6604

##### Charge

2

##### Retention Time

66.01

##### Predicted Retention Time

-

##### Area

0

##### Parts Per Million

1.2

##### Fragmentation mode

ETHCD

##### Originating file

01 D:\separate\_stitch\_analyses\xle-disambiguation\20210325\_F59\_3ug\_DENOVO\_12.csv

#### Meta Information from PEAKS

##### Scan Identifier

F3:11194

##### Original sequence

T

F

P

A

V

L

Q

S

S

G

L

Y

##### Posttranslational Modifications

##### Source File

D:\separate\_stitch\_analyses\xle-disambiguation\raw\20210323\_F1\_UM1\_Peng0013\_SA\_F59\_ingel\_3ug\_chymo.raw

##### Fraction

3

##### Scan Feature

-

##### De Novo Score

96

##### ConfidenceScore

96

### m/z

641.8383

##### Mass

1281.6604

##### Charge

2

##### Retention Time

63.34

##### Predicted Retention Time

-

##### Area

0

##### Parts Per Million

1.3

##### Fragmentation mode

HCD

##### Originating file

01 D:\separate\_stitch\_analyses\xle-disambiguation\20210325\_F59\_3ug\_DENOVO\_12.csv
