## Supplementary material for "A handle on mass coincidence errors in *de novo* sequencing of antibodies by bottom-up proteomics": Combined_046.html

Details Combined\_046 | Stitch OverviewUndefined

### Read Combined\_046

#### Sequence (length=7)

FYYCAKD

#### Spectrum 4867? Spectrum 4867 The raw spectrum of this peptide as annotated by Hecklib. The fragments are coloured according to ion type (see legend). Any peaks with a star '\*' as text can be hovered over to see the full details, first the ion type second the mass shift type. By hovering over the amino acids in the peptide or ions in the legend the corresponding peaks are highlighted. By toggling the 'Unassigned' label you can turn the background (unassigned) peaks on or off in the plot. By updating the slider in the Ion legend you can update the spectrum to only show the top X% of the peaks with labels. The top X% means any peak that is within X% of the highest intensity. By dragging in the spectrum you can zoom in to a specific part of the spectrum and use 'Zoom Out' to get back to the original zoom level. The annotation of the spectrum is based on the given sequence in the peptides file and is done with different software so inconsistencies are likely. The peaks are annotated based on the given sequence, with 20 ppm tolerance.

Copy Data

##### Spectrum 4867 (TSV)

###### Preview

```
Loading example...
```

*Click on the button to copy the data to your clipboard.*

Mz MinMz MaxIntensity Max

WidthHeightPeptide font sizePeptide stroke widthSpectrum font sizeSpectrum stroke widthCompact peptide

Ion legend

wxyz

abcd

OtherUnassignedIon^Charge^_Position_Show for top:%

FYYCAKD

02.24e+44.49e+46.73e+48.97e+4

Zoom Out

y^+1^_1_y^+1^_2_y^+1^_2_y^+1^_2_a^+1^_2_b^+1^_2_y^+1^_3_y^+1^_3_y^+1^_4_\*y^+1^_4_y^+1^_5_y^+1^_5_y^+1^_6_

0776155323293105

Fragment Matches Table

Show background peaks

| Position | Ion type | Intensity | mz Theoretical | mz Error (Th) | mz Error (ppm) | Charge | Series Number |
| --- | --- | --- | --- | --- | --- | --- | --- |
| - | - | 8.867E+04 | 120.1 | - | - | 0 | - |
| - | - | 6222 | 121.1 | - | - | 0 | - |
| - | - | 457 | 127.1 | - | - | 0 | - |
| - | - | 1074 | 127.1 | - | - | 0 | - |
| - | - | 803.1 | 128.1 | - | - | 0 | - |
| - | - | 1112 | 128.1 | - | - | 0 | - |
| - | - | 938.7 | 129.1 | - | - | 0 | - |
| - | - | 5.456E+04 | 129.1 | - | - | 0 | - |
| - | - | 1478 | 130.1 | - | - | 0 | - |
| - | - | 1457 | 130.1 | - | - | 0 | - |
| - | - | 1821 | 130.1 | - | - | 0 | - |
| - | - | 2859 | 130.1 | - | - | 0 | - |
| - | - | 709 | 131.1 | - | - | 0 | - |
| - | - | 872.1 | 133.1 | - | - | 0 | - |
| - | - | 2255 | 134 | - | - | 0 | - |
| 7 | y | 4747 | 134 | 0.000275 | 2.052 | +1 | 1 |
| - | - | 950.8 | 134.1 | - | - | 0 | - |
| - | - | 8.885E+04 | 136.1 | - | - | 0 | - |
| - | - | 7062 | 137.1 | - | - | 0 | - |
| - | - | 954.9 | 139.1 | - | - | 0 | - |
| - | - | 1032 | 140.1 | - | - | 0 | - |
| - | - | 866.5 | 141.1 | - | - | 0 | - |
| - | - | 1843 | 141.1 | - | - | 0 | - |
| - | - | 602.5 | 142.1 | - | - | 0 | - |
| - | - | 566.9 | 142.1 | - | - | 0 | - |
| - | - | 567.3 | 143.1 | - | - | 0 | - |
| - | - | 1755 | 144.1 | - | - | 0 | - |
| - | - | 463.1 | 145 | - | - | 0 | - |
| - | - | 1403 | 145.1 | - | - | 0 | - |
| - | - | 1160 | 146.1 | - | - | 0 | - |
| - | - | 882.7 | 147 | - | - | 0 | - |
| - | - | 1135 | 147.1 | - | - | 0 | - |
| - | - | 1140 | 149 | - | - | 0 | - |
| - | - | 621.8 | 149 | - | - | 0 | - |
| - | - | 3580 | 152.1 | - | - | 0 | - |
| - | - | 683.8 | 153.1 | - | - | 0 | - |
| - | - | 675.9 | 155.1 | - | - | 0 | - |
| - | - | 2055 | 155.1 | - | - | 0 | - |
| - | - | 570.1 | 157.1 | - | - | 0 | - |
| - | - | 487.6 | 157.1 | - | - | 0 | - |
| - | - | 2171 | 157.1 | - | - | 0 | - |
| - | - | 638.2 | 158.1 | - | - | 0 | - |
| - | - | 1690 | 159.1 | - | - | 0 | - |
| - | - | 1896 | 159.1 | - | - | 0 | - |
| - | - | 8390 | 163.1 | - | - | 0 | - |
| - | - | 1466 | 165.1 | - | - | 0 | - |
| - | - | 421.2 | 166 | - | - | 0 | - |
| - | - | 769.2 | 166.1 | - | - | 0 | - |
| - | - | 538.2 | 167.1 | - | - | 0 | - |
| - | - | 983.6 | 169.1 | - | - | 0 | - |
| - | - | 1654 | 169.1 | - | - | 0 | - |
| - | - | 548.8 | 171.1 | - | - | 0 | - |
| - | - | 731 | 171.1 | - | - | 0 | - |
| - | - | 2843 | 171.1 | - | - | 0 | - |
| - | - | 624.4 | 173.1 | - | - | 0 | - |
| - | - | 1316 | 173.5 | - | - | 0 | - |
| - | - | 938.3 | 175.1 | - | - | 0 | - |
| - | - | 1097 | 181.1 | - | - | 0 | - |
| - | - | 729.1 | 181.1 | - | - | 0 | - |
| - | - | 559.4 | 182.1 | - | - | 0 | - |
| - | - | 1331 | 182.1 | - | - | 0 | - |
| - | - | 1457 | 183.1 | - | - | 0 | - |
| - | - | 926.4 | 183.1 | - | - | 0 | - |
| - | - | 624.4 | 185.1 | - | - | 0 | - |
| - | - | 1450 | 185.1 | - | - | 0 | - |
| - | - | 1.738E+04 | 185.2 | - | - | 0 | - |
| - | - | 1869 | 186.1 | - | - | 0 | - |
| - | - | 1482 | 186.2 | - | - | 0 | - |
| - | - | 979.8 | 187.1 | - | - | 0 | - |
| - | - | 728.5 | 187.1 | - | - | 0 | - |
| - | - | 629 | 188.1 | - | - | 0 | - |
| - | - | 574.6 | 195.1 | - | - | 0 | - |
| - | - | 483.1 | 198.1 | - | - | 0 | - |
| - | - | 1148 | 199.1 | - | - | 0 | - |
| - | - | 846.2 | 199.1 | - | - | 0 | - |
| - | - | 1.452E+04 | 200.1 | - | - | 0 | - |
| - | - | 1041 | 201.1 | - | - | 0 | - |
| - | - | 1186 | 201.1 | - | - | 0 | - |
| - | - | 1776 | 202.1 | - | - | 0 | - |
| - | - | 1.085E+04 | 203.1 | - | - | 0 | - |
| - | - | 883.3 | 204.1 | - | - | 0 | - |
| - | - | 2187 | 205.1 | - | - | 0 | - |
| - | - | 1161 | 207.1 | - | - | 0 | - |
| - | - | 1049 | 210.1 | - | - | 0 | - |
| - | - | 880.1 | 211.1 | - | - | 0 | - |
| - | - | 1193 | 211.1 | - | - | 0 | - |
| - | - | 8089 | 213.2 | - | - | 0 | - |
| - | - | 1029 | 214.2 | - | - | 0 | - |
| - | - | 910.9 | 215.1 | - | - | 0 | - |
| - | - | 2289 | 215.1 | - | - | 0 | - |
| - | - | 1038 | 216.1 | - | - | 0 | - |
| - | - | 685.1 | 216.1 | - | - | 0 | - |
| - | - | 654.2 | 218.2 | - | - | 0 | - |
| - | - | 777.6 | 224.1 | - | - | 0 | - |
| - | - | 649.1 | 224.1 | - | - | 0 | - |
| - | - | 825.1 | 225.1 | - | - | 0 | - |
| - | - | 833.7 | 226.1 | - | - | 0 | - |
| - | - | 9416 | 226.2 | - | - | 0 | - |
| - | - | 840.8 | 227.1 | - | - | 0 | - |
| - | - | 1196 | 227.2 | - | - | 0 | - |
| - | - | 1613 | 227.2 | - | - | 0 | - |
| - | - | 693.9 | 228.1 | - | - | 0 | - |
| - | - | 1110 | 233.1 | - | - | 0 | - |
| - | - | 5712 | 233.1 | - | - | 0 | - |
| - | - | 1672 | 233.1 | - | - | 0 | - |
| - | - | 3729 | 233.2 | - | - | 0 | - |
| - | - | 564.1 | 234.1 | - | - | 0 | - |
| - | - | 617.6 | 234.2 | - | - | 0 | - |
| - | - | 1029 | 239.2 | - | - | 0 | - |
| - | - | 2419 | 240.1 | - | - | 0 | - |
| - | - | 603.4 | 243.1 | - | - | 0 | - |
| 6 | y | 1151 | 244.1 | 0.0005204 | 2.132 | +1 | 2 |
| 6 | y | 2893 | 245.1 | 0.0001137 | 0.4638 | +1 | 2 |
| - | - | 879.9 | 246.2 | - | - | 0 | - |
| - | - | 641 | 249.1 | - | - | 0 | - |
| - | - | 677 | 249.2 | - | - | 0 | - |
| - | - | 711.9 | 251.2 | - | - | 0 | - |
| - | - | 1062 | 252.1 | - | - | 0 | - |
| - | - | 561.4 | 252.1 | - | - | 0 | - |
| - | - | 519.5 | 252.2 | - | - | 0 | - |
| - | - | 3316 | 253.2 | - | - | 0 | - |
| - | - | 800.9 | 254.1 | - | - | 0 | - |
| - | - | 996.5 | 256.2 | - | - | 0 | - |
| - | - | 527.9 | 261.1 | - | - | 0 | - |
| - | - | 709.1 | 261.2 | - | - | 0 | - |
| 6 | y | 1.396E+04 | 262.1 | 8.435E-05 | 0.3218 | +1 | 2 |
| - | - | 1416 | 263.1 | - | - | 0 | - |
| - | - | 597.1 | 268.3 | - | - | 0 | - |
| - | - | 766.4 | 270.2 | - | - | 0 | - |
| - | - | 704.4 | 271.2 | - | - | 0 | - |
| - | - | 575.7 | 272.1 | - | - | 0 | - |
| - | - | 690.2 | 274.1 | - | - | 0 | - |
| - | - | 1463 | 279.2 | - | - | 0 | - |
| - | - | 1061 | 281.1 | - | - | 0 | - |
| - | - | 586.2 | 282.1 | - | - | 0 | - |
| 2 | a | 4.836E+04 | 283.1 | 0.0001523 | 0.5379 | +1 | 2 |
| - | - | 8895 | 284.1 | - | - | 0 | - |
| - | - | 873.7 | 285.2 | - | - | 0 | - |
| - | - | 1950 | 287.2 | - | - | 0 | - |
| - | - | 1.002E+04 | 297.1 | - | - | 0 | - |
| - | - | 594.1 | 297.1 | - | - | 0 | - |
| - | - | 2285 | 297.2 | - | - | 0 | - |
| - | - | 1786 | 298.1 | - | - | 0 | - |
| - | - | 6997 | 299.1 | - | - | 0 | - |
| - | - | 4467 | 299.2 | - | - | 0 | - |
| - | - | 932.3 | 300.2 | - | - | 0 | - |
| - | - | 6224 | 304.1 | - | - | 0 | - |
| - | - | 543.3 | 304.2 | - | - | 0 | - |
| - | - | 996.1 | 305.1 | - | - | 0 | - |
| - | - | 728.6 | 306.1 | - | - | 0 | - |
| - | - | 1135 | 309.1 | - | - | 0 | - |
| 2 | b | 1.016E+04 | 311.1 | 0.0002633 | 0.8463 | +1 | 2 |
| - | - | 1534 | 312.1 | - | - | 0 | - |
| 5 | y | 856.4 | 315.2 | 0.0007079 | 2.246 | +1 | 3 |
| - | - | 1205 | 316.2 | - | - | 0 | - |
| - | - | 941.2 | 323.2 | - | - | 0 | - |
| - | - | 6004 | 325.1 | - | - | 0 | - |
| - | - | 595.1 | 326.1 | - | - | 0 | - |
| - | - | 3787 | 327.1 | - | - | 0 | - |
| 5 | y | 8414 | 333.2 | 0.0001105 | 0.3315 | +1 | 3 |
| - | - | 2065 | 334.2 | - | - | 0 | - |
| - | - | 891.4 | 338.2 | - | - | 0 | - |
| - | - | 727.2 | 340.2 | - | - | 0 | - |
| - | - | 842.6 | 340.2 | - | - | 0 | - |
| - | - | 2447 | 340.3 | - | - | 0 | - |
| - | - | 686.4 | 349.8 | - | - | 0 | - |
| - | - | 2495 | 361.2 | - | - | 0 | - |
| - | - | 6691 | 368.2 | - | - | 0 | - |
| - | - | 660.2 | 394.2 | - | - | 0 | - |
| - | - | 3677 | 396.1 | - | - | 0 | - |
| - | - | 1411 | 396.2 | - | - | 0 | - |
| - | - | 671.2 | 397.1 | - | - | 0 | - |
| - | - | 1587 | 398.2 | - | - | 0 | - |
| - | - | 659.5 | 413.7 | - | - | 0 | - |
| - | - | 675.2 | 431.2 | - | - | 0 | - |
| - | - | 645 | 440.2 | - | - | 0 | - |
| - | - | 609.6 | 441.3 | - | - | 0 | - |
| - | - | 605.2 | 442.2 | - | - | 0 | - |
| - | - | 1034 | 442.3 | - | - | 0 | - |
| - | - | 601.2 | 466.2 | - | - | 0 | - |
| - | - | 646.6 | 466.7 | - | - | 0 | - |
| - | - | 1314 | 467.3 | - | - | 0 | - |
| - | - | 1824 | 470.3 | - | - | 0 | - |
| - | - | 994 | 472.3 | - | - | 0 | - |
| 4 | y | 1268 | 476.2 | 0.005305 | 11.14 | +1 | 4 |
| - | - | 634.1 | 481.2 | - | - | 0 | - |
| - | - | 1308 | 482.2 | - | - | 0 | - |
| - | - | 1240 | 482.3 | - | - | 0 | - |
| 0 | Precursor | 1359 | 484.2 | 0.00364 | 7.517 | +2 | -1 |
| - | - | 855.3 | 484.3 | - | - | 0 | - |
| - | - | 3700 | 484.3 | - | - | 0 | - |
| - | - | 887.4 | 484.7 | - | - | 0 | - |
| - | - | 1163 | 484.8 | - | - | 0 | - |
| - | - | 1486 | 485.3 | - | - | 0 | - |
| - | - | 2423 | 485.4 | - | - | 0 | - |
| - | - | 1979 | 488.1 | - | - | 0 | - |
| - | - | 958.8 | 488.3 | - | - | 0 | - |
| - | - | 1228 | 488.8 | - | - | 0 | - |
| - | - | 2556 | 489.8 | - | - | 0 | - |
| - | - | 1184 | 490.3 | - | - | 0 | - |
| - | - | 1219 | 490.8 | - | - | 0 | - |
| 4 | y | 1.018E+04 | 494.2 | 0.004395 | 8.893 | +1 | 4 |
| - | - | 750.6 | 494.2 | - | - | 0 | - |
| - | - | 2687 | 495.2 | - | - | 0 | - |
| - | - | 594.5 | 509.3 | - | - | 0 | - |
| - | - | 2648 | 513.4 | - | - | 0 | - |
| - | - | 666.9 | 514.4 | - | - | 0 | - |
| - | - | 582.6 | 515.2 | - | - | 0 | - |
| - | - | 2203 | 524.2 | - | - | 0 | - |
| - | - | 663.4 | 525.2 | - | - | 0 | - |
| - | - | 1040 | 539.3 | - | - | 0 | - |
| - | - | 5592 | 553.3 | - | - | 0 | - |
| - | - | 1216 | 554.3 | - | - | 0 | - |
| - | - | 799 | 559.2 | - | - | 0 | - |
| - | - | 1677 | 563.8 | - | - | 0 | - |
| - | - | 1053 | 564.3 | - | - | 0 | - |
| - | - | 798.8 | 564.8 | - | - | 0 | - |
| - | - | 9107 | 570.3 | - | - | 0 | - |
| - | - | 3649 | 571.3 | - | - | 0 | - |
| - | - | 1715 | 577.8 | - | - | 0 | - |
| - | - | 2241 | 578.3 | - | - | 0 | - |
| - | - | 1129 | 578.8 | - | - | 0 | - |
| - | - | 3324 | 605.3 | - | - | 0 | - |
| - | - | 1523 | 606.3 | - | - | 0 | - |
| - | - | 1417 | 627.3 | - | - | 0 | - |
| - | - | 1215 | 627.8 | - | - | 0 | - |
| 3 | y | 1986 | 639.2 | 0.006011 | 9.404 | +1 | 5 |
| - | - | 1003 | 645.3 | - | - | 0 | - |
| - | - | 1525 | 645.8 | - | - | 0 | - |
| - | - | 741.7 | 646.3 | - | - | 0 | - |
| - | - | 1016 | 651.4 | - | - | 0 | - |
| - | - | 2187 | 656.4 | - | - | 0 | - |
| 3 | y | 3.846E+04 | 657.3 | 0.00504 | 7.669 | +1 | 5 |
| - | - | 1.467E+04 | 658.3 | - | - | 0 | - |
| - | - | 3660 | 659.3 | - | - | 0 | - |
| - | - | 1554 | 666.4 | - | - | 0 | - |
| - | - | 1555 | 668.4 | - | - | 0 | - |
| - | - | 996.4 | 669.4 | - | - | 0 | - |
| - | - | 861.8 | 686.4 | - | - | 0 | - |
| - | - | 1509 | 708.4 | - | - | 0 | - |
| - | - | 884.9 | 716.3 | - | - | 0 | - |
| - | - | 1263 | 734.3 | - | - | 0 | - |
| - | - | 1263 | 739.4 | - | - | 0 | - |
| - | - | 1020 | 753.4 | - | - | 0 | - |
| - | - | 9665 | 755.4 | - | - | 0 | - |
| - | - | 3873 | 756.4 | - | - | 0 | - |
| - | - | 1316 | 757.4 | - | - | 0 | - |
| - | - | 4668 | 765.5 | - | - | 0 | - |
| - | - | 1445 | 766.5 | - | - | 0 | - |
| - | - | 710.1 | 787.4 | - | - | 0 | - |
| - | - | 3132 | 805.4 | - | - | 0 | - |
| - | - | 1447 | 806.4 | - | - | 0 | - |
| 2 | y | 1.206E+04 | 820.3 | 0.005991 | 7.303 | +1 | 6 |
| - | - | 4965 | 821.3 | - | - | 0 | - |
| - | - | 1383 | 822.3 | - | - | 0 | - |
| - | - | 774.1 | 868.5 | - | - | 0 | - |
| - | - | 663.7 | 869.5 | - | - | 0 | - |
| - | - | 734.4 | 928.4 | - | - | 0 | - |
| - | - | 671.5 | 948.4 | - | - | 0 | - |
| - | - | 819 | 993.5 | - | - | 0 | - |
| - | - | 951.7 | 1127 | - | - | 0 | - |
| - | - | 735.6 | 1475 | - | - | 0 | - |
| - | - | 614.3 | 2001 | - | - | 0 | - |
| - | - | 910.9 | 3075 | - | - | 0 | - |

m/z Charge Intensity FragmentType MassShift Position
120.08106994628906 0 88672.24
121.0843734741211 0 6221.8843
127.05068969726562 0 457.04758
127.0868911743164 0 1074.1975
128.07093811035156 0 803.0853
128.10736083984375 0 1112.0532
129.06622314453125 0 938.6894
129.10250854492188 0 54555.816
130.0502471923828 0 1478.2421
130.0654296875 0 1457.3661
130.0865936279297 0 1820.8209
130.10577392578125 0 2858.5703
131.1182098388672 0 709.0263
133.06106567382812 0 872.06805
134.0272979736328 0 2254.7915
134.04505920410156 0 4747.4014 y 6
134.0603790283203 0 950.81287
136.07598876953125 0 88851.93
137.07931518554688 0 7061.7114
139.0865478515625 0 954.853
140.0821075439453 0 1031.704
141.0662078857422 0 866.45996
141.10240173339844 0 1843.4025
142.0982208251953 0 602.49567
142.1228790283203 0 566.85455
143.1180419921875 0 567.3042
144.10211181640625 0 1755.265
145.04977416992188 0 463.05075
145.06103515625 0 1402.5822
146.06027221679688 0 1159.8779
147.04425048828125 0 882.69006
147.11305236816406 0 1134.883
148.95458984375 0 1140.0046
149.02330017089844 0 621.798
152.07081604003906 0 3579.9705
153.06625366210938 0 683.7537
155.0819091796875 0 675.8513
155.11807250976562 0 2055.3003
157.06138610839844 0 570.0783
157.09765625 0 487.63193
157.1338653564453 0 2171.2734
158.0922088623047 0 638.213
159.07669067382812 0 1690.0724
159.09193420410156 0 1895.9065
163.07162475585938 0 8390.466
165.10240173339844 0 1466.0321
165.97996520996094 0 421.17926
166.08631896972656 0 769.2268
167.117919921875 0 538.15295
169.06048583984375 0 983.6351
169.09719848632812 0 1653.6404
171.07693481445312 0 548.77936
171.11297607421875 0 731.01276
171.1495819091797 0 2843.3591
173.0557098388672 0 624.4256
173.45162963867188 0 1315.6924
175.08665466308594 0 938.3361
181.09710693359375 0 1097.2299
181.13323974609375 0 729.08716
182.0819091796875 0 559.39984
182.12905883789062 0 1331.3176
183.11288452148438 0 1457.1013
183.1493682861328 0 926.3528
185.0925750732422 0 624.4034
185.12869262695312 0 1449.5376
185.16506958007812 0 17377.094
186.0875701904297 0 1868.7733
186.16835021972656 0 1481.7291
187.10755920410156 0 979.78705
187.14390563964844 0 728.47003
188.1397705078125 0 628.9554
195.1128387451172 0 574.6275
198.08755493164062 0 483.05026
199.1078643798828 0 1148.1282
199.1432342529297 0 846.1851
200.1395263671875 0 14515.98
201.1234130859375 0 1041.1771
201.14271545410156 0 1186.4261
202.10781860351562 0 1775.7555
203.1028289794922 0 10845.023
204.1063690185547 0 883.28784
205.06439208984375 0 2186.8457
207.1125030517578 0 1160.879
210.1235809326172 0 1049.3956
211.10797119140625 0 880.14325
211.1444091796875 0 1193.2286
213.16006469726562 0 8088.9136
214.1631317138672 0 1029.3536
215.10256958007812 0 910.91504
215.13900756835938 0 2288.7153
216.06594848632812 0 1037.7726
216.09764099121094 0 685.13043
218.15072631835938 0 654.2245
224.10374450683594 0 777.5917
224.1393585205078 0 649.0524
225.0983123779297 0 825.0888
226.11843872070312 0 833.6949
226.1551513671875 0 9416.3
227.10231018066406 0 840.77405
227.15895080566406 0 1195.6644
227.17579650878906 0 1613.3705
228.13462829589844 0 693.891
233.05953979492188 0 1109.7246
233.0922393798828 0 5712.359
233.1287841796875 0 1672.2393
233.1651153564453 0 3728.5708
234.05868530273438 0 564.1196
234.16986083984375 0 617.5794
239.1505584716797 0 1029.0938
240.13433837890625 0 2418.5942
243.0980224609375 0 603.3523
244.128662109375 0 1151.0913 y Water loss 5
245.11331176757812 0 2893.255 y Ammonia loss 5
246.18125915527344 0 879.92566
249.09823608398438 0 641.0147
249.1600341796875 0 677.0481
251.15061950683594 0 711.9028
252.0692596435547 0 1061.7678
252.1348114013672 0 561.35706
252.16976928710938 0 519.4558
253.16610717773438 0 3315.9158
254.1498565673828 0 800.86194
256.1767883300781 0 996.50494
261.12518310546875 0 527.8695
261.1600036621094 0 709.07806
262.13983154296875 0 13959.191 y 5
263.14251708984375 0 1416.2683
268.3064270019531 0 597.0604
270.18017578125 0 766.43445
271.1753845214844 0 704.4128
272.1239929199219 0 575.665
274.1405029296875 0 690.1519
279.1816711425781 0 1463.4497
281.1486511230469 0 1061.4998
282.14447021484375 0 586.2136
283.1442565917969 0 48356.395 a 1
284.1474304199219 0 8895.255
285.152099609375 0 873.71155
287.2077941894531 0 1949.8606
297.090576171875 0 10018.527
297.10748291015625 0 594.06604
297.19219970703125 0 2285.4902
298.093994140625 0 1786.0701
299.1392517089844 0 6996.85
299.1717529296875 0 4467.284
300.1756591796875 0 932.3426
304.12939453125 0 6224.115
304.162353515625 0 543.32733
305.131591796875 0 996.08057
306.14404296875 0 728.57983
309.14453125 0 1135.3164
311.1392822265625 0 10163.01 b 1
312.1430969238281 0 1533.5759
315.16558837890625 0 856.4212 y Water loss 4
316.18609619140625 0 1205.2295
323.2081298828125 0 941.1743
325.0855407714844 0 6003.951
326.0849609375 0 595.065
327.1334228515625 0 3786.9377
333.1769714355469 0 8414.142 y 4
334.1808776855469 0 2065.1968
338.24383544921875 0 891.351
340.18682861328125 0 727.1781
340.23602294921875 0 842.561
340.2591247558594 0 2446.956
349.7813720703125 0 686.4043
361.1534118652344 0 2494.8032
368.22900390625 0 6690.883
394.2095947265625 0 660.18
396.1224670410156 0 3677.172
396.15484619140625 0 1411.2223
397.1250305175781 0 671.2253
398.2410583496094 0 1586.6312
413.6688232421875 0 659.5029
431.2129211425781 0 675.1699
440.2480773925781 0 645.0108
441.2534484863281 0 609.6175
442.2350769042969 0 605.18207
442.3125915527344 0 1033.6575
466.2208557128906 0 601.2056
466.7201232910156 0 646.6137
467.3230285644531 0 1313.981
470.3078308105469 0 1824.254
472.2752685546875 0 994.0286
476.18017578125 0 1268.452 y Water loss 3
481.2487487792969 0 634.10284
482.2372131347656 0 1307.6692
482.2735290527344 0 1239.7472
484.195556640625 0 1358.8116 Precursor
484.28118896484375 0 855.3123
484.3497619628906 0 3700.4707
484.69830322265625 0 887.39923
484.7860107421875 0 1162.6425
485.287353515625 0 1486.1882
485.3548278808594 0 2423.4197
488.14971923828125 0 1979.3756
488.2989807128906 0 958.8236
488.7767333984375 0 1227.8932
489.7837829589844 0 2555.7688
490.2859191894531 0 1183.9985
490.7861633300781 0 1218.8794
494.191650390625 0 10181.65 y 3
494.2322692871094 0 750.58466
495.1944885253906 0 2686.7927
509.2817077636719 0 594.4997
513.3507080078125 0 2648.0203
514.3551635742188 0 666.9203
515.2371826171875 0 582.61426
524.2178955078125 0 2202.845
525.2222290039062 0 663.4015
539.329833984375 0 1039.7183
553.3089599609375 0 5591.8228
554.3092651367188 0 1215.7039
559.1849975585938 0 799.015
563.7704467773438 0 1676.7913
564.2742309570312 0 1053.352
564.7737426757812 0 798.7647
570.3243408203125 0 9107.097
571.3271484375 0 3648.9338
577.7682495117188 0 1714.677
578.2703247070312 0 2240.5474
578.7711791992188 0 1129.0245
605.3064575195312 0 3323.757
606.3119506835938 0 1522.5336
627.3067016601562 0 1417.1707
627.8070068359375 0 1214.8329
639.2427978515625 0 1985.9663 y Water loss 2
645.2998657226562 0 1002.85645
645.8016967773438 0 1525.0896
646.3056640625 0 741.70526
651.36328125 0 1015.7081
656.3590087890625 0 2187.2095
657.2543334960938 0 38462.086 y 2
658.2572021484375 0 14672.323
659.2548828125 0 3660.3474
666.3927001953125 0 1554.0048
668.3739624023438 0 1555.3535
669.3718872070312 0 996.38354
686.3837890625 0 861.829
708.4019775390625 0 1508.9525
716.33740234375 0 884.9435
734.3497924804688 0 1263.0399
739.4091796875 0 1262.6199
753.4137573242188 0 1019.7018
755.4038696289062 0 9664.521
756.4058227539062 0 3872.7559
757.411865234375 0 1315.8837
765.4611206054688 0 4667.7817
766.4631958007812 0 1445.2979
787.3742065429688 0 710.07605
805.3850708007812 0 3131.586
806.3883666992188 0 1446.8859
820.3167114257812 0 12057.215 y 1
821.3192138671875 0 4964.715
822.3204345703125 0 1382.8408
868.483642578125 0 774.1197
869.4947509765625 0 663.7295
928.4335327148438 0 734.4375
948.4242553710938 0 671.53656
993.5210571289062 0 819.0153
1126.5428466796875 0 951.6682
1475.026611328125 0 735.57874
2000.7052001953125 0 614.3314
3074.559814453125 0 910.93005

Spectrum Details

|  |  |
| --- | --- |
| Matched peaks? Matched peaks The total absolute number of peaks matched. Additionally in brackets the total fraction of peaks matched and the total number of peaks is shown. | 14 (5.30% of 264) |
| FDR? FDR The false discovery rate estimated for this peptide. It is calculated by matching all theoretical fragments with a non-integer shift with the raw peaks for this spectrum. This is done with 40 different shifts. The resulting percentage is the average number of annotated peaks over the number of annotated peaks with the correct spectrum. | 3.40% |
| Satellite FDR? Satellite FDR See the FDR for details on its calculation. This satellite ion specific FDR only contains the satellite ions (d/w) for I/L/J positions. | - |
| PSM Score? PSM Score The PSM Score as given by Hecklib to this annotated spectrum. It is shown with three significant figures. | 185 |

#### Spectrum 4945? Spectrum 4945 The raw spectrum of this peptide as annotated by Hecklib. The fragments are coloured according to ion type (see legend). Any peaks with a star '\*' as text can be hovered over to see the full details, first the ion type second the mass shift type. By hovering over the amino acids in the peptide or ions in the legend the corresponding peaks are highlighted. By toggling the 'Unassigned' label you can turn the background (unassigned) peaks on or off in the plot. By updating the slider in the Ion legend you can update the spectrum to only show the top X% of the peaks with labels. The top X% means any peak that is within X% of the highest intensity. By dragging in the spectrum you can zoom in to a specific part of the spectrum and use 'Zoom Out' to get back to the original zoom level. The annotation of the spectrum is based on the given sequence in the peptides file and is done with different software so inconsistencies are likely. The peaks are annotated based on the given sequence, with 20 ppm tolerance.

Copy Data

##### Spectrum 4945 (TSV)

###### Preview

```
Loading example...
```

*Click on the button to copy the data to your clipboard.*

Mz MinMz MaxIntensity Max

WidthHeightPeptide font sizePeptide stroke widthSpectrum font sizeSpectrum stroke widthCompact peptide

Ion legend

wxyz

abcd

OtherUnassignedIon^Charge^_Position_Show for top:%

FYYCAKD

02.21e+44.43e+46.64e+48.85e+4

Zoom Out

y^+1^_1_y^+1^_2_y^+1^_2_y^+1^_2_a^+1^_2_b^+1^_2_y^+1^_3_y^+1^_3_y^+2^_6_y^+1^_4_\*y^+1^_4_y^+1^_5_y^+1^_5_y^+1^_6_

043687213081744

Fragment Matches Table

Show background peaks

| Position | Ion type | Intensity | mz Theoretical | mz Error (Th) | mz Error (ppm) | Charge | Series Number |
| --- | --- | --- | --- | --- | --- | --- | --- |
| - | - | 8.763E+04 | 120.1 | - | - | 0 | - |
| - | - | 747 | 121.1 | - | - | 0 | - |
| - | - | 6816 | 121.1 | - | - | 0 | - |
| - | - | 398.8 | 122.1 | - | - | 0 | - |
| - | - | 690.2 | 123.1 | - | - | 0 | - |
| - | - | 774.2 | 127.1 | - | - | 0 | - |
| - | - | 727.6 | 128.1 | - | - | 0 | - |
| - | - | 1196 | 128.1 | - | - | 0 | - |
| - | - | 357.9 | 128.6 | - | - | 0 | - |
| - | - | 779.8 | 129.1 | - | - | 0 | - |
| - | - | 5.329E+04 | 129.1 | - | - | 0 | - |
| - | - | 885.8 | 130.1 | - | - | 0 | - |
| - | - | 1539 | 130.1 | - | - | 0 | - |
| - | - | 1156 | 130.1 | - | - | 0 | - |
| - | - | 3275 | 130.1 | - | - | 0 | - |
| - | - | 1068 | 131.1 | - | - | 0 | - |
| - | - | 623.7 | 133.1 | - | - | 0 | - |
| - | - | 548.5 | 133.1 | - | - | 0 | - |
| - | - | 1549 | 134 | - | - | 0 | - |
| 7 | y | 3923 | 134 | 0.000275 | 2.052 | +1 | 1 |
| - | - | 8.134E+04 | 136.1 | - | - | 0 | - |
| - | - | 7696 | 137.1 | - | - | 0 | - |
| - | - | 630.5 | 138.1 | - | - | 0 | - |
| - | - | 1054 | 139.1 | - | - | 0 | - |
| - | - | 1674 | 141.1 | - | - | 0 | - |
| - | - | 432.5 | 143.1 | - | - | 0 | - |
| - | - | 476.9 | 143.1 | - | - | 0 | - |
| - | - | 1875 | 144.1 | - | - | 0 | - |
| - | - | 677.1 | 145.1 | - | - | 0 | - |
| - | - | 690.4 | 146.1 | - | - | 0 | - |
| - | - | 626.1 | 147 | - | - | 0 | - |
| - | - | 542.5 | 147.1 | - | - | 0 | - |
| - | - | 1212 | 147.1 | - | - | 0 | - |
| - | - | 727.5 | 149 | - | - | 0 | - |
| - | - | 1961 | 152.1 | - | - | 0 | - |
| - | - | 1423 | 155.1 | - | - | 0 | - |
| - | - | 528.4 | 156.1 | - | - | 0 | - |
| - | - | 1256 | 157.1 | - | - | 0 | - |
| - | - | 463.5 | 158.1 | - | - | 0 | - |
| - | - | 1778 | 159.1 | - | - | 0 | - |
| - | - | 677.7 | 160.1 | - | - | 0 | - |
| - | - | 517.6 | 162.9 | - | - | 0 | - |
| - | - | 1426 | 163.1 | - | - | 0 | - |
| - | - | 1526 | 165.1 | - | - | 0 | - |
| - | - | 2085 | 166.1 | - | - | 0 | - |
| - | - | 490.2 | 167.1 | - | - | 0 | - |
| - | - | 916.1 | 169.1 | - | - | 0 | - |
| - | - | 1169 | 169.1 | - | - | 0 | - |
| - | - | 1144 | 171.1 | - | - | 0 | - |
| - | - | 2025 | 171.1 | - | - | 0 | - |
| - | - | 673 | 175.1 | - | - | 0 | - |
| - | - | 461.2 | 178.1 | - | - | 0 | - |
| - | - | 625.5 | 179.1 | - | - | 0 | - |
| - | - | 669.3 | 180.1 | - | - | 0 | - |
| - | - | 535.4 | 180.1 | - | - | 0 | - |
| - | - | 1113 | 181.1 | - | - | 0 | - |
| - | - | 831.7 | 181.1 | - | - | 0 | - |
| - | - | 1011 | 182.1 | - | - | 0 | - |
| - | - | 1694 | 183.1 | - | - | 0 | - |
| - | - | 1271 | 183.1 | - | - | 0 | - |
| - | - | 755 | 185.1 | - | - | 0 | - |
| - | - | 772.9 | 185.1 | - | - | 0 | - |
| - | - | 1.98E+04 | 185.2 | - | - | 0 | - |
| - | - | 2022 | 186.1 | - | - | 0 | - |
| - | - | 2004 | 186.2 | - | - | 0 | - |
| - | - | 1547 | 187.1 | - | - | 0 | - |
| - | - | 937.3 | 187.1 | - | - | 0 | - |
| - | - | 545.2 | 189.1 | - | - | 0 | - |
| - | - | 818.7 | 191.1 | - | - | 0 | - |
| - | - | 604 | 193.1 | - | - | 0 | - |
| - | - | 712.8 | 195.1 | - | - | 0 | - |
| - | - | 621.9 | 197.1 | - | - | 0 | - |
| - | - | 511.3 | 199.1 | - | - | 0 | - |
| - | - | 1084 | 199.1 | - | - | 0 | - |
| - | - | 971.8 | 199.1 | - | - | 0 | - |
| - | - | 1.365E+04 | 200.1 | - | - | 0 | - |
| - | - | 986.4 | 200.1 | - | - | 0 | - |
| - | - | 1342 | 201.1 | - | - | 0 | - |
| - | - | 1604 | 202.1 | - | - | 0 | - |
| - | - | 1.021E+04 | 203.1 | - | - | 0 | - |
| - | - | 838.2 | 204.1 | - | - | 0 | - |
| - | - | 2620 | 205.1 | - | - | 0 | - |
| - | - | 751.6 | 205.1 | - | - | 0 | - |
| - | - | 619.4 | 208.1 | - | - | 0 | - |
| - | - | 899.3 | 209.1 | - | - | 0 | - |
| - | - | 1674 | 210.1 | - | - | 0 | - |
| - | - | 1198 | 211.1 | - | - | 0 | - |
| - | - | 8007 | 213.2 | - | - | 0 | - |
| - | - | 938.2 | 214.2 | - | - | 0 | - |
| - | - | 1271 | 215.1 | - | - | 0 | - |
| - | - | 664.9 | 216.1 | - | - | 0 | - |
| - | - | 570.2 | 223.1 | - | - | 0 | - |
| - | - | 1305 | 224.1 | - | - | 0 | - |
| - | - | 747.2 | 226.1 | - | - | 0 | - |
| - | - | 1.1E+04 | 226.2 | - | - | 0 | - |
| - | - | 1293 | 227.1 | - | - | 0 | - |
| - | - | 1009 | 227.2 | - | - | 0 | - |
| - | - | 763.1 | 227.2 | - | - | 0 | - |
| - | - | 997.8 | 228.1 | - | - | 0 | - |
| - | - | 1349 | 233.1 | - | - | 0 | - |
| - | - | 4712 | 233.1 | - | - | 0 | - |
| - | - | 6183 | 233.2 | - | - | 0 | - |
| - | - | 694 | 234.1 | - | - | 0 | - |
| - | - | 645.5 | 234.1 | - | - | 0 | - |
| - | - | 1476 | 239.2 | - | - | 0 | - |
| - | - | 836.1 | 240.1 | - | - | 0 | - |
| 6 | y | 864.2 | 244.1 | 0.0004867 | 1.994 | +1 | 2 |
| 6 | y | 2779 | 245.1 | 0.00019 | 0.775 | +1 | 2 |
| - | - | 628.1 | 247.1 | - | - | 0 | - |
| - | - | 844.9 | 249.2 | - | - | 0 | - |
| - | - | 575.3 | 251.1 | - | - | 0 | - |
| - | - | 822.3 | 251.1 | - | - | 0 | - |
| - | - | 1168 | 252.1 | - | - | 0 | - |
| - | - | 854.8 | 253.2 | - | - | 0 | - |
| - | - | 603.4 | 254.1 | - | - | 0 | - |
| - | - | 2134 | 261.2 | - | - | 0 | - |
| - | - | 999.3 | 262.1 | - | - | 0 | - |
| 6 | y | 1.405E+04 | 262.1 | 0.0001454 | 0.5546 | +1 | 2 |
| - | - | 1325 | 263.1 | - | - | 0 | - |
| - | - | 462.5 | 265.1 | - | - | 0 | - |
| - | - | 577.2 | 269.2 | - | - | 0 | - |
| - | - | 758.6 | 271.2 | - | - | 0 | - |
| - | - | 1376 | 279.2 | - | - | 0 | - |
| - | - | 545.8 | 282.9 | - | - | 0 | - |
| 2 | a | 4.888E+04 | 283.1 | 0.0001523 | 0.5379 | +1 | 2 |
| - | - | 7777 | 284.1 | - | - | 0 | - |
| - | - | 637.2 | 284.7 | - | - | 0 | - |
| - | - | 877.5 | 285.2 | - | - | 0 | - |
| - | - | 1058 | 287.2 | - | - | 0 | - |
| - | - | 7328 | 297.1 | - | - | 0 | - |
| - | - | 912.6 | 297.1 | - | - | 0 | - |
| - | - | 2844 | 297.2 | - | - | 0 | - |
| - | - | 1750 | 298.1 | - | - | 0 | - |
| - | - | 6109 | 299.1 | - | - | 0 | - |
| - | - | 4639 | 299.2 | - | - | 0 | - |
| - | - | 1186 | 300.1 | - | - | 0 | - |
| - | - | 5967 | 304.1 | - | - | 0 | - |
| - | - | 1158 | 305.1 | - | - | 0 | - |
| 2 | b | 8742 | 311.1 | 0.0003854 | 1.239 | +1 | 2 |
| - | - | 2402 | 312.1 | - | - | 0 | - |
| 5 | y | 1046 | 315.2 | 0.001337 | 4.241 | +1 | 3 |
| - | - | 1712 | 323.2 | - | - | 0 | - |
| - | - | 5824 | 325.1 | - | - | 0 | - |
| - | - | 935 | 326.1 | - | - | 0 | - |
| - | - | 3188 | 327.1 | - | - | 0 | - |
| - | - | 794.3 | 327.2 | - | - | 0 | - |
| 5 | y | 1.009E+04 | 333.2 | 0.0002325 | 0.6979 | +1 | 3 |
| - | - | 1417 | 334.2 | - | - | 0 | - |
| - | - | 1506 | 340.2 | - | - | 0 | - |
| - | - | 888.7 | 341.2 | - | - | 0 | - |
| - | - | 1364 | 354.7 | - | - | 0 | - |
| - | - | 2313 | 361.2 | - | - | 0 | - |
| - | - | 769.7 | 363.1 | - | - | 0 | - |
| - | - | 642.4 | 364.1 | - | - | 0 | - |
| - | - | 6398 | 368.2 | - | - | 0 | - |
| - | - | 981.8 | 369.2 | - | - | 0 | - |
| - | - | 1048 | 376.2 | - | - | 0 | - |
| - | - | 2630 | 396.1 | - | - | 0 | - |
| - | - | 1794 | 396.2 | - | - | 0 | - |
| - | - | 662.5 | 397.1 | - | - | 0 | - |
| - | - | 1551 | 398.2 | - | - | 0 | - |
| 2 | y | 1215 | 410.7 | 0.002422 | 5.899 | +2 | 6 |
| - | - | 599.6 | 431.8 | - | - | 0 | - |
| - | - | 739.5 | 464.3 | - | - | 0 | - |
| - | - | 642.9 | 468.7 | - | - | 0 | - |
| - | - | 1399 | 472.3 | - | - | 0 | - |
| 4 | y | 1071 | 476.2 | 0.002284 | 4.796 | +1 | 4 |
| - | - | 877.3 | 479.3 | - | - | 0 | - |
| - | - | 824.6 | 479.8 | - | - | 0 | - |
| - | - | 977.3 | 482.2 | - | - | 0 | - |
| - | - | 1528 | 482.3 | - | - | 0 | - |
| 0 | Precursor | 952.5 | 484.2 | 0.001168 | 2.412 | +2 | -1 |
| - | - | 633.4 | 484.3 | - | - | 0 | - |
| - | - | 723.2 | 484.3 | - | - | 0 | - |
| - | - | 748.8 | 484.3 | - | - | 0 | - |
| - | - | 803.1 | 484.7 | - | - | 0 | - |
| - | - | 1574 | 484.8 | - | - | 0 | - |
| - | - | 681.1 | 485.2 | - | - | 0 | - |
| - | - | 1450 | 485.4 | - | - | 0 | - |
| - | - | 762.5 | 487.8 | - | - | 0 | - |
| - | - | 2305 | 488.1 | - | - | 0 | - |
| - | - | 669.7 | 488.3 | - | - | 0 | - |
| - | - | 865.5 | 489.8 | - | - | 0 | - |
| - | - | 991.7 | 490.3 | - | - | 0 | - |
| 4 | y | 1.15E+04 | 494.2 | 0.004304 | 8.708 | +1 | 4 |
| - | - | 1593 | 495.2 | - | - | 0 | - |
| - | - | 1079 | 496.2 | - | - | 0 | - |
| - | - | 1940 | 521.8 | - | - | 0 | - |
| - | - | 2110 | 524.2 | - | - | 0 | - |
| - | - | 665.7 | 529.7 | - | - | 0 | - |
| - | - | 966 | 542.2 | - | - | 0 | - |
| - | - | 796.6 | 543.3 | - | - | 0 | - |
| - | - | 5119 | 553.3 | - | - | 0 | - |
| - | - | 2159 | 554.3 | - | - | 0 | - |
| - | - | 701.4 | 555.3 | - | - | 0 | - |
| - | - | 780.8 | 559.2 | - | - | 0 | - |
| - | - | 1099 | 563.8 | - | - | 0 | - |
| - | - | 1.205E+04 | 570.3 | - | - | 0 | - |
| - | - | 2722 | 571.3 | - | - | 0 | - |
| - | - | 3674 | 577.8 | - | - | 0 | - |
| - | - | 2106 | 578.3 | - | - | 0 | - |
| - | - | 836.5 | 578.8 | - | - | 0 | - |
| - | - | 788.1 | 593.3 | - | - | 0 | - |
| - | - | 665.3 | 594.3 | - | - | 0 | - |
| - | - | 1082 | 605.3 | - | - | 0 | - |
| - | - | 676.5 | 606.3 | - | - | 0 | - |
| - | - | 857.1 | 627.3 | - | - | 0 | - |
| 3 | y | 1850 | 639.2 | 0.005889 | 9.213 | +1 | 5 |
| - | - | 1387 | 645.3 | - | - | 0 | - |
| - | - | 852.6 | 645.8 | - | - | 0 | - |
| - | - | 979.1 | 656.4 | - | - | 0 | - |
| 3 | y | 4.103E+04 | 657.3 | 0.005163 | 7.855 | +1 | 5 |
| - | - | 1.283E+04 | 658.3 | - | - | 0 | - |
| - | - | 3701 | 659.3 | - | - | 0 | - |
| - | - | 1034 | 666.4 | - | - | 0 | - |
| - | - | 668.8 | 687.3 | - | - | 0 | - |
| - | - | 1491 | 708.4 | - | - | 0 | - |
| - | - | 1046 | 709.4 | - | - | 0 | - |
| - | - | 1199 | 720.4 | - | - | 0 | - |
| - | - | 752 | 738.4 | - | - | 0 | - |
| - | - | 1.084E+04 | 755.4 | - | - | 0 | - |
| - | - | 3948 | 756.4 | - | - | 0 | - |
| - | - | 622.5 | 757.4 | - | - | 0 | - |
| - | - | 4564 | 765.5 | - | - | 0 | - |
| - | - | 1995 | 766.5 | - | - | 0 | - |
| - | - | 800.7 | 779.4 | - | - | 0 | - |
| 2 | y | 1.001E+04 | 820.3 | 0.005259 | 6.411 | +1 | 6 |
| - | - | 3985 | 821.3 | - | - | 0 | - |
| - | - | 2408 | 822.3 | - | - | 0 | - |
| - | - | 1185 | 868.5 | - | - | 0 | - |
| - | - | 655.3 | 1033 | - | - | 0 | - |
| - | - | 668.7 | 1727 | - | - | 0 | - |

m/z Charge Intensity FragmentType MassShift Position
120.08108520507812 0 87632.79
121.07978057861328 0 747.02594
121.08442687988281 0 6815.744
122.07152557373047 0 398.80902
123.11690521240234 0 690.2123
127.08685302734375 0 774.2151
128.07090759277344 0 727.6483
128.10743713378906 0 1196.4961
128.615478515625 0 357.94263
129.06582641601562 0 779.7688
129.10252380371094 0 53287.79
130.05030822753906 0 885.8341
130.06536865234375 0 1538.5217
130.08670043945312 0 1155.8997
130.10592651367188 0 3275.113
131.11830139160156 0 1068.3815
133.0610809326172 0 623.6747
133.08602905273438 0 548.49634
134.02734375 0 1549.1957
134.04505920410156 0 3922.803 y 6
136.0760040283203 0 81339.29
137.079345703125 0 7695.636
138.06668090820312 0 630.4757
139.08677673339844 0 1054.0068
141.10244750976562 0 1674.2866
143.0819854736328 0 432.4893
143.11831665039062 0 476.90292
144.1021270751953 0 1874.5553
145.06117248535156 0 677.0554
146.06016540527344 0 690.4122
147.04429626464844 0 626.1483
147.0652618408203 0 542.5372
147.1131591796875 0 1212.1724
149.02317810058594 0 727.5276
152.07102966308594 0 1961.463
155.1179962158203 0 1422.5267
156.0769805908203 0 528.37616
157.13380432128906 0 1255.9882
158.09373474121094 0 463.48807
159.0918426513672 0 1777.506
160.07582092285156 0 677.69293
162.85301208496094 0 517.6474
163.0716094970703 0 1426.3156
165.10260009765625 0 1526.3296
166.053466796875 0 2085.3374
167.08155822753906 0 490.2196
169.06089782714844 0 916.1046
169.09732055664062 0 1168.6101
171.1130828857422 0 1144.3348
171.14939880371094 0 2025.003
175.0865020751953 0 673.0017
178.13482666015625 0 461.24002
179.117431640625 0 625.5103
180.06564331054688 0 669.2931
180.1134796142578 0 535.3871
181.0975341796875 0 1113.2296
181.1337127685547 0 831.6763
182.1293487548828 0 1010.9015
183.11289978027344 0 1693.5905
183.14942932128906 0 1270.5045
185.09275817871094 0 754.9983
185.12939453125 0 772.88184
185.16510009765625 0 19801.836
186.08763122558594 0 2021.865
186.168212890625 0 2004.2393
187.1077117919922 0 1546.8647
187.14398193359375 0 937.27716
189.10244750976562 0 545.1555
191.08221435546875 0 818.70746
193.09788513183594 0 604.0423
195.1129913330078 0 712.79834
197.1289520263672 0 621.8967
199.09344482421875 0 511.29425
199.1082305908203 0 1084.4384
199.14453125 0 971.75885
200.13954162597656 0 13648.763
200.14967346191406 0 986.40704
201.14273071289062 0 1341.5751
202.1074676513672 0 1603.8918
203.10279846191406 0 10207.969
204.10658264160156 0 838.151
205.0643768310547 0 2619.508
205.0963134765625 0 751.5756
208.1079559326172 0 619.4161
209.0919952392578 0 899.2755
210.12400817871094 0 1674.467
211.1080780029297 0 1198.2072
213.16000366210938 0 8007.0156
214.1643829345703 0 938.2495
215.13914489746094 0 1271.067
216.06581115722656 0 664.942
223.10888671875 0 570.23
224.13951110839844 0 1305.2822
226.1187286376953 0 747.2308
226.15524291992188 0 11000.644
227.1024627685547 0 1292.7253
227.15853881835938 0 1008.84283
227.17640686035156 0 763.0997
228.13458251953125 0 997.797
233.0586700439453 0 1348.7413
233.09242248535156 0 4711.8184
233.1650390625 0 6183.494
234.09620666503906 0 694.0298
234.12423706054688 0 645.51086
239.15054321289062 0 1476.4977
240.13478088378906 0 836.1478
244.12966918945312 0 864.22754 y Water loss 5
245.11338806152344 0 2778.6763 y Ammonia loss 5
247.1445770263672 0 628.12524
249.15966796875 0 844.93475
251.0699920654297 0 575.3006
251.1497344970703 0 822.30334
252.0687713623047 0 1167.5736
253.16697692871094 0 854.76025
254.11595153808594 0 603.3868
261.1597595214844 0 2133.7808
262.125 0 999.3498
262.139892578125 0 14046.951 y 5
263.1424560546875 0 1325.3612
265.079345703125 0 462.51416
269.197021484375 0 577.23114
271.177490234375 0 758.5664
279.181396484375 0 1375.5309
282.9248962402344 0 545.8181
283.1442565917969 0 48875.938 a 1
284.14752197265625 0 7777.113
284.6825866699219 0 637.2215
285.15155029296875 0 877.5365
287.2074890136719 0 1057.6427
297.0904235839844 0 7327.9326
297.107666015625 0 912.6411
297.1925354003906 0 2843.5305
298.09503173828125 0 1750.1301
299.13922119140625 0 6109.1562
299.171630859375 0 4638.72
300.142578125 0 1186.2606
304.12945556640625 0 5966.9004
305.13232421875 0 1157.7845
311.139404296875 0 8741.856 b 1
312.1427307128906 0 2402.3882
315.1676330566406 0 1045.5485 y Water loss 4
323.20849609375 0 1711.6978
325.08551025390625 0 5823.9556
326.0872802734375 0 935.0071
327.13397216796875 0 3188.093
327.16741943359375 0 794.2882
333.1770935058594 0 10089.227 y 4
334.181884765625 0 1416.581
340.2341613769531 0 1505.6599
341.18255615234375 0 888.67444
354.7054138183594 0 1363.5674
361.1534118652344 0 2312.5664
363.1385803222656 0 769.7244
364.1252136230469 0 642.3619
368.22900390625 0 6398.3027
369.2329406738281 0 981.829
376.1980285644531 0 1047.6891
396.12237548828125 0 2630.174
396.15435791015625 0 1793.9744
397.1291809082031 0 662.5453
398.23895263671875 0 1551.1487
410.6625671386719 0 1215.0134 y 1
431.7622985839844 0 599.6409
464.26043701171875 0 739.50256
468.7446594238281 0 642.94055
472.27734375 0 1399.01
476.1831970214844 0 1071.0826 y Water loss 3
479.2623291015625 0 877.3118
479.7519836425781 0 824.55536
482.2376708984375 0 977.26935
482.271484375 0 1527.8058
484.1980285644531 0 952.4506 Precursor
484.2829895019531 0 633.42633
484.31744384765625 0 723.221
484.349853515625 0 748.82166
484.6989440917969 0 803.1093
484.7879333496094 0 1573.5062
485.2000427246094 0 681.1082
485.3592529296875 0 1450.1124
487.7865295410156 0 762.52313
488.1485900878906 0 2304.9583
488.3006591796875 0 669.67395
489.7838134765625 0 865.5379
490.2829895019531 0 991.7097
494.1917419433594 0 11503.34 y 3
495.1947326660156 0 1592.644
496.1897888183594 0 1079.0173
521.7835693359375 0 1939.5627
524.21630859375 0 2110.1355
529.7202758789062 0 665.665
542.2289428710938 0 966.0112
543.303466796875 0 796.5703
553.309326171875 0 5119.204
554.3106689453125 0 2158.7908
555.3147583007812 0 701.3769
559.1871948242188 0 780.7846
563.7682495117188 0 1098.5525
570.324462890625 0 12053.727
571.3273315429688 0 2722.2314
577.76806640625 0 3674.3245
578.2699584960938 0 2105.9844
578.7728881835938 0 836.45874
593.3056030273438 0 788.1375
594.30908203125 0 665.33246
605.30419921875 0 1082.0149
606.3173217773438 0 676.4554
627.3009033203125 0 857.12427
639.242919921875 0 1849.6938 y Water loss 2
645.3012084960938 0 1387.2126
645.8026733398438 0 852.58246
656.3597412109375 0 979.11676
657.2542114257812 0 41028.008 y 2
658.2572631835938 0 12832.375
659.2559814453125 0 3701.3198
666.3933715820312 0 1033.7197
687.2774047851562 0 668.83734
708.4020385742188 0 1491.1613
709.4043579101562 0 1046.3412
720.3609619140625 0 1198.8918
738.3832397460938 0 752.0037
755.4034423828125 0 10843.102
756.406005859375 0 3948.38
757.4110717773438 0 622.52234
765.4603271484375 0 4563.6543
766.461669921875 0 1995.212
779.3860473632812 0 800.69885
820.3174438476562 0 10014.426 y 1
821.320068359375 0 3985.1042
822.3189086914062 0 2408.1016
868.4837646484375 0 1184.9481
1032.89794921875 0 655.3235
1726.834228515625 0 668.734

Spectrum Details

|  |  |
| --- | --- |
| Matched peaks? Matched peaks The total absolute number of peaks matched. Additionally in brackets the total fraction of peaks matched and the total number of peaks is shown. | 15 (6.47% of 232) |
| FDR? FDR The false discovery rate estimated for this peptide. It is calculated by matching all theoretical fragments with a non-integer shift with the raw peaks for this spectrum. This is done with 40 different shifts. The resulting percentage is the average number of annotated peaks over the number of annotated peaks with the correct spectrum. | 2.38% |
| Satellite FDR? Satellite FDR See the FDR for details on its calculation. This satellite ion specific FDR only contains the satellite ions (d/w) for I/L/J positions. | - |
| PSM Score? PSM Score The PSM Score as given by Hecklib to this annotated spectrum. It is shown with three significant figures. | 185 |

#### Reverse Lookup? Reverse Lookup All places where this read could be placed.

| Group | Segment | Template | Template Part | Read Part | Score | Unique |
| --- | --- | --- | --- | --- | --- | --- |
| Homo sapiens Heavy Chain | IGHV | IGHV3-9 | [93..99] | [0..7] | 47 | False |
| Homo sapiens Heavy Chain | IGHV | IGHV3-43 | [93..99] | [0..7] | 47 | False |

| Recombined | Template Part | Read Part | Score | Unique |
| --- | --- | --- | --- | --- |
| REC-0-1 | [92..99] | [0..7] | 56 | True |

#### Meta Information from Multiple reads

##### Number of combined reads

2

##### Intensity

1

##### TotalArea

0

#### Positional Score

Copy Data

##### Positional Score (TSV)

###### Preview

```
Loading example...
```

*Click on the button to copy the data to your clipboard.*

000123456

Label Value
"0" 0
"1" 0
"2" 0
"3" 0
"4" 0
"5" 0
"6" 0

#### Meta Information from PEAKS

##### Scan Identifier

F2:4867

##### Original sequence

F

Y

Y

C

+58.01

A

K

D

##### Posttranslational Modifications

Carboxymethyl

##### Source File

D:\separate\_stitch\_analyses\xle-disambiguation\raw\20210323\_F1\_UM1\_Peng0013\_SA\_F59\_ingel\_3ug\_TL.raw

##### Fraction

2

##### Scan Feature

-

##### De Novo Score

98

##### ConfidenceScore

98

### m/z

484.1977

##### Mass

966.3793

##### Charge

2

##### Retention Time

26.16

##### Predicted Retention Time

-

##### Area

0

##### Parts Per Million

1.6

##### Fragmentation mode

HCD

##### Originating file

01 D:\separate\_stitch\_analyses\xle-disambiguation\20210325\_F59\_3ug\_DENOVO\_12.csv

#### Meta Information from PEAKS

##### Scan Identifier

F2:4945

##### Original sequence

F

Y

Y

C

+58.01

A

K

D

##### Posttranslational Modifications

Carboxymethyl

##### Source File

D:\separate\_stitch\_analyses\xle-disambiguation\raw\20210323\_F1\_UM1\_Peng0013\_SA\_F59\_ingel\_3ug\_TL.raw

##### Fraction

2

##### Scan Feature

-

##### De Novo Score

98

##### ConfidenceScore

98

### m/z

484.1978

##### Mass

966.3793

##### Charge

2

##### Retention Time

26.61

##### Predicted Retention Time

-

##### Area

0

##### Parts Per Million

1.9

##### Fragmentation mode

HCD

##### Originating file

01 D:\separate\_stitch\_analyses\xle-disambiguation\20210325\_F59\_3ug\_DENOVO\_12.csv
