## Supplementary material for "A handle on mass coincidence errors in *de novo* sequencing of antibodies by bottom-up proteomics": Combined_047.html

Details Combined\_047 | Stitch OverviewUndefined

### Read Combined\_047

#### Sequence (length=26)

VAWKADSSVPKAGVETTTPSKQSNNK

#### Spectrum 3914? Spectrum 3914 The raw spectrum of this peptide as annotated by Hecklib. The fragments are coloured according to ion type (see legend). Any peaks with a star '\*' as text can be hovered over to see the full details, first the ion type second the mass shift type. By hovering over the amino acids in the peptide or ions in the legend the corresponding peaks are highlighted. By toggling the 'Unassigned' label you can turn the background (unassigned) peaks on or off in the plot. By updating the slider in the Ion legend you can update the spectrum to only show the top X% of the peaks with labels. The top X% means any peak that is within X% of the highest intensity. By dragging in the spectrum you can zoom in to a specific part of the spectrum and use 'Zoom Out' to get back to the original zoom level. The annotation of the spectrum is based on the given sequence in the peptides file and is done with different software so inconsistencies are likely. The peaks are annotated based on the given sequence, with 20 ppm tolerance.

Copy Data

##### Spectrum 3914 (TSV)

###### Preview

```
Loading example...
```

*Click on the button to copy the data to your clipboard.*

Mz MinMz MaxIntensity Max

WidthHeightPeptide font sizePeptide stroke widthSpectrum font sizeSpectrum stroke widthCompact peptide

Ion legend

wxyz

abcd

OtherUnassignedIon^Charge^_Position_Show for top:%

VAWKADSSVPKAGVETTTPSKQSNNK

02.05e+54.09e+56.14e+58.18e+5

Zoom Out

y^+1^_1_a^+1^_2_y^+1^_1_b^+1^_2_y^+2^_4_y^+3^_6_b^+2^_4_y^+1^_2_y^+1^_2_b^+2^_6_b^+1^_3_y^+1^_3_y^+1^_3_y^+2^_7_b^+2^_8_b^+3^_12_b^+2^_8_y^+2^_8_y^+2^_8_y^+1^_4_y^+1^_4_y^+2^_8_y^+1^_4_b^+2^_9_b^+1^_4_y^+2^_9_y^+2^_10_b^+1^_5_y^+1^_5_y^+1^_5_b^+2^_11_y^+1^_5_y^+3^_17_y^+2^_11_b^+3^_18_b^+2^_12_b^+2^_12_b^+2^_12_b^+2^_13_y^+3^_19_y^+2^_12_b^+1^_6_y^+3^_20_b^+2^_14_b^+2^_14_y^+1^_6_y^+1^_6_y^+2^_13_b^+3^_21_y^+2^_13_y^+1^_6_b^+1^_7_y^+3^_22_y^+2^_14_y^+3^_22_b^+1^_7_b^+2^_15_y^+2^_15_b^+3^_23_b^+3^_23_y^+2^_15_y^+3^_23_y^+1^_7_y^+3^_23_b^+2^_16_b^+2^_16_b^+2^_16_b^+1^_8_y^+2^_16_y^+2^_16_b^+1^_8_y^+3^_24_y^+3^_24_y^+3^_24_b^+2^_17_b^+2^_17_b^+3^_25_b^+3^_25_b^+2^_17_y^+3^_25_y^+3^_25_y^+1^_8_y^+1^_8_y^+1^_8_\*\*b^+2^_18_b^+2^_18_\*b^+2^_18_y^+2^_18_y^+2^_18_b^+2^_19_y^+2^_19_y^+2^_19_y^+1^_9_y^+1^_9_y^+2^_19_y^+1^_9_y^+2^_20_y^+2^_20_b^+1^_10_b^+1^_10_y^+2^_20_b^+1^_10_y^+2^_21_y^+2^_21_y^+1^_10_y^+1^_10_y^+2^_21_y^+1^_10_y^+2^_22_y^+2^_22_y^+2^_22_b^+2^_22_b^+2^_22_b^+1^_11_b^+2^_23_b^+2^_23_y^+2^_23_y^+1^_11_y^+1^_11_y^+1^_11_b^+1^_12_b^+2^_24_b^+2^_24_b^+1^_12_b^+1^_13_b^+2^_25_b^+1^_13_y^+1^_12_y^+1^_12_y^+1^_12_b^+1^_14_y^+1^_13_y^+1^_13_y^+1^_14_y^+1^_14_y^+1^_14_b^+1^_15_y^+1^_15_y^+1^_15_y^+1^_15_b^+1^_16_y^+1^_16_

047795314301907

Fragment Matches Table

Show background peaks

| Position | Ion type | Intensity | mz Theoretical | mz Error (Th) | mz Error (ppm) | Charge | Series Number |
| --- | --- | --- | --- | --- | --- | --- | --- |
| - | - | 438.9 | 120.6 | - | - | 0 | - |
| - | - | 616.7 | 126.1 | - | - | 0 | - |
| - | - | 633.7 | 126.1 | - | - | 0 | - |
| - | - | 1055 | 127.1 | - | - | 0 | - |
| - | - | 381.8 | 128.1 | - | - | 0 | - |
| - | - | 428 | 128.2 | - | - | 0 | - |
| - | - | 379.7 | 128.3 | - | - | 0 | - |
| - | - | 369.4 | 128.7 | - | - | 0 | - |
| - | - | 1071 | 129.1 | - | - | 0 | - |
| - | - | 1.395E+05 | 129.1 | - | - | 0 | - |
| - | - | 3440 | 130.1 | - | - | 0 | - |
| 26 | y | 4625 | 130.1 | 0.0002928 | 2.251 | +1 | 1 |
| - | - | 1082 | 130.1 | - | - | 0 | - |
| - | - | 8844 | 130.1 | - | - | 0 | - |
| - | - | 852.8 | 132.1 | - | - | 0 | - |
| - | - | 9456 | 132.1 | - | - | 0 | - |
| - | - | 375.9 | 132.2 | - | - | 0 | - |
| - | - | 942.9 | 133.1 | - | - | 0 | - |
| - | - | 1060 | 136.1 | - | - | 0 | - |
| - | - | 375.9 | 137.9 | - | - | 0 | - |
| - | - | 1870 | 138.1 | - | - | 0 | - |
| - | - | 1564 | 139.1 | - | - | 0 | - |
| - | - | 560.5 | 140.1 | - | - | 0 | - |
| - | - | 882.4 | 140.1 | - | - | 0 | - |
| - | - | 596.3 | 140.5 | - | - | 0 | - |
| - | - | 1124 | 141.1 | - | - | 0 | - |
| - | - | 2171 | 141.1 | - | - | 0 | - |
| - | - | 754.7 | 142.1 | - | - | 0 | - |
| - | - | 416.9 | 143 | - | - | 0 | - |
| 2 | a | 1.332E+04 | 143.1 | 0.0003203 | 2.238 | +1 | 2 |
| - | - | 1143 | 144.1 | - | - | 0 | - |
| - | - | 480.4 | 144.5 | - | - | 0 | - |
| - | - | 759.8 | 146.1 | - | - | 0 | - |
| - | - | 912.6 | 147.1 | - | - | 0 | - |
| 26 | y | 5666 | 147.1 | 0.000355 | 2.413 | +1 | 1 |
| - | - | 420 | 148.1 | - | - | 0 | - |
| - | - | 513.3 | 148.9 | - | - | 0 | - |
| - | - | 514.7 | 153.1 | - | - | 0 | - |
| - | - | 3625 | 153.1 | - | - | 0 | - |
| - | - | 1683 | 155.1 | - | - | 0 | - |
| - | - | 6328 | 155.1 | - | - | 0 | - |
| - | - | 590.6 | 155.2 | - | - | 0 | - |
| - | - | 2537 | 157.1 | - | - | 0 | - |
| - | - | 771.2 | 157.1 | - | - | 0 | - |
| - | - | 8868 | 157.1 | - | - | 0 | - |
| - | - | 663.4 | 158.1 | - | - | 0 | - |
| - | - | 966 | 158.1 | - | - | 0 | - |
| - | - | 1.512E+05 | 159.1 | - | - | 0 | - |
| - | - | 1006 | 160.1 | - | - | 0 | - |
| - | - | 801.9 | 160.1 | - | - | 0 | - |
| - | - | 1.677E+04 | 160.1 | - | - | 0 | - |
| - | - | 1016 | 161.1 | - | - | 0 | - |
| - | - | 526.3 | 163.1 | - | - | 0 | - |
| - | - | 3103 | 167.1 | - | - | 0 | - |
| - | - | 2170 | 167.1 | - | - | 0 | - |
| - | - | 511.9 | 168.1 | - | - | 0 | - |
| - | - | 1050 | 169.1 | - | - | 0 | - |
| - | - | 882.9 | 169.1 | - | - | 0 | - |
| - | - | 1.693E+04 | 169.1 | - | - | 0 | - |
| - | - | 1.653E+04 | 170.1 | - | - | 0 | - |
| - | - | 603.7 | 170.1 | - | - | 0 | - |
| - | - | 1814 | 170.1 | - | - | 0 | - |
| - | - | 2037 | 171.1 | - | - | 0 | - |
| - | - | 836.8 | 171.1 | - | - | 0 | - |
| 2 | b | 1.51E+04 | 171.1 | 0.000355 | 2.075 | +1 | 2 |
| - | - | 1644 | 172.1 | - | - | 0 | - |
| - | - | 1352 | 172.1 | - | - | 0 | - |
| - | - | 672.5 | 173.1 | - | - | 0 | - |
| - | - | 557.2 | 173.1 | - | - | 0 | - |
| - | - | 1468 | 173.5 | - | - | 0 | - |
| - | - | 1323 | 175.1 | - | - | 0 | - |
| - | - | 603.6 | 180.1 | - | - | 0 | - |
| - | - | 2064 | 181.1 | - | - | 0 | - |
| - | - | 697 | 181.1 | - | - | 0 | - |
| - | - | 5862 | 182.1 | - | - | 0 | - |
| - | - | 3213 | 183.1 | - | - | 0 | - |
| - | - | 690.4 | 183.1 | - | - | 0 | - |
| - | - | 3370 | 183.1 | - | - | 0 | - |
| - | - | 676.6 | 184.1 | - | - | 0 | - |
| - | - | 3420 | 184.1 | - | - | 0 | - |
| - | - | 1197 | 185.1 | - | - | 0 | - |
| - | - | 1.592E+04 | 185.1 | - | - | 0 | - |
| - | - | 3185 | 185.1 | - | - | 0 | - |
| - | - | 1431 | 186.1 | - | - | 0 | - |
| - | - | 458.6 | 186.1 | - | - | 0 | - |
| - | - | 3077 | 187.1 | - | - | 0 | - |
| - | - | 1.228E+04 | 187.1 | - | - | 0 | - |
| - | - | 1602 | 188.1 | - | - | 0 | - |
| - | - | 583.6 | 193.1 | - | - | 0 | - |
| - | - | 543.8 | 194.1 | - | - | 0 | - |
| - | - | 665.5 | 195 | - | - | 0 | - |
| - | - | 3038 | 195.1 | - | - | 0 | - |
| - | - | 3321 | 195.1 | - | - | 0 | - |
| - | - | 1106 | 196.1 | - | - | 0 | - |
| - | - | 1.101E+04 | 197.1 | - | - | 0 | - |
| - | - | 1564 | 198.1 | - | - | 0 | - |
| - | - | 3664 | 198.1 | - | - | 0 | - |
| - | - | 674.8 | 198.1 | - | - | 0 | - |
| - | - | 1008 | 199.1 | - | - | 0 | - |
| - | - | 4051 | 199.1 | - | - | 0 | - |
| - | - | 3287 | 200.1 | - | - | 0 | - |
| - | - | 979.7 | 201.1 | - | - | 0 | - |
| - | - | 3496 | 202.1 | - | - | 0 | - |
| - | - | 2444 | 202.1 | - | - | 0 | - |
| - | - | 828.6 | 203.1 | - | - | 0 | - |
| - | - | 2116 | 203.1 | - | - | 0 | - |
| - | - | 4278 | 204.1 | - | - | 0 | - |
| - | - | 3250 | 205.1 | - | - | 0 | - |
| - | - | 622.5 | 205.1 | - | - | 0 | - |
| - | - | 1436 | 208.1 | - | - | 0 | - |
| - | - | 3647 | 209.1 | - | - | 0 | - |
| - | - | 532 | 210.1 | - | - | 0 | - |
| - | - | 2760 | 210.1 | - | - | 0 | - |
| - | - | 724.8 | 210.2 | - | - | 0 | - |
| - | - | 967.1 | 212.1 | - | - | 0 | - |
| - | - | 1147 | 212.1 | - | - | 0 | - |
| - | - | 3372 | 213.1 | - | - | 0 | - |
| - | - | 1820 | 213.1 | - | - | 0 | - |
| - | - | 3160 | 214.2 | - | - | 0 | - |
| - | - | 2044 | 216.1 | - | - | 0 | - |
| - | - | 1.087E+04 | 216.1 | - | - | 0 | - |
| - | - | 991.8 | 216.1 | - | - | 0 | - |
| - | - | 601.5 | 217.1 | - | - | 0 | - |
| - | - | 1021 | 221.1 | - | - | 0 | - |
| - | - | 4917 | 222.1 | - | - | 0 | - |
| 23 | y | 1627 | 223.1 | 0.002325 | 10.42 | +2 | 4 |
| - | - | 937.3 | 224.1 | - | - | 0 | - |
| - | - | 951.8 | 225.1 | - | - | 0 | - |
| - | - | 662.6 | 226.1 | - | - | 0 | - |
| - | - | 9145 | 226.1 | - | - | 0 | - |
| - | - | 567.8 | 227.1 | - | - | 0 | - |
| - | - | 6413 | 228.1 | - | - | 0 | - |
| - | - | 2133 | 228.2 | - | - | 0 | - |
| - | - | 990.8 | 229.1 | - | - | 0 | - |
| - | - | 2303 | 229.1 | - | - | 0 | - |
| - | - | 989.2 | 229.1 | - | - | 0 | - |
| - | - | 719.9 | 230.1 | - | - | 0 | - |
| - | - | 5514 | 230.1 | - | - | 0 | - |
| - | - | 1571 | 231.1 | - | - | 0 | - |
| - | - | 590.1 | 231.1 | - | - | 0 | - |
| - | - | 853.8 | 233.2 | - | - | 0 | - |
| - | - | 1010 | 236.1 | - | - | 0 | - |
| - | - | 653.2 | 236.1 | - | - | 0 | - |
| - | - | 753.1 | 237.1 | - | - | 0 | - |
| - | - | 1347 | 238.1 | - | - | 0 | - |
| - | - | 876.9 | 238.2 | - | - | 0 | - |
| - | - | 2065 | 239.2 | - | - | 0 | - |
| 21 | y | 4459 | 240.1 | 0.001762 | 7.339 | +3 | 6 |
| - | - | 724.5 | 241.1 | - | - | 0 | - |
| - | - | 718.9 | 241.1 | - | - | 0 | - |
| - | - | 915.3 | 242.1 | - | - | 0 | - |
| 4 | b | 1383 | 243.1 | 0.001854 | 7.623 | +2 | 4 |
| - | - | 719.6 | 244.1 | - | - | 0 | - |
| 25 | y | 5006 | 244.1 | 0.0005325 | 2.181 | +1 | 2 |
| - | - | 768 | 249.1 | - | - | 0 | - |
| - | - | 1176 | 250.1 | - | - | 0 | - |
| - | - | 534.9 | 250.1 | - | - | 0 | - |
| - | - | 642.5 | 250.2 | - | - | 0 | - |
| - | - | 661 | 251.1 | - | - | 0 | - |
| - | - | 1386 | 252.1 | - | - | 0 | - |
| - | - | 778 | 252.1 | - | - | 0 | - |
| - | - | 543.1 | 253.2 | - | - | 0 | - |
| - | - | 1003 | 254.1 | - | - | 0 | - |
| - | - | 4234 | 254.1 | - | - | 0 | - |
| - | - | 483.2 | 255.1 | - | - | 0 | - |
| - | - | 543.5 | 255.1 | - | - | 0 | - |
| - | - | 2293 | 256.1 | - | - | 0 | - |
| - | - | 892.5 | 256.1 | - | - | 0 | - |
| - | - | 1378 | 256.1 | - | - | 0 | - |
| - | - | 766.7 | 256.2 | - | - | 0 | - |
| - | - | 3696 | 257.2 | - | - | 0 | - |
| - | - | 642.9 | 258.1 | - | - | 0 | - |
| - | - | 9732 | 258.1 | - | - | 0 | - |
| - | - | 728.5 | 258.1 | - | - | 0 | - |
| - | - | 673.1 | 258.1 | - | - | 0 | - |
| - | - | 1352 | 259.1 | - | - | 0 | - |
| 25 | y | 1.266E+04 | 261.2 | 0.0003048 | 1.167 | +1 | 2 |
| - | - | 1241 | 262.2 | - | - | 0 | - |
| - | - | 3083 | 266.2 | - | - | 0 | - |
| - | - | 953.5 | 267.1 | - | - | 0 | - |
| - | - | 495.3 | 267.2 | - | - | 0 | - |
| - | - | 573.2 | 268.1 | - | - | 0 | - |
| - | - | 3890 | 268.1 | - | - | 0 | - |
| - | - | 788.9 | 268.2 | - | - | 0 | - |
| - | - | 599.1 | 269.2 | - | - | 0 | - |
| - | - | 838.7 | 270.1 | - | - | 0 | - |
| - | - | 971 | 270.1 | - | - | 0 | - |
| - | - | 6725 | 270.2 | - | - | 0 | - |
| - | - | 3186 | 272.1 | - | - | 0 | - |
| - | - | 1392 | 274.1 | - | - | 0 | - |
| - | - | 2544 | 278.1 | - | - | 0 | - |
| - | - | 676.3 | 282.1 | - | - | 0 | - |
| - | - | 832.3 | 282.1 | - | - | 0 | - |
| - | - | 524.5 | 282.2 | - | - | 0 | - |
| - | - | 1455 | 283.2 | - | - | 0 | - |
| - | - | 1666 | 284.1 | - | - | 0 | - |
| - | - | 3708 | 284.2 | - | - | 0 | - |
| - | - | 846.7 | 284.2 | - | - | 0 | - |
| - | - | 1938 | 285.2 | - | - | 0 | - |
| - | - | 5141 | 286.1 | - | - | 0 | - |
| - | - | 1161 | 287.1 | - | - | 0 | - |
| - | - | 717.7 | 287.2 | - | - | 0 | - |
| - | - | 631.5 | 290.1 | - | - | 0 | - |
| - | - | 602.7 | 292.1 | - | - | 0 | - |
| - | - | 958.2 | 294.1 | - | - | 0 | - |
| - | - | 819.7 | 295.1 | - | - | 0 | - |
| - | - | 2773 | 295.2 | - | - | 0 | - |
| - | - | 3731 | 296.1 | - | - | 0 | - |
| - | - | 656.6 | 296.2 | - | - | 0 | - |
| - | - | 2207 | 297.2 | - | - | 0 | - |
| - | - | 694.3 | 297.2 | - | - | 0 | - |
| - | - | 3.604E+04 | 298.2 | - | - | 0 | - |
| - | - | 7252 | 299.2 | - | - | 0 | - |
| - | - | 984.6 | 299.2 | - | - | 0 | - |
| - | - | 896.5 | 300.1 | - | - | 0 | - |
| - | - | 563.4 | 300.2 | - | - | 0 | - |
| - | - | 1421 | 301.2 | - | - | 0 | - |
| - | - | 840.7 | 303.2 | - | - | 0 | - |
| - | - | 745.2 | 306.1 | - | - | 0 | - |
| - | - | 817 | 307.1 | - | - | 0 | - |
| - | - | 660.3 | 307.2 | - | - | 0 | - |
| - | - | 1613 | 309.2 | - | - | 0 | - |
| - | - | 1191 | 309.2 | - | - | 0 | - |
| - | - | 1237 | 311.2 | - | - | 0 | - |
| - | - | 1503 | 311.2 | - | - | 0 | - |
| - | - | 1985 | 312.1 | - | - | 0 | - |
| - | - | 1195 | 312.2 | - | - | 0 | - |
| - | - | 4117 | 312.2 | - | - | 0 | - |
| - | - | 1641 | 313.1 | - | - | 0 | - |
| - | - | 1449 | 313.2 | - | - | 0 | - |
| - | - | 2.183E+04 | 313.2 | - | - | 0 | - |
| - | - | 1787 | 314.1 | - | - | 0 | - |
| - | - | 2533 | 314.2 | - | - | 0 | - |
| - | - | 1233 | 315.1 | - | - | 0 | - |
| - | - | 3489 | 315.2 | - | - | 0 | - |
| - | - | 2.925E+04 | 315.2 | - | - | 0 | - |
| - | - | 766.7 | 316.1 | - | - | 0 | - |
| - | - | 6281 | 316.2 | - | - | 0 | - |
| - | - | 862.4 | 317.1 | - | - | 0 | - |
| - | - | 943.8 | 317.2 | - | - | 0 | - |
| - | - | 649.7 | 319.2 | - | - | 0 | - |
| - | - | 697.8 | 321.2 | - | - | 0 | - |
| - | - | 5672 | 323.1 | - | - | 0 | - |
| - | - | 656.1 | 323.2 | - | - | 0 | - |
| - | - | 3188 | 325.1 | - | - | 0 | - |
| - | - | 811.3 | 325.2 | - | - | 0 | - |
| - | - | 730.6 | 325.2 | - | - | 0 | - |
| - | - | 1.284E+04 | 325.2 | - | - | 0 | - |
| - | - | 1547 | 326.2 | - | - | 0 | - |
| - | - | 2893 | 326.2 | - | - | 0 | - |
| 6 | b | 945.6 | 327.2 | 0.006139 | 18.76 | +2 | 6 |
| - | - | 2218 | 327.2 | - | - | 0 | - |
| - | - | 3110 | 328.2 | - | - | 0 | - |
| - | - | 1370 | 329.2 | - | - | 0 | - |
| - | - | 680.7 | 329.2 | - | - | 0 | - |
| - | - | 1274 | 330.1 | - | - | 0 | - |
| - | - | 670.6 | 330.2 | - | - | 0 | - |
| - | - | 2725 | 330.2 | - | - | 0 | - |
| - | - | 910.3 | 336.2 | - | - | 0 | - |
| - | - | 794.3 | 338.2 | - | - | 0 | - |
| - | - | 1112 | 339.2 | - | - | 0 | - |
| - | - | 974 | 339.2 | - | - | 0 | - |
| - | - | 1223 | 340.2 | - | - | 0 | - |
| - | - | 2516 | 341.1 | - | - | 0 | - |
| - | - | 5306 | 343.1 | - | - | 0 | - |
| - | - | 1646 | 343.2 | - | - | 0 | - |
| - | - | 1441 | 343.2 | - | - | 0 | - |
| - | - | 630.5 | 343.9 | - | - | 0 | - |
| - | - | 670.8 | 344.1 | - | - | 0 | - |
| - | - | 2430 | 344.2 | - | - | 0 | - |
| - | - | 819.5 | 347.2 | - | - | 0 | - |
| - | - | 883.2 | 351.2 | - | - | 0 | - |
| - | - | 1173 | 352.2 | - | - | 0 | - |
| - | - | 1114 | 353.1 | - | - | 0 | - |
| - | - | 981.2 | 353.2 | - | - | 0 | - |
| - | - | 2007 | 354.2 | - | - | 0 | - |
| - | - | 6512 | 356.2 | - | - | 0 | - |
| 3 | b | 1.633E+04 | 357.2 | 0.0004488 | 1.256 | +1 | 3 |
| - | - | 941 | 357.2 | - | - | 0 | - |
| 24 | y | 2985 | 358.2 | 0.0007111 | 1.985 | +1 | 3 |
| - | - | 2585 | 358.2 | - | - | 0 | - |
| - | - | 2503 | 361.1 | - | - | 0 | - |
| - | - | 723.9 | 364.1 | - | - | 0 | - |
| - | - | 571.2 | 364.2 | - | - | 0 | - |
| - | - | 1038 | 365.2 | - | - | 0 | - |
| - | - | 857.8 | 368.2 | - | - | 0 | - |
| - | - | 708.6 | 368.2 | - | - | 0 | - |
| - | - | 487 | 368.8 | - | - | 0 | - |
| - | - | 2826 | 369.1 | - | - | 0 | - |
| - | - | 478.9 | 369.2 | - | - | 0 | - |
| - | - | 741.2 | 369.2 | - | - | 0 | - |
| - | - | 516.3 | 370.2 | - | - | 0 | - |
| - | - | 970.7 | 371.2 | - | - | 0 | - |
| - | - | 1159 | 371.2 | - | - | 0 | - |
| - | - | 874.6 | 372.2 | - | - | 0 | - |
| - | - | 1697 | 374.2 | - | - | 0 | - |
| 24 | y | 7566 | 375.2 | 0.0004071 | 1.085 | +1 | 3 |
| - | - | 1173 | 376.2 | - | - | 0 | - |
| - | - | 2322 | 378.3 | - | - | 0 | - |
| - | - | 2436 | 379.2 | - | - | 0 | - |
| - | - | 638.6 | 380.2 | - | - | 0 | - |
| - | - | 587.1 | 380.2 | - | - | 0 | - |
| - | - | 1550 | 382.1 | - | - | 0 | - |
| - | - | 753.9 | 382.2 | - | - | 0 | - |
| - | - | 1046 | 382.2 | - | - | 0 | - |
| - | - | 917.4 | 383.2 | - | - | 0 | - |
| - | - | 1743 | 384.2 | - | - | 0 | - |
| - | - | 1304 | 384.2 | - | - | 0 | - |
| - | - | 6835 | 386.2 | - | - | 0 | - |
| - | - | 1502 | 387.2 | - | - | 0 | - |
| - | - | 1308 | 387.2 | - | - | 0 | - |
| - | - | 1175 | 387.2 | - | - | 0 | - |
| - | - | 1627 | 392.2 | - | - | 0 | - |
| - | - | 762.3 | 392.2 | - | - | 0 | - |
| - | - | 743.6 | 396.2 | - | - | 0 | - |
| - | - | 1166 | 396.2 | - | - | 0 | - |
| - | - | 2723 | 396.3 | - | - | 0 | - |
| - | - | 2116 | 397.2 | - | - | 0 | - |
| - | - | 1145 | 397.3 | - | - | 0 | - |
| - | - | 2100 | 402.2 | - | - | 0 | - |
| 20 | y | 644.9 | 403.2 | 0.001683 | 4.175 | +2 | 7 |
| - | - | 861.5 | 405.2 | - | - | 0 | - |
| - | - | 842.3 | 405.2 | - | - | 0 | - |
| - | - | 2035 | 406.2 | - | - | 0 | - |
| - | - | 886.9 | 410.1 | - | - | 0 | - |
| - | - | 2557 | 410.2 | - | - | 0 | - |
| - | - | 734.8 | 412.2 | - | - | 0 | - |
| - | - | 781.2 | 412.2 | - | - | 0 | - |
| - | - | 1591 | 412.3 | - | - | 0 | - |
| - | - | 600.9 | 413.3 | - | - | 0 | - |
| 8 | b | 856.1 | 414.2 | 0.001292 | 3.119 | +2 | 8 |
| 12 | b | 3518 | 414.2 | 0.007654 | 18.48 | +3 | 12 |
| - | - | 2489 | 415.2 | - | - | 0 | - |
| - | - | 698.7 | 415.2 | - | - | 0 | - |
| - | - | 1431 | 416.2 | - | - | 0 | - |
| - | - | 1175 | 422.2 | - | - | 0 | - |
| 8 | b | 1041 | 423.2 | 0.005089 | 12.02 | +2 | 8 |
| - | - | 5766 | 423.2 | - | - | 0 | - |
| - | - | 896.7 | 424.2 | - | - | 0 | - |
| - | - | 2665 | 424.2 | - | - | 0 | - |
| - | - | 1325 | 426.2 | - | - | 0 | - |
| - | - | 1004 | 426.2 | - | - | 0 | - |
| - | - | 1737 | 427.2 | - | - | 0 | - |
| - | - | 1298 | 427.2 | - | - | 0 | - |
| - | - | 836.2 | 428.1 | - | - | 0 | - |
| - | - | 1218 | 428.2 | - | - | 0 | - |
| - | - | 979.6 | 430.2 | - | - | 0 | - |
| - | - | 668 | 431.2 | - | - | 0 | - |
| - | - | 778.1 | 431.3 | - | - | 0 | - |
| - | - | 656 | 438.2 | - | - | 0 | - |
| - | - | 1061 | 439.2 | - | - | 0 | - |
| - | - | 2149 | 440.2 | - | - | 0 | - |
| - | - | 2051 | 440.2 | - | - | 0 | - |
| - | - | 634.1 | 441.2 | - | - | 0 | - |
| - | - | 1590 | 441.2 | - | - | 0 | - |
| - | - | 9158 | 441.2 | - | - | 0 | - |
| - | - | 929 | 441.3 | - | - | 0 | - |
| - | - | 713.2 | 442.2 | - | - | 0 | - |
| - | - | 1680 | 442.2 | - | - | 0 | - |
| 19 | y | 743.5 | 442.7 | 0.007857 | 17.75 | +2 | 8 |
| 19 | y | 1044 | 443.2 | 0.0007222 | 1.629 | +2 | 8 |
| - | - | 1140 | 443.7 | - | - | 0 | - |
| - | - | 1247 | 444.2 | - | - | 0 | - |
| 23 | y | 1753 | 444.2 | 0.000855 | 1.925 | +1 | 4 |
| - | - | 920 | 445.2 | - | - | 0 | - |
| 23 | y | 2514 | 445.2 | 0.0001158 | 0.2601 | +1 | 4 |
| 19 | y | 6108 | 451.7 | 0.001079 | 2.388 | +2 | 8 |
| - | - | 673 | 452.2 | - | - | 0 | - |
| - | - | 2133 | 452.2 | - | - | 0 | - |
| - | - | 1171 | 452.7 | - | - | 0 | - |
| - | - | 2027 | 453.2 | - | - | 0 | - |
| - | - | 936.1 | 453.2 | - | - | 0 | - |
| - | - | 2384 | 453.3 | - | - | 0 | - |
| - | - | 658.3 | 454.2 | - | - | 0 | - |
| - | - | 935.3 | 455.3 | - | - | 0 | - |
| - | - | 2734 | 457.3 | - | - | 0 | - |
| - | - | 1596 | 458.2 | - | - | 0 | - |
| - | - | 1564 | 458.2 | - | - | 0 | - |
| - | - | 714.4 | 458.3 | - | - | 0 | - |
| - | - | 1035 | 459.2 | - | - | 0 | - |
| - | - | 1124 | 461.2 | - | - | 0 | - |
| 23 | y | 1.754E+04 | 462.2 | 0.001063 | 2.3 | +1 | 4 |
| - | - | 897.9 | 462.7 | - | - | 0 | - |
| - | - | 5154 | 463.2 | - | - | 0 | - |
| 9 | b | 1018 | 464.2 | 0.005473 | 11.79 | +2 | 9 |
| - | - | 1444 | 465.3 | - | - | 0 | - |
| - | - | 1169 | 468.2 | - | - | 0 | - |
| - | - | 879.3 | 470.2 | - | - | 0 | - |
| - | - | 3990 | 471.2 | - | - | 0 | - |
| - | - | 1090 | 472.2 | - | - | 0 | - |
| - | - | 695.4 | 476.2 | - | - | 0 | - |
| - | - | 2087 | 483.3 | - | - | 0 | - |
| 4 | b | 7000 | 485.3 | 0.003511 | 7.234 | +1 | 4 |
| - | - | 1637 | 486.2 | - | - | 0 | - |
| - | - | 1919 | 486.3 | - | - | 0 | - |
| - | - | 782.8 | 487.2 | - | - | 0 | - |
| - | - | 6671 | 489.2 | - | - | 0 | - |
| - | - | 1254 | 490.2 | - | - | 0 | - |
| - | - | 833.9 | 493.2 | - | - | 0 | - |
| - | - | 1154 | 493.2 | - | - | 0 | - |
| - | - | 754.4 | 499.2 | - | - | 0 | - |
| - | - | 6117 | 501.2 | - | - | 0 | - |
| 18 | y | 2464 | 502.3 | 0.003931 | 7.826 | +2 | 9 |
| - | - | 1072 | 502.8 | - | - | 0 | - |
| - | - | 889.6 | 503.2 | - | - | 0 | - |
| - | - | 869.1 | 507.3 | - | - | 0 | - |
| - | - | 1698 | 510.3 | - | - | 0 | - |
| - | - | 762.8 | 513.3 | - | - | 0 | - |
| - | - | 769.2 | 517.2 | - | - | 0 | - |
| - | - | 882.5 | 520.2 | - | - | 0 | - |
| - | - | 1539 | 521.2 | - | - | 0 | - |
| - | - | 997.6 | 523.3 | - | - | 0 | - |
| - | - | 1386 | 524.3 | - | - | 0 | - |
| - | - | 1837 | 524.4 | - | - | 0 | - |
| - | - | 599.8 | 526.8 | - | - | 0 | - |
| - | - | 713.9 | 527.3 | - | - | 0 | - |
| - | - | 3988 | 528.3 | - | - | 0 | - |
| - | - | 1326 | 528.3 | - | - | 0 | - |
| - | - | 882.9 | 529.3 | - | - | 0 | - |
| - | - | 832 | 529.3 | - | - | 0 | - |
| - | - | 1027 | 535.8 | - | - | 0 | - |
| - | - | 1205 | 537.2 | - | - | 0 | - |
| - | - | 2567 | 538.2 | - | - | 0 | - |
| - | - | 6218 | 538.3 | - | - | 0 | - |
| - | - | 1102 | 539.2 | - | - | 0 | - |
| - | - | 1746 | 539.3 | - | - | 0 | - |
| - | - | 908.3 | 540.2 | - | - | 0 | - |
| - | - | 2042 | 540.3 | - | - | 0 | - |
| - | - | 1010 | 541.3 | - | - | 0 | - |
| - | - | 1101 | 542.3 | - | - | 0 | - |
| 17 | y | 608.4 | 544.3 | 0.008026 | 14.75 | +2 | 10 |
| - | - | 1096 | 552.3 | - | - | 0 | - |
| - | - | 6356 | 552.4 | - | - | 0 | - |
| - | - | 1629 | 553.4 | - | - | 0 | - |
| - | - | 678.9 | 553.8 | - | - | 0 | - |
| - | - | 685.5 | 554.3 | - | - | 0 | - |
| - | - | 4411 | 555.3 | - | - | 0 | - |
| - | - | 2647 | 556.2 | - | - | 0 | - |
| 5 | b | 1.623E+04 | 556.3 | 0.0007573 | 1.361 | +1 | 5 |
| - | - | 3368 | 557.3 | - | - | 0 | - |
| - | - | 4802 | 557.3 | - | - | 0 | - |
| - | - | 2055 | 558.3 | - | - | 0 | - |
| - | - | 859.7 | 562.3 | - | - | 0 | - |
| - | - | 661.5 | 564.3 | - | - | 0 | - |
| - | - | 1173 | 564.8 | - | - | 0 | - |
| - | - | 822.1 | 565.3 | - | - | 0 | - |
| - | - | 1459 | 568.3 | - | - | 0 | - |
| - | - | 848.5 | 570.3 | - | - | 0 | - |
| - | - | 2406 | 570.3 | - | - | 0 | - |
| - | - | 963.2 | 571.3 | - | - | 0 | - |
| 22 | y | 7758 | 572.3 | 0.001756 | 3.069 | +1 | 5 |
| 22 | y | 9885 | 573.3 | 0.001688 | 2.945 | +1 | 5 |
| - | - | 1730 | 574.3 | - | - | 0 | - |
| - | - | 766.5 | 584.3 | - | - | 0 | - |
| 11 | b | 999.9 | 585.3 | 0.009796 | 16.74 | +2 | 11 |
| - | - | 804.2 | 585.8 | - | - | 0 | - |
| - | - | 2144 | 586.3 | - | - | 0 | - |
| - | - | 1172 | 588.3 | - | - | 0 | - |
| - | - | 604.2 | 589.3 | - | - | 0 | - |
| 22 | y | 1.176E+04 | 590.3 | 0.000652 | 1.105 | +1 | 5 |
| - | - | 3697 | 591.3 | - | - | 0 | - |
| - | - | 741.5 | 592.3 | - | - | 0 | - |
| 10 | y | 1085 | 596.3 | 0.005863 | 9.832 | +3 | 17 |
| - | - | 1048 | 597.3 | - | - | 0 | - |
| 16 | y | 5239 | 603.3 | 0.0006367 | 1.055 | +2 | 11 |
| - | - | 2857 | 603.8 | - | - | 0 | - |
| 18 | b | 852.3 | 604.3 | 0.001402 | 2.32 | +3 | 18 |
| - | - | 1037 | 606.8 | - | - | 0 | - |
| 12 | b | 2361 | 611.8 | 6.155E-05 | 0.1006 | +2 | 12 |
| 12 | b | 1534 | 612.3 | 0.009152 | 14.95 | +2 | 12 |
| - | - | 687.9 | 614.3 | - | - | 0 | - |
| - | - | 1032 | 614.8 | - | - | 0 | - |
| 12 | b | 4287 | 620.8 | 0.0003944 | 0.6353 | +2 | 12 |
| - | - | 2124 | 621.3 | - | - | 0 | - |
| - | - | 1073 | 621.8 | - | - | 0 | - |
| - | - | 1953 | 624.3 | - | - | 0 | - |
| - | - | 1262 | 625.3 | - | - | 0 | - |
| - | - | 592.7 | 626.8 | - | - | 0 | - |
| - | - | 1206 | 627.3 | - | - | 0 | - |
| - | - | 647.4 | 628.4 | - | - | 0 | - |
| - | - | 1842 | 639.4 | - | - | 0 | - |
| - | - | 1169 | 640.3 | - | - | 0 | - |
| - | - | 835.8 | 641.3 | - | - | 0 | - |
| - | - | 5362 | 642.3 | - | - | 0 | - |
| - | - | 1232 | 643.3 | - | - | 0 | - |
| - | - | 683.8 | 644.3 | - | - | 0 | - |
| 13 | b | 3075 | 649.3 | 0.001182 | 1.821 | +2 | 13 |
| - | - | 3325 | 649.8 | - | - | 0 | - |
| - | - | 770.4 | 650.4 | - | - | 0 | - |
| - | - | 1228 | 651.3 | - | - | 0 | - |
| - | - | 633 | 655.2 | - | - | 0 | - |
| - | - | 1512 | 657.3 | - | - | 0 | - |
| - | - | 1333 | 657.4 | - | - | 0 | - |
| - | - | 2099 | 658.3 | - | - | 0 | - |
| 8 | y | 943.2 | 658.3 | 0.008752 | 13.29 | +3 | 19 |
| - | - | 751.6 | 663.4 | - | - | 0 | - |
| - | - | 3365 | 667.3 | - | - | 0 | - |
| 15 | y | 1937 | 667.8 | 0.001496 | 2.24 | +2 | 12 |
| - | - | 3125 | 668.3 | - | - | 0 | - |
| - | - | 1124 | 668.8 | - | - | 0 | - |
| - | - | 1183 | 669.3 | - | - | 0 | - |
| 6 | b | 1.304E+04 | 671.4 | 0.0005476 | 0.8157 | +1 | 6 |
| - | - | 4460 | 672.4 | - | - | 0 | - |
| - | - | 5108 | 675.3 | - | - | 0 | - |
| - | - | 1697 | 676.3 | - | - | 0 | - |
| - | - | 1036 | 677.4 | - | - | 0 | - |
| - | - | 1764 | 678.4 | - | - | 0 | - |
| - | - | 992.6 | 678.9 | - | - | 0 | - |
| - | - | 3720 | 681.4 | - | - | 0 | - |
| - | - | 1520 | 682.4 | - | - | 0 | - |
| - | - | 964.9 | 683.3 | - | - | 0 | - |
| - | - | 1421 | 684.3 | - | - | 0 | - |
| - | - | 3779 | 684.9 | - | - | 0 | - |
| - | - | 1.052E+04 | 685.4 | - | - | 0 | - |
| - | - | 757.3 | 685.9 | - | - | 0 | - |
| - | - | 3436 | 686.4 | - | - | 0 | - |
| 7 | y | 2482 | 687.4 | 0.00656 | 9.543 | +3 | 20 |
| - | - | 699 | 688.3 | - | - | 0 | - |
| - | - | 1041 | 688.4 | - | - | 0 | - |
| 14 | b | 892.4 | 690.4 | 0.01176 | 17.03 | +2 | 14 |
| - | - | 727.4 | 695.3 | - | - | 0 | - |
| 14 | b | 2409 | 698.9 | 0.0004996 | 0.7148 | +2 | 14 |
| - | - | 1703 | 699.4 | - | - | 0 | - |
| 21 | y | 1780 | 700.4 | 0.002447 | 3.495 | +1 | 6 |
| 21 | y | 3749 | 701.4 | 0.0009637 | 1.374 | +1 | 6 |
| - | - | 2170 | 702.4 | - | - | 0 | - |
| - | - | 683 | 705.4 | - | - | 0 | - |
| 14 | y | 1021 | 708.4 | 0.001502 | 2.12 | +2 | 13 |
| - | - | 641.4 | 710.9 | - | - | 0 | - |
| 21 | b | 755.1 | 714.4 | 0.01004 | 14.05 | +3 | 21 |
| - | - | 940.7 | 716.4 | - | - | 0 | - |
| 14 | y | 842.5 | 717.4 | 0.004948 | 6.897 | +2 | 13 |
| 21 | y | 8032 | 718.4 | 0.0003545 | 0.4935 | +1 | 6 |
| - | - | 2660 | 719.4 | - | - | 0 | - |
| - | - | 692.3 | 720.4 | - | - | 0 | - |
| - | - | 937 | 726.4 | - | - | 0 | - |
| - | - | 1521 | 727.9 | - | - | 0 | - |
| - | - | 1017 | 728.4 | - | - | 0 | - |
| - | - | 795.5 | 728.9 | - | - | 0 | - |
| - | - | 840.5 | 729.4 | - | - | 0 | - |
| - | - | 5926 | 738.4 | - | - | 0 | - |
| - | - | 1947 | 739.3 | - | - | 0 | - |
| - | - | 1164 | 739.4 | - | - | 0 | - |
| 7 | b | 1463 | 740.4 | 0.009174 | 12.39 | +1 | 7 |
| - | - | 1616 | 742.4 | - | - | 0 | - |
| 5 | y | 1081 | 743.4 | 0.005229 | 7.034 | +3 | 22 |
| - | - | 1838 | 744.4 | - | - | 0 | - |
| - | - | 1030 | 745.4 | - | - | 0 | - |
| 13 | y | 2247 | 745.9 | 0.0009625 | 1.29 | +2 | 14 |
| - | - | 1943 | 746.4 | - | - | 0 | - |
| - | - | 728.8 | 746.9 | - | - | 0 | - |
| - | - | 869.7 | 747.3 | - | - | 0 | - |
| 5 | y | 862.1 | 749.4 | 0.007058 | 9.418 | +3 | 22 |
| - | - | 856.1 | 749.9 | - | - | 0 | - |
| - | - | 1399 | 752.4 | - | - | 0 | - |
| - | - | 7568 | 756.4 | - | - | 0 | - |
| - | - | 3690 | 757.4 | - | - | 0 | - |
| 7 | b | 6014 | 758.4 | 0.0001354 | 0.1786 | +1 | 7 |
| - | - | 2226 | 759.4 | - | - | 0 | - |
| - | - | 936.5 | 760.4 | - | - | 0 | - |
| - | - | 813.3 | 761.4 | - | - | 0 | - |
| 15 | b | 4192 | 763.4 | 0.0003502 | 0.4587 | +2 | 15 |
| - | - | 4389 | 763.9 | - | - | 0 | - |
| - | - | 1865 | 764.4 | - | - | 0 | - |
| - | - | 1063 | 768.4 | - | - | 0 | - |
| - | - | 2735 | 770.4 | - | - | 0 | - |
| - | - | 1095 | 771.4 | - | - | 0 | - |
| - | - | 900.5 | 771.7 | - | - | 0 | - |
| 12 | y | 1056 | 772.4 | 0.007035 | 9.108 | +2 | 15 |
| - | - | 747.3 | 772.8 | - | - | 0 | - |
| - | - | 799.2 | 773.4 | - | - | 0 | - |
| - | - | 3779 | 774.4 | - | - | 0 | - |
| - | - | 1120 | 775.4 | - | - | 0 | - |
| - | - | 1180 | 775.9 | - | - | 0 | - |
| - | - | 982.6 | 776.4 | - | - | 0 | - |
| - | - | 1195 | 779.4 | - | - | 0 | - |
| 23 | b | 667.2 | 780.1 | 0.00493 | 6.32 | +3 | 23 |
| 23 | b | 1487 | 780.4 | 0.001115 | 1.429 | +3 | 23 |
| - | - | 740.8 | 780.9 | - | - | 0 | - |
| 12 | y | 5070 | 781.4 | 0.0001712 | 0.2191 | +2 | 15 |
| - | - | 2671 | 781.9 | - | - | 0 | - |
| - | - | 1115 | 782.4 | - | - | 0 | - |
| - | - | 961.3 | 782.5 | - | - | 0 | - |
| - | - | 1622 | 783.4 | - | - | 0 | - |
| 4 | y | 2687 | 786.4 | 0.003174 | 4.036 | +3 | 23 |
| - | - | 1949 | 786.7 | - | - | 0 | - |
| 20 | y | 2151 | 787.4 | 0.001883 | 2.392 | +1 | 7 |
| - | - | 4018 | 788.4 | - | - | 0 | - |
| - | - | 1287 | 789.4 | - | - | 0 | - |
| - | - | 928.4 | 790.4 | - | - | 0 | - |
| - | - | 668.7 | 791.9 | - | - | 0 | - |
| 4 | y | 7502 | 792.1 | 0.0007534 | 0.9512 | +3 | 23 |
| - | - | 1.175E+04 | 792.4 | - | - | 0 | - |
| - | - | 6747 | 792.7 | - | - | 0 | - |
| - | - | 1754 | 793.1 | - | - | 0 | - |
| - | - | 865.1 | 793.4 | - | - | 0 | - |
| - | - | 1145 | 795.4 | - | - | 0 | - |
| - | - | 921.8 | 799.1 | - | - | 0 | - |
| - | - | 937.6 | 799.4 | - | - | 0 | - |
| - | - | 1132 | 799.9 | - | - | 0 | - |
| 16 | b | 911.1 | 804.9 | 0.001662 | 2.064 | +2 | 16 |
| 16 | b | 6096 | 805.4 | 0.002757 | 3.423 | +2 | 16 |
| - | - | 973.1 | 805.9 | - | - | 0 | - |
| - | - | 2961 | 806.4 | - | - | 0 | - |
| - | - | 1034 | 807.4 | - | - | 0 | - |
| - | - | 932.7 | 809.4 | - | - | 0 | - |
| - | - | 845.4 | 810.4 | - | - | 0 | - |
| - | - | 917.2 | 812.4 | - | - | 0 | - |
| - | - | 2427 | 813.4 | - | - | 0 | - |
| 16 | b | 3297 | 813.9 | 0.0002026 | 0.249 | +2 | 16 |
| - | - | 4748 | 814.4 | - | - | 0 | - |
| - | - | 1569 | 814.9 | - | - | 0 | - |
| - | - | 1822 | 815.4 | - | - | 0 | - |
| - | - | 702.9 | 815.9 | - | - | 0 | - |
| - | - | 733.6 | 818.9 | - | - | 0 | - |
| - | - | 861.2 | 820.9 | - | - | 0 | - |
| - | - | 773.7 | 824.4 | - | - | 0 | - |
| - | - | 801.7 | 826.9 | - | - | 0 | - |
| - | - | 605.4 | 827.1 | - | - | 0 | - |
| 8 | b | 4716 | 827.4 | 0.0005763 | 0.6966 | +1 | 8 |
| - | - | 4178 | 828.4 | - | - | 0 | - |
| - | - | 849.3 | 828.9 | - | - | 0 | - |
| - | - | 1981 | 829.4 | - | - | 0 | - |
| - | - | 1920 | 829.7 | - | - | 0 | - |
| - | - | 1698 | 829.9 | - | - | 0 | - |
| - | - | 2096 | 830.4 | - | - | 0 | - |
| - | - | 1049 | 830.9 | - | - | 0 | - |
| - | - | 1415 | 831.4 | - | - | 0 | - |
| - | - | 1183 | 833.5 | - | - | 0 | - |
| - | - | 957.7 | 834.4 | - | - | 0 | - |
| - | - | 1747 | 836.3 | - | - | 0 | - |
| 11 | y | 1385 | 836.4 | 0.005078 | 6.071 | +2 | 16 |
| - | - | 869.3 | 836.8 | - | - | 0 | - |
| 11 | y | 1963 | 836.9 | 0.003952 | 4.722 | +2 | 16 |
| - | - | 682 | 837 | - | - | 0 | - |
| - | - | 2095 | 837.4 | - | - | 0 | - |
| - | - | 2465 | 838 | - | - | 0 | - |
| - | - | 936 | 838.4 | - | - | 0 | - |
| - | - | 1642 | 839.4 | - | - | 0 | - |
| - | - | 792.8 | 841.4 | - | - | 0 | - |
| - | - | 1223 | 842.4 | - | - | 0 | - |
| - | - | 826.4 | 842.8 | - | - | 0 | - |
| - | - | 1001 | 843.4 | - | - | 0 | - |
| 8 | b | 1.691E+04 | 845.4 | 0.009062 | 10.72 | +1 | 8 |
| - | - | 5598 | 845.9 | - | - | 0 | - |
| - | - | 8777 | 846.4 | - | - | 0 | - |
| - | - | 1352 | 846.9 | - | - | 0 | - |
| - | - | 2524 | 847.4 | - | - | 0 | - |
| 3 | y | 1480 | 848.1 | 0.001655 | 1.951 | +3 | 24 |
| 3 | y | 4230 | 848.4 | 0.005179 | 6.104 | +3 | 24 |
| - | - | 1666 | 848.8 | - | - | 0 | - |
| - | - | 1610 | 849.1 | - | - | 0 | - |
| - | - | 1357 | 849.4 | - | - | 0 | - |
| - | - | 1034 | 852.9 | - | - | 0 | - |
| - | - | 980.3 | 853.4 | - | - | 0 | - |
| 3 | y | 1.111E+04 | 854.1 | 0.0006219 | 0.7282 | +3 | 24 |
| - | - | 1.494E+04 | 854.4 | - | - | 0 | - |
| - | - | 1.206E+04 | 854.8 | - | - | 0 | - |
| - | - | 9486 | 855.1 | - | - | 0 | - |
| 17 | b | 4324 | 855.4 | 0.0002224 | 0.26 | +2 | 17 |
| 17 | b | 1068 | 855.9 | 0.007177 | 8.385 | +2 | 17 |
| 25 | b | 2061 | 856.4 | 0.01183 | 13.82 | +3 | 25 |
| - | - | 1239 | 856.8 | - | - | 0 | - |
| - | - | 1549 | 857.1 | - | - | 0 | - |
| - | - | 2585 | 857.4 | - | - | 0 | - |
| - | - | 767.9 | 857.6 | - | - | 0 | - |
| 25 | b | 1115 | 862.1 | 0.01647 | 19.1 | +3 | 25 |
| - | - | 3057 | 862.4 | - | - | 0 | - |
| - | - | 936.6 | 862.6 | - | - | 0 | - |
| - | - | 2052 | 862.8 | - | - | 0 | - |
| - | - | 1423 | 863.1 | - | - | 0 | - |
| - | - | 2413 | 863.4 | - | - | 0 | - |
| 17 | b | 3529 | 864.5 | 0.0009215 | 1.066 | +2 | 17 |
| - | - | 2874 | 865 | - | - | 0 | - |
| - | - | 2701 | 865.5 | - | - | 0 | - |
| - | - | 2985 | 866.5 | - | - | 0 | - |
| - | - | 2590 | 867.4 | - | - | 0 | - |
| - | - | 2323 | 868.1 | - | - | 0 | - |
| - | - | 4826 | 868.4 | - | - | 0 | - |
| - | - | 1208 | 868.8 | - | - | 0 | - |
| - | - | 994.5 | 869.4 | - | - | 0 | - |
| - | - | 5484 | 871.4 | - | - | 0 | - |
| 2 | y | 1136 | 871.8 | 0.0008258 | 0.9473 | +3 | 25 |
| - | - | 3591 | 872.4 | - | - | 0 | - |
| - | - | 1144 | 873.1 | - | - | 0 | - |
| - | - | 2037 | 873.4 | - | - | 0 | - |
| - | - | 1234 | 875.4 | - | - | 0 | - |
| 2 | y | 3481 | 877.8 | 0.001455 | 1.657 | +3 | 25 |
| - | - | 5374 | 878.1 | - | - | 0 | - |
| - | - | 3446 | 878.5 | - | - | 0 | - |
| - | - | 1995 | 878.8 | - | - | 0 | - |
| - | - | 1518 | 879.5 | - | - | 0 | - |
| - | - | 1091 | 881.1 | - | - | 0 | - |
| - | - | 879.3 | 882.5 | - | - | 0 | - |
| - | - | 1230 | 883.5 | - | - | 0 | - |
| 19 | y | 8302 | 884.5 | 0.00944 | 10.67 | +1 | 8 |
| 19 | y | 1.405E+04 | 885.4 | 0.004123 | 4.657 | +1 | 8 |
| - | - | 7197 | 886.4 | - | - | 0 | - |
| - | - | 1622 | 886.8 | - | - | 0 | - |
| - | - | 1077 | 887.1 | - | - | 0 | - |
| - | - | 2880 | 887.5 | - | - | 0 | - |
| - | - | 1975 | 888.5 | - | - | 0 | - |
| - | - | 1560 | 890.5 | - | - | 0 | - |
| - | - | 1261 | 890.8 | - | - | 0 | - |
| - | - | 1446 | 891.1 | - | - | 0 | - |
| - | - | 1813 | 891.5 | - | - | 0 | - |
| - | - | 1217 | 891.8 | - | - | 0 | - |
| - | - | 1141 | 892.5 | - | - | 0 | - |
| - | - | 2466 | 893.5 | - | - | 0 | - |
| - | - | 1252 | 895.5 | - | - | 0 | - |
| - | - | 1269 | 895.8 | - | - | 0 | - |
| - | - | 974.1 | 896 | - | - | 0 | - |
| - | - | 2266 | 896.1 | - | - | 0 | - |
| - | - | 4642 | 896.5 | - | - | 0 | - |
| - | - | 4217 | 896.8 | - | - | 0 | - |
| - | - | 1484 | 897 | - | - | 0 | - |
| - | - | 2278 | 897.1 | - | - | 0 | - |
| - | - | 5403 | 897.5 | - | - | 0 | - |
| - | - | 1801 | 898 | - | - | 0 | - |
| - | - | 847.5 | 898.5 | - | - | 0 | - |
| - | - | 1825 | 898.8 | - | - | 0 | - |
| - | - | 3868 | 899.1 | - | - | 0 | - |
| - | - | 5854 | 899.5 | - | - | 0 | - |
| - | - | 2189 | 899.8 | - | - | 0 | - |
| - | - | 2945 | 900.1 | - | - | 0 | - |
| - | - | 3306 | 900.5 | - | - | 0 | - |
| - | - | 2933 | 900.8 | - | - | 0 | - |
| - | - | 979 | 901 | - | - | 0 | - |
| - | - | 1548 | 901.1 | - | - | 0 | - |
| - | - | 2905 | 901.5 | - | - | 0 | - |
| 19 | y | 7.063E+04 | 902.5 | 0.0005842 | 0.6474 | +1 | 8 |
| - | - | 3.358E+04 | 903.5 | - | - | 0 | - |
| - | - | 1.151E+04 | 904.5 | - | - | 0 | - |
| 0 | Precursor | 1.471E+04 | 904.8 | 0.0006776 | 0.7489 | +3 | -1 |
| 0 | Precursor | 3.461E+04 | 905.1 | 0.004447 | 4.913 | +3 | -1 |
| - | - | 2.994E+04 | 905.5 | - | - | 0 | - |
| - | - | 2.261E+04 | 905.8 | - | - | 0 | - |
| 18 | b | 3018 | 906 | 0.0006752 | 0.7452 | +2 | 18 |
| - | - | 1.309E+04 | 906.1 | - | - | 0 | - |
| 18 | b | 8373 | 906.5 | 0.00824 | 9.09 | +2 | 18 |
| - | - | 2001 | 906.8 | - | - | 0 | - |
| - | - | 2282 | 907 | - | - | 0 | - |
| - | - | 1603 | 909.5 | - | - | 0 | - |
| - | - | 3930 | 909.8 | - | - | 0 | - |
| - | - | 1.433E+04 | 910.1 | - | - | 0 | - |
| - | - | 2.314E+04 | 910.5 | - | - | 0 | - |
| 0 | Precursor | 5.191E+05 | 910.8 | 0.0006226 | 0.6836 | +3 | -1 |
| - | - | 8.102E+05 | 911.1 | - | - | 0 | - |
| - | - | 6.503E+05 | 911.5 | - | - | 0 | - |
| - | - | 3.726E+05 | 911.8 | - | - | 0 | - |
| - | - | 1.467E+05 | 912.1 | - | - | 0 | - |
| - | - | 4.574E+04 | 912.5 | - | - | 0 | - |
| - | - | 1120 | 913.5 | - | - | 0 | - |
| - | - | 883.9 | 914.5 | - | - | 0 | - |
| 18 | b | 7653 | 915 | 0.002473 | 2.703 | +2 | 18 |
| - | - | 8933 | 915.5 | - | - | 0 | - |
| - | - | 2775 | 916 | - | - | 0 | - |
| - | - | 1768 | 916.5 | - | - | 0 | - |
| - | - | 810.4 | 917 | - | - | 0 | - |
| - | - | 1060 | 930.7 | - | - | 0 | - |
| 9 | y | 2808 | 935 | 0.005802 | 6.205 | +2 | 18 |
| - | - | 1929 | 935.5 | - | - | 0 | - |
| - | - | 1221 | 936 | - | - | 0 | - |
| - | - | 1075 | 936.5 | - | - | 0 | - |
| - | - | 1084 | 941.9 | - | - | 0 | - |
| - | - | 2814 | 942.5 | - | - | 0 | - |
| 9 | y | 1.276E+04 | 943.5 | 0.001735 | 1.839 | +2 | 18 |
| - | - | 1.282E+04 | 944 | - | - | 0 | - |
| - | - | 6681 | 944.5 | - | - | 0 | - |
| - | - | 2327 | 945 | - | - | 0 | - |
| - | - | 896 | 945.5 | - | - | 0 | - |
| 19 | b | 1052 | 954.5 | 0.001271 | 1.331 | +2 | 19 |
| - | - | 870.4 | 958.5 | - | - | 0 | - |
| - | - | 2248 | 959.5 | - | - | 0 | - |
| - | - | 1195 | 966.5 | - | - | 0 | - |
| - | - | 868.6 | 967.5 | - | - | 0 | - |
| - | - | 1752 | 968.5 | - | - | 0 | - |
| - | - | 1445 | 969.5 | - | - | 0 | - |
| - | - | 1514 | 970.5 | - | - | 0 | - |
| - | - | 1392 | 972.5 | - | - | 0 | - |
| 8 | y | 974.2 | 978 | 0.002213 | 2.263 | +2 | 19 |
| 8 | y | 1661 | 978.5 | 0.001812 | 1.852 | +2 | 19 |
| - | - | 1171 | 979 | - | - | 0 | - |
| - | - | 1147 | 979.5 | - | - | 0 | - |
| - | - | 2728 | 984.5 | - | - | 0 | - |
| 18 | y | 2909 | 985.5 | 0.01151 | 11.67 | +1 | 9 |
| 18 | y | 4192 | 986.5 | 0.00686 | 6.954 | +1 | 9 |
| 8 | y | 6706 | 987 | 0.0002934 | 0.2973 | +2 | 19 |
| - | - | 6734 | 987.5 | - | - | 0 | - |
| - | - | 3103 | 988 | - | - | 0 | - |
| - | - | 1575 | 988.5 | - | - | 0 | - |
| - | - | 1359 | 994 | - | - | 0 | - |
| - | - | 1199 | 994.5 | - | - | 0 | - |
| - | - | 1533 | 999.5 | - | - | 0 | - |
| - | - | 1198 | 1001 | - | - | 0 | - |
| 18 | y | 2.029E+04 | 1004 | 0.001063 | 1.059 | +1 | 9 |
| - | - | 9443 | 1005 | - | - | 0 | - |
| - | - | 3112 | 1006 | - | - | 0 | - |
| - | - | 1274 | 1007 | - | - | 0 | - |
| - | - | 1533 | 1008 | - | - | 0 | - |
| - | - | 1160 | 1013 | - | - | 0 | - |
| - | - | 3544 | 1014 | - | - | 0 | - |
| 7 | y | 1652 | 1022 | 0.0004662 | 0.4564 | +2 | 20 |
| 7 | y | 2518 | 1022 | 0.004902 | 4.796 | +2 | 20 |
| - | - | 2802 | 1023 | - | - | 0 | - |
| 10 | b | 3605 | 1024 | 0.0002331 | 0.2278 | +1 | 10 |
| 10 | b | 1885 | 1025 | 0.01545 | 15.08 | +1 | 10 |
| - | - | 1135 | 1026 | - | - | 0 | - |
| 7 | y | 1.299E+04 | 1031 | 0.000477 | 0.4629 | +2 | 20 |
| - | - | 1.318E+04 | 1031 | - | - | 0 | - |
| - | - | 9821 | 1032 | - | - | 0 | - |
| - | - | 3117 | 1032 | - | - | 0 | - |
| - | - | 1204 | 1033 | - | - | 0 | - |
| - | - | 1643 | 1041 | - | - | 0 | - |
| 10 | b | 1.575E+04 | 1042 | 0.0001167 | 0.112 | +1 | 10 |
| - | - | 9687 | 1043 | - | - | 0 | - |
| - | - | 4390 | 1044 | - | - | 0 | - |
| - | - | 971.4 | 1045 | - | - | 0 | - |
| - | - | 1439 | 1052 | - | - | 0 | - |
| - | - | 1553 | 1053 | - | - | 0 | - |
| - | - | 1512 | 1058 | - | - | 0 | - |
| - | - | 5058 | 1059 | - | - | 0 | - |
| - | - | 3340 | 1060 | - | - | 0 | - |
| - | - | 1582 | 1061 | - | - | 0 | - |
| - | - | 1146 | 1070 | - | - | 0 | - |
| - | - | 2184 | 1071 | - | - | 0 | - |
| - | - | 1466 | 1071 | - | - | 0 | - |
| - | - | 3241 | 1072 | - | - | 0 | - |
| - | - | 1089 | 1072 | - | - | 0 | - |
| - | - | 1211 | 1073 | - | - | 0 | - |
| - | - | 1119 | 1078 | - | - | 0 | - |
| 6 | y | 1605 | 1079 | 0.004495 | 4.166 | +2 | 21 |
| 6 | y | 3864 | 1080 | 0.006872 | 6.366 | +2 | 21 |
| - | - | 4247 | 1080 | - | - | 0 | - |
| - | - | 1987 | 1081 | - | - | 0 | - |
| 17 | y | 878.4 | 1087 | 0.004293 | 3.951 | +1 | 10 |
| 17 | y | 3598 | 1088 | 0.008559 | 7.87 | +1 | 10 |
| 6 | y | 1.413E+04 | 1088 | 0.0004212 | 0.3871 | +2 | 21 |
| - | - | 2.145E+04 | 1089 | - | - | 0 | - |
| - | - | 1.328E+04 | 1089 | - | - | 0 | - |
| - | - | 5959 | 1090 | - | - | 0 | - |
| - | - | 1584 | 1090 | - | - | 0 | - |
| - | - | 950.1 | 1094 | - | - | 0 | - |
| - | - | 905.2 | 1099 | - | - | 0 | - |
| - | - | 1741 | 1101 | - | - | 0 | - |
| - | - | 1213 | 1102 | - | - | 0 | - |
| - | - | 1175 | 1102 | - | - | 0 | - |
| - | - | 853.3 | 1103 | - | - | 0 | - |
| 17 | y | 1.642E+04 | 1105 | 0.0003201 | 0.2898 | +1 | 10 |
| - | - | 1.015E+04 | 1106 | - | - | 0 | - |
| - | - | 1541 | 1106 | - | - | 0 | - |
| - | - | 5302 | 1107 | - | - | 0 | - |
| - | - | 1862 | 1107 | - | - | 0 | - |
| - | - | 2156 | 1108 | - | - | 0 | - |
| 5 | y | 4400 | 1115 | 0.002295 | 2.059 | +2 | 22 |
| 5 | y | 8697 | 1115 | 0.005527 | 4.957 | +2 | 22 |
| - | - | 6316 | 1116 | - | - | 0 | - |
| - | - | 3748 | 1116 | - | - | 0 | - |
| - | - | 1294 | 1117 | - | - | 0 | - |
| - | - | 1469 | 1123 | - | - | 0 | - |
| - | - | 1260 | 1123 | - | - | 0 | - |
| 5 | y | 3.714E+04 | 1124 | 0.0001869 | 0.1664 | +2 | 22 |
| - | - | 4.217E+04 | 1124 | - | - | 0 | - |
| - | - | 3.065E+04 | 1125 | - | - | 0 | - |
| - | - | 1.461E+04 | 1125 | - | - | 0 | - |
| - | - | 5036 | 1126 | - | - | 0 | - |
| 22 | b | 1302 | 1126 | 0.01113 | 9.884 | +2 | 22 |
| - | - | 951.7 | 1127 | - | - | 0 | - |
| - | - | 916.7 | 1128 | - | - | 0 | - |
| 22 | b | 1693 | 1135 | 0.004217 | 3.715 | +2 | 22 |
| - | - | 1487 | 1136 | - | - | 0 | - |
| - | - | 1305 | 1143 | - | - | 0 | - |
| - | - | 1759 | 1144 | - | - | 0 | - |
| - | - | 1069 | 1151 | - | - | 0 | - |
| - | - | 1016 | 1162 | - | - | 0 | - |
| - | - | 1005 | 1166 | - | - | 0 | - |
| 11 | b | 5025 | 1170 | 0.01329 | 11.36 | +1 | 11 |
| 23 | b | 1775 | 1170 | 0.008558 | 7.314 | +2 | 23 |
| - | - | 3345 | 1171 | - | - | 0 | - |
| - | - | 2098 | 1172 | - | - | 0 | - |
| 23 | b | 2734 | 1179 | 0.0003224 | 0.2736 | +2 | 23 |
| 4 | y | 2663 | 1179 | 0.009315 | 7.9 | +2 | 23 |
| - | - | 2413 | 1180 | - | - | 0 | - |
| - | - | 1999 | 1180 | - | - | 0 | - |
| 16 | y | 1.23E+04 | 1188 | 0.01142 | 9.619 | +1 | 11 |
| - | - | 1.215E+04 | 1188 | - | - | 0 | - |
| 16 | y | 9901 | 1189 | 0.02179 | 18.34 | +1 | 11 |
| - | - | 4423 | 1189 | - | - | 0 | - |
| - | - | 2505 | 1190 | - | - | 0 | - |
| 16 | y | 1.86E+04 | 1206 | 4.869E-06 | 0.004039 | +1 | 11 |
| - | - | 1.165E+04 | 1207 | - | - | 0 | - |
| - | - | 5973 | 1208 | - | - | 0 | - |
| - | - | 1102 | 1209 | - | - | 0 | - |
| 12 | b | 1111 | 1223 | 0.0002909 | 0.2379 | +1 | 12 |
| - | - | 1054 | 1224 | - | - | 0 | - |
| 24 | b | 1658 | 1227 | 0.01462 | 11.92 | +2 | 24 |
| 24 | b | 1965 | 1236 | 0.003476 | 2.813 | +2 | 24 |
| - | - | 2118 | 1236 | - | - | 0 | - |
| - | - | 1582 | 1237 | - | - | 0 | - |
| 12 | b | 1846 | 1241 | 0.0003575 | 0.2882 | +1 | 12 |
| - | - | 1797 | 1242 | - | - | 0 | - |
| - | - | 1528 | 1271 | - | - | 0 | - |
| - | - | 1592 | 1273 | - | - | 0 | - |
| - | - | 1131 | 1273 | - | - | 0 | - |
| 13 | b | 2331 | 1281 | 0.006869 | 5.363 | +1 | 13 |
| - | - | 3367 | 1281 | - | - | 0 | - |
| - | - | 2283 | 1282 | - | - | 0 | - |
| 25 | b | 1450 | 1293 | 0.0008915 | 0.6897 | +2 | 25 |
| - | - | 2014 | 1293 | - | - | 0 | - |
| - | - | 1746 | 1294 | - | - | 0 | - |
| - | - | 1191 | 1294 | - | - | 0 | - |
| 13 | b | 2662 | 1298 | 0.003691 | 2.845 | +1 | 13 |
| - | - | 2261 | 1299 | - | - | 0 | - |
| - | - | 893.8 | 1300 | - | - | 0 | - |
| - | - | 1817 | 1302 | - | - | 0 | - |
| - | - | 1051 | 1302 | - | - | 0 | - |
| - | - | 1520 | 1303 | - | - | 0 | - |
| 15 | y | 4494 | 1317 | 0.008626 | 6.551 | +1 | 12 |
| 15 | y | 5166 | 1318 | 0.01045 | 7.931 | +1 | 12 |
| - | - | 2194 | 1319 | - | - | 0 | - |
| - | - | 1196 | 1320 | - | - | 0 | - |
| - | - | 1254 | 1327 | - | - | 0 | - |
| - | - | 1220 | 1328 | - | - | 0 | - |
| 15 | y | 1.958E+04 | 1335 | 0.001451 | 1.087 | +1 | 12 |
| - | - | 1.453E+04 | 1336 | - | - | 0 | - |
| - | - | 5398 | 1337 | - | - | 0 | - |
| - | - | 1860 | 1338 | - | - | 0 | - |
| - | - | 2672 | 1345 | - | - | 0 | - |
| - | - | 1575 | 1346 | - | - | 0 | - |
| - | - | 1030 | 1356 | - | - | 0 | - |
| - | - | 892.4 | 1363 | - | - | 0 | - |
| - | - | 977.8 | 1373 | - | - | 0 | - |
| 14 | b | 2081 | 1397 | 0.006129 | 4.388 | +1 | 14 |
| - | - | 1666 | 1398 | - | - | 0 | - |
| - | - | 1276 | 1399 | - | - | 0 | - |
| 14 | y | 1497 | 1417 | 0.001729 | 1.22 | +1 | 13 |
| - | - | 1179 | 1430 | - | - | 0 | - |
| 14 | y | 4495 | 1434 | 0.007731 | 5.392 | +1 | 13 |
| - | - | 3664 | 1435 | - | - | 0 | - |
| - | - | 1837 | 1436 | - | - | 0 | - |
| - | - | 1065 | 1447 | - | - | 0 | - |
| - | - | 1037 | 1455 | - | - | 0 | - |
| - | - | 1036 | 1457 | - | - | 0 | - |
| 13 | y | 2492 | 1473 | 0.009324 | 6.331 | +1 | 14 |
| 13 | y | 3479 | 1474 | 0.0131 | 8.89 | +1 | 14 |
| - | - | 2761 | 1475 | - | - | 0 | - |
| - | - | 986.9 | 1476 | - | - | 0 | - |
| 13 | y | 1.347E+04 | 1491 | 0.002827 | 1.896 | +1 | 14 |
| - | - | 1.247E+04 | 1492 | - | - | 0 | - |
| - | - | 5050 | 1493 | - | - | 0 | - |
| - | - | 1423 | 1494 | - | - | 0 | - |
| 15 | b | 2317 | 1526 | 0.008683 | 5.691 | +1 | 15 |
| - | - | 1257 | 1527 | - | - | 0 | - |
| - | - | 839.2 | 1528 | - | - | 0 | - |
| 12 | y | 1495 | 1544 | 0.004681 | 3.032 | +1 | 15 |
| 12 | y | 3188 | 1545 | 0.004675 | 3.026 | +1 | 15 |
| - | - | 1413 | 1546 | - | - | 0 | - |
| 12 | y | 1.244E+04 | 1562 | 0.004174 | 2.673 | +1 | 15 |
| - | - | 8992 | 1563 | - | - | 0 | - |
| - | - | 4699 | 1564 | - | - | 0 | - |
| - | - | 1034 | 1565 | - | - | 0 | - |
| 16 | b | 1253 | 1627 | 0.008266 | 5.081 | +1 | 16 |
| - | - | 998.7 | 1657 | - | - | 0 | - |
| - | - | 965.5 | 1675 | - | - | 0 | - |
| 11 | y | 2301 | 1690 | 0.004289 | 2.538 | +1 | 16 |
| - | - | 1830 | 1691 | - | - | 0 | - |
| - | - | 1188 | 1830 | - | - | 0 | - |
| - | - | 905.6 | 1888 | - | - | 0 | - |

m/z Charge Intensity FragmentType MassShift Position
120.60368347167969 0 438.90475
126.05531311035156 0 616.7077
126.09142303466797 0 633.7434
127.0867919921875 0 1054.8436
128.07037353515625 0 381.75256
128.24725341796875 0 427.95865
128.3443145751953 0 379.74326
128.68746948242188 0 369.39108
129.06593322753906 0 1070.7305
129.10260009765625 0 139473.78
130.06539916992188 0 3440.0417
130.0865478515625 0 4624.908 y Ammonia loss 25
130.1001434326172 0 1081.7997
130.10594177246094 0 8844.285
132.07594299316406 0 852.77124
132.08111572265625 0 9455.808
132.21456909179688 0 375.9034
133.0847930908203 0 942.911
136.07586669921875 0 1059.5193
137.9406280517578 0 375.9279
138.12808227539062 0 1870.132
139.08694458007812 0 1563.6873
140.0712127685547 0 560.4541
140.1070098876953 0 882.4023
140.45883178710938 0 596.28174
141.06610107421875 0 1123.7599
141.1026153564453 0 2170.7917
142.06565856933594 0 754.7409
143.02481079101562 0 416.88068
143.1182098388672 0 13318.606 a 1
144.12127685546875 0 1143.0612
144.49880981445312 0 480.41544
146.12905883789062 0 759.7938
147.07644653320312 0 912.618
147.1131591796875 0 5665.715 y 25
148.11663818359375 0 420.003
148.94764709472656 0 513.32056
153.06619262695312 0 514.7289
153.10255432128906 0 3625.1704
155.0819091796875 0 1683.4241
155.11822509765625 0 6328.0034
155.1547393798828 0 590.61664
157.0611114501953 0 2537.2766
157.09068298339844 0 771.17084
157.09747314453125 0 8867.587
158.06028747558594 0 663.4109
158.1009521484375 0 965.97217
159.09207153320312 0 151150.34
160.07608032226562 0 1005.64514
160.08924865722656 0 801.8893
160.0953826904297 0 16767.453
161.0989227294922 0 1016.4984
163.0872344970703 0 526.30896
167.08177185058594 0 3102.7112
167.1182861328125 0 2169.8186
168.0657958984375 0 511.88077
169.0760040283203 0 1049.544
169.09732055664062 0 882.94257
169.13389587402344 0 16934.4
170.0603790283203 0 16532.502
170.12966918945312 0 603.6799
170.1373291015625 0 1814.1525
171.06370544433594 0 2037.1962
171.076904296875 0 836.7545
171.1131591796875 0 15095.773 b 1
172.11659240722656 0 1644.4846
172.144775390625 0 1352.2563
173.09251403808594 0 672.52576
173.1483154296875 0 557.2134
173.45266723632812 0 1468.1466
175.10816955566406 0 1323.15
180.11343383789062 0 603.63776
181.097412109375 0 2063.987
181.13385009765625 0 696.978
182.12918090820312 0 5861.668
183.11314392089844 0 3213.1206
183.13270568847656 0 690.4433
183.14962768554688 0 3369.9155
184.07229614257812 0 676.56195
184.1084747314453 0 3419.599
185.05662536621094 0 1197.4572
185.09242248535156 0 15924.908
185.12884521484375 0 3184.8252
186.09596252441406 0 1430.6289
186.1236114501953 0 458.5708
187.07171630859375 0 3077.135
187.08692932128906 0 12276.449
188.09051513671875 0 1602.3348
193.0970916748047 0 583.5728
194.1292266845703 0 543.82745
195.04063415527344 0 665.4855
195.0767364501953 0 3038.0032
195.11328125 0 3320.6243
196.10829162597656 0 1105.5146
197.12875366210938 0 11013.221
198.08749389648438 0 1563.5939
198.1240997314453 0 3663.5342
198.13291931152344 0 674.84796
199.07142639160156 0 1007.64087
199.1080322265625 0 4050.5938
200.13975524902344 0 3286.845
201.12376403808594 0 979.6701
202.08255004882812 0 3496.3672
202.11871337890625 0 2444.3372
203.06674194335938 0 828.57526
203.10287475585938 0 2115.844
204.1134796142578 0 4277.7163
205.09754943847656 0 3250.1467
205.1165771484375 0 622.4847
208.1085205078125 0 1435.845
209.09226989746094 0 3647.0522
210.09652709960938 0 531.9933
210.12371826171875 0 2760.0908
210.15957641601562 0 724.77783
212.06698608398438 0 967.1202
212.13934326171875 0 1147.303
213.08726501464844 0 3372.1038
213.12368774414062 0 1819.6992
214.15524291992188 0 3160.4106
216.09788513183594 0 2044.4503
216.1347198486328 0 10869.877
216.14535522460938 0 991.7532
217.1385040283203 0 601.49084
221.0915985107422 0 1021.2258
222.12413024902344 0 4917.2705
223.1080322265625 0 1627.027 y Ammonia loss 22
224.10321044921875 0 937.3449
225.1384735107422 0 951.7592
226.0830841064453 0 662.55585
226.11895751953125 0 9144.824
227.10348510742188 0 567.8468
228.1345672607422 0 6413.479
228.17117309570312 0 2132.7026
229.09347534179688 0 990.75244
229.11868286132812 0 2302.527
229.13861083984375 0 989.17914
230.07870483398438 0 719.93134
230.12911987304688 0 5513.554
231.09812927246094 0 1570.7468
231.13272094726562 0 590.0571
233.16171264648438 0 853.8096
236.103271484375 0 1010.10834
236.13906860351562 0 653.21075
237.08653259277344 0 753.0685
238.11875915527344 0 1347.1556
238.15545654296875 0 876.8893
239.15078735351562 0 2065.4607
240.1346893310547 0 4458.6 y 20
241.1177978515625 0 724.5149
241.13775634765625 0 718.8791
242.1139373779297 0 915.3435
243.14532470703125 0 1382.7596 b 3
244.09396362304688 0 719.62897
244.1297149658203 0 5005.9507 y Ammonia loss 24
249.1228790283203 0 767.9846
250.1194610595703 0 1175.7308
250.1390380859375 0 534.9476
250.1541290283203 0 642.4639
251.10137939453125 0 660.9968
252.09814453125 0 1385.6426
252.134765625 0 778.03577
253.16647338867188 0 543.11224
254.07752990722656 0 1003.11676
254.1139678955078 0 4233.8086
255.06166076660156 0 483.1919
255.11843872070312 0 543.54047
256.0931701660156 0 2293.01
256.12994384765625 0 892.5357
256.1446838378906 0 1377.883
256.16571044921875 0 766.69037
257.1612243652344 0 3695.5227
258.10931396484375 0 642.8722
258.1240539550781 0 9732.333
258.14105224609375 0 728.46747
258.145263671875 0 673.1042
259.1274108886719 0 1351.6033
261.1560363769531 0 12661.557 y 24
262.158447265625 0 1240.5293
266.15008544921875 0 3083.2737
267.145751953125 0 953.4768
267.1697692871094 0 495.25864
268.09320068359375 0 573.245
268.12945556640625 0 3889.832
268.1646728515625 0 788.8945
269.1612854003906 0 599.10345
270.1083984375 0 838.6943
270.1446533203125 0 970.9923
270.1604309082031 0 6724.711
272.08843994140625 0 3186.017
274.10333251953125 0 1392.4318
278.11370849609375 0 2543.5837
282.1082763671875 0 676.33466
282.1450500488281 0 832.32874
282.18267822265625 0 524.46423
283.1769104003906 0 1455.4791
284.1249694824219 0 1666.3114
284.16082763671875 0 3707.8003
284.17694091796875 0 846.6587
285.1566467285156 0 1938.0188
286.1402282714844 0 5140.867
287.14434814453125 0 1161.3132
287.17315673828125 0 717.67377
290.09832763671875 0 631.482
292.128662109375 0 602.6891
294.14532470703125 0 958.2035
295.1039123535156 0 819.6862
295.1766357421875 0 2772.9617
296.1242980957031 0 3730.9978
296.17822265625 0 656.57983
297.1561279296875 0 2207.4119
297.1923522949219 0 694.29834
298.15557861328125 0 36041.195
299.1588439941406 0 7252.4307
299.20855712890625 0 984.6469
300.12017822265625 0 896.4592
300.1603088378906 0 563.419
301.1873779296875 0 1420.6837
303.1670227050781 0 840.72504
306.1083068847656 0 745.24133
307.1413879394531 0 817.009
307.1770324707031 0 660.2636
309.1565246582031 0 1613.1436
309.1926574707031 0 1190.905
311.1714172363281 0 1237.2802
311.20849609375 0 1502.5594
312.1311950683594 0 1984.9231
312.1553649902344 0 1194.7806
312.1717529296875 0 4116.6973
313.1142578125 0 1641.2833
313.153564453125 0 1448.8641
313.1875 0 21833.057
314.1354675292969 0 1786.6458
314.190673828125 0 2532.6357
315.1310119628906 0 1232.8075
315.1661376953125 0 3489.257
315.1823425292969 0 29246.42
316.128173828125 0 766.7093
316.1854553222656 0 6281.4863
317.1097717285156 0 862.4416
317.18829345703125 0 943.791
319.17669677734375 0 649.7328
321.15594482421875 0 697.8394
323.1355895996094 0 5671.631
323.1705017089844 0 656.0505
325.1151428222656 0 3187.5198
325.15045166015625 0 811.27795
325.1883239746094 0 730.6357
325.2240905761719 0 12835.361
326.1833801269531 0 1546.8772
326.2274169921875 0 2893.3
327.16778564453125 0 945.62256 b Water loss 5
327.20281982421875 0 2218.127
328.23480224609375 0 3110.1584
329.1827087402344 0 1370.1842
329.2366638183594 0 680.7408
330.1417236328125 0 1274.1847
330.16607666015625 0 670.55115
330.214599609375 0 2725.167
336.1694641113281 0 910.2693
338.21905517578125 0 794.317
339.1644592285156 0 1112.358
339.2023620605469 0 974.0483
340.16400146484375 0 1223.3942
341.14599609375 0 2516.394
343.1254577636719 0 5306.062
343.1615905761719 0 1645.8419
343.1979064941406 0 1441.2106
343.8589782714844 0 630.47363
344.1283874511719 0 670.76166
344.1938171386719 0 2429.9153
347.1714172363281 0 819.5141
351.1661682128906 0 883.225
352.2001647949219 0 1172.5653
353.1457214355469 0 1113.5721
353.1824035644531 0 981.17413
354.1778564453125 0 2007.3788
356.2298278808594 0 6512.4834
357.19256591796875 0 16333.69 b 2
357.2314453125 0 941.0211
358.1728210449219 0 2984.874 y Ammonia loss 23
358.1966857910156 0 2585.2737
361.135986328125 0 2502.9465
364.12384033203125 0 723.8807
364.1995544433594 0 571.1734
365.1826171875 0 1038.4034
368.1894836425781 0 857.7677
368.21038818359375 0 708.62555
368.82452392578125 0 486.97943
369.1406555175781 0 2825.6597
369.1653747558594 0 478.8863
369.213134765625 0 741.19086
370.2433166503906 0 516.30383
371.15673828125 0 970.70746
371.1923522949219 0 1159.1172
372.1888122558594 0 874.5895
374.21832275390625 0 1697.3165
375.1990661621094 0 7565.5015 y 23
376.20135498046875 0 1173.0192
378.2503356933594 0 2322.363
379.162109375 0 2436.3398
380.1647033691406 0 638.5655
380.22882080078125 0 587.07416
382.136474609375 0 1550.283
382.17364501953125 0 753.90137
382.2086486816406 0 1046.3854
383.19232177734375 0 917.42334
384.1888732910156 0 1742.5345
384.2257385253906 0 1304.3947
386.2190856933594 0 6834.6978
387.15234375 0 1501.9779
387.1884765625 0 1308.0142
387.2224426269531 0 1174.9716
392.1561279296875 0 1627.2279
392.1921081542969 0 762.2847
396.19256591796875 0 743.5575
396.22857666015625 0 1165.7249
396.26123046875 0 2723.276
397.17291259765625 0 2115.5745
397.2643127441406 0 1145.3599
402.19927978515625 0 2100.375
403.2100830078125 0 644.9062 y 19
405.1974792480469 0 861.52576
405.2254333496094 0 842.3387
406.20928955078125 0 2034.7316
410.13299560546875 0 886.94183
410.167236328125 0 2557.4678
412.1861267089844 0 734.76526
412.2207946777344 0 781.18054
412.2561340332031 0 1590.9144
413.2584228515625 0 600.8548
414.2072448730469 0 856.09546 b Water loss 7
414.2353210449219 0 3517.9097 b 11
415.1835021972656 0 2489.0015
415.23590087890625 0 698.65894
416.18902587890625 0 1431.4282
422.1678161621094 0 1174.7585
423.2061462402344 0 1041.0096 b 7
423.2359313964844 0 5766.482
424.18829345703125 0 896.65546
424.2197265625 0 2664.6448
426.1729431152344 0 1324.7023
426.2016296386719 0 1003.94666
427.1580810546875 0 1736.534
427.1946105957031 0 1297.7538
428.1435546875 0 836.164
428.1776428222656 0 1217.8428
430.19476318359375 0 979.5957
431.22906494140625 0 667.9832
431.2601318359375 0 778.0578
438.21466064453125 0 655.9669
439.2438659667969 0 1061.3466
440.1778259277344 0 2149.4854
440.2150573730469 0 2050.8262
441.17950439453125 0 634.0604
441.2113952636719 0 1590.3442
441.2464599609375 0 9158.277
441.2792053222656 0 928.99335
442.19384765625 0 713.21484
442.24951171875 0 1679.6359
442.74072265625 0 743.4727 y Water loss 18
443.2241516113281 0 1044.2706 y Ammonia loss 18
443.7261962890625 0 1140.1982
444.1859436035156 0 1246.5177
444.2209777832031 0 1753.0294 y Water loss 22
445.170166015625 0 919.9605
445.2042541503906 0 2513.6694 y Ammonia loss 22
451.7392272949219 0 6107.6533 y 18
452.1802062988281 0 673.0169
452.24078369140625 0 2132.5303
452.7408752441406 0 1171.294
453.21044921875 0 2026.7798
453.2486572265625 0 936.0845
453.28424072265625 0 2383.763
454.2131652832031 0 658.2727
455.2998962402344 0 935.3227
457.2555236816406 0 2734.1233
458.1888732910156 0 1595.677
458.2285461425781 0 1564.1642
458.261962890625 0 714.4362
459.22418212890625 0 1034.6387
461.2373046875 0 1124.0896
462.23175048828125 0 17535.25 y 22
462.7355651855469 0 897.94366
463.2342834472656 0 5154.177
464.2376403808594 0 1018.1377 b Ammonia loss 8
465.2817077636719 0 1443.5596
468.2098693847656 0 1169.383
470.2250061035156 0 879.25916
471.2213134765625 0 3990.0518
472.22271728515625 0 1089.6316
476.1789245605469 0 695.3889
483.2935485839844 0 2087.0647
485.2835693359375 0 6999.7676 b 3
486.2201843261719 0 1637.1973
486.2881774902344 0 1918.9396
487.22119140625 0 782.7659
489.2319030761719 0 6670.521
490.2333068847656 0 1253.79
493.20477294921875 0 833.8539
493.2398986816406 0 1153.8735
499.2135009765625 0 754.3562
501.2463684082031 0 6117.0996
502.258056640625 0 2464.3008 y 17
502.76287841796875 0 1072.3982
503.1889953613281 0 889.6216
507.25408935546875 0 869.0516
510.2679138183594 0 1698.4658
513.2718505859375 0 762.8248
517.2281494140625 0 769.18604
520.2149658203125 0 882.52734
521.2010498046875 0 1539.1884
523.2510986328125 0 997.56934
524.2853393554688 0 1385.5547
524.3562622070312 0 1836.9747
526.7796630859375 0 599.84015
527.2536010742188 0 713.914
528.2785034179688 0 3988.0303
528.3262329101562 0 1325.694
529.2778930664062 0 882.9499
529.3320922851562 0 832.0379
535.7841186523438 0 1027.206
537.2425537109375 0 1204.9463
538.2257690429688 0 2567.1191
538.3145751953125 0 6218.43
539.2466430664062 0 1101.9352
539.31640625 0 1746.1799
540.2489013671875 0 908.3004
540.3169555664062 0 2041.6304
541.2631225585938 0 1010.189
542.295654296875 0 1100.7367
544.2805786132812 0 608.41876 y Ammonia loss 16
552.31201171875 0 1095.8162
552.351318359375 0 6356.138
553.35546875 0 1629.3329
553.801513671875 0 678.93555
554.2647705078125 0 685.5177
555.2536010742188 0 4411.305
556.2415161132812 0 2647.0503
556.324951171875 0 16233.375 b 4
557.2564697265625 0 3368.4417
557.3282470703125 0 4801.935
558.3287353515625 0 2054.5818
562.302978515625 0 859.66766
564.2974243164062 0 661.5075
564.7972412109375 0 1173.293
565.297607421875 0 822.064
568.3096313476562 0 1458.528
570.2755126953125 0 848.4693
570.3255004882812 0 2406.4978
571.271240234375 0 963.21106
572.2804565429688 0 7757.939 y Water loss 21
573.264404296875 0 9884.751 y Ammonia loss 21
574.2688598632812 0 1729.7682
584.3443603515625 0 766.54565
585.3095092773438 0 999.90247 b 10
585.8131103515625 0 804.2187
586.3191528320312 0 2144.1357
588.2791137695312 0 1171.9181
589.2791137695312 0 604.17535
590.2899169921875 0 11763.201 y 21
591.2921752929688 0 3696.991
592.298095703125 0 741.54944
596.3087768554688 0 1085.345 y 9
597.3011474609375 0 1047.7
603.310302734375 0 5238.8535 y 15
603.812255859375 0 2856.6545
604.3173828125 0 852.2727 b Water loss 17
606.8391723632812 0 1037.3706
611.8326416015625 0 2361.134 b Water loss 11
612.333740234375 0 1534.2045 b Ammonia loss 11
614.328857421875 0 687.89417
614.8316040039062 0 1031.5586
620.8382568359375 0 4287.4 b 11
621.3389282226562 0 2124.236
621.8397827148438 0 1072.9246
624.3056030273438 0 1953.1421
625.288330078125 0 1262.0331
626.8303833007812 0 592.6685
627.3483276367188 0 1206.3878
628.3565673828125 0 647.3554
639.3823852539062 0 1842.3542
640.2763061523438 0 1169.3113
641.3451538085938 0 835.8371
642.3186645507812 0 5362.293
643.328857421875 0 1232.304
644.3363037109375 0 683.77563
649.347412109375 0 3075.0085 b 12
649.8494262695312 0 3324.8267
650.3523559570312 0 770.41895
651.3475952148438 0 1227.6515
655.21875 0 632.99744
657.2999267578125 0 1512.1678
657.3584594726562 0 1332.5233
658.2926025390625 0 2098.799
658.3568725585938 0 943.1994 y 7
663.3766479492188 0 751.5742
667.341552734375 0 3365.1003
667.8324584960938 0 1937.4066 y 14
668.3384399414062 0 3124.5386
668.8404541015625 0 1123.5319
669.3489990234375 0 1182.527
671.3516845703125 0 13039.951 b 5
672.3551025390625 0 4460.456
675.3102416992188 0 5107.5225
676.3119506835938 0 1696.9762
677.3638305664062 0 1035.8931
678.352783203125 0 1764.1342
678.8548583984375 0 992.6278
681.3934326171875 0 3719.9087
682.4004516601562 0 1519.7198
683.344970703125 0 964.9492
684.332275390625 0 1421.1038
684.8857421875 0 3778.6704
685.3526000976562 0 10521.417
685.8888549804688 0 757.2811
686.3555297851562 0 3436.1943
687.3653564453125 0 2482.3103 y 6
688.2960815429688 0 699.0326
688.3661499023438 0 1041.1084
690.3812866210938 0 892.4146 b Ammonia loss 13
695.3392944335938 0 727.3979
698.88330078125 0 2408.5073 b 13
699.3841552734375 0 1703.435
700.3712158203125 0 1779.7344 y Water loss 20
701.358642578125 0 3749.003 y Ammonia loss 20
702.3690795898438 0 2169.821
705.3632202148438 0 682.96313
708.3613891601562 0 1020.67773 y Water loss 13
710.8698120117188 0 641.4106
714.369384765625 0 755.1141 b 20
716.3641967773438 0 940.74524
717.3701171875 0 842.47345 y 13
718.3845825195312 0 8031.8677 y 20
719.38818359375 0 2660.0012
720.3878784179688 0 692.2593
726.3717651367188 0 937.0286
727.8733520507812 0 1520.8226
728.3859252929688 0 1016.85486
728.8743286132812 0 795.4616
729.3722534179688 0 840.54926
738.3773193359375 0 5926.4424
739.3362426757812 0 1946.8466
739.3954467773438 0 1164.2288
740.3817749023438 0 1463.3303 b Water loss 6
742.3741455078125 0 1616.1271
743.3713989257812 0 1080.6086 y Water loss 4
744.3854370117188 0 1837.5073
745.392578125 0 1030.3857
745.8749389648438 0 2246.7388 y 12
746.36376953125 0 1943.0496
746.8818359375 0 728.785
747.3490600585938 0 869.659
749.38720703125 0 862.0575 y 4
749.9130249023438 0 856.09784
752.388671875 0 1399.363
756.3782348632812 0 7568.453
757.3534545898438 0 3689.8628
758.38330078125 0 6014.207 b 6
759.3876342773438 0 2226.284
760.3903198242188 0 936.4585
761.3900756835938 0 813.3448
763.4037475585938 0 4192.4775 b 14
763.9050903320312 0 4389.4917
764.4102172851562 0 1864.9291
768.4255981445312 0 1062.7441
770.39599609375 0 2734.8357
771.3961791992188 0 1094.7457
771.7355346679688 0 900.4521
772.3821411132812 0 1056.2157 y Water loss 11
772.7935791015625 0 747.2935
773.3931274414062 0 799.22455
774.3753662109375 0 3779.1838
775.3779296875 0 1119.5569
775.904296875 0 1179.5947
776.4015502929688 0 982.6354
779.393310546875 0 1195.0597
780.0776977539062 0 667.1597 b Water loss 22
780.399658203125 0 1487.4208 b Ammonia loss 22
780.8963012695312 0 740.8436
781.394287109375 0 5069.7812 y 11
781.8956909179688 0 2670.7344
782.3846435546875 0 1115.0397
782.4506225585938 0 961.3208
783.4404907226562 0 1622.1459
786.4061279296875 0 2686.921 y Ammonia loss 3
786.7384643554688 0 1949.4553
787.40380859375 0 2151.1436 y Water loss 19
788.4073486328125 0 4018.2734
789.4202270507812 0 1287.1934
790.4019775390625 0 928.3521
791.9267578125 0 668.68976
792.0792236328125 0 7501.7964 y 3
792.412353515625 0 11750.339
792.7467651367188 0 6746.585
793.0799560546875 0 1753.5994
793.414306640625 0 865.1175
795.4201049804688 0 1144.9779
799.1339721679688 0 921.79486
799.4085083007812 0 937.6342
799.9291381835938 0 1132.3456
804.92431640625 0 911.0639 b Water loss 15
805.4174194335938 0 6095.956 b Ammonia loss 15
805.9205322265625 0 973.1384
806.4207763671875 0 2961.1028
807.4241943359375 0 1034.4692
809.3939208984375 0 932.6957
810.3843994140625 0 845.3742
812.4141235351562 0 917.2349
813.4154663085938 0 2427.1306
813.927734375 0 3297.2964 b 15
814.42724609375 0 4747.7393
814.9301147460938 0 1568.8236
815.4157104492188 0 1821.5739
815.9232788085938 0 702.9116
818.9224853515625 0 733.61896
820.9088134765625 0 861.221
824.4180297851562 0 773.73303
826.9267578125 0 801.718
827.0689086914062 0 605.3656
827.404052734375 0 4715.5786 b Water loss 7
828.4095458984375 0 4177.708
828.9287719726562 0 849.3065
829.4219970703125 0 1981.4297
829.68701171875 0 1920.0996
829.932373046875 0 1697.5844
830.4246215820312 0 2095.8264
830.9234008789062 0 1049.0774
831.4281005859375 0 1415.3385
833.5029296875 0 1182.8116
834.429443359375 0 957.7195
836.2968139648438 0 1747.2434
836.4315795898438 0 1385.4142 y Water loss 10
836.849853515625 0 869.2857
836.9326171875 0 1963.1733 y Ammonia loss 10
837.0459594726562 0 681.95386
837.4313354492188 0 2095.3555
837.9867553710938 0 2465.0122
838.435791015625 0 936.0216
839.429443359375 0 1641.5587
841.4386596679688 0 792.7822
842.4417724609375 0 1222.8081
842.757568359375 0 826.3925
843.427978515625 0 1000.97394
845.4242553710938 0 16909.88 b 7
845.9432373046875 0 5598.2817
846.4255981445312 0 8777.222
846.9448852539062 0 1351.5751
847.428466796875 0 2524.2751
848.0997314453125 0 1479.722 y Water loss 2
848.4345703125 0 4230.044 y Ammonia loss 2
848.7689819335938 0 1665.6729
849.0985717773438 0 1609.9178
849.4341430664062 0 1356.8383
852.9210815429688 0 1033.8495
853.4358520507812 0 980.2532
854.1055297851562 0 11110.17 y 2
854.4393920898438 0 14938.326
854.7736206054688 0 12056.296
855.1078491210938 0 9486.339
855.4467163085938 0 4324.047 b Water loss 16
855.9456787109375 0 1067.6449 b Ammonia loss 16
856.4412231445312 0 2061.1936 b Ammonia loss 24
856.758056640625 0 1238.9064
857.0970458984375 0 1549.4552
857.428955078125 0 2585.2153
857.5592651367188 0 767.94666
862.0884399414062 0 1114.7435 b 24
862.4403076171875 0 3057.4153
862.5761108398438 0 936.6349
862.7682495117188 0 2051.5295
863.1054077148438 0 1422.6882
863.4358520507812 0 2412.8086
864.4526977539062 0 3528.653 b 16
864.953369140625 0 2873.7556
865.4591674804688 0 2700.8608
866.4692993164062 0 2984.6477
867.4419555664062 0 2590.259
868.1099243164062 0 2323.2908
868.43408203125 0 4825.7876
868.7776489257812 0 1207.5842
869.4402465820312 0 994.46484
871.4317016601562 0 5484.014
871.78125 0 1135.8658 y Water loss 1
872.4426879882812 0 3591.1475
873.1085815429688 0 1143.8915
873.4403076171875 0 2036.7365
875.437744140625 0 1234.1433
877.785400390625 0 3481.3818 y 1
878.1183471679688 0 5374.315
878.4535522460938 0 3445.6833
878.7833862304688 0 1995.411
879.4591674804688 0 1518.081
881.1168212890625 0 1090.5245
882.4577026367188 0 879.2937
883.490966796875 0 1230.1614
884.4678955078125 0 8301.792 y Water loss 18
885.4465942382812 0 14047.758 y Ammonia loss 18
886.4495239257812 0 7197.3135
886.7869873046875 0 1622.4307
887.1199951171875 0 1076.7167
887.4577026367188 0 2880.0295
888.4618530273438 0 1974.5016
890.4634399414062 0 1560.0371
890.7931518554688 0 1260.5652
891.1309204101562 0 1445.9283
891.4593505859375 0 1813.3525
891.7935791015625 0 1216.8251
892.4721069335938 0 1141.3494
893.4646606445312 0 2466.487
895.4786987304688 0 1251.8865
895.7978515625 0 1268.7512
895.9796752929688 0 974.112
896.1371459960938 0 2265.6484
896.46337890625 0 4642.467
896.794189453125 0 4216.9688
896.9633178710938 0 1484.4437
897.129150390625 0 2278.229
897.4636840820312 0 5402.9233
897.9657592773438 0 1801.2207
898.4737548828125 0 847.52185
898.7953491210938 0 1824.8882
899.1317138671875 0 3868.4653
899.4638061523438 0 5853.6206
899.7991943359375 0 2189.4036
900.1328735351562 0 2945.3062
900.4635009765625 0 3306.4644
900.8021240234375 0 2933.4734
900.9728393554688 0 978.9746
901.1354370117188 0 1547.958
901.47021484375 0 2905.4937
902.4696044921875 0 70632.195 y 18
903.4724731445312 0 33583.05
904.4722290039062 0 11506.679
904.8025512695312 0 14708.996 Precursor Water loss
905.1356811523438 0 34605.34 Precursor Ammonia loss
905.4707641601562 0 29941.713
905.8023071289062 0 22606.92
905.9710083007812 0 3018.2915 b Water loss 17
906.1360473632812 0 13092.8
906.4705810546875 0 8373.032 b Ammonia loss 17
906.8026123046875 0 2000.7177
906.9694213867188 0 2281.9258
909.4657592773438 0 1602.8417
909.79052734375 0 3929.8523
910.133544921875 0 14325.233
910.4679565429688 0 23144.604
910.807373046875 0 519132.88 Precursor
911.14208984375 0 810172.3
911.4764404296875 0 650265.7
911.8099365234375 0 372612.06
912.1439819335938 0 146664.16
912.4777221679688 0 45741.535
913.4844970703125 0 1120.0776
914.4874877929688 0 883.94006
914.9780883789062 0 7653.491 b 17
915.4771728515625 0 8933.02
915.9768676757812 0 2775.1377
916.4774780273438 0 1768.3961
916.986572265625 0 810.3502
930.66015625 0 1059.6823
934.9950561523438 0 2808.1074 y Ammonia loss 8
935.49755859375 0 1929.0552
935.9953002929688 0 1221.1168
936.4892578125 0 1074.7625
941.8602905273438 0 1084.0079
942.478271484375 0 2814.229
943.5007934570312 0 12761.862 y 8
944.0046997070312 0 12820.306
944.5044555664062 0 6680.975
945.007568359375 0 2327.4458
945.4939575195312 0 895.97955
954.4979858398438 0 1052.4777 b Water loss 18
958.4721069335938 0 870.3555
959.4874877929688 0 2248.3188
966.517822265625 0 1194.6145
967.5101928710938 0 868.58325
968.5010986328125 0 1752.1272
969.4624633789062 0 1444.8125
970.483154296875 0 1514.0044
972.5031127929688 0 1392.1345
978.0110473632812 0 974.2043 y Water loss 7
978.507080078125 0 1660.6622 y Ammonia loss 7
979.0061645507812 0 1171.3734
979.5059814453125 0 1147.2865
984.533447265625 0 2727.9653
985.5176391601562 0 2908.7832 y Water loss 17
986.4970092773438 0 4192.4487 y Ammonia loss 17
987.0182495117188 0 6706.267 y 7
987.516357421875 0 6733.9336
988.0230102539062 0 3102.5232
988.512451171875 0 1574.6298
993.9910888671875 0 1359.1737
994.4981079101562 0 1199.1891
999.5223388671875 0 1533.1227
1000.5265502929688 0 1198.1549
1003.5177612304688 0 20291.818 y 17
1004.5196533203125 0 9443.199
1005.5174560546875 0 3112.3662
1006.518310546875 0 1273.6803
1007.5213012695312 0 1533.3574
1013.0172729492188 0 1159.5841
1013.5343627929688 0 3543.762
1021.52880859375 0 1651.8416 y Water loss 6
1022.0261840820312 0 2518.2207 y Ammonia loss 6
1022.528076171875 0 2801.6755
1023.5255737304688 0 3604.9072 b Water loss 9
1024.5252685546875 0 1884.6301 b Ammonia loss 9
1025.5283203125 0 1134.7057
1030.5350341796875 0 12986.143 y 6
1031.0360107421875 0 13179.95
1031.537841796875 0 9820.597
1032.03857421875 0 3117.1462
1032.53759765625 0 1204.4897
1040.5709228515625 0 1642.869
1041.5362548828125 0 15752.132 b 9
1042.538330078125 0 9687.2705
1043.539794921875 0 4389.891
1044.5484619140625 0 971.395
1051.5172119140625 0 1439.4094
1052.5406494140625 0 1552.8154
1057.540771484375 0 1511.9653
1058.560302734375 0 5058.336
1059.5552978515625 0 3340.1782
1060.5352783203125 0 1582.1118
1069.53466796875 0 1146.2654
1070.547119140625 0 2183.8596
1071.040283203125 0 1465.9781
1071.5565185546875 0 3241.497
1072.06005859375 0 1088.9493
1072.554931640625 0 1211.0549
1077.5234375 0 1118.5239
1079.0472412109375 0 1605.131 y Water loss 5
1079.5416259765625 0 3863.5989 y Ammonia loss 5
1080.0372314453125 0 4246.9707
1080.5447998046875 0 1986.798
1086.55810546875 0 878.3532 y Water loss 16
1087.54638671875 0 3597.6619 y Ammonia loss 16
1088.047607421875 0 14125.56 y 5
1088.548583984375 0 21452.914
1089.0504150390625 0 13281.195
1089.5513916015625 0 5959.2363
1090.056884765625 0 1584.1794
1094.0577392578125 0 950.104
1098.547119140625 0 905.2326
1100.5645751953125 0 1740.7068
1101.5615234375 0 1212.5479
1102.052734375 0 1175.2056
1102.556884765625 0 853.33563
1104.564697265625 0 16424.938 y 16
1105.565185546875 0 10153.108
1106.0546875 0 1541.3209
1106.5643310546875 0 5302.0024
1107.0494384765625 0 1862.096
1107.555908203125 0 2156.299
1114.5635986328125 0 4400.4126 y Water loss 4
1115.058837890625 0 8697.291 y Ammonia loss 4
1115.5599365234375 0 6316.115
1116.0594482421875 0 3748.4949
1116.55322265625 0 1294.482
1122.562255859375 0 1469.0082
1123.0621337890625 0 1260.2511
1123.5667724609375 0 37139.754 y 4
1124.0677490234375 0 42168.406
1124.569091796875 0 30649.426
1125.071044921875 0 14606.327
1125.5740966796875 0 5036.0195
1126.078369140625 0 1302.2195 b Water loss 21
1127.093994140625 0 951.69806
1127.579345703125 0 916.74664
1135.0989990234375 0 1692.9437 b 21
1135.5906982421875 0 1486.7305
1142.574951171875 0 1304.6425
1143.57421875 0 1758.723
1151.0855712890625 0 1068.8162
1161.5906982421875 0 1015.82416
1165.614501953125 0 1004.875
1169.6180419921875 0 5024.6616 b 10
1170.1060791015625 0 1774.5146 b Ammonia loss 22
1170.614501953125 0 3345.0208
1171.6119384765625 0 2097.8547
1178.6104736328125 0 2733.7554 b 22
1179.110107421875 0 2662.7244 y Ammonia loss 3
1179.6051025390625 0 2412.9707
1180.1103515625 0 1998.5795
1187.6129150390625 0 12297.647 y Water loss 15
1188.1151123046875 0 12151.476
1188.6072998046875 0 9901.4795 y Ammonia loss 15
1189.1168212890625 0 4423.4575
1189.59814453125 0 2505.4492
1205.612060546875 0 18601.69 y 15
1206.6153564453125 0 11649.581
1207.612548828125 0 5972.65
1208.6151123046875 0 1101.7991
1222.6575927734375 0 1111.2483 b Water loss 11
1223.6663818359375 0 1054.4811
1226.6416015625 0 1658.3654 b Water loss 23
1235.6287841796875 0 1965.0834 b 23
1236.1292724609375 0 2118.1882
1236.641845703125 0 1581.597
1240.6680908203125 0 1845.9337 b 11
1241.668212890625 0 1797.4587
1270.65966796875 0 1527.5228
1272.640625 0 1591.7728
1273.1448974609375 0 1131.4717
1280.656494140625 0 2331.0774 b Ammonia loss 12
1281.15380859375 0 3367.0432
1281.6505126953125 0 2283.484
1292.65283203125 0 1450.2119 b 24
1293.1541748046875 0 2014.3519
1293.65869140625 0 1745.6995
1294.157958984375 0 1191.1825
1297.693603515625 0 2661.5125 b 12
1298.6878662109375 0 2261.1775
1299.6768798828125 0 893.76624
1301.6497802734375 0 1816.732
1302.1605224609375 0 1050.9946
1302.6409912109375 0 1519.85
1316.6527099609375 0 4493.5903 y Water loss 14
1317.6385498046875 0 5166.2427 y Ammonia loss 14
1318.6436767578125 0 2194.3442
1319.6376953125 0 1195.6136
1326.6583251953125 0 1253.5496
1327.6678466796875 0 1220.2931
1334.6531982421875 0 19581.605 y 14
1335.6568603515625 0 14531.973
1336.6600341796875 0 5397.696
1337.6661376953125 0 1859.6956
1344.6614990234375 0 2671.9429
1345.66015625 0 1574.7642
1355.7005615234375 0 1029.683
1362.6517333984375 0 892.3995
1372.7025146484375 0 977.8122
1396.752197265625 0 2081.2693 b 13
1397.753662109375 0 1665.883
1398.748779296875 0 1276.011
1416.6982421875 0 1496.7751 y Ammonia loss 13
1429.7393798828125 0 1178.5665
1433.71533203125 0 4494.7305 y 13
1434.7237548828125 0 3664.054
1435.7244873046875 0 1836.9568
1446.7244873046875 0 1065.2623
1454.743408203125 0 1036.6746
1456.72265625 0 1035.6594
1472.7432861328125 0 2492.4304 y Water loss 12
1473.7310791015625 0 3478.874 y Ammonia loss 12
1474.719482421875 0 2760.6777
1475.721435546875 0 986.9294
1490.74169921875 0 13473.124 y 12
1491.7437744140625 0 12472.123
1492.749267578125 0 5050.1333
1493.7457275390625 0 1423.4841
1525.792236328125 0 2316.5452 b 14
1526.783447265625 0 1256.725
1527.7825927734375 0 839.21735
1543.7757568359375 0 1494.7214 y Water loss 11
1544.759765625 0 3187.944 y Ammonia loss 11
1545.762939453125 0 1412.7068
1561.7774658203125 0 12443.011 y 11
1562.7825927734375 0 8991.614
1563.7862548828125 0 4699.324
1564.78173828125 0 1033.6738
1626.84033203125 0 1253.13 b 15
1656.845947265625 0 998.7082
1674.86962890625 0 965.5453
1689.872314453125 0 2300.51 y 10
1690.8773193359375 0 1829.6133
1829.952880859375 0 1188.103
1887.96630859375 0 905.60724

Spectrum Details

|  |  |
| --- | --- |
| Matched peaks? Matched peaks The total absolute number of peaks matched. Additionally in brackets the total fraction of peaks matched and the total number of peaks is shown. | 145 (14.98% of 968) |
| FDR? FDR The false discovery rate estimated for this peptide. It is calculated by matching all theoretical fragments with a non-integer shift with the raw peaks for this spectrum. This is done with 40 different shifts. The resulting percentage is the average number of annotated peaks over the number of annotated peaks with the correct spectrum. | 0.34% |
| Satellite FDR? Satellite FDR See the FDR for details on its calculation. This satellite ion specific FDR only contains the satellite ions (d/w) for I/L/J positions. | - |
| PSM Score? PSM Score The PSM Score as given by Hecklib to this annotated spectrum. It is shown with three significant figures. | 830 |

#### Spectrum 3895? Spectrum 3895 The raw spectrum of this peptide as annotated by Hecklib. The fragments are coloured according to ion type (see legend). Any peaks with a star '\*' as text can be hovered over to see the full details, first the ion type second the mass shift type. By hovering over the amino acids in the peptide or ions in the legend the corresponding peaks are highlighted. By toggling the 'Unassigned' label you can turn the background (unassigned) peaks on or off in the plot. By updating the slider in the Ion legend you can update the spectrum to only show the top X% of the peaks with labels. The top X% means any peak that is within X% of the highest intensity. By dragging in the spectrum you can zoom in to a specific part of the spectrum and use 'Zoom Out' to get back to the original zoom level. The annotation of the spectrum is based on the given sequence in the peptides file and is done with different software so inconsistencies are likely. The peaks are annotated based on the given sequence, with 20 ppm tolerance.

Copy Data

##### Spectrum 3895 (TSV)

###### Preview

```
Loading example...
```

*Click on the button to copy the data to your clipboard.*

Mz MinMz MaxIntensity Max

WidthHeightPeptide font sizePeptide stroke widthSpectrum font sizeSpectrum stroke widthCompact peptide

Ion legend

wxyz

abcd

OtherUnassignedIon^Charge^_Position_Show for top:%

VAWKADSSVPKAGVETTTPSKQSNNK

01.52e+53.04e+54.56e+56.08e+5

Zoom Out

y^+1^_1_a^+1^_2_y^+1^_1_a^+2^_3_b^+1^_2_y^+2^_4_y^+3^_6_b^+2^_4_y^+1^_2_y^+1^_2_b^+1^_3_y^+1^_3_y^+1^_3_b^+2^_8_b^+3^_12_y^+2^_8_y^+1^_4_y^+1^_4_y^+2^_8_y^+1^_4_b^+2^_9_b^+1^_4_y^+2^_9_b^+1^_5_y^+1^_5_y^+1^_5_y^+1^_5_y^+2^_11_b^+2^_12_b^+2^_12_b^+2^_13_y^+2^_12_y^+2^_12_b^+1^_6_y^+3^_20_b^+2^_14_y^+1^_6_y^+1^_6_y^+2^_13_y^+1^_6_y^+3^_22_y^+2^_14_b^+1^_7_b^+2^_15_y^+2^_15_b^+3^_23_y^+2^_15_b^+3^_23_y^+3^_23_y^+1^_7_y^+3^_23_b^+2^_16_b^+2^_16_b^+2^_16_b^+1^_8_y^+2^_16_b^+1^_8_y^+3^_24_y^+3^_24_y^+3^_24_b^+2^_17_b^+3^_25_b^+2^_17_y^+3^_25_y^+1^_8_y^+1^_8_y^+1^_8_\*\*b^+2^_18_b^+2^_18_\*b^+2^_18_y^+2^_18_y^+2^_18_y^+2^_18_y^+1^_9_y^+1^_9_y^+2^_19_y^+1^_9_y^+2^_20_y^+2^_20_b^+1^_10_b^+1^_10_y^+2^_20_b^+1^_10_b^+2^_21_y^+2^_21_y^+2^_21_y^+1^_10_y^+1^_10_y^+2^_21_y^+1^_10_y^+2^_22_y^+2^_22_y^+2^_22_b^+2^_22_b^+1^_11_b^+2^_23_y^+2^_23_y^+1^_11_y^+1^_11_b^+2^_24_b^+2^_24_b^+1^_12_y^+2^_24_b^+1^_13_b^+2^_25_y^+1^_12_y^+1^_12_y^+1^_12_\*b^+1^_14_y^+1^_13_y^+1^_13_y^+1^_14_y^+1^_14_y^+1^_14_b^+1^_15_y^+1^_15_y^+1^_15_y^+1^_15_y^+1^_16_

045290413561808

Fragment Matches Table

Show background peaks

| Position | Ion type | Intensity | mz Theoretical | mz Error (Th) | mz Error (ppm) | Charge | Series Number |
| --- | --- | --- | --- | --- | --- | --- | --- |
| - | - | 394.8 | 122.9 | - | - | 0 | - |
| - | - | 585.7 | 126.1 | - | - | 0 | - |
| - | - | 742.8 | 127.1 | - | - | 0 | - |
| - | - | 382.1 | 128.1 | - | - | 0 | - |
| - | - | 1235 | 129.1 | - | - | 0 | - |
| - | - | 9.794E+04 | 129.1 | - | - | 0 | - |
| - | - | 2831 | 130.1 | - | - | 0 | - |
| 26 | y | 3172 | 130.1 | 0.0004759 | 3.658 | +1 | 1 |
| - | - | 836.9 | 130.1 | - | - | 0 | - |
| - | - | 6770 | 130.1 | - | - | 0 | - |
| - | - | 6251 | 132.1 | - | - | 0 | - |
| - | - | 644.2 | 136.1 | - | - | 0 | - |
| - | - | 456 | 137.5 | - | - | 0 | - |
| - | - | 1412 | 138.1 | - | - | 0 | - |
| - | - | 1586 | 139.1 | - | - | 0 | - |
| - | - | 1228 | 141.1 | - | - | 0 | - |
| - | - | 1284 | 141.1 | - | - | 0 | - |
| 2 | a | 8602 | 143.1 | 0.0005034 | 3.517 | +1 | 2 |
| - | - | 552.8 | 146.1 | - | - | 0 | - |
| - | - | 507 | 147.1 | - | - | 0 | - |
| 26 | y | 3755 | 147.1 | 0.0004161 | 2.828 | +1 | 1 |
| - | - | 808.3 | 148.9 | - | - | 0 | - |
| - | - | 2352 | 153.1 | - | - | 0 | - |
| - | - | 561 | 154.1 | - | - | 0 | - |
| - | - | 423 | 154.2 | - | - | 0 | - |
| - | - | 965.5 | 155.1 | - | - | 0 | - |
| - | - | 4323 | 155.1 | - | - | 0 | - |
| - | - | 631.5 | 155.2 | - | - | 0 | - |
| - | - | 604.7 | 156.1 | - | - | 0 | - |
| - | - | 1262 | 157.1 | - | - | 0 | - |
| - | - | 7590 | 157.1 | - | - | 0 | - |
| - | - | 728.5 | 158.1 | - | - | 0 | - |
| - | - | 580.6 | 158.1 | - | - | 0 | - |
| - | - | 1.079E+05 | 159.1 | - | - | 0 | - |
| - | - | 989.6 | 160.1 | - | - | 0 | - |
| - | - | 1.078E+04 | 160.1 | - | - | 0 | - |
| - | - | 827.4 | 161.1 | - | - | 0 | - |
| 3 | a | 839.9 | 165.1 | 0.0004217 | 2.554 | +2 | 3 |
| - | - | 592.8 | 165.8 | - | - | 0 | - |
| - | - | 549.5 | 167.1 | - | - | 0 | - |
| - | - | 2461 | 167.1 | - | - | 0 | - |
| - | - | 1314 | 167.1 | - | - | 0 | - |
| - | - | 924.7 | 169.1 | - | - | 0 | - |
| - | - | 1.258E+04 | 169.1 | - | - | 0 | - |
| - | - | 1.321E+04 | 170.1 | - | - | 0 | - |
| - | - | 1662 | 170.1 | - | - | 0 | - |
| - | - | 1426 | 171.1 | - | - | 0 | - |
| - | - | 646.9 | 171.1 | - | - | 0 | - |
| 2 | b | 1.256E+04 | 171.1 | 0.0004466 | 2.61 | +1 | 2 |
| - | - | 1021 | 171.1 | - | - | 0 | - |
| - | - | 943.8 | 172.1 | - | - | 0 | - |
| - | - | 594.6 | 172.1 | - | - | 0 | - |
| - | - | 1577 | 173.4 | - | - | 0 | - |
| - | - | 396.5 | 174.1 | - | - | 0 | - |
| - | - | 786.6 | 175.1 | - | - | 0 | - |
| - | - | 1042 | 175.1 | - | - | 0 | - |
| - | - | 483.8 | 180.1 | - | - | 0 | - |
| - | - | 1500 | 181.1 | - | - | 0 | - |
| - | - | 1294 | 181.1 | - | - | 0 | - |
| - | - | 3822 | 182.1 | - | - | 0 | - |
| - | - | 421.3 | 182.7 | - | - | 0 | - |
| - | - | 2031 | 183.1 | - | - | 0 | - |
| - | - | 644.2 | 183.1 | - | - | 0 | - |
| - | - | 1959 | 183.1 | - | - | 0 | - |
| - | - | 578 | 184.1 | - | - | 0 | - |
| - | - | 2494 | 184.1 | - | - | 0 | - |
| - | - | 1061 | 185.1 | - | - | 0 | - |
| - | - | 1.107E+04 | 185.1 | - | - | 0 | - |
| - | - | 2302 | 185.1 | - | - | 0 | - |
| - | - | 1198 | 186.1 | - | - | 0 | - |
| - | - | 2610 | 187.1 | - | - | 0 | - |
| - | - | 8889 | 187.1 | - | - | 0 | - |
| - | - | 1416 | 188.1 | - | - | 0 | - |
| - | - | 601.4 | 193.1 | - | - | 0 | - |
| - | - | 1642 | 195.1 | - | - | 0 | - |
| - | - | 1925 | 195.1 | - | - | 0 | - |
| - | - | 717.3 | 196.1 | - | - | 0 | - |
| - | - | 8011 | 197.1 | - | - | 0 | - |
| - | - | 1323 | 198.1 | - | - | 0 | - |
| - | - | 2818 | 198.1 | - | - | 0 | - |
| - | - | 628.7 | 199.1 | - | - | 0 | - |
| - | - | 3046 | 199.1 | - | - | 0 | - |
| - | - | 2362 | 200.1 | - | - | 0 | - |
| - | - | 1120 | 201.1 | - | - | 0 | - |
| - | - | 2426 | 202.1 | - | - | 0 | - |
| - | - | 2484 | 202.1 | - | - | 0 | - |
| - | - | 713.3 | 203.1 | - | - | 0 | - |
| - | - | 1719 | 203.1 | - | - | 0 | - |
| - | - | 3178 | 204.1 | - | - | 0 | - |
| - | - | 2168 | 205.1 | - | - | 0 | - |
| - | - | 738.3 | 205.1 | - | - | 0 | - |
| - | - | 524.9 | 206.1 | - | - | 0 | - |
| - | - | 1405 | 208.1 | - | - | 0 | - |
| - | - | 2827 | 209.1 | - | - | 0 | - |
| - | - | 1958 | 210.1 | - | - | 0 | - |
| - | - | 590.5 | 212.1 | - | - | 0 | - |
| - | - | 1248 | 212.1 | - | - | 0 | - |
| - | - | 2855 | 213.1 | - | - | 0 | - |
| - | - | 1229 | 213.1 | - | - | 0 | - |
| - | - | 482.4 | 213.1 | - | - | 0 | - |
| - | - | 2360 | 214.2 | - | - | 0 | - |
| - | - | 1075 | 216.1 | - | - | 0 | - |
| - | - | 8342 | 216.1 | - | - | 0 | - |
| - | - | 543.4 | 217.1 | - | - | 0 | - |
| - | - | 692.2 | 221.1 | - | - | 0 | - |
| - | - | 3585 | 222.1 | - | - | 0 | - |
| 23 | y | 657.4 | 223.1 | 0.003683 | 16.51 | +2 | 4 |
| - | - | 704.2 | 223.1 | - | - | 0 | - |
| - | - | 595.4 | 224.1 | - | - | 0 | - |
| - | - | 690.2 | 224.1 | - | - | 0 | - |
| - | - | 581.4 | 226.1 | - | - | 0 | - |
| - | - | 6848 | 226.1 | - | - | 0 | - |
| - | - | 533.4 | 227.1 | - | - | 0 | - |
| - | - | 5055 | 228.1 | - | - | 0 | - |
| - | - | 1944 | 228.2 | - | - | 0 | - |
| - | - | 1013 | 229.1 | - | - | 0 | - |
| - | - | 2110 | 229.1 | - | - | 0 | - |
| - | - | 3452 | 230.1 | - | - | 0 | - |
| - | - | 1037 | 231.1 | - | - | 0 | - |
| - | - | 690.4 | 231.1 | - | - | 0 | - |
| - | - | 825 | 236.1 | - | - | 0 | - |
| - | - | 739.9 | 236.1 | - | - | 0 | - |
| - | - | 1631 | 238.1 | - | - | 0 | - |
| - | - | 754.8 | 238.2 | - | - | 0 | - |
| - | - | 1240 | 239.2 | - | - | 0 | - |
| 21 | y | 3631 | 240.1 | 0.001884 | 7.847 | +3 | 6 |
| - | - | 1228 | 242.1 | - | - | 0 | - |
| 4 | b | 1305 | 243.1 | 0.001945 | 8 | +2 | 4 |
| 25 | y | 4803 | 244.1 | 0.0006088 | 2.494 | +1 | 2 |
| - | - | 926.3 | 249.1 | - | - | 0 | - |
| - | - | 601.4 | 250.1 | - | - | 0 | - |
| - | - | 851.9 | 250.2 | - | - | 0 | - |
| - | - | 802.9 | 252.1 | - | - | 0 | - |
| - | - | 782.5 | 254.1 | - | - | 0 | - |
| - | - | 3780 | 254.1 | - | - | 0 | - |
| - | - | 1623 | 256.1 | - | - | 0 | - |
| - | - | 680.4 | 256.1 | - | - | 0 | - |
| - | - | 868.2 | 256.1 | - | - | 0 | - |
| - | - | 619.8 | 256.2 | - | - | 0 | - |
| - | - | 3039 | 257.2 | - | - | 0 | - |
| - | - | 7328 | 258.1 | - | - | 0 | - |
| - | - | 1443 | 259.1 | - | - | 0 | - |
| 25 | y | 7346 | 261.2 | 0.0004879 | 1.868 | +1 | 2 |
| - | - | 607.4 | 262.2 | - | - | 0 | - |
| - | - | 1826 | 266.2 | - | - | 0 | - |
| - | - | 917.5 | 267.1 | - | - | 0 | - |
| - | - | 2554 | 268.1 | - | - | 0 | - |
| - | - | 1003 | 269.2 | - | - | 0 | - |
| - | - | 699.9 | 270.1 | - | - | 0 | - |
| - | - | 5336 | 270.2 | - | - | 0 | - |
| - | - | 662.3 | 271.1 | - | - | 0 | - |
| - | - | 709.7 | 271.2 | - | - | 0 | - |
| - | - | 2031 | 272.1 | - | - | 0 | - |
| - | - | 689.8 | 272.1 | - | - | 0 | - |
| - | - | 1236 | 274.1 | - | - | 0 | - |
| - | - | 1920 | 278.1 | - | - | 0 | - |
| - | - | 506.8 | 281.8 | - | - | 0 | - |
| - | - | 650.2 | 282.2 | - | - | 0 | - |
| - | - | 1185 | 284.1 | - | - | 0 | - |
| - | - | 591.2 | 284.1 | - | - | 0 | - |
| - | - | 2500 | 284.2 | - | - | 0 | - |
| - | - | 1260 | 285.2 | - | - | 0 | - |
| - | - | 3288 | 286.1 | - | - | 0 | - |
| - | - | 604.5 | 286.2 | - | - | 0 | - |
| - | - | 752.1 | 290.1 | - | - | 0 | - |
| - | - | 650.3 | 294.1 | - | - | 0 | - |
| - | - | 714 | 295.1 | - | - | 0 | - |
| - | - | 939.8 | 295.1 | - | - | 0 | - |
| - | - | 2085 | 295.2 | - | - | 0 | - |
| - | - | 2241 | 296.1 | - | - | 0 | - |
| - | - | 1197 | 297.2 | - | - | 0 | - |
| - | - | 2.849E+04 | 298.2 | - | - | 0 | - |
| - | - | 4452 | 299.2 | - | - | 0 | - |
| - | - | 537.5 | 300.2 | - | - | 0 | - |
| - | - | 655.5 | 301.2 | - | - | 0 | - |
| - | - | 865 | 306.1 | - | - | 0 | - |
| - | - | 695.7 | 306.1 | - | - | 0 | - |
| - | - | 900.3 | 309.2 | - | - | 0 | - |
| - | - | 718.7 | 309.2 | - | - | 0 | - |
| - | - | 868 | 311.2 | - | - | 0 | - |
| - | - | 993.8 | 311.2 | - | - | 0 | - |
| - | - | 608.1 | 312.1 | - | - | 0 | - |
| - | - | 916.2 | 312.2 | - | - | 0 | - |
| - | - | 3050 | 312.2 | - | - | 0 | - |
| - | - | 1568 | 313.1 | - | - | 0 | - |
| - | - | 1205 | 313.2 | - | - | 0 | - |
| - | - | 1.677E+04 | 313.2 | - | - | 0 | - |
| - | - | 2064 | 314.1 | - | - | 0 | - |
| - | - | 2763 | 314.2 | - | - | 0 | - |
| - | - | 669.7 | 315.1 | - | - | 0 | - |
| - | - | 1826 | 315.2 | - | - | 0 | - |
| - | - | 2.157E+04 | 315.2 | - | - | 0 | - |
| - | - | 625.6 | 316.1 | - | - | 0 | - |
| - | - | 3955 | 316.2 | - | - | 0 | - |
| - | - | 594.3 | 317.1 | - | - | 0 | - |
| - | - | 2778 | 323.1 | - | - | 0 | - |
| - | - | 1083 | 324.1 | - | - | 0 | - |
| - | - | 2854 | 325.1 | - | - | 0 | - |
| - | - | 736.6 | 325.2 | - | - | 0 | - |
| - | - | 8906 | 325.2 | - | - | 0 | - |
| - | - | 1132 | 326.2 | - | - | 0 | - |
| - | - | 1419 | 326.2 | - | - | 0 | - |
| - | - | 819 | 327.2 | - | - | 0 | - |
| - | - | 1336 | 327.2 | - | - | 0 | - |
| - | - | 2728 | 328.2 | - | - | 0 | - |
| - | - | 1161 | 329.2 | - | - | 0 | - |
| - | - | 1025 | 330.1 | - | - | 0 | - |
| - | - | 738.2 | 330.2 | - | - | 0 | - |
| - | - | 2294 | 330.2 | - | - | 0 | - |
| - | - | 604.7 | 338.2 | - | - | 0 | - |
| - | - | 1197 | 339.2 | - | - | 0 | - |
| - | - | 1055 | 339.2 | - | - | 0 | - |
| - | - | 512.6 | 340.2 | - | - | 0 | - |
| - | - | 1750 | 341.1 | - | - | 0 | - |
| - | - | 4252 | 343.1 | - | - | 0 | - |
| - | - | 1106 | 343.2 | - | - | 0 | - |
| - | - | 890.8 | 343.2 | - | - | 0 | - |
| - | - | 2870 | 344.2 | - | - | 0 | - |
| - | - | 1673 | 347.2 | - | - | 0 | - |
| - | - | 736.4 | 351.1 | - | - | 0 | - |
| - | - | 1093 | 353.1 | - | - | 0 | - |
| - | - | 720.2 | 353.2 | - | - | 0 | - |
| - | - | 1466 | 354.2 | - | - | 0 | - |
| - | - | 4995 | 356.2 | - | - | 0 | - |
| 3 | b | 1.132E+04 | 357.2 | 0.000754 | 2.111 | +1 | 3 |
| - | - | 928.3 | 357.2 | - | - | 0 | - |
| 24 | y | 1472 | 358.2 | 0.001474 | 4.115 | +1 | 3 |
| - | - | 1570 | 358.2 | - | - | 0 | - |
| - | - | 1685 | 361.1 | - | - | 0 | - |
| - | - | 794.8 | 368.2 | - | - | 0 | - |
| - | - | 1787 | 369.1 | - | - | 0 | - |
| - | - | 1548 | 369.2 | - | - | 0 | - |
| - | - | 935.1 | 371.2 | - | - | 0 | - |
| - | - | 1240 | 371.2 | - | - | 0 | - |
| - | - | 1109 | 374.2 | - | - | 0 | - |
| 24 | y | 5682 | 375.2 | 0.0003461 | 0.9224 | +1 | 3 |
| - | - | 632.9 | 375.2 | - | - | 0 | - |
| - | - | 830 | 376.2 | - | - | 0 | - |
| - | - | 1739 | 378.3 | - | - | 0 | - |
| - | - | 898.2 | 379.2 | - | - | 0 | - |
| - | - | 2226 | 384.2 | - | - | 0 | - |
| - | - | 1422 | 384.2 | - | - | 0 | - |
| - | - | 4789 | 386.2 | - | - | 0 | - |
| - | - | 779.8 | 387.2 | - | - | 0 | - |
| - | - | 1333 | 387.2 | - | - | 0 | - |
| - | - | 875.5 | 387.2 | - | - | 0 | - |
| - | - | 809.3 | 392.2 | - | - | 0 | - |
| - | - | 761.3 | 395.2 | - | - | 0 | - |
| - | - | 2980 | 396.3 | - | - | 0 | - |
| - | - | 1884 | 397.2 | - | - | 0 | - |
| - | - | 723.3 | 397.3 | - | - | 0 | - |
| - | - | 629.7 | 398.2 | - | - | 0 | - |
| - | - | 1622 | 402.2 | - | - | 0 | - |
| - | - | 665.3 | 410.1 | - | - | 0 | - |
| - | - | 2819 | 410.2 | - | - | 0 | - |
| - | - | 1000 | 412.2 | - | - | 0 | - |
| - | - | 930.9 | 412.3 | - | - | 0 | - |
| 8 | b | 1024 | 414.2 | 0.0002643 | 0.6381 | +2 | 8 |
| 12 | b | 1740 | 414.2 | 0.008173 | 19.73 | +3 | 12 |
| - | - | 556.4 | 415.1 | - | - | 0 | - |
| - | - | 2017 | 415.2 | - | - | 0 | - |
| - | - | 921.9 | 422.2 | - | - | 0 | - |
| - | - | 935.9 | 423.2 | - | - | 0 | - |
| - | - | 4222 | 423.2 | - | - | 0 | - |
| - | - | 1934 | 424.2 | - | - | 0 | - |
| - | - | 1092 | 427.2 | - | - | 0 | - |
| - | - | 643.5 | 427.2 | - | - | 0 | - |
| - | - | 1034 | 428.2 | - | - | 0 | - |
| - | - | 682.3 | 430.2 | - | - | 0 | - |
| - | - | 855.7 | 436.2 | - | - | 0 | - |
| - | - | 760.7 | 439.2 | - | - | 0 | - |
| - | - | 2060 | 440.2 | - | - | 0 | - |
| - | - | 1270 | 440.2 | - | - | 0 | - |
| - | - | 841.6 | 441.2 | - | - | 0 | - |
| - | - | 5893 | 441.2 | - | - | 0 | - |
| - | - | 1584 | 442.2 | - | - | 0 | - |
| 19 | y | 1062 | 443.2 | 0.0003255 | 0.7343 | +2 | 8 |
| - | - | 650 | 443.7 | - | - | 0 | - |
| - | - | 1293 | 444.2 | - | - | 0 | - |
| 23 | y | 1250 | 444.2 | 0.001525 | 3.434 | +1 | 4 |
| 23 | y | 1608 | 445.2 | 0.00155 | 3.482 | +1 | 4 |
| 19 | y | 4930 | 451.7 | 0.0009264 | 2.051 | +2 | 8 |
| - | - | 2005 | 452.2 | - | - | 0 | - |
| - | - | 2013 | 453.2 | - | - | 0 | - |
| - | - | 2136 | 453.3 | - | - | 0 | - |
| - | - | 956.9 | 457.3 | - | - | 0 | - |
| - | - | 1549 | 458.2 | - | - | 0 | - |
| - | - | 1405 | 458.2 | - | - | 0 | - |
| 23 | y | 1.469E+04 | 462.2 | 0.001429 | 3.092 | +1 | 4 |
| - | - | 2974 | 463.2 | - | - | 0 | - |
| 9 | b | 789.2 | 464.2 | 0.003703 | 7.976 | +2 | 9 |
| - | - | 1250 | 468.2 | - | - | 0 | - |
| - | - | 1421 | 470.2 | - | - | 0 | - |
| - | - | 2619 | 471.2 | - | - | 0 | - |
| - | - | 651 | 472.2 | - | - | 0 | - |
| - | - | 701.4 | 483.3 | - | - | 0 | - |
| 4 | b | 4879 | 485.3 | 0.0029 | 5.977 | +1 | 4 |
| - | - | 1162 | 486.2 | - | - | 0 | - |
| - | - | 1965 | 486.3 | - | - | 0 | - |
| - | - | 3789 | 489.2 | - | - | 0 | - |
| - | - | 1748 | 490.2 | - | - | 0 | - |
| - | - | 2347 | 501.2 | - | - | 0 | - |
| 18 | y | 2178 | 502.3 | 0.001825 | 3.634 | +2 | 9 |
| - | - | 710.2 | 510.2 | - | - | 0 | - |
| - | - | 941.4 | 510.3 | - | - | 0 | - |
| - | - | 1271 | 521.2 | - | - | 0 | - |
| - | - | 1578 | 524.3 | - | - | 0 | - |
| - | - | 964.5 | 524.4 | - | - | 0 | - |
| - | - | 583.3 | 525.3 | - | - | 0 | - |
| - | - | 2155 | 528.3 | - | - | 0 | - |
| - | - | 1226 | 528.3 | - | - | 0 | - |
| - | - | 905.9 | 529.3 | - | - | 0 | - |
| - | - | 1061 | 535.8 | - | - | 0 | - |
| - | - | 1221 | 537.2 | - | - | 0 | - |
| - | - | 1573 | 538.2 | - | - | 0 | - |
| - | - | 4862 | 538.3 | - | - | 0 | - |
| - | - | 1070 | 539.2 | - | - | 0 | - |
| - | - | 839 | 539.3 | - | - | 0 | - |
| - | - | 711.2 | 540.3 | - | - | 0 | - |
| - | - | 1669 | 542.3 | - | - | 0 | - |
| - | - | 649.8 | 552.3 | - | - | 0 | - |
| - | - | 4280 | 552.4 | - | - | 0 | - |
| - | - | 1278 | 553.4 | - | - | 0 | - |
| - | - | 2368 | 555.3 | - | - | 0 | - |
| - | - | 2303 | 556.2 | - | - | 0 | - |
| 5 | b | 1.083E+04 | 556.3 | 0.001185 | 2.129 | +1 | 5 |
| - | - | 2749 | 557.3 | - | - | 0 | - |
| - | - | 4416 | 557.3 | - | - | 0 | - |
| - | - | 1002 | 558.3 | - | - | 0 | - |
| - | - | 850.5 | 564.8 | - | - | 0 | - |
| - | - | 753.7 | 568.3 | - | - | 0 | - |
| - | - | 1276 | 570.3 | - | - | 0 | - |
| - | - | 578 | 571.3 | - | - | 0 | - |
| 22 | y | 5102 | 572.3 | 0.001695 | 2.962 | +1 | 5 |
| 22 | y | 6874 | 573.3 | 0.002604 | 4.542 | +1 | 5 |
| - | - | 1814 | 574.3 | - | - | 0 | - |
| - | - | 1700 | 586.3 | - | - | 0 | - |
| 22 | y | 9919 | 590.3 | 0.0007131 | 1.208 | +1 | 5 |
| - | - | 2913 | 591.3 | - | - | 0 | - |
| - | - | 838.2 | 592.3 | - | - | 0 | - |
| - | - | 672.8 | 600.3 | - | - | 0 | - |
| 16 | y | 3124 | 603.3 | 0.001064 | 1.763 | +2 | 11 |
| - | - | 1223 | 603.8 | - | - | 0 | - |
| - | - | 978.8 | 606.3 | - | - | 0 | - |
| - | - | 828.4 | 607.3 | - | - | 0 | - |
| 12 | b | 1254 | 612.3 | 0.009885 | 16.14 | +2 | 12 |
| - | - | 1154 | 612.8 | - | - | 0 | - |
| 12 | b | 2046 | 620.8 | 0.001615 | 2.602 | +2 | 12 |
| - | - | 1634 | 621.3 | - | - | 0 | - |
| - | - | 997.6 | 621.8 | - | - | 0 | - |
| - | - | 603.2 | 624.3 | - | - | 0 | - |
| - | - | 965.7 | 625.3 | - | - | 0 | - |
| - | - | 962.3 | 639.4 | - | - | 0 | - |
| - | - | 3267 | 642.3 | - | - | 0 | - |
| - | - | 1101 | 643.3 | - | - | 0 | - |
| 13 | b | 2535 | 649.3 | 0.001076 | 1.657 | +2 | 13 |
| - | - | 2002 | 649.8 | - | - | 0 | - |
| - | - | 1074 | 650 | - | - | 0 | - |
| - | - | 1092 | 650.4 | - | - | 0 | - |
| - | - | 1144 | 657.3 | - | - | 0 | - |
| - | - | 989.2 | 657.4 | - | - | 0 | - |
| - | - | 785.1 | 658.3 | - | - | 0 | - |
| 15 | y | 863.2 | 659.3 | 0.007385 | 11.2 | +2 | 12 |
| - | - | 2717 | 667.3 | - | - | 0 | - |
| 15 | y | 1691 | 667.8 | 0.002778 | 4.159 | +2 | 12 |
| - | - | 3356 | 668.3 | - | - | 0 | - |
| - | - | 715.9 | 668.8 | - | - | 0 | - |
| - | - | 1631 | 669.3 | - | - | 0 | - |
| - | - | 1010 | 670.4 | - | - | 0 | - |
| 6 | b | 9191 | 671.4 | 0.001402 | 2.088 | +1 | 6 |
| - | - | 3575 | 672.4 | - | - | 0 | - |
| - | - | 3278 | 675.3 | - | - | 0 | - |
| - | - | 2192 | 676.3 | - | - | 0 | - |
| - | - | 802.5 | 677.3 | - | - | 0 | - |
| - | - | 3460 | 681.4 | - | - | 0 | - |
| - | - | 643.5 | 681.6 | - | - | 0 | - |
| - | - | 1574 | 682.4 | - | - | 0 | - |
| - | - | 1946 | 684.9 | - | - | 0 | - |
| - | - | 5752 | 685.4 | - | - | 0 | - |
| - | - | 2944 | 686.4 | - | - | 0 | - |
| 7 | y | 2405 | 687.4 | 0.00601 | 8.744 | +3 | 20 |
| - | - | 703 | 697.4 | - | - | 0 | - |
| 14 | b | 2501 | 698.9 | 0.001171 | 1.675 | +2 | 14 |
| - | - | 1214 | 699.4 | - | - | 0 | - |
| - | - | 957 | 699.9 | - | - | 0 | - |
| 21 | y | 1210 | 700.4 | 0.004266 | 6.092 | +1 | 6 |
| 21 | y | 2763 | 701.4 | 0.002551 | 3.637 | +1 | 6 |
| - | - | 1040 | 702.4 | - | - | 0 | - |
| 14 | y | 970 | 717.4 | 0.006779 | 9.449 | +2 | 13 |
| - | - | 594 | 717.9 | - | - | 0 | - |
| 21 | y | 5459 | 718.4 | 0.001636 | 2.278 | +1 | 6 |
| - | - | 1795 | 719.4 | - | - | 0 | - |
| - | - | 860.2 | 719.9 | - | - | 0 | - |
| - | - | 632.6 | 720.4 | - | - | 0 | - |
| - | - | 690.2 | 722.3 | - | - | 0 | - |
| - | - | 767.2 | 726.4 | - | - | 0 | - |
| - | - | 841.5 | 728.9 | - | - | 0 | - |
| - | - | 1072 | 729.4 | - | - | 0 | - |
| - | - | 849.4 | 732.4 | - | - | 0 | - |
| - | - | 3358 | 738.4 | - | - | 0 | - |
| - | - | 962.8 | 739.4 | - | - | 0 | - |
| - | - | 954.4 | 740.3 | - | - | 0 | - |
| - | - | 1002 | 742.4 | - | - | 0 | - |
| 5 | y | 645.8 | 743.4 | 0.00236 | 3.175 | +3 | 22 |
| - | - | 2346 | 744.4 | - | - | 0 | - |
| - | - | 1060 | 745.4 | - | - | 0 | - |
| 13 | y | 1939 | 745.9 | 0.0003803 | 0.5099 | +2 | 14 |
| - | - | 1672 | 746.4 | - | - | 0 | - |
| - | - | 651.8 | 746.9 | - | - | 0 | - |
| - | - | 773.5 | 748.4 | - | - | 0 | - |
| - | - | 823.6 | 749.2 | - | - | 0 | - |
| - | - | 5092 | 756.4 | - | - | 0 | - |
| - | - | 1342 | 757.3 | - | - | 0 | - |
| - | - | 1751 | 757.4 | - | - | 0 | - |
| 7 | b | 4373 | 758.4 | 0.001112 | 1.466 | +1 | 7 |
| - | - | 2469 | 759.4 | - | - | 0 | - |
| 15 | b | 3430 | 763.4 | 0.001542 | 2.02 | +2 | 15 |
| - | - | 3434 | 763.9 | - | - | 0 | - |
| - | - | 725 | 764.4 | - | - | 0 | - |
| - | - | 656.1 | 766.9 | - | - | 0 | - |
| - | - | 1212 | 770.4 | - | - | 0 | - |
| 12 | y | 1081 | 772.4 | 0.0001673 | 0.2166 | +2 | 15 |
| - | - | 691.8 | 774.1 | - | - | 0 | - |
| - | - | 1097 | 774.2 | - | - | 0 | - |
| - | - | 2626 | 774.4 | - | - | 0 | - |
| - | - | 1676 | 775.4 | - | - | 0 | - |
| 23 | b | 895 | 780.4 | 0.004989 | 6.392 | +3 | 23 |
| - | - | 709.5 | 781.1 | - | - | 0 | - |
| 12 | y | 2488 | 781.4 | 0.0005002 | 0.6401 | +2 | 15 |
| - | - | 1691 | 781.9 | - | - | 0 | - |
| - | - | 1794 | 782.4 | - | - | 0 | - |
| 23 | b | 1041 | 786.1 | 0.001827 | 2.324 | +3 | 23 |
| 4 | y | 1226 | 786.4 | 0.001587 | 2.018 | +3 | 23 |
| - | - | 1307 | 786.7 | - | - | 0 | - |
| - | - | 1253 | 787.1 | - | - | 0 | - |
| 20 | y | 1430 | 787.4 | 0.003915 | 4.972 | +1 | 7 |
| - | - | 2611 | 788.4 | - | - | 0 | - |
| - | - | 929.6 | 789.4 | - | - | 0 | - |
| - | - | 919.7 | 789.6 | - | - | 0 | - |
| 4 | y | 6977 | 792.1 | 0.001425 | 1.799 | +3 | 23 |
| - | - | 5371 | 792.4 | - | - | 0 | - |
| - | - | 6191 | 792.7 | - | - | 0 | - |
| - | - | 2837 | 793.1 | - | - | 0 | - |
| - | - | 973.2 | 794.9 | - | - | 0 | - |
| 16 | b | 1066 | 804.9 | 0.0001359 | 0.1688 | +2 | 16 |
| 16 | b | 5152 | 805.4 | 0.004222 | 5.242 | +2 | 16 |
| - | - | 2213 | 806.4 | - | - | 0 | - |
| - | - | 1120 | 807.4 | - | - | 0 | - |
| - | - | 1115 | 809.4 | - | - | 0 | - |
| - | - | 797.4 | 810.4 | - | - | 0 | - |
| - | - | 2450 | 813.4 | - | - | 0 | - |
| 16 | b | 1277 | 813.9 | 0.012 | 14.75 | +2 | 16 |
| - | - | 2615 | 814.4 | - | - | 0 | - |
| - | - | 1245 | 814.9 | - | - | 0 | - |
| - | - | 1037 | 815.4 | - | - | 0 | - |
| - | - | 1210 | 818.9 | - | - | 0 | - |
| - | - | 973.5 | 821.4 | - | - | 0 | - |
| - | - | 764.2 | 824.4 | - | - | 0 | - |
| 8 | b | 3487 | 827.4 | 0.00217 | 2.623 | +1 | 8 |
| - | - | 853.8 | 828.2 | - | - | 0 | - |
| - | - | 2955 | 828.4 | - | - | 0 | - |
| - | - | 1014 | 828.9 | - | - | 0 | - |
| - | - | 1140 | 829.4 | - | - | 0 | - |
| - | - | 1537 | 829.9 | - | - | 0 | - |
| - | - | 792 | 830.2 | - | - | 0 | - |
| - | - | 1840 | 830.4 | - | - | 0 | - |
| - | - | 1674 | 830.9 | - | - | 0 | - |
| - | - | 1109 | 831.4 | - | - | 0 | - |
| - | - | 1452 | 833.4 | - | - | 0 | - |
| 11 | y | 2063 | 836.9 | 0.004074 | 4.868 | +2 | 16 |
| - | - | 891.4 | 837.4 | - | - | 0 | - |
| - | - | 721.5 | 837.9 | - | - | 0 | - |
| - | - | 831.4 | 838.4 | - | - | 0 | - |
| - | - | 729.5 | 841.4 | - | - | 0 | - |
| - | - | 979.3 | 842.4 | - | - | 0 | - |
| - | - | 889.7 | 843.4 | - | - | 0 | - |
| - | - | 802.7 | 843.7 | - | - | 0 | - |
| - | - | 636 | 844.4 | - | - | 0 | - |
| 8 | b | 1.197E+04 | 845.4 | 0.007536 | 8.914 | +1 | 8 |
| - | - | 4759 | 845.9 | - | - | 0 | - |
| - | - | 6865 | 846.4 | - | - | 0 | - |
| - | - | 1126 | 847 | - | - | 0 | - |
| - | - | 1518 | 847.4 | - | - | 0 | - |
| 3 | y | 949.6 | 848.1 | 0.0005424 | 0.6396 | +3 | 24 |
| 3 | y | 2650 | 848.4 | 0.002676 | 3.155 | +3 | 24 |
| - | - | 1463 | 848.8 | - | - | 0 | - |
| - | - | 1828 | 849.1 | - | - | 0 | - |
| - | - | 1436 | 849.4 | - | - | 0 | - |
| - | - | 2498 | 852.6 | - | - | 0 | - |
| 3 | y | 8050 | 854.1 | 0.001537 | 1.8 | +3 | 24 |
| - | - | 1.045E+04 | 854.4 | - | - | 0 | - |
| - | - | 1.004E+04 | 854.8 | - | - | 0 | - |
| - | - | 3962 | 855.1 | - | - | 0 | - |
| 17 | b | 4027 | 855.4 | 0.002219 | 2.594 | +2 | 17 |
| 25 | b | 2075 | 856.4 | 0.01617 | 18.88 | +3 | 25 |
| - | - | 846.9 | 856.8 | - | - | 0 | - |
| - | - | 1157 | 857.4 | - | - | 0 | - |
| - | - | 772 | 858.4 | - | - | 0 | - |
| - | - | 1340 | 862.4 | - | - | 0 | - |
| - | - | 1449 | 862.8 | - | - | 0 | - |
| - | - | 1556 | 863.1 | - | - | 0 | - |
| 17 | b | 3372 | 864.5 | 0.00263 | 3.043 | +2 | 17 |
| - | - | 2307 | 865 | - | - | 0 | - |
| - | - | 1706 | 865.5 | - | - | 0 | - |
| - | - | 1991 | 866.5 | - | - | 0 | - |
| - | - | 2288 | 867.4 | - | - | 0 | - |
| - | - | 826.8 | 868.1 | - | - | 0 | - |
| - | - | 3429 | 868.4 | - | - | 0 | - |
| - | - | 1180 | 868.8 | - | - | 0 | - |
| - | - | 856.1 | 869.4 | - | - | 0 | - |
| - | - | 1035 | 870.5 | - | - | 0 | - |
| - | - | 818.4 | 870.9 | - | - | 0 | - |
| - | - | 2939 | 871.4 | - | - | 0 | - |
| - | - | 2509 | 872.4 | - | - | 0 | - |
| - | - | 1049 | 873.4 | - | - | 0 | - |
| 2 | y | 1624 | 877.8 | 0.0001729 | 0.197 | +3 | 25 |
| - | - | 3104 | 878.1 | - | - | 0 | - |
| - | - | 1023 | 878.5 | - | - | 0 | - |
| - | - | 826.6 | 880 | - | - | 0 | - |
| - | - | 983.1 | 881.5 | - | - | 0 | - |
| - | - | 1210 | 883.5 | - | - | 0 | - |
| 19 | y | 6800 | 884.5 | 0.01145 | 12.95 | +1 | 8 |
| 19 | y | 1.183E+04 | 885.4 | 0.005771 | 6.518 | +1 | 8 |
| - | - | 5540 | 886.5 | - | - | 0 | - |
| - | - | 1064 | 886.8 | - | - | 0 | - |
| - | - | 934.3 | 887.1 | - | - | 0 | - |
| - | - | 2172 | 887.5 | - | - | 0 | - |
| - | - | 927.8 | 891.1 | - | - | 0 | - |
| - | - | 847 | 892.5 | - | - | 0 | - |
| - | - | 1472 | 895.8 | - | - | 0 | - |
| - | - | 2204 | 896.1 | - | - | 0 | - |
| - | - | 5032 | 896.5 | - | - | 0 | - |
| - | - | 3018 | 896.8 | - | - | 0 | - |
| - | - | 1019 | 897 | - | - | 0 | - |
| - | - | 2175 | 897.1 | - | - | 0 | - |
| - | - | 2520 | 897.5 | - | - | 0 | - |
| - | - | 1075 | 898.8 | - | - | 0 | - |
| - | - | 2725 | 899.1 | - | - | 0 | - |
| - | - | 5324 | 899.5 | - | - | 0 | - |
| - | - | 3038 | 899.8 | - | - | 0 | - |
| - | - | 1697 | 900.1 | - | - | 0 | - |
| - | - | 2340 | 900.5 | - | - | 0 | - |
| - | - | 1492 | 900.8 | - | - | 0 | - |
| - | - | 1344 | 901.1 | - | - | 0 | - |
| - | - | 1547 | 901.5 | - | - | 0 | - |
| 19 | y | 4.974E+04 | 902.5 | 0.001317 | 1.459 | +1 | 8 |
| - | - | 2.238E+04 | 903.5 | - | - | 0 | - |
| - | - | 6723 | 904.5 | - | - | 0 | - |
| 0 | Precursor | 9052 | 904.8 | 5.485E-05 | 0.06062 | +3 | -1 |
| 0 | Precursor | 1.707E+04 | 905.1 | 0.005424 | 5.992 | +3 | -1 |
| - | - | 2.062E+04 | 905.5 | - | - | 0 | - |
| - | - | 1.65E+04 | 905.8 | - | - | 0 | - |
| 18 | b | 2035 | 906 | 0.0004921 | 0.5431 | +2 | 18 |
| - | - | 9612 | 906.1 | - | - | 0 | - |
| 18 | b | 6566 | 906.5 | 0.009339 | 10.3 | +2 | 18 |
| - | - | 2448 | 906.8 | - | - | 0 | - |
| - | - | 1269 | 907 | - | - | 0 | - |
| - | - | 795.7 | 907.5 | - | - | 0 | - |
| - | - | 1189 | 909.5 | - | - | 0 | - |
| - | - | 4440 | 909.8 | - | - | 0 | - |
| - | - | 1.204E+04 | 910.1 | - | - | 0 | - |
| - | - | 1.762E+04 | 910.5 | - | - | 0 | - |
| 0 | Precursor | 3.774E+05 | 910.8 | 0.0009889 | 1.086 | +3 | -1 |
| - | - | 6.016E+05 | 911.1 | - | - | 0 | - |
| - | - | 4.866E+05 | 911.5 | - | - | 0 | - |
| - | - | 2.88E+05 | 911.8 | - | - | 0 | - |
| - | - | 1.181E+05 | 912.1 | - | - | 0 | - |
| - | - | 3.345E+04 | 912.5 | - | - | 0 | - |
| - | - | 917.6 | 913.5 | - | - | 0 | - |
| - | - | 1119 | 914.5 | - | - | 0 | - |
| 18 | b | 5246 | 915 | 0.001008 | 1.102 | +2 | 18 |
| - | - | 4545 | 915.5 | - | - | 0 | - |
| - | - | 3210 | 916 | - | - | 0 | - |
| - | - | 1107 | 916.5 | - | - | 0 | - |
| 9 | y | 846.9 | 934.5 | 0.003045 | 3.258 | +2 | 18 |
| 9 | y | 1471 | 935 | 0.004703 | 5.03 | +2 | 18 |
| - | - | 1356 | 935.5 | - | - | 0 | - |
| - | - | 773.9 | 941.5 | - | - | 0 | - |
| - | - | 1971 | 942.5 | - | - | 0 | - |
| 9 | y | 9225 | 943.5 | 0.001125 | 1.192 | +2 | 18 |
| - | - | 9258 | 944 | - | - | 0 | - |
| - | - | 4669 | 944.5 | - | - | 0 | - |
| - | - | 1988 | 945 | - | - | 0 | - |
| - | - | 763.3 | 945.5 | - | - | 0 | - |
| - | - | 833.3 | 949 | - | - | 0 | - |
| - | - | 1891 | 959.5 | - | - | 0 | - |
| - | - | 973.9 | 969.5 | - | - | 0 | - |
| - | - | 1227 | 970.5 | - | - | 0 | - |
| - | - | 1565 | 984.5 | - | - | 0 | - |
| 18 | y | 1783 | 985.5 | 0.00705 | 7.153 | +1 | 9 |
| 18 | y | 2937 | 986.5 | 0.01247 | 12.65 | +1 | 9 |
| 8 | y | 3823 | 987 | 0.002026 | 2.053 | +2 | 19 |
| - | - | 4372 | 987.5 | - | - | 0 | - |
| - | - | 2425 | 988 | - | - | 0 | - |
| - | - | 1394 | 988.5 | - | - | 0 | - |
| - | - | 883.2 | 993.5 | - | - | 0 | - |
| - | - | 1191 | 999.5 | - | - | 0 | - |
| - | - | 943.6 | 1001 | - | - | 0 | - |
| 18 | y | 1.441E+04 | 1004 | 0.001612 | 1.606 | +1 | 9 |
| - | - | 8420 | 1005 | - | - | 0 | - |
| - | - | 3118 | 1006 | - | - | 0 | - |
| - | - | 1280 | 1007 | - | - | 0 | - |
| - | - | 877.7 | 1008 | - | - | 0 | - |
| - | - | 1926 | 1014 | - | - | 0 | - |
| - | - | 1235 | 1015 | - | - | 0 | - |
| 7 | y | 1650 | 1022 | 0.003135 | 3.069 | +2 | 20 |
| 7 | y | 2501 | 1022 | 0.002948 | 2.885 | +2 | 20 |
| - | - | 1204 | 1023 | - | - | 0 | - |
| - | - | 1010 | 1023 | - | - | 0 | - |
| 10 | b | 2511 | 1024 | 5.002E-05 | 0.04887 | +1 | 10 |
| 10 | b | 1571 | 1025 | 0.0152 | 14.84 | +1 | 10 |
| - | - | 927.8 | 1026 | - | - | 0 | - |
| 7 | y | 1.022E+04 | 1031 | 0.0005991 | 0.5814 | +2 | 20 |
| - | - | 1.138E+04 | 1031 | - | - | 0 | - |
| - | - | 5982 | 1032 | - | - | 0 | - |
| - | - | 2081 | 1032 | - | - | 0 | - |
| - | - | 1103 | 1033 | - | - | 0 | - |
| 10 | b | 1.161E+04 | 1042 | 0.001226 | 1.177 | +1 | 10 |
| - | - | 6778 | 1043 | - | - | 0 | - |
| - | - | 2186 | 1044 | - | - | 0 | - |
| - | - | 1713 | 1051 | - | - | 0 | - |
| - | - | 1227 | 1052 | - | - | 0 | - |
| - | - | 3088 | 1059 | - | - | 0 | - |
| - | - | 1647 | 1060 | - | - | 0 | - |
| - | - | 883.3 | 1061 | - | - | 0 | - |
| - | - | 1654 | 1071 | - | - | 0 | - |
| 21 | b | 1052 | 1071 | 0.006899 | 6.442 | +2 | 21 |
| - | - | 3448 | 1072 | - | - | 0 | - |
| 6 | y | 1876 | 1079 | 0.002664 | 2.469 | +2 | 21 |
| 6 | y | 3233 | 1080 | 0.005163 | 4.783 | +2 | 21 |
| - | - | 1086 | 1080 | - | - | 0 | - |
| - | - | 1385 | 1081 | - | - | 0 | - |
| 17 | y | 1018 | 1087 | 0.004049 | 3.726 | +1 | 10 |
| 17 | y | 1543 | 1088 | 0.0121 | 11.12 | +1 | 10 |
| 6 | y | 1.138E+04 | 1088 | 0.0005553 | 0.5104 | +2 | 21 |
| - | - | 1.187E+04 | 1089 | - | - | 0 | - |
| - | - | 8470 | 1089 | - | - | 0 | - |
| - | - | 3559 | 1090 | - | - | 0 | - |
| - | - | 990.2 | 1090 | - | - | 0 | - |
| - | - | 1130 | 1100 | - | - | 0 | - |
| - | - | 998.1 | 1102 | - | - | 0 | - |
| 17 | y | 1.033E+04 | 1105 | 0.002273 | 2.058 | +1 | 10 |
| - | - | 7146 | 1106 | - | - | 0 | - |
| - | - | 1767 | 1106 | - | - | 0 | - |
| - | - | 4552 | 1107 | - | - | 0 | - |
| - | - | 1376 | 1108 | - | - | 0 | - |
| 5 | y | 4182 | 1115 | 0.001441 | 1.293 | +2 | 22 |
| 5 | y | 5314 | 1115 | 0.006626 | 5.942 | +2 | 22 |
| - | - | 4933 | 1116 | - | - | 0 | - |
| - | - | 3270 | 1116 | - | - | 0 | - |
| - | - | 1972 | 1117 | - | - | 0 | - |
| 5 | y | 2.806E+04 | 1124 | 0.0009193 | 0.8182 | +2 | 22 |
| - | - | 3.249E+04 | 1124 | - | - | 0 | - |
| - | - | 2.122E+04 | 1125 | - | - | 0 | - |
| - | - | 9390 | 1125 | - | - | 0 | - |
| - | - | 2305 | 1126 | - | - | 0 | - |
| 22 | b | 911.8 | 1126 | 0.007346 | 6.524 | +2 | 22 |
| - | - | 823.3 | 1128 | - | - | 0 | - |
| - | - | 1999 | 1136 | - | - | 0 | - |
| - | - | 1397 | 1136 | - | - | 0 | - |
| - | - | 1033 | 1141 | - | - | 0 | - |
| - | - | 943 | 1143 | - | - | 0 | - |
| - | - | 1444 | 1144 | - | - | 0 | - |
| 11 | b | 3750 | 1170 | 0.01085 | 9.277 | +1 | 11 |
| - | - | 2523 | 1171 | - | - | 0 | - |
| - | - | 872.9 | 1172 | - | - | 0 | - |
| - | - | 817.2 | 1173 | - | - | 0 | - |
| 23 | b | 1263 | 1179 | 0.0005666 | 0.4807 | +2 | 23 |
| 4 | y | 2428 | 1179 | 0.007362 | 6.244 | +2 | 23 |
| - | - | 2644 | 1180 | - | - | 0 | - |
| 16 | y | 7642 | 1188 | 0.01179 | 9.928 | +1 | 11 |
| - | - | 9907 | 1188 | - | - | 0 | - |
| - | - | 8757 | 1189 | - | - | 0 | - |
| - | - | 3186 | 1189 | - | - | 0 | - |
| - | - | 2515 | 1190 | - | - | 0 | - |
| 16 | y | 1.381E+04 | 1206 | 0.0008594 | 0.7128 | +1 | 11 |
| - | - | 9617 | 1207 | - | - | 0 | - |
| - | - | 3895 | 1208 | - | - | 0 | - |
| - | - | 1195 | 1209 | - | - | 0 | - |
| 24 | b | 1110 | 1227 | 0.008765 | 7.145 | +2 | 24 |
| - | - | 1031 | 1228 | - | - | 0 | - |
| 24 | b | 1098 | 1236 | 0.005802 | 4.695 | +2 | 24 |
| - | - | 947.6 | 1237 | - | - | 0 | - |
| 12 | b | 933.8 | 1241 | 0.007804 | 6.29 | +1 | 12 |
| - | - | 1208 | 1242 | - | - | 0 | - |
| - | - | 781.6 | 1271 | - | - | 0 | - |
| 3 | y | 1217 | 1272 | 0.003838 | 3.017 | +2 | 24 |
| - | - | 1064 | 1274 | - | - | 0 | - |
| 13 | b | 1368 | 1281 | 0.003695 | 2.885 | +1 | 13 |
| - | - | 3466 | 1281 | - | - | 0 | - |
| - | - | 2449 | 1282 | - | - | 0 | - |
| - | - | 960.3 | 1282 | - | - | 0 | - |
| 25 | b | 1196 | 1293 | 0.002282 | 1.766 | +2 | 25 |
| - | - | 1042 | 1293 | - | - | 0 | - |
| - | - | 1184 | 1294 | - | - | 0 | - |
| - | - | 1046 | 1299 | - | - | 0 | - |
| - | - | 1718 | 1302 | - | - | 0 | - |
| - | - | 949.5 | 1302 | - | - | 0 | - |
| - | - | 928.7 | 1309 | - | - | 0 | - |
| 15 | y | 2443 | 1317 | 0.004231 | 3.214 | +1 | 12 |
| 15 | y | 3003 | 1318 | 0.008985 | 6.819 | +1 | 12 |
| - | - | 1170 | 1319 | - | - | 0 | - |
| - | - | 1278 | 1320 | - | - | 0 | - |
| - | - | 1277 | 1327 | - | - | 0 | - |
| - | - | 834.3 | 1328 | - | - | 0 | - |
| 15 | y | 1.266E+04 | 1335 | 0.0003519 | 0.2637 | +1 | 12 |
| - | - | 8732 | 1336 | - | - | 0 | - |
| - | - | 3165 | 1337 | - | - | 0 | - |
| - | - | 950.5 | 1338 | - | - | 0 | - |
| - | - | 1233 | 1345 | - | - | 0 | - |
| - | - | 1618 | 1346 | - | - | 0 | - |
| 0 | Precursor | 844.2 | 1357 | 0.01456 | 10.73 | +2 | -1 |
| 14 | b | 1896 | 1397 | 0.0062 | 4.439 | +1 | 14 |
| - | - | 1382 | 1398 | - | - | 0 | - |
| 14 | y | 818.9 | 1417 | 0.01426 | 10.07 | +1 | 13 |
| 14 | y | 3130 | 1434 | 0.001139 | 0.7943 | +1 | 13 |
| - | - | 2648 | 1435 | - | - | 0 | - |
| - | - | 1495 | 1436 | - | - | 0 | - |
| 13 | y | 2578 | 1473 | 0.002733 | 1.855 | +1 | 14 |
| 13 | y | 2587 | 1474 | 0.01335 | 9.056 | +1 | 14 |
| - | - | 1910 | 1475 | - | - | 0 | - |
| 13 | y | 9713 | 1491 | 0.0006299 | 0.4226 | +1 | 14 |
| - | - | 9154 | 1492 | - | - | 0 | - |
| - | - | 4472 | 1493 | - | - | 0 | - |
| - | - | 778.4 | 1494 | - | - | 0 | - |
| 15 | b | 1155 | 1526 | 0.009049 | 5.931 | +1 | 15 |
| - | - | 1501 | 1527 | - | - | 0 | - |
| 12 | y | 969.4 | 1544 | 0.01286 | 8.33 | +1 | 15 |
| 12 | y | 1549 | 1545 | 0.01212 | 7.846 | +1 | 15 |
| - | - | 1286 | 1546 | - | - | 0 | - |
| - | - | 1018 | 1547 | - | - | 0 | - |
| 12 | y | 7892 | 1562 | 0.001977 | 1.266 | +1 | 15 |
| - | - | 5827 | 1563 | - | - | 0 | - |
| - | - | 4112 | 1564 | - | - | 0 | - |
| - | - | 832.1 | 1565 | - | - | 0 | - |
| 11 | y | 1186 | 1690 | 0.001115 | 0.6598 | +1 | 16 |
| - | - | 1267 | 1691 | - | - | 0 | - |
| - | - | 708.6 | 1731 | - | - | 0 | - |
| - | - | 876.4 | 1790 | - | - | 0 | - |

m/z Charge Intensity FragmentType MassShift Position
122.90099334716797 0 394.7826
126.09188079833984 0 585.7288
127.08702087402344 0 742.828
128.07127380371094 0 382.11203
129.0662841796875 0 1235.0264
129.10269165039062 0 97938.02
130.06553649902344 0 2831.4358
130.08673095703125 0 3171.7126 y Ammonia loss 25
130.10035705566406 0 836.85693
130.10604858398438 0 6770.424
132.08123779296875 0 6250.847
136.07608032226562 0 644.2146
137.52474975585938 0 455.97794
138.12832641601562 0 1412.4059
139.0869903564453 0 1586.0336
141.06637573242188 0 1227.5122
141.1025390625 0 1284.3914
143.11839294433594 0 8602.23 a 1
146.1295166015625 0 552.7886
147.07659912109375 0 507.0232
147.11322021484375 0 3754.9397 y 25
148.94786071777344 0 808.27454
153.10275268554688 0 2352.0554
154.10549926757812 0 560.97815
154.2359619140625 0 422.95087
155.0817413330078 0 965.478
155.1183624267578 0 4323.474
155.15447998046875 0 631.5085
156.12179565429688 0 604.6711
157.06121826171875 0 1261.824
157.09764099121094 0 7590.014
158.06092834472656 0 728.45624
158.1008758544922 0 580.55084
159.09217834472656 0 107941.76
160.08843994140625 0 989.5526
160.09556579589844 0 10783.796
161.0987091064453 0 827.3645
165.1026611328125 0 839.8825 a 2
165.80584716796875 0 592.8201
167.07496643066406 0 549.47455
167.08206176757812 0 2461.4995
167.11834716796875 0 1313.7986
169.097412109375 0 924.70013
169.13401794433594 0 12577.12
170.0605010986328 0 13214.766
170.1374053955078 0 1661.7482
171.06396484375 0 1426.4229
171.0767364501953 0 646.9201
171.11325073242188 0 12563.71 b 1
171.12091064453125 0 1020.57623
172.11666870117188 0 943.8331
172.14520263671875 0 594.6111
173.43862915039062 0 1576.5789
174.14077758789062 0 396.53802
175.07196044921875 0 786.63403
175.10806274414062 0 1041.895
180.1132049560547 0 483.79663
181.09776306152344 0 1500.231
181.13406372070312 0 1294.2906
182.12925720214844 0 3822.1804
182.65451049804688 0 421.26422
183.11328125 0 2030.7485
183.1324462890625 0 644.2423
183.1495819091797 0 1959.0703
184.07229614257812 0 578.01666
184.10855102539062 0 2494.1528
185.05575561523438 0 1060.6041
185.09255981445312 0 11067.445
185.12896728515625 0 2301.598
186.09597778320312 0 1197.7476
187.07174682617188 0 2610.1145
187.0870819091797 0 8888.627
188.0903778076172 0 1415.6428
193.13380432128906 0 601.3917
195.07705688476562 0 1641.6509
195.11326599121094 0 1924.5691
196.1078338623047 0 717.29315
197.12892150878906 0 8011.0874
198.08763122558594 0 1323.4934
198.1242218017578 0 2817.872
199.0714874267578 0 628.7221
199.10804748535156 0 3046.125
200.13992309570312 0 2362.2969
201.1239471435547 0 1119.73
202.0826416015625 0 2426.1333
202.11903381347656 0 2484.4976
203.0670623779297 0 713.25903
203.10284423828125 0 1719.1742
204.1138916015625 0 3177.5735
205.09771728515625 0 2168.0388
205.1165313720703 0 738.31354
206.1005096435547 0 524.87604
208.10861206054688 0 1404.5133
209.09237670898438 0 2826.7954
210.1240234375 0 1957.6764
212.06671142578125 0 590.51465
212.1396484375 0 1247.9648
213.0875701904297 0 2854.7341
213.12364196777344 0 1229.2842
213.14219665527344 0 482.35226
214.15603637695312 0 2359.5547
216.09811401367188 0 1075.0643
216.1348114013672 0 8341.558
217.1386260986328 0 543.413
221.0924072265625 0 692.17413
222.12428283691406 0 3584.5332
223.10939025878906 0 657.4011 y Ammonia loss 22
223.12762451171875 0 704.156
224.10415649414062 0 595.3643
224.1399383544922 0 690.23016
226.0824432373047 0 581.40393
226.11924743652344 0 6847.6177
227.1033935546875 0 533.42303
228.1348419189453 0 5054.632
228.1713104248047 0 1943.7726
229.0943145751953 0 1012.78424
229.11900329589844 0 2109.9487
230.1293487548828 0 3452.01
231.09808349609375 0 1036.935
231.13291931152344 0 690.4164
236.10403442382812 0 825.00934
236.13958740234375 0 739.9487
238.1195831298828 0 1630.6023
238.1549835205078 0 754.78235
239.1513671875 0 1240.0448
240.1348114013672 0 3630.711 y 20
242.11390686035156 0 1228.2427
243.14523315429688 0 1304.6769 b 3
244.12979125976562 0 4802.625 y Ammonia loss 24
249.1241455078125 0 926.289
250.117919921875 0 601.43665
250.15554809570312 0 851.8601
252.0980987548828 0 802.927
254.07704162597656 0 782.5142
254.11399841308594 0 3779.784
256.09332275390625 0 1623.3049
256.1297607421875 0 680.38464
256.14434814453125 0 868.2278
256.1656494140625 0 619.7612
257.1613464355469 0 3039.2476
258.1242980957031 0 7328.17
259.12701416015625 0 1442.5497
261.1562194824219 0 7345.9307 y 24
262.1614990234375 0 607.4359
266.15057373046875 0 1826.3335
267.1463317871094 0 917.4965
268.12969970703125 0 2554.3474
269.1613464355469 0 1002.98364
270.10882568359375 0 699.9469
270.1605529785156 0 5336.167
271.14166259765625 0 662.26013
271.16302490234375 0 709.73987
272.08868408203125 0 2031.1738
272.12493896484375 0 689.82275
274.10333251953125 0 1235.8506
278.11407470703125 0 1919.7688
281.7962646484375 0 506.83636
282.18145751953125 0 650.1831
284.1244812011719 0 1185.059
284.14312744140625 0 591.1728
284.16119384765625 0 2500.0908
285.15704345703125 0 1260.408
286.140380859375 0 3288.485
286.17645263671875 0 604.5149
290.09832763671875 0 752.0735
294.1442565917969 0 650.27563
295.10205078125 0 713.973
295.14019775390625 0 939.753
295.1772766113281 0 2085.0793
296.1247253417969 0 2241.0115
297.1564636230469 0 1197.1874
298.1558532714844 0 28486.074
299.1591491699219 0 4452.4746
300.1607971191406 0 537.5308
301.1870422363281 0 655.4588
306.1097412109375 0 864.977
306.1449890136719 0 695.70557
309.15643310546875 0 900.3023
309.19384765625 0 718.6869
311.1719970703125 0 867.9513
311.20831298828125 0 993.7941
312.1337890625 0 608.1347
312.153564453125 0 916.199
312.1715087890625 0 3050.3115
313.1156005859375 0 1567.8925
313.1541748046875 0 1204.7448
313.1879577636719 0 16769.004
314.1352844238281 0 2063.5964
314.1904296875 0 2763.0806
315.132568359375 0 669.6811
315.1666564941406 0 1826.3099
315.1824951171875 0 21572.984
316.1272888183594 0 625.631
316.185791015625 0 3954.6912
317.1093444824219 0 594.3062
323.13604736328125 0 2778.253
324.1385192871094 0 1083.4908
325.11541748046875 0 2853.6665
325.1889343261719 0 736.6264
325.2243347167969 0 8906.378
326.1836853027344 0 1131.9833
326.2275085449219 0 1419.1461
327.1671142578125 0 819.03516
327.2035827636719 0 1336.3577
328.23492431640625 0 2727.9822
329.1813659667969 0 1160.8975
330.1412048339844 0 1024.5217
330.16552734375 0 738.22675
330.21429443359375 0 2293.973
338.2195739746094 0 604.6689
339.1663513183594 0 1197.4563
339.2041320800781 0 1055.1014
340.1639404296875 0 512.5844
341.14630126953125 0 1749.7903
343.125732421875 0 4252.1216
343.1613464355469 0 1106.0079
343.198974609375 0 890.75934
344.1937255859375 0 2870.2063
347.1721496582031 0 1673.0312
351.13006591796875 0 736.3716
353.1453552246094 0 1092.8564
353.1820068359375 0 720.18066
354.1785888671875 0 1466.0331
356.2298583984375 0 4994.501
357.19287109375 0 11319.475 b 2
357.2323303222656 0 928.2819
358.173583984375 0 1471.6815 y Ammonia loss 23
358.19635009765625 0 1570.0537
361.1361389160156 0 1684.8169
368.1911926269531 0 794.7554
369.14105224609375 0 1786.6433
369.1766052246094 0 1548.3031
371.15557861328125 0 935.11456
371.1928405761719 0 1239.808
374.2197265625 0 1108.8175
375.1990051269531 0 5681.8984 y 23
375.2223205566406 0 632.87573
376.2021484375 0 830.00134
378.25030517578125 0 1738.8235
379.1605529785156 0 898.15405
384.1892395019531 0 2226.115
384.22509765625 0 1421.8568
386.2194519042969 0 4789.0957
387.1527404785156 0 779.8134
387.1881408691406 0 1333.4484
387.22216796875 0 875.49603
392.157958984375 0 809.33466
395.1927490234375 0 761.29877
396.26165771484375 0 2980.0217
397.1737365722656 0 1884.4136
397.2607116699219 0 723.2625
398.1744689941406 0 629.7123
402.19927978515625 0 1621.9421
410.13482666015625 0 665.34985
410.16815185546875 0 2818.537
412.2232360839844 0 1000.4784
412.25494384765625 0 930.9425
414.2056884765625 0 1023.8155 b Water loss 7
414.23583984375 0 1740.0128 b 11
415.14849853515625 0 556.42523
415.1832580566406 0 2017.2434
422.1690368652344 0 921.9487
423.20208740234375 0 935.93396
423.2359313964844 0 4221.5586
424.2209777832031 0 1934.4
427.1572265625 0 1092.0071
427.1972961425781 0 643.48474
428.1777648925781 0 1034.2019
430.23065185546875 0 682.311
436.1839904785156 0 855.70935
439.2450866699219 0 760.6799
440.17913818359375 0 2060.042
440.2139892578125 0 1270.1145
441.2127990722656 0 841.64154
441.24676513671875 0 5892.717
442.2488708496094 0 1583.5198
443.22454833984375 0 1061.7125 y Ammonia loss 18
443.7239074707031 0 650.00275
444.1855163574219 0 1292.5461
444.2185974121094 0 1250.3478 y Water loss 22
445.2056884765625 0 1607.6107 y Ammonia loss 22
451.73907470703125 0 4929.7183 y 18
452.24176025390625 0 2004.9266
453.20977783203125 0 2012.965
453.2832336425781 0 2136.444
457.25341796875 0 956.8584
458.1900939941406 0 1549.179
458.22711181640625 0 1404.6469
462.23211669921875 0 14694.055 y 22
463.23492431640625 0 2973.935
464.2358703613281 0 789.1862 b Ammonia loss 8
468.2105712890625 0 1250.2352
470.2256774902344 0 1421.071
471.2213134765625 0 2619.2163
472.2251892089844 0 651.0282
483.2948913574219 0 701.4055
485.2841796875 0 4878.8184 b 3
486.2204895019531 0 1161.8306
486.2890625 0 1965.289
489.23175048828125 0 3788.8357
490.2348937988281 0 1747.6342
501.24615478515625 0 2346.9912
502.2601623535156 0 2177.6575 y 17
510.23162841796875 0 710.241
510.26910400390625 0 941.4233
521.2000122070312 0 1270.7081
524.28515625 0 1578.3698
524.3587646484375 0 964.51526
525.2704467773438 0 583.28864
528.279296875 0 2154.622
528.3287353515625 0 1225.7206
529.2723999023438 0 905.91833
535.78515625 0 1061.4854
537.2401123046875 0 1221.4147
538.2275390625 0 1573.0732
538.3145751953125 0 4861.883
539.248046875 0 1070.4092
539.3158569335938 0 838.9545
540.3135375976562 0 711.2231
542.293701171875 0 1669.187
552.313232421875 0 649.8419
552.3522338867188 0 4279.928
553.35498046875 0 1278.297
555.2527465820312 0 2367.6692
556.2380981445312 0 2302.9912
556.3253784179688 0 10825.367 b 4
557.25634765625 0 2749.3801
557.3284301757812 0 4415.8003
558.3279418945312 0 1001.7016
564.7974853515625 0 850.4954
568.31201171875 0 753.71924
570.3245239257812 0 1275.7048
571.271484375 0 578.002
572.2803955078125 0 5102.4604 y Water loss 21
573.2653198242188 0 6874.31 y Ammonia loss 21
574.26806640625 0 1814.2179
586.3220825195312 0 1699.85
590.2899780273438 0 9918.831 y 21
591.2938232421875 0 2912.5818
592.2949829101562 0 838.23016
600.3286743164062 0 672.8439
603.3107299804688 0 3123.7004 y 15
603.8150024414062 0 1223.1875
606.2938842773438 0 978.75543
607.2879028320312 0 828.3736
612.33447265625 0 1253.7415 b Ammonia loss 11
612.836181640625 0 1154.2434
620.8394775390625 0 2045.5112 b 11
621.3402709960938 0 1634.4318
621.8447265625 0 997.6037
624.30126953125 0 603.1826
625.3057250976562 0 965.67847
639.3844604492188 0 962.2862
642.3218383789062 0 3267.1782
643.3268432617188 0 1100.9851
649.3496704101562 0 2534.738 b 12
649.8487548828125 0 2001.7637
650.0199584960938 0 1074.4186
650.3549194335938 0 1091.767
657.2987670898438 0 1144.0865
657.3515014648438 0 989.1685
658.2877197265625 0 785.0531
659.310302734375 0 863.2178 y Ammonia loss 14
667.3408203125 0 2716.9272
667.833740234375 0 1690.985 y 14
668.3386840820312 0 3356.3723
668.8390502929688 0 715.92267
669.3472900390625 0 1630.5106
670.355712890625 0 1009.6445
671.3525390625 0 9191.068 b 5
672.3557739257812 0 3575.1714
675.310546875 0 3277.5479
676.3115234375 0 2191.907
677.3150634765625 0 802.5236
681.3944091796875 0 3459.8943
681.579345703125 0 643.5259
682.4015502929688 0 1573.6987
684.8861083984375 0 1945.8708
685.3534545898438 0 5752.001
686.3560791015625 0 2944.2837
687.3648071289062 0 2405.221 y 6
697.3512573242188 0 703.0275
698.8839721679688 0 2500.981 b 13
699.3893432617188 0 1214.031
699.888427734375 0 956.9672
700.3779296875 0 1210.0615 y Water loss 20
701.3602294921875 0 2762.7534 y Ammonia loss 20
702.378173828125 0 1040.1926
717.3719482421875 0 970.0369 y 13
717.869384765625 0 593.9824
718.3858642578125 0 5458.93 y 20
719.3861694335938 0 1795.1968
719.8746948242188 0 860.2471
720.374267578125 0 632.5838
722.308349609375 0 690.17896
726.3853759765625 0 767.1662
728.876220703125 0 841.48016
729.37353515625 0 1072.2825
732.390625 0 849.39954
738.3775024414062 0 3357.537
739.3905639648438 0 962.77527
740.322998046875 0 954.418
742.373291015625 0 1001.90906
743.374267578125 0 645.82855 y Water loss 4
744.3837890625 0 2345.5854
745.383056640625 0 1059.634
745.8762817382812 0 1939.4086 y 12
746.3560791015625 0 1672.3486
746.8788452148438 0 651.83295
748.3925170898438 0 773.49506
749.217041015625 0 823.5666
756.3764038085938 0 5092.12
757.3433837890625 0 1342.2682
757.3959350585938 0 1751.3257
758.38427734375 0 4372.7812 b 6
759.3825073242188 0 2468.5588
763.4056396484375 0 3430.3374 b 14
763.9053955078125 0 3433.5667
764.424560546875 0 724.97687
766.9436645507812 0 656.07764
770.395263671875 0 1212.1389
772.3893432617188 0 1081.034 y Water loss 11
774.1092529296875 0 691.76135
774.1797485351562 0 1097.01
774.3773803710938 0 2625.5122
775.3815307617188 0 1675.8949
780.40576171875 0 895.00867 b Ammonia loss 22
781.0753784179688 0 709.46313
781.3949584960938 0 2488.0063 y 11
781.89794921875 0 1690.6415
782.4392700195312 0 1793.9331
786.074462890625 0 1041.2881 b 22
786.404541015625 0 1226.2997 y Ammonia loss 3
786.7372436523438 0 1306.5784
787.0694580078125 0 1252.5165
787.4096069335938 0 1430.0443 y Water loss 19
788.4058227539062 0 2610.8396
789.408447265625 0 929.58795
789.5578002929688 0 919.709
792.0798950195312 0 6976.598 y 3
792.4138793945312 0 5371.3013
792.7476196289062 0 6190.6904
793.08154296875 0 2837.2861
794.9171752929688 0 973.2462
804.9227905273438 0 1066.221 b Water loss 15
805.4188842773438 0 5151.577 b Ammonia loss 15
806.4232177734375 0 2213.1038
807.4210815429688 0 1119.672
809.3981323242188 0 1115.364
810.3960571289062 0 797.3511
813.4182739257812 0 2449.8904
813.93994140625 0 1277.1892 b 15
814.431396484375 0 2614.8306
814.9306030273438 0 1244.751
815.4182739257812 0 1036.8065
818.9149169921875 0 1210.2274
821.419921875 0 973.5282
824.4242553710938 0 764.169
827.4067993164062 0 3487.3906 b Water loss 7
828.1653442382812 0 853.772
828.4078979492188 0 2955.0095
828.9296264648438 0 1014.4282
829.4266967773438 0 1139.8892
829.9277954101562 0 1537.2164
830.1875610351562 0 791.96375
830.4301147460938 0 1839.5178
830.928955078125 0 1673.6449
831.4166259765625 0 1109.3484
833.4310302734375 0 1451.673
836.9327392578125 0 2062.606 y Ammonia loss 10
837.433349609375 0 891.36206
837.9434204101562 0 721.4702
838.4397583007812 0 831.37695
841.4398803710938 0 729.4993
842.4398193359375 0 979.28546
843.4469604492188 0 889.7201
843.6688842773438 0 802.69684
844.4327392578125 0 635.9948
845.4227294921875 0 11968.855 b 7
845.9447021484375 0 4758.953
846.42724609375 0 6864.8506
846.9561157226562 0 1125.9491
847.4241943359375 0 1518.2666
848.1019287109375 0 949.6397 y Water loss 2
848.4320678710938 0 2649.9612 y Ammonia loss 2
848.769287109375 0 1463.0603
849.0999145507812 0 1827.8583
849.4351806640625 0 1436.3772
852.6276245117188 0 2498.0298
854.1064453125 0 8049.622 y 2
854.4398193359375 0 10445.773
854.7752075195312 0 10038.077
855.107421875 0 3961.8552
855.4442749023438 0 4026.7551 b Water loss 16
856.445556640625 0 2074.782 b Ammonia loss 24
856.7681884765625 0 846.8677
857.4053955078125 0 1156.745
858.4042358398438 0 771.99475
862.435546875 0 1339.5997
862.7763061523438 0 1448.6887
863.0996704101562 0 1555.8021
864.4544067382812 0 3372.045 b 16
864.9531860351562 0 2306.7893
865.4537353515625 0 1705.983
866.4688720703125 0 1990.9224
867.4472045898438 0 2287.511
868.1046142578125 0 826.8212
868.434326171875 0 3428.7087
868.7785034179688 0 1179.5623
869.4375610351562 0 856.13245
870.4526977539062 0 1034.8475
870.9430541992188 0 818.4242
871.434326171875 0 2938.9692
872.4441528320312 0 2509.2585
873.4301147460938 0 1048.9358
877.7841186523438 0 1624.1742 y 1
878.1212768554688 0 3103.9314
878.4550170898438 0 1022.5612
879.95263671875 0 826.5519
881.455078125 0 983.1057
883.4789428710938 0 1209.7108
884.4699096679688 0 6799.9097 y Water loss 18
885.4482421875 0 11834.832 y Ammonia loss 18
886.4505004882812 0 5540.406
886.7858276367188 0 1064.4233
887.1195068359375 0 934.31866
887.4554443359375 0 2172.068
891.1295776367188 0 927.8201
892.4589233398438 0 846.9834
895.7997436523438 0 1472.2518
896.132568359375 0 2204.0757
896.466552734375 0 5031.952
896.7974853515625 0 3017.844
896.9684448242188 0 1019.15497
897.13818359375 0 2174.5183
897.4708251953125 0 2519.6006
898.8078002929688 0 1074.6407
899.1293334960938 0 2725.283
899.46630859375 0 5324.0405
899.7965698242188 0 3038.4258
900.13330078125 0 1697.303
900.47119140625 0 2339.9294
900.7984008789062 0 1492.2964
901.122802734375 0 1343.7573
901.4720458984375 0 1546.5474
902.4703369140625 0 49737.914 y 18
903.4729614257812 0 22381.477
904.4755249023438 0 6723.2793
904.8032836914062 0 9052.44 Precursor Water loss
905.1366577148438 0 17074.44 Precursor Ammonia loss
905.4692993164062 0 20617.977
905.8030395507812 0 16497.623
905.9708251953125 0 2035.4836 b Water loss 17
906.1361083984375 0 9612.024
906.4716796875 0 6565.908 b Ammonia loss 17
906.8050537109375 0 2448.0105
906.9690551757812 0 1269.1108
907.4524536132812 0 795.7472
909.4522094726562 0 1188.7356
909.7926635742188 0 4440.462
910.1331787109375 0 12041.898
910.4688720703125 0 17616.8
910.8077392578125 0 377402.12 Precursor
911.1426391601562 0 601610.3
911.4768676757812 0 486648.16
911.8104248046875 0 288035.53
912.1446533203125 0 118090.25
912.4774169921875 0 33447.805
913.47509765625 0 917.61444
914.4746704101562 0 1118.5065
914.9766235351562 0 5245.971 b 17
915.4784545898438 0 4544.9346
915.979736328125 0 3209.7483
916.4840698242188 0 1106.7896
934.4942016601562 0 846.89496 y Water loss 8
934.9939575195312 0 1470.742 y Ammonia loss 8
935.4945678710938 0 1356.0137
941.5020141601562 0 773.8546
942.4804077148438 0 1971.217
943.5014038085938 0 9224.833 y 8
944.0048828125 0 9257.7
944.5054931640625 0 4668.683
945.0090942382812 0 1988.4956
945.4697265625 0 763.3375
949.0076904296875 0 833.2619
959.491455078125 0 1891.1631
969.4710083007812 0 973.8868
970.4818115234375 0 1226.8661
984.5350952148438 0 1564.9879
985.51318359375 0 1782.6409 y Water loss 17
986.5026245117188 0 2936.902 y Ammonia loss 17
987.0205688476562 0 3822.908 y 7
987.5155639648438 0 4372.355
988.0240478515625 0 2424.6858
988.5223999023438 0 1393.9463
993.5018310546875 0 883.24835
999.5250244140625 0 1190.7947
1000.5228881835938 0 943.6052
1003.518310546875 0 14413.724 y 17
1004.5194091796875 0 8420.29
1005.5216064453125 0 3118.0415
1006.5166015625 0 1279.7222
1007.5013427734375 0 877.71423
1013.5354614257812 0 1926.3972
1014.5377197265625 0 1235.4335
1021.5324096679688 0 1650.4949 y Water loss 6
1022.0242309570312 0 2501.1426 y Ammonia loss 6
1022.5333251953125 0 1203.6885
1023.030517578125 0 1010.18274
1023.5257568359375 0 2510.6323 b Water loss 9
1024.5250244140625 0 1570.5293 b Ammonia loss 9
1025.5224609375 0 927.81366
1030.53515625 0 10221.59 y 6
1031.0372314453125 0 11381.661
1031.5374755859375 0 5982.3677
1032.0362548828125 0 2080.5454
1032.5338134765625 0 1103.0519
1041.53759765625 0 11607.173 b 9
1042.5372314453125 0 6778.3247
1043.5419921875 0 2186.377
1051.0196533203125 0 1713.4467
1051.517578125 0 1226.653
1058.5628662109375 0 3088.188
1059.5489501953125 0 1646.6249
1060.5313720703125 0 883.28265
1070.54638671875 0 1653.7748
1071.05859375 0 1052.3363 b 20
1071.5611572265625 0 3447.537
1079.04541015625 0 1875.8829 y Water loss 5
1079.5399169921875 0 3233.2246 y Ammonia loss 5
1080.044189453125 0 1085.611
1080.5379638671875 0 1384.9269
1086.557861328125 0 1018.17596 y Water loss 16
1087.5499267578125 0 1543.3846 y Ammonia loss 16
1088.048583984375 0 11381.996 y 5
1088.55029296875 0 11869.527
1089.0511474609375 0 8470.443
1089.5538330078125 0 3559.4263
1090.0504150390625 0 990.1892
1099.5543212890625 0 1130.006
1102.0440673828125 0 998.06476
1104.566650390625 0 10328.135 y 16
1105.5665283203125 0 7146.431
1106.0577392578125 0 1767.3384
1106.56005859375 0 4551.804
1107.5750732421875 0 1375.6104
1114.562744140625 0 4181.558 y Water loss 4
1115.0599365234375 0 5314.4707 y Ammonia loss 4
1115.558837890625 0 4933.1763
1116.061767578125 0 3270.4634
1116.563232421875 0 1971.9406
1123.5675048828125 0 28056.82 y 4
1124.0693359375 0 32492.795
1124.5697021484375 0 21223.936
1125.072509765625 0 9389.66
1125.5718994140625 0 2305.4575
1126.0821533203125 0 911.79584 b Water loss 21
1127.5814208984375 0 823.26306
1135.5928955078125 0 1999.1251
1136.1009521484375 0 1397.4675
1140.591064453125 0 1033.4572
1142.5638427734375 0 943.0051
1143.596435546875 0 1444.1409
1169.6204833984375 0 3749.7822 b 10
1170.6148681640625 0 2522.8403
1171.60498046875 0 872.88654
1172.5955810546875 0 817.16833
1178.6102294921875 0 1263.0717 b 22
1179.108154296875 0 2427.5325 y Ammonia loss 3
1179.60888671875 0 2643.5557
1187.61328125 0 7642.4717 y Water loss 15
1188.11572265625 0 9907.336
1188.61083984375 0 8756.933
1189.1214599609375 0 3185.963
1189.6031494140625 0 2514.6267
1205.6129150390625 0 13806.445 y 15
1206.61474609375 0 9617.031
1207.6143798828125 0 3894.9204
1208.6077880859375 0 1194.613
1226.6357421875 0 1110.3541 b Water loss 23
1227.6390380859375 0 1031.4103
1235.6380615234375 0 1097.7272 b 23
1236.635498046875 0 947.5909
1240.66064453125 0 933.79584 b 11
1241.6646728515625 0 1208.3395
1270.6705322265625 0 781.6116
1272.144287109375 0 1216.6423 y Ammonia loss 2
1273.633544921875 0 1064.2032
1280.65966796875 0 1368.3512 b Ammonia loss 12
1281.15771484375 0 3465.9988
1281.6578369140625 0 2448.5151
1282.1590576171875 0 960.29504
1292.656005859375 0 1196.4025 b 24
1293.155517578125 0 1041.5591
1294.1488037109375 0 1183.9158
1298.657958984375 0 1045.5924
1301.6461181640625 0 1718.1078
1302.171142578125 0 949.51385
1308.6397705078125 0 928.71204
1316.6483154296875 0 2443.3667 y Water loss 14
1317.6370849609375 0 3003.3342 y Ammonia loss 14
1318.6385498046875 0 1169.7832
1319.641357421875 0 1278.288
1326.6605224609375 0 1277.2574
1327.6639404296875 0 834.29364
1334.654296875 0 12659.149 y 14
1335.6583251953125 0 8732.424
1336.6585693359375 0 3165.4663
1337.652587890625 0 950.51874
1344.6573486328125 0 1232.9315
1345.65673828125 0 1617.5895
1356.6866455078125 0 844.1905 Precursor Water loss
1396.7645263671875 0 1895.765 b 13
1397.7615966796875 0 1381.5917
1416.6822509765625 0 818.8921 y Ammonia loss 13
1433.721923828125 0 3130.3286 y 13
1434.721923828125 0 2648.1519
1435.71875 0 1494.8334
1472.7366943359375 0 2578.0625 y Water loss 12
1473.7313232421875 0 2587.063 y Ammonia loss 12
1474.7196044921875 0 1910.4507
1490.743896484375 0 9712.881 y 12
1491.748291015625 0 9153.73
1492.7486572265625 0 4471.9673
1493.7388916015625 0 778.432
1525.7918701171875 0 1154.6005 b 14
1526.7923583984375 0 1500.7866
1543.783935546875 0 969.3817 y Water loss 11
1544.7672119140625 0 1548.6519 y Ammonia loss 11
1545.7650146484375 0 1286.0592
1546.75146484375 0 1017.832
1561.7796630859375 0 7892.3022 y 11
1562.7850341796875 0 5827.135
1563.7841796875 0 4112.0723
1564.7723388671875 0 832.0534
1689.87548828125 0 1186.3474 y 10
1690.873779296875 0 1266.8025
1730.8779296875 0 708.63666
1789.921875 0 876.3959

Spectrum Details

|  |  |
| --- | --- |
| Matched peaks? Matched peaks The total absolute number of peaks matched. Additionally in brackets the total fraction of peaks matched and the total number of peaks is shown. | 123 (16.64% of 739) |
| FDR? FDR The false discovery rate estimated for this peptide. It is calculated by matching all theoretical fragments with a non-integer shift with the raw peaks for this spectrum. This is done with 40 different shifts. The resulting percentage is the average number of annotated peaks over the number of annotated peaks with the correct spectrum. | 0.21% |
| Satellite FDR? Satellite FDR See the FDR for details on its calculation. This satellite ion specific FDR only contains the satellite ions (d/w) for I/L/J positions. | - |
| PSM Score? PSM Score The PSM Score as given by Hecklib to this annotated spectrum. It is shown with three significant figures. | 738 |

#### Reverse Lookup? Reverse Lookup All places where this read could be placed.

| Group | Segment | Template | Template Part | Read Part | Score | Unique |
| --- | --- | --- | --- | --- | --- | --- |
| Homo sapiens Light Chain | IGLC | IGLC2 | [39..65] | [0..26] | 196 | False |
| Homo sapiens Light Chain | IGLC | IGLC3 | [37..63] | [0..26] | 196 | False |

| Recombined | Template Part | Read Part | Score | Unique |
| --- | --- | --- | --- | --- |
| REC-0-1\_002 | [150..176] | [0..26] | 196 | True |

#### Meta Information from Multiple reads

##### Number of combined reads

2

##### Intensity

0.5735

##### TotalArea

1.267E+07

#### Positional Score

Copy Data

##### Positional Score (TSV)

###### Preview

```
Loading example...
```

*Click on the button to copy the data to your clipboard.*

00024681012141618202224

Label Value
"0" 0
"" 0
"2" 0
"" 0
"4" 0
"" 0
"6" 0
"" 0
"8" 0
"" 0
"10" 0
"" 0
"12" 0
"" 0
"14" 0
"" 0
"16" 0
"" 0
"18" 0
"" 0
"20" 0
"" 0
"22" 0
"" 0
"24" 0
"" 0

#### Meta Information from PEAKS

##### Scan Identifier

F2:3914

##### Original sequence

V

A

W

K

A

D

S

S

V

P

K

A

G

V

E

T

T

T

P

S

K

Q

S

N

N

K

##### Posttranslational Modifications

##### Source File

D:\separate\_stitch\_analyses\xle-disambiguation\raw\20210323\_F1\_UM1\_Peng0013\_SA\_F59\_ingel\_3ug\_TL.raw

##### Fraction

2

##### Scan Feature

F2:17434

##### De Novo Score

98

##### ConfidenceScore

98

### m/z

910.8077

##### Mass

2729.3982

##### Charge

3

##### Retention Time

20.71

##### Predicted Retention Time

-

##### Area

6.336E+06

##### Parts Per Million

1.2

##### Fragmentation mode

HCD

##### Originating file

01 D:\separate\_stitch\_analyses\xle-disambiguation\20210325\_F59\_3ug\_DENOVO\_12.csv

#### Meta Information from PEAKS

##### Scan Identifier

F2:3895

##### Original sequence

V

A

W

K

A

D

S

S

V

P

K

A

G

V

E

T

T

T

P

S

K

Q

S

N

N

K

##### Posttranslational Modifications

##### Source File

D:\separate\_stitch\_analyses\xle-disambiguation\raw\20210323\_F1\_UM1\_Peng0013\_SA\_F59\_ingel\_3ug\_TL.raw

##### Fraction

2

##### Scan Feature

F2:17434

##### De Novo Score

98

##### ConfidenceScore

98

### m/z

910.8077

##### Mass

2729.3982

##### Charge

3

##### Retention Time

20.71

##### Predicted Retention Time

-

##### Area

6.336E+06

##### Parts Per Million

1.2

##### Fragmentation mode

HCD

##### Originating file

01 D:\separate\_stitch\_analyses\xle-disambiguation\20210325\_F59\_3ug\_DENOVO\_12.csv
