## Supplementary material for "A handle on mass coincidence errors in *de novo* sequencing of antibodies by bottom-up proteomics": Combined_048.html

Details Combined\_048 | Stitch OverviewUndefined

### Read Combined\_048

#### Sequence (length=13)

VSNKALPAPIEKT

#### Spectrum 4088? Spectrum 4088 The raw spectrum of this peptide as annotated by Hecklib. The fragments are coloured according to ion type (see legend). Any peaks with a star '\*' as text can be hovered over to see the full details, first the ion type second the mass shift type. By hovering over the amino acids in the peptide or ions in the legend the corresponding peaks are highlighted. By toggling the 'Unassigned' label you can turn the background (unassigned) peaks on or off in the plot. By updating the slider in the Ion legend you can update the spectrum to only show the top X% of the peaks with labels. The top X% means any peak that is within X% of the highest intensity. By dragging in the spectrum you can zoom in to a specific part of the spectrum and use 'Zoom Out' to get back to the original zoom level. The annotation of the spectrum is based on the given sequence in the peptides file and is done with different software so inconsistencies are likely. The peaks are annotated based on the given sequence, with 20 ppm tolerance.

Copy Data

##### Spectrum 4088 (TSV)

###### Preview

```
Loading example...
```

*Click on the button to copy the data to your clipboard.*

Mz MinMz MaxIntensity Max

WidthHeightPeptide font sizePeptide stroke widthSpectrum font sizeSpectrum stroke widthCompact peptide

Ion legend

wxyz

abcd

OtherUnassignedIon^Charge^_Position_Show for top:%

VSNKALPAPIEKT

01.62e+53.24e+54.86e+56.48e+5

Zoom Out

y^+1^_1_a^+1^_2_d^+1^_2_a^+1^_2_b^+1^_2_b^+1^_2_y^+3^_6_y^+1^_2_y^+1^_2_y^+2^_4_b^+2^_5_y^+1^_2_b^+2^_5_a^+1^_3_a^+1^_3_b^+1^_3_b^+1^_3_y^+3^_8_y^+2^_5_y^+2^_5_y^+2^_5_b^+2^_6_b^+2^_6_b^+1^_3_b^+2^_6_y^+2^_6_b^+3^_10_b^+2^_7_b^+2^_7_b^+2^_7_y^+1^_3_y^+1^_3_y^+2^_7_b^+3^_11_y^+1^_3_y^+2^_7_b^+2^_8_b^+2^_8_b^+2^_8_b^+1^_4_b^+1^_4_y^+2^_8_b^+1^_4_b^+2^_9_y^+2^_8_b^+2^_9_\*\*\*y^+2^_9_y^+1^_4_b^+1^_5_b^+2^_10_y^+1^_4_b^+1^_5_y^+2^_10_y^+2^_10_b^+2^_11_y^+1^_5_y^+2^_11_y^+2^_11_y^+1^_5_y^+2^_11_b^+1^_6_b^+1^_6_b^+1^_6_y^+2^_12_y^+2^_12_y^+2^_12_y^+1^_6_\*b^+1^_7_y^+1^_7_y^+1^_7_b^+1^_8_b^+1^_8_b^+1^_8_y^+1^_8_y^+1^_8_y^+1^_9_y^+1^_9_y^+1^_9_b^+1^_10_y^+1^_10_b^+1^_11_

0776155223293105

Fragment Matches Table

Show background peaks

| Position | Ion type | Intensity | mz Theoretical | mz Error (Th) | mz Error (ppm) | Charge | Series Number |
| --- | --- | --- | --- | --- | --- | --- | --- |
| 13 | y | 3.625E+04 | 120.1 | 0.0005891 | 4.906 | +1 | 1 |
| - | - | 1670 | 120.1 | - | - | 0 | - |
| - | - | 1799 | 121.1 | - | - | 0 | - |
| - | - | 372.6 | 121.2 | - | - | 0 | - |
| - | - | 416.5 | 122.1 | - | - | 0 | - |
| - | - | 919.2 | 123.1 | - | - | 0 | - |
| - | - | 717.6 | 124.1 | - | - | 0 | - |
| - | - | 1209 | 125.1 | - | - | 0 | - |
| - | - | 805.2 | 126.1 | - | - | 0 | - |
| - | - | 843.5 | 126.1 | - | - | 0 | - |
| - | - | 2298 | 127.1 | - | - | 0 | - |
| - | - | 887.6 | 127.1 | - | - | 0 | - |
| - | - | 417.7 | 127.7 | - | - | 0 | - |
| - | - | 698.8 | 128.1 | - | - | 0 | - |
| - | - | 2322 | 129.1 | - | - | 0 | - |
| - | - | 4.936E+05 | 129.1 | - | - | 0 | - |
| - | - | 2091 | 130.1 | - | - | 0 | - |
| - | - | 4439 | 130.1 | - | - | 0 | - |
| - | - | 3336 | 130.1 | - | - | 0 | - |
| - | - | 2.936E+04 | 130.1 | - | - | 0 | - |
| - | - | 544.8 | 131.1 | - | - | 0 | - |
| - | - | 449 | 131.1 | - | - | 0 | - |
| - | - | 689.2 | 131.1 | - | - | 0 | - |
| - | - | 1.361E+04 | 132.1 | - | - | 0 | - |
| - | - | 362.9 | 133.1 | - | - | 0 | - |
| - | - | 778.1 | 133.1 | - | - | 0 | - |
| - | - | 393.6 | 133.9 | - | - | 0 | - |
| - | - | 389.9 | 135.1 | - | - | 0 | - |
| - | - | 861.7 | 136.1 | - | - | 0 | - |
| - | - | 1486 | 137.1 | - | - | 0 | - |
| - | - | 2845 | 138.1 | - | - | 0 | - |
| - | - | 9759 | 139.1 | - | - | 0 | - |
| - | - | 774.8 | 140.1 | - | - | 0 | - |
| - | - | 1.013E+04 | 140.1 | - | - | 0 | - |
| 2 | a | 1.298E+05 | 141.1 | 0.00062 | 4.394 | +1 | 2 |
| - | - | 1168 | 141.1 | - | - | 0 | - |
| - | - | 1437 | 142.1 | - | - | 0 | - |
| - | - | 1160 | 142.1 | - | - | 0 | - |
| - | - | 1.114E+04 | 142.1 | - | - | 0 | - |
| - | - | 752.7 | 142.1 | - | - | 0 | - |
| - | - | 465 | 143 | - | - | 0 | - |
| - | - | 467.8 | 143.1 | - | - | 0 | - |
| 2 | d | 5103 | 143.1 | 0.0006102 | 4.264 | +1 | 2 |
| - | - | 757.2 | 146.1 | - | - | 0 | - |
| - | - | 661.4 | 146.1 | - | - | 0 | - |
| - | - | 1.323E+04 | 147.1 | - | - | 0 | - |
| - | - | 753.1 | 148.1 | - | - | 0 | - |
| - | - | 1575 | 151.1 | - | - | 0 | - |
| - | - | 550.8 | 152.1 | - | - | 0 | - |
| - | - | 792.3 | 152.1 | - | - | 0 | - |
| - | - | 7847 | 152.1 | - | - | 0 | - |
| - | - | 3134 | 153.1 | - | - | 0 | - |
| - | - | 818.4 | 153.1 | - | - | 0 | - |
| - | - | 891.9 | 154.1 | - | - | 0 | - |
| - | - | 548.2 | 155.1 | - | - | 0 | - |
| - | - | 2.219E+04 | 155.1 | - | - | 0 | - |
| - | - | 1086 | 156.1 | - | - | 0 | - |
| - | - | 1042 | 156.1 | - | - | 0 | - |
| - | - | 2506 | 156.1 | - | - | 0 | - |
| - | - | 7643 | 157.1 | - | - | 0 | - |
| - | - | 1754 | 157.1 | - | - | 0 | - |
| - | - | 2.131E+04 | 157.1 | - | - | 0 | - |
| - | - | 1285 | 158.1 | - | - | 0 | - |
| 2 | a | 6.203E+04 | 159.1 | 0.0006144 | 3.862 | +1 | 2 |
| - | - | 4180 | 160.1 | - | - | 0 | - |
| - | - | 707.8 | 163.1 | - | - | 0 | - |
| - | - | 6318 | 165.1 | - | - | 0 | - |
| - | - | 701 | 165.1 | - | - | 0 | - |
| - | - | 2678 | 167 | - | - | 0 | - |
| - | - | 4309 | 167.1 | - | - | 0 | - |
| - | - | 4177 | 167.1 | - | - | 0 | - |
| - | - | 677.3 | 168 | - | - | 0 | - |
| - | - | 730.8 | 168.1 | - | - | 0 | - |
| - | - | 879.4 | 168.1 | - | - | 0 | - |
| 2 | b | 1.446E+05 | 169.1 | 0.0006853 | 4.052 | +1 | 2 |
| - | - | 7373 | 169.1 | - | - | 0 | - |
| - | - | 1045 | 169.2 | - | - | 0 | - |
| - | - | 583.4 | 170.1 | - | - | 0 | - |
| - | - | 779.5 | 170.1 | - | - | 0 | - |
| - | - | 1.303E+04 | 170.1 | - | - | 0 | - |
| - | - | 749.1 | 170.1 | - | - | 0 | - |
| - | - | 965.6 | 171.1 | - | - | 0 | - |
| - | - | 5115 | 171.1 | - | - | 0 | - |
| - | - | 2362 | 172.1 | - | - | 0 | - |
| - | - | 857.9 | 173.1 | - | - | 0 | - |
| - | - | 1683 | 173.5 | - | - | 0 | - |
| - | - | 3336 | 174.1 | - | - | 0 | - |
| - | - | 643.6 | 175.3 | - | - | 0 | - |
| - | - | 845.1 | 177.1 | - | - | 0 | - |
| - | - | 773.6 | 179.1 | - | - | 0 | - |
| - | - | 1030 | 179.1 | - | - | 0 | - |
| - | - | 1462 | 180.1 | - | - | 0 | - |
| - | - | 537.4 | 180.9 | - | - | 0 | - |
| - | - | 3619 | 181.1 | - | - | 0 | - |
| - | - | 2.004E+04 | 181.1 | - | - | 0 | - |
| - | - | 1.643E+04 | 182.1 | - | - | 0 | - |
| - | - | 2326 | 182.1 | - | - | 0 | - |
| - | - | 3704 | 183.1 | - | - | 0 | - |
| - | - | 2.908E+05 | 183.1 | - | - | 0 | - |
| - | - | 5822 | 184.1 | - | - | 0 | - |
| - | - | 6833 | 184.1 | - | - | 0 | - |
| - | - | 3.091E+04 | 184.2 | - | - | 0 | - |
| - | - | 5359 | 185.1 | - | - | 0 | - |
| - | - | 2.957E+04 | 185.1 | - | - | 0 | - |
| - | - | 1549 | 185.2 | - | - | 0 | - |
| - | - | 7408 | 185.2 | - | - | 0 | - |
| - | - | 1119 | 186.1 | - | - | 0 | - |
| - | - | 1628 | 186.1 | - | - | 0 | - |
| - | - | 2557 | 186.1 | - | - | 0 | - |
| - | - | 1289 | 186.2 | - | - | 0 | - |
| - | - | 676.1 | 186.2 | - | - | 0 | - |
| 2 | b | 6.093E+04 | 187.1 | 0.0007407 | 3.959 | +1 | 2 |
| - | - | 2718 | 187.1 | - | - | 0 | - |
| - | - | 5147 | 188.1 | - | - | 0 | - |
| - | - | 1639 | 191.1 | - | - | 0 | - |
| - | - | 591.5 | 191.1 | - | - | 0 | - |
| - | - | 1401 | 193.1 | - | - | 0 | - |
| - | - | 1559 | 193.1 | - | - | 0 | - |
| - | - | 1623 | 194.1 | - | - | 0 | - |
| - | - | 1.304E+04 | 195.1 | - | - | 0 | - |
| - | - | 741.3 | 195.1 | - | - | 0 | - |
| - | - | 609 | 196.1 | - | - | 0 | - |
| - | - | 1096 | 196.1 | - | - | 0 | - |
| - | - | 4179 | 197.1 | - | - | 0 | - |
| - | - | 4028 | 197.1 | - | - | 0 | - |
| - | - | 1.311E+04 | 197.2 | - | - | 0 | - |
| - | - | 5897 | 198.1 | - | - | 0 | - |
| - | - | 1679 | 198.2 | - | - | 0 | - |
| - | - | 2769 | 199.1 | - | - | 0 | - |
| - | - | 620.3 | 199.1 | - | - | 0 | - |
| - | - | 554.1 | 200.1 | - | - | 0 | - |
| - | - | 5647 | 200.1 | - | - | 0 | - |
| - | - | 1170 | 201.1 | - | - | 0 | - |
| - | - | 1529 | 201.1 | - | - | 0 | - |
| - | - | 949.3 | 201.1 | - | - | 0 | - |
| - | - | 2.858E+04 | 202.1 | - | - | 0 | - |
| - | - | 1023 | 202.2 | - | - | 0 | - |
| - | - | 883.2 | 203.1 | - | - | 0 | - |
| - | - | 1746 | 203.1 | - | - | 0 | - |
| - | - | 1563 | 204.1 | - | - | 0 | - |
| - | - | 1085 | 205.1 | - | - | 0 | - |
| - | - | 1109 | 207.1 | - | - | 0 | - |
| - | - | 748.1 | 207.1 | - | - | 0 | - |
| - | - | 1.266E+04 | 208.1 | - | - | 0 | - |
| - | - | 591.8 | 208.1 | - | - | 0 | - |
| - | - | 2.5E+04 | 209.1 | - | - | 0 | - |
| - | - | 1107 | 209.1 | - | - | 0 | - |
| - | - | 1620 | 209.1 | - | - | 0 | - |
| - | - | 3588 | 209.2 | - | - | 0 | - |
| - | - | 2007 | 210.1 | - | - | 0 | - |
| - | - | 5432 | 210.1 | - | - | 0 | - |
| - | - | 990.8 | 210.1 | - | - | 0 | - |
| - | - | 2214 | 211.1 | - | - | 0 | - |
| - | - | 1.6E+05 | 211.1 | - | - | 0 | - |
| - | - | 2093 | 212.1 | - | - | 0 | - |
| - | - | 878.2 | 212.1 | - | - | 0 | - |
| - | - | 1.903E+04 | 212.1 | - | - | 0 | - |
| - | - | 861.6 | 213.1 | - | - | 0 | - |
| - | - | 1.047E+04 | 213.1 | - | - | 0 | - |
| - | - | 870.2 | 213.2 | - | - | 0 | - |
| - | - | 4608 | 213.2 | - | - | 0 | - |
| - | - | 1200 | 214.1 | - | - | 0 | - |
| - | - | 4579 | 214.1 | - | - | 0 | - |
| 8 | y | 811.2 | 214.1 | 0.002271 | 10.6 | +3 | 6 |
| - | - | 909 | 214.6 | - | - | 0 | - |
| - | - | 1.728E+04 | 215.1 | - | - | 0 | - |
| - | - | 1160 | 216.1 | - | - | 0 | - |
| - | - | 516.3 | 216.6 | - | - | 0 | - |
| - | - | 1416 | 217.2 | - | - | 0 | - |
| - | - | 645.6 | 218.1 | - | - | 0 | - |
| - | - | 2988 | 218.2 | - | - | 0 | - |
| - | - | 1.281E+04 | 219.1 | - | - | 0 | - |
| - | - | 1156 | 219.6 | - | - | 0 | - |
| - | - | 1042 | 220.1 | - | - | 0 | - |
| - | - | 999.3 | 221.1 | - | - | 0 | - |
| - | - | 801.9 | 221.1 | - | - | 0 | - |
| - | - | 4.202E+04 | 222.1 | - | - | 0 | - |
| - | - | 4362 | 223.1 | - | - | 0 | - |
| - | - | 5332 | 223.1 | - | - | 0 | - |
| - | - | 5160 | 223.2 | - | - | 0 | - |
| - | - | 1265 | 224.1 | - | - | 0 | - |
| - | - | 1435 | 225.1 | - | - | 0 | - |
| - | - | 7089 | 225.1 | - | - | 0 | - |
| - | - | 552.1 | 225.2 | - | - | 0 | - |
| - | - | 7387 | 225.6 | - | - | 0 | - |
| - | - | 6.291E+04 | 226.1 | - | - | 0 | - |
| - | - | 2397 | 226.1 | - | - | 0 | - |
| - | - | 1615 | 226.2 | - | - | 0 | - |
| - | - | 1828 | 227.1 | - | - | 0 | - |
| - | - | 5407 | 227.1 | - | - | 0 | - |
| - | - | 7021 | 227.2 | - | - | 0 | - |
| - | - | 1633 | 228.1 | - | - | 0 | - |
| - | - | 891.6 | 228.1 | - | - | 0 | - |
| - | - | 8306 | 228.1 | - | - | 0 | - |
| - | - | 2128 | 228.2 | - | - | 0 | - |
| - | - | 1820 | 229.1 | - | - | 0 | - |
| - | - | 2106 | 229.1 | - | - | 0 | - |
| - | - | 1049 | 229.1 | - | - | 0 | - |
| - | - | 1230 | 229.2 | - | - | 0 | - |
| - | - | 661.7 | 230.1 | - | - | 0 | - |
| 12 | y | 2.507E+04 | 230.1 | 0.0008694 | 3.778 | +1 | 2 |
| 12 | y | 2.117E+04 | 231.1 | 0.0007863 | 3.402 | +1 | 2 |
| - | - | 2574 | 231.2 | - | - | 0 | - |
| - | - | 2165 | 232.1 | - | - | 0 | - |
| - | - | 2604 | 234.1 | - | - | 0 | - |
| - | - | 1026 | 234.6 | - | - | 0 | - |
| - | - | 698.6 | 235.1 | - | - | 0 | - |
| - | - | 1222 | 235.6 | - | - | 0 | - |
| - | - | 835 | 236.1 | - | - | 0 | - |
| - | - | 1576 | 236.1 | - | - | 0 | - |
| - | - | 1907 | 236.2 | - | - | 0 | - |
| - | - | 1.393E+04 | 236.6 | - | - | 0 | - |
| - | - | 880.9 | 237.1 | - | - | 0 | - |
| 10 | y | 835.5 | 237.1 | 0.003518 | 14.83 | +2 | 4 |
| - | - | 3417 | 237.1 | - | - | 0 | - |
| - | - | 3646 | 237.2 | - | - | 0 | - |
| - | - | 2164 | 238.1 | - | - | 0 | - |
| - | - | 2196 | 239.1 | - | - | 0 | - |
| - | - | 2221 | 239.2 | - | - | 0 | - |
| - | - | 3.913E+05 | 240.1 | - | - | 0 | - |
| - | - | 932.4 | 240.6 | - | - | 0 | - |
| - | - | 1063 | 240.6 | - | - | 0 | - |
| - | - | 1187 | 241 | - | - | 0 | - |
| - | - | 6233 | 241.1 | - | - | 0 | - |
| - | - | 4.286E+04 | 241.1 | - | - | 0 | - |
| - | - | 579.2 | 241.6 | - | - | 0 | - |
| 5 | b | 2806 | 241.6 | 0.0005872 | 2.43 | +2 | 5 |
| - | - | 3883 | 242.1 | - | - | 0 | - |
| - | - | 1.823E+04 | 243.1 | - | - | 0 | - |
| - | - | 4.841E+04 | 243.1 | - | - | 0 | - |
| - | - | 1665 | 243.2 | - | - | 0 | - |
| - | - | 1.516E+04 | 243.7 | - | - | 0 | - |
| - | - | 1540 | 244.1 | - | - | 0 | - |
| - | - | 7736 | 244.2 | - | - | 0 | - |
| - | - | 1158 | 244.2 | - | - | 0 | - |
| - | - | 636.2 | 245.2 | - | - | 0 | - |
| - | - | 536 | 247.1 | - | - | 0 | - |
| 12 | y | 1.494E+05 | 248.2 | 0.0009706 | 3.911 | +1 | 2 |
| - | - | 1282 | 249.1 | - | - | 0 | - |
| - | - | 1.595E+04 | 249.2 | - | - | 0 | - |
| - | - | 1.63E+04 | 249.6 | - | - | 0 | - |
| - | - | 970.1 | 249.7 | - | - | 0 | - |
| - | - | 934.1 | 250.1 | - | - | 0 | - |
| - | - | 3877 | 250.1 | - | - | 0 | - |
| - | - | 980.4 | 250.2 | - | - | 0 | - |
| 5 | b | 6899 | 250.6 | 0.0007217 | 2.879 | +2 | 5 |
| - | - | 2944 | 251.1 | - | - | 0 | - |
| - | - | 6354 | 252.1 | - | - | 0 | - |
| - | - | 5265 | 252.2 | - | - | 0 | - |
| - | - | 4168 | 253.1 | - | - | 0 | - |
| - | - | 3841 | 253.1 | - | - | 0 | - |
| - | - | 727.6 | 253.6 | - | - | 0 | - |
| - | - | 719.9 | 254.1 | - | - | 0 | - |
| - | - | 5104 | 254.2 | - | - | 0 | - |
| - | - | 3060 | 254.2 | - | - | 0 | - |
| - | - | 1246 | 254.7 | - | - | 0 | - |
| - | - | 883.8 | 255.1 | - | - | 0 | - |
| 3 | a | 2148 | 255.1 | 0.003642 | 14.27 | +1 | 3 |
| - | - | 4205 | 255.2 | - | - | 0 | - |
| - | - | 806 | 255.6 | - | - | 0 | - |
| 3 | a | 1853 | 256.1 | 0.00125 | 4.879 | +1 | 3 |
| - | - | 756.2 | 256.2 | - | - | 0 | - |
| - | - | 632 | 257.7 | - | - | 0 | - |
| - | - | 9.073E+04 | 258.1 | - | - | 0 | - |
| - | - | 648.2 | 259.1 | - | - | 0 | - |
| - | - | 1150 | 259.1 | - | - | 0 | - |
| - | - | 9994 | 259.1 | - | - | 0 | - |
| - | - | 797.1 | 260.2 | - | - | 0 | - |
| - | - | 2495 | 261.1 | - | - | 0 | - |
| - | - | 6518 | 262.1 | - | - | 0 | - |
| - | - | 1866 | 262.6 | - | - | 0 | - |
| - | - | 1210 | 262.7 | - | - | 0 | - |
| - | - | 2916 | 263.2 | - | - | 0 | - |
| - | - | 1.47E+04 | 263.7 | - | - | 0 | - |
| - | - | 771 | 264.1 | - | - | 0 | - |
| - | - | 4038 | 264.2 | - | - | 0 | - |
| - | - | 1.042E+04 | 264.2 | - | - | 0 | - |
| - | - | 1691 | 265.1 | - | - | 0 | - |
| - | - | 4794 | 265.2 | - | - | 0 | - |
| - | - | 924 | 265.2 | - | - | 0 | - |
| - | - | 935 | 266.1 | - | - | 0 | - |
| - | - | 1.975E+04 | 266.2 | - | - | 0 | - |
| - | - | 892.2 | 267.1 | - | - | 0 | - |
| - | - | 2271 | 267.2 | - | - | 0 | - |
| - | - | 639.1 | 267.2 | - | - | 0 | - |
| - | - | 4912 | 268.1 | - | - | 0 | - |
| - | - | 671.4 | 268.2 | - | - | 0 | - |
| - | - | 2226 | 268.2 | - | - | 0 | - |
| - | - | 767.7 | 268.2 | - | - | 0 | - |
| - | - | 3392 | 269.2 | - | - | 0 | - |
| - | - | 1673 | 269.2 | - | - | 0 | - |
| - | - | 2342 | 270.1 | - | - | 0 | - |
| - | - | 916.3 | 270.2 | - | - | 0 | - |
| - | - | 923.7 | 270.7 | - | - | 0 | - |
| - | - | 6301 | 270.7 | - | - | 0 | - |
| - | - | 3328 | 271.1 | - | - | 0 | - |
| - | - | 845.7 | 271.2 | - | - | 0 | - |
| - | - | 3066 | 271.2 | - | - | 0 | - |
| - | - | 3792 | 271.7 | - | - | 0 | - |
| - | - | 1747 | 272.2 | - | - | 0 | - |
| - | - | 2512 | 275.7 | - | - | 0 | - |
| - | - | 1.106E+04 | 276.2 | - | - | 0 | - |
| - | - | 874.2 | 276.2 | - | - | 0 | - |
| - | - | 3641 | 276.7 | - | - | 0 | - |
| - | - | 5356 | 277.1 | - | - | 0 | - |
| - | - | 2678 | 277.2 | - | - | 0 | - |
| - | - | 979.9 | 278.1 | - | - | 0 | - |
| - | - | 885.9 | 278.1 | - | - | 0 | - |
| - | - | 819.1 | 278.2 | - | - | 0 | - |
| - | - | 4742 | 278.2 | - | - | 0 | - |
| - | - | 1.309E+04 | 279.1 | - | - | 0 | - |
| - | - | 557.2 | 279.2 | - | - | 0 | - |
| - | - | 6841 | 280.1 | - | - | 0 | - |
| - | - | 2033 | 280.1 | - | - | 0 | - |
| - | - | 2964 | 280.2 | - | - | 0 | - |
| - | - | 557 | 281.1 | - | - | 0 | - |
| - | - | 671.1 | 281.2 | - | - | 0 | - |
| - | - | 1562 | 281.7 | - | - | 0 | - |
| - | - | 688.3 | 282.1 | - | - | 0 | - |
| - | - | 875.5 | 282.2 | - | - | 0 | - |
| - | - | 1.941E+04 | 282.2 | - | - | 0 | - |
| 3 | b | 1.336E+04 | 283.1 | 0.001123 | 3.967 | +1 | 3 |
| - | - | 2889 | 283.2 | - | - | 0 | - |
| - | - | 3901 | 283.7 | - | - | 0 | - |
| 3 | b | 844.7 | 284.1 | 0.002246 | 7.904 | +1 | 3 |
| - | - | 1826 | 284.1 | - | - | 0 | - |
| - | - | 1126 | 284.2 | - | - | 0 | - |
| 6 | y | 734.8 | 284.2 | 0.002994 | 10.54 | +3 | 8 |
| - | - | 2575 | 284.2 | - | - | 0 | - |
| - | - | 5512 | 284.2 | - | - | 0 | - |
| - | - | 1.155E+04 | 284.7 | - | - | 0 | - |
| - | - | 2996 | 285.2 | - | - | 0 | - |
| 9 | y | 4.943E+04 | 285.2 | 0.003323 | 11.65 | +2 | 5 |
| - | - | 7505 | 285.2 | - | - | 0 | - |
| 9 | y | 1.172E+04 | 285.7 | 0.0006343 | 2.22 | +2 | 5 |
| - | - | 1.099E+04 | 285.7 | - | - | 0 | - |
| - | - | 685.1 | 286.1 | - | - | 0 | - |
| - | - | 3338 | 286.2 | - | - | 0 | - |
| - | - | 1472 | 286.2 | - | - | 0 | - |
| - | - | 1114 | 286.2 | - | - | 0 | - |
| - | - | 795 | 287.2 | - | - | 0 | - |
| - | - | 1233 | 289.7 | - | - | 0 | - |
| - | - | 4213 | 290.2 | - | - | 0 | - |
| - | - | 2473 | 290.7 | - | - | 0 | - |
| - | - | 776.7 | 290.7 | - | - | 0 | - |
| - | - | 965.9 | 292.2 | - | - | 0 | - |
| - | - | 1.827E+04 | 292.2 | - | - | 0 | - |
| - | - | 5271 | 292.7 | - | - | 0 | - |
| - | - | 840.5 | 293.1 | - | - | 0 | - |
| - | - | 445.6 | 293.2 | - | - | 0 | - |
| - | - | 9.077E+04 | 293.2 | - | - | 0 | - |
| - | - | 2.639E+04 | 293.7 | - | - | 0 | - |
| 9 | y | 4.45E+05 | 294.2 | 0.001154 | 3.922 | +2 | 5 |
| - | - | 1.34E+05 | 294.7 | - | - | 0 | - |
| - | - | 2.105E+04 | 295.1 | - | - | 0 | - |
| - | - | 3.25E+04 | 295.2 | - | - | 0 | - |
| - | - | 4710 | 295.2 | - | - | 0 | - |
| - | - | 4864 | 295.7 | - | - | 0 | - |
| - | - | 2984 | 296.1 | - | - | 0 | - |
| - | - | 1.727E+04 | 296.2 | - | - | 0 | - |
| - | - | 2073 | 296.2 | - | - | 0 | - |
| - | - | 917.7 | 296.2 | - | - | 0 | - |
| - | - | 1.17E+04 | 297.2 | - | - | 0 | - |
| - | - | 3740 | 297.2 | - | - | 0 | - |
| - | - | 7381 | 297.2 | - | - | 0 | - |
| - | - | 1668 | 297.7 | - | - | 0 | - |
| - | - | 2030 | 297.7 | - | - | 0 | - |
| - | - | 1766 | 298.1 | - | - | 0 | - |
| - | - | 1615 | 298.2 | - | - | 0 | - |
| 6 | b | 4861 | 298.2 | 0.0009594 | 3.217 | +2 | 6 |
| 6 | b | 3196 | 298.7 | 0.001383 | 4.631 | +2 | 6 |
| - | - | 3.57E+04 | 299.2 | - | - | 0 | - |
| - | - | 1.168E+04 | 299.7 | - | - | 0 | - |
| - | - | 2818 | 300.2 | - | - | 0 | - |
| - | - | 1393 | 300.2 | - | - | 0 | - |
| - | - | 927.9 | 300.7 | - | - | 0 | - |
| 3 | b | 1.642E+04 | 301.2 | 0.001148 | 3.813 | +1 | 3 |
| - | - | 1081 | 301.2 | - | - | 0 | - |
| - | - | 1758 | 302.2 | - | - | 0 | - |
| - | - | 1679 | 305.1 | - | - | 0 | - |
| - | - | 940.7 | 305.2 | - | - | 0 | - |
| - | - | 1.964E+04 | 306.2 | - | - | 0 | - |
| - | - | 4330 | 306.2 | - | - | 0 | - |
| - | - | 4929 | 306.7 | - | - | 0 | - |
| - | - | 1212 | 306.7 | - | - | 0 | - |
| - | - | 991.1 | 307.1 | - | - | 0 | - |
| 6 | b | 1.968E+04 | 307.2 | 0.000804 | 2.617 | +2 | 6 |
| - | - | 6981 | 307.7 | - | - | 0 | - |
| - | - | 923.2 | 308.2 | - | - | 0 | - |
| - | - | 3021 | 309.2 | - | - | 0 | - |
| - | - | 902.3 | 309.7 | - | - | 0 | - |
| - | - | 1222 | 311.1 | - | - | 0 | - |
| - | - | 718.6 | 311.2 | - | - | 0 | - |
| - | - | 1668 | 311.2 | - | - | 0 | - |
| - | - | 1.623E+04 | 312.2 | - | - | 0 | - |
| - | - | 8440 | 312.2 | - | - | 0 | - |
| - | - | 1713 | 312.7 | - | - | 0 | - |
| - | - | 3.92E+04 | 313.2 | - | - | 0 | - |
| - | - | 2199 | 313.2 | - | - | 0 | - |
| - | - | 2189 | 313.2 | - | - | 0 | - |
| - | - | 1.37E+04 | 313.2 | - | - | 0 | - |
| - | - | 4925 | 314.2 | - | - | 0 | - |
| - | - | 2.271E+04 | 314.2 | - | - | 0 | - |
| - | - | 1907 | 314.2 | - | - | 0 | - |
| - | - | 2318 | 315.2 | - | - | 0 | - |
| - | - | 2977 | 315.2 | - | - | 0 | - |
| - | - | 4394 | 315.2 | - | - | 0 | - |
| - | - | 1.91E+04 | 318.2 | - | - | 0 | - |
| - | - | 1367 | 318.7 | - | - | 0 | - |
| - | - | 3625 | 319.2 | - | - | 0 | - |
| - | - | 1916 | 319.2 | - | - | 0 | - |
| - | - | 1408 | 319.7 | - | - | 0 | - |
| - | - | 947.9 | 320.2 | - | - | 0 | - |
| - | - | 6104 | 320.2 | - | - | 0 | - |
| - | - | 1.541E+04 | 320.7 | - | - | 0 | - |
| - | - | 1359 | 321.2 | - | - | 0 | - |
| - | - | 5634 | 321.2 | - | - | 0 | - |
| - | - | 786 | 321.2 | - | - | 0 | - |
| - | - | 3518 | 322.2 | - | - | 0 | - |
| - | - | 7102 | 322.2 | - | - | 0 | - |
| - | - | 9836 | 323.1 | - | - | 0 | - |
| - | - | 7390 | 323.2 | - | - | 0 | - |
| - | - | 7099 | 323.2 | - | - | 0 | - |
| - | - | 1247 | 324.1 | - | - | 0 | - |
| - | - | 2489 | 324.2 | - | - | 0 | - |
| - | - | 720.7 | 324.2 | - | - | 0 | - |
| - | - | 773.9 | 324.2 | - | - | 0 | - |
| - | - | 2963 | 325.2 | - | - | 0 | - |
| - | - | 6452 | 325.2 | - | - | 0 | - |
| - | - | 1905 | 325.2 | - | - | 0 | - |
| - | - | 6436 | 325.7 | - | - | 0 | - |
| - | - | 2153 | 326.2 | - | - | 0 | - |
| - | - | 898.8 | 326.7 | - | - | 0 | - |
| - | - | 2584 | 327.2 | - | - | 0 | - |
| - | - | 691.1 | 327.2 | - | - | 0 | - |
| - | - | 1.565E+04 | 327.7 | - | - | 0 | - |
| - | - | 5029 | 328.2 | - | - | 0 | - |
| - | - | 2216 | 328.7 | - | - | 0 | - |
| 8 | y | 2454 | 329.7 | 0.001121 | 3.4 | +2 | 6 |
| - | - | 3.875E+04 | 330.2 | - | - | 0 | - |
| - | - | 5895 | 331.2 | - | - | 0 | - |
| 10 | b | 643.2 | 331.2 | 0.003005 | 9.074 | +3 | 10 |
| - | - | 3116 | 331.2 | - | - | 0 | - |
| - | - | 668.6 | 332.2 | - | - | 0 | - |
| - | - | 1824 | 332.7 | - | - | 0 | - |
| - | - | 4057 | 332.7 | - | - | 0 | - |
| - | - | 1.084E+04 | 333.2 | - | - | 0 | - |
| - | - | 3982 | 333.7 | - | - | 0 | - |
| - | - | 783 | 334.2 | - | - | 0 | - |
| - | - | 1.79E+04 | 334.7 | - | - | 0 | - |
| - | - | 5359 | 335.2 | - | - | 0 | - |
| - | - | 739.8 | 335.2 | - | - | 0 | - |
| - | - | 2127 | 335.7 | - | - | 0 | - |
| - | - | 917.3 | 336.2 | - | - | 0 | - |
| - | - | 998.1 | 336.2 | - | - | 0 | - |
| - | - | 614.1 | 337.2 | - | - | 0 | - |
| - | - | 1608 | 337.2 | - | - | 0 | - |
| - | - | 608.3 | 338.2 | - | - | 0 | - |
| - | - | 3168 | 338.2 | - | - | 0 | - |
| - | - | 945.4 | 338.2 | - | - | 0 | - |
| - | - | 1924 | 338.7 | - | - | 0 | - |
| - | - | 3603 | 339.2 | - | - | 0 | - |
| - | - | 2776 | 339.2 | - | - | 0 | - |
| - | - | 1170 | 339.2 | - | - | 0 | - |
| - | - | 1.92E+05 | 340.2 | - | - | 0 | - |
| - | - | 1234 | 341.1 | - | - | 0 | - |
| - | - | 3.782E+04 | 341.2 | - | - | 0 | - |
| - | - | 4939 | 341.2 | - | - | 0 | - |
| - | - | 8392 | 341.7 | - | - | 0 | - |
| - | - | 1.181E+05 | 341.7 | - | - | 0 | - |
| - | - | 2276 | 342.1 | - | - | 0 | - |
| - | - | 4388 | 342.2 | - | - | 0 | - |
| - | - | 3541 | 342.2 | - | - | 0 | - |
| - | - | 4.06E+04 | 342.2 | - | - | 0 | - |
| - | - | 9779 | 342.7 | - | - | 0 | - |
| - | - | 844.9 | 343.2 | - | - | 0 | - |
| - | - | 975.8 | 343.2 | - | - | 0 | - |
| - | - | 975.7 | 344.2 | - | - | 0 | - |
| 7 | b | 8200 | 346.7 | 0.001311 | 3.781 | +2 | 7 |
| 7 | b | 1.522E+04 | 347.2 | 0.002192 | 6.315 | +2 | 7 |
| - | - | 4409 | 347.7 | - | - | 0 | - |
| - | - | 3683 | 348.2 | - | - | 0 | - |
| - | - | 1096 | 349.2 | - | - | 0 | - |
| - | - | 2937 | 349.2 | - | - | 0 | - |
| - | - | 858.5 | 349.2 | - | - | 0 | - |
| - | - | 1069 | 350.2 | - | - | 0 | - |
| - | - | 2340 | 351.2 | - | - | 0 | - |
| - | - | 3581 | 351.2 | - | - | 0 | - |
| - | - | 1.243E+04 | 351.2 | - | - | 0 | - |
| - | - | 924.1 | 352.2 | - | - | 0 | - |
| - | - | 2038 | 352.2 | - | - | 0 | - |
| - | - | 691.4 | 353.1 | - | - | 0 | - |
| - | - | 783.8 | 353.2 | - | - | 0 | - |
| - | - | 1.322E+04 | 353.2 | - | - | 0 | - |
| - | - | 954.4 | 354.2 | - | - | 0 | - |
| - | - | 2477 | 354.2 | - | - | 0 | - |
| - | - | 3119 | 354.7 | - | - | 0 | - |
| - | - | 872.2 | 355.2 | - | - | 0 | - |
| - | - | 709.8 | 355.2 | - | - | 0 | - |
| 7 | b | 1.276E+05 | 355.7 | 0.001247 | 3.506 | +2 | 7 |
| - | - | 5721 | 356.2 | - | - | 0 | - |
| - | - | 4.579E+04 | 356.2 | - | - | 0 | - |
| - | - | 1.115E+04 | 356.7 | - | - | 0 | - |
| - | - | 819 | 357.2 | - | - | 0 | - |
| - | - | 5417 | 357.2 | - | - | 0 | - |
| - | - | 2789 | 358.2 | - | - | 0 | - |
| - | - | 2729 | 358.2 | - | - | 0 | - |
| 11 | y | 9.249E+04 | 359.2 | 0.001276 | 3.551 | +1 | 3 |
| - | - | 637.5 | 359.7 | - | - | 0 | - |
| - | - | 1390 | 359.7 | - | - | 0 | - |
| 11 | y | 1264 | 360.2 | 0.0004143 | 1.15 | +1 | 3 |
| - | - | 1.521E+04 | 360.2 | - | - | 0 | - |
| - | - | 617.5 | 360.7 | - | - | 0 | - |
| - | - | 2767 | 361.2 | - | - | 0 | - |
| - | - | 2143 | 363.2 | - | - | 0 | - |
| - | - | 741.7 | 364.2 | - | - | 0 | - |
| - | - | 1628 | 364.2 | - | - | 0 | - |
| - | - | 5316 | 365.2 | - | - | 0 | - |
| - | - | 2685 | 365.2 | - | - | 0 | - |
| - | - | 2310 | 365.3 | - | - | 0 | - |
| - | - | 2.069E+04 | 366.2 | - | - | 0 | - |
| - | - | 2146 | 366.2 | - | - | 0 | - |
| - | - | 1072 | 366.3 | - | - | 0 | - |
| - | - | 893.3 | 367.2 | - | - | 0 | - |
| - | - | 2794 | 367.2 | - | - | 0 | - |
| - | - | 747.1 | 367.3 | - | - | 0 | - |
| - | - | 5549 | 368.2 | - | - | 0 | - |
| - | - | 1.309E+04 | 368.2 | - | - | 0 | - |
| - | - | 9860 | 368.7 | - | - | 0 | - |
| 7 | y | 9930 | 369.2 | 0.003764 | 10.19 | +2 | 7 |
| - | - | 3584 | 369.7 | - | - | 0 | - |
| - | - | 792.3 | 370.2 | - | - | 0 | - |
| - | - | 1775 | 370.2 | - | - | 0 | - |
| - | - | 801.6 | 371.2 | - | - | 0 | - |
| - | - | 2.007E+04 | 371.2 | - | - | 0 | - |
| - | - | 2040 | 372.2 | - | - | 0 | - |
| - | - | 3746 | 372.2 | - | - | 0 | - |
| - | - | 6240 | 373.2 | - | - | 0 | - |
| - | - | 5080 | 373.7 | - | - | 0 | - |
| 11 | b | 2616 | 374.2 | 0.001138 | 3.041 | +3 | 11 |
| - | - | 946.4 | 375.2 | - | - | 0 | - |
| - | - | 756.3 | 375.2 | - | - | 0 | - |
| - | - | 1342 | 376.2 | - | - | 0 | - |
| - | - | 835 | 376.2 | - | - | 0 | - |
| 11 | y | 1.247E+05 | 377.2 | 0.001423 | 3.771 | +1 | 3 |
| - | - | 6.88E+04 | 377.2 | - | - | 0 | - |
| - | - | 391.7 | 377.7 | - | - | 0 | - |
| - | - | 3.122E+04 | 377.7 | - | - | 0 | - |
| 7 | y | 1.095E+05 | 378.2 | 0.0006179 | 1.634 | +2 | 7 |
| - | - | 4.587E+04 | 378.7 | - | - | 0 | - |
| - | - | 1.65E+04 | 379.2 | - | - | 0 | - |
| - | - | 3141 | 379.7 | - | - | 0 | - |
| - | - | 4507 | 380.2 | - | - | 0 | - |
| - | - | 1.043E+04 | 381.2 | - | - | 0 | - |
| 8 | b | 5123 | 382.2 | 0.0002173 | 0.5684 | +2 | 8 |
| - | - | 2617 | 382.2 | - | - | 0 | - |
| - | - | 1142 | 382.3 | - | - | 0 | - |
| 8 | b | 7153 | 382.7 | 0.003014 | 7.876 | +2 | 8 |
| - | - | 4.727E+04 | 383.2 | - | - | 0 | - |
| - | - | 3980 | 383.2 | - | - | 0 | - |
| - | - | 1453 | 383.7 | - | - | 0 | - |
| - | - | 1.682E+04 | 384.2 | - | - | 0 | - |
| - | - | 3630 | 384.2 | - | - | 0 | - |
| - | - | 1012 | 384.3 | - | - | 0 | - |
| - | - | 751.8 | 384.3 | - | - | 0 | - |
| - | - | 3065 | 385.2 | - | - | 0 | - |
| - | - | 1813 | 385.2 | - | - | 0 | - |
| - | - | 1119 | 385.3 | - | - | 0 | - |
| - | - | 1307 | 385.5 | - | - | 0 | - |
| - | - | 2103 | 385.9 | - | - | 0 | - |
| - | - | 948.2 | 386.2 | - | - | 0 | - |
| - | - | 2882 | 386.2 | - | - | 0 | - |
| - | - | 2843 | 387.2 | - | - | 0 | - |
| - | - | 763.2 | 387.2 | - | - | 0 | - |
| - | - | 4846 | 389.2 | - | - | 0 | - |
| - | - | 1735 | 390.2 | - | - | 0 | - |
| - | - | 952.2 | 390.2 | - | - | 0 | - |
| 8 | b | 3.802E+04 | 391.2 | 0.001458 | 3.728 | +2 | 8 |
| - | - | 1.403E+04 | 391.7 | - | - | 0 | - |
| - | - | 2686 | 392.2 | - | - | 0 | - |
| - | - | 4604 | 392.2 | - | - | 0 | - |
| - | - | 1085 | 392.3 | - | - | 0 | - |
| - | - | 2600 | 393.2 | - | - | 0 | - |
| - | - | 2298 | 393.2 | - | - | 0 | - |
| - | - | 2812 | 394.2 | - | - | 0 | - |
| - | - | 8668 | 394.2 | - | - | 0 | - |
| - | - | 1190 | 395.2 | - | - | 0 | - |
| - | - | 3269 | 396.2 | - | - | 0 | - |
| - | - | 619.1 | 396.3 | - | - | 0 | - |
| - | - | 1516 | 398.2 | - | - | 0 | - |
| - | - | 5163 | 398.2 | - | - | 0 | - |
| - | - | 995.2 | 399.2 | - | - | 0 | - |
| - | - | 3460 | 399.2 | - | - | 0 | - |
| - | - | 1.369E+04 | 399.3 | - | - | 0 | - |
| - | - | 2570 | 400.3 | - | - | 0 | - |
| - | - | 7.224E+04 | 401.2 | - | - | 0 | - |
| - | - | 1.275E+04 | 402.2 | - | - | 0 | - |
| - | - | 2079 | 403.2 | - | - | 0 | - |
| - | - | 1234 | 404.2 | - | - | 0 | - |
| - | - | 852.4 | 404.7 | - | - | 0 | - |
| - | - | 691 | 406.2 | - | - | 0 | - |
| - | - | 1993 | 408.2 | - | - | 0 | - |
| - | - | 8119 | 409.2 | - | - | 0 | - |
| - | - | 2020 | 409.3 | - | - | 0 | - |
| - | - | 3403 | 410.2 | - | - | 0 | - |
| - | - | 8676 | 410.2 | - | - | 0 | - |
| - | - | 6471 | 410.3 | - | - | 0 | - |
| - | - | 6168 | 411.2 | - | - | 0 | - |
| 4 | b | 3.272E+04 | 411.2 | 0.002867 | 6.971 | +1 | 4 |
| - | - | 1243 | 411.3 | - | - | 0 | - |
| - | - | 1072 | 412.2 | - | - | 0 | - |
| 4 | b | 1.37E+04 | 412.2 | 0.004145 | 10.06 | +1 | 4 |
| - | - | 7042 | 412.3 | - | - | 0 | - |
| - | - | 2711 | 413.2 | - | - | 0 | - |
| - | - | 854.1 | 413.3 | - | - | 0 | - |
| - | - | 1098 | 413.3 | - | - | 0 | - |
| - | - | 1268 | 414.2 | - | - | 0 | - |
| - | - | 2014 | 415.2 | - | - | 0 | - |
| - | - | 1258 | 417.2 | - | - | 0 | - |
| - | - | 906.7 | 419.2 | - | - | 0 | - |
| - | - | 2556 | 420.2 | - | - | 0 | - |
| - | - | 710.9 | 420.3 | - | - | 0 | - |
| - | - | 772.8 | 421.2 | - | - | 0 | - |
| - | - | 2049 | 422.2 | - | - | 0 | - |
| - | - | 1560 | 422.2 | - | - | 0 | - |
| - | - | 1072 | 424.2 | - | - | 0 | - |
| - | - | 745.3 | 424.3 | - | - | 0 | - |
| - | - | 813.5 | 425.2 | - | - | 0 | - |
| - | - | 4254 | 425.3 | - | - | 0 | - |
| 6 | y | 1341 | 425.8 | 0.007844 | 18.42 | +2 | 8 |
| - | - | 2057 | 426.2 | - | - | 0 | - |
| - | - | 1090 | 426.3 | - | - | 0 | - |
| - | - | 3.131E+04 | 427.2 | - | - | 0 | - |
| - | - | 3.063E+04 | 427.3 | - | - | 0 | - |
| - | - | 1012 | 427.3 | - | - | 0 | - |
| - | - | 4607 | 428.2 | - | - | 0 | - |
| - | - | 7504 | 428.3 | - | - | 0 | - |
| - | - | 6576 | 429.2 | - | - | 0 | - |
| 4 | b | 6.858E+04 | 429.2 | 0.001705 | 3.973 | +1 | 4 |
| - | - | 1142 | 430.2 | - | - | 0 | - |
| - | - | 1.594E+04 | 430.3 | - | - | 0 | - |
| 9 | b | 3071 | 431.2 | 0.00715 | 16.58 | +2 | 9 |
| - | - | 978.4 | 434.2 | - | - | 0 | - |
| - | - | 908.4 | 434.2 | - | - | 0 | - |
| 6 | y | 2276 | 434.8 | 0.002012 | 4.629 | +2 | 8 |
| - | - | 2256 | 435.2 | - | - | 0 | - |
| - | - | 1935 | 435.3 | - | - | 0 | - |
| - | - | 1786 | 436.3 | - | - | 0 | - |
| - | - | 3795 | 437.2 | - | - | 0 | - |
| - | - | 5079 | 437.3 | - | - | 0 | - |
| - | - | 4194 | 438.2 | - | - | 0 | - |
| - | - | 1879 | 438.3 | - | - | 0 | - |
| - | - | 8541 | 439.2 | - | - | 0 | - |
| - | - | 919.4 | 439.3 | - | - | 0 | - |
| 9 | b | 7953 | 439.8 | 0.001688 | 3.838 | +2 | 9 |
| - | - | 3277 | 440.3 | - | - | 0 | - |
| - | - | 1639 | 440.8 | - | - | 0 | - |
| - | - | 1286 | 441.2 | - | - | 0 | - |
| - | - | 827.8 | 441.3 | - | - | 0 | - |
| - | - | 5784 | 443.3 | - | - | 0 | - |
| - | - | 2540 | 444.3 | - | - | 0 | - |
| - | - | 957.1 | 446.2 | - | - | 0 | - |
| - | - | 1545 | 446.3 | - | - | 0 | - |
| - | - | 2918 | 447.2 | - | - | 0 | - |
| - | - | 627.9 | 447.3 | - | - | 0 | - |
| - | - | 1543 | 448.2 | - | - | 0 | - |
| - | - | 902 | 449.2 | - | - | 0 | - |
| - | - | 1462 | 450.2 | - | - | 0 | - |
| - | - | 6802 | 450.3 | - | - | 0 | - |
| 0 | Precursor | 1327 | 450.6 | 0.00304 | 6.746 | +3 | -1 |
| 0 | Precursor | 933.5 | 450.9 | 0.006211 | 13.77 | +3 | -1 |
| - | - | 4136 | 451.3 | - | - | 0 | - |
| - | - | 677 | 452.2 | - | - | 0 | - |
| - | - | 2891 | 452.3 | - | - | 0 | - |
| - | - | 5094 | 453.2 | - | - | 0 | - |
| - | - | 3619 | 453.3 | - | - | 0 | - |
| - | - | 2139 | 453.3 | - | - | 0 | - |
| - | - | 3861 | 454.3 | - | - | 0 | - |
| - | - | 3537 | 455.2 | - | - | 0 | - |
| - | - | 1.812E+04 | 455.3 | - | - | 0 | - |
| - | - | 984.4 | 456.2 | - | - | 0 | - |
| - | - | 3674 | 456.3 | - | - | 0 | - |
| 0 | Precursor | 1228 | 456.6 | 0.00202 | 4.425 | +3 | -1 |
| - | - | 1952 | 456.9 | - | - | 0 | - |
| - | - | 2201 | 457.3 | - | - | 0 | - |
| - | - | 814.8 | 461.3 | - | - | 0 | - |
| - | - | 2678 | 462.3 | - | - | 0 | - |
| - | - | 5059 | 463.2 | - | - | 0 | - |
| - | - | 895 | 463.3 | - | - | 0 | - |
| - | - | 1296 | 463.3 | - | - | 0 | - |
| - | - | 8183 | 464.3 | - | - | 0 | - |
| - | - | 978.1 | 464.3 | - | - | 0 | - |
| - | - | 1.305E+04 | 465.2 | - | - | 0 | - |
| - | - | 2821 | 466.2 | - | - | 0 | - |
| - | - | 3395 | 467.3 | - | - | 0 | - |
| - | - | 1214 | 467.3 | - | - | 0 | - |
| - | - | 1.223E+05 | 468.3 | - | - | 0 | - |
| - | - | 3.933E+04 | 469.3 | - | - | 0 | - |
| 5 | y | 1.072E+04 | 470.3 | 0.001316 | 2.799 | +2 | 9 |
| - | - | 1832 | 471.3 | - | - | 0 | - |
| 10 | y | 2.407E+04 | 472.3 | 0.006476 | 13.71 | +1 | 4 |
| - | - | 4982 | 473.3 | - | - | 0 | - |
| - | - | 734.7 | 474.3 | - | - | 0 | - |
| - | - | 8409 | 479.3 | - | - | 0 | - |
| - | - | 1403 | 479.3 | - | - | 0 | - |
| - | - | 5552 | 480.3 | - | - | 0 | - |
| - | - | 6371 | 480.3 | - | - | 0 | - |
| - | - | 6691 | 481.2 | - | - | 0 | - |
| - | - | 1731 | 481.3 | - | - | 0 | - |
| - | - | 2.401E+04 | 481.3 | - | - | 0 | - |
| 5 | b | 1.065E+05 | 482.3 | 0.00189 | 3.918 | +1 | 5 |
| - | - | 5431 | 482.3 | - | - | 0 | - |
| - | - | 2E+04 | 483.3 | - | - | 0 | - |
| - | - | 3151 | 484.3 | - | - | 0 | - |
| - | - | 2795 | 484.3 | - | - | 0 | - |
| - | - | 3395 | 485.3 | - | - | 0 | - |
| - | - | 6.732E+04 | 486.3 | - | - | 0 | - |
| 10 | b | 1.703E+04 | 487.3 | 0.007724 | 15.85 | +2 | 10 |
| - | - | 3270 | 488.3 | - | - | 0 | - |
| 10 | y | 8.66E+04 | 490.3 | 0.001862 | 3.797 | +1 | 4 |
| - | - | 984.6 | 491.2 | - | - | 0 | - |
| - | - | 2.365E+04 | 491.3 | - | - | 0 | - |
| - | - | 3568 | 492.3 | - | - | 0 | - |
| - | - | 1124 | 493.2 | - | - | 0 | - |
| - | - | 1096 | 495.3 | - | - | 0 | - |
| - | - | 594.5 | 496.3 | - | - | 0 | - |
| - | - | 8728 | 496.3 | - | - | 0 | - |
| - | - | 3741 | 496.3 | - | - | 0 | - |
| - | - | 1.53E+04 | 497.3 | - | - | 0 | - |
| - | - | 3760 | 497.3 | - | - | 0 | - |
| - | - | 5.225E+04 | 498.3 | - | - | 0 | - |
| - | - | 4936 | 498.3 | - | - | 0 | - |
| - | - | 1.12E+04 | 499.3 | - | - | 0 | - |
| - | - | 1473 | 499.3 | - | - | 0 | - |
| 5 | b | 1.717E+05 | 500.3 | 0.001365 | 2.729 | +1 | 5 |
| - | - | 4.109E+04 | 501.3 | - | - | 0 | - |
| - | - | 8192 | 502.3 | - | - | 0 | - |
| - | - | 822.5 | 503.3 | - | - | 0 | - |
| - | - | 1394 | 506.3 | - | - | 0 | - |
| - | - | 1226 | 507.3 | - | - | 0 | - |
| - | - | 4273 | 507.3 | - | - | 0 | - |
| - | - | 3.394E+04 | 508.3 | - | - | 0 | - |
| - | - | 1062 | 509.2 | - | - | 0 | - |
| - | - | 8304 | 509.3 | - | - | 0 | - |
| - | - | 2080 | 509.3 | - | - | 0 | - |
| - | - | 5826 | 510.3 | - | - | 0 | - |
| - | - | 632.5 | 510.3 | - | - | 0 | - |
| - | - | 994.8 | 512.3 | - | - | 0 | - |
| - | - | 8.601E+04 | 514.3 | - | - | 0 | - |
| - | - | 2.455E+04 | 515.3 | - | - | 0 | - |
| - | - | 4051 | 516.3 | - | - | 0 | - |
| - | - | 708 | 517.3 | - | - | 0 | - |
| - | - | 1099 | 521.3 | - | - | 0 | - |
| - | - | 1224 | 522.3 | - | - | 0 | - |
| - | - | 1223 | 523.3 | - | - | 0 | - |
| - | - | 8318 | 524.3 | - | - | 0 | - |
| - | - | 1.605E+04 | 524.3 | - | - | 0 | - |
| - | - | 3107 | 525.3 | - | - | 0 | - |
| 4 | y | 897.4 | 525.8 | 0.006255 | 11.9 | +2 | 10 |
| - | - | 4697 | 526.3 | - | - | 0 | - |
| - | - | 3.81E+04 | 526.3 | - | - | 0 | - |
| - | - | 773.6 | 527.3 | - | - | 0 | - |
| - | - | 9706 | 527.3 | - | - | 0 | - |
| - | - | 1350 | 528.3 | - | - | 0 | - |
| - | - | 5673 | 533.3 | - | - | 0 | - |
| - | - | 3420 | 534.3 | - | - | 0 | - |
| 4 | y | 6602 | 534.3 | 0.0004271 | 0.7994 | +2 | 10 |
| - | - | 2607 | 534.8 | - | - | 0 | - |
| - | - | 835 | 535.3 | - | - | 0 | - |
| - | - | 1142 | 535.3 | - | - | 0 | - |
| - | - | 979.9 | 536.3 | - | - | 0 | - |
| - | - | 1250 | 537.3 | - | - | 0 | - |
| - | - | 845.1 | 538.3 | - | - | 0 | - |
| - | - | 1.842E+04 | 539.3 | - | - | 0 | - |
| - | - | 1.297E+04 | 540.3 | - | - | 0 | - |
| - | - | 4432 | 541.3 | - | - | 0 | - |
| - | - | 4881 | 542.3 | - | - | 0 | - |
| - | - | 8930 | 542.3 | - | - | 0 | - |
| - | - | 1279 | 543.3 | - | - | 0 | - |
| - | - | 3846 | 543.3 | - | - | 0 | - |
| - | - | 1185 | 548.3 | - | - | 0 | - |
| - | - | 4858 | 550.3 | - | - | 0 | - |
| - | - | 2150 | 551.3 | - | - | 0 | - |
| - | - | 3923 | 551.3 | - | - | 0 | - |
| - | - | 3387 | 552.3 | - | - | 0 | - |
| 11 | b | 5184 | 552.3 | 0.008305 | 15.04 | +2 | 11 |
| - | - | 2901 | 553.3 | - | - | 0 | - |
| - | - | 795 | 554.3 | - | - | 0 | - |
| - | - | 1794 | 554.3 | - | - | 0 | - |
| - | - | 1925 | 557.3 | - | - | 0 | - |
| - | - | 1032 | 558.3 | - | - | 0 | - |
| - | - | 1540 | 560.3 | - | - | 0 | - |
| - | - | 870.3 | 561.3 | - | - | 0 | - |
| - | - | 1564 | 562.3 | - | - | 0 | - |
| - | - | 2540 | 566.3 | - | - | 0 | - |
| - | - | 643.9 | 567.3 | - | - | 0 | - |
| - | - | 7195 | 567.4 | - | - | 0 | - |
| - | - | 1.553E+04 | 568.3 | - | - | 0 | - |
| 9 | y | 4.322E+04 | 569.3 | 0.003425 | 6.015 | +1 | 5 |
| - | - | 1.332E+04 | 570.3 | - | - | 0 | - |
| - | - | 6104 | 571.3 | - | - | 0 | - |
| - | - | 1711 | 572.3 | - | - | 0 | - |
| - | - | 2613 | 576.3 | - | - | 0 | - |
| - | - | 2122 | 577.3 | - | - | 0 | - |
| - | - | 4460 | 577.8 | - | - | 0 | - |
| - | - | 1594 | 578.3 | - | - | 0 | - |
| - | - | 8492 | 578.3 | - | - | 0 | - |
| - | - | 1521 | 579.3 | - | - | 0 | - |
| - | - | 7117 | 579.3 | - | - | 0 | - |
| - | - | 2107 | 580.3 | - | - | 0 | - |
| - | - | 1298 | 580.4 | - | - | 0 | - |
| 3 | y | 946 | 582.3 | 0.004524 | 7.768 | +2 | 11 |
| 3 | y | 2192 | 582.8 | 0.003468 | 5.951 | +2 | 11 |
| - | - | 1.299E+04 | 583.4 | - | - | 0 | - |
| - | - | 3993 | 584.4 | - | - | 0 | - |
| - | - | 1095 | 584.8 | - | - | 0 | - |
| - | - | 2736 | 585.3 | - | - | 0 | - |
| - | - | 3.614E+04 | 585.4 | - | - | 0 | - |
| - | - | 1610 | 586.3 | - | - | 0 | - |
| - | - | 1.047E+04 | 586.4 | - | - | 0 | - |
| 9 | y | 6.415E+05 | 587.3 | 0.002137 | 3.639 | +1 | 5 |
| - | - | 1.994E+05 | 588.3 | - | - | 0 | - |
| - | - | 4.36E+04 | 589.3 | - | - | 0 | - |
| - | - | 6401 | 590.4 | - | - | 0 | - |
| 3 | y | 2.117E+04 | 591.3 | 0.002035 | 3.441 | +2 | 11 |
| - | - | 1.199E+04 | 591.9 | - | - | 0 | - |
| - | - | 4800 | 592.4 | - | - | 0 | - |
| - | - | 1480 | 592.9 | - | - | 0 | - |
| - | - | 4176 | 593.3 | - | - | 0 | - |
| - | - | 3805 | 594.3 | - | - | 0 | - |
| 6 | b | 4.644E+04 | 595.4 | 0.001993 | 3.348 | +1 | 6 |
| 6 | b | 3.032E+04 | 596.3 | 0.006747 | 11.31 | +1 | 6 |
| - | - | 1.012E+05 | 597.3 | - | - | 0 | - |
| - | - | 3.226E+04 | 598.3 | - | - | 0 | - |
| - | - | 6750 | 599.3 | - | - | 0 | - |
| - | - | 832.8 | 600.3 | - | - | 0 | - |
| - | - | 899.1 | 604.3 | - | - | 0 | - |
| - | - | 1169 | 605.3 | - | - | 0 | - |
| - | - | 916.9 | 606.3 | - | - | 0 | - |
| - | - | 1068 | 608.8 | - | - | 0 | - |
| - | - | 905.8 | 610.3 | - | - | 0 | - |
| - | - | 6.071E+04 | 611.4 | - | - | 0 | - |
| - | - | 1.857E+04 | 612.4 | - | - | 0 | - |
| 6 | b | 3.432E+05 | 613.4 | 0.001926 | 3.141 | +1 | 6 |
| - | - | 1.129E+05 | 614.4 | - | - | 0 | - |
| - | - | 2.299E+04 | 615.4 | - | - | 0 | - |
| - | - | 3349 | 616.4 | - | - | 0 | - |
| - | - | 1077 | 617.9 | - | - | 0 | - |
| - | - | 847.4 | 618.4 | - | - | 0 | - |
| - | - | 2922 | 621.4 | - | - | 0 | - |
| - | - | 4697 | 622.4 | - | - | 0 | - |
| - | - | 6786 | 623.4 | - | - | 0 | - |
| - | - | 2012 | 624.4 | - | - | 0 | - |
| 2 | y | 4560 | 625.9 | 0.002167 | 3.463 | +2 | 12 |
| 2 | y | 6821 | 626.4 | 0.00546 | 8.717 | +2 | 12 |
| - | - | 3435 | 626.9 | - | - | 0 | - |
| - | - | 1088 | 627.3 | - | - | 0 | - |
| - | - | 825.4 | 627.4 | - | - | 0 | - |
| - | - | 676.6 | 627.8 | - | - | 0 | - |
| - | - | 712.9 | 632.4 | - | - | 0 | - |
| - | - | 1515 | 633.9 | - | - | 0 | - |
| - | - | 700.4 | 634.4 | - | - | 0 | - |
| 2 | y | 7.324E+04 | 634.9 | 0.002256 | 3.553 | +2 | 12 |
| - | - | 4.996E+04 | 635.4 | - | - | 0 | - |
| - | - | 1.803E+04 | 635.9 | - | - | 0 | - |
| - | - | 3.627E+04 | 636.4 | - | - | 0 | - |
| - | - | 1038 | 636.9 | - | - | 0 | - |
| - | - | 1.316E+04 | 637.4 | - | - | 0 | - |
| - | - | 3141 | 638.4 | - | - | 0 | - |
| - | - | 4.798E+04 | 639.4 | - | - | 0 | - |
| - | - | 2686 | 639.9 | - | - | 0 | - |
| - | - | 2.466E+04 | 640.4 | - | - | 0 | - |
| - | - | 6230 | 641.4 | - | - | 0 | - |
| - | - | 1586 | 642.4 | - | - | 0 | - |
| - | - | 4091 | 647.4 | - | - | 0 | - |
| - | - | 1579 | 648.4 | - | - | 0 | - |
| - | - | 5651 | 650.4 | - | - | 0 | - |
| - | - | 4691 | 651.4 | - | - | 0 | - |
| - | - | 1620 | 652.4 | - | - | 0 | - |
| - | - | 1.282E+04 | 654.4 | - | - | 0 | - |
| - | - | 4368 | 655.4 | - | - | 0 | - |
| - | - | 2620 | 656.4 | - | - | 0 | - |
| - | - | 1028 | 657.4 | - | - | 0 | - |
| 8 | y | 7.342E+04 | 658.4 | 0.002194 | 3.333 | +1 | 6 |
| - | - | 2.595E+04 | 659.4 | - | - | 0 | - |
| - | - | 6178 | 660.4 | - | - | 0 | - |
| - | - | 1410 | 661.4 | - | - | 0 | - |
| - | - | 6325 | 664.4 | - | - | 0 | - |
| - | - | 6938 | 665.4 | - | - | 0 | - |
| - | - | 2756 | 666.4 | - | - | 0 | - |
| - | - | 1990 | 667.4 | - | - | 0 | - |
| - | - | 4.291E+04 | 668.4 | - | - | 0 | - |
| - | - | 1.552E+04 | 669.4 | - | - | 0 | - |
| - | - | 3031 | 670.4 | - | - | 0 | - |
| - | - | 1608 | 671.4 | - | - | 0 | - |
| - | - | 753 | 675.3 | - | - | 0 | - |
| - | - | 6.076E+04 | 682.4 | - | - | 0 | - |
| - | - | 2.1E+04 | 683.4 | - | - | 0 | - |
| 0 | Precursor | 1.754E+04 | 684.4 | 0.004853 | 7.091 | +2 | -1 |
| - | - | 5442 | 685.4 | - | - | 0 | - |
| - | - | 892.3 | 686.4 | - | - | 0 | - |
| - | - | 1544 | 691.5 | - | - | 0 | - |
| - | - | 1148 | 692.4 | - | - | 0 | - |
| - | - | 1450 | 693.4 | - | - | 0 | - |
| - | - | 1796 | 701.4 | - | - | 0 | - |
| - | - | 2326 | 710.4 | - | - | 0 | - |
| 7 | b | 5474 | 710.4 | 0.002263 | 3.186 | +1 | 7 |
| - | - | 932.9 | 711.4 | - | - | 0 | - |
| - | - | 1364 | 711.4 | - | - | 0 | - |
| - | - | 1909 | 712.4 | - | - | 0 | - |
| - | - | 937.1 | 713.4 | - | - | 0 | - |
| - | - | 853 | 714.4 | - | - | 0 | - |
| - | - | 1119 | 719.4 | - | - | 0 | - |
| - | - | 4059 | 736.4 | - | - | 0 | - |
| 7 | y | 8353 | 737.4 | 0.005039 | 6.833 | +1 | 7 |
| - | - | 4229 | 738.4 | - | - | 0 | - |
| - | - | 815.6 | 739.4 | - | - | 0 | - |
| - | - | 1644 | 746.4 | - | - | 0 | - |
| - | - | 722.5 | 747.4 | - | - | 0 | - |
| - | - | 2751 | 749.5 | - | - | 0 | - |
| - | - | 1157 | 750.5 | - | - | 0 | - |
| - | - | 971.3 | 751.4 | - | - | 0 | - |
| - | - | 2.273E+04 | 753.5 | - | - | 0 | - |
| - | - | 7846 | 754.5 | - | - | 0 | - |
| 7 | y | 1.632E+05 | 755.4 | 0.002348 | 3.108 | +1 | 7 |
| - | - | 7.173E+04 | 756.4 | - | - | 0 | - |
| - | - | 1.748E+04 | 757.4 | - | - | 0 | - |
| - | - | 2796 | 758.4 | - | - | 0 | - |
| - | - | 776.3 | 759.4 | - | - | 0 | - |
| 8 | b | 8384 | 763.4 | 0.001471 | 1.927 | +1 | 8 |
| 8 | b | 1.057E+04 | 764.4 | 0.007323 | 9.58 | +1 | 8 |
| - | - | 4451 | 765.4 | - | - | 0 | - |
| - | - | 1264 | 766.4 | - | - | 0 | - |
| - | - | 1827 | 779.4 | - | - | 0 | - |
| - | - | 1460 | 780.4 | - | - | 0 | - |
| 8 | b | 1.636E+05 | 781.5 | 0.002259 | 2.89 | +1 | 8 |
| - | - | 6.554E+04 | 782.5 | - | - | 0 | - |
| - | - | 1.989E+04 | 783.5 | - | - | 0 | - |
| - | - | 2613 | 784.5 | - | - | 0 | - |
| - | - | 1682 | 805.5 | - | - | 0 | - |
| - | - | 1125 | 806.5 | - | - | 0 | - |
| - | - | 1073 | 811.4 | - | - | 0 | - |
| - | - | 2030 | 820.5 | - | - | 0 | - |
| - | - | 954.4 | 821.5 | - | - | 0 | - |
| - | - | 1048 | 838.5 | - | - | 0 | - |
| 6 | y | 1576 | 850.5 | 7.644E-05 | 0.08987 | +1 | 8 |
| - | - | 938.7 | 864.5 | - | - | 0 | - |
| 6 | y | 1.9E+04 | 868.5 | 0.002573 | 2.963 | +1 | 8 |
| - | - | 9420 | 869.5 | - | - | 0 | - |
| - | - | 2767 | 870.5 | - | - | 0 | - |
| - | - | 3618 | 892.5 | - | - | 0 | - |
| - | - | 1057 | 893.5 | - | - | 0 | - |
| - | - | 801.5 | 894.5 | - | - | 0 | - |
| 5 | y | 827.3 | 921.5 | 0.002513 | 2.727 | +1 | 9 |
| 5 | y | 741.2 | 922.5 | 0.016 | 17.34 | +1 | 9 |
| - | - | 1539 | 934.5 | - | - | 0 | - |
| - | - | 1448 | 935.5 | - | - | 0 | - |
| 5 | y | 2.063E+04 | 939.6 | 0.002203 | 2.344 | +1 | 9 |
| - | - | 1.172E+04 | 940.6 | - | - | 0 | - |
| - | - | 2273 | 941.6 | - | - | 0 | - |
| 10 | b | 1928 | 991.6 | 0.0009022 | 0.9098 | +1 | 10 |
| - | - | 979.9 | 1005 | - | - | 0 | - |
| - | - | 6710 | 1022 | - | - | 0 | - |
| - | - | 3232 | 1023 | - | - | 0 | - |
| - | - | 795 | 1024 | - | - | 0 | - |
| 4 | y | 2044 | 1068 | 0.001417 | 1.327 | +1 | 10 |
| - | - | 1486 | 1069 | - | - | 0 | - |
| 11 | b | 2634 | 1121 | 0.000267 | 0.2382 | +1 | 11 |
| - | - | 1485 | 1122 | - | - | 0 | - |
| - | - | 758.9 | 3074 | - | - | 0 | - |

m/z Charge Intensity FragmentType MassShift Position
120.06610870361328 0 36248.72 y 12
120.0814208984375 0 1669.8197
121.06956481933594 0 1799.1577
121.17300415039062 0 372.5958
122.09676361083984 0 416.5073
123.09239959716797 0 919.2051
124.07626342773438 0 717.627
125.1080551147461 0 1208.9702
126.0556640625 0 805.163
126.0919189453125 0 843.48425
127.08727264404297 0 2297.6704
127.12348175048828 0 887.56555
127.6581039428711 0 417.71857
128.07144165039062 0 698.7652
129.0663604736328 0 2322.3928
129.10287475585938 0 493643.3
130.0504150390625 0 2091.182
130.0868377685547 0 4438.775
130.09999084472656 0 3336.4766
130.10614013671875 0 29357.918
131.10696411132812 0 544.75494
131.10992431640625 0 448.95966
131.11863708496094 0 689.20685
132.07733154296875 0 13606.536
133.06178283691406 0 362.92288
133.0806884765625 0 778.1255
133.89453125 0 393.58807
135.0564727783203 0 389.8988
136.0764923095703 0 861.7236
137.10801696777344 0 1486.0813
138.12841796875 0 2845.4946
139.0872344970703 0 9758.732
140.0904998779297 0 774.7802
140.14398193359375 0 10127.788
141.1028594970703 0 129818.56 a Water loss 1
141.1472930908203 0 1167.8195
142.08689880371094 0 1437.0813
142.09983825683594 0 1159.5204
142.10626220703125 0 11142.803
142.12350463867188 0 752.7011
143.04627990722656 0 464.97653
143.1080780029297 0 467.77505
143.11849975585938 0 5103.4585 d 1
146.060791015625 0 757.23474
146.12942504882812 0 661.41113
147.11346435546875 0 13229.975
148.1167449951172 0 753.0577
151.0870361328125 0 1574.9467
152.0710906982422 0 550.845
152.10740661621094 0 792.3017
152.14402770996094 0 7847.267
153.1029815673828 0 3133.7441
153.14744567871094 0 818.4089
154.08657836914062 0 891.9087
155.08279418945312 0 548.17847
155.11851501464844 0 22192.6
156.07728576660156 0 1085.6033
156.11415100097656 0 1042.1381
156.1219024658203 0 2506.0637
157.0614471435547 0 7642.8
157.09774780273438 0 1753.5355
157.13414001464844 0 21314.512
158.13735961914062 0 1284.7666
159.11341857910156 0 62034.477 a 1
160.11683654785156 0 4179.515
163.0873565673828 0 707.82336
165.10284423828125 0 6318.445
165.13917541503906 0 701.02014
167.04579162597656 0 2678.096
167.08224487304688 0 4308.574
167.1185760498047 0 4177.1094
168.0493927001953 0 677.2812
168.0779266357422 0 730.7972
168.11415100097656 0 879.35114
169.09783935546875 0 144613.5 b Water loss 1
169.13417053222656 0 7373.027
169.1708221435547 0 1045.3394
170.08206176757812 0 583.4397
170.09449768066406 0 779.5122
170.10122680664062 0 13034.192
170.13742065429688 0 749.0544
171.1051788330078 0 965.6331
171.1134490966797 0 5115.316
172.14520263671875 0 2362.384
173.12908935546875 0 857.8929
173.45147705078125 0 1682.6189
174.08802795410156 0 3335.7212
175.26463317871094 0 643.55804
177.1031036376953 0 845.056
179.08213806152344 0 773.58923
179.1183624267578 0 1030.148
180.11361694335938 0 1461.9937
180.91270446777344 0 537.4028
181.09788513183594 0 3619.3706
181.13429260253906 0 20035.469
182.12950134277344 0 16430.22
182.1378936767578 0 2326.1084
183.11363220214844 0 3704.1765
183.1499481201172 0 290835.66
184.0723876953125 0 5822.1504
184.1088104248047 0 6833.0195
184.1533660888672 0 30911.967
185.056396484375 0 5359.331
185.129150390625 0 29569.293
185.1560821533203 0 1549.2949
185.16552734375 0 7407.7124
186.08750915527344 0 1119.4309
186.12428283691406 0 1628.3215
186.13291931152344 0 2557.2732
186.16091918945312 0 1289.3737
186.1691131591797 0 676.1363
187.10845947265625 0 60933.098 b 1
187.14500427246094 0 2717.5693
188.11178588867188 0 5146.8296
191.0820770263672 0 1639.3518
191.1187744140625 0 591.52924
193.0977783203125 0 1400.6982
193.13404846191406 0 1559.312
194.12899780273438 0 1623.1973
195.11351013183594 0 13041.145
195.14898681640625 0 741.32904
196.10813903808594 0 608.9945
196.1176300048828 0 1095.9457
197.0928955078125 0 4178.7124
197.12924194335938 0 4028.2227
197.16561889648438 0 13113.09
198.12437438964844 0 5897.2783
198.16879272460938 0 1678.9736
199.1084442138672 0 2768.5732
199.14491271972656 0 620.2906
200.11203002929688 0 554.1294
200.14019775390625 0 5647.4106
201.08799743652344 0 1170.2765
201.0990447998047 0 1528.8341
201.12405395507812 0 949.30945
202.0829315185547 0 28579.13
202.15585327148438 0 1023.2802
203.0675048828125 0 883.22705
203.08644104003906 0 1746.1814
204.13494873046875 0 1562.9111
205.09788513183594 0 1085.0175
207.11337280273438 0 1108.8234
207.1497344970703 0 748.1007
208.10877990722656 0 12662.792
208.14515686035156 0 591.769
209.0928192138672 0 25001.898
209.11131286621094 0 1107.2997
209.12916564941406 0 1620.2004
209.16563415527344 0 3588.1812
210.0960235595703 0 2006.9897
210.1245574951172 0 5431.6377
210.13523864746094 0 990.7777
211.1082763671875 0 2214.2795
211.14488220214844 0 160046.27
212.06729125976562 0 2093.2544
212.12857055664062 0 878.1976
212.14830017089844 0 19028.143
213.08799743652344 0 861.6218
213.12413024902344 0 10465.223
213.15029907226562 0 870.19727
213.16061401367188 0 4608.256
214.10812377929688 0 1199.7178
214.11935424804688 0 4579.2715
214.1292724609375 0 811.22534 y Water loss 7
214.6205596923828 0 909.0201
215.13975524902344 0 17276.291
216.14364624023438 0 1160.0078
216.63441467285156 0 516.3242
217.1669464111328 0 1416.4885
218.1212921142578 0 645.5945
218.1507568359375 0 2988.4004
219.1096649169922 0 12810.069
219.62158203125 0 1156.2333
220.11233520507812 0 1042.0884
221.059814453125 0 999.2805
221.0930633544922 0 801.9358
222.12452697753906 0 42018.48
223.10833740234375 0 4362.3267
223.1278533935547 0 5331.7993
223.1812286376953 0 5160.4287
224.14048767089844 0 1265.305
225.1241912841797 0 1434.9794
225.13551330566406 0 7088.5464
225.1613311767578 0 552.08563
225.64013671875 0 7387.225
226.1195068359375 0 62911.547
226.140869140625 0 2397.3228
226.1556854248047 0 1614.6498
227.10345458984375 0 1828.3422
227.1234893798828 0 5406.939
227.1762237548828 0 7021.214
228.09893798828125 0 1632.7185
228.12342834472656 0 891.5557
228.13523864746094 0 8306.049
228.17152404785156 0 2127.873
229.09393310546875 0 1819.5012
229.11911010742188 0 2105.859
229.1387939453125 0 1048.7784
229.1549530029297 0 1229.9791
230.07806396484375 0 661.68005
230.15078735351562 0 25073.506 y Water loss 11
231.1347198486328 0 21174.193 y Ammonia loss 11
231.1543426513672 0 2573.813
232.1382598876953 0 2164.934
234.1246795654297 0 2604.041
234.64454650878906 0 1026.1848
235.14686584472656 0 698.58264
235.6406707763672 0 1222.4806
236.10357666015625 0 835.0308
236.14036560058594 0 1575.7637
236.1767120361328 0 1907.4381
236.64849853515625 0 13927.229
237.12384033203125 0 880.86255
237.137451171875 0 835.52313 y Ammonia loss 9
237.14984130859375 0 3416.5142
237.1607208251953 0 3646.0554
238.1195068359375 0 2163.807
239.1399383544922 0 2195.7593
239.17640686035156 0 2220.8884
240.13519287109375 0 391294.5
240.63223266601562 0 932.43195
240.64527893066406 0 1062.5452
241.03152465820312 0 1187.4741
241.11924743652344 0 6232.7573
241.13845825195312 0 42859.223
241.62794494628906 0 579.17505
241.6403045654297 0 2805.8333 b Water loss 4
242.1407470703125 0 3882.6104
243.13441467285156 0 18231.531
243.1461944580078 0 48412.637
243.18275451660156 0 1664.8387
243.65090942382812 0 15158.14
244.13809204101562 0 1540.3679
244.15066528320312 0 7735.9624
244.16554260253906 0 1158.2529
245.15159606933594 0 636.2462
247.14453125 0 535.9787
248.1614532470703 0 149356.9 y 11
249.13511657714844 0 1282.1057
249.16465759277344 0 15949.311
249.63807678222656 0 16303.688
249.65469360351562 0 970.1083
250.11935424804688 0 934.05304
250.1397247314453 0 3876.91
250.16697692871094 0 980.35156
250.64572143554688 0 6899.339 b 4
251.14898681640625 0 2943.5752
252.135009765625 0 6353.6475
252.17166137695312 0 5265.168
253.11871337890625 0 4168.442
253.13075256347656 0 3841.2893
253.64065551757812 0 727.58417
254.11328125 0 719.86707
254.1508026123047 0 5104.221
254.1874542236328 0 3059.692
254.6500244140625 0 1245.902
255.11058044433594 0 883.77124
255.1415252685547 0 2148.0576 a Water loss 2
255.1712646484375 0 4204.9424
255.64244079589844 0 806.0032
256.13043212890625 0 1852.7996 a Ammonia loss 2
256.1751403808594 0 756.176
257.6529235839844 0 631.97327
258.14569091796875 0 90734.445
259.0929260253906 0 648.2354
259.1308288574219 0 1150.3522
259.1491394042969 0 9994.137
260.1982421875 0 797.1494
261.1352233886719 0 2494.6252
262.11962890625 0 6518.3193
262.6461181640625 0 1866.2943
262.6637268066406 0 1210.076
263.1741638183594 0 2916.0703
263.653564453125 0 14699.277
264.0990295410156 0 770.9744
264.1553039550781 0 4038.4712
264.1716003417969 0 10423.382
265.1304016113281 0 1691.3104
265.1556701660156 0 4793.528
265.1741638183594 0 924.0451
266.11419677734375 0 935.0152
266.15081787109375 0 19753.629
267.1432189941406 0 892.15686
267.1551513671875 0 2271.23
267.16986083984375 0 639.07623
268.1300964355469 0 4911.5776
268.1511535644531 0 671.36017
268.1665954589844 0 2225.687
268.2021484375 0 767.7038
269.1617736816406 0 3391.9277
269.1988220214844 0 1673.0868
270.1453857421875 0 2342.2654
270.1818542480469 0 916.3135
270.66375732421875 0 923.6982
270.6795959472656 0 6300.909
271.1409912109375 0 3328.3403
271.1579895019531 0 845.73987
271.172607421875 0 3066.2366
271.6703796386719 0 3792.0986
272.1611328125 0 1746.7181
275.6716003417969 0 2511.7195
276.1564025878906 0 11056.793
276.1722412109375 0 874.2185
276.6595458984375 0 3640.9905
277.1306457519531 0 5355.898
277.1587219238281 0 2678.4868
278.1142578125 0 979.8664
278.1490478515625 0 885.9225
278.163818359375 0 819.0621
278.1875 0 4742.3687
279.1461486816406 0 13093.162
279.18377685546875 0 557.1737
280.130126953125 0 6840.731
280.1485900878906 0 2033.4888
280.166748046875 0 2963.5945
281.1313171386719 0 556.97394
281.16259765625 0 671.06134
281.65386962890625 0 1562.2761
282.10955810546875 0 688.26874
282.1561584472656 0 875.5142
282.1822509765625 0 19406.566
283.1412048339844 0 13356.281 b Water loss 2
283.18487548828125 0 2888.7227
283.66949462890625 0 3901.1794
284.1263427734375 0 844.7076 b Ammonia loss 2
284.1443786621094 0 1825.5201
284.1612548828125 0 1125.7789
284.16961669921875 0 734.78107 y Water loss 5
284.1848449707031 0 2575.3086
284.19818115234375 0 5512.042
284.6774597167969 0 11548.149
285.1560974121094 0 2995.6086
285.171630859375 0 49431.33 y Water loss 8
285.22955322265625 0 7504.5063
285.66094970703125 0 11719.151 y Ammonia loss 8
285.67474365234375 0 10988.487
286.14361572265625 0 685.06104
286.1625671386719 0 3337.526
286.1885070800781 0 1472.3428
286.23291015625 0 1113.6415
287.1733703613281 0 794.9693
289.6695861816406 0 1233.2583
290.1671447753906 0 4213.356
290.65899658203125 0 2473.1416
290.6721496582031 0 776.6709
292.16632080078125 0 965.91907
292.1828918457031 0 18273.521
292.68414306640625 0 5270.779
293.12640380859375 0 840.4745
293.175048828125 0 445.61453
293.1907043457031 0 90769.336
293.6921691894531 0 26390.498
294.17474365234375 0 444973.97 y 8
294.67620849609375 0 133960.73
295.14111328125 0 21053.576
295.177490234375 0 32497.127
295.2139587402344 0 4709.721
295.67852783203125 0 4863.611
296.1438293457031 0 2983.838
296.1728515625 0 17272.484
296.1973876953125 0 2072.8845
296.21478271484375 0 917.6702
297.15667724609375 0 11695.088
297.1752624511719 0 3740.1719
297.19354248046875 0 7380.9937
297.66693115234375 0 1668.1217
297.6852722167969 0 2030.1412
298.140625 0 1766.0878
298.16094970703125 0 1615.0728
298.1827087402344 0 4861.325 b Water loss 5
298.6751403808594 0 3195.878 b Ammonia loss 5
299.17254638671875 0 35702.113
299.6741638183594 0 11684.089
300.1698303222656 0 2817.7737
300.1932678222656 0 1393.3845
300.6763000488281 0 927.87146
301.15179443359375 0 16416.73 b 2
301.1705322265625 0 1081.2365
302.15472412109375 0 1758.2368
305.12579345703125 0 1678.8792
305.1611328125 0 940.7
306.1803283691406 0 19642.422
306.19873046875 0 4330.489
306.68170166015625 0 4929.3374
306.6994934082031 0 1212.0154
307.14129638671875 0 991.0811
307.1878356933594 0 19676.84 b 5
307.6892395019531 0 6981.186
308.193603515625 0 923.2085
309.156982421875 0 3021.129
309.68572998046875 0 902.2538
311.1365051269531 0 1221.8219
311.173095703125 0 718.58136
311.2099304199219 0 1667.7146
312.1678466796875 0 16228.291
312.19342041015625 0 8440.217
312.6808776855469 0 1712.5907
313.15185546875 0 39204.52
313.16851806640625 0 2198.5693
313.19390869140625 0 2189.2898
313.2245788574219 0 13696.672
314.1545104980469 0 4924.5327
314.18353271484375 0 22707.89
314.22442626953125 0 1906.8777
315.1674499511719 0 2317.993
315.1863708496094 0 2976.7937
315.2041320800781 0 4393.995
318.17852783203125 0 19099.92
318.69085693359375 0 1367.251
319.18524169921875 0 3624.624
319.2069396972656 0 1916.0585
319.68865966796875 0 1407.5876
320.1746826171875 0 947.87427
320.1961975097656 0 6103.523
320.6940612792969 0 15410.614
321.1565856933594 0 1358.598
321.1949462890625 0 5633.6333
321.22869873046875 0 786.0116
322.1527404785156 0 3517.5676
322.1774597167969 0 7102.4976
323.1362609863281 0 9836.143
323.1735534667969 0 7389.8594
323.2091369628906 0 7099.0664
324.13653564453125 0 1246.9567
324.157958984375 0 2488.6963
324.18060302734375 0 720.65027
324.21441650390625 0 773.9301
325.1518859863281 0 2963.315
325.1880798339844 0 6451.543
325.224365234375 0 1904.6576
325.68597412109375 0 6435.7173
326.1863708496094 0 2153.0862
326.67974853515625 0 898.77246
327.1673889160156 0 2584.3364
327.18804931640625 0 691.0836
327.69976806640625 0 15654.861
328.201171875 0 5029.492
328.70623779296875 0 2216.4705
329.6932678222656 0 2453.9604 y 7
330.1784362792969 0 38748.594
331.1813049316406 0 5894.6646
331.1996765136719 0 643.2079 b 9
331.23541259765625 0 3115.9658
332.1825866699219 0 668.5791
332.693115234375 0 1823.8401
332.7125549316406 0 4057.2422
333.2035827636719 0 10840.121
333.7052001953125 0 3981.6133
334.2098083496094 0 782.99023
334.691162109375 0 17896.365
335.1924743652344 0 5358.802
335.20526123046875 0 739.7989
335.6939697265625 0 2127.4514
336.16943359375 0 917.28107
336.1917419433594 0 998.1272
337.15228271484375 0 614.12537
337.1899108886719 0 1607.5225
338.1783447265625 0 608.34357
338.1955261230469 0 3168.2834
338.2171325683594 0 945.42944
338.6974182128906 0 1923.9608
339.16766357421875 0 3603.056
339.2041320800781 0 2775.6367
339.2389221191406 0 1169.6099
340.1881103515625 0 192005.02
341.1460266113281 0 1233.7705
341.18853759765625 0 37816.6
341.21923828125 0 4939.307
341.69879150390625 0 8392.254
341.7173767089844 0 118085.945
342.0803527832031 0 2275.7434
342.16790771484375 0 4387.818
342.1965026855469 0 3541.068
342.2190856933594 0 40603.684
342.72015380859375 0 9779.072
343.16778564453125 0 844.919
343.2220153808594 0 975.8122
344.1939697265625 0 975.71265
346.7094421386719 0 8199.519 b Water loss 6
347.20233154296875 0 15224.649 b Ammonia loss 6
347.7027282714844 0 4409.349
348.1675109863281 0 3683.325
349.1513366699219 0 1096.1864
349.1888732910156 0 2936.9165
349.2243957519531 0 858.46686
350.18524169921875 0 1069.4138
351.16815185546875 0 2340.0154
351.2037048339844 0 3580.8125
351.24029541015625 0 12433.705
352.2060241699219 0 924.08704
352.24346923828125 0 2038.1178
353.1468505859375 0 691.4496
353.18292236328125 0 783.8435
353.2195129394531 0 13217.343
354.1785888671875 0 954.4339
354.22271728515625 0 2476.6123
354.725341796875 0 3118.7178
355.1611633300781 0 872.19934
355.2013854980469 0 709.77484
355.71466064453125 0 127636.92 b 6
356.1934509277344 0 5720.9155
356.2161865234375 0 45792.39
356.7171630859375 0 11153.02
357.19464111328125 0 818.9942
357.21636962890625 0 5417.1523
358.1734619140625 0 2788.5684
358.19873046875 0 2729.4028
359.19378662109375 0 92494.69 y Water loss 10
359.6925964355469 0 637.51794
359.7181091308594 0 1389.7172
360.17694091796875 0 1264.3434 y Ammonia loss 10
360.1972351074219 0 15211.618
360.70843505859375 0 617.4644
361.1986389160156 0 2766.909
363.2039794921875 0 2143.2886
364.20538330078125 0 741.7222
364.23504638671875 0 1628.4133
365.19415283203125 0 5316.16
365.2197265625 0 2684.5376
365.25628662109375 0 2310.091
366.1783142089844 0 20687.951
366.21673583984375 0 2145.817
366.2530212402344 0 1071.6328
367.15948486328125 0 893.3064
367.18121337890625 0 2793.985
367.2718200683594 0 747.11536
368.1942138671875 0 5549.061
368.2306213378906 0 13093.267
368.7227478027344 0 9860.3
369.2170104980469 0 9930.014 y Water loss 6
369.7150573730469 0 3583.8833
370.1730041503906 0 792.32776
370.2132263183594 0 1775.0017
371.20428466796875 0 801.61005
371.230224609375 0 20070.818
372.1893310546875 0 2040.066
372.2325744628906 0 3746.1487
373.22076416015625 0 6240.214
373.714599609375 0 5079.866
374.218017578125 0 2615.728 b 10
375.1812438964844 0 946.4462
375.2039489746094 0 756.30316
376.16204833984375 0 1341.9869
376.20111083984375 0 835.0402
377.2044982910156 0 124691.67 y 10
377.2356262207031 0 68799.05
377.7149658203125 0 391.70157
377.7373962402344 0 31224.357
378.2191467285156 0 109523.18 y 6
378.7212219238281 0 45869.305
379.2337646484375 0 16500.6
379.7239685058594 0 3141.0205
380.2367248535156 0 4507.477
381.2148742675781 0 10427.717
382.2264709472656 0 5122.6943 b Water loss 7
382.2476806640625 0 2617.4648
382.281982421875 0 1141.8414
382.7217102050781 0 7152.821 b Ammonia loss 7
383.20513916015625 0 47269.81
383.226318359375 0 3979.9966
383.7212219238281 0 1453.3619
384.1888427734375 0 16824.658
384.2091979980469 0 3629.9778
384.26226806640625 0 1012.094
384.2975769042969 0 751.8021
385.1912536621094 0 3064.56
385.2197265625 0 1812.862
385.2622375488281 0 1119.3029
385.5296936035156 0 1307.0675
385.864501953125 0 2102.6492
386.2010803222656 0 948.16846
386.2410583496094 0 2882.1301
387.1886901855469 0 2842.6948
387.241943359375 0 763.183
389.2409362792969 0 4846.391
390.2228698730469 0 1735.0167
390.2478332519531 0 952.1772
391.2334289550781 0 38020.004 b 7
391.73486328125 0 14032.448
392.19403076171875 0 2685.8167
392.23516845703125 0 4603.9683
392.2650451660156 0 1084.7672
393.189697265625 0 2600.0132
393.2200622558594 0 2298.4866
394.174072265625 0 2812.1492
394.2103576660156 0 8668.237
395.21435546875 0 1190.2457
396.22515869140625 0 3269.4775
396.2562255859375 0 619.10004
398.2048034667969 0 1516.2604
398.2413635253906 0 5162.8184
399.2072448730469 0 995.2402
399.2386169433594 0 3460.1016
399.27276611328125 0 13694.709
400.2762145996094 0 2570.3362
401.2157897949219 0 72237.35
402.2185974121094 0 12746.571
403.22064208984375 0 2078.9414
404.2398681640625 0 1234.1096
404.7412414550781 0 852.4482
406.211181640625 0 690.98236
408.2258605957031 0 1992.8018
409.2210693359375 0 8119.497
409.2568664550781 0 2019.7058
410.204833984375 0 3402.812
410.2412414550781 0 8675.958
410.27734375 0 6470.97
411.19842529296875 0 6167.9673
411.232177734375 0 32723.682 b Water loss 3
411.28533935546875 0 1242.7885
412.1841735839844 0 1072.3124
412.22320556640625 0 13697.471 b Ammonia loss 3
412.29351806640625 0 7041.9414
413.22393798828125 0 2710.567
413.2529602050781 0 854.1124
413.2967224121094 0 1097.5005
414.2348937988281 0 1268.1378
415.2325439453125 0 2013.803
417.22393798828125 0 1257.9539
419.24188232421875 0 906.72534
420.2254333496094 0 2556.4153
420.2579040527344 0 710.93945
421.22686767578125 0 772.83765
422.2052001953125 0 2048.509
422.24188232421875 0 1560.044
424.2220458984375 0 1072.1572
424.2588195800781 0 745.3239
425.2203063964844 0 813.54645
425.2530212402344 0 4253.51
425.76312255859375 0 1341.3685 y Water loss 5
426.23681640625 0 2056.7412
426.2660827636719 0 1089.9467
427.2316589355469 0 31311.24
427.2680969238281 0 30628.252
427.3009338378906 0 1011.6307
428.2343444824219 0 4607.1987
428.2688293457031 0 7503.514
429.2118835449219 0 6576.0776
429.247314453125 0 68583.484 b 3
430.2146301269531 0 1142.4836
430.25006103515625 0 15941.891
431.2522277832031 0 3070.9746 b Ammonia loss 8
434.2054443359375 0 978.3574
434.2412109375 0 908.3824
434.7625732421875 0 2276.358 y 5
435.2367248535156 0 2255.719
435.2649841308594 0 1935.2883
436.2591857910156 0 1786.3359
437.21484375 0 3794.51
437.2513427734375 0 5079
438.23614501953125 0 4194.411
438.27197265625 0 1879.0186
439.2321472167969 0 8541.17
439.2663269042969 0 919.4473
439.7600402832031 0 7953.1455 b 8
440.2597961425781 0 3276.6792
440.76336669921875 0 1639.2001
441.2477722167969 0 1286.4945
441.2823486328125 0 827.7529
443.2628479003906 0 5783.8433
444.2605895996094 0 2540.018
446.205322265625 0 957.05493
446.2757873535156 0 1545.1075
447.23712158203125 0 2917.629
447.2724609375 0 627.8553
448.22076416015625 0 1543.4445
449.22259521484375 0 902.0019
450.2373352050781 0 1462.1511
450.2729797363281 0 6802.0225
450.6007995605469 0 1327.4286 Precursor Water loss
450.9319763183594 0 933.5469 Precursor Ammonia loss
451.2698974609375 0 4135.852
452.2222595214844 0 676.9931
452.2607727050781 0 2890.798
453.247314453125 0 5094.095
453.28302001953125 0 3618.6257
453.3204040527344 0 2139.372
454.2723693847656 0 3860.754
455.2262878417969 0 3536.5752
455.2628479003906 0 18116.355
456.2297668457031 0 984.3656
456.2650451660156 0 3673.5884
456.6033020019531 0 1227.9269 Precursor
456.9369201660156 0 1951.9855
457.2735290527344 0 2200.7695
461.25555419921875 0 814.7852
462.2725830078125 0 2677.6792
463.23187255859375 0 5059.459
463.267333984375 0 894.9908
463.3050537109375 0 1295.5468
464.2635803222656 0 8183.1255
464.3236999511719 0 978.1377
465.24786376953125 0 13053.402
466.249755859375 0 2821.079
467.26165771484375 0 3394.9675
467.2992858886719 0 1213.5941
468.28350830078125 0 122290.57
469.28411865234375 0 39333.316
470.2778015136719 0 10719.287 y 4
471.28594970703125 0 1831.6726
472.2830505371094 0 24066.262 y Water loss 9
473.28631591796875 0 4981.86
474.27777099609375 0 734.7225
479.2630615234375 0 8409.105
479.2991638183594 0 1402.6724
480.25689697265625 0 5552.118
480.2854309082031 0 6371.293
481.2430419921875 0 6691.3857
481.28277587890625 0 1730.9884
481.31524658203125 0 24008.18
482.2740478515625 0 106504.54 b Water loss 4
482.314697265625 0 5431.1777
483.2759704589844 0 19996.197
484.2619934082031 0 3151.3174
484.2799987792969 0 2794.9387
485.2742614746094 0 3394.7043
486.299072265625 0 67324.4
487.3028259277344 0 17031.951 b Water loss 9
488.304931640625 0 3270.293
490.28900146484375 0 86595.88 y 9
491.22900390625 0 984.64557
491.2927551269531 0 23645.248
492.2953796386719 0 3568.0037
493.24322509765625 0 1123.5647
495.2962646484375 0 1096.176
496.2538757324219 0 594.53894
496.2894287109375 0 8728.117
496.32666015625 0 3740.6555
497.2740478515625 0 15303.332
497.3092041015625 0 3760.4353
498.26934814453125 0 52249.29
498.3059387207031 0 4936.2524
499.2718505859375 0 11200.432
499.302490234375 0 1473.4614
500.2840881347656 0 171678.64 b 4
501.2870178222656 0 41085.527
502.2895812988281 0 8192.326
503.2908020019531 0 822.47144
506.2737731933594 0 1394.1439
507.2578125 0 1225.9281
507.29486083984375 0 4272.7544
508.27972412109375 0 33944.617
509.23675537109375 0 1061.9675
509.2803955078125 0 8304.057
509.31097412109375 0 2079.726
510.2720947265625 0 5825.83
510.3118896484375 0 632.5434
512.3244018554688 0 994.76794
514.30029296875 0 86010.805
515.3025512695312 0 24550.746
516.3045043945312 0 4051.226
517.310791015625 0 708.02716
521.3108520507812 0 1098.9136
522.3059692382812 0 1224.1965
523.287841796875 0 1222.7527
524.2835083007812 0 8318.451
524.321533203125 0 16045.246
525.3356323242188 0 3107.1814
525.819580078125 0 897.4083 y Ammonia loss 3
526.2622680664062 0 4697.0156
526.3003540039062 0 38097.707
527.263427734375 0 773.60675
527.3034057617188 0 9705.764
528.30859375 0 1349.9216
533.309326171875 0 5673.3037
534.26904296875 0 3419.5417
534.3270263671875 0 6602.152 y 3
534.829833984375 0 2607.3203
535.2864990234375 0 835.0331
535.3309326171875 0 1141.592
536.2821655273438 0 979.8647
537.3040161132812 0 1250.4116
538.302001953125 0 845.13086
539.320556640625 0 18415.076
540.3178100585938 0 12969.926
541.3234252929688 0 4432.3457
542.2942504882812 0 4881.4316
542.33203125 0 8929.677
543.2913208007812 0 1279.261
543.3312377929688 0 3845.6482
548.3190307617188 0 1184.9083
550.3369750976562 0 4857.857
551.291748046875 0 2150.1406
551.3263549804688 0 3923.375
552.278564453125 0 3387.049
552.3167114257812 0 5184.317 b Ammonia loss 10
553.3135986328125 0 2901.338
554.2882080078125 0 795.03827
554.3292846679688 0 1793.7335
557.3352661132812 0 1925.3079
558.3345947265625 0 1032.1667
560.3203735351562 0 1539.941
561.3115234375 0 870.31683
562.303466796875 0 1564.1245
566.3313598632812 0 2539.8252
567.3245239257812 0 643.8684
567.36328125 0 7195.081
568.3485717773438 0 15533.914
569.332763671875 0 43223.336 y Water loss 8
570.3355102539062 0 13324.835
571.3260498046875 0 6103.763
572.324462890625 0 1710.8892
576.3170776367188 0 2613.0513
577.3466186523438 0 2121.7095
577.7910766601562 0 4459.506
578.2901611328125 0 1593.9042
578.3323364257812 0 8491.727
579.2880249023438 0 1521.2175
579.3297729492188 0 7116.5044
580.3131713867188 0 2106.77
580.3838500976562 0 1298.1427
582.3382568359375 0 945.962 y Water loss 2
582.8382568359375 0 2192.0994 y Ammonia loss 2
583.3576049804688 0 12990.509
584.3579711914062 0 3992.5237
584.8482055664062 0 1094.5664
585.3316040039062 0 2735.521
585.373779296875 0 36135.035
586.3323974609375 0 1609.9153
586.377197265625 0 10469.691
587.342041015625 0 641545.8 y 8
588.3448486328125 0 199439.94
589.3473510742188 0 43598.21
590.3501586914062 0 6401.3853
591.35009765625 0 21170.127 y 2
591.8516235351562 0 11991.059
592.3532104492188 0 4800.081
592.85595703125 0 1479.6215
593.3446655273438 0 4175.735
594.3289184570312 0 3805.3335
595.3582153320312 0 46444.867 b Water loss 5
596.3469848632812 0 30315.912 b Ammonia loss 5
597.3381958007812 0 101157.83
598.3408813476562 0 32261.777
599.3430786132812 0 6749.8394
600.3429565429688 0 832.79395
604.3486938476562 0 899.13965
605.3492431640625 0 1168.6409
606.32275390625 0 916.8975
608.8397827148438 0 1068.3645
610.3361206054688 0 905.7758
611.3531494140625 0 60706.836
612.3562622070312 0 18573.986
613.3687133789062 0 343171 b 5
614.3717651367188 0 112895.81
615.3742065429688 0 22993.312
616.3775024414062 0 3348.7112
617.8505249023438 0 1076.5215
618.3619995117188 0 847.4137
621.366943359375 0 2921.8713
622.3616333007812 0 4697.1
623.3543701171875 0 6785.9634
624.3565673828125 0 2012.393
625.8609619140625 0 4560.289 y Water loss 1
626.3562622070312 0 6821.133 y Ammonia loss 1
626.858154296875 0 3434.9531
627.2824096679688 0 1087.8076
627.3548583984375 0 825.35754
627.7838134765625 0 676.59375
632.354736328125 0 712.9404
633.875732421875 0 1515.3297
634.3795776367188 0 700.3766
634.8663330078125 0 73235.71 y 1
635.3674926757812 0 49960.344
635.8685302734375 0 18030.904
636.373291015625 0 36268.28
636.8695068359375 0 1038.4572
637.3748779296875 0 13160.453
638.376220703125 0 3140.5952
639.3843994140625 0 47978.66
639.8583984375 0 2686.2378
640.3811645507812 0 24663.828
641.3795776367188 0 6229.9526
642.3836059570312 0 1586.3333
647.35400390625 0 4091.2776
648.3604736328125 0 1579.1351
650.364013671875 0 5651.2627
651.35302734375 0 4690.892
652.3563842773438 0 1619.6936
654.392578125 0 12823.688
655.3955688476562 0 4368.065
656.4088134765625 0 2619.9897
657.4141235351562 0 1028.2793
658.3792114257812 0 73421.63 y 7
659.3821411132812 0 25952.133
660.384521484375 0 6178.133
661.3883056640625 0 1410.1808
664.379638671875 0 6324.9014
665.3658447265625 0 6938.2104
666.3634643554688 0 2755.5547
667.3818969726562 0 1990.362
668.37451171875 0 42905.652
669.3779296875 0 15516.6455
670.3807373046875 0 3031.154
671.3812866210938 0 1607.8171
675.3446655273438 0 753.01855
682.3905639648438 0 60761.996
683.3935546875 0 20997.094
684.4031372070312 0 17542.295 Precursor
685.4070434570312 0 5441.893
686.4113159179688 0 892.2701
691.45361328125 0 1544.2505
692.3851928710938 0 1148.392
693.3619995117188 0 1450.115
701.397705078125 0 1795.5793
710.38427734375 0 2326.092
710.4218139648438 0 5474.163 b 6
711.3812866210938 0 932.94147
711.4300537109375 0 1363.9529
712.3662109375 0 1909.3033
713.3657836914062 0 937.05725
714.37109375 0 853.0272
719.408203125 0 1118.5826
736.440185546875 0 4059.198
737.4242553710938 0 8353.471 y Water loss 6
738.4242553710938 0 4228.9673
739.4257202148438 0 815.59106
746.4238891601562 0 1644.066
747.4268798828125 0 722.4748
749.4578857421875 0 2750.501
750.4620971679688 0 1157.4327
751.4484252929688 0 971.32294
753.4637451171875 0 22727.355
754.4672241210938 0 7845.914
755.43212890625 0 163204.97 y 6
756.4351196289062 0 71725.27
757.4380493164062 0 17484.314
758.4392700195312 0 2795.5662
759.447509765625 0 776.3282
763.4475708007812 0 8383.632 b Water loss 7
764.4374389648438 0 10571.137 b Ammonia loss 7
765.4375610351562 0 4451.355
766.4352416992188 0 1264.2028
779.445068359375 0 1827.4072
780.4457397460938 0 1459.8696
781.4589233398438 0 163574.02 b 7
782.4620361328125 0 65538.65
783.4644165039062 0 19887.312
784.467529296875 0 2612.6606
805.4964599609375 0 1681.6614
806.50244140625 0 1125.0443
811.4329223632812 0 1073.1614
820.4957885742188 0 2029.9756
821.4990844726562 0 954.4296
838.5000610351562 0 1048.2982
850.5033569335938 0 1575.8491 y Water loss 5
864.5376586914062 0 938.70514
868.5164184570312 0 19001.729 y 5
869.5188598632812 0 9420.304
870.5225219726562 0 2767.08
892.52685546875 0 3617.5576
893.5303344726562 0 1057.4493
894.5354614257812 0 801.48004
921.5429077148438 0 827.2677 y Water loss 4
922.5404052734375 0 741.1581 y Ammonia loss 4
934.5420532226562 0 1539.2839
935.5374755859375 0 1448.1936
939.5531616210938 0 20630.084 y 4
940.5555419921875 0 11723.794
941.5584716796875 0 2273.4111
991.5925903320312 0 1927.515 b 9
1004.5440063476562 0 979.8617
1021.5706176757812 0 6710.1265
1022.5743408203125 0 3232.327
1023.5786743164062 0 795.0375
1067.6473388671875 0 2044.4534 y 3
1068.65234375 0 1486.4257
1120.6363525390625 0 2634.4814 b 10
1121.6453857421875 0 1485.2242
3074.051513671875 0 758.9378

Spectrum Details

|  |  |
| --- | --- |
| Matched peaks? Matched peaks The total absolute number of peaks matched. Additionally in brackets the total fraction of peaks matched and the total number of peaks is shown. | 85 (8.76% of 970) |
| FDR? FDR The false discovery rate estimated for this peptide. It is calculated by matching all theoretical fragments with a non-integer shift with the raw peaks for this spectrum. This is done with 40 different shifts. The resulting percentage is the average number of annotated peaks over the number of annotated peaks with the correct spectrum. | 0.08% |
| Satellite FDR? Satellite FDR See the FDR for details on its calculation. This satellite ion specific FDR only contains the satellite ions (d/w) for I/L/J positions. | - |
| PSM Score? PSM Score The PSM Score as given by Hecklib to this annotated spectrum. It is shown with three significant figures. | 649 |

#### Spectrum 4150? Spectrum 4150 The raw spectrum of this peptide as annotated by Hecklib. The fragments are coloured according to ion type (see legend). Any peaks with a star '\*' as text can be hovered over to see the full details, first the ion type second the mass shift type. By hovering over the amino acids in the peptide or ions in the legend the corresponding peaks are highlighted. By toggling the 'Unassigned' label you can turn the background (unassigned) peaks on or off in the plot. By updating the slider in the Ion legend you can update the spectrum to only show the top X% of the peaks with labels. The top X% means any peak that is within X% of the highest intensity. By dragging in the spectrum you can zoom in to a specific part of the spectrum and use 'Zoom Out' to get back to the original zoom level. The annotation of the spectrum is based on the given sequence in the peptides file and is done with different software so inconsistencies are likely. The peaks are annotated based on the given sequence, with 20 ppm tolerance.

Copy Data

##### Spectrum 4150 (TSV)

###### Preview

```
Loading example...
```

*Click on the button to copy the data to your clipboard.*

Mz MinMz MaxIntensity Max

WidthHeightPeptide font sizePeptide stroke widthSpectrum font sizeSpectrum stroke widthCompact peptide

Ion legend

wxyz

abcd

OtherUnassignedIon^Charge^_Position_Show for top:%

VSNKALPAPIEKT

02.22e+44.45e+46.67e+48.89e+4

Zoom Out

y^+1^_1_w^+1^_2_z^+2^_3_y^+1^_2_y^+1^_2_z^+1^_2_y^+1^_2_y^+2^_5_y^+2^_5_c^+1^_3_w^+1^_3_c^+2^_6_c^+1^_3_c^+2^_7_y^+1^_3_z^+1^_3_y^+1^_3_y^+2^_7_c^+2^_8_c^+1^_4_w^+1^_4_c^+1^_4_w^+1^_4_z^+1^_4_y^+1^_4_c^+1^_5_c^+1^_5_z^+2^_10_y^+2^_10_w^+1^_5_w^+2^_11_c^+2^_11_z^+2^_11_y^+1^_5_y^+2^_11_c^+1^_6_c^+1^_6_w^+2^_12_c^+2^_12_z^+1^_6_y^+2^_12_z^+2^_12_c^+2^_12_y^+2^_12_y^+1^_6_z^+1^_6_y^+1^_6_w^+1^_7_c^+1^_7_y^+1^_7_c^+1^_8_c^+1^_8_w^+1^_8_z^+1^_8_z^+1^_8_c^+1^_9_z^+1^_9_y^+1^_9_c^+1^_10_w^+1^_10_c^+1^_10_y^+1^_10_z^+1^_10_y^+1^_10_w^+1^_11_c^+1^_11_z^+1^_11_w^+1^_12_z^+1^_12_c^+1^_12_

0645128919342578

Fragment Matches Table

Show background peaks

| Position | Ion type | Intensity | mz Theoretical | mz Error (Th) | mz Error (ppm) | Charge | Series Number |
| --- | --- | --- | --- | --- | --- | --- | --- |
| 13 | y | 1929 | 120.1 | 7.029E-05 | 0.5854 | +1 | 1 |
| - | - | 1466 | 120.1 | - | - | 0 | - |
| - | - | 850.3 | 127.1 | - | - | 0 | - |
| - | - | 1004 | 128.1 | - | - | 0 | - |
| - | - | 2278 | 129.1 | - | - | 0 | - |
| - | - | 4101 | 129.1 | - | - | 0 | - |
| - | - | 725.1 | 131.1 | - | - | 0 | - |
| - | - | 3050 | 132.1 | - | - | 0 | - |
| - | - | 1517 | 136.1 | - | - | 0 | - |
| - | - | 418.5 | 141.1 | - | - | 0 | - |
| - | - | 4989 | 141.1 | - | - | 0 | - |
| - | - | 1869 | 141.1 | - | - | 0 | - |
| - | - | 474.9 | 141.4 | - | - | 0 | - |
| - | - | 835.3 | 142.1 | - | - | 0 | - |
| - | - | 1631 | 147.1 | - | - | 0 | - |
| - | - | 645.8 | 148.9 | - | - | 0 | - |
| - | - | 427.1 | 154.6 | - | - | 0 | - |
| - | - | 486.3 | 158.1 | - | - | 0 | - |
| - | - | 589.8 | 158.7 | - | - | 0 | - |
| - | - | 2069 | 159.1 | - | - | 0 | - |
| - | - | 976.7 | 165.1 | - | - | 0 | - |
| - | - | 2525 | 169.1 | - | - | 0 | - |
| - | - | 439.4 | 171.7 | - | - | 0 | - |
| - | - | 3095 | 173.4 | - | - | 0 | - |
| 12 | w | 2820 | 174.1 | 0.0002249 | 1.292 | +1 | 2 |
| - | - | 515.5 | 176.3 | - | - | 0 | - |
| - | - | 465.1 | 178.3 | - | - | 0 | - |
| 11 | z | 1059 | 181.1 | 0.001507 | 8.319 | +2 | 3 |
| - | - | 1.33E+04 | 183.1 | - | - | 0 | - |
| - | - | 822.1 | 183.1 | - | - | 0 | - |
| - | - | 1492 | 184.1 | - | - | 0 | - |
| - | - | 764.5 | 186.1 | - | - | 0 | - |
| - | - | 2055 | 187.1 | - | - | 0 | - |
| - | - | 762.6 | 187.1 | - | - | 0 | - |
| - | - | 459.4 | 189.9 | - | - | 0 | - |
| - | - | 669 | 191.1 | - | - | 0 | - |
| - | - | 518.6 | 194.2 | - | - | 0 | - |
| - | - | 492.6 | 199.1 | - | - | 0 | - |
| - | - | 635.9 | 201.1 | - | - | 0 | - |
| - | - | 1647 | 202.1 | - | - | 0 | - |
| - | - | 6398 | 209.1 | - | - | 0 | - |
| - | - | 1333 | 211.1 | - | - | 0 | - |
| - | - | 4134 | 213.1 | - | - | 0 | - |
| - | - | 1009 | 215.1 | - | - | 0 | - |
| - | - | 538 | 217.2 | - | - | 0 | - |
| - | - | 867.1 | 219.1 | - | - | 0 | - |
| - | - | 640.7 | 225.1 | - | - | 0 | - |
| - | - | 833.4 | 226.1 | - | - | 0 | - |
| - | - | 564.3 | 227.1 | - | - | 0 | - |
| 12 | y | 1173 | 230.1 | 0.0004886 | 2.123 | +1 | 2 |
| 12 | y | 7625 | 231.1 | 8.441E-05 | 0.3652 | +1 | 2 |
| 12 | z | 1152 | 232.1 | 0.000316 | 1.361 | +1 | 2 |
| - | - | 1992 | 240.1 | - | - | 0 | - |
| - | - | 1094 | 242.1 | - | - | 0 | - |
| - | - | 1601 | 243.1 | - | - | 0 | - |
| - | - | 2481 | 244.1 | - | - | 0 | - |
| - | - | 510.6 | 244.1 | - | - | 0 | - |
| 12 | y | 1509 | 248.2 | 0.0005586 | 2.251 | +1 | 2 |
| - | - | 575.1 | 249.2 | - | - | 0 | - |
| - | - | 798.2 | 251.1 | - | - | 0 | - |
| - | - | 1622 | 258.1 | - | - | 0 | - |
| - | - | 7433 | 260.1 | - | - | 0 | - |
| - | - | 693.2 | 273.2 | - | - | 0 | - |
| - | - | 2260 | 274.1 | - | - | 0 | - |
| - | - | 1456 | 274.2 | - | - | 0 | - |
| - | - | 749.8 | 275.1 | - | - | 0 | - |
| - | - | 2874 | 279.1 | - | - | 0 | - |
| - | - | 3857 | 284.2 | - | - | 0 | - |
| 9 | y | 639.7 | 285.2 | 0.0008882 | 3.115 | +2 | 5 |
| - | - | 2929 | 293.2 | - | - | 0 | - |
| - | - | 1113 | 293.7 | - | - | 0 | - |
| 9 | y | 1.765E+04 | 294.2 | 8.553E-05 | 0.2907 | +2 | 5 |
| - | - | 1171 | 294.2 | - | - | 0 | - |
| - | - | 5576 | 294.7 | - | - | 0 | - |
| - | - | 1577 | 295.2 | - | - | 0 | - |
| - | - | 591.1 | 295.7 | - | - | 0 | - |
| - | - | 1370 | 299.2 | - | - | 0 | - |
| 3 | c | 2407 | 301.2 | 0.0004463 | 1.482 | +1 | 3 |
| 11 | w | 2.542E+04 | 302.2 | 0.0002173 | 0.7192 | +1 | 3 |
| - | - | 3669 | 303.2 | - | - | 0 | - |
| - | - | 615.5 | 306.2 | - | - | 0 | - |
| 6 | c | 925 | 307.2 | 0.0005903 | 1.922 | +2 | 6 |
| - | - | 1388 | 310.1 | - | - | 0 | - |
| - | - | 998.5 | 313.2 | - | - | 0 | - |
| - | - | 1113 | 314.1 | - | - | 0 | - |
| 3 | c | 1.346E+04 | 318.2 | 0.0002644 | 0.8309 | +1 | 3 |
| - | - | 628.1 | 318.7 | - | - | 0 | - |
| - | - | 3051 | 319.1 | - | - | 0 | - |
| - | - | 1713 | 319.2 | - | - | 0 | - |
| - | - | 753.7 | 327.7 | - | - | 0 | - |
| - | - | 819.8 | 328.2 | - | - | 0 | - |
| - | - | 747.8 | 330.2 | - | - | 0 | - |
| - | - | 829.7 | 334.7 | - | - | 0 | - |
| - | - | 543.3 | 336.5 | - | - | 0 | - |
| - | - | 1699 | 337.2 | - | - | 0 | - |
| - | - | 2889 | 338.1 | - | - | 0 | - |
| - | - | 1942 | 340.2 | - | - | 0 | - |
| - | - | 2736 | 341.7 | - | - | 0 | - |
| - | - | 1571 | 342.2 | - | - | 0 | - |
| - | - | 507.7 | 343.6 | - | - | 0 | - |
| - | - | 680.8 | 347.2 | - | - | 0 | - |
| - | - | 1394 | 349.1 | - | - | 0 | - |
| - | - | 540.1 | 352.2 | - | - | 0 | - |
| - | - | 8293 | 355.2 | - | - | 0 | - |
| - | - | 1028 | 355.2 | - | - | 0 | - |
| 7 | c | 7052 | 355.7 | 2.628E-05 | 0.07389 | +2 | 7 |
| - | - | 3231 | 356.2 | - | - | 0 | - |
| - | - | 751.3 | 356.7 | - | - | 0 | - |
| - | - | 821.8 | 357.2 | - | - | 0 | - |
| - | - | 1296 | 358.2 | - | - | 0 | - |
| 11 | y | 1046 | 359.2 | 0.001062 | 2.957 | +1 | 3 |
| 11 | z | 2477 | 361.2 | 9.658E-05 | 0.2674 | +1 | 3 |
| - | - | 4084 | 366.1 | - | - | 0 | - |
| - | - | 3678 | 366.7 | - | - | 0 | - |
| - | - | 1473 | 367.2 | - | - | 0 | - |
| - | - | 599.5 | 368.7 | - | - | 0 | - |
| - | - | 693.4 | 372.2 | - | - | 0 | - |
| - | - | 780.3 | 374.3 | - | - | 0 | - |
| 11 | y | 2950 | 377.2 | 0.0001338 | 0.3548 | +1 | 3 |
| - | - | 2578 | 377.2 | - | - | 0 | - |
| - | - | 1365 | 377.7 | - | - | 0 | - |
| 7 | y | 4910 | 378.2 | 0.0002366 | 0.6255 | +2 | 7 |
| - | - | 3429 | 378.7 | - | - | 0 | - |
| - | - | 558.9 | 379.2 | - | - | 0 | - |
| - | - | 4692 | 380.7 | - | - | 0 | - |
| - | - | 1146 | 381.2 | - | - | 0 | - |
| - | - | 789.3 | 381.7 | - | - | 0 | - |
| - | - | 709.3 | 388.7 | - | - | 0 | - |
| 8 | c | 2121 | 391.2 | 0.0004818 | 1.232 | +2 | 8 |
| - | - | 731.5 | 391.7 | - | - | 0 | - |
| - | - | 662.9 | 394.2 | - | - | 0 | - |
| - | - | 611.3 | 394.7 | - | - | 0 | - |
| - | - | 740.8 | 402.2 | - | - | 0 | - |
| - | - | 1203 | 402.3 | - | - | 0 | - |
| - | - | 3108 | 402.7 | - | - | 0 | - |
| - | - | 4367 | 403.2 | - | - | 0 | - |
| - | - | 2191 | 403.7 | - | - | 0 | - |
| - | - | 755.3 | 404.7 | - | - | 0 | - |
| - | - | 5350 | 409.2 | - | - | 0 | - |
| - | - | 2557 | 409.7 | - | - | 0 | - |
| - | - | 1247 | 410.2 | - | - | 0 | - |
| - | - | 582.4 | 410.7 | - | - | 0 | - |
| - | - | 643.7 | 411.2 | - | - | 0 | - |
| - | - | 915.1 | 412.2 | - | - | 0 | - |
| - | - | 1219 | 415.2 | - | - | 0 | - |
| - | - | 619.5 | 415.7 | - | - | 0 | - |
| - | - | 1.985E+04 | 418.2 | - | - | 0 | - |
| - | - | 1.117E+04 | 418.7 | - | - | 0 | - |
| - | - | 2513 | 419.2 | - | - | 0 | - |
| - | - | 1013 | 419.7 | - | - | 0 | - |
| - | - | 819.4 | 421.2 | - | - | 0 | - |
| - | - | 5251 | 424.2 | - | - | 0 | - |
| - | - | 1903 | 424.7 | - | - | 0 | - |
| 4 | c | 3816 | 429.2 | 0.0003624 | 0.8444 | +1 | 4 |
| - | - | 698 | 430.3 | - | - | 0 | - |
| - | - | 1241 | 431.2 | - | - | 0 | - |
| - | - | 6933 | 433.2 | - | - | 0 | - |
| - | - | 3071 | 433.7 | - | - | 0 | - |
| - | - | 549.2 | 434.2 | - | - | 0 | - |
| - | - | 1463 | 437.2 | - | - | 0 | - |
| - | - | 4687 | 439.2 | - | - | 0 | - |
| - | - | 698.7 | 440.2 | - | - | 0 | - |
| 10 | w | 3920 | 445.2 | 0.0002933 | 0.6587 | +1 | 4 |
| - | - | 2388 | 446.2 | - | - | 0 | - |
| - | - | 917.5 | 446.2 | - | - | 0 | - |
| 4 | c | 2.428E+04 | 446.3 | 0.0004247 | 0.9516 | +1 | 4 |
| - | - | 1301 | 446.7 | - | - | 0 | - |
| - | - | 5103 | 447.3 | - | - | 0 | - |
| - | - | 3209 | 451.9 | - | - | 0 | - |
| - | - | 6684 | 455.2 | - | - | 0 | - |
| - | - | 2385 | 455.7 | - | - | 0 | - |
| - | - | 1651 | 456.2 | - | - | 0 | - |
| - | - | 1442 | 456.2 | - | - | 0 | - |
| - | - | 7491 | 457.2 | - | - | 0 | - |
| - | - | 2025 | 457.3 | - | - | 0 | - |
| 10 | w | 1163 | 459.2 | 0.000848 | 1.847 | +1 | 4 |
| - | - | 3125 | 468.3 | - | - | 0 | - |
| 10 | z | 613.7 | 474.3 | 0.004194 | 8.844 | +1 | 4 |
| - | - | 785.2 | 481.3 | - | - | 0 | - |
| - | - | 2058 | 482.3 | - | - | 0 | - |
| - | - | 937.7 | 483.2 | - | - | 0 | - |
| 10 | y | 2755 | 490.3 | 0.0006411 | 1.308 | +1 | 4 |
| - | - | 741.7 | 491.3 | - | - | 0 | - |
| - | - | 7291 | 495.3 | - | - | 0 | - |
| - | - | 1848 | 496.2 | - | - | 0 | - |
| - | - | 1149 | 497.3 | - | - | 0 | - |
| - | - | 2672 | 498.3 | - | - | 0 | - |
| - | - | 1103 | 498.8 | - | - | 0 | - |
| 5 | c | 5440 | 500.3 | 8.189E-06 | 0.01637 | +1 | 5 |
| - | - | 1668 | 501.3 | - | - | 0 | - |
| - | - | 869.3 | 503.2 | - | - | 0 | - |
| - | - | 660.3 | 503.8 | - | - | 0 | - |
| - | - | 677 | 508.3 | - | - | 0 | - |
| - | - | 6886 | 508.8 | - | - | 0 | - |
| - | - | 1634 | 510.2 | - | - | 0 | - |
| - | - | 899 | 511.2 | - | - | 0 | - |
| - | - | 8639 | 512.2 | - | - | 0 | - |
| - | - | 2690 | 513.2 | - | - | 0 | - |
| - | - | 1640 | 514.3 | - | - | 0 | - |
| - | - | 835 | 515.3 | - | - | 0 | - |
| 5 | c | 2.687E+04 | 517.3 | 0.0003592 | 0.6944 | +1 | 5 |
| - | - | 5218 | 518.3 | - | - | 0 | - |
| - | - | 1220 | 519.3 | - | - | 0 | - |
| 4 | z | 4075 | 526.3 | 0.0006478 | 1.231 | +2 | 10 |
| - | - | 2666 | 526.8 | - | - | 0 | - |
| 4 | y | 2830 | 534.3 | 0.001587 | 2.97 | +2 | 10 |
| - | - | 1106 | 534.8 | - | - | 0 | - |
| - | - | 7309 | 539.3 | - | - | 0 | - |
| - | - | 615 | 541.3 | - | - | 0 | - |
| - | - | 1217 | 542.8 | - | - | 0 | - |
| - | - | 1566 | 543.3 | - | - | 0 | - |
| 9 | w | 1846 | 544.3 | 0.0006354 | 1.167 | +1 | 5 |
| - | - | 991.2 | 545.3 | - | - | 0 | - |
| - | - | 6143 | 555.3 | - | - | 0 | - |
| - | - | 2815 | 555.8 | - | - | 0 | - |
| - | - | 873.1 | 556.3 | - | - | 0 | - |
| - | - | 3744 | 556.8 | - | - | 0 | - |
| - | - | 638.5 | 558.3 | - | - | 0 | - |
| - | - | 744.9 | 560.3 | - | - | 0 | - |
| 3 | w | 8560 | 561.3 | 0.0003328 | 0.5929 | +2 | 11 |
| - | - | 4826 | 561.8 | - | - | 0 | - |
| - | - | 1420 | 562.3 | - | - | 0 | - |
| - | - | 730.5 | 562.8 | - | - | 0 | - |
| 11 | c | 1048 | 569.3 | 0.003901 | 6.852 | +2 | 11 |
| - | - | 1519 | 570.3 | - | - | 0 | - |
| - | - | 1120 | 574.8 | - | - | 0 | - |
| - | - | 789.6 | 576.8 | - | - | 0 | - |
| - | - | 821 | 578.3 | - | - | 0 | - |
| 3 | z | 4998 | 583.3 | 0.0005935 | 1.017 | +2 | 11 |
| - | - | 4315 | 583.8 | - | - | 0 | - |
| - | - | 1218 | 584.3 | - | - | 0 | - |
| - | - | 1462 | 584.8 | - | - | 0 | - |
| - | - | 2792 | 585.3 | - | - | 0 | - |
| - | - | 997.9 | 585.4 | - | - | 0 | - |
| - | - | 1010 | 586.3 | - | - | 0 | - |
| - | - | 2747 | 586.4 | - | - | 0 | - |
| 9 | y | 2.348E+04 | 587.3 | 0.0002454 | 0.4178 | +1 | 5 |
| - | - | 1168 | 587.4 | - | - | 0 | - |
| - | - | 7273 | 588.3 | - | - | 0 | - |
| - | - | 1411 | 589.3 | - | - | 0 | - |
| - | - | 630.2 | 590.8 | - | - | 0 | - |
| 3 | y | 3922 | 591.3 | 0.0004067 | 0.6877 | +2 | 11 |
| - | - | 2707 | 591.8 | - | - | 0 | - |
| - | - | 3100 | 592.2 | - | - | 0 | - |
| - | - | 1187 | 592.4 | - | - | 0 | - |
| - | - | 1075 | 593.2 | - | - | 0 | - |
| - | - | 1613 | 595.4 | - | - | 0 | - |
| - | - | 829.8 | 595.8 | - | - | 0 | - |
| - | - | 900.9 | 596.3 | - | - | 0 | - |
| - | - | 3048 | 597.3 | - | - | 0 | - |
| - | - | 818.2 | 597.8 | - | - | 0 | - |
| - | - | 1056 | 598.3 | - | - | 0 | - |
| - | - | 3306 | 598.3 | - | - | 0 | - |
| - | - | 1356 | 598.8 | - | - | 0 | - |
| - | - | 926.8 | 599.3 | - | - | 0 | - |
| - | - | 1089 | 611.4 | - | - | 0 | - |
| - | - | 912.8 | 611.9 | - | - | 0 | - |
| 6 | c | 1712 | 612.4 | 0.002644 | 4.318 | +1 | 6 |
| - | - | 1523 | 612.9 | - | - | 0 | - |
| - | - | 992.5 | 613.3 | - | - | 0 | - |
| 6 | c | 1.18E+04 | 613.4 | 0.0007057 | 1.15 | +1 | 6 |
| - | - | 2002 | 613.8 | - | - | 0 | - |
| - | - | 3626 | 614.4 | - | - | 0 | - |
| 2 | w | 831.2 | 618.4 | 0.0007807 | 1.263 | +2 | 12 |
| - | - | 942.3 | 619.4 | - | - | 0 | - |
| - | - | 958 | 620.9 | - | - | 0 | - |
| - | - | 1009 | 624.8 | - | - | 0 | - |
| 12 | c | 814.6 | 624.9 | 0.005349 | 8.56 | +2 | 12 |
| 8 | z | 794.2 | 625.3 | 0.01064 | 17.01 | +1 | 6 |
| - | - | 889.9 | 625.4 | - | - | 0 | - |
| 2 | y | 2247 | 625.9 | 0.009735 | 15.55 | +2 | 12 |
| - | - | 5085 | 626.4 | - | - | 0 | - |
| 2 | z | 1956 | 626.9 | 0.003153 | 5.029 | +2 | 12 |
| - | - | 3452 | 626.9 | - | - | 0 | - |
| - | - | 3863 | 627.4 | - | - | 0 | - |
| - | - | 1173 | 627.9 | - | - | 0 | - |
| - | - | 785.1 | 632.4 | - | - | 0 | - |
| - | - | 1463 | 632.8 | - | - | 0 | - |
| - | - | 1364 | 632.9 | - | - | 0 | - |
| 12 | c | 7.381E+04 | 633.4 | 0.0001313 | 0.2073 | +2 | 12 |
| - | - | 2321 | 633.8 | - | - | 0 | - |
| - | - | 4.593E+04 | 633.9 | - | - | 0 | - |
| - | - | 1.883E+04 | 634.4 | - | - | 0 | - |
| 2 | y | 8799 | 634.9 | 0.004087 | 6.437 | +2 | 12 |
| - | - | 5043 | 635.4 | - | - | 0 | - |
| - | - | 1574 | 635.9 | - | - | 0 | - |
| - | - | 922.1 | 639.4 | - | - | 0 | - |
| - | - | 983 | 639.9 | - | - | 0 | - |
| 8 | y | 5076 | 640.4 | 0.01197 | 18.69 | +1 | 6 |
| - | - | 2639 | 640.9 | - | - | 0 | - |
| - | - | 1572 | 641.4 | - | - | 0 | - |
| 8 | z | 3.729E+04 | 642.4 | 0.004534 | 7.058 | +1 | 6 |
| - | - | 1240 | 642.8 | - | - | 0 | - |
| - | - | 1.074E+04 | 643.4 | - | - | 0 | - |
| - | - | 2645 | 644.4 | - | - | 0 | - |
| - | - | 5776 | 648.4 | - | - | 0 | - |
| - | - | 3588 | 648.9 | - | - | 0 | - |
| - | - | 1817 | 648.9 | - | - | 0 | - |
| - | - | 2720 | 649.4 | - | - | 0 | - |
| - | - | 2205 | 650.3 | - | - | 0 | - |
| - | - | 749.9 | 651.3 | - | - | 0 | - |
| - | - | 2755 | 653.9 | - | - | 0 | - |
| - | - | 2704 | 654.4 | - | - | 0 | - |
| - | - | 934.2 | 654.9 | - | - | 0 | - |
| - | - | 3626 | 655.4 | - | - | 0 | - |
| - | - | 2550 | 655.9 | - | - | 0 | - |
| - | - | 2248 | 656.3 | - | - | 0 | - |
| - | - | 1.381E+04 | 656.9 | - | - | 0 | - |
| - | - | 6438 | 657.4 | - | - | 0 | - |
| - | - | 3424 | 657.9 | - | - | 0 | - |
| - | - | 1.001E+04 | 658.3 | - | - | 0 | - |
| 8 | y | 4981 | 658.4 | 0.0007903 | 1.2 | +1 | 6 |
| - | - | 927.7 | 658.8 | - | - | 0 | - |
| - | - | 2009 | 659.3 | - | - | 0 | - |
| - | - | 1090 | 659.4 | - | - | 0 | - |
| - | - | 1021 | 661.9 | - | - | 0 | - |
| - | - | 8282 | 662.4 | - | - | 0 | - |
| - | - | 6852 | 662.9 | - | - | 0 | - |
| - | - | 4313 | 663.4 | - | - | 0 | - |
| - | - | 1578 | 663.9 | - | - | 0 | - |
| - | - | 910.5 | 664.4 | - | - | 0 | - |
| - | - | 6106 | 665.3 | - | - | 0 | - |
| - | - | 4044 | 667.4 | - | - | 0 | - |
| - | - | 2381 | 667.9 | - | - | 0 | - |
| - | - | 2636 | 668.4 | - | - | 0 | - |
| - | - | 1.342E+04 | 668.9 | - | - | 0 | - |
| - | - | 9709 | 669.4 | - | - | 0 | - |
| - | - | 3322 | 669.9 | - | - | 0 | - |
| - | - | 2682 | 670.4 | - | - | 0 | - |
| - | - | 1207 | 670.9 | - | - | 0 | - |
| - | - | 730 | 675.4 | - | - | 0 | - |
| - | - | 1995 | 675.9 | - | - | 0 | - |
| - | - | 3.428E+04 | 676.4 | - | - | 0 | - |
| - | - | 2.157E+04 | 676.9 | - | - | 0 | - |
| - | - | 9011 | 677.4 | - | - | 0 | - |
| - | - | 826.3 | 677.8 | - | - | 0 | - |
| - | - | 1813 | 677.9 | - | - | 0 | - |
| - | - | 2.01E+04 | 678.3 | - | - | 0 | - |
| - | - | 2141 | 682.4 | - | - | 0 | - |
| - | - | 1858 | 683.4 | - | - | 0 | - |
| - | - | 1755 | 683.9 | - | - | 0 | - |
| - | - | 6.183E+04 | 684.4 | - | - | 0 | - |
| - | - | 8.804E+04 | 684.9 | - | - | 0 | - |
| - | - | 5.247E+04 | 685.4 | - | - | 0 | - |
| - | - | 2.156E+04 | 685.9 | - | - | 0 | - |
| - | - | 1.627E+04 | 686.3 | - | - | 0 | - |
| - | - | 4204 | 686.4 | - | - | 0 | - |
| - | - | 7054 | 686.9 | - | - | 0 | - |
| - | - | 770.6 | 686.9 | - | - | 0 | - |
| - | - | 735.9 | 701.4 | - | - | 0 | - |
| 7 | w | 841.5 | 712.4 | 0.001578 | 2.215 | +1 | 7 |
| - | - | 1213 | 713.4 | - | - | 0 | - |
| - | - | 7045 | 715.4 | - | - | 0 | - |
| 7 | c | 4.67E+04 | 727.4 | 6.702E-05 | 0.09214 | +1 | 7 |
| - | - | 952.6 | 728.4 | - | - | 0 | - |
| - | - | 1.627E+04 | 728.4 | - | - | 0 | - |
| - | - | 4166 | 729.5 | - | - | 0 | - |
| - | - | 1166 | 731.4 | - | - | 0 | - |
| - | - | 760.2 | 747.4 | - | - | 0 | - |
| - | - | 1253 | 749.3 | - | - | 0 | - |
| - | - | 5338 | 750.3 | - | - | 0 | - |
| - | - | 2373 | 751.3 | - | - | 0 | - |
| - | - | 2026 | 754.5 | - | - | 0 | - |
| 7 | y | 1.076E+04 | 755.4 | 0.0003335 | 0.4415 | +1 | 7 |
| - | - | 4418 | 756.4 | - | - | 0 | - |
| - | - | 1886 | 757.4 | - | - | 0 | - |
| 8 | c | 1004 | 780.5 | 0.002998 | 3.841 | +1 | 8 |
| 8 | c | 6923 | 781.5 | 0.001526 | 1.953 | +1 | 8 |
| - | - | 2124 | 782.3 | - | - | 0 | - |
| - | - | 2658 | 782.5 | - | - | 0 | - |
| - | - | 2153 | 783.3 | - | - | 0 | - |
| - | - | 1688 | 783.4 | - | - | 0 | - |
| - | - | 1210 | 784.3 | - | - | 0 | - |
| - | - | 1004 | 784.4 | - | - | 0 | - |
| - | - | 1668 | 808.4 | - | - | 0 | - |
| 6 | w | 7939 | 809.4 | 0.0003434 | 0.4243 | +1 | 8 |
| - | - | 3702 | 810.4 | - | - | 0 | - |
| - | - | 2080 | 813.4 | - | - | 0 | - |
| - | - | 6111 | 814.4 | - | - | 0 | - |
| - | - | 4051 | 821.4 | - | - | 0 | - |
| - | - | 1188 | 821.5 | - | - | 0 | - |
| - | - | 3985 | 822.4 | - | - | 0 | - |
| - | - | 2047 | 822.5 | - | - | 0 | - |
| - | - | 1744 | 823.4 | - | - | 0 | - |
| - | - | 955.3 | 823.5 | - | - | 0 | - |
| - | - | 1167 | 824.5 | - | - | 0 | - |
| - | - | 969.7 | 825.5 | - | - | 0 | - |
| 6 | z | 791 | 834.5 | 0.005614 | 6.727 | +1 | 8 |
| - | - | 5368 | 837.3 | - | - | 0 | - |
| - | - | 1499 | 838.3 | - | - | 0 | - |
| - | - | 1450 | 839.4 | - | - | 0 | - |
| - | - | 2058 | 851.5 | - | - | 0 | - |
| 6 | z | 1.675E+04 | 852.5 | 0.0009726 | 1.141 | +1 | 8 |
| - | - | 9403 | 853.5 | - | - | 0 | - |
| - | - | 1664 | 854.3 | - | - | 0 | - |
| - | - | 2709 | 854.5 | - | - | 0 | - |
| - | - | 776.9 | 866.4 | - | - | 0 | - |
| - | - | 1.47E+04 | 867.5 | - | - | 0 | - |
| - | - | 6266 | 868.5 | - | - | 0 | - |
| - | - | 876.7 | 869.5 | - | - | 0 | - |
| - | - | 840.9 | 877.4 | - | - | 0 | - |
| - | - | 8742 | 878.4 | - | - | 0 | - |
| - | - | 2749 | 892.4 | - | - | 0 | - |
| - | - | 808.4 | 893.4 | - | - | 0 | - |
| - | - | 1166 | 894.5 | - | - | 0 | - |
| - | - | 2.014E+04 | 895.4 | - | - | 0 | - |
| 9 | c | 4949 | 895.5 | 0.001493 | 1.667 | +1 | 9 |
| - | - | 1.215E+04 | 896.4 | - | - | 0 | - |
| - | - | 2889 | 896.5 | - | - | 0 | - |
| - | - | 3589 | 897.4 | - | - | 0 | - |
| - | - | 827.8 | 898.4 | - | - | 0 | - |
| - | - | 6214 | 909.4 | - | - | 0 | - |
| - | - | 3.552E+04 | 910.4 | - | - | 0 | - |
| - | - | 1.875E+04 | 911.4 | - | - | 0 | - |
| - | - | 6775 | 912.4 | - | - | 0 | - |
| - | - | 1624 | 913.4 | - | - | 0 | - |
| - | - | 3090 | 914.5 | - | - | 0 | - |
| - | - | 6539 | 915.5 | - | - | 0 | - |
| - | - | 1361 | 918.5 | - | - | 0 | - |
| 5 | z | 1.4E+04 | 923.5 | 0.0001748 | 0.1892 | +1 | 9 |
| - | - | 8343 | 924.5 | - | - | 0 | - |
| - | - | 2412 | 925.5 | - | - | 0 | - |
| 5 | y | 1.544E+04 | 939.6 | 0.0002998 | 0.3191 | +1 | 9 |
| - | - | 9096 | 940.6 | - | - | 0 | - |
| - | - | 2266 | 941.6 | - | - | 0 | - |
| - | - | 906.6 | 942.6 | - | - | 0 | - |
| - | - | 2312 | 949.6 | - | - | 0 | - |
| - | - | 1403 | 950.6 | - | - | 0 | - |
| - | - | 981.3 | 951.6 | - | - | 0 | - |
| - | - | 950.6 | 964.6 | - | - | 0 | - |
| - | - | 2330 | 965.6 | - | - | 0 | - |
| - | - | 1001 | 966.6 | - | - | 0 | - |
| - | - | 954.7 | 989.6 | - | - | 0 | - |
| 10 | c | 872.6 | 991.6 | 0.007887 | 7.954 | +1 | 10 |
| 4 | w | 773.9 | 993.6 | 0.001709 | 1.72 | +1 | 10 |
| - | - | 2817 | 995.6 | - | - | 0 | - |
| - | - | 1810 | 996.6 | - | - | 0 | - |
| - | - | 7707 | 1007 | - | - | 0 | - |
| - | - | 5255 | 1008 | - | - | 0 | - |
| 10 | c | 6.044E+04 | 1009 | 0.0001685 | 0.1671 | +1 | 10 |
| - | - | 3.132E+04 | 1010 | - | - | 0 | - |
| - | - | 9802 | 1011 | - | - | 0 | - |
| - | - | 2651 | 1012 | - | - | 0 | - |
| - | - | 838.9 | 1016 | - | - | 0 | - |
| - | - | 1.074E+04 | 1017 | - | - | 0 | - |
| - | - | 787.1 | 1023 | - | - | 0 | - |
| - | - | 1239 | 1032 | - | - | 0 | - |
| - | - | 1351 | 1033 | - | - | 0 | - |
| - | - | 760.1 | 1035 | - | - | 0 | - |
| 4 | y | 859.9 | 1051 | 0.005505 | 5.24 | +1 | 10 |
| 4 | z | 2.464E+04 | 1052 | 0.0001227 | 0.1167 | +1 | 10 |
| - | - | 2.694E+04 | 1053 | - | - | 0 | - |
| - | - | 1.159E+04 | 1054 | - | - | 0 | - |
| - | - | 5299 | 1055 | - | - | 0 | - |
| - | - | 1051 | 1056 | - | - | 0 | - |
| - | - | 2584 | 1064 | - | - | 0 | - |
| - | - | 1607 | 1065 | - | - | 0 | - |
| - | - | 1138 | 1067 | - | - | 0 | - |
| 4 | y | 4461 | 1068 | 0.0006583 | 0.6166 | +1 | 10 |
| - | - | 1998 | 1069 | - | - | 0 | - |
| - | - | 2379 | 1078 | - | - | 0 | - |
| - | - | 5851 | 1079 | - | - | 0 | - |
| - | - | 1543 | 1094 | - | - | 0 | - |
| - | - | 1008 | 1095 | - | - | 0 | - |
| - | - | 767.6 | 1096 | - | - | 0 | - |
| - | - | 1584 | 1108 | - | - | 0 | - |
| - | - | 1571 | 1110 | - | - | 0 | - |
| - | - | 934.3 | 1111 | - | - | 0 | - |
| 3 | w | 3247 | 1122 | 0.0002516 | 0.2243 | +1 | 11 |
| - | - | 1872 | 1123 | - | - | 0 | - |
| - | - | 881.4 | 1124 | - | - | 0 | - |
| - | - | 764.9 | 1136 | - | - | 0 | - |
| - | - | 937.7 | 1137 | - | - | 0 | - |
| 11 | c | 8.092E+04 | 1138 | 0.0004032 | 0.3544 | +1 | 11 |
| - | - | 4.613E+04 | 1139 | - | - | 0 | - |
| - | - | 1.986E+04 | 1140 | - | - | 0 | - |
| - | - | 4620 | 1141 | - | - | 0 | - |
| - | - | 918.4 | 1150 | - | - | 0 | - |
| 3 | z | 3159 | 1166 | 0.001017 | 0.8726 | +1 | 11 |
| - | - | 2.022E+04 | 1167 | - | - | 0 | - |
| - | - | 1.047E+04 | 1168 | - | - | 0 | - |
| - | - | 5155 | 1169 | - | - | 0 | - |
| - | - | 1141 | 1170 | - | - | 0 | - |
| - | - | 4509 | 1194 | - | - | 0 | - |
| - | - | 2184 | 1222 | - | - | 0 | - |
| - | - | 3208 | 1223 | - | - | 0 | - |
| - | - | 1309 | 1224 | - | - | 0 | - |
| 2 | w | 745.9 | 1236 | 0.003833 | 3.102 | +1 | 12 |
| - | - | 732.2 | 1239 | - | - | 0 | - |
| - | - | 1444 | 1250 | - | - | 0 | - |
| - | - | 801.9 | 1251 | - | - | 0 | - |
| 2 | z | 905.6 | 1253 | 0.00915 | 7.304 | +1 | 12 |
| - | - | 5359 | 1254 | - | - | 0 | - |
| - | - | 3355 | 1255 | - | - | 0 | - |
| - | - | 1247 | 1256 | - | - | 0 | - |
| 12 | c | 8022 | 1266 | 0.0007618 | 0.6018 | +1 | 12 |
| - | - | 1.653E+04 | 1267 | - | - | 0 | - |
| - | - | 9481 | 1268 | - | - | 0 | - |
| - | - | 4112 | 1269 | - | - | 0 | - |
| - | - | 1095 | 1270 | - | - | 0 | - |
| - | - | 2276 | 1284 | - | - | 0 | - |
| - | - | 1995 | 1285 | - | - | 0 | - |
| - | - | 2290 | 1308 | - | - | 0 | - |
| - | - | 1332 | 1311 | - | - | 0 | - |
| - | - | 5126 | 1314 | - | - | 0 | - |
| - | - | 1012 | 1315 | - | - | 0 | - |
| - | - | 4235 | 1324 | - | - | 0 | - |
| - | - | 1.923E+04 | 1325 | - | - | 0 | - |
| - | - | 1.172E+04 | 1326 | - | - | 0 | - |
| - | - | 4695 | 1327 | - | - | 0 | - |
| - | - | 1614 | 1328 | - | - | 0 | - |
| - | - | 893.7 | 1340 | - | - | 0 | - |
| - | - | 1755 | 1341 | - | - | 0 | - |
| - | - | 2348 | 1342 | - | - | 0 | - |
| - | - | 1477 | 1343 | - | - | 0 | - |
| - | - | 977.3 | 1351 | - | - | 0 | - |
| - | - | 6327 | 1352 | - | - | 0 | - |
| - | - | 4.065E+04 | 1353 | - | - | 0 | - |
| - | - | 2.581E+04 | 1354 | - | - | 0 | - |
| - | - | 1.086E+04 | 1355 | - | - | 0 | - |
| - | - | 2129 | 1356 | - | - | 0 | - |
| - | - | 2206 | 1356 | - | - | 0 | - |
| - | - | 1.821E+04 | 1357 | - | - | 0 | - |
| - | - | 1392 | 1358 | - | - | 0 | - |
| - | - | 643.7 | 1367 | - | - | 0 | - |
| - | - | 5466 | 1368 | - | - | 0 | - |
| - | - | 1.831E+04 | 1369 | - | - | 0 | - |
| - | - | 5.824E+04 | 1370 | - | - | 0 | - |
| - | - | 3.628E+04 | 1371 | - | - | 0 | - |
| - | - | 1.446E+04 | 1372 | - | - | 0 | - |
| - | - | 3525 | 1373 | - | - | 0 | - |
| - | - | 1475 | 1373 | - | - | 0 | - |
| - | - | 1.029E+04 | 1374 | - | - | 0 | - |
| - | - | 625.4 | 2553 | - | - | 0 | - |

m/z Charge Intensity FragmentType MassShift Position
120.06558990478516 0 1929.2662 y 12
120.08097839355469 0 1466.0469
127.08676147460938 0 850.32684
128.09463500976562 0 1003.9737
129.06597900390625 0 2278.116
129.10231018066406 0 4101.3027
131.11801147460938 0 725.0529
132.0768280029297 0 3049.5576
136.0758056640625 0 1516.8254
141.07394409179688 0 418.4718
141.07852172851562 0 4988.9897
141.10250854492188 0 1868.7363
141.35951232910156 0 474.93903
142.08212280273438 0 835.2562
147.07655334472656 0 1630.9146
148.94784545898438 0 645.79095
154.57676696777344 0 427.0541
158.06961059570312 0 486.27805
158.7045440673828 0 589.76605
159.1126708984375 0 2069.2966
165.10255432128906 0 976.737
169.0972137451172 0 2524.929
171.70907592773438 0 439.39032
173.44021606445312 0 3094.6565
174.07630920410156 0 2820.459 w 11
176.25608825683594 0 515.4544
178.28317260742188 0 465.12845
181.09732055664062 0 1059.4907 z 10
183.11293029785156 0 13297.954
183.1495361328125 0 822.1009
184.11648559570312 0 1491.9169
186.10011291503906 0 764.52814
187.10765075683594 0 2055.2432
187.14434814453125 0 762.5805
189.87057495117188 0 459.35492
191.11782836914062 0 668.9592
194.18392944335938 0 518.58984
199.1078338623047 0 492.63174
201.12356567382812 0 635.8958
202.0819854736328 0 1646.8698
209.09202575683594 0 6398.288
211.14407348632812 0 1333.1094
213.1234588623047 0 4134.418
215.13966369628906 0 1009.10455
217.1659393310547 0 538.0249
219.1091766357422 0 867.0928
225.1232452392578 0 640.739
226.1188201904297 0 833.39734
227.1023406982422 0 564.3072
230.14942932128906 0 1172.8085 y Water loss 11
231.13401794433594 0 7624.732 y Ammonia loss 11
232.14207458496094 0 1151.6787 z 11
240.13449096679688 0 1991.9828
242.1135711669922 0 1093.5892
243.145751953125 0 1600.9779
244.12918090820312 0 2481.433
244.1438446044922 0 510.56757
248.16104125976562 0 1508.9961 y 11
249.16090393066406 0 575.10944
251.1028594970703 0 798.15015
258.1444091796875 0 1621.8247
260.1240234375 0 7432.7334
273.15679931640625 0 693.1967
274.13983154296875 0 2259.6758
274.16265869140625 0 1455.7489
275.14459228515625 0 749.7852
279.0973815917969 0 2873.8242
284.16046142578125 0 3856.6492
285.16741943359375 0 639.6937 y Water loss 8
293.18975830078125 0 2928.727
293.6904296875 0 1112.9547
294.1736755371094 0 17648.43 y 8
294.1925354003906 0 1170.5236
294.6752014160156 0 5576.284
295.1769714355469 0 1577.334
295.67730712890625 0 591.11816
299.1707763671875 0 1369.9242
301.1510925292969 0 2407.2065 c Ammonia loss 2
302.1712646484375 0 25422.262 w 10
303.17462158203125 0 3668.7283
306.17974853515625 0 615.4798
307.1876220703125 0 924.9959 c Ammonia loss 5
310.1398010253906 0 1388.0886
313.1510314941406 0 998.46295
314.1356201171875 0 1112.8218
318.1774597167969 0 13460.542 c 2
318.66558837890625 0 628.08905
319.1479797363281 0 3051.2397
319.1811828613281 0 1713.0745
327.69970703125 0 753.6803
328.18524169921875 0 819.79956
330.1781005859375 0 747.77747
334.6898193359375 0 829.68536
336.5360107421875 0 543.3315
337.1507568359375 0 1699.3193
338.1346740722656 0 2889.1797
340.186767578125 0 1942.24
341.716552734375 0 2736.1584
342.2176208496094 0 1571.1394
343.6495056152344 0 507.74954
347.20166015625 0 680.78064
349.1181335449219 0 1394.2273
352.16259765625 0 540.05194
355.161376953125 0 8292.519
355.19915771484375 0 1027.7465
355.71343994140625 0 7051.8516 c Ammonia loss 6
356.215087890625 0 3230.938
356.71588134765625 0 751.2914
357.19097900390625 0 821.8476
358.2449035644531 0 1295.9456
359.1935729980469 0 1046.0034 y Water loss 10
361.1844482421875 0 2477.2559 z 10
366.14471435546875 0 4083.535
366.6658935546875 0 3678.3103
367.1653747558594 0 1472.6924
368.71923828125 0 599.45386
372.15496826171875 0 693.39777
374.2514953613281 0 780.3313
377.20294189453125 0 2949.7732 y 10
377.2347106933594 0 2578.257
377.7349548339844 0 1365.4202
378.2182922363281 0 4909.8633 y 6
378.719970703125 0 3429.2322
379.22308349609375 0 558.90393
380.6637268066406 0 4691.9688
381.1654968261719 0 1146.384
381.6649169921875 0 789.2626
388.68170166015625 0 709.3165
391.2324523925781 0 2121.4714 c Ammonia loss 7
391.73480224609375 0 731.5286
394.1693115234375 0 662.9106
394.6620178222656 0 611.2625
402.17547607421875 0 740.7668
402.2607116699219 0 1203.3091
402.67803955078125 0 3108.0234
403.1761779785156 0 4366.6606
403.6761474609375 0 2190.8708
404.7127380371094 0 755.28937
409.17449951171875 0 5349.5015
409.672119140625 0 2556.6
410.17071533203125 0 1246.6188
410.67333984375 0 582.4335
411.2241516113281 0 643.65717
412.2215881347656 0 915.0546
415.173828125 0 1219.3121
415.6669006347656 0 619.53125
418.18035888671875 0 19846.762
418.6818542480469 0 11168.096
419.1832580566406 0 2512.866
419.68365478515625 0 1012.73785
421.2102966308594 0 819.4008
424.180419921875 0 5251.235
424.680419921875 0 1902.851
429.2459716796875 0 3816.3489 c Ammonia loss 3
430.2515869140625 0 698.0145
431.1873779296875 0 1241.0878
433.1856689453125 0 6932.734
433.68719482421875 0 3070.902
434.1897888183594 0 549.2456
437.1867980957031 0 1463.0658
439.2191162109375 0 4686.6294
440.192626953125 0 698.6851
445.2295837402344 0 3919.8123 w 9
446.19512939453125 0 2388.3748
446.23602294921875 0 917.5179
446.2725830078125 0 24283.678 c 3
446.6955871582031 0 1300.89
447.275146484375 0 5103.1084
451.89813232421875 0 3208.877
455.200927734375 0 6684.1416
455.7015380859375 0 2384.9583
456.2007141113281 0 1650.654
456.2453918457031 0 1441.7728
457.22967529296875 0 7490.88
457.33819580078125 0 2024.7195
459.24578857421875 0 1162.9325 w 9
468.2818908691406 0 3125.4653
474.26422119140625 0 613.6899 z 9
481.3150634765625 0 785.2279
482.2718200683594 0 2057.5996
483.17938232421875 0 937.68225
490.28778076171875 0 2755.149 y 9
491.2903137207031 0 741.6606
495.2564392089844 0 7291.45
496.1623229980469 0 1847.5121
497.3210754394531 0 1149.4354
498.28582763671875 0 2671.533
498.78857421875 0 1103.1976
500.28271484375 0 5440.18 c Ammonia loss 4
501.2877197265625 0 1668.2958
503.2225341796875 0 869.3248
503.7742919921875 0 660.28424
508.2615051269531 0 677.00055
508.76177978515625 0 6885.675
510.17822265625 0 1633.8926
511.1825256347656 0 898.9691
512.1932373046875 0 8638.696
513.196533203125 0 2690.032
514.2997436523438 0 1640.4076
515.30078125 0 834.95404
517.3096313476562 0 26866.977 c 4
518.3124389648438 0 5218.059
519.3137817382812 0 1219.7965
526.3165893554688 0 4074.5713 z 3
526.8189086914062 0 2665.6172
534.3281860351562 0 2830.165 y 3
534.8265380859375 0 1105.7288
539.294677734375 0 7308.889
541.3289794921875 0 614.9508
542.79296875 0 1217.3982
543.3233032226562 0 1565.9762
544.29833984375 0 1845.6415 w 8
545.302490234375 0 991.1854
555.306884765625 0 6142.587
555.809326171875 0 2815.1057
556.3114013671875 0 873.06006
556.7908935546875 0 3743.9805
558.2731323242188 0 638.5322
560.2899169921875 0 744.8533
561.3322143554688 0 8560.409 w 2
561.8329467773438 0 4826.1543
562.3355102539062 0 1419.8385
562.835205078125 0 730.4836
569.3310546875 0 1048.0343 c 10
570.336181640625 0 1519.09
574.811767578125 0 1120.0042
576.84228515625 0 789.56323
578.2910766601562 0 821.0378
583.3392944335938 0 4998.1597 z 2
583.841064453125 0 4314.6426
584.3477783203125 0 1218.2994
584.845458984375 0 1462.2633
585.3233642578125 0 2792.4011
585.370361328125 0 997.8698
586.3279418945312 0 1009.59033
586.378662109375 0 2747.1772
587.3401489257812 0 23480.764 y 8
587.392578125 0 1168.4081
588.3430786132812 0 7272.8574
589.3468017578125 0 1410.9861
590.842529296875 0 630.19745
591.34765625 0 3922.19 y 2
591.8499755859375 0 2707.422
592.2307739257812 0 3100.1738
592.3523559570312 0 1186.9663
593.2328491210938 0 1075.0201
595.3564453125 0 1612.7614
595.8110961914062 0 829.7812
596.3494262695312 0 900.9277
597.3387451171875 0 3047.9146
597.808349609375 0 818.24963
598.2836303710938 0 1056.4159
598.3464965820312 0 3306.2148
598.816162109375 0 1355.5853
599.3215942382812 0 926.82434
611.3502807617188 0 1088.9539
611.8822021484375 0 912.8464
612.380126953125 0 1711.5048 c Water loss 5
612.8870239257812 0 1522.8528
613.3156127929688 0 992.4562
613.3674926757812 0 11797.983 c Ammonia loss 5
613.8294067382812 0 2002.0662
614.3701171875 0 3625.6184
618.3541259765625 0 831.2107 w 1
619.3544311523438 0 942.2527
620.8596801757812 0 957.9546
624.8236694335938 0 1009.2487
624.87451171875 0 814.56836 c Ammonia loss 11
625.3211059570312 0 794.16864 z Ammonia loss 7
625.3756713867188 0 889.93463
625.8490600585938 0 2246.5679 y Water loss 1
626.3789672851562 0 5084.5625
626.8515625 0 1956.1787 z 1
626.8811645507812 0 3451.575
627.365234375 0 3863.1455
627.8615112304688 0 1172.9391
632.377197265625 0 785.1346
632.8245239257812 0 1463.0818
632.8804321289062 0 1364.1791
633.382568359375 0 73810.08 c 11
633.8298950195312 0 2320.5261
633.884033203125 0 45929.57
634.3834228515625 0 18831.48
634.8681640625 0 8799.063 y 1
635.3658447265625 0 5042.6597
635.8677368164062 0 1574.1892
639.3844604492188 0 922.0983
639.8550415039062 0 982.9829
640.37841796875 0 5075.5557 y Water loss 7
640.8751220703125 0 2638.841
641.380859375 0 1572.166
642.353759765625 0 37286.188 z 7
642.8414306640625 0 1240.3417
643.3617553710938 0 10744.213
644.364990234375 0 2644.858
648.3570556640625 0 5776.0757
648.8596801757812 0 3588.4758
648.8956298828125 0 1816.7545
649.3643188476562 0 2720.2964
650.3071899414062 0 2204.98
651.345703125 0 749.9466
653.90185546875 0 2754.7705
654.3925170898438 0 2703.8818
654.8839721679688 0 934.23236
655.392822265625 0 3626.435
655.885009765625 0 2550.209
656.3458251953125 0 2247.688
656.8699340820312 0 13808.944
657.3731689453125 0 6437.816
657.8736572265625 0 3423.9714
658.3189086914062 0 10012.466
658.3778076171875 0 4981.112 y 7
658.8217163085938 0 927.7424
659.3231201171875 0 2009.4813
659.3795166015625 0 1089.7302
661.8942260742188 0 1021.06885
662.3917236328125 0 8281.67
662.8934326171875 0 6851.8774
663.3900146484375 0 4313.4883
663.87890625 0 1578.0127
664.375732421875 0 910.51917
665.3378295898438 0 6105.5645
667.3842163085938 0 4043.583
667.8853759765625 0 2380.984
668.3798828125 0 2636.111
668.8778076171875 0 13417.597
669.3792114257812 0 9709.041
669.8803100585938 0 3321.5815
670.3827514648438 0 2682.1208
670.8865966796875 0 1207.3071
675.3806762695312 0 729.9925
675.8870849609375 0 1995.4733
676.389404296875 0 34283.457
676.8904418945312 0 21568.25
677.3919067382812 0 9011.105
677.83740234375 0 826.31714
677.8927001953125 0 1812.8555
678.3397827148438 0 20096.92
682.3881225585938 0 2141.4792
683.3907470703125 0 1858.0619
683.8934326171875 0 1754.9988
684.3984375 0 61825.055
684.9014282226562 0 88040.6
685.4028930664062 0 52473.074
685.90478515625 0 21563.17
686.3489379882812 0 16274.139
686.4083251953125 0 4204.1826
686.8524169921875 0 7053.8423
686.9053955078125 0 770.5626
701.3930053710938 0 735.9323
712.38916015625 0 841.5251 w 6
713.3899536132812 0 1213.3263
715.3861694335938 0 7044.7485
727.4461669921875 0 46702.816 c 6
728.3749389648438 0 952.6267
728.44921875 0 16267.79
729.4515380859375 0 4166.301
731.4071044921875 0 1165.8846
747.4071044921875 0 760.2038
749.3342895507812 0 1252.5253
750.3361206054688 0 5338.3145
751.3404541015625 0 2373.2795
754.4685668945312 0 2026.487
755.4301147460938 0 10762.3955 y 6
756.4332275390625 0 4417.6235
757.4345703125 0 1885.5054
780.4756469726562 0 1004.2506 c Water loss 7
781.4581909179688 0 6923.251 c Ammonia loss 7
782.3421630859375 0 2124.4802
782.4594116210938 0 2657.6665
783.34423828125 0 2153.225
783.4285888671875 0 1687.6748
784.3462524414062 0 1209.6537
784.4271240234375 0 1003.91046
808.4205322265625 0 1667.5256
809.4400024414062 0 7939.032 w 5
810.443359375 0 3702.075
813.4134521484375 0 2079.7166
814.4212646484375 0 6111.1855
821.3738403320312 0 4050.8806
821.49951171875 0 1187.8783
822.3793334960938 0 3984.7705
822.5068359375 0 2046.5571
823.3834838867188 0 1744.2018
823.509765625 0 955.279
824.5009765625 0 1166.6562
825.5088500976562 0 969.7007
834.4789428710938 0 790.9794 z Water loss 5
837.3442993164062 0 5367.636
838.3477783203125 0 1499.1333
839.354248046875 0 1450.2341
851.5166015625 0 2058.4517
852.49609375 0 16745.504 z 5
853.49853515625 0 9402.538
854.3351440429688 0 1664.1609
854.5021362304688 0 2709.4602
866.3699340820312 0 776.86017
867.4698486328125 0 14697.441
868.4780883789062 0 6266.149
869.4712524414062 0 876.73663
877.4443969726562 0 840.94714
878.4495849609375 0 8741.784
892.3870849609375 0 2748.9868
893.3900756835938 0 808.353
894.4765014648438 0 1166.1807
895.3738403320312 0 20142.09
895.5344848632812 0 4949.285 c 8
896.376953125 0 12146.33
896.5379028320312 0 2888.6272
897.3790283203125 0 3588.8464
898.3837890625 0 827.83844
909.3902587890625 0 6214.498
910.3986206054688 0 35522.02
911.4009399414062 0 18746.736
912.4021606445312 0 6774.903
913.3984985351562 0 1623.6478
914.4639282226562 0 3089.5825
915.4691162109375 0 6538.845
918.5297241210938 0 1361.1157
923.5324096679688 0 13996.497 z 4
924.535400390625 0 8343.438
925.5379638671875 0 2411.921
939.5506591796875 0 15437.083 y 4
940.5542602539062 0 9095.697
941.55810546875 0 2265.9038
942.55908203125 0 906.59717
949.5955200195312 0 2311.72
950.5982666015625 0 1403.4312
951.5969848632812 0 981.30426
964.6055908203125 0 950.5864
965.6160888671875 0 2329.7559
966.61474609375 0 1000.9836
989.5615234375 0 954.6865
991.6013793945312 0 872.5737 c Ammonia loss 9
993.559814453125 0 773.8958 w 3
995.564453125 0 2817.0288
996.5669555664062 0 1810.2943
1006.5458374023438 0 7706.6157
1007.5587158203125 0 5255.294
1008.619873046875 0 60439.324 c 9
1009.6228637695312 0 31323.693
1010.6258544921875 0 9801.715
1011.6297607421875 0 2651.4878
1015.5012817382812 0 838.9214
1016.5153198242188 0 10739.556
1022.56591796875 0 787.1208
1032.4989013671875 0 1239.1914
1033.4951171875 0 1350.667
1034.50244140625 0 760.0504
1050.6248779296875 0 859.91864 y Ammonia loss 3
1051.6270751953125 0 24641.617 z 3
1052.6334228515625 0 26935.21
1053.63623046875 0 11593.628
1054.6395263671875 0 5298.6284
1055.645751953125 0 1050.7755
1063.6397705078125 0 2583.7021
1064.64111328125 0 1607.237
1066.6317138671875 0 1138.3433
1067.645263671875 0 4461.056 y 3
1068.652587890625 0 1998.2887
1077.58837890625 0 2379.2102
1078.5897216796875 0 5850.847
1093.64697265625 0 1542.8701
1094.65625 0 1008.1113
1095.64013671875 0 767.55054
1107.6044921875 0 1583.5536
1109.61669921875 0 1570.6927
1110.6063232421875 0 934.34375
1121.65673828125 0 3246.7053 w 2
1122.6595458984375 0 1871.5597
1123.6640625 0 881.3725
1135.6414794921875 0 764.94727
1136.649169921875 0 937.6785
1137.6622314453125 0 80918.54 c 10
1138.6654052734375 0 46128.04
1139.6688232421875 0 19858.693
1140.672607421875 0 4619.872
1149.7169189453125 0 918.37067
1165.671142578125 0 3158.562 z 2
1166.677490234375 0 20218.146
1167.6802978515625 0 10474.937
1168.6824951171875 0 5154.9067
1169.6875 0 1140.7196
1193.6175537109375 0 4508.9136
1221.7470703125 0 2183.8687
1222.748291015625 0 3207.9746
1223.7506103515625 0 1309.2493
1235.7032470703125 0 745.9313 w 1
1238.76953125 0 732.2448
1249.7366943359375 0 1444.177
1250.7470703125 0 801.9489
1252.7113037109375 0 905.5714 z 1
1253.7093505859375 0 5359.447
1254.7098388671875 0 3354.662
1255.7081298828125 0 1246.9587
1265.7568359375 0 8022.263 c 11
1266.7626953125 0 16531.434
1267.76708984375 0 9480.971
1268.7623291015625 0 4112.2837
1269.7626953125 0 1094.5404
1283.67333984375 0 2276.4258
1284.681396484375 0 1994.9308
1307.7611083984375 0 2289.5278
1310.7625732421875 0 1332.347
1313.6619873046875 0 5126.195
1314.663330078125 0 1011.9381
1323.775390625 0 4235.1387
1324.782470703125 0 19234.69
1325.785400390625 0 11720.273
1326.78662109375 0 4694.6167
1327.781005859375 0 1613.7367
1339.6441650390625 0 893.7374
1340.80078125 0 1754.8875
1341.80419921875 0 2348.4902
1342.7996826171875 0 1477.2131
1350.75439453125 0 977.3444
1351.7701416015625 0 6327.0874
1352.776611328125 0 40649.965
1353.77978515625 0 25810.297
1354.782470703125 0 10860.495
1355.663330078125 0 2128.836
1355.7952880859375 0 2205.7388
1356.67822265625 0 18206.81
1357.679443359375 0 1391.9457
1366.73779296875 0 643.7305
1367.78759765625 0 5466.357
1368.79541015625 0 18307.48
1369.8028564453125 0 58242.19
1370.805419921875 0 36279.242
1371.806884765625 0 14464.2295
1372.6981201171875 0 3525.0034
1372.8388671875 0 1474.9519
1373.7044677734375 0 10294.373
2552.69873046875 0 625.38916

Spectrum Details

|  |  |
| --- | --- |
| Matched peaks? Matched peaks The total absolute number of peaks matched. Additionally in brackets the total fraction of peaks matched and the total number of peaks is shown. | 70 (13.11% of 534) |
| FDR? FDR The false discovery rate estimated for this peptide. It is calculated by matching all theoretical fragments with a non-integer shift with the raw peaks for this spectrum. This is done with 40 different shifts. The resulting percentage is the average number of annotated peaks over the number of annotated peaks with the correct spectrum. | 1.97% |
| Satellite FDR? Satellite FDR See the FDR for details on its calculation. This satellite ion specific FDR only contains the satellite ions (d/w) for I/L/J positions. | 1.59% |
| PSM Score? PSM Score The PSM Score as given by Hecklib to this annotated spectrum. It is shown with three significant figures. | 553 |

#### Spectrum 4179? Spectrum 4179 The raw spectrum of this peptide as annotated by Hecklib. The fragments are coloured according to ion type (see legend). Any peaks with a star '\*' as text can be hovered over to see the full details, first the ion type second the mass shift type. By hovering over the amino acids in the peptide or ions in the legend the corresponding peaks are highlighted. By toggling the 'Unassigned' label you can turn the background (unassigned) peaks on or off in the plot. By updating the slider in the Ion legend you can update the spectrum to only show the top X% of the peaks with labels. The top X% means any peak that is within X% of the highest intensity. By dragging in the spectrum you can zoom in to a specific part of the spectrum and use 'Zoom Out' to get back to the original zoom level. The annotation of the spectrum is based on the given sequence in the peptides file and is done with different software so inconsistencies are likely. The peaks are annotated based on the given sequence, with 20 ppm tolerance.

Copy Data

##### Spectrum 4179 (TSV)

###### Preview

```
Loading example...
```

*Click on the button to copy the data to your clipboard.*

Mz MinMz MaxIntensity Max

WidthHeightPeptide font sizePeptide stroke widthSpectrum font sizeSpectrum stroke widthCompact peptide

Ion legend

wxyz

abcd

OtherUnassignedIon^Charge^_Position_Show for top:%

VSNKALPAPIEKT

09.59e+31.92e+42.88e+43.83e+4

Zoom Out

y^+1^_1_w^+1^_2_y^+1^_2_y^+1^_2_y^+2^_5_c^+1^_3_w^+1^_3_c^+2^_6_c^+1^_3_c^+2^_7_z^+1^_3_y^+2^_7_y^+1^_3_y^+2^_7_c^+2^_8_c^+1^_4_w^+1^_4_c^+1^_4_z^+1^_4_y^+1^_4_c^+1^_5_c^+1^_5_z^+2^_10_y^+2^_10_w^+1^_5_w^+2^_11_z^+2^_11_y^+1^_5_y^+2^_11_c^+1^_6_c^+1^_6_y^+2^_12_z^+2^_12_c^+2^_12_y^+2^_12_y^+1^_6_z^+1^_6_y^+1^_6_c^+1^_7_y^+1^_7_c^+1^_8_w^+1^_8_z^+1^_8_c^+1^_9_z^+1^_9_y^+1^_9_c^+1^_10_z^+1^_10_y^+1^_10_w^+1^_11_c^+1^_11_z^+1^_11_c^+1^_12_

035871610741432

Fragment Matches Table

Show background peaks

| Position | Ion type | Intensity | mz Theoretical | mz Error (Th) | mz Error (ppm) | Charge | Series Number |
| --- | --- | --- | --- | --- | --- | --- | --- |
| 13 | y | 809 | 120.1 | 0.0002349 | 1.956 | +1 | 1 |
| - | - | 457 | 125.9 | - | - | 0 | - |
| - | - | 395.6 | 127.4 | - | - | 0 | - |
| - | - | 892 | 129.1 | - | - | 0 | - |
| - | - | 1353 | 132.1 | - | - | 0 | - |
| - | - | 2185 | 141.1 | - | - | 0 | - |
| - | - | 732.8 | 141.1 | - | - | 0 | - |
| - | - | 466.3 | 147.7 | - | - | 0 | - |
| - | - | 739 | 148.9 | - | - | 0 | - |
| - | - | 1168 | 159.1 | - | - | 0 | - |
| - | - | 508 | 165.1 | - | - | 0 | - |
| - | - | 498.1 | 167.1 | - | - | 0 | - |
| - | - | 1209 | 169.1 | - | - | 0 | - |
| - | - | 836.7 | 173.1 | - | - | 0 | - |
| - | - | 2094 | 173.5 | - | - | 0 | - |
| 12 | w | 1566 | 174.1 | 3.977E-06 | 0.02285 | +1 | 2 |
| - | - | 5003 | 183.1 | - | - | 0 | - |
| - | - | 2407 | 187.1 | - | - | 0 | - |
| - | - | 506.5 | 190 | - | - | 0 | - |
| - | - | 792.6 | 202.1 | - | - | 0 | - |
| - | - | 1313 | 215.1 | - | - | 0 | - |
| 12 | y | 538.4 | 230.1 | 0.00174 | 7.56 | +1 | 2 |
| - | - | 488.8 | 236.8 | - | - | 0 | - |
| - | - | 755.3 | 239.1 | - | - | 0 | - |
| - | - | 1177 | 240.1 | - | - | 0 | - |
| - | - | 671.5 | 243.1 | - | - | 0 | - |
| 12 | y | 1402 | 248.2 | 0.0004179 | 1.684 | +1 | 2 |
| - | - | 598.5 | 258.1 | - | - | 0 | - |
| - | - | 494.5 | 262.3 | - | - | 0 | - |
| - | - | 616.4 | 283.1 | - | - | 0 | - |
| - | - | 1358 | 284.2 | - | - | 0 | - |
| - | - | 1599 | 293.2 | - | - | 0 | - |
| - | - | 804.8 | 293.7 | - | - | 0 | - |
| 9 | y | 7929 | 294.2 | 0.0001586 | 0.5392 | +2 | 5 |
| - | - | 1776 | 294.7 | - | - | 0 | - |
| 3 | c | 1016 | 301.2 | 0.0007134 | 2.369 | +1 | 3 |
| - | - | 982 | 302.2 | - | - | 0 | - |
| 11 | w | 1.091E+04 | 302.2 | 0.000271 | 0.8967 | +1 | 3 |
| - | - | 2322 | 303.2 | - | - | 0 | - |
| - | - | 539.9 | 306.7 | - | - | 0 | - |
| 6 | c | 635.2 | 307.2 | 0.001943 | 6.324 | +2 | 6 |
| - | - | 3349 | 309.2 | - | - | 0 | - |
| - | - | 773.3 | 310.2 | - | - | 0 | - |
| 3 | c | 5503 | 318.2 | 0.0002239 | 0.7037 | +1 | 3 |
| - | - | 619.7 | 330.2 | - | - | 0 | - |
| - | - | 709.4 | 334.7 | - | - | 0 | - |
| - | - | 1450 | 340.2 | - | - | 0 | - |
| - | - | 1194 | 341.7 | - | - | 0 | - |
| 7 | c | 3046 | 355.7 | 0.0004315 | 1.213 | +2 | 7 |
| - | - | 798.9 | 356.2 | - | - | 0 | - |
| 11 | z | 1130 | 361.2 | 0.0004833 | 1.338 | +1 | 3 |
| 7 | y | 682.6 | 369.2 | 1.035E-05 | 0.02804 | +2 | 7 |
| 11 | y | 1419 | 377.2 | 0.0001949 | 0.5166 | +1 | 3 |
| - | - | 1803 | 377.2 | - | - | 0 | - |
| 7 | y | 2186 | 378.2 | 0.0002822 | 0.7462 | +2 | 7 |
| - | - | 1166 | 378.7 | - | - | 0 | - |
| 8 | c | 1018 | 391.2 | 0.001654 | 4.229 | +2 | 8 |
| - | - | 627.8 | 391.7 | - | - | 0 | - |
| 4 | c | 1542 | 429.2 | 0.0007667 | 1.786 | +1 | 4 |
| 10 | w | 2757 | 445.2 | 7.964E-05 | 0.1789 | +1 | 4 |
| 4 | c | 9726 | 446.3 | 3.31E-05 | 0.07418 | +1 | 4 |
| - | - | 1937 | 447.3 | - | - | 0 | - |
| - | - | 841.7 | 455.2 | - | - | 0 | - |
| - | - | 961.3 | 456.2 | - | - | 0 | - |
| 10 | z | 657.4 | 456.3 | 0.0008624 | 1.89 | +1 | 4 |
| - | - | 1018 | 456.9 | - | - | 0 | - |
| - | - | 877.8 | 457.2 | - | - | 0 | - |
| - | - | 1572 | 468.3 | - | - | 0 | - |
| 10 | y | 825.8 | 490.3 | 0.001007 | 2.054 | +1 | 4 |
| - | - | 1573 | 498.3 | - | - | 0 | - |
| - | - | 668.9 | 498.8 | - | - | 0 | - |
| 5 | c | 1900 | 500.3 | 0.0006327 | 1.265 | +1 | 5 |
| - | - | 565.5 | 501.3 | - | - | 0 | - |
| 5 | c | 1.057E+04 | 517.3 | 0.0001291 | 0.2495 | +1 | 5 |
| - | - | 2395 | 518.3 | - | - | 0 | - |
| 4 | z | 1237 | 526.3 | 0.001991 | 3.782 | +2 | 10 |
| - | - | 1544 | 526.8 | - | - | 0 | - |
| 4 | y | 1109 | 534.3 | 0.0004271 | 0.7994 | +2 | 10 |
| - | - | 740.2 | 534.8 | - | - | 0 | - |
| - | - | 990 | 543.3 | - | - | 0 | - |
| 9 | w | 701.1 | 544.3 | 0.0003913 | 0.7189 | +1 | 5 |
| - | - | 666.5 | 554.3 | - | - | 0 | - |
| - | - | 1846 | 555.3 | - | - | 0 | - |
| - | - | 1198 | 555.8 | - | - | 0 | - |
| - | - | 707 | 556.3 | - | - | 0 | - |
| 3 | w | 4356 | 561.3 | 0.0002165 | 0.3857 | +2 | 11 |
| - | - | 1943 | 561.8 | - | - | 0 | - |
| - | - | 937.5 | 562.3 | - | - | 0 | - |
| 3 | z | 2312 | 583.3 | 0.001387 | 2.378 | +2 | 11 |
| - | - | 1374 | 583.8 | - | - | 0 | - |
| - | - | 862.8 | 584.3 | - | - | 0 | - |
| - | - | 823.8 | 586.4 | - | - | 0 | - |
| 9 | y | 1.195E+04 | 587.3 | 0.0001208 | 0.2058 | +1 | 5 |
| - | - | 3507 | 588.3 | - | - | 0 | - |
| 3 | y | 2521 | 591.3 | 0.0008339 | 1.41 | +2 | 11 |
| - | - | 1373 | 591.8 | - | - | 0 | - |
| - | - | 611.9 | 597.3 | - | - | 0 | - |
| - | - | 863.7 | 597.8 | - | - | 0 | - |
| - | - | 1126 | 598.3 | - | - | 0 | - |
| - | - | 651.8 | 599.3 | - | - | 0 | - |
| - | - | 741.8 | 608.3 | - | - | 0 | - |
| - | - | 703.5 | 611.3 | - | - | 0 | - |
| 6 | c | 770.2 | 612.4 | 0.001201 | 1.961 | +1 | 6 |
| 6 | c | 5408 | 613.4 | 0.0002784 | 0.4539 | +1 | 6 |
| - | - | 2265 | 614.4 | - | - | 0 | - |
| - | - | 751.8 | 618.8 | - | - | 0 | - |
| 2 | y | 1590 | 625.9 | 0.006988 | 11.17 | +2 | 12 |
| - | - | 2208 | 626.4 | - | - | 0 | - |
| 2 | z | 2006 | 626.9 | 0.003866 | 6.168 | +2 | 12 |
| - | - | 1490 | 627.4 | - | - | 0 | - |
| - | - | 913.8 | 627.9 | - | - | 0 | - |
| 12 | c | 2.876E+04 | 633.4 | 9.211E-06 | 0.01454 | +2 | 12 |
| - | - | 1.736E+04 | 633.9 | - | - | 0 | - |
| - | - | 9096 | 634.4 | - | - | 0 | - |
| 2 | y | 3329 | 634.9 | 0.006284 | 9.898 | +2 | 12 |
| - | - | 2035 | 635.4 | - | - | 0 | - |
| - | - | 809.2 | 639.4 | - | - | 0 | - |
| 8 | y | 2133 | 640.4 | 0.009036 | 14.11 | +1 | 6 |
| - | - | 1666 | 640.9 | - | - | 0 | - |
| - | - | 637.2 | 641.9 | - | - | 0 | - |
| 8 | z | 1.382E+04 | 642.4 | 0.0005052 | 0.7865 | +1 | 6 |
| - | - | 4053 | 643.4 | - | - | 0 | - |
| - | - | 794.1 | 644.4 | - | - | 0 | - |
| - | - | 2664 | 648.4 | - | - | 0 | - |
| - | - | 1824 | 648.9 | - | - | 0 | - |
| - | - | 628.4 | 649.4 | - | - | 0 | - |
| - | - | 923.4 | 653.9 | - | - | 0 | - |
| - | - | 1295 | 654.4 | - | - | 0 | - |
| - | - | 1828 | 655.4 | - | - | 0 | - |
| - | - | 1040 | 655.9 | - | - | 0 | - |
| - | - | 5703 | 656.9 | - | - | 0 | - |
| - | - | 3895 | 657.4 | - | - | 0 | - |
| - | - | 1462 | 657.9 | - | - | 0 | - |
| 8 | y | 2264 | 658.4 | 0.0009123 | 1.386 | +1 | 6 |
| - | - | 628.5 | 661.9 | - | - | 0 | - |
| - | - | 3347 | 662.4 | - | - | 0 | - |
| - | - | 3460 | 662.9 | - | - | 0 | - |
| - | - | 1651 | 663.4 | - | - | 0 | - |
| - | - | 1763 | 667.4 | - | - | 0 | - |
| - | - | 1269 | 667.9 | - | - | 0 | - |
| - | - | 1137 | 668.4 | - | - | 0 | - |
| - | - | 6708 | 668.9 | - | - | 0 | - |
| - | - | 4842 | 669.4 | - | - | 0 | - |
| - | - | 1599 | 669.9 | - | - | 0 | - |
| - | - | 906.6 | 670.4 | - | - | 0 | - |
| - | - | 762.9 | 670.9 | - | - | 0 | - |
| - | - | 706.8 | 675.9 | - | - | 0 | - |
| - | - | 1.373E+04 | 676.4 | - | - | 0 | - |
| - | - | 8046 | 676.9 | - | - | 0 | - |
| - | - | 4184 | 677.4 | - | - | 0 | - |
| - | - | 840.6 | 677.9 | - | - | 0 | - |
| - | - | 2.536E+04 | 684.4 | - | - | 0 | - |
| - | - | 3.797E+04 | 684.9 | - | - | 0 | - |
| - | - | 2.513E+04 | 685.4 | - | - | 0 | - |
| - | - | 1.033E+04 | 685.9 | - | - | 0 | - |
| - | - | 2335 | 686.4 | - | - | 0 | - |
| - | - | 1365 | 695.3 | - | - | 0 | - |
| - | - | 608.4 | 696.3 | - | - | 0 | - |
| - | - | 829.7 | 697.3 | - | - | 0 | - |
| 7 | c | 2.171E+04 | 727.4 | 5.989E-06 | 0.008233 | +1 | 7 |
| - | - | 7759 | 728.4 | - | - | 0 | - |
| - | - | 1799 | 729.4 | - | - | 0 | - |
| - | - | 750.2 | 754.5 | - | - | 0 | - |
| 7 | y | 5077 | 755.4 | 0.0001504 | 0.1991 | +1 | 7 |
| - | - | 2068 | 756.4 | - | - | 0 | - |
| 8 | c | 3276 | 781.5 | 0.001099 | 1.406 | +1 | 8 |
| - | - | 1675 | 782.5 | - | - | 0 | - |
| 6 | w | 3187 | 809.4 | 0.0001448 | 0.1789 | +1 | 8 |
| - | - | 1758 | 810.4 | - | - | 0 | - |
| - | - | 647.4 | 821.5 | - | - | 0 | - |
| 6 | z | 7736 | 852.5 | 0.0009116 | 1.069 | +1 | 8 |
| - | - | 3674 | 853.5 | - | - | 0 | - |
| - | - | 1804 | 854.5 | - | - | 0 | - |
| - | - | 1995 | 861.3 | - | - | 0 | - |
| - | - | 782.9 | 862.3 | - | - | 0 | - |
| - | - | 6005 | 867.5 | - | - | 0 | - |
| - | - | 2999 | 868.5 | - | - | 0 | - |
| - | - | 1132 | 878.3 | - | - | 0 | - |
| - | - | 1375 | 879.3 | - | - | 0 | - |
| - | - | 730.3 | 880.3 | - | - | 0 | - |
| - | - | 627.7 | 895.4 | - | - | 0 | - |
| 9 | c | 2503 | 895.5 | 0.002836 | 3.166 | +1 | 9 |
| - | - | 964.1 | 896.5 | - | - | 0 | - |
| - | - | 1610 | 903.9 | - | - | 0 | - |
| - | - | 870.8 | 906.4 | - | - | 0 | - |
| - | - | 2465 | 912.4 | - | - | 0 | - |
| - | - | 1753 | 913.4 | - | - | 0 | - |
| - | - | 951.2 | 913.9 | - | - | 0 | - |
| - | - | 1012 | 914.4 | - | - | 0 | - |
| - | - | 794.6 | 914.9 | - | - | 0 | - |
| - | - | 633.7 | 915.4 | - | - | 0 | - |
| 5 | z | 6163 | 923.5 | 0.0003135 | 0.3395 | +1 | 9 |
| - | - | 3709 | 924.5 | - | - | 0 | - |
| - | - | 650.6 | 925.5 | - | - | 0 | - |
| 5 | y | 5768 | 939.6 | 0.0003608 | 0.384 | +1 | 9 |
| - | - | 2951 | 940.6 | - | - | 0 | - |
| - | - | 692.2 | 941.6 | - | - | 0 | - |
| - | - | 688.3 | 949.6 | - | - | 0 | - |
| - | - | 817.1 | 950.6 | - | - | 0 | - |
| - | - | 1649 | 965.6 | - | - | 0 | - |
| - | - | 1045 | 995.6 | - | - | 0 | - |
| 10 | c | 2.688E+04 | 1009 | 0.0006249 | 0.6196 | +1 | 10 |
| - | - | 1.356E+04 | 1010 | - | - | 0 | - |
| - | - | 5190 | 1011 | - | - | 0 | - |
| - | - | 1094 | 1012 | - | - | 0 | - |
| 4 | z | 8572 | 1052 | 0.0001214 | 0.1155 | +1 | 10 |
| - | - | 1.174E+04 | 1053 | - | - | 0 | - |
| - | - | 5597 | 1054 | - | - | 0 | - |
| - | - | 1790 | 1055 | - | - | 0 | - |
| - | - | 927.8 | 1064 | - | - | 0 | - |
| - | - | 825.5 | 1065 | - | - | 0 | - |
| 4 | y | 1671 | 1068 | 0.002123 | 1.989 | +1 | 10 |
| - | - | 690.3 | 1111 | - | - | 0 | - |
| 3 | w | 2130 | 1122 | 0.002556 | 2.279 | +1 | 11 |
| - | - | 921.6 | 1123 | - | - | 0 | - |
| 11 | c | 3.165E+04 | 1138 | 0.0004513 | 0.3966 | +1 | 11 |
| - | - | 2.02E+04 | 1139 | - | - | 0 | - |
| - | - | 8265 | 1140 | - | - | 0 | - |
| - | - | 1495 | 1141 | - | - | 0 | - |
| 3 | z | 945.7 | 1166 | 0.001912 | 1.641 | +1 | 11 |
| - | - | 7530 | 1167 | - | - | 0 | - |
| - | - | 4570 | 1168 | - | - | 0 | - |
| - | - | 1647 | 1169 | - | - | 0 | - |
| - | - | 667.7 | 1222 | - | - | 0 | - |
| - | - | 1466 | 1223 | - | - | 0 | - |
| - | - | 723.2 | 1224 | - | - | 0 | - |
| - | - | 2348 | 1254 | - | - | 0 | - |
| - | - | 1730 | 1255 | - | - | 0 | - |
| 12 | c | 3201 | 1266 | 0.0008838 | 0.6983 | +1 | 12 |
| - | - | 8003 | 1267 | - | - | 0 | - |
| - | - | 5014 | 1268 | - | - | 0 | - |
| - | - | 1403 | 1269 | - | - | 0 | - |
| - | - | 719.3 | 1311 | - | - | 0 | - |
| - | - | 1554 | 1324 | - | - | 0 | - |
| - | - | 8218 | 1325 | - | - | 0 | - |
| - | - | 6310 | 1326 | - | - | 0 | - |
| - | - | 1393 | 1327 | - | - | 0 | - |
| - | - | 671.1 | 1341 | - | - | 0 | - |
| - | - | 835.5 | 1342 | - | - | 0 | - |
| - | - | 686.9 | 1344 | - | - | 0 | - |
| - | - | 2077 | 1352 | - | - | 0 | - |
| - | - | 1.904E+04 | 1353 | - | - | 0 | - |
| - | - | 1.244E+04 | 1354 | - | - | 0 | - |
| - | - | 5626 | 1355 | - | - | 0 | - |
| - | - | 1707 | 1356 | - | - | 0 | - |
| - | - | 2708 | 1368 | - | - | 0 | - |
| - | - | 7899 | 1369 | - | - | 0 | - |
| - | - | 2.495E+04 | 1370 | - | - | 0 | - |
| - | - | 1.812E+04 | 1371 | - | - | 0 | - |
| - | - | 6170 | 1372 | - | - | 0 | - |
| - | - | 2115 | 1373 | - | - | 0 | - |
| - | - | 628.8 | 1417 | - | - | 0 | - |

m/z Charge Intensity FragmentType MassShift Position
120.0652847290039 0 808.9923 y 12
125.94856262207031 0 457.03592
127.40185546875 0 395.5588
129.10214233398438 0 892.04114
132.07652282714844 0 1352.9822
141.07815551757812 0 2184.8457
141.10208129882812 0 732.8367
147.74185180664062 0 466.33893
148.94769287109375 0 738.9749
159.11312866210938 0 1168.015
165.10162353515625 0 508.03378
167.14085388183594 0 498.07065
169.09693908691406 0 1208.9421
173.0919952392578 0 836.7463
173.45098876953125 0 2093.6262
174.07608032226562 0 1566.2511 w 11
183.11264038085938 0 5003.182
187.10745239257812 0 2406.8074
190.02499389648438 0 506.4668
202.08155822753906 0 792.55444
215.13900756835938 0 1312.8119
230.14817810058594 0 538.3544 y Water loss 11
236.76234436035156 0 488.81604
239.0883331298828 0 755.2875
240.13436889648438 0 1177.1816
243.14454650878906 0 671.5457
248.16006469726562 0 1401.7686 y 11
258.14453125 0 598.49036
262.3204650878906 0 494.54837
283.13946533203125 0 616.4491
284.1605529785156 0 1357.6448
293.1896057128906 0 1599.3225
293.69134521484375 0 804.7537
294.1734313964844 0 7928.5186 y 8
294.6746520996094 0 1775.7449
301.1499328613281 0 1016.047 c Ammonia loss 2
302.15313720703125 0 981.99615
302.1707763671875 0 10912.41 w 10
303.17401123046875 0 2321.6973
306.68365478515625 0 539.9199
307.1850891113281 0 635.1793 c Ammonia loss 5
309.2033996582031 0 3348.8374
310.20892333984375 0 773.3255
318.1769714355469 0 5503.111 c 2
330.1773681640625 0 619.7054
334.6900939941406 0 709.36505
340.1874084472656 0 1449.9447
341.71539306640625 0 1194.3114
355.7129821777344 0 3045.877 c Ammonia loss 6
356.21484375 0 798.9047
361.1838684082031 0 1129.6237 z 10
369.2132568359375 0 682.5836 y Water loss 6
377.202880859375 0 1418.6339 y 10
377.2342224121094 0 1802.6582
378.21881103515625 0 2185.7341 y 6
378.7200622558594 0 1166.016
391.2303161621094 0 1017.6936 c Ammonia loss 7
391.73199462890625 0 627.81305
429.2448425292969 0 1541.5026 c Ammonia loss 3
445.2293701171875 0 2757.371 w 9
446.2721252441406 0 9726.233 c 3
447.2745666503906 0 1936.9144
455.24920654296875 0 841.6679
456.21343994140625 0 961.3292
456.2569885253906 0 657.4334 z Water loss 9
456.8570556640625 0 1018.3947
457.2073669433594 0 877.7906
468.2806701660156 0 1572.2819
490.28814697265625 0 825.7935 y 9
498.2848205566406 0 1572.9498
498.78656005859375 0 668.9485
500.2833557128906 0 1900.1244 c Ammonia loss 4
501.2886047363281 0 565.53046
517.3091430664062 0 10566.684 c 4
518.3123779296875 0 2395.2722
526.3152465820312 0 1237.4348 z 3
526.8178100585938 0 1543.8064
534.3270263671875 0 1108.6726 y 3
534.828857421875 0 740.178
543.3250122070312 0 989.964
544.298095703125 0 701.09735 w 8
554.3336791992188 0 666.51227
555.3057861328125 0 1846.0608
555.8095703125 0 1198.2972
556.3106079101562 0 707.01904
561.3316650390625 0 4355.7646 w 2
561.8344116210938 0 1943.3302
562.33642578125 0 937.5398
583.340087890625 0 2312.4553 z 2
583.842041015625 0 1373.6136
584.34619140625 0 862.80963
586.378173828125 0 823.81793
587.3397827148438 0 11949.798 y 8
588.3419799804688 0 3506.673
591.3472290039062 0 2521.387 y 2
591.8492431640625 0 1373.4829
597.3387451171875 0 611.90533
597.8133544921875 0 863.74493
598.3487548828125 0 1126.1606
599.3245239257812 0 651.84265
608.2930908203125 0 741.7765
611.3493041992188 0 703.46497
612.3839721679688 0 770.1867 c Water loss 5
613.3670654296875 0 5408.3765 c Ammonia loss 5
614.3690185546875 0 2264.7712
618.8482666015625 0 751.84467
625.851806640625 0 1590.259 y Water loss 1
626.3778076171875 0 2207.6824
626.8585815429688 0 2005.7279 z 1
627.3651733398438 0 1490.3481
627.8580932617188 0 913.7801
633.3824462890625 0 28755.115 c 11
633.8834228515625 0 17357.979
634.3838500976562 0 9096.091
634.870361328125 0 3328.553 y 1
635.3652954101562 0 2034.806
639.3778686523438 0 809.2481
640.37548828125 0 2132.6465 y Water loss 7
640.8773193359375 0 1666.3544
641.8845825195312 0 637.218
642.3577880859375 0 13815.657 z 7
643.3616943359375 0 4052.6611
644.364990234375 0 794.14026
648.3583984375 0 2663.712
648.8895263671875 0 1824.0448
649.360595703125 0 628.38257
653.9013671875 0 923.3575
654.38916015625 0 1295.2715
655.39306640625 0 1828.0586
655.8931274414062 0 1039.5952
656.8707885742188 0 5703.157
657.3731079101562 0 3894.602
657.8753051757812 0 1461.5787
658.3779296875 0 2264.2769 y 7
661.8873291015625 0 628.4502
662.3916015625 0 3346.5574
662.8922729492188 0 3460.3613
663.3869018554688 0 1650.9099
667.384033203125 0 1763.454
667.886474609375 0 1269.1787
668.3807373046875 0 1137.3392
668.877685546875 0 6708.334
669.3788452148438 0 4841.581
669.879150390625 0 1599.4731
670.3868408203125 0 906.5965
670.8865966796875 0 762.8978
675.899658203125 0 706.83234
676.3892211914062 0 13729.236
676.890380859375 0 8045.939
677.3909912109375 0 4184.4077
677.8899536132812 0 840.5602
684.3980712890625 0 25356.955
684.9012451171875 0 37969.832
685.4025268554688 0 25134.701
685.9041137695312 0 10327.365
686.40380859375 0 2334.897
695.3304443359375 0 1365.4812
696.3301391601562 0 608.36993
697.3469848632812 0 829.6671
727.4461059570312 0 21707.105 c 6
728.4483642578125 0 7759.04
729.449951171875 0 1798.674
754.476806640625 0 750.17694
755.429931640625 0 5077.057 y 6
756.4339599609375 0 2068.0527
781.457763671875 0 3276.3293 c Ammonia loss 7
782.4590454101562 0 1675.4008
809.4404907226562 0 3187.4253 w 5
810.4413452148438 0 1757.7683
821.504638671875 0 647.4055
852.4960327148438 0 7736.368 z 5
853.4979858398438 0 3674.478
854.5025634765625 0 1804.2936
861.3079833984375 0 1995.0801
862.3118286132812 0 782.94586
867.4692993164062 0 6005.278
868.4789428710938 0 2998.9785
878.3167114257812 0 1132.118
879.3182983398438 0 1375.0575
880.3269653320312 0 730.33386
895.4146118164062 0 627.7416
895.5331420898438 0 2502.9358 c 8
896.5447998046875 0 964.14264
903.9248046875 0 1610.4016
906.4208984375 0 870.8152
912.44091796875 0 2465.1775
913.4417724609375 0 1753.0488
913.9368286132812 0 951.24896
914.4235229492188 0 1012.4606
914.9224853515625 0 794.57135
915.4220581054688 0 633.73334
923.5319213867188 0 6162.8 z 4
924.5343627929688 0 3708.7888
925.5408325195312 0 650.6473
939.5505981445312 0 5767.8496 y 4
940.5537719726562 0 2950.8489
941.5629272460938 0 692.2099
949.59912109375 0 688.349
950.6012573242188 0 817.097
965.61474609375 0 1649.1567
995.5654296875 0 1045.4025
1008.6206665039062 0 26877.04 c 9
1009.6234741210938 0 13561.313
1010.6251831054688 0 5190.379
1011.6264038085938 0 1093.7318
1051.6273193359375 0 8572.49 z 3
1052.6331787109375 0 11735.876
1053.6373291015625 0 5596.5083
1054.6409912109375 0 1790.4174
1063.63330078125 0 927.8265
1064.6483154296875 0 825.4579
1067.643798828125 0 1671.1624 y 3
1110.6031494140625 0 690.3181
1121.6539306640625 0 2129.6667 w 2
1122.6654052734375 0 921.60565
1137.6630859375 0 31653.979 c 10
1138.666015625 0 20197.79
1139.6693115234375 0 8265.029
1140.667724609375 0 1495.0315
1165.668212890625 0 945.6504 z 2
1166.6778564453125 0 7530.4375
1167.68017578125 0 4570.165
1168.6820068359375 0 1646.9384
1221.748779296875 0 667.67883
1222.7452392578125 0 1466.0669
1223.7684326171875 0 723.217
1253.708740234375 0 2347.8545
1254.7039794921875 0 1730.32
1265.7567138671875 0 3200.6863 c 11
1266.763671875 0 8003.059
1267.765380859375 0 5014.0522
1268.76904296875 0 1403.1438
1310.7623291015625 0 719.26843
1323.778076171875 0 1553.8065
1324.7823486328125 0 8217.541
1325.78369140625 0 6309.8
1326.7933349609375 0 1393.2933
1340.7958984375 0 671.14734
1341.8223876953125 0 835.5401
1343.8201904296875 0 686.9047
1351.77099609375 0 2077.1653
1352.77734375 0 19039.045
1353.7806396484375 0 12440.111
1354.7840576171875 0 5626.1465
1355.7867431640625 0 1706.6176
1367.786376953125 0 2707.989
1368.7967529296875 0 7898.7554
1369.8045654296875 0 24954.71
1370.8076171875 0 18122.049
1371.80810546875 0 6169.6914
1372.8089599609375 0 2114.7998
1417.4544677734375 0 628.75165

Spectrum Details

|  |  |
| --- | --- |
| Matched peaks? Matched peaks The total absolute number of peaks matched. Additionally in brackets the total fraction of peaks matched and the total number of peaks is shown. | 53 (21.03% of 252) |
| FDR? FDR The false discovery rate estimated for this peptide. It is calculated by matching all theoretical fragments with a non-integer shift with the raw peaks for this spectrum. This is done with 40 different shifts. The resulting percentage is the average number of annotated peaks over the number of annotated peaks with the correct spectrum. | 1.03% |
| Satellite FDR? Satellite FDR See the FDR for details on its calculation. This satellite ion specific FDR only contains the satellite ions (d/w) for I/L/J positions. | 0.00% |
| PSM Score? PSM Score The PSM Score as given by Hecklib to this annotated spectrum. It is shown with three significant figures. | 378 |

#### Reverse Lookup? Reverse Lookup All places where this read could be placed.

| Group | Segment | Template | Template Part | Read Part | Score | Unique |
| --- | --- | --- | --- | --- | --- | --- |
| Homo sapiens Heavy Chain | IGHC | IGHG1 | [205..218] | [0..13] | 104 | False |
| Homo sapiens Heavy Chain | IGHC | IGHG3 | [252..265] | [0..13] | 104 | False |
| Homo sapiens Heavy Chain | IGHC | IGHG2 | [201..214] | [0..13] | 95 | False |

| Recombined | Template Part | Read Part | Score | Unique |
| --- | --- | --- | --- | --- |
| REC-0-1 | [330..343] | [0..13] | 104 | True |

#### Meta Information from Multiple reads

##### Number of combined reads

3

##### Intensity

0.6104

##### TotalArea

1.109E+08

##### Changes to the peptide sequence

VSNKALPAPIEKT

J→ISupport for Isoleucine based on side chain ions (2 for I 0 for L) (Position: 10)

J→LSupport for Leucine based on side chain ions (1 for L 0 for I) (Position: 6)

L→JNo support for either Leucine or Isoleucine based on side chain ions (Position: 10)

L→JNo support for either Leucine or Isoleucine based on side chain ions (Position: 6)

#### Positional Score

Copy Data

##### Positional Score (TSV)

###### Preview

```
Loading example...
```

*Click on the button to copy the data to your clipboard.*

100123456789101112

Label Value
"0" 0.333
"1" 0.327
"2" 0.323
"3" 0.333
"4" 0.333
"5" 0.323
"6" 0.32
"7" 0.32
"8" 0.32
"9" 0.323
"10" 0.327
"11" 0.323
"12" 0.32

#### Meta Information from PEAKS

##### Scan Identifier

F2:4088

##### Original sequence

V

S

N

K

A

L

P

A

P

L

E

K

T

##### Posttranslational Modifications

##### Source File

D:\separate\_stitch\_analyses\xle-disambiguation\raw\20210323\_F1\_UM1\_Peng0013\_SA\_F59\_ingel\_3ug\_TL.raw

##### Fraction

2

##### Scan Feature

F2:2491

##### De Novo Score

99

##### ConfidenceScore

99

### m/z

456.6018

##### Mass

1366.782

##### Charge

3

##### Retention Time

21.84

##### Predicted Retention Time

-

##### Area

5.522E+07

##### Parts Per Million

1.2

##### Fragmentation mode

HCD

##### Originating file

01 D:\separate\_stitch\_analyses\xle-disambiguation\20210325\_F59\_3ug\_DENOVO\_12.csv

#### Meta Information from PEAKS

##### Scan Identifier

F1:4179

##### Original sequence

V

S

N

K

A

L

P

A

P

L

E

K

T

##### Posttranslational Modifications

##### Source File

D:\separate\_stitch\_analyses\xle-disambiguation\raw\20210323\_F1\_UM1\_Peng0013\_SA\_F59\_ingel\_3ug\_ELA.raw

##### Fraction

1

##### Scan Feature

F1:2457

##### De Novo Score

98

##### ConfidenceScore

98

### m/z

456.6023

##### Mass

1366.782

##### Charge

3

##### Retention Time

22.01

##### Predicted Retention Time

-

##### Area

4.356E+05

##### Parts Per Million

2.2

##### Fragmentation mode

ETHCD

##### Originating file

01 D:\separate\_stitch\_analyses\xle-disambiguation\20210325\_F59\_3ug\_DENOVO\_12.csv

#### Meta Information from PEAKS

##### Scan Identifier

F2:4150

##### Original sequence

V

S

N

K

A

L

P

A

P

L

E

K

T

##### Posttranslational Modifications

##### Source File

D:\separate\_stitch\_analyses\xle-disambiguation\raw\20210323\_F1\_UM1\_Peng0013\_SA\_F59\_ingel\_3ug\_TL.raw

##### Fraction

2

##### Scan Feature

F2:2491

##### De Novo Score

97

##### ConfidenceScore

97

### m/z

456.6018

##### Mass

1366.782

##### Charge

3

##### Retention Time

21.84

##### Predicted Retention Time

-

##### Area

5.522E+07

##### Parts Per Million

1.2

##### Fragmentation mode

ETHCD

##### Originating file

01 D:\separate\_stitch\_analyses\xle-disambiguation\20210325\_F59\_3ug\_DENOVO\_12.csv
