## Supplementary material for "A handle on mass coincidence errors in *de novo* sequencing of antibodies by bottom-up proteomics": Combined_049.html

Details Combined\_049 | Stitch OverviewUndefined

### Read Combined\_049

#### Sequence (length=13)

VTJFPPSSEEJQA

#### Spectrum 9705? Spectrum 9705 The raw spectrum of this peptide as annotated by Hecklib. The fragments are coloured according to ion type (see legend). Any peaks with a star '\*' as text can be hovered over to see the full details, first the ion type second the mass shift type. By hovering over the amino acids in the peptide or ions in the legend the corresponding peaks are highlighted. By toggling the 'Unassigned' label you can turn the background (unassigned) peaks on or off in the plot. By updating the slider in the Ion legend you can update the spectrum to only show the top X% of the peaks with labels. The top X% means any peak that is within X% of the highest intensity. By dragging in the spectrum you can zoom in to a specific part of the spectrum and use 'Zoom Out' to get back to the original zoom level. The annotation of the spectrum is based on the given sequence in the peptides file and is done with different software so inconsistencies are likely. The peaks are annotated based on the given sequence, with 20 ppm tolerance.

Copy Data

##### Spectrum 9705 (TSV)

###### Preview

```
Loading example...
```

*Click on the button to copy the data to your clipboard.*

Mz MinMz MaxIntensity Max

WidthHeightPeptide font sizePeptide stroke widthSpectrum font sizeSpectrum stroke widthCompact peptide

Ion legend

wxyz

abcd

OtherUnassignedIon^Charge^_Position_Show for top:%

VTJFPPSSEEJQA

04.12e+48.24e+41.24e+51.65e+5

Zoom Out

y^+1^_2_y^+1^_2_y^+2^_5_y^+1^_3_y^+1^_3_y^+1^_4_w^+1^_6_y^+1^_6_c^+1^_6_y^+1^_6_c^+1^_7_y^+1^_7_c^+1^_7_y^+1^_7_c^+1^_8_y^+1^_8_y^+1^_9_y^+1^_9_z^+1^_9_y^+1^_9_c^+1^_9_y^+1^_10_z^+1^_10_c^+1^_10_z^+1^_11_z^+1^_11_c^+1^_11_z^+1^_12_z^+1^_12_w^+1^_12_y^+1^_12_z^+1^_12_y^+1^_12_c^+1^_12_c^+1^_12_

0790158023703160

Fragment Matches Table

Show background peaks

| Position | Ion type | Intensity | mz Theoretical | mz Error (Th) | mz Error (ppm) | Charge | Series Number |
| --- | --- | --- | --- | --- | --- | --- | --- |
| - | - | 1243 | 120.1 | - | - | 0 | - |
| - | - | 726.3 | 149 | - | - | 0 | - |
| - | - | 429.4 | 149 | - | - | 0 | - |
| - | - | 620 | 173.1 | - | - | 0 | - |
| - | - | 3299 | 173.4 | - | - | 0 | - |
| - | - | 2071 | 183.1 | - | - | 0 | - |
| - | - | 8114 | 200.1 | - | - | 0 | - |
| 12 | y | 6778 | 201.1 | 0.0001443 | 0.7178 | +1 | 2 |
| - | - | 3517 | 201.1 | - | - | 0 | - |
| - | - | 536.7 | 202.1 | - | - | 0 | - |
| - | - | 555.9 | 204.5 | - | - | 0 | - |
| - | - | 1386 | 215.1 | - | - | 0 | - |
| 12 | y | 1.671E+04 | 218.1 | 0.0001761 | 0.8072 | +1 | 2 |
| - | - | 1201 | 219.1 | - | - | 0 | - |
| - | - | 732.7 | 225.1 | - | - | 0 | - |
| - | - | 1916 | 233.2 | - | - | 0 | - |
| - | - | 568.2 | 236.7 | - | - | 0 | - |
| - | - | 5264 | 242.2 | - | - | 0 | - |
| - | - | 494.7 | 243.2 | - | - | 0 | - |
| - | - | 963.4 | 251.2 | - | - | 0 | - |
| - | - | 1701 | 261.2 | - | - | 0 | - |
| - | - | 678.9 | 267.1 | - | - | 0 | - |
| - | - | 838.9 | 269.2 | - | - | 0 | - |
| - | - | 1939 | 282.1 | - | - | 0 | - |
| 9 | y | 2896 | 295.1 | 0.0008035 | 2.722 | +2 | 5 |
| - | - | 841 | 296.1 | - | - | 0 | - |
| - | - | 2.835E+04 | 296.2 | - | - | 0 | - |
| - | - | 3492 | 297.2 | - | - | 0 | - |
| - | - | 736.8 | 303.2 | - | - | 0 | - |
| 11 | y | 1103 | 314.2 | 0.001133 | 3.606 | +1 | 3 |
| - | - | 1.208E+04 | 314.2 | - | - | 0 | - |
| - | - | 1869 | 315.2 | - | - | 0 | - |
| 11 | y | 2813 | 331.2 | 0.0003101 | 0.9362 | +1 | 3 |
| - | - | 796.9 | 346.1 | - | - | 0 | - |
| - | - | 1599 | 346.2 | - | - | 0 | - |
| - | - | 1410 | 348.2 | - | - | 0 | - |
| - | - | 833.2 | 353.2 | - | - | 0 | - |
| - | - | 667.7 | 358.3 | - | - | 0 | - |
| - | - | 810.9 | 360.2 | - | - | 0 | - |
| - | - | 4263 | 362.2 | - | - | 0 | - |
| - | - | 665.7 | 363.2 | - | - | 0 | - |
| - | - | 1298 | 369.2 | - | - | 0 | - |
| - | - | 1422 | 370.7 | - | - | 0 | - |
| - | - | 779.1 | 371.2 | - | - | 0 | - |
| - | - | 563.9 | 390.2 | - | - | 0 | - |
| - | - | 758.7 | 408.2 | - | - | 0 | - |
| - | - | 920.4 | 411.3 | - | - | 0 | - |
| - | - | 556.5 | 415.2 | - | - | 0 | - |
| - | - | 606.9 | 415.3 | - | - | 0 | - |
| - | - | 1616 | 416.3 | - | - | 0 | - |
| - | - | 599.6 | 422.5 | - | - | 0 | - |
| - | - | 724 | 424.2 | - | - | 0 | - |
| - | - | 1.592E+04 | 433.3 | - | - | 0 | - |
| - | - | 4420 | 434.3 | - | - | 0 | - |
| - | - | 586.9 | 434.7 | - | - | 0 | - |
| - | - | 740.1 | 435.3 | - | - | 0 | - |
| - | - | 1348 | 440.2 | - | - | 0 | - |
| - | - | 1308 | 443.3 | - | - | 0 | - |
| - | - | 885.7 | 459.2 | - | - | 0 | - |
| 10 | y | 697 | 460.2 | 0.0009909 | 2.153 | +1 | 4 |
| - | - | 1.799E+04 | 461.3 | - | - | 0 | - |
| - | - | 4305 | 462.3 | - | - | 0 | - |
| - | - | 863.6 | 463.3 | - | - | 0 | - |
| - | - | 716.4 | 480.2 | - | - | 0 | - |
| - | - | 1762 | 498.2 | - | - | 0 | - |
| - | - | 801.9 | 530.2 | - | - | 0 | - |
| - | - | 623.2 | 530.3 | - | - | 0 | - |
| - | - | 1031 | 530.3 | - | - | 0 | - |
| - | - | 1062 | 540.3 | - | - | 0 | - |
| - | - | 1438 | 544.3 | - | - | 0 | - |
| - | - | 1698 | 544.8 | - | - | 0 | - |
| - | - | 714.6 | 545.3 | - | - | 0 | - |
| - | - | 3324 | 558.3 | - | - | 0 | - |
| - | - | 888.2 | 559.3 | - | - | 0 | - |
| - | - | 945.2 | 587.3 | - | - | 0 | - |
| - | - | 865 | 599.3 | - | - | 0 | - |
| - | - | 1188 | 600.8 | - | - | 0 | - |
| - | - | 1235 | 601.3 | - | - | 0 | - |
| - | - | 577.9 | 603.3 | - | - | 0 | - |
| - | - | 1.198E+04 | 627.3 | - | - | 0 | - |
| - | - | 3906 | 628.3 | - | - | 0 | - |
| - | - | 1543 | 628.4 | - | - | 0 | - |
| - | - | 935.6 | 629.4 | - | - | 0 | - |
| 8 | w | 666.8 | 643.3 | 0.002942 | 4.573 | +1 | 6 |
| - | - | 1239 | 643.3 | - | - | 0 | - |
| - | - | 842.9 | 644.3 | - | - | 0 | - |
| - | - | 640 | 645.3 | - | - | 0 | - |
| - | - | 1030 | 645.4 | - | - | 0 | - |
| - | - | 971.9 | 655.4 | - | - | 0 | - |
| - | - | 793.5 | 656.4 | - | - | 0 | - |
| - | - | 1448 | 657.4 | - | - | 0 | - |
| - | - | 1226 | 658.3 | - | - | 0 | - |
| 8 | y | 902.6 | 659.3 | 0.002617 | 3.97 | +1 | 6 |
| - | - | 3277 | 671.4 | - | - | 0 | - |
| 6 | c | 4285 | 672.4 | 0.0008191 | 1.218 | +1 | 6 |
| - | - | 1850 | 673.4 | - | - | 0 | - |
| - | - | 642.3 | 674.3 | - | - | 0 | - |
| 8 | y | 1199 | 676.3 | 0.002572 | 3.803 | +1 | 6 |
| - | - | 720.5 | 688.6 | - | - | 0 | - |
| - | - | 678.9 | 708.3 | - | - | 0 | - |
| - | - | 927.8 | 712.4 | - | - | 0 | - |
| - | - | 1344 | 715.4 | - | - | 0 | - |
| - | - | 7103 | 716.4 | - | - | 0 | - |
| - | - | 3089 | 717.4 | - | - | 0 | - |
| - | - | 707 | 722.3 | - | - | 0 | - |
| - | - | 9726 | 740.3 | - | - | 0 | - |
| - | - | 1226 | 740.4 | - | - | 0 | - |
| - | - | 3069 | 741.4 | - | - | 0 | - |
| 7 | c | 595.8 | 741.4 | 0.01313 | 17.71 | +1 | 7 |
| - | - | 1119 | 742.3 | - | - | 0 | - |
| - | - | 2840 | 742.4 | - | - | 0 | - |
| - | - | 1921 | 743.4 | - | - | 0 | - |
| - | - | 2274 | 744.4 | - | - | 0 | - |
| - | - | 1224 | 745.3 | - | - | 0 | - |
| 7 | y | 744 | 746.3 | 0.001792 | 2.401 | +1 | 7 |
| - | - | 2573 | 757.4 | - | - | 0 | - |
| - | - | 1.691E+04 | 758.4 | - | - | 0 | - |
| 7 | c | 1.955E+04 | 759.4 | 0.001231 | 1.621 | +1 | 7 |
| - | - | 7677 | 760.4 | - | - | 0 | - |
| - | - | 1531 | 761.4 | - | - | 0 | - |
| 7 | y | 2369 | 763.3 | 9.871E-05 | 0.1293 | +1 | 7 |
| - | - | 1895 | 771.4 | - | - | 0 | - |
| - | - | 878.1 | 772.4 | - | - | 0 | - |
| - | - | 922 | 774.3 | - | - | 0 | - |
| - | - | 592.7 | 785.4 | - | - | 0 | - |
| - | - | 2112 | 803.5 | - | - | 0 | - |
| - | - | 1370 | 829.4 | - | - | 0 | - |
| - | - | 1066 | 830.4 | - | - | 0 | - |
| - | - | 1561 | 839.4 | - | - | 0 | - |
| - | - | 1492 | 840.4 | - | - | 0 | - |
| - | - | 1.148E+04 | 845.5 | - | - | 0 | - |
| 8 | c | 5.424E+04 | 846.5 | 6.553E-05 | 0.07742 | +1 | 8 |
| - | - | 2.537E+04 | 847.5 | - | - | 0 | - |
| - | - | 5502 | 848.5 | - | - | 0 | - |
| - | - | 1380 | 850.4 | - | - | 0 | - |
| - | - | 984 | 851.4 | - | - | 0 | - |
| 6 | y | 8820 | 860.4 | 0.0006042 | 0.7023 | +1 | 8 |
| - | - | 3690 | 861.4 | - | - | 0 | - |
| - | - | 1.709E+04 | 868.4 | - | - | 0 | - |
| - | - | 7917 | 869.4 | - | - | 0 | - |
| - | - | 2294 | 870.4 | - | - | 0 | - |
| - | - | 934.6 | 871.4 | - | - | 0 | - |
| - | - | 1868 | 887.4 | - | - | 0 | - |
| - | - | 1167 | 888.4 | - | - | 0 | - |
| - | - | 1949 | 931.5 | - | - | 0 | - |
| - | - | 907.7 | 932.5 | - | - | 0 | - |
| - | - | 2521 | 938.4 | - | - | 0 | - |
| 5 | y | 2969 | 939.4 | 0.0007624 | 0.8116 | +1 | 9 |
| 5 | y | 5423 | 940.4 | 0.006311 | 6.711 | +1 | 9 |
| 5 | z | 3006 | 941.4 | 0.0002324 | 0.2469 | +1 | 9 |
| - | - | 834.4 | 942.4 | - | - | 0 | - |
| - | - | 1.977E+04 | 956.4 | - | - | 0 | - |
| 5 | y | 9.036E+04 | 957.5 | 8.648E-05 | 0.09032 | +1 | 9 |
| - | - | 3.901E+04 | 958.5 | - | - | 0 | - |
| - | - | 1.141E+04 | 959.5 | - | - | 0 | - |
| - | - | 1337 | 960.5 | - | - | 0 | - |
| - | - | 5965 | 974.5 | - | - | 0 | - |
| 9 | c | 6.645E+04 | 975.5 | 0.0005022 | 0.5148 | +1 | 9 |
| - | - | 3.587E+04 | 976.5 | - | - | 0 | - |
| - | - | 9130 | 977.5 | - | - | 0 | - |
| - | - | 1124 | 978.5 | - | - | 0 | - |
| - | - | 1200 | 985.4 | - | - | 0 | - |
| - | - | 903.6 | 988.5 | - | - | 0 | - |
| - | - | 924.2 | 1000 | - | - | 0 | - |
| - | - | 1018 | 1002 | - | - | 0 | - |
| - | - | 862.7 | 1015 | - | - | 0 | - |
| - | - | 1429 | 1016 | - | - | 0 | - |
| - | - | 1076 | 1017 | - | - | 0 | - |
| - | - | 751.1 | 1053 | - | - | 0 | - |
| - | - | 726.4 | 1054 | - | - | 0 | - |
| - | - | 1836 | 1061 | - | - | 0 | - |
| - | - | 2353 | 1062 | - | - | 0 | - |
| - | - | 1709 | 1063 | - | - | 0 | - |
| - | - | 796.2 | 1066 | - | - | 0 | - |
| - | - | 1942 | 1070 | - | - | 0 | - |
| - | - | 916.8 | 1071 | - | - | 0 | - |
| - | - | 1372 | 1086 | - | - | 0 | - |
| 4 | y | 1572 | 1087 | 0.006996 | 6.439 | +1 | 10 |
| - | - | 1.48E+04 | 1088 | - | - | 0 | - |
| 4 | z | 2.301E+04 | 1089 | 0.01052 | 9.661 | +1 | 10 |
| - | - | 1.429E+04 | 1090 | - | - | 0 | - |
| - | - | 4206 | 1091 | - | - | 0 | - |
| - | - | 629.5 | 1102 | - | - | 0 | - |
| - | - | 8727 | 1104 | - | - | 0 | - |
| 10 | c | 7.223E+04 | 1105 | 0.01041 | 9.428 | +1 | 10 |
| - | - | 4.497E+04 | 1106 | - | - | 0 | - |
| - | - | 1.635E+04 | 1107 | - | - | 0 | - |
| - | - | 2041 | 1108 | - | - | 0 | - |
| - | - | 726.1 | 1120 | - | - | 0 | - |
| - | - | 750.8 | 1130 | - | - | 0 | - |
| - | - | 1360 | 1131 | - | - | 0 | - |
| - | - | 1005 | 1132 | - | - | 0 | - |
| - | - | 1773 | 1156 | - | - | 0 | - |
| - | - | 1176 | 1157 | - | - | 0 | - |
| - | - | 950.6 | 1158 | - | - | 0 | - |
| - | - | 6060 | 1174 | - | - | 0 | - |
| - | - | 3790 | 1175 | - | - | 0 | - |
| - | - | 1314 | 1176 | - | - | 0 | - |
| - | - | 2658 | 1183 | - | - | 0 | - |
| 3 | z | 1921 | 1184 | 0.02161 | 18.26 | +1 | 11 |
| - | - | 2163 | 1185 | - | - | 0 | - |
| - | - | 991 | 1186 | - | - | 0 | - |
| - | - | 8458 | 1201 | - | - | 0 | - |
| 3 | z | 1.713E+04 | 1202 | 0.00885 | 7.365 | +1 | 11 |
| - | - | 1.213E+04 | 1203 | - | - | 0 | - |
| - | - | 4209 | 1204 | - | - | 0 | - |
| - | - | 818.6 | 1214 | - | - | 0 | - |
| - | - | 1142 | 1215 | - | - | 0 | - |
| - | - | 833.9 | 1216 | - | - | 0 | - |
| - | - | 1207 | 1217 | - | - | 0 | - |
| 11 | c | 5.016E+04 | 1218 | 0.002925 | 2.402 | +1 | 11 |
| - | - | 3.441E+04 | 1219 | - | - | 0 | - |
| - | - | 1.243E+04 | 1220 | - | - | 0 | - |
| - | - | 1646 | 1221 | - | - | 0 | - |
| - | - | 708.2 | 1231 | - | - | 0 | - |
| - | - | 3490 | 1244 | - | - | 0 | - |
| - | - | 2279 | 1245 | - | - | 0 | - |
| - | - | 1086 | 1246 | - | - | 0 | - |
| - | - | 1615 | 1247 | - | - | 0 | - |
| - | - | 1480 | 1248 | - | - | 0 | - |
| - | - | 647.8 | 1255 | - | - | 0 | - |
| - | - | 1570 | 1283 | - | - | 0 | - |
| - | - | 808.3 | 1284 | - | - | 0 | - |
| 2 | z | 2.109E+04 | 1285 | 0.0005541 | 0.4313 | +1 | 12 |
| 2 | z | 1.566E+04 | 1286 | 0.01897 | 14.76 | +1 | 12 |
| - | - | 4908 | 1287 | - | - | 0 | - |
| 2 | w | 1407 | 1288 | 0.02041 | 15.85 | +1 | 12 |
| 2 | y | 5322 | 1301 | 0.0216 | 16.61 | +1 | 12 |
| - | - | 5251 | 1302 | - | - | 0 | - |
| 2 | z | 2.24E+04 | 1303 | 0.002553 | 1.96 | +1 | 12 |
| - | - | 1.636E+04 | 1304 | - | - | 0 | - |
| - | - | 5515 | 1305 | - | - | 0 | - |
| - | - | 1660 | 1306 | - | - | 0 | - |
| - | - | 1114 | 1313 | - | - | 0 | - |
| - | - | 1697 | 1314 | - | - | 0 | - |
| - | - | 1176 | 1315 | - | - | 0 | - |
| - | - | 932.1 | 1317 | - | - | 0 | - |
| - | - | 2459 | 1318 | - | - | 0 | - |
| 2 | y | 2918 | 1319 | 0.004818 | 3.654 | +1 | 12 |
| 12 | c | 753.4 | 1329 | 0.01436 | 10.81 | +1 | 12 |
| - | - | 942.6 | 1330 | - | - | 0 | - |
| - | - | 1975 | 1331 | - | - | 0 | - |
| - | - | 1833 | 1332 | - | - | 0 | - |
| - | - | 629.5 | 1333 | - | - | 0 | - |
| - | - | 1377 | 1341 | - | - | 0 | - |
| - | - | 680.7 | 1342 | - | - | 0 | - |
| - | - | 1546 | 1344 | - | - | 0 | - |
| - | - | 1441 | 1345 | - | - | 0 | - |
| 12 | c | 1.383E+05 | 1346 | 0.001444 | 1.073 | +1 | 12 |
| - | - | 1.107E+05 | 1347 | - | - | 0 | - |
| - | - | 4.888E+04 | 1348 | - | - | 0 | - |
| - | - | 9761 | 1349 | - | - | 0 | - |
| - | - | 834.3 | 1350 | - | - | 0 | - |
| - | - | 3553 | 1356 | - | - | 0 | - |
| - | - | 3330 | 1357 | - | - | 0 | - |
| - | - | 2076 | 1358 | - | - | 0 | - |
| - | - | 796.6 | 1361 | - | - | 0 | - |
| - | - | 2938 | 1363 | - | - | 0 | - |
| - | - | 2864 | 1364 | - | - | 0 | - |
| - | - | 4468 | 1373 | - | - | 0 | - |
| - | - | 1.504E+04 | 1374 | - | - | 0 | - |
| - | - | 1.257E+04 | 1375 | - | - | 0 | - |
| - | - | 4712 | 1376 | - | - | 0 | - |
| - | - | 1090 | 1377 | - | - | 0 | - |
| - | - | 7497 | 1384 | - | - | 0 | - |
| - | - | 5726 | 1385 | - | - | 0 | - |
| - | - | 2382 | 1386 | - | - | 0 | - |
| - | - | 4085 | 1391 | - | - | 0 | - |
| - | - | 4022 | 1392 | - | - | 0 | - |
| - | - | 1606 | 1393 | - | - | 0 | - |
| - | - | 6556 | 1401 | - | - | 0 | - |
| - | - | 2.476E+04 | 1402 | - | - | 0 | - |
| - | - | 1.817E+04 | 1403 | - | - | 0 | - |
| - | - | 6983 | 1404 | - | - | 0 | - |
| - | - | 1912 | 1405 | - | - | 0 | - |
| - | - | 3946 | 1417 | - | - | 0 | - |
| - | - | 4.619E+04 | 1418 | - | - | 0 | - |
| - | - | 1.632E+05 | 1419 | - | - | 0 | - |
| - | - | 1.199E+05 | 1420 | - | - | 0 | - |
| - | - | 1683 | 1421 | - | - | 0 | - |
| - | - | 4.665E+04 | 1421 | - | - | 0 | - |
| - | - | 7909 | 1422 | - | - | 0 | - |
| - | - | 1326 | 2106 | - | - | 0 | - |
| - | - | 899.5 | 2125 | - | - | 0 | - |
| - | - | 1075 | 2126 | - | - | 0 | - |
| - | - | 948.5 | 2128 | - | - | 0 | - |
| - | - | 1719 | 2132 | - | - | 0 | - |
| - | - | 841.7 | 3030 | - | - | 0 | - |
| - | - | 765.2 | 3129 | - | - | 0 | - |

m/z Charge Intensity FragmentType MassShift Position
120.08099365234375 0 1242.912
148.95486450195312 0 726.3482
149.00137329101562 0 429.37207
173.12893676757812 0 619.9807
173.439453125 0 3299.178
183.1131134033203 0 2071.4534
200.10317993164062 0 8114.364
201.08712768554688 0 6777.6455 y Ammonia loss 11
201.1234588623047 0 3516.796
202.09046936035156 0 536.7207
204.5411376953125 0 555.9052
215.13919067382812 0 1386.2675
218.11370849609375 0 16713.057 y 11
219.11767578125 0 1201.3771
225.12423706054688 0 732.69104
233.16481018066406 0 1915.9062
236.72930908203125 0 568.23663
242.1503143310547 0 5263.957
243.1539764404297 0 494.7392
251.1758270263672 0 963.4073
261.15966796875 0 1700.8304
267.14971923828125 0 678.8807
269.1854248046875 0 838.86646
282.1451721191406 0 1939.3591
295.14422607421875 0 2895.8203 y 8
296.1482238769531 0 841.02484
296.19720458984375 0 28345.385
297.1999816894531 0 3491.572
303.17144775390625 0 736.81525
314.17218017578125 0 1102.5485 y Ammonia loss 10
314.2077941894531 0 12077.394
315.2109680175781 0 1868.816
331.1979064941406 0 2812.5112 y 10
346.12451171875 0 796.93134
346.1756896972656 0 1598.6599
348.19171142578125 0 1410.4783
353.18182373046875 0 833.2109
358.2587585449219 0 667.6586
360.2287292480469 0 810.86694
362.2073059082031 0 4263.2095
363.20953369140625 0 665.6934
369.1770324707031 0 1298.0613
370.6767883300781 0 1421.6306
371.1933898925781 0 779.1245
390.20361328125 0 563.9375
408.2283020019531 0 758.71564
411.2618103027344 0 920.42206
415.1525573730469 0 556.4585
415.2720642089844 0 606.8918
416.25445556640625 0 1615.6565
422.4696350097656 0 599.61194
424.2211608886719 0 723.9861
433.2816162109375 0 15922.739
434.2850646972656 0 4420.257
434.70526123046875 0 586.92145
435.2876892089844 0 740.0727
440.2179870605469 0 1347.9586
443.2648010253906 0 1308.4775
459.207763671875 0 885.65247
460.2411804199219 0 696.9905 y 9
461.276611328125 0 17986.812
462.27911376953125 0 4305.484
463.2823181152344 0 863.5699
480.21002197265625 0 716.3762
498.22015380859375 0 1761.769
530.2116088867188 0 801.89856
530.2733764648438 0 623.20056
530.3335571289062 0 1031.1326
540.3194580078125 0 1062.073
544.2694702148438 0 1438.2141
544.7699584960938 0 1697.7622
545.274169921875 0 714.6368
558.3297729492188 0 3323.775
559.3294677734375 0 888.2063
587.2678833007812 0 945.245
599.2666625976562 0 865.04266
600.8118286132812 0 1187.7642
601.313232421875 0 1235.2654
603.2838745117188 0 577.8691
627.2623291015625 0 11981.262
628.266845703125 0 3905.523
628.3954467773438 0 1543.4996
629.4010009765625 0 935.6118
643.2904052734375 0 666.7747 w 7
643.3470458984375 0 1239.2318
644.3457641601562 0 842.8678
645.2962646484375 0 640.0465
645.3560180664062 0 1029.8483
655.3827514648438 0 971.9394
656.38818359375 0 793.52716
657.3941040039062 0 1448.3359
658.2794189453125 0 1225.8687
659.28564453125 0 902.5816 y Ammonia loss 7
671.40087890625 0 3276.692
672.4071044921875 0 4285.009 c 5
673.4116821289062 0 1849.7844
674.3014526367188 0 642.295
676.3173828125 0 1199.4006 y 7
688.6207885742188 0 720.47675
708.3228149414062 0 678.9353
712.3512573242188 0 927.75073
715.4267578125 0 1344.1866
716.4347534179688 0 7102.853
717.4359130859375 0 3089.493
722.3281860351562 0 707.00903
740.346435546875 0 9725.925
740.4217529296875 0 1226.1908
741.3506469726562 0 3068.5728
741.416259765625 0 595.8203 c Water loss 6
742.3485717773438 0 1118.6077
742.4149780273438 0 2839.6558
743.41943359375 0 1920.9696
744.4276123046875 0 2273.5112
745.3117065429688 0 1224.0631
746.3220825195312 0 743.9907 y Ammonia loss 6
757.3731689453125 0 2572.8616
758.4326782226562 0 16909.863
759.438720703125 0 19551.734 c 6
760.44287109375 0 7676.546
761.447998046875 0 1531.3911
763.3467407226562 0 2369.3484 y 6
771.351318359375 0 1895.4822
772.3526611328125 0 878.09
774.3268432617188 0 922.019
785.4442138671875 0 592.7461
803.4651489257812 0 2111.93
829.445556640625 0 1369.5527
830.4456787109375 0 1066.374
839.414306640625 0 1560.5657
840.412353515625 0 1491.8108
845.4650268554688 0 11475.758
846.4720458984375 0 54236.285 c 7
847.4754028320312 0 25373.941
848.4783325195312 0 5502.299
850.3936767578125 0 1379.6744
851.3982543945312 0 984.0353
860.4002075195312 0 8820.352 y 5
861.4026489257812 0 3690.0295
868.4053344726562 0 17093.117
869.4066772460938 0 7917.3
870.4107055664062 0 2293.508
871.404052734375 0 934.6436
887.416259765625 0 1867.989
888.4212646484375 0 1166.9221
931.4971923828125 0 1948.8478
932.5028686523438 0 907.7036
938.436279296875 0 2520.923
939.4410400390625 0 2969.1638 y Water loss 4
940.43212890625 0 5423.165 y Ammonia loss 4
941.4334106445312 0 3006.237 z 4
942.4327392578125 0 834.4374
956.445068359375 0 19765.887
957.4524536132812 0 90357.95 y 4
958.4566040039062 0 39006.01
959.4609375 0 11414.398
960.4616088867188 0 1337.13
974.506591796875 0 5964.5063
975.5150756835938 0 66452.65 c 8
976.517578125 0 35872.906
977.5210571289062 0 9129.82
978.5211791992188 0 1124.1272
985.440673828125 0 1199.601
988.4638671875 0 903.6438
1000.4873046875 0 924.23975
1001.5095825195312 0 1018.16846
1015.4721069335938 0 862.7101
1016.4762573242188 0 1429.4402
1017.4788818359375 0 1075.9772
1053.0416259765625 0 751.08594
1053.54296875 0 726.3598
1060.5433349609375 0 1836.3315
1061.549560546875 0 2353.3154
1062.5477294921875 0 1709.4768
1066.0650634765625 0 796.1569
1069.515869140625 0 1942.0518
1070.51708984375 0 916.84106
1085.51123046875 0 1372.2297
1086.5172119140625 0 1572.3733 y Water loss 3
1087.5302734375 0 14801.737
1088.5125732421875 0 23007.805 z 3
1089.5111083984375 0 14291.624
1090.5091552734375 0 4205.8755
1101.5830078125 0 629.48505
1103.5198974609375 0 8726.522
1104.5467529296875 0 72225.14 c 9
1105.550537109375 0 44971.062
1106.5560302734375 0 16352.981
1107.561279296875 0 2041.4714
1120.489013671875 0 726.13586
1129.5626220703125 0 750.7655
1130.5640869140625 0 1359.6814
1131.5574951171875 0 1004.818
1155.6123046875 0 1772.9197
1156.6207275390625 0 1175.6758
1157.6219482421875 0 950.6357
1173.626708984375 0 6060.174
1174.6300048828125 0 3789.8223
1175.6328125 0 1314.332
1182.60302734375 0 2657.5845
1183.59716796875 0 1920.9272 z Water loss 2
1184.5924072265625 0 2162.6572
1185.59814453125 0 990.9871
1200.6143798828125 0 8458.449
1201.594970703125 0 17128.984 z 2
1202.595458984375 0 12134.387
1203.5933837890625 0 4209.1846
1213.5771484375 0 818.62854
1214.5877685546875 0 1141.5085
1215.607666015625 0 833.9024
1216.6038818359375 0 1206.8643
1217.6383056640625 0 50157.516 c 10
1218.641357421875 0 34410.25
1219.64404296875 0 12429.569
1220.6488037109375 0 1645.7974
1230.6138916015625 0 708.2335
1243.6590576171875 0 3490.0757
1244.6639404296875 0 2279.1365
1245.548583984375 0 1086.0441
1246.5731201171875 0 1614.6704
1247.5784912109375 0 1479.6724
1255.3880615234375 0 647.7671
1282.602783203125 0 1569.6443
1283.653076171875 0 808.27075
1284.6226806640625 0 21089.525 z Water loss 1
1285.626220703125 0 15660.227 z Ammonia loss 1
1286.629150390625 0 4907.566
1287.6307373046875 0 1407.2637 w 1
1300.620361328125 0 5321.9375 y Water loss 1
1301.658447265625 0 5251.0234
1302.6363525390625 0 22398.537 z 1
1303.640380859375 0 16357.06
1304.6429443359375 0 5515.28
1305.64208984375 0 1660.0105
1312.660400390625 0 1113.9154
1313.6715087890625 0 1697.0752
1314.682861328125 0 1176.2422
1316.6689453125 0 932.0647
1317.6419677734375 0 2459.2415
1318.647705078125 0 2917.7244 y 1
1328.6876220703125 0 753.3711 c Ammonia loss 11
1329.6732177734375 0 942.5674
1330.669189453125 0 1974.5896
1331.6761474609375 0 1833.1418
1332.6986083984375 0 629.50964
1340.685546875 0 1376.5598
1341.676025390625 0 680.68256
1343.6754150390625 0 1546.4387
1344.693115234375 0 1440.5798
1345.6983642578125 0 138308.9 c 11
1346.70166015625 0 110722.625
1347.7041015625 0 48883.16
1348.7056884765625 0 9760.794
1349.7060546875 0 834.2985
1355.69970703125 0 3552.8513
1356.7144775390625 0 3329.9248
1357.72021484375 0 2076.06
1360.6766357421875 0 796.5649
1362.6617431640625 0 2938.0867
1363.6634521484375 0 2863.8164
1372.7186279296875 0 4467.959
1373.70947265625 0 15037.11
1374.71044921875 0 12568.027
1375.71044921875 0 4711.7075
1376.7191162109375 0 1089.9536
1383.6876220703125 0 7496.5127
1384.6954345703125 0 5725.7427
1385.6954345703125 0 2381.782
1390.732177734375 0 4084.8308
1391.735595703125 0 4022.2073
1392.7281494140625 0 1605.9531
1400.7164306640625 0 6555.9014
1401.7054443359375 0 24756.121
1402.705810546875 0 18165.92
1403.705322265625 0 6983.185
1404.70458984375 0 1912.3954
1416.7115478515625 0 3945.8262
1417.7188720703125 0 46187.426
1418.7265625 0 163245.56
1419.7293701171875 0 119881.805
1420.5545654296875 0 1682.5941
1420.732177734375 0 46654.57
1421.7337646484375 0 7908.8325
2106.083984375 0 1326.0448
2125.1044921875 0 899.4635
2126.08447265625 0 1074.5953
2128.0927734375 0 948.5303
2132.12744140625 0 1719.4259
3030.3046875 0 841.7482
3128.6865234375 0 765.1989

Spectrum Details

|  |  |
| --- | --- |
| Matched peaks? Matched peaks The total absolute number of peaks matched. Additionally in brackets the total fraction of peaks matched and the total number of peaks is shown. | 35 (12.11% of 289) |
| FDR? FDR The false discovery rate estimated for this peptide. It is calculated by matching all theoretical fragments with a non-integer shift with the raw peaks for this spectrum. This is done with 40 different shifts. The resulting percentage is the average number of annotated peaks over the number of annotated peaks with the correct spectrum. | 4.56% |
| Satellite FDR? Satellite FDR See the FDR for details on its calculation. This satellite ion specific FDR only contains the satellite ions (d/w) for I/L/J positions. | ∞ |
| PSM Score? PSM Score The PSM Score as given by Hecklib to this annotated spectrum. It is shown with three significant figures. | 404 |

#### Spectrum 10021? Spectrum 10021 The raw spectrum of this peptide as annotated by Hecklib. The fragments are coloured according to ion type (see legend). Any peaks with a star '\*' as text can be hovered over to see the full details, first the ion type second the mass shift type. By hovering over the amino acids in the peptide or ions in the legend the corresponding peaks are highlighted. By toggling the 'Unassigned' label you can turn the background (unassigned) peaks on or off in the plot. By updating the slider in the Ion legend you can update the spectrum to only show the top X% of the peaks with labels. The top X% means any peak that is within X% of the highest intensity. By dragging in the spectrum you can zoom in to a specific part of the spectrum and use 'Zoom Out' to get back to the original zoom level. The annotation of the spectrum is based on the given sequence in the peptides file and is done with different software so inconsistencies are likely. The peaks are annotated based on the given sequence, with 20 ppm tolerance.

Copy Data

##### Spectrum 10021 (TSV)

###### Preview

```
Loading example...
```

*Click on the button to copy the data to your clipboard.*

Mz MinMz MaxIntensity Max

WidthHeightPeptide font sizePeptide stroke widthSpectrum font sizeSpectrum stroke widthCompact peptide

Ion legend

wxyz

abcd

OtherUnassignedIon^Charge^_Position_Show for top:%

VTJFPPSSEEJQA

04.15e+48.30e+41.24e+51.66e+5

Zoom Out

y^+1^_2_y^+1^_2_y^+2^_5_y^+1^_3_y^+1^_3_y^+1^_4_y^+2^_9_y^+1^_5_w^+1^_6_y^+1^_6_c^+1^_6_y^+1^_6_c^+1^_7_y^+1^_7_c^+1^_7_y^+1^_7_y^+1^_8_c^+1^_8_y^+1^_8_y^+1^_9_y^+1^_9_z^+1^_9_y^+1^_9_c^+1^_9_y^+1^_10_z^+1^_10_c^+1^_10_z^+1^_11_c^+1^_11_z^+1^_12_z^+1^_12_y^+1^_12_z^+1^_12_y^+1^_12_c^+1^_12_c^+1^_12_c^+1^_12_

0538107716152153

Fragment Matches Table

Show background peaks

| Position | Ion type | Intensity | mz Theoretical | mz Error (Th) | mz Error (ppm) | Charge | Series Number |
| --- | --- | --- | --- | --- | --- | --- | --- |
| - | - | 1404 | 120.1 | - | - | 0 | - |
| - | - | 1109 | 133.1 | - | - | 0 | - |
| - | - | 362.6 | 134.7 | - | - | 0 | - |
| - | - | 395.5 | 137.8 | - | - | 0 | - |
| - | - | 489.9 | 143.9 | - | - | 0 | - |
| - | - | 856.5 | 149 | - | - | 0 | - |
| - | - | 550.2 | 165.3 | - | - | 0 | - |
| - | - | 460 | 165.5 | - | - | 0 | - |
| - | - | 432.8 | 171.6 | - | - | 0 | - |
| - | - | 3321 | 173.5 | - | - | 0 | - |
| - | - | 1832 | 183.1 | - | - | 0 | - |
| - | - | 485 | 197.1 | - | - | 0 | - |
| - | - | 7645 | 200.1 | - | - | 0 | - |
| 12 | y | 6140 | 201.1 | 0.0001443 | 0.7178 | +1 | 2 |
| - | - | 806.6 | 201.1 | - | - | 0 | - |
| - | - | 3642 | 201.1 | - | - | 0 | - |
| - | - | 592.8 | 213.1 | - | - | 0 | - |
| - | - | 554.5 | 215 | - | - | 0 | - |
| - | - | 1.121E+04 | 215.1 | - | - | 0 | - |
| - | - | 1246 | 216.1 | - | - | 0 | - |
| 12 | y | 1.527E+04 | 218.1 | 0.0001608 | 0.7372 | +1 | 2 |
| - | - | 525.9 | 218.3 | - | - | 0 | - |
| - | - | 1475 | 219.1 | - | - | 0 | - |
| - | - | 525.7 | 222.5 | - | - | 0 | - |
| - | - | 589.3 | 225.1 | - | - | 0 | - |
| - | - | 1354 | 233.2 | - | - | 0 | - |
| - | - | 5204 | 242.2 | - | - | 0 | - |
| - | - | 827.1 | 261.2 | - | - | 0 | - |
| - | - | 504.5 | 268.4 | - | - | 0 | - |
| - | - | 461.5 | 272.9 | - | - | 0 | - |
| - | - | 449.1 | 275.4 | - | - | 0 | - |
| - | - | 1016 | 280.2 | - | - | 0 | - |
| - | - | 1721 | 282.1 | - | - | 0 | - |
| - | - | 566.5 | 292.1 | - | - | 0 | - |
| 9 | y | 915.9 | 295.1 | 0.0001931 | 0.6543 | +2 | 5 |
| - | - | 2.755E+04 | 296.2 | - | - | 0 | - |
| - | - | 509.6 | 297.1 | - | - | 0 | - |
| - | - | 5306 | 297.2 | - | - | 0 | - |
| - | - | 682.6 | 303.2 | - | - | 0 | - |
| 11 | y | 1370 | 314.2 | 0.001011 | 3.217 | +1 | 3 |
| - | - | 1.296E+04 | 314.2 | - | - | 0 | - |
| - | - | 1235 | 315.2 | - | - | 0 | - |
| - | - | 589.8 | 330.2 | - | - | 0 | - |
| 11 | y | 2158 | 331.2 | 0.0002185 | 0.6598 | +1 | 3 |
| - | - | 750.3 | 332.2 | - | - | 0 | - |
| - | - | 1060 | 346.1 | - | - | 0 | - |
| - | - | 593 | 346.2 | - | - | 0 | - |
| - | - | 856.5 | 348.2 | - | - | 0 | - |
| - | - | 640.2 | 349.2 | - | - | 0 | - |
| - | - | 1100 | 353.2 | - | - | 0 | - |
| - | - | 3764 | 362.2 | - | - | 0 | - |
| - | - | 691.9 | 363.2 | - | - | 0 | - |
| - | - | 698.4 | 369.2 | - | - | 0 | - |
| - | - | 579.5 | 370.7 | - | - | 0 | - |
| - | - | 812.1 | 371.2 | - | - | 0 | - |
| - | - | 701.3 | 398.2 | - | - | 0 | - |
| - | - | 911.6 | 411.3 | - | - | 0 | - |
| - | - | 578.1 | 415.2 | - | - | 0 | - |
| - | - | 823 | 415.3 | - | - | 0 | - |
| - | - | 1444 | 416.3 | - | - | 0 | - |
| - | - | 589.3 | 432.2 | - | - | 0 | - |
| - | - | 1.644E+04 | 433.3 | - | - | 0 | - |
| - | - | 4417 | 434.3 | - | - | 0 | - |
| - | - | 623.9 | 434.7 | - | - | 0 | - |
| - | - | 822 | 435.3 | - | - | 0 | - |
| - | - | 1289 | 441.2 | - | - | 0 | - |
| - | - | 2189 | 443.3 | - | - | 0 | - |
| - | - | 603.6 | 454.2 | - | - | 0 | - |
| - | - | 755.4 | 459.2 | - | - | 0 | - |
| 10 | y | 697 | 460.2 | 0.0002585 | 0.5616 | +1 | 4 |
| - | - | 1.654E+04 | 461.3 | - | - | 0 | - |
| - | - | 4891 | 462.3 | - | - | 0 | - |
| 5 | y | 835.5 | 470.2 | 0.001364 | 2.902 | +2 | 9 |
| - | - | 759.4 | 480.2 | - | - | 0 | - |
| - | - | 598.9 | 480.2 | - | - | 0 | - |
| - | - | 1689 | 498.2 | - | - | 0 | - |
| - | - | 1421 | 530.3 | - | - | 0 | - |
| - | - | 549.4 | 530.8 | - | - | 0 | - |
| - | - | 825.8 | 540.3 | - | - | 0 | - |
| - | - | 1456 | 544.3 | - | - | 0 | - |
| - | - | 1257 | 544.8 | - | - | 0 | - |
| - | - | 1034 | 545.3 | - | - | 0 | - |
| - | - | 3519 | 558.3 | - | - | 0 | - |
| - | - | 1166 | 559.3 | - | - | 0 | - |
| - | - | 924.7 | 587.3 | - | - | 0 | - |
| 9 | y | 672 | 589.3 | 0.001533 | 2.601 | +1 | 5 |
| - | - | 699.8 | 600.8 | - | - | 0 | - |
| - | - | 778.4 | 609.3 | - | - | 0 | - |
| - | - | 1.09E+04 | 627.3 | - | - | 0 | - |
| - | - | 3279 | 628.3 | - | - | 0 | - |
| - | - | 903.2 | 628.4 | - | - | 0 | - |
| - | - | 752.3 | 629.4 | - | - | 0 | - |
| 8 | w | 936 | 643.3 | 0.001538 | 2.391 | +1 | 6 |
| - | - | 1124 | 643.3 | - | - | 0 | - |
| - | - | 1185 | 644.3 | - | - | 0 | - |
| - | - | 708.4 | 645.3 | - | - | 0 | - |
| - | - | 752.4 | 645.4 | - | - | 0 | - |
| - | - | 601.7 | 646.3 | - | - | 0 | - |
| - | - | 997.3 | 655.4 | - | - | 0 | - |
| - | - | 738.7 | 656.4 | - | - | 0 | - |
| - | - | 780.6 | 657.4 | - | - | 0 | - |
| - | - | 851.5 | 658.3 | - | - | 0 | - |
| 8 | y | 1019 | 659.3 | 0.00396 | 6.007 | +1 | 6 |
| - | - | 747 | 660.3 | - | - | 0 | - |
| - | - | 742.2 | 663.4 | - | - | 0 | - |
| - | - | 3152 | 671.4 | - | - | 0 | - |
| 6 | c | 4971 | 672.4 | 0.000758 | 1.127 | +1 | 6 |
| - | - | 2090 | 673.4 | - | - | 0 | - |
| - | - | 736 | 674.4 | - | - | 0 | - |
| 8 | y | 1493 | 676.3 | 0.001961 | 2.9 | +1 | 6 |
| - | - | 877 | 677.3 | - | - | 0 | - |
| - | - | 616.3 | 708.4 | - | - | 0 | - |
| - | - | 639.2 | 712.3 | - | - | 0 | - |
| - | - | 799.1 | 714.4 | - | - | 0 | - |
| - | - | 714.1 | 715.4 | - | - | 0 | - |
| - | - | 6571 | 716.4 | - | - | 0 | - |
| - | - | 1932 | 717.4 | - | - | 0 | - |
| - | - | 614.4 | 718.4 | - | - | 0 | - |
| - | - | 1029 | 722.3 | - | - | 0 | - |
| - | - | 1.026E+04 | 740.3 | - | - | 0 | - |
| - | - | 913.2 | 740.4 | - | - | 0 | - |
| - | - | 3511 | 741.3 | - | - | 0 | - |
| 7 | c | 867.3 | 741.4 | 0.01014 | 13.67 | +1 | 7 |
| - | - | 836.5 | 742.3 | - | - | 0 | - |
| - | - | 3139 | 742.4 | - | - | 0 | - |
| - | - | 1383 | 743.4 | - | - | 0 | - |
| - | - | 1381 | 744.4 | - | - | 0 | - |
| - | - | 842.5 | 745.3 | - | - | 0 | - |
| - | - | 751 | 745.4 | - | - | 0 | - |
| 7 | y | 899.9 | 746.3 | 0.002358 | 3.16 | +1 | 7 |
| - | - | 2819 | 757.4 | - | - | 0 | - |
| - | - | 904.1 | 758.4 | - | - | 0 | - |
| - | - | 1.608E+04 | 758.4 | - | - | 0 | - |
| 7 | c | 1.951E+04 | 759.4 | 0.001231 | 1.621 | +1 | 7 |
| - | - | 6948 | 760.4 | - | - | 0 | - |
| - | - | 1692 | 761.4 | - | - | 0 | - |
| 7 | y | 1987 | 763.3 | 8.44E-05 | 0.1106 | +1 | 7 |
| - | - | 2784 | 771.4 | - | - | 0 | - |
| - | - | 957.5 | 772.4 | - | - | 0 | - |
| - | - | 1484 | 774.3 | - | - | 0 | - |
| - | - | 783.2 | 775.3 | - | - | 0 | - |
| - | - | 1946 | 803.5 | - | - | 0 | - |
| - | - | 1428 | 829.5 | - | - | 0 | - |
| - | - | 808.1 | 830.5 | - | - | 0 | - |
| - | - | 1380 | 839.4 | - | - | 0 | - |
| - | - | 895.9 | 840.4 | - | - | 0 | - |
| - | - | 764.2 | 841.4 | - | - | 0 | - |
| 6 | y | 689 | 843.4 | 0.0006177 | 0.7324 | +1 | 8 |
| - | - | 9981 | 845.5 | - | - | 0 | - |
| 8 | c | 5.109E+04 | 846.5 | 6.553E-05 | 0.07742 | +1 | 8 |
| - | - | 2.261E+04 | 847.5 | - | - | 0 | - |
| - | - | 826.9 | 848.4 | - | - | 0 | - |
| - | - | 5468 | 848.5 | - | - | 0 | - |
| 6 | y | 7992 | 860.4 | 0.001154 | 1.341 | +1 | 8 |
| - | - | 4182 | 861.4 | - | - | 0 | - |
| - | - | 1277 | 862.4 | - | - | 0 | - |
| - | - | 1.559E+04 | 868.4 | - | - | 0 | - |
| - | - | 6751 | 869.4 | - | - | 0 | - |
| - | - | 1939 | 870.4 | - | - | 0 | - |
| - | - | 2762 | 887.4 | - | - | 0 | - |
| - | - | 1279 | 888.4 | - | - | 0 | - |
| - | - | 1754 | 931.5 | - | - | 0 | - |
| - | - | 766.4 | 932.5 | - | - | 0 | - |
| - | - | 2070 | 938.4 | - | - | 0 | - |
| 5 | y | 3183 | 939.4 | 0.0018 | 1.916 | +1 | 9 |
| 5 | y | 5738 | 940.4 | 0.007593 | 8.074 | +1 | 9 |
| 5 | z | 2268 | 941.4 | 0.002819 | 2.995 | +1 | 9 |
| - | - | 1.863E+04 | 956.4 | - | - | 0 | - |
| 5 | y | 9.091E+04 | 957.5 | 0.0002085 | 0.2178 | +1 | 9 |
| - | - | 4.003E+04 | 958.5 | - | - | 0 | - |
| - | - | 1.077E+04 | 959.5 | - | - | 0 | - |
| - | - | 1435 | 960.5 | - | - | 0 | - |
| - | - | 631.3 | 964 | - | - | 0 | - |
| - | - | 6445 | 974.5 | - | - | 0 | - |
| 9 | c | 5.838E+04 | 975.5 | 0.0006853 | 0.7025 | +1 | 9 |
| - | - | 3.317E+04 | 976.5 | - | - | 0 | - |
| - | - | 9405 | 977.5 | - | - | 0 | - |
| - | - | 921.1 | 978.5 | - | - | 0 | - |
| - | - | 1119 | 985.4 | - | - | 0 | - |
| - | - | 817.2 | 1001 | - | - | 0 | - |
| - | - | 713.3 | 1002 | - | - | 0 | - |
| - | - | 1362 | 1009 | - | - | 0 | - |
| - | - | 845.2 | 1010 | - | - | 0 | - |
| - | - | 769.1 | 1015 | - | - | 0 | - |
| - | - | 1505 | 1016 | - | - | 0 | - |
| - | - | 704.4 | 1026 | - | - | 0 | - |
| - | - | 742.4 | 1029 | - | - | 0 | - |
| - | - | 920.5 | 1033 | - | - | 0 | - |
| - | - | 1150 | 1033 | - | - | 0 | - |
| - | - | 704.9 | 1047 | - | - | 0 | - |
| - | - | 817.4 | 1054 | - | - | 0 | - |
| - | - | 920.7 | 1055 | - | - | 0 | - |
| - | - | 884.3 | 1057 | - | - | 0 | - |
| - | - | 736.8 | 1057 | - | - | 0 | - |
| - | - | 931.3 | 1058 | - | - | 0 | - |
| - | - | 1729 | 1061 | - | - | 0 | - |
| - | - | 2508 | 1062 | - | - | 0 | - |
| - | - | 698.2 | 1062 | - | - | 0 | - |
| - | - | 1080 | 1063 | - | - | 0 | - |
| - | - | 1175 | 1070 | - | - | 0 | - |
| - | - | 773.8 | 1071 | - | - | 0 | - |
| - | - | 885 | 1086 | - | - | 0 | - |
| 4 | y | 900.6 | 1087 | 0.009193 | 8.461 | +1 | 10 |
| - | - | 1.652E+04 | 1088 | - | - | 0 | - |
| 4 | z | 2.415E+04 | 1089 | 0.01039 | 9.549 | +1 | 10 |
| - | - | 1.444E+04 | 1090 | - | - | 0 | - |
| - | - | 4076 | 1091 | - | - | 0 | - |
| - | - | 1509 | 1092 | - | - | 0 | - |
| - | - | 1108 | 1103 | - | - | 0 | - |
| - | - | 8038 | 1104 | - | - | 0 | - |
| 10 | c | 6.985E+04 | 1105 | 0.01054 | 9.538 | +1 | 10 |
| - | - | 4.213E+04 | 1106 | - | - | 0 | - |
| - | - | 1.389E+04 | 1107 | - | - | 0 | - |
| - | - | 944.6 | 1108 | - | - | 0 | - |
| - | - | 1272 | 1130 | - | - | 0 | - |
| - | - | 1237 | 1131 | - | - | 0 | - |
| - | - | 993.6 | 1132 | - | - | 0 | - |
| - | - | 1547 | 1156 | - | - | 0 | - |
| - | - | 888 | 1157 | - | - | 0 | - |
| - | - | 702.1 | 1165 | - | - | 0 | - |
| - | - | 882.5 | 1170 | - | - | 0 | - |
| - | - | 5019 | 1174 | - | - | 0 | - |
| - | - | 3154 | 1175 | - | - | 0 | - |
| - | - | 1639 | 1176 | - | - | 0 | - |
| - | - | 2948 | 1183 | - | - | 0 | - |
| - | - | 2350 | 1184 | - | - | 0 | - |
| - | - | 1415 | 1185 | - | - | 0 | - |
| - | - | 1574 | 1186 | - | - | 0 | - |
| - | - | 9290 | 1201 | - | - | 0 | - |
| 3 | z | 1.706E+04 | 1202 | 0.01031 | 8.584 | +1 | 11 |
| - | - | 1.065E+04 | 1203 | - | - | 0 | - |
| - | - | 3111 | 1204 | - | - | 0 | - |
| - | - | 1178 | 1214 | - | - | 0 | - |
| - | - | 769.1 | 1215 | - | - | 0 | - |
| - | - | 812.5 | 1216 | - | - | 0 | - |
| - | - | 1365 | 1217 | - | - | 0 | - |
| 11 | c | 4.711E+04 | 1218 | 0.002192 | 1.801 | +1 | 11 |
| - | - | 3.189E+04 | 1219 | - | - | 0 | - |
| - | - | 1.091E+04 | 1220 | - | - | 0 | - |
| - | - | 1765 | 1221 | - | - | 0 | - |
| - | - | 846.1 | 1224 | - | - | 0 | - |
| - | - | 965.5 | 1231 | - | - | 0 | - |
| - | - | 889.3 | 1232 | - | - | 0 | - |
| - | - | 3461 | 1244 | - | - | 0 | - |
| - | - | 2101 | 1245 | - | - | 0 | - |
| - | - | 887.1 | 1246 | - | - | 0 | - |
| - | - | 2331 | 1247 | - | - | 0 | - |
| - | - | 992 | 1248 | - | - | 0 | - |
| - | - | 641.5 | 1255 | - | - | 0 | - |
| - | - | 999.2 | 1267 | - | - | 0 | - |
| - | - | 627.4 | 1278 | - | - | 0 | - |
| - | - | 1784 | 1283 | - | - | 0 | - |
| - | - | 1586 | 1284 | - | - | 0 | - |
| 2 | z | 1.674E+04 | 1285 | 0.0006762 | 0.5263 | +1 | 12 |
| 2 | z | 1.376E+04 | 1286 | 0.01909 | 14.85 | +1 | 12 |
| - | - | 4878 | 1287 | - | - | 0 | - |
| - | - | 680.8 | 1288 | - | - | 0 | - |
| - | - | 861.2 | 1294 | - | - | 0 | - |
| - | - | 841.9 | 1295 | - | - | 0 | - |
| - | - | 859.5 | 1296 | - | - | 0 | - |
| 2 | y | 5207 | 1301 | 0.02428 | 18.67 | +1 | 12 |
| - | - | 5884 | 1302 | - | - | 0 | - |
| 2 | z | 2.244E+04 | 1303 | 0.002675 | 2.054 | +1 | 12 |
| - | - | 1.69E+04 | 1304 | - | - | 0 | - |
| - | - | 5750 | 1305 | - | - | 0 | - |
| - | - | 1424 | 1306 | - | - | 0 | - |
| - | - | 1087 | 1310 | - | - | 0 | - |
| - | - | 1987 | 1311 | - | - | 0 | - |
| - | - | 1753 | 1312 | - | - | 0 | - |
| - | - | 1858 | 1313 | - | - | 0 | - |
| - | - | 1860 | 1314 | - | - | 0 | - |
| - | - | 964.7 | 1315 | - | - | 0 | - |
| - | - | 903.5 | 1317 | - | - | 0 | - |
| - | - | 2557 | 1318 | - | - | 0 | - |
| 2 | y | 2913 | 1319 | 0.001645 | 1.247 | +1 | 12 |
| - | - | 985.1 | 1320 | - | - | 0 | - |
| - | - | 807.7 | 1326 | - | - | 0 | - |
| - | - | 989.4 | 1327 | - | - | 0 | - |
| 12 | c | 760.1 | 1328 | 0.01871 | 14.09 | +1 | 12 |
| 12 | c | 1238 | 1329 | 0.00179 | 1.347 | +1 | 12 |
| - | - | 1372 | 1330 | - | - | 0 | - |
| - | - | 2726 | 1331 | - | - | 0 | - |
| - | - | 2550 | 1332 | - | - | 0 | - |
| - | - | 821 | 1333 | - | - | 0 | - |
| - | - | 1124 | 1341 | - | - | 0 | - |
| - | - | 1176 | 1342 | - | - | 0 | - |
| - | - | 1428 | 1343 | - | - | 0 | - |
| - | - | 828.2 | 1344 | - | - | 0 | - |
| - | - | 1425 | 1345 | - | - | 0 | - |
| 12 | c | 1.354E+05 | 1346 | 0.001322 | 0.9822 | +1 | 12 |
| - | - | 1.053E+05 | 1347 | - | - | 0 | - |
| - | - | 4.676E+04 | 1348 | - | - | 0 | - |
| - | - | 9643 | 1349 | - | - | 0 | - |
| - | - | 665.9 | 1350 | - | - | 0 | - |
| - | - | 953.5 | 1355 | - | - | 0 | - |
| - | - | 2848 | 1356 | - | - | 0 | - |
| - | - | 3577 | 1357 | - | - | 0 | - |
| - | - | 1844 | 1358 | - | - | 0 | - |
| - | - | 2212 | 1360 | - | - | 0 | - |
| - | - | 2105 | 1361 | - | - | 0 | - |
| - | - | 898.2 | 1362 | - | - | 0 | - |
| - | - | 3262 | 1363 | - | - | 0 | - |
| - | - | 2506 | 1364 | - | - | 0 | - |
| - | - | 1090 | 1365 | - | - | 0 | - |
| - | - | 2220 | 1368 | - | - | 0 | - |
| - | - | 910.4 | 1372 | - | - | 0 | - |
| - | - | 4189 | 1373 | - | - | 0 | - |
| - | - | 1.37E+04 | 1374 | - | - | 0 | - |
| - | - | 1.195E+04 | 1375 | - | - | 0 | - |
| - | - | 4565 | 1376 | - | - | 0 | - |
| - | - | 1797 | 1377 | - | - | 0 | - |
| - | - | 2742 | 1383 | - | - | 0 | - |
| - | - | 9732 | 1384 | - | - | 0 | - |
| - | - | 7621 | 1385 | - | - | 0 | - |
| - | - | 5087 | 1386 | - | - | 0 | - |
| - | - | 1836 | 1387 | - | - | 0 | - |
| - | - | 851.1 | 1388 | - | - | 0 | - |
| - | - | 4620 | 1391 | - | - | 0 | - |
| - | - | 3117 | 1392 | - | - | 0 | - |
| - | - | 1090 | 1393 | - | - | 0 | - |
| - | - | 2632 | 1399 | - | - | 0 | - |
| - | - | 3178 | 1400 | - | - | 0 | - |
| - | - | 8120 | 1401 | - | - | 0 | - |
| - | - | 2.589E+04 | 1402 | - | - | 0 | - |
| - | - | 2.032E+04 | 1403 | - | - | 0 | - |
| - | - | 8097 | 1404 | - | - | 0 | - |
| - | - | 1558 | 1405 | - | - | 0 | - |
| - | - | 2749 | 1416 | - | - | 0 | - |
| - | - | 5087 | 1417 | - | - | 0 | - |
| - | - | 4.369E+04 | 1418 | - | - | 0 | - |
| - | - | 1.643E+05 | 1419 | - | - | 0 | - |
| - | - | 1.108E+05 | 1420 | - | - | 0 | - |
| - | - | 4.631E+04 | 1421 | - | - | 0 | - |
| - | - | 6717 | 1422 | - | - | 0 | - |
| - | - | 949.3 | 1451 | - | - | 0 | - |
| - | - | 744.5 | 1475 | - | - | 0 | - |
| - | - | 1150 | 2048 | - | - | 0 | - |
| - | - | 1057 | 2051 | - | - | 0 | - |
| - | - | 1201 | 2065 | - | - | 0 | - |
| - | - | 930.2 | 2066 | - | - | 0 | - |
| - | - | 830.3 | 2068 | - | - | 0 | - |
| - | - | 777.3 | 2096 | - | - | 0 | - |
| - | - | 872.5 | 2097 | - | - | 0 | - |
| - | - | 1116 | 2106 | - | - | 0 | - |
| - | - | 1571 | 2107 | - | - | 0 | - |
| - | - | 2002 | 2108 | - | - | 0 | - |
| - | - | 1235 | 2109 | - | - | 0 | - |
| - | - | 854 | 2112 | - | - | 0 | - |
| - | - | 806.4 | 2114 | - | - | 0 | - |
| - | - | 1022 | 2123 | - | - | 0 | - |
| - | - | 2474 | 2124 | - | - | 0 | - |
| - | - | 2187 | 2125 | - | - | 0 | - |
| - | - | 880.8 | 2126 | - | - | 0 | - |
| - | - | 1088 | 2127 | - | - | 0 | - |
| - | - | 942.3 | 2129 | - | - | 0 | - |
| - | - | 1053 | 2130 | - | - | 0 | - |
| - | - | 1115 | 2131 | - | - | 0 | - |
| - | - | 987.7 | 2132 | - | - | 0 | - |

m/z Charge Intensity FragmentType MassShift Position
120.08100128173828 0 1404.2943
133.08631896972656 0 1108.913
134.65811157226562 0 362.60837
137.82333374023438 0 395.48895
143.93951416015625 0 489.92178
148.9537811279297 0 856.5399
165.31629943847656 0 550.17267
165.48171997070312 0 460.01657
171.62667846679688 0 432.77795
173.450927734375 0 3321.1082
183.11322021484375 0 1831.559
197.1291046142578 0 484.9847
200.10299682617188 0 7645.3613
201.08712768554688 0 6140.1514 y Ammonia loss 11
201.10606384277344 0 806.62897
201.12356567382812 0 3642.0137
213.12289428710938 0 592.774
214.9657440185547 0 554.4767
215.1392059326172 0 11206.886
216.14205932617188 0 1246.1469
218.1136932373047 0 15265.172 y 11
218.33050537109375 0 525.8868
219.11724853515625 0 1475.0989
222.47862243652344 0 525.7362
225.1234588623047 0 589.32056
233.1654815673828 0 1353.5171
242.15016174316406 0 5203.6562
261.1590576171875 0 827.0522
268.36669921875 0 504.52884
272.8739318847656 0 461.53967
275.4154357910156 0 449.1424
280.1659851074219 0 1016.2249
282.1451721191406 0 1721.2339
292.0840148925781 0 566.5111
295.14483642578125 0 915.89526 y 8
296.1971740722656 0 27553.018
297.1347351074219 0 509.5954
297.2004699707031 0 5305.833
303.1700439453125 0 682.6472
314.17205810546875 0 1370.0332 y Ammonia loss 10
314.2079162597656 0 12955.48
315.21087646484375 0 1235.0442
330.1833801269531 0 589.8141
331.19781494140625 0 2158.0918 y 10
332.19903564453125 0 750.312
346.1250915527344 0 1060.3457
346.1771240234375 0 593.0276
348.1920166015625 0 856.493
349.1938171386719 0 640.1771
353.1821594238281 0 1100.0078
362.2075500488281 0 3763.5046
363.21026611328125 0 691.86615
369.1757507324219 0 698.37445
370.67547607421875 0 579.51996
371.1929016113281 0 812.0827
398.20257568359375 0 701.2834
411.26092529296875 0 911.6471
415.2320251464844 0 578.05743
415.2680358886719 0 822.987
416.2553405761719 0 1443.7944
432.166748046875 0 589.30383
433.2816162109375 0 16441.193
434.2846984863281 0 4416.5312
434.70843505859375 0 623.8644
435.2861633300781 0 821.98096
441.2332458496094 0 1289.2861
443.265869140625 0 2189.416
454.2279052734375 0 603.63275
459.2120666503906 0 755.40424
460.2404479980469 0 696.9532 y 9
461.27655029296875 0 16538.957
462.2801513671875 0 4891.2544
470.2231750488281 0 835.52826 y Water loss 4
480.208740234375 0 759.44696
480.24761962890625 0 598.9461
498.2202453613281 0 1688.6348
530.3353271484375 0 1420.5348
530.7734985351562 0 549.37225
540.3182983398438 0 825.7568
544.2697143554688 0 1456.0809
544.7713012695312 0 1256.8956
545.269775390625 0 1034.05
558.3292846679688 0 3518.7676
559.3347778320312 0 1166.2058
587.2648315429688 0 924.7142
589.28125 0 672.0164 y 8
600.81396484375 0 699.7555
609.2535400390625 0 778.4297
627.262451171875 0 10899.953
628.2655639648438 0 3279.1567
628.3920288085938 0 903.19867
629.4036865234375 0 752.3049
643.2918090820312 0 935.98694 w 7
643.3480224609375 0 1123.7312
644.2911376953125 0 1184.5718
645.29638671875 0 708.4404
645.3607177734375 0 752.3597
646.2972412109375 0 601.6725
655.3822631835938 0 997.33997
656.38720703125 0 738.7359
657.3951416015625 0 780.6159
658.2822875976562 0 851.5314
659.2843017578125 0 1018.85266 y Ammonia loss 7
660.31787109375 0 746.97845
663.4196166992188 0 742.20886
671.4011840820312 0 3151.6194
672.4071655273438 0 4970.7676 c 5
673.410888671875 0 2089.799
674.4267578125 0 736.0209
676.3167724609375 0 1493.4733 y 7
677.322265625 0 876.9885
708.4248046875 0 616.3055
712.3497314453125 0 639.22375
714.4180297851562 0 799.0974
715.4230346679688 0 714.0618
716.4352416992188 0 6570.961
717.4368896484375 0 1931.8933
718.436279296875 0 614.3904
722.3334350585938 0 1029.4707
740.3463134765625 0 10263.61
740.4179077148438 0 913.2489
741.3497924804688 0 3510.9841
741.4192504882812 0 867.2868 c Water loss 6
742.3466796875 0 836.4629
742.4130249023438 0 3138.964
743.41650390625 0 1383.0739
744.426025390625 0 1380.8849
745.31103515625 0 842.4603
745.4314575195312 0 751.0416
746.3179321289062 0 899.8524 y Ammonia loss 6
757.3714599609375 0 2818.573
758.37548828125 0 904.0669
758.4329833984375 0 16075.147
759.438720703125 0 19511.717 c 6
760.4422607421875 0 6947.7344
761.4451293945312 0 1692.0205
763.346923828125 0 1987.469 y 6
771.3529052734375 0 2784.3862
772.3515625 0 957.5184
774.331787109375 0 1483.9506
775.337646484375 0 783.2013
803.4686889648438 0 1945.6124
829.4508056640625 0 1428.1326
830.4511108398438 0 808.1379
839.4146728515625 0 1380.2367
840.4116821289062 0 895.8608
841.4159545898438 0 764.21906
843.3724365234375 0 688.97437 y Ammonia loss 5
845.4649658203125 0 9981.005
846.4720458984375 0 51087.74 c 7
847.4754638671875 0 22614.15
848.4006958007812 0 826.8731
848.478271484375 0 5468.258
860.4007568359375 0 7991.666 y 5
861.4026489257812 0 4182.2466
862.406005859375 0 1276.652
868.405517578125 0 15585.726
869.4085083007812 0 6750.8037
870.4122314453125 0 1939.4062
887.4142456054688 0 2762.0615
888.413818359375 0 1278.554
931.5001220703125 0 1753.5371
932.502685546875 0 766.4317
938.43310546875 0 2069.6606
939.4400024414062 0 3183.294 y Water loss 4
940.4334106445312 0 5738.0747 y Ammonia loss 4
941.4364624023438 0 2267.6592 z 4
956.4456787109375 0 18631.082
957.4525756835938 0 90909.26 y 4
958.4566650390625 0 40032.37
959.4627075195312 0 10770.547
960.4740600585938 0 1434.708
964.0252685546875 0 631.3393
974.5079345703125 0 6445.162
975.5152587890625 0 58384.66 c 8
976.51806640625 0 33172.47
977.5213623046875 0 9405.121
978.5283203125 0 921.10406
985.4430541992188 0 1118.8158
1000.5477905273438 0 817.1762
1001.5150146484375 0 713.3231
1009.0477294921875 0 1362.2568
1009.5382690429688 0 845.2135
1015.4751586914062 0 769.06384
1016.4812622070312 0 1505.1841
1025.5533447265625 0 704.3557
1029.4810791015625 0 742.36914
1032.55078125 0 920.4769
1033.052734375 0 1150.4832
1047.486328125 0 704.862
1053.5499267578125 0 817.40784
1054.540771484375 0 920.72876
1056.518310546875 0 884.2888
1057.0439453125 0 736.76544
1057.5394287109375 0 931.25684
1060.5435791015625 0 1729.2035
1061.544677734375 0 2508.127
1062.0660400390625 0 698.1938
1062.55908203125 0 1080.1489
1069.5181884765625 0 1174.557
1070.5135498046875 0 773.75433
1085.51171875 0 885.04175
1086.5194091796875 0 900.584 y Water loss 3
1087.5296630859375 0 16523.908
1088.512451171875 0 24150.44 z 3
1089.5107421875 0 14437.414
1090.5118408203125 0 4076.1558
1091.5162353515625 0 1508.8392
1102.5419921875 0 1107.9843
1103.5211181640625 0 8037.954
1104.546630859375 0 69852.1 c 9
1105.5506591796875 0 42127.445
1106.55517578125 0 13892.697
1107.5528564453125 0 944.6097
1129.563720703125 0 1272.2957
1130.56298828125 0 1237.376
1131.568359375 0 993.6349
1155.61669921875 0 1546.5687
1156.6162109375 0 887.9939
1164.59033203125 0 702.07983
1169.5179443359375 0 882.4997
1173.6273193359375 0 5018.931
1174.6328125 0 3153.8848
1175.6329345703125 0 1639.3265
1182.603515625 0 2947.9714
1183.6051025390625 0 2349.8774
1184.5994873046875 0 1415.3301
1185.5927734375 0 1574.4945
1200.6153564453125 0 9290.278
1201.596435546875 0 17055.074 z 2
1202.59375 0 10652.397
1203.600341796875 0 3110.8103
1213.5845947265625 0 1178.0011
1214.595458984375 0 769.10706
1215.6007080078125 0 812.4812
1216.6033935546875 0 1364.6923
1217.6390380859375 0 47108.484 c 10
1218.6417236328125 0 31886.744
1219.64404296875 0 10914.462
1220.6451416015625 0 1765.3795
1223.6143798828125 0 846.0919
1230.60009765625 0 965.52167
1231.5999755859375 0 889.294
1243.656005859375 0 3461.0366
1244.65576171875 0 2100.7778
1245.6458740234375 0 887.1324
1246.58154296875 0 2331.4373
1247.5823974609375 0 992.03546
1254.6024169921875 0 641.51746
1266.6182861328125 0 999.2271
1277.6553955078125 0 627.41833
1282.6072998046875 0 1784.1754
1283.6212158203125 0 1585.5366
1284.62255859375 0 16743.867 z Water loss 1
1285.6263427734375 0 13755.4795 z Ammonia loss 1
1286.6322021484375 0 4878.238
1287.6448974609375 0 680.8484
1293.6357421875 0 861.1913
1294.6461181640625 0 841.8839
1295.65478515625 0 859.47107
1300.61767578125 0 5207.259 y Water loss 1
1301.6630859375 0 5883.568
1302.636474609375 0 22436.775 z 1
1303.6402587890625 0 16902.182
1304.6446533203125 0 5750.1978
1305.6461181640625 0 1424.4159
1309.6480712890625 0 1087.3617
1310.6497802734375 0 1987.402
1311.6566162109375 0 1753.117
1312.66259765625 0 1858.4554
1313.658935546875 0 1859.9506
1314.689697265625 0 964.727
1316.674560546875 0 903.54803
1317.639892578125 0 2556.79
1318.65087890625 0 2913.3247 y 1
1319.659423828125 0 985.08545
1325.64404296875 0 807.7108
1326.65283203125 0 989.381
1327.6705322265625 0 760.0813 c Water loss 11
1328.675048828125 0 1238.3945 c Ammonia loss 11
1329.6893310546875 0 1371.5648
1330.6693115234375 0 2725.7686
1331.6737060546875 0 2550.4312
1332.6943359375 0 820.97363
1340.6839599609375 0 1123.7771
1341.640869140625 0 1175.5032
1342.661376953125 0 1428.4686
1343.6788330078125 0 828.2348
1344.6859130859375 0 1425.1877
1345.698486328125 0 135417.34 c 11
1346.7021484375 0 105336.09
1347.7047119140625 0 46764.043
1348.706298828125 0 9642.893
1349.677490234375 0 665.91797
1354.692138671875 0 953.4659
1355.6943359375 0 2848.4688
1356.710205078125 0 3576.7546
1357.642822265625 0 1843.9684
1359.650390625 0 2212.3523
1360.6551513671875 0 2105.3887
1361.6624755859375 0 898.152
1362.6661376953125 0 3262.04
1363.6622314453125 0 2505.8535
1364.66015625 0 1089.5159
1367.656982421875 0 2220.3994
1371.7081298828125 0 910.3639
1372.719970703125 0 4188.6406
1373.70947265625 0 13703.272
1374.7122802734375 0 11949.204
1375.7127685546875 0 4565.1147
1376.7149658203125 0 1796.707
1382.6529541015625 0 2742.3499
1383.6763916015625 0 9732.443
1384.67822265625 0 7620.7847
1385.679443359375 0 5087.384
1386.674560546875 0 1836.356
1387.6507568359375 0 851.06256
1390.728759765625 0 4619.949
1391.73388671875 0 3116.967
1392.741455078125 0 1090.1268
1398.645263671875 0 2632.4263
1399.6507568359375 0 3177.9224
1400.677001953125 0 8120.363
1401.6966552734375 0 25890.6
1402.701904296875 0 20315.521
1403.69775390625 0 8097.4106
1404.7008056640625 0 1557.9657
1415.654052734375 0 2749.3
1416.663818359375 0 5086.5225
1417.7154541015625 0 43687.188
1418.7266845703125 0 164270.89
1419.728515625 0 110811.375
1420.731201171875 0 46313.45
1421.7344970703125 0 6716.683
1450.723876953125 0 949.31146
1475.4407958984375 0 744.45123
2048.08203125 0 1149.5576
2051.087646484375 0 1056.8584
2065.101806640625 0 1200.5698
2066.10107421875 0 930.20575
2068.070068359375 0 830.2505
2096.07763671875 0 777.2822
2097.10205078125 0 872.4599
2106.10400390625 0 1115.791
2107.094482421875 0 1571.4991
2108.093017578125 0 2002.1976
2109.053955078125 0 1235.4446
2112.08154296875 0 854.03937
2114.05859375 0 806.434
2123.087890625 0 1021.67255
2124.103759765625 0 2473.9607
2125.1015625 0 2186.8718
2126.099365234375 0 880.8001
2127.064453125 0 1087.6586
2129.09375 0 942.3049
2130.084716796875 0 1052.7491
2131.0712890625 0 1114.6885
2132.076171875 0 987.7264

Spectrum Details

|  |  |
| --- | --- |
| Matched peaks? Matched peaks The total absolute number of peaks matched. Additionally in brackets the total fraction of peaks matched and the total number of peaks is shown. | 37 (10.34% of 358) |
| FDR? FDR The false discovery rate estimated for this peptide. It is calculated by matching all theoretical fragments with a non-integer shift with the raw peaks for this spectrum. This is done with 40 different shifts. The resulting percentage is the average number of annotated peaks over the number of annotated peaks with the correct spectrum. | 4.76% |
| Satellite FDR? Satellite FDR See the FDR for details on its calculation. This satellite ion specific FDR only contains the satellite ions (d/w) for I/L/J positions. | ∞ |
| PSM Score? PSM Score The PSM Score as given by Hecklib to this annotated spectrum. It is shown with three significant figures. | 440 |

#### Spectrum 10080? Spectrum 10080 The raw spectrum of this peptide as annotated by Hecklib. The fragments are coloured according to ion type (see legend). Any peaks with a star '\*' as text can be hovered over to see the full details, first the ion type second the mass shift type. By hovering over the amino acids in the peptide or ions in the legend the corresponding peaks are highlighted. By toggling the 'Unassigned' label you can turn the background (unassigned) peaks on or off in the plot. By updating the slider in the Ion legend you can update the spectrum to only show the top X% of the peaks with labels. The top X% means any peak that is within X% of the highest intensity. By dragging in the spectrum you can zoom in to a specific part of the spectrum and use 'Zoom Out' to get back to the original zoom level. The annotation of the spectrum is based on the given sequence in the peptides file and is done with different software so inconsistencies are likely. The peaks are annotated based on the given sequence, with 20 ppm tolerance.

Copy Data

##### Spectrum 10080 (TSV)

###### Preview

```
Loading example...
```

*Click on the button to copy the data to your clipboard.*

Mz MinMz MaxIntensity Max

WidthHeightPeptide font sizePeptide stroke widthSpectrum font sizeSpectrum stroke widthCompact peptide

Ion legend

wxyz

abcd

OtherUnassignedIon^Charge^_Position_Show for top:%

VTJFPPSSEEJQA

02.72e+45.43e+48.15e+41.09e+5

Zoom Out

y^+1^_2_y^+1^_2_y^+1^_3_y^+1^_3_y^+1^_4_w^+1^_6_c^+1^_6_y^+1^_6_y^+1^_7_c^+1^_7_y^+1^_7_c^+1^_8_y^+1^_8_y^+1^_9_y^+1^_9_z^+1^_9_y^+1^_9_c^+1^_9_z^+1^_10_c^+1^_10_z^+1^_11_z^+1^_11_c^+1^_11_z^+1^_12_z^+1^_12_z^+1^_12_y^+1^_12_c^+1^_12_c^+1^_12_

0538107716152153

Fragment Matches Table

Show background peaks

| Position | Ion type | Intensity | mz Theoretical | mz Error (Th) | mz Error (ppm) | Charge | Series Number |
| --- | --- | --- | --- | --- | --- | --- | --- |
| - | - | 1097 | 120.1 | - | - | 0 | - |
| - | - | 452.2 | 124.2 | - | - | 0 | - |
| - | - | 375.2 | 130.2 | - | - | 0 | - |
| - | - | 954.8 | 133.1 | - | - | 0 | - |
| - | - | 384.2 | 134.7 | - | - | 0 | - |
| - | - | 374 | 135.9 | - | - | 0 | - |
| - | - | 465.8 | 142.2 | - | - | 0 | - |
| - | - | 410.8 | 150.9 | - | - | 0 | - |
| - | - | 482.2 | 154.7 | - | - | 0 | - |
| - | - | 603.1 | 173.1 | - | - | 0 | - |
| - | - | 1648 | 183.1 | - | - | 0 | - |
| - | - | 449.5 | 187 | - | - | 0 | - |
| - | - | 492.1 | 191 | - | - | 0 | - |
| - | - | 5240 | 200.1 | - | - | 0 | - |
| 12 | y | 4336 | 201.1 | 0.0001443 | 0.7178 | +1 | 2 |
| - | - | 2615 | 201.1 | - | - | 0 | - |
| - | - | 1596 | 213.1 | - | - | 0 | - |
| - | - | 1.237E+04 | 215.1 | - | - | 0 | - |
| - | - | 1116 | 216.1 | - | - | 0 | - |
| 12 | y | 1.173E+04 | 218.1 | 8.45E-05 | 0.3874 | +1 | 2 |
| - | - | 1029 | 227.1 | - | - | 0 | - |
| - | - | 1562 | 233.2 | - | - | 0 | - |
| - | - | 3171 | 242.2 | - | - | 0 | - |
| - | - | 599.3 | 243.1 | - | - | 0 | - |
| - | - | 697.9 | 261.2 | - | - | 0 | - |
| - | - | 611.3 | 267.1 | - | - | 0 | - |
| - | - | 621.9 | 280.2 | - | - | 0 | - |
| - | - | 557 | 282.1 | - | - | 0 | - |
| - | - | 630.9 | 283.2 | - | - | 0 | - |
| - | - | 2.038E+04 | 296.2 | - | - | 0 | - |
| - | - | 3320 | 297.2 | - | - | 0 | - |
| - | - | 623.6 | 303.2 | - | - | 0 | - |
| 11 | y | 1313 | 314.2 | 0.002293 | 7.297 | +1 | 3 |
| - | - | 8457 | 314.2 | - | - | 0 | - |
| - | - | 1433 | 315.2 | - | - | 0 | - |
| - | - | 622.9 | 330.2 | - | - | 0 | - |
| 11 | y | 1814 | 331.2 | 0.0002185 | 0.6598 | +1 | 3 |
| - | - | 629.2 | 346.1 | - | - | 0 | - |
| - | - | 1287 | 348.2 | - | - | 0 | - |
| - | - | 792.1 | 353.2 | - | - | 0 | - |
| - | - | 2602 | 362.2 | - | - | 0 | - |
| - | - | 870.7 | 371.2 | - | - | 0 | - |
| - | - | 552.6 | 395.2 | - | - | 0 | - |
| - | - | 678.4 | 398.2 | - | - | 0 | - |
| - | - | 752.1 | 411.3 | - | - | 0 | - |
| - | - | 997 | 416.3 | - | - | 0 | - |
| - | - | 1247 | 427.3 | - | - | 0 | - |
| - | - | 1.077E+04 | 433.3 | - | - | 0 | - |
| - | - | 2456 | 434.3 | - | - | 0 | - |
| - | - | 751.4 | 441.2 | - | - | 0 | - |
| 10 | y | 696.5 | 442.2 | 0.006734 | 15.23 | +1 | 4 |
| - | - | 1365 | 443.3 | - | - | 0 | - |
| - | - | 1.3E+04 | 461.3 | - | - | 0 | - |
| - | - | 2905 | 462.3 | - | - | 0 | - |
| - | - | 700.3 | 480.2 | - | - | 0 | - |
| - | - | 1497 | 498.2 | - | - | 0 | - |
| - | - | 637.1 | 540.3 | - | - | 0 | - |
| - | - | 1356 | 544.3 | - | - | 0 | - |
| - | - | 1871 | 544.8 | - | - | 0 | - |
| - | - | 2439 | 558.3 | - | - | 0 | - |
| - | - | 1245 | 559.3 | - | - | 0 | - |
| - | - | 618.3 | 562.2 | - | - | 0 | - |
| - | - | 867.7 | 587.3 | - | - | 0 | - |
| - | - | 731.5 | 600.8 | - | - | 0 | - |
| - | - | 575.2 | 625.5 | - | - | 0 | - |
| - | - | 7646 | 627.3 | - | - | 0 | - |
| - | - | 2123 | 628.3 | - | - | 0 | - |
| - | - | 1456 | 628.4 | - | - | 0 | - |
| - | - | 862.6 | 629.4 | - | - | 0 | - |
| 8 | w | 646.3 | 643.3 | 0.001965 | 3.055 | +1 | 6 |
| - | - | 1112 | 643.3 | - | - | 0 | - |
| - | - | 682.9 | 644.3 | - | - | 0 | - |
| - | - | 845.4 | 645.4 | - | - | 0 | - |
| - | - | 610.7 | 649.4 | - | - | 0 | - |
| - | - | 803.6 | 655.4 | - | - | 0 | - |
| - | - | 850 | 657.4 | - | - | 0 | - |
| - | - | 634.9 | 658.3 | - | - | 0 | - |
| - | - | 2607 | 671.4 | - | - | 0 | - |
| 6 | c | 2371 | 672.4 | 0.00204 | 3.034 | +1 | 6 |
| - | - | 1604 | 673.4 | - | - | 0 | - |
| - | - | 614.2 | 674.3 | - | - | 0 | - |
| 8 | y | 727.8 | 676.3 | 0.006095 | 9.012 | +1 | 6 |
| - | - | 772.9 | 708.4 | - | - | 0 | - |
| - | - | 1050 | 710.3 | - | - | 0 | - |
| - | - | 843.1 | 714.4 | - | - | 0 | - |
| - | - | 1022 | 715.4 | - | - | 0 | - |
| - | - | 3812 | 716.4 | - | - | 0 | - |
| - | - | 1582 | 717.4 | - | - | 0 | - |
| - | - | 848.4 | 722.3 | - | - | 0 | - |
| - | - | 5919 | 740.3 | - | - | 0 | - |
| - | - | 2783 | 741.4 | - | - | 0 | - |
| - | - | 2208 | 742.4 | - | - | 0 | - |
| - | - | 1192 | 744.4 | - | - | 0 | - |
| - | - | 706.7 | 745.3 | - | - | 0 | - |
| 7 | y | 857.7 | 746.3 | 0.005776 | 7.74 | +1 | 7 |
| - | - | 2386 | 757.4 | - | - | 0 | - |
| - | - | 832.7 | 758.4 | - | - | 0 | - |
| - | - | 9883 | 758.4 | - | - | 0 | - |
| 7 | c | 1.314E+04 | 759.4 | 0.00117 | 1.541 | +1 | 7 |
| - | - | 5394 | 760.4 | - | - | 0 | - |
| - | - | 1091 | 761.4 | - | - | 0 | - |
| 7 | y | 2003 | 763.3 | 9.871E-05 | 0.1293 | +1 | 7 |
| - | - | 1070 | 771.4 | - | - | 0 | - |
| - | - | 717.8 | 772.4 | - | - | 0 | - |
| - | - | 1017 | 774.3 | - | - | 0 | - |
| - | - | 1252 | 803.5 | - | - | 0 | - |
| - | - | 1197 | 829.4 | - | - | 0 | - |
| - | - | 804.8 | 830.5 | - | - | 0 | - |
| - | - | 1174 | 839.4 | - | - | 0 | - |
| - | - | 799.3 | 840.4 | - | - | 0 | - |
| - | - | 7626 | 845.5 | - | - | 0 | - |
| 8 | c | 3.373E+04 | 846.5 | 0.0001266 | 0.1495 | +1 | 8 |
| - | - | 1.557E+04 | 847.5 | - | - | 0 | - |
| - | - | 4052 | 848.5 | - | - | 0 | - |
| - | - | 780 | 850.4 | - | - | 0 | - |
| 6 | y | 5877 | 860.4 | 0.00213 | 2.476 | +1 | 8 |
| - | - | 3293 | 861.4 | - | - | 0 | - |
| - | - | 650 | 867.4 | - | - | 0 | - |
| - | - | 1.124E+04 | 868.4 | - | - | 0 | - |
| - | - | 4941 | 869.4 | - | - | 0 | - |
| - | - | 1578 | 870.4 | - | - | 0 | - |
| - | - | 1670 | 887.4 | - | - | 0 | - |
| - | - | 828.6 | 888.4 | - | - | 0 | - |
| - | - | 1488 | 931.5 | - | - | 0 | - |
| - | - | 1218 | 932.5 | - | - | 0 | - |
| - | - | 1994 | 938.4 | - | - | 0 | - |
| 5 | y | 2585 | 939.4 | 0.0002131 | 0.2268 | +1 | 9 |
| 5 | y | 3010 | 940.4 | 0.01241 | 13.2 | +1 | 9 |
| 5 | z | 2477 | 941.4 | 0.002063 | 2.192 | +1 | 9 |
| - | - | 770.9 | 951.5 | - | - | 0 | - |
| - | - | 1.235E+04 | 956.4 | - | - | 0 | - |
| 5 | y | 6.313E+04 | 957.5 | 0.0003306 | 0.3453 | +1 | 9 |
| - | - | 2.893E+04 | 958.5 | - | - | 0 | - |
| - | - | 7634 | 959.5 | - | - | 0 | - |
| - | - | 948.7 | 960.5 | - | - | 0 | - |
| - | - | 4696 | 974.5 | - | - | 0 | - |
| 9 | c | 3.997E+04 | 975.5 | 0.001052 | 1.078 | +1 | 9 |
| - | - | 2.256E+04 | 976.5 | - | - | 0 | - |
| - | - | 5829 | 977.5 | - | - | 0 | - |
| - | - | 697.3 | 978.5 | - | - | 0 | - |
| - | - | 899.3 | 985.4 | - | - | 0 | - |
| - | - | 1158 | 1001 | - | - | 0 | - |
| - | - | 978.6 | 1002 | - | - | 0 | - |
| - | - | 693.6 | 1005 | - | - | 0 | - |
| - | - | 897.1 | 1009 | - | - | 0 | - |
| - | - | 1156 | 1010 | - | - | 0 | - |
| - | - | 829.7 | 1010 | - | - | 0 | - |
| - | - | 673.8 | 1019 | - | - | 0 | - |
| - | - | 778 | 1025 | - | - | 0 | - |
| - | - | 720.6 | 1026 | - | - | 0 | - |
| - | - | 1027 | 1032 | - | - | 0 | - |
| - | - | 1189 | 1033 | - | - | 0 | - |
| - | - | 811.9 | 1033 | - | - | 0 | - |
| - | - | 1864 | 1034 | - | - | 0 | - |
| - | - | 908.5 | 1040 | - | - | 0 | - |
| - | - | 786 | 1042 | - | - | 0 | - |
| - | - | 781.6 | 1047 | - | - | 0 | - |
| - | - | 979.1 | 1048 | - | - | 0 | - |
| - | - | 1016 | 1048 | - | - | 0 | - |
| - | - | 1619 | 1049 | - | - | 0 | - |
| - | - | 1423 | 1054 | - | - | 0 | - |
| - | - | 986 | 1054 | - | - | 0 | - |
| - | - | 1471 | 1055 | - | - | 0 | - |
| - | - | 1011 | 1056 | - | - | 0 | - |
| - | - | 1433 | 1056 | - | - | 0 | - |
| - | - | 1188 | 1057 | - | - | 0 | - |
| - | - | 2551 | 1057 | - | - | 0 | - |
| - | - | 1926 | 1058 | - | - | 0 | - |
| - | - | 1384 | 1061 | - | - | 0 | - |
| - | - | 1730 | 1062 | - | - | 0 | - |
| - | - | 1062 | 1062 | - | - | 0 | - |
| - | - | 1567 | 1063 | - | - | 0 | - |
| - | - | 1246 | 1063 | - | - | 0 | - |
| - | - | 1027 | 1064 | - | - | 0 | - |
| - | - | 1037 | 1065 | - | - | 0 | - |
| - | - | 1627 | 1065 | - | - | 0 | - |
| - | - | 2398 | 1066 | - | - | 0 | - |
| - | - | 755 | 1085 | - | - | 0 | - |
| - | - | 1398 | 1086 | - | - | 0 | - |
| - | - | 1.034E+04 | 1088 | - | - | 0 | - |
| 4 | z | 1.612E+04 | 1089 | 0.01186 | 10.89 | +1 | 10 |
| - | - | 1.066E+04 | 1090 | - | - | 0 | - |
| - | - | 2802 | 1091 | - | - | 0 | - |
| - | - | 1149 | 1092 | - | - | 0 | - |
| - | - | 5457 | 1104 | - | - | 0 | - |
| 10 | c | 4.797E+04 | 1105 | 0.008949 | 8.102 | +1 | 10 |
| - | - | 2.943E+04 | 1106 | - | - | 0 | - |
| - | - | 1.004E+04 | 1107 | - | - | 0 | - |
| - | - | 1449 | 1108 | - | - | 0 | - |
| - | - | 790.9 | 1156 | - | - | 0 | - |
| - | - | 786 | 1157 | - | - | 0 | - |
| - | - | 3292 | 1174 | - | - | 0 | - |
| - | - | 2090 | 1175 | - | - | 0 | - |
| - | - | 899.8 | 1176 | - | - | 0 | - |
| - | - | 1541 | 1183 | - | - | 0 | - |
| 3 | z | 1533 | 1184 | 0.01258 | 10.63 | +1 | 11 |
| - | - | 1068 | 1185 | - | - | 0 | - |
| - | - | 673.6 | 1186 | - | - | 0 | - |
| - | - | 4913 | 1201 | - | - | 0 | - |
| 3 | z | 1.244E+04 | 1202 | 0.007751 | 6.451 | +1 | 11 |
| - | - | 8460 | 1203 | - | - | 0 | - |
| - | - | 2980 | 1204 | - | - | 0 | - |
| - | - | 794.8 | 1205 | - | - | 0 | - |
| - | - | 1029 | 1214 | - | - | 0 | - |
| - | - | 720.6 | 1215 | - | - | 0 | - |
| - | - | 640.7 | 1216 | - | - | 0 | - |
| - | - | 694.7 | 1217 | - | - | 0 | - |
| 11 | c | 2.96E+04 | 1218 | 0.00207 | 1.7 | +1 | 11 |
| - | - | 1.962E+04 | 1219 | - | - | 0 | - |
| - | - | 8147 | 1220 | - | - | 0 | - |
| - | - | 756.1 | 1229 | - | - | 0 | - |
| - | - | 3128 | 1244 | - | - | 0 | - |
| - | - | 1068 | 1245 | - | - | 0 | - |
| - | - | 1067 | 1246 | - | - | 0 | - |
| - | - | 1954 | 1247 | - | - | 0 | - |
| - | - | 1219 | 1248 | - | - | 0 | - |
| - | - | 1109 | 1261 | - | - | 0 | - |
| - | - | 1171 | 1262 | - | - | 0 | - |
| - | - | 789.1 | 1268 | - | - | 0 | - |
| - | - | 821.6 | 1269 | - | - | 0 | - |
| - | - | 679 | 1276 | - | - | 0 | - |
| - | - | 1143 | 1278 | - | - | 0 | - |
| - | - | 1103 | 1283 | - | - | 0 | - |
| 2 | z | 1.263E+04 | 1285 | 0.001399 | 1.089 | +1 | 12 |
| 2 | z | 8491 | 1286 | 0.0208 | 16.18 | +1 | 12 |
| - | - | 4060 | 1287 | - | - | 0 | - |
| - | - | 920.8 | 1292 | - | - | 0 | - |
| - | - | 3650 | 1293 | - | - | 0 | - |
| - | - | 2742 | 1294 | - | - | 0 | - |
| - | - | 1447 | 1295 | - | - | 0 | - |
| - | - | 671.3 | 1296 | - | - | 0 | - |
| - | - | 2783 | 1301 | - | - | 0 | - |
| - | - | 3818 | 1302 | - | - | 0 | - |
| 2 | z | 1.6E+04 | 1303 | 0.002675 | 2.054 | +1 | 12 |
| - | - | 9852 | 1304 | - | - | 0 | - |
| - | - | 4679 | 1305 | - | - | 0 | - |
| - | - | 935.8 | 1306 | - | - | 0 | - |
| - | - | 1140 | 1309 | - | - | 0 | - |
| - | - | 2277 | 1310 | - | - | 0 | - |
| - | - | 2438 | 1311 | - | - | 0 | - |
| - | - | 949.2 | 1312 | - | - | 0 | - |
| - | - | 1488 | 1313 | - | - | 0 | - |
| - | - | 1166 | 1314 | - | - | 0 | - |
| - | - | 1181 | 1315 | - | - | 0 | - |
| - | - | 767 | 1316 | - | - | 0 | - |
| - | - | 680.7 | 1317 | - | - | 0 | - |
| - | - | 1709 | 1318 | - | - | 0 | - |
| 2 | y | 2050 | 1319 | 0.0141 | 10.69 | +1 | 12 |
| - | - | 658.3 | 1320 | - | - | 0 | - |
| - | - | 1009 | 1327 | - | - | 0 | - |
| 12 | c | 934.7 | 1329 | 0.02018 | 15.19 | +1 | 12 |
| - | - | 915.8 | 1330 | - | - | 0 | - |
| - | - | 1641 | 1331 | - | - | 0 | - |
| - | - | 1110 | 1332 | - | - | 0 | - |
| - | - | 824.3 | 1340 | - | - | 0 | - |
| - | - | 1127 | 1341 | - | - | 0 | - |
| - | - | 716.3 | 1343 | - | - | 0 | - |
| - | - | 1459 | 1344 | - | - | 0 | - |
| 12 | c | 8.865E+04 | 1346 | 0.0007114 | 0.5286 | +1 | 12 |
| - | - | 7.373E+04 | 1347 | - | - | 0 | - |
| - | - | 3.24E+04 | 1348 | - | - | 0 | - |
| - | - | 6714 | 1349 | - | - | 0 | - |
| - | - | 2128 | 1356 | - | - | 0 | - |
| - | - | 1580 | 1357 | - | - | 0 | - |
| - | - | 3342 | 1358 | - | - | 0 | - |
| - | - | 2873 | 1359 | - | - | 0 | - |
| - | - | 2571 | 1360 | - | - | 0 | - |
| - | - | 2342 | 1361 | - | - | 0 | - |
| - | - | 725.6 | 1362 | - | - | 0 | - |
| - | - | 2276 | 1363 | - | - | 0 | - |
| - | - | 1446 | 1364 | - | - | 0 | - |
| - | - | 795.6 | 1365 | - | - | 0 | - |
| - | - | 1182 | 1369 | - | - | 0 | - |
| - | - | 873.3 | 1372 | - | - | 0 | - |
| - | - | 2505 | 1373 | - | - | 0 | - |
| - | - | 1.067E+04 | 1374 | - | - | 0 | - |
| - | - | 8102 | 1375 | - | - | 0 | - |
| - | - | 3475 | 1376 | - | - | 0 | - |
| - | - | 1880 | 1383 | - | - | 0 | - |
| - | - | 9862 | 1384 | - | - | 0 | - |
| - | - | 8397 | 1385 | - | - | 0 | - |
| - | - | 4510 | 1386 | - | - | 0 | - |
| - | - | 2100 | 1387 | - | - | 0 | - |
| - | - | 945.6 | 1388 | - | - | 0 | - |
| - | - | 3277 | 1391 | - | - | 0 | - |
| - | - | 2214 | 1392 | - | - | 0 | - |
| - | - | 931.8 | 1393 | - | - | 0 | - |
| - | - | 3852 | 1399 | - | - | 0 | - |
| - | - | 6427 | 1400 | - | - | 0 | - |
| - | - | 8288 | 1401 | - | - | 0 | - |
| - | - | 1.846E+04 | 1402 | - | - | 0 | - |
| - | - | 1.527E+04 | 1403 | - | - | 0 | - |
| - | - | 5603 | 1404 | - | - | 0 | - |
| - | - | 1029 | 1405 | - | - | 0 | - |
| - | - | 5246 | 1416 | - | - | 0 | - |
| - | - | 1.27E+04 | 1417 | - | - | 0 | - |
| - | - | 3.243E+04 | 1418 | - | - | 0 | - |
| - | - | 1.076E+05 | 1419 | - | - | 0 | - |
| - | - | 7.895E+04 | 1420 | - | - | 0 | - |
| - | - | 3.117E+04 | 1421 | - | - | 0 | - |
| - | - | 5634 | 1422 | - | - | 0 | - |
| - | - | 1079 | 1991 | - | - | 0 | - |
| - | - | 728.8 | 1992 | - | - | 0 | - |
| - | - | 799.7 | 2017 | - | - | 0 | - |
| - | - | 940 | 2018 | - | - | 0 | - |
| - | - | 873.5 | 2022 | - | - | 0 | - |
| - | - | 820.4 | 2034 | - | - | 0 | - |
| - | - | 1055 | 2048 | - | - | 0 | - |
| - | - | 1291 | 2049 | - | - | 0 | - |
| - | - | 1360 | 2050 | - | - | 0 | - |
| - | - | 1014 | 2063 | - | - | 0 | - |
| - | - | 1379 | 2064 | - | - | 0 | - |
| - | - | 1377 | 2065 | - | - | 0 | - |
| - | - | 1333 | 2066 | - | - | 0 | - |
| - | - | 979.3 | 2067 | - | - | 0 | - |
| - | - | 1792 | 2079 | - | - | 0 | - |
| - | - | 1415 | 2081 | - | - | 0 | - |
| - | - | 1056 | 2082 | - | - | 0 | - |
| - | - | 1212 | 2093 | - | - | 0 | - |
| - | - | 936.4 | 2094 | - | - | 0 | - |
| - | - | 1797 | 2095 | - | - | 0 | - |
| - | - | 2185 | 2096 | - | - | 0 | - |
| - | - | 2364 | 2097 | - | - | 0 | - |
| - | - | 2513 | 2106 | - | - | 0 | - |
| - | - | 2433 | 2107 | - | - | 0 | - |
| - | - | 2379 | 2108 | - | - | 0 | - |
| - | - | 2026 | 2109 | - | - | 0 | - |
| - | - | 1160 | 2110 | - | - | 0 | - |
| - | - | 2726 | 2111 | - | - | 0 | - |
| - | - | 3314 | 2112 | - | - | 0 | - |
| - | - | 3474 | 2113 | - | - | 0 | - |
| - | - | 3553 | 2114 | - | - | 0 | - |
| - | - | 1069 | 2115 | - | - | 0 | - |
| - | - | 1260 | 2123 | - | - | 0 | - |
| - | - | 3214 | 2124 | - | - | 0 | - |
| - | - | 2691 | 2125 | - | - | 0 | - |
| - | - | 1855 | 2126 | - | - | 0 | - |
| - | - | 2868 | 2127 | - | - | 0 | - |
| - | - | 2084 | 2128 | - | - | 0 | - |
| - | - | 2513 | 2129 | - | - | 0 | - |
| - | - | 2239 | 2130 | - | - | 0 | - |
| - | - | 2119 | 2131 | - | - | 0 | - |
| - | - | 1257 | 2132 | - | - | 0 | - |

m/z Charge Intensity FragmentType MassShift Position
120.08082580566406 0 1097.2898
124.2164306640625 0 452.2316
130.1639404296875 0 375.20227
133.08609008789062 0 954.8115
134.72589111328125 0 384.19888
135.9489288330078 0 374.04532
142.24591064453125 0 465.7664
150.8905792236328 0 410.79007
154.65621948242188 0 482.2319
173.1285858154297 0 603.10785
183.11306762695312 0 1647.9255
186.99639892578125 0 449.4981
191.04592895507812 0 492.0626
200.10299682617188 0 5239.528
201.08712768554688 0 4335.632 y Ammonia loss 11
201.12344360351562 0 2615.489
213.12374877929688 0 1596.4723
215.13905334472656 0 12368.42
216.143310546875 0 1116.2626
218.11361694335938 0 11730.924 y 11
227.10317993164062 0 1029.1945
233.16477966308594 0 1562.3367
242.15016174316406 0 3171.0444
243.13328552246094 0 599.25104
261.16021728515625 0 697.9014
267.14996337890625 0 611.33936
280.1658630371094 0 621.9168
282.14532470703125 0 556.9737
283.1733703613281 0 630.8614
296.1971130371094 0 20381.08
297.2002868652344 0 3320.2476
303.1698913574219 0 623.5947
314.17333984375 0 1312.5669 y Ammonia loss 10
314.20770263671875 0 8457.242
315.2110595703125 0 1433.3909
330.1817626953125 0 622.9046
331.19781494140625 0 1813.6385 y 10
346.12396240234375 0 629.2092
348.1925048828125 0 1286.5051
353.1820068359375 0 792.1313
362.20806884765625 0 2601.9214
371.1935119628906 0 870.71783
395.2334899902344 0 552.5801
398.2028503417969 0 678.44604
411.26165771484375 0 752.13007
416.2541198730469 0 996.96796
427.30377197265625 0 1246.898
433.2817077636719 0 10771.622
434.2844543457031 0 2456.462
441.234375 0 751.4279
442.2363586425781 0 696.5232 y Water loss 9
443.2665710449219 0 1364.6299
461.2763366699219 0 12996.708
462.27984619140625 0 2905.0706
480.208984375 0 700.25836
498.2200012207031 0 1496.9547
540.3199462890625 0 637.1353
544.2681274414062 0 1355.5309
544.7708129882812 0 1870.735
558.3287963867188 0 2438.7952
559.3320922851562 0 1244.769
562.1858520507812 0 618.3074
587.263916015625 0 867.7036
600.814697265625 0 731.5223
625.4769287109375 0 575.2462
627.2626953125 0 7646.409
628.2657470703125 0 2123.4468
628.3958129882812 0 1456.2244
629.40234375 0 862.64166
643.2913818359375 0 646.2732 w 7
643.3463134765625 0 1111.673
644.2911376953125 0 682.9206
645.3563842773438 0 845.4275
649.4326171875 0 610.6805
655.3821411132812 0 803.56866
657.3973999023438 0 850.04395
658.2789916992188 0 634.8624
671.4006958007812 0 2606.8726
672.4058837890625 0 2371.4316 c 5
673.4118041992188 0 1604.1641
674.2987670898438 0 614.1829
676.3087158203125 0 727.84045 y 7
708.4052734375 0 772.8592
710.3416748046875 0 1050.4667
714.4227905273438 0 843.09
715.4266357421875 0 1022.4908
716.4349365234375 0 3812.2144
717.4363403320312 0 1581.5829
722.3355102539062 0 848.40216
740.3465576171875 0 5918.7715
741.350341796875 0 2782.5417
742.416015625 0 2208.43
744.4237060546875 0 1192.4766
745.3129272460938 0 706.71826
746.3145141601562 0 857.7216 y Ammonia loss 6
757.3734130859375 0 2386.2295
758.36572265625 0 832.7155
758.432373046875 0 9882.782
759.4387817382812 0 13137.647 c 6
760.4428100585938 0 5393.785
761.4432373046875 0 1091.4757
763.3467407226562 0 2002.7518 y 6
771.3541259765625 0 1069.5377
772.3521118164062 0 717.81775
774.3291625976562 0 1016.5322
803.4690551757812 0 1252.4363
829.4486694335938 0 1196.7627
830.4537963867188 0 804.784
839.4137573242188 0 1173.77
840.4226684570312 0 799.3098
845.46533203125 0 7626.026
846.4721069335938 0 33728.656 c 7
847.4756469726562 0 15574.684
848.4774169921875 0 4052.1353
850.3931884765625 0 779.9572
860.4017333984375 0 5876.6895 y 5
861.404052734375 0 3293.2986
867.4064331054688 0 650.02014
868.4059448242188 0 11236.661
869.4078979492188 0 4940.525
870.41259765625 0 1578.4932
887.4165649414062 0 1669.6725
888.4227905273438 0 828.56146
931.501708984375 0 1487.8243
932.5009765625 0 1218.251
938.43701171875 0 1993.6007
939.4415893554688 0 2585.168 y Water loss 4
940.438232421875 0 3009.7246 y Ammonia loss 4
941.4315795898438 0 2477.4922 z 4
951.45263671875 0 770.90485
956.4452514648438 0 12346.432
957.4526977539062 0 63126.98 y 4
958.4573974609375 0 28926.338
959.460205078125 0 7634.309
960.4622802734375 0 948.6961
974.5086059570312 0 4696.296
975.515625 0 39971.598 c 8
976.518798828125 0 22559.812
977.5214233398438 0 5829.249
978.5391845703125 0 697.2815
985.44384765625 0 899.3304
1000.5399169921875 0 1158.0355
1001.5370483398438 0 978.5717
1004.5152587890625 0 693.56854
1009.052001953125 0 897.111
1009.5547485351562 0 1156.4109
1010.4574584960938 0 829.6893
1019.0234375 0 673.76495
1024.5462646484375 0 778.0147
1026.05224609375 0 720.6227
1032.0458984375 0 1026.8474
1032.5445556640625 0 1189.2874
1033.0438232421875 0 811.8978
1033.5499267578125 0 1863.9551
1040.0430908203125 0 908.49414
1042.044189453125 0 786.0376
1047.0308837890625 0 781.6177
1047.5303955078125 0 979.1157
1048.0350341796875 0 1016.38354
1048.5352783203125 0 1619.2109
1053.5487060546875 0 1423.2793
1054.052978515625 0 985.9851
1054.5426025390625 0 1470.7009
1055.5361328125 0 1011.0755
1056.0350341796875 0 1432.832
1056.5458984375 0 1188.1588
1057.042236328125 0 2550.7666
1057.5408935546875 0 1926.4578
1060.5394287109375 0 1384.304
1061.548095703125 0 1729.6206
1062.0618896484375 0 1061.5623
1062.5567626953125 0 1566.7205
1063.0350341796875 0 1245.6077
1064.0396728515625 0 1026.9523
1064.54541015625 0 1037.4823
1065.04345703125 0 1626.5532
1065.5447998046875 0 2398.1626
1084.529052734375 0 754.996
1085.5133056640625 0 1398.3567
1087.5306396484375 0 10340.962
1088.513916015625 0 16116.96 z 3
1089.5103759765625 0 10658.084
1090.5093994140625 0 2802.191
1091.515625 0 1148.5521
1103.521240234375 0 5456.6377
1104.5482177734375 0 47972.99 c 9
1105.5506591796875 0 29433.55
1106.5543212890625 0 10036.147
1107.5577392578125 0 1449.1956
1155.62158203125 0 790.89136
1156.6197509765625 0 786.0307
1173.6302490234375 0 3291.5254
1174.634033203125 0 2089.507
1175.6427001953125 0 899.7956
1182.60400390625 0 1541.0769
1183.588134765625 0 1532.5463 z Water loss 2
1184.60986328125 0 1068.4878
1185.6082763671875 0 673.6234
1200.6134033203125 0 4913.0205
1201.5938720703125 0 12441.791 z 2
1202.5965576171875 0 8459.551
1203.5946044921875 0 2979.9304
1204.5966796875 0 794.7543
1213.5849609375 0 1029.3955
1214.5859375 0 720.6198
1215.6024169921875 0 640.70404
1216.6231689453125 0 694.6798
1217.63916015625 0 29595.998 c 10
1218.6419677734375 0 19619.174
1219.6451416015625 0 8146.6426
1228.6435546875 0 756.065
1243.6597900390625 0 3128.0127
1244.6673583984375 0 1067.8788
1245.643798828125 0 1067.4446
1246.576171875 0 1953.784
1247.572998046875 0 1218.54
1260.645263671875 0 1109.4987
1261.650390625 0 1170.573
1267.630859375 0 789.1362
1268.6126708984375 0 821.62787
1275.6431884765625 0 679.0374
1277.632568359375 0 1142.8008
1282.598876953125 0 1103.4031
1284.6246337890625 0 12628.42 z Water loss 1
1285.6280517578125 0 8491.015 z Ammonia loss 1
1286.63134765625 0 4059.8345
1291.665283203125 0 920.7909
1292.646240234375 0 3649.839
1293.645263671875 0 2741.7454
1294.6514892578125 0 1447.0364
1295.6435546875 0 671.3212
1300.6136474609375 0 2783.1794
1301.6539306640625 0 3817.547
1302.636474609375 0 15996.7705 z 1
1303.6414794921875 0 9852.283
1304.6422119140625 0 4679.277
1305.657470703125 0 935.7921
1308.64306640625 0 1140.0823
1309.6650390625 0 2276.9973
1310.663330078125 0 2437.952
1311.656005859375 0 949.1887
1312.65234375 0 1488.4984
1313.6693115234375 0 1165.7467
1314.663818359375 0 1180.8889
1315.5726318359375 0 766.9794
1316.61328125 0 680.6682
1317.6466064453125 0 1709.1987
1318.638427734375 0 2049.7937 y 1
1319.6571044921875 0 658.34033
1326.635009765625 0 1008.73224
1328.653076171875 0 934.69934 c Ammonia loss 11
1329.669189453125 0 915.8133
1330.670166015625 0 1641.1898
1331.6729736328125 0 1109.9927
1339.6114501953125 0 824.2888
1340.6287841796875 0 1126.9701
1342.6429443359375 0 716.30695
1343.6656494140625 0 1459.2648
1345.6990966796875 0 88649.734 c 11
1346.702880859375 0 73729.5
1347.705078125 0 32402.473
1348.7030029296875 0 6714.164
1355.693603515625 0 2127.9805
1356.712890625 0 1579.6644
1357.6361083984375 0 3341.6763
1358.6365966796875 0 2872.787
1359.6396484375 0 2570.9604
1360.6507568359375 0 2341.6
1361.6640625 0 725.63916
1362.6668701171875 0 2275.9604
1363.6661376953125 0 1446.443
1364.657958984375 0 795.55066
1368.6575927734375 0 1182.2562
1371.68408203125 0 873.3336
1372.720703125 0 2504.6682
1373.7110595703125 0 10665.178
1374.7139892578125 0 8101.9453
1375.70703125 0 3474.8484
1382.6522216796875 0 1879.9783
1383.670654296875 0 9862.002
1384.670166015625 0 8396.948
1385.672119140625 0 4509.51
1386.6678466796875 0 2100.4968
1387.667724609375 0 945.6315
1390.7291259765625 0 3276.8708
1391.736328125 0 2213.563
1392.747314453125 0 931.7908
1398.6392822265625 0 3852.48
1399.64697265625 0 6427.048
1400.6767578125 0 8288.223
1401.69482421875 0 18462.887
1402.6983642578125 0 15266.531
1403.6995849609375 0 5603.1436
1404.7130126953125 0 1029.4746
1415.6446533203125 0 5246.278
1416.650634765625 0 12695.066
1417.7115478515625 0 32431.664
1418.72607421875 0 107568.88
1419.7286376953125 0 78945.8
1420.7315673828125 0 31170.46
1421.7332763671875 0 5633.518
1991.0523681640625 0 1078.8901
1992.0498046875 0 728.83746
2017.0689697265625 0 799.68567
2018.09716796875 0 940.0291
2022.0322265625 0 873.52167
2034.07666015625 0 820.3942
2048.070068359375 0 1055.1981
2049.063232421875 0 1290.955
2050.076904296875 0 1359.9258
2063.087890625 0 1013.808
2064.10498046875 0 1378.9039
2065.087646484375 0 1376.9745
2066.096435546875 0 1332.6816
2067.099609375 0 979.29144
2079.077392578125 0 1791.6825
2081.078125 0 1415.0214
2082.074462890625 0 1056.0377
2093.062744140625 0 1212.1727
2094.060791015625 0 936.44507
2095.06787109375 0 1796.7538
2096.071044921875 0 2185.3145
2097.075927734375 0 2363.703
2106.084716796875 0 2512.8506
2107.095703125 0 2432.5002
2108.086669921875 0 2378.7727
2109.067138671875 0 2026.4971
2110.10302734375 0 1160.122
2111.06591796875 0 2725.9312
2112.0634765625 0 3313.5642
2113.078125 0 3473.6104
2114.082763671875 0 3553.0232
2115.09521484375 0 1069.1489
2123.086181640625 0 1260.0355
2124.10693359375 0 3214.2964
2125.11083984375 0 2691.0696
2126.09423828125 0 1855.4796
2127.088623046875 0 2868.456
2128.0810546875 0 2083.8235
2129.08203125 0 2513.0464
2130.086669921875 0 2239.2954
2131.0810546875 0 2118.7622
2132.10888671875 0 1257.3246

Spectrum Details

|  |  |
| --- | --- |
| Matched peaks? Matched peaks The total absolute number of peaks matched. Additionally in brackets the total fraction of peaks matched and the total number of peaks is shown. | 29 (8.45% of 343) |
| FDR? FDR The false discovery rate estimated for this peptide. It is calculated by matching all theoretical fragments with a non-integer shift with the raw peaks for this spectrum. This is done with 40 different shifts. The resulting percentage is the average number of annotated peaks over the number of annotated peaks with the correct spectrum. | 4.76% |
| Satellite FDR? Satellite FDR See the FDR for details on its calculation. This satellite ion specific FDR only contains the satellite ions (d/w) for I/L/J positions. | ∞ |
| PSM Score? PSM Score The PSM Score as given by Hecklib to this annotated spectrum. It is shown with three significant figures. | 336 |

#### Spectrum 10143? Spectrum 10143 The raw spectrum of this peptide as annotated by Hecklib. The fragments are coloured according to ion type (see legend). Any peaks with a star '\*' as text can be hovered over to see the full details, first the ion type second the mass shift type. By hovering over the amino acids in the peptide or ions in the legend the corresponding peaks are highlighted. By toggling the 'Unassigned' label you can turn the background (unassigned) peaks on or off in the plot. By updating the slider in the Ion legend you can update the spectrum to only show the top X% of the peaks with labels. The top X% means any peak that is within X% of the highest intensity. By dragging in the spectrum you can zoom in to a specific part of the spectrum and use 'Zoom Out' to get back to the original zoom level. The annotation of the spectrum is based on the given sequence in the peptides file and is done with different software so inconsistencies are likely. The peaks are annotated based on the given sequence, with 20 ppm tolerance.

Copy Data

##### Spectrum 10143 (TSV)

###### Preview

```
Loading example...
```

*Click on the button to copy the data to your clipboard.*

Mz MinMz MaxIntensity Max

WidthHeightPeptide font sizePeptide stroke widthSpectrum font sizeSpectrum stroke widthCompact peptide

Ion legend

wxyz

abcd

OtherUnassignedIon^Charge^_Position_Show for top:%

VTJFPPSSEEJQA

01.95e+43.89e+45.84e+47.79e+4

Zoom Out

y^+1^_2_y^+1^_2_y^+1^_3_y^+1^_3_c^+1^_6_c^+1^_7_y^+1^_7_c^+1^_8_y^+1^_8_y^+1^_9_y^+1^_9_z^+1^_9_y^+1^_9_c^+1^_9_y^+1^_10_z^+1^_10_c^+1^_10_z^+1^_11_z^+1^_11_c^+1^_11_z^+1^_12_z^+1^_12_w^+1^_12_y^+1^_12_z^+1^_12_y^+1^_12_c^+1^_12_

0538107716152153

Fragment Matches Table

Show background peaks

| Position | Ion type | Intensity | mz Theoretical | mz Error (Th) | mz Error (ppm) | Charge | Series Number |
| --- | --- | --- | --- | --- | --- | --- | --- |
| - | - | 796.4 | 120.1 | - | - | 0 | - |
| - | - | 491 | 130.1 | - | - | 0 | - |
| - | - | 344.4 | 132.3 | - | - | 0 | - |
| - | - | 1133 | 133.1 | - | - | 0 | - |
| - | - | 404.5 | 137 | - | - | 0 | - |
| - | - | 447.6 | 140.4 | - | - | 0 | - |
| - | - | 463.8 | 148.8 | - | - | 0 | - |
| - | - | 585.6 | 148.9 | - | - | 0 | - |
| - | - | 662.4 | 148.9 | - | - | 0 | - |
| - | - | 975.2 | 148.9 | - | - | 0 | - |
| - | - | 1029 | 148.9 | - | - | 0 | - |
| - | - | 1350 | 148.9 | - | - | 0 | - |
| - | - | 2967 | 148.9 | - | - | 0 | - |
| - | - | 6271 | 149 | - | - | 0 | - |
| - | - | 3580 | 149 | - | - | 0 | - |
| - | - | 1462 | 149 | - | - | 0 | - |
| - | - | 1025 | 149 | - | - | 0 | - |
| - | - | 943.9 | 149 | - | - | 0 | - |
| - | - | 781.5 | 149 | - | - | 0 | - |
| - | - | 475.3 | 149 | - | - | 0 | - |
| - | - | 446.9 | 159.7 | - | - | 0 | - |
| - | - | 396.4 | 163.1 | - | - | 0 | - |
| - | - | 546.5 | 173.1 | - | - | 0 | - |
| - | - | 670 | 173.4 | - | - | 0 | - |
| - | - | 1257 | 183.1 | - | - | 0 | - |
| - | - | 420.6 | 198.9 | - | - | 0 | - |
| - | - | 3149 | 200.1 | - | - | 0 | - |
| 12 | y | 3648 | 201.1 | 0.0001138 | 0.5661 | +1 | 2 |
| - | - | 1933 | 201.1 | - | - | 0 | - |
| - | - | 449.2 | 203.2 | - | - | 0 | - |
| - | - | 505.6 | 210 | - | - | 0 | - |
| - | - | 587.2 | 213.1 | - | - | 0 | - |
| - | - | 8775 | 215.1 | - | - | 0 | - |
| - | - | 770.3 | 216.1 | - | - | 0 | - |
| 12 | y | 7173 | 218.1 | 0.0001913 | 0.8771 | +1 | 2 |
| - | - | 849.3 | 227.1 | - | - | 0 | - |
| - | - | 1054 | 233.2 | - | - | 0 | - |
| - | - | 522 | 233.4 | - | - | 0 | - |
| - | - | 2300 | 242.2 | - | - | 0 | - |
| - | - | 540.3 | 271.5 | - | - | 0 | - |
| - | - | 1166 | 282.1 | - | - | 0 | - |
| - | - | 590.5 | 293.5 | - | - | 0 | - |
| - | - | 1.73E+04 | 296.2 | - | - | 0 | - |
| - | - | 2108 | 297.2 | - | - | 0 | - |
| - | - | 564.2 | 299.2 | - | - | 0 | - |
| 11 | y | 989 | 314.2 | 0.001011 | 3.217 | +1 | 3 |
| - | - | 5341 | 314.2 | - | - | 0 | - |
| 11 | y | 1081 | 331.2 | 0.000188 | 0.5677 | +1 | 3 |
| - | - | 503.4 | 333.5 | - | - | 0 | - |
| - | - | 518.2 | 342.6 | - | - | 0 | - |
| - | - | 660.5 | 346.1 | - | - | 0 | - |
| - | - | 743.3 | 346.2 | - | - | 0 | - |
| - | - | 498 | 354.2 | - | - | 0 | - |
| - | - | 1305 | 362.2 | - | - | 0 | - |
| - | - | 528.6 | 371.2 | - | - | 0 | - |
| - | - | 556.3 | 382.2 | - | - | 0 | - |
| - | - | 1066 | 416.3 | - | - | 0 | - |
| - | - | 7209 | 433.3 | - | - | 0 | - |
| - | - | 1403 | 434.3 | - | - | 0 | - |
| - | - | 1326 | 441.2 | - | - | 0 | - |
| - | - | 1174 | 443.3 | - | - | 0 | - |
| - | - | 7573 | 461.3 | - | - | 0 | - |
| - | - | 2207 | 462.3 | - | - | 0 | - |
| - | - | 1119 | 498.2 | - | - | 0 | - |
| - | - | 614.7 | 528.8 | - | - | 0 | - |
| - | - | 574.5 | 530.3 | - | - | 0 | - |
| - | - | 1498 | 544.3 | - | - | 0 | - |
| - | - | 745.6 | 544.8 | - | - | 0 | - |
| - | - | 866.1 | 558.3 | - | - | 0 | - |
| - | - | 685 | 600.8 | - | - | 0 | - |
| - | - | 4959 | 627.3 | - | - | 0 | - |
| - | - | 2278 | 628.3 | - | - | 0 | - |
| - | - | 1054 | 657.4 | - | - | 0 | - |
| - | - | 660.3 | 658.3 | - | - | 0 | - |
| - | - | 1776 | 671.4 | - | - | 0 | - |
| 6 | c | 2208 | 672.4 | 0.0005237 | 0.7789 | +1 | 6 |
| - | - | 790.2 | 673.4 | - | - | 0 | - |
| - | - | 587.4 | 691.4 | - | - | 0 | - |
| - | - | 3005 | 716.4 | - | - | 0 | - |
| - | - | 4361 | 740.3 | - | - | 0 | - |
| - | - | 1378 | 741.3 | - | - | 0 | - |
| - | - | 1462 | 742.4 | - | - | 0 | - |
| - | - | 924.3 | 743.4 | - | - | 0 | - |
| - | - | 764.8 | 744.4 | - | - | 0 | - |
| - | - | 1169 | 745.3 | - | - | 0 | - |
| - | - | 1416 | 757.4 | - | - | 0 | - |
| - | - | 6743 | 758.4 | - | - | 0 | - |
| 7 | c | 1.168E+04 | 759.4 | 0.0009871 | 1.3 | +1 | 7 |
| - | - | 3229 | 760.4 | - | - | 0 | - |
| 7 | y | 1215 | 763.3 | 0.000526 | 0.689 | +1 | 7 |
| - | - | 1003 | 771.4 | - | - | 0 | - |
| - | - | 1030 | 803.5 | - | - | 0 | - |
| - | - | 999.9 | 829.4 | - | - | 0 | - |
| - | - | 651.8 | 833.3 | - | - | 0 | - |
| - | - | 4122 | 845.5 | - | - | 0 | - |
| 8 | c | 2.637E+04 | 846.5 | 0.0005538 | 0.6543 | +1 | 8 |
| - | - | 1.123E+04 | 847.5 | - | - | 0 | - |
| - | - | 3598 | 848.5 | - | - | 0 | - |
| 6 | y | 4159 | 860.4 | 0.001703 | 1.979 | +1 | 8 |
| - | - | 1863 | 861.4 | - | - | 0 | - |
| - | - | 8196 | 868.4 | - | - | 0 | - |
| - | - | 3434 | 869.4 | - | - | 0 | - |
| - | - | 793.3 | 872 | - | - | 0 | - |
| - | - | 1441 | 887.4 | - | - | 0 | - |
| - | - | 794.4 | 931.5 | - | - | 0 | - |
| - | - | 1988 | 938.4 | - | - | 0 | - |
| 5 | y | 1531 | 939.4 | 0.001983 | 2.111 | +1 | 9 |
| 5 | y | 3014 | 940.4 | 0.003747 | 3.985 | +1 | 9 |
| 5 | z | 957.9 | 941.4 | 0.005139 | 5.458 | +1 | 9 |
| - | - | 9714 | 956.4 | - | - | 0 | - |
| 5 | y | 4.425E+04 | 957.5 | 0.0004527 | 0.4728 | +1 | 9 |
| - | - | 2.106E+04 | 958.5 | - | - | 0 | - |
| - | - | 4920 | 959.5 | - | - | 0 | - |
| - | - | 650.6 | 960.5 | - | - | 0 | - |
| - | - | 2995 | 974.5 | - | - | 0 | - |
| 9 | c | 2.712E+04 | 975.5 | 0.0009905 | 1.015 | +1 | 9 |
| - | - | 1.673E+04 | 976.5 | - | - | 0 | - |
| - | - | 3886 | 977.5 | - | - | 0 | - |
| - | - | 856.8 | 1009 | - | - | 0 | - |
| - | - | 1423 | 1009 | - | - | 0 | - |
| - | - | 745.4 | 1015 | - | - | 0 | - |
| - | - | 843.5 | 1016 | - | - | 0 | - |
| - | - | 785 | 1020 | - | - | 0 | - |
| - | - | 727.5 | 1025 | - | - | 0 | - |
| - | - | 1141 | 1026 | - | - | 0 | - |
| - | - | 1070 | 1032 | - | - | 0 | - |
| - | - | 1790 | 1032 | - | - | 0 | - |
| - | - | 1197 | 1033 | - | - | 0 | - |
| - | - | 1594 | 1049 | - | - | 0 | - |
| - | - | 902.6 | 1054 | - | - | 0 | - |
| - | - | 917 | 1054 | - | - | 0 | - |
| - | - | 1197 | 1055 | - | - | 0 | - |
| - | - | 842.1 | 1055 | - | - | 0 | - |
| - | - | 1181 | 1056 | - | - | 0 | - |
| - | - | 1539 | 1057 | - | - | 0 | - |
| - | - | 3136 | 1057 | - | - | 0 | - |
| - | - | 1571 | 1058 | - | - | 0 | - |
| - | - | 683.1 | 1061 | - | - | 0 | - |
| - | - | 1624 | 1062 | - | - | 0 | - |
| - | - | 917.3 | 1063 | - | - | 0 | - |
| - | - | 1004 | 1064 | - | - | 0 | - |
| - | - | 1326 | 1065 | - | - | 0 | - |
| - | - | 755.3 | 1065 | - | - | 0 | - |
| - | - | 2329 | 1066 | - | - | 0 | - |
| - | - | 1012 | 1066 | - | - | 0 | - |
| 4 | y | 676.8 | 1087 | 0.009681 | 8.91 | +1 | 10 |
| - | - | 8653 | 1088 | - | - | 0 | - |
| 4 | z | 1.216E+04 | 1089 | 0.00954 | 8.764 | +1 | 10 |
| - | - | 8627 | 1090 | - | - | 0 | - |
| - | - | 2371 | 1091 | - | - | 0 | - |
| - | - | 3488 | 1104 | - | - | 0 | - |
| 10 | c | 3.762E+04 | 1105 | 0.009437 | 8.544 | +1 | 10 |
| - | - | 2.102E+04 | 1106 | - | - | 0 | - |
| - | - | 7231 | 1107 | - | - | 0 | - |
| - | - | 724.6 | 1131 | - | - | 0 | - |
| - | - | 2178 | 1174 | - | - | 0 | - |
| - | - | 2212 | 1175 | - | - | 0 | - |
| - | - | 1104 | 1183 | - | - | 0 | - |
| 3 | z | 934.7 | 1184 | 0.01905 | 16.09 | +1 | 11 |
| - | - | 1306 | 1185 | - | - | 0 | - |
| - | - | 799.3 | 1186 | - | - | 0 | - |
| - | - | 3779 | 1201 | - | - | 0 | - |
| 3 | z | 8388 | 1202 | 0.009704 | 8.076 | +1 | 11 |
| - | - | 6098 | 1203 | - | - | 0 | - |
| - | - | 2293 | 1204 | - | - | 0 | - |
| - | - | 868.1 | 1205 | - | - | 0 | - |
| 11 | c | 2.193E+04 | 1218 | 0.001948 | 1.6 | +1 | 11 |
| - | - | 1.629E+04 | 1219 | - | - | 0 | - |
| - | - | 5738 | 1220 | - | - | 0 | - |
| - | - | 725.6 | 1221 | - | - | 0 | - |
| - | - | 1784 | 1244 | - | - | 0 | - |
| - | - | 792.4 | 1245 | - | - | 0 | - |
| - | - | 748.1 | 1247 | - | - | 0 | - |
| - | - | 773.6 | 1277 | - | - | 0 | - |
| - | - | 673.4 | 1284 | - | - | 0 | - |
| 2 | z | 9465 | 1285 | 0.001277 | 0.994 | +1 | 12 |
| 2 | z | 6755 | 1286 | 0.0186 | 14.47 | +1 | 12 |
| - | - | 2391 | 1287 | - | - | 0 | - |
| 2 | w | 805.6 | 1288 | 0.01162 | 9.027 | +1 | 12 |
| - | - | 898.1 | 1293 | - | - | 0 | - |
| - | - | 810.8 | 1294 | - | - | 0 | - |
| 2 | y | 2501 | 1301 | 0.02038 | 15.67 | +1 | 12 |
| - | - | 3033 | 1302 | - | - | 0 | - |
| 2 | z | 1.201E+04 | 1303 | 0.001699 | 1.304 | +1 | 12 |
| - | - | 9139 | 1304 | - | - | 0 | - |
| - | - | 3430 | 1305 | - | - | 0 | - |
| - | - | 738.8 | 1309 | - | - | 0 | - |
| - | - | 994.3 | 1311 | - | - | 0 | - |
| - | - | 1065 | 1312 | - | - | 0 | - |
| - | - | 729.4 | 1313 | - | - | 0 | - |
| - | - | 910.8 | 1314 | - | - | 0 | - |
| - | - | 749.9 | 1315 | - | - | 0 | - |
| - | - | 1512 | 1318 | - | - | 0 | - |
| 2 | y | 1069 | 1319 | 0.009708 | 7.362 | +1 | 12 |
| - | - | 827.1 | 1320 | - | - | 0 | - |
| - | - | 1136 | 1331 | - | - | 0 | - |
| - | - | 1428 | 1332 | - | - | 0 | - |
| - | - | 923.7 | 1336 | - | - | 0 | - |
| - | - | 639.9 | 1342 | - | - | 0 | - |
| - | - | 686.8 | 1344 | - | - | 0 | - |
| - | - | 811.2 | 1345 | - | - | 0 | - |
| 12 | c | 6.56E+04 | 1346 | 0.0005893 | 0.4379 | +1 | 12 |
| - | - | 5.255E+04 | 1347 | - | - | 0 | - |
| - | - | 2.214E+04 | 1348 | - | - | 0 | - |
| - | - | 4367 | 1349 | - | - | 0 | - |
| - | - | 753.5 | 1350 | - | - | 0 | - |
| - | - | 990.6 | 1352 | - | - | 0 | - |
| - | - | 770.2 | 1353 | - | - | 0 | - |
| - | - | 1590 | 1356 | - | - | 0 | - |
| - | - | 1500 | 1357 | - | - | 0 | - |
| - | - | 1578 | 1358 | - | - | 0 | - |
| - | - | 800.4 | 1359 | - | - | 0 | - |
| - | - | 1108 | 1360 | - | - | 0 | - |
| - | - | 914 | 1361 | - | - | 0 | - |
| - | - | 1855 | 1363 | - | - | 0 | - |
| - | - | 1386 | 1364 | - | - | 0 | - |
| - | - | 1018 | 1366 | - | - | 0 | - |
| - | - | 1725 | 1373 | - | - | 0 | - |
| - | - | 7025 | 1374 | - | - | 0 | - |
| - | - | 4723 | 1375 | - | - | 0 | - |
| - | - | 2031 | 1376 | - | - | 0 | - |
| - | - | 1320 | 1382 | - | - | 0 | - |
| - | - | 2371 | 1383 | - | - | 0 | - |
| - | - | 6469 | 1384 | - | - | 0 | - |
| - | - | 4571 | 1385 | - | - | 0 | - |
| - | - | 2447 | 1386 | - | - | 0 | - |
| - | - | 2408 | 1391 | - | - | 0 | - |
| - | - | 2063 | 1392 | - | - | 0 | - |
| - | - | 2404 | 1399 | - | - | 0 | - |
| - | - | 3906 | 1400 | - | - | 0 | - |
| - | - | 6078 | 1401 | - | - | 0 | - |
| - | - | 1.427E+04 | 1402 | - | - | 0 | - |
| - | - | 9795 | 1403 | - | - | 0 | - |
| - | - | 3526 | 1404 | - | - | 0 | - |
| - | - | 874.5 | 1405 | - | - | 0 | - |
| - | - | 4247 | 1416 | - | - | 0 | - |
| - | - | 7462 | 1417 | - | - | 0 | - |
| - | - | 2.216E+04 | 1418 | - | - | 0 | - |
| - | - | 7.708E+04 | 1419 | - | - | 0 | - |
| - | - | 6.071E+04 | 1420 | - | - | 0 | - |
| - | - | 2.233E+04 | 1421 | - | - | 0 | - |
| - | - | 3369 | 1422 | - | - | 0 | - |
| - | - | 997.8 | 2023 | - | - | 0 | - |
| - | - | 835.9 | 2049 | - | - | 0 | - |
| - | - | 1111 | 2051 | - | - | 0 | - |
| - | - | 1282 | 2064 | - | - | 0 | - |
| - | - | 1085 | 2065 | - | - | 0 | - |
| - | - | 899.8 | 2080 | - | - | 0 | - |
| - | - | 2274 | 2095 | - | - | 0 | - |
| - | - | 2171 | 2096 | - | - | 0 | - |
| - | - | 1387 | 2097 | - | - | 0 | - |
| - | - | 1006 | 2099 | - | - | 0 | - |
| - | - | 1594 | 2106 | - | - | 0 | - |
| - | - | 2502 | 2107 | - | - | 0 | - |
| - | - | 1819 | 2108 | - | - | 0 | - |
| - | - | 1502 | 2109 | - | - | 0 | - |
| - | - | 1008 | 2110 | - | - | 0 | - |
| - | - | 2214 | 2111 | - | - | 0 | - |
| - | - | 2677 | 2112 | - | - | 0 | - |
| - | - | 2637 | 2113 | - | - | 0 | - |
| - | - | 2400 | 2114 | - | - | 0 | - |
| - | - | 1711 | 2115 | - | - | 0 | - |
| - | - | 2464 | 2124 | - | - | 0 | - |
| - | - | 1942 | 2125 | - | - | 0 | - |
| - | - | 1820 | 2126 | - | - | 0 | - |
| - | - | 1240 | 2127 | - | - | 0 | - |
| - | - | 1752 | 2128 | - | - | 0 | - |
| - | - | 1423 | 2129 | - | - | 0 | - |
| - | - | 1919 | 2130 | - | - | 0 | - |
| - | - | 1892 | 2131 | - | - | 0 | - |
| - | - | 2012 | 2132 | - | - | 0 | - |

m/z Charge Intensity FragmentType MassShift Position
120.08081817626953 0 796.37604
130.06137084960938 0 491.04623
132.31480407714844 0 344.41373
133.08615112304688 0 1132.7566
137.04734802246094 0 404.5083
140.38284301757812 0 447.5762
148.77987670898438 0 463.75415
148.90191650390625 0 585.5582
148.90953063964844 0 662.4272
148.91690063476562 0 975.163
148.924072265625 0 1029.0544
148.93128967285156 0 1349.7333
148.9386444091797 0 2967.134
148.955078125 0 6270.9785
148.96292114257812 0 3579.9653
148.97071838378906 0 1462.2386
148.97787475585938 0 1024.9763
148.98507690429688 0 943.93884
148.9921875 0 781.5395
148.9994659423828 0 475.28174
159.73863220214844 0 446.86694
163.0570526123047 0 396.44128
173.12896728515625 0 546.47845
173.44093322753906 0 669.9908
183.1136932373047 0 1257.0554
198.8501739501953 0 420.61603
200.1030731201172 0 3148.8877
201.08709716796875 0 3648.341 y Ammonia loss 11
201.12353515625 0 1933.2498
203.24049377441406 0 449.2486
210.00997924804688 0 505.62616
213.12307739257812 0 587.2184
215.13917541503906 0 8774.592
216.14268493652344 0 770.2623
218.1137237548828 0 7173.1543 y 11
227.10293579101562 0 849.29956
233.1649627685547 0 1053.9055
233.4415283203125 0 522.0417
242.15025329589844 0 2299.5354
271.4621276855469 0 540.277
282.1453552246094 0 1166.445
293.4715270996094 0 590.54333
296.1971740722656 0 17296.66
297.1999206542969 0 2108.1943
299.171875 0 564.15094
314.17205810546875 0 989.0218 y Ammonia loss 10
314.2076721191406 0 5340.766
331.1977844238281 0 1081.3225 y 10
333.53277587890625 0 503.43048
342.5884704589844 0 518.21655
346.1250915527344 0 660.5146
346.1760559082031 0 743.3013
354.1617431640625 0 498.02774
362.20758056640625 0 1305.1807
371.1940002441406 0 528.60364
382.1971130371094 0 556.2569
416.2541198730469 0 1065.8708
433.28155517578125 0 7209.301
434.2848205566406 0 1403.2731
441.23590087890625 0 1326.4703
443.2659606933594 0 1173.8262
461.2766418457031 0 7572.9307
462.2784423828125 0 2206.9355
498.220458984375 0 1118.8489
528.7857666015625 0 614.6789
530.3397216796875 0 574.4964
544.2709350585938 0 1498.0138
544.7681274414062 0 745.56226
558.33154296875 0 866.0765
600.81201171875 0 685.00793
627.26220703125 0 4958.922
628.2661743164062 0 2278.4521
657.3909301757812 0 1053.5106
658.2786865234375 0 660.25073
671.4000854492188 0 1775.9329
672.408447265625 0 2208.4724 c 5
673.4071044921875 0 790.2101
691.4151000976562 0 587.4054
716.4346923828125 0 3004.994
740.3475341796875 0 4361.4067
741.349609375 0 1377.9718
742.4176025390625 0 1462.3176
743.4168090820312 0 924.26886
744.4271850585938 0 764.8409
745.3167724609375 0 1169.1786
757.373779296875 0 1416.0155
758.4337158203125 0 6742.7344
759.43896484375 0 11683.686 c 6
760.44189453125 0 3228.6523
763.3463134765625 0 1215.0886 y 6
771.3526611328125 0 1003.3226
803.4669189453125 0 1029.6034
829.445556640625 0 999.9234
833.337646484375 0 651.76416
845.46484375 0 4121.821
846.4725341796875 0 26371.941 c 7
847.4752197265625 0 11233.369
848.4781494140625 0 3598.2932
860.4013061523438 0 4158.823 y 5
861.40087890625 0 1863.2589
868.4060668945312 0 8196.073
869.407470703125 0 3433.6025
871.9627685546875 0 793.2895
887.4161376953125 0 1440.7367
931.4994506835938 0 794.38086
938.4329833984375 0 1987.6913
939.4398193359375 0 1530.9672 y Water loss 4
940.4295654296875 0 3013.91 y Ammonia loss 4
941.4387817382812 0 957.9006 z 4
956.4454345703125 0 9714.289
957.4528198242188 0 44247.277 y 4
958.4566650390625 0 21063.598
959.46044921875 0 4920.243
960.4657592773438 0 650.5943
974.50732421875 0 2994.764
975.5155639648438 0 27123.383 c 8
976.5180053710938 0 16733.496
977.5224609375 0 3885.608
1008.5390014648438 0 856.8383
1009.048583984375 0 1422.546
1015.4725341796875 0 745.38434
1016.4699096679688 0 843.5114
1019.53125 0 785.0256
1025.0426025390625 0 727.5394
1025.551513671875 0 1140.946
1031.5416259765625 0 1070.2985
1032.04443359375 0 1790.3823
1032.5501708984375 0 1197.1786
1048.5430908203125 0 1594.0447
1053.5465087890625 0 902.6136
1054.0467529296875 0 917.04346
1054.5440673828125 0 1196.7938
1055.030517578125 0 842.13727
1056.043701171875 0 1180.9545
1056.5386962890625 0 1539.0896
1057.042236328125 0 3135.6748
1057.5478515625 0 1570.8138
1060.541259765625 0 683.13135
1061.54931640625 0 1624.1731
1062.5570068359375 0 917.3361
1064.0380859375 0 1004.4665
1064.5455322265625 0 1325.8949
1065.0452880859375 0 755.27167
1065.546630859375 0 2328.644
1066.056640625 0 1012.4449
1086.5198974609375 0 676.80804 y Water loss 3
1087.5318603515625 0 8653.015
1088.5115966796875 0 12163.316 z 3
1089.510498046875 0 8626.567
1090.507568359375 0 2370.7246
1103.518798828125 0 3487.7986
1104.5477294921875 0 37624.04 c 9
1105.5511474609375 0 21021.002
1106.554931640625 0 7230.9834
1130.5548095703125 0 724.59515
1173.62939453125 0 2178.3667
1174.6295166015625 0 2211.5
1182.605712890625 0 1103.6266
1183.5946044921875 0 934.65393 z Water loss 2
1184.5941162109375 0 1305.5275
1185.5821533203125 0 799.2624
1200.616455078125 0 3778.8352
1201.5958251953125 0 8388.403 z 2
1202.5958251953125 0 6098.4214
1203.5989990234375 0 2293.2156
1204.5804443359375 0 868.11914
1217.6392822265625 0 21930.035 c 10
1218.6424560546875 0 16293.692
1219.646484375 0 5737.8145
1220.6431884765625 0 725.6087
1243.6529541015625 0 1784.2473
1244.6756591796875 0 792.444
1246.574951171875 0 748.1315
1276.6097412109375 0 773.57513
1283.6572265625 0 673.38403
1284.62451171875 0 9465.324 z Water loss 1
1285.6258544921875 0 6755.233 z Ammonia loss 1
1286.6285400390625 0 2391.1055
1287.6219482421875 0 805.5863 w 1
1292.6529541015625 0 898.1121
1293.6474609375 0 810.8006
1300.62158203125 0 2500.5642 y Water loss 1
1301.6549072265625 0 3032.5598
1302.635498046875 0 12010.283 z 1
1303.6400146484375 0 9138.546
1304.6427001953125 0 3430.0867
1308.6553955078125 0 738.8226
1310.6668701171875 0 994.28644
1311.655029296875 0 1064.6892
1312.650390625 0 729.4273
1313.666015625 0 910.8096
1314.65966796875 0 749.87756
1317.638427734375 0 1511.9567
1318.6622314453125 0 1069.1578 y 1
1319.669189453125 0 827.1363
1330.6719970703125 0 1135.8713
1331.669921875 0 1427.737
1335.683349609375 0 923.69836
1341.6448974609375 0 639.8934
1343.6849365234375 0 686.81744
1344.679931640625 0 811.1896
1345.69921875 0 65596.22 c 11
1346.7030029296875 0 52550.207
1347.7047119140625 0 22139.834
1348.7076416015625 0 4367.0796
1349.711669921875 0 753.4663
1351.68994140625 0 990.5939
1352.6917724609375 0 770.2252
1355.6854248046875 0 1589.5061
1356.7105712890625 0 1499.9663
1357.6529541015625 0 1578.01
1358.624755859375 0 800.3974
1359.6494140625 0 1108.2046
1360.648193359375 0 913.9763
1362.6685791015625 0 1854.7452
1363.6773681640625 0 1385.8545
1365.6773681640625 0 1018.3259
1372.726806640625 0 1725.1656
1373.70849609375 0 7025.31
1374.715576171875 0 4723.3003
1375.7119140625 0 2031.3271
1381.6622314453125 0 1319.6985
1382.658203125 0 2371.0718
1383.67578125 0 6468.5205
1384.680908203125 0 4570.514
1385.6776123046875 0 2447.191
1390.7352294921875 0 2408.0486
1391.73095703125 0 2063.2815
1398.6507568359375 0 2403.9568
1399.655029296875 0 3905.8237
1400.6868896484375 0 6078.1104
1401.697265625 0 14274.263
1402.6949462890625 0 9794.914
1403.7049560546875 0 3525.7556
1404.707763671875 0 874.48193
1415.651611328125 0 4246.933
1416.6591796875 0 7461.9785
1417.7154541015625 0 22155.824
1418.7271728515625 0 77080.48
1419.72998046875 0 60712.992
1420.7320556640625 0 22334.686
1421.7269287109375 0 3368.7336
2023.0601806640625 0 997.76324
2049.083740234375 0 835.93726
2051.070556640625 0 1111.4694
2064.081787109375 0 1282.1606
2065.062255859375 0 1085.4281
2080.06640625 0 899.78485
2095.06298828125 0 2274.0203
2096.06591796875 0 2170.7366
2097.079833984375 0 1387.2489
2099.084716796875 0 1006.05383
2106.075927734375 0 1594.0482
2107.096923828125 0 2501.51
2108.095458984375 0 1818.7651
2109.070068359375 0 1502.224
2110.071533203125 0 1008.09265
2111.0712890625 0 2214.0771
2112.06396484375 0 2677.1726
2113.069091796875 0 2637.2847
2114.08251953125 0 2399.9336
2115.079833984375 0 1711.2135
2124.105224609375 0 2463.7236
2125.106689453125 0 1942.2334
2126.094970703125 0 1820.496
2127.094970703125 0 1240.4042
2128.07373046875 0 1752.3069
2129.071044921875 0 1423.0245
2130.093017578125 0 1919.4921
2131.0908203125 0 1891.6952
2132.0869140625 0 2011.5192

Spectrum Details

|  |  |
| --- | --- |
| Matched peaks? Matched peaks The total absolute number of peaks matched. Additionally in brackets the total fraction of peaks matched and the total number of peaks is shown. | 27 (9.96% of 271) |
| FDR? FDR The false discovery rate estimated for this peptide. It is calculated by matching all theoretical fragments with a non-integer shift with the raw peaks for this spectrum. This is done with 40 different shifts. The resulting percentage is the average number of annotated peaks over the number of annotated peaks with the correct spectrum. | 4.23% |
| Satellite FDR? Satellite FDR See the FDR for details on its calculation. This satellite ion specific FDR only contains the satellite ions (d/w) for I/L/J positions. | ∞ |
| PSM Score? PSM Score The PSM Score as given by Hecklib to this annotated spectrum. It is shown with three significant figures. | 302 |

#### Spectrum 9831? Spectrum 9831 The raw spectrum of this peptide as annotated by Hecklib. The fragments are coloured according to ion type (see legend). Any peaks with a star '\*' as text can be hovered over to see the full details, first the ion type second the mass shift type. By hovering over the amino acids in the peptide or ions in the legend the corresponding peaks are highlighted. By toggling the 'Unassigned' label you can turn the background (unassigned) peaks on or off in the plot. By updating the slider in the Ion legend you can update the spectrum to only show the top X% of the peaks with labels. The top X% means any peak that is within X% of the highest intensity. By dragging in the spectrum you can zoom in to a specific part of the spectrum and use 'Zoom Out' to get back to the original zoom level. The annotation of the spectrum is based on the given sequence in the peptides file and is done with different software so inconsistencies are likely. The peaks are annotated based on the given sequence, with 20 ppm tolerance.

Copy Data

##### Spectrum 9831 (TSV)

###### Preview

```
Loading example...
```

*Click on the button to copy the data to your clipboard.*

Mz MinMz MaxIntensity Max

WidthHeightPeptide font sizePeptide stroke widthSpectrum font sizeSpectrum stroke widthCompact peptide

Ion legend

wxyz

abcd

OtherUnassignedIon^Charge^_Position_Show for top:%

VTJFPPSSEEJQA

07.25e+41.45e+52.17e+52.90e+5

Zoom Out

y^+1^_2_y^+1^_2_y^+2^_5_y^+1^_3_y^+1^_3_z^+2^_7_y^+1^_4_y^+1^_4_y^+2^_9_z^+1^_5_y^+1^_5_w^+1^_6_y^+1^_6_c^+1^_6_y^+1^_6_c^+1^_7_z^+1^_7_c^+1^_7_y^+1^_7_y^+1^_8_z^+1^_8_c^+1^_8_y^+1^_8_w^+1^_9_z^+1^_9_y^+1^_9_y^+1^_9_z^+1^_9_y^+1^_9_c^+1^_9_z^+1^_10_y^+1^_10_z^+1^_10_c^+1^_10_w^+1^_11_z^+1^_11_c^+1^_11_z^+1^_12_z^+1^_12_w^+1^_12_y^+1^_12_z^+1^_12_c^+1^_12_c^+1^_12_

0764152922933057

Fragment Matches Table

Show background peaks

| Position | Ion type | Intensity | mz Theoretical | mz Error (Th) | mz Error (ppm) | Charge | Series Number |
| --- | --- | --- | --- | --- | --- | --- | --- |
| - | - | 3366 | 120.1 | - | - | 0 | - |
| - | - | 395.8 | 129.1 | - | - | 0 | - |
| - | - | 4247 | 129.1 | - | - | 0 | - |
| - | - | 1314 | 131.1 | - | - | 0 | - |
| - | - | 369.7 | 132.1 | - | - | 0 | - |
| - | - | 581.1 | 133.1 | - | - | 0 | - |
| - | - | 4163 | 134 | - | - | 0 | - |
| - | - | 443.4 | 136 | - | - | 0 | - |
| - | - | 879.7 | 136.1 | - | - | 0 | - |
| - | - | 450.3 | 148.9 | - | - | 0 | - |
| - | - | 454.6 | 148.9 | - | - | 0 | - |
| - | - | 445.7 | 148.9 | - | - | 0 | - |
| - | - | 574.2 | 148.9 | - | - | 0 | - |
| - | - | 689.2 | 148.9 | - | - | 0 | - |
| - | - | 1281 | 148.9 | - | - | 0 | - |
| - | - | 1227 | 148.9 | - | - | 0 | - |
| - | - | 1996 | 148.9 | - | - | 0 | - |
| - | - | 3279 | 148.9 | - | - | 0 | - |
| - | - | 3650 | 149 | - | - | 0 | - |
| - | - | 2249 | 149 | - | - | 0 | - |
| - | - | 1228 | 149 | - | - | 0 | - |
| - | - | 948.4 | 149 | - | - | 0 | - |
| - | - | 547.5 | 149 | - | - | 0 | - |
| - | - | 531.4 | 149 | - | - | 0 | - |
| - | - | 425.9 | 161.1 | - | - | 0 | - |
| - | - | 538.1 | 163.5 | - | - | 0 | - |
| - | - | 1783 | 167.1 | - | - | 0 | - |
| - | - | 3130 | 169.1 | - | - | 0 | - |
| - | - | 536 | 169.1 | - | - | 0 | - |
| - | - | 1678 | 173.1 | - | - | 0 | - |
| - | - | 914.4 | 173.4 | - | - | 0 | - |
| - | - | 451.8 | 173.6 | - | - | 0 | - |
| - | - | 763.5 | 177.1 | - | - | 0 | - |
| - | - | 1979 | 183.1 | - | - | 0 | - |
| - | - | 551.8 | 197.1 | - | - | 0 | - |
| - | - | 1128 | 197.1 | - | - | 0 | - |
| - | - | 459.6 | 198.4 | - | - | 0 | - |
| - | - | 2518 | 199.1 | - | - | 0 | - |
| - | - | 1.069E+04 | 200.1 | - | - | 0 | - |
| - | - | 1054 | 200.1 | - | - | 0 | - |
| 12 | y | 1.093E+04 | 201.1 | 0.0001901 | 0.9455 | +1 | 2 |
| - | - | 698 | 201.1 | - | - | 0 | - |
| - | - | 5867 | 201.1 | - | - | 0 | - |
| - | - | 1294 | 202.1 | - | - | 0 | - |
| - | - | 474.8 | 202.1 | - | - | 0 | - |
| - | - | 1137 | 209.1 | - | - | 0 | - |
| - | - | 1.123E+04 | 213.1 | - | - | 0 | - |
| - | - | 831.6 | 214.1 | - | - | 0 | - |
| - | - | 2.87E+05 | 215.1 | - | - | 0 | - |
| - | - | 641.9 | 216.1 | - | - | 0 | - |
| - | - | 2.943E+04 | 216.1 | - | - | 0 | - |
| - | - | 2232 | 217.1 | - | - | 0 | - |
| 12 | y | 2.487E+04 | 218.1 | 0.0002676 | 1.227 | +1 | 2 |
| - | - | 2456 | 219.1 | - | - | 0 | - |
| - | - | 1842 | 225.1 | - | - | 0 | - |
| - | - | 1125 | 225.1 | - | - | 0 | - |
| - | - | 2263 | 226.1 | - | - | 0 | - |
| - | - | 1.241E+04 | 227.1 | - | - | 0 | - |
| - | - | 1716 | 228.1 | - | - | 0 | - |
| - | - | 3169 | 228.2 | - | - | 0 | - |
| - | - | 1427 | 229.1 | - | - | 0 | - |
| - | - | 1643 | 231.1 | - | - | 0 | - |
| - | - | 2987 | 233.2 | - | - | 0 | - |
| - | - | 7938 | 242.2 | - | - | 0 | - |
| - | - | 647.6 | 243.1 | - | - | 0 | - |
| - | - | 818.8 | 243.2 | - | - | 0 | - |
| - | - | 964 | 245.1 | - | - | 0 | - |
| - | - | 787.5 | 245.1 | - | - | 0 | - |
| - | - | 1.26E+04 | 247.1 | - | - | 0 | - |
| - | - | 2881 | 247.1 | - | - | 0 | - |
| - | - | 908.8 | 247.1 | - | - | 0 | - |
| - | - | 1086 | 248.1 | - | - | 0 | - |
| - | - | 832.6 | 251.2 | - | - | 0 | - |
| - | - | 628.9 | 259.1 | - | - | 0 | - |
| - | - | 832.1 | 259.1 | - | - | 0 | - |
| - | - | 2059 | 261.2 | - | - | 0 | - |
| - | - | 556.2 | 272.5 | - | - | 0 | - |
| - | - | 1.613E+04 | 275.1 | - | - | 0 | - |
| - | - | 788 | 275.1 | - | - | 0 | - |
| - | - | 716.9 | 279.1 | - | - | 0 | - |
| - | - | 864.4 | 280.2 | - | - | 0 | - |
| - | - | 1897 | 282.1 | - | - | 0 | - |
| 9 | y | 945.9 | 295.1 | 0.0007729 | 2.619 | +2 | 5 |
| - | - | 4.435E+04 | 296.2 | - | - | 0 | - |
| - | - | 6814 | 297.2 | - | - | 0 | - |
| - | - | 1192 | 298.2 | - | - | 0 | - |
| - | - | 661.4 | 298.2 | - | - | 0 | - |
| - | - | 933.2 | 299.2 | - | - | 0 | - |
| - | - | 1295 | 303.2 | - | - | 0 | - |
| 11 | y | 2364 | 314.2 | 0.0006141 | 1.955 | +1 | 3 |
| - | - | 1.863E+04 | 314.2 | - | - | 0 | - |
| - | - | 3570 | 315.2 | - | - | 0 | - |
| - | - | 2911 | 324.2 | - | - | 0 | - |
| - | - | 1249 | 325.1 | - | - | 0 | - |
| - | - | 3027 | 326.2 | - | - | 0 | - |
| - | - | 1680 | 330.2 | - | - | 0 | - |
| - | - | 532.5 | 330.6 | - | - | 0 | - |
| 11 | y | 4571 | 331.2 | 0.0003711 | 1.121 | +1 | 3 |
| - | - | 653.5 | 332.2 | - | - | 0 | - |
| - | - | 928.4 | 343.2 | - | - | 0 | - |
| - | - | 2340 | 344.2 | - | - | 0 | - |
| - | - | 629.3 | 347.2 | - | - | 0 | - |
| - | - | 1412 | 348.2 | - | - | 0 | - |
| - | - | 1224 | 353.2 | - | - | 0 | - |
| - | - | 633.3 | 360.2 | - | - | 0 | - |
| - | - | 4897 | 362.2 | - | - | 0 | - |
| - | - | 937.8 | 363.2 | - | - | 0 | - |
| - | - | 1143 | 369.2 | - | - | 0 | - |
| - | - | 1306 | 371.2 | - | - | 0 | - |
| - | - | 673.6 | 373.1 | - | - | 0 | - |
| 7 | z | 8355 | 374.2 | 0.006315 | 16.88 | +2 | 7 |
| - | - | 1820 | 375.2 | - | - | 0 | - |
| - | - | 1117 | 388.1 | - | - | 0 | - |
| - | - | 1049 | 391.2 | - | - | 0 | - |
| - | - | 704.1 | 393.3 | - | - | 0 | - |
| - | - | 763.3 | 395.2 | - | - | 0 | - |
| - | - | 762.2 | 398.2 | - | - | 0 | - |
| - | - | 1533 | 407.2 | - | - | 0 | - |
| - | - | 1517 | 411.3 | - | - | 0 | - |
| - | - | 650.8 | 415.3 | - | - | 0 | - |
| - | - | 2281 | 416.3 | - | - | 0 | - |
| - | - | 6793 | 423.2 | - | - | 0 | - |
| - | - | 1650 | 424.2 | - | - | 0 | - |
| - | - | 9102 | 427.3 | - | - | 0 | - |
| - | - | 2063 | 428.3 | - | - | 0 | - |
| - | - | 2.592E+04 | 433.3 | - | - | 0 | - |
| - | - | 7482 | 434.3 | - | - | 0 | - |
| - | - | 744.3 | 434.7 | - | - | 0 | - |
| - | - | 655.7 | 435.3 | - | - | 0 | - |
| - | - | 2242 | 439.2 | - | - | 0 | - |
| - | - | 899.3 | 440.2 | - | - | 0 | - |
| - | - | 2.348E+04 | 441.2 | - | - | 0 | - |
| 10 | y | 6875 | 442.2 | 0.008443 | 19.09 | +1 | 4 |
| - | - | 1199 | 443.2 | - | - | 0 | - |
| - | - | 2388 | 443.3 | - | - | 0 | - |
| - | - | 917.2 | 443.3 | - | - | 0 | - |
| - | - | 1116 | 451.2 | - | - | 0 | - |
| - | - | 1424 | 455.2 | - | - | 0 | - |
| - | - | 716.1 | 456.2 | - | - | 0 | - |
| - | - | 600 | 457.2 | - | - | 0 | - |
| - | - | 1142 | 459.2 | - | - | 0 | - |
| - | - | 665.1 | 459.3 | - | - | 0 | - |
| 10 | y | 761.5 | 460.2 | 0.001876 | 4.076 | +1 | 4 |
| - | - | 2.655E+04 | 461.3 | - | - | 0 | - |
| - | - | 7645 | 462.3 | - | - | 0 | - |
| - | - | 627.4 | 463.3 | - | - | 0 | - |
| 5 | y | 1771 | 470.2 | 0.0008939 | 1.901 | +2 | 9 |
| - | - | 1904 | 476.3 | - | - | 0 | - |
| - | - | 768.4 | 477.3 | - | - | 0 | - |
| - | - | 6227 | 479.2 | - | - | 0 | - |
| - | - | 735.5 | 479.7 | - | - | 0 | - |
| - | - | 1804 | 480.2 | - | - | 0 | - |
| - | - | 628.5 | 482.2 | - | - | 0 | - |
| - | - | 2270 | 492.3 | - | - | 0 | - |
| - | - | 3042 | 498.2 | - | - | 0 | - |
| - | - | 893.8 | 499.2 | - | - | 0 | - |
| - | - | 1793 | 502.3 | - | - | 0 | - |
| - | - | 880.2 | 505.7 | - | - | 0 | - |
| - | - | 862 | 506.7 | - | - | 0 | - |
| - | - | 656.6 | 507.2 | - | - | 0 | - |
| - | - | 1671 | 518.3 | - | - | 0 | - |
| - | - | 1.326E+04 | 519.3 | - | - | 0 | - |
| - | - | 1217 | 519.7 | - | - | 0 | - |
| - | - | 1361 | 520.2 | - | - | 0 | - |
| - | - | 6961 | 520.3 | - | - | 0 | - |
| - | - | 2541 | 520.7 | - | - | 0 | - |
| - | - | 2795 | 521.3 | - | - | 0 | - |
| - | - | 682.6 | 529.2 | - | - | 0 | - |
| - | - | 1407 | 530.2 | - | - | 0 | - |
| - | - | 785 | 530.3 | - | - | 0 | - |
| - | - | 2267 | 530.3 | - | - | 0 | - |
| - | - | 758.8 | 530.8 | - | - | 0 | - |
| - | - | 1518 | 540.3 | - | - | 0 | - |
| - | - | 1096 | 542.3 | - | - | 0 | - |
| - | - | 2307 | 544.3 | - | - | 0 | - |
| - | - | 2592 | 544.8 | - | - | 0 | - |
| - | - | 732.7 | 546.2 | - | - | 0 | - |
| - | - | 1522 | 554.3 | - | - | 0 | - |
| 9 | z | 644 | 555.3 | 0.009507 | 17.12 | +1 | 5 |
| - | - | 4596 | 558.3 | - | - | 0 | - |
| - | - | 1717 | 559.3 | - | - | 0 | - |
| - | - | 1759 | 568.2 | - | - | 0 | - |
| - | - | 793.4 | 569.2 | - | - | 0 | - |
| - | - | 5702 | 570.3 | - | - | 0 | - |
| 9 | y | 2629 | 571.3 | 0.009462 | 16.56 | +1 | 5 |
| - | - | 1849 | 584.8 | - | - | 0 | - |
| - | - | 1567 | 585.3 | - | - | 0 | - |
| - | - | 1144 | 585.8 | - | - | 0 | - |
| - | - | 2841 | 586.3 | - | - | 0 | - |
| - | - | 2183 | 587.3 | - | - | 0 | - |
| - | - | 5079 | 588.3 | - | - | 0 | - |
| - | - | 3484 | 589.3 | - | - | 0 | - |
| - | - | 2134 | 591.3 | - | - | 0 | - |
| - | - | 2456 | 599.3 | - | - | 0 | - |
| - | - | 682.5 | 600.3 | - | - | 0 | - |
| - | - | 1938 | 600.8 | - | - | 0 | - |
| - | - | 1394 | 601.3 | - | - | 0 | - |
| - | - | 688.1 | 602.3 | - | - | 0 | - |
| - | - | 713.6 | 603.3 | - | - | 0 | - |
| - | - | 1312 | 604.3 | - | - | 0 | - |
| - | - | 1252 | 605.3 | - | - | 0 | - |
| - | - | 2429 | 607.3 | - | - | 0 | - |
| - | - | 1166 | 609.3 | - | - | 0 | - |
| - | - | 1.236E+04 | 617.3 | - | - | 0 | - |
| - | - | 3681 | 618.3 | - | - | 0 | - |
| - | - | 1722 | 619.3 | - | - | 0 | - |
| - | - | 1.613E+04 | 627.3 | - | - | 0 | - |
| - | - | 5278 | 628.3 | - | - | 0 | - |
| - | - | 1603 | 628.4 | - | - | 0 | - |
| - | - | 1336 | 629.3 | - | - | 0 | - |
| - | - | 1019 | 629.4 | - | - | 0 | - |
| - | - | 809.7 | 633.3 | - | - | 0 | - |
| - | - | 1.495E+04 | 634.3 | - | - | 0 | - |
| - | - | 9432 | 635.3 | - | - | 0 | - |
| - | - | 3580 | 636.3 | - | - | 0 | - |
| 8 | w | 1734 | 643.3 | 0.003735 | 5.807 | +1 | 6 |
| - | - | 1070 | 643.3 | - | - | 0 | - |
| - | - | 1382 | 644.3 | - | - | 0 | - |
| - | - | 734.1 | 645.3 | - | - | 0 | - |
| - | - | 1482 | 655.4 | - | - | 0 | - |
| - | - | 692.8 | 656.4 | - | - | 0 | - |
| - | - | 1657 | 658.3 | - | - | 0 | - |
| 8 | y | 1120 | 659.3 | 0.002862 | 4.34 | +1 | 6 |
| - | - | 1874 | 663.4 | - | - | 0 | - |
| - | - | 1104 | 667.3 | - | - | 0 | - |
| - | - | 2279 | 669.3 | - | - | 0 | - |
| - | - | 939 | 670.3 | - | - | 0 | - |
| - | - | 3677 | 671.4 | - | - | 0 | - |
| 6 | c | 3970 | 672.4 | 0.0002087 | 0.3104 | +1 | 6 |
| - | - | 1059 | 673.4 | - | - | 0 | - |
| - | - | 980.4 | 674.3 | - | - | 0 | - |
| 8 | y | 2496 | 676.3 | 0.0009238 | 1.366 | +1 | 6 |
| - | - | 757 | 677.3 | - | - | 0 | - |
| - | - | 636.3 | 682.3 | - | - | 0 | - |
| - | - | 800 | 685.3 | - | - | 0 | - |
| - | - | 1243 | 686.3 | - | - | 0 | - |
| - | - | 959.4 | 690.3 | - | - | 0 | - |
| - | - | 3944 | 691.3 | - | - | 0 | - |
| - | - | 2297 | 691.8 | - | - | 0 | - |
| - | - | 930.8 | 692.3 | - | - | 0 | - |
| - | - | 821.7 | 692.8 | - | - | 0 | - |
| - | - | 718 | 697.4 | - | - | 0 | - |
| - | - | 778.8 | 705.4 | - | - | 0 | - |
| - | - | 1304 | 709.3 | - | - | 0 | - |
| - | - | 774.9 | 710.8 | - | - | 0 | - |
| - | - | 721 | 712.4 | - | - | 0 | - |
| - | - | 1095 | 715.4 | - | - | 0 | - |
| - | - | 951.3 | 716.4 | - | - | 0 | - |
| - | - | 6664 | 716.4 | - | - | 0 | - |
| - | - | 3073 | 717.4 | - | - | 0 | - |
| - | - | 801.3 | 720.3 | - | - | 0 | - |
| - | - | 1077 | 722.3 | - | - | 0 | - |
| - | - | 712.6 | 723.3 | - | - | 0 | - |
| - | - | 1196 | 724.4 | - | - | 0 | - |
| - | - | 1515 | 728.4 | - | - | 0 | - |
| - | - | 1.362E+04 | 740.3 | - | - | 0 | - |
| - | - | 1524 | 740.4 | - | - | 0 | - |
| - | - | 4972 | 741.3 | - | - | 0 | - |
| 7 | c | 951.1 | 741.4 | 0.009343 | 12.6 | +1 | 7 |
| - | - | 718.2 | 742.4 | - | - | 0 | - |
| - | - | 3491 | 742.4 | - | - | 0 | - |
| - | - | 1871 | 743.4 | - | - | 0 | - |
| - | - | 2585 | 744.4 | - | - | 0 | - |
| - | - | 1154 | 745.3 | - | - | 0 | - |
| - | - | 743.8 | 745.4 | - | - | 0 | - |
| 7 | z | 728.1 | 747.3 | 0.003977 | 5.322 | +1 | 7 |
| - | - | 760 | 750.3 | - | - | 0 | - |
| - | - | 5121 | 752.4 | - | - | 0 | - |
| - | - | 2860 | 753.4 | - | - | 0 | - |
| - | - | 704 | 754.4 | - | - | 0 | - |
| - | - | 3679 | 757.4 | - | - | 0 | - |
| - | - | 1.616E+04 | 758.4 | - | - | 0 | - |
| 7 | c | 2.049E+04 | 759.4 | 0.00117 | 1.541 | +1 | 7 |
| - | - | 7067 | 760.4 | - | - | 0 | - |
| - | - | 1696 | 761.4 | - | - | 0 | - |
| - | - | 1254 | 762.3 | - | - | 0 | - |
| 7 | y | 2647 | 763.3 | 0.001976 | 2.589 | +1 | 7 |
| - | - | 1133 | 764.4 | - | - | 0 | - |
| - | - | 977.2 | 766.3 | - | - | 0 | - |
| - | - | 1150 | 767.4 | - | - | 0 | - |
| - | - | 2971 | 768.4 | - | - | 0 | - |
| - | - | 2136 | 769.4 | - | - | 0 | - |
| - | - | 1043 | 770.4 | - | - | 0 | - |
| - | - | 3993 | 771.4 | - | - | 0 | - |
| - | - | 1289 | 772.4 | - | - | 0 | - |
| - | - | 1146 | 774.3 | - | - | 0 | - |
| - | - | 894.8 | 779.4 | - | - | 0 | - |
| - | - | 1.131E+04 | 780.4 | - | - | 0 | - |
| - | - | 6068 | 781.4 | - | - | 0 | - |
| - | - | 2274 | 782.4 | - | - | 0 | - |
| - | - | 1176 | 784.4 | - | - | 0 | - |
| - | - | 1677 | 785.4 | - | - | 0 | - |
| - | - | 890.8 | 786.4 | - | - | 0 | - |
| - | - | 4437 | 794.3 | - | - | 0 | - |
| - | - | 2837 | 795.3 | - | - | 0 | - |
| - | - | 1.086E+04 | 796.4 | - | - | 0 | - |
| - | - | 1.628E+04 | 797.4 | - | - | 0 | - |
| - | - | 7354 | 798.4 | - | - | 0 | - |
| - | - | 2259 | 799.4 | - | - | 0 | - |
| - | - | 2207 | 803.5 | - | - | 0 | - |
| - | - | 929 | 804.5 | - | - | 0 | - |
| - | - | 5111 | 812.4 | - | - | 0 | - |
| - | - | 4524 | 813.4 | - | - | 0 | - |
| - | - | 2769 | 814.4 | - | - | 0 | - |
| - | - | 1576 | 815.4 | - | - | 0 | - |
| - | - | 1778 | 820.4 | - | - | 0 | - |
| - | - | 940.3 | 825.4 | - | - | 0 | - |
| - | - | 909.4 | 827.5 | - | - | 0 | - |
| - | - | 668.8 | 828.4 | - | - | 0 | - |
| - | - | 1395 | 829.4 | - | - | 0 | - |
| - | - | 1615 | 829.4 | - | - | 0 | - |
| - | - | 1778 | 830.4 | - | - | 0 | - |
| - | - | 1292 | 830.5 | - | - | 0 | - |
| - | - | 934.6 | 836.4 | - | - | 0 | - |
| - | - | 826.9 | 837.4 | - | - | 0 | - |
| - | - | 725.1 | 838.4 | - | - | 0 | - |
| - | - | 1799 | 839.4 | - | - | 0 | - |
| - | - | 2345 | 840.4 | - | - | 0 | - |
| 6 | y | 727.5 | 842.4 | 0.00647 | 7.681 | +1 | 8 |
| - | - | 5506 | 843.4 | - | - | 0 | - |
| 6 | z | 1940 | 844.4 | 0.00407 | 4.82 | +1 | 8 |
| - | - | 982.9 | 845.4 | - | - | 0 | - |
| - | - | 1.113E+04 | 845.5 | - | - | 0 | - |
| 8 | c | 5.567E+04 | 846.5 | 0.0001176 | 0.1389 | +1 | 8 |
| - | - | 2.524E+04 | 847.5 | - | - | 0 | - |
| - | - | 5216 | 848.5 | - | - | 0 | - |
| - | - | 1135 | 850.4 | - | - | 0 | - |
| - | - | 638.2 | 851.4 | - | - | 0 | - |
| - | - | 951.8 | 852.4 | - | - | 0 | - |
| - | - | 824.2 | 859.4 | - | - | 0 | - |
| 6 | y | 1.131E+04 | 860.4 | 0.0008484 | 0.986 | +1 | 8 |
| - | - | 6276 | 861.4 | - | - | 0 | - |
| - | - | 939 | 862.4 | - | - | 0 | - |
| - | - | 1021 | 867.4 | - | - | 0 | - |
| - | - | 2.517E+04 | 868.4 | - | - | 0 | - |
| - | - | 1.085E+04 | 869.4 | - | - | 0 | - |
| - | - | 3360 | 870.4 | - | - | 0 | - |
| - | - | 885.8 | 877.4 | - | - | 0 | - |
| - | - | 1133 | 881.4 | - | - | 0 | - |
| - | - | 1852 | 883.4 | - | - | 0 | - |
| - | - | 768.5 | 884.4 | - | - | 0 | - |
| - | - | 744.9 | 885.4 | - | - | 0 | - |
| - | - | 3713 | 887.4 | - | - | 0 | - |
| - | - | 2072 | 888.4 | - | - | 0 | - |
| - | - | 956.4 | 889.4 | - | - | 0 | - |
| - | - | 768.4 | 891.4 | - | - | 0 | - |
| - | - | 1871 | 893.4 | - | - | 0 | - |
| - | - | 1146 | 894.4 | - | - | 0 | - |
| - | - | 3541 | 895.4 | - | - | 0 | - |
| - | - | 1086 | 896.4 | - | - | 0 | - |
| - | - | 950.5 | 897.4 | - | - | 0 | - |
| - | - | 879.5 | 899.4 | - | - | 0 | - |
| - | - | 2171 | 900.4 | - | - | 0 | - |
| - | - | 1156 | 901.4 | - | - | 0 | - |
| - | - | 1004 | 902.4 | - | - | 0 | - |
| - | - | 1176 | 903.4 | - | - | 0 | - |
| - | - | 1110 | 904.4 | - | - | 0 | - |
| - | - | 738 | 911.4 | - | - | 0 | - |
| - | - | 842.9 | 912.4 | - | - | 0 | - |
| - | - | 1103 | 913.4 | - | - | 0 | - |
| 5 | w | 1264 | 914.4 | 0.0007817 | 0.8549 | +1 | 9 |
| - | - | 5737 | 915.4 | - | - | 0 | - |
| - | - | 5231 | 916.4 | - | - | 0 | - |
| - | - | 2768 | 917.4 | - | - | 0 | - |
| - | - | 2479 | 918.4 | - | - | 0 | - |
| - | - | 826.1 | 919.4 | - | - | 0 | - |
| - | - | 957.6 | 922.4 | - | - | 0 | - |
| 5 | z | 1831 | 923.4 | 0.01518 | 16.44 | +1 | 9 |
| - | - | 2779 | 925.4 | - | - | 0 | - |
| - | - | 2087 | 926.4 | - | - | 0 | - |
| - | - | 3479 | 927.4 | - | - | 0 | - |
| - | - | 2245 | 928.4 | - | - | 0 | - |
| - | - | 764.3 | 929.4 | - | - | 0 | - |
| - | - | 2043 | 931.5 | - | - | 0 | - |
| - | - | 2925 | 932.4 | - | - | 0 | - |
| - | - | 4498 | 933.4 | - | - | 0 | - |
| - | - | 2846 | 934.4 | - | - | 0 | - |
| - | - | 1678 | 935.4 | - | - | 0 | - |
| - | - | 2924 | 938.4 | - | - | 0 | - |
| 5 | y | 4637 | 939.4 | 0.001312 | 1.396 | +1 | 9 |
| 5 | y | 9008 | 940.4 | 0.007043 | 7.489 | +1 | 9 |
| 5 | z | 9249 | 941.4 | 0.01659 | 17.62 | +1 | 9 |
| - | - | 6810 | 942.4 | - | - | 0 | - |
| - | - | 2.186E+04 | 943.4 | - | - | 0 | - |
| - | - | 1.05E+04 | 944.4 | - | - | 0 | - |
| - | - | 6176 | 945.4 | - | - | 0 | - |
| - | - | 2396 | 946.4 | - | - | 0 | - |
| - | - | 637.2 | 947.4 | - | - | 0 | - |
| - | - | 1247 | 948.4 | - | - | 0 | - |
| - | - | 1011 | 949.4 | - | - | 0 | - |
| - | - | 3597 | 950.4 | - | - | 0 | - |
| - | - | 2521 | 951.5 | - | - | 0 | - |
| - | - | 1070 | 955.4 | - | - | 0 | - |
| - | - | 2.273E+04 | 956.4 | - | - | 0 | - |
| 5 | y | 1.191E+05 | 957.5 | 0.0001577 | 0.1647 | +1 | 9 |
| - | - | 5.801E+04 | 958.5 | - | - | 0 | - |
| - | - | 1.576E+04 | 959.5 | - | - | 0 | - |
| - | - | 4591 | 960.4 | - | - | 0 | - |
| - | - | 6927 | 961.4 | - | - | 0 | - |
| - | - | 3514 | 962.4 | - | - | 0 | - |
| - | - | 973.7 | 963.4 | - | - | 0 | - |
| - | - | 6215 | 974.5 | - | - | 0 | - |
| - | - | 2841 | 975.4 | - | - | 0 | - |
| 9 | c | 6.622E+04 | 975.5 | 0.0005633 | 0.5774 | +1 | 9 |
| - | - | 1343 | 976.4 | - | - | 0 | - |
| - | - | 3.406E+04 | 976.5 | - | - | 0 | - |
| - | - | 1743 | 977.4 | - | - | 0 | - |
| - | - | 8717 | 977.5 | - | - | 0 | - |
| - | - | 2190 | 978.4 | - | - | 0 | - |
| - | - | 2600 | 979.5 | - | - | 0 | - |
| - | - | 1302 | 980.5 | - | - | 0 | - |
| - | - | 750 | 981.5 | - | - | 0 | - |
| - | - | 1279 | 984.5 | - | - | 0 | - |
| - | - | 1519 | 985.4 | - | - | 0 | - |
| - | - | 792.7 | 986.5 | - | - | 0 | - |
| - | - | 1164 | 987.5 | - | - | 0 | - |
| - | - | 899 | 988.5 | - | - | 0 | - |
| - | - | 761.5 | 989.5 | - | - | 0 | - |
| - | - | 1110 | 990.5 | - | - | 0 | - |
| - | - | 1275 | 991.5 | - | - | 0 | - |
| - | - | 1043 | 992.5 | - | - | 0 | - |
| - | - | 1931 | 993.5 | - | - | 0 | - |
| - | - | 4894 | 994.5 | - | - | 0 | - |
| - | - | 1.278E+04 | 995.5 | - | - | 0 | - |
| - | - | 6306 | 996.5 | - | - | 0 | - |
| - | - | 2456 | 997.5 | - | - | 0 | - |
| - | - | 803 | 999.5 | - | - | 0 | - |
| - | - | 1976 | 1000 | - | - | 0 | - |
| - | - | 903.4 | 1001 | - | - | 0 | - |
| - | - | 1004 | 1003 | - | - | 0 | - |
| - | - | 1111 | 1006 | - | - | 0 | - |
| - | - | 2201 | 1008 | - | - | 0 | - |
| - | - | 2221 | 1009 | - | - | 0 | - |
| - | - | 6491 | 1010 | - | - | 0 | - |
| - | - | 1.248E+04 | 1011 | - | - | 0 | - |
| - | - | 7262 | 1012 | - | - | 0 | - |
| - | - | 3328 | 1013 | - | - | 0 | - |
| - | - | 1117 | 1014 | - | - | 0 | - |
| - | - | 3278 | 1015 | - | - | 0 | - |
| - | - | 2435 | 1016 | - | - | 0 | - |
| - | - | 1372 | 1017 | - | - | 0 | - |
| - | - | 638.3 | 1022 | - | - | 0 | - |
| - | - | 896.1 | 1025 | - | - | 0 | - |
| - | - | 3719 | 1026 | - | - | 0 | - |
| - | - | 3267 | 1027 | - | - | 0 | - |
| - | - | 6394 | 1028 | - | - | 0 | - |
| - | - | 5614 | 1029 | - | - | 0 | - |
| - | - | 2852 | 1030 | - | - | 0 | - |
| - | - | 932.2 | 1031 | - | - | 0 | - |
| - | - | 1891 | 1040 | - | - | 0 | - |
| - | - | 1232 | 1042 | - | - | 0 | - |
| - | - | 1694 | 1042 | - | - | 0 | - |
| - | - | 1216 | 1044 | - | - | 0 | - |
| - | - | 3020 | 1046 | - | - | 0 | - |
| - | - | 1947 | 1047 | - | - | 0 | - |
| - | - | 776.7 | 1048 | - | - | 0 | - |
| - | - | 835.6 | 1054 | - | - | 0 | - |
| - | - | 1919 | 1056 | - | - | 0 | - |
| - | - | 1.056E+04 | 1057 | - | - | 0 | - |
| - | - | 7145 | 1059 | - | - | 0 | - |
| - | - | 3305 | 1060 | - | - | 0 | - |
| - | - | 2778 | 1061 | - | - | 0 | - |
| - | - | 2864 | 1062 | - | - | 0 | - |
| - | - | 1699 | 1062 | - | - | 0 | - |
| - | - | 848.9 | 1063 | - | - | 0 | - |
| - | - | 1086 | 1064 | - | - | 0 | - |
| - | - | 879.6 | 1069 | - | - | 0 | - |
| - | - | 2790 | 1070 | - | - | 0 | - |
| 4 | z | 1549 | 1070 | 0.01852 | 17.3 | +1 | 10 |
| - | - | 1770 | 1072 | - | - | 0 | - |
| - | - | 3355 | 1073 | - | - | 0 | - |
| - | - | 3331 | 1075 | - | - | 0 | - |
| - | - | 2106 | 1076 | - | - | 0 | - |
| - | - | 963.1 | 1077 | - | - | 0 | - |
| - | - | 2377 | 1078 | - | - | 0 | - |
| - | - | 1411 | 1079 | - | - | 0 | - |
| - | - | 1516 | 1080 | - | - | 0 | - |
| - | - | 825.3 | 1081 | - | - | 0 | - |
| - | - | 940.3 | 1084 | - | - | 0 | - |
| - | - | 753.6 | 1085 | - | - | 0 | - |
| - | - | 1643 | 1086 | - | - | 0 | - |
| 4 | y | 2093 | 1087 | 0.008949 | 8.236 | +1 | 10 |
| - | - | 2.319E+04 | 1088 | - | - | 0 | - |
| 4 | z | 2.814E+04 | 1089 | 0.01174 | 10.78 | +1 | 10 |
| - | - | 1.737E+04 | 1090 | - | - | 0 | - |
| - | - | 6371 | 1091 | - | - | 0 | - |
| - | - | 6443 | 1092 | - | - | 0 | - |
| - | - | 3426 | 1093 | - | - | 0 | - |
| - | - | 3030 | 1094 | - | - | 0 | - |
| - | - | 2817 | 1095 | - | - | 0 | - |
| - | - | 1928 | 1096 | - | - | 0 | - |
| - | - | 841.6 | 1097 | - | - | 0 | - |
| - | - | 839.5 | 1098 | - | - | 0 | - |
| - | - | 1510 | 1102 | - | - | 0 | - |
| - | - | 8445 | 1104 | - | - | 0 | - |
| 10 | c | 7.872E+04 | 1105 | 0.01151 | 10.42 | +1 | 10 |
| - | - | 4.64E+04 | 1106 | - | - | 0 | - |
| - | - | 1.555E+04 | 1107 | - | - | 0 | - |
| - | - | 3047 | 1108 | - | - | 0 | - |
| - | - | 2360 | 1109 | - | - | 0 | - |
| - | - | 4290 | 1110 | - | - | 0 | - |
| - | - | 4274 | 1111 | - | - | 0 | - |
| - | - | 2158 | 1112 | - | - | 0 | - |
| - | - | 822.3 | 1113 | - | - | 0 | - |
| - | - | 898.7 | 1114 | - | - | 0 | - |
| - | - | 1139 | 1117 | - | - | 0 | - |
| - | - | 663.4 | 1118 | - | - | 0 | - |
| - | - | 1452 | 1124 | - | - | 0 | - |
| - | - | 913.7 | 1125 | - | - | 0 | - |
| - | - | 2492 | 1126 | - | - | 0 | - |
| - | - | 3207 | 1127 | - | - | 0 | - |
| - | - | 3136 | 1128 | - | - | 0 | - |
| - | - | 1821 | 1129 | - | - | 0 | - |
| - | - | 1686 | 1130 | - | - | 0 | - |
| - | - | 1739 | 1131 | - | - | 0 | - |
| - | - | 929.6 | 1132 | - | - | 0 | - |
| - | - | 1217 | 1140 | - | - | 0 | - |
| - | - | 1394 | 1141 | - | - | 0 | - |
| - | - | 3323 | 1142 | - | - | 0 | - |
| - | - | 1769 | 1143 | - | - | 0 | - |
| - | - | 2413 | 1144 | - | - | 0 | - |
| - | - | 1964 | 1145 | - | - | 0 | - |
| - | - | 910.7 | 1146 | - | - | 0 | - |
| - | - | 2021 | 1149 | - | - | 0 | - |
| - | - | 1721 | 1150 | - | - | 0 | - |
| - | - | 3573 | 1152 | - | - | 0 | - |
| - | - | 3794 | 1153 | - | - | 0 | - |
| - | - | 2843 | 1154 | - | - | 0 | - |
| - | - | 985.2 | 1155 | - | - | 0 | - |
| - | - | 1043 | 1156 | - | - | 0 | - |
| - | - | 1610 | 1157 | - | - | 0 | - |
| - | - | 1823 | 1158 | - | - | 0 | - |
| 3 | w | 2256 | 1159 | 0.008519 | 7.353 | +1 | 11 |
| - | - | 1453 | 1160 | - | - | 0 | - |
| - | - | 1922 | 1164 | - | - | 0 | - |
| - | - | 1206 | 1165 | - | - | 0 | - |
| - | - | 1058 | 1166 | - | - | 0 | - |
| - | - | 1.178E+04 | 1168 | - | - | 0 | - |
| - | - | 1.607E+04 | 1169 | - | - | 0 | - |
| - | - | 3.432E+04 | 1170 | - | - | 0 | - |
| - | - | 2.335E+04 | 1171 | - | - | 0 | - |
| - | - | 7671 | 1172 | - | - | 0 | - |
| - | - | 1751 | 1173 | - | - | 0 | - |
| - | - | 6201 | 1174 | - | - | 0 | - |
| - | - | 3721 | 1175 | - | - | 0 | - |
| - | - | 1909 | 1176 | - | - | 0 | - |
| - | - | 3686 | 1183 | - | - | 0 | - |
| - | - | 1202 | 1183 | - | - | 0 | - |
| - | - | 1834 | 1184 | - | - | 0 | - |
| - | - | 1702 | 1184 | - | - | 0 | - |
| - | - | 1974 | 1185 | - | - | 0 | - |
| - | - | 9191 | 1186 | - | - | 0 | - |
| - | - | 7453 | 1187 | - | - | 0 | - |
| - | - | 3555 | 1188 | - | - | 0 | - |
| - | - | 1773 | 1189 | - | - | 0 | - |
| - | - | 1195 | 1191 | - | - | 0 | - |
| - | - | 879.8 | 1195 | - | - | 0 | - |
| - | - | 1.489E+04 | 1201 | - | - | 0 | - |
| 3 | z | 1.853E+04 | 1202 | 0.01044 | 8.686 | +1 | 11 |
| - | - | 1.26E+04 | 1203 | - | - | 0 | - |
| - | - | 4421 | 1204 | - | - | 0 | - |
| - | - | 2847 | 1204 | - | - | 0 | - |
| - | - | 7717 | 1205 | - | - | 0 | - |
| - | - | 4479 | 1206 | - | - | 0 | - |
| - | - | 2975 | 1207 | - | - | 0 | - |
| - | - | 5685 | 1208 | - | - | 0 | - |
| - | - | 3887 | 1209 | - | - | 0 | - |
| - | - | 1451 | 1210 | - | - | 0 | - |
| - | - | 842 | 1216 | - | - | 0 | - |
| - | - | 2035 | 1217 | - | - | 0 | - |
| 11 | c | 4.94E+04 | 1218 | 0.003657 | 3.004 | +1 | 11 |
| - | - | 3.279E+04 | 1219 | - | - | 0 | - |
| - | - | 1.214E+04 | 1220 | - | - | 0 | - |
| - | - | 2215 | 1221 | - | - | 0 | - |
| - | - | 2123 | 1222 | - | - | 0 | - |
| - | - | 2747 | 1223 | - | - | 0 | - |
| - | - | 8821 | 1224 | - | - | 0 | - |
| - | - | 3584 | 1225 | - | - | 0 | - |
| - | - | 2150 | 1226 | - | - | 0 | - |
| - | - | 1542 | 1231 | - | - | 0 | - |
| - | - | 1021 | 1232 | - | - | 0 | - |
| - | - | 910.4 | 1233 | - | - | 0 | - |
| - | - | 812 | 1234 | - | - | 0 | - |
| - | - | 2100 | 1239 | - | - | 0 | - |
| - | - | 3784 | 1240 | - | - | 0 | - |
| - | - | 4202 | 1241 | - | - | 0 | - |
| - | - | 3333 | 1242 | - | - | 0 | - |
| - | - | 1059 | 1243 | - | - | 0 | - |
| - | - | 3567 | 1244 | - | - | 0 | - |
| - | - | 1464 | 1245 | - | - | 0 | - |
| - | - | 842.7 | 1246 | - | - | 0 | - |
| - | - | 2823 | 1247 | - | - | 0 | - |
| - | - | 1978 | 1248 | - | - | 0 | - |
| - | - | 916.1 | 1249 | - | - | 0 | - |
| - | - | 895.8 | 1253 | - | - | 0 | - |
| - | - | 914 | 1255 | - | - | 0 | - |
| - | - | 2393 | 1256 | - | - | 0 | - |
| - | - | 1786 | 1257 | - | - | 0 | - |
| - | - | 1219 | 1258 | - | - | 0 | - |
| - | - | 8437 | 1261 | - | - | 0 | - |
| - | - | 7627 | 1262 | - | - | 0 | - |
| - | - | 4516 | 1263 | - | - | 0 | - |
| - | - | 1641 | 1264 | - | - | 0 | - |
| - | - | 1793 | 1267 | - | - | 0 | - |
| - | - | 1246 | 1268 | - | - | 0 | - |
| - | - | 1047 | 1269 | - | - | 0 | - |
| - | - | 3505 | 1274 | - | - | 0 | - |
| - | - | 8663 | 1275 | - | - | 0 | - |
| - | - | 9914 | 1276 | - | - | 0 | - |
| - | - | 2.308E+04 | 1277 | - | - | 0 | - |
| - | - | 1.56E+04 | 1278 | - | - | 0 | - |
| - | - | 8598 | 1279 | - | - | 0 | - |
| - | - | 4472 | 1280 | - | - | 0 | - |
| - | - | 1680 | 1281 | - | - | 0 | - |
| - | - | 734.5 | 1282 | - | - | 0 | - |
| - | - | 3557 | 1283 | - | - | 0 | - |
| - | - | 5531 | 1284 | - | - | 0 | - |
| 2 | z | 2.288E+04 | 1285 | 0.003728 | 2.902 | +1 | 12 |
| 2 | z | 1.519E+04 | 1286 | 0.01787 | 13.9 | +1 | 12 |
| - | - | 5822 | 1287 | - | - | 0 | - |
| 2 | w | 1288 | 1288 | 0.009182 | 7.131 | +1 | 12 |
| - | - | 1094 | 1290 | - | - | 0 | - |
| - | - | 8101 | 1291 | - | - | 0 | - |
| - | - | 1.083E+04 | 1292 | - | - | 0 | - |
| - | - | 1.673E+04 | 1293 | - | - | 0 | - |
| - | - | 1.339E+04 | 1294 | - | - | 0 | - |
| - | - | 7733 | 1295 | - | - | 0 | - |
| - | - | 2836 | 1296 | - | - | 0 | - |
| - | - | 2303 | 1297 | - | - | 0 | - |
| - | - | 941.6 | 1298 | - | - | 0 | - |
| - | - | 2824 | 1300 | - | - | 0 | - |
| - | - | 8533 | 1301 | - | - | 0 | - |
| 2 | y | 6437 | 1302 | 0.004397 | 3.378 | +1 | 12 |
| 2 | z | 2.385E+04 | 1303 | 0.0008441 | 0.648 | +1 | 12 |
| - | - | 1.78E+04 | 1304 | - | - | 0 | - |
| - | - | 6639 | 1305 | - | - | 0 | - |
| - | - | 1906 | 1306 | - | - | 0 | - |
| - | - | 851.2 | 1307 | - | - | 0 | - |
| - | - | 5965 | 1308 | - | - | 0 | - |
| - | - | 1.914E+04 | 1309 | - | - | 0 | - |
| - | - | 2.29E+04 | 1310 | - | - | 0 | - |
| - | - | 6.753E+04 | 1311 | - | - | 0 | - |
| - | - | 4.876E+04 | 1312 | - | - | 0 | - |
| - | - | 1.965E+04 | 1313 | - | - | 0 | - |
| - | - | 5206 | 1314 | - | - | 0 | - |
| - | - | 1656 | 1315 | - | - | 0 | - |
| - | - | 3676 | 1316 | - | - | 0 | - |
| - | - | 3270 | 1317 | - | - | 0 | - |
| - | - | 1.53E+04 | 1318 | - | - | 0 | - |
| - | - | 1.097E+04 | 1319 | - | - | 0 | - |
| - | - | 5399 | 1320 | - | - | 0 | - |
| - | - | 1559 | 1321 | - | - | 0 | - |
| - | - | 2622 | 1323 | - | - | 0 | - |
| - | - | 8961 | 1324 | - | - | 0 | - |
| - | - | 1.356E+04 | 1325 | - | - | 0 | - |
| - | - | 2.878E+04 | 1326 | - | - | 0 | - |
| - | - | 1.958E+04 | 1327 | - | - | 0 | - |
| - | - | 1.44E+04 | 1328 | - | - | 0 | - |
| 12 | c | 8582 | 1329 | 0.006023 | 4.533 | +1 | 12 |
| - | - | 2495 | 1330 | - | - | 0 | - |
| - | - | 3623 | 1331 | - | - | 0 | - |
| - | - | 3236 | 1332 | - | - | 0 | - |
| - | - | 754 | 1333 | - | - | 0 | - |
| - | - | 946.9 | 1336 | - | - | 0 | - |
| - | - | 1386 | 1337 | - | - | 0 | - |
| - | - | 1647 | 1338 | - | - | 0 | - |
| - | - | 4516 | 1339 | - | - | 0 | - |
| - | - | 6925 | 1340 | - | - | 0 | - |
| - | - | 7589 | 1341 | - | - | 0 | - |
| - | - | 1.994E+04 | 1342 | - | - | 0 | - |
| - | - | 1.648E+04 | 1343 | - | - | 0 | - |
| - | - | 8990 | 1344 | - | - | 0 | - |
| - | - | 4790 | 1345 | - | - | 0 | - |
| 12 | c | 1.44E+05 | 1346 | 0.001932 | 1.436 | +1 | 12 |
| - | - | 1.114E+05 | 1347 | - | - | 0 | - |
| - | - | 5.015E+04 | 1348 | - | - | 0 | - |
| - | - | 1.077E+04 | 1349 | - | - | 0 | - |
| - | - | 1117 | 1351 | - | - | 0 | - |
| - | - | 1292 | 1352 | - | - | 0 | - |
| - | - | 1578 | 1353 | - | - | 0 | - |
| - | - | 3060 | 1354 | - | - | 0 | - |
| - | - | 6273 | 1355 | - | - | 0 | - |
| - | - | 5912 | 1356 | - | - | 0 | - |
| - | - | 1.009E+04 | 1357 | - | - | 0 | - |
| - | - | 1.803E+04 | 1358 | - | - | 0 | - |
| - | - | 1.45E+04 | 1359 | - | - | 0 | - |
| - | - | 4.913E+04 | 1360 | - | - | 0 | - |
| - | - | 3.761E+04 | 1361 | - | - | 0 | - |
| - | - | 1.767E+04 | 1362 | - | - | 0 | - |
| - | - | 8055 | 1363 | - | - | 0 | - |
| - | - | 1727 | 1364 | - | - | 0 | - |
| - | - | 3130 | 1365 | - | - | 0 | - |
| - | - | 4523 | 1366 | - | - | 0 | - |
| - | - | 9861 | 1367 | - | - | 0 | - |
| - | - | 3.502E+04 | 1368 | - | - | 0 | - |
| - | - | 2.765E+04 | 1369 | - | - | 0 | - |
| - | - | 1.486E+04 | 1370 | - | - | 0 | - |
| - | - | 7229 | 1371 | - | - | 0 | - |
| - | - | 3870 | 1372 | - | - | 0 | - |
| - | - | 6571 | 1373 | - | - | 0 | - |
| - | - | 1.733E+04 | 1374 | - | - | 0 | - |
| - | - | 1.434E+04 | 1375 | - | - | 0 | - |
| - | - | 6538 | 1376 | - | - | 0 | - |
| - | - | 837.3 | 1377 | - | - | 0 | - |
| - | - | 2280 | 1381 | - | - | 0 | - |
| - | - | 1.293E+04 | 1382 | - | - | 0 | - |
| - | - | 8.865E+04 | 1383 | - | - | 0 | - |
| - | - | 1.385E+05 | 1384 | - | - | 0 | - |
| - | - | 1.042E+05 | 1385 | - | - | 0 | - |
| - | - | 8.317E+04 | 1386 | - | - | 0 | - |
| - | - | 4.207E+04 | 1387 | - | - | 0 | - |
| - | - | 1.746E+04 | 1388 | - | - | 0 | - |
| - | - | 8210 | 1389 | - | - | 0 | - |
| - | - | 1918 | 1390 | - | - | 0 | - |
| - | - | 6492 | 1391 | - | - | 0 | - |
| - | - | 4500 | 1392 | - | - | 0 | - |
| - | - | 1740 | 1393 | - | - | 0 | - |
| - | - | 961.8 | 1396 | - | - | 0 | - |
| - | - | 4622 | 1398 | - | - | 0 | - |
| - | - | 3.472E+04 | 1399 | - | - | 0 | - |
| - | - | 5.123E+04 | 1400 | - | - | 0 | - |
| - | - | 1.406E+05 | 1401 | - | - | 0 | - |
| - | - | 2.021E+05 | 1402 | - | - | 0 | - |
| - | - | 1.272E+05 | 1403 | - | - | 0 | - |
| - | - | 5.008E+04 | 1404 | - | - | 0 | - |
| - | - | 1.137E+04 | 1405 | - | - | 0 | - |
| - | - | 2604 | 1415 | - | - | 0 | - |
| - | - | 2.614E+04 | 1416 | - | - | 0 | - |
| - | - | 4.834E+04 | 1417 | - | - | 0 | - |
| - | - | 7.814E+04 | 1418 | - | - | 0 | - |
| - | - | 1.814E+05 | 1419 | - | - | 0 | - |
| - | - | 1.235E+05 | 1420 | - | - | 0 | - |
| - | - | 5.003E+04 | 1421 | - | - | 0 | - |
| - | - | 1.19E+04 | 1422 | - | - | 0 | - |
| - | - | 4536 | 1422 | - | - | 0 | - |
| - | - | 897.4 | 2125 | - | - | 0 | - |
| - | - | 745 | 3027 | - | - | 0 | - |

m/z Charge Intensity FragmentType MassShift Position
120.08098602294922 0 3366.4912
129.0660400390625 0 395.83612
129.1024932861328 0 4246.5103
131.1183624267578 0 1313.9191
132.13699340820312 0 369.7276
133.0865020751953 0 581.0547
134.02732849121094 0 4163.204
136.0226593017578 0 443.39706
136.07615661621094 0 879.684
148.88502502441406 0 450.29266
148.89157104492188 0 454.58362
148.89892578125 0 445.7497
148.9059295654297 0 574.1895
148.91299438476562 0 689.1992
148.9206085205078 0 1281.4055
148.9276123046875 0 1226.9752
148.93478393554688 0 1995.9474
148.94264221191406 0 3279.265
148.95921325683594 0 3649.6218
148.96701049804688 0 2248.6077
148.97378540039062 0 1227.926
148.99639892578125 0 948.4372
149.0037078857422 0 547.54865
149.01101684570312 0 531.40094
161.09085083007812 0 425.8567
163.52561950683594 0 538.08875
167.11801147460938 0 1782.8988
169.1337890625 0 3130.2114
169.140625 0 536.0042
173.1285400390625 0 1677.6766
173.44993591308594 0 914.37616
173.63111877441406 0 451.84195
177.11244201660156 0 763.5364
183.11300659179688 0 1978.6531
197.0915985107422 0 551.75726
197.12852478027344 0 1128.3047
198.3815155029297 0 459.5698
199.10784912109375 0 2518.4355
200.10308837890625 0 10691.113
200.11317443847656 0 1053.7725
201.08717346191406 0 10934.492 y Ammonia loss 11
201.10633850097656 0 698.01324
201.12355041503906 0 5866.6567
202.090576171875 0 1294.0562
202.12673950195312 0 474.75635
209.09237670898438 0 1137.1892
213.12355041503906 0 11233.477
214.12705993652344 0 831.6275
215.13931274414062 0 287002.9
216.102783203125 0 641.8784
216.142578125 0 29433.38
217.14434814453125 0 2231.8035
218.11380004882812 0 24868.95 y 11
219.11715698242188 0 2455.9033
225.08714294433594 0 1841.7277
225.12380981445312 0 1125.2145
226.0951385498047 0 2263.1672
227.1029052734375 0 12406.418
228.10601806640625 0 1715.8667
228.17088317871094 0 3169.0876
229.11863708496094 0 1427.2052
231.1341552734375 0 1643.472
233.16505432128906 0 2987.4985
242.1502227783203 0 7938.4946
243.11276245117188 0 647.5645
243.15240478515625 0 818.7686
245.11428833007812 0 963.99725
245.1288299560547 0 787.517
247.1113739013672 0 12604.231
247.12879943847656 0 2880.6797
247.14303588867188 0 908.8477
248.11427307128906 0 1086.4971
251.17578125 0 832.55597
259.09356689453125 0 628.8805
259.107177734375 0 832.1358
261.1592102050781 0 2059.2705
272.54736328125 0 556.2077
275.106201171875 0 16131.78
275.12310791015625 0 788.0283
279.0987854003906 0 716.87256
280.1663818359375 0 864.4197
282.14501953125 0 1897.2186
295.1442565917969 0 945.8843 y 8
296.197265625 0 44350.18
297.20050048828125 0 6813.6523
298.1767578125 0 1191.5834
298.2027893066406 0 661.4242
299.1712341308594 0 933.2024
303.1700744628906 0 1295.2139
314.1716613769531 0 2363.9873 y Ammonia loss 10
314.20794677734375 0 18627.29
315.2113342285156 0 3570.0432
324.1559143066406 0 2911.3962
325.1189270019531 0 1248.7025
326.171875 0 3027.3904
330.1812744140625 0 1680.199
330.61767578125 0 532.4736
331.1979675292969 0 4570.731 y 10
332.2017517089844 0 653.46106
343.1976623535156 0 928.36774
344.18231201171875 0 2339.5947
347.18084716796875 0 629.30475
348.1914367675781 0 1411.8585
353.1824035644531 0 1223.7748
360.2237548828125 0 633.3283
362.2078857421875 0 4896.551
363.2104187011719 0 937.7726
369.17578125 0 1142.6405
371.1941223144531 0 1306.3922
373.138427734375 0 673.56335
374.17401123046875 0 8355.329 z 6
375.17718505859375 0 1820.0055
388.14996337890625 0 1116.7759
391.2018127441406 0 1048.9619
393.2503967285156 0 704.1258
395.2287902832031 0 763.3162
398.24249267578125 0 762.23895
407.19232177734375 0 1533.0349
411.2605895996094 0 1517.3768
415.27117919921875 0 650.8182
416.2547607421875 0 2281.2886
423.2245788574219 0 6792.7856
424.2252197265625 0 1650.379
427.3031921386719 0 9101.55
428.3067932128906 0 2063.1682
433.2815246582031 0 25918.611
434.2845458984375 0 7482.1045
434.7069396972656 0 744.2628
435.2897644042969 0 655.7287
439.2193603515625 0 2242.371
440.22088623046875 0 899.30164
441.2350158691406 0 23484.723
442.2380676269531 0 6874.6045 y Water loss 9
443.2357482910156 0 1198.8644
443.26617431640625 0 2387.5679
443.2978515625 0 917.1717
451.2203674316406 0 1115.515
455.2142333984375 0 1423.7986
456.2169494628906 0 716.1465
457.2231750488281 0 599.9996
459.2095642089844 0 1141.6288
459.29400634765625 0 665.1332
460.2420654296875 0 761.5064 y 9
461.27655029296875 0 26553.916
462.2796936035156 0 7645.0503
463.284423828125 0 627.4422
470.2254333496094 0 1770.506 y Water loss 4
476.29107666015625 0 1904.3615
477.29058837890625 0 768.41125
479.21429443359375 0 6226.9023
479.748291015625 0 735.52637
480.2144470214844 0 1804.4285
482.22418212890625 0 628.4661
492.2861633300781 0 2270.1548
498.21990966796875 0 3041.9675
499.2254943847656 0 893.8458
502.27020263671875 0 1793.4926
505.73504638671875 0 880.17126
506.74658203125 0 862.0015
507.2443542480469 0 656.62915
518.2662353515625 0 1670.7784
519.29638671875 0 13255.045
519.7343139648438 0 1216.8418
520.2369995117188 0 1360.5208
520.282958984375 0 6961.2725
520.7400512695312 0 2540.9907
521.2846069335938 0 2794.638
529.24072265625 0 682.6006
530.208251953125 0 1407.1614
530.2730102539062 0 785.031
530.33349609375 0 2266.6218
530.7726440429688 0 758.8259
540.3199462890625 0 1518.153
542.3307495117188 0 1095.517
544.26904296875 0 2307.1816
544.7717895507812 0 2591.5024
546.243896484375 0 732.6891
554.2620849609375 0 1522.0999
555.2630004882812 0 643.95935 z Water loss 8
558.328857421875 0 4596.5
559.3314208984375 0 1717.255
568.2427978515625 0 1759.3628
569.2451171875 0 793.38104
570.2567138671875 0 5702.2065
571.2627563476562 0 2628.6958 y Water loss 8
584.75927734375 0 1849.2931
585.2626342773438 0 1566.8391
585.7603149414062 0 1143.781
586.25146484375 0 2840.862
587.2642211914062 0 2183.3494
588.3035278320312 0 5079.0225
589.3017578125 0 3483.6665
591.3156127929688 0 2134.2173
599.28271484375 0 2455.9778
600.2844848632812 0 682.5473
600.810546875 0 1938.236
601.3138427734375 0 1394.2733
602.2845458984375 0 688.0536
603.2894897460938 0 713.61523
604.2999877929688 0 1312.1416
605.2992553710938 0 1252.1355
607.3109741210938 0 2429.1775
609.2520751953125 0 1165.9602
617.2965698242188 0 12360.311
618.299072265625 0 3680.7942
619.2969970703125 0 1722.0746
627.2625732421875 0 16132.141
628.265625 0 5277.547
628.3939819335938 0 1603.1832
629.2674560546875 0 1336.4427
629.3978881835938 0 1018.93494
633.317626953125 0 809.7184
634.3236694335938 0 14947.215
635.3154296875 0 9432.307
636.3138427734375 0 3579.8167
643.2896118164062 0 1733.7234 w 7
643.3464965820312 0 1069.6935
644.2900390625 0 1382.262
645.2893676757812 0 734.0763
655.3834838867188 0 1481.5801
656.3867797851562 0 692.8478
658.2806396484375 0 1656.5477
659.285400390625 0 1119.6327 y Ammonia loss 7
663.4175415039062 0 1874.4448
667.3107299804688 0 1103.8977
669.3223266601562 0 2279.2715
670.3286743164062 0 939.01935
671.400390625 0 3676.6362
672.40771484375 0 3970.1765 c 5
673.414794921875 0 1058.6792
674.2977294921875 0 980.3709
676.3157348632812 0 2496.202 y 7
677.3182373046875 0 756.9861
682.3112182617188 0 636.29443
685.3195190429688 0 800.02454
686.321044921875 0 1242.5012
690.3336791992188 0 959.4001
691.3222045898438 0 3943.9934
691.8224487304688 0 2296.508
692.3233642578125 0 930.8161
692.8256225585938 0 821.66187
697.4158325195312 0 718.0193
705.3931884765625 0 778.76013
709.2882080078125 0 1303.6833
710.7923583984375 0 774.9211
712.3553466796875 0 721.0298
715.4217529296875 0 1095.3492
716.3642578125 0 951.2868
716.4338989257812 0 6663.882
717.4373168945312 0 3072.8567
720.3411865234375 0 801.31696
722.3348388671875 0 1076.9762
723.3417358398438 0 712.6185
724.40771484375 0 1195.8789
728.366455078125 0 1514.949
740.34619140625 0 13615.393
740.4185180664062 0 1523.9952
741.3489379882812 0 4972.045
741.4200439453125 0 951.13226 c Water loss 6
742.3573608398438 0 718.21814
742.413818359375 0 3491.4338
743.4176635742188 0 1870.7482
744.424560546875 0 2584.956
745.31201171875 0 1153.9602
745.4332885742188 0 743.8075
747.3320922851562 0 728.0583 z 6
750.3452758789062 0 760.02954
752.3650512695312 0 5120.7256
753.3685302734375 0 2859.6504
754.37451171875 0 703.98816
757.373779296875 0 3678.5027
758.4326171875 0 16157.472
759.4387817382812 0 20490.635 c 6
760.442138671875 0 7066.8413
761.4448852539062 0 1695.9727
762.3462524414062 0 1253.9395
763.3488159179688 0 2646.8494 y 6
764.3567504882812 0 1132.9803
766.3485717773438 0 977.2259
767.3565673828125 0 1150.1316
768.3604736328125 0 2970.7493
769.3630981445312 0 2135.7637
770.3667602539062 0 1042.7938
771.3544311523438 0 3992.555
772.3524780273438 0 1289.1691
774.3309936523438 0 1146.2985
779.356201171875 0 894.7911
780.3599853515625 0 11308.884
781.3637084960938 0 6068.3496
782.3651123046875 0 2273.968
784.3593139648438 0 1176.1168
785.3681030273438 0 1676.6025
786.3695678710938 0 890.8282
794.336669921875 0 4436.814
795.3433837890625 0 2837.4062
796.3538818359375 0 10857.414
797.3837280273438 0 16279.962
798.385986328125 0 7353.7437
799.3848266601562 0 2259.0222
803.4671020507812 0 2207.3125
804.471923828125 0 928.9823
812.3546752929688 0 5111.231
813.372314453125 0 4524.2456
814.3713989257812 0 2768.645
815.3799438476562 0 1576.2091
820.3781127929688 0 1778.3544
825.3744506835938 0 940.3207
827.456298828125 0 909.4249
828.3524169921875 0 668.75146
829.3699951171875 0 1394.8475
829.4490356445312 0 1615.3308
830.3697509765625 0 1777.857
830.4530029296875 0 1292.2855
836.369384765625 0 934.63763
837.3754272460938 0 826.88776
838.3843994140625 0 725.14557
839.4111328125 0 1798.666
840.4124145507812 0 2345.0789
842.382568359375 0 727.45996 y Water loss 5
843.3900146484375 0 5505.779
844.3849487304688 0 1939.5739 z 5
845.3768920898438 0 982.8505
845.4647827148438 0 11132.283
846.4718627929688 0 55672.03 c 7
847.4752807617188 0 25241.54
848.4788208007812 0 5216.155
850.3927001953125 0 1134.6406
851.3950805664062 0 638.16174
852.3882446289062 0 951.8406
859.4089965820312 0 824.1625
860.4004516601562 0 11307.785 y 5
861.4041137695312 0 6275.7275
862.4116821289062 0 938.9687
867.3960571289062 0 1020.71893
868.4052734375 0 25173.408
869.4085083007812 0 10854.6045
870.4117431640625 0 3359.75
877.4174194335938 0 885.7718
881.387939453125 0 1132.7406
883.4012451171875 0 1851.9469
884.4149780273438 0 768.4967
885.4176635742188 0 744.8836
887.4198608398438 0 3713.2983
888.4232788085938 0 2071.921
889.4244384765625 0 956.36206
891.3993530273438 0 768.437
893.4032592773438 0 1871.2253
894.4038696289062 0 1145.842
895.419921875 0 3540.9275
896.4194946289062 0 1085.6368
897.4273681640625 0 950.46814
899.4209594726562 0 879.4648
900.434814453125 0 2171.028
901.4420776367188 0 1156.1543
902.4368286132812 0 1004.4894
903.4263305664062 0 1176.1262
904.4255981445312 0 1109.666
911.4149780273438 0 738.04236
912.4077758789062 0 842.85815
913.4127807617188 0 1102.681
914.4109497070312 0 1264.2555 w 4
915.4264526367188 0 5736.6567
916.4331665039062 0 5231.2334
917.4375610351562 0 2768.1091
918.4461059570312 0 2478.9976
919.4451293945312 0 826.098
922.4334106445312 0 957.5664
923.4078979492188 0 1830.5406 z Water loss 4
925.408203125 0 2778.6897
926.4193115234375 0 2086.6663
927.4251708984375 0 3478.8872
928.4270629882812 0 2244.5537
929.426513671875 0 764.2761
931.502197265625 0 2043.1824
932.4280395507812 0 2924.5884
933.4361572265625 0 4498.2188
934.4467163085938 0 2845.6643
935.4466552734375 0 1677.6969
938.4317626953125 0 2923.8123
939.4404907226562 0 4636.734 y Water loss 4
940.432861328125 0 9008.44 y Ammonia loss 4
941.4170532226562 0 9249.28 z 4
942.4152221679688 0 6809.8716
943.4223022460938 0 21858.152
944.4263916015625 0 10502.8
945.430908203125 0 6176.3076
946.4443359375 0 2395.8042
947.4287719726562 0 637.19666
948.4257202148438 0 1247.0948
949.4281005859375 0 1011.4249
950.4443359375 0 3596.9475
951.451416015625 0 2521.055
955.43115234375 0 1070.4017
956.4445190429688 0 22731.947
957.4522094726562 0 119088.625 y 4
958.4564819335938 0 58008.24
959.4527587890625 0 15762.916
960.4308471679688 0 4591.392
961.4306640625 0 6926.747
962.4364624023438 0 3513.749
963.4422607421875 0 973.66846
974.5074462890625 0 6215.1025
975.4227294921875 0 2840.7485
975.51513671875 0 66222.21 c 8
976.4219360351562 0 1343.419
976.5181884765625 0 34058.31
977.4229125976562 0 1743.3107
977.5206909179688 0 8716.652
978.4359130859375 0 2189.5847
979.463623046875 0 2599.5642
980.4571533203125 0 1302.4749
981.4783935546875 0 749.99084
984.4561767578125 0 1278.9996
985.4489135742188 0 1518.9167
986.4994506835938 0 792.6829
987.4835815429688 0 1163.5317
988.468505859375 0 898.95264
989.4742431640625 0 761.51276
990.4691772460938 0 1109.9808
991.4585571289062 0 1274.8578
992.4768676757812 0 1042.795
993.45751953125 0 1930.5571
994.4649047851562 0 4893.809
995.4722290039062 0 12783.241
996.4771118164062 0 6306.108
997.4761352539062 0 2456.22
999.4554443359375 0 803.03064
1000.4945068359375 0 1975.7306
1001.4876708984375 0 903.44476
1002.5108642578125 0 1003.5767
1006.454833984375 0 1110.5096
1008.47021484375 0 2200.619
1009.4685668945312 0 2220.849
1010.4699096679688 0 6491.381
1011.4686889648438 0 12475.852
1012.4716186523438 0 7262.3716
1013.4776000976562 0 3328.1992
1014.4865112304688 0 1116.6066
1015.4767456054688 0 3277.6118
1016.47607421875 0 2435.419
1017.471435546875 0 1371.5892
1022.4578857421875 0 638.3472
1025.4442138671875 0 896.07837
1026.4664306640625 0 3719.15
1027.471923828125 0 3266.8264
1028.4749755859375 0 6394.1245
1029.47705078125 0 5614.324
1030.4796142578125 0 2852.4026
1031.48046875 0 932.2351
1040.4764404296875 0 1891.243
1041.503173828125 0 1232.1788
1042.4884033203125 0 1693.7843
1044.4940185546875 0 1215.5112
1046.475341796875 0 3019.9006
1047.4854736328125 0 1946.7938
1048.488037109375 0 776.7367
1054.4681396484375 0 835.56256
1056.491455078125 0 1918.9221
1057.4993896484375 0 10560.121
1058.5018310546875 0 7144.775
1059.50439453125 0 3304.6133
1060.5361328125 0 2777.614
1061.5477294921875 0 2863.786
1062.4664306640625 0 1698.851
1063.4766845703125 0 848.9439
1064.4847412109375 0 1086.0494
1068.5234375 0 879.6197
1069.52001953125 0 2790.2507
1070.510009765625 0 1548.7222 z Water loss 3
1072.4913330078125 0 1770.1796
1073.494384765625 0 3355.0994
1074.5009765625 0 3330.9165
1075.5023193359375 0 2106.2612
1076.504150390625 0 963.0519
1077.5235595703125 0 2376.6382
1078.5335693359375 0 1411.046
1079.54248046875 0 1516.4421
1081.4786376953125 0 825.3136
1083.53857421875 0 940.3431
1084.5191650390625 0 753.5825
1085.51611328125 0 1643.4385
1086.5191650390625 0 2093.0952 y Water loss 3
1087.5294189453125 0 23192.158
1088.5137939453125 0 28141.94 z 3
1089.509521484375 0 17369.898
1090.5074462890625 0 6371.399
1091.5072021484375 0 6443.376
1092.512451171875 0 3426.4546
1093.532470703125 0 3029.94
1094.5345458984375 0 2817.045
1095.54150390625 0 1928.1969
1096.5467529296875 0 841.57745
1097.53125 0 839.50165
1101.5537109375 0 1510.1505
1103.5191650390625 0 8444.865
1104.545654296875 0 78717.984 c 9
1105.54931640625 0 46396.64
1106.5531005859375 0 15546.058
1107.5452880859375 0 3046.5989
1108.53759765625 0 2360.343
1109.545166015625 0 4290.359
1110.53515625 0 4274.0874
1111.53369140625 0 2157.6821
1112.53759765625 0 822.3096
1113.5460205078125 0 898.70026
1116.5841064453125 0 1139.2578
1117.5728759765625 0 663.4434
1123.5135498046875 0 1452.2039
1124.5093994140625 0 913.7321
1125.531982421875 0 2491.9573
1126.5443115234375 0 3206.7412
1127.54541015625 0 3136.3647
1128.5401611328125 0 1820.8285
1129.5552978515625 0 1686.361
1130.550048828125 0 1739.1654
1131.5513916015625 0 929.649
1139.5203857421875 0 1217.417
1140.51220703125 0 1393.843
1141.5250244140625 0 3322.7595
1142.5281982421875 0 1768.762
1143.540283203125 0 2413.4072
1144.547119140625 0 1964.141
1145.5306396484375 0 910.71185
1149.4940185546875 0 2021.0931
1150.4991455078125 0 1720.579
1151.505859375 0 3573.399
1152.51416015625 0 3793.537
1153.51904296875 0 2842.8408
1154.5185546875 0 985.2161
1155.633056640625 0 1043.3561
1156.6162109375 0 1610.2578
1157.5106201171875 0 1822.9938
1158.5228271484375 0 2256.481 w 2
1159.5279541015625 0 1452.639
1163.5631103515625 0 1922.2354
1164.5797119140625 0 1205.5072
1165.5855712890625 0 1057.6417
1167.502685546875 0 11777.378
1168.509033203125 0 16066.695
1169.5172119140625 0 34321.348
1170.521240234375 0 23347.404
1171.5238037109375 0 7671.383
1172.5233154296875 0 1751.1141
1173.6265869140625 0 6200.7695
1174.6282958984375 0 3720.9744
1175.635009765625 0 1909.0582
1182.6051025390625 0 3686.4888
1183.4871826171875 0 1201.7405
1183.611328125 0 1833.7354
1184.49169921875 0 1702.1154
1184.6107177734375 0 1973.767
1185.5120849609375 0 9191.114
1186.51708984375 0 7453.085
1187.5223388671875 0 3554.5928
1188.52880859375 0 1773.425
1190.623779296875 0 1194.5264
1194.620849609375 0 879.79193
1200.6143798828125 0 14893.587
1201.5965576171875 0 18527.828 z 2
1202.5948486328125 0 12600.609
1203.521728515625 0 4421.234
1203.601318359375 0 2847.0608
1204.5308837890625 0 7716.579
1205.535400390625 0 4479.2915
1206.607421875 0 2974.7407
1207.6239013671875 0 5684.869
1208.6258544921875 0 3886.586
1209.632568359375 0 1450.7479
1215.5994873046875 0 841.98065
1216.61279296875 0 2035.4003
1217.6375732421875 0 49400.445 c 10
1218.64013671875 0 32789.918
1219.6429443359375 0 12137.246
1220.633056640625 0 2214.948
1221.61572265625 0 2122.9236
1222.6142578125 0 2747.362
1223.616943359375 0 8820.772
1224.62060546875 0 3583.973
1225.6158447265625 0 2150.0027
1230.5953369140625 0 1542.0244
1231.6016845703125 0 1020.8314
1232.605712890625 0 910.3914
1233.614990234375 0 812.02234
1238.5985107421875 0 2099.8735
1239.616455078125 0 3784.3816
1240.6260986328125 0 4201.5986
1241.62255859375 0 3333.079
1242.6190185546875 0 1059.2108
1243.6500244140625 0 3566.666
1244.671142578125 0 1464.3668
1245.685546875 0 842.6771
1246.5814208984375 0 2822.9526
1247.5826416015625 0 1977.7567
1248.63916015625 0 916.0865
1252.615966796875 0 895.84406
1254.589599609375 0 913.97723
1255.587158203125 0 2392.832
1256.5855712890625 0 1785.5929
1257.601318359375 0 1218.5808
1260.6070556640625 0 8436.684
1261.6123046875 0 7627.0933
1262.61279296875 0 4515.733
1263.6314697265625 0 1641.2964
1266.6280517578125 0 1792.6455
1267.6038818359375 0 1245.7699
1268.6307373046875 0 1046.6898
1273.6007080078125 0 3505.1
1274.620849609375 0 8662.874
1275.633056640625 0 9913.544
1276.629150390625 0 23082.406
1277.635009765625 0 15603.1875
1278.6290283203125 0 8597.553
1279.6285400390625 0 4471.921
1280.6365966796875 0 1679.6381
1281.656494140625 0 734.4947
1282.5997314453125 0 3556.678
1283.6031494140625 0 5531.0254
1284.6195068359375 0 22875.734 z Water loss 1
1285.6251220703125 0 15190.828 z Ammonia loss 1
1286.626708984375 0 5821.9893
1287.6195068359375 0 1288.3826 w 1
1289.634765625 0 1093.946
1290.6168212890625 0 8101.141
1291.6319580078125 0 10826.674
1292.638916015625 0 16734.932
1293.6473388671875 0 13388.8955
1294.6446533203125 0 7733.4
1295.6260986328125 0 2835.8062
1296.6265869140625 0 2303.0007
1297.6029052734375 0 941.5682
1299.599609375 0 2824.1438
1300.6141357421875 0 8532.986
1301.63037109375 0 6437.426 y Ammonia loss 1
1302.6346435546875 0 23853.266 z 1
1303.6373291015625 0 17802.562
1304.6414794921875 0 6638.9023
1305.646240234375 0 1906.2452
1306.6009521484375 0 851.2412
1307.637451171875 0 5965.1865
1308.6341552734375 0 19136.768
1309.6478271484375 0 22900.963
1310.6510009765625 0 67534.125
1311.6534423828125 0 48758.09
1312.654541015625 0 19645.012
1313.654052734375 0 5206.1445
1314.6380615234375 0 1655.9641
1315.5960693359375 0 3676.3418
1316.5985107421875 0 3270.0159
1317.606201171875 0 15299.944
1318.6142578125 0 10966.774
1319.6171875 0 5398.8877
1320.616943359375 0 1559.2922
1322.6300048828125 0 2621.893
1323.6331787109375 0 8960.761
1324.634521484375 0 13563.984
1325.6405029296875 0 28782.998
1326.645263671875 0 19581.25
1327.65576171875 0 14395.84
1328.667236328125 0 8581.619 c Ammonia loss 11
1329.6708984375 0 2495.3164
1330.6629638671875 0 3622.8262
1331.670654296875 0 3236.4558
1332.6728515625 0 754.0109
1335.64453125 0 946.9189
1336.62451171875 0 1385.9474
1337.6307373046875 0 1647.3843
1338.6427001953125 0 4515.9214
1339.639404296875 0 6924.591
1340.635498046875 0 7588.789
1341.6370849609375 0 19936.602
1342.6375732421875 0 16481.19
1343.6435546875 0 8990.096
1344.6502685546875 0 4790.468
1345.6978759765625 0 143991.66 c 11
1346.701416015625 0 111375.1
1347.7034912109375 0 50150.66
1348.7039794921875 0 10767.72
1350.6639404296875 0 1117.4581
1351.6400146484375 0 1292.3618
1352.6458740234375 0 1578.1982
1353.6444091796875 0 3059.7556
1354.6435546875 0 6273.114
1355.6529541015625 0 5912.447
1356.6522216796875 0 10093.666
1357.6383056640625 0 18032.3
1358.636962890625 0 14496.587
1359.64697265625 0 49131.6
1360.6507568359375 0 37607.496
1361.652099609375 0 17670.53
1362.6561279296875 0 8054.8916
1363.667236328125 0 1726.8838
1364.637939453125 0 3130.4546
1365.6353759765625 0 4522.657
1366.6416015625 0 9860.784
1367.6533203125 0 35021.457
1368.6546630859375 0 27648.207
1369.6553955078125 0 14864.63
1370.6531982421875 0 7228.6147
1371.64697265625 0 3870.291
1372.6729736328125 0 6570.966
1373.6922607421875 0 17330.37
1374.699462890625 0 14336.08
1375.6990966796875 0 6538.154
1376.6878662109375 0 837.2789
1380.63232421875 0 2280.3408
1381.634765625 0 12930.146
1382.64111328125 0 88650.09
1383.6485595703125 0 138530.53
1384.651123046875 0 104226.33
1385.6578369140625 0 83165.336
1386.6627197265625 0 42073.26
1387.6632080078125 0 17464.299
1388.66357421875 0 8209.883
1389.652099609375 0 1918.0502
1390.693603515625 0 6491.8047
1391.6917724609375 0 4499.7197
1392.6900634765625 0 1740.4116
1395.625244140625 0 961.84155
1397.6351318359375 0 4622.0986
1398.635498046875 0 34719.65
1399.64111328125 0 51229.402
1400.6505126953125 0 140641.67
1401.66015625 0 202060.66
1402.6640625 0 127195.55
1403.6644287109375 0 50083.016
1404.665771484375 0 11373.341
1414.6463623046875 0 2604.2205
1415.6400146484375 0 26138.818
1416.645751953125 0 48340.445
1417.669921875 0 78141.05
1418.7088623046875 0 181399.14
1419.71728515625 0 123518.2
1420.7200927734375 0 50030.9
1421.584716796875 0 11903.6
1421.7381591796875 0 4535.999
2125.0830078125 0 897.3904
3027.109619140625 0 744.99756

Spectrum Details

|  |  |
| --- | --- |
| Matched peaks? Matched peaks The total absolute number of peaks matched. Additionally in brackets the total fraction of peaks matched and the total number of peaks is shown. | 44 (5.97% of 737) |
| FDR? FDR The false discovery rate estimated for this peptide. It is calculated by matching all theoretical fragments with a non-integer shift with the raw peaks for this spectrum. This is done with 40 different shifts. The resulting percentage is the average number of annotated peaks over the number of annotated peaks with the correct spectrum. | 6.01% |
| Satellite FDR? Satellite FDR See the FDR for details on its calculation. This satellite ion specific FDR only contains the satellite ions (d/w) for I/L/J positions. | 7.14% |
| PSM Score? PSM Score The PSM Score as given by Hecklib to this annotated spectrum. It is shown with three significant figures. | 495 |

#### Spectrum 9903? Spectrum 9903 The raw spectrum of this peptide as annotated by Hecklib. The fragments are coloured according to ion type (see legend). Any peaks with a star '\*' as text can be hovered over to see the full details, first the ion type second the mass shift type. By hovering over the amino acids in the peptide or ions in the legend the corresponding peaks are highlighted. By toggling the 'Unassigned' label you can turn the background (unassigned) peaks on or off in the plot. By updating the slider in the Ion legend you can update the spectrum to only show the top X% of the peaks with labels. The top X% means any peak that is within X% of the highest intensity. By dragging in the spectrum you can zoom in to a specific part of the spectrum and use 'Zoom Out' to get back to the original zoom level. The annotation of the spectrum is based on the given sequence in the peptides file and is done with different software so inconsistencies are likely. The peaks are annotated based on the given sequence, with 20 ppm tolerance.

Copy Data

##### Spectrum 9903 (TSV)

###### Preview

```
Loading example...
```

*Click on the button to copy the data to your clipboard.*

Mz MinMz MaxIntensity Max

WidthHeightPeptide font sizePeptide stroke widthSpectrum font sizeSpectrum stroke widthCompact peptide

Ion legend

wxyz

abcd

OtherUnassignedIon^Charge^_Position_Show for top:%

VTJFPPSSEEJQA

06.43e+41.29e+51.93e+52.57e+5

Zoom Out

a^+1^_2_a^+1^_2_b^+1^_2_y^+1^_2_b^+1^_2_y^+1^_2_a^+1^_3_a^+1^_3_y^+2^_5_b^+1^_3_y^+1^_3_b^+1^_3_y^+1^_3_y^+2^_7_y^+1^_4_b^+1^_4_y^+1^_4_b^+1^_4_y^+2^_9_b^+2^_9_b^+2^_9_b^+2^_10_b^+1^_5_b^+2^_10_b^+1^_5_y^+1^_5_b^+2^_11_b^+1^_6_b^+1^_6_y^+1^_6_y^+1^_6_b^+1^_7_b^+1^_7_y^+1^_7_y^+1^_7_b^+1^_8_b^+1^_8_y^+1^_8_y^+1^_8_y^+1^_8_y^+1^_9_y^+1^_9_y^+1^_9_b^+1^_10_y^+1^_10_b^+1^_10_y^+1^_10_b^+1^_11_y^+1^_11_b^+1^_11_y^+1^_11_

0765152922943059

Fragment Matches Table

Show background peaks

| Position | Ion type | Intensity | mz Theoretical | mz Error (Th) | mz Error (ppm) | Charge | Series Number |
| --- | --- | --- | --- | --- | --- | --- | --- |
| - | - | 3.22E+04 | 120.1 | - | - | 0 | - |
| - | - | 367.9 | 120.3 | - | - | 0 | - |
| - | - | 2587 | 121.1 | - | - | 0 | - |
| - | - | 8051 | 126.1 | - | - | 0 | - |
| - | - | 431.4 | 127.1 | - | - | 0 | - |
| - | - | 462.7 | 127.1 | - | - | 0 | - |
| - | - | 699.5 | 127.1 | - | - | 0 | - |
| - | - | 8861 | 128.1 | - | - | 0 | - |
| - | - | 4680 | 129.1 | - | - | 0 | - |
| - | - | 734.1 | 129.1 | - | - | 0 | - |
| - | - | 1.289E+04 | 129.1 | - | - | 0 | - |
| - | - | 1017 | 129.1 | - | - | 0 | - |
| - | - | 3131 | 130.1 | - | - | 0 | - |
| - | - | 587.9 | 130.1 | - | - | 0 | - |
| - | - | 1650 | 130.1 | - | - | 0 | - |
| - | - | 904.2 | 130.1 | - | - | 0 | - |
| - | - | 393.9 | 130.8 | - | - | 0 | - |
| - | - | 657 | 131 | - | - | 0 | - |
| - | - | 9471 | 131.1 | - | - | 0 | - |
| - | - | 738.4 | 132.1 | - | - | 0 | - |
| - | - | 538.3 | 133.1 | - | - | 0 | - |
| - | - | 499 | 133.1 | - | - | 0 | - |
| - | - | 2775 | 134 | - | - | 0 | - |
| - | - | 4437 | 136.1 | - | - | 0 | - |
| - | - | 1778 | 138.1 | - | - | 0 | - |
| - | - | 7082 | 138.1 | - | - | 0 | - |
| - | - | 1732 | 139.1 | - | - | 0 | - |
| - | - | 370 | 139.1 | - | - | 0 | - |
| - | - | 702.4 | 139.1 | - | - | 0 | - |
| - | - | 758.2 | 141.1 | - | - | 0 | - |
| - | - | 827.1 | 141.1 | - | - | 0 | - |
| - | - | 524.7 | 143.1 | - | - | 0 | - |
| - | - | 774.4 | 143.1 | - | - | 0 | - |
| - | - | 436.9 | 144.1 | - | - | 0 | - |
| - | - | 473.2 | 145.1 | - | - | 0 | - |
| - | - | 693.6 | 147 | - | - | 0 | - |
| - | - | 3449 | 147.1 | - | - | 0 | - |
| - | - | 660.8 | 148.1 | - | - | 0 | - |
| - | - | 445.5 | 148.8 | - | - | 0 | - |
| - | - | 385.3 | 150.2 | - | - | 0 | - |
| - | - | 460.9 | 151.1 | - | - | 0 | - |
| - | - | 1125 | 152.1 | - | - | 0 | - |
| - | - | 782 | 153.1 | - | - | 0 | - |
| - | - | 584.2 | 153.1 | - | - | 0 | - |
| - | - | 384 | 153.6 | - | - | 0 | - |
| - | - | 3904 | 154.1 | - | - | 0 | - |
| - | - | 1794 | 154.1 | - | - | 0 | - |
| - | - | 8333 | 155.1 | - | - | 0 | - |
| 2 | a | 2630 | 155.1 | 0.0002898 | 1.868 | +1 | 2 |
| - | - | 1090 | 156.1 | - | - | 0 | - |
| - | - | 593.4 | 156.1 | - | - | 0 | - |
| - | - | 2621 | 157.1 | - | - | 0 | - |
| - | - | 7788 | 157.1 | - | - | 0 | - |
| - | - | 921 | 159.1 | - | - | 0 | - |
| - | - | 421.4 | 161.7 | - | - | 0 | - |
| - | - | 459.1 | 162.1 | - | - | 0 | - |
| - | - | 1353 | 165.1 | - | - | 0 | - |
| - | - | 1430 | 166.1 | - | - | 0 | - |
| - | - | 495.4 | 167 | - | - | 0 | - |
| - | - | 5851 | 167.1 | - | - | 0 | - |
| - | - | 7108 | 167.1 | - | - | 0 | - |
| - | - | 882.9 | 169.1 | - | - | 0 | - |
| - | - | 1.06E+04 | 169.1 | - | - | 0 | - |
| - | - | 998 | 170.1 | - | - | 0 | - |
| - | - | 1446 | 171.1 | - | - | 0 | - |
| - | - | 524.3 | 171.1 | - | - | 0 | - |
| - | - | 3331 | 172.1 | - | - | 0 | - |
| 2 | a | 2.442E+04 | 173.1 | 0.0002842 | 1.641 | +1 | 2 |
| - | - | 482 | 173.3 | - | - | 0 | - |
| - | - | 1012 | 174.1 | - | - | 0 | - |
| - | - | 1684 | 174.1 | - | - | 0 | - |
| - | - | 852.7 | 174.1 | - | - | 0 | - |
| - | - | 3088 | 175.1 | - | - | 0 | - |
| - | - | 517.2 | 178.1 | - | - | 0 | - |
| - | - | 534 | 180.1 | - | - | 0 | - |
| - | - | 499.9 | 181.1 | - | - | 0 | - |
| - | - | 3207 | 181.1 | - | - | 0 | - |
| - | - | 898.2 | 181.1 | - | - | 0 | - |
| - | - | 3853 | 183.1 | - | - | 0 | - |
| 2 | b | 8.226E+04 | 183.1 | 0.0003398 | 1.855 | +1 | 2 |
| - | - | 1621 | 183.1 | - | - | 0 | - |
| - | - | 658.1 | 184.1 | - | - | 0 | - |
| - | - | 7865 | 184.1 | - | - | 0 | - |
| - | - | 1.305E+04 | 185.1 | - | - | 0 | - |
| - | - | 534.9 | 185.1 | - | - | 0 | - |
| - | - | 479.9 | 185.2 | - | - | 0 | - |
| - | - | 802.3 | 186.1 | - | - | 0 | - |
| - | - | 1.353E+04 | 187.1 | - | - | 0 | - |
| - | - | 1092 | 188.1 | - | - | 0 | - |
| - | - | 6156 | 189.1 | - | - | 0 | - |
| - | - | 560.1 | 189.1 | - | - | 0 | - |
| - | - | 1616 | 195.1 | - | - | 0 | - |
| - | - | 1.346E+04 | 195.1 | - | - | 0 | - |
| - | - | 745.2 | 196.1 | - | - | 0 | - |
| - | - | 727.9 | 197.1 | - | - | 0 | - |
| - | - | 1.77E+04 | 197.1 | - | - | 0 | - |
| - | - | 1470 | 198.1 | - | - | 0 | - |
| - | - | 6477 | 199.1 | - | - | 0 | - |
| - | - | 4984 | 199.1 | - | - | 0 | - |
| - | - | 1.033E+05 | 200.1 | - | - | 0 | - |
| 12 | y | 8.791E+04 | 201.1 | 0.0003275 | 1.628 | +1 | 2 |
| - | - | 8125 | 201.1 | - | - | 0 | - |
| 2 | b | 4.583E+04 | 201.1 | 0.0002273 | 1.13 | +1 | 2 |
| - | - | 811.8 | 201.1 | - | - | 0 | - |
| - | - | 7402 | 202.1 | - | - | 0 | - |
| - | - | 4210 | 202.1 | - | - | 0 | - |
| - | - | 1061 | 203.1 | - | - | 0 | - |
| - | - | 2608 | 203.1 | - | - | 0 | - |
| - | - | 643.3 | 207.1 | - | - | 0 | - |
| - | - | 1413 | 208.1 | - | - | 0 | - |
| - | - | 1037 | 208.1 | - | - | 0 | - |
| - | - | 4833 | 209.1 | - | - | 0 | - |
| - | - | 760.7 | 211.1 | - | - | 0 | - |
| - | - | 1278 | 211.1 | - | - | 0 | - |
| - | - | 3234 | 212.1 | - | - | 0 | - |
| - | - | 1791 | 213.1 | - | - | 0 | - |
| - | - | 4757 | 213.1 | - | - | 0 | - |
| - | - | 1256 | 214.2 | - | - | 0 | - |
| - | - | 1.252E+05 | 215.1 | - | - | 0 | - |
| - | - | 867.4 | 216.1 | - | - | 0 | - |
| - | - | 1.375E+04 | 216.1 | - | - | 0 | - |
| - | - | 1.683E+04 | 217.1 | - | - | 0 | - |
| - | - | 7684 | 217.1 | - | - | 0 | - |
| - | - | 1218 | 217.1 | - | - | 0 | - |
| - | - | 1442 | 218.1 | - | - | 0 | - |
| 12 | y | 6.918E+04 | 218.1 | 0.0003286 | 1.507 | +1 | 2 |
| - | - | 6600 | 219.1 | - | - | 0 | - |
| - | - | 5069 | 219.1 | - | - | 0 | - |
| - | - | 673.4 | 224.1 | - | - | 0 | - |
| - | - | 702.4 | 225.1 | - | - | 0 | - |
| - | - | 1.005E+04 | 225.1 | - | - | 0 | - |
| - | - | 696.9 | 226.1 | - | - | 0 | - |
| - | - | 2331 | 226.1 | - | - | 0 | - |
| - | - | 685.3 | 226.2 | - | - | 0 | - |
| - | - | 1.005E+04 | 227.1 | - | - | 0 | - |
| - | - | 937.5 | 228.1 | - | - | 0 | - |
| - | - | 2769 | 228.2 | - | - | 0 | - |
| - | - | 1186 | 229.1 | - | - | 0 | - |
| - | - | 1597 | 229.2 | - | - | 0 | - |
| - | - | 589.7 | 229.2 | - | - | 0 | - |
| - | - | 1495 | 231.1 | - | - | 0 | - |
| - | - | 2.625E+04 | 233.2 | - | - | 0 | - |
| - | - | 3671 | 234.2 | - | - | 0 | - |
| - | - | 4925 | 236.1 | - | - | 0 | - |
| - | - | 652.3 | 237.1 | - | - | 0 | - |
| - | - | 635.5 | 240.1 | - | - | 0 | - |
| - | - | 525.2 | 240.5 | - | - | 0 | - |
| - | - | 6002 | 241.1 | - | - | 0 | - |
| - | - | 674.6 | 242.1 | - | - | 0 | - |
| - | - | 915.2 | 242.1 | - | - | 0 | - |
| - | - | 3.338E+04 | 242.2 | - | - | 0 | - |
| - | - | 4135 | 243.1 | - | - | 0 | - |
| - | - | 4516 | 243.2 | - | - | 0 | - |
| - | - | 926.1 | 244.1 | - | - | 0 | - |
| - | - | 5407 | 244.1 | - | - | 0 | - |
| - | - | 6260 | 245.1 | - | - | 0 | - |
| - | - | 772 | 246.1 | - | - | 0 | - |
| - | - | 1.941E+04 | 247.1 | - | - | 0 | - |
| - | - | 4594 | 247.1 | - | - | 0 | - |
| - | - | 1560 | 248.1 | - | - | 0 | - |
| - | - | 776.3 | 249.1 | - | - | 0 | - |
| - | - | 602 | 249.1 | - | - | 0 | - |
| - | - | 658.7 | 250.1 | - | - | 0 | - |
| - | - | 1184 | 251.1 | - | - | 0 | - |
| - | - | 790.4 | 251.2 | - | - | 0 | - |
| - | - | 1.32E+04 | 251.2 | - | - | 0 | - |
| - | - | 1950 | 252.1 | - | - | 0 | - |
| - | - | 1497 | 252.2 | - | - | 0 | - |
| - | - | 531.8 | 253.2 | - | - | 0 | - |
| - | - | 1.498E+04 | 254.1 | - | - | 0 | - |
| - | - | 3060 | 254.2 | - | - | 0 | - |
| - | - | 1951 | 255.1 | - | - | 0 | - |
| - | - | 1.127E+04 | 259.1 | - | - | 0 | - |
| - | - | 670.1 | 259.1 | - | - | 0 | - |
| - | - | 1165 | 260.1 | - | - | 0 | - |
| - | - | 988.5 | 260.2 | - | - | 0 | - |
| - | - | 9302 | 261.2 | - | - | 0 | - |
| - | - | 754.8 | 262.1 | - | - | 0 | - |
| - | - | 910.4 | 262.2 | - | - | 0 | - |
| - | - | 1205 | 263.1 | - | - | 0 | - |
| - | - | 701 | 264.1 | - | - | 0 | - |
| - | - | 6434 | 264.1 | - | - | 0 | - |
| - | - | 613.8 | 264.2 | - | - | 0 | - |
| - | - | 916.5 | 265.1 | - | - | 0 | - |
| - | - | 863.3 | 267.1 | - | - | 0 | - |
| - | - | 1971 | 268.1 | - | - | 0 | - |
| 3 | a | 1961 | 268.2 | 0.0005611 | 2.092 | +1 | 3 |
| - | - | 3082 | 269.2 | - | - | 0 | - |
| - | - | 1039 | 270.1 | - | - | 0 | - |
| - | - | 1578 | 270.2 | - | - | 0 | - |
| - | - | 9543 | 272.1 | - | - | 0 | - |
| - | - | 1657 | 273.1 | - | - | 0 | - |
| - | - | 1.203E+04 | 275.1 | - | - | 0 | - |
| - | - | 3933 | 275.2 | - | - | 0 | - |
| - | - | 764.5 | 276.1 | - | - | 0 | - |
| - | - | 622 | 277.2 | - | - | 0 | - |
| - | - | 707.9 | 279.1 | - | - | 0 | - |
| - | - | 9531 | 280.2 | - | - | 0 | - |
| - | - | 1163 | 281.2 | - | - | 0 | - |
| - | - | 1740 | 282.1 | - | - | 0 | - |
| - | - | 3.545E+04 | 282.1 | - | - | 0 | - |
| - | - | 565.6 | 283.1 | - | - | 0 | - |
| - | - | 4616 | 283.1 | - | - | 0 | - |
| - | - | 5376 | 285.2 | - | - | 0 | - |
| - | - | 1449 | 285.2 | - | - | 0 | - |
| - | - | 7199 | 286.1 | - | - | 0 | - |
| - | - | 1167 | 286.2 | - | - | 0 | - |
| 3 | a | 1101 | 286.2 | 0.0006776 | 2.367 | +1 | 3 |
| - | - | 2626 | 294.2 | - | - | 0 | - |
| 9 | y | 810.9 | 295.1 | 0.0002236 | 0.7577 | +2 | 5 |
| - | - | 1128 | 296.1 | - | - | 0 | - |
| 3 | b | 2.304E+05 | 296.2 | 0.000489 | 1.651 | +1 | 3 |
| - | - | 852.4 | 297.1 | - | - | 0 | - |
| - | - | 3.637E+04 | 297.2 | - | - | 0 | - |
| - | - | 1.041E+04 | 298.2 | - | - | 0 | - |
| - | - | 1479 | 298.2 | - | - | 0 | - |
| - | - | 1396 | 299.1 | - | - | 0 | - |
| - | - | 787.9 | 299.1 | - | - | 0 | - |
| - | - | 1972 | 299.2 | - | - | 0 | - |
| - | - | 1280 | 301.2 | - | - | 0 | - |
| - | - | 9514 | 303.2 | - | - | 0 | - |
| - | - | 3885 | 304.1 | - | - | 0 | - |
| - | - | 2095 | 304.2 | - | - | 0 | - |
| - | - | 660.4 | 305.1 | - | - | 0 | - |
| - | - | 1565 | 306.1 | - | - | 0 | - |
| - | - | 1295 | 308.2 | - | - | 0 | - |
| - | - | 1540 | 310.1 | - | - | 0 | - |
| - | - | 812 | 310.1 | - | - | 0 | - |
| - | - | 739.9 | 311.1 | - | - | 0 | - |
| - | - | 1099 | 312.2 | - | - | 0 | - |
| - | - | 1129 | 313.2 | - | - | 0 | - |
| - | - | 1740 | 314.1 | - | - | 0 | - |
| 11 | y | 1.086E+04 | 314.2 | 0.0006751 | 2.149 | +1 | 3 |
| 3 | b | 5.264E+04 | 314.2 | 0.0004834 | 1.539 | +1 | 3 |
| - | - | 1885 | 315.2 | - | - | 0 | - |
| - | - | 9108 | 315.2 | - | - | 0 | - |
| - | - | 2231 | 317.2 | - | - | 0 | - |
| - | - | 712.6 | 319.1 | - | - | 0 | - |
| - | - | 1034 | 321.2 | - | - | 0 | - |
| - | - | 1576 | 323.2 | - | - | 0 | - |
| - | - | 2218 | 324.2 | - | - | 0 | - |
| - | - | 673.8 | 324.2 | - | - | 0 | - |
| - | - | 785.3 | 325.2 | - | - | 0 | - |
| - | - | 1900 | 326.2 | - | - | 0 | - |
| - | - | 6362 | 328.1 | - | - | 0 | - |
| - | - | 982.7 | 328.2 | - | - | 0 | - |
| - | - | 821.7 | 329.1 | - | - | 0 | - |
| - | - | 6509 | 330.2 | - | - | 0 | - |
| - | - | 998.1 | 331.2 | - | - | 0 | - |
| 11 | y | 1.307E+04 | 331.2 | 0.0004627 | 1.397 | +1 | 3 |
| - | - | 2169 | 332.2 | - | - | 0 | - |
| - | - | 593.5 | 333.1 | - | - | 0 | - |
| - | - | 3264 | 333.2 | - | - | 0 | - |
| - | - | 1137 | 334.2 | - | - | 0 | - |
| - | - | 2440 | 334.2 | - | - | 0 | - |
| - | - | 1388 | 336.2 | - | - | 0 | - |
| - | - | 2363 | 337.2 | - | - | 0 | - |
| - | - | 1324 | 338.1 | - | - | 0 | - |
| - | - | 2411 | 339.2 | - | - | 0 | - |
| - | - | 743.8 | 340.2 | - | - | 0 | - |
| - | - | 7882 | 341.2 | - | - | 0 | - |
| - | - | 2051 | 342.2 | - | - | 0 | - |
| - | - | 1648 | 343.2 | - | - | 0 | - |
| - | - | 2117 | 344.2 | - | - | 0 | - |
| - | - | 1.896E+04 | 346.1 | - | - | 0 | - |
| - | - | 8755 | 346.2 | - | - | 0 | - |
| - | - | 2524 | 347.1 | - | - | 0 | - |
| - | - | 2154 | 347.2 | - | - | 0 | - |
| - | - | 6822 | 348.2 | - | - | 0 | - |
| - | - | 1536 | 349.2 | - | - | 0 | - |
| - | - | 1.64E+04 | 351.2 | - | - | 0 | - |
| - | - | 2431 | 352.2 | - | - | 0 | - |
| - | - | 6026 | 353.2 | - | - | 0 | - |
| - | - | 3330 | 354.2 | - | - | 0 | - |
| - | - | 914.6 | 354.2 | - | - | 0 | - |
| - | - | 3966 | 355.2 | - | - | 0 | - |
| - | - | 1691 | 356.1 | - | - | 0 | - |
| - | - | 1589 | 360.2 | - | - | 0 | - |
| - | - | 2.067E+04 | 362.2 | - | - | 0 | - |
| - | - | 4836 | 363.2 | - | - | 0 | - |
| - | - | 4356 | 365.1 | - | - | 0 | - |
| - | - | 1475 | 365.3 | - | - | 0 | - |
| - | - | 1238 | 367.2 | - | - | 0 | - |
| - | - | 2.267E+04 | 369.2 | - | - | 0 | - |
| - | - | 4373 | 370.2 | - | - | 0 | - |
| - | - | 1188 | 370.7 | - | - | 0 | - |
| - | - | 756.2 | 371.2 | - | - | 0 | - |
| - | - | 4240 | 371.2 | - | - | 0 | - |
| - | - | 2426 | 372.2 | - | - | 0 | - |
| 7 | y | 7645 | 373.2 | 0.0004046 | 1.084 | +2 | 7 |
| - | - | 3639 | 374.2 | - | - | 0 | - |
| - | - | 929.4 | 375.2 | - | - | 0 | - |
| - | - | 775.1 | 377.2 | - | - | 0 | - |
| - | - | 729 | 379.1 | - | - | 0 | - |
| - | - | 619.2 | 379.2 | - | - | 0 | - |
| - | - | 860.1 | 379.2 | - | - | 0 | - |
| - | - | 1076 | 381.2 | - | - | 0 | - |
| - | - | 671.2 | 382.2 | - | - | 0 | - |
| - | - | 1.152E+04 | 383.2 | - | - | 0 | - |
| - | - | 768.9 | 383.2 | - | - | 0 | - |
| - | - | 576.3 | 383.2 | - | - | 0 | - |
| - | - | 1738 | 383.3 | - | - | 0 | - |
| - | - | 1628 | 384.2 | - | - | 0 | - |
| - | - | 2809 | 386.2 | - | - | 0 | - |
| - | - | 1272 | 390.2 | - | - | 0 | - |
| - | - | 1355 | 393.2 | - | - | 0 | - |
| - | - | 3855 | 393.3 | - | - | 0 | - |
| - | - | 1192 | 394.3 | - | - | 0 | - |
| - | - | 1530 | 395.2 | - | - | 0 | - |
| - | - | 1431 | 395.2 | - | - | 0 | - |
| - | - | 2205 | 397.1 | - | - | 0 | - |
| - | - | 4998 | 398.2 | - | - | 0 | - |
| - | - | 1061 | 399.2 | - | - | 0 | - |
| - | - | 1646 | 400.2 | - | - | 0 | - |
| - | - | 1.815E+04 | 401.2 | - | - | 0 | - |
| - | - | 775.3 | 401.2 | - | - | 0 | - |
| - | - | 3051 | 402.2 | - | - | 0 | - |
| - | - | 957.9 | 407.2 | - | - | 0 | - |
| - | - | 902.2 | 411.2 | - | - | 0 | - |
| - | - | 7546 | 411.3 | - | - | 0 | - |
| - | - | 1334 | 412.3 | - | - | 0 | - |
| - | - | 1147 | 413.2 | - | - | 0 | - |
| - | - | 3719 | 415.1 | - | - | 0 | - |
| - | - | 2647 | 415.3 | - | - | 0 | - |
| - | - | 810.2 | 416.2 | - | - | 0 | - |
| - | - | 8882 | 416.3 | - | - | 0 | - |
| - | - | 2993 | 417.3 | - | - | 0 | - |
| - | - | 695 | 421.2 | - | - | 0 | - |
| - | - | 1357 | 423.2 | - | - | 0 | - |
| - | - | 2020 | 423.2 | - | - | 0 | - |
| - | - | 1077 | 424.2 | - | - | 0 | - |
| - | - | 995.7 | 425.2 | - | - | 0 | - |
| - | - | 1393 | 427.2 | - | - | 0 | - |
| - | - | 2216 | 429.2 | - | - | 0 | - |
| - | - | 802.1 | 431.2 | - | - | 0 | - |
| - | - | 7107 | 433.2 | - | - | 0 | - |
| - | - | 7.033E+04 | 433.3 | - | - | 0 | - |
| - | - | 1550 | 434.2 | - | - | 0 | - |
| - | - | 3369 | 434.2 | - | - | 0 | - |
| - | - | 1.575E+04 | 434.3 | - | - | 0 | - |
| - | - | 1348 | 434.7 | - | - | 0 | - |
| - | - | 1584 | 435.2 | - | - | 0 | - |
| - | - | 2167 | 435.3 | - | - | 0 | - |
| - | - | 1466 | 440.2 | - | - | 0 | - |
| - | - | 1897 | 441.2 | - | - | 0 | - |
| - | - | 4866 | 441.2 | - | - | 0 | - |
| 10 | y | 1599 | 442.2 | 0.005025 | 11.36 | +1 | 4 |
| - | - | 1733 | 443.1 | - | - | 0 | - |
| - | - | 960.2 | 443.2 | - | - | 0 | - |
| - | - | 3516 | 443.2 | - | - | 0 | - |
| 4 | b | 7564 | 443.3 | 0.0004955 | 1.118 | +1 | 4 |
| - | - | 756.6 | 444.2 | - | - | 0 | - |
| - | - | 2220 | 444.3 | - | - | 0 | - |
| - | - | 995.3 | 445.2 | - | - | 0 | - |
| - | - | 715.7 | 446.2 | - | - | 0 | - |
| - | - | 1171 | 450.2 | - | - | 0 | - |
| - | - | 1067 | 451.2 | - | - | 0 | - |
| - | - | 5976 | 452.2 | - | - | 0 | - |
| - | - | 1013 | 453.2 | - | - | 0 | - |
| - | - | 670.9 | 458.2 | - | - | 0 | - |
| - | - | 5772 | 459.2 | - | - | 0 | - |
| - | - | 1419 | 459.3 | - | - | 0 | - |
| - | - | 1166 | 460.2 | - | - | 0 | - |
| 10 | y | 2308 | 460.2 | 0.001235 | 2.683 | +1 | 4 |
| 4 | b | 6.389E+04 | 461.3 | 0.000612 | 1.327 | +1 | 4 |
| - | - | 6228 | 462.2 | - | - | 0 | - |
| - | - | 1.758E+04 | 462.3 | - | - | 0 | - |
| - | - | 1046 | 463.2 | - | - | 0 | - |
| - | - | 2281 | 463.3 | - | - | 0 | - |
| - | - | 870.2 | 464.3 | - | - | 0 | - |
| - | - | 1630 | 466.2 | - | - | 0 | - |
| - | - | 2421 | 468.2 | - | - | 0 | - |
| - | - | 1123 | 468.2 | - | - | 0 | - |
| - | - | 1769 | 469.2 | - | - | 0 | - |
| 5 | y | 2.136E+04 | 470.2 | 0.0005887 | 1.252 | +2 | 9 |
| 9 | b | 905.5 | 470.7 | 0.002256 | 4.792 | +2 | 9 |
| - | - | 3429 | 471.2 | - | - | 0 | - |
| - | - | 1576 | 476.2 | - | - | 0 | - |
| - | - | 1288 | 479.2 | - | - | 0 | - |
| 9 | b | 722 | 479.7 | 0.00351 | 7.316 | +2 | 9 |
| - | - | 1.955E+04 | 480.2 | - | - | 0 | - |
| - | - | 4113 | 481.2 | - | - | 0 | - |
| - | - | 1718 | 482.2 | - | - | 0 | - |
| - | - | 989.4 | 482.3 | - | - | 0 | - |
| - | - | 3611 | 484.2 | - | - | 0 | - |
| - | - | 988.1 | 485.2 | - | - | 0 | - |
| - | - | 3392 | 494.2 | - | - | 0 | - |
| - | - | 1582 | 495.2 | - | - | 0 | - |
| - | - | 5.861E+04 | 498.2 | - | - | 0 | - |
| - | - | 1.481E+04 | 499.2 | - | - | 0 | - |
| - | - | 4038 | 500.2 | - | - | 0 | - |
| - | - | 997 | 500.8 | - | - | 0 | - |
| - | - | 7267 | 502.2 | - | - | 0 | - |
| - | - | 816.5 | 502.3 | - | - | 0 | - |
| - | - | 1838 | 503.2 | - | - | 0 | - |
| - | - | 1217 | 508.3 | - | - | 0 | - |
| - | - | 1464 | 510.2 | - | - | 0 | - |
| - | - | 7049 | 512.2 | - | - | 0 | - |
| - | - | 889.9 | 512.3 | - | - | 0 | - |
| - | - | 1467 | 513.2 | - | - | 0 | - |
| - | - | 816.4 | 514.3 | - | - | 0 | - |
| - | - | 1896 | 515.2 | - | - | 0 | - |
| - | - | 1192 | 516.2 | - | - | 0 | - |
| - | - | 764.1 | 525.8 | - | - | 0 | - |
| - | - | 1335 | 526.3 | - | - | 0 | - |
| - | - | 1583 | 528.2 | - | - | 0 | - |
| - | - | 2.413E+04 | 530.2 | - | - | 0 | - |
| - | - | 6313 | 530.3 | - | - | 0 | - |
| - | - | 810.3 | 530.8 | - | - | 0 | - |
| - | - | 5547 | 531.2 | - | - | 0 | - |
| - | - | 1455 | 531.3 | - | - | 0 | - |
| - | - | 1107 | 532.2 | - | - | 0 | - |
| 10 | b | 638.8 | 535.3 | 0.007393 | 13.81 | +2 | 10 |
| - | - | 787.1 | 535.8 | - | - | 0 | - |
| 5 | b | 4734 | 540.3 | 0.0006186 | 1.145 | +1 | 5 |
| - | - | 1115 | 541.3 | - | - | 0 | - |
| - | - | 691.4 | 542.3 | - | - | 0 | - |
| 10 | b | 5720 | 544.3 | 0.0002092 | 0.3843 | +2 | 10 |
| - | - | 4934 | 544.8 | - | - | 0 | - |
| - | - | 1872 | 545.3 | - | - | 0 | - |
| - | - | 3148 | 546.2 | - | - | 0 | - |
| - | - | 799.8 | 547.2 | - | - | 0 | - |
| - | - | 943.8 | 551.3 | - | - | 0 | - |
| - | - | 683.9 | 554.3 | - | - | 0 | - |
| 5 | b | 1.12E+04 | 558.3 | 0.0004299 | 0.77 | +1 | 5 |
| - | - | 4227 | 559.3 | - | - | 0 | - |
| - | - | 988.7 | 560.3 | - | - | 0 | - |
| - | - | 1513 | 563.2 | - | - | 0 | - |
| - | - | 859 | 569.3 | - | - | 0 | - |
| - | - | 1764 | 570.3 | - | - | 0 | - |
| - | - | 904.6 | 573.2 | - | - | 0 | - |
| - | - | 762.8 | 579.3 | - | - | 0 | - |
| - | - | 6010 | 581.3 | - | - | 0 | - |
| - | - | 1925 | 582.3 | - | - | 0 | - |
| - | - | 768.8 | 583.3 | - | - | 0 | - |
| - | - | 755.6 | 584.8 | - | - | 0 | - |
| - | - | 4394 | 587.3 | - | - | 0 | - |
| - | - | 1559 | 588.3 | - | - | 0 | - |
| 9 | y | 1570 | 589.3 | 0.004693 | 7.964 | +1 | 5 |
| - | - | 4593 | 591.2 | - | - | 0 | - |
| - | - | 886.5 | 592.2 | - | - | 0 | - |
| - | - | 1232 | 595.3 | - | - | 0 | - |
| - | - | 3569 | 597.3 | - | - | 0 | - |
| - | - | 1550 | 598.3 | - | - | 0 | - |
| - | - | 3.549E+04 | 599.3 | - | - | 0 | - |
| - | - | 737.9 | 599.8 | - | - | 0 | - |
| - | - | 1.073E+04 | 600.3 | - | - | 0 | - |
| - | - | 966.8 | 600.3 | - | - | 0 | - |
| 11 | b | 4956 | 600.8 | 0.0008497 | 1.414 | +2 | 11 |
| - | - | 1441 | 601.3 | - | - | 0 | - |
| - | - | 2101 | 601.3 | - | - | 0 | - |
| - | - | 1028 | 601.8 | - | - | 0 | - |
| - | - | 886.6 | 607.3 | - | - | 0 | - |
| - | - | 1.235E+04 | 609.3 | - | - | 0 | - |
| - | - | 4066 | 610.3 | - | - | 0 | - |
| - | - | 4996 | 615.3 | - | - | 0 | - |
| - | - | 1447 | 616.3 | - | - | 0 | - |
| - | - | 6683 | 617.3 | - | - | 0 | - |
| - | - | 1764 | 618.3 | - | - | 0 | - |
| - | - | 2978 | 625.3 | - | - | 0 | - |
| - | - | 1758 | 626.3 | - | - | 0 | - |
| - | - | 1.378E+05 | 627.3 | - | - | 0 | - |
| - | - | 3.981E+04 | 628.3 | - | - | 0 | - |
| - | - | 7412 | 629.3 | - | - | 0 | - |
| - | - | 803.8 | 629.3 | - | - | 0 | - |
| - | - | 1938 | 634.3 | - | - | 0 | - |
| 6 | b | 1047 | 637.4 | 0.001303 | 2.044 | +1 | 6 |
| - | - | 686.8 | 642.3 | - | - | 0 | - |
| - | - | 1.091E+04 | 643.3 | - | - | 0 | - |
| - | - | 8760 | 644.3 | - | - | 0 | - |
| - | - | 4767 | 645.3 | - | - | 0 | - |
| - | - | 1401 | 646.3 | - | - | 0 | - |
| 6 | b | 3944 | 655.4 | 0.0005761 | 0.879 | +1 | 6 |
| - | - | 1898 | 656.4 | - | - | 0 | - |
| 8 | y | 1359 | 659.3 | 0.001458 | 2.211 | +1 | 6 |
| - | - | 2784 | 660.3 | - | - | 0 | - |
| - | - | 1080 | 661.3 | - | - | 0 | - |
| - | - | 905.3 | 663.3 | - | - | 0 | - |
| - | - | 2452 | 674.3 | - | - | 0 | - |
| 8 | y | 4487 | 676.3 | 0.003426 | 5.066 | +1 | 6 |
| - | - | 1315 | 677.3 | - | - | 0 | - |
| - | - | 1015 | 680.3 | - | - | 0 | - |
| - | - | 720.5 | 686.3 | - | - | 0 | - |
| - | - | 657.5 | 686.4 | - | - | 0 | - |
| - | - | 1050 | 691.3 | - | - | 0 | - |
| - | - | 4537 | 694.3 | - | - | 0 | - |
| - | - | 1648 | 695.3 | - | - | 0 | - |
| - | - | 1640 | 698.3 | - | - | 0 | - |
| - | - | 2538 | 704.3 | - | - | 0 | - |
| - | - | 1571 | 705.3 | - | - | 0 | - |
| - | - | 806.7 | 708.3 | - | - | 0 | - |
| - | - | 798.7 | 708.4 | - | - | 0 | - |
| - | - | 709.8 | 708.7 | - | - | 0 | - |
| - | - | 3.354E+04 | 712.4 | - | - | 0 | - |
| - | - | 1.352E+04 | 713.4 | - | - | 0 | - |
| - | - | 2202 | 714.4 | - | - | 0 | - |
| - | - | 1018 | 714.4 | - | - | 0 | - |
| - | - | 8331 | 722.3 | - | - | 0 | - |
| - | - | 2922 | 723.3 | - | - | 0 | - |
| 7 | b | 2819 | 724.4 | 0.0004334 | 0.5983 | +1 | 7 |
| - | - | 1539 | 725.4 | - | - | 0 | - |
| - | - | 3430 | 726.3 | - | - | 0 | - |
| - | - | 1371 | 727.3 | - | - | 0 | - |
| - | - | 1054 | 728.3 | - | - | 0 | - |
| - | - | 1573 | 729.3 | - | - | 0 | - |
| - | - | 890.9 | 735.3 | - | - | 0 | - |
| - | - | 1653 | 739.4 | - | - | 0 | - |
| - | - | 7.43E+04 | 740.3 | - | - | 0 | - |
| - | - | 2.717E+04 | 741.3 | - | - | 0 | - |
| - | - | 5944 | 742.4 | - | - | 0 | - |
| 7 | b | 1.175E+04 | 742.4 | 0.001209 | 1.628 | +1 | 7 |
| - | - | 1442 | 743.4 | - | - | 0 | - |
| - | - | 4955 | 743.4 | - | - | 0 | - |
| - | - | 1432 | 744.4 | - | - | 0 | - |
| 7 | y | 2634 | 746.3 | 0.01357 | 18.19 | +1 | 7 |
| - | - | 886.5 | 747.3 | - | - | 0 | - |
| - | - | 1676 | 752.4 | - | - | 0 | - |
| - | - | 1883 | 753.3 | - | - | 0 | - |
| - | - | 1031 | 754.3 | - | - | 0 | - |
| - | - | 1321 | 756.3 | - | - | 0 | - |
| - | - | 2.097E+04 | 757.4 | - | - | 0 | - |
| - | - | 1.12E+04 | 758.4 | - | - | 0 | - |
| - | - | 2714 | 759.4 | - | - | 0 | - |
| 7 | y | 1557 | 763.3 | 0.002404 | 3.149 | +1 | 7 |
| - | - | 1270 | 768.4 | - | - | 0 | - |
| - | - | 637.3 | 769.4 | - | - | 0 | - |
| - | - | 1.403E+04 | 771.4 | - | - | 0 | - |
| - | - | 6590 | 772.4 | - | - | 0 | - |
| - | - | 999.6 | 773.4 | - | - | 0 | - |
| - | - | 1.022E+04 | 774.3 | - | - | 0 | - |
| - | - | 5726 | 775.3 | - | - | 0 | - |
| - | - | 1477 | 776.3 | - | - | 0 | - |
| - | - | 2050 | 780.4 | - | - | 0 | - |
| - | - | 1443 | 781.4 | - | - | 0 | - |
| - | - | 855.8 | 788.4 | - | - | 0 | - |
| - | - | 1358 | 791.4 | - | - | 0 | - |
| - | - | 1006 | 794.3 | - | - | 0 | - |
| - | - | 901.9 | 795.3 | - | - | 0 | - |
| - | - | 738.6 | 795.4 | - | - | 0 | - |
| - | - | 2692 | 796.4 | - | - | 0 | - |
| - | - | 1638 | 797.4 | - | - | 0 | - |
| - | - | 1253 | 801.4 | - | - | 0 | - |
| - | - | 1438 | 809.4 | - | - | 0 | - |
| 8 | b | 4588 | 811.4 | 0.006339 | 7.812 | +1 | 8 |
| - | - | 1118 | 812.3 | - | - | 0 | - |
| - | - | 2102 | 812.4 | - | - | 0 | - |
| - | - | 777.2 | 821.4 | - | - | 0 | - |
| - | - | 1075 | 822.4 | - | - | 0 | - |
| - | - | 899.6 | 823.4 | - | - | 0 | - |
| - | - | 1079 | 823.4 | - | - | 0 | - |
| - | - | 987.3 | 825.4 | - | - | 0 | - |
| - | - | 782 | 826.4 | - | - | 0 | - |
| - | - | 2266 | 827.4 | - | - | 0 | - |
| - | - | 838.3 | 828.4 | - | - | 0 | - |
| 8 | b | 5999 | 829.4 | 0.0002474 | 0.2983 | +1 | 8 |
| - | - | 3246 | 830.4 | - | - | 0 | - |
| - | - | 729.9 | 832.4 | - | - | 0 | - |
| - | - | 933.9 | 833.4 | - | - | 0 | - |
| - | - | 8434 | 839.4 | - | - | 0 | - |
| - | - | 7128 | 840.4 | - | - | 0 | - |
| - | - | 4951 | 841.4 | - | - | 0 | - |
| 6 | y | 3582 | 842.4 | 0.006042 | 7.172 | +1 | 8 |
| 6 | y | 5731 | 843.4 | 0.006768 | 8.024 | +1 | 8 |
| - | - | 1756 | 844.4 | - | - | 0 | - |
| - | - | 815.8 | 845.4 | - | - | 0 | - |
| - | - | 7645 | 850.4 | - | - | 0 | - |
| - | - | 4560 | 851.4 | - | - | 0 | - |
| - | - | 1608 | 852.4 | - | - | 0 | - |
| - | - | 836.4 | 853.4 | - | - | 0 | - |
| - | - | 4564 | 859.4 | - | - | 0 | - |
| 6 | y | 2.393E+04 | 860.4 | 0.001398 | 1.624 | +1 | 8 |
| - | - | 1.118E+04 | 861.4 | - | - | 0 | - |
| - | - | 2021 | 862.4 | - | - | 0 | - |
| - | - | 8.844E+04 | 868.4 | - | - | 0 | - |
| - | - | 4.018E+04 | 869.4 | - | - | 0 | - |
| - | - | 1125 | 869.5 | - | - | 0 | - |
| - | - | 9538 | 870.4 | - | - | 0 | - |
| - | - | 837 | 871.4 | - | - | 0 | - |
| - | - | 2323 | 886.4 | - | - | 0 | - |
| - | - | 1.595E+04 | 887.4 | - | - | 0 | - |
| - | - | 6686 | 888.4 | - | - | 0 | - |
| - | - | 1892 | 889.4 | - | - | 0 | - |
| - | - | 1293 | 895.4 | - | - | 0 | - |
| - | - | 894.9 | 896.4 | - | - | 0 | - |
| - | - | 1679 | 904.4 | - | - | 0 | - |
| - | - | 939 | 905.4 | - | - | 0 | - |
| - | - | 1068 | 912.5 | - | - | 0 | - |
| - | - | 712.7 | 913.5 | - | - | 0 | - |
| - | - | 1041 | 915.4 | - | - | 0 | - |
| - | - | 962.3 | 916.4 | - | - | 0 | - |
| - | - | 2028 | 921.4 | - | - | 0 | - |
| - | - | 3993 | 922.4 | - | - | 0 | - |
| - | - | 2473 | 923.4 | - | - | 0 | - |
| - | - | 958.1 | 926.4 | - | - | 0 | - |
| - | - | 809.1 | 927.4 | - | - | 0 | - |
| - | - | 1387 | 930.5 | - | - | 0 | - |
| 5 | y | 1.163E+04 | 939.4 | 0.0005793 | 0.6166 | +1 | 9 |
| 5 | y | 2.996E+04 | 940.4 | 0.006128 | 6.516 | +1 | 9 |
| - | - | 1.571E+04 | 941.4 | - | - | 0 | - |
| - | - | 5554 | 942.4 | - | - | 0 | - |
| - | - | 5129 | 943.4 | - | - | 0 | - |
| - | - | 2821 | 944.4 | - | - | 0 | - |
| - | - | 1115 | 956.4 | - | - | 0 | - |
| 5 | y | 2.547E+05 | 957.5 | 0.0002187 | 0.2284 | +1 | 9 |
| - | - | 1.279E+05 | 958.5 | - | - | 0 | - |
| - | - | 3.832E+04 | 959.5 | - | - | 0 | - |
| - | - | 4832 | 960.5 | - | - | 0 | - |
| - | - | 920.7 | 970.5 | - | - | 0 | - |
| - | - | 1010 | 971.5 | - | - | 0 | - |
| - | - | 832 | 972.5 | - | - | 0 | - |
| - | - | 950.1 | 977.4 | - | - | 0 | - |
| - | - | 1376 | 986.5 | - | - | 0 | - |
| - | - | 1915 | 987.5 | - | - | 0 | - |
| - | - | 4430 | 988.5 | - | - | 0 | - |
| - | - | 2059 | 989.5 | - | - | 0 | - |
| - | - | 738.4 | 990.5 | - | - | 0 | - |
| - | - | 3014 | 1000 | - | - | 0 | - |
| - | - | 1744 | 1001 | - | - | 0 | - |
| - | - | 5403 | 1015 | - | - | 0 | - |
| - | - | 3681 | 1016 | - | - | 0 | - |
| - | - | 754.6 | 1017 | - | - | 0 | - |
| - | - | 1979 | 1036 | - | - | 0 | - |
| - | - | 1389 | 1037 | - | - | 0 | - |
| - | - | 784.7 | 1044 | - | - | 0 | - |
| - | - | 805.6 | 1051 | - | - | 0 | - |
| - | - | 1171 | 1052 | - | - | 0 | - |
| - | - | 861.1 | 1053 | - | - | 0 | - |
| - | - | 4208 | 1054 | - | - | 0 | - |
| - | - | 1972 | 1055 | - | - | 0 | - |
| - | - | 906 | 1056 | - | - | 0 | - |
| - | - | 2582 | 1060 | - | - | 0 | - |
| - | - | 1243 | 1061 | - | - | 0 | - |
| 10 | b | 6114 | 1070 | 0.0008877 | 0.83 | +1 | 10 |
| - | - | 4585 | 1071 | - | - | 0 | - |
| - | - | 1897 | 1072 | - | - | 0 | - |
| - | - | 1585 | 1084 | - | - | 0 | - |
| - | - | 1031 | 1085 | - | - | 0 | - |
| - | - | 1555 | 1086 | - | - | 0 | - |
| 4 | y | 1655 | 1087 | 0.005042 | 4.641 | +1 | 10 |
| 10 | b | 6.289E+04 | 1088 | 0.001687 | 1.551 | +1 | 10 |
| - | - | 4.028E+04 | 1089 | - | - | 0 | - |
| - | - | 1201 | 1089 | - | - | 0 | - |
| - | - | 1.277E+04 | 1090 | - | - | 0 | - |
| - | - | 1045 | 1091 | - | - | 0 | - |
| - | - | 2722 | 1102 | - | - | 0 | - |
| - | - | 2019 | 1103 | - | - | 0 | - |
| - | - | 1461 | 1104 | - | - | 0 | - |
| 4 | y | 1.275E+04 | 1105 | 9.297E-05 | 0.08417 | +1 | 10 |
| - | - | 7268 | 1106 | - | - | 0 | - |
| - | - | 1152 | 1107 | - | - | 0 | - |
| - | - | 975.6 | 1129 | - | - | 0 | - |
| - | - | 1012 | 1130 | - | - | 0 | - |
| - | - | 845.9 | 1152 | - | - | 0 | - |
| - | - | 931.4 | 1157 | - | - | 0 | - |
| - | - | 1718 | 1165 | - | - | 0 | - |
| - | - | 1108 | 1166 | - | - | 0 | - |
| - | - | 1881 | 1167 | - | - | 0 | - |
| - | - | 2184 | 1169 | - | - | 0 | - |
| - | - | 5574 | 1170 | - | - | 0 | - |
| - | - | 3863 | 1171 | - | - | 0 | - |
| - | - | 1453 | 1172 | - | - | 0 | - |
| - | - | 2217 | 1173 | - | - | 0 | - |
| - | - | 2313 | 1174 | - | - | 0 | - |
| 11 | b | 7784 | 1183 | 0.0002349 | 0.1986 | +1 | 11 |
| - | - | 5126 | 1184 | - | - | 0 | - |
| - | - | 1882 | 1185 | - | - | 0 | - |
| - | - | 1605 | 1187 | - | - | 0 | - |
| - | - | 742.3 | 1199 | - | - | 0 | - |
| 3 | y | 659.9 | 1200 | 0.01448 | 12.07 | +1 | 11 |
| 11 | b | 3.425E+04 | 1201 | 0.001156 | 0.9629 | +1 | 11 |
| - | - | 2.26E+04 | 1202 | - | - | 0 | - |
| - | - | 8321 | 1203 | - | - | 0 | - |
| - | - | 1046 | 1204 | - | - | 0 | - |
| 3 | y | 1370 | 1218 | 0.002943 | 2.417 | +1 | 11 |
| - | - | 1220 | 1219 | - | - | 0 | - |
| - | - | 801.4 | 1636 | - | - | 0 | - |
| - | - | 660.5 | 1639 | - | - | 0 | - |
| - | - | 684.9 | 3028 | - | - | 0 | - |

m/z Charge Intensity FragmentType MassShift Position
120.08109283447266 0 32196.588
120.30674743652344 0 367.89954
121.08442687988281 0 2586.5388
126.05529022216797 0 8050.862
127.05120086669922 0 431.4291
127.05818176269531 0 462.65085
127.0869369506836 0 699.46436
128.10726928710938 0 8860.769
129.06613159179688 0 4680.088
129.09759521484375 0 734.1126
129.10255432128906 0 12885.548
129.1106414794922 0 1016.9301
130.0500946044922 0 3130.9043
130.05514526367188 0 587.8896
130.0654296875 0 1649.9899
130.10569763183594 0 904.2431
130.7958526611328 0 393.9099
131.0450897216797 0 656.9628
131.11814880371094 0 9471.412
132.10206604003906 0 738.39325
133.06126403808594 0 538.2849
133.08599853515625 0 499.0352
134.02732849121094 0 2775.4214
136.0759735107422 0 4436.703
138.0917205810547 0 1778.166
138.12799072265625 0 7081.7695
139.08680725097656 0 1731.5503
139.1232452392578 0 369.99136
139.13124084472656 0 702.4119
141.06614685058594 0 758.15125
141.10272216796875 0 827.0707
143.08126831054688 0 524.6894
143.1181640625 0 774.44037
144.06570434570312 0 436.9309
145.0614776611328 0 473.15308
147.04437255859375 0 693.6284
147.07669067382812 0 3449.2183
148.06057739257812 0 660.7673
148.75462341308594 0 445.50174
150.23709106445312 0 385.2839
151.12364196777344 0 460.86713
152.0710906982422 0 1124.544
153.06626892089844 0 782.01935
153.1024932861328 0 584.174
153.6389923095703 0 384.01587
154.08651733398438 0 3904.079
154.09788513183594 0 1793.7449
155.081787109375 0 8333.279
155.11817932128906 0 2629.682 a Water loss 1
156.06581115722656 0 1090.0558
156.08489990234375 0 593.4145
157.06106567382812 0 2620.8777
157.0974578857422 0 7788.387
159.09176635742188 0 920.9616
161.69638061523438 0 421.38165
162.0546417236328 0 459.09323
165.102783203125 0 1352.5183
166.08651733398438 0 1429.6161
167.04486083984375 0 495.4031
167.0818634033203 0 5851.207
167.1182098388672 0 7108.3423
169.0974884033203 0 882.8706
169.1338348388672 0 10601.785
170.13710021972656 0 998.02814
171.07672119140625 0 1445.7543
171.1128692626953 0 524.34296
172.11204528808594 0 3330.68
173.1287384033203 0 24421.432 a 1
173.25296020507812 0 482.0247
174.0551300048828 0 1012.2993
174.12765502929688 0 1684.2512
174.13331604003906 0 852.656
175.07162475585938 0 3088.4841
178.05044555664062 0 517.2127
180.11256408691406 0 534.02844
181.06109619140625 0 499.87518
181.0974578857422 0 3207.3687
181.1342010498047 0 898.2217
183.0767364501953 0 3852.7622
183.11314392089844 0 82262.73 b Water loss 1
183.1492156982422 0 1621.0049
184.10861206054688 0 658.0818
184.11651611328125 0 7864.9526
185.09237670898438 0 13049.962
185.12875366210938 0 534.9324
185.16510009765625 0 479.88962
186.09564208984375 0 802.2647
187.14442443847656 0 13534.124
188.1477508544922 0 1091.5939
189.08737182617188 0 6155.762
189.1234130859375 0 560.0954
195.07664489746094 0 1615.8931
195.11306762695312 0 13460.527
196.11672973632812 0 745.18445
197.09249877929688 0 727.8966
197.12872314453125 0 17700.746
198.1322784423828 0 1469.9814
199.07150268554688 0 6477.1245
199.10804748535156 0 4983.521
200.10325622558594 0 103251.13
201.08731079101562 0 87911.81 y Ammonia loss 11
201.10638427734375 0 8124.75
201.12359619140625 0 45827.082 b 1
201.13534545898438 0 811.7972
202.09054565429688 0 7402.069
202.12696838378906 0 4210.178
203.09239196777344 0 1060.7544
203.10313415527344 0 2607.6628
207.11326599121094 0 643.33405
208.09678649902344 0 1413.2761
208.1077880859375 0 1037.1631
209.09222412109375 0 4832.8013
211.10800170898438 0 760.66003
211.14468383789062 0 1277.5142
212.13951110839844 0 3234.3003
213.08731079101562 0 1791.2891
213.12364196777344 0 4757.017
214.15548706054688 0 1255.692
215.13934326171875 0 125223.88
216.09837341308594 0 867.4191
216.14263916015625 0 13750.883
217.0821990966797 0 16827.424
217.13372802734375 0 7683.927
217.14459228515625 0 1217.8201
218.08668518066406 0 1442.1279
218.11386108398438 0 69184.09 y 11
219.11712646484375 0 6600.4434
219.1495819091797 0 5068.613
224.1029815673828 0 673.40875
225.0872802734375 0 702.40515
225.12367248535156 0 10050.058
226.09596252441406 0 696.86646
226.1188507080078 0 2331.1096
226.1554412841797 0 685.2746
227.10293579101562 0 10049.304
228.10691833496094 0 937.4917
228.1708984375 0 2768.9077
229.1187286376953 0 1185.5953
229.15504455566406 0 1597.2117
229.17396545410156 0 589.6533
231.0979766845703 0 1494.7622
233.1652069091797 0 26250.723
234.1687469482422 0 3671.457
236.1034393310547 0 4924.6035
237.12332153320312 0 652.3238
240.13455200195312 0 635.5393
240.52085876464844 0 525.1721
241.08224487304688 0 6002.1973
242.08604431152344 0 674.5605
242.1136474609375 0 915.1839
242.15029907226562 0 33376.08
243.13433837890625 0 4135.324
243.15335083007812 0 4515.8955
244.09678649902344 0 926.14325
244.1295928955078 0 5407.4873
245.1290740966797 0 6259.6025
246.13221740722656 0 771.967
247.11135864257812 0 19412.305
247.1443634033203 0 4593.7627
248.1148681640625 0 1560.4666
249.10879516601562 0 776.3041
249.1234893798828 0 602.0206
250.08323669433594 0 658.65546
251.10232543945312 0 1183.9656
251.15191650390625 0 790.39264
251.17578125 0 13197.063
252.1345977783203 0 1950.1427
252.1803436279297 0 1497.1564
253.15513610839844 0 531.8475
254.11392211914062 0 14981.189
254.1502685546875 0 3059.9207
255.11709594726562 0 1950.7505
259.09271240234375 0 11269.192
259.10675048828125 0 670.05524
260.0963134765625 0 1165.2491
260.1601867675781 0 988.45953
261.1596374511719 0 9302.301
262.1416931152344 0 754.8158
262.1636657714844 0 910.3702
263.1394958496094 0 1204.7291
264.097412109375 0 701.0465
264.1343688964844 0 6434.0747
264.16778564453125 0 613.8137
265.13775634765625 0 916.50604
267.1496276855469 0 863.319
268.09326171875 0 1970.9169
268.2025146484375 0 1960.7435 a Water loss 2
269.18634033203125 0 3082.1526
270.14501953125 0 1038.6967
270.182373046875 0 1578.436
272.1243591308594 0 9543.194
273.126220703125 0 1657.3138
275.1061706542969 0 12027.468
275.1756591796875 0 3932.711
276.1080322265625 0 764.5383
277.1559753417969 0 621.97943
279.0986022949219 0 707.91693
280.1658020019531 0 9531.304
281.1693115234375 0 1162.5354
282.1086120605469 0 1739.5964
282.1451416015625 0 35450.02
283.1111755371094 0 565.57153
283.1479187011719 0 4616.2866
285.1601257324219 0 5375.6396
285.1966552734375 0 1448.894
286.1036682128906 0 7198.6787
286.1628723144531 0 1166.9589
286.21319580078125 0 1100.9133 a 2
294.18206787109375 0 2626.4846
295.1448059082031 0 810.9495 y 8
296.089599609375 0 1128.1771
296.1973571777344 0 230363.83 b Water loss 2
297.1454162597656 0 852.35297
297.2005310058594 0 36371.19
298.1763916015625 0 10409.012
298.2021179199219 0 1478.8698
299.0878601074219 0 1396.037
299.1402587890625 0 787.87915
299.1768798828125 0 1972.4725
301.1558532714844 0 1280.0779
303.1708679199219 0 9513.583
304.1141052246094 0 3885.4407
304.1727294921875 0 2094.6133
305.11907958984375 0 660.41626
306.1455993652344 0 1565.3906
308.1604309082031 0 1294.9077
310.10406494140625 0 1540.0964
310.1370544433594 0 811.98505
311.1392822265625 0 739.8943
312.1922607421875 0 1098.8251
313.19036865234375 0 1129.0366
314.1352844238281 0 1739.6553
314.1717224121094 0 10860.584 y Ammonia loss 10
314.2079162597656 0 52642.555 b 2
315.173583984375 0 1885.3046
315.2109069824219 0 9107.733
317.1864929199219 0 2230.6223
319.1402587890625 0 712.6334
321.15582275390625 0 1033.7645
323.17144775390625 0 1575.5757
324.15545654296875 0 2217.6946
324.1922912597656 0 673.8222
325.18829345703125 0 785.2607
326.171142578125 0 1899.7705
328.11431884765625 0 6361.9136
328.1651611328125 0 982.7191
329.1173095703125 0 821.71375
330.1820068359375 0 6509.226
331.1783142089844 0 998.09515
331.19805908203125 0 13074.216 y 10
332.20147705078125 0 2168.625
333.13702392578125 0 593.54395
333.1566162109375 0 3263.683
334.1588439941406 0 1137.3586
334.2132873535156 0 2440.0493
336.1564025878906 0 1387.6658
337.1511535644531 0 2363.474
338.1337585449219 0 1324.312
339.16644287109375 0 2410.9575
340.188232421875 0 743.83655
341.183349609375 0 7881.584
342.1849670410156 0 2051.1003
343.1981201171875 0 1648.2434
344.1968994140625 0 2117.3955
346.12493896484375 0 18956.951
346.176513671875 0 8755.35
347.1292724609375 0 2524.3784
347.1802978515625 0 2153.714
348.19207763671875 0 6822.3765
349.1946105957031 0 1535.672
351.16680908203125 0 16402.26
352.169189453125 0 2430.5237
353.1820983886719 0 6025.7197
354.166015625 0 3330.2341
354.187255859375 0 914.62317
355.16204833984375 0 3965.8083
356.1442565917969 0 1691.2578
360.22723388671875 0 1589.3773
362.2076416015625 0 20673.531
363.209716796875 0 4836.2437
365.14605712890625 0 4356.038
365.2555847167969 0 1475.1827
367.23486328125 0 1237.6584
369.1771545410156 0 22673.646
370.1807861328125 0 4372.787
370.6771240234375 0 1188.4535
371.1535949707031 0 756.1508
371.19268798828125 0 4240.3677
372.1767578125 0 2425.734
373.17218017578125 0 7645.3047 y Water loss 6
374.17437744140625 0 3639.4453
375.1744689941406 0 929.39185
377.2181396484375 0 775.07025
379.12689208984375 0 728.99756
379.1628723144531 0 619.1563
379.1934814453125 0 860.12665
381.2148742675781 0 1076.0793
382.21826171875 0 671.19653
383.15655517578125 0 11520.124
383.1872863769531 0 768.9394
383.195068359375 0 576.2746
383.26513671875 0 1738.0944
384.1585388183594 0 1627.5477
386.2045593261719 0 2809.0273
390.20355224609375 0 1271.8635
393.1761779785156 0 1354.9895
393.2503356933594 0 3855.1763
394.254150390625 0 1192.3513
395.1932067871094 0 1530.3885
395.22943115234375 0 1430.6348
397.13629150390625 0 2205.4995
398.24395751953125 0 4997.5444
399.24810791015625 0 1060.694
400.2235412597656 0 1645.8965
401.1670837402344 0 18149.65
401.21612548828125 0 775.2531
402.1703186035156 0 3050.8677
407.1927185058594 0 957.90656
411.15484619140625 0 902.2393
411.26080322265625 0 7546.27
412.2636413574219 0 1333.9688
413.2043151855469 0 1146.8712
415.1466979980469 0 3719.3984
415.2712097167969 0 2647.4648
416.192626953125 0 810.185
416.2550048828125 0 8882.429
417.2565612792969 0 2993.4233
421.1727294921875 0 694.9658
423.1887512207031 0 1357.1677
423.2241516113281 0 2019.7368
424.220947265625 0 1077.4895
425.20172119140625 0 995.6956
427.23455810546875 0 1392.7202
429.2138977050781 0 2216.0503
431.21697998046875 0 802.09143
433.1571044921875 0 7107.3633
433.2815856933594 0 70334.195
434.16192626953125 0 1549.8258
434.2044982910156 0 3368.5125
434.2846984863281 0 15746.498
434.7069091796875 0 1348.3076
435.206787109375 0 1584.3397
435.2863464355469 0 2166.6956
440.21429443359375 0 1465.6732
441.19732666015625 0 1897.1631
441.2352600097656 0 4866.1665
442.2346496582031 0 1599.2396 y Water loss 9
443.1413879394531 0 1732.8702
443.1762390136719 0 960.17957
443.2300109863281 0 3515.8503
443.2657775878906 0 7564.438 b Water loss 3
444.1836853027344 0 756.55493
444.2686462402344 0 2220.1543
445.2431640625 0 995.349
446.2456359863281 0 715.6922
450.2349548339844 0 1170.7354
451.21923828125 0 1067.3463
452.2146301269531 0 5975.7783
453.21771240234375 0 1013.0533
458.15069580078125 0 670.9108
459.2091979980469 0 5772.12
459.26129150390625 0 1418.8822
460.209228515625 0 1166.4644
460.2414245605469 0 2308.4622 y 9
461.2764587402344 0 63886.125 b 3
462.198974609375 0 6227.6133
462.2795104980469 0 17581.648
463.2005615234375 0 1045.747
463.2815246582031 0 2280.7307
464.2519226074219 0 870.2377
466.1942138671875 0 1630.4719
468.20928955078125 0 2421.1235
468.2436218261719 0 1122.946
469.2452087402344 0 1768.6974
470.2251281738281 0 21363.23 y Water loss 4
470.7401123046875 0 905.4545 b Water loss 8
471.2291259765625 0 3429.3098
476.1774597167969 0 1576.3423
479.21295166015625 0 1288.403
479.744140625 0 721.9935 b 8
480.2097473144531 0 19550.992
481.213134765625 0 4112.807
482.2236633300781 0 1717.9498
482.2629699707031 0 989.42413
484.2043151855469 0 3611.024
485.20916748046875 0 988.1064
494.18896484375 0 3392.405
495.19317626953125 0 1581.8293
498.22015380859375 0 58610.43
499.22314453125 0 14806.423
500.23052978515625 0 4037.9324
500.75347900390625 0 996.9716
502.21478271484375 0 7267.404
502.2696838378906 0 816.53564
503.21868896484375 0 1837.6097
508.3153381347656 0 1216.638
510.2193603515625 0 1463.9789
512.1990356445312 0 7048.795
512.3236694335938 0 889.8653
513.2030639648438 0 1467.4164
514.30029296875 0 816.43884
515.2474365234375 0 1896.321
516.2476196289062 0 1192.1217
525.75732421875 0 764.08655
526.2622680664062 0 1335.4701
528.2321166992188 0 1582.8806
530.2098388671875 0 24126.092
530.333740234375 0 6313.0244
530.7726440429688 0 810.32904
531.2123413085938 0 5546.6343
531.33642578125 0 1454.9702
532.2130126953125 0 1107.2317
535.2710571289062 0 638.7949 b Water loss 9
535.76708984375 0 787.12555
540.3186645507812 0 4733.9634 b Water loss 4
541.3226318359375 0 1114.5542
542.2593383789062 0 691.3817
544.2687377929688 0 5720.1353 b 9
544.770751953125 0 4933.658
545.2714233398438 0 1871.5397
546.2404174804688 0 3147.6167
547.2448120117188 0 799.75244
551.2808227539062 0 943.8485
554.2531127929688 0 683.94257
558.3290405273438 0 11202.674 b 4
559.3317260742188 0 4227.081
560.3333740234375 0 988.6974
563.2468872070312 0 1512.6439
569.251708984375 0 858.95764
570.25537109375 0 1764.2026
573.2272338867188 0 904.5902
579.2809448242188 0 762.7738
581.258056640625 0 6010.109
582.252197265625 0 1925.0646
583.2578735351562 0 768.84766
584.7572021484375 0 755.61566
587.2675170898438 0 4393.711
588.26953125 0 1558.5394
589.2874755859375 0 1569.6031 y 8
591.2417602539062 0 4593.2026
592.2420043945312 0 886.45325
595.34716796875 0 1232.0402
597.2894287109375 0 3569.1655
598.2904052734375 0 1549.8073
599.2680053710938 0 35488.914
599.7987670898438 0 737.92065
600.2705688476562 0 10728.091
600.3184204101562 0 966.7894
600.8118286132812 0 4956.4453 b 10
601.2684326171875 0 1440.6249
601.3140869140625 0 2100.65
601.811767578125 0 1028.1991
607.2703857421875 0 886.6312
609.2525634765625 0 12352.984
610.2543334960938 0 4066.2793
615.299560546875 0 4995.5264
616.3032836914062 0 1447.1292
617.2969360351562 0 6682.549
618.2988891601562 0 1764.1222
625.2839965820312 0 2977.6978
626.2852783203125 0 1757.9268
627.262451171875 0 137758.05
628.2657470703125 0 39808.887
629.2687377929688 0 7411.931
629.3262939453125 0 803.78577
634.3206787109375 0 1938.0751
637.3695068359375 0 1046.9314 b Water loss 5
642.3173828125 0 686.75226
643.29345703125 0 10910.48
644.2921752929688 0 8760.188
645.2899780273438 0 4766.7397
646.2943115234375 0 1400.9027
655.3807983398438 0 3944.3652 b 5
656.384521484375 0 1898.4921
659.2868041992188 0 1358.5543 y Ammonia loss 7
660.32177734375 0 2783.9119
661.3236083984375 0 1079.952
663.31298828125 0 905.29565
674.2991943359375 0 2452.2986
676.3182373046875 0 4487.039 y 7
677.316162109375 0 1314.6882
680.3214721679688 0 1015.2081
686.306640625 0 720.54913
686.380126953125 0 657.47284
691.3248901367188 0 1050.0856
694.3409423828125 0 4536.5205
695.3353271484375 0 1648.0668
698.3336791992188 0 1639.502
704.326171875 0 2537.5845
705.322509765625 0 1571.098
708.315185546875 0 806.67834
708.3794555664062 0 798.67786
708.695556640625 0 709.75665
712.3515014648438 0 33542.383
713.35546875 0 13519.533
714.35498046875 0 2201.658
714.4154663085938 0 1017.9977
722.3361206054688 0 8331.232
723.3399047851562 0 2922.3167
724.4024047851562 0 2818.8115 b Water loss 6
725.4080810546875 0 1539.4277
726.33056640625 0 3430.1064
727.3303833007812 0 1371.1199
728.3267211914062 0 1054.2721
729.3174438476562 0 1573.4766
735.3390502929688 0 890.882
739.3622436523438 0 1652.9137
740.346435546875 0 74304.375
741.3494262695312 0 27168.252
742.3500366210938 0 5943.97
742.4146118164062 0 11748.489 b 6
743.353271484375 0 1442.389
743.4176635742188 0 4955.3657
744.4192504882812 0 1431.6733
746.3338623046875 0 2634.4048 y Ammonia loss 6
747.33349609375 0 886.4929
752.3649291992188 0 1675.7094
753.3488159179688 0 1883.4353
754.3345336914062 0 1030.991
756.3211059570312 0 1321.4923
757.3729248046875 0 20970.25
758.375 0 11202.099
759.3761596679688 0 2714.133
763.3492431640625 0 1557.3231 y 6
768.3590087890625 0 1269.9783
769.3658447265625 0 637.3048
771.3516235351562 0 14033.789
772.3536987304688 0 6590.0093
773.3583374023438 0 999.5978
774.3307495117188 0 10219.015
775.334228515625 0 5726.0083
776.33349609375 0 1476.775
780.3575439453125 0 2050.126
781.3641357421875 0 1442.9796
788.3875122070312 0 855.78534
791.3609619140625 0 1358.0212
794.3375244140625 0 1006.48944
795.3353271484375 0 901.88165
795.439697265625 0 738.6335
796.3504638671875 0 2692.159
797.36962890625 0 1637.9031
801.44921875 0 1253.2645
809.37109375 0 1437.514
811.4285278320312 0 4587.9956 b Water loss 7
812.3485717773438 0 1117.6135
812.4332275390625 0 2102.23
821.4005126953125 0 777.2193
822.4058227539062 0 1074.7571
823.3751831054688 0 899.5782
823.4490966796875 0 1078.6476
825.3790893554688 0 987.3381
826.3720092773438 0 781.9962
827.3798217773438 0 2266.2715
828.3805541992188 0 838.3067
829.4456787109375 0 5999.3535 b 7
830.4492797851562 0 3246.453
832.3798217773438 0 729.94824
833.3809204101562 0 933.925
839.41455078125 0 8433.551
840.4130249023438 0 7127.9814
841.4092407226562 0 4950.7363
842.3950805664062 0 3581.9763 y Water loss 5
843.3798217773438 0 5731.2705 y Ammonia loss 5
844.3858642578125 0 1756.266
845.3961181640625 0 815.7663
850.3934936523438 0 7645.3516
851.3892211914062 0 4559.656
852.390869140625 0 1607.904
853.4105224609375 0 836.4335
859.4197387695312 0 4563.5107
860.4010009765625 0 23930.982 y 5
861.4036865234375 0 11182.354
862.4058837890625 0 2020.9011
868.4047241210938 0 88440.57
869.4068603515625 0 40184.96
869.50732421875 0 1124.761
870.4080200195312 0 9538.388
871.4095458984375 0 836.9682
886.4146728515625 0 2322.6978
887.4142456054688 0 15945.46
888.4159545898438 0 6686.2817
889.4197998046875 0 1891.5604
895.4168090820312 0 1293.3317
896.4478759765625 0 894.8651
904.4364624023438 0 1679.1272
905.4317626953125 0 939.04
912.4649658203125 0 1067.5825
913.4791870117188 0 712.66583
915.4290771484375 0 1041.24
916.4274291992188 0 962.28766
921.4326171875 0 2028.4556
922.4439697265625 0 3993.382
923.43408203125 0 2473.4294
926.4253540039062 0 958.1171
927.426025390625 0 809.09106
930.4880981445312 0 1387.3115
939.4412231445312 0 11630.844 y Water loss 4
940.4319458007812 0 29957.852 y Ammonia loss 4
941.4310913085938 0 15712.821
942.4338989257812 0 5554.401
943.4231567382812 0 5128.914
944.427734375 0 2821.3948
956.4489135742188 0 1115.0983
957.4521484375 0 254650.14 y 4
958.4566650390625 0 127949.46
959.4605102539062 0 38322.695
960.468994140625 0 4831.6846
970.456787109375 0 920.7165
971.4554443359375 0 1009.79333
972.4634399414062 0 832.0323
977.4209594726562 0 950.0827
986.4778442382812 0 1376.4155
987.4821166992188 0 1915.0706
988.4647216796875 0 4430.001
989.4688720703125 0 2058.7349
990.4727783203125 0 738.42114
1000.4974365234375 0 3013.9429
1001.4945678710938 0 1743.8008
1015.4719848632812 0 5402.855
1016.4752807617188 0 3681.4143
1017.4845581054688 0 754.6299
1035.5364990234375 0 1978.9167
1036.53466796875 0 1389.4441
1043.5164794921875 0 784.65656
1050.5145263671875 0 805.62756
1051.5013427734375 0 1171.1437
1052.5113525390625 0 861.06274
1053.5438232421875 0 4208.353
1054.5489501953125 0 1971.8586
1055.5445556640625 0 905.9628
1059.537841796875 0 2581.7332
1060.5396728515625 0 1243.2812
1069.5191650390625 0 6114.369 b Water loss 9
1070.5216064453125 0 4584.7
1071.5269775390625 0 1896.806
1083.538330078125 0 1585.2891
1084.5369873046875 0 1031.349
1085.519775390625 0 1554.9374
1086.5152587890625 0 1655.1646 y Water loss 3
1087.5289306640625 0 62893.96 b 9
1088.53173828125 0 40276.094
1088.674072265625 0 1201.3835
1089.5347900390625 0 12774.033
1090.5369873046875 0 1044.5619
1101.5457763671875 0 2721.8184
1102.5474853515625 0 2018.5349
1103.5567626953125 0 1460.9502
1104.5208740234375 0 12749.202 y 3
1105.5230712890625 0 7267.8228
1106.5262451171875 0 1152.1202
1128.547119140625 0 975.5556
1129.5545654296875 0 1012.3571
1151.502197265625 0 845.9343
1156.6053466796875 0 931.4313
1164.5848388671875 0 1718.3021
1165.591552734375 0 1107.6694
1167.4991455078125 0 1880.9298
1168.5035400390625 0 2184.3992
1169.5162353515625 0 5574.3647
1170.520263671875 0 3863.437
1171.5220947265625 0 1453.1609
1172.6185302734375 0 2216.7134
1173.6224365234375 0 2312.9905
1182.6038818359375 0 7783.8955 b Water loss 10
1183.604736328125 0 5125.8457
1184.611083984375 0 1882.3424
1186.5186767578125 0 1604.6361
1198.5980224609375 0 742.3263
1199.6087646484375 0 659.8776 y Water loss 2
1200.613525390625 0 34250.504 b 10
1201.6158447265625 0 22602.521
1202.6199951171875 0 8320.622
1203.6307373046875 0 1046.3857
1217.6077880859375 0 1370.1919 y 2
1218.607421875 0 1219.723
1635.6297607421875 0 801.4468
1638.6280517578125 0 660.49603
3028.255615234375 0 684.86444

Spectrum Details

|  |  |
| --- | --- |
| Matched peaks? Matched peaks The total absolute number of peaks matched. Additionally in brackets the total fraction of peaks matched and the total number of peaks is shown. | 51 (7.52% of 678) |
| FDR? FDR The false discovery rate estimated for this peptide. It is calculated by matching all theoretical fragments with a non-integer shift with the raw peaks for this spectrum. This is done with 40 different shifts. The resulting percentage is the average number of annotated peaks over the number of annotated peaks with the correct spectrum. | 1.49% |
| Satellite FDR? Satellite FDR See the FDR for details on its calculation. This satellite ion specific FDR only contains the satellite ions (d/w) for I/L/J positions. | - |
| PSM Score? PSM Score The PSM Score as given by Hecklib to this annotated spectrum. It is shown with three significant figures. | 704 |

#### Reverse Lookup? Reverse Lookup All places where this read could be placed.

| Group | Segment | Template | Template Part | Read Part | Score | Unique |
| --- | --- | --- | --- | --- | --- | --- |
| Homo sapiens Light Chain | IGLC | IGLC2 | [8..21] | [0..13] | 104 | False |
| Homo sapiens Light Chain | IGLC | IGLC3 | [6..19] | [0..13] | 104 | False |
| Homo sapiens Light Chain | IGLC | IGLC6 | [8..21] | [0..13] | 104 | False |
| Homo sapiens Light Chain | IGLC | IGLC7 | [8..21] | [0..13] | 104 | False |

| Recombined | Template Part | Read Part | Score | Unique |
| --- | --- | --- | --- | --- |
| REC-0-1\_002 | [119..132] | [0..13] | 104 | True |

#### Meta Information from Multiple reads

##### Number of combined reads

6

##### Intensity

0.7242

##### TotalArea

3.605E+08

##### Changes to the peptide sequence

VTJFPPSSEEJQA

L→JNo support for either Leucine or Isoleucine based on side chain ions (Position: 3)

J→LSupport for Leucine based on side chain ions (1 for L 0 for I) (Position: 3)

L→JNo support for either Leucine or Isoleucine based on side chain ions (Position: 11)

L→JNo support for either Leucine or Isoleucine based on side chain ions (Position: 3)

#### Positional Score

Copy Data

##### Positional Score (TSV)

###### Preview

```
Loading example...
```

*Click on the button to copy the data to your clipboard.*

100123456789101112

Label Value
"0" 0.608
"1" 0.59
"2" 0.647
"3" 0.647
"4" 0.657
"5" 0.653
"6" 0.652
"7" 0.655
"8" 0.653
"9" 0.662
"10" 0.667
"11" 0.65
"12" 0.657

#### Meta Information from PEAKS

##### Scan Identifier

F1:9705

##### Original sequence

V

T

L

F

P

P

S

S

E

E

L

Q

A

##### Posttranslational Modifications

##### Source File

D:\separate\_stitch\_analyses\xle-disambiguation\raw\20210323\_F1\_UM1\_Peng0013\_SA\_F59\_ingel\_3ug\_ELA.raw

##### Fraction

1

##### Scan Feature

F1:13275

##### De Novo Score

98

##### ConfidenceScore

98

### m/z

709.3657

##### Mass

1416.7136

##### Charge

2

##### Retention Time

54.13

##### Predicted Retention Time

-

##### Area

6.009E+07

##### Parts Per Million

2.3

##### Fragmentation mode

ETHCD

##### Originating file

01 D:\separate\_stitch\_analyses\xle-disambiguation\20210325\_F59\_3ug\_DENOVO\_12.csv

#### Meta Information from PEAKS

##### Scan Identifier

F1:10021

##### Original sequence

V

T

L

F

P

P

S

S

E

E

L

Q

A

##### Posttranslational Modifications

##### Source File

D:\separate\_stitch\_analyses\xle-disambiguation\raw\20210323\_F1\_UM1\_Peng0013\_SA\_F59\_ingel\_3ug\_ELA.raw

##### Fraction

1

##### Scan Feature

F1:13275

##### De Novo Score

98

##### ConfidenceScore

98

### m/z

709.3657

##### Mass

1416.7136

##### Charge

2

##### Retention Time

54.13

##### Predicted Retention Time

-

##### Area

6.009E+07

##### Parts Per Million

2.3

##### Fragmentation mode

ETHCD

##### Originating file

01 D:\separate\_stitch\_analyses\xle-disambiguation\20210325\_F59\_3ug\_DENOVO\_12.csv

#### Meta Information from PEAKS

##### Scan Identifier

F1:10080

##### Original sequence

V

T

L

F

P

P

S

S

E

E

L

Q

A

##### Posttranslational Modifications

##### Source File

D:\separate\_stitch\_analyses\xle-disambiguation\raw\20210323\_F1\_UM1\_Peng0013\_SA\_F59\_ingel\_3ug\_ELA.raw

##### Fraction

1

##### Scan Feature

F1:13275

##### De Novo Score

98

##### ConfidenceScore

98

### m/z

709.3657

##### Mass

1416.7136

##### Charge

2

##### Retention Time

54.13

##### Predicted Retention Time

-

##### Area

6.009E+07

##### Parts Per Million

2.3

##### Fragmentation mode

ETHCD

##### Originating file

01 D:\separate\_stitch\_analyses\xle-disambiguation\20210325\_F59\_3ug\_DENOVO\_12.csv

#### Meta Information from PEAKS

##### Scan Identifier

F1:10143

##### Original sequence

V

T

L

F

P

P

S

S

E

E

L

Q

A

##### Posttranslational Modifications

##### Source File

D:\separate\_stitch\_analyses\xle-disambiguation\raw\20210323\_F1\_UM1\_Peng0013\_SA\_F59\_ingel\_3ug\_ELA.raw

##### Fraction

1

##### Scan Feature

F1:13275

##### De Novo Score

98

##### ConfidenceScore

98

### m/z

709.3657

##### Mass

1416.7136

##### Charge

2

##### Retention Time

54.13

##### Predicted Retention Time

-

##### Area

6.009E+07

##### Parts Per Million

2.3

##### Fragmentation mode

ETHCD

##### Originating file

01 D:\separate\_stitch\_analyses\xle-disambiguation\20210325\_F59\_3ug\_DENOVO\_12.csv

#### Meta Information from PEAKS

##### Scan Identifier

F1:9831

##### Original sequence

V

T

L

F

P

P

S

S

E

E

L

Q

A

##### Posttranslational Modifications

##### Source File

D:\separate\_stitch\_analyses\xle-disambiguation\raw\20210323\_F1\_UM1\_Peng0013\_SA\_F59\_ingel\_3ug\_ELA.raw

##### Fraction

1

##### Scan Feature

F1:13275

##### De Novo Score

95

##### ConfidenceScore

95

### m/z

709.3657

##### Mass

1416.7136

##### Charge

2

##### Retention Time

54.13

##### Predicted Retention Time

-

##### Area

6.009E+07

##### Parts Per Million

2.3

##### Fragmentation mode

ETHCD

##### Originating file

01 D:\separate\_stitch\_analyses\xle-disambiguation\20210325\_F59\_3ug\_DENOVO\_12.csv

#### Meta Information from PEAKS

##### Scan Identifier

F1:9903

##### Original sequence

V

T

L

F

P

P

S

S

E

E

L

Q

A

##### Posttranslational Modifications

##### Source File

D:\separate\_stitch\_analyses\xle-disambiguation\raw\20210323\_F1\_UM1\_Peng0013\_SA\_F59\_ingel\_3ug\_ELA.raw

##### Fraction

1

##### Scan Feature

F1:13275

##### De Novo Score

95

##### ConfidenceScore

95

### m/z

709.3657

##### Mass

1416.7136

##### Charge

2

##### Retention Time

54.13

##### Predicted Retention Time

-

##### Area

6.009E+07

##### Parts Per Million

2.3

##### Fragmentation mode

HCD

##### Originating file

01 D:\separate\_stitch\_analyses\xle-disambiguation\20210325\_F59\_3ug\_DENOVO\_12.csv
