## Supplementary material for "A handle on mass coincidence errors in *de novo* sequencing of antibodies by bottom-up proteomics": Combined_050.html

Details Combined\_050 | Stitch OverviewUndefined

### Read Combined\_050

#### Sequence (length=12)

SNKALPAPJEKT

#### Spectrum 3813? Spectrum 3813 The raw spectrum of this peptide as annotated by Hecklib. The fragments are coloured according to ion type (see legend). Any peaks with a star '\*' as text can be hovered over to see the full details, first the ion type second the mass shift type. By hovering over the amino acids in the peptide or ions in the legend the corresponding peaks are highlighted. By toggling the 'Unassigned' label you can turn the background (unassigned) peaks on or off in the plot. By updating the slider in the Ion legend you can update the spectrum to only show the top X% of the peaks with labels. The top X% means any peak that is within X% of the highest intensity. By dragging in the spectrum you can zoom in to a specific part of the spectrum and use 'Zoom Out' to get back to the original zoom level. The annotation of the spectrum is based on the given sequence in the peptides file and is done with different software so inconsistencies are likely. The peaks are annotated based on the given sequence, with 20 ppm tolerance.

Copy Data

##### Spectrum 3813 (TSV)

###### Preview

```
Loading example...
```

*Click on the button to copy the data to your clipboard.*

Mz MinMz MaxIntensity Max

WidthHeightPeptide font sizePeptide stroke widthSpectrum font sizeSpectrum stroke widthCompact peptide

Ion legend

wxyz

abcd

OtherUnassignedIon^Charge^_Position_Show for top:%

SNKALPAPJEKT

07.97e+41.59e+52.39e+53.19e+5

Zoom Out

y^+1^_1_c^+2^_3_w^+1^_2_c^+1^_2_c^+1^_2_w^+2^_4_y^+1^_2_z^+1^_2_z^+3^_7_y^+1^_2_c^+2^_5_y^+2^_5_y^+2^_5_z^+2^_5_y^+2^_5_w^+1^_3_c^+2^_6_z^+2^_6_c^+1^_3_c^+1^_3_c^+3^_10_c^+2^_7_z^+1^_3_c^+1^_3_y^+1^_3_z^+1^_3_y^+1^_3_y^+2^_7_c^+2^_8_c^+1^_4_c^+1^_4_c^+1^_4_y^+2^_8_w^+1^_4_w^+1^_4_z^+2^_9_y^+2^_9_y^+1^_4_z^+1^_4_y^+1^_4_c^+2^_10_c^+1^_5_z^+2^_10_z^+2^_10_c^+2^_10_y^+2^_10_z^+2^_10_y^+2^_10_w^+1^_5_w^+2^_11_y^+1^_5_z^+1^_5_z^+2^_11_c^+2^_11_z^+2^_11_c^+2^_11_y^+1^_5_y^+2^_11_c^+1^_6_z^+1^_6_c^+1^_6_y^+1^_6_z^+1^_6_y^+1^_6_c^+1^_7_c^+1^_7_w^+1^_7_y^+1^_7_c^+1^_8_w^+1^_8_y^+1^_8_z^+1^_8_c^+1^_9_c^+1^_9_z^+1^_9_y^+1^_9_w^+1^_10_c^+1^_10_c^+1^_10_y^+1^_10_z^+1^_10_y^+1^_10_w^+1^_11_z^+1^_11_c^+1^_11_y^+1^_11_

0792158323753167

Fragment Matches Table

Show background peaks

| Position | Ion type | Intensity | mz Theoretical | mz Error (Th) | mz Error (ppm) | Charge | Series Number |
| --- | --- | --- | --- | --- | --- | --- | --- |
| - | - | 536.6 | 120.1 | - | - | 0 | - |
| 12 | y | 8303 | 120.1 | 0.0003449 | 2.873 | +1 | 1 |
| - | - | 2101 | 127.1 | - | - | 0 | - |
| - | - | 961.4 | 128.1 | - | - | 0 | - |
| - | - | 2795 | 128.1 | - | - | 0 | - |
| - | - | 436.6 | 128.1 | - | - | 0 | - |
| - | - | 4.162E+04 | 129.1 | - | - | 0 | - |
| - | - | 2622 | 130.1 | - | - | 0 | - |
| - | - | 2813 | 131.1 | - | - | 0 | - |
| - | - | 541.4 | 132.1 | - | - | 0 | - |
| - | - | 1348 | 133.1 | - | - | 0 | - |
| - | - | 478.2 | 134.2 | - | - | 0 | - |
| - | - | 569.6 | 141 | - | - | 0 | - |
| - | - | 2.153E+04 | 141.1 | - | - | 0 | - |
| - | - | 5821 | 141.1 | - | - | 0 | - |
| - | - | 605.2 | 142.1 | - | - | 0 | - |
| - | - | 1818 | 142.1 | - | - | 0 | - |
| - | - | 534.6 | 146.1 | - | - | 0 | - |
| - | - | 7954 | 146.1 | - | - | 0 | - |
| - | - | 472.4 | 147.1 | - | - | 0 | - |
| - | - | 1602 | 155.1 | - | - | 0 | - |
| - | - | 2281 | 156.1 | - | - | 0 | - |
| - | - | 3240 | 157.1 | - | - | 0 | - |
| - | - | 3057 | 157.1 | - | - | 0 | - |
| - | - | 2335 | 158.1 | - | - | 0 | - |
| 3 | c | 3210 | 165.1 | 0.002418 | 14.64 | +2 | 3 |
| - | - | 1105 | 167 | - | - | 0 | - |
| - | - | 8868 | 169.1 | - | - | 0 | - |
| - | - | 434.3 | 169.5 | - | - | 0 | - |
| - | - | 971.7 | 170.1 | - | - | 0 | - |
| - | - | 700.7 | 170.1 | - | - | 0 | - |
| 11 | w | 8648 | 174.1 | 0.0003928 | 2.256 | +1 | 2 |
| - | - | 1638 | 174.1 | - | - | 0 | - |
| - | - | 743.7 | 175.1 | - | - | 0 | - |
| - | - | 791.8 | 177.1 | - | - | 0 | - |
| - | - | 3.81E+04 | 183.1 | - | - | 0 | - |
| - | - | 7163 | 183.1 | - | - | 0 | - |
| - | - | 654.7 | 184.1 | - | - | 0 | - |
| - | - | 4061 | 184.1 | - | - | 0 | - |
| - | - | 994.8 | 184.2 | - | - | 0 | - |
| - | - | 2190 | 185.1 | - | - | 0 | - |
| - | - | 5030 | 185.1 | - | - | 0 | - |
| - | - | 1743 | 185.1 | - | - | 0 | - |
| - | - | 1437 | 187.1 | - | - | 0 | - |
| - | - | 990.5 | 188.1 | - | - | 0 | - |
| - | - | 667.8 | 197.2 | - | - | 0 | - |
| - | - | 1441 | 199.1 | - | - | 0 | - |
| - | - | 785.7 | 200.1 | - | - | 0 | - |
| - | - | 784.2 | 201.1 | - | - | 0 | - |
| - | - | 867.5 | 201.1 | - | - | 0 | - |
| 2 | c | 1.121E+04 | 202.1 | 0.0003177 | 1.572 | +1 | 2 |
| - | - | 1688 | 202.1 | - | - | 0 | - |
| - | - | 1261 | 203.1 | - | - | 0 | - |
| - | - | 487.4 | 206.6 | - | - | 0 | - |
| - | - | 1095 | 208.1 | - | - | 0 | - |
| - | - | 6339 | 211.1 | - | - | 0 | - |
| - | - | 883.2 | 212.1 | - | - | 0 | - |
| - | - | 554.6 | 212.6 | - | - | 0 | - |
| - | - | 632.2 | 215.1 | - | - | 0 | - |
| - | - | 954.4 | 216.1 | - | - | 0 | - |
| - | - | 6078 | 217.2 | - | - | 0 | - |
| - | - | 557 | 217.7 | - | - | 0 | - |
| - | - | 581.5 | 218.2 | - | - | 0 | - |
| 2 | c | 6701 | 219.1 | 0.0005326 | 2.431 | +1 | 2 |
| - | - | 668.4 | 219.1 | - | - | 0 | - |
| - | - | 524.2 | 222.1 | - | - | 0 | - |
| - | - | 2833 | 225.1 | - | - | 0 | - |
| - | - | 548.6 | 225.5 | - | - | 0 | - |
| - | - | 6301 | 226.1 | - | - | 0 | - |
| - | - | 1094 | 227.1 | - | - | 0 | - |
| - | - | 4078 | 229.1 | - | - | 0 | - |
| 9 | w | 687.4 | 230.1 | 0.004237 | 18.41 | +2 | 4 |
| 11 | y | 958.7 | 230.1 | 6.138E-05 | 0.2667 | +1 | 2 |
| 11 | z | 3444 | 232.1 | 0.0004076 | 1.756 | +1 | 2 |
| - | - | 1280 | 235.1 | - | - | 0 | - |
| - | - | 782.5 | 235.6 | - | - | 0 | - |
| - | - | 665.3 | 239.1 | - | - | 0 | - |
| - | - | 1.694E+04 | 240.1 | - | - | 0 | - |
| - | - | 616.6 | 240.6 | - | - | 0 | - |
| 6 | z | 1121 | 241.1 | 0.0007599 | 3.152 | +3 | 7 |
| - | - | 3539 | 242.1 | - | - | 0 | - |
| - | - | 3253 | 243.1 | - | - | 0 | - |
| - | - | 2514 | 243.1 | - | - | 0 | - |
| - | - | 8811 | 243.7 | - | - | 0 | - |
| - | - | 2349 | 244.2 | - | - | 0 | - |
| - | - | 961.5 | 245.1 | - | - | 0 | - |
| 11 | y | 1.292E+04 | 248.2 | 0.0004823 | 1.944 | +1 | 2 |
| - | - | 1779 | 249.2 | - | - | 0 | - |
| - | - | 9138 | 249.6 | - | - | 0 | - |
| - | - | 1786 | 250.1 | - | - | 0 | - |
| - | - | 1007 | 253.2 | - | - | 0 | - |
| 5 | c | 1630 | 257.7 | 0.0004345 | 1.686 | +2 | 5 |
| - | - | 5960 | 258.1 | - | - | 0 | - |
| - | - | 524.2 | 259.1 | - | - | 0 | - |
| - | - | 1525 | 259.2 | - | - | 0 | - |
| - | - | 698.8 | 262.6 | - | - | 0 | - |
| - | - | 1268 | 268.2 | - | - | 0 | - |
| - | - | 978.7 | 270.7 | - | - | 0 | - |
| - | - | 657 | 271.2 | - | - | 0 | - |
| - | - | 1954 | 276.2 | - | - | 0 | - |
| - | - | 871.8 | 282.2 | - | - | 0 | - |
| - | - | 1757 | 283.7 | - | - | 0 | - |
| - | - | 1.497E+04 | 284.2 | - | - | 0 | - |
| - | - | 2942 | 285.2 | - | - | 0 | - |
| 8 | y | 6337 | 285.2 | 9.477E-05 | 0.3323 | +2 | 5 |
| 8 | y | 2495 | 285.7 | 0.001563 | 5.472 | +2 | 5 |
| - | - | 1071 | 285.7 | - | - | 0 | - |
| 8 | z | 1072 | 286.2 | 0.001691 | 5.91 | +2 | 5 |
| - | - | 829.4 | 288.7 | - | - | 0 | - |
| - | - | 2307 | 289.2 | - | - | 0 | - |
| - | - | 750.5 | 290.1 | - | - | 0 | - |
| - | - | 3.029E+04 | 292.2 | - | - | 0 | - |
| - | - | 8271 | 292.7 | - | - | 0 | - |
| - | - | 1844 | 293.2 | - | - | 0 | - |
| 8 | y | 1.086E+05 | 294.2 | 0.0006043 | 2.054 | +2 | 5 |
| - | - | 3.609E+04 | 294.7 | - | - | 0 | - |
| - | - | 4055 | 295.1 | - | - | 0 | - |
| - | - | 5860 | 295.2 | - | - | 0 | - |
| - | - | 865.3 | 295.7 | - | - | 0 | - |
| - | - | 3333 | 297.2 | - | - | 0 | - |
| - | - | 2091 | 297.7 | - | - | 0 | - |
| - | - | 1540 | 298.2 | - | - | 0 | - |
| - | - | 573.9 | 298.7 | - | - | 0 | - |
| 10 | w | 1.179E+05 | 302.2 | 0.0007056 | 2.335 | +1 | 3 |
| - | - | 1061 | 303.1 | - | - | 0 | - |
| - | - | 1.671E+04 | 303.2 | - | - | 0 | - |
| - | - | 2200 | 304.2 | - | - | 0 | - |
| 6 | c | 5.059E+04 | 306.2 | 0.0006639 | 2.168 | +2 | 6 |
| - | - | 1.692E+04 | 306.7 | - | - | 0 | - |
| - | - | 5043 | 307.2 | - | - | 0 | - |
| - | - | 6664 | 312.2 | - | - | 0 | - |
| - | - | 620.8 | 312.2 | - | - | 0 | - |
| - | - | 1.164E+04 | 313.2 | - | - | 0 | - |
| 7 | z | 998.1 | 313.2 | 0.0006252 | 1.996 | +2 | 6 |
| - | - | 3174 | 313.2 | - | - | 0 | - |
| - | - | 2709 | 314.2 | - | - | 0 | - |
| - | - | 990.5 | 316.2 | - | - | 0 | - |
| - | - | 4142 | 319.2 | - | - | 0 | - |
| - | - | 632.7 | 319.7 | - | - | 0 | - |
| - | - | 6742 | 327.2 | - | - | 0 | - |
| - | - | 2.245E+04 | 327.7 | - | - | 0 | - |
| - | - | 8340 | 328.2 | - | - | 0 | - |
| - | - | 2152 | 328.7 | - | - | 0 | - |
| 3 | c | 818.3 | 329.2 | 0.001987 | 6.036 | +1 | 3 |
| 3 | c | 2.34E+04 | 330.2 | 0.0006916 | 2.095 | +1 | 3 |
| - | - | 3834 | 331.2 | - | - | 0 | - |
| - | - | 778.7 | 331.2 | - | - | 0 | - |
| - | - | 976.9 | 332.2 | - | - | 0 | - |
| - | - | 1204 | 332.7 | - | - | 0 | - |
| - | - | 2520 | 333.2 | - | - | 0 | - |
| - | - | 1202 | 333.7 | - | - | 0 | - |
| - | - | 668.8 | 334.2 | - | - | 0 | - |
| - | - | 918.5 | 334.2 | - | - | 0 | - |
| - | - | 1.501E+04 | 340.2 | - | - | 0 | - |
| 10 | c | 2921 | 341.2 | 0.00453 | 13.28 | +3 | 10 |
| 7 | c | 1.262E+04 | 341.7 | 0.0006922 | 2.026 | +2 | 7 |
| - | - | 4422 | 342.2 | - | - | 0 | - |
| - | - | 1578 | 342.7 | - | - | 0 | - |
| 10 | z | 645.1 | 343.2 | 0.002055 | 5.989 | +1 | 3 |
| - | - | 1278 | 344.2 | - | - | 0 | - |
| 3 | c | 3.695E+04 | 347.2 | 0.0007539 | 2.171 | +1 | 3 |
| - | - | 5864 | 348.2 | - | - | 0 | - |
| - | - | 1045 | 353.2 | - | - | 0 | - |
| 10 | y | 7561 | 359.2 | 0.0004821 | 1.342 | +1 | 3 |
| - | - | 1023 | 360.2 | - | - | 0 | - |
| 10 | z | 8451 | 361.2 | 0.0005849 | 1.619 | +1 | 3 |
| - | - | 1492 | 362.2 | - | - | 0 | - |
| - | - | 1085 | 366.2 | - | - | 0 | - |
| - | - | 984.4 | 368.2 | - | - | 0 | - |
| - | - | 1885 | 371.2 | - | - | 0 | - |
| - | - | 1045 | 373.2 | - | - | 0 | - |
| - | - | 579.3 | 373.3 | - | - | 0 | - |
| - | - | 1434 | 374.2 | - | - | 0 | - |
| 10 | y | 1.757E+04 | 377.2 | 0.0005986 | 1.587 | +1 | 3 |
| 6 | y | 1.684E+04 | 378.2 | 9.91E-05 | 0.262 | +2 | 7 |
| - | - | 5905 | 378.7 | - | - | 0 | - |
| - | - | 824.8 | 379.2 | - | - | 0 | - |
| - | - | 682.8 | 379.7 | - | - | 0 | - |
| - | - | 712.8 | 380.2 | - | - | 0 | - |
| - | - | 717.2 | 381.2 | - | - | 0 | - |
| - | - | 1.072E+04 | 383.2 | - | - | 0 | - |
| - | - | 2231 | 384.2 | - | - | 0 | - |
| - | - | 1161 | 384.2 | - | - | 0 | - |
| 8 | c | 4171 | 390.2 | 0.0007995 | 2.049 | +2 | 8 |
| - | - | 734.8 | 390.7 | - | - | 0 | - |
| - | - | 942.6 | 391.2 | - | - | 0 | - |
| - | - | 803.9 | 400.2 | - | - | 0 | - |
| 4 | c | 1336 | 400.2 | 0.00106 | 2.649 | +1 | 4 |
| 4 | c | 4.164E+04 | 401.2 | 0.0009008 | 2.245 | +1 | 4 |
| - | - | 8097 | 402.2 | - | - | 0 | - |
| - | - | 1265 | 403.2 | - | - | 0 | - |
| - | - | 837.3 | 410.3 | - | - | 0 | - |
| - | - | 1520 | 411.2 | - | - | 0 | - |
| - | - | 839.4 | 415.3 | - | - | 0 | - |
| 4 | c | 9.114E+04 | 418.2 | 0.000902 | 2.157 | +1 | 4 |
| - | - | 1.941E+04 | 419.2 | - | - | 0 | - |
| - | - | 2392 | 420.2 | - | - | 0 | - |
| - | - | 649.3 | 423.2 | - | - | 0 | - |
| - | - | 824.7 | 425.2 | - | - | 0 | - |
| - | - | 3841 | 427.2 | - | - | 0 | - |
| - | - | 875.3 | 427.3 | - | - | 0 | - |
| - | - | 1266 | 428.2 | - | - | 0 | - |
| 5 | y | 1852 | 434.8 | 0.0007307 | 1.681 | +2 | 8 |
| - | - | 1218 | 435.3 | - | - | 0 | - |
| - | - | 678.9 | 439.2 | - | - | 0 | - |
| - | - | 822.9 | 443.3 | - | - | 0 | - |
| - | - | 680.1 | 443.3 | - | - | 0 | - |
| - | - | 5130 | 444.3 | - | - | 0 | - |
| 9 | w | 2.493E+04 | 445.2 | 0.001087 | 2.441 | +1 | 4 |
| - | - | 5165 | 446.2 | - | - | 0 | - |
| - | - | 1223 | 447.2 | - | - | 0 | - |
| - | - | 1525 | 455.3 | - | - | 0 | - |
| 9 | w | 2147 | 459.2 | 0.001458 | 3.176 | +1 | 4 |
| 4 | z | 1105 | 462.3 | 0.007527 | 16.28 | +2 | 9 |
| - | - | 1.624E+04 | 468.3 | - | - | 0 | - |
| - | - | 4813 | 469.3 | - | - | 0 | - |
| 4 | y | 1603 | 470.3 | 0.003322 | 7.065 | +2 | 9 |
| - | - | 745.7 | 471.3 | - | - | 0 | - |
| 9 | y | 1517 | 472.3 | 0.001898 | 4.019 | +1 | 4 |
| 9 | z | 2193 | 474.3 | 0.0007494 | 1.58 | +1 | 4 |
| - | - | 1939 | 475.8 | - | - | 0 | - |
| - | - | 1864 | 476.3 | - | - | 0 | - |
| - | - | 832.4 | 479.3 | - | - | 0 | - |
| - | - | 1476 | 480.3 | - | - | 0 | - |
| - | - | 1090 | 481.2 | - | - | 0 | - |
| - | - | 6075 | 481.3 | - | - | 0 | - |
| - | - | 1927 | 482.3 | - | - | 0 | - |
| - | - | 1967 | 485.3 | - | - | 0 | - |
| - | - | 8810 | 486.3 | - | - | 0 | - |
| - | - | 4225 | 487.3 | - | - | 0 | - |
| - | - | 1302 | 488.3 | - | - | 0 | - |
| - | - | 722.2 | 489.3 | - | - | 0 | - |
| - | - | 891.2 | 489.8 | - | - | 0 | - |
| 9 | y | 2.205E+04 | 490.3 | 0.00116 | 2.366 | +1 | 4 |
| - | - | 1188 | 490.8 | - | - | 0 | - |
| - | - | 5302 | 491.3 | - | - | 0 | - |
| - | - | 1197 | 492.3 | - | - | 0 | - |
| - | - | 2657 | 496.3 | - | - | 0 | - |
| - | - | 3903 | 497.3 | - | - | 0 | - |
| - | - | 1426 | 497.3 | - | - | 0 | - |
| - | - | 3.013E+04 | 498.3 | - | - | 0 | - |
| - | - | 1.621E+04 | 498.8 | - | - | 0 | - |
| - | - | 7184 | 499.3 | - | - | 0 | - |
| - | - | 1453 | 499.8 | - | - | 0 | - |
| - | - | 1232 | 500.3 | - | - | 0 | - |
| - | - | 3757 | 504.8 | - | - | 0 | - |
| - | - | 1501 | 505.3 | - | - | 0 | - |
| - | - | 1023 | 505.8 | - | - | 0 | - |
| - | - | 1136 | 508.3 | - | - | 0 | - |
| 10 | c | 1508 | 511.3 | 0.006804 | 13.31 | +2 | 10 |
| - | - | 1240 | 511.8 | - | - | 0 | - |
| - | - | 739.8 | 511.9 | - | - | 0 | - |
| - | - | 1187 | 513.3 | - | - | 0 | - |
| 5 | c | 7.231E+04 | 514.3 | 0.0008823 | 1.715 | +1 | 5 |
| - | - | 1.865E+04 | 515.3 | - | - | 0 | - |
| - | - | 1243 | 516.3 | - | - | 0 | - |
| - | - | 3247 | 516.3 | - | - | 0 | - |
| 3 | z | 759.8 | 517.3 | 0.00269 | 5.199 | +2 | 10 |
| 3 | z | 1050 | 517.8 | 0.006767 | 13.07 | +2 | 10 |
| - | - | 1146 | 518.3 | - | - | 0 | - |
| - | - | 1228 | 519.3 | - | - | 0 | - |
| 10 | c | 937.3 | 519.8 | 0.00308 | 5.926 | +2 | 10 |
| - | - | 632.6 | 520.3 | - | - | 0 | - |
| - | - | 910.9 | 523.3 | - | - | 0 | - |
| - | - | 1280 | 524.3 | - | - | 0 | - |
| - | - | 1379 | 524.8 | - | - | 0 | - |
| 3 | y | 2109 | 525.8 | 0.0009467 | 1.801 | +2 | 10 |
| 3 | z | 3.844E+04 | 526.3 | 0.0001595 | 0.3031 | +2 | 10 |
| - | - | 2.373E+04 | 526.8 | - | - | 0 | - |
| - | - | 1.108E+04 | 527.3 | - | - | 0 | - |
| - | - | 3320 | 527.8 | - | - | 0 | - |
| - | - | 9171 | 528.3 | - | - | 0 | - |
| - | - | 2046 | 529.3 | - | - | 0 | - |
| - | - | 1532 | 532.3 | - | - | 0 | - |
| - | - | 2072 | 532.8 | - | - | 0 | - |
| - | - | 5184 | 533.8 | - | - | 0 | - |
| 3 | y | 2.151E+04 | 534.3 | 0.0003663 | 0.6856 | +2 | 10 |
| - | - | 1.082E+04 | 534.8 | - | - | 0 | - |
| - | - | 5142 | 535.3 | - | - | 0 | - |
| - | - | 1303 | 539.3 | - | - | 0 | - |
| - | - | 1.068E+04 | 540.3 | - | - | 0 | - |
| - | - | 3981 | 541.3 | - | - | 0 | - |
| - | - | 1130 | 542.3 | - | - | 0 | - |
| 8 | w | 7616 | 544.3 | 0.001002 | 1.84 | +1 | 5 |
| - | - | 3527 | 545.3 | - | - | 0 | - |
| - | - | 1682 | 547.8 | - | - | 0 | - |
| - | - | 1803 | 548.3 | - | - | 0 | - |
| - | - | 3606 | 548.8 | - | - | 0 | - |
| - | - | 1355 | 549.3 | - | - | 0 | - |
| - | - | 662.1 | 549.8 | - | - | 0 | - |
| - | - | 1244 | 551.3 | - | - | 0 | - |
| - | - | 1724 | 552.3 | - | - | 0 | - |
| - | - | 949.5 | 552.3 | - | - | 0 | - |
| - | - | 2646 | 553.8 | - | - | 0 | - |
| - | - | 8760 | 554.3 | - | - | 0 | - |
| - | - | 4083 | 554.8 | - | - | 0 | - |
| - | - | 1.31E+04 | 555.3 | - | - | 0 | - |
| - | - | 8730 | 555.8 | - | - | 0 | - |
| - | - | 3030 | 556.3 | - | - | 0 | - |
| - | - | 831.2 | 556.8 | - | - | 0 | - |
| - | - | 1887 | 557.3 | - | - | 0 | - |
| - | - | 2163 | 558.3 | - | - | 0 | - |
| - | - | 812 | 560.3 | - | - | 0 | - |
| - | - | 757.1 | 560.8 | - | - | 0 | - |
| 2 | w | 1.505E+04 | 561.3 | 0.00076 | 1.354 | +2 | 11 |
| - | - | 1.179E+04 | 561.8 | - | - | 0 | - |
| - | - | 5733 | 562.3 | - | - | 0 | - |
| - | - | 6900 | 562.8 | - | - | 0 | - |
| - | - | 4735 | 562.8 | - | - | 0 | - |
| - | - | 5903 | 563.3 | - | - | 0 | - |
| - | - | 1767 | 563.4 | - | - | 0 | - |
| - | - | 2534 | 563.8 | - | - | 0 | - |
| - | - | 1269 | 564.6 | - | - | 0 | - |
| - | - | 1673 | 565 | - | - | 0 | - |
| - | - | 902.7 | 565.6 | - | - | 0 | - |
| - | - | 707.1 | 566.8 | - | - | 0 | - |
| - | - | 3875 | 567.3 | - | - | 0 | - |
| - | - | 4304 | 567.8 | - | - | 0 | - |
| - | - | 1610 | 568.3 | - | - | 0 | - |
| - | - | 1864 | 568.8 | - | - | 0 | - |
| - | - | 1.032E+04 | 569.3 | - | - | 0 | - |
| 8 | y | 3451 | 570.3 | 0.005798 | 10.17 | +1 | 5 |
| - | - | 1887 | 570.3 | - | - | 0 | - |
| 8 | z | 2086 | 571.3 | 0.003894 | 6.815 | +1 | 5 |
| - | - | 1208 | 571.8 | - | - | 0 | - |
| - | - | 741 | 573.9 | - | - | 0 | - |
| 2 | z | 726.7 | 574.3 | 0.001542 | 2.685 | +2 | 11 |
| 11 | c | 4078 | 575.3 | 0.001592 | 2.768 | +2 | 11 |
| - | - | 5205 | 575.8 | - | - | 0 | - |
| - | - | 1.572E+04 | 576.3 | - | - | 0 | - |
| - | - | 1.475E+04 | 576.8 | - | - | 0 | - |
| - | - | 5046 | 577.3 | - | - | 0 | - |
| - | - | 876.8 | 577.8 | - | - | 0 | - |
| - | - | 1066 | 577.9 | - | - | 0 | - |
| - | - | 944.7 | 578.3 | - | - | 0 | - |
| - | - | 737.6 | 579.3 | - | - | 0 | - |
| - | - | 761.2 | 579.8 | - | - | 0 | - |
| - | - | 1226 | 580.4 | - | - | 0 | - |
| 2 | z | 1.28E+04 | 583.3 | 0.00334 | 5.726 | +2 | 11 |
| 11 | c | 2.926E+05 | 583.8 | 0.000952 | 1.631 | +2 | 11 |
| - | - | 1.85E+05 | 584.4 | - | - | 0 | - |
| - | - | 5.477E+04 | 584.9 | - | - | 0 | - |
| - | - | 2.644E+04 | 585.3 | - | - | 0 | - |
| - | - | 1.528E+04 | 585.4 | - | - | 0 | - |
| - | - | 9852 | 585.8 | - | - | 0 | - |
| - | - | 7109 | 586.3 | - | - | 0 | - |
| 8 | y | 1.27E+05 | 587.3 | 0.0011 | 1.873 | +1 | 5 |
| - | - | 4.085E+04 | 588.3 | - | - | 0 | - |
| - | - | 8925 | 589.3 | - | - | 0 | - |
| - | - | 1198 | 589.8 | - | - | 0 | - |
| - | - | 1694 | 590.4 | - | - | 0 | - |
| - | - | 2.209E+04 | 590.8 | - | - | 0 | - |
| 2 | y | 1.223E+04 | 591.3 | 0.001872 | 3.165 | +2 | 11 |
| - | - | 4479 | 591.8 | - | - | 0 | - |
| - | - | 1907 | 592.3 | - | - | 0 | - |
| - | - | 712.3 | 592.9 | - | - | 0 | - |
| - | - | 4842 | 595.4 | - | - | 0 | - |
| - | - | 7888 | 595.9 | - | - | 0 | - |
| - | - | 7051 | 596.4 | - | - | 0 | - |
| - | - | 1927 | 596.9 | - | - | 0 | - |
| - | - | 2.804E+04 | 597.3 | - | - | 0 | - |
| - | - | 1.69E+04 | 597.8 | - | - | 0 | - |
| - | - | 9108 | 598.3 | - | - | 0 | - |
| - | - | 8.562E+04 | 598.8 | - | - | 0 | - |
| - | - | 4.379E+04 | 599.3 | - | - | 0 | - |
| - | - | 2.304E+04 | 599.8 | - | - | 0 | - |
| - | - | 6041 | 600.3 | - | - | 0 | - |
| - | - | 1815 | 600.8 | - | - | 0 | - |
| - | - | 3752 | 603.4 | - | - | 0 | - |
| - | - | 2880 | 603.9 | - | - | 0 | - |
| - | - | 8026 | 604.4 | - | - | 0 | - |
| - | - | 1.768E+04 | 604.9 | - | - | 0 | - |
| - | - | 1.007E+04 | 605.4 | - | - | 0 | - |
| - | - | 1.565E+04 | 605.9 | - | - | 0 | - |
| - | - | 8755 | 606.4 | - | - | 0 | - |
| - | - | 3889 | 606.8 | - | - | 0 | - |
| - | - | 3.964E+04 | 607.3 | - | - | 0 | - |
| - | - | 2.442E+04 | 607.8 | - | - | 0 | - |
| - | - | 9799 | 608.3 | - | - | 0 | - |
| - | - | 2284 | 608.8 | - | - | 0 | - |
| 6 | c | 2009 | 611.4 | 0.001097 | 1.794 | +1 | 6 |
| - | - | 1743 | 612.3 | - | - | 0 | - |
| - | - | 1.246E+04 | 612.9 | - | - | 0 | - |
| - | - | 1.272E+04 | 613.4 | - | - | 0 | - |
| - | - | 8374 | 613.8 | - | - | 0 | - |
| - | - | 5731 | 614.3 | - | - | 0 | - |
| - | - | 2233 | 614.8 | - | - | 0 | - |
| - | - | 2342 | 615.3 | - | - | 0 | - |
| - | - | 779.7 | 616.3 | - | - | 0 | - |
| - | - | 3270 | 617.9 | - | - | 0 | - |
| - | - | 1.674E+04 | 618.4 | - | - | 0 | - |
| - | - | 1.267E+04 | 618.9 | - | - | 0 | - |
| - | - | 4420 | 619.4 | - | - | 0 | - |
| - | - | 2064 | 619.9 | - | - | 0 | - |
| - | - | 759 | 620.4 | - | - | 0 | - |
| - | - | 6726 | 620.9 | - | - | 0 | - |
| - | - | 4815 | 621.4 | - | - | 0 | - |
| - | - | 1415 | 621.9 | - | - | 0 | - |
| - | - | 2296 | 623.4 | - | - | 0 | - |
| 7 | z | 993.9 | 624.3 | 0.004505 | 7.216 | +1 | 6 |
| - | - | 1188 | 625.9 | - | - | 0 | - |
| - | - | 1.289E+04 | 626.4 | - | - | 0 | - |
| - | - | 9.366E+04 | 626.9 | - | - | 0 | - |
| - | - | 6.621E+04 | 627.4 | - | - | 0 | - |
| - | - | 841.2 | 627.8 | - | - | 0 | - |
| - | - | 2.865E+04 | 627.9 | - | - | 0 | - |
| 6 | c | 1.329E+05 | 628.4 | 0.0005488 | 0.8734 | +1 | 6 |
| - | - | 1273 | 628.9 | - | - | 0 | - |
| - | - | 4.543E+04 | 629.4 | - | - | 0 | - |
| - | - | 8830 | 630.4 | - | - | 0 | - |
| - | - | 920.7 | 631.4 | - | - | 0 | - |
| - | - | 838.4 | 633.3 | - | - | 0 | - |
| - | - | 1663 | 633.8 | - | - | 0 | - |
| - | - | 1633 | 634.7 | - | - | 0 | - |
| - | - | 2.352E+05 | 634.9 | - | - | 0 | - |
| - | - | 3.156E+05 | 635.4 | - | - | 0 | - |
| - | - | 1.692E+05 | 635.9 | - | - | 0 | - |
| - | - | 6.608E+04 | 636.4 | - | - | 0 | - |
| - | - | 1.801E+04 | 636.9 | - | - | 0 | - |
| - | - | 3951 | 637.4 | - | - | 0 | - |
| 7 | y | 1077 | 640.4 | 0.0002466 | 0.385 | +1 | 6 |
| 7 | z | 1.07E+05 | 642.4 | 0.001143 | 1.779 | +1 | 6 |
| - | - | 4.183E+04 | 643.4 | - | - | 0 | - |
| - | - | 9866 | 644.4 | - | - | 0 | - |
| - | - | 1441 | 645.4 | - | - | 0 | - |
| - | - | 846.7 | 651.4 | - | - | 0 | - |
| - | - | 3249 | 653.4 | - | - | 0 | - |
| - | - | 6508 | 654.4 | - | - | 0 | - |
| - | - | 7433 | 655.4 | - | - | 0 | - |
| - | - | 3425 | 656.4 | - | - | 0 | - |
| - | - | 1316 | 657.4 | - | - | 0 | - |
| 7 | y | 1.934E+04 | 658.4 | 0.00134 | 2.035 | +1 | 6 |
| - | - | 7724 | 659.4 | - | - | 0 | - |
| - | - | 1872 | 660.4 | - | - | 0 | - |
| - | - | 1456 | 664.4 | - | - | 0 | - |
| - | - | 4020 | 665.4 | - | - | 0 | - |
| - | - | 2125 | 666.4 | - | - | 0 | - |
| - | - | 1018 | 667.4 | - | - | 0 | - |
| - | - | 1624 | 674.3 | - | - | 0 | - |
| 7 | c | 851.3 | 681.4 | 0.009765 | 14.33 | +1 | 7 |
| 7 | c | 4.754E+04 | 682.4 | 0.001459 | 2.138 | +1 | 7 |
| - | - | 1.833E+04 | 683.4 | - | - | 0 | - |
| - | - | 3701 | 684.4 | - | - | 0 | - |
| - | - | 928.1 | 707.5 | - | - | 0 | - |
| 6 | w | 4143 | 712.4 | 0.00109 | 1.53 | +1 | 7 |
| - | - | 2743 | 713.4 | - | - | 0 | - |
| - | - | 2225 | 725.4 | - | - | 0 | - |
| - | - | 4711 | 726.4 | - | - | 0 | - |
| - | - | 1204 | 727.4 | - | - | 0 | - |
| - | - | 3496 | 750.5 | - | - | 0 | - |
| - | - | 2806 | 751.5 | - | - | 0 | - |
| - | - | 4180 | 752.5 | - | - | 0 | - |
| - | - | 1188 | 753.5 | - | - | 0 | - |
| - | - | 1041 | 754.4 | - | - | 0 | - |
| 6 | y | 3.471E+04 | 755.4 | 0.001432 | 1.896 | +1 | 7 |
| - | - | 1.415E+04 | 756.4 | - | - | 0 | - |
| - | - | 4472 | 757.4 | - | - | 0 | - |
| - | - | 942 | 758.4 | - | - | 0 | - |
| - | - | 859.6 | 766.5 | - | - | 0 | - |
| - | - | 1247 | 778.5 | - | - | 0 | - |
| - | - | 4412 | 780.5 | - | - | 0 | - |
| - | - | 1889 | 781.5 | - | - | 0 | - |
| - | - | 3306 | 783.4 | - | - | 0 | - |
| - | - | 3832 | 784.4 | - | - | 0 | - |
| 8 | c | 3.193E+04 | 796.5 | 0.0003929 | 0.4932 | +1 | 8 |
| - | - | 1.478E+04 | 797.5 | - | - | 0 | - |
| - | - | 3651 | 798.5 | - | - | 0 | - |
| 5 | w | 4.173E+04 | 809.4 | 0.001488 | 1.838 | +1 | 8 |
| - | - | 1.82E+04 | 810.4 | - | - | 0 | - |
| - | - | 4360 | 811.4 | - | - | 0 | - |
| - | - | 1125 | 812.5 | - | - | 0 | - |
| - | - | 799.6 | 820.5 | - | - | 0 | - |
| - | - | 3790 | 821.5 | - | - | 0 | - |
| - | - | 7988 | 822.5 | - | - | 0 | - |
| - | - | 4711 | 823.5 | - | - | 0 | - |
| - | - | 904.4 | 824.5 | - | - | 0 | - |
| - | - | 764.5 | 831.8 | - | - | 0 | - |
| - | - | 2676 | 837.5 | - | - | 0 | - |
| - | - | 974.5 | 838.5 | - | - | 0 | - |
| - | - | 720.3 | 838.9 | - | - | 0 | - |
| - | - | 990.6 | 839.5 | - | - | 0 | - |
| - | - | 2426 | 847 | - | - | 0 | - |
| - | - | 4287 | 847.4 | - | - | 0 | - |
| - | - | 4261 | 847.9 | - | - | 0 | - |
| - | - | 2455 | 848.4 | - | - | 0 | - |
| - | - | 3841 | 849.5 | - | - | 0 | - |
| - | - | 2056 | 850.5 | - | - | 0 | - |
| 5 | y | 2412 | 851.5 | 0.01338 | 15.71 | +1 | 8 |
| 5 | z | 1.198E+05 | 852.5 | 0.001034 | 1.213 | +1 | 8 |
| - | - | 6.014E+04 | 853.5 | - | - | 0 | - |
| - | - | 1.982E+04 | 854.5 | - | - | 0 | - |
| - | - | 3666 | 855.5 | - | - | 0 | - |
| - | - | 1745 | 865.5 | - | - | 0 | - |
| - | - | 8000 | 866.5 | - | - | 0 | - |
| - | - | 4.777E+04 | 867.5 | - | - | 0 | - |
| - | - | 3141 | 867.5 | - | - | 0 | - |
| - | - | 2.071E+04 | 868.5 | - | - | 0 | - |
| - | - | 6509 | 869.5 | - | - | 0 | - |
| - | - | 1799 | 870.5 | - | - | 0 | - |
| - | - | 697.1 | 871.5 | - | - | 0 | - |
| - | - | 1066 | 884.5 | - | - | 0 | - |
| 9 | c | 1954 | 892.5 | 0.004279 | 4.795 | +1 | 9 |
| - | - | 2111 | 893.5 | - | - | 0 | - |
| - | - | 2047 | 894.5 | - | - | 0 | - |
| - | - | 3015 | 908.5 | - | - | 0 | - |
| 9 | c | 2.522E+05 | 909.6 | 0.0009846 | 1.083 | +1 | 9 |
| - | - | 1.277E+05 | 910.6 | - | - | 0 | - |
| - | - | 3.781E+04 | 911.6 | - | - | 0 | - |
| - | - | 7200 | 912.6 | - | - | 0 | - |
| - | - | 1464 | 913.6 | - | - | 0 | - |
| 4 | z | 5.56E+04 | 923.5 | 0.0009682 | 1.048 | +1 | 9 |
| - | - | 3.306E+04 | 924.5 | - | - | 0 | - |
| - | - | 1.085E+04 | 925.5 | - | - | 0 | - |
| - | - | 2127 | 926.5 | - | - | 0 | - |
| - | - | 1158 | 938.5 | - | - | 0 | - |
| 4 | y | 3.473E+04 | 939.6 | 0.0004937 | 0.5254 | +1 | 9 |
| - | - | 1.931E+04 | 940.6 | - | - | 0 | - |
| - | - | 6217 | 941.6 | - | - | 0 | - |
| - | - | 1275 | 942.6 | - | - | 0 | - |
| - | - | 7495 | 949.6 | - | - | 0 | - |
| - | - | 6539 | 950.6 | - | - | 0 | - |
| - | - | 1830 | 951.6 | - | - | 0 | - |
| - | - | 1217 | 965.6 | - | - | 0 | - |
| - | - | 2702 | 978.6 | - | - | 0 | - |
| - | - | 1746 | 979.6 | - | - | 0 | - |
| - | - | 723.4 | 980.6 | - | - | 0 | - |
| - | - | 855.3 | 992.6 | - | - | 0 | - |
| 3 | w | 7761 | 993.6 | 0.001587 | 1.597 | +1 | 10 |
| - | - | 6320 | 994.6 | - | - | 0 | - |
| - | - | 6010 | 995.6 | - | - | 0 | - |
| - | - | 2688 | 996.6 | - | - | 0 | - |
| - | - | 1683 | 1007 | - | - | 0 | - |
| - | - | 3532 | 1008 | - | - | 0 | - |
| - | - | 2381 | 1009 | - | - | 0 | - |
| - | - | 1122 | 1010 | - | - | 0 | - |
| 10 | c | 1006 | 1022 | 0.001237 | 1.211 | +1 | 10 |
| - | - | 1042 | 1023 | - | - | 0 | - |
| - | - | 3065 | 1024 | - | - | 0 | - |
| - | - | 797.8 | 1025 | - | - | 0 | - |
| 10 | c | 3.022E+05 | 1039 | 0.001726 | 1.662 | +1 | 10 |
| - | - | 1.84E+05 | 1040 | - | - | 0 | - |
| - | - | 6.097E+04 | 1041 | - | - | 0 | - |
| - | - | 1.569E+04 | 1042 | - | - | 0 | - |
| - | - | 2087 | 1043 | - | - | 0 | - |
| 3 | y | 6103 | 1051 | 0.001965 | 1.87 | +1 | 10 |
| 3 | z | 4.509E+04 | 1052 | 0.0007318 | 0.6958 | +1 | 10 |
| - | - | 7.424E+04 | 1053 | - | - | 0 | - |
| - | - | 3.884E+04 | 1054 | - | - | 0 | - |
| - | - | 1.261E+04 | 1055 | - | - | 0 | - |
| - | - | 2759 | 1056 | - | - | 0 | - |
| - | - | 9545 | 1064 | - | - | 0 | - |
| - | - | 5195 | 1065 | - | - | 0 | - |
| - | - | 3979 | 1066 | - | - | 0 | - |
| - | - | 3412 | 1067 | - | - | 0 | - |
| 3 | y | 5540 | 1068 | 0.003466 | 3.246 | +1 | 10 |
| - | - | 4560 | 1069 | - | - | 0 | - |
| - | - | 1714 | 1070 | - | - | 0 | - |
| - | - | 2107 | 1095 | - | - | 0 | - |
| - | - | 947 | 1096 | - | - | 0 | - |
| - | - | 725.1 | 1107 | - | - | 0 | - |
| - | - | 1880 | 1108 | - | - | 0 | - |
| - | - | 1715 | 1109 | - | - | 0 | - |
| - | - | 2044 | 1110 | - | - | 0 | - |
| - | - | 2102 | 1111 | - | - | 0 | - |
| - | - | 902.5 | 1112 | - | - | 0 | - |
| 2 | w | 2984 | 1122 | 0.002327 | 2.074 | +1 | 11 |
| - | - | 7812 | 1123 | - | - | 0 | - |
| - | - | 1.031E+04 | 1124 | - | - | 0 | - |
| - | - | 5134 | 1125 | - | - | 0 | - |
| - | - | 1640 | 1126 | - | - | 0 | - |
| - | - | 1107 | 1140 | - | - | 0 | - |
| - | - | 4980 | 1151 | - | - | 0 | - |
| - | - | 6574 | 1152 | - | - | 0 | - |
| - | - | 4785 | 1153 | - | - | 0 | - |
| - | - | 1785 | 1154 | - | - | 0 | - |
| 2 | z | 3212 | 1166 | 0.001017 | 0.8726 | +1 | 11 |
| 11 | c | 4.529E+04 | 1167 | 0.006322 | 5.419 | +1 | 11 |
| - | - | 5.255E+04 | 1168 | - | - | 0 | - |
| - | - | 2.876E+04 | 1169 | - | - | 0 | - |
| - | - | 9943 | 1170 | - | - | 0 | - |
| - | - | 3258 | 1171 | - | - | 0 | - |
| - | - | 950.1 | 1172 | - | - | 0 | - |
| 2 | y | 2183 | 1182 | 0.02027 | 17.15 | +1 | 11 |
| - | - | 1645 | 1183 | - | - | 0 | - |
| - | - | 863.3 | 1195 | - | - | 0 | - |
| - | - | 1723 | 1197 | - | - | 0 | - |
| - | - | 5606 | 1198 | - | - | 0 | - |
| - | - | 3573 | 1199 | - | - | 0 | - |
| - | - | 1797 | 1200 | - | - | 0 | - |
| - | - | 1110 | 1207 | - | - | 0 | - |
| - | - | 2907 | 1208 | - | - | 0 | - |
| - | - | 2338 | 1209 | - | - | 0 | - |
| - | - | 861.9 | 1211 | - | - | 0 | - |
| - | - | 3447 | 1212 | - | - | 0 | - |
| - | - | 2514 | 1213 | - | - | 0 | - |
| - | - | 987.3 | 1214 | - | - | 0 | - |
| - | - | 6758 | 1225 | - | - | 0 | - |
| - | - | 2.829E+04 | 1226 | - | - | 0 | - |
| - | - | 2.034E+04 | 1227 | - | - | 0 | - |
| - | - | 6185 | 1228 | - | - | 0 | - |
| - | - | 1881 | 1229 | - | - | 0 | - |
| - | - | 1.051E+04 | 1236 | - | - | 0 | - |
| - | - | 6934 | 1237 | - | - | 0 | - |
| - | - | 2999 | 1238 | - | - | 0 | - |
| - | - | 862.2 | 1239 | - | - | 0 | - |
| - | - | 1891 | 1242 | - | - | 0 | - |
| - | - | 3421 | 1243 | - | - | 0 | - |
| - | - | 1485 | 1244 | - | - | 0 | - |
| - | - | 1.243E+04 | 1253 | - | - | 0 | - |
| - | - | 1.09E+05 | 1254 | - | - | 0 | - |
| - | - | 7.13E+04 | 1255 | - | - | 0 | - |
| - | - | 2.695E+04 | 1256 | - | - | 0 | - |
| - | - | 7426 | 1257 | - | - | 0 | - |
| - | - | 1989 | 1258 | - | - | 0 | - |
| - | - | 8931 | 1269 | - | - | 0 | - |
| - | - | 4.353E+04 | 1270 | - | - | 0 | - |
| - | - | 1.368E+05 | 1271 | - | - | 0 | - |
| - | - | 8.73E+04 | 1272 | - | - | 0 | - |
| - | - | 3.525E+04 | 1273 | - | - | 0 | - |
| - | - | 1.028E+04 | 1274 | - | - | 0 | - |
| - | - | 1763 | 1275 | - | - | 0 | - |
| - | - | 687.3 | 1694 | - | - | 0 | - |
| - | - | 778 | 3040 | - | - | 0 | - |
| - | - | 651.2 | 3136 | - | - | 0 | - |

m/z Charge Intensity FragmentType MassShift Position
120.06146240234375 0 536.6035
120.06586456298828 0 8302.878 y 11
127.08699035644531 0 2101.2944
128.0711212158203 0 961.40454
128.09474182128906 0 2795.0593
128.10780334472656 0 436.5901
129.1025848388672 0 41616.41
130.1059112548828 0 2622.0981
131.11830139160156 0 2813.3323
132.07716369628906 0 541.3844
133.08619689941406 0 1348.359
134.18702697753906 0 478.2439
141.04234313964844 0 569.62823
141.0787353515625 0 21526.877
141.10256958007812 0 5821
142.07501220703125 0 605.184
142.08216857910156 0 1817.8613
146.1234130859375 0 534.60144
146.12911987304688 0 7953.5464
147.1326141357422 0 472.4358
155.11834716796875 0 1602.4434
156.06585693359375 0 2281.094
157.06105041503906 0 3240.2598
157.1338348388672 0 3056.8584
158.06906127929688 0 2335.1272
165.10264587402344 0 3210.436 c Water loss 2
167.04541015625 0 1104.8455
169.09751892089844 0 8868.336
169.51943969726562 0 434.29877
170.10142517089844 0 971.7349
170.11761474609375 0 700.6643
174.07647705078125 0 8647.826 w 10
174.08740234375 0 1638.2722
175.07943725585938 0 743.6792
177.11231994628906 0 791.81085
183.1132049560547 0 38097.47
183.14955139160156 0 7162.837
184.07188415527344 0 654.66296
184.1167755126953 0 4061.2263
184.15249633789062 0 994.7605
185.05606079101562 0 2189.7625
185.07986450195312 0 5030.4365
185.1289520263672 0 1743.2689
187.14459228515625 0 1437.4409
188.092529296875 0 990.4678
197.1522216796875 0 667.79913
199.1080780029297 0 1441.4464
200.1397705078125 0 785.65454
201.1243438720703 0 784.169
201.14781188964844 0 867.4547
202.08255004882812 0 11205.993 c Ammonia loss 1
202.1067352294922 0 1688.4132
203.08651733398438 0 1261.4498
206.59632873535156 0 487.4382
208.1087646484375 0 1095.3757
211.14450073242188 0 6339.053
212.14833068847656 0 883.1837
212.63128662109375 0 554.587
215.13938903808594 0 632.2061
216.1470489501953 0 954.4251
217.16627502441406 0 6077.5283
217.72879028320312 0 556.993
218.16983032226562 0 581.46454
219.10931396484375 0 6701.047 c 1
219.12205505371094 0 668.39465
222.12326049804688 0 524.1924
225.12368774414062 0 2833.1191
225.474609375 0 548.6199
226.11895751953125 0 6301.139
227.1265106201172 0 1093.8884
229.11868286132812 0 4077.932
230.1218719482422 0 687.41693 w 8
230.1498565673828 0 958.72 y Water loss 10
232.1421661376953 0 3444.1746 z 10
235.1426239013672 0 1279.8131
235.6417694091797 0 782.4996
239.13951110839844 0 665.34906
240.13470458984375 0 16941.254
240.6329345703125 0 616.621
241.13758850097656 0 1120.55 z Water loss 5
242.126708984375 0 3538.7095
243.13421630859375 0 3253.4597
243.14588928222656 0 2513.6714
243.6553497314453 0 8811.044
244.15682983398438 0 2348.7825
245.12466430664062 0 961.54626
248.1609649658203 0 12915.908 y 10
249.16397094726562 0 1778.6941
249.63760375976562 0 9138.47
250.1388397216797 0 1786.4962
253.21531677246094 0 1006.7403
257.65325927734375 0 1630.3782 c Ammonia loss 4
258.14520263671875 0 5960.364
259.14276123046875 0 524.2097
259.1768798828125 0 1524.5747
262.6451721191406 0 698.7804
268.1785583496094 0 1267.7588
270.6601867675781 0 978.7157
271.16009521484375 0 657.0299
276.1517639160156 0 1953.9879
282.18096923828125 0 871.78876
283.6688537597656 0 1757.4316
284.160888671875 0 14971.954
285.1551818847656 0 2942.1672
285.168212890625 0 6336.5015 y Water loss 7
285.65875244140625 0 2494.735 y Ammonia loss 7
285.67266845703125 0 1071.2278
286.16253662109375 0 1072.3523 z 7
288.6608581542969 0 829.4494
289.16351318359375 0 2306.5776
290.1120910644531 0 750.45337
292.182373046875 0 30289.379
292.6837158203125 0 8271.448
293.1849060058594 0 1844.1232
294.1741943359375 0 108588.57 y 7
294.67572021484375 0 36087.37
295.14056396484375 0 4055.4739
295.1766357421875 0 5859.9087
295.67767333984375 0 865.2521
297.174560546875 0 3333.2979
297.6669616699219 0 2091.205
298.1675720214844 0 1540.1902
298.669189453125 0 573.9251
302.1717529296875 0 117892.266 w 9
303.1191101074219 0 1060.9045
303.17486572265625 0 16707.35
304.1766052246094 0 2199.6055
306.17987060546875 0 50593.387 c Ammonia loss 5
306.6813049316406 0 16924.484
307.1824951171875 0 5043.415
312.1673889160156 0 6663.9624
312.2135925292969 0 620.8175
313.1512451171875 0 11638.564
313.1701354980469 0 998.1025 z Ammonia loss 6
313.224365234375 0 3174.4756
314.1553649902344 0 2709.3135
316.1627502441406 0 990.48596
319.1877136230469 0 4141.93
319.68902587890625 0 632.68945
327.2019958496094 0 6741.9507
327.7009582519531 0 22453.973
328.20233154296875 0 8339.553
328.70361328125 0 2151.834
329.1911926269531 0 818.2874 c Water loss 2
330.1778869628906 0 23400.975 c Ammonia loss 2
331.1811218261719 0 3834.483
331.2347412109375 0 778.6625
332.16888427734375 0 976.90344
332.6927795410156 0 1204.2214
333.1851501464844 0 2519.707
333.6864318847656 0 1201.9066
334.1560363769531 0 668.8473
334.18609619140625 0 918.5438
340.1874084472656 0 15005.641
341.1895446777344 0 2920.6687 c Ammonia loss 9
341.6984558105469 0 12618.683 c Ammonia loss 6
342.19952392578125 0 4421.684
342.7015686035156 0 1578.2607
343.17584228515625 0 645.10205 z Water loss 9
344.2282409667969 0 1277.9419
347.2044982910156 0 36953.73 c 2
348.20751953125 0 5864.274
353.2191467285156 0 1044.5469
359.1929931640625 0 7561.368 y Water loss 9
360.1978454589844 0 1022.6448
361.1849365234375 0 8451.329 z 9
362.1876525878906 0 1491.6283
366.177978515625 0 1085.0128
368.230712890625 0 984.3558
371.22979736328125 0 1885.01
373.22015380859375 0 1044.5573
373.33282470703125 0 579.28143
374.2279052734375 0 1433.734
377.20367431640625 0 17567.123 y 9
378.2186279296875 0 16837.727 y 5
378.7205810546875 0 5905.416
379.2216491699219 0 824.80194
379.7236022949219 0 682.8387
380.1904296875 0 712.77563
381.2158203125 0 717.1574
383.2044677734375 0 10722.072
384.1884460449219 0 2230.686
384.21075439453125 0 1160.8804
390.2249450683594 0 4171.4146 c Ammonia loss 7
390.72625732421875 0 734.7548
391.2288818359375 0 942.57104
400.2053527832031 0 803.9412
400.2313537597656 0 1335.758 c Water loss 3
401.2152099609375 0 41638.402 c Ammonia loss 3
402.2179260253906 0 8096.839
403.2200622558594 0 1265.205
410.2782897949219 0 837.27344
411.2250671386719 0 1519.7567
415.25927734375 0 839.4116
418.24176025390625 0 91137.66 c 3
419.2445068359375 0 19408.736
420.24627685546875 0 2392.4805
423.22222900390625 0 649.3323
425.21368408203125 0 824.7228
427.2309265136719 0 3840.8835
427.26458740234375 0 875.34436
428.2349548339844 0 1265.9542
434.76129150390625 0 1851.7457 y 4
435.2621154785156 0 1218.3978
439.2297668457031 0 678.92804
443.2640380859375 0 822.93146
443.2957458496094 0 680.0815
444.2581481933594 0 5130.0137
445.2303771972656 0 24930.783 w 8
446.2339172363281 0 5165.056
447.2359924316406 0 1222.6753
455.2625427246094 0 1525.3779
459.24639892578125 0 2147.0127 w 8
462.27728271484375 0 1104.9517 z 3
468.2828369140625 0 16238.028
469.28466796875 0 4812.869
470.2824401855469 0 1603.4518 y 3
471.2919006347656 0 745.68964
472.2784729003906 0 1517.1407 y Water loss 8
474.2691650390625 0 2192.9414 z 8
475.80694580078125 0 1938.6887
476.3064880371094 0 1864.1389
479.2607116699219 0 832.39026
480.2584228515625 0 1475.5686
481.2403564453125 0 1090.4813
481.31475830078125 0 6074.7256
482.3167724609375 0 1926.5685
485.3086853027344 0 1967.0785
486.30078125 0 8809.557
487.3090515136719 0 4224.7827
488.31793212890625 0 1301.8381
489.2796325683594 0 722.1698
489.7741394042969 0 891.1652
490.2882995605469 0 22049.502 y 8
490.81005859375 0 1188.0713
491.2914123535156 0 5301.656
492.29412841796875 0 1197.3146
496.2884521484375 0 2656.6448
497.2738342285156 0 3902.97
497.3226623535156 0 1426.326
498.2848815917969 0 30132.607
498.78857421875 0 16210.399
499.2861022949219 0 7184.0933
499.7908020019531 0 1452.5854
500.2755432128906 0 1231.5364
504.7904052734375 0 3757.4775
505.29156494140625 0 1501.3157
505.7932434082031 0 1022.69336
508.2796325683594 0 1135.521
511.2806701660156 0 1508.2688 c Ammonia loss 9
511.8019104003906 0 1240.3022
511.9454040527344 0 739.7543
513.2899169921875 0 1186.6655
514.2992553710938 0 72311.34 c Ammonia loss 4
515.3018798828125 0 18652.686
516.2655029296875 0 1242.7712
516.3052368164062 0 3247.0457
517.3092651367188 0 759.7585 z Water loss 2
517.8107299804688 0 1049.8274 z Ammonia loss 2
518.3179321289062 0 1145.7666
519.2955322265625 0 1227.7955
519.7976684570312 0 937.3227 c 9
520.2969970703125 0 632.5501
523.2991333007812 0 910.8552
524.3092651367188 0 1279.5165
524.8045043945312 0 1379.1884
525.8123779296875 0 2109.1733 y Ammonia loss 2
526.3170776367188 0 38443.17 z 2
526.8187255859375 0 23730.83
527.3215942382812 0 11079.08
527.8225708007812 0 3320.0125
528.3026123046875 0 9171.013
529.30517578125 0 2046.4377
532.3359375 0 1531.7399
532.8338623046875 0 2072.0208
533.813720703125 0 5183.5044
534.3262329101562 0 21511.38 y 2
534.8271484375 0 10819.133
535.3280029296875 0 5141.901
539.3200073242188 0 1303.2478
540.3160400390625 0 10676.997
541.3186645507812 0 3981.2717
542.3141479492188 0 1129.6422
544.2987060546875 0 7615.5723 w 7
545.303955078125 0 3527.49
547.8242797851562 0 1681.6304
548.3206176757812 0 1802.6815
548.8175659179688 0 3605.918
549.3192749023438 0 1354.5253
549.8154907226562 0 662.0662
551.2945556640625 0 1244.3724
552.2782592773438 0 1723.7273
552.3212280273438 0 949.4907
553.8338012695312 0 2646.1665
554.3364868164062 0 8760.373
554.8383178710938 0 4082.8647
555.3084716796875 0 13102.043
555.8103637695312 0 8730.391
556.3101806640625 0 3030.2358
556.809326171875 0 831.1621
557.3411865234375 0 1886.8992
558.3483276367188 0 2162.7458
560.3214721679688 0 811.97186
560.8233642578125 0 757.0591
561.3326416015625 0 15053.793 w 1
561.8356323242188 0 11788.302
562.3383178710938 0 5733.328
562.8135375976562 0 6899.907
562.849853515625 0 4735.359
563.315673828125 0 5902.5454
563.3538818359375 0 1766.6819
563.8171997070312 0 2533.6924
564.6307983398438 0 1269.2413
564.9630126953125 0 1672.7838
565.6337890625 0 902.6877
566.829833984375 0 707.10016
567.3340454101562 0 3875.3906
567.8327026367188 0 4303.92
568.3292236328125 0 1610.0525
568.8289794921875 0 1863.6218
569.3065795898438 0 10320.367
570.3075561523438 0 3450.7869 y Ammonia loss 7
570.3421020507812 0 1886.8707
571.3250732421875 0 2086.19 z 7
571.8297119140625 0 1208.3112
573.8548583984375 0 741.00934
574.3349609375 0 726.66156 z Water loss 1
575.3365478515625 0 4078.0356 c Ammonia loss 10
575.835205078125 0 5204.9873
576.342529296875 0 15717.663
576.8446044921875 0 14754.682
577.3473510742188 0 5046.0547
577.8126220703125 0 876.7517
577.8572387695312 0 1065.638
578.32568359375 0 944.66754
579.3099975585938 0 737.64484
579.8110961914062 0 761.1784
580.3589477539062 0 1225.8915
583.342041015625 0 12804.481 z 1
583.8491821289062 0 292561.25 c 10
584.3506469726562 0 184988.17
584.8515625 0 54767.77
585.3292846679688 0 26440.842
585.3553466796875 0 15277.263
585.8318481445312 0 9852.455
586.3322143554688 0 7108.5864
587.3410034179688 0 126954.75 y 7
588.343994140625 0 40853.39
589.3465576171875 0 8925.483
589.8206787109375 0 1198.274
590.3506469726562 0 1693.9169
590.84521484375 0 22085.041
591.34619140625 0 12226.756 y 1
591.845703125 0 4478.6113
592.3487548828125 0 1906.8479
592.8562622070312 0 712.33356
595.362060546875 0 4842.3965
595.8569946289062 0 7888.1787
596.3577880859375 0 7051.3364
596.8574829101562 0 1927.252
597.3491821289062 0 28037.34
597.8494873046875 0 16897.344
598.339599609375 0 9108.355
598.824951171875 0 85621.516
599.3269653320312 0 43790.664
599.82861328125 0 23039.887
600.331298828125 0 6040.744
600.8316040039062 0 1815.3076
603.3612060546875 0 3752.291
603.8606567382812 0 2879.6885
604.3663330078125 0 8025.5835
604.8521118164062 0 17683.098
605.35205078125 0 10069.255
605.8567504882812 0 15646.399
606.3577880859375 0 8754.608
606.8436279296875 0 3888.916
607.3378295898438 0 39642.3
607.839111328125 0 24421.113
608.3411254882812 0 9799.365
608.8430786132812 0 2284.3467
611.3522338867188 0 2008.8944 c Ammonia loss 5
612.3494873046875 0 1742.608
612.8578491210938 0 12463.652
613.3599243164062 0 12716.3
613.8473510742188 0 8373.957
614.3447875976562 0 5731.244
614.8468627929688 0 2232.6836
615.3369750976562 0 2341.7637
616.3389892578125 0 779.665
617.8529663085938 0 3270.4233
618.353271484375 0 16742.508
618.8551635742188 0 12667.676
619.3565673828125 0 4420.287
619.8560180664062 0 2064.474
620.3604736328125 0 758.9934
620.850341796875 0 6726.258
621.3521118164062 0 4815.3916
621.8507080078125 0 1414.8778
623.3541259765625 0 2296.12
624.3522338867188 0 993.85474 z Water loss 6
625.86083984375 0 1187.8917
626.36376953125 0 12890.646
626.8568725585938 0 93657.2
627.3577880859375 0 66209.21
627.7686767578125 0 841.2205
627.8588256835938 0 28647.2
628.3782348632812 0 132933.84 c 5
628.8656005859375 0 1273.4406
629.3817749023438 0 45431.414
630.3843994140625 0 8829.676
631.3831787109375 0 920.7432
633.2871704101562 0 838.42224
633.7910766601562 0 1663.165
634.7415771484375 0 1633.1498
634.8652954101562 0 235229.38
635.367919921875 0 315598.94
635.869384765625 0 169160.38
636.3712768554688 0 66079.805
636.872802734375 0 18010.25
637.3751220703125 0 3950.9722
640.36669921875 0 1077.3435 y Water loss 6
642.3594360351562 0 107033.45 z 6
643.3624267578125 0 41826.527
644.3641967773438 0 9865.548
645.3636474609375 0 1441.0259
651.3599243164062 0 846.67456
653.3992309570312 0 3249.1846
654.3953857421875 0 6508.0264
655.4005126953125 0 7432.9785
656.4056396484375 0 3424.9668
657.4148559570312 0 1316.3881
658.3783569335938 0 19338.988 y 6
659.3812866210938 0 7723.87
660.3834838867188 0 1872.2596
664.3781127929688 0 1455.6395
665.3634033203125 0 4020.1394
666.3692626953125 0 2124.6582
667.3716430664062 0 1017.93964
674.3489990234375 0 1624.3005
681.3944702148438 0 851.2644 c Water loss 6
682.3897094726562 0 47542.688 c Ammonia loss 6
683.3929443359375 0 18329.3
684.3956298828125 0 3700.9512
707.4564819335938 0 928.10266
712.388671875 0 4142.7017 w 5
713.3947143554688 0 2742.7212
725.4312744140625 0 2225.2852
726.4373779296875 0 4711.182
727.443359375 0 1203.5515
750.4649047851562 0 3495.7263
751.4691162109375 0 2805.6716
752.4591064453125 0 4180.2627
753.4662475585938 0 1187.9983
754.4163818359375 0 1041.485
755.4312133789062 0 34714.594 y 5
756.4346313476562 0 14149.194
757.4387817382812 0 4471.5527
758.4434814453125 0 941.9626
766.4873657226562 0 859.59454
778.4967651367188 0 1246.6366
780.4732666015625 0 4412.0967
781.4759521484375 0 1889.1266
783.4261474609375 0 3306.0396
784.4312133789062 0 3832.449
796.4679565429688 0 31933.092 c 7
797.470458984375 0 14776.744
798.4738159179688 0 3651.1575
809.4418334960938 0 41725.14 w 4
810.4451293945312 0 18202.957
811.4466552734375 0 4360.3623
812.4515380859375 0 1124.5199
820.4950561523438 0 799.62805
821.5045776367188 0 3789.7239
822.5079956054688 0 7987.925
823.5111694335938 0 4710.5957
824.51708984375 0 904.40283
831.8004760742188 0 764.45685
837.5215454101562 0 2676.409
838.5233154296875 0 974.4831
838.9364624023438 0 720.28326
839.5281372070312 0 990.64557
846.950927734375 0 2425.9946
847.4475708007812 0 4287.1045
847.9483642578125 0 4261.419
848.4487915039062 0 2454.9126
849.5211791992188 0 3841.457
850.5245971679688 0 2055.503
851.5006713867188 0 2411.71 y Ammonia loss 4
852.4961547851562 0 119832.65 z 4
853.4993286132812 0 60136.8
854.5022583007812 0 19815.44
855.5050659179688 0 3665.8022
865.5348510742188 0 1745.4749
866.5465087890625 0 8000.3574
867.4705200195312 0 47771.2
867.54443359375 0 3141.051
868.4810180664062 0 20707.371
869.5049438476562 0 6509.1826
870.5123901367188 0 1799.0702
871.52685546875 0 697.1214
884.481201171875 0 1066.2355
892.5293579101562 0 1954.0503 c Ammonia loss 8
893.5305786132812 0 2111.231
894.53466796875 0 2047.0579
908.5441284179688 0 3014.8914
909.5526123046875 0 252173.38 c 8
910.555419921875 0 127743.73
911.5584716796875 0 37808.34
912.5614013671875 0 7199.8057
913.5653686523438 0 1463.717
923.533203125 0 55596.15 z 3
924.5360107421875 0 33061.734
925.5399169921875 0 10846.677
926.5416259765625 0 2126.9473
938.5337524414062 0 1157.6671
939.5514526367188 0 34733.58 y 3
940.5548706054688 0 19314.87
941.557373046875 0 6216.5537
942.5648803710938 0 1275.0986
949.5963134765625 0 7495.1836
950.6002197265625 0 6538.9697
951.6000366210938 0 1829.9338
965.6107177734375 0 1217.037
978.5652465820312 0 2701.8748
979.5752563476562 0 1746.0507
980.5657958984375 0 723.3874
992.55712890625 0 855.3074
993.5631103515625 0 7761.417 w 2
994.570068359375 0 6319.852
995.5717163085938 0 6009.591
996.580810546875 0 2688.364
1006.5816040039062 0 1682.9253
1007.5768432617188 0 3531.9312
1008.5824584960938 0 2381.0862
1009.5907592773438 0 1121.553
1021.5689086914062 0 1006.0809 c Ammonia loss 9
1022.5775756835938 0 1041.6123
1023.5874633789062 0 3064.7527
1024.5830078125 0 797.82025
1038.595947265625 0 302212.7 c 9
1039.598388671875 0 183997.03
1040.6011962890625 0 60972.176
1041.6046142578125 0 15692.011
1042.6080322265625 0 2086.5117
1050.621337890625 0 6102.5547 y Ammonia loss 2
1051.6279296875 0 45092.86 z 2
1052.6346435546875 0 74243.25
1053.63818359375 0 38837.3
1054.6407470703125 0 12614.203
1055.644287109375 0 2759.3933
1063.6392822265625 0 9545.334
1064.643798828125 0 5194.839
1065.644287109375 0 3978.8577
1066.62744140625 0 3412.489
1067.6424560546875 0 5539.529 y 2
1068.64794921875 0 4559.645
1069.65185546875 0 1713.8433
1094.606201171875 0 2106.8022
1095.6014404296875 0 947.0444
1106.6591796875 0 725.06464
1107.6116943359375 0 1879.564
1108.6490478515625 0 1715.3965
1109.6146240234375 0 2043.5327
1110.6190185546875 0 2102.1702
1111.6141357421875 0 902.4794
1121.6588134765625 0 2984.0703 w 1
1122.673583984375 0 7812.3584
1123.6806640625 0 10307.504
1124.6868896484375 0 5133.83
1125.695556640625 0 1640.0151
1139.708251953125 0 1107.0463
1150.6719970703125 0 4980.2983
1151.676025390625 0 6573.901
1152.6785888671875 0 4784.5986
1153.6864013671875 0 1784.8093
1165.671142578125 0 3211.6472 z 1
1166.682861328125 0 45289.047 c 10
1167.692138671875 0 52554.582
1168.69580078125 0 28761.773
1169.689697265625 0 9942.782
1170.6834716796875 0 3258.3574
1171.6744384765625 0 950.1112
1181.6685791015625 0 2183.4304 y 1
1182.671875 0 1645.2305
1194.63525390625 0 863.35
1196.64306640625 0 1723.232
1197.6519775390625 0 5605.847
1198.6531982421875 0 3573.0078
1199.66259765625 0 1797.3514
1206.72265625 0 1109.8799
1207.7127685546875 0 2906.948
1208.705078125 0 2337.6694
1210.703125 0 861.89905
1211.7005615234375 0 3447.204
1212.703857421875 0 2513.9756
1213.6903076171875 0 987.2665
1224.711669921875 0 6757.8154
1225.71533203125 0 28294.408
1226.718017578125 0 20342.36
1227.7208251953125 0 6185.337
1228.7239990234375 0 1880.9855
1235.698974609375 0 10508.517
1236.700439453125 0 6934.42
1237.7021484375 0 2998.8342
1238.705810546875 0 862.2479
1241.7322998046875 0 1890.5405
1242.7401123046875 0 3420.6343
1243.739501953125 0 1484.6733
1252.706787109375 0 12434.995
1253.7098388671875 0 109049.81
1254.712890625 0 71297.89
1255.7156982421875 0 26945.953
1256.717529296875 0 7425.9155
1257.7177734375 0 1988.7354
1268.72119140625 0 8930.576
1269.7276611328125 0 43532.453
1270.7359619140625 0 136813.75
1271.738525390625 0 87303.766
1272.7415771484375 0 35250.918
1273.7445068359375 0 10284.366
1274.746826171875 0 1763.1782
1693.897216796875 0 687.29553
3040.048583984375 0 778.00226
3135.632568359375 0 651.18677

Spectrum Details

|  |  |
| --- | --- |
| Matched peaks? Matched peaks The total absolute number of peaks matched. Additionally in brackets the total fraction of peaks matched and the total number of peaks is shown. | 86 (13.78% of 624) |
| FDR? FDR The false discovery rate estimated for this peptide. It is calculated by matching all theoretical fragments with a non-integer shift with the raw peaks for this spectrum. This is done with 40 different shifts. The resulting percentage is the average number of annotated peaks over the number of annotated peaks with the correct spectrum. | 0.03% |
| Satellite FDR? Satellite FDR See the FDR for details on its calculation. This satellite ion specific FDR only contains the satellite ions (d/w) for I/L/J positions. | 0.00% |
| PSM Score? PSM Score The PSM Score as given by Hecklib to this annotated spectrum. It is shown with three significant figures. | 670 |

#### Spectrum 3842? Spectrum 3842 The raw spectrum of this peptide as annotated by Hecklib. The fragments are coloured according to ion type (see legend). Any peaks with a star '\*' as text can be hovered over to see the full details, first the ion type second the mass shift type. By hovering over the amino acids in the peptide or ions in the legend the corresponding peaks are highlighted. By toggling the 'Unassigned' label you can turn the background (unassigned) peaks on or off in the plot. By updating the slider in the Ion legend you can update the spectrum to only show the top X% of the peaks with labels. The top X% means any peak that is within X% of the highest intensity. By dragging in the spectrum you can zoom in to a specific part of the spectrum and use 'Zoom Out' to get back to the original zoom level. The annotation of the spectrum is based on the given sequence in the peptides file and is done with different software so inconsistencies are likely. The peaks are annotated based on the given sequence, with 20 ppm tolerance.

Copy Data

##### Spectrum 3842 (TSV)

###### Preview

```
Loading example...
```

*Click on the button to copy the data to your clipboard.*

Mz MinMz MaxIntensity Max

WidthHeightPeptide font sizePeptide stroke widthSpectrum font sizeSpectrum stroke widthCompact peptide

Ion legend

wxyz

abcd

OtherUnassignedIon^Charge^_Position_Show for top:%

SNKALPAPJEKT

02.00e+53.99e+55.99e+57.99e+5

Zoom Out

c^+1^_2_y^+1^_2_c^+1^_3_y^+1^_3_y^+1^_3_c^+1^_4_c^+2^_10_c^+1^_5_y^+2^_10_c^+2^_11_y^+1^_5_y^+2^_11_c^+1^_6_c^+1^_6_z^+1^_6_y^+1^_6_c^+1^_7_y^+1^_7_y^+1^_7_c^+1^_8_w^+1^_8_z^+1^_8_y^+1^_8_c^+1^_9_c^+1^_9_z^+1^_9_y^+1^_9_w^+1^_10_c^+1^_10_z^+1^_10_c^+1^_10_z^+1^_10_y^+1^_10_w^+1^_11_z^+1^_11_c^+1^_11_y^+1^_11_

03296579861315

Fragment Matches Table

Show background peaks

| Position | Ion type | Intensity | mz Theoretical | mz Error (Th) | mz Error (ppm) | Charge | Series Number |
| --- | --- | --- | --- | --- | --- | --- | --- |
| - | - | 712 | 127.2 | - | - | 0 | - |
| - | - | 777.5 | 128.4 | - | - | 0 | - |
| - | - | 2.719E+04 | 129.1 | - | - | 0 | - |
| - | - | 1430 | 130.1 | - | - | 0 | - |
| - | - | 690.8 | 134.1 | - | - | 0 | - |
| - | - | 1853 | 149 | - | - | 0 | - |
| - | - | 929.4 | 155.1 | - | - | 0 | - |
| - | - | 850.6 | 162.3 | - | - | 0 | - |
| - | - | 945.9 | 170.1 | - | - | 0 | - |
| - | - | 862.4 | 170.9 | - | - | 0 | - |
| - | - | 3351 | 173.5 | - | - | 0 | - |
| - | - | 856.8 | 175.2 | - | - | 0 | - |
| - | - | 968 | 185.1 | - | - | 0 | - |
| - | - | 1078 | 198.6 | - | - | 0 | - |
| 2 | c | 4498 | 202.1 | 0.0002774 | 1.372 | +1 | 2 |
| - | - | 3640 | 226.1 | - | - | 0 | - |
| - | - | 1045 | 231.3 | - | - | 0 | - |
| - | - | 1065 | 234.5 | - | - | 0 | - |
| - | - | 999.8 | 235.3 | - | - | 0 | - |
| - | - | 6126 | 240.1 | - | - | 0 | - |
| - | - | 1449 | 243.1 | - | - | 0 | - |
| 11 | y | 1.32E+04 | 248.2 | 5.947E-06 | 0.02396 | +1 | 2 |
| - | - | 1459 | 249.2 | - | - | 0 | - |
| - | - | 2744 | 258.1 | - | - | 0 | - |
| - | - | 890.6 | 265.5 | - | - | 0 | - |
| - | - | 1353 | 275.3 | - | - | 0 | - |
| - | - | 1129 | 295.1 | - | - | 0 | - |
| - | - | 1026 | 296.1 | - | - | 0 | - |
| - | - | 3544 | 312.2 | - | - | 0 | - |
| - | - | 5890 | 313.2 | - | - | 0 | - |
| - | - | 2809 | 313.2 | - | - | 0 | - |
| - | - | 2112 | 314.2 | - | - | 0 | - |
| 3 | c | 1.321E+04 | 330.2 | 0.0001423 | 0.431 | +1 | 3 |
| - | - | 2027 | 331.2 | - | - | 0 | - |
| 10 | y | 1201 | 359.2 | 2.436E-05 | 0.06782 | +1 | 3 |
| 10 | y | 6107 | 377.2 | 0.0005916 | 1.568 | +1 | 3 |
| - | - | 8938 | 383.2 | - | - | 0 | - |
| - | - | 3243 | 384.2 | - | - | 0 | - |
| - | - | 2250 | 390.7 | - | - | 0 | - |
| 4 | c | 2.419E+04 | 401.2 | 0.0001379 | 0.3437 | +1 | 4 |
| - | - | 1280 | 401.7 | - | - | 0 | - |
| - | - | 4874 | 402.2 | - | - | 0 | - |
| - | - | 1876 | 455.3 | - | - | 0 | - |
| - | - | 6770 | 468.3 | - | - | 0 | - |
| - | - | 1974 | 469.3 | - | - | 0 | - |
| - | - | 5009 | 486.3 | - | - | 0 | - |
| - | - | 1576 | 496.3 | - | - | 0 | - |
| - | - | 6162 | 497.3 | - | - | 0 | - |
| - | - | 2329 | 498.3 | - | - | 0 | - |
| - | - | 3025 | 508.3 | - | - | 0 | - |
| 10 | c | 2281 | 511.3 | 0.0006424 | 1.256 | +2 | 10 |
| 5 | c | 7.735E+04 | 514.3 | 9.429E-05 | 0.1833 | +1 | 5 |
| - | - | 2.118E+04 | 515.3 | - | - | 0 | - |
| - | - | 3070 | 516.3 | - | - | 0 | - |
| 3 | y | 1.855E+04 | 534.3 | 0.0003051 | 0.571 | +2 | 10 |
| - | - | 8157 | 534.8 | - | - | 0 | - |
| - | - | 3886 | 535.3 | - | - | 0 | - |
| - | - | 1667 | 539.3 | - | - | 0 | - |
| 11 | c | 5085 | 575.3 | 0.001104 | 1.919 | +2 | 11 |
| - | - | 3386 | 575.8 | - | - | 0 | - |
| - | - | 2738 | 584.3 | - | - | 0 | - |
| - | - | 1990 | 584.8 | - | - | 0 | - |
| - | - | 3492 | 586.3 | - | - | 0 | - |
| 8 | y | 2.23E+04 | 587.3 | 0.000426 | 0.7253 | +1 | 5 |
| - | - | 5527 | 588.3 | - | - | 0 | - |
| 2 | y | 1815 | 591.3 | 0.003011 | 5.092 | +2 | 11 |
| - | - | 1567 | 596.8 | - | - | 0 | - |
| - | - | 2053 | 603.9 | - | - | 0 | - |
| - | - | 2244 | 608.3 | - | - | 0 | - |
| 6 | c | 2787 | 611.4 | 1.695E-06 | 0.002773 | +1 | 6 |
| - | - | 1185 | 612.3 | - | - | 0 | - |
| - | - | 2271 | 617.3 | - | - | 0 | - |
| - | - | 3084 | 617.8 | - | - | 0 | - |
| - | - | 1657 | 618.4 | - | - | 0 | - |
| - | - | 1816 | 625.8 | - | - | 0 | - |
| - | - | 2E+04 | 625.9 | - | - | 0 | - |
| - | - | 2.271E+04 | 626.4 | - | - | 0 | - |
| - | - | 1.554E+04 | 626.9 | - | - | 0 | - |
| - | - | 8929 | 627.4 | - | - | 0 | - |
| 6 | c | 5.853E+04 | 628.4 | 0.0003667 | 0.5836 | +1 | 6 |
| - | - | 1.684E+04 | 629.4 | - | - | 0 | - |
| - | - | 3679 | 630.4 | - | - | 0 | - |
| - | - | 6.633E+05 | 634.9 | - | - | 0 | - |
| - | - | 4.62E+05 | 635.4 | - | - | 0 | - |
| - | - | 1.851E+05 | 635.9 | - | - | 0 | - |
| - | - | 4.113E+04 | 636.4 | - | - | 0 | - |
| - | - | 5619 | 637.4 | - | - | 0 | - |
| 7 | z | 5.966E+04 | 642.4 | 0.0002611 | 0.4064 | +1 | 6 |
| - | - | 2.465E+04 | 643.4 | - | - | 0 | - |
| - | - | 4278 | 644.4 | - | - | 0 | - |
| - | - | 1102 | 645.4 | - | - | 0 | - |
| - | - | 2.83E+04 | 655.4 | - | - | 0 | - |
| - | - | 2.609E+04 | 656.4 | - | - | 0 | - |
| - | - | 7672 | 657.4 | - | - | 0 | - |
| 7 | y | 2210 | 658.4 | 0.002139 | 3.25 | +1 | 6 |
| - | - | 1358 | 659.4 | - | - | 0 | - |
| 7 | c | 1.497E+04 | 682.4 | 0.0001892 | 0.2773 | +1 | 7 |
| - | - | 5434 | 683.4 | - | - | 0 | - |
| 6 | y | 2665 | 737.4 | 0.001431 | 1.94 | +1 | 7 |
| - | - | 2252 | 749.5 | - | - | 0 | - |
| - | - | 1239 | 749.7 | - | - | 0 | - |
| - | - | 6843 | 751.5 | - | - | 0 | - |
| - | - | 2150 | 752.5 | - | - | 0 | - |
| - | - | 8269 | 754.4 | - | - | 0 | - |
| 6 | y | 1.235E+05 | 755.4 | 2.837E-05 | 0.03755 | +1 | 7 |
| - | - | 4.773E+04 | 756.4 | - | - | 0 | - |
| - | - | 2047 | 756.5 | - | - | 0 | - |
| - | - | 1.07E+04 | 757.4 | - | - | 0 | - |
| - | - | 1967 | 780.5 | - | - | 0 | - |
| - | - | 2066 | 782.4 | - | - | 0 | - |
| - | - | 8969 | 783.4 | - | - | 0 | - |
| - | - | 3350 | 784.4 | - | - | 0 | - |
| - | - | 1499 | 795.5 | - | - | 0 | - |
| 8 | c | 7592 | 796.5 | 0.001133 | 1.423 | +1 | 8 |
| - | - | 4977 | 797.5 | - | - | 0 | - |
| 5 | w | 5327 | 809.4 | 0.005516 | 6.814 | +1 | 8 |
| - | - | 3147 | 810.4 | - | - | 0 | - |
| - | - | 1236 | 811.9 | - | - | 0 | - |
| - | - | 1934 | 820.5 | - | - | 0 | - |
| - | - | 2474 | 822.5 | - | - | 0 | - |
| - | - | 1302 | 850.5 | - | - | 0 | - |
| 5 | z | 2.155E+05 | 852.5 | 0.0004312 | 0.5058 | +1 | 8 |
| - | - | 1.373E+05 | 853.5 | - | - | 0 | - |
| - | - | 4.148E+04 | 854.5 | - | - | 0 | - |
| - | - | 6318 | 855.5 | - | - | 0 | - |
| - | - | 5915 | 867.5 | - | - | 0 | - |
| 5 | y | 4.342E+04 | 868.5 | 0.0005395 | 0.6212 | +1 | 8 |
| - | - | 2.061E+04 | 869.5 | - | - | 0 | - |
| - | - | 4295 | 870.5 | - | - | 0 | - |
| - | - | 1894 | 884.5 | - | - | 0 | - |
| - | - | 3240 | 890.5 | - | - | 0 | - |
| 9 | c | 1.306E+04 | 892.5 | 0.00135 | 1.512 | +1 | 9 |
| - | - | 6035 | 893.5 | - | - | 0 | - |
| - | - | 3980 | 906.6 | - | - | 0 | - |
| - | - | 1.035E+05 | 908.5 | - | - | 0 | - |
| 9 | c | 2.538E+05 | 909.6 | 0.0009685 | 1.065 | +1 | 9 |
| - | - | 1.121E+05 | 910.6 | - | - | 0 | - |
| - | - | 2.861E+04 | 911.6 | - | - | 0 | - |
| - | - | 2454 | 912.6 | - | - | 0 | - |
| 4 | z | 2.534E+04 | 923.5 | 0.0004799 | 0.5197 | +1 | 9 |
| - | - | 4.605E+04 | 924.5 | - | - | 0 | - |
| - | - | 1.85E+04 | 925.5 | - | - | 0 | - |
| - | - | 3690 | 926.5 | - | - | 0 | - |
| - | - | 5060 | 933.6 | - | - | 0 | - |
| - | - | 2329 | 934.6 | - | - | 0 | - |
| - | - | 8842 | 938.5 | - | - | 0 | - |
| 4 | y | 8.836E+04 | 939.6 | 0.0008491 | 0.9037 | +1 | 9 |
| - | - | 4.48E+04 | 940.6 | - | - | 0 | - |
| - | - | 1.153E+04 | 941.6 | - | - | 0 | - |
| - | - | 6103 | 950.6 | - | - | 0 | - |
| - | - | 2221 | 951.6 | - | - | 0 | - |
| - | - | 1.032E+04 | 979.6 | - | - | 0 | - |
| - | - | 5084 | 980.6 | - | - | 0 | - |
| - | - | 3182 | 981.5 | - | - | 0 | - |
| - | - | 1914 | 982.5 | - | - | 0 | - |
| - | - | 4099 | 989.6 | - | - | 0 | - |
| - | - | 1968 | 990.6 | - | - | 0 | - |
| 3 | w | 3280 | 993.6 | 0.004699 | 4.73 | +1 | 10 |
| - | - | 4569 | 994.6 | - | - | 0 | - |
| - | - | 4.163E+04 | 995.6 | - | - | 0 | - |
| - | - | 2.147E+04 | 996.6 | - | - | 0 | - |
| - | - | 6852 | 997.6 | - | - | 0 | - |
| - | - | 1352 | 1005 | - | - | 0 | - |
| - | - | 3454 | 1008 | - | - | 0 | - |
| - | - | 3799 | 1009 | - | - | 0 | - |
| - | - | 3167 | 1010 | - | - | 0 | - |
| - | - | 1891 | 1021 | - | - | 0 | - |
| 10 | c | 2.071E+04 | 1022 | 0.0001057 | 0.1035 | +1 | 10 |
| - | - | 1.243E+04 | 1023 | - | - | 0 | - |
| - | - | 4353 | 1024 | - | - | 0 | - |
| 3 | z | 1609 | 1035 | 0.009337 | 9.024 | +1 | 10 |
| - | - | 9169 | 1038 | - | - | 0 | - |
| 10 | c | 1.692E+05 | 1039 | 0.0002617 | 0.2519 | +1 | 10 |
| - | - | 9.737E+04 | 1040 | - | - | 0 | - |
| - | - | 2.995E+04 | 1041 | - | - | 0 | - |
| - | - | 3678 | 1042 | - | - | 0 | - |
| 3 | z | 2.283E+05 | 1052 | 6.534E-07 | 0.0006213 | +1 | 10 |
| - | - | 1.355E+05 | 1053 | - | - | 0 | - |
| - | - | 4.241E+04 | 1054 | - | - | 0 | - |
| - | - | 4919 | 1055 | - | - | 0 | - |
| - | - | 1.385E+04 | 1065 | - | - | 0 | - |
| - | - | 6979 | 1066 | - | - | 0 | - |
| - | - | 3281 | 1067 | - | - | 0 | - |
| 3 | y | 2.409E+04 | 1068 | 0.0006583 | 0.6166 | +1 | 10 |
| - | - | 1.23E+04 | 1069 | - | - | 0 | - |
| - | - | 2210 | 1070 | - | - | 0 | - |
| - | - | 1463 | 1080 | - | - | 0 | - |
| - | - | 3819 | 1094 | - | - | 0 | - |
| - | - | 2299 | 1095 | - | - | 0 | - |
| - | - | 6.608E+04 | 1110 | - | - | 0 | - |
| - | - | 4.155E+04 | 1111 | - | - | 0 | - |
| - | - | 1.416E+04 | 1112 | - | - | 0 | - |
| - | - | 1408 | 1113 | - | - | 0 | - |
| 2 | w | 2636 | 1122 | 7.466E-06 | 0.006656 | +1 | 11 |
| - | - | 3777 | 1123 | - | - | 0 | - |
| - | - | 1.117E+04 | 1124 | - | - | 0 | - |
| - | - | 7876 | 1125 | - | - | 0 | - |
| - | - | 2917 | 1126 | - | - | 0 | - |
| - | - | 1.583E+04 | 1152 | - | - | 0 | - |
| - | - | 1.792E+04 | 1153 | - | - | 0 | - |
| - | - | 8576 | 1154 | - | - | 0 | - |
| - | - | 1530 | 1155 | - | - | 0 | - |
| 2 | z | 5.693E+04 | 1166 | 0.0005697 | 0.4887 | +1 | 11 |
| 11 | c | 1.578E+05 | 1167 | 0.004125 | 3.536 | +1 | 11 |
| - | - | 8.883E+04 | 1168 | - | - | 0 | - |
| - | - | 4.654E+04 | 1169 | - | - | 0 | - |
| - | - | 2.714E+04 | 1170 | - | - | 0 | - |
| - | - | 8474 | 1171 | - | - | 0 | - |
| - | - | 1834 | 1181 | - | - | 0 | - |
| 2 | y | 2061 | 1182 | 0.01368 | 11.58 | +1 | 11 |
| - | - | 4516 | 1197 | - | - | 0 | - |
| - | - | 4029 | 1198 | - | - | 0 | - |
| - | - | 3716 | 1199 | - | - | 0 | - |
| - | - | 3352 | 1200 | - | - | 0 | - |
| - | - | 3675 | 1208 | - | - | 0 | - |
| - | - | 1910 | 1209 | - | - | 0 | - |
| - | - | 4682 | 1211 | - | - | 0 | - |
| - | - | 2263 | 1212 | - | - | 0 | - |
| - | - | 2335 | 1213 | - | - | 0 | - |
| - | - | 9759 | 1214 | - | - | 0 | - |
| - | - | 4929 | 1215 | - | - | 0 | - |
| - | - | 2041 | 1216 | - | - | 0 | - |
| - | - | 1.093E+05 | 1225 | - | - | 0 | - |
| - | - | 6.882E+04 | 1226 | - | - | 0 | - |
| - | - | 2.871E+04 | 1227 | - | - | 0 | - |
| - | - | 3474 | 1228 | - | - | 0 | - |
| - | - | 7063 | 1235 | - | - | 0 | - |
| - | - | 4729 | 1236 | - | - | 0 | - |
| - | - | 3230 | 1237 | - | - | 0 | - |
| - | - | 1.753E+04 | 1242 | - | - | 0 | - |
| - | - | 1.366E+04 | 1243 | - | - | 0 | - |
| - | - | 4777 | 1244 | - | - | 0 | - |
| - | - | 1.739E+04 | 1252 | - | - | 0 | - |
| - | - | 5.484E+04 | 1253 | - | - | 0 | - |
| - | - | 3.557E+04 | 1254 | - | - | 0 | - |
| - | - | 1.447E+04 | 1255 | - | - | 0 | - |
| - | - | 1683 | 1256 | - | - | 0 | - |
| - | - | 3.469E+05 | 1269 | - | - | 0 | - |
| - | - | 7.908E+05 | 1270 | - | - | 0 | - |
| - | - | 4.856E+05 | 1271 | - | - | 0 | - |
| - | - | 1.608E+05 | 1272 | - | - | 0 | - |
| - | - | 2.128E+04 | 1273 | - | - | 0 | - |
| - | - | 1937 | 1302 | - | - | 0 | - |

m/z Charge Intensity FragmentType MassShift Position
127.18523406982422 0 711.9971
128.37229919433594 0 777.45465
129.10220336914062 0 27191.574
130.1055450439453 0 1430.4669
134.1193389892578 0 690.78015
148.95372009277344 0 1853.2083
155.11904907226562 0 929.44037
162.27772521972656 0 850.5744
170.10887145996094 0 945.91724
170.90415954589844 0 862.4199
173.45204162597656 0 3350.524
175.187744140625 0 856.7668
185.0557861328125 0 967.9584
198.5738525390625 0 1077.663
202.0819549560547 0 4498.1665 c Ammonia loss 1
226.11842346191406 0 3639.702
231.2921600341797 0 1045.2992
234.4995880126953 0 1065.0631
235.2653350830078 0 999.83453
240.1342315673828 0 6126.0815
243.14547729492188 0 1448.6016
248.1604766845703 0 13201.801 y 10
249.16375732421875 0 1459.4644
258.14471435546875 0 2743.5881
265.4803161621094 0 890.5532
275.3067932128906 0 1352.6493
295.1389465332031 0 1129.2863
296.1421203613281 0 1026.3586
312.16619873046875 0 3543.8513
313.15069580078125 0 5890.2866
313.22369384765625 0 2809.3594
314.1535339355469 0 2111.7122
330.1773376464844 0 13212.92 c Ammonia loss 2
331.18121337890625 0 2027.2784
359.1925354003906 0 1200.9485 y Water loss 9
377.2024841308594 0 6107.16 y 9
383.2033996582031 0 8937.706
384.18719482421875 0 3243.0408
390.7252197265625 0 2249.583
401.2144470214844 0 24187.748 c Ammonia loss 3
401.6744384765625 0 1279.6046
402.21649169921875 0 4873.952
455.2602844238281 0 1876.3082
468.2822265625 0 6769.7637
469.2808837890625 0 1974.434
486.3034973144531 0 5009.0356
496.287841796875 0 1576.1553
497.27252197265625 0 6161.791
498.2740478515625 0 2328.8586
508.27685546875 0 3025.286
511.2881164550781 0 2280.5308 c Ammonia loss 9
514.2982788085938 0 77349.37 c Ammonia loss 4
515.3012084960938 0 21178.736
516.3045043945312 0 3070.126
534.326904296875 0 18553.348 y 2
534.8280029296875 0 8156.8965
535.3303833007812 0 3885.7493
539.3187866210938 0 1667.0018
575.3360595703125 0 5084.6367 c Ammonia loss 10
575.8368530273438 0 3385.6082
584.340087890625 0 2737.894
584.8424072265625 0 1989.569
586.33154296875 0 3492.388
587.3394775390625 0 22295.47 y 7
588.3432006835938 0 5527.1714
591.35107421875 0 1815.0568 y 1
596.8295288085938 0 1566.9971
603.8651733398438 0 2052.7903
608.3414916992188 0 2244.0146
611.3511352539062 0 2787.011 c Ammonia loss 5
612.3490600585938 0 1185.4924
617.3472900390625 0 2270.574
617.84619140625 0 3083.8542
618.35009765625 0 1656.7528
625.8057250976562 0 1816.3484
625.859130859375 0 19995.627
626.3551025390625 0 22706.844
626.8540649414062 0 15535.235
627.365234375 0 8929.374
628.3773193359375 0 58531.25 c 5
629.380615234375 0 16841.955
630.38134765625 0 3678.5784
634.8640747070312 0 663324.4
635.3656005859375 0 461966.22
635.8668823242188 0 185077.95
636.3690185546875 0 41131.492
637.3781127929688 0 5619.0234
642.3580322265625 0 59662.184 z 6
643.362060546875 0 24647.914
644.3671264648438 0 4278.3457
645.3648681640625 0 1101.9128
655.4009399414062 0 28297.156
656.4067993164062 0 26088.465
657.4112548828125 0 7671.5635
658.3748779296875 0 2210.2551 y 6
659.3796997070312 0 1357.9657
682.3880615234375 0 14969.247 c Ammonia loss 6
683.390625 0 5433.722
737.4177856445312 0 2665.2788 y Water loss 5
749.4506225585938 0 2251.6367
749.6599731445312 0 1239.3313
751.4710693359375 0 6843.4463
752.4734497070312 0 2150.0146
754.4215698242188 0 8268.879
755.4298095703125 0 123483.69 y 5
756.4327392578125 0 47733.047
756.5111083984375 0 2046.6647
757.43505859375 0 10696.04
780.4686279296875 0 1966.5397
782.4161376953125 0 2065.676
783.4254150390625 0 8969.484
784.4259033203125 0 3349.8064
795.4610595703125 0 1499.241
796.4664306640625 0 7591.8096 c 7
797.4702758789062 0 4976.8945
809.4458618164062 0 5327.349 w 4
810.4473266601562 0 3147.2275
811.9288330078125 0 1235.5092
820.4960327148438 0 1933.5305
822.5070190429688 0 2474.0105
850.5249633789062 0 1302.3553
852.4946899414062 0 215520.42 z 4
853.4990844726562 0 137300.58
854.5025634765625 0 41478.656
855.5054931640625 0 6317.8105
867.4765625 0 5915.45
868.5133056640625 0 43420.023 y 4
869.5167236328125 0 20613.842
870.52197265625 0 4294.5156
884.4811401367188 0 1893.7358
890.5357055664062 0 3240.1484
892.5264282226562 0 13063.392 c Ammonia loss 8
893.5292358398438 0 6035.3823
906.5889282226562 0 3980.372
908.5438232421875 0 103459.91
909.5506591796875 0 253804.69 c 8
910.553466796875 0 112090.01
911.5565185546875 0 28611.354
912.5576171875 0 2453.9119
923.53271484375 0 25335.04 z 3
924.5380859375 0 46051.59
925.5419921875 0 18498.398
926.5438232421875 0 3690.0725
933.575927734375 0 5060.277
934.5789794921875 0 2328.665
938.5418701171875 0 8842.305
939.5501098632812 0 88359.5 y 3
940.5537109375 0 44804.207
941.5556030273438 0 11525.878
950.6016845703125 0 6103.0933
951.60791015625 0 2221.206
979.6070556640625 0 10324.271
980.6069946289062 0 5083.97
981.53173828125 0 3181.7886
982.5264282226562 0 1913.7745
989.626708984375 0 4098.9727
990.6321411132812 0 1968.4829
993.5662231445312 0 3280.2446 w 2
994.57568359375 0 4569.1357
995.565185546875 0 41631.52
996.5677490234375 0 21465.45
997.5723876953125 0 6851.8027
1004.5471801757812 0 1352.1442
1007.6298217773438 0 3453.5422
1008.5841674804688 0 3799.4065
1009.59521484375 0 3167.0186
1020.6316528320312 0 1890.7906
1021.5675659179688 0 20714.572 c Ammonia loss 9
1022.570556640625 0 12430.179
1023.5729370117188 0 4353.465
1034.6099853515625 0 1608.9948 z Ammonia loss 2
1037.5869140625 0 9169.26
1038.594482421875 0 169180.11 c 9
1039.5966796875 0 97367.945
1040.599609375 0 29953.295
1041.6002197265625 0 3677.765
1051.627197265625 0 228346.16 z 2
1052.630126953125 0 135492.14
1053.63330078125 0 42412.957
1054.6361083984375 0 4919.3286
1064.64697265625 0 13847.063
1065.647705078125 0 6978.7993
1066.6434326171875 0 3280.652
1067.645263671875 0 24087.773 y 2
1068.6485595703125 0 12297.792
1069.6470947265625 0 2210.0085
1079.672607421875 0 1462.6036
1093.6463623046875 0 3819.1138
1094.623291015625 0 2298.6523
1109.6072998046875 0 66079.87
1110.6099853515625 0 41552.355
1111.61279296875 0 14158.813
1112.6190185546875 0 1407.9183
1121.656494140625 0 2636.3347 w 1
1122.6707763671875 0 3777.454
1123.6820068359375 0 11172.178
1124.6854248046875 0 7876.1885
1125.6873779296875 0 2917.25
1151.6759033203125 0 15833.455
1152.6822509765625 0 17919.684
1153.6845703125 0 8575.977
1154.678466796875 0 1530.0352
1165.6695556640625 0 56934.01 z 1
1166.68505859375 0 157829 c 10
1167.689208984375 0 88829.41
1168.6668701171875 0 46544.594
1169.654541015625 0 27135.506
1170.65380859375 0 8474.184
1180.6884765625 0 1834.4934
1181.6751708984375 0 2060.6526 y 1
1196.641357421875 0 4515.889
1197.656005859375 0 4029.1946
1198.665283203125 0 3715.8857
1199.662109375 0 3351.743
1207.706787109375 0 3674.9016
1208.7125244140625 0 1910.4423
1210.6871337890625 0 4682.1343
1211.691650390625 0 2263.211
1212.689453125 0 2334.8079
1213.6646728515625 0 9758.641
1214.6673583984375 0 4928.751
1215.6710205078125 0 2040.9448
1224.7060546875 0 109270.5
1225.709228515625 0 68821.7
1226.711669921875 0 28711.508
1227.7109375 0 3473.9653
1234.68896484375 0 7062.763
1235.697021484375 0 4729.278
1236.6881103515625 0 3229.9438
1241.732421875 0 17530.334
1242.73486328125 0 13664.735
1243.734375 0 4777.253
1251.71728515625 0 17392.828
1252.70458984375 0 54844.445
1253.704833984375 0 35567.47
1254.7060546875 0 14470.358
1255.7109375 0 1683.4524
1268.7205810546875 0 346879.5
1269.7266845703125 0 790769.56
1270.7293701171875 0 485622
1271.73291015625 0 160799.17
1272.736083984375 0 21283.785
1301.71630859375 0 1937.3086

Spectrum Details

|  |  |
| --- | --- |
| Matched peaks? Matched peaks The total absolute number of peaks matched. Additionally in brackets the total fraction of peaks matched and the total number of peaks is shown. | 37 (15.23% of 243) |
| FDR? FDR The false discovery rate estimated for this peptide. It is calculated by matching all theoretical fragments with a non-integer shift with the raw peaks for this spectrum. This is done with 40 different shifts. The resulting percentage is the average number of annotated peaks over the number of annotated peaks with the correct spectrum. | 0.19% |
| Satellite FDR? Satellite FDR See the FDR for details on its calculation. This satellite ion specific FDR only contains the satellite ions (d/w) for I/L/J positions. | 0.00% |
| PSM Score? PSM Score The PSM Score as given by Hecklib to this annotated spectrum. It is shown with three significant figures. | 358 |

#### Reverse Lookup? Reverse Lookup All places where this read could be placed.

| Group | Segment | Template | Template Part | Read Part | Score | Unique |
| --- | --- | --- | --- | --- | --- | --- |
| Homo sapiens Heavy Chain | IGHC | IGHG1 | [206..218] | [0..12] | 96 | False |
| Homo sapiens Heavy Chain | IGHC | IGHG3 | [253..265] | [0..12] | 96 | False |
| Homo sapiens Heavy Chain | IGHC | IGHG2 | [202..214] | [0..12] | 87 | False |

| Recombined | Template Part | Read Part | Score | Unique |
| --- | --- | --- | --- | --- |
| REC-0-1 | [331..343] | [0..12] | 96 | True |

#### Meta Information from Multiple reads

##### Number of combined reads

2

##### Intensity

0.6866

##### TotalArea

7.987E+07

##### Changes to the peptide sequence

SNKALPAPJEKT

I→JNo support for either Leucine or Isoleucine based on side chain ions (Position: 9)

J→LSupport for Leucine based on side chain ions (1 for L 0 for I) (Position: 5)

L→ISupport for Isoleucine based on side chain ions (3 for I 0 for L) (Position: 9)

L→JEqual support for both Leucine and Isoleucine based on side chain ions (1 ions for both) (Position: 5)

#### Positional Score

Copy Data

##### Positional Score (TSV)

###### Preview

```
Loading example...
```

*Click on the button to copy the data to your clipboard.*

0001234567891011

Label Value
"0" 0
"1" 0
"2" 0
"3" 0
"4" 0
"5" 0
"6" 0
"7" 0
"8" 0
"9" 0
"10" 0
"11" 0

#### Meta Information from PEAKS

##### Scan Identifier

F1:3813

##### Original sequence

S

N

K

A

L

P

A

P

L

E

K

T

##### Posttranslational Modifications

##### Source File

D:\separate\_stitch\_analyses\xle-disambiguation\raw\20210323\_F1\_UM1\_Peng0013\_SA\_F59\_ingel\_3ug\_ELA.raw

##### Fraction

1

##### Scan Feature

F1:1355

##### De Novo Score

98

##### ConfidenceScore

98

### m/z

423.5797

##### Mass

1267.7136

##### Charge

3

##### Retention Time

20

##### Predicted Retention Time

-

##### Area

6.043E+07

##### Parts Per Million

2.8

##### Fragmentation mode

ETHCD

##### Originating file

01 D:\separate\_stitch\_analyses\xle-disambiguation\20210325\_F59\_3ug\_DENOVO\_12.csv

#### Meta Information from PEAKS

##### Scan Identifier

F1:3842

##### Original sequence

S

N

K

A

L

P

A

P

L

E

K

T

##### Posttranslational Modifications

##### Source File

D:\separate\_stitch\_analyses\xle-disambiguation\raw\20210323\_F1\_UM1\_Peng0013\_SA\_F59\_ingel\_3ug\_ELA.raw

##### Fraction

1

##### Scan Feature

F1:9926

##### De Novo Score

98

##### ConfidenceScore

98

### m/z

634.8644

##### Mass

1267.7136

##### Charge

2

##### Retention Time

20

##### Predicted Retention Time

-

##### Area

1.944E+07

##### Parts Per Million

0.5

##### Fragmentation mode

ETHCD

##### Originating file

01 D:\separate\_stitch\_analyses\xle-disambiguation\20210325\_F59\_3ug\_DENOVO\_12.csv
