## Supplementary material for "A handle on mass coincidence errors in *de novo* sequencing of antibodies by bottom-up proteomics": Combined_051.html

Details Combined\_051 | Stitch OverviewUndefined

### Read Combined\_051

#### Sequence (length=19)

JTGTSTVGVGRGVLGDQKN

#### Spectrum 4926? Spectrum 4926 The raw spectrum of this peptide as annotated by Hecklib. The fragments are coloured according to ion type (see legend). Any peaks with a star '\*' as text can be hovered over to see the full details, first the ion type second the mass shift type. By hovering over the amino acids in the peptide or ions in the legend the corresponding peaks are highlighted. By toggling the 'Unassigned' label you can turn the background (unassigned) peaks on or off in the plot. By updating the slider in the Ion legend you can update the spectrum to only show the top X% of the peaks with labels. The top X% means any peak that is within X% of the highest intensity. By dragging in the spectrum you can zoom in to a specific part of the spectrum and use 'Zoom Out' to get back to the original zoom level. The annotation of the spectrum is based on the given sequence in the peptides file and is done with different software so inconsistencies are likely. The peaks are annotated based on the given sequence, with 20 ppm tolerance.

Copy Data

##### Spectrum 4926 (TSV)

###### Preview

```
Loading example...
```

*Click on the button to copy the data to your clipboard.*

Mz MinMz MaxIntensity Max

WidthHeightPeptide font sizePeptide stroke widthSpectrum font sizeSpectrum stroke widthCompact peptide

Ion legend

wxyz

abcd

OtherUnassignedIon^Charge^_Position_Show for top:%

JTGTSTVGVGRGVLGDQKN

02.09e+54.18e+56.26e+58.35e+5

Zoom Out

y^+2^_2_y^+1^_1_w^+1^_2_y^+2^_3_z^+1^_2_y^+1^_2_z^+1^_2_y^+1^_2_c^+1^_3_w^+1^_3_y^+2^_6_w^+3^_11_y^+1^_3_z^+1^_3_y^+2^_7_y^+1^_3_c^+1^_4_c^+3^_14_w^+1^_4_w^+3^_14_c^+1^_5_c^+1^_5_y^+1^_4_z^+1^_4_y^+3^_15_y^+2^_9_y^+3^_15_c^+3^_16_y^+1^_4_y^+2^_10_z^+2^_10_c^+2^_11_y^+2^_10_y^+3^_16_y^+3^_16_c^+3^_17_z^+3^_17_y^+3^_17_y^+3^_17_z^+3^_17_c^+2^_12_y^+1^_5_z^+1^_5_y^+3^_17_y^+1^_5_z^+3^_18_z^+3^_18_y^+2^_11_c^+3^_18_y^+3^_18_y^+3^_18_z^+3^_18_c^+1^_6_y^+3^_18_z^+2^_12_y^+2^_12_z^+2^_12_c^+2^_13_y^+2^_12_w^+1^_6_y^+2^_13_z^+2^_13_c^+2^_14_y^+1^_6_z^+1^_6_y^+1^_6_c^+1^_7_c^+2^_15_c^+2^_15_z^+2^_14_y^+2^_14_y^+2^_14_z^+2^_14_y^+2^_14_z^+2^_15_w^+2^_15_c^+1^_8_y^+2^_15_c^+2^_16_c^+2^_16_z^+1^_7_z^+1^_7_w^+1^_7_y^+2^_15_c^+2^_16_y^+1^_7_z^+1^_7_y^+1^_7_z^+2^_16_z^+2^_16_w^+2^_16_y^+2^_16_y^+2^_16_z^+2^_16_y^+2^_16_c^+2^_17_c^+2^_17_z^+2^_17_z^+2^_17_c^+2^_17_y^+2^_17_y^+2^_17_z^+2^_17_y^+2^_17_y^+1^_8_c^+1^_9_z^+2^_18_z^+2^_18_w^+2^_18_c^+2^_18_y^+2^_18_y^+2^_18_z^+2^_18_c^+1^_10_y^+2^_18_c^+1^_10_z^+1^_9_y^+1^_9_z^+1^_10_c^+1^_11_y^+1^_10_c^+1^_11_c^+1^_12_c^+1^_12_c^+1^_12_z^+1^_11_y^+1^_11_y^+1^_12_z^+1^_12_c^+1^_13_y^+1^_12_c^+1^_13_z^+1^_13_c^+1^_14_c^+1^_14_c^+1^_15_c^+1^_15_y^+1^_14_z^+1^_14_y^+1^_14_c^+1^_16_y^+1^_15_c^+1^_16_z^+1^_16_c^+1^_17_z^+1^_17_z^+1^_18_c^+1^_18_

047194214131884

Fragment Matches Table

Show background peaks

| Position | Ion type | Intensity | mz Theoretical | mz Error (Th) | mz Error (ppm) | Charge | Series Number |
| --- | --- | --- | --- | --- | --- | --- | --- |
| - | - | 1059 | 120.1 | - | - | 0 | - |
| - | - | 927.9 | 120.1 | - | - | 0 | - |
| - | - | 429.7 | 125.1 | - | - | 0 | - |
| - | - | 3163 | 125.1 | - | - | 0 | - |
| - | - | 2.56E+04 | 129.1 | - | - | 0 | - |
| - | - | 1399 | 130.1 | - | - | 0 | - |
| 18 | y | 2451 | 131.1 | 0.0004509 | 3.44 | +2 | 2 |
| - | - | 690.7 | 132.1 | - | - | 0 | - |
| - | - | 1161 | 133.1 | - | - | 0 | - |
| 19 | y | 1.517E+04 | 133.1 | 0.000526 | 3.953 | +1 | 1 |
| - | - | 372.3 | 138.9 | - | - | 0 | - |
| - | - | 3.237E+04 | 142.1 | - | - | 0 | - |
| - | - | 352.8 | 143.1 | - | - | 0 | - |
| - | - | 2872 | 143.1 | - | - | 0 | - |
| - | - | 731 | 148.9 | - | - | 0 | - |
| - | - | 465.6 | 149.4 | - | - | 0 | - |
| - | - | 431.7 | 150.7 | - | - | 0 | - |
| - | - | 6389 | 152.1 | - | - | 0 | - |
| - | - | 983 | 153.1 | - | - | 0 | - |
| - | - | 5435 | 157.1 | - | - | 0 | - |
| - | - | 7397 | 159.1 | - | - | 0 | - |
| - | - | 632.6 | 159.1 | - | - | 0 | - |
| - | - | 576 | 160.1 | - | - | 0 | - |
| - | - | 1234 | 165.1 | - | - | 0 | - |
| - | - | 463.2 | 167 | - | - | 0 | - |
| - | - | 485.6 | 167.8 | - | - | 0 | - |
| - | - | 1.04E+04 | 169.1 | - | - | 0 | - |
| - | - | 732.2 | 170.1 | - | - | 0 | - |
| - | - | 1135 | 171.1 | - | - | 0 | - |
| - | - | 2452 | 173.4 | - | - | 0 | - |
| - | - | 5495 | 183.1 | - | - | 0 | - |
| - | - | 622.8 | 185.1 | - | - | 0 | - |
| 18 | w | 1103 | 187.1 | 0.0008408 | 4.495 | +1 | 2 |
| - | - | 2.358E+05 | 187.1 | - | - | 0 | - |
| - | - | 2.156E+04 | 188.1 | - | - | 0 | - |
| - | - | 573.7 | 189.1 | - | - | 0 | - |
| - | - | 780.8 | 189.1 | - | - | 0 | - |
| 17 | y | 2595 | 195.1 | 0.002595 | 13.3 | +2 | 3 |
| - | - | 1.341E+04 | 197.1 | - | - | 0 | - |
| - | - | 465.2 | 198.1 | - | - | 0 | - |
| - | - | 1285 | 198.1 | - | - | 0 | - |
| - | - | 569.9 | 199.1 | - | - | 0 | - |
| - | - | 2404 | 201.1 | - | - | 0 | - |
| - | - | 3972 | 201.1 | - | - | 0 | - |
| - | - | 1319 | 213.1 | - | - | 0 | - |
| - | - | 1312 | 214.1 | - | - | 0 | - |
| - | - | 3.467E+05 | 215.1 | - | - | 0 | - |
| - | - | 3.703E+04 | 216.1 | - | - | 0 | - |
| - | - | 3456 | 217.1 | - | - | 0 | - |
| - | - | 1485 | 228.1 | - | - | 0 | - |
| 18 | z | 2230 | 228.1 | 0.0009002 | 3.946 | +1 | 2 |
| - | - | 1721 | 232.1 | - | - | 0 | - |
| - | - | 570.2 | 235.1 | - | - | 0 | - |
| - | - | 2479 | 239.2 | - | - | 0 | - |
| - | - | 7513 | 240.1 | - | - | 0 | - |
| - | - | 1247 | 241.1 | - | - | 0 | - |
| - | - | 4925 | 242.1 | - | - | 0 | - |
| - | - | 1050 | 244.1 | - | - | 0 | - |
| 18 | y | 1142 | 244.1 | 0.0003188 | 1.306 | +1 | 2 |
| 18 | z | 7.539E+04 | 245.1 | 0.0008404 | 3.428 | +1 | 2 |
| - | - | 1.023E+04 | 246.1 | - | - | 0 | - |
| - | - | 1107 | 247.1 | - | - | 0 | - |
| - | - | 3.385E+04 | 254.2 | - | - | 0 | - |
| - | - | 5142 | 255.2 | - | - | 0 | - |
| - | - | 3733 | 257.2 | - | - | 0 | - |
| - | - | 7146 | 260.1 | - | - | 0 | - |
| - | - | 1021 | 261.1 | - | - | 0 | - |
| 18 | y | 1.926E+04 | 261.2 | 0.0007625 | 2.92 | +1 | 2 |
| - | - | 1917 | 262.2 | - | - | 0 | - |
| - | - | 2054 | 263.1 | - | - | 0 | - |
| - | - | 531.2 | 263.6 | - | - | 0 | - |
| - | - | 510.7 | 264.3 | - | - | 0 | - |
| - | - | 802.6 | 270.1 | - | - | 0 | - |
| - | - | 1087 | 271.2 | - | - | 0 | - |
| - | - | 6664 | 272.2 | - | - | 0 | - |
| - | - | 855.1 | 274.2 | - | - | 0 | - |
| - | - | 1251 | 284.2 | - | - | 0 | - |
| - | - | 1303 | 285.2 | - | - | 0 | - |
| - | - | 833.2 | 286.1 | - | - | 0 | - |
| 3 | c | 1319 | 289.2 | 0.0003157 | 1.092 | +1 | 3 |
| - | - | 808.1 | 298.1 | - | - | 0 | - |
| - | - | 636.9 | 298.2 | - | - | 0 | - |
| - | - | 617.5 | 299.2 | - | - | 0 | - |
| - | - | 1792 | 301.1 | - | - | 0 | - |
| - | - | 2532 | 302.1 | - | - | 0 | - |
| - | - | 2249 | 311.1 | - | - | 0 | - |
| - | - | 1426 | 312.2 | - | - | 0 | - |
| 17 | w | 2.7E+04 | 315.2 | 0.001154 | 3.66 | +1 | 3 |
| - | - | 3310 | 316.2 | - | - | 0 | - |
| - | - | 1458 | 316.2 | - | - | 0 | - |
| - | - | 615.7 | 317.2 | - | - | 0 | - |
| - | - | 644.1 | 319.6 | - | - | 0 | - |
| - | - | 571.4 | 327.2 | - | - | 0 | - |
| - | - | 4290 | 327.2 | - | - | 0 | - |
| - | - | 731.1 | 328.2 | - | - | 0 | - |
| - | - | 8869 | 328.2 | - | - | 0 | - |
| - | - | 2756 | 328.2 | - | - | 0 | - |
| - | - | 6883 | 329.1 | - | - | 0 | - |
| - | - | 2236 | 329.2 | - | - | 0 | - |
| - | - | 4187 | 329.2 | - | - | 0 | - |
| - | - | 1602 | 330.1 | - | - | 0 | - |
| - | - | 869.6 | 337.2 | - | - | 0 | - |
| 14 | y | 2606 | 337.7 | 0.001408 | 4.17 | +2 | 6 |
| - | - | 805.4 | 338.2 | - | - | 0 | - |
| - | - | 8859 | 345.2 | - | - | 0 | - |
| - | - | 1756 | 346.2 | - | - | 0 | - |
| - | - | 4354 | 347.2 | - | - | 0 | - |
| - | - | 997.5 | 355.2 | - | - | 0 | - |
| - | - | 1.097E+04 | 355.2 | - | - | 0 | - |
| - | - | 2068 | 356.2 | - | - | 0 | - |
| - | - | 767.4 | 359.2 | - | - | 0 | - |
| - | - | 848.9 | 364.2 | - | - | 0 | - |
| 9 | w | 7857 | 371.2 | 0.004437 | 11.95 | +3 | 11 |
| 17 | y | 1.337E+04 | 372.2 | 0.001266 | 3.401 | +1 | 3 |
| 17 | z | 6.53E+04 | 373.2 | 0.002199 | 5.893 | +1 | 3 |
| - | - | 1.543E+04 | 374.2 | - | - | 0 | - |
| - | - | 2366 | 375.2 | - | - | 0 | - |
| - | - | 9150 | 386.2 | - | - | 0 | - |
| 13 | y | 2235 | 387.2 | 0.0028 | 7.231 | +2 | 7 |
| 17 | y | 6.123E+04 | 389.2 | 0.001359 | 3.491 | +1 | 3 |
| - | - | 8469 | 390.2 | - | - | 0 | - |
| 4 | c | 2026 | 390.2 | 0.006165 | 15.8 | +1 | 4 |
| - | - | 717.6 | 391.2 | - | - | 0 | - |
| - | - | 1028 | 394.2 | - | - | 0 | - |
| - | - | 1100 | 396.2 | - | - | 0 | - |
| - | - | 663.4 | 398.2 | - | - | 0 | - |
| - | - | 1135 | 400.2 | - | - | 0 | - |
| - | - | 952 | 400.2 | - | - | 0 | - |
| - | - | 783.2 | 406.2 | - | - | 0 | - |
| - | - | 2316 | 410.2 | - | - | 0 | - |
| - | - | 986.5 | 411.2 | - | - | 0 | - |
| - | - | 3089 | 412.2 | - | - | 0 | - |
| - | - | 1473 | 414.2 | - | - | 0 | - |
| - | - | 2282 | 414.2 | - | - | 0 | - |
| - | - | 3013 | 415.2 | - | - | 0 | - |
| - | - | 805.4 | 416.2 | - | - | 0 | - |
| - | - | 723.3 | 416.2 | - | - | 0 | - |
| - | - | 1783 | 418.2 | - | - | 0 | - |
| - | - | 5580 | 424.2 | - | - | 0 | - |
| - | - | 1440 | 425.2 | - | - | 0 | - |
| - | - | 6161 | 428.2 | - | - | 0 | - |
| - | - | 4664 | 429.2 | - | - | 0 | - |
| - | - | 4167 | 430.2 | - | - | 0 | - |
| - | - | 1753 | 432.2 | - | - | 0 | - |
| 14 | c | 1199 | 433.3 | 0.006425 | 14.83 | +3 | 14 |
| - | - | 2830 | 434.3 | - | - | 0 | - |
| - | - | 1068 | 435.3 | - | - | 0 | - |
| - | - | 1536 | 436.2 | - | - | 0 | - |
| - | - | 667.5 | 436.7 | - | - | 0 | - |
| - | - | 571.9 | 437.8 | - | - | 0 | - |
| - | - | 1.169E+04 | 442.2 | - | - | 0 | - |
| 16 | w | 2920 | 443.2 | 0.008311 | 18.75 | +1 | 4 |
| - | - | 8.003E+04 | 444.2 | - | - | 0 | - |
| - | - | 1.846E+04 | 445.2 | - | - | 0 | - |
| - | - | 2850 | 446.2 | - | - | 0 | - |
| - | - | 825.9 | 447.2 | - | - | 0 | - |
| - | - | 1882 | 448.2 | - | - | 0 | - |
| 6 | w | 1155 | 456.3 | 0.006605 | 14.48 | +3 | 14 |
| 5 | c | 999.6 | 459.3 | 0.001181 | 2.571 | +1 | 5 |
| - | - | 7306 | 460.2 | - | - | 0 | - |
| - | - | 1753 | 461.2 | - | - | 0 | - |
| - | - | 1492 | 463.3 | - | - | 0 | - |
| - | - | 1005 | 464.2 | - | - | 0 | - |
| - | - | 801.7 | 474.2 | - | - | 0 | - |
| - | - | 971.9 | 475.3 | - | - | 0 | - |
| - | - | 3840 | 476.3 | - | - | 0 | - |
| 5 | c | 1.004E+05 | 477.3 | 0.001816 | 3.805 | +1 | 5 |
| - | - | 2.39E+04 | 478.3 | - | - | 0 | - |
| - | - | 3898 | 479.3 | - | - | 0 | - |
| - | - | 951.7 | 480.2 | - | - | 0 | - |
| - | - | 807.6 | 484.3 | - | - | 0 | - |
| - | - | 1228 | 485.2 | - | - | 0 | - |
| 16 | y | 846.6 | 486.2 | 0.005213 | 10.72 | +1 | 4 |
| 16 | z | 5.201E+04 | 488.2 | 0.001868 | 3.825 | +1 | 4 |
| - | - | 3.597E+04 | 489.2 | - | - | 0 | - |
| - | - | 1738 | 489.8 | - | - | 0 | - |
| - | - | 8185 | 490.2 | - | - | 0 | - |
| 5 | y | 795 | 490.3 | 0.003893 | 7.94 | +3 | 15 |
| - | - | 674.8 | 490.3 | - | - | 0 | - |
| - | - | 1095 | 491.2 | - | - | 0 | - |
| 11 | y | 2486 | 493.8 | 0.003021 | 6.118 | +2 | 9 |
| - | - | 1609 | 494.3 | - | - | 0 | - |
| 5 | y | 727.1 | 496.3 | 0.001653 | 3.33 | +3 | 15 |
| 16 | c | 693.1 | 496.6 | 0.001506 | 3.032 | +3 | 16 |
| - | - | 668.9 | 497.3 | - | - | 0 | - |
| - | - | 1763 | 501.3 | - | - | 0 | - |
| - | - | 700.4 | 501.7 | - | - | 0 | - |
| - | - | 4458 | 503.2 | - | - | 0 | - |
| 16 | y | 2.312E+04 | 504.2 | 0.001515 | 3.005 | +1 | 4 |
| - | - | 5636 | 505.2 | - | - | 0 | - |
| - | - | 812.9 | 506.2 | - | - | 0 | - |
| - | - | 804 | 507.3 | - | - | 0 | - |
| - | - | 889.1 | 510.7 | - | - | 0 | - |
| - | - | 3059 | 511.3 | - | - | 0 | - |
| - | - | 879.5 | 512.3 | - | - | 0 | - |
| 10 | y | 833.4 | 513.8 | 0.006693 | 13.03 | +2 | 10 |
| 10 | z | 841.2 | 514.3 | 0.002963 | 5.762 | +2 | 10 |
| - | - | 1086 | 514.8 | - | - | 0 | - |
| 11 | c | 4075 | 515.3 | 0.0001416 | 0.2748 | +2 | 11 |
| - | - | 1745 | 515.8 | - | - | 0 | - |
| - | - | 2753 | 516.3 | - | - | 0 | - |
| - | - | 1929 | 517.9 | - | - | 0 | - |
| - | - | 1943 | 518.3 | - | - | 0 | - |
| 10 | y | 2.418E+04 | 522.3 | 0.001902 | 3.642 | +2 | 10 |
| - | - | 1.084E+04 | 522.8 | - | - | 0 | - |
| - | - | 5257 | 523.3 | - | - | 0 | - |
| 4 | y | 2316 | 523.9 | 0.002699 | 5.151 | +3 | 16 |
| - | - | 1636 | 524.3 | - | - | 0 | - |
| - | - | 6529 | 525.3 | - | - | 0 | - |
| - | - | 1265 | 525.8 | - | - | 0 | - |
| - | - | 1263 | 526.3 | - | - | 0 | - |
| - | - | 1701 | 526.3 | - | - | 0 | - |
| - | - | 844.6 | 527.3 | - | - | 0 | - |
| - | - | 799.8 | 527.8 | - | - | 0 | - |
| - | - | 956.3 | 528.3 | - | - | 0 | - |
| - | - | 924.1 | 528.6 | - | - | 0 | - |
| - | - | 6387 | 529.3 | - | - | 0 | - |
| 4 | y | 1067 | 530 | 0.00583 | 11 | +3 | 16 |
| - | - | 975.9 | 531.3 | - | - | 0 | - |
| 17 | c | 1659 | 533.3 | 0.001095 | 2.054 | +3 | 17 |
| - | - | 3687 | 534.3 | - | - | 0 | - |
| - | - | 1948 | 534.8 | - | - | 0 | - |
| - | - | 6294 | 535.3 | - | - | 0 | - |
| - | - | 1490 | 536.3 | - | - | 0 | - |
| - | - | 1254 | 536.8 | - | - | 0 | - |
| - | - | 4935 | 537 | - | - | 0 | - |
| - | - | 3527 | 537.3 | - | - | 0 | - |
| 3 | z | 2825 | 537.6 | 0.004492 | 8.355 | +3 | 17 |
| - | - | 946.2 | 538 | - | - | 0 | - |
| - | - | 4051 | 542.3 | - | - | 0 | - |
| 3 | y | 1.153E+04 | 543 | 0.001404 | 2.586 | +3 | 17 |
| 3 | y | 1.323E+04 | 543.3 | 0.002012 | 3.703 | +3 | 17 |
| 3 | z | 4809 | 543.6 | 0.006036 | 11.1 | +3 | 17 |
| 12 | c | 2116 | 543.8 | 0.001617 | 2.973 | +2 | 12 |
| - | - | 1430 | 544 | - | - | 0 | - |
| 15 | y | 1166 | 544.2 | 0.001809 | 3.324 | +1 | 5 |
| - | - | 1709 | 544.3 | - | - | 0 | - |
| 15 | z | 6357 | 545.2 | 0.001553 | 2.848 | +1 | 5 |
| - | - | 738.8 | 545.3 | - | - | 0 | - |
| - | - | 4.725E+04 | 546.3 | - | - | 0 | - |
| - | - | 1171 | 546.8 | - | - | 0 | - |
| - | - | 1.162E+04 | 547.3 | - | - | 0 | - |
| - | - | 2259 | 548.3 | - | - | 0 | - |
| 3 | y | 1717 | 549 | 0.001422 | 2.591 | +3 | 17 |
| - | - | 1031 | 549.3 | - | - | 0 | - |
| - | - | 1249 | 549.6 | - | - | 0 | - |
| - | - | 687.5 | 550.3 | - | - | 0 | - |
| - | - | 1932 | 551.3 | - | - | 0 | - |
| - | - | 1486 | 551.8 | - | - | 0 | - |
| - | - | 637.7 | 553 | - | - | 0 | - |
| - | - | 1263 | 559.3 | - | - | 0 | - |
| - | - | 1908 | 560.3 | - | - | 0 | - |
| 15 | y | 2.698E+04 | 561.3 | 0.002604 | 4.639 | +1 | 5 |
| - | - | 659.6 | 562 | - | - | 0 | - |
| - | - | 8276 | 562.3 | - | - | 0 | - |
| - | - | 1055 | 563.3 | - | - | 0 | - |
| - | - | 1800 | 564.6 | - | - | 0 | - |
| - | - | 2381 | 565 | - | - | 0 | - |
| - | - | 1919 | 565.3 | - | - | 0 | - |
| - | - | 2959 | 570.3 | - | - | 0 | - |
| - | - | 4460 | 570.6 | - | - | 0 | - |
| - | - | 6587 | 571 | - | - | 0 | - |
| 2 | z | 5666 | 571.3 | 0.003207 | 5.613 | +3 | 18 |
| 2 | z | 3008 | 571.6 | 0.00473 | 8.275 | +3 | 18 |
| 9 | y | 9502 | 571.8 | 0.002302 | 4.026 | +2 | 11 |
| - | - | 1658 | 572 | - | - | 0 | - |
| - | - | 6953 | 572.3 | - | - | 0 | - |
| - | - | 3299 | 572.8 | - | - | 0 | - |
| - | - | 668.1 | 576 | - | - | 0 | - |
| 18 | c | 3248 | 576.3 | 0.003746 | 6.501 | +3 | 18 |
| 2 | y | 3.122E+04 | 576.6 | 0.002682 | 4.652 | +3 | 18 |
| 2 | y | 3.436E+04 | 577 | 0.007929 | 13.74 | +3 | 18 |
| 2 | z | 1.733E+04 | 577.3 | 0.006643 | 11.51 | +3 | 18 |
| - | - | 5659 | 577.6 | - | - | 0 | - |
| - | - | 1999 | 578 | - | - | 0 | - |
| 6 | c | 3.163E+04 | 578.3 | 0.001806 | 3.123 | +1 | 6 |
| - | - | 1.01E+04 | 579.3 | - | - | 0 | - |
| - | - | 4098 | 579.8 | - | - | 0 | - |
| - | - | 1884 | 580.3 | - | - | 0 | - |
| - | - | 1515 | 582 | - | - | 0 | - |
| - | - | 1742 | 582.3 | - | - | 0 | - |
| 2 | y | 2.733E+04 | 582.6 | 0.002151 | 3.693 | +3 | 18 |
| - | - | 2.354E+04 | 583 | - | - | 0 | - |
| 8 | z | 1.428E+04 | 583.3 | 0.0004995 | 0.8563 | +2 | 12 |
| - | - | 5074 | 583.6 | - | - | 0 | - |
| - | - | 933.3 | 583.8 | - | - | 0 | - |
| - | - | 3442 | 584.3 | - | - | 0 | - |
| - | - | 2862 | 584.8 | - | - | 0 | - |
| - | - | 921.9 | 585.3 | - | - | 0 | - |
| - | - | 1997 | 586.3 | - | - | 0 | - |
| - | - | 1950 | 588 | - | - | 0 | - |
| - | - | 1053 | 588.3 | - | - | 0 | - |
| - | - | 4044 | 588.3 | - | - | 0 | - |
| - | - | 951.7 | 588.7 | - | - | 0 | - |
| - | - | 896.5 | 591.3 | - | - | 0 | - |
| 8 | y | 3469 | 591.8 | 0.002708 | 4.576 | +2 | 12 |
| 8 | z | 7704 | 592.3 | 0.002092 | 3.531 | +2 | 12 |
| - | - | 5123 | 592.8 | - | - | 0 | - |
| 13 | c | 1.056E+04 | 593.3 | 2.581E-06 | 0.00435 | +2 | 13 |
| - | - | 1709 | 593.7 | - | - | 0 | - |
| - | - | 6168 | 593.8 | - | - | 0 | - |
| - | - | 1876 | 594.3 | - | - | 0 | - |
| - | - | 1069 | 594.7 | - | - | 0 | - |
| - | - | 1669 | 598.3 | - | - | 0 | - |
| - | - | 1416 | 599.3 | - | - | 0 | - |
| - | - | 5696 | 599.7 | - | - | 0 | - |
| - | - | 2263 | 599.8 | - | - | 0 | - |
| - | - | 6225 | 600 | - | - | 0 | - |
| 8 | y | 9.037E+04 | 600.3 | 0.002068 | 3.445 | +2 | 12 |
| - | - | 5.268E+04 | 600.8 | - | - | 0 | - |
| - | - | 2.184E+04 | 601.3 | - | - | 0 | - |
| - | - | 5649 | 601.8 | - | - | 0 | - |
| - | - | 1131 | 602.3 | - | - | 0 | - |
| - | - | 4671 | 602.3 | - | - | 0 | - |
| - | - | 4577 | 602.7 | - | - | 0 | - |
| - | - | 2676 | 603 | - | - | 0 | - |
| - | - | 1102 | 603.3 | - | - | 0 | - |
| - | - | 1627 | 605.3 | - | - | 0 | - |
| - | - | 2980 | 605.7 | - | - | 0 | - |
| - | - | 3419 | 606 | - | - | 0 | - |
| - | - | 3323 | 606.3 | - | - | 0 | - |
| - | - | 2.006E+04 | 608.3 | - | - | 0 | - |
| - | - | 2.64E+04 | 608.7 | - | - | 0 | - |
| - | - | 1.997E+04 | 609 | - | - | 0 | - |
| - | - | 7152 | 609.3 | - | - | 0 | - |
| - | - | 2549 | 609.7 | - | - | 0 | - |
| - | - | 1055 | 610 | - | - | 0 | - |
| - | - | 1667 | 610.8 | - | - | 0 | - |
| - | - | 1681 | 611.3 | - | - | 0 | - |
| - | - | 2973 | 613.7 | - | - | 0 | - |
| - | - | 3254 | 613.8 | - | - | 0 | - |
| - | - | 3914 | 614 | - | - | 0 | - |
| - | - | 1.836E+05 | 614.3 | - | - | 0 | - |
| - | - | 1.919E+05 | 614.7 | - | - | 0 | - |
| - | - | 1.113E+05 | 615 | - | - | 0 | - |
| 14 | w | 3.934E+04 | 615.3 | 0.002049 | 3.33 | +1 | 6 |
| - | - | 4.039E+04 | 615.3 | - | - | 0 | - |
| - | - | 1.347E+04 | 615.7 | - | - | 0 | - |
| - | - | 1.077E+04 | 616.3 | - | - | 0 | - |
| - | - | 3011 | 617.3 | - | - | 0 | - |
| - | - | 2212 | 618.8 | - | - | 0 | - |
| - | - | 1648 | 619.3 | - | - | 0 | - |
| - | - | 1303 | 619.4 | - | - | 0 | - |
| - | - | 1.006E+04 | 619.7 | - | - | 0 | - |
| - | - | 1000 | 619.8 | - | - | 0 | - |
| - | - | 1329 | 619.9 | - | - | 0 | - |
| - | - | 7720 | 620 | - | - | 0 | - |
| - | - | 2.748E+05 | 620.3 | - | - | 0 | - |
| - | - | 2.651E+05 | 620.7 | - | - | 0 | - |
| - | - | 1.605E+05 | 621 | - | - | 0 | - |
| - | - | 5.767E+04 | 621.3 | - | - | 0 | - |
| - | - | 1.615E+04 | 621.7 | - | - | 0 | - |
| - | - | 5449 | 624.3 | - | - | 0 | - |
| - | - | 1567 | 625.3 | - | - | 0 | - |
| - | - | 1282 | 628.8 | - | - | 0 | - |
| - | - | 7195 | 632.4 | - | - | 0 | - |
| - | - | 9377 | 633.4 | - | - | 0 | - |
| - | - | 1.645E+04 | 634.4 | - | - | 0 | - |
| - | - | 1877 | 634.9 | - | - | 0 | - |
| - | - | 5270 | 635.4 | - | - | 0 | - |
| - | - | 1128 | 636.4 | - | - | 0 | - |
| - | - | 1121 | 639.4 | - | - | 0 | - |
| - | - | 706 | 640.3 | - | - | 0 | - |
| - | - | 2437 | 640.9 | - | - | 0 | - |
| 7 | y | 2950 | 641.3 | 0.001705 | 2.658 | +2 | 13 |
| 7 | z | 5583 | 641.9 | 0.003896 | 6.069 | +2 | 13 |
| - | - | 1.655E+04 | 642.3 | - | - | 0 | - |
| - | - | 4545 | 643.4 | - | - | 0 | - |
| - | - | 989.9 | 644.3 | - | - | 0 | - |
| 14 | c | 5.003E+04 | 649.9 | 0.009559 | 14.71 | +2 | 14 |
| - | - | 3.332E+04 | 650.4 | - | - | 0 | - |
| - | - | 1.472E+04 | 650.9 | - | - | 0 | - |
| - | - | 5459 | 651.4 | - | - | 0 | - |
| - | - | 2357 | 655.4 | - | - | 0 | - |
| - | - | 1509 | 656.4 | - | - | 0 | - |
| 14 | y | 1226 | 657.3 | 0.001852 | 2.817 | +1 | 6 |
| 14 | z | 4.683E+04 | 658.3 | 0.002389 | 3.628 | +1 | 6 |
| - | - | 2.401E+04 | 659.3 | - | - | 0 | - |
| - | - | 1.658E+04 | 660.4 | - | - | 0 | - |
| - | - | 5420 | 661.4 | - | - | 0 | - |
| - | - | 3993 | 663.4 | - | - | 0 | - |
| - | - | 2765 | 663.9 | - | - | 0 | - |
| - | - | 1836 | 664.4 | - | - | 0 | - |
| - | - | 2078 | 669.4 | - | - | 0 | - |
| - | - | 1307 | 669.9 | - | - | 0 | - |
| - | - | 2118 | 670.4 | - | - | 0 | - |
| - | - | 1848 | 670.9 | - | - | 0 | - |
| - | - | 864.3 | 671.4 | - | - | 0 | - |
| - | - | 1674 | 672.4 | - | - | 0 | - |
| 14 | y | 3.122E+04 | 674.3 | 0.002219 | 3.291 | +1 | 6 |
| - | - | 1.106E+04 | 675.4 | - | - | 0 | - |
| - | - | 1.321E+04 | 676.4 | - | - | 0 | - |
| 7 | c | 4.237E+04 | 677.4 | 0.001874 | 2.766 | +1 | 7 |
| 15 | c | 2050 | 677.9 | 0.01302 | 19.21 | +2 | 15 |
| 15 | c | 1.55E+04 | 678.4 | 0.001072 | 1.58 | +2 | 15 |
| - | - | 827.7 | 678.9 | - | - | 0 | - |
| - | - | 3055 | 679.4 | - | - | 0 | - |
| - | - | 1299 | 679.9 | - | - | 0 | - |
| - | - | 1124 | 680.4 | - | - | 0 | - |
| 6 | z | 1276 | 683.9 | 0.007308 | 10.69 | +2 | 14 |
| - | - | 2093 | 684.4 | - | - | 0 | - |
| - | - | 2541 | 685.4 | - | - | 0 | - |
| - | - | 1447 | 686.4 | - | - | 0 | - |
| - | - | 1386 | 687.4 | - | - | 0 | - |
| - | - | 1254 | 688.4 | - | - | 0 | - |
| - | - | 5046 | 690.4 | - | - | 0 | - |
| - | - | 804.7 | 690.9 | - | - | 0 | - |
| 6 | y | 1.457E+04 | 691.4 | 0.0005737 | 0.8298 | +2 | 14 |
| 6 | y | 4.864E+04 | 691.9 | 0.001913 | 2.765 | +2 | 14 |
| 6 | z | 4.954E+04 | 692.4 | 0.0009913 | 1.432 | +2 | 14 |
| - | - | 3.729E+04 | 692.9 | - | - | 0 | - |
| - | - | 1.562E+04 | 693.4 | - | - | 0 | - |
| - | - | 4662 | 693.9 | - | - | 0 | - |
| - | - | 1202 | 698.4 | - | - | 0 | - |
| - | - | 6597 | 699.4 | - | - | 0 | - |
| - | - | 5420 | 699.9 | - | - | 0 | - |
| 6 | y | 7.734E+04 | 700.4 | 0.001822 | 2.602 | +2 | 14 |
| - | - | 5.136E+04 | 700.9 | - | - | 0 | - |
| - | - | 1.181E+04 | 701.3 | - | - | 0 | - |
| - | - | 2.169E+04 | 701.4 | - | - | 0 | - |
| - | - | 6104 | 701.9 | - | - | 0 | - |
| - | - | 5015 | 702.3 | - | - | 0 | - |
| - | - | 1041 | 702.4 | - | - | 0 | - |
| - | - | 1835 | 703.4 | - | - | 0 | - |
| - | - | 5157 | 704.4 | - | - | 0 | - |
| - | - | 2768 | 704.9 | - | - | 0 | - |
| - | - | 3443 | 705.4 | - | - | 0 | - |
| - | - | 870.3 | 705.9 | - | - | 0 | - |
| - | - | 2724 | 706.9 | - | - | 0 | - |
| - | - | 2515 | 707.4 | - | - | 0 | - |
| - | - | 3360 | 707.9 | - | - | 0 | - |
| - | - | 1952 | 708.4 | - | - | 0 | - |
| - | - | 937.4 | 708.9 | - | - | 0 | - |
| - | - | 3200 | 713.4 | - | - | 0 | - |
| - | - | 3984 | 713.9 | - | - | 0 | - |
| - | - | 3427 | 714.3 | - | - | 0 | - |
| - | - | 2408 | 714.4 | - | - | 0 | - |
| - | - | 2123 | 714.9 | - | - | 0 | - |
| - | - | 838 | 715.4 | - | - | 0 | - |
| - | - | 4453 | 717.4 | - | - | 0 | - |
| - | - | 1888 | 717.9 | - | - | 0 | - |
| - | - | 2285 | 718.4 | - | - | 0 | - |
| - | - | 1012 | 719.4 | - | - | 0 | - |
| - | - | 971.4 | 720.4 | - | - | 0 | - |
| - | - | 1783 | 721.4 | - | - | 0 | - |
| - | - | 1119 | 721.9 | - | - | 0 | - |
| - | - | 3003 | 725.9 | - | - | 0 | - |
| - | - | 2313 | 726.4 | - | - | 0 | - |
| 5 | z | 9585 | 726.9 | 0.007593 | 10.45 | +2 | 15 |
| 5 | w | 5702 | 727.4 | 0.00716 | 9.843 | +2 | 15 |
| - | - | 3631 | 727.9 | - | - | 0 | - |
| - | - | 3416 | 728.4 | - | - | 0 | - |
| - | - | 1704 | 728.9 | - | - | 0 | - |
| - | - | 1025 | 729.4 | - | - | 0 | - |
| - | - | 1035 | 732.4 | - | - | 0 | - |
| - | - | 3743 | 733.4 | - | - | 0 | - |
| 8 | c | 5.748E+04 | 734.4 | 0.002199 | 2.995 | +1 | 8 |
| 5 | y | 7023 | 734.9 | 0.004332 | 5.895 | +2 | 15 |
| 16 | c | 3.132E+04 | 735.4 | 0.003485 | 4.739 | +2 | 16 |
| 16 | c | 1.336E+05 | 735.9 | 0.003306 | 4.492 | +2 | 16 |
| - | - | 1.481E+05 | 736.4 | - | - | 0 | - |
| - | - | 7.588E+04 | 736.9 | - | - | 0 | - |
| - | - | 3.058E+04 | 737.4 | - | - | 0 | - |
| - | - | 7821 | 737.9 | - | - | 0 | - |
| - | - | 1260 | 738.4 | - | - | 0 | - |
| 13 | z | 1170 | 739.4 | 0.006856 | 9.273 | +1 | 7 |
| 13 | z | 1149 | 740.4 | 0.01411 | 19.06 | +1 | 7 |
| - | - | 1385 | 741.4 | - | - | 0 | - |
| 13 | w | 3950 | 742.4 | 0.003592 | 4.839 | +1 | 7 |
| - | - | 1778 | 742.9 | - | - | 0 | - |
| - | - | 1.121E+04 | 743.4 | - | - | 0 | - |
| 5 | y | 1.658E+05 | 743.9 | 0.002532 | 3.403 | +2 | 15 |
| 16 | c | 1.31E+05 | 744.4 | 0.006143 | 8.252 | +2 | 16 |
| - | - | 5.856E+04 | 744.9 | - | - | 0 | - |
| - | - | 1.618E+04 | 745.4 | - | - | 0 | - |
| - | - | 4562 | 745.9 | - | - | 0 | - |
| - | - | 2701 | 747.9 | - | - | 0 | - |
| - | - | 3943 | 748.4 | - | - | 0 | - |
| - | - | 3024 | 748.9 | - | - | 0 | - |
| - | - | 901.9 | 749.4 | - | - | 0 | - |
| - | - | 1302 | 751.4 | - | - | 0 | - |
| - | - | 996.6 | 754.4 | - | - | 0 | - |
| 13 | y | 906.4 | 756.4 | 0.00552 | 7.298 | +1 | 7 |
| - | - | 1.087E+04 | 756.9 | - | - | 0 | - |
| 13 | z | 5.238E+04 | 757.4 | 0.004348 | 5.741 | +1 | 7 |
| - | - | 6969 | 757.9 | - | - | 0 | - |
| - | - | 2.4E+04 | 758.4 | - | - | 0 | - |
| - | - | 1629 | 758.9 | - | - | 0 | - |
| - | - | 5264 | 759.4 | - | - | 0 | - |
| - | - | 1282 | 760.4 | - | - | 0 | - |
| - | - | 1187 | 763.4 | - | - | 0 | - |
| - | - | 3037 | 764.4 | - | - | 0 | - |
| - | - | 2684 | 764.9 | - | - | 0 | - |
| - | - | 2040 | 765.4 | - | - | 0 | - |
| - | - | 4039 | 765.9 | - | - | 0 | - |
| - | - | 2982 | 766.4 | - | - | 0 | - |
| - | - | 1519 | 766.9 | - | - | 0 | - |
| - | - | 1154 | 769.4 | - | - | 0 | - |
| - | - | 6770 | 770.4 | - | - | 0 | - |
| - | - | 3376 | 771.4 | - | - | 0 | - |
| - | - | 3012 | 772.4 | - | - | 0 | - |
| 13 | y | 1.83E+04 | 773.4 | 0.002836 | 3.667 | +1 | 7 |
| - | - | 6374 | 774.4 | - | - | 0 | - |
| - | - | 1326 | 775.4 | - | - | 0 | - |
| - | - | 1095 | 776.9 | - | - | 0 | - |
| 4 | z | 1615 | 777.4 | 0.002065 | 2.656 | +2 | 16 |
| 4 | z | 2027 | 777.9 | 0.01518 | 19.52 | +2 | 16 |
| - | - | 4302 | 778.4 | - | - | 0 | - |
| 4 | w | 4569 | 778.9 | 0.006628 | 8.509 | +2 | 16 |
| - | - | 3543 | 779.4 | - | - | 0 | - |
| - | - | 1678 | 779.9 | - | - | 0 | - |
| - | - | 1937 | 780.4 | - | - | 0 | - |
| - | - | 1195 | 781.4 | - | - | 0 | - |
| - | - | 1332 | 782.4 | - | - | 0 | - |
| - | - | 2228 | 783.4 | - | - | 0 | - |
| 4 | y | 7751 | 785.4 | 0.004184 | 5.328 | +2 | 16 |
| 4 | y | 2.434E+04 | 785.9 | 0.003503 | 4.457 | +2 | 16 |
| 4 | z | 5.626E+04 | 786.4 | 0.00252 | 3.204 | +2 | 16 |
| - | - | 5.818E+04 | 786.9 | - | - | 0 | - |
| - | - | 3.315E+04 | 787.4 | - | - | 0 | - |
| - | - | 1.155E+04 | 787.9 | - | - | 0 | - |
| - | - | 3734 | 788.4 | - | - | 0 | - |
| - | - | 1.182E+04 | 789.5 | - | - | 0 | - |
| - | - | 1.163E+04 | 790.5 | - | - | 0 | - |
| - | - | 5007 | 790.9 | - | - | 0 | - |
| - | - | 3069 | 791.5 | - | - | 0 | - |
| - | - | 3395 | 791.9 | - | - | 0 | - |
| - | - | 1338 | 792.4 | - | - | 0 | - |
| - | - | 4363 | 792.9 | - | - | 0 | - |
| - | - | 5884 | 793.4 | - | - | 0 | - |
| - | - | 5481 | 793.9 | - | - | 0 | - |
| 4 | y | 6.622E+04 | 794.4 | 0.002496 | 3.142 | +2 | 16 |
| - | - | 5.495E+04 | 794.9 | - | - | 0 | - |
| - | - | 2.324E+04 | 795.4 | - | - | 0 | - |
| - | - | 8150 | 795.9 | - | - | 0 | - |
| - | - | 1730 | 796.4 | - | - | 0 | - |
| - | - | 832.9 | 796.9 | - | - | 0 | - |
| - | - | 1099 | 797.4 | - | - | 0 | - |
| - | - | 5200 | 798.4 | - | - | 0 | - |
| - | - | 2689 | 798.9 | - | - | 0 | - |
| 17 | c | 3085 | 799.4 | 0.01391 | 17.4 | +2 | 17 |
| 17 | c | 5671 | 799.9 | 0.002135 | 2.669 | +2 | 17 |
| - | - | 5883 | 800.4 | - | - | 0 | - |
| - | - | 4987 | 800.9 | - | - | 0 | - |
| - | - | 4888 | 801.4 | - | - | 0 | - |
| - | - | 2444 | 801.9 | - | - | 0 | - |
| - | - | 5907 | 804.9 | - | - | 0 | - |
| - | - | 9887 | 805.4 | - | - | 0 | - |
| 3 | z | 8185 | 805.9 | 0.003601 | 4.468 | +2 | 17 |
| 3 | z | 4001 | 806.4 | 0.01373 | 17.03 | +2 | 17 |
| - | - | 1588 | 806.9 | - | - | 0 | - |
| - | - | 7958 | 807.4 | - | - | 0 | - |
| - | - | 5930 | 807.9 | - | - | 0 | - |
| 17 | c | 5372 | 808.4 | 0.002595 | 3.21 | +2 | 17 |
| - | - | 2947 | 808.9 | - | - | 0 | - |
| - | - | 7114 | 811.4 | - | - | 0 | - |
| - | - | 3400 | 812.4 | - | - | 0 | - |
| - | - | 1566 | 812.9 | - | - | 0 | - |
| - | - | 2580 | 813.4 | - | - | 0 | - |
| 3 | y | 4.592E+04 | 813.9 | 0.002723 | 3.345 | +2 | 17 |
| 3 | y | 7.739E+04 | 814.4 | 0.007114 | 8.735 | +2 | 17 |
| 3 | z | 7.177E+04 | 814.9 | 0.003812 | 4.677 | +2 | 17 |
| - | - | 9.777E+04 | 815.4 | - | - | 0 | - |
| - | - | 2.384E+04 | 815.9 | - | - | 0 | - |
| - | - | 3.351E+04 | 816.4 | - | - | 0 | - |
| - | - | 3368 | 816.9 | - | - | 0 | - |
| - | - | 7733 | 817.4 | - | - | 0 | - |
| - | - | 1586 | 818.4 | - | - | 0 | - |
| - | - | 1136 | 821.5 | - | - | 0 | - |
| - | - | 1.579E+04 | 821.9 | - | - | 0 | - |
| - | - | 1.53E+04 | 822.4 | - | - | 0 | - |
| 3 | y | 8.268E+05 | 822.9 | 0.003056 | 3.713 | +2 | 17 |
| - | - | 7.207E+05 | 823.4 | - | - | 0 | - |
| - | - | 3.634E+05 | 823.9 | - | - | 0 | - |
| - | - | 1.151E+05 | 824.4 | - | - | 0 | - |
| - | - | 2.703E+04 | 824.9 | - | - | 0 | - |
| - | - | 1035 | 829.9 | - | - | 0 | - |
| 12 | y | 2.162E+04 | 830.4 | 0.00304 | 3.66 | +1 | 8 |
| - | - | 8437 | 831.4 | - | - | 0 | - |
| - | - | 1.432E+04 | 832.5 | - | - | 0 | - |
| 9 | c | 2.752E+04 | 833.5 | 0.001351 | 1.621 | +1 | 9 |
| - | - | 1.04E+04 | 834.5 | - | - | 0 | - |
| - | - | 2992 | 835.5 | - | - | 0 | - |
| - | - | 1860 | 836 | - | - | 0 | - |
| - | - | 1454 | 836.4 | - | - | 0 | - |
| - | - | 1844 | 836.9 | - | - | 0 | - |
| - | - | 3191 | 837.4 | - | - | 0 | - |
| - | - | 2358 | 837.9 | - | - | 0 | - |
| - | - | 1702 | 839.5 | - | - | 0 | - |
| - | - | 6568 | 843.5 | - | - | 0 | - |
| - | - | 7034 | 844 | - | - | 0 | - |
| - | - | 4952 | 844.5 | - | - | 0 | - |
| - | - | 2317 | 844.9 | - | - | 0 | - |
| - | - | 5143 | 846.5 | - | - | 0 | - |
| - | - | 4712 | 847.5 | - | - | 0 | - |
| - | - | 2084 | 848.5 | - | - | 0 | - |
| - | - | 1029 | 849.5 | - | - | 0 | - |
| - | - | 8074 | 850.5 | - | - | 0 | - |
| - | - | 6444 | 851 | - | - | 0 | - |
| - | - | 5271 | 851.5 | - | - | 0 | - |
| - | - | 3109 | 852 | - | - | 0 | - |
| - | - | 1549 | 852.4 | - | - | 0 | - |
| - | - | 1996 | 855 | - | - | 0 | - |
| - | - | 7263 | 855.5 | - | - | 0 | - |
| - | - | 7154 | 856 | - | - | 0 | - |
| 2 | z | 7128 | 856.4 | 0.002101 | 2.453 | +2 | 18 |
| 2 | z | 8835 | 856.9 | 0.01461 | 17.05 | +2 | 18 |
| - | - | 6378 | 857.5 | - | - | 0 | - |
| 2 | w | 6742 | 857.9 | 0.008068 | 9.403 | +2 | 18 |
| - | - | 6600 | 858.4 | - | - | 0 | - |
| - | - | 3252 | 858.9 | - | - | 0 | - |
| - | - | 1543 | 863.5 | - | - | 0 | - |
| 18 | c | 5810 | 864 | 0.00311 | 3.6 | +2 | 18 |
| 2 | y | 5.624E+04 | 864.5 | 0.002748 | 3.179 | +2 | 18 |
| 2 | y | 9.036E+04 | 864.9 | 0.008421 | 9.736 | +2 | 18 |
| 2 | z | 1.008E+05 | 865.5 | 0.003776 | 4.363 | +2 | 18 |
| - | - | 6.404E+04 | 866 | - | - | 0 | - |
| - | - | 2.906E+04 | 866.5 | - | - | 0 | - |
| - | - | 1.019E+04 | 867 | - | - | 0 | - |
| - | - | 2279 | 867.5 | - | - | 0 | - |
| - | - | 5825 | 869.5 | - | - | 0 | - |
| - | - | 4805 | 869.9 | - | - | 0 | - |
| - | - | 4368 | 870.4 | - | - | 0 | - |
| - | - | 1709 | 871 | - | - | 0 | - |
| - | - | 7066 | 871.5 | - | - | 0 | - |
| - | - | 9828 | 872 | - | - | 0 | - |
| 10 | c | 3.357E+05 | 872.5 | 0.008214 | 9.414 | +1 | 10 |
| - | - | 3.179E+05 | 873 | - | - | 0 | - |
| 2 | y | 4.572E+05 | 873.5 | 0.009429 | 10.79 | +2 | 18 |
| - | - | 3.968E+05 | 874 | - | - | 0 | - |
| - | - | 2.11E+05 | 874.5 | - | - | 0 | - |
| - | - | 6.775E+04 | 875 | - | - | 0 | - |
| - | - | 1.875E+04 | 875.5 | - | - | 0 | - |
| - | - | 1287 | 876.5 | - | - | 0 | - |
| - | - | 1.958E+04 | 878.5 | - | - | 0 | - |
| - | - | 2.029E+04 | 879 | - | - | 0 | - |
| - | - | 1.217E+04 | 879.5 | - | - | 0 | - |
| - | - | 9169 | 880 | - | - | 0 | - |
| - | - | 4875 | 880.5 | - | - | 0 | - |
| - | - | 1026 | 881 | - | - | 0 | - |
| - | - | 1263 | 885.4 | - | - | 0 | - |
| - | - | 2360 | 886 | - | - | 0 | - |
| - | - | 3954 | 886.5 | - | - | 0 | - |
| - | - | 3883 | 887 | - | - | 0 | - |
| - | - | 2495 | 887.5 | - | - | 0 | - |
| - | - | 1614 | 888.5 | - | - | 0 | - |
| - | - | 4876 | 889.5 | - | - | 0 | - |
| 10 | c | 8.732E+04 | 890.5 | 0.002837 | 3.186 | +1 | 10 |
| - | - | 4.331E+04 | 891.5 | - | - | 0 | - |
| - | - | 1333 | 892 | - | - | 0 | - |
| - | - | 1.182E+04 | 892.5 | - | - | 0 | - |
| - | - | 1606 | 893 | - | - | 0 | - |
| - | - | 3951 | 893.5 | - | - | 0 | - |
| - | - | 7014 | 894 | - | - | 0 | - |
| - | - | 6471 | 894.5 | - | - | 0 | - |
| - | - | 2540 | 895 | - | - | 0 | - |
| - | - | 2295 | 895.5 | - | - | 0 | - |
| - | - | 3993 | 899.5 | - | - | 0 | - |
| - | - | 1.488E+04 | 900 | - | - | 0 | - |
| - | - | 7.525E+04 | 900.5 | - | - | 0 | - |
| - | - | 1.039E+05 | 901 | - | - | 0 | - |
| - | - | 7.784E+04 | 901.5 | - | - | 0 | - |
| - | - | 3.89E+04 | 902 | - | - | 0 | - |
| - | - | 1.244E+04 | 902.5 | - | - | 0 | - |
| - | - | 2478 | 903 | - | - | 0 | - |
| - | - | 1275 | 906.5 | - | - | 0 | - |
| - | - | 6.29E+04 | 907.5 | - | - | 0 | - |
| - | - | 7.915E+04 | 908 | - | - | 0 | - |
| - | - | 5.82E+04 | 908.5 | - | - | 0 | - |
| - | - | 4.371E+04 | 909 | - | - | 0 | - |
| - | - | 2.73E+04 | 909.5 | - | - | 0 | - |
| - | - | 1.373E+04 | 910 | - | - | 0 | - |
| - | - | 5380 | 910.5 | - | - | 0 | - |
| - | - | 6619 | 911.5 | - | - | 0 | - |
| - | - | 3780 | 912.5 | - | - | 0 | - |
| - | - | 6631 | 913 | - | - | 0 | - |
| - | - | 9764 | 913.5 | - | - | 0 | - |
| - | - | 5677 | 914 | - | - | 0 | - |
| - | - | 6913 | 914.5 | - | - | 0 | - |
| - | - | 5148 | 915 | - | - | 0 | - |
| - | - | 3913 | 915.5 | - | - | 0 | - |
| - | - | 2682 | 916.5 | - | - | 0 | - |
| - | - | 1099 | 917.5 | - | - | 0 | - |
| - | - | 1527 | 920.5 | - | - | 0 | - |
| - | - | 5586 | 921 | - | - | 0 | - |
| - | - | 2.498E+04 | 921.5 | - | - | 0 | - |
| - | - | 1.695E+05 | 922 | - | - | 0 | - |
| - | - | 1.633E+05 | 922.5 | - | - | 0 | - |
| - | - | 9.13E+04 | 923 | - | - | 0 | - |
| - | - | 1411 | 923.4 | - | - | 0 | - |
| - | - | 3.305E+04 | 923.5 | - | - | 0 | - |
| - | - | 9657 | 924 | - | - | 0 | - |
| - | - | 9380 | 926.5 | - | - | 0 | - |
| - | - | 4689 | 927.5 | - | - | 0 | - |
| - | - | 1516 | 928.5 | - | - | 0 | - |
| - | - | 1172 | 928.5 | - | - | 0 | - |
| - | - | 6783 | 929 | - | - | 0 | - |
| - | - | 1.379E+04 | 929.5 | - | - | 0 | - |
| - | - | 1.869E+05 | 930 | - | - | 0 | - |
| - | - | 2.014E+05 | 930.5 | - | - | 0 | - |
| - | - | 1.267E+05 | 931 | - | - | 0 | - |
| - | - | 4.85E+04 | 931.5 | - | - | 0 | - |
| - | - | 1.288E+04 | 932 | - | - | 0 | - |
| - | - | 1709 | 932.5 | - | - | 0 | - |
| - | - | 3203 | 939.5 | - | - | 0 | - |
| - | - | 1658 | 940.5 | - | - | 0 | - |
| 11 | z | 6.176E+04 | 970.5 | 0.002928 | 3.017 | +1 | 9 |
| - | - | 4.351E+04 | 971.5 | - | - | 0 | - |
| - | - | 1.6E+04 | 972.5 | - | - | 0 | - |
| - | - | 4165 | 973.5 | - | - | 0 | - |
| - | - | 2095 | 983.6 | - | - | 0 | - |
| - | - | 4413 | 985.5 | - | - | 0 | - |
| 11 | y | 1.105E+04 | 986.5 | 0.006238 | 6.323 | +1 | 9 |
| - | - | 4977 | 987.5 | - | - | 0 | - |
| - | - | 1434 | 988.6 | - | - | 0 | - |
| - | - | 1107 | 998.6 | - | - | 0 | - |
| - | - | 1054 | 1002 | - | - | 0 | - |
| - | - | 1.19E+04 | 1003 | - | - | 0 | - |
| - | - | 6026 | 1004 | - | - | 0 | - |
| - | - | 1871 | 1005 | - | - | 0 | - |
| - | - | 1079 | 1011 | - | - | 0 | - |
| - | - | 2184 | 1012 | - | - | 0 | - |
| - | - | 2192 | 1013 | - | - | 0 | - |
| 10 | z | 1.516E+04 | 1028 | 0.004536 | 4.414 | +1 | 10 |
| - | - | 3.385E+04 | 1029 | - | - | 0 | - |
| 11 | c | 1.726E+04 | 1030 | 0.01283 | 12.46 | +1 | 11 |
| - | - | 5643 | 1031 | - | - | 0 | - |
| - | - | 1012 | 1032 | - | - | 0 | - |
| - | - | 1330 | 1039 | - | - | 0 | - |
| - | - | 1134 | 1042 | - | - | 0 | - |
| - | - | 5072 | 1043 | - | - | 0 | - |
| 10 | y | 2.773E+04 | 1044 | 0.003146 | 3.014 | +1 | 10 |
| - | - | 1.703E+04 | 1045 | - | - | 0 | - |
| - | - | 3.321E+04 | 1046 | - | - | 0 | - |
| 11 | c | 4.147E+04 | 1047 | 0.0005418 | 0.5177 | +1 | 11 |
| - | - | 1.847E+04 | 1048 | - | - | 0 | - |
| - | - | 5556 | 1049 | - | - | 0 | - |
| - | - | 1101 | 1051 | - | - | 0 | - |
| - | - | 2064 | 1056 | - | - | 0 | - |
| - | - | 8751 | 1060 | - | - | 0 | - |
| - | - | 6232 | 1061 | - | - | 0 | - |
| - | - | 2457 | 1062 | - | - | 0 | - |
| - | - | 5776 | 1068 | - | - | 0 | - |
| - | - | 3131 | 1069 | - | - | 0 | - |
| - | - | 1566 | 1070 | - | - | 0 | - |
| - | - | 2136 | 1071 | - | - | 0 | - |
| - | - | 1445 | 1072 | - | - | 0 | - |
| - | - | 2823 | 1073 | - | - | 0 | - |
| - | - | 1362 | 1074 | - | - | 0 | - |
| - | - | 3316 | 1083 | - | - | 0 | - |
| - | - | 1171 | 1084 | - | - | 0 | - |
| - | - | 8512 | 1085 | - | - | 0 | - |
| 12 | c | 4967 | 1086 | 0.009784 | 9.013 | +1 | 12 |
| 12 | c | 2520 | 1087 | 0.01905 | 17.54 | +1 | 12 |
| - | - | 1000 | 1088 | - | - | 0 | - |
| - | - | 1477 | 1098 | - | - | 0 | - |
| - | - | 3963 | 1099 | - | - | 0 | - |
| - | - | 2169 | 1100 | - | - | 0 | - |
| - | - | 1083 | 1101 | - | - | 0 | - |
| - | - | 1733 | 1102 | - | - | 0 | - |
| - | - | 4540 | 1103 | - | - | 0 | - |
| 12 | c | 7.139E+04 | 1104 | 0.003614 | 3.275 | +1 | 12 |
| - | - | 4.189E+04 | 1105 | - | - | 0 | - |
| - | - | 1.456E+04 | 1106 | - | - | 0 | - |
| - | - | 2758 | 1107 | - | - | 0 | - |
| 9 | z | 3.701E+04 | 1127 | 0.003627 | 3.219 | +1 | 11 |
| - | - | 2.721E+04 | 1128 | - | - | 0 | - |
| - | - | 1.137E+04 | 1129 | - | - | 0 | - |
| - | - | 2485 | 1130 | - | - | 0 | - |
| - | - | 2403 | 1140 | - | - | 0 | - |
| - | - | 1949 | 1141 | - | - | 0 | - |
| - | - | 9019 | 1142 | - | - | 0 | - |
| 9 | y | 1.79E+04 | 1143 | 0.003824 | 3.346 | +1 | 11 |
| - | - | 8753 | 1144 | - | - | 0 | - |
| - | - | 3642 | 1145 | - | - | 0 | - |
| - | - | 1437 | 1158 | - | - | 0 | - |
| - | - | 7074 | 1159 | - | - | 0 | - |
| - | - | 4996 | 1160 | - | - | 0 | - |
| - | - | 1872 | 1161 | - | - | 0 | - |
| 8 | y | 2022 | 1183 | 0.002683 | 2.269 | +1 | 12 |
| 8 | z | 2.936E+04 | 1184 | 0.003769 | 3.185 | +1 | 12 |
| - | - | 4.271E+04 | 1185 | - | - | 0 | - |
| 13 | c | 2.701E+04 | 1186 | 0.01213 | 10.23 | +1 | 13 |
| - | - | 1.002E+04 | 1187 | - | - | 0 | - |
| - | - | 3230 | 1188 | - | - | 0 | - |
| - | - | 8166 | 1199 | - | - | 0 | - |
| 8 | y | 3.659E+04 | 1200 | 0.002746 | 2.289 | +1 | 12 |
| - | - | 2.22E+04 | 1201 | - | - | 0 | - |
| - | - | 1.244E+04 | 1202 | - | - | 0 | - |
| 13 | c | 7.352E+04 | 1203 | 0.002583 | 2.148 | +1 | 13 |
| - | - | 4.512E+04 | 1204 | - | - | 0 | - |
| - | - | 1.543E+04 | 1205 | - | - | 0 | - |
| - | - | 3180 | 1206 | - | - | 0 | - |
| - | - | 1642 | 1227 | - | - | 0 | - |
| - | - | 1881 | 1228 | - | - | 0 | - |
| - | - | 838.3 | 1229 | - | - | 0 | - |
| - | - | 1043 | 1239 | - | - | 0 | - |
| - | - | 1204 | 1243 | - | - | 0 | - |
| - | - | 1.681E+04 | 1257 | - | - | 0 | - |
| - | - | 9863 | 1258 | - | - | 0 | - |
| - | - | 5440 | 1259 | - | - | 0 | - |
| - | - | 1595 | 1260 | - | - | 0 | - |
| - | - | 1004 | 1261 | - | - | 0 | - |
| - | - | 5476 | 1272 | - | - | 0 | - |
| - | - | 3917 | 1273 | - | - | 0 | - |
| - | - | 2663 | 1274 | - | - | 0 | - |
| 7 | z | 1.858E+04 | 1283 | 0.002372 | 1.849 | +1 | 13 |
| - | - | 1.412E+04 | 1284 | - | - | 0 | - |
| - | - | 6255 | 1285 | - | - | 0 | - |
| - | - | 2069 | 1286 | - | - | 0 | - |
| - | - | 4211 | 1298 | - | - | 0 | - |
| 14 | c | 7047 | 1299 | 0.02002 | 15.41 | +1 | 14 |
| - | - | 6273 | 1300 | - | - | 0 | - |
| - | - | 2184 | 1301 | - | - | 0 | - |
| - | - | 1073 | 1313 | - | - | 0 | - |
| - | - | 1285 | 1314 | - | - | 0 | - |
| - | - | 2.45E+04 | 1315 | - | - | 0 | - |
| 14 | c | 2.606E+04 | 1316 | 0.0009145 | 0.695 | +1 | 14 |
| - | - | 1.296E+04 | 1317 | - | - | 0 | - |
| - | - | 4938 | 1318 | - | - | 0 | - |
| - | - | 1287 | 1319 | - | - | 0 | - |
| - | - | 2954 | 1326 | - | - | 0 | - |
| - | - | 1987 | 1327 | - | - | 0 | - |
| - | - | 893.1 | 1328 | - | - | 0 | - |
| - | - | 8633 | 1329 | - | - | 0 | - |
| - | - | 9589 | 1330 | - | - | 0 | - |
| - | - | 4805 | 1331 | - | - | 0 | - |
| - | - | 1353 | 1332 | - | - | 0 | - |
| - | - | 3341 | 1340 | - | - | 0 | - |
| - | - | 2213 | 1341 | - | - | 0 | - |
| - | - | 1175 | 1342 | - | - | 0 | - |
| - | - | 968.6 | 1355 | - | - | 0 | - |
| 15 | c | 1247 | 1356 | 0.02098 | 15.47 | +1 | 15 |
| - | - | 7222 | 1358 | - | - | 0 | - |
| - | - | 4461 | 1359 | - | - | 0 | - |
| - | - | 2766 | 1360 | - | - | 0 | - |
| - | - | 1129 | 1368 | - | - | 0 | - |
| - | - | 2698 | 1371 | - | - | 0 | - |
| - | - | 1.786E+04 | 1372 | - | - | 0 | - |
| 15 | c | 1.157E+05 | 1373 | 0.002036 | 1.483 | +1 | 15 |
| - | - | 8.545E+04 | 1374 | - | - | 0 | - |
| - | - | 3.691E+04 | 1375 | - | - | 0 | - |
| - | - | 9271 | 1376 | - | - | 0 | - |
| - | - | 1320 | 1377 | - | - | 0 | - |
| 6 | y | 1759 | 1383 | 0.0007381 | 0.5338 | +1 | 14 |
| 6 | z | 4.535E+04 | 1384 | 0.002667 | 1.928 | +1 | 14 |
| - | - | 4.076E+04 | 1385 | - | - | 0 | - |
| - | - | 2.208E+04 | 1386 | - | - | 0 | - |
| - | - | 8468 | 1387 | - | - | 0 | - |
| - | - | 1662 | 1388 | - | - | 0 | - |
| - | - | 1102 | 1399 | - | - | 0 | - |
| 6 | y | 3846 | 1400 | 0.001286 | 0.9189 | +1 | 14 |
| - | - | 2639 | 1401 | - | - | 0 | - |
| - | - | 1278 | 1402 | - | - | 0 | - |
| - | - | 3281 | 1444 | - | - | 0 | - |
| - | - | 1844 | 1445 | - | - | 0 | - |
| - | - | 1176 | 1446 | - | - | 0 | - |
| 16 | c | 4682 | 1471 | 0.008002 | 5.44 | +1 | 16 |
| - | - | 1.177E+04 | 1472 | - | - | 0 | - |
| - | - | 8226 | 1473 | - | - | 0 | - |
| - | - | 3445 | 1474 | - | - | 0 | - |
| - | - | 2241 | 1486 | - | - | 0 | - |
| 5 | y | 5314 | 1487 | 0.0006209 | 0.4176 | +1 | 15 |
| 16 | c | 6.957E+04 | 1488 | 0.001094 | 0.7352 | +1 | 16 |
| - | - | 5.661E+04 | 1489 | - | - | 0 | - |
| - | - | 2.313E+04 | 1490 | - | - | 0 | - |
| - | - | 5605 | 1491 | - | - | 0 | - |
| - | - | 2791 | 1514 | - | - | 0 | - |
| - | - | 1883 | 1515 | - | - | 0 | - |
| - | - | 1034 | 1516 | - | - | 0 | - |
| 4 | z | 2130 | 1572 | 0.0194 | 12.34 | +1 | 16 |
| - | - | 4035 | 1573 | - | - | 0 | - |
| - | - | 2840 | 1574 | - | - | 0 | - |
| - | - | 1083 | 1575 | - | - | 0 | - |
| - | - | 2434 | 1614 | - | - | 0 | - |
| - | - | 4356 | 1615 | - | - | 0 | - |
| 17 | c | 5.824E+04 | 1616 | 0.001354 | 0.8381 | +1 | 17 |
| - | - | 5.46E+04 | 1617 | - | - | 0 | - |
| - | - | 644.3 | 1617 | - | - | 0 | - |
| - | - | 2.531E+04 | 1618 | - | - | 0 | - |
| - | - | 7614 | 1619 | - | - | 0 | - |
| - | - | 1286 | 1620 | - | - | 0 | - |
| 3 | z | 1259 | 1629 | 0.01234 | 7.574 | +1 | 17 |
| - | - | 7640 | 1630 | - | - | 0 | - |
| - | - | 6384 | 1631 | - | - | 0 | - |
| - | - | 3078 | 1632 | - | - | 0 | - |
| - | - | 1071 | 1701 | - | - | 0 | - |
| - | - | 1063 | 1714 | - | - | 0 | - |
| 2 | z | 2693 | 1730 | 0.005307 | 3.068 | +1 | 18 |
| - | - | 8016 | 1731 | - | - | 0 | - |
| - | - | 4484 | 1732 | - | - | 0 | - |
| - | - | 2092 | 1733 | - | - | 0 | - |
| 18 | c | 4958 | 1744 | 0.0008354 | 0.479 | +1 | 18 |
| - | - | 7442 | 1745 | - | - | 0 | - |
| - | - | 5399 | 1746 | - | - | 0 | - |
| - | - | 2010 | 1747 | - | - | 0 | - |
| - | - | 1067 | 1748 | - | - | 0 | - |
| - | - | 2449 | 1759 | - | - | 0 | - |
| - | - | 1701 | 1760 | - | - | 0 | - |
| - | - | 7825 | 1801 | - | - | 0 | - |
| - | - | 9161 | 1802 | - | - | 0 | - |
| - | - | 5051 | 1803 | - | - | 0 | - |
| - | - | 2103 | 1804 | - | - | 0 | - |
| - | - | 3198 | 1816 | - | - | 0 | - |
| - | - | 2607 | 1817 | - | - | 0 | - |
| - | - | 1933 | 1818 | - | - | 0 | - |
| - | - | 1188 | 1832 | - | - | 0 | - |
| - | - | 5250 | 1833 | - | - | 0 | - |
| - | - | 4835 | 1834 | - | - | 0 | - |
| - | - | 2712 | 1835 | - | - | 0 | - |
| - | - | 1375 | 1836 | - | - | 0 | - |
| - | - | 3103 | 1843 | - | - | 0 | - |
| - | - | 1.349E+04 | 1844 | - | - | 0 | - |
| - | - | 1.309E+04 | 1845 | - | - | 0 | - |
| - | - | 6726 | 1846 | - | - | 0 | - |
| - | - | 2653 | 1847 | - | - | 0 | - |
| - | - | 934.3 | 1848 | - | - | 0 | - |
| - | - | 2226 | 1859 | - | - | 0 | - |
| - | - | 9779 | 1860 | - | - | 0 | - |
| - | - | 2.939E+04 | 1861 | - | - | 0 | - |
| - | - | 2.769E+04 | 1862 | - | - | 0 | - |
| - | - | 1.59E+04 | 1863 | - | - | 0 | - |
| - | - | 5950 | 1864 | - | - | 0 | - |
| - | - | 1346 | 1865 | - | - | 0 | - |

m/z Charge Intensity FragmentType MassShift Position
120.06623840332031 0 1059.1293
120.08157348632812 0 927.86523
125.10359954833984 0 429.74777
125.10785675048828 0 3163.0476
129.10275268554688 0 25595.307
130.10626220703125 0 1399.2051
131.0819549560547 0 2450.7751 y 17
132.1026611328125 0 690.6961
133.05613708496094 0 1160.7559
133.06129455566406 0 15173.013 y 18
138.9042205810547 0 372.32028
142.12315368652344 0 32374.727
143.1210174560547 0 352.7633
143.12657165527344 0 2872.4944
148.94728088378906 0 731.0148
149.3524932861328 0 465.56342
150.70407104492188 0 431.6821
152.10751342773438 0 6389.2266
153.11090087890625 0 982.9703
157.13409423828125 0 5435.14
159.0769805908203 0 7396.799
159.14991760253906 0 632.55005
160.0802459716797 0 576.0116
165.10293579101562 0 1234.034
167.00999450683594 0 463.16595
167.77346801757812 0 485.59198
169.1341552734375 0 10399.814
170.1377716064453 0 732.236
171.1134796142578 0 1135.4336
173.44046020507812 0 2452.309
183.11349487304688 0 5495.03
185.12908935546875 0 622.7525
187.07217407226562 0 1102.9967 w 17
187.14474487304688 0 235802.75
188.14816284179688 0 21559.125
189.0876922607422 0 573.69415
189.14952087402344 0 780.80096
195.11338806152344 0 2595.152 y 16
197.1291046142578 0 13408.486
198.11331176757812 0 465.17526
198.13255310058594 0 1284.7498
199.1333465576172 0 569.9029
201.1239471435547 0 2403.65
201.147705078125 0 3971.8357
213.12399291992188 0 1319.0751
214.13185119628906 0 1311.8608
215.1396942138672 0 346694.22
216.14305114746094 0 37028.074
217.14523315429688 0 3455.6104
228.0986785888672 0 1484.8219
228.11135864257812 0 2229.5923 z Ammonia loss 17
232.13009643554688 0 1721.4526
235.10940551757812 0 570.18066
239.15127563476562 0 2478.5696
240.13504028320312 0 7512.746
241.1416015625 0 1247.2725
242.11415100097656 0 4925.3403
244.0935516357422 0 1049.8743
244.12950134277344 0 1142.3486 y Ammonia loss 17
245.13784790039062 0 75387.6 z 17
246.14260864257812 0 10225.928
247.1485137939453 0 1106.6589
254.15069580078125 0 33854.03
255.15403747558594 0 5142.07
257.1614990234375 0 3733.2915
260.1248779296875 0 7146.2524
261.1289978027344 0 1020.83044
261.156494140625 0 19258.18 y 17
262.1598205566406 0 1916.6066
263.1028137207031 0 2054.2605
263.6232604980469 0 531.2458
264.34637451171875 0 510.74152
270.1456604003906 0 802.56024
271.1660461425781 0 1087.4563
272.1612854003906 0 6663.6807
274.1885070800781 0 855.07324
284.1612548828125 0 1250.895
285.1934509277344 0 1302.8904
286.1401062011719 0 833.18427
289.1873474121094 0 1318.5187 c 2
298.14190673828125 0 808.1075
298.1773376464844 0 636.89685
299.1775207519531 0 617.4554
301.1153869628906 0 1791.5238
302.1220703125 0 2532.0938
311.136474609375 0 2248.7166
312.1803894042969 0 1426.4214
315.1674499511719 0 27004.332 w 16
316.17022705078125 0 3310.161
316.18817138671875 0 1457.7227
317.19000244140625 0 615.7243
319.5775451660156 0 644.14545
327.1688232421875 0 571.4087
327.2039794921875 0 4290.0117
328.1673278808594 0 731.11255
328.1878662109375 0 8868.57
328.21075439453125 0 2755.7393
329.1466369628906 0 6883.0503
329.1856689453125 0 2235.6492
329.20733642578125 0 4186.9434
330.1498107910156 0 1602.0288
337.1886901855469 0 869.58875
337.6784362792969 0 2606.096 y 13
338.1781005859375 0 805.3505
345.21435546875 0 8858.561
346.2176818847656 0 1755.57
347.1572570800781 0 4354.464
355.161865234375 0 997.5262
355.198486328125 0 10974.44
356.2008972167969 0 2067.9155
359.1948547363281 0 767.41077
364.1865539550781 0 848.85724
371.2044372558594 0 7857.4585 w 8
372.18902587890625 0 13370.827 y Ammonia loss 16
373.1977844238281 0 65300.035 z 16
374.2011413574219 0 15430.853
375.20489501953125 0 2365.8618
386.2048645019531 0 9150.176
387.20843505859375 0 2235.488 y 12
389.2156677246094 0 61231.33 y 16
390.2182922363281 0 8469.479
390.2408752441406 0 2026.0222 c 3
391.2395324707031 0 717.636
394.1737060546875 0 1028.3379
396.2234802246094 0 1099.5022
398.20703125 0 663.35516
400.2195129394531 0 1134.8824
400.2467041015625 0 951.96204
406.2087707519531 0 783.15186
410.2045593261719 0 2315.5476
411.2027893066406 0 986.45325
412.18426513671875 0 3089.3594
414.2024230957031 0 1473.4608
414.2359313964844 0 2282.204
415.2434387207031 0 3012.9507
416.2129211425781 0 805.43353
416.2450866699219 0 723.2546
418.2308654785156 0 1783.1235
424.2203674316406 0 5579.954
425.2250671386719 0 1440.295
428.21563720703125 0 6161.202
429.2126770019531 0 4664.142
430.1947021484375 0 4167.4507
432.248046875 0 1752.8553
433.2513122558594 0 1199.0742 c Water loss 13
434.26226806640625 0 2829.825
435.2669372558594 0 1067.6316
436.2249755859375 0 1536.424
436.7271423339844 0 667.46564
437.8472900390625 0 571.94867
442.2311096191406 0 11690.87
443.2331848144531 0 2920.473 w 15
444.23443603515625 0 80028.01
445.2374572753906 0 18463.25
446.2306823730469 0 2850.3289
447.22955322265625 0 825.87726
448.20440673828125 0 1881.9105
456.2593688964844 0 1155.0443 w 5
459.2573547363281 0 999.5954 c Water loss 4
460.24224853515625 0 7305.949
461.24493408203125 0 1752.8513
463.2508544921875 0 1491.7684
464.21551513671875 0 1005.0011
474.1709899902344 0 801.7072
475.2533874511719 0 971.8636
476.2603454589844 0 3840.3376
477.2685546875 0 100356.45 c 4
478.2713928222656 0 23904.834
479.27362060546875 0 3898.2861
480.2442626953125 0 951.6648
484.263916015625 0 807.56134
485.2372741699219 0 1228.4653
486.23590087890625 0 846.6455 y Water loss 15
488.2243957519531 0 52011.277 z 15
489.23052978515625 0 35965.945
489.7854309082031 0 1737.8656
490.2331848144531 0 8185.35
490.2709655761719 0 794.9598 y Water loss 4
490.28839111328125 0 674.8159
491.2351989746094 0 1095.3983
493.7755432128906 0 2486.0186 y 10
494.27545166015625 0 1609.3866
496.2722473144531 0 727.14636 y 4
496.6092224121094 0 693.14014 c 15
497.27923583984375 0 668.89014
501.2561340332031 0 1762.7155
501.72698974609375 0 700.3901
503.23681640625 0 4457.9624
504.2427673339844 0 23119.754 y 15
505.2458801269531 0 5636.058
506.2445983886719 0 812.90936
507.2535400390625 0 803.97473
510.73272705078125 0 889.1341
511.2524719238281 0 3058.9915
512.2574462890625 0 879.5155
513.7766723632812 0 833.42957 y Ammonia loss 9
514.27685546875 0 841.1771 z 9
514.7844848632812 0 1086.2285
515.2881469726562 0 4074.6968 c Ammonia loss 10
515.790771484375 0 1744.6069
516.291748046875 0 2752.5415
517.9484252929688 0 1929.0593
518.2839965820312 0 1943.2014
522.28515625 0 24178.635 y 9
522.78662109375 0 10838.363
523.288818359375 0 5257.089
523.9523315429688 0 2316.4443 y Water loss 3
524.2886962890625 0 1636.1747
525.2684326171875 0 6529.4683
525.7921142578125 0 1264.6399
526.2689819335938 0 1262.5635
526.3111572265625 0 1700.5891
527.274658203125 0 844.5976
527.79150390625 0 799.79486
528.2872924804688 0 956.33136
528.61669921875 0 924.1163
529.2633056640625 0 6386.658
529.958984375 0 1066.8324 y 3
531.2794799804688 0 975.90643
533.2944946289062 0 1658.828 c Water loss 16
534.302734375 0 3687.0396
534.802490234375 0 1947.7805
535.30908203125 0 6293.574
536.3124389648438 0 1490.3538
536.7884521484375 0 1253.531
536.954833984375 0 4935.405
537.2919311523438 0 3527.265
537.6217041015625 0 2824.906 z Water loss 2
537.9600830078125 0 946.2242
542.2960205078125 0 4050.8923
542.9581909179688 0 11527.713 y Water loss 2
543.2868041992188 0 13231.898 y Ammonia loss 2
543.6267700195312 0 4809.452 z 2
543.8003540039062 0 2115.5347 c Ammonia loss 11
543.9620361328125 0 1430.2943
544.2379760742188 0 1165.9044 y Ammonia loss 14
544.2866821289062 0 1709.4764
545.2455444335938 0 6356.9517 z 14
545.2847900390625 0 738.7954
546.25390625 0 47248.63
546.77392578125 0 1170.9376
547.2611083984375 0 11618.282
548.2653198242188 0 2258.8806
548.9617309570312 0 1717.081 y 2
549.2929077148438 0 1031.0869
549.6328735351562 0 1248.676
550.3031005859375 0 687.5387
551.2606811523438 0 1932.3925
551.7588500976562 0 1485.7793
552.9755249023438 0 637.6739
559.301513671875 0 1263.43
560.2606811523438 0 1908.0272
561.2653198242188 0 26978.688 y 14
561.9671630859375 0 659.61127
562.2680053710938 0 8275.687
563.267822265625 0 1055.0597
564.6382446289062 0 1800.0475
564.9674072265625 0 2381.179
565.2991333007812 0 1918.9465
570.32421875 0 2959.1318
570.6392822265625 0 4460.455
570.9725952148438 0 6587.206
571.302978515625 0 5666.284 z Water loss 1
571.6325073242188 0 3008.3596 z Ammonia loss 1
571.8197631835938 0 9501.705 y 8
571.9710693359375 0 1657.8872
572.3203125 0 6953.067
572.8229370117188 0 3299.3784
575.9710083007812 0 668.138
576.3159790039062 0 3247.7214 c Ammonia loss 17
576.6420288085938 0 31221.414 y Water loss 1
576.9752807617188 0 34360.42 y Ammonia loss 1
577.3099365234375 0 17334.732 z 1
577.6439819335938 0 5659.494
577.9771118164062 0 1999.324
578.3162231445312 0 31630.469 c 5
579.324462890625 0 10098.623
579.8384399414062 0 4098.2104
580.3167114257812 0 1883.6708
581.9728393554688 0 1515.4257
582.3095703125 0 1742.1072
582.64501953125 0 27327.695 y 1
582.9791870117188 0 23540.373
583.3130493164062 0 14277.575 z Water loss 7
583.6466674804688 0 5074.455
583.8400268554688 0 933.2946
584.3313598632812 0 3441.9111
584.83154296875 0 2861.896
585.3080444335938 0 921.93207
586.288330078125 0 1997.2566
587.9816284179688 0 1949.918
588.263427734375 0 1052.811
588.3158569335938 0 4044.1812
588.6549072265625 0 951.74963
591.3370361328125 0 896.54407
591.817626953125 0 3469.18 y Ammonia loss 7
592.3209228515625 0 7703.853 z 7
592.8233642578125 0 5122.7305
593.3329467773438 0 10563.315 c Ammonia loss 12
593.6568603515625 0 1709.3236
593.8357543945312 0 6168.4814
594.3379516601562 0 1875.8193
594.6570434570312 0 1069.4231
598.3257446289062 0 1669.3066
599.3258056640625 0 1415.6658
599.6610717773438 0 5696.042
599.826904296875 0 2262.992
599.9928588867188 0 6224.794
600.3302612304688 0 90372.88 y 7
600.8319091796875 0 52679.68
601.3324584960938 0 21841.092
601.8341674804688 0 5648.883
602.2688598632812 0 1130.9396
602.3256225585938 0 4671.2
602.6626586914062 0 4576.8477
602.997802734375 0 2676.2139
603.3219604492188 0 1101.5428
605.3292846679688 0 1626.8894
605.6622314453125 0 2980.0952
605.9971923828125 0 3419.4927
606.330078125 0 3323.304
608.332763671875 0 20058.508
608.665771484375 0 26401.531
608.9993286132812 0 19970.451
609.3319091796875 0 7152.0044
609.6665649414062 0 2548.6956
609.9929809570312 0 1054.543
610.7901611328125 0 1666.8729
611.2948608398438 0 1681.282
613.663818359375 0 2973.2026
613.8241577148438 0 3254.4485
613.99658203125 0 3913.7693
614.3360595703125 0 183610
614.6702270507812 0 191943.72
615.0044555664062 0 111343.24
615.2753295898438 0 39337.2 w 13
615.3385009765625 0 40385.63
615.671142578125 0 13469.928
616.2786254882812 0 10773.686
617.2803955078125 0 3011.2551
618.7943725585938 0 2212.061
619.2968139648438 0 1648.4663
619.3510131835938 0 1303.3927
619.6674194335938 0 10063.549
619.7959594726562 0 1000.4957
619.8588256835938 0 1329.1486
620.0027465820312 0 7720.0547
620.3394775390625 0 274761.2
620.6739501953125 0 265071.7
621.0079345703125 0 160488.17
621.3419189453125 0 57671.645
621.6759643554688 0 16153.526
624.337646484375 0 5448.926
625.3389892578125 0 1567.33
628.835205078125 0 1281.7976
632.3638305664062 0 7194.644
633.3699951171875 0 9376.739
634.3780517578125 0 16447.076
634.861328125 0 1877.307
635.380126953125 0 5269.5015
636.3817138671875 0 1127.9707
639.3947143554688 0 1121.39
640.3382568359375 0 706.0467
640.8723754882812 0 2436.8916
641.350830078125 0 2949.5679 y Ammonia loss 6
641.85693359375 0 5582.5674 z 6
642.3494873046875 0 16554.115
643.35400390625 0 4544.988
644.3489990234375 0 989.9102
649.8654174804688 0 50032.582 c Ammonia loss 13
650.3670654296875 0 33323.367
650.8683471679688 0 14718.555
651.3694458007812 0 5459.0854
655.353759765625 0 2357.2292
656.3536376953125 0 1508.6011
657.3220825195312 0 1226.0453 y Ammonia loss 13
658.3304443359375 0 46832.17 z 13
659.336669921875 0 24009.148
660.3541259765625 0 16583.957
661.3590698242188 0 5419.957
663.3579711914062 0 3992.7874
663.8606567382812 0 2764.7864
664.36279296875 0 1835.9198
669.367431640625 0 2077.777
669.8771362304688 0 1306.8224
670.3632202148438 0 2118.0657
670.8561401367188 0 1848.467
671.3651123046875 0 864.32367
672.3797607421875 0 1673.6272
674.3489990234375 0 31221.473 y 13
675.3523559570312 0 11059.556
676.374755859375 0 13209.326
677.3847045898438 0 42372.19 c 6
677.8806762695312 0 2050.3855 c Water loss 14
678.3867797851562 0 15502.072 c Ammonia loss 14
678.881591796875 0 827.6804
679.3861694335938 0 3054.664
679.8601684570312 0 1298.7264
680.3909301757812 0 1123.86
683.8709106445312 0 1275.8911 z Ammonia loss 5
684.3685302734375 0 2092.9883
685.3552856445312 0 2540.739
686.3694458007812 0 1447.0748
687.3825073242188 0 1386.0618
688.3818359375 0 1253.5504
690.3933715820312 0 5045.828
690.8693237304688 0 804.7369
691.3815307617188 0 14573.753 y Water loss 5
691.8748779296875 0 48642.21 y Ammonia loss 5
692.3778686523438 0 49538.434 z 5
692.8800048828125 0 37287.13
693.3814697265625 0 15616.187
693.88427734375 0 4662.4443
698.386962890625 0 1201.9181
699.37158203125 0 6597.4463
699.8843383789062 0 5419.823
700.3880615234375 0 77336.74 y 5
700.8899536132812 0 51359.094
701.3338012695312 0 11813.872
701.3916015625 0 21693.969
701.89208984375 0 6103.6143
702.3394165039062 0 5015.1377
702.3975219726562 0 1041.1895
703.3565063476562 0 1835.0464
704.3710327148438 0 5156.785
704.8751220703125 0 2767.634
705.3778686523438 0 3442.955
705.8824462890625 0 870.26807
706.8845825195312 0 2724.2502
707.3839721679688 0 2515.142
707.8646850585938 0 3359.794
708.365234375 0 1952.4824
708.8775634765625 0 937.3933
713.4044189453125 0 3200.0151
713.8923950195312 0 3984.2708
714.3466796875 0 3427.3286
714.3992919921875 0 2407.8674
714.8892822265625 0 2122.6196
715.4086303710938 0 838.0043
717.3809814453125 0 4453.3047
717.8872680664062 0 1887.7117
718.3893432617188 0 2284.972
719.3880615234375 0 1011.7948
720.388671875 0 971.4063
721.4039916992188 0 1782.545
721.9005126953125 0 1119.4503
725.8934326171875 0 3002.5002
726.3955078125 0 2312.686
726.8952026367188 0 9584.751 z Water loss 4
727.398681640625 0 5702.4834 w 4
727.8975830078125 0 3630.714
728.3946533203125 0 3416.3562
728.8981323242188 0 1703.796
729.3893432617188 0 1024.5022
732.3932495117188 0 1034.6003
733.3983764648438 0 3743.0635
734.406494140625 0 57479.91 c 7
734.8926391601562 0 7023.217 y Water loss 4
735.4036865234375 0 31324.312 c Water loss 15
735.8958740234375 0 133559.12 c Ammonia loss 15
736.3980712890625 0 148115.75
736.8995361328125 0 75880.125
737.4016723632812 0 30575.438
737.9028930664062 0 7821.254
738.4021606445312 0 1259.7396
739.3927612304688 0 1169.9342 z Water loss 12
740.384033203125 0 1149.154 z Ammonia loss 12
741.3816528320312 0 1384.9265
742.3765869140625 0 3950.3667 w 12
742.8970947265625 0 1777.766
743.3963012695312 0 11206.059
743.90478515625 0 165779.4 y 4
744.4063110351562 0 130953.74 c 15
744.9076538085938 0 58563.633
745.4091796875 0 16183.193
745.9111938476562 0 4561.874
747.9051513671875 0 2700.6553
748.4033203125 0 3942.6296
748.9016723632812 0 3023.6287
749.4080810546875 0 901.87933
751.4027099609375 0 1302.0044
754.4208374023438 0 996.57367
756.3941650390625 0 906.3612 y Ammonia loss 12
756.91259765625 0 10870.595
757.4008178710938 0 52376.305 z 12
757.9089965820312 0 6968.7617
758.4033813476562 0 24004.066
758.8982543945312 0 1628.8163
759.4065551757812 0 5263.774
760.4086303710938 0 1281.7759
763.4109497070312 0 1187.2479
764.4197998046875 0 3037.2505
764.91162109375 0 2684.4668
765.4015502929688 0 2039.6329
765.91650390625 0 4039.3643
766.4195556640625 0 2981.6445
766.91748046875 0 1518.5342
769.418212890625 0 1153.7146
770.4307250976562 0 6769.8647
771.4352416992188 0 3375.5264
772.4253540039062 0 3012.2673
773.4180297851562 0 18300.24 y 12
774.4219360351562 0 6373.511
775.4181518554688 0 1326.168
776.9119873046875 0 1095.1729
777.4135131835938 0 1615.1898 z Water loss 3
777.9186401367188 0 2027.3424 z Ammonia loss 3
778.4135131835938 0 4301.988
778.91162109375 0 4568.879 w 3
779.4111938476562 0 3543.4246
779.9148559570312 0 1678.0709
780.4225463867188 0 1936.8602
781.426513671875 0 1194.5675
782.4191284179688 0 1331.7637
783.4097900390625 0 2228.128
785.4166259765625 0 7751.12 y Water loss 3
785.9163208007812 0 24342.88 y Ammonia loss 3
786.4192504882812 0 56260.8 z 3
786.920654296875 0 58175.12
787.4229125976562 0 33154.594
787.9244384765625 0 11554.807
788.4331665039062 0 3734.1062
789.4609985351562 0 11816.86
790.4671020507812 0 11628.676
790.9259033203125 0 5007.018
791.4710693359375 0 3068.5183
791.9276123046875 0 3395.396
792.420654296875 0 1338.0624
792.927490234375 0 4363.299
793.4269409179688 0 5883.803
793.9243774414062 0 5481.445
794.4285888671875 0 66215.73 y 3
794.9297485351562 0 54949.605
795.4315185546875 0 23243.838
795.9327392578125 0 8149.5723
796.42431640625 0 1729.625
796.9268188476562 0 832.9485
797.428955078125 0 1098.7795
798.4298706054688 0 5199.8496
798.9263305664062 0 2688.6182
799.4225463867188 0 3084.636 c Water loss 16
799.9306030273438 0 5671.258 c Ammonia loss 16
800.4325561523438 0 5883.0483
800.9306640625 0 4987.0854
801.4249267578125 0 4887.8574
801.9220581054688 0 2444.4243
804.9282836914062 0 5907.4473
805.427734375 0 9886.547
805.92578125 0 8185.162 z Water loss 2
806.4279174804688 0 4000.7827 z Ammonia loss 2
806.9326171875 0 1587.7234
807.4345703125 0 7957.982
807.9344482421875 0 5929.975
808.4391479492188 0 5372.1357 c 16
808.9407958984375 0 2946.96
811.4456176757812 0 7114.218
812.4457397460938 0 3400.0984
812.9301147460938 0 1565.901
813.43603515625 0 2579.9211
813.9342651367188 0 45916.594 y Water loss 2
814.4306640625 0 77388.625 y Ammonia loss 2
814.9312744140625 0 71774.88 z 2
815.4309692382812 0 97767.38
815.9335327148438 0 23835.691
816.4351196289062 0 33508.188
816.9390869140625 0 3368.3694
817.4407958984375 0 7732.646
818.4385986328125 0 1585.7874
821.4684448242188 0 1136.437
821.93310546875 0 15785.432
822.4345703125 0 15304.54
822.9398803710938 0 826757.1 y 2
823.4412841796875 0 720669.75
823.9423217773438 0 363381.25
824.4435424804688 0 115055.35
824.9446411132812 0 27032.588
829.9292602539062 0 1035.3915
830.439697265625 0 21617.758 y 11
831.4434814453125 0 8436.885
832.464599609375 0 14323.159
833.4740600585938 0 27521.81 c 8
834.476806640625 0 10402.805
835.4814453125 0 2991.5166
835.9503784179688 0 1859.9043
836.4326171875 0 1453.6285
836.9366455078125 0 1843.8179
837.4326782226562 0 3191.4675
837.9281616210938 0 2357.623
839.483154296875 0 1701.5222
843.4522094726562 0 6567.972
843.9546508789062 0 7034.397
844.451904296875 0 4951.636
844.9445190429688 0 2317.0627
846.482666015625 0 5142.977
847.4876708984375 0 4712.3667
848.4968872070312 0 2083.7197
849.50390625 0 1029.2593
850.48388671875 0 8074.4834
850.9810791015625 0 6444.0522
851.4732055664062 0 5271.1826
851.955078125 0 3109.1628
852.4487915039062 0 1549.4983
854.973388671875 0 1995.9064
855.4550170898438 0 7263.252
855.954345703125 0 7153.73
856.4481201171875 0 7127.555 z Water loss 1
856.95263671875 0 8834.956 z Ammonia loss 1
857.45068359375 0 6377.8423
857.9476318359375 0 6741.8906 w 1
858.446044921875 0 6599.9985
858.9483032226562 0 3252.465
863.458984375 0 1543.4602
863.9728393554688 0 5809.514 c Ammonia loss 17
864.4581298828125 0 56237.938 y Water loss 1
864.955810546875 0 90361.48 y Ammonia loss 1
865.455078125 0 100763.62 z 1
865.9563598632812 0 64036.016
866.4575805664062 0 29063.496
866.95703125 0 10192.543
867.4578857421875 0 2278.8552
869.4500732421875 0 5825.046
869.9490966796875 0 4804.701
870.448486328125 0 4368.011
870.9678344726562 0 1708.8884
871.4810180664062 0 7065.9385
871.9839477539062 0 9827.765
872.4918212890625 0 335749.62 c Water loss 9
872.9931640625 0 317913.03
873.4700927734375 0 457183.1 y 1
873.96728515625 0 396817.4
874.467529296875 0 211015.48
874.9674682617188 0 67745.13
875.4706420898438 0 18753.75
876.4854125976562 0 1286.7894
878.4562377929688 0 19583.32
878.95849609375 0 20290.754
879.4597778320312 0 12174.796
879.9635620117188 0 9168.907
880.4637451171875 0 4875.412
880.9566650390625 0 1025.6948
885.4488525390625 0 1263.4938
886.0022583007812 0 2360.1157
886.4820556640625 0 3953.8835
886.9815673828125 0 3882.7156
887.480224609375 0 2495.2786
888.4743041992188 0 1613.6888
889.4854736328125 0 4875.9165
890.4970092773438 0 87320.71 c 9
891.4999389648438 0 43305.42
891.986328125 0 1332.6807
892.4993896484375 0 11823.733
892.9812622070312 0 1606.1095
893.4871826171875 0 3951.1802
893.9744262695312 0 7014.1426
894.4735717773438 0 6470.6533
894.971923828125 0 2539.9272
895.4816284179688 0 2294.5227
899.4913940429688 0 3992.591
899.99609375 0 14884.588
900.4985961914062 0 75247.68
900.9961547851562 0 103873.375
901.4955444335938 0 77842.25
901.9962768554688 0 38902.96
902.494140625 0 12437.424
902.9920043945312 0 2478.0713
906.4771728515625 0 1274.6099
907.4788818359375 0 62897.242
907.9857788085938 0 79147.31
908.4901123046875 0 58201.203
908.9929809570312 0 43712.32
909.4957885742188 0 27296.258
909.9981079101562 0 13727.914
910.5078125 0 5380.1147
911.509765625 0 6619.067
912.4983520507812 0 3779.8416
912.9911499023438 0 6630.8677
913.492919921875 0 9764.129
913.9945068359375 0 5676.626
914.484130859375 0 6913.309
914.9876098632812 0 5148.2866
915.485595703125 0 3913.019
916.4871826171875 0 2682.4617
917.4775390625 0 1099.1527
920.4934692382812 0 1527.2357
920.9910278320312 0 5585.9185
921.5004272460938 0 24977.781
921.9976196289062 0 169533.9
922.4984741210938 0 163305.44
922.9989013671875 0 91303.41
923.4081420898438 0 1410.5731
923.5003051757812 0 33051.49
924.001953125 0 9656.52
926.5347290039062 0 9380.28
927.5363159179688 0 4688.611
928.485107421875 0 1516.2355
928.5422973632812 0 1171.6133
928.9950561523438 0 6782.642
929.500732421875 0 13791.537
930.0055541992188 0 186886.6
930.50732421875 0 201413.72
931.0084838867188 0 126699.92
931.5098266601562 0 48495.883
932.0108032226562 0 12878.846
932.5042114257812 0 1708.5559
939.5048828125 0 3202.796
940.508056640625 0 1657.7971
970.52197265625 0 61761.56 z 10
971.5260009765625 0 43513.848
972.5277709960938 0 16004.704
973.5258178710938 0 4164.8555
983.5574340820312 0 2095.1733
985.5391845703125 0 4412.904
986.5440063476562 0 11052.425 y 10
987.5455322265625 0 4977.4165
988.5559692382812 0 1434.1086
998.5651245117188 0 1106.6249
1001.5655517578125 0 1053.9983
1002.583984375 0 11904.267
1003.5891723632812 0 6025.6255
1004.5873413085938 0 1871.2998
1010.5808715820312 0 1078.6123
1011.562744140625 0 2184.3188
1012.5626831054688 0 2191.9172
1027.5450439453125 0 15162.143 z 9
1028.551513671875 0 33854.15
1029.555908203125 0 17258.453 c Ammonia loss 10
1030.5606689453125 0 5642.5933
1031.570068359375 0 1012.18774
1038.571533203125 0 1329.925
1041.5950927734375 0 1133.9523
1042.5567626953125 0 5071.8735
1043.5623779296875 0 27731.682 y 9
1044.5655517578125 0 17034.584
1045.5877685546875 0 33206.125
1046.5958251953125 0 41465.035 c 10
1047.5999755859375 0 18473.05
1048.6046142578125 0 5555.639
1050.5941162109375 0 1101.3588
1055.5711669921875 0 2064.193
1059.604248046875 0 8751.197
1060.6114501953125 0 6232.223
1061.6107177734375 0 2457.2185
1067.5999755859375 0 5776.3027
1068.6021728515625 0 3130.9314
1069.5985107421875 0 1566.4823
1070.5511474609375 0 2135.9297
1071.546142578125 0 1444.5012
1072.5712890625 0 2822.7634
1073.5748291015625 0 1362.4299
1082.617431640625 0 3315.8665
1083.6124267578125 0 1170.5015
1084.6142578125 0 8511.917
1085.615966796875 0 4966.7437 c Water loss 11
1086.6092529296875 0 2520.178 c Ammonia loss 11
1087.599609375 0 1000.36584
1097.62109375 0 1476.7275
1098.6011962890625 0 3963.2031
1099.60107421875 0 2169.2388
1100.5919189453125 0 1082.5887
1101.603271484375 0 1732.5765
1102.6116943359375 0 4540.285
1103.620361328125 0 71388.16 c 11
1104.6231689453125 0 41889.176
1105.6253662109375 0 14561.7705
1106.6263427734375 0 2757.8923
1126.612548828125 0 37009.918 z 8
1127.615234375 0 27206.67
1128.619140625 0 11365.33
1129.620361328125 0 2484.9307
1139.6361083984375 0 2402.6445
1140.6263427734375 0 1949.378
1141.625244140625 0 9018.937
1142.6314697265625 0 17899.018 y 8
1143.63427734375 0 8753.465
1144.6392822265625 0 3642.206
1157.671630859375 0 1436.5713
1158.67529296875 0 7074.229
1159.67626953125 0 4996.203
1160.6732177734375 0 1872.2748
1182.625244140625 0 2022.3843 y Ammonia loss 7
1183.6341552734375 0 29355.736 z 7
1184.6397705078125 0 42706.22
1185.646484375 0 27011.775 c Ammonia loss 12
1186.64990234375 0 10016.526
1187.6580810546875 0 3230.4883
1198.6466064453125 0 8166.4854
1199.65185546875 0 36592.33 y 7
1200.655517578125 0 22204.764
1201.6651611328125 0 12439.345
1202.687744140625 0 73517.66 c 12
1203.6912841796875 0 45118.613
1204.693603515625 0 15430.844
1205.7000732421875 0 3180.0752
1226.648193359375 0 1642.1265
1227.654541015625 0 1880.8665
1228.6881103515625 0 838.29315
1238.719482421875 0 1043.3838
1242.690185546875 0 1204.33
1256.6627197265625 0 16806.916
1257.6658935546875 0 9862.973
1258.6759033203125 0 5439.955
1259.6856689453125 0 1594.7401
1260.6910400390625 0 1004.4786
1271.757568359375 0 5476.11
1272.7545166015625 0 3917.0989
1273.75927734375 0 2663.4536
1282.701171875 0 18581.965 z 6
1283.7037353515625 0 14119.3125
1284.704833984375 0 6254.8013
1285.712158203125 0 2068.7217
1297.7149658203125 0 4210.852
1298.72265625 0 7047.357 c Ammonia loss 13
1299.727294921875 0 6273.2397
1300.7298583984375 0 2183.845
1312.73681640625 0 1072.7959
1313.7415771484375 0 1285.4661
1314.76318359375 0 24501.523
1315.768310546875 0 26064.004 c 13
1316.7720947265625 0 12956.327
1317.7750244140625 0 4937.582
1318.7716064453125 0 1287.3286
1325.7186279296875 0 2954.1846
1326.71240234375 0 1986.8774
1327.7105712890625 0 893.06006
1328.779296875 0 8632.739
1329.78564453125 0 9588.7
1330.7879638671875 0 4804.7236
1331.7928466796875 0 1353.1969
1339.7579345703125 0 3341.0862
1340.7490234375 0 2212.7307
1341.760986328125 0 1174.8573
1354.7509765625 0 968.6425
1355.7431640625 0 1246.5894 c Ammonia loss 14
1357.7125244140625 0 7221.5234
1358.714599609375 0 4460.8267
1359.717041015625 0 2766.2622
1367.713134765625 0 1128.7344
1370.7718505859375 0 2698.304
1371.78369140625 0 17861.463
1372.792724609375 0 115744.97 c 14
1373.79638671875 0 85447.24
1374.798828125 0 36911.645
1375.800048828125 0 9270.91
1376.7978515625 0 1319.5342
1382.7379150390625 0 1759.4248 y Ammonia loss 5
1383.7491455078125 0 45347.613 z 5
1384.74951171875 0 40757.004
1385.7523193359375 0 22083.688
1386.7552490234375 0 8468.483
1387.7547607421875 0 1661.9084
1398.759521484375 0 1101.7655
1399.763916015625 0 3845.803 y 5
1400.7628173828125 0 2639.0208
1401.7708740234375 0 1277.8639
1443.807861328125 0 3281.489
1444.8062744140625 0 1843.9847
1445.8206787109375 0 1175.86
1470.7830810546875 0 4681.745 c Ammonia loss 15
1471.7882080078125 0 11769.675
1472.7918701171875 0 8226.274
1473.7930908203125 0 3445.0356
1485.7940673828125 0 2241.434
1486.7978515625 0 5314.473 y 4
1487.8187255859375 0 69572.26 c 15
1488.822021484375 0 56613.402
1489.8238525390625 0 23131.896
1490.82666015625 0 5604.8623
1513.8172607421875 0 2790.5664
1514.8240966796875 0 1882.9546
1515.8291015625 0 1033.7117
1571.8455810546875 0 2129.7864 z 3
1572.8450927734375 0 4034.8984
1573.840087890625 0 2839.7231
1574.8616943359375 0 1083.3386
1613.859130859375 0 2433.9155
1614.864013671875 0 4355.582
1615.8775634765625 0 58238.703 c 16
1616.88037109375 0 54602.29
1617.0665283203125 0 644.3
1617.883056640625 0 25309.97
1618.885009765625 0 7614.3506
1619.8857421875 0 1285.6694
1628.8599853515625 0 1259.1879 z 2
1629.8555908203125 0 7640.3
1630.8529052734375 0 6383.76
1631.8597412109375 0 3077.7068
1700.935302734375 0 1070.7179
1713.90625 0 1062.7466
1729.900634765625 0 2693.232 z 1
1730.90576171875 0 8016.0547
1731.9112548828125 0 4483.5317
1732.913818359375 0 2092.0554
1743.9703369140625 0 4957.7046 c 17
1744.9781494140625 0 7442.336
1745.9776611328125 0 5398.604
1746.979248046875 0 2010.0037
1747.9794921875 0 1067.1849
1758.927490234375 0 2449.0845
1759.9205322265625 0 1700.641
1800.96728515625 0 7824.813
1801.972412109375 0 9161.251
1802.975830078125 0 5050.8457
1803.974365234375 0 2102.6829
1815.9993896484375 0 3198.4312
1816.994384765625 0 2607.4453
1817.993896484375 0 1932.9922
1832.0302734375 0 1187.7258
1833.021240234375 0 5249.7686
1834.024658203125 0 4835.054
1835.0296630859375 0 2711.8562
1836.0455322265625 0 1375.0135
1842.991943359375 0 3102.63
1843.990234375 0 13494.537
1844.993896484375 0 13089.482
1845.9991455078125 0 6726.0576
1846.994873046875 0 2652.5237
1847.9937744140625 0 934.2606
1859.0008544921875 0 2225.8582
1860.00634765625 0 9778.583
1861.0157470703125 0 29386.777
1862.0184326171875 0 27685.36
1863.0201416015625 0 15902.551
1864.02294921875 0 5950.4053
1865.0048828125 0 1346.246

Spectrum Details

|  |  |
| --- | --- |
| Matched peaks? Matched peaks The total absolute number of peaks matched. Additionally in brackets the total fraction of peaks matched and the total number of peaks is shown. | 148 (16.02% of 924) |
| FDR? FDR The false discovery rate estimated for this peptide. It is calculated by matching all theoretical fragments with a non-integer shift with the raw peaks for this spectrum. This is done with 40 different shifts. The resulting percentage is the average number of annotated peaks over the number of annotated peaks with the correct spectrum. | 0.55% |
| Satellite FDR? Satellite FDR See the FDR for details on its calculation. This satellite ion specific FDR only contains the satellite ions (d/w) for I/L/J positions. | 9.52% |
| PSM Score? PSM Score The PSM Score as given by Hecklib to this annotated spectrum. It is shown with three significant figures. | 904 |

#### Spectrum 4992? Spectrum 4992 The raw spectrum of this peptide as annotated by Hecklib. The fragments are coloured according to ion type (see legend). Any peaks with a star '\*' as text can be hovered over to see the full details, first the ion type second the mass shift type. By hovering over the amino acids in the peptide or ions in the legend the corresponding peaks are highlighted. By toggling the 'Unassigned' label you can turn the background (unassigned) peaks on or off in the plot. By updating the slider in the Ion legend you can update the spectrum to only show the top X% of the peaks with labels. The top X% means any peak that is within X% of the highest intensity. By dragging in the spectrum you can zoom in to a specific part of the spectrum and use 'Zoom Out' to get back to the original zoom level. The annotation of the spectrum is based on the given sequence in the peptides file and is done with different software so inconsistencies are likely. The peaks are annotated based on the given sequence, with 20 ppm tolerance.

Copy Data

##### Spectrum 4992 (TSV)

###### Preview

```
Loading example...
```

*Click on the button to copy the data to your clipboard.*

Mz MinMz MaxIntensity Max

WidthHeightPeptide font sizePeptide stroke widthSpectrum font sizeSpectrum stroke widthCompact peptide

Ion legend

wxyz

abcd

OtherUnassignedIon^Charge^_Position_Show for top:%

JTGTSTVGVGRGVLGDQKN

04.81e+39.62e+31.44e+41.92e+4

Zoom Out

y^+2^_2_z^+1^_2_y^+1^_2_w^+1^_3_z^+1^_3_c^+3^_14_c^+1^_5_z^+1^_4_y^+1^_4_y^+1^_5_c^+1^_6_y^+3^_18_y^+2^_12_w^+1^_6_c^+2^_14_z^+1^_6_y^+1^_6_c^+1^_7_y^+2^_14_z^+2^_14_y^+2^_14_c^+1^_8_c^+2^_16_c^+2^_16_y^+2^_15_c^+2^_16_z^+1^_7_y^+1^_7_y^+2^_16_z^+2^_16_y^+2^_16_y^+2^_17_y^+2^_17_z^+2^_17_y^+2^_17_y^+1^_8_c^+1^_9_y^+2^_18_y^+2^_18_z^+2^_18_c^+1^_10_c^+1^_10_z^+1^_9_z^+1^_10_c^+1^_11_y^+1^_10_c^+1^_11_c^+1^_12_z^+1^_11_y^+1^_11_z^+1^_12_c^+1^_13_y^+1^_12_c^+1^_13_z^+1^_13_c^+1^_14_c^+1^_15_z^+1^_14_y^+1^_14_c^+1^_16_y^+1^_15_c^+1^_16_z^+1^_16_c^+1^_17_z^+1^_18_c^+1^_18_

0776155223283104

Fragment Matches Table

Show background peaks

| Position | Ion type | Intensity | mz Theoretical | mz Error (Th) | mz Error (ppm) | Charge | Series Number |
| --- | --- | --- | --- | --- | --- | --- | --- |
| - | - | 983 | 120.1 | - | - | 0 | - |
| - | - | 554.2 | 120.1 | - | - | 0 | - |
| - | - | 410.5 | 127.9 | - | - | 0 | - |
| - | - | 623.7 | 129.1 | - | - | 0 | - |
| 18 | y | 353.7 | 131.1 | 0.0002373 | 1.81 | +2 | 2 |
| - | - | 427.5 | 144 | - | - | 0 | - |
| - | - | 433.3 | 165 | - | - | 0 | - |
| - | - | 4292 | 173.4 | - | - | 0 | - |
| - | - | 2629 | 187.1 | - | - | 0 | - |
| - | - | 3512 | 215.1 | - | - | 0 | - |
| - | - | 772.1 | 235.1 | - | - | 0 | - |
| 18 | z | 4578 | 245.1 | 0.0001667 | 0.6801 | +1 | 2 |
| - | - | 619.4 | 246.1 | - | - | 0 | - |
| - | - | 585.2 | 247.6 | - | - | 0 | - |
| 18 | y | 769.4 | 261.2 | 0.001922 | 7.36 | +1 | 2 |
| - | - | 1389 | 263.1 | - | - | 0 | - |
| - | - | 551.2 | 264.1 | - | - | 0 | - |
| 17 | w | 1750 | 315.2 | 0.0002381 | 0.7556 | +1 | 3 |
| - | - | 483 | 315.4 | - | - | 0 | - |
| - | - | 1237 | 316.2 | - | - | 0 | - |
| 17 | z | 2627 | 373.2 | 0.0004904 | 1.314 | +1 | 3 |
| - | - | 560.2 | 399 | - | - | 0 | - |
| - | - | 674.2 | 415 | - | - | 0 | - |
| - | - | 1501 | 436.2 | - | - | 0 | - |
| 14 | c | 1035 | 439.3 | 0.0007307 | 1.664 | +3 | 14 |
| - | - | 3359 | 444.2 | - | - | 0 | - |
| - | - | 700.1 | 445.2 | - | - | 0 | - |
| - | - | 845.8 | 459.8 | - | - | 0 | - |
| 5 | c | 4655 | 477.3 | 1.553E-05 | 0.03255 | +1 | 5 |
| - | - | 1320 | 478.3 | - | - | 0 | - |
| 16 | z | 2734 | 488.2 | 5.493E-05 | 0.1125 | +1 | 4 |
| - | - | 1750 | 489.2 | - | - | 0 | - |
| - | - | 5699 | 489.8 | - | - | 0 | - |
| - | - | 2532 | 490.3 | - | - | 0 | - |
| - | - | 1210 | 490.8 | - | - | 0 | - |
| - | - | 780.5 | 491.3 | - | - | 0 | - |
| 16 | y | 1374 | 504.2 | 0.00121 | 2.4 | +1 | 4 |
| - | - | 1135 | 510.7 | - | - | 0 | - |
| - | - | 795.6 | 537.3 | - | - | 0 | - |
| - | - | 866.2 | 543.8 | - | - | 0 | - |
| - | - | 2998 | 546.3 | - | - | 0 | - |
| - | - | 880.7 | 546.8 | - | - | 0 | - |
| - | - | 1250 | 551.3 | - | - | 0 | - |
| - | - | 1116 | 551.8 | - | - | 0 | - |
| - | - | 724.2 | 560.8 | - | - | 0 | - |
| 15 | y | 1488 | 561.3 | 0.00297 | 5.292 | +1 | 5 |
| - | - | 2227 | 575.3 | - | - | 0 | - |
| - | - | 1741 | 577.3 | - | - | 0 | - |
| 6 | c | 1018 | 578.3 | 0.0007685 | 1.329 | +1 | 6 |
| - | - | 827.6 | 579.3 | - | - | 0 | - |
| 2 | y | 599.6 | 582.6 | 0.002518 | 4.321 | +3 | 18 |
| 8 | y | 1501 | 600.3 | 0.0009837 | 1.639 | +2 | 12 |
| - | - | 730.5 | 601.8 | - | - | 0 | - |
| - | - | 1976 | 610.8 | - | - | 0 | - |
| - | - | 901.9 | 611.3 | - | - | 0 | - |
| - | - | 2962 | 614.3 | - | - | 0 | - |
| - | - | 2351 | 614.7 | - | - | 0 | - |
| - | - | 1071 | 615 | - | - | 0 | - |
| 14 | w | 2209 | 615.3 | 0.001317 | 2.14 | +1 | 6 |
| - | - | 2135 | 615.3 | - | - | 0 | - |
| - | - | 746.7 | 616.3 | - | - | 0 | - |
| - | - | 1526 | 618.8 | - | - | 0 | - |
| - | - | 2973 | 619.3 | - | - | 0 | - |
| - | - | 966.2 | 619.3 | - | - | 0 | - |
| - | - | 3494 | 619.9 | - | - | 0 | - |
| - | - | 830.4 | 620.3 | - | - | 0 | - |
| - | - | 2387 | 620.4 | - | - | 0 | - |
| - | - | 1751 | 620.7 | - | - | 0 | - |
| - | - | 989.2 | 620.8 | - | - | 0 | - |
| - | - | 1305 | 620.9 | - | - | 0 | - |
| - | - | 909 | 621.3 | - | - | 0 | - |
| - | - | 702 | 642.4 | - | - | 0 | - |
| 14 | c | 1319 | 649.9 | 0.01151 | 17.71 | +2 | 14 |
| - | - | 607.3 | 650.4 | - | - | 0 | - |
| 14 | z | 1850 | 658.3 | 0.0001139 | 0.173 | +1 | 6 |
| - | - | 1029 | 659.3 | - | - | 0 | - |
| - | - | 827.2 | 672.4 | - | - | 0 | - |
| 14 | y | 718.2 | 674.3 | 0.003884 | 5.76 | +1 | 6 |
| - | - | 812.4 | 676.4 | - | - | 0 | - |
| 7 | c | 1624 | 677.4 | 0.000873 | 1.289 | +1 | 7 |
| - | - | 1029 | 687.4 | - | - | 0 | - |
| 6 | y | 2873 | 691.9 | 8.206E-05 | 0.1186 | +2 | 14 |
| 6 | z | 2229 | 692.4 | 4.628E-05 | 0.06684 | +2 | 14 |
| - | - | 1979 | 692.9 | - | - | 0 | - |
| - | - | 879.7 | 693.4 | - | - | 0 | - |
| - | - | 963.9 | 696.4 | - | - | 0 | - |
| - | - | 1912 | 697.4 | - | - | 0 | - |
| - | - | 661.6 | 699.4 | - | - | 0 | - |
| 6 | y | 1407 | 700.4 | 0.0001742 | 0.2487 | +2 | 14 |
| - | - | 988.5 | 700.9 | - | - | 0 | - |
| - | - | 692.6 | 713.9 | - | - | 0 | - |
| 8 | c | 3383 | 734.4 | 0.001768 | 2.407 | +1 | 8 |
| 16 | c | 1544 | 735.4 | 0.006781 | 9.221 | +2 | 16 |
| 16 | c | 5526 | 735.9 | 0.006907 | 9.385 | +2 | 16 |
| - | - | 6366 | 736.4 | - | - | 0 | - |
| - | - | 2488 | 736.9 | - | - | 0 | - |
| - | - | 1441 | 737.4 | - | - | 0 | - |
| - | - | 665.1 | 738.3 | - | - | 0 | - |
| - | - | 2753 | 741.4 | - | - | 0 | - |
| - | - | 1485 | 743.4 | - | - | 0 | - |
| 5 | y | 2701 | 743.9 | 3.189E-05 | 0.04287 | +2 | 15 |
| 16 | c | 2532 | 744.4 | 0.01035 | 13.91 | +2 | 16 |
| - | - | 1087 | 744.9 | - | - | 0 | - |
| - | - | 609.9 | 746.4 | - | - | 0 | - |
| 13 | z | 1877 | 757.4 | 0.001541 | 2.034 | +1 | 7 |
| - | - | 920.5 | 758.4 | - | - | 0 | - |
| - | - | 718.1 | 770.4 | - | - | 0 | - |
| 13 | y | 960.4 | 773.4 | 0.002535 | 3.278 | +1 | 7 |
| - | - | 937.3 | 781.3 | - | - | 0 | - |
| 4 | y | 998.3 | 785.9 | 0.003211 | 4.086 | +2 | 16 |
| 4 | z | 2326 | 786.4 | 0.001814 | 2.306 | +2 | 16 |
| - | - | 3341 | 786.9 | - | - | 0 | - |
| - | - | 1593 | 787.4 | - | - | 0 | - |
| - | - | 881.2 | 787.9 | - | - | 0 | - |
| - | - | 990.9 | 789.5 | - | - | 0 | - |
| 4 | y | 611.5 | 794.4 | 0.006049 | 7.614 | +2 | 16 |
| - | - | 667.3 | 807.4 | - | - | 0 | - |
| 3 | y | 753.8 | 813.9 | 0.004554 | 5.595 | +2 | 17 |
| 3 | y | 1475 | 814.4 | 0.005893 | 7.236 | +2 | 17 |
| 3 | z | 2095 | 814.9 | 0.002835 | 3.479 | +2 | 17 |
| - | - | 3037 | 815.4 | - | - | 0 | - |
| - | - | 2395 | 816.4 | - | - | 0 | - |
| - | - | 764.2 | 820.4 | - | - | 0 | - |
| 3 | y | 8332 | 822.9 | 5.709E-05 | 0.06937 | +2 | 17 |
| - | - | 8685 | 823.4 | - | - | 0 | - |
| - | - | 4301 | 823.9 | - | - | 0 | - |
| - | - | 683.6 | 824.4 | - | - | 0 | - |
| - | - | 706.3 | 824.5 | - | - | 0 | - |
| - | - | 807.8 | 825.5 | - | - | 0 | - |
| 12 | y | 1169 | 830.4 | 0.00404 | 4.865 | +1 | 8 |
| - | - | 1208 | 832.5 | - | - | 0 | - |
| 9 | c | 1660 | 833.5 | 0.001151 | 1.381 | +1 | 9 |
| - | - | 683.4 | 834.5 | - | - | 0 | - |
| - | - | 999.3 | 838.4 | - | - | 0 | - |
| - | - | 1199 | 839.4 | - | - | 0 | - |
| 2 | y | 854.8 | 864.5 | 0.01257 | 14.54 | +2 | 18 |
| 2 | y | 1762 | 864.9 | 0.000612 | 0.7076 | +2 | 18 |
| 2 | z | 3702 | 865.5 | 0.001229 | 1.42 | +2 | 18 |
| - | - | 2125 | 866 | - | - | 0 | - |
| - | - | 1506 | 866.5 | - | - | 0 | - |
| - | - | 865.8 | 871.4 | - | - | 0 | - |
| 10 | c | 1.905E+04 | 872.5 | 0.00504 | 5.776 | +1 | 10 |
| - | - | 1.214E+04 | 873 | - | - | 0 | - |
| - | - | 8902 | 873.5 | - | - | 0 | - |
| - | - | 4379 | 874 | - | - | 0 | - |
| - | - | 1776 | 874.5 | - | - | 0 | - |
| - | - | 696.9 | 879.5 | - | - | 0 | - |
| 10 | c | 3980 | 890.5 | 0.0006421 | 0.7211 | +1 | 10 |
| - | - | 2176 | 891.5 | - | - | 0 | - |
| - | - | 908 | 892.5 | - | - | 0 | - |
| - | - | 3343 | 900.5 | - | - | 0 | - |
| - | - | 5855 | 901 | - | - | 0 | - |
| - | - | 2503 | 901.5 | - | - | 0 | - |
| - | - | 1702 | 902 | - | - | 0 | - |
| - | - | 1382 | 905.4 | - | - | 0 | - |
| - | - | 3304 | 907.5 | - | - | 0 | - |
| - | - | 4115 | 908 | - | - | 0 | - |
| - | - | 1501 | 908.5 | - | - | 0 | - |
| - | - | 1557 | 909 | - | - | 0 | - |
| - | - | 1180 | 909.5 | - | - | 0 | - |
| - | - | 820 | 913 | - | - | 0 | - |
| - | - | 809 | 921.5 | - | - | 0 | - |
| - | - | 7749 | 922 | - | - | 0 | - |
| - | - | 8820 | 922.5 | - | - | 0 | - |
| - | - | 4372 | 923 | - | - | 0 | - |
| - | - | 1748 | 923.4 | - | - | 0 | - |
| - | - | 1672 | 923.5 | - | - | 0 | - |
| - | - | 847 | 924 | - | - | 0 | - |
| - | - | 815.9 | 929.5 | - | - | 0 | - |
| - | - | 9821 | 930 | - | - | 0 | - |
| - | - | 9456 | 930.5 | - | - | 0 | - |
| - | - | 822.8 | 930.9 | - | - | 0 | - |
| - | - | 6418 | 931 | - | - | 0 | - |
| - | - | 923.5 | 931.4 | - | - | 0 | - |
| - | - | 1736 | 931.5 | - | - | 0 | - |
| - | - | 627.5 | 931.9 | - | - | 0 | - |
| - | - | 779.9 | 939.6 | - | - | 0 | - |
| - | - | 1064 | 940.6 | - | - | 0 | - |
| - | - | 749.7 | 959.5 | - | - | 0 | - |
| - | - | 1971 | 960.5 | - | - | 0 | - |
| - | - | 845.4 | 961.5 | - | - | 0 | - |
| 11 | z | 3262 | 970.5 | 0.001466 | 1.511 | +1 | 9 |
| - | - | 1952 | 971.5 | - | - | 0 | - |
| - | - | 1166 | 972.5 | - | - | 0 | - |
| - | - | 1021 | 976.5 | - | - | 0 | - |
| - | - | 947.5 | 977.5 | - | - | 0 | - |
| - | - | 694.6 | 979.6 | - | - | 0 | - |
| - | - | 809.4 | 1003 | - | - | 0 | - |
| 10 | z | 1509 | 1028 | 0.001934 | 1.882 | +1 | 10 |
| - | - | 1491 | 1029 | - | - | 0 | - |
| 11 | c | 1676 | 1030 | 0.02015 | 19.57 | +1 | 11 |
| - | - | 917.7 | 1042 | - | - | 0 | - |
| 10 | y | 918.2 | 1044 | 0.002169 | 2.079 | +1 | 10 |
| - | - | 2100 | 1046 | - | - | 0 | - |
| 11 | c | 2701 | 1047 | 0.001045 | 0.9986 | +1 | 11 |
| - | - | 716.2 | 1048 | - | - | 0 | - |
| - | - | 705.7 | 1093 | - | - | 0 | - |
| 12 | c | 4166 | 1104 | 7.416E-05 | 0.0672 | +1 | 12 |
| - | - | 2229 | 1105 | - | - | 0 | - |
| - | - | 908 | 1106 | - | - | 0 | - |
| 9 | z | 3153 | 1127 | 0.0009412 | 0.8354 | +1 | 11 |
| - | - | 1614 | 1128 | - | - | 0 | - |
| - | - | 3076 | 1137 | - | - | 0 | - |
| - | - | 1624 | 1138 | - | - | 0 | - |
| - | - | 1080 | 1139 | - | - | 0 | - |
| - | - | 1090 | 1142 | - | - | 0 | - |
| 9 | y | 841.8 | 1143 | 0.000693 | 0.6065 | +1 | 11 |
| - | - | 736 | 1144 | - | - | 0 | - |
| - | - | 1110 | 1155 | - | - | 0 | - |
| - | - | 900.7 | 1156 | - | - | 0 | - |
| - | - | 784.1 | 1178 | - | - | 0 | - |
| - | - | 809 | 1180 | - | - | 0 | - |
| - | - | 1076 | 1180 | - | - | 0 | - |
| - | - | 859.2 | 1181 | - | - | 0 | - |
| 8 | z | 1542 | 1184 | 0.003066 | 2.591 | +1 | 12 |
| - | - | 2575 | 1185 | - | - | 0 | - |
| 13 | c | 1323 | 1186 | 0.01567 | 13.21 | +1 | 13 |
| - | - | 841.2 | 1187 | - | - | 0 | - |
| 8 | y | 2452 | 1200 | 0.0007925 | 0.6606 | +1 | 12 |
| - | - | 1806 | 1201 | - | - | 0 | - |
| - | - | 924.1 | 1202 | - | - | 0 | - |
| 13 | c | 4420 | 1203 | 0.001118 | 0.9298 | +1 | 13 |
| - | - | 2380 | 1204 | - | - | 0 | - |
| - | - | 2229 | 1221 | - | - | 0 | - |
| - | - | 1882 | 1222 | - | - | 0 | - |
| - | - | 6382 | 1223 | - | - | 0 | - |
| - | - | 4766 | 1224 | - | - | 0 | - |
| - | - | 1949 | 1225 | - | - | 0 | - |
| - | - | 903.1 | 1227 | - | - | 0 | - |
| - | - | 690.3 | 1232 | - | - | 0 | - |
| - | - | 670.8 | 1237 | - | - | 0 | - |
| - | - | 3358 | 1238 | - | - | 0 | - |
| - | - | 5910 | 1239 | - | - | 0 | - |
| - | - | 983.7 | 1239 | - | - | 0 | - |
| - | - | 6834 | 1240 | - | - | 0 | - |
| - | - | 3255 | 1240 | - | - | 0 | - |
| - | - | 4551 | 1241 | - | - | 0 | - |
| - | - | 1976 | 1241 | - | - | 0 | - |
| - | - | 1709 | 1242 | - | - | 0 | - |
| - | - | 1207 | 1242 | - | - | 0 | - |
| - | - | 1096 | 1243 | - | - | 0 | - |
| 7 | z | 1440 | 1283 | 0.0005575 | 0.4347 | +1 | 13 |
| - | - | 863.7 | 1284 | - | - | 0 | - |
| - | - | 1930 | 1315 | - | - | 0 | - |
| 14 | c | 1980 | 1316 | 0.002135 | 1.623 | +1 | 14 |
| - | - | 1322 | 1317 | - | - | 0 | - |
| - | - | 1083 | 1329 | - | - | 0 | - |
| - | - | 782.2 | 1330 | - | - | 0 | - |
| - | - | 2820 | 1372 | - | - | 0 | - |
| 15 | c | 8915 | 1373 | 0.002725 | 1.985 | +1 | 15 |
| - | - | 6441 | 1374 | - | - | 0 | - |
| - | - | 2576 | 1375 | - | - | 0 | - |
| 6 | z | 2852 | 1384 | 0.003802 | 2.748 | +1 | 14 |
| - | - | 4944 | 1385 | - | - | 0 | - |
| - | - | 3435 | 1386 | - | - | 0 | - |
| - | - | 1082 | 1387 | - | - | 0 | - |
| 6 | y | 758.7 | 1400 | 0.01239 | 8.855 | +1 | 14 |
| 16 | c | 1409 | 1471 | 0.02082 | 14.15 | +1 | 16 |
| - | - | 3544 | 1472 | - | - | 0 | - |
| - | - | 3119 | 1473 | - | - | 0 | - |
| - | - | 1664 | 1474 | - | - | 0 | - |
| 5 | y | 1727 | 1487 | 0.004405 | 2.963 | +1 | 15 |
| 16 | c | 5930 | 1488 | 0.001836 | 1.234 | +1 | 16 |
| - | - | 5000 | 1489 | - | - | 0 | - |
| - | - | 1907 | 1490 | - | - | 0 | - |
| - | - | 891.6 | 1491 | - | - | 0 | - |
| - | - | 1130 | 1514 | - | - | 0 | - |
| - | - | 1007 | 1515 | - | - | 0 | - |
| - | - | 708.6 | 1552 | - | - | 0 | - |
| 4 | z | 719.6 | 1572 | 0.01549 | 9.855 | +1 | 16 |
| - | - | 2475 | 1573 | - | - | 0 | - |
| - | - | 1712 | 1574 | - | - | 0 | - |
| - | - | 889.4 | 1575 | - | - | 0 | - |
| - | - | 707.6 | 1615 | - | - | 0 | - |
| 17 | c | 7628 | 1616 | 0.001942 | 1.202 | +1 | 17 |
| - | - | 6963 | 1617 | - | - | 0 | - |
| - | - | 2654 | 1618 | - | - | 0 | - |
| - | - | 1019 | 1619 | - | - | 0 | - |
| - | - | 4321 | 1630 | - | - | 0 | - |
| - | - | 2655 | 1631 | - | - | 0 | - |
| - | - | 1338 | 1632 | - | - | 0 | - |
| - | - | 777.8 | 1686 | - | - | 0 | - |
| 2 | z | 1261 | 1730 | 0.01263 | 7.302 | +1 | 18 |
| - | - | 3633 | 1731 | - | - | 0 | - |
| - | - | 3349 | 1732 | - | - | 0 | - |
| - | - | 1990 | 1733 | - | - | 0 | - |
| 18 | c | 2773 | 1744 | 0.003033 | 1.739 | +1 | 18 |
| - | - | 5019 | 1745 | - | - | 0 | - |
| - | - | 2414 | 1746 | - | - | 0 | - |
| - | - | 1384 | 1747 | - | - | 0 | - |
| - | - | 2714 | 1801 | - | - | 0 | - |
| - | - | 2609 | 1802 | - | - | 0 | - |
| - | - | 1340 | 1803 | - | - | 0 | - |
| - | - | 1040 | 1804 | - | - | 0 | - |
| - | - | 1653 | 1816 | - | - | 0 | - |
| - | - | 1639 | 1817 | - | - | 0 | - |
| - | - | 1242 | 1818 | - | - | 0 | - |
| - | - | 1357 | 1832 | - | - | 0 | - |
| - | - | 3631 | 1833 | - | - | 0 | - |
| - | - | 2530 | 1834 | - | - | 0 | - |
| - | - | 1711 | 1835 | - | - | 0 | - |
| - | - | 1080 | 1836 | - | - | 0 | - |
| - | - | 2358 | 1843 | - | - | 0 | - |
| - | - | 8758 | 1844 | - | - | 0 | - |
| - | - | 7926 | 1845 | - | - | 0 | - |
| - | - | 4367 | 1846 | - | - | 0 | - |
| - | - | 2087 | 1847 | - | - | 0 | - |
| - | - | 2834 | 1859 | - | - | 0 | - |
| - | - | 5542 | 1860 | - | - | 0 | - |
| - | - | 1.904E+04 | 1861 | - | - | 0 | - |
| - | - | 1.782E+04 | 1862 | - | - | 0 | - |
| - | - | 7917 | 1863 | - | - | 0 | - |
| - | - | 2912 | 1864 | - | - | 0 | - |
| - | - | 711.6 | 2231 | - | - | 0 | - |
| - | - | 970.6 | 3073 | - | - | 0 | - |

m/z Charge Intensity FragmentType MassShift Position
120.06553649902344 0 982.97363
120.080810546875 0 554.1805
127.91344451904297 0 410.5056
129.10227966308594 0 623.69025
131.0817413330078 0 353.6525 y 17
144.01231384277344 0 427.5227
165.04957580566406 0 433.31097
173.44036865234375 0 4292.062
187.1439971923828 0 2628.9207
215.13897705078125 0 3512.4634
235.1077117919922 0 772.1136
245.1368408203125 0 4577.603 z 17
246.14305114746094 0 619.382
247.5884246826172 0 585.23065
261.15765380859375 0 769.4271 y 17
263.1029357910156 0 1389.2073
264.1058349609375 0 551.21124
315.1665344238281 0 1750.1167 w 16
315.39666748046875 0 483.03195
316.18731689453125 0 1236.9203
373.1960754394531 0 2627.322 z 16
399.001708984375 0 560.23126
414.9939880371094 0 674.1519
436.22564697265625 0 1501.3995
439.2605285644531 0 1034.7533 c 13
444.2325439453125 0 3358.7642
445.23590087890625 0 700.0558
459.77459716796875 0 845.7858
477.2667541503906 0 4654.6274 c 4
478.2679138183594 0 1320.4419
488.22247314453125 0 2734.3372 z 15
489.2300109863281 0 1749.9111
489.7846984863281 0 5699.473
490.2858581542969 0 2532.1538
490.7916259765625 0 1210.2207
491.294189453125 0 780.536
504.2424621582031 0 1374.0002 y 15
510.72894287109375 0 1134.7578
537.26025390625 0 795.62964
543.8134765625 0 866.1799
546.2562866210938 0 2998.2163
546.7655029296875 0 880.72955
551.2590942382812 0 1249.969
551.7599487304688 0 1115.6288
560.7654418945312 0 724.1994
561.2656860351562 0 1487.8612 y 14
575.302734375 0 2227.4817
577.3158569335938 0 1741.0029
578.315185546875 0 1017.54645 c 5
579.3198852539062 0 827.62415
582.6453857421875 0 599.55743 y 1
600.3272094726562 0 1500.8712 y 7
601.784423828125 0 730.52545
610.7844848632812 0 1975.6276
611.2951049804688 0 901.91724
614.3350830078125 0 2962.0854
614.6668701171875 0 2350.7993
615.0045776367188 0 1070.9033
615.2745971679688 0 2209.0872 w 13
615.34423828125 0 2135.1968
616.2725830078125 0 746.73
618.7913208007812 0 1525.6617
619.2913818359375 0 2973.434
619.3427734375 0 966.18994
619.860595703125 0 3493.9875
620.2930297851562 0 830.4284
620.3554077148438 0 2386.68
620.6714477539062 0 1751.019
620.8007202148438 0 989.1595
620.8614501953125 0 1304.8693
621.3037109375 0 908.9966
642.3502807617188 0 701.988
649.8634643554688 0 1318.5858 c Ammonia loss 13
650.37109375 0 607.317
658.3279418945312 0 1850.1085 z 13
659.3355102539062 0 1028.8212
672.36962890625 0 827.1565
674.3428955078125 0 718.2189 y 13
676.3660278320312 0 812.42883
677.3819580078125 0 1623.7107 c 6
687.37939453125 0 1029.268
691.873046875 0 2872.6836 y Ammonia loss 5
692.3768310546875 0 2229.0256 z 5
692.8761596679688 0 1978.9388
693.3777465820312 0 879.7075
696.4296875 0 963.94885
697.4386596679688 0 1912.353
699.3767700195312 0 661.6104
700.3864135742188 0 1407.1348 y 5
700.8876953125 0 988.5026
713.886474609375 0 692.63837
734.4025268554688 0 3383.4072 c 7
735.400390625 0 1543.6885 c Water loss 15
735.8922729492188 0 5526.124 c Ammonia loss 15
736.3952026367188 0 6366.1323
736.8969116210938 0 2487.6287
737.3970947265625 0 1440.7189
738.348388671875 0 665.07465
741.3876953125 0 2752.5654
743.3946533203125 0 1485.1086
743.9022216796875 0 2700.911 y 4
744.402099609375 0 2531.549 c 15
744.9095458984375 0 1087.0659
746.3536376953125 0 609.8559
757.3980102539062 0 1876.5242 z 12
758.3986206054688 0 920.5445
770.4310302734375 0 718.05524
773.4126586914062 0 960.3766 y 12
781.3470458984375 0 937.3114
785.9096069335938 0 998.26855 y Ammonia loss 3
786.4149169921875 0 2326.0234 z 3
786.9190673828125 0 3341.1003
787.4204711914062 0 1593.1211
787.9238891601562 0 881.222
789.452880859375 0 990.89374
794.4200439453125 0 611.516 y 3
807.4349365234375 0 667.27374
813.9360961914062 0 753.7964 y Water loss 2
814.429443359375 0 1474.5945 y Ammonia loss 2
814.9302978515625 0 2094.8562 z 2
815.4271240234375 0 3037.2712
816.4317626953125 0 2394.773
820.3594360351562 0 764.1501
822.936767578125 0 8332.075 y 2
823.4384765625 0 8685.061
823.9396362304688 0 4301.1367
824.4464111328125 0 683.6431
824.5245971679688 0 706.25574
825.5335693359375 0 807.82135
830.4326171875 0 1169.086 y 11
832.4646606445312 0 1208.2418
833.4715576171875 0 1659.579 c 8
834.4716186523438 0 683.42126
838.3734741210938 0 999.28125
839.3743896484375 0 1199.3094
864.4428100585938 0 854.8176 y Water loss 1
864.94677734375 0 1761.5791 y Ammonia loss 1
865.4500732421875 0 3702.1484 z 1
865.95361328125 0 2125.1917
866.4512329101562 0 1505.729
871.442626953125 0 865.8343
872.4886474609375 0 19052.076 c Water loss 9
872.9898071289062 0 12144.104
873.4793701171875 0 8901.501
873.9708251953125 0 4379.152
874.4657592773438 0 1775.6482
879.4557495117188 0 696.9471
890.4935302734375 0 3980.303 c 9
891.4965209960938 0 2176.1077
892.497314453125 0 908.0204
900.4957275390625 0 3342.8413
900.9918823242188 0 5854.9526
901.4939575195312 0 2502.5361
901.9913330078125 0 1701.7346
905.394287109375 0 1381.8973
907.4734497070312 0 3304.0107
907.9823608398438 0 4115.305
908.4911499023438 0 1501.0438
908.989501953125 0 1556.5533
909.49609375 0 1179.5715
912.9935913085938 0 820.02057
921.4933471679688 0 809.0376
921.9937133789062 0 7749.2446
922.4956665039062 0 8819.898
922.9960327148438 0 4371.7715
923.397705078125 0 1748.0867
923.5015869140625 0 1672.2838
923.9913330078125 0 847.0156
929.4879150390625 0 815.9179
930.0016479492188 0 9820.983
930.5034790039062 0 9455.81
930.9192504882812 0 822.8152
931.0064697265625 0 6417.936
931.4203491210938 0 923.46295
931.5100708007812 0 1735.6619
931.9093627929688 0 627.45514
939.5598754882812 0 779.91223
940.55859375 0 1064.3524
959.4828491210938 0 749.7374
960.4705810546875 0 1971.2128
961.4671630859375 0 845.4034
970.517578125 0 3261.5032 z 10
971.5216674804688 0 1951.758
972.529052734375 0 1165.7167
976.4778442382812 0 1020.978
977.485107421875 0 947.4606
979.5638427734375 0 694.6105
1002.5811157226562 0 809.3806
1027.53857421875 0 1509.4148 z 9
1028.5504150390625 0 1491.3854
1029.548583984375 0 1676.2247 c Ammonia loss 10
1041.6114501953125 0 917.7314
1043.5614013671875 0 918.17017 y 9
1045.5875244140625 0 2100.1577
1046.59423828125 0 2700.5303 c 10
1047.594970703125 0 716.2499
1092.5203857421875 0 705.66064
1103.6168212890625 0 4165.5557 c 11
1104.6185302734375 0 2229.3877
1105.61767578125 0 907.97107
1126.60986328125 0 3152.8438 z 8
1127.614501953125 0 1614.0536
1136.5494384765625 0 3075.8027
1137.5479736328125 0 1624.0509
1138.5609130859375 0 1080.0833
1141.613525390625 0 1089.5786
1142.626953125 0 841.814 y 8
1143.625732421875 0 735.9933
1154.6920166015625 0 1110.0466
1155.6922607421875 0 900.6776
1177.5780029296875 0 784.1477
1179.5653076171875 0 808.98883
1179.7003173828125 0 1075.8383
1180.6927490234375 0 859.2238
1183.6273193359375 0 1541.8615 z 7
1184.634033203125 0 2575.4763
1185.6429443359375 0 1322.7834 c Ammonia loss 12
1186.63818359375 0 841.2226
1199.64990234375 0 2452.1448 y 7
1200.654541015625 0 1805.8004
1201.65087890625 0 924.1498
1202.686279296875 0 4419.758 c 12
1203.6864013671875 0 2380.2522
1220.558837890625 0 2228.8674
1221.5655517578125 0 1881.5543
1222.56103515625 0 6382.3926
1223.563232421875 0 4765.7466
1224.5633544921875 0 1949.0647
1226.540771484375 0 903.0851
1231.519775390625 0 690.3207
1236.5787353515625 0 670.8373
1237.5902099609375 0 3357.5752
1238.5802001953125 0 5910.395
1238.72119140625 0 983.6891
1239.5843505859375 0 6834.154
1239.7225341796875 0 3255.0417
1240.58642578125 0 4550.8325
1240.7247314453125 0 1976.0981
1241.5838623046875 0 1709.2817
1241.7269287109375 0 1207.0189
1242.708984375 0 1095.5966
1282.6982421875 0 1440.0487 z 6
1283.701416015625 0 863.6587
1314.7596435546875 0 1930.4241
1315.76708984375 0 1980.2683 c 13
1316.76220703125 0 1322.2849
1328.7930908203125 0 1082.928
1329.7733154296875 0 782.1749
1371.782470703125 0 2819.9775
1372.7879638671875 0 8914.635 c 14
1373.791259765625 0 6440.5156
1374.7952880859375 0 2576.146
1383.74267578125 0 2852.343 z 5
1384.746826171875 0 4943.7744
1385.7542724609375 0 3435.1956
1386.7581787109375 0 1081.5847
1399.7528076171875 0 758.6698 y 5
1470.770263671875 0 1409.1821 c Ammonia loss 15
1471.7861328125 0 3544.0063
1472.789306640625 0 3119.15
1473.7891845703125 0 1663.5598
1486.8016357421875 0 1726.6057 y 4
1487.8157958984375 0 5929.9688 c 15
1488.818603515625 0 5000.0083
1489.818603515625 0 1906.6617
1490.8311767578125 0 891.5525
1513.8236083984375 0 1130.4473
1514.80810546875 0 1006.9606
1551.81982421875 0 708.60376
1571.8416748046875 0 719.55414 z 3
1572.8343505859375 0 2474.6636
1573.83740234375 0 1712.2834
1574.8309326171875 0 889.40234
1614.8494873046875 0 707.6079
1615.874267578125 0 7628.497 c 16
1616.8756103515625 0 6963.277
1617.87548828125 0 2653.954
1618.8741455078125 0 1018.8289
1629.849853515625 0 4321.425
1630.85107421875 0 2654.6362
1631.85791015625 0 1338.3918
1685.8857421875 0 777.79987
1729.907958984375 0 1260.9078 z 1
1730.906982421875 0 3633.0366
1731.90625 0 3349.3713
1732.902587890625 0 1989.5933
1743.9681396484375 0 2773.1897 c 17
1744.973388671875 0 5019.458
1745.974853515625 0 2414.459
1746.979248046875 0 1384.1812
1800.981201171875 0 2714.2754
1801.9730224609375 0 2609.2976
1802.978759765625 0 1339.7808
1803.96728515625 0 1039.7869
1815.9942626953125 0 1653.4805
1816.9771728515625 0 1639.3181
1817.99853515625 0 1242.3481
1831.999755859375 0 1357.2734
1833.0133056640625 0 3631.2454
1834.0189208984375 0 2529.9336
1835.0216064453125 0 1710.578
1836.025146484375 0 1079.9246
1842.978271484375 0 2357.784
1843.9879150390625 0 8758.2295
1844.98974609375 0 7926.1797
1845.9932861328125 0 4366.7856
1846.9979248046875 0 2086.5078
1858.9954833984375 0 2834.2407
1860.001220703125 0 5542.3984
1861.0107421875 0 19039.041
1862.0135498046875 0 17824.281
1863.015380859375 0 7916.7856
1864.02099609375 0 2912.2273
2231.306884765625 0 711.63947
3072.9169921875 0 970.5996

Spectrum Details

|  |  |
| --- | --- |
| Matched peaks? Matched peaks The total absolute number of peaks matched. Additionally in brackets the total fraction of peaks matched and the total number of peaks is shown. | 66 (20.95% of 315) |
| FDR? FDR The false discovery rate estimated for this peptide. It is calculated by matching all theoretical fragments with a non-integer shift with the raw peaks for this spectrum. This is done with 40 different shifts. The resulting percentage is the average number of annotated peaks over the number of annotated peaks with the correct spectrum. | 1.55% |
| Satellite FDR? Satellite FDR See the FDR for details on its calculation. This satellite ion specific FDR only contains the satellite ions (d/w) for I/L/J positions. | 0.00% |
| PSM Score? PSM Score The PSM Score as given by Hecklib to this annotated spectrum. It is shown with three significant figures. | 494 |

#### Reverse Lookup? Reverse Lookup All places where this read could be placed.

| Group | Segment | Template | Template Part | Read Part | Score | Unique |
| --- | --- | --- | --- | --- | --- | --- |
| Decoy | Decoy | THER | [232..251] | [0..19] | 152 | True |

| Recombined | Template Part | Read Part | Score | Unique |
| --- | --- | --- | --- | --- |
| THER | [232..251] | [0..19] | 152 | True |

#### Meta Information from Multiple reads

##### Number of combined reads

2

##### Intensity

0.6694

##### TotalArea

5.302E+07

##### Changes to the peptide sequence

JTGTSTVGVGRGVLGDQKN

L→JNo support for either Leucine or Isoleucine based on side chain ions (Position: 1)

#### Positional Score

Copy Data

##### Positional Score (TSV)

###### Preview

```
Loading example...
```

*Click on the button to copy the data to your clipboard.*

000123456789101112131415161718

Label Value
"0" 0
"1" 0
"2" 0
"3" 0
"4" 0
"5" 0
"6" 0
"7" 0
"8" 0
"9" 0
"10" 0
"11" 0
"12" 0
"13" 0
"14" 0
"15" 0
"16" 0
"17" 0
"18" 0

#### Meta Information from PEAKS

##### Scan Identifier

F2:4926

##### Original sequence

L

T

G

T

S

T

V

G

V

G

R

G

V

L

G

D

Q

K

N

##### Posttranslational Modifications

##### Source File

D:\separate\_stitch\_analyses\xle-disambiguation\raw\20210323\_F1\_UM1\_Peng0013\_SA\_F59\_ingel\_3ug\_TL.raw

##### Fraction

2

##### Scan Feature

F2:9036

##### De Novo Score

99

##### ConfidenceScore

99

### m/z

620.3391

##### Mass

1857.9907

##### Charge

3

##### Retention Time

26.53

##### Predicted Retention Time

-

##### Area

2.651E+07

##### Parts Per Million

2.6

##### Fragmentation mode

ETHCD

##### Originating file

01 D:\separate\_stitch\_analyses\xle-disambiguation\20210325\_F59\_3ug\_DENOVO\_12.csv

#### Meta Information from PEAKS

##### Scan Identifier

F2:4992

##### Original sequence

L

T

G

T

S

T

V

G

V

G

R

G

V

L

G

D

Q

K

N

##### Posttranslational Modifications

##### Source File

D:\separate\_stitch\_analyses\xle-disambiguation\raw\20210323\_F1\_UM1\_Peng0013\_SA\_F59\_ingel\_3ug\_TL.raw

##### Fraction

2

##### Scan Feature

F2:9036

##### De Novo Score

98

##### ConfidenceScore

98

### m/z

620.3391

##### Mass

1857.9907

##### Charge

3

##### Retention Time

26.53

##### Predicted Retention Time

-

##### Area

2.651E+07

##### Parts Per Million

2.6

##### Fragmentation mode

ETHCD

##### Originating file

01 D:\separate\_stitch\_analyses\xle-disambiguation\20210325\_F59\_3ug\_DENOVO\_12.csv
