## Supplementary material for "A handle on mass coincidence errors in *de novo* sequencing of antibodies by bottom-up proteomics": Combined_052.html

Details Combined\_052 | Stitch OverviewUndefined

### Read Combined\_052

#### Sequence (length=10)

QVSLQDKTGF

#### Spectrum 5529? Spectrum 5529 The raw spectrum of this peptide as annotated by Hecklib. The fragments are coloured according to ion type (see legend). Any peaks with a star '\*' as text can be hovered over to see the full details, first the ion type second the mass shift type. By hovering over the amino acids in the peptide or ions in the legend the corresponding peaks are highlighted. By toggling the 'Unassigned' label you can turn the background (unassigned) peaks on or off in the plot. By updating the slider in the Ion legend you can update the spectrum to only show the top X% of the peaks with labels. The top X% means any peak that is within X% of the highest intensity. By dragging in the spectrum you can zoom in to a specific part of the spectrum and use 'Zoom Out' to get back to the original zoom level. The annotation of the spectrum is based on the given sequence in the peptides file and is done with different software so inconsistencies are likely. The peaks are annotated based on the given sequence, with 20 ppm tolerance.

Copy Data

##### Spectrum 5529 (TSV)

###### Preview

```
Loading example...
```

*Click on the button to copy the data to your clipboard.*

Mz MinMz MaxIntensity Max

WidthHeightPeptide font sizePeptide stroke widthSpectrum font sizeSpectrum stroke widthCompact peptide

Ion legend

wxyz

abcd

OtherUnassignedIon^Charge^_Position_Show for top:%

QVSLQDKTGF

08.27e+41.65e+52.48e+53.31e+5

Zoom Out

y^+1^_1_a^+1^_2_d^+1^_2_a^+1^_2_b^+1^_2_y^+1^_2_b^+1^_2_a^+1^_3_a^+1^_3_y^+2^_5_a^+1^_3_b^+1^_3_b^+1^_3_y^+1^_3_b^+1^_3_y^+1^_3_b^+2^_6_b^+2^_6_y^+2^_6_y^+2^_6_y^+2^_6_b^+2^_7_y^+2^_7_y^+2^_7_b^+1^_4_b^+1^_4_b^+1^_4_y^+1^_4_y^+2^_8_y^+2^_8_b^+2^_8_b^+2^_8_y^+2^_8_b^+2^_8_y^+1^_4_b^+2^_9_b^+2^_9_b^+2^_9_y^+2^_9_b^+1^_5_b^+1^_5_y^+1^_5_y^+1^_5_\*\*b^+1^_5_\*y^+1^_5_b^+1^_6_b^+1^_6_b^+1^_6_y^+1^_6_y^+1^_6_y^+1^_6_b^+1^_7_b^+1^_7_y^+1^_7_y^+1^_7_b^+1^_7_y^+1^_7_y^+1^_8_y^+1^_8_b^+1^_8_b^+1^_8_y^+1^_8_b^+1^_8_b^+1^_9_b^+1^_9_b^+1^_9_y^+1^_9_y^+1^_9_

0827165424823309

Fragment Matches Table

Show background peaks

| Position | Ion type | Intensity | mz Theoretical | mz Error (Th) | mz Error (ppm) | Charge | Series Number |
| --- | --- | --- | --- | --- | --- | --- | --- |
| - | - | 6.004E+04 | 120.1 | - | - | 0 | - |
| - | - | 4710 | 121.1 | - | - | 0 | - |
| - | - | 428.3 | 121.9 | - | - | 0 | - |
| - | - | 439.5 | 122.3 | - | - | 0 | - |
| - | - | 808.6 | 123.1 | - | - | 0 | - |
| - | - | 492.9 | 124.1 | - | - | 0 | - |
| - | - | 346 | 124.5 | - | - | 0 | - |
| - | - | 790.2 | 126.1 | - | - | 0 | - |
| - | - | 764.1 | 127.1 | - | - | 0 | - |
| - | - | 2185 | 127.1 | - | - | 0 | - |
| - | - | 395.7 | 127.6 | - | - | 0 | - |
| - | - | 869.8 | 128.1 | - | - | 0 | - |
| - | - | 3384 | 128.1 | - | - | 0 | - |
| - | - | 1.55E+04 | 129.1 | - | - | 0 | - |
| - | - | 1.166E+05 | 129.1 | - | - | 0 | - |
| - | - | 616.3 | 130 | - | - | 0 | - |
| - | - | 678.2 | 130.1 | - | - | 0 | - |
| - | - | 447 | 130.1 | - | - | 0 | - |
| - | - | 1096 | 130.1 | - | - | 0 | - |
| - | - | 7380 | 130.1 | - | - | 0 | - |
| - | - | 884 | 131 | - | - | 0 | - |
| - | - | 3994 | 131.1 | - | - | 0 | - |
| - | - | 6803 | 131.1 | - | - | 0 | - |
| - | - | 540.7 | 132.1 | - | - | 0 | - |
| - | - | 964.6 | 133.1 | - | - | 0 | - |
| - | - | 1101 | 133.1 | - | - | 0 | - |
| - | - | 5250 | 136.1 | - | - | 0 | - |
| - | - | 573.9 | 137.1 | - | - | 0 | - |
| - | - | 3.565E+04 | 138.1 | - | - | 0 | - |
| - | - | 477.7 | 139.1 | - | - | 0 | - |
| - | - | 1791 | 139.1 | - | - | 0 | - |
| - | - | 3225 | 139.1 | - | - | 0 | - |
| - | - | 1036 | 140.1 | - | - | 0 | - |
| - | - | 2448 | 141.1 | - | - | 0 | - |
| - | - | 4712 | 141.1 | - | - | 0 | - |
| - | - | 456.6 | 144.9 | - | - | 0 | - |
| - | - | 479 | 146.1 | - | - | 0 | - |
| - | - | 455.8 | 147.1 | - | - | 0 | - |
| - | - | 468.5 | 148.6 | - | - | 0 | - |
| - | - | 459 | 148.8 | - | - | 0 | - |
| - | - | 542.1 | 148.8 | - | - | 0 | - |
| - | - | 540.8 | 148.9 | - | - | 0 | - |
| - | - | 530 | 148.9 | - | - | 0 | - |
| - | - | 460 | 148.9 | - | - | 0 | - |
| - | - | 579.3 | 148.9 | - | - | 0 | - |
| - | - | 738.6 | 148.9 | - | - | 0 | - |
| - | - | 1021 | 148.9 | - | - | 0 | - |
| - | - | 1192 | 148.9 | - | - | 0 | - |
| - | - | 1072 | 148.9 | - | - | 0 | - |
| - | - | 1454 | 148.9 | - | - | 0 | - |
| - | - | 3401 | 148.9 | - | - | 0 | - |
| - | - | 5732 | 149 | - | - | 0 | - |
| - | - | 3551 | 149 | - | - | 0 | - |
| - | - | 1081 | 149 | - | - | 0 | - |
| - | - | 1167 | 149 | - | - | 0 | - |
| - | - | 833.6 | 149 | - | - | 0 | - |
| - | - | 998 | 149 | - | - | 0 | - |
| - | - | 916.4 | 149 | - | - | 0 | - |
| - | - | 611.6 | 149 | - | - | 0 | - |
| - | - | 493.1 | 149 | - | - | 0 | - |
| - | - | 451.6 | 149 | - | - | 0 | - |
| - | - | 458.4 | 149 | - | - | 0 | - |
| - | - | 1296 | 149.1 | - | - | 0 | - |
| - | - | 4523 | 151.1 | - | - | 0 | - |
| - | - | 443.9 | 151.1 | - | - | 0 | - |
| - | - | 446.5 | 152.1 | - | - | 0 | - |
| - | - | 598.5 | 153.1 | - | - | 0 | - |
| - | - | 1572 | 154.1 | - | - | 0 | - |
| - | - | 1049 | 155.1 | - | - | 0 | - |
| - | - | 1412 | 155.1 | - | - | 0 | - |
| - | - | 4.563E+04 | 155.1 | - | - | 0 | - |
| - | - | 3650 | 156.1 | - | - | 0 | - |
| - | - | 484.2 | 157.1 | - | - | 0 | - |
| - | - | 877.2 | 157.1 | - | - | 0 | - |
| - | - | 450 | 157.1 | - | - | 0 | - |
| - | - | 1.827E+04 | 159.1 | - | - | 0 | - |
| - | - | 1787 | 159.1 | - | - | 0 | - |
| - | - | 755.4 | 160.1 | - | - | 0 | - |
| - | - | 574.1 | 164.8 | - | - | 0 | - |
| - | - | 2192 | 165.1 | - | - | 0 | - |
| - | - | 550.4 | 165.1 | - | - | 0 | - |
| - | - | 1741 | 165.1 | - | - | 0 | - |
| 10 | y | 8.373E+04 | 166.1 | 0.0003843 | 2.314 | +1 | 1 |
| - | - | 938.7 | 167.1 | - | - | 0 | - |
| - | - | 7686 | 167.1 | - | - | 0 | - |
| - | - | 5520 | 167.1 | - | - | 0 | - |
| - | - | 4119 | 168.1 | - | - | 0 | - |
| - | - | 1029 | 168.1 | - | - | 0 | - |
| - | - | 1749 | 169.1 | - | - | 0 | - |
| - | - | 551 | 172.1 | - | - | 0 | - |
| - | - | 1.067E+05 | 173.1 | - | - | 0 | - |
| - | - | 704.4 | 174.1 | - | - | 0 | - |
| - | - | 8860 | 174.1 | - | - | 0 | - |
| - | - | 635.2 | 175.1 | - | - | 0 | - |
| - | - | 1089 | 177.1 | - | - | 0 | - |
| - | - | 1.837E+04 | 177.1 | - | - | 0 | - |
| - | - | 1068 | 177.1 | - | - | 0 | - |
| - | - | 1771 | 178.1 | - | - | 0 | - |
| - | - | 693.6 | 179.1 | - | - | 0 | - |
| - | - | 1064 | 180.1 | - | - | 0 | - |
| - | - | 550.8 | 180.1 | - | - | 0 | - |
| - | - | 4833 | 181.1 | - | - | 0 | - |
| - | - | 804.7 | 181.1 | - | - | 0 | - |
| - | - | 3184 | 182.1 | - | - | 0 | - |
| - | - | 1.411E+05 | 182.1 | - | - | 0 | - |
| 2 | a | 1.928E+05 | 183.1 | 0.0004313 | 2.355 | +1 | 2 |
| - | - | 1.338E+04 | 183.1 | - | - | 0 | - |
| - | - | 1699 | 183.1 | - | - | 0 | - |
| - | - | 653.2 | 184.1 | - | - | 0 | - |
| - | - | 2523 | 184.1 | - | - | 0 | - |
| - | - | 1.832E+04 | 184.1 | - | - | 0 | - |
| - | - | 709.4 | 184.1 | - | - | 0 | - |
| - | - | 1079 | 184.1 | - | - | 0 | - |
| - | - | 951.7 | 185.1 | - | - | 0 | - |
| - | - | 941.7 | 185.1 | - | - | 0 | - |
| - | - | 8876 | 185.1 | - | - | 0 | - |
| 2 | d | 949.6 | 186.1 | 0.003311 | 17.79 | +1 | 2 |
| - | - | 5877 | 187.1 | - | - | 0 | - |
| - | - | 682.1 | 188.1 | - | - | 0 | - |
| - | - | 593 | 191.1 | - | - | 0 | - |
| - | - | 1096 | 193.1 | - | - | 0 | - |
| - | - | 6351 | 194.1 | - | - | 0 | - |
| - | - | 1453 | 195.1 | - | - | 0 | - |
| - | - | 1266 | 196.1 | - | - | 0 | - |
| - | - | 1858 | 197.1 | - | - | 0 | - |
| - | - | 1403 | 197.1 | - | - | 0 | - |
| - | - | 8545 | 198.1 | - | - | 0 | - |
| - | - | 2558 | 198.1 | - | - | 0 | - |
| - | - | 4001 | 199.1 | - | - | 0 | - |
| - | - | 1447 | 199.1 | - | - | 0 | - |
| 2 | a | 2.296E+05 | 200.1 | 0.000402 | 2.009 | +1 | 2 |
| - | - | 3.807E+04 | 201.1 | - | - | 0 | - |
| - | - | 2.013E+04 | 201.1 | - | - | 0 | - |
| - | - | 2979 | 202.1 | - | - | 0 | - |
| - | - | 572.6 | 202.1 | - | - | 0 | - |
| - | - | 1089 | 202.2 | - | - | 0 | - |
| - | - | 2631 | 205.1 | - | - | 0 | - |
| - | - | 605.8 | 207.1 | - | - | 0 | - |
| - | - | 1472 | 208.1 | - | - | 0 | - |
| - | - | 1702 | 208.1 | - | - | 0 | - |
| - | - | 1517 | 209.1 | - | - | 0 | - |
| - | - | 1.371E+04 | 209.1 | - | - | 0 | - |
| - | - | 1532 | 210.1 | - | - | 0 | - |
| - | - | 5.742E+04 | 210.1 | - | - | 0 | - |
| 2 | b | 3.167E+04 | 211.1 | 0.0003897 | 1.846 | +1 | 2 |
| - | - | 5716 | 211.1 | - | - | 0 | - |
| - | - | 631.3 | 211.1 | - | - | 0 | - |
| - | - | 488.3 | 211.1 | - | - | 0 | - |
| - | - | 2864 | 212.1 | - | - | 0 | - |
| - | - | 950.6 | 212.1 | - | - | 0 | - |
| - | - | 4304 | 212.1 | - | - | 0 | - |
| - | - | 1382 | 213.1 | - | - | 0 | - |
| - | - | 684.4 | 213.1 | - | - | 0 | - |
| - | - | 645.2 | 214.1 | - | - | 0 | - |
| - | - | 558.8 | 214.2 | - | - | 0 | - |
| - | - | 1312 | 215.1 | - | - | 0 | - |
| - | - | 3813 | 216.1 | - | - | 0 | - |
| - | - | 583 | 216.5 | - | - | 0 | - |
| - | - | 549.3 | 217.1 | - | - | 0 | - |
| - | - | 2765 | 218.2 | - | - | 0 | - |
| - | - | 529.1 | 218.8 | - | - | 0 | - |
| - | - | 560 | 219.2 | - | - | 0 | - |
| - | - | 1613 | 222.1 | - | - | 0 | - |
| 9 | y | 5.711E+04 | 223.1 | 0.0003897 | 1.747 | +1 | 2 |
| - | - | 6019 | 224.1 | - | - | 0 | - |
| - | - | 693.2 | 225.1 | - | - | 0 | - |
| - | - | 4329 | 225.1 | - | - | 0 | - |
| - | - | 2.047E+04 | 226.1 | - | - | 0 | - |
| - | - | 4.289E+04 | 226.1 | - | - | 0 | - |
| - | - | 1.456E+04 | 227.1 | - | - | 0 | - |
| - | - | 1987 | 227.1 | - | - | 0 | - |
| - | - | 2621 | 227.1 | - | - | 0 | - |
| - | - | 4932 | 227.1 | - | - | 0 | - |
| - | - | 911.6 | 228.1 | - | - | 0 | - |
| 2 | b | 7.667E+04 | 228.1 | 0.0003909 | 1.714 | +1 | 2 |
| - | - | 1543 | 229.1 | - | - | 0 | - |
| - | - | 8442 | 229.1 | - | - | 0 | - |
| - | - | 5068 | 230.2 | - | - | 0 | - |
| - | - | 779.6 | 231.2 | - | - | 0 | - |
| - | - | 627.3 | 233.1 | - | - | 0 | - |
| - | - | 796.3 | 234.1 | - | - | 0 | - |
| - | - | 535 | 236.1 | - | - | 0 | - |
| - | - | 764 | 238.2 | - | - | 0 | - |
| - | - | 1236 | 239.2 | - | - | 0 | - |
| - | - | 3730 | 240.1 | - | - | 0 | - |
| - | - | 608.1 | 241.1 | - | - | 0 | - |
| - | - | 1.65E+04 | 242.2 | - | - | 0 | - |
| - | - | 2844 | 243.1 | - | - | 0 | - |
| - | - | 585.6 | 243.1 | - | - | 0 | - |
| - | - | 1796 | 243.2 | - | - | 0 | - |
| - | - | 3.395E+04 | 244.1 | - | - | 0 | - |
| - | - | 3.861E+04 | 244.1 | - | - | 0 | - |
| - | - | 3042 | 245.1 | - | - | 0 | - |
| - | - | 3819 | 245.1 | - | - | 0 | - |
| - | - | 811 | 248.2 | - | - | 0 | - |
| - | - | 568.1 | 250.1 | - | - | 0 | - |
| - | - | 796.2 | 251.2 | - | - | 0 | - |
| - | - | 2762 | 252.1 | - | - | 0 | - |
| - | - | 1170 | 253.1 | - | - | 0 | - |
| - | - | 579.4 | 254.1 | - | - | 0 | - |
| - | - | 1805 | 254.1 | - | - | 0 | - |
| - | - | 801.2 | 254.2 | - | - | 0 | - |
| - | - | 639.3 | 255.1 | - | - | 0 | - |
| - | - | 2871 | 258.1 | - | - | 0 | - |
| - | - | 1471 | 261.1 | - | - | 0 | - |
| - | - | 1271 | 261.1 | - | - | 0 | - |
| - | - | 597.7 | 261.8 | - | - | 0 | - |
| - | - | 735.2 | 262.1 | - | - | 0 | - |
| - | - | 551.9 | 264.1 | - | - | 0 | - |
| - | - | 1574 | 265.1 | - | - | 0 | - |
| - | - | 2136 | 266.1 | - | - | 0 | - |
| - | - | 1604 | 266.2 | - | - | 0 | - |
| - | - | 2206 | 267.1 | - | - | 0 | - |
| - | - | 1034 | 268.1 | - | - | 0 | - |
| 3 | a | 2225 | 269.2 | 0.0002243 | 0.8332 | +1 | 3 |
| 3 | a | 5475 | 270.1 | 0.0002175 | 0.8051 | +1 | 3 |
| - | - | 910.9 | 276.1 | - | - | 0 | - |
| - | - | 1673 | 278.1 | - | - | 0 | - |
| - | - | 9421 | 279.1 | - | - | 0 | - |
| - | - | 1.176E+04 | 280.1 | - | - | 0 | - |
| - | - | 1106 | 280.1 | - | - | 0 | - |
| - | - | 1585 | 281.1 | - | - | 0 | - |
| - | - | 1337 | 282.1 | - | - | 0 | - |
| - | - | 3461 | 282.2 | - | - | 0 | - |
| - | - | 1.099E+04 | 283.1 | - | - | 0 | - |
| 6 | y | 1988 | 284.1 | 0.001265 | 4.451 | +2 | 5 |
| - | - | 684.4 | 284.2 | - | - | 0 | - |
| - | - | 809.2 | 286.1 | - | - | 0 | - |
| 3 | a | 2519 | 287.2 | 0.0003102 | 1.08 | +1 | 3 |
| - | - | 688.7 | 288.2 | - | - | 0 | - |
| - | - | 1898 | 290.1 | - | - | 0 | - |
| - | - | 1163 | 291.1 | - | - | 0 | - |
| - | - | 2036 | 292.1 | - | - | 0 | - |
| - | - | 3009 | 294.1 | - | - | 0 | - |
| - | - | 641.8 | 296.1 | - | - | 0 | - |
| 3 | b | 4.507E+04 | 297.2 | 0.0005184 | 1.745 | +1 | 3 |
| 3 | b | 3.312E+04 | 298.1 | 0.00042 | 1.409 | +1 | 3 |
| - | - | 6139 | 298.2 | - | - | 0 | - |
| - | - | 3539 | 299.1 | - | - | 0 | - |
| - | - | 1204 | 300.2 | - | - | 0 | - |
| - | - | 837.6 | 300.2 | - | - | 0 | - |
| - | - | 2385 | 301.2 | - | - | 0 | - |
| - | - | 1096 | 305.2 | - | - | 0 | - |
| - | - | 1882 | 305.2 | - | - | 0 | - |
| 8 | y | 4399 | 306.1 | 0.0004006 | 1.308 | +1 | 3 |
| - | - | 5610 | 309.2 | - | - | 0 | - |
| - | - | 2620 | 310.1 | - | - | 0 | - |
| - | - | 1091 | 310.2 | - | - | 0 | - |
| - | - | 1034 | 311.1 | - | - | 0 | - |
| - | - | 2105 | 311.2 | - | - | 0 | - |
| 3 | b | 1.934E+04 | 315.2 | 0.0006043 | 1.918 | +1 | 3 |
| - | - | 2919 | 316.2 | - | - | 0 | - |
| - | - | 1488 | 317.2 | - | - | 0 | - |
| - | - | 1034 | 317.2 | - | - | 0 | - |
| - | - | 3928 | 319.1 | - | - | 0 | - |
| - | - | 586.1 | 322.1 | - | - | 0 | - |
| 8 | y | 2.327E+04 | 324.2 | 0.0005781 | 1.783 | +1 | 3 |
| - | - | 3701 | 325.2 | - | - | 0 | - |
| 6 | b | 7111 | 327.2 | 0.0007569 | 2.314 | +2 | 6 |
| - | - | 660.6 | 327.2 | - | - | 0 | - |
| - | - | 1137 | 328.2 | - | - | 0 | - |
| - | - | 1001 | 328.2 | - | - | 0 | - |
| - | - | 909.7 | 329.1 | - | - | 0 | - |
| - | - | 1.944E+04 | 329.2 | - | - | 0 | - |
| - | - | 3010 | 330.2 | - | - | 0 | - |
| 6 | b | 4467 | 336.2 | 0.004495 | 13.37 | +2 | 6 |
| - | - | 2.132E+04 | 337.2 | - | - | 0 | - |
| - | - | 658.9 | 338.1 | - | - | 0 | - |
| - | - | 4221 | 338.2 | - | - | 0 | - |
| 5 | y | 6160 | 339.2 | 0.0003907 | 1.152 | +2 | 6 |
| 5 | y | 1757 | 339.7 | 0.004477 | 13.18 | +2 | 6 |
| - | - | 3374 | 340.2 | - | - | 0 | - |
| - | - | 558.4 | 341.2 | - | - | 0 | - |
| - | - | 1.227E+04 | 345.2 | - | - | 0 | - |
| - | - | 2660 | 346.2 | - | - | 0 | - |
| 5 | y | 9969 | 348.2 | 0.0002353 | 0.6759 | +2 | 6 |
| - | - | 2621 | 348.7 | - | - | 0 | - |
| - | - | 577.3 | 348.7 | - | - | 0 | - |
| - | - | 1073 | 349.2 | - | - | 0 | - |
| - | - | 1149 | 349.2 | - | - | 0 | - |
| - | - | 814.9 | 352.2 | - | - | 0 | - |
| - | - | 648.3 | 352.2 | - | - | 0 | - |
| - | - | 2.813E+04 | 354.2 | - | - | 0 | - |
| - | - | 1.19E+04 | 355.2 | - | - | 0 | - |
| - | - | 3792 | 355.2 | - | - | 0 | - |
| - | - | 1612 | 356.2 | - | - | 0 | - |
| - | - | 548.8 | 356.2 | - | - | 0 | - |
| - | - | 8861 | 357.2 | - | - | 0 | - |
| - | - | 1517 | 358.2 | - | - | 0 | - |
| - | - | 617 | 359.2 | - | - | 0 | - |
| - | - | 3511 | 364.2 | - | - | 0 | - |
| - | - | 2.418E+04 | 365.2 | - | - | 0 | - |
| - | - | 1296 | 366.2 | - | - | 0 | - |
| - | - | 2769 | 366.2 | - | - | 0 | - |
| - | - | 1693 | 367.2 | - | - | 0 | - |
| - | - | 2.419E+04 | 372.2 | - | - | 0 | - |
| - | - | 910.7 | 373.2 | - | - | 0 | - |
| - | - | 2796 | 373.2 | - | - | 0 | - |
| - | - | 626.6 | 373.5 | - | - | 0 | - |
| - | - | 1057 | 373.7 | - | - | 0 | - |
| - | - | 1005 | 374.2 | - | - | 0 | - |
| - | - | 602.3 | 375.2 | - | - | 0 | - |
| - | - | 1488 | 376.2 | - | - | 0 | - |
| - | - | 989.7 | 380.2 | - | - | 0 | - |
| - | - | 1728 | 380.2 | - | - | 0 | - |
| - | - | 698.9 | 382.2 | - | - | 0 | - |
| - | - | 1056 | 382.2 | - | - | 0 | - |
| - | - | 6614 | 382.2 | - | - | 0 | - |
| - | - | 650.9 | 382.7 | - | - | 0 | - |
| - | - | 2.688E+04 | 383.2 | - | - | 0 | - |
| - | - | 2543 | 384.2 | - | - | 0 | - |
| - | - | 4327 | 384.2 | - | - | 0 | - |
| - | - | 1959 | 385.2 | - | - | 0 | - |
| - | - | 757.5 | 385.2 | - | - | 0 | - |
| - | - | 749.9 | 389.2 | - | - | 0 | - |
| 7 | b | 573.7 | 391.7 | 0.00326 | 8.322 | +2 | 7 |
| - | - | 1.008E+04 | 392.2 | - | - | 0 | - |
| - | - | 3603 | 393.2 | - | - | 0 | - |
| - | - | 2.205E+04 | 393.2 | - | - | 0 | - |
| - | - | 1.091E+04 | 394.2 | - | - | 0 | - |
| - | - | 4252 | 394.2 | - | - | 0 | - |
| - | - | 2232 | 395.2 | - | - | 0 | - |
| 4 | y | 4145 | 396.2 | 0.0007748 | 1.955 | +2 | 7 |
| - | - | 2589 | 396.7 | - | - | 0 | - |
| - | - | 2062 | 398.2 | - | - | 0 | - |
| - | - | 1129 | 400.3 | - | - | 0 | - |
| - | - | 1140 | 401.7 | - | - | 0 | - |
| - | - | 7043 | 402.2 | - | - | 0 | - |
| - | - | 1206 | 403.2 | - | - | 0 | - |
| 4 | y | 1.543E+04 | 404.7 | 0.0005923 | 1.463 | +2 | 7 |
| - | - | 5893 | 405.2 | - | - | 0 | - |
| - | - | 2153 | 405.7 | - | - | 0 | - |
| - | - | 654.9 | 406.2 | - | - | 0 | - |
| - | - | 677.2 | 408.2 | - | - | 0 | - |
| - | - | 823.6 | 409.3 | - | - | 0 | - |
| - | - | 1051 | 410.2 | - | - | 0 | - |
| 4 | b | 2.03E+04 | 410.2 | 0.0004693 | 1.144 | +1 | 4 |
| - | - | 1.514E+04 | 410.7 | - | - | 0 | - |
| - | - | 3068 | 411.2 | - | - | 0 | - |
| 4 | b | 2.683E+04 | 411.2 | 0.0002394 | 0.5821 | +1 | 4 |
| - | - | 647.5 | 411.7 | - | - | 0 | - |
| - | - | 3007 | 412.2 | - | - | 0 | - |
| - | - | 4263 | 412.2 | - | - | 0 | - |
| - | - | 679.1 | 413.2 | - | - | 0 | - |
| - | - | 756.7 | 413.2 | - | - | 0 | - |
| - | - | 1327 | 418.7 | - | - | 0 | - |
| - | - | 2659 | 419.2 | - | - | 0 | - |
| - | - | 1037 | 419.2 | - | - | 0 | - |
| - | - | 1567 | 419.7 | - | - | 0 | - |
| - | - | 6559 | 420.2 | - | - | 0 | - |
| - | - | 2953 | 423.7 | - | - | 0 | - |
| - | - | 2540 | 424.2 | - | - | 0 | - |
| - | - | 1171 | 426.2 | - | - | 0 | - |
| - | - | 2887 | 427.2 | - | - | 0 | - |
| - | - | 1402 | 427.2 | - | - | 0 | - |
| - | - | 4032 | 427.3 | - | - | 0 | - |
| - | - | 3884 | 427.7 | - | - | 0 | - |
| - | - | 715 | 428.2 | - | - | 0 | - |
| - | - | 2462 | 428.2 | - | - | 0 | - |
| 4 | b | 9723 | 428.3 | 0.0002077 | 0.485 | +1 | 4 |
| - | - | 848.1 | 428.7 | - | - | 0 | - |
| - | - | 1626 | 429.2 | - | - | 0 | - |
| - | - | 2683 | 429.3 | - | - | 0 | - |
| - | - | 1197 | 430.2 | - | - | 0 | - |
| - | - | 1.035E+04 | 432.7 | - | - | 0 | - |
| - | - | 5393 | 433.2 | - | - | 0 | - |
| - | - | 2536 | 433.7 | - | - | 0 | - |
| 7 | y | 3793 | 434.2 | 0.0002862 | 0.6591 | +1 | 4 |
| - | - | 1166 | 435.2 | - | - | 0 | - |
| - | - | 588.4 | 436.2 | - | - | 0 | - |
| - | - | 3964 | 437.2 | - | - | 0 | - |
| - | - | 6717 | 438.2 | - | - | 0 | - |
| - | - | 677.5 | 438.7 | - | - | 0 | - |
| - | - | 855.2 | 439.2 | - | - | 0 | - |
| 3 | y | 1730 | 439.2 | 0.00283 | 6.444 | +2 | 8 |
| 3 | y | 6913 | 439.7 | 0.001362 | 3.098 | +2 | 8 |
| - | - | 3447 | 440.2 | - | - | 0 | - |
| 8 | b | 1.783E+04 | 441.7 | 0.0005421 | 1.227 | +2 | 8 |
| 8 | b | 7815 | 442.2 | 0.008778 | 19.85 | +2 | 8 |
| - | - | 2665 | 442.7 | - | - | 0 | - |
| - | - | 2.013E+04 | 444.2 | - | - | 0 | - |
| - | - | 3655 | 445.2 | - | - | 0 | - |
| - | - | 1037 | 445.2 | - | - | 0 | - |
| - | - | 1217 | 445.3 | - | - | 0 | - |
| - | - | 715.4 | 446.2 | - | - | 0 | - |
| - | - | 621.1 | 447.7 | - | - | 0 | - |
| 3 | y | 1.247E+04 | 448.2 | 0.001058 | 2.359 | +2 | 8 |
| - | - | 7413 | 448.7 | - | - | 0 | - |
| - | - | 2662 | 449.2 | - | - | 0 | - |
| - | - | 2171 | 450.2 | - | - | 0 | - |
| 8 | b | 2313 | 450.7 | 0.0005088 | 1.129 | +2 | 8 |
| - | - | 922.3 | 451.2 | - | - | 0 | - |
| 7 | y | 4.86E+04 | 452.3 | 0.0004942 | 1.093 | +1 | 4 |
| - | - | 1094 | 452.7 | - | - | 0 | - |
| - | - | 1.202E+04 | 453.3 | - | - | 0 | - |
| - | - | 1510 | 454.3 | - | - | 0 | - |
| - | - | 7736 | 455.2 | - | - | 0 | - |
| - | - | 1.838E+04 | 456.2 | - | - | 0 | - |
| - | - | 4338 | 457.2 | - | - | 0 | - |
| - | - | 3882 | 461.2 | - | - | 0 | - |
| - | - | 3944 | 461.7 | - | - | 0 | - |
| - | - | 2500 | 462.2 | - | - | 0 | - |
| - | - | 888.9 | 467.2 | - | - | 0 | - |
| - | - | 986.6 | 467.3 | - | - | 0 | - |
| - | - | 1457 | 469.2 | - | - | 0 | - |
| 9 | b | 1.2E+04 | 470.2 | 0.0004304 | 0.9153 | +2 | 9 |
| 9 | b | 7222 | 470.7 | 0.007934 | 16.85 | +2 | 9 |
| - | - | 2377 | 471.2 | - | - | 0 | - |
| - | - | 1.162E+04 | 473.2 | - | - | 0 | - |
| - | - | 3648 | 474.2 | - | - | 0 | - |
| 9 | b | 9749 | 479.3 | 0.0008854 | 1.847 | +2 | 9 |
| - | - | 3135 | 479.8 | - | - | 0 | - |
| - | - | 1351 | 480.3 | - | - | 0 | - |
| - | - | 4634 | 485.3 | - | - | 0 | - |
| - | - | 782.9 | 486.3 | - | - | 0 | - |
| - | - | 3802 | 495.2 | - | - | 0 | - |
| - | - | 797.4 | 495.3 | - | - | 0 | - |
| - | - | 854.9 | 496.2 | - | - | 0 | - |
| 2 | y | 2112 | 497.8 | 0.0008472 | 1.702 | +2 | 9 |
| - | - | 1050 | 498.3 | - | - | 0 | - |
| - | - | 2493 | 510.3 | - | - | 0 | - |
| - | - | 3344 | 512.2 | - | - | 0 | - |
| - | - | 7072 | 513.2 | - | - | 0 | - |
| - | - | 1950 | 514.2 | - | - | 0 | - |
| - | - | 801.8 | 520.8 | - | - | 0 | - |
| - | - | 3123 | 521.3 | - | - | 0 | - |
| - | - | 1694 | 521.8 | - | - | 0 | - |
| - | - | 2847 | 522.3 | - | - | 0 | - |
| - | - | 1328 | 522.7 | - | - | 0 | - |
| - | - | 928.1 | 522.8 | - | - | 0 | - |
| - | - | 1164 | 524.3 | - | - | 0 | - |
| - | - | 2792 | 529.8 | - | - | 0 | - |
| - | - | 6754 | 530.3 | - | - | 0 | - |
| - | - | 2003 | 530.8 | - | - | 0 | - |
| - | - | 3984 | 531.3 | - | - | 0 | - |
| - | - | 2004 | 532.2 | - | - | 0 | - |
| - | - | 2696 | 534.3 | - | - | 0 | - |
| - | - | 988.8 | 534.8 | - | - | 0 | - |
| - | - | 1078 | 535.3 | - | - | 0 | - |
| - | - | 2182 | 535.8 | - | - | 0 | - |
| - | - | 835.1 | 537.3 | - | - | 0 | - |
| 5 | b | 1.675E+04 | 538.3 | 0.0002109 | 0.3918 | +1 | 5 |
| - | - | 1100 | 538.8 | - | - | 0 | - |
| 5 | b | 9566 | 539.3 | 0.003317 | 6.151 | +1 | 5 |
| - | - | 2079 | 540.3 | - | - | 0 | - |
| - | - | 998.6 | 542.3 | - | - | 0 | - |
| - | - | 917 | 542.7 | - | - | 0 | - |
| - | - | 728.1 | 543.3 | - | - | 0 | - |
| - | - | 1.788E+04 | 543.8 | - | - | 0 | - |
| - | - | 1.585E+04 | 544.3 | - | - | 0 | - |
| - | - | 6952 | 544.8 | - | - | 0 | - |
| - | - | 2866 | 545.3 | - | - | 0 | - |
| 6 | y | 9273 | 549.3 | 0.0004123 | 0.7506 | +1 | 5 |
| 6 | y | 1065 | 550.3 | 0.007425 | 13.49 | +1 | 5 |
| - | - | 2177 | 550.3 | - | - | 0 | - |
| - | - | 1039 | 551.3 | - | - | 0 | - |
| - | - | 3234 | 551.7 | - | - | 0 | - |
| - | - | 2294 | 552.2 | - | - | 0 | - |
| - | - | 1022 | 552.3 | - | - | 0 | - |
| 0 | Precursor | 3.276E+05 | 552.8 | 0.0005529 | 1 | +2 | -1 |
| 0 | Precursor | 2.088E+05 | 553.3 | 0.0094 | 16.99 | +2 | -1 |
| - | - | 7.48E+04 | 553.8 | - | - | 0 | - |
| - | - | 8894 | 554.3 | - | - | 0 | - |
| - | - | 1230 | 555.3 | - | - | 0 | - |
| 5 | b | 2989 | 556.3 | 0.0003884 | 0.6982 | +1 | 5 |
| 0 | Precursor | 4.833E+04 | 561.8 | 0.0002755 | 0.4903 | +2 | -1 |
| - | - | 3.379E+04 | 562.3 | - | - | 0 | - |
| - | - | 1.01E+04 | 562.8 | - | - | 0 | - |
| - | - | 2641 | 563.3 | - | - | 0 | - |
| 6 | y | 1.072E+05 | 567.3 | 0.0001015 | 0.1789 | +1 | 5 |
| - | - | 3.228E+04 | 568.3 | - | - | 0 | - |
| - | - | 4914 | 569.3 | - | - | 0 | - |
| - | - | 1.578E+04 | 572.3 | - | - | 0 | - |
| - | - | 4945 | 573.3 | - | - | 0 | - |
| - | - | 1019 | 574.3 | - | - | 0 | - |
| - | - | 5913 | 586.3 | - | - | 0 | - |
| - | - | 2611 | 587.3 | - | - | 0 | - |
| - | - | 1096 | 590.3 | - | - | 0 | - |
| - | - | 1005 | 593.3 | - | - | 0 | - |
| - | - | 940.1 | 594.3 | - | - | 0 | - |
| - | - | 947.4 | 600.3 | - | - | 0 | - |
| - | - | 871.9 | 608.3 | - | - | 0 | - |
| - | - | 8680 | 611.3 | - | - | 0 | - |
| - | - | 2963 | 612.3 | - | - | 0 | - |
| - | - | 1031 | 612.9 | - | - | 0 | - |
| - | - | 965.9 | 615.1 | - | - | 0 | - |
| - | - | 2249 | 618.3 | - | - | 0 | - |
| - | - | 2104 | 618.5 | - | - | 0 | - |
| - | - | 2171 | 618.7 | - | - | 0 | - |
| - | - | 695.4 | 618.9 | - | - | 0 | - |
| - | - | 1181 | 620.3 | - | - | 0 | - |
| - | - | 1622 | 625.3 | - | - | 0 | - |
| - | - | 967.6 | 628.3 | - | - | 0 | - |
| - | - | 2395 | 629.3 | - | - | 0 | - |
| - | - | 1143 | 632.3 | - | - | 0 | - |
| - | - | 1496 | 633.3 | - | - | 0 | - |
| - | - | 1220 | 635.3 | - | - | 0 | - |
| - | - | 1532 | 636.3 | - | - | 0 | - |
| - | - | 1.078E+04 | 637.3 | - | - | 0 | - |
| - | - | 3739 | 638.3 | - | - | 0 | - |
| - | - | 772.9 | 638.8 | - | - | 0 | - |
| - | - | 1882 | 642.3 | - | - | 0 | - |
| - | - | 2796 | 643.3 | - | - | 0 | - |
| - | - | 1622 | 644.3 | - | - | 0 | - |
| - | - | 2439 | 645.4 | - | - | 0 | - |
| - | - | 808.5 | 646.4 | - | - | 0 | - |
| - | - | 3891 | 649.3 | - | - | 0 | - |
| - | - | 1715 | 650.3 | - | - | 0 | - |
| 6 | b | 2.503E+04 | 653.3 | 0.0003064 | 0.469 | +1 | 6 |
| 6 | b | 9965 | 654.3 | 0.01293 | 19.77 | +1 | 6 |
| - | - | 8154 | 655.3 | - | - | 0 | - |
| - | - | 3374 | 656.3 | - | - | 0 | - |
| - | - | 3055 | 659.3 | - | - | 0 | - |
| - | - | 5372 | 660.3 | - | - | 0 | - |
| - | - | 1592 | 661.3 | - | - | 0 | - |
| - | - | 787.7 | 661.4 | - | - | 0 | - |
| - | - | 7013 | 670.4 | - | - | 0 | - |
| 6 | b | 3600 | 671.3 | 0.009273 | 13.81 | +1 | 6 |
| - | - | 1551 | 672.3 | - | - | 0 | - |
| - | - | 3.145E+04 | 673.4 | - | - | 0 | - |
| - | - | 1.185E+04 | 674.4 | - | - | 0 | - |
| - | - | 1682 | 675.4 | - | - | 0 | - |
| 5 | y | 3.275E+04 | 677.3 | 6.229E-05 | 0.09196 | +1 | 6 |
| 5 | y | 7.501E+04 | 678.3 | 0.001948 | 2.871 | +1 | 6 |
| - | - | 2.642E+04 | 679.3 | - | - | 0 | - |
| - | - | 5262 | 680.3 | - | - | 0 | - |
| - | - | 1023 | 686.4 | - | - | 0 | - |
| - | - | 786 | 694 | - | - | 0 | - |
| - | - | 888.4 | 694.4 | - | - | 0 | - |
| 5 | y | 1.013E+05 | 695.3 | 4.351E-06 | 0.006257 | +1 | 6 |
| - | - | 3.563E+04 | 696.3 | - | - | 0 | - |
| - | - | 7691 | 697.3 | - | - | 0 | - |
| - | - | 1063 | 705.3 | - | - | 0 | - |
| - | - | 1022 | 706.3 | - | - | 0 | - |
| - | - | 4564 | 712.4 | - | - | 0 | - |
| - | - | 2611 | 713.4 | - | - | 0 | - |
| - | - | 881.3 | 714.4 | - | - | 0 | - |
| - | - | 1.257E+04 | 730.4 | - | - | 0 | - |
| - | - | 4382 | 731.4 | - | - | 0 | - |
| - | - | 1069 | 732.4 | - | - | 0 | - |
| - | - | 632.3 | 736.4 | - | - | 0 | - |
| - | - | 1248 | 748.4 | - | - | 0 | - |
| - | - | 680.9 | 755.4 | - | - | 0 | - |
| - | - | 1779 | 763.4 | - | - | 0 | - |
| - | - | 1846 | 764.4 | - | - | 0 | - |
| - | - | 1464 | 765.4 | - | - | 0 | - |
| - | - | 743.1 | 765.7 | - | - | 0 | - |
| - | - | 2706 | 772.4 | - | - | 0 | - |
| - | - | 996.8 | 772.9 | - | - | 0 | - |
| - | - | 2668 | 773.4 | - | - | 0 | - |
| - | - | 1552 | 774.4 | - | - | 0 | - |
| 7 | b | 1.807E+04 | 781.4 | 0.0002962 | 0.3791 | +1 | 7 |
| 7 | b | 9208 | 782.4 | 0.01367 | 17.48 | +1 | 7 |
| - | - | 2313 | 783.4 | - | - | 0 | - |
| - | - | 794.6 | 783.9 | - | - | 0 | - |
| - | - | 670.5 | 784.9 | - | - | 0 | - |
| 4 | y | 1990 | 790.4 | 0.001726 | 2.184 | +1 | 7 |
| 4 | y | 1498 | 791.4 | 0.01334 | 16.86 | +1 | 7 |
| - | - | 983.9 | 794.4 | - | - | 0 | - |
| 7 | b | 2731 | 799.4 | 0.001095 | 1.37 | +1 | 7 |
| - | - | 1238 | 800.4 | - | - | 0 | - |
| 4 | y | 3.126E+04 | 808.4 | 0.0005722 | 0.7078 | +1 | 7 |
| - | - | 1.43E+04 | 809.4 | - | - | 0 | - |
| - | - | 2825 | 810.4 | - | - | 0 | - |
| - | - | 629.5 | 811.4 | - | - | 0 | - |
| - | - | 799.3 | 818.4 | - | - | 0 | - |
| - | - | 1230 | 820.4 | - | - | 0 | - |
| - | - | 3172 | 821.4 | - | - | 0 | - |
| - | - | 1954 | 822.4 | - | - | 0 | - |
| - | - | 1430 | 829.4 | - | - | 0 | - |
| - | - | 874.7 | 830.4 | - | - | 0 | - |
| - | - | 842.9 | 833.4 | - | - | 0 | - |
| - | - | 1212 | 838.4 | - | - | 0 | - |
| - | - | 755.8 | 839.4 | - | - | 0 | - |
| - | - | 2871 | 846.4 | - | - | 0 | - |
| - | - | 2921 | 847.4 | - | - | 0 | - |
| - | - | 1603 | 848.4 | - | - | 0 | - |
| - | - | 1827 | 849.4 | - | - | 0 | - |
| - | - | 867.3 | 850.4 | - | - | 0 | - |
| - | - | 1635 | 855.5 | - | - | 0 | - |
| - | - | 1862 | 859.4 | - | - | 0 | - |
| - | - | 1905 | 860.4 | - | - | 0 | - |
| - | - | 775 | 861.4 | - | - | 0 | - |
| - | - | 1383 | 864.5 | - | - | 0 | - |
| - | - | 5908 | 865.4 | - | - | 0 | - |
| - | - | 2133 | 866.4 | - | - | 0 | - |
| - | - | 1413 | 867.4 | - | - | 0 | - |
| 3 | y | 9456 | 877.4 | 0.001589 | 1.811 | +1 | 8 |
| 3 | y | 8457 | 878.4 | 0.009146 | 10.41 | +1 | 8 |
| - | - | 3405 | 879.4 | - | - | 0 | - |
| 8 | b | 1982 | 882.5 | 0.0004262 | 0.4829 | +1 | 8 |
| 8 | b | 1.805E+04 | 883.5 | 6.891E-05 | 0.078 | +1 | 8 |
| - | - | 8773 | 884.5 | - | - | 0 | - |
| - | - | 1951 | 885.5 | - | - | 0 | - |
| 3 | y | 1.757E+05 | 895.5 | 0.0009234 | 1.031 | +1 | 8 |
| - | - | 9.034E+04 | 896.5 | - | - | 0 | - |
| - | - | 2.287E+04 | 897.5 | - | - | 0 | - |
| - | - | 2099 | 898.5 | - | - | 0 | - |
| 8 | b | 3775 | 900.5 | 0.00141 | 1.566 | +1 | 8 |
| - | - | 3087 | 901.5 | - | - | 0 | - |
| - | - | 680.8 | 902.5 | - | - | 0 | - |
| - | - | 1298 | 905.4 | - | - | 0 | - |
| - | - | 4608 | 922.5 | - | - | 0 | - |
| - | - | 3765 | 923.5 | - | - | 0 | - |
| - | - | 1569 | 924.5 | - | - | 0 | - |
| 9 | b | 2605 | 939.5 | 0.00179 | 1.905 | +1 | 9 |
| 9 | b | 2.058E+04 | 940.5 | 0.0004145 | 0.4407 | +1 | 9 |
| - | - | 9137 | 941.5 | - | - | 0 | - |
| - | - | 2825 | 942.5 | - | - | 0 | - |
| 9 | b | 7873 | 957.5 | 0.0007185 | 0.7503 | +1 | 9 |
| - | - | 3346 | 958.5 | - | - | 0 | - |
| 2 | y | 671.8 | 976.5 | 0.009953 | 10.19 | +1 | 9 |
| 2 | y | 9745 | 994.5 | 0.001222 | 1.229 | +1 | 9 |
| - | - | 5446 | 995.5 | - | - | 0 | - |
| - | - | 1534 | 996.5 | - | - | 0 | - |
| - | - | 918.7 | 1004 | - | - | 0 | - |
| - | - | 699.7 | 1477 | - | - | 0 | - |
| - | - | 769.6 | 1857 | - | - | 0 | - |
| - | - | 712.9 | 2721 | - | - | 0 | - |
| - | - | 651.2 | 3276 | - | - | 0 | - |

m/z Charge Intensity FragmentType MassShift Position
120.08116912841797 0 60040.887
121.08450317382812 0 4709.5435
121.85179901123047 0 428.2754
122.27020263671875 0 439.45615
123.09195709228516 0 808.56165
124.07574462890625 0 492.88553
124.53675079345703 0 345.99783
126.09161376953125 0 790.18756
127.0870590209961 0 764.1126
127.12342071533203 0 2184.8074
127.64633178710938 0 395.70953
128.08213806152344 0 869.84235
128.10731506347656 0 3383.9302
129.06625366210938 0 15499.607
129.10263061523438 0 116586.37
130.0499725341797 0 616.277
130.06973266601562 0 678.1771
130.08680725097656 0 447.03052
130.10025024414062 0 1096.1588
130.1060028076172 0 7379.681
131.04954528808594 0 883.96857
131.0818634033203 0 3993.57
131.11825561523438 0 6803.049
132.10238647460938 0 540.7015
133.06130981445312 0 964.5853
133.08615112304688 0 1100.59
136.0760498046875 0 5250.092
137.10784912109375 0 573.89417
138.09173583984375 0 35651.57
139.05064392089844 0 477.68234
139.0869598388672 0 1790.558
139.0950927734375 0 3224.5806
140.10740661621094 0 1035.9944
141.06617736816406 0 2447.889
141.1026153564453 0 4712.087
144.8901824951172 0 456.64297
146.06044006347656 0 479.02435
147.1132354736328 0 455.80682
148.58975219726562 0 468.48297
148.80255126953125 0 459.0251
148.83853149414062 0 542.11066
148.8749237060547 0 540.7516
148.88198852539062 0 529.9636
148.8895263671875 0 459.99146
148.89697265625 0 579.3212
148.90371704101562 0 738.6449
148.91127014160156 0 1020.91895
148.9182586669922 0 1191.6826
148.9259033203125 0 1072.3574
148.93276977539062 0 1454.1614
148.94046020507812 0 3401.112
148.95706176757812 0 5731.7246
148.96493530273438 0 3551.2969
148.97210693359375 0 1081.1824
148.9795379638672 0 1167.0605
148.9870147705078 0 833.6493
148.9940948486328 0 998.01044
149.0010986328125 0 916.4385
149.00814819335938 0 611.6157
149.02291870117188 0 493.12634
149.03013610839844 0 451.60867
149.0453338623047 0 458.38242
149.0599365234375 0 1296.0615
151.08706665039062 0 4523.1226
151.09707641601562 0 443.9178
152.14315795898438 0 446.4657
153.10250854492188 0 598.53186
154.09774780273438 0 1572.2789
155.0708770751953 0 1048.8964
155.08200073242188 0 1411.9535
155.11827087402344 0 45625.13
156.12161254882812 0 3649.7954
157.06080627441406 0 484.2273
157.09718322753906 0 877.2224
157.13397216796875 0 450.01242
159.07676696777344 0 18272.611
159.1130828857422 0 1787.0511
160.08062744140625 0 755.3583
164.80514526367188 0 574.1435
165.05526733398438 0 2192.3057
165.06607055664062 0 550.3631
165.10267639160156 0 1740.7611
166.08663940429688 0 83734.11 y 9
167.0829315185547 0 938.74054
167.09010314941406 0 7685.6655
167.1182098388672 0 5519.8623
168.1134796142578 0 4119.4604
168.1210479736328 0 1028.578
169.09751892089844 0 1749.0896
172.07229614257812 0 550.9863
173.12887573242188 0 106741.91
174.12498474121094 0 704.4376
174.13226318359375 0 8859.519
175.13502502441406 0 635.24884
177.08750915527344 0 1089.4673
177.10263061523438 0 18373.594
177.11146545410156 0 1068.2903
178.1060791015625 0 1771.4174
179.11788940429688 0 693.6376
180.07691955566406 0 1063.5723
180.11386108398438 0 550.7607
181.06124877929688 0 4833.0894
181.09754943847656 0 804.70483
182.0815887451172 0 3184.2454
182.12925720214844 0 141126.78
183.1132354736328 0 192813.89 a Ammonia loss 1
183.1325225830078 0 13383.062
183.1494140625 0 1698.7247
184.1003875732422 0 653.1555
184.10894775390625 0 2523.075
184.11659240722656 0 18324.54
184.13613891601562 0 709.40027
184.14462280273438 0 1079.3958
185.09237670898438 0 951.66956
185.1192626953125 0 941.6522
185.12881469726562 0 8876.442
186.12701416015625 0 949.62946 d 1
187.10806274414062 0 5877.2344
188.11196899414062 0 682.1169
191.0828857421875 0 592.9569
193.09762573242188 0 1096.0286
194.12921142578125 0 6351.0205
195.1132049560547 0 1452.6252
196.108154296875 0 1265.9913
197.09234619140625 0 1857.9143
197.12864685058594 0 1402.9999
198.08763122558594 0 8544.816
198.12393188476562 0 2557.6082
199.07167053222656 0 4001.4885
199.10791015625 0 1447.3593
200.13975524902344 0 229634.06 a 1
201.12368774414062 0 38070.844
201.14305114746094 0 20132.928
202.12725830078125 0 2979.2593
202.14512634277344 0 572.57513
202.15528869628906 0 1088.7135
205.0975799560547 0 2631.2117
207.112548828125 0 605.8319
208.09730529785156 0 1472.2034
208.1083984375 0 1702.0057
209.055908203125 0 1516.6971
209.09242248535156 0 13705.699
210.09519958496094 0 1532.3003
210.12408447265625 0 57422.312
211.1081085205078 0 31668.39 b Ammonia loss 1
211.12730407714844 0 5715.727
211.13906860351562 0 631.34827
211.14315795898438 0 488.31427
212.11148071289062 0 2863.5955
212.1294403076172 0 950.60925
212.13973999023438 0 4304.2646
213.1240234375 0 1382.3392
213.14418029785156 0 684.42
214.1191864013672 0 645.21844
214.156005859375 0 558.8026
215.13949584960938 0 1312.4388
216.09817504882812 0 3812.5312
216.5135498046875 0 582.9889
217.09689331054688 0 549.34216
218.1501007080078 0 2764.7603
218.75128173828125 0 529.11707
219.15377807617188 0 560.02655
222.12408447265625 0 1613.285
223.1081085205078 0 57112.87 y 8
224.111328125 0 6018.53
225.11244201660156 0 693.2301
225.1238555908203 0 4328.674
226.0826873779297 0 20472.342
226.1190185546875 0 42888.633
227.066650390625 0 14561.701
227.08633422851562 0 1987.382
227.10284423828125 0 2621.1892
227.1225128173828 0 4931.511
228.07009887695312 0 911.5862
228.13465881347656 0 76665.81 b 1
229.11862182617188 0 1543.2971
229.13796997070312 0 8441.502
230.15037536621094 0 5068.247
231.15155029296875 0 779.55536
233.12953186035156 0 627.34894
234.12420654296875 0 796.3327
236.10440063476562 0 535.02374
238.15472412109375 0 763.9689
239.1511993408203 0 1235.6262
240.13470458984375 0 3730.3145
241.13787841796875 0 608.1327
242.150390625 0 16503.59
243.10916137695312 0 2844.0178
243.13290405273438 0 585.5936
243.15411376953125 0 1795.6455
244.09320068359375 0 33948.555
244.12962341308594 0 38610.23
245.09617614746094 0 3041.5884
245.13294982910156 0 3818.8784
248.16004943847656 0 810.9716
250.11795043945312 0 568.1227
251.15061950683594 0 796.2072
252.13475036621094 0 2761.9224
253.11862182617188 0 1170.1316
254.0768585205078 0 579.4406
254.1139678955078 0 1805.366
254.18682861328125 0 801.18195
255.14524841308594 0 639.2836
258.1453857421875 0 2871.0654
261.11968994140625 0 1471.2058
261.1330261230469 0 1271.2188
261.8021545410156 0 597.716
262.138916015625 0 735.16785
264.1345520019531 0 551.92065
265.1298522949219 0 1573.9542
266.1138916015625 0 2135.6418
266.15032958984375 0 1604.4576
267.1455383300781 0 2206.397
268.12908935546875 0 1034.2601
269.1610412597656 0 2224.8455 a Water loss 2
270.1450500488281 0 5474.8296 a Ammonia loss 2
276.1351318359375 0 910.9082
278.14990234375 0 1673.0219
279.1457214355469 0 9420.973
280.12969970703125 0 11762.712
280.1475524902344 0 1105.788
281.1322326660156 0 1585.2114
282.1455383300781 0 1336.7366
282.1816101074219 0 3460.5225
283.1404724121094 0 10989.69
284.1435546875 0 1987.6002 y 5
284.15985107421875 0 684.35156
286.1399230957031 0 809.24445
287.17169189453125 0 2518.6055 a 2
288.1748046875 0 688.6856
290.11505126953125 0 1897.8638
291.1454772949219 0 1163.3456
292.1294860839844 0 2036.2223
294.1450500488281 0 3009.2976
296.1230163574219 0 641.8481
297.15625 0 45070.883 b Water loss 2
298.1401672363281 0 33121.45 b Ammonia loss 2
298.1593933105469 0 6139.172
299.1427917480469 0 3539.4543
300.1566467285156 0 1203.886
300.1918640136719 0 837.5715
301.1518249511719 0 2384.741
305.1605224609375 0 1095.7139
305.18206787109375 0 1881.5361
306.1452331542969 0 4399.2744 y Water loss 7
309.15625 0 5609.8037
310.1400451660156 0 2620.1575
310.1587829589844 0 1091.0194
311.1368103027344 0 1033.7642
311.1714782714844 0 2105.4233
315.1669006347656 0 19335.967 b 2
316.1703796386719 0 2919.0022
317.1817321777344 0 1488.1614
317.21820068359375 0 1034.3796
319.140380859375 0 3928.0984
322.1395568847656 0 586.1274
324.1559753417969 0 23270.463 y 7
325.15875244140625 0 3701.2605
327.16705322265625 0 7110.9634 b Water loss 5
327.1990661621094 0 660.5824
328.15081787109375 0 1137.2087
328.1707763671875 0 1000.90643
329.1486511230469 0 909.6781
329.1825866699219 0 19438.123
330.18560791015625 0 3010.0635
336.1670837402344 0 4467.348 b 5
337.151123046875 0 21322.184
338.135009765625 0 658.9496
338.15460205078125 0 4220.896
339.16668701171875 0 6159.811 y Water loss 4
339.66278076171875 0 1756.7549 y Ammonia loss 4
340.1515197753906 0 3374.4182
341.1545104980469 0 558.42566
345.1773986816406 0 12274.969
346.17999267578125 0 2659.6768
348.17181396484375 0 9968.633 y 4
348.67340087890625 0 2620.563
348.7229309082031 0 577.27136
349.1515808105469 0 1072.598
349.17449951171875 0 1148.7213
352.197998046875 0 814.9161
352.2345275878906 0 648.2926
354.17742919921875 0 28134.875
355.1613464355469 0 11898.261
355.1816711425781 0 3792.426
356.16436767578125 0 1611.6616
356.1914367675781 0 548.8299
357.1769714355469 0 8860.555
358.17889404296875 0 1516.8196
359.1665344238281 0 617.0035
364.2347106933594 0 3510.5815
365.21868896484375 0 24183.885
366.1781921386719 0 1295.9521
366.2215270996094 0 2768.6257
367.161376953125 0 1693.3591
372.18798828125 0 24191.299
373.16864013671875 0 910.6563
373.19146728515625 0 2796.2761
373.48333740234375 0 626.57495
373.6940002441406 0 1057.4572
374.1969909667969 0 1004.6519
375.1772766113281 0 602.33606
376.1630554199219 0 1487.9929
380.1921081542969 0 989.65765
380.2284240722656 0 1727.852
382.1744689941406 0 698.9086
382.21331787109375 0 1055.6979
382.2451477050781 0 6613.869
382.7064208984375 0 650.8996
383.2292785644531 0 26884.361
384.18780517578125 0 2543.3945
384.23248291015625 0 4327.2104
385.1725158691406 0 1958.8854
385.23480224609375 0 757.4506
389.2189636230469 0 749.8868
391.70904541015625 0 573.70306 b Ammonia loss 6
392.2297668457031 0 10079.584
393.18743896484375 0 3603.2158
393.2138671875 0 22048.814
394.17266845703125 0 10905.383
394.2161865234375 0 4252.2695
395.1752624511719 0 2231.5337
396.20111083984375 0 4144.7505 y Ammonia loss 3
396.7032165527344 0 2588.815
398.2035217285156 0 2061.666
400.2551574707031 0 1129.031
401.7142333984375 0 1139.9694
402.1997985839844 0 7043.194
403.20355224609375 0 1205.8192
404.7142028808594 0 15426.861 y 3
405.21551513671875 0 5893.287
405.7180480957031 0 2153.477
406.24554443359375 0 654.94916
408.2246398925781 0 677.1842
409.2549133300781 0 823.56586
410.2087097167969 0 1050.578
410.2402648925781 0 20304.383 b Water loss 3
410.71978759765625 0 15139.916
411.1990966796875 0 3068.3345
411.22357177734375 0 26826.94 b Ammonia loss 3
411.720703125 0 647.5219
412.183349609375 0 3007.331
412.2267761230469 0 4263.4946
413.18621826171875 0 679.11304
413.2274169921875 0 756.71204
418.7370300292969 0 1327.1958
419.2042236328125 0 2659.3438
419.23260498046875 0 1037.4938
419.72540283203125 0 1566.7229
420.18817138671875 0 6558.926
423.727783203125 0 2952.6985
424.22552490234375 0 2539.5083
426.1975402832031 0 1171.0345
427.18328857421875 0 2886.9233
427.23095703125 0 1402.0616
427.2669982910156 0 4031.7185
427.74066162109375 0 3884.2297
428.18695068359375 0 714.976
428.2153015136719 0 2461.7627
428.2501525878906 0 9722.507 b 3
428.73516845703125 0 848.05927
429.2135009765625 0 1625.9879
429.25262451171875 0 2682.9185
430.2264099121094 0 1196.928
432.73272705078125 0 10346.379
433.23101806640625 0 5393.4956
433.73004150390625 0 2536.43
434.2400817871094 0 3792.9622 y Water loss 6
435.2334289550781 0 1166.12
436.23748779296875 0 588.40826
437.2147521972656 0 3963.5032
438.19903564453125 0 6717.2725
438.731201171875 0 677.46094
439.1980895996094 0 855.2497
439.2271728515625 0 1729.9886 y Water loss 2
439.71771240234375 0 6913.3765 y Ammonia loss 2
440.21954345703125 0 3447.4438
441.7381591796875 0 17831.518 b Water loss 7
442.2384033203125 0 7815.0103 b Ammonia loss 7
442.7368469238281 0 2665.1296
444.20965576171875 0 20131.238
445.21136474609375 0 3655.4678
445.24273681640625 0 1037.2396
445.2772216796875 0 1216.7207
446.2143859863281 0 715.36
447.74298095703125 0 621.062
448.2306823730469 0 12465.828 y 2
448.7318420410156 0 7412.7666
449.234619140625 0 2662.041
450.23504638671875 0 2171.3743
450.743408203125 0 2313.1104 b 7
451.24053955078125 0 922.3114
452.2508544921875 0 48595.703 y 6
452.7309875488281 0 1093.964
453.2538146972656 0 12024.441
454.2568359375 0 1510.1388
455.2252502441406 0 7735.8984
456.2095642089844 0 18377.986
457.2127685546875 0 4338.372
461.24261474609375 0 3882.42
461.7398376464844 0 3944.2317
462.2373962402344 0 2499.7085
467.2257080078125 0 888.86646
467.25982666015625 0 986.6288
469.2054443359375 0 1457.1494
470.248779296875 0 12000.798 b Water loss 8
470.748291015625 0 7221.6226 b Ammonia loss 8
471.2481994628906 0 2377.111
473.2361755371094 0 11624.253
474.2384338378906 0 3647.7866
479.2545166015625 0 9749.131 b 8
479.7555236816406 0 3134.534
480.25543212890625 0 1351.2357
485.27252197265625 0 4634.0176
486.2737731933594 0 782.92194
495.2195739746094 0 3801.8562
495.25555419921875 0 797.3695
496.22027587890625 0 854.8592
497.7646789550781 0 2112.128 y 1
498.26605224609375 0 1050.3146
510.3036193847656 0 2493.1099
512.2468872070312 0 3344.1274
513.2313842773438 0 7071.7817
514.2340087890625 0 1950.2163
520.7799072265625 0 801.8047
521.2725219726562 0 3122.8254
521.77001953125 0 1693.7754
522.2657470703125 0 2846.7625
522.6859130859375 0 1327.6288
522.762451171875 0 928.06226
524.2832641601562 0 1163.5615
529.7860717773438 0 2791.7627
530.2576904296875 0 6754.3076
530.7783813476562 0 2003.167
531.257080078125 0 3983.587
532.2439575195312 0 2004.2112
534.304443359375 0 2695.6042
534.7759399414062 0 988.7713
535.2711181640625 0 1077.607
535.7685546875 0 2182.2222
537.2666625976562 0 835.0746
538.298583984375 0 16751.258 b Water loss 4
538.7913208007812 0 1100.386
539.2857055664062 0 9566.254 b Ammonia loss 4
540.2869873046875 0 2078.5208
542.2935791015625 0 998.60187
542.684326171875 0 916.9643
543.3011474609375 0 728.1152
543.7828979492188 0 17878.574
544.280029296875 0 15850.805
544.77978515625 0 6951.7637
545.2777099609375 0 2866.465
549.2671508789062 0 9272.722 y Water loss 5
550.2581787109375 0 1064.6465 y Ammonia loss 5
550.2972412109375 0 2177.118
551.282958984375 0 1039.2264
551.6887817382812 0 3233.5054
552.1905517578125 0 2294.3196
552.2877197265625 0 1021.5971
552.7883911132812 0 327599.16 Precursor Water loss
553.2892456054688 0 208824.03 Precursor Ammonia loss
553.79052734375 0 74801.9
554.2913818359375 0 8894.002
555.2811279296875 0 1229.767
556.309326171875 0 2989.1555 b 4
561.7933959960938 0 48334.645 Precursor
562.294921875 0 33785.14
562.7960815429688 0 10098.235
563.2850952148438 0 2640.9802
567.2774047851562 0 107215.02 y 5
568.2803344726562 0 32277.61
569.2835693359375 0 4914.476
572.30419921875 0 15776.352
573.306640625 0 4944.9287
574.3071899414062 0 1019.45325
586.3196411132812 0 5912.594
587.3251342773438 0 2610.9995
590.3151245117188 0 1095.85
593.299072265625 0 1005.0858
594.2899780273438 0 940.14954
600.3305053710938 0 947.38354
608.3054809570312 0 871.9256
611.314453125 0 8679.9795
612.3153686523438 0 2963.3247
612.9467163085938 0 1030.7427
615.1420288085938 0 965.9261
618.34521484375 0 2248.9114
618.54833984375 0 2104.3562
618.7488403320312 0 2171.3718
618.9442138671875 0 695.4249
620.3042602539062 0 1180.6465
625.3319702148438 0 1621.5776
628.3397216796875 0 967.5802
629.326171875 0 2394.675
632.3016357421875 0 1143.4377
633.3046264648438 0 1496.2239
635.3121337890625 0 1220.084
636.3028564453125 0 1532.2034
637.3303833007812 0 10781.826
638.3321533203125 0 3739.289
638.759033203125 0 772.85156
642.2908325195312 0 1882.4941
643.3410034179688 0 2795.8096
644.3428344726562 0 1621.5496
645.3554077148438 0 2438.9792
646.3548583984375 0 808.4956
649.3310546875 0 3891.4204
650.3301391601562 0 1715.4004
653.3256225585938 0 25033.748 b Water loss 5
654.322265625 0 9964.718 b Ammonia loss 5
655.337890625 0 8153.927
656.3387451171875 0 3374.0571
659.3148193359375 0 3054.8943
660.3011474609375 0 5372.4067
661.2969970703125 0 1591.9431
661.3584594726562 0 787.7163
670.3516845703125 0 7013.472
671.3451538085938 0 3600.1128 b 5
672.3493041992188 0 1550.8618
673.3517456054688 0 31445.66
674.3544311523438 0 11845.162
675.355224609375 0 1681.7239
677.3253784179688 0 32749.125 y Water loss 4
678.311279296875 0 75007.68 y Ammonia loss 4
679.3133544921875 0 26416.732
680.3160400390625 0 5262.3296
686.3504638671875 0 1022.788
693.9898071289062 0 786.02264
694.357177734375 0 888.3653
695.3358764648438 0 101265.7 y 4
696.3389282226562 0 35628.87
697.341552734375 0 7690.526
705.3226928710938 0 1063.1008
706.3099975585938 0 1022.3738
712.361572265625 0 4564.205
713.360595703125 0 2610.7166
714.3582763671875 0 881.2977
730.3729858398438 0 12570.942
731.3749389648438 0 4382.336
732.3721313476562 0 1068.6425
736.3934936523438 0 632.2535
748.3833618164062 0 1248.177
755.3682861328125 0 680.8887
763.410400390625 0 1779.4084
764.39892578125 0 1846.4618
765.3978271484375 0 1464.4805
765.6724853515625 0 743.13837
772.4171142578125 0 2706.2305
772.929931640625 0 996.8067
773.3828125 0 2667.8184
774.3857421875 0 1552.0466
781.4199829101562 0 18067.654 b Water loss 6
782.41796875 0 9208.409 b Ammonia loss 6
783.4203491210938 0 2313.1912
783.8648681640625 0 794.58307
784.8677978515625 0 670.4975
790.4076538085938 0 1990.4331 y Water loss 3
791.40673828125 0 1497.5286 y Ammonia loss 3
794.3947143554688 0 983.93774
799.4297485351562 0 2730.9414 b 6
800.4326782226562 0 1237.623
808.4193725585938 0 31263.445 y 3
809.4227294921875 0 14304.782
810.4246215820312 0 2824.901
811.4357299804688 0 629.46295
818.4002075195312 0 799.2543
820.4282836914062 0 1229.7041
821.4154663085938 0 3172.4138
822.4212036132812 0 1954.0327
829.4420166015625 0 1430.1711
830.428466796875 0 874.7136
833.4171752929688 0 842.8699
838.4422607421875 0 1212.3771
839.4363403320312 0 755.8247
846.4471435546875 0 2871.2168
847.4365844726562 0 2921.3677
848.4334716796875 0 1602.752
849.4414672851562 0 1826.8468
850.444580078125 0 867.3085
855.45654296875 0 1634.9581
859.4331665039062 0 1862.0336
860.4268188476562 0 1904.5631
861.417236328125 0 774.99817
864.4574584960938 0 1382.6201
865.4417724609375 0 5908.1353
866.4385986328125 0 2132.7803
867.4456787109375 0 1413.3505
877.4398193359375 0 9455.997 y Water loss 2
878.4345703125 0 8456.87 y Ammonia loss 2
879.4340209960938 0 3404.8188
882.4683837890625 0 1981.829 b Water loss 7
883.451904296875 0 18050.73 b Ammonia loss 7
884.4550170898438 0 8772.902
885.45849609375 0 1950.5728
895.4510498046875 0 175711.75 y 2
896.4542236328125 0 90344.086
897.4567260742188 0 22869.895
898.4564819335938 0 2099.0466
900.4771118164062 0 3775.4617 b 7
901.48046875 0 3087.4673
902.479736328125 0 680.8209
905.4423828125 0 1297.6754
922.4630737304688 0 4608.057
923.4581909179688 0 3764.7344
924.4570922851562 0 1568.935
939.4912109375 0 2605.1936 b Water loss 8
940.4730224609375 0 20584.318 b Ammonia loss 8
941.475341796875 0 9136.648
942.4794921875 0 2824.6519
957.499267578125 0 7873.2026 b 8
958.5001831054688 0 3345.7832
976.519775390625 0 671.7738 y Water loss 1
994.5191650390625 0 9745.403 y 1
995.5213623046875 0 5445.691
996.5216674804688 0 1534.3379
1004.4990844726562 0 918.6765
1477.17919921875 0 699.68726
1857.2464599609375 0 769.59106
2721.11669921875 0 712.8871
3275.909423828125 0 651.2166

Spectrum Details

|  |  |
| --- | --- |
| Matched peaks? Matched peaks The total absolute number of peaks matched. Additionally in brackets the total fraction of peaks matched and the total number of peaks is shown. | 71 (11.45% of 620) |
| FDR? FDR The false discovery rate estimated for this peptide. It is calculated by matching all theoretical fragments with a non-integer shift with the raw peaks for this spectrum. This is done with 40 different shifts. The resulting percentage is the average number of annotated peaks over the number of annotated peaks with the correct spectrum. | 0.17% |
| Satellite FDR? Satellite FDR See the FDR for details on its calculation. This satellite ion specific FDR only contains the satellite ions (d/w) for I/L/J positions. | - |
| PSM Score? PSM Score The PSM Score as given by Hecklib to this annotated spectrum. It is shown with three significant figures. | 896 |

#### Spectrum 5655? Spectrum 5655 The raw spectrum of this peptide as annotated by Hecklib. The fragments are coloured according to ion type (see legend). Any peaks with a star '\*' as text can be hovered over to see the full details, first the ion type second the mass shift type. By hovering over the amino acids in the peptide or ions in the legend the corresponding peaks are highlighted. By toggling the 'Unassigned' label you can turn the background (unassigned) peaks on or off in the plot. By updating the slider in the Ion legend you can update the spectrum to only show the top X% of the peaks with labels. The top X% means any peak that is within X% of the highest intensity. By dragging in the spectrum you can zoom in to a specific part of the spectrum and use 'Zoom Out' to get back to the original zoom level. The annotation of the spectrum is based on the given sequence in the peptides file and is done with different software so inconsistencies are likely. The peaks are annotated based on the given sequence, with 20 ppm tolerance.

Copy Data

##### Spectrum 5655 (TSV)

###### Preview

```
Loading example...
```

*Click on the button to copy the data to your clipboard.*

Mz MinMz MaxIntensity Max

WidthHeightPeptide font sizePeptide stroke widthSpectrum font sizeSpectrum stroke widthCompact peptide

Ion legend

wxyz

abcd

OtherUnassignedIon^Charge^_Position_Show for top:%

QVSLQDKTGF

04.59e+49.18e+41.38e+51.84e+5

Zoom Out

y^+1^_1_a^+1^_2_d^+1^_2_a^+1^_2_b^+1^_2_y^+1^_2_b^+1^_2_a^+1^_3_a^+1^_3_y^+2^_5_a^+1^_3_b^+1^_3_b^+1^_3_y^+1^_3_b^+1^_3_y^+1^_3_b^+2^_6_b^+2^_6_y^+2^_6_y^+2^_6_y^+2^_7_y^+2^_7_b^+1^_4_b^+1^_4_b^+1^_4_y^+1^_4_y^+2^_8_y^+2^_8_b^+2^_8_b^+2^_8_y^+2^_8_y^+1^_4_b^+2^_9_b^+2^_9_b^+2^_9_y^+2^_9_b^+1^_5_b^+1^_5_y^+1^_5_\*\*b^+1^_5_\*y^+1^_5_b^+1^_6_b^+1^_6_b^+1^_6_y^+1^_6_y^+1^_6_y^+1^_6_b^+1^_7_b^+1^_7_y^+1^_7_b^+1^_7_y^+1^_7_y^+1^_8_y^+1^_8_b^+1^_8_y^+1^_8_b^+1^_8_b^+1^_9_b^+1^_9_b^+1^_9_y^+1^_9_

0779155923383118

Fragment Matches Table

Show background peaks

| Position | Ion type | Intensity | mz Theoretical | mz Error (Th) | mz Error (ppm) | Charge | Series Number |
| --- | --- | --- | --- | --- | --- | --- | --- |
| - | - | 4.104E+04 | 120.1 | - | - | 0 | - |
| - | - | 396.5 | 120.2 | - | - | 0 | - |
| - | - | 2546 | 121.1 | - | - | 0 | - |
| - | - | 336.6 | 122.4 | - | - | 0 | - |
| - | - | 464.9 | 125.1 | - | - | 0 | - |
| - | - | 690.5 | 126.1 | - | - | 0 | - |
| - | - | 386 | 126.7 | - | - | 0 | - |
| - | - | 487.5 | 127.1 | - | - | 0 | - |
| - | - | 1633 | 127.1 | - | - | 0 | - |
| - | - | 420.3 | 128.1 | - | - | 0 | - |
| - | - | 2982 | 128.1 | - | - | 0 | - |
| - | - | 8227 | 129.1 | - | - | 0 | - |
| - | - | 7.047E+04 | 129.1 | - | - | 0 | - |
| - | - | 451.4 | 130 | - | - | 0 | - |
| - | - | 755.9 | 130.1 | - | - | 0 | - |
| - | - | 4431 | 130.1 | - | - | 0 | - |
| - | - | 1057 | 131 | - | - | 0 | - |
| - | - | 2931 | 131.1 | - | - | 0 | - |
| - | - | 3947 | 131.1 | - | - | 0 | - |
| - | - | 886.1 | 132.1 | - | - | 0 | - |
| - | - | 1049 | 133.1 | - | - | 0 | - |
| - | - | 1211 | 133.1 | - | - | 0 | - |
| - | - | 3890 | 136.1 | - | - | 0 | - |
| - | - | 406.5 | 137.1 | - | - | 0 | - |
| - | - | 2.102E+04 | 138.1 | - | - | 0 | - |
| - | - | 388.9 | 139.1 | - | - | 0 | - |
| - | - | 1604 | 139.1 | - | - | 0 | - |
| - | - | 1656 | 139.1 | - | - | 0 | - |
| - | - | 465.5 | 140.1 | - | - | 0 | - |
| - | - | 1320 | 141.1 | - | - | 0 | - |
| - | - | 3647 | 141.1 | - | - | 0 | - |
| - | - | 589.4 | 148.9 | - | - | 0 | - |
| - | - | 689.3 | 149.1 | - | - | 0 | - |
| - | - | 471.3 | 150.1 | - | - | 0 | - |
| - | - | 3089 | 151.1 | - | - | 0 | - |
| - | - | 697.4 | 153.1 | - | - | 0 | - |
| - | - | 1240 | 154.1 | - | - | 0 | - |
| - | - | 1300 | 155.1 | - | - | 0 | - |
| - | - | 2.566E+04 | 155.1 | - | - | 0 | - |
| - | - | 496.9 | 156.1 | - | - | 0 | - |
| - | - | 1783 | 156.1 | - | - | 0 | - |
| - | - | 1756 | 156.1 | - | - | 0 | - |
| - | - | 507.3 | 157.1 | - | - | 0 | - |
| - | - | 628.2 | 157.1 | - | - | 0 | - |
| - | - | 522.8 | 157.1 | - | - | 0 | - |
| - | - | 1.081E+04 | 159.1 | - | - | 0 | - |
| - | - | 502 | 159.1 | - | - | 0 | - |
| - | - | 1190 | 159.1 | - | - | 0 | - |
| - | - | 939.1 | 165.1 | - | - | 0 | - |
| - | - | 1254 | 165.1 | - | - | 0 | - |
| 10 | y | 5.294E+04 | 166.1 | 0.0001402 | 0.8442 | +1 | 1 |
| - | - | 991.3 | 167.1 | - | - | 0 | - |
| - | - | 5522 | 167.1 | - | - | 0 | - |
| - | - | 4852 | 167.1 | - | - | 0 | - |
| - | - | 3474 | 168.1 | - | - | 0 | - |
| - | - | 1718 | 169.1 | - | - | 0 | - |
| - | - | 722.1 | 169.1 | - | - | 0 | - |
| - | - | 581 | 171.1 | - | - | 0 | - |
| - | - | 7.195E+04 | 173.1 | - | - | 0 | - |
| - | - | 1540 | 173.5 | - | - | 0 | - |
| - | - | 781.7 | 174.1 | - | - | 0 | - |
| - | - | 5643 | 174.1 | - | - | 0 | - |
| - | - | 1226 | 175.1 | - | - | 0 | - |
| - | - | 442.3 | 177.1 | - | - | 0 | - |
| - | - | 1.061E+04 | 177.1 | - | - | 0 | - |
| - | - | 380.6 | 177.1 | - | - | 0 | - |
| - | - | 1058 | 178.1 | - | - | 0 | - |
| - | - | 3076 | 181.1 | - | - | 0 | - |
| - | - | 1578 | 182.1 | - | - | 0 | - |
| - | - | 8.129E+04 | 182.1 | - | - | 0 | - |
| 2 | a | 1.199E+05 | 183.1 | 0.0001261 | 0.6889 | +1 | 2 |
| - | - | 8123 | 183.1 | - | - | 0 | - |
| - | - | 587.3 | 184.1 | - | - | 0 | - |
| - | - | 2056 | 184.1 | - | - | 0 | - |
| - | - | 1.093E+04 | 184.1 | - | - | 0 | - |
| - | - | 607.8 | 184.1 | - | - | 0 | - |
| - | - | 3762 | 185.1 | - | - | 0 | - |
| 2 | d | 790.3 | 186.1 | 6.084E-05 | 0.3269 | +1 | 2 |
| - | - | 4974 | 187.1 | - | - | 0 | - |
| - | - | 1530 | 187.1 | - | - | 0 | - |
| - | - | 1246 | 188.1 | - | - | 0 | - |
| - | - | 4912 | 194.1 | - | - | 0 | - |
| - | - | 1283 | 195.1 | - | - | 0 | - |
| - | - | 553.8 | 196.1 | - | - | 0 | - |
| - | - | 757.1 | 197.1 | - | - | 0 | - |
| - | - | 1335 | 197.1 | - | - | 0 | - |
| - | - | 4930 | 198.1 | - | - | 0 | - |
| - | - | 1702 | 198.1 | - | - | 0 | - |
| - | - | 2659 | 199.1 | - | - | 0 | - |
| - | - | 1050 | 199.1 | - | - | 0 | - |
| 2 | a | 1.269E+05 | 200.1 | 9.681E-05 | 0.4837 | +1 | 2 |
| - | - | 4.622E+04 | 201.1 | - | - | 0 | - |
| - | - | 1.184E+04 | 201.1 | - | - | 0 | - |
| - | - | 3252 | 202.1 | - | - | 0 | - |
| - | - | 2099 | 205.1 | - | - | 0 | - |
| - | - | 830 | 208.1 | - | - | 0 | - |
| - | - | 830.8 | 208.1 | - | - | 0 | - |
| - | - | 982.6 | 209.1 | - | - | 0 | - |
| - | - | 8630 | 209.1 | - | - | 0 | - |
| - | - | 3.363E+04 | 210.1 | - | - | 0 | - |
| 2 | b | 2.253E+04 | 211.1 | 2.353E-05 | 0.1115 | +1 | 2 |
| - | - | 4122 | 211.1 | - | - | 0 | - |
| - | - | 560.6 | 211.1 | - | - | 0 | - |
| - | - | 927.8 | 211.1 | - | - | 0 | - |
| - | - | 1537 | 212.1 | - | - | 0 | - |
| - | - | 3196 | 212.1 | - | - | 0 | - |
| - | - | 851.3 | 213.1 | - | - | 0 | - |
| - | - | 625.7 | 213.1 | - | - | 0 | - |
| - | - | 853.7 | 215.1 | - | - | 0 | - |
| - | - | 2425 | 216.1 | - | - | 0 | - |
| - | - | 1995 | 218.1 | - | - | 0 | - |
| - | - | 1397 | 222.1 | - | - | 0 | - |
| 9 | y | 3.62E+04 | 223.1 | 8.457E-05 | 0.379 | +1 | 2 |
| - | - | 3506 | 224.1 | - | - | 0 | - |
| - | - | 3258 | 225.1 | - | - | 0 | - |
| - | - | 1.146E+04 | 226.1 | - | - | 0 | - |
| - | - | 2.798E+04 | 226.1 | - | - | 0 | - |
| - | - | 7928 | 227.1 | - | - | 0 | - |
| - | - | 1203 | 227.1 | - | - | 0 | - |
| - | - | 1807 | 227.1 | - | - | 0 | - |
| - | - | 2989 | 227.1 | - | - | 0 | - |
| 2 | b | 4.448E+04 | 228.1 | 8.576E-05 | 0.3759 | +1 | 2 |
| - | - | 5955 | 229.1 | - | - | 0 | - |
| - | - | 4720 | 229.1 | - | - | 0 | - |
| - | - | 3119 | 230.1 | - | - | 0 | - |
| - | - | 1196 | 234.1 | - | - | 0 | - |
| - | - | 695.6 | 238.1 | - | - | 0 | - |
| - | - | 1350 | 239.2 | - | - | 0 | - |
| - | - | 3000 | 240.1 | - | - | 0 | - |
| - | - | 1.082E+04 | 242.1 | - | - | 0 | - |
| - | - | 404.3 | 242.2 | - | - | 0 | - |
| - | - | 1502 | 243.1 | - | - | 0 | - |
| - | - | 810.7 | 243.2 | - | - | 0 | - |
| - | - | 863.9 | 244.1 | - | - | 0 | - |
| - | - | 2.153E+04 | 244.1 | - | - | 0 | - |
| - | - | 2.502E+04 | 244.1 | - | - | 0 | - |
| - | - | 1922 | 245.1 | - | - | 0 | - |
| - | - | 2125 | 245.1 | - | - | 0 | - |
| - | - | 761.8 | 249.1 | - | - | 0 | - |
| - | - | 587.9 | 250.2 | - | - | 0 | - |
| - | - | 961.7 | 251.2 | - | - | 0 | - |
| - | - | 1669 | 252.1 | - | - | 0 | - |
| - | - | 925.7 | 253.1 | - | - | 0 | - |
| - | - | 1861 | 258.1 | - | - | 0 | - |
| - | - | 1157 | 261.1 | - | - | 0 | - |
| - | - | 646.8 | 262.1 | - | - | 0 | - |
| - | - | 905.8 | 265.1 | - | - | 0 | - |
| - | - | 1377 | 266.1 | - | - | 0 | - |
| - | - | 1526 | 267.1 | - | - | 0 | - |
| - | - | 694.6 | 268.1 | - | - | 0 | - |
| 3 | a | 1127 | 269.2 | 0.0005387 | 2.001 | +1 | 3 |
| 3 | a | 4346 | 270.1 | 0.0003013 | 1.115 | +1 | 3 |
| - | - | 1831 | 279.1 | - | - | 0 | - |
| - | - | 4316 | 279.1 | - | - | 0 | - |
| - | - | 7105 | 280.1 | - | - | 0 | - |
| - | - | 952.8 | 280.1 | - | - | 0 | - |
| - | - | 728.1 | 281.1 | - | - | 0 | - |
| - | - | 2307 | 282.2 | - | - | 0 | - |
| - | - | 6682 | 283.1 | - | - | 0 | - |
| 6 | y | 1237 | 284.1 | 0.001478 | 5.203 | +2 | 5 |
| 3 | a | 2315 | 287.2 | 5.05E-06 | 0.01759 | +1 | 3 |
| - | - | 809.5 | 290.1 | - | - | 0 | - |
| - | - | 1053 | 291.1 | - | - | 0 | - |
| - | - | 1437 | 292.1 | - | - | 0 | - |
| - | - | 714.1 | 294.1 | - | - | 0 | - |
| 3 | b | 2.537E+04 | 297.2 | 4.032E-07 | 0.001357 | +1 | 3 |
| 3 | b | 2.337E+04 | 298.1 | 6.823E-05 | 0.2289 | +1 | 3 |
| - | - | 2992 | 298.2 | - | - | 0 | - |
| - | - | 3755 | 299.1 | - | - | 0 | - |
| - | - | 687.8 | 300.2 | - | - | 0 | - |
| - | - | 996.5 | 305.2 | - | - | 0 | - |
| 8 | y | 2815 | 306.1 | 0.0001792 | 0.5855 | +1 | 3 |
| - | - | 3937 | 309.2 | - | - | 0 | - |
| - | - | 1611 | 310.1 | - | - | 0 | - |
| - | - | 751 | 310.2 | - | - | 0 | - |
| - | - | 1661 | 311.2 | - | - | 0 | - |
| 3 | b | 1.083E+04 | 315.2 | 2.451E-05 | 0.07777 | +1 | 3 |
| - | - | 1749 | 316.1 | - | - | 0 | - |
| - | - | 1790 | 316.2 | - | - | 0 | - |
| - | - | 1319 | 317.2 | - | - | 0 | - |
| - | - | 2539 | 319.1 | - | - | 0 | - |
| 8 | y | 1.318E+04 | 324.2 | 5.929E-05 | 0.1829 | +1 | 3 |
| - | - | 1803 | 325.2 | - | - | 0 | - |
| 6 | b | 4052 | 327.2 | 0.0001281 | 0.3915 | +2 | 6 |
| - | - | 897.4 | 328.2 | - | - | 0 | - |
| - | - | 771.1 | 328.2 | - | - | 0 | - |
| - | - | 649.1 | 329.1 | - | - | 0 | - |
| - | - | 1.229E+04 | 329.2 | - | - | 0 | - |
| - | - | 1558 | 330.2 | - | - | 0 | - |
| 6 | b | 2653 | 336.2 | 0.005044 | 15 | +2 | 6 |
| - | - | 1.271E+04 | 337.2 | - | - | 0 | - |
| - | - | 728.3 | 337.2 | - | - | 0 | - |
| - | - | 2425 | 338.2 | - | - | 0 | - |
| 5 | y | 4830 | 339.2 | 5.503E-05 | 0.1622 | +2 | 6 |
| - | - | 1030 | 339.7 | - | - | 0 | - |
| - | - | 1674 | 340.2 | - | - | 0 | - |
| - | - | 9956 | 345.2 | - | - | 0 | - |
| - | - | 1934 | 346.2 | - | - | 0 | - |
| 5 | y | 7203 | 348.2 | 0.0005581 | 1.603 | +2 | 6 |
| - | - | 2633 | 348.7 | - | - | 0 | - |
| - | - | 1.737E+04 | 354.2 | - | - | 0 | - |
| - | - | 6939 | 355.2 | - | - | 0 | - |
| - | - | 1788 | 355.2 | - | - | 0 | - |
| - | - | 978.8 | 356.2 | - | - | 0 | - |
| - | - | 6362 | 357.2 | - | - | 0 | - |
| - | - | 2782 | 364.2 | - | - | 0 | - |
| - | - | 1.397E+04 | 365.2 | - | - | 0 | - |
| - | - | 782.1 | 366.2 | - | - | 0 | - |
| - | - | 2919 | 366.2 | - | - | 0 | - |
| - | - | 1426 | 367.2 | - | - | 0 | - |
| - | - | 987.9 | 371.2 | - | - | 0 | - |
| - | - | 1.43E+04 | 372.2 | - | - | 0 | - |
| - | - | 2411 | 373.2 | - | - | 0 | - |
| - | - | 741.3 | 376.2 | - | - | 0 | - |
| - | - | 965.9 | 380.2 | - | - | 0 | - |
| - | - | 602.3 | 382.2 | - | - | 0 | - |
| - | - | 3460 | 382.2 | - | - | 0 | - |
| - | - | 1.682E+04 | 383.2 | - | - | 0 | - |
| - | - | 1486 | 384.2 | - | - | 0 | - |
| - | - | 2418 | 384.2 | - | - | 0 | - |
| - | - | 6593 | 392.2 | - | - | 0 | - |
| - | - | 1619 | 393.2 | - | - | 0 | - |
| - | - | 1.364E+04 | 393.2 | - | - | 0 | - |
| - | - | 6968 | 394.2 | - | - | 0 | - |
| - | - | 2548 | 394.2 | - | - | 0 | - |
| - | - | 1258 | 395.2 | - | - | 0 | - |
| 4 | y | 1825 | 396.2 | 0.0005306 | 1.339 | +2 | 7 |
| - | - | 1344 | 396.7 | - | - | 0 | - |
| - | - | 967.4 | 400.3 | - | - | 0 | - |
| - | - | 3413 | 402.2 | - | - | 0 | - |
| 4 | y | 8715 | 404.7 | 0.0001707 | 0.4217 | +2 | 7 |
| - | - | 5706 | 405.2 | - | - | 0 | - |
| - | - | 1010 | 405.7 | - | - | 0 | - |
| - | - | 1130 | 410.2 | - | - | 0 | - |
| 4 | b | 1.078E+04 | 410.2 | 1.154E-05 | 0.02814 | +1 | 4 |
| - | - | 7865 | 410.7 | - | - | 0 | - |
| - | - | 1315 | 411.2 | - | - | 0 | - |
| 4 | b | 1.834E+04 | 411.2 | 0.0007277 | 1.77 | +1 | 4 |
| - | - | 1100 | 411.7 | - | - | 0 | - |
| - | - | 737.8 | 412.2 | - | - | 0 | - |
| - | - | 2650 | 412.2 | - | - | 0 | - |
| - | - | 2948 | 412.2 | - | - | 0 | - |
| - | - | 620.2 | 418.2 | - | - | 0 | - |
| - | - | 1213 | 419.2 | - | - | 0 | - |
| - | - | 594 | 419.7 | - | - | 0 | - |
| - | - | 2702 | 420.2 | - | - | 0 | - |
| - | - | 1656 | 423.7 | - | - | 0 | - |
| - | - | 1658 | 424.2 | - | - | 0 | - |
| - | - | 1075 | 426.2 | - | - | 0 | - |
| - | - | 1446 | 427.2 | - | - | 0 | - |
| - | - | 1066 | 427.2 | - | - | 0 | - |
| - | - | 2170 | 427.3 | - | - | 0 | - |
| - | - | 1772 | 427.7 | - | - | 0 | - |
| - | - | 753.9 | 428.2 | - | - | 0 | - |
| - | - | 1515 | 428.2 | - | - | 0 | - |
| 4 | b | 5550 | 428.3 | 0.001001 | 2.338 | +1 | 4 |
| - | - | 721 | 429.3 | - | - | 0 | - |
| - | - | 871.1 | 430.7 | - | - | 0 | - |
| - | - | 6253 | 432.7 | - | - | 0 | - |
| - | - | 2471 | 433.2 | - | - | 0 | - |
| - | - | 786.8 | 433.7 | - | - | 0 | - |
| 7 | y | 2957 | 434.2 | 1.897E-05 | 0.04369 | +1 | 4 |
| - | - | 1618 | 435.2 | - | - | 0 | - |
| - | - | 1239 | 435.7 | - | - | 0 | - |
| - | - | 2836 | 437.2 | - | - | 0 | - |
| - | - | 4346 | 438.2 | - | - | 0 | - |
| - | - | 1201 | 439.2 | - | - | 0 | - |
| 3 | y | 1110 | 439.2 | 0.002983 | 6.791 | +2 | 8 |
| 3 | y | 4704 | 439.7 | 0.00185 | 4.208 | +2 | 8 |
| - | - | 1759 | 440.2 | - | - | 0 | - |
| - | - | 935.9 | 440.7 | - | - | 0 | - |
| 8 | b | 7027 | 441.7 | 0.0001759 | 0.3983 | +2 | 8 |
| 8 | b | 4201 | 442.2 | 0.008504 | 19.23 | +2 | 8 |
| - | - | 874.6 | 442.7 | - | - | 0 | - |
| - | - | 1.32E+04 | 444.2 | - | - | 0 | - |
| - | - | 2246 | 445.2 | - | - | 0 | - |
| 3 | y | 8692 | 448.2 | 5.046E-05 | 0.1126 | +2 | 8 |
| - | - | 1173 | 448.3 | - | - | 0 | - |
| - | - | 4549 | 448.7 | - | - | 0 | - |
| - | - | 1028 | 450.2 | - | - | 0 | - |
| 7 | y | 3.126E+04 | 452.3 | 0.0002382 | 0.5267 | +1 | 4 |
| - | - | 5960 | 453.3 | - | - | 0 | - |
| - | - | 950.9 | 454.3 | - | - | 0 | - |
| - | - | 4332 | 455.2 | - | - | 0 | - |
| - | - | 8507 | 456.2 | - | - | 0 | - |
| - | - | 723.3 | 457.2 | - | - | 0 | - |
| - | - | 2846 | 457.2 | - | - | 0 | - |
| - | - | 2387 | 461.2 | - | - | 0 | - |
| - | - | 3516 | 461.7 | - | - | 0 | - |
| - | - | 1907 | 462.2 | - | - | 0 | - |
| - | - | 770.4 | 468.2 | - | - | 0 | - |
| 9 | b | 5310 | 470.2 | 0.0005462 | 1.161 | +2 | 9 |
| 9 | b | 2859 | 470.7 | 0.007965 | 16.92 | +2 | 9 |
| - | - | 1829 | 471.2 | - | - | 0 | - |
| - | - | 5580 | 473.2 | - | - | 0 | - |
| - | - | 1563 | 474.2 | - | - | 0 | - |
| 9 | b | 6067 | 479.3 | 0.0002133 | 0.445 | +2 | 9 |
| - | - | 2484 | 479.8 | - | - | 0 | - |
| - | - | 1589 | 480.3 | - | - | 0 | - |
| - | - | 1109 | 485.2 | - | - | 0 | - |
| - | - | 3212 | 485.3 | - | - | 0 | - |
| - | - | 690.4 | 486.2 | - | - | 0 | - |
| - | - | 1692 | 495.2 | - | - | 0 | - |
| 2 | y | 2064 | 497.8 | 0.001136 | 2.283 | +2 | 9 |
| - | - | 1364 | 498.3 | - | - | 0 | - |
| - | - | 1915 | 510.3 | - | - | 0 | - |
| - | - | 2221 | 512.2 | - | - | 0 | - |
| - | - | 4487 | 513.2 | - | - | 0 | - |
| - | - | 1659 | 514.2 | - | - | 0 | - |
| - | - | 983.6 | 521.3 | - | - | 0 | - |
| - | - | 1395 | 522.3 | - | - | 0 | - |
| - | - | 1050 | 522.7 | - | - | 0 | - |
| - | - | 873.6 | 522.8 | - | - | 0 | - |
| - | - | 1241 | 524.3 | - | - | 0 | - |
| - | - | 1093 | 529.8 | - | - | 0 | - |
| - | - | 4322 | 530.3 | - | - | 0 | - |
| - | - | 742.1 | 530.8 | - | - | 0 | - |
| - | - | 1641 | 531.3 | - | - | 0 | - |
| - | - | 1115 | 532.2 | - | - | 0 | - |
| - | - | 3895 | 534.3 | - | - | 0 | - |
| - | - | 1093 | 534.8 | - | - | 0 | - |
| - | - | 1138 | 535.3 | - | - | 0 | - |
| - | - | 941.1 | 535.3 | - | - | 0 | - |
| - | - | 1313 | 535.8 | - | - | 0 | - |
| 5 | b | 1.022E+04 | 538.3 | 0.0005826 | 1.082 | +1 | 5 |
| 5 | b | 5171 | 539.3 | 0.003317 | 6.151 | +1 | 5 |
| - | - | 802.1 | 542.7 | - | - | 0 | - |
| - | - | 2680 | 543.3 | - | - | 0 | - |
| - | - | 1.194E+04 | 543.8 | - | - | 0 | - |
| - | - | 9841 | 544.3 | - | - | 0 | - |
| - | - | 3893 | 544.8 | - | - | 0 | - |
| - | - | 1726 | 545.3 | - | - | 0 | - |
| - | - | 650.5 | 545.8 | - | - | 0 | - |
| 6 | y | 5594 | 549.3 | 4.605E-05 | 0.08384 | +1 | 5 |
| - | - | 1182 | 550.3 | - | - | 0 | - |
| - | - | 5973 | 551.7 | - | - | 0 | - |
| - | - | 3168 | 552.2 | - | - | 0 | - |
| 0 | Precursor | 1.818E+05 | 552.8 | 0.0006068 | 1.098 | +2 | -1 |
| 0 | Precursor | 1.104E+05 | 553.3 | 0.008423 | 15.22 | +2 | -1 |
| - | - | 4.116E+04 | 553.8 | - | - | 0 | - |
| - | - | 7517 | 554.3 | - | - | 0 | - |
| - | - | 1384 | 555.3 | - | - | 0 | - |
| 5 | b | 1809 | 556.3 | 0.001443 | 2.593 | +1 | 5 |
| - | - | 653.7 | 558.3 | - | - | 0 | - |
| 0 | Precursor | 2.671E+04 | 561.8 | 0.0008232 | 1.465 | +2 | -1 |
| - | - | 1.7E+04 | 562.3 | - | - | 0 | - |
| - | - | 687.4 | 562.7 | - | - | 0 | - |
| - | - | 5932 | 562.8 | - | - | 0 | - |
| - | - | 1098 | 563.3 | - | - | 0 | - |
| 6 | y | 7.441E+04 | 567.3 | 0.0005699 | 1.005 | +1 | 5 |
| - | - | 1.955E+04 | 568.3 | - | - | 0 | - |
| - | - | 3662 | 569.3 | - | - | 0 | - |
| - | - | 1.054E+04 | 572.3 | - | - | 0 | - |
| - | - | 2234 | 573.3 | - | - | 0 | - |
| - | - | 790.9 | 574.3 | - | - | 0 | - |
| - | - | 873.5 | 583.3 | - | - | 0 | - |
| - | - | 4819 | 586.3 | - | - | 0 | - |
| - | - | 1025 | 587.3 | - | - | 0 | - |
| - | - | 788.4 | 597.9 | - | - | 0 | - |
| - | - | 639.9 | 599.6 | - | - | 0 | - |
| - | - | 609.7 | 601.8 | - | - | 0 | - |
| - | - | 640.6 | 606.3 | - | - | 0 | - |
| - | - | 959.5 | 608.3 | - | - | 0 | - |
| - | - | 5617 | 611.3 | - | - | 0 | - |
| - | - | 1724 | 612.3 | - | - | 0 | - |
| - | - | 977 | 625.3 | - | - | 0 | - |
| - | - | 623.4 | 626.3 | - | - | 0 | - |
| - | - | 1242 | 629.3 | - | - | 0 | - |
| - | - | 647.4 | 632.6 | - | - | 0 | - |
| - | - | 912.2 | 632.9 | - | - | 0 | - |
| - | - | 2167 | 636.3 | - | - | 0 | - |
| - | - | 5201 | 637.3 | - | - | 0 | - |
| - | - | 1724 | 638.3 | - | - | 0 | - |
| - | - | 2043 | 641.9 | - | - | 0 | - |
| - | - | 899.5 | 642.3 | - | - | 0 | - |
| - | - | 1981 | 642.4 | - | - | 0 | - |
| - | - | 1685 | 643.3 | - | - | 0 | - |
| - | - | 1700 | 645.4 | - | - | 0 | - |
| - | - | 6678 | 649.3 | - | - | 0 | - |
| - | - | 2437 | 650.3 | - | - | 0 | - |
| 6 | b | 1.443E+04 | 653.3 | 0.001097 | 1.68 | +1 | 6 |
| 6 | b | 5587 | 654.3 | 0.01068 | 16.32 | +1 | 6 |
| - | - | 5192 | 655.3 | - | - | 0 | - |
| - | - | 2244 | 656.3 | - | - | 0 | - |
| - | - | 3083 | 659.3 | - | - | 0 | - |
| - | - | 3840 | 660.3 | - | - | 0 | - |
| - | - | 1728 | 661.3 | - | - | 0 | - |
| - | - | 4604 | 670.4 | - | - | 0 | - |
| 6 | b | 1826 | 671.3 | 0.007686 | 11.45 | +1 | 6 |
| - | - | 1.971E+04 | 673.4 | - | - | 0 | - |
| - | - | 7843 | 674.4 | - | - | 0 | - |
| - | - | 1022 | 675.4 | - | - | 0 | - |
| 5 | y | 1.928E+04 | 677.3 | 0.001036 | 1.53 | +1 | 6 |
| 5 | y | 4.55E+04 | 678.3 | 0.0005438 | 0.8017 | +1 | 6 |
| - | - | 1.616E+04 | 679.3 | - | - | 0 | - |
| - | - | 2247 | 680.3 | - | - | 0 | - |
| - | - | 940.6 | 686.3 | - | - | 0 | - |
| 5 | y | 6.21E+04 | 695.3 | 0.0009199 | 1.323 | +1 | 6 |
| - | - | 2.59E+04 | 696.3 | - | - | 0 | - |
| - | - | 4744 | 697.3 | - | - | 0 | - |
| - | - | 1497 | 711.9 | - | - | 0 | - |
| - | - | 3214 | 712.4 | - | - | 0 | - |
| - | - | 1215 | 713.4 | - | - | 0 | - |
| - | - | 680 | 713.5 | - | - | 0 | - |
| - | - | 8230 | 730.4 | - | - | 0 | - |
| - | - | 2633 | 731.4 | - | - | 0 | - |
| - | - | 1662 | 748.4 | - | - | 0 | - |
| - | - | 1024 | 755.4 | - | - | 0 | - |
| - | - | 674.7 | 763.4 | - | - | 0 | - |
| - | - | 891.2 | 764.4 | - | - | 0 | - |
| - | - | 684.9 | 765.4 | - | - | 0 | - |
| - | - | 964.5 | 772.4 | - | - | 0 | - |
| - | - | 1807 | 773.4 | - | - | 0 | - |
| - | - | 898.4 | 774.4 | - | - | 0 | - |
| 7 | b | 1.024E+04 | 781.4 | 0.001029 | 1.316 | +1 | 7 |
| 7 | b | 5040 | 782.4 | 0.01392 | 17.79 | +1 | 7 |
| - | - | 1067 | 783.4 | - | - | 0 | - |
| 4 | y | 1171 | 790.4 | 0.005572 | 7.049 | +1 | 7 |
| 7 | b | 1455 | 799.4 | 0.001889 | 2.363 | +1 | 7 |
| 4 | y | 1.979E+04 | 808.4 | 0.001366 | 1.689 | +1 | 7 |
| - | - | 8573 | 809.4 | - | - | 0 | - |
| - | - | 2836 | 810.4 | - | - | 0 | - |
| - | - | 2545 | 821.4 | - | - | 0 | - |
| - | - | 1216 | 822.4 | - | - | 0 | - |
| - | - | 1034 | 839.4 | - | - | 0 | - |
| - | - | 1592 | 846.4 | - | - | 0 | - |
| - | - | 1716 | 847.4 | - | - | 0 | - |
| - | - | 784.9 | 848.4 | - | - | 0 | - |
| - | - | 1053 | 849.4 | - | - | 0 | - |
| - | - | 786.9 | 855.5 | - | - | 0 | - |
| - | - | 1544 | 859.4 | - | - | 0 | - |
| - | - | 908.8 | 860.4 | - | - | 0 | - |
| - | - | 663.4 | 864.5 | - | - | 0 | - |
| - | - | 4030 | 865.4 | - | - | 0 | - |
| - | - | 1471 | 866.4 | - | - | 0 | - |
| 3 | y | 6013 | 877.4 | 0.002627 | 2.994 | +1 | 8 |
| 3 | y | 4767 | 878.4 | 0.008048 | 9.161 | +1 | 8 |
| - | - | 1627 | 879.4 | - | - | 0 | - |
| 8 | b | 1.114E+04 | 883.5 | 0.0006182 | 0.6998 | +1 | 8 |
| - | - | 4808 | 884.5 | - | - | 0 | - |
| 3 | y | 1.131E+05 | 895.5 | 0.002266 | 2.531 | +1 | 8 |
| - | - | 5.309E+04 | 896.5 | - | - | 0 | - |
| - | - | 1.379E+04 | 897.5 | - | - | 0 | - |
| - | - | 772.3 | 898.5 | - | - | 0 | - |
| 8 | b | 2919 | 900.5 | 0.002265 | 2.515 | +1 | 8 |
| - | - | 1761 | 901.5 | - | - | 0 | - |
| - | - | 718.6 | 902.5 | - | - | 0 | - |
| - | - | 2722 | 922.5 | - | - | 0 | - |
| - | - | 2897 | 923.5 | - | - | 0 | - |
| - | - | 937.1 | 924.5 | - | - | 0 | - |
| 9 | b | 1832 | 939.5 | 0.006206 | 6.606 | +1 | 9 |
| 9 | b | 1.223E+04 | 940.5 | 0.0009028 | 0.9599 | +1 | 9 |
| - | - | 7245 | 941.5 | - | - | 0 | - |
| - | - | 1914 | 942.5 | - | - | 0 | - |
| 9 | b | 3474 | 957.5 | 0.004564 | 4.766 | +1 | 9 |
| - | - | 2177 | 958.5 | - | - | 0 | - |
| - | - | 987.1 | 959.5 | - | - | 0 | - |
| 2 | y | 6405 | 994.5 | 0.002199 | 2.211 | +1 | 9 |
| - | - | 2569 | 995.5 | - | - | 0 | - |
| - | - | 647 | 1712 | - | - | 0 | - |
| - | - | 828.6 | 2886 | - | - | 0 | - |
| - | - | 830.6 | 3087 | - | - | 0 | - |

m/z Charge Intensity FragmentType MassShift Position
120.08097839355469 0 41038.445
120.2103500366211 0 396.5366
121.08429718017578 0 2545.8594
122.44593811035156 0 336.64847
125.10733795166016 0 464.9029
126.09142303466797 0 690.4737
126.6981430053711 0 386.02765
127.08714294433594 0 487.4773
127.12320709228516 0 1632.7842
128.07066345214844 0 420.29025
128.107177734375 0 2981.941
129.06602478027344 0 8227.337
129.10243225097656 0 70467.89
130.04995727539062 0 451.3657
130.10009765625 0 755.8686
130.10572814941406 0 4431.445
131.04920959472656 0 1056.7667
131.0816650390625 0 2931.277
131.11807250976562 0 3946.649
132.10227966308594 0 886.0639
133.0610809326172 0 1049.0758
133.0861053466797 0 1210.8982
136.07582092285156 0 3890.4546
137.07919311523438 0 406.45377
138.0915069580078 0 21021.377
139.05010986328125 0 388.90808
139.0867462158203 0 1604.0583
139.09487915039062 0 1655.6877
140.1072998046875 0 465.4892
141.06600952148438 0 1320.294
141.10238647460938 0 3646.8752
148.94747924804688 0 589.4145
149.0599822998047 0 689.31494
150.0911865234375 0 471.32346
151.08689880371094 0 3089.4138
153.10208129882812 0 697.3817
154.09774780273438 0 1239.6897
155.08172607421875 0 1300.1377
155.11802673339844 0 25658.852
156.07643127441406 0 496.8988
156.10206604003906 0 1782.9536
156.12124633789062 0 1756.183
157.06044006347656 0 507.3002
157.0972137451172 0 628.15985
157.13352966308594 0 522.84753
159.0765380859375 0 10808.762
159.09144592285156 0 501.9635
159.11282348632812 0 1190.2192
165.05474853515625 0 939.0559
165.10256958007812 0 1254.0507
166.08639526367188 0 52943.55 y 9
167.08262634277344 0 991.3103
167.08984375 0 5521.7534
167.11793518066406 0 4852.2515
168.1133575439453 0 3473.6777
169.09722900390625 0 1717.9204
169.13368225097656 0 722.1104
171.11305236816406 0 581.0282
173.12857055664062 0 71953.25
173.4518280029297 0 1539.8889
174.0552978515625 0 781.65533
174.1319580078125 0 5643.146
175.08645629882812 0 1225.7274
177.08798217773438 0 442.33978
177.1023406982422 0 10614.82
177.11093139648438 0 380.5701
178.10577392578125 0 1057.578
181.06094360351562 0 3076.15
182.08155822753906 0 1577.8917
182.12892150878906 0 81292.13
183.11293029785156 0 119906.61 a Ammonia loss 1
183.13233947753906 0 8122.538
184.09767150878906 0 587.2814
184.1081085205078 0 2055.766
184.11631774902344 0 10932.106
184.1442108154297 0 607.771
185.12860107421875 0 3762.356
186.12376403808594 0 790.3274 d 1
187.10791015625 0 4974.438
187.14447021484375 0 1530.0243
188.07054138183594 0 1246.3453
194.12887573242188 0 4911.7285
195.11265563964844 0 1283.3336
196.10787963867188 0 553.83105
197.09152221679688 0 757.1113
197.1282501220703 0 1334.7208
198.0873260498047 0 4929.6274
198.1236572265625 0 1702.0803
199.07131958007812 0 2658.6716
199.10772705078125 0 1049.8236
200.1394500732422 0 126861.54 a 1
201.12342834472656 0 46221.113
201.14271545410156 0 11842.241
202.1268768310547 0 3252.023
205.09707641601562 0 2098.8723
208.09654235839844 0 829.95667
208.10763549804688 0 830.7584
209.055419921875 0 982.5759
209.09214782714844 0 8630.13
210.123779296875 0 33627.914
211.1077423095703 0 22529.572 b Ammonia loss 1
211.12693786621094 0 4121.6553
211.1375732421875 0 560.6197
211.14390563964844 0 927.80255
212.11099243164062 0 1537.4303
212.13929748535156 0 3195.5688
213.12376403808594 0 851.336
213.14283752441406 0 625.7055
215.13943481445312 0 853.71436
216.09796142578125 0 2424.922
218.14996337890625 0 1994.585
222.12376403808594 0 1397.0941
223.10780334472656 0 36204.92 y 8
224.1109161376953 0 3506.2437
225.12353515625 0 3258.3018
226.0823211669922 0 11455.077
226.11865234375 0 27976.467
227.06631469726562 0 7928.3613
227.0862579345703 0 1202.9406
227.10240173339844 0 1806.512
227.12222290039062 0 2988.6953
228.1343536376953 0 44484.465 b 1
229.1183624267578 0 5955.429
229.13772583007812 0 4719.906
230.14991760253906 0 3118.8203
234.12326049804688 0 1196.4703
238.1185302734375 0 695.58
239.1503143310547 0 1349.6937
240.13421630859375 0 3000.2852
242.14993286132812 0 10816.553
242.1627197265625 0 404.32056
243.1089630126953 0 1501.7147
243.15406799316406 0 810.68854
244.0786895751953 0 863.9011
244.0928497314453 0 21530.34
244.1291961669922 0 25015.398
245.09642028808594 0 1922.3788
245.13233947753906 0 2125.0686
249.12271118164062 0 761.82587
250.15505981445312 0 587.9262
251.15016174316406 0 961.74207
252.13436889648438 0 1669.4584
253.117431640625 0 925.70636
258.1446533203125 0 1860.5697
261.1197204589844 0 1157.3787
262.1193542480469 0 646.75867
265.1296081542969 0 905.83167
266.113525390625 0 1377.0131
267.1455993652344 0 1526.2252
268.1298828125 0 694.56177
269.1602783203125 0 1127.2013 a Water loss 2
270.14453125 0 4346.149 a Ammonia loss 2
279.1323547363281 0 1831.3037
279.1454162597656 0 4315.6143
280.1292724609375 0 7104.7407
280.1464538574219 0 952.8187
281.1328125 0 728.1461
282.181396484375 0 2306.9463
283.1401062011719 0 6682.4175
284.1437683105469 0 1236.5116 y 5
287.17138671875 0 2315.4119 a 2
290.1144104003906 0 809.5412
291.14422607421875 0 1053.4417
292.1297912597656 0 1437.3903
294.1440124511719 0 714.1205
297.1557312011719 0 25371.709 b Water loss 2
298.1396789550781 0 23366.285 b Ammonia loss 2
298.15850830078125 0 2992.4077
299.1424865722656 0 3754.7898
300.154541015625 0 687.7924
305.1812438964844 0 996.5039
306.1446533203125 0 2815.0593 y Water loss 7
309.15570068359375 0 3937.3186
310.13983154296875 0 1611.0183
310.1593017578125 0 750.95215
311.17181396484375 0 1661.4764
315.16632080078125 0 10831.883 b 2
316.1497497558594 0 1748.7477
316.1701965332031 0 1789.6431
317.182861328125 0 1319.1719
319.1396484375 0 2538.6482
324.15545654296875 0 13182.594 y 7
325.1580505371094 0 1802.9891
327.1661682128906 0 4051.6052 b Water loss 5
328.150390625 0 897.3728
328.1704406738281 0 771.09216
329.1488037109375 0 649.05225
329.1819763183594 0 12287.9
330.18414306640625 0 1558.4523
336.1665344238281 0 2653.3438 b 5
337.15057373046875 0 12712.938
337.17041015625 0 728.2784
338.15325927734375 0 2425.1736
339.1663513183594 0 4830.005 y Water loss 4
339.6659851074219 0 1030.3231
340.1505126953125 0 1673.8475
345.1768493652344 0 9955.713
346.18011474609375 0 1934.419
348.1710205078125 0 7202.919 y 4
348.6725769042969 0 2632.5715
354.17694091796875 0 17369.629
355.1608581542969 0 6938.9917
355.18084716796875 0 1788.1997
356.1636657714844 0 978.79205
357.1764831542969 0 6361.6387
364.23382568359375 0 2781.602
365.21807861328125 0 13965.454
366.1992492675781 0 782.1233
366.2218933105469 0 2919.3003
367.1613464355469 0 1426.357
371.2031555175781 0 987.8982
372.18743896484375 0 14300.174
373.1896057128906 0 2410.5557
376.16021728515625 0 741.273
380.2305603027344 0 965.9452
382.2179870605469 0 602.3377
382.2444763183594 0 3459.888
383.2284240722656 0 16818.748
384.1870422363281 0 1485.9097
384.2320861816406 0 2418.335
392.2288513183594 0 6593.1367
393.1859436035156 0 1618.9453
393.21319580078125 0 13641.6045
394.1718444824219 0 6967.7812
394.21539306640625 0 2548.1782
395.17431640625 0 1258.3843
396.20086669921875 0 1824.9933 y Ammonia loss 3
396.70263671875 0 1343.5789
400.2568359375 0 967.4422
402.1995544433594 0 3412.9585
404.71343994140625 0 8714.714 y 3
405.2147216796875 0 5705.783
405.7156066894531 0 1010.01465
410.2086181640625 0 1129.9714
410.23980712890625 0 10784.21 b Water loss 3
410.7190856933594 0 7865.0176
411.2019348144531 0 1315.4532
411.22308349609375 0 18338.975 b Ammonia loss 3
411.72314453125 0 1100.0676
412.1548767089844 0 737.7971
412.1830749511719 0 2650.3179
412.2254943847656 0 2947.9307
418.2087707519531 0 620.2171
419.2038269042969 0 1212.5603
419.72369384765625 0 593.9559
420.1867980957031 0 2701.8337
423.7284851074219 0 1656.446
424.2253112792969 0 1658.4396
426.19677734375 0 1075.3597
427.1832580566406 0 1445.8088
427.23248291015625 0 1066.1451
427.267333984375 0 2170.0942
427.74072265625 0 1771.9629
428.1597595214844 0 753.8719
428.2145080566406 0 1515.4805
428.2493591308594 0 5549.6553 b 3
429.2550354003906 0 721.0443
430.7116394042969 0 871.0982
432.732177734375 0 6252.8223
433.2279052734375 0 2470.984
433.7276916503906 0 786.7516
434.2397766113281 0 2957.295 y Water loss 6
435.23406982421875 0 1618.1213
435.7265930175781 0 1238.6516
437.2144775390625 0 2835.5244
438.1982116699219 0 4346.239
439.1971130371094 0 1201.2499
439.2273254394531 0 1109.6499 y Water loss 2
439.71820068359375 0 4704.097 y Ammonia loss 2
440.2191467285156 0 1759.0669
440.7184143066406 0 935.93427
441.73779296875 0 7027.174 b Water loss 7
442.2381286621094 0 4200.656 b Ammonia loss 7
442.73699951171875 0 874.56854
444.2088928222656 0 13196.139
445.2115783691406 0 2246.0684
448.22967529296875 0 8692.436 y 2
448.26275634765625 0 1173.3237
448.7314147949219 0 4548.717
450.234375 0 1028.0715
452.2501220703125 0 31255.883 y 6
453.25341796875 0 5959.5073
454.2578430175781 0 950.90906
455.22418212890625 0 4331.6406
456.20904541015625 0 8506.9
457.17578125 0 723.27435
457.2107849121094 0 2846.4155
461.24139404296875 0 2387.2964
461.73870849609375 0 3515.5913
462.23736572265625 0 1906.5619
468.2466735839844 0 770.43176
470.247802734375 0 5309.8926 b Water loss 8
470.7483215332031 0 2859.0166 b Ammonia loss 8
471.24517822265625 0 1828.7953
473.23529052734375 0 5580.131
474.23712158203125 0 1563.0062
479.25341796875 0 6067.3984 b 8
479.754638671875 0 2483.793
480.25421142578125 0 1589.07
485.2367858886719 0 1109.2106
485.2726745605469 0 3212.351
486.2349548339844 0 690.4086
495.2188720703125 0 1691.8153
497.7626953125 0 2063.809 y 1
498.2630920410156 0 1364.4346
510.3026428222656 0 1914.864
512.2469482421875 0 2221.2979
513.2303466796875 0 4486.9634
514.232666015625 0 1658.6049
521.2692260742188 0 983.59875
522.2684326171875 0 1395.0417
522.685791015625 0 1050.2794
522.76123046875 0 873.5921
524.2820434570312 0 1241.4414
529.7791748046875 0 1093.2136
530.256591796875 0 4322.2217
530.7755126953125 0 742.1039
531.2578735351562 0 1641.293
532.2430419921875 0 1115.1705
534.3027954101562 0 3894.5703
534.7785034179688 0 1092.8721
535.268798828125 0 1138.4233
535.310791015625 0 941.09534
535.7735595703125 0 1312.8723
538.2977905273438 0 10224.109 b Water loss 4
539.2857055664062 0 5170.852 b Ammonia loss 4
542.6848754882812 0 802.1213
543.301025390625 0 2680.4148
543.7821044921875 0 11936.649
544.2793579101562 0 9841.461
544.7776489257812 0 3893.2344
545.274169921875 0 1725.5457
545.763916015625 0 650.4544
549.2667846679688 0 5593.522 y Water loss 5
550.2625732421875 0 1181.7993
551.6881713867188 0 5972.7124
552.18994140625 0 3168.1484
552.7872314453125 0 181823.92 Precursor Water loss
553.2882690429688 0 110375.19 Precursor Ammonia loss
553.7896118164062 0 41158.77
554.2911376953125 0 7517.067
555.27978515625 0 1383.7473
556.3074951171875 0 1809.1404 b 4
558.2523193359375 0 653.66956
561.7922973632812 0 26708.635 Precursor
562.2935791015625 0 16999.871
562.6538696289062 0 687.37494
562.7953491210938 0 5931.7935
563.291259765625 0 1097.8713
567.2767333984375 0 74414.664 y 5
568.2797241210938 0 19552.523
569.281982421875 0 3661.7593
572.3033447265625 0 10535.349
573.3079833984375 0 2233.9275
574.3104858398438 0 790.9325
583.2706909179688 0 873.48425
586.3189697265625 0 4818.724
587.3223266601562 0 1024.902
597.9470825195312 0 788.44244
599.5799560546875 0 639.92487
601.78564453125 0 609.6987
606.2868041992188 0 640.6317
608.30322265625 0 959.5062
611.314208984375 0 5617.227
612.3182983398438 0 1723.8367
625.3302001953125 0 976.9676
626.3251342773438 0 623.4038
629.325439453125 0 1241.7179
632.6143798828125 0 647.42126
632.8663940429688 0 912.1833
636.3001098632812 0 2167.2134
637.3289794921875 0 5201.285
638.3262939453125 0 1723.5425
641.8541870117188 0 2042.5955
642.2920532226562 0 899.5413
642.357177734375 0 1980.6064
643.3419189453125 0 1685.189
645.3558959960938 0 1700.1968
649.3297119140625 0 6678.37
650.3311767578125 0 2437.3857
653.32421875 0 14425.559 b Water loss 5
654.3200073242188 0 5587.4243 b Ammonia loss 5
655.338134765625 0 5191.5347
656.3346557617188 0 2244.235
659.3135375976562 0 3083.4282
660.3005981445312 0 3839.6284
661.3001708984375 0 1728.4329
670.3515014648438 0 4603.775
671.3435668945312 0 1825.7509 b 5
673.3505859375 0 19708.37
674.3535766601562 0 7842.8726
675.357177734375 0 1022.2504
677.3242797851562 0 19278.318 y Water loss 4
678.3098754882812 0 45498.367 y Ammonia loss 4
679.31201171875 0 16159.849
680.3118286132812 0 2246.8218
686.3427124023438 0 940.63184
695.3349609375 0 62097.043 y 4
696.337890625 0 25898.602
697.3392333984375 0 4744.2227
711.8506469726562 0 1497.1343
712.3591918945312 0 3213.9104
713.3576049804688 0 1214.9889
713.4730224609375 0 679.97363
730.3715209960938 0 8229.915
731.3758544921875 0 2633.3708
748.3770141601562 0 1661.7717
755.364501953125 0 1023.7949
763.412353515625 0 674.7345
764.3942260742188 0 891.2326
765.3888549804688 0 684.8922
772.414794921875 0 964.4954
773.3798217773438 0 1806.8992
774.3851318359375 0 898.4257
781.4192504882812 0 10238.523 b Water loss 6
782.418212890625 0 5039.8384 b Ammonia loss 6
783.4216918945312 0 1067.3136
790.40380859375 0 1170.6033 y Water loss 3
799.428955078125 0 1455.1321 b 6
808.4185791015625 0 19791.602 y 3
809.4212646484375 0 8573.446
810.423583984375 0 2835.7085
821.4136352539062 0 2545.0464
822.4166259765625 0 1215.5386
839.4327392578125 0 1033.8188
846.4453735351562 0 1592.1633
847.4363403320312 0 1715.7866
848.4242553710938 0 784.8692
849.44580078125 0 1053.142
855.4554443359375 0 786.94336
859.4270629882812 0 1543.6753
860.4227905273438 0 908.7671
864.4523315429688 0 663.39496
865.4408569335938 0 4030.0613
866.4392700195312 0 1471.382
877.4387817382812 0 6013.189 y Water loss 2
878.4334716796875 0 4766.9077 y Ammonia loss 2
879.4329223632812 0 1627.4672
883.4513549804688 0 11135.846 b Ammonia loss 7
884.4523315429688 0 4808.356
895.44970703125 0 113142.36 y 2
896.4525146484375 0 53093.6
897.4553833007812 0 13793.896
898.458984375 0 772.3009
900.4762573242188 0 2918.853 b 7
901.47314453125 0 1761.4036
902.4700317382812 0 718.59875
922.4625244140625 0 2721.7998
923.4553833007812 0 2897.2053
924.4518432617188 0 937.1225
939.4832153320312 0 1832.1586 b Water loss 8
940.4725341796875 0 12226.186 b Ammonia loss 8
941.474609375 0 7245.441
942.4803466796875 0 1913.9521
957.4954223632812 0 3474.07 b 8
958.4896850585938 0 2177.2788
959.5023193359375 0 987.10864
994.5181884765625 0 6404.813 y 1
995.52001953125 0 2569.4287
1711.5938720703125 0 647.03876
2885.630859375 0 828.56195
3086.826904296875 0 830.62335

Spectrum Details

|  |  |
| --- | --- |
| Matched peaks? Matched peaks The total absolute number of peaks matched. Additionally in brackets the total fraction of peaks matched and the total number of peaks is shown. | 64 (13.85% of 462) |
| FDR? FDR The false discovery rate estimated for this peptide. It is calculated by matching all theoretical fragments with a non-integer shift with the raw peaks for this spectrum. This is done with 40 different shifts. The resulting percentage is the average number of annotated peaks over the number of annotated peaks with the correct spectrum. | 0.11% |
| Satellite FDR? Satellite FDR See the FDR for details on its calculation. This satellite ion specific FDR only contains the satellite ions (d/w) for I/L/J positions. | - |
| PSM Score? PSM Score The PSM Score as given by Hecklib to this annotated spectrum. It is shown with three significant figures. | 784 |

#### Spectrum 5387? Spectrum 5387 The raw spectrum of this peptide as annotated by Hecklib. The fragments are coloured according to ion type (see legend). Any peaks with a star '\*' as text can be hovered over to see the full details, first the ion type second the mass shift type. By hovering over the amino acids in the peptide or ions in the legend the corresponding peaks are highlighted. By toggling the 'Unassigned' label you can turn the background (unassigned) peaks on or off in the plot. By updating the slider in the Ion legend you can update the spectrum to only show the top X% of the peaks with labels. The top X% means any peak that is within X% of the highest intensity. By dragging in the spectrum you can zoom in to a specific part of the spectrum and use 'Zoom Out' to get back to the original zoom level. The annotation of the spectrum is based on the given sequence in the peptides file and is done with different software so inconsistencies are likely. The peaks are annotated based on the given sequence, with 20 ppm tolerance.

Copy Data

##### Spectrum 5387 (TSV)

###### Preview

```
Loading example...
```

*Click on the button to copy the data to your clipboard.*

Mz MinMz MaxIntensity Max

WidthHeightPeptide font sizePeptide stroke widthSpectrum font sizeSpectrum stroke widthCompact peptide

Ion legend

wxyz

abcd

OtherUnassignedIon^Charge^_Position_Show for top:%

QVSLQDKTGF

04.76e+69.51e+61.43e+71.90e+7

Zoom Out

y^+1^_1_a^+1^_2_d^+1^_2_a^+1^_2_b^+1^_2_y^+1^_2_b^+1^_2_a^+1^_3_a^+1^_3_y^+2^_5_a^+1^_3_b^+1^_3_b^+1^_3_y^+1^_3_b^+1^_3_y^+1^_3_b^+2^_6_b^+2^_6_y^+2^_6_y^+2^_6_y^+2^_6_b^+2^_7_b^+2^_7_y^+2^_7_y^+2^_7_y^+2^_7_b^+1^_4_b^+1^_4_b^+1^_4_y^+1^_4_y^+2^_8_y^+2^_8_b^+2^_8_y^+2^_8_b^+2^_8_y^+1^_4_b^+2^_9_b^+2^_9_b^+2^_9_y^+2^_9_b^+1^_5_b^+1^_5_y^+1^_5_\*\*b^+1^_5_\*y^+1^_5_b^+1^_6_b^+1^_6_y^+1^_6_y^+1^_6_y^+1^_6_b^+1^_7_b^+1^_7_y^+1^_7_y^+1^_7_b^+1^_7_y^+1^_7_y^+1^_8_y^+1^_8_b^+1^_8_b^+1^_8_y^+1^_8_b^+1^_8_b^+1^_9_b^+1^_9_b^+1^_9_y^+1^_9_y^+1^_9_y^+1^_9_

0840167925193359

Fragment Matches Table

Show background peaks

| Position | Ion type | Intensity | mz Theoretical | mz Error (Th) | mz Error (ppm) | Charge | Series Number |
| --- | --- | --- | --- | --- | --- | --- | --- |
| - | - | 3.222E+06 | 120.1 | - | - | 0 | - |
| - | - | 2.809E+04 | 121.1 | - | - | 0 | - |
| - | - | 2.253E+05 | 121.1 | - | - | 0 | - |
| - | - | 2.669E+04 | 123.1 | - | - | 0 | - |
| - | - | 1.6E+04 | 124 | - | - | 0 | - |
| - | - | 2.071E+04 | 124.1 | - | - | 0 | - |
| - | - | 1.886E+04 | 124.7 | - | - | 0 | - |
| - | - | 3.518E+04 | 126.1 | - | - | 0 | - |
| - | - | 2.81E+04 | 127.1 | - | - | 0 | - |
| - | - | 1.402E+05 | 127.1 | - | - | 0 | - |
| - | - | 1.655E+05 | 128.1 | - | - | 0 | - |
| - | - | 8.17E+05 | 129.1 | - | - | 0 | - |
| - | - | 6.184E+06 | 129.1 | - | - | 0 | - |
| - | - | 3.541E+04 | 130.1 | - | - | 0 | - |
| - | - | 3.538E+04 | 130.1 | - | - | 0 | - |
| - | - | 2.716E+04 | 130.1 | - | - | 0 | - |
| - | - | 3.88E+05 | 130.1 | - | - | 0 | - |
| - | - | 4.515E+04 | 131 | - | - | 0 | - |
| - | - | 2.839E+05 | 131.1 | - | - | 0 | - |
| - | - | 4.026E+05 | 131.1 | - | - | 0 | - |
| - | - | 2.246E+04 | 132.1 | - | - | 0 | - |
| - | - | 2.131E+04 | 132.2 | - | - | 0 | - |
| - | - | 6.709E+04 | 133.1 | - | - | 0 | - |
| - | - | 2.091E+04 | 137.1 | - | - | 0 | - |
| - | - | 2.006E+06 | 138.1 | - | - | 0 | - |
| - | - | 2.089E+04 | 139.1 | - | - | 0 | - |
| - | - | 1.322E+05 | 139.1 | - | - | 0 | - |
| - | - | 1.729E+05 | 139.1 | - | - | 0 | - |
| - | - | 4.078E+04 | 140.1 | - | - | 0 | - |
| - | - | 9.866E+04 | 141.1 | - | - | 0 | - |
| - | - | 2.382E+05 | 141.1 | - | - | 0 | - |
| - | - | 1.897E+04 | 143.1 | - | - | 0 | - |
| - | - | 3.49E+04 | 147.1 | - | - | 0 | - |
| - | - | 1.825E+04 | 147.6 | - | - | 0 | - |
| - | - | 1.744E+04 | 148.6 | - | - | 0 | - |
| - | - | 3.21E+04 | 149 | - | - | 0 | - |
| - | - | 4.691E+04 | 149.1 | - | - | 0 | - |
| - | - | 3.189E+04 | 149.1 | - | - | 0 | - |
| - | - | 2.861E+05 | 151.1 | - | - | 0 | - |
| - | - | 1.275E+05 | 154.1 | - | - | 0 | - |
| - | - | 2.73E+04 | 154.1 | - | - | 0 | - |
| - | - | 8.827E+04 | 155.1 | - | - | 0 | - |
| - | - | 2.58E+06 | 155.1 | - | - | 0 | - |
| - | - | 2.751E+04 | 156.1 | - | - | 0 | - |
| - | - | 3.969E+04 | 156.1 | - | - | 0 | - |
| - | - | 2.032E+05 | 156.1 | - | - | 0 | - |
| - | - | 3.835E+04 | 157.1 | - | - | 0 | - |
| - | - | 2.374E+04 | 157.1 | - | - | 0 | - |
| - | - | 9.723E+05 | 159.1 | - | - | 0 | - |
| - | - | 1.104E+05 | 159.1 | - | - | 0 | - |
| - | - | 3.443E+04 | 160.1 | - | - | 0 | - |
| - | - | 2.159E+04 | 160.8 | - | - | 0 | - |
| - | - | 6.258E+04 | 165.1 | - | - | 0 | - |
| 10 | y | 4.46E+06 | 166.1 | 0.0003843 | 2.314 | +1 | 1 |
| - | - | 4.347E+04 | 167.1 | - | - | 0 | - |
| - | - | 4.168E+05 | 167.1 | - | - | 0 | - |
| - | - | 3.75E+05 | 167.1 | - | - | 0 | - |
| - | - | 3.048E+05 | 168.1 | - | - | 0 | - |
| - | - | 3.079E+04 | 168.1 | - | - | 0 | - |
| - | - | 1.977E+04 | 169.1 | - | - | 0 | - |
| - | - | 6.407E+04 | 169.1 | - | - | 0 | - |
| - | - | 2.787E+04 | 169.1 | - | - | 0 | - |
| - | - | 3.231E+04 | 169.1 | - | - | 0 | - |
| - | - | 3.622E+04 | 171.1 | - | - | 0 | - |
| - | - | 4.378E+04 | 172.1 | - | - | 0 | - |
| - | - | 5.977E+06 | 173.1 | - | - | 0 | - |
| - | - | 1.139E+05 | 173.4 | - | - | 0 | - |
| - | - | 3.546E+04 | 174.1 | - | - | 0 | - |
| - | - | 4.378E+05 | 174.1 | - | - | 0 | - |
| - | - | 5.034E+04 | 177.1 | - | - | 0 | - |
| - | - | 9.754E+05 | 177.1 | - | - | 0 | - |
| - | - | 2.345E+04 | 178.1 | - | - | 0 | - |
| - | - | 1.128E+05 | 178.1 | - | - | 0 | - |
| - | - | 4.767E+04 | 180.1 | - | - | 0 | - |
| - | - | 3.166E+04 | 180.1 | - | - | 0 | - |
| - | - | 2.582E+05 | 181.1 | - | - | 0 | - |
| - | - | 2.313E+04 | 181.1 | - | - | 0 | - |
| - | - | 2.683E+04 | 182.1 | - | - | 0 | - |
| - | - | 7.999E+06 | 182.1 | - | - | 0 | - |
| 2 | a | 1.133E+07 | 183.1 | 0.0004161 | 2.272 | +1 | 2 |
| - | - | 8.114E+05 | 183.1 | - | - | 0 | - |
| - | - | 5.218E+04 | 184.1 | - | - | 0 | - |
| - | - | 1.727E+05 | 184.1 | - | - | 0 | - |
| - | - | 9.896E+05 | 184.1 | - | - | 0 | - |
| - | - | 6.752E+04 | 184.1 | - | - | 0 | - |
| - | - | 5.168E+04 | 185.1 | - | - | 0 | - |
| - | - | 4.385E+05 | 185.1 | - | - | 0 | - |
| 2 | d | 4.638E+04 | 186.1 | 0.003265 | 17.54 | +1 | 2 |
| - | - | 2.949E+05 | 187.1 | - | - | 0 | - |
| - | - | 3.535E+04 | 188.1 | - | - | 0 | - |
| - | - | 4.507E+04 | 191.1 | - | - | 0 | - |
| - | - | 2.808E+04 | 191.6 | - | - | 0 | - |
| - | - | 7.406E+04 | 193.1 | - | - | 0 | - |
| - | - | 4.833E+05 | 194.1 | - | - | 0 | - |
| - | - | 6.731E+04 | 195.1 | - | - | 0 | - |
| - | - | 4.268E+04 | 195.1 | - | - | 0 | - |
| - | - | 5.943E+04 | 196.1 | - | - | 0 | - |
| - | - | 9.553E+04 | 197.1 | - | - | 0 | - |
| - | - | 3.027E+04 | 197.1 | - | - | 0 | - |
| - | - | 4.859E+05 | 198.1 | - | - | 0 | - |
| - | - | 1.665E+05 | 198.1 | - | - | 0 | - |
| - | - | 2.168E+05 | 199.1 | - | - | 0 | - |
| - | - | 3.912E+04 | 199.1 | - | - | 0 | - |
| - | - | 2.654E+04 | 199.1 | - | - | 0 | - |
| 2 | a | 1.247E+07 | 200.1 | 0.0003715 | 1.856 | +1 | 2 |
| - | - | 2.116E+06 | 201.1 | - | - | 0 | - |
| - | - | 1.175E+06 | 201.1 | - | - | 0 | - |
| - | - | 2.042E+05 | 202.1 | - | - | 0 | - |
| - | - | 4.829E+04 | 202.1 | - | - | 0 | - |
| - | - | 6.224E+04 | 202.2 | - | - | 0 | - |
| - | - | 2.984E+04 | 204.1 | - | - | 0 | - |
| - | - | 3.04E+04 | 204.1 | - | - | 0 | - |
| - | - | 1.193E+05 | 205.1 | - | - | 0 | - |
| - | - | 8.936E+04 | 208.1 | - | - | 0 | - |
| - | - | 8.984E+04 | 208.1 | - | - | 0 | - |
| - | - | 8.254E+04 | 209.1 | - | - | 0 | - |
| - | - | 7.358E+05 | 209.1 | - | - | 0 | - |
| - | - | 6.472E+04 | 210.1 | - | - | 0 | - |
| - | - | 3.474E+06 | 210.1 | - | - | 0 | - |
| 2 | b | 1.822E+06 | 211.1 | 0.000344 | 1.629 | +1 | 2 |
| - | - | 3.667E+05 | 211.1 | - | - | 0 | - |
| - | - | 1.546E+05 | 212.1 | - | - | 0 | - |
| - | - | 1.693E+05 | 212.1 | - | - | 0 | - |
| - | - | 5.646E+04 | 213.1 | - | - | 0 | - |
| - | - | 3.476E+04 | 214.2 | - | - | 0 | - |
| - | - | 4.432E+04 | 215.1 | - | - | 0 | - |
| - | - | 2.133E+05 | 216.1 | - | - | 0 | - |
| - | - | 3.315E+04 | 217.1 | - | - | 0 | - |
| - | - | 1.826E+05 | 218.2 | - | - | 0 | - |
| - | - | 2.356E+04 | 222.1 | - | - | 0 | - |
| 9 | y | 3.104E+06 | 223.1 | 0.0003897 | 1.747 | +1 | 2 |
| - | - | 3.233E+05 | 224.1 | - | - | 0 | - |
| - | - | 4.443E+04 | 225.1 | - | - | 0 | - |
| - | - | 3.102E+04 | 225.1 | - | - | 0 | - |
| - | - | 1.843E+05 | 225.1 | - | - | 0 | - |
| - | - | 1.113E+06 | 226.1 | - | - | 0 | - |
| - | - | 2.471E+06 | 226.1 | - | - | 0 | - |
| - | - | 7.708E+05 | 227.1 | - | - | 0 | - |
| - | - | 1.036E+05 | 227.1 | - | - | 0 | - |
| - | - | 1.508E+05 | 227.1 | - | - | 0 | - |
| - | - | 2.881E+05 | 227.1 | - | - | 0 | - |
| - | - | 5.912E+04 | 228.1 | - | - | 0 | - |
| 2 | b | 4.505E+06 | 228.1 | 0.0003909 | 1.714 | +1 | 2 |
| - | - | 5.112E+04 | 229.1 | - | - | 0 | - |
| - | - | 4.645E+05 | 229.1 | - | - | 0 | - |
| - | - | 1.948E+04 | 230.1 | - | - | 0 | - |
| - | - | 3.098E+05 | 230.2 | - | - | 0 | - |
| - | - | 6.444E+04 | 231.2 | - | - | 0 | - |
| - | - | 4.245E+04 | 234.1 | - | - | 0 | - |
| - | - | 5.342E+04 | 236.1 | - | - | 0 | - |
| - | - | 2.361E+04 | 237.4 | - | - | 0 | - |
| - | - | 2.817E+04 | 238.1 | - | - | 0 | - |
| - | - | 7.8E+04 | 240.1 | - | - | 0 | - |
| - | - | 9.782E+05 | 242.2 | - | - | 0 | - |
| - | - | 1.841E+05 | 243.1 | - | - | 0 | - |
| - | - | 1.088E+05 | 243.2 | - | - | 0 | - |
| - | - | 1.927E+06 | 244.1 | - | - | 0 | - |
| - | - | 2.162E+06 | 244.1 | - | - | 0 | - |
| - | - | 1.604E+05 | 245.1 | - | - | 0 | - |
| - | - | 2.005E+05 | 245.1 | - | - | 0 | - |
| - | - | 7.427E+04 | 248.1 | - | - | 0 | - |
| - | - | 3.43E+04 | 248.1 | - | - | 0 | - |
| - | - | 2.441E+04 | 250.2 | - | - | 0 | - |
| - | - | 2.727E+04 | 251.2 | - | - | 0 | - |
| - | - | 1.515E+05 | 252.1 | - | - | 0 | - |
| - | - | 7.466E+04 | 253.1 | - | - | 0 | - |
| - | - | 9.634E+04 | 254.1 | - | - | 0 | - |
| - | - | 4.246E+04 | 254.2 | - | - | 0 | - |
| - | - | 1.372E+05 | 258.1 | - | - | 0 | - |
| - | - | 1.134E+05 | 261.1 | - | - | 0 | - |
| - | - | 2.972E+04 | 262.1 | - | - | 0 | - |
| - | - | 5.115E+04 | 262.1 | - | - | 0 | - |
| - | - | 5.221E+04 | 264.1 | - | - | 0 | - |
| - | - | 5.383E+04 | 265.1 | - | - | 0 | - |
| - | - | 9.397E+04 | 266.1 | - | - | 0 | - |
| - | - | 8.68E+04 | 266.1 | - | - | 0 | - |
| - | - | 1.743E+05 | 267.1 | - | - | 0 | - |
| - | - | 4.203E+04 | 268.1 | - | - | 0 | - |
| 3 | a | 1.023E+05 | 269.2 | 0.0003769 | 1.4 | +1 | 3 |
| 3 | a | 3.159E+05 | 270.1 | 0.0004311 | 1.596 | +1 | 3 |
| - | - | 4.243E+04 | 270.2 | - | - | 0 | - |
| - | - | 3.438E+04 | 271.1 | - | - | 0 | - |
| - | - | 4.747E+04 | 274.1 | - | - | 0 | - |
| - | - | 9.348E+04 | 278.2 | - | - | 0 | - |
| - | - | 5.326E+05 | 279.1 | - | - | 0 | - |
| - | - | 6.221E+05 | 280.1 | - | - | 0 | - |
| - | - | 7.436E+04 | 280.1 | - | - | 0 | - |
| - | - | 7.218E+04 | 281.1 | - | - | 0 | - |
| - | - | 7.648E+04 | 282.1 | - | - | 0 | - |
| - | - | 1.579E+05 | 282.2 | - | - | 0 | - |
| - | - | 6.252E+05 | 283.1 | - | - | 0 | - |
| - | - | 2.795E+04 | 283.2 | - | - | 0 | - |
| - | - | 2.297E+04 | 284.1 | - | - | 0 | - |
| 6 | y | 4.395E+04 | 284.1 | 0.002302 | 8.103 | +2 | 5 |
| - | - | 3.537E+04 | 284.2 | - | - | 0 | - |
| - | - | 2.885E+04 | 285.2 | - | - | 0 | - |
| 3 | a | 1.382E+05 | 287.2 | 0.0003713 | 1.293 | +1 | 3 |
| - | - | 4.724E+04 | 290.1 | - | - | 0 | - |
| - | - | 8.599E+04 | 291.1 | - | - | 0 | - |
| - | - | 1.516E+05 | 292.1 | - | - | 0 | - |
| - | - | 2.947E+04 | 293.1 | - | - | 0 | - |
| - | - | 4.286E+04 | 293.2 | - | - | 0 | - |
| - | - | 1.226E+05 | 294.1 | - | - | 0 | - |
| - | - | 3.708E+04 | 296.2 | - | - | 0 | - |
| 3 | b | 2.507E+06 | 297.2 | 0.0005489 | 1.847 | +1 | 3 |
| 3 | b | 1.842E+06 | 298.1 | 0.0005116 | 1.716 | +1 | 3 |
| - | - | 3.297E+05 | 298.2 | - | - | 0 | - |
| - | - | 2.487E+05 | 299.1 | - | - | 0 | - |
| - | - | 3.419E+04 | 299.2 | - | - | 0 | - |
| - | - | 2.791E+04 | 301.1 | - | - | 0 | - |
| - | - | 1.163E+05 | 301.2 | - | - | 0 | - |
| - | - | 4.335E+04 | 301.2 | - | - | 0 | - |
| - | - | 2.712E+04 | 305.2 | - | - | 0 | - |
| - | - | 1.404E+05 | 305.2 | - | - | 0 | - |
| 8 | y | 2.66E+05 | 306.1 | 0.0005532 | 1.807 | +1 | 3 |
| - | - | 3.409E+04 | 306.2 | - | - | 0 | - |
| - | - | 3.389E+04 | 307.1 | - | - | 0 | - |
| - | - | 4.017E+05 | 309.2 | - | - | 0 | - |
| - | - | 1.461E+05 | 310.1 | - | - | 0 | - |
| - | - | 4.296E+04 | 310.2 | - | - | 0 | - |
| - | - | 2.334E+04 | 311.1 | - | - | 0 | - |
| - | - | 1.601E+05 | 311.2 | - | - | 0 | - |
| 3 | b | 1.017E+06 | 315.2 | 0.0005128 | 1.627 | +1 | 3 |
| - | - | 1.615E+05 | 316.2 | - | - | 0 | - |
| - | - | 7.826E+04 | 317.2 | - | - | 0 | - |
| - | - | 2.037E+05 | 319.1 | - | - | 0 | - |
| - | - | 2.875E+04 | 322.1 | - | - | 0 | - |
| - | - | 2.799E+04 | 323.2 | - | - | 0 | - |
| 8 | y | 1.249E+06 | 324.2 | 0.0006391 | 1.972 | +1 | 3 |
| - | - | 2.096E+05 | 325.2 | - | - | 0 | - |
| 6 | b | 3.605E+05 | 327.2 | 0.0006349 | 1.94 | +2 | 6 |
| - | - | 8.458E+04 | 328.2 | - | - | 0 | - |
| - | - | 9.117E+04 | 328.2 | - | - | 0 | - |
| - | - | 6.047E+04 | 329.1 | - | - | 0 | - |
| - | - | 1.035E+06 | 329.2 | - | - | 0 | - |
| - | - | 1.712E+05 | 330.2 | - | - | 0 | - |
| 6 | b | 2.947E+05 | 336.2 | 0.004342 | 12.92 | +2 | 6 |
| - | - | 1.238E+06 | 337.2 | - | - | 0 | - |
| - | - | 3.705E+04 | 338.1 | - | - | 0 | - |
| - | - | 2.144E+05 | 338.2 | - | - | 0 | - |
| 5 | y | 4.278E+05 | 339.2 | 0.0005128 | 1.512 | +2 | 6 |
| 5 | y | 9.95E+04 | 339.7 | 0.005636 | 16.59 | +2 | 6 |
| - | - | 1.615E+05 | 340.2 | - | - | 0 | - |
| - | - | 8.408E+05 | 345.2 | - | - | 0 | - |
| - | - | 1.309E+05 | 346.2 | - | - | 0 | - |
| - | - | 5.405E+04 | 347.2 | - | - | 0 | - |
| 5 | y | 4.858E+05 | 348.2 | 0.000449 | 1.289 | +2 | 6 |
| - | - | 4.077E+04 | 348.2 | - | - | 0 | - |
| - | - | 1.846E+05 | 348.7 | - | - | 0 | - |
| - | - | 3.107E+04 | 349.2 | - | - | 0 | - |
| - | - | 3.34E+04 | 352.2 | - | - | 0 | - |
| - | - | 1.582E+06 | 354.2 | - | - | 0 | - |
| - | - | 7.268E+05 | 355.2 | - | - | 0 | - |
| - | - | 1.793E+05 | 355.2 | - | - | 0 | - |
| - | - | 1.273E+05 | 356.2 | - | - | 0 | - |
| - | - | 5.205E+05 | 357.2 | - | - | 0 | - |
| - | - | 4.1E+04 | 357.2 | - | - | 0 | - |
| - | - | 9.412E+04 | 358.2 | - | - | 0 | - |
| - | - | 2.032E+05 | 364.2 | - | - | 0 | - |
| - | - | 1.305E+06 | 365.2 | - | - | 0 | - |
| - | - | 4.1E+04 | 366.2 | - | - | 0 | - |
| - | - | 4.951E+04 | 366.2 | - | - | 0 | - |
| - | - | 2.413E+05 | 366.2 | - | - | 0 | - |
| - | - | 8.251E+04 | 367.2 | - | - | 0 | - |
| - | - | 1.217E+06 | 372.2 | - | - | 0 | - |
| - | - | 4.637E+04 | 373.2 | - | - | 0 | - |
| - | - | 2.431E+05 | 373.2 | - | - | 0 | - |
| - | - | 4.015E+04 | 373.7 | - | - | 0 | - |
| - | - | 6.764E+04 | 374.2 | - | - | 0 | - |
| - | - | 4.649E+04 | 375.2 | - | - | 0 | - |
| - | - | 9.742E+04 | 376.2 | - | - | 0 | - |
| - | - | 9.437E+04 | 380.2 | - | - | 0 | - |
| - | - | 3.506E+04 | 381.2 | - | - | 0 | - |
| - | - | 5.331E+04 | 382.2 | - | - | 0 | - |
| - | - | 3.613E+04 | 382.2 | - | - | 0 | - |
| - | - | 4.94E+05 | 382.2 | - | - | 0 | - |
| - | - | 2.979E+04 | 382.7 | - | - | 0 | - |
| - | - | 3.683E+04 | 383.2 | - | - | 0 | - |
| - | - | 1.478E+06 | 383.2 | - | - | 0 | - |
| - | - | 1.697E+05 | 384.2 | - | - | 0 | - |
| - | - | 2.82E+05 | 384.2 | - | - | 0 | - |
| - | - | 9.509E+04 | 385.2 | - | - | 0 | - |
| - | - | 4.277E+04 | 389.2 | - | - | 0 | - |
| - | - | 3.605E+04 | 391.2 | - | - | 0 | - |
| 7 | b | 5.367E+04 | 391.2 | 0.0007913 | 2.023 | +2 | 7 |
| 7 | b | 3.194E+04 | 391.7 | 0.003321 | 8.478 | +2 | 7 |
| - | - | 2.064E+04 | 392.2 | - | - | 0 | - |
| - | - | 6.16E+05 | 392.2 | - | - | 0 | - |
| - | - | 2.007E+05 | 393.2 | - | - | 0 | - |
| - | - | 1.153E+06 | 393.2 | - | - | 0 | - |
| - | - | 6.778E+05 | 394.2 | - | - | 0 | - |
| - | - | 2.788E+05 | 394.2 | - | - | 0 | - |
| - | - | 1.086E+05 | 395.2 | - | - | 0 | - |
| 4 | y | 4.954E+04 | 395.7 | 0.002121 | 5.36 | +2 | 7 |
| 4 | y | 2.097E+05 | 396.2 | 0.0005001 | 1.262 | +2 | 7 |
| - | - | 1.406E+05 | 396.7 | - | - | 0 | - |
| - | - | 2.646E+04 | 398.2 | - | - | 0 | - |
| - | - | 8.117E+04 | 400.3 | - | - | 0 | - |
| - | - | 5.255E+04 | 401.7 | - | - | 0 | - |
| - | - | 2.182E+04 | 402.2 | - | - | 0 | - |
| - | - | 3.471E+05 | 402.2 | - | - | 0 | - |
| - | - | 5.718E+04 | 403.2 | - | - | 0 | - |
| 4 | y | 7.129E+05 | 404.7 | 0.0006228 | 1.539 | +2 | 7 |
| - | - | 2.989E+05 | 405.2 | - | - | 0 | - |
| - | - | 1.143E+05 | 405.7 | - | - | 0 | - |
| - | - | 4.367E+04 | 406.2 | - | - | 0 | - |
| - | - | 2.977E+04 | 408.2 | - | - | 0 | - |
| - | - | 2.851E+04 | 408.2 | - | - | 0 | - |
| - | - | 4.558E+04 | 409.3 | - | - | 0 | - |
| - | - | 7.497E+04 | 410.2 | - | - | 0 | - |
| 4 | b | 1.047E+06 | 410.2 | 0.0005609 | 1.367 | +1 | 4 |
| - | - | 7.473E+05 | 410.7 | - | - | 0 | - |
| - | - | 1.572E+05 | 411.2 | - | - | 0 | - |
| 4 | b | 1.486E+06 | 411.2 | 0.0001268 | 0.3084 | +1 | 4 |
| - | - | 7.161E+04 | 411.7 | - | - | 0 | - |
| - | - | 2.12E+05 | 412.2 | - | - | 0 | - |
| - | - | 2.277E+05 | 412.2 | - | - | 0 | - |
| - | - | 3.309E+04 | 413.2 | - | - | 0 | - |
| - | - | 6.441E+04 | 418.7 | - | - | 0 | - |
| - | - | 1.57E+05 | 419.2 | - | - | 0 | - |
| - | - | 4.072E+04 | 419.2 | - | - | 0 | - |
| - | - | 9.186E+04 | 419.7 | - | - | 0 | - |
| - | - | 3.292E+05 | 420.2 | - | - | 0 | - |
| - | - | 4.23E+04 | 420.2 | - | - | 0 | - |
| - | - | 1.007E+05 | 421.2 | - | - | 0 | - |
| - | - | 1.858E+05 | 423.7 | - | - | 0 | - |
| - | - | 7.303E+04 | 424.2 | - | - | 0 | - |
| - | - | 1.195E+05 | 426.2 | - | - | 0 | - |
| - | - | 1.344E+05 | 427.2 | - | - | 0 | - |
| - | - | 9.508E+04 | 427.2 | - | - | 0 | - |
| - | - | 2.384E+05 | 427.3 | - | - | 0 | - |
| - | - | 2.18E+05 | 427.7 | - | - | 0 | - |
| - | - | 1.596E+05 | 428.2 | - | - | 0 | - |
| 4 | b | 5.305E+05 | 428.3 | 0.0007875 | 1.839 | +1 | 4 |
| - | - | 8.714E+04 | 429.2 | - | - | 0 | - |
| - | - | 1.165E+05 | 429.3 | - | - | 0 | - |
| - | - | 2.778E+04 | 430.7 | - | - | 0 | - |
| - | - | 4.421E+04 | 431.2 | - | - | 0 | - |
| - | - | 4.33E+04 | 432.2 | - | - | 0 | - |
| - | - | 5.048E+05 | 432.7 | - | - | 0 | - |
| - | - | 4.24E+05 | 433.2 | - | - | 0 | - |
| - | - | 1.407E+05 | 433.7 | - | - | 0 | - |
| 7 | y | 2.721E+05 | 434.2 | 0.0002936 | 0.6762 | +1 | 4 |
| - | - | 3.924E+04 | 435.2 | - | - | 0 | - |
| - | - | 3.57E+04 | 436.7 | - | - | 0 | - |
| - | - | 2.417E+05 | 437.2 | - | - | 0 | - |
| - | - | 3.652E+04 | 437.2 | - | - | 0 | - |
| - | - | 4.019E+05 | 438.2 | - | - | 0 | - |
| - | - | 5.566E+04 | 439.2 | - | - | 0 | - |
| 3 | y | 8.535E+04 | 439.2 | 0.001945 | 4.429 | +2 | 8 |
| 3 | y | 3.513E+05 | 439.7 | 0.001393 | 3.167 | +2 | 8 |
| - | - | 2.181E+05 | 440.2 | - | - | 0 | - |
| - | - | 6.562E+04 | 440.7 | - | - | 0 | - |
| 8 | b | 8.687E+05 | 441.7 | 0.0007863 | 1.78 | +2 | 8 |
| - | - | 4.794E+05 | 442.2 | - | - | 0 | - |
| - | - | 1.414E+05 | 442.7 | - | - | 0 | - |
| - | - | 2.747E+04 | 443.2 | - | - | 0 | - |
| - | - | 1.043E+06 | 444.2 | - | - | 0 | - |
| - | - | 1.989E+05 | 445.2 | - | - | 0 | - |
| - | - | 5.032E+04 | 445.2 | - | - | 0 | - |
| - | - | 6.363E+04 | 445.3 | - | - | 0 | - |
| 3 | y | 7.932E+05 | 448.2 | 0.0009965 | 2.223 | +2 | 8 |
| - | - | 4.464E+05 | 448.7 | - | - | 0 | - |
| - | - | 7.615E+04 | 449.2 | - | - | 0 | - |
| - | - | 7.563E+04 | 450.2 | - | - | 0 | - |
| 8 | b | 1.287E+05 | 450.7 | 0.0007835 | 1.738 | +2 | 8 |
| - | - | 7.018E+04 | 451.2 | - | - | 0 | - |
| 7 | y | 2.616E+06 | 452.3 | 0.0005858 | 1.295 | +1 | 4 |
| - | - | 5.138E+04 | 452.7 | - | - | 0 | - |
| - | - | 7.055E+05 | 453.3 | - | - | 0 | - |
| - | - | 1.019E+05 | 454.3 | - | - | 0 | - |
| - | - | 3.619E+05 | 455.2 | - | - | 0 | - |
| - | - | 9.158E+05 | 456.2 | - | - | 0 | - |
| - | - | 2.13E+05 | 457.2 | - | - | 0 | - |
| - | - | 4.943E+04 | 457.2 | - | - | 0 | - |
| - | - | 2.514E+05 | 461.2 | - | - | 0 | - |
| - | - | 2.395E+05 | 461.7 | - | - | 0 | - |
| - | - | 2.124E+05 | 462.2 | - | - | 0 | - |
| - | - | 5.332E+04 | 462.7 | - | - | 0 | - |
| - | - | 2.919E+04 | 467.2 | - | - | 0 | - |
| - | - | 4.015E+04 | 467.3 | - | - | 0 | - |
| - | - | 5.297E+04 | 468.2 | - | - | 0 | - |
| - | - | 4.912E+04 | 469.2 | - | - | 0 | - |
| - | - | 4.471E+04 | 470.2 | - | - | 0 | - |
| 9 | b | 6.607E+05 | 470.2 | 0.0004609 | 0.9802 | +2 | 9 |
| 9 | b | 3.657E+05 | 470.7 | 0.009033 | 19.19 | +2 | 9 |
| - | - | 1.622E+05 | 471.2 | - | - | 0 | - |
| - | - | 5.956E+05 | 473.2 | - | - | 0 | - |
| - | - | 1.377E+05 | 474.2 | - | - | 0 | - |
| - | - | 5.03E+04 | 477.2 | - | - | 0 | - |
| 9 | b | 5.811E+05 | 479.3 | 0.0007328 | 1.529 | +2 | 9 |
| - | - | 2.758E+05 | 479.8 | - | - | 0 | - |
| - | - | 1.12E+05 | 480.3 | - | - | 0 | - |
| - | - | 4.983E+04 | 485.2 | - | - | 0 | - |
| - | - | 2.35E+05 | 485.3 | - | - | 0 | - |
| - | - | 3.095E+04 | 486.2 | - | - | 0 | - |
| - | - | 3.887E+04 | 486.3 | - | - | 0 | - |
| - | - | 6.567E+04 | 494.2 | - | - | 0 | - |
| - | - | 1.704E+05 | 495.2 | - | - | 0 | - |
| - | - | 7.376E+04 | 496.2 | - | - | 0 | - |
| 2 | y | 1.359E+05 | 497.8 | 0.0006946 | 1.395 | +2 | 9 |
| - | - | 6.046E+04 | 498.3 | - | - | 0 | - |
| - | - | 6.697E+04 | 502.8 | - | - | 0 | - |
| - | - | 5.041E+04 | 503.3 | - | - | 0 | - |
| - | - | 5.925E+04 | 505.2 | - | - | 0 | - |
| - | - | 2.859E+04 | 506.3 | - | - | 0 | - |
| - | - | 1.992E+05 | 510.3 | - | - | 0 | - |
| - | - | 4.881E+04 | 511.3 | - | - | 0 | - |
| - | - | 1.669E+05 | 512.2 | - | - | 0 | - |
| - | - | 4.705E+05 | 513.2 | - | - | 0 | - |
| - | - | 3.31E+04 | 513.8 | - | - | 0 | - |
| - | - | 1.198E+05 | 514.2 | - | - | 0 | - |
| - | - | 7.148E+04 | 520.3 | - | - | 0 | - |
| - | - | 1.855E+05 | 521.3 | - | - | 0 | - |
| - | - | 3.518E+04 | 521.8 | - | - | 0 | - |
| - | - | 1.328E+05 | 522.3 | - | - | 0 | - |
| - | - | 8.835E+04 | 522.8 | - | - | 0 | - |
| - | - | 4.16E+04 | 523.3 | - | - | 0 | - |
| - | - | 6.017E+04 | 524.3 | - | - | 0 | - |
| - | - | 3.516E+04 | 527.3 | - | - | 0 | - |
| - | - | 1.621E+05 | 529.8 | - | - | 0 | - |
| - | - | 3.872E+05 | 530.3 | - | - | 0 | - |
| - | - | 4.887E+04 | 530.8 | - | - | 0 | - |
| - | - | 2.109E+05 | 531.3 | - | - | 0 | - |
| - | - | 1.186E+05 | 532.2 | - | - | 0 | - |
| - | - | 8.118E+04 | 534.8 | - | - | 0 | - |
| - | - | 1.406E+05 | 535.3 | - | - | 0 | - |
| - | - | 8.589E+04 | 535.8 | - | - | 0 | - |
| - | - | 4.705E+04 | 537.3 | - | - | 0 | - |
| 5 | b | 1E+06 | 538.3 | 0.000455 | 0.8453 | +1 | 5 |
| - | - | 3.707E+04 | 538.8 | - | - | 0 | - |
| 5 | b | 5.232E+05 | 539.3 | 0.004171 | 7.735 | +1 | 5 |
| - | - | 9.752E+04 | 540.3 | - | - | 0 | - |
| - | - | 5.705E+04 | 543.3 | - | - | 0 | - |
| - | - | 9.574E+05 | 543.8 | - | - | 0 | - |
| - | - | 9.211E+05 | 544.3 | - | - | 0 | - |
| - | - | 3.974E+05 | 544.8 | - | - | 0 | - |
| - | - | 1.449E+05 | 545.3 | - | - | 0 | - |
| - | - | 4.569E+04 | 548.3 | - | - | 0 | - |
| 6 | y | 4.784E+05 | 549.3 | 0.0005343 | 0.9728 | +1 | 5 |
| - | - | 9.524E+04 | 550.3 | - | - | 0 | - |
| - | - | 8.181E+04 | 550.3 | - | - | 0 | - |
| - | - | 4.921E+04 | 551.3 | - | - | 0 | - |
| 0 | Precursor | 1.883E+07 | 552.8 | 0.0007971 | 1.442 | +2 | -1 |
| 0 | Precursor | 1.241E+07 | 553.3 | 0.009644 | 17.43 | +2 | -1 |
| - | - | 4.167E+06 | 553.8 | - | - | 0 | - |
| - | - | 5.526E+05 | 554.3 | - | - | 0 | - |
| - | - | 8.714E+04 | 555.3 | - | - | 0 | - |
| - | - | 3.765E+04 | 555.3 | - | - | 0 | - |
| 5 | b | 1.569E+05 | 556.3 | 0.0002663 | 0.4787 | +1 | 5 |
| - | - | 7.388E+04 | 557.3 | - | - | 0 | - |
| - | - | 4.946E+04 | 561.7 | - | - | 0 | - |
| 0 | Precursor | 2.625E+06 | 561.8 | 0.0004586 | 0.8162 | +2 | -1 |
| - | - | 1.804E+06 | 562.3 | - | - | 0 | - |
| - | - | 5.989E+05 | 562.8 | - | - | 0 | - |
| 6 | y | 6.223E+06 | 567.3 | 0.0003456 | 0.6093 | +1 | 5 |
| - | - | 1.731E+06 | 568.3 | - | - | 0 | - |
| - | - | 2.965E+05 | 569.3 | - | - | 0 | - |
| - | - | 9.67E+05 | 572.3 | - | - | 0 | - |
| - | - | 2.579E+05 | 573.3 | - | - | 0 | - |
| - | - | 3.149E+04 | 574.3 | - | - | 0 | - |
| - | - | 3.36E+04 | 576.3 | - | - | 0 | - |
| - | - | 3.086E+04 | 577.3 | - | - | 0 | - |
| - | - | 4.37E+04 | 581.3 | - | - | 0 | - |
| - | - | 3.768E+05 | 586.3 | - | - | 0 | - |
| - | - | 1.23E+05 | 587.3 | - | - | 0 | - |
| - | - | 6.675E+04 | 593.3 | - | - | 0 | - |
| - | - | 3.884E+04 | 594.3 | - | - | 0 | - |
| - | - | 2.818E+04 | 608.3 | - | - | 0 | - |
| - | - | 5.612E+05 | 611.3 | - | - | 0 | - |
| - | - | 1.669E+05 | 612.3 | - | - | 0 | - |
| - | - | 3.327E+04 | 613.3 | - | - | 0 | - |
| - | - | 4.357E+04 | 619.3 | - | - | 0 | - |
| - | - | 6.803E+04 | 620.3 | - | - | 0 | - |
| - | - | 5.069E+04 | 625.3 | - | - | 0 | - |
| - | - | 6.864E+04 | 627.3 | - | - | 0 | - |
| - | - | 3.682E+04 | 628.3 | - | - | 0 | - |
| - | - | 1.467E+05 | 629.3 | - | - | 0 | - |
| - | - | 1.03E+05 | 632.3 | - | - | 0 | - |
| - | - | 6.36E+04 | 633.3 | - | - | 0 | - |
| - | - | 9.137E+04 | 635.3 | - | - | 0 | - |
| - | - | 1.557E+05 | 636.3 | - | - | 0 | - |
| - | - | 5.568E+05 | 637.3 | - | - | 0 | - |
| - | - | 2.293E+05 | 638.3 | - | - | 0 | - |
| - | - | 3.251E+04 | 639.3 | - | - | 0 | - |
| - | - | 7.202E+04 | 642.3 | - | - | 0 | - |
| - | - | 1.727E+05 | 643.3 | - | - | 0 | - |
| - | - | 5.831E+04 | 644.3 | - | - | 0 | - |
| - | - | 1.46E+05 | 645.4 | - | - | 0 | - |
| - | - | 5.311E+04 | 646.4 | - | - | 0 | - |
| - | - | 6.224E+04 | 649.3 | - | - | 0 | - |
| - | - | 4.422E+04 | 651.3 | - | - | 0 | - |
| 6 | b | 1.314E+06 | 653.3 | 0.0004895 | 0.7493 | +1 | 6 |
| - | - | 5.68E+05 | 654.3 | - | - | 0 | - |
| - | - | 4.776E+05 | 655.3 | - | - | 0 | - |
| - | - | 1.527E+05 | 656.3 | - | - | 0 | - |
| - | - | 4.576E+04 | 657.3 | - | - | 0 | - |
| - | - | 1.821E+05 | 659.3 | - | - | 0 | - |
| - | - | 3.016E+05 | 660.3 | - | - | 0 | - |
| - | - | 8.356E+04 | 661.3 | - | - | 0 | - |
| - | - | 3.247E+05 | 670.4 | - | - | 0 | - |
| 6 | b | 2.166E+05 | 671.3 | 0.009578 | 14.27 | +1 | 6 |
| - | - | 6.532E+04 | 672.3 | - | - | 0 | - |
| - | - | 1.894E+06 | 673.4 | - | - | 0 | - |
| - | - | 6.397E+05 | 674.4 | - | - | 0 | - |
| - | - | 1.369E+05 | 675.4 | - | - | 0 | - |
| 5 | y | 1.921E+06 | 677.3 | 0.0002454 | 0.3623 | +1 | 6 |
| 5 | y | 4.157E+06 | 678.3 | 0.002314 | 3.411 | +1 | 6 |
| - | - | 1.523E+06 | 679.3 | - | - | 0 | - |
| - | - | 2.748E+05 | 680.3 | - | - | 0 | - |
| - | - | 4.377E+04 | 686.3 | - | - | 0 | - |
| - | - | 3.954E+04 | 694.4 | - | - | 0 | - |
| 5 | y | 5.841E+06 | 695.3 | 0.0003008 | 0.4326 | +1 | 6 |
| - | - | 2.171E+06 | 696.3 | - | - | 0 | - |
| - | - | 4.076E+05 | 697.3 | - | - | 0 | - |
| - | - | 4.046E+04 | 705.3 | - | - | 0 | - |
| - | - | 5.977E+04 | 706.3 | - | - | 0 | - |
| - | - | 2.569E+05 | 712.4 | - | - | 0 | - |
| - | - | 1.234E+05 | 713.4 | - | - | 0 | - |
| - | - | 7.98E+05 | 730.4 | - | - | 0 | - |
| - | - | 3.082E+05 | 731.4 | - | - | 0 | - |
| - | - | 4.803E+04 | 732.4 | - | - | 0 | - |
| - | - | 3.638E+04 | 737.4 | - | - | 0 | - |
| - | - | 3.746E+04 | 742.2 | - | - | 0 | - |
| - | - | 9.258E+04 | 748.4 | - | - | 0 | - |
| - | - | 3.215E+04 | 754.4 | - | - | 0 | - |
| - | - | 1.099E+05 | 763.4 | - | - | 0 | - |
| - | - | 1.777E+05 | 764.4 | - | - | 0 | - |
| - | - | 3.603E+04 | 765.4 | - | - | 0 | - |
| - | - | 7.565E+04 | 772.4 | - | - | 0 | - |
| - | - | 1.962E+05 | 773.4 | - | - | 0 | - |
| - | - | 6.94E+04 | 774.4 | - | - | 0 | - |
| 7 | b | 9.404E+05 | 781.4 | 0.000131 | 0.1677 | +1 | 7 |
| 7 | b | 4.307E+05 | 782.4 | 0.01447 | 18.49 | +1 | 7 |
| - | - | 1.453E+05 | 783.4 | - | - | 0 | - |
| 4 | y | 1.236E+05 | 790.4 | 0.000532 | 0.6731 | +1 | 7 |
| 4 | y | 9.241E+04 | 791.4 | 0.01572 | 19.87 | +1 | 7 |
| 7 | b | 1.855E+05 | 799.4 | 0.0004239 | 0.5303 | +1 | 7 |
| - | - | 6.932E+04 | 800.4 | - | - | 0 | - |
| 4 | y | 1.792E+06 | 808.4 | 8.396E-05 | 0.1039 | +1 | 7 |
| - | - | 8.358E+05 | 809.4 | - | - | 0 | - |
| - | - | 2.281E+04 | 809.5 | - | - | 0 | - |
| - | - | 1.81E+05 | 810.4 | - | - | 0 | - |
| - | - | 3.703E+04 | 819.4 | - | - | 0 | - |
| - | - | 7.384E+04 | 820.4 | - | - | 0 | - |
| - | - | 2.057E+05 | 821.4 | - | - | 0 | - |
| - | - | 9.487E+04 | 822.4 | - | - | 0 | - |
| - | - | 3.029E+04 | 828.4 | - | - | 0 | - |
| - | - | 6.692E+04 | 829.4 | - | - | 0 | - |
| - | - | 3.201E+04 | 830.4 | - | - | 0 | - |
| - | - | 4.724E+04 | 833.4 | - | - | 0 | - |
| - | - | 4.05E+04 | 834.4 | - | - | 0 | - |
| - | - | 5.142E+04 | 838.4 | - | - | 0 | - |
| - | - | 1.167E+05 | 839.4 | - | - | 0 | - |
| - | - | 1.834E+05 | 846.4 | - | - | 0 | - |
| - | - | 1.822E+05 | 847.4 | - | - | 0 | - |
| - | - | 5.306E+04 | 848.4 | - | - | 0 | - |
| - | - | 7.026E+04 | 849.4 | - | - | 0 | - |
| - | - | 3.174E+04 | 850.4 | - | - | 0 | - |
| - | - | 3.521E+04 | 851.4 | - | - | 0 | - |
| - | - | 7.554E+04 | 855.5 | - | - | 0 | - |
| - | - | 1.234E+05 | 859.4 | - | - | 0 | - |
| - | - | 7.84E+04 | 860.4 | - | - | 0 | - |
| - | - | 5.33E+04 | 864.5 | - | - | 0 | - |
| - | - | 3.181E+05 | 865.4 | - | - | 0 | - |
| - | - | 1.798E+05 | 866.4 | - | - | 0 | - |
| - | - | 6.921E+04 | 867.4 | - | - | 0 | - |
| 3 | y | 5.577E+05 | 877.4 | 5.876E-05 | 0.06697 | +1 | 8 |
| 3 | y | 4.941E+05 | 878.4 | 0.01226 | 13.96 | +1 | 8 |
| - | - | 1.557E+05 | 879.4 | - | - | 0 | - |
| - | - | 4.572E+04 | 880.4 | - | - | 0 | - |
| 8 | b | 7.584E+04 | 882.5 | 0.0004283 | 0.4854 | +1 | 8 |
| 8 | b | 9.006E+05 | 883.5 | 0.0001142 | 0.1293 | +1 | 8 |
| - | - | 4.815E+05 | 884.5 | - | - | 0 | - |
| - | - | 1.107E+05 | 885.5 | - | - | 0 | - |
| 3 | y | 9.831E+06 | 895.5 | 0.0004351 | 0.4859 | +1 | 8 |
| - | - | 5.192E+06 | 896.5 | - | - | 0 | - |
| - | - | 1.34E+06 | 897.5 | - | - | 0 | - |
| - | - | 5.061E+04 | 898.5 | - | - | 0 | - |
| 8 | b | 2.727E+05 | 900.5 | 0.000495 | 0.5497 | +1 | 8 |
| - | - | 1.268E+05 | 901.5 | - | - | 0 | - |
| - | - | 6.112E+04 | 905.4 | - | - | 0 | - |
| - | - | 3.817E+04 | 906.4 | - | - | 0 | - |
| - | - | 2.642E+05 | 922.5 | - | - | 0 | - |
| - | - | 1.516E+05 | 923.5 | - | - | 0 | - |
| - | - | 6.844E+04 | 924.5 | - | - | 0 | - |
| 9 | b | 1.001E+05 | 939.5 | 0.0004468 | 0.4756 | +1 | 9 |
| 9 | b | 1.17E+06 | 940.5 | 0.0005621 | 0.5977 | +1 | 9 |
| - | - | 6.842E+05 | 941.5 | - | - | 0 | - |
| - | - | 1.467E+05 | 942.5 | - | - | 0 | - |
| 9 | b | 2.841E+05 | 957.5 | 0.0001081 | 0.1129 | +1 | 9 |
| - | - | 2.145E+05 | 958.5 | - | - | 0 | - |
| - | - | 7.155E+04 | 959.5 | - | - | 0 | - |
| 2 | y | 4.95E+04 | 976.5 | 0.0009113 | 0.9332 | +1 | 9 |
| 2 | y | 3.608E+04 | 977.5 | 0.01062 | 10.86 | +1 | 9 |
| - | - | 3.133E+04 | 978.5 | - | - | 0 | - |
| 2 | y | 5.134E+05 | 994.5 | 6.242E-05 | 0.06276 | +1 | 9 |
| - | - | 2.87E+05 | 995.5 | - | - | 0 | - |
| - | - | 1.032E+05 | 996.5 | - | - | 0 | - |
| - | - | 2.806E+04 | 996.6 | - | - | 0 | - |
| - | - | 5.212E+04 | 1004 | - | - | 0 | - |
| - | - | 3.757E+04 | 1006 | - | - | 0 | - |
| - | - | 2.707E+04 | 1202 | - | - | 0 | - |
| - | - | 2.848E+04 | 2436 | - | - | 0 | - |
| - | - | 3.266E+04 | 2563 | - | - | 0 | - |
| - | - | 4.239E+04 | 3325 | - | - | 0 | - |

m/z Charge Intensity FragmentType MassShift Position
120.08116149902344 0 3222297
121.07965850830078 0 28087.8
121.08443450927734 0 225294.9
123.09191131591797 0 26691.28
123.99446868896484 0 16004.316
124.07624816894531 0 20706.195
124.73091125488281 0 18863.133
126.09144592285156 0 35181.59
127.08677673339844 0 28098.207
127.12334442138672 0 140160.12
128.10739135742188 0 165517.34
129.0662384033203 0 816950.25
129.1026153564453 0 6183557.5
130.05059814453125 0 35412.934
130.0696258544922 0 35379.027
130.1000518798828 0 27155.797
130.10595703125 0 387972
131.04966735839844 0 45148.94
131.0818634033203 0 283869.22
131.11819458007812 0 402629.53
132.12156677246094 0 22462.893
132.22549438476562 0 21305.348
133.06117248535156 0 67089.664
137.10716247558594 0 20906.031
138.0917205810547 0 2006384.8
139.050537109375 0 20888.887
139.0871124267578 0 132202.84
139.09510803222656 0 172865.9
140.10736083984375 0 40783.49
141.0663299560547 0 98655.664
141.10260009765625 0 238215.23
143.11814880371094 0 18973.268
147.1131134033203 0 34903.355
147.60401916503906 0 18254.602
148.58932495117188 0 17438.691
148.95518493652344 0 32098.69
149.0602264404297 0 46913.04
149.10789489746094 0 31893.234
151.08694458007812 0 286067.47
154.09791564941406 0 127476.984
154.13429260253906 0 27303.28
155.08197021484375 0 88273.94
155.1182403564453 0 2580011.8
156.1014404296875 0 27507.79
156.11508178710938 0 39686.395
156.12155151367188 0 203184.66
157.06117248535156 0 38352.574
157.09825134277344 0 23736.387
159.07675170898438 0 972285.06
159.1131134033203 0 110403.914
160.07992553710938 0 34434.24
160.77137756347656 0 21591.123
165.1024627685547 0 62575.51
166.08663940429688 0 4459715.5 y 9
167.08213806152344 0 43468.953
167.08998107910156 0 416823.06
167.11822509765625 0 375007.38
168.113525390625 0 304813.94
168.12118530273438 0 30785.287
169.0605926513672 0 19771.52
169.09771728515625 0 64069.246
169.11639404296875 0 27871.887
169.13392639160156 0 32310.559
171.0762481689453 0 36221.88
172.1083984375 0 43783.9
173.1288604736328 0 5977340.5
173.4395294189453 0 113860.72
174.1249542236328 0 35456.043
174.1322479248047 0 437791.88
177.08795166015625 0 50336.19
177.1026153564453 0 975397.1
178.09803771972656 0 23445.31
178.10604858398438 0 112809.695
180.07749938964844 0 47673.484
180.11390686035156 0 31661.883
181.06121826171875 0 258169.39
181.0982666015625 0 23132.432
182.06517028808594 0 26830.996
182.12925720214844 0 7999250
183.11322021484375 0 11325795 a Ammonia loss 1
183.1325225830078 0 811398.6
184.09774780273438 0 52176.75
184.10867309570312 0 172732.7
184.11660766601562 0 989588.3
184.14480590820312 0 67515.15
185.1195068359375 0 51680.02
185.12884521484375 0 438504.06
186.12696838378906 0 46376.04 d 1
187.1081085205078 0 294907.44
188.11077880859375 0 35347.453
191.08212280273438 0 45072.812
191.6269073486328 0 28080.395
193.097412109375 0 74062.586
194.12913513183594 0 483252.25
195.11305236816406 0 67313.76
195.13279724121094 0 42676.504
196.10829162597656 0 59434.09
197.09231567382812 0 95533.09
197.12864685058594 0 30274.713
198.08763122558594 0 485918.4
198.12416076660156 0 166509.38
199.07168579101562 0 216839.1
199.09104919433594 0 39121.23
199.10777282714844 0 26541.514
200.1397247314453 0 12471779 a 1
201.12367248535156 0 2115862.8
201.1430206298828 0 1174660.1
202.12692260742188 0 204245.12
202.1450958251953 0 48292.016
202.15557861328125 0 62244.305
204.1134490966797 0 29843.277
204.1335906982422 0 30402.607
205.0973358154297 0 119283.65
208.0970458984375 0 89363.86
208.10853576660156 0 89842.234
209.05569458007812 0 82543.05
209.09243774414062 0 735796.3
210.09596252441406 0 64724.645
210.1240692138672 0 3474062.5
211.10806274414062 0 1822419.5 b Ammonia loss 1
211.12738037109375 0 366742.66
212.11146545410156 0 154604.7
212.1398468017578 0 169319.58
213.1241912841797 0 56455.51
214.1554718017578 0 34755.402
215.10276794433594 0 44322.62
216.09828186035156 0 213255.11
217.1015625 0 33153.504
218.1503448486328 0 182603.31
222.1236572265625 0 23561.332
223.1081085205078 0 3103808.5 y 8
224.1112823486328 0 323328.28
225.09861755371094 0 44425.996
225.112548828125 0 31019.053
225.12371826171875 0 184304.38
226.08265686035156 0 1113103.8
226.1189727783203 0 2471035.8
227.06661987304688 0 770770.2
227.08636474609375 0 103615.56
227.1028289794922 0 150834.86
227.1224365234375 0 288137.2
228.0700225830078 0 59121.54
228.13465881347656 0 4504762.5 b 1
229.1182861328125 0 51115.613
229.137939453125 0 464510.12
230.13963317871094 0 19479.076
230.1503143310547 0 309793.8
231.1507568359375 0 64436.594
234.12445068359375 0 42451.617
236.10342407226562 0 53421.5
237.4227752685547 0 23605.057
238.1193389892578 0 28173.934
240.13479614257812 0 77999.02
242.15037536621094 0 978225
243.1092529296875 0 184140.66
243.1533203125 0 108797.266
244.09320068359375 0 1927350.1
244.12962341308594 0 2162334.2
245.0962371826172 0 160443.16
245.13279724121094 0 200529.69
248.1033172607422 0 74269.54
248.13995361328125 0 34304.152
250.15480041503906 0 24405.328
251.15016174316406 0 27273.223
252.13475036621094 0 151548.64
253.1186065673828 0 74660.88
254.11415100097656 0 96339.555
254.1865997314453 0 42459.348
258.1448669433594 0 137221.69
261.1202392578125 0 113371.44
262.1206359863281 0 29720.06
262.13916015625 0 51146.477
264.13433837890625 0 52207.984
265.1307067871094 0 53830.414
266.11383056640625 0 93971.37
266.1499328613281 0 86796.07
267.1456604003906 0 174336.84
268.13037109375 0 42032.18
269.16119384765625 0 102296.08 a Water loss 2
270.145263671875 0 315933.44 a Ammonia loss 2
270.1607360839844 0 42428.258
271.1496887207031 0 34380.836
274.1184387207031 0 47469.71
278.1502685546875 0 93483.33
279.1456298828125 0 532554.75
280.1296691894531 0 622114.94
280.1477355957031 0 74363.305
281.1326599121094 0 72179.15
282.14520263671875 0 76484.25
282.1817626953125 0 157923.36
283.1405334472656 0 625156.25
283.17822265625 0 27948.709
284.1243896484375 0 22972.572
284.14459228515625 0 43945.54 y 5
284.16064453125 0 35367.598
285.1916809082031 0 28846.416
287.1717529296875 0 138198.4 a 2
290.1137390136719 0 47239.523
291.1457824707031 0 85991.266
292.12982177734375 0 151615.25
293.13372802734375 0 29465.836
293.1604309082031 0 42857.574
294.1455383300781 0 122617.766
296.17254638671875 0 37082.08
297.1562805175781 0 2507414.5 b Water loss 2
298.1402587890625 0 1842157.9 b Ammonia loss 2
298.15936279296875 0 329685.94
299.1432800292969 0 248660.47
299.1734924316406 0 34185.35
301.1329650878906 0 27912.201
301.1515197753906 0 116320.375
301.18701171875 0 43350.03
305.1651611328125 0 27122.643
305.1825256347656 0 140360.97
306.1453857421875 0 265959.84 y Water loss 7
306.18408203125 0 34093.438
307.1478576660156 0 33885.418
309.1562805175781 0 401677.16
310.1399841308594 0 146129.31
310.1589050292969 0 42956
311.13970947265625 0 23338.45
311.17218017578125 0 160098.8
315.16680908203125 0 1017328.8 b 2
316.1696472167969 0 161507.97
317.1824035644531 0 78262.08
319.1407775878906 0 203700.16
322.14019775390625 0 28752.328
323.172607421875 0 27986.006
324.1560363769531 0 1249387.2 y 7
325.15869140625 0 209624.98
327.16693115234375 0 360515.47 b Water loss 5
328.1504821777344 0 84580.89
328.1735534667969 0 91167.4
329.1492004394531 0 60466.617
329.18255615234375 0 1035237.4
330.18572998046875 0 171190.28
336.167236328125 0 294693.4 b 5
337.1512451171875 0 1238372.9
338.13458251953125 0 37050.52
338.1548767089844 0 214355
339.16680908203125 0 427848.78 y Water loss 4
339.6639404296875 0 99496.125 y Ammonia loss 4
340.1513671875 0 161485.39
345.17730712890625 0 840785.25
346.1803283691406 0 130869.94
347.20941162109375 0 54049.152
348.1720275878906 0 485755.8 y 4
348.1937255859375 0 40772.453
348.6737365722656 0 184598.58
349.1509704589844 0 31071.285
352.2352600097656 0 33402.035
354.1775817871094 0 1581910.2
355.161376953125 0 726753.8
355.1816101074219 0 179297.8
356.1645812988281 0 127290.71
357.17718505859375 0 520494.28
357.2000427246094 0 41002.02
358.17901611328125 0 94118.84
364.23486328125 0 203223.98
365.2186584472656 0 1304917.9
366.1783447265625 0 41004.867
366.20062255859375 0 49511.527
366.22222900390625 0 241267.55
367.1613464355469 0 82505.28
372.1880798339844 0 1216860.2
373.1685791015625 0 46374.43
373.1913757324219 0 243078.95
373.6966247558594 0 40149.15
374.20013427734375 0 67644.64
375.17645263671875 0 46488.777
376.1615295410156 0 97418.22
380.228759765625 0 94372.92
381.1767578125 0 35060.387
382.17279052734375 0 53305.64
382.2123718261719 0 36131.906
382.2452087402344 0 493998.34
382.7042236328125 0 29790.45
383.15594482421875 0 36830.363
383.2293701171875 0 1478187.6
384.188232421875 0 169728.23
384.2322692871094 0 281993.53
385.1722412109375 0 95086.14
389.2186279296875 0 42774.785
391.16015625 0 36045.746
391.2145690917969 0 53669.586 b Water loss 6
391.7091064453125 0 31943.178 b Ammonia loss 6
392.2035827636719 0 20642.184
392.2298889160156 0 615982.5
393.18682861328125 0 200686.75
393.2138977050781 0 1152501
394.1727600097656 0 677797.5
394.21673583984375 0 278776.25
395.17559814453125 0 108637.75
395.71044921875 0 49537.773 y Water loss 3
396.2008361816406 0 209741.34 y Ammonia loss 3
396.7026672363281 0 140607.69
398.2031555175781 0 26464.959
400.256103515625 0 81169.41
401.7160949707031 0 52551.684
402.1755676269531 0 21820.186
402.19989013671875 0 347052.25
403.2008972167969 0 57178.12
404.7142333984375 0 712867 y 3
405.2155456542969 0 298902.56
405.71807861328125 0 114286.47
406.2456970214844 0 43667.047
408.1890869140625 0 29772.629
408.22296142578125 0 28511.816
409.25567626953125 0 45575.457
410.20526123046875 0 74965.04
410.2403564453125 0 1046545.4 b Water loss 3
410.71978759765625 0 747250.5
411.19866943359375 0 157185.5
411.22393798828125 0 1486354 b Ammonia loss 3
411.72161865234375 0 71608.89
412.18316650390625 0 211984.67
412.22698974609375 0 227724.61
413.1855163574219 0 33085.035
418.7347717285156 0 64406.52
419.2040710449219 0 157024.7
419.2279357910156 0 40719.125
419.72564697265625 0 91860.305
420.1883850097656 0 329192.16
420.2164306640625 0 42298.504
421.1902770996094 0 100746.71
423.72821044921875 0 185844.11
424.2250061035156 0 73026.266
426.19976806640625 0 119509.445
427.1831359863281 0 134384.5
427.23126220703125 0 95081.43
427.2669677734375 0 238439.6
427.7406921386719 0 217976.5
428.21453857421875 0 159594.39
428.24957275390625 0 530521 b 3
429.2125549316406 0 87140.734
429.2532958984375 0 116523.93
430.71539306640625 0 27780.764
431.2128601074219 0 44208.883
432.2268981933594 0 43299.598
432.73297119140625 0 504844.16
433.2306823730469 0 423980.38
433.72796630859375 0 140731.3
434.239501953125 0 272052.8 y Water loss 6
435.24822998046875 0 39239.37
436.7436218261719 0 35703.44
437.2149963378906 0 241732.86
437.24688720703125 0 36523.832
438.19891357421875 0 401864.66
439.1964111328125 0 55657.594
439.2262878417969 0 85352.72 y Water loss 2
439.7177429199219 0 351336.53 y Ammonia loss 2
440.2191467285156 0 218053.72
440.7174987792969 0 65616.89
441.7384033203125 0 868661 b Water loss 7
442.2389831542969 0 479440.94
442.7392578125 0 141379.33
443.2315368652344 0 27465.197
444.2096862792969 0 1042905
445.2124328613281 0 198882.72
445.2411804199219 0 50319.977
445.2773742675781 0 63631.34
448.2306213378906 0 793191.4 y 2
448.73199462890625 0 446423.62
449.2319641113281 0 76151.51
450.23492431640625 0 75627.23
450.7436828613281 0 128685.82 b 7
451.2408752441406 0 70176.836
452.2509460449219 0 2616143.2 y 6
452.7301330566406 0 51378.44
453.2540283203125 0 705466.06
454.2568359375 0 101856
455.2255859375 0 361947.84
456.2098388671875 0 915752.75
457.2113952636719 0 212999.61
457.24462890625 0 49429.14
461.242919921875 0 251374.95
461.73773193359375 0 239473.67
462.2377624511719 0 212359.34
462.7377624511719 0 53320.332
467.2264404296875 0 29190.361
467.2594299316406 0 40151.914
468.2485046386719 0 52973.87
469.2029113769531 0 49121.992
470.20806884765625 0 44709.598
470.2488098144531 0 660667.1 b Water loss 8
470.7493896484375 0 365737.62 b Ammonia loss 8
471.2481384277344 0 162196.31
473.2361755371094 0 595555.3
474.23846435546875 0 137653.22
477.2103576660156 0 50299.324
479.2543640136719 0 581080.75 b 8
479.7553405761719 0 275764.72
480.2569274902344 0 112046.32
485.2359619140625 0 49830.16
485.2724609375 0 235033.6
486.2342224121094 0 30948.205
486.27618408203125 0 38869.9
494.23681640625 0 65666.62
495.2203063964844 0 170385.38
496.220458984375 0 73757.07
497.7645263671875 0 135923.06 y 1
498.26617431640625 0 60455.242
502.7568664550781 0 66971.04
503.2627868652344 0 50413.676
505.24072265625 0 59247.95
506.26995849609375 0 28585.818
510.3040771484375 0 199238.73
511.30889892578125 0 48805.27
512.2472534179688 0 166873.4
513.2314453125 0 470544.94
513.7537841796875 0 33097.797
514.2329711914062 0 119756.664
520.286376953125 0 71477.85
521.2726440429688 0 185473.72
521.7733154296875 0 35179.543
522.2645874023438 0 132846.48
522.7646484375 0 88349.35
523.2536010742188 0 41595.316
524.2852172851562 0 60167.965
527.2579345703125 0 35156.945
529.7852783203125 0 162058.78
530.2572631835938 0 387174.88
530.7772216796875 0 48868.703
531.2578735351562 0 210885.77
532.2451782226562 0 118582.76
534.7772216796875 0 81179.88
535.2738647460938 0 140574.4
535.7726440429688 0 85890.75
537.2694702148438 0 47052.293
538.298828125 0 1000096.2 b Water loss 4
538.791259765625 0 37067.445
539.2865600585938 0 523184.12 b Ammonia loss 4
540.2886962890625 0 97516.766
543.2789916992188 0 57048.86
543.7828369140625 0 957373.75
544.2805786132812 0 921073.06
544.7802124023438 0 397358.9
545.2777099609375 0 144944.58
548.269287109375 0 45685.19
549.2672729492188 0 478410.06 y Water loss 5
550.2662963867188 0 95241.16
550.3030395507812 0 81811.44
551.2791137695312 0 49212.383
552.7886352539062 0 18834736 Precursor Water loss
553.2894897460938 0 12414413 Precursor Ammonia loss
553.790771484375 0 4166939
554.2919921875 0 552625.2
555.281982421875 0 87135.99
555.328125 0 37652.67
556.3092041015625 0 156854.6 b 4
557.3126220703125 0 73881.984
561.7400512695312 0 49458.95
561.7935791015625 0 2625159.5 Precursor
562.294921875 0 1804422.9
562.796142578125 0 598914.3
567.2776489257812 0 6222890.5 y 5
568.280517578125 0 1731307.1
569.2839965820312 0 296476.75
572.3042602539062 0 967030.6
573.306884765625 0 257935.66
574.3091430664062 0 31494.078
576.2802734375 0 33601.805
577.2655029296875 0 30856.78
581.3404541015625 0 43701.2
586.31982421875 0 376774.88
587.3217163085938 0 122982.58
593.3031616210938 0 66751.664
594.2871704101562 0 38835.195
608.302978515625 0 28177.357
611.3153686523438 0 561241.3
612.31787109375 0 166912.62
613.3173217773438 0 33273.773
619.32080078125 0 43569.484
620.3043823242188 0 68030.93
625.3273315429688 0 50694.008
627.3452758789062 0 68643.45
628.3313598632812 0 36824.805
629.3263549804688 0 146724.31
632.3043212890625 0 103043.82
633.3016357421875 0 63600.516
635.3150024414062 0 91369.46
636.3007202148438 0 155651.14
637.3306274414062 0 556791.75
638.3303833007812 0 229281.31
639.3271484375 0 32510.871
642.2892456054688 0 72020.27
643.3423461914062 0 172724.1
644.341796875 0 58313.215
645.3562622070312 0 145967.11
646.359619140625 0 53105.14
649.3319702148438 0 62244.984
651.3094482421875 0 44224.68
653.3258056640625 0 1314123.6 b Water loss 5
654.3231811523438 0 568019.56
655.3386840820312 0 477570.78
656.3369750976562 0 152729.81
657.3429565429688 0 45757.73
659.3143920898438 0 182148.77
660.301025390625 0 301566.06
661.302978515625 0 83558.2
670.3515014648438 0 324662.53
671.345458984375 0 216555.36 b 5
672.3419799804688 0 65321.684
673.3518676757812 0 1894035.1
674.3545532226562 0 639658.56
675.3562622070312 0 136894.66
677.3255615234375 0 1920593 y Water loss 4
678.3116455078125 0 4156703.2 y Ammonia loss 4
679.313232421875 0 1522973.4
680.31494140625 0 274829.4
686.3465576171875 0 43769.746
694.35693359375 0 39535.152
695.336181640625 0 5840915 y 4
696.3391723632812 0 2171103.8
697.3411865234375 0 407573.75
705.3226928710938 0 40463.277
706.3086547851562 0 59765.16
712.3618774414062 0 256930.16
713.3585205078125 0 123373.23
730.373046875 0 797984.94
731.3760375976562 0 308222.1
732.3765869140625 0 48027.176
737.4000854492188 0 36376.625
742.2089233398438 0 37460.67
748.3814697265625 0 92577.93
754.4105224609375 0 32151.682
763.4085693359375 0 109910.97
764.3976440429688 0 177710.62
765.4010009765625 0 36028.38
772.4164428710938 0 75653.375
773.38525390625 0 196238.72
774.3844604492188 0 69400.914
781.42041015625 0 940440.5 b Water loss 6
782.4187622070312 0 430662.25 b Ammonia loss 6
783.4199829101562 0 145325.56
790.409912109375 0 123590.69 y Water loss 3
791.4091186523438 0 92406.04 y Ammonia loss 3
799.430419921875 0 185526.92 b 6
800.4327392578125 0 69317.82
808.4198608398438 0 1792263.4 y 3
809.422607421875 0 835781.75
809.5094604492188 0 22810.762
810.4269409179688 0 181031.5
819.392578125 0 37025.473
820.4295043945312 0 73836.414
821.4190673828125 0 205660.98
822.4170532226562 0 94873.74
828.435302734375 0 30287.916
829.439208984375 0 66924.15
830.4151000976562 0 32013.365
833.416259765625 0 47243.324
834.416259765625 0 40496.48
838.4428100585938 0 51416.9
839.4332275390625 0 116706.78
846.4462280273438 0 183369.52
847.4371948242188 0 182156
848.4292602539062 0 53055.547
849.4403686523438 0 70259.69
850.4451293945312 0 31740.455
851.4318237304688 0 35206.086
855.456298828125 0 75544.53
859.4320068359375 0 123368.57
860.4212036132812 0 78397.03
864.4564208984375 0 53300.055
865.442138671875 0 318068.1
866.4397583007812 0 179842.72
867.4368896484375 0 69213.875
877.4414672851562 0 557721.1 y Water loss 2
878.4376831054688 0 494133.44 y Ammonia loss 2
879.4354248046875 0 155739.4
880.4323120117188 0 45720.02
882.467529296875 0 75838.055 b Water loss 7
883.4520874023438 0 900566.6 b Ammonia loss 7
884.4547729492188 0 481524.56
885.4577026367188 0 110702.15
895.4515380859375 0 9830807 y 2
896.4546508789062 0 5191705.5
897.4572143554688 0 1339776.1
898.4630737304688 0 50614.785
900.47802734375 0 272736 b 7
901.4804077148438 0 126830.59
905.4356689453125 0 61115.71
906.4291381835938 0 38172.473
922.46337890625 0 264230.72
923.459716796875 0 151642.72
924.4598388671875 0 68439.74
939.4898681640625 0 100087.8 b Water loss 8
940.4739990234375 0 1169734.9 b Ammonia loss 8
941.4771118164062 0 684240.5
942.4775390625 0 146720.9
957.4998779296875 0 284060.9 b 8
958.49755859375 0 214491.11
959.4982299804688 0 71548.29
976.5089111328125 0 49496 y Water loss 1
977.5044555664062 0 36084.695 y Ammonia loss 1
978.5055541992188 0 31333.875
994.5203247070312 0 513375.72 y 1
995.5231323242188 0 287004.5
996.5260620117188 0 103181.38
996.6495971679688 0 28060.326
1004.4993896484375 0 52120.668
1005.508056640625 0 37571.59
1201.6138916015625 0 27072.191
2435.52392578125 0 28484.018
2562.904296875 0 32661.168
3325.306884765625 0 42389.59

Spectrum Details

|  |  |
| --- | --- |
| Matched peaks? Matched peaks The total absolute number of peaks matched. Additionally in brackets the total fraction of peaks matched and the total number of peaks is shown. | 71 (11.72% of 606) |
| FDR? FDR The false discovery rate estimated for this peptide. It is calculated by matching all theoretical fragments with a non-integer shift with the raw peaks for this spectrum. This is done with 40 different shifts. The resulting percentage is the average number of annotated peaks over the number of annotated peaks with the correct spectrum. | 0.07% |
| Satellite FDR? Satellite FDR See the FDR for details on its calculation. This satellite ion specific FDR only contains the satellite ions (d/w) for I/L/J positions. | - |
| PSM Score? PSM Score The PSM Score as given by Hecklib to this annotated spectrum. It is shown with three significant figures. | 867 |

#### Spectrum 5588? Spectrum 5588 The raw spectrum of this peptide as annotated by Hecklib. The fragments are coloured according to ion type (see legend). Any peaks with a star '\*' as text can be hovered over to see the full details, first the ion type second the mass shift type. By hovering over the amino acids in the peptide or ions in the legend the corresponding peaks are highlighted. By toggling the 'Unassigned' label you can turn the background (unassigned) peaks on or off in the plot. By updating the slider in the Ion legend you can update the spectrum to only show the top X% of the peaks with labels. The top X% means any peak that is within X% of the highest intensity. By dragging in the spectrum you can zoom in to a specific part of the spectrum and use 'Zoom Out' to get back to the original zoom level. The annotation of the spectrum is based on the given sequence in the peptides file and is done with different software so inconsistencies are likely. The peaks are annotated based on the given sequence, with 20 ppm tolerance.

Copy Data

##### Spectrum 5588 (TSV)

###### Preview

```
Loading example...
```

*Click on the button to copy the data to your clipboard.*

Mz MinMz MaxIntensity Max

WidthHeightPeptide font sizePeptide stroke widthSpectrum font sizeSpectrum stroke widthCompact peptide

Ion legend

wxyz

abcd

OtherUnassignedIon^Charge^_Position_Show for top:%

QVSLQDKTGF

04.65e+49.30e+41.40e+51.86e+5

Zoom Out

y^+1^_1_a^+1^_2_d^+1^_2_a^+1^_2_b^+1^_2_y^+1^_2_b^+1^_2_a^+1^_3_a^+1^_3_y^+2^_5_a^+1^_3_b^+1^_3_b^+1^_3_y^+1^_3_b^+1^_3_y^+1^_3_b^+2^_6_b^+2^_6_y^+2^_6_y^+2^_6_y^+2^_6_y^+2^_7_y^+2^_7_b^+1^_4_b^+1^_4_b^+1^_4_y^+1^_4_y^+2^_8_y^+2^_8_b^+2^_8_y^+2^_8_b^+2^_8_y^+1^_4_b^+2^_9_b^+2^_9_b^+2^_9_y^+2^_9_y^+2^_9_b^+1^_5_b^+1^_5_y^+1^_5_y^+1^_5_\*\*b^+1^_5_\*y^+1^_5_b^+1^_6_b^+1^_6_y^+1^_6_y^+1^_6_y^+1^_6_b^+1^_7_b^+1^_7_y^+1^_7_y^+1^_7_b^+1^_7_y^+1^_7_y^+1^_8_y^+1^_8_b^+1^_8_b^+1^_8_y^+1^_8_b^+1^_8_b^+1^_9_b^+1^_9_b^+1^_9_y^+1^_9_

0855170925643419

Fragment Matches Table

Show background peaks

| Position | Ion type | Intensity | mz Theoretical | mz Error (Th) | mz Error (ppm) | Charge | Series Number |
| --- | --- | --- | --- | --- | --- | --- | --- |
| - | - | 4.003E+04 | 120.1 | - | - | 0 | - |
| - | - | 3185 | 121.1 | - | - | 0 | - |
| - | - | 872.4 | 123 | - | - | 0 | - |
| - | - | 430.9 | 123.1 | - | - | 0 | - |
| - | - | 682.9 | 125.1 | - | - | 0 | - |
| - | - | 521.9 | 126.1 | - | - | 0 | - |
| - | - | 373.7 | 126.4 | - | - | 0 | - |
| - | - | 731.5 | 127.1 | - | - | 0 | - |
| - | - | 1540 | 127.1 | - | - | 0 | - |
| - | - | 2192 | 128.1 | - | - | 0 | - |
| - | - | 875.5 | 129 | - | - | 0 | - |
| - | - | 8299 | 129.1 | - | - | 0 | - |
| - | - | 7.22E+04 | 129.1 | - | - | 0 | - |
| - | - | 677.9 | 130.1 | - | - | 0 | - |
| - | - | 480.8 | 130.1 | - | - | 0 | - |
| - | - | 701.7 | 130.1 | - | - | 0 | - |
| - | - | 3861 | 130.1 | - | - | 0 | - |
| - | - | 1234 | 131 | - | - | 0 | - |
| - | - | 2606 | 131.1 | - | - | 0 | - |
| - | - | 2760 | 131.1 | - | - | 0 | - |
| - | - | 418.8 | 132.1 | - | - | 0 | - |
| - | - | 612.2 | 132.1 | - | - | 0 | - |
| - | - | 817.2 | 133.1 | - | - | 0 | - |
| - | - | 4069 | 133.1 | - | - | 0 | - |
| - | - | 9872 | 136.1 | - | - | 0 | - |
| - | - | 691.3 | 137.1 | - | - | 0 | - |
| - | - | 1.834E+04 | 138.1 | - | - | 0 | - |
| - | - | 515.8 | 139.1 | - | - | 0 | - |
| - | - | 1904 | 139.1 | - | - | 0 | - |
| - | - | 1486 | 139.1 | - | - | 0 | - |
| - | - | 689.8 | 140.1 | - | - | 0 | - |
| - | - | 1579 | 141.1 | - | - | 0 | - |
| - | - | 3725 | 141.1 | - | - | 0 | - |
| - | - | 680 | 146.1 | - | - | 0 | - |
| - | - | 566.2 | 147.1 | - | - | 0 | - |
| - | - | 937.2 | 148.9 | - | - | 0 | - |
| - | - | 2895 | 151.1 | - | - | 0 | - |
| - | - | 493.9 | 151.1 | - | - | 0 | - |
| - | - | 1014 | 154.1 | - | - | 0 | - |
| - | - | 1495 | 155.1 | - | - | 0 | - |
| - | - | 1305 | 155.1 | - | - | 0 | - |
| - | - | 2.478E+04 | 155.1 | - | - | 0 | - |
| - | - | 2296 | 156.1 | - | - | 0 | - |
| - | - | 438.6 | 157.1 | - | - | 0 | - |
| - | - | 841.3 | 157.1 | - | - | 0 | - |
| - | - | 565.9 | 157.1 | - | - | 0 | - |
| - | - | 9502 | 159.1 | - | - | 0 | - |
| - | - | 968.9 | 159.1 | - | - | 0 | - |
| - | - | 894.1 | 159.1 | - | - | 0 | - |
| - | - | 1100 | 159.1 | - | - | 0 | - |
| - | - | 5609 | 165.1 | - | - | 0 | - |
| - | - | 736.3 | 165.1 | - | - | 0 | - |
| - | - | 881.3 | 166.1 | - | - | 0 | - |
| 10 | y | 4.903E+04 | 166.1 | 0.0002775 | 1.671 | +1 | 1 |
| - | - | 881.9 | 167.1 | - | - | 0 | - |
| - | - | 4812 | 167.1 | - | - | 0 | - |
| - | - | 3264 | 167.1 | - | - | 0 | - |
| - | - | 2626 | 168.1 | - | - | 0 | - |
| - | - | 1699 | 169.1 | - | - | 0 | - |
| - | - | 570.3 | 169.1 | - | - | 0 | - |
| - | - | 624 | 171.1 | - | - | 0 | - |
| - | - | 818.5 | 172.1 | - | - | 0 | - |
| - | - | 6.279E+04 | 173.1 | - | - | 0 | - |
| - | - | 2430 | 173.4 | - | - | 0 | - |
| - | - | 4170 | 174.1 | - | - | 0 | - |
| - | - | 681.2 | 177.1 | - | - | 0 | - |
| - | - | 808.4 | 177.1 | - | - | 0 | - |
| - | - | 1.059E+04 | 177.1 | - | - | 0 | - |
| - | - | 892.5 | 177.1 | - | - | 0 | - |
| - | - | 1148 | 178.1 | - | - | 0 | - |
| - | - | 3265 | 181.1 | - | - | 0 | - |
| - | - | 566.7 | 181.1 | - | - | 0 | - |
| - | - | 499.8 | 182.1 | - | - | 0 | - |
| - | - | 9134 | 182.1 | - | - | 0 | - |
| - | - | 8.295E+04 | 182.1 | - | - | 0 | - |
| 2 | a | 1.157E+05 | 183.1 | 0.0002787 | 1.522 | +1 | 2 |
| - | - | 7546 | 183.1 | - | - | 0 | - |
| - | - | 2006 | 183.1 | - | - | 0 | - |
| - | - | 444.3 | 183.9 | - | - | 0 | - |
| - | - | 1741 | 184.1 | - | - | 0 | - |
| - | - | 1.044E+04 | 184.1 | - | - | 0 | - |
| - | - | 638 | 185.1 | - | - | 0 | - |
| - | - | 3918 | 185.1 | - | - | 0 | - |
| 2 | d | 770.3 | 186.1 | 0.0004881 | 2.622 | +1 | 2 |
| - | - | 3716 | 187.1 | - | - | 0 | - |
| - | - | 2115 | 188.1 | - | - | 0 | - |
| - | - | 3649 | 194.1 | - | - | 0 | - |
| - | - | 1121 | 195.1 | - | - | 0 | - |
| - | - | 756.1 | 197.1 | - | - | 0 | - |
| - | - | 560.8 | 197.1 | - | - | 0 | - |
| - | - | 549.3 | 197.2 | - | - | 0 | - |
| - | - | 4205 | 198.1 | - | - | 0 | - |
| - | - | 2743 | 198.1 | - | - | 0 | - |
| - | - | 3216 | 199.1 | - | - | 0 | - |
| - | - | 789.1 | 199.1 | - | - | 0 | - |
| 2 | a | 1.313E+05 | 200.1 | 0.0002647 | 1.322 | +1 | 2 |
| - | - | 584.5 | 201.1 | - | - | 0 | - |
| - | - | 2.35E+04 | 201.1 | - | - | 0 | - |
| - | - | 1.31E+04 | 201.1 | - | - | 0 | - |
| - | - | 1947 | 202.1 | - | - | 0 | - |
| - | - | 674.4 | 202.1 | - | - | 0 | - |
| - | - | 1065 | 202.2 | - | - | 0 | - |
| - | - | 1499 | 205.1 | - | - | 0 | - |
| - | - | 876.2 | 207.1 | - | - | 0 | - |
| - | - | 587.8 | 208.1 | - | - | 0 | - |
| - | - | 955.7 | 208.1 | - | - | 0 | - |
| - | - | 741.4 | 209.1 | - | - | 0 | - |
| - | - | 7236 | 209.1 | - | - | 0 | - |
| - | - | 3.535E+04 | 210.1 | - | - | 0 | - |
| 2 | b | 1.948E+04 | 211.1 | 0.0002524 | 1.196 | +1 | 2 |
| - | - | 3629 | 211.1 | - | - | 0 | - |
| - | - | 905.2 | 211.1 | - | - | 0 | - |
| - | - | 2034 | 212.1 | - | - | 0 | - |
| - | - | 2232 | 212.1 | - | - | 0 | - |
| - | - | 1316 | 213.1 | - | - | 0 | - |
| - | - | 702.1 | 214.1 | - | - | 0 | - |
| - | - | 1028 | 215.1 | - | - | 0 | - |
| - | - | 2096 | 216.1 | - | - | 0 | - |
| - | - | 1141 | 216.1 | - | - | 0 | - |
| - | - | 830.1 | 217.1 | - | - | 0 | - |
| - | - | 644.5 | 217.1 | - | - | 0 | - |
| - | - | 529.4 | 217.7 | - | - | 0 | - |
| - | - | 2070 | 218.2 | - | - | 0 | - |
| - | - | 819.6 | 222.1 | - | - | 0 | - |
| 9 | y | 3.453E+04 | 223.1 | 0.0002524 | 1.131 | +1 | 2 |
| - | - | 3071 | 224.1 | - | - | 0 | - |
| - | - | 2629 | 225.1 | - | - | 0 | - |
| - | - | 1.062E+04 | 226.1 | - | - | 0 | - |
| - | - | 2.693E+04 | 226.1 | - | - | 0 | - |
| - | - | 9438 | 227.1 | - | - | 0 | - |
| - | - | 974.8 | 227.1 | - | - | 0 | - |
| - | - | 966.5 | 227.1 | - | - | 0 | - |
| - | - | 3068 | 227.1 | - | - | 0 | - |
| - | - | 1084 | 228.1 | - | - | 0 | - |
| 2 | b | 4.559E+04 | 228.1 | 0.0002689 | 1.179 | +1 | 2 |
| - | - | 2416 | 229.1 | - | - | 0 | - |
| - | - | 5109 | 229.1 | - | - | 0 | - |
| - | - | 578.6 | 230.1 | - | - | 0 | - |
| - | - | 3186 | 230.2 | - | - | 0 | - |
| - | - | 697.3 | 234.1 | - | - | 0 | - |
| - | - | 585.3 | 239.1 | - | - | 0 | - |
| - | - | 1746 | 239.2 | - | - | 0 | - |
| - | - | 2626 | 240.1 | - | - | 0 | - |
| - | - | 1.031E+04 | 242.2 | - | - | 0 | - |
| - | - | 1690 | 243.1 | - | - | 0 | - |
| - | - | 1653 | 243.2 | - | - | 0 | - |
| - | - | 2.072E+04 | 244.1 | - | - | 0 | - |
| - | - | 2.265E+04 | 244.1 | - | - | 0 | - |
| - | - | 3283 | 245.1 | - | - | 0 | - |
| - | - | 1603 | 245.1 | - | - | 0 | - |
| - | - | 2225 | 245.1 | - | - | 0 | - |
| - | - | 701.9 | 248.2 | - | - | 0 | - |
| - | - | 1853 | 252.1 | - | - | 0 | - |
| - | - | 986 | 253.1 | - | - | 0 | - |
| - | - | 958.4 | 254.1 | - | - | 0 | - |
| - | - | 1252 | 258.1 | - | - | 0 | - |
| - | - | 1062 | 261.1 | - | - | 0 | - |
| - | - | 646.6 | 261.1 | - | - | 0 | - |
| - | - | 1089 | 266.1 | - | - | 0 | - |
| - | - | 1212 | 266.2 | - | - | 0 | - |
| - | - | 1515 | 267.1 | - | - | 0 | - |
| 3 | a | 1114 | 269.2 | 0.0001938 | 0.7199 | +1 | 3 |
| 3 | a | 3308 | 270.1 | 0.0002785 | 1.031 | +1 | 3 |
| - | - | 767.9 | 278.1 | - | - | 0 | - |
| - | - | 3641 | 279.1 | - | - | 0 | - |
| - | - | 6206 | 280.1 | - | - | 0 | - |
| - | - | 731.9 | 280.1 | - | - | 0 | - |
| - | - | 810.8 | 281.1 | - | - | 0 | - |
| - | - | 2006 | 282.2 | - | - | 0 | - |
| - | - | 6983 | 283.1 | - | - | 0 | - |
| 6 | y | 605.9 | 284.1 | 0.0006239 | 2.196 | +2 | 5 |
| 3 | a | 1176 | 287.2 | 0.001216 | 4.233 | +1 | 3 |
| - | - | 703.8 | 291.1 | - | - | 0 | - |
| - | - | 954.5 | 292.1 | - | - | 0 | - |
| - | - | 671.5 | 293.2 | - | - | 0 | - |
| - | - | 1257 | 294.1 | - | - | 0 | - |
| 3 | b | 2.5E+04 | 297.2 | 0.0003658 | 1.231 | +1 | 3 |
| 3 | b | 1.982E+04 | 298.1 | 0.0002369 | 0.7947 | +1 | 3 |
| - | - | 2779 | 298.2 | - | - | 0 | - |
| - | - | 3429 | 299.1 | - | - | 0 | - |
| - | - | 1385 | 300.2 | - | - | 0 | - |
| - | - | 1445 | 301.2 | - | - | 0 | - |
| - | - | 698.6 | 301.2 | - | - | 0 | - |
| - | - | 1240 | 305.2 | - | - | 0 | - |
| - | - | 1366 | 305.2 | - | - | 0 | - |
| 8 | y | 1923 | 306.1 | 0.0003701 | 1.209 | +1 | 3 |
| - | - | 3551 | 309.2 | - | - | 0 | - |
| - | - | 1256 | 310.1 | - | - | 0 | - |
| - | - | 1798 | 311.2 | - | - | 0 | - |
| - | - | 1510 | 312.2 | - | - | 0 | - |
| 3 | b | 1.121E+04 | 315.2 | 0.0002381 | 0.7556 | +1 | 3 |
| - | - | 719.6 | 316.2 | - | - | 0 | - |
| - | - | 2627 | 319.1 | - | - | 0 | - |
| - | - | 675.2 | 323.2 | - | - | 0 | - |
| 8 | y | 1.417E+04 | 324.2 | 0.0004255 | 1.313 | +1 | 3 |
| - | - | 2038 | 325.2 | - | - | 0 | - |
| 6 | b | 4970 | 327.2 | 0.0001771 | 0.5413 | +2 | 6 |
| - | - | 1.161E+04 | 329.2 | - | - | 0 | - |
| - | - | 1983 | 330.2 | - | - | 0 | - |
| 6 | b | 3424 | 336.2 | 0.004678 | 13.92 | +2 | 6 |
| - | - | 1.2E+04 | 337.2 | - | - | 0 | - |
| - | - | 526.7 | 337.2 | - | - | 0 | - |
| - | - | 711.5 | 338.1 | - | - | 0 | - |
| - | - | 2525 | 338.2 | - | - | 0 | - |
| 5 | y | 4019 | 339.2 | 0.0006959 | 2.052 | +2 | 6 |
| 5 | y | 581.2 | 339.7 | 0.001242 | 3.656 | +2 | 6 |
| - | - | 2574 | 340.2 | - | - | 0 | - |
| - | - | 6565 | 345.2 | - | - | 0 | - |
| - | - | 869.8 | 346.2 | - | - | 0 | - |
| 5 | y | 4962 | 348.2 | 0.0001004 | 0.2883 | +2 | 6 |
| - | - | 1543 | 348.7 | - | - | 0 | - |
| - | - | 1.779E+04 | 354.2 | - | - | 0 | - |
| - | - | 6486 | 355.2 | - | - | 0 | - |
| - | - | 2074 | 355.2 | - | - | 0 | - |
| - | - | 1400 | 356.2 | - | - | 0 | - |
| - | - | 5150 | 357.2 | - | - | 0 | - |
| - | - | 2845 | 358.2 | - | - | 0 | - |
| - | - | 2045 | 364.2 | - | - | 0 | - |
| - | - | 1.308E+04 | 365.2 | - | - | 0 | - |
| - | - | 907.8 | 365.2 | - | - | 0 | - |
| - | - | 814.2 | 366.2 | - | - | 0 | - |
| - | - | 2531 | 366.2 | - | - | 0 | - |
| - | - | 817.6 | 367.2 | - | - | 0 | - |
| - | - | 1.174E+04 | 372.2 | - | - | 0 | - |
| - | - | 2459 | 373.2 | - | - | 0 | - |
| - | - | 3366 | 382.2 | - | - | 0 | - |
| - | - | 1.549E+04 | 383.2 | - | - | 0 | - |
| - | - | 1961 | 384.2 | - | - | 0 | - |
| - | - | 2469 | 384.2 | - | - | 0 | - |
| - | - | 2086 | 385.2 | - | - | 0 | - |
| - | - | 848.6 | 392.2 | - | - | 0 | - |
| - | - | 6426 | 392.2 | - | - | 0 | - |
| - | - | 1828 | 393.2 | - | - | 0 | - |
| - | - | 9980 | 393.2 | - | - | 0 | - |
| - | - | 6283 | 394.2 | - | - | 0 | - |
| - | - | 2425 | 394.2 | - | - | 0 | - |
| - | - | 889.7 | 395.2 | - | - | 0 | - |
| 4 | y | 2443 | 396.2 | 0.001538 | 3.881 | +2 | 7 |
| - | - | 896.4 | 396.7 | - | - | 0 | - |
| - | - | 664.9 | 398.2 | - | - | 0 | - |
| - | - | 3819 | 402.2 | - | - | 0 | - |
| - | - | 635.6 | 403.2 | - | - | 0 | - |
| 4 | y | 7704 | 404.7 | 0.0004397 | 1.086 | +2 | 7 |
| - | - | 4523 | 405.2 | - | - | 0 | - |
| - | - | 1109 | 410.2 | - | - | 0 | - |
| 4 | b | 1E+04 | 410.2 | 0.0002862 | 0.6976 | +1 | 4 |
| - | - | 7276 | 410.7 | - | - | 0 | - |
| - | - | 1729 | 411.2 | - | - | 0 | - |
| 4 | b | 1.458E+04 | 411.2 | 0.0003615 | 0.879 | +1 | 4 |
| - | - | 2395 | 412.2 | - | - | 0 | - |
| - | - | 3301 | 412.2 | - | - | 0 | - |
| - | - | 1509 | 419.2 | - | - | 0 | - |
| - | - | 1236 | 419.7 | - | - | 0 | - |
| - | - | 4143 | 420.2 | - | - | 0 | - |
| - | - | 1269 | 420.2 | - | - | 0 | - |
| - | - | 1606 | 423.7 | - | - | 0 | - |
| - | - | 1268 | 424.2 | - | - | 0 | - |
| - | - | 714.9 | 425.2 | - | - | 0 | - |
| - | - | 763.7 | 426.2 | - | - | 0 | - |
| - | - | 1337 | 427.2 | - | - | 0 | - |
| - | - | 1018 | 427.2 | - | - | 0 | - |
| - | - | 2893 | 427.3 | - | - | 0 | - |
| - | - | 1815 | 427.7 | - | - | 0 | - |
| - | - | 1989 | 428.2 | - | - | 0 | - |
| 4 | b | 5474 | 428.3 | 0.000696 | 1.625 | +1 | 4 |
| - | - | 1239 | 429.2 | - | - | 0 | - |
| - | - | 665.2 | 429.3 | - | - | 0 | - |
| - | - | 5726 | 432.7 | - | - | 0 | - |
| - | - | 4524 | 433.2 | - | - | 0 | - |
| - | - | 1426 | 433.7 | - | - | 0 | - |
| 7 | y | 2853 | 434.2 | 0.0004767 | 1.098 | +1 | 4 |
| - | - | 709.1 | 435.2 | - | - | 0 | - |
| - | - | 775.6 | 436 | - | - | 0 | - |
| - | - | 2389 | 437.2 | - | - | 0 | - |
| - | - | 3824 | 438.2 | - | - | 0 | - |
| - | - | 1105 | 439.2 | - | - | 0 | - |
| 3 | y | 902.2 | 439.2 | 0.001701 | 3.873 | +2 | 8 |
| 3 | y | 3570 | 439.7 | 0.0009654 | 2.195 | +2 | 8 |
| - | - | 1508 | 440.2 | - | - | 0 | - |
| 8 | b | 8852 | 441.7 | 0.0004506 | 1.02 | +2 | 8 |
| - | - | 3356 | 442.2 | - | - | 0 | - |
| - | - | 757.1 | 442.7 | - | - | 0 | - |
| - | - | 9988 | 444.2 | - | - | 0 | - |
| - | - | 2284 | 445.2 | - | - | 0 | - |
| 3 | y | 8383 | 448.2 | 5.046E-05 | 0.1126 | +2 | 8 |
| - | - | 3684 | 448.7 | - | - | 0 | - |
| - | - | 984.4 | 449.2 | - | - | 0 | - |
| - | - | 931.8 | 450.2 | - | - | 0 | - |
| 8 | b | 1168 | 450.7 | 0.0006508 | 1.444 | +2 | 8 |
| 7 | y | 2.888E+04 | 452.3 | 0.000128 | 0.2831 | +1 | 4 |
| - | - | 6042 | 453.3 | - | - | 0 | - |
| - | - | 880.7 | 454.3 | - | - | 0 | - |
| - | - | 3366 | 455.2 | - | - | 0 | - |
| - | - | 9125 | 456.2 | - | - | 0 | - |
| - | - | 1360 | 457.2 | - | - | 0 | - |
| - | - | 2812 | 461.2 | - | - | 0 | - |
| - | - | 2285 | 461.7 | - | - | 0 | - |
| - | - | 2286 | 462.2 | - | - | 0 | - |
| - | - | 770.5 | 463.2 | - | - | 0 | - |
| - | - | 727.3 | 467.2 | - | - | 0 | - |
| - | - | 713.9 | 468.2 | - | - | 0 | - |
| 9 | b | 6956 | 470.2 | 5.788E-05 | 0.1231 | +2 | 9 |
| 9 | b | 4719 | 470.7 | 0.008575 | 18.22 | +2 | 9 |
| - | - | 1378 | 471.2 | - | - | 0 | - |
| - | - | 5634 | 473.2 | - | - | 0 | - |
| - | - | 906.4 | 474.2 | - | - | 0 | - |
| 9 | b | 4826 | 479.3 | 0.000336 | 0.7012 | +2 | 9 |
| - | - | 2690 | 479.8 | - | - | 0 | - |
| - | - | 843.4 | 480.3 | - | - | 0 | - |
| - | - | 2519 | 485.3 | - | - | 0 | - |
| 2 | y | 700.8 | 489.3 | 0.00251 | 5.131 | +2 | 9 |
| - | - | 1079 | 494.2 | - | - | 0 | - |
| - | - | 1545 | 495.2 | - | - | 0 | - |
| - | - | 940.9 | 496.2 | - | - | 0 | - |
| 2 | y | 1498 | 497.8 | 0.0008008 | 1.609 | +2 | 9 |
| - | - | 1214 | 498.3 | - | - | 0 | - |
| - | - | 1951 | 510.3 | - | - | 0 | - |
| - | - | 2706 | 512.2 | - | - | 0 | - |
| - | - | 4698 | 513.2 | - | - | 0 | - |
| - | - | 1017 | 513.7 | - | - | 0 | - |
| - | - | 1555 | 514.2 | - | - | 0 | - |
| - | - | 2125 | 521.3 | - | - | 0 | - |
| - | - | 1665 | 522.3 | - | - | 0 | - |
| - | - | 817.4 | 524.3 | - | - | 0 | - |
| - | - | 1743 | 529.8 | - | - | 0 | - |
| - | - | 4239 | 530.3 | - | - | 0 | - |
| - | - | 2213 | 531.3 | - | - | 0 | - |
| - | - | 3327 | 534.3 | - | - | 0 | - |
| - | - | 2098 | 534.8 | - | - | 0 | - |
| - | - | 1466 | 535.3 | - | - | 0 | - |
| - | - | 1077 | 537.3 | - | - | 0 | - |
| 5 | b | 1.155E+04 | 538.3 | 9.429E-05 | 0.1752 | +1 | 5 |
| 5 | b | 4907 | 539.3 | 0.003622 | 6.716 | +1 | 5 |
| - | - | 2786 | 539.8 | - | - | 0 | - |
| - | - | 1020 | 540.3 | - | - | 0 | - |
| - | - | 5399 | 543.3 | - | - | 0 | - |
| - | - | 8988 | 543.8 | - | - | 0 | - |
| - | - | 7842 | 544.3 | - | - | 0 | - |
| - | - | 4210 | 544.8 | - | - | 0 | - |
| - | - | 1133 | 545.3 | - | - | 0 | - |
| 6 | y | 4687 | 549.3 | 0.0002902 | 0.5283 | +1 | 5 |
| 6 | y | 1071 | 550.3 | 0.01023 | 18.6 | +1 | 5 |
| - | - | 895.3 | 550.3 | - | - | 0 | - |
| - | - | 5696 | 551.7 | - | - | 0 | - |
| - | - | 3179 | 552.2 | - | - | 0 | - |
| 0 | Precursor | 1.842E+05 | 552.8 | 3.596E-06 | 0.006506 | +2 | -1 |
| 0 | Precursor | 1.167E+05 | 553.3 | 0.008911 | 16.11 | +2 | -1 |
| - | - | 4.244E+04 | 553.8 | - | - | 0 | - |
| - | - | 6374 | 554.3 | - | - | 0 | - |
| - | - | 1283 | 555.3 | - | - | 0 | - |
| 5 | b | 1090 | 556.3 | 0.001914 | 3.441 | +1 | 5 |
| - | - | 778.5 | 557.1 | - | - | 0 | - |
| - | - | 688.6 | 557.3 | - | - | 0 | - |
| - | - | 896.4 | 560.8 | - | - | 0 | - |
| - | - | 720.5 | 561.3 | - | - | 0 | - |
| - | - | 892.4 | 561.6 | - | - | 0 | - |
| 0 | Precursor | 2.365E+04 | 561.8 | 9.076E-05 | 0.1616 | +2 | -1 |
| - | - | 1.449E+04 | 562.3 | - | - | 0 | - |
| - | - | 3651 | 562.8 | - | - | 0 | - |
| - | - | 781.9 | 563.3 | - | - | 0 | - |
| 6 | y | 6.599E+04 | 567.3 | 8.162E-05 | 0.1439 | +1 | 5 |
| - | - | 1.915E+04 | 568.3 | - | - | 0 | - |
| - | - | 1056 | 568.3 | - | - | 0 | - |
| - | - | 3218 | 569.3 | - | - | 0 | - |
| - | - | 8722 | 572.3 | - | - | 0 | - |
| - | - | 3175 | 573.3 | - | - | 0 | - |
| - | - | 740.5 | 574.3 | - | - | 0 | - |
| - | - | 4177 | 586.3 | - | - | 0 | - |
| - | - | 1072 | 587.3 | - | - | 0 | - |
| - | - | 760.4 | 593.3 | - | - | 0 | - |
| - | - | 5597 | 611.3 | - | - | 0 | - |
| - | - | 1663 | 612.3 | - | - | 0 | - |
| - | - | 676 | 618.5 | - | - | 0 | - |
| - | - | 806.8 | 627.3 | - | - | 0 | - |
| - | - | 809.8 | 628.3 | - | - | 0 | - |
| - | - | 727.2 | 629.3 | - | - | 0 | - |
| - | - | 968.4 | 633.3 | - | - | 0 | - |
| - | - | 744.6 | 635.3 | - | - | 0 | - |
| - | - | 1331 | 636.3 | - | - | 0 | - |
| - | - | 5109 | 637.3 | - | - | 0 | - |
| - | - | 2297 | 638.3 | - | - | 0 | - |
| - | - | 8523 | 641.9 | - | - | 0 | - |
| - | - | 627.2 | 642.3 | - | - | 0 | - |
| - | - | 3606 | 642.4 | - | - | 0 | - |
| - | - | 1562 | 642.9 | - | - | 0 | - |
| - | - | 1610 | 643.3 | - | - | 0 | - |
| - | - | 1575 | 645.4 | - | - | 0 | - |
| - | - | 4496 | 649.3 | - | - | 0 | - |
| - | - | 1498 | 650.3 | - | - | 0 | - |
| - | - | 809.8 | 652.3 | - | - | 0 | - |
| 6 | b | 1.394E+04 | 653.3 | 0.0001208 | 0.1849 | +1 | 6 |
| - | - | 4980 | 654.3 | - | - | 0 | - |
| - | - | 5155 | 655.3 | - | - | 0 | - |
| - | - | 1806 | 656.3 | - | - | 0 | - |
| - | - | 2276 | 659.3 | - | - | 0 | - |
| - | - | 3069 | 660.3 | - | - | 0 | - |
| - | - | 3970 | 670.4 | - | - | 0 | - |
| 6 | b | 1628 | 671.3 | 0.013 | 19.36 | +1 | 6 |
| - | - | 1.949E+04 | 673.4 | - | - | 0 | - |
| - | - | 6398 | 674.4 | - | - | 0 | - |
| - | - | 1278 | 675.4 | - | - | 0 | - |
| 5 | y | 1.91E+04 | 677.3 | 0.0003039 | 0.4487 | +1 | 6 |
| 5 | y | 4.19E+04 | 678.3 | 0.001581 | 2.331 | +1 | 6 |
| - | - | 1.464E+04 | 679.3 | - | - | 0 | - |
| - | - | 2509 | 680.3 | - | - | 0 | - |
| - | - | 854.7 | 685.3 | - | - | 0 | - |
| - | - | 969.4 | 694.3 | - | - | 0 | - |
| 5 | y | 5.67E+04 | 695.3 | 0.0004316 | 0.6207 | +1 | 6 |
| - | - | 2.176E+04 | 696.3 | - | - | 0 | - |
| - | - | 3953 | 697.3 | - | - | 0 | - |
| - | - | 2971 | 712.4 | - | - | 0 | - |
| - | - | 1163 | 713.4 | - | - | 0 | - |
| - | - | 7670 | 730.4 | - | - | 0 | - |
| - | - | 2400 | 731.4 | - | - | 0 | - |
| - | - | 1218 | 755.4 | - | - | 0 | - |
| - | - | 1579 | 755.9 | - | - | 0 | - |
| - | - | 752.9 | 765.4 | - | - | 0 | - |
| - | - | 1566 | 773.4 | - | - | 0 | - |
| - | - | 840.8 | 774.4 | - | - | 0 | - |
| 7 | b | 1.086E+04 | 781.4 | 0.0006014 | 0.7696 | +1 | 7 |
| 7 | b | 4954 | 782.4 | 0.01294 | 16.54 | +1 | 7 |
| - | - | 991.8 | 783.4 | - | - | 0 | - |
| 4 | y | 909.3 | 790.4 | 0.006513 | 8.241 | +1 | 7 |
| 4 | y | 742.1 | 791.4 | 0.0134 | 16.94 | +1 | 7 |
| 7 | b | 1537 | 799.4 | 0.002079 | 2.6 | +1 | 7 |
| 4 | y | 1.695E+04 | 808.4 | 0.001122 | 1.387 | +1 | 7 |
| - | - | 8451 | 809.4 | - | - | 0 | - |
| - | - | 2169 | 810.4 | - | - | 0 | - |
| - | - | 642.8 | 820.4 | - | - | 0 | - |
| - | - | 1774 | 821.4 | - | - | 0 | - |
| - | - | 855.5 | 822.4 | - | - | 0 | - |
| - | - | 1090 | 829.4 | - | - | 0 | - |
| - | - | 864 | 839.4 | - | - | 0 | - |
| - | - | 1161 | 846.4 | - | - | 0 | - |
| - | - | 2015 | 847.4 | - | - | 0 | - |
| - | - | 917.2 | 860.4 | - | - | 0 | - |
| - | - | 777.9 | 864.5 | - | - | 0 | - |
| - | - | 2609 | 865.4 | - | - | 0 | - |
| - | - | 2130 | 866.4 | - | - | 0 | - |
| - | - | 651.6 | 867.4 | - | - | 0 | - |
| 3 | y | 6386 | 877.4 | 0.001284 | 1.463 | +1 | 8 |
| 3 | y | 4716 | 878.4 | 0.01031 | 11.73 | +1 | 8 |
| - | - | 1564 | 879.4 | - | - | 0 | - |
| 8 | b | 770.9 | 882.5 | 0.002379 | 2.696 | +1 | 8 |
| 8 | b | 9526 | 883.5 | 0.0003131 | 0.3543 | +1 | 8 |
| - | - | 4275 | 884.5 | - | - | 0 | - |
| - | - | 945.9 | 885.5 | - | - | 0 | - |
| 3 | y | 9.488E+04 | 895.5 | 0.001534 | 1.713 | +1 | 8 |
| - | - | 4.652E+04 | 896.5 | - | - | 0 | - |
| - | - | 1.391E+04 | 897.5 | - | - | 0 | - |
| - | - | 1795 | 898.5 | - | - | 0 | - |
| 8 | b | 2337 | 900.5 | 0.001899 | 2.109 | +1 | 8 |
| - | - | 1064 | 901.5 | - | - | 0 | - |
| - | - | 767.2 | 902.5 | - | - | 0 | - |
| - | - | 655.8 | 904.5 | - | - | 0 | - |
| - | - | 3047 | 922.5 | - | - | 0 | - |
| - | - | 1932 | 923.5 | - | - | 0 | - |
| - | - | 1058 | 924.5 | - | - | 0 | - |
| 9 | b | 1299 | 939.5 | 0.001201 | 1.278 | +1 | 9 |
| 9 | b | 1.08E+04 | 940.5 | 0.0006586 | 0.7003 | +1 | 9 |
| - | - | 5713 | 941.5 | - | - | 0 | - |
| - | - | 1734 | 942.5 | - | - | 0 | - |
| 9 | b | 3066 | 957.5 | 0.0002912 | 0.3041 | +1 | 9 |
| - | - | 1936 | 958.5 | - | - | 0 | - |
| - | - | 957 | 959.5 | - | - | 0 | - |
| 2 | y | 5231 | 994.5 | 0.002016 | 2.027 | +1 | 9 |
| - | - | 2828 | 995.5 | - | - | 0 | - |
| - | - | 615.8 | 1246 | - | - | 0 | - |
| - | - | 678.2 | 1929 | - | - | 0 | - |
| - | - | 750.4 | 3085 | - | - | 0 | - |
| - | - | 675.4 | 3385 | - | - | 0 | - |

m/z Charge Intensity FragmentType MassShift Position
120.0810775756836 0 40027.035
121.08441162109375 0 3185.033
123.04462432861328 0 872.4338
123.09208679199219 0 430.93054
125.06002044677734 0 682.9163
126.09192657470703 0 521.89056
126.4386215209961 0 373.6906
127.08684539794922 0 731.4747
127.12326049804688 0 1540.2268
128.10726928710938 0 2191.9731
129.0183868408203 0 875.52435
129.06617736816406 0 8299.481
129.10255432128906 0 72199.016
130.0504150390625 0 677.8924
130.0865020751953 0 480.77304
130.10079956054688 0 701.7023
130.1059112548828 0 3861.0586
131.04949951171875 0 1234.1394
131.08180236816406 0 2606.499
131.1182098388672 0 2760.3826
132.0850372314453 0 418.8361
132.10202026367188 0 612.19446
133.06097412109375 0 817.1836
133.08624267578125 0 4068.5654
136.0760040283203 0 9871.502
137.07916259765625 0 691.2729
138.09161376953125 0 18344.732
139.05064392089844 0 515.78796
139.08694458007812 0 1903.7375
139.094970703125 0 1485.6311
140.1068878173828 0 689.80896
141.0663299560547 0 1578.8596
141.10255432128906 0 3725.321
146.0601806640625 0 680.04926
147.076904296875 0 566.2104
148.9473419189453 0 937.1795
151.08682250976562 0 2894.7612
151.09690856933594 0 493.8752
154.09739685058594 0 1014.17645
155.0704345703125 0 1495.3877
155.08181762695312 0 1304.5908
155.11817932128906 0 24783.773
156.12155151367188 0 2295.5034
157.06101989746094 0 438.57938
157.0972900390625 0 841.2772
157.1336669921875 0 565.8894
159.07664489746094 0 9502.316
159.08387756347656 0 968.92017
159.0918426513672 0 894.10077
159.11322021484375 0 1099.936
165.05487060546875 0 5609.0425
165.1027374267578 0 736.28326
166.05860900878906 0 881.2648
166.08653259277344 0 49025.07 y 9
167.08255004882812 0 881.88885
167.08981323242188 0 4811.925
167.1180419921875 0 3263.9717
168.11343383789062 0 2625.5947
169.09730529785156 0 1698.8978
169.1337890625 0 570.30725
171.1125030517578 0 624.0298
172.10800170898438 0 818.45404
173.12875366210938 0 62786.695
173.4395751953125 0 2430.3618
174.13218688964844 0 4170.4546
177.08779907226562 0 681.245
177.094482421875 0 808.3539
177.1024627685547 0 10591.599
177.1112823486328 0 892.45154
178.10589599609375 0 1147.8131
181.06105041503906 0 3265.218
181.09796142578125 0 566.67804
182.0641326904297 0 499.81223
182.08151245117188 0 9134.117
182.1291046142578 0 82954.27
183.1130828857422 0 115724.72 a Ammonia loss 1
183.1324462890625 0 7545.623
183.1492156982422 0 2006.2332
183.94032287597656 0 444.31625
184.1086883544922 0 1740.808
184.11651611328125 0 10443.91
185.08108520507812 0 637.96844
185.1287384033203 0 3917.6765
186.1241912841797 0 770.3012 d 1
187.10806274414062 0 3716.4817
188.07089233398438 0 2115.0728
194.12913513183594 0 3649.1921
195.1128692626953 0 1120.6217
197.0927734375 0 756.12384
197.12890625 0 560.7521
197.16506958007812 0 549.2739
198.08753967285156 0 4205.0815
198.12379455566406 0 2743.3933
199.071533203125 0 3216.4963
199.10842895507812 0 789.1134
200.13961791992188 0 131276.84 a 1
201.08761596679688 0 584.488
201.1235809326172 0 23504.67
201.14300537109375 0 13096.066
202.12680053710938 0 1946.9797
202.14385986328125 0 674.4181
202.15487670898438 0 1064.8201
205.09783935546875 0 1499.1268
207.1132049560547 0 876.2048
208.09706115722656 0 587.7691
208.1078338623047 0 955.7392
209.05665588378906 0 741.4414
209.09231567382812 0 7236.0938
210.12396240234375 0 35354.35
211.10797119140625 0 19480.129 b Ammonia loss 1
211.1273956298828 0 3628.6255
211.14450073242188 0 905.1713
212.11154174804688 0 2033.6636
212.139404296875 0 2232.2932
213.1239013671875 0 1316.3862
214.1186981201172 0 702.10046
215.1393280029297 0 1028.4022
216.09805297851562 0 2095.7405
216.1343536376953 0 1140.8367
217.08221435546875 0 830.0866
217.10009765625 0 644.5466
217.69415283203125 0 529.415
218.15003967285156 0 2070.3044
222.12379455566406 0 819.5649
223.10797119140625 0 34530.8 y 8
224.1112518310547 0 3071.1287
225.1237030029297 0 2629.1003
226.08241271972656 0 10624.876
226.1188507080078 0 26931.76
227.06651306152344 0 9437.998
227.0855712890625 0 974.7786
227.10269165039062 0 966.53894
227.12222290039062 0 3068.14
228.07029724121094 0 1084.4785
228.13453674316406 0 45592.71 b 1
229.1184539794922 0 2415.636
229.13775634765625 0 5109.398
230.1227569580078 0 578.6133
230.1504364013672 0 3186.2766
234.12399291992188 0 697.32965
239.11476135253906 0 585.2876
239.15037536621094 0 1746.0925
240.1343231201172 0 2626.2915
242.1502685546875 0 10309.136
243.10911560058594 0 1690.1526
243.15362548828125 0 1652.9984
244.09303283691406 0 20719.564
244.12948608398438 0 22647.205
245.07705688476562 0 3282.7388
245.09579467773438 0 1603.4766
245.1322021484375 0 2225.2659
248.16065979003906 0 701.9038
252.1343994140625 0 1852.7552
253.11831665039062 0 986.0251
254.11341857910156 0 958.37067
258.1451721191406 0 1251.9323
261.1193542480469 0 1061.6775
261.13372802734375 0 646.5851
266.1132507324219 0 1089.1425
266.15020751953125 0 1211.8955
267.1456604003906 0 1515.1598
269.1610107421875 0 1113.5267 a Water loss 2
270.1451110839844 0 3308.423 a Ammonia loss 2
278.14947509765625 0 767.85626
279.1455993652344 0 3640.5515
280.12945556640625 0 6205.6943
280.1468505859375 0 731.8537
281.1319885253906 0 810.7553
282.1809997558594 0 2005.894
283.1403503417969 0 6983.2085
284.1429138183594 0 605.902 y 5
287.170166015625 0 1176.2842 a 2
291.1458435058594 0 703.7929
292.1301574707031 0 954.52905
293.16082763671875 0 671.4767
294.14410400390625 0 1256.6598
297.1560974121094 0 25003.947 b Water loss 2
298.1399841308594 0 19821.361 b Ammonia loss 2
298.1587219238281 0 2779.2297
299.14276123046875 0 3428.6523
300.15594482421875 0 1385.2987
301.150634765625 0 1445.149
301.18780517578125 0 698.5668
305.1613464355469 0 1239.7607
305.1819763183594 0 1366.4229
306.14520263671875 0 1923.2998 y Water loss 7
309.1560974121094 0 3551.3062
310.1396789550781 0 1255.7721
311.17181396484375 0 1797.9371
312.15545654296875 0 1510.0991
315.1665344238281 0 11212.085 b 2
316.1680908203125 0 719.59357
319.14056396484375 0 2626.5823
323.1728515625 0 675.1935
324.15582275390625 0 14170.76 y 7
325.1583557128906 0 2037.9141
327.1664733886719 0 4970.0703 b Water loss 5
329.1821594238281 0 11613.154
330.18572998046875 0 1983.3032
336.1669006347656 0 3424.3562 b 5
337.1510314941406 0 12000.863
337.1715393066406 0 526.71985
338.1340637207031 0 711.5468
338.154541015625 0 2524.7407
339.1669921875 0 4019.1135 y Water loss 4
339.6595458984375 0 581.16547 y Ammonia loss 4
340.1507263183594 0 2573.9739
345.177001953125 0 6565.076
346.1781311035156 0 869.83344
348.1714782714844 0 4962.0645 y 4
348.67266845703125 0 1542.7552
354.17718505859375 0 17790.137
355.16131591796875 0 6486.4326
355.18115234375 0 2074.176
356.16424560546875 0 1400.4479
357.17694091796875 0 5150.3535
358.16064453125 0 2844.7336
364.23382568359375 0 2044.628
365.21844482421875 0 13083.554
365.241455078125 0 907.8469
366.1989440917969 0 814.1997
366.2215576171875 0 2531.0288
367.16180419921875 0 817.58
372.1880187988281 0 11740.702
373.1910095214844 0 2459.2944
382.2447814941406 0 3365.5518
383.2289123535156 0 15489.611
384.1880798339844 0 1961.3307
384.2323303222656 0 2469.1418
385.1717224121094 0 2086.3955
392.19671630859375 0 848.64264
392.22967529296875 0 6426.4956
393.1868896484375 0 1828.257
393.2135925292969 0 9980.238
394.1725158691406 0 6283.2783
394.2156982421875 0 2424.5078
395.1761474609375 0 889.6855
396.2018737792969 0 2443.4712 y Ammonia loss 3
396.70318603515625 0 896.4453
398.2023620605469 0 664.9264
402.1988830566406 0 3818.5664
403.2021179199219 0 635.6147
404.71405029296875 0 7703.769 y 3
405.21527099609375 0 4523.194
410.2084655761719 0 1109.0254
410.2400817871094 0 10000.618 b Water loss 3
410.71978759765625 0 7276.3433
411.1965637207031 0 1729.4164
411.22344970703125 0 14575.428 b Ammonia loss 3
412.1830139160156 0 2395.3196
412.22698974609375 0 3300.9653
419.2049255371094 0 1509.4027
419.723876953125 0 1235.7572
420.1870422363281 0 4143.3115
420.21490478515625 0 1269.2268
423.72705078125 0 1606.3131
424.22589111328125 0 1267.6102
425.2137145996094 0 714.8746
426.1985168457031 0 763.73584
427.1826171875 0 1336.8354
427.2327575683594 0 1017.54126
427.2672119140625 0 2892.9248
427.74078369140625 0 1815.2778
428.2144470214844 0 1988.6735
428.2496643066406 0 5473.791 b 3
429.21173095703125 0 1238.6415
429.2524719238281 0 665.2056
432.7325744628906 0 5726.385
433.2301940917969 0 4524.222
433.7301025390625 0 1425.6088
434.23931884765625 0 2852.5295 y Water loss 6
435.2403259277344 0 709.14307
436.0353088378906 0 775.5581
437.21490478515625 0 2388.6592
438.1983337402344 0 3824.4172
439.1959228515625 0 1104.582
439.2260437011719 0 902.15857 y Water loss 2
439.7173156738281 0 3570.0315 y Ammonia loss 2
440.21820068359375 0 1508.4244
441.7380676269531 0 8851.746 b Water loss 7
442.23931884765625 0 3356.2478
442.73809814453125 0 757.0997
444.20928955078125 0 9988.329
445.2115783691406 0 2284.112
448.22967529296875 0 8383.494 y 2
448.73175048828125 0 3684.2207
449.2325744628906 0 984.42914
450.23291015625 0 931.8442
450.74224853515625 0 1168.4313 b 7
452.25048828125 0 28875.227 y 6
453.25372314453125 0 6042.2437
454.2580261230469 0 880.70917
455.2254333496094 0 3366.4705
456.2093200683594 0 9125.172
457.21142578125 0 1360.3293
461.24169921875 0 2811.8699
461.7378234863281 0 2285.4587
462.2381591796875 0 2286.1128
463.2314147949219 0 770.4555
467.22735595703125 0 727.2856
468.2483825683594 0 713.89526
470.248291015625 0 6956.174 b Water loss 8
470.7489318847656 0 4719.135 b Ammonia loss 8
471.2492980957031 0 1378.0765
473.23583984375 0 5634.3804
474.24090576171875 0 906.4395
479.25396728515625 0 4825.9307 b 8
479.7549743652344 0 2689.7715
480.25341796875 0 843.3584
485.27203369140625 0 2519.2664
489.248046875 0 700.79626 y Ammonia loss 1
494.23382568359375 0 1079.4218
495.2204284667969 0 1544.5005
496.2186584472656 0 940.862
497.7630310058594 0 1497.5707 y 1
498.26507568359375 0 1214.1227
510.3041687011719 0 1950.6143
512.2460327148438 0 2705.9004
513.2308349609375 0 4697.956
513.6815795898438 0 1016.59924
514.2328491210938 0 1554.5122
521.2719116210938 0 2124.8264
522.2655029296875 0 1665.1543
524.2831420898438 0 817.37537
529.7852783203125 0 1742.7988
530.2568969726562 0 4239.109
531.256103515625 0 2212.6514
534.30322265625 0 3326.873
534.7769165039062 0 2098.0535
535.2708129882812 0 1466.1985
537.2686157226562 0 1077.3484
538.2982788085938 0 11545.086 b Water loss 4
539.2860107421875 0 4907.399 b Ammonia loss 4
539.810302734375 0 2786.1892
540.3134155273438 0 1020.25476
543.301025390625 0 5399.0254
543.7822875976562 0 8987.86
544.2816162109375 0 7841.823
544.7797241210938 0 4209.7173
545.2684326171875 0 1132.979
549.2670288085938 0 4686.9614 y Water loss 5
550.260986328125 0 1070.804 y Ammonia loss 5
550.3041381835938 0 895.3286
551.6892700195312 0 5696.226
552.1903076171875 0 3178.8257
552.787841796875 0 184204.72 Precursor Water loss
553.2887573242188 0 116672.46 Precursor Ammonia loss
553.7896728515625 0 42437.62
554.2908935546875 0 6373.5654
555.2771606445312 0 1282.6583
556.3108520507812 0 1089.5122 b 4
557.06005859375 0 778.53107
557.306640625 0 688.6111
560.8172607421875 0 896.4109
561.3253784179688 0 720.52515
561.5703735351562 0 892.43304
561.7930297851562 0 23654.438 Precursor
562.2947998046875 0 14488.413
562.7952880859375 0 3650.7378
563.2850341796875 0 781.85547
567.2772216796875 0 65989.44 y 5
568.280029296875 0 19146.379
568.3270263671875 0 1055.8936
569.2840576171875 0 3218.1428
572.3031005859375 0 8721.65
573.3058471679688 0 3174.9321
574.305419921875 0 740.53235
586.3192138671875 0 4177.156
587.3213500976562 0 1072.4542
593.303466796875 0 760.4203
611.3145751953125 0 5596.9805
612.3168334960938 0 1662.5967
618.5404052734375 0 676.0111
627.3436889648438 0 806.8358
628.3340454101562 0 809.7513
629.3276977539062 0 727.24445
633.3065795898438 0 968.4261
635.3130493164062 0 744.6172
636.3002319335938 0 1330.7139
637.3305053710938 0 5108.631
638.3284912109375 0 2296.7654
641.8546142578125 0 8522.885
642.2936401367188 0 627.2068
642.3553466796875 0 3606.199
642.8568115234375 0 1561.9514
643.3414306640625 0 1610.0323
645.35791015625 0 1574.5005
649.3292846679688 0 4496.2695
650.3314819335938 0 1498.095
652.3455200195312 0 809.7876
653.3251953125 0 13936.007 b Water loss 5
654.3229370117188 0 4979.9683
655.3380126953125 0 5155.476
656.3377685546875 0 1805.6671
659.3146362304688 0 2275.8223
660.301025390625 0 3068.9238
670.3515625 0 3969.7876
671.348876953125 0 1628.496 b 5
673.3511352539062 0 19494.691
674.35400390625 0 6397.6587
675.354248046875 0 1277.9288
677.3250122070312 0 19104.66 y Water loss 4
678.3109130859375 0 41895.094 y Ammonia loss 4
679.3128051757812 0 14635.623
680.31591796875 0 2509.0815
685.3302612304688 0 854.6539
694.3397827148438 0 969.3802
695.33544921875 0 56701.03 y 4
696.3372802734375 0 21762.293
697.3380737304688 0 3952.5508
712.3591918945312 0 2970.6387
713.3588256835938 0 1162.9424
730.3726806640625 0 7670.438
731.3764038085938 0 2399.9575
755.3657836914062 0 1217.871
755.8690795898438 0 1578.6101
765.4000854492188 0 752.9062
773.3828735351562 0 1565.8982
774.3836669921875 0 840.7603
781.419677734375 0 10861.023 b Water loss 6
782.417236328125 0 4953.9307 b Ammonia loss 6
783.4163818359375 0 991.7968
790.4158935546875 0 909.30963 y Water loss 3
791.4067993164062 0 742.1271 y Ammonia loss 3
799.4329223632812 0 1536.9944 b 6
808.4188232421875 0 16947.771 y 3
809.4213256835938 0 8451.494
810.4239501953125 0 2168.5193
820.4312133789062 0 642.82385
821.4168701171875 0 1774.2715
822.4254150390625 0 855.4788
829.4275512695312 0 1089.5815
839.4425048828125 0 863.959
846.4453735351562 0 1161.0801
847.4371337890625 0 2014.6636
860.420654296875 0 917.2299
864.45751953125 0 777.91156
865.4400634765625 0 2608.9485
866.4334106445312 0 2129.6172
867.4342041015625 0 651.60645
877.4401245117188 0 6385.833 y Water loss 2
878.4357299804688 0 4715.8438 y Ammonia loss 2
879.4403686523438 0 1564.474
882.4703369140625 0 770.8862 b Water loss 7
883.45166015625 0 9525.735 b Ammonia loss 7
884.4537353515625 0 4274.7944
885.461669921875 0 945.86237
895.450439453125 0 94884.234 y 2
896.4530639648438 0 46521.793
897.4556274414062 0 13907.363
898.4559936523438 0 1794.8003
900.4766235351562 0 2336.8054 b 7
901.4815673828125 0 1064.0574
902.478271484375 0 767.1762
904.458251953125 0 655.83435
922.4656982421875 0 3047.2239
923.4614868164062 0 1932.3903
924.453369140625 0 1058.1719
939.4882202148438 0 1298.8369 b Water loss 8
940.4727783203125 0 10798.89 b Ammonia loss 8
941.475341796875 0 5712.8525
942.4779663085938 0 1733.7296
957.4996948242188 0 3066.0393 b 8
958.4954833984375 0 1935.7294
959.4839477539062 0 956.9504
994.5183715820312 0 5230.936 y 1
995.5215454101562 0 2827.9368
1245.9473876953125 0 615.8028
1928.7252197265625 0 678.2045
3085.17822265625 0 750.38617
3384.837158203125 0 675.38806

Spectrum Details

|  |  |
| --- | --- |
| Matched peaks? Matched peaks The total absolute number of peaks matched. Additionally in brackets the total fraction of peaks matched and the total number of peaks is shown. | 68 (14.44% of 471) |
| FDR? FDR The false discovery rate estimated for this peptide. It is calculated by matching all theoretical fragments with a non-integer shift with the raw peaks for this spectrum. This is done with 40 different shifts. The resulting percentage is the average number of annotated peaks over the number of annotated peaks with the correct spectrum. | 0.04% |
| Satellite FDR? Satellite FDR See the FDR for details on its calculation. This satellite ion specific FDR only contains the satellite ions (d/w) for I/L/J positions. | - |
| PSM Score? PSM Score The PSM Score as given by Hecklib to this annotated spectrum. It is shown with three significant figures. | 838 |

#### Spectrum 5464? Spectrum 5464 The raw spectrum of this peptide as annotated by Hecklib. The fragments are coloured according to ion type (see legend). Any peaks with a star '\*' as text can be hovered over to see the full details, first the ion type second the mass shift type. By hovering over the amino acids in the peptide or ions in the legend the corresponding peaks are highlighted. By toggling the 'Unassigned' label you can turn the background (unassigned) peaks on or off in the plot. By updating the slider in the Ion legend you can update the spectrum to only show the top X% of the peaks with labels. The top X% means any peak that is within X% of the highest intensity. By dragging in the spectrum you can zoom in to a specific part of the spectrum and use 'Zoom Out' to get back to the original zoom level. The annotation of the spectrum is based on the given sequence in the peptides file and is done with different software so inconsistencies are likely. The peaks are annotated based on the given sequence, with 20 ppm tolerance.

Copy Data

##### Spectrum 5464 (TSV)

###### Preview

```
Loading example...
```

*Click on the button to copy the data to your clipboard.*

Mz MinMz MaxIntensity Max

WidthHeightPeptide font sizePeptide stroke widthSpectrum font sizeSpectrum stroke widthCompact peptide

Ion legend

wxyz

abcd

OtherUnassignedIon^Charge^_Position_Show for top:%

QVSLQDKTGF

01.44e+52.89e+54.33e+55.78e+5

Zoom Out

y^+1^_1_y^+1^_2_c^+1^_2_c^+1^_3_y^+1^_3_y^+2^_6_y^+2^_6_y^+2^_7_z^+2^_7_c^+2^_7_y^+2^_7_c^+1^_4_z^+1^_4_y^+2^_8_y^+2^_8_z^+2^_8_c^+1^_4_y^+2^_8_c^+2^_8_y^+1^_4_c^+2^_9_y^+2^_9_y^+1^_5_z^+1^_5_c^+1^_5_y^+1^_5_c^+1^_5_w^+1^_6_c^+1^_6_c^+1^_6_y^+1^_6_y^+1^_6_z^+1^_6_c^+1^_6_y^+1^_6_w^+1^_7_z^+1^_7_c^+1^_7_c^+1^_7_y^+1^_7_c^+1^_7_z^+1^_8_w^+1^_8_y^+1^_8_y^+1^_8_z^+1^_8_y^+1^_8_c^+1^_8_c^+1^_8_c^+1^_9_c^+1^_9_c^+1^_9_z^+1^_9_y^+1^_9_

0779155823373116

Fragment Matches Table

Show background peaks

| Position | Ion type | Intensity | mz Theoretical | mz Error (Th) | mz Error (ppm) | Charge | Series Number |
| --- | --- | --- | --- | --- | --- | --- | --- |
| - | - | 7153 | 120.1 | - | - | 0 | - |
| - | - | 853.4 | 121.1 | - | - | 0 | - |
| - | - | 741.2 | 126.4 | - | - | 0 | - |
| - | - | 2582 | 129.1 | - | - | 0 | - |
| - | - | 932.2 | 131.1 | - | - | 0 | - |
| - | - | 868.9 | 135.6 | - | - | 0 | - |
| - | - | 914.6 | 147.6 | - | - | 0 | - |
| - | - | 2101 | 155.1 | - | - | 0 | - |
| - | - | 842.2 | 155.3 | - | - | 0 | - |
| - | - | 2147 | 159.1 | - | - | 0 | - |
| - | - | 1823 | 166.1 | - | - | 0 | - |
| 10 | y | 2.94E+04 | 166.1 | 0.0002775 | 1.671 | +1 | 1 |
| - | - | 1813 | 167.1 | - | - | 0 | - |
| - | - | 1.01E+04 | 173.1 | - | - | 0 | - |
| - | - | 2079 | 173.5 | - | - | 0 | - |
| - | - | 966.9 | 174.1 | - | - | 0 | - |
| - | - | 2493 | 177.1 | - | - | 0 | - |
| - | - | 5263 | 182.1 | - | - | 0 | - |
| - | - | 1.507E+04 | 183.1 | - | - | 0 | - |
| - | - | 942.8 | 184.1 | - | - | 0 | - |
| - | - | 867.1 | 184.5 | - | - | 0 | - |
| - | - | 886.2 | 190.9 | - | - | 0 | - |
| - | - | 965.8 | 198.9 | - | - | 0 | - |
| - | - | 7.193E+04 | 200.1 | - | - | 0 | - |
| - | - | 8498 | 201.1 | - | - | 0 | - |
| - | - | 5763 | 201.1 | - | - | 0 | - |
| - | - | 1.293E+04 | 210.1 | - | - | 0 | - |
| - | - | 1.037E+04 | 211.1 | - | - | 0 | - |
| - | - | 1024 | 212.1 | - | - | 0 | - |
| - | - | 1345 | 223.1 | - | - | 0 | - |
| 9 | y | 2.647E+04 | 223.1 | 0.0003134 | 1.405 | +1 | 2 |
| - | - | 953 | 226.1 | - | - | 0 | - |
| - | - | 905.6 | 227.6 | - | - | 0 | - |
| 2 | c | 5.998E+04 | 228.1 | 0.0002841 | 1.245 | +1 | 2 |
| - | - | 6109 | 229.1 | - | - | 0 | - |
| - | - | 3224 | 242.2 | - | - | 0 | - |
| - | - | 1891 | 244.1 | - | - | 0 | - |
| - | - | 3128 | 244.1 | - | - | 0 | - |
| - | - | 1734 | 270.1 | - | - | 0 | - |
| - | - | 1691 | 280.1 | - | - | 0 | - |
| - | - | 6583 | 297.2 | - | - | 0 | - |
| - | - | 9567 | 298.1 | - | - | 0 | - |
| - | - | 1999 | 299.1 | - | - | 0 | - |
| - | - | 1225 | 300.2 | - | - | 0 | - |
| 3 | c | 7724 | 315.2 | 0.0006043 | 1.918 | +1 | 3 |
| - | - | 1051 | 316.2 | - | - | 0 | - |
| 8 | y | 1.284E+04 | 324.2 | 0.0004255 | 1.313 | +1 | 3 |
| - | - | 2161 | 325.2 | - | - | 0 | - |
| - | - | 7553 | 329.2 | - | - | 0 | - |
| - | - | 1382 | 330.2 | - | - | 0 | - |
| - | - | 1359 | 337.1 | - | - | 0 | - |
| 5 | y | 1804 | 339.2 | 0.001215 | 3.581 | +2 | 6 |
| - | - | 1612 | 345.2 | - | - | 0 | - |
| 5 | y | 7489 | 348.2 | 0.0004184 | 1.202 | +2 | 6 |
| - | - | 3489 | 348.7 | - | - | 0 | - |
| - | - | 1424 | 357.2 | - | - | 0 | - |
| - | - | 8925 | 365.2 | - | - | 0 | - |
| - | - | 2739 | 366.2 | - | - | 0 | - |
| - | - | 4802 | 372.2 | - | - | 0 | - |
| - | - | 2369 | 376.2 | - | - | 0 | - |
| - | - | 1382 | 380.2 | - | - | 0 | - |
| - | - | 1.772E+04 | 383.2 | - | - | 0 | - |
| - | - | 3984 | 384.2 | - | - | 0 | - |
| - | - | 1257 | 392.2 | - | - | 0 | - |
| - | - | 1.781E+04 | 393.2 | - | - | 0 | - |
| - | - | 3314 | 394.2 | - | - | 0 | - |
| 4 | y | 3934 | 396.2 | 0.000378 | 0.9542 | +2 | 7 |
| 4 | z | 1915 | 396.7 | 0.00207 | 5.217 | +2 | 7 |
| - | - | 1054 | 398.2 | - | - | 0 | - |
| 7 | c | 993.2 | 400.2 | 0.00146 | 3.648 | +2 | 7 |
| - | - | 1480 | 402.2 | - | - | 0 | - |
| 4 | y | 1.188E+04 | 404.7 | 0.0005617 | 1.388 | +2 | 7 |
| - | - | 4721 | 405.2 | - | - | 0 | - |
| - | - | 1348 | 405.7 | - | - | 0 | - |
| - | - | 7017 | 410.2 | - | - | 0 | - |
| - | - | 2741 | 410.7 | - | - | 0 | - |
| - | - | 1.804E+04 | 411.2 | - | - | 0 | - |
| - | - | 4274 | 412.2 | - | - | 0 | - |
| - | - | 1333 | 419.2 | - | - | 0 | - |
| - | - | 1098 | 421 | - | - | 0 | - |
| 4 | c | 7473 | 428.3 | 0.0004027 | 0.9403 | +1 | 4 |
| - | - | 1446 | 429.3 | - | - | 0 | - |
| - | - | 1003 | 430.7 | - | - | 0 | - |
| - | - | 3518 | 432.7 | - | - | 0 | - |
| - | - | 3965 | 433.2 | - | - | 0 | - |
| - | - | 1061 | 433.7 | - | - | 0 | - |
| 7 | z | 3.969E+04 | 436.2 | 0.0007246 | 1.661 | +1 | 4 |
| - | - | 3.193E+04 | 437.2 | - | - | 0 | - |
| - | - | 6282 | 438.2 | - | - | 0 | - |
| 3 | y | 1810 | 439.2 | 0.001137 | 2.588 | +2 | 8 |
| 3 | y | 7150 | 439.7 | 0.0009654 | 2.195 | +2 | 8 |
| 3 | z | 2598 | 440.2 | 0.003435 | 7.804 | +2 | 8 |
| - | - | 5521 | 441.7 | - | - | 0 | - |
| - | - | 3606 | 442.2 | - | - | 0 | - |
| - | - | 1391 | 442.7 | - | - | 0 | - |
| - | - | 4687 | 443.3 | - | - | 0 | - |
| - | - | 4390 | 444.2 | - | - | 0 | - |
| - | - | 1072 | 444.3 | - | - | 0 | - |
| 4 | c | 1389 | 445.3 | 0.0006785 | 1.524 | +1 | 4 |
| 3 | y | 1.479E+04 | 448.2 | 0.0007524 | 1.679 | +2 | 8 |
| - | - | 8025 | 448.7 | - | - | 0 | - |
| - | - | 1743 | 449.2 | - | - | 0 | - |
| 8 | c | 1678 | 450.7 | 0.0006614 | 1.467 | +2 | 8 |
| - | - | 3592 | 451.2 | - | - | 0 | - |
| 7 | y | 2.477E+04 | 452.3 | 0.0004027 | 0.8904 | +1 | 4 |
| - | - | 5513 | 453.3 | - | - | 0 | - |
| - | - | 1489 | 455.2 | - | - | 0 | - |
| - | - | 1385 | 457.2 | - | - | 0 | - |
| - | - | 6224 | 461.7 | - | - | 0 | - |
| - | - | 2748 | 462.2 | - | - | 0 | - |
| - | - | 1066 | 465 | - | - | 0 | - |
| - | - | 1373 | 465.3 | - | - | 0 | - |
| - | - | 5596 | 470.2 | - | - | 0 | - |
| - | - | 3465 | 470.7 | - | - | 0 | - |
| - | - | 1882 | 471.2 | - | - | 0 | - |
| - | - | 2714 | 473.2 | - | - | 0 | - |
| 9 | c | 5183 | 479.3 | 0.0006107 | 1.274 | +2 | 9 |
| - | - | 2114 | 479.8 | - | - | 0 | - |
| - | - | 2892 | 487.3 | - | - | 0 | - |
| 2 | y | 2125 | 497.8 | 0.0005725 | 1.15 | +2 | 9 |
| - | - | 1207 | 498.3 | - | - | 0 | - |
| - | - | 1.267E+04 | 507.3 | - | - | 0 | - |
| - | - | 7130 | 508.3 | - | - | 0 | - |
| - | - | 1731 | 509.3 | - | - | 0 | - |
| - | - | 1308 | 516.3 | - | - | 0 | - |
| - | - | 1845 | 521.3 | - | - | 0 | - |
| - | - | 1748 | 522.3 | - | - | 0 | - |
| - | - | 3661 | 529.3 | - | - | 0 | - |
| - | - | 1567 | 529.8 | - | - | 0 | - |
| - | - | 1618 | 530.3 | - | - | 0 | - |
| - | - | 5488 | 530.3 | - | - | 0 | - |
| - | - | 1415 | 530.8 | - | - | 0 | - |
| - | - | 1690 | 531.3 | - | - | 0 | - |
| - | - | 5032 | 538.3 | - | - | 0 | - |
| - | - | 7559 | 539.3 | - | - | 0 | - |
| - | - | 1986 | 540.3 | - | - | 0 | - |
| - | - | 1.152E+04 | 543.8 | - | - | 0 | - |
| - | - | 1.372E+04 | 544.3 | - | - | 0 | - |
| - | - | 5240 | 544.8 | - | - | 0 | - |
| - | - | 2746 | 545.3 | - | - | 0 | - |
| 6 | y | 2326 | 549.3 | 0.0008695 | 1.583 | +1 | 5 |
| 6 | z | 1.069E+04 | 551.3 | 2.672E-05 | 0.04848 | +1 | 5 |
| - | - | 5.562E+04 | 552.3 | - | - | 0 | - |
| - | - | 3.015E+05 | 552.8 | - | - | 0 | - |
| - | - | 1.966E+05 | 553.3 | - | - | 0 | - |
| - | - | 6.064E+04 | 553.8 | - | - | 0 | - |
| - | - | 5489 | 554.3 | - | - | 0 | - |
| 5 | c | 3787 | 556.3 | 0.0007713 | 1.386 | +1 | 5 |
| - | - | 6.601E+04 | 561.8 | - | - | 0 | - |
| - | - | 3.95E+04 | 562.3 | - | - | 0 | - |
| - | - | 1.49E+04 | 562.8 | - | - | 0 | - |
| - | - | 1442 | 563.3 | - | - | 0 | - |
| 6 | y | 6.388E+04 | 567.3 | 0.0002236 | 0.3941 | +1 | 5 |
| - | - | 2.009E+04 | 568.3 | - | - | 0 | - |
| - | - | 4136 | 569.3 | - | - | 0 | - |
| - | - | 2.262E+04 | 572.3 | - | - | 0 | - |
| 5 | c | 5.854E+04 | 573.3 | 0.0002208 | 0.3851 | +1 | 5 |
| - | - | 1.661E+04 | 574.3 | - | - | 0 | - |
| - | - | 2244 | 575.3 | - | - | 0 | - |
| 5 | w | 5395 | 621.3 | 0.001866 | 3.003 | +1 | 6 |
| - | - | 2100 | 622.3 | - | - | 0 | - |
| - | - | 2333 | 631.3 | - | - | 0 | - |
| - | - | 1555 | 632.3 | - | - | 0 | - |
| - | - | 9598 | 644.3 | - | - | 0 | - |
| - | - | 1.683E+04 | 645.4 | - | - | 0 | - |
| - | - | 4263 | 646.4 | - | - | 0 | - |
| - | - | 4238 | 653.3 | - | - | 0 | - |
| - | - | 4468 | 654.3 | - | - | 0 | - |
| - | - | 2019 | 655.3 | - | - | 0 | - |
| 6 | c | 2020 | 670.4 | 0.001279 | 1.908 | +1 | 6 |
| 6 | c | 2437 | 671.3 | 0.002071 | 3.085 | +1 | 6 |
| - | - | 1198 | 672.3 | - | - | 0 | - |
| - | - | 7171 | 673.4 | - | - | 0 | - |
| - | - | 2224 | 674.4 | - | - | 0 | - |
| - | - | 1774 | 675.3 | - | - | 0 | - |
| 5 | y | 1.139E+04 | 677.3 | 0.0007337 | 1.083 | +1 | 6 |
| 5 | y | 2.835E+04 | 678.3 | 0.00152 | 2.241 | +1 | 6 |
| 5 | z | 1.04E+05 | 679.3 | 0.000104 | 0.1531 | +1 | 6 |
| - | - | 5.925E+04 | 680.3 | - | - | 0 | - |
| - | - | 1358 | 680.4 | - | - | 0 | - |
| - | - | 1.444E+04 | 681.3 | - | - | 0 | - |
| - | - | 2719 | 682.3 | - | - | 0 | - |
| - | - | 4.22E+04 | 687.4 | - | - | 0 | - |
| 6 | c | 3.242E+05 | 688.4 | 0.000302 | 0.4387 | +1 | 6 |
| - | - | 1.084E+05 | 689.4 | - | - | 0 | - |
| - | - | 1938 | 689.4 | - | - | 0 | - |
| - | - | 2.095E+04 | 690.4 | - | - | 0 | - |
| - | - | 1313 | 691.4 | - | - | 0 | - |
| - | - | 4464 | 694.3 | - | - | 0 | - |
| 5 | y | 7.01E+04 | 695.3 | 0.0002398 | 0.3449 | +1 | 6 |
| - | - | 2.431E+04 | 696.3 | - | - | 0 | - |
| - | - | 6553 | 697.3 | - | - | 0 | - |
| - | - | 1397 | 721.4 | - | - | 0 | - |
| - | - | 3334 | 730.4 | - | - | 0 | - |
| - | - | 1665 | 731.4 | - | - | 0 | - |
| - | - | 3149 | 734.4 | - | - | 0 | - |
| - | - | 1545 | 735.4 | - | - | 0 | - |
| - | - | 1696 | 739.8 | - | - | 0 | - |
| - | - | 1307 | 744.4 | - | - | 0 | - |
| 4 | w | 7.846E+04 | 749.3 | 0.001211 | 1.616 | +1 | 7 |
| - | - | 2.847E+04 | 750.4 | - | - | 0 | - |
| - | - | 7077 | 751.4 | - | - | 0 | - |
| - | - | 1.462E+04 | 757.4 | - | - | 0 | - |
| - | - | 5092 | 758.4 | - | - | 0 | - |
| - | - | 1397 | 759.4 | - | - | 0 | - |
| - | - | 1308 | 764.4 | - | - | 0 | - |
| - | - | 3526 | 765.9 | - | - | 0 | - |
| - | - | 3594 | 766.4 | - | - | 0 | - |
| - | - | 1268 | 771.5 | - | - | 0 | - |
| - | - | 1.547E+04 | 772.4 | - | - | 0 | - |
| - | - | 5911 | 773.4 | - | - | 0 | - |
| - | - | 2218 | 774.5 | - | - | 0 | - |
| - | - | 1164 | 775.9 | - | - | 0 | - |
| - | - | 7423 | 781.4 | - | - | 0 | - |
| - | - | 4201 | 782.4 | - | - | 0 | - |
| - | - | 1365 | 783.9 | - | - | 0 | - |
| 4 | z | 3.356E+05 | 792.4 | 0.0005127 | 0.647 | +1 | 7 |
| - | - | 2.379E+05 | 793.4 | - | - | 0 | - |
| - | - | 7.144E+04 | 794.4 | - | - | 0 | - |
| - | - | 1.002E+04 | 795.4 | - | - | 0 | - |
| 7 | c | 1738 | 798.4 | 0.009633 | 12.07 | +1 | 7 |
| 7 | c | 8771 | 799.4 | 0.0007358 | 0.9203 | +1 | 7 |
| - | - | 4415 | 800.4 | - | - | 0 | - |
| - | - | 5583 | 801.4 | - | - | 0 | - |
| - | - | 1736 | 802.5 | - | - | 0 | - |
| - | - | 1.168E+04 | 807.4 | - | - | 0 | - |
| 4 | y | 3.678E+04 | 808.4 | 0.0008164 | 1.01 | +1 | 7 |
| - | - | 1.407E+04 | 809.4 | - | - | 0 | - |
| - | - | 4802 | 810.4 | - | - | 0 | - |
| - | - | 7.284E+04 | 815.4 | - | - | 0 | - |
| 7 | c | 2.132E+05 | 816.5 | 0.0002396 | 0.2935 | +1 | 7 |
| - | - | 8.872E+04 | 817.5 | - | - | 0 | - |
| - | - | 1.731E+04 | 818.5 | - | - | 0 | - |
| - | - | 1967 | 819.5 | - | - | 0 | - |
| - | - | 2804 | 821.4 | - | - | 0 | - |
| - | - | 2.313E+04 | 823.4 | - | - | 0 | - |
| - | - | 1.158E+04 | 824.4 | - | - | 0 | - |
| - | - | 3291 | 825.4 | - | - | 0 | - |
| - | - | 3550 | 831.9 | - | - | 0 | - |
| - | - | 1013 | 832.4 | - | - | 0 | - |
| - | - | 1713 | 832.9 | - | - | 0 | - |
| - | - | 1842 | 835.4 | - | - | 0 | - |
| - | - | 1865 | 836.4 | - | - | 0 | - |
| - | - | 1375 | 837.4 | - | - | 0 | - |
| - | - | 2864 | 840.4 | - | - | 0 | - |
| - | - | 1520 | 840.9 | - | - | 0 | - |
| - | - | 3221 | 850.5 | - | - | 0 | - |
| - | - | 1897 | 851.5 | - | - | 0 | - |
| - | - | 8347 | 855.5 | - | - | 0 | - |
| - | - | 4782 | 856.5 | - | - | 0 | - |
| - | - | 3028 | 859.5 | - | - | 0 | - |
| 3 | z | 1242 | 861.4 | 0.001847 | 2.144 | +1 | 8 |
| 3 | w | 1767 | 862.4 | 0.0005778 | 0.67 | +1 | 8 |
| - | - | 1169 | 863.4 | - | - | 0 | - |
| - | - | 3103 | 865.4 | - | - | 0 | - |
| - | - | 2865 | 866.4 | - | - | 0 | - |
| - | - | 1625 | 872.5 | - | - | 0 | - |
| - | - | 8391 | 873.5 | - | - | 0 | - |
| - | - | 5979 | 874.5 | - | - | 0 | - |
| - | - | 1696 | 875.5 | - | - | 0 | - |
| 3 | y | 6200 | 877.4 | 0.003542 | 4.037 | +1 | 8 |
| 3 | y | 5604 | 878.4 | 0.009696 | 11.04 | +1 | 8 |
| 3 | z | 1.048E+05 | 879.4 | 0.0002836 | 0.3225 | +1 | 8 |
| - | - | 6.391E+04 | 880.4 | - | - | 0 | - |
| - | - | 2.402E+04 | 881.4 | - | - | 0 | - |
| - | - | 3141 | 882.4 | - | - | 0 | - |
| - | - | 1.232E+04 | 883.5 | - | - | 0 | - |
| - | - | 7836 | 884.5 | - | - | 0 | - |
| - | - | 1688 | 885.5 | - | - | 0 | - |
| - | - | 2060 | 886.5 | - | - | 0 | - |
| - | - | 2.789E+04 | 894.4 | - | - | 0 | - |
| 3 | y | 2.252E+05 | 895.5 | 0.0003131 | 0.3496 | +1 | 8 |
| - | - | 1.124E+05 | 896.5 | - | - | 0 | - |
| - | - | 2.713E+04 | 897.5 | - | - | 0 | - |
| - | - | 3794 | 898.5 | - | - | 0 | - |
| 8 | c | 3801 | 900.5 | 0.0008612 | 0.9564 | +1 | 8 |
| - | - | 2211 | 901.5 | - | - | 0 | - |
| 8 | c | 4.823E+04 | 917.5 | 0.0001886 | 0.2056 | +1 | 8 |
| - | - | 2.237E+04 | 918.5 | - | - | 0 | - |
| - | - | 5715 | 919.5 | - | - | 0 | - |
| - | - | 3038 | 922.5 | - | - | 0 | - |
| - | - | 2205 | 923.5 | - | - | 0 | - |
| - | - | 7.835E+04 | 930.5 | - | - | 0 | - |
| - | - | 4.054E+04 | 931.5 | - | - | 0 | - |
| - | - | 9923 | 932.5 | - | - | 0 | - |
| - | - | 1.817E+04 | 940.5 | - | - | 0 | - |
| - | - | 9398 | 941.5 | - | - | 0 | - |
| - | - | 1499 | 942.5 | - | - | 0 | - |
| 9 | c | 1796 | 956.5 | 0.003073 | 3.212 | +1 | 9 |
| 9 | c | 8302 | 957.5 | 0.003981 | 4.158 | +1 | 9 |
| - | - | 6361 | 958.5 | - | - | 0 | - |
| - | - | 2600 | 959.5 | - | - | 0 | - |
| - | - | 1513 | 969.4 | - | - | 0 | - |
| - | - | 1618 | 970.4 | - | - | 0 | - |
| 9 | c | 4.657E+05 | 974.5 | 0.0001372 | 0.1408 | +1 | 9 |
| - | - | 2.336E+05 | 975.5 | - | - | 0 | - |
| - | - | 1427 | 975.8 | - | - | 0 | - |
| - | - | 6.609E+04 | 976.5 | - | - | 0 | - |
| - | - | 5779 | 977.5 | - | - | 0 | - |
| 2 | z | 2.402E+04 | 978.5 | 0.0001982 | 0.2026 | +1 | 9 |
| - | - | 1.549E+04 | 979.5 | - | - | 0 | - |
| - | - | 1993 | 980.5 | - | - | 0 | - |
| - | - | 1344 | 992.5 | - | - | 0 | - |
| 2 | y | 7449 | 994.5 | 0.002443 | 2.456 | +1 | 9 |
| - | - | 6930 | 995.5 | - | - | 0 | - |
| - | - | 1319 | 996.5 | - | - | 0 | - |
| - | - | 1722 | 1008 | - | - | 0 | - |
| - | - | 1844 | 1010 | - | - | 0 | - |
| - | - | 2904 | 1017 | - | - | 0 | - |
| - | - | 2155 | 1018 | - | - | 0 | - |
| - | - | 1462 | 1023 | - | - | 0 | - |
| - | - | 3410 | 1024 | - | - | 0 | - |
| - | - | 2651 | 1034 | - | - | 0 | - |
| - | - | 6408 | 1035 | - | - | 0 | - |
| - | - | 4470 | 1036 | - | - | 0 | - |
| - | - | 3268 | 1037 | - | - | 0 | - |
| - | - | 3.63E+04 | 1049 | - | - | 0 | - |
| - | - | 2.151E+04 | 1050 | - | - | 0 | - |
| - | - | 3.537E+04 | 1051 | - | - | 0 | - |
| - | - | 1.993E+04 | 1052 | - | - | 0 | - |
| - | - | 7956 | 1053 | - | - | 0 | - |
| - | - | 2079 | 1054 | - | - | 0 | - |
| - | - | 4858 | 1062 | - | - | 0 | - |
| - | - | 1.167E+04 | 1063 | - | - | 0 | - |
| - | - | 4.492E+04 | 1064 | - | - | 0 | - |
| - | - | 2.626E+04 | 1065 | - | - | 0 | - |
| - | - | 8393 | 1066 | - | - | 0 | - |
| - | - | 9027 | 1068 | - | - | 0 | - |
| - | - | 3760 | 1069 | - | - | 0 | - |
| - | - | 1464 | 1070 | - | - | 0 | - |
| - | - | 1525 | 1074 | - | - | 0 | - |
| - | - | 1.309E+04 | 1078 | - | - | 0 | - |
| - | - | 6.04E+04 | 1079 | - | - | 0 | - |
| - | - | 5.422E+04 | 1080 | - | - | 0 | - |
| - | - | 2.329E+04 | 1081 | - | - | 0 | - |
| - | - | 5182 | 1082 | - | - | 0 | - |
| - | - | 1925 | 1088 | - | - | 0 | - |
| - | - | 4792 | 1089 | - | - | 0 | - |
| - | - | 1777 | 1090 | - | - | 0 | - |
| - | - | 2.651E+04 | 1096 | - | - | 0 | - |
| - | - | 1.636E+04 | 1097 | - | - | 0 | - |
| - | - | 5215 | 1098 | - | - | 0 | - |
| - | - | 2.53E+04 | 1106 | - | - | 0 | - |
| - | - | 5.722E+05 | 1107 | - | - | 0 | - |
| - | - | 3.462E+05 | 1108 | - | - | 0 | - |
| - | - | 2527 | 1108 | - | - | 0 | - |
| - | - | 1.202E+05 | 1109 | - | - | 0 | - |
| - | - | 1.182E+04 | 1110 | - | - | 0 | - |
| - | - | 1332 | 1116 | - | - | 0 | - |
| - | - | 2.513E+05 | 1123 | - | - | 0 | - |
| - | - | 4.831E+05 | 1124 | - | - | 0 | - |
| - | - | 2.549E+05 | 1125 | - | - | 0 | - |
| - | - | 7.412E+04 | 1126 | - | - | 0 | - |
| - | - | 7749 | 1127 | - | - | 0 | - |
| - | - | 2852 | 1139 | - | - | 0 | - |
| - | - | 1787 | 1149 | - | - | 0 | - |
| - | - | 3604 | 1260 | - | - | 0 | - |
| - | - | 1653 | 1261 | - | - | 0 | - |
| - | - | 1815 | 1335 | - | - | 0 | - |
| - | - | 3071 | 1375 | - | - | 0 | - |
| - | - | 1670 | 1376 | - | - | 0 | - |
| - | - | 1611 | 1532 | - | - | 0 | - |
| - | - | 1412 | 1533 | - | - | 0 | - |
| - | - | 1511 | 1534 | - | - | 0 | - |
| - | - | 1878 | 1551 | - | - | 0 | - |
| - | - | 1442 | 1621 | - | - | 0 | - |
| - | - | 1316 | 1622 | - | - | 0 | - |
| - | - | 2147 | 1623 | - | - | 0 | - |
| - | - | 3760 | 1635 | - | - | 0 | - |
| - | - | 2319 | 1636 | - | - | 0 | - |
| - | - | 1556 | 1637 | - | - | 0 | - |
| - | - | 8044 | 1664 | - | - | 0 | - |
| - | - | 5612 | 1665 | - | - | 0 | - |
| - | - | 4575 | 1666 | - | - | 0 | - |
| - | - | 1509 | 1667 | - | - | 0 | - |
| - | - | 7112 | 1681 | - | - | 0 | - |
| - | - | 5933 | 1682 | - | - | 0 | - |
| - | - | 3253 | 1683 | - | - | 0 | - |
| - | - | 2178 | 1684 | - | - | 0 | - |
| - | - | 1679 | 3085 | - | - | 0 | - |

m/z Charge Intensity FragmentType MassShift Position
120.08106231689453 0 7153.4233
121.08455657958984 0 853.39307
126.37870788574219 0 741.2034
129.10250854492188 0 2582.2598
131.11842346191406 0 932.20264
135.61940002441406 0 868.8969
147.62835693359375 0 914.5641
155.1183624267578 0 2101.4248
155.29466247558594 0 842.202
159.07669067382812 0 2146.5234
166.07888793945312 0 1823.2631
166.08653259277344 0 29404.758 y 9
167.0901336669922 0 1812.7344
173.1287078857422 0 10098.681
173.45318603515625 0 2079.4333
174.13211059570312 0 966.8658
177.10243225097656 0 2493.323
182.12913513183594 0 5263.34
183.113037109375 0 15065.847
184.1170654296875 0 942.7614
184.5094757080078 0 867.0577
190.8750457763672 0 886.2453
198.9008331298828 0 965.77
200.13958740234375 0 71931.69
201.1236572265625 0 8497.603
201.14291381835938 0 5762.7217
210.12399291992188 0 12928.122
211.10789489746094 0 10367.411
212.11138916015625 0 1023.9181
223.09571838378906 0 1345.2592
223.1080322265625 0 26469.64 y 8
226.11813354492188 0 953.0114
227.63168334960938 0 905.57227
228.13455200195312 0 59976.844 c Ammonia loss 1
229.13787841796875 0 6108.8013
242.1504669189453 0 3223.7983
244.09249877929688 0 1891.1552
244.1291961669922 0 3128.2568
270.1462707519531 0 1733.8192
280.12921142578125 0 1691.4739
297.156005859375 0 6582.88
298.1401672363281 0 9567.107
299.1436767578125 0 1999.1532
300.156005859375 0 1224.9558
315.1669006347656 0 7724.162 c Ammonia loss 2
316.17120361328125 0 1050.9684
324.15582275390625 0 12841.439 y 7
325.1595458984375 0 2161.0945
329.1822814941406 0 7552.844
330.1876525878906 0 1381.6964
337.1498718261719 0 1358.7379
339.1675109863281 0 1803.619 y Water loss 4
345.17730712890625 0 1611.7295
348.1719970703125 0 7488.998 y 4
348.6727294921875 0 3489.0894
357.1790466308594 0 1423.5167
365.2185974121094 0 8924.772
366.22296142578125 0 2738.6123
372.18768310546875 0 4801.6494
376.19769287109375 0 2368.8054
380.1928405761719 0 1381.8114
383.22906494140625 0 17721.49
384.2327575683594 0 3983.5632
392.23040771484375 0 1256.7877
393.2137145996094 0 17813.494
394.2158508300781 0 3314.2412
396.2007141113281 0 3934.4458 y Ammonia loss 3
396.7021789550781 0 1915.3386 z 3
398.2070007324219 0 1054.4664
400.22052001953125 0 993.18085 c Ammonia loss 6
402.1980895996094 0 1480.1779
404.71417236328125 0 11876.12 y 3
405.21551513671875 0 4721.398
405.7162170410156 0 1347.7201
410.2407531738281 0 7017.4077
410.7192687988281 0 2741.386
411.22406005859375 0 18040.314
412.22705078125 0 4273.9297
419.2237854003906 0 1332.8533
421.0416564941406 0 1097.6013
428.2507629394531 0 7472.547 c Ammonia loss 3
429.252685546875 0 1446.4436
430.7140808105469 0 1002.7667
432.7334899902344 0 3517.8306
433.2269287109375 0 3965.0078
433.72979736328125 0 1061.0886
436.23236083984375 0 39691.793 z 6
437.23846435546875 0 31928.73
438.24072265625 0 6282.4717
439.22320556640625 0 1810.4216 y Water loss 2
439.7173156738281 0 7150.414 y Ammonia loss 2
440.2168273925781 0 2598.4192 z 2
441.73870849609375 0 5520.5376
442.2379455566406 0 3606.4004
442.73907470703125 0 1390.7102
443.2745361328125 0 4686.5015
444.208984375 0 4389.6387
444.27978515625 0 1071.7367
445.277587890625 0 1388.7262 c 3
448.2303771972656 0 14787.839 y 2
448.7319030761719 0 8025.389
449.2324523925781 0 1743.2347
450.7435607910156 0 1677.8767 c Ammonia loss 7
451.24310302734375 0 3592.1187
452.2507629394531 0 24765.268 y 6
453.2541809082031 0 5512.725
455.2264099121094 0 1489.232
457.24407958984375 0 1384.6647
461.7361755371094 0 6224.278
462.2375183105469 0 2747.8135
465.0129089355469 0 1065.7601
465.2528381347656 0 1373.3175
470.2485046386719 0 5596.164
470.74859619140625 0 3465.285
471.24798583984375 0 1882.3986
473.237060546875 0 2713.5552
479.2542419433594 0 5182.5703 c Ammonia loss 8
479.75616455078125 0 2114.1138
487.2875061035156 0 2891.6467
497.764404296875 0 2125.2378 y 1
498.2652893066406 0 1207.4094
507.2695007324219 0 12667.221
508.274169921875 0 7130.34
509.27691650390625 0 1730.7642
516.2661743164062 0 1308.2404
521.2734985351562 0 1845.2473
522.265869140625 0 1747.951
529.3231201171875 0 3660.501
529.7840576171875 0 1566.7788
530.2603149414062 0 1617.5227
530.3064575195312 0 5487.6753
530.77490234375 0 1414.9048
531.3077392578125 0 1690.1929
538.2992553710938 0 5031.84
539.2847900390625 0 7559.1284
540.2854614257812 0 1986.3014
543.7835083007812 0 11524.093
544.281005859375 0 13720.281
544.77783203125 0 5239.6587
545.2765502929688 0 2745.6538
549.265869140625 0 2325.781 y Water loss 5
551.2586059570312 0 10693.931 z 5
552.266845703125 0 55619.02
552.7884521484375 0 301502.16
553.2889404296875 0 196591.72
553.7910766601562 0 60642.133
554.284423828125 0 5488.711
556.3081665039062 0 3787.082 c Ammonia loss 4
561.79345703125 0 66008.305
562.294921875 0 39504.793
562.7965087890625 0 14899.451
563.29833984375 0 1442.3969
567.2775268554688 0 63877.664 y 5
568.2803344726562 0 20088.361
569.2843017578125 0 4135.6597
572.3268432617188 0 22620.01
573.3352661132812 0 58536.668 c 4
574.337158203125 0 16614.81
575.3383178710938 0 2244.3564
621.2897338867188 0 5394.5415 w 4
622.2918701171875 0 2100.42
631.2955322265625 0 2333.3997
632.2996215820312 0 1555.1884
644.349609375 0 9597.876
645.3565063476562 0 16834.156
646.359130859375 0 4263.237
653.3262939453125 0 4237.768
654.3140869140625 0 4468.2925
655.3195190429688 0 2018.7001
670.3505859375 0 2020.0278 c Water loss 5
671.3379516601562 0 2437.193 c Ammonia loss 5
672.3389282226562 0 1198.1833
673.3521118164062 0 7171.386
674.3544311523438 0 2223.502
675.3284912109375 0 1773.5927
677.3260498046875 0 11392.922 y Water loss 4
678.3108520507812 0 28352.906 y Ammonia loss 4
679.3172607421875 0 104016.45 z 4
680.3221435546875 0 59245.184
680.3905029296875 0 1357.8541
681.3261108398438 0 14444.832
682.3316040039062 0 2719.2876
687.3552856445312 0 42196.223
688.3627319335938 0 324196.84 c 5
689.3655395507812 0 108386.78
689.4375610351562 0 1938.087
690.3678588867188 0 20950.043
691.3704223632812 0 1312.9178
694.3269653320312 0 4463.8604
695.3361206054688 0 70095.33 y 4
696.3392944335938 0 24305.275
697.3411254882812 0 6553.4927
721.3668212890625 0 1397.2135
730.3723754882812 0 3334.0845
731.3792724609375 0 1664.5381
734.3718872070312 0 3149.3958
735.3775634765625 0 1545.2351
739.8453369140625 0 1695.6954
744.4197387695312 0 1307.3274
749.34765625 0 78462.91 w 3
750.3505249023438 0 28470.328
751.3544921875 0 7076.5815
757.4303588867188 0 14622.768
758.4314575195312 0 5092.113
759.439453125 0 1397.245
764.3899536132812 0 1307.6346
765.8779907226562 0 3525.6758
766.3772583007812 0 3594.4534
771.455078125 0 1267.8918
772.4442138671875 0 15472.52
773.4465942382812 0 5910.9385
774.45556640625 0 2217.5818
775.8568115234375 0 1164.0742
781.4201049804688 0 7423.033
782.4169311523438 0 4200.7583
783.8717651367188 0 1365.0287
792.4017333984375 0 335603.3 z 3
793.4060668945312 0 237922.19
794.4098510742188 0 71435.25
795.4143676757812 0 10016.342
798.4371948242188 0 1738.3087 c Water loss 6
799.4315795898438 0 8770.957 c Ammonia loss 6
800.434814453125 0 4415.3926
801.4451904296875 0 5583.422
802.4523315429688 0 1736.1996
807.4116821289062 0 11680.774
808.4191284179688 0 36784.594 y 3
809.4228515625 0 14070.615
810.42138671875 0 4801.971
815.4498901367188 0 72842.39
816.4571533203125 0 213197.4 c 6
817.4602661132812 0 88722.02
818.463134765625 0 17308.357
819.4692993164062 0 1967.3505
821.4115600585938 0 2803.8452
823.3709106445312 0 23131.977
824.377685546875 0 11577.716
825.3819580078125 0 3290.5378
831.8953247070312 0 3550.0232
832.3953247070312 0 1012.67303
832.89501953125 0 1712.8596
835.4422607421875 0 1842.2759
836.3764038085938 0 1865.0588
837.3851928710938 0 1375.2722
840.4056396484375 0 2863.7417
840.911865234375 0 1520.4696
850.4547119140625 0 3221.219
851.4569702148438 0 1897.1648
855.4788208007812 0 8347.461
856.4827880859375 0 4782.192
859.4708862304688 0 3027.631
861.4208374023438 0 1242.2083 z Water loss 2
862.429931640625 0 1766.5536 w 2
863.4305419921875 0 1168.6508
865.4426879882812 0 3103.37
866.4408569335938 0 2865.3313
872.4794921875 0 1624.7511
873.489990234375 0 8391.085
874.493896484375 0 5978.62
875.4994506835938 0 1696.2181
877.4378662109375 0 6199.534 y Water loss 2
878.4351196289062 0 5603.6294 y Ammonia loss 2
879.4335327148438 0 104788.2 z 2
880.4375610351562 0 63912.457
881.4414672851562 0 24024.945
882.4446411132812 0 3140.6018
883.4529418945312 0 12318.72
884.4555053710938 0 7836.349
885.461181640625 0 1687.6029
886.5247802734375 0 2060.4429
894.4439086914062 0 27887.598
895.45166015625 0 225151.34 y 2
896.4549560546875 0 112433.305
897.4571533203125 0 27133.346
898.4594116210938 0 3793.7275
900.4776611328125 0 3801.0518 c Ammonia loss 7
901.4868774414062 0 2210.8389
917.5048828125 0 48233.043 c 7
918.50732421875 0 22366.398
919.5094604492188 0 5715.3213
922.461669921875 0 3037.814
923.4525146484375 0 2204.7017
930.5126342773438 0 78352.76
931.5158081054688 0 40539.023
932.5189819335938 0 9922.521
940.4747314453125 0 18170.977
941.4766235351562 0 9397.756
942.4771118164062 0 1499.1562
956.51904296875 0 1796.2007 c Water loss 8
957.5039672851562 0 8302.04 c Ammonia loss 8
958.5010986328125 0 6360.5957
959.5059204101562 0 2600.123
969.4378051757812 0 1512.8737
970.4493408203125 0 1618.2668
974.5266723632812 0 465732.4 c 8
975.5294799804688 0 233602.45
975.7687377929688 0 1427.486
976.5317993164062 0 66092.27
977.5338134765625 0 5778.8467
978.50146484375 0 24023.361 z 1
979.504638671875 0 15485.495
980.5078735351562 0 1992.7206
992.5082397460938 0 1343.9866
994.5179443359375 0 7448.5703 y 1
995.52197265625 0 6929.536
996.5338745117188 0 1319.2573
1007.5358276367188 0 1721.997
1009.5682373046875 0 1843.6012
1016.5018920898438 0 2904.403
1017.5052490234375 0 2154.6511
1022.50830078125 0 1462.0238
1023.5079345703125 0 3410.3997
1033.5909423828125 0 2650.602
1034.572265625 0 6407.916
1035.5137939453125 0 4470.0615
1036.5228271484375 0 3268.4966
1048.531982421875 0 36302.9
1049.5352783203125 0 21511.693
1050.514404296875 0 35365.324
1051.5133056640625 0 19932.74
1052.5341796875 0 7956.296
1053.54931640625 0 2078.8218
1061.580810546875 0 4857.995
1062.5767822265625 0 11672.102
1063.5662841796875 0 44923.324
1064.5667724609375 0 26263.342
1065.5684814453125 0 8392.871
1067.5255126953125 0 9026.721
1068.5245361328125 0 3759.9194
1069.5347900390625 0 1464.3209
1073.52294921875 0 1525.1831
1077.5806884765625 0 13086.498
1078.5672607421875 0 60404.383
1079.5802001953125 0 54215.29
1080.5885009765625 0 23291.5
1081.598876953125 0 5181.7734
1087.5655517578125 0 1925.3197
1088.55810546875 0 4792.1543
1089.5537109375 0 1777.2297
1095.592041015625 0 26506.547
1096.59423828125 0 16362.818
1097.59375 0 5215.439
1105.5755615234375 0 25304.012
1106.560302734375 0 572227.1
1107.5634765625 0 346247.03
1107.84619140625 0 2527.1907
1108.5654296875 0 120201.336
1109.5675048828125 0 11824.529
1115.5152587890625 0 1331.5966
1122.5787353515625 0 251308.25
1123.5853271484375 0 483139.47
1124.5882568359375 0 254918.84
1125.5919189453125 0 74124.72
1126.5948486328125 0 7748.6367
1138.5447998046875 0 2852.2922
1148.5457763671875 0 1786.5046
1259.6226806640625 0 3603.9346
1260.622802734375 0 1652.8599
1334.62548828125 0 1814.8604
1374.64453125 0 3071.3167
1375.6527099609375 0 1669.7019
1531.7574462890625 0 1611.1852
1532.754150390625 0 1411.6771
1533.754638671875 0 1511.1151
1550.7025146484375 0 1877.875
1620.7835693359375 0 1441.5497
1621.7784423828125 0 1316.4823
1622.7786865234375 0 2147.396
1634.7930908203125 0 3759.707
1635.810546875 0 2319.1523
1636.8111572265625 0 1555.8969
1663.7894287109375 0 8044.141
1664.7940673828125 0 5612.1924
1665.7916259765625 0 4574.714
1666.7957763671875 0 1509.0847
1680.8126220703125 0 7112.4595
1681.8150634765625 0 5933.369
1682.7977294921875 0 3252.967
1683.7999267578125 0 2178.4167
3085.364990234375 0 1678.817

Spectrum Details

|  |  |
| --- | --- |
| Matched peaks? Matched peaks The total absolute number of peaks matched. Additionally in brackets the total fraction of peaks matched and the total number of peaks is shown. | 54 (14.21% of 380) |
| FDR? FDR The false discovery rate estimated for this peptide. It is calculated by matching all theoretical fragments with a non-integer shift with the raw peaks for this spectrum. This is done with 40 different shifts. The resulting percentage is the average number of annotated peaks over the number of annotated peaks with the correct spectrum. | 0.71% |
| Satellite FDR? Satellite FDR See the FDR for details on its calculation. This satellite ion specific FDR only contains the satellite ions (d/w) for I/L/J positions. | 11.90% |
| PSM Score? PSM Score The PSM Score as given by Hecklib to this annotated spectrum. It is shown with three significant figures. | 572 |

#### Spectrum 5767? Spectrum 5767 The raw spectrum of this peptide as annotated by Hecklib. The fragments are coloured according to ion type (see legend). Any peaks with a star '\*' as text can be hovered over to see the full details, first the ion type second the mass shift type. By hovering over the amino acids in the peptide or ions in the legend the corresponding peaks are highlighted. By toggling the 'Unassigned' label you can turn the background (unassigned) peaks on or off in the plot. By updating the slider in the Ion legend you can update the spectrum to only show the top X% of the peaks with labels. The top X% means any peak that is within X% of the highest intensity. By dragging in the spectrum you can zoom in to a specific part of the spectrum and use 'Zoom Out' to get back to the original zoom level. The annotation of the spectrum is based on the given sequence in the peptides file and is done with different software so inconsistencies are likely. The peaks are annotated based on the given sequence, with 20 ppm tolerance.

Copy Data

##### Spectrum 5767 (TSV)

###### Preview

```
Loading example...
```

*Click on the button to copy the data to your clipboard.*

Mz MinMz MaxIntensity Max

WidthHeightPeptide font sizePeptide stroke widthSpectrum font sizeSpectrum stroke widthCompact peptide

Ion legend

wxyz

abcd

OtherUnassignedIon^Charge^_Position_Show for top:%

QVSLQDKTGF

01.96e+43.92e+45.89e+47.85e+4

Zoom Out

y^+1^_1_y^+1^_2_c^+1^_2_c^+1^_3_y^+1^_3_y^+2^_6_y^+2^_7_c^+1^_4_z^+1^_4_y^+2^_8_z^+2^_8_y^+2^_8_c^+2^_8_y^+1^_4_c^+2^_9_z^+1^_5_y^+1^_5_c^+1^_5_w^+1^_6_y^+1^_6_y^+1^_6_z^+1^_6_c^+1^_6_y^+1^_6_w^+1^_7_z^+1^_7_c^+1^_7_y^+1^_7_c^+1^_7_y^+1^_8_y^+1^_8_z^+1^_8_y^+1^_8_c^+1^_8_c^+1^_8_c^+1^_9_c^+1^_9_c^+1^_9_y^+1^_9_z^+1^_9_y^+1^_9_

0779155723363114

Fragment Matches Table

Show background peaks

| Position | Ion type | Intensity | mz Theoretical | mz Error (Th) | mz Error (ppm) | Charge | Series Number |
| --- | --- | --- | --- | --- | --- | --- | --- |
| - | - | 1339 | 120.1 | - | - | 0 | - |
| - | - | 518.4 | 129.1 | - | - | 0 | - |
| - | - | 433.1 | 139.4 | - | - | 0 | - |
| - | - | 830.7 | 149 | - | - | 0 | - |
| - | - | 457.3 | 155.1 | - | - | 0 | - |
| - | - | 446.8 | 157.3 | - | - | 0 | - |
| 10 | y | 4043 | 166.1 | 3.34E-05 | 0.2011 | +1 | 1 |
| - | - | 1372 | 173.1 | - | - | 0 | - |
| - | - | 975.3 | 177.1 | - | - | 0 | - |
| - | - | 1356 | 182.1 | - | - | 0 | - |
| - | - | 2638 | 183.1 | - | - | 0 | - |
| - | - | 465.9 | 197.2 | - | - | 0 | - |
| - | - | 1.194E+04 | 200.1 | - | - | 0 | - |
| - | - | 2906 | 201.1 | - | - | 0 | - |
| - | - | 803.9 | 201.1 | - | - | 0 | - |
| - | - | 2201 | 210.1 | - | - | 0 | - |
| - | - | 2361 | 211.1 | - | - | 0 | - |
| - | - | 546 | 217.6 | - | - | 0 | - |
| 9 | y | 3676 | 223.1 | 0.0002206 | 0.9888 | +1 | 2 |
| 2 | c | 8546 | 228.1 | 0.0001431 | 0.6274 | +1 | 2 |
| - | - | 524.2 | 234.4 | - | - | 0 | - |
| - | - | 555.1 | 238 | - | - | 0 | - |
| - | - | 596.3 | 258.6 | - | - | 0 | - |
| - | - | 748.9 | 297.2 | - | - | 0 | - |
| - | - | 2335 | 298.1 | - | - | 0 | - |
| 3 | c | 1468 | 315.2 | 3.653E-05 | 0.1159 | +1 | 3 |
| 8 | y | 1513 | 324.2 | 0.0005171 | 1.595 | +1 | 3 |
| - | - | 797.6 | 329.2 | - | - | 0 | - |
| 5 | y | 947.5 | 348.2 | 0.0002835 | 0.8142 | +2 | 6 |
| - | - | 1074 | 365.2 | - | - | 0 | - |
| - | - | 2061 | 380.2 | - | - | 0 | - |
| - | - | 569.5 | 381.8 | - | - | 0 | - |
| - | - | 1977 | 383.2 | - | - | 0 | - |
| - | - | 720.3 | 392.2 | - | - | 0 | - |
| - | - | 3026 | 393.2 | - | - | 0 | - |
| - | - | 1773 | 398.2 | - | - | 0 | - |
| 4 | y | 1172 | 404.7 | 0.001172 | 2.896 | +2 | 7 |
| - | - | 1045 | 410.2 | - | - | 0 | - |
| - | - | 2775 | 411.2 | - | - | 0 | - |
| - | - | 662.6 | 412.2 | - | - | 0 | - |
| - | - | 506.7 | 415.1 | - | - | 0 | - |
| - | - | 1461 | 417.1 | - | - | 0 | - |
| - | - | 1752 | 419.2 | - | - | 0 | - |
| 4 | c | 944 | 428.3 | 0.001105 | 2.579 | +1 | 4 |
| - | - | 804.8 | 430.2 | - | - | 0 | - |
| 7 | z | 9574 | 436.2 | 0.000435 | 0.9972 | +1 | 4 |
| - | - | 6236 | 437.2 | - | - | 0 | - |
| - | - | 1512 | 438.2 | - | - | 0 | - |
| 3 | y | 712.1 | 439.7 | 0.0004384 | 0.997 | +2 | 8 |
| 3 | z | 630.5 | 440.2 | 0.002581 | 5.863 | +2 | 8 |
| - | - | 940.6 | 441.7 | - | - | 0 | - |
| - | - | 822.7 | 442.2 | - | - | 0 | - |
| - | - | 746.2 | 444.2 | - | - | 0 | - |
| 3 | y | 1546 | 448.2 | 0.0007829 | 1.747 | +2 | 8 |
| - | - | 920.5 | 448.7 | - | - | 0 | - |
| 8 | c | 675.4 | 450.7 | 0.0001931 | 0.4284 | +2 | 8 |
| 7 | y | 3822 | 452.3 | 0.000696 | 1.539 | +1 | 4 |
| - | - | 1208 | 470.2 | - | - | 0 | - |
| 9 | c | 1201 | 479.3 | 0.0002445 | 0.5101 | +2 | 9 |
| - | - | 633.4 | 507.1 | - | - | 0 | - |
| - | - | 3447 | 507.3 | - | - | 0 | - |
| - | - | 1064 | 508.3 | - | - | 0 | - |
| - | - | 1087 | 530.3 | - | - | 0 | - |
| - | - | 788.3 | 534.3 | - | - | 0 | - |
| - | - | 989.3 | 539.3 | - | - | 0 | - |
| - | - | 719.9 | 542.7 | - | - | 0 | - |
| - | - | 925.9 | 543.8 | - | - | 0 | - |
| - | - | 664.1 | 544.1 | - | - | 0 | - |
| - | - | 1010 | 544.3 | - | - | 0 | - |
| - | - | 745.2 | 544.8 | - | - | 0 | - |
| 6 | z | 2753 | 551.3 | 0.0003395 | 0.6158 | +1 | 5 |
| - | - | 1611 | 551.7 | - | - | 0 | - |
| - | - | 1169 | 552.2 | - | - | 0 | - |
| - | - | 1.297E+04 | 552.3 | - | - | 0 | - |
| - | - | 3.261E+04 | 552.8 | - | - | 0 | - |
| - | - | 3.097E+04 | 553.3 | - | - | 0 | - |
| - | - | 1.388E+04 | 553.8 | - | - | 0 | - |
| - | - | 1625 | 554.3 | - | - | 0 | - |
| - | - | 7640 | 561.8 | - | - | 0 | - |
| - | - | 791 | 562.1 | - | - | 0 | - |
| - | - | 933.1 | 562.2 | - | - | 0 | - |
| - | - | 5186 | 562.3 | - | - | 0 | - |
| - | - | 831.6 | 562.3 | - | - | 0 | - |
| - | - | 2180 | 562.8 | - | - | 0 | - |
| - | - | 905.6 | 563.3 | - | - | 0 | - |
| 6 | y | 1.037E+04 | 567.3 | 0.000753 | 1.327 | +1 | 5 |
| - | - | 3217 | 568.3 | - | - | 0 | - |
| - | - | 3358 | 572.3 | - | - | 0 | - |
| 5 | c | 9608 | 573.3 | 0.001014 | 1.769 | +1 | 5 |
| - | - | 2941 | 574.3 | - | - | 0 | - |
| 5 | w | 928.6 | 621.3 | 8.723E-05 | 0.1404 | +1 | 6 |
| - | - | 892.6 | 631.3 | - | - | 0 | - |
| - | - | 676.6 | 634.3 | - | - | 0 | - |
| - | - | 1875 | 644.3 | - | - | 0 | - |
| - | - | 2073 | 645.4 | - | - | 0 | - |
| - | - | 653.6 | 646.4 | - | - | 0 | - |
| - | - | 817.4 | 649.3 | - | - | 0 | - |
| - | - | 739.2 | 654.3 | - | - | 0 | - |
| - | - | 760.2 | 671.8 | - | - | 0 | - |
| - | - | 1034 | 673.4 | - | - | 0 | - |
| 5 | y | 1442 | 677.3 | 0.0007947 | 1.173 | +1 | 6 |
| 5 | y | 4144 | 678.3 | 0.0001165 | 0.1718 | +1 | 6 |
| 5 | z | 1.622E+04 | 679.3 | 0.0008726 | 1.284 | +1 | 6 |
| - | - | 9802 | 680.3 | - | - | 0 | - |
| - | - | 3275 | 681.3 | - | - | 0 | - |
| - | - | 5749 | 687.4 | - | - | 0 | - |
| 6 | c | 4.131E+04 | 688.4 | 0.001407 | 2.044 | +1 | 6 |
| - | - | 2.017E+04 | 689.4 | - | - | 0 | - |
| - | - | 4361 | 690.4 | - | - | 0 | - |
| 5 | y | 9275 | 695.3 | 0.001103 | 1.586 | +1 | 6 |
| - | - | 5287 | 696.3 | - | - | 0 | - |
| - | - | 1091 | 697.3 | - | - | 0 | - |
| - | - | 1362 | 724.4 | - | - | 0 | - |
| 4 | w | 1.278E+04 | 749.3 | 0.0006203 | 0.8278 | +1 | 7 |
| - | - | 5692 | 750.3 | - | - | 0 | - |
| - | - | 1733 | 751.3 | - | - | 0 | - |
| - | - | 2584 | 757.4 | - | - | 0 | - |
| - | - | 718.3 | 758.4 | - | - | 0 | - |
| - | - | 1137 | 758.9 | - | - | 0 | - |
| - | - | 611.8 | 761.4 | - | - | 0 | - |
| - | - | 2635 | 772.4 | - | - | 0 | - |
| - | - | 878.6 | 773.4 | - | - | 0 | - |
| - | - | 619.6 | 779.4 | - | - | 0 | - |
| - | - | 827 | 779.9 | - | - | 0 | - |
| 4 | z | 4.792E+04 | 792.4 | 0.0009522 | 1.202 | +1 | 7 |
| - | - | 4.228E+04 | 793.4 | - | - | 0 | - |
| - | - | 1.557E+04 | 794.4 | - | - | 0 | - |
| - | - | 3693 | 794.9 | - | - | 0 | - |
| - | - | 5024 | 795.4 | - | - | 0 | - |
| - | - | 1439 | 795.9 | - | - | 0 | - |
| - | - | 720.3 | 796.4 | - | - | 0 | - |
| 7 | c | 891.5 | 799.4 | 0.008542 | 10.68 | +1 | 7 |
| - | - | 827.7 | 800.4 | - | - | 0 | - |
| - | - | 770.6 | 801.3 | - | - | 0 | - |
| - | - | 1870 | 807.4 | - | - | 0 | - |
| 4 | y | 4464 | 808.4 | 0.002403 | 2.973 | +1 | 7 |
| - | - | 3576 | 809.4 | - | - | 0 | - |
| - | - | 1.104E+04 | 815.4 | - | - | 0 | - |
| 7 | c | 3.2E+04 | 816.5 | 0.00323 | 3.957 | +1 | 7 |
| - | - | 1.837E+04 | 817.5 | - | - | 0 | - |
| - | - | 5657 | 818.5 | - | - | 0 | - |
| - | - | 2734 | 823.4 | - | - | 0 | - |
| - | - | 2267 | 824.4 | - | - | 0 | - |
| - | - | 1710 | 832.4 | - | - | 0 | - |
| - | - | 611 | 841.4 | - | - | 0 | - |
| - | - | 3635 | 844.4 | - | - | 0 | - |
| - | - | 1592 | 844.9 | - | - | 0 | - |
| - | - | 606.9 | 850.5 | - | - | 0 | - |
| - | - | 1126 | 855.5 | - | - | 0 | - |
| - | - | 1399 | 856.5 | - | - | 0 | - |
| - | - | 667.2 | 870.7 | - | - | 0 | - |
| - | - | 679.4 | 872.5 | - | - | 0 | - |
| - | - | 1156 | 873.5 | - | - | 0 | - |
| - | - | 1145 | 874.5 | - | - | 0 | - |
| - | - | 572.3 | 876.4 | - | - | 0 | - |
| 3 | y | 1408 | 877.4 | 0.001035 | 1.18 | +1 | 8 |
| 3 | y | 1110 | 878.4 | 0.003836 | 4.367 | +1 | 8 |
| 3 | z | 1.301E+04 | 879.4 | 0.0009982 | 1.135 | +1 | 8 |
| - | - | 1.262E+04 | 880.4 | - | - | 0 | - |
| - | - | 5911 | 881.4 | - | - | 0 | - |
| - | - | 1967 | 882.4 | - | - | 0 | - |
| - | - | 1400 | 883.4 | - | - | 0 | - |
| - | - | 1175 | 884.5 | - | - | 0 | - |
| - | - | 4205 | 894.4 | - | - | 0 | - |
| 3 | y | 3.08E+04 | 895.5 | 0.003182 | 3.553 | +1 | 8 |
| - | - | 1.999E+04 | 896.4 | - | - | 0 | - |
| - | - | 6668 | 897.4 | - | - | 0 | - |
| 8 | c | 713 | 900.5 | 0.002692 | 2.99 | +1 | 8 |
| - | - | 683 | 902.5 | - | - | 0 | - |
| 8 | c | 6340 | 917.5 | 0.001837 | 2.002 | +1 | 8 |
| - | - | 5699 | 918.5 | - | - | 0 | - |
| - | - | 1492 | 919.5 | - | - | 0 | - |
| - | - | 672.5 | 923.4 | - | - | 0 | - |
| - | - | 9729 | 930.5 | - | - | 0 | - |
| - | - | 7164 | 931.5 | - | - | 0 | - |
| - | - | 2570 | 932.5 | - | - | 0 | - |
| - | - | 703.6 | 938.3 | - | - | 0 | - |
| - | - | 1926 | 940.5 | - | - | 0 | - |
| - | - | 1442 | 941.5 | - | - | 0 | - |
| - | - | 647.5 | 947.5 | - | - | 0 | - |
| - | - | 1077 | 956.3 | - | - | 0 | - |
| 9 | c | 804.3 | 956.5 | 0.004984 | 5.211 | +1 | 9 |
| 9 | c | 2017 | 957.5 | 0.00154 | 1.608 | +1 | 9 |
| - | - | 1753 | 958.5 | - | - | 0 | - |
| - | - | 727.4 | 968.4 | - | - | 0 | - |
| - | - | 1080 | 969.4 | - | - | 0 | - |
| 9 | c | 6.224E+04 | 974.5 | 0.001328 | 1.362 | +1 | 9 |
| - | - | 4.999E+04 | 975.5 | - | - | 0 | - |
| 2 | y | 1.821E+04 | 976.5 | 0.008976 | 9.192 | +1 | 9 |
| - | - | 1594 | 977.5 | - | - | 0 | - |
| 2 | z | 3370 | 978.5 | 0.001175 | 1.201 | +1 | 9 |
| - | - | 4773 | 979.5 | - | - | 0 | - |
| - | - | 1512 | 980.5 | - | - | 0 | - |
| - | - | 2181 | 985.4 | - | - | 0 | - |
| - | - | 1075 | 986.4 | - | - | 0 | - |
| - | - | 655.8 | 987.4 | - | - | 0 | - |
| - | - | 914.4 | 993.5 | - | - | 0 | - |
| 2 | y | 1003 | 994.5 | 0.001952 | 1.962 | +1 | 9 |
| - | - | 640.5 | 995.5 | - | - | 0 | - |
| - | - | 842.7 | 1034 | - | - | 0 | - |
| - | - | 1034 | 1035 | - | - | 0 | - |
| - | - | 954.3 | 1036 | - | - | 0 | - |
| - | - | 4871 | 1049 | - | - | 0 | - |
| - | - | 4295 | 1050 | - | - | 0 | - |
| - | - | 6068 | 1051 | - | - | 0 | - |
| - | - | 2970 | 1052 | - | - | 0 | - |
| - | - | 718.7 | 1052 | - | - | 0 | - |
| - | - | 642.9 | 1062 | - | - | 0 | - |
| - | - | 2038 | 1063 | - | - | 0 | - |
| - | - | 5728 | 1064 | - | - | 0 | - |
| - | - | 4327 | 1065 | - | - | 0 | - |
| - | - | 2081 | 1066 | - | - | 0 | - |
| - | - | 1023 | 1068 | - | - | 0 | - |
| - | - | 858 | 1069 | - | - | 0 | - |
| - | - | 1542 | 1078 | - | - | 0 | - |
| - | - | 8462 | 1079 | - | - | 0 | - |
| - | - | 7969 | 1080 | - | - | 0 | - |
| - | - | 4802 | 1081 | - | - | 0 | - |
| - | - | 2040 | 1082 | - | - | 0 | - |
| - | - | 658.9 | 1084 | - | - | 0 | - |
| - | - | 945.5 | 1089 | - | - | 0 | - |
| - | - | 2123 | 1096 | - | - | 0 | - |
| - | - | 3357 | 1097 | - | - | 0 | - |
| - | - | 1550 | 1103 | - | - | 0 | - |
| - | - | 1082 | 1104 | - | - | 0 | - |
| - | - | 3605 | 1106 | - | - | 0 | - |
| - | - | 7.77E+04 | 1107 | - | - | 0 | - |
| - | - | 6.931E+04 | 1108 | - | - | 0 | - |
| - | - | 2.809E+04 | 1109 | - | - | 0 | - |
| - | - | 3751 | 1110 | - | - | 0 | - |
| - | - | 917 | 1121 | - | - | 0 | - |
| - | - | 3.269E+04 | 1123 | - | - | 0 | - |
| - | - | 7.253E+04 | 1124 | - | - | 0 | - |
| - | - | 5.199E+04 | 1125 | - | - | 0 | - |
| - | - | 1.94E+04 | 1126 | - | - | 0 | - |
| - | - | 3243 | 1127 | - | - | 0 | - |
| - | - | 668.5 | 1537 | - | - | 0 | - |
| - | - | 784.4 | 1574 | - | - | 0 | - |
| - | - | 888.6 | 1575 | - | - | 0 | - |
| - | - | 2011 | 1589 | - | - | 0 | - |
| - | - | 2070 | 1590 | - | - | 0 | - |
| - | - | 3119 | 1591 | - | - | 0 | - |
| - | - | 2006 | 1592 | - | - | 0 | - |
| - | - | 1133 | 1593 | - | - | 0 | - |
| - | - | 1544 | 1645 | - | - | 0 | - |
| - | - | 2004 | 1646 | - | - | 0 | - |
| - | - | 908.9 | 1647 | - | - | 0 | - |
| - | - | 796.5 | 1648 | - | - | 0 | - |
| - | - | 773.1 | 1663 | - | - | 0 | - |
| - | - | 1675 | 1664 | - | - | 0 | - |
| - | - | 1594 | 1665 | - | - | 0 | - |
| - | - | 929.1 | 1666 | - | - | 0 | - |
| - | - | 1051 | 1668 | - | - | 0 | - |
| - | - | 3633 | 1681 | - | - | 0 | - |
| - | - | 2366 | 1682 | - | - | 0 | - |
| - | - | 2260 | 1683 | - | - | 0 | - |
| - | - | 848.6 | 1685 | - | - | 0 | - |
| - | - | 934.4 | 1688 | - | - | 0 | - |
| - | - | 6301 | 1689 | - | - | 0 | - |
| - | - | 2589 | 1690 | - | - | 0 | - |
| - | - | 689.3 | 2092 | - | - | 0 | - |
| - | - | 744.8 | 2349 | - | - | 0 | - |
| - | - | 766.9 | 3083 | - | - | 0 | - |

m/z Charge Intensity FragmentType MassShift Position
120.08067321777344 0 1338.8196
129.10231018066406 0 518.4401
139.40374755859375 0 433.1093
148.9547882080078 0 830.67474
155.1182403564453 0 457.33484
157.3001708984375 0 446.7538
166.08628845214844 0 4043.2056 y 9
173.12831115722656 0 1371.5865
177.10220336914062 0 975.3097
182.12892150878906 0 1355.5334
183.11285400390625 0 2638.4539
197.1985321044922 0 465.8873
200.1391143798828 0 11944.057
201.12289428710938 0 2906.0024
201.1431121826172 0 803.93604
210.1236114501953 0 2200.8928
211.10736083984375 0 2360.9556
217.572265625 0 546.0095
223.1074981689453 0 3676.4404 y 8
228.13412475585938 0 8545.526 c Ammonia loss 1
234.35342407226562 0 524.23615
237.95993041992188 0 555.0828
258.6334228515625 0 596.28595
297.1570739746094 0 748.85284
298.1395263671875 0 2335.315
315.166259765625 0 1468.3539 c Ammonia loss 2
324.1559143066406 0 1513.021 y 7
329.1817321777344 0 797.62085
348.1712951660156 0 947.5271 y 4
365.21826171875 0 1073.5504
380.191650390625 0 2061.397
381.82586669921875 0 569.46497
383.228515625 0 1977.389
392.2299499511719 0 720.3193
393.2129211425781 0 3026.4822
398.20379638671875 0 1772.5623
404.71478271484375 0 1171.6581 y 3
410.2398986816406 0 1044.6624
411.2234191894531 0 2774.9321
412.2248229980469 0 662.5726
415.1174621582031 0 506.71515
417.0616455078125 0 1461.4122
419.21728515625 0 1751.5635
428.25146484375 0 944.0395 c Ammonia loss 3
430.23089599609375 0 804.8115
436.231201171875 0 9573.524 z 6
437.23724365234375 0 6235.5576
438.2415771484375 0 1512.0315
439.7159118652344 0 712.09674 y Ammonia loss 2
440.2176818847656 0 630.5019 z 2
441.7386474609375 0 940.58813
442.2380676269531 0 822.743
444.20928955078125 0 746.16724
448.23040771484375 0 1546.1294 y 2
448.7286682128906 0 920.51447
450.7427062988281 0 675.4102 c Ammonia loss 7
452.2496643066406 0 3822.468 y 6
470.2489318847656 0 1208.0574
479.2538757324219 0 1200.5122 c Ammonia loss 8
507.12677001953125 0 633.37463
507.2685241699219 0 3447.3992
508.27362060546875 0 1064.0845
530.3040161132812 0 1086.5908
534.3018188476562 0 788.3457
539.28369140625 0 989.3244
542.6842041015625 0 719.8652
543.782470703125 0 925.9399
544.1041259765625 0 664.06635
544.2769165039062 0 1010.32715
544.7731323242188 0 745.1857
551.2582397460938 0 2752.5012 z 5
551.688720703125 0 1610.5337
552.1889038085938 0 1168.6848
552.2657470703125 0 12965.125
552.787353515625 0 32610.28
553.28466796875 0 30971.873
553.7852172851562 0 13879.132
554.282470703125 0 1625.4688
561.7925415039062 0 7639.5015
562.1156005859375 0 791.0178
562.2442016601562 0 933.13464
562.2916259765625 0 5186.213
562.3411865234375 0 831.6024
562.7909545898438 0 2179.8218
563.2813110351562 0 905.6271
567.2765502929688 0 10372.209 y 5
568.2783813476562 0 3216.7158
572.3259887695312 0 3357.9688
573.33447265625 0 9608.186 c 4
574.3308715820312 0 2941.4753
621.2877807617188 0 928.5834 w 4
631.2920532226562 0 892.55554
634.3182373046875 0 676.62134
644.3479614257812 0 1874.6343
645.3557739257812 0 2073.1357
646.35595703125 0 653.58057
649.3321533203125 0 817.43823
654.3125610351562 0 739.20514
671.8113403320312 0 760.15314
673.3529052734375 0 1034.2822
677.3261108398438 0 1441.6654 y Water loss 4
678.3094482421875 0 4144.4033 y Ammonia loss 4
679.3162841796875 0 16221.888 z 4
680.3141479492188 0 9801.784
681.31396484375 0 3274.983
687.3534545898438 0 5748.958
688.3610229492188 0 41307.805 c 5
689.3572998046875 0 20174.18
690.3573608398438 0 4361.4272
695.3347778320312 0 9275.033 y 4
696.327880859375 0 5287.336
697.3287353515625 0 1090.6377
724.3648071289062 0 1361.9371
749.3458251953125 0 12777.457 w 3
750.343994140625 0 5691.6455
751.3433227539062 0 1732.53
757.4291381835938 0 2584.0413
758.4177856445312 0 718.3095
758.864501953125 0 1137.4088
761.3649291992188 0 611.79553
772.4420776367188 0 2634.5186
773.435546875 0 878.64777
779.3513793945312 0 619.5596
779.857421875 0 827.014
792.4002685546875 0 47923.934 z 3
793.39990234375 0 42283.81
794.39990234375 0 15569.521
794.8759765625 0 3692.7373
795.3854370117188 0 5023.5684
795.876708984375 0 1438.5918
796.38037109375 0 720.2738
799.4223022460938 0 891.45087 c Ammonia loss 6
800.4203491210938 0 827.654
801.2691040039062 0 770.5757
807.4075927734375 0 1869.9191
808.4175415039062 0 4463.6445 y 3
809.4107055664062 0 3576.0732
815.4486083984375 0 11043.187
816.4541625976562 0 31999.934 c 6
817.4508666992188 0 18367.12
818.453857421875 0 5656.5957
823.3692626953125 0 2734.4917
824.3685913085938 0 2266.6296
832.4429321289062 0 1710.1251
841.3804321289062 0 611.0184
844.4000854492188 0 3634.6802
844.9073486328125 0 1591.905
850.4525146484375 0 606.8558
855.4801025390625 0 1126.46
856.4754028320312 0 1399.0037
870.7141723632812 0 667.2103
872.4808959960938 0 679.3936
873.4857788085938 0 1155.5546
874.4942626953125 0 1144.953
876.4479370117188 0 572.3494
877.4424438476562 0 1408.1993 y Water loss 2
878.4292602539062 0 1110.2542 y Ammonia loss 2
879.4322509765625 0 13008.954 z 2
880.4298706054688 0 12615.73
881.4293212890625 0 5910.5615
882.4324340820312 0 1966.814
883.4479370117188 0 1399.9498
884.451171875 0 1174.9622
894.4435424804688 0 4205.1177
895.4487915039062 0 30799.354 y 2
896.4457397460938 0 19985.395
897.4469604492188 0 6667.7256
900.475830078125 0 712.9691 c Ammonia loss 7
902.4826049804688 0 683.01605
917.5032348632812 0 6339.7837 c 7
918.4995727539062 0 5698.7627
919.4926147460938 0 1491.6423
923.447509765625 0 672.47144
930.510986328125 0 9728.658
931.509521484375 0 7164.237
932.5090942382812 0 2569.5999
938.3234252929688 0 703.61255
940.4718017578125 0 1925.7393
941.4668579101562 0 1442.489
947.4697875976562 0 647.51056
956.3313598632812 0 1076.6637
956.510986328125 0 804.3383 c Water loss 8
957.5015258789062 0 2017.4187 c Ammonia loss 8
958.496337890625 0 1752.7983
968.4323120117188 0 727.39484
969.4354248046875 0 1080.2441
974.5252075195312 0 62243.21 c 8
975.52001953125 0 49991.79
976.518798828125 0 18212.314 y Water loss 1
977.5228881835938 0 1594.1993
978.50048828125 0 3369.6125 z 1
979.4915771484375 0 4773.436
980.49267578125 0 1511.5474
985.4010009765625 0 2181.322
986.4015502929688 0 1074.7161
987.40478515625 0 655.7873
993.4735107421875 0 914.36304
994.5223388671875 0 1002.64105 y 1
995.501953125 0 640.4859
1033.5947265625 0 842.68304
1034.581298828125 0 1033.7526
1035.531494140625 0 954.25195
1048.5318603515625 0 4871.3403
1049.5267333984375 0 4294.6787
1050.5145263671875 0 6068.1406
1051.507080078125 0 2969.85
1052.4849853515625 0 718.66156
1061.5833740234375 0 642.85406
1062.5736083984375 0 2038.2263
1063.562744140625 0 5727.9453
1064.5604248046875 0 4327.2173
1065.5621337890625 0 2081.0618
1067.534912109375 0 1022.69855
1068.5162353515625 0 857.9546
1077.58056640625 0 1542.2837
1078.565673828125 0 8461.954
1079.578369140625 0 7969.4614
1080.578125 0 4801.609
1081.5888671875 0 2040.3447
1083.6435546875 0 658.9434
1088.540283203125 0 945.45905
1095.587890625 0 2122.636
1096.5845947265625 0 3357.4133
1103.3739013671875 0 1549.8083
1104.369140625 0 1081.5234
1105.5684814453125 0 3605.1382
1106.5587158203125 0 77704.8
1107.5548095703125 0 69312.73
1108.5555419921875 0 28088.42
1109.5548095703125 0 3751.3071
1121.3828125 0 916.97876
1122.5772705078125 0 32687.186
1123.5799560546875 0 72533.445
1124.579345703125 0 51990.047
1125.5804443359375 0 19401.156
1126.5765380859375 0 3243.0361
1537.1658935546875 0 668.4779
1573.742431640625 0 784.39417
1574.7291259765625 0 888.5897
1588.740234375 0 2010.842
1589.7471923828125 0 2069.7566
1590.757080078125 0 3119.1897
1591.7525634765625 0 2005.7197
1592.7650146484375 0 1133.208
1644.726806640625 0 1544.3279
1645.734619140625 0 2004.346
1646.734619140625 0 908.86255
1647.7513427734375 0 796.46387
1662.748291015625 0 773.0528
1663.7642822265625 0 1674.5436
1664.7535400390625 0 1594.2382
1665.7608642578125 0 929.0672
1667.76220703125 0 1050.5161
1680.772216796875 0 3633.4436
1681.7706298828125 0 2365.9602
1682.7672119140625 0 2260.0037
1684.8001708984375 0 848.55664
1687.756591796875 0 934.4267
1688.7957763671875 0 6300.595
1689.7994384765625 0 2588.919
2092.025146484375 0 689.2743
2349.037353515625 0 744.8168
3083.20751953125 0 766.8791

Spectrum Details

|  |  |
| --- | --- |
| Matched peaks? Matched peaks The total absolute number of peaks matched. Additionally in brackets the total fraction of peaks matched and the total number of peaks is shown. | 41 (15.59% of 263) |
| FDR? FDR The false discovery rate estimated for this peptide. It is calculated by matching all theoretical fragments with a non-integer shift with the raw peaks for this spectrum. This is done with 40 different shifts. The resulting percentage is the average number of annotated peaks over the number of annotated peaks with the correct spectrum. | 0.81% |
| Satellite FDR? Satellite FDR See the FDR for details on its calculation. This satellite ion specific FDR only contains the satellite ions (d/w) for I/L/J positions. | 4.76% |
| PSM Score? PSM Score The PSM Score as given by Hecklib to this annotated spectrum. It is shown with three significant figures. | 450 |

#### Reverse Lookup? Reverse Lookup All places where this read could be placed.

| Group | Segment | Template | Template Part | Read Part | Score | Unique |
| --- | --- | --- | --- | --- | --- | --- |
| Decoy | Decoy | TRYP | [22..32] | [0..10] | 39 | True |

| Recombined | Template Part | Read Part | Score | Unique |
| --- | --- | --- | --- | --- |
| TRYP | [22..32] | [0..10] | 39 | True |

#### Meta Information from Multiple reads

##### Number of combined reads

6

##### Intensity

0.9406

##### TotalArea

1.879E+09

##### Changes to the peptide sequence

QVSLQDKTGF

J→LSupport for Leucine based on side chain ions (1 for L 0 for I) (Position: 4)

L→JNo support for either Leucine or Isoleucine based on side chain ions (Position: 4)

#### Positional Score

Copy Data

##### Positional Score (TSV)

###### Preview

```
Loading example...
```

*Click on the button to copy the data to your clipboard.*

100123456789

Label Value
"0" 0.598
"1" 0.623
"2" 0.655
"3" 0.655
"4" 0.628
"5" 0.658
"6" 0.66
"7" 0.66
"8" 0.667
"9" 0.667

#### Meta Information from PEAKS

##### Scan Identifier

F3:5529

##### Original sequence

Q

V

S

L

Q

D

K

T

G

F

##### Posttranslational Modifications

##### Source File

D:\separate\_stitch\_analyses\xle-disambiguation\raw\20210323\_F1\_UM1\_Peng0013\_SA\_F59\_ingel\_3ug\_chymo.raw

##### Fraction

3

##### Scan Feature

F3:6793

##### De Novo Score

98

##### ConfidenceScore

98

### m/z

561.7946

##### Mass

1121.5718

##### Charge

2

##### Retention Time

29.79

##### Predicted Retention Time

-

##### Area

6.263E+08

##### Parts Per Million

2.5

##### Fragmentation mode

HCD

##### Originating file

01 D:\separate\_stitch\_analyses\xle-disambiguation\20210325\_F59\_3ug\_DENOVO\_12.csv

#### Meta Information from PEAKS

##### Scan Identifier

F3:5655

##### Original sequence

Q

V

S

L

Q

D

K

T

G

F

##### Posttranslational Modifications

##### Source File

D:\separate\_stitch\_analyses\xle-disambiguation\raw\20210323\_F1\_UM1\_Peng0013\_SA\_F59\_ingel\_3ug\_chymo.raw

##### Fraction

3

##### Scan Feature

-

##### De Novo Score

98

##### ConfidenceScore

98

### m/z

561.7938

##### Mass

1121.5718

##### Charge

2

##### Retention Time

31.29

##### Predicted Retention Time

-

##### Area

0

##### Parts Per Million

1.2

##### Fragmentation mode

HCD

##### Originating file

01 D:\separate\_stitch\_analyses\xle-disambiguation\20210325\_F59\_3ug\_DENOVO\_12.csv

#### Meta Information from PEAKS

##### Scan Identifier

F3:5387

##### Original sequence

Q

V

S

L

Q

D

K

T

G

F

##### Posttranslational Modifications

##### Source File

D:\separate\_stitch\_analyses\xle-disambiguation\raw\20210323\_F1\_UM1\_Peng0013\_SA\_F59\_ingel\_3ug\_chymo.raw

##### Fraction

3

##### Scan Feature

F3:6793

##### De Novo Score

98

##### ConfidenceScore

98

### m/z

561.7946

##### Mass

1121.5718

##### Charge

2

##### Retention Time

29.79

##### Predicted Retention Time

-

##### Area

6.263E+08

##### Parts Per Million

2.5

##### Fragmentation mode

HCD

##### Originating file

01 D:\separate\_stitch\_analyses\xle-disambiguation\20210325\_F59\_3ug\_DENOVO\_12.csv

#### Meta Information from PEAKS

##### Scan Identifier

F3:5588

##### Original sequence

Q

V

S

L

Q

D

K

T

G

F

##### Posttranslational Modifications

##### Source File

D:\separate\_stitch\_analyses\xle-disambiguation\raw\20210323\_F1\_UM1\_Peng0013\_SA\_F59\_ingel\_3ug\_chymo.raw

##### Fraction

3

##### Scan Feature

-

##### De Novo Score

98

##### ConfidenceScore

98

### m/z

561.7938

##### Mass

1121.5718

##### Charge

2

##### Retention Time

30.94

##### Predicted Retention Time

-

##### Area

0

##### Parts Per Million

1.1

##### Fragmentation mode

HCD

##### Originating file

01 D:\separate\_stitch\_analyses\xle-disambiguation\20210325\_F59\_3ug\_DENOVO\_12.csv

#### Meta Information from PEAKS

##### Scan Identifier

F3:5464

##### Original sequence

Q

V

S

L

Q

D

K

T

G

F

##### Posttranslational Modifications

##### Source File

D:\separate\_stitch\_analyses\xle-disambiguation\raw\20210323\_F1\_UM1\_Peng0013\_SA\_F59\_ingel\_3ug\_chymo.raw

##### Fraction

3

##### Scan Feature

F3:6793

##### De Novo Score

97

##### ConfidenceScore

97

### m/z

561.7946

##### Mass

1121.5718

##### Charge

2

##### Retention Time

29.79

##### Predicted Retention Time

-

##### Area

6.263E+08

##### Parts Per Million

2.5

##### Fragmentation mode

ETHCD

##### Originating file

01 D:\separate\_stitch\_analyses\xle-disambiguation\20210325\_F59\_3ug\_DENOVO\_12.csv

#### Meta Information from PEAKS

##### Scan Identifier

F3:5767

##### Original sequence

Q

V

S

L

Q

D

K

T

G

F

##### Posttranslational Modifications

##### Source File

D:\separate\_stitch\_analyses\xle-disambiguation\raw\20210323\_F1\_UM1\_Peng0013\_SA\_F59\_ingel\_3ug\_chymo.raw

##### Fraction

3

##### Scan Feature

-

##### De Novo Score

95

##### ConfidenceScore

95

### m/z

561.7933

##### Mass

1121.5718

##### Charge

2

##### Retention Time

31.91

##### Predicted Retention Time

-

##### Area

0

##### Parts Per Million

0.2

##### Fragmentation mode

ETHCD

##### Originating file

01 D:\separate\_stitch\_analyses\xle-disambiguation\20210325\_F59\_3ug\_DENOVO\_12.csv
