## Supplementary material for "A handle on mass coincidence errors in *de novo* sequencing of antibodies by bottom-up proteomics": Combined_053.html

Details Combined\_053 | Stitch OverviewUndefined

### Read Combined\_053

#### Sequence (length=15)

VJHQDWLDGKEYKCK

#### Spectrum 4849? Spectrum 4849 The raw spectrum of this peptide as annotated by Hecklib. The fragments are coloured according to ion type (see legend). Any peaks with a star '\*' as text can be hovered over to see the full details, first the ion type second the mass shift type. By hovering over the amino acids in the peptide or ions in the legend the corresponding peaks are highlighted. By toggling the 'Unassigned' label you can turn the background (unassigned) peaks on or off in the plot. By updating the slider in the Ion legend you can update the spectrum to only show the top X% of the peaks with labels. The top X% means any peak that is within X% of the highest intensity. By dragging in the spectrum you can zoom in to a specific part of the spectrum and use 'Zoom Out' to get back to the original zoom level. The annotation of the spectrum is based on the given sequence in the peptides file and is done with different software so inconsistencies are likely. The peaks are annotated based on the given sequence, with 20 ppm tolerance.

Copy Data

##### Spectrum 4849 (TSV)

###### Preview

```
Loading example...
```

*Click on the button to copy the data to your clipboard.*

Mz MinMz MaxIntensity Max

WidthHeightPeptide font sizePeptide stroke widthSpectrum font sizeSpectrum stroke widthCompact peptide

Ion legend

wxyz

abcd

OtherUnassignedIon^Charge^_Position_Show for top:%

VJHQDWLDGKEYKCK

05.33e+31.07e+41.60e+42.13e+4

Zoom Out

y^+1^_2_c^+1^_3_y^+2^_6_y^+1^_3_y^+2^_7_c^+1^_4_c^+1^_4_y^+3^_13_z^+3^_13_y^+3^_13_z^+1^_4_y^+1^_4_y^+3^_14_c^+1^_5_w^+1^_5_z^+1^_5_y^+2^_11_y^+1^_5_y^+2^_12_c^+1^_6_c^+2^_13_c^+2^_13_z^+1^_6_z^+2^_13_y^+2^_13_y^+1^_6_w^+2^_14_c^+1^_7_y^+1^_7_c^+2^_14_y^+1^_7_z^+1^_7_y^+2^_14_z^+2^_14_c^+1^_7_c^+1^_8_z^+1^_8_c^+1^_8_y^+1^_8_c^+1^_9_z^+1^_9_y^+1^_9_c^+1^_10_z^+1^_10_y^+1^_10_c^+1^_11_z^+1^_11_c^+1^_12_z^+1^_12_c^+1^_13_c^+1^_14_

0837167425113348

Fragment Matches Table

Show background peaks

| Position | Ion type | Intensity | mz Theoretical | mz Error (Th) | mz Error (ppm) | Charge | Series Number |
| --- | --- | --- | --- | --- | --- | --- | --- |
| - | - | 389.1 | 121.8 | - | - | 0 | - |
| - | - | 377.9 | 128.6 | - | - | 0 | - |
| - | - | 1036 | 129.1 | - | - | 0 | - |
| - | - | 420.7 | 138.1 | - | - | 0 | - |
| - | - | 400.4 | 144.9 | - | - | 0 | - |
| - | - | 417.2 | 153 | - | - | 0 | - |
| - | - | 1304 | 155.1 | - | - | 0 | - |
| - | - | 776.8 | 177 | - | - | 0 | - |
| - | - | 434.3 | 179.9 | - | - | 0 | - |
| - | - | 1189 | 185.2 | - | - | 0 | - |
| - | - | 921.9 | 213.2 | - | - | 0 | - |
| - | - | 510.1 | 237.4 | - | - | 0 | - |
| - | - | 805.7 | 239.1 | - | - | 0 | - |
| - | - | 969.5 | 266.1 | - | - | 0 | - |
| - | - | 1868 | 283.2 | - | - | 0 | - |
| - | - | 535.2 | 299.1 | - | - | 0 | - |
| 14 | y | 841.2 | 308.1 | 0.004669 | 15.15 | +1 | 2 |
| - | - | 596.5 | 332.2 | - | - | 0 | - |
| - | - | 553.9 | 339.2 | - | - | 0 | - |
| - | - | 3273 | 350.2 | - | - | 0 | - |
| - | - | 1656 | 355.1 | - | - | 0 | - |
| 3 | c | 991.9 | 367.2 | 0.0002812 | 0.7658 | +1 | 3 |
| - | - | 581.3 | 401.2 | - | - | 0 | - |
| - | - | 587.6 | 402.2 | - | - | 0 | - |
| - | - | 571.8 | 415.1 | - | - | 0 | - |
| 10 | y | 3893 | 420.2 | 0.000516 | 1.228 | +2 | 6 |
| - | - | 1142 | 421.2 | - | - | 0 | - |
| 13 | y | 966.7 | 436.2 | 0.005425 | 12.44 | +1 | 3 |
| 9 | y | 1853 | 457.2 | 0.002885 | 6.311 | +2 | 7 |
| - | - | 1223 | 457.7 | - | - | 0 | - |
| 4 | c | 1172 | 478.3 | 0.0005408 | 1.131 | +1 | 4 |
| 4 | c | 5090 | 495.3 | 4.018E-05 | 0.08113 | +1 | 4 |
| - | - | 1299 | 496.3 | - | - | 0 | - |
| 3 | y | 874.3 | 564.3 | 0.006381 | 11.31 | +3 | 13 |
| 3 | z | 579.2 | 564.6 | 0.0003342 | 0.5919 | +3 | 13 |
| - | - | 2361 | 566.3 | - | - | 0 | - |
| - | - | 906.1 | 567.3 | - | - | 0 | - |
| 3 | y | 2480 | 569.9 | 0.0006785 | 1.19 | +3 | 13 |
| - | - | 2014 | 570.3 | - | - | 0 | - |
| - | - | 2352 | 570.6 | - | - | 0 | - |
| 12 | z | 3325 | 583.3 | 0.004527 | 7.762 | +1 | 4 |
| - | - | 1930 | 584.3 | - | - | 0 | - |
| 12 | y | 1698 | 599.3 | 0.004453 | 7.43 | +1 | 4 |
| 2 | y | 754.6 | 607.6 | 0.002434 | 4.006 | +3 | 14 |
| 5 | c | 4734 | 610.3 | 0.0008408 | 1.378 | +1 | 5 |
| - | - | 1210 | 611.3 | - | - | 0 | - |
| - | - | 700.1 | 631.8 | - | - | 0 | - |
| - | - | 2673 | 635 | - | - | 0 | - |
| - | - | 1071 | 635.3 | - | - | 0 | - |
| - | - | 823.2 | 635.6 | - | - | 0 | - |
| - | - | 881.7 | 639.3 | - | - | 0 | - |
| - | - | 980 | 640.3 | - | - | 0 | - |
| - | - | 1.483E+04 | 640.6 | - | - | 0 | - |
| - | - | 1.215E+04 | 641 | - | - | 0 | - |
| - | - | 1.029E+04 | 641.3 | - | - | 0 | - |
| - | - | 4453 | 641.4 | - | - | 0 | - |
| - | - | 3734 | 641.7 | - | - | 0 | - |
| - | - | 1746 | 642 | - | - | 0 | - |
| 11 | w | 2591 | 653.3 | 0.004519 | 6.917 | +1 | 5 |
| - | - | 1074 | 654.3 | - | - | 0 | - |
| - | - | 647.7 | 677.4 | - | - | 0 | - |
| - | - | 744.9 | 678.4 | - | - | 0 | - |
| 11 | z | 4377 | 712.3 | 0.00464 | 6.514 | +1 | 5 |
| 5 | y | 2639 | 713.3 | 0.0109 | 15.28 | +2 | 11 |
| 11 | y | 866.3 | 728.3 | 0.003467 | 4.76 | +1 | 5 |
| - | - | 627 | 734.4 | - | - | 0 | - |
| 4 | y | 751.3 | 776.9 | 0.002284 | 2.939 | +2 | 12 |
| 6 | c | 4018 | 796.4 | 0.001084 | 1.361 | +1 | 6 |
| - | - | 1763 | 797.4 | - | - | 0 | - |
| 13 | c | 557.7 | 806.9 | 0.0036 | 4.461 | +2 | 13 |
| 13 | c | 5143 | 815.4 | 0.0001518 | 0.1862 | +2 | 13 |
| - | - | 3644 | 815.9 | - | - | 0 | - |
| - | - | 2936 | 816.4 | - | - | 0 | - |
| - | - | 888 | 831.5 | - | - | 0 | - |
| 10 | z | 1600 | 840.4 | 0.004937 | 5.875 | +1 | 6 |
| - | - | 3124 | 841.4 | - | - | 0 | - |
| - | - | 1980 | 842.4 | - | - | 0 | - |
| 3 | z | 2059 | 846.4 | 0.000255 | 0.3013 | +2 | 13 |
| - | - | 992.5 | 846.9 | - | - | 0 | - |
| - | - | 1496 | 847.4 | - | - | 0 | - |
| 3 | y | 1535 | 854.4 | 0.003269 | 3.827 | +2 | 13 |
| - | - | 1940 | 854.9 | - | - | 0 | - |
| 10 | y | 781.2 | 856.4 | 0.01097 | 12.8 | +1 | 6 |
| 2 | w | 978.2 | 881.4 | 0.001838 | 2.085 | +2 | 14 |
| - | - | 732.7 | 881.9 | - | - | 0 | - |
| 7 | c | 1475 | 892.5 | 0.002049 | 2.295 | +1 | 7 |
| 9 | y | 1508 | 895.4 | 0.00325 | 3.629 | +1 | 7 |
| 14 | c | 6604 | 895.9 | 0.0008345 | 0.9314 | +2 | 14 |
| 9 | y | 6570 | 896.4 | 0.009012 | 10.05 | +1 | 7 |
| - | - | 4202 | 896.9 | - | - | 0 | - |
| 9 | z | 3532 | 897.4 | 0.001133 | 1.262 | +1 | 7 |
| - | - | 1348 | 898.4 | - | - | 0 | - |
| 2 | y | 983.8 | 902.4 | 0.01735 | 19.23 | +2 | 14 |
| 2 | z | 1368 | 902.9 | 0.002025 | 2.242 | +2 | 14 |
| - | - | 625 | 903.9 | - | - | 0 | - |
| 7 | c | 5612 | 909.5 | 0.0001553 | 0.1707 | +1 | 7 |
| - | - | 2623 | 910.5 | - | - | 0 | - |
| - | - | 963.7 | 911.5 | - | - | 0 | - |
| - | - | 769.7 | 916.5 | - | - | 0 | - |
| - | - | 721.3 | 922.4 | - | - | 0 | - |
| - | - | 906.8 | 924.4 | - | - | 0 | - |
| - | - | 789.4 | 924.9 | - | - | 0 | - |
| - | - | 4067 | 931 | - | - | 0 | - |
| - | - | 1.138E+04 | 931.5 | - | - | 0 | - |
| - | - | 9064 | 932 | - | - | 0 | - |
| - | - | 6631 | 932.5 | - | - | 0 | - |
| - | - | 1839 | 933 | - | - | 0 | - |
| - | - | 785.2 | 933.5 | - | - | 0 | - |
| - | - | 1781 | 938 | - | - | 0 | - |
| - | - | 3021 | 938.5 | - | - | 0 | - |
| - | - | 3444 | 939 | - | - | 0 | - |
| - | - | 2383 | 939.5 | - | - | 0 | - |
| - | - | 645.8 | 940 | - | - | 0 | - |
| - | - | 1232 | 943.5 | - | - | 0 | - |
| - | - | 1606 | 952 | - | - | 0 | - |
| - | - | 5058 | 952.5 | - | - | 0 | - |
| - | - | 3700 | 953 | - | - | 0 | - |
| - | - | 3261 | 953.5 | - | - | 0 | - |
| - | - | 2363 | 955.5 | - | - | 0 | - |
| - | - | 938.5 | 958.9 | - | - | 0 | - |
| - | - | 996.5 | 960 | - | - | 0 | - |
| - | - | 1.491E+04 | 960.5 | - | - | 0 | - |
| - | - | 1.734E+04 | 961 | - | - | 0 | - |
| - | - | 1.438E+04 | 961.5 | - | - | 0 | - |
| - | - | 7607 | 962 | - | - | 0 | - |
| - | - | 2333 | 962.5 | - | - | 0 | - |
| - | - | 1641 | 963 | - | - | 0 | - |
| - | - | 1260 | 966.5 | - | - | 0 | - |
| - | - | 703.7 | 967.5 | - | - | 0 | - |
| 8 | c | 1116 | 1007 | 0.001587 | 1.575 | +1 | 8 |
| - | - | 719.9 | 1009 | - | - | 0 | - |
| 8 | z | 3530 | 1012 | 0.003662 | 3.617 | +1 | 8 |
| - | - | 2056 | 1013 | - | - | 0 | - |
| - | - | 803.6 | 1014 | - | - | 0 | - |
| - | - | 841.8 | 1024 | - | - | 0 | - |
| 8 | c | 2024 | 1025 | 0.0001818 | 0.1775 | +1 | 8 |
| - | - | 1598 | 1026 | - | - | 0 | - |
| 8 | y | 1519 | 1028 | 0.0009013 | 0.8763 | +1 | 8 |
| - | - | 872.5 | 1067 | - | - | 0 | - |
| - | - | 2223 | 1081 | - | - | 0 | - |
| 9 | c | 3565 | 1082 | 0.001992 | 1.842 | +1 | 9 |
| - | - | 1973 | 1083 | - | - | 0 | - |
| 7 | z | 3258 | 1126 | 0.003009 | 2.673 | +1 | 9 |
| - | - | 1700 | 1127 | - | - | 0 | - |
| 7 | y | 754.7 | 1142 | 0.005253 | 4.602 | +1 | 9 |
| 10 | c | 9349 | 1210 | 0.0009452 | 0.7814 | +1 | 10 |
| - | - | 5831 | 1211 | - | - | 0 | - |
| - | - | 1194 | 1212 | - | - | 0 | - |
| - | - | 1043 | 1213 | - | - | 0 | - |
| - | - | 783.9 | 1262 | - | - | 0 | - |
| - | - | 2427 | 1281 | - | - | 0 | - |
| - | - | 1073 | 1282 | - | - | 0 | - |
| - | - | 836.9 | 1296 | - | - | 0 | - |
| 6 | z | 3176 | 1312 | 0.003098 | 2.362 | +1 | 10 |
| - | - | 2894 | 1313 | - | - | 0 | - |
| - | - | 1208 | 1314 | - | - | 0 | - |
| 6 | y | 3279 | 1328 | 0.0005167 | 0.3892 | +1 | 10 |
| - | - | 2385 | 1329 | - | - | 0 | - |
| - | - | 753.3 | 1338 | - | - | 0 | - |
| 11 | c | 5909 | 1339 | 0.0007105 | 0.5307 | +1 | 11 |
| - | - | 4353 | 1340 | - | - | 0 | - |
| - | - | 1683 | 1341 | - | - | 0 | - |
| - | - | 2225 | 1383 | - | - | 0 | - |
| - | - | 2172 | 1384 | - | - | 0 | - |
| - | - | 873.5 | 1385 | - | - | 0 | - |
| 5 | z | 4961 | 1427 | 0.005993 | 4.201 | +1 | 11 |
| - | - | 5843 | 1428 | - | - | 0 | - |
| - | - | 2873 | 1429 | - | - | 0 | - |
| - | - | 788.9 | 1430 | - | - | 0 | - |
| - | - | 1200 | 1444 | - | - | 0 | - |
| 12 | c | 4205 | 1502 | 0.001217 | 0.8102 | +1 | 12 |
| - | - | 3274 | 1503 | - | - | 0 | - |
| - | - | 1341 | 1504 | - | - | 0 | - |
| - | - | 736.4 | 1546 | - | - | 0 | - |
| 4 | z | 4513 | 1555 | 0.003536 | 2.274 | +1 | 12 |
| - | - | 4181 | 1556 | - | - | 0 | - |
| - | - | 1800 | 1557 | - | - | 0 | - |
| - | - | 747.9 | 1587 | - | - | 0 | - |
| 13 | c | 1671 | 1630 | 0.00243 | 1.491 | +1 | 13 |
| - | - | 1797 | 1631 | - | - | 0 | - |
| - | - | 1056 | 1632 | - | - | 0 | - |
| - | - | 1296 | 1693 | - | - | 0 | - |
| - | - | 1654 | 1694 | - | - | 0 | - |
| - | - | 1271 | 1695 | - | - | 0 | - |
| - | - | 713.5 | 1779 | - | - | 0 | - |
| 14 | c | 1886 | 1791 | 0.001701 | 0.95 | +1 | 14 |
| - | - | 2951 | 1792 | - | - | 0 | - |
| - | - | 1591 | 1793 | - | - | 0 | - |
| - | - | 797.7 | 1794 | - | - | 0 | - |
| - | - | 1749 | 1806 | - | - | 0 | - |
| - | - | 1763 | 1807 | - | - | 0 | - |
| - | - | 1394 | 1808 | - | - | 0 | - |
| - | - | 1223 | 1861 | - | - | 0 | - |
| - | - | 2661 | 1862 | - | - | 0 | - |
| - | - | 5582 | 1863 | - | - | 0 | - |
| - | - | 3608 | 1864 | - | - | 0 | - |
| - | - | 2340 | 1865 | - | - | 0 | - |
| - | - | 1618 | 1876 | - | - | 0 | - |
| - | - | 3066 | 1877 | - | - | 0 | - |
| - | - | 2192 | 1878 | - | - | 0 | - |
| - | - | 1168 | 1879 | - | - | 0 | - |
| - | - | 1278 | 1893 | - | - | 0 | - |
| - | - | 2682 | 1894 | - | - | 0 | - |
| - | - | 2465 | 1895 | - | - | 0 | - |
| - | - | 1268 | 1896 | - | - | 0 | - |
| - | - | 769.5 | 1903 | - | - | 0 | - |
| - | - | 2173 | 1904 | - | - | 0 | - |
| - | - | 5181 | 1905 | - | - | 0 | - |
| - | - | 5312 | 1906 | - | - | 0 | - |
| - | - | 3548 | 1907 | - | - | 0 | - |
| - | - | 1386 | 1908 | - | - | 0 | - |
| - | - | 2643 | 1920 | - | - | 0 | - |
| - | - | 8102 | 1921 | - | - | 0 | - |
| - | - | 2.112E+04 | 1922 | - | - | 0 | - |
| - | - | 2.098E+04 | 1923 | - | - | 0 | - |
| - | - | 1.115E+04 | 1924 | - | - | 0 | - |
| - | - | 6112 | 1925 | - | - | 0 | - |
| - | - | 2273 | 1926 | - | - | 0 | - |
| - | - | 670.7 | 1985 | - | - | 0 | - |
| - | - | 832.3 | 1996 | - | - | 0 | - |
| - | - | 696.5 | 3315 | - | - | 0 | - |

m/z Charge Intensity FragmentType MassShift Position
121.78231811523438 0 389.13104
128.60977172851562 0 377.90097
129.10226440429688 0 1036.0054
138.09478759765625 0 420.67358
144.94308471679688 0 400.37982
153.00559997558594 0 417.20062
155.09243774414062 0 1304.222
177.0399169921875 0 776.7686
179.87063598632812 0 434.28568
185.1652069091797 0 1189.2262
213.15977478027344 0 921.9013
237.3798370361328 0 510.10294
239.09402465820312 0 805.7203
266.1239013671875 0 969.4654
283.151123046875 0 1867.6481
299.06134033203125 0 535.24646
308.1273193359375 0 841.2144 y 13
332.2090148925781 0 596.48236
339.2093200683594 0 553.89624
350.2181091308594 0 3272.9238
355.0691223144531 0 1655.9308
367.24493408203125 0 991.93866 c 2
401.2395935058594 0 581.3365
402.2437744140625 0 587.59015
415.0824279785156 0 571.83673
420.2037658691406 0 3893.2896 y Ammonia loss 9
421.2061767578125 0 1142.417
436.2215270996094 0 966.6995 y 12
457.22540283203125 0 1852.8936 y 8
457.72674560546875 0 1222.9286
478.2767028808594 0 1172.4912 c Ammonia loss 3
495.3038330078125 0 5089.907 c 3
496.3065490722656 0 1298.5094
564.2637329101562 0 874.3209 y Ammonia loss 2
564.5936279296875 0 579.23474 z 2
566.316650390625 0 2361.199
567.3213500976562 0 906.05316
569.9321899414062 0 2479.923 y 2
570.2662963867188 0 2013.6685
570.599365234375 0 2352.4
583.2670288085938 0 3325.0142 z 11
584.270751953125 0 1929.6725
599.2858276367188 0 1697.554 y 11
607.6251220703125 0 754.644 y 1
610.3298950195312 0 4733.7715 c 4
611.33251953125 0 1210.1125
631.8309936523438 0 700.06067
634.9789428710938 0 2672.921
635.3133544921875 0 1070.8448
635.6448364257812 0 823.2189
639.272705078125 0 881.67346
640.3299560546875 0 980.0273
640.6488037109375 0 14826.485
640.9829711914062 0 12145.932
641.3182373046875 0 10293.677
641.3973999023438 0 4453.127
641.651611328125 0 3733.563
641.9843139648438 0 1745.7258
653.2963256835938 0 2590.9685 w 10
654.298828125 0 1073.6362
677.3673095703125 0 647.7307
678.374755859375 0 744.8599
712.3095092773438 0 4376.701 z 10
713.312744140625 0 2638.7456 y Ammonia loss 4
728.3294067382812 0 866.27985 y 10
734.3624267578125 0 626.96954
776.858642578125 0 751.3094 y Water loss 3
796.4111328125 0 4017.902 c 5
797.4114990234375 0 1762.9385
806.9131469726562 0 557.6833 c Ammonia loss 12
815.4229736328125 0 5142.877 c 12
815.9237060546875 0 3644.1155
816.4270629882812 0 2936.1638
831.4500122070312 0 888.04114
840.4041748046875 0 1600.1155 z 9
841.4109497070312 0 3123.575
842.4129638671875 0 1979.8137
846.3860473632812 0 2058.8657 z 2
846.8832397460938 0 992.53937
847.3873901367188 0 1495.9619
854.3923950195312 0 1535.1399 y 2
854.897705078125 0 1940.0546
856.4168701171875 0 781.2109 y 9
881.3991088867188 0 978.18115 w 1
881.9024047851562 0 732.7093
892.4655151367188 0 1475.1793 c Ammonia loss 6
895.4354858398438 0 1507.7845 y Water loss 8
895.9315795898438 0 6603.8804 c 13
896.4317626953125 0 6570.257 y Ammonia loss 8
896.9339599609375 0 4202.448
897.429443359375 0 3532.0994 z 8
898.4331665039062 0 1348.2898
902.4417724609375 0 983.76685 y Ammonia loss 1
902.9303588867188 0 1368.171 z 1
903.9370727539062 0 624.972
909.4939575195312 0 5612.213 c 6
910.4953002929688 0 2623.2908
911.5051879882812 0 963.7497
916.4620361328125 0 769.7296
922.4498901367188 0 721.2971
924.4315795898438 0 906.76465
924.93212890625 0 789.3916
930.96337890625 0 4067.1333
931.467041015625 0 11376.168
931.9671630859375 0 9064.132
932.4688720703125 0 6630.7817
932.9688110351562 0 1838.5009
933.4638061523438 0 785.2477
937.9689331054688 0 1780.8604
938.4666137695312 0 3021.0947
938.9641723632812 0 3443.5864
939.46630859375 0 2382.6462
939.964599609375 0 645.8328
943.4660034179688 0 1231.844
951.968994140625 0 1605.8557
952.4624633789062 0 5058.074
952.9617309570312 0 3700.1187
953.463134765625 0 3260.8162
955.4530029296875 0 2362.7688
958.933349609375 0 938.4635
959.9606323242188 0 996.4918
960.4699096679688 0 14905.335
960.9720458984375 0 17343.754
961.4743041992188 0 14377.852
961.97265625 0 7606.717
962.4712524414062 0 2333.022
962.9577026367188 0 1640.9233
966.4943237304688 0 1260.1342
967.484619140625 0 703.72156
1007.492919921875 0 1115.7328 c Ammonia loss 7
1008.5010986328125 0 719.94446
1012.453857421875 0 3530.0286 z 7
1013.4592895507812 0 2056.4521
1014.4595336914062 0 803.57385
1023.517333984375 0 841.7549
1024.5208740234375 0 2024.0535 c 7
1025.527099609375 0 1597.9376
1028.475341796875 0 1518.6554 y 7
1066.880126953125 0 872.5435
1080.5355224609375 0 2223.102
1081.54052734375 0 3565.489 c 8
1082.548095703125 0 1973.4778
1125.53857421875 0 3257.5308 z 6
1126.5445556640625 0 1700.4506
1141.5550537109375 0 754.6867 y 6
1209.638427734375 0 9348.597 c 9
1210.64013671875 0 5830.744
1211.6414794921875 0 1194.2886
1212.6484375 0 1043.0647
1261.548583984375 0 783.862
1280.6466064453125 0 2427.4922
1281.6690673828125 0 1072.6833
1295.6602783203125 0 836.88086
1311.6177978515625 0 3175.7603 z 5
1312.6195068359375 0 2893.7397
1313.621337890625 0 1207.9468
1327.64013671875 0 3278.589 y 5
1328.6416015625 0 2385.3083
1337.669189453125 0 753.3256
1338.6807861328125 0 5909.2266 c 10
1339.6810302734375 0 4353.317
1340.6810302734375 0 1682.8329
1382.648681640625 0 2224.9304
1383.657958984375 0 2171.9358
1384.6544189453125 0 873.458
1426.641845703125 0 4961.389 z 4
1427.6451416015625 0 5843.472
1428.647216796875 0 2873.0408
1429.6534423828125 0 788.8648
1443.6715087890625 0 1200.2743
1501.7421875 0 4205.035 c 11
1502.7467041015625 0 3274.0234
1503.7464599609375 0 1341.3092
1545.7740478515625 0 736.37445
1554.702880859375 0 4513.338 z 3
1555.705322265625 0 4181.003
1556.7127685546875 0 1800.0453
1586.8275146484375 0 747.9273
1629.8359375 0 1670.7507 c 12
1630.83984375 0 1796.6125
1631.837646484375 0 1055.6622
1692.7652587890625 0 1295.9175
1693.7740478515625 0 1654.197
1694.7779541015625 0 1271.466
1778.8270263671875 0 713.47754
1790.8592529296875 0 1886.1907 c 13
1791.856201171875 0 2950.5234
1792.8631591796875 0 1591.2296
1793.8687744140625 0 797.7097
1805.84619140625 0 1749.1301
1806.8599853515625 0 1762.5193
1807.8592529296875 0 1393.6208
1860.91650390625 0 1223.4005
1861.9232177734375 0 2661.3594
1862.9315185546875 0 5581.5693
1863.9306640625 0 3607.813
1864.93896484375 0 2339.7424
1875.937744140625 0 1617.8517
1876.936767578125 0 3065.653
1877.9298095703125 0 2192.2534
1878.940185546875 0 1167.7544
1892.94287109375 0 1277.7465
1893.9539794921875 0 2682.339
1894.9512939453125 0 2465.0447
1895.95654296875 0 1268.1052
1902.9169921875 0 769.4743
1903.934326171875 0 2172.568
1904.924560546875 0 5181.17
1905.92626953125 0 5311.7656
1906.9251708984375 0 3547.737
1907.9190673828125 0 1386.3762
1919.9298095703125 0 2643.0544
1920.9384765625 0 8101.5566
1921.946044921875 0 21115.875
1922.9520263671875 0 20983.158
1923.95263671875 0 11151.653
1924.9517822265625 0 6112.387
1925.95361328125 0 2273.238
1985.14697265625 0 670.7368
1996.2213134765625 0 832.34753
3314.72119140625 0 696.4885

Spectrum Details

|  |  |
| --- | --- |
| Matched peaks? Matched peaks The total absolute number of peaks matched. Additionally in brackets the total fraction of peaks matched and the total number of peaks is shown. | 51 (23.08% of 221) |
| FDR? FDR The false discovery rate estimated for this peptide. It is calculated by matching all theoretical fragments with a non-integer shift with the raw peaks for this spectrum. This is done with 40 different shifts. The resulting percentage is the average number of annotated peaks over the number of annotated peaks with the correct spectrum. | 0.37% |
| Satellite FDR? Satellite FDR See the FDR for details on its calculation. This satellite ion specific FDR only contains the satellite ions (d/w) for I/L/J positions. | 0.00% |
| PSM Score? PSM Score The PSM Score as given by Hecklib to this annotated spectrum. It is shown with three significant figures. | 415 |

#### Spectrum 4847? Spectrum 4847 The raw spectrum of this peptide as annotated by Hecklib. The fragments are coloured according to ion type (see legend). Any peaks with a star '\*' as text can be hovered over to see the full details, first the ion type second the mass shift type. By hovering over the amino acids in the peptide or ions in the legend the corresponding peaks are highlighted. By toggling the 'Unassigned' label you can turn the background (unassigned) peaks on or off in the plot. By updating the slider in the Ion legend you can update the spectrum to only show the top X% of the peaks with labels. The top X% means any peak that is within X% of the highest intensity. By dragging in the spectrum you can zoom in to a specific part of the spectrum and use 'Zoom Out' to get back to the original zoom level. The annotation of the spectrum is based on the given sequence in the peptides file and is done with different software so inconsistencies are likely. The peaks are annotated based on the given sequence, with 20 ppm tolerance.

Copy Data

##### Spectrum 4847 (TSV)

###### Preview

```
Loading example...
```

*Click on the button to copy the data to your clipboard.*

Mz MinMz MaxIntensity Max

WidthHeightPeptide font sizePeptide stroke widthSpectrum font sizeSpectrum stroke widthCompact peptide

Ion legend

wxyz

abcd

OtherUnassignedIon^Charge^_Position_Show for top:%

VJHQDWLDGKEYKCK

02.06e+44.13e+46.19e+48.25e+4

Zoom Out

y^+1^_1_y^+1^_1_d^+1^_2_b^+3^_4_a^+1^_2_b^+1^_2_y^+4^_7_y^+1^_2_b^+2^_5_y^+2^_4_y^+3^_7_y^+1^_2_b^+4^_11_y^+3^_8_b^+1^_3_y^+2^_5_y^+2^_5_b^+4^_12_b^+2^_6_b^+2^_6_b^+2^_6_y^+1^_3_y^+2^_6_y^+2^_6_y^+1^_3_y^+3^_10_y^+3^_10_b^+2^_7_y^+2^_7_y^+2^_7_y^+2^_7_\*\*b^+1^_4_\*b^+2^_8_y^+2^_8_y^+2^_8_y^+2^_8_y^+3^_12_y^+3^_12_y^+2^_9_y^+2^_9_y^+3^_13_y^+3^_13_y^+3^_13_y^+2^_9_b^+1^_5_b^+1^_5_b^+1^_5_y^+1^_4_y^+3^_14_y^+3^_14_y^+3^_14_b^+2^_11_y^+2^_10_y^+2^_10_y^+1^_5_y^+1^_5_y^+2^_11_y^+1^_5_b^+1^_6_b^+1^_6_y^+2^_12_y^+2^_12_b^+1^_6_y^+2^_12_y^+1^_6_b^+1^_7_y^+1^_7_y^+1^_8_y^+1^_8_y^+1^_8_y^+1^_9_y^+1^_11_

0797159423913188

Fragment Matches Table

Show background peaks

| Position | Ion type | Intensity | mz Theoretical | mz Error (Th) | mz Error (ppm) | Charge | Series Number |
| --- | --- | --- | --- | --- | --- | --- | --- |
| - | - | 1317 | 120.1 | - | - | 0 | - |
| - | - | 4931 | 120.1 | - | - | 0 | - |
| - | - | 607 | 121.1 | - | - | 0 | - |
| - | - | 450.8 | 122.1 | - | - | 0 | - |
| - | - | 1456 | 127.1 | - | - | 0 | - |
| - | - | 718.4 | 127.1 | - | - | 0 | - |
| - | - | 455.5 | 128.1 | - | - | 0 | - |
| - | - | 672.3 | 129.1 | - | - | 0 | - |
| - | - | 7.071E+04 | 129.1 | - | - | 0 | - |
| - | - | 963.6 | 130.1 | - | - | 0 | - |
| - | - | 2.205E+04 | 130.1 | - | - | 0 | - |
| 15 | y | 7693 | 130.1 | 0.0003233 | 2.485 | +1 | 1 |
| - | - | 739.8 | 130.1 | - | - | 0 | - |
| - | - | 5041 | 130.1 | - | - | 0 | - |
| - | - | 2013 | 131.1 | - | - | 0 | - |
| - | - | 920.4 | 131.1 | - | - | 0 | - |
| - | - | 4834 | 132.1 | - | - | 0 | - |
| - | - | 635.6 | 133.1 | - | - | 0 | - |
| - | - | 1868 | 134 | - | - | 0 | - |
| - | - | 727.1 | 134 | - | - | 0 | - |
| - | - | 6037 | 136.1 | - | - | 0 | - |
| - | - | 732.7 | 138.1 | - | - | 0 | - |
| - | - | 772.7 | 139.1 | - | - | 0 | - |
| - | - | 493.4 | 140.1 | - | - | 0 | - |
| - | - | 1760 | 141.1 | - | - | 0 | - |
| - | - | 930.8 | 143 | - | - | 0 | - |
| - | - | 420.1 | 144.1 | - | - | 0 | - |
| - | - | 1690 | 145.1 | - | - | 0 | - |
| - | - | 493.9 | 147 | - | - | 0 | - |
| - | - | 436 | 147.1 | - | - | 0 | - |
| 15 | y | 7055 | 147.1 | 0.0003398 | 2.31 | +1 | 1 |
| - | - | 850.8 | 149 | - | - | 0 | - |
| - | - | 509.9 | 150.1 | - | - | 0 | - |
| - | - | 687.8 | 151.1 | - | - | 0 | - |
| - | - | 738 | 152.1 | - | - | 0 | - |
| - | - | 924.9 | 153.1 | - | - | 0 | - |
| - | - | 514.4 | 155.1 | - | - | 0 | - |
| - | - | 4615 | 155.1 | - | - | 0 | - |
| - | - | 553.2 | 156.1 | - | - | 0 | - |
| - | - | 540.2 | 157.1 | - | - | 0 | - |
| 2 | d | 1712 | 157.1 | 0.000341 | 2.17 | +1 | 2 |
| - | - | 650.3 | 158.1 | - | - | 0 | - |
| - | - | 1066 | 158.1 | - | - | 0 | - |
| - | - | 4.462E+04 | 159.1 | - | - | 0 | - |
| 4 | b | 5269 | 160.1 | 0.001837 | 11.47 | +3 | 4 |
| - | - | 689 | 163.1 | - | - | 0 | - |
| - | - | 530.5 | 165.1 | - | - | 0 | - |
| - | - | 1067 | 166.1 | - | - | 0 | - |
| - | - | 6059 | 166.1 | - | - | 0 | - |
| - | - | 680.7 | 168.1 | - | - | 0 | - |
| - | - | 941 | 169.1 | - | - | 0 | - |
| - | - | 1923 | 169.1 | - | - | 0 | - |
| - | - | 3339 | 170.1 | - | - | 0 | - |
| - | - | 747.3 | 171.1 | - | - | 0 | - |
| - | - | 1420 | 173.1 | - | - | 0 | - |
| - | - | 883 | 173.1 | - | - | 0 | - |
| - | - | 504.7 | 176.4 | - | - | 0 | - |
| - | - | 2539 | 178.1 | - | - | 0 | - |
| - | - | 656.2 | 178.9 | - | - | 0 | - |
| - | - | 633.3 | 180 | - | - | 0 | - |
| - | - | 1029 | 180.1 | - | - | 0 | - |
| - | - | 693.2 | 181.1 | - | - | 0 | - |
| - | - | 876.9 | 181.1 | - | - | 0 | - |
| - | - | 1418 | 183.1 | - | - | 0 | - |
| - | - | 649 | 185.1 | - | - | 0 | - |
| 2 | a | 1.131E+04 | 185.2 | 0.0003519 | 1.901 | +1 | 2 |
| - | - | 6813 | 186.1 | - | - | 0 | - |
| - | - | 1141 | 186.2 | - | - | 0 | - |
| - | - | 4573 | 187.1 | - | - | 0 | - |
| - | - | 535.9 | 187.1 | - | - | 0 | - |
| - | - | 813.7 | 187.1 | - | - | 0 | - |
| - | - | 802.6 | 188.1 | - | - | 0 | - |
| - | - | 1746 | 195.1 | - | - | 0 | - |
| - | - | 492.9 | 197.3 | - | - | 0 | - |
| - | - | 753.9 | 198.1 | - | - | 0 | - |
| - | - | 2108 | 198.1 | - | - | 0 | - |
| - | - | 1033 | 199.1 | - | - | 0 | - |
| - | - | 1026 | 199.1 | - | - | 0 | - |
| - | - | 1105 | 201.1 | - | - | 0 | - |
| - | - | 1481 | 205.1 | - | - | 0 | - |
| - | - | 552.3 | 209.1 | - | - | 0 | - |
| - | - | 584.9 | 210.1 | - | - | 0 | - |
| - | - | 658.6 | 211.1 | - | - | 0 | - |
| - | - | 843 | 213.1 | - | - | 0 | - |
| - | - | 632 | 213.1 | - | - | 0 | - |
| 2 | b | 3220 | 213.2 | 0.0003256 | 1.528 | +1 | 2 |
| - | - | 2350 | 215.1 | - | - | 0 | - |
| - | - | 488.2 | 215.6 | - | - | 0 | - |
| - | - | 1052 | 216.1 | - | - | 0 | - |
| - | - | 651.2 | 216.1 | - | - | 0 | - |
| - | - | 714.3 | 221.1 | - | - | 0 | - |
| - | - | 1050 | 223.1 | - | - | 0 | - |
| - | - | 3534 | 223.2 | - | - | 0 | - |
| - | - | 841.9 | 225 | - | - | 0 | - |
| - | - | 1198 | 226.1 | - | - | 0 | - |
| - | - | 532.9 | 226.1 | - | - | 0 | - |
| - | - | 1637 | 226.2 | - | - | 0 | - |
| - | - | 1436 | 227.1 | - | - | 0 | - |
| - | - | 1710 | 227.1 | - | - | 0 | - |
| 9 | y | 1273 | 229.1 | 0.00087 | 3.797 | +4 | 7 |
| - | - | 1486 | 232.1 | - | - | 0 | - |
| - | - | 1095 | 233.1 | - | - | 0 | - |
| - | - | 1886 | 233.2 | - | - | 0 | - |
| - | - | 5353 | 234.1 | - | - | 0 | - |
| - | - | 728.1 | 235.1 | - | - | 0 | - |
| - | - | 742.1 | 238.1 | - | - | 0 | - |
| - | - | 1480 | 240.1 | - | - | 0 | - |
| - | - | 779.8 | 241.1 | - | - | 0 | - |
| - | - | 560.9 | 243.1 | - | - | 0 | - |
| - | - | 4101 | 244.1 | - | - | 0 | - |
| - | - | 1719 | 245.1 | - | - | 0 | - |
| - | - | 770.1 | 247.1 | - | - | 0 | - |
| - | - | 1594 | 247.1 | - | - | 0 | - |
| - | - | 1839 | 248.1 | - | - | 0 | - |
| - | - | 3726 | 249.1 | - | - | 0 | - |
| - | - | 663.1 | 249.2 | - | - | 0 | - |
| - | - | 642.3 | 250.1 | - | - | 0 | - |
| - | - | 1.776E+04 | 251.2 | - | - | 0 | - |
| - | - | 2068 | 252.2 | - | - | 0 | - |
| - | - | 1094 | 252.6 | - | - | 0 | - |
| - | - | 775.3 | 254.7 | - | - | 0 | - |
| - | - | 916.3 | 255.1 | - | - | 0 | - |
| - | - | 593.2 | 256.1 | - | - | 0 | - |
| - | - | 632.3 | 257.1 | - | - | 0 | - |
| - | - | 1013 | 261.1 | - | - | 0 | - |
| - | - | 616.5 | 261.1 | - | - | 0 | - |
| - | - | 1156 | 261.2 | - | - | 0 | - |
| - | - | 677.2 | 263.1 | - | - | 0 | - |
| - | - | 1348 | 265.1 | - | - | 0 | - |
| - | - | 1265 | 265.1 | - | - | 0 | - |
| - | - | 889 | 265.2 | - | - | 0 | - |
| - | - | 8839 | 266.1 | - | - | 0 | - |
| - | - | 1168 | 267.1 | - | - | 0 | - |
| - | - | 8764 | 270.1 | - | - | 0 | - |
| - | - | 914.4 | 270.2 | - | - | 0 | - |
| - | - | 1726 | 270.6 | - | - | 0 | - |
| - | - | 3231 | 272.1 | - | - | 0 | - |
| - | - | 2120 | 272.2 | - | - | 0 | - |
| - | - | 1444 | 273.1 | - | - | 0 | - |
| - | - | 2649 | 274.1 | - | - | 0 | - |
| - | - | 1049 | 279.1 | - | - | 0 | - |
| - | - | 7392 | 283.1 | - | - | 0 | - |
| - | - | 624.8 | 284.1 | - | - | 0 | - |
| - | - | 1564 | 284.1 | - | - | 0 | - |
| - | - | 5489 | 290.1 | - | - | 0 | - |
| 14 | y | 1123 | 291.1 | 0.003877 | 13.32 | +1 | 2 |
| - | - | 1085 | 291.1 | - | - | 0 | - |
| - | - | 4301 | 292.2 | - | - | 0 | - |
| 5 | b | 5926 | 297.2 | 0.0004879 | 1.642 | +2 | 5 |
| - | - | 918.5 | 297.7 | - | - | 0 | - |
| - | - | 1447 | 299.1 | - | - | 0 | - |
| - | - | 500.7 | 300.1 | - | - | 0 | - |
| 12 | y | 3369 | 300.1 | 0.001928 | 6.423 | +2 | 4 |
| - | - | 544.8 | 300.2 | - | - | 0 | - |
| - | - | 776.5 | 300.6 | - | - | 0 | - |
| - | - | 2475 | 301.2 | - | - | 0 | - |
| - | - | 1012 | 302.1 | - | - | 0 | - |
| - | - | 958.6 | 302.2 | - | - | 0 | - |
| 9 | y | 2608 | 305.2 | 0.001419 | 4.651 | +3 | 7 |
| - | - | 745.4 | 305.5 | - | - | 0 | - |
| - | - | 770 | 306.2 | - | - | 0 | - |
| 14 | y | 1.875E+04 | 308.1 | 0.004089 | 13.27 | +1 | 2 |
| - | - | 1036 | 308.1 | - | - | 0 | - |
| - | - | 3457 | 309.1 | - | - | 0 | - |
| - | - | 713.5 | 309.2 | - | - | 0 | - |
| - | - | 828.3 | 310.1 | - | - | 0 | - |
| - | - | 684.5 | 311.2 | - | - | 0 | - |
| - | - | 2579 | 315.2 | - | - | 0 | - |
| - | - | 791.1 | 318.2 | - | - | 0 | - |
| - | - | 711.8 | 319.2 | - | - | 0 | - |
| - | - | 1056 | 323.2 | - | - | 0 | - |
| 11 | b | 2443 | 326.7 | 0.001952 | 5.976 | +4 | 11 |
| - | - | 1058 | 327.2 | - | - | 0 | - |
| - | - | 2342 | 332.2 | - | - | 0 | - |
| - | - | 755.1 | 333.2 | - | - | 0 | - |
| - | - | 753.7 | 333.2 | - | - | 0 | - |
| - | - | 631.9 | 340.2 | - | - | 0 | - |
| - | - | 782.3 | 340.7 | - | - | 0 | - |
| 8 | y | 1732 | 343.5 | 0.001265 | 3.684 | +3 | 8 |
| - | - | 868 | 343.8 | - | - | 0 | - |
| - | - | 1582 | 345.1 | - | - | 0 | - |
| - | - | 4404 | 346.1 | - | - | 0 | - |
| - | - | 653 | 347.1 | - | - | 0 | - |
| 3 | b | 1.07E+04 | 350.2 | 0.0002669 | 0.7621 | +1 | 3 |
| - | - | 1855 | 351.2 | - | - | 0 | - |
| - | - | 624.2 | 354.2 | - | - | 0 | - |
| - | - | 692.5 | 355.1 | - | - | 0 | - |
| 11 | y | 1025 | 355.7 | 0.002347 | 6.6 | +2 | 5 |
| - | - | 855.1 | 358.7 | - | - | 0 | - |
| - | - | 729.5 | 360 | - | - | 0 | - |
| - | - | 676.3 | 362.2 | - | - | 0 | - |
| - | - | 5890 | 363.1 | - | - | 0 | - |
| - | - | 917.5 | 364.1 | - | - | 0 | - |
| 11 | y | 631.1 | 364.7 | 0.006409 | 17.58 | +2 | 5 |
| - | - | 724.4 | 367.1 | - | - | 0 | - |
| - | - | 1366 | 367.2 | - | - | 0 | - |
| 12 | b | 4974 | 367.7 | 0.006933 | 18.86 | +4 | 12 |
| - | - | 1615 | 368.2 | - | - | 0 | - |
| - | - | 769.7 | 368.7 | - | - | 0 | - |
| - | - | 2.639E+04 | 376.2 | - | - | 0 | - |
| - | - | 9990 | 376.7 | - | - | 0 | - |
| - | - | 3616 | 377.2 | - | - | 0 | - |
| - | - | 668.9 | 378.2 | - | - | 0 | - |
| - | - | 3061 | 379.2 | - | - | 0 | - |
| - | - | 1.526E+04 | 381.2 | - | - | 0 | - |
| 6 | b | 4767 | 381.2 | 1.877E-05 | 0.04925 | +2 | 6 |
| - | - | 1943 | 381.5 | - | - | 0 | - |
| 6 | b | 1117 | 381.7 | 0.006638 | 17.39 | +2 | 6 |
| - | - | 1080 | 381.9 | - | - | 0 | - |
| - | - | 2126 | 382.2 | - | - | 0 | - |
| - | - | 761.7 | 385.1 | - | - | 0 | - |
| - | - | 1635 | 389.2 | - | - | 0 | - |
| 6 | b | 1.278E+04 | 390.2 | 0.0002906 | 0.7448 | +2 | 6 |
| - | - | 4947 | 390.7 | - | - | 0 | - |
| - | - | 830.9 | 391.2 | - | - | 0 | - |
| - | - | 923.6 | 391.2 | - | - | 0 | - |
| - | - | 644.3 | 392.7 | - | - | 0 | - |
| - | - | 736.4 | 395.2 | - | - | 0 | - |
| - | - | 726.1 | 398.2 | - | - | 0 | - |
| - | - | 1006 | 402.2 | - | - | 0 | - |
| - | - | 848.5 | 403.2 | - | - | 0 | - |
| - | - | 717.6 | 407.2 | - | - | 0 | - |
| - | - | 5548 | 412.2 | - | - | 0 | - |
| - | - | 1238 | 413.1 | - | - | 0 | - |
| - | - | 730.7 | 413.2 | - | - | 0 | - |
| - | - | 886.5 | 414.2 | - | - | 0 | - |
| - | - | 927.1 | 415.2 | - | - | 0 | - |
| - | - | 1798 | 418.2 | - | - | 0 | - |
| - | - | 1095 | 419.2 | - | - | 0 | - |
| 13 | y | 1244 | 419.2 | 0.007379 | 17.6 | +1 | 3 |
| 10 | y | 909.2 | 419.7 | 0.001459 | 3.475 | +2 | 6 |
| - | - | 1698 | 421.2 | - | - | 0 | - |
| - | - | 833 | 421.7 | - | - | 0 | - |
| - | - | 866 | 423.7 | - | - | 0 | - |
| - | - | 1017 | 424.2 | - | - | 0 | - |
| - | - | 672.3 | 424.7 | - | - | 0 | - |
| 10 | y | 1775 | 428.7 | 0.001675 | 3.907 | +2 | 6 |
| - | - | 1242 | 429.2 | - | - | 0 | - |
| - | - | 1201 | 430.2 | - | - | 0 | - |
| - | - | 6001 | 430.2 | - | - | 0 | - |
| - | - | 710.7 | 431.2 | - | - | 0 | - |
| - | - | 677.4 | 432.2 | - | - | 0 | - |
| - | - | 3934 | 432.7 | - | - | 0 | - |
| - | - | 1819 | 433.2 | - | - | 0 | - |
| - | - | 698 | 433.7 | - | - | 0 | - |
| - | - | 1595 | 435.2 | - | - | 0 | - |
| - | - | 929.6 | 436.2 | - | - | 0 | - |
| 13 | y | 5687 | 436.2 | 0.00399 | 9.147 | +1 | 3 |
| - | - | 4901 | 437.2 | - | - | 0 | - |
| - | - | 659.8 | 437.2 | - | - | 0 | - |
| 6 | y | 1200 | 437.5 | 0.0001528 | 0.3493 | +3 | 10 |
| - | - | 1071 | 437.9 | - | - | 0 | - |
| - | - | 1280 | 438.2 | - | - | 0 | - |
| - | - | 897.6 | 440.3 | - | - | 0 | - |
| - | - | 793.8 | 441.3 | - | - | 0 | - |
| - | - | 1005 | 441.7 | - | - | 0 | - |
| 6 | y | 676.5 | 443.2 | 0.001108 | 2.5 | +3 | 10 |
| 7 | b | 1895 | 446.7 | 0.0003595 | 0.8048 | +2 | 7 |
| - | - | 876.9 | 447.2 | - | - | 0 | - |
| 9 | y | 2374 | 448.2 | 0.001692 | 3.776 | +2 | 7 |
| 9 | y | 929.8 | 448.7 | 0.003675 | 8.191 | +2 | 7 |
| - | - | 1300 | 450.2 | - | - | 0 | - |
| - | - | 734.3 | 450.7 | - | - | 0 | - |
| - | - | 1969 | 453.2 | - | - | 0 | - |
| 9 | y | 2.261E+04 | 457.2 | 0.001939 | 4.242 | +2 | 7 |
| - | - | 1.116E+04 | 457.7 | - | - | 0 | - |
| - | - | 2860 | 458.2 | - | - | 0 | - |
| - | - | 1394 | 458.7 | - | - | 0 | - |
| - | - | 1539 | 460.2 | - | - | 0 | - |
| - | - | 693.3 | 468.3 | - | - | 0 | - |
| - | - | 1198 | 468.7 | - | - | 0 | - |
| - | - | 881.9 | 471.2 | - | - | 0 | - |
| 0 | Precursor | 2152 | 476.2 | 0.0001934 | 0.406 | +4 | -1 |
| 0 | Precursor | 1141 | 476.5 | 0.002155 | 4.522 | +4 | -1 |
| - | - | 1231 | 476.7 | - | - | 0 | - |
| - | - | 863.2 | 477 | - | - | 0 | - |
| - | - | 820.6 | 477.2 | - | - | 0 | - |
| - | - | 1355 | 478.2 | - | - | 0 | - |
| 4 | b | 3632 | 478.3 | 0.0006493 | 1.358 | +1 | 4 |
| - | - | 739.7 | 479.3 | - | - | 0 | - |
| - | - | 703 | 479.8 | - | - | 0 | - |
| - | - | 1021 | 480.5 | - | - | 0 | - |
| 0 | Precursor | 1.128E+04 | 480.7 | 8.796E-05 | 0.183 | +4 | -1 |
| - | - | 1.069E+04 | 481 | - | - | 0 | - |
| - | - | 4176 | 481.2 | - | - | 0 | - |
| - | - | 859.9 | 481.3 | - | - | 0 | - |
| - | - | 1418 | 481.5 | - | - | 0 | - |
| - | - | 1438 | 481.7 | - | - | 0 | - |
| - | - | 1978 | 483.2 | - | - | 0 | - |
| - | - | 706.7 | 483.7 | - | - | 0 | - |
| - | - | 620.1 | 486.2 | - | - | 0 | - |
| - | - | 639.4 | 487.8 | - | - | 0 | - |
| - | - | 1920 | 488.8 | - | - | 0 | - |
| - | - | 4308 | 489.8 | - | - | 0 | - |
| - | - | 1442 | 490.3 | - | - | 0 | - |
| - | - | 1183 | 491.3 | - | - | 0 | - |
| - | - | 7296 | 494.2 | - | - | 0 | - |
| 8 | b | 1585 | 495.2 | 0.00626 | 12.64 | +2 | 8 |
| - | - | 1044 | 496.2 | - | - | 0 | - |
| - | - | 675.7 | 496.7 | - | - | 0 | - |
| - | - | 2291 | 497.2 | - | - | 0 | - |
| - | - | 688.1 | 497.7 | - | - | 0 | - |
| - | - | 634.1 | 504.2 | - | - | 0 | - |
| 8 | y | 1.355E+04 | 505.7 | 0.002377 | 4.7 | +2 | 8 |
| 8 | y | 1.024E+04 | 506.2 | 0.003418 | 6.752 | +2 | 8 |
| - | - | 5955 | 506.7 | - | - | 0 | - |
| - | - | 1390 | 507.2 | - | - | 0 | - |
| - | - | 728.8 | 512.2 | - | - | 0 | - |
| - | - | 687.6 | 514.2 | - | - | 0 | - |
| - | - | 2814 | 514.2 | - | - | 0 | - |
| 8 | y | 8.169E+04 | 514.7 | 0.002014 | 3.912 | +2 | 8 |
| - | - | 3.891E+04 | 515.2 | - | - | 0 | - |
| - | - | 1279 | 515.3 | - | - | 0 | - |
| - | - | 1.963E+04 | 515.7 | - | - | 0 | - |
| - | - | 5077 | 516.2 | - | - | 0 | - |
| - | - | 1634 | 516.7 | - | - | 0 | - |
| 4 | y | 1053 | 518.2 | 0.002748 | 5.302 | +3 | 12 |
| 4 | y | 1236 | 518.6 | 0.004513 | 8.704 | +3 | 12 |
| - | - | 887.4 | 518.9 | - | - | 0 | - |
| - | - | 689.1 | 522.2 | - | - | 0 | - |
| - | - | 670.2 | 523.2 | - | - | 0 | - |
| - | - | 693.4 | 530.2 | - | - | 0 | - |
| - | - | 721.4 | 532.2 | - | - | 0 | - |
| - | - | 3699 | 539.2 | - | - | 0 | - |
| - | - | 953.2 | 540.2 | - | - | 0 | - |
| - | - | 930.6 | 543.3 | - | - | 0 | - |
| - | - | 911.2 | 549.2 | - | - | 0 | - |
| - | - | 729.9 | 549.3 | - | - | 0 | - |
| - | - | 803.9 | 553.8 | - | - | 0 | - |
| - | - | 700.7 | 558.3 | - | - | 0 | - |
| - | - | 1080 | 558.6 | - | - | 0 | - |
| 7 | y | 6858 | 562.3 | 0.002447 | 4.353 | +2 | 9 |
| 7 | y | 2455 | 562.8 | 0.007071 | 12.56 | +2 | 9 |
| - | - | 1846 | 563.3 | - | - | 0 | - |
| 3 | y | 2933 | 563.9 | 0.001429 | 2.535 | +3 | 13 |
| - | - | 1462 | 564.2 | - | - | 0 | - |
| 3 | y | 2114 | 564.3 | 0.004977 | 8.821 | +3 | 13 |
| - | - | 2957 | 564.6 | - | - | 0 | - |
| - | - | 1176 | 565.3 | - | - | 0 | - |
| - | - | 4463 | 567.2 | - | - | 0 | - |
| - | - | 1160 | 568.2 | - | - | 0 | - |
| - | - | 1079 | 569.6 | - | - | 0 | - |
| 3 | y | 1.036E+04 | 569.9 | 0.0009226 | 1.619 | +3 | 13 |
| - | - | 7691 | 570.3 | - | - | 0 | - |
| - | - | 5312 | 570.6 | - | - | 0 | - |
| - | - | 931.6 | 570.9 | - | - | 0 | - |
| 7 | y | 2.536E+04 | 571.3 | 0.002603 | 4.556 | +2 | 9 |
| - | - | 1.531E+04 | 571.8 | - | - | 0 | - |
| - | - | 7887 | 572.3 | - | - | 0 | - |
| - | - | 2976 | 572.8 | - | - | 0 | - |
| - | - | 1309 | 575.2 | - | - | 0 | - |
| 5 | b | 1704 | 575.3 | 0.001239 | 2.153 | +1 | 5 |
| 5 | b | 1591 | 576.3 | 0.002086 | 3.621 | +1 | 5 |
| - | - | 917.4 | 577.3 | - | - | 0 | - |
| - | - | 1162 | 581.3 | - | - | 0 | - |
| - | - | 2432 | 582.2 | - | - | 0 | - |
| - | - | 1016 | 592.3 | - | - | 0 | - |
| - | - | 3770 | 593.3 | - | - | 0 | - |
| 5 | b | 2.167E+04 | 593.3 | 1.246E-05 | 0.021 | +1 | 5 |
| - | - | 1177 | 594.3 | - | - | 0 | - |
| - | - | 7222 | 594.3 | - | - | 0 | - |
| - | - | 1692 | 595.3 | - | - | 0 | - |
| - | - | 1486 | 595.9 | - | - | 0 | - |
| - | - | 1046 | 598 | - | - | 0 | - |
| 12 | y | 1.055E+04 | 599.3 | 0.004514 | 7.532 | +1 | 4 |
| - | - | 1930 | 600.3 | - | - | 0 | - |
| - | - | 1511 | 601.3 | - | - | 0 | - |
| 2 | y | 4315 | 601.6 | 0.002148 | 3.57 | +3 | 14 |
| 2 | y | 3975 | 602 | 0.003587 | 5.96 | +3 | 14 |
| - | - | 3785 | 602.3 | - | - | 0 | - |
| - | - | 866.2 | 602.6 | - | - | 0 | - |
| - | - | 801.1 | 606.3 | - | - | 0 | - |
| - | - | 1079 | 607.3 | - | - | 0 | - |
| 2 | y | 1.37E+04 | 607.6 | 0.001153 | 1.897 | +3 | 14 |
| - | - | 1.354E+04 | 608 | - | - | 0 | - |
| - | - | 8043 | 608.3 | - | - | 0 | - |
| - | - | 2463 | 608.6 | - | - | 0 | - |
| - | - | 1325 | 609 | - | - | 0 | - |
| - | - | 764.4 | 635.3 | - | - | 0 | - |
| - | - | 2672 | 650.3 | - | - | 0 | - |
| - | - | 744.2 | 651.3 | - | - | 0 | - |
| 11 | b | 2743 | 652.3 | 0.005844 | 8.959 | +2 | 11 |
| 6 | y | 696 | 655.8 | 0.001167 | 1.779 | +2 | 10 |
| - | - | 1572 | 656.4 | - | - | 0 | - |
| - | - | 1816 | 662.3 | - | - | 0 | - |
| 6 | y | 3044 | 664.3 | 0.002281 | 3.434 | +2 | 10 |
| - | - | 2207 | 664.8 | - | - | 0 | - |
| - | - | 975.1 | 665.3 | - | - | 0 | - |
| - | - | 911.2 | 666.3 | - | - | 0 | - |
| - | - | 7617 | 680.3 | - | - | 0 | - |
| - | - | 2547 | 681.3 | - | - | 0 | - |
| - | - | 716.9 | 682.3 | - | - | 0 | - |
| - | - | 869.2 | 688.3 | - | - | 0 | - |
| - | - | 1002 | 708.3 | - | - | 0 | - |
| 11 | y | 4405 | 710.3 | 0.005353 | 7.536 | +1 | 5 |
| 11 | y | 2167 | 711.3 | 0.01411 | 19.84 | +1 | 5 |
| - | - | 2559 | 721.4 | - | - | 0 | - |
| 5 | y | 1312 | 721.8 | 0.00556 | 7.702 | +2 | 11 |
| - | - | 2007 | 722.3 | - | - | 0 | - |
| 11 | y | 1.317E+04 | 728.3 | 0.005176 | 7.106 | +1 | 5 |
| - | - | 4406 | 729.3 | - | - | 0 | - |
| - | - | 1417 | 730.3 | - | - | 0 | - |
| - | - | 683.6 | 733.4 | - | - | 0 | - |
| - | - | 879.6 | 734.4 | - | - | 0 | - |
| - | - | 923.2 | 735.4 | - | - | 0 | - |
| - | - | 5341 | 751.4 | - | - | 0 | - |
| - | - | 2673 | 752.4 | - | - | 0 | - |
| 6 | b | 675.6 | 761.4 | 0.003774 | 4.957 | +1 | 6 |
| 6 | b | 1767 | 762.4 | 0.004744 | 6.222 | +1 | 6 |
| - | - | 4109 | 763.4 | - | - | 0 | - |
| - | - | 1544 | 764.4 | - | - | 0 | - |
| - | - | 681.5 | 765.4 | - | - | 0 | - |
| - | - | 1109 | 768.3 | - | - | 0 | - |
| - | - | 1872 | 772.4 | - | - | 0 | - |
| - | - | 1081 | 773.4 | - | - | 0 | - |
| 4 | y | 1152 | 776.9 | 0.003687 | 4.747 | +2 | 12 |
| 4 | y | 1956 | 777.4 | 0.001127 | 1.45 | +2 | 12 |
| - | - | 1649 | 777.4 | - | - | 0 | - |
| - | - | 1695 | 777.9 | - | - | 0 | - |
| 6 | b | 2.72E+04 | 779.4 | 0.0005651 | 0.7251 | +1 | 6 |
| - | - | 949.1 | 780.3 | - | - | 0 | - |
| - | - | 1.154E+04 | 780.4 | - | - | 0 | - |
| - | - | 2530 | 781.4 | - | - | 0 | - |
| - | - | 908.1 | 785.4 | - | - | 0 | - |
| 4 | y | 1547 | 785.9 | 0.00427 | 5.434 | +2 | 12 |
| - | - | 1416 | 786.4 | - | - | 0 | - |
| - | - | 1332 | 786.9 | - | - | 0 | - |
| - | - | 1672 | 793.4 | - | - | 0 | - |
| - | - | 1144 | 795.3 | - | - | 0 | - |
| 10 | y | 2054 | 856.4 | 0.006022 | 7.032 | +1 | 6 |
| - | - | 2087 | 882.4 | - | - | 0 | - |
| - | - | 962.9 | 883.4 | - | - | 0 | - |
| 7 | b | 1019 | 892.5 | 0.005703 | 6.39 | +1 | 7 |
| - | - | 2408 | 900.4 | - | - | 0 | - |
| - | - | 902.2 | 901.4 | - | - | 0 | - |
| - | - | 971.8 | 908.4 | - | - | 0 | - |
| 9 | y | 4094 | 913.4 | 0.005819 | 6.37 | +1 | 7 |
| - | - | 2401 | 914.4 | - | - | 0 | - |
| - | - | 685.1 | 918.3 | - | - | 0 | - |
| 8 | y | 1571 | 1010 | 0.003154 | 3.121 | +1 | 8 |
| 8 | y | 1392 | 1011 | 0.0005014 | 0.4957 | +1 | 8 |
| 8 | y | 7735 | 1028 | 0.006517 | 6.336 | +1 | 8 |
| - | - | 4360 | 1029 | - | - | 0 | - |
| - | - | 1803 | 1030 | - | - | 0 | - |
| 7 | y | 712.8 | 1142 | 0.007084 | 6.206 | +1 | 9 |
| - | - | 1135 | 1143 | - | - | 0 | - |
| - | - | 587.4 | 1259 | - | - | 0 | - |
| 5 | y | 651.5 | 1443 | 0.01214 | 8.418 | +1 | 11 |
| - | - | 678.8 | 2664 | - | - | 0 | - |
| - | - | 1137 | 3075 | - | - | 0 | - |
| - | - | 900.5 | 3075 | - | - | 0 | - |
| - | - | 741.3 | 3156 | - | - | 0 | - |

m/z Charge Intensity FragmentType MassShift Position
120.06591033935547 0 1316.9354
120.0811538696289 0 4931.254
121.08451080322266 0 607.04706
122.07191467285156 0 450.78162
127.0506591796875 0 1456.1376
127.08694458007812 0 718.44476
128.10728454589844 0 455.54352
129.06614685058594 0 672.2846
129.10260009765625 0 70705.48
130.05029296875 0 963.5676
130.06546020507812 0 22054.105
130.08657836914062 0 7693.12 y Ammonia loss 14
130.10081481933594 0 739.83594
130.10594177246094 0 5041.3604
131.06895446777344 0 2013.2899
131.11834716796875 0 920.4011
132.08108520507812 0 4834.024
133.0609893798828 0 635.5568
134.02743530273438 0 1868.0472
134.0450439453125 0 727.1184
136.07603454589844 0 6036.8193
138.06654357910156 0 732.74396
139.0870361328125 0 772.714
140.08250427246094 0 493.3695
141.10263061523438 0 1759.926
143.04539489746094 0 930.7621
144.08114624023438 0 420.0595
145.06117248535156 0 1689.5302
147.044189453125 0 493.90823
147.07679748535156 0 435.98984
147.11314392089844 0 7055.048 y 14
149.0238494873047 0 850.7924
150.0551300048828 0 509.9324
151.0872802734375 0 687.7892
152.07102966308594 0 738.02844
153.1027069091797 0 924.89435
155.08164978027344 0 514.39056
155.09307861328125 0 4615.3193
156.07699584960938 0 553.17487
157.0974578857422 0 540.1786
157.13388061523438 0 1711.6875 d 1
158.060302734375 0 650.3371
158.0843505859375 0 1065.9296
159.09205627441406 0 44622.324
160.09542846679688 0 5269.463 b 3
163.07192993164062 0 688.9579
165.10263061523438 0 530.52527
166.0535888671875 0 1067.1866
166.06143188476562 0 6058.793
168.08154296875 0 680.67377
169.07679748535156 0 941.0149
169.0973663330078 0 1922.5624
170.06036376953125 0 3339.3782
171.11305236816406 0 747.31244
173.0561065673828 0 1420.4327
173.12901306152344 0 883.01776
176.41934204101562 0 504.67258
178.13418579101562 0 2538.94
178.87728881835938 0 656.1824
180.03321838378906 0 633.3031
180.11329650878906 0 1029.1439
181.0609893798828 0 693.2167
181.09742736816406 0 876.93256
183.1130828857422 0 1418.3125
185.12918090820312 0 649.018
185.16519165039062 0 11309.484 a 1
186.12403869628906 0 6813.387
186.16856384277344 0 1141.2365
187.08685302734375 0 4573.2207
187.10797119140625 0 535.936
187.127685546875 0 813.7419
188.09056091308594 0 802.57574
195.11294555664062 0 1745.7107
197.3196258544922 0 492.87262
198.087646484375 0 753.8714
198.12744140625 0 2108.3435
199.07106018066406 0 1032.9974
199.10800170898438 0 1025.9103
201.1239013671875 0 1104.6924
205.09730529785156 0 1480.6357
209.0942840576172 0 552.29535
210.12637329101562 0 584.883
211.10768127441406 0 658.586
213.12356567382812 0 843.00885
213.1383514404297 0 631.9769
213.1600799560547 0 3220.0542 b 1
215.1392059326172 0 2350.0017
215.59573364257812 0 488.2219
216.09872436523438 0 1051.693
216.13381958007812 0 651.1745
221.1034698486328 0 714.2844
223.10794067382812 0 1049.9551
223.15573120117188 0 3533.8591
225.04261779785156 0 841.9497
226.0833740234375 0 1198.0039
226.11978149414062 0 532.92523
226.1552734375 0 1636.6735
227.0670166015625 0 1436.4043
227.08511352539062 0 1710.2009
229.11865234375 0 1272.5262 y 8
232.14450073242188 0 1485.5483
233.14022827148438 0 1094.8517
233.16482543945312 0 1886.0476
234.12416076660156 0 5353.0493
235.12789916992188 0 728.11847
238.11878967285156 0 742.058
240.13479614257812 0 1480.4963
241.11903381347656 0 779.77527
243.1100616455078 0 560.92316
244.0931396484375 0 4101.257
245.0958709716797 0 1719.4951
247.1073760986328 0 770.10406
247.14442443847656 0 1593.5537
248.11444091796875 0 1839.4482
249.0988311767578 0 3726.4812
249.15966796875 0 663.06287
250.10186767578125 0 642.3291
251.15060424804688 0 17764.465
252.1540069580078 0 2068.2424
252.6035919189453 0 1093.8347
254.65530395507812 0 775.3084
255.14599609375 0 916.2565
256.1080017089844 0 593.16425
257.1285400390625 0 632.2746
261.1199035644531 0 1013.29236
261.13433837890625 0 616.521
261.1609802246094 0 1155.6565
263.1042785644531 0 677.1684
265.11767578125 0 1347.8401
265.1304626464844 0 1264.8696
265.1549072265625 0 888.97455
266.1248474121094 0 8838.847
267.12841796875 0 1168.3234
270.1219482421875 0 8763.919
270.16180419921875 0 914.4264
270.62274169921875 0 1725.5435
272.1064147949219 0 3230.9756
272.1758728027344 0 2120.3464
273.0903015136719 0 1443.9928
274.11859130859375 0 2649.0793
279.1460876464844 0 1048.6034
283.1404113769531 0 7392.484
284.10357666015625 0 624.82367
284.1436462402344 0 1564.1934
290.1170959472656 0 5489.373
291.1015625 0 1123.3435 y Ammonia loss 13
291.1201171875 0 1084.632
292.16607666015625 0 4301.231
297.1562194824219 0 5926.2783 b 4
297.6575622558594 0 918.46716
299.06170654296875 0 1447.2462
300.1302185058594 0 500.70438
300.1468505859375 0 3368.551 y 11
300.1689453125 0 544.8353
300.6484069824219 0 776.503
301.15106201171875 0 2475.1838
302.1129455566406 0 1011.9695
302.1513671875 0 958.6475
305.1531982421875 0 2608.4114 y 8
305.4876708984375 0 745.36633
306.23089599609375 0 769.9576
308.1278991699219 0 18751.008 y 13
308.1472473144531 0 1035.7466
309.13079833984375 0 3456.9753
309.1666564941406 0 713.4736
310.1239318847656 0 828.29486
311.16400146484375 0 684.5078
315.1666259765625 0 2578.5432
318.15179443359375 0 791.1008
319.1518859863281 0 711.75916
323.20843505859375 0 1056.4836
326.66424560546875 0 2442.6458 b Water loss 10
327.1656494140625 0 1057.5854
332.2085876464844 0 2342.3918
333.1773376464844 0 755.09955
333.2117919921875 0 753.655
340.1614685058594 0 631.86523
340.6615295410156 0 782.3254
343.49566650390625 0 1731.7816 y 7
343.8298645019531 0 868.0132
345.13031005859375 0 1582.1628
346.114501953125 0 4403.6753
347.1156921386719 0 653.013
350.21893310546875 0 10697.48 b 2
351.2214050292969 0 1855.1875
354.216796875 0 624.23303
355.0699768066406 0 692.5202
355.6624450683594 0 1025.1077 y Water loss 10
358.6802673339844 0 855.1198
360.02789306640625 0 729.48
362.186279296875 0 676.28345
363.1411437988281 0 5889.6826
364.1231689453125 0 917.4933
364.6636657714844 0 631.1495 y 10
367.1397705078125 0 724.3734
367.24560546875 0 1365.6278
367.6849670410156 0 4974.275 b Ammonia loss 11
368.1855163574219 0 1615.3956
368.6871643066406 0 769.6558
376.197998046875 0 26393.73
376.69952392578125 0 9989.719
377.2008972167969 0 3615.6575
378.2181396484375 0 668.8977
379.20953369140625 0 3060.797
381.15185546875 0 15263.276
381.19012451171875 0 4767.1807 b Water loss 5
381.5241394042969 0 1943.345
381.6887512207031 0 1117.0775 b Ammonia loss 5
381.8585510253906 0 1080.0679
382.1544189453125 0 2126.5
385.1484680175781 0 761.6888
389.2096862792969 0 1635.4232
390.1956787109375 0 12784.035 b 5
390.6964416503906 0 4946.9995
391.1728820800781 0 830.9392
391.1985778808594 0 923.56085
392.73248291015625 0 644.2975
395.157470703125 0 736.43427
398.1778869628906 0 726.0859
402.1749572753906 0 1005.5243
403.2005310058594 0 848.4791
407.20501708984375 0 717.56824
412.1832580566406 0 5548.399
413.1422424316406 0 1237.6719
413.16796875 0 730.71564
414.2359619140625 0 886.4892
415.1968994140625 0 927.0646
418.2132873535156 0 1798.0448
419.1662902832031 0 1094.6414
419.1930236816406 0 1243.5353 y Ammonia loss 12
419.7108154296875 0 909.24475 y Water loss 9
421.2086486816406 0 1697.9423
421.73565673828125 0 833.0022
423.73187255859375 0 865.954
424.2272644042969 0 1016.6138
424.731201171875 0 672.30426
428.71588134765625 0 1774.5441 y 9
429.2152404785156 0 1242.2325
430.1666259765625 0 1200.8873
430.19403076171875 0 6000.805
431.19805908203125 0 710.72
432.20709228515625 0 677.3616
432.74066162109375 0 3933.722
433.2431335449219 0 1818.8461
433.742431640625 0 698.01984
435.1818542480469 0 1594.7906
436.1875305175781 0 929.59515
436.22296142578125 0 5687.206 y 12
437.1781005859375 0 4901.207
437.22412109375 0 659.81067
437.5426940917969 0 1199.7609 y Ammonia loss 5
437.876953125 0 1070.8756
438.1777648925781 0 1279.5068
440.25067138671875 0 897.60297
441.25860595703125 0 793.83374
441.6874084472656 0 1005.3138
443.2169494628906 0 676.49567 y 5
446.737060546875 0 1895.0789 b 6
447.2386779785156 0 876.9004
448.2213134765625 0 2374.366 y Water loss 8
448.71868896484375 0 929.79236 y Ammonia loss 8
450.23590087890625 0 1300.43
450.6913757324219 0 734.268
453.1803894042969 0 1969.3513
457.2263488769531 0 22607.893 y 8
457.72796630859375 0 11158.971
458.228271484375 0 2859.9224
458.7269592285156 0 1394.4232
460.24560546875 0 1538.9756
468.2515869140625 0 693.2561
468.7435607910156 0 1198.2561
471.19256591796875 0 881.93854
476.23675537109375 0 2151.6655 Precursor Water loss
476.485107421875 0 1141.248 Precursor Ammonia loss
476.7376403808594 0 1231.414
476.9853210449219 0 863.1531
477.2080078125 0 820.5518
478.2313537597656 0 1354.795
478.27789306640625 0 3632.045 b 3
479.3130187988281 0 739.7107
479.7517395019531 0 702.95636
480.4944152832031 0 1020.6281
480.739501953125 0 11284.28 Precursor
480.989501953125 0 10686.814
481.2397155761719 0 4175.781
481.2749938964844 0 859.9385
481.48980712890625 0 1418.3923
481.741943359375 0 1437.97
483.2237854003906 0 1977.9056
483.7270812988281 0 706.71515
486.2389221191406 0 620.08386
487.76947021484375 0 639.4471
488.7761535644531 0 1919.5071
489.78387451171875 0 4308.3726
490.28497314453125 0 1442.4836
491.27557373046875 0 1183.1453
494.23577880859375 0 7295.78
495.2393493652344 0 1584.9174 b Water loss 7
496.23895263671875 0 1044.219
496.7261962890625 0 675.65924
497.224853515625 0 2291.352
497.72088623046875 0 688.0756
504.1974182128906 0 634.0931
505.7341003417969 0 13552.58 y Water loss 7
506.2319030761719 0 10242.658 y Ammonia loss 7
506.7326965332031 0 5955.0103
507.2305908203125 0 1389.8728
512.2350463867188 0 728.82886
514.185546875 0 687.62555
514.2477416992188 0 2814.439
514.73974609375 0 81692.96 y 7
515.2410278320312 0 38906.574
515.2855834960938 0 1279.4072
515.7410278320312 0 19634.445
516.2421264648438 0 5076.8843
516.7432250976562 0 1633.5942
518.2402954101562 0 1052.6364 y Water loss 3
518.5755615234375 0 1235.7849 y Ammonia loss 3
518.9076538085938 0 887.4341
522.2250366210938 0 689.14886
523.246826171875 0 670.16406
530.2210083007812 0 693.4346
532.1912231445312 0 721.4377
539.2357177734375 0 3699.4563
540.2379760742188 0 953.2437
543.2681274414062 0 930.6046
549.2225341796875 0 911.2187
549.30908203125 0 729.8537
553.766357421875 0 803.8793
558.2548217773438 0 700.70825
558.587646484375 0 1079.5154
562.2760620117188 0 6857.871 y Water loss 6
562.777587890625 0 2454.9668 y Ammonia loss 6
563.2750244140625 0 1845.8447
563.9279174804688 0 2933.0837 y Water loss 2
564.21142578125 0 1461.6979
564.2623291015625 0 2114.456 y Ammonia loss 2
564.5953369140625 0 2956.7786
565.2639770507812 0 1175.9635
567.2314453125 0 4462.856
568.2352905273438 0 1159.5936
569.6043701171875 0 1079.3126
569.9319458007812 0 10362.738 y 2
570.2664794921875 0 7690.92
570.599853515625 0 5311.748
570.9337768554688 0 931.57184
571.2811889648438 0 25363.965 y 6
571.7833251953125 0 15309.663
572.2835083007812 0 7887.152
572.7836303710938 0 2976.0942
575.2459106445312 0 1308.9673
575.2948608398438 0 1704.1575 b Water loss 4
576.2797241210938 0 1590.5297 b Ammonia loss 4
577.2879028320312 0 917.3653
581.2748413085938 0 1162.3705
582.2225952148438 0 2432.1697
592.2725219726562 0 1016.25916
593.255615234375 0 3769.7634
593.30419921875 0 21670.477 b 4
594.2597045898438 0 1177.2374
594.3069458007812 0 7221.768
595.3088989257812 0 1691.5459
595.9445190429688 0 1486.0831
597.9592895507812 0 1046.4733
599.2857666015625 0 10554.131 y 11
600.2901000976562 0 1929.848
601.291015625 0 1511.0698
601.6218872070312 0 4314.8267 y Water loss 1
601.9556274414062 0 3975.4421 y Ammonia loss 1
602.2899169921875 0 3785.0708
602.6234741210938 0 866.164
606.3255615234375 0 801.07446
607.2949829101562 0 1079.0226
607.6264038085938 0 13697.749 y 1
607.9606323242188 0 13539.668
608.293701171875 0 8042.905
608.6278686523438 0 2463.2039
608.9598999023438 0 1324.8086
635.297607421875 0 764.421
650.3268432617188 0 2671.737
651.326904296875 0 744.24725
652.3192749023438 0 2743.3137 b Water loss 10
655.8113403320312 0 696.031 y Ammonia loss 5
656.3594360351562 0 1571.9395
662.3126220703125 0 1815.5435
664.3211669921875 0 3043.978 y 5
664.82275390625 0 2206.6526
665.325439453125 0 975.1273
666.298828125 0 911.16205
680.3145751953125 0 7616.632
681.3171997070312 0 2547.303
682.322021484375 0 716.9208
688.331298828125 0 869.1949
708.314453125 0 1002.0582
710.3169555664062 0 4405.35 y Water loss 10
711.3204345703125 0 2166.7168 y Ammonia loss 10
721.3510131835938 0 2559.4922
721.8313598632812 0 1311.905 y 4
722.3475341796875 0 2006.9719
728.3276977539062 0 13172.152 y 10
729.330810546875 0 4405.6406
730.331787109375 0 1416.6882
733.3760375976562 0 683.641
734.3666381835938 0 879.6286
735.4059448242188 0 923.1603
751.388427734375 0 5340.975
752.3912963867188 0 2672.6533
761.376708984375 0 675.5855 b Water loss 5
762.3616943359375 0 1767.3822 b Ammonia loss 5
763.4085693359375 0 4109.3296
764.409423828125 0 1543.8368
765.4158325195312 0 681.47473
768.3424072265625 0 1108.5005
772.3606567382812 0 1872.1448
773.3590698242188 0 1080.655
776.8572387695312 0 1151.9526 y Water loss 3
777.351806640625 0 1955.6954 y Ammonia loss 3
777.4168701171875 0 1649.1351
777.8584594726562 0 1694.9438
779.3829345703125 0 27204.873 b 5
780.31103515625 0 949.0946
780.3858032226562 0 11537.361
781.3886108398438 0 2530.2761
785.3527221679688 0 908.12103
785.8619384765625 0 1546.553 y 3
786.3619995117188 0 1416.3148
786.8663940429688 0 1331.9729
793.3970336914062 0 1672.4031
795.3411254882812 0 1143.8883
856.4218139648438 0 2054.3174 y 9
882.3659057617188 0 2087.1372
883.3619995117188 0 962.9371
892.4732666015625 0 1019.1921 b 6
900.3753662109375 0 2408.2075
901.3838500976562 0 902.16833
908.4254150390625 0 971.82764
913.4434814453125 0 4093.8623 y 8
914.44482421875 0 2400.8833
918.3285522460938 0 685.0513
1010.4625244140625 0 1571.0397 y Water loss 7
1011.4501953125 0 1391.8501 y Ammonia loss 7
1028.4697265625 0 7735.4536 y 7
1029.4747314453125 0 4359.5503
1030.4732666015625 0 1803.4409
1141.55322265625 0 712.7529 y 6
1142.5562744140625 0 1135.2753
1259.26904296875 0 587.44135
1442.6544189453125 0 651.50305 y 4
2664.394775390625 0 678.84357
3074.638427734375 0 1137.3306
3075.224609375 0 900.53973
3156.4013671875 0 741.3296

Spectrum Details

|  |  |
| --- | --- |
| Matched peaks? Matched peaks The total absolute number of peaks matched. Additionally in brackets the total fraction of peaks matched and the total number of peaks is shown. | 75 (16.59% of 452) |
| FDR? FDR The false discovery rate estimated for this peptide. It is calculated by matching all theoretical fragments with a non-integer shift with the raw peaks for this spectrum. This is done with 40 different shifts. The resulting percentage is the average number of annotated peaks over the number of annotated peaks with the correct spectrum. | 0.44% |
| Satellite FDR? Satellite FDR See the FDR for details on its calculation. This satellite ion specific FDR only contains the satellite ions (d/w) for I/L/J positions. | 4.76% |
| PSM Score? PSM Score The PSM Score as given by Hecklib to this annotated spectrum. It is shown with three significant figures. | 353 |

#### Spectrum 4845? Spectrum 4845 The raw spectrum of this peptide as annotated by Hecklib. The fragments are coloured according to ion type (see legend). Any peaks with a star '\*' as text can be hovered over to see the full details, first the ion type second the mass shift type. By hovering over the amino acids in the peptide or ions in the legend the corresponding peaks are highlighted. By toggling the 'Unassigned' label you can turn the background (unassigned) peaks on or off in the plot. By updating the slider in the Ion legend you can update the spectrum to only show the top X% of the peaks with labels. The top X% means any peak that is within X% of the highest intensity. By dragging in the spectrum you can zoom in to a specific part of the spectrum and use 'Zoom Out' to get back to the original zoom level. The annotation of the spectrum is based on the given sequence in the peptides file and is done with different software so inconsistencies are likely. The peaks are annotated based on the given sequence, with 20 ppm tolerance.

Copy Data

##### Spectrum 4845 (TSV)

###### Preview

```
Loading example...
```

*Click on the button to copy the data to your clipboard.*

Mz MinMz MaxIntensity Max

WidthHeightPeptide font sizePeptide stroke widthSpectrum font sizeSpectrum stroke widthCompact peptide

Ion legend

wxyz

abcd

OtherUnassignedIon^Charge^_Position_Show for top:%

VJHQDWLDGKEYKCK

04.06e+38.12e+31.22e+41.62e+4

Zoom Out

y^+1^_1_c^+2^_5_y^+2^_4_y^+3^_7_y^+1^_2_w^+2^_5_y^+2^_5_c^+3^_9_w^+1^_3_c^+1^_3_c^+4^_12_c^+2^_6_w^+2^_6_y^+2^_6_y^+2^_6_y^+1^_3_c^+2^_7_z^+2^_7_y^+2^_7_c^+1^_4_w^+3^_12_c^+3^_12_y^+2^_8_y^+2^_8_z^+2^_8_y^+2^_8_y^+3^_12_c^+2^_9_c^+2^_9_w^+2^_9_c^+3^_13_z^+2^_9_z^+3^_13_y^+3^_13_y^+2^_9_z^+1^_4_w^+3^_14_c^+1^_5_c^+3^_14_y^+1^_4_c^+2^_10_c^+1^_5_w^+1^_5_z^+2^_10_y^+2^_10_c^+2^_11_z^+1^_5_y^+2^_11_z^+2^_11_y^+1^_5_c^+2^_12_z^+2^_12_c^+1^_6_c^+1^_6_c^+2^_13_z^+2^_13_c^+1^_7_c^+2^_14_y^+1^_7_z^+1^_7_c^+1^_7_c^+1^_8_c^+1^_8_c^+1^_9_

0777155323303107

Fragment Matches Table

Show background peaks

| Position | Ion type | Intensity | mz Theoretical | mz Error (Th) | mz Error (ppm) | Charge | Series Number |
| --- | --- | --- | --- | --- | --- | --- | --- |
| - | - | 818.7 | 120.1 | - | - | 0 | - |
| - | - | 467.3 | 123.1 | - | - | 0 | - |
| - | - | 576.7 | 125.1 | - | - | 0 | - |
| - | - | 612.9 | 126.1 | - | - | 0 | - |
| - | - | 757.7 | 129.1 | - | - | 0 | - |
| - | - | 3028 | 129.1 | - | - | 0 | - |
| - | - | 6357 | 130.1 | - | - | 0 | - |
| - | - | 723.8 | 131.1 | - | - | 0 | - |
| - | - | 831 | 133.1 | - | - | 0 | - |
| - | - | 813 | 133.1 | - | - | 0 | - |
| - | - | 497.5 | 136.1 | - | - | 0 | - |
| - | - | 707.8 | 140.1 | - | - | 0 | - |
| - | - | 457.7 | 142.1 | - | - | 0 | - |
| - | - | 397.5 | 142.8 | - | - | 0 | - |
| - | - | 571.4 | 147.1 | - | - | 0 | - |
| 15 | y | 1121 | 147.1 | 0.0002024 | 1.376 | +1 | 1 |
| - | - | 1325 | 149 | - | - | 0 | - |
| - | - | 2288 | 155.1 | - | - | 0 | - |
| - | - | 591.5 | 156.1 | - | - | 0 | - |
| - | - | 1797 | 159.1 | - | - | 0 | - |
| - | - | 924.6 | 165.1 | - | - | 0 | - |
| - | - | 691 | 167.1 | - | - | 0 | - |
| - | - | 762.9 | 170.1 | - | - | 0 | - |
| - | - | 436.7 | 173 | - | - | 0 | - |
| - | - | 652.6 | 173.4 | - | - | 0 | - |
| - | - | 2484 | 173.5 | - | - | 0 | - |
| - | - | 735.9 | 173.5 | - | - | 0 | - |
| - | - | 500.9 | 175.1 | - | - | 0 | - |
| - | - | 524.3 | 176.4 | - | - | 0 | - |
| - | - | 427.1 | 179.4 | - | - | 0 | - |
| - | - | 2119 | 185.2 | - | - | 0 | - |
| - | - | 1201 | 190.1 | - | - | 0 | - |
| - | - | 2940 | 199.2 | - | - | 0 | - |
| - | - | 1766 | 201.1 | - | - | 0 | - |
| - | - | 948.9 | 213.2 | - | - | 0 | - |
| - | - | 882.7 | 214 | - | - | 0 | - |
| - | - | 909.5 | 215.1 | - | - | 0 | - |
| - | - | 656.3 | 218 | - | - | 0 | - |
| - | - | 1718 | 221.1 | - | - | 0 | - |
| - | - | 1415 | 225 | - | - | 0 | - |
| - | - | 1068 | 233.2 | - | - | 0 | - |
| - | - | 854.1 | 235.1 | - | - | 0 | - |
| - | - | 2273 | 239.1 | - | - | 0 | - |
| - | - | 613.4 | 239.2 | - | - | 0 | - |
| - | - | 1263 | 251.2 | - | - | 0 | - |
| - | - | 907.3 | 255 | - | - | 0 | - |
| - | - | 608.9 | 261.2 | - | - | 0 | - |
| - | - | 698.1 | 263.1 | - | - | 0 | - |
| - | - | 758.5 | 269 | - | - | 0 | - |
| - | - | 947 | 275 | - | - | 0 | - |
| - | - | 515.4 | 280.8 | - | - | 0 | - |
| - | - | 794.6 | 284.2 | - | - | 0 | - |
| - | - | 4795 | 295.1 | - | - | 0 | - |
| - | - | 1360 | 296.1 | - | - | 0 | - |
| 5 | c | 678.7 | 297.2 | 0.0007328 | 2.466 | +2 | 5 |
| - | - | 1667 | 299.1 | - | - | 0 | - |
| 12 | y | 826.8 | 300.1 | 0.002019 | 6.728 | +2 | 4 |
| - | - | 641.5 | 302.2 | - | - | 0 | - |
| - | - | 822.7 | 303.2 | - | - | 0 | - |
| 9 | y | 598.6 | 305.2 | 0.001511 | 4.951 | +3 | 7 |
| 14 | y | 2013 | 308.1 | 0.004303 | 13.97 | +1 | 2 |
| - | - | 983.3 | 313.1 | - | - | 0 | - |
| - | - | 810.6 | 318.2 | - | - | 0 | - |
| - | - | 979.3 | 318.2 | - | - | 0 | - |
| - | - | 1235 | 326.7 | - | - | 0 | - |
| 11 | w | 8421 | 327.2 | 0.001961 | 5.994 | +2 | 5 |
| - | - | 3495 | 327.7 | - | - | 0 | - |
| - | - | 747 | 328.2 | - | - | 0 | - |
| - | - | 1094 | 329.2 | - | - | 0 | - |
| - | - | 1776 | 340.3 | - | - | 0 | - |
| - | - | 1238 | 341.2 | - | - | 0 | - |
| - | - | 2150 | 350.2 | - | - | 0 | - |
| - | - | 2865 | 351.2 | - | - | 0 | - |
| 11 | y | 595.4 | 356.2 | 0.003783 | 10.62 | +2 | 5 |
| - | - | 2168 | 359 | - | - | 0 | - |
| - | - | 703.5 | 360 | - | - | 0 | - |
| 9 | c | 742.6 | 361.2 | 0.006906 | 19.12 | +3 | 9 |
| 13 | w | 1218 | 362.1 | 0.003546 | 9.791 | +1 | 3 |
| - | - | 932.3 | 367.1 | - | - | 0 | - |
| 3 | c | 4202 | 367.2 | 9.813E-05 | 0.2672 | +1 | 3 |
| - | - | 2828 | 367.3 | - | - | 0 | - |
| - | - | 533.6 | 367.3 | - | - | 0 | - |
| - | - | 2478 | 367.7 | - | - | 0 | - |
| - | - | 1167 | 368.2 | - | - | 0 | - |
| - | - | 781.1 | 368.3 | - | - | 0 | - |
| - | - | 562.3 | 369.2 | - | - | 0 | - |
| - | - | 774.4 | 375.2 | - | - | 0 | - |
| 12 | c | 3117 | 376.2 | 0.007086 | 18.84 | +4 | 12 |
| - | - | 1687 | 376.7 | - | - | 0 | - |
| - | - | 689 | 383.3 | - | - | 0 | - |
| - | - | 779.9 | 384.1 | - | - | 0 | - |
| - | - | 878.7 | 384.2 | - | - | 0 | - |
| - | - | 1994 | 384.2 | - | - | 0 | - |
| - | - | 1146 | 384.2 | - | - | 0 | - |
| - | - | 1161 | 384.6 | - | - | 0 | - |
| - | - | 4440 | 384.8 | - | - | 0 | - |
| - | - | 1205 | 384.9 | - | - | 0 | - |
| - | - | 656.8 | 385.2 | - | - | 0 | - |
| - | - | 1485 | 385.2 | - | - | 0 | - |
| - | - | 926.2 | 385.2 | - | - | 0 | - |
| - | - | 2131 | 385.3 | - | - | 0 | - |
| - | - | 717.5 | 385.3 | - | - | 0 | - |
| - | - | 2073 | 386.2 | - | - | 0 | - |
| 6 | c | 3746 | 390.2 | 0.0002906 | 0.7448 | +2 | 6 |
| - | - | 1547 | 390.7 | - | - | 0 | - |
| - | - | 1525 | 391.2 | - | - | 0 | - |
| 10 | w | 1405 | 391.7 | 0.002109 | 5.384 | +2 | 6 |
| - | - | 613.6 | 399.7 | - | - | 0 | - |
| - | - | 670.6 | 402.2 | - | - | 0 | - |
| 10 | y | 4152 | 420.2 | 0.000516 | 1.228 | +2 | 6 |
| - | - | 871.6 | 421.2 | - | - | 0 | - |
| - | - | 642.8 | 424.2 | - | - | 0 | - |
| 10 | y | 1440 | 428.7 | 0.001522 | 3.551 | +2 | 6 |
| - | - | 848.2 | 429.2 | - | - | 0 | - |
| - | - | 3517 | 432.7 | - | - | 0 | - |
| - | - | 2338 | 433.2 | - | - | 0 | - |
| - | - | 622.2 | 433.7 | - | - | 0 | - |
| 13 | y | 3536 | 436.2 | 0.003746 | 8.587 | +1 | 3 |
| - | - | 676.7 | 439.3 | - | - | 0 | - |
| - | - | 887.3 | 439.9 | - | - | 0 | - |
| - | - | 587.9 | 441.7 | - | - | 0 | - |
| 7 | c | 3807 | 446.7 | 0.0008917 | 1.996 | +2 | 7 |
| - | - | 974 | 447.2 | - | - | 0 | - |
| - | - | 2010 | 448.3 | - | - | 0 | - |
| - | - | 733.4 | 448.8 | - | - | 0 | - |
| 9 | z | 1935 | 449.2 | 0.0003731 | 0.8306 | +2 | 7 |
| - | - | 579.7 | 456.1 | - | - | 0 | - |
| 9 | y | 3766 | 457.2 | 0.001939 | 4.242 | +2 | 7 |
| - | - | 2027 | 457.7 | - | - | 0 | - |
| - | - | 875.6 | 458.2 | - | - | 0 | - |
| - | - | 1366 | 461.2 | - | - | 0 | - |
| - | - | 899.7 | 466.2 | - | - | 0 | - |
| - | - | 994.9 | 466.5 | - | - | 0 | - |
| - | - | 1136 | 473 | - | - | 0 | - |
| - | - | 1482 | 473.2 | - | - | 0 | - |
| - | - | 684.3 | 473.5 | - | - | 0 | - |
| - | - | 1036 | 476.5 | - | - | 0 | - |
| - | - | 713.3 | 476.7 | - | - | 0 | - |
| - | - | 611.4 | 477 | - | - | 0 | - |
| - | - | 751.5 | 477.2 | - | - | 0 | - |
| - | - | 717.1 | 477.7 | - | - | 0 | - |
| - | - | 5986 | 478.2 | - | - | 0 | - |
| - | - | 3536 | 478.7 | - | - | 0 | - |
| - | - | 624.5 | 479.2 | - | - | 0 | - |
| - | - | 7023 | 480.7 | - | - | 0 | - |
| - | - | 1.095E+04 | 481 | - | - | 0 | - |
| - | - | 6448 | 481.2 | - | - | 0 | - |
| - | - | 3441 | 481.5 | - | - | 0 | - |
| - | - | 1187 | 481.7 | - | - | 0 | - |
| - | - | 830.2 | 482 | - | - | 0 | - |
| - | - | 1042 | 484.8 | - | - | 0 | - |
| - | - | 609.6 | 485.3 | - | - | 0 | - |
| - | - | 754.7 | 486.2 | - | - | 0 | - |
| - | - | 765.6 | 494.2 | - | - | 0 | - |
| 4 | c | 6168 | 495.3 | 0.000559 | 1.129 | +1 | 4 |
| - | - | 1941 | 496.3 | - | - | 0 | - |
| 4 | w | 3584 | 499.6 | 0.000539 | 1.079 | +3 | 12 |
| - | - | 1407 | 499.9 | - | - | 0 | - |
| 12 | c | 1534 | 501.3 | 0.002444 | 4.876 | +3 | 12 |
| - | - | 934 | 501.6 | - | - | 0 | - |
| 8 | y | 1174 | 505.7 | 0.001217 | 2.407 | +2 | 8 |
| 8 | y | 899 | 506.2 | 0.003357 | 6.631 | +2 | 8 |
| 8 | z | 4119 | 506.7 | 0.002051 | 4.048 | +2 | 8 |
| - | - | 2706 | 507.2 | - | - | 0 | - |
| - | - | 1132 | 507.7 | - | - | 0 | - |
| - | - | 808.6 | 511.6 | - | - | 0 | - |
| - | - | 678.8 | 512.3 | - | - | 0 | - |
| - | - | 1345 | 512.6 | - | - | 0 | - |
| - | - | 1496 | 512.9 | - | - | 0 | - |
| - | - | 1179 | 513.3 | - | - | 0 | - |
| 8 | y | 1.07E+04 | 514.7 | 0.001831 | 3.556 | +2 | 8 |
| - | - | 6991 | 515.2 | - | - | 0 | - |
| - | - | 1966 | 515.7 | - | - | 0 | - |
| - | - | 589.9 | 519.7 | - | - | 0 | - |
| 4 | y | 930.1 | 524.2 | 0.0009331 | 1.78 | +3 | 12 |
| 9 | c | 728 | 532.3 | 0.002465 | 4.631 | +2 | 9 |
| - | - | 960 | 540.9 | - | - | 0 | - |
| 9 | c | 6489 | 541.3 | 0.001103 | 2.038 | +2 | 9 |
| 7 | w | 1099 | 541.7 | 0.007235 | 13.35 | +2 | 9 |
| - | - | 2510 | 541.8 | - | - | 0 | - |
| - | - | 1549 | 542.2 | - | - | 0 | - |
| - | - | 1169 | 542.3 | - | - | 0 | - |
| 13 | c | 4960 | 544 | 0.0008089 | 1.487 | +3 | 13 |
| - | - | 5448 | 544.3 | - | - | 0 | - |
| - | - | 1878 | 544.6 | - | - | 0 | - |
| - | - | 1311 | 545 | - | - | 0 | - |
| - | - | 615.1 | 545.6 | - | - | 0 | - |
| - | - | 1135 | 550.3 | - | - | 0 | - |
| - | - | 1116 | 552.6 | - | - | 0 | - |
| - | - | 1003 | 553 | - | - | 0 | - |
| 7 | z | 1225 | 563.3 | 0.001144 | 2.031 | +2 | 9 |
| 3 | z | 821.6 | 564.6 | 0.001921 | 3.403 | +3 | 13 |
| - | - | 973.7 | 564.9 | - | - | 0 | - |
| - | - | 749.8 | 567.3 | - | - | 0 | - |
| 3 | y | 1101 | 569.9 | 0.002265 | 3.975 | +3 | 13 |
| - | - | 1668 | 570.3 | - | - | 0 | - |
| 7 | y | 4194 | 571.3 | 0.002725 | 4.77 | +2 | 9 |
| - | - | 2547 | 571.8 | - | - | 0 | - |
| - | - | 875.7 | 572.3 | - | - | 0 | - |
| - | - | 2052 | 576.4 | - | - | 0 | - |
| - | - | 1177 | 576.9 | - | - | 0 | - |
| - | - | 1551 | 578.3 | - | - | 0 | - |
| 12 | z | 5806 | 583.3 | 0.004222 | 7.239 | +1 | 4 |
| - | - | 2184 | 584.3 | - | - | 0 | - |
| - | - | 691.6 | 585.3 | - | - | 0 | - |
| - | - | 978 | 585.4 | - | - | 0 | - |
| - | - | 685.9 | 586.4 | - | - | 0 | - |
| 2 | w | 1778 | 587.9 | 0.0001107 | 0.1883 | +3 | 14 |
| - | - | 1005 | 588.3 | - | - | 0 | - |
| - | - | 1628 | 588.6 | - | - | 0 | - |
| 5 | c | 2663 | 593.3 | 0.0005618 | 0.9469 | +1 | 5 |
| - | - | 879.7 | 597.3 | - | - | 0 | - |
| 14 | c | 7935 | 597.6 | 0.0006828 | 1.143 | +3 | 14 |
| - | - | 4306 | 598 | - | - | 0 | - |
| - | - | 3336 | 598.3 | - | - | 0 | - |
| - | - | 1571 | 598.6 | - | - | 0 | - |
| 12 | y | 2140 | 599.3 | 0.002499 | 4.171 | +1 | 4 |
| - | - | 596.8 | 601.5 | - | - | 0 | - |
| 10 | c | 1.608E+04 | 605.3 | 0.0005575 | 0.921 | +2 | 10 |
| - | - | 9784 | 605.8 | - | - | 0 | - |
| - | - | 3858 | 606.3 | - | - | 0 | - |
| - | - | 1429 | 606.8 | - | - | 0 | - |
| - | - | 1440 | 610.3 | - | - | 0 | - |
| 5 | c | 4669 | 610.3 | 0.0001357 | 0.2224 | +1 | 5 |
| - | - | 1400 | 611.3 | - | - | 0 | - |
| - | - | 1025 | 615.6 | - | - | 0 | - |
| - | - | 1006 | 616 | - | - | 0 | - |
| - | - | 879.2 | 616.3 | - | - | 0 | - |
| - | - | 991.5 | 616.6 | - | - | 0 | - |
| - | - | 646.4 | 620.7 | - | - | 0 | - |
| - | - | 6292 | 621.3 | - | - | 0 | - |
| - | - | 5685 | 621.6 | - | - | 0 | - |
| - | - | 4088 | 622 | - | - | 0 | - |
| - | - | 2567 | 622.3 | - | - | 0 | - |
| - | - | 1152 | 622.7 | - | - | 0 | - |
| - | - | 953.5 | 626.3 | - | - | 0 | - |
| - | - | 960.6 | 630.6 | - | - | 0 | - |
| - | - | 799.5 | 635.3 | - | - | 0 | - |
| - | - | 9396 | 635.6 | - | - | 0 | - |
| - | - | 7816 | 636 | - | - | 0 | - |
| - | - | 4850 | 636.3 | - | - | 0 | - |
| - | - | 1002 | 637 | - | - | 0 | - |
| - | - | 665.4 | 637.3 | - | - | 0 | - |
| - | - | 2712 | 640.6 | - | - | 0 | - |
| - | - | 5378 | 641 | - | - | 0 | - |
| - | - | 9433 | 641.3 | - | - | 0 | - |
| - | - | 6316 | 641.7 | - | - | 0 | - |
| - | - | 3589 | 642 | - | - | 0 | - |
| - | - | 1238 | 642.3 | - | - | 0 | - |
| - | - | 848.3 | 642.7 | - | - | 0 | - |
| - | - | 983.2 | 652.3 | - | - | 0 | - |
| 11 | w | 1060 | 653.3 | 0.003665 | 5.61 | +1 | 5 |
| 6 | z | 1013 | 656.3 | 0.004638 | 7.067 | +2 | 10 |
| - | - | 658.9 | 657.3 | - | - | 0 | - |
| 6 | y | 963 | 664.3 | 0.00547 | 8.234 | +2 | 10 |
| 11 | c | 5035 | 669.8 | 0.0016 | 2.388 | +2 | 11 |
| - | - | 2952 | 670.3 | - | - | 0 | - |
| - | - | 2699 | 670.8 | - | - | 0 | - |
| - | - | 1550 | 680.3 | - | - | 0 | - |
| - | - | 1096 | 681.3 | - | - | 0 | - |
| - | - | 1265 | 685.4 | - | - | 0 | - |
| 11 | z | 6091 | 712.3 | 0.003541 | 4.972 | +1 | 5 |
| 5 | y | 2928 | 713.3 | 0.00614 | 8.608 | +2 | 11 |
| 5 | z | 1673 | 713.8 | 0.001264 | 1.77 | +2 | 11 |
| - | - | 4604 | 714.3 | - | - | 0 | - |
| - | - | 1472 | 714.8 | - | - | 0 | - |
| - | - | 705 | 715.3 | - | - | 0 | - |
| 11 | y | 691.3 | 728.3 | 0.002551 | 3.503 | +1 | 5 |
| - | - | 794.1 | 729.4 | - | - | 0 | - |
| - | - | 827.6 | 729.9 | - | - | 0 | - |
| - | - | 826.3 | 748.4 | - | - | 0 | - |
| - | - | 1021 | 749.3 | - | - | 0 | - |
| 12 | c | 3105 | 751.4 | 0.003627 | 4.827 | +2 | 12 |
| - | - | 3159 | 751.9 | - | - | 0 | - |
| - | - | 1361 | 752.4 | - | - | 0 | - |
| - | - | 868.7 | 752.9 | - | - | 0 | - |
| - | - | 970.6 | 763.4 | - | - | 0 | - |
| - | - | 2154 | 768.5 | - | - | 0 | - |
| - | - | 723.1 | 768.9 | - | - | 0 | - |
| - | - | 1242 | 769.4 | - | - | 0 | - |
| - | - | 821.5 | 769.5 | - | - | 0 | - |
| 4 | z | 1139 | 777.9 | 0.0006364 | 0.8182 | +2 | 12 |
| - | - | 3657 | 778.4 | - | - | 0 | - |
| - | - | 3181 | 778.9 | - | - | 0 | - |
| 6 | c | 5977 | 779.4 | 0.004044 | 5.189 | +1 | 6 |
| - | - | 3132 | 780.4 | - | - | 0 | - |
| - | - | 1244 | 781.4 | - | - | 0 | - |
| - | - | 872.3 | 781.8 | - | - | 0 | - |
| - | - | 821.2 | 793.4 | - | - | 0 | - |
| 6 | c | 1424 | 796.4 | 0.0008081 | 1.015 | +1 | 6 |
| - | - | 1716 | 797.4 | - | - | 0 | - |
| 13 | c | 1306 | 815.4 | 0.002227 | 2.731 | +2 | 13 |
| - | - | 1399 | 815.9 | - | - | 0 | - |
| - | - | 1529 | 816.4 | - | - | 0 | - |
| - | - | 1042 | 828.4 | - | - | 0 | - |
| - | - | 871 | 828.9 | - | - | 0 | - |
| - | - | 2230 | 841.4 | - | - | 0 | - |
| - | - | 827.4 | 842.4 | - | - | 0 | - |
| 3 | z | 940.3 | 846.4 | 0.004628 | 5.468 | +2 | 13 |
| - | - | 3398 | 846.9 | - | - | 0 | - |
| - | - | 2758 | 847.4 | - | - | 0 | - |
| - | - | 956 | 866.9 | - | - | 0 | - |
| - | - | 753.7 | 867.4 | - | - | 0 | - |
| - | - | 1291 | 888.4 | - | - | 0 | - |
| 7 | c | 2616 | 892.5 | 0.0008889 | 0.996 | +1 | 7 |
| - | - | 1539 | 893.5 | - | - | 0 | - |
| - | - | 946.7 | 894.5 | - | - | 0 | - |
| 14 | c | 1789 | 895.9 | 0.001323 | 1.476 | +2 | 14 |
| 9 | y | 3984 | 896.4 | 0.009317 | 10.39 | +1 | 7 |
| - | - | 1415 | 896.9 | - | - | 0 | - |
| 9 | z | 1366 | 897.4 | 0.006436 | 7.171 | +1 | 7 |
| - | - | 1688 | 903.4 | - | - | 0 | - |
| - | - | 1554 | 903.9 | - | - | 0 | - |
| 7 | c | 4160 | 909.5 | 3.32E-05 | 0.03651 | +1 | 7 |
| - | - | 3076 | 910.5 | - | - | 0 | - |
| - | - | 766.4 | 911.5 | - | - | 0 | - |
| - | - | 914.7 | 915 | - | - | 0 | - |
| - | - | 803.2 | 924 | - | - | 0 | - |
| - | - | 820.7 | 931.5 | - | - | 0 | - |
| - | - | 1014 | 932 | - | - | 0 | - |
| - | - | 1494 | 932.5 | - | - | 0 | - |
| - | - | 754 | 953 | - | - | 0 | - |
| - | - | 1058 | 953.5 | - | - | 0 | - |
| - | - | 884.4 | 954 | - | - | 0 | - |
| - | - | 1481 | 961.5 | - | - | 0 | - |
| - | - | 1996 | 966.5 | - | - | 0 | - |
| - | - | 1468 | 967.5 | - | - | 0 | - |
| 8 | c | 685.5 | 1007 | 0.00177 | 1.757 | +1 | 8 |
| - | - | 1233 | 1013 | - | - | 0 | - |
| - | - | 718.3 | 1014 | - | - | 0 | - |
| 8 | c | 759.3 | 1025 | 0.004457 | 4.35 | +1 | 8 |
| - | - | 844.1 | 1026 | - | - | 0 | - |
| 9 | c | 1301 | 1082 | 0.0005274 | 0.4876 | +1 | 9 |
| - | - | 2230 | 1083 | - | - | 0 | - |
| - | - | 1061 | 1127 | - | - | 0 | - |
| - | - | 1120 | 1211 | - | - | 0 | - |
| - | - | 1139 | 1212 | - | - | 0 | - |
| - | - | 684 | 1324 | - | - | 0 | - |
| - | - | 768.4 | 1340 | - | - | 0 | - |
| - | - | 681.7 | 2895 | - | - | 0 | - |
| - | - | 751.5 | 3076 | - | - | 0 | - |

m/z Charge Intensity FragmentType MassShift Position
120.08074951171875 0 818.7028
123.11710357666016 0 467.3193
125.09623718261719 0 576.6883
126.10317993164062 0 612.895
129.05494689941406 0 757.7054
129.1024169921875 0 3027.941
130.0653839111328 0 6356.527
131.0690155029297 0 723.8479
133.0607452392578 0 830.99347
133.08619689941406 0 812.95215
136.07586669921875 0 497.53577
140.0819549560547 0 707.7791
142.09751892089844 0 457.738
142.8414764404297 0 397.533
147.07687377929688 0 571.4017
147.11300659179688 0 1121.2848 y 14
149.04531860351562 0 1325.2103
155.0929412841797 0 2287.9421
156.07701110839844 0 591.46027
159.09193420410156 0 1797.1953
165.1023406982422 0 924.59576
167.05532836914062 0 691.00446
170.0604705810547 0 762.94714
173.0135498046875 0 436.67358
173.43524169921875 0 652.61176
173.45037841796875 0 2483.733
173.45680236816406 0 735.9155
175.13287353515625 0 500.89194
176.38796997070312 0 524.2704
179.42469787597656 0 427.09998
185.1650848388672 0 2119.3618
190.0825653076172 0 1200.6759
199.1696319580078 0 2940.2568
201.12342834472656 0 1765.8469
213.15953063964844 0 948.93774
214.040283203125 0 882.6964
215.13922119140625 0 909.4873
218.0354461669922 0 656.288
221.08448791503906 0 1717.9929
225.0435333251953 0 1414.6942
233.16502380371094 0 1067.7207
235.11769104003906 0 854.0853
239.0951385498047 0 2272.7976
239.2359161376953 0 613.42706
251.1509246826172 0 1262.8478
255.0303955078125 0 907.31146
261.1593322753906 0 608.8652
263.1484680175781 0 698.08435
269.044921875 0 758.54297
275.03448486328125 0 946.9648
280.83184814453125 0 515.42944
284.18524169921875 0 794.6396
295.1034851074219 0 4795.4946
296.103515625 0 1360.0114
297.1549987792969 0 678.711 c Ammonia loss 4
299.0611572265625 0 1666.6069
300.1467590332031 0 826.83124 y 11
302.19720458984375 0 641.50867
303.1922607421875 0 822.65125
305.1531066894531 0 598.5649 y 8
308.127685546875 0 2012.9117 y 13
313.1142272949219 0 983.3045
318.1779479980469 0 810.60895
318.21527099609375 0 979.33704
326.66229248046875 0 1235.4907
327.152099609375 0 8420.805 w 10
327.6537780761719 0 3495.1326
328.1521301269531 0 746.9523
329.2186584472656 0 1094.1428
340.2601623535156 0 1776.0117
341.181640625 0 1237.7341
350.2189025878906 0 2150.4944
351.16510009765625 0 2864.7314
356.16058349609375 0 595.38635 y Ammonia loss 10
359.0286865234375 0 2168.024
360.03155517578125 0 703.54767
361.1925964355469 0 742.617 c 8
362.1390075683594 0 1217.6501 w 12
367.06793212890625 0 932.28955
367.2451171875 0 4201.558 c 2
367.26934814453125 0 2827.6829
367.2926330566406 0 533.5571
367.6850891113281 0 2477.7078
368.18780517578125 0 1167.0581
368.2734069824219 0 781.0627
369.17633056640625 0 562.2678
375.2137756347656 0 774.4238
376.1983947753906 0 3116.8628 c 11
376.6994323730469 0 1686.7135
383.2948303222656 0 689.03296
384.1051025390625 0 779.9022
384.1628112792969 0 878.6982
384.19110107421875 0 1994.3384
384.2492980957031 0 1145.6862
384.5820617675781 0 1160.9691
384.77239990234375 0 4439.8213
384.91693115234375 0 1204.6906
385.1512145996094 0 656.8307
385.20086669921875 0 1485.2839
385.24957275390625 0 926.1778
385.276123046875 0 2130.547
385.3097839355469 0 717.49664
386.201416015625 0 2073.166
390.1956787109375 0 3745.526 c Ammonia loss 5
390.6981201171875 0 1546.7854
391.19940185546875 0 1524.8364
391.6732482910156 0 1405.2815 w 9
399.7428894042969 0 613.6457
402.2125244140625 0 670.61975
420.2037658691406 0 4151.5044 y Ammonia loss 9
421.2088623046875 0 871.5795
424.2294921875 0 642.8196
428.7160339355469 0 1440.2666 y 9
429.2152404785156 0 848.23615
432.740478515625 0 3516.5278
433.2414245605469 0 2338.3296
433.743408203125 0 622.2424
436.22320556640625 0 3535.924 y 12
439.2623596191406 0 676.6805
439.8818359375 0 887.34106
441.70611572265625 0 587.9237
446.7383117675781 0 3806.5632 c Ammonia loss 6
447.2397155761719 0 973.9846
448.2663879394531 0 2010.0624
448.7657165527344 0 733.43036
449.21929931640625 0 1935.2186 z 8
456.1108093261719 0 579.6647
457.2263488769531 0 3765.943 y 8
457.7272033691406 0 2026.5411
458.2305603027344 0 875.5838
461.2386169433594 0 1365.9417
466.2384338378906 0 899.7163
466.4858093261719 0 994.8775
472.9801330566406 0 1135.7593
473.2296447753906 0 1481.7544
473.4785461425781 0 684.3019
476.4857482910156 0 1036.3916
476.732421875 0 713.3335
476.98468017578125 0 611.41174
477.228515625 0 751.49817
477.6959533691406 0 717.07874
478.2315979003906 0 5985.995
478.7330322265625 0 3536.0286
479.234375 0 624.53534
480.7396545410156 0 7023.0483
480.99041748046875 0 10952.449
481.2414245605469 0 6448.3657
481.49176025390625 0 3441.3306
481.7424621582031 0 1186.6069
481.9931335449219 0 830.16406
484.78765869140625 0 1042.143
485.29046630859375 0 609.6307
486.24713134765625 0 754.7143
494.2371520996094 0 765.58405
495.3043518066406 0 6168.4253 c 3
496.30706787109375 0 1941.2463
499.5633544921875 0 3584.0562 w 3
499.8971862792969 0 1406.9294
501.2550964355469 0 1534.3265 c 11
501.590576171875 0 933.96063
505.7352600097656 0 1174.0256 y Water loss 7
506.2318420410156 0 898.9619 y Ammonia loss 7
506.7303466796875 0 4119.17 z 7
507.2331848144531 0 2706.0413
507.7322998046875 0 1131.5037
511.5981750488281 0 808.64166
512.2633056640625 0 678.7924
512.6038208007812 0 1345.2954
512.9383544921875 0 1496.3866
513.2666015625 0 1178.9084
514.7399291992188 0 10703.329 y 7
515.2415161132812 0 6991.354
515.741943359375 0 1966.3438
519.7498168945312 0 589.88824
524.2474975585938 0 930.1498 y 3
532.2671508789062 0 727.96814 c Water loss 8
540.912109375 0 959.9872
541.2760009765625 0 6488.5806 c 8
541.7398071289062 0 1098.7563 w 6
541.7777099609375 0 2510.343
542.2452392578125 0 1548.9958
542.2825317382812 0 1168.6185
543.9517822265625 0 4959.782 c 12
544.2855224609375 0 5447.956
544.6179809570312 0 1878.0098
544.9528198242188 0 1311.0444
545.5794067382812 0 615.10126
550.25390625 0 1134.9532
552.6240844726562 0 1116.3519
552.9578857421875 0 1002.5623
563.2755737304688 0 1225.0315 z 6
564.59521484375 0 821.5973 z 2
564.9251098632812 0 973.70465
567.2969970703125 0 749.8016
569.9306030273438 0 1100.7742 y 2
570.2648315429688 0 1668.3881
571.2810668945312 0 4194.172 y 6
571.783203125 0 2547.3438
572.2816162109375 0 875.6893
576.3685913085938 0 2052.007
576.8695678710938 0 1176.6324
578.3431396484375 0 1550.6683
583.267333984375 0 5805.786 z 11
584.2709350585938 0 2183.9595
585.267578125 0 691.56555
585.3746948242188 0 977.95605
586.3841552734375 0 685.92084
587.936279296875 0 1778.4072 w 1
588.26953125 0 1004.92285
588.6069946289062 0 1627.6464
593.3047485351562 0 2662.606 c Ammonia loss 4
597.29833984375 0 879.6727
597.6233520507812 0 7935.1987 c 13
597.957275390625 0 4305.6006
598.2896118164062 0 3335.8599
598.62646484375 0 1570.8173
599.2877807617188 0 2140.15 y 11
601.4605102539062 0 596.7792
605.3229370117188 0 16080.099 c 9
605.8244018554688 0 9784.146
606.326171875 0 3857.8105
606.82763671875 0 1429.3966
610.281494140625 0 1439.6176
610.3308715820312 0 4668.875 c 4
611.3335571289062 0 1400.434
615.6396484375 0 1024.5211
615.9733276367188 0 1006.4936
616.3118286132812 0 879.23346
616.6471557617188 0 991.4868
620.6506958007812 0 646.43713
621.3134765625 0 6291.966
621.6491088867188 0 5684.7256
621.9817504882812 0 4087.879
622.3187866210938 0 2567.4902
622.6530151367188 0 1152.4385
626.3145751953125 0 953.5058
630.6438598632812 0 960.58234
635.3129272460938 0 799.48474
635.64599609375 0 9396.389
635.9810791015625 0 7815.654
636.314208984375 0 4850.2983
636.9805297851562 0 1002.44543
637.3052978515625 0 665.4129
640.649658203125 0 2711.9116
640.9841918945312 0 5378.057
641.3204956054688 0 9432.848
641.65478515625 0 6316.251
641.98974609375 0 3589.4644
642.3226928710938 0 1238.1356
642.65283203125 0 848.2765
652.319091796875 0 983.2188
653.2971801757812 0 1059.9614 w 10
656.3094482421875 0 1013.3725 z 5
657.3136596679688 0 658.93225
664.3289184570312 0 963.0368 y 5
669.8452758789062 0 5035.0244 c 10
670.3455810546875 0 2951.7134
670.8472900390625 0 2699.0947
680.3129272460938 0 1549.501
681.32275390625 0 1095.7297
685.3949584960938 0 1264.5477
712.3106079101562 0 6090.595 z 10
713.3175048828125 0 2927.9763 y Ammonia loss 4
713.8262939453125 0 1673.1425 z 4
714.3264770507812 0 4604.3467
714.8296508789062 0 1472.1481
715.3296508789062 0 705.02966
728.330322265625 0 691.3343 y 10
729.3812866210938 0 794.1445
729.8695068359375 0 827.55646
748.3508911132812 0 826.3319
749.3461303710938 0 1020.50256
751.3789672851562 0 3105.2334 c 11
751.8768920898438 0 3158.5725
752.378662109375 0 1361.2539
752.8814086914062 0 868.68616
763.4060668945312 0 970.62335
768.5377807617188 0 2153.7764
768.9041748046875 0 723.12317
769.4052124023438 0 1242.1466
769.53759765625 0 821.5499
777.8574829101562 0 1138.9169 z 3
778.358642578125 0 3657.29
778.86181640625 0 3180.6047
779.3794555664062 0 5976.838 c Ammonia loss 5
780.3862915039062 0 3132.1123
781.3864135742188 0 1243.611
781.8416748046875 0 872.33185
793.4136352539062 0 821.2248
796.4092407226562 0 1424.0155 c 5
797.41162109375 0 1716.4939
815.425048828125 0 1305.6542 c 12
815.9273071289062 0 1399.2374
816.4284057617188 0 1529.3663
828.42822265625 0 1041.8801
828.9331665039062 0 871.0098
841.4102172851562 0 2230.2886
842.4066772460938 0 827.4199
846.3909301757812 0 940.2971 z 2
846.8887939453125 0 3398.1938
847.3890991210938 0 2758.2454
866.92822265625 0 956.03973
867.4270629882812 0 753.715
888.4244995117188 0 1290.5951
892.4666748046875 0 2616.2869 c Ammonia loss 6
893.471435546875 0 1538.7993
894.4806518554688 0 946.7
895.9310913085938 0 1788.8352 c 13
896.4320678710938 0 3983.8188 y Ammonia loss 8
896.9360961914062 0 1414.5095
897.43701171875 0 1365.9497 z 8
903.4301147460938 0 1688.1456
903.9312744140625 0 1553.8456
909.4940795898438 0 4160.3003 c 6
910.4967651367188 0 3076.158
911.508056640625 0 766.435
914.9578247070312 0 914.7105
923.9647216796875 0 803.20447
931.4738159179688 0 820.6891
931.9647827148438 0 1013.60767
932.4712524414062 0 1493.9048
952.9664916992188 0 754.0091
953.4689331054688 0 1058.3346
953.9623413085938 0 884.3786
961.4798583984375 0 1480.8877
966.4921264648438 0 1995.7091
967.4965209960938 0 1468.3008
1007.4962768554688 0 685.4789 c Ammonia loss 7
1013.4608154296875 0 1232.7466
1014.4661254882812 0 718.25745
1024.5255126953125 0 759.2841 c 7
1025.5260009765625 0 844.12775
1081.5419921875 0 1301.4783 c 8
1082.550537109375 0 2229.8599
1126.5430908203125 0 1060.8356
1210.638671875 0 1120.344
1211.6448974609375 0 1138.8754
1323.970947265625 0 684.04517
1339.682373046875 0 768.44836
2894.969482421875 0 681.6612
3075.775634765625 0 751.51135

Spectrum Details

|  |  |
| --- | --- |
| Matched peaks? Matched peaks The total absolute number of peaks matched. Additionally in brackets the total fraction of peaks matched and the total number of peaks is shown. | 64 (18.77% of 341) |
| FDR? FDR The false discovery rate estimated for this peptide. It is calculated by matching all theoretical fragments with a non-integer shift with the raw peaks for this spectrum. This is done with 40 different shifts. The resulting percentage is the average number of annotated peaks over the number of annotated peaks with the correct spectrum. | 2.79% |
| Satellite FDR? Satellite FDR See the FDR for details on its calculation. This satellite ion specific FDR only contains the satellite ions (d/w) for I/L/J positions. | 1.19% |
| PSM Score? PSM Score The PSM Score as given by Hecklib to this annotated spectrum. It is shown with three significant figures. | 201 |

#### Reverse Lookup? Reverse Lookup All places where this read could be placed.

| Group | Segment | Template | Template Part | Read Part | Score | Unique |
| --- | --- | --- | --- | --- | --- | --- |
| Homo sapiens Heavy Chain | IGHC | IGHG1 | [190..205] | [0..15] | 115 | False |
| Homo sapiens Heavy Chain | IGHC | IGHG3 | [237..252] | [0..15] | 115 | False |
| Homo sapiens Heavy Chain | IGHC | IGHG2 | [186..201] | [0..15] | 106 | False |
| Homo sapiens Heavy Chain | IGHC | IGHG4 | [187..202] | [0..15] | 115 | False |

| Recombined | Template Part | Read Part | Score | Unique |
| --- | --- | --- | --- | --- |
| REC-0-1 | [315..330] | [0..15] | 120 | True |

#### Meta Information from Multiple reads

##### Number of combined reads

3

##### Intensity

0.3615

##### TotalArea

9.552E+05

##### Changes to the peptide sequence

VJHQDWLDGKEYKCK

J→LSupport for Leucine based on side chain ions (1 for L 0 for I) (Position: 7)

I→JEqual support for both Leucine and Isoleucine based on side chain ions (1 ions for both) (Position: 2)

L→ISupport for Isoleucine based on side chain ions (1 for I 0 for L) (Position: 2)

L→JNo support for either Leucine or Isoleucine based on side chain ions (Position: 7)

#### Positional Score

Copy Data

##### Positional Score (TSV)

###### Preview

```
Loading example...
```

*Click on the button to copy the data to your clipboard.*

1001234567891011121314

Label Value
"0" 0.283
"1" 0.31
"2" 0.317
"3" 0.317
"4" 0.323
"5" 0.303
"6" 0.327
"7" 0.323
"8" 0.32
"9" 0.323
"10" 0.327
"11" 0.33
"12" 0.327
"13" 0.323
"14" 0.317

#### Meta Information from PEAKS

##### Scan Identifier

F2:4849

##### Original sequence

V

L

H

Q

D

W

L

D

G

K

E

Y

K

C

+58.01

K

##### Posttranslational Modifications

Carboxymethyl

##### Source File

D:\separate\_stitch\_analyses\xle-disambiguation\raw\20210323\_F1\_UM1\_Peng0013\_SA\_F59\_ingel\_3ug\_TL.raw

##### Fraction

2

##### Scan Feature

F2:9992

##### De Novo Score

99

##### ConfidenceScore

99

### m/z

640.65

##### Mass

1918.9248

##### Charge

3

##### Retention Time

26.03

##### Predicted Retention Time

-

##### Area

2.632E+05

##### Parts Per Million

1.8

##### Fragmentation mode

ETHCD

##### Originating file

01 D:\separate\_stitch\_analyses\xle-disambiguation\20210325\_F59\_3ug\_DENOVO\_12.csv

#### Meta Information from PEAKS

##### Scan Identifier

F2:4847

##### Original sequence

V

L

H

Q

D

W

L

D

G

K

E

Y

K

C

+58.01

K

##### Posttranslational Modifications

Carboxymethyl

##### Source File

D:\separate\_stitch\_analyses\xle-disambiguation\raw\20210323\_F1\_UM1\_Peng0013\_SA\_F59\_ingel\_3ug\_TL.raw

##### Fraction

2

##### Scan Feature

F2:3312

##### De Novo Score

98

##### ConfidenceScore

98

### m/z

480.7397

##### Mass

1918.9248

##### Charge

4

##### Retention Time

26.03

##### Predicted Retention Time

-

##### Area

5.636E+05

##### Parts Per Million

2.5

##### Fragmentation mode

HCD

##### Originating file

01 D:\separate\_stitch\_analyses\xle-disambiguation\20210325\_F59\_3ug\_DENOVO\_12.csv

#### Meta Information from PEAKS

##### Scan Identifier

F2:4845

##### Original sequence

V

L

H

Q

D

W

L

D

G

K

E

Y

K

C

+58.01

K

##### Posttranslational Modifications

Carboxymethyl

##### Source File

D:\separate\_stitch\_analyses\xle-disambiguation\raw\20210323\_F1\_UM1\_Peng0013\_SA\_F59\_ingel\_3ug\_TL.raw

##### Fraction

2

##### Scan Feature

F2:564

##### De Novo Score

95

##### ConfidenceScore

95

### m/z

384.7927

##### Mass

1918.9248

##### Charge

5

##### Retention Time

26.03

##### Predicted Retention Time

-

##### Area

1.284E+05

##### Parts Per Million

1.3

##### Fragmentation mode

ETHCD

##### Originating file

01 D:\separate\_stitch\_analyses\xle-disambiguation\20210325\_F59\_3ug\_DENOVO\_12.csv
