## Supplementary material for "A handle on mass coincidence errors in *de novo* sequencing of antibodies by bottom-up proteomics": Combined_054.html

Details Combined\_054 | Stitch OverviewUndefined

### Read Combined\_054

#### Sequence (length=9)

QVYTJPPSR

#### Spectrum 4836? Spectrum 4836 The raw spectrum of this peptide as annotated by Hecklib. The fragments are coloured according to ion type (see legend). Any peaks with a star '\*' as text can be hovered over to see the full details, first the ion type second the mass shift type. By hovering over the amino acids in the peptide or ions in the legend the corresponding peaks are highlighted. By toggling the 'Unassigned' label you can turn the background (unassigned) peaks on or off in the plot. By updating the slider in the Ion legend you can update the spectrum to only show the top X% of the peaks with labels. The top X% means any peak that is within X% of the highest intensity. By dragging in the spectrum you can zoom in to a specific part of the spectrum and use 'Zoom Out' to get back to the original zoom level. The annotation of the spectrum is based on the given sequence in the peptides file and is done with different software so inconsistencies are likely. The peaks are annotated based on the given sequence, with 20 ppm tolerance.

Copy Data

##### Spectrum 4836 (TSV)

###### Preview

```
Loading example...
```

*Click on the button to copy the data to your clipboard.*

Mz MinMz MaxIntensity Max

WidthHeightPeptide font sizePeptide stroke widthSpectrum font sizeSpectrum stroke widthCompact peptide

Ion legend

wxyz

abcd

OtherUnassignedIon^Charge^_Position_Show for top:%

QVYTJPPSR

01.09e+42.18e+43.27e+44.36e+4

Zoom Out

y^+1^_1_y^+1^_1_a^+1^_2_a^+1^_2_b^+1^_2_b^+1^_2_y^+2^_4_y^+1^_2_y^+1^_2_y^+1^_3_y^+1^_3_a^+1^_3_y^+1^_3_b^+1^_3_b^+1^_3_y^+1^_4_y^+1^_4_y^+1^_4_b^+1^_4_b^+1^_4_b^+1^_4_\*\*y^+1^_5_b^+1^_5_y^+1^_6_y^+1^_7_y^+1^_7_y^+1^_8_

0778155523333110

Fragment Matches Table

Show background peaks

| Position | Ion type | Intensity | mz Theoretical | mz Error (Th) | mz Error (ppm) | Charge | Series Number |
| --- | --- | --- | --- | --- | --- | --- | --- |
| - | - | 1179 | 120.1 | - | - | 0 | - |
| - | - | 611.7 | 126.1 | - | - | 0 | - |
| - | - | 1157 | 129.1 | - | - | 0 | - |
| - | - | 1603 | 129.1 | - | - | 0 | - |
| - | - | 699.6 | 130.1 | - | - | 0 | - |
| - | - | 455.5 | 133.1 | - | - | 0 | - |
| - | - | 1.419E+04 | 136.1 | - | - | 0 | - |
| - | - | 1331 | 137.1 | - | - | 0 | - |
| - | - | 2054 | 138.1 | - | - | 0 | - |
| - | - | 395.5 | 147 | - | - | 0 | - |
| - | - | 444.2 | 147.1 | - | - | 0 | - |
| - | - | 845.7 | 148.9 | - | - | 0 | - |
| - | - | 369 | 149.4 | - | - | 0 | - |
| - | - | 447.7 | 151.2 | - | - | 0 | - |
| - | - | 509.3 | 155.1 | - | - | 0 | - |
| - | - | 2892 | 155.1 | - | - | 0 | - |
| - | - | 910 | 157.1 | - | - | 0 | - |
| - | - | 534.6 | 157.1 | - | - | 0 | - |
| - | - | 587.5 | 157.1 | - | - | 0 | - |
| 9 | y | 501.4 | 158.1 | 3.468E-05 | 0.2194 | +1 | 1 |
| - | - | 535.2 | 160.1 | - | - | 0 | - |
| - | - | 637.4 | 169.1 | - | - | 0 | - |
| 9 | y | 4340 | 175.1 | 0.00028 | 1.599 | +1 | 1 |
| - | - | 1.358E+04 | 182.1 | - | - | 0 | - |
| 2 | a | 1.233E+04 | 183.1 | 0.0001872 | 1.022 | +1 | 2 |
| - | - | 1311 | 183.1 | - | - | 0 | - |
| - | - | 1425 | 184.1 | - | - | 0 | - |
| - | - | 1641 | 185.1 | - | - | 0 | - |
| - | - | 2518 | 187.1 | - | - | 0 | - |
| - | - | 955 | 192.1 | - | - | 0 | - |
| - | - | 568.1 | 195.1 | - | - | 0 | - |
| 2 | a | 1.628E+04 | 200.1 | 0.0001578 | 0.7887 | +1 | 2 |
| - | - | 1536 | 201.1 | - | - | 0 | - |
| - | - | 917.5 | 202.1 | - | - | 0 | - |
| - | - | 772.6 | 202.1 | - | - | 0 | - |
| - | - | 483.3 | 202.8 | - | - | 0 | - |
| - | - | 665.6 | 203.1 | - | - | 0 | - |
| - | - | 465.6 | 204.2 | - | - | 0 | - |
| - | - | 6502 | 210.1 | - | - | 0 | - |
| 2 | b | 1705 | 211.1 | 0.0002066 | 0.9788 | +1 | 2 |
| - | - | 756.9 | 211.1 | - | - | 0 | - |
| - | - | 740 | 215.1 | - | - | 0 | - |
| - | - | 482 | 219.1 | - | - | 0 | - |
| - | - | 560.8 | 226.1 | - | - | 0 | - |
| 2 | b | 8173 | 228.1 | 0.0002536 | 1.112 | +1 | 2 |
| 6 | y | 1893 | 228.6 | 0.0002182 | 0.9545 | +2 | 4 |
| - | - | 666.8 | 229.1 | - | - | 0 | - |
| - | - | 822.2 | 230.1 | - | - | 0 | - |
| - | - | 1951 | 237.1 | - | - | 0 | - |
| - | - | 676.7 | 244.2 | - | - | 0 | - |
| 8 | y | 880.7 | 245.1 | 0.0002097 | 0.8554 | +1 | 2 |
| - | - | 3602 | 247.1 | - | - | 0 | - |
| 8 | y | 1021 | 262.2 | 0.0002951 | 1.126 | +1 | 2 |
| - | - | 2056 | 265.1 | - | - | 0 | - |
| - | - | 997.3 | 282.1 | - | - | 0 | - |
| - | - | 4025 | 299.2 | - | - | 0 | - |
| - | - | 1403 | 300.2 | - | - | 0 | - |
| - | - | 694.6 | 307 | - | - | 0 | - |
| - | - | 808 | 309.2 | - | - | 0 | - |
| - | - | 545.6 | 337.1 | - | - | 0 | - |
| 7 | y | 604.2 | 341.2 | 0.000919 | 2.693 | +1 | 3 |
| 7 | y | 3786 | 342.2 | 1.027E-05 | 0.03001 | +1 | 3 |
| 3 | a | 768.3 | 346.2 | 0.001144 | 3.304 | +1 | 3 |
| 7 | y | 4545 | 359.2 | 0.0001311 | 0.3651 | +1 | 3 |
| - | - | 741.1 | 360.2 | - | - | 0 | - |
| - | - | 552.8 | 370.9 | - | - | 0 | - |
| - | - | 1543 | 373.2 | - | - | 0 | - |
| 3 | b | 1267 | 374.2 | 3.423E-05 | 0.09147 | +1 | 3 |
| - | - | 594.2 | 388.9 | - | - | 0 | - |
| 3 | b | 1660 | 391.2 | 5.614E-05 | 0.1435 | +1 | 3 |
| - | - | 2421 | 396.2 | - | - | 0 | - |
| - | - | 2871 | 418.2 | - | - | 0 | - |
| - | - | 2207 | 418.7 | - | - | 0 | - |
| - | - | 905.5 | 422.2 | - | - | 0 | - |
| - | - | 800.6 | 430.2 | - | - | 0 | - |
| 6 | y | 1078 | 438.2 | 0.002078 | 4.741 | +1 | 4 |
| 6 | y | 6260 | 439.2 | 0.0006927 | 1.577 | +1 | 4 |
| - | - | 1328 | 440.2 | - | - | 0 | - |
| 6 | y | 4.312E+04 | 456.3 | 0.0001751 | 0.3837 | +1 | 4 |
| - | - | 7913 | 457.2 | - | - | 0 | - |
| - | - | 1.006E+04 | 457.3 | - | - | 0 | - |
| - | - | 2456 | 458.2 | - | - | 0 | - |
| - | - | 707.1 | 458.3 | - | - | 0 | - |
| 4 | b | 1351 | 474.2 | 0.0001836 | 0.3871 | +1 | 4 |
| 4 | b | 4587 | 475.2 | 8.524E-05 | 0.1794 | +1 | 4 |
| - | - | 1322 | 476.2 | - | - | 0 | - |
| 4 | b | 834.1 | 492.2 | 0.0009817 | 1.994 | +1 | 4 |
| - | - | 1632 | 513.3 | - | - | 0 | - |
| 0 | Precursor | 1.637E+04 | 521.8 | 0.0004098 | 0.7854 | +2 | -1 |
| 0 | Precursor | 1.32E+04 | 522.3 | 0.009352 | 17.91 | +2 | -1 |
| - | - | 2541 | 522.8 | - | - | 0 | - |
| - | - | 865.7 | 531.3 | - | - | 0 | - |
| - | - | 1155 | 532.3 | - | - | 0 | - |
| - | - | 960.5 | 542.3 | - | - | 0 | - |
| 5 | y | 7102 | 569.3 | 0.0001182 | 0.2075 | +1 | 5 |
| - | - | 1772 | 570.3 | - | - | 0 | - |
| - | - | 1789 | 570.3 | - | - | 0 | - |
| - | - | 868.2 | 571.3 | - | - | 0 | - |
| 5 | b | 1583 | 587.3 | 0.0004758 | 0.8102 | +1 | 5 |
| - | - | 840.9 | 588.3 | - | - | 0 | - |
| - | - | 977.8 | 641.4 | - | - | 0 | - |
| 4 | y | 9600 | 670.4 | 0.0006165 | 0.9196 | +1 | 6 |
| - | - | 3920 | 671.4 | - | - | 0 | - |
| - | - | 673.3 | 672.4 | - | - | 0 | - |
| - | - | 890.1 | 673.9 | - | - | 0 | - |
| 3 | y | 1005 | 816.4 | 0.009296 | 11.39 | +1 | 7 |
| 3 | y | 2.262E+04 | 833.5 | 0.001323 | 1.587 | +1 | 7 |
| - | - | 1.219E+04 | 834.5 | - | - | 0 | - |
| - | - | 2534 | 835.5 | - | - | 0 | - |
| - | - | 558.7 | 848 | - | - | 0 | - |
| 2 | y | 913.7 | 932.5 | 0.004551 | 4.881 | +1 | 8 |
| - | - | 543.2 | 1053 | - | - | 0 | - |
| - | - | 538.7 | 1318 | - | - | 0 | - |
| - | - | 585.5 | 1799 | - | - | 0 | - |
| - | - | 619.4 | 1882 | - | - | 0 | - |
| - | - | 655.5 | 2362 | - | - | 0 | - |
| - | - | 830.5 | 3079 | - | - | 0 | - |
| - | - | 983.9 | 3080 | - | - | 0 | - |

m/z Charge Intensity FragmentType MassShift Position
120.08113861083984 0 1179.0267
126.05535125732422 0 611.67206
129.066162109375 0 1156.6813
129.1024932861328 0 1603.1626
130.08648681640625 0 699.59656
133.06088256835938 0 455.51547
136.07594299316406 0 14193.282
137.07916259765625 0 1331.3685
138.0915985107422 0 2054.482
147.04412841796875 0 395.48282
147.1127166748047 0 444.23093
148.94711303710938 0 845.67584
149.40432739257812 0 369.02112
151.2476348876953 0 447.68008
155.11209106445312 0 509.27667
155.11827087402344 0 2892.135
157.09730529785156 0 910.02356
157.1082000732422 0 534.5957
157.1336669921875 0 587.5134
158.09243774414062 0 501.35104 y Ammonia loss 8
160.07501220703125 0 535.2125
169.13343811035156 0 637.3736
175.11923217773438 0 4339.628 y 8
182.12905883789062 0 13583.9375
183.1129913330078 0 12328.822 a Ammonia loss 1
183.13226318359375 0 1311.0833
184.116455078125 0 1424.5184
185.09217834472656 0 1640.9816
187.14439392089844 0 2518.494
192.10198974609375 0 954.97974
195.1128692626953 0 568.0702
200.13951110839844 0 16279.623 a 1
201.14297485351562 0 1535.763
202.08578491210938 0 917.4588
202.1188201904297 0 772.60474
202.8184051513672 0 483.2714
203.102783203125 0 665.56213
204.17556762695312 0 465.61893
210.1239471435547 0 6502.371
211.10792541503906 0 1704.7283 b Ammonia loss 1
211.12689208984375 0 756.8856
215.13912963867188 0 740.0368
219.11370849609375 0 482.0433
226.1183319091797 0 560.84125
228.134521484375 0 8173.051 b 1
228.63211059570312 0 1892.7787 y 5
229.13607788085938 0 666.7781
230.081787109375 0 822.168
237.12330627441406 0 1950.5668
244.16534423828125 0 676.73236
245.1242218017578 0 880.67505 y Ammonia loss 7
247.1078643798828 0 3602.416
262.1512756347656 0 1020.7245 y 7
265.1186218261719 0 2056.2168
282.14508056640625 0 997.3385
299.1715393066406 0 4024.7854
300.1559753417969 0 1403.1711
306.98687744140625 0 694.6284
309.2032470703125 0 808.0476
337.149658203125 0 545.61017
341.1922607421875 0 604.22186 y Water loss 6
342.17718505859375 0 3785.6885 y Ammonia loss 6
346.1772766113281 0 768.3156 a Ammonia loss 2
359.20361328125 0 4545.1777 y 6
360.1900329589844 0 741.09265
370.8825988769531 0 552.7832
373.1866149902344 0 1543.4541
374.17108154296875 0 1266.9077 b Ammonia loss 2
388.8914489746094 0 594.1888
391.1975402832031 0 1660.4133 b 2
396.22381591796875 0 2420.7795
418.23455810546875 0 2871.458
418.736572265625 0 2206.7366
422.20635986328125 0 905.47974
430.2084045410156 0 800.5566
438.2438659667969 0 1077.8542 y Water loss 5
439.23065185546875 0 6259.8145 y Ammonia loss 5
440.2340087890625 0 1328.221
456.2566833496094 0 43119.99 y 5
457.2086486816406 0 7912.8496
457.25933837890625 0 10061.452
458.2119445800781 0 2456.0498
458.26031494140625 0 707.11615
474.2348937988281 0 1351.4297 b Water loss 3
475.21881103515625 0 4586.657 b Ammonia loss 3
476.222412109375 0 1322.2015
492.2442932128906 0 834.0709 b 3
513.2754516601562 0 1632.1687
521.7872314453125 0 16370.624 Precursor Water loss
522.2890014648438 0 13195.649 Precursor Ammonia loss
522.7896728515625 0 2541.3342
531.2955932617188 0 865.69354
532.308349609375 0 1155.1108
542.2944946289062 0 960.5225
569.3404541015625 0 7102.089 y 4
570.2941284179688 0 1771.8505
570.3447875976562 0 1789.2427
571.2960205078125 0 868.21643
587.3182983398438 0 1583.4661 b Water loss 4
588.318115234375 0 840.93695
641.4006958007812 0 977.7912
670.3876342773438 0 9599.905 y 3
671.3912963867188 0 3920.17
672.3866577148438 0 673.28973
673.8688354492188 0 890.11566
816.434326171875 0 1005.1845 y Ammonia loss 2
833.4502563476562 0 22616.275 y 2
834.4530639648438 0 12191.159
835.456298828125 0 2534.0142
847.9529418945312 0 558.7438
932.5154418945312 0 913.73 y 1
1053.2713623046875 0 543.23834
1317.75830078125 0 538.7068
1799.4970703125 0 585.53186
1882.330810546875 0 619.4022
2361.5458984375 0 655.4557
3078.85400390625 0 830.5012
3079.539794921875 0 983.9179

Spectrum Details

|  |  |
| --- | --- |
| Matched peaks? Matched peaks The total absolute number of peaks matched. Additionally in brackets the total fraction of peaks matched and the total number of peaks is shown. | 29 (24.58% of 118) |
| FDR? FDR The false discovery rate estimated for this peptide. It is calculated by matching all theoretical fragments with a non-integer shift with the raw peaks for this spectrum. This is done with 40 different shifts. The resulting percentage is the average number of annotated peaks over the number of annotated peaks with the correct spectrum. | 0.00% |
| Satellite FDR? Satellite FDR See the FDR for details on its calculation. This satellite ion specific FDR only contains the satellite ions (d/w) for I/L/J positions. | - |
| PSM Score? PSM Score The PSM Score as given by Hecklib to this annotated spectrum. It is shown with three significant figures. | 383 |

#### Spectrum 4893? Spectrum 4893 The raw spectrum of this peptide as annotated by Hecklib. The fragments are coloured according to ion type (see legend). Any peaks with a star '\*' as text can be hovered over to see the full details, first the ion type second the mass shift type. By hovering over the amino acids in the peptide or ions in the legend the corresponding peaks are highlighted. By toggling the 'Unassigned' label you can turn the background (unassigned) peaks on or off in the plot. By updating the slider in the Ion legend you can update the spectrum to only show the top X% of the peaks with labels. The top X% means any peak that is within X% of the highest intensity. By dragging in the spectrum you can zoom in to a specific part of the spectrum and use 'Zoom Out' to get back to the original zoom level. The annotation of the spectrum is based on the given sequence in the peptides file and is done with different software so inconsistencies are likely. The peaks are annotated based on the given sequence, with 20 ppm tolerance.

Copy Data

##### Spectrum 4893 (TSV)

###### Preview

```
Loading example...
```

*Click on the button to copy the data to your clipboard.*

Mz MinMz MaxIntensity Max

WidthHeightPeptide font sizePeptide stroke widthSpectrum font sizeSpectrum stroke widthCompact peptide

Ion legend

wxyz

abcd

OtherUnassignedIon^Charge^_Position_Show for top:%

QVYTJPPSR

09.15e+31.83e+42.75e+43.66e+4

Zoom Out

y^+1^_1_y^+1^_1_a^+1^_2_a^+1^_2_b^+1^_2_b^+1^_2_y^+2^_4_y^+1^_2_y^+1^_2_y^+1^_3_a^+1^_3_y^+1^_3_b^+1^_3_b^+1^_3_y^+1^_4_y^+1^_4_y^+1^_4_b^+1^_4_b^+1^_4_\*\*\*y^+1^_5_b^+1^_5_y^+1^_6_y^+1^_7_y^+1^_7_y^+1^_7_y^+1^_8_

047995914381917

Fragment Matches Table

Show background peaks

| Position | Ion type | Intensity | mz Theoretical | mz Error (Th) | mz Error (ppm) | Charge | Series Number |
| --- | --- | --- | --- | --- | --- | --- | --- |
| - | - | 841.8 | 120.1 | - | - | 0 | - |
| - | - | 374.8 | 120.5 | - | - | 0 | - |
| - | - | 339.6 | 120.8 | - | - | 0 | - |
| - | - | 993.5 | 129.1 | - | - | 0 | - |
| - | - | 1574 | 129.1 | - | - | 0 | - |
| - | - | 3640 | 133.1 | - | - | 0 | - |
| - | - | 503.4 | 133.1 | - | - | 0 | - |
| - | - | 909.2 | 136.1 | - | - | 0 | - |
| - | - | 1.32E+04 | 136.1 | - | - | 0 | - |
| - | - | 1255 | 137.1 | - | - | 0 | - |
| - | - | 2390 | 138.1 | - | - | 0 | - |
| - | - | 403.4 | 138.5 | - | - | 0 | - |
| - | - | 431.8 | 146.2 | - | - | 0 | - |
| - | - | 1868 | 155.1 | - | - | 0 | - |
| 9 | y | 513.2 | 158.1 | 1.942E-05 | 0.1228 | +1 | 1 |
| - | - | 1635 | 173.4 | - | - | 0 | - |
| 9 | y | 4912 | 175.1 | 0.0001167 | 0.6665 | +1 | 1 |
| - | - | 1.299E+04 | 182.1 | - | - | 0 | - |
| 2 | a | 1.087E+04 | 183.1 | 5.696E-05 | 0.3111 | +1 | 2 |
| - | - | 1095 | 183.1 | - | - | 0 | - |
| - | - | 1346 | 184.1 | - | - | 0 | - |
| - | - | 1157 | 185.1 | - | - | 0 | - |
| - | - | 661.1 | 185.2 | - | - | 0 | - |
| - | - | 777.7 | 187.1 | - | - | 0 | - |
| - | - | 1453 | 187.1 | - | - | 0 | - |
| - | - | 561.5 | 192.1 | - | - | 0 | - |
| - | - | 549.7 | 193.6 | - | - | 0 | - |
| - | - | 619 | 195.1 | - | - | 0 | - |
| 2 | a | 1.527E+04 | 200.1 | 0.0002084 | 1.041 | +1 | 2 |
| - | - | 500.9 | 200.7 | - | - | 0 | - |
| - | - | 976.8 | 201.1 | - | - | 0 | - |
| - | - | 507.4 | 202 | - | - | 0 | - |
| - | - | 744 | 202.1 | - | - | 0 | - |
| - | - | 692.5 | 204.1 | - | - | 0 | - |
| - | - | 576.1 | 209.1 | - | - | 0 | - |
| - | - | 6893 | 210.1 | - | - | 0 | - |
| 2 | b | 1311 | 211.1 | 8.328E-05 | 0.3945 | +1 | 2 |
| - | - | 697.3 | 215.1 | - | - | 0 | - |
| 2 | b | 5175 | 228.1 | 0.0001736 | 0.7611 | +1 | 2 |
| 6 | y | 1472 | 228.6 | 0.0002335 | 1.021 | +2 | 4 |
| - | - | 764.9 | 229.1 | - | - | 0 | - |
| - | - | 1536 | 230.1 | - | - | 0 | - |
| - | - | 468.8 | 232.5 | - | - | 0 | - |
| - | - | 605 | 235.1 | - | - | 0 | - |
| - | - | 1834 | 237.1 | - | - | 0 | - |
| 8 | y | 992.4 | 245.1 | 0.0001486 | 0.6064 | +1 | 2 |
| - | - | 3212 | 247.1 | - | - | 0 | - |
| - | - | 785.6 | 248.1 | - | - | 0 | - |
| - | - | 563.6 | 253.4 | - | - | 0 | - |
| 8 | y | 1194 | 262.2 | 1.012E-05 | 0.0386 | +1 | 2 |
| - | - | 2465 | 265.1 | - | - | 0 | - |
| - | - | 714 | 278.2 | - | - | 0 | - |
| - | - | 909.9 | 282.1 | - | - | 0 | - |
| - | - | 3057 | 299.2 | - | - | 0 | - |
| - | - | 1563 | 300.2 | - | - | 0 | - |
| - | - | 759.7 | 307 | - | - | 0 | - |
| - | - | 650.6 | 309.2 | - | - | 0 | - |
| - | - | 569 | 321.3 | - | - | 0 | - |
| - | - | 1062 | 326 | - | - | 0 | - |
| - | - | 582.8 | 338.4 | - | - | 0 | - |
| 7 | y | 3184 | 342.2 | 0.000407 | 1.189 | +1 | 3 |
| 3 | a | 856.6 | 346.2 | 0.0001683 | 0.4863 | +1 | 3 |
| - | - | 502.7 | 348.1 | - | - | 0 | - |
| - | - | 628.2 | 358.8 | - | - | 0 | - |
| 7 | y | 3586 | 359.2 | 0.0006194 | 1.724 | +1 | 3 |
| - | - | 1609 | 373.2 | - | - | 0 | - |
| 3 | b | 1454 | 374.2 | 0.000393 | 1.05 | +1 | 3 |
| 3 | b | 1691 | 391.2 | 0.0006665 | 1.704 | +1 | 3 |
| - | - | 3222 | 396.2 | - | - | 0 | - |
| - | - | 673.4 | 397.2 | - | - | 0 | - |
| - | - | 717.6 | 414.9 | - | - | 0 | - |
| - | - | 1315 | 418.2 | - | - | 0 | - |
| - | - | 669 | 421 | - | - | 0 | - |
| - | - | 642 | 422.2 | - | - | 0 | - |
| - | - | 694 | 424.3 | - | - | 0 | - |
| 6 | y | 1358 | 438.2 | 0.00101 | 2.304 | +1 | 4 |
| 6 | y | 6090 | 439.2 | 0.0002839 | 0.6463 | +1 | 4 |
| - | - | 1215 | 440.2 | - | - | 0 | - |
| 6 | y | 3.625E+04 | 456.3 | 0.0001911 | 0.4189 | +1 | 4 |
| - | - | 5742 | 457.2 | - | - | 0 | - |
| - | - | 8732 | 457.3 | - | - | 0 | - |
| - | - | 1489 | 458.2 | - | - | 0 | - |
| - | - | 1106 | 458.3 | - | - | 0 | - |
| - | - | 626.9 | 459.2 | - | - | 0 | - |
| 4 | b | 1809 | 474.2 | 0.0008845 | 1.865 | +1 | 4 |
| 4 | b | 4463 | 475.2 | 0.0004515 | 0.95 | +1 | 4 |
| - | - | 1125 | 476.2 | - | - | 0 | - |
| - | - | 603.8 | 479.9 | - | - | 0 | - |
| - | - | 784.4 | 513.3 | - | - | 0 | - |
| 0 | Precursor | 1.623E+04 | 521.8 | 0.000715 | 1.37 | +2 | -1 |
| 0 | Precursor | 9436 | 522.3 | 0.008742 | 16.74 | +2 | -1 |
| - | - | 3382 | 522.8 | - | - | 0 | - |
| - | - | 788 | 523.3 | - | - | 0 | - |
| - | - | 1197 | 530.3 | - | - | 0 | - |
| 0 | Precursor | 915.7 | 530.8 | 0.002213 | 4.169 | +2 | -1 |
| 5 | y | 6445 | 569.3 | 0.0006675 | 1.172 | +1 | 5 |
| - | - | 1578 | 570.3 | - | - | 0 | - |
| - | - | 1787 | 570.3 | - | - | 0 | - |
| - | - | 1077 | 571.3 | - | - | 0 | - |
| 5 | b | 1311 | 587.3 | 0.001025 | 1.745 | +1 | 5 |
| 4 | y | 8622 | 670.4 | 0.001044 | 1.557 | +1 | 6 |
| - | - | 3584 | 671.4 | - | - | 0 | - |
| - | - | 672.7 | 672.4 | - | - | 0 | - |
| 3 | y | 1010 | 815.4 | 0.003576 | 4.385 | +1 | 7 |
| 3 | y | 946.7 | 816.4 | 0.008503 | 10.41 | +1 | 7 |
| 3 | y | 2.148E+04 | 833.5 | 0.001445 | 1.734 | +1 | 7 |
| - | - | 1.091E+04 | 834.5 | - | - | 0 | - |
| - | - | 2821 | 835.5 | - | - | 0 | - |
| 2 | y | 958.5 | 932.5 | 0.003147 | 3.375 | +1 | 8 |
| - | - | 1016 | 933.5 | - | - | 0 | - |
| - | - | 716.6 | 1898 | - | - | 0 | - |

m/z Charge Intensity FragmentType MassShift Position
120.08082580566406 0 841.83307
120.49706268310547 0 374.77405
120.82332611083984 0 339.55936
129.06593322753906 0 993.48224
129.10218811035156 0 1573.6309
133.06076049804688 0 3640.0796
133.08612060546875 0 503.38757
136.07008361816406 0 909.2017
136.07571411132812 0 13200.469
137.07925415039062 0 1255.0632
138.0914306640625 0 2389.9578
138.49798583984375 0 403.36954
146.15780639648438 0 431.80685
155.11778259277344 0 1867.9344
158.09242248535156 0 513.20197 y Ammonia loss 8
173.43887329101562 0 1634.8132
175.11883544921875 0 4912.1733 y 8
182.128662109375 0 12986.225
183.1127471923828 0 10865.272 a Ammonia loss 1
183.13238525390625 0 1094.6251
184.11666870117188 0 1345.5768
185.092041015625 0 1157.3252
185.1651611328125 0 661.08527
187.07119750976562 0 777.67303
187.14401245117188 0 1453.4297
192.1018524169922 0 561.453
193.61953735351562 0 549.66675
195.11277770996094 0 619.00165
200.13914489746094 0 15268.987 a 1
200.7345733642578 0 500.9025
201.14291381835938 0 976.798
201.9652099609375 0 507.43744
202.11880493164062 0 744.0167
204.09707641601562 0 692.4549
209.09173583984375 0 576.104
210.12355041503906 0 6892.7485
211.10763549804688 0 1310.8463 b Ammonia loss 1
215.13868713378906 0 697.3482
228.13409423828125 0 5175.362 b 1
228.6321258544922 0 1472.5 y 5
229.1332550048828 0 764.9004
230.0809783935547 0 1536.4862
232.54995727539062 0 468.80377
235.14430236816406 0 605.0355
237.12301635742188 0 1833.7725
245.12428283691406 0 992.3792 y Ammonia loss 7
247.10743713378906 0 3211.5435
248.11087036132812 0 785.6115
253.3817138671875 0 563.5566
262.1509704589844 0 1193.8679 y 7
265.1176452636719 0 2465.221
278.15936279296875 0 713.9543
282.1451110839844 0 909.9307
299.1710205078125 0 3056.9788
300.15478515625 0 1562.5496
306.98486328125 0 759.679
309.2031555175781 0 650.5563
321.2824401855469 0 569.03986
326.0094909667969 0 1062.0325
338.4468688964844 0 582.77655
342.1767883300781 0 3183.96 y Ammonia loss 6
346.17596435546875 0 856.62805 a Ammonia loss 2
348.0930480957031 0 502.66556
358.81683349609375 0 628.21466
359.203125 0 3586.4575 y 6
373.18670654296875 0 1608.5096
374.170654296875 0 1453.7045 b Ammonia loss 2
391.1969299316406 0 1690.9952 b 2
396.2234191894531 0 3221.9775
397.22625732421875 0 673.3595
414.9211120605469 0 717.63385
418.2334899902344 0 1315.4978
420.9871520996094 0 668.98145
422.2043762207031 0 641.9692
424.2514953613281 0 694.0445
438.24493408203125 0 1358.4307 y Water loss 5
439.22967529296875 0 6090.0376 y Ammonia loss 5
440.2340087890625 0 1214.8838
456.2563171386719 0 36252.957 y 5
457.20806884765625 0 5741.68
457.2596130371094 0 8731.974
458.2106018066406 0 1489.0525
458.2610778808594 0 1105.9746
459.2104187011719 0 626.89594
474.23382568359375 0 1809.3899 b Water loss 3
475.21917724609375 0 4463.381 b Ammonia loss 3
476.2204895019531 0 1124.8137
479.8655700683594 0 603.7675
513.2794799804688 0 784.35895
521.7869262695312 0 16229.469 Precursor Water loss
522.2883911132812 0 9436.295 Precursor Ammonia loss
522.7894287109375 0 3382.1614
523.295166015625 0 787.9655
530.3152465820312 0 1196.8357
530.7907104492188 0 915.691 Precursor
569.3399047851562 0 6445.217 y 4
570.291748046875 0 1577.7183
570.3427124023438 0 1786.9191
571.2935180664062 0 1076.569
587.3177490234375 0 1310.8748 b Water loss 4
670.38720703125 0 8622.246 y 3
671.388916015625 0 3583.7668
672.3916625976562 0 672.6547
815.4374389648438 0 1010.4532 y Water loss 2
816.4335327148438 0 946.72754 y Ammonia loss 2
833.4501342773438 0 21478.518 y 2
834.4527587890625 0 10914.798
835.4581909179688 0 2820.615
932.516845703125 0 958.487 y 1
933.512939453125 0 1015.5391
1898.206298828125 0 716.5927

Spectrum Details

|  |  |
| --- | --- |
| Matched peaks? Matched peaks The total absolute number of peaks matched. Additionally in brackets the total fraction of peaks matched and the total number of peaks is shown. | 29 (26.13% of 111) |
| FDR? FDR The false discovery rate estimated for this peptide. It is calculated by matching all theoretical fragments with a non-integer shift with the raw peaks for this spectrum. This is done with 40 different shifts. The resulting percentage is the average number of annotated peaks over the number of annotated peaks with the correct spectrum. | 0.33% |
| Satellite FDR? Satellite FDR See the FDR for details on its calculation. This satellite ion specific FDR only contains the satellite ions (d/w) for I/L/J positions. | - |
| PSM Score? PSM Score The PSM Score as given by Hecklib to this annotated spectrum. It is shown with three significant figures. | 363 |

#### Spectrum 4660? Spectrum 4660 The raw spectrum of this peptide as annotated by Hecklib. The fragments are coloured according to ion type (see legend). Any peaks with a star '\*' as text can be hovered over to see the full details, first the ion type second the mass shift type. By hovering over the amino acids in the peptide or ions in the legend the corresponding peaks are highlighted. By toggling the 'Unassigned' label you can turn the background (unassigned) peaks on or off in the plot. By updating the slider in the Ion legend you can update the spectrum to only show the top X% of the peaks with labels. The top X% means any peak that is within X% of the highest intensity. By dragging in the spectrum you can zoom in to a specific part of the spectrum and use 'Zoom Out' to get back to the original zoom level. The annotation of the spectrum is based on the given sequence in the peptides file and is done with different software so inconsistencies are likely. The peaks are annotated based on the given sequence, with 20 ppm tolerance.

Copy Data

##### Spectrum 4660 (TSV)

###### Preview

```
Loading example...
```

*Click on the button to copy the data to your clipboard.*

Mz MinMz MaxIntensity Max

WidthHeightPeptide font sizePeptide stroke widthSpectrum font sizeSpectrum stroke widthCompact peptide

Ion legend

wxyz

abcd

OtherUnassignedIon^Charge^_Position_Show for top:%

QVYTJPPSR

01.31e+42.62e+43.92e+45.23e+4

Zoom Out

y^+1^_1_a^+1^_2_a^+1^_2_b^+1^_2_b^+1^_2_y^+2^_4_y^+1^_2_y^+1^_2_y^+1^_3_y^+1^_3_a^+1^_3_y^+1^_3_b^+1^_3_b^+1^_3_y^+1^_4_y^+1^_4_y^+1^_4_b^+1^_4_b^+1^_4_\*\*\*y^+1^_5_y^+1^_5_b^+1^_5_y^+1^_6_y^+1^_7_y^+1^_7_y^+1^_8_

0778155523333110

Fragment Matches Table

Show background peaks

| Position | Ion type | Intensity | mz Theoretical | mz Error (Th) | mz Error (ppm) | Charge | Series Number |
| --- | --- | --- | --- | --- | --- | --- | --- |
| - | - | 983.1 | 120.1 | - | - | 0 | - |
| - | - | 369.7 | 121.3 | - | - | 0 | - |
| - | - | 754.3 | 126.1 | - | - | 0 | - |
| - | - | 358 | 128.3 | - | - | 0 | - |
| - | - | 1272 | 129.1 | - | - | 0 | - |
| - | - | 1329 | 129.1 | - | - | 0 | - |
| - | - | 512 | 129.8 | - | - | 0 | - |
| - | - | 1.827E+04 | 136.1 | - | - | 0 | - |
| - | - | 1322 | 137.1 | - | - | 0 | - |
| - | - | 2550 | 138.1 | - | - | 0 | - |
| - | - | 456.4 | 148.7 | - | - | 0 | - |
| - | - | 432.5 | 148.9 | - | - | 0 | - |
| - | - | 538.9 | 148.9 | - | - | 0 | - |
| - | - | 783.4 | 148.9 | - | - | 0 | - |
| - | - | 711.4 | 148.9 | - | - | 0 | - |
| - | - | 1203 | 148.9 | - | - | 0 | - |
| - | - | 1259 | 148.9 | - | - | 0 | - |
| - | - | 2643 | 148.9 | - | - | 0 | - |
| - | - | 4344 | 148.9 | - | - | 0 | - |
| - | - | 3868 | 149 | - | - | 0 | - |
| - | - | 2142 | 149 | - | - | 0 | - |
| - | - | 1350 | 149 | - | - | 0 | - |
| - | - | 1139 | 149 | - | - | 0 | - |
| - | - | 635.2 | 149 | - | - | 0 | - |
| - | - | 808.7 | 149 | - | - | 0 | - |
| - | - | 630.9 | 149 | - | - | 0 | - |
| - | - | 418.5 | 149 | - | - | 0 | - |
| - | - | 600.3 | 149 | - | - | 0 | - |
| - | - | 495 | 149.1 | - | - | 0 | - |
| - | - | 488.6 | 149.1 | - | - | 0 | - |
| - | - | 3498 | 155.1 | - | - | 0 | - |
| - | - | 596 | 157.1 | - | - | 0 | - |
| 9 | y | 4501 | 175.1 | 6.639E-05 | 0.3791 | +1 | 1 |
| - | - | 1.612E+04 | 182.1 | - | - | 0 | - |
| 2 | a | 1.447E+04 | 183.1 | 3.459E-05 | 0.1889 | +1 | 2 |
| - | - | 1001 | 183.1 | - | - | 0 | - |
| - | - | 1131 | 184.1 | - | - | 0 | - |
| - | - | 1203 | 185.1 | - | - | 0 | - |
| - | - | 1943 | 187.1 | - | - | 0 | - |
| - | - | 894.9 | 192.1 | - | - | 0 | - |
| - | - | 443.1 | 194.9 | - | - | 0 | - |
| - | - | 951.5 | 195.1 | - | - | 0 | - |
| 2 | a | 1.87E+04 | 200.1 | 5.262E-06 | 0.02629 | +1 | 2 |
| - | - | 1012 | 201.1 | - | - | 0 | - |
| - | - | 1350 | 202.1 | - | - | 0 | - |
| - | - | 1124 | 203.1 | - | - | 0 | - |
| - | - | 9393 | 210.1 | - | - | 0 | - |
| 2 | b | 1177 | 211.1 | 0.0001761 | 0.8343 | +1 | 2 |
| - | - | 812.4 | 211.1 | - | - | 0 | - |
| - | - | 932 | 215.1 | - | - | 0 | - |
| 2 | b | 7903 | 228.1 | 3.998E-05 | 0.1753 | +1 | 2 |
| 6 | y | 2506 | 228.6 | 0.0001632 | 0.714 | +2 | 4 |
| - | - | 810.8 | 229.1 | - | - | 0 | - |
| - | - | 1737 | 230.1 | - | - | 0 | - |
| - | - | 542.5 | 235.1 | - | - | 0 | - |
| - | - | 2790 | 237.1 | - | - | 0 | - |
| - | - | 855.3 | 238.1 | - | - | 0 | - |
| 8 | y | 1335 | 245.1 | 0.0002249 | 0.9176 | +1 | 2 |
| - | - | 5236 | 247.1 | - | - | 0 | - |
| - | - | 774.5 | 248.1 | - | - | 0 | - |
| 8 | y | 847 | 262.2 | 0.0001932 | 0.7371 | +1 | 2 |
| - | - | 2475 | 265.1 | - | - | 0 | - |
| - | - | 1318 | 282.1 | - | - | 0 | - |
| - | - | 2909 | 299.2 | - | - | 0 | - |
| - | - | 1611 | 300.2 | - | - | 0 | - |
| - | - | 512.2 | 317.8 | - | - | 0 | - |
| - | - | 607.6 | 325.2 | - | - | 0 | - |
| 7 | y | 826.1 | 341.2 | 0.0002712 | 0.7948 | +1 | 3 |
| 7 | y | 4651 | 342.2 | 0.000407 | 1.189 | +1 | 3 |
| 3 | a | 632.9 | 346.2 | 0.0007177 | 2.073 | +1 | 3 |
| 7 | y | 4935 | 359.2 | 0.0006194 | 1.724 | +1 | 3 |
| - | - | 689.7 | 360.2 | - | - | 0 | - |
| - | - | 2485 | 373.2 | - | - | 0 | - |
| 3 | b | 1881 | 374.2 | 0.001644 | 4.394 | +1 | 3 |
| 3 | b | 1259 | 391.2 | 2.562E-05 | 0.06549 | +1 | 3 |
| - | - | 583.7 | 391.8 | - | - | 0 | - |
| - | - | 2932 | 396.2 | - | - | 0 | - |
| - | - | 571.2 | 397.2 | - | - | 0 | - |
| - | - | 936.7 | 421.2 | - | - | 0 | - |
| - | - | 581.2 | 422.2 | - | - | 0 | - |
| - | - | 1069 | 430.2 | - | - | 0 | - |
| - | - | 561.4 | 438 | - | - | 0 | - |
| 6 | y | 1078 | 438.2 | 0.000155 | 0.3537 | +1 | 4 |
| 6 | y | 7402 | 439.2 | 0.0002229 | 0.5074 | +1 | 4 |
| - | - | 1004 | 440.2 | - | - | 0 | - |
| - | - | 870.4 | 447.2 | - | - | 0 | - |
| 6 | y | 5.178E+04 | 456.3 | 0.0003437 | 0.7534 | +1 | 4 |
| - | - | 381.7 | 457.2 | - | - | 0 | - |
| - | - | 1.103E+04 | 457.2 | - | - | 0 | - |
| - | - | 1.076E+04 | 457.3 | - | - | 0 | - |
| - | - | 3504 | 458.2 | - | - | 0 | - |
| - | - | 1705 | 458.3 | - | - | 0 | - |
| 4 | b | 2113 | 474.2 | 0.0007625 | 1.608 | +1 | 4 |
| 4 | b | 4938 | 475.2 | 0.0002505 | 0.527 | +1 | 4 |
| - | - | 1192 | 476.2 | - | - | 0 | - |
| - | - | 1279 | 512.8 | - | - | 0 | - |
| - | - | 713.2 | 513.3 | - | - | 0 | - |
| - | - | 691.1 | 513.8 | - | - | 0 | - |
| 0 | Precursor | 2.293E+04 | 521.8 | 0.0008981 | 1.721 | +2 | -1 |
| 0 | Precursor | 1.48E+04 | 522.3 | 0.008498 | 16.27 | +2 | -1 |
| - | - | 3646 | 522.8 | - | - | 0 | - |
| 0 | Precursor | 1197 | 530.8 | 0.001481 | 2.79 | +2 | -1 |
| - | - | 927.1 | 531.3 | - | - | 0 | - |
| - | - | 1228 | 542.3 | - | - | 0 | - |
| - | - | 699.8 | 543.3 | - | - | 0 | - |
| 5 | y | 645.6 | 551.3 | 7.055E-05 | 0.128 | +1 | 5 |
| - | - | 889.4 | 562.3 | - | - | 0 | - |
| 5 | y | 8976 | 569.3 | 0.001156 | 2.03 | +1 | 5 |
| - | - | 3169 | 570.3 | - | - | 0 | - |
| - | - | 1887 | 570.3 | - | - | 0 | - |
| 5 | b | 1910 | 587.3 | 1.243E-05 | 0.02117 | +1 | 5 |
| - | - | 600.7 | 657.1 | - | - | 0 | - |
| 4 | y | 1.341E+04 | 670.4 | 0.001654 | 2.467 | +1 | 6 |
| - | - | 4883 | 671.4 | - | - | 0 | - |
| - | - | 662.9 | 691.6 | - | - | 0 | - |
| - | - | 656.1 | 713.9 | - | - | 0 | - |
| 3 | y | 1737 | 815.4 | 0.001928 | 2.364 | +1 | 7 |
| - | - | 784.9 | 816.4 | - | - | 0 | - |
| 3 | y | 2.849E+04 | 833.5 | 0.002177 | 2.613 | +1 | 7 |
| - | - | 1.418E+04 | 834.5 | - | - | 0 | - |
| - | - | 2954 | 835.5 | - | - | 0 | - |
| 2 | y | 1217 | 932.5 | 0.004246 | 4.553 | +1 | 8 |
| - | - | 823.2 | 3080 | - | - | 0 | - |

m/z Charge Intensity FragmentType MassShift Position
120.08094787597656 0 983.09796
121.253662109375 0 369.72354
126.05530548095703 0 754.2678
128.33554077148438 0 357.96857
129.0658721923828 0 1272.0278
129.1025390625 0 1328.8706
129.77694702148438 0 511.99142
136.07583618164062 0 18270.406
137.07923889160156 0 1321.7982
138.09149169921875 0 2549.6035
148.6630859375 0 456.42767
148.8939208984375 0 432.47137
148.90093994140625 0 538.86566
148.9081573486328 0 783.44116
148.91506958007812 0 711.4471
148.92214965820312 0 1202.8589
148.92950439453125 0 1259.2714
148.93661499023438 0 2642.9858
148.94439697265625 0 4344.2705
148.96090698242188 0 3867.6138
148.96868896484375 0 2142.4597
148.97567749023438 0 1350.1726
148.98291015625 0 1139.3047
148.9903564453125 0 635.2064
148.99728393554688 0 808.7052
149.00425720214844 0 630.92755
149.01173400878906 0 418.508
149.01918029785156 0 600.2794
149.0551300048828 0 494.97226
149.1057891845703 0 488.57254
155.1179962158203 0 3498.256
157.0970916748047 0 596.02704
175.1190185546875 0 4501.0327 y 8
182.1288299560547 0 16119.858
183.1128387451172 0 14473.57 a Ammonia loss 1
183.1321563720703 0 1000.7369
184.11648559570312 0 1130.8555
185.09231567382812 0 1202.9535
187.1439208984375 0 1942.9668
192.1020050048828 0 894.88776
194.87054443359375 0 443.08292
195.11300659179688 0 951.53705
200.1393585205078 0 18695.303 a 1
201.14320373535156 0 1012.1766
202.11849975585938 0 1349.9076
203.10272216796875 0 1123.7189
210.1236572265625 0 9392.812
211.10789489746094 0 1177.3833 b Ammonia loss 1
211.12745666503906 0 812.37665
215.13832092285156 0 932.04974
228.13430786132812 0 7903.39 b 1
228.63172912597656 0 2505.5476 y 5
229.13731384277344 0 810.8371
230.08099365234375 0 1736.5143
235.14552307128906 0 542.51044
237.12315368652344 0 2789.723
238.12644958496094 0 855.30365
245.12420654296875 0 1334.8119 y Ammonia loss 7
247.10760498046875 0 5235.846
248.111083984375 0 774.52594
262.1507873535156 0 846.9712 y 7
265.1178894042969 0 2474.5847
282.1444091796875 0 1318.1881
299.1714172363281 0 2908.74
300.15533447265625 0 1611.2003
317.84649658203125 0 512.1554
325.17059326171875 0 607.58105
341.1934509277344 0 826.0806 y Water loss 6
342.1767883300781 0 4651.43 y Ammonia loss 6
346.1754150390625 0 632.8808 a Ammonia loss 2
359.203125 0 4934.6113 y 6
360.20855712890625 0 689.68317
373.1869201660156 0 2484.5066
374.1694030761719 0 1881.348 b Ammonia loss 2
391.19757080078125 0 1258.971 b 2
391.772705078125 0 583.739
396.2236633300781 0 2931.8425
397.2285461425781 0 571.1557
421.21759033203125 0 936.748
422.199951171875 0 581.1765
430.2071228027344 0 1068.81
437.95782470703125 0 561.4151
438.24578857421875 0 1078.2928 y Water loss 5
439.229736328125 0 7401.9062 y Ammonia loss 5
440.2317810058594 0 1003.7932
447.2238464355469 0 870.44727
456.25616455078125 0 51783.25 y 5
457.1749267578125 0 381.74338
457.2078857421875 0 11028.811
457.2589416503906 0 10760.527
458.2107849121094 0 3504.2585
458.2620849609375 0 1705.0968
474.23394775390625 0 2112.9119 b Water loss 3
475.2184753417969 0 4938.107 b Ammonia loss 3
476.2251281738281 0 1191.7646
512.7826538085938 0 1278.8202
513.279296875 0 713.1778
513.7737426757812 0 691.142
521.7867431640625 0 22925.691 Precursor Water loss
522.2881469726562 0 14799.466 Precursor Ammonia loss
522.7886352539062 0 3645.9746
530.7914428710938 0 1197.0148 Precursor
531.2945556640625 0 927.08624
542.2970581054688 0 1227.5471
543.2953491210938 0 699.80756
551.330078125 0 645.57275 y Water loss 4
562.296875 0 889.35297
569.3394165039062 0 8975.526 y 4
570.2911376953125 0 3169.2808
570.3424682617188 0 1886.9523
587.3187866210938 0 1910.4259 b Water loss 4
657.102294921875 0 600.666
670.3865966796875 0 13409.081 y 3
671.3897705078125 0 4882.9443
691.6048583984375 0 662.8535
713.8897094726562 0 656.12964
815.4390869140625 0 1737.126 y Water loss 2
816.443603515625 0 784.9202
833.4494018554688 0 28492.215 y 2
834.4518432617188 0 14176.606
835.4552612304688 0 2953.5188
932.5157470703125 0 1217.055 y 1
3079.583984375 0 823.21295

Spectrum Details

|  |  |
| --- | --- |
| Matched peaks? Matched peaks The total absolute number of peaks matched. Additionally in brackets the total fraction of peaks matched and the total number of peaks is shown. | 29 (23.58% of 123) |
| FDR? FDR The false discovery rate estimated for this peptide. It is calculated by matching all theoretical fragments with a non-integer shift with the raw peaks for this spectrum. This is done with 40 different shifts. The resulting percentage is the average number of annotated peaks over the number of annotated peaks with the correct spectrum. | 0.16% |
| Satellite FDR? Satellite FDR See the FDR for details on its calculation. This satellite ion specific FDR only contains the satellite ions (d/w) for I/L/J positions. | - |
| PSM Score? PSM Score The PSM Score as given by Hecklib to this annotated spectrum. It is shown with three significant figures. | 363 |

#### Reverse Lookup? Reverse Lookup All places where this read could be placed.

| Group | Segment | Template | Template Part | Read Part | Score | Unique |
| --- | --- | --- | --- | --- | --- | --- |
| Homo sapiens Heavy Chain | IGHC | IGHG1 | [229..238] | [0..9] | 72 | False |
| Homo sapiens Heavy Chain | IGHC | IGHG3 | [276..285] | [0..9] | 72 | False |
| Homo sapiens Heavy Chain | IGHC | IGHG2 | [225..234] | [0..9] | 72 | False |

| Recombined | Template Part | Read Part | Score | Unique |
| --- | --- | --- | --- | --- |
| REC-0-1 | [354..363] | [0..9] | 72 | True |

#### Meta Information from Multiple reads

##### Number of combined reads

3

##### Intensity

0.6056

##### TotalArea

1.148E+06

##### Changes to the peptide sequence

QVYTJPPSR

L→JNo support for either Leucine or Isoleucine based on side chain ions (Position: 5)

#### Positional Score

Copy Data

##### Positional Score (TSV)

###### Preview

```
Loading example...
```

*Click on the button to copy the data to your clipboard.*

10012345678

Label Value
"0" 0.287
"1" 0.31
"2" 0.33
"3" 0.33
"4" 0.333
"5" 0.33
"6" 0.323
"7" 0.327
"8" 0.33

#### Meta Information from PEAKS

##### Scan Identifier

F4:4836

##### Original sequence

Q

V

Y

T

L

P

P

S

R

##### Posttranslational Modifications

##### Source File

D:\separate\_stitch\_analyses\xle-disambiguation\raw\20210323\_F1\_UM1\_Peng0013\_SA\_F59\_ingel\_3ug\_tryp.raw

##### Fraction

4

##### Scan Feature

-

##### De Novo Score

98

##### ConfidenceScore

96

### m/z

530.7934

##### Mass

1059.5713

##### Charge

2

##### Retention Time

25.86

##### Predicted Retention Time

26.03

##### Area

0

##### Parts Per Million

0.9

##### Fragmentation mode

HCD

##### Originating file

01 D:\separate\_stitch\_analyses\xle-disambiguation\20210325\_F59\_3ug\_DENOVO\_12.csv

#### Meta Information from PEAKS

##### Scan Identifier

F4:4893

##### Original sequence

Q

V

Y

T

L

P

P

S

R

##### Posttranslational Modifications

##### Source File

D:\separate\_stitch\_analyses\xle-disambiguation\raw\20210323\_F1\_UM1\_Peng0013\_SA\_F59\_ingel\_3ug\_tryp.raw

##### Fraction

4

##### Scan Feature

F4:3899

##### De Novo Score

98

##### ConfidenceScore

95

### m/z

530.7932

##### Mass

1059.5713

##### Charge

2

##### Retention Time

25.93

##### Predicted Retention Time

26.03

##### Area

3.732E+05

##### Parts Per Million

0.6

##### Fragmentation mode

HCD

##### Originating file

01 D:\separate\_stitch\_analyses\xle-disambiguation\20210325\_F59\_3ug\_DENOVO\_12.csv

#### Meta Information from PEAKS

##### Scan Identifier

F4:4660

##### Original sequence

Q

V

Y

T

L

P

P

S

R

##### Posttranslational Modifications

##### Source File

D:\separate\_stitch\_analyses\xle-disambiguation\raw\20210323\_F1\_UM1\_Peng0013\_SA\_F59\_ingel\_3ug\_tryp.raw

##### Fraction

4

##### Scan Feature

F4:3898

##### De Novo Score

97

##### ConfidenceScore

96

### m/z

530.7932

##### Mass

1059.5713

##### Charge

2

##### Retention Time

25.11

##### Predicted Retention Time

26.03

##### Area

7.743E+05

##### Parts Per Million

0.6

##### Fragmentation mode

HCD

##### Originating file

01 D:\separate\_stitch\_analyses\xle-disambiguation\20210325\_F59\_3ug\_DENOVO\_12.csv
