## Supplementary material for "A handle on mass coincidence errors in *de novo* sequencing of antibodies by bottom-up proteomics": Combined_055.html

Details Combined\_055 | Stitch OverviewUndefined

### Read Combined\_055

#### Sequence (length=14)

JFPPSSEEJQANKA

#### Spectrum 6428? Spectrum 6428 The raw spectrum of this peptide as annotated by Hecklib. The fragments are coloured according to ion type (see legend). Any peaks with a star '\*' as text can be hovered over to see the full details, first the ion type second the mass shift type. By hovering over the amino acids in the peptide or ions in the legend the corresponding peaks are highlighted. By toggling the 'Unassigned' label you can turn the background (unassigned) peaks on or off in the plot. By updating the slider in the Ion legend you can update the spectrum to only show the top X% of the peaks with labels. The top X% means any peak that is within X% of the highest intensity. By dragging in the spectrum you can zoom in to a specific part of the spectrum and use 'Zoom Out' to get back to the original zoom level. The annotation of the spectrum is based on the given sequence in the peptides file and is done with different software so inconsistencies are likely. The peaks are annotated based on the given sequence, with 20 ppm tolerance.

Copy Data

##### Spectrum 6428 (TSV)

###### Preview

```
Loading example...
```

*Click on the button to copy the data to your clipboard.*

Mz MinMz MaxIntensity Max

WidthHeightPeptide font sizePeptide stroke widthSpectrum font sizeSpectrum stroke widthCompact peptide

Ion legend

wxyz

abcd

OtherUnassignedIon^Charge^_Position_Show for top:%

JFPPSSEEJQANKA

02.17e+64.35e+66.52e+68.70e+6

Zoom Out

y^+1^_2_y^+2^_6_y^+1^_3_y^+1^_4_z^+1^_5_y^+1^_5_w^+1^_6_y^+2^_11_c^+2^_11_z^+2^_12_y^+2^_12_y^+2^_12_z^+2^_12_c^+1^_6_y^+2^_12_y^+1^_6_w^+1^_7_z^+2^_13_y^+2^_13_c^+1^_7_y^+1^_7_w^+1^_8_z^+1^_8_y^+1^_8_c^+1^_8_y^+1^_8_c^+1^_8_w^+1^_9_y^+1^_9_y^+1^_9_z^+1^_9_y^+1^_9_c^+1^_9_z^+1^_10_y^+1^_10_c^+1^_10_c^+1^_10_y^+1^_11_y^+1^_11_c^+1^_11_c^+1^_11_y^+1^_12_c^+1^_12_c^+1^_12_z^+1^_13_c^+1^_13_

0773154523183090

Fragment Matches Table

Show background peaks

| Position | Ion type | Intensity | mz Theoretical | mz Error (Th) | mz Error (ppm) | Charge | Series Number |
| --- | --- | --- | --- | --- | --- | --- | --- |
| - | - | 4.185E+04 | 120.1 | - | - | 0 | - |
| - | - | 4863 | 120.1 | - | - | 0 | - |
| - | - | 6.708E+04 | 129.1 | - | - | 0 | - |
| - | - | 4160 | 153.5 | - | - | 0 | - |
| - | - | 4517 | 154 | - | - | 0 | - |
| - | - | 4884 | 175.8 | - | - | 0 | - |
| - | - | 5775 | 188.6 | - | - | 0 | - |
| - | - | 5537 | 207.9 | - | - | 0 | - |
| - | - | 5503 | 209.8 | - | - | 0 | - |
| 13 | y | 4.775E+04 | 218.1 | 0.0003659 | 1.677 | +1 | 2 |
| - | - | 5478 | 222.4 | - | - | 0 | - |
| - | - | 3.208E+04 | 226.1 | - | - | 0 | - |
| - | - | 3.91E+05 | 233.2 | - | - | 0 | - |
| - | - | 4.412E+04 | 234.2 | - | - | 0 | - |
| - | - | 2.527E+04 | 243.1 | - | - | 0 | - |
| - | - | 1.052E+04 | 245.1 | - | - | 0 | - |
| - | - | 5374 | 246.7 | - | - | 0 | - |
| - | - | 4.174E+05 | 261.2 | - | - | 0 | - |
| - | - | 6.253E+04 | 262.2 | - | - | 0 | - |
| - | - | 1.821E+04 | 282.1 | - | - | 0 | - |
| - | - | 5813 | 288.9 | - | - | 0 | - |
| - | - | 1.99E+04 | 297.2 | - | - | 0 | - |
| 9 | y | 1.433E+04 | 314.2 | 0.006167 | 19.63 | +2 | 6 |
| 12 | y | 3.427E+04 | 332.2 | 0.0004529 | 1.364 | +1 | 3 |
| - | - | 6.464E+04 | 358.2 | - | - | 0 | - |
| - | - | 1.513E+04 | 359.2 | - | - | 0 | - |
| - | - | 6788 | 400.8 | - | - | 0 | - |
| 11 | y | 6.209E+04 | 403.2 | 0.0006622 | 1.642 | +1 | 4 |
| - | - | 1.118E+04 | 404.2 | - | - | 0 | - |
| - | - | 6206 | 408.2 | - | - | 0 | - |
| - | - | 9177 | 425.2 | - | - | 0 | - |
| - | - | 9632 | 434.7 | - | - | 0 | - |
| - | - | 1.282E+04 | 455.3 | - | - | 0 | - |
| - | - | 1.131E+04 | 470.2 | - | - | 0 | - |
| - | - | 1.524E+04 | 498.2 | - | - | 0 | - |
| 10 | z | 9.843E+04 | 515.3 | 0.0006952 | 1.349 | +1 | 5 |
| - | - | 1.138E+05 | 516.3 | - | - | 0 | - |
| - | - | 2.686E+04 | 517.3 | - | - | 0 | - |
| 10 | y | 4.512E+04 | 531.3 | 0.001075 | 2.024 | +1 | 5 |
| - | - | 9616 | 532.3 | - | - | 0 | - |
| - | - | 7440 | 574.3 | - | - | 0 | - |
| - | - | 2.666E+04 | 582.3 | - | - | 0 | - |
| - | - | 5.269E+04 | 582.8 | - | - | 0 | - |
| - | - | 3.287E+04 | 583.3 | - | - | 0 | - |
| - | - | 1.267E+04 | 583.8 | - | - | 0 | - |
| 9 | w | 1.197E+04 | 585.3 | 0.0003343 | 0.5711 | +1 | 6 |
| 4 | y | 5.414E+04 | 587.3 | 0.0005619 | 0.9568 | +2 | 11 |
| - | - | 3.485E+04 | 587.8 | - | - | 0 | - |
| - | - | 1.123E+04 | 588.3 | - | - | 0 | - |
| - | - | 3.356E+04 | 591.3 | - | - | 0 | - |
| - | - | 2.18E+04 | 591.8 | - | - | 0 | - |
| 11 | c | 2.156E+04 | 600.3 | 0.0004854 | 0.8086 | +2 | 11 |
| - | - | 1.259E+04 | 600.8 | - | - | 0 | - |
| - | - | 1.037E+04 | 613.8 | - | - | 0 | - |
| - | - | 8210 | 617.8 | - | - | 0 | - |
| - | - | 1.905E+04 | 618.3 | - | - | 0 | - |
| 3 | z | 8488 | 618.8 | 0.006049 | 9.775 | +2 | 12 |
| - | - | 8051 | 620.3 | - | - | 0 | - |
| 3 | y | 6.539E+04 | 626.8 | 0.0007636 | 1.218 | +2 | 12 |
| - | - | 1.12E+04 | 627.3 | - | - | 0 | - |
| 3 | y | 1.492E+05 | 627.3 | 0.002896 | 4.617 | +2 | 12 |
| 3 | z | 8.544E+04 | 627.8 | 8.255E-05 | 0.1315 | +2 | 12 |
| - | - | 3.363E+04 | 628.3 | - | - | 0 | - |
| 6 | c | 3.631E+05 | 628.3 | 0.009779 | 15.56 | +1 | 6 |
| - | - | 1.527E+04 | 628.8 | - | - | 0 | - |
| - | - | 2.349E+05 | 629.4 | - | - | 0 | - |
| - | - | 5.96E+04 | 630.4 | - | - | 0 | - |
| - | - | 3.287E+04 | 634.8 | - | - | 0 | - |
| - | - | 4.089E+04 | 635.3 | - | - | 0 | - |
| 3 | y | 8.612E+06 | 635.8 | 0.001035 | 1.629 | +2 | 12 |
| - | - | 5.747E+06 | 636.3 | - | - | 0 | - |
| - | - | 2.037E+06 | 636.8 | - | - | 0 | - |
| - | - | 2.277E+05 | 637.3 | - | - | 0 | - |
| - | - | 7545 | 643.4 | - | - | 0 | - |
| 9 | y | 4.942E+04 | 644.4 | 0.0006903 | 1.071 | +1 | 6 |
| - | - | 1.461E+04 | 645.4 | - | - | 0 | - |
| - | - | 7023 | 671.7 | - | - | 0 | - |
| 8 | w | 1.173E+04 | 698.4 | 0.003099 | 4.438 | +1 | 7 |
| 2 | z | 1.635E+04 | 701.3 | 0.008551 | 12.19 | +2 | 13 |
| - | - | 8988 | 702.3 | - | - | 0 | - |
| 2 | y | 1.244E+05 | 709.4 | 0.000764 | 1.077 | +2 | 13 |
| - | - | 8.726E+04 | 709.9 | - | - | 0 | - |
| - | - | 4.598E+04 | 710.4 | - | - | 0 | - |
| - | - | 6612 | 730.9 | - | - | 0 | - |
| - | - | 3.464E+04 | 740.3 | - | - | 0 | - |
| - | - | 1.111E+04 | 741.3 | - | - | 0 | - |
| 7 | c | 4.848E+05 | 757.4 | 0.009178 | 12.12 | +1 | 7 |
| - | - | 1.079E+04 | 757.9 | - | - | 0 | - |
| - | - | 5.601E+05 | 758.4 | - | - | 0 | - |
| - | - | 1.834E+05 | 759.4 | - | - | 0 | - |
| - | - | 2.919E+04 | 760.4 | - | - | 0 | - |
| - | - | 2.192E+05 | 765.9 | - | - | 0 | - |
| - | - | 2.054E+05 | 766.4 | - | - | 0 | - |
| - | - | 9.164E+04 | 766.9 | - | - | 0 | - |
| - | - | 1.514E+04 | 767.4 | - | - | 0 | - |
| 8 | y | 7.33E+04 | 773.4 | 0.0003336 | 0.4313 | +1 | 7 |
| - | - | 2.807E+04 | 774.4 | - | - | 0 | - |
| 7 | w | 9982 | 827.4 | 0.003517 | 4.251 | +1 | 8 |
| - | - | 8.35E+04 | 868.4 | - | - | 0 | - |
| 7 | z | 2.914E+04 | 869.4 | 0.004799 | 5.52 | +1 | 8 |
| 7 | y | 1.723E+04 | 885.4 | 0.003149 | 3.557 | +1 | 8 |
| 8 | c | 3.326E+05 | 886.4 | 0.00931 | 10.5 | +1 | 8 |
| - | - | 8.886E+05 | 887.4 | - | - | 0 | - |
| - | - | 3.638E+05 | 888.4 | - | - | 0 | - |
| - | - | 8.3E+04 | 889.5 | - | - | 0 | - |
| - | - | 6.021E+04 | 901.5 | - | - | 0 | - |
| 7 | y | 6.229E+04 | 902.5 | 0.001976 | 2.19 | +1 | 8 |
| - | - | 6.022E+04 | 903.4 | - | - | 0 | - |
| 8 | c | 1.245E+05 | 904.4 | 0.0008814 | 0.9745 | +1 | 8 |
| - | - | 5.717E+04 | 905.4 | - | - | 0 | - |
| - | - | 1.039E+04 | 906.4 | - | - | 0 | - |
| - | - | 4.19E+04 | 939.4 | - | - | 0 | - |
| - | - | 3.67E+04 | 940.4 | - | - | 0 | - |
| - | - | 9526 | 941.4 | - | - | 0 | - |
| 6 | w | 3.265E+04 | 956.5 | 0.001311 | 1.37 | +1 | 9 |
| - | - | 1.218E+04 | 957.5 | - | - | 0 | - |
| - | - | 7107 | 958.5 | - | - | 0 | - |
| 6 | y | 1.891E+04 | 971.5 | 0.0152 | 15.65 | +1 | 9 |
| 6 | y | 2.234E+04 | 972.5 | 0.001517 | 1.559 | +1 | 9 |
| 6 | z | 7.876E+04 | 973.5 | 0.00242 | 2.485 | +1 | 9 |
| - | - | 1.249E+06 | 974.5 | - | - | 0 | - |
| - | - | 5.937E+05 | 975.5 | - | - | 0 | - |
| - | - | 1.556E+05 | 976.5 | - | - | 0 | - |
| - | - | 8663 | 977.5 | - | - | 0 | - |
| - | - | 2.368E+05 | 988.5 | - | - | 0 | - |
| 6 | y | 3.78E+05 | 989.5 | 0.001229 | 1.242 | +1 | 9 |
| - | - | 1.483E+05 | 990.5 | - | - | 0 | - |
| - | - | 3.42E+04 | 991.5 | - | - | 0 | - |
| - | - | 1.655E+04 | 1001 | - | - | 0 | - |
| - | - | 1.295E+04 | 1001 | - | - | 0 | - |
| - | - | 2.074E+05 | 1017 | - | - | 0 | - |
| 9 | c | 3.516E+05 | 1018 | 0.0009682 | 0.9516 | +1 | 9 |
| - | - | 1.859E+05 | 1019 | - | - | 0 | - |
| - | - | 3.559E+04 | 1020 | - | - | 0 | - |
| - | - | 1.083E+04 | 1036 | - | - | 0 | - |
| - | - | 7.215E+04 | 1053 | - | - | 0 | - |
| - | - | 3.575E+04 | 1054 | - | - | 0 | - |
| - | - | 8753 | 1055 | - | - | 0 | - |
| 5 | z | 3.165E+04 | 1061 | 0.0001763 | 0.1662 | +1 | 10 |
| - | - | 4.358E+05 | 1062 | - | - | 0 | - |
| - | - | 2.276E+05 | 1063 | - | - | 0 | - |
| - | - | 6.951E+04 | 1064 | - | - | 0 | - |
| - | - | 8923 | 1074 | - | - | 0 | - |
| - | - | 3.279E+04 | 1076 | - | - | 0 | - |
| 5 | y | 1.345E+05 | 1077 | 0.0003593 | 0.3338 | +1 | 10 |
| - | - | 6.935E+04 | 1078 | - | - | 0 | - |
| - | - | 1.475E+04 | 1079 | - | - | 0 | - |
| - | - | 1.18E+04 | 1085 | - | - | 0 | - |
| - | - | 7.307E+04 | 1102 | - | - | 0 | - |
| - | - | 5.528E+04 | 1103 | - | - | 0 | - |
| - | - | 2.614E+04 | 1104 | - | - | 0 | - |
| 10 | c | 2.387E+04 | 1129 | 0.002648 | 2.346 | +1 | 10 |
| - | - | 1.093E+04 | 1130 | - | - | 0 | - |
| - | - | 9864 | 1131 | - | - | 0 | - |
| - | - | 2.077E+05 | 1145 | - | - | 0 | - |
| 10 | c | 1.493E+06 | 1146 | 0.0002687 | 0.2346 | +1 | 10 |
| - | - | 9.609E+05 | 1147 | - | - | 0 | - |
| - | - | 2.69E+05 | 1148 | - | - | 0 | - |
| - | - | 2.869E+04 | 1149 | - | - | 0 | - |
| 4 | y | 1.387E+04 | 1157 | 0.005042 | 4.36 | +1 | 11 |
| - | - | 8718 | 1172 | - | - | 0 | - |
| - | - | 9.417E+04 | 1173 | - | - | 0 | - |
| 4 | y | 8.373E+05 | 1174 | 0.000954 | 0.8129 | +1 | 11 |
| - | - | 4.96E+05 | 1175 | - | - | 0 | - |
| - | - | 1.444E+05 | 1176 | - | - | 0 | - |
| - | - | 1.833E+04 | 1177 | - | - | 0 | - |
| - | - | 1.21E+04 | 1182 | - | - | 0 | - |
| 11 | c | 1.584E+04 | 1200 | 0.007732 | 6.446 | +1 | 11 |
| - | - | 9007 | 1201 | - | - | 0 | - |
| - | - | 1.891E+04 | 1202 | - | - | 0 | - |
| - | - | 1.091E+04 | 1215 | - | - | 0 | - |
| - | - | 1.79E+05 | 1216 | - | - | 0 | - |
| 11 | c | 2.326E+05 | 1217 | 0.002421 | 1.99 | +1 | 11 |
| - | - | 1.07E+05 | 1218 | - | - | 0 | - |
| - | - | 2.381E+04 | 1219 | - | - | 0 | - |
| - | - | 4.104E+04 | 1270 | - | - | 0 | - |
| 3 | y | 1.298E+06 | 1271 | 0.0002962 | 0.2331 | +1 | 12 |
| - | - | 8.832E+05 | 1272 | - | - | 0 | - |
| - | - | 2.824E+05 | 1273 | - | - | 0 | - |
| - | - | 6.167E+04 | 1274 | - | - | 0 | - |
| - | - | 2.093E+05 | 1287 | - | - | 0 | - |
| - | - | 1.642E+05 | 1288 | - | - | 0 | - |
| - | - | 8.227E+04 | 1289 | - | - | 0 | - |
| - | - | 2.941E+04 | 1299 | - | - | 0 | - |
| - | - | 2.549E+04 | 1300 | - | - | 0 | - |
| 12 | c | 1.625E+04 | 1314 | 0.001221 | 0.9298 | +1 | 12 |
| - | - | 1.205E+04 | 1315 | - | - | 0 | - |
| - | - | 8.43E+04 | 1330 | - | - | 0 | - |
| 12 | c | 6.712E+05 | 1331 | 0.0009151 | 0.6877 | +1 | 12 |
| - | - | 4.698E+05 | 1332 | - | - | 0 | - |
| - | - | 1.719E+05 | 1333 | - | - | 0 | - |
| - | - | 1.849E+04 | 1334 | - | - | 0 | - |
| - | - | 7258 | 1358 | - | - | 0 | - |
| 2 | z | 2.347E+05 | 1402 | 0.0003034 | 0.2165 | +1 | 13 |
| - | - | 1.874E+05 | 1403 | - | - | 0 | - |
| - | - | 7.43E+04 | 1404 | - | - | 0 | - |
| - | - | 1.299E+04 | 1405 | - | - | 0 | - |
| - | - | 2.198E+05 | 1415 | - | - | 0 | - |
| - | - | 1.822E+05 | 1416 | - | - | 0 | - |
| - | - | 7.431E+04 | 1417 | - | - | 0 | - |
| - | - | 9486 | 1418 | - | - | 0 | - |
| - | - | 6729 | 1420 | - | - | 0 | - |
| - | - | 8650 | 1423 | - | - | 0 | - |
| - | - | 9249 | 1428 | - | - | 0 | - |
| - | - | 8153 | 1430 | - | - | 0 | - |
| - | - | 3.017E+04 | 1431 | - | - | 0 | - |
| - | - | 3.01E+04 | 1432 | - | - | 0 | - |
| - | - | 1.392E+04 | 1433 | - | - | 0 | - |
| - | - | 5.001E+04 | 1443 | - | - | 0 | - |
| - | - | 4.71E+04 | 1444 | - | - | 0 | - |
| - | - | 4.607E+04 | 1445 | - | - | 0 | - |
| - | - | 1.101E+04 | 1446 | - | - | 0 | - |
| - | - | 8489 | 1447 | - | - | 0 | - |
| - | - | 1.177E+04 | 1457 | - | - | 0 | - |
| - | - | 1.207E+04 | 1458 | - | - | 0 | - |
| 13 | c | 1.513E+06 | 1459 | 0.00164 | 1.124 | +1 | 13 |
| - | - | 1.358E+06 | 1460 | - | - | 0 | - |
| - | - | 6.367E+05 | 1461 | - | - | 0 | - |
| - | - | 1.446E+05 | 1462 | - | - | 0 | - |
| - | - | 1.656E+04 | 1463 | - | - | 0 | - |
| - | - | 1.995E+04 | 1472 | - | - | 0 | - |
| - | - | 8.183E+04 | 1473 | - | - | 0 | - |
| - | - | 7.583E+04 | 1474 | - | - | 0 | - |
| - | - | 2.772E+04 | 1475 | - | - | 0 | - |
| - | - | 7.587E+04 | 1476 | - | - | 0 | - |
| - | - | 5.079E+04 | 1477 | - | - | 0 | - |
| - | - | 2.196E+04 | 1478 | - | - | 0 | - |
| - | - | 8996 | 1485 | - | - | 0 | - |
| - | - | 6.281E+04 | 1486 | - | - | 0 | - |
| - | - | 9.859E+05 | 1487 | - | - | 0 | - |
| - | - | 8.062E+05 | 1488 | - | - | 0 | - |
| - | - | 3.744E+05 | 1489 | - | - | 0 | - |
| - | - | 7.318E+04 | 1490 | - | - | 0 | - |
| - | - | 9349 | 1491 | - | - | 0 | - |
| - | - | 1.175E+04 | 1498 | - | - | 0 | - |
| - | - | 8562 | 1501 | - | - | 0 | - |
| - | - | 4.823E+04 | 1504 | - | - | 0 | - |
| - | - | 4.77E+04 | 1505 | - | - | 0 | - |
| - | - | 2.213E+04 | 1506 | - | - | 0 | - |
| - | - | 1.632E+04 | 1513 | - | - | 0 | - |
| - | - | 9.131E+04 | 1514 | - | - | 0 | - |
| - | - | 1.665E+06 | 1515 | - | - | 0 | - |
| - | - | 1.358E+06 | 1516 | - | - | 0 | - |
| - | - | 6.408E+05 | 1517 | - | - | 0 | - |
| - | - | 1.153E+05 | 1518 | - | - | 0 | - |
| - | - | 2.168E+04 | 1529 | - | - | 0 | - |
| - | - | 8.3E+04 | 1530 | - | - | 0 | - |
| - | - | 2.59E+06 | 1531 | - | - | 0 | - |
| - | - | 7.926E+06 | 1532 | - | - | 0 | - |
| - | - | 5.994E+06 | 1533 | - | - | 0 | - |
| - | - | 2.366E+06 | 1534 | - | - | 0 | - |
| - | - | 3.591E+05 | 1535 | - | - | 0 | - |
| - | - | 1.23E+04 | 1564 | - | - | 0 | - |
| - | - | 1.036E+04 | 1565 | - | - | 0 | - |
| - | - | 7443 | 3060 | - | - | 0 | - |

m/z Charge Intensity FragmentType MassShift Position
120.08110809326172 0 41850.605
120.0848159790039 0 4863.4985
129.1024627685547 0 67075.28
153.51133728027344 0 4160.386
154.0150146484375 0 4516.5703
175.8019256591797 0 4884.433
188.5968017578125 0 5774.856
207.8646240234375 0 5537.3667
209.7594757080078 0 5503.304
218.15028381347656 0 47750.348 y 12
222.3973846435547 0 5478.088
226.11904907226562 0 32079.936
233.1652374267578 0 391046.9
234.16842651367188 0 44116.984
243.14553833007812 0 25272.064
245.12892150878906 0 10516.4
246.70513916015625 0 5374.3765
261.16009521484375 0 417375.66
262.1632385253906 0 62526.883
282.14544677734375 0 18213.408
288.86749267578125 0 5813.1387
297.1562194824219 0 19897.094
314.1828308105469 0 14331.321 y Ammonia loss 8
332.19329833984375 0 34267.86 y 11
358.2127990722656 0 64642.652
359.21630859375 0 15126.258
400.8192443847656 0 6788.351
403.2306213378906 0 62085.1 y 10
404.2343444824219 0 11178.489
408.19097900390625 0 6205.9756
425.21490478515625 0 9176.783
434.7060852050781 0 9631.609
455.2653503417969 0 12823.457
470.2230529785156 0 11305.524
498.22064208984375 0 15239.704
515.2705078125 0 98427.305 z 9
516.277587890625 0 113770.52
517.2796020507812 0 26857.998
531.2896118164062 0 45120.13 y 9
532.2911376953125 0 9615.95
574.2689208984375 0 7439.9365
582.2886962890625 0 26660.768
582.7830810546875 0 52691
583.2821655273438 0 32867.402
583.782958984375 0 12666.308
585.2987670898438 0 11974.806 w 8
587.29150390625 0 54136.223 y 3
587.79248046875 0 34845.062
588.2931518554688 0 11226.324
591.29443359375 0 33559.5
591.7953491210938 0 21802.242
600.30029296875 0 21558.605 c Ammonia loss 10
600.801025390625 0 12585.167
613.80859375 0 10365.439
617.80810546875 0 8210.49
618.2984619140625 0 19046.162
618.796630859375 0 8487.941 z Water loss 2
620.3043823242188 0 8051.171
626.8128051757812 0 65392.66 y Water loss 2
627.2645874023438 0 11204.123
627.3069458007812 0 149184.83 y Ammonia loss 2
627.8080444335938 0 85436.79 z 2
628.3065185546875 0 33634.96
628.3551025390625 0 363082.3 c Water loss 5
628.809326171875 0 15270.892
629.360107421875 0 234921.44
630.3636474609375 0 59602.11
634.8110961914062 0 32871.77
635.312744140625 0 40885.51
635.818359375 0 8611731 y 2
636.31982421875 0 5746895
636.8206787109375 0 2037255.4
637.32177734375 0 227748.1
643.3624877929688 0 7544.901
644.373291015625 0 49416.105 y 8
645.375 0 14608.392
671.6641235351562 0 7022.882
698.3800659179688 0 11734.542 w 7
701.3336181640625 0 16352.443 z 1
702.337646484375 0 8987.743
709.352294921875 0 124373.42 y 1
709.8538208007812 0 87259.33
710.35400390625 0 45983.914
730.9102783203125 0 6612.0347
740.345947265625 0 34641.973
741.346435546875 0 11111.727
757.3970947265625 0 484805.03 c Water loss 6
757.8892822265625 0 10791.389
758.4033203125 0 560127.7
759.40673828125 0 183419.83
760.4100341796875 0 29185.357
765.8944091796875 0 219245.64
766.3956298828125 0 205390.62
766.8973388671875 0 91644.75
767.3976440429688 0 15135.906
773.41552734375 0 73295.35 y 7
774.4188842773438 0 28069.59
827.4222412109375 0 9982.446 w 6
868.4052124023438 0 83498.19
869.40771484375 0 29142.51 z Ammonia loss 6
885.4343872070312 0 17231.85 y Ammonia loss 6
886.4398193359375 0 332572.94 c Water loss 7
887.4464111328125 0 888646.8
888.4497680664062 0 363818.3
889.4530029296875 0 83002.914
901.4500732421875 0 60206.51
902.455810546875 0 62285.906 y 6
903.4420166015625 0 60224.684
904.4419555664062 0 124509.47 c 7
905.4447631835938 0 57165.473
906.4443969726562 0 10390.902
939.443115234375 0 41898.65
940.4454956054688 0 36704.62
941.44482421875 0 9525.963
956.467041015625 0 32653.994 w 5
957.4575805664062 0 12183.65
958.4725952148438 0 7107.0894
971.4640502929688 0 18905.701 y Water loss 5
972.4647827148438 0 22337.496 y Ammonia loss 5
973.4735107421875 0 78760.375 z 5
974.4794311523438 0 1248697.1
975.4827270507812 0 593742.9
976.4849853515625 0 155611.1
977.4833374023438 0 8662.641
988.4828491210938 0 236837.67
989.4885864257812 0 378041.3 y 5
990.4921875 0 148294.3
991.494140625 0 34196.004
1000.5042114257812 0 16554.287
1001.4949340820312 0 12945.099
1016.5156860351562 0 207378.1
1017.524169921875 0 351596.66 c 8
1018.52783203125 0 185934.77
1019.531005859375 0 35590.38
1036.4658203125 0 10828.085
1053.4862060546875 0 72146.06
1054.48779296875 0 35747.9
1055.4931640625 0 8753.477
1060.5032958984375 0 31645.502 z 4
1061.511474609375 0 435786.2
1062.514404296875 0 227561.95
1063.51806640625 0 69512.28
1073.5509033203125 0 8923.272
1075.517822265625 0 32790.582
1076.521484375 0 134507.92 y 4
1077.5262451171875 0 69345.59
1078.5289306640625 0 14747.859
1084.5250244140625 0 11800.052
1101.5706787109375 0 73067.77
1102.5736083984375 0 55282.32
1103.5753173828125 0 26135.088
1128.559814453125 0 23870.314 c Ammonia loss 9
1129.568603515625 0 10931.917
1130.5709228515625 0 9864.3125
1144.5767822265625 0 207689.75
1145.583984375 0 1492712.9 c 9
1146.587158203125 0 960944.7
1147.590087890625 0 269017.34
1148.5928955078125 0 28689.24
1156.5531005859375 0 13872.57 y Ammonia loss 3
1171.571533203125 0 8717.706
1172.56982421875 0 94174.2
1173.5755615234375 0 837264.8 y 3
1174.577880859375 0 495955.6
1175.580322265625 0 144353.42
1176.585205078125 0 18329.248
1181.57861328125 0 12099.303
1199.5865478515625 0 15836.443 c Ammonia loss 10
1200.584716796875 0 9007.462
1201.576416015625 0 18909.414
1214.6246337890625 0 10912.778
1215.6141357421875 0 178978.61
1216.618408203125 0 232588.22 c 10
1217.6209716796875 0 107008.69
1218.6275634765625 0 23813.215
1269.61767578125 0 41044.934
1270.6270751953125 0 1298177.5 y 2
1271.6295166015625 0 883234.2
1272.632080078125 0 282357.34
1273.633056640625 0 61673.75
1286.6490478515625 0 209279.78
1287.6531982421875 0 164241.69
1288.6552734375 0 82267.01
1298.621826171875 0 29408.64
1299.62548828125 0 25493.107
1313.635986328125 0 16249.133 c Ammonia loss 11
1314.6480712890625 0 12046.696
1329.655517578125 0 84300.33
1330.662841796875 0 671240.6 c 11
1331.665771484375 0 469788.9
1332.6685791015625 0 171908.73
1333.676025390625 0 18490.46
1357.7044677734375 0 7257.643
1401.6767578125 0 234704.33 z 1
1402.67919921875 0 187365.11
1403.681640625 0 74301.22
1404.6953125 0 12987.193
1414.7435302734375 0 219844.95
1415.744873046875 0 182164.4
1416.7471923828125 0 74307.016
1417.7320556640625 0 9486.189
1419.922607421875 0 6728.957
1422.5882568359375 0 8650.41
1427.740966796875 0 9248.65
1429.7156982421875 0 8153.3193
1430.7064208984375 0 30168.354
1431.70849609375 0 30102.13
1432.7099609375 0 13915.326
1442.7431640625 0 50009.855
1443.732666015625 0 47104.723
1444.7353515625 0 46068.17
1445.734619140625 0 11005.403
1446.7078857421875 0 8488.855
1456.74658203125 0 11765.636
1457.7283935546875 0 12070.847
1458.757080078125 0 1513058.5 c 12
1459.75927734375 0 1357899.5
1460.7603759765625 0 636692.6
1461.7586669921875 0 144637.77
1462.7469482421875 0 16555.564
1471.7235107421875 0 19949.172
1472.7427978515625 0 81827.52
1473.7513427734375 0 75834.766
1474.7564697265625 0 27724.846
1475.7286376953125 0 75870.89
1476.729736328125 0 50793.16
1477.7318115234375 0 21955.68
1484.7501220703125 0 8995.946
1485.7744140625 0 62813.17
1486.7659912109375 0 985865.8
1487.76904296875 0 806229
1488.7684326171875 0 374375.94
1489.7642822265625 0 73180.78
1490.74853515625 0 9349.483
1497.7486572265625 0 11753.105
1500.7435302734375 0 8562.185
1503.7916259765625 0 48227.58
1504.792236328125 0 47703.973
1505.794677734375 0 22126.28
1512.7432861328125 0 16319.597
1513.7713623046875 0 91313.36
1514.7611083984375 0 1665266.1
1515.7633056640625 0 1358071.2
1516.76513671875 0 640824.7
1517.765625 0 115304.92
1528.764892578125 0 21676.432
1529.7674560546875 0 83004.8
1530.778564453125 0 2589893
1531.786376953125 0 7926433
1532.7891845703125 0 5994169
1533.7913818359375 0 2366480
1534.79345703125 0 359105.62
1563.7984619140625 0 12296.377
1564.76220703125 0 10361.409
3059.637451171875 0 7442.7446

Spectrum Details

|  |  |
| --- | --- |
| Matched peaks? Matched peaks The total absolute number of peaks matched. Additionally in brackets the total fraction of peaks matched and the total number of peaks is shown. | 46 (18.04% of 255) |
| FDR? FDR The false discovery rate estimated for this peptide. It is calculated by matching all theoretical fragments with a non-integer shift with the raw peaks for this spectrum. This is done with 40 different shifts. The resulting percentage is the average number of annotated peaks over the number of annotated peaks with the correct spectrum. | 0.10% |
| Satellite FDR? Satellite FDR See the FDR for details on its calculation. This satellite ion specific FDR only contains the satellite ions (d/w) for I/L/J positions. | 0.00% |
| PSM Score? PSM Score The PSM Score as given by Hecklib to this annotated spectrum. It is shown with three significant figures. | 401 |

#### Spectrum 6165? Spectrum 6165 The raw spectrum of this peptide as annotated by Hecklib. The fragments are coloured according to ion type (see legend). Any peaks with a star '\*' as text can be hovered over to see the full details, first the ion type second the mass shift type. By hovering over the amino acids in the peptide or ions in the legend the corresponding peaks are highlighted. By toggling the 'Unassigned' label you can turn the background (unassigned) peaks on or off in the plot. By updating the slider in the Ion legend you can update the spectrum to only show the top X% of the peaks with labels. The top X% means any peak that is within X% of the highest intensity. By dragging in the spectrum you can zoom in to a specific part of the spectrum and use 'Zoom Out' to get back to the original zoom level. The annotation of the spectrum is based on the given sequence in the peptides file and is done with different software so inconsistencies are likely. The peaks are annotated based on the given sequence, with 20 ppm tolerance.

Copy Data

##### Spectrum 6165 (TSV)

###### Preview

```
Loading example...
```

*Click on the button to copy the data to your clipboard.*

Mz MinMz MaxIntensity Max

WidthHeightPeptide font sizePeptide stroke widthSpectrum font sizeSpectrum stroke widthCompact peptide

Ion legend

wxyz

abcd

OtherUnassignedIon^Charge^_Position_Show for top:%

JFPPSSEEJQANKA

04.69e+39.38e+31.41e+41.88e+4

Zoom Out

w^+1^_2_z^+1^_2_y^+1^_2_y^+2^_4_z^+1^_2_y^+1^_2_w^+1^_3_y^+2^_6_y^+1^_3_z^+2^_7_y^+1^_4_y^+2^_7_y^+1^_4_w^+1^_5_c^+2^_9_y^+1^_5_z^+1^_5_y^+1^_5_c^+1^_5_w^+1^_6_y^+2^_11_c^+2^_11_c^+2^_11_y^+2^_12_y^+2^_12_c^+1^_6_y^+2^_12_y^+1^_6_c^+1^_6_c^+2^_12_w^+1^_7_z^+2^_13_c^+2^_13_c^+1^_7_y^+1^_7_c^+1^_7_w^+1^_8_c^+1^_8_y^+1^_8_c^+1^_8_z^+1^_9_y^+1^_9_c^+1^_9_z^+1^_10_y^+1^_10_c^+1^_10_y^+1^_11_c^+1^_11_y^+1^_12_c^+1^_12_c^+1^_12_z^+1^_13_c^+1^_13_

0768153723053073

Fragment Matches Table

Show background peaks

| Position | Ion type | Intensity | mz Theoretical | mz Error (Th) | mz Error (ppm) | Charge | Series Number |
| --- | --- | --- | --- | --- | --- | --- | --- |
| - | - | 636.1 | 120.1 | - | - | 0 | - |
| - | - | 1895 | 120.1 | - | - | 0 | - |
| - | - | 568.2 | 126.1 | - | - | 0 | - |
| - | - | 1029 | 128.1 | - | - | 0 | - |
| - | - | 522.2 | 129.1 | - | - | 0 | - |
| - | - | 2360 | 129.1 | - | - | 0 | - |
| - | - | 369.5 | 129.8 | - | - | 0 | - |
| - | - | 1620 | 131.1 | - | - | 0 | - |
| - | - | 691.6 | 136.1 | - | - | 0 | - |
| - | - | 664.7 | 140.1 | - | - | 0 | - |
| - | - | 447.4 | 142.1 | - | - | 0 | - |
| 13 | w | 1215 | 144.1 | 0.0001052 | 0.7301 | +1 | 2 |
| - | - | 757.3 | 148.9 | - | - | 0 | - |
| - | - | 467.1 | 158.1 | - | - | 0 | - |
| - | - | 2184 | 159.1 | - | - | 0 | - |
| - | - | 549.3 | 160.1 | - | - | 0 | - |
| - | - | 561.1 | 165.1 | - | - | 0 | - |
| - | - | 417.3 | 165.6 | - | - | 0 | - |
| - | - | 464.7 | 166.1 | - | - | 0 | - |
| - | - | 478.3 | 171.1 | - | - | 0 | - |
| - | - | 702.9 | 173.1 | - | - | 0 | - |
| - | - | 702.6 | 173.4 | - | - | 0 | - |
| - | - | 2447 | 173.4 | - | - | 0 | - |
| - | - | 1657 | 173.5 | - | - | 0 | - |
| - | - | 458.6 | 173.5 | - | - | 0 | - |
| - | - | 2619 | 183.1 | - | - | 0 | - |
| - | - | 576 | 185.1 | - | - | 0 | - |
| 13 | z | 502.5 | 185.1 | 0.0001984 | 1.072 | +1 | 2 |
| - | - | 4556 | 187.1 | - | - | 0 | - |
| - | - | 1115 | 187.1 | - | - | 0 | - |
| - | - | 640.8 | 188.1 | - | - | 0 | - |
| - | - | 510.5 | 193.6 | - | - | 0 | - |
| - | - | 467.1 | 197.1 | - | - | 0 | - |
| 13 | y | 851.9 | 201.1 | 4.424E-05 | 0.22 | +1 | 2 |
| 11 | y | 632.2 | 202.1 | 0.003117 | 15.42 | +2 | 4 |
| 13 | z | 1982 | 202.1 | 0.0003192 | 1.579 | +1 | 2 |
| - | - | 1065 | 203.1 | - | - | 0 | - |
| - | - | 1840 | 212.1 | - | - | 0 | - |
| - | - | 543.9 | 213.4 | - | - | 0 | - |
| - | - | 482.1 | 215.1 | - | - | 0 | - |
| - | - | 1670 | 215.1 | - | - | 0 | - |
| - | - | 773.6 | 216.1 | - | - | 0 | - |
| 13 | y | 2327 | 218.1 | 0.0004428 | 2.03 | +1 | 2 |
| - | - | 958.1 | 219.1 | - | - | 0 | - |
| - | - | 2414 | 219.1 | - | - | 0 | - |
| - | - | 1345 | 220.1 | - | - | 0 | - |
| - | - | 893.5 | 221.1 | - | - | 0 | - |
| - | - | 523 | 221.8 | - | - | 0 | - |
| - | - | 474.1 | 227.5 | - | - | 0 | - |
| - | - | 997.5 | 229.1 | - | - | 0 | - |
| - | - | 702.3 | 229.2 | - | - | 0 | - |
| - | - | 645.3 | 232.1 | - | - | 0 | - |
| - | - | 792.6 | 233.2 | - | - | 0 | - |
| - | - | 4793 | 233.2 | - | - | 0 | - |
| - | - | 896 | 234.2 | - | - | 0 | - |
| - | - | 468.3 | 236.1 | - | - | 0 | - |
| - | - | 565.9 | 243.1 | - | - | 0 | - |
| - | - | 503.7 | 244 | - | - | 0 | - |
| - | - | 3086 | 261.2 | - | - | 0 | - |
| - | - | 578.7 | 270.1 | - | - | 0 | - |
| 12 | w | 2104 | 272.2 | 0.0005095 | 1.872 | +1 | 3 |
| - | - | 604.2 | 274.1 | - | - | 0 | - |
| - | - | 1326 | 283.1 | - | - | 0 | - |
| - | - | 1048 | 284.1 | - | - | 0 | - |
| - | - | 546.1 | 289.4 | - | - | 0 | - |
| - | - | 916.3 | 299.1 | - | - | 0 | - |
| - | - | 1.116E+04 | 299.2 | - | - | 0 | - |
| - | - | 1877 | 300.1 | - | - | 0 | - |
| - | - | 1656 | 300.2 | - | - | 0 | - |
| - | - | 1119 | 300.2 | - | - | 0 | - |
| - | - | 754 | 301.1 | - | - | 0 | - |
| - | - | 1565 | 301.2 | - | - | 0 | - |
| - | - | 747.3 | 302.2 | - | - | 0 | - |
| - | - | 655.5 | 311.1 | - | - | 0 | - |
| - | - | 569.6 | 311.3 | - | - | 0 | - |
| - | - | 1853 | 312.1 | - | - | 0 | - |
| - | - | 834.8 | 313.1 | - | - | 0 | - |
| - | - | 1076 | 322.2 | - | - | 0 | - |
| 9 | y | 631.9 | 322.7 | 0.001798 | 5.571 | +2 | 6 |
| - | - | 719.6 | 323.2 | - | - | 0 | - |
| - | - | 699.3 | 324.2 | - | - | 0 | - |
| - | - | 546.6 | 325.2 | - | - | 0 | - |
| - | - | 987.8 | 327.1 | - | - | 0 | - |
| - | - | 1194 | 328.1 | - | - | 0 | - |
| - | - | 606.5 | 329.1 | - | - | 0 | - |
| 12 | y | 1020 | 332.2 | 0.0007067 | 2.127 | +1 | 3 |
| - | - | 947.5 | 332.2 | - | - | 0 | - |
| - | - | 1208 | 343.2 | - | - | 0 | - |
| - | - | 746.8 | 357.2 | - | - | 0 | - |
| - | - | 1678 | 358.2 | - | - | 0 | - |
| - | - | 1403 | 359.2 | - | - | 0 | - |
| - | - | 788.9 | 360.2 | - | - | 0 | - |
| - | - | 917 | 371.2 | - | - | 0 | - |
| 8 | z | 581.9 | 379.2 | 0.00583 | 15.37 | +2 | 7 |
| 11 | y | 6105 | 386.2 | 0.0006513 | 1.686 | +1 | 4 |
| 8 | y | 1603 | 387.2 | 0.004387 | 11.33 | +2 | 7 |
| - | - | 1560 | 392.2 | - | - | 0 | - |
| - | - | 1210 | 393.2 | - | - | 0 | - |
| - | - | 835.2 | 395.2 | - | - | 0 | - |
| - | - | 563.5 | 401.3 | - | - | 0 | - |
| - | - | 1129 | 402.2 | - | - | 0 | - |
| 11 | y | 1476 | 403.2 | 0.002023 | 5.018 | +1 | 4 |
| - | - | 623.4 | 404.2 | - | - | 0 | - |
| - | - | 952 | 430.2 | - | - | 0 | - |
| - | - | 1004 | 449.3 | - | - | 0 | - |
| - | - | 6258 | 453.3 | - | - | 0 | - |
| - | - | 1019 | 454.3 | - | - | 0 | - |
| 10 | w | 1.076E+04 | 457.2 | 0.0001072 | 0.2345 | +1 | 5 |
| - | - | 2636 | 458.2 | - | - | 0 | - |
| - | - | 708.4 | 459.2 | - | - | 0 | - |
| - | - | 2073 | 467.3 | - | - | 0 | - |
| - | - | 700.1 | 468.2 | - | - | 0 | - |
| - | - | 597.8 | 473.3 | - | - | 0 | - |
| - | - | 588.5 | 476.2 | - | - | 0 | - |
| - | - | 3243 | 488.3 | - | - | 0 | - |
| - | - | 677.2 | 488.3 | - | - | 0 | - |
| - | - | 914.1 | 489.3 | - | - | 0 | - |
| - | - | 636.3 | 492.3 | - | - | 0 | - |
| - | - | 848.3 | 494.2 | - | - | 0 | - |
| - | - | 657.9 | 494.3 | - | - | 0 | - |
| - | - | 790.3 | 495.3 | - | - | 0 | - |
| 9 | c | 1884 | 509.3 | 0.002719 | 5.338 | +2 | 9 |
| - | - | 936.1 | 510.2 | - | - | 0 | - |
| - | - | 3906 | 510.3 | - | - | 0 | - |
| - | - | 3620 | 510.3 | - | - | 0 | - |
| - | - | 961.4 | 510.4 | - | - | 0 | - |
| - | - | 756.9 | 511.3 | - | - | 0 | - |
| - | - | 732.7 | 511.3 | - | - | 0 | - |
| - | - | 770.2 | 511.4 | - | - | 0 | - |
| - | - | 2456 | 512.2 | - | - | 0 | - |
| - | - | 2044 | 512.3 | - | - | 0 | - |
| 10 | y | 1549 | 514.3 | 0.002344 | 4.558 | +1 | 5 |
| 10 | z | 892.7 | 515.3 | 0.002343 | 4.547 | +1 | 5 |
| 10 | y | 2948 | 531.3 | 0.0006949 | 1.308 | +1 | 5 |
| - | - | 586.1 | 545 | - | - | 0 | - |
| 5 | c | 1861 | 559.3 | 0.002021 | 3.614 | +1 | 5 |
| - | - | 995 | 560.3 | - | - | 0 | - |
| - | - | 1069 | 570.3 | - | - | 0 | - |
| - | - | 650.7 | 582.8 | - | - | 0 | - |
| 9 | w | 1.368E+04 | 585.3 | 0.001494 | 2.552 | +1 | 6 |
| - | - | 4137 | 586.3 | - | - | 0 | - |
| 4 | y | 1218 | 587.3 | 0.006726 | 11.45 | +2 | 11 |
| - | - | 784.1 | 591.3 | - | - | 0 | - |
| 11 | c | 2546 | 599.8 | 0.003229 | 5.383 | +2 | 11 |
| 11 | c | 2239 | 600.3 | 0.004214 | 7.02 | +2 | 11 |
| - | - | 1093 | 600.8 | - | - | 0 | - |
| - | - | 965.1 | 610.3 | - | - | 0 | - |
| - | - | 1048 | 614.3 | - | - | 0 | - |
| - | - | 823.8 | 614.8 | - | - | 0 | - |
| - | - | 1917 | 616.3 | - | - | 0 | - |
| - | - | 668.3 | 617.3 | - | - | 0 | - |
| - | - | 665.7 | 618.3 | - | - | 0 | - |
| 3 | y | 1312 | 626.8 | 0.004852 | 7.74 | +2 | 12 |
| - | - | 2598 | 627.3 | - | - | 0 | - |
| 3 | y | 1555 | 627.3 | 0.003629 | 5.785 | +2 | 12 |
| 6 | c | 2110 | 628.3 | 0.008314 | 13.23 | +1 | 6 |
| - | - | 1744 | 635.3 | - | - | 0 | - |
| 3 | y | 1.737E+04 | 635.8 | 0.001406 | 2.211 | +2 | 12 |
| - | - | 1.14E+04 | 636.3 | - | - | 0 | - |
| - | - | 5366 | 636.8 | - | - | 0 | - |
| - | - | 1925 | 637.3 | - | - | 0 | - |
| 9 | y | 1507 | 644.4 | 0.002545 | 3.949 | +1 | 6 |
| 6 | c | 9447 | 646.4 | 0.001579 | 2.443 | +1 | 6 |
| - | - | 3496 | 647.4 | - | - | 0 | - |
| 12 | c | 1234 | 657.3 | 0.003883 | 5.907 | +2 | 12 |
| - | - | 780.9 | 657.8 | - | - | 0 | - |
| - | - | 784.4 | 660.3 | - | - | 0 | - |
| - | - | 4711 | 673.4 | - | - | 0 | - |
| - | - | 787.5 | 674 | - | - | 0 | - |
| - | - | 1458 | 679.3 | - | - | 0 | - |
| - | - | 1220 | 679.7 | - | - | 0 | - |
| - | - | 1581 | 680 | - | - | 0 | - |
| - | - | 1425 | 680.3 | - | - | 0 | - |
| 8 | w | 6965 | 698.4 | 0.002062 | 2.952 | +1 | 7 |
| - | - | 3887 | 699.4 | - | - | 0 | - |
| 2 | z | 2211 | 701.3 | 0.006659 | 9.494 | +2 | 13 |
| - | - | 972.5 | 701.8 | - | - | 0 | - |
| - | - | 1794 | 702.3 | - | - | 0 | - |
| - | - | 823.7 | 708.4 | - | - | 0 | - |
| - | - | 1073 | 714.3 | - | - | 0 | - |
| - | - | 1021 | 714.9 | - | - | 0 | - |
| - | - | 768.4 | 725.4 | - | - | 0 | - |
| 13 | c | 1.041E+04 | 729.9 | 0.001773 | 2.429 | +2 | 13 |
| - | - | 1.017E+04 | 730.4 | - | - | 0 | - |
| - | - | 6540 | 730.9 | - | - | 0 | - |
| - | - | 2492 | 731.4 | - | - | 0 | - |
| - | - | 620.2 | 733.8 | - | - | 0 | - |
| - | - | 959.9 | 736.4 | - | - | 0 | - |
| - | - | 1011 | 736.9 | - | - | 0 | - |
| - | - | 859.9 | 738.4 | - | - | 0 | - |
| - | - | 1493 | 740.3 | - | - | 0 | - |
| - | - | 2319 | 743.9 | - | - | 0 | - |
| - | - | 2773 | 744.4 | - | - | 0 | - |
| - | - | 1310 | 744.9 | - | - | 0 | - |
| - | - | 723.3 | 752.4 | - | - | 0 | - |
| 7 | c | 9316 | 757.4 | 0.00631 | 8.331 | +1 | 7 |
| - | - | 1423 | 757.9 | - | - | 0 | - |
| - | - | 4541 | 758.4 | - | - | 0 | - |
| - | - | 1455 | 759.4 | - | - | 0 | - |
| - | - | 599.6 | 763.9 | - | - | 0 | - |
| - | - | 1039 | 764.4 | - | - | 0 | - |
| - | - | 7033 | 765.9 | - | - | 0 | - |
| - | - | 776.6 | 766.3 | - | - | 0 | - |
| - | - | 1.397E+04 | 766.4 | - | - | 0 | - |
| - | - | 1.011E+04 | 766.9 | - | - | 0 | - |
| - | - | 892.1 | 767.1 | - | - | 0 | - |
| - | - | 4855 | 767.4 | - | - | 0 | - |
| - | - | 897.4 | 767.9 | - | - | 0 | - |
| 8 | y | 982.8 | 773.4 | 0.0007651 | 0.9892 | +1 | 7 |
| 7 | c | 1.733E+04 | 775.4 | 0.001386 | 1.788 | +1 | 7 |
| - | - | 6565 | 776.4 | - | - | 0 | - |
| - | - | 1463 | 777.4 | - | - | 0 | - |
| - | - | 805.6 | 801.4 | - | - | 0 | - |
| - | - | 931.5 | 809.4 | - | - | 0 | - |
| - | - | 972.5 | 812.4 | - | - | 0 | - |
| - | - | 762.7 | 817.8 | - | - | 0 | - |
| 7 | w | 1921 | 827.4 | 0.002174 | 2.628 | +1 | 8 |
| - | - | 1774 | 828.4 | - | - | 0 | - |
| - | - | 960.4 | 861.4 | - | - | 0 | - |
| - | - | 995.3 | 862.4 | - | - | 0 | - |
| - | - | 825.8 | 868.4 | - | - | 0 | - |
| - | - | 774.1 | 876.9 | - | - | 0 | - |
| 8 | c | 1.213E+04 | 886.4 | 0.007784 | 8.781 | +1 | 8 |
| - | - | 6233 | 887.4 | - | - | 0 | - |
| - | - | 1814 | 888.4 | - | - | 0 | - |
| - | - | 1117 | 892.4 | - | - | 0 | - |
| 7 | y | 2949 | 902.5 | 0.01547 | 17.14 | +1 | 8 |
| - | - | 964 | 903.4 | - | - | 0 | - |
| 8 | c | 1.858E+04 | 904.4 | 0.002048 | 2.265 | +1 | 8 |
| - | - | 694 | 904.9 | - | - | 0 | - |
| - | - | 8377 | 905.4 | - | - | 0 | - |
| - | - | 2882 | 906.4 | - | - | 0 | - |
| - | - | 792 | 907.4 | - | - | 0 | - |
| - | - | 803.9 | 909.5 | - | - | 0 | - |
| - | - | 734.1 | 918.4 | - | - | 0 | - |
| 6 | z | 9496 | 973.5 | 0.00344 | 3.534 | +1 | 9 |
| - | - | 6754 | 974.5 | - | - | 0 | - |
| - | - | 1737 | 975.5 | - | - | 0 | - |
| - | - | 1120 | 981.5 | - | - | 0 | - |
| - | - | 1190 | 982.5 | - | - | 0 | - |
| 6 | y | 5270 | 989.5 | 0.00306 | 3.092 | +1 | 9 |
| - | - | 2535 | 990.5 | - | - | 0 | - |
| - | - | 723.5 | 997 | - | - | 0 | - |
| - | - | 1196 | 997.5 | - | - | 0 | - |
| - | - | 1007 | 998.5 | - | - | 0 | - |
| - | - | 1226 | 1001 | - | - | 0 | - |
| - | - | 2405 | 1002 | - | - | 0 | - |
| - | - | 1634 | 1003 | - | - | 0 | - |
| - | - | 929.4 | 1003 | - | - | 0 | - |
| - | - | 1951 | 1003 | - | - | 0 | - |
| - | - | 1050 | 1004 | - | - | 0 | - |
| - | - | 1500 | 1007 | - | - | 0 | - |
| - | - | 1498 | 1010 | - | - | 0 | - |
| - | - | 1610 | 1011 | - | - | 0 | - |
| - | - | 1640 | 1011 | - | - | 0 | - |
| - | - | 2039 | 1012 | - | - | 0 | - |
| - | - | 1313 | 1012 | - | - | 0 | - |
| - | - | 1338 | 1013 | - | - | 0 | - |
| - | - | 814.7 | 1013 | - | - | 0 | - |
| - | - | 833 | 1015 | - | - | 0 | - |
| - | - | 814.6 | 1016 | - | - | 0 | - |
| - | - | 834.2 | 1017 | - | - | 0 | - |
| 9 | c | 1.046E+04 | 1018 | 0.002067 | 2.031 | +1 | 9 |
| - | - | 7424 | 1019 | - | - | 0 | - |
| - | - | 1808 | 1019 | - | - | 0 | - |
| - | - | 4688 | 1020 | - | - | 0 | - |
| - | - | 2208 | 1020 | - | - | 0 | - |
| - | - | 3142 | 1020 | - | - | 0 | - |
| - | - | 1079 | 1021 | - | - | 0 | - |
| - | - | 2150 | 1021 | - | - | 0 | - |
| - | - | 872.2 | 1022 | - | - | 0 | - |
| - | - | 1207 | 1022 | - | - | 0 | - |
| - | - | 1337 | 1022 | - | - | 0 | - |
| - | - | 1318 | 1023 | - | - | 0 | - |
| - | - | 1026 | 1023 | - | - | 0 | - |
| - | - | 1703 | 1023 | - | - | 0 | - |
| - | - | 903.8 | 1024 | - | - | 0 | - |
| 5 | z | 2531 | 1061 | 0.003242 | 3.057 | +1 | 10 |
| - | - | 1693 | 1062 | - | - | 0 | - |
| 5 | y | 1130 | 1077 | 0.002204 | 2.047 | +1 | 10 |
| - | - | 1097 | 1078 | - | - | 0 | - |
| - | - | 1063 | 1113 | - | - | 0 | - |
| - | - | 769.3 | 1140 | - | - | 0 | - |
| 10 | c | 6274 | 1146 | 0.004614 | 4.028 | +1 | 10 |
| - | - | 4417 | 1147 | - | - | 0 | - |
| - | - | 1480 | 1148 | - | - | 0 | - |
| 4 | y | 2280 | 1174 | 0.001243 | 1.059 | +1 | 11 |
| - | - | 760.4 | 1175 | - | - | 0 | - |
| - | - | 710.7 | 1197 | - | - | 0 | - |
| - | - | 1248 | 1197 | - | - | 0 | - |
| 11 | c | 1.382E+04 | 1217 | 0.002543 | 2.09 | +1 | 11 |
| - | - | 8771 | 1218 | - | - | 0 | - |
| - | - | 3144 | 1219 | - | - | 0 | - |
| - | - | 1069 | 1220 | - | - | 0 | - |
| - | - | 880.8 | 1226 | - | - | 0 | - |
| 3 | y | 1640 | 1271 | 0.00762 | 5.997 | +1 | 12 |
| - | - | 798.1 | 1272 | - | - | 0 | - |
| - | - | 963.7 | 1283 | - | - | 0 | - |
| 12 | c | 883.7 | 1313 | 0.02416 | 18.41 | +1 | 12 |
| 12 | c | 1.838E+04 | 1331 | 0.004577 | 3.44 | +1 | 12 |
| - | - | 1.382E+04 | 1332 | - | - | 0 | - |
| - | - | 6842 | 1333 | - | - | 0 | - |
| - | - | 1946 | 1334 | - | - | 0 | - |
| 2 | z | 2627 | 1402 | 0.005308 | 3.787 | +1 | 13 |
| - | - | 8155 | 1403 | - | - | 0 | - |
| - | - | 6371 | 1404 | - | - | 0 | - |
| - | - | 1947 | 1405 | - | - | 0 | - |
| - | - | 983.7 | 1416 | - | - | 0 | - |
| - | - | 1099 | 1443 | - | - | 0 | - |
| - | - | 757.8 | 1445 | - | - | 0 | - |
| 13 | c | 1649 | 1459 | 0.009574 | 6.563 | +1 | 13 |
| - | - | 4082 | 1460 | - | - | 0 | - |
| - | - | 2732 | 1461 | - | - | 0 | - |
| - | - | 1217 | 1462 | - | - | 0 | - |
| - | - | 928.2 | 1487 | - | - | 0 | - |
| - | - | 3955 | 1488 | - | - | 0 | - |
| - | - | 3555 | 1489 | - | - | 0 | - |
| - | - | 1407 | 1490 | - | - | 0 | - |
| - | - | 748 | 1491 | - | - | 0 | - |
| - | - | 1088 | 1502 | - | - | 0 | - |
| - | - | 974.1 | 1503 | - | - | 0 | - |
| - | - | 1217 | 1504 | - | - | 0 | - |
| - | - | 1465 | 1505 | - | - | 0 | - |
| - | - | 853.6 | 1511 | - | - | 0 | - |
| - | - | 1214 | 1512 | - | - | 0 | - |
| - | - | 1782 | 1515 | - | - | 0 | - |
| - | - | 9305 | 1516 | - | - | 0 | - |
| - | - | 7074 | 1517 | - | - | 0 | - |
| - | - | 3251 | 1518 | - | - | 0 | - |
| - | - | 1096 | 1519 | - | - | 0 | - |
| - | - | 981.9 | 1525 | - | - | 0 | - |
| - | - | 1659 | 1525 | - | - | 0 | - |
| - | - | 936.2 | 1526 | - | - | 0 | - |
| - | - | 1163 | 1526 | - | - | 0 | - |
| - | - | 1432 | 1530 | - | - | 0 | - |
| - | - | 775.9 | 1531 | - | - | 0 | - |
| - | - | 4474 | 1532 | - | - | 0 | - |
| - | - | 1.123E+04 | 1533 | - | - | 0 | - |
| - | - | 8220 | 1534 | - | - | 0 | - |
| - | - | 1454 | 1534 | - | - | 0 | - |
| - | - | 3679 | 1535 | - | - | 0 | - |
| - | - | 1048 | 1536 | - | - | 0 | - |
| - | - | 686.8 | 1796 | - | - | 0 | - |
| - | - | 812.2 | 2037 | - | - | 0 | - |
| - | - | 783 | 2038 | - | - | 0 | - |
| - | - | 816.1 | 2040 | - | - | 0 | - |
| - | - | 847 | 2488 | - | - | 0 | - |
| - | - | 856.1 | 3043 | - | - | 0 | - |

m/z Charge Intensity FragmentType MassShift Position
120.06511688232422 0 636.1286
120.08055877685547 0 1894.9867
126.05491638183594 0 568.1615
128.0943145751953 0 1028.5577
129.05462646484375 0 522.191
129.10202026367188 0 2359.755
129.7900848388672 0 369.4679
131.117431640625 0 1620.4604
136.07542419433594 0 691.5757
140.0817413330078 0 664.6743
142.09671020507812 0 447.3723
144.06541442871094 0 1214.8364 w 12
148.94728088378906 0 757.3419
158.0921173095703 0 467.06238
159.11245727539062 0 2183.622
160.11624145507812 0 549.323
165.1018829345703 0 561.0824
165.61183166503906 0 417.27066
166.06118774414062 0 464.73618
171.07632446289062 0 478.32773
173.12879943847656 0 702.89246
173.4287872314453 0 702.56757
173.43881225585938 0 2447.208
173.4536895751953 0 1656.8025
173.46563720703125 0 458.62363
183.1122589111328 0 2619.1084
185.09130859375 0 575.9617
185.10484313964844 0 502.48135 z Ammonia loss 12
187.10733032226562 0 4556.027
187.14402770996094 0 1114.8221
188.1110076904297 0 640.7783
193.61538696289062 0 510.4538
197.1287841796875 0 467.05963
201.1234130859375 0 851.8609 y Ammonia loss 12
202.12173461914062 0 632.19214 y 10
202.13087463378906 0 1981.9236 z 12
203.10220336914062 0 1065.3157
212.13873291015625 0 1840.2611
213.35122680664062 0 543.87714
215.1017608642578 0 482.06573
215.1384735107422 0 1669.823
216.1420440673828 0 773.61444
218.14947509765625 0 2327.106 y 12
219.09698486328125 0 958.1064
219.13348388671875 0 2414.393
220.1373748779297 0 1345.2743
221.13894653320312 0 893.53204
221.78599548339844 0 522.9559
227.4549560546875 0 474.08786
229.10885620117188 0 997.48395
229.154052734375 0 702.2742
232.12913513183594 0 645.28906
233.15139770507812 0 792.58795
233.1643829345703 0 4792.995
234.1676483154297 0 895.9834
236.07528686523438 0 468.33218
243.1331787109375 0 565.90216
243.9796905517578 0 503.66977
261.15899658203125 0 3086.4658
270.1451721191406 0 578.65356
272.15997314453125 0 2104.4697 w 11
274.1385498046875 0 604.15497
283.14495849609375 0 1325.5972
284.1465148925781 0 1047.6636
289.39697265625 0 546.1233
299.1401672363281 0 916.27673
299.1708984375 0 11163.618
300.1419677734375 0 1877.2328
300.1741027832031 0 1656.0297
300.1919250488281 0 1119.2184
301.1442565917969 0 754.01984
301.19439697265625 0 1564.8452
302.1960144042969 0 747.289
311.1405334472656 0 655.4689
311.259033203125 0 569.632
312.14068603515625 0 1852.8612
313.1444091796875 0 834.7969
322.1601867675781 0 1076.1194
322.6881408691406 0 631.869 y 8
323.16326904296875 0 719.62823
324.1658935546875 0 699.3339
325.1864318847656 0 546.59796
327.13433837890625 0 987.8014
328.1365051269531 0 1194.3846
329.13970947265625 0 606.4868
332.192138671875 0 1020.0372 y 11
332.2152404785156 0 947.46893
343.197509765625 0 1207.959
357.1756591796875 0 746.81824
358.18048095703125 0 1678.459
359.1814270019531 0 1403.2843
360.2139587402344 0 788.94324
371.2017822265625 0 916.9841
379.20770263671875 0 581.91534 z 7
386.2027587890625 0 6105.2715 y Ammonia loss 10
387.20684814453125 0 1602.9226 y 7
392.1842041015625 0 1560.0486
393.1863098144531 0 1209.6162
395.2276306152344 0 835.19336
401.33648681640625 0 563.49536
402.2420349121094 0 1129.031
403.2279357910156 0 1475.8069 y 10
404.23614501953125 0 623.4008
430.2313537597656 0 952.0261
449.3115234375 0 1004.3765
453.3427429199219 0 6257.8335
454.34844970703125 0 1018.9537
457.2406311035156 0 10758.611 w 9
458.2423400878906 0 2635.6443
459.24725341796875 0 708.3937
467.3226623535156 0 2073.3276
468.2416076660156 0 700.0932
473.2977600097656 0 597.7859
476.23040771484375 0 588.54987
488.2726135253906 0 3243.1558
488.3102111816406 0 677.16
489.27252197265625 0 914.08496
492.2539978027344 0 636.298
494.22381591796875 0 848.3349
494.26171875 0 657.85095
495.302978515625 0 790.29565
509.26348876953125 0 1884.309 c 8
510.2132263183594 0 936.0989
510.2536315917969 0 3905.591
510.28863525390625 0 3619.8594
510.3761901855469 0 961.44116
511.2590026855469 0 756.86896
511.2923583984375 0 732.6669
511.3923645019531 0 770.19446
512.2317504882812 0 2455.827
512.2704467773438 0 2044.1167
514.2596435546875 0 1549.3146 y Ammonia loss 9
515.2721557617188 0 892.7398 z 9
531.287841796875 0 2948.1204 y 9
545.0243530273438 0 586.05304
559.3218383789062 0 1861.1565 c 4
560.3272094726562 0 995.00903
570.323486328125 0 1068.6166
582.78173828125 0 650.68134
585.297607421875 0 13683.75 w 8
586.3005981445312 0 4137.0024
587.2976684570312 0 1218.3524 y 3
591.2910766601562 0 784.07745
599.8055419921875 0 2545.6096 c Water loss 10
600.3049926757812 0 2239.1492 c Ammonia loss 10
600.80859375 0 1092.9978
610.334228515625 0 965.0541
614.312744140625 0 1047.9739
614.812744140625 0 823.83014
616.32958984375 0 1916.5212
617.3338012695312 0 668.2912
618.3004150390625 0 665.70654
626.8071899414062 0 1311.8391 y Water loss 2
627.2601318359375 0 2598.3816
627.3076782226562 0 1554.9438 y Ammonia loss 2
628.3536376953125 0 2110.47 c Water loss 5
635.31201171875 0 1743.806
635.81591796875 0 17368.12 y 2
636.3170776367188 0 11397.165
636.8190307617188 0 5366.0317
637.32177734375 0 1924.9099
644.3700561523438 0 1507.1182 y 8
646.3543090820312 0 9447.472 c 5
647.3580322265625 0 3496.1907
657.318359375 0 1233.7676 c Ammonia loss 11
657.8147583007812 0 780.9194
660.3355712890625 0 784.3614
673.3522338867188 0 4710.694
673.99365234375 0 787.4531
679.3345947265625 0 1458.4574
679.667724609375 0 1219.8829
679.9994506835938 0 1581.2852
680.3301391601562 0 1424.8104
698.381103515625 0 6964.7036 w 7
699.383056640625 0 3886.7554
701.3355102539062 0 2210.759 z 1
701.842529296875 0 972.4888
702.3397216796875 0 1794.2325
708.3736572265625 0 823.69806
714.3389282226562 0 1073.4192
714.8822631835938 0 1020.6005
725.3538208007812 0 768.3872
729.8812255859375 0 10408.3 c 12
730.3824462890625 0 10168.22
730.8847045898438 0 6539.8086
731.3848266601562 0 2491.9648
733.8490600585938 0 620.22314
736.3533935546875 0 959.8567
736.8634643554688 0 1010.7914
738.3656616210938 0 859.8503
740.3455810546875 0 1493.2887
743.8858642578125 0 2319.3486
744.3858032226562 0 2772.8938
744.8882446289062 0 1309.8729
752.3522338867188 0 723.273
757.3942260742188 0 9315.705 c Water loss 6
757.893310546875 0 1423.2319
758.3948974609375 0 4540.8833
759.3960571289062 0 1455.1755
763.9171142578125 0 599.61237
764.423828125 0 1039.4889
765.890625 0 7033.204
766.3212280273438 0 776.5806
766.3939208984375 0 13969.863
766.8963623046875 0 10106.067
767.1091918945312 0 892.1487
767.3980102539062 0 4855.0796
767.9058227539062 0 897.3924
773.4144287109375 0 982.8491 y 7
775.3970947265625 0 17333.66 c 6
776.3995361328125 0 6565.139
777.4048461914062 0 1463.4578
801.4121704101562 0 805.57477
809.3800048828125 0 931.5071
812.38330078125 0 972.5321
817.8133544921875 0 762.6629
827.423583984375 0 1920.5823 w 6
828.4286499023438 0 1774.2622
861.3902587890625 0 960.42114
862.3981323242188 0 995.26
868.4058227539062 0 825.7956
876.8848876953125 0 774.115
886.4382934570312 0 12127.503 c Water loss 7
887.4367065429688 0 6232.7437
888.4354248046875 0 1813.5817
892.4140014648438 0 1116.878
902.4423217773438 0 2949.2595 y 6
903.4456176757812 0 963.971
904.4390258789062 0 18580.428 c 7
904.9235229492188 0 694.0493
905.4408569335938 0 8376.723
906.4443359375 0 2882.4946
907.4465942382812 0 792.01825
909.4909057617188 0 803.9064
918.4264526367188 0 734.07806
973.4676513671875 0 9495.525 z 5
974.4727172851562 0 6754.1133
975.473388671875 0 1736.7217
981.49365234375 0 1119.7197
982.4865112304688 0 1189.6953
989.4867553710938 0 5270.152 y 5
990.493896484375 0 2534.7517
996.9825439453125 0 723.48596
997.4605102539062 0 1196.3936
998.4817504882812 0 1007.1049
1001.4764404296875 0 1226.3717
1002.4644165039062 0 2405.2188
1002.7939453125 0 1633.753
1003.1292724609375 0 929.4099
1003.471923828125 0 1951.0293
1004.4586791992188 0 1049.8221
1007.4641723632812 0 1500.0754
1009.9949951171875 0 1497.7329
1010.9898681640625 0 1609.5262
1011.4827270507812 0 1639.7766
1011.9872436523438 0 2038.7761
1012.4764404296875 0 1313.4724
1012.9796752929688 0 1337.9182
1013.4927978515625 0 814.7429
1015.484619140625 0 833.0239
1016.466552734375 0 814.55804
1016.7936401367188 0 834.2318
1017.5230712890625 0 10461.317 c 8
1018.5218505859375 0 7424.3545
1019.0018920898438 0 1808.3783
1019.516845703125 0 4688.45
1019.99951171875 0 2208.2388
1020.4956665039062 0 3142.1846
1020.9951171875 0 1078.9476
1021.4893188476562 0 2150.2239
1022.0089111328125 0 872.1826
1022.1348266601562 0 1207.2932
1022.4605102539062 0 1337.351
1022.80126953125 0 1317.8623
1023.133056640625 0 1025.7805
1023.499267578125 0 1703.0662
1023.9644165039062 0 903.8238
1060.4998779296875 0 2531.2117 z 4
1061.5003662109375 0 1693.2603
1076.5240478515625 0 1129.8009 y 4
1077.528076171875 0 1097.2838
1112.5433349609375 0 1063.4917
1139.511962890625 0 769.3022
1145.5791015625 0 6273.508 c 9
1146.581787109375 0 4416.5547
1147.58935546875 0 1479.691
1173.5733642578125 0 2280.2178 y 3
1174.573974609375 0 760.3745
1196.534912109375 0 710.72797
1197.0447998046875 0 1248.3628
1216.6182861328125 0 13815.428 c 10
1217.62109375 0 8771.385
1218.6229248046875 0 3143.735
1219.62646484375 0 1069.3484
1226.04052734375 0 880.8243
1270.6197509765625 0 1639.988 y 2
1271.63525390625 0 798.10236
1282.5823974609375 0 963.7467
1312.6290283203125 0 883.7445 c Water loss 11
1330.6591796875 0 18378.672 c 11
1331.661376953125 0 13818.6045
1332.6641845703125 0 6842.086
1333.670654296875 0 1945.5063
1401.6717529296875 0 2627.2593 z 1
1402.677978515625 0 8155.1367
1403.6824951171875 0 6371.469
1404.6842041015625 0 1947.3402
1415.746826171875 0 983.72186
1442.7391357421875 0 1099.1252
1444.6875 0 757.781
1458.7491455078125 0 1649.4758 c 12
1459.7574462890625 0 4082.463
1460.7640380859375 0 2732.2646
1461.7664794921875 0 1216.9946
1486.7601318359375 0 928.16473
1487.7677001953125 0 3955.183
1488.771240234375 0 3555.107
1489.7601318359375 0 1407.0867
1490.732666015625 0 748.0189
1501.714599609375 0 1088.3484
1502.685546875 0 974.11774
1503.6951904296875 0 1216.9341
1505.1986083984375 0 1465.0796
1510.7098388671875 0 853.5887
1512.201904296875 0 1213.6777
1514.7640380859375 0 1782.3673
1515.76171875 0 9304.817
1516.7655029296875 0 7074.376
1517.7645263671875 0 3250.6875
1518.7647705078125 0 1096.397
1524.6966552734375 0 981.8856
1525.191650390625 0 1658.89
1525.6846923828125 0 936.164
1526.187744140625 0 1162.5298
1529.7655029296875 0 1432.428
1530.76611328125 0 775.85156
1531.77880859375 0 4474.027
1532.788330078125 0 11232.2295
1533.7891845703125 0 8219.612
1534.20361328125 0 1454.473
1534.7882080078125 0 3679.1475
1535.80126953125 0 1047.6414
1796.3516845703125 0 686.833
2036.9962158203125 0 812.20245
2037.9788818359375 0 783.00525
2040.0015869140625 0 816.07153
2488.145751953125 0 847.02106
3042.91162109375 0 856.07886

Spectrum Details

|  |  |
| --- | --- |
| Matched peaks? Matched peaks The total absolute number of peaks matched. Additionally in brackets the total fraction of peaks matched and the total number of peaks is shown. | 53 (15.23% of 348) |
| FDR? FDR The false discovery rate estimated for this peptide. It is calculated by matching all theoretical fragments with a non-integer shift with the raw peaks for this spectrum. This is done with 40 different shifts. The resulting percentage is the average number of annotated peaks over the number of annotated peaks with the correct spectrum. | 0.58% |
| Satellite FDR? Satellite FDR See the FDR for details on its calculation. This satellite ion specific FDR only contains the satellite ions (d/w) for I/L/J positions. | 2.38% |
| PSM Score? PSM Score The PSM Score as given by Hecklib to this annotated spectrum. It is shown with three significant figures. | 436 |

#### Spectrum 6286? Spectrum 6286 The raw spectrum of this peptide as annotated by Hecklib. The fragments are coloured according to ion type (see legend). Any peaks with a star '\*' as text can be hovered over to see the full details, first the ion type second the mass shift type. By hovering over the amino acids in the peptide or ions in the legend the corresponding peaks are highlighted. By toggling the 'Unassigned' label you can turn the background (unassigned) peaks on or off in the plot. By updating the slider in the Ion legend you can update the spectrum to only show the top X% of the peaks with labels. The top X% means any peak that is within X% of the highest intensity. By dragging in the spectrum you can zoom in to a specific part of the spectrum and use 'Zoom Out' to get back to the original zoom level. The annotation of the spectrum is based on the given sequence in the peptides file and is done with different software so inconsistencies are likely. The peaks are annotated based on the given sequence, with 20 ppm tolerance.

Copy Data

##### Spectrum 6286 (TSV)

###### Preview

```
Loading example...
```

*Click on the button to copy the data to your clipboard.*

Mz MinMz MaxIntensity Max

WidthHeightPeptide font sizePeptide stroke widthSpectrum font sizeSpectrum stroke widthCompact peptide

Ion legend

wxyz

abcd

OtherUnassignedIon^Charge^_Position_Show for top:%

JFPPSSEEJQANKA

02.50e+44.99e+47.49e+49.99e+4

Zoom Out

w^+1^_2_z^+1^_2_y^+1^_2_z^+1^_2_y^+1^_2_y^+2^_5_w^+1^_3_z^+2^_6_y^+2^_6_y^+1^_3_z^+1^_3_y^+2^_6_y^+1^_3_z^+1^_4_y^+1^_4_y^+2^_7_y^+1^_4_y^+3^_12_w^+1^_5_c^+1^_4_y^+3^_13_c^+2^_9_y^+1^_5_z^+1^_5_y^+1^_5_c^+1^_5_w^+1^_6_y^+2^_11_c^+2^_11_c^+2^_11_z^+2^_12_y^+2^_12_y^+2^_12_z^+2^_12_c^+1^_6_y^+2^_12_y^+1^_6_c^+1^_6_c^+2^_12_c^+2^_12_c^+2^_12_z^+2^_13_z^+2^_13_w^+1^_7_z^+2^_13_c^+2^_13_c^+2^_13_z^+1^_7_c^+1^_7_y^+1^_7_c^+1^_7_w^+1^_8_z^+1^_8_c^+1^_8_y^+1^_8_c^+1^_8_w^+1^_9_y^+1^_9_z^+1^_9_y^+1^_9_c^+1^_9_z^+1^_10_y^+1^_10_c^+1^_10_c^+1^_10_y^+1^_11_c^+1^_11_c^+1^_11_c^+1^_11_w^+1^_12_z^+1^_12_y^+1^_12_c^+1^_12_c^+1^_12_z^+1^_13_y^+1^_13_z^+1^_13_y^+1^_13_c^+1^_13_

0789157823663155

Fragment Matches Table

Show background peaks

| Position | Ion type | Intensity | mz Theoretical | mz Error (Th) | mz Error (ppm) | Charge | Series Number |
| --- | --- | --- | --- | --- | --- | --- | --- |
| - | - | 4491 | 120.1 | - | - | 0 | - |
| - | - | 464.8 | 121.1 | - | - | 0 | - |
| - | - | 323.3 | 122.6 | - | - | 0 | - |
| - | - | 2667 | 126.1 | - | - | 0 | - |
| - | - | 1111 | 127.1 | - | - | 0 | - |
| - | - | 3134 | 128.1 | - | - | 0 | - |
| - | - | 725.7 | 129.1 | - | - | 0 | - |
| - | - | 8843 | 129.1 | - | - | 0 | - |
| - | - | 538.6 | 130 | - | - | 0 | - |
| - | - | 665.7 | 130.1 | - | - | 0 | - |
| - | - | 603.5 | 131.1 | - | - | 0 | - |
| - | - | 442.1 | 131.1 | - | - | 0 | - |
| - | - | 526.7 | 131.1 | - | - | 0 | - |
| - | - | 7522 | 131.1 | - | - | 0 | - |
| - | - | 601.1 | 132.1 | - | - | 0 | - |
| - | - | 555 | 136.1 | - | - | 0 | - |
| - | - | 556.1 | 140.1 | - | - | 0 | - |
| - | - | 529.5 | 140.1 | - | - | 0 | - |
| - | - | 622.2 | 141.1 | - | - | 0 | - |
| 13 | w | 6335 | 144.1 | 0.0002 | 1.388 | +1 | 2 |
| - | - | 456.9 | 145 | - | - | 0 | - |
| - | - | 464.3 | 152.4 | - | - | 0 | - |
| - | - | 723.7 | 155.1 | - | - | 0 | - |
| - | - | 490.3 | 155.3 | - | - | 0 | - |
| - | - | 1965 | 156.1 | - | - | 0 | - |
| - | - | 2990 | 165.1 | - | - | 0 | - |
| - | - | 623.5 | 166.1 | - | - | 0 | - |
| - | - | 1421 | 169.1 | - | - | 0 | - |
| - | - | 686.2 | 171.1 | - | - | 0 | - |
| - | - | 798.6 | 171.1 | - | - | 0 | - |
| - | - | 866 | 173.1 | - | - | 0 | - |
| - | - | 1403 | 173.4 | - | - | 0 | - |
| - | - | 531.9 | 181.1 | - | - | 0 | - |
| - | - | 8854 | 183.1 | - | - | 0 | - |
| - | - | 848.2 | 184.1 | - | - | 0 | - |
| 13 | z | 2757 | 185.1 | 3.052E-05 | 0.1649 | +1 | 2 |
| - | - | 594.1 | 189.1 | - | - | 0 | - |
| - | - | 477.9 | 194.6 | - | - | 0 | - |
| - | - | 568.9 | 197.1 | - | - | 0 | - |
| - | - | 540.5 | 198.1 | - | - | 0 | - |
| - | - | 865.5 | 199.1 | - | - | 0 | - |
| - | - | 1027 | 199.1 | - | - | 0 | - |
| 13 | y | 1970 | 201.1 | 0.0003036 | 1.51 | +1 | 2 |
| - | - | 1244 | 202.1 | - | - | 0 | - |
| - | - | 1010 | 202.1 | - | - | 0 | - |
| 13 | z | 1.455E+04 | 202.1 | 9.275E-05 | 0.4588 | +1 | 2 |
| - | - | 669.9 | 203.1 | - | - | 0 | - |
| - | - | 989.8 | 210.1 | - | - | 0 | - |
| - | - | 645.5 | 211.1 | - | - | 0 | - |
| - | - | 1.25E+04 | 212.1 | - | - | 0 | - |
| - | - | 1417 | 213.1 | - | - | 0 | - |
| - | - | 573 | 215.1 | - | - | 0 | - |
| - | - | 742.9 | 216.1 | - | - | 0 | - |
| - | - | 1797 | 217.1 | - | - | 0 | - |
| 13 | y | 9228 | 218.1 | 0.0001065 | 0.4881 | +1 | 2 |
| - | - | 972.2 | 219.1 | - | - | 0 | - |
| - | - | 573.5 | 219.2 | - | - | 0 | - |
| - | - | 588.8 | 228.1 | - | - | 0 | - |
| - | - | 5745 | 229.1 | - | - | 0 | - |
| - | - | 529.9 | 230.1 | - | - | 0 | - |
| - | - | 987.9 | 230.1 | - | - | 0 | - |
| - | - | 798 | 232.2 | - | - | 0 | - |
| - | - | 1113 | 233.2 | - | - | 0 | - |
| - | - | 2.557E+04 | 233.2 | - | - | 0 | - |
| - | - | 3191 | 234.2 | - | - | 0 | - |
| - | - | 995.3 | 241.1 | - | - | 0 | - |
| - | - | 1746 | 242.1 | - | - | 0 | - |
| - | - | 2232 | 243.1 | - | - | 0 | - |
| - | - | 644 | 245.1 | - | - | 0 | - |
| - | - | 2140 | 247.1 | - | - | 0 | - |
| - | - | 718.4 | 254.2 | - | - | 0 | - |
| - | - | 699.4 | 257.1 | - | - | 0 | - |
| - | - | 2388 | 260.2 | - | - | 0 | - |
| - | - | 1.662E+04 | 261.2 | - | - | 0 | - |
| - | - | 2966 | 262.2 | - | - | 0 | - |
| - | - | 891.5 | 265.2 | - | - | 0 | - |
| 10 | y | 2265 | 266.1 | 1.213E-05 | 0.04557 | +2 | 5 |
| - | - | 2164 | 270.1 | - | - | 0 | - |
| 12 | w | 9065 | 272.2 | 7.035E-05 | 0.2585 | +1 | 3 |
| - | - | 1234 | 273.2 | - | - | 0 | - |
| - | - | 898 | 281.1 | - | - | 0 | - |
| - | - | 2488 | 282.1 | - | - | 0 | - |
| - | - | 660.4 | 282.2 | - | - | 0 | - |
| - | - | 635.4 | 288.2 | - | - | 0 | - |
| - | - | 1415 | 297.2 | - | - | 0 | - |
| - | - | 5.468E+04 | 299.2 | - | - | 0 | - |
| - | - | 7492 | 300.2 | - | - | 0 | - |
| - | - | 841.6 | 301.2 | - | - | 0 | - |
| - | - | 1614 | 305.2 | - | - | 0 | - |
| 9 | z | 587.5 | 306.2 | 0.002995 | 9.783 | +2 | 6 |
| - | - | 664.9 | 311.3 | - | - | 0 | - |
| 9 | y | 2097 | 314.2 | 0.003329 | 10.59 | +2 | 6 |
| 12 | y | 754.4 | 315.2 | 0.0009215 | 2.924 | +1 | 3 |
| 12 | z | 696 | 316.2 | 0.001507 | 4.767 | +1 | 3 |
| 9 | y | 2486 | 322.7 | 0.0004911 | 1.522 | +2 | 6 |
| - | - | 834.7 | 323.2 | - | - | 0 | - |
| - | - | 1432 | 325.2 | - | - | 0 | - |
| 12 | y | 4836 | 332.2 | 0.0001478 | 0.4448 | +1 | 3 |
| - | - | 1421 | 342.2 | - | - | 0 | - |
| - | - | 6966 | 343.2 | - | - | 0 | - |
| - | - | 999.5 | 344.2 | - | - | 0 | - |
| - | - | 1123 | 351.2 | - | - | 0 | - |
| - | - | 743.7 | 352.2 | - | - | 0 | - |
| - | - | 3094 | 358.2 | - | - | 0 | - |
| - | - | 798.9 | 360.2 | - | - | 0 | - |
| - | - | 1450 | 368.2 | - | - | 0 | - |
| - | - | 830.6 | 369.2 | - | - | 0 | - |
| - | - | 1408 | 369.2 | - | - | 0 | - |
| 11 | z | 958.5 | 370.2 | 0.00332 | 8.969 | +1 | 4 |
| - | - | 2435 | 370.7 | - | - | 0 | - |
| - | - | 536.1 | 371.7 | - | - | 0 | - |
| - | - | 986.5 | 377.2 | - | - | 0 | - |
| 11 | y | 2.96E+04 | 386.2 | 0.0002642 | 0.6842 | +1 | 4 |
| 8 | y | 8431 | 387.2 | 0.003319 | 8.571 | +2 | 7 |
| - | - | 1083 | 388.2 | - | - | 0 | - |
| - | - | 723.2 | 394.2 | - | - | 0 | - |
| - | - | 630.6 | 394.5 | - | - | 0 | - |
| - | - | 592.4 | 396 | - | - | 0 | - |
| - | - | 680.6 | 401.3 | - | - | 0 | - |
| - | - | 523.1 | 403.2 | - | - | 0 | - |
| 11 | y | 1.432E+04 | 403.2 | 0.0005401 | 1.339 | +1 | 4 |
| - | - | 527.8 | 403.7 | - | - | 0 | - |
| - | - | 2719 | 404.2 | - | - | 0 | - |
| - | - | 699.8 | 412.2 | - | - | 0 | - |
| - | - | 879.7 | 413.2 | - | - | 0 | - |
| - | - | 665.1 | 415.2 | - | - | 0 | - |
| 3 | y | 649.6 | 418.5 | 0.002588 | 6.183 | +3 | 12 |
| - | - | 859.9 | 425.2 | - | - | 0 | - |
| - | - | 928.3 | 428.3 | - | - | 0 | - |
| - | - | 937.9 | 429.3 | - | - | 0 | - |
| - | - | 698 | 430.3 | - | - | 0 | - |
| - | - | 593.8 | 431.2 | - | - | 0 | - |
| - | - | 1177 | 434.7 | - | - | 0 | - |
| - | - | 917 | 444.2 | - | - | 0 | - |
| - | - | 587.7 | 444.7 | - | - | 0 | - |
| - | - | 592.3 | 449.1 | - | - | 0 | - |
| - | - | 1714 | 449.1 | - | - | 0 | - |
| - | - | 980.6 | 455.3 | - | - | 0 | - |
| - | - | 1000 | 456.3 | - | - | 0 | - |
| 10 | w | 5.17E+04 | 457.2 | 0.000626 | 1.369 | +1 | 5 |
| - | - | 1.013E+04 | 458.2 | - | - | 0 | - |
| - | - | 1753 | 459.2 | - | - | 0 | - |
| - | - | 572.5 | 470.2 | - | - | 0 | - |
| - | - | 685.8 | 471.2 | - | - | 0 | - |
| - | - | 2029 | 471.3 | - | - | 0 | - |
| - | - | 818.7 | 472.3 | - | - | 0 | - |
| 4 | c | 1266 | 472.3 | 0.001992 | 4.218 | +1 | 4 |
| 2 | y | 1321 | 473.2 | 0.001184 | 2.501 | +3 | 13 |
| - | - | 1544 | 474.3 | - | - | 0 | - |
| - | - | 778.6 | 475.2 | - | - | 0 | - |
| - | - | 716.2 | 475.7 | - | - | 0 | - |
| - | - | 771 | 480.2 | - | - | 0 | - |
| - | - | 2458 | 488.3 | - | - | 0 | - |
| - | - | 737.2 | 490.3 | - | - | 0 | - |
| - | - | 804.9 | 493.2 | - | - | 0 | - |
| - | - | 817.4 | 497.3 | - | - | 0 | - |
| - | - | 2281 | 498.2 | - | - | 0 | - |
| 9 | c | 707.8 | 500.3 | 0.005279 | 10.55 | +2 | 9 |
| - | - | 1018 | 500.8 | - | - | 0 | - |
| - | - | 1257 | 501.3 | - | - | 0 | - |
| - | - | 708.1 | 509.2 | - | - | 0 | - |
| - | - | 1731 | 510.3 | - | - | 0 | - |
| - | - | 2965 | 511.3 | - | - | 0 | - |
| - | - | 715.2 | 511.8 | - | - | 0 | - |
| - | - | 916.4 | 512.3 | - | - | 0 | - |
| - | - | 1853 | 513.3 | - | - | 0 | - |
| 10 | y | 7030 | 514.3 | 0.001013 | 1.97 | +1 | 5 |
| 10 | z | 5454 | 515.3 | 0.0007697 | 1.494 | +1 | 5 |
| - | - | 1603 | 516.3 | - | - | 0 | - |
| 10 | y | 1.48E+04 | 531.3 | 0.0002817 | 0.5302 | +1 | 5 |
| - | - | 3260 | 532.3 | - | - | 0 | - |
| - | - | 608.8 | 533.3 | - | - | 0 | - |
| - | - | 1435 | 542.3 | - | - | 0 | - |
| - | - | 1192 | 557.3 | - | - | 0 | - |
| - | - | 546 | 559.3 | - | - | 0 | - |
| 5 | c | 1.425E+04 | 559.3 | 0.0004813 | 0.8604 | +1 | 5 |
| - | - | 5139 | 560.3 | - | - | 0 | - |
| - | - | 1177 | 561.3 | - | - | 0 | - |
| - | - | 5280 | 570.3 | - | - | 0 | - |
| - | - | 685.5 | 571.3 | - | - | 0 | - |
| - | - | 1725 | 571.3 | - | - | 0 | - |
| - | - | 1083 | 577.8 | - | - | 0 | - |
| - | - | 1030 | 578.3 | - | - | 0 | - |
| 9 | w | 7.302E+04 | 585.3 | 0.0004592 | 0.7845 | +1 | 6 |
| - | - | 2.033E+04 | 586.3 | - | - | 0 | - |
| 4 | y | 6674 | 587.3 | 0.007581 | 12.91 | +2 | 11 |
| - | - | 2183 | 587.8 | - | - | 0 | - |
| - | - | 1429 | 588.3 | - | - | 0 | - |
| - | - | 1967 | 591.3 | - | - | 0 | - |
| - | - | 1197 | 591.8 | - | - | 0 | - |
| - | - | 780.6 | 592.3 | - | - | 0 | - |
| - | - | 1305 | 599.3 | - | - | 0 | - |
| 11 | c | 1.156E+04 | 599.8 | 0.001825 | 3.042 | +2 | 11 |
| 11 | c | 7842 | 600.3 | 0.007022 | 11.7 | +2 | 11 |
| - | - | 3042 | 600.8 | - | - | 0 | - |
| - | - | 1354 | 609.3 | - | - | 0 | - |
| - | - | 645.1 | 610.2 | - | - | 0 | - |
| - | - | 2185 | 613.8 | - | - | 0 | - |
| - | - | 8451 | 614.3 | - | - | 0 | - |
| - | - | 5366 | 614.8 | - | - | 0 | - |
| - | - | 2452 | 615.3 | - | - | 0 | - |
| - | - | 1435 | 616.3 | - | - | 0 | - |
| - | - | 950.6 | 618.3 | - | - | 0 | - |
| 3 | z | 778.8 | 618.8 | 0.005988 | 9.676 | +2 | 12 |
| 3 | y | 2220 | 626.8 | 0.001067 | 1.703 | +2 | 12 |
| - | - | 1.155E+04 | 627.3 | - | - | 0 | - |
| 3 | y | 3747 | 627.3 | 0.005704 | 9.093 | +2 | 12 |
| 3 | z | 4214 | 627.8 | 0.0001616 | 0.2574 | +2 | 12 |
| - | - | 2625 | 628.3 | - | - | 0 | - |
| - | - | 1200 | 628.3 | - | - | 0 | - |
| 6 | c | 1.073E+04 | 628.3 | 0.008925 | 14.2 | +1 | 6 |
| - | - | 828.4 | 628.8 | - | - | 0 | - |
| - | - | 3650 | 629.4 | - | - | 0 | - |
| - | - | 599.5 | 630.3 | - | - | 0 | - |
| - | - | 869.6 | 634.8 | - | - | 0 | - |
| - | - | 1.003E+04 | 635.3 | - | - | 0 | - |
| 3 | y | 9.188E+04 | 635.8 | 0.000242 | 0.3806 | +2 | 12 |
| - | - | 6.633E+04 | 636.3 | - | - | 0 | - |
| - | - | 2.681E+04 | 636.8 | - | - | 0 | - |
| - | - | 9041 | 637.3 | - | - | 0 | - |
| - | - | 1613 | 637.8 | - | - | 0 | - |
| 9 | y | 7307 | 644.4 | 0.0006903 | 1.071 | +1 | 6 |
| - | - | 2649 | 645.4 | - | - | 0 | - |
| 6 | c | 5.054E+04 | 646.4 | 0.0007404 | 1.146 | +1 | 6 |
| - | - | 1027 | 647.3 | - | - | 0 | - |
| - | - | 2.077E+04 | 647.4 | - | - | 0 | - |
| - | - | 4057 | 648.4 | - | - | 0 | - |
| 12 | c | 6987 | 656.8 | 0.01181 | 17.99 | +2 | 12 |
| 12 | c | 3623 | 657.3 | 0.001258 | 1.914 | +2 | 12 |
| - | - | 886 | 657.8 | - | - | 0 | - |
| 12 | c | 730.2 | 665.8 | 0.002875 | 4.318 | +2 | 12 |
| - | - | 2844 | 672.4 | - | - | 0 | - |
| - | - | 2805 | 673.4 | - | - | 0 | - |
| - | - | 736.6 | 680.7 | - | - | 0 | - |
| - | - | 696.2 | 681 | - | - | 0 | - |
| - | - | 1190 | 691.8 | - | - | 0 | - |
| 2 | z | 2132 | 692.3 | 9.451E-05 | 0.1365 | +2 | 13 |
| 2 | z | 2496 | 692.8 | 0.003747 | 5.409 | +2 | 13 |
| - | - | 1645 | 693.3 | - | - | 0 | - |
| 8 | w | 3.269E+04 | 698.4 | 0.0006847 | 0.9805 | +1 | 7 |
| - | - | 1.408E+04 | 699.4 | - | - | 0 | - |
| - | - | 2856 | 700.4 | - | - | 0 | - |
| 2 | z | 1.814E+04 | 701.3 | 0.003729 | 5.317 | +2 | 13 |
| - | - | 7521 | 701.8 | - | - | 0 | - |
| - | - | 6190 | 702.3 | - | - | 0 | - |
| - | - | 1822 | 702.8 | - | - | 0 | - |
| - | - | 915.3 | 703.3 | - | - | 0 | - |
| - | - | 2270 | 707.9 | - | - | 0 | - |
| - | - | 1572 | 708.4 | - | - | 0 | - |
| - | - | 1060 | 713.4 | - | - | 0 | - |
| - | - | 1734 | 714.3 | - | - | 0 | - |
| - | - | 3860 | 714.4 | - | - | 0 | - |
| - | - | 3888 | 714.9 | - | - | 0 | - |
| - | - | 2649 | 715.4 | - | - | 0 | - |
| - | - | 3047 | 715.9 | - | - | 0 | - |
| - | - | 1578 | 716.4 | - | - | 0 | - |
| 13 | c | 631.4 | 721.4 | 0.001614 | 2.238 | +2 | 13 |
| - | - | 894.7 | 721.9 | - | - | 0 | - |
| - | - | 1530 | 722.4 | - | - | 0 | - |
| - | - | 691.6 | 722.9 | - | - | 0 | - |
| - | - | 608.4 | 727.8 | - | - | 0 | - |
| - | - | 955.1 | 728.3 | - | - | 0 | - |
| 13 | c | 5.182E+04 | 729.9 | 5.846E-05 | 0.08009 | +2 | 13 |
| - | - | 5.473E+04 | 730.4 | - | - | 0 | - |
| - | - | 3.277E+04 | 730.9 | - | - | 0 | - |
| - | - | 1113 | 731 | - | - | 0 | - |
| - | - | 1.125E+04 | 731.4 | - | - | 0 | - |
| - | - | 2544 | 731.9 | - | - | 0 | - |
| - | - | 4733 | 736.4 | - | - | 0 | - |
| - | - | 3442 | 736.9 | - | - | 0 | - |
| - | - | 2214 | 737.4 | - | - | 0 | - |
| - | - | 1205 | 737.9 | - | - | 0 | - |
| - | - | 3093 | 738.4 | - | - | 0 | - |
| - | - | 2452 | 738.9 | - | - | 0 | - |
| 8 | z | 1086 | 739.4 | 0.01268 | 17.14 | +1 | 7 |
| - | - | 5445 | 740.3 | - | - | 0 | - |
| - | - | 2085 | 741.3 | - | - | 0 | - |
| - | - | 1317 | 743.4 | - | - | 0 | - |
| - | - | 1.634E+04 | 743.9 | - | - | 0 | - |
| - | - | 1.159E+04 | 744.4 | - | - | 0 | - |
| - | - | 7364 | 744.9 | - | - | 0 | - |
| - | - | 2300 | 745.4 | - | - | 0 | - |
| - | - | 746.9 | 745.9 | - | - | 0 | - |
| 7 | c | 4.564E+04 | 757.4 | 0.008629 | 11.39 | +1 | 7 |
| - | - | 3801 | 757.9 | - | - | 0 | - |
| - | - | 2.044E+04 | 758.4 | - | - | 0 | - |
| - | - | 1193 | 758.9 | - | - | 0 | - |
| - | - | 5746 | 759.4 | - | - | 0 | - |
| - | - | 652.3 | 759.9 | - | - | 0 | - |
| - | - | 917.4 | 760.4 | - | - | 0 | - |
| - | - | 1953 | 765.4 | - | - | 0 | - |
| - | - | 3.795E+04 | 765.9 | - | - | 0 | - |
| - | - | 6.825E+04 | 766.4 | - | - | 0 | - |
| - | - | 4.646E+04 | 766.9 | - | - | 0 | - |
| - | - | 2.152E+04 | 767.4 | - | - | 0 | - |
| - | - | 7045 | 767.9 | - | - | 0 | - |
| - | - | 1767 | 768.4 | - | - | 0 | - |
| 8 | y | 4294 | 773.4 | 0.0005777 | 0.747 | +1 | 7 |
| - | - | 1543 | 774.4 | - | - | 0 | - |
| 7 | c | 8.829E+04 | 775.4 | 0.0006278 | 0.8097 | +1 | 7 |
| - | - | 4.07E+04 | 776.4 | - | - | 0 | - |
| - | - | 1.049E+04 | 777.4 | - | - | 0 | - |
| - | - | 1443 | 778.4 | - | - | 0 | - |
| - | - | 2433 | 801.4 | - | - | 0 | - |
| - | - | 2066 | 802.4 | - | - | 0 | - |
| - | - | 1876 | 814.4 | - | - | 0 | - |
| - | - | 1675 | 815.4 | - | - | 0 | - |
| 7 | w | 1.361E+04 | 827.4 | 0.000328 | 0.3964 | +1 | 8 |
| - | - | 7042 | 828.4 | - | - | 0 | - |
| - | - | 2056 | 829.4 | - | - | 0 | - |
| - | - | 820.9 | 843.4 | - | - | 0 | - |
| - | - | 747.3 | 844.4 | - | - | 0 | - |
| - | - | 855 | 860.4 | - | - | 0 | - |
| - | - | 751.1 | 860.7 | - | - | 0 | - |
| - | - | 652.5 | 861.4 | - | - | 0 | - |
| - | - | 2530 | 868.4 | - | - | 0 | - |
| 7 | z | 1131 | 869.4 | 0.002846 | 3.273 | +1 | 8 |
| - | - | 1021 | 870.4 | - | - | 0 | - |
| - | - | 1239 | 871.4 | - | - | 0 | - |
| 8 | c | 6.362E+04 | 886.4 | 0.009127 | 10.3 | +1 | 8 |
| - | - | 3.112E+04 | 887.4 | - | - | 0 | - |
| - | - | 1.083E+04 | 888.4 | - | - | 0 | - |
| - | - | 2478 | 889.4 | - | - | 0 | - |
| - | - | 723.6 | 890.4 | - | - | 0 | - |
| - | - | 1574 | 901.5 | - | - | 0 | - |
| 7 | y | 1.321E+04 | 902.5 | 0.01138 | 12.61 | +1 | 8 |
| - | - | 6401 | 903.4 | - | - | 0 | - |
| 8 | c | 9.001E+04 | 904.4 | 0.0008814 | 0.9745 | +1 | 8 |
| - | - | 4.684E+04 | 905.4 | - | - | 0 | - |
| - | - | 1.513E+04 | 906.4 | - | - | 0 | - |
| - | - | 2212 | 907.4 | - | - | 0 | - |
| - | - | 887.5 | 916.5 | - | - | 0 | - |
| - | - | 4126 | 917.4 | - | - | 0 | - |
| - | - | 2588 | 918.4 | - | - | 0 | - |
| - | - | 1052 | 930.5 | - | - | 0 | - |
| - | - | 687.6 | 931.5 | - | - | 0 | - |
| - | - | 1384 | 935.4 | - | - | 0 | - |
| - | - | 1513 | 939.4 | - | - | 0 | - |
| - | - | 744.3 | 941.5 | - | - | 0 | - |
| - | - | 854.2 | 942.5 | - | - | 0 | - |
| - | - | 708 | 946.5 | - | - | 0 | - |
| - | - | 1513 | 954.5 | - | - | 0 | - |
| 6 | w | 1228 | 956.5 | 0.002897 | 3.029 | +1 | 9 |
| - | - | 1080 | 961.4 | - | - | 0 | - |
| - | - | 770.1 | 964.5 | - | - | 0 | - |
| - | - | 1960 | 971.5 | - | - | 0 | - |
| 6 | y | 2083 | 972.5 | 0.003671 | 3.775 | +1 | 9 |
| 6 | z | 5.291E+04 | 973.5 | 0.001199 | 1.232 | +1 | 9 |
| - | - | 2.898E+04 | 974.5 | - | - | 0 | - |
| - | - | 8671 | 975.5 | - | - | 0 | - |
| - | - | 2422 | 976.5 | - | - | 0 | - |
| - | - | 847.3 | 977.5 | - | - | 0 | - |
| - | - | 1184 | 984.4 | - | - | 0 | - |
| - | - | 1938 | 985.4 | - | - | 0 | - |
| - | - | 734 | 987.4 | - | - | 0 | - |
| - | - | 1291 | 988.5 | - | - | 0 | - |
| 6 | y | 1.443E+04 | 989.5 | 0.003243 | 3.277 | +1 | 9 |
| - | - | 7278 | 990.5 | - | - | 0 | - |
| - | - | 2753 | 991.5 | - | - | 0 | - |
| - | - | 1583 | 1000 | - | - | 0 | - |
| - | - | 1446 | 1001 | - | - | 0 | - |
| - | - | 1479 | 1003 | - | - | 0 | - |
| - | - | 865 | 1003 | - | - | 0 | - |
| - | - | 2239 | 1004 | - | - | 0 | - |
| - | - | 2999 | 1005 | - | - | 0 | - |
| - | - | 2632 | 1006 | - | - | 0 | - |
| - | - | 749.2 | 1007 | - | - | 0 | - |
| - | - | 5989 | 1007 | - | - | 0 | - |
| - | - | 707.9 | 1011 | - | - | 0 | - |
| - | - | 1023 | 1013 | - | - | 0 | - |
| - | - | 927.3 | 1016 | - | - | 0 | - |
| - | - | 1902 | 1016 | - | - | 0 | - |
| - | - | 1020 | 1017 | - | - | 0 | - |
| 9 | c | 6.042E+04 | 1018 | 0.0005577 | 0.548 | +1 | 9 |
| - | - | 3.598E+04 | 1019 | - | - | 0 | - |
| - | - | 1.253E+04 | 1020 | - | - | 0 | - |
| - | - | 3582 | 1021 | - | - | 0 | - |
| - | - | 1179 | 1021 | - | - | 0 | - |
| - | - | 1436 | 1022 | - | - | 0 | - |
| - | - | 1885 | 1022 | - | - | 0 | - |
| - | - | 823.1 | 1023 | - | - | 0 | - |
| - | - | 735.1 | 1044 | - | - | 0 | - |
| - | - | 2003 | 1053 | - | - | 0 | - |
| - | - | 1012 | 1054 | - | - | 0 | - |
| - | - | 1031 | 1058 | - | - | 0 | - |
| 5 | z | 1.108E+04 | 1061 | 0.001275 | 1.202 | +1 | 10 |
| - | - | 8889 | 1062 | - | - | 0 | - |
| - | - | 2703 | 1063 | - | - | 0 | - |
| - | - | 910.7 | 1071 | - | - | 0 | - |
| 5 | y | 6784 | 1077 | 6.899E-06 | 0.006409 | +1 | 10 |
| - | - | 3809 | 1078 | - | - | 0 | - |
| - | - | 1020 | 1079 | - | - | 0 | - |
| - | - | 2314 | 1102 | - | - | 0 | - |
| - | - | 1721 | 1103 | - | - | 0 | - |
| 10 | c | 1536 | 1129 | 0.001991 | 1.764 | +1 | 10 |
| - | - | 819.2 | 1130 | - | - | 0 | - |
| - | - | 824.8 | 1132 | - | - | 0 | - |
| 10 | c | 3.502E+04 | 1146 | 0.000757 | 0.6608 | +1 | 10 |
| - | - | 2.273E+04 | 1147 | - | - | 0 | - |
| - | - | 8058 | 1148 | - | - | 0 | - |
| - | - | 1463 | 1149 | - | - | 0 | - |
| - | - | 752.4 | 1172 | - | - | 0 | - |
| 4 | y | 1.1E+04 | 1174 | 0.001442 | 1.229 | +1 | 11 |
| - | - | 6345 | 1175 | - | - | 0 | - |
| - | - | 3202 | 1176 | - | - | 0 | - |
| - | - | 979.7 | 1197 | - | - | 0 | - |
| - | - | 809.3 | 1197 | - | - | 0 | - |
| 11 | c | 1180 | 1199 | 0.002355 | 1.964 | +1 | 11 |
| 11 | c | 1721 | 1200 | 0.00362 | 3.018 | +1 | 11 |
| - | - | 1522 | 1201 | - | - | 0 | - |
| - | - | 1410 | 1216 | - | - | 0 | - |
| 11 | c | 7.255E+04 | 1217 | 0.0005084 | 0.4179 | +1 | 11 |
| - | - | 4.887E+04 | 1218 | - | - | 0 | - |
| - | - | 1.931E+04 | 1219 | - | - | 0 | - |
| - | - | 5196 | 1220 | - | - | 0 | - |
| - | - | 1018 | 1221 | - | - | 0 | - |
| 3 | w | 3989 | 1228 | 0.02005 | 16.33 | +1 | 12 |
| - | - | 2834 | 1229 | - | - | 0 | - |
| - | - | 1401 | 1230 | - | - | 0 | - |
| - | - | 748.7 | 1231 | - | - | 0 | - |
| 3 | z | 1015 | 1255 | 0.008052 | 6.418 | +1 | 12 |
| - | - | 979.2 | 1270 | - | - | 0 | - |
| 3 | y | 5682 | 1271 | 0.001413 | 1.112 | +1 | 12 |
| - | - | 5157 | 1272 | - | - | 0 | - |
| - | - | 3370 | 1273 | - | - | 0 | - |
| - | - | 2017 | 1274 | - | - | 0 | - |
| - | - | 2474 | 1287 | - | - | 0 | - |
| - | - | 3248 | 1288 | - | - | 0 | - |
| - | - | 961.6 | 1290 | - | - | 0 | - |
| - | - | 3681 | 1300 | - | - | 0 | - |
| - | - | 1740 | 1301 | - | - | 0 | - |
| 12 | c | 1207 | 1313 | 0.02502 | 19.06 | +1 | 12 |
| - | - | 4207 | 1330 | - | - | 0 | - |
| 12 | c | 9.889E+04 | 1331 | 0.0003047 | 0.229 | +1 | 12 |
| - | - | 7.456E+04 | 1332 | - | - | 0 | - |
| - | - | 3.678E+04 | 1333 | - | - | 0 | - |
| - | - | 1.001E+04 | 1334 | - | - | 0 | - |
| - | - | 1227 | 1335 | - | - | 0 | - |
| - | - | 772.3 | 1359 | - | - | 0 | - |
| 2 | z | 1237 | 1385 | 0.02429 | 17.54 | +1 | 13 |
| - | - | 1032 | 1386 | - | - | 0 | - |
| - | - | 923.7 | 1387 | - | - | 0 | - |
| 2 | y | 774.3 | 1401 | 0.003493 | 2.494 | +1 | 13 |
| 2 | z | 1.195E+04 | 1402 | 6.279E-05 | 0.04479 | +1 | 13 |
| - | - | 3.433E+04 | 1403 | - | - | 0 | - |
| - | - | 2.605E+04 | 1404 | - | - | 0 | - |
| - | - | 1.178E+04 | 1405 | - | - | 0 | - |
| - | - | 3052 | 1406 | - | - | 0 | - |
| - | - | 4111 | 1415 | - | - | 0 | - |
| - | - | 4764 | 1416 | - | - | 0 | - |
| - | - | 2152 | 1417 | - | - | 0 | - |
| 2 | y | 1599 | 1418 | 0.009659 | 6.813 | +1 | 13 |
| - | - | 1479 | 1419 | - | - | 0 | - |
| - | - | 1048 | 1420 | - | - | 0 | - |
| - | - | 4261 | 1443 | - | - | 0 | - |
| - | - | 3202 | 1444 | - | - | 0 | - |
| - | - | 2268 | 1445 | - | - | 0 | - |
| 13 | c | 5229 | 1459 | 0.004081 | 2.798 | +1 | 13 |
| - | - | 1.962E+04 | 1460 | - | - | 0 | - |
| - | - | 1.313E+04 | 1461 | - | - | 0 | - |
| - | - | 7582 | 1462 | - | - | 0 | - |
| - | - | 2380 | 1463 | - | - | 0 | - |
| - | - | 1414 | 1470 | - | - | 0 | - |
| - | - | 2851 | 1471 | - | - | 0 | - |
| - | - | 1973 | 1472 | - | - | 0 | - |
| - | - | 817.9 | 1473 | - | - | 0 | - |
| - | - | 860.3 | 1474 | - | - | 0 | - |
| - | - | 3968 | 1487 | - | - | 0 | - |
| - | - | 1.866E+04 | 1488 | - | - | 0 | - |
| - | - | 1.492E+04 | 1489 | - | - | 0 | - |
| - | - | 7449 | 1490 | - | - | 0 | - |
| - | - | 1234 | 1491 | - | - | 0 | - |
| - | - | 875.2 | 1492 | - | - | 0 | - |
| - | - | 774.9 | 1497 | - | - | 0 | - |
| - | - | 1668 | 1499 | - | - | 0 | - |
| - | - | 1367 | 1500 | - | - | 0 | - |
| - | - | 849.7 | 1504 | - | - | 0 | - |
| - | - | 1711 | 1505 | - | - | 0 | - |
| - | - | 1033 | 1506 | - | - | 0 | - |
| - | - | 1520 | 1514 | - | - | 0 | - |
| - | - | 4943 | 1515 | - | - | 0 | - |
| - | - | 3.781E+04 | 1516 | - | - | 0 | - |
| - | - | 3.538E+04 | 1517 | - | - | 0 | - |
| - | - | 1.782E+04 | 1518 | - | - | 0 | - |
| - | - | 5379 | 1519 | - | - | 0 | - |
| - | - | 1863 | 1520 | - | - | 0 | - |
| - | - | 894.6 | 1525 | - | - | 0 | - |
| - | - | 862.1 | 1530 | - | - | 0 | - |
| - | - | 3203 | 1531 | - | - | 0 | - |
| - | - | 1.565E+04 | 1532 | - | - | 0 | - |
| - | - | 5.162E+04 | 1533 | - | - | 0 | - |
| - | - | 3.999E+04 | 1534 | - | - | 0 | - |
| - | - | 806.3 | 1534 | - | - | 0 | - |
| - | - | 1.879E+04 | 1535 | - | - | 0 | - |
| - | - | 6117 | 1536 | - | - | 0 | - |
| - | - | 1515 | 1537 | - | - | 0 | - |
| - | - | 899.9 | 2023 | - | - | 0 | - |
| - | - | 899.2 | 3124 | - | - | 0 | - |

m/z Charge Intensity FragmentType MassShift Position
120.08096313476562 0 4491.2812
121.0843505859375 0 464.76395
122.58422088623047 0 323.32074
126.05514526367188 0 2666.7302
127.0868148803711 0 1111.3179
128.09463500976562 0 3133.685
129.0662078857422 0 725.7426
129.10240173339844 0 8842.633
130.04995727539062 0 538.5906
130.10577392578125 0 665.6732
131.05795288085938 0 603.47437
131.0701904296875 0 442.12466
131.1133575439453 0 526.6879
131.11807250976562 0 7522.2695
132.12130737304688 0 601.0655
136.0760040283203 0 554.95624
140.08221435546875 0 556.09
140.10765075683594 0 529.4833
141.10256958007812 0 622.17255
144.0657196044922 0 6334.8354 w 12
145.0107879638672 0 456.854
152.38462829589844 0 464.26956
155.09396362304688 0 723.718
155.29153442382812 0 490.2882
156.05313110351562 0 1965.377
165.1024169921875 0 2990.0845
166.08636474609375 0 623.4871
169.06112670898438 0 1420.8005
171.07681274414062 0 686.177
171.1132049560547 0 798.59216
173.1287078857422 0 866.00037
173.43885803222656 0 1403.3453
181.09747314453125 0 531.91656
183.11297607421875 0 8853.601
184.11639404296875 0 848.1757
185.1046142578125 0 2756.844 z Ammonia loss 12
189.0872344970703 0 594.0897
194.58619689941406 0 477.91693
197.12823486328125 0 568.9053
198.0999755859375 0 540.52496
199.07247924804688 0 865.4791
199.107666015625 0 1026.7197
201.12367248535156 0 1969.7826 y Ammonia loss 12
202.0859375 0 1244.1161
202.1227264404297 0 1009.88696
202.13128662109375 0 14553.0625 z 12
203.1366729736328 0 669.85876
210.1000518798828 0 989.7616
211.1075439453125 0 645.47095
212.13951110839844 0 12496.99
213.14276123046875 0 1416.6564
215.13882446289062 0 573.0134
216.0979766845703 0 742.854
217.14553833007812 0 1796.581
218.1500244140625 0 9228.078 y 12
219.13421630859375 0 972.24506
219.15380859375 0 573.5057
228.13409423828125 0 588.7578
229.1097869873047 0 5744.7886
230.08462524414062 0 529.94775
230.11573791503906 0 987.8516
232.15708923339844 0 798.041
233.15155029296875 0 1112.8638
233.16506958007812 0 25573.092
234.16844177246094 0 3190.5388
241.0828094482422 0 995.25037
242.14988708496094 0 1746.1882
243.14535522460938 0 2232.281
245.14157104492188 0 644.04266
247.11123657226562 0 2140.4895
254.15060424804688 0 718.3905
257.1420593261719 0 699.3707
260.1521301269531 0 2387.5793
261.1597900390625 0 16619.002
262.16314697265625 0 2965.667
265.15435791015625 0 891.5432
266.1479187011719 0 2265.0122 y 9
270.1446838378906 0 2163.6917
272.1605529785156 0 9064.759 w 11
273.1646728515625 0 1234.4752
281.1382141113281 0 897.9755
282.1449890136719 0 2488.2253
282.1610412597656 0 660.39307
288.1548767089844 0 635.4397
297.156005859375 0 1414.6082
299.1716613769531 0 54678.86
300.1748962402344 0 7492.3247
301.17767333984375 0 841.5625
305.21002197265625 0 1614.3489
306.164306640625 0 587.49066 z Ammonia loss 8
311.25848388671875 0 664.91187
314.17999267578125 0 2096.572 y Ammonia loss 8
315.1653747558594 0 754.39215 y Ammonia loss 11
316.1756286621094 0 696.00946 z 11
322.6904296875 0 2486.264 y 8
323.1903076171875 0 834.737
325.1875915527344 0 1431.779
332.1929931640625 0 4836.3125 y 11
342.19384765625 0 1421.2734
343.1978759765625 0 6966.0146
344.2004089355469 0 999.5348
351.166748046875 0 1122.7722
352.17340087890625 0 743.73914
358.21258544921875 0 3093.773
360.2151184082031 0 798.9342
368.1927795410156 0 1450.1506
369.1768798828125 0 830.55493
369.2032775878906 0 1407.9952
370.1813659667969 0 958.4683 z Ammonia loss 10
370.67828369140625 0 2435.0044
371.6816711425781 0 536.05896
377.177734375 0 986.50214
386.20367431640625 0 29600.654 y Ammonia loss 10
387.2079162597656 0 8430.878 y 7
388.21124267578125 0 1083.4252
394.18194580078125 0 723.2031
394.53009033203125 0 630.55334
396.0284729003906 0 592.4453
401.26031494140625 0 680.5856
403.2030944824219 0 523.1087
403.2304992675781 0 14324.429 y 10
403.6854248046875 0 527.8406
404.232421875 0 2718.766
412.2117614746094 0 699.8416
413.2177734375 0 879.7439
415.2318115234375 0 665.062
418.5410461425781 0 649.5855 y Ammonia loss 2
425.2162170410156 0 859.9301
428.2774658203125 0 928.3021
429.2825927734375 0 937.93286
430.2935791015625 0 697.9543
431.222412109375 0 593.8198
434.7087707519531 0 1176.7914
444.2118225097656 0 916.9664
444.71307373046875 0 587.70685
449.0679016113281 0 592.317
449.1261901855469 0 1713.9164
455.26776123046875 0 980.5948
456.27008056640625 0 1000.05914
457.24114990234375 0 51702.016 w 9
458.2444152832031 0 10131.013
459.2450256347656 0 1752.8225
470.2250671386719 0 572.48285
471.2225341796875 0 685.8425
471.2569274902344 0 2029.0991
472.25946044921875 0 818.65625
472.2938232421875 0 1266.3124 c 3
473.235595703125 0 1321.3525 y 1
474.30731201171875 0 1543.7222
475.1771545410156 0 778.64923
475.67529296875 0 716.1711
480.2102966308594 0 771.0482
488.2734069824219 0 2457.8352
490.2608642578125 0 737.23785
493.1866455078125 0 804.90015
497.2623291015625 0 817.41376
498.2194519042969 0 2280.9207
500.2556457519531 0 707.7825 c Water loss 8
500.75250244140625 0 1018.19867
501.2535400390625 0 1256.6129
509.2320861816406 0 708.114
510.25750732421875 0 1730.7975
511.26129150390625 0 2965.4517
511.7667541503906 0 715.2203
512.3180541992188 0 916.38245
513.2780151367188 0 1852.5034
514.2630004882812 0 7030.1553 y Ammonia loss 9
515.26904296875 0 5454.4365 z 9
516.2750854492188 0 1603.457
531.288818359375 0 14797.374 y 9
532.2914428710938 0 3260.2297
533.2918701171875 0 608.803
542.2962646484375 0 1434.617
557.3211669921875 0 1191.829
559.2838745117188 0 545.9939
559.3243408203125 0 14248.503 c 4
560.3274536132812 0 5138.5796
561.3323974609375 0 1177.1438
570.3251953125 0 5280.1655
571.283203125 0 685.5034
571.329833984375 0 1725.0347
577.7999267578125 0 1082.6519
578.30126953125 0 1030.208
585.299560546875 0 73024.14 w 8
586.302490234375 0 20328.629
587.2985229492188 0 6673.6865 y 3
587.7923583984375 0 2182.5342
588.297119140625 0 1429.2164
591.2923583984375 0 1966.7832
591.794677734375 0 1196.5748
592.2935180664062 0 780.6415
599.3053588867188 0 1304.6327
599.8069458007812 0 11555.526 c Water loss 10
600.3078002929688 0 7841.6855 c Ammonia loss 10
600.8084106445312 0 3042.439
609.2522583007812 0 1353.7264
610.2490234375 0 645.1115
613.8102416992188 0 2185.0198
614.314453125 0 8450.94
614.8162841796875 0 5365.7134
615.315673828125 0 2452.2456
616.3344116210938 0 1435.4702
618.2992553710938 0 950.5631
618.7966918945312 0 778.8135 z Water loss 2
626.8109741210938 0 2219.606 y Water loss 2
627.2621459960938 0 11546.35
627.3097534179688 0 3746.511 y Ammonia loss 2
627.8078002929688 0 4213.6206 z 2
628.2642211914062 0 2624.7344
628.3109130859375 0 1199.6407
628.354248046875 0 10734.885 c Water loss 5
628.8072509765625 0 828.39087
629.3551025390625 0 3650.4102
630.34130859375 0 599.5088
634.8072509765625 0 869.55414
635.3139038085938 0 10030.411
635.8175659179688 0 91877.27 y 2
636.3192749023438 0 66330.78
636.8206787109375 0 26808.42
637.322265625 0 9040.52
637.8241577148438 0 1612.8773
644.373291015625 0 7307.162 y 8
645.37451171875 0 2649.0046
646.3566284179688 0 50539.176 c 5
647.2975463867188 0 1026.9261
647.3594360351562 0 20772.64
648.3618774414062 0 4057.325
656.8184204101562 0 6986.669 c Water loss 11
657.3209838867188 0 3623.1152 c Ammonia loss 11
657.8211059570312 0 885.9871
665.8326416015625 0 730.2447 c 11
672.3720092773438 0 2843.889
673.3576049804688 0 2804.9514
680.654296875 0 736.6302
680.9906005859375 0 696.1566
691.8352661132812 0 1189.6792
692.3367919921875 0 2132.138 z Water loss 1
692.8326416015625 0 2495.8901 z Ammonia loss 1
693.3324584960938 0 1644.749
698.3838500976562 0 32693.072 w 7
699.3856201171875 0 14077.379
700.3875122070312 0 2855.98
701.3384399414062 0 18141.459 z 1
701.8438720703125 0 7520.913
702.3422241210938 0 6190.418
702.8466796875 0 1821.9841
703.3468017578125 0 915.33746
707.8749389648438 0 2270.4617
708.375732421875 0 1571.6106
713.4016723632812 0 1060.4132
714.335205078125 0 1734.1992
714.38134765625 0 3859.509
714.8787231445312 0 3887.8313
715.379638671875 0 2648.9272
715.8573608398438 0 3046.7317
716.357666015625 0 1578.2695
721.371337890625 0 631.4265 c Ammonia loss 12
721.8721923828125 0 894.6936
722.3644409179688 0 1530.4773
722.8717041015625 0 691.60236
727.8419799804688 0 608.39
728.3468627929688 0 955.0889
729.883056640625 0 51818.348 c 12
730.3849487304688 0 54733.39
730.8861694335938 0 32767.627
730.96142578125 0 1112.64
731.3865966796875 0 11249.171
731.8883056640625 0 2544.097
736.3585815429688 0 4733.143
736.8704223632812 0 3441.97
737.3646850585938 0 2213.9272
737.8809814453125 0 1204.5051
738.3695068359375 0 3093.236
738.870361328125 0 2451.5083
739.3732299804688 0 1085.7815 z Water loss 7
740.3487548828125 0 5444.8574
741.3497924804688 0 2084.5405
743.3925170898438 0 1317.428
743.8873901367188 0 16336.808
744.3895874023438 0 11586.38
744.888916015625 0 7364.1323
745.3908081054688 0 2299.7478
745.8896484375 0 746.93964
757.3965454101562 0 45640.605 c Water loss 6
757.8896484375 0 3801.206
758.3966064453125 0 20435.074
758.8920288085938 0 1193.0615
759.3956909179688 0 5746.472
759.8900146484375 0 652.25726
760.3890380859375 0 917.38513
765.3889770507812 0 1952.5659
765.89404296875 0 37954.5
766.3967895507812 0 68250.57
766.898681640625 0 46458.07
767.399658203125 0 21521.582
767.8997192382812 0 7045.322
768.3993530273438 0 1767.354
773.415771484375 0 4294.3853 y 7
774.4193115234375 0 1542.9364
775.3991088867188 0 88293.234 c 6
776.4019165039062 0 40700.062
777.4047241210938 0 10493.05
778.4074096679688 0 1442.9952
801.4125366210938 0 2432.792
802.4158325195312 0 2066.3174
814.4124145507812 0 1876.1433
815.4194946289062 0 1675.2599
827.4260864257812 0 13611.256 w 6
828.4290161132812 0 7042.2534
829.4290161132812 0 2055.603
843.4313354492188 0 820.8997
844.4359741210938 0 747.2714
860.4376831054688 0 855.00116
860.7301635742188 0 751.1125
861.419189453125 0 652.53265
868.4082641601562 0 2530.3284
869.40966796875 0 1130.6273 z Ammonia loss 6
870.4130249023438 0 1020.66016
871.403076171875 0 1239.0616
886.4396362304688 0 63615.53 c Water loss 7
887.4375 0 31121.139
888.4383544921875 0 10830.518
889.4373168945312 0 2478.0706
890.43017578125 0 723.56586
901.457763671875 0 1573.8716
902.4464111328125 0 13205.324 y 6
903.4480590820312 0 6400.729
904.4419555664062 0 90012.16 c 7
905.4442138671875 0 46843.477
906.4468994140625 0 15130.585
907.4486083984375 0 2211.8826
916.4569702148438 0 887.45465
917.408935546875 0 4126.342
918.4180908203125 0 2587.7034
930.4669799804688 0 1051.6633
931.4520874023438 0 687.641
935.433837890625 0 1383.8219
939.443603515625 0 1512.9585
941.45166015625 0 744.30927
942.4574584960938 0 854.15594
946.4810791015625 0 707.9744
954.455078125 0 1512.8693
956.4654541015625 0 1228.1553 w 5
961.4356689453125 0 1079.7937
964.486083984375 0 770.0544
971.45654296875 0 1960.29
972.4595947265625 0 2082.639 y Ammonia loss 5
973.4722900390625 0 52905.84 z 5
974.475341796875 0 28981.832
975.4772338867188 0 8671.382
976.4822387695312 0 2422.2063
977.4705200195312 0 847.2614
984.3621215820312 0 1184.4958
985.3710327148438 0 1938.1467
987.44140625 0 733.9785
988.4862670898438 0 1291.2274
989.486572265625 0 14425.138 y 5
990.4906616210938 0 7277.778
991.4888305664062 0 2753.3862
1000.4988403320312 0 1583.1868
1001.49462890625 0 1445.5558
1002.7987670898438 0 1478.8082
1003.1329956054688 0 864.9989
1004.4536743164062 0 2239.347
1005.4529418945312 0 2998.9448
1006.4441528320312 0 2632.4094
1007.1307373046875 0 749.2083
1007.453857421875 0 5988.7603
1011.4745483398438 0 707.92053
1013.48583984375 0 1023.47424
1015.516357421875 0 927.2799
1016.4708862304688 0 1902.282
1016.8008422851562 0 1019.5125
1017.5256958007812 0 60422.18 c 8
1018.5280151367188 0 35981.773
1019.531005859375 0 12526.908
1020.5264282226562 0 3582.4712
1021.4864501953125 0 1178.7083
1022.3668212890625 0 1436.2186
1022.4688720703125 0 1884.9517
1023.3507690429688 0 823.0712
1043.5419921875 0 735.1179
1053.4805908203125 0 2002.9718
1054.4832763671875 0 1011.8113
1058.4869384765625 0 1031.4528
1060.50439453125 0 11084.007 z 4
1061.508056640625 0 8889.423
1062.5106201171875 0 2703.197
1070.503662109375 0 910.6616
1076.5218505859375 0 6783.6914 y 4
1077.5262451171875 0 3808.8127
1078.5299072265625 0 1019.8504
1101.5657958984375 0 2313.6553
1102.5679931640625 0 1721.1564
1128.55517578125 0 1536.2267 c Ammonia loss 9
1129.5518798828125 0 819.1983
1131.5633544921875 0 824.79803
1145.58447265625 0 35020.254 c 9
1146.5869140625 0 22729.666
1147.5902099609375 0 8058.3237
1148.5941162109375 0 1462.9158
1171.5609130859375 0 752.3558
1173.5760498046875 0 11002.796 y 3
1174.5782470703125 0 6344.994
1175.576904296875 0 3201.939
1196.5302734375 0 979.7053
1197.038818359375 0 809.3033
1198.60791015625 0 1179.5891 c Water loss 10
1199.597900390625 0 1721.1339 c Ammonia loss 10
1200.573486328125 0 1521.8573
1215.60986328125 0 1409.6967
1216.621337890625 0 72550.68 c 10
1217.6236572265625 0 48874.496
1218.626953125 0 19314.7
1219.6290283203125 0 5196.3135
1220.630126953125 0 1017.61163
1227.605224609375 0 3989.3633 w 2
1228.6046142578125 0 2834.1885
1229.5908203125 0 1401.0122
1230.6146240234375 0 748.65466
1254.61669921875 0 1015.4741 z 2
1269.622314453125 0 979.16705
1270.6287841796875 0 5682.308 y 2
1271.6322021484375 0 5157.4443
1272.638427734375 0 3369.7017
1273.6365966796875 0 2017.2461
1286.6527099609375 0 2473.6406
1287.65087890625 0 3247.9763
1290.0701904296875 0 961.5885
1299.6295166015625 0 3680.6648
1300.6275634765625 0 1739.931
1312.628173828125 0 1207.1237 c Water loss 11
1329.656005859375 0 4206.9097
1330.6634521484375 0 98890.51 c 11
1331.6666259765625 0 74555.11
1332.6680908203125 0 36779.4
1333.6707763671875 0 10013.831
1334.6829833984375 0 1226.6154
1358.669921875 0 772.31
1384.6748046875 0 1237.1449 z Ammonia loss 1
1385.6607666015625 0 1032.0914
1386.6588134765625 0 923.65594
1400.6727294921875 0 774.2759 y Ammonia loss 1
1401.6771240234375 0 11946.309 z 1
1402.6827392578125 0 34327.523
1403.6864013671875 0 26052.543
1404.6905517578125 0 11776.807
1405.69091796875 0 3052.1143
1414.738525390625 0 4111.084
1415.7451171875 0 4764.3193
1416.7467041015625 0 2152.453
1417.7054443359375 0 1599.2028 y 1
1418.7030029296875 0 1478.571
1419.7208251953125 0 1048.3175
1442.7401123046875 0 4260.599
1443.739501953125 0 3201.9512
1444.7503662109375 0 2267.9255
1458.754638671875 0 5228.9824 c 12
1459.763671875 0 19615.023
1460.767578125 0 13125.985
1461.7686767578125 0 7581.62
1462.7734375 0 2380.275
1469.7572021484375 0 1414.0905
1470.7598876953125 0 2850.5154
1471.76318359375 0 1972.8363
1472.7607421875 0 817.8623
1473.758544921875 0 860.27344
1486.767578125 0 3967.8203
1487.7724609375 0 18661.256
1488.7757568359375 0 14919.64
1489.7786865234375 0 7448.5156
1490.780517578125 0 1234.0775
1491.7965087890625 0 875.1616
1496.7154541015625 0 774.9329
1498.7418212890625 0 1667.5271
1499.73828125 0 1367.1156
1503.6973876953125 0 849.72235
1504.794677734375 0 1710.6467
1505.7989501953125 0 1032.5377
1513.76220703125 0 1520.4734
1514.7635498046875 0 4943.489
1515.7677001953125 0 37812.676
1516.7711181640625 0 35382.973
1517.7728271484375 0 17820.75
1518.7747802734375 0 5378.873
1519.7685546875 0 1863.0096
1525.1942138671875 0 894.5669
1529.7572021484375 0 862.11115
1530.774658203125 0 3202.6558
1531.78466796875 0 15651.658
1532.7926025390625 0 51621.74
1533.794921875 0 39987.062
1534.1680908203125 0 806.3072
1534.798583984375 0 18794.348
1535.7967529296875 0 6116.524
1536.8095703125 0 1515.3524
2022.9771728515625 0 899.9327
3123.9716796875 0 899.1572

Spectrum Details

|  |  |
| --- | --- |
| Matched peaks? Matched peaks The total absolute number of peaks matched. Additionally in brackets the total fraction of peaks matched and the total number of peaks is shown. | 79 (15.86% of 498) |
| FDR? FDR The false discovery rate estimated for this peptide. It is calculated by matching all theoretical fragments with a non-integer shift with the raw peaks for this spectrum. This is done with 40 different shifts. The resulting percentage is the average number of annotated peaks over the number of annotated peaks with the correct spectrum. | 0.54% |
| Satellite FDR? Satellite FDR See the FDR for details on its calculation. This satellite ion specific FDR only contains the satellite ions (d/w) for I/L/J positions. | 2.38% |
| PSM Score? PSM Score The PSM Score as given by Hecklib to this annotated spectrum. It is shown with three significant figures. | 706 |

#### Spectrum 6136? Spectrum 6136 The raw spectrum of this peptide as annotated by Hecklib. The fragments are coloured according to ion type (see legend). Any peaks with a star '\*' as text can be hovered over to see the full details, first the ion type second the mass shift type. By hovering over the amino acids in the peptide or ions in the legend the corresponding peaks are highlighted. By toggling the 'Unassigned' label you can turn the background (unassigned) peaks on or off in the plot. By updating the slider in the Ion legend you can update the spectrum to only show the top X% of the peaks with labels. The top X% means any peak that is within X% of the highest intensity. By dragging in the spectrum you can zoom in to a specific part of the spectrum and use 'Zoom Out' to get back to the original zoom level. The annotation of the spectrum is based on the given sequence in the peptides file and is done with different software so inconsistencies are likely. The peaks are annotated based on the given sequence, with 20 ppm tolerance.

Copy Data

##### Spectrum 6136 (TSV)

###### Preview

```
Loading example...
```

*Click on the button to copy the data to your clipboard.*

Mz MinMz MaxIntensity Max

WidthHeightPeptide font sizePeptide stroke widthSpectrum font sizeSpectrum stroke widthCompact peptide

Ion legend

wxyz

abcd

OtherUnassignedIon^Charge^_Position_Show for top:%

JFPPSSEEJQANKA

01.31e+52.63e+53.94e+55.25e+5

Zoom Out

y^+1^_2_y^+1^_2_y^+2^_6_y^+1^_3_y^+1^_4_z^+1^_5_z^+1^_5_y^+1^_5_w^+1^_6_y^+2^_11_c^+2^_11_y^+2^_12_y^+2^_12_z^+2^_12_c^+1^_6_y^+2^_12_y^+1^_6_w^+1^_7_z^+2^_13_y^+2^_13_c^+2^_13_c^+1^_7_y^+1^_7_w^+1^_8_z^+1^_8_y^+1^_8_c^+1^_8_y^+1^_8_c^+1^_8_w^+1^_9_y^+1^_9_y^+1^_9_z^+1^_9_y^+1^_9_c^+1^_9_z^+1^_10_y^+1^_10_c^+1^_10_c^+1^_10_y^+1^_11_c^+1^_11_y^+1^_12_y^+1^_12_c^+1^_12_c^+1^_12_c^+1^_12_z^+1^_13_c^+1^_13_

0775155023253100

Fragment Matches Table

Show background peaks

| Position | Ion type | Intensity | mz Theoretical | mz Error (Th) | mz Error (ppm) | Charge | Series Number |
| --- | --- | --- | --- | --- | --- | --- | --- |
| - | - | 2401 | 120.1 | - | - | 0 | - |
| - | - | 4640 | 129.1 | - | - | 0 | - |
| - | - | 511.5 | 140.1 | - | - | 0 | - |
| - | - | 509.1 | 152.3 | - | - | 0 | - |
| - | - | 1836 | 159.1 | - | - | 0 | - |
| - | - | 566.5 | 168.9 | - | - | 0 | - |
| - | - | 1291 | 173.4 | - | - | 0 | - |
| - | - | 569.4 | 174.1 | - | - | 0 | - |
| - | - | 459.2 | 174.8 | - | - | 0 | - |
| - | - | 4502 | 187.1 | - | - | 0 | - |
| - | - | 510.8 | 189.1 | - | - | 0 | - |
| 13 | y | 515.4 | 201.1 | 0.0006729 | 3.346 | +1 | 2 |
| - | - | 2415 | 215.1 | - | - | 0 | - |
| - | - | 614.7 | 216.1 | - | - | 0 | - |
| - | - | 874 | 217.1 | - | - | 0 | - |
| 13 | y | 2371 | 218.1 | 0.0002896 | 1.327 | +1 | 2 |
| - | - | 5547 | 219.1 | - | - | 0 | - |
| - | - | 711.5 | 220.1 | - | - | 0 | - |
| - | - | 1638 | 226.1 | - | - | 0 | - |
| - | - | 2.372E+04 | 233.2 | - | - | 0 | - |
| - | - | 2977 | 234.2 | - | - | 0 | - |
| - | - | 527 | 236.1 | - | - | 0 | - |
| - | - | 1229 | 243.1 | - | - | 0 | - |
| - | - | 600.3 | 246.9 | - | - | 0 | - |
| - | - | 2.082E+04 | 261.2 | - | - | 0 | - |
| - | - | 4235 | 262.2 | - | - | 0 | - |
| - | - | 795.2 | 282.1 | - | - | 0 | - |
| - | - | 1922 | 297.2 | - | - | 0 | - |
| - | - | 5349 | 300.2 | - | - | 0 | - |
| - | - | 643 | 301.2 | - | - | 0 | - |
| 9 | y | 633.1 | 314.2 | 0.00574 | 18.27 | +2 | 6 |
| - | - | 1066 | 314.2 | - | - | 0 | - |
| 12 | y | 1608 | 332.2 | 0.0006666 | 2.007 | +1 | 3 |
| - | - | 4564 | 332.2 | - | - | 0 | - |
| - | - | 3533 | 358.2 | - | - | 0 | - |
| - | - | 1078 | 360.2 | - | - | 0 | - |
| - | - | 8372 | 401.2 | - | - | 0 | - |
| - | - | 1133 | 402.2 | - | - | 0 | - |
| 11 | y | 2686 | 403.2 | 0.0009979 | 2.475 | +1 | 4 |
| - | - | 615.7 | 414.8 | - | - | 0 | - |
| - | - | 621.2 | 434.7 | - | - | 0 | - |
| - | - | 751.2 | 455.3 | - | - | 0 | - |
| - | - | 658 | 482.3 | - | - | 0 | - |
| - | - | 2045 | 488.3 | - | - | 0 | - |
| - | - | 767.8 | 498.2 | - | - | 0 | - |
| 10 | z | 2188 | 498.2 | 0.008049 | 16.15 | +1 | 5 |
| - | - | 3222 | 500.3 | - | - | 0 | - |
| - | - | 1187 | 501.3 | - | - | 0 | - |
| 10 | z | 4694 | 515.3 | 0.0006342 | 1.231 | +1 | 5 |
| - | - | 8137 | 516.3 | - | - | 0 | - |
| - | - | 1934 | 517.3 | - | - | 0 | - |
| - | - | 1080 | 518.3 | - | - | 0 | - |
| - | - | 1.64E+04 | 518.3 | - | - | 0 | - |
| - | - | 4645 | 519.3 | - | - | 0 | - |
| 10 | y | 1766 | 531.3 | 0.001014 | 1.909 | +1 | 5 |
| - | - | 2403 | 542.3 | - | - | 0 | - |
| - | - | 662.5 | 550.3 | - | - | 0 | - |
| - | - | 743 | 582.2 | - | - | 0 | - |
| - | - | 1865 | 582.3 | - | - | 0 | - |
| - | - | 3093 | 582.8 | - | - | 0 | - |
| - | - | 892.6 | 583.3 | - | - | 0 | - |
| 9 | w | 913.4 | 585.3 | 0.0006423 | 1.097 | +1 | 6 |
| 4 | y | 3711 | 587.3 | 0.001294 | 2.204 | +2 | 11 |
| - | - | 2247 | 587.8 | - | - | 0 | - |
| - | - | 2082 | 591.3 | - | - | 0 | - |
| 11 | c | 1266 | 600.3 | 0.0009127 | 1.52 | +2 | 11 |
| - | - | 3065 | 616.3 | - | - | 0 | - |
| - | - | 740.2 | 617.3 | - | - | 0 | - |
| - | - | 811.7 | 618.3 | - | - | 0 | - |
| 3 | y | 5357 | 626.8 | 0.000335 | 0.5345 | +2 | 12 |
| - | - | 930.6 | 627.3 | - | - | 0 | - |
| 3 | y | 9644 | 627.3 | 0.003507 | 5.59 | +2 | 12 |
| 3 | z | 5139 | 627.8 | 0.0004668 | 0.7435 | +2 | 12 |
| - | - | 2703 | 628.3 | - | - | 0 | - |
| 6 | c | 2.011E+04 | 628.3 | 0.009657 | 15.37 | +1 | 6 |
| - | - | 647.4 | 628.8 | - | - | 0 | - |
| - | - | 1.569E+04 | 629.4 | - | - | 0 | - |
| - | - | 4239 | 630.4 | - | - | 0 | - |
| - | - | 752.4 | 631.4 | - | - | 0 | - |
| - | - | 1571 | 634.8 | - | - | 0 | - |
| - | - | 1189 | 635.3 | - | - | 0 | - |
| 3 | y | 5.198E+05 | 635.8 | 0.0009134 | 1.437 | +2 | 12 |
| - | - | 3.298E+05 | 636.3 | - | - | 0 | - |
| - | - | 1.255E+05 | 636.8 | - | - | 0 | - |
| - | - | 1.444E+04 | 637.3 | - | - | 0 | - |
| 9 | y | 2434 | 644.4 | 0.0004462 | 0.6925 | +1 | 6 |
| - | - | 5973 | 673.4 | - | - | 0 | - |
| - | - | 2021 | 674.4 | - | - | 0 | - |
| 8 | w | 993.6 | 698.4 | 0.0008411 | 1.204 | +1 | 7 |
| 2 | z | 1140 | 701.3 | 0.01032 | 14.72 | +2 | 13 |
| 2 | y | 6673 | 709.4 | 0.0009471 | 1.335 | +2 | 13 |
| - | - | 6064 | 709.9 | - | - | 0 | - |
| - | - | 2201 | 710.4 | - | - | 0 | - |
| 13 | c | 766.8 | 721.4 | 0.0009429 | 1.307 | +2 | 13 |
| - | - | 1460 | 740.3 | - | - | 0 | - |
| - | - | 765.5 | 743.5 | - | - | 0 | - |
| 7 | c | 2.949E+04 | 757.4 | 0.009056 | 11.96 | +1 | 7 |
| - | - | 3.67E+04 | 758.4 | - | - | 0 | - |
| - | - | 1.244E+04 | 759.4 | - | - | 0 | - |
| - | - | 2011 | 760.4 | - | - | 0 | - |
| - | - | 1.344E+04 | 765.9 | - | - | 0 | - |
| - | - | 1.159E+04 | 766.4 | - | - | 0 | - |
| - | - | 6253 | 766.9 | - | - | 0 | - |
| - | - | 874.4 | 767.4 | - | - | 0 | - |
| 8 | y | 4088 | 773.4 | 0.001005 | 1.299 | +1 | 7 |
| - | - | 1449 | 774.4 | - | - | 0 | - |
| 7 | w | 1145 | 827.4 | 0.002052 | 2.48 | +1 | 8 |
| - | - | 1127 | 862.4 | - | - | 0 | - |
| - | - | 4261 | 868.4 | - | - | 0 | - |
| 7 | z | 2301 | 869.4 | 0.003517 | 4.045 | +1 | 8 |
| 7 | y | 1107 | 885.4 | 0.005164 | 5.832 | +1 | 8 |
| 8 | c | 2.443E+04 | 886.4 | 0.009798 | 11.05 | +1 | 8 |
| - | - | 5.722E+04 | 887.4 | - | - | 0 | - |
| - | - | 2.04E+04 | 888.5 | - | - | 0 | - |
| - | - | 5021 | 889.5 | - | - | 0 | - |
| - | - | 880.5 | 890.4 | - | - | 0 | - |
| - | - | 3402 | 901.5 | - | - | 0 | - |
| 7 | y | 3802 | 902.5 | 0.001549 | 1.717 | +1 | 8 |
| - | - | 2608 | 903.4 | - | - | 0 | - |
| 8 | c | 8751 | 904.4 | 0.0006983 | 0.7721 | +1 | 8 |
| - | - | 3302 | 905.4 | - | - | 0 | - |
| - | - | 874.1 | 906.5 | - | - | 0 | - |
| - | - | 584.5 | 923.4 | - | - | 0 | - |
| - | - | 2961 | 939.4 | - | - | 0 | - |
| - | - | 2159 | 940.4 | - | - | 0 | - |
| 6 | w | 2500 | 956.5 | 0.0009478 | 0.9909 | +1 | 9 |
| - | - | 1938 | 957.5 | - | - | 0 | - |
| 6 | y | 838.2 | 971.5 | 0.005191 | 5.343 | +1 | 9 |
| 6 | y | 799.9 | 972.5 | 0.002981 | 3.066 | +1 | 9 |
| 6 | z | 4700 | 973.5 | 0.001626 | 1.67 | +1 | 9 |
| - | - | 7.694E+04 | 974.5 | - | - | 0 | - |
| - | - | 3.776E+04 | 975.5 | - | - | 0 | - |
| - | - | 9242 | 976.5 | - | - | 0 | - |
| - | - | 1.356E+04 | 988.5 | - | - | 0 | - |
| 6 | y | 2.217E+04 | 989.5 | 0.0009847 | 0.9952 | +1 | 9 |
| - | - | 8984 | 990.5 | - | - | 0 | - |
| - | - | 2320 | 991.5 | - | - | 0 | - |
| - | - | 738.9 | 999.5 | - | - | 0 | - |
| - | - | 1084 | 1001 | - | - | 0 | - |
| - | - | 821.6 | 1002 | - | - | 0 | - |
| - | - | 904.1 | 1015 | - | - | 0 | - |
| - | - | 792.6 | 1016 | - | - | 0 | - |
| - | - | 1.372E+04 | 1017 | - | - | 0 | - |
| 9 | c | 2.299E+04 | 1018 | 0.001334 | 1.311 | +1 | 9 |
| - | - | 1.224E+04 | 1019 | - | - | 0 | - |
| - | - | 2304 | 1020 | - | - | 0 | - |
| - | - | 892.3 | 1021 | - | - | 0 | - |
| - | - | 1364 | 1021 | - | - | 0 | - |
| - | - | 1938 | 1022 | - | - | 0 | - |
| - | - | 2277 | 1022 | - | - | 0 | - |
| - | - | 1220 | 1023 | - | - | 0 | - |
| - | - | 4351 | 1053 | - | - | 0 | - |
| - | - | 2789 | 1054 | - | - | 0 | - |
| 5 | z | 2088 | 1061 | 0.001885 | 1.778 | +1 | 10 |
| - | - | 2.858E+04 | 1062 | - | - | 0 | - |
| - | - | 1.535E+04 | 1063 | - | - | 0 | - |
| - | - | 3595 | 1064 | - | - | 0 | - |
| - | - | 820.8 | 1065 | - | - | 0 | - |
| - | - | 1792 | 1076 | - | - | 0 | - |
| 5 | y | 9033 | 1077 | 0.0002372 | 0.2204 | +1 | 10 |
| - | - | 3367 | 1078 | - | - | 0 | - |
| - | - | 1080 | 1079 | - | - | 0 | - |
| - | - | 4331 | 1102 | - | - | 0 | - |
| - | - | 2395 | 1103 | - | - | 0 | - |
| - | - | 1175 | 1104 | - | - | 0 | - |
| - | - | 721.8 | 1113 | - | - | 0 | - |
| 10 | c | 1491 | 1129 | 0.004554 | 4.035 | +1 | 10 |
| - | - | 1006 | 1130 | - | - | 0 | - |
| - | - | 1.317E+04 | 1145 | - | - | 0 | - |
| 10 | c | 8.967E+04 | 1146 | 0.0002687 | 0.2346 | +1 | 10 |
| - | - | 5.684E+04 | 1147 | - | - | 0 | - |
| - | - | 1.936E+04 | 1148 | - | - | 0 | - |
| - | - | 1691 | 1149 | - | - | 0 | - |
| - | - | 6701 | 1173 | - | - | 0 | - |
| 4 | y | 4.655E+04 | 1174 | 0.000954 | 0.8129 | +1 | 11 |
| - | - | 2.594E+04 | 1175 | - | - | 0 | - |
| - | - | 8692 | 1176 | - | - | 0 | - |
| - | - | 912.1 | 1177 | - | - | 0 | - |
| - | - | 1081 | 1182 | - | - | 0 | - |
| - | - | 1040 | 1183 | - | - | 0 | - |
| - | - | 846.9 | 1184 | - | - | 0 | - |
| - | - | 697 | 1197 | - | - | 0 | - |
| - | - | 695.9 | 1202 | - | - | 0 | - |
| - | - | 1.209E+04 | 1216 | - | - | 0 | - |
| 11 | c | 1.117E+04 | 1217 | 0.00291 | 2.391 | +1 | 11 |
| - | - | 4913 | 1218 | - | - | 0 | - |
| - | - | 1616 | 1219 | - | - | 0 | - |
| - | - | 956.8 | 1227 | - | - | 0 | - |
| 3 | y | 905.4 | 1254 | 0.009163 | 7.309 | +1 | 12 |
| - | - | 1990 | 1270 | - | - | 0 | - |
| 3 | y | 8.551E+04 | 1271 | 0.0002962 | 0.2331 | +1 | 12 |
| - | - | 5.612E+04 | 1272 | - | - | 0 | - |
| - | - | 1.927E+04 | 1273 | - | - | 0 | - |
| - | - | 2102 | 1274 | - | - | 0 | - |
| - | - | 1.221E+04 | 1287 | - | - | 0 | - |
| - | - | 1.091E+04 | 1288 | - | - | 0 | - |
| - | - | 789.9 | 1288 | - | - | 0 | - |
| - | - | 4095 | 1289 | - | - | 0 | - |
| - | - | 756.2 | 1290 | - | - | 0 | - |
| - | - | 1533 | 1299 | - | - | 0 | - |
| - | - | 1178 | 1300 | - | - | 0 | - |
| - | - | 716.7 | 1305 | - | - | 0 | - |
| 12 | c | 778.3 | 1313 | 0.01977 | 15.06 | +1 | 12 |
| 12 | c | 991.4 | 1314 | 0.003541 | 2.695 | +1 | 12 |
| - | - | 5210 | 1330 | - | - | 0 | - |
| 12 | c | 4.225E+04 | 1331 | 0.0005489 | 0.4125 | +1 | 12 |
| - | - | 2.723E+04 | 1332 | - | - | 0 | - |
| - | - | 1.142E+04 | 1333 | - | - | 0 | - |
| - | - | 1373 | 1334 | - | - | 0 | - |
| - | - | 973.6 | 1359 | - | - | 0 | - |
| - | - | 1124 | 1360 | - | - | 0 | - |
| 2 | z | 1.57E+04 | 1402 | 0.0009138 | 0.6519 | +1 | 13 |
| - | - | 1.197E+04 | 1403 | - | - | 0 | - |
| - | - | 3794 | 1404 | - | - | 0 | - |
| - | - | 1193 | 1405 | - | - | 0 | - |
| - | - | 1.303E+04 | 1415 | - | - | 0 | - |
| - | - | 1.119E+04 | 1416 | - | - | 0 | - |
| - | - | 4246 | 1417 | - | - | 0 | - |
| - | - | 1236 | 1424 | - | - | 0 | - |
| - | - | 2621 | 1431 | - | - | 0 | - |
| - | - | 1486 | 1432 | - | - | 0 | - |
| - | - | 842.9 | 1433 | - | - | 0 | - |
| - | - | 2395 | 1443 | - | - | 0 | - |
| - | - | 3100 | 1444 | - | - | 0 | - |
| - | - | 2459 | 1445 | - | - | 0 | - |
| - | - | 745 | 1445 | - | - | 0 | - |
| - | - | 801.1 | 1448 | - | - | 0 | - |
| - | - | 867.5 | 1457 | - | - | 0 | - |
| - | - | 1236 | 1458 | - | - | 0 | - |
| 13 | c | 9.15E+04 | 1459 | 0.002006 | 1.375 | +1 | 13 |
| - | - | 7.757E+04 | 1460 | - | - | 0 | - |
| - | - | 3.788E+04 | 1461 | - | - | 0 | - |
| - | - | 6743 | 1462 | - | - | 0 | - |
| - | - | 1218 | 1463 | - | - | 0 | - |
| - | - | 1519 | 1472 | - | - | 0 | - |
| - | - | 4444 | 1473 | - | - | 0 | - |
| - | - | 4389 | 1474 | - | - | 0 | - |
| - | - | 2167 | 1475 | - | - | 0 | - |
| - | - | 4451 | 1476 | - | - | 0 | - |
| - | - | 4379 | 1477 | - | - | 0 | - |
| - | - | 1702 | 1478 | - | - | 0 | - |
| - | - | 4600 | 1486 | - | - | 0 | - |
| - | - | 5.93E+04 | 1487 | - | - | 0 | - |
| - | - | 4.844E+04 | 1488 | - | - | 0 | - |
| - | - | 1760 | 1488 | - | - | 0 | - |
| - | - | 2.326E+04 | 1489 | - | - | 0 | - |
| - | - | 4323 | 1490 | - | - | 0 | - |
| - | - | 900.4 | 1495 | - | - | 0 | - |
| - | - | 1026 | 1497 | - | - | 0 | - |
| - | - | 855.7 | 1498 | - | - | 0 | - |
| - | - | 672.4 | 1499 | - | - | 0 | - |
| - | - | 841.6 | 1500 | - | - | 0 | - |
| - | - | 1536 | 1501 | - | - | 0 | - |
| - | - | 1467 | 1502 | - | - | 0 | - |
| - | - | 1002 | 1502 | - | - | 0 | - |
| - | - | 1947 | 1503 | - | - | 0 | - |
| - | - | 2601 | 1503 | - | - | 0 | - |
| - | - | 1938 | 1504 | - | - | 0 | - |
| - | - | 2087 | 1504 | - | - | 0 | - |
| - | - | 990.5 | 1504 | - | - | 0 | - |
| - | - | 2633 | 1505 | - | - | 0 | - |
| - | - | 774.6 | 1505 | - | - | 0 | - |
| - | - | 1493 | 1506 | - | - | 0 | - |
| - | - | 670.4 | 1507 | - | - | 0 | - |
| - | - | 1003 | 1510 | - | - | 0 | - |
| - | - | 1733 | 1510 | - | - | 0 | - |
| - | - | 1971 | 1511 | - | - | 0 | - |
| - | - | 1679 | 1511 | - | - | 0 | - |
| - | - | 1829 | 1512 | - | - | 0 | - |
| - | - | 679.3 | 1512 | - | - | 0 | - |
| - | - | 813 | 1513 | - | - | 0 | - |
| - | - | 5664 | 1514 | - | - | 0 | - |
| - | - | 1.006E+05 | 1515 | - | - | 0 | - |
| - | - | 8.961E+04 | 1516 | - | - | 0 | - |
| - | - | 4.038E+04 | 1517 | - | - | 0 | - |
| - | - | 5670 | 1518 | - | - | 0 | - |
| - | - | 911.7 | 1520 | - | - | 0 | - |
| - | - | 1136 | 1521 | - | - | 0 | - |
| - | - | 1066 | 1522 | - | - | 0 | - |
| - | - | 985.7 | 1523 | - | - | 0 | - |
| - | - | 4614 | 1524 | - | - | 0 | - |
| - | - | 7007 | 1524 | - | - | 0 | - |
| - | - | 5408 | 1525 | - | - | 0 | - |
| - | - | 5306 | 1525 | - | - | 0 | - |
| - | - | 3459 | 1526 | - | - | 0 | - |
| - | - | 2003 | 1526 | - | - | 0 | - |
| - | - | 2260 | 1529 | - | - | 0 | - |
| - | - | 1969 | 1529 | - | - | 0 | - |
| - | - | 9652 | 1530 | - | - | 0 | - |
| - | - | 2111 | 1530 | - | - | 0 | - |
| - | - | 1.683E+05 | 1531 | - | - | 0 | - |
| - | - | 4.772E+05 | 1532 | - | - | 0 | - |
| - | - | 3.557E+05 | 1533 | - | - | 0 | - |
| - | - | 1.189E+04 | 1533 | - | - | 0 | - |
| - | - | 1.504E+05 | 1534 | - | - | 0 | - |
| - | - | 5005 | 1534 | - | - | 0 | - |
| - | - | 2.101E+04 | 1535 | - | - | 0 | - |
| - | - | 888.4 | 1881 | - | - | 0 | - |
| - | - | 692.1 | 2002 | - | - | 0 | - |
| - | - | 674.8 | 2024 | - | - | 0 | - |
| - | - | 764.1 | 2295 | - | - | 0 | - |
| - | - | 1028 | 2297 | - | - | 0 | - |
| - | - | 853.7 | 2302 | - | - | 0 | - |
| - | - | 1513 | 2391 | - | - | 0 | - |
| - | - | 1860 | 2392 | - | - | 0 | - |
| - | - | 1730 | 2393 | - | - | 0 | - |
| - | - | 1079 | 2577 | - | - | 0 | - |
| - | - | 852.6 | 2704 | - | - | 0 | - |
| - | - | 756.3 | 2705 | - | - | 0 | - |
| - | - | 926.1 | 2707 | - | - | 0 | - |
| - | - | 1069 | 3005 | - | - | 0 | - |
| - | - | 1626 | 3006 | - | - | 0 | - |
| - | - | 858 | 3020 | - | - | 0 | - |
| - | - | 957.2 | 3021 | - | - | 0 | - |
| - | - | 843.5 | 3042 | - | - | 0 | - |
| - | - | 1010 | 3048 | - | - | 0 | - |
| - | - | 1053 | 3049 | - | - | 0 | - |
| - | - | 1058 | 3050 | - | - | 0 | - |
| - | - | 795.7 | 3057 | - | - | 0 | - |
| - | - | 744.1 | 3058 | - | - | 0 | - |
| - | - | 874.4 | 3060 | - | - | 0 | - |
| - | - | 2437 | 3063 | - | - | 0 | - |
| - | - | 4031 | 3064 | - | - | 0 | - |
| - | - | 3908 | 3065 | - | - | 0 | - |
| - | - | 4000 | 3066 | - | - | 0 | - |
| - | - | 1743 | 3067 | - | - | 0 | - |
| - | - | 1007 | 3068 | - | - | 0 | - |
| - | - | 900.1 | 3069 | - | - | 0 | - |

m/z Charge Intensity FragmentType MassShift Position
120.08106994628906 0 2400.9126
129.10250854492188 0 4639.638
140.14593505859375 0 511.4879
152.31886291503906 0 509.0744
159.11294555664062 0 1835.824
168.8926544189453 0 566.53076
173.43795776367188 0 1290.9144
174.13584899902344 0 569.4419
174.8147735595703 0 459.19846
187.1079864501953 0 4502.1025
189.05206298828125 0 510.75635
201.12269592285156 0 515.35675 y Ammonia loss 12
215.13909912109375 0 2414.8645
216.14230346679688 0 614.71533
217.13343811035156 0 873.96436
218.15020751953125 0 2370.701 y 12
219.13427734375 0 5546.8384
220.13723754882812 0 711.50006
226.1192169189453 0 1638.465
233.1652069091797 0 23724.66
234.1685791015625 0 2977.0566
236.11317443847656 0 526.97144
243.14573669433594 0 1228.6138
246.9043426513672 0 600.27905
261.16009521484375 0 20818.975
262.163330078125 0 4235.203
282.1455383300781 0 795.16095
297.1564025878906 0 1921.6637
300.1922912597656 0 5349.118
301.1954345703125 0 643.00085
314.1824035644531 0 633.1116 y Ammonia loss 8
314.20751953125 0 1066.17
332.1935119628906 0 1607.8383 y 11
332.2184753417969 0 4564.01
358.2133483886719 0 3532.9265
360.2143249511719 0 1077.5889
401.2399597167969 0 8371.8955
402.24346923828125 0 1132.5363
403.23095703125 0 2685.5159 y 10
414.8130187988281 0 615.7413
434.7046813964844 0 621.183
455.2657775878906 0 751.1529
482.2969665527344 0 658.0116
488.27349853515625 0 2045.0787
498.2161865234375 0 767.79205
498.2513122558594 0 2188.0173 z Ammonia loss 9
500.3091125488281 0 3222.2742
501.312255859375 0 1187.1066
515.2704467773438 0 4694.2866 z 9
516.27734375 0 8136.8774
517.2800903320312 0 1933.7902
518.27734375 0 1080.1794
518.3192138671875 0 16397.844
519.3221435546875 0 4645.2188
531.28955078125 0 1765.7047 y 9
542.2772827148438 0 2402.6716
550.3074951171875 0 662.5341
582.2365112304688 0 742.95105
582.2901611328125 0 1864.8074
582.7838134765625 0 3093.0732
583.2866821289062 0 892.5576
585.2997436523438 0 913.4031 w 8
587.292236328125 0 3711.121 y 3
587.79345703125 0 2247.1501
591.2944946289062 0 2081.7612
600.2998657226562 0 1266.0026 c Ammonia loss 10
616.3311157226562 0 3064.784
617.3353271484375 0 740.17596
618.298095703125 0 811.70886
626.8117065429688 0 5356.727 y Water loss 2
627.2634887695312 0 930.6379
627.3075561523438 0 9644.367 y Ammonia loss 2
627.8074951171875 0 5139.406 z 2
628.30615234375 0 2703.3877
628.35498046875 0 20105.771 c Water loss 5
628.8135986328125 0 647.4289
629.3601684570312 0 15693.177
630.3640747070312 0 4239.2983
631.3682861328125 0 752.3565
634.8115234375 0 1570.6932
635.3092651367188 0 1188.9316
635.8182373046875 0 519804.5 y 2
636.31982421875 0 329800.84
636.82080078125 0 125483.67
637.322265625 0 14437.395
644.373046875 0 2434.2551 y 8
673.3538208007812 0 5973.1035
674.355712890625 0 2020.5767
698.38232421875 0 993.6303 w 7
701.3318481445312 0 1140.3451 z 1
709.3524780273438 0 6672.645 y 1
709.8536987304688 0 6063.92
710.3558959960938 0 2201.2324
721.3706665039062 0 766.77844 c Ammonia loss 12
740.34765625 0 1460.1812
743.4891967773438 0 765.4913
757.39697265625 0 29485.969 c Water loss 6
758.4033813476562 0 36702.77
759.4064331054688 0 12442.55
760.4095458984375 0 2011.4281
765.8944702148438 0 13444.55
766.3954467773438 0 11588.21
766.8968505859375 0 6252.546
767.4014892578125 0 874.4422
773.4161987304688 0 4087.9602 y 7
774.4179077148438 0 1448.9312
827.4237060546875 0 1144.611 w 6
862.4057006835938 0 1127.393
868.40576171875 0 4260.8164
869.4089965820312 0 2300.528 z Ammonia loss 6
885.4364013671875 0 1107.3818 y Ammonia loss 6
886.4403076171875 0 24428.674 c Water loss 7
887.4466552734375 0 57215.695
888.450439453125 0 20399.31
889.4539184570312 0 5021.1445
890.4451904296875 0 880.501
901.4501342773438 0 3401.5925
902.4562377929688 0 3801.7288 y 6
903.4424438476562 0 2608.1282
904.4417724609375 0 8750.937 c 7
905.4441528320312 0 3302.2158
906.4508666992188 0 874.1435
923.415283203125 0 584.48956
939.4410400390625 0 2961.4182
940.443603515625 0 2158.6572
956.4692993164062 0 2500.2107 w 5
957.4686279296875 0 1938.3604
971.4740600585938 0 838.21967 y Water loss 5
972.4662475585938 0 799.89685 y Ammonia loss 5
973.4727172851562 0 4700.3066 z 5
974.4795532226562 0 76939.1
975.4832153320312 0 37755.586
976.4846801757812 0 9241.874
988.4830322265625 0 13564.091
989.4888305664062 0 22170.562 y 5
990.4921875 0 8984.04
991.4942626953125 0 2320.1929
999.475830078125 0 738.9064
1000.5011596679688 0 1084.1786
1002.4854125976562 0 821.57416
1015.4873046875 0 904.09607
1016.1370239257812 0 792.6273
1016.5160522460938 0 13716.459
1017.5238037109375 0 22992.771 c 8
1018.527587890625 0 12241.639
1019.5291137695312 0 2303.8843
1021.129150390625 0 892.2901
1021.4655151367188 0 1364.1111
1021.7998046875 0 1937.9355
1022.1366577148438 0 2276.5698
1022.8071899414062 0 1219.6003
1053.4853515625 0 4351.4473
1054.4873046875 0 2789.3223
1060.5050048828125 0 2087.556 z 4
1061.51171875 0 28579.893
1062.513916015625 0 15346.79
1063.5174560546875 0 3595.385
1064.51904296875 0 820.82446
1075.5159912109375 0 1791.9813
1076.5216064453125 0 9033.329 y 4
1077.5245361328125 0 3367.4702
1078.52685546875 0 1080.2003
1101.5701904296875 0 4330.628
1102.575439453125 0 2395.0833
1103.5806884765625 0 1175.3774
1112.5570068359375 0 721.7608
1128.5526123046875 0 1490.6118 c Ammonia loss 9
1129.56005859375 0 1006.1639
1144.57666015625 0 13170.273
1145.583984375 0 89670.05 c 9
1146.5872802734375 0 56843.766
1147.589599609375 0 19364.299
1148.5899658203125 0 1691.0737
1172.568603515625 0 6701.037
1173.5755615234375 0 46553.668 y 3
1174.5775146484375 0 25941.91
1175.580810546875 0 8691.804
1176.5849609375 0 912.1274
1181.5771484375 0 1081.3481
1182.5860595703125 0 1039.6091
1183.5802001953125 0 846.8527
1196.532958984375 0 697.00146
1201.575927734375 0 695.8544
1215.613037109375 0 12094.294
1216.617919921875 0 11166.155 c 10
1217.6219482421875 0 4912.5093
1218.6287841796875 0 1616.4712
1227.0435791015625 0 956.77057
1253.6099853515625 0 905.3652 y Ammonia loss 2
1269.6182861328125 0 1989.9264
1270.6270751953125 0 85509.05 y 2
1271.6295166015625 0 56119.316
1272.63330078125 0 19271.28
1273.6287841796875 0 2102.1333
1286.6490478515625 0 12212.723
1287.65234375 0 10908.527
1288.0736083984375 0 789.8892
1288.6558837890625 0 4095.3337
1289.658203125 0 756.2336
1298.619384765625 0 1532.7291
1299.61865234375 0 1177.9579
1304.5731201171875 0 716.6972
1312.6334228515625 0 778.3447 c Water loss 11
1313.6336669921875 0 991.44366 c Ammonia loss 11
1329.6519775390625 0 5210.364
1330.6632080078125 0 42247.2 c 11
1331.666259765625 0 27225.98
1332.66796875 0 11422.452
1333.6676025390625 0 1372.5305
1359.1124267578125 0 973.58026
1360.10400390625 0 1124.151
1401.6761474609375 0 15703.112 z 1
1402.67919921875 0 11970.814
1403.6795654296875 0 3793.7312
1404.681640625 0 1192.8646
1414.743896484375 0 13033.68
1415.7452392578125 0 11190.139
1416.744140625 0 4245.9585
1423.64404296875 0 1236.3522
1430.709228515625 0 2620.705
1431.700439453125 0 1486.4218
1432.69482421875 0 842.90643
1442.73974609375 0 2395.26
1443.7271728515625 0 3100.197
1444.7314453125 0 2458.8381
1445.1402587890625 0 744.98175
1447.6737060546875 0 801.09186
1456.728271484375 0 867.5083
1457.738037109375 0 1236.4199
1458.7567138671875 0 91498.54 c 12
1459.7596435546875 0 77571.16
1460.7613525390625 0 37877.94
1461.7586669921875 0 6743.1333
1462.7421875 0 1217.546
1471.7076416015625 0 1519.3772
1472.7445068359375 0 4444.2188
1473.745361328125 0 4388.688
1474.7479248046875 0 2166.8271
1475.722900390625 0 4450.9756
1476.7269287109375 0 4379.3477
1477.7288818359375 0 1701.5847
1485.7803955078125 0 4599.687
1486.7657470703125 0 59303.78
1487.7685546875 0 48442.305
1488.1978759765625 0 1760.2087
1488.7684326171875 0 23255.598
1489.75634765625 0 4323.4443
1495.1988525390625 0 900.38684
1496.7542724609375 0 1025.9252
1497.7335205078125 0 855.74475
1498.769287109375 0 672.4368
1500.225341796875 0 841.5999
1500.7447509765625 0 1536.3318
1501.6982421875 0 1467.1019
1502.1990966796875 0 1001.58484
1502.7022705078125 0 1946.8463
1503.1947021484375 0 2600.879
1503.7186279296875 0 1937.6953
1503.7880859375 0 2086.6428
1504.214599609375 0 990.54974
1504.7652587890625 0 2632.9417
1505.2252197265625 0 774.556
1505.7977294921875 0 1493.4408
1507.2203369140625 0 670.4143
1509.7012939453125 0 1002.69946
1510.1923828125 0 1732.7434
1510.69970703125 0 1971.2946
1511.2012939453125 0 1679.1252
1511.7022705078125 0 1828.971
1512.2366943359375 0 679.2538
1512.7518310546875 0 813.02246
1513.7684326171875 0 5664.084
1514.7607421875 0 100553.22
1515.76318359375 0 89610.516
1516.765380859375 0 40377.19
1517.76904296875 0 5669.7217
1519.722412109375 0 911.65155
1520.74658203125 0 1136.0533
1521.7633056640625 0 1066.3258
1523.1947021484375 0 985.65076
1523.688232421875 0 4614.276
1524.1900634765625 0 7006.7676
1524.6976318359375 0 5408.351
1525.1966552734375 0 5305.613
1525.7093505859375 0 3458.6943
1526.2025146484375 0 2003.289
1528.75830078125 0 2260.3032
1529.215576171875 0 1969.005
1529.763427734375 0 9651.68
1530.2930908203125 0 2110.787
1530.7783203125 0 168266.52
1531.7855224609375 0 477153.88
1532.7884521484375 0 355671.75
1533.1932373046875 0 11894.589
1533.79052734375 0 150429.22
1534.2310791015625 0 5005.099
1534.7904052734375 0 21009.762
1880.8214111328125 0 888.42224
2001.9532470703125 0 692.072
2023.971923828125 0 674.75226
2295.044921875 0 764.1373
2297.177001953125 0 1027.7166
2302.056884765625 0 853.6663
2391.038330078125 0 1512.687
2392.043212890625 0 1860.46
2393.059814453125 0 1730.1051
2577.12451171875 0 1079.1559
2704.17626953125 0 852.5924
2705.181884765625 0 756.2638
2707.198486328125 0 926.1145
3005.34423828125 0 1068.557
3006.40185546875 0 1625.8496
3020.3837890625 0 858.01086
3021.370361328125 0 957.1768
3041.52197265625 0 843.54095
3048.378173828125 0 1010.1913
3049.33935546875 0 1053.4374
3050.348876953125 0 1057.9307
3057.484130859375 0 795.729
3058.49560546875 0 744.1378
3059.5078125 0 874.39075
3063.415283203125 0 2437.4175
3064.404541015625 0 4030.9758
3065.400390625 0 3907.9485
3066.4140625 0 4000.3328
3067.44140625 0 1743.2626
3068.4326171875 0 1006.8534
3069.4716796875 0 900.1323

Spectrum Details

|  |  |
| --- | --- |
| Matched peaks? Matched peaks The total absolute number of peaks matched. Additionally in brackets the total fraction of peaks matched and the total number of peaks is shown. | 48 (14.63% of 328) |
| FDR? FDR The false discovery rate estimated for this peptide. It is calculated by matching all theoretical fragments with a non-integer shift with the raw peaks for this spectrum. This is done with 40 different shifts. The resulting percentage is the average number of annotated peaks over the number of annotated peaks with the correct spectrum. | 0.40% |
| Satellite FDR? Satellite FDR See the FDR for details on its calculation. This satellite ion specific FDR only contains the satellite ions (d/w) for I/L/J positions. | 0.00% |
| PSM Score? PSM Score The PSM Score as given by Hecklib to this annotated spectrum. It is shown with three significant figures. | 436 |

#### Spectrum 6225? Spectrum 6225 The raw spectrum of this peptide as annotated by Hecklib. The fragments are coloured according to ion type (see legend). Any peaks with a star '\*' as text can be hovered over to see the full details, first the ion type second the mass shift type. By hovering over the amino acids in the peptide or ions in the legend the corresponding peaks are highlighted. By toggling the 'Unassigned' label you can turn the background (unassigned) peaks on or off in the plot. By updating the slider in the Ion legend you can update the spectrum to only show the top X% of the peaks with labels. The top X% means any peak that is within X% of the highest intensity. By dragging in the spectrum you can zoom in to a specific part of the spectrum and use 'Zoom Out' to get back to the original zoom level. The annotation of the spectrum is based on the given sequence in the peptides file and is done with different software so inconsistencies are likely. The peaks are annotated based on the given sequence, with 20 ppm tolerance.

Copy Data

##### Spectrum 6225 (TSV)

###### Preview

```
Loading example...
```

*Click on the button to copy the data to your clipboard.*

Mz MinMz MaxIntensity Max

WidthHeightPeptide font sizePeptide stroke widthSpectrum font sizeSpectrum stroke widthCompact peptide

Ion legend

wxyz

abcd

OtherUnassignedIon^Charge^_Position_Show for top:%

JFPPSSEEJQANKA

01.35e+42.71e+44.06e+45.42e+4

Zoom Out

w^+1^_2_z^+1^_2_y^+1^_2_y^+2^_4_z^+1^_2_y^+1^_2_y^+2^_5_w^+1^_3_y^+2^_6_y^+2^_6_y^+1^_3_y^+1^_4_y^+2^_7_y^+1^_4_z^+3^_12_w^+1^_5_c^+1^_4_z^+2^_9_y^+1^_5_z^+1^_5_y^+2^_10_y^+1^_5_c^+1^_5_w^+1^_6_y^+2^_11_c^+2^_11_c^+2^_11_z^+2^_12_y^+2^_12_y^+2^_12_z^+2^_12_c^+1^_6_y^+2^_12_y^+1^_6_c^+1^_6_c^+2^_12_c^+2^_12_z^+2^_13_z^+2^_13_w^+1^_7_z^+2^_13_c^+2^_13_c^+2^_13_y^+1^_7_c^+1^_7_y^+1^_7_c^+1^_7_w^+1^_8_y^+1^_8_c^+1^_8_y^+1^_8_c^+1^_8_y^+1^_9_z^+1^_9_y^+1^_9_c^+1^_9_z^+1^_10_y^+1^_10_c^+1^_10_y^+1^_11_c^+1^_11_c^+1^_11_w^+1^_12_z^+1^_12_y^+1^_12_c^+1^_12_z^+1^_13_c^+1^_13_

038877611641552

Fragment Matches Table

Show background peaks

| Position | Ion type | Intensity | mz Theoretical | mz Error (Th) | mz Error (ppm) | Charge | Series Number |
| --- | --- | --- | --- | --- | --- | --- | --- |
| - | - | 3081 | 120.1 | - | - | 0 | - |
| - | - | 1285 | 126.1 | - | - | 0 | - |
| - | - | 923.3 | 127.1 | - | - | 0 | - |
| - | - | 1728 | 128.1 | - | - | 0 | - |
| - | - | 485.5 | 129.1 | - | - | 0 | - |
| - | - | 4019 | 129.1 | - | - | 0 | - |
| - | - | 444 | 130.1 | - | - | 0 | - |
| - | - | 4423 | 131.1 | - | - | 0 | - |
| - | - | 3377 | 132.1 | - | - | 0 | - |
| - | - | 428.7 | 133.1 | - | - | 0 | - |
| - | - | 372 | 134.6 | - | - | 0 | - |
| - | - | 907.2 | 136.1 | - | - | 0 | - |
| - | - | 494.8 | 140.6 | - | - | 0 | - |
| 13 | w | 4480 | 144.1 | 9.318E-05 | 0.6468 | +1 | 2 |
| - | - | 1540 | 145.1 | - | - | 0 | - |
| - | - | 490 | 145.1 | - | - | 0 | - |
| - | - | 573.9 | 155.1 | - | - | 0 | - |
| - | - | 1460 | 165.1 | - | - | 0 | - |
| - | - | 516.5 | 169.1 | - | - | 0 | - |
| - | - | 640.5 | 171.1 | - | - | 0 | - |
| - | - | 597.8 | 173.1 | - | - | 0 | - |
| - | - | 3048 | 173.4 | - | - | 0 | - |
| - | - | 5408 | 183.1 | - | - | 0 | - |
| 13 | z | 1749 | 185.1 | 1.526E-05 | 0.08242 | +1 | 2 |
| - | - | 583.3 | 186.1 | - | - | 0 | - |
| - | - | 655.1 | 187.1 | - | - | 0 | - |
| - | - | 546.1 | 187.1 | - | - | 0 | - |
| - | - | 675.2 | 187.1 | - | - | 0 | - |
| 13 | y | 1684 | 201.1 | 1.373E-05 | 0.06824 | +1 | 2 |
| - | - | 776.4 | 202.1 | - | - | 0 | - |
| 11 | y | 799 | 202.1 | 0.003788 | 18.74 | +2 | 4 |
| 13 | z | 8303 | 202.1 | 4.697E-05 | 0.2324 | +1 | 2 |
| - | - | 630.2 | 203.1 | - | - | 0 | - |
| - | - | 638.2 | 211.1 | - | - | 0 | - |
| - | - | 7477 | 212.1 | - | - | 0 | - |
| - | - | 1010 | 213.1 | - | - | 0 | - |
| - | - | 1280 | 215.1 | - | - | 0 | - |
| - | - | 480.4 | 216.1 | - | - | 0 | - |
| - | - | 526.6 | 217.1 | - | - | 0 | - |
| 13 | y | 5998 | 218.1 | 4.612E-05 | 0.2114 | +1 | 2 |
| - | - | 718.3 | 219.2 | - | - | 0 | - |
| - | - | 473 | 224.1 | - | - | 0 | - |
| - | - | 2757 | 229.1 | - | - | 0 | - |
| - | - | 793.4 | 229.2 | - | - | 0 | - |
| - | - | 773.1 | 230.1 | - | - | 0 | - |
| - | - | 3994 | 232.1 | - | - | 0 | - |
| - | - | 8270 | 233.1 | - | - | 0 | - |
| - | - | 1.447E+04 | 233.2 | - | - | 0 | - |
| - | - | 1176 | 234.2 | - | - | 0 | - |
| - | - | 1119 | 234.2 | - | - | 0 | - |
| - | - | 556.3 | 238.2 | - | - | 0 | - |
| - | - | 1146 | 243.1 | - | - | 0 | - |
| - | - | 606.9 | 247.1 | - | - | 0 | - |
| - | - | 988.9 | 251.2 | - | - | 0 | - |
| - | - | 783 | 260.2 | - | - | 0 | - |
| - | - | 7115 | 261.2 | - | - | 0 | - |
| - | - | 1111 | 262.2 | - | - | 0 | - |
| 10 | y | 942.7 | 266.1 | 7.942E-05 | 0.2984 | +2 | 5 |
| - | - | 600.7 | 266.6 | - | - | 0 | - |
| - | - | 1369 | 270.1 | - | - | 0 | - |
| 12 | w | 5599 | 272.2 | 8.224E-05 | 0.3022 | +1 | 3 |
| - | - | 583.7 | 281.1 | - | - | 0 | - |
| - | - | 1476 | 282.1 | - | - | 0 | - |
| - | - | 1705 | 283.1 | - | - | 0 | - |
| - | - | 656.5 | 297.2 | - | - | 0 | - |
| - | - | 577.5 | 298.1 | - | - | 0 | - |
| - | - | 2.976E+04 | 299.2 | - | - | 0 | - |
| - | - | 3790 | 300.2 | - | - | 0 | - |
| - | - | 691.5 | 302.2 | - | - | 0 | - |
| - | - | 625.8 | 307.5 | - | - | 0 | - |
| - | - | 666.5 | 311.2 | - | - | 0 | - |
| - | - | 595.2 | 311.3 | - | - | 0 | - |
| 9 | y | 878.4 | 314.2 | 0.001437 | 4.573 | +2 | 6 |
| 9 | y | 1382 | 322.7 | 0.0002165 | 0.6708 | +2 | 6 |
| - | - | 967.2 | 325.2 | - | - | 0 | - |
| 12 | y | 2309 | 332.2 | 6.585E-05 | 0.1982 | +1 | 3 |
| - | - | 959.4 | 342.2 | - | - | 0 | - |
| - | - | 3809 | 343.2 | - | - | 0 | - |
| - | - | 2453 | 358.2 | - | - | 0 | - |
| - | - | 703.8 | 369.2 | - | - | 0 | - |
| - | - | 1158 | 370.7 | - | - | 0 | - |
| 11 | y | 1.648E+04 | 386.2 | 2.01E-05 | 0.05204 | +1 | 4 |
| 8 | y | 4077 | 387.2 | 0.003746 | 9.675 | +2 | 7 |
| - | - | 1192 | 395.2 | - | - | 0 | - |
| - | - | 887.8 | 402.2 | - | - | 0 | - |
| - | - | 2700 | 403.2 | - | - | 0 | - |
| 11 | y | 7905 | 403.2 | 7.026E-05 | 0.1743 | +1 | 4 |
| - | - | 583.8 | 404.2 | - | - | 0 | - |
| - | - | 1047 | 404.2 | - | - | 0 | - |
| 3 | z | 659.2 | 418.9 | 0.002116 | 5.052 | +3 | 12 |
| - | - | 656.9 | 428.3 | - | - | 0 | - |
| - | - | 896.3 | 434.7 | - | - | 0 | - |
| - | - | 590.7 | 439.1 | - | - | 0 | - |
| - | - | 748.2 | 444.2 | - | - | 0 | - |
| - | - | 1573 | 449.1 | - | - | 0 | - |
| - | - | 751.3 | 453.3 | - | - | 0 | - |
| - | - | 854.1 | 455.3 | - | - | 0 | - |
| 10 | w | 2.866E+04 | 457.2 | 0.0002293 | 0.5015 | +1 | 5 |
| - | - | 6041 | 458.2 | - | - | 0 | - |
| - | - | 1180 | 459.2 | - | - | 0 | - |
| - | - | 764.1 | 465.2 | - | - | 0 | - |
| - | - | 737.6 | 471.3 | - | - | 0 | - |
| 4 | c | 720.4 | 472.3 | 0.00361 | 7.643 | +1 | 4 |
| - | - | 776.5 | 474.3 | - | - | 0 | - |
| 6 | z | 1138 | 487.2 | 0.001177 | 2.416 | +2 | 9 |
| - | - | 924.9 | 488.2 | - | - | 0 | - |
| - | - | 2205 | 488.3 | - | - | 0 | - |
| - | - | 942.4 | 494.2 | - | - | 0 | - |
| - | - | 1523 | 496.2 | - | - | 0 | - |
| - | - | 674.1 | 497.3 | - | - | 0 | - |
| - | - | 2178 | 498.2 | - | - | 0 | - |
| - | - | 711.2 | 510.3 | - | - | 0 | - |
| - | - | 1054 | 511.3 | - | - | 0 | - |
| - | - | 833.7 | 511.3 | - | - | 0 | - |
| - | - | 646.3 | 511.8 | - | - | 0 | - |
| - | - | 776.6 | 512.2 | - | - | 0 | - |
| - | - | 1129 | 512.3 | - | - | 0 | - |
| - | - | 1264 | 512.3 | - | - | 0 | - |
| - | - | 1398 | 513.3 | - | - | 0 | - |
| - | - | 2538 | 514.2 | - | - | 0 | - |
| 10 | y | 3247 | 514.3 | 0.0009519 | 1.851 | +1 | 5 |
| - | - | 996 | 515.2 | - | - | 0 | - |
| 10 | z | 2635 | 515.3 | 0.0007697 | 1.494 | +1 | 5 |
| - | - | 985.6 | 516.3 | - | - | 0 | - |
| 5 | y | 2363 | 530.3 | 0.006535 | 12.32 | +2 | 10 |
| - | - | 3160 | 531.2 | - | - | 0 | - |
| 10 | y | 8315 | 531.3 | 3.753E-05 | 0.07063 | +1 | 5 |
| - | - | 1349 | 532.3 | - | - | 0 | - |
| - | - | 2067 | 532.3 | - | - | 0 | - |
| - | - | 1217 | 542.3 | - | - | 0 | - |
| - | - | 731.1 | 543.3 | - | - | 0 | - |
| - | - | 1088 | 557.3 | - | - | 0 | - |
| 5 | c | 7020 | 559.3 | 0.0001291 | 0.2308 | +1 | 5 |
| - | - | 2723 | 560.3 | - | - | 0 | - |
| - | - | 671.5 | 561.3 | - | - | 0 | - |
| - | - | 2192 | 570.3 | - | - | 0 | - |
| - | - | 788.9 | 571.3 | - | - | 0 | - |
| 9 | w | 4.048E+04 | 585.3 | 3.192E-05 | 0.05454 | +1 | 6 |
| - | - | 1.196E+04 | 586.3 | - | - | 0 | - |
| 4 | y | 4090 | 587.3 | 0.007825 | 13.32 | +2 | 11 |
| - | - | 881.9 | 591.3 | - | - | 0 | - |
| - | - | 1203 | 599.3 | - | - | 0 | - |
| 11 | c | 7095 | 599.8 | 0.001886 | 3.144 | +2 | 11 |
| 11 | c | 4456 | 600.3 | 0.007327 | 12.21 | +2 | 11 |
| - | - | 2443 | 600.8 | - | - | 0 | - |
| - | - | 790.6 | 601.3 | - | - | 0 | - |
| - | - | 796.9 | 602.3 | - | - | 0 | - |
| - | - | 2089 | 613.8 | - | - | 0 | - |
| - | - | 3790 | 614.3 | - | - | 0 | - |
| - | - | 2247 | 614.8 | - | - | 0 | - |
| - | - | 1100 | 615.3 | - | - | 0 | - |
| - | - | 1422 | 616.3 | - | - | 0 | - |
| - | - | 1131 | 618.3 | - | - | 0 | - |
| - | - | 600.3 | 619 | - | - | 0 | - |
| 3 | z | 625 | 619.3 | 0.002573 | 4.155 | +2 | 12 |
| 3 | y | 1268 | 626.8 | 0.002106 | 3.36 | +2 | 12 |
| - | - | 5297 | 627.3 | - | - | 0 | - |
| 3 | y | 2004 | 627.3 | 0.006009 | 9.579 | +2 | 12 |
| 3 | z | 1244 | 627.8 | 0.0006499 | 1.035 | +2 | 12 |
| - | - | 1669 | 628.3 | - | - | 0 | - |
| 6 | c | 5166 | 628.3 | 0.00862 | 13.72 | +1 | 6 |
| - | - | 765.4 | 629.3 | - | - | 0 | - |
| - | - | 1724 | 629.4 | - | - | 0 | - |
| - | - | 4729 | 635.3 | - | - | 0 | - |
| 3 | y | 5.174E+04 | 635.8 | 0.0002463 | 0.3874 | +2 | 12 |
| - | - | 3.553E+04 | 636.3 | - | - | 0 | - |
| - | - | 896.2 | 636.4 | - | - | 0 | - |
| - | - | 1.436E+04 | 636.8 | - | - | 0 | - |
| - | - | 4881 | 637.3 | - | - | 0 | - |
| - | - | 1350 | 637.8 | - | - | 0 | - |
| 9 | y | 3947 | 644.4 | 0.0009576 | 1.486 | +1 | 6 |
| - | - | 1353 | 645.4 | - | - | 0 | - |
| 6 | c | 2.943E+04 | 646.4 | 0.0002522 | 0.3901 | +1 | 6 |
| - | - | 9385 | 647.4 | - | - | 0 | - |
| - | - | 1893 | 648.4 | - | - | 0 | - |
| - | - | 575.7 | 649.3 | - | - | 0 | - |
| 12 | c | 2089 | 656.8 | 0.01096 | 16.69 | +2 | 12 |
| 12 | c | 1966 | 657.3 | 0.0003897 | 0.5929 | +2 | 12 |
| - | - | 1192 | 657.8 | - | - | 0 | - |
| - | - | 1753 | 672.4 | - | - | 0 | - |
| - | - | 4187 | 673.4 | - | - | 0 | - |
| - | - | 810.5 | 674.4 | - | - | 0 | - |
| 2 | z | 916.6 | 692.3 | 0.0006438 | 0.9299 | +2 | 13 |
| 2 | z | 1157 | 692.8 | 0.002771 | 3.999 | +2 | 13 |
| - | - | 698 | 693.3 | - | - | 0 | - |
| - | - | 1011 | 696.3 | - | - | 0 | - |
| 8 | w | 1.92E+04 | 698.4 | 0.0001087 | 0.1557 | +1 | 7 |
| - | - | 832.4 | 698.9 | - | - | 0 | - |
| - | - | 614.4 | 699.3 | - | - | 0 | - |
| - | - | 6648 | 699.4 | - | - | 0 | - |
| - | - | 2284 | 700.4 | - | - | 0 | - |
| 2 | z | 1.015E+04 | 701.3 | 0.003912 | 5.578 | +2 | 13 |
| - | - | 4626 | 701.8 | - | - | 0 | - |
| - | - | 3596 | 702.3 | - | - | 0 | - |
| - | - | 1427 | 702.9 | - | - | 0 | - |
| - | - | 983.3 | 703.3 | - | - | 0 | - |
| - | - | 747.4 | 707.9 | - | - | 0 | - |
| - | - | 1133 | 708.4 | - | - | 0 | - |
| - | - | 2533 | 714.3 | - | - | 0 | - |
| - | - | 1877 | 714.4 | - | - | 0 | - |
| - | - | 966.3 | 714.9 | - | - | 0 | - |
| - | - | 955.3 | 715.3 | - | - | 0 | - |
| - | - | 2213 | 715.4 | - | - | 0 | - |
| - | - | 1389 | 715.9 | - | - | 0 | - |
| 13 | c | 999.7 | 721.4 | 0.003818 | 5.293 | +2 | 13 |
| - | - | 815.5 | 721.9 | - | - | 0 | - |
| 13 | c | 2.667E+04 | 729.9 | 0.0001857 | 0.2544 | +2 | 13 |
| - | - | 2.732E+04 | 730.4 | - | - | 0 | - |
| - | - | 1.648E+04 | 730.9 | - | - | 0 | - |
| - | - | 7719 | 731.4 | - | - | 0 | - |
| - | - | 1581 | 731.9 | - | - | 0 | - |
| - | - | 1808 | 732.4 | - | - | 0 | - |
| - | - | 1563 | 736.4 | - | - | 0 | - |
| - | - | 1572 | 736.9 | - | - | 0 | - |
| - | - | 1508 | 737.4 | - | - | 0 | - |
| - | - | 555.8 | 737.9 | - | - | 0 | - |
| - | - | 2418 | 738.4 | - | - | 0 | - |
| - | - | 1159 | 738.9 | - | - | 0 | - |
| - | - | 2709 | 740.3 | - | - | 0 | - |
| - | - | 1427 | 741.4 | - | - | 0 | - |
| - | - | 8545 | 743.9 | - | - | 0 | - |
| - | - | 5897 | 744.4 | - | - | 0 | - |
| - | - | 4057 | 744.9 | - | - | 0 | - |
| - | - | 674.6 | 745.4 | - | - | 0 | - |
| - | - | 937.4 | 745.9 | - | - | 0 | - |
| 8 | y | 1073 | 755.4 | 0.004123 | 5.458 | +1 | 7 |
| 7 | c | 2.428E+04 | 757.4 | 0.00808 | 10.67 | +1 | 7 |
| - | - | 1540 | 757.9 | - | - | 0 | - |
| - | - | 1.309E+04 | 758.4 | - | - | 0 | - |
| - | - | 3140 | 759.4 | - | - | 0 | - |
| - | - | 741.5 | 760.4 | - | - | 0 | - |
| - | - | 1475 | 765.4 | - | - | 0 | - |
| - | - | 1162 | 765.8 | - | - | 0 | - |
| - | - | 2.318E+04 | 765.9 | - | - | 0 | - |
| - | - | 3.982E+04 | 766.4 | - | - | 0 | - |
| - | - | 2.323E+04 | 766.9 | - | - | 0 | - |
| - | - | 1.211E+04 | 767.4 | - | - | 0 | - |
| - | - | 1779 | 767.8 | - | - | 0 | - |
| - | - | 3860 | 767.9 | - | - | 0 | - |
| - | - | 902.7 | 768.3 | - | - | 0 | - |
| - | - | 1683 | 768.4 | - | - | 0 | - |
| 8 | y | 2677 | 773.4 | 0.001192 | 1.542 | +1 | 7 |
| - | - | 1267 | 774.4 | - | - | 0 | - |
| 7 | c | 4.489E+04 | 775.4 | 0.0002006 | 0.2587 | +1 | 7 |
| - | - | 2.096E+04 | 776.4 | - | - | 0 | - |
| - | - | 6355 | 777.4 | - | - | 0 | - |
| - | - | 1167 | 778.4 | - | - | 0 | - |
| - | - | 2119 | 786.4 | - | - | 0 | - |
| - | - | 2184 | 787.4 | - | - | 0 | - |
| - | - | 1116 | 788.4 | - | - | 0 | - |
| - | - | 756.6 | 797.4 | - | - | 0 | - |
| - | - | 2163 | 801.4 | - | - | 0 | - |
| - | - | 873.2 | 802.4 | - | - | 0 | - |
| - | - | 2138 | 803.4 | - | - | 0 | - |
| - | - | 1451 | 804.4 | - | - | 0 | - |
| - | - | 1146 | 814.4 | - | - | 0 | - |
| - | - | 1122 | 815.4 | - | - | 0 | - |
| - | - | 804.3 | 816.4 | - | - | 0 | - |
| 7 | w | 6765 | 827.4 | 0.0007096 | 0.8576 | +1 | 8 |
| - | - | 3064 | 828.4 | - | - | 0 | - |
| - | - | 898.1 | 829.4 | - | - | 0 | - |
| - | - | 805 | 830.4 | - | - | 0 | - |
| - | - | 1914 | 832.4 | - | - | 0 | - |
| - | - | 936.5 | 861.4 | - | - | 0 | - |
| - | - | 1418 | 868.4 | - | - | 0 | - |
| - | - | 1243 | 869.4 | - | - | 0 | - |
| - | - | 706 | 872.4 | - | - | 0 | - |
| - | - | 768.2 | 874.4 | - | - | 0 | - |
| 7 | y | 752.8 | 885.4 | 0.00448 | 5.06 | +1 | 8 |
| 8 | c | 3.365E+04 | 886.4 | 0.008516 | 9.608 | +1 | 8 |
| - | - | 1.778E+04 | 887.4 | - | - | 0 | - |
| - | - | 5919 | 888.4 | - | - | 0 | - |
| - | - | 1730 | 889.4 | - | - | 0 | - |
| - | - | 1137 | 890.4 | - | - | 0 | - |
| - | - | 776.6 | 891.4 | - | - | 0 | - |
| - | - | 882.5 | 901.5 | - | - | 0 | - |
| 7 | y | 8019 | 902.5 | 0.01144 | 12.67 | +1 | 8 |
| - | - | 4169 | 903.4 | - | - | 0 | - |
| 8 | c | 5.296E+04 | 904.4 | 0.0001562 | 0.1727 | +1 | 8 |
| - | - | 2.576E+04 | 905.4 | - | - | 0 | - |
| - | - | 5888 | 906.4 | - | - | 0 | - |
| - | - | 1497 | 907.5 | - | - | 0 | - |
| - | - | 752.7 | 915.5 | - | - | 0 | - |
| - | - | 2261 | 917.4 | - | - | 0 | - |
| - | - | 792.3 | 918.4 | - | - | 0 | - |
| - | - | 715.5 | 919.4 | - | - | 0 | - |
| - | - | 772.8 | 929.5 | - | - | 0 | - |
| - | - | 1051 | 930.5 | - | - | 0 | - |
| - | - | 979.1 | 931.5 | - | - | 0 | - |
| - | - | 2005 | 944.5 | - | - | 0 | - |
| - | - | 2284 | 945.5 | - | - | 0 | - |
| - | - | 1.189E+04 | 946.5 | - | - | 0 | - |
| - | - | 4799 | 947.5 | - | - | 0 | - |
| - | - | 1939 | 948.5 | - | - | 0 | - |
| - | - | 690 | 960.5 | - | - | 0 | - |
| - | - | 6561 | 962.5 | - | - | 0 | - |
| - | - | 7260 | 963.5 | - | - | 0 | - |
| - | - | 3356 | 964.5 | - | - | 0 | - |
| - | - | 1183 | 965.5 | - | - | 0 | - |
| - | - | 927.5 | 971.5 | - | - | 0 | - |
| 6 | y | 1051 | 972.5 | 0.006906 | 7.102 | +1 | 9 |
| 6 | z | 3.285E+04 | 973.5 | 0.0005885 | 0.6045 | +1 | 9 |
| - | - | 1.534E+04 | 974.5 | - | - | 0 | - |
| - | - | 5368 | 975.5 | - | - | 0 | - |
| - | - | 1011 | 976.5 | - | - | 0 | - |
| - | - | 1009 | 978.5 | - | - | 0 | - |
| - | - | 920.1 | 983.5 | - | - | 0 | - |
| - | - | 831.9 | 984.4 | - | - | 0 | - |
| - | - | 1071 | 985.5 | - | - | 0 | - |
| - | - | 950.9 | 988.5 | - | - | 0 | - |
| 6 | y | 8681 | 989.5 | 0.002205 | 2.229 | +1 | 9 |
| - | - | 4076 | 990.5 | - | - | 0 | - |
| - | - | 1577 | 991.5 | - | - | 0 | - |
| - | - | 1485 | 997.5 | - | - | 0 | - |
| - | - | 968.8 | 1000 | - | - | 0 | - |
| - | - | 2585 | 1002 | - | - | 0 | - |
| - | - | 1033 | 1003 | - | - | 0 | - |
| - | - | 1062 | 1003 | - | - | 0 | - |
| - | - | 1662 | 1003 | - | - | 0 | - |
| - | - | 1973 | 1004 | - | - | 0 | - |
| - | - | 1758 | 1005 | - | - | 0 | - |
| - | - | 943.9 | 1006 | - | - | 0 | - |
| - | - | 733.4 | 1007 | - | - | 0 | - |
| - | - | 851.6 | 1012 | - | - | 0 | - |
| - | - | 1039 | 1012 | - | - | 0 | - |
| - | - | 927.2 | 1013 | - | - | 0 | - |
| 9 | c | 3.2E+04 | 1018 | 0.0009682 | 0.9516 | +1 | 9 |
| - | - | 1.962E+04 | 1019 | - | - | 0 | - |
| - | - | 1.057E+04 | 1020 | - | - | 0 | - |
| - | - | 4283 | 1021 | - | - | 0 | - |
| - | - | 844 | 1021 | - | - | 0 | - |
| - | - | 802.7 | 1022 | - | - | 0 | - |
| - | - | 1176 | 1022 | - | - | 0 | - |
| - | - | 1656 | 1022 | - | - | 0 | - |
| - | - | 1098 | 1022 | - | - | 0 | - |
| - | - | 1230 | 1023 | - | - | 0 | - |
| - | - | 1056 | 1024 | - | - | 0 | - |
| - | - | 845.7 | 1053 | - | - | 0 | - |
| - | - | 969.1 | 1054 | - | - | 0 | - |
| 5 | z | 6513 | 1061 | 0.0009224 | 0.8697 | +1 | 10 |
| - | - | 4449 | 1062 | - | - | 0 | - |
| - | - | 2095 | 1063 | - | - | 0 | - |
| - | - | 777.3 | 1071 | - | - | 0 | - |
| 5 | y | 3148 | 1077 | 0.0002372 | 0.2204 | +1 | 10 |
| - | - | 1821 | 1078 | - | - | 0 | - |
| - | - | 872.5 | 1143 | - | - | 0 | - |
| 10 | c | 1.944E+04 | 1146 | 0.0008299 | 0.7245 | +1 | 10 |
| - | - | 1.257E+04 | 1147 | - | - | 0 | - |
| - | - | 4622 | 1148 | - | - | 0 | - |
| 4 | y | 6569 | 1174 | 0.0007098 | 0.6049 | +1 | 11 |
| - | - | 3857 | 1175 | - | - | 0 | - |
| - | - | 1599 | 1176 | - | - | 0 | - |
| - | - | 925.3 | 1177 | - | - | 0 | - |
| 11 | c | 747.1 | 1199 | 8.68E-05 | 0.07242 | +1 | 11 |
| - | - | 739.4 | 1201 | - | - | 0 | - |
| - | - | 1383 | 1216 | - | - | 0 | - |
| 11 | c | 3.79E+04 | 1217 | 0.0004681 | 0.3848 | +1 | 11 |
| - | - | 2.562E+04 | 1218 | - | - | 0 | - |
| - | - | 1.168E+04 | 1219 | - | - | 0 | - |
| - | - | 2974 | 1220 | - | - | 0 | - |
| 3 | w | 2076 | 1228 | 0.01651 | 13.45 | +1 | 12 |
| - | - | 1574 | 1229 | - | - | 0 | - |
| 3 | z | 735.2 | 1255 | 0.005244 | 4.18 | +1 | 12 |
| 3 | y | 3802 | 1271 | 0.001029 | 0.8095 | +1 | 12 |
| - | - | 3072 | 1272 | - | - | 0 | - |
| - | - | 2617 | 1273 | - | - | 0 | - |
| - | - | 796 | 1282 | - | - | 0 | - |
| - | - | 1516 | 1287 | - | - | 0 | - |
| - | - | 1479 | 1288 | - | - | 0 | - |
| - | - | 1242 | 1289 | - | - | 0 | - |
| - | - | 772.2 | 1290 | - | - | 0 | - |
| - | - | 1814 | 1300 | - | - | 0 | - |
| - | - | 2190 | 1330 | - | - | 0 | - |
| 12 | c | 5.365E+04 | 1331 | 0.00177 | 1.33 | +1 | 12 |
| - | - | 4.164E+04 | 1332 | - | - | 0 | - |
| - | - | 1.918E+04 | 1333 | - | - | 0 | - |
| - | - | 5407 | 1334 | - | - | 0 | - |
| - | - | 1013 | 1387 | - | - | 0 | - |
| - | - | 825.9 | 1395 | - | - | 0 | - |
| 2 | z | 7628 | 1402 | 0.0009138 | 0.6519 | +1 | 13 |
| - | - | 1.873E+04 | 1403 | - | - | 0 | - |
| - | - | 1.414E+04 | 1404 | - | - | 0 | - |
| - | - | 5574 | 1405 | - | - | 0 | - |
| - | - | 2143 | 1406 | - | - | 0 | - |
| - | - | 1380 | 1415 | - | - | 0 | - |
| - | - | 2870 | 1416 | - | - | 0 | - |
| - | - | 1188 | 1417 | - | - | 0 | - |
| - | - | 2731 | 1443 | - | - | 0 | - |
| - | - | 2163 | 1444 | - | - | 0 | - |
| 13 | c | 3293 | 1459 | 0.0006795 | 0.4658 | +1 | 13 |
| - | - | 1.033E+04 | 1460 | - | - | 0 | - |
| - | - | 7690 | 1461 | - | - | 0 | - |
| - | - | 3386 | 1462 | - | - | 0 | - |
| - | - | 1146 | 1470 | - | - | 0 | - |
| - | - | 1391 | 1471 | - | - | 0 | - |
| - | - | 910.3 | 1472 | - | - | 0 | - |
| - | - | 923.3 | 1474 | - | - | 0 | - |
| - | - | 918.4 | 1475 | - | - | 0 | - |
| - | - | 753 | 1476 | - | - | 0 | - |
| - | - | 1942 | 1487 | - | - | 0 | - |
| - | - | 1.117E+04 | 1488 | - | - | 0 | - |
| - | - | 7490 | 1489 | - | - | 0 | - |
| - | - | 4400 | 1490 | - | - | 0 | - |
| - | - | 792.6 | 1491 | - | - | 0 | - |
| - | - | 1013 | 1499 | - | - | 0 | - |
| - | - | 1758 | 1504 | - | - | 0 | - |
| - | - | 2864 | 1515 | - | - | 0 | - |
| - | - | 2.441E+04 | 1516 | - | - | 0 | - |
| - | - | 1.906E+04 | 1517 | - | - | 0 | - |
| - | - | 9910 | 1518 | - | - | 0 | - |
| - | - | 2756 | 1519 | - | - | 0 | - |
| - | - | 988.1 | 1525 | - | - | 0 | - |
| - | - | 1054 | 1525 | - | - | 0 | - |
| - | - | 2498 | 1531 | - | - | 0 | - |
| - | - | 9708 | 1532 | - | - | 0 | - |
| - | - | 2.798E+04 | 1533 | - | - | 0 | - |
| - | - | 988.3 | 1533 | - | - | 0 | - |
| - | - | 2.088E+04 | 1534 | - | - | 0 | - |
| - | - | 1204 | 1534 | - | - | 0 | - |
| - | - | 1.2E+04 | 1535 | - | - | 0 | - |
| - | - | 3006 | 1536 | - | - | 0 | - |
| - | - | 1177 | 1537 | - | - | 0 | - |

m/z Charge Intensity FragmentType MassShift Position
120.08087158203125 0 3081.4385
126.05513763427734 0 1285.4749
127.08686065673828 0 923.2778
128.09449768066406 0 1728.4916
129.06578063964844 0 485.50485
129.10231018066406 0 4018.5603
130.0501708984375 0 443.99377
131.11801147460938 0 4422.519
132.10198974609375 0 3376.8127
133.1053924560547 0 428.6981
134.64027404785156 0 371.99768
136.07577514648438 0 907.1761
140.6073760986328 0 494.8027
144.06561279296875 0 4480.09 w 12
145.0606231689453 0 1539.8018
145.09706115722656 0 490.00354
155.09326171875 0 573.896
165.102294921875 0 1460.2642
169.06106567382812 0 516.49695
171.0762176513672 0 640.50726
173.1288299560547 0 597.7759
173.44041442871094 0 3047.5264
183.1128387451172 0 5407.5176
185.10462951660156 0 1748.7886 z Ammonia loss 12
186.08749389648438 0 583.3235
187.0713348388672 0 655.06915
187.10800170898438 0 546.0911
187.14382934570312 0 675.2425
201.12338256835938 0 1683.7905 y Ammonia loss 12
202.08567810058594 0 776.42535
202.12240600585938 0 799.01166 y 10
202.13124084472656 0 8303.131 z 12
203.1024932861328 0 630.23083
211.10684204101562 0 638.2443
212.13934326171875 0 7476.506
213.14320373535156 0 1010.48895
215.1392059326172 0 1280.1937
216.0977325439453 0 480.37097
217.0814208984375 0 526.59973
218.14987182617188 0 5998.4463 y 12
219.1534423828125 0 718.3164
224.0748748779297 0 473.03668
229.1095428466797 0 2756.926
229.15484619140625 0 793.4084
230.11585998535156 0 773.076
232.09312438964844 0 3994.3286
233.14967346191406 0 8270.428
233.16481018066406 0 14465.861
234.15341186523438 0 1175.6251
234.16831970214844 0 1118.6969
238.1556396484375 0 556.3382
243.14581298828125 0 1146.0576
247.12852478027344 0 606.8523
251.15028381347656 0 988.9322
260.1520690917969 0 782.96124
261.1596374511719 0 7115.269
262.1629333496094 0 1110.67
266.1478271484375 0 942.73566 y 9
266.6492919921875 0 600.7359
270.14453125 0 1368.9182
272.160400390625 0 5598.5454 w 11
281.12274169921875 0 583.67816
282.14495849609375 0 1475.6539
283.14068603515625 0 1704.9552
297.15643310546875 0 656.49585
298.1277770996094 0 577.5162
299.1714782714844 0 29762.197
300.1741638183594 0 3789.9453
302.17144775390625 0 691.5178
307.463623046875 0 625.82196
311.1606140136719 0 666.46094
311.2589111328125 0 595.1504
314.1781005859375 0 878.41364 y Ammonia loss 8
322.6901550292969 0 1381.995 y 8
325.1873474121094 0 967.2226
332.1927795410156 0 2308.88 y 11
342.19451904296875 0 959.3607
343.19769287109375 0 3809.433
358.2114562988281 0 2453.3843
369.2003479003906 0 703.8376
370.6790771484375 0 1158.1864
386.20343017578125 0 16476.488 y Ammonia loss 10
387.2074890136719 0 4077.3455 y 7
395.2275695800781 0 1192.2738
402.185791015625 0 887.8498
403.1929626464844 0 2700.4663
403.2298889160156 0 7904.9736 y 10
404.1939697265625 0 583.7783
404.23272705078125 0 1047.3507
418.8722839355469 0 659.16974 z 2
428.2773132324219 0 656.89325
434.7062683105469 0 896.26
439.1081848144531 0 590.6583
444.2109375 0 748.1841
449.126220703125 0 1572.5114
453.3433837890625 0 751.32733
455.26507568359375 0 854.13043
457.2407531738281 0 28662.102 w 9
458.24383544921875 0 6041.2266
459.24786376953125 0 1180.372
465.24151611328125 0 764.14935
471.2554931640625 0 737.57916
472.2954406738281 0 720.3644 c 3
474.306640625 0 776.5182
487.2380065917969 0 1137.5834 z 5
488.2392272949219 0 924.86505
488.27423095703125 0 2204.5994
494.200439453125 0 942.3577
496.2154846191406 0 1522.962
497.2980651855469 0 674.1056
498.2182312011719 0 2178.1729
510.3311462402344 0 711.2383
511.26214599609375 0 1054.0062
511.3270263671875 0 833.73346
511.814697265625 0 646.29626
512.2083129882812 0 776.61334
512.2658081054688 0 1128.9642
512.3173217773438 0 1263.7571
513.2784423828125 0 1398.3041
514.2247314453125 0 2538.4612
514.262939453125 0 3246.696 y Ammonia loss 9
515.2325439453125 0 996.0028
515.26904296875 0 2635.2722 z 9
516.317138671875 0 985.59216
530.2447509765625 0 2362.57 y Ammonia loss 4
531.2481689453125 0 3159.7903
531.28857421875 0 8315.459 y 9
532.2527465820312 0 1349.296
532.2919921875 0 2067.3083
542.2965698242188 0 1216.8645
543.2517700195312 0 731.06946
557.3187255859375 0 1088.213
559.32373046875 0 7020.222 c 4
560.3273315429688 0 2723.3867
561.3307495117188 0 671.48694
570.3253173828125 0 2192.288
571.3280639648438 0 788.9061
585.2991333007812 0 40480.656 w 8
586.3019409179688 0 11957.695
587.2987670898438 0 4089.576 y 3
591.2911376953125 0 881.87683
599.3020629882812 0 1202.8425
599.806884765625 0 7094.553 c Water loss 10
600.30810546875 0 4455.8945 c Ammonia loss 10
600.8104248046875 0 2442.8657
601.3099365234375 0 790.627
602.258544921875 0 796.92914
613.809814453125 0 2089.4194
614.3145751953125 0 3789.6863
614.8170166015625 0 2247.4873
615.3154296875 0 1099.631
616.3312377929688 0 1422.3518
618.2894897460938 0 1131.3075
618.9619140625 0 600.31226
619.2921142578125 0 624.9862 z Ammonia loss 2
626.8141479492188 0 1267.5356 y Water loss 2
627.2612915039062 0 5296.571
627.31005859375 0 2004.2535 y Ammonia loss 2
627.8073120117188 0 1244.1058 z 2
628.26416015625 0 1668.9436
628.3539428710938 0 5166.047 c Water loss 5
629.264404296875 0 765.3537
629.3545532226562 0 1723.7175
635.3137817382812 0 4728.5894
635.8170776367188 0 51735.367 y 2
636.3187255859375 0 35531.133
636.381103515625 0 896.2399
636.8203125 0 14362.818
637.32177734375 0 4880.555
637.8263549804688 0 1349.9568
644.3716430664062 0 3947.0479 y 8
645.3762817382812 0 1353.036
646.3561401367188 0 29432.965 c 5
647.3588256835938 0 9384.868
648.360595703125 0 1892.505
649.3153076171875 0 575.6768
656.8192749023438 0 2088.663 c Water loss 11
657.3226318359375 0 1965.8668 c Ammonia loss 11
657.8208618164062 0 1192.338
672.3701782226562 0 1752.508
673.3555908203125 0 4187.3745
674.3514404296875 0 810.4801
692.3362426757812 0 916.6284 z Water loss 1
692.8316650390625 0 1157.3092 z Ammonia loss 1
693.3385620117188 0 698.04626
696.324462890625 0 1010.9147
698.383056640625 0 19204.389 w 7
698.8860473632812 0 832.4444
699.3326416015625 0 614.4406
699.3851318359375 0 6647.7305
700.386962890625 0 2284.3535
701.3382568359375 0 10149.009 z 1
701.843505859375 0 4626.4106
702.3407592773438 0 3595.7478
702.8513793945312 0 1427.0283
703.3445434570312 0 983.2982
707.8784790039062 0 747.41736
708.3778076171875 0 1133.0952
714.3385620117188 0 2533.3843
714.386962890625 0 1876.7815
714.883056640625 0 966.2853
715.3379516601562 0 955.2587
715.3945922851562 0 2212.9197
715.857177734375 0 1389.3671
721.3659057617188 0 999.66547 c Ammonia loss 12
721.8731689453125 0 815.46094
729.8828125 0 26673.594 c 12
730.3849487304688 0 27318.59
730.8858642578125 0 16476.377
731.3824462890625 0 7719.425
731.8869018554688 0 1580.9779
732.3778686523438 0 1807.9122
736.3538818359375 0 1563.3384
736.8609619140625 0 1571.5796
737.3724365234375 0 1508.3798
737.8871459960938 0 555.8235
738.3649291992188 0 2418.3054
738.8699951171875 0 1159.3558
740.3465576171875 0 2709.4146
741.3515625 0 1427.3501
743.8873291015625 0 8545.419
744.3878784179688 0 5896.6157
744.8871459960938 0 4056.7153
745.3773193359375 0 674.63696
745.8818969726562 0 937.36383
755.4087524414062 0 1072.6642 y Water loss 7
757.39599609375 0 24282.398 c Water loss 6
757.8873901367188 0 1539.7966
758.3953857421875 0 13092.347
759.3952026367188 0 3140.009
760.3887329101562 0 741.48755
765.3916015625 0 1474.9033
765.7658081054688 0 1162.4208
765.8933715820312 0 23178.564
766.3958740234375 0 39818.96
766.8975830078125 0 23233.627
767.399169921875 0 12114.129
767.7564697265625 0 1779.4932
767.898681640625 0 3860.1426
768.2562255859375 0 902.73303
768.3986206054688 0 1682.6625
773.4140014648438 0 2676.542 y 7
774.42041015625 0 1266.8707
775.398681640625 0 44886.34 c 6
776.4015502929688 0 20964.588
777.4046630859375 0 6354.7695
778.4017333984375 0 1167.4507
786.3998413085938 0 2119.365
787.4027099609375 0 2183.8477
788.410888671875 0 1115.7864
797.3775634765625 0 756.5772
801.4124755859375 0 2162.7053
802.4295654296875 0 873.17017
803.3805541992188 0 2137.9155
804.3787841796875 0 1451.1885
814.4173583984375 0 1145.6862
815.3950805664062 0 1122.4309
816.3898315429688 0 804.27277
827.425048828125 0 6764.6245 w 6
828.42919921875 0 3064.348
829.4342651367188 0 898.0758
830.4261474609375 0 804.99805
832.4140014648438 0 1914.0894
861.43408203125 0 936.5419
868.4103393554688 0 1418.3298
869.3948974609375 0 1243.4054
872.4046020507812 0 705.9741
874.4363403320312 0 768.17834
885.4267578125 0 752.76447 y Ammonia loss 6
886.4390258789062 0 33648.727 c Water loss 7
887.4368286132812 0 17776.53
888.438232421875 0 5918.974
889.4229736328125 0 1730.149
890.4190063476562 0 1137.2129
891.4170532226562 0 776.6096
901.4542846679688 0 882.53143
902.4463500976562 0 8018.645 y 6
903.4447631835938 0 4169.1304
904.44091796875 0 52959.273 c 7
905.443115234375 0 25757.094
906.446044921875 0 5888.2417
907.4503784179688 0 1497.1799
915.46875 0 752.7496
917.4091796875 0 2260.5361
918.4194946289062 0 792.2774
919.448974609375 0 715.5427
929.4583129882812 0 772.81805
930.4574584960938 0 1051.2428
931.4600219726562 0 979.07477
944.4624633789062 0 2004.624
945.4673461914062 0 2284.095
946.4827270507812 0 11894.986
947.486328125 0 4799.365
948.4917602539062 0 1938.5875
960.4661254882812 0 689.9944
962.4791870117188 0 6561.0454
963.4807739257812 0 7260.287
964.485107421875 0 3355.5237
965.4846801757812 0 1182.6056
971.4595947265625 0 927.48596
972.4563598632812 0 1051.2208 y Ammonia loss 5
973.4716796875 0 32851.75 z 5
974.4749755859375 0 15340.123
975.4772338867188 0 5367.9653
976.4747924804688 0 1010.9901
978.4700317382812 0 1008.83954
983.4791870117188 0 920.10364
984.4488525390625 0 831.8931
985.4701538085938 0 1071.4458
988.4762573242188 0 950.9421
989.4876098632812 0 8681.029 y 5
990.4904174804688 0 4075.712
991.4943237304688 0 1577.1611
997.4655151367188 0 1485.4529
1000.4862670898438 0 968.7724
1002.458251953125 0 2584.987
1002.7933959960938 0 1032.7115
1003.1285400390625 0 1062.4279
1003.480712890625 0 1662.214
1004.4671630859375 0 1972.5981
1005.4681396484375 0 1758.0831
1006.4391479492188 0 943.92816
1007.4459838867188 0 733.3507
1011.9664916992188 0 851.56323
1012.4909057617188 0 1038.914
1012.9794311523438 0 927.1965
1017.524169921875 0 32000.494 c 8
1018.5245361328125 0 19618.639
1019.5188598632812 0 10573.734
1020.507080078125 0 4283.056
1020.99169921875 0 843.9532
1021.51123046875 0 802.73224
1022.00146484375 0 1176.336
1022.1342163085938 0 1655.9432
1022.4901733398438 0 1097.5931
1023.4323120117188 0 1230.2273
1024.4464111328125 0 1056.4822
1053.488525390625 0 845.7266
1054.4903564453125 0 969.09607
1060.502197265625 0 6513.0923 z 4
1061.5054931640625 0 4448.635
1062.5120849609375 0 2094.9077
1070.51318359375 0 777.2836
1076.5216064453125 0 3148.4739 y 4
1077.5257568359375 0 1820.659
1142.5537109375 0 872.471
1145.5828857421875 0 19444.637 c 9
1146.5853271484375 0 12567.856
1147.5899658203125 0 4621.883
1173.5753173828125 0 6568.865 y 3
1174.5726318359375 0 3856.9858
1175.587646484375 0 1599.0375
1176.5772705078125 0 925.33514
1198.6103515625 0 747.06604 c Water loss 10
1200.5704345703125 0 739.42957
1215.6182861328125 0 1382.857
1216.620361328125 0 37895.543 c 10
1217.623046875 0 25623.25
1218.6256103515625 0 11678.381
1219.63037109375 0 2974.1826
1227.6016845703125 0 2076.346 w 2
1228.592041015625 0 1574.1738
1254.6138916015625 0 735.1732 z 2
1270.6263427734375 0 3801.9478 y 2
1271.630859375 0 3072.01
1272.63525390625 0 2617.0256
1281.5875244140625 0 795.99603
1286.6461181640625 0 1516.2833
1287.6517333984375 0 1479.1774
1288.6546630859375 0 1242.2124
1290.07763671875 0 772.24896
1299.626708984375 0 1813.591
1329.6512451171875 0 2190.477
1330.6619873046875 0 53649.918 c 11
1331.664794921875 0 41640.734
1332.6676025390625 0 19184.129
1333.668701171875 0 5407.216
1386.6676025390625 0 1013.22424
1395.12353515625 0 825.886
1401.6761474609375 0 7628.4746 z 1
1402.681884765625 0 18727.746
1403.686279296875 0 14142.985
1404.68603515625 0 5573.739
1405.68212890625 0 2142.79
1414.74169921875 0 1379.9889
1415.744873046875 0 2869.9973
1416.7518310546875 0 1187.6144
1442.735595703125 0 2730.6587
1443.74609375 0 2163.3457
1458.7593994140625 0 3293.4758 c 12
1459.7642822265625 0 10331.451
1460.7659912109375 0 7689.5005
1461.7645263671875 0 3386.3372
1469.760986328125 0 1146.3574
1470.762939453125 0 1391.4386
1471.7637939453125 0 910.3266
1473.751953125 0 923.313
1474.75830078125 0 918.4211
1476.499267578125 0 752.98224
1486.7601318359375 0 1941.5366
1487.7698974609375 0 11169.494
1488.773681640625 0 7490.102
1489.7740478515625 0 4400.117
1490.7730712890625 0 792.6049
1498.745361328125 0 1013.4614
1504.1865234375 0 1758.0238
1514.7662353515625 0 2864.2993
1515.76611328125 0 24412.984
1516.769287109375 0 19064.428
1517.7716064453125 0 9909.585
1518.7720947265625 0 2755.5122
1524.693115234375 0 988.09393
1525.2127685546875 0 1054.1494
1530.7755126953125 0 2498.1677
1531.7833251953125 0 9707.799
1532.7913818359375 0 27977.074
1533.1651611328125 0 988.27997
1533.7950439453125 0 20881.709
1534.1837158203125 0 1203.5739
1534.796630859375 0 11995.784
1535.796630859375 0 3006.0862
1536.796875 0 1176.9496

Spectrum Details

|  |  |
| --- | --- |
| Matched peaks? Matched peaks The total absolute number of peaks matched. Additionally in brackets the total fraction of peaks matched and the total number of peaks is shown. | 68 (16.11% of 422) |
| FDR? FDR The false discovery rate estimated for this peptide. It is calculated by matching all theoretical fragments with a non-integer shift with the raw peaks for this spectrum. This is done with 40 different shifts. The resulting percentage is the average number of annotated peaks over the number of annotated peaks with the correct spectrum. | 0.46% |
| Satellite FDR? Satellite FDR See the FDR for details on its calculation. This satellite ion specific FDR only contains the satellite ions (d/w) for I/L/J positions. | 2.38% |
| PSM Score? PSM Score The PSM Score as given by Hecklib to this annotated spectrum. It is shown with three significant figures. | 528 |

#### Spectrum 6410? Spectrum 6410 The raw spectrum of this peptide as annotated by Hecklib. The fragments are coloured according to ion type (see legend). Any peaks with a star '\*' as text can be hovered over to see the full details, first the ion type second the mass shift type. By hovering over the amino acids in the peptide or ions in the legend the corresponding peaks are highlighted. By toggling the 'Unassigned' label you can turn the background (unassigned) peaks on or off in the plot. By updating the slider in the Ion legend you can update the spectrum to only show the top X% of the peaks with labels. The top X% means any peak that is within X% of the highest intensity. By dragging in the spectrum you can zoom in to a specific part of the spectrum and use 'Zoom Out' to get back to the original zoom level. The annotation of the spectrum is based on the given sequence in the peptides file and is done with different software so inconsistencies are likely. The peaks are annotated based on the given sequence, with 20 ppm tolerance.

Copy Data

##### Spectrum 6410 (TSV)

###### Preview

```
Loading example...
```

*Click on the button to copy the data to your clipboard.*

Mz MinMz MaxIntensity Max

WidthHeightPeptide font sizePeptide stroke widthSpectrum font sizeSpectrum stroke widthCompact peptide

Ion legend

wxyz

abcd

OtherUnassignedIon^Charge^_Position_Show for top:%

JFPPSSEEJQANKA

04.49e+48.98e+41.35e+51.80e+5

Zoom Out

a^+2^_3_y^+1^_2_y^+2^_4_y^+1^_2_b^+2^_4_a^+1^_2_y^+2^_5_b^+1^_2_y^+2^_5_b^+2^_6_y^+2^_6_b^+2^_6_y^+2^_6_y^+1^_3_b^+1^_3_b^+2^_7_b^+3^_10_b^+2^_7_y^+1^_4_y^+2^_7_b^+3^_11_y^+1^_4_y^+3^_12_y^+3^_12_b^+2^_8_y^+2^_8_b^+2^_8_b^+1^_4_b^+3^_13_y^+2^_9_b^+2^_9_\*y^+1^_5_b^+1^_5_y^+1^_5_b^+1^_5_b^+2^_10_y^+2^_11_y^+2^_11_y^+2^_11_b^+2^_11_b^+2^_11_b^+1^_6_y^+2^_12_y^+2^_12_b^+1^_6_y^+2^_12_y^+1^_6_y^+2^_13_b^+2^_13_b^+1^_7_y^+1^_7_y^+1^_7_\*b^+1^_7_y^+1^_7_b^+1^_8_y^+1^_8_y^+1^_8_b^+1^_8_y^+1^_8_y^+1^_9_y^+1^_9_b^+1^_9_y^+1^_9_b^+1^_9_y^+1^_10_y^+1^_10_y^+1^_10_b^+1^_10_y^+1^_11_y^+1^_11_b^+1^_11_b^+1^_12_

0768153623043072

Fragment Matches Table

Show background peaks

| Position | Ion type | Intensity | mz Theoretical | mz Error (Th) | mz Error (ppm) | Charge | Series Number |
| --- | --- | --- | --- | --- | --- | --- | --- |
| - | - | 1.344E+05 | 120.1 | - | - | 0 | - |
| - | - | 1.217E+04 | 121.1 | - | - | 0 | - |
| - | - | 420.4 | 122.1 | - | - | 0 | - |
| - | - | 8443 | 126.1 | - | - | 0 | - |
| - | - | 703.1 | 126.1 | - | - | 0 | - |
| - | - | 904.4 | 127.1 | - | - | 0 | - |
| - | - | 735.6 | 127.1 | - | - | 0 | - |
| - | - | 1154 | 127.1 | - | - | 0 | - |
| - | - | 446.1 | 128.1 | - | - | 0 | - |
| - | - | 603.9 | 129.1 | - | - | 0 | - |
| - | - | 7289 | 129.1 | - | - | 0 | - |
| - | - | 1.176E+05 | 129.1 | - | - | 0 | - |
| - | - | 9522 | 130.1 | - | - | 0 | - |
| - | - | 538.6 | 130.1 | - | - | 0 | - |
| - | - | 5286 | 130.1 | - | - | 0 | - |
| - | - | 549.5 | 130.1 | - | - | 0 | - |
| - | - | 861.2 | 130.1 | - | - | 0 | - |
| - | - | 1061 | 130.1 | - | - | 0 | - |
| - | - | 7173 | 130.1 | - | - | 0 | - |
| - | - | 396.4 | 131 | - | - | 0 | - |
| - | - | 508.4 | 131 | - | - | 0 | - |
| - | - | 606 | 131.1 | - | - | 0 | - |
| - | - | 419.3 | 131.1 | - | - | 0 | - |
| - | - | 7044 | 131.1 | - | - | 0 | - |
| - | - | 438 | 132.1 | - | - | 0 | - |
| - | - | 1047 | 132.1 | - | - | 0 | - |
| - | - | 1273 | 134 | - | - | 0 | - |
| - | - | 467.8 | 134 | - | - | 0 | - |
| - | - | 3578 | 136.1 | - | - | 0 | - |
| - | - | 3607 | 138.1 | - | - | 0 | - |
| - | - | 2977 | 139.1 | - | - | 0 | - |
| - | - | 1176 | 140.1 | - | - | 0 | - |
| - | - | 825.8 | 140.1 | - | - | 0 | - |
| - | - | 3829 | 141.1 | - | - | 0 | - |
| - | - | 1389 | 141.1 | - | - | 0 | - |
| - | - | 519.7 | 142.1 | - | - | 0 | - |
| - | - | 638.8 | 142.1 | - | - | 0 | - |
| - | - | 791.6 | 145.1 | - | - | 0 | - |
| - | - | 1540 | 146.1 | - | - | 0 | - |
| - | - | 3399 | 147.1 | - | - | 0 | - |
| - | - | 2656 | 147.1 | - | - | 0 | - |
| - | - | 457.3 | 148.1 | - | - | 0 | - |
| - | - | 455.7 | 148.1 | - | - | 0 | - |
| - | - | 462.5 | 149 | - | - | 0 | - |
| - | - | 483.2 | 152 | - | - | 0 | - |
| - | - | 1148 | 152.1 | - | - | 0 | - |
| - | - | 671.2 | 153.1 | - | - | 0 | - |
| - | - | 961.5 | 153.1 | - | - | 0 | - |
| - | - | 370.8 | 153.1 | - | - | 0 | - |
| - | - | 2897 | 154.1 | - | - | 0 | - |
| - | - | 5277 | 154.1 | - | - | 0 | - |
| - | - | 7678 | 155.1 | - | - | 0 | - |
| - | - | 930.5 | 155.1 | - | - | 0 | - |
| - | - | 2608 | 157.1 | - | - | 0 | - |
| - | - | 8717 | 157.1 | - | - | 0 | - |
| - | - | 1308 | 158.1 | - | - | 0 | - |
| - | - | 642.1 | 158.1 | - | - | 0 | - |
| - | - | 829.7 | 159.1 | - | - | 0 | - |
| - | - | 2099 | 163.1 | - | - | 0 | - |
| - | - | 1100 | 163.1 | - | - | 0 | - |
| - | - | 652.9 | 164.1 | - | - | 0 | - |
| - | - | 1381 | 165.1 | - | - | 0 | - |
| 3 | a | 759.1 | 165.6 | 0.0005818 | 3.513 | +2 | 3 |
| - | - | 1142 | 166.1 | - | - | 0 | - |
| - | - | 1167 | 166.1 | - | - | 0 | - |
| - | - | 5608 | 167.1 | - | - | 0 | - |
| - | - | 6497 | 167.1 | - | - | 0 | - |
| - | - | 2174 | 169.1 | - | - | 0 | - |
| - | - | 1420 | 169.1 | - | - | 0 | - |
| - | - | 416.2 | 169.5 | - | - | 0 | - |
| - | - | 613.9 | 170.1 | - | - | 0 | - |
| - | - | 1909 | 171.1 | - | - | 0 | - |
| - | - | 1145 | 172.1 | - | - | 0 | - |
| - | - | 2364 | 172.1 | - | - | 0 | - |
| - | - | 868.9 | 172.1 | - | - | 0 | - |
| - | - | 778.2 | 173.1 | - | - | 0 | - |
| - | - | 1369 | 173.1 | - | - | 0 | - |
| - | - | 3581 | 173.5 | - | - | 0 | - |
| - | - | 454.3 | 174.1 | - | - | 0 | - |
| - | - | 3348 | 175.1 | - | - | 0 | - |
| - | - | 601.7 | 175.1 | - | - | 0 | - |
| - | - | 552.2 | 175.4 | - | - | 0 | - |
| - | - | 668 | 177.1 | - | - | 0 | - |
| - | - | 506.3 | 177.3 | - | - | 0 | - |
| - | - | 1401 | 180.1 | - | - | 0 | - |
| - | - | 447 | 181.1 | - | - | 0 | - |
| - | - | 2736 | 181.1 | - | - | 0 | - |
| - | - | 795.8 | 181.1 | - | - | 0 | - |
| - | - | 8152 | 182.1 | - | - | 0 | - |
| - | - | 842.7 | 182.1 | - | - | 0 | - |
| - | - | 5169 | 183.1 | - | - | 0 | - |
| - | - | 541.8 | 183.1 | - | - | 0 | - |
| - | - | 2240 | 183.1 | - | - | 0 | - |
| - | - | 1194 | 183.1 | - | - | 0 | - |
| - | - | 494.6 | 184.1 | - | - | 0 | - |
| - | - | 5629 | 184.1 | - | - | 0 | - |
| - | - | 1.204E+04 | 185.1 | - | - | 0 | - |
| - | - | 4970 | 186.1 | - | - | 0 | - |
| - | - | 1281 | 186.1 | - | - | 0 | - |
| - | - | 1221 | 186.1 | - | - | 0 | - |
| - | - | 591.1 | 187.1 | - | - | 0 | - |
| - | - | 1755 | 187.1 | - | - | 0 | - |
| - | - | 1269 | 187.1 | - | - | 0 | - |
| - | - | 4442 | 189.1 | - | - | 0 | - |
| - | - | 1664 | 191.1 | - | - | 0 | - |
| - | - | 518.7 | 193.1 | - | - | 0 | - |
| - | - | 1521 | 195.1 | - | - | 0 | - |
| - | - | 1.436E+04 | 195.1 | - | - | 0 | - |
| - | - | 1112 | 196.1 | - | - | 0 | - |
| - | - | 459.9 | 196.9 | - | - | 0 | - |
| - | - | 1.085E+04 | 197.1 | - | - | 0 | - |
| - | - | 467 | 197.2 | - | - | 0 | - |
| - | - | 897.7 | 198.1 | - | - | 0 | - |
| - | - | 6492 | 198.1 | - | - | 0 | - |
| - | - | 8698 | 199.1 | - | - | 0 | - |
| - | - | 816.7 | 199.1 | - | - | 0 | - |
| - | - | 4772 | 199.1 | - | - | 0 | - |
| - | - | 1393 | 199.1 | - | - | 0 | - |
| - | - | 746 | 200.1 | - | - | 0 | - |
| - | - | 8383 | 200.1 | - | - | 0 | - |
| - | - | 616.4 | 200.1 | - | - | 0 | - |
| - | - | 1956 | 200.1 | - | - | 0 | - |
| - | - | 703.3 | 201.1 | - | - | 0 | - |
| - | - | 994.2 | 201.1 | - | - | 0 | - |
| 13 | y | 1.078E+04 | 201.1 | 0.0003036 | 1.51 | +1 | 2 |
| - | - | 591.8 | 202.1 | - | - | 0 | - |
| 11 | y | 1905 | 202.1 | 0.0001871 | 0.9258 | +2 | 4 |
| - | - | 1274 | 203.1 | - | - | 0 | - |
| - | - | 1105 | 204.1 | - | - | 0 | - |
| - | - | 642.4 | 204.6 | - | - | 0 | - |
| - | - | 586.5 | 205.1 | - | - | 0 | - |
| - | - | 566.1 | 205.1 | - | - | 0 | - |
| - | - | 563 | 206.1 | - | - | 0 | - |
| - | - | 491.2 | 207.1 | - | - | 0 | - |
| - | - | 1132 | 207.1 | - | - | 0 | - |
| - | - | 862.5 | 208.1 | - | - | 0 | - |
| - | - | 6967 | 208.1 | - | - | 0 | - |
| - | - | 2.913E+04 | 209.1 | - | - | 0 | - |
| - | - | 669.3 | 209.1 | - | - | 0 | - |
| - | - | 2001 | 210.1 | - | - | 0 | - |
| - | - | 989.3 | 211.1 | - | - | 0 | - |
| - | - | 2186 | 212.1 | - | - | 0 | - |
| - | - | 1854 | 213.1 | - | - | 0 | - |
| - | - | 1867 | 213.1 | - | - | 0 | - |
| - | - | 705.6 | 214.1 | - | - | 0 | - |
| - | - | 571.3 | 215.1 | - | - | 0 | - |
| - | - | 5783 | 215.1 | - | - | 0 | - |
| - | - | 512.3 | 216.1 | - | - | 0 | - |
| - | - | 599.4 | 216.1 | - | - | 0 | - |
| - | - | 1.277E+04 | 217.1 | - | - | 0 | - |
| - | - | 3229 | 217.1 | - | - | 0 | - |
| - | - | 1105 | 218.1 | - | - | 0 | - |
| 13 | y | 5.094E+04 | 218.1 | 0.0003811 | 1.747 | +1 | 2 |
| - | - | 4843 | 219.2 | - | - | 0 | - |
| - | - | 1601 | 221.1 | - | - | 0 | - |
| - | - | 652 | 221.1 | - | - | 0 | - |
| - | - | 1463 | 221.6 | - | - | 0 | - |
| - | - | 574.3 | 223.1 | - | - | 0 | - |
| - | - | 586.3 | 223.2 | - | - | 0 | - |
| - | - | 1004 | 224.1 | - | - | 0 | - |
| - | - | 654.7 | 224.1 | - | - | 0 | - |
| - | - | 8560 | 225.1 | - | - | 0 | - |
| - | - | 5036 | 225.1 | - | - | 0 | - |
| - | - | 9.305E+04 | 226.1 | - | - | 0 | - |
| - | - | 3778 | 227.1 | - | - | 0 | - |
| - | - | 8160 | 227.1 | - | - | 0 | - |
| - | - | 761 | 228.1 | - | - | 0 | - |
| 4 | b | 5655 | 228.1 | 0.001727 | 7.571 | +2 | 4 |
| - | - | 548.8 | 229.1 | - | - | 0 | - |
| - | - | 1207 | 229.1 | - | - | 0 | - |
| - | - | 1154 | 229.2 | - | - | 0 | - |
| - | - | 1336 | 231.1 | - | - | 0 | - |
| 2 | a | 1.777E+05 | 233.2 | 0.0004587 | 1.967 | +1 | 2 |
| - | - | 1124 | 234.1 | - | - | 0 | - |
| - | - | 2653 | 234.1 | - | - | 0 | - |
| - | - | 2.778E+04 | 234.2 | - | - | 0 | - |
| - | - | 2241 | 235.2 | - | - | 0 | - |
| - | - | 4858 | 236.1 | - | - | 0 | - |
| - | - | 566.5 | 236.1 | - | - | 0 | - |
| - | - | 854 | 236.1 | - | - | 0 | - |
| - | - | 609.6 | 237.1 | - | - | 0 | - |
| - | - | 739.9 | 238.1 | - | - | 0 | - |
| - | - | 848.6 | 239.2 | - | - | 0 | - |
| - | - | 1505 | 240.1 | - | - | 0 | - |
| - | - | 1172 | 240.1 | - | - | 0 | - |
| - | - | 1457 | 240.6 | - | - | 0 | - |
| - | - | 8883 | 241.1 | - | - | 0 | - |
| - | - | 571.2 | 242.1 | - | - | 0 | - |
| - | - | 630.1 | 242.1 | - | - | 0 | - |
| - | - | 2.059E+04 | 242.2 | - | - | 0 | - |
| - | - | 1692 | 243.1 | - | - | 0 | - |
| - | - | 2.529E+04 | 243.1 | - | - | 0 | - |
| - | - | 1164 | 244.1 | - | - | 0 | - |
| - | - | 5293 | 244.1 | - | - | 0 | - |
| - | - | 1915 | 244.1 | - | - | 0 | - |
| - | - | 3278 | 245.1 | - | - | 0 | - |
| - | - | 1705 | 248.6 | - | - | 0 | - |
| - | - | 834.2 | 248.7 | - | - | 0 | - |
| - | - | 7828 | 249.1 | - | - | 0 | - |
| - | - | 1466 | 249.2 | - | - | 0 | - |
| - | - | 1549 | 249.6 | - | - | 0 | - |
| - | - | 3510 | 251.2 | - | - | 0 | - |
| - | - | 2018 | 252.1 | - | - | 0 | - |
| - | - | 2197 | 252.1 | - | - | 0 | - |
| - | - | 1.528E+04 | 254.1 | - | - | 0 | - |
| - | - | 2355 | 254.2 | - | - | 0 | - |
| - | - | 1680 | 255.1 | - | - | 0 | - |
| - | - | 506.5 | 257.1 | - | - | 0 | - |
| - | - | 1.506E+04 | 257.1 | - | - | 0 | - |
| 10 | y | 2075 | 257.6 | 0.0008735 | 3.39 | +2 | 5 |
| - | - | 2670 | 257.6 | - | - | 0 | - |
| - | - | 697.5 | 258.1 | - | - | 0 | - |
| - | - | 8957 | 259.1 | - | - | 0 | - |
| - | - | 807.8 | 260.1 | - | - | 0 | - |
| 2 | b | 7.345E+04 | 261.2 | 0.0002493 | 0.9546 | +1 | 2 |
| - | - | 1252 | 262.1 | - | - | 0 | - |
| - | - | 1.261E+04 | 262.2 | - | - | 0 | - |
| - | - | 1077 | 263.2 | - | - | 0 | - |
| - | - | 1.163E+04 | 264.1 | - | - | 0 | - |
| - | - | 1822 | 265.1 | - | - | 0 | - |
| - | - | 572.7 | 265.2 | - | - | 0 | - |
| - | - | 833.5 | 266.1 | - | - | 0 | - |
| - | - | 790.4 | 266.1 | - | - | 0 | - |
| 10 | y | 1.338E+04 | 266.1 | 0.0002258 | 0.8482 | +2 | 5 |
| - | - | 3522 | 266.6 | - | - | 0 | - |
| - | - | 667.3 | 268.1 | - | - | 0 | - |
| - | - | 1218 | 269.2 | - | - | 0 | - |
| - | - | 1212 | 269.7 | - | - | 0 | - |
| - | - | 788 | 270.1 | - | - | 0 | - |
| - | - | 871.9 | 270.1 | - | - | 0 | - |
| - | - | 1033 | 272.1 | - | - | 0 | - |
| - | - | 6077 | 272.1 | - | - | 0 | - |
| - | - | 1244 | 273.1 | - | - | 0 | - |
| - | - | 1123 | 277.2 | - | - | 0 | - |
| - | - | 628 | 278.1 | - | - | 0 | - |
| - | - | 1188 | 278.2 | - | - | 0 | - |
| - | - | 5205 | 279.1 | - | - | 0 | - |
| - | - | 8653 | 280.1 | - | - | 0 | - |
| - | - | 1016 | 280.1 | - | - | 0 | - |
| - | - | 926.1 | 281.1 | - | - | 0 | - |
| - | - | 1148 | 282.1 | - | - | 0 | - |
| - | - | 2385 | 282.1 | - | - | 0 | - |
| - | - | 3.356E+04 | 282.1 | - | - | 0 | - |
| - | - | 828.9 | 282.6 | - | - | 0 | - |
| - | - | 4101 | 283.1 | - | - | 0 | - |
| - | - | 1107 | 285.2 | - | - | 0 | - |
| - | - | 4781 | 286.1 | - | - | 0 | - |
| - | - | 1027 | 292.2 | - | - | 0 | - |
| - | - | 1043 | 296.1 | - | - | 0 | - |
| - | - | 4352 | 296.1 | - | - | 0 | - |
| - | - | 5087 | 296.2 | - | - | 0 | - |
| - | - | 1138 | 296.6 | - | - | 0 | - |
| - | - | 4663 | 297.1 | - | - | 0 | - |
| - | - | 3.046E+04 | 297.2 | - | - | 0 | - |
| - | - | 1131 | 298.1 | - | - | 0 | - |
| - | - | 4208 | 298.1 | - | - | 0 | - |
| - | - | 3541 | 298.2 | - | - | 0 | - |
| - | - | 2377 | 299.2 | - | - | 0 | - |
| - | - | 783.1 | 300.1 | - | - | 0 | - |
| - | - | 1872 | 300.2 | - | - | 0 | - |
| - | - | 2519 | 304.1 | - | - | 0 | - |
| - | - | 698.6 | 305.2 | - | - | 0 | - |
| - | - | 1792 | 306.1 | - | - | 0 | - |
| 6 | b | 2119 | 306.2 | 0.0007625 | 2.491 | +2 | 6 |
| - | - | 1218 | 307.1 | - | - | 0 | - |
| - | - | 2339 | 308.2 | - | - | 0 | - |
| - | - | 1894 | 310.1 | - | - | 0 | - |
| - | - | 588.4 | 313.2 | - | - | 0 | - |
| - | - | 3830 | 313.2 | - | - | 0 | - |
| - | - | 2385 | 314.1 | - | - | 0 | - |
| - | - | 4874 | 314.1 | - | - | 0 | - |
| 9 | y | 1.159E+04 | 314.2 | 0.00458 | 14.58 | +2 | 6 |
| - | - | 1862 | 314.6 | - | - | 0 | - |
| - | - | 1614 | 314.7 | - | - | 0 | - |
| - | - | 1165 | 315.1 | - | - | 0 | - |
| - | - | 822 | 315.1 | - | - | 0 | - |
| 6 | b | 1.467E+04 | 315.2 | 0.001468 | 4.658 | +2 | 6 |
| - | - | 1239 | 315.2 | - | - | 0 | - |
| - | - | 2343 | 316.2 | - | - | 0 | - |
| - | - | 814.1 | 319.1 | - | - | 0 | - |
| - | - | 638.3 | 321.2 | - | - | 0 | - |
| 9 | y | 1.546E+04 | 322.7 | 0.0004911 | 1.522 | +2 | 6 |
| - | - | 1320 | 323.2 | - | - | 0 | - |
| - | - | 4918 | 323.2 | - | - | 0 | - |
| - | - | 1419 | 323.7 | - | - | 0 | - |
| - | - | 769 | 324.2 | - | - | 0 | - |
| - | - | 635.9 | 324.2 | - | - | 0 | - |
| - | - | 1886 | 325.2 | - | - | 0 | - |
| - | - | 1132 | 325.2 | - | - | 0 | - |
| - | - | 1570 | 326.2 | - | - | 0 | - |
| - | - | 4906 | 328.1 | - | - | 0 | - |
| - | - | 677.9 | 329.1 | - | - | 0 | - |
| - | - | 1413 | 329.7 | - | - | 0 | - |
| - | - | 733.8 | 330.2 | - | - | 0 | - |
| 12 | y | 3.918E+04 | 332.2 | 0.000514 | 1.547 | +1 | 3 |
| - | - | 6550 | 333.2 | - | - | 0 | - |
| - | - | 5709 | 333.2 | - | - | 0 | - |
| - | - | 867.5 | 334.2 | - | - | 0 | - |
| - | - | 1492 | 336.2 | - | - | 0 | - |
| - | - | 1490 | 337.2 | - | - | 0 | - |
| - | - | 633.1 | 337.2 | - | - | 0 | - |
| - | - | 1097 | 338.1 | - | - | 0 | - |
| - | - | 4136 | 338.7 | - | - | 0 | - |
| - | - | 3822 | 339.2 | - | - | 0 | - |
| - | - | 1461 | 340.2 | - | - | 0 | - |
| - | - | 6352 | 341.2 | - | - | 0 | - |
| - | - | 1801 | 342.2 | - | - | 0 | - |
| - | - | 3419 | 343.2 | - | - | 0 | - |
| - | - | 1720 | 343.2 | - | - | 0 | - |
| - | - | 570.7 | 344.2 | - | - | 0 | - |
| - | - | 614.2 | 344.2 | - | - | 0 | - |
| - | - | 8626 | 346.1 | - | - | 0 | - |
| - | - | 1830 | 347.1 | - | - | 0 | - |
| - | - | 2441 | 347.7 | - | - | 0 | - |
| - | - | 1206 | 348.2 | - | - | 0 | - |
| - | - | 557 | 349.1 | - | - | 0 | - |
| - | - | 1418 | 350.2 | - | - | 0 | - |
| - | - | 2.78E+04 | 351.2 | - | - | 0 | - |
| - | - | 5601 | 352.2 | - | - | 0 | - |
| - | - | 8247 | 353.2 | - | - | 0 | - |
| - | - | 2282 | 354.2 | - | - | 0 | - |
| - | - | 1391 | 354.2 | - | - | 0 | - |
| - | - | 3421 | 355.2 | - | - | 0 | - |
| - | - | 1051 | 356.1 | - | - | 0 | - |
| - | - | 737.7 | 356.7 | - | - | 0 | - |
| 3 | b | 4395 | 358.2 | 0.0002198 | 0.6137 | +1 | 3 |
| - | - | 970 | 359.2 | - | - | 0 | - |
| - | - | 2345 | 361.7 | - | - | 0 | - |
| - | - | 1555 | 362.2 | - | - | 0 | - |
| - | - | 851.9 | 363.2 | - | - | 0 | - |
| - | - | 3325 | 365.1 | - | - | 0 | - |
| - | - | 1075 | 365.2 | - | - | 0 | - |
| - | - | 680.1 | 365.7 | - | - | 0 | - |
| - | - | 8867 | 368.2 | - | - | 0 | - |
| - | - | 2.814E+04 | 369.2 | - | - | 0 | - |
| - | - | 5112 | 370.2 | - | - | 0 | - |
| 7 | b | 1.109E+04 | 370.7 | 0.006374 | 17.19 | +2 | 7 |
| 10 | b | 2985 | 371.2 | 0.001517 | 4.087 | +3 | 10 |
| - | - | 1610 | 372.2 | - | - | 0 | - |
| - | - | 5152 | 373.2 | - | - | 0 | - |
| - | - | 1410 | 374.2 | - | - | 0 | - |
| 7 | b | 1543 | 379.7 | 0.001588 | 4.183 | +2 | 7 |
| - | - | 601.2 | 380.2 | - | - | 0 | - |
| - | - | 7753 | 383.2 | - | - | 0 | - |
| - | - | 994.9 | 384.2 | - | - | 0 | - |
| - | - | 3517 | 385.2 | - | - | 0 | - |
| 11 | y | 2.223E+04 | 386.2 | 0.0004779 | 1.237 | +1 | 4 |
| - | - | 603.4 | 387.2 | - | - | 0 | - |
| 8 | y | 4058 | 387.2 | 0.00338 | 8.729 | +2 | 7 |
| - | - | 1370 | 390.2 | - | - | 0 | - |
| - | - | 770.3 | 392.2 | - | - | 0 | - |
| - | - | 1530 | 393.2 | - | - | 0 | - |
| 11 | b | 986.4 | 394.5 | 0.0003859 | 0.9781 | +3 | 11 |
| - | - | 1216 | 395.2 | - | - | 0 | - |
| - | - | 581.9 | 396.2 | - | - | 0 | - |
| - | - | 1063 | 397.1 | - | - | 0 | - |
| - | - | 1.194E+04 | 401.2 | - | - | 0 | - |
| - | - | 3333 | 402.2 | - | - | 0 | - |
| 11 | y | 9.132E+04 | 403.2 | 0.0006011 | 1.491 | +1 | 4 |
| - | - | 2.16E+04 | 404.2 | - | - | 0 | - |
| - | - | 726.5 | 405.2 | - | - | 0 | - |
| - | - | 2727 | 405.2 | - | - | 0 | - |
| - | - | 5800 | 407.2 | - | - | 0 | - |
| - | - | 1.508E+04 | 408.2 | - | - | 0 | - |
| - | - | 3236 | 409.2 | - | - | 0 | - |
| - | - | 1140 | 410.2 | - | - | 0 | - |
| - | - | 804.1 | 412.5 | - | - | 0 | - |
| - | - | 771 | 413.2 | - | - | 0 | - |
| - | - | 1774 | 414.2 | - | - | 0 | - |
| - | - | 3569 | 415.1 | - | - | 0 | - |
| - | - | 682.7 | 415.2 | - | - | 0 | - |
| 3 | y | 959.7 | 418.2 | 0.003048 | 7.288 | +3 | 12 |
| 3 | y | 1615 | 418.5 | 0.004083 | 9.756 | +3 | 12 |
| - | - | 1232 | 418.9 | - | - | 0 | - |
| - | - | 727.5 | 421.2 | - | - | 0 | - |
| - | - | 8030 | 424.2 | - | - | 0 | - |
| - | - | 1029 | 424.5 | - | - | 0 | - |
| - | - | 1156 | 425.1 | - | - | 0 | - |
| - | - | 7320 | 425.2 | - | - | 0 | - |
| - | - | 2164 | 425.7 | - | - | 0 | - |
| - | - | 3588 | 426.2 | - | - | 0 | - |
| - | - | 708.4 | 426.7 | - | - | 0 | - |
| - | - | 2286 | 427.2 | - | - | 0 | - |
| - | - | 1399 | 427.3 | - | - | 0 | - |
| - | - | 1436 | 429.2 | - | - | 0 | - |
| - | - | 4154 | 433.2 | - | - | 0 | - |
| - | - | 1098 | 434.2 | - | - | 0 | - |
| - | - | 3476 | 434.2 | - | - | 0 | - |
| - | - | 4389 | 434.7 | - | - | 0 | - |
| 8 | b | 4232 | 435.2 | 0.002023 | 4.649 | +2 | 8 |
| - | - | 1088 | 435.7 | - | - | 0 | - |
| - | - | 840.3 | 440.2 | - | - | 0 | - |
| - | - | 1482 | 441.2 | - | - | 0 | - |
| - | - | 5799 | 442.2 | - | - | 0 | - |
| - | - | 836 | 443.1 | - | - | 0 | - |
| 7 | y | 3151 | 443.2 | 0.006024 | 13.59 | +2 | 8 |
| 8 | b | 2479 | 444.2 | 0.002234 | 5.029 | +2 | 8 |
| - | - | 2372 | 444.7 | - | - | 0 | - |
| - | - | 703.6 | 448.2 | - | - | 0 | - |
| - | - | 2000 | 450.2 | - | - | 0 | - |
| - | - | 786.3 | 451.2 | - | - | 0 | - |
| - | - | 4747 | 452.2 | - | - | 0 | - |
| - | - | 1164 | 453.2 | - | - | 0 | - |
| - | - | 731.4 | 454.2 | - | - | 0 | - |
| - | - | 1078 | 454.3 | - | - | 0 | - |
| 4 | b | 3856 | 455.3 | 0.0005566 | 1.223 | +1 | 4 |
| - | - | 1131 | 456.3 | - | - | 0 | - |
| - | - | 1728 | 459.2 | - | - | 0 | - |
| - | - | 1194 | 461.2 | - | - | 0 | - |
| - | - | 7696 | 462.2 | - | - | 0 | - |
| - | - | 1849 | 463.2 | - | - | 0 | - |
| - | - | 830.5 | 464.2 | - | - | 0 | - |
| - | - | 1244 | 464.3 | - | - | 0 | - |
| - | - | 1131 | 466.2 | - | - | 0 | - |
| - | - | 561.8 | 467.2 | - | - | 0 | - |
| - | - | 2496 | 468.2 | - | - | 0 | - |
| - | - | 695.7 | 469.2 | - | - | 0 | - |
| - | - | 2.089E+04 | 470.2 | - | - | 0 | - |
| - | - | 1664 | 470.7 | - | - | 0 | - |
| - | - | 5164 | 471.2 | - | - | 0 | - |
| 13 | b | 742.8 | 475.6 | 0.008243 | 17.33 | +3 | 13 |
| - | - | 1329 | 476.2 | - | - | 0 | - |
| - | - | 738.7 | 477.3 | - | - | 0 | - |
| - | - | 2583 | 478.2 | - | - | 0 | - |
| - | - | 2775 | 479.2 | - | - | 0 | - |
| - | - | 2.017E+04 | 480.2 | - | - | 0 | - |
| - | - | 4673 | 481.2 | - | - | 0 | - |
| - | - | 2061 | 482.2 | - | - | 0 | - |
| - | - | 1034 | 482.3 | - | - | 0 | - |
| - | - | 2748 | 484.2 | - | - | 0 | - |
| - | - | 1365 | 485.2 | - | - | 0 | - |
| 6 | y | 822.6 | 486.2 | 0.001854 | 3.812 | +2 | 9 |
| - | - | 981.6 | 488.3 | - | - | 0 | - |
| - | - | 704.1 | 489.3 | - | - | 0 | - |
| - | - | 772.9 | 492.2 | - | - | 0 | - |
| - | - | 782.7 | 493.2 | - | - | 0 | - |
| - | - | 2211 | 494.2 | - | - | 0 | - |
| - | - | 1566 | 494.2 | - | - | 0 | - |
| - | - | 636.7 | 495.5 | - | - | 0 | - |
| - | - | 7499 | 496.3 | - | - | 0 | - |
| - | - | 1.452E+04 | 497.2 | - | - | 0 | - |
| - | - | 6.876E+04 | 498.2 | - | - | 0 | - |
| - | - | 1.81E+04 | 499.2 | - | - | 0 | - |
| - | - | 4892 | 500.2 | - | - | 0 | - |
| 9 | b | 1611 | 500.8 | 0.0006744 | 1.347 | +2 | 9 |
| - | - | 1550 | 501.3 | - | - | 0 | - |
| - | - | 3595 | 502.2 | - | - | 0 | - |
| - | - | 1365 | 503.2 | - | - | 0 | - |
| - | - | 935.3 | 503.3 | - | - | 0 | - |
| 0 | Precursor | 1243 | 505.3 | 0.009583 | 18.97 | +3 | -1 |
| - | - | 636.7 | 506.3 | - | - | 0 | - |
| - | - | 676.5 | 507.7 | - | - | 0 | - |
| - | - | 1854 | 509.3 | - | - | 0 | - |
| - | - | 1182 | 510.2 | - | - | 0 | - |
| - | - | 1255 | 511.3 | - | - | 0 | - |
| - | - | 813.9 | 511.8 | - | - | 0 | - |
| - | - | 4900 | 512.2 | - | - | 0 | - |
| - | - | 1234 | 513.2 | - | - | 0 | - |
| - | - | 2.669E+04 | 513.3 | - | - | 0 | - |
| 10 | y | 6.784E+04 | 514.3 | 0.0009519 | 1.851 | +1 | 5 |
| - | - | 1.798E+04 | 515.3 | - | - | 0 | - |
| - | - | 3238 | 516.3 | - | - | 0 | - |
| - | - | 1069 | 518.2 | - | - | 0 | - |
| - | - | 1722 | 520.3 | - | - | 0 | - |
| - | - | 898.4 | 520.3 | - | - | 0 | - |
| 5 | b | 1323 | 524.3 | 0.0003078 | 0.5871 | +1 | 5 |
| - | - | 695.3 | 526.3 | - | - | 0 | - |
| - | - | 1.272E+04 | 530.2 | - | - | 0 | - |
| - | - | 3156 | 531.2 | - | - | 0 | - |
| 10 | y | 7.735E+04 | 531.3 | 0.0004037 | 0.7599 | +1 | 5 |
| - | - | 2.095E+04 | 532.3 | - | - | 0 | - |
| - | - | 3800 | 533.3 | - | - | 0 | - |
| - | - | 1241 | 534.3 | - | - | 0 | - |
| - | - | 2988 | 538.3 | - | - | 0 | - |
| - | - | 858.4 | 539.3 | - | - | 0 | - |
| - | - | 2486 | 540.3 | - | - | 0 | - |
| - | - | 955.7 | 540.8 | - | - | 0 | - |
| - | - | 966.7 | 541.3 | - | - | 0 | - |
| - | - | 1210 | 542.3 | - | - | 0 | - |
| 5 | b | 2216 | 542.3 | 0.0004801 | 0.8852 | +1 | 5 |
| - | - | 1035 | 543.3 | - | - | 0 | - |
| - | - | 1077 | 543.3 | - | - | 0 | - |
| - | - | 1205 | 546.2 | - | - | 0 | - |
| - | - | 667.4 | 554.3 | - | - | 0 | - |
| - | - | 2320 | 555.3 | - | - | 0 | - |
| - | - | 823.7 | 556.2 | - | - | 0 | - |
| 10 | b | 1037 | 556.3 | 0.004063 | 7.305 | +2 | 10 |
| - | - | 657.3 | 557.3 | - | - | 0 | - |
| - | - | 645.4 | 560.8 | - | - | 0 | - |
| - | - | 745.2 | 562.3 | - | - | 0 | - |
| - | - | 1125 | 563.2 | - | - | 0 | - |
| - | - | 993.2 | 570.3 | - | - | 0 | - |
| - | - | 2177 | 573.2 | - | - | 0 | - |
| - | - | 876.1 | 573.3 | - | - | 0 | - |
| - | - | 1951 | 573.8 | - | - | 0 | - |
| - | - | 1591 | 574.3 | - | - | 0 | - |
| - | - | 951.6 | 574.8 | - | - | 0 | - |
| 4 | y | 1540 | 578.3 | 0.0009917 | 1.715 | +2 | 11 |
| 4 | y | 5089 | 578.8 | 0.001813 | 3.132 | +2 | 11 |
| - | - | 3038 | 579.3 | - | - | 0 | - |
| - | - | 5086 | 581.3 | - | - | 0 | - |
| - | - | 887.5 | 582.2 | - | - | 0 | - |
| - | - | 1923 | 582.3 | - | - | 0 | - |
| - | - | 5075 | 582.8 | - | - | 0 | - |
| - | - | 4400 | 583.3 | - | - | 0 | - |
| - | - | 1345 | 583.8 | - | - | 0 | - |
| - | - | 800.8 | 584.3 | - | - | 0 | - |
| 4 | y | 1.428E+04 | 587.3 | 0.0003788 | 0.645 | +2 | 11 |
| - | - | 8429 | 587.8 | - | - | 0 | - |
| - | - | 3083 | 588.3 | - | - | 0 | - |
| - | - | 877.8 | 588.8 | - | - | 0 | - |
| - | - | 4333 | 591.2 | - | - | 0 | - |
| 11 | b | 1959 | 591.3 | 0.001917 | 3.242 | +2 | 11 |
| 11 | b | 1764 | 591.8 | 0.004488 | 7.584 | +2 | 11 |
| - | - | 1233 | 592.2 | - | - | 0 | - |
| - | - | 2096 | 592.3 | - | - | 0 | - |
| - | - | 752.5 | 593.2 | - | - | 0 | - |
| - | - | 2068 | 593.3 | - | - | 0 | - |
| - | - | 1569 | 597.3 | - | - | 0 | - |
| - | - | 2.92E+04 | 599.3 | - | - | 0 | - |
| - | - | 6485 | 600.3 | - | - | 0 | - |
| - | - | 1591 | 600.8 | - | - | 0 | - |
| - | - | 2055 | 601.3 | - | - | 0 | - |
| - | - | 2358 | 601.3 | - | - | 0 | - |
| - | - | 1166 | 602.3 | - | - | 0 | - |
| - | - | 2388 | 607.8 | - | - | 0 | - |
| - | - | 699.5 | 608.3 | - | - | 0 | - |
| - | - | 1525 | 608.8 | - | - | 0 | - |
| - | - | 1.725E+04 | 609.3 | - | - | 0 | - |
| - | - | 4481 | 609.3 | - | - | 0 | - |
| - | - | 1040 | 609.8 | - | - | 0 | - |
| - | - | 4307 | 610.3 | - | - | 0 | - |
| - | - | 1576 | 610.3 | - | - | 0 | - |
| - | - | 701.8 | 610.8 | - | - | 0 | - |
| - | - | 865.9 | 611.3 | - | - | 0 | - |
| 6 | b | 7720 | 611.3 | 0.002368 | 3.873 | +1 | 6 |
| - | - | 800.6 | 612.3 | - | - | 0 | - |
| - | - | 3065 | 612.3 | - | - | 0 | - |
| - | - | 839.5 | 612.8 | - | - | 0 | - |
| - | - | 803.5 | 613.8 | - | - | 0 | - |
| - | - | 2296 | 615.3 | - | - | 0 | - |
| - | - | 996.9 | 616.3 | - | - | 0 | - |
| - | - | 2370 | 617.8 | - | - | 0 | - |
| - | - | 6803 | 618.3 | - | - | 0 | - |
| - | - | 5507 | 618.8 | - | - | 0 | - |
| - | - | 2347 | 619.3 | - | - | 0 | - |
| - | - | 985.6 | 619.4 | - | - | 0 | - |
| - | - | 966.8 | 619.8 | - | - | 0 | - |
| - | - | 1702 | 625.3 | - | - | 0 | - |
| - | - | 817.5 | 626.3 | - | - | 0 | - |
| 3 | y | 1.082E+04 | 626.8 | 0.0003364 | 0.5366 | +2 | 12 |
| - | - | 1.61E+05 | 627.3 | - | - | 0 | - |
| 3 | y | 1.929E+04 | 627.3 | 0.003934 | 6.271 | +2 | 12 |
| - | - | 2.119E+04 | 627.8 | - | - | 0 | - |
| - | - | 5.509E+04 | 628.3 | - | - | 0 | - |
| - | - | 2680 | 628.8 | - | - | 0 | - |
| - | - | 1.267E+04 | 629.3 | - | - | 0 | - |
| 6 | b | 8716 | 629.3 | 0.000312 | 0.4958 | +1 | 6 |
| - | - | 1766 | 630.3 | - | - | 0 | - |
| - | - | 2828 | 630.3 | - | - | 0 | - |
| - | - | 886.9 | 631.3 | - | - | 0 | - |
| - | - | 797.7 | 632.8 | - | - | 0 | - |
| - | - | 2180 | 633.3 | - | - | 0 | - |
| - | - | 1043 | 633.9 | - | - | 0 | - |
| - | - | 1639 | 634.8 | - | - | 0 | - |
| - | - | 1026 | 635.3 | - | - | 0 | - |
| 3 | y | 1.604E+05 | 635.8 | 0.000303 | 0.4766 | +2 | 12 |
| - | - | 1.065E+05 | 636.3 | - | - | 0 | - |
| - | - | 4.849E+04 | 636.8 | - | - | 0 | - |
| - | - | 1.275E+04 | 637.3 | - | - | 0 | - |
| - | - | 1963 | 637.8 | - | - | 0 | - |
| - | - | 703.1 | 643 | - | - | 0 | - |
| - | - | 5503 | 643.3 | - | - | 0 | - |
| - | - | 2140 | 644.3 | - | - | 0 | - |
| 9 | y | 3.085E+04 | 644.4 | 0.0003241 | 0.503 | +1 | 6 |
| - | - | 1897 | 645.3 | - | - | 0 | - |
| - | - | 1.106E+04 | 645.4 | - | - | 0 | - |
| - | - | 2072 | 646.4 | - | - | 0 | - |
| - | - | 1055 | 649.3 | - | - | 0 | - |
| - | - | 725.1 | 656.3 | - | - | 0 | - |
| - | - | 1135 | 660.3 | - | - | 0 | - |
| - | - | 1367 | 667.3 | - | - | 0 | - |
| - | - | 2589 | 668.9 | - | - | 0 | - |
| - | - | 1385 | 676.3 | - | - | 0 | - |
| - | - | 911.9 | 678.3 | - | - | 0 | - |
| - | - | 623.7 | 680.3 | - | - | 0 | - |
| - | - | 695.1 | 682.4 | - | - | 0 | - |
| - | - | 3115 | 682.9 | - | - | 0 | - |
| - | - | 588.4 | 683 | - | - | 0 | - |
| - | - | 645.8 | 683.3 | - | - | 0 | - |
| - | - | 2110 | 683.4 | - | - | 0 | - |
| - | - | 819 | 684.4 | - | - | 0 | - |
| - | - | 703.4 | 686.3 | - | - | 0 | - |
| - | - | 5254 | 694.3 | - | - | 0 | - |
| - | - | 2369 | 695.3 | - | - | 0 | - |
| - | - | 746.9 | 696.3 | - | - | 0 | - |
| - | - | 3758 | 704.3 | - | - | 0 | - |
| - | - | 1495 | 705.3 | - | - | 0 | - |
| - | - | 726.4 | 706.3 | - | - | 0 | - |
| - | - | 740.9 | 707.3 | - | - | 0 | - |
| - | - | 916 | 708.3 | - | - | 0 | - |
| 2 | y | 975.5 | 709.4 | 0.004487 | 6.326 | +2 | 13 |
| - | - | 1028 | 709.8 | - | - | 0 | - |
| 13 | b | 2.187E+04 | 712.4 | 0.01282 | 17.99 | +2 | 13 |
| - | - | 8806 | 713.4 | - | - | 0 | - |
| - | - | 2602 | 714.4 | - | - | 0 | - |
| - | - | 1870 | 721.4 | - | - | 0 | - |
| - | - | 1.053E+04 | 722.3 | - | - | 0 | - |
| - | - | 4118 | 723.3 | - | - | 0 | - |
| - | - | 1087 | 724.3 | - | - | 0 | - |
| - | - | 3136 | 730.4 | - | - | 0 | - |
| - | - | 1947 | 731.4 | - | - | 0 | - |
| - | - | 710.6 | 736.3 | - | - | 0 | - |
| - | - | 1003 | 738.3 | - | - | 0 | - |
| - | - | 2846 | 738.4 | - | - | 0 | - |
| - | - | 3521 | 739.4 | - | - | 0 | - |
| 7 | b | 5.075E+04 | 740.4 | 0.01371 | 18.52 | +1 | 7 |
| - | - | 2.36E+04 | 741.4 | - | - | 0 | - |
| - | - | 6243 | 742.4 | - | - | 0 | - |
| - | - | 1806 | 743.4 | - | - | 0 | - |
| - | - | 987.8 | 746.3 | - | - | 0 | - |
| - | - | 1147 | 753.3 | - | - | 0 | - |
| - | - | 1183 | 755.3 | - | - | 0 | - |
| 8 | y | 2658 | 755.4 | 0.0002171 | 0.2874 | +1 | 7 |
| 8 | y | 3856 | 756.4 | 0.003506 | 4.635 | +1 | 7 |
| 0 | Precursor | 1.397E+04 | 757.4 | 0.006692 | 8.836 | +2 | -1 |
| 7 | b | 2.449E+04 | 758.4 | 0.0007487 | 0.9872 | +1 | 7 |
| - | - | 9164 | 759.4 | - | - | 0 | - |
| - | - | 2694 | 760.4 | - | - | 0 | - |
| - | - | 1661 | 770.4 | - | - | 0 | - |
| - | - | 4184 | 771.4 | - | - | 0 | - |
| - | - | 1835 | 772.4 | - | - | 0 | - |
| 8 | y | 1.39E+04 | 773.4 | 0.0004556 | 0.5891 | +1 | 7 |
| - | - | 2359 | 774.3 | - | - | 0 | - |
| - | - | 4056 | 774.4 | - | - | 0 | - |
| - | - | 1240 | 775.3 | - | - | 0 | - |
| - | - | 1581 | 775.4 | - | - | 0 | - |
| - | - | 891.6 | 806.4 | - | - | 0 | - |
| - | - | 886.5 | 832.4 | - | - | 0 | - |
| - | - | 1076 | 833.4 | - | - | 0 | - |
| - | - | 1393 | 840.4 | - | - | 0 | - |
| - | - | 1240 | 841.4 | - | - | 0 | - |
| - | - | 3499 | 842.4 | - | - | 0 | - |
| - | - | 1040 | 843.4 | - | - | 0 | - |
| - | - | 4621 | 850.4 | - | - | 0 | - |
| - | - | 3542 | 851.4 | - | - | 0 | - |
| - | - | 1339 | 852.4 | - | - | 0 | - |
| - | - | 1619 | 853.4 | - | - | 0 | - |
| - | - | 1025 | 854.4 | - | - | 0 | - |
| - | - | 7004 | 859.4 | - | - | 0 | - |
| - | - | 2992 | 860.4 | - | - | 0 | - |
| - | - | 992.6 | 867.4 | - | - | 0 | - |
| - | - | 2.558E+04 | 868.4 | - | - | 0 | - |
| 8 | b | 1.675E+04 | 869.4 | 0.002595 | 2.985 | +1 | 8 |
| - | - | 6160 | 870.4 | - | - | 0 | - |
| - | - | 1234 | 871.4 | - | - | 0 | - |
| 7 | y | 902.4 | 884.4 | 0.002886 | 3.263 | +1 | 8 |
| 7 | y | 934.6 | 885.4 | 0.004248 | 4.798 | +1 | 8 |
| 8 | b | 5.317E+04 | 887.4 | 0.0004015 | 0.4525 | +1 | 8 |
| - | - | 2.722E+04 | 888.4 | - | - | 0 | - |
| - | - | 8102 | 889.4 | - | - | 0 | - |
| - | - | 1843 | 890.4 | - | - | 0 | - |
| - | - | 1865 | 894.4 | - | - | 0 | - |
| 7 | y | 2772 | 902.5 | 0.0009387 | 1.04 | +1 | 8 |
| - | - | 1019 | 903.5 | - | - | 0 | - |
| - | - | 867.5 | 904.4 | - | - | 0 | - |
| - | - | 3467 | 911.4 | - | - | 0 | - |
| - | - | 1373 | 912.5 | - | - | 0 | - |
| - | - | 2180 | 921.4 | - | - | 0 | - |
| - | - | 3491 | 922.4 | - | - | 0 | - |
| - | - | 1209 | 923.4 | - | - | 0 | - |
| - | - | 757.1 | 924.4 | - | - | 0 | - |
| - | - | 1.415E+04 | 939.4 | - | - | 0 | - |
| - | - | 7570 | 940.4 | - | - | 0 | - |
| - | - | 1485 | 941.4 | - | - | 0 | - |
| - | - | 1991 | 956.5 | - | - | 0 | - |
| - | - | 1013 | 957.5 | - | - | 0 | - |
| 6 | y | 1045 | 971.5 | 0.01337 | 13.76 | +1 | 9 |
| 6 | y | 2503 | 972.5 | 0.01439 | 14.8 | +1 | 9 |
| - | - | 826.7 | 973.5 | - | - | 0 | - |
| 9 | b | 1221 | 982.5 | 0.001208 | 1.229 | +1 | 9 |
| 6 | y | 3507 | 989.5 | 0.002205 | 2.229 | +1 | 9 |
| - | - | 1619 | 990.5 | - | - | 0 | - |
| 9 | b | 8291 | 1000 | 0.00103 | 1.03 | +1 | 9 |
| - | - | 3811 | 1002 | - | - | 0 | - |
| - | - | 2147 | 1003 | - | - | 0 | - |
| - | - | 729.6 | 1025 | - | - | 0 | - |
| - | - | 824.2 | 1035 | - | - | 0 | - |
| - | - | 4050 | 1036 | - | - | 0 | - |
| - | - | 2545 | 1037 | - | - | 0 | - |
| - | - | 892.6 | 1038 | - | - | 0 | - |
| - | - | 792.6 | 1041 | - | - | 0 | - |
| - | - | 1.232E+04 | 1053 | - | - | 0 | - |
| - | - | 7183 | 1054 | - | - | 0 | - |
| - | - | 2925 | 1055 | - | - | 0 | - |
| - | - | 909 | 1056 | - | - | 0 | - |
| 5 | y | 802.6 | 1059 | 0.01482 | 14 | +1 | 10 |
| 5 | y | 3663 | 1059 | 0.001898 | 1.791 | +1 | 10 |
| - | - | 1748 | 1061 | - | - | 0 | - |
| - | - | 872.5 | 1069 | - | - | 0 | - |
| - | - | 1044 | 1071 | - | - | 0 | - |
| 5 | y | 7040 | 1077 | 6.899E-06 | 0.006409 | +1 | 10 |
| - | - | 3237 | 1078 | - | - | 0 | - |
| - | - | 1038 | 1085 | - | - | 0 | - |
| 10 | b | 2315 | 1129 | 0.001014 | 0.8987 | +1 | 10 |
| - | - | 779.3 | 1130 | - | - | 0 | - |
| - | - | 1162 | 1139 | - | - | 0 | - |
| 4 | y | 2595 | 1157 | 0.0002814 | 0.2433 | +1 | 11 |
| - | - | 1238 | 1158 | - | - | 0 | - |
| 4 | y | 6481 | 1174 | 0.000755 | 0.6433 | +1 | 11 |
| - | - | 4296 | 1175 | - | - | 0 | - |
| - | - | 1422 | 1176 | - | - | 0 | - |
| 11 | b | 841.3 | 1200 | 0.001629 | 1.358 | +1 | 11 |
| - | - | 803.5 | 1201 | - | - | 0 | - |
| 12 | b | 1026 | 1314 | 7.647E-07 | 0.0005821 | +1 | 12 |
| - | - | 625.7 | 1846 | - | - | 0 | - |
| - | - | 647 | 2598 | - | - | 0 | - |
| - | - | 759.3 | 3042 | - | - | 0 | - |

m/z Charge Intensity FragmentType MassShift Position
120.0811538696289 0 134386.98
121.08446502685547 0 12167.6455
122.07176971435547 0 420.39276
126.05529022216797 0 8443.211
126.06643676757812 0 703.0876
127.05062866210938 0 904.3971
127.05857849121094 0 735.5728
127.08711242675781 0 1154.2228
128.0824432373047 0 446.05145
129.0552520751953 0 603.8538
129.06617736816406 0 7289.077
129.10260009765625 0 117605.93
130.05020141601562 0 9521.916
130.061279296875 0 538.57
130.0654296875 0 5286.033
130.0699005126953 0 549.4604
130.0863800048828 0 861.2213
130.1002960205078 0 1061.3718
130.10594177246094 0 7172.901
130.9914093017578 0 396.39124
131.0454864501953 0 508.3631
131.06863403320312 0 605.98785
131.10720825195312 0 419.26736
131.11825561523438 0 7044.457
132.0813446044922 0 438.01065
132.10218811035156 0 1047.2004
134.02728271484375 0 1272.5586
134.04470825195312 0 467.7746
136.07595825195312 0 3578.2378
138.12802124023438 0 3606.6
139.08694458007812 0 2977.2146
140.08241271972656 0 1175.7703
140.09030151367188 0 825.7872
141.06619262695312 0 3828.7402
141.1025390625 0 1388.8771
142.06982421875 0 519.679
142.09783935546875 0 638.83594
145.06130981445312 0 791.61774
146.06028747558594 0 1540.0706
147.07681274414062 0 3398.5085
147.11317443847656 0 2655.8284
148.06072998046875 0 457.3076
148.07566833496094 0 455.692
149.02371215820312 0 462.46747
152.03480529785156 0 483.18567
152.0708770751953 0 1147.9363
153.06631469726562 0 671.1502
153.10269165039062 0 961.4622
153.1363525390625 0 370.75555
154.08670043945312 0 2896.681
154.09786987304688 0 5276.822
155.08180236816406 0 7678.264
155.11839294433594 0 930.48376
157.06103515625 0 2607.634
157.09750366210938 0 8716.866
158.09262084960938 0 1307.8351
158.10040283203125 0 642.10614
159.11337280273438 0 829.73334
163.07179260253906 0 2098.5627
163.0866241455078 0 1099.5641
164.07000732421875 0 652.8872
165.10263061523438 0 1380.5391
165.61302185058594 0 759.06537 a 2
166.06141662597656 0 1142.4594
166.0867462158203 0 1166.7537
167.0818328857422 0 5608.372
167.1182403564453 0 6497.425
169.06103515625 0 2173.92
169.0977020263672 0 1419.6505
169.48538208007812 0 416.24683
170.0595703125 0 613.86285
171.07679748535156 0 1909.297
172.07205200195312 0 1145.2693
172.10838317871094 0 2363.9294
172.1138458251953 0 868.9006
173.0922088623047 0 778.20856
173.12869262695312 0 1369.0298
173.45071411132812 0 3581.3748
174.05569458007812 0 454.30228
175.07174682617188 0 3347.83
175.08609008789062 0 601.6962
175.3733673095703 0 552.22217
177.102294921875 0 667.9935
177.30799865722656 0 506.2768
180.1136016845703 0 1400.5125
181.06143188476562 0 447.00998
181.09750366210938 0 2736.051
181.13389587402344 0 795.80005
182.09274291992188 0 8152.385
182.12901306152344 0 842.7188
183.07672119140625 0 5168.504
183.09642028808594 0 541.7592
183.11326599121094 0 2240.1147
183.14967346191406 0 1194.3082
184.08038330078125 0 494.62717
184.1083526611328 0 5629.0864
185.09239196777344 0 12036.355
186.08773803710938 0 4969.7505
186.09571838378906 0 1280.5619
186.12445068359375 0 1220.5272
187.07229614257812 0 591.09344
187.1079559326172 0 1755.1333
187.14480590820312 0 1268.7976
189.08740234375 0 4441.6025
191.0819091796875 0 1663.9333
193.09674072265625 0 518.6609
195.0766143798828 0 1521.205
195.11309814453125 0 14358.578
196.11720275878906 0 1112.1614
196.88400268554688 0 459.8706
197.1288299560547 0 10848.478
197.1971435546875 0 467.01163
198.08782958984375 0 897.6736
198.12400817871094 0 6491.8345
199.0716552734375 0 8698.042
199.08203125 0 816.7438
199.1082000732422 0 4771.6294
199.11917114257812 0 1392.6414
200.0743408203125 0 746.04926
200.10325622558594 0 8382.717
200.1127471923828 0 616.42206
200.13946533203125 0 1956.022
201.0876922607422 0 703.2699
201.10743713378906 0 994.18243
201.12367248535156 0 10778.666 y Ammonia loss 12
202.1083221435547 0 591.7629
202.11880493164062 0 1904.9742 y 10
203.10279846191406 0 1273.8245
204.10598754882812 0 1105.1947
204.5975799560547 0 642.43066
205.09970092773438 0 586.47217
205.1186981201172 0 566.1011
206.0924835205078 0 563.0035
207.09767150878906 0 491.1592
207.11309814453125 0 1132.32
208.09791564941406 0 862.5357
208.1083221435547 0 6967.1123
209.09242248535156 0 29131.398
209.1292724609375 0 669.32916
210.09597778320312 0 2000.7834
211.1448974609375 0 989.25006
212.13967895507812 0 2185.8303
213.08738708496094 0 1854.026
213.12350463867188 0 1867.0029
214.138427734375 0 705.6291
215.10293579101562 0 571.3003
215.13934326171875 0 5782.956
216.13333129882812 0 512.33344
216.1439971923828 0 599.4126
217.08224487304688 0 12768.478
217.13363647460938 0 3229.239
218.0850830078125 0 1104.7252
218.15029907226562 0 50939.004 y 12
219.15350341796875 0 4842.775
221.05868530273438 0 1600.5068
221.10340881347656 0 652.0461
221.62403869628906 0 1463.0413
223.0711669921875 0 574.3329
223.1564178466797 0 586.26886
224.1038360595703 0 1003.5556
224.1396026611328 0 654.7488
225.12362670898438 0 8559.564
225.13516235351562 0 5035.9316
226.11898803710938 0 93049.38
227.1029815673828 0 3777.5933
227.1222686767578 0 8159.625
228.12469482421875 0 760.99884
228.13455200195312 0 5655.214 b 3
229.11924743652344 0 548.84033
229.13510131835938 0 1207.1085
229.1549835205078 0 1153.7025
231.09796142578125 0 1336.0475
233.16529846191406 0 177737.77 a 1
234.08750915527344 0 1123.6499
234.12396240234375 0 2652.8206
234.16859436035156 0 27777
235.17178344726562 0 2240.5867
236.10330200195312 0 4858.157
236.12808227539062 0 566.4851
236.14059448242188 0 853.99384
237.1238555908203 0 609.5734
238.0818634033203 0 739.8914
239.15017700195312 0 848.5627
240.097900390625 0 1504.6528
240.13450622558594 0 1171.8301
240.608642578125 0 1457.1871
241.0823516845703 0 8882.812
242.08627319335938 0 571.1593
242.11354064941406 0 630.10614
242.15036010742188 0 20589.064
243.13380432128906 0 1691.7539
243.1456298828125 0 25288.674
244.09323120117188 0 1164.445
244.13037109375 0 5292.714
244.1490478515625 0 1914.609
245.12924194335938 0 3277.5227
248.62945556640625 0 1705.0618
248.650634765625 0 834.2442
249.121826171875 0 7828.358
249.17178344726562 0 1465.7589
249.62298583984375 0 1549.4398
251.15089416503906 0 3509.9956
252.09817504882812 0 2018.1128
252.13470458984375 0 2196.851
254.1138458251953 0 15276.1
254.15011596679688 0 2355.491
255.11660766601562 0 1680.4644
257.1283264160156 0 506.5003
257.14300537109375 0 15060.267
257.6337585449219 0 2074.7449 y Ammonia loss 9
257.6449890136719 0 2669.6824
258.1095886230469 0 697.52
259.0927734375 0 8957.16
260.0961608886719 0 807.78876
261.1600036621094 0 73447.664 b 1
262.118896484375 0 1252.0991
262.1632995605469 0 12612.121
263.1661071777344 0 1076.5769
264.1345520019531 0 11631.833
265.13775634765625 0 1821.5763
265.1529846191406 0 572.74896
266.1250305175781 0 833.5422
266.13104248046875 0 790.3774
266.14813232421875 0 13382.01 y 9
266.64959716796875 0 3522.0474
268.0933532714844 0 667.2823
269.1607971191406 0 1217.5468
269.6529846191406 0 1211.5945
270.10845947265625 0 788.01874
270.14434814453125 0 871.9015
272.1083679199219 0 1033.3883
272.1242980957031 0 6077.001
273.1230773925781 0 1244.2753
277.1661682128906 0 1122.5566
278.1499938964844 0 628.0121
278.16748046875 0 1188.2975
279.1455383300781 0 5204.7417
280.1296081542969 0 8653.49
280.14739990234375 0 1016.4451
281.13336181640625 0 926.0672
282.10980224609375 0 1148.1785
282.1277160644531 0 2385.392
282.1452331542969 0 33561.176
282.6277770996094 0 828.90314
283.1479187011719 0 4100.774
285.1966552734375 0 1106.9802
286.1037902832031 0 4780.822
292.1656799316406 0 1026.6782
296.0875549316406 0 1042.9135
296.1358337402344 0 4351.679
296.1722717285156 0 5087.3022
296.6263122558594 0 1138.3745
297.1199951171875 0 4662.662
297.1564025878906 0 30461.682
298.12359619140625 0 1130.9349
298.1405334472656 0 4207.9634
298.15985107421875 0 3540.979
299.1715393066406 0 2377.252
300.1188659667969 0 783.0571
300.15570068359375 0 1871.585
304.11492919921875 0 2519.3364
305.17242431640625 0 698.63696
306.14471435546875 0 1791.7032
306.1637878417969 0 2118.7896 b Water loss 5
307.1409912109375 0 1217.9763
308.1609802246094 0 2339.0652
310.10406494140625 0 1893.5117
313.1672058105469 0 588.3906
313.1877136230469 0 3830.348
314.13311767578125 0 2384.8606
314.14654541015625 0 4874.397
314.1812438964844 0 11593.388 y Ammonia loss 8
314.63604736328125 0 1862.1428
314.67877197265625 0 1613.8313
315.1317443847656 0 1165.2069
315.14996337890625 0 822.0277
315.1668395996094 0 14666.372 b 5
315.18505859375 0 1239.4576
316.16986083984375 0 2343.3904
319.14093017578125 0 814.06335
321.15478515625 0 638.30225
322.6904296875 0 15459.967 y 8
323.1716003417969 0 1320.3566
323.19183349609375 0 4917.5425
323.69244384765625 0 1418.6343
324.1549987792969 0 769.0445
324.1759338378906 0 635.86084
325.15191650390625 0 1885.5217
325.1879577636719 0 1131.8578
326.1706237792969 0 1569.8835
328.11431884765625 0 4905.724
329.116943359375 0 677.93427
329.66436767578125 0 1412.9352
330.16558837890625 0 733.7618
332.193359375 0 39177.402 y 11
333.1564025878906 0 6550.3936
333.1963195800781 0 5708.875
334.1589050292969 0 867.5005
336.1560974121094 0 1492.0126
337.1512756347656 0 1489.7246
337.1897277832031 0 633.1441
338.13446044921875 0 1096.7467
338.66937255859375 0 4135.567
339.1679992675781 0 3821.9138
340.17236328125 0 1461.2162
341.1829528808594 0 6351.626
342.183349609375 0 1801.4708
343.1617431640625 0 3418.6655
343.1875 0 1719.7003
344.16552734375 0 570.7196
344.1910400390625 0 614.1624
346.12469482421875 0 8625.973
347.1295166015625 0 1829.6132
347.6749267578125 0 2440.822
348.1756286621094 0 1205.7284
349.1133728027344 0 556.97156
350.1825256347656 0 1417.7761
351.16668701171875 0 27798.467
352.1697082519531 0 5601.0713
353.18212890625 0 8247.056
354.1662292480469 0 2281.69
354.1868896484375 0 1390.7767
355.1617126464844 0 3420.6316
356.1451110839844 0 1050.9325
356.6800231933594 0 737.7022
358.2127380371094 0 4394.9697 b 2
359.216064453125 0 969.98334
361.6746826171875 0 2345.199
362.1819763183594 0 1555.1592
363.1647033691406 0 851.88965
365.1460266113281 0 3324.9248
365.18212890625 0 1075.4169
365.68994140625 0 680.1179
368.19317626953125 0 8866.614
369.1772766113281 0 28135.654
370.18023681640625 0 5111.9976
370.6779479980469 0 11089.042 b Water loss 6
371.1802062988281 0 2984.5864 b Ammonia loss 9
372.17718505859375 0 1609.5608
373.1722717285156 0 5151.9087
374.1756896972656 0 1409.662
379.6911926269531 0 1542.5527 b 6
380.1925048828125 0 601.20734
383.15679931640625 0 7753.074
384.1604919433594 0 994.9496
385.2196350097656 0 3516.6921
386.2038879394531 0 22234.332 y Ammonia loss 10
387.1825866699219 0 603.40326
387.2078552246094 0 4058.4917 y 7
390.1776123046875 0 1369.7228
392.1943664550781 0 770.3366
393.17718505859375 0 1530.274
394.53314208984375 0 986.35095 b Water loss 10
395.1938171386719 0 1215.9757
396.1881408691406 0 581.8873
397.1357116699219 0 1062.6042
401.1672668457031 0 11943.948
402.17083740234375 0 3332.9905
403.2305603027344 0 91315.89 y 10
404.2335205078125 0 21595.635
405.160400390625 0 726.5453
405.23504638671875 0 2727.0198
407.20404052734375 0 5800.0977
408.1884460449219 0 15080.653
409.1921081542969 0 3235.7312
410.2046203613281 0 1140.1533
412.532958984375 0 804.06866
413.20477294921875 0 770.95435
414.1990966796875 0 1774.0198
415.146484375 0 3568.8604
415.2005920410156 0 682.6776
418.2135009765625 0 959.71545 y Water loss 2
418.54254150390625 0 1614.688 y Ammonia loss 2
418.8751220703125 0 1231.7717
421.20709228515625 0 727.4633
424.2294006347656 0 8030.1396
424.54656982421875 0 1028.935
425.1313781738281 0 1155.7942
425.21502685546875 0 7319.8633
425.7015686035156 0 2163.5498
426.2003173828125 0 3587.5508
426.7046203613281 0 708.36444
427.2307434082031 0 2285.8489
427.2718200683594 0 1398.7467
429.2129211425781 0 1435.5927
433.15692138671875 0 4154.101
434.16253662109375 0 1098.0769
434.2043151855469 0 3476.4995
434.70703125 0 4389.3228
435.2076416015625 0 4232.375 b Water loss 7
435.7076110839844 0 1087.7101
440.21630859375 0 840.27875
441.1985168457031 0 1482.4058
442.2410583496094 0 5799.1567
443.1419982910156 0 836.0367
443.22528076171875 0 3151.463 y Ammonia loss 6
444.213134765625 0 2479.207 b 7
444.71240234375 0 2372.1938
448.1852722167969 0 703.6169
450.19879150390625 0 2000.0188
451.2286682128906 0 786.3368
452.2146301269531 0 4746.8984
453.2159729003906 0 1163.8694
454.2293701171875 0 731.44507
454.2647705078125 0 1077.638
455.2658386230469 0 3855.6255 b 3
456.26971435546875 0 1130.5477
459.2097473144531 0 1727.9032
461.21820068359375 0 1194.0751
462.19940185546875 0 7696.0537
463.20196533203125 0 1848.985
464.2154235839844 0 830.45483
464.25299072265625 0 1244.0499
466.1941223144531 0 1130.531
467.1849060058594 0 561.7591
468.2095031738281 0 2496.3167
469.24176025390625 0 695.6741
470.22528076171875 0 20892.32
470.72698974609375 0 1663.8202
471.228515625 0 5164.0093
475.5816345214844 0 742.82367 b Ammonia loss 12
476.1783752441406 0 1329.0238
477.26007080078125 0 738.68384
478.24072265625 0 2582.7087
479.2267150878906 0 2774.6343
480.20989990234375 0 20165.584
481.21282958984375 0 4673.0215
482.22210693359375 0 2061.1086
482.2601318359375 0 1034.1832
484.2049865722656 0 2748.1882
485.2102966308594 0 1365.1589
486.2451171875 0 822.638 y Water loss 5
488.2794189453125 0 981.59906
489.2766418457031 0 704.0794
492.2131652832031 0 772.9382
493.20452880859375 0 782.7151
494.1887512207031 0 2210.7114
494.2356872558594 0 1565.9457
495.4945373535156 0 636.70447
496.2510070800781 0 7499.321
497.2364196777344 0 14524.542
498.2203369140625 0 68758.45
499.22344970703125 0 18102.178
500.22711181640625 0 4891.976
500.75225830078125 0 1610.716 b 8
501.2532958984375 0 1549.5703
502.2152099609375 0 3594.8555
503.2189636230469 0 1364.9309
503.2634582519531 0 935.2651
505.2463684082031 0 1243.2429 Precursor Ammonia loss
506.2621154785156 0 636.66705
507.74224853515625 0 676.48663
509.2568359375 0 1853.8379
510.2178039550781 0 1182.3536
511.26300048828125 0 1255.0964
511.7602844238281 0 813.8652
512.19970703125 0 4899.6465
513.2054443359375 0 1234.3982
513.2785034179688 0 26688.354
514.262939453125 0 67835.414 y Ammonia loss 9
515.2654418945312 0 17983.066
516.2666625976562 0 3237.8545
518.2432250976562 0 1069.0248
520.2501831054688 0 1722.3633
520.2894897460938 0 898.4051
524.2864379882812 0 1322.6023 b Water loss 4
526.3137817382812 0 695.2579
530.2098999023438 0 12722.454
531.21337890625 0 3156.4
531.2889404296875 0 77354.4 y 9
532.2914428710938 0 20952.14
533.2930297851562 0 3799.6453
534.2952270507812 0 1240.7024
538.2994995117188 0 2987.5845
539.3019409179688 0 858.4211
540.3104858398438 0 2486.3865
540.814697265625 0 955.7117
541.26953125 0 966.7315
542.2570190429688 0 1209.8226
542.2977905273438 0 2216.017 b 4
543.2613525390625 0 1035.2285
543.3029174804688 0 1076.8225
546.2420043945312 0 1204.8198
554.2578735351562 0 667.3708
555.325927734375 0 2319.584
556.2252197265625 0 823.65405
556.2730102539062 0 1037.2979 b Ammonia loss 9
557.2759399414062 0 657.26483
560.7662353515625 0 645.35693
562.2655639648438 0 745.153
563.2462768554688 0 1124.9677
570.2694091796875 0 993.1775
573.2304077148438 0 2176.8884
573.3334350585938 0 876.14575
573.7759399414062 0 1951.0757
574.2744140625 0 1591.4001
574.7740478515625 0 951.581
578.28466796875 0 1539.7869 y Water loss 3
578.7794799804688 0 5088.7065 y Ammonia loss 3
579.28125 0 3038.454
581.2579956054688 0 5085.8657
582.2478637695312 0 887.48846
582.28955078125 0 1922.8708
582.7810668945312 0 5075.2153
583.28173828125 0 4400.117
583.783935546875 0 1344.9563
584.3189697265625 0 800.8213
587.2913208007812 0 14275.407 y 3
587.7925415039062 0 8428.797
588.2930908203125 0 3083.113
588.7943725585938 0 877.78687
591.2410278320312 0 4332.517
591.2935791015625 0 1959.2932 b Water loss 10
591.7919921875 0 1764.4998 b Ammonia loss 10
592.2438354492188 0 1233.2886
592.2991943359375 0 2096.1108
593.2474975585938 0 752.5333
593.307373046875 0 2068.1584
597.2874755859375 0 1568.6812
599.2677612304688 0 29201.895
600.27294921875 0 6485.325
600.7972412109375 0 1591.1465
601.2728881835938 0 2055.3738
601.335205078125 0 2357.7314
602.3341674804688 0 1166.1232
607.796630859375 0 2387.7104
608.278076171875 0 699.5162
608.7993774414062 0 1525.1973
609.252197265625 0 17246.072
609.33740234375 0 4481.23
609.7945556640625 0 1039.8245
610.2550659179688 0 4307.3916
610.3380126953125 0 1576.3716
610.793212890625 0 701.8281
611.2587890625 0 865.92065
611.31640625 0 7720.354 b Water loss 5
612.2630004882812 0 800.5853
612.3182373046875 0 3064.5225
612.81640625 0 839.5245
613.8064575195312 0 803.4935
615.2998657226562 0 2295.9426
616.2984619140625 0 996.8997
617.806396484375 0 2369.8308
618.3018798828125 0 6802.8413
618.7993774414062 0 5507.2866
619.2969970703125 0 2346.6628
619.3508911132812 0 985.56085
619.796630859375 0 966.84045
625.2842407226562 0 1702.4695
626.284423828125 0 817.4921
626.8123779296875 0 10815.874 y Water loss 2
627.26220703125 0 160982.73
627.3079833984375 0 19294.12 y Ammonia loss 2
627.8069458007812 0 21191.664
628.2653198242188 0 55091.246
628.81005859375 0 2680.2925
629.267578125 0 12668.175
629.3296508789062 0 8715.594 b 5
630.270263671875 0 1766.2993
630.3329467773438 0 2827.516
631.3331909179688 0 886.9424
632.8482055664062 0 797.72516
633.3482666015625 0 2180.0513
633.8518676757812 0 1043.451
634.8103637695312 0 1638.6023
635.314697265625 0 1025.866
635.817626953125 0 160399.73 y 2
636.3189697265625 0 106537.46
636.8206176757812 0 48488.773
637.321533203125 0 12751.14
637.8228149414062 0 1963.1047
642.9552612304688 0 703.12054
643.296142578125 0 5503.3647
644.2970581054688 0 2139.679
644.3729248046875 0 30854.252 y 8
645.2908325195312 0 1896.7314
645.3753662109375 0 11062.821
646.3780517578125 0 2072.492
649.3318481445312 0 1054.6301
656.3270874023438 0 725.1156
660.321533203125 0 1134.5531
667.3438110351562 0 1366.6562
668.8870239257812 0 2589.2764
676.3281860351562 0 1385.2411
678.3321533203125 0 911.93665
680.3169555664062 0 623.735
682.390380859375 0 695.09534
682.886474609375 0 3114.7527
682.9611206054688 0 588.4331
683.29833984375 0 645.78564
683.3861694335938 0 2109.8323
684.3683471679688 0 819.0085
686.3175048828125 0 703.38556
694.3411865234375 0 5254.0254
695.3408813476562 0 2369.4968
696.3494873046875 0 746.9262
704.324462890625 0 3758.2473
705.330322265625 0 1495.0642
706.3322143554688 0 726.3854
707.3323364257812 0 740.869
708.3353881835938 0 916.0465
709.3560180664062 0 975.4587 y 1
709.8494873046875 0 1027.7815
712.3516235351562 0 21866.514 b Water loss 12
713.3545532226562 0 8805.985
714.3575439453125 0 2602.2717
721.3504638671875 0 1869.6597
722.337890625 0 10530.06
723.3421630859375 0 4118.1543
724.3414916992188 0 1087.1226
730.3779296875 0 3135.9456
731.37744140625 0 1947.3021
736.3220825195312 0 710.60565
738.3047485351562 0 1003.25903
738.378173828125 0 2845.7124
739.3645629882812 0 3521.3499
740.34765625 0 50749.008 b Water loss 6
741.3504028320312 0 23596.36
742.353271484375 0 6243.079
743.3557739257812 0 1806.2385
746.3385009765625 0 987.7731
753.342529296875 0 1147.4855
755.3308715820312 0 1182.697
755.4048461914062 0 2658.2275 y Water loss 7
756.3921508789062 0 3856.3572 y Ammonia loss 7
757.3735961914062 0 13965.565 Precursor Ammonia loss
758.3726806640625 0 24491.234 b 6
759.375732421875 0 9163.6455
760.3779907226562 0 2693.723
770.3682861328125 0 1661.1743
771.3535766601562 0 4184.2114
772.3538208007812 0 1834.8
773.4156494140625 0 13899.44 y 7
774.3330078125 0 2359.3115
774.4176635742188 0 4055.6418
775.3388671875 0 1240.2595
775.4188232421875 0 1580.8807
806.366943359375 0 891.6038
832.3868408203125 0 886.5287
833.376220703125 0 1076.0797
840.4115600585938 0 1393.4095
841.4092407226562 0 1239.7151
842.3936157226562 0 3498.9302
843.3978881835938 0 1039.8317
850.394775390625 0 4621.407
851.3931884765625 0 3542.047
852.3997192382812 0 1339.4404
853.4273681640625 0 1619.1112
854.4307250976562 0 1025.0624
859.4199829101562 0 7003.7617
860.4235229492188 0 2992.0645
867.4306640625 0 992.5627
868.405029296875 0 25575.867
869.4065551757812 0 16753.613 b Water loss 7
870.4088134765625 0 6159.7573
871.4132080078125 0 1234.3538
884.4443359375 0 902.3844 y Water loss 6
885.4354858398438 0 934.6312 y Ammonia loss 6
887.4141235351562 0 53168.133 b 7
888.41748046875 0 27215.459
889.419677734375 0 8102.368
890.4248657226562 0 1843.174
894.4227294921875 0 1864.9371
902.4568481445312 0 2772.4966 y 6
903.462890625 0 1018.87103
904.4086303710938 0 867.4582
911.4470825195312 0 3466.9702
912.4500122070312 0 1373.1437
921.43115234375 0 2180.4854
922.4182739257812 0 3491.2336
923.41015625 0 1209.0457
924.42236328125 0 757.11804
939.4409790039062 0 14152.52
940.444580078125 0 7569.5977
941.4432373046875 0 1485.1241
956.4588623046875 0 1991.0939
957.45751953125 0 1012.81995
971.4658813476562 0 1044.9904 y Water loss 5
972.4776611328125 0 2503.2954 y Ammonia loss 5
973.4866333007812 0 826.72833
982.48681640625 0 1221.4175 b Water loss 8
989.4876098632812 0 3507.3613 y 5
990.4967651367188 0 1619.4612
1000.49755859375 0 8290.566 b 8
1001.5004272460938 0 3811.3733
1002.5036010742188 0 2147.0337
1025.4920654296875 0 729.6357
1035.4730224609375 0 824.2379
1036.462158203125 0 4049.501
1037.46240234375 0 2544.9019
1038.4703369140625 0 892.6243
1041.4830322265625 0 792.64026
1053.4837646484375 0 12318.216
1054.4849853515625 0 7183.229
1055.487548828125 0 2925.4563
1056.4825439453125 0 909.01886
1058.4964599609375 0 802.64044 y Water loss 4
1059.4971923828125 0 3662.964 y Ammonia loss 4
1060.500732421875 0 1747.7035
1069.495849609375 0 872.5462
1071.4964599609375 0 1044.0131
1076.5218505859375 0 7039.832 y 4
1077.521728515625 0 3236.7925
1084.5196533203125 0 1037.8247
1128.55615234375 0 2315.4531 b 9
1129.5584716796875 0 779.31287
1138.5386962890625 0 1161.8705
1156.54833984375 0 2595.293 y Ammonia loss 3
1157.556640625 0 1237.6382
1173.5738525390625 0 6480.561 y 3
1174.5745849609375 0 4295.863
1175.5755615234375 0 1422.0497
1199.5926513671875 0 841.2807 b 10
1200.5875244140625 0 803.5227
1313.63720703125 0 1025.9606 b 11
1846.379150390625 0 625.7066
2598.447998046875 0 647.0441
3041.58447265625 0 759.27594

Spectrum Details

|  |  |
| --- | --- |
| Matched peaks? Matched peaks The total absolute number of peaks matched. Additionally in brackets the total fraction of peaks matched and the total number of peaks is shown. | 74 (10.31% of 718) |
| FDR? FDR The false discovery rate estimated for this peptide. It is calculated by matching all theoretical fragments with a non-integer shift with the raw peaks for this spectrum. This is done with 40 different shifts. The resulting percentage is the average number of annotated peaks over the number of annotated peaks with the correct spectrum. | 0.35% |
| Satellite FDR? Satellite FDR See the FDR for details on its calculation. This satellite ion specific FDR only contains the satellite ions (d/w) for I/L/J positions. | - |
| PSM Score? PSM Score The PSM Score as given by Hecklib to this annotated spectrum. It is shown with three significant figures. | 636 |

#### Spectrum 6346? Spectrum 6346 The raw spectrum of this peptide as annotated by Hecklib. The fragments are coloured according to ion type (see legend). Any peaks with a star '\*' as text can be hovered over to see the full details, first the ion type second the mass shift type. By hovering over the amino acids in the peptide or ions in the legend the corresponding peaks are highlighted. By toggling the 'Unassigned' label you can turn the background (unassigned) peaks on or off in the plot. By updating the slider in the Ion legend you can update the spectrum to only show the top X% of the peaks with labels. The top X% means any peak that is within X% of the highest intensity. By dragging in the spectrum you can zoom in to a specific part of the spectrum and use 'Zoom Out' to get back to the original zoom level. The annotation of the spectrum is based on the given sequence in the peptides file and is done with different software so inconsistencies are likely. The peaks are annotated based on the given sequence, with 20 ppm tolerance.

Copy Data

##### Spectrum 6346 (TSV)

###### Preview

```
Loading example...
```

*Click on the button to copy the data to your clipboard.*

Mz MinMz MaxIntensity Max

WidthHeightPeptide font sizePeptide stroke widthSpectrum font sizeSpectrum stroke widthCompact peptide

Ion legend

wxyz

abcd

OtherUnassignedIon^Charge^_Position_Show for top:%

JFPPSSEEJQANKA

05.22e+41.04e+51.57e+52.09e+5

Zoom Out

y^+1^_2_y^+2^_4_y^+1^_2_b^+2^_4_a^+1^_2_y^+2^_5_b^+1^_2_y^+2^_5_b^+2^_6_y^+2^_6_b^+2^_6_y^+2^_6_y^+1^_3_b^+1^_3_b^+2^_7_b^+3^_10_y^+2^_7_b^+2^_7_y^+1^_4_y^+2^_7_b^+3^_11_b^+3^_11_y^+1^_4_y^+3^_12_y^+3^_12_b^+2^_8_y^+2^_8_b^+2^_8_b^+1^_4_b^+2^_9_y^+1^_5_b^+1^_5_y^+1^_5_b^+1^_5_b^+2^_10_y^+2^_11_y^+2^_11_y^+2^_11_b^+2^_11_b^+2^_11_b^+1^_6_y^+2^_12_y^+2^_12_b^+1^_6_y^+2^_12_y^+1^_6_y^+2^_13_b^+2^_13_b^+1^_7_y^+1^_7_y^+1^_7_\*b^+1^_7_y^+1^_7_b^+1^_8_y^+1^_8_y^+1^_8_b^+1^_8_y^+1^_8_y^+1^_9_y^+1^_9_b^+1^_9_y^+1^_9_b^+1^_9_y^+1^_10_y^+1^_10_y^+1^_10_b^+1^_10_y^+1^_11_y^+1^_11_b^+1^_11_b^+1^_11_

0855171025643419

Fragment Matches Table

Show background peaks

| Position | Ion type | Intensity | mz Theoretical | mz Error (Th) | mz Error (ppm) | Charge | Series Number |
| --- | --- | --- | --- | --- | --- | --- | --- |
| - | - | 1.454E+05 | 120.1 | - | - | 0 | - |
| - | - | 605.7 | 121.1 | - | - | 0 | - |
| - | - | 1.291E+04 | 121.1 | - | - | 0 | - |
| - | - | 598.8 | 125.1 | - | - | 0 | - |
| - | - | 760.7 | 126.1 | - | - | 0 | - |
| - | - | 8389 | 126.1 | - | - | 0 | - |
| - | - | 620.1 | 127.1 | - | - | 0 | - |
| - | - | 457 | 127.1 | - | - | 0 | - |
| - | - | 1430 | 127.1 | - | - | 0 | - |
| - | - | 414.5 | 128.1 | - | - | 0 | - |
| - | - | 7360 | 129.1 | - | - | 0 | - |
| - | - | 1.238E+05 | 129.1 | - | - | 0 | - |
| - | - | 9201 | 130.1 | - | - | 0 | - |
| - | - | 4747 | 130.1 | - | - | 0 | - |
| - | - | 521.1 | 130.1 | - | - | 0 | - |
| - | - | 882.8 | 130.1 | - | - | 0 | - |
| - | - | 7801 | 130.1 | - | - | 0 | - |
| - | - | 381.2 | 130.3 | - | - | 0 | - |
| - | - | 7242 | 131.1 | - | - | 0 | - |
| - | - | 513.1 | 132.1 | - | - | 0 | - |
| - | - | 792.8 | 132.1 | - | - | 0 | - |
| - | - | 1837 | 132.1 | - | - | 0 | - |
| - | - | 595.4 | 132.1 | - | - | 0 | - |
| - | - | 1004 | 134 | - | - | 0 | - |
| - | - | 393.3 | 134.5 | - | - | 0 | - |
| - | - | 3686 | 136.1 | - | - | 0 | - |
| - | - | 4084 | 138.1 | - | - | 0 | - |
| - | - | 549.3 | 139.1 | - | - | 0 | - |
| - | - | 3928 | 139.1 | - | - | 0 | - |
| - | - | 419.6 | 139.1 | - | - | 0 | - |
| - | - | 565.4 | 140.1 | - | - | 0 | - |
| - | - | 353.2 | 141.1 | - | - | 0 | - |
| - | - | 3826 | 141.1 | - | - | 0 | - |
| - | - | 940.6 | 141.1 | - | - | 0 | - |
| - | - | 443.8 | 145.1 | - | - | 0 | - |
| - | - | 1309 | 146.1 | - | - | 0 | - |
| - | - | 3128 | 147.1 | - | - | 0 | - |
| - | - | 3212 | 147.1 | - | - | 0 | - |
| - | - | 555.4 | 148.1 | - | - | 0 | - |
| - | - | 505.8 | 150.1 | - | - | 0 | - |
| - | - | 591.7 | 152.1 | - | - | 0 | - |
| - | - | 869 | 152.1 | - | - | 0 | - |
| - | - | 1270 | 153.1 | - | - | 0 | - |
| - | - | 921.2 | 153.1 | - | - | 0 | - |
| - | - | 4081 | 154.1 | - | - | 0 | - |
| - | - | 6110 | 154.1 | - | - | 0 | - |
| - | - | 9243 | 155.1 | - | - | 0 | - |
| - | - | 562.6 | 155.1 | - | - | 0 | - |
| - | - | 554.7 | 155.1 | - | - | 0 | - |
| - | - | 1643 | 155.1 | - | - | 0 | - |
| - | - | 501.8 | 156.1 | - | - | 0 | - |
| - | - | 824.7 | 156.1 | - | - | 0 | - |
| - | - | 2889 | 157.1 | - | - | 0 | - |
| - | - | 1.12E+04 | 157.1 | - | - | 0 | - |
| - | - | 552.7 | 157.1 | - | - | 0 | - |
| - | - | 881.8 | 157.1 | - | - | 0 | - |
| - | - | 1269 | 158.1 | - | - | 0 | - |
| - | - | 758.5 | 158.1 | - | - | 0 | - |
| - | - | 821.9 | 159.1 | - | - | 0 | - |
| - | - | 1739 | 163.1 | - | - | 0 | - |
| - | - | 1232 | 163.1 | - | - | 0 | - |
| - | - | 2009 | 165.1 | - | - | 0 | - |
| - | - | 1683 | 166.1 | - | - | 0 | - |
| - | - | 556.3 | 166.1 | - | - | 0 | - |
| - | - | 7116 | 167.1 | - | - | 0 | - |
| - | - | 6379 | 167.1 | - | - | 0 | - |
| - | - | 534.7 | 168.1 | - | - | 0 | - |
| - | - | 610.7 | 168.1 | - | - | 0 | - |
| - | - | 843.7 | 168.1 | - | - | 0 | - |
| - | - | 2481 | 169.1 | - | - | 0 | - |
| - | - | 1293 | 169.1 | - | - | 0 | - |
| - | - | 579.5 | 170 | - | - | 0 | - |
| - | - | 2570 | 171.1 | - | - | 0 | - |
| - | - | 1346 | 172.1 | - | - | 0 | - |
| - | - | 3054 | 172.1 | - | - | 0 | - |
| - | - | 744 | 173.1 | - | - | 0 | - |
| - | - | 1835 | 173.1 | - | - | 0 | - |
| - | - | 882.8 | 174.1 | - | - | 0 | - |
| - | - | 507.7 | 174.1 | - | - | 0 | - |
| - | - | 3138 | 175.1 | - | - | 0 | - |
| - | - | 543.4 | 175.1 | - | - | 0 | - |
| - | - | 467.6 | 176.6 | - | - | 0 | - |
| - | - | 1482 | 180.1 | - | - | 0 | - |
| - | - | 1715 | 181.1 | - | - | 0 | - |
| - | - | 9238 | 182.1 | - | - | 0 | - |
| - | - | 577.6 | 182.1 | - | - | 0 | - |
| - | - | 6786 | 183.1 | - | - | 0 | - |
| - | - | 663.7 | 183.1 | - | - | 0 | - |
| - | - | 1235 | 183.1 | - | - | 0 | - |
| - | - | 962 | 183.1 | - | - | 0 | - |
| - | - | 848.1 | 184.1 | - | - | 0 | - |
| - | - | 7325 | 184.1 | - | - | 0 | - |
| - | - | 1.383E+04 | 185.1 | - | - | 0 | - |
| - | - | 6755 | 186.1 | - | - | 0 | - |
| - | - | 1280 | 186.1 | - | - | 0 | - |
| - | - | 1156 | 186.1 | - | - | 0 | - |
| - | - | 732.3 | 187.1 | - | - | 0 | - |
| - | - | 1351 | 187.1 | - | - | 0 | - |
| - | - | 985.5 | 187.1 | - | - | 0 | - |
| - | - | 5793 | 189.1 | - | - | 0 | - |
| - | - | 1631 | 191.1 | - | - | 0 | - |
| - | - | 591.8 | 191.1 | - | - | 0 | - |
| - | - | 630.2 | 193.1 | - | - | 0 | - |
| - | - | 1713 | 195.1 | - | - | 0 | - |
| - | - | 1.446E+04 | 195.1 | - | - | 0 | - |
| - | - | 1500 | 196.1 | - | - | 0 | - |
| - | - | 1.456E+04 | 197.1 | - | - | 0 | - |
| - | - | 826.9 | 198.1 | - | - | 0 | - |
| - | - | 6868 | 198.1 | - | - | 0 | - |
| - | - | 8903 | 199.1 | - | - | 0 | - |
| - | - | 4439 | 199.1 | - | - | 0 | - |
| - | - | 1469 | 199.1 | - | - | 0 | - |
| - | - | 625.7 | 199.1 | - | - | 0 | - |
| - | - | 1269 | 200.1 | - | - | 0 | - |
| - | - | 9035 | 200.1 | - | - | 0 | - |
| - | - | 1975 | 200.1 | - | - | 0 | - |
| - | - | 523.1 | 200.5 | - | - | 0 | - |
| - | - | 1664 | 201.1 | - | - | 0 | - |
| 13 | y | 1.393E+04 | 201.1 | 0.0002579 | 1.282 | +1 | 2 |
| 11 | y | 2051 | 202.1 | 0.0002024 | 1.001 | +2 | 4 |
| - | - | 1469 | 203.1 | - | - | 0 | - |
| - | - | 1229 | 207.1 | - | - | 0 | - |
| - | - | 1362 | 208.1 | - | - | 0 | - |
| - | - | 1.013E+04 | 208.1 | - | - | 0 | - |
| - | - | 2.737E+04 | 209.1 | - | - | 0 | - |
| - | - | 1185 | 209.1 | - | - | 0 | - |
| - | - | 2480 | 210.1 | - | - | 0 | - |
| - | - | 537.8 | 210.8 | - | - | 0 | - |
| - | - | 1692 | 211.1 | - | - | 0 | - |
| - | - | 3205 | 212.1 | - | - | 0 | - |
| - | - | 1909 | 213.1 | - | - | 0 | - |
| - | - | 1515 | 213.1 | - | - | 0 | - |
| - | - | 665.6 | 214.1 | - | - | 0 | - |
| - | - | 869.7 | 214.1 | - | - | 0 | - |
| - | - | 7043 | 215.1 | - | - | 0 | - |
| - | - | 613.8 | 216.1 | - | - | 0 | - |
| - | - | 944.3 | 216.1 | - | - | 0 | - |
| - | - | 1.367E+04 | 217.1 | - | - | 0 | - |
| - | - | 4543 | 217.1 | - | - | 0 | - |
| - | - | 1418 | 218.1 | - | - | 0 | - |
| 13 | y | 5.434E+04 | 218.1 | 0.0003506 | 1.607 | +1 | 2 |
| - | - | 4125 | 219.2 | - | - | 0 | - |
| - | - | 1543 | 221.1 | - | - | 0 | - |
| - | - | 856 | 221.1 | - | - | 0 | - |
| - | - | 759.6 | 221.6 | - | - | 0 | - |
| - | - | 663.3 | 223.1 | - | - | 0 | - |
| - | - | 611.3 | 223.2 | - | - | 0 | - |
| - | - | 1129 | 224.1 | - | - | 0 | - |
| - | - | 8372 | 225.1 | - | - | 0 | - |
| - | - | 4997 | 225.1 | - | - | 0 | - |
| - | - | 1.032E+05 | 226.1 | - | - | 0 | - |
| - | - | 3689 | 227.1 | - | - | 0 | - |
| - | - | 1.177E+04 | 227.1 | - | - | 0 | - |
| - | - | 577.1 | 227.1 | - | - | 0 | - |
| - | - | 1009 | 228.1 | - | - | 0 | - |
| 4 | b | 7506 | 228.1 | 0.001697 | 7.437 | +2 | 4 |
| - | - | 928.7 | 229.1 | - | - | 0 | - |
| - | - | 820 | 229.2 | - | - | 0 | - |
| - | - | 1926 | 231.1 | - | - | 0 | - |
| 2 | a | 2.067E+05 | 233.2 | 0.0004587 | 1.967 | +1 | 2 |
| - | - | 1025 | 234.1 | - | - | 0 | - |
| - | - | 1776 | 234.1 | - | - | 0 | - |
| - | - | 3.142E+04 | 234.2 | - | - | 0 | - |
| - | - | 806.8 | 235.1 | - | - | 0 | - |
| - | - | 2375 | 235.2 | - | - | 0 | - |
| - | - | 6201 | 236.1 | - | - | 0 | - |
| - | - | 680.5 | 236.1 | - | - | 0 | - |
| - | - | 871.7 | 237.1 | - | - | 0 | - |
| - | - | 592.9 | 238.1 | - | - | 0 | - |
| - | - | 660.1 | 238.1 | - | - | 0 | - |
| - | - | 811.3 | 239.2 | - | - | 0 | - |
| - | - | 865.4 | 240.1 | - | - | 0 | - |
| - | - | 673.5 | 240.1 | - | - | 0 | - |
| - | - | 1023 | 240.6 | - | - | 0 | - |
| - | - | 8470 | 241.1 | - | - | 0 | - |
| - | - | 764.1 | 242.1 | - | - | 0 | - |
| - | - | 2.006E+04 | 242.2 | - | - | 0 | - |
| - | - | 2199 | 243.1 | - | - | 0 | - |
| - | - | 2.83E+04 | 243.1 | - | - | 0 | - |
| - | - | 5397 | 244.1 | - | - | 0 | - |
| - | - | 3353 | 244.1 | - | - | 0 | - |
| - | - | 3531 | 245.1 | - | - | 0 | - |
| - | - | 933.2 | 247.1 | - | - | 0 | - |
| - | - | 1565 | 248.6 | - | - | 0 | - |
| - | - | 1256 | 248.7 | - | - | 0 | - |
| - | - | 6467 | 249.1 | - | - | 0 | - |
| - | - | 925.3 | 249.6 | - | - | 0 | - |
| - | - | 1125 | 250.2 | - | - | 0 | - |
| - | - | 3705 | 251.2 | - | - | 0 | - |
| - | - | 1585 | 252.1 | - | - | 0 | - |
| - | - | 725.7 | 252.1 | - | - | 0 | - |
| - | - | 2678 | 252.1 | - | - | 0 | - |
| - | - | 846.7 | 253.1 | - | - | 0 | - |
| - | - | 1.733E+04 | 254.1 | - | - | 0 | - |
| - | - | 2129 | 254.1 | - | - | 0 | - |
| - | - | 2238 | 255.1 | - | - | 0 | - |
| - | - | 2.043E+04 | 257.1 | - | - | 0 | - |
| 10 | y | 2435 | 257.6 | 0.0008124 | 3.153 | +2 | 5 |
| - | - | 3185 | 257.6 | - | - | 0 | - |
| - | - | 730.1 | 258.1 | - | - | 0 | - |
| - | - | 1.027E+04 | 259.1 | - | - | 0 | - |
| - | - | 845.4 | 260.1 | - | - | 0 | - |
| 2 | b | 8.289E+04 | 261.2 | 0.0002188 | 0.8378 | +1 | 2 |
| - | - | 1504 | 262.1 | - | - | 0 | - |
| - | - | 1.293E+04 | 262.2 | - | - | 0 | - |
| - | - | 1550 | 263.2 | - | - | 0 | - |
| - | - | 1.2E+04 | 264.1 | - | - | 0 | - |
| - | - | 1654 | 265.1 | - | - | 0 | - |
| 10 | y | 1.344E+04 | 266.1 | 0.0002258 | 0.8482 | +2 | 5 |
| - | - | 3377 | 266.6 | - | - | 0 | - |
| - | - | 703 | 267.2 | - | - | 0 | - |
| - | - | 1861 | 268.1 | - | - | 0 | - |
| - | - | 894.1 | 268.1 | - | - | 0 | - |
| - | - | 739.2 | 269.1 | - | - | 0 | - |
| - | - | 1398 | 269.2 | - | - | 0 | - |
| - | - | 1364 | 269.7 | - | - | 0 | - |
| - | - | 714.9 | 270.1 | - | - | 0 | - |
| - | - | 870.5 | 272.1 | - | - | 0 | - |
| - | - | 7073 | 272.1 | - | - | 0 | - |
| - | - | 1686 | 273.1 | - | - | 0 | - |
| - | - | 786.3 | 278.2 | - | - | 0 | - |
| - | - | 606 | 278.2 | - | - | 0 | - |
| - | - | 6572 | 279.1 | - | - | 0 | - |
| - | - | 9072 | 280.1 | - | - | 0 | - |
| - | - | 1025 | 280.1 | - | - | 0 | - |
| - | - | 840.4 | 281.1 | - | - | 0 | - |
| - | - | 1060 | 281.1 | - | - | 0 | - |
| - | - | 1366 | 282.1 | - | - | 0 | - |
| - | - | 2171 | 282.1 | - | - | 0 | - |
| - | - | 3.47E+04 | 282.1 | - | - | 0 | - |
| - | - | 751.9 | 282.6 | - | - | 0 | - |
| - | - | 4811 | 283.1 | - | - | 0 | - |
| - | - | 740.6 | 285.2 | - | - | 0 | - |
| - | - | 5672 | 286.1 | - | - | 0 | - |
| - | - | 558.8 | 291.7 | - | - | 0 | - |
| - | - | 551.9 | 292.1 | - | - | 0 | - |
| - | - | 588 | 292.2 | - | - | 0 | - |
| - | - | 711.7 | 292.7 | - | - | 0 | - |
| - | - | 1905 | 296.1 | - | - | 0 | - |
| - | - | 4905 | 296.1 | - | - | 0 | - |
| - | - | 5294 | 296.2 | - | - | 0 | - |
| - | - | 1081 | 296.6 | - | - | 0 | - |
| - | - | 6349 | 297.1 | - | - | 0 | - |
| - | - | 3.346E+04 | 297.2 | - | - | 0 | - |
| - | - | 1030 | 298.1 | - | - | 0 | - |
| - | - | 4817 | 298.1 | - | - | 0 | - |
| - | - | 4553 | 298.2 | - | - | 0 | - |
| - | - | 1069 | 299.1 | - | - | 0 | - |
| - | - | 2316 | 299.2 | - | - | 0 | - |
| - | - | 968.1 | 300.1 | - | - | 0 | - |
| - | - | 627.9 | 300.1 | - | - | 0 | - |
| - | - | 2227 | 300.2 | - | - | 0 | - |
| - | - | 3658 | 304.1 | - | - | 0 | - |
| - | - | 1612 | 305.1 | - | - | 0 | - |
| - | - | 829.7 | 305.7 | - | - | 0 | - |
| - | - | 1256 | 306.1 | - | - | 0 | - |
| 6 | b | 1165 | 306.2 | 0.0003353 | 1.095 | +2 | 6 |
| - | - | 2766 | 308.2 | - | - | 0 | - |
| - | - | 794.5 | 308.2 | - | - | 0 | - |
| - | - | 1103 | 310.1 | - | - | 0 | - |
| - | - | 3561 | 313.2 | - | - | 0 | - |
| - | - | 4401 | 314.1 | - | - | 0 | - |
| - | - | 3802 | 314.1 | - | - | 0 | - |
| 9 | y | 1.474E+04 | 314.2 | 0.004275 | 13.61 | +2 | 6 |
| - | - | 1477 | 314.6 | - | - | 0 | - |
| - | - | 2437 | 314.7 | - | - | 0 | - |
| - | - | 758 | 315.1 | - | - | 0 | - |
| - | - | 620.5 | 315.1 | - | - | 0 | - |
| 6 | b | 1.483E+04 | 315.2 | 0.001529 | 4.852 | +2 | 6 |
| - | - | 1555 | 315.2 | - | - | 0 | - |
| - | - | 2328 | 316.2 | - | - | 0 | - |
| - | - | 553.8 | 319.1 | - | - | 0 | - |
| - | - | 1162 | 321.2 | - | - | 0 | - |
| - | - | 893.6 | 322.2 | - | - | 0 | - |
| 9 | y | 1.804E+04 | 322.7 | 0.0003691 | 1.144 | +2 | 6 |
| - | - | 1564 | 323.2 | - | - | 0 | - |
| - | - | 5567 | 323.2 | - | - | 0 | - |
| - | - | 572.2 | 323.7 | - | - | 0 | - |
| - | - | 2002 | 324.2 | - | - | 0 | - |
| - | - | 1384 | 325.2 | - | - | 0 | - |
| - | - | 799.9 | 325.2 | - | - | 0 | - |
| - | - | 1445 | 326.2 | - | - | 0 | - |
| - | - | 5640 | 328.1 | - | - | 0 | - |
| - | - | 641.9 | 328.7 | - | - | 0 | - |
| - | - | 709.4 | 329.1 | - | - | 0 | - |
| - | - | 1915 | 329.7 | - | - | 0 | - |
| - | - | 794.5 | 330.2 | - | - | 0 | - |
| 12 | y | 4.512E+04 | 332.2 | 0.0004529 | 1.364 | +1 | 3 |
| - | - | 6974 | 333.2 | - | - | 0 | - |
| - | - | 760.6 | 333.2 | - | - | 0 | - |
| - | - | 6259 | 333.2 | - | - | 0 | - |
| - | - | 1354 | 334.2 | - | - | 0 | - |
| - | - | 686.7 | 334.2 | - | - | 0 | - |
| - | - | 1424 | 336.2 | - | - | 0 | - |
| - | - | 1920 | 337.2 | - | - | 0 | - |
| - | - | 952.1 | 338.1 | - | - | 0 | - |
| - | - | 5081 | 338.7 | - | - | 0 | - |
| - | - | 4151 | 339.2 | - | - | 0 | - |
| - | - | 6288 | 341.2 | - | - | 0 | - |
| - | - | 2435 | 342.2 | - | - | 0 | - |
| - | - | 3689 | 343.2 | - | - | 0 | - |
| - | - | 2308 | 343.2 | - | - | 0 | - |
| - | - | 9463 | 346.1 | - | - | 0 | - |
| - | - | 1110 | 347.1 | - | - | 0 | - |
| - | - | 3271 | 347.7 | - | - | 0 | - |
| - | - | 1066 | 348.2 | - | - | 0 | - |
| - | - | 945.2 | 350.2 | - | - | 0 | - |
| - | - | 2.804E+04 | 351.2 | - | - | 0 | - |
| - | - | 5607 | 352.2 | - | - | 0 | - |
| - | - | 9079 | 353.2 | - | - | 0 | - |
| - | - | 2153 | 354.2 | - | - | 0 | - |
| - | - | 1035 | 354.2 | - | - | 0 | - |
| - | - | 3729 | 355.2 | - | - | 0 | - |
| - | - | 1169 | 356.1 | - | - | 0 | - |
| 3 | b | 4775 | 358.2 | 0.0002198 | 0.6137 | +1 | 3 |
| - | - | 1186 | 359.2 | - | - | 0 | - |
| - | - | 1046 | 360.2 | - | - | 0 | - |
| - | - | 609.8 | 361.2 | - | - | 0 | - |
| - | - | 2112 | 361.7 | - | - | 0 | - |
| - | - | 933.8 | 362.2 | - | - | 0 | - |
| - | - | 840.7 | 363.2 | - | - | 0 | - |
| - | - | 4814 | 365.1 | - | - | 0 | - |
| - | - | 728.5 | 365.2 | - | - | 0 | - |
| - | - | 687.6 | 366.1 | - | - | 0 | - |
| - | - | 1267 | 367.2 | - | - | 0 | - |
| - | - | 1.07E+04 | 368.2 | - | - | 0 | - |
| - | - | 3.147E+04 | 369.2 | - | - | 0 | - |
| - | - | 882.5 | 369.7 | - | - | 0 | - |
| - | - | 6747 | 370.2 | - | - | 0 | - |
| 7 | b | 1.388E+04 | 370.7 | 0.006282 | 16.95 | +2 | 7 |
| 10 | b | 3432 | 371.2 | 0.0007411 | 1.997 | +3 | 10 |
| - | - | 3301 | 371.2 | - | - | 0 | - |
| - | - | 1047 | 371.7 | - | - | 0 | - |
| - | - | 1655 | 372.2 | - | - | 0 | - |
| - | - | 5363 | 373.2 | - | - | 0 | - |
| - | - | 862.7 | 374.2 | - | - | 0 | - |
| - | - | 938.6 | 377.2 | - | - | 0 | - |
| 8 | y | 784.7 | 378.7 | 0.001563 | 4.128 | +2 | 7 |
| - | - | 1154 | 379.2 | - | - | 0 | - |
| 7 | b | 1574 | 379.7 | 0.0002151 | 0.5666 | +2 | 7 |
| - | - | 977.1 | 380.2 | - | - | 0 | - |
| - | - | 863.8 | 381.2 | - | - | 0 | - |
| - | - | 7555 | 383.2 | - | - | 0 | - |
| - | - | 1806 | 384.2 | - | - | 0 | - |
| - | - | 4430 | 385.2 | - | - | 0 | - |
| 11 | y | 2.618E+04 | 386.2 | 0.0003558 | 0.9213 | +1 | 4 |
| 8 | y | 4921 | 387.2 | 0.003502 | 9.044 | +2 | 7 |
| - | - | 706.3 | 388.5 | - | - | 0 | - |
| - | - | 647 | 392.2 | - | - | 0 | - |
| - | - | 1284 | 393.2 | - | - | 0 | - |
| - | - | 765.4 | 394.2 | - | - | 0 | - |
| 11 | b | 700 | 394.5 | 0.0008742 | 2.216 | +3 | 11 |
| 11 | b | 894.6 | 394.9 | 0.00487 | 12.33 | +3 | 11 |
| - | - | 582.2 | 395.2 | - | - | 0 | - |
| - | - | 875.7 | 395.2 | - | - | 0 | - |
| - | - | 642.2 | 396.2 | - | - | 0 | - |
| - | - | 581.8 | 396.7 | - | - | 0 | - |
| - | - | 1194 | 397.1 | - | - | 0 | - |
| - | - | 1.399E+04 | 401.2 | - | - | 0 | - |
| - | - | 2771 | 402.2 | - | - | 0 | - |
| 11 | y | 1.124E+05 | 403.2 | 0.000418 | 1.037 | +1 | 4 |
| - | - | 2.035E+04 | 404.2 | - | - | 0 | - |
| - | - | 916.7 | 405.2 | - | - | 0 | - |
| - | - | 3779 | 405.2 | - | - | 0 | - |
| - | - | 7022 | 407.2 | - | - | 0 | - |
| - | - | 1.571E+04 | 408.2 | - | - | 0 | - |
| - | - | 2915 | 409.2 | - | - | 0 | - |
| - | - | 1753 | 410.2 | - | - | 0 | - |
| - | - | 676.7 | 411.2 | - | - | 0 | - |
| - | - | 595.8 | 411.2 | - | - | 0 | - |
| - | - | 520.4 | 413.2 | - | - | 0 | - |
| - | - | 1395 | 414.2 | - | - | 0 | - |
| - | - | 2167 | 415.1 | - | - | 0 | - |
| 3 | y | 1190 | 418.2 | 0.001583 | 3.785 | +3 | 12 |
| 3 | y | 1526 | 418.5 | 0.002435 | 5.818 | +3 | 12 |
| - | - | 1052 | 418.9 | - | - | 0 | - |
| - | - | 617 | 423.2 | - | - | 0 | - |
| - | - | 8000 | 424.2 | - | - | 0 | - |
| - | - | 924.7 | 424.5 | - | - | 0 | - |
| - | - | 1.038E+04 | 425.2 | - | - | 0 | - |
| - | - | 995.8 | 425.7 | - | - | 0 | - |
| - | - | 2431 | 426.2 | - | - | 0 | - |
| - | - | 811.5 | 426.2 | - | - | 0 | - |
| - | - | 3572 | 427.2 | - | - | 0 | - |
| - | - | 1483 | 427.3 | - | - | 0 | - |
| - | - | 640.6 | 428.2 | - | - | 0 | - |
| - | - | 977.5 | 429.2 | - | - | 0 | - |
| - | - | 779.9 | 430.2 | - | - | 0 | - |
| - | - | 4527 | 433.2 | - | - | 0 | - |
| - | - | 741 | 434.2 | - | - | 0 | - |
| - | - | 3077 | 434.2 | - | - | 0 | - |
| - | - | 5763 | 434.7 | - | - | 0 | - |
| 8 | b | 4686 | 435.2 | 0.002176 | 4.999 | +2 | 8 |
| - | - | 926.1 | 435.7 | - | - | 0 | - |
| - | - | 684.7 | 440.2 | - | - | 0 | - |
| - | - | 1250 | 441.2 | - | - | 0 | - |
| - | - | 6307 | 442.2 | - | - | 0 | - |
| - | - | 1340 | 443.1 | - | - | 0 | - |
| - | - | 682.3 | 443.2 | - | - | 0 | - |
| 7 | y | 3773 | 443.2 | 0.006054 | 13.66 | +2 | 8 |
| 8 | b | 1889 | 444.2 | 0.001135 | 2.556 | +2 | 8 |
| - | - | 1391 | 444.7 | - | - | 0 | - |
| - | - | 889.8 | 445.2 | - | - | 0 | - |
| - | - | 786.5 | 448.2 | - | - | 0 | - |
| - | - | 2174 | 450.2 | - | - | 0 | - |
| - | - | 5535 | 452.2 | - | - | 0 | - |
| - | - | 1819 | 453.2 | - | - | 0 | - |
| - | - | 843.9 | 454.3 | - | - | 0 | - |
| 4 | b | 5792 | 455.3 | 0.001106 | 2.429 | +1 | 4 |
| - | - | 1393 | 456.3 | - | - | 0 | - |
| - | - | 2061 | 459.2 | - | - | 0 | - |
| - | - | 809.2 | 460.3 | - | - | 0 | - |
| - | - | 1759 | 461.2 | - | - | 0 | - |
| - | - | 764.3 | 461.7 | - | - | 0 | - |
| - | - | 6490 | 462.2 | - | - | 0 | - |
| - | - | 1927 | 463.2 | - | - | 0 | - |
| - | - | 996.2 | 464.2 | - | - | 0 | - |
| - | - | 1128 | 466.2 | - | - | 0 | - |
| - | - | 3408 | 468.2 | - | - | 0 | - |
| - | - | 2.327E+04 | 470.2 | - | - | 0 | - |
| - | - | 2055 | 470.7 | - | - | 0 | - |
| - | - | 5291 | 471.2 | - | - | 0 | - |
| - | - | 1208 | 476.2 | - | - | 0 | - |
| - | - | 2515 | 478.2 | - | - | 0 | - |
| - | - | 3172 | 479.2 | - | - | 0 | - |
| - | - | 2.018E+04 | 480.2 | - | - | 0 | - |
| - | - | 5548 | 481.2 | - | - | 0 | - |
| - | - | 2066 | 482.2 | - | - | 0 | - |
| - | - | 872.6 | 482.3 | - | - | 0 | - |
| - | - | 3093 | 484.2 | - | - | 0 | - |
| - | - | 904.5 | 487.3 | - | - | 0 | - |
| - | - | 1146 | 488.3 | - | - | 0 | - |
| - | - | 2265 | 494.2 | - | - | 0 | - |
| - | - | 1577 | 494.2 | - | - | 0 | - |
| - | - | 8507 | 496.3 | - | - | 0 | - |
| - | - | 1.646E+04 | 497.2 | - | - | 0 | - |
| - | - | 7.458E+04 | 498.2 | - | - | 0 | - |
| - | - | 1.882E+04 | 499.2 | - | - | 0 | - |
| - | - | 3847 | 500.2 | - | - | 0 | - |
| 9 | b | 1700 | 500.8 | 5.797E-05 | 0.1158 | +2 | 9 |
| - | - | 1022 | 501.3 | - | - | 0 | - |
| - | - | 941.9 | 501.8 | - | - | 0 | - |
| - | - | 4439 | 502.2 | - | - | 0 | - |
| - | - | 1402 | 503.2 | - | - | 0 | - |
| - | - | 1237 | 503.3 | - | - | 0 | - |
| - | - | 824.8 | 511.3 | - | - | 0 | - |
| - | - | 5339 | 512.2 | - | - | 0 | - |
| - | - | 1315 | 512.2 | - | - | 0 | - |
| - | - | 2198 | 513.2 | - | - | 0 | - |
| - | - | 2.885E+04 | 513.3 | - | - | 0 | - |
| 10 | y | 7.331E+04 | 514.3 | 0.0008908 | 1.732 | +1 | 5 |
| - | - | 2.025E+04 | 515.3 | - | - | 0 | - |
| - | - | 2700 | 516.3 | - | - | 0 | - |
| - | - | 854.1 | 521.3 | - | - | 0 | - |
| 5 | b | 1967 | 524.3 | 0.0007908 | 1.508 | +1 | 5 |
| - | - | 1868 | 528.2 | - | - | 0 | - |
| - | - | 1.856E+04 | 530.2 | - | - | 0 | - |
| - | - | 3797 | 531.2 | - | - | 0 | - |
| 10 | y | 9.127E+04 | 531.3 | 0.0001596 | 0.3004 | +1 | 5 |
| - | - | 897.5 | 532.2 | - | - | 0 | - |
| - | - | 2.428E+04 | 532.3 | - | - | 0 | - |
| - | - | 4170 | 533.3 | - | - | 0 | - |
| - | - | 1717 | 534.3 | - | - | 0 | - |
| - | - | 1037 | 535.2 | - | - | 0 | - |
| - | - | 3366 | 538.3 | - | - | 0 | - |
| - | - | 1102 | 539.3 | - | - | 0 | - |
| - | - | 645.2 | 539.8 | - | - | 0 | - |
| - | - | 821.6 | 539.8 | - | - | 0 | - |
| - | - | 1919 | 540.3 | - | - | 0 | - |
| - | - | 1036 | 542.3 | - | - | 0 | - |
| 5 | b | 3705 | 542.3 | 0.0007242 | 1.335 | +1 | 5 |
| - | - | 1497 | 543.3 | - | - | 0 | - |
| - | - | 708.2 | 543.8 | - | - | 0 | - |
| - | - | 976 | 545.2 | - | - | 0 | - |
| - | - | 1058 | 546.2 | - | - | 0 | - |
| - | - | 759.3 | 549 | - | - | 0 | - |
| - | - | 1112 | 553.3 | - | - | 0 | - |
| - | - | 2186 | 555.3 | - | - | 0 | - |
| 10 | b | 1166 | 556.3 | 0.001378 | 2.477 | +2 | 10 |
| - | - | 1243 | 556.3 | - | - | 0 | - |
| - | - | 1513 | 563.2 | - | - | 0 | - |
| - | - | 1134 | 565.3 | - | - | 0 | - |
| - | - | 1843 | 569.8 | - | - | 0 | - |
| - | - | 1248 | 570.3 | - | - | 0 | - |
| - | - | 930.1 | 571.3 | - | - | 0 | - |
| - | - | 1740 | 573.2 | - | - | 0 | - |
| - | - | 1402 | 573.8 | - | - | 0 | - |
| - | - | 870.3 | 574.3 | - | - | 0 | - |
| 4 | y | 2051 | 578.3 | 0.0006865 | 1.187 | +2 | 11 |
| 4 | y | 5301 | 578.8 | 0.002423 | 4.186 | +2 | 11 |
| - | - | 2825 | 579.3 | - | - | 0 | - |
| - | - | 1105 | 579.8 | - | - | 0 | - |
| - | - | 846.2 | 580.3 | - | - | 0 | - |
| - | - | 5902 | 581.3 | - | - | 0 | - |
| - | - | 1689 | 582.3 | - | - | 0 | - |
| - | - | 6329 | 582.8 | - | - | 0 | - |
| - | - | 3103 | 583.3 | - | - | 0 | - |
| - | - | 1202 | 583.8 | - | - | 0 | - |
| - | - | 1090 | 584.3 | - | - | 0 | - |
| 4 | y | 1.608E+04 | 587.3 | 0.0004147 | 0.706 | +2 | 11 |
| - | - | 1.076E+04 | 587.8 | - | - | 0 | - |
| - | - | 4280 | 588.3 | - | - | 0 | - |
| - | - | 5756 | 591.2 | - | - | 0 | - |
| 11 | b | 1248 | 591.3 | 0.002527 | 4.274 | +2 | 11 |
| 11 | b | 2383 | 591.8 | 0.006747 | 11.4 | +2 | 11 |
| - | - | 2348 | 592.2 | - | - | 0 | - |
| - | - | 1621 | 592.3 | - | - | 0 | - |
| - | - | 1715 | 593.3 | - | - | 0 | - |
| - | - | 763.1 | 594.3 | - | - | 0 | - |
| - | - | 1576 | 597.3 | - | - | 0 | - |
| - | - | 3.03E+04 | 599.3 | - | - | 0 | - |
| - | - | 706.7 | 599.8 | - | - | 0 | - |
| - | - | 8026 | 600.3 | - | - | 0 | - |
| - | - | 2502 | 600.8 | - | - | 0 | - |
| - | - | 1106 | 601.3 | - | - | 0 | - |
| - | - | 2765 | 601.3 | - | - | 0 | - |
| - | - | 807.7 | 607.3 | - | - | 0 | - |
| - | - | 728 | 608.3 | - | - | 0 | - |
| - | - | 848.6 | 608.8 | - | - | 0 | - |
| - | - | 1.977E+04 | 609.3 | - | - | 0 | - |
| - | - | 5384 | 609.3 | - | - | 0 | - |
| - | - | 1456 | 609.8 | - | - | 0 | - |
| - | - | 5388 | 610.3 | - | - | 0 | - |
| - | - | 1066 | 610.3 | - | - | 0 | - |
| - | - | 1107 | 611.3 | - | - | 0 | - |
| 6 | b | 7266 | 611.3 | 0.003406 | 5.571 | +1 | 6 |
| - | - | 3290 | 612.3 | - | - | 0 | - |
| - | - | 827.1 | 613.3 | - | - | 0 | - |
| - | - | 2696 | 615.3 | - | - | 0 | - |
| - | - | 3099 | 617.8 | - | - | 0 | - |
| - | - | 8185 | 618.3 | - | - | 0 | - |
| - | - | 5375 | 618.8 | - | - | 0 | - |
| - | - | 2656 | 619.3 | - | - | 0 | - |
| - | - | 804.5 | 622.3 | - | - | 0 | - |
| - | - | 2510 | 625.3 | - | - | 0 | - |
| - | - | 951.3 | 626.3 | - | - | 0 | - |
| 3 | y | 1.4E+04 | 626.8 | 2.986E-05 | 0.04764 | +2 | 12 |
| - | - | 1.802E+05 | 627.3 | - | - | 0 | - |
| 3 | y | 1.93E+04 | 627.3 | 0.003812 | 6.077 | +2 | 12 |
| - | - | 2.293E+04 | 627.8 | - | - | 0 | - |
| - | - | 5.789E+04 | 628.3 | - | - | 0 | - |
| - | - | 2152 | 628.8 | - | - | 0 | - |
| - | - | 1.35E+04 | 629.3 | - | - | 0 | - |
| 6 | b | 9931 | 629.3 | 5.421E-05 | 0.08613 | +1 | 6 |
| - | - | 2053 | 630.3 | - | - | 0 | - |
| - | - | 3291 | 630.3 | - | - | 0 | - |
| - | - | 808.2 | 631.3 | - | - | 0 | - |
| - | - | 826.5 | 632.8 | - | - | 0 | - |
| - | - | 2365 | 633.4 | - | - | 0 | - |
| - | - | 1160 | 634.8 | - | - | 0 | - |
| - | - | 855.7 | 635.3 | - | - | 0 | - |
| 3 | y | 1.73E+05 | 635.8 | 5.889E-05 | 0.09262 | +2 | 12 |
| - | - | 1.277E+05 | 636.3 | - | - | 0 | - |
| - | - | 5.389E+04 | 636.8 | - | - | 0 | - |
| - | - | 1.647E+04 | 637.3 | - | - | 0 | - |
| - | - | 2508 | 637.8 | - | - | 0 | - |
| - | - | 5879 | 643.3 | - | - | 0 | - |
| - | - | 3190 | 644.3 | - | - | 0 | - |
| 9 | y | 3.095E+04 | 644.4 | 1.896E-05 | 0.02942 | +1 | 6 |
| - | - | 1632 | 645.3 | - | - | 0 | - |
| - | - | 1.214E+04 | 645.4 | - | - | 0 | - |
| - | - | 2587 | 646.4 | - | - | 0 | - |
| - | - | 764.7 | 649.3 | - | - | 0 | - |
| - | - | 1054 | 658.3 | - | - | 0 | - |
| - | - | 1757 | 660.3 | - | - | 0 | - |
| - | - | 822.1 | 661.3 | - | - | 0 | - |
| - | - | 890.9 | 662.3 | - | - | 0 | - |
| - | - | 816.3 | 664.8 | - | - | 0 | - |
| - | - | 1237 | 667.3 | - | - | 0 | - |
| - | - | 779.8 | 674.3 | - | - | 0 | - |
| - | - | 773.9 | 676.3 | - | - | 0 | - |
| - | - | 765 | 677.3 | - | - | 0 | - |
| - | - | 941 | 680.3 | - | - | 0 | - |
| - | - | 668.5 | 682.4 | - | - | 0 | - |
| - | - | 1762 | 682.6 | - | - | 0 | - |
| - | - | 779.6 | 682.9 | - | - | 0 | - |
| - | - | 1597 | 683 | - | - | 0 | - |
| - | - | 1427 | 683.4 | - | - | 0 | - |
| - | - | 682.2 | 684.3 | - | - | 0 | - |
| - | - | 1283 | 684.4 | - | - | 0 | - |
| - | - | 1137 | 686.3 | - | - | 0 | - |
| - | - | 6423 | 694.3 | - | - | 0 | - |
| - | - | 2152 | 695.3 | - | - | 0 | - |
| - | - | 774.4 | 696.4 | - | - | 0 | - |
| - | - | 4110 | 704.3 | - | - | 0 | - |
| - | - | 2012 | 705.3 | - | - | 0 | - |
| 2 | y | 1204 | 709.4 | 2.942E-05 | 0.04147 | +2 | 13 |
| - | - | 990.6 | 709.8 | - | - | 0 | - |
| 13 | b | 2.403E+04 | 712.4 | 0.01282 | 17.99 | +2 | 13 |
| - | - | 1.021E+04 | 713.4 | - | - | 0 | - |
| - | - | 3468 | 714.4 | - | - | 0 | - |
| - | - | 751.5 | 715.4 | - | - | 0 | - |
| - | - | 1356 | 721.4 | - | - | 0 | - |
| - | - | 1.239E+04 | 722.3 | - | - | 0 | - |
| - | - | 5732 | 723.3 | - | - | 0 | - |
| - | - | 1573 | 724.3 | - | - | 0 | - |
| - | - | 676.7 | 729.3 | - | - | 0 | - |
| - | - | 4511 | 730.4 | - | - | 0 | - |
| - | - | 2142 | 731.4 | - | - | 0 | - |
| - | - | 940.8 | 738.3 | - | - | 0 | - |
| - | - | 2408 | 738.4 | - | - | 0 | - |
| - | - | 4470 | 739.4 | - | - | 0 | - |
| 7 | b | 6.241E+04 | 740.4 | 0.01426 | 19.26 | +1 | 7 |
| - | - | 2.379E+04 | 741.4 | - | - | 0 | - |
| - | - | 6263 | 742.4 | - | - | 0 | - |
| - | - | 1058 | 743.4 | - | - | 0 | - |
| - | - | 707.3 | 746.3 | - | - | 0 | - |
| - | - | 1557 | 753.3 | - | - | 0 | - |
| - | - | 989.1 | 754.3 | - | - | 0 | - |
| 8 | y | 3282 | 755.4 | 0.0006444 | 0.853 | +1 | 7 |
| 8 | y | 4538 | 756.4 | 0.004849 | 6.411 | +1 | 7 |
| 0 | Precursor | 1.55E+04 | 757.4 | 0.006448 | 8.514 | +2 | -1 |
| 7 | b | 2.785E+04 | 758.4 | 0.0006266 | 0.8263 | +1 | 7 |
| - | - | 1.188E+04 | 759.4 | - | - | 0 | - |
| - | - | 3071 | 760.4 | - | - | 0 | - |
| - | - | 1830 | 770.4 | - | - | 0 | - |
| - | - | 4644 | 771.4 | - | - | 0 | - |
| - | - | 2200 | 772.4 | - | - | 0 | - |
| 8 | y | 1.496E+04 | 773.4 | 0.0001505 | 0.1946 | +1 | 7 |
| - | - | 858.1 | 774.3 | - | - | 0 | - |
| - | - | 3443 | 774.3 | - | - | 0 | - |
| - | - | 6803 | 774.4 | - | - | 0 | - |
| - | - | 1312 | 775.3 | - | - | 0 | - |
| - | - | 1467 | 775.4 | - | - | 0 | - |
| - | - | 776.9 | 791.4 | - | - | 0 | - |
| - | - | 774.8 | 807.4 | - | - | 0 | - |
| - | - | 923 | 810.4 | - | - | 0 | - |
| - | - | 845.6 | 815.4 | - | - | 0 | - |
| - | - | 1158 | 833.4 | - | - | 0 | - |
| - | - | 2412 | 840.4 | - | - | 0 | - |
| - | - | 2077 | 841.4 | - | - | 0 | - |
| - | - | 3028 | 842.4 | - | - | 0 | - |
| - | - | 1125 | 843.4 | - | - | 0 | - |
| - | - | 4258 | 850.4 | - | - | 0 | - |
| - | - | 3871 | 851.4 | - | - | 0 | - |
| - | - | 1741 | 852.4 | - | - | 0 | - |
| - | - | 2025 | 853.4 | - | - | 0 | - |
| - | - | 799.2 | 854.4 | - | - | 0 | - |
| - | - | 6472 | 859.4 | - | - | 0 | - |
| - | - | 4258 | 860.4 | - | - | 0 | - |
| - | - | 924.6 | 861.4 | - | - | 0 | - |
| - | - | 2.883E+04 | 868.4 | - | - | 0 | - |
| 8 | b | 1.961E+04 | 869.4 | 0.002107 | 2.423 | +1 | 8 |
| - | - | 7534 | 870.4 | - | - | 0 | - |
| - | - | 1202 | 871.4 | - | - | 0 | - |
| 7 | y | 823.9 | 884.4 | 0.01491 | 16.86 | +1 | 8 |
| 7 | y | 1530 | 885.4 | 0.0008911 | 1.006 | +1 | 8 |
| - | - | 1231 | 886.4 | - | - | 0 | - |
| 8 | b | 5.647E+04 | 887.4 | 0.0008288 | 0.9339 | +1 | 8 |
| - | - | 3.014E+04 | 888.4 | - | - | 0 | - |
| - | - | 9178 | 889.4 | - | - | 0 | - |
| - | - | 1244 | 890.4 | - | - | 0 | - |
| - | - | 691.4 | 893.4 | - | - | 0 | - |
| - | - | 1972 | 894.4 | - | - | 0 | - |
| 7 | y | 2822 | 902.5 | 0.003929 | 4.354 | +1 | 8 |
| - | - | 1650 | 903.5 | - | - | 0 | - |
| - | - | 2707 | 911.4 | - | - | 0 | - |
| - | - | 1681 | 912.4 | - | - | 0 | - |
| - | - | 830 | 914.4 | - | - | 0 | - |
| - | - | 3404 | 921.4 | - | - | 0 | - |
| - | - | 3951 | 922.4 | - | - | 0 | - |
| - | - | 2060 | 923.4 | - | - | 0 | - |
| - | - | 1.557E+04 | 939.4 | - | - | 0 | - |
| - | - | 8731 | 940.4 | - | - | 0 | - |
| - | - | 2390 | 941.4 | - | - | 0 | - |
| - | - | 2112 | 956.5 | - | - | 0 | - |
| - | - | 1936 | 957.5 | - | - | 0 | - |
| 6 | y | 968.8 | 971.5 | 0.002566 | 2.641 | +1 | 9 |
| 6 | y | 1780 | 972.5 | 0.01287 | 13.23 | +1 | 9 |
| - | - | 892.5 | 973.5 | - | - | 0 | - |
| 9 | b | 1212 | 982.5 | 0.0002314 | 0.2355 | +1 | 9 |
| - | - | 922 | 983.5 | - | - | 0 | - |
| 6 | y | 4475 | 989.5 | 0.001534 | 1.55 | +1 | 9 |
| - | - | 2382 | 990.5 | - | - | 0 | - |
| 9 | b | 9218 | 1000 | 0.002007 | 2.006 | +1 | 9 |
| - | - | 5093 | 1002 | - | - | 0 | - |
| - | - | 2029 | 1003 | - | - | 0 | - |
| - | - | 1264 | 1025 | - | - | 0 | - |
| - | - | 1258 | 1035 | - | - | 0 | - |
| - | - | 3678 | 1036 | - | - | 0 | - |
| - | - | 1672 | 1037 | - | - | 0 | - |
| - | - | 1338 | 1041 | - | - | 0 | - |
| - | - | 661.7 | 1042 | - | - | 0 | - |
| - | - | 1.216E+04 | 1053 | - | - | 0 | - |
| - | - | 7636 | 1054 | - | - | 0 | - |
| - | - | 2472 | 1055 | - | - | 0 | - |
| 5 | y | 818 | 1059 | 0.01216 | 11.49 | +1 | 10 |
| 5 | y | 2877 | 1059 | 0.002752 | 2.598 | +1 | 10 |
| - | - | 1244 | 1061 | - | - | 0 | - |
| - | - | 876.8 | 1062 | - | - | 0 | - |
| - | - | 706.3 | 1069 | - | - | 0 | - |
| - | - | 994.9 | 1070 | - | - | 0 | - |
| - | - | 1395 | 1072 | - | - | 0 | - |
| 5 | y | 7045 | 1077 | 0.0002372 | 0.2204 | +1 | 10 |
| - | - | 3933 | 1078 | - | - | 0 | - |
| - | - | 1092 | 1079 | - | - | 0 | - |
| - | - | 1067 | 1087 | - | - | 0 | - |
| - | - | 776 | 1087 | - | - | 0 | - |
| 10 | b | 2327 | 1129 | 0.0005259 | 0.466 | +1 | 10 |
| - | - | 1438 | 1139 | - | - | 0 | - |
| 4 | y | 2610 | 1157 | 0.001258 | 1.088 | +1 | 11 |
| - | - | 1483 | 1158 | - | - | 0 | - |
| 4 | y | 8220 | 1174 | 0.001976 | 1.683 | +1 | 11 |
| - | - | 5547 | 1175 | - | - | 0 | - |
| - | - | 2507 | 1176 | - | - | 0 | - |
| 11 | b | 890.6 | 1182 | 0.005469 | 4.628 | +1 | 11 |
| 11 | b | 854.2 | 1200 | 0.002155 | 1.797 | +1 | 11 |
| - | - | 696.2 | 1231 | - | - | 0 | - |
| - | - | 703.8 | 1407 | - | - | 0 | - |
| - | - | 777.7 | 1445 | - | - | 0 | - |
| - | - | 773.1 | 3385 | - | - | 0 | - |

m/z Charge Intensity FragmentType MassShift Position
120.0811538696289 0 145422.47
121.07905578613281 0 605.7041
121.08446502685547 0 12914.43
125.07119750976562 0 598.8418
126.05068969726562 0 760.6836
126.0552978515625 0 8389.017
127.05075073242188 0 620.0845
127.058349609375 0 457.00134
127.08686828613281 0 1429.5737
128.08262634277344 0 414.47348
129.06619262695312 0 7360.0664
129.1026153564453 0 123818.66
130.05020141601562 0 9201.426
130.0654296875 0 4746.737
130.0865478515625 0 521.1093
130.10020446777344 0 882.76337
130.10594177246094 0 7800.826
130.26695251464844 0 381.21237
131.11819458007812 0 7242.42
132.0764617919922 0 513.0869
132.08145141601562 0 792.7909
132.10220336914062 0 1837.1488
132.12176513671875 0 595.3846
134.02743530273438 0 1004.36224
134.48849487304688 0 393.34073
136.07598876953125 0 3685.9927
138.12802124023438 0 4083.7385
139.05072021484375 0 549.3062
139.0869140625 0 3928.3997
139.10546875 0 419.55518
140.07106018066406 0 565.3695
141.06216430664062 0 353.15073
141.0662078857422 0 3826.2961
141.10264587402344 0 940.59937
145.06092834472656 0 443.82904
146.0604705810547 0 1308.7864
147.07681274414062 0 3128.2727
147.11312866210938 0 3212.072
148.07574462890625 0 555.4255
150.12022399902344 0 505.81607
152.0706329345703 0 591.72064
152.1078338623047 0 869.00494
153.06602478027344 0 1270.2379
153.1027069091797 0 921.1679
154.08660888671875 0 4081.1357
154.09786987304688 0 6110.0005
155.0818328857422 0 9242.997
155.0933380126953 0 562.55023
155.1007537841797 0 554.7292
155.1183319091797 0 1642.5521
156.0781707763672 0 501.77576
156.08523559570312 0 824.7215
157.0610809326172 0 2889.1775
157.0975341796875 0 11200.782
157.10580444335938 0 552.74115
157.10855102539062 0 881.778
158.09315490722656 0 1269.1494
158.10023498535156 0 758.4523
159.11302185058594 0 821.94147
163.07167053222656 0 1738.861
163.08724975585938 0 1231.5046
165.1023712158203 0 2009.279
166.06146240234375 0 1682.8113
166.08682250976562 0 556.3212
167.0818634033203 0 7115.8286
167.11822509765625 0 6378.578
168.06549072265625 0 534.742
168.0849609375 0 610.67883
168.12176513671875 0 843.73395
169.06100463867188 0 2481.04
169.09716796875 0 1292.8224
170.0452117919922 0 579.4572
171.0768585205078 0 2569.515
172.07196044921875 0 1345.7185
172.108642578125 0 3053.908
173.09246826171875 0 744.0418
173.12875366210938 0 1835.4355
174.05581665039062 0 882.75055
174.13302612304688 0 507.7441
175.07168579101562 0 3138.4976
175.0873565673828 0 543.3549
176.6092071533203 0 467.55002
180.1136932373047 0 1482.0453
181.0976104736328 0 1714.5226
182.0927276611328 0 9237.6455
182.1290283203125 0 577.58936
183.0767822265625 0 6786.098
183.09584045410156 0 663.71497
183.1131591796875 0 1234.865
183.14927673339844 0 962.0351
184.09951782226562 0 848.0889
184.10842895507812 0 7325.441
185.09242248535156 0 13830.796
186.08767700195312 0 6755.2876
186.0960693359375 0 1280.3248
186.12417602539062 0 1156.0288
187.071533203125 0 732.29535
187.1078643798828 0 1351.4768
187.1447296142578 0 985.4928
189.08729553222656 0 5793.3877
191.08172607421875 0 1631.3856
191.1025848388672 0 591.8287
193.0973663330078 0 630.2081
195.07672119140625 0 1712.6705
195.11314392089844 0 14463.659
196.11672973632812 0 1499.5894
197.12875366210938 0 14559.348
198.08731079101562 0 826.9136
198.12403869628906 0 6867.5234
199.0716094970703 0 8902.573
199.10806274414062 0 4439.391
199.11903381347656 0 1469.1552
199.12794494628906 0 625.7308
200.07521057128906 0 1268.9805
200.10321044921875 0 9035.382
200.1395263671875 0 1975.3595
200.52708435058594 0 523.149
201.08718872070312 0 1664.3535
201.12362670898438 0 13928.906 y Ammonia loss 12
202.1188201904297 0 2050.888 y 10
203.10308837890625 0 1469.0658
207.11309814453125 0 1228.5426
208.09727478027344 0 1361.6824
208.10836791992188 0 10126.403
209.09239196777344 0 27366.643
209.11114501953125 0 1184.6946
210.09573364257812 0 2480.1467
210.7667694091797 0 537.84906
211.14447021484375 0 1691.927
212.13961791992188 0 3205.0557
213.08729553222656 0 1908.5248
213.1236572265625 0 1514.6942
214.11985778808594 0 665.6457
214.13919067382812 0 869.68555
215.1393280029297 0 7043.329
216.09861755371094 0 613.80634
216.14337158203125 0 944.3456
217.08221435546875 0 13672.514
217.1338653564453 0 4543.2554
218.0856475830078 0 1417.7872
218.1502685546875 0 54340.387 y 12
219.1530303955078 0 4125.182
221.05946350097656 0 1542.7029
221.10440063476562 0 856.0181
221.62374877929688 0 759.6255
223.108642578125 0 663.35
223.15518188476562 0 611.26855
224.1029510498047 0 1129.3195
225.1235809326172 0 8371.86
225.13519287109375 0 4996.9287
226.11898803710938 0 103199.38
227.10292053222656 0 3689.2725
227.12229919433594 0 11765.199
227.14149475097656 0 577.1195
228.12338256835938 0 1009.387
228.13458251953125 0 7505.9507 b 3
229.13514709472656 0 928.691
229.15499877929688 0 819.9804
231.09799194335938 0 1925.7064
233.16529846191406 0 206690.97 a 1
234.08755493164062 0 1025.226
234.12435913085938 0 1775.696
234.1685333251953 0 31415.66
235.1083221435547 0 806.78265
235.17185974121094 0 2375.0305
236.1034698486328 0 6201.1074
236.13970947265625 0 680.50287
237.107177734375 0 871.6939
238.08151245117188 0 592.8705
238.11973571777344 0 660.0645
239.15077209472656 0 811.2837
240.09814453125 0 865.39294
240.13504028320312 0 673.477
240.6082763671875 0 1022.6141
241.08229064941406 0 8469.516
242.08737182617188 0 764.1037
242.1502685546875 0 20062.355
243.13412475585938 0 2199.1167
243.1456298828125 0 28298.803
244.13009643554688 0 5396.7944
244.149169921875 0 3353.1365
245.12939453125 0 3530.7239
247.11029052734375 0 933.19934
248.62986755371094 0 1564.8314
248.65048217773438 0 1255.7153
249.1217041015625 0 6466.851
249.6234130859375 0 925.33716
250.15489196777344 0 1125.4541
251.15052795410156 0 3704.505
252.09767150878906 0 1584.5704
252.11093139648438 0 725.7041
252.1345977783203 0 2678.0225
253.13043212890625 0 846.71533
254.11386108398438 0 17326.68
254.14999389648438 0 2128.632
255.11669921875 0 2237.6567
257.1429748535156 0 20428.465
257.6338195800781 0 2435.3547 y Ammonia loss 9
257.6446533203125 0 3184.7166
258.1087646484375 0 730.0965
259.0928039550781 0 10266.02
260.095947265625 0 845.3819
261.15997314453125 0 82893.47 b 1
262.1188049316406 0 1503.5927
262.1632995605469 0 12933.525
263.16546630859375 0 1549.8002
264.134521484375 0 11995.76
265.1374816894531 0 1653.829
266.14813232421875 0 13442.759 y 9
266.6497802734375 0 3377.4424
267.1506652832031 0 703.03516
268.0932312011719 0 1860.8649
268.1413269042969 0 894.07965
269.12530517578125 0 739.15643
269.1609191894531 0 1398.4286
269.6526794433594 0 1363.683
270.14471435546875 0 714.94763
272.1086730957031 0 870.51807
272.1242370605469 0 7072.6816
273.12249755859375 0 1685.9
278.1500244140625 0 786.32465
278.1668701171875 0 606.0419
279.14581298828125 0 6572.023
280.1295166015625 0 9071.886
280.1471252441406 0 1025.1394
281.1151123046875 0 840.4471
281.1329345703125 0 1059.5104
282.1086120605469 0 1366.3634
282.1275329589844 0 2170.5283
282.1451721191406 0 34700.83
282.6293640136719 0 751.8619
283.1477355957031 0 4811.006
285.195556640625 0 740.59985
286.1037902832031 0 5671.9155
291.7032775878906 0 558.7581
292.1319885253906 0 551.9312
292.1662902832031 0 587.99976
292.66876220703125 0 711.7457
296.12200927734375 0 1904.7196
296.13580322265625 0 4905.315
296.1722106933594 0 5293.5806
296.62548828125 0 1080.6613
297.11968994140625 0 6348.917
297.15625 0 33458.277
298.12225341796875 0 1030.2173
298.1402893066406 0 4817.4995
298.1595458984375 0 4552.799
299.1423645019531 0 1068.6333
299.1716003417969 0 2315.7795
300.1197204589844 0 968.0742
300.13775634765625 0 627.8593
300.156005859375 0 2227.2585
304.1142578125 0 3657.5044
305.1298522949219 0 1612.0675
305.6646423339844 0 829.72095
306.1456298828125 0 1255.5725
306.1633605957031 0 1164.8662 b Water loss 5
308.16119384765625 0 2765.958
308.1984558105469 0 794.53687
310.1047058105469 0 1102.8553
313.1876525878906 0 3561.148
314.13421630859375 0 4400.9683
314.1477966308594 0 3801.9082
314.1809387207031 0 14736.072 y Ammonia loss 8
314.6374816894531 0 1476.9359
314.6782531738281 0 2437.1191
315.13299560546875 0 758.01605
315.1493835449219 0 620.46045
315.1667785644531 0 14829.26 b 5
315.185302734375 0 1554.9597
316.1697082519531 0 2327.7993
319.1395263671875 0 553.7869
321.156494140625 0 1161.969
322.1766662597656 0 893.60406
322.6903076171875 0 18035.947 y 8
323.1716613769531 0 1564.4901
323.1914978027344 0 5567.1475
323.69232177734375 0 572.2133
324.1559143066406 0 2001.9244
325.1517333984375 0 1383.5396
325.18609619140625 0 799.85583
326.1712341308594 0 1444.5522
328.114501953125 0 5640.0454
328.6653747558594 0 641.8933
329.1184997558594 0 709.42725
329.66448974609375 0 1914.9905
330.165771484375 0 794.51013
332.19329833984375 0 45122.7 y 11
333.15625 0 6973.63
333.1775817871094 0 760.62775
333.19647216796875 0 6259.066
334.1585998535156 0 1353.5391
334.1998596191406 0 686.7178
336.1553955078125 0 1424.0314
337.15118408203125 0 1919.7733
338.134765625 0 952.06085
338.66888427734375 0 5081.0884
339.1680603027344 0 4151.2266
341.182373046875 0 6287.5264
342.18414306640625 0 2435.4766
343.1612854003906 0 3689.3813
343.18707275390625 0 2308.2915
346.1249084472656 0 9462.767
347.1319580078125 0 1109.8486
347.6744689941406 0 3271.0183
348.1763610839844 0 1066.233
350.1840515136719 0 945.1672
351.16668701171875 0 28039.162
352.1692810058594 0 5606.5923
353.1822509765625 0 9079.3955
354.1661376953125 0 2152.746
354.1878356933594 0 1035.1617
355.16217041015625 0 3728.5205
356.1454162597656 0 1169.3541
358.2127380371094 0 4774.995 b 2
359.2164001464844 0 1185.8282
360.2147521972656 0 1046.276
361.17633056640625 0 609.8025
361.6753234863281 0 2112.0896
362.18194580078125 0 933.791
363.1689453125 0 840.6858
365.1459045410156 0 4814.388
365.1839904785156 0 728.4702
366.1446533203125 0 687.59625
367.2087707519531 0 1267.1478
368.1930847167969 0 10698.698
369.1771545410156 0 31465.518
369.693603515625 0 882.4773
370.1803283691406 0 6747.1445
370.67803955078125 0 13880.78 b Water loss 6
371.1824645996094 0 3431.9023 b Ammonia loss 9
371.18951416015625 0 3300.7886
371.6827697753906 0 1047.278
372.1764221191406 0 1654.6903
373.1719055175781 0 5363.01
374.1759338378906 0 862.68695
377.1841125488281 0 938.5667
378.69952392578125 0 784.65424 y Ammonia loss 7
379.18988037109375 0 1154.1765
379.6898193359375 0 1573.942 b 6
380.1909484863281 0 977.0606
381.15203857421875 0 863.76807
383.156494140625 0 7555.416
384.1595764160156 0 1806.0436
385.219482421875 0 4430.051
386.2037658691406 0 26179.402 y Ammonia loss 10
387.2077331542969 0 4921.237 y 7
388.52801513671875 0 706.3072
392.1938171386719 0 647.02954
393.17669677734375 0 1284.0347
394.1801452636719 0 765.3895
394.53363037109375 0 700.0008 b Water loss 10
394.8656311035156 0 894.63855 b Ammonia loss 10
395.1927490234375 0 582.2022
395.2301330566406 0 875.6675
396.188720703125 0 642.22485
396.7104187011719 0 581.77905
397.133544921875 0 1193.5369
401.1671142578125 0 13991.461
402.1708984375 0 2770.8552
403.2303771972656 0 112390.72 y 10
404.2332763671875 0 20351.14
405.161865234375 0 916.66956
405.2349548339844 0 3778.8955
407.20367431640625 0 7022.337
408.1883239746094 0 15710.057
409.1917419433594 0 2915.0398
410.20184326171875 0 1753.3927
411.1525573730469 0 676.73895
411.2029113769531 0 595.7613
413.20758056640625 0 520.37683
414.1992492675781 0 1395.0536
415.14605712890625 0 2167.359
418.2120361328125 0 1190.0697 y Water loss 2
418.5408935546875 0 1526.1696 y Ammonia loss 2
418.8728942871094 0 1052.2994
423.1882019042969 0 617.0348
424.2289733886719 0 7999.6074
424.5486755371094 0 924.7053
425.21527099609375 0 10379.289
425.7020263671875 0 995.829
426.1993713378906 0 2431.0012
426.22442626953125 0 811.5169
427.23028564453125 0 3571.6719
427.2713317871094 0 1483.2563
428.23736572265625 0 640.5797
429.2135925292969 0 977.5488
430.21356201171875 0 779.90796
433.15716552734375 0 4527.1177
434.1674499511719 0 741.04785
434.204833984375 0 3077.108
434.70721435546875 0 5763.368
435.2077941894531 0 4686.4956 b Water loss 7
435.7098083496094 0 926.11096
440.2125244140625 0 684.6527
441.1986083984375 0 1249.791
442.2405700683594 0 6307.435
443.1419677734375 0 1339.7202
443.175537109375 0 682.2754
443.2253112792969 0 3772.697 y Ammonia loss 6
444.2120361328125 0 1889.0217 b 7
444.7120666503906 0 1391.2952
445.214599609375 0 889.75354
448.1838684082031 0 786.5149
450.198974609375 0 2173.5945
452.2147521972656 0 5534.7847
453.2158508300781 0 1819.3628
454.2687072753906 0 843.898
455.2663879394531 0 5792.212 b 3
456.27001953125 0 1392.8218
459.2091064453125 0 2061.4937
460.25299072265625 0 809.16656
461.21820068359375 0 1758.792
461.7183532714844 0 764.2946
462.1993103027344 0 6490.094
463.2013244628906 0 1926.9082
464.2134704589844 0 996.20026
466.1944885253906 0 1128.4224
468.20916748046875 0 3407.677
470.2252197265625 0 23269.504
470.726318359375 0 2054.94
471.2294921875 0 5291.2905
476.1805419921875 0 1208.4067
478.2406311035156 0 2515.4207
479.2250061035156 0 3171.709
480.20965576171875 0 20179.832
481.2122802734375 0 5547.754
482.223876953125 0 2065.9827
482.2609558105469 0 872.5561
484.2047424316406 0 3092.9204
487.2518005371094 0 904.5445
488.2758483886719 0 1146.0535
494.1893005371094 0 2265.4531
494.23590087890625 0 1576.6874
496.2518310546875 0 8506.802
497.23651123046875 0 16463.285
498.22021484375 0 74577.14
499.2229919433594 0 18819.36
500.2284851074219 0 3846.9866
500.75299072265625 0 1699.5557 b 8
501.2564392089844 0 1022.38464
501.75469970703125 0 941.91345
502.21527099609375 0 4438.7676
503.21881103515625 0 1401.933
503.2613830566406 0 1236.8292
511.26300048828125 0 824.829
512.1990966796875 0 5339.3555
512.2398071289062 0 1314.8914
513.202392578125 0 2198.1907
513.2782592773438 0 28849.246
514.2628784179688 0 73309.73 y Ammonia loss 9
515.2647705078125 0 20252.54
516.2653198242188 0 2700.3674
521.2739868164062 0 854.14374
524.2875366210938 0 1966.5266 b Water loss 4
528.23095703125 0 1868.1254
530.2095947265625 0 18555.77
531.2128295898438 0 3796.6052
531.2886962890625 0 91272.43 y 9
532.2213134765625 0 897.532
532.29150390625 0 24278.906
533.2940063476562 0 4170.141
534.2584838867188 0 1716.9
535.2493286132812 0 1036.6185
538.2982177734375 0 3365.6904
539.2989501953125 0 1101.6996
539.7592163085938 0 645.17706
539.8115844726562 0 821.5556
540.311767578125 0 1919.076
542.2572631835938 0 1036.3727
542.2980346679688 0 3704.8855 b 4
543.3007202148438 0 1496.5167
543.7623291015625 0 708.1683
545.2376708984375 0 975.9682
546.2394409179688 0 1058.182
548.9900512695312 0 759.3361
553.261474609375 0 1111.627
555.3246459960938 0 2186.444
556.2703247070312 0 1165.8875 b Ammonia loss 9
556.3212280273438 0 1242.8331
563.24560546875 0 1513.1125
565.2674560546875 0 1133.9119
569.7756958007812 0 1843.3092
570.267578125 0 1247.938
571.271484375 0 930.0964
573.2310791015625 0 1740.2118
573.7798461914062 0 1402.0161
574.274169921875 0 870.2847
578.2849731445312 0 2050.8542 y Water loss 3
578.7800903320312 0 5300.598 y Ammonia loss 3
579.2813110351562 0 2824.927
579.7804565429688 0 1104.7833
580.2762451171875 0 846.2265
581.2581787109375 0 5901.857
582.2926635742188 0 1688.6649
582.783447265625 0 6329.2446
583.2822265625 0 3102.7056
583.7848510742188 0 1202.1229
584.3184204101562 0 1089.9624
587.29052734375 0 16080.158 y 3
587.792236328125 0 10756.938
588.2938232421875 0 4280.4136
591.2413940429688 0 5756.266
591.29296875 0 1247.5591 b Water loss 10
591.7942504882812 0 2383.4766 b Ammonia loss 10
592.2437744140625 0 2347.535
592.3054809570312 0 1620.6052
593.3043823242188 0 1714.9758
594.311767578125 0 763.0596
597.2890014648438 0 1575.7755
599.2673950195312 0 30298.129
599.79833984375 0 706.7355
600.2717895507812 0 8026.2666
600.7986450195312 0 2501.7188
601.2738647460938 0 1106.0609
601.3338012695312 0 2764.6929
607.2725830078125 0 807.7391
608.283203125 0 728.0084
608.7994995117188 0 848.6348
609.2515869140625 0 19770.559
609.3368530273438 0 5383.8804
609.787353515625 0 1455.5078
610.2556762695312 0 5388.4346
610.3375854492188 0 1066.428
611.2568359375 0 1107.1536
611.3153686523438 0 7265.97 b Water loss 5
612.3173217773438 0 3290.0808
613.3129272460938 0 827.06555
615.298095703125 0 2696.1582
617.8056640625 0 3098.96
618.3011474609375 0 8184.6475
618.7991943359375 0 5374.63
619.300537109375 0 2656.0825
622.2771606445312 0 804.5189
625.2835693359375 0 2510.0896
626.2828979492188 0 951.267
626.81201171875 0 14002.035 y Water loss 2
627.2620849609375 0 180193.22
627.307861328125 0 19300.936 y Ammonia loss 2
627.8065185546875 0 22930.027
628.2650146484375 0 57893.824
628.80908203125 0 2152.0757
629.2674560546875 0 13496.236
629.3292846679688 0 9931.042 b 5
630.2695922851562 0 2053.1284
630.3333740234375 0 3290.5664
631.3345336914062 0 808.19855
632.8377075195312 0 826.4587
633.3511962890625 0 2364.635
634.8086547851562 0 1160.195
635.3120727539062 0 855.6989
635.8173828125 0 173034.23 y 2
636.31884765625 0 127654.65
636.8203125 0 53890.652
637.3212280273438 0 16466.932
637.8226928710938 0 2508.206
643.2944946289062 0 5879.2056
644.295166015625 0 3190.451
644.3726196289062 0 30946.467 y 8
645.2872314453125 0 1632.3987
645.3753662109375 0 12144.048
646.3782958984375 0 2587.356
649.3292846679688 0 764.67865
658.3019409179688 0 1054.4791
660.3198852539062 0 1757.4677
661.3135986328125 0 822.1282
662.3226928710938 0 890.9315
664.8294067382812 0 816.2744
667.3438720703125 0 1236.5824
674.2980346679688 0 779.7538
676.3305053710938 0 773.924
677.3374633789062 0 764.9506
680.31640625 0 940.9736
682.38916015625 0 668.5467
682.6318359375 0 1762.4158
682.8851928710938 0 779.63214
682.9661254882812 0 1597.0619
683.3875732421875 0 1426.8933
684.2998046875 0 682.2394
684.3682861328125 0 1282.9211
686.3132934570312 0 1136.9186
694.3411865234375 0 6422.98
695.3397216796875 0 2151.7664
696.3509521484375 0 774.4229
704.325439453125 0 4110.428
705.3272094726562 0 2011.8845
709.3515014648438 0 1204.3993 y 1
709.8489379882812 0 990.5592
712.3516235351562 0 24031.543 b Water loss 12
713.3544311523438 0 10214.924
714.3557739257812 0 3467.5208
715.360595703125 0 751.4875
721.3548583984375 0 1355.9177
722.3384399414062 0 12393.826
723.340576171875 0 5731.709
724.3419799804688 0 1572.5051
729.3099365234375 0 676.7208
730.3772583007812 0 4510.542
731.3787841796875 0 2141.6064
738.3035888671875 0 940.8151
738.379150390625 0 2407.54
739.364990234375 0 4469.5063
740.3471069335938 0 62412.168 b Water loss 6
741.350341796875 0 23788.8
742.3526611328125 0 6263.222
743.3583984375 0 1058.0574
746.339111328125 0 707.2587
753.3421630859375 0 1557.1147
754.3392333984375 0 989.1083
755.4052734375 0 3281.973 y Water loss 7
756.3934936523438 0 4537.709 y Ammonia loss 7
757.3738403320312 0 15501.175 Precursor Ammonia loss
758.37255859375 0 27849.701 b 6
759.375244140625 0 11883.652
760.3773193359375 0 3071.3542
770.3668212890625 0 1830.0575
771.3551025390625 0 4644.4893
772.3517456054688 0 2200.394
773.4153442382812 0 14962.454 y 7
774.2575073242188 0 858.05316
774.33154296875 0 3442.7588
774.4178466796875 0 6803.2363
775.3400268554688 0 1311.9696
775.4186401367188 0 1467.0656
791.376220703125 0 776.89374
807.3677978515625 0 774.811
810.3785400390625 0 923.0111
815.3757934570312 0 845.6043
833.3859252929688 0 1158.1849
840.4099731445312 0 2412.3386
841.4122314453125 0 2077.2625
842.3916015625 0 3028.2773
843.3946533203125 0 1124.9066
850.3936767578125 0 4257.7114
851.391357421875 0 3870.8022
852.3947143554688 0 1741.238
853.424072265625 0 2025.4188
854.42724609375 0 799.1566
859.4179077148438 0 6471.6733
860.4220581054688 0 4257.6807
861.4271240234375 0 924.6349
868.4044799804688 0 28831.861
869.4060668945312 0 19608.441 b Water loss 7
870.4087524414062 0 7533.708
871.4111938476562 0 1201.608
884.4323120117188 0 823.92065 y Water loss 6
885.43212890625 0 1529.8208 y Ammonia loss 6
886.423095703125 0 1231.0133
887.4136962890625 0 56465.18 b 7
888.4168701171875 0 30140.768
889.4191284179688 0 9178.204
890.421630859375 0 1243.8291
893.4188232421875 0 691.43274
894.4261474609375 0 1972.3828
902.453857421875 0 2822.4744 y 6
903.4572143554688 0 1649.6556
911.44287109375 0 2706.9692
912.446044921875 0 1681.0873
914.4271850585938 0 830.0345
921.4315795898438 0 3404.222
922.4241943359375 0 3951.2144
923.4210815429688 0 2060.3508
939.4405517578125 0 15572.15
940.4427490234375 0 8730.563
941.4463500976562 0 2389.6135
956.456787109375 0 2111.716
957.4576416015625 0 1935.8489
971.4766845703125 0 968.8208 y Water loss 5
972.4761352539062 0 1780.0338 y Ammonia loss 5
973.4810791015625 0 892.4538
982.48779296875 0 1211.5172 b Water loss 8
983.4812622070312 0 922.03973
989.48828125 0 4475.19 y 5
990.4891357421875 0 2382.1743
1000.49658203125 0 9217.562 b 8
1001.5016479492188 0 5092.99
1002.5034790039062 0 2029.3489
1025.4820556640625 0 1263.9407
1035.470703125 0 1258.1162
1036.460693359375 0 3677.6338
1037.459228515625 0 1671.5349
1041.4849853515625 0 1338.3832
1042.4949951171875 0 661.6601
1053.4835205078125 0 12159.972
1054.48193359375 0 7635.7695
1055.4844970703125 0 2471.5112
1058.5234375 0 817.9502 y Water loss 4
1059.498046875 0 2876.905 y Ammonia loss 4
1060.50537109375 0 1243.9535
1061.503173828125 0 876.791
1068.5025634765625 0 706.2748
1070.499755859375 0 994.92267
1071.5037841796875 0 1394.5663
1076.5216064453125 0 7045.393 y 4
1077.5234375 0 3933.3066
1078.5220947265625 0 1092.0206
1086.50537109375 0 1067.4548
1087.4908447265625 0 776.0267
1128.556640625 0 2326.9219 b 9
1138.53662109375 0 1437.8553
1156.54931640625 0 2610.2668 y Ammonia loss 3
1157.552490234375 0 1482.7758
1173.5726318359375 0 8219.537 y 3
1174.575439453125 0 5546.6494
1175.57666015625 0 2506.9456
1181.5782470703125 0 890.55536 b Water loss 10
1199.596435546875 0 854.183 b 10
1231.1268310546875 0 696.21173
1407.18701171875 0 703.7514
1445.350830078125 0 777.7126
3385.295166015625 0 773.10065

Spectrum Details

|  |  |
| --- | --- |
| Matched peaks? Matched peaks The total absolute number of peaks matched. Additionally in brackets the total fraction of peaks matched and the total number of peaks is shown. | 72 (10.13% of 711) |
| FDR? FDR The false discovery rate estimated for this peptide. It is calculated by matching all theoretical fragments with a non-integer shift with the raw peaks for this spectrum. This is done with 40 different shifts. The resulting percentage is the average number of annotated peaks over the number of annotated peaks with the correct spectrum. | 0.36% |
| Satellite FDR? Satellite FDR See the FDR for details on its calculation. This satellite ion specific FDR only contains the satellite ions (d/w) for I/L/J positions. | - |
| PSM Score? PSM Score The PSM Score as given by Hecklib to this annotated spectrum. It is shown with three significant figures. | 636 |

#### Spectrum 6472? Spectrum 6472 The raw spectrum of this peptide as annotated by Hecklib. The fragments are coloured according to ion type (see legend). Any peaks with a star '\*' as text can be hovered over to see the full details, first the ion type second the mass shift type. By hovering over the amino acids in the peptide or ions in the legend the corresponding peaks are highlighted. By toggling the 'Unassigned' label you can turn the background (unassigned) peaks on or off in the plot. By updating the slider in the Ion legend you can update the spectrum to only show the top X% of the peaks with labels. The top X% means any peak that is within X% of the highest intensity. By dragging in the spectrum you can zoom in to a specific part of the spectrum and use 'Zoom Out' to get back to the original zoom level. The annotation of the spectrum is based on the given sequence in the peptides file and is done with different software so inconsistencies are likely. The peaks are annotated based on the given sequence, with 20 ppm tolerance.

Copy Data

##### Spectrum 6472 (TSV)

###### Preview

```
Loading example...
```

*Click on the button to copy the data to your clipboard.*

Mz MinMz MaxIntensity Max

WidthHeightPeptide font sizePeptide stroke widthSpectrum font sizeSpectrum stroke widthCompact peptide

Ion legend

wxyz

abcd

OtherUnassignedIon^Charge^_Position_Show for top:%

JFPPSSEEJQANKA

03.85e+47.70e+41.15e+51.54e+5

Zoom Out

y^+1^_2_y^+2^_4_y^+1^_2_b^+2^_4_a^+1^_2_y^+2^_5_b^+1^_2_y^+2^_5_b^+2^_6_y^+2^_6_b^+2^_6_y^+2^_6_b^+3^_9_y^+1^_3_b^+1^_3_b^+2^_7_b^+3^_10_b^+2^_7_y^+1^_4_y^+2^_7_y^+1^_4_y^+3^_12_y^+3^_12_b^+2^_8_y^+2^_8_b^+2^_8_b^+1^_4_b^+2^_9_y^+1^_5_b^+1^_5_y^+1^_5_y^+2^_10_b^+1^_5_b^+2^_10_y^+2^_11_y^+2^_11_b^+2^_11_b^+2^_11_b^+1^_6_y^+2^_12_y^+2^_12_b^+1^_6_y^+2^_12_y^+1^_6_b^+2^_13_b^+1^_7_y^+1^_7_y^+1^_7_\*b^+1^_7_y^+1^_7_b^+1^_8_y^+1^_8_b^+1^_8_y^+1^_8_y^+1^_9_b^+1^_9_y^+1^_9_b^+1^_9_y^+1^_10_y^+1^_10_y^+1^_10_b^+1^_10_y^+1^_11_y^+1^_11_y^+1^_11_

0768153623043072

Fragment Matches Table

Show background peaks

| Position | Ion type | Intensity | mz Theoretical | mz Error (Th) | mz Error (ppm) | Charge | Series Number |
| --- | --- | --- | --- | --- | --- | --- | --- |
| - | - | 1.138E+05 | 120.1 | - | - | 0 | - |
| - | - | 1.093E+04 | 121.1 | - | - | 0 | - |
| - | - | 348 | 122.1 | - | - | 0 | - |
| - | - | 447.3 | 125.1 | - | - | 0 | - |
| - | - | 414.8 | 126 | - | - | 0 | - |
| - | - | 6852 | 126.1 | - | - | 0 | - |
| - | - | 1014 | 127.1 | - | - | 0 | - |
| - | - | 454.3 | 127.1 | - | - | 0 | - |
| - | - | 1247 | 127.1 | - | - | 0 | - |
| - | - | 462.2 | 128.5 | - | - | 0 | - |
| - | - | 438.6 | 129.1 | - | - | 0 | - |
| - | - | 6095 | 129.1 | - | - | 0 | - |
| - | - | 9.771E+04 | 129.1 | - | - | 0 | - |
| - | - | 7682 | 130.1 | - | - | 0 | - |
| - | - | 4172 | 130.1 | - | - | 0 | - |
| - | - | 969.2 | 130.1 | - | - | 0 | - |
| - | - | 1102 | 130.1 | - | - | 0 | - |
| - | - | 7024 | 130.1 | - | - | 0 | - |
| - | - | 588.5 | 131.1 | - | - | 0 | - |
| - | - | 5980 | 131.1 | - | - | 0 | - |
| - | - | 1289 | 132.1 | - | - | 0 | - |
| - | - | 398.6 | 133.4 | - | - | 0 | - |
| - | - | 1253 | 134 | - | - | 0 | - |
| - | - | 5900 | 136.1 | - | - | 0 | - |
| - | - | 1983 | 137.1 | - | - | 0 | - |
| - | - | 3014 | 138.1 | - | - | 0 | - |
| - | - | 1023 | 139.1 | - | - | 0 | - |
| - | - | 3110 | 139.1 | - | - | 0 | - |
| - | - | 1131 | 140.1 | - | - | 0 | - |
| - | - | 480.3 | 140.1 | - | - | 0 | - |
| - | - | 2875 | 141.1 | - | - | 0 | - |
| - | - | 1852 | 141.1 | - | - | 0 | - |
| - | - | 846.7 | 142.1 | - | - | 0 | - |
| - | - | 573.6 | 143.1 | - | - | 0 | - |
| - | - | 499.1 | 143.1 | - | - | 0 | - |
| - | - | 1935 | 145.1 | - | - | 0 | - |
| - | - | 741 | 146.1 | - | - | 0 | - |
| - | - | 2659 | 147.1 | - | - | 0 | - |
| - | - | 2020 | 147.1 | - | - | 0 | - |
| - | - | 488.3 | 148.1 | - | - | 0 | - |
| - | - | 2258 | 152.1 | - | - | 0 | - |
| - | - | 472.6 | 153.1 | - | - | 0 | - |
| - | - | 596.9 | 154.1 | - | - | 0 | - |
| - | - | 3116 | 154.1 | - | - | 0 | - |
| - | - | 5407 | 154.1 | - | - | 0 | - |
| - | - | 471.8 | 155 | - | - | 0 | - |
| - | - | 5678 | 155.1 | - | - | 0 | - |
| - | - | 1722 | 155.1 | - | - | 0 | - |
| - | - | 2577 | 157.1 | - | - | 0 | - |
| - | - | 8072 | 157.1 | - | - | 0 | - |
| - | - | 512 | 157.1 | - | - | 0 | - |
| - | - | 967.4 | 158.1 | - | - | 0 | - |
| - | - | 689.9 | 158.1 | - | - | 0 | - |
| - | - | 787.6 | 159.1 | - | - | 0 | - |
| - | - | 920.1 | 160.1 | - | - | 0 | - |
| - | - | 1703 | 163.1 | - | - | 0 | - |
| - | - | 936 | 163.1 | - | - | 0 | - |
| - | - | 1395 | 165.1 | - | - | 0 | - |
| - | - | 958.6 | 166.1 | - | - | 0 | - |
| - | - | 5682 | 167.1 | - | - | 0 | - |
| - | - | 5984 | 167.1 | - | - | 0 | - |
| - | - | 2173 | 169.1 | - | - | 0 | - |
| - | - | 1464 | 169.1 | - | - | 0 | - |
| - | - | 667.2 | 170.1 | - | - | 0 | - |
| - | - | 1292 | 171.1 | - | - | 0 | - |
| - | - | 677.8 | 171.1 | - | - | 0 | - |
| - | - | 1368 | 172.1 | - | - | 0 | - |
| - | - | 2913 | 172.1 | - | - | 0 | - |
| - | - | 1356 | 173.1 | - | - | 0 | - |
| - | - | 567.6 | 174.1 | - | - | 0 | - |
| - | - | 539.1 | 174.1 | - | - | 0 | - |
| - | - | 2659 | 175.1 | - | - | 0 | - |
| - | - | 575.5 | 179.1 | - | - | 0 | - |
| - | - | 1033 | 180.1 | - | - | 0 | - |
| - | - | 623.6 | 181.1 | - | - | 0 | - |
| - | - | 1824 | 181.1 | - | - | 0 | - |
| - | - | 7772 | 182.1 | - | - | 0 | - |
| - | - | 891.4 | 182.1 | - | - | 0 | - |
| - | - | 5684 | 183.1 | - | - | 0 | - |
| - | - | 564.3 | 183.1 | - | - | 0 | - |
| - | - | 1006 | 183.1 | - | - | 0 | - |
| - | - | 1092 | 183.1 | - | - | 0 | - |
| - | - | 5269 | 184.1 | - | - | 0 | - |
| - | - | 1.063E+04 | 185.1 | - | - | 0 | - |
| - | - | 731.2 | 185.1 | - | - | 0 | - |
| - | - | 661.4 | 185.2 | - | - | 0 | - |
| - | - | 3758 | 186.1 | - | - | 0 | - |
| - | - | 679.8 | 186.1 | - | - | 0 | - |
| - | - | 1796 | 186.1 | - | - | 0 | - |
| - | - | 776.7 | 187.1 | - | - | 0 | - |
| - | - | 993.4 | 187.1 | - | - | 0 | - |
| - | - | 1298 | 187.1 | - | - | 0 | - |
| - | - | 3550 | 189.1 | - | - | 0 | - |
| - | - | 1290 | 191.1 | - | - | 0 | - |
| - | - | 1500 | 195.1 | - | - | 0 | - |
| - | - | 1.188E+04 | 195.1 | - | - | 0 | - |
| - | - | 1027 | 196.1 | - | - | 0 | - |
| - | - | 1.074E+04 | 197.1 | - | - | 0 | - |
| - | - | 4716 | 198.1 | - | - | 0 | - |
| - | - | 635.6 | 198.1 | - | - | 0 | - |
| - | - | 7285 | 199.1 | - | - | 0 | - |
| - | - | 3542 | 199.1 | - | - | 0 | - |
| - | - | 1230 | 199.1 | - | - | 0 | - |
| - | - | 581.3 | 200.1 | - | - | 0 | - |
| - | - | 6122 | 200.1 | - | - | 0 | - |
| - | - | 770.1 | 200.1 | - | - | 0 | - |
| - | - | 2200 | 200.1 | - | - | 0 | - |
| - | - | 617.4 | 201.1 | - | - | 0 | - |
| - | - | 910.7 | 201.1 | - | - | 0 | - |
| 13 | y | 1.092E+04 | 201.1 | 0.0003036 | 1.51 | +1 | 2 |
| - | - | 859.5 | 201.1 | - | - | 0 | - |
| - | - | 684.4 | 202.1 | - | - | 0 | - |
| 11 | y | 2026 | 202.1 | 0.0003397 | 1.681 | +2 | 4 |
| - | - | 940.4 | 203.1 | - | - | 0 | - |
| - | - | 456.4 | 203.1 | - | - | 0 | - |
| - | - | 1139 | 204.1 | - | - | 0 | - |
| - | - | 1402 | 207.1 | - | - | 0 | - |
| - | - | 1118 | 208.1 | - | - | 0 | - |
| - | - | 6565 | 208.1 | - | - | 0 | - |
| - | - | 2.256E+04 | 209.1 | - | - | 0 | - |
| - | - | 1665 | 210.1 | - | - | 0 | - |
| - | - | 660.9 | 210.1 | - | - | 0 | - |
| - | - | 989.4 | 211.1 | - | - | 0 | - |
| - | - | 1116 | 211.1 | - | - | 0 | - |
| - | - | 1592 | 212.1 | - | - | 0 | - |
| - | - | 1509 | 213.1 | - | - | 0 | - |
| - | - | 858.1 | 213.1 | - | - | 0 | - |
| - | - | 1959 | 213.1 | - | - | 0 | - |
| - | - | 531 | 213.1 | - | - | 0 | - |
| - | - | 804.6 | 214.1 | - | - | 0 | - |
| - | - | 626.7 | 214.2 | - | - | 0 | - |
| - | - | 545.5 | 215.1 | - | - | 0 | - |
| - | - | 5655 | 215.1 | - | - | 0 | - |
| - | - | 9325 | 217.1 | - | - | 0 | - |
| - | - | 3661 | 217.1 | - | - | 0 | - |
| - | - | 566.6 | 218.1 | - | - | 0 | - |
| 13 | y | 4.344E+04 | 218.1 | 0.0004116 | 1.887 | +1 | 2 |
| - | - | 4158 | 219.2 | - | - | 0 | - |
| - | - | 1475 | 221.1 | - | - | 0 | - |
| - | - | 778.9 | 221.1 | - | - | 0 | - |
| - | - | 991.4 | 221.6 | - | - | 0 | - |
| - | - | 702.4 | 223.1 | - | - | 0 | - |
| - | - | 1040 | 223.2 | - | - | 0 | - |
| - | - | 7271 | 225.1 | - | - | 0 | - |
| - | - | 3730 | 225.1 | - | - | 0 | - |
| - | - | 7.931E+04 | 226.1 | - | - | 0 | - |
| - | - | 4021 | 227.1 | - | - | 0 | - |
| - | - | 8393 | 227.1 | - | - | 0 | - |
| - | - | 540.9 | 228.1 | - | - | 0 | - |
| 4 | b | 6279 | 228.1 | 0.001605 | 7.036 | +2 | 4 |
| - | - | 671.9 | 229.1 | - | - | 0 | - |
| - | - | 1233 | 229.2 | - | - | 0 | - |
| - | - | 1625 | 231.1 | - | - | 0 | - |
| 2 | a | 1.525E+05 | 233.2 | 0.0005045 | 2.164 | +1 | 2 |
| - | - | 779.5 | 234.1 | - | - | 0 | - |
| - | - | 2012 | 234.1 | - | - | 0 | - |
| - | - | 1076 | 234.1 | - | - | 0 | - |
| - | - | 2.167E+04 | 234.2 | - | - | 0 | - |
| - | - | 1909 | 235.2 | - | - | 0 | - |
| - | - | 635.6 | 235.6 | - | - | 0 | - |
| - | - | 3394 | 236.1 | - | - | 0 | - |
| - | - | 1007 | 236.1 | - | - | 0 | - |
| - | - | 654.7 | 239.1 | - | - | 0 | - |
| - | - | 585.3 | 239.2 | - | - | 0 | - |
| - | - | 1504 | 240.1 | - | - | 0 | - |
| - | - | 1392 | 240.1 | - | - | 0 | - |
| - | - | 7045 | 241.1 | - | - | 0 | - |
| - | - | 631.8 | 242.1 | - | - | 0 | - |
| - | - | 1.656E+04 | 242.2 | - | - | 0 | - |
| - | - | 1544 | 243.1 | - | - | 0 | - |
| - | - | 2.014E+04 | 243.1 | - | - | 0 | - |
| - | - | 748.5 | 244.1 | - | - | 0 | - |
| - | - | 5960 | 244.1 | - | - | 0 | - |
| - | - | 2332 | 244.1 | - | - | 0 | - |
| - | - | 770.4 | 245.1 | - | - | 0 | - |
| - | - | 2884 | 245.1 | - | - | 0 | - |
| - | - | 1554 | 248.6 | - | - | 0 | - |
| - | - | 6614 | 249.1 | - | - | 0 | - |
| - | - | 935.8 | 249.6 | - | - | 0 | - |
| - | - | 721.7 | 250.2 | - | - | 0 | - |
| - | - | 3351 | 251.2 | - | - | 0 | - |
| - | - | 1449 | 252.1 | - | - | 0 | - |
| - | - | 2637 | 252.1 | - | - | 0 | - |
| - | - | 1.221E+04 | 254.1 | - | - | 0 | - |
| - | - | 1954 | 254.2 | - | - | 0 | - |
| - | - | 1083 | 255.1 | - | - | 0 | - |
| - | - | 1192 | 255.1 | - | - | 0 | - |
| - | - | 700.4 | 257.1 | - | - | 0 | - |
| - | - | 1.233E+04 | 257.1 | - | - | 0 | - |
| 10 | y | 2371 | 257.6 | 0.0003547 | 1.377 | +2 | 5 |
| - | - | 2274 | 257.6 | - | - | 0 | - |
| - | - | 706.9 | 258.1 | - | - | 0 | - |
| - | - | 2308 | 258.1 | - | - | 0 | - |
| - | - | 8181 | 259.1 | - | - | 0 | - |
| - | - | 717.2 | 260.1 | - | - | 0 | - |
| 2 | b | 6.133E+04 | 261.2 | 0.0003103 | 1.188 | +1 | 2 |
| - | - | 1098 | 262.1 | - | - | 0 | - |
| - | - | 1.117E+04 | 262.2 | - | - | 0 | - |
| - | - | 604.6 | 263.2 | - | - | 0 | - |
| - | - | 8799 | 264.1 | - | - | 0 | - |
| - | - | 1078 | 265.1 | - | - | 0 | - |
| 10 | y | 9191 | 266.1 | 0.0004089 | 1.536 | +2 | 5 |
| - | - | 2692 | 266.6 | - | - | 0 | - |
| - | - | 1511 | 268.1 | - | - | 0 | - |
| - | - | 583.5 | 268.2 | - | - | 0 | - |
| - | - | 1407 | 269.2 | - | - | 0 | - |
| - | - | 587.7 | 269.7 | - | - | 0 | - |
| - | - | 5479 | 272.1 | - | - | 0 | - |
| - | - | 809.6 | 273.1 | - | - | 0 | - |
| - | - | 1120 | 278.2 | - | - | 0 | - |
| - | - | 986.6 | 278.2 | - | - | 0 | - |
| - | - | 757.7 | 279.1 | - | - | 0 | - |
| - | - | 5021 | 279.1 | - | - | 0 | - |
| - | - | 7107 | 280.1 | - | - | 0 | - |
| - | - | 550.4 | 281.1 | - | - | 0 | - |
| - | - | 762.9 | 281.1 | - | - | 0 | - |
| - | - | 1061 | 282.1 | - | - | 0 | - |
| - | - | 934.8 | 282.1 | - | - | 0 | - |
| - | - | 2.76E+04 | 282.1 | - | - | 0 | - |
| - | - | 638 | 282.6 | - | - | 0 | - |
| - | - | 4502 | 283.1 | - | - | 0 | - |
| - | - | 833.1 | 285.2 | - | - | 0 | - |
| - | - | 3883 | 286.1 | - | - | 0 | - |
| - | - | 633.6 | 292.2 | - | - | 0 | - |
| - | - | 691.8 | 296.1 | - | - | 0 | - |
| - | - | 3605 | 296.1 | - | - | 0 | - |
| - | - | 3750 | 296.2 | - | - | 0 | - |
| - | - | 4253 | 297.1 | - | - | 0 | - |
| - | - | 2.701E+04 | 297.2 | - | - | 0 | - |
| - | - | 893.2 | 298.1 | - | - | 0 | - |
| - | - | 988.3 | 298.1 | - | - | 0 | - |
| - | - | 3704 | 298.1 | - | - | 0 | - |
| - | - | 3769 | 298.2 | - | - | 0 | - |
| - | - | 875.6 | 299.1 | - | - | 0 | - |
| - | - | 2864 | 299.2 | - | - | 0 | - |
| - | - | 677.4 | 300.1 | - | - | 0 | - |
| - | - | 1100 | 300.2 | - | - | 0 | - |
| - | - | 3202 | 304.1 | - | - | 0 | - |
| - | - | 1222 | 305.1 | - | - | 0 | - |
| - | - | 682.1 | 305.2 | - | - | 0 | - |
| - | - | 1240 | 306.1 | - | - | 0 | - |
| 6 | b | 1056 | 306.2 | 0.001342 | 4.384 | +2 | 6 |
| - | - | 1241 | 307.1 | - | - | 0 | - |
| - | - | 2654 | 308.2 | - | - | 0 | - |
| - | - | 1404 | 310.1 | - | - | 0 | - |
| - | - | 718.4 | 313.2 | - | - | 0 | - |
| - | - | 2964 | 313.2 | - | - | 0 | - |
| - | - | 2824 | 314.1 | - | - | 0 | - |
| - | - | 3259 | 314.1 | - | - | 0 | - |
| 9 | y | 9078 | 314.2 | 0.004 | 12.73 | +2 | 6 |
| - | - | 1444 | 314.6 | - | - | 0 | - |
| - | - | 1645 | 314.7 | - | - | 0 | - |
| - | - | 1150 | 315.1 | - | - | 0 | - |
| 6 | b | 1.105E+04 | 315.2 | 0.001377 | 4.368 | +2 | 6 |
| - | - | 1299 | 315.2 | - | - | 0 | - |
| - | - | 674.4 | 316.1 | - | - | 0 | - |
| - | - | 2023 | 316.2 | - | - | 0 | - |
| - | - | 549.8 | 319.1 | - | - | 0 | - |
| - | - | 1332 | 321.2 | - | - | 0 | - |
| 9 | y | 1.297E+04 | 322.7 | 0.0005827 | 1.806 | +2 | 6 |
| - | - | 1174 | 323.2 | - | - | 0 | - |
| - | - | 3173 | 323.2 | - | - | 0 | - |
| - | - | 581.7 | 323.7 | - | - | 0 | - |
| - | - | 1426 | 325.2 | - | - | 0 | - |
| - | - | 809.5 | 327.2 | - | - | 0 | - |
| - | - | 2440 | 328.1 | - | - | 0 | - |
| 9 | b | 649.6 | 328.2 | 0.002091 | 6.372 | +3 | 9 |
| - | - | 976 | 329.7 | - | - | 0 | - |
| - | - | 1308 | 330.2 | - | - | 0 | - |
| 12 | y | 3.153E+04 | 332.2 | 0.000575 | 1.731 | +1 | 3 |
| - | - | 5128 | 333.2 | - | - | 0 | - |
| - | - | 1052 | 333.2 | - | - | 0 | - |
| - | - | 4534 | 333.2 | - | - | 0 | - |
| - | - | 1011 | 336.2 | - | - | 0 | - |
| - | - | 793.6 | 337.2 | - | - | 0 | - |
| - | - | 1023 | 338.1 | - | - | 0 | - |
| - | - | 4222 | 338.7 | - | - | 0 | - |
| - | - | 3517 | 339.2 | - | - | 0 | - |
| - | - | 767 | 340.2 | - | - | 0 | - |
| - | - | 4702 | 341.2 | - | - | 0 | - |
| - | - | 1565 | 342.2 | - | - | 0 | - |
| - | - | 1608 | 343.2 | - | - | 0 | - |
| - | - | 1622 | 343.2 | - | - | 0 | - |
| - | - | 956.6 | 344.2 | - | - | 0 | - |
| - | - | 585.3 | 344.2 | - | - | 0 | - |
| - | - | 6756 | 346.1 | - | - | 0 | - |
| - | - | 1897 | 347.1 | - | - | 0 | - |
| - | - | 2712 | 347.7 | - | - | 0 | - |
| - | - | 1011 | 348.2 | - | - | 0 | - |
| - | - | 717.9 | 350.2 | - | - | 0 | - |
| - | - | 2.237E+04 | 351.2 | - | - | 0 | - |
| - | - | 4331 | 352.2 | - | - | 0 | - |
| - | - | 6353 | 353.2 | - | - | 0 | - |
| - | - | 1501 | 354.2 | - | - | 0 | - |
| - | - | 923.4 | 354.2 | - | - | 0 | - |
| - | - | 2493 | 355.2 | - | - | 0 | - |
| 3 | b | 3444 | 358.2 | 0.0002198 | 0.6137 | +1 | 3 |
| - | - | 1580 | 361.7 | - | - | 0 | - |
| - | - | 1000 | 362.2 | - | - | 0 | - |
| - | - | 4099 | 365.1 | - | - | 0 | - |
| - | - | 727.6 | 366.7 | - | - | 0 | - |
| - | - | 6927 | 368.2 | - | - | 0 | - |
| - | - | 2.551E+04 | 369.2 | - | - | 0 | - |
| - | - | 4931 | 370.2 | - | - | 0 | - |
| 7 | b | 8984 | 370.7 | 0.006069 | 16.37 | +2 | 7 |
| 10 | b | 2419 | 371.2 | 0.0007847 | 2.114 | +3 | 10 |
| - | - | 1437 | 372.2 | - | - | 0 | - |
| - | - | 3724 | 373.2 | - | - | 0 | - |
| - | - | 879.6 | 374.2 | - | - | 0 | - |
| - | - | 627.2 | 375.1 | - | - | 0 | - |
| - | - | 726.7 | 375.2 | - | - | 0 | - |
| - | - | 1545 | 375.7 | - | - | 0 | - |
| - | - | 587.2 | 377.2 | - | - | 0 | - |
| - | - | 1450 | 379.2 | - | - | 0 | - |
| 7 | b | 2016 | 379.7 | 0.00107 | 2.817 | +2 | 7 |
| - | - | 686.4 | 380.2 | - | - | 0 | - |
| - | - | 786.3 | 381.2 | - | - | 0 | - |
| - | - | 1377 | 382.7 | - | - | 0 | - |
| - | - | 5682 | 383.2 | - | - | 0 | - |
| - | - | 960.8 | 384.2 | - | - | 0 | - |
| - | - | 3357 | 385.2 | - | - | 0 | - |
| 11 | y | 1.753E+04 | 386.2 | 0.0005694 | 1.474 | +1 | 4 |
| 8 | y | 3925 | 387.2 | 0.003563 | 9.202 | +2 | 7 |
| - | - | 664.3 | 389.2 | - | - | 0 | - |
| - | - | 1537 | 397.1 | - | - | 0 | - |
| - | - | 1.069E+04 | 401.2 | - | - | 0 | - |
| - | - | 2187 | 402.2 | - | - | 0 | - |
| 11 | y | 7.976E+04 | 403.2 | 0.0006316 | 1.566 | +1 | 4 |
| - | - | 1.371E+04 | 404.2 | - | - | 0 | - |
| - | - | 1925 | 405.2 | - | - | 0 | - |
| - | - | 4274 | 407.2 | - | - | 0 | - |
| - | - | 1.152E+04 | 408.2 | - | - | 0 | - |
| - | - | 2715 | 409.2 | - | - | 0 | - |
| - | - | 1446 | 410.2 | - | - | 0 | - |
| - | - | 700.5 | 411.2 | - | - | 0 | - |
| - | - | 836.2 | 413.2 | - | - | 0 | - |
| - | - | 1264 | 414.2 | - | - | 0 | - |
| - | - | 2691 | 415.1 | - | - | 0 | - |
| - | - | 820.8 | 416.1 | - | - | 0 | - |
| 3 | y | 1092 | 418.2 | 0.002956 | 7.069 | +3 | 12 |
| 3 | y | 1259 | 418.5 | 0.001733 | 4.141 | +3 | 12 |
| - | - | 832 | 418.9 | - | - | 0 | - |
| - | - | 786.3 | 423.2 | - | - | 0 | - |
| - | - | 5223 | 424.2 | - | - | 0 | - |
| - | - | 1054 | 424.5 | - | - | 0 | - |
| - | - | 7000 | 425.2 | - | - | 0 | - |
| - | - | 1011 | 425.7 | - | - | 0 | - |
| - | - | 2296 | 426.2 | - | - | 0 | - |
| - | - | 1302 | 427.2 | - | - | 0 | - |
| - | - | 1269 | 429.2 | - | - | 0 | - |
| - | - | 3743 | 433.2 | - | - | 0 | - |
| - | - | 2730 | 434.2 | - | - | 0 | - |
| - | - | 5359 | 434.7 | - | - | 0 | - |
| 8 | b | 3585 | 435.2 | 0.001962 | 4.509 | +2 | 8 |
| - | - | 1115 | 435.7 | - | - | 0 | - |
| - | - | 989 | 437.2 | - | - | 0 | - |
| - | - | 906.8 | 437.7 | - | - | 0 | - |
| - | - | 804.9 | 440.2 | - | - | 0 | - |
| - | - | 1237 | 441.2 | - | - | 0 | - |
| - | - | 4977 | 442.2 | - | - | 0 | - |
| - | - | 686.7 | 443.1 | - | - | 0 | - |
| 7 | y | 2777 | 443.2 | 0.006878 | 15.52 | +2 | 8 |
| 8 | b | 1966 | 444.2 | 0.001318 | 2.968 | +2 | 8 |
| - | - | 1942 | 444.7 | - | - | 0 | - |
| - | - | 2434 | 450.2 | - | - | 0 | - |
| - | - | 1189 | 451.2 | - | - | 0 | - |
| - | - | 4252 | 452.2 | - | - | 0 | - |
| - | - | 1461 | 453.2 | - | - | 0 | - |
| - | - | 790.4 | 454.3 | - | - | 0 | - |
| 4 | b | 2217 | 455.3 | 0.001045 | 2.295 | +1 | 4 |
| - | - | 1921 | 459.2 | - | - | 0 | - |
| - | - | 706.2 | 459.7 | - | - | 0 | - |
| - | - | 912.3 | 461.2 | - | - | 0 | - |
| - | - | 5003 | 462.2 | - | - | 0 | - |
| - | - | 1101 | 463.2 | - | - | 0 | - |
| - | - | 953.2 | 464.2 | - | - | 0 | - |
| - | - | 1122 | 466.2 | - | - | 0 | - |
| - | - | 2250 | 468.2 | - | - | 0 | - |
| - | - | 607 | 469.2 | - | - | 0 | - |
| - | - | 2.021E+04 | 470.2 | - | - | 0 | - |
| - | - | 912 | 470.7 | - | - | 0 | - |
| - | - | 4002 | 471.2 | - | - | 0 | - |
| - | - | 718 | 474.2 | - | - | 0 | - |
| - | - | 1360 | 476.2 | - | - | 0 | - |
| - | - | 642.8 | 477.7 | - | - | 0 | - |
| - | - | 2212 | 478.2 | - | - | 0 | - |
| - | - | 2239 | 479.2 | - | - | 0 | - |
| - | - | 1.575E+04 | 480.2 | - | - | 0 | - |
| - | - | 4631 | 481.2 | - | - | 0 | - |
| - | - | 1397 | 482.2 | - | - | 0 | - |
| - | - | 1409 | 482.3 | - | - | 0 | - |
| - | - | 2493 | 484.2 | - | - | 0 | - |
| - | - | 1188 | 488.3 | - | - | 0 | - |
| - | - | 1857 | 494.2 | - | - | 0 | - |
| - | - | 2257 | 494.2 | - | - | 0 | - |
| - | - | 758.3 | 495.2 | - | - | 0 | - |
| - | - | 882.3 | 495.2 | - | - | 0 | - |
| - | - | 5080 | 496.3 | - | - | 0 | - |
| - | - | 1.126E+04 | 497.2 | - | - | 0 | - |
| - | - | 5.567E+04 | 498.2 | - | - | 0 | - |
| - | - | 1.399E+04 | 499.2 | - | - | 0 | - |
| - | - | 2630 | 500.2 | - | - | 0 | - |
| 9 | b | 964.2 | 500.8 | 0.001675 | 3.346 | +2 | 9 |
| - | - | 697.1 | 501.7 | - | - | 0 | - |
| - | - | 3019 | 502.2 | - | - | 0 | - |
| - | - | 928.6 | 508.2 | - | - | 0 | - |
| - | - | 3321 | 510.2 | - | - | 0 | - |
| - | - | 1903 | 510.7 | - | - | 0 | - |
| - | - | 2203 | 511.3 | - | - | 0 | - |
| - | - | 830.3 | 511.8 | - | - | 0 | - |
| - | - | 3743 | 512.2 | - | - | 0 | - |
| - | - | 1314 | 512.2 | - | - | 0 | - |
| - | - | 648.2 | 512.3 | - | - | 0 | - |
| - | - | 1221 | 513.2 | - | - | 0 | - |
| - | - | 2.48E+04 | 513.3 | - | - | 0 | - |
| 10 | y | 5.151E+04 | 514.3 | 0.001074 | 2.088 | +1 | 5 |
| - | - | 1.337E+04 | 515.3 | - | - | 0 | - |
| - | - | 1938 | 516.3 | - | - | 0 | - |
| 5 | b | 1096 | 524.3 | 0.0007298 | 1.392 | +1 | 5 |
| - | - | 1.206E+04 | 530.2 | - | - | 0 | - |
| - | - | 3608 | 531.2 | - | - | 0 | - |
| 10 | y | 6.534E+04 | 531.3 | 0.0004037 | 0.7599 | +1 | 5 |
| - | - | 843 | 532.2 | - | - | 0 | - |
| - | - | 1.628E+04 | 532.3 | - | - | 0 | - |
| - | - | 3146 | 533.3 | - | - | 0 | - |
| - | - | 828.6 | 538.3 | - | - | 0 | - |
| - | - | 2865 | 538.3 | - | - | 0 | - |
| 5 | y | 838.1 | 538.8 | 0.002048 | 3.801 | +2 | 10 |
| - | - | 679.4 | 539.3 | - | - | 0 | - |
| - | - | 876.7 | 542.3 | - | - | 0 | - |
| 5 | b | 2462 | 542.3 | 0.0007852 | 1.448 | +1 | 5 |
| - | - | 884.5 | 543.3 | - | - | 0 | - |
| - | - | 1466 | 546.2 | - | - | 0 | - |
| - | - | 658.9 | 549.5 | - | - | 0 | - |
| - | - | 2044 | 555.3 | - | - | 0 | - |
| 10 | b | 1471 | 556.3 | 0.0015 | 2.696 | +2 | 10 |
| - | - | 702 | 560.3 | - | - | 0 | - |
| - | - | 1128 | 563.2 | - | - | 0 | - |
| - | - | 918.1 | 569.3 | - | - | 0 | - |
| - | - | 1158 | 569.8 | - | - | 0 | - |
| - | - | 1122 | 571.3 | - | - | 0 | - |
| - | - | 1252 | 573.2 | - | - | 0 | - |
| - | - | 844.8 | 573.8 | - | - | 0 | - |
| - | - | 1409 | 574.3 | - | - | 0 | - |
| 4 | y | 4096 | 578.8 | 0.003644 | 6.295 | +2 | 11 |
| - | - | 2238 | 579.3 | - | - | 0 | - |
| - | - | 4779 | 581.3 | - | - | 0 | - |
| - | - | 1454 | 582.3 | - | - | 0 | - |
| - | - | 5151 | 582.8 | - | - | 0 | - |
| - | - | 2807 | 583.3 | - | - | 0 | - |
| - | - | 754 | 583.3 | - | - | 0 | - |
| - | - | 1056 | 583.8 | - | - | 0 | - |
| 4 | y | 9484 | 587.3 | 0.0007809 | 1.33 | +2 | 11 |
| - | - | 7135 | 587.8 | - | - | 0 | - |
| - | - | 3436 | 588.3 | - | - | 0 | - |
| - | - | 887.3 | 588.8 | - | - | 0 | - |
| - | - | 4402 | 591.2 | - | - | 0 | - |
| 11 | b | 1872 | 591.3 | 0.001135 | 1.919 | +2 | 11 |
| 11 | b | 1658 | 591.8 | 0.004855 | 8.203 | +2 | 11 |
| - | - | 1149 | 592.2 | - | - | 0 | - |
| - | - | 672.9 | 592.3 | - | - | 0 | - |
| - | - | 1625 | 593.3 | - | - | 0 | - |
| - | - | 1112 | 597.3 | - | - | 0 | - |
| - | - | 2.461E+04 | 599.3 | - | - | 0 | - |
| - | - | 5680 | 600.3 | - | - | 0 | - |
| - | - | 2313 | 600.8 | - | - | 0 | - |
| - | - | 1120 | 601.3 | - | - | 0 | - |
| - | - | 1340 | 601.3 | - | - | 0 | - |
| - | - | 836.5 | 607.3 | - | - | 0 | - |
| - | - | 1.442E+04 | 609.3 | - | - | 0 | - |
| - | - | 2831 | 609.3 | - | - | 0 | - |
| - | - | 864.6 | 609.8 | - | - | 0 | - |
| - | - | 3960 | 610.3 | - | - | 0 | - |
| - | - | 1286 | 610.3 | - | - | 0 | - |
| - | - | 1087 | 611.3 | - | - | 0 | - |
| 6 | b | 6719 | 611.3 | 0.003345 | 5.471 | +1 | 6 |
| - | - | 1966 | 612.3 | - | - | 0 | - |
| - | - | 1032 | 613.3 | - | - | 0 | - |
| - | - | 2859 | 615.3 | - | - | 0 | - |
| - | - | 840.5 | 617.3 | - | - | 0 | - |
| - | - | 1874 | 617.8 | - | - | 0 | - |
| - | - | 5904 | 618.3 | - | - | 0 | - |
| - | - | 4033 | 618.8 | - | - | 0 | - |
| - | - | 1482 | 619.3 | - | - | 0 | - |
| - | - | 821.7 | 620.3 | - | - | 0 | - |
| - | - | 1558 | 625.3 | - | - | 0 | - |
| - | - | 1442 | 626.3 | - | - | 0 | - |
| 3 | y | 9457 | 626.8 | 0.0003364 | 0.5366 | +2 | 12 |
| - | - | 1.324E+05 | 627.3 | - | - | 0 | - |
| 3 | y | 1.482E+04 | 627.3 | 0.004178 | 6.661 | +2 | 12 |
| - | - | 1.654E+04 | 627.8 | - | - | 0 | - |
| - | - | 4.278E+04 | 628.3 | - | - | 0 | - |
| - | - | 1598 | 628.8 | - | - | 0 | - |
| - | - | 1.046E+04 | 629.3 | - | - | 0 | - |
| 6 | b | 6864 | 629.3 | 0.0006782 | 1.078 | +1 | 6 |
| - | - | 1508 | 630.3 | - | - | 0 | - |
| - | - | 2543 | 630.3 | - | - | 0 | - |
| - | - | 669 | 633.4 | - | - | 0 | - |
| - | - | 1457 | 633.9 | - | - | 0 | - |
| 3 | y | 1.375E+05 | 635.8 | 0.000242 | 0.3806 | +2 | 12 |
| - | - | 8.625E+04 | 636.3 | - | - | 0 | - |
| - | - | 3.957E+04 | 636.8 | - | - | 0 | - |
| - | - | 1.312E+04 | 637.3 | - | - | 0 | - |
| - | - | 2349 | 637.8 | - | - | 0 | - |
| - | - | 4255 | 643.3 | - | - | 0 | - |
| - | - | 2697 | 644.3 | - | - | 0 | - |
| 9 | y | 2.487E+04 | 644.4 | 0.0002021 | 0.3136 | +1 | 6 |
| - | - | 2093 | 645.3 | - | - | 0 | - |
| - | - | 9458 | 645.4 | - | - | 0 | - |
| - | - | 1796 | 646.4 | - | - | 0 | - |
| - | - | 1150 | 658.3 | - | - | 0 | - |
| - | - | 983.4 | 660.3 | - | - | 0 | - |
| - | - | 1744 | 676.3 | - | - | 0 | - |
| - | - | 1797 | 683 | - | - | 0 | - |
| - | - | 1093 | 683.3 | - | - | 0 | - |
| - | - | 1411 | 683.4 | - | - | 0 | - |
| - | - | 1025 | 683.9 | - | - | 0 | - |
| - | - | 1522 | 684.4 | - | - | 0 | - |
| - | - | 4043 | 694.3 | - | - | 0 | - |
| - | - | 1602 | 695.3 | - | - | 0 | - |
| - | - | 1247 | 701.3 | - | - | 0 | - |
| - | - | 3742 | 704.3 | - | - | 0 | - |
| - | - | 819.3 | 705.3 | - | - | 0 | - |
| - | - | 987.8 | 706.3 | - | - | 0 | - |
| 13 | b | 1.725E+04 | 712.4 | 0.013 | 18.25 | +2 | 13 |
| - | - | 7762 | 713.4 | - | - | 0 | - |
| - | - | 2425 | 714.4 | - | - | 0 | - |
| - | - | 1020 | 721.4 | - | - | 0 | - |
| - | - | 7317 | 722.3 | - | - | 0 | - |
| - | - | 3364 | 723.3 | - | - | 0 | - |
| - | - | 832.3 | 724.3 | - | - | 0 | - |
| - | - | 2452 | 730.4 | - | - | 0 | - |
| - | - | 1173 | 731.4 | - | - | 0 | - |
| - | - | 1549 | 738.4 | - | - | 0 | - |
| - | - | 3309 | 739.4 | - | - | 0 | - |
| 7 | b | 4.581E+04 | 740.4 | 0.01383 | 18.68 | +1 | 7 |
| - | - | 1.558E+04 | 741.3 | - | - | 0 | - |
| - | - | 4896 | 742.4 | - | - | 0 | - |
| - | - | 1619 | 743.4 | - | - | 0 | - |
| - | - | 1418 | 753.3 | - | - | 0 | - |
| - | - | 1028 | 754.3 | - | - | 0 | - |
| - | - | 785.9 | 755.3 | - | - | 0 | - |
| 8 | y | 3176 | 755.4 | 2.703E-05 | 0.03579 | +1 | 7 |
| 8 | y | 2826 | 756.4 | 0.005093 | 6.733 | +1 | 7 |
| 0 | Precursor | 1.061E+04 | 757.4 | 0.006021 | 7.949 | +2 | -1 |
| 7 | b | 1.965E+04 | 758.4 | 0.0009318 | 1.229 | +1 | 7 |
| - | - | 8464 | 759.4 | - | - | 0 | - |
| - | - | 1860 | 760.4 | - | - | 0 | - |
| - | - | 1542 | 770.4 | - | - | 0 | - |
| - | - | 3117 | 771.4 | - | - | 0 | - |
| - | - | 1712 | 772.4 | - | - | 0 | - |
| 8 | y | 1.119E+04 | 773.4 | 0.0001505 | 0.1946 | +1 | 7 |
| - | - | 2970 | 774.3 | - | - | 0 | - |
| - | - | 3784 | 774.4 | - | - | 0 | - |
| - | - | 1020 | 775.3 | - | - | 0 | - |
| - | - | 1287 | 775.4 | - | - | 0 | - |
| - | - | 806.4 | 834.4 | - | - | 0 | - |
| - | - | 1211 | 840.4 | - | - | 0 | - |
| - | - | 1417 | 841.4 | - | - | 0 | - |
| - | - | 2518 | 842.4 | - | - | 0 | - |
| - | - | 945.9 | 843.4 | - | - | 0 | - |
| - | - | 3416 | 850.4 | - | - | 0 | - |
| - | - | 3213 | 851.4 | - | - | 0 | - |
| - | - | 850.8 | 852.4 | - | - | 0 | - |
| - | - | 1193 | 853.4 | - | - | 0 | - |
| - | - | 3979 | 859.4 | - | - | 0 | - |
| - | - | 3106 | 860.4 | - | - | 0 | - |
| - | - | 2.147E+04 | 868.4 | - | - | 0 | - |
| 8 | b | 1.482E+04 | 869.4 | 0.002045 | 2.353 | +1 | 8 |
| - | - | 4964 | 870.4 | - | - | 0 | - |
| - | - | 1536 | 871.4 | - | - | 0 | - |
| 7 | y | 1076 | 884.4 | 0.002851 | 3.224 | +1 | 8 |
| 8 | b | 4.045E+04 | 887.4 | 0.0003405 | 0.3837 | +1 | 8 |
| - | - | 2.228E+04 | 888.4 | - | - | 0 | - |
| - | - | 6570 | 889.4 | - | - | 0 | - |
| - | - | 1804 | 890.4 | - | - | 0 | - |
| - | - | 836.6 | 891.4 | - | - | 0 | - |
| - | - | 1033 | 892.4 | - | - | 0 | - |
| - | - | 1006 | 894.4 | - | - | 0 | - |
| - | - | 796 | 895.4 | - | - | 0 | - |
| 7 | y | 2627 | 902.5 | 0.001488 | 1.649 | +1 | 8 |
| - | - | 760.3 | 903.5 | - | - | 0 | - |
| - | - | 2325 | 911.4 | - | - | 0 | - |
| - | - | 1919 | 921.4 | - | - | 0 | - |
| - | - | 2445 | 922.4 | - | - | 0 | - |
| - | - | 1446 | 923.4 | - | - | 0 | - |
| - | - | 9478 | 939.4 | - | - | 0 | - |
| - | - | 5415 | 940.4 | - | - | 0 | - |
| - | - | 1177 | 941.4 | - | - | 0 | - |
| - | - | 1723 | 956.5 | - | - | 0 | - |
| - | - | 1240 | 957.5 | - | - | 0 | - |
| - | - | 1095 | 958.5 | - | - | 0 | - |
| 6 | y | 1772 | 972.5 | 0.01476 | 15.18 | +1 | 9 |
| - | - | 716 | 973.5 | - | - | 0 | - |
| 9 | b | 1231 | 982.5 | 0.005541 | 5.64 | +1 | 9 |
| - | - | 1041 | 983.5 | - | - | 0 | - |
| 6 | y | 2092 | 989.5 | 0.0003133 | 0.3167 | +1 | 9 |
| - | - | 2364 | 990.5 | - | - | 0 | - |
| 9 | b | 6283 | 1000 | 0.001214 | 1.213 | +1 | 9 |
| - | - | 4438 | 1002 | - | - | 0 | - |
| - | - | 776.2 | 1003 | - | - | 0 | - |
| - | - | 698 | 1018 | - | - | 0 | - |
| - | - | 1348 | 1025 | - | - | 0 | - |
| - | - | 1005 | 1035 | - | - | 0 | - |
| - | - | 2895 | 1036 | - | - | 0 | - |
| - | - | 1533 | 1037 | - | - | 0 | - |
| - | - | 1075 | 1041 | - | - | 0 | - |
| - | - | 1.008E+04 | 1053 | - | - | 0 | - |
| - | - | 7101 | 1054 | - | - | 0 | - |
| - | - | 2270 | 1055 | - | - | 0 | - |
| 5 | y | 1124 | 1059 | 0.0006589 | 0.6225 | +1 | 10 |
| 5 | y | 2385 | 1059 | 0.003595 | 3.393 | +1 | 10 |
| - | - | 1212 | 1060 | - | - | 0 | - |
| - | - | 1024 | 1072 | - | - | 0 | - |
| 5 | y | 5581 | 1077 | 0.0001152 | 0.107 | +1 | 10 |
| - | - | 2475 | 1078 | - | - | 0 | - |
| - | - | 1046 | 1079 | - | - | 0 | - |
| 10 | b | 1095 | 1129 | 0.001258 | 1.115 | +1 | 10 |
| - | - | 808.7 | 1130 | - | - | 0 | - |
| - | - | 1048 | 1139 | - | - | 0 | - |
| 4 | y | 848.7 | 1156 | 0.006758 | 5.848 | +1 | 11 |
| 4 | y | 2000 | 1157 | 0.002892 | 2.501 | +1 | 11 |
| - | - | 1653 | 1158 | - | - | 0 | - |
| 4 | y | 7281 | 1174 | 0.001487 | 1.267 | +1 | 11 |
| - | - | 3842 | 1175 | - | - | 0 | - |
| - | - | 1338 | 1176 | - | - | 0 | - |
| - | - | 709.1 | 2192 | - | - | 0 | - |
| - | - | 636.2 | 3042 | - | - | 0 | - |

m/z Charge Intensity FragmentType MassShift Position
120.0811767578125 0 113825.945
121.08451843261719 0 10925.296
122.0878677368164 0 347.9741
125.07164764404297 0 447.29175
125.98983764648438 0 414.76
126.05533599853516 0 6852.2207
127.05076599121094 0 1013.8888
127.05840301513672 0 454.26663
127.08721923828125 0 1247.3428
128.546630859375 0 462.23505
129.0547637939453 0 438.56732
129.0662384033203 0 6094.676
129.10264587402344 0 97707.9
130.0502166748047 0 7682.303
130.06546020507812 0 4172.346
130.08676147460938 0 969.24066
130.100341796875 0 1102.464
130.10595703125 0 7024.422
131.06895446777344 0 588.4824
131.1182403564453 0 5979.5063
132.1023712158203 0 1289.2167
133.4365234375 0 398.58713
134.02749633789062 0 1253.4938
136.07603454589844 0 5900.329
137.07945251464844 0 1982.9315
138.12808227539062 0 3014.3538
139.05078125 0 1023.0867
139.0869903564453 0 3110.0706
140.0823516845703 0 1131.4253
140.10789489746094 0 480.26263
141.0662078857422 0 2875.3162
141.10263061523438 0 1851.9346
142.09799194335938 0 846.7069
143.08187866210938 0 573.57275
143.11842346191406 0 499.12973
145.06101989746094 0 1934.5121
146.0606231689453 0 740.9688
147.07669067382812 0 2659.3147
147.11312866210938 0 2020.169
148.06056213378906 0 488.33276
152.07095336914062 0 2257.6853
153.0659637451172 0 472.61252
154.05062866210938 0 596.90265
154.0866241455078 0 3116.0806
154.09791564941406 0 5406.9004
155.04559326171875 0 471.80432
155.0818634033203 0 5678.4375
155.11822509765625 0 1721.718
157.061279296875 0 2576.8196
157.09754943847656 0 8072.1855
157.10867309570312 0 512.03925
158.09291076660156 0 967.44257
158.1002655029297 0 689.9302
159.11306762695312 0 787.59296
160.09544372558594 0 920.12616
163.07168579101562 0 1703.0693
163.08749389648438 0 936.0244
165.1027374267578 0 1395.1892
166.06143188476562 0 958.61786
167.08180236816406 0 5682.321
167.11825561523438 0 5983.5425
169.0611572265625 0 2173.1477
169.09756469726562 0 1463.9832
170.06057739257812 0 667.1588
171.0767822265625 0 1291.6377
171.14968872070312 0 677.81396
172.0722198486328 0 1367.71
172.10879516601562 0 2913.1895
173.12876892089844 0 1356.0664
174.05540466308594 0 567.57294
174.13502502441406 0 539.12213
175.0716552734375 0 2658.6895
179.11814880371094 0 575.54565
180.11343383789062 0 1032.6486
181.06106567382812 0 623.63525
181.0974578857422 0 1823.9271
182.0928192138672 0 7772.352
182.12911987304688 0 891.43756
183.0767364501953 0 5683.674
183.09664916992188 0 564.29663
183.1131591796875 0 1005.7918
183.14947509765625 0 1091.7435
184.10841369628906 0 5269.4478
185.09242248535156 0 10631.373
185.11195373535156 0 731.19904
185.16488647460938 0 661.4372
186.0877685546875 0 3757.8125
186.09568786621094 0 679.77325
186.12380981445312 0 1795.861
187.0721893310547 0 776.723
187.10826110839844 0 993.4209
187.14439392089844 0 1297.944
189.08743286132812 0 3549.769
191.08216857910156 0 1289.5911
195.0765838623047 0 1500.2874
195.1131134033203 0 11876.274
196.1165771484375 0 1027.4763
197.12879943847656 0 10743.725
198.12403869628906 0 4716.2856
198.13092041015625 0 635.58966
199.0717315673828 0 7284.9956
199.10813903808594 0 3542.3296
199.11920166015625 0 1229.8348
200.07530212402344 0 581.25104
200.10317993164062 0 6121.943
200.1118927001953 0 770.0818
200.1398468017578 0 2200.346
201.0872344970703 0 617.36084
201.10707092285156 0 910.67487
201.12367248535156 0 10918.261 y Ammonia loss 12
201.13363647460938 0 859.5232
202.10916137695312 0 684.3553
202.11895751953125 0 2026.3295 y 10
203.10279846191406 0 940.41315
203.14112854003906 0 456.44623
204.10626220703125 0 1139.208
207.11322021484375 0 1401.9763
208.09715270996094 0 1118.2723
208.10845947265625 0 6565.0757
209.09246826171875 0 22559.307
210.09597778320312 0 1664.5682
210.12432861328125 0 660.8965
211.10797119140625 0 989.38544
211.14422607421875 0 1116.2307
212.13987731933594 0 1592.2852
213.0873260498047 0 1508.6321
213.11268615722656 0 858.10583
213.12388610839844 0 1958.9113
213.14251708984375 0 531.0284
214.08241271972656 0 804.61755
214.15591430664062 0 626.7118
215.103515625 0 545.4895
215.13937377929688 0 5655.374
217.08233642578125 0 9324.54
217.13377380371094 0 3660.62
218.0865936279297 0 566.59076
218.15032958984375 0 43440.293 y 12
219.1534881591797 0 4157.8364
221.0591278076172 0 1474.9175
221.1041717529297 0 778.8821
221.6237335205078 0 991.3523
223.10830688476562 0 702.41766
223.156005859375 0 1039.9384
225.1236572265625 0 7270.527
225.1352081298828 0 3730.473
226.11903381347656 0 79305.15
227.10301208496094 0 4020.6003
227.12234497070312 0 8392.556
228.1237030029297 0 540.85333
228.13467407226562 0 6278.9097 b 3
229.13552856445312 0 671.93243
229.155029296875 0 1233.1019
231.0980987548828 0 1625.2042
233.16534423828125 0 152472.67 a 1
234.087158203125 0 779.52057
234.124267578125 0 2011.9541
234.14683532714844 0 1075.5017
234.16856384277344 0 21669.898
235.17178344726562 0 1909.035
235.615966796875 0 635.58826
236.1033477783203 0 3394.427
236.13986206054688 0 1007.0234
239.11473083496094 0 654.74445
239.15025329589844 0 585.2858
240.09776306152344 0 1504.1191
240.13507080078125 0 1391.5789
241.08236694335938 0 7044.8257
242.08560180664062 0 631.81635
242.15036010742188 0 16559.738
243.13397216796875 0 1543.5502
243.14566040039062 0 20144.854
244.09356689453125 0 748.5087
244.1300811767578 0 5959.686
244.14881896972656 0 2331.6956
245.07728576660156 0 770.38275
245.12913513183594 0 2883.5754
248.6297149658203 0 1553.7528
249.12176513671875 0 6614.361
249.6232452392578 0 935.7732
250.15478515625 0 721.704
251.15090942382812 0 3350.5803
252.09815979003906 0 1448.7565
252.13455200195312 0 2636.8757
254.11395263671875 0 12206.204
254.150634765625 0 1954.0468
255.09820556640625 0 1083.1327
255.11737060546875 0 1191.7346
257.1285095214844 0 700.44745
257.14300537109375 0 12326.419
257.63427734375 0 2371.2004 y Ammonia loss 9
257.64532470703125 0 2274.0796
258.108642578125 0 706.86554
258.1453857421875 0 2307.986
259.0927734375 0 8181.176
260.0964660644531 0 717.1869
261.1600646972656 0 61333.89 b 1
262.1189270019531 0 1097.6628
262.1634826660156 0 11167.144
263.1641845703125 0 604.62634
264.13458251953125 0 8798.56
265.1379089355469 0 1078.2428
266.1483154296875 0 9190.956 y 9
266.64984130859375 0 2692.2651
268.09283447265625 0 1511.147
268.1669006347656 0 583.4699
269.1607360839844 0 1407.2871
269.6531066894531 0 587.6787
272.1243591308594 0 5478.895
273.12445068359375 0 809.64825
278.15057373046875 0 1120.0367
278.1676025390625 0 986.61084
279.110107421875 0 757.68567
279.1458435058594 0 5020.6416
280.1295471191406 0 7107.0044
281.1145935058594 0 550.4289
281.13226318359375 0 762.92365
282.1084289550781 0 1061.4735
282.12908935546875 0 934.8023
282.145263671875 0 27596.719
282.6293029785156 0 637.95557
283.1477966308594 0 4502.2485
285.1981506347656 0 833.08044
286.1039733886719 0 3883.072
292.16571044921875 0 633.5944
296.08734130859375 0 691.761
296.1357727050781 0 3604.9744
296.172119140625 0 3749.6392
297.1199035644531 0 4253.4175
297.1564025878906 0 27010.18
298.10394287109375 0 893.1608
298.12335205078125 0 988.32635
298.140380859375 0 3704.1511
298.1600036621094 0 3769.0022
299.14398193359375 0 875.63837
299.171630859375 0 2863.5823
300.11834716796875 0 677.42285
300.15704345703125 0 1099.625
304.1142272949219 0 3202.152
305.13055419921875 0 1221.8121
305.160400390625 0 682.0578
306.1455383300781 0 1239.6235
306.16436767578125 0 1056.0574 b Water loss 5
307.1402587890625 0 1241.2148
308.1606140136719 0 2653.6997
310.1041259765625 0 1403.7085
313.1624755859375 0 718.4174
313.1878662109375 0 2964.1401
314.1340637207031 0 2824.0608
314.1471252441406 0 3259.054
314.1806640625 0 9077.587 y Ammonia loss 8
314.6365051269531 0 1443.8604
314.6783752441406 0 1644.8918
315.1480712890625 0 1150.4209
315.16693115234375 0 11049.072 b 5
315.1871337890625 0 1299.4846
316.1499328613281 0 674.37274
316.16949462890625 0 2023.1957
319.1401672363281 0 549.782
321.1563415527344 0 1331.8618
322.6905212402344 0 12969.082 y 8
323.17254638671875 0 1173.8213
323.19189453125 0 3172.7366
323.6938171386719 0 581.7035
325.1514892578125 0 1426.3813
327.16754150390625 0 809.5375
328.1142272949219 0 2440.4673
328.16961669921875 0 649.63354 b Water loss 8
329.6641540527344 0 975.9782
330.16558837890625 0 1307.7554
332.19342041015625 0 31533.488 y 11
333.1561279296875 0 5128.2695
333.17767333984375 0 1051.8452
333.1966552734375 0 4533.5947
336.1561279296875 0 1011.3385
337.1514587402344 0 793.6412
338.1343994140625 0 1023.2523
338.6689758300781 0 4222.2466
339.1685485839844 0 3516.6667
340.1701354980469 0 766.9865
341.1829528808594 0 4702.4863
342.1820068359375 0 1565.0258
343.1614074707031 0 1608.0975
343.18896484375 0 1621.9597
344.1637268066406 0 956.5865
344.1927795410156 0 585.3254
346.1249084472656 0 6755.877
347.129638671875 0 1896.5667
347.6745300292969 0 2712.2876
348.17840576171875 0 1010.5514
350.181396484375 0 717.87476
351.1667785644531 0 22371.422
352.1698913574219 0 4330.5615
353.18218994140625 0 6352.7407
354.1660461425781 0 1501.4491
354.1876525878906 0 923.43317
355.1628112792969 0 2493.2979
358.2127380371094 0 3444.3677 b 2
361.67681884765625 0 1579.9392
362.175048828125 0 1000.1049
365.1459045410156 0 4099.434
366.6632995605469 0 727.56525
368.1929016113281 0 6927.1665
369.17724609375 0 25514.336
370.1802673339844 0 4931.461
370.6782531738281 0 8984.074 b Water loss 6
371.1809387207031 0 2418.6992 b Ammonia loss 9
372.1774597167969 0 1436.6802
373.1726379394531 0 3723.8428
374.1756286621094 0 879.6158
375.1402893066406 0 627.1666
375.1710205078125 0 726.7275
375.66595458984375 0 1545.473
377.1766357421875 0 587.2156
379.1924133300781 0 1450.3092
379.690673828125 0 2015.8038 b 6
380.1939392089844 0 686.3591
381.1525573730469 0 786.34845
382.66168212890625 0 1376.6763
383.15679931640625 0 5681.5747
384.1581115722656 0 960.8074
385.2198181152344 0 3357.1558
386.2039794921875 0 17527.889 y Ammonia loss 10
387.2076721191406 0 3925.462 y 7
389.19000244140625 0 664.2563
397.13623046875 0 1536.6841
401.16741943359375 0 10693.964
402.16998291015625 0 2187.1306
403.2305908203125 0 79760.734 y 10
404.2333984375 0 13708.739
405.2357177734375 0 1925.1846
407.2042236328125 0 4274.394
408.18853759765625 0 11522.907
409.191162109375 0 2714.6255
410.20416259765625 0 1445.7219
411.2033386230469 0 700.4639
413.2047424316406 0 836.16974
414.1980895996094 0 1264.217
415.1474304199219 0 2691.0056
416.14910888671875 0 820.7675
418.2134094238281 0 1092.088 y Water loss 2
418.5401916503906 0 1258.8196 y Ammonia loss 2
418.8747863769531 0 832.01215
423.1867980957031 0 786.3299
424.2281799316406 0 5223.487
424.5496520996094 0 1053.7096
425.21533203125 0 6999.697
425.7015380859375 0 1010.50037
426.2001647949219 0 2296.4604
427.2315673828125 0 1301.9567
429.21563720703125 0 1268.645
433.157470703125 0 3743.3994
434.2047119140625 0 2729.6738
434.70635986328125 0 5358.962
435.20758056640625 0 3585.3054 b Water loss 7
435.7076721191406 0 1115.1904
437.20501708984375 0 988.95703
437.7106628417969 0 906.7886
440.2155456542969 0 804.9363
441.199951171875 0 1236.7164
442.2398986816406 0 4977.2036
443.141845703125 0 686.6973
443.22613525390625 0 2776.7378 y Ammonia loss 6
444.21221923828125 0 1965.5991 b 7
444.7121887207031 0 1942.0222
450.1990966796875 0 2433.6038
451.23095703125 0 1188.6915
452.2140808105469 0 4251.954
453.2166748046875 0 1460.7433
454.26641845703125 0 790.4072
455.2663269042969 0 2217.4636 b 3
459.208251953125 0 1921.2833
459.7220153808594 0 706.22034
461.2201232910156 0 912.3456
462.1991882324219 0 5002.933
463.2037658691406 0 1101.4987
464.21307373046875 0 953.1521
466.1916809082031 0 1121.9119
468.20947265625 0 2249.9458
469.2393798828125 0 606.95605
470.22528076171875 0 20208.316
470.7281188964844 0 911.9595
471.2293395996094 0 4002.388
474.2312316894531 0 717.9875
476.1774597167969 0 1360.0372
477.7445983886719 0 642.78186
478.24151611328125 0 2212.0115
479.2266845703125 0 2238.9385
480.2099304199219 0 15753.987
481.2132263183594 0 4630.7725
482.22064208984375 0 1396.729
482.25982666015625 0 1408.8105
484.2039794921875 0 2492.6956
488.2765197753906 0 1188.1697
494.1895446777344 0 1856.5039
494.2340393066406 0 2256.6208
495.1907653808594 0 758.29315
495.2344970703125 0 882.2849
496.2520751953125 0 5080.263
497.23638916015625 0 11259.265
498.22039794921875 0 55665.438
499.22332763671875 0 13990.274
500.2278137207031 0 2629.6619
500.7546081542969 0 964.15466 b 8
501.7286071777344 0 697.14575
502.21649169921875 0 3019.377
508.2489013671875 0 928.5592
510.24407958984375 0 3321.2705
510.7449645996094 0 1902.7135
511.2525939941406 0 2202.8384
511.7541198730469 0 830.2903
512.1990356445312 0 3742.6768
512.2355346679688 0 1314.4242
512.3478393554688 0 648.2454
513.206298828125 0 1221.2927
513.2785034179688 0 24804.531
514.2630615234375 0 51506.293 y Ammonia loss 9
515.2650146484375 0 13372.261
516.2652587890625 0 1938.3419
524.2874755859375 0 1096.0116 b Water loss 4
530.2099609375 0 12059.913
531.2134399414062 0 3607.505
531.2889404296875 0 65340.65 y 9
532.2173461914062 0 843.01117
532.2913818359375 0 16282.023
533.2932739257812 0 3146.4233
538.259033203125 0 828.6079
538.2993774414062 0 2865.0183
538.7625122070312 0 838.1182 y 4
539.302001953125 0 679.3597
542.2576293945312 0 876.7385
542.298095703125 0 2462.0178 b 4
543.302001953125 0 884.5469
546.2389526367188 0 1465.8611
549.4910888671875 0 658.89825
555.3252563476562 0 2043.6488
556.2704467773438 0 1470.6833 b Ammonia loss 9
560.271240234375 0 702.03094
563.2468872070312 0 1128.0338
569.2572631835938 0 918.0602
569.7766723632812 0 1157.8501
571.2772216796875 0 1122.183
573.230224609375 0 1251.739
573.7744750976562 0 844.8112
574.2711181640625 0 1409.3008
578.7813110351562 0 4095.9368 y Ammonia loss 3
579.2798461914062 0 2238.0457
581.2578125 0 4779.4263
582.292236328125 0 1453.6366
582.78271484375 0 5150.705
583.2811279296875 0 2807.049
583.326416015625 0 753.9866
583.7830810546875 0 1055.5673
587.2901611328125 0 9483.599 y 3
587.7930297851562 0 7134.838
588.2940063476562 0 3435.668
588.791015625 0 887.29736
591.2412719726562 0 4402.173
591.296630859375 0 1872.2478 b Water loss 10
591.7923583984375 0 1657.5178 b Ammonia loss 10
592.242919921875 0 1149.2501
592.3034057617188 0 672.93744
593.3059692382812 0 1625.0747
597.285888671875 0 1111.5897
599.2677001953125 0 24610.107
600.2723388671875 0 5680.1504
600.80029296875 0 2313.0981
601.2794189453125 0 1120.0154
601.3350219726562 0 1339.8871
607.3009033203125 0 836.4684
609.2517700195312 0 14422.982
609.3365478515625 0 2831.048
609.7887573242188 0 864.639
610.2545776367188 0 3960.052
610.33935546875 0 1286.3596
611.258056640625 0 1086.7743
611.3154296875 0 6718.742 b Water loss 5
612.31494140625 0 1966.0366
613.3182373046875 0 1032.0961
615.298095703125 0 2858.946
617.2925415039062 0 840.52814
617.8068237304688 0 1874.3845
618.3002319335938 0 5904.0957
618.7980346679688 0 4032.6697
619.2978515625 0 1482.1619
620.3045654296875 0 821.7022
625.2862548828125 0 1558.1833
626.2825927734375 0 1441.8721
626.8123779296875 0 9456.873 y Water loss 2
627.26220703125 0 132425.56
627.3082275390625 0 14820.957 y Ammonia loss 2
627.8067016601562 0 16537.184
628.265380859375 0 42784.81
628.8112182617188 0 1597.78
629.26708984375 0 10460.206
629.3300170898438 0 6863.749 b 5
630.2688598632812 0 1507.5686
630.333251953125 0 2542.903
633.352783203125 0 668.9657
633.852294921875 0 1457.2703
635.8175659179688 0 137454.08 y 2
636.319091796875 0 86250.8
636.8202514648438 0 39574.26
637.3214111328125 0 13115.36
637.8212280273438 0 2348.8833
643.2949829101562 0 4255.2656
644.2960815429688 0 2696.6045
644.372802734375 0 24865.46 y 8
645.287841796875 0 2092.621
645.3763427734375 0 9458.413
646.379150390625 0 1795.6698
658.3047485351562 0 1149.6617
660.3226318359375 0 983.41327
676.3281860351562 0 1744.4097
682.9649047851562 0 1797.262
683.3017578125 0 1092.7175
683.3865966796875 0 1410.5085
683.8866577148438 0 1025.2076
684.3687133789062 0 1522.177
694.3427734375 0 4043.3518
695.3402709960938 0 1601.7563
701.2711181640625 0 1247.2546
704.3242797851562 0 3741.5388
705.3302612304688 0 819.2917
706.33349609375 0 987.8016
712.3514404296875 0 17247.9 b Water loss 12
713.3553466796875 0 7762.0083
714.356689453125 0 2424.9023
721.35302734375 0 1019.59143
722.337890625 0 7317.4805
723.34228515625 0 3364.1267
724.3463745117188 0 832.31573
730.37646484375 0 2451.8264
731.3811645507812 0 1172.798
738.3773803710938 0 1549.0795
739.366455078125 0 3309.2334
740.3475341796875 0 45809.156 b Water loss 6
741.3499755859375 0 15584.679
742.3538818359375 0 4895.816
743.3548583984375 0 1618.8027
753.3441162109375 0 1417.6823
754.3359985351562 0 1027.9231
755.3341674804688 0 785.91736
755.4046020507812 0 3175.9055 y Water loss 7
756.3937377929688 0 2826.2668 y Ammonia loss 7
757.374267578125 0 10611.298 Precursor Ammonia loss
758.3728637695312 0 19653.7 b 6
759.375732421875 0 8464.094
760.3805541992188 0 1859.713
770.3689575195312 0 1541.6353
771.3551025390625 0 3116.677
772.3529663085938 0 1712.2092
773.4153442382812 0 11193.714 y 7
774.3330078125 0 2969.8125
774.4193725585938 0 3784.0005
775.3385620117188 0 1019.6962
775.4174194335938 0 1286.9557
834.3660278320312 0 806.41345
840.4117431640625 0 1210.6709
841.4100952148438 0 1417.3206
842.3944091796875 0 2518.4658
843.3919067382812 0 945.9248
850.3941650390625 0 3415.9434
851.3920288085938 0 3212.6296
852.3869018554688 0 850.7595
853.4324951171875 0 1192.9199
859.4169921875 0 3978.727
860.4193115234375 0 3106.085
868.4049072265625 0 21467.88
869.406005859375 0 14817.931 b Water loss 7
870.4083251953125 0 4964.1855
871.4110107421875 0 1536.2511
884.4500732421875 0 1075.7413 y Water loss 6
887.4141845703125 0 40452.785 b 7
888.417724609375 0 22277.785
889.4207763671875 0 6570.143
890.42333984375 0 1803.5769
891.3981323242188 0 836.6416
892.3873291015625 0 1033.1423
894.420166015625 0 1005.6416
895.4307861328125 0 795.969
902.456298828125 0 2626.935 y 6
903.457763671875 0 760.28754
911.4448852539062 0 2324.6655
921.4285888671875 0 1918.6766
922.4208374023438 0 2444.5051
923.4186401367188 0 1445.5549
939.4402465820312 0 9478.186
940.4435424804688 0 5415.374
941.4476318359375 0 1177.4242
956.4590454101562 0 1723.3513
957.47216796875 0 1240.1827
958.4553833007812 0 1095.0505
972.47802734375 0 1772.052 y Ammonia loss 5
973.4910888671875 0 716.0176
982.4824829101562 0 1231.3271 b Water loss 8
983.4879150390625 0 1040.821
989.489501953125 0 2091.5505 y 5
990.4889526367188 0 2364.1448
1000.4973754882812 0 6282.7446 b 8
1001.5000610351562 0 4437.9805
1002.5056762695312 0 776.2159
1018.4506225585938 0 698.01843
1025.483154296875 0 1347.7786
1035.469970703125 0 1004.96954
1036.45947265625 0 2894.5986
1037.46435546875 0 1532.9739
1041.480224609375 0 1074.8735
1053.4840087890625 0 10077.229
1054.484375 0 7101.0435
1055.4835205078125 0 2269.5642
1058.5106201171875 0 1124.3148 y Water loss 4
1059.49169921875 0 2385.3853 y Ammonia loss 4
1060.4969482421875 0 1212.0133
1071.5040283203125 0 1024.2147
1076.521728515625 0 5581.1143 y 4
1077.5224609375 0 2474.9268
1078.5252685546875 0 1046.0374
1128.555908203125 0 1095.161 b 9
1129.5648193359375 0 808.73376
1138.5406494140625 0 1047.8387
1155.57080078125 0 848.73914 y Water loss 3
1156.545166015625 0 1999.8416 y Ammonia loss 3
1157.5582275390625 0 1653.4907
1173.5731201171875 0 7281.4478 y 3
1174.5760498046875 0 3841.9475
1175.5726318359375 0 1337.717
2192.373291015625 0 709.10675
3041.540283203125 0 636.2201

Spectrum Details

|  |  |
| --- | --- |
| Matched peaks? Matched peaks The total absolute number of peaks matched. Additionally in brackets the total fraction of peaks matched and the total number of peaks is shown. | 66 (10.51% of 628) |
| FDR? FDR The false discovery rate estimated for this peptide. It is calculated by matching all theoretical fragments with a non-integer shift with the raw peaks for this spectrum. This is done with 40 different shifts. The resulting percentage is the average number of annotated peaks over the number of annotated peaks with the correct spectrum. | 0.36% |
| Satellite FDR? Satellite FDR See the FDR for details on its calculation. This satellite ion specific FDR only contains the satellite ions (d/w) for I/L/J positions. | - |
| PSM Score? PSM Score The PSM Score as given by Hecklib to this annotated spectrum. It is shown with three significant figures. | 569 |

#### Reverse Lookup? Reverse Lookup All places where this read could be placed.

| Group | Segment | Template | Template Part | Read Part | Score | Unique |
| --- | --- | --- | --- | --- | --- | --- |
| Homo sapiens Light Chain | IGLC | IGLC2 | [10..24] | [0..14] | 112 | False |
| Homo sapiens Light Chain | IGLC | IGLC3 | [8..22] | [0..14] | 112 | False |
| Homo sapiens Light Chain | IGLC | IGLC6 | [10..24] | [0..14] | 112 | False |
| Homo sapiens Light Chain | IGLC | IGLC7 | [10..24] | [0..14] | 112 | False |

| Recombined | Template Part | Read Part | Score | Unique |
| --- | --- | --- | --- | --- |
| REC-0-1\_002 | [121..135] | [0..14] | 112 | True |

#### Meta Information from Multiple reads

##### Number of combined reads

8

##### Intensity

0.7531

##### TotalArea

2.821E+09

##### Changes to the peptide sequence

JFPPSSEEJQANKA

L→JNo support for either Leucine or Isoleucine based on side chain ions (Position: 9)

J→LSupport for Leucine based on side chain ions (1 for L 0 for I) (Position: 9)

L→JEqual support for both Leucine and Isoleucine based on side chain ions (1 ions for both) (Position: 9)

L→JNo support for either Leucine or Isoleucine based on side chain ions (Position: 1)

#### Positional Score

Copy Data

##### Positional Score (TSV)

###### Preview

```
Loading example...
```

*Click on the button to copy the data to your clipboard.*

10012345678910111213

Label Value
"0" 0.75
"1" 0.73
"2" 0.734
"3" 0.742
"4" 0.749
"5" 0.746
"6" 0.746
"7" 0.749
"8" 0.75
"9" 0.738
"10" 0.738
"11" 0.731
"12" 0.647
"13" 0.691

#### Meta Information from PEAKS

##### Scan Identifier

F1:6428

##### Original sequence

L

F

P

P

S

S

E

E

L

Q

A

N

K

A

##### Posttranslational Modifications

##### Source File

D:\separate\_stitch\_analyses\xle-disambiguation\raw\20210323\_F1\_UM1\_Peng0013\_SA\_F59\_ingel\_3ug\_ELA.raw

##### Fraction

1

##### Scan Feature

F1:15397

##### De Novo Score

98

##### ConfidenceScore

98

### m/z

765.8953

##### Mass

1529.7725

##### Charge

2

##### Retention Time

34.15

##### Predicted Retention Time

-

##### Area

1.295E+09

##### Parts Per Million

2.3

##### Fragmentation mode

ETHCD

##### Originating file

01 D:\separate\_stitch\_analyses\xle-disambiguation\20210325\_F59\_3ug\_DENOVO\_12.csv

#### Meta Information from PEAKS

##### Scan Identifier

F1:6165

##### Original sequence

L

F

P

P

S

S

E

E

L

Q

A

N

K

A

##### Posttranslational Modifications

##### Source File

D:\separate\_stitch\_analyses\xle-disambiguation\raw\20210323\_F1\_UM1\_Peng0013\_SA\_F59\_ingel\_3ug\_ELA.raw

##### Fraction

1

##### Scan Feature

F1:4455

##### De Novo Score

98

##### ConfidenceScore

98

### m/z

510.9326

##### Mass

1529.7725

##### Charge

3

##### Retention Time

34.15

##### Predicted Retention Time

-

##### Area

3.838E+07

##### Parts Per Million

2.3

##### Fragmentation mode

ETHCD

##### Originating file

01 D:\separate\_stitch\_analyses\xle-disambiguation\20210325\_F59\_3ug\_DENOVO\_12.csv

#### Meta Information from PEAKS

##### Scan Identifier

F1:6286

##### Original sequence

L

F

P

P

S

S

E

E

L

Q

A

N

K

A

##### Posttranslational Modifications

##### Source File

D:\separate\_stitch\_analyses\xle-disambiguation\raw\20210323\_F1\_UM1\_Peng0013\_SA\_F59\_ingel\_3ug\_ELA.raw

##### Fraction

1

##### Scan Feature

F1:4455

##### De Novo Score

98

##### ConfidenceScore

98

### m/z

510.9326

##### Mass

1529.7725

##### Charge

3

##### Retention Time

34.15

##### Predicted Retention Time

-

##### Area

3.838E+07

##### Parts Per Million

2.3

##### Fragmentation mode

ETHCD

##### Originating file

01 D:\separate\_stitch\_analyses\xle-disambiguation\20210325\_F59\_3ug\_DENOVO\_12.csv

#### Meta Information from PEAKS

##### Scan Identifier

F1:6136

##### Original sequence

L

F

P

P

S

S

E

E

L

Q

A

N

K

A

##### Posttranslational Modifications

##### Source File

D:\separate\_stitch\_analyses\xle-disambiguation\raw\20210323\_F1\_UM1\_Peng0013\_SA\_F59\_ingel\_3ug\_ELA.raw

##### Fraction

1

##### Scan Feature

F1:15397

##### De Novo Score

98

##### ConfidenceScore

98

### m/z

765.8953

##### Mass

1529.7725

##### Charge

2

##### Retention Time

34.15

##### Predicted Retention Time

-

##### Area

1.295E+09

##### Parts Per Million

2.3

##### Fragmentation mode

ETHCD

##### Originating file

01 D:\separate\_stitch\_analyses\xle-disambiguation\20210325\_F59\_3ug\_DENOVO\_12.csv

#### Meta Information from PEAKS

##### Scan Identifier

F1:6225

##### Original sequence

L

F

P

P

S

S

E

E

L

Q

A

N

K

A

##### Posttranslational Modifications

##### Source File

D:\separate\_stitch\_analyses\xle-disambiguation\raw\20210323\_F1\_UM1\_Peng0013\_SA\_F59\_ingel\_3ug\_ELA.raw

##### Fraction

1

##### Scan Feature

F1:4455

##### De Novo Score

98

##### ConfidenceScore

98

### m/z

510.9326

##### Mass

1529.7725

##### Charge

3

##### Retention Time

34.15

##### Predicted Retention Time

-

##### Area

3.838E+07

##### Parts Per Million

2.3

##### Fragmentation mode

ETHCD

##### Originating file

01 D:\separate\_stitch\_analyses\xle-disambiguation\20210325\_F59\_3ug\_DENOVO\_12.csv

#### Meta Information from PEAKS

##### Scan Identifier

F1:6410

##### Original sequence

L

F

P

P

S

S

E

E

L

Q

A

N

K

A

##### Posttranslational Modifications

##### Source File

D:\separate\_stitch\_analyses\xle-disambiguation\raw\20210323\_F1\_UM1\_Peng0013\_SA\_F59\_ingel\_3ug\_ELA.raw

##### Fraction

1

##### Scan Feature

F1:4455

##### De Novo Score

97

##### ConfidenceScore

97

### m/z

510.9326

##### Mass

1529.7725

##### Charge

3

##### Retention Time

34.15

##### Predicted Retention Time

-

##### Area

3.838E+07

##### Parts Per Million

2.3

##### Fragmentation mode

HCD

##### Originating file

01 D:\separate\_stitch\_analyses\xle-disambiguation\20210325\_F59\_3ug\_DENOVO\_12.csv

#### Meta Information from PEAKS

##### Scan Identifier

F1:6346

##### Original sequence

L

F

P

P

S

S

E

E

L

Q

A

N

K

A

##### Posttranslational Modifications

##### Source File

D:\separate\_stitch\_analyses\xle-disambiguation\raw\20210323\_F1\_UM1\_Peng0013\_SA\_F59\_ingel\_3ug\_ELA.raw

##### Fraction

1

##### Scan Feature

F1:4455

##### De Novo Score

96

##### ConfidenceScore

96

### m/z

510.9326

##### Mass

1529.7725

##### Charge

3

##### Retention Time

34.15

##### Predicted Retention Time

-

##### Area

3.838E+07

##### Parts Per Million

2.3

##### Fragmentation mode

HCD

##### Originating file

01 D:\separate\_stitch\_analyses\xle-disambiguation\20210325\_F59\_3ug\_DENOVO\_12.csv

#### Meta Information from PEAKS

##### Scan Identifier

F1:6472

##### Original sequence

L

F

P

P

S

S

E

E

L

Q

A

N

K

A

##### Posttranslational Modifications

##### Source File

D:\separate\_stitch\_analyses\xle-disambiguation\raw\20210323\_F1\_UM1\_Peng0013\_SA\_F59\_ingel\_3ug\_ELA.raw

##### Fraction

1

##### Scan Feature

F1:4455

##### De Novo Score

96

##### ConfidenceScore

96

### m/z

510.9326

##### Mass

1529.7725

##### Charge

3

##### Retention Time

34.15

##### Predicted Retention Time

-

##### Area

3.838E+07

##### Parts Per Million

2.3

##### Fragmentation mode

HCD

##### Originating file

01 D:\separate\_stitch\_analyses\xle-disambiguation\20210325\_F59\_3ug\_DENOVO\_12.csv
