## Supplementary material for "A handle on mass coincidence errors in *de novo* sequencing of antibodies by bottom-up proteomics": Combined_056.html

Details Combined\_056 | Stitch OverviewUndefined

### Read Combined\_056

#### Sequence (length=15)

JHQDWLDGKEYKCKK

#### Spectrum 5173? Spectrum 5173 The raw spectrum of this peptide as annotated by Hecklib. The fragments are coloured according to ion type (see legend). Any peaks with a star '\*' as text can be hovered over to see the full details, first the ion type second the mass shift type. By hovering over the amino acids in the peptide or ions in the legend the corresponding peaks are highlighted. By toggling the 'Unassigned' label you can turn the background (unassigned) peaks on or off in the plot. By updating the slider in the Ion legend you can update the spectrum to only show the top X% of the peaks with labels. The top X% means any peak that is within X% of the highest intensity. By dragging in the spectrum you can zoom in to a specific part of the spectrum and use 'Zoom Out' to get back to the original zoom level. The annotation of the spectrum is based on the given sequence in the peptides file and is done with different software so inconsistencies are likely. The peaks are annotated based on the given sequence, with 20 ppm tolerance.

Copy Data

##### Spectrum 5173 (TSV)

###### Preview

```
Loading example...
```

*Click on the button to copy the data to your clipboard.*

Mz MinMz MaxIntensity Max

WidthHeightPeptide font sizePeptide stroke widthSpectrum font sizeSpectrum stroke widthCompact peptide

Ion legend

wxyz

abcd

OtherUnassignedIon^Charge^_Position_Show for top:%

JHQDWLDGKEYKCKK

04.72e+49.44e+41.42e+51.89e+5

Zoom Out

c^+1^_2_z^+1^_2_y^+1^_2_c^+1^_3_c^+1^_3_c^+1^_4_c^+1^_4_y^+2^_7_y^+2^_7_w^+1^_4_y^+2^_8_c^+3^_14_z^+1^_4_c^+3^_14_y^+2^_9_c^+2^_10_y^+1^_4_y^+3^_14_y^+3^_14_z^+3^_14_y^+3^_14_c^+1^_5_c^+1^_5_c^+2^_12_z^+1^_5_y^+1^_5_c^+1^_6_c^+1^_6_y^+2^_12_w^+1^_6_w^+2^_13_c^+2^_13_z^+2^_13_y^+2^_13_y^+2^_13_z^+2^_13_y^+2^_13_z^+1^_6_z^+1^_6_c^+2^_14_c^+1^_7_c^+2^_14_y^+1^_6_c^+1^_7_z^+2^_14_y^+2^_14_c^+1^_8_w^+1^_7_c^+1^_8_y^+1^_7_z^+1^_7_y^+1^_7_z^+1^_8_c^+1^_9_y^+1^_8_c^+1^_9_z^+1^_9_y^+1^_9_c^+1^_10_y^+1^_10_z^+1^_10_y^+1^_10_c^+1^_11_y^+1^_11_y^+1^_11_z^+1^_11_c^+1^_12_c^+1^_12_c^+1^_12_z^+1^_12_y^+1^_12_z^+1^_12_y^+1^_12_w^+1^_13_c^+1^_13_z^+1^_13_c^+1^_14_

0769153823073075

Fragment Matches Table

Show background peaks

| Position | Ion type | Intensity | mz Theoretical | mz Error (Th) | mz Error (ppm) | Charge | Series Number |
| --- | --- | --- | --- | --- | --- | --- | --- |
| - | - | 424 | 122.8 | - | - | 0 | - |
| - | - | 2126 | 129.1 | - | - | 0 | - |
| - | - | 387.4 | 136.1 | - | - | 0 | - |
| - | - | 594.8 | 149 | - | - | 0 | - |
| - | - | 1108 | 155.1 | - | - | 0 | - |
| - | - | 3939 | 166.1 | - | - | 0 | - |
| - | - | 515.2 | 170.8 | - | - | 0 | - |
| - | - | 485.4 | 171.1 | - | - | 0 | - |
| - | - | 1802 | 173.1 | - | - | 0 | - |
| - | - | 586.7 | 173.1 | - | - | 0 | - |
| - | - | 577.4 | 178.1 | - | - | 0 | - |
| - | - | 3705 | 185.2 | - | - | 0 | - |
| - | - | 470.3 | 190 | - | - | 0 | - |
| - | - | 686.2 | 201.1 | - | - | 0 | - |
| - | - | 568.5 | 204.1 | - | - | 0 | - |
| - | - | 1664 | 205.1 | - | - | 0 | - |
| - | - | 907.2 | 207.2 | - | - | 0 | - |
| - | - | 481 | 214 | - | - | 0 | - |
| - | - | 619.4 | 217.1 | - | - | 0 | - |
| - | - | 3739 | 223.2 | - | - | 0 | - |
| - | - | 665.5 | 224.2 | - | - | 0 | - |
| - | - | 607.2 | 230.2 | - | - | 0 | - |
| - | - | 5915 | 234.1 | - | - | 0 | - |
| - | - | 735.3 | 239.1 | - | - | 0 | - |
| - | - | 592.9 | 242.1 | - | - | 0 | - |
| - | - | 527.7 | 248.1 | - | - | 0 | - |
| - | - | 1411 | 250.1 | - | - | 0 | - |
| - | - | 2.763E+04 | 251.2 | - | - | 0 | - |
| - | - | 3512 | 252.2 | - | - | 0 | - |
| - | - | 1089 | 260.1 | - | - | 0 | - |
| - | - | 575.6 | 263.6 | - | - | 0 | - |
| - | - | 551.3 | 264.1 | - | - | 0 | - |
| - | - | 1025 | 266.1 | - | - | 0 | - |
| 2 | c | 1.651E+04 | 268.2 | 0.0001351 | 0.5039 | +1 | 2 |
| - | - | 1834 | 269.2 | - | - | 0 | - |
| - | - | 495.9 | 277.2 | - | - | 0 | - |
| - | - | 1499 | 283.2 | - | - | 0 | - |
| - | - | 1577 | 294.2 | - | - | 0 | - |
| - | - | 665.3 | 299.2 | - | - | 0 | - |
| - | - | 520.7 | 311.1 | - | - | 0 | - |
| 14 | z | 2.981E+04 | 317.2 | 0.004402 | 13.88 | +1 | 2 |
| - | - | 6433 | 318.2 | - | - | 0 | - |
| - | - | 991 | 319.2 | - | - | 0 | - |
| 14 | y | 2974 | 333.2 | 0.004724 | 14.18 | +1 | 2 |
| - | - | 856.6 | 342 | - | - | 0 | - |
| - | - | 610.5 | 343 | - | - | 0 | - |
| - | - | 2867 | 352.2 | - | - | 0 | - |
| - | - | 2373 | 353.2 | - | - | 0 | - |
| - | - | 2632 | 355.1 | - | - | 0 | - |
| - | - | 2021 | 356.1 | - | - | 0 | - |
| - | - | 1629 | 357.1 | - | - | 0 | - |
| - | - | 2512 | 359 | - | - | 0 | - |
| - | - | 2329 | 360 | - | - | 0 | - |
| - | - | 1254 | 361 | - | - | 0 | - |
| - | - | 562.8 | 370.1 | - | - | 0 | - |
| 3 | c | 4384 | 379.2 | 0.0001201 | 0.3167 | +1 | 3 |
| - | - | 868.2 | 380.2 | - | - | 0 | - |
| - | - | 646.6 | 383.2 | - | - | 0 | - |
| - | - | 2.438E+04 | 387.2 | - | - | 0 | - |
| - | - | 5002 | 388.2 | - | - | 0 | - |
| - | - | 642.7 | 393.1 | - | - | 0 | - |
| - | - | 4863 | 395.2 | - | - | 0 | - |
| 3 | c | 3.982E+04 | 396.2 | 2.735E-05 | 0.06902 | +1 | 3 |
| - | - | 7042 | 397.2 | - | - | 0 | - |
| - | - | 831.5 | 398.2 | - | - | 0 | - |
| - | - | 1308 | 401.2 | - | - | 0 | - |
| - | - | 3115 | 429.1 | - | - | 0 | - |
| - | - | 5854 | 430.1 | - | - | 0 | - |
| - | - | 5026 | 431.1 | - | - | 0 | - |
| - | - | 1435 | 432.1 | - | - | 0 | - |
| - | - | 802.7 | 451.2 | - | - | 0 | - |
| - | - | 7199 | 467.2 | - | - | 0 | - |
| - | - | 3286 | 468.3 | - | - | 0 | - |
| - | - | 834.1 | 469.3 | - | - | 0 | - |
| - | - | 585.9 | 474.3 | - | - | 0 | - |
| 4 | c | 4946 | 494.2 | 0.00445 | 9.003 | +1 | 4 |
| - | - | 1422 | 495.2 | - | - | 0 | - |
| - | - | 979.1 | 496.3 | - | - | 0 | - |
| - | - | 792.8 | 497.8 | - | - | 0 | - |
| 4 | c | 3.284E+04 | 511.3 | 3.768E-05 | 0.0737 | +1 | 4 |
| - | - | 8310 | 512.3 | - | - | 0 | - |
| 9 | y | 1029 | 513.3 | 0.00935 | 18.22 | +2 | 7 |
| - | - | 5747 | 515.3 | - | - | 0 | - |
| - | - | 2397 | 516.3 | - | - | 0 | - |
| 9 | y | 729.4 | 521.8 | 0.0008728 | 1.673 | +2 | 7 |
| - | - | 2301 | 547.3 | - | - | 0 | - |
| 12 | w | 1009 | 548.2 | 0.006916 | 12.61 | +1 | 4 |
| 8 | y | 5308 | 550.3 | 0.004769 | 8.666 | +2 | 8 |
| - | - | 3400 | 550.8 | - | - | 0 | - |
| - | - | 1425 | 551.3 | - | - | 0 | - |
| - | - | 926.9 | 551.8 | - | - | 0 | - |
| - | - | 829.2 | 555.3 | - | - | 0 | - |
| - | - | 683.9 | 590.3 | - | - | 0 | - |
| - | - | 1119 | 597 | - | - | 0 | - |
| - | - | 2036 | 597.3 | - | - | 0 | - |
| 14 | c | 3672 | 601.6 | 0.0006828 | 1.135 | +3 | 14 |
| - | - | 3126 | 602 | - | - | 0 | - |
| - | - | 1962 | 602.3 | - | - | 0 | - |
| - | - | 733.8 | 602.6 | - | - | 0 | - |
| 12 | z | 2.787E+04 | 606.3 | 0.008869 | 14.63 | +1 | 4 |
| 14 | c | 1.214E+04 | 607.3 | 0.009165 | 15.09 | +3 | 14 |
| 7 | y | 2663 | 607.8 | 0.003287 | 5.408 | +2 | 9 |
| - | - | 3989 | 608.3 | - | - | 0 | - |
| - | - | 641.8 | 608.8 | - | - | 0 | - |
| 10 | c | 1205 | 611.8 | 0.000174 | 0.2845 | +2 | 10 |
| - | - | 1057 | 612.3 | - | - | 0 | - |
| - | - | 3050 | 619.2 | - | - | 0 | - |
| - | - | 5379 | 622.2 | - | - | 0 | - |
| 12 | y | 1.084E+04 | 622.3 | 0.008978 | 14.43 | +1 | 4 |
| - | - | 4622 | 623.2 | - | - | 0 | - |
| - | - | 2920 | 623.3 | - | - | 0 | - |
| - | - | 794.3 | 624.2 | - | - | 0 | - |
| - | - | 1023 | 624.3 | - | - | 0 | - |
| - | - | 968 | 625.3 | - | - | 0 | - |
| - | - | 859.6 | 625.7 | - | - | 0 | - |
| 2 | y | 1609 | 626 | 2.092E-05 | 0.03342 | +3 | 14 |
| 2 | y | 779.4 | 626.3 | 0.01186 | 18.94 | +3 | 14 |
| 2 | z | 921.8 | 626.6 | 0.0006569 | 1.048 | +3 | 14 |
| - | - | 1355 | 628.6 | - | - | 0 | - |
| - | - | 2770 | 629 | - | - | 0 | - |
| - | - | 1948 | 629.3 | - | - | 0 | - |
| - | - | 1660 | 629.6 | - | - | 0 | - |
| - | - | 806.5 | 631.6 | - | - | 0 | - |
| 2 | y | 1.194E+04 | 632 | 0.003073 | 4.863 | +3 | 14 |
| - | - | 9726 | 632.3 | - | - | 0 | - |
| - | - | 5275 | 632.6 | - | - | 0 | - |
| - | - | 3434 | 633 | - | - | 0 | - |
| - | - | 843 | 633.3 | - | - | 0 | - |
| - | - | 981.6 | 634.3 | - | - | 0 | - |
| - | - | 1.541E+04 | 634.6 | - | - | 0 | - |
| - | - | 1.806E+04 | 635 | - | - | 0 | - |
| - | - | 8994 | 635.3 | - | - | 0 | - |
| - | - | 4242 | 635.6 | - | - | 0 | - |
| - | - | 1659 | 636 | - | - | 0 | - |
| - | - | 1817 | 637.2 | - | - | 0 | - |
| - | - | 845.3 | 637.3 | - | - | 0 | - |
| - | - | 945.7 | 637.4 | - | - | 0 | - |
| - | - | 891.8 | 638.3 | - | - | 0 | - |
| - | - | 1089 | 640.6 | - | - | 0 | - |
| - | - | 871.4 | 641.3 | - | - | 0 | - |
| - | - | 847.5 | 650.3 | - | - | 0 | - |
| - | - | 739.5 | 650.6 | - | - | 0 | - |
| - | - | 618.8 | 651.3 | - | - | 0 | - |
| - | - | 1271 | 653.3 | - | - | 0 | - |
| - | - | 2074 | 654.3 | - | - | 0 | - |
| - | - | 2723 | 658 | - | - | 0 | - |
| - | - | 3787 | 658.3 | - | - | 0 | - |
| - | - | 2126 | 658.6 | - | - | 0 | - |
| - | - | 1400 | 659 | - | - | 0 | - |
| - | - | 1.215E+04 | 663.7 | - | - | 0 | - |
| - | - | 1.373E+04 | 664 | - | - | 0 | - |
| - | - | 8834 | 664.3 | - | - | 0 | - |
| - | - | 7277 | 664.7 | - | - | 0 | - |
| - | - | 3268 | 665 | - | - | 0 | - |
| - | - | 1054 | 668.3 | - | - | 0 | - |
| - | - | 867.4 | 668.3 | - | - | 0 | - |
| - | - | 1958 | 669 | - | - | 0 | - |
| - | - | 1.136E+04 | 669.3 | - | - | 0 | - |
| - | - | 1.575E+05 | 669.7 | - | - | 0 | - |
| - | - | 1.869E+05 | 670 | - | - | 0 | - |
| - | - | 1.233E+05 | 670.3 | - | - | 0 | - |
| - | - | 4.748E+04 | 670.7 | - | - | 0 | - |
| - | - | 1.69E+04 | 671 | - | - | 0 | - |
| - | - | 1320 | 678.4 | - | - | 0 | - |
| 5 | c | 2559 | 680.3 | 0.001259 | 1.851 | +1 | 5 |
| - | - | 893.9 | 681.3 | - | - | 0 | - |
| 5 | c | 2.657E+04 | 697.3 | 0.0002042 | 0.2929 | +1 | 5 |
| - | - | 9485 | 698.3 | - | - | 0 | - |
| - | - | 1958 | 699.3 | - | - | 0 | - |
| - | - | 922.9 | 711.3 | - | - | 0 | - |
| - | - | 980.7 | 712.3 | - | - | 0 | - |
| 12 | c | 3358 | 757.4 | 0.004063 | 5.365 | +2 | 12 |
| - | - | 2390 | 757.9 | - | - | 0 | - |
| - | - | 1098 | 758.4 | - | - | 0 | - |
| - | - | 2394 | 766.4 | - | - | 0 | - |
| - | - | 1887 | 767.4 | - | - | 0 | - |
| - | - | 954 | 768.4 | - | - | 0 | - |
| 11 | z | 2.488E+04 | 769.4 | 0.008721 | 11.34 | +1 | 5 |
| - | - | 1.293E+04 | 770.4 | - | - | 0 | - |
| - | - | 4175 | 771.4 | - | - | 0 | - |
| - | - | 994.9 | 772.4 | - | - | 0 | - |
| - | - | 792.2 | 777.4 | - | - | 0 | - |
| 11 | y | 5697 | 785.4 | 0.008769 | 11.16 | +1 | 5 |
| - | - | 2847 | 786.4 | - | - | 0 | - |
| - | - | 902.7 | 787.4 | - | - | 0 | - |
| 6 | c | 3392 | 793.4 | 0.001241 | 1.564 | +1 | 6 |
| - | - | 1293 | 794.4 | - | - | 0 | - |
| - | - | 1194 | 807.4 | - | - | 0 | - |
| - | - | 720.9 | 808.4 | - | - | 0 | - |
| - | - | 2000 | 809.4 | - | - | 0 | - |
| 6 | c | 3.371E+04 | 810.4 | 0.0002228 | 0.2749 | +1 | 6 |
| - | - | 1.418E+04 | 811.4 | - | - | 0 | - |
| - | - | 4071 | 812.4 | - | - | 0 | - |
| 4 | y | 4163 | 814.9 | 0.005124 | 6.288 | +2 | 12 |
| - | - | 3539 | 815.4 | - | - | 0 | - |
| - | - | 1845 | 815.9 | - | - | 0 | - |
| - | - | 641.9 | 816.4 | - | - | 0 | - |
| - | - | 1032 | 817.4 | - | - | 0 | - |
| - | - | 1546 | 818.4 | - | - | 0 | - |
| - | - | 720.2 | 818.9 | - | - | 0 | - |
| - | - | 1788 | 826.4 | - | - | 0 | - |
| - | - | 1334 | 826.9 | - | - | 0 | - |
| 10 | w | 3988 | 839.4 | 0.009446 | 11.25 | +1 | 6 |
| - | - | 1229 | 840.4 | - | - | 0 | - |
| 3 | w | 1129 | 841.9 | 0.000519 | 0.6165 | +2 | 13 |
| - | - | 1514 | 842.9 | - | - | 0 | - |
| - | - | 1633 | 843.4 | - | - | 0 | - |
| - | - | 685.9 | 843.9 | - | - | 0 | - |
| 13 | c | 2880 | 846.4 | 0.000502 | 0.5931 | +2 | 13 |
| - | - | 1635 | 846.9 | - | - | 0 | - |
| - | - | 757.5 | 847.4 | - | - | 0 | - |
| - | - | 867.3 | 860.4 | - | - | 0 | - |
| - | - | 886.7 | 861.4 | - | - | 0 | - |
| 3 | z | 983.8 | 861.9 | 0.004204 | 4.878 | +2 | 13 |
| - | - | 9481 | 867.4 | - | - | 0 | - |
| - | - | 4171 | 868.4 | - | - | 0 | - |
| 3 | y | 2901 | 869.9 | 0.005937 | 6.825 | +2 | 13 |
| 3 | y | 9273 | 870.4 | 0.0001021 | 0.1173 | +2 | 13 |
| 3 | z | 8889 | 870.9 | 0.005764 | 6.618 | +2 | 13 |
| - | - | 5556 | 871.4 | - | - | 0 | - |
| - | - | 2706 | 871.9 | - | - | 0 | - |
| - | - | 875.8 | 872.4 | - | - | 0 | - |
| - | - | 726.7 | 873.9 | - | - | 0 | - |
| - | - | 635 | 874.4 | - | - | 0 | - |
| 3 | y | 1.396E+04 | 878.9 | 0.004139 | 4.709 | +2 | 13 |
| - | - | 1.561E+04 | 879.4 | - | - | 0 | - |
| - | - | 9617 | 879.9 | - | - | 0 | - |
| 10 | z | 3814 | 880.4 | 0.01589 | 18.05 | +1 | 6 |
| - | - | 1631 | 880.9 | - | - | 0 | - |
| - | - | 4117 | 881.4 | - | - | 0 | - |
| - | - | 4103 | 882.4 | - | - | 0 | - |
| - | - | 1849 | 883.4 | - | - | 0 | - |
| - | - | 1894 | 888.4 | - | - | 0 | - |
| - | - | 3621 | 888.9 | - | - | 0 | - |
| - | - | 1875 | 889.4 | - | - | 0 | - |
| - | - | 974.6 | 889.9 | - | - | 0 | - |
| - | - | 969.3 | 892.5 | - | - | 0 | - |
| - | - | 1043 | 893.5 | - | - | 0 | - |
| - | - | 988.5 | 893.9 | - | - | 0 | - |
| - | - | 2169 | 894.4 | - | - | 0 | - |
| - | - | 823.3 | 894.9 | - | - | 0 | - |
| - | - | 732.6 | 896 | - | - | 0 | - |
| 10 | z | 2.528E+04 | 898.4 | 0.008956 | 9.969 | +1 | 6 |
| - | - | 1.272E+04 | 899.4 | - | - | 0 | - |
| - | - | 5294 | 900.4 | - | - | 0 | - |
| - | - | 1195 | 901.4 | - | - | 0 | - |
| 14 | c | 2837 | 901.9 | 0.0006304 | 0.6989 | +2 | 14 |
| - | - | 2720 | 902.4 | - | - | 0 | - |
| - | - | 1204 | 902.9 | - | - | 0 | - |
| - | - | 955.2 | 903.4 | - | - | 0 | - |
| - | - | 977.4 | 903.9 | - | - | 0 | - |
| 7 | c | 4860 | 908.4 | 0.003351 | 3.688 | +1 | 7 |
| - | - | 1905 | 909 | - | - | 0 | - |
| - | - | 3152 | 909.4 | - | - | 0 | - |
| - | - | 2143 | 909.9 | - | - | 0 | - |
| 14 | c | 2.352E+04 | 910.4 | 0.002085 | 2.29 | +2 | 14 |
| - | - | 2.353E+04 | 910.9 | - | - | 0 | - |
| - | - | 1.29E+04 | 911.4 | - | - | 0 | - |
| - | - | 5613 | 911.9 | - | - | 0 | - |
| - | - | 1762 | 912.4 | - | - | 0 | - |
| 10 | y | 4717 | 914.4 | 0.008149 | 8.911 | +1 | 6 |
| - | - | 1601 | 915.4 | - | - | 0 | - |
| - | - | 1416 | 916 | - | - | 0 | - |
| - | - | 2416 | 917.4 | - | - | 0 | - |
| - | - | 2567 | 917.9 | - | - | 0 | - |
| - | - | 788.6 | 918.4 | - | - | 0 | - |
| - | - | 691.4 | 920.5 | - | - | 0 | - |
| - | - | 1008 | 923 | - | - | 0 | - |
| - | - | 728.1 | 923.5 | - | - | 0 | - |
| - | - | 1043 | 924 | - | - | 0 | - |
| - | - | 5737 | 924.4 | - | - | 0 | - |
| 7 | c | 1.903E+04 | 925.5 | 0.001043 | 1.127 | +1 | 7 |
| - | - | 7854 | 926.5 | - | - | 0 | - |
| - | - | 1901 | 927.5 | - | - | 0 | - |
| - | - | 1720 | 929 | - | - | 0 | - |
| - | - | 1054 | 930 | - | - | 0 | - |
| - | - | 867.8 | 930.5 | - | - | 0 | - |
| - | - | 795.3 | 931 | - | - | 0 | - |
| - | - | 843.1 | 931.4 | - | - | 0 | - |
| - | - | 1136 | 932 | - | - | 0 | - |
| - | - | 2327 | 935.5 | - | - | 0 | - |
| - | - | 974.9 | 936 | - | - | 0 | - |
| - | - | 1562 | 936.5 | - | - | 0 | - |
| - | - | 1885 | 937 | - | - | 0 | - |
| - | - | 1136 | 937.5 | - | - | 0 | - |
| - | - | 7352 | 938 | - | - | 0 | - |
| - | - | 1.107E+04 | 938.5 | - | - | 0 | - |
| - | - | 8170 | 939 | - | - | 0 | - |
| 2 | z | 1.449E+04 | 939.4 | 0.006956 | 7.404 | +2 | 14 |
| - | - | 1.101E+04 | 939.9 | - | - | 0 | - |
| - | - | 9550 | 940.4 | - | - | 0 | - |
| - | - | 2581 | 940.9 | - | - | 0 | - |
| - | - | 745 | 943 | - | - | 0 | - |
| - | - | 1040 | 943.5 | - | - | 0 | - |
| - | - | 902.1 | 944 | - | - | 0 | - |
| - | - | 873.5 | 944.5 | - | - | 0 | - |
| - | - | 1090 | 945.5 | - | - | 0 | - |
| - | - | 790.5 | 946 | - | - | 0 | - |
| - | - | 3036 | 946.5 | - | - | 0 | - |
| - | - | 1291 | 947 | - | - | 0 | - |
| 2 | y | 1257 | 947.4 | 0.002965 | 3.129 | +2 | 14 |
| - | - | 823.8 | 948.4 | - | - | 0 | - |
| - | - | 3400 | 950.5 | - | - | 0 | - |
| - | - | 4661 | 951 | - | - | 0 | - |
| - | - | 3571 | 951.5 | - | - | 0 | - |
| - | - | 3320 | 952 | - | - | 0 | - |
| - | - | 5247 | 952.5 | - | - | 0 | - |
| - | - | 3404 | 953 | - | - | 0 | - |
| - | - | 1788 | 953.5 | - | - | 0 | - |
| - | - | 2012 | 954 | - | - | 0 | - |
| - | - | 884.9 | 954.5 | - | - | 0 | - |
| - | - | 833.8 | 959 | - | - | 0 | - |
| - | - | 4122 | 959.5 | - | - | 0 | - |
| - | - | 4.319E+04 | 960 | - | - | 0 | - |
| - | - | 4.57E+04 | 960.5 | - | - | 0 | - |
| - | - | 3.114E+04 | 961 | - | - | 0 | - |
| - | - | 1.265E+04 | 961.5 | - | - | 0 | - |
| - | - | 3641 | 962 | - | - | 0 | - |
| - | - | 3101 | 965 | - | - | 0 | - |
| 8 | c | 2470 | 965.4 | 0.01662 | 17.21 | +1 | 8 |
| - | - | 3991 | 966 | - | - | 0 | - |
| - | - | 4448 | 966.5 | - | - | 0 | - |
| - | - | 2217 | 967 | - | - | 0 | - |
| - | - | 2615 | 967.5 | - | - | 0 | - |
| - | - | 6088 | 968 | - | - | 0 | - |
| 9 | w | 8419 | 968.4 | 0.01449 | 14.96 | +1 | 7 |
| - | - | 4002 | 969 | - | - | 0 | - |
| - | - | 2854 | 969.5 | - | - | 0 | - |
| - | - | 1280 | 972.5 | - | - | 0 | - |
| - | - | 2076 | 973 | - | - | 0 | - |
| - | - | 3087 | 973.5 | - | - | 0 | - |
| - | - | 1.11E+04 | 974 | - | - | 0 | - |
| - | - | 3.573E+04 | 974.5 | - | - | 0 | - |
| - | - | 8.612E+04 | 975 | - | - | 0 | - |
| - | - | 7.68E+04 | 975.5 | - | - | 0 | - |
| - | - | 4.981E+04 | 976 | - | - | 0 | - |
| - | - | 1.843E+04 | 976.5 | - | - | 0 | - |
| - | - | 6147 | 977 | - | - | 0 | - |
| - | - | 772.8 | 978.5 | - | - | 0 | - |
| - | - | 750.3 | 980.5 | - | - | 0 | - |
| - | - | 2.129E+04 | 981.5 | - | - | 0 | - |
| - | - | 1.739E+04 | 982 | - | - | 0 | - |
| 8 | c | 4.102E+04 | 982.5 | 0.0038 | 3.867 | +1 | 8 |
| - | - | 7679 | 983 | - | - | 0 | - |
| - | - | 1.772E+04 | 983.5 | - | - | 0 | - |
| - | - | 4100 | 984.5 | - | - | 0 | - |
| - | - | 1036 | 985.5 | - | - | 0 | - |
| - | - | 2724 | 986.5 | - | - | 0 | - |
| - | - | 2450 | 987 | - | - | 0 | - |
| - | - | 2839 | 987.5 | - | - | 0 | - |
| - | - | 811 | 988 | - | - | 0 | - |
| - | - | 890.2 | 991 | - | - | 0 | - |
| - | - | 1066 | 994 | - | - | 0 | - |
| - | - | 1641 | 995 | - | - | 0 | - |
| - | - | 7507 | 995.5 | - | - | 0 | - |
| - | - | 2.829E+04 | 996 | - | - | 0 | - |
| - | - | 2.89E+04 | 996.5 | - | - | 0 | - |
| - | - | 1.556E+04 | 997 | - | - | 0 | - |
| - | - | 9575 | 997.5 | - | - | 0 | - |
| - | - | 2428 | 998 | - | - | 0 | - |
| - | - | 905.4 | 1002 | - | - | 0 | - |
| - | - | 1660 | 1003 | - | - | 0 | - |
| - | - | 7935 | 1003 | - | - | 0 | - |
| - | - | 7.35E+04 | 1004 | - | - | 0 | - |
| - | - | 1.422E+05 | 1004 | - | - | 0 | - |
| - | - | 1.058E+05 | 1005 | - | - | 0 | - |
| - | - | 6.134E+04 | 1005 | - | - | 0 | - |
| - | - | 2.296E+04 | 1006 | - | - | 0 | - |
| - | - | 6239 | 1006 | - | - | 0 | - |
| - | - | 852.3 | 1012 | - | - | 0 | - |
| 9 | y | 1179 | 1025 | 0.002948 | 2.877 | +1 | 7 |
| 9 | z | 2.16E+04 | 1027 | 0.009315 | 9.074 | +1 | 7 |
| - | - | 2.182E+04 | 1028 | - | - | 0 | - |
| - | - | 1.027E+04 | 1029 | - | - | 0 | - |
| - | - | 2937 | 1030 | - | - | 0 | - |
| - | - | 1140 | 1040 | - | - | 0 | - |
| - | - | 1723 | 1042 | - | - | 0 | - |
| 9 | y | 7419 | 1043 | 0.007775 | 7.458 | +1 | 7 |
| - | - | 4392 | 1044 | - | - | 0 | - |
| - | - | 1760 | 1045 | - | - | 0 | - |
| - | - | 1947 | 1067 | - | - | 0 | - |
| - | - | 1686 | 1068 | - | - | 0 | - |
| 8 | z | 1.105E+04 | 1084 | 0.00905 | 8.352 | +1 | 8 |
| - | - | 1.29E+04 | 1085 | - | - | 0 | - |
| - | - | 6293 | 1086 | - | - | 0 | - |
| - | - | 2058 | 1087 | - | - | 0 | - |
| - | - | 846.6 | 1088 | - | - | 0 | - |
| 9 | c | 877.2 | 1093 | 0.0008868 | 0.8117 | +1 | 9 |
| - | - | 2744 | 1099 | - | - | 0 | - |
| 8 | y | 7225 | 1100 | 0.01129 | 10.27 | +1 | 8 |
| - | - | 3691 | 1101 | - | - | 0 | - |
| - | - | 1183 | 1102 | - | - | 0 | - |
| - | - | 1619 | 1108 | - | - | 0 | - |
| - | - | 1030 | 1109 | - | - | 0 | - |
| - | - | 2560 | 1110 | - | - | 0 | - |
| 9 | c | 3.112E+04 | 1111 | 0.0005872 | 0.5287 | +1 | 9 |
| - | - | 2.142E+04 | 1112 | - | - | 0 | - |
| - | - | 6874 | 1113 | - | - | 0 | - |
| - | - | 944.8 | 1114 | - | - | 0 | - |
| - | - | 2442 | 1137 | - | - | 0 | - |
| - | - | 9623 | 1142 | - | - | 0 | - |
| - | - | 6243 | 1143 | - | - | 0 | - |
| - | - | 3006 | 1144 | - | - | 0 | - |
| - | - | 1812 | 1155 | - | - | 0 | - |
| - | - | 1064 | 1156 | - | - | 0 | - |
| - | - | 2790 | 1196 | - | - | 0 | - |
| - | - | 2462 | 1197 | - | - | 0 | - |
| - | - | 819.7 | 1198 | - | - | 0 | - |
| 7 | z | 2.598E+04 | 1199 | 0.008649 | 7.216 | +1 | 9 |
| - | - | 1.721E+04 | 1200 | - | - | 0 | - |
| - | - | 8906 | 1201 | - | - | 0 | - |
| - | - | 2573 | 1202 | - | - | 0 | - |
| - | - | 1149 | 1214 | - | - | 0 | - |
| 7 | y | 6214 | 1215 | 0.008208 | 6.758 | +1 | 9 |
| - | - | 3711 | 1216 | - | - | 0 | - |
| - | - | 1279 | 1217 | - | - | 0 | - |
| - | - | 2675 | 1239 | - | - | 0 | - |
| 10 | c | 2.993E+04 | 1240 | 0.0005778 | 0.4661 | +1 | 10 |
| - | - | 1.904E+04 | 1241 | - | - | 0 | - |
| - | - | 6990 | 1242 | - | - | 0 | - |
| - | - | 2272 | 1243 | - | - | 0 | - |
| - | - | 3219 | 1268 | - | - | 0 | - |
| - | - | 2050 | 1269 | - | - | 0 | - |
| - | - | 1928 | 1270 | - | - | 0 | - |
| 6 | y | 1266 | 1311 | 0.01521 | 11.6 | +1 | 10 |
| 6 | z | 1.334E+04 | 1312 | 0.00824 | 6.282 | +1 | 10 |
| - | - | 1.123E+04 | 1313 | - | - | 0 | - |
| - | - | 5777 | 1314 | - | - | 0 | - |
| - | - | 2124 | 1315 | - | - | 0 | - |
| - | - | 727.3 | 1323 | - | - | 0 | - |
| - | - | 859 | 1327 | - | - | 0 | - |
| 6 | y | 2765 | 1328 | 0.01207 | 9.093 | +1 | 10 |
| - | - | 1804 | 1329 | - | - | 0 | - |
| - | - | 1121 | 1330 | - | - | 0 | - |
| - | - | 824.3 | 1337 | - | - | 0 | - |
| - | - | 800.5 | 1339 | - | - | 0 | - |
| - | - | 818.4 | 1342 | - | - | 0 | - |
| - | - | 5810 | 1359 | - | - | 0 | - |
| - | - | 3695 | 1360 | - | - | 0 | - |
| - | - | 1013 | 1361 | - | - | 0 | - |
| - | - | 741.3 | 1390 | - | - | 0 | - |
| - | - | 4383 | 1402 | - | - | 0 | - |
| 11 | c | 3.294E+04 | 1403 | 0.001162 | 0.8285 | +1 | 11 |
| - | - | 2.511E+04 | 1404 | - | - | 0 | - |
| - | - | 1.135E+04 | 1405 | - | - | 0 | - |
| - | - | 3485 | 1406 | - | - | 0 | - |
| - | - | 1383 | 1407 | - | - | 0 | - |
| - | - | 1296 | 1442 | - | - | 0 | - |
| - | - | 1029 | 1482 | - | - | 0 | - |
| - | - | 2982 | 1487 | - | - | 0 | - |
| - | - | 2278 | 1488 | - | - | 0 | - |
| 5 | y | 873.8 | 1496 | 0.02833 | 18.94 | +1 | 11 |
| 5 | y | 2594 | 1497 | 0.007674 | 5.127 | +1 | 11 |
| 5 | z | 1.312E+04 | 1498 | 0.007719 | 5.154 | +1 | 11 |
| - | - | 1.545E+04 | 1499 | - | - | 0 | - |
| - | - | 6951 | 1500 | - | - | 0 | - |
| - | - | 2659 | 1501 | - | - | 0 | - |
| - | - | 1287 | 1502 | - | - | 0 | - |
| 12 | c | 1954 | 1513 | 0.02819 | 18.63 | +1 | 12 |
| 12 | c | 7014 | 1514 | 0.006832 | 4.513 | +1 | 12 |
| - | - | 5491 | 1515 | - | - | 0 | - |
| - | - | 3294 | 1516 | - | - | 0 | - |
| - | - | 1211 | 1517 | - | - | 0 | - |
| - | - | 811.4 | 1522 | - | - | 0 | - |
| - | - | 825.2 | 1523 | - | - | 0 | - |
| - | - | 1638 | 1525 | - | - | 0 | - |
| - | - | 1621 | 1526 | - | - | 0 | - |
| - | - | 1014 | 1527 | - | - | 0 | - |
| - | - | 880.3 | 1529 | - | - | 0 | - |
| - | - | 3980 | 1530 | - | - | 0 | - |
| 12 | c | 2.069E+04 | 1531 | 0.001521 | 0.9934 | +1 | 12 |
| - | - | 1.922E+04 | 1532 | - | - | 0 | - |
| - | - | 8509 | 1533 | - | - | 0 | - |
| - | - | 3013 | 1534 | - | - | 0 | - |
| - | - | 831.8 | 1540 | - | - | 0 | - |
| - | - | 969.5 | 1551 | - | - | 0 | - |
| - | - | 1535 | 1552 | - | - | 0 | - |
| - | - | 1033 | 1553 | - | - | 0 | - |
| - | - | 1599 | 1554 | - | - | 0 | - |
| - | - | 1324 | 1555 | - | - | 0 | - |
| - | - | 1452 | 1556 | - | - | 0 | - |
| - | - | 2978 | 1557 | - | - | 0 | - |
| - | - | 2654 | 1558 | - | - | 0 | - |
| - | - | 905.7 | 1559 | - | - | 0 | - |
| - | - | 7392 | 1569 | - | - | 0 | - |
| - | - | 4604 | 1570 | - | - | 0 | - |
| - | - | 3525 | 1571 | - | - | 0 | - |
| - | - | 1688 | 1572 | - | - | 0 | - |
| 4 | z | 724.1 | 1596 | 0.001155 | 0.7239 | +1 | 12 |
| - | - | 947.3 | 1600 | - | - | 0 | - |
| 4 | y | 1767 | 1612 | 0.008527 | 5.29 | +1 | 12 |
| 4 | z | 2.524E+04 | 1613 | 0.01049 | 6.506 | +1 | 12 |
| - | - | 3.224E+04 | 1614 | - | - | 0 | - |
| - | - | 2.037E+04 | 1615 | - | - | 0 | - |
| - | - | 8610 | 1616 | - | - | 0 | - |
| - | - | 3514 | 1617 | - | - | 0 | - |
| - | - | 1147 | 1618 | - | - | 0 | - |
| - | - | 1780 | 1628 | - | - | 0 | - |
| 4 | y | 6935 | 1629 | 0.01152 | 7.071 | +1 | 12 |
| - | - | 5633 | 1630 | - | - | 0 | - |
| - | - | 3577 | 1631 | - | - | 0 | - |
| - | - | 1602 | 1632 | - | - | 0 | - |
| - | - | 1911 | 1633 | - | - | 0 | - |
| - | - | 1005 | 1634 | - | - | 0 | - |
| - | - | 1323 | 1654 | - | - | 0 | - |
| - | - | 977.3 | 1667 | - | - | 0 | - |
| 3 | w | 1019 | 1683 | 0.002184 | 1.298 | +1 | 13 |
| - | - | 1174 | 1684 | - | - | 0 | - |
| - | - | 1282 | 1685 | - | - | 0 | - |
| - | - | 4083 | 1691 | - | - | 0 | - |
| 13 | c | 3.733E+04 | 1692 | 0.006301 | 3.724 | +1 | 13 |
| - | - | 3.193E+04 | 1693 | - | - | 0 | - |
| - | - | 1.945E+04 | 1694 | - | - | 0 | - |
| - | - | 6820 | 1695 | - | - | 0 | - |
| - | - | 2019 | 1696 | - | - | 0 | - |
| - | - | 1140 | 1697 | - | - | 0 | - |
| 3 | z | 1.35E+04 | 1741 | 0.01145 | 6.579 | +1 | 13 |
| - | - | 2.146E+04 | 1742 | - | - | 0 | - |
| - | - | 1.444E+04 | 1743 | - | - | 0 | - |
| - | - | 7331 | 1744 | - | - | 0 | - |
| - | - | 2442 | 1745 | - | - | 0 | - |
| - | - | 1378 | 1758 | - | - | 0 | - |
| - | - | 1023 | 1759 | - | - | 0 | - |
| - | - | 2641 | 1777 | - | - | 0 | - |
| - | - | 1900 | 1778 | - | - | 0 | - |
| - | - | 1407 | 1779 | - | - | 0 | - |
| 14 | c | 3375 | 1820 | 0.008246 | 4.531 | +1 | 14 |
| - | - | 6777 | 1821 | - | - | 0 | - |
| - | - | 4432 | 1822 | - | - | 0 | - |
| - | - | 3361 | 1823 | - | - | 0 | - |
| - | - | 897.5 | 1848 | - | - | 0 | - |
| - | - | 2296 | 1861 | - | - | 0 | - |
| - | - | 1708 | 1862 | - | - | 0 | - |
| - | - | 1245 | 1875 | - | - | 0 | - |
| - | - | 3413 | 1876 | - | - | 0 | - |
| - | - | 4206 | 1877 | - | - | 0 | - |
| - | - | 3726 | 1878 | - | - | 0 | - |
| - | - | 8006 | 1879 | - | - | 0 | - |
| - | - | 8425 | 1880 | - | - | 0 | - |
| - | - | 4936 | 1881 | - | - | 0 | - |
| - | - | 2476 | 1882 | - | - | 0 | - |
| - | - | 1296 | 1890 | - | - | 0 | - |
| - | - | 1705 | 1891 | - | - | 0 | - |
| - | - | 2104 | 1892 | - | - | 0 | - |
| - | - | 2055 | 1893 | - | - | 0 | - |
| - | - | 1355 | 1901 | - | - | 0 | - |
| - | - | 1001 | 1903 | - | - | 0 | - |
| - | - | 1431 | 1904 | - | - | 0 | - |
| - | - | 1766 | 1905 | - | - | 0 | - |
| - | - | 3184 | 1917 | - | - | 0 | - |
| - | - | 7426 | 1918 | - | - | 0 | - |
| - | - | 9848 | 1919 | - | - | 0 | - |
| - | - | 1.204E+04 | 1920 | - | - | 0 | - |
| - | - | 9471 | 1921 | - | - | 0 | - |
| - | - | 7585 | 1922 | - | - | 0 | - |
| - | - | 4443 | 1923 | - | - | 0 | - |
| - | - | 1628 | 1924 | - | - | 0 | - |
| - | - | 2553 | 1932 | - | - | 0 | - |
| - | - | 5597 | 1933 | - | - | 0 | - |
| - | - | 3925 | 1934 | - | - | 0 | - |
| - | - | 1867 | 1935 | - | - | 0 | - |
| - | - | 2184 | 1936 | - | - | 0 | - |
| - | - | 1140 | 1937 | - | - | 0 | - |
| - | - | 1569 | 1947 | - | - | 0 | - |
| - | - | 4147 | 1948 | - | - | 0 | - |
| - | - | 1.948E+04 | 1949 | - | - | 0 | - |
| - | - | 2.906E+04 | 1950 | - | - | 0 | - |
| - | - | 2.256E+04 | 1951 | - | - | 0 | - |
| - | - | 1.198E+04 | 1952 | - | - | 0 | - |
| - | - | 4806 | 1953 | - | - | 0 | - |
| - | - | 1630 | 1954 | - | - | 0 | - |
| - | - | 1662 | 1962 | - | - | 0 | - |
| - | - | 7266 | 1963 | - | - | 0 | - |
| - | - | 1.581E+04 | 1964 | - | - | 0 | - |
| - | - | 1.311E+04 | 1965 | - | - | 0 | - |
| - | - | 9092 | 1966 | - | - | 0 | - |
| - | - | 2580 | 1967 | - | - | 0 | - |
| - | - | 1499 | 1974 | - | - | 0 | - |
| - | - | 1610 | 1975 | - | - | 0 | - |
| - | - | 6842 | 1980 | - | - | 0 | - |
| - | - | 1.791E+04 | 1981 | - | - | 0 | - |
| - | - | 1.744E+04 | 1982 | - | - | 0 | - |
| - | - | 9248 | 1983 | - | - | 0 | - |
| - | - | 4092 | 1984 | - | - | 0 | - |
| - | - | 993.3 | 1985 | - | - | 0 | - |
| - | - | 1292 | 1990 | - | - | 0 | - |
| - | - | 1.214E+04 | 1991 | - | - | 0 | - |
| - | - | 3.549E+04 | 1992 | - | - | 0 | - |
| - | - | 3.254E+04 | 1993 | - | - | 0 | - |
| - | - | 1.922E+04 | 1994 | - | - | 0 | - |
| - | - | 9448 | 1995 | - | - | 0 | - |
| - | - | 2835 | 1996 | - | - | 0 | - |
| - | - | 1687 | 2006 | - | - | 0 | - |
| - | - | 9165 | 2007 | - | - | 0 | - |
| - | - | 4.207E+04 | 2008 | - | - | 0 | - |
| - | - | 1.411E+05 | 2009 | - | - | 0 | - |
| - | - | 1.291E+05 | 2010 | - | - | 0 | - |
| - | - | 8.324E+04 | 2011 | - | - | 0 | - |
| - | - | 3.561E+04 | 2012 | - | - | 0 | - |
| - | - | 1.178E+04 | 2013 | - | - | 0 | - |
| - | - | 1488 | 2211 | - | - | 0 | - |
| - | - | 1029 | 2213 | - | - | 0 | - |
| - | - | 923.1 | 3044 | - | - | 0 | - |
| - | - | 1077 | 3045 | - | - | 0 | - |

m/z Charge Intensity FragmentType MassShift Position
122.76618194580078 0 423.99423
129.102294921875 0 2126.071
136.07603454589844 0 387.3844
149.0448760986328 0 594.8163
155.0929718017578 0 1107.9855
166.06112670898438 0 3939.419
170.77484130859375 0 515.234
171.07635498046875 0 485.35712
173.0919189453125 0 1802.0122
173.12779235839844 0 586.73975
178.13429260253906 0 577.3931
185.16488647460938 0 3705.107
190.02943420410156 0 470.33463
201.1239776611328 0 686.19055
204.13470458984375 0 568.52716
205.1182098388672 0 1664.3701
207.1607208251953 0 907.1769
214.0253143310547 0 480.99585
217.1182403564453 0 619.44257
223.1552734375 0 3738.9543
224.15939331054688 0 665.4793
230.1864013671875 0 607.2183
234.12376403808594 0 5914.6167
239.09437561035156 0 735.268
242.1494903564453 0 592.9291
248.11276245117188 0 527.71783
250.14215087890625 0 1410.5587
251.15023803710938 0 27634.842
252.15382385253906 0 3512.3865
260.1228332519531 0 1089.0496
263.5672607421875 0 575.5914
264.1180419921875 0 551.34235
266.1243591308594 0 1025.2748
268.1766662597656 0 16508.783 c 1
269.17999267578125 0 1833.6509
277.1537780761719 0 495.85602
283.152099609375 0 1498.6022
294.1919860839844 0 1577.0101
299.18408203125 0 665.2928
311.14166259765625 0 520.7455
317.19464111328125 0 29813.514 z 13
318.19976806640625 0 6433.157
319.2019958496094 0 990.95544
333.2130432128906 0 2973.5125 y 13
342.0182189941406 0 856.56964
343.0179748535156 0 610.51685
352.2212829589844 0 2867.2983
353.2283020019531 0 2373.0251
355.0698547363281 0 2632.2512
356.0709533691406 0 2021.1926
357.06805419921875 0 1628.5546
359.0284118652344 0 2511.8384
360.0293884277344 0 2329.1008
361.0254821777344 0 1253.533
370.1250915527344 0 562.8267
379.2087097167969 0 4384.2686 c Ammonia loss 2
380.2112731933594 0 868.2495
383.2275695800781 0 646.6435
387.2238464355469 0 24375.611
388.2268371582031 0 5002.4146
393.1458740234375 0 642.69073
395.2277526855469 0 4862.6157
396.2353515625 0 39818.453 c 2
397.23846435546875 0 7041.8125
398.2407531738281 0 831.46466
401.240478515625 0 1307.5391
429.0892028808594 0 3115.1367
430.0894470214844 0 5853.604
431.08740234375 0 5025.6626
432.08782958984375 0 1435.4006
451.230712890625 0 802.6582
467.248779296875 0 7199.08
468.2549743652344 0 3286.0562
469.2578125 0 834.078
474.2735595703125 0 585.9008
494.2313232421875 0 4946.4307 c Ammonia loss 3
495.234619140625 0 1422.2124
496.2511291503906 0 979.07104
497.7558288574219 0 792.84344
511.2623596191406 0 32841.684 c 3
512.265380859375 0 8310.122
513.26611328125 0 1028.8099 y Ammonia loss 8
515.3192749023438 0 5747.052
516.3229370117188 0 2396.8462
521.7691650390625 0 729.402 y 8
547.2908935546875 0 2301.0474
548.2406005859375 0 1008.54376 w 11
550.2760009765625 0 5307.753 y 7
550.7781982421875 0 3400.017
551.2811279296875 0 1424.6464
551.7782592773438 0 926.90216
555.2701416015625 0 829.18097
590.284423828125 0 683.8962
596.9518432617188 0 1118.914
597.282470703125 0 2036.1782
601.6233520507812 0 3672.0017 c Ammonia loss 13
601.9556274414062 0 3126.3867
602.2905883789062 0 1962.282
602.6263427734375 0 733.8331
606.3043212890625 0 27867.096 z 11
607.3087158203125 0 12139.064 c 13
607.7909545898438 0 2662.7788 y 6
608.30322265625 0 3988.7507
608.7948608398438 0 641.8215
611.7960205078125 0 1205.3334 c Ammonia loss 9
612.297607421875 0 1057.029
619.2296752929688 0 3050.1536
622.2415771484375 0 5379.491
622.3229370117188 0 10840.356 y 11
623.2433471679688 0 4622.054
623.32568359375 0 2920.099
624.2467041015625 0 794.25867
624.32275390625 0 1023.4453
625.3131103515625 0 967.9753
625.6502685546875 0 859.6421
625.96435546875 0 1608.7615 y Water loss 1
626.30419921875 0 779.44104 y Ammonia loss 1
626.6276245117188 0 921.8051 z 1
628.6441040039062 0 1354.5278
628.9744262695312 0 2769.9583
629.3078002929688 0 1947.991
629.6435546875 0 1659.7249
631.6380615234375 0 806.4592
631.9647827148438 0 11940.189 y 1
632.2998046875 0 9726.242
632.6331176757812 0 5275.059
632.9686279296875 0 3433.7556
633.3059692382812 0 842.9735
634.3169555664062 0 981.61804
634.6453857421875 0 15406.377
634.9796142578125 0 18058.074
635.3125610351562 0 8994.094
635.6467895507812 0 4242.236
635.9807739257812 0 1659.3127
637.2396850585938 0 1816.9883
637.3026733398438 0 845.33716
637.3554077148438 0 945.6924
638.3048706054688 0 891.7906
640.6455688476562 0 1089.0941
641.3202514648438 0 871.3853
650.3245849609375 0 847.5256
650.6493530273438 0 739.52435
651.3021850585938 0 618.78284
653.326171875 0 1271.1093
654.3336791992188 0 2074.068
657.9799194335938 0 2723.0288
658.3140869140625 0 3787.3445
658.6471557617188 0 2126.415
658.9857788085938 0 1400.3765
663.655029296875 0 12147.366
663.9884033203125 0 13734.93
664.3229370117188 0 8834.236
664.6560668945312 0 7276.5913
664.98828125 0 3268.1343
668.2843627929688 0 1053.8186
668.3469848632812 0 867.35754
668.9871826171875 0 1957.7803
669.3308715820312 0 11362.984
669.659912109375 0 157469.94
669.9940185546875 0 186891.03
670.328125 0 123309.44
670.6616821289062 0 47481.277
670.9952392578125 0 16895.25
678.378662109375 0 1320.227
680.3163452148438 0 2558.9736 c Ammonia loss 4
681.3169555664062 0 893.8614
697.3414306640625 0 26567.95 c 4
698.3447265625 0 9485.096
699.3482055664062 0 1958.1746
711.30029296875 0 922.9253
712.3036499023438 0 980.70703
757.3712768554688 0 3357.8857 c Ammonia loss 11
757.8740234375 0 2389.534
758.3753662109375 0 1098.4266
766.409912109375 0 2393.6218
767.4157104492188 0 1886.8905
768.4198608398438 0 953.966
769.3677978515625 0 24881.215 z 10
770.3712768554688 0 12929.934
771.370361328125 0 4174.9717
772.3712158203125 0 994.8836
777.3624877929688 0 792.19696
785.386474609375 0 5696.6094 y 10
786.3892822265625 0 2846.9856
787.3839721679688 0 902.654
793.400390625 0 3391.951 c Ammonia loss 5
794.4000854492188 0 1293.1921
807.4266967773438 0 1194.2538
808.4262084960938 0 720.9373
809.4136962890625 0 2000.3248
810.4254760742188 0 33712.594 c 5
811.4281616210938 0 14181.062
812.4301147460938 0 4070.7708
814.88427734375 0 4163.486 y 3
815.3869018554688 0 3538.8408
815.8873901367188 0 1844.9285
816.3872680664062 0 641.9005
817.39208984375 0 1031.5643
818.385986328125 0 1546.1444
818.8851928710938 0 720.15717
826.395263671875 0 1788.0463
826.8920288085938 0 1334.432
839.3963623046875 0 3988.2703 w 9
840.3998413085938 0 1229.3833
841.8952026367188 0 1129.0387 w 2
842.9412841796875 0 1514.2433
843.4435424804688 0 1633.2933
843.9495239257812 0 685.86145
846.397705078125 0 2880.3767 c 12
846.897216796875 0 1634.5084
847.39892578125 0 757.49066
860.4367065429688 0 867.28546
861.4005737304688 0 886.73816
861.8998413085938 0 983.814 z Water loss 2
867.4247436523438 0 9480.773
868.4263305664062 0 4170.8286
869.907470703125 0 2900.6536 y Water loss 2
870.405517578125 0 9273.37 y Ammonia loss 2
870.903564453125 0 8888.56 z 2
871.4049682617188 0 5556.245
871.9085083007812 0 2705.871
872.4112548828125 0 875.78204
873.9170532226562 0 726.73566
874.4358520507812 0 634.95917
878.91455078125 0 13964.713 y 2
879.4158935546875 0 15610.455
879.9171752929688 0 9617.305
880.4244384765625 0 3813.862 z Water loss 9
880.9182739257812 0 1630.8234
881.4408569335938 0 4116.8623
882.4446411132812 0 4103.245
883.4481811523438 0 1849.4363
888.4356079101562 0 1894.1714
888.9391479492188 0 3620.9412
889.4365844726562 0 1874.6069
889.9376220703125 0 974.64764
892.485595703125 0 969.2635
893.4807739257812 0 1042.7921
893.9472045898438 0 988.53186
894.44873046875 0 2168.754
894.9476928710938 0 823.27875
895.9532470703125 0 732.5949
898.41015625 0 25281.072 z 9
899.412841796875 0 12719.776
900.4126586914062 0 5294.3237
901.4156494140625 0 1194.6886
901.9330444335938 0 2836.541 c Ammonia loss 13
902.435546875 0 2720.116
902.9322509765625 0 1203.9412
903.432861328125 0 955.1959
903.92529296875 0 977.36945
908.429443359375 0 4860.3604 c Ammonia loss 6
908.9539184570312 0 1904.8368
909.4338989257812 0 3152.1775
909.946533203125 0 2142.9211
910.443603515625 0 23517.05 c 13
910.9445190429688 0 23526.977
911.4444580078125 0 12902.458
911.9437255859375 0 5613.272
912.4491577148438 0 1762.2828
914.4296875 0 4716.952 y 9
915.4283447265625 0 1600.9352
915.9678955078125 0 1416.0117
917.4417114257812 0 2416.484
917.9407348632812 0 2566.821
918.4443969726562 0 788.56604
920.4680786132812 0 691.42535
922.9754028320312 0 1008.3321
923.4608764648438 0 728.0542
923.9530639648438 0 1043.364
924.448974609375 0 5736.801
925.4515991210938 0 19029.682 c 6
926.4551391601562 0 7853.626
927.4609985351562 0 1900.6895
928.9739379882812 0 1720.2865
929.9788208007812 0 1054.4998
930.4737548828125 0 867.79663
930.9653930664062 0 795.3013
931.4436645507812 0 843.1094
931.958740234375 0 1136.1527
935.5169067382812 0 2327.4226
935.98876953125 0 974.85706
936.502685546875 0 1562.2958
936.9857177734375 0 1884.7472
937.49365234375 0 1136.3523
937.9736328125 0 7352.038
938.466064453125 0 11068.025
938.9599609375 0 8170.482
939.4457397460938 0 14490.324 z 1
939.939208984375 0 11011.317
940.4426879882812 0 9549.814
940.9422607421875 0 2581.061
942.978759765625 0 744.96246
943.4757690429688 0 1039.5342
943.9747314453125 0 902.0936
944.4785766601562 0 873.52356
945.4619140625 0 1089.7997
945.9619750976562 0 790.52423
946.46337890625 0 3035.9956
946.9513549804688 0 1290.9855
947.4511108398438 0 1256.7351 y 1
948.4462890625 0 823.7588
950.4779052734375 0 3400.0256
950.9788818359375 0 4660.551
951.4758911132812 0 3571.2488
951.9837036132812 0 3320.2732
952.4736328125 0 5246.5176
952.9722900390625 0 3403.8005
953.4675903320312 0 1788.2543
953.95361328125 0 2011.9932
954.4539794921875 0 884.89734
958.9896240234375 0 833.75385
959.4874267578125 0 4121.851
959.9779663085938 0 43191.188
960.4797973632812 0 45703.613
960.9807739257812 0 31137.865
961.4800415039062 0 12653.576
961.980712890625 0 3641.3445
964.9758911132812 0 3100.9814
965.4641723632812 0 2469.8274 c Ammonia loss 7
965.9761352539062 0 3991.3442
966.4743041992188 0 4447.7627
966.9730224609375 0 2216.8608
967.4739990234375 0 2615.3623
967.9611206054688 0 6087.6987
968.462890625 0 8419 w 8
968.9624633789062 0 4002.171
969.4639282226562 0 2853.8833
972.4777221679688 0 1280.3923
972.9784545898438 0 2076.1606
973.478759765625 0 3087.1025
973.9784545898438 0 11098.476
974.469970703125 0 35731.56
974.9784545898438 0 86121.75
975.4808349609375 0 76796.47
975.9832153320312 0 49805.23
976.4835205078125 0 18429.148
976.9859619140625 0 6146.6616
978.4810180664062 0 772.8034
980.4773559570312 0 750.25494
981.4783325195312 0 21285.1
981.9837036132812 0 17392.049
982.4779052734375 0 41015.793 c 7
982.9847412109375 0 7678.829
983.4793090820312 0 17723.553
984.4816284179688 0 4100.239
985.48388671875 0 1036.4368
986.4808349609375 0 2723.8906
986.9761352539062 0 2450.1733
987.4771118164062 0 2839.2285
987.9761962890625 0 810.97144
990.973876953125 0 890.16943
993.9552001953125 0 1065.8005
994.97021484375 0 1641.2235
995.4844360351562 0 7506.8105
995.9773559570312 0 28286.908
996.4790649414062 0 28895.33
996.97802734375 0 15563.8
997.4801025390625 0 9574.566
997.9784545898438 0 2428.2666
1002.4596557617188 0 905.4144
1002.9759521484375 0 1659.7518
1003.4894409179688 0 7934.847
1003.9866943359375 0 73502.59
1004.4883422851562 0 142202.08
1004.9898071289062 0 105813.73
1005.4905395507812 0 61341.082
1005.9903564453125 0 22955.166
1006.4913330078125 0 6239.398
1012.4583740234375 0 852.27734
1024.519287109375 0 1178.8713 y Water loss 8
1026.5047607421875 0 21603.453 z 8
1027.5101318359375 0 21817.83
1028.510986328125 0 10272.609
1029.5167236328125 0 2936.6917
1039.5294189453125 0 1139.7137
1041.517822265625 0 1723.2808
1042.5250244140625 0 7419.2964 y 8
1043.526611328125 0 4392.1597
1044.5283203125 0 1760.4395
1066.552734375 0 1947.2838
1067.552490234375 0 1686.1287
1083.5264892578125 0 11048.243 z 7
1084.531005859375 0 12896.072
1085.534912109375 0 6292.5435
1086.5350341796875 0 2058.3943
1088.197265625 0 846.5899
1092.5576171875 0 877.15576 c Water loss 8
1098.53662109375 0 2744.1978
1099.54296875 0 7225.1772 y 7
1100.547119140625 0 3690.6433
1101.5465087890625 0 1182.8202
1107.567626953125 0 1619.0458
1108.54931640625 0 1029.5406
1109.57568359375 0 2560.1343
1110.5684814453125 0 31117.082 c 8
1111.5712890625 0 21416.746
1112.5721435546875 0 6874.0044
1113.5733642578125 0 944.77496
1136.58447265625 0 2442.319
1141.5545654296875 0 9623.061
1142.55859375 0 6242.7935
1143.5587158203125 0 3006.4656
1154.565673828125 0 1812.2106
1155.567138671875 0 1064.3531
1195.600341796875 0 2789.856
1196.60595703125 0 2461.5337
1197.5826416015625 0 819.71985
1198.5538330078125 0 25979.625 z 6
1199.5574951171875 0 17210.295
1200.55859375 0 8905.585
1201.55908203125 0 2573.0964
1213.5703125 0 1149.4595
1214.572998046875 0 6213.574 y 6
1215.5762939453125 0 3710.729
1216.5728759765625 0 1278.5952
1238.6116943359375 0 2674.6204
1239.611083984375 0 29928.354 c 9
1240.6134033203125 0 19039.625
1241.6158447265625 0 6990.077
1242.617919921875 0 2271.8577
1267.6412353515625 0 3219.0066
1268.6407470703125 0 2049.6216
1269.592529296875 0 1927.6873
1310.6539306640625 0 1265.9402 y Ammonia loss 5
1311.6383056640625 0 13342.307 z 5
1312.639892578125 0 11233.751
1313.6427001953125 0 5777.123
1314.6475830078125 0 2123.6968
1322.64892578125 0 727.2813
1326.647705078125 0 859.00494
1327.6531982421875 0 2764.7634 y 5
1328.6519775390625 0 1803.9578
1329.64990234375 0 1121.3956
1336.6236572265625 0 824.2615
1338.70751953125 0 800.4547
1341.60693359375 0 818.44574
1358.660400390625 0 5810.319
1359.6632080078125 0 3695.4136
1360.6702880859375 0 1012.5561
1390.1793212890625 0 741.3357
1401.6776123046875 0 4383.2627
1402.673828125 0 32940.23 c 10
1403.677490234375 0 25113.094
1404.67919921875 0 11346.031
1405.681640625 0 3485.0918
1406.70556640625 0 1383.4741
1441.6478271484375 0 1296.1715
1481.71923828125 0 1028.5215
1486.754150390625 0 2982.3074
1487.7635498046875 0 2278.1748
1495.7056884765625 0 873.8254 y Water loss 4
1496.7257080078125 0 2594.1802 y Ammonia loss 4
1497.7181396484375 0 13115.314 z 4
1498.7183837890625 0 15454.323
1499.720703125 0 6951.296
1500.7215576171875 0 2659.0715
1501.730712890625 0 1286.7971
1512.731201171875 0 1953.9166 c Water loss 11
1513.736572265625 0 7013.906 c Ammonia loss 11
1514.7391357421875 0 5491.0415
1515.745849609375 0 3294.23
1516.74853515625 0 1210.8739
1521.748291015625 0 811.4396
1522.744873046875 0 825.1593
1524.7486572265625 0 1637.807
1525.73681640625 0 1621.1167
1526.737060546875 0 1013.95685
1528.767333984375 0 880.26263
1529.77197265625 0 3980.0225
1530.7684326171875 0 20692.514 c 11
1531.77197265625 0 19224.902
1532.772705078125 0 8509.472
1533.772216796875 0 3013.3853
1539.6695556640625 0 831.8042
1550.7520751953125 0 969.4571
1551.7496337890625 0 1535.434
1552.7257080078125 0 1033.036
1553.72705078125 0 1599.4424
1554.73291015625 0 1324.489
1555.7464599609375 0 1451.8602
1556.759765625 0 2977.9482
1557.7867431640625 0 2653.6687
1558.791259765625 0 905.71246
1568.75146484375 0 7392.486
1569.7520751953125 0 4603.584
1570.757080078125 0 3525.223
1571.75634765625 0 1687.9061
1595.72509765625 0 724.0804 z Ammonia loss 3
1599.74609375 0 947.3101
1611.7364501953125 0 1766.7395 y Ammonia loss 3
1612.7423095703125 0 25236.002 z 3
1613.7457275390625 0 32244.58
1614.7486572265625 0 20372.797
1615.7530517578125 0 8609.598
1616.7552490234375 0 3514.1023
1617.7626953125 0 1147.3966
1627.7479248046875 0 1780.3811
1628.760009765625 0 6935.0034 y 3
1629.7615966796875 0 5632.756
1630.7608642578125 0 3577.3977
1631.7723388671875 0 1602.0258
1632.77294921875 0 1911.4229
1633.77587890625 0 1005.28094
1653.7923583984375 0 1323.359
1666.8494873046875 0 977.3175
1682.7799072265625 0 1019.16846 w 2
1683.7742919921875 0 1173.69
1684.794189453125 0 1281.6742
1690.7880859375 0 4083.0813
1691.7828369140625 0 37333.438 c 12
1692.7861328125 0 31929.469
1693.787109375 0 19453.352
1694.7916259765625 0 6819.7397
1695.7900390625 0 2018.635
1696.804443359375 0 1139.6847
1740.7999267578125 0 13499.243 z 2
1741.80419921875 0 21456.018
1742.809326171875 0 14444.899
1743.811279296875 0 7330.5396
1744.809326171875 0 2442.3374
1757.8212890625 0 1377.791
1758.846435546875 0 1022.7373
1776.871826171875 0 2640.5154
1777.8779296875 0 1899.9705
1778.88134765625 0 1407.2341
1819.8758544921875 0 3375.4314 c 13
1820.8792724609375 0 6776.872
1821.8857421875 0 4432.2593
1822.8870849609375 0 3360.7366
1847.9375 0 897.547
1860.93603515625 0 2295.7295
1861.94091796875 0 1708.2078
1874.9447021484375 0 1244.8004
1875.944091796875 0 3412.86
1876.94287109375 0 4206.281
1877.9178466796875 0 3726.1277
1878.8701171875 0 8005.501
1879.8699951171875 0 8424.566
1880.87255859375 0 4936.0605
1881.872314453125 0 2476.0535
1889.9569091796875 0 1296.4124
1890.9683837890625 0 1704.9482
1891.96240234375 0 2104.4622
1892.950927734375 0 2054.7083
1900.9666748046875 0 1354.731
1902.9105224609375 0 1000.7037
1903.9454345703125 0 1431.4912
1904.9439697265625 0 1766.3926
1916.9847412109375 0 3183.9382
1917.989013671875 0 7426.4717
1918.9742431640625 0 9848.474
1919.9599609375 0 12039.142
1920.9608154296875 0 9471.269
1921.9591064453125 0 7585.158
1922.9674072265625 0 4443.2837
1923.9647216796875 0 1628.0518
1931.94091796875 0 2553.3718
1932.9461669921875 0 5597.026
1933.9405517578125 0 3925.4526
1934.952392578125 0 1867.4907
1935.9384765625 0 2183.6829
1936.9393310546875 0 1140.0077
1946.947265625 0 1569.1318
1947.9517822265625 0 4147.1807
1948.95458984375 0 19478.08
1949.959716796875 0 29060.783
1950.9610595703125 0 22558.926
1951.9658203125 0 11976.93
1952.9635009765625 0 4806.2495
1953.9505615234375 0 1630.28
1961.9759521484375 0 1661.665
1962.964599609375 0 7266.2314
1963.9627685546875 0 15811.809
1964.9615478515625 0 13105.0205
1965.9619140625 0 9092.351
1966.962646484375 0 2580.085
1973.9588623046875 0 1498.8123
1974.9251708984375 0 1610.2881
1979.9727783203125 0 6842.105
1980.978759765625 0 17910.455
1981.982421875 0 17436.277
1982.9844970703125 0 9247.919
1983.986572265625 0 4092.2415
1984.99755859375 0 993.33026
1989.954833984375 0 1292.2416
1990.9583740234375 0 12135.193
1991.95166015625 0 35486.83
1992.9542236328125 0 32536.643
1993.95458984375 0 19220.084
1994.9544677734375 0 9448.27
1995.95751953125 0 2834.9082
2005.9737548828125 0 1686.6721
2006.9619140625 0 9165.265
2007.9713134765625 0 42073.68
2008.975341796875 0 141078.03
2009.9766845703125 0 129065.58
2010.97998046875 0 83236.57
2011.9808349609375 0 35614.043
2012.9786376953125 0 11784.555
2211.047607421875 0 1487.5735
2213.0556640625 0 1029.1849
3044.3212890625 0 923.08813
3044.94873046875 0 1076.677

Spectrum Details

|  |  |
| --- | --- |
| Matched peaks? Matched peaks The total absolute number of peaks matched. Additionally in brackets the total fraction of peaks matched and the total number of peaks is shown. | 77 (12.75% of 604) |
| FDR? FDR The false discovery rate estimated for this peptide. It is calculated by matching all theoretical fragments with a non-integer shift with the raw peaks for this spectrum. This is done with 40 different shifts. The resulting percentage is the average number of annotated peaks over the number of annotated peaks with the correct spectrum. | 0.99% |
| Satellite FDR? Satellite FDR See the FDR for details on its calculation. This satellite ion specific FDR only contains the satellite ions (d/w) for I/L/J positions. | - |
| PSM Score? PSM Score The PSM Score as given by Hecklib to this annotated spectrum. It is shown with three significant figures. | 671 |

#### Spectrum 5138? Spectrum 5138 The raw spectrum of this peptide as annotated by Hecklib. The fragments are coloured according to ion type (see legend). Any peaks with a star '\*' as text can be hovered over to see the full details, first the ion type second the mass shift type. By hovering over the amino acids in the peptide or ions in the legend the corresponding peaks are highlighted. By toggling the 'Unassigned' label you can turn the background (unassigned) peaks on or off in the plot. By updating the slider in the Ion legend you can update the spectrum to only show the top X% of the peaks with labels. The top X% means any peak that is within X% of the highest intensity. By dragging in the spectrum you can zoom in to a specific part of the spectrum and use 'Zoom Out' to get back to the original zoom level. The annotation of the spectrum is based on the given sequence in the peptides file and is done with different software so inconsistencies are likely. The peaks are annotated based on the given sequence, with 20 ppm tolerance.

Copy Data

##### Spectrum 5138 (TSV)

###### Preview

```
Loading example...
```

*Click on the button to copy the data to your clipboard.*

Mz MinMz MaxIntensity Max

WidthHeightPeptide font sizePeptide stroke widthSpectrum font sizeSpectrum stroke widthCompact peptide

Ion legend

wxyz

abcd

OtherUnassignedIon^Charge^_Position_Show for top:%

JHQDWLDGKEYKCKK

01.94e+43.87e+45.81e+47.75e+4

Zoom Out

y^+4^_5_z^+2^_3_c^+1^_2_y^+2^_4_z^+1^_2_y^+1^_2_c^+2^_5_y^+3^_8_c^+1^_3_y^+2^_5_c^+1^_3_c^+2^_6_y^+3^_9_w^+2^_6_z^+3^_10_y^+3^_10_y^+2^_6_z^+2^_6_y^+4^_14_c^+1^_4_y^+3^_11_c^+3^_12_c^+3^_12_c^+1^_4_y^+2^_7_z^+2^_7_y^+2^_7_z^+2^_8_w^+1^_4_y^+2^_8_c^+2^_9_w^+2^_9_y^+3^_13_y^+3^_13_z^+3^_13_y^+3^_13_z^+2^_9_y^+2^_9_y^+2^_9_z^+2^_9_c^+3^_14_c^+3^_14_z^+1^_4_c^+3^_14_y^+2^_9_c^+2^_10_c^+2^_10_y^+1^_4_y^+3^_14_z^+3^_14_w^+2^_10_y^+2^_10_y^+2^_10_z^+2^_10_y^+2^_10_c^+1^_5_c^+1^_5_c^+1^_5_c^+2^_11_y^+2^_11_y^+2^_11_z^+2^_11_c^+2^_12_c^+2^_12_c^+2^_12_z^+1^_5_w^+2^_12_y^+1^_5_c^+1^_6_y^+2^_12_z^+2^_12_c^+1^_6_y^+2^_12_w^+1^_6_w^+2^_13_c^+2^_13_y^+2^_13_z^+2^_13_y^+1^_6_z^+1^_6_c^+2^_14_c^+1^_7_c^+2^_14_y^+1^_6_c^+1^_7_z^+2^_14_y^+2^_14_c^+1^_8_c^+1^_8_z^+1^_7_z^+1^_8_y^+1^_8_c^+1^_9_z^+1^_9_c^+1^_10_y^+1^_10_z^+1^_10_c^+1^_11_z^+1^_11_c^+1^_12_c^+1^_12_z^+1^_12_w^+1^_13_c^+1^_13_z^+1^_13_z^+1^_14_

0780156123413122

Fragment Matches Table

Show background peaks

| Position | Ion type | Intensity | mz Theoretical | mz Error (Th) | mz Error (ppm) | Charge | Series Number |
| --- | --- | --- | --- | --- | --- | --- | --- |
| - | - | 406.6 | 126.1 | - | - | 0 | - |
| - | - | 463.6 | 129.1 | - | - | 0 | - |
| - | - | 1951 | 129.1 | - | - | 0 | - |
| - | - | 2203 | 130.1 | - | - | 0 | - |
| - | - | 530.5 | 131.1 | - | - | 0 | - |
| - | - | 793.3 | 131.1 | - | - | 0 | - |
| - | - | 514.1 | 133.1 | - | - | 0 | - |
| - | - | 654.9 | 136.1 | - | - | 0 | - |
| - | - | 465.7 | 140.1 | - | - | 0 | - |
| - | - | 497.9 | 141.2 | - | - | 0 | - |
| - | - | 456.8 | 143.1 | - | - | 0 | - |
| - | - | 420.7 | 146.1 | - | - | 0 | - |
| - | - | 430.2 | 148.5 | - | - | 0 | - |
| - | - | 1068 | 149 | - | - | 0 | - |
| - | - | 382.8 | 152 | - | - | 0 | - |
| - | - | 3626 | 155.1 | - | - | 0 | - |
| - | - | 1585 | 159.1 | - | - | 0 | - |
| - | - | 3579 | 166.1 | - | - | 0 | - |
| - | - | 686.2 | 168.1 | - | - | 0 | - |
| - | - | 4204 | 171.1 | - | - | 0 | - |
| - | - | 1.586E+04 | 173.1 | - | - | 0 | - |
| - | - | 578.3 | 173.1 | - | - | 0 | - |
| - | - | 755 | 173.5 | - | - | 0 | - |
| - | - | 643.3 | 174.1 | - | - | 0 | - |
| - | - | 1415 | 174.1 | - | - | 0 | - |
| - | - | 573.8 | 175.1 | - | - | 0 | - |
| - | - | 1685 | 183.1 | - | - | 0 | - |
| - | - | 429.2 | 183.5 | - | - | 0 | - |
| - | - | 3935 | 185.2 | - | - | 0 | - |
| - | - | 2232 | 189.1 | - | - | 0 | - |
| - | - | 2572 | 191.1 | - | - | 0 | - |
| - | - | 513.9 | 191.1 | - | - | 0 | - |
| 11 | y | 484.2 | 197.1 | 0.001262 | 6.401 | +4 | 5 |
| - | - | 711.9 | 201.1 | - | - | 0 | - |
| - | - | 5627 | 201.1 | - | - | 0 | - |
| - | - | 1958 | 202.1 | - | - | 0 | - |
| - | - | 1291 | 202.1 | - | - | 0 | - |
| - | - | 3951 | 203.1 | - | - | 0 | - |
| - | - | 1586 | 205.1 | - | - | 0 | - |
| - | - | 4437 | 215.1 | - | - | 0 | - |
| - | - | 1027 | 217.1 | - | - | 0 | - |
| - | - | 634.5 | 218.1 | - | - | 0 | - |
| - | - | 500.9 | 220.9 | - | - | 0 | - |
| - | - | 3939 | 223.2 | - | - | 0 | - |
| 13 | z | 1.564E+04 | 231.1 | 0.001974 | 8.541 | +2 | 3 |
| - | - | 805.3 | 231.1 | - | - | 0 | - |
| - | - | 827.2 | 233.1 | - | - | 0 | - |
| - | - | 3852 | 234.1 | - | - | 0 | - |
| - | - | 620.3 | 239.1 | - | - | 0 | - |
| - | - | 867.3 | 250.1 | - | - | 0 | - |
| - | - | 627 | 250.2 | - | - | 0 | - |
| - | - | 1.682E+04 | 251.2 | - | - | 0 | - |
| - | - | 2441 | 252.2 | - | - | 0 | - |
| - | - | 1581 | 260.1 | - | - | 0 | - |
| - | - | 1044 | 261.1 | - | - | 0 | - |
| 2 | c | 2066 | 268.2 | 0.0002572 | 0.9591 | +1 | 2 |
| - | - | 695 | 270.1 | - | - | 0 | - |
| - | - | 1907 | 274.1 | - | - | 0 | - |
| - | - | 1563 | 281.1 | - | - | 0 | - |
| - | - | 609.5 | 282 | - | - | 0 | - |
| - | - | 1223 | 282.2 | - | - | 0 | - |
| - | - | 1659 | 283.2 | - | - | 0 | - |
| - | - | 4086 | 284.2 | - | - | 0 | - |
| - | - | 1864 | 286.2 | - | - | 0 | - |
| - | - | 976.3 | 288.2 | - | - | 0 | - |
| - | - | 3300 | 294.2 | - | - | 0 | - |
| - | - | 1.361E+04 | 296.2 | - | - | 0 | - |
| - | - | 2727 | 297.2 | - | - | 0 | - |
| - | - | 7772 | 299.1 | - | - | 0 | - |
| - | - | 1106 | 299.2 | - | - | 0 | - |
| - | - | 845.4 | 300.1 | - | - | 0 | - |
| - | - | 473.9 | 300.1 | - | - | 0 | - |
| - | - | 6857 | 302.2 | - | - | 0 | - |
| - | - | 1046 | 303.2 | - | - | 0 | - |
| 12 | y | 587.5 | 311.7 | 0.00477 | 15.31 | +2 | 4 |
| - | - | 2211 | 312.2 | - | - | 0 | - |
| - | - | 5362 | 314.2 | - | - | 0 | - |
| 14 | z | 1042 | 317.2 | 0.003761 | 11.86 | +1 | 2 |
| - | - | 1268 | 318.2 | - | - | 0 | - |
| - | - | 3626 | 326.7 | - | - | 0 | - |
| - | - | 2203 | 327.2 | - | - | 0 | - |
| - | - | 596.1 | 327.7 | - | - | 0 | - |
| - | - | 6966 | 330.2 | - | - | 0 | - |
| - | - | 1729 | 331.2 | - | - | 0 | - |
| - | - | 694.9 | 331.7 | - | - | 0 | - |
| 14 | y | 595.8 | 333.2 | 0.004693 | 14.09 | +1 | 2 |
| 5 | c | 2409 | 340.7 | 0.0004145 | 1.217 | +2 | 5 |
| - | - | 1154 | 341.2 | - | - | 0 | - |
| - | - | 537.2 | 351.2 | - | - | 0 | - |
| - | - | 1086 | 352.2 | - | - | 0 | - |
| - | - | 3536 | 355.2 | - | - | 0 | - |
| - | - | 1123 | 359 | - | - | 0 | - |
| 8 | y | 1260 | 367.2 | 0.002563 | 6.98 | +3 | 8 |
| - | - | 634.2 | 367.5 | - | - | 0 | - |
| - | - | 1009 | 367.7 | - | - | 0 | - |
| - | - | 584.4 | 369.4 | - | - | 0 | - |
| - | - | 555 | 371.2 | - | - | 0 | - |
| - | - | 4248 | 373.2 | - | - | 0 | - |
| - | - | 1138 | 373.2 | - | - | 0 | - |
| - | - | 774.4 | 378.2 | - | - | 0 | - |
| 3 | c | 7307 | 379.2 | 0.0004253 | 1.121 | +1 | 3 |
| - | - | 676.7 | 380.2 | - | - | 0 | - |
| - | - | 1175 | 383.2 | - | - | 0 | - |
| - | - | 2277 | 383.2 | - | - | 0 | - |
| - | - | 1299 | 384.2 | - | - | 0 | - |
| - | - | 3196 | 387.2 | - | - | 0 | - |
| - | - | 869.3 | 388.2 | - | - | 0 | - |
| - | - | 728.5 | 389.2 | - | - | 0 | - |
| - | - | 567.1 | 390 | - | - | 0 | - |
| 11 | y | 1982 | 393.2 | 0.004727 | 12.02 | +2 | 5 |
| 3 | c | 1.035E+04 | 396.2 | 3.171E-06 | 0.008002 | +1 | 3 |
| 6 | c | 634.9 | 397.2 | 0.003147 | 7.922 | +2 | 6 |
| - | - | 1649 | 397.2 | - | - | 0 | - |
| - | - | 743.1 | 399.2 | - | - | 0 | - |
| - | - | 515.2 | 401.2 | - | - | 0 | - |
| - | - | 1.148E+04 | 401.2 | - | - | 0 | - |
| - | - | 3312 | 402.2 | - | - | 0 | - |
| 7 | y | 1093 | 405.5 | 0.001249 | 3.081 | +3 | 9 |
| - | - | 1192 | 407.7 | - | - | 0 | - |
| - | - | 754.1 | 409.2 | - | - | 0 | - |
| - | - | 2850 | 415 | - | - | 0 | - |
| - | - | 695.8 | 415.2 | - | - | 0 | - |
| - | - | 813.2 | 416 | - | - | 0 | - |
| - | - | 931.6 | 417.2 | - | - | 0 | - |
| 10 | w | 2.396E+04 | 420.2 | 0.004394 | 10.46 | +2 | 6 |
| - | - | 9205 | 420.7 | - | - | 0 | - |
| - | - | 4103 | 421.2 | - | - | 0 | - |
| - | - | 1228 | 421.7 | - | - | 0 | - |
| - | - | 675.3 | 430.1 | - | - | 0 | - |
| - | - | 1180 | 431.2 | - | - | 0 | - |
| 6 | z | 719.5 | 432.2 | 0.007323 | 16.94 | +3 | 10 |
| - | - | 1153 | 435.2 | - | - | 0 | - |
| - | - | 919.1 | 435.7 | - | - | 0 | - |
| - | - | 969.1 | 436.2 | - | - | 0 | - |
| - | - | 698.1 | 442.2 | - | - | 0 | - |
| 6 | y | 1536 | 443.2 | 0.003127 | 7.056 | +3 | 10 |
| - | - | 1513 | 444.2 | - | - | 0 | - |
| - | - | 1168 | 445.7 | - | - | 0 | - |
| - | - | 1304 | 448.3 | - | - | 0 | - |
| 10 | y | 1276 | 448.7 | 0.003071 | 6.844 | +2 | 6 |
| - | - | 1312 | 449.2 | - | - | 0 | - |
| 10 | z | 2270 | 449.7 | 0.004152 | 9.232 | +2 | 6 |
| - | - | 1076 | 450.2 | - | - | 0 | - |
| - | - | 602.7 | 454.3 | - | - | 0 | - |
| - | - | 5573 | 458.2 | - | - | 0 | - |
| - | - | 2997 | 458.7 | - | - | 0 | - |
| - | - | 1689 | 459.2 | - | - | 0 | - |
| - | - | 626.1 | 459.7 | - | - | 0 | - |
| - | - | 1204 | 460.2 | - | - | 0 | - |
| - | - | 4031 | 460.7 | - | - | 0 | - |
| - | - | 2062 | 461.2 | - | - | 0 | - |
| - | - | 1487 | 462.2 | - | - | 0 | - |
| - | - | 900.6 | 466.2 | - | - | 0 | - |
| - | - | 4299 | 466.3 | - | - | 0 | - |
| - | - | 2069 | 467.2 | - | - | 0 | - |
| - | - | 1129 | 468.3 | - | - | 0 | - |
| - | - | 1230 | 468.8 | - | - | 0 | - |
| - | - | 885.5 | 474 | - | - | 0 | - |
| 2 | y | 1055 | 474.2 | 0.002788 | 5.879 | +4 | 14 |
| - | - | 1001 | 474.5 | - | - | 0 | - |
| - | - | 678.1 | 475.3 | - | - | 0 | - |
| - | - | 797.3 | 476.2 | - | - | 0 | - |
| - | - | 733.8 | 476.8 | - | - | 0 | - |
| - | - | 978.4 | 483.3 | - | - | 0 | - |
| - | - | 1921 | 484.2 | - | - | 0 | - |
| - | - | 2.567E+04 | 484.3 | - | - | 0 | - |
| - | - | 1713 | 484.7 | - | - | 0 | - |
| - | - | 1603 | 485.2 | - | - | 0 | - |
| - | - | 7584 | 485.3 | - | - | 0 | - |
| - | - | 912.2 | 485.7 | - | - | 0 | - |
| - | - | 807.3 | 486.2 | - | - | 0 | - |
| - | - | 1707 | 486.3 | - | - | 0 | - |
| - | - | 888.5 | 487.3 | - | - | 0 | - |
| - | - | 3053 | 488.3 | - | - | 0 | - |
| - | - | 639.6 | 489.8 | - | - | 0 | - |
| - | - | 4379 | 493.7 | - | - | 0 | - |
| 4 | c | 7441 | 494.2 | 0.0003111 | 0.6295 | +1 | 4 |
| - | - | 1751 | 494.7 | - | - | 0 | - |
| - | - | 1594 | 495.2 | - | - | 0 | - |
| - | - | 1029 | 496.2 | - | - | 0 | - |
| - | - | 1843 | 496.8 | - | - | 0 | - |
| - | - | 813.9 | 497.3 | - | - | 0 | - |
| - | - | 1037 | 498 | - | - | 0 | - |
| - | - | 999 | 498.2 | - | - | 0 | - |
| - | - | 1111 | 498.5 | - | - | 0 | - |
| 5 | y | 985.5 | 499.2 | 0.006299 | 12.62 | +3 | 11 |
| - | - | 4138 | 501.3 | - | - | 0 | - |
| - | - | 7077 | 502.2 | - | - | 0 | - |
| - | - | 3.49E+04 | 502.3 | - | - | 0 | - |
| - | - | 1.508E+04 | 502.5 | - | - | 0 | - |
| - | - | 1.916E+04 | 502.8 | - | - | 0 | - |
| - | - | 8044 | 503 | - | - | 0 | - |
| - | - | 3399 | 503.1 | - | - | 0 | - |
| - | - | 7966 | 503.2 | - | - | 0 | - |
| - | - | 8230 | 503.3 | - | - | 0 | - |
| - | - | 697.1 | 503.5 | - | - | 0 | - |
| - | - | 1939 | 503.8 | - | - | 0 | - |
| - | - | 3422 | 503.8 | - | - | 0 | - |
| - | - | 1271 | 504.1 | - | - | 0 | - |
| - | - | 2080 | 504.2 | - | - | 0 | - |
| - | - | 1718 | 504.3 | - | - | 0 | - |
| 12 | c | 748.8 | 504.9 | 0.002925 | 5.792 | +3 | 12 |
| 12 | c | 742.1 | 505.3 | 0.004178 | 8.27 | +3 | 12 |
| - | - | 636.6 | 506.7 | - | - | 0 | - |
| 4 | c | 1.87E+04 | 511.3 | 0.0001149 | 0.2248 | +1 | 4 |
| - | - | 4726 | 512.3 | - | - | 0 | - |
| 9 | y | 918.3 | 512.8 | 0.006577 | 12.83 | +2 | 7 |
| - | - | 1145 | 513.3 | - | - | 0 | - |
| 9 | z | 1079 | 513.8 | 0.00384 | 7.474 | +2 | 7 |
| - | - | 852.8 | 514.3 | - | - | 0 | - |
| - | - | 4649 | 515.3 | - | - | 0 | - |
| - | - | 1877 | 516.3 | - | - | 0 | - |
| 9 | y | 5932 | 521.8 | 0.004474 | 8.575 | +2 | 7 |
| - | - | 2206 | 522.3 | - | - | 0 | - |
| - | - | 981.7 | 522.8 | - | - | 0 | - |
| - | - | 661.4 | 533.8 | - | - | 0 | - |
| - | - | 1212 | 538.3 | - | - | 0 | - |
| 8 | z | 2711 | 542.3 | 0.005539 | 10.21 | +2 | 8 |
| - | - | 1593 | 542.8 | - | - | 0 | - |
| - | - | 1423 | 543.3 | - | - | 0 | - |
| - | - | 1151 | 546.3 | - | - | 0 | - |
| - | - | 691.9 | 546.8 | - | - | 0 | - |
| - | - | 2204 | 547.3 | - | - | 0 | - |
| 12 | w | 1051 | 548.2 | 0.009479 | 17.29 | +1 | 4 |
| - | - | 921.3 | 548.3 | - | - | 0 | - |
| - | - | 2271 | 549.3 | - | - | 0 | - |
| 8 | y | 5480 | 550.3 | 0.006173 | 11.22 | +2 | 8 |
| - | - | 3953 | 550.8 | - | - | 0 | - |
| - | - | 804.7 | 553.8 | - | - | 0 | - |
| - | - | 1742 | 554.3 | - | - | 0 | - |
| - | - | 1575 | 554.8 | - | - | 0 | - |
| - | - | 7619 | 555.3 | - | - | 0 | - |
| 9 | c | 2.339E+04 | 555.8 | 0.001622 | 2.919 | +2 | 9 |
| - | - | 1.249E+04 | 556.3 | - | - | 0 | - |
| - | - | 5139 | 556.8 | - | - | 0 | - |
| - | - | 1475 | 557.3 | - | - | 0 | - |
| - | - | 755.2 | 563.8 | - | - | 0 | - |
| - | - | 1884 | 564.2 | - | - | 0 | - |
| - | - | 3267 | 568.8 | - | - | 0 | - |
| - | - | 2277 | 569.3 | - | - | 0 | - |
| - | - | 2255 | 570.3 | - | - | 0 | - |
| - | - | 2119 | 570.7 | - | - | 0 | - |
| - | - | 1.459E+04 | 571.3 | - | - | 0 | - |
| - | - | 8541 | 571.8 | - | - | 0 | - |
| - | - | 3535 | 572.3 | - | - | 0 | - |
| 7 | w | 5806 | 577.3 | 0.005473 | 9.481 | +2 | 9 |
| - | - | 3285 | 577.8 | - | - | 0 | - |
| - | - | 2011 | 578.3 | - | - | 0 | - |
| - | - | 648.3 | 578.8 | - | - | 0 | - |
| 3 | y | 974.1 | 580.3 | 0.003433 | 5.917 | +3 | 13 |
| 3 | y | 929.8 | 580.6 | 0.005964 | 10.27 | +3 | 13 |
| 3 | z | 821.8 | 580.9 | 0.003457 | 5.951 | +3 | 13 |
| 3 | y | 991.1 | 586.3 | 0.002994 | 5.106 | +3 | 13 |
| - | - | 914.7 | 586.6 | - | - | 0 | - |
| 7 | z | 671.2 | 590.8 | 0.005742 | 9.72 | +2 | 9 |
| - | - | 700.7 | 593.3 | - | - | 0 | - |
| - | - | 666.9 | 598.3 | - | - | 0 | - |
| 7 | y | 1610 | 598.8 | 0.003986 | 6.656 | +2 | 9 |
| 7 | y | 5038 | 599.3 | 0.006692 | 11.17 | +2 | 9 |
| 7 | z | 1.326E+04 | 599.8 | 0.002531 | 4.219 | +2 | 9 |
| - | - | 8192 | 600.3 | - | - | 0 | - |
| - | - | 988.8 | 600.4 | - | - | 0 | - |
| - | - | 3729 | 600.8 | - | - | 0 | - |
| 14 | c | 1034 | 601.3 | 0.0006195 | 1.03 | +3 | 14 |
| 14 | c | 1289 | 601.6 | 0.002186 | 3.633 | +3 | 14 |
| - | - | 915.1 | 602 | - | - | 0 | - |
| - | - | 861 | 602.3 | - | - | 0 | - |
| - | - | 755.8 | 603 | - | - | 0 | - |
| - | - | 1101 | 603.3 | - | - | 0 | - |
| 12 | z | 1.497E+04 | 606.3 | 0.009846 | 16.24 | +1 | 4 |
| - | - | 1101 | 606.6 | - | - | 0 | - |
| - | - | 5523 | 607 | - | - | 0 | - |
| 14 | c | 1.909E+04 | 607.3 | 0.0009248 | 1.523 | +3 | 14 |
| - | - | 1.167E+04 | 607.6 | - | - | 0 | - |
| 7 | y | 1.809E+04 | 607.8 | 0.003592 | 5.91 | +2 | 9 |
| - | - | 4592 | 608 | - | - | 0 | - |
| - | - | 1.235E+04 | 608.3 | - | - | 0 | - |
| - | - | 4667 | 608.8 | - | - | 0 | - |
| - | - | 1668 | 609.3 | - | - | 0 | - |
| - | - | 1092 | 611.6 | - | - | 0 | - |
| 10 | c | 2799 | 611.8 | 0.005319 | 8.694 | +2 | 10 |
| - | - | 768.1 | 612 | - | - | 0 | - |
| - | - | 1788 | 612.3 | - | - | 0 | - |
| - | - | 2641 | 612.8 | - | - | 0 | - |
| - | - | 2228 | 613.3 | - | - | 0 | - |
| - | - | 888.7 | 613.8 | - | - | 0 | - |
| - | - | 660.2 | 615.3 | - | - | 0 | - |
| - | - | 8432 | 619.8 | - | - | 0 | - |
| 10 | c | 2.336E+04 | 620.3 | 0.002298 | 3.705 | +2 | 10 |
| - | - | 1.166E+04 | 620.8 | - | - | 0 | - |
| - | - | 7690 | 621.3 | - | - | 0 | - |
| - | - | 680.9 | 621.6 | - | - | 0 | - |
| - | - | 1154 | 621.8 | - | - | 0 | - |
| 12 | y | 8840 | 622.3 | 0.009832 | 15.8 | +1 | 4 |
| - | - | 2525 | 623.3 | - | - | 0 | - |
| - | - | 1200 | 624.3 | - | - | 0 | - |
| - | - | 1021 | 624.7 | - | - | 0 | - |
| - | - | 839.2 | 625.3 | - | - | 0 | - |
| - | - | 1168 | 625.7 | - | - | 0 | - |
| 2 | y | 2024 | 626 | 0.01247 | 19.92 | +3 | 14 |
| - | - | 5367 | 626.3 | - | - | 0 | - |
| 2 | z | 5054 | 626.6 | 0.01076 | 17.17 | +3 | 14 |
| - | - | 2419 | 627 | - | - | 0 | - |
| - | - | 708.3 | 627.3 | - | - | 0 | - |
| - | - | 670.6 | 628.1 | - | - | 0 | - |
| - | - | 882.8 | 630.6 | - | - | 0 | - |
| - | - | 1036 | 631.3 | - | - | 0 | - |
| - | - | 1493 | 631.6 | - | - | 0 | - |
| - | - | 928.5 | 632.6 | - | - | 0 | - |
| - | - | 5650 | 634.3 | - | - | 0 | - |
| - | - | 2172 | 634.6 | - | - | 0 | - |
| 6 | w | 1.019E+04 | 634.8 | 0.0004513 | 0.711 | +2 | 10 |
| - | - | 734.8 | 635 | - | - | 0 | - |
| - | - | 6621 | 635.3 | - | - | 0 | - |
| - | - | 2497 | 635.6 | - | - | 0 | - |
| - | - | 2164 | 635.8 | - | - | 0 | - |
| - | - | 1343 | 636 | - | - | 0 | - |
| - | - | 767.1 | 636.3 | - | - | 0 | - |
| - | - | 7008 | 640 | - | - | 0 | - |
| - | - | 2.834E+04 | 640.3 | - | - | 0 | - |
| - | - | 2.44E+04 | 640.7 | - | - | 0 | - |
| - | - | 1.411E+04 | 641 | - | - | 0 | - |
| - | - | 7867 | 641.3 | - | - | 0 | - |
| - | - | 2316 | 641.7 | - | - | 0 | - |
| - | - | 966.1 | 642.3 | - | - | 0 | - |
| - | - | 713 | 643.3 | - | - | 0 | - |
| - | - | 759.7 | 644.3 | - | - | 0 | - |
| - | - | 1609 | 644.7 | - | - | 0 | - |
| - | - | 706.1 | 645.3 | - | - | 0 | - |
| - | - | 1756 | 645.6 | - | - | 0 | - |
| - | - | 2098 | 646 | - | - | 0 | - |
| - | - | 894.3 | 646.3 | - | - | 0 | - |
| - | - | 1824 | 649.3 | - | - | 0 | - |
| - | - | 6673 | 649.6 | - | - | 0 | - |
| - | - | 2.09E+04 | 650 | - | - | 0 | - |
| - | - | 2.952E+04 | 650.3 | - | - | 0 | - |
| - | - | 2.439E+04 | 650.7 | - | - | 0 | - |
| - | - | 1.62E+04 | 651 | - | - | 0 | - |
| - | - | 4421 | 651.3 | - | - | 0 | - |
| - | - | 1792 | 651.7 | - | - | 0 | - |
| - | - | 2426 | 652.3 | - | - | 0 | - |
| - | - | 1374 | 653.3 | - | - | 0 | - |
| - | - | 1866 | 654.3 | - | - | 0 | - |
| - | - | 3627 | 654.7 | - | - | 0 | - |
| - | - | 4958 | 655 | - | - | 0 | - |
| 6 | y | 2864 | 655.3 | 0.00372 | 5.677 | +2 | 10 |
| - | - | 1686 | 655.7 | - | - | 0 | - |
| 6 | y | 2382 | 655.8 | 0.007629 | 11.63 | +2 | 10 |
| 6 | z | 5638 | 656.3 | 0.002998 | 4.567 | +2 | 10 |
| - | - | 2519 | 656.8 | - | - | 0 | - |
| - | - | 1929 | 657.3 | - | - | 0 | - |
| - | - | 1488 | 657.8 | - | - | 0 | - |
| - | - | 834.9 | 658 | - | - | 0 | - |
| - | - | 1558 | 658.3 | - | - | 0 | - |
| - | - | 817.6 | 663.3 | - | - | 0 | - |
| - | - | 1465 | 663.7 | - | - | 0 | - |
| - | - | 3118 | 663.8 | - | - | 0 | - |
| - | - | 3231 | 664 | - | - | 0 | - |
| 6 | y | 1.071E+04 | 664.3 | 0.005096 | 7.672 | +2 | 10 |
| - | - | 3385 | 664.7 | - | - | 0 | - |
| - | - | 4897 | 664.8 | - | - | 0 | - |
| - | - | 2209 | 665 | - | - | 0 | - |
| - | - | 3904 | 665.3 | - | - | 0 | - |
| - | - | 1360 | 669 | - | - | 0 | - |
| - | - | 1.609E+04 | 669.3 | - | - | 0 | - |
| - | - | 6.254E+04 | 669.7 | - | - | 0 | - |
| - | - | 7.673E+04 | 670 | - | - | 0 | - |
| - | - | 5.05E+04 | 670.3 | - | - | 0 | - |
| - | - | 3.024E+04 | 670.7 | - | - | 0 | - |
| - | - | 1.205E+04 | 671 | - | - | 0 | - |
| - | - | 5316 | 671.3 | - | - | 0 | - |
| - | - | 1768 | 671.7 | - | - | 0 | - |
| - | - | 701.1 | 672 | - | - | 0 | - |
| - | - | 849.3 | 672.3 | - | - | 0 | - |
| - | - | 2069 | 674.3 | - | - | 0 | - |
| - | - | 740.3 | 675.3 | - | - | 0 | - |
| - | - | 933.9 | 678.3 | - | - | 0 | - |
| 5 | c | 964.4 | 679.3 | 0.008835 | 13 | +1 | 5 |
| - | - | 2640 | 679.8 | - | - | 0 | - |
| 5 | c | 8441 | 680.3 | 0.0007711 | 1.134 | +1 | 5 |
| - | - | 2719 | 681.3 | - | - | 0 | - |
| - | - | 2183 | 690.3 | - | - | 0 | - |
| 5 | c | 2.85E+04 | 697.3 | 0.0002653 | 0.3804 | +1 | 5 |
| - | - | 1.066E+04 | 698.3 | - | - | 0 | - |
| - | - | 1736 | 699.4 | - | - | 0 | - |
| - | - | 1209 | 700.4 | - | - | 0 | - |
| - | - | 8415 | 701.3 | - | - | 0 | - |
| 11 | c | 2.036E+04 | 701.8 | 0.001884 | 2.685 | +2 | 11 |
| - | - | 1.291E+04 | 702.3 | - | - | 0 | - |
| - | - | 3233 | 702.8 | - | - | 0 | - |
| - | - | 1852 | 703.4 | - | - | 0 | - |
| - | - | 1050 | 704.4 | - | - | 0 | - |
| - | - | 760 | 706.3 | - | - | 0 | - |
| - | - | 5924 | 716.4 | - | - | 0 | - |
| - | - | 2194 | 717.4 | - | - | 0 | - |
| - | - | 1059 | 721.3 | - | - | 0 | - |
| - | - | 1093 | 722.3 | - | - | 0 | - |
| - | - | 1301 | 727.4 | - | - | 0 | - |
| 5 | y | 904.3 | 748.4 | 0.01213 | 16.2 | +2 | 11 |
| 5 | y | 3519 | 748.9 | 0.00496 | 6.623 | +2 | 11 |
| 5 | z | 8071 | 749.4 | 0.002981 | 3.978 | +2 | 11 |
| - | - | 4358 | 749.9 | - | - | 0 | - |
| - | - | 2006 | 750.4 | - | - | 0 | - |
| - | - | 1084 | 750.9 | - | - | 0 | - |
| - | - | 1676 | 754.9 | - | - | 0 | - |
| - | - | 2325 | 755.9 | - | - | 0 | - |
| 12 | c | 1348 | 756.9 | 0.009858 | 13.03 | +2 | 12 |
| 12 | c | 5292 | 757.4 | 2.588E-05 | 0.03418 | +2 | 12 |
| - | - | 2892 | 757.9 | - | - | 0 | - |
| - | - | 1068 | 758.4 | - | - | 0 | - |
| - | - | 748.2 | 758.9 | - | - | 0 | - |
| - | - | 735.7 | 761.4 | - | - | 0 | - |
| - | - | 769.6 | 763.4 | - | - | 0 | - |
| - | - | 1755 | 765.4 | - | - | 0 | - |
| 12 | c | 4645 | 765.9 | 0.001827 | 2.385 | +2 | 12 |
| - | - | 3887 | 766.4 | - | - | 0 | - |
| - | - | 764 | 767.3 | - | - | 0 | - |
| - | - | 933.2 | 767.4 | - | - | 0 | - |
| 11 | z | 2.388E+04 | 769.4 | 0.009271 | 12.05 | +1 | 5 |
| - | - | 1.196E+04 | 770.4 | - | - | 0 | - |
| - | - | 889.3 | 770.9 | - | - | 0 | - |
| - | - | 5452 | 771.4 | - | - | 0 | - |
| - | - | 1425 | 772.4 | - | - | 0 | - |
| - | - | 886.1 | 777.4 | - | - | 0 | - |
| - | - | 829.6 | 777.9 | - | - | 0 | - |
| - | - | 783.1 | 778.9 | - | - | 0 | - |
| 4 | w | 1034 | 784.4 | 0.00569 | 7.254 | +2 | 12 |
| - | - | 4343 | 784.9 | - | - | 0 | - |
| 11 | y | 1.028E+04 | 785.4 | 0.009989 | 12.72 | +1 | 5 |
| - | - | 975.4 | 785.9 | - | - | 0 | - |
| - | - | 3748 | 786.4 | - | - | 0 | - |
| - | - | 2050 | 787.4 | - | - | 0 | - |
| 6 | c | 1808 | 793.4 | 0.002665 | 3.359 | +1 | 6 |
| - | - | 3666 | 803.4 | - | - | 0 | - |
| 4 | y | 5819 | 806.4 | 0.005892 | 7.307 | +2 | 12 |
| 4 | z | 1.197E+04 | 806.9 | 0.002354 | 2.917 | +2 | 12 |
| - | - | 1.186E+04 | 807.4 | - | - | 0 | - |
| - | - | 5199 | 807.9 | - | - | 0 | - |
| - | - | 1745 | 808.4 | - | - | 0 | - |
| - | - | 906.3 | 809.4 | - | - | 0 | - |
| 6 | c | 4.76E+04 | 810.4 | 0.0009552 | 1.179 | +1 | 6 |
| - | - | 2.184E+04 | 811.4 | - | - | 0 | - |
| - | - | 4911 | 812.4 | - | - | 0 | - |
| - | - | 4271 | 814.4 | - | - | 0 | - |
| 4 | y | 1903 | 814.9 | 0.005856 | 7.187 | +2 | 12 |
| - | - | 3100 | 815.4 | - | - | 0 | - |
| - | - | 951.4 | 816.4 | - | - | 0 | - |
| - | - | 1180 | 819.4 | - | - | 0 | - |
| - | - | 1538 | 831.4 | - | - | 0 | - |
| - | - | 1098 | 832.4 | - | - | 0 | - |
| 10 | w | 2635 | 839.4 | 0.008347 | 9.944 | +1 | 6 |
| - | - | 2266 | 840.4 | - | - | 0 | - |
| - | - | 1186 | 841.4 | - | - | 0 | - |
| 3 | w | 1729 | 841.9 | 0.0009463 | 1.124 | +2 | 13 |
| - | - | 1237 | 842.4 | - | - | 0 | - |
| - | - | 960.4 | 843 | - | - | 0 | - |
| - | - | 1203 | 843.4 | - | - | 0 | - |
| - | - | 4032 | 845.9 | - | - | 0 | - |
| 13 | c | 1.477E+04 | 846.4 | 0.0002304 | 0.2722 | +2 | 13 |
| - | - | 9149 | 846.9 | - | - | 0 | - |
| - | - | 6204 | 847.4 | - | - | 0 | - |
| - | - | 2336 | 847.9 | - | - | 0 | - |
| - | - | 1388 | 848.4 | - | - | 0 | - |
| - | - | 1253 | 848.9 | - | - | 0 | - |
| - | - | 835.4 | 849.4 | - | - | 0 | - |
| - | - | 1.052E+04 | 867.4 | - | - | 0 | - |
| - | - | 5409 | 868.4 | - | - | 0 | - |
| - | - | 1507 | 869.4 | - | - | 0 | - |
| 3 | y | 7624 | 870.4 | 0.006938 | 7.971 | +2 | 13 |
| 3 | z | 1.334E+04 | 870.9 | 0.00143 | 1.642 | +2 | 13 |
| - | - | 8507 | 871.4 | - | - | 0 | - |
| - | - | 5729 | 871.9 | - | - | 0 | - |
| - | - | 2252 | 872.4 | - | - | 0 | - |
| - | - | 801.1 | 872.9 | - | - | 0 | - |
| - | - | 781.2 | 875.4 | - | - | 0 | - |
| - | - | 818.6 | 879.4 | - | - | 0 | - |
| - | - | 1238 | 880.5 | - | - | 0 | - |
| - | - | 3384 | 881.4 | - | - | 0 | - |
| - | - | 1611 | 881.9 | - | - | 0 | - |
| - | - | 1926 | 882.4 | - | - | 0 | - |
| - | - | 4210 | 888.4 | - | - | 0 | - |
| - | - | 1953 | 888.9 | - | - | 0 | - |
| - | - | 1469 | 889.4 | - | - | 0 | - |
| - | - | 1179 | 890.5 | - | - | 0 | - |
| 10 | y | 722.2 | 896.4 | 0.01383 | 15.43 | +1 | 6 |
| 10 | z | 2.84E+04 | 898.4 | 0.009017 | 10.04 | +1 | 6 |
| - | - | 1.864E+04 | 899.4 | - | - | 0 | - |
| - | - | 9737 | 900.4 | - | - | 0 | - |
| - | - | 2422 | 901.4 | - | - | 0 | - |
| 14 | c | 788.4 | 901.9 | 0.009236 | 10.24 | +2 | 14 |
| 7 | c | 1397 | 908.4 | 0.004815 | 5.301 | +1 | 7 |
| - | - | 1740 | 910 | - | - | 0 | - |
| 14 | c | 9963 | 910.4 | 0.002085 | 2.29 | +2 | 14 |
| - | - | 6390 | 910.9 | - | - | 0 | - |
| - | - | 4144 | 911.4 | - | - | 0 | - |
| - | - | 2369 | 911.9 | - | - | 0 | - |
| 10 | y | 2770 | 914.4 | 0.00998 | 10.91 | +1 | 6 |
| - | - | 3597 | 915.4 | - | - | 0 | - |
| - | - | 2853 | 916.4 | - | - | 0 | - |
| - | - | 1009 | 917.4 | - | - | 0 | - |
| - | - | 1778 | 924.5 | - | - | 0 | - |
| 7 | c | 8234 | 925.5 | 0.001399 | 1.511 | +1 | 7 |
| - | - | 4253 | 926.5 | - | - | 0 | - |
| - | - | 968.5 | 927.5 | - | - | 0 | - |
| - | - | 898.2 | 930 | - | - | 0 | - |
| - | - | 2891 | 930.5 | - | - | 0 | - |
| - | - | 2344 | 931 | - | - | 0 | - |
| - | - | 1652 | 931.5 | - | - | 0 | - |
| - | - | 900.9 | 932 | - | - | 0 | - |
| - | - | 3923 | 932.5 | - | - | 0 | - |
| - | - | 2071 | 933.5 | - | - | 0 | - |
| - | - | 952.1 | 935.5 | - | - | 0 | - |
| - | - | 1952 | 937.5 | - | - | 0 | - |
| - | - | 1031 | 938 | - | - | 0 | - |
| - | - | 7715 | 938.5 | - | - | 0 | - |
| - | - | 2065 | 939 | - | - | 0 | - |
| 2 | z | 8550 | 939.4 | 0.003111 | 3.311 | +2 | 14 |
| - | - | 1.045E+04 | 939.9 | - | - | 0 | - |
| - | - | 6112 | 940.4 | - | - | 0 | - |
| - | - | 2276 | 940.9 | - | - | 0 | - |
| - | - | 3390 | 945.5 | - | - | 0 | - |
| - | - | 2375 | 946.5 | - | - | 0 | - |
| 2 | y | 826.8 | 947.4 | 0.009496 | 10.02 | +2 | 14 |
| - | - | 1811 | 952 | - | - | 0 | - |
| - | - | 2717 | 952.5 | - | - | 0 | - |
| - | - | 1830 | 953 | - | - | 0 | - |
| - | - | 1587 | 953.5 | - | - | 0 | - |
| - | - | 841.4 | 958 | - | - | 0 | - |
| - | - | 1134 | 959.5 | - | - | 0 | - |
| - | - | 4091 | 960 | - | - | 0 | - |
| - | - | 5960 | 960.5 | - | - | 0 | - |
| - | - | 3313 | 961 | - | - | 0 | - |
| - | - | 1620 | 961.5 | - | - | 0 | - |
| - | - | 1497 | 962.5 | - | - | 0 | - |
| - | - | 974 | 963.5 | - | - | 0 | - |
| 8 | c | 904.1 | 965.4 | 0.01906 | 19.74 | +1 | 8 |
| - | - | 4705 | 966.5 | - | - | 0 | - |
| - | - | 4274 | 967 | - | - | 0 | - |
| - | - | 3330 | 967.5 | - | - | 0 | - |
| - | - | 1828 | 968 | - | - | 0 | - |
| - | - | 1757 | 968.5 | - | - | 0 | - |
| - | - | 1914 | 973.5 | - | - | 0 | - |
| - | - | 975.8 | 974 | - | - | 0 | - |
| - | - | 6982 | 974.5 | - | - | 0 | - |
| - | - | 1.726E+04 | 975 | - | - | 0 | - |
| - | - | 1.517E+04 | 975.5 | - | - | 0 | - |
| - | - | 8880 | 976 | - | - | 0 | - |
| - | - | 4619 | 976.5 | - | - | 0 | - |
| - | - | 1836 | 977 | - | - | 0 | - |
| - | - | 1522 | 977.5 | - | - | 0 | - |
| - | - | 8964 | 981.5 | - | - | 0 | - |
| - | - | 8443 | 982 | - | - | 0 | - |
| 8 | c | 3.215E+04 | 982.5 | 0.00618 | 6.29 | +1 | 8 |
| - | - | 6647 | 983 | - | - | 0 | - |
| - | - | 1.284E+04 | 983.5 | - | - | 0 | - |
| - | - | 1278 | 984 | - | - | 0 | - |
| - | - | 3547 | 984.5 | - | - | 0 | - |
| - | - | 1211 | 987 | - | - | 0 | - |
| - | - | 1552 | 987.5 | - | - | 0 | - |
| - | - | 961.7 | 988 | - | - | 0 | - |
| - | - | 5624 | 988.5 | - | - | 0 | - |
| - | - | 3906 | 989.5 | - | - | 0 | - |
| - | - | 2128 | 990.5 | - | - | 0 | - |
| - | - | 2727 | 995.5 | - | - | 0 | - |
| - | - | 1.338E+04 | 996 | - | - | 0 | - |
| - | - | 2.655E+04 | 996.5 | - | - | 0 | - |
| - | - | 2.29E+04 | 997 | - | - | 0 | - |
| - | - | 1.314E+04 | 997.5 | - | - | 0 | - |
| - | - | 6466 | 998 | - | - | 0 | - |
| - | - | 2448 | 998.5 | - | - | 0 | - |
| - | - | 1049 | 999 | - | - | 0 | - |
| - | - | 1640 | 1003 | - | - | 0 | - |
| - | - | 5454 | 1004 | - | - | 0 | - |
| - | - | 1.85E+04 | 1004 | - | - | 0 | - |
| - | - | 2.731E+04 | 1005 | - | - | 0 | - |
| - | - | 2.754E+04 | 1005 | - | - | 0 | - |
| - | - | 1.49E+04 | 1006 | - | - | 0 | - |
| - | - | 9964 | 1006 | - | - | 0 | - |
| - | - | 3120 | 1007 | - | - | 0 | - |
| - | - | 3533 | 1007 | - | - | 0 | - |
| - | - | 2075 | 1008 | - | - | 0 | - |
| - | - | 983.8 | 1008 | - | - | 0 | - |
| 9 | z | 3663 | 1027 | 0.00785 | 7.647 | +1 | 7 |
| - | - | 8202 | 1028 | - | - | 0 | - |
| - | - | 3737 | 1029 | - | - | 0 | - |
| - | - | 1906 | 1030 | - | - | 0 | - |
| - | - | 1664 | 1067 | - | - | 0 | - |
| - | - | 1171 | 1068 | - | - | 0 | - |
| 8 | z | 1243 | 1084 | 0.009904 | 9.141 | +1 | 8 |
| - | - | 3486 | 1085 | - | - | 0 | - |
| - | - | 1329 | 1086 | - | - | 0 | - |
| 8 | y | 1740 | 1100 | 0.01422 | 12.94 | +1 | 8 |
| - | - | 1463 | 1101 | - | - | 0 | - |
| - | - | 8483 | 1110 | - | - | 0 | - |
| 9 | c | 2.855E+04 | 1111 | 0.001122 | 1.01 | +1 | 9 |
| - | - | 1.843E+04 | 1112 | - | - | 0 | - |
| - | - | 6195 | 1113 | - | - | 0 | - |
| - | - | 1395 | 1114 | - | - | 0 | - |
| - | - | 1271 | 1141 | - | - | 0 | - |
| - | - | 1150 | 1142 | - | - | 0 | - |
| - | - | 1373 | 1143 | - | - | 0 | - |
| - | - | 1723 | 1196 | - | - | 0 | - |
| - | - | 3321 | 1197 | - | - | 0 | - |
| - | - | 1653 | 1198 | - | - | 0 | - |
| 7 | z | 6401 | 1199 | 0.008563 | 7.144 | +1 | 9 |
| - | - | 1.329E+04 | 1200 | - | - | 0 | - |
| - | - | 6478 | 1201 | - | - | 0 | - |
| - | - | 2967 | 1202 | - | - | 0 | - |
| - | - | 1556 | 1203 | - | - | 0 | - |
| - | - | 2333 | 1239 | - | - | 0 | - |
| 10 | c | 6808 | 1240 | 0.004793 | 3.867 | +1 | 10 |
| - | - | 5907 | 1241 | - | - | 0 | - |
| - | - | 3319 | 1242 | - | - | 0 | - |
| - | - | 1052 | 1243 | - | - | 0 | - |
| 6 | y | 1005 | 1311 | 0.01692 | 12.91 | +1 | 10 |
| 6 | z | 3968 | 1312 | 0.006286 | 4.792 | +1 | 10 |
| - | - | 6573 | 1313 | - | - | 0 | - |
| - | - | 2972 | 1314 | - | - | 0 | - |
| - | - | 1666 | 1315 | - | - | 0 | - |
| - | - | 893.2 | 1317 | - | - | 0 | - |
| - | - | 1588 | 1359 | - | - | 0 | - |
| - | - | 2018 | 1360 | - | - | 0 | - |
| - | - | 1220 | 1361 | - | - | 0 | - |
| - | - | 984.8 | 1387 | - | - | 0 | - |
| - | - | 1656 | 1402 | - | - | 0 | - |
| 11 | c | 4077 | 1403 | 0.005063 | 3.61 | +1 | 11 |
| - | - | 3712 | 1404 | - | - | 0 | - |
| - | - | 1906 | 1405 | - | - | 0 | - |
| - | - | 1125 | 1440 | - | - | 0 | - |
| 5 | z | 2758 | 1498 | 0.001192 | 0.7958 | +1 | 11 |
| - | - | 3232 | 1499 | - | - | 0 | - |
| - | - | 2459 | 1500 | - | - | 0 | - |
| - | - | 989.5 | 1501 | - | - | 0 | - |
| 12 | c | 1013 | 1514 | 0.008671 | 5.728 | +1 | 12 |
| - | - | 1246 | 1515 | - | - | 0 | - |
| - | - | 756.6 | 1530 | - | - | 0 | - |
| 12 | c | 2094 | 1531 | 0.002863 | 1.871 | +1 | 12 |
| - | - | 2226 | 1532 | - | - | 0 | - |
| - | - | 922.1 | 1533 | - | - | 0 | - |
| - | - | 1473 | 1555 | - | - | 0 | - |
| - | - | 1158 | 1569 | - | - | 0 | - |
| 4 | z | 3301 | 1613 | 0.008783 | 5.446 | +1 | 12 |
| - | - | 6950 | 1614 | - | - | 0 | - |
| - | - | 4052 | 1615 | - | - | 0 | - |
| - | - | 1707 | 1616 | - | - | 0 | - |
| 3 | w | 890 | 1683 | 0.01869 | 11.11 | +1 | 13 |
| - | - | 874.9 | 1691 | - | - | 0 | - |
| 13 | c | 2986 | 1692 | 0.002639 | 1.56 | +1 | 13 |
| - | - | 3526 | 1693 | - | - | 0 | - |
| - | - | 2046 | 1694 | - | - | 0 | - |
| - | - | 959.4 | 1695 | - | - | 0 | - |
| 3 | z | 3041 | 1741 | 0.007102 | 4.08 | +1 | 13 |
| - | - | 4417 | 1742 | - | - | 0 | - |
| - | - | 3009 | 1743 | - | - | 0 | - |
| - | - | 897.4 | 1744 | - | - | 0 | - |
| - | - | 818.7 | 1745 | - | - | 0 | - |
| - | - | 967.1 | 1777 | - | - | 0 | - |
| - | - | 1286 | 1778 | - | - | 0 | - |
| - | - | 1001 | 1822 | - | - | 0 | - |
| 2 | z | 1149 | 1878 | 0.03132 | 16.68 | +1 | 14 |
| - | - | 2000 | 1879 | - | - | 0 | - |
| - | - | 1325 | 1880 | - | - | 0 | - |
| - | - | 1160 | 1904 | - | - | 0 | - |
| - | - | 865.5 | 1919 | - | - | 0 | - |
| - | - | 803.1 | 1920 | - | - | 0 | - |
| - | - | 1345 | 1921 | - | - | 0 | - |
| - | - | 812.3 | 1922 | - | - | 0 | - |
| - | - | 948.7 | 1923 | - | - | 0 | - |
| - | - | 1044 | 1933 | - | - | 0 | - |
| - | - | 950.4 | 1949 | - | - | 0 | - |
| - | - | 1724 | 1950 | - | - | 0 | - |
| - | - | 2414 | 1951 | - | - | 0 | - |
| - | - | 1805 | 1952 | - | - | 0 | - |
| - | - | 931.2 | 1964 | - | - | 0 | - |
| - | - | 1467 | 1965 | - | - | 0 | - |
| - | - | 804.4 | 1981 | - | - | 0 | - |
| - | - | 2160 | 1992 | - | - | 0 | - |
| - | - | 2774 | 1993 | - | - | 0 | - |
| - | - | 1822 | 1994 | - | - | 0 | - |
| - | - | 1219 | 1995 | - | - | 0 | - |
| - | - | 825.9 | 1996 | - | - | 0 | - |
| - | - | 2235 | 2008 | - | - | 0 | - |
| - | - | 3396 | 2009 | - | - | 0 | - |
| - | - | 3285 | 2010 | - | - | 0 | - |
| - | - | 2457 | 2011 | - | - | 0 | - |
| - | - | 853.7 | 2012 | - | - | 0 | - |
| - | - | 697.7 | 3091 | - | - | 0 | - |

m/z Charge Intensity FragmentType MassShift Position
126.09109497070312 0 406.60056
129.06573486328125 0 463.58682
129.10226440429688 0 1951.3761
130.06517028808594 0 2203.0537
131.06895446777344 0 530.546
131.11814880371094 0 793.326
133.0612030029297 0 514.0798
136.0756072998047 0 654.92224
140.08212280273438 0 465.74033
141.2064208984375 0 497.89572
143.0821533203125 0 456.81866
146.09268188476562 0 420.6979
148.54071044921875 0 430.17416
149.04489135742188 0 1067.6699
152.0164337158203 0 382.76074
155.0928497314453 0 3626.2412
159.09152221679688 0 1584.9064
166.0611572265625 0 3578.6772
168.13818359375 0 686.2072
171.07644653320312 0 4204.3706
173.092041015625 0 15859.181
173.1285858154297 0 578.33356
173.4540557861328 0 755.00104
174.08750915527344 0 643.2625
174.09542846679688 0 1414.5194
175.07130432128906 0 573.8351
183.1126708984375 0 1685.4214
183.5270233154297 0 429.17285
185.16488647460938 0 3934.9998
189.08709716796875 0 2232.3694
191.10250854492188 0 2572.4185
191.11111450195312 0 513.9003
197.10552978515625 0 484.2049 y 10
201.08702087402344 0 711.9261
201.1233367919922 0 5627.0166
202.08212280273438 0 1957.6812
202.12689208984375 0 1291.4786
203.1024169921875 0 3950.5818
205.11842346191406 0 1586.2957
215.13888549804688 0 4436.963
217.11842346191406 0 1026.722
218.1496124267578 0 634.5107
220.87364196777344 0 500.91187
223.15538024902344 0 3939.3638
231.09750366210938 0 15644.382 z Ammonia loss 12
231.110595703125 0 805.2627
233.13995361328125 0 827.1591
234.1238250732422 0 3852.4
239.14002990722656 0 620.3447
250.1429901123047 0 867.31866
250.16632080078125 0 627.02673
251.15013122558594 0 16818.988
252.15354919433594 0 2441.244
260.1239013671875 0 1580.6086
261.1184387207031 0 1044.2118
268.1765441894531 0 2065.5508 c 1
270.1444091796875 0 695.01245
274.1185302734375 0 1907.3405
281.05096435546875 0 1563.3718
282.0486145019531 0 609.54785
282.1816101074219 0 1223.229
283.1501159667969 0 1659.3202
284.1605529785156 0 4086.1587
286.1761474609375 0 1863.6233
288.1549987792969 0 976.3171
294.1810607910156 0 3300.2449
296.1967468261719 0 13613.618
297.2001953125 0 2726.6255
299.0617370605469 0 7772.309
299.2074279785156 0 1106.1381
300.06268310546875 0 845.3826
300.118896484375 0 473.85773
302.17095947265625 0 6856.5703
303.1751403808594 0 1045.7972
311.6648254394531 0 587.4724 y 11
312.1915588378906 0 2211.2175
314.20758056640625 0 5361.6743
317.1952819824219 0 1041.9816 z 13
318.2024841308594 0 1267.8016
326.6636047363281 0 3626.4004
327.166015625 0 2203.3997
327.6670227050781 0 596.0953
330.2024230957031 0 6966.277
331.2055969238281 0 1728.6462
331.6559143066406 0 694.91644
333.21307373046875 0 595.75195 y 13
340.6607666015625 0 2408.5168 c Ammonia loss 4
341.16131591796875 0 1153.7565
351.2135314941406 0 537.1776
352.2214660644531 0 1085.9371
355.19720458984375 0 3536.123
359.02825927734375 0 1122.7637
367.1870422363281 0 1259.9733 y 7
367.5184020996094 0 634.21704
367.68035888671875 0 1008.62384
369.37359619140625 0 584.43396
371.1936340332031 0 554.99084
373.20819091796875 0 4248.169
373.2428283691406 0 1137.8456
378.20196533203125 0 774.3588
379.2084045410156 0 7307.4106 c Ammonia loss 2
380.2112121582031 0 676.72864
383.2055969238281 0 1174.9147
383.2288513183594 0 2276.859
384.2306823730469 0 1299.2808
387.2236633300781 0 3196.4302
388.2260437011719 0 869.25684
389.2029113769531 0 728.49884
390.03411865234375 0 567.13916
393.196533203125 0 1982.1477 y 10
396.2353820800781 0 10353.725 c 2
397.20635986328125 0 634.93823 c Ammonia loss 5
397.23858642578125 0 1649.4327
399.22418212890625 0 743.0514
401.20849609375 0 515.21893
401.23944091796875 0 11481.88
402.24249267578125 0 3311.851
405.5306701660156 0 1092.9863 y 6
407.7058410644531 0 1192.483
409.2156066894531 0 754.08545
415.03643798828125 0 2850.399
415.2424621582031 0 695.8071
416.03631591796875 0 813.2059
417.2331848144531 0 931.6232
420.2021484375 0 23961.518 w 9
420.70330810546875 0 9204.778
421.203857421875 0 4102.8306
421.7036437988281 0 1227.9888
430.09027099609375 0 675.2597
431.2271728515625 0 1179.8835
432.2041931152344 0 719.5401 z Ammonia loss 5
435.2267761230469 0 1152.9131
435.7237854003906 0 919.0593
436.221435546875 0 969.10754
442.2274475097656 0 698.1174
443.2234802246094 0 1535.5515 y 5
444.2307434082031 0 1512.8284
445.7308654785156 0 1167.6
448.255615234375 0 1303.7225
448.7142028808594 0 1275.841 y Water loss 9
449.2242736816406 0 1311.9338
449.71734619140625 0 2270.4573 z 9
450.2206115722656 0 1076.2683
454.25933837890625 0 602.7008
458.2297058105469 0 5573.0034
458.72943115234375 0 2996.7705
459.227783203125 0 1689.1328
459.7287292480469 0 626.10864
460.23809814453125 0 1203.6758
460.73492431640625 0 4030.8677
461.2368469238281 0 2062.3599
462.2016296386719 0 1486.8613
466.2340087890625 0 900.5933
466.2661437988281 0 4299.166
467.24658203125 0 2068.8906
468.2593688964844 0 1128.556
468.76531982421875 0 1230.1718
473.9807434082031 0 885.4684
474.2304992675781 0 1054.6458 y 1
474.4765930175781 0 1001.3098
475.2972717285156 0 678.1365
476.22857666015625 0 797.279
476.7522277832031 0 733.83356
483.2613830566406 0 978.3782
484.1984558105469 0 1921.09
484.27679443359375 0 25668.379
484.74859619140625 0 1713.1445
485.2344665527344 0 1602.6631
485.28009033203125 0 7583.812
485.72796630859375 0 912.1904
486.2210998535156 0 807.2859
486.29144287109375 0 1707.1985
487.2878723144531 0 888.5239
488.2931213378906 0 3053.2195
489.80914306640625 0 639.58124
493.747314453125 0 4379.2695
494.236083984375 0 7441.264 c Ammonia loss 3
494.7445068359375 0 1750.6686
495.2376403808594 0 1594.1589
496.2489318847656 0 1029.1119
496.7749938964844 0 1842.8354
497.27685546875 0 813.8697
497.9958190917969 0 1037.0665
498.243408203125 0 998.99744
498.4973449707031 0 1111.14
499.24322509765625 0 985.50916 y Water loss 4
501.2723693847656 0 4137.658
502.2496337890625 0 7076.8726
502.2875061035156 0 34904.96
502.4990234375 0 15076.093
502.75042724609375 0 19155.72
502.9995422363281 0 8044.343
503.1077575683594 0 3399.0496
503.24993896484375 0 7965.7007
503.2911071777344 0 8229.559
503.4997253417969 0 697.14935
503.751220703125 0 1939.0771
503.8067626953125 0 3421.5068
504.10760498046875 0 1270.8776
504.2425537109375 0 2079.9019
504.303466796875 0 1717.6727
504.9217224121094 0 748.7796 c Water loss 11
505.24847412109375 0 742.09467 c Ammonia loss 11
506.7303771972656 0 636.6272
511.26220703125 0 18695.564 c 3
512.2649536132812 0 4726.203
512.7581787109375 0 918.266 y Water loss 8
513.2673950195312 0 1144.6506
513.7568359375 0 1079.3527 z 8
514.2578735351562 0 852.84796
515.31884765625 0 4649.348
516.32275390625 0 1876.7639
521.7655639648438 0 5932.47 y 8
522.2679443359375 0 2205.954
522.7664184570312 0 981.66626
533.7794189453125 0 661.44836
538.2713012695312 0 1212.4546
542.265869140625 0 2710.8582 z 7
542.7684326171875 0 1592.971
543.2647094726562 0 1422.7286
546.2620239257812 0 1150.6692
546.7691650390625 0 691.92487
547.2862548828125 0 2204.0303
548.238037109375 0 1051.1112 w 11
548.2908325195312 0 921.28485
549.2685546875 0 2271.4038
550.2745971679688 0 5480.231 y 7
550.7772216796875 0 3953.1921
553.7962646484375 0 804.72534
554.2919921875 0 1742.0668
554.7862548828125 0 1575.1877
555.2957153320312 0 7619.166
555.789794921875 0 23387.992 c 8
556.2904052734375 0 12489.27
556.7913208007812 0 5139.4365
557.2898559570312 0 1474.6005
563.7567749023438 0 755.2449
564.2449951171875 0 1883.6986
568.7962036132812 0 3267.4978
569.2960815429688 0 2277.0571
570.274658203125 0 2254.6184
570.7489013671875 0 2118.599
571.281005859375 0 14588.708
571.7822265625 0 8540.623
572.2837524414062 0 3534.521
577.2805786132812 0 5805.5767 w 6
577.7835693359375 0 3284.6099
578.2813110351562 0 2010.7898
578.7865600585938 0 648.2949
580.2745971679688 0 974.08545 y Water loss 2
580.6119995117188 0 929.75055 y Ammonia loss 2
580.9454345703125 0 821.82806 z 2
586.2845458984375 0 991.07367 y 2
586.6137084960938 0 914.7279
590.7853393554688 0 671.2487 z Water loss 6
593.28173828125 0 700.6694
598.29931640625 0 666.9434
598.7849731445312 0 1610.386 y Water loss 6
599.2876586914062 0 5038.171 y Ammonia loss 6
599.7823486328125 0 13261.001 z 6
600.282958984375 0 8192.083
600.370361328125 0 988.7656
600.7842407226562 0 3729.0383
601.29541015625 0 1033.6278 c Water loss 13
601.626220703125 0 1288.6227 c Ammonia loss 13
601.95947265625 0 915.0582
602.2894287109375 0 860.954
602.96044921875 0 755.7553
603.2838134765625 0 1101.3872
606.3033447265625 0 14967.534 z 11
606.6341552734375 0 1101.4635
606.9686279296875 0 5523.1196
607.3004760742188 0 19089.695 c 13
607.6325073242188 0 11672.876
607.7906494140625 0 18092.312 y 6
607.9671630859375 0 4592.1577
608.2943115234375 0 12346.413
608.7918701171875 0 4667.387
609.2977294921875 0 1667.92
611.6329956054688 0 1091.8156
611.801513671875 0 2799.2915 c Ammonia loss 9
611.9664916992188 0 768.0686
612.300537109375 0 1787.7391
612.7994995117188 0 2640.9863
613.301513671875 0 2227.8943
613.8025512695312 0 888.6641
615.304931640625 0 660.1508
619.8172607421875 0 8431.539
620.311767578125 0 23362.94 c 9
620.8118286132812 0 11655.192
621.3124389648438 0 7690.0513
621.640625 0 680.8887
621.8137817382812 0 1153.757
622.3220825195312 0 8840.026 y 11
623.324951171875 0 2524.7576
624.3241577148438 0 1200.476
624.6661376953125 0 1021.0726
625.3153686523438 0 839.19885
625.6508178710938 0 1167.5996
625.976806640625 0 2023.5334 y Water loss 1
626.3064575195312 0 5367.1147
626.6390380859375 0 5053.955 z 1
626.9714965820312 0 2418.8877
627.3056030273438 0 708.2503
628.0895385742188 0 670.6019
630.6359252929688 0 882.84564
631.3121337890625 0 1036.4504
631.6375122070312 0 1493.37
632.6373901367188 0 928.5009
634.3048095703125 0 5649.9004
634.6488647460938 0 2172.082
634.799072265625 0 10185.909 w 5
634.9905395507812 0 734.80475
635.3024291992188 0 6620.857
635.6449584960938 0 2496.5889
635.7982788085938 0 2163.5623
635.9805908203125 0 1342.8943
636.3198852539062 0 767.09204
639.9918212890625 0 7007.9546
640.32275390625 0 28343.3
640.6560668945312 0 24396.576
640.990234375 0 14105.006
641.3238525390625 0 7867.265
641.6569213867188 0 2315.7952
642.3109741210938 0 966.11615
643.327880859375 0 712.9662
644.3153686523438 0 759.74414
644.6568603515625 0 1608.9702
645.30322265625 0 706.11145
645.6459350585938 0 1756.1012
645.9788818359375 0 2098.46
646.3189697265625 0 894.3345
649.3250122070312 0 1823.778
649.6488037109375 0 6672.84
649.9769897460938 0 20899.318
650.3191528320312 0 29516.717
650.6533203125 0 24392.229
650.9912719726562 0 16197.562
651.3253173828125 0 4420.571
651.6528930664062 0 1791.8711
652.3187866210938 0 2425.8816
653.3248291015625 0 1374.3262
654.3283081054688 0 1865.7709
654.66064453125 0 3627.1406
654.9945678710938 0 4957.919
655.3272705078125 0 2863.6858 y Water loss 5
655.6552124023438 0 1686.036
655.8306274414062 0 2382.2112 y Ammonia loss 5
656.3239135742188 0 5637.837 z 5
656.8257446289062 0 2518.7346
657.3257446289062 0 1929.1234
657.83251953125 0 1488.1135
657.9901123046875 0 834.85803
658.3199462890625 0 1557.9224
663.3306884765625 0 817.644
663.6561889648438 0 1465.0227
663.8387451171875 0 3118.291
663.9917602539062 0 3230.6968
664.3311767578125 0 10712.327 y 5
664.6570434570312 0 3385.0178
664.8342895507812 0 4897.402
664.9864501953125 0 2209.0867
665.330322265625 0 3903.9724
668.9910888671875 0 1359.5153
669.3309936523438 0 16090.358
669.6622314453125 0 62539.586
669.99609375 0 76726.45
670.3297119140625 0 50499.133
670.6634521484375 0 30237.807
670.9970092773438 0 12054.1875
671.331298828125 0 5315.6284
671.6610717773438 0 1768.1077
671.993408203125 0 701.14795
672.3253784179688 0 849.2835
674.324462890625 0 2069.0542
675.3224487304688 0 740.3002
678.329833984375 0 933.90686
679.3399047851562 0 964.3546 c Water loss 4
679.83544921875 0 2639.7065
680.3158569335938 0 8441.075 c Ammonia loss 4
681.3197631835938 0 2719.3623
690.338623046875 0 2182.8926
697.3413696289062 0 28497.326 c 4
698.3436279296875 0 10658.44
699.3501586914062 0 1736.1047
700.3590698242188 0 1208.9698
701.3485717773438 0 8415.495
701.843017578125 0 20356.639 c 10
702.3432006835938 0 12910.455
702.8462524414062 0 3232.6348
703.3516235351562 0 1852.125
704.367431640625 0 1049.954
706.330078125 0 760.04565
716.4047241210938 0 5923.7656
717.4019165039062 0 2193.6807
721.3306884765625 0 1058.5259
722.3336791992188 0 1093.0537
727.3593139648438 0 1300.9418
748.3585205078125 0 904.3232 y Water loss 4
748.8676147460938 0 3519.061 y Ammonia loss 4
749.3635864257812 0 8070.5513 z 4
749.8650512695312 0 4358.1533
750.364013671875 0 2005.994
750.8634643554688 0 1083.662
754.8832397460938 0 1675.8846
755.8861694335938 0 2324.8896
756.8734741210938 0 1347.8329 c Water loss 11
757.3753662109375 0 5291.9326 c Ammonia loss 11
757.8734130859375 0 2892.3184
758.3768920898438 0 1067.6951
758.8883666992188 0 748.22107
761.3841552734375 0 735.69183
763.3626098632812 0 769.6261
765.3980102539062 0 1754.6587
765.8904418945312 0 4645.457 c 11
766.3952026367188 0 3887.2798
767.3423461914062 0 764.04614
767.4146118164062 0 933.1793
769.3672485351562 0 23878.006 z 10
770.3703002929688 0 11958.124
770.8643188476562 0 889.25586
771.3717651367188 0 5452.316
772.3726806640625 0 1425.2323
777.3729858398438 0 886.1436
777.8656005859375 0 829.604
778.8929443359375 0 783.05585
784.3869018554688 0 1034.4523 w 3
784.88134765625 0 4342.7583
785.38525390625 0 10279.752 y 10
785.8768310546875 0 975.3527
786.3875732421875 0 3747.806
787.3958129882812 0 2050.474
793.396484375 0 1808.3899 c Ammonia loss 5
803.4234008789062 0 3665.8972
806.3820190429688 0 5818.9106 y Ammonia loss 3
806.877685546875 0 11969.599 z 3
807.37890625 0 11855.051
807.8798828125 0 5199.001
808.3784790039062 0 1745.4343
809.40380859375 0 906.2855
810.4247436523438 0 47600.383 c 5
811.427734375 0 21836.562
812.4301147460938 0 4910.82
814.4033813476562 0 4271.3076
814.883544921875 0 1903.4044 y 3
815.3975219726562 0 3099.7747
816.4005737304688 0 951.35834
819.3981323242188 0 1179.9558
831.4324340820312 0 1538.4344
832.4246826171875 0 1098.4548
839.3974609375 0 2635.15 w 9
840.4039916992188 0 2266.439
841.3939208984375 0 1185.6873
841.8956298828125 0 1728.8823 w 2
842.3995361328125 0 1236.9426
842.954833984375 0 960.38275
843.4442138671875 0 1202.8392
845.9014282226562 0 4032.41
846.3984375 0 14765.792 c 12
846.8978881835938 0 9149.43
847.3997192382812 0 6203.5913
847.9015502929688 0 2335.6528
848.4028930664062 0 1388.2792
848.91455078125 0 1253.4238
849.4083251953125 0 835.35406
867.423095703125 0 10515.282
868.425537109375 0 5409.3784
869.4261474609375 0 1506.6763
870.412353515625 0 7624.36 y Ammonia loss 2
870.9078979492188 0 13336.855 z 2
871.4105224609375 0 8507.328
871.9104614257812 0 5729.0195
872.41015625 0 2251.852
872.9049072265625 0 801.0729
875.4428100585938 0 781.1559
879.414306640625 0 818.5918
880.4534912109375 0 1237.7417
881.4429931640625 0 3384.45
881.9441528320312 0 1611.142
882.4420776367188 0 1926.458
888.4415893554688 0 4209.9023
888.9380493164062 0 1953.0066
889.4410400390625 0 1468.7488
890.45166015625 0 1179.4901
896.4411010742188 0 722.18365 y Water loss 9
898.4100952148438 0 28403.158 z 9
899.4144897460938 0 18640.547
900.4168090820312 0 9737.048
901.4212036132812 0 2421.8123
901.941650390625 0 788.3524 c Ammonia loss 13
908.430908203125 0 1397.3812 c Ammonia loss 6
909.953369140625 0 1739.7374
910.443603515625 0 9962.977 c 13
910.9439697265625 0 6389.5024
911.4450073242188 0 4144.224
911.9490356445312 0 2368.541
914.4278564453125 0 2770.2883 y 9
915.4424438476562 0 3597.3777
916.44580078125 0 2853.2334
917.4330444335938 0 1009.07526
924.4550170898438 0 1777.6388
925.4540405273438 0 8233.557 c 6
926.4559326171875 0 4253.4756
927.4502563476562 0 968.5489
929.9647216796875 0 898.1852
930.4733276367188 0 2890.748
930.9712524414062 0 2343.9697
931.47314453125 0 1652.426
931.9607543945312 0 900.89874
932.478515625 0 3922.9075
933.4721069335938 0 2071.3567
935.4592895507812 0 952.1073
937.4793701171875 0 1951.7544
937.9774169921875 0 1030.8329
938.47119140625 0 7714.7134
938.9649047851562 0 2065.0542
939.44189453125 0 8549.872 z 1
939.9395751953125 0 10450.536
940.4415283203125 0 6112.045
940.9424438476562 0 2276.4563
945.4832153320312 0 3389.62
946.4864501953125 0 2374.51
947.4576416015625 0 826.8025 y 1
951.9696044921875 0 1811.0608
952.47998046875 0 2717.1155
952.9769287109375 0 1829.7842
953.4738159179688 0 1586.9427
957.9816284179688 0 841.43945
959.4812622070312 0 1134.2035
959.9818725585938 0 4091.2085
960.482177734375 0 5960.2397
960.9830932617188 0 3313.3955
961.4840698242188 0 1619.8655
962.4752197265625 0 1496.5023
963.458984375 0 974.0482
965.4666137695312 0 904.1274 c Ammonia loss 7
966.4774169921875 0 4704.795
966.9745483398438 0 4274.449
967.4760131835938 0 3330.3425
967.9749145507812 0 1828.044
968.4774169921875 0 1757.201
973.4767456054688 0 1913.6522
973.9791259765625 0 975.77563
974.4864501953125 0 6981.9165
974.9841918945312 0 17255.467
975.486083984375 0 15174.228
975.9874877929688 0 8879.531
976.4889526367188 0 4618.543
976.9871215820312 0 1835.547
977.4835815429688 0 1521.6512
981.4878540039062 0 8964.123
981.9888916015625 0 8442.982
982.4802856445312 0 32154.723 c 7
982.98828125 0 6647.166
983.4786987304688 0 12835.899
983.9878540039062 0 1278.3842
984.4855346679688 0 3546.686
986.9781494140625 0 1210.9789
987.4667358398438 0 1552.3253
987.985107421875 0 961.70905
988.4779663085938 0 5623.9224
989.4763793945312 0 3906.0483
990.4743041992188 0 2127.5896
995.4892578125 0 2726.6675
995.9857788085938 0 13379.845
996.4830322265625 0 26551.271
996.9833984375 0 22903.252
997.484130859375 0 13138.199
997.9850463867188 0 6465.7705
998.483642578125 0 2448.2366
998.9881591796875 0 1049.316
1003.4879150390625 0 1639.9623
1003.9901123046875 0 5454.01
1004.494873046875 0 18501.664
1004.995361328125 0 27311.098
1005.4966430664062 0 27536.016
1005.9960327148438 0 14898.782
1006.4965209960938 0 9964.143
1006.9957885742188 0 3119.6719
1007.4849853515625 0 3532.9849
1007.6144409179688 0 2075.4697
1008.4760131835938 0 983.79114
1026.5062255859375 0 3662.784 z 8
1027.5113525390625 0 8202.398
1028.5155029296875 0 3736.5862
1029.514892578125 0 1905.8313
1066.55224609375 0 1664.2449
1067.5614013671875 0 1171.409
1083.525634765625 0 1243.4727 z 7
1084.5382080078125 0 3486.2969
1085.5384521484375 0 1329.02
1099.5400390625 0 1740.4167 y 7
1100.5499267578125 0 1463.4869
1109.583984375 0 8483.255
1110.5701904296875 0 28548.607 c 8
1111.5733642578125 0 18431.355
1112.5755615234375 0 6194.756
1113.574462890625 0 1394.6581
1140.56103515625 0 1270.6984
1141.55712890625 0 1150.1807
1142.5623779296875 0 1372.5121
1195.6082763671875 0 1722.7136
1196.6031494140625 0 3320.9185
1197.5936279296875 0 1653.064
1198.571044921875 0 6401.2065 z 6
1199.56298828125 0 13285.48
1200.56396484375 0 6478.359
1201.5667724609375 0 2967.4514
1202.561279296875 0 1555.7585
1238.6214599609375 0 2333.4385
1239.616455078125 0 6808.4346 c 9
1240.6185302734375 0 5907.3247
1241.6180419921875 0 3319.1719
1242.6185302734375 0 1051.6244
1310.6556396484375 0 1005.36786 y Ammonia loss 5
1311.65283203125 0 3968.4895 z 5
1312.64697265625 0 6573.119
1313.652099609375 0 2971.5286
1314.6583251953125 0 1666.0679
1316.6678466796875 0 893.23346
1358.6710205078125 0 1588.4241
1359.670166015625 0 2018.0115
1360.6689453125 0 1219.799
1386.6517333984375 0 984.765
1401.683349609375 0 1656.2604
1402.6800537109375 0 4077.0845 c 10
1403.68115234375 0 3712.3481
1404.6827392578125 0 1906.3762
1439.71875 0 1125.291
1497.72705078125 0 2758.1213 z 4
1498.725341796875 0 3231.6016
1499.725830078125 0 2459.3635
1500.72509765625 0 989.4506
1513.7520751953125 0 1013.43005 c Ammonia loss 11
1514.7598876953125 0 1246.4716
1529.7802734375 0 756.56396
1530.76708984375 0 2094.0623 c 11
1531.7763671875 0 2226.4058
1532.78125 0 922.06995
1554.740966796875 0 1472.5391
1568.7464599609375 0 1158.0262
1612.7440185546875 0 3300.6406 z 3
1613.7509765625 0 6949.603
1614.75390625 0 4051.6787
1615.754150390625 0 1707.4375
1682.80078125 0 890.0386 w 2
1690.7962646484375 0 874.948
1691.7864990234375 0 2985.9976 c 12
1692.791748046875 0 3525.6965
1693.7921142578125 0 2045.9524
1694.7978515625 0 959.3737
1740.8184814453125 0 3040.8467 z 2
1741.8121337890625 0 4417.0693
1742.81494140625 0 3009.3704
1743.8128662109375 0 897.40485
1744.7899169921875 0 818.6576
1776.87255859375 0 967.054
1777.8831787109375 0 1285.8857
1821.892578125 0 1000.75726
1877.901611328125 0 1148.8789 z 1
1878.8719482421875 0 2000.3616
1879.865966796875 0 1325.2571
1903.9453125 0 1160.2526
1918.98583984375 0 865.5171
1919.98388671875 0 803.13306
1920.9752197265625 0 1344.5906
1921.9522705078125 0 812.256
1922.9705810546875 0 948.6618
1932.9193115234375 0 1044.3871
1948.968505859375 0 950.3836
1949.9599609375 0 1723.5684
1950.975341796875 0 2413.8071
1951.977783203125 0 1805.3833
1963.9873046875 0 931.1571
1964.972412109375 0 1466.941
1980.9669189453125 0 804.3612
1991.9580078125 0 2160.3396
1992.9609375 0 2773.883
1993.9599609375 0 1822.3881
1994.9532470703125 0 1219.3656
1995.9664306640625 0 825.9374
2007.96435546875 0 2235.0837
2008.9788818359375 0 3396.1099
2009.9876708984375 0 3284.9233
2010.9765625 0 2457.1812
2011.9796142578125 0 853.71954
3090.9775390625 0 697.70825

Spectrum Details

|  |  |
| --- | --- |
| Matched peaks? Matched peaks The total absolute number of peaks matched. Additionally in brackets the total fraction of peaks matched and the total number of peaks is shown. | 106 (15.43% of 687) |
| FDR? FDR The false discovery rate estimated for this peptide. It is calculated by matching all theoretical fragments with a non-integer shift with the raw peaks for this spectrum. This is done with 40 different shifts. The resulting percentage is the average number of annotated peaks over the number of annotated peaks with the correct spectrum. | 0.40% |
| Satellite FDR? Satellite FDR See the FDR for details on its calculation. This satellite ion specific FDR only contains the satellite ions (d/w) for I/L/J positions. | 0.00% |
| PSM Score? PSM Score The PSM Score as given by Hecklib to this annotated spectrum. It is shown with three significant figures. | 487 |

#### Spectrum 4177? Spectrum 4177 The raw spectrum of this peptide as annotated by Hecklib. The fragments are coloured according to ion type (see legend). Any peaks with a star '\*' as text can be hovered over to see the full details, first the ion type second the mass shift type. By hovering over the amino acids in the peptide or ions in the legend the corresponding peaks are highlighted. By toggling the 'Unassigned' label you can turn the background (unassigned) peaks on or off in the plot. By updating the slider in the Ion legend you can update the spectrum to only show the top X% of the peaks with labels. The top X% means any peak that is within X% of the highest intensity. By dragging in the spectrum you can zoom in to a specific part of the spectrum and use 'Zoom Out' to get back to the original zoom level. The annotation of the spectrum is based on the given sequence in the peptides file and is done with different software so inconsistencies are likely. The peaks are annotated based on the given sequence, with 20 ppm tolerance.

Copy Data

##### Spectrum 4177 (TSV)

###### Preview

```
Loading example...
```

*Click on the button to copy the data to your clipboard.*

Mz MinMz MaxIntensity Max

WidthHeightPeptide font sizePeptide stroke widthSpectrum font sizeSpectrum stroke widthCompact peptide

Ion legend

wxyz

abcd

OtherUnassignedIon^Charge^_Position_Show for top:%

JHQDWLDGKEYKCKK

01.69e+43.37e+45.06e+46.75e+4

Zoom Out

c^+4^_6_c^+1^_2_z^+1^_2_y^+1^_2_c^+2^_5_y^+3^_8_c^+1^_3_y^+2^_5_c^+1^_3_y^+3^_9_w^+2^_6_w^+2^_7_c^+1^_4_y^+3^_11_y^+3^_11_z^+2^_7_c^+1^_4_y^+2^_7_y^+2^_7_z^+2^_8_y^+2^_8_c^+2^_9_c^+2^_9_c^+2^_9_w^+2^_9_y^+3^_13_y^+3^_13_z^+1^_4_y^+2^_9_y^+2^_9_z^+2^_9_z^+1^_4_c^+3^_14_y^+2^_9_c^+3^_14_y^+1^_4_c^+2^_10_y^+3^_14_w^+2^_10_y^+3^_14_y^+2^_10_z^+2^_10_y^+2^_10_c^+1^_5_c^+2^_11_c^+1^_5_z^+2^_11_y^+2^_11_y^+2^_11_z^+2^_11_c^+2^_12_y^+1^_5_z^+1^_5_c^+2^_12_y^+1^_5_c^+1^_6_z^+2^_12_y^+2^_12_c^+1^_6_w^+1^_6_w^+2^_13_c^+2^_13_y^+2^_13_z^+2^_13_z^+1^_6_y^+1^_6_z^+1^_6_c^+2^_14_y^+1^_6_c^+2^_14_c^+1^_7_c^+1^_7_z^+2^_14_c^+1^_8_z^+1^_7_z^+1^_8_c^+1^_9_z^+1^_9_y^+1^_9_c^+1^_10_z^+1^_10_c^+1^_11_z^+1^_11_c^+1^_12_c^+1^_12_z^+1^_12_c^+1^_13_z^+1^_13_

0512102515372049

Fragment Matches Table

Show background peaks

| Position | Ion type | Intensity | mz Theoretical | mz Error (Th) | mz Error (ppm) | Charge | Series Number |
| --- | --- | --- | --- | --- | --- | --- | --- |
| - | - | 361.3 | 126.1 | - | - | 0 | - |
| - | - | 1534 | 129.1 | - | - | 0 | - |
| - | - | 571.6 | 136.1 | - | - | 0 | - |
| - | - | 443.4 | 137.1 | - | - | 0 | - |
| - | - | 612.2 | 146.1 | - | - | 0 | - |
| - | - | 532.6 | 149 | - | - | 0 | - |
| - | - | 474.5 | 149.1 | - | - | 0 | - |
| - | - | 2880 | 155.1 | - | - | 0 | - |
| - | - | 429.4 | 163.1 | - | - | 0 | - |
| - | - | 3185 | 166.1 | - | - | 0 | - |
| - | - | 607.9 | 171.9 | - | - | 0 | - |
| - | - | 558.4 | 173.1 | - | - | 0 | - |
| - | - | 1177 | 173.4 | - | - | 0 | - |
| - | - | 1280 | 175.1 | - | - | 0 | - |
| - | - | 889.9 | 181.2 | - | - | 0 | - |
| - | - | 1029 | 183.1 | - | - | 0 | - |
| - | - | 3050 | 185.2 | - | - | 0 | - |
| - | - | 513.5 | 187.1 | - | - | 0 | - |
| 6 | c | 2840 | 203.1 | 0.000282 | 1.389 | +4 | 6 |
| - | - | 1409 | 205.1 | - | - | 0 | - |
| - | - | 4627 | 209.2 | - | - | 0 | - |
| - | - | 533.3 | 210.2 | - | - | 0 | - |
| - | - | 1106 | 213.1 | - | - | 0 | - |
| - | - | 1825 | 215.1 | - | - | 0 | - |
| - | - | 665.9 | 216.1 | - | - | 0 | - |
| - | - | 2056 | 223.2 | - | - | 0 | - |
| - | - | 800.6 | 230.1 | - | - | 0 | - |
| - | - | 3199 | 234.1 | - | - | 0 | - |
| - | - | 633.8 | 235.1 | - | - | 0 | - |
| - | - | 1.089E+04 | 237.2 | - | - | 0 | - |
| - | - | 2111 | 238.2 | - | - | 0 | - |
| - | - | 1.232E+04 | 251.2 | - | - | 0 | - |
| - | - | 1633 | 252.2 | - | - | 0 | - |
| 2 | c | 937.8 | 268.2 | 0.0004752 | 1.772 | +1 | 2 |
| - | - | 631.3 | 278.1 | - | - | 0 | - |
| - | - | 662.3 | 282.2 | - | - | 0 | - |
| - | - | 536.7 | 290.8 | - | - | 0 | - |
| - | - | 1412 | 299.1 | - | - | 0 | - |
| - | - | 693.2 | 300.1 | - | - | 0 | - |
| - | - | 895.3 | 301.1 | - | - | 0 | - |
| - | - | 772 | 309.2 | - | - | 0 | - |
| 14 | z | 813.3 | 317.2 | 0.004188 | 13.2 | +1 | 2 |
| - | - | 767.5 | 318.2 | - | - | 0 | - |
| 14 | y | 598.3 | 333.2 | 0.005975 | 17.93 | +1 | 2 |
| - | - | 2270 | 334.7 | - | - | 0 | - |
| - | - | 1042 | 335.2 | - | - | 0 | - |
| 5 | c | 1484 | 348.7 | 0.002449 | 7.025 | +2 | 5 |
| 8 | y | 1213 | 367.2 | 0.003387 | 9.224 | +3 | 8 |
| - | - | 751.9 | 376.1 | - | - | 0 | - |
| 3 | c | 3321 | 379.2 | 0.0001811 | 0.4776 | +1 | 3 |
| - | - | 595.1 | 385.2 | - | - | 0 | - |
| - | - | 2838 | 387.2 | - | - | 0 | - |
| - | - | 661.9 | 389.5 | - | - | 0 | - |
| 11 | y | 673.3 | 393.2 | 0.004757 | 12.1 | +2 | 5 |
| - | - | 672.6 | 394.2 | - | - | 0 | - |
| - | - | 589.1 | 396.2 | - | - | 0 | - |
| 3 | c | 3705 | 396.2 | 0.0002473 | 0.6242 | +1 | 3 |
| - | - | 3197 | 401 | - | - | 0 | - |
| - | - | 2370 | 402 | - | - | 0 | - |
| - | - | 1681 | 403 | - | - | 0 | - |
| 7 | y | 1035 | 405.5 | 0.001829 | 4.511 | +3 | 9 |
| - | - | 3930 | 419 | - | - | 0 | - |
| - | - | 2045 | 420 | - | - | 0 | - |
| 10 | w | 1.024E+04 | 420.2 | 0.004119 | 9.803 | +2 | 6 |
| - | - | 5774 | 420.7 | - | - | 0 | - |
| - | - | 1580 | 421 | - | - | 0 | - |
| - | - | 2341 | 421.2 | - | - | 0 | - |
| - | - | 733.3 | 421.7 | - | - | 0 | - |
| - | - | 843 | 431.2 | - | - | 0 | - |
| - | - | 580.4 | 449.1 | - | - | 0 | - |
| - | - | 1422 | 460.7 | - | - | 0 | - |
| - | - | 1095 | 461.2 | - | - | 0 | - |
| - | - | 689.3 | 462.2 | - | - | 0 | - |
| - | - | 794.8 | 468.3 | - | - | 0 | - |
| - | - | 1522 | 468.8 | - | - | 0 | - |
| - | - | 647.3 | 472.2 | - | - | 0 | - |
| - | - | 599.6 | 474.3 | - | - | 0 | - |
| - | - | 843.1 | 479.2 | - | - | 0 | - |
| - | - | 1671 | 484.3 | - | - | 0 | - |
| 9 | w | 1240 | 484.7 | 0.006708 | 13.84 | +2 | 7 |
| - | - | 1041 | 484.8 | - | - | 0 | - |
| - | - | 6045 | 489.1 | - | - | 0 | - |
| - | - | 3694 | 490.1 | - | - | 0 | - |
| - | - | 2643 | 491.1 | - | - | 0 | - |
| 4 | c | 1.689E+04 | 494.2 | 0.0003297 | 0.6671 | +1 | 4 |
| - | - | 4674 | 495.2 | - | - | 0 | - |
| - | - | 677.8 | 495.8 | - | - | 0 | - |
| - | - | 1511 | 496.2 | - | - | 0 | - |
| - | - | 2232 | 496.8 | - | - | 0 | - |
| - | - | 974.2 | 497.3 | - | - | 0 | - |
| - | - | 732.1 | 502 | - | - | 0 | - |
| - | - | 2588 | 504.2 | - | - | 0 | - |
| 5 | y | 2977 | 504.6 | 0.0008189 | 1.623 | +3 | 11 |
| 5 | y | 2284 | 504.9 | 0.004977 | 9.858 | +3 | 11 |
| 9 | z | 885.6 | 505.2 | 0.007716 | 15.27 | +2 | 7 |
| - | - | 7482 | 506.2 | - | - | 0 | - |
| - | - | 9711 | 506.3 | - | - | 0 | - |
| - | - | 596.2 | 506.3 | - | - | 0 | - |
| - | - | 985.5 | 506.3 | - | - | 0 | - |
| - | - | 1.014E+04 | 506.5 | - | - | 0 | - |
| - | - | 595.2 | 506.6 | - | - | 0 | - |
| - | - | 7627 | 506.7 | - | - | 0 | - |
| - | - | 5857 | 507 | - | - | 0 | - |
| - | - | 880.5 | 507.2 | - | - | 0 | - |
| - | - | 1897 | 507.3 | - | - | 0 | - |
| - | - | 808.7 | 507.5 | - | - | 0 | - |
| - | - | 1982 | 507.9 | - | - | 0 | - |
| 4 | c | 1.373E+04 | 511.3 | 0.0003429 | 0.6706 | +1 | 4 |
| - | - | 4390 | 512.3 | - | - | 0 | - |
| 9 | y | 751 | 512.8 | 0.002121 | 4.137 | +2 | 7 |
| - | - | 563.1 | 513.7 | - | - | 0 | - |
| - | - | 2611 | 515.3 | - | - | 0 | - |
| - | - | 1128 | 516.3 | - | - | 0 | - |
| - | - | 783.4 | 518.8 | - | - | 0 | - |
| 9 | y | 3394 | 521.8 | 0.004535 | 8.691 | +2 | 7 |
| - | - | 2029 | 522.3 | - | - | 0 | - |
| - | - | 660 | 522.8 | - | - | 0 | - |
| - | - | 794.4 | 533.8 | - | - | 0 | - |
| - | - | 937.3 | 534.3 | - | - | 0 | - |
| - | - | 1305 | 537.3 | - | - | 0 | - |
| 8 | z | 1454 | 542.3 | 0.004501 | 8.3 | +2 | 8 |
| - | - | 1165 | 542.8 | - | - | 0 | - |
| - | - | 1350 | 547.3 | - | - | 0 | - |
| - | - | 788.8 | 548.2 | - | - | 0 | - |
| 8 | y | 3398 | 550.3 | 0.003304 | 6.004 | +2 | 8 |
| - | - | 2953 | 550.8 | - | - | 0 | - |
| - | - | 650.8 | 551.3 | - | - | 0 | - |
| - | - | 2059 | 552.3 | - | - | 0 | - |
| - | - | 1396 | 553.3 | - | - | 0 | - |
| - | - | 1116 | 554.3 | - | - | 0 | - |
| 9 | c | 1758 | 554.8 | 0.006534 | 11.78 | +2 | 9 |
| 9 | c | 1798 | 555.3 | 0.00476 | 8.573 | +2 | 9 |
| - | - | 782.2 | 555.8 | - | - | 0 | - |
| - | - | 3681 | 563.3 | - | - | 0 | - |
| 9 | c | 1.86E+04 | 563.8 | 0.003143 | 5.575 | +2 | 9 |
| - | - | 1003 | 564.2 | - | - | 0 | - |
| - | - | 9200 | 564.3 | - | - | 0 | - |
| - | - | 3128 | 564.8 | - | - | 0 | - |
| - | - | 1180 | 565.3 | - | - | 0 | - |
| - | - | 1651 | 570.3 | - | - | 0 | - |
| - | - | 2588 | 570.3 | - | - | 0 | - |
| - | - | 1800 | 570.7 | - | - | 0 | - |
| - | - | 801 | 571.2 | - | - | 0 | - |
| - | - | 1.12E+04 | 571.3 | - | - | 0 | - |
| - | - | 586.6 | 571.3 | - | - | 0 | - |
| - | - | 6449 | 571.8 | - | - | 0 | - |
| - | - | 1809 | 572.3 | - | - | 0 | - |
| - | - | 1222 | 572.8 | - | - | 0 | - |
| - | - | 702.9 | 576.8 | - | - | 0 | - |
| 7 | w | 2252 | 577.3 | 0.001811 | 3.138 | +2 | 9 |
| - | - | 1451 | 577.8 | - | - | 0 | - |
| - | - | 862.8 | 578.8 | - | - | 0 | - |
| 3 | y | 952.3 | 585.6 | 0.003355 | 5.73 | +3 | 13 |
| 3 | y | 1180 | 585.9 | 0.002868 | 4.895 | +3 | 13 |
| - | - | 888.1 | 586.6 | - | - | 0 | - |
| - | - | 689.1 | 588.3 | - | - | 0 | - |
| 12 | z | 715.1 | 589.3 | 0.00234 | 3.97 | +1 | 4 |
| - | - | 632.1 | 592.3 | - | - | 0 | - |
| 7 | y | 1490 | 598.8 | 0.003375 | 5.637 | +2 | 9 |
| 7 | y | 2556 | 599.3 | 0.005044 | 8.417 | +2 | 9 |
| 7 | z | 1.127E+04 | 599.8 | 0.003812 | 6.356 | +2 | 9 |
| - | - | 7178 | 600.3 | - | - | 0 | - |
| - | - | 3279 | 600.8 | - | - | 0 | - |
| - | - | 656.3 | 600.9 | - | - | 0 | - |
| - | - | 1300 | 601.3 | - | - | 0 | - |
| - | - | 714.1 | 602.3 | - | - | 0 | - |
| 12 | z | 9765 | 606.3 | 0.008869 | 14.63 | +1 | 4 |
| - | - | 718.7 | 606.8 | - | - | 0 | - |
| 14 | c | 1151 | 607 | 0.008917 | 14.69 | +3 | 14 |
| - | - | 5446 | 607.3 | - | - | 0 | - |
| - | - | 1381 | 607.6 | - | - | 0 | - |
| 7 | y | 1.46E+04 | 607.8 | 0.003958 | 6.512 | +2 | 9 |
| - | - | 8632 | 608.3 | - | - | 0 | - |
| - | - | 4167 | 608.8 | - | - | 0 | - |
| - | - | 1223 | 609.3 | - | - | 0 | - |
| - | - | 982.8 | 611.6 | - | - | 0 | - |
| - | - | 749.9 | 612 | - | - | 0 | - |
| - | - | 2455 | 612.3 | - | - | 0 | - |
| 14 | c | 8572 | 612.6 | 0.0004536 | 0.7405 | +3 | 14 |
| - | - | 1992 | 612.8 | - | - | 0 | - |
| - | - | 8124 | 613 | - | - | 0 | - |
| - | - | 6473 | 613.3 | - | - | 0 | - |
| - | - | 1248 | 613.6 | - | - | 0 | - |
| - | - | 1002 | 615.3 | - | - | 0 | - |
| - | - | 1005 | 615.8 | - | - | 0 | - |
| - | - | 1841 | 621 | - | - | 0 | - |
| - | - | 3179 | 621.3 | - | - | 0 | - |
| - | - | 2127 | 621.6 | - | - | 0 | - |
| - | - | 984.9 | 622 | - | - | 0 | - |
| 12 | y | 6535 | 622.3 | 0.009588 | 15.41 | +1 | 4 |
| - | - | 2732 | 623.3 | - | - | 0 | - |
| - | - | 1732 | 624.3 | - | - | 0 | - |
| - | - | 894.3 | 625.3 | - | - | 0 | - |
| - | - | 2631 | 626.3 | - | - | 0 | - |
| - | - | 2928 | 626.6 | - | - | 0 | - |
| - | - | 673.6 | 626.8 | - | - | 0 | - |
| - | - | 2464 | 627 | - | - | 0 | - |
| - | - | 986.4 | 627.3 | - | - | 0 | - |
| - | - | 3792 | 627.8 | - | - | 0 | - |
| 10 | c | 1.451E+04 | 628.3 | 0.00327 | 5.205 | +2 | 10 |
| - | - | 1015 | 628.4 | - | - | 0 | - |
| - | - | 1.079E+04 | 628.8 | - | - | 0 | - |
| - | - | 4327 | 629.3 | - | - | 0 | - |
| - | - | 1492 | 629.8 | - | - | 0 | - |
| - | - | 715.2 | 630.6 | - | - | 0 | - |
| - | - | 762 | 631 | - | - | 0 | - |
| 2 | y | 1599 | 631.3 | 0.0106 | 16.79 | +3 | 14 |
| - | - | 2141 | 631.6 | - | - | 0 | - |
| - | - | 1007 | 632 | - | - | 0 | - |
| - | - | 1974 | 634.3 | - | - | 0 | - |
| 6 | w | 3659 | 634.8 | 0.002893 | 4.557 | +2 | 10 |
| - | - | 3228 | 635.3 | - | - | 0 | - |
| - | - | 1150 | 635.8 | - | - | 0 | - |
| - | - | 773.5 | 636.3 | - | - | 0 | - |
| 2 | y | 1224 | 637.3 | 0.0008499 | 1.334 | +3 | 14 |
| - | - | 713.9 | 637.4 | - | - | 0 | - |
| - | - | 1191 | 640 | - | - | 0 | - |
| - | - | 1236 | 640.3 | - | - | 0 | - |
| - | - | 1166 | 640.6 | - | - | 0 | - |
| - | - | 729.4 | 641 | - | - | 0 | - |
| - | - | 1059 | 641.3 | - | - | 0 | - |
| - | - | 4435 | 645.3 | - | - | 0 | - |
| - | - | 1.891E+04 | 645.7 | - | - | 0 | - |
| - | - | 2.259E+04 | 646 | - | - | 0 | - |
| - | - | 1.029E+04 | 646.3 | - | - | 0 | - |
| - | - | 3436 | 646.7 | - | - | 0 | - |
| - | - | 2603 | 647 | - | - | 0 | - |
| - | - | 1489 | 649.7 | - | - | 0 | - |
| - | - | 1210 | 650 | - | - | 0 | - |
| - | - | 771.7 | 650.3 | - | - | 0 | - |
| - | - | 1495 | 651.3 | - | - | 0 | - |
| - | - | 983.5 | 651.6 | - | - | 0 | - |
| - | - | 842.3 | 654.7 | - | - | 0 | - |
| - | - | 1931 | 655 | - | - | 0 | - |
| - | - | 9875 | 655.3 | - | - | 0 | - |
| - | - | 1.525E+04 | 655.6 | - | - | 0 | - |
| 6 | y | 1505 | 655.8 | 0.007079 | 10.79 | +2 | 10 |
| - | - | 1.519E+04 | 656 | - | - | 0 | - |
| 6 | z | 1.201E+04 | 656.3 | 0.005195 | 7.915 | +2 | 10 |
| - | - | 5419 | 656.7 | - | - | 0 | - |
| - | - | 1789 | 656.8 | - | - | 0 | - |
| - | - | 1256 | 657 | - | - | 0 | - |
| - | - | 1467 | 657.3 | - | - | 0 | - |
| - | - | 1448 | 660 | - | - | 0 | - |
| - | - | 2595 | 660.3 | - | - | 0 | - |
| - | - | 849.4 | 663.8 | - | - | 0 | - |
| 6 | y | 8496 | 664.3 | 0.003693 | 5.558 | +2 | 10 |
| - | - | 4599 | 664.8 | - | - | 0 | - |
| - | - | 1773 | 665.3 | - | - | 0 | - |
| - | - | 863.1 | 665.8 | - | - | 0 | - |
| - | - | 971.1 | 669 | - | - | 0 | - |
| - | - | 3353 | 669.3 | - | - | 0 | - |
| - | - | 4973 | 669.7 | - | - | 0 | - |
| - | - | 754.5 | 669.8 | - | - | 0 | - |
| - | - | 3573 | 670 | - | - | 0 | - |
| - | - | 2554 | 670.3 | - | - | 0 | - |
| - | - | 898.1 | 673.7 | - | - | 0 | - |
| - | - | 1173 | 674 | - | - | 0 | - |
| - | - | 831.6 | 674.3 | - | - | 0 | - |
| - | - | 1476 | 674.4 | - | - | 0 | - |
| - | - | 6917 | 674.7 | - | - | 0 | - |
| - | - | 4.251E+04 | 675 | - | - | 0 | - |
| - | - | 6.679E+04 | 675.3 | - | - | 0 | - |
| - | - | 4.98E+04 | 675.7 | - | - | 0 | - |
| - | - | 3.003E+04 | 676 | - | - | 0 | - |
| - | - | 1.206E+04 | 676.3 | - | - | 0 | - |
| - | - | 4959 | 676.7 | - | - | 0 | - |
| - | - | 896 | 677 | - | - | 0 | - |
| - | - | 1295 | 682.4 | - | - | 0 | - |
| - | - | 1165 | 682.4 | - | - | 0 | - |
| - | - | 1631 | 687.8 | - | - | 0 | - |
| - | - | 853.7 | 688.3 | - | - | 0 | - |
| - | - | 768.1 | 688.8 | - | - | 0 | - |
| - | - | 4619 | 690.8 | - | - | 0 | - |
| - | - | 1.456E+04 | 691.3 | - | - | 0 | - |
| - | - | 9123 | 691.8 | - | - | 0 | - |
| - | - | 7999 | 692.3 | - | - | 0 | - |
| - | - | 2335 | 692.8 | - | - | 0 | - |
| - | - | 970.3 | 693.3 | - | - | 0 | - |
| 5 | c | 4850 | 696.3 | 0.00479 | 6.879 | +1 | 5 |
| - | - | 1564 | 697.3 | - | - | 0 | - |
| - | - | 3287 | 709.3 | - | - | 0 | - |
| 11 | c | 1.378E+04 | 709.8 | 0.003588 | 5.055 | +2 | 11 |
| - | - | 1.017E+04 | 710.3 | - | - | 0 | - |
| - | - | 3932 | 710.8 | - | - | 0 | - |
| - | - | 1737 | 711.3 | - | - | 0 | - |
| - | - | 850.3 | 711.8 | - | - | 0 | - |
| 5 | c | 1.304E+04 | 713.3 | 0.00473 | 6.631 | +1 | 5 |
| - | - | 4868 | 714.3 | - | - | 0 | - |
| - | - | 1489 | 715.3 | - | - | 0 | - |
| - | - | 688.7 | 726.4 | - | - | 0 | - |
| - | - | 983.5 | 726.9 | - | - | 0 | - |
| - | - | 1137 | 727.4 | - | - | 0 | - |
| - | - | 1463 | 733.4 | - | - | 0 | - |
| - | - | 839.6 | 733.9 | - | - | 0 | - |
| - | - | 953.9 | 734.4 | - | - | 0 | - |
| - | - | 811.6 | 742.4 | - | - | 0 | - |
| 5 | z | 1218 | 748.8 | 0.004612 | 6.159 | +2 | 11 |
| - | - | 705.5 | 753.4 | - | - | 0 | - |
| - | - | 1.188E+04 | 755.4 | - | - | 0 | - |
| - | - | 7987 | 755.9 | - | - | 0 | - |
| 5 | y | 7905 | 756.4 | 0.002915 | 3.855 | +2 | 11 |
| - | - | 3694 | 756.4 | - | - | 0 | - |
| 5 | y | 6782 | 756.9 | 0.00764 | 10.09 | +2 | 11 |
| 5 | z | 2619 | 757.4 | 0.002812 | 3.713 | +2 | 11 |
| - | - | 1231 | 757.9 | - | - | 0 | - |
| 12 | c | 751.6 | 765.4 | 0.01052 | 13.74 | +2 | 12 |
| 11 | y | 796.2 | 768.4 | 0.01133 | 14.75 | +1 | 5 |
| 11 | z | 1.625E+04 | 769.4 | 0.009515 | 12.37 | +1 | 5 |
| - | - | 7016 | 770.4 | - | - | 0 | - |
| - | - | 2698 | 771.4 | - | - | 0 | - |
| - | - | 1041 | 772.4 | - | - | 0 | - |
| 12 | c | 4039 | 773.9 | 0.003836 | 4.957 | +2 | 12 |
| - | - | 1930 | 774.4 | - | - | 0 | - |
| - | - | 1164 | 774.9 | - | - | 0 | - |
| - | - | 1212 | 782.4 | - | - | 0 | - |
| - | - | 732.7 | 783.4 | - | - | 0 | - |
| 11 | y | 4838 | 785.4 | 0.01011 | 12.87 | +1 | 5 |
| - | - | 1606 | 786.4 | - | - | 0 | - |
| - | - | 811.9 | 787.4 | - | - | 0 | - |
| - | - | 1905 | 789.5 | - | - | 0 | - |
| - | - | 1052 | 790.5 | - | - | 0 | - |
| - | - | 1148 | 792.9 | - | - | 0 | - |
| 6 | c | 1010 | 809.4 | 0.007457 | 9.213 | +1 | 6 |
| - | - | 1369 | 810.4 | - | - | 0 | - |
| - | - | 1065 | 812.9 | - | - | 0 | - |
| 4 | z | 7962 | 814.9 | 0.001626 | 1.996 | +2 | 12 |
| - | - | 6514 | 815.4 | - | - | 0 | - |
| - | - | 3768 | 815.9 | - | - | 0 | - |
| - | - | 2534 | 816.4 | - | - | 0 | - |
| 4 | y | 1219 | 822.9 | 0.0006732 | 0.8181 | +2 | 12 |
| - | - | 942.8 | 823.4 | - | - | 0 | - |
| - | - | 712.7 | 823.9 | - | - | 0 | - |
| - | - | 1236 | 825.4 | - | - | 0 | - |
| 6 | c | 3.338E+04 | 826.4 | 0.005016 | 6.07 | +1 | 6 |
| - | - | 859.6 | 827.3 | - | - | 0 | - |
| - | - | 1.372E+04 | 827.4 | - | - | 0 | - |
| - | - | 3843 | 828.4 | - | - | 0 | - |
| - | - | 804.7 | 829.4 | - | - | 0 | - |
| 10 | w | 1705 | 839.4 | 0.007492 | 8.926 | +1 | 6 |
| - | - | 1171 | 840.4 | - | - | 0 | - |
| 3 | w | 1308 | 849.9 | 0.001927 | 2.268 | +2 | 13 |
| - | - | 2425 | 853.9 | - | - | 0 | - |
| 13 | c | 8981 | 854.4 | 0.001385 | 1.621 | +2 | 13 |
| - | - | 8580 | 854.9 | - | - | 0 | - |
| - | - | 4717 | 855.4 | - | - | 0 | - |
| - | - | 1156 | 855.9 | - | - | 0 | - |
| - | - | 783.6 | 876.4 | - | - | 0 | - |
| 3 | y | 1251 | 878.4 | 0.007238 | 8.24 | +2 | 13 |
| 3 | z | 6207 | 878.9 | 0.0007025 | 0.7993 | +2 | 13 |
| - | - | 4751 | 879.4 | - | - | 0 | - |
| - | - | 2745 | 879.9 | - | - | 0 | - |
| 10 | z | 1254 | 880.4 | 0.003518 | 3.996 | +1 | 6 |
| - | - | 8672 | 883.4 | - | - | 0 | - |
| - | - | 4152 | 884.4 | - | - | 0 | - |
| - | - | 1490 | 885.4 | - | - | 0 | - |
| - | - | 969 | 887.4 | - | - | 0 | - |
| - | - | 1623 | 889.4 | - | - | 0 | - |
| 10 | y | 3947 | 896.4 | 0.00858 | 9.572 | +1 | 6 |
| - | - | 2845 | 896.9 | - | - | 0 | - |
| - | - | 2590 | 897.4 | - | - | 0 | - |
| 10 | z | 2.096E+04 | 898.4 | 0.008101 | 9.018 | +1 | 6 |
| - | - | 1.333E+04 | 899.4 | - | - | 0 | - |
| - | - | 4513 | 900.4 | - | - | 0 | - |
| - | - | 1775 | 901.4 | - | - | 0 | - |
| - | - | 1132 | 902 | - | - | 0 | - |
| - | - | 868.5 | 903.4 | - | - | 0 | - |
| 14 | c | 823.1 | 909.9 | 0.01057 | 11.62 | +2 | 14 |
| - | - | 962.3 | 910.4 | - | - | 0 | - |
| 10 | y | 2243 | 914.4 | 0.01047 | 11.45 | +1 | 6 |
| - | - | 1153 | 915.4 | - | - | 0 | - |
| - | - | 1688 | 917.9 | - | - | 0 | - |
| 14 | c | 3966 | 918.4 | 0.002854 | 3.107 | +2 | 14 |
| - | - | 4819 | 918.9 | - | - | 0 | - |
| - | - | 3249 | 919.4 | - | - | 0 | - |
| - | - | 1033 | 919.9 | - | - | 0 | - |
| 7 | c | 662.4 | 924.4 | 0.009933 | 10.74 | +1 | 7 |
| - | - | 3098 | 939 | - | - | 0 | - |
| - | - | 1824 | 939.5 | - | - | 0 | - |
| - | - | 1052 | 940 | - | - | 0 | - |
| - | - | 1245 | 940.4 | - | - | 0 | - |
| 7 | c | 5555 | 941.4 | 0.004624 | 4.911 | +1 | 7 |
| - | - | 2709 | 942.5 | - | - | 0 | - |
| - | - | 740.8 | 945.5 | - | - | 0 | - |
| - | - | 2220 | 946 | - | - | 0 | - |
| - | - | 2682 | 946.5 | - | - | 0 | - |
| - | - | 2028 | 947 | - | - | 0 | - |
| 2 | z | 3147 | 947.4 | 0.00451 | 4.76 | +2 | 14 |
| - | - | 4548 | 947.9 | - | - | 0 | - |
| - | - | 3372 | 948.4 | - | - | 0 | - |
| - | - | 1155 | 948.9 | - | - | 0 | - |
| - | - | 2321 | 954.5 | - | - | 0 | - |
| - | - | 1084 | 959.5 | - | - | 0 | - |
| - | - | 1614 | 960 | - | - | 0 | - |
| - | - | 1135 | 960.5 | - | - | 0 | - |
| - | - | 939.7 | 961 | - | - | 0 | - |
| - | - | 1188 | 961.5 | - | - | 0 | - |
| - | - | 3357 | 968 | - | - | 0 | - |
| - | - | 4740 | 968.5 | - | - | 0 | - |
| - | - | 2171 | 969 | - | - | 0 | - |
| - | - | 1234 | 969.5 | - | - | 0 | - |
| - | - | 1003 | 973.5 | - | - | 0 | - |
| - | - | 774.3 | 974 | - | - | 0 | - |
| - | - | 3794 | 974.5 | - | - | 0 | - |
| - | - | 3586 | 975 | - | - | 0 | - |
| - | - | 3001 | 975.5 | - | - | 0 | - |
| - | - | 1552 | 976 | - | - | 0 | - |
| - | - | 699.2 | 980.5 | - | - | 0 | - |
| - | - | 1432 | 981.5 | - | - | 0 | - |
| - | - | 2125 | 982 | - | - | 0 | - |
| - | - | 4204 | 982.5 | - | - | 0 | - |
| - | - | 1.064E+04 | 983 | - | - | 0 | - |
| - | - | 1.356E+04 | 983.5 | - | - | 0 | - |
| - | - | 7797 | 984 | - | - | 0 | - |
| - | - | 4613 | 984.5 | - | - | 0 | - |
| - | - | 1877 | 985 | - | - | 0 | - |
| - | - | 1548 | 989.5 | - | - | 0 | - |
| - | - | 4694 | 990 | - | - | 0 | - |
| - | - | 4746 | 990.5 | - | - | 0 | - |
| - | - | 3459 | 991 | - | - | 0 | - |
| - | - | 1595 | 991.5 | - | - | 0 | - |
| - | - | 904.7 | 992 | - | - | 0 | - |
| - | - | 4296 | 992.6 | - | - | 0 | - |
| - | - | 3584 | 993.6 | - | - | 0 | - |
| - | - | 1175 | 994.6 | - | - | 0 | - |
| - | - | 2182 | 995.5 | - | - | 0 | - |
| - | - | 2722 | 996 | - | - | 0 | - |
| - | - | 1648 | 996.5 | - | - | 0 | - |
| - | - | 3819 | 997.5 | - | - | 0 | - |
| 8 | c | 1.577E+04 | 998.5 | 0.006597 | 6.608 | +1 | 8 |
| - | - | 8562 | 999.5 | - | - | 0 | - |
| - | - | 2784 | 1000 | - | - | 0 | - |
| - | - | 3805 | 1003 | - | - | 0 | - |
| - | - | 1.548E+04 | 1004 | - | - | 0 | - |
| - | - | 2.249E+04 | 1004 | - | - | 0 | - |
| - | - | 1.674E+04 | 1005 | - | - | 0 | - |
| - | - | 1.045E+04 | 1005 | - | - | 0 | - |
| - | - | 3402 | 1006 | - | - | 0 | - |
| - | - | 3087 | 1006 | - | - | 0 | - |
| - | - | 1755 | 1011 | - | - | 0 | - |
| - | - | 1864 | 1011 | - | - | 0 | - |
| - | - | 1949 | 1011 | - | - | 0 | - |
| - | - | 1092 | 1012 | - | - | 0 | - |
| - | - | 3570 | 1012 | - | - | 0 | - |
| - | - | 1.321E+04 | 1012 | - | - | 0 | - |
| - | - | 2.093E+04 | 1013 | - | - | 0 | - |
| - | - | 2.246E+04 | 1013 | - | - | 0 | - |
| - | - | 1.346E+04 | 1014 | - | - | 0 | - |
| - | - | 6763 | 1014 | - | - | 0 | - |
| - | - | 3779 | 1015 | - | - | 0 | - |
| - | - | 1262 | 1015 | - | - | 0 | - |
| 9 | z | 1280 | 1027 | 0.01005 | 9.787 | +1 | 7 |
| - | - | 5247 | 1028 | - | - | 0 | - |
| - | - | 3849 | 1029 | - | - | 0 | - |
| 8 | z | 1854 | 1084 | 0.003401 | 3.139 | +1 | 8 |
| - | - | 3200 | 1085 | - | - | 0 | - |
| - | - | 1837 | 1086 | - | - | 0 | - |
| - | - | 1537 | 1111 | - | - | 0 | - |
| - | - | 3270 | 1126 | - | - | 0 | - |
| 9 | c | 1.445E+04 | 1127 | 0.006849 | 6.08 | +1 | 9 |
| - | - | 1.05E+04 | 1128 | - | - | 0 | - |
| - | - | 3670 | 1129 | - | - | 0 | - |
| - | - | 1089 | 1130 | - | - | 0 | - |
| - | - | 959.3 | 1141 | - | - | 0 | - |
| - | - | 1846 | 1142 | - | - | 0 | - |
| - | - | 1091 | 1143 | - | - | 0 | - |
| 7 | z | 3097 | 1199 | 0.004046 | 3.376 | +1 | 9 |
| - | - | 9020 | 1200 | - | - | 0 | - |
| - | - | 4972 | 1201 | - | - | 0 | - |
| - | - | 2564 | 1202 | - | - | 0 | - |
| - | - | 1212 | 1213 | - | - | 0 | - |
| - | - | 1056 | 1214 | - | - | 0 | - |
| 7 | y | 1222 | 1215 | 0.004058 | 3.341 | +1 | 9 |
| - | - | 913.6 | 1240 | - | - | 0 | - |
| - | - | 1152 | 1255 | - | - | 0 | - |
| 10 | c | 4225 | 1256 | 0.009666 | 7.699 | +1 | 10 |
| - | - | 3715 | 1257 | - | - | 0 | - |
| - | - | 1577 | 1258 | - | - | 0 | - |
| 6 | z | 1227 | 1312 | 0.01215 | 9.26 | +1 | 10 |
| - | - | 4128 | 1313 | - | - | 0 | - |
| - | - | 1897 | 1314 | - | - | 0 | - |
| - | - | 1017 | 1375 | - | - | 0 | - |
| - | - | 1958 | 1376 | - | - | 0 | - |
| - | - | 948 | 1377 | - | - | 0 | - |
| - | - | 2476 | 1382 | - | - | 0 | - |
| - | - | 3206 | 1383 | - | - | 0 | - |
| - | - | 1845 | 1384 | - | - | 0 | - |
| 11 | c | 1794 | 1419 | 0.0103 | 7.262 | +1 | 11 |
| - | - | 3395 | 1420 | - | - | 0 | - |
| - | - | 1981 | 1421 | - | - | 0 | - |
| - | - | 1140 | 1513 | - | - | 0 | - |
| 5 | z | 1767 | 1514 | 0.01217 | 8.039 | +1 | 11 |
| - | - | 1391 | 1515 | - | - | 0 | - |
| 12 | c | 867.3 | 1530 | 0.001471 | 0.9613 | +1 | 12 |
| - | - | 766.7 | 1545 | - | - | 0 | - |
| 12 | c | 1860 | 1547 | 0.0004227 | 0.2733 | +1 | 12 |
| - | - | 1447 | 1548 | - | - | 0 | - |
| 4 | z | 1013 | 1629 | 0.002559 | 1.571 | +1 | 12 |
| - | - | 3926 | 1630 | - | - | 0 | - |
| - | - | 2761 | 1631 | - | - | 0 | - |
| - | - | 1436 | 1632 | - | - | 0 | - |
| - | - | 1004 | 1707 | - | - | 0 | - |
| 13 | c | 2118 | 1708 | 0.006751 | 3.953 | +1 | 13 |
| - | - | 2201 | 1709 | - | - | 0 | - |
| - | - | 1238 | 1710 | - | - | 0 | - |
| - | - | 930.2 | 1711 | - | - | 0 | - |
| 3 | z | 2167 | 1757 | 0.0007203 | 0.41 | +1 | 13 |
| - | - | 2608 | 1758 | - | - | 0 | - |
| - | - | 2738 | 1759 | - | - | 0 | - |
| - | - | 1864 | 1760 | - | - | 0 | - |
| - | - | 969.6 | 1837 | - | - | 0 | - |
| - | - | 807.8 | 1895 | - | - | 0 | - |
| - | - | 787.1 | 1896 | - | - | 0 | - |
| - | - | 726.1 | 1936 | - | - | 0 | - |
| - | - | 790.4 | 1937 | - | - | 0 | - |
| - | - | 1260 | 1938 | - | - | 0 | - |
| - | - | 1156 | 1949 | - | - | 0 | - |
| - | - | 1771 | 1966 | - | - | 0 | - |
| - | - | 1373 | 1967 | - | - | 0 | - |
| - | - | 1398 | 1968 | - | - | 0 | - |
| - | - | 743.3 | 1982 | - | - | 0 | - |
| - | - | 699.3 | 2007 | - | - | 0 | - |
| - | - | 1930 | 2008 | - | - | 0 | - |
| - | - | 2956 | 2009 | - | - | 0 | - |
| - | - | 2028 | 2010 | - | - | 0 | - |
| - | - | 1213 | 2011 | - | - | 0 | - |
| - | - | 922.2 | 2024 | - | - | 0 | - |
| - | - | 2965 | 2025 | - | - | 0 | - |
| - | - | 2828 | 2026 | - | - | 0 | - |
| - | - | 2114 | 2027 | - | - | 0 | - |
| - | - | 1354 | 2028 | - | - | 0 | - |
| - | - | 946.5 | 2029 | - | - | 0 | - |

m/z Charge Intensity FragmentType MassShift Position
126.07347869873047 0 361.33362
129.10214233398438 0 1534.0074
136.07545471191406 0 571.6359
137.1147003173828 0 443.3905
146.092529296875 0 612.2398
149.04483032226562 0 532.648
149.0934600830078 0 474.54825
155.09280395507812 0 2879.6836
163.0718994140625 0 429.39572
166.06117248535156 0 3184.621
171.87277221679688 0 607.88055
173.09234619140625 0 558.37115
173.43861389160156 0 1176.7358
175.0865936279297 0 1279.7109
181.169677734375 0 889.9325
183.11288452148438 0 1029.3505
185.16490173339844 0 3050.3398
187.1077880859375 0 513.4907
203.1024627685547 0 2839.5005 c Ammonia loss 5
205.11849975585938 0 1408.587
209.16481018066406 0 4627.392
210.1687469482422 0 533.33795
213.08731079101562 0 1105.8872
215.1385498046875 0 1825.1399
216.1432342529297 0 665.9248
223.1552734375 0 2055.5264
230.11399841308594 0 800.6032
234.12376403808594 0 3199.4778
235.12701416015625 0 633.8166
237.15988159179688 0 10893.929
238.1632843017578 0 2110.7124
251.15029907226562 0 12318.047
252.15403747558594 0 1633.105
268.1772766113281 0 937.7697 c 1
278.1492004394531 0 631.2514
282.18133544921875 0 662.2719
290.8166198730469 0 536.6838
299.0619201660156 0 1411.5126
300.0614013671875 0 693.2294
301.0591125488281 0 895.3038
309.2038269042969 0 772.046
317.1948547363281 0 813.3437 z 13
318.2027893066406 0 767.54724
333.2117919921875 0 598.2901 y 13
334.6611328125 0 2270.4863
335.1625061035156 0 1041.744
348.65863037109375 0 1483.5472 c Ammonia loss 4
367.18621826171875 0 1212.8185 y 7
376.1369934082031 0 751.93036
379.2086486816406 0 3321.3286 c Ammonia loss 2
385.2293701171875 0 595.0829
387.2236633300781 0 2838.387
389.4936218261719 0 661.85034
393.1965026855469 0 673.26294 y 10
394.1986999511719 0 672.56366
396.1923828125 0 589.05457
396.2356262207031 0 3705.1987 c 2
400.9852600097656 0 3196.5232
401.9850158691406 0 2369.6672
402.9835205078125 0 1680.6588
405.53009033203125 0 1034.5402 y 6
418.9955139160156 0 3929.7483
419.9953308105469 0 2045.0903
420.2024230957031 0 10239.581 w 9
420.7036437988281 0 5774.3457
420.9939270019531 0 1579.7695
421.2043151855469 0 2340.777
421.7023620605469 0 733.30365
431.2286376953125 0 842.9678
449.1242370605469 0 580.42896
460.7371826171875 0 1422.2771
461.23956298828125 0 1094.852
462.20294189453125 0 689.2884
468.2659912109375 0 794.7841
468.76519775390625 0 1522.0383
472.2498779296875 0 647.3441
474.2786865234375 0 599.5666
479.22552490234375 0 843.05206
484.2503967285156 0 1670.731
484.72113037109375 0 1240.3713 w 8
484.7557373046875 0 1041.2365
489.0555419921875 0 6045.3433
490.05670166015625 0 3694.287
491.05596923828125 0 2643.0022
494.2354431152344 0 16891.998 c Ammonia loss 3
495.2379150390625 0 4674.4185
495.765625 0 677.81934
496.2491455078125 0 1511.2825
496.7753601074219 0 2232.046
497.2807922363281 0 974.18726
501.9952697753906 0 732.1358
504.2444763183594 0 2587.8833
504.5787048339844 0 2976.6877 y Water loss 4
504.9125061035156 0 2283.8064 y Ammonia loss 4
505.23968505859375 0 885.5709 z Ammonia loss 8
506.22576904296875 0 7482.3516
506.2599792480469 0 9711.444
506.2978210449219 0 596.18524
506.3053283691406 0 985.50433
506.4967956542969 0 10137.353
506.58203125 0 595.2167
506.7455749511719 0 7627.084
506.99749755859375 0 5856.5723
507.2283935546875 0 880.50604
507.26544189453125 0 1897.3162
507.4990539550781 0 808.73956
507.9436950683594 0 1981.5173
511.2626647949219 0 13726.505 c 3
512.2650756835938 0 4390.3594
512.7626342773438 0 750.98254 y Water loss 8
513.7391967773438 0 563.1052
515.3191528320312 0 2611.0618
516.3240356445312 0 1127.8441
518.7636108398438 0 783.36426
521.7655029296875 0 3393.754 y 8
522.2679443359375 0 2029.2672
522.7625122070312 0 659.98737
533.7755126953125 0 794.4218
534.2723388671875 0 937.2899
537.2789306640625 0 1305.2755
542.2669067382812 0 1453.8483 z 7
542.7686157226562 0 1164.5635
547.2913818359375 0 1350.3455
548.2365112304688 0 788.7583
550.2774658203125 0 3398.2031 y 7
550.7772827148438 0 2953.436
551.2784423828125 0 650.7871
552.2642822265625 0 2058.9446
553.3092041015625 0 1396.3977
554.2925415039062 0 1115.6663
554.784423828125 0 1757.7463 c Water loss 8
555.274658203125 0 1798.4146 c Ammonia loss 8
555.7786254882812 0 782.1653
563.2936401367188 0 3681.0056
563.7863159179688 0 18598.623 c 8
564.2423095703125 0 1002.94525
564.28662109375 0 9199.603
564.7899780273438 0 3127.5933
565.2909545898438 0 1179.6255
570.271240234375 0 1651.1904
570.3251953125 0 2587.8726
570.7474975585938 0 1800.2463
571.2378540039062 0 801.0018
571.2811889648438 0 11199.263
571.3282470703125 0 586.59534
571.7828369140625 0 6449.241
572.2833862304688 0 1809.1696
572.7882080078125 0 1221.8738
576.7948608398438 0 702.9299
577.2842407226562 0 2252.3354 w 6
577.7861938476562 0 1451.083
578.7888793945312 0 862.7991
585.6046752929688 0 952.3397 y Water loss 2
585.9389038085938 0 1179.9572 y Ammonia loss 2
586.605224609375 0 888.13995
588.2811279296875 0 689.1335
589.2843017578125 0 715.082 z Ammonia loss 11
592.2789306640625 0 632.1427
598.7855834960938 0 1489.8608 y Water loss 6
599.2860107421875 0 2555.5564 y Ammonia loss 6
599.7810668945312 0 11268.865 z 6
600.2817993164062 0 7178.2783
600.7836303710938 0 3279.364
600.946044921875 0 656.3029
601.2881469726562 0 1300.3518
602.281494140625 0 714.11676
606.3043212890625 0 9764.931 z 11
606.802001953125 0 718.6946
606.9629516601562 0 1150.678 c Ammonia loss 13
607.3043823242188 0 5446.4004
607.6373901367188 0 1380.8583
607.790283203125 0 14603.6455 y 6
608.2924194335938 0 8631.625
608.7926025390625 0 4166.534
609.2933349609375 0 1223.0465
611.6434936523438 0 982.7674
611.9729614257812 0 749.8975
612.3006591796875 0 2455.0432
612.6300048828125 0 8572.437 c 13
612.8018798828125 0 1992.269
612.96435546875 0 8124.035
613.2996826171875 0 6473.0713
613.626953125 0 1248.0867
615.3087768554688 0 1002.30585
615.8062744140625 0 1005.0374
620.970458984375 0 1841.2736
621.30517578125 0 3178.9482
621.6390991210938 0 2127.1218
621.9733276367188 0 984.85596
622.3223266601562 0 6535.2354 y 11
623.3262939453125 0 2732.0452
624.3273315429688 0 1732.3273
625.323486328125 0 894.2535
626.3063354492188 0 2631.4236
626.6386108398438 0 2928.0762
626.8162231445312 0 673.61676
626.9722290039062 0 2464.2708
627.3096923828125 0 986.3698
627.8145141601562 0 3792.3318
628.3077392578125 0 14507.603 c 9
628.373291015625 0 1014.53754
628.8087158203125 0 10792.915
629.3106689453125 0 4326.8193
629.8115234375 0 1492.4473
630.644287109375 0 715.2203
630.9874877929688 0 761.9954
631.304931640625 0 1599.4928 y Water loss 1
631.6432495117188 0 2141.4548
631.9744262695312 0 1006.83374
634.3140258789062 0 1974.2659
634.796630859375 0 3659.0564 w 5
635.2984619140625 0 3227.6328
635.7980346679688 0 1150.266
636.298828125 0 773.52405
637.2987060546875 0 1223.9413 y 1
637.3576049804688 0 713.87695
639.9813232421875 0 1191.266
640.3174438476562 0 1235.8137
640.6446533203125 0 1166.1428
640.9784545898438 0 729.3959
641.2938842773438 0 1058.9891
645.3257446289062 0 4434.6445
645.654052734375 0 18905.797
645.9876098632812 0 22589.617
646.3215942382812 0 10291.691
646.654296875 0 3436.0627
646.9912109375 0 2602.7
649.6516723632812 0 1488.8146
649.9853515625 0 1210.4845
650.313232421875 0 771.74817
651.3091430664062 0 1495.3027
651.645263671875 0 983.5415
654.6575927734375 0 842.29034
654.9815063476562 0 1930.8018
655.3101806640625 0 9875.088
655.6488647460938 0 15246.446
655.830078125 0 1504.8109 y Ammonia loss 5
655.9871826171875 0 15185.532
656.3217163085938 0 12007.328 z 5
656.6578979492188 0 5418.756
656.825927734375 0 1789.4877
656.9901123046875 0 1256.4634
657.3241577148438 0 1467.095
659.9940185546875 0 1448.3074
660.3260498046875 0 2594.8672
663.8386840820312 0 849.4388
664.3325805664062 0 8496.269 y 5
664.8330078125 0 4598.7285
665.3375854492188 0 1772.5422
665.8331909179688 0 863.0801
668.9827880859375 0 971.1036
669.3245849609375 0 3353.2954
669.6557006835938 0 4973.2783
669.8477783203125 0 754.45374
669.9908447265625 0 3573.2415
670.3192749023438 0 2553.956
673.700439453125 0 898.1199
674.0374755859375 0 1172.7463
674.2920532226562 0 831.5517
674.3643798828125 0 1475.5791
674.6617431640625 0 6917.4985
674.9931640625 0 42507.55
675.3270874023438 0 66786.79
675.6610717773438 0 49798.965
675.9952392578125 0 30032.396
676.3294677734375 0 12059.622
676.6632080078125 0 4959.0293
676.999755859375 0 896.04706
682.36376953125 0 1295.0171
682.4263916015625 0 1165.4188
687.8298950195312 0 1630.563
688.3367919921875 0 853.7323
688.8382568359375 0 768.0746
690.8436279296875 0 4618.9033
691.3394165039062 0 14561.068
691.8406372070312 0 9122.79
692.3402099609375 0 7998.6357
692.842041015625 0 2334.9446
693.3421630859375 0 970.2751
696.3098754882812 0 4849.9116 c Ammonia loss 4
697.3126831054688 0 1563.9563
709.3474731445312 0 3287.1492
709.8397216796875 0 13778.02 c 10
710.3407592773438 0 10173.368
710.84228515625 0 3931.846
711.3432006835938 0 1736.836
711.8474731445312 0 850.26715
713.3363647460938 0 13035.009 c 4
714.3394775390625 0 4868.3057
715.3411865234375 0 1488.8157
726.3594970703125 0 688.7266
726.8617553710938 0 983.5173
727.3611450195312 0 1137.0452
733.3516845703125 0 1462.8151
733.8599853515625 0 839.6316
734.3587036132812 0 953.9037
742.3543701171875 0 811.6426
748.8529052734375 0 1217.8374 z Ammonia loss 4
753.3980102539062 0 705.5011
755.4022216796875 0 11876.299
755.86474609375 0 7987.2695
756.3627319335938 0 7904.709 y Water loss 4
756.415283203125 0 3693.6873
756.8652954101562 0 6782.216 y Ammonia loss 4
757.3643798828125 0 2619.1924 z 4
757.8595581054688 0 1230.9998
765.380859375 0 751.6413 c Ammonia loss 11
768.3573608398438 0 796.16974 y Ammonia loss 10
769.3670043945312 0 16252.907 z 10
770.3711547851562 0 7016.0244
771.3729248046875 0 2698.3608
772.3688354492188 0 1041.4989
773.887451171875 0 4038.7205 c 11
774.3881225585938 0 1929.8727
774.8970947265625 0 1164.2384
782.4039916992188 0 1211.5598
783.4095458984375 0 732.7007
785.3851318359375 0 4838.368 y 10
786.386962890625 0 1606.3389
787.3860473632812 0 811.8828
789.4595947265625 0 1905.3884
790.466796875 0 1051.6417
792.8762817382812 0 1147.7583
809.3966064453125 0 1009.908 c Ammonia loss 5
810.39404296875 0 1369.1033
812.8853759765625 0 1065.3116
814.8734130859375 0 7962.0703 z 3
815.3740234375 0 6513.5273
815.87548828125 0 3767.5354
816.3753662109375 0 2534.1436
822.8837280273438 0 1218.613 y 3
823.3761596679688 0 942.8231
823.8871459960938 0 712.7417
825.4093017578125 0 1235.8445
826.4207153320312 0 33384.395 c 5
827.34130859375 0 859.56433
827.4229125976562 0 13717.731
828.4269409179688 0 3843.4116
829.4334106445312 0 804.65027
839.3983154296875 0 1705.1974 w 9
840.4005737304688 0 1170.9424
849.8877563476562 0 1307.964 w 2
853.8999633789062 0 2424.9277
854.3945922851562 0 8980.778 c 12
854.8956909179688 0 8580.182
855.3961181640625 0 4716.793
855.8938598632812 0 1155.5985
876.4189453125 0 783.63513
878.4076538085938 0 1250.8497 y Ammonia loss 2
878.9036254882812 0 6207.2144 z 2
879.40625 0 4750.64
879.90380859375 0 2744.9778
880.405029296875 0 1253.922 z Water loss 9
883.417236328125 0 8671.946
884.4209594726562 0 4152.271
885.4261474609375 0 1489.6852
887.4132690429688 0 968.99567
889.4341430664062 0 1622.7915
896.4358520507812 0 3946.5586 y Water loss 9
896.9369506835938 0 2844.5525
897.4348754882812 0 2590.1296
898.4110107421875 0 20957.549 z 9
899.4146118164062 0 13328.726
900.4160766601562 0 4513.236
901.4130249023438 0 1775.3582
901.9846801757812 0 1131.9624
903.42529296875 0 868.5265
909.93798828125 0 823.0556 c Ammonia loss 13
910.4364013671875 0 962.27527
914.4273681640625 0 2242.6953 y 9
915.4304809570312 0 1152.6223
917.944091796875 0 1687.9244
918.4435424804688 0 3965.5186 c 13
918.9435424804688 0 4818.6235
919.4415893554688 0 3249.194
919.9465942382812 0 1032.5801
924.426025390625 0 662.432 c Ammonia loss 6
938.9675903320312 0 3098.1062
939.4700317382812 0 1824.1417
939.9639282226562 0 1052.4963
940.44482421875 0 1244.9447
941.447265625 0 5555.4727 c 6
942.4514770507812 0 2709.2375
945.4609375 0 740.79443
945.9711303710938 0 2220.1562
946.475341796875 0 2682.0664
946.9683227539062 0 2027.8883
947.4382934570312 0 3147.4216 z 1
947.9351196289062 0 4547.6523
948.4361572265625 0 3371.668
948.9319458007812 0 1154.768
954.45849609375 0 2320.8127
959.472412109375 0 1083.8323
959.9718627929688 0 1614.2465
960.4683227539062 0 1134.6123
960.9744873046875 0 939.67487
961.4735107421875 0 1188.3992
967.9772338867188 0 3356.909
968.4786376953125 0 4740.1045
968.97998046875 0 2170.9194
969.4862670898438 0 1233.9498
973.4655151367188 0 1002.6925
973.9801025390625 0 774.3317
974.4773559570312 0 3793.5432
974.9742431640625 0 3585.921
975.4752197265625 0 3000.7705
975.9776000976562 0 1551.5634
980.4928588867188 0 699.20667
981.47998046875 0 1431.6791
981.9884033203125 0 2125.2292
982.482421875 0 4203.5605
982.9816284179688 0 10637.963
983.4819946289062 0 13559.841
983.9830322265625 0 7796.932
984.483154296875 0 4612.828
984.9857788085938 0 1877.1224
989.4873657226562 0 1548.012
989.9887084960938 0 4694.0903
990.4835205078125 0 4746.345
990.988525390625 0 3458.6284
991.48193359375 0 1594.6696
991.9861450195312 0 904.7362
992.5643310546875 0 4295.839
993.5658569335938 0 3584.3564
994.5736694335938 0 1174.8525
995.4774780273438 0 2181.5842
995.9781494140625 0 2722.1677
996.4769287109375 0 1648.332
997.4804077148438 0 3819.141
998.470703125 0 15771.828 c 7
999.4744262695312 0 8562.172
1000.4789428710938 0 2783.6538
1003.4888916015625 0 3805.321
1003.9846801757812 0 15476.277
1004.4822998046875 0 22490.393
1004.9818115234375 0 16744.781
1005.483154296875 0 10448.41
1005.9795532226562 0 3402.006
1006.4893798828125 0 3087.3286
1010.5706787109375 0 1754.5947
1011.0532836914062 0 1863.7634
1011.4767456054688 0 1948.6614
1011.58203125 0 1091.8951
1011.985107421875 0 3570.4446
1012.4856567382812 0 13213.426
1012.990966796875 0 20929.822
1013.49072265625 0 22464.145
1013.9935913085938 0 13455.653
1014.490234375 0 6763.201
1014.9993896484375 0 3778.921
1015.4834594726562 0 1261.6014
1026.5040283203125 0 1279.7551 z 8
1027.511962890625 0 5246.6284
1028.513427734375 0 3848.523
1083.5389404296875 0 1853.5491 z 7
1084.5367431640625 0 3200.3484
1085.5413818359375 0 1836.7595
1110.53857421875 0 1536.5007
1125.5771484375 0 3269.5542
1126.56591796875 0 14448.34 c 8
1127.5679931640625 0 10498.724
1128.5682373046875 0 3669.6147
1129.5760498046875 0 1089.2297
1140.550537109375 0 959.3487
1141.5548095703125 0 1846.1753
1142.553466796875 0 1091.1606
1198.5665283203125 0 3096.7104 z 6
1199.563720703125 0 9020.075
1200.564208984375 0 4972.333
1201.5623779296875 0 2564.2737
1212.5894775390625 0 1212.3579
1213.5977783203125 0 1056.0087
1214.5771484375 0 1221.5645 y 6
1239.6031494140625 0 913.5753
1254.625244140625 0 1151.6998
1255.611328125 0 4225.439 c 9
1256.612060546875 0 3714.5557
1257.6136474609375 0 1577.207
1311.65869140625 0 1226.5057 z 5
1312.6451416015625 0 4127.8794
1313.6451416015625 0 1896.9061
1374.666015625 0 1016.67236
1375.669189453125 0 1958.1177
1376.6636962890625 0 947.9532
1381.67578125 0 2475.7695
1382.674072265625 0 3206.098
1383.681640625 0 1845.3151
1418.67529296875 0 1793.5513 c 10
1419.674560546875 0 3395.119
1420.674560546875 0 1980.5077
1512.7421875 0 1139.5676
1513.72802734375 0 1766.8716 z 4
1514.7337646484375 0 1390.5342
1529.73193359375 0 867.2545 c Ammonia loss 11
1545.095947265625 0 766.70544
1546.7603759765625 0 1860.0028 c 11
1547.7703857421875 0 1446.7576
1628.745361328125 0 1013.06366 z 3
1629.74169921875 0 3925.8264
1630.7470703125 0 2761.2576
1631.7470703125 0 1435.9146
1706.7894287109375 0 1004.3336
1707.785888671875 0 2117.9414 c 12
1708.7825927734375 0 2201.1758
1709.7845458984375 0 1237.9071
1710.7999267578125 0 930.2334
1756.8006591796875 0 2166.546 z 2
1757.801513671875 0 2607.6096
1758.8067626953125 0 2738.3364
1759.8157958984375 0 1863.7861
1836.8988037109375 0 969.63135
1894.8624267578125 0 807.8451
1895.852783203125 0 787.0602
1935.96875 0 726.14703
1936.9710693359375 0 790.39667
1937.966064453125 0 1260.2993
1948.9515380859375 0 1155.7363
1965.9525146484375 0 1770.6362
1966.9622802734375 0 1372.6354
1967.9449462890625 0 1397.6184
1981.9610595703125 0 743.3261
2006.9613037109375 0 699.3178
2007.9554443359375 0 1929.6282
2008.9576416015625 0 2956.3486
2009.9490966796875 0 2028.4061
2010.97021484375 0 1212.9578
2023.9696044921875 0 922.1865
2024.97705078125 0 2965.3381
2025.974609375 0 2828.1785
2026.9786376953125 0 2113.796
2027.96826171875 0 1354.24
2029.014892578125 0 946.53107

Spectrum Details

|  |  |
| --- | --- |
| Matched peaks? Matched peaks The total absolute number of peaks matched. Additionally in brackets the total fraction of peaks matched and the total number of peaks is shown. | 88 (16.54% of 532) |
| FDR? FDR The false discovery rate estimated for this peptide. It is calculated by matching all theoretical fragments with a non-integer shift with the raw peaks for this spectrum. This is done with 40 different shifts. The resulting percentage is the average number of annotated peaks over the number of annotated peaks with the correct spectrum. | 0.62% |
| Satellite FDR? Satellite FDR See the FDR for details on its calculation. This satellite ion specific FDR only contains the satellite ions (d/w) for I/L/J positions. | 0.00% |
| PSM Score? PSM Score The PSM Score as given by Hecklib to this annotated spectrum. It is shown with three significant figures. | 470 |

#### Spectrum 5273? Spectrum 5273 The raw spectrum of this peptide as annotated by Hecklib. The fragments are coloured according to ion type (see legend). Any peaks with a star '\*' as text can be hovered over to see the full details, first the ion type second the mass shift type. By hovering over the amino acids in the peptide or ions in the legend the corresponding peaks are highlighted. By toggling the 'Unassigned' label you can turn the background (unassigned) peaks on or off in the plot. By updating the slider in the Ion legend you can update the spectrum to only show the top X% of the peaks with labels. The top X% means any peak that is within X% of the highest intensity. By dragging in the spectrum you can zoom in to a specific part of the spectrum and use 'Zoom Out' to get back to the original zoom level. The annotation of the spectrum is based on the given sequence in the peptides file and is done with different software so inconsistencies are likely. The peaks are annotated based on the given sequence, with 20 ppm tolerance.

Copy Data

##### Spectrum 5273 (TSV)

###### Preview

```
Loading example...
```

*Click on the button to copy the data to your clipboard.*

Mz MinMz MaxIntensity Max

WidthHeightPeptide font sizePeptide stroke widthSpectrum font sizeSpectrum stroke widthCompact peptide

Ion legend

wxyz

abcd

OtherUnassignedIon^Charge^_Position_Show for top:%

JHQDWLDGKEYKCKK

04.36e+48.72e+41.31e+51.74e+5

Zoom Out

y^+4^_3_c^+2^_4_c^+1^_2_z^+3^_6_z^+1^_2_c^+3^_7_z^+2^_4_y^+2^_4_z^+1^_2_y^+1^_2_c^+2^_5_y^+3^_7_y^+3^_7_c^+5^_14_y^+3^_8_z^+3^_8_y^+3^_8_y^+4^_11_y^+4^_11_z^+4^_11_c^+1^_3_y^+2^_5_z^+2^_5_y^+2^_5_c^+1^_3_c^+2^_6_z^+4^_12_y^+3^_9_y^+3^_9_z^+3^_9_y^+3^_9_w^+2^_6_w^+3^_10_z^+3^_10_y^+3^_10_y^+3^_10_y^+4^_13_y^+3^_10_y^+2^_6_y^+2^_6_z^+2^_6_c^+4^_14_c^+2^_7_c^+4^_14_y^+2^_6_c^+2^_7_z^+4^_14_z^+4^_14_y^+4^_14_y^+4^_14_z^+4^_14_y^+4^_14_z^+1^_3_c^+2^_8_c^+2^_8_w^+2^_7_c^+2^_8_c^+1^_4_z^+3^_11_c^+1^_4_y^+3^_11_y^+3^_11_z^+3^_11_c^+3^_12_c^+3^_12_c^+1^_4_y^+2^_7_z^+2^_7_y^+2^_7_w^+3^_12_y^+3^_12_y^+2^_8_y^+2^_8_z^+2^_8_y^+3^_12_w^+1^_4_y^+2^_8_c^+2^_9_w^+3^_13_c^+3^_13_w^+2^_9_y^+3^_13_y^+3^_13_z^+3^_13_y^+3^_13_z^+1^_4_z^+2^_9_z^+2^_9_y^+2^_9_y^+2^_9_z^+2^_9_c^+3^_14_c^+3^_14_z^+1^_4_c^+3^_14_y^+2^_9_c^+2^_10_c^+2^_10_c^+2^_10_y^+1^_4_z^+3^_14_y^+3^_14_w^+2^_10_z^+2^_10_y^+2^_10_y^+2^_10_z^+2^_10_y^+2^_10_c^+1^_5_c^+1^_5_c^+2^_11_c^+2^_11_c^+1^_5_c^+2^_11_y^+2^_11_z^+2^_11_c^+2^_12_c^+2^_12_y^+1^_5_z^+1^_5_w^+2^_12_y^+1^_5_c^+1^_6_y^+2^_12_z^+2^_12_c^+1^_6_y^+2^_12_w^+1^_6_w^+2^_13_c^+2^_13_z^+2^_13_z^+2^_13_y^+2^_13_y^+1^_6_z^+1^_6_c^+1^_7_c^+2^_14_y^+1^_6_c^+1^_7_z^+2^_14_c^+1^_8_c^+1^_8_z^+1^_7_z^+1^_8_y^+1^_8_c^+1^_9_z^+1^_9_c^+1^_10_z^+1^_10_c^+1^_11_z^+1^_11_

047594914241899

Fragment Matches Table

Show background peaks

| Position | Ion type | Intensity | mz Theoretical | mz Error (Th) | mz Error (ppm) | Charge | Series Number |
| --- | --- | --- | --- | --- | --- | --- | --- |
| 13 | y | 636 | 120.1 | 0.0004713 | 3.926 | +4 | 3 |
| - | - | 501.8 | 120.1 | - | - | 0 | - |
| - | - | 355.1 | 122 | - | - | 0 | - |
| - | - | 627 | 127.1 | - | - | 0 | - |
| - | - | 455.9 | 127.1 | - | - | 0 | - |
| - | - | 472.1 | 128.1 | - | - | 0 | - |
| - | - | 359.1 | 129 | - | - | 0 | - |
| - | - | 5536 | 129 | - | - | 0 | - |
| - | - | 531.8 | 129.1 | - | - | 0 | - |
| - | - | 1.455E+04 | 129.1 | - | - | 0 | - |
| - | - | 4.229E+04 | 130.1 | - | - | 0 | - |
| - | - | 482.6 | 130.1 | - | - | 0 | - |
| - | - | 3544 | 131.1 | - | - | 0 | - |
| - | - | 1571 | 131.1 | - | - | 0 | - |
| - | - | 656 | 132.1 | - | - | 0 | - |
| - | - | 3576 | 132.1 | - | - | 0 | - |
| - | - | 687.4 | 132.1 | - | - | 0 | - |
| - | - | 3729 | 133.1 | - | - | 0 | - |
| - | - | 2493 | 133.1 | - | - | 0 | - |
| - | - | 3862 | 136.1 | - | - | 0 | - |
| - | - | 397.8 | 139 | - | - | 0 | - |
| - | - | 3374 | 139 | - | - | 0 | - |
| - | - | 548.4 | 144.1 | - | - | 0 | - |
| - | - | 468.1 | 146.1 | - | - | 0 | - |
| - | - | 1402 | 146.1 | - | - | 0 | - |
| - | - | 666.4 | 147 | - | - | 0 | - |
| - | - | 962 | 147.1 | - | - | 0 | - |
| - | - | 1.559E+04 | 155.1 | - | - | 0 | - |
| - | - | 700.7 | 156.1 | - | - | 0 | - |
| - | - | 1214 | 156.1 | - | - | 0 | - |
| - | - | 815.5 | 156.1 | - | - | 0 | - |
| - | - | 1.329E+04 | 157 | - | - | 0 | - |
| - | - | 742.4 | 157.1 | - | - | 0 | - |
| - | - | 611.4 | 158.1 | - | - | 0 | - |
| - | - | 1.756E+04 | 159.1 | - | - | 0 | - |
| - | - | 554.6 | 159.1 | - | - | 0 | - |
| - | - | 1739 | 160.1 | - | - | 0 | - |
| - | - | 525 | 164.1 | - | - | 0 | - |
| - | - | 1853 | 164.1 | - | - | 0 | - |
| - | - | 1.298E+04 | 166.1 | - | - | 0 | - |
| - | - | 747.8 | 167.1 | - | - | 0 | - |
| - | - | 1217 | 168.1 | - | - | 0 | - |
| - | - | 9040 | 170.1 | - | - | 0 | - |
| - | - | 940.1 | 171.1 | - | - | 0 | - |
| - | - | 548.1 | 171.1 | - | - | 0 | - |
| - | - | 2998 | 172.1 | - | - | 0 | - |
| - | - | 2582 | 173.1 | - | - | 0 | - |
| - | - | 599.2 | 173.1 | - | - | 0 | - |
| - | - | 981.8 | 173.4 | - | - | 0 | - |
| - | - | 1722 | 174.1 | - | - | 0 | - |
| - | - | 1894 | 177.1 | - | - | 0 | - |
| - | - | 1233 | 178.1 | - | - | 0 | - |
| - | - | 5444 | 183.1 | - | - | 0 | - |
| - | - | 660.5 | 183.1 | - | - | 0 | - |
| - | - | 783.3 | 183.1 | - | - | 0 | - |
| - | - | 718.5 | 183.1 | - | - | 0 | - |
| - | - | 953.4 | 185.1 | - | - | 0 | - |
| - | - | 1.605E+04 | 185.1 | - | - | 0 | - |
| - | - | 936.8 | 185.1 | - | - | 0 | - |
| - | - | 1.228E+04 | 185.2 | - | - | 0 | - |
| - | - | 505.1 | 186.1 | - | - | 0 | - |
| - | - | 473.3 | 186.1 | - | - | 0 | - |
| - | - | 3167 | 186.1 | - | - | 0 | - |
| - | - | 1177 | 186.2 | - | - | 0 | - |
| - | - | 3813 | 187.1 | - | - | 0 | - |
| - | - | 1278 | 187.1 | - | - | 0 | - |
| - | - | 499.7 | 189.1 | - | - | 0 | - |
| - | - | 7604 | 190.1 | - | - | 0 | - |
| - | - | 916.7 | 192.1 | - | - | 0 | - |
| - | - | 2467 | 198.1 | - | - | 0 | - |
| - | - | 2882 | 199.2 | - | - | 0 | - |
| - | - | 2179 | 201.1 | - | - | 0 | - |
| - | - | 570.7 | 202.1 | - | - | 0 | - |
| - | - | 1537 | 203.1 | - | - | 0 | - |
| - | - | 958.7 | 203.2 | - | - | 0 | - |
| - | - | 7068 | 205.1 | - | - | 0 | - |
| - | - | 1074 | 207.2 | - | - | 0 | - |
| - | - | 574.7 | 209.2 | - | - | 0 | - |
| - | - | 1606 | 210.1 | - | - | 0 | - |
| - | - | 1078 | 211.6 | - | - | 0 | - |
| - | - | 8734 | 213.1 | - | - | 0 | - |
| - | - | 659.5 | 215.1 | - | - | 0 | - |
| - | - | 1.219E+04 | 217 | - | - | 0 | - |
| - | - | 600.1 | 221.1 | - | - | 0 | - |
| - | - | 1605 | 223.1 | - | - | 0 | - |
| - | - | 1.216E+04 | 223.2 | - | - | 0 | - |
| - | - | 1367 | 224.2 | - | - | 0 | - |
| - | - | 499.7 | 225.1 | - | - | 0 | - |
| - | - | 722.6 | 226.2 | - | - | 0 | - |
| - | - | 882.7 | 227.1 | - | - | 0 | - |
| - | - | 543.3 | 227.1 | - | - | 0 | - |
| - | - | 551.9 | 229.1 | - | - | 0 | - |
| - | - | 5765 | 229.1 | - | - | 0 | - |
| - | - | 676.7 | 229.2 | - | - | 0 | - |
| - | - | 690 | 230.1 | - | - | 0 | - |
| - | - | 3107 | 233.1 | - | - | 0 | - |
| - | - | 1.258E+04 | 234.1 | - | - | 0 | - |
| - | - | 1902 | 235.1 | - | - | 0 | - |
| - | - | 1609 | 235.2 | - | - | 0 | - |
| - | - | 620.4 | 236.1 | - | - | 0 | - |
| - | - | 2017 | 236.6 | - | - | 0 | - |
| - | - | 703 | 241.1 | - | - | 0 | - |
| - | - | 990.3 | 244.1 | - | - | 0 | - |
| - | - | 733.2 | 245.1 | - | - | 0 | - |
| - | - | 833.9 | 246.1 | - | - | 0 | - |
| - | - | 1313 | 246.1 | - | - | 0 | - |
| - | - | 731.1 | 247.1 | - | - | 0 | - |
| 4 | c | 2906 | 247.6 | 0.0003546 | 1.432 | +2 | 4 |
| - | - | 1990 | 250.1 | - | - | 0 | - |
| - | - | 1533 | 250.1 | - | - | 0 | - |
| - | - | 2145 | 250.2 | - | - | 0 | - |
| - | - | 5.376E+04 | 251.2 | - | - | 0 | - |
| - | - | 7401 | 252.2 | - | - | 0 | - |
| - | - | 651.4 | 252.6 | - | - | 0 | - |
| - | - | 1178 | 255.2 | - | - | 0 | - |
| - | - | 720.2 | 256.6 | - | - | 0 | - |
| - | - | 2.212E+04 | 259.2 | - | - | 0 | - |
| - | - | 1674 | 261.1 | - | - | 0 | - |
| - | - | 1752 | 264.2 | - | - | 0 | - |
| - | - | 1765 | 265.1 | - | - | 0 | - |
| - | - | 767.1 | 265.6 | - | - | 0 | - |
| - | - | 1503 | 266.1 | - | - | 0 | - |
| 2 | c | 1.009E+04 | 268.2 | 0.0004142 | 1.544 | +1 | 2 |
| - | - | 2114 | 269.1 | - | - | 0 | - |
| - | - | 1692 | 269.2 | - | - | 0 | - |
| - | - | 6347 | 270.1 | - | - | 0 | - |
| - | - | 671.4 | 270.2 | - | - | 0 | - |
| - | - | 831.3 | 270.6 | - | - | 0 | - |
| - | - | 1962 | 272.1 | - | - | 0 | - |
| - | - | 1464 | 272.2 | - | - | 0 | - |
| - | - | 7656 | 273.1 | - | - | 0 | - |
| - | - | 1500 | 274.1 | - | - | 0 | - |
| - | - | 1.014E+04 | 274.1 | - | - | 0 | - |
| - | - | 2823 | 274.7 | - | - | 0 | - |
| - | - | 917 | 275.2 | - | - | 0 | - |
| - | - | 895.4 | 279.2 | - | - | 0 | - |
| - | - | 659.8 | 280.2 | - | - | 0 | - |
| - | - | 5185 | 282.2 | - | - | 0 | - |
| - | - | 1219 | 283.1 | - | - | 0 | - |
| - | - | 1099 | 284.1 | - | - | 0 | - |
| - | - | 845 | 285.2 | - | - | 0 | - |
| - | - | 977.4 | 286.1 | - | - | 0 | - |
| - | - | 976 | 288.2 | - | - | 0 | - |
| - | - | 3505 | 290.1 | - | - | 0 | - |
| - | - | 687.5 | 292.2 | - | - | 0 | - |
| 10 | z | 1057 | 294.1 | 0.005451 | 18.53 | +3 | 6 |
| - | - | 1903 | 294.2 | - | - | 0 | - |
| - | - | 1349 | 295.1 | - | - | 0 | - |
| - | - | 918.8 | 296.2 | - | - | 0 | - |
| - | - | 854.7 | 297.1 | - | - | 0 | - |
| - | - | 686.6 | 297.2 | - | - | 0 | - |
| 14 | z | 963.9 | 300.2 | 0.001046 | 3.485 | +1 | 2 |
| - | - | 1243 | 301.2 | - | - | 0 | - |
| - | - | 1380 | 302.1 | - | - | 0 | - |
| 7 | c | 6899 | 303.2 | 7.267E-05 | 0.2397 | +3 | 7 |
| 12 | z | 2399 | 303.7 | 0.005174 | 17.04 | +2 | 4 |
| - | - | 1070 | 304.2 | - | - | 0 | - |
| - | - | 772.8 | 307.1 | - | - | 0 | - |
| - | - | 1475 | 309.1 | - | - | 0 | - |
| - | - | 694.3 | 309.6 | - | - | 0 | - |
| 12 | y | 6768 | 311.7 | 0.003885 | 12.47 | +2 | 4 |
| - | - | 1863 | 312.2 | - | - | 0 | - |
| - | - | 773.2 | 315.2 | - | - | 0 | - |
| 14 | z | 2993 | 317.2 | 0.003731 | 11.76 | +1 | 2 |
| - | - | 2402 | 317.7 | - | - | 0 | - |
| - | - | 7405 | 318.2 | - | - | 0 | - |
| - | - | 1102 | 318.2 | - | - | 0 | - |
| - | - | 2310 | 318.7 | - | - | 0 | - |
| - | - | 666.5 | 319.2 | - | - | 0 | - |
| - | - | 1241 | 323.1 | - | - | 0 | - |
| - | - | 3.289E+04 | 326.7 | - | - | 0 | - |
| - | - | 1.337E+04 | 327.2 | - | - | 0 | - |
| - | - | 3032 | 327.7 | - | - | 0 | - |
| - | - | 3462 | 329.2 | - | - | 0 | - |
| - | - | 815.9 | 329.2 | - | - | 0 | - |
| - | - | 4070 | 331.7 | - | - | 0 | - |
| - | - | 1915 | 332.2 | - | - | 0 | - |
| - | - | 669.1 | 332.2 | - | - | 0 | - |
| 14 | y | 2719 | 333.2 | 0.004053 | 12.16 | +1 | 2 |
| - | - | 1062 | 337.8 | - | - | 0 | - |
| - | - | 625.9 | 338.2 | - | - | 0 | - |
| - | - | 700.1 | 338.7 | - | - | 0 | - |
| - | - | 1544 | 339.7 | - | - | 0 | - |
| - | - | 901.5 | 340.2 | - | - | 0 | - |
| 5 | c | 1.36E+04 | 340.7 | 0.0007146 | 2.098 | +2 | 5 |
| - | - | 6605 | 341.2 | - | - | 0 | - |
| - | - | 1358 | 341.7 | - | - | 0 | - |
| 9 | y | 711.4 | 342.5 | 0.002357 | 6.881 | +3 | 7 |
| 9 | y | 563.2 | 348.2 | 0.002092 | 6.008 | +3 | 7 |
| - | - | 2159 | 349.2 | - | - | 0 | - |
| - | - | 1523 | 350.2 | - | - | 0 | - |
| - | - | 634 | 350.5 | - | - | 0 | - |
| - | - | 678.5 | 350.8 | - | - | 0 | - |
| - | - | 1341 | 351.2 | - | - | 0 | - |
| - | - | 1181 | 353.7 | - | - | 0 | - |
| - | - | 649.9 | 359.2 | - | - | 0 | - |
| - | - | 1034 | 360.7 | - | - | 0 | - |
| 14 | c | 1211 | 361.2 | 0.001756 | 4.863 | +5 | 14 |
| - | - | 662.5 | 361.2 | - | - | 0 | - |
| - | - | 3926 | 361.2 | - | - | 0 | - |
| 8 | y | 1285 | 361.5 | 0.003184 | 8.807 | +3 | 8 |
| 8 | z | 1098 | 361.9 | 0.001532 | 4.233 | +3 | 8 |
| - | - | 1077 | 362.2 | - | - | 0 | - |
| - | - | 2072 | 363.2 | - | - | 0 | - |
| - | - | 1228 | 364.2 | - | - | 0 | - |
| - | - | 680.7 | 364.7 | - | - | 0 | - |
| - | - | 1350 | 365.9 | - | - | 0 | - |
| - | - | 628.8 | 366.2 | - | - | 0 | - |
| 8 | y | 2.619E+04 | 367.2 | 0.002838 | 7.728 | +3 | 8 |
| - | - | 1.618E+04 | 367.5 | - | - | 0 | - |
| - | - | 1479 | 367.7 | - | - | 0 | - |
| - | - | 4920 | 367.9 | - | - | 0 | - |
| - | - | 545.8 | 368.1 | - | - | 0 | - |
| - | - | 2514 | 368.2 | - | - | 0 | - |
| - | - | 1081 | 369.2 | - | - | 0 | - |
| - | - | 1126 | 371.2 | - | - | 0 | - |
| - | - | 1805 | 374.2 | - | - | 0 | - |
| 5 | y | 5628 | 374.7 | 0.00452 | 12.06 | +4 | 11 |
| 5 | y | 3242 | 374.9 | 0.00125 | 3.334 | +4 | 11 |
| 5 | z | 2789 | 375.2 | 0.006956 | 18.54 | +4 | 11 |
| - | - | 724.4 | 375.4 | - | - | 0 | - |
| - | - | 699.7 | 375.7 | - | - | 0 | - |
| - | - | 1398 | 376.2 | - | - | 0 | - |
| - | - | 2584 | 378.2 | - | - | 0 | - |
| - | - | 859.5 | 378.2 | - | - | 0 | - |
| 3 | c | 1.503E+04 | 379.2 | 0.0003072 | 0.81 | +1 | 3 |
| - | - | 512.3 | 379.2 | - | - | 0 | - |
| - | - | 833.6 | 379.5 | - | - | 0 | - |
| - | - | 1545 | 379.7 | - | - | 0 | - |
| - | - | 2595 | 380.2 | - | - | 0 | - |
| - | - | 646 | 380.2 | - | - | 0 | - |
| - | - | 2048 | 381.2 | - | - | 0 | - |
| - | - | 758.8 | 381.5 | - | - | 0 | - |
| - | - | 1.131E+04 | 383.2 | - | - | 0 | - |
| - | - | 1033 | 383.2 | - | - | 0 | - |
| - | - | 6575 | 383.7 | - | - | 0 | - |
| - | - | 1997 | 384.2 | - | - | 0 | - |
| - | - | 1283 | 384.2 | - | - | 0 | - |
| 11 | y | 1108 | 384.7 | 0.004575 | 11.89 | +2 | 5 |
| 11 | z | 6479 | 385.2 | 0.002353 | 6.109 | +2 | 5 |
| - | - | 2.012E+04 | 385.2 | - | - | 0 | - |
| - | - | 8743 | 385.5 | - | - | 0 | - |
| - | - | 4603 | 385.9 | - | - | 0 | - |
| - | - | 1683 | 386.2 | - | - | 0 | - |
| - | - | 2622 | 386.2 | - | - | 0 | - |
| - | - | 768.6 | 386.5 | - | - | 0 | - |
| - | - | 3.096E+04 | 387.2 | - | - | 0 | - |
| - | - | 901.1 | 388.2 | - | - | 0 | - |
| - | - | 6585 | 388.2 | - | - | 0 | - |
| - | - | 962.7 | 388.7 | - | - | 0 | - |
| - | - | 3506 | 389.2 | - | - | 0 | - |
| - | - | 988.9 | 389.2 | - | - | 0 | - |
| - | - | 826.3 | 390.2 | - | - | 0 | - |
| 11 | y | 2.18E+04 | 393.2 | 0.004055 | 10.31 | +2 | 5 |
| - | - | 1.012E+04 | 393.7 | - | - | 0 | - |
| - | - | 2922 | 394.2 | - | - | 0 | - |
| - | - | 1084 | 394.7 | - | - | 0 | - |
| - | - | 1043 | 395.2 | - | - | 0 | - |
| - | - | 1648 | 395.4 | - | - | 0 | - |
| - | - | 1228 | 395.6 | - | - | 0 | - |
| 3 | c | 2.445E+04 | 396.2 | 0.0006746 | 1.702 | +1 | 3 |
| 6 | c | 1.015E+04 | 397.2 | 0.0006748 | 1.699 | +2 | 6 |
| - | - | 5810 | 397.2 | - | - | 0 | - |
| - | - | 5158 | 397.7 | - | - | 0 | - |
| - | - | 2461 | 398.2 | - | - | 0 | - |
| - | - | 1028 | 398.2 | - | - | 0 | - |
| - | - | 2612 | 398.6 | - | - | 0 | - |
| - | - | 3290 | 398.8 | - | - | 0 | - |
| - | - | 5541 | 399 | - | - | 0 | - |
| - | - | 2785 | 399.2 | - | - | 0 | - |
| - | - | 713 | 399.4 | - | - | 0 | - |
| 4 | z | 691.5 | 399.4 | 0.002723 | 6.818 | +4 | 12 |
| 7 | y | 1492 | 399.5 | 0.003496 | 8.75 | +3 | 9 |
| 7 | y | 709.4 | 399.9 | 0.001209 | 3.024 | +3 | 9 |
| 7 | z | 1706 | 400.2 | 0.0005725 | 1.431 | +3 | 9 |
| - | - | 1269 | 401.2 | - | - | 0 | - |
| - | - | 2225 | 401.2 | - | - | 0 | - |
| - | - | 2857 | 401.3 | - | - | 0 | - |
| - | - | 678.6 | 402 | - | - | 0 | - |
| - | - | 4105 | 402.2 | - | - | 0 | - |
| - | - | 832.4 | 402.2 | - | - | 0 | - |
| - | - | 2537 | 402.3 | - | - | 0 | - |
| - | - | 6052 | 402.4 | - | - | 0 | - |
| - | - | 2149 | 402.6 | - | - | 0 | - |
| - | - | 1347 | 402.8 | - | - | 0 | - |
| - | - | 962.5 | 403.2 | - | - | 0 | - |
| - | - | 942.9 | 403.2 | - | - | 0 | - |
| - | - | 824.1 | 403.3 | - | - | 0 | - |
| - | - | 714.2 | 403.6 | - | - | 0 | - |
| - | - | 3067 | 404.2 | - | - | 0 | - |
| - | - | 1853 | 404.7 | - | - | 0 | - |
| 7 | y | 8987 | 405.5 | 0.002196 | 5.414 | +3 | 9 |
| - | - | 8297 | 405.9 | - | - | 0 | - |
| - | - | 1858 | 406.2 | - | - | 0 | - |
| - | - | 754.7 | 409.1 | - | - | 0 | - |
| - | - | 671 | 409.2 | - | - | 0 | - |
| - | - | 734.9 | 413.9 | - | - | 0 | - |
| - | - | 1376 | 418.2 | - | - | 0 | - |
| 10 | w | 1.443E+05 | 420.2 | 0.003722 | 8.858 | +2 | 6 |
| - | - | 6.483E+04 | 420.7 | - | - | 0 | - |
| - | - | 2.045E+04 | 421.2 | - | - | 0 | - |
| - | - | 4679 | 421.7 | - | - | 0 | - |
| - | - | 935.8 | 422.2 | - | - | 0 | - |
| - | - | 819.4 | 423.2 | - | - | 0 | - |
| 6 | w | 1.302E+04 | 423.5 | 0.002147 | 5.068 | +3 | 10 |
| - | - | 7600 | 423.9 | - | - | 0 | - |
| - | - | 3177 | 424.2 | - | - | 0 | - |
| - | - | 796.6 | 424.5 | - | - | 0 | - |
| - | - | 1213 | 430.2 | - | - | 0 | - |
| - | - | 1022 | 431.2 | - | - | 0 | - |
| 6 | z | 895.5 | 432.2 | 0.004943 | 11.44 | +3 | 10 |
| - | - | 830.6 | 434.2 | - | - | 0 | - |
| - | - | 1032 | 435.2 | - | - | 0 | - |
| - | - | 1294 | 435.2 | - | - | 0 | - |
| - | - | 1107 | 436.2 | - | - | 0 | - |
| 6 | y | 1543 | 437.2 | 0.002047 | 4.682 | +3 | 10 |
| 6 | y | 2186 | 437.6 | 0.003291 | 7.521 | +3 | 10 |
| - | - | 932.7 | 438.7 | - | - | 0 | - |
| - | - | 747.3 | 439.2 | - | - | 0 | - |
| - | - | 706.5 | 439.7 | - | - | 0 | - |
| 3 | y | 776.5 | 440 | 0.002532 | 5.755 | +4 | 13 |
| - | - | 1504 | 440.7 | - | - | 0 | - |
| - | - | 806.9 | 441.2 | - | - | 0 | - |
| - | - | 1140 | 442.9 | - | - | 0 | - |
| - | - | 1229 | 443.2 | - | - | 0 | - |
| 6 | y | 1.963E+04 | 443.2 | 0.001968 | 4.44 | +3 | 10 |
| - | - | 1.424E+04 | 443.6 | - | - | 0 | - |
| - | - | 3615 | 443.9 | - | - | 0 | - |
| - | - | 1794 | 444.2 | - | - | 0 | - |
| - | - | 1194 | 445.7 | - | - | 0 | - |
| - | - | 1688 | 446.2 | - | - | 0 | - |
| - | - | 713.4 | 446.7 | - | - | 0 | - |
| - | - | 2288 | 447.2 | - | - | 0 | - |
| - | - | 1941 | 448.2 | - | - | 0 | - |
| 10 | y | 2642 | 448.7 | 0.003498 | 7.796 | +2 | 6 |
| 10 | y | 1084 | 449.2 | 0.005379 | 11.97 | +2 | 6 |
| 10 | z | 2738 | 449.7 | 0.002226 | 4.951 | +2 | 6 |
| - | - | 1175 | 450.2 | - | - | 0 | - |
| 14 | c | 805.5 | 451.2 | 0.003148 | 6.977 | +4 | 14 |
| - | - | 871.2 | 451.9 | - | - | 0 | - |
| - | - | 865.1 | 452.2 | - | - | 0 | - |
| - | - | 1728 | 452.2 | - | - | 0 | - |
| - | - | 1034 | 452.5 | - | - | 0 | - |
| - | - | 865.6 | 452.7 | - | - | 0 | - |
| - | - | 685.9 | 453 | - | - | 0 | - |
| - | - | 885.9 | 453.6 | - | - | 0 | - |
| - | - | 853.1 | 454.6 | - | - | 0 | - |
| 7 | c | 1.269E+04 | 454.7 | 0.001089 | 2.394 | +2 | 7 |
| - | - | 1214 | 455 | - | - | 0 | - |
| - | - | 6545 | 455.2 | - | - | 0 | - |
| 14 | c | 3692 | 455.7 | 0.004986 | 10.94 | +4 | 14 |
| - | - | 1563 | 456 | - | - | 0 | - |
| - | - | 2314 | 456.2 | - | - | 0 | - |
| - | - | 639.4 | 456.7 | - | - | 0 | - |
| 10 | y | 3438 | 457.7 | 0.003379 | 7.382 | +2 | 6 |
| - | - | 1949 | 458.2 | - | - | 0 | - |
| - | - | 625.8 | 459 | - | - | 0 | - |
| - | - | 830.7 | 459.2 | - | - | 0 | - |
| - | - | 796.7 | 459.5 | - | - | 0 | - |
| - | - | 1588 | 459.7 | - | - | 0 | - |
| - | - | 1185 | 460 | - | - | 0 | - |
| - | - | 841.4 | 460.2 | - | - | 0 | - |
| - | - | 1.929E+04 | 460.7 | - | - | 0 | - |
| - | - | 9533 | 461.2 | - | - | 0 | - |
| - | - | 1027 | 461.6 | - | - | 0 | - |
| - | - | 3615 | 461.7 | - | - | 0 | - |
| - | - | 2820 | 462.2 | - | - | 0 | - |
| - | - | 1301 | 462.2 | - | - | 0 | - |
| - | - | 1954 | 462.7 | - | - | 0 | - |
| 7 | c | 1102 | 463.2 | 0.000528 | 1.14 | +2 | 7 |
| - | - | 2438 | 465.5 | - | - | 0 | - |
| 2 | z | 1861 | 465.7 | 0.00109 | 2.34 | +4 | 14 |
| 2 | z | 1348 | 466 | 0.008949 | 19.21 | +4 | 14 |
| - | - | 3374 | 466.2 | - | - | 0 | - |
| - | - | 2672 | 467.3 | - | - | 0 | - |
| - | - | 8385 | 468.3 | - | - | 0 | - |
| - | - | 930 | 468.3 | - | - | 0 | - |
| - | - | 5177 | 468.8 | - | - | 0 | - |
| - | - | 2619 | 469.2 | - | - | 0 | - |
| - | - | 899.8 | 469.5 | - | - | 0 | - |
| 2 | y | 2185 | 469.7 | 0.002292 | 4.879 | +4 | 14 |
| 2 | y | 7205 | 470 | 0.008235 | 17.52 | +4 | 14 |
| 2 | z | 7842 | 470.2 | 0.007103 | 15.11 | +4 | 14 |
| - | - | 1164 | 470.3 | - | - | 0 | - |
| - | - | 4129 | 470.5 | - | - | 0 | - |
| - | - | 1053 | 470.7 | - | - | 0 | - |
| - | - | 732.3 | 473 | - | - | 0 | - |
| - | - | 1672 | 473.2 | - | - | 0 | - |
| - | - | 779.8 | 473.5 | - | - | 0 | - |
| - | - | 2470 | 473.7 | - | - | 0 | - |
| - | - | 901 | 473.9 | - | - | 0 | - |
| - | - | 2410 | 474 | - | - | 0 | - |
| 2 | y | 7819 | 474.2 | 0.0005994 | 1.264 | +4 | 14 |
| - | - | 7247 | 474.5 | - | - | 0 | - |
| - | - | 4101 | 474.7 | - | - | 0 | - |
| - | - | 2260 | 476 | - | - | 0 | - |
| - | - | 854.7 | 476.2 | - | - | 0 | - |
| - | - | 1827 | 476.5 | - | - | 0 | - |
| - | - | 2227 | 476.7 | - | - | 0 | - |
| - | - | 1184 | 477 | - | - | 0 | - |
| - | - | 3250 | 477.2 | - | - | 0 | - |
| - | - | 993.6 | 477.5 | - | - | 0 | - |
| 13 | z | 1903 | 478.2 | 0.0009191 | 1.922 | +1 | 3 |
| - | - | 923.6 | 479.2 | - | - | 0 | - |
| - | - | 986.5 | 479.6 | - | - | 0 | - |
| - | - | 1353 | 479.7 | - | - | 0 | - |
| - | - | 1033 | 479.9 | - | - | 0 | - |
| - | - | 1672 | 480.2 | - | - | 0 | - |
| - | - | 3.646E+04 | 480.5 | - | - | 0 | - |
| - | - | 3.744E+04 | 480.7 | - | - | 0 | - |
| - | - | 1.958E+04 | 481 | - | - | 0 | - |
| - | - | 1.072E+04 | 481.2 | - | - | 0 | - |
| - | - | 3256 | 481.5 | - | - | 0 | - |
| - | - | 1163 | 481.7 | - | - | 0 | - |
| 8 | c | 4991 | 482.7 | 0.0005532 | 1.146 | +2 | 8 |
| 8 | c | 7675 | 483.2 | 0.002564 | 5.306 | +2 | 8 |
| - | - | 1602 | 483.5 | - | - | 0 | - |
| - | - | 2469 | 483.7 | - | - | 0 | - |
| - | - | 1665 | 484 | - | - | 0 | - |
| - | - | 1.108E+04 | 484.2 | - | - | 0 | - |
| - | - | 3764 | 484.5 | - | - | 0 | - |
| - | - | 759.4 | 484.6 | - | - | 0 | - |
| 9 | w | 5579 | 484.7 | 0.002955 | 6.096 | +2 | 7 |
| - | - | 2852 | 484.8 | - | - | 0 | - |
| - | - | 1561 | 485 | - | - | 0 | - |
| - | - | 3037 | 485.2 | - | - | 0 | - |
| - | - | 1607 | 485.7 | - | - | 0 | - |
| - | - | 969.2 | 487.3 | - | - | 0 | - |
| - | - | 3299 | 487.5 | - | - | 0 | - |
| - | - | 7.966E+04 | 487.7 | - | - | 0 | - |
| - | - | 7.516E+04 | 488 | - | - | 0 | - |
| - | - | 4.501E+04 | 488.2 | - | - | 0 | - |
| - | - | 2.186E+04 | 488.5 | - | - | 0 | - |
| - | - | 1.183E+04 | 488.7 | - | - | 0 | - |
| - | - | 4145 | 489 | - | - | 0 | - |
| - | - | 2522 | 489.2 | - | - | 0 | - |
| - | - | 1282 | 491.2 | - | - | 0 | - |
| 8 | c | 7289 | 491.7 | 0.001008 | 2.05 | +2 | 8 |
| - | - | 770.6 | 492 | - | - | 0 | - |
| - | - | 4909 | 492.2 | - | - | 0 | - |
| - | - | 2925 | 492.7 | - | - | 0 | - |
| 4 | c | 791.8 | 493.3 | 0.002393 | 4.852 | +1 | 4 |
| - | - | 2403 | 493.6 | - | - | 0 | - |
| 5 | z | 1650 | 493.9 | 0.00907 | 18.36 | +3 | 11 |
| 4 | c | 3.412E+04 | 494.2 | 0.003076 | 6.224 | +1 | 4 |
| - | - | 812.9 | 494.5 | - | - | 0 | - |
| - | - | 8559 | 495.2 | - | - | 0 | - |
| - | - | 784.5 | 495.8 | - | - | 0 | - |
| - | - | 887.9 | 496.2 | - | - | 0 | - |
| - | - | 1178 | 496.3 | - | - | 0 | - |
| - | - | 4140 | 496.8 | - | - | 0 | - |
| - | - | 3515 | 497.3 | - | - | 0 | - |
| - | - | 2706 | 497.8 | - | - | 0 | - |
| - | - | 4346 | 498.2 | - | - | 0 | - |
| - | - | 3653 | 498.5 | - | - | 0 | - |
| - | - | 3112 | 498.7 | - | - | 0 | - |
| - | - | 970 | 499 | - | - | 0 | - |
| 5 | y | 8597 | 499.2 | 0.008465 | 16.96 | +3 | 11 |
| - | - | 884.7 | 499.5 | - | - | 0 | - |
| 5 | y | 9583 | 499.6 | 0.002578 | 5.16 | +3 | 11 |
| - | - | 1748 | 499.7 | - | - | 0 | - |
| 5 | z | 7693 | 499.9 | 0.003131 | 6.264 | +3 | 11 |
| - | - | 5981 | 500.2 | - | - | 0 | - |
| - | - | 1947 | 500.6 | - | - | 0 | - |
| - | - | 1013 | 500.9 | - | - | 0 | - |
| - | - | 1548 | 502 | - | - | 0 | - |
| - | - | 4072 | 502.2 | - | - | 0 | - |
| - | - | 1.209E+05 | 502.5 | - | - | 0 | - |
| - | - | 1.728E+05 | 502.7 | - | - | 0 | - |
| - | - | 1.27E+05 | 503 | - | - | 0 | - |
| - | - | 5.97E+04 | 503.2 | - | - | 0 | - |
| - | - | 2.197E+04 | 503.5 | - | - | 0 | - |
| - | - | 8123 | 503.7 | - | - | 0 | - |
| - | - | 2801 | 504 | - | - | 0 | - |
| - | - | 1586 | 504.2 | - | - | 0 | - |
| 12 | c | 6470 | 505.3 | 0.002347 | 4.646 | +3 | 12 |
| - | - | 6724 | 505.6 | - | - | 0 | - |
| - | - | 3024 | 505.7 | - | - | 0 | - |
| - | - | 2631 | 505.9 | - | - | 0 | - |
| - | - | 5658 | 506.3 | - | - | 0 | - |
| - | - | 806.7 | 506.6 | - | - | 0 | - |
| - | - | 1726 | 506.7 | - | - | 0 | - |
| - | - | 899.9 | 506.9 | - | - | 0 | - |
| - | - | 1773 | 507.3 | - | - | 0 | - |
| - | - | 948.2 | 510.3 | - | - | 0 | - |
| - | - | 938.4 | 510.6 | - | - | 0 | - |
| 12 | c | 4.44E+04 | 510.9 | 0.0009084 | 1.778 | +3 | 12 |
| 4 | c | 7.929E+04 | 511.3 | 0.0008006 | 1.566 | +1 | 4 |
| - | - | 1.618E+04 | 511.6 | - | - | 0 | - |
| - | - | 6261 | 511.9 | - | - | 0 | - |
| - | - | 1.278E+04 | 512.3 | - | - | 0 | - |
| 9 | y | 1839 | 512.8 | 0.005051 | 9.85 | +2 | 7 |
| - | - | 1522 | 513.2 | - | - | 0 | - |
| - | - | 2072 | 513.3 | - | - | 0 | - |
| 9 | z | 1.995E+04 | 513.8 | 0.004023 | 7.831 | +2 | 7 |
| - | - | 1.7E+04 | 514.3 | - | - | 0 | - |
| - | - | 1565 | 514.6 | - | - | 0 | - |
| - | - | 7038 | 514.8 | - | - | 0 | - |
| - | - | 1273 | 514.9 | - | - | 0 | - |
| - | - | 1162 | 515.3 | - | - | 0 | - |
| - | - | 7144 | 515.3 | - | - | 0 | - |
| - | - | 877.4 | 515.8 | - | - | 0 | - |
| - | - | 2026 | 516.3 | - | - | 0 | - |
| - | - | 992.6 | 517.3 | - | - | 0 | - |
| - | - | 1064 | 517.6 | - | - | 0 | - |
| - | - | 1211 | 518.3 | - | - | 0 | - |
| - | - | 1592 | 519.9 | - | - | 0 | - |
| - | - | 1694 | 520.3 | - | - | 0 | - |
| - | - | 1760 | 521.3 | - | - | 0 | - |
| 9 | y | 1.592E+04 | 521.8 | 0.003558 | 6.82 | +2 | 7 |
| - | - | 9707 | 522.3 | - | - | 0 | - |
| - | - | 3606 | 522.8 | - | - | 0 | - |
| 4 | w | 1257 | 523.3 | 0.009058 | 17.31 | +3 | 12 |
| - | - | 2117 | 523.6 | - | - | 0 | - |
| - | - | 1852 | 523.9 | - | - | 0 | - |
| - | - | 1203 | 525.3 | - | - | 0 | - |
| - | - | 1152 | 525.4 | - | - | 0 | - |
| - | - | 3171 | 526.3 | - | - | 0 | - |
| - | - | 1434 | 526.8 | - | - | 0 | - |
| - | - | 761.9 | 528.3 | - | - | 0 | - |
| - | - | 1010 | 529.3 | - | - | 0 | - |
| - | - | 848 | 529.6 | - | - | 0 | - |
| - | - | 1343 | 530.3 | - | - | 0 | - |
| - | - | 1168 | 530.6 | - | - | 0 | - |
| - | - | 991.2 | 533.8 | - | - | 0 | - |
| - | - | 2177 | 534.3 | - | - | 0 | - |
| - | - | 1114 | 534.3 | - | - | 0 | - |
| - | - | 2383 | 535 | - | - | 0 | - |
| - | - | 3548 | 535.3 | - | - | 0 | - |
| - | - | 5358 | 535.6 | - | - | 0 | - |
| - | - | 1.285E+04 | 535.9 | - | - | 0 | - |
| - | - | 1.259E+04 | 536.3 | - | - | 0 | - |
| - | - | 6707 | 536.6 | - | - | 0 | - |
| - | - | 4310 | 536.9 | - | - | 0 | - |
| - | - | 3288 | 537.3 | - | - | 0 | - |
| 4 | y | 795.3 | 537.9 | 0.002703 | 5.025 | +3 | 12 |
| - | - | 948.1 | 538.3 | - | - | 0 | - |
| - | - | 844.8 | 538.6 | - | - | 0 | - |
| - | - | 833.7 | 539.2 | - | - | 0 | - |
| 8 | y | 2655 | 541.3 | 0.005834 | 10.78 | +2 | 8 |
| 8 | y | 1612 | 541.8 | 0.006919 | 12.77 | +2 | 8 |
| 8 | z | 1.358E+04 | 542.3 | 0.004257 | 7.85 | +2 | 8 |
| - | - | 6972 | 542.8 | - | - | 0 | - |
| - | - | 3493 | 543.3 | - | - | 0 | - |
| 4 | y | 3348 | 543.6 | 0.002098 | 3.859 | +3 | 12 |
| - | - | 1364 | 543.8 | - | - | 0 | - |
| - | - | 4077 | 543.9 | - | - | 0 | - |
| - | - | 1696 | 544.3 | - | - | 0 | - |
| - | - | 2392 | 546.3 | - | - | 0 | - |
| - | - | 2391 | 546.8 | - | - | 0 | - |
| - | - | 4438 | 547.3 | - | - | 0 | - |
| - | - | 2043 | 547.8 | - | - | 0 | - |
| 12 | w | 1992 | 548.2 | 0.00832 | 15.17 | +1 | 4 |
| - | - | 1793 | 548.3 | - | - | 0 | - |
| - | - | 1103 | 548.8 | - | - | 0 | - |
| - | - | 2714 | 549.6 | - | - | 0 | - |
| - | - | 1197 | 549.8 | - | - | 0 | - |
| - | - | 1987 | 549.9 | - | - | 0 | - |
| 8 | y | 3.442E+04 | 550.3 | 0.004158 | 7.557 | +2 | 8 |
| - | - | 1193 | 550.6 | - | - | 0 | - |
| - | - | 2.095E+04 | 550.8 | - | - | 0 | - |
| - | - | 5788 | 551.3 | - | - | 0 | - |
| - | - | 1118 | 551.6 | - | - | 0 | - |
| - | - | 1502 | 551.8 | - | - | 0 | - |
| - | - | 2384 | 552.3 | - | - | 0 | - |
| - | - | 1252 | 554.3 | - | - | 0 | - |
| - | - | 1957 | 554.8 | - | - | 0 | - |
| - | - | 6201 | 555.3 | - | - | 0 | - |
| 9 | c | 1.4E+05 | 555.8 | 0.0002796 | 0.5031 | +2 | 9 |
| - | - | 9.141E+04 | 556.3 | - | - | 0 | - |
| - | - | 2.892E+04 | 556.8 | - | - | 0 | - |
| - | - | 1099 | 556.9 | - | - | 0 | - |
| - | - | 7323 | 557.3 | - | - | 0 | - |
| - | - | 1001 | 557.6 | - | - | 0 | - |
| - | - | 1769 | 557.8 | - | - | 0 | - |
| - | - | 783.3 | 560.9 | - | - | 0 | - |
| - | - | 1687 | 561.3 | - | - | 0 | - |
| 3 | w | 5.271E+04 | 561.6 | 0.002263 | 4.029 | +3 | 13 |
| - | - | 4.797E+04 | 561.9 | - | - | 0 | - |
| - | - | 2.494E+04 | 562.3 | - | - | 0 | - |
| - | - | 7428 | 562.6 | - | - | 0 | - |
| - | - | 1918 | 562.6 | - | - | 0 | - |
| - | - | 3769 | 562.9 | - | - | 0 | - |
| - | - | 2219 | 563.3 | - | - | 0 | - |
| - | - | 1165 | 563.3 | - | - | 0 | - |
| - | - | 7296 | 563.8 | - | - | 0 | - |
| - | - | 999.1 | 563.9 | - | - | 0 | - |
| - | - | 4383 | 564.3 | - | - | 0 | - |
| - | - | 1069 | 564.3 | - | - | 0 | - |
| 13 | c | 2.725E+04 | 564.6 | 0.0009495 | 1.682 | +3 | 13 |
| - | - | 1673 | 564.8 | - | - | 0 | - |
| - | - | 2.445E+04 | 564.9 | - | - | 0 | - |
| - | - | 1.358E+04 | 565.3 | - | - | 0 | - |
| - | - | 7145 | 565.6 | - | - | 0 | - |
| - | - | 3279 | 565.9 | - | - | 0 | - |
| - | - | 4654 | 566.3 | - | - | 0 | - |
| - | - | 4019 | 566.6 | - | - | 0 | - |
| - | - | 2604 | 566.9 | - | - | 0 | - |
| - | - | 2706 | 567.2 | - | - | 0 | - |
| - | - | 2579 | 567.3 | - | - | 0 | - |
| - | - | 908.8 | 568.2 | - | - | 0 | - |
| - | - | 1658 | 568.3 | - | - | 0 | - |
| - | - | 5113 | 568.8 | - | - | 0 | - |
| - | - | 2168 | 569.3 | - | - | 0 | - |
| - | - | 1014 | 569.8 | - | - | 0 | - |
| - | - | 796.5 | 569.9 | - | - | 0 | - |
| - | - | 3557 | 570.3 | - | - | 0 | - |
| - | - | 4351 | 570.8 | - | - | 0 | - |
| - | - | 2096 | 571.3 | - | - | 0 | - |
| - | - | 2566 | 571.8 | - | - | 0 | - |
| - | - | 972.2 | 572.3 | - | - | 0 | - |
| - | - | 1864 | 573 | - | - | 0 | - |
| - | - | 1807 | 573.3 | - | - | 0 | - |
| - | - | 805.1 | 574 | - | - | 0 | - |
| - | - | 1109 | 574.3 | - | - | 0 | - |
| 7 | w | 2406 | 577.3 | 0.005351 | 9.27 | +2 | 9 |
| - | - | 918.4 | 577.3 | - | - | 0 | - |
| - | - | 1.286E+04 | 577.8 | - | - | 0 | - |
| - | - | 1630 | 578 | - | - | 0 | - |
| - | - | 8608 | 578.3 | - | - | 0 | - |
| - | - | 9505 | 578.8 | - | - | 0 | - |
| - | - | 3878 | 579.3 | - | - | 0 | - |
| - | - | 1466 | 579.8 | - | - | 0 | - |
| 3 | y | 1175 | 580.3 | 0.003739 | 6.443 | +3 | 13 |
| 3 | y | 2344 | 580.6 | 0.002119 | 3.649 | +3 | 13 |
| 3 | z | 1.349E+04 | 580.9 | 0.002768 | 4.766 | +3 | 13 |
| - | - | 1.197E+04 | 581.3 | - | - | 0 | - |
| - | - | 5006 | 581.6 | - | - | 0 | - |
| - | - | 2090 | 581.9 | - | - | 0 | - |
| - | - | 4503 | 582.3 | - | - | 0 | - |
| - | - | 1570 | 583.3 | - | - | 0 | - |
| - | - | 962.8 | 585.8 | - | - | 0 | - |
| 3 | y | 7628 | 586.3 | 0.001218 | 2.077 | +3 | 13 |
| - | - | 5568 | 586.6 | - | - | 0 | - |
| - | - | 2449 | 586.9 | - | - | 0 | - |
| - | - | 3020 | 587.3 | - | - | 0 | - |
| - | - | 1965 | 587.6 | - | - | 0 | - |
| - | - | 1016 | 588 | - | - | 0 | - |
| - | - | 7575 | 588.3 | - | - | 0 | - |
| - | - | 6053 | 588.6 | - | - | 0 | - |
| - | - | 2639 | 589 | - | - | 0 | - |
| 12 | z | 841.5 | 589.3 | 0.00234 | 3.97 | +1 | 4 |
| 7 | z | 2303 | 590.8 | 0.004095 | 6.931 | +2 | 9 |
| 7 | z | 2913 | 591.3 | 0.009279 | 15.69 | +2 | 9 |
| - | - | 1087 | 592 | - | - | 0 | - |
| - | - | 1645 | 592.3 | - | - | 0 | - |
| - | - | 8990 | 592.6 | - | - | 0 | - |
| - | - | 8127 | 593 | - | - | 0 | - |
| - | - | 3515 | 593.3 | - | - | 0 | - |
| - | - | 1217 | 593.6 | - | - | 0 | - |
| - | - | 1078 | 597 | - | - | 0 | - |
| - | - | 1758 | 597.3 | - | - | 0 | - |
| - | - | 2756 | 598.3 | - | - | 0 | - |
| 7 | y | 2716 | 598.8 | 0.007001 | 11.69 | +2 | 9 |
| 7 | y | 4095 | 599.3 | 0.003579 | 5.972 | +2 | 9 |
| 7 | z | 4.241E+04 | 599.8 | 0.003324 | 5.542 | +2 | 9 |
| - | - | 4.448E+04 | 600.3 | - | - | 0 | - |
| - | - | 2.494E+04 | 600.8 | - | - | 0 | - |
| 14 | c | 8928 | 601.3 | 0.009225 | 15.34 | +3 | 14 |
| 14 | c | 2762 | 601.6 | 0.00481 | 7.996 | +3 | 14 |
| - | - | 2152 | 601.8 | - | - | 0 | - |
| - | - | 2339 | 602 | - | - | 0 | - |
| - | - | 1759 | 602.3 | - | - | 0 | - |
| - | - | 2667 | 602.6 | - | - | 0 | - |
| - | - | 1160 | 602.8 | - | - | 0 | - |
| - | - | 7676 | 602.9 | - | - | 0 | - |
| - | - | 9119 | 603.3 | - | - | 0 | - |
| - | - | 2875 | 603.6 | - | - | 0 | - |
| - | - | 1306 | 604 | - | - | 0 | - |
| - | - | 1021 | 604.3 | - | - | 0 | - |
| 12 | z | 3.034E+04 | 606.3 | 0.007771 | 12.82 | +1 | 4 |
| - | - | 2084 | 606.6 | - | - | 0 | - |
| - | - | 9579 | 607 | - | - | 0 | - |
| 14 | c | 5.026E+04 | 607.3 | 0.0006807 | 1.121 | +3 | 14 |
| - | - | 3.887E+04 | 607.6 | - | - | 0 | - |
| 7 | y | 4.638E+04 | 607.8 | 0.003714 | 6.11 | +2 | 9 |
| - | - | 2.202E+04 | 608 | - | - | 0 | - |
| - | - | 3.874E+04 | 608.3 | - | - | 0 | - |
| - | - | 2172 | 608.6 | - | - | 0 | - |
| - | - | 1.189E+04 | 608.8 | - | - | 0 | - |
| - | - | 4466 | 609.3 | - | - | 0 | - |
| - | - | 5381 | 610.3 | - | - | 0 | - |
| 10 | c | 2703 | 611.3 | 0.007312 | 11.96 | +2 | 10 |
| 10 | c | 1.061E+04 | 611.8 | 0.002756 | 4.504 | +2 | 10 |
| - | - | 6237 | 612 | - | - | 0 | - |
| - | - | 1.739E+04 | 612.3 | - | - | 0 | - |
| - | - | 3212 | 612.6 | - | - | 0 | - |
| - | - | 3909 | 612.8 | - | - | 0 | - |
| - | - | 2048 | 613 | - | - | 0 | - |
| - | - | 2753 | 613.3 | - | - | 0 | - |
| - | - | 1824 | 615.3 | - | - | 0 | - |
| - | - | 888.6 | 616.3 | - | - | 0 | - |
| - | - | 968 | 618.3 | - | - | 0 | - |
| - | - | 1079 | 619.8 | - | - | 0 | - |
| - | - | 897.4 | 620 | - | - | 0 | - |
| 10 | c | 5.594E+04 | 620.3 | 0.001139 | 1.836 | +2 | 10 |
| - | - | 5639 | 620.6 | - | - | 0 | - |
| - | - | 3.649E+04 | 620.8 | - | - | 0 | - |
| - | - | 1.051E+04 | 621 | - | - | 0 | - |
| - | - | 1334 | 621.2 | - | - | 0 | - |
| - | - | 2.262E+04 | 621.3 | - | - | 0 | - |
| - | - | 5646 | 621.7 | - | - | 0 | - |
| - | - | 4684 | 621.8 | - | - | 0 | - |
| - | - | 1821 | 622 | - | - | 0 | - |
| - | - | 1034 | 622.2 | - | - | 0 | - |
| 12 | y | 1.668E+04 | 622.3 | 0.009039 | 14.52 | +1 | 4 |
| - | - | 4676 | 623.3 | - | - | 0 | - |
| - | - | 3694 | 625.7 | - | - | 0 | - |
| - | - | 5818 | 626 | - | - | 0 | - |
| - | - | 7835 | 626.3 | - | - | 0 | - |
| 2 | z | 1.543E+04 | 626.6 | 0.003188 | 5.088 | +3 | 14 |
| - | - | 1155 | 626.8 | - | - | 0 | - |
| - | - | 1.549E+04 | 627 | - | - | 0 | - |
| - | - | 9644 | 627.3 | - | - | 0 | - |
| - | - | 4956 | 627.6 | - | - | 0 | - |
| - | - | 863 | 627.8 | - | - | 0 | - |
| - | - | 3237 | 628 | - | - | 0 | - |
| - | - | 1139 | 628.3 | - | - | 0 | - |
| - | - | 955.4 | 630 | - | - | 0 | - |
| - | - | 1641 | 630.3 | - | - | 0 | - |
| - | - | 6128 | 630.6 | - | - | 0 | - |
| - | - | 7368 | 631 | - | - | 0 | - |
| - | - | 4260 | 631.3 | - | - | 0 | - |
| - | - | 1334 | 631.6 | - | - | 0 | - |
| 2 | y | 3100 | 632 | 8.267E-05 | 0.1308 | +3 | 14 |
| - | - | 4079 | 632.3 | - | - | 0 | - |
| - | - | 1044 | 632.6 | - | - | 0 | - |
| - | - | 1021 | 633.3 | - | - | 0 | - |
| - | - | 2497 | 634.3 | - | - | 0 | - |
| - | - | 2981 | 634.6 | - | - | 0 | - |
| 6 | w | 2.604E+04 | 634.8 | 0.003625 | 5.711 | +2 | 10 |
| - | - | 3992 | 635 | - | - | 0 | - |
| - | - | 1.855E+04 | 635.3 | - | - | 0 | - |
| - | - | 2633 | 635.7 | - | - | 0 | - |
| - | - | 6731 | 635.8 | - | - | 0 | - |
| - | - | 2759 | 636 | - | - | 0 | - |
| - | - | 2278 | 636.3 | - | - | 0 | - |
| - | - | 1246 | 636.8 | - | - | 0 | - |
| - | - | 1047 | 637.4 | - | - | 0 | - |
| - | - | 1054 | 638.3 | - | - | 0 | - |
| - | - | 1532 | 639 | - | - | 0 | - |
| - | - | 1568 | 639.3 | - | - | 0 | - |
| - | - | 1054 | 639.7 | - | - | 0 | - |
| - | - | 1.868E+04 | 640.3 | - | - | 0 | - |
| - | - | 2.423E+04 | 640.7 | - | - | 0 | - |
| - | - | 1099 | 640.8 | - | - | 0 | - |
| - | - | 1.709E+04 | 641 | - | - | 0 | - |
| - | - | 1.068E+04 | 641.3 | - | - | 0 | - |
| - | - | 3836 | 641.7 | - | - | 0 | - |
| - | - | 1325 | 642 | - | - | 0 | - |
| - | - | 5385 | 643.8 | - | - | 0 | - |
| - | - | 5581 | 644.3 | - | - | 0 | - |
| - | - | 6271 | 644.7 | - | - | 0 | - |
| - | - | 1714 | 644.8 | - | - | 0 | - |
| - | - | 1.084E+04 | 645 | - | - | 0 | - |
| - | - | 6128 | 645.3 | - | - | 0 | - |
| - | - | 6917 | 645.7 | - | - | 0 | - |
| - | - | 4754 | 646 | - | - | 0 | - |
| - | - | 4065 | 646.3 | - | - | 0 | - |
| - | - | 2243 | 646.7 | - | - | 0 | - |
| 6 | z | 1302 | 647.3 | 0.004124 | 6.371 | +2 | 10 |
| - | - | 2599 | 649.3 | - | - | 0 | - |
| - | - | 1370 | 649.7 | - | - | 0 | - |
| - | - | 1.619E+04 | 650 | - | - | 0 | - |
| - | - | 7.619E+04 | 650.3 | - | - | 0 | - |
| - | - | 8.224E+04 | 650.7 | - | - | 0 | - |
| - | - | 1791 | 650.8 | - | - | 0 | - |
| - | - | 4.694E+04 | 651 | - | - | 0 | - |
| - | - | 2.535E+04 | 651.3 | - | - | 0 | - |
| - | - | 1.036E+04 | 651.7 | - | - | 0 | - |
| - | - | 1182 | 651.8 | - | - | 0 | - |
| - | - | 2109 | 652 | - | - | 0 | - |
| - | - | 7035 | 652.3 | - | - | 0 | - |
| - | - | 3286 | 653.3 | - | - | 0 | - |
| - | - | 2117 | 654.3 | - | - | 0 | - |
| - | - | 1373 | 654.7 | - | - | 0 | - |
| - | - | 3967 | 655 | - | - | 0 | - |
| 6 | y | 7242 | 655.3 | 0.005796 | 8.844 | +2 | 10 |
| - | - | 4128 | 655.7 | - | - | 0 | - |
| 6 | y | 1998 | 655.8 | 0.0002435 | 0.3713 | +2 | 10 |
| - | - | 3882 | 656 | - | - | 0 | - |
| 6 | z | 2.319E+04 | 656.3 | 0.003669 | 5.59 | +2 | 10 |
| - | - | 1.927E+04 | 656.8 | - | - | 0 | - |
| - | - | 1015 | 657 | - | - | 0 | - |
| - | - | 1.224E+04 | 657.3 | - | - | 0 | - |
| - | - | 4874 | 657.8 | - | - | 0 | - |
| - | - | 2162 | 658.3 | - | - | 0 | - |
| - | - | 2432 | 658.7 | - | - | 0 | - |
| - | - | 3655 | 659 | - | - | 0 | - |
| - | - | 3923 | 659.3 | - | - | 0 | - |
| - | - | 2472 | 659.7 | - | - | 0 | - |
| - | - | 2356 | 663.3 | - | - | 0 | - |
| - | - | 2691 | 663.8 | - | - | 0 | - |
| - | - | 2393 | 664 | - | - | 0 | - |
| 6 | y | 3.523E+04 | 664.3 | 0.005341 | 8.039 | +2 | 10 |
| - | - | 9.192E+04 | 664.7 | - | - | 0 | - |
| - | - | 1.278E+04 | 664.8 | - | - | 0 | - |
| - | - | 1.001E+05 | 665 | - | - | 0 | - |
| - | - | 5.363E+04 | 665.3 | - | - | 0 | - |
| - | - | 2.294E+04 | 665.7 | - | - | 0 | - |
| - | - | 3387 | 665.8 | - | - | 0 | - |
| - | - | 1.04E+04 | 666 | - | - | 0 | - |
| - | - | 6255 | 666.3 | - | - | 0 | - |
| - | - | 1.926E+04 | 669.7 | - | - | 0 | - |
| - | - | 4.211E+04 | 670 | - | - | 0 | - |
| - | - | 6.876E+04 | 670.3 | - | - | 0 | - |
| - | - | 5.274E+04 | 670.7 | - | - | 0 | - |
| - | - | 2.945E+04 | 671 | - | - | 0 | - |
| - | - | 1.388E+04 | 671.3 | - | - | 0 | - |
| - | - | 6006 | 671.7 | - | - | 0 | - |
| - | - | 2775 | 671.8 | - | - | 0 | - |
| - | - | 1794 | 672 | - | - | 0 | - |
| - | - | 2921 | 672.3 | - | - | 0 | - |
| - | - | 1705 | 672.8 | - | - | 0 | - |
| 5 | c | 2108 | 679.3 | 0.01024 | 15.07 | +1 | 5 |
| - | - | 3.043E+04 | 679.8 | - | - | 0 | - |
| 5 | c | 4.95E+04 | 680.3 | 0.006081 | 8.939 | +1 | 5 |
| - | - | 1.012E+04 | 680.8 | - | - | 0 | - |
| - | - | 2.222E+04 | 681.3 | - | - | 0 | - |
| - | - | 5177 | 682.3 | - | - | 0 | - |
| - | - | 1984 | 683.3 | - | - | 0 | - |
| - | - | 1092 | 683.8 | - | - | 0 | - |
| - | - | 1149 | 685.3 | - | - | 0 | - |
| - | - | 1.364E+04 | 691.3 | - | - | 0 | - |
| - | - | 1.053E+04 | 691.8 | - | - | 0 | - |
| - | - | 5534 | 692.3 | - | - | 0 | - |
| 11 | c | 2242 | 692.8 | 0.003321 | 4.794 | +2 | 11 |
| 11 | c | 1112 | 693.3 | 0.005759 | 8.307 | +2 | 11 |
| - | - | 2997 | 693.8 | - | - | 0 | - |
| - | - | 2134 | 694.3 | - | - | 0 | - |
| 5 | c | 2.735E+04 | 697.3 | 0.0008334 | 1.195 | +1 | 5 |
| - | - | 1.355E+04 | 698.3 | - | - | 0 | - |
| - | - | 2830 | 699.3 | - | - | 0 | - |
| - | - | 1717 | 701.3 | - | - | 0 | - |
| 11 | c | 4.351E+04 | 701.8 | 0.0006635 | 0.9454 | +2 | 11 |
| - | - | 3.617E+04 | 702.3 | - | - | 0 | - |
| - | - | 1.484E+04 | 702.8 | - | - | 0 | - |
| - | - | 4933 | 703.3 | - | - | 0 | - |
| - | - | 2052 | 703.9 | - | - | 0 | - |
| - | - | 3419 | 705.4 | - | - | 0 | - |
| - | - | 2391 | 705.9 | - | - | 0 | - |
| - | - | 2079 | 711.3 | - | - | 0 | - |
| - | - | 1103 | 711.4 | - | - | 0 | - |
| - | - | 2097 | 714.4 | - | - | 0 | - |
| - | - | 1089 | 719.3 | - | - | 0 | - |
| - | - | 980.4 | 719.9 | - | - | 0 | - |
| - | - | 4275 | 720.4 | - | - | 0 | - |
| - | - | 2811 | 720.9 | - | - | 0 | - |
| - | - | 3718 | 721.3 | - | - | 0 | - |
| - | - | 2447 | 722.4 | - | - | 0 | - |
| - | - | 1335 | 723.4 | - | - | 0 | - |
| - | - | 2331 | 726.9 | - | - | 0 | - |
| - | - | 1023 | 727.4 | - | - | 0 | - |
| - | - | 1470 | 728.4 | - | - | 0 | - |
| - | - | 962.1 | 741.4 | - | - | 0 | - |
| - | - | 3035 | 742.3 | - | - | 0 | - |
| - | - | 2461 | 742.9 | - | - | 0 | - |
| - | - | 1091 | 743.4 | - | - | 0 | - |
| - | - | 3794 | 744.4 | - | - | 0 | - |
| - | - | 2593 | 744.9 | - | - | 0 | - |
| - | - | 1056 | 745.4 | - | - | 0 | - |
| 5 | y | 2430 | 748.9 | 0.006393 | 8.537 | +2 | 11 |
| 5 | z | 1.229E+04 | 749.4 | 0.003775 | 5.037 | +2 | 11 |
| - | - | 1.559E+04 | 749.9 | - | - | 0 | - |
| - | - | 1.106E+04 | 750.4 | - | - | 0 | - |
| - | - | 5235 | 750.9 | - | - | 0 | - |
| - | - | 1919 | 751.4 | - | - | 0 | - |
| - | - | 1262 | 752.4 | - | - | 0 | - |
| 12 | c | 3574 | 757.4 | 0.002538 | 3.351 | +2 | 12 |
| - | - | 4411 | 757.9 | - | - | 0 | - |
| - | - | 3207 | 758.4 | - | - | 0 | - |
| - | - | 1142 | 758.9 | - | - | 0 | - |
| - | - | 2278 | 762.9 | - | - | 0 | - |
| - | - | 1202 | 763.4 | - | - | 0 | - |
| - | - | 1262 | 763.9 | - | - | 0 | - |
| - | - | 2428 | 765.4 | - | - | 0 | - |
| 12 | c | 4211 | 765.9 | 0.001095 | 1.429 | +2 | 12 |
| - | - | 9597 | 766.4 | - | - | 0 | - |
| - | - | 5364 | 766.9 | - | - | 0 | - |
| - | - | 3087 | 767.4 | - | - | 0 | - |
| 11 | y | 2884 | 768.4 | 0.009929 | 12.92 | +1 | 5 |
| 11 | z | 1.691E+04 | 769.4 | 0.009026 | 11.73 | +1 | 5 |
| - | - | 1.24E+04 | 770.4 | - | - | 0 | - |
| - | - | 1183 | 770.9 | - | - | 0 | - |
| - | - | 4932 | 771.4 | - | - | 0 | - |
| - | - | 1789 | 776.4 | - | - | 0 | - |
| - | - | 1883 | 777.4 | - | - | 0 | - |
| - | - | 3591 | 777.9 | - | - | 0 | - |
| - | - | 2588 | 778.4 | - | - | 0 | - |
| - | - | 1218 | 778.9 | - | - | 0 | - |
| - | - | 1228 | 779.4 | - | - | 0 | - |
| 4 | w | 2276 | 784.4 | 0.001295 | 1.651 | +2 | 12 |
| - | - | 1721 | 784.9 | - | - | 0 | - |
| 11 | y | 8142 | 785.4 | 0.009196 | 11.71 | +1 | 5 |
| - | - | 2858 | 785.9 | - | - | 0 | - |
| - | - | 3146 | 786.4 | - | - | 0 | - |
| - | - | 1987 | 787.4 | - | - | 0 | - |
| 6 | c | 2.049E+04 | 793.4 | 0.0006305 | 0.7947 | +1 | 6 |
| - | - | 1.275E+04 | 794.4 | - | - | 0 | - |
| - | - | 3965 | 795.4 | - | - | 0 | - |
| - | - | 1385 | 796.4 | - | - | 0 | - |
| - | - | 1269 | 802.4 | - | - | 0 | - |
| - | - | 3071 | 802.9 | - | - | 0 | - |
| - | - | 2808 | 803.4 | - | - | 0 | - |
| - | - | 3211 | 803.9 | - | - | 0 | - |
| - | - | 3374 | 804.4 | - | - | 0 | - |
| - | - | 1693 | 804.9 | - | - | 0 | - |
| - | - | 1736 | 805.4 | - | - | 0 | - |
| 4 | y | 2541 | 806.4 | 0.001559 | 1.933 | +2 | 12 |
| 4 | z | 9965 | 806.9 | 0.003391 | 4.203 | +2 | 12 |
| - | - | 2.272E+04 | 807.4 | - | - | 0 | - |
| - | - | 1.721E+04 | 807.9 | - | - | 0 | - |
| - | - | 7468 | 808.4 | - | - | 0 | - |
| - | - | 3355 | 808.9 | - | - | 0 | - |
| - | - | 1915 | 809.4 | - | - | 0 | - |
| 6 | c | 8.342E+04 | 810.4 | 0.0003265 | 0.4029 | +1 | 6 |
| - | - | 4.455E+04 | 811.4 | - | - | 0 | - |
| - | - | 1.236E+04 | 812.4 | - | - | 0 | - |
| - | - | 1150 | 813.4 | - | - | 0 | - |
| 4 | y | 2344 | 814.9 | 0.003964 | 4.865 | +2 | 12 |
| - | - | 1550 | 815.4 | - | - | 0 | - |
| - | - | 2241 | 816.9 | - | - | 0 | - |
| - | - | 4239 | 817.4 | - | - | 0 | - |
| - | - | 2822 | 817.9 | - | - | 0 | - |
| - | - | 1719 | 818.4 | - | - | 0 | - |
| - | - | 1094 | 823.9 | - | - | 0 | - |
| - | - | 2192 | 824.4 | - | - | 0 | - |
| - | - | 1111 | 824.9 | - | - | 0 | - |
| - | - | 2867 | 826.9 | - | - | 0 | - |
| - | - | 2683 | 827.4 | - | - | 0 | - |
| - | - | 1924 | 828.4 | - | - | 0 | - |
| - | - | 1808 | 838.4 | - | - | 0 | - |
| - | - | 1347 | 838.9 | - | - | 0 | - |
| 10 | w | 2793 | 839.4 | 0.003647 | 4.345 | +1 | 6 |
| - | - | 3225 | 840.4 | - | - | 0 | - |
| - | - | 3361 | 841.4 | - | - | 0 | - |
| 3 | w | 6986 | 841.9 | 0.005158 | 6.126 | +2 | 13 |
| - | - | 5695 | 842.4 | - | - | 0 | - |
| - | - | 1346 | 842.9 | - | - | 0 | - |
| - | - | 1075 | 843.4 | - | - | 0 | - |
| - | - | 1201 | 845.4 | - | - | 0 | - |
| 13 | c | 8232 | 846.4 | 0.001662 | 1.963 | +2 | 13 |
| - | - | 9975 | 846.9 | - | - | 0 | - |
| - | - | 6671 | 847.4 | - | - | 0 | - |
| - | - | 3965 | 847.9 | - | - | 0 | - |
| - | - | 1850 | 848.4 | - | - | 0 | - |
| - | - | 1028 | 848.9 | - | - | 0 | - |
| - | - | 1641 | 859.4 | - | - | 0 | - |
| - | - | 1664 | 859.9 | - | - | 0 | - |
| - | - | 2241 | 860.4 | - | - | 0 | - |
| - | - | 1144 | 861.4 | - | - | 0 | - |
| 3 | z | 1271 | 862.4 | 0.005802 | 6.728 | +2 | 13 |
| - | - | 1040 | 865.4 | - | - | 0 | - |
| - | - | 1180 | 866.4 | - | - | 0 | - |
| - | - | 2325 | 867.4 | - | - | 0 | - |
| 3 | z | 1.078E+04 | 870.9 | 0.003993 | 4.585 | +2 | 13 |
| - | - | 3.708E+04 | 871.4 | - | - | 0 | - |
| - | - | 3.461E+04 | 871.9 | - | - | 0 | - |
| - | - | 1.341E+04 | 872.4 | - | - | 0 | - |
| - | - | 5685 | 872.9 | - | - | 0 | - |
| - | - | 2964 | 873.4 | - | - | 0 | - |
| 3 | y | 2349 | 878.9 | 0.003041 | 3.459 | +2 | 13 |
| - | - | 2832 | 879.4 | - | - | 0 | - |
| - | - | 945 | 879.9 | - | - | 0 | - |
| - | - | 2653 | 880.4 | - | - | 0 | - |
| - | - | 2141 | 880.9 | - | - | 0 | - |
| - | - | 6825 | 881.4 | - | - | 0 | - |
| - | - | 2195 | 881.9 | - | - | 0 | - |
| - | - | 4560 | 882.4 | - | - | 0 | - |
| - | - | 1253 | 883.5 | - | - | 0 | - |
| - | - | 2042 | 888.4 | - | - | 0 | - |
| - | - | 9715 | 888.9 | - | - | 0 | - |
| - | - | 8702 | 889.4 | - | - | 0 | - |
| - | - | 5176 | 889.9 | - | - | 0 | - |
| - | - | 2022 | 890.4 | - | - | 0 | - |
| - | - | 1008 | 895.9 | - | - | 0 | - |
| 10 | y | 1243 | 896.4 | 0.002904 | 3.24 | +1 | 6 |
| 10 | z | 1.311E+04 | 898.4 | 0.00859 | 9.561 | +1 | 6 |
| - | - | 1.855E+04 | 899.4 | - | - | 0 | - |
| - | - | 7782 | 900.4 | - | - | 0 | - |
| - | - | 2307 | 901.4 | - | - | 0 | - |
| - | - | 1718 | 902.4 | - | - | 0 | - |
| - | - | 980.5 | 902.9 | - | - | 0 | - |
| - | - | 1111 | 903.4 | - | - | 0 | - |
| 7 | c | 9180 | 908.4 | 0.0007871 | 0.8664 | +1 | 7 |
| - | - | 5298 | 909.4 | - | - | 0 | - |
| 14 | c | 6018 | 910.4 | 0.007273 | 7.989 | +2 | 14 |
| - | - | 4684 | 910.9 | - | - | 0 | - |
| - | - | 3236 | 911.4 | - | - | 0 | - |
| - | - | 1004 | 911.9 | - | - | 0 | - |
| 10 | y | 2892 | 914.4 | 0.009492 | 10.38 | +1 | 6 |
| - | - | 988 | 915.4 | - | - | 0 | - |
| - | - | 1553 | 923.5 | - | - | 0 | - |
| 7 | c | 4974 | 925.5 | 0.0001883 | 0.2035 | +1 | 7 |
| - | - | 4154 | 926.5 | - | - | 0 | - |
| - | - | 1227 | 927.5 | - | - | 0 | - |
| - | - | 2274 | 931 | - | - | 0 | - |
| - | - | 2174 | 931.5 | - | - | 0 | - |
| - | - | 2805 | 932 | - | - | 0 | - |
| - | - | 5138 | 938.5 | - | - | 0 | - |
| - | - | 1262 | 939 | - | - | 0 | - |
| 2 | z | 4412 | 939.4 | 0.01825 | 19.42 | +2 | 14 |
| - | - | 1.427E+04 | 939.9 | - | - | 0 | - |
| - | - | 1.415E+04 | 940.4 | - | - | 0 | - |
| - | - | 6609 | 940.9 | - | - | 0 | - |
| - | - | 3882 | 941.4 | - | - | 0 | - |
| - | - | 989.7 | 945.5 | - | - | 0 | - |
| - | - | 995.3 | 946 | - | - | 0 | - |
| - | - | 1473 | 946.5 | - | - | 0 | - |
| - | - | 3384 | 952.5 | - | - | 0 | - |
| - | - | 2660 | 953 | - | - | 0 | - |
| - | - | 2626 | 953.5 | - | - | 0 | - |
| - | - | 1449 | 954 | - | - | 0 | - |
| - | - | 1182 | 954.5 | - | - | 0 | - |
| - | - | 1332 | 958.5 | - | - | 0 | - |
| - | - | 1259 | 960.5 | - | - | 0 | - |
| - | - | 2443 | 961 | - | - | 0 | - |
| - | - | 1763 | 961.5 | - | - | 0 | - |
| - | - | 1292 | 962 | - | - | 0 | - |
| - | - | 1038 | 964 | - | - | 0 | - |
| 8 | c | 1054 | 964.5 | 0.007895 | 8.185 | +1 | 8 |
| - | - | 3518 | 966.5 | - | - | 0 | - |
| - | - | 2405 | 967 | - | - | 0 | - |
| - | - | 6714 | 967.5 | - | - | 0 | - |
| - | - | 4940 | 968 | - | - | 0 | - |
| - | - | 3764 | 968.5 | - | - | 0 | - |
| - | - | 973.6 | 969.5 | - | - | 0 | - |
| - | - | 2116 | 974.5 | - | - | 0 | - |
| - | - | 4590 | 975 | - | - | 0 | - |
| - | - | 7399 | 975.5 | - | - | 0 | - |
| - | - | 5638 | 976 | - | - | 0 | - |
| - | - | 3127 | 976.5 | - | - | 0 | - |
| - | - | 1956 | 977 | - | - | 0 | - |
| - | - | 933.8 | 981.5 | - | - | 0 | - |
| 8 | c | 1.928E+04 | 982.5 | 0.0007479 | 0.7612 | +1 | 8 |
| - | - | 1701 | 983 | - | - | 0 | - |
| - | - | 1.406E+04 | 983.5 | - | - | 0 | - |
| - | - | 5623 | 984.5 | - | - | 0 | - |
| - | - | 1553 | 985.5 | - | - | 0 | - |
| - | - | 1286 | 991 | - | - | 0 | - |
| - | - | 9439 | 996.5 | - | - | 0 | - |
| - | - | 8346 | 997 | - | - | 0 | - |
| - | - | 7048 | 997.5 | - | - | 0 | - |
| - | - | 3092 | 998 | - | - | 0 | - |
| - | - | 5237 | 1005 | - | - | 0 | - |
| - | - | 7956 | 1005 | - | - | 0 | - |
| - | - | 5393 | 1006 | - | - | 0 | - |
| - | - | 1409 | 1010 | - | - | 0 | - |
| 9 | z | 2920 | 1027 | 0.009315 | 9.074 | +1 | 7 |
| - | - | 1.076E+04 | 1028 | - | - | 0 | - |
| - | - | 5126 | 1029 | - | - | 0 | - |
| - | - | 2507 | 1030 | - | - | 0 | - |
| - | - | 1699 | 1052 | - | - | 0 | - |
| - | - | 1811 | 1067 | - | - | 0 | - |
| 8 | z | 1479 | 1084 | 0.009294 | 8.577 | +1 | 8 |
| - | - | 2844 | 1085 | - | - | 0 | - |
| - | - | 1849 | 1086 | - | - | 0 | - |
| - | - | 4014 | 1095 | - | - | 0 | - |
| - | - | 2915 | 1096 | - | - | 0 | - |
| 8 | y | 1354 | 1100 | 0.007632 | 6.941 | +1 | 8 |
| 9 | c | 1.784E+04 | 1111 | 0.0008313 | 0.7486 | +1 | 9 |
| - | - | 1.635E+04 | 1112 | - | - | 0 | - |
| - | - | 7988 | 1113 | - | - | 0 | - |
| - | - | 2423 | 1114 | - | - | 0 | - |
| - | - | 1384 | 1137 | - | - | 0 | - |
| - | - | 1155 | 1141 | - | - | 0 | - |
| - | - | 881.6 | 1142 | - | - | 0 | - |
| - | - | 1113 | 1182 | - | - | 0 | - |
| - | - | 1153 | 1197 | - | - | 0 | - |
| 7 | z | 1093 | 1199 | 0.002668 | 2.226 | +1 | 9 |
| - | - | 7305 | 1200 | - | - | 0 | - |
| - | - | 4866 | 1201 | - | - | 0 | - |
| - | - | 1190 | 1202 | - | - | 0 | - |
| - | - | 1158 | 1224 | - | - | 0 | - |
| - | - | 1373 | 1225 | - | - | 0 | - |
| - | - | 930.5 | 1226 | - | - | 0 | - |
| 10 | c | 4532 | 1240 | 0.002165 | 1.746 | +1 | 10 |
| - | - | 3291 | 1241 | - | - | 0 | - |
| - | - | 2733 | 1242 | - | - | 0 | - |
| - | - | 786 | 1243 | - | - | 0 | - |
| - | - | 1122 | 1265 | - | - | 0 | - |
| - | - | 1209 | 1266 | - | - | 0 | - |
| - | - | 1187 | 1267 | - | - | 0 | - |
| 6 | z | 1620 | 1312 | 0.01581 | 12.05 | +1 | 10 |
| - | - | 5098 | 1313 | - | - | 0 | - |
| - | - | 4850 | 1314 | - | - | 0 | - |
| - | - | 2175 | 1315 | - | - | 0 | - |
| - | - | 985.8 | 1345 | - | - | 0 | - |
| - | - | 1174 | 1359 | - | - | 0 | - |
| - | - | 4171 | 1360 | - | - | 0 | - |
| - | - | 2740 | 1361 | - | - | 0 | - |
| - | - | 1344 | 1362 | - | - | 0 | - |
| - | - | 1022 | 1388 | - | - | 0 | - |
| 11 | c | 2576 | 1403 | 0.003481 | 2.482 | +1 | 11 |
| - | - | 3499 | 1404 | - | - | 0 | - |
| - | - | 2251 | 1405 | - | - | 0 | - |
| 5 | z | 1562 | 1498 | 0.002959 | 1.975 | +1 | 11 |
| - | - | 1236 | 1499 | - | - | 0 | - |
| - | - | 1014 | 1516 | - | - | 0 | - |
| - | - | 880.5 | 1533 | - | - | 0 | - |
| - | - | 1211 | 1614 | - | - | 0 | - |
| - | - | 1034 | 1615 | - | - | 0 | - |
| - | - | 930.9 | 1693 | - | - | 0 | - |
| - | - | 852.6 | 1694 | - | - | 0 | - |
| - | - | 1664 | 1743 | - | - | 0 | - |
| - | - | 1040 | 1744 | - | - | 0 | - |
| - | - | 799.5 | 1880 | - | - | 0 | - |

m/z Charge Intensity FragmentType MassShift Position
120.05758666992188 0 636.0194 y Ammonia loss 12
120.06596374511719 0 501.75964
121.96086120605469 0 355.08963
127.08695983886719 0 626.9649
127.1121826171875 0 455.94553
128.09487915039062 0 472.08057
129.0144500732422 0 359.1314
129.01858520507812 0 5535.9834
129.06642150878906 0 531.76855
129.1025848388672 0 14550.411
130.06546020507812 0 42292.766
130.10609436035156 0 482.55603
131.06883239746094 0 3543.5276
131.11822509765625 0 1570.7872
132.0660858154297 0 656.04895
132.0811767578125 0 3575.8425
132.10231018066406 0 687.44775
133.0611114501953 0 3728.846
133.08615112304688 0 2492.8354
136.07606506347656 0 3861.9011
138.99818420410156 0 397.80753
139.00291442871094 0 3374.3057
144.08131408691406 0 548.39606
146.06063842773438 0 468.09042
146.0928192138672 0 1402.0101
147.04483032226562 0 666.4442
147.0764617919922 0 961.9766
155.09307861328125 0 15594.27
156.06613159179688 0 700.7218
156.09625244140625 0 1214.465
156.1383514404297 0 815.4525
157.01348876953125 0 13292.084
157.13394165039062 0 742.4111
158.06088256835938 0 611.43005
159.09202575683594 0 17559.293
159.11363220214844 0 554.5662
160.09536743164062 0 1738.9287
164.0712890625 0 524.99304
164.082275390625 0 1853.0112
166.0614471435547 0 12981.335
167.0650634765625 0 747.80115
168.1384735107422 0 1216.9459
170.0604248046875 0 9039.727
171.06869506835938 0 940.1232
171.13845825195312 0 548.1371
172.0719451904297 0 2998.266
173.05613708496094 0 2581.7998
173.09251403808594 0 599.24445
173.43910217285156 0 981.842
174.07667541503906 0 1721.5656
177.1124267578125 0 1893.8234
178.1343231201172 0 1232.5046
183.1130828857422 0 5444.428
183.12246704101562 0 660.5209
183.1259765625 0 783.34406
183.1498565673828 0 718.5292
185.07179260253906 0 953.447
185.0812225341797 0 16045.137
185.09141540527344 0 936.83246
185.1652069091797 0 12283.735
186.05227661132812 0 505.081
186.07858276367188 0 473.3156
186.1239471435547 0 3166.6375
186.17298889160156 0 1177.2518
187.08702087402344 0 3813.125
187.10818481445312 0 1277.8904
189.09901428222656 0 499.65738
190.08267211914062 0 7604.463
192.0765838623047 0 916.7029
198.12803649902344 0 2466.6177
199.1697235107422 0 2881.6167
201.12342834472656 0 2179.2686
202.11874389648438 0 570.7031
203.10264587402344 0 1536.6478
203.15078735351562 0 958.7173
205.11868286132812 0 7067.971
207.16055297851562 0 1073.6611
209.17083740234375 0 574.6763
210.1279754638672 0 1606.3247
211.6295928955078 0 1078.028
213.07615661621094 0 8733.914
215.13966369628906 0 659.5292
217.03468322753906 0 12189.625
221.13865661621094 0 600.06635
223.1192626953125 0 1605.0193
223.1557159423828 0 12161.679
224.15875244140625 0 1366.7909
225.12457275390625 0 499.71808
226.15524291992188 0 722.6314
227.0669403076172 0 882.70807
227.1378936767578 0 543.28217
229.0825653076172 0 551.8717
229.11874389648438 0 5765.1006
229.1550750732422 0 676.6632
230.1225128173828 0 690.0164
233.13987731933594 0 3106.5542
234.1241455078125 0 12581.605
235.1276092529297 0 1901.8564
235.1542510986328 0 1608.7094
236.13827514648438 0 620.38477
236.63352966308594 0 2016.9585
241.1432647705078 0 702.9943
244.09266662597656 0 990.34674
245.14755249023438 0 733.22504
246.1099395751953 0 833.87335
246.12350463867188 0 1313.4419
247.0756072998047 0 731.0777
247.6211700439453 0 2905.741 c Ammonia loss 3
250.10800170898438 0 1989.7836
250.14273071289062 0 1533.4504
250.166748046875 0 2144.6934
251.15072631835938 0 53764.938
252.15399169921875 0 7401.4395
252.60440063476562 0 651.40405
255.15020751953125 0 1178.3503
256.6317443847656 0 720.21246
259.1542663574219 0 22118.025
261.1195983886719 0 1673.6459
264.17059326171875 0 1751.6957
265.1436767578125 0 1764.6904
265.6354064941406 0 767.1106
266.1250915527344 0 1503.4799
268.1772155761719 0 10086.178 c 1
269.1383972167969 0 2113.928
269.1806945800781 0 1691.9806
270.1219787597656 0 6347.0513
270.1811828613281 0 671.3652
270.623291015625 0 831.28705
272.1065979003906 0 1962.1548
272.1763000488281 0 1463.6418
273.09686279296875 0 7656.1294
274.1194763183594 0 1500.0686
274.1495361328125 0 10138.159
274.65142822265625 0 2823.1765
275.1517639160156 0 916.9837
279.18084716796875 0 895.4426
280.1504821777344 0 659.7584
282.18170166015625 0 5184.8184
283.1390380859375 0 1218.699
284.119873046875 0 1099.3882
285.15716552734375 0 845.0375
286.1405334472656 0 977.4091
288.1556091308594 0 976.0321
290.11724853515625 0 3504.5881
292.1651916503906 0 687.459
294.146484375 0 1057.3167 z Water loss 9
294.193115234375 0 1902.7219
295.1041259765625 0 1348.8508
296.1972351074219 0 918.7582
297.1280212402344 0 854.6516
297.19390869140625 0 686.55237
300.17144775390625 0 963.91315 z Ammonia loss 13
301.151611328125 0 1243.2802
302.14404296875 0 1379.8049
303.15228271484375 0 6899.4287 c Water loss 6
303.6550598144531 0 2398.6738 z 11
304.155029296875 0 1070.1533
307.14324951171875 0 772.82
309.1455078125 0 1475.3777
309.6473693847656 0 694.3095
311.66571044921875 0 6768.3784 y 11
312.1667785644531 0 1863.0514
315.1663818359375 0 773.24
317.1953125 0 2992.9146 z 13
317.65869140625 0 2401.6658
318.15130615234375 0 7404.714
318.201416015625 0 1101.6602
318.65301513671875 0 2309.5857
319.15521240234375 0 666.5207
323.14190673828125 0 1241.1482
326.6643981933594 0 32886.605
327.16595458984375 0 13368.976
327.66741943359375 0 3032.3403
329.1606140136719 0 3461.509
329.1913146972656 0 815.9119
331.6564025878906 0 4070.0513
332.1532897949219 0 1914.6992
332.1934814453125 0 669.0614
333.2137145996094 0 2719.4565 y 13
337.84857177734375 0 1061.8289
338.1824645996094 0 625.94366
338.672607421875 0 700.0501
339.6950988769531 0 1543.969
340.19891357421875 0 901.5344
340.6618957519531 0 13602.115 c Ammonia loss 4
341.16339111328125 0 6605.0874
341.6640319824219 0 1358.2141
342.50457763671875 0 711.35785 y Ammonia loss 8
348.18035888671875 0 563.22925 y 8
349.18914794921875 0 2159.2126
350.1796875 0 1523.431
350.51239013671875 0 633.96704
350.8472900390625 0 678.4986
351.2151794433594 0 1340.737
353.66839599609375 0 1180.5498
359.1502685546875 0 649.86017
360.6944885253906 0 1033.5314
361.17877197265625 0 1210.801 c Water loss 13
361.2015380859375 0 662.46106
361.2225646972656 0 3925.7515
361.51727294921875 0 1285.2737 y Ammonia loss 7
361.8515625 0 1097.8098 z 7
362.1825866699219 0 1076.878
363.2021179199219 0 2072.4675
364.19647216796875 0 1228.4958
364.69598388671875 0 680.6779
365.8548889160156 0 1350.4161
366.1907653808594 0 628.82324
367.186767578125 0 26193.21 y 7
367.5210266113281 0 16179.134
367.6825866699219 0 1478.6715
367.85491943359375 0 4919.9644
368.1105651855469 0 545.78864
368.1873779296875 0 2514.2244
369.2126159667969 0 1081.1816
371.1920471191406 0 1125.6875
374.20123291015625 0 1805.4581
374.6934814453125 0 5627.943 y Water loss 4
374.9337158203125 0 3241.5735 y Ammonia loss 4
375.1938781738281 0 2788.7478 z 4
375.43511962890625 0 724.44055
375.696533203125 0 699.66626
376.19281005859375 0 1398.1438
378.2030029296875 0 2583.5337
378.2268371582031 0 859.49774
379.2091369628906 0 15032.437 c Ammonia loss 2
379.2356262207031 0 512.2632
379.521484375 0 833.59064
379.742431640625 0 1545.2107
380.2118835449219 0 2595.2048
380.2388000488281 0 646.041
381.1516418457031 0 2047.6696
381.5211486816406 0 758.7638
383.2060546875 0 11307.13
383.2305603027344 0 1032.9191
383.7076110839844 0 6574.551
384.2079162597656 0 1997.2726
384.23333740234375 0 1282.7334
384.68341064453125 0 1107.8982 y Ammonia loss 10
385.1895446777344 0 6479.289 z 10
385.24371337890625 0 20119.582
385.5257873535156 0 8742.589
385.8596496582031 0 4603.3647
386.1949157714844 0 1682.7983
386.24688720703125 0 2621.9312
386.52581787109375 0 768.6065
387.2243347167969 0 30955.139
388.1991271972656 0 901.1135
388.2266540527344 0 6584.9546
388.6913757324219 0 962.7128
389.18701171875 0 3506.2427
389.22857666015625 0 988.92346
390.19085693359375 0 826.33075
393.19720458984375 0 21797.791 y 10
393.6990966796875 0 10117.23
394.1983947753906 0 2922.2712
394.7002258300781 0 1083.8578
395.19366455078125 0 1042.8258
395.39300537109375 0 1648.0558
395.5929870605469 0 1227.8346
396.2360534667969 0 24454.24 c 2
397.2038879394531 0 10145.3955 c Ammonia loss 5
397.2395324707031 0 5810.442
397.7057800292969 0 5158.1514
398.2063903808594 0 2460.5923
398.2364807128906 0 1028.0302
398.5968322753906 0 2612.0642
398.7958984375 0 3289.568
398.9953308105469 0 5541.1978
399.1959228515625 0 2784.9531
399.3965759277344 0 712.96564
399.43829345703125 0 691.49866 z Water loss 3
399.52490234375 0 1492.42 y Water loss 6
399.8551940917969 0 709.4186 y Ammonia loss 6
400.1917724609375 0 1706.2328 z 6
401.1882019042969 0 1268.5728
401.23895263671875 0 2225.481
401.2854919433594 0 2856.506
401.958251953125 0 678.5705
402.21246337890625 0 4104.653
402.2405090332031 0 832.4026
402.27081298828125 0 2537.4922
402.4000244140625 0 6052.0967
402.59967041015625 0 2148.6594
402.8015441894531 0 1346.5549
403.188720703125 0 962.5288
403.2159729003906 0 942.9414
403.27996826171875 0 824.14667
403.56878662109375 0 714.1652
404.21673583984375 0 3066.7966
404.71875 0 1852.8208
405.52972412109375 0 8986.564 y 6
405.8640441894531 0 8297.216
406.19781494140625 0 1858.0844
409.1461486816406 0 754.73846
409.22039794921875 0 670.98755
413.8849792480469 0 734.90625
418.2144470214844 0 1375.7266
420.20281982421875 0 144316.3 w 9
420.7043151855469 0 64826.344
421.20458984375 0 20449.195
421.7051696777344 0 4679.376
422.20550537109375 0 935.8279
423.2247314453125 0 819.44586
423.5332946777344 0 13016.1 w 5
423.8679504394531 0 7600.375
424.2012634277344 0 3177.1929
424.5339050292969 0 796.5669
430.19696044921875 0 1213.3815
431.22943115234375 0 1021.5161
432.2065734863281 0 895.52374 z Ammonia loss 5
434.20684814453125 0 830.6285
435.1655578613281 0 1031.6415
435.22412109375 0 1293.8329
436.2272033691406 0 1107.1873
437.2210388183594 0 1542.7264 y Water loss 5
437.55438232421875 0 2185.5586 y Ammonia loss 5
438.7422790527344 0 932.69934
439.24359130859375 0 747.30743
439.74176025390625 0 706.4689
439.96551513671875 0 776.5101 y 2
440.71746826171875 0 1504.3539
441.2207946777344 0 806.9237
442.8941345214844 0 1139.8531
443.192138671875 0 1228.5214
443.2246398925781 0 19626.184 y 5
443.558837890625 0 14236.358
443.8928527832031 0 3614.9143
444.2270202636719 0 1793.7074
445.71136474609375 0 1194.0708
446.2121887207031 0 1687.731
446.74420166015625 0 713.3842
447.2204895019531 0 2287.5017
448.22198486328125 0 1941.2892
448.7137756347656 0 2641.6873 y Water loss 9
449.21466064453125 0 1084.0813 y Ammonia loss 9
449.7109680175781 0 2738.0037 z 9
450.20709228515625 0 1174.8706
451.22698974609375 0 805.4702 c Water loss 13
451.8853454589844 0 871.2036
452.19036865234375 0 865.09064
452.2250671386719 0 1727.7583
452.4727478027344 0 1034.4181
452.7202453613281 0 865.56323
452.97369384765625 0 685.89307
453.5728454589844 0 885.9088
454.570556640625 0 853.08984
454.7177734375 0 12688.802 c Ammonia loss 6
454.98870849609375 0 1214.4412
455.2188720703125 0 6545.3657
455.72149658203125 0 3691.7673 c 13
455.96612548828125 0 1563.4349
456.21832275390625 0 2313.8308
456.7312927246094 0 639.38995
457.71917724609375 0 3438.4207 y 9
458.22137451171875 0 1949.4143
458.9821472167969 0 625.8471
459.2175598144531 0 830.7413
459.47076416015625 0 796.6809
459.724365234375 0 1588.249
459.9728088378906 0 1184.6799
460.2254333496094 0 841.4088
460.7359619140625 0 19293.74
461.23681640625 0 9533.181
461.5516662597656 0 1027.4572
461.7366027832031 0 3614.9236
462.20147705078125 0 2820.1682
462.2358703613281 0 1301.2511
462.7345275878906 0 1953.6655
463.22943115234375 0 1102.3883 c 6
465.4668884277344 0 2438.2937
465.71929931640625 0 1860.8789 z Water loss 1
465.975341796875 0 1347.9421 z Ammonia loss 1
466.2409362792969 0 3373.9033
467.2508239746094 0 2671.5293
468.2636413574219 0 8384.915
468.3190612792969 0 929.9776
468.7655944824219 0 5177.2607
469.2364501953125 0 2619.3926
469.4820861816406 0 899.79224
469.7227783203125 0 2185.11 y Water loss 1
469.97930908203125 0 7204.913 y Ammonia loss 1
470.2301330566406 0 7842.4434 z 1
470.3324890136719 0 1163.8291
470.47998046875 0 4129.2856
470.7267150878906 0 1053.3877
472.9721374511719 0 732.31946
473.2349548339844 0 1672.1667
473.48272705078125 0 779.7624
473.7339782714844 0 2470.2686
473.90350341796875 0 901.04114
473.98046875 0 2410.242
474.22711181640625 0 7819.3096 y 1
474.47735595703125 0 7246.5547
474.7272033691406 0 4101.339
475.9867248535156 0 2260.4019
476.2461853027344 0 854.70593
476.4885559082031 0 1826.8402
476.7376403808594 0 2226.6562
476.9823303222656 0 1183.792
477.2289733886719 0 3249.8726
477.4798583984375 0 993.6256
478.2191467285156 0 1902.6409 z 12
479.22674560546875 0 923.6093
479.5701599121094 0 986.53314
479.74957275390625 0 1352.6852
479.90301513671875 0 1033.1985
480.2466735839844 0 1671.9574
480.4934387207031 0 36458.555
480.74407958984375 0 37438.625
480.9941101074219 0 19583.895
481.2449035644531 0 10722.224
481.4949035644531 0 3256.0786
481.7441101074219 0 1163.482
482.7359619140625 0 4991.3623 c Water loss 7
483.22998046875 0 7675.138 c Ammonia loss 7
483.4842834472656 0 1602.36
483.7318420410156 0 2468.9807
483.98883056640625 0 1664.5211
484.24969482421875 0 11076.222
484.48773193359375 0 3763.5923
484.5745849609375 0 759.44055
484.7248840332031 0 5578.508 w 8
484.75433349609375 0 2851.996
484.9862365722656 0 1561.364
485.2286682128906 0 3036.5825
485.7259826660156 0 1607.4164
487.29144287109375 0 969.2484
487.47918701171875 0 3299.2278
487.7295227050781 0 79661.19
487.9828186035156 0 75161.69
488.23419189453125 0 45014.445
488.4842224121094 0 21857.377
488.7453308105469 0 11834.211
488.9949951171875 0 4144.575
489.2475891113281 0 2521.7239
491.2456970214844 0 1282.0369
491.74169921875 0 7289.324 c 7
491.9961242675781 0 770.6002
492.2417907714844 0 4909.4434
492.7423400878906 0 2925.1404
493.254150390625 0 791.8311 c Water loss 3
493.56512451171875 0 2402.561
493.90087890625 0 1650.4556 z Water loss 4
494.2326965332031 0 34121.6 c Ammonia loss 3
494.4875793457031 0 812.87415
495.2371520996094 0 8558.74
495.76666259765625 0 784.4785
496.2309875488281 0 887.9125
496.2620544433594 0 1177.6381
496.7750244140625 0 4140.287
497.2764892578125 0 3515.41
497.7552795410156 0 2706.0884
498.24658203125 0 4345.5083
498.49371337890625 0 3652.6956
498.7457580566406 0 3111.8604
498.9961242675781 0 969.9971
499.2410583496094 0 8596.66 y Water loss 4
499.4936828613281 0 884.68915
499.574951171875 0 9583.404 y Ammonia loss 4
499.74468994140625 0 1747.6268
499.91033935546875 0 7692.622 z 4
500.2451171875 0 5981.0107
500.5769348144531 0 1946.754
500.91064453125 0 1013.2421
501.9929504394531 0 1548.3102
502.2493591308594 0 4072.0273
502.4974365234375 0 120858.12
502.7483215332031 0 172756.2
502.99896240234375 0 126976.625
503.2494812011719 0 59700.01
503.5001220703125 0 21970.03
503.7497253417969 0 8123.051
504.0008239746094 0 2801.104
504.2481994628906 0 1585.7329
505.25030517578125 0 6469.969 c Ammonia loss 11
505.5850524902344 0 6724.3
505.74053955078125 0 3023.8843
505.9179992675781 0 2631.0613
506.2596130371094 0 5657.548
506.5916442871094 0 806.6595
506.7334289550781 0 1726.162
506.9320068359375 0 899.8937
507.26544189453125 0 1772.9486
510.26068115234375 0 948.1784
510.5951843261719 0 938.3962
510.9290771484375 0 44399.336 c 11
511.26312255859375 0 79290.92 c 3
511.5971984863281 0 16178.947
511.9314880371094 0 6261.291
512.2659301757812 0 12781.556
512.7597045898438 0 1839.0457 y Water loss 8
513.2328491210938 0 1521.5796
513.2683715820312 0 2072.1975
513.7566528320312 0 19952.924 z 8
514.2575073242188 0 17000.133
514.5803833007812 0 1565.1874
514.7603149414062 0 7038.226
514.9116821289062 0 1272.7313
515.2577514648438 0 1161.9734
515.3197021484375 0 7144.465
515.7651977539062 0 877.3606
516.323486328125 0 2025.9633
517.2808227539062 0 992.62146
517.5963134765625 0 1063.6268
518.2881469726562 0 1211.2523
519.93603515625 0 1592.4443
520.266845703125 0 1694.4055
521.2770385742188 0 1760.335
521.7664794921875 0 15915.946 y 8
522.2677001953125 0 9707.311
522.76806640625 0 3605.5713
523.265625 0 1257.3237 w 3
523.591064453125 0 2116.5928
523.9259643554688 0 1852.2637
525.2731323242188 0 1203.31
525.35107421875 0 1151.6924
526.2553100585938 0 3171.1646
526.75341796875 0 1434.4385
528.2548828125 0 761.92535
529.28173828125 0 1009.61224
529.618896484375 0 848.00055
530.27880859375 0 1342.535
530.6096801757812 0 1167.6279
533.7716674804688 0 991.16187
534.2691650390625 0 2176.524
534.3111572265625 0 1114.2476
534.95166015625 0 2383.2139
535.2835083007812 0 3548.4783
535.6165771484375 0 5358.49
535.9474487304688 0 12853.473
536.2827758789062 0 12591.944
536.617919921875 0 6706.8926
536.9493408203125 0 4309.517
537.2789306640625 0 3287.8896
537.9225463867188 0 795.2517 y Ammonia loss 3
538.2755737304688 0 948.0903
538.58984375 0 844.79083
539.2377319335938 0 833.715
541.2696533203125 0 2654.999 y Water loss 7
541.7744140625 0 1612.043 y Ammonia loss 7
542.2671508789062 0 13577.449 z 7
542.7684936523438 0 6971.616
543.271728515625 0 3493.2083
543.59326171875 0 3347.757 y 3
543.7683715820312 0 1364.1069
543.927734375 0 4077.49
544.262939453125 0 1695.8259
546.2619018554688 0 2391.562
546.764404296875 0 2390.733
547.2864379882812 0 4437.969
547.7753295898438 0 2043.2793
548.2391967773438 0 1992.3945 w 11
548.2900390625 0 1792.9792
548.7501220703125 0 1102.9207
549.608154296875 0 2713.9604
549.7605590820312 0 1197.3949
549.93994140625 0 1987.3206
550.276611328125 0 34418.785 y 7
550.59814453125 0 1192.7112
550.7784423828125 0 20948.047
551.2774658203125 0 5787.7617
551.5958862304688 0 1118.341
551.7786865234375 0 1501.6724
552.2619018554688 0 2384.268
554.2930297851562 0 1252.0198
554.7877197265625 0 1957.2595
555.2700805664062 0 6201.128
555.7884521484375 0 140021.27 c 8
556.2901000976562 0 91405.91
556.791015625 0 28922.146
556.9360961914062 0 1098.7402
557.291259765625 0 7323.2524
557.61376953125 0 1001.3587
557.7904663085938 0 1768.8838
560.9307250976562 0 783.3074
561.2692260742188 0 1687.0513
561.5966186523438 0 52709.48 w 2
561.9309692382812 0 47974.664
562.2648315429688 0 24935.314
562.5980224609375 0 7427.66
562.635009765625 0 1917.5764
562.9320678710938 0 3768.8442
563.2664184570312 0 2219.027
563.3126831054688 0 1165.1864
563.7816162109375 0 7296.3857
563.9264526367188 0 999.1099
564.279296875 0 4383.0566
564.3247680664062 0 1068.8641
564.6002807617188 0 27246.66 c 12
564.781982421875 0 1673.0232
564.934326171875 0 24452.035
565.2693481445312 0 13584.148
565.6027221679688 0 7144.8706
565.9356079101562 0 3278.5068
566.2752075195312 0 4653.927
566.6077270507812 0 4019.1604
566.9425659179688 0 2604.458
567.2302856445312 0 2706.2693
567.2798461914062 0 2578.9397
568.2335205078125 0 908.83606
568.2830810546875 0 1658.2361
568.7957763671875 0 5112.6406
569.2950439453125 0 2168.1296
569.7769775390625 0 1014.2404
569.9351806640625 0 796.5186
570.2761840820312 0 3557.2812
570.7772216796875 0 4351.181
571.2794799804688 0 2096.3872
571.7813110351562 0 2566.468
572.2809448242188 0 972.22784
572.9553833007812 0 1863.6174
573.2881469726562 0 1806.925
573.9573974609375 0 805.11035
574.287841796875 0 1108.8834
577.2807006835938 0 2405.7678 w 6
577.3304443359375 0 918.4301
577.7857055664062 0 12856.072
577.9584350585938 0 1629.8088
578.2861328125 0 8608.285
578.78759765625 0 9505.242
579.2874145507812 0 3878.1409
579.7852783203125 0 1466.0671
580.2742919921875 0 1175.2875 y Water loss 2
580.608154296875 0 2344.0874 y Ammonia loss 2
580.939208984375 0 13485.001 z 2
581.2738647460938 0 11969.251
581.6078491210938 0 5006.3936
581.943359375 0 2089.9004
582.2742309570312 0 4503.0264
583.2810668945312 0 1570.4255
585.7825927734375 0 962.8078
586.2803344726562 0 7628.4775 y 2
586.6142578125 0 5568.45
586.9484252929688 0 2449.1184
587.2837524414062 0 3019.8838
587.6240234375 0 1965.4609
587.9584350585938 0 1015.586
588.2838134765625 0 7575.0996
588.6168212890625 0 6052.798
588.9503784179688 0 2638.8884
589.2843017578125 0 841.5392 z Ammonia loss 11
590.78369140625 0 2303.1274 z Water loss 6
591.2808837890625 0 2913.3987 z Ammonia loss 6
591.9570922851562 0 1086.8082
592.290283203125 0 1644.7747
592.6272583007812 0 8990.195
592.9602661132812 0 8127.323
593.2913818359375 0 3515.02
593.6300048828125 0 1217.2185
596.9635009765625 0 1077.769
597.2886962890625 0 1758.235
598.302734375 0 2756.0342
598.7959594726562 0 2715.9834 y Water loss 6
599.2845458984375 0 4094.632 y Ammonia loss 6
599.7815551757812 0 42414.773 z 6
600.2835693359375 0 44482.273
600.7850341796875 0 24936.926
601.2868041992188 0 8928.211 c Water loss 13
601.6288452148438 0 2761.7017 c Ammonia loss 13
601.7861938476562 0 2152.3513
601.9573974609375 0 2339.1013
602.290283203125 0 1758.5911
602.619873046875 0 2667.324
602.794677734375 0 1159.8376
602.94775390625 0 7676.2637
603.2828369140625 0 9119.121
603.6170043945312 0 2875.2195
603.9521484375 0 1306.2698
604.282470703125 0 1020.8088
606.305419921875 0 30344.152 z 11
606.6343383789062 0 2084.1772
606.9683837890625 0 9579.167
607.3002319335938 0 50255.734 c 13
607.6328735351562 0 38871.41
607.79052734375 0 46384.145 y 6
607.9661254882812 0 22020.025
608.2947387695312 0 38738.543
608.6328735351562 0 2172.195
608.7926635742188 0 11893.814
609.2975463867188 0 4466.42
610.2854614257812 0 5381.4346
611.296875 0 2702.9702 c Water loss 9
611.7989501953125 0 10606.145 c Ammonia loss 9
611.9610595703125 0 6236.532
612.2997436523438 0 17392.428
612.63134765625 0 3211.8345
612.8046875 0 3908.802
612.9602661132812 0 2047.8882
613.3021240234375 0 2752.7202
615.303955078125 0 1823.9857
616.30224609375 0 888.5668
618.3338012695312 0 967.96356
619.8154296875 0 1078.9512
619.9900512695312 0 897.3969
620.3106079101562 0 55941.645 c 9
620.6387939453125 0 5639.427
620.812255859375 0 36494.9
620.9789428710938 0 10509.714
621.24169921875 0 1334.3423
621.3137817382812 0 22624.424
621.6505126953125 0 5646.419
621.814453125 0 4683.7266
621.9745483398438 0 1821.1934
622.242431640625 0 1034.0645
622.3228759765625 0 16681.861 y 11
623.3262939453125 0 4676.039
625.6504516601562 0 3694.0818
625.9833984375 0 5818.0547
626.3092041015625 0 7835.419
626.6314697265625 0 15433.0205 z 1
626.8190307617188 0 1154.9362
626.9631958007812 0 15489.257
627.2982788085938 0 9643.708
627.6300048828125 0 4956.0146
627.8202514648438 0 862.983
627.9640502929688 0 3237.1973
628.3048706054688 0 1138.8868
629.9913330078125 0 955.4099
630.3109130859375 0 1641.1028
630.6437377929688 0 6128.075
630.9796752929688 0 7367.8965
631.3060302734375 0 4259.502
631.6474609375 0 1333.922
631.9677734375 0 3100.0657 y 1
632.3016357421875 0 4079.1472
632.6358642578125 0 1043.9908
633.3131713867188 0 1020.9955
634.3215942382812 0 2497.4636
634.6484985351562 0 2981.186
634.7958984375 0 26038.627 w 5
634.98193359375 0 3991.5276
635.2999267578125 0 18554.025
635.654296875 0 2633.1636
635.7977905273438 0 6730.92
635.988525390625 0 2759.184
636.33740234375 0 2278.2817
636.8042602539062 0 1245.5125
637.3553466796875 0 1046.561
638.2701416015625 0 1054.1337
638.9849853515625 0 1531.7965
639.3214721679688 0 1568.1465
639.6607666015625 0 1053.6548
640.322265625 0 18684.191
640.6561889648438 0 24232.91
640.816650390625 0 1099.2549
640.990234375 0 17090.816
641.3241577148438 0 10684.982
641.6563720703125 0 3836.1497
641.9948120117188 0 1324.6866
643.8240966796875 0 5385.0176
644.3229370117188 0 5581.282
644.652587890625 0 6271.3574
644.8248291015625 0 1713.9276
644.98583984375 0 10835.033
645.3202514648438 0 6128.2295
645.650146484375 0 6916.503
645.9824829101562 0 4754.3574
646.3167114257812 0 4064.5422
646.6500854492188 0 2242.812
647.3175048828125 0 1301.5648 z Water loss 5
649.3215942382812 0 2599.487
649.6534423828125 0 1369.691
649.9813232421875 0 16186.448
650.3230590820312 0 76188.26
650.6586303710938 0 82243.33
650.82470703125 0 1790.7579
650.9927978515625 0 46941.5
651.326416015625 0 25347.416
651.66162109375 0 10361.13
651.8107299804688 0 1181.9093
651.9955444335938 0 2108.7446
652.3206787109375 0 7034.6094
653.323974609375 0 3286.1936
654.3294067382812 0 2117.2485
654.6580810546875 0 1373.2484
654.9932250976562 0 3967.4985
655.3251953125 0 7242.326 y Water loss 5
655.6581420898438 0 4127.7305
655.8232421875 0 1997.9546 y Ammonia loss 5
655.9929809570312 0 3881.8518
656.3232421875 0 23193.262 z 5
656.8246459960938 0 19268.996
656.994140625 0 1014.524
657.3264770507812 0 12236.996
657.831787109375 0 4873.8755
658.332763671875 0 2162.094
658.6502075195312 0 2431.7473
658.9864501953125 0 3654.8984
659.3192138671875 0 3922.5168
659.6527709960938 0 2472.0647
663.2909545898438 0 2355.6174
663.82666015625 0 2690.5852
663.9911499023438 0 2393.1694
664.3309326171875 0 35233.184 y 5
664.6568603515625 0 91918.5
664.8333740234375 0 12783.948
664.990966796875 0 100141.58
665.3264770507812 0 53634.023
665.6585693359375 0 22941.39
665.8351440429688 0 3387.1367
665.9925537109375 0 10398.263
666.327392578125 0 6255.231
669.6602172851562 0 19257.414
669.9954833984375 0 42111.98
670.3306884765625 0 68758.125
670.6654663085938 0 52743.555
670.999755859375 0 29454.63
671.3325805664062 0 13881.6455
671.6676635742188 0 6005.785
671.8247680664062 0 2774.8833
672.0036010742188 0 1793.6306
672.31494140625 0 2920.8894
672.8106689453125 0 1704.9093
679.34130859375 0 2107.8276 c Water loss 4
679.8351440429688 0 30431.402
680.3211669921875 0 49501.324 c Ammonia loss 4
680.8384399414062 0 10124.786
681.3214721679688 0 22223.547
682.3275756835938 0 5177.4824
683.3295288085938 0 1984.3473
683.8358764648438 0 1092.2682
685.3243408203125 0 1148.5508
691.3373413085938 0 13642.617
691.8385620117188 0 10528.337
692.3389282226562 0 5534.2153
692.8391723632812 0 2242.2092 c Water loss 10
693.3336181640625 0 1112.3965 c Ammonia loss 10
693.8339233398438 0 2996.8413
694.337158203125 0 2133.9724
697.3424682617188 0 27354.271 c 4
698.3468627929688 0 13549.24
699.3455810546875 0 2830.1172
701.3442993164062 0 1717.2139
701.841796875 0 43510.457 c 10
702.3434448242188 0 36170.88
702.8458251953125 0 14840.766
703.3477172851562 0 4932.589
703.8529663085938 0 2051.6794
705.3558959960938 0 3418.653
705.8533325195312 0 2391.178
711.307861328125 0 2078.9126
711.3677368164062 0 1102.6022
714.377197265625 0 2096.9014
719.330810546875 0 1088.9775
719.8541259765625 0 980.4417
720.3560791015625 0 4275.292
720.8605346679688 0 2811.3577
721.3480224609375 0 3717.783
722.3579711914062 0 2447.0352
723.3654174804688 0 1334.7301
726.8587036132812 0 2331.3845
727.3576049804688 0 1023.082
728.3655395507812 0 1470.0737
741.3665771484375 0 962.1426
742.3475952148438 0 3034.5957
742.850341796875 0 2461.2217
743.3502807617188 0 1091.2218
744.3857421875 0 3794.3179
744.8867797851562 0 2593.4087
745.39111328125 0 1055.512
748.8562622070312 0 2429.8345 y Ammonia loss 4
749.36279296875 0 12293.692 z 4
749.8650512695312 0 15591.488
750.367431640625 0 11061.844
750.8692626953125 0 5234.7285
751.3658447265625 0 1918.6252
752.3788452148438 0 1262.1588
757.372802734375 0 3574.1558 c Ammonia loss 11
757.874755859375 0 4411.344
758.378173828125 0 3207.0757
758.884033203125 0 1142.4734
762.864501953125 0 2278.0896
763.3673095703125 0 1202.1371
763.8621826171875 0 1262.3126
765.4025268554688 0 2427.918
765.8897094726562 0 4211.079 c 11
766.3985595703125 0 9597.262
766.89306640625 0 5364.419
767.4059448242188 0 3086.8599
768.3587646484375 0 2884.1777 y Ammonia loss 10
769.3674926757812 0 16910.809 z 10
770.3724975585938 0 12399.53
770.8724365234375 0 1182.6306
771.3723754882812 0 4931.8394
776.3776245117188 0 1788.623
777.3712158203125 0 1883.1284
777.8704833984375 0 3591.2227
778.37060546875 0 2587.8755
778.87841796875 0 1217.778
779.4012451171875 0 1227.683
784.3825073242188 0 2276.0283 w 3
784.8782958984375 0 1720.627
785.3860473632812 0 8141.744 y 10
785.8827514648438 0 2857.793
786.3870849609375 0 3145.679
787.3978881835938 0 1987.0693
793.3997802734375 0 20491.764 c Ammonia loss 5
794.4033203125 0 12749.156
795.408203125 0 3965.0205
796.415283203125 0 1384.6147
802.4306640625 0 1268.7397
802.9241943359375 0 3070.6755
803.4207763671875 0 2808.3926
803.9170532226562 0 3210.9468
804.423828125 0 3374.2317
804.92724609375 0 1692.5034
805.4188232421875 0 1736.408
806.377685546875 0 2540.6797 y Ammonia loss 3
806.8766479492188 0 9965.494 z 3
807.3793334960938 0 22723.957
807.8812866210938 0 17208.229
808.3825073242188 0 7468.149
808.8839111328125 0 3354.8682
809.388427734375 0 1914.657
810.426025390625 0 83415.65 c 5
811.4291381835938 0 44547.5
812.4324951171875 0 12357.347
813.4405517578125 0 1150.382
814.8854370117188 0 2344.183 y 3
815.3840942382812 0 1550.114
816.8905029296875 0 2240.6538
817.3916625976562 0 4239.1934
817.8920288085938 0 2822.3894
818.394775390625 0 1718.9397
823.9066162109375 0 1094.1908
824.4011840820312 0 2191.999
824.8953247070312 0 1110.8109
826.8995971679688 0 2866.684
827.401123046875 0 2683.1973
828.40283203125 0 1924.4172
838.39013671875 0 1808.4094
838.8877563476562 0 1347.0714
839.4021606445312 0 2792.7925 w 9
840.4041137695312 0 3224.7332
841.4026489257812 0 3361.3796
841.8998413085938 0 6985.6016 w 2
842.4015502929688 0 5694.899
842.904052734375 0 1345.5706
843.40478515625 0 1075.493
845.4014282226562 0 1201.3958
846.3965454101562 0 8232.214 c 12
846.8987426757812 0 9975.086
847.4008178710938 0 6670.925
847.9011840820312 0 3965.2761
848.4021606445312 0 1849.8213
848.9033203125 0 1027.854
859.43408203125 0 1641.3131
859.9343872070312 0 1663.842
860.4293823242188 0 2241.0833
861.431640625 0 1143.6014
862.40185546875 0 1271.3329 z Ammonia loss 2
865.42626953125 0 1040.1251
866.4297485351562 0 1179.9349
867.4313354492188 0 2325.397
870.9053344726562 0 10779.23 z 2
871.4088745117188 0 37081.895
871.9102172851562 0 34612.38
872.4110107421875 0 13414.424
872.9127807617188 0 5685.1465
873.4158325195312 0 2964.152
878.9156494140625 0 2349.0579 y 2
879.417724609375 0 2831.5227
879.9281616210938 0 945.00146
880.4266967773438 0 2652.822
880.9268798828125 0 2140.693
881.4390869140625 0 6824.541
881.9403686523438 0 2194.7368
882.4436645507812 0 4560.0747
883.4527587890625 0 1252.9178
888.43896484375 0 2041.8623
888.93994140625 0 9715.381
889.4429931640625 0 8702.417
889.9412841796875 0 5175.885
890.440673828125 0 2021.6831
895.9317626953125 0 1008.4755
896.43017578125 0 1243.3213 y Water loss 9
898.4105224609375 0 13114.591 z 9
899.4166870117188 0 18554.852
900.4188842773438 0 7782.1694
901.4213256835938 0 2307.0144
902.4287109375 0 1718.2924
902.9319458007812 0 980.5149
903.4357299804688 0 1110.7648
908.4268798828125 0 9179.587 c Ammonia loss 6
909.4314575195312 0 5297.6514
910.4384155273438 0 6018.475 c 13
910.9391479492188 0 4683.9463
911.4449462890625 0 3236.1716
911.9487915039062 0 1003.53534
914.4283447265625 0 2891.9504 y 9
915.4353637695312 0 987.99084
923.45751953125 0 1552.9186
925.4524536132812 0 4973.632 c 6
926.4581298828125 0 4154.1914
927.4589233398438 0 1227.3495
930.9683227539062 0 2273.8354
931.4697265625 0 2174.2988
931.97314453125 0 2805.4006
938.4661865234375 0 5138.204
938.9701538085938 0 1262.3203
939.45703125 0 4411.716 z 1
939.9392700195312 0 14274.62
940.4407958984375 0 14149.3955
940.94091796875 0 6608.9897
941.4423828125 0 3881.7332
945.48388671875 0 989.7325
945.9796752929688 0 995.336
946.4853515625 0 1472.6549
952.4735107421875 0 3383.8274
952.9771728515625 0 2660.365
953.4769287109375 0 2626.02
953.9772338867188 0 1449.2927
954.477294921875 0 1181.8936
958.4784545898438 0 1331.5752
960.4813232421875 0 1258.8881
960.9813232421875 0 2443.2212
961.4818115234375 0 1762.5104
961.9840698242188 0 1292.4161
963.9682006835938 0 1038.1694
964.471435546875 0 1054.104 c Water loss 7
966.4595947265625 0 3518.4858
966.9776000976562 0 2404.899
967.47216796875 0 6714.213
967.9798583984375 0 4939.7124
968.4828491210938 0 3763.5774
969.5014038085938 0 973.602
974.4776000976562 0 2115.7056
974.984130859375 0 4590.31
975.4845581054688 0 7398.686
975.9871215820312 0 5638.419
976.4886474609375 0 3126.763
976.9937133789062 0 1955.925
981.4724731445312 0 933.8108
982.474853515625 0 19284.729 c 7
982.9824829101562 0 1701.0848
983.4802856445312 0 14056.991
984.4832763671875 0 5623.1235
985.4885864257812 0 1552.6821
990.9825439453125 0 1286.4524
996.48193359375 0 9439.403
996.9832763671875 0 8345.689
997.4843139648438 0 7048.2744
997.9853515625 0 3091.9583
1004.9917602539062 0 5236.772
1005.4951171875 0 7956.2563
1005.9972534179688 0 5392.916
1010.4658813476562 0 1408.5436
1026.5047607421875 0 2919.747 z 8
1027.512451171875 0 10763.558
1028.5157470703125 0 5126.0483
1029.5162353515625 0 2507.1375
1051.5546875 0 1699.234
1066.553466796875 0 1811.1691
1083.5262451171875 0 1478.6597 z 7
1084.536376953125 0 2844.02
1085.532958984375 0 1849.4253
1094.5469970703125 0 4013.5251
1095.552734375 0 2914.5833
1099.546630859375 0 1354.493 y 7
1110.5682373046875 0 17838.56 c 8
1111.5723876953125 0 16347.527
1112.578125 0 7987.928
1113.5838623046875 0 2422.5833
1136.52685546875 0 1384.2124
1140.54736328125 0 1155.1368
1141.557861328125 0 881.5934
1181.6036376953125 0 1113.3319
1196.6021728515625 0 1152.7286
1198.559814453125 0 1092.6464 z 6
1199.5609130859375 0 7304.872
1200.5615234375 0 4866.3457
1201.5758056640625 0 1190.005
1223.586669921875 0 1157.9645
1224.61083984375 0 1373.1641
1225.602783203125 0 930.53284
1239.6094970703125 0 4532.145 c 9
1240.615478515625 0 3290.5183
1241.6226806640625 0 2732.8494
1242.6221923828125 0 785.95685
1264.6092529296875 0 1122.0659
1265.6163330078125 0 1208.7954
1266.6142578125 0 1186.6646
1311.6307373046875 0 1620.393 z 5
1312.641845703125 0 5097.7134
1313.6458740234375 0 4849.952
1314.642333984375 0 2175.2727
1344.6502685546875 0 985.7899
1358.656494140625 0 1174.2015
1359.6658935546875 0 4171.3267
1360.6689453125 0 2740.3586
1361.6771240234375 0 1343.7861
1387.6583251953125 0 1022.34216
1402.6715087890625 0 2576.0215 c 10
1403.6766357421875 0 3498.895
1404.6873779296875 0 2251.2412
1497.722900390625 0 1561.7681 z 4
1498.72119140625 0 1236.3064
1515.7564697265625 0 1014.3819
1532.789306640625 0 880.4684
1613.74658203125 0 1210.7501
1614.734375 0 1034.0631
1692.7918701171875 0 930.90796
1693.7952880859375 0 852.61707
1742.812744140625 0 1664.011
1743.8006591796875 0 1040.21
1879.8873291015625 0 799.4868

Spectrum Details

|  |  |
| --- | --- |
| Matched peaks? Matched peaks The total absolute number of peaks matched. Additionally in brackets the total fraction of peaks matched and the total number of peaks is shown. | 151 (13.64% of 1107) |
| FDR? FDR The false discovery rate estimated for this peptide. It is calculated by matching all theoretical fragments with a non-integer shift with the raw peaks for this spectrum. This is done with 40 different shifts. The resulting percentage is the average number of annotated peaks over the number of annotated peaks with the correct spectrum. | 1.12% |
| Satellite FDR? Satellite FDR See the FDR for details on its calculation. This satellite ion specific FDR only contains the satellite ions (d/w) for I/L/J positions. | 2.38% |
| PSM Score? PSM Score The PSM Score as given by Hecklib to this annotated spectrum. It is shown with three significant figures. | 452 |

#### Spectrum 5486? Spectrum 5486 The raw spectrum of this peptide as annotated by Hecklib. The fragments are coloured according to ion type (see legend). Any peaks with a star '\*' as text can be hovered over to see the full details, first the ion type second the mass shift type. By hovering over the amino acids in the peptide or ions in the legend the corresponding peaks are highlighted. By toggling the 'Unassigned' label you can turn the background (unassigned) peaks on or off in the plot. By updating the slider in the Ion legend you can update the spectrum to only show the top X% of the peaks with labels. The top X% means any peak that is within X% of the highest intensity. By dragging in the spectrum you can zoom in to a specific part of the spectrum and use 'Zoom Out' to get back to the original zoom level. The annotation of the spectrum is based on the given sequence in the peptides file and is done with different software so inconsistencies are likely. The peaks are annotated based on the given sequence, with 20 ppm tolerance.

Copy Data

##### Spectrum 5486 (TSV)

###### Preview

```
Loading example...
```

*Click on the button to copy the data to your clipboard.*

Mz MinMz MaxIntensity Max

WidthHeightPeptide font sizePeptide stroke widthSpectrum font sizeSpectrum stroke widthCompact peptide

Ion legend

wxyz

abcd

OtherUnassignedIon^Charge^_Position_Show for top:%

JHQDWLDGKEYKCKK

08.74e+31.75e+42.62e+43.50e+4

Zoom Out

w^+2^_2_c^+1^_2_c^+2^_5_y^+3^_8_c^+1^_3_y^+2^_5_c^+1^_3_c^+3^_10_w^+2^_6_y^+3^_10_z^+2^_6_y^+4^_14_c^+1^_4_c^+1^_4_y^+2^_7_w^+1^_4_y^+2^_8_c^+2^_9_w^+2^_9_y^+3^_13_y^+2^_9_z^+2^_9_z^+1^_4_c^+3^_14_y^+2^_9_c^+2^_10_c^+2^_10_c^+2^_10_z^+3^_14_y^+1^_4_y^+3^_14_y^+3^_14_y^+3^_14_w^+2^_10_z^+2^_10_y^+2^_10_z^+2^_10_y^+2^_10_c^+1^_5_c^+1^_5_c^+1^_5_c^+2^_11_y^+2^_11_z^+2^_11_c^+2^_12_c^+2^_12_y^+1^_5_z^+1^_5_y^+1^_5_c^+1^_6_y^+2^_12_z^+2^_12_c^+1^_6_y^+2^_12_w^+2^_13_c^+2^_13_y^+2^_13_z^+2^_13_y^+2^_13_z^+1^_6_c^+1^_7_c^+2^_14_c^+1^_7_z^+2^_14_y^+2^_14_c^+1^_8_c^+1^_8_w^+1^_7_c^+1^_8_z^+1^_7_c^+1^_9_z^+1^_9_c^+1^_10_z^+1^_10_c^+1^_11_z^+1^_11_c^+1^_12_c^+1^_12_z^+1^_12_c^+1^_13_z^+1^_13_

0508101615242032

Fragment Matches Table

Show background peaks

| Position | Ion type | Intensity | mz Theoretical | mz Error (Th) | mz Error (ppm) | Charge | Series Number |
| --- | --- | --- | --- | --- | --- | --- | --- |
| - | - | 391.3 | 120.1 | - | - | 0 | - |
| - | - | 355.7 | 120.9 | - | - | 0 | - |
| - | - | 391.5 | 129.1 | - | - | 0 | - |
| - | - | 2291 | 129.1 | - | - | 0 | - |
| - | - | 1969 | 130.1 | - | - | 0 | - |
| 14 | w | 431 | 130.1 | 0.0004984 | 3.832 | +2 | 2 |
| - | - | 506.5 | 133.1 | - | - | 0 | - |
| - | - | 1297 | 133.1 | - | - | 0 | - |
| - | - | 679.8 | 136.1 | - | - | 0 | - |
| - | - | 411 | 138.8 | - | - | 0 | - |
| - | - | 540.1 | 140.1 | - | - | 0 | - |
| - | - | 559.3 | 147.1 | - | - | 0 | - |
| - | - | 866.4 | 149 | - | - | 0 | - |
| - | - | 1114 | 149 | - | - | 0 | - |
| - | - | 2091 | 155.1 | - | - | 0 | - |
| - | - | 459.6 | 156.1 | - | - | 0 | - |
| - | - | 464.2 | 156.6 | - | - | 0 | - |
| - | - | 900.4 | 159.1 | - | - | 0 | - |
| - | - | 452.2 | 165.1 | - | - | 0 | - |
| - | - | 1509 | 166.1 | - | - | 0 | - |
| - | - | 596.5 | 167.1 | - | - | 0 | - |
| - | - | 480.7 | 168.1 | - | - | 0 | - |
| - | - | 1054 | 171.1 | - | - | 0 | - |
| - | - | 998.5 | 173.1 | - | - | 0 | - |
| - | - | 3785 | 173.5 | - | - | 0 | - |
| - | - | 638.8 | 174.1 | - | - | 0 | - |
| - | - | 508.7 | 175.1 | - | - | 0 | - |
| - | - | 628.7 | 177.1 | - | - | 0 | - |
| - | - | 945.1 | 183.1 | - | - | 0 | - |
| - | - | 2631 | 185.2 | - | - | 0 | - |
| - | - | 973.1 | 187.1 | - | - | 0 | - |
| - | - | 1372 | 191.1 | - | - | 0 | - |
| - | - | 1248 | 199.1 | - | - | 0 | - |
| - | - | 645.8 | 201.1 | - | - | 0 | - |
| - | - | 1361 | 202.1 | - | - | 0 | - |
| - | - | 2466 | 203.1 | - | - | 0 | - |
| - | - | 561.9 | 205.1 | - | - | 0 | - |
| - | - | 1085 | 207.1 | - | - | 0 | - |
| - | - | 1825 | 209.1 | - | - | 0 | - |
| - | - | 725.5 | 213.1 | - | - | 0 | - |
| - | - | 7861 | 215.1 | - | - | 0 | - |
| - | - | 667.9 | 216.1 | - | - | 0 | - |
| - | - | 638.4 | 217.1 | - | - | 0 | - |
| - | - | 1172 | 217.2 | - | - | 0 | - |
| - | - | 1738 | 218.1 | - | - | 0 | - |
| - | - | 1785 | 223.2 | - | - | 0 | - |
| - | - | 1670 | 227.1 | - | - | 0 | - |
| - | - | 639.8 | 229.1 | - | - | 0 | - |
| - | - | 762.2 | 229.2 | - | - | 0 | - |
| - | - | 640.8 | 230.1 | - | - | 0 | - |
| - | - | 924.4 | 231.2 | - | - | 0 | - |
| - | - | 2669 | 234.1 | - | - | 0 | - |
| - | - | 816.2 | 239.1 | - | - | 0 | - |
| - | - | 808.9 | 242.2 | - | - | 0 | - |
| - | - | 9716 | 251.1 | - | - | 0 | - |
| - | - | 1745 | 252.2 | - | - | 0 | - |
| - | - | 730.7 | 261.1 | - | - | 0 | - |
| 2 | c | 1200 | 268.2 | 0.0001351 | 0.5039 | +1 | 2 |
| - | - | 550.9 | 275.1 | - | - | 0 | - |
| - | - | 1952 | 281.1 | - | - | 0 | - |
| - | - | 495.2 | 282.1 | - | - | 0 | - |
| - | - | 868.7 | 282.2 | - | - | 0 | - |
| - | - | 1184 | 283.2 | - | - | 0 | - |
| - | - | 536.4 | 293.6 | - | - | 0 | - |
| - | - | 8037 | 299.1 | - | - | 0 | - |
| - | - | 903.7 | 299.1 | - | - | 0 | - |
| - | - | 673.7 | 299.2 | - | - | 0 | - |
| - | - | 1576 | 300.1 | - | - | 0 | - |
| - | - | 682.7 | 300.1 | - | - | 0 | - |
| - | - | 1201 | 302.2 | - | - | 0 | - |
| - | - | 562.6 | 310.1 | - | - | 0 | - |
| - | - | 1910 | 326.7 | - | - | 0 | - |
| - | - | 508.9 | 329.1 | - | - | 0 | - |
| - | - | 928.1 | 332.2 | - | - | 0 | - |
| - | - | 1379 | 338.1 | - | - | 0 | - |
| - | - | 1497 | 340.2 | - | - | 0 | - |
| 5 | c | 1088 | 340.7 | 0.0002619 | 0.7689 | +2 | 5 |
| - | - | 767.4 | 354.1 | - | - | 0 | - |
| - | - | 1596 | 359 | - | - | 0 | - |
| - | - | 1201 | 367.2 | - | - | 0 | - |
| 8 | y | 606.3 | 367.2 | 0.005462 | 14.88 | +3 | 8 |
| - | - | 556.5 | 367.7 | - | - | 0 | - |
| 3 | c | 4688 | 379.2 | 0.0005779 | 1.524 | +1 | 3 |
| - | - | 1006 | 380.2 | - | - | 0 | - |
| - | - | 2073 | 387.2 | - | - | 0 | - |
| 11 | y | 1340 | 393.2 | 0.004269 | 10.86 | +2 | 5 |
| - | - | 1250 | 393.7 | - | - | 0 | - |
| 3 | c | 4479 | 396.2 | 0.0006377 | 1.609 | +1 | 3 |
| - | - | 1157 | 397.2 | - | - | 0 | - |
| - | - | 664.8 | 407.7 | - | - | 0 | - |
| 10 | c | 645.6 | 408.2 | 0.001677 | 4.108 | +3 | 10 |
| - | - | 2058 | 415 | - | - | 0 | - |
| - | - | 1314 | 416 | - | - | 0 | - |
| 10 | w | 9720 | 420.2 | 0.004516 | 10.75 | +2 | 6 |
| - | - | 5197 | 420.7 | - | - | 0 | - |
| - | - | 1450 | 421.2 | - | - | 0 | - |
| - | - | 1036 | 421.2 | - | - | 0 | - |
| - | - | 732.3 | 431.2 | - | - | 0 | - |
| - | - | 703.2 | 435.7 | - | - | 0 | - |
| 6 | y | 614.1 | 443.2 | 0.008773 | 19.79 | +3 | 10 |
| - | - | 1282 | 444.2 | - | - | 0 | - |
| - | - | 805.8 | 444.3 | - | - | 0 | - |
| - | - | 761.4 | 449.2 | - | - | 0 | - |
| 10 | z | 1663 | 449.7 | 0.00641 | 14.25 | +2 | 6 |
| - | - | 7528 | 457.3 | - | - | 0 | - |
| - | - | 604.8 | 457.7 | - | - | 0 | - |
| - | - | 3160 | 458.2 | - | - | 0 | - |
| - | - | 2861 | 458.3 | - | - | 0 | - |
| - | - | 1958 | 458.7 | - | - | 0 | - |
| - | - | 4610 | 459.3 | - | - | 0 | - |
| - | - | 1147 | 460.3 | - | - | 0 | - |
| - | - | 1755 | 460.7 | - | - | 0 | - |
| - | - | 1010 | 461.2 | - | - | 0 | - |
| - | - | 989.4 | 466.2 | - | - | 0 | - |
| - | - | 791.7 | 467.2 | - | - | 0 | - |
| - | - | 1381 | 467.2 | - | - | 0 | - |
| - | - | 1195 | 468.3 | - | - | 0 | - |
| - | - | 690.5 | 468.8 | - | - | 0 | - |
| - | - | 1711 | 469.2 | - | - | 0 | - |
| - | - | 704.8 | 470.2 | - | - | 0 | - |
| - | - | 809 | 471.3 | - | - | 0 | - |
| - | - | 1007 | 472.3 | - | - | 0 | - |
| 2 | y | 1161 | 474.2 | 0.0005078 | 1.071 | +4 | 14 |
| - | - | 1132 | 474.5 | - | - | 0 | - |
| - | - | 712.9 | 476.5 | - | - | 0 | - |
| - | - | 1415 | 483.2 | - | - | 0 | - |
| - | - | 1647 | 484.2 | - | - | 0 | - |
| - | - | 903 | 484.7 | - | - | 0 | - |
| - | - | 3054 | 485.2 | - | - | 0 | - |
| - | - | 4256 | 486.3 | - | - | 0 | - |
| - | - | 789.4 | 487.2 | - | - | 0 | - |
| - | - | 712.1 | 487.3 | - | - | 0 | - |
| - | - | 2156 | 488.3 | - | - | 0 | - |
| - | - | 3913 | 493.7 | - | - | 0 | - |
| 4 | c | 4393 | 494.2 | 0.003271 | 6.619 | +1 | 4 |
| - | - | 1439 | 494.7 | - | - | 0 | - |
| - | - | 676.9 | 495.2 | - | - | 0 | - |
| - | - | 729.6 | 496.3 | - | - | 0 | - |
| - | - | 766.6 | 498.2 | - | - | 0 | - |
| - | - | 2490 | 501.2 | - | - | 0 | - |
| - | - | 1683 | 501.3 | - | - | 0 | - |
| - | - | 1428 | 501.7 | - | - | 0 | - |
| - | - | 5016 | 502.3 | - | - | 0 | - |
| - | - | 5088 | 502.5 | - | - | 0 | - |
| - | - | 1.213E+04 | 502.8 | - | - | 0 | - |
| - | - | 4070 | 503 | - | - | 0 | - |
| - | - | 4363 | 503.1 | - | - | 0 | - |
| - | - | 4859 | 503.3 | - | - | 0 | - |
| - | - | 643.4 | 503.5 | - | - | 0 | - |
| - | - | 897.9 | 503.8 | - | - | 0 | - |
| - | - | 1573 | 504.1 | - | - | 0 | - |
| - | - | 2052 | 504.3 | - | - | 0 | - |
| - | - | 2527 | 504.3 | - | - | 0 | - |
| - | - | 606.1 | 504.3 | - | - | 0 | - |
| - | - | 849.2 | 506.3 | - | - | 0 | - |
| 4 | c | 1.302E+04 | 511.3 | 0.000298 | 0.5829 | +1 | 4 |
| - | - | 2716 | 512.3 | - | - | 0 | - |
| - | - | 1754 | 515.3 | - | - | 0 | - |
| - | - | 651.9 | 516.3 | - | - | 0 | - |
| - | - | 1789 | 519.3 | - | - | 0 | - |
| 9 | y | 2304 | 521.8 | 0.005267 | 10.1 | +2 | 7 |
| - | - | 1059 | 522.3 | - | - | 0 | - |
| - | - | 988.4 | 522.8 | - | - | 0 | - |
| - | - | 764.5 | 528.3 | - | - | 0 | - |
| - | - | 1111 | 530.8 | - | - | 0 | - |
| - | - | 644.1 | 535.3 | - | - | 0 | - |
| - | - | 1175 | 547.3 | - | - | 0 | - |
| 12 | w | 730.8 | 548.2 | 0.008991 | 16.4 | +1 | 4 |
| 8 | y | 2865 | 550.3 | 0.006905 | 12.55 | +2 | 8 |
| - | - | 1998 | 550.8 | - | - | 0 | - |
| - | - | 811.3 | 553.8 | - | - | 0 | - |
| - | - | 895.4 | 554.8 | - | - | 0 | - |
| - | - | 3113 | 555.3 | - | - | 0 | - |
| 9 | c | 9628 | 555.8 | 0.0006458 | 1.162 | +2 | 9 |
| - | - | 4587 | 556.3 | - | - | 0 | - |
| - | - | 1883 | 556.8 | - | - | 0 | - |
| - | - | 866.4 | 563.8 | - | - | 0 | - |
| - | - | 1700 | 564.3 | - | - | 0 | - |
| - | - | 1024 | 568.8 | - | - | 0 | - |
| - | - | 683.7 | 569.3 | - | - | 0 | - |
| - | - | 833.8 | 570 | - | - | 0 | - |
| - | - | 1551 | 570.3 | - | - | 0 | - |
| - | - | 3371 | 571.3 | - | - | 0 | - |
| - | - | 2179 | 571.8 | - | - | 0 | - |
| - | - | 1194 | 572.3 | - | - | 0 | - |
| 7 | w | 1847 | 577.3 | 0.006511 | 11.28 | +2 | 9 |
| - | - | 1467 | 577.8 | - | - | 0 | - |
| - | - | 1604 | 578.3 | - | - | 0 | - |
| 3 | y | 999.6 | 580.3 | 0.003006 | 5.181 | +3 | 13 |
| - | - | 646 | 581.3 | - | - | 0 | - |
| - | - | 646.3 | 584.3 | - | - | 0 | - |
| - | - | 679.8 | 586.6 | - | - | 0 | - |
| - | - | 734.4 | 588 | - | - | 0 | - |
| - | - | 723.3 | 593 | - | - | 0 | - |
| - | - | 1896 | 597.3 | - | - | 0 | - |
| - | - | 738.2 | 597.6 | - | - | 0 | - |
| - | - | 906.4 | 597.8 | - | - | 0 | - |
| - | - | 841.4 | 598 | - | - | 0 | - |
| - | - | 575.9 | 598.8 | - | - | 0 | - |
| 7 | y | 2036 | 599.3 | 0.007241 | 12.08 | +2 | 9 |
| 7 | z | 5694 | 599.8 | 0.003263 | 5.44 | +2 | 9 |
| - | - | 3838 | 600.3 | - | - | 0 | - |
| - | - | 1193 | 600.8 | - | - | 0 | - |
| - | - | 751.6 | 601.4 | - | - | 0 | - |
| - | - | 1150 | 602 | - | - | 0 | - |
| - | - | 1184 | 603 | - | - | 0 | - |
| - | - | 882 | 603.3 | - | - | 0 | - |
| - | - | 668.6 | 604.8 | - | - | 0 | - |
| - | - | 759.3 | 605 | - | - | 0 | - |
| 12 | z | 7670 | 606.3 | 0.009602 | 15.84 | +1 | 4 |
| - | - | 2470 | 607 | - | - | 0 | - |
| 14 | c | 1.11E+04 | 607.3 | 0.0004366 | 0.7189 | +3 | 14 |
| - | - | 6795 | 607.6 | - | - | 0 | - |
| 7 | y | 8386 | 607.8 | 0.003714 | 6.11 | +2 | 9 |
| - | - | 2550 | 608 | - | - | 0 | - |
| - | - | 6072 | 608.3 | - | - | 0 | - |
| - | - | 2608 | 608.8 | - | - | 0 | - |
| - | - | 1246 | 609.3 | - | - | 0 | - |
| 10 | c | 818.9 | 611.3 | 0.002027 | 3.315 | +2 | 10 |
| - | - | 798 | 611.6 | - | - | 0 | - |
| 10 | c | 926.6 | 611.8 | 0.005929 | 9.692 | +2 | 10 |
| - | - | 981.3 | 612 | - | - | 0 | - |
| - | - | 1635 | 612.3 | - | - | 0 | - |
| - | - | 1069 | 612.8 | - | - | 0 | - |
| - | - | 1145 | 613.3 | - | - | 0 | - |
| - | - | 808.8 | 614.6 | - | - | 0 | - |
| - | - | 803.6 | 616.3 | - | - | 0 | - |
| - | - | 790.2 | 617.6 | - | - | 0 | - |
| - | - | 3518 | 619.8 | - | - | 0 | - |
| 10 | c | 1.018E+04 | 620.3 | 0.001383 | 2.229 | +2 | 10 |
| - | - | 5449 | 620.8 | - | - | 0 | - |
| 2 | z | 859.6 | 621 | 0.008905 | 14.34 | +3 | 14 |
| - | - | 2840 | 621.3 | - | - | 0 | - |
| - | - | 705.9 | 621.6 | - | - | 0 | - |
| - | - | 1385 | 621.8 | - | - | 0 | - |
| 12 | y | 3757 | 622.3 | 0.008734 | 14.03 | +1 | 4 |
| - | - | 735 | 623.3 | - | - | 0 | - |
| - | - | 1079 | 625.6 | - | - | 0 | - |
| 2 | y | 990.8 | 626 | 0.00112 | 1.789 | +3 | 14 |
| 2 | y | 3034 | 626.3 | 0.01039 | 16.6 | +3 | 14 |
| - | - | 2115 | 626.6 | - | - | 0 | - |
| - | - | 1490 | 627 | - | - | 0 | - |
| - | - | 990.4 | 628 | - | - | 0 | - |
| - | - | 831.8 | 631.3 | - | - | 0 | - |
| - | - | 945 | 631.6 | - | - | 0 | - |
| 2 | y | 1575 | 632 | 0.002646 | 4.187 | +3 | 14 |
| - | - | 1009 | 632.3 | - | - | 0 | - |
| - | - | 677.5 | 633 | - | - | 0 | - |
| - | - | 2805 | 634.3 | - | - | 0 | - |
| - | - | 1475 | 634.6 | - | - | 0 | - |
| 6 | w | 6588 | 634.8 | 0.001489 | 2.346 | +2 | 10 |
| - | - | 1232 | 635 | - | - | 0 | - |
| - | - | 4930 | 635.3 | - | - | 0 | - |
| - | - | 1038 | 635.6 | - | - | 0 | - |
| - | - | 2676 | 635.8 | - | - | 0 | - |
| - | - | 2722 | 636.3 | - | - | 0 | - |
| - | - | 824 | 636.6 | - | - | 0 | - |
| - | - | 956.3 | 637 | - | - | 0 | - |
| - | - | 1474 | 637.3 | - | - | 0 | - |
| - | - | 793.2 | 639.3 | - | - | 0 | - |
| - | - | 3154 | 640 | - | - | 0 | - |
| - | - | 1.211E+04 | 640.3 | - | - | 0 | - |
| - | - | 8351 | 640.7 | - | - | 0 | - |
| - | - | 7869 | 641 | - | - | 0 | - |
| - | - | 2038 | 641.3 | - | - | 0 | - |
| - | - | 1097 | 641.7 | - | - | 0 | - |
| - | - | 806.9 | 642.6 | - | - | 0 | - |
| - | - | 891.7 | 644.3 | - | - | 0 | - |
| - | - | 1251 | 644.6 | - | - | 0 | - |
| - | - | 1685 | 645 | - | - | 0 | - |
| - | - | 1629 | 645.3 | - | - | 0 | - |
| - | - | 2355 | 645.6 | - | - | 0 | - |
| - | - | 4051 | 646 | - | - | 0 | - |
| - | - | 3580 | 646.3 | - | - | 0 | - |
| - | - | 1105 | 646.6 | - | - | 0 | - |
| - | - | 985.5 | 647 | - | - | 0 | - |
| 6 | z | 1096 | 647.3 | 0.01011 | 15.61 | +2 | 10 |
| - | - | 918.2 | 647.6 | - | - | 0 | - |
| - | - | 1575 | 648.3 | - | - | 0 | - |
| - | - | 941.9 | 648.6 | - | - | 0 | - |
| - | - | 753.4 | 649 | - | - | 0 | - |
| - | - | 1000 | 649.3 | - | - | 0 | - |
| - | - | 2996 | 649.6 | - | - | 0 | - |
| - | - | 9374 | 650 | - | - | 0 | - |
| - | - | 1.599E+04 | 650.3 | - | - | 0 | - |
| - | - | 9455 | 650.6 | - | - | 0 | - |
| - | - | 6115 | 651 | - | - | 0 | - |
| - | - | 4743 | 651.3 | - | - | 0 | - |
| - | - | 3146 | 651.6 | - | - | 0 | - |
| - | - | 1771 | 652 | - | - | 0 | - |
| - | - | 857.9 | 652.3 | - | - | 0 | - |
| - | - | 746.7 | 652.6 | - | - | 0 | - |
| - | - | 833.3 | 653 | - | - | 0 | - |
| - | - | 950.7 | 653.3 | - | - | 0 | - |
| - | - | 910.4 | 653.6 | - | - | 0 | - |
| - | - | 1165 | 654.3 | - | - | 0 | - |
| - | - | 1761 | 655 | - | - | 0 | - |
| - | - | 818.2 | 655.3 | - | - | 0 | - |
| - | - | 861.6 | 655.6 | - | - | 0 | - |
| 6 | y | 863.8 | 655.8 | 0.002624 | 4.001 | +2 | 10 |
| 6 | z | 3395 | 656.3 | 0.00666 | 10.15 | +2 | 10 |
| - | - | 2876 | 656.6 | - | - | 0 | - |
| - | - | 1036 | 656.8 | - | - | 0 | - |
| - | - | 3497 | 657 | - | - | 0 | - |
| - | - | 3032 | 657.3 | - | - | 0 | - |
| - | - | 1142 | 657.6 | - | - | 0 | - |
| - | - | 1131 | 658 | - | - | 0 | - |
| - | - | 969.1 | 658.3 | - | - | 0 | - |
| - | - | 1845 | 659 | - | - | 0 | - |
| - | - | 903.6 | 660 | - | - | 0 | - |
| - | - | 1579 | 662.3 | - | - | 0 | - |
| - | - | 2997 | 662.6 | - | - | 0 | - |
| - | - | 898.1 | 663 | - | - | 0 | - |
| - | - | 1238 | 664 | - | - | 0 | - |
| 6 | y | 6172 | 664.3 | 0.008576 | 12.91 | +2 | 10 |
| - | - | 1658 | 664.6 | - | - | 0 | - |
| - | - | 1999 | 664.8 | - | - | 0 | - |
| - | - | 1842 | 665 | - | - | 0 | - |
| - | - | 3099 | 665.3 | - | - | 0 | - |
| - | - | 1151 | 665.7 | - | - | 0 | - |
| - | - | 924.5 | 668.3 | - | - | 0 | - |
| - | - | 1672 | 668.7 | - | - | 0 | - |
| - | - | 985.5 | 669 | - | - | 0 | - |
| - | - | 4484 | 669.3 | - | - | 0 | - |
| - | - | 2.478E+04 | 669.7 | - | - | 0 | - |
| - | - | 3.46E+04 | 670 | - | - | 0 | - |
| - | - | 2.594E+04 | 670.3 | - | - | 0 | - |
| - | - | 1.249E+04 | 670.7 | - | - | 0 | - |
| - | - | 6833 | 671 | - | - | 0 | - |
| - | - | 3326 | 671.3 | - | - | 0 | - |
| - | - | 1863 | 671.7 | - | - | 0 | - |
| - | - | 861.2 | 675.3 | - | - | 0 | - |
| 5 | c | 742.9 | 679.3 | 0.008346 | 12.29 | +1 | 5 |
| 5 | c | 4231 | 680.3 | 0.0001444 | 0.2122 | +1 | 5 |
| - | - | 1201 | 681.3 | - | - | 0 | - |
| - | - | 1062 | 682.3 | - | - | 0 | - |
| - | - | 767.9 | 683.3 | - | - | 0 | - |
| - | - | 1958 | 690.3 | - | - | 0 | - |
| - | - | 1686 | 691.3 | - | - | 0 | - |
| - | - | 1052 | 691.8 | - | - | 0 | - |
| - | - | 831.4 | 695.9 | - | - | 0 | - |
| 5 | c | 1.247E+04 | 697.3 | 0.0005094 | 0.7305 | +1 | 5 |
| - | - | 5679 | 698.3 | - | - | 0 | - |
| - | - | 1835 | 699.3 | - | - | 0 | - |
| - | - | 1314 | 699.8 | - | - | 0 | - |
| - | - | 1119 | 700.3 | - | - | 0 | - |
| - | - | 2255 | 701.3 | - | - | 0 | - |
| 11 | c | 9892 | 701.8 | 0.002311 | 3.293 | +2 | 11 |
| - | - | 6106 | 702.3 | - | - | 0 | - |
| - | - | 2242 | 702.8 | - | - | 0 | - |
| - | - | 851.4 | 703.3 | - | - | 0 | - |
| - | - | 838.5 | 704.4 | - | - | 0 | - |
| - | - | 752.1 | 711.3 | - | - | 0 | - |
| - | - | 906.1 | 713.3 | - | - | 0 | - |
| - | - | 673.7 | 714.3 | - | - | 0 | - |
| - | - | 4366 | 716.4 | - | - | 0 | - |
| - | - | 1434 | 717.4 | - | - | 0 | - |
| - | - | 773.8 | 726.4 | - | - | 0 | - |
| 5 | y | 1667 | 748.9 | 0.005875 | 7.845 | +2 | 11 |
| 5 | z | 4065 | 749.4 | 0.00292 | 3.897 | +2 | 11 |
| - | - | 4253 | 749.9 | - | - | 0 | - |
| - | - | 1937 | 750.4 | - | - | 0 | - |
| - | - | 876.1 | 750.9 | - | - | 0 | - |
| - | - | 1076 | 754.4 | - | - | 0 | - |
| - | - | 2931 | 754.9 | - | - | 0 | - |
| - | - | 1584 | 755.4 | - | - | 0 | - |
| 12 | c | 2604 | 757.4 | 2.588E-05 | 0.03418 | +2 | 12 |
| - | - | 1428 | 757.9 | - | - | 0 | - |
| - | - | 955.8 | 758.4 | - | - | 0 | - |
| - | - | 780.9 | 761.4 | - | - | 0 | - |
| 12 | c | 1450 | 765.9 | 0.002559 | 3.342 | +2 | 12 |
| - | - | 1421 | 766.4 | - | - | 0 | - |
| 11 | y | 955.8 | 768.4 | 0.01078 | 14.03 | +1 | 5 |
| 11 | z | 9780 | 769.4 | 0.009332 | 12.13 | +1 | 5 |
| - | - | 5536 | 770.4 | - | - | 0 | - |
| - | - | 1916 | 771.4 | - | - | 0 | - |
| - | - | 1675 | 784.9 | - | - | 0 | - |
| 11 | y | 4172 | 785.4 | 0.01054 | 13.42 | +1 | 5 |
| - | - | 1748 | 786.4 | - | - | 0 | - |
| - | - | 976.6 | 787.4 | - | - | 0 | - |
| - | - | 746 | 789.4 | - | - | 0 | - |
| 6 | c | 1370 | 793.4 | 0.004191 | 5.283 | +1 | 6 |
| - | - | 805.3 | 798.4 | - | - | 0 | - |
| - | - | 2110 | 803.4 | - | - | 0 | - |
| - | - | 1032 | 804.4 | - | - | 0 | - |
| 4 | y | 1853 | 806.4 | 0.01047 | 12.98 | +2 | 12 |
| 4 | z | 5686 | 806.9 | 0.001499 | 1.858 | +2 | 12 |
| - | - | 6753 | 807.4 | - | - | 0 | - |
| - | - | 3698 | 807.9 | - | - | 0 | - |
| 6 | c | 2.253E+04 | 810.4 | 0.0002838 | 0.3502 | +1 | 6 |
| - | - | 9656 | 811.4 | - | - | 0 | - |
| - | - | 2380 | 812.4 | - | - | 0 | - |
| - | - | 2641 | 814.4 | - | - | 0 | - |
| 4 | y | 1569 | 814.9 | 0.005368 | 6.587 | +2 | 12 |
| - | - | 2064 | 815.4 | - | - | 0 | - |
| - | - | 1813 | 831.4 | - | - | 0 | - |
| - | - | 1375 | 839.4 | - | - | 0 | - |
| - | - | 1234 | 841.4 | - | - | 0 | - |
| 3 | w | 1356 | 841.9 | 0.0008853 | 1.052 | +2 | 13 |
| - | - | 1776 | 842.4 | - | - | 0 | - |
| - | - | 816.7 | 843.4 | - | - | 0 | - |
| - | - | 1234 | 845.9 | - | - | 0 | - |
| 13 | c | 5188 | 846.4 | 0.0007462 | 0.8816 | +2 | 13 |
| - | - | 4776 | 846.9 | - | - | 0 | - |
| - | - | 3087 | 847.4 | - | - | 0 | - |
| - | - | 923.7 | 849.4 | - | - | 0 | - |
| - | - | 738.3 | 862.9 | - | - | 0 | - |
| - | - | 3302 | 867.4 | - | - | 0 | - |
| - | - | 923.1 | 868.4 | - | - | 0 | - |
| 3 | y | 3101 | 870.4 | 0.006816 | 7.831 | +2 | 13 |
| 3 | z | 6127 | 870.9 | 0.001003 | 1.151 | +2 | 13 |
| - | - | 5658 | 871.4 | - | - | 0 | - |
| - | - | 2264 | 871.9 | - | - | 0 | - |
| - | - | 807 | 872.4 | - | - | 0 | - |
| - | - | 854.2 | 874.4 | - | - | 0 | - |
| - | - | 877.4 | 878.4 | - | - | 0 | - |
| 3 | y | 714.6 | 878.9 | 0.001271 | 1.446 | +2 | 13 |
| - | - | 1141 | 879.4 | - | - | 0 | - |
| - | - | 1777 | 881.4 | - | - | 0 | - |
| - | - | 794.2 | 882.4 | - | - | 0 | - |
| - | - | 1703 | 888.4 | - | - | 0 | - |
| - | - | 1044 | 888.9 | - | - | 0 | - |
| 10 | z | 1.175E+04 | 898.4 | 0.008346 | 9.289 | +1 | 6 |
| - | - | 9452 | 899.4 | - | - | 0 | - |
| - | - | 4287 | 900.4 | - | - | 0 | - |
| - | - | 1465 | 901.4 | - | - | 0 | - |
| - | - | 856.5 | 902.4 | - | - | 0 | - |
| - | - | 710.9 | 904.9 | - | - | 0 | - |
| 7 | c | 1220 | 908.4 | 0.0005557 | 0.6117 | +1 | 7 |
| 14 | c | 3642 | 910.4 | 0.003855 | 4.234 | +2 | 14 |
| - | - | 3465 | 910.9 | - | - | 0 | - |
| - | - | 2841 | 911.4 | - | - | 0 | - |
| - | - | 1060 | 911.9 | - | - | 0 | - |
| - | - | 803.7 | 912.4 | - | - | 0 | - |
| - | - | 954 | 914.4 | - | - | 0 | - |
| - | - | 1518 | 915.4 | - | - | 0 | - |
| - | - | 934.6 | 916.4 | - | - | 0 | - |
| - | - | 697.2 | 918.9 | - | - | 0 | - |
| 7 | c | 2968 | 925.5 | 0.001226 | 1.325 | +1 | 7 |
| - | - | 2025 | 926.5 | - | - | 0 | - |
| - | - | 819.2 | 926.9 | - | - | 0 | - |
| - | - | 841.1 | 930.5 | - | - | 0 | - |
| - | - | 1645 | 931 | - | - | 0 | - |
| - | - | 1658 | 931.5 | - | - | 0 | - |
| - | - | 2419 | 932.5 | - | - | 0 | - |
| - | - | 969.6 | 932.9 | - | - | 0 | - |
| - | - | 949.9 | 933.5 | - | - | 0 | - |
| - | - | 847.3 | 933.9 | - | - | 0 | - |
| - | - | 968.5 | 937.5 | - | - | 0 | - |
| - | - | 3962 | 938.5 | - | - | 0 | - |
| - | - | 1869 | 939 | - | - | 0 | - |
| 2 | z | 3594 | 939.4 | 0.01257 | 13.38 | +2 | 14 |
| - | - | 3935 | 939.9 | - | - | 0 | - |
| - | - | 2616 | 940.4 | - | - | 0 | - |
| - | - | 1695 | 940.9 | - | - | 0 | - |
| - | - | 1171 | 941.5 | - | - | 0 | - |
| - | - | 828.5 | 942.4 | - | - | 0 | - |
| - | - | 841 | 944.5 | - | - | 0 | - |
| - | - | 2261 | 945.5 | - | - | 0 | - |
| - | - | 750 | 946 | - | - | 0 | - |
| - | - | 1783 | 946.5 | - | - | 0 | - |
| - | - | 891.8 | 947 | - | - | 0 | - |
| 2 | y | 1060 | 947.4 | 0.01603 | 16.92 | +2 | 14 |
| - | - | 869.3 | 951.4 | - | - | 0 | - |
| - | - | 1182 | 952 | - | - | 0 | - |
| - | - | 998.5 | 953 | - | - | 0 | - |
| - | - | 1052 | 953.5 | - | - | 0 | - |
| - | - | 885.1 | 954 | - | - | 0 | - |
| - | - | 1805 | 955.5 | - | - | 0 | - |
| - | - | 1675 | 960 | - | - | 0 | - |
| - | - | 4008 | 960.5 | - | - | 0 | - |
| - | - | 1090 | 961 | - | - | 0 | - |
| - | - | 1263 | 961.5 | - | - | 0 | - |
| - | - | 1615 | 962.5 | - | - | 0 | - |
| - | - | 1038 | 963 | - | - | 0 | - |
| - | - | 1552 | 963.5 | - | - | 0 | - |
| 8 | c | 2009 | 964.5 | 0.004862 | 5.041 | +1 | 8 |
| 8 | c | 744.9 | 965.4 | 0.006667 | 6.906 | +1 | 8 |
| - | - | 1156 | 966 | - | - | 0 | - |
| - | - | 1138 | 966.5 | - | - | 0 | - |
| - | - | 1473 | 967 | - | - | 0 | - |
| - | - | 3352 | 967.5 | - | - | 0 | - |
| - | - | 1448 | 968 | - | - | 0 | - |
| 9 | w | 3053 | 968.4 | 0.01589 | 16.41 | +1 | 7 |
| - | - | 2077 | 969 | - | - | 0 | - |
| - | - | 1642 | 969.5 | - | - | 0 | - |
| - | - | 1036 | 971 | - | - | 0 | - |
| - | - | 896.3 | 971.5 | - | - | 0 | - |
| - | - | 1512 | 972.5 | - | - | 0 | - |
| - | - | 1046 | 973 | - | - | 0 | - |
| - | - | 811.4 | 973.5 | - | - | 0 | - |
| - | - | 2500 | 974 | - | - | 0 | - |
| - | - | 2051 | 974.5 | - | - | 0 | - |
| - | - | 8173 | 975 | - | - | 0 | - |
| - | - | 7717 | 975.5 | - | - | 0 | - |
| - | - | 4115 | 976 | - | - | 0 | - |
| - | - | 4206 | 976.5 | - | - | 0 | - |
| - | - | 2551 | 977 | - | - | 0 | - |
| - | - | 2529 | 977.5 | - | - | 0 | - |
| - | - | 767.6 | 978 | - | - | 0 | - |
| - | - | 1510 | 978.5 | - | - | 0 | - |
| - | - | 1503 | 980.5 | - | - | 0 | - |
| - | - | 4198 | 981.5 | - | - | 0 | - |
| - | - | 3001 | 982 | - | - | 0 | - |
| 8 | c | 1.341E+04 | 982.5 | 0.003983 | 4.054 | +1 | 8 |
| - | - | 2636 | 983 | - | - | 0 | - |
| - | - | 7118 | 983.5 | - | - | 0 | - |
| - | - | 2864 | 984.5 | - | - | 0 | - |
| - | - | 3005 | 985 | - | - | 0 | - |
| - | - | 5265 | 985.5 | - | - | 0 | - |
| - | - | 2567 | 986 | - | - | 0 | - |
| - | - | 2591 | 986.5 | - | - | 0 | - |
| - | - | 2837 | 987.5 | - | - | 0 | - |
| - | - | 1011 | 988 | - | - | 0 | - |
| - | - | 3378 | 988.5 | - | - | 0 | - |
| - | - | 2806 | 989.5 | - | - | 0 | - |
| - | - | 914.9 | 990 | - | - | 0 | - |
| - | - | 2212 | 993.5 | - | - | 0 | - |
| - | - | 3067 | 994 | - | - | 0 | - |
| - | - | 2432 | 994.5 | - | - | 0 | - |
| - | - | 1650 | 995 | - | - | 0 | - |
| - | - | 2009 | 995.5 | - | - | 0 | - |
| - | - | 4944 | 996 | - | - | 0 | - |
| - | - | 1.3E+04 | 996.5 | - | - | 0 | - |
| - | - | 1.052E+04 | 997 | - | - | 0 | - |
| - | - | 6334 | 997.5 | - | - | 0 | - |
| - | - | 2947 | 998 | - | - | 0 | - |
| - | - | 1506 | 998.5 | - | - | 0 | - |
| - | - | 1278 | 999 | - | - | 0 | - |
| - | - | 1366 | 999.5 | - | - | 0 | - |
| - | - | 2454 | 1002 | - | - | 0 | - |
| - | - | 1224 | 1003 | - | - | 0 | - |
| - | - | 2120 | 1003 | - | - | 0 | - |
| - | - | 3616 | 1004 | - | - | 0 | - |
| - | - | 9742 | 1004 | - | - | 0 | - |
| - | - | 1.481E+04 | 1005 | - | - | 0 | - |
| - | - | 1.084E+04 | 1005 | - | - | 0 | - |
| - | - | 6460 | 1006 | - | - | 0 | - |
| - | - | 4351 | 1006 | - | - | 0 | - |
| - | - | 1552 | 1007 | - | - | 0 | - |
| - | - | 1892 | 1007 | - | - | 0 | - |
| - | - | 1576 | 1008 | - | - | 0 | - |
| 9 | z | 1267 | 1027 | 0.006507 | 6.339 | +1 | 7 |
| - | - | 3915 | 1028 | - | - | 0 | - |
| - | - | 1999 | 1029 | - | - | 0 | - |
| - | - | 999.8 | 1085 | - | - | 0 | - |
| - | - | 2688 | 1110 | - | - | 0 | - |
| 9 | c | 1.308E+04 | 1111 | 0.001244 | 1.12 | +1 | 9 |
| - | - | 7726 | 1112 | - | - | 0 | - |
| - | - | 3216 | 1113 | - | - | 0 | - |
| - | - | 1103 | 1142 | - | - | 0 | - |
| - | - | 864.8 | 1196 | - | - | 0 | - |
| - | - | 1901 | 1197 | - | - | 0 | - |
| - | - | 850.2 | 1198 | - | - | 0 | - |
| 7 | z | 2277 | 1199 | 0.004535 | 3.783 | +1 | 9 |
| - | - | 6041 | 1200 | - | - | 0 | - |
| - | - | 3016 | 1201 | - | - | 0 | - |
| - | - | 1798 | 1202 | - | - | 0 | - |
| 10 | c | 2992 | 1240 | 0.004427 | 3.571 | +1 | 10 |
| - | - | 3190 | 1241 | - | - | 0 | - |
| - | - | 1641 | 1242 | - | - | 0 | - |
| 6 | z | 1377 | 1312 | 0.002503 | 1.908 | +1 | 10 |
| - | - | 3012 | 1313 | - | - | 0 | - |
| - | - | 1572 | 1314 | - | - | 0 | - |
| - | - | 1042 | 1359 | - | - | 0 | - |
| 11 | c | 2059 | 1403 | 0.008237 | 5.873 | +1 | 11 |
| - | - | 1656 | 1404 | - | - | 0 | - |
| - | - | 867.2 | 1405 | - | - | 0 | - |
| 5 | z | 1175 | 1498 | 0.004912 | 3.279 | +1 | 11 |
| - | - | 1949 | 1499 | - | - | 0 | - |
| - | - | 1986 | 1500 | - | - | 0 | - |
| - | - | 847.7 | 1511 | - | - | 0 | - |
| 12 | c | 879.4 | 1514 | 0.002804 | 1.852 | +1 | 12 |
| 12 | c | 1009 | 1531 | 0.01191 | 7.778 | +1 | 12 |
| - | - | 784.2 | 1532 | - | - | 0 | - |
| 4 | z | 1624 | 1613 | 0.0002498 | 0.1549 | +1 | 12 |
| - | - | 3158 | 1614 | - | - | 0 | - |
| - | - | 1932 | 1615 | - | - | 0 | - |
| - | - | 1095 | 1616 | - | - | 0 | - |
| 13 | c | 1581 | 1692 | 0.006517 | 3.852 | +1 | 13 |
| - | - | 1686 | 1693 | - | - | 0 | - |
| - | - | 907.3 | 1694 | - | - | 0 | - |
| 3 | z | 1789 | 1741 | 0.005027 | 2.888 | +1 | 13 |
| - | - | 2365 | 1742 | - | - | 0 | - |
| - | - | 1283 | 1743 | - | - | 0 | - |
| - | - | 1080 | 1744 | - | - | 0 | - |
| - | - | 762.2 | 1761 | - | - | 0 | - |
| - | - | 1203 | 1821 | - | - | 0 | - |
| - | - | 981.8 | 1921 | - | - | 0 | - |
| - | - | 901.9 | 1951 | - | - | 0 | - |
| - | - | 900.2 | 1952 | - | - | 0 | - |
| - | - | 1028 | 1992 | - | - | 0 | - |
| - | - | 1115 | 1993 | - | - | 0 | - |
| - | - | 1223 | 2009 | - | - | 0 | - |
| - | - | 1191 | 2010 | - | - | 0 | - |
| - | - | 1310 | 2011 | - | - | 0 | - |
| - | - | 898 | 2012 | - | - | 0 | - |

m/z Charge Intensity FragmentType MassShift Position
120.06556701660156 0 391.27307
120.87273406982422 0 355.701
129.06585693359375 0 391.53955
129.10218811035156 0 2290.9841
130.06507873535156 0 1968.6411
130.06982421875 0 431.01505 w 13
133.0603485107422 0 506.4843
133.08592224121094 0 1297.4045
136.0753936767578 0 679.8111
138.76266479492188 0 411.01904
140.08218383789062 0 540.14764
147.0767822265625 0 559.3007
148.95376586914062 0 866.3606
149.04478454589844 0 1113.6504
155.0924530029297 0 2090.5374
156.07638549804688 0 459.58148
156.62660217285156 0 464.18396
159.09170532226562 0 900.3568
165.0775146484375 0 452.18518
166.06077575683594 0 1508.6202
167.05508422851562 0 596.4502
168.13803100585938 0 480.67642
171.14898681640625 0 1054.4928
173.0917510986328 0 998.5471
173.45086669921875 0 3785.4575
174.08746337890625 0 638.7856
175.07090759277344 0 508.6932
177.1115264892578 0 628.73224
183.11270141601562 0 945.1224
185.16465759277344 0 2630.9092
187.10745239257812 0 973.11194
191.10252380371094 0 1372.0096
199.1438751220703 0 1247.6875
201.12294006347656 0 645.82965
202.0822296142578 0 1361.1685
203.10231018066406 0 2465.9783
205.11766052246094 0 561.8966
207.0977783203125 0 1084.7858
209.09181213378906 0 1824.969
213.12310791015625 0 725.4658
215.13877868652344 0 7861.232
216.1422576904297 0 667.90753
217.08213806152344 0 638.4095
217.154541015625 0 1171.7046
218.1135711669922 0 1738.4081
223.1551055908203 0 1784.77
227.1026153564453 0 1670.086
229.11810302734375 0 639.76953
229.15419006347656 0 762.1632
230.1127471923828 0 640.82007
231.1695556640625 0 924.433
234.1232452392578 0 2668.581
239.0950927734375 0 816.1555
242.15008544921875 0 808.888
251.14996337890625 0 9715.968
252.15328979492188 0 1744.9205
261.11883544921875 0 730.7437
268.1766662597656 0 1200.2195 c 1
275.1235656738281 0 550.9427
281.0507507324219 0 1951.5717
282.1247253417969 0 495.19824
282.1817626953125 0 868.73175
283.151611328125 0 1183.7795
293.6220397949219 0 536.43976
299.06146240234375 0 8037.0654
299.13421630859375 0 903.67096
299.2097473144531 0 673.729
300.06353759765625 0 1575.8436
300.131103515625 0 682.6548
302.1708984375 0 1201.2701
310.14093017578125 0 562.62146
326.66387939453125 0 1909.7942
329.1457214355469 0 508.91525
332.1820983886719 0 928.1316
338.1349182128906 0 1378.7478
340.1866149902344 0 1497.3679
340.6609191894531 0 1088.4316 c Ammonia loss 4
354.1296081542969 0 767.38354
359.02801513671875 0 1595.9742
367.1612548828125 0 1201.3864
367.18414306640625 0 606.31946 y 7
367.679443359375 0 556.48737
379.208251953125 0 4687.8906 c Ammonia loss 2
380.2107849121094 0 1006.0929
387.22308349609375 0 2072.8242
393.1969909667969 0 1340.2903 y 10
393.6980895996094 0 1249.5319
396.2347412109375 0 4478.558 c 2
397.2387390136719 0 1157.4868
407.7070617675781 0 664.8267
408.1982116699219 0 645.6269 c Ammonia loss 9
415.03662109375 0 2058.436
416.03594970703125 0 1313.8425
420.2020263671875 0 9719.635 w 9
420.7034606933594 0 5197.1895
421.2015380859375 0 1450.4354
421.232421875 0 1035.5292
431.2247009277344 0 732.2945
435.7214050292969 0 703.24243
443.21783447265625 0 614.14685 y 5
444.2330017089844 0 1282.4883
444.2794494628906 0 805.8473
449.221435546875 0 761.4403
449.7196044921875 0 1663.3267 z 9
457.2657775878906 0 7528.402
457.71307373046875 0 604.7822
458.2300720214844 0 3160.1313
458.2692565917969 0 2860.853
458.7288513183594 0 1957.8936
459.2798156738281 0 4610.07
460.2834167480469 0 1147.3802
460.73370361328125 0 1755.2731
461.2373962402344 0 1010.4957
466.2401428222656 0 989.44196
467.21356201171875 0 791.66406
467.2477111816406 0 1380.6913
468.2630310058594 0 1194.6349
468.7653503417969 0 690.45386
469.2274475097656 0 1711.0638
470.2325439453125 0 704.7638
471.2677001953125 0 809.0045
472.30059814453125 0 1007.10864
474.2272033691406 0 1160.6962 y 1
474.47705078125 0 1132.0208
476.4849853515625 0 712.86017
483.2076110839844 0 1415.0824
484.24169921875 0 1647.0364
484.74835205078125 0 903.03577
485.2355651855469 0 3054.1177
486.2919921875 0 4256.223
487.2403259277344 0 789.391
487.28460693359375 0 712.1308
488.2933654785156 0 2155.5469
493.74664306640625 0 3913.3555
494.2390441894531 0 4393.3076 c Ammonia loss 3
494.74774169921875 0 1438.5985
495.2404479980469 0 676.92505
496.2533874511719 0 729.55096
498.2425537109375 0 766.5653
501.2185974121094 0 2489.9185
501.3037109375 0 1682.9049
501.74798583984375 0 1427.8234
502.2681884765625 0 5016.0015
502.4976806640625 0 5087.558
502.75030517578125 0 12128.0205
502.9975891113281 0 4069.7715
503.10723876953125 0 4363.409
503.25140380859375 0 4858.6396
503.505859375 0 643.41077
503.7688903808594 0 897.937
504.1068115234375 0 1572.8892
504.2630615234375 0 2051.7983
504.3019104003906 0 2527.2075
504.3371276855469 0 606.07477
506.26171875 0 849.24915
511.26202392578125 0 13016.724 c 3
512.2645263671875 0 2715.6816
515.317626953125 0 1753.9855
516.3256225585938 0 651.8749
519.2962036132812 0 1788.9868
521.7647705078125 0 2303.9263 y 8
522.2681884765625 0 1059.0822
522.7671508789062 0 988.407
528.2685546875 0 764.4627
530.775634765625 0 1111.4442
535.2879638671875 0 644.1455
547.2901000976562 0 1174.9323
548.238525390625 0 730.77185 w 11
550.2738647460938 0 2865.1501 y 7
550.7781982421875 0 1997.9092
553.7952880859375 0 811.3269
554.7932739257812 0 895.43146
555.2954711914062 0 3113.298
555.788818359375 0 9628.201 c 8
556.2889404296875 0 4586.7563
556.7893676757812 0 1882.8138
563.7691650390625 0 866.3883
564.2723388671875 0 1699.6967
568.7979125976562 0 1024.2936
569.2920532226562 0 683.7429
569.9581909179688 0 833.8179
570.275146484375 0 1551.3372
571.2814331054688 0 3371.3755
571.7810668945312 0 2178.9734
572.2798461914062 0 1194.3928
577.279541015625 0 1846.9847 w 6
577.7847290039062 0 1466.6042
578.2835693359375 0 1603.5209
580.2750244140625 0 999.6131 y Water loss 2
581.2813110351562 0 646.01154
584.3497924804688 0 646.2586
586.6096801757812 0 679.81287
587.955322265625 0 734.4381
592.9505004882812 0 723.28455
597.2898559570312 0 1895.6478
597.6278076171875 0 738.18646
597.79833984375 0 906.394
597.9638671875 0 841.37897
598.8082275390625 0 575.9135
599.2882080078125 0 2036.24 y Ammonia loss 6
599.7816162109375 0 5694.5 z 6
600.2816772460938 0 3837.8364
600.7813720703125 0 1192.7432
601.3746948242188 0 751.5646
601.9583740234375 0 1149.6735
602.9502563476562 0 1183.6194
603.28173828125 0 882.00934
604.7919921875 0 668.59467
604.9688110351562 0 759.3325
606.3035888671875 0 7669.7656 z 11
606.96875 0 2470.2937
607.2999877929688 0 11096.0625 c 13
607.6326293945312 0 6795.0674
607.79052734375 0 8385.553 y 6
607.9652709960938 0 2549.919
608.2944946289062 0 6072.4155
608.7921142578125 0 2608.3604
609.2969360351562 0 1246.4242
611.3062133789062 0 818.949 c Water loss 9
611.6296997070312 0 797.9818
611.8021240234375 0 926.61597 c Ammonia loss 9
611.9601440429688 0 981.32654
612.2973022460938 0 1634.6154
612.80126953125 0 1068.539
613.3017578125 0 1144.8386
614.6343994140625 0 808.79865
616.2883911132812 0 803.5777
617.6256103515625 0 790.248
619.816162109375 0 3517.8677
620.3108520507812 0 10178.277 c 9
620.8113403320312 0 5449.389
620.961669921875 0 859.57855 z Ammonia loss 1
621.30859375 0 2839.7683
621.6270141601562 0 705.8816
621.8135375976562 0 1384.6783
622.3231811523438 0 3756.849 y 11
623.3312377929688 0 734.9612
625.6481323242188 0 1078.5847
625.9654541015625 0 990.7586 y Water loss 1
626.302734375 0 3034.2744 y Ammonia loss 1
626.6409301757812 0 2114.9985
626.9685668945312 0 1489.9799
627.9668579101562 0 990.40234
631.29052734375 0 831.8153
631.6358642578125 0 945.0205
631.9652099609375 0 1574.7574 y 1
632.3060913085938 0 1008.83124
632.9696655273438 0 677.516
634.308837890625 0 2804.9277
634.6473999023438 0 1474.998
634.7980346679688 0 6587.553 w 5
634.9805297851562 0 1232.1327
635.30029296875 0 4929.6685
635.6486206054688 0 1038.3282
635.8005981445312 0 2675.7634
636.2991943359375 0 2722.4224
636.6300659179688 0 823.9996
636.9671630859375 0 956.2529
637.3082885742188 0 1473.8228
639.3140869140625 0 793.17633
639.9901733398438 0 3153.687
640.3213500976562 0 12107.139
640.6549072265625 0 8351.111
640.9876098632812 0 7869.1216
641.3176879882812 0 2038.3225
641.6513671875 0 1096.642
642.634521484375 0 806.8629
644.30712890625 0 891.6533
644.6409301757812 0 1250.6705
644.9698486328125 0 1684.7615
645.3055419921875 0 1628.6504
645.644287109375 0 2354.8164
645.978759765625 0 4050.5303
646.3128051757812 0 3580.0544
646.6461181640625 0 1104.7566
646.97802734375 0 985.48645
647.3115234375 0 1096.4182 z Water loss 5
647.6348876953125 0 918.18146
648.3101806640625 0 1575.2369
648.64453125 0 941.91327
648.9752197265625 0 753.4167
649.319091796875 0 999.99994
649.6446533203125 0 2995.5327
649.9758911132812 0 9374.028
650.317138671875 0 15990.368
650.6478271484375 0 9454.504
650.9865112304688 0 6114.955
651.3169555664062 0 4743.3003
651.6477661132812 0 3146.203
651.9778442382812 0 1771.4348
652.310791015625 0 857.93695
652.6355590820312 0 746.66235
652.973876953125 0 833.3028
653.3170776367188 0 950.6602
653.6326293945312 0 910.3975
654.3209838867188 0 1165.2721
654.9945678710938 0 1761.0667
655.3172607421875 0 818.23895
655.6454467773438 0 861.6461
655.8256225585938 0 863.82074 y Ammonia loss 5
656.3202514648438 0 3395.1143 z 5
656.64208984375 0 2875.5623
656.82275390625 0 1036.077
656.9754028320312 0 3496.6775
657.3103637695312 0 3031.7356
657.6439819335938 0 1141.8193
657.9775390625 0 1130.9186
658.313232421875 0 969.10516
658.9866943359375 0 1845.1344
659.9880981445312 0 903.5524
662.3038330078125 0 1579.3835
662.6438598632812 0 2997.264
662.9737548828125 0 898.12317
663.99169921875 0 1237.9705
664.3276977539062 0 6172.135 y 5
664.6497802734375 0 1658.3889
664.8328857421875 0 1999.1013
664.9837036132812 0 1841.9741
665.3251953125 0 3098.9082
665.6548461914062 0 1151.4531
668.314697265625 0 924.4666
668.6517333984375 0 1671.6886
668.9871215820312 0 985.53076
669.3289794921875 0 4483.5195
669.6609497070312 0 24780.031
669.994873046875 0 34604.324
670.3292846679688 0 25944.35
670.6622314453125 0 12492.585
670.9961547851562 0 6832.668
671.327880859375 0 3325.8672
671.6608276367188 0 1863.4089
675.32763671875 0 861.19293
679.3394165039062 0 742.8631 c Water loss 4
680.31494140625 0 4230.6963 c Ammonia loss 4
681.3152465820312 0 1200.94
682.3278198242188 0 1061.5521
683.3389282226562 0 767.87006
690.3427124023438 0 1957.521
691.3314819335938 0 1685.9758
691.8297729492188 0 1052.247
695.9256591796875 0 831.3684
697.3411254882812 0 12470.636 c 4
698.3448486328125 0 5678.6797
699.344970703125 0 1834.5577
699.8394775390625 0 1314.4036
700.3436889648438 0 1119.0765
701.349365234375 0 2254.544
701.8434448242188 0 9891.948 c 10
702.3429565429688 0 6105.5986
702.8450317382812 0 2242.4092
703.3489379882812 0 851.4459
704.3668212890625 0 838.4674
711.333984375 0 752.12427
713.3421020507812 0 906.07776
714.3309326171875 0 673.6699
716.4039306640625 0 4365.6846
717.4066772460938 0 1433.9082
726.3633422851562 0 773.80695
748.8685302734375 0 1667.4498 y Ammonia loss 4
749.3636474609375 0 4065.3423 z 4
749.8627319335938 0 4252.556
750.3646850585938 0 1937.2479
750.8645629882812 0 876.07336
754.3675537109375 0 1075.762
754.8677978515625 0 2930.7747
755.361572265625 0 1583.5372
757.3753662109375 0 2604.4043 c Ammonia loss 11
757.8726806640625 0 1428.0114
758.3706665039062 0 955.81256
761.3798217773438 0 780.896
765.8911743164062 0 1450.4042 c 11
766.3952026367188 0 1421.4498
768.35791015625 0 955.7861 y Ammonia loss 10
769.3671875 0 9779.986 z 10
770.3671264648438 0 5535.64
771.3682250976562 0 1916.3751
784.8814086914062 0 1674.7032
785.3847045898438 0 4172.457 y 10
786.3798828125 0 1748.016
787.3991088867188 0 976.6186
789.3598022460938 0 746.0398
793.3949584960938 0 1370.1093 c Ammonia loss 5
798.3836059570312 0 805.3499
803.427490234375 0 2110.1746
804.4265747070312 0 1031.5677
806.3865966796875 0 1852.8704 y Ammonia loss 3
806.8785400390625 0 5686.471 z 3
807.3784790039062 0 6752.75
807.8765869140625 0 3697.5144
810.4254150390625 0 22533.541 c 5
811.4277954101562 0 9656.378
812.427490234375 0 2380.1147
814.404296875 0 2641.2605
814.884033203125 0 1569.2904 y 3
815.3960571289062 0 2063.7473
831.4290161132812 0 1813.4193
839.386474609375 0 1374.5149
841.4016723632812 0 1233.6898
841.8955688476562 0 1356.3379 w 2
842.3987426757812 0 1776.2565
843.384521484375 0 816.7225
845.9004516601562 0 1234.0966
846.3974609375 0 5188.095 c 12
846.8978271484375 0 4776.2866
847.3984375 0 3087.023
849.4197998046875 0 923.6654
862.9015502929688 0 738.30963
867.4227294921875 0 3301.6736
868.423583984375 0 923.1416
870.4122314453125 0 3100.6382 y Ammonia loss 2
870.9083251953125 0 6127.353 z 2
871.407958984375 0 5657.823
871.9082641601562 0 2264.0066
872.4139404296875 0 807.00433
874.4221801757812 0 854.208
878.4029541015625 0 877.42316
878.9174194335938 0 714.5733 y 2
879.40673828125 0 1140.688
881.435302734375 0 1777.4713
882.4406127929688 0 794.23566
888.4365844726562 0 1702.7692
888.9429321289062 0 1043.8746
898.4107666015625 0 11747.983 z 9
899.4153442382812 0 9452.157
900.419189453125 0 4287.4956
901.417236328125 0 1465.0975
902.4296875 0 856.4594
904.9269409179688 0 710.9197
908.425537109375 0 1219.8319 c Ammonia loss 6
910.4418334960938 0 3642.0393 c 13
910.9437866210938 0 3464.9717
911.4448852539062 0 2841.4868
911.9464111328125 0 1059.7092
912.4419555664062 0 803.6869
914.4185180664062 0 953.9699
915.4423217773438 0 1518.4584
916.4434814453125 0 934.616
918.94140625 0 697.23724
925.451416015625 0 2968.33 c 6
926.4512939453125 0 2024.772
926.9381713867188 0 819.2112
930.4559326171875 0 841.1073
930.9676513671875 0 1645.4845
931.470458984375 0 1657.798
932.4775390625 0 2418.5923
932.9388427734375 0 969.55084
933.4772338867188 0 949.8515
933.9342041015625 0 847.3048
937.4628295898438 0 968.49884
938.47265625 0 3962.3938
938.9635009765625 0 1868.8748
939.4513549804688 0 3593.8115 z 1
939.93994140625 0 3935.206
940.4467163085938 0 2616.166
940.9415283203125 0 1694.6768
941.45263671875 0 1170.9266
942.4296264648438 0 828.5095
944.4557495117188 0 841.0485
945.4834594726562 0 2260.9404
945.9640502929688 0 749.9685
946.4697265625 0 1783.2595
946.9552612304688 0 891.83026
947.4641723632812 0 1060.3229 y 1
951.4442749023438 0 869.2545
951.9617919921875 0 1181.9166
952.9780883789062 0 998.49335
953.4666137695312 0 1051.7534
953.9714965820312 0 885.11566
955.4583129882812 0 1804.9519
959.9746704101562 0 1675.3127
960.4766235351562 0 4008.3215
960.9741821289062 0 1090.4003
961.4713134765625 0 1263.442
962.4589233398438 0 1615.168
962.9608154296875 0 1038.4152
963.4622802734375 0 1552.4998
964.4586791992188 0 2008.9587 c Water loss 7
965.4542236328125 0 744.8636 c Ammonia loss 7
965.9667358398438 0 1156.3606
966.4737548828125 0 1138.4415
966.9750366210938 0 1473.0988
967.4730224609375 0 3352.0376
967.968994140625 0 1448.1543
968.4642944335938 0 3052.9978 w 8
968.961669921875 0 2076.7993
969.4657592773438 0 1642.4225
970.9640502929688 0 1036.1735
971.4586181640625 0 896.3433
972.45947265625 0 1511.5249
972.968505859375 0 1046.3524
973.47314453125 0 811.3849
973.9794921875 0 2499.8267
974.4777221679688 0 2051.2314
974.98388671875 0 8173.057
975.4830932617188 0 7717.3057
975.982666015625 0 4115.315
976.475830078125 0 4205.5933
976.97314453125 0 2551.4722
977.4712524414062 0 2529.4246
977.9609985351562 0 767.58453
978.4532470703125 0 1509.9487
980.4677124023438 0 1502.7832
981.4835815429688 0 4198.235
981.9861450195312 0 3000.5632
982.4780883789062 0 13411.7705 c 7
982.988525390625 0 2636.3623
983.4786376953125 0 7118.0396
984.4686889648438 0 2863.9805
984.9600219726562 0 3004.9834
985.4721069335938 0 5265.455
985.9658203125 0 2566.7327
986.4701538085938 0 2590.5054
987.4696655273438 0 2836.9856
987.965576171875 0 1011.0646
988.4776611328125 0 3377.9014
989.4801025390625 0 2805.9973
989.955322265625 0 914.8982
993.4598999023438 0 2212.26
993.96826171875 0 3066.9668
994.4660034179688 0 2432.2166
994.9683837890625 0 1649.7999
995.474365234375 0 2008.7794
995.9822998046875 0 4944.09
996.481689453125 0 12997.2705
996.9804077148438 0 10520.452
997.4849853515625 0 6333.703
997.9804077148438 0 2946.981
998.4768676757812 0 1506.4501
998.9577026367188 0 1277.5452
999.4560546875 0 1365.6715
1002.4686279296875 0 2454.0469
1002.9644775390625 0 1223.7083
1003.4862670898438 0 2119.8218
1003.9832763671875 0 3616.404
1004.4949340820312 0 9742.365
1004.9935302734375 0 14808.939
1005.4943237304688 0 10841
1005.99609375 0 6460.3438
1006.4937133789062 0 4351.4053
1006.987060546875 0 1552.3519
1007.4805908203125 0 1891.7803
1008.449462890625 0 1575.742
1026.507568359375 0 1266.9149 z 8
1027.5098876953125 0 3914.654
1028.5087890625 0 1999.4427
1084.529296875 0 999.7715
1109.5849609375 0 2688.3367
1110.5703125 0 13081.935 c 8
1111.5716552734375 0 7726.045
1112.571044921875 0 3216.1174
1141.5494384765625 0 1102.7635
1195.5933837890625 0 864.84247
1196.599609375 0 1900.6547
1197.59716796875 0 850.18024
1198.5670166015625 0 2277.1 z 6
1199.563232421875 0 6041.077
1200.568359375 0 3016.3533
1201.5679931640625 0 1798.198
1239.6160888671875 0 2992.0237 c 9
1240.61669921875 0 3190.0938
1241.6260986328125 0 1640.5123
1311.64404296875 0 1376.5424 z 5
1312.647216796875 0 3011.5647
1313.6517333984375 0 1572.1185
1358.6685791015625 0 1041.703
1402.6832275390625 0 2059.4595 c 10
1403.679931640625 0 1655.6554
1404.68408203125 0 867.2152
1497.720947265625 0 1174.6929 z 4
1498.7197265625 0 1949.2772
1499.7227783203125 0 1985.899
1510.751953125 0 847.711
1513.7406005859375 0 879.35657 c Ammonia loss 11
1530.7818603515625 0 1009.4373 c 11
1531.772705078125 0 784.19543
1612.7530517578125 0 1624.3616 z 3
1613.7548828125 0 3157.7585
1614.7564697265625 0 1932.3634
1615.7750244140625 0 1094.5487
1691.795654296875 0 1581.262 c 12
1692.7950439453125 0 1685.9648
1693.787841796875 0 907.3124
1740.81640625 0 1788.9966 z 2
1741.806396484375 0 2364.6877
1742.8101806640625 0 1283.0938
1743.832275390625 0 1079.98
1760.843017578125 0 762.1617
1820.8837890625 0 1203.2698
1920.959228515625 0 981.82556
1951.004150390625 0 901.8732
1951.955810546875 0 900.1642
1991.953857421875 0 1027.6595
1992.96044921875 0 1114.741
2008.9647216796875 0 1223.2427
2009.974365234375 0 1191.3364
2010.98095703125 0 1310.3871
2011.9896240234375 0 897.9803

Spectrum Details

|  |  |
| --- | --- |
| Matched peaks? Matched peaks The total absolute number of peaks matched. Additionally in brackets the total fraction of peaks matched and the total number of peaks is shown. | 81 (13.57% of 597) |
| FDR? FDR The false discovery rate estimated for this peptide. It is calculated by matching all theoretical fragments with a non-integer shift with the raw peaks for this spectrum. This is done with 40 different shifts. The resulting percentage is the average number of annotated peaks over the number of annotated peaks with the correct spectrum. | 0.38% |
| Satellite FDR? Satellite FDR See the FDR for details on its calculation. This satellite ion specific FDR only contains the satellite ions (d/w) for I/L/J positions. | 0.00% |
| PSM Score? PSM Score The PSM Score as given by Hecklib to this annotated spectrum. It is shown with three significant figures. | 385 |

#### Spectrum 5469? Spectrum 5469 The raw spectrum of this peptide as annotated by Hecklib. The fragments are coloured according to ion type (see legend). Any peaks with a star '\*' as text can be hovered over to see the full details, first the ion type second the mass shift type. By hovering over the amino acids in the peptide or ions in the legend the corresponding peaks are highlighted. By toggling the 'Unassigned' label you can turn the background (unassigned) peaks on or off in the plot. By updating the slider in the Ion legend you can update the spectrum to only show the top X% of the peaks with labels. The top X% means any peak that is within X% of the highest intensity. By dragging in the spectrum you can zoom in to a specific part of the spectrum and use 'Zoom Out' to get back to the original zoom level. The annotation of the spectrum is based on the given sequence in the peptides file and is done with different software so inconsistencies are likely. The peaks are annotated based on the given sequence, with 20 ppm tolerance.

Copy Data

##### Spectrum 5469 (TSV)

###### Preview

```
Loading example...
```

*Click on the button to copy the data to your clipboard.*

Mz MinMz MaxIntensity Max

WidthHeightPeptide font sizePeptide stroke widthSpectrum font sizeSpectrum stroke widthCompact peptide

Ion legend

wxyz

abcd

OtherUnassignedIon^Charge^_Position_Show for top:%

JHQDWLDGKEYKCKK

01.01e+42.01e+43.02e+44.03e+4

Zoom Out

c^+4^_6_y^+1^_1_c^+1^_2_y^+1^_2_c^+2^_5_y^+4^_11_c^+1^_3_y^+2^_5_c^+1^_3_w^+2^_6_z^+2^_6_c^+1^_4_y^+3^_11_c^+1^_4_y^+2^_7_z^+2^_8_z^+2^_8_w^+1^_4_y^+2^_8_c^+2^_9_w^+2^_9_y^+3^_13_z^+2^_9_y^+2^_9_y^+2^_9_z^+2^_9_c^+3^_14_y^+1^_4_z^+1^_4_c^+3^_14_y^+2^_9_c^+2^_10_c^+2^_10_c^+2^_10_y^+1^_4_y^+3^_14_z^+3^_14_y^+3^_14_w^+2^_10_z^+2^_10_z^+2^_10_y^+2^_10_z^+2^_10_y^+2^_10_c^+1^_5_c^+1^_5_c^+2^_11_y^+2^_11_z^+2^_11_z^+1^_5_c^+2^_12_c^+2^_12_y^+1^_5_z^+1^_5_y^+1^_5_c^+1^_6_y^+2^_12_z^+2^_12_c^+1^_6_y^+2^_12_c^+2^_13_y^+2^_13_z^+2^_13_z^+1^_6_c^+2^_14_c^+1^_7_c^+2^_14_y^+1^_6_c^+1^_7_z^+2^_14_y^+2^_14_c^+1^_8_c^+1^_8_w^+1^_7_c^+1^_8_z^+1^_7_c^+1^_9_z^+1^_9_c^+1^_10_z^+1^_10_c^+1^_11_z^+1^_11_z^+1^_12_c^+1^_13_z^+1^_13_

0770153923093079

Fragment Matches Table

Show background peaks

| Position | Ion type | Intensity | mz Theoretical | mz Error (Th) | mz Error (ppm) | Charge | Series Number |
| --- | --- | --- | --- | --- | --- | --- | --- |
| - | - | 429.1 | 126.1 | - | - | 0 | - |
| - | - | 3547 | 129.1 | - | - | 0 | - |
| - | - | 2247 | 130.1 | - | - | 0 | - |
| - | - | 382.7 | 131.4 | - | - | 0 | - |
| - | - | 429 | 133.1 | - | - | 0 | - |
| - | - | 1276 | 133.1 | - | - | 0 | - |
| - | - | 722.9 | 136.1 | - | - | 0 | - |
| - | - | 436.3 | 140.1 | - | - | 0 | - |
| - | - | 464.6 | 144.5 | - | - | 0 | - |
| - | - | 803.9 | 146.1 | - | - | 0 | - |
| - | - | 1067 | 148.9 | - | - | 0 | - |
| - | - | 488.3 | 149 | - | - | 0 | - |
| - | - | 1245 | 149 | - | - | 0 | - |
| - | - | 3161 | 155.1 | - | - | 0 | - |
| - | - | 454.8 | 156.1 | - | - | 0 | - |
| - | - | 516.7 | 156.9 | - | - | 0 | - |
| - | - | 775.7 | 159.1 | - | - | 0 | - |
| - | - | 733 | 162.1 | - | - | 0 | - |
| - | - | 462.7 | 164.1 | - | - | 0 | - |
| - | - | 605.1 | 165.1 | - | - | 0 | - |
| - | - | 3310 | 166.1 | - | - | 0 | - |
| - | - | 1607 | 168.1 | - | - | 0 | - |
| - | - | 1194 | 173.1 | - | - | 0 | - |
| - | - | 4354 | 173.5 | - | - | 0 | - |
| - | - | 556.3 | 174.1 | - | - | 0 | - |
| - | - | 489.2 | 176.4 | - | - | 0 | - |
| - | - | 1103 | 177.1 | - | - | 0 | - |
| - | - | 1499 | 183.1 | - | - | 0 | - |
| - | - | 455.9 | 184.7 | - | - | 0 | - |
| - | - | 2797 | 185.2 | - | - | 0 | - |
| - | - | 1079 | 187.1 | - | - | 0 | - |
| - | - | 1753 | 191.1 | - | - | 0 | - |
| 6 | c | 1179 | 199.1 | 0.003032 | 15.23 | +4 | 6 |
| - | - | 1624 | 202.1 | - | - | 0 | - |
| - | - | 1548 | 203.1 | - | - | 0 | - |
| - | - | 687.4 | 204.1 | - | - | 0 | - |
| 15 | y | 869.2 | 205.1 | 0.003892 | 18.98 | +1 | 1 |
| - | - | 454.5 | 208 | - | - | 0 | - |
| - | - | 914.6 | 209.1 | - | - | 0 | - |
| - | - | 712.8 | 213.1 | - | - | 0 | - |
| - | - | 9141 | 215.1 | - | - | 0 | - |
| - | - | 974.2 | 217.2 | - | - | 0 | - |
| - | - | 600.6 | 217.3 | - | - | 0 | - |
| - | - | 1131 | 218.1 | - | - | 0 | - |
| - | - | 524 | 218.2 | - | - | 0 | - |
| - | - | 683.4 | 221.1 | - | - | 0 | - |
| - | - | 2524 | 223.2 | - | - | 0 | - |
| - | - | 3345 | 227.1 | - | - | 0 | - |
| - | - | 566.2 | 229.2 | - | - | 0 | - |
| - | - | 597.2 | 230.2 | - | - | 0 | - |
| - | - | 791.6 | 231.2 | - | - | 0 | - |
| - | - | 2981 | 234.1 | - | - | 0 | - |
| - | - | 589.5 | 235.1 | - | - | 0 | - |
| - | - | 668.6 | 242.2 | - | - | 0 | - |
| - | - | 1208 | 246.2 | - | - | 0 | - |
| - | - | 1.497E+04 | 251.2 | - | - | 0 | - |
| - | - | 1632 | 252.2 | - | - | 0 | - |
| - | - | 901.7 | 261.1 | - | - | 0 | - |
| 2 | c | 1510 | 268.2 | 0.0003226 | 1.203 | +1 | 2 |
| - | - | 763.2 | 274.1 | - | - | 0 | - |
| - | - | 944 | 275.1 | - | - | 0 | - |
| - | - | 1052 | 281.1 | - | - | 0 | - |
| - | - | 1120 | 283.2 | - | - | 0 | - |
| - | - | 7005 | 299.1 | - | - | 0 | - |
| - | - | 1118 | 300.1 | - | - | 0 | - |
| - | - | 1712 | 300.2 | - | - | 0 | - |
| - | - | 779.4 | 302.2 | - | - | 0 | - |
| - | - | 797.9 | 317.1 | - | - | 0 | - |
| - | - | 703.7 | 318.2 | - | - | 0 | - |
| - | - | 551.3 | 319.2 | - | - | 0 | - |
| - | - | 2006 | 326.7 | - | - | 0 | - |
| 14 | y | 1108 | 333.2 | 0.002832 | 8.499 | +1 | 2 |
| - | - | 1268 | 338.1 | - | - | 0 | - |
| - | - | 1976 | 340.2 | - | - | 0 | - |
| 5 | c | 674.5 | 340.7 | 0.002485 | 7.294 | +2 | 5 |
| - | - | 499.2 | 341.2 | - | - | 0 | - |
| - | - | 740 | 349.7 | - | - | 0 | - |
| - | - | 803.7 | 359 | - | - | 0 | - |
| 5 | y | 664 | 374.7 | 0.005221 | 13.94 | +4 | 11 |
| - | - | 746.9 | 378.2 | - | - | 0 | - |
| 3 | c | 5220 | 379.2 | 0.0005513 | 1.454 | +1 | 3 |
| - | - | 1452 | 380.2 | - | - | 0 | - |
| - | - | 612.4 | 381.2 | - | - | 0 | - |
| - | - | 735 | 383.2 | - | - | 0 | - |
| - | - | 629 | 386.2 | - | - | 0 | - |
| - | - | 2036 | 387.2 | - | - | 0 | - |
| 11 | y | 1448 | 393.2 | 0.003597 | 9.149 | +2 | 5 |
| 3 | c | 5743 | 396.2 | 0.0005525 | 1.394 | +1 | 3 |
| - | - | 1024 | 397.2 | - | - | 0 | - |
| - | - | 695.4 | 403.5 | - | - | 0 | - |
| - | - | 749.6 | 407.2 | - | - | 0 | - |
| - | - | 3123 | 415 | - | - | 0 | - |
| - | - | 721.6 | 416 | - | - | 0 | - |
| 10 | w | 1.004E+04 | 420.2 | 0.003661 | 8.713 | +2 | 6 |
| - | - | 392.3 | 420.2 | - | - | 0 | - |
| - | - | 4436 | 420.7 | - | - | 0 | - |
| - | - | 1817 | 421.2 | - | - | 0 | - |
| - | - | 782.2 | 428.3 | - | - | 0 | - |
| - | - | 632.2 | 435.7 | - | - | 0 | - |
| - | - | 620.5 | 445.2 | - | - | 0 | - |
| - | - | 768.8 | 447.9 | - | - | 0 | - |
| 10 | z | 2013 | 449.7 | 0.005373 | 11.95 | +2 | 6 |
| - | - | 6557 | 457.3 | - | - | 0 | - |
| - | - | 3101 | 458.2 | - | - | 0 | - |
| - | - | 3064 | 458.3 | - | - | 0 | - |
| - | - | 1983 | 458.7 | - | - | 0 | - |
| - | - | 5585 | 459.3 | - | - | 0 | - |
| - | - | 1322 | 460.3 | - | - | 0 | - |
| - | - | 1556 | 460.7 | - | - | 0 | - |
| - | - | 763.2 | 461.2 | - | - | 0 | - |
| - | - | 1287 | 467.2 | - | - | 0 | - |
| - | - | 880.1 | 468.3 | - | - | 0 | - |
| - | - | 1194 | 469.2 | - | - | 0 | - |
| - | - | 1008 | 472.3 | - | - | 0 | - |
| - | - | 773.3 | 478.3 | - | - | 0 | - |
| - | - | 587.1 | 481.2 | - | - | 0 | - |
| - | - | 1150 | 484.2 | - | - | 0 | - |
| - | - | 2587 | 485.2 | - | - | 0 | - |
| - | - | 785.3 | 485.3 | - | - | 0 | - |
| - | - | 5698 | 486.3 | - | - | 0 | - |
| - | - | 1262 | 487.2 | - | - | 0 | - |
| - | - | 1047 | 488.3 | - | - | 0 | - |
| - | - | 911.3 | 489.2 | - | - | 0 | - |
| - | - | 698.5 | 492.5 | - | - | 0 | - |
| - | - | 3482 | 493.7 | - | - | 0 | - |
| 4 | c | 6577 | 494.2 | 0.001776 | 3.593 | +1 | 4 |
| - | - | 1288 | 494.7 | - | - | 0 | - |
| - | - | 1260 | 495.2 | - | - | 0 | - |
| - | - | 611.7 | 496.3 | - | - | 0 | - |
| - | - | 784.4 | 498.3 | - | - | 0 | - |
| - | - | 1195 | 498.7 | - | - | 0 | - |
| 5 | y | 1408 | 499.2 | 0.004712 | 9.438 | +3 | 11 |
| - | - | 2511 | 501.2 | - | - | 0 | - |
| - | - | 1309 | 501.8 | - | - | 0 | - |
| - | - | 4983 | 502.3 | - | - | 0 | - |
| - | - | 5398 | 502.5 | - | - | 0 | - |
| - | - | 1.223E+04 | 502.8 | - | - | 0 | - |
| - | - | 3321 | 503 | - | - | 0 | - |
| - | - | 3808 | 503.1 | - | - | 0 | - |
| - | - | 7091 | 503.2 | - | - | 0 | - |
| - | - | 832.1 | 503.3 | - | - | 0 | - |
| - | - | 1245 | 503.8 | - | - | 0 | - |
| - | - | 1327 | 504.1 | - | - | 0 | - |
| - | - | 2047 | 504.3 | - | - | 0 | - |
| - | - | 3121 | 504.3 | - | - | 0 | - |
| - | - | 971.3 | 504.3 | - | - | 0 | - |
| - | - | 853.5 | 505.6 | - | - | 0 | - |
| - | - | 986.8 | 506.3 | - | - | 0 | - |
| - | - | 642.2 | 506.7 | - | - | 0 | - |
| - | - | 934.1 | 507.3 | - | - | 0 | - |
| 4 | c | 1.442E+04 | 511.3 | 0.001136 | 2.223 | +1 | 4 |
| - | - | 992.6 | 511.3 | - | - | 0 | - |
| - | - | 3840 | 512.3 | - | - | 0 | - |
| - | - | 928.4 | 513.3 | - | - | 0 | - |
| - | - | 1950 | 515.3 | - | - | 0 | - |
| - | - | 762.4 | 519.3 | - | - | 0 | - |
| - | - | 1505 | 519.3 | - | - | 0 | - |
| 9 | y | 2901 | 521.8 | 0.003314 | 6.352 | +2 | 7 |
| - | - | 792.2 | 522.3 | - | - | 0 | - |
| - | - | 835 | 522.8 | - | - | 0 | - |
| - | - | 826.8 | 527.8 | - | - | 0 | - |
| - | - | 835.9 | 529.3 | - | - | 0 | - |
| 8 | z | 663.7 | 533.3 | 0.009167 | 17.19 | +2 | 8 |
| - | - | 1639 | 535.3 | - | - | 0 | - |
| - | - | 637.2 | 536.3 | - | - | 0 | - |
| 8 | z | 1472 | 542.3 | 0.005294 | 9.763 | +2 | 8 |
| - | - | 825.4 | 543.3 | - | - | 0 | - |
| - | - | 935.6 | 547.3 | - | - | 0 | - |
| - | - | 588.9 | 547.8 | - | - | 0 | - |
| 12 | w | 1124 | 548.2 | 0.009723 | 17.74 | +1 | 4 |
| - | - | 799.4 | 548.8 | - | - | 0 | - |
| - | - | 624.6 | 549.3 | - | - | 0 | - |
| - | - | 1493 | 550.2 | - | - | 0 | - |
| 8 | y | 2180 | 550.3 | 0.002388 | 4.34 | +2 | 8 |
| - | - | 1574 | 550.8 | - | - | 0 | - |
| - | - | 871.4 | 552.3 | - | - | 0 | - |
| - | - | 1190 | 555.3 | - | - | 0 | - |
| - | - | 1977 | 555.3 | - | - | 0 | - |
| 9 | c | 8855 | 555.8 | 0.001378 | 2.48 | +2 | 9 |
| - | - | 4498 | 556.3 | - | - | 0 | - |
| - | - | 1207 | 556.8 | - | - | 0 | - |
| - | - | 634.5 | 563.3 | - | - | 0 | - |
| - | - | 5212 | 564.3 | - | - | 0 | - |
| - | - | 2539 | 564.8 | - | - | 0 | - |
| - | - | 898.6 | 565.3 | - | - | 0 | - |
| - | - | 815.5 | 565.8 | - | - | 0 | - |
| - | - | 868.2 | 568.8 | - | - | 0 | - |
| - | - | 2521 | 571.3 | - | - | 0 | - |
| - | - | 2272 | 571.8 | - | - | 0 | - |
| - | - | 2066 | 572.3 | - | - | 0 | - |
| - | - | 788.1 | 573.3 | - | - | 0 | - |
| 7 | w | 2485 | 577.3 | 0.00407 | 7.05 | +2 | 9 |
| - | - | 1909 | 577.8 | - | - | 0 | - |
| - | - | 1272 | 578.3 | - | - | 0 | - |
| 3 | y | 1262 | 580.3 | 0.003433 | 5.917 | +3 | 13 |
| - | - | 953 | 590.6 | - | - | 0 | - |
| 7 | z | 657.6 | 590.8 | 0.006648 | 11.25 | +2 | 9 |
| - | - | 1541 | 591.3 | - | - | 0 | - |
| - | - | 794.8 | 592.8 | - | - | 0 | - |
| - | - | 1069 | 596.3 | - | - | 0 | - |
| - | - | 2120 | 597.3 | - | - | 0 | - |
| - | - | 937.3 | 597.6 | - | - | 0 | - |
| - | - | 876.3 | 598 | - | - | 0 | - |
| - | - | 763.7 | 598.3 | - | - | 0 | - |
| 7 | y | 1054 | 598.8 | 0.001911 | 3.191 | +2 | 9 |
| - | - | 756.5 | 599 | - | - | 0 | - |
| 7 | y | 2932 | 599.3 | 0.007791 | 13 | +2 | 9 |
| 7 | z | 6086 | 599.8 | 0.002286 | 3.812 | +2 | 9 |
| - | - | 3560 | 600.3 | - | - | 0 | - |
| - | - | 1697 | 600.8 | - | - | 0 | - |
| 14 | c | 848.7 | 601.3 | 0.001657 | 2.756 | +3 | 14 |
| - | - | 750.6 | 601.4 | - | - | 0 | - |
| - | - | 771.5 | 602 | - | - | 0 | - |
| - | - | 903.7 | 602.3 | - | - | 0 | - |
| - | - | 786.2 | 602.9 | - | - | 0 | - |
| - | - | 983.7 | 605 | - | - | 0 | - |
| 12 | y | 1061 | 605.3 | 0.008674 | 14.33 | +1 | 4 |
| - | - | 810.7 | 605.6 | - | - | 0 | - |
| 12 | z | 6026 | 606.3 | 0.009052 | 14.93 | +1 | 4 |
| - | - | 740 | 606.6 | - | - | 0 | - |
| - | - | 2072 | 607 | - | - | 0 | - |
| 14 | c | 9289 | 607.3 | 0.002146 | 3.533 | +3 | 14 |
| - | - | 4762 | 607.6 | - | - | 0 | - |
| 7 | y | 7885 | 607.8 | 0.00347 | 5.709 | +2 | 9 |
| - | - | 3238 | 608 | - | - | 0 | - |
| - | - | 7591 | 608.3 | - | - | 0 | - |
| - | - | 1081 | 608.6 | - | - | 0 | - |
| - | - | 1815 | 608.8 | - | - | 0 | - |
| - | - | 600.2 | 609 | - | - | 0 | - |
| - | - | 994.3 | 609.3 | - | - | 0 | - |
| 10 | c | 1670 | 611.3 | 0.003772 | 6.17 | +2 | 10 |
| 10 | c | 846.7 | 611.8 | 0.01051 | 17.17 | +2 | 10 |
| - | - | 1006 | 612 | - | - | 0 | - |
| - | - | 2691 | 612.3 | - | - | 0 | - |
| - | - | 944.2 | 612.6 | - | - | 0 | - |
| - | - | 1811 | 612.8 | - | - | 0 | - |
| - | - | 1039 | 613.6 | - | - | 0 | - |
| - | - | 924.7 | 615.3 | - | - | 0 | - |
| - | - | 1146 | 615.6 | - | - | 0 | - |
| - | - | 816.3 | 616.6 | - | - | 0 | - |
| - | - | 813 | 617.3 | - | - | 0 | - |
| - | - | 1005 | 619.3 | - | - | 0 | - |
| - | - | 3956 | 619.8 | - | - | 0 | - |
| 10 | c | 1.437E+04 | 620.3 | 0.001749 | 2.82 | +2 | 10 |
| - | - | 809.7 | 620.6 | - | - | 0 | - |
| - | - | 7589 | 620.8 | - | - | 0 | - |
| - | - | 1309 | 621 | - | - | 0 | - |
| - | - | 4003 | 621.3 | - | - | 0 | - |
| - | - | 1316 | 621.8 | - | - | 0 | - |
| - | - | 941.1 | 622 | - | - | 0 | - |
| 12 | y | 3935 | 622.3 | 0.01166 | 18.74 | +1 | 4 |
| - | - | 809.2 | 622.6 | - | - | 0 | - |
| - | - | 1096 | 623.3 | - | - | 0 | - |
| - | - | 1147 | 623.6 | - | - | 0 | - |
| - | - | 1215 | 625.6 | - | - | 0 | - |
| 2 | y | 2688 | 626 | 0.01082 | 17.29 | +3 | 14 |
| - | - | 3055 | 626.3 | - | - | 0 | - |
| 2 | z | 4301 | 626.6 | 0.01143 | 18.24 | +3 | 14 |
| - | - | 705.5 | 626.8 | - | - | 0 | - |
| - | - | 1479 | 627 | - | - | 0 | - |
| - | - | 2992 | 627.3 | - | - | 0 | - |
| - | - | 922.2 | 627.6 | - | - | 0 | - |
| - | - | 1179 | 628 | - | - | 0 | - |
| - | - | 1580 | 629.3 | - | - | 0 | - |
| - | - | 588.3 | 629.6 | - | - | 0 | - |
| - | - | 813.8 | 629.8 | - | - | 0 | - |
| - | - | 666.6 | 630 | - | - | 0 | - |
| - | - | 990.2 | 630.3 | - | - | 0 | - |
| - | - | 1051 | 630.6 | - | - | 0 | - |
| - | - | 1746 | 631 | - | - | 0 | - |
| - | - | 2692 | 631.3 | - | - | 0 | - |
| - | - | 2171 | 631.6 | - | - | 0 | - |
| 2 | y | 2728 | 632 | 0.0002658 | 0.4206 | +3 | 14 |
| - | - | 2480 | 632.3 | - | - | 0 | - |
| - | - | 1563 | 633 | - | - | 0 | - |
| - | - | 959.3 | 633.3 | - | - | 0 | - |
| - | - | 829.9 | 634 | - | - | 0 | - |
| - | - | 2712 | 634.3 | - | - | 0 | - |
| - | - | 1083 | 634.6 | - | - | 0 | - |
| 6 | w | 5770 | 634.8 | 0.001123 | 1.769 | +2 | 10 |
| - | - | 1008 | 635 | - | - | 0 | - |
| - | - | 5546 | 635.3 | - | - | 0 | - |
| - | - | 1953 | 635.6 | - | - | 0 | - |
| - | - | 2351 | 635.8 | - | - | 0 | - |
| - | - | 1960 | 636 | - | - | 0 | - |
| - | - | 2764 | 636.3 | - | - | 0 | - |
| - | - | 2578 | 636.6 | - | - | 0 | - |
| - | - | 2574 | 637 | - | - | 0 | - |
| - | - | 2334 | 637.3 | - | - | 0 | - |
| - | - | 2279 | 637.6 | - | - | 0 | - |
| - | - | 2277 | 638 | - | - | 0 | - |
| - | - | 1672 | 638.3 | - | - | 0 | - |
| - | - | 1156 | 638.6 | - | - | 0 | - |
| - | - | 898.9 | 639.3 | - | - | 0 | - |
| - | - | 2029 | 640 | - | - | 0 | - |
| - | - | 1.432E+04 | 640.3 | - | - | 0 | - |
| - | - | 1.451E+04 | 640.7 | - | - | 0 | - |
| - | - | 8352 | 641 | - | - | 0 | - |
| - | - | 5193 | 641.3 | - | - | 0 | - |
| - | - | 2953 | 641.6 | - | - | 0 | - |
| - | - | 2281 | 642 | - | - | 0 | - |
| - | - | 3022 | 642.3 | - | - | 0 | - |
| - | - | 3390 | 642.6 | - | - | 0 | - |
| - | - | 3334 | 643 | - | - | 0 | - |
| - | - | 1059 | 643.3 | - | - | 0 | - |
| - | - | 1529 | 644 | - | - | 0 | - |
| - | - | 2079 | 644.3 | - | - | 0 | - |
| - | - | 1772 | 644.6 | - | - | 0 | - |
| - | - | 2164 | 645 | - | - | 0 | - |
| - | - | 3764 | 645.3 | - | - | 0 | - |
| - | - | 6958 | 645.6 | - | - | 0 | - |
| - | - | 9487 | 646 | - | - | 0 | - |
| - | - | 6740 | 646.3 | - | - | 0 | - |
| - | - | 3883 | 646.6 | - | - | 0 | - |
| - | - | 2074 | 647 | - | - | 0 | - |
| 6 | z | 1954 | 647.3 | 0.01188 | 18.35 | +2 | 10 |
| - | - | 1204 | 647.6 | - | - | 0 | - |
| 6 | z | 1309 | 647.8 | 0.0009536 | 1.472 | +2 | 10 |
| - | - | 2782 | 648 | - | - | 0 | - |
| - | - | 2601 | 648.3 | - | - | 0 | - |
| - | - | 1663 | 648.6 | - | - | 0 | - |
| - | - | 2039 | 649 | - | - | 0 | - |
| - | - | 1745 | 649.3 | - | - | 0 | - |
| - | - | 4058 | 649.6 | - | - | 0 | - |
| - | - | 9012 | 650 | - | - | 0 | - |
| - | - | 1.679E+04 | 650.3 | - | - | 0 | - |
| - | - | 1.376E+04 | 650.7 | - | - | 0 | - |
| - | - | 1.153E+04 | 651 | - | - | 0 | - |
| - | - | 8593 | 651.3 | - | - | 0 | - |
| - | - | 9160 | 651.6 | - | - | 0 | - |
| - | - | 4219 | 652 | - | - | 0 | - |
| - | - | 4661 | 652.3 | - | - | 0 | - |
| - | - | 2680 | 652.6 | - | - | 0 | - |
| - | - | 915.3 | 653 | - | - | 0 | - |
| - | - | 2366 | 653.3 | - | - | 0 | - |
| - | - | 1578 | 653.6 | - | - | 0 | - |
| - | - | 986.2 | 654 | - | - | 0 | - |
| - | - | 2058 | 654.7 | - | - | 0 | - |
| - | - | 1749 | 655 | - | - | 0 | - |
| 6 | y | 1162 | 655.3 | 0.007383 | 11.27 | +2 | 10 |
| 6 | z | 8038 | 656.3 | 0.01227 | 18.7 | +2 | 10 |
| - | - | 6234 | 656.6 | - | - | 0 | - |
| - | - | 1581 | 656.8 | - | - | 0 | - |
| - | - | 8488 | 657 | - | - | 0 | - |
| - | - | 7984 | 657.3 | - | - | 0 | - |
| - | - | 4384 | 657.6 | - | - | 0 | - |
| - | - | 3268 | 658 | - | - | 0 | - |
| - | - | 782 | 658.3 | - | - | 0 | - |
| - | - | 1270 | 658.6 | - | - | 0 | - |
| - | - | 2306 | 659 | - | - | 0 | - |
| - | - | 2103 | 659.3 | - | - | 0 | - |
| - | - | 1636 | 659.6 | - | - | 0 | - |
| - | - | 1102 | 660 | - | - | 0 | - |
| - | - | 1253 | 660.3 | - | - | 0 | - |
| - | - | 900.3 | 660.6 | - | - | 0 | - |
| - | - | 1033 | 661.3 | - | - | 0 | - |
| - | - | 1913 | 662 | - | - | 0 | - |
| - | - | 4823 | 662.3 | - | - | 0 | - |
| - | - | 7749 | 662.6 | - | - | 0 | - |
| - | - | 5990 | 663 | - | - | 0 | - |
| - | - | 3630 | 663.3 | - | - | 0 | - |
| - | - | 2631 | 663.6 | - | - | 0 | - |
| - | - | 1215 | 663.8 | - | - | 0 | - |
| - | - | 2332 | 664 | - | - | 0 | - |
| 6 | y | 5953 | 664.3 | 0.008453 | 12.72 | +2 | 10 |
| - | - | 3590 | 664.7 | - | - | 0 | - |
| - | - | 3098 | 664.8 | - | - | 0 | - |
| - | - | 3693 | 665 | - | - | 0 | - |
| - | - | 2457 | 665.3 | - | - | 0 | - |
| - | - | 2428 | 665.7 | - | - | 0 | - |
| - | - | 1626 | 666.3 | - | - | 0 | - |
| - | - | 1203 | 668 | - | - | 0 | - |
| - | - | 3456 | 668.3 | - | - | 0 | - |
| - | - | 3250 | 668.6 | - | - | 0 | - |
| - | - | 2975 | 669 | - | - | 0 | - |
| - | - | 7585 | 669.3 | - | - | 0 | - |
| - | - | 2.342E+04 | 669.7 | - | - | 0 | - |
| - | - | 3.987E+04 | 670 | - | - | 0 | - |
| - | - | 2.813E+04 | 670.3 | - | - | 0 | - |
| - | - | 1.309E+04 | 670.7 | - | - | 0 | - |
| - | - | 7452 | 671 | - | - | 0 | - |
| - | - | 4112 | 671.3 | - | - | 0 | - |
| - | - | 1456 | 671.7 | - | - | 0 | - |
| - | - | 1010 | 672 | - | - | 0 | - |
| - | - | 1018 | 673.4 | - | - | 0 | - |
| - | - | 945.5 | 674.3 | - | - | 0 | - |
| 5 | c | 4108 | 680.3 | 0.001443 | 2.12 | +1 | 5 |
| - | - | 2170 | 681.3 | - | - | 0 | - |
| - | - | 3119 | 682.3 | - | - | 0 | - |
| - | - | 1579 | 683.3 | - | - | 0 | - |
| - | - | 1575 | 690.3 | - | - | 0 | - |
| - | - | 1295 | 691.3 | - | - | 0 | - |
| - | - | 938.5 | 691.8 | - | - | 0 | - |
| - | - | 1029 | 694.8 | - | - | 0 | - |
| - | - | 807.8 | 695.9 | - | - | 0 | - |
| 5 | c | 1.609E+04 | 697.3 | 0.0004061 | 0.5824 | +1 | 5 |
| - | - | 8286 | 698.3 | - | - | 0 | - |
| - | - | 2423 | 699.3 | - | - | 0 | - |
| - | - | 1064 | 699.8 | - | - | 0 | - |
| - | - | 1673 | 700.3 | - | - | 0 | - |
| - | - | 2941 | 701.3 | - | - | 0 | - |
| 11 | c | 8676 | 701.8 | 0.00225 | 3.206 | +2 | 11 |
| - | - | 9048 | 702.3 | - | - | 0 | - |
| - | - | 2541 | 702.8 | - | - | 0 | - |
| - | - | 1112 | 703.3 | - | - | 0 | - |
| - | - | 782.5 | 708.8 | - | - | 0 | - |
| - | - | 822.9 | 711.3 | - | - | 0 | - |
| - | - | 1104 | 712.3 | - | - | 0 | - |
| - | - | 1206 | 712.8 | - | - | 0 | - |
| - | - | 1583 | 713.3 | - | - | 0 | - |
| - | - | 1026 | 714.3 | - | - | 0 | - |
| - | - | 3957 | 716.4 | - | - | 0 | - |
| - | - | 1399 | 717.4 | - | - | 0 | - |
| - | - | 1154 | 721.3 | - | - | 0 | - |
| - | - | 879.9 | 721.8 | - | - | 0 | - |
| - | - | 1165 | 722.3 | - | - | 0 | - |
| - | - | 1215 | 723.3 | - | - | 0 | - |
| - | - | 882.1 | 735.3 | - | - | 0 | - |
| - | - | 894.8 | 742.8 | - | - | 0 | - |
| - | - | 950 | 746.3 | - | - | 0 | - |
| 5 | y | 904.7 | 748.9 | 0.005814 | 7.764 | +2 | 11 |
| 5 | z | 6938 | 749.4 | 0.00292 | 3.897 | +2 | 11 |
| - | - | 3182 | 749.9 | - | - | 0 | - |
| - | - | 2309 | 750.4 | - | - | 0 | - |
| 11 | z | 1164 | 752.3 | 0.01472 | 19.56 | +1 | 5 |
| - | - | 1028 | 754.4 | - | - | 0 | - |
| - | - | 2184 | 754.9 | - | - | 0 | - |
| - | - | 1780 | 755.4 | - | - | 0 | - |
| 12 | c | 2002 | 757.4 | 0.002354 | 3.109 | +2 | 12 |
| - | - | 1922 | 757.9 | - | - | 0 | - |
| - | - | 1683 | 758.4 | - | - | 0 | - |
| 12 | c | 1652 | 765.9 | 0.0006673 | 0.8713 | +2 | 12 |
| - | - | 2057 | 766.4 | - | - | 0 | - |
| 11 | y | 885.1 | 768.4 | 0.00938 | 12.21 | +1 | 5 |
| 11 | z | 1.001E+04 | 769.4 | 0.008355 | 10.86 | +1 | 5 |
| - | - | 5913 | 770.4 | - | - | 0 | - |
| - | - | 3041 | 771.4 | - | - | 0 | - |
| - | - | 2278 | 784.9 | - | - | 0 | - |
| 11 | y | 4847 | 785.4 | 0.009867 | 12.56 | +1 | 5 |
| - | - | 1224 | 786.4 | - | - | 0 | - |
| - | - | 815.6 | 788.4 | - | - | 0 | - |
| 6 | c | 1620 | 793.4 | 0.002401 | 3.026 | +1 | 6 |
| - | - | 955.8 | 797.4 | - | - | 0 | - |
| - | - | 2417 | 803.4 | - | - | 0 | - |
| 4 | y | 2732 | 806.4 | 0.009493 | 11.77 | +2 | 12 |
| 4 | z | 7939 | 806.9 | 0.0008888 | 1.102 | +2 | 12 |
| - | - | 6023 | 807.4 | - | - | 0 | - |
| - | - | 3494 | 807.9 | - | - | 0 | - |
| - | - | 1054 | 808.4 | - | - | 0 | - |
| - | - | 813.4 | 809.4 | - | - | 0 | - |
| 6 | c | 2.586E+04 | 810.4 | 0.0005707 | 0.7041 | +1 | 6 |
| - | - | 1475 | 811.4 | - | - | 0 | - |
| - | - | 9387 | 811.4 | - | - | 0 | - |
| - | - | 1206 | 812.4 | - | - | 0 | - |
| - | - | 2539 | 812.4 | - | - | 0 | - |
| - | - | 2330 | 814.4 | - | - | 0 | - |
| 4 | y | 1865 | 814.9 | 0.0004852 | 0.5955 | +2 | 12 |
| - | - | 1410 | 815.4 | - | - | 0 | - |
| - | - | 1050 | 826.4 | - | - | 0 | - |
| - | - | 1028 | 827.4 | - | - | 0 | - |
| - | - | 1841 | 831.4 | - | - | 0 | - |
| - | - | 1778 | 831.8 | - | - | 0 | - |
| - | - | 1065 | 832.4 | - | - | 0 | - |
| - | - | 3705 | 837.5 | - | - | 0 | - |
| - | - | 4048 | 837.8 | - | - | 0 | - |
| - | - | 1292 | 838.1 | - | - | 0 | - |
| - | - | 1076 | 838.5 | - | - | 0 | - |
| - | - | 805 | 839.4 | - | - | 0 | - |
| - | - | 1683 | 840.4 | - | - | 0 | - |
| - | - | 2245 | 842.4 | - | - | 0 | - |
| - | - | 842.6 | 843.4 | - | - | 0 | - |
| - | - | 823.7 | 845.4 | - | - | 0 | - |
| - | - | 2258 | 845.9 | - | - | 0 | - |
| 13 | c | 6433 | 846.4 | 0.001146 | 1.354 | +2 | 13 |
| - | - | 4051 | 846.9 | - | - | 0 | - |
| - | - | 2267 | 847.4 | - | - | 0 | - |
| - | - | 1177 | 847.9 | - | - | 0 | - |
| - | - | 1257 | 848.9 | - | - | 0 | - |
| - | - | 1154 | 851 | - | - | 0 | - |
| - | - | 896.3 | 854.4 | - | - | 0 | - |
| - | - | 742.1 | 860.4 | - | - | 0 | - |
| - | - | 1050 | 862.9 | - | - | 0 | - |
| - | - | 3045 | 867.4 | - | - | 0 | - |
| - | - | 1179 | 868.4 | - | - | 0 | - |
| - | - | 938.6 | 869.4 | - | - | 0 | - |
| 3 | y | 2528 | 870.4 | 0.004497 | 5.166 | +2 | 13 |
| 3 | z | 8373 | 870.9 | 0.004177 | 4.796 | +2 | 13 |
| - | - | 5106 | 871.4 | - | - | 0 | - |
| - | - | 2828 | 871.9 | - | - | 0 | - |
| - | - | 2173 | 872.4 | - | - | 0 | - |
| - | - | 945.5 | 873.4 | - | - | 0 | - |
| - | - | 829.7 | 878.4 | - | - | 0 | - |
| - | - | 883.1 | 878.9 | - | - | 0 | - |
| - | - | 4131 | 881.4 | - | - | 0 | - |
| - | - | 1257 | 883.4 | - | - | 0 | - |
| - | - | 1192 | 888.4 | - | - | 0 | - |
| - | - | 903.4 | 888.9 | - | - | 0 | - |
| - | - | 1075 | 889.9 | - | - | 0 | - |
| 10 | z | 1.061E+04 | 898.4 | 0.006881 | 7.659 | +1 | 6 |
| - | - | 9753 | 899.4 | - | - | 0 | - |
| - | - | 4902 | 900.4 | - | - | 0 | - |
| 14 | c | 1176 | 901.4 | 0.01749 | 19.41 | +2 | 14 |
| - | - | 956.7 | 902.4 | - | - | 0 | - |
| 7 | c | 1195 | 908.4 | 0.004767 | 5.248 | +1 | 7 |
| 14 | c | 4009 | 910.4 | 0.00117 | 1.285 | +2 | 14 |
| - | - | 4575 | 910.9 | - | - | 0 | - |
| - | - | 2201 | 911.4 | - | - | 0 | - |
| - | - | 1265 | 911.9 | - | - | 0 | - |
| - | - | 824.7 | 913.4 | - | - | 0 | - |
| 10 | y | 1476 | 914.4 | 0.004426 | 4.84 | +1 | 6 |
| - | - | 2245 | 915.4 | - | - | 0 | - |
| - | - | 1061 | 917.9 | - | - | 0 | - |
| - | - | 949.1 | 918.9 | - | - | 0 | - |
| - | - | 1174 | 919.4 | - | - | 0 | - |
| - | - | 1069 | 922.4 | - | - | 0 | - |
| - | - | 1311 | 923.4 | - | - | 0 | - |
| - | - | 1870 | 924.5 | - | - | 0 | - |
| 7 | c | 4895 | 925.5 | 0.0002493 | 0.2694 | +1 | 7 |
| - | - | 2575 | 926.5 | - | - | 0 | - |
| - | - | 1177 | 927.5 | - | - | 0 | - |
| - | - | 841.8 | 930.5 | - | - | 0 | - |
| - | - | 1940 | 931 | - | - | 0 | - |
| - | - | 1574 | 931.5 | - | - | 0 | - |
| - | - | 3264 | 932.5 | - | - | 0 | - |
| - | - | 1606 | 938 | - | - | 0 | - |
| - | - | 4692 | 938.5 | - | - | 0 | - |
| - | - | 2204 | 939 | - | - | 0 | - |
| 2 | z | 6187 | 939.4 | 0.01794 | 19.1 | +2 | 14 |
| - | - | 6547 | 939.9 | - | - | 0 | - |
| - | - | 4636 | 940.4 | - | - | 0 | - |
| - | - | 2141 | 940.9 | - | - | 0 | - |
| - | - | 957.2 | 941.5 | - | - | 0 | - |
| - | - | 1651 | 942.5 | - | - | 0 | - |
| - | - | 986.9 | 943.4 | - | - | 0 | - |
| - | - | 1161 | 943.9 | - | - | 0 | - |
| - | - | 1643 | 945.5 | - | - | 0 | - |
| - | - | 1967 | 946.5 | - | - | 0 | - |
| - | - | 1759 | 947 | - | - | 0 | - |
| 2 | y | 3090 | 947.4 | 0.01023 | 10.8 | +2 | 14 |
| - | - | 1093 | 948 | - | - | 0 | - |
| - | - | 1163 | 948.5 | - | - | 0 | - |
| - | - | 1192 | 949.5 | - | - | 0 | - |
| - | - | 910.2 | 950.4 | - | - | 0 | - |
| - | - | 1583 | 953 | - | - | 0 | - |
| - | - | 1635 | 953.5 | - | - | 0 | - |
| - | - | 1640 | 954 | - | - | 0 | - |
| - | - | 2654 | 954.5 | - | - | 0 | - |
| - | - | 2192 | 955 | - | - | 0 | - |
| - | - | 3027 | 955.5 | - | - | 0 | - |
| - | - | 2781 | 956 | - | - | 0 | - |
| - | - | 1680 | 957 | - | - | 0 | - |
| - | - | 2140 | 957.4 | - | - | 0 | - |
| - | - | 1906 | 959.5 | - | - | 0 | - |
| - | - | 3017 | 960 | - | - | 0 | - |
| - | - | 3951 | 960.5 | - | - | 0 | - |
| - | - | 1835 | 961 | - | - | 0 | - |
| - | - | 1256 | 961.5 | - | - | 0 | - |
| - | - | 1848 | 962 | - | - | 0 | - |
| - | - | 2572 | 962.5 | - | - | 0 | - |
| - | - | 2678 | 963 | - | - | 0 | - |
| - | - | 3090 | 963.5 | - | - | 0 | - |
| - | - | 2718 | 964 | - | - | 0 | - |
| 8 | c | 1776 | 964.5 | 0.01078 | 11.18 | +1 | 8 |
| - | - | 2013 | 965 | - | - | 0 | - |
| 8 | c | 1835 | 965.4 | 0.0124 | 12.85 | +1 | 8 |
| - | - | 1288 | 966 | - | - | 0 | - |
| - | - | 3069 | 966.5 | - | - | 0 | - |
| - | - | 2462 | 967 | - | - | 0 | - |
| - | - | 3848 | 967.5 | - | - | 0 | - |
| - | - | 4422 | 968 | - | - | 0 | - |
| 9 | w | 6473 | 968.4 | 0.01718 | 17.73 | +1 | 7 |
| - | - | 3685 | 969 | - | - | 0 | - |
| - | - | 3417 | 969.5 | - | - | 0 | - |
| - | - | 1166 | 970 | - | - | 0 | - |
| - | - | 3053 | 970.5 | - | - | 0 | - |
| - | - | 1791 | 971 | - | - | 0 | - |
| - | - | 2639 | 971.5 | - | - | 0 | - |
| - | - | 1559 | 972 | - | - | 0 | - |
| - | - | 3248 | 972.5 | - | - | 0 | - |
| - | - | 1243 | 973 | - | - | 0 | - |
| - | - | 2396 | 973.5 | - | - | 0 | - |
| - | - | 2450 | 974 | - | - | 0 | - |
| - | - | 2729 | 974.5 | - | - | 0 | - |
| - | - | 7929 | 975 | - | - | 0 | - |
| - | - | 8510 | 975.5 | - | - | 0 | - |
| - | - | 6552 | 976 | - | - | 0 | - |
| - | - | 7535 | 976.5 | - | - | 0 | - |
| - | - | 6614 | 977 | - | - | 0 | - |
| - | - | 5787 | 977.5 | - | - | 0 | - |
| - | - | 3933 | 978 | - | - | 0 | - |
| - | - | 2824 | 978.5 | - | - | 0 | - |
| - | - | 1652 | 979 | - | - | 0 | - |
| - | - | 1891 | 980 | - | - | 0 | - |
| - | - | 1599 | 980.5 | - | - | 0 | - |
| - | - | 1683 | 981 | - | - | 0 | - |
| - | - | 4472 | 981.5 | - | - | 0 | - |
| - | - | 3343 | 982 | - | - | 0 | - |
| 8 | c | 1.362E+04 | 982.5 | 0.005081 | 5.172 | +1 | 8 |
| - | - | 2981 | 983 | - | - | 0 | - |
| - | - | 7362 | 983.5 | - | - | 0 | - |
| - | - | 1241 | 984 | - | - | 0 | - |
| - | - | 5619 | 984.5 | - | - | 0 | - |
| - | - | 7816 | 985 | - | - | 0 | - |
| - | - | 8586 | 985.5 | - | - | 0 | - |
| - | - | 3986 | 986 | - | - | 0 | - |
| - | - | 3708 | 986.5 | - | - | 0 | - |
| - | - | 2497 | 987 | - | - | 0 | - |
| - | - | 2131 | 987.5 | - | - | 0 | - |
| - | - | 2670 | 988 | - | - | 0 | - |
| - | - | 4839 | 988.5 | - | - | 0 | - |
| - | - | 2282 | 989 | - | - | 0 | - |
| - | - | 4611 | 989.5 | - | - | 0 | - |
| - | - | 2040 | 990.5 | - | - | 0 | - |
| - | - | 1218 | 991.5 | - | - | 0 | - |
| - | - | 2293 | 993 | - | - | 0 | - |
| - | - | 7632 | 993.5 | - | - | 0 | - |
| - | - | 7738 | 994 | - | - | 0 | - |
| - | - | 5726 | 994.5 | - | - | 0 | - |
| - | - | 4081 | 995 | - | - | 0 | - |
| - | - | 2559 | 995.5 | - | - | 0 | - |
| - | - | 8644 | 996 | - | - | 0 | - |
| - | - | 1.668E+04 | 996.5 | - | - | 0 | - |
| - | - | 1.247E+04 | 997 | - | - | 0 | - |
| - | - | 1.006E+04 | 997.5 | - | - | 0 | - |
| - | - | 4813 | 998 | - | - | 0 | - |
| - | - | 4273 | 998.5 | - | - | 0 | - |
| - | - | 3169 | 999 | - | - | 0 | - |
| - | - | 2388 | 999.5 | - | - | 0 | - |
| - | - | 3680 | 1002 | - | - | 0 | - |
| - | - | 2904 | 1003 | - | - | 0 | - |
| - | - | 3811 | 1003 | - | - | 0 | - |
| - | - | 4263 | 1004 | - | - | 0 | - |
| - | - | 9629 | 1004 | - | - | 0 | - |
| - | - | 1.549E+04 | 1005 | - | - | 0 | - |
| - | - | 1.279E+04 | 1005 | - | - | 0 | - |
| - | - | 7256 | 1006 | - | - | 0 | - |
| - | - | 4463 | 1006 | - | - | 0 | - |
| - | - | 2404 | 1007 | - | - | 0 | - |
| - | - | 1972 | 1007 | - | - | 0 | - |
| - | - | 1082 | 1008 | - | - | 0 | - |
| 9 | z | 1728 | 1027 | 0.01041 | 10.14 | +1 | 7 |
| - | - | 3422 | 1028 | - | - | 0 | - |
| - | - | 1486 | 1029 | - | - | 0 | - |
| - | - | 1056 | 1030 | - | - | 0 | - |
| - | - | 1099 | 1067 | - | - | 0 | - |
| - | - | 1734 | 1085 | - | - | 0 | - |
| - | - | 3770 | 1110 | - | - | 0 | - |
| 9 | c | 1.597E+04 | 1111 | 0.002465 | 2.219 | +1 | 9 |
| - | - | 9736 | 1112 | - | - | 0 | - |
| - | - | 2894 | 1113 | - | - | 0 | - |
| - | - | 872.2 | 1197 | - | - | 0 | - |
| 7 | z | 2667 | 1199 | 0.008929 | 7.45 | +1 | 9 |
| - | - | 5582 | 1200 | - | - | 0 | - |
| - | - | 3534 | 1201 | - | - | 0 | - |
| - | - | 1157 | 1202 | - | - | 0 | - |
| - | - | 999.6 | 1216 | - | - | 0 | - |
| - | - | 933.4 | 1226 | - | - | 0 | - |
| - | - | 1252 | 1239 | - | - | 0 | - |
| 10 | c | 4533 | 1240 | 0.005404 | 4.359 | +1 | 10 |
| - | - | 3391 | 1241 | - | - | 0 | - |
| - | - | 2213 | 1242 | - | - | 0 | - |
| 6 | z | 1927 | 1312 | 0.007263 | 5.537 | +1 | 10 |
| - | - | 3269 | 1313 | - | - | 0 | - |
| - | - | 1705 | 1314 | - | - | 0 | - |
| - | - | 949 | 1359 | - | - | 0 | - |
| - | - | 1035 | 1402 | - | - | 0 | - |
| 11 | c | 1716 | 1403 | 0.004209 | 3.001 | +1 | 11 |
| - | - | 2215 | 1404 | - | - | 0 | - |
| - | - | 1162 | 1405 | - | - | 0 | - |
| - | - | 833.4 | 1406 | - | - | 0 | - |
| - | - | 873.5 | 1475 | - | - | 0 | - |
| 5 | z | 1300 | 1498 | 0.003877 | 2.589 | +1 | 11 |
| - | - | 2331 | 1499 | - | - | 0 | - |
| - | - | 1366 | 1500 | - | - | 0 | - |
| - | - | 1205 | 1515 | - | - | 0 | - |
| 4 | z | 1060 | 1613 | 0.009272 | 5.749 | +1 | 12 |
| - | - | 3363 | 1614 | - | - | 0 | - |
| - | - | 1959 | 1615 | - | - | 0 | - |
| - | - | 1114 | 1616 | - | - | 0 | - |
| 13 | c | 859 | 1692 | 0.0157 | 9.28 | +1 | 13 |
| - | - | 2296 | 1693 | - | - | 0 | - |
| - | - | 807.1 | 1695 | - | - | 0 | - |
| 3 | z | 1327 | 1741 | 0.001609 | 0.9242 | +1 | 13 |
| - | - | 2392 | 1742 | - | - | 0 | - |
| - | - | 2280 | 1743 | - | - | 0 | - |
| - | - | 1665 | 1744 | - | - | 0 | - |
| - | - | 981.5 | 1821 | - | - | 0 | - |
| - | - | 966.3 | 1879 | - | - | 0 | - |
| - | - | 772.4 | 1904 | - | - | 0 | - |
| - | - | 2050 | 1950 | - | - | 0 | - |
| - | - | 1777 | 1951 | - | - | 0 | - |
| - | - | 1136 | 1953 | - | - | 0 | - |
| - | - | 1186 | 1954 | - | - | 0 | - |
| - | - | 1051 | 1970 | - | - | 0 | - |
| - | - | 1034 | 1971 | - | - | 0 | - |
| - | - | 924.5 | 1972 | - | - | 0 | - |
| - | - | 820.8 | 1989 | - | - | 0 | - |
| - | - | 880.1 | 1992 | - | - | 0 | - |
| - | - | 1064 | 1993 | - | - | 0 | - |
| - | - | 739.5 | 1994 | - | - | 0 | - |
| - | - | 876.2 | 2008 | - | - | 0 | - |
| - | - | 1784 | 2009 | - | - | 0 | - |
| - | - | 1541 | 2010 | - | - | 0 | - |
| - | - | 1622 | 2011 | - | - | 0 | - |
| - | - | 831.7 | 3048 | - | - | 0 | - |
| - | - | 727.3 | 3048 | - | - | 0 | - |

m/z Charge Intensity FragmentType MassShift Position
126.06696319580078 0 429.1314
129.1025390625 0 3546.7751
130.06536865234375 0 2246.5876
131.36260986328125 0 382.7321
133.06076049804688 0 428.97433
133.08616638183594 0 1275.9016
136.0760040283203 0 722.9231
140.0823516845703 0 436.26303
144.4639892578125 0 464.5679
146.09286499023438 0 803.916
148.9470977783203 0 1067.3966
149.02391052246094 0 488.25885
149.04502868652344 0 1245.1785
155.09300231933594 0 3161.189
156.07757568359375 0 454.82562
156.91632080078125 0 516.742
159.0921173095703 0 775.71063
162.05499267578125 0 733.0351
164.08236694335938 0 462.73465
165.10272216796875 0 605.0873
166.0614013671875 0 3310.0044
168.1388397216797 0 1607.4196
173.0924530029297 0 1193.985
173.45025634765625 0 4353.9297
174.08802795410156 0 556.33997
176.35350036621094 0 489.20627
177.1128387451172 0 1103.0203
183.11312866210938 0 1499.1802
184.65748596191406 0 455.9182
185.1650848388672 0 2797.3633
187.10804748535156 0 1078.7114
191.10292053222656 0 1753.4258
199.1082763671875 0 1179.3348 c Ammonia loss 5
202.08273315429688 0 1624.3937
203.1028289794922 0 1548.4617
204.13497924804688 0 687.3815
205.11891174316406 0 869.20746 y 14
207.96127319335938 0 454.48993
209.09259033203125 0 914.5517
213.12332153320312 0 712.76514
215.13938903808594 0 9140.86
217.1551055908203 0 974.1633
217.33831787109375 0 600.599
218.1143341064453 0 1130.7343
218.1586151123047 0 524.0056
221.08445739746094 0 683.43634
223.15562438964844 0 2523.9343
227.10289001464844 0 3345.1282
229.15542602539062 0 566.1804
230.1611328125 0 597.2449
231.17076110839844 0 791.56586
234.12416076660156 0 2980.9216
235.1292724609375 0 589.529
242.1505584716797 0 668.6425
246.1813201904297 0 1208.0283
251.15069580078125 0 14972.269
252.15379333496094 0 1632.2151
261.1196594238281 0 901.73334
268.1771240234375 0 1510.22 c 1
274.1178894042969 0 763.1935
275.12396240234375 0 944.0164
281.0514831542969 0 1051.641
283.1519470214844 0 1120.4614
299.0622863769531 0 7005.028
300.0633850097656 0 1117.5931
300.1923828125 0 1711.5189
302.1716613769531 0 779.3627
317.1464538574219 0 797.9454
318.2040100097656 0 703.69586
319.1643981933594 0 551.3285
326.66461181640625 0 2006.183
333.2149353027344 0 1108.0691 y 13
338.1352233886719 0 1267.5216
340.1874084472656 0 1975.6989
340.6636657714844 0 674.474 c Ammonia loss 4
341.16259765625 0 499.15244
349.6809387207031 0 739.9987
359.0278625488281 0 803.71985
374.6941833496094 0 663.9974 y Water loss 4
378.20184326171875 0 746.9471
379.2093811035156 0 5220.039 c Ammonia loss 2
380.21221923828125 0 1451.7769
381.1521911621094 0 612.4363
383.207275390625 0 735.031
386.15582275390625 0 629.0261
387.2236022949219 0 2035.6477
393.1976623535156 0 1448.1337 y 10
396.2359313964844 0 5742.912 c 2
397.23968505859375 0 1023.83026
403.49945068359375 0 695.3849
407.1970520019531 0 749.55096
415.0380554199219 0 3122.9534
416.0382385253906 0 721.6178
420.202880859375 0 10043.66 w 9
420.2326354980469 0 392.25357
420.7045593261719 0 4435.549
421.20452880859375 0 1816.5947
428.2875061035156 0 782.2116
435.721923828125 0 632.23834
445.1958923339844 0 620.47797
447.92401123046875 0 768.75586
449.71856689453125 0 2012.9232 z 9
457.2664489746094 0 6556.659
458.23065185546875 0 3101.1702
458.27032470703125 0 3063.9128
458.7304992675781 0 1983.1766
459.2804870605469 0 5585.3857
460.28594970703125 0 1322.4258
460.735595703125 0 1555.598
461.2358093261719 0 763.23425
467.2488098144531 0 1287.2363
468.2628479003906 0 880.1143
469.2305603027344 0 1194.4655
472.3017883300781 0 1007.8528
478.2501220703125 0 773.3347
481.2317810058594 0 587.073
484.2444763183594 0 1149.8817
485.23193359375 0 2587.3252
485.3110656738281 0 785.2585
486.29302978515625 0 5697.8833
487.2413330078125 0 1261.6718
488.294189453125 0 1046.6472
489.23419189453125 0 911.25977
492.48846435546875 0 698.49866
493.74993896484375 0 3482.1807
494.237548828125 0 6577.309 c Ammonia loss 3
494.7453918457031 0 1287.922
495.2372741699219 0 1259.8745
496.2513427734375 0 611.6936
498.2567138671875 0 784.4034
498.74700927734375 0 1195.3539
499.24481201171875 0 1407.8893 y Water loss 4
501.21978759765625 0 2510.6284
501.75067138671875 0 1308.9888
502.2720947265625 0 4983.119
502.49853515625 0 5397.505
502.7514953613281 0 12232.711
502.9986267089844 0 3320.7556
503.1084289550781 0 3807.956
503.2498474121094 0 7091.476
503.3219299316406 0 832.0844
503.75738525390625 0 1245.1764
504.1068115234375 0 1326.5104
504.2640380859375 0 2046.7585
504.3029479980469 0 3120.8828
504.339599609375 0 971.32874
505.5876159667969 0 853.4808
506.2618103027344 0 986.8057
506.7278137207031 0 642.2198
507.2644958496094 0 934.0696
511.2634582519531 0 14418.312 c 3
511.3028259277344 0 992.57214
512.2652587890625 0 3839.9854
513.2696533203125 0 928.44794
515.3193969726562 0 1950.4215
519.2750854492188 0 762.4114
519.2984619140625 0 1504.8818
521.7667236328125 0 2901.4937 y 8
522.2697143554688 0 792.2058
522.7700805664062 0 834.95154
527.7677001953125 0 826.83734
529.2651977539062 0 835.9175
533.2569580078125 0 663.6565 z Water loss 7
535.2921142578125 0 1638.8097
536.2870483398438 0 637.23425
542.26611328125 0 1472.1093 z 7
543.2703857421875 0 825.42334
547.2897338867188 0 935.5733
547.7724609375 0 588.9025
548.23779296875 0 1124.3337 w 11
548.75390625 0 799.429
549.2625732421875 0 624.59235
550.2457275390625 0 1492.6919
550.2783813476562 0 2180.268 y 7
550.7778930664062 0 1574.4719
552.2601928710938 0 871.41626
555.26220703125 0 1190.0326
555.3007202148438 0 1976.553
555.78955078125 0 8854.62 c 8
556.2899780273438 0 4497.599
556.79296875 0 1207.255
563.2689819335938 0 634.5386
564.274169921875 0 5211.5405
564.7720336914062 0 2539.4048
565.2757568359375 0 898.5838
565.7708129882812 0 815.45953
568.7974853515625 0 868.16547
571.280517578125 0 2520.8352
571.781982421875 0 2271.7087
572.2784423828125 0 2066.3823
573.2723999023438 0 788.06085
577.281982421875 0 2485.3943 w 6
577.7830810546875 0 1908.8378
578.29052734375 0 1271.9021
580.2745971679688 0 1262.1891 y Water loss 2
590.627685546875 0 952.9637
590.77294921875 0 657.628 z Water loss 6
591.2853393554688 0 1541.4532
592.781005859375 0 794.82855
596.2890625 0 1069.4542
597.29443359375 0 2119.7512
597.6299438476562 0 937.344
597.9612426757812 0 876.32153
598.302001953125 0 763.74866
598.7870483398438 0 1053.9481 y Water loss 6
598.9632568359375 0 756.50085
599.2887573242188 0 2931.6772 y Ammonia loss 6
599.7825927734375 0 6085.6787 z 6
600.2833862304688 0 3560.1963
600.7843627929688 0 1697.3167
601.2943725585938 0 848.71814 c Water loss 13
601.3784790039062 0 750.601
601.9605102539062 0 771.5225
602.28955078125 0 903.6969
602.9476928710938 0 786.19696
604.967041015625 0 983.6616
605.2966918945312 0 1060.7782 y Ammonia loss 11
605.6286010742188 0 810.7398
606.3041381835938 0 6026.014 z 11
606.6345825195312 0 739.9537
606.9677124023438 0 2072.033
607.3016967773438 0 9289.015 c 13
607.63134765625 0 4762.2344
607.790771484375 0 7885.145 y 6
607.96533203125 0 3237.581
608.2941284179688 0 7591.4824
608.6292724609375 0 1080.9915
608.7928466796875 0 1815.4921
608.9572143554688 0 600.1674
609.2903442382812 0 994.3051
611.3004150390625 0 1669.5487 c Water loss 9
611.8067016601562 0 846.67957 c Ammonia loss 9
611.9697265625 0 1006.2181
612.3029174804688 0 2690.9531
612.62939453125 0 944.24023
612.8035888671875 0 1810.6959
613.6179809570312 0 1038.6357
615.3059692382812 0 924.73627
615.6278686523438 0 1146.1222
616.6298217773438 0 816.3006
617.289306640625 0 813.02686
619.2993774414062 0 1005.07074
619.8177490234375 0 3956.427
620.3112182617188 0 14370.472 c 9
620.6378784179688 0 809.7109
620.81298828125 0 7588.552
620.97412109375 0 1308.8417
621.3108520507812 0 4002.89
621.815185546875 0 1315.5679
621.9694213867188 0 941.0856
622.3202514648438 0 3934.5166 y 11
622.632080078125 0 809.2049
623.3282470703125 0 1096.1742
623.6314086914062 0 1147.0582
625.6424560546875 0 1214.828
625.9751586914062 0 2688.05 y Water loss 1
626.3096313476562 0 3055.0623
626.6397094726562 0 4301.158 z 1
626.8064575195312 0 705.5129
626.9635620117188 0 1478.5542
627.2955322265625 0 2992.218
627.6316528320312 0 922.16705
627.968017578125 0 1179.285
629.3013916015625 0 1580.382
629.6351928710938 0 588.26935
629.8084106445312 0 813.756
629.9671630859375 0 666.5975
630.3191528320312 0 990.21533
630.6367797851562 0 1051.0934
630.971923828125 0 1746.2678
631.3033447265625 0 2692.0522
631.63623046875 0 2171.4668
631.9675903320312 0 2728.0698 y 1
632.3018798828125 0 2480.2104
632.9760131835938 0 1563.321
633.2997436523438 0 959.31195
633.9700927734375 0 829.8621
634.3041381835938 0 2712.116
634.6401977539062 0 1083.4717
634.7984008789062 0 5769.729 w 5
634.9862060546875 0 1007.80273
635.30078125 0 5545.6807
635.6396484375 0 1952.9454
635.7984008789062 0 2351.4282
635.9678344726562 0 1959.6447
636.300537109375 0 2763.523
636.6375122070312 0 2577.5225
636.971435546875 0 2573.6738
637.304931640625 0 2334.3796
637.6392822265625 0 2278.809
637.9712524414062 0 2277.095
638.304931640625 0 1672.3558
638.6310424804688 0 1156.042
639.3097534179688 0 898.9095
639.9910278320312 0 2029.0936
640.3219604492188 0 14322.057
640.6553955078125 0 14505.254
640.9892578125 0 8351.79
641.318359375 0 5193.23
641.6472778320312 0 2953.399
641.9767456054688 0 2281.4216
642.3082885742188 0 3021.9956
642.6423950195312 0 3390.2605
642.971923828125 0 3333.673
643.3094482421875 0 1059.3903
643.9677124023438 0 1528.9645
644.3046264648438 0 2078.531
644.6392822265625 0 1771.7131
644.979736328125 0 2164.0454
645.3099975585938 0 3764.2598
645.6437377929688 0 6958.028
645.9791259765625 0 9486.685
646.3139038085938 0 6739.9976
646.64892578125 0 3882.501
646.9811401367188 0 2073.7832
647.3097534179688 0 1953.8578 z Water loss 5
647.641357421875 0 1203.8486
647.8126831054688 0 1308.955 z Ammonia loss 5
647.9752807617188 0 2782.354
648.3069458007812 0 2601.2017
648.6405029296875 0 1663.0297
648.9747314453125 0 2038.6437
649.3184204101562 0 1744.8223
649.6451416015625 0 4057.8186
649.9765014648438 0 9011.854
650.3167724609375 0 16794.932
650.651611328125 0 13759.524
650.98291015625 0 11534.715
651.3116455078125 0 8592.568
651.64697265625 0 9160.456
651.9798583984375 0 4219.214
652.3124389648438 0 4661.412
652.64697265625 0 2679.8662
652.9735717773438 0 915.2756
653.3095703125 0 2365.6794
653.6432495117188 0 1577.7937
653.9761962890625 0 986.1621
654.6559448242188 0 2057.8408
654.98681640625 0 1748.6329
655.3236083984375 0 1161.7655 y Water loss 5
656.3146362304688 0 8037.91 z 5
656.6419677734375 0 6234.261
656.8232421875 0 1580.8119
656.9771728515625 0 8487.53
657.3121337890625 0 7984.1587
657.645751953125 0 4384.451
657.9764404296875 0 3268.342
658.3062744140625 0 782.0361
658.6426391601562 0 1270.3695
658.9785766601562 0 2306.4653
659.3088989257812 0 2102.7583
659.6470336914062 0 1635.7239
659.9761352539062 0 1101.5845
660.313720703125 0 1252.8002
660.642333984375 0 900.3483
661.29931640625 0 1032.9286
661.9743041992188 0 1913.0192
662.3079223632812 0 4822.9824
662.6438598632812 0 7749.092
662.9783325195312 0 5989.6655
663.3121948242188 0 3629.6025
663.64697265625 0 2630.8337
663.8384399414062 0 1215.1914
663.9801025390625 0 2332.2805
664.3278198242188 0 5952.7427 y 5
664.6561889648438 0 3589.965
664.834228515625 0 3097.8691
664.9848022460938 0 3692.7112
665.326171875 0 2457.115
665.656494140625 0 2428.217
666.3010864257812 0 1626.1565
667.9782104492188 0 1202.5867
668.3158569335938 0 3456.2107
668.6488647460938 0 3250.3906
668.9835205078125 0 2974.5598
669.3280639648438 0 7584.7485
669.6621704101562 0 23424.328
669.9962768554688 0 39868.695
670.3289184570312 0 28128.09
670.6632690429688 0 13087.583
670.9964599609375 0 7452.457
671.32666015625 0 4111.7827
671.66162109375 0 1455.9744
671.9865112304688 0 1009.7412
673.354736328125 0 1018.3715
674.3204345703125 0 945.4875
680.3165283203125 0 4108.1133 c Ammonia loss 4
681.3197021484375 0 2169.8054
682.3348388671875 0 3119.351
683.3397216796875 0 1579.4911
690.3402709960938 0 1574.8376
691.3336791992188 0 1295.1154
691.83251953125 0 938.4561
694.83154296875 0 1029.0991
695.925048828125 0 807.7752
697.342041015625 0 16093.482 c 4
698.3452758789062 0 8286.149
699.3430786132812 0 2423.381
699.8427124023438 0 1063.6605
700.3427124023438 0 1672.6176
701.3478393554688 0 2941.0818
701.8433837890625 0 8676.398 c 10
702.3428955078125 0 9047.619
702.8487548828125 0 2541.4167
703.348388671875 0 1111.7992
708.8365478515625 0 782.54596
711.31396484375 0 822.8694
712.3341064453125 0 1104.4948
712.8434448242188 0 1206.343
713.334716796875 0 1582.9999
714.3328247070312 0 1026.1919
716.4048461914062 0 3956.706
717.4093017578125 0 1398.546
721.3370971679688 0 1154.1843
721.8363647460938 0 879.87897
722.345458984375 0 1165.2985
723.3413696289062 0 1215.2375
735.3375244140625 0 882.0807
742.837890625 0 894.79724
746.3402099609375 0 950.03613
748.8684692382812 0 904.6935 y Ammonia loss 4
749.3636474609375 0 6938.42 z 4
749.86328125 0 3182.4172
750.366943359375 0 2309.143
752.3646850585938 0 1163.7163 z Ammonia loss 10
754.3682250976562 0 1028.2517
754.8633422851562 0 2183.566
755.3624877929688 0 1780.1958
757.3729858398438 0 2002.169 c Ammonia loss 11
757.8780517578125 0 1921.5259
758.3751831054688 0 1682.6105
765.8892822265625 0 1651.6523 c 11
766.3958740234375 0 2057.1794
768.3593139648438 0 885.13214 y Ammonia loss 10
769.3681640625 0 10007.624 z 10
770.3693237304688 0 5912.7266
771.364501953125 0 3040.9739
784.8784790039062 0 2277.9104
785.3853759765625 0 4846.577 y 10
786.3881225585938 0 1224.3898
788.3511352539062 0 815.59235
793.4015502929688 0 1619.6975 c Ammonia loss 5
797.3780517578125 0 955.799
803.42529296875 0 2416.8203
806.3856201171875 0 2732.4304 y Ammonia loss 3
806.879150390625 0 7939.0806 z 3
807.3801879882812 0 6023.4126
807.8794555664062 0 3494.0972
808.3761596679688 0 1053.7723
809.4092407226562 0 813.36237
810.42626953125 0 25861.494 c 5
811.3624267578125 0 1474.9111
811.4283447265625 0 9386.82
812.36962890625 0 1206.0066
812.4313354492188 0 2539.0378
814.4069213867188 0 2329.9321
814.888916015625 0 1865.2001 y 3
815.396484375 0 1409.7549
826.4139404296875 0 1050.0525
827.38916015625 0 1027.9353
831.4468994140625 0 1840.5083
831.7852783203125 0 1778.064
832.4435424804688 0 1064.5905
837.455322265625 0 3705.071
837.7884521484375 0 4047.7117
838.12255859375 0 1292.3043
838.4600830078125 0 1076.4196
839.3885498046875 0 804.99615
840.4005126953125 0 1682.7268
842.4051513671875 0 2244.7007
843.3998413085938 0 842.5665
845.3863525390625 0 823.69147
845.9020385742188 0 2258.3088
846.3993530273438 0 6433.274 c 12
846.8984375 0 4051.304
847.3997192382812 0 2267.0212
847.9038696289062 0 1176.5222
848.914306640625 0 1257.2903
850.9686279296875 0 1153.5817
854.3948974609375 0 896.3109
860.41552734375 0 742.12164
862.8931884765625 0 1049.8579
867.4256591796875 0 3044.6519
868.4288330078125 0 1179.3993
869.43359375 0 938.62006
870.409912109375 0 2527.6125 y Ammonia loss 2
870.9051513671875 0 8372.79 z 2
871.4086303710938 0 5106.2627
871.9085693359375 0 2827.9583
872.4104614257812 0 2172.7314
873.4202880859375 0 945.49023
878.4033813476562 0 829.73834
878.8971557617188 0 883.07227
881.4369506835938 0 4130.6436
883.4124145507812 0 1257.4559
888.4348754882812 0 1191.5004
888.9468383789062 0 903.44556
889.90087890625 0 1074.9478
898.4122314453125 0 10613.153 z 9
899.4158325195312 0 9752.856
900.4212646484375 0 4902.4854
901.4229125976562 0 1176.2518 c Water loss 13
902.4386596679688 0 956.7333
908.4213256835938 0 1195.1694 c Ammonia loss 6
910.4445190429688 0 4009.468 c 13
910.9445190429688 0 4575.4053
911.4409790039062 0 2201.172
911.9481201171875 0 1264.5817
913.4296875 0 824.7024
914.4334106445312 0 1476.1993 y 9
915.4432373046875 0 2244.99
917.9276123046875 0 1061.2623
918.9452514648438 0 949.0944
919.44384765625 0 1173.9178
922.4402465820312 0 1068.9565
923.4425659179688 0 1310.9194
924.4527587890625 0 1869.8911
925.452392578125 0 4895.0376 c 6
926.4501342773438 0 2574.511
927.4535522460938 0 1176.9163
930.4576416015625 0 841.84546
930.9761962890625 0 1939.8994
931.4744262695312 0 1574.4056
932.4783325195312 0 3263.888
937.9619750976562 0 1605.8823
938.4710083007812 0 4692.218
938.9580078125 0 2203.8188
939.4567260742188 0 6186.984 z 1
939.94287109375 0 6546.937
940.4462280273438 0 4636.498
940.9430541992188 0 2140.5032
941.4524536132812 0 957.2423
942.4513549804688 0 1651.37
943.431884765625 0 986.931
943.936279296875 0 1161.1434
945.479248046875 0 1642.6774
946.4766845703125 0 1967.1074
946.9603881835938 0 1759.1664
947.4583740234375 0 3090.3188 y 1
947.951171875 0 1092.8984
948.466552734375 0 1162.9661
949.4563598632812 0 1191.76
950.448974609375 0 910.2247
952.9703369140625 0 1582.941
953.4674072265625 0 1634.971
953.9649658203125 0 1639.886
954.4502563476562 0 2654.0732
954.9580078125 0 2192.2996
955.4570922851562 0 3026.988
955.9555053710938 0 2781.3254
956.95556640625 0 1680.3667
957.442138671875 0 2139.9783
959.4534912109375 0 1905.9117
959.974853515625 0 3017.2385
960.4719848632812 0 3950.8303
960.9708251953125 0 1835.3704
961.4734497070312 0 1256.1558
961.9666748046875 0 1847.9072
962.45556640625 0 2571.9277
962.9544677734375 0 2677.9216
963.4569091796875 0 3090.2417
963.9570922851562 0 2717.8325
964.4527587890625 0 1776.1301 c Water loss 7
964.9583740234375 0 2013.4207
965.4599609375 0 1834.6611 c Ammonia loss 7
965.9641723632812 0 1288.3523
966.4705200195312 0 3068.717
966.9686279296875 0 2461.7542
967.47119140625 0 3847.6812
967.9617919921875 0 4421.558
968.465576171875 0 6473.0737 w 8
968.9663696289062 0 3684.5044
969.4641723632812 0 3416.791
969.963623046875 0 1166.4797
970.4536743164062 0 3053.3906
970.953857421875 0 1791.1864
971.4591674804688 0 2639.224
971.9558715820312 0 1559.1453
972.4622192382812 0 3248.051
972.956787109375 0 1243.3829
973.4642333984375 0 2396.3967
973.9701538085938 0 2450.394
974.4829711914062 0 2728.6587
974.981689453125 0 7929.46
975.4811401367188 0 8510.458
975.9788208007812 0 6552.1885
976.4705200195312 0 7534.8516
976.9646606445312 0 6613.8057
977.4656372070312 0 5787.3135
977.9677124023438 0 3933.4348
978.4713134765625 0 2823.6365
978.9625854492188 0 1652.4308
979.9625244140625 0 1890.7487
980.4629516601562 0 1599.1273
980.9637451171875 0 1682.5204
981.4813842773438 0 4471.577
981.9873046875 0 3342.7383
982.4791870117188 0 13616.299 c 7
982.9828491210938 0 2981.248
983.4791870117188 0 7362.4224
983.9710083007812 0 1240.9252
984.4650268554688 0 5619.368
984.9603271484375 0 7816.0723
985.46435546875 0 8586.444
985.9611206054688 0 3985.7659
986.4661865234375 0 3708.1643
986.9666748046875 0 2497.347
987.4678344726562 0 2131.3403
987.9650268554688 0 2670.248
988.476318359375 0 4839.17
988.968505859375 0 2281.771
989.4764404296875 0 4610.793
990.4683837890625 0 2039.6858
991.4663696289062 0 1218.2069
992.962646484375 0 2293.0398
993.46435546875 0 7631.888
993.9653930664062 0 7737.924
994.4656372070312 0 5726.4536
994.9635620117188 0 4081.3914
995.4717407226562 0 2558.6482
995.98095703125 0 8644.338
996.4811401367188 0 16681.256
996.9788818359375 0 12470.8545
997.4786376953125 0 10060.2295
997.9778442382812 0 4812.8
998.4763793945312 0 4273.4937
998.974853515625 0 3169.0027
999.4666748046875 0 2387.6428
1002.4686279296875 0 3680.2273
1002.9747314453125 0 2903.7727
1003.4829711914062 0 3810.8892
1003.9869995117188 0 4263.2275
1004.4913940429688 0 9628.706
1004.9913940429688 0 15493.572
1005.494873046875 0 12786.95
1005.9927368164062 0 7255.55
1006.4944458007812 0 4463.1826
1006.98388671875 0 2403.8071
1007.4776611328125 0 1971.7418
1008.4650268554688 0 1082.2314
1026.503662109375 0 1727.5255 z 8
1027.5107421875 0 3421.8267
1028.514892578125 0 1485.8641
1029.515380859375 0 1055.519
1066.55810546875 0 1099.0559
1084.5380859375 0 1733.9109
1109.58251953125 0 3770.2295
1110.571533203125 0 15973.993 c 8
1111.57421875 0 9735.782
1112.566162109375 0 2893.744
1196.59765625 0 872.2358
1198.5714111328125 0 2666.9797 z 6
1199.5662841796875 0 5581.515
1200.5660400390625 0 3533.778
1201.5640869140625 0 1156.7554
1215.5894775390625 0 999.56085
1225.613525390625 0 933.41296
1238.6182861328125 0 1251.6587
1239.6170654296875 0 4533.187 c 9
1240.61572265625 0 3391.013
1241.6202392578125 0 2213.1663
1311.65380859375 0 1927.1901 z 5
1312.6488037109375 0 3268.853
1313.649169921875 0 1705.3224
1358.6761474609375 0 949.01807
1401.6640625 0 1034.7388
1402.67919921875 0 1716.4648 c 10
1403.6822509765625 0 2214.7695
1404.680908203125 0 1161.7673
1405.6842041015625 0 833.44147
1474.68115234375 0 873.4744
1497.729736328125 0 1299.689 z 4
1498.721923828125 0 2330.9062
1499.7296142578125 0 1366.2976
1514.74853515625 0 1204.7748
1612.7435302734375 0 1059.8646 z 3
1613.7520751953125 0 3362.7239
1614.7550048828125 0 1959.0723
1615.74658203125 0 1113.6248
1691.7734375 0 858.97986 c 12
1692.8023681640625 0 2296.3696
1694.7911376953125 0 807.1461
1740.81298828125 0 1326.5327 z 2
1741.8167724609375 0 2391.7593
1742.8077392578125 0 2279.9697
1743.818115234375 0 1665.075
1820.8814697265625 0 981.527
1878.895751953125 0 966.2675
1903.914794921875 0 772.411
1949.9620361328125 0 2050.3223
1950.940185546875 0 1777.081
1952.9256591796875 0 1135.8945
1953.9234619140625 0 1186.0474
1969.9090576171875 0 1051.486
1970.91552734375 0 1034.0211
1971.9010009765625 0 924.5437
1988.9339599609375 0 820.80334
1991.91455078125 0 880.08954
1992.976318359375 0 1063.9285
1993.9427490234375 0 739.4827
2007.982666015625 0 876.17786
2008.9696044921875 0 1784.3684
2009.97900390625 0 1540.6205
2010.984619140625 0 1621.7006
3047.835205078125 0 831.72375
3048.419921875 0 727.2502

Spectrum Details

|  |  |
| --- | --- |
| Matched peaks? Matched peaks The total absolute number of peaks matched. Additionally in brackets the total fraction of peaks matched and the total number of peaks is shown. | 85 (12.04% of 706) |
| FDR? FDR The false discovery rate estimated for this peptide. It is calculated by matching all theoretical fragments with a non-integer shift with the raw peaks for this spectrum. This is done with 40 different shifts. The resulting percentage is the average number of annotated peaks over the number of annotated peaks with the correct spectrum. | 0.59% |
| Satellite FDR? Satellite FDR See the FDR for details on its calculation. This satellite ion specific FDR only contains the satellite ions (d/w) for I/L/J positions. | 0.00% |
| PSM Score? PSM Score The PSM Score as given by Hecklib to this annotated spectrum. It is shown with three significant figures. | 419 |

#### Spectrum 5830? Spectrum 5830 The raw spectrum of this peptide as annotated by Hecklib. The fragments are coloured according to ion type (see legend). Any peaks with a star '\*' as text can be hovered over to see the full details, first the ion type second the mass shift type. By hovering over the amino acids in the peptide or ions in the legend the corresponding peaks are highlighted. By toggling the 'Unassigned' label you can turn the background (unassigned) peaks on or off in the plot. By updating the slider in the Ion legend you can update the spectrum to only show the top X% of the peaks with labels. The top X% means any peak that is within X% of the highest intensity. By dragging in the spectrum you can zoom in to a specific part of the spectrum and use 'Zoom Out' to get back to the original zoom level. The annotation of the spectrum is based on the given sequence in the peptides file and is done with different software so inconsistencies are likely. The peaks are annotated based on the given sequence, with 20 ppm tolerance.

Copy Data

##### Spectrum 5830 (TSV)

###### Preview

```
Loading example...
```

*Click on the button to copy the data to your clipboard.*

Mz MinMz MaxIntensity Max

WidthHeightPeptide font sizePeptide stroke widthSpectrum font sizeSpectrum stroke widthCompact peptide

Ion legend

wxyz

abcd

OtherUnassignedIon^Charge^_Position_Show for top:%

JHQDWLDGKEYKCKK

04.37e+38.74e+31.31e+41.75e+4

Zoom Out

c^+3^_3_c^+1^_3_c^+1^_3_w^+2^_6_z^+2^_6_y^+2^_6_c^+1^_4_c^+1^_4_y^+2^_7_z^+2^_8_y^+2^_8_c^+2^_9_w^+2^_9_y^+2^_9_z^+2^_9_z^+1^_4_c^+3^_14_y^+2^_9_c^+2^_10_y^+1^_4_y^+3^_14_w^+2^_10_z^+2^_10_y^+2^_10_c^+1^_5_c^+1^_5_c^+2^_11_z^+2^_11_c^+2^_12_z^+1^_5_y^+1^_5_y^+2^_12_z^+2^_12_c^+1^_6_w^+1^_6_c^+2^_13_y^+2^_13_z^+2^_13_z^+1^_6_c^+2^_14_c^+1^_7_z^+2^_14_c^+1^_8_z^+1^_7_c^+1^_9_z^+1^_9_c^+1^_10_z^+1^_10_y^+1^_11_z^+1^_11_c^+1^_13_z^+1^_13_

0508101515232030

Fragment Matches Table

Show background peaks

| Position | Ion type | Intensity | mz Theoretical | mz Error (Th) | mz Error (ppm) | Charge | Series Number |
| --- | --- | --- | --- | --- | --- | --- | --- |
| - | - | 601.7 | 129.1 | - | - | 0 | - |
| - | - | 989.1 | 129.1 | - | - | 0 | - |
| - | - | 1076 | 130.1 | - | - | 0 | - |
| - | - | 395.4 | 131.1 | - | - | 0 | - |
| - | - | 830 | 132.1 | - | - | 0 | - |
| 3 | c | 475.7 | 132.7 | 0.001274 | 9.599 | +3 | 3 |
| - | - | 1552 | 133.1 | - | - | 0 | - |
| - | - | 422.8 | 133.1 | - | - | 0 | - |
| - | - | 432.8 | 134 | - | - | 0 | - |
| - | - | 458.3 | 140.1 | - | - | 0 | - |
| - | - | 444.5 | 145.3 | - | - | 0 | - |
| - | - | 368.5 | 147.9 | - | - | 0 | - |
| - | - | 390 | 148.5 | - | - | 0 | - |
| - | - | 669.8 | 148.9 | - | - | 0 | - |
| - | - | 1258 | 149 | - | - | 0 | - |
| - | - | 1005 | 155.1 | - | - | 0 | - |
| - | - | 689.6 | 156.1 | - | - | 0 | - |
| - | - | 444.6 | 162.3 | - | - | 0 | - |
| - | - | 953.9 | 166.1 | - | - | 0 | - |
| - | - | 788.8 | 174.1 | - | - | 0 | - |
| - | - | 1359 | 177.1 | - | - | 0 | - |
| - | - | 2059 | 185.2 | - | - | 0 | - |
| - | - | 699.5 | 187.1 | - | - | 0 | - |
| - | - | 1571 | 191.1 | - | - | 0 | - |
| - | - | 538.3 | 194.3 | - | - | 0 | - |
| - | - | 839.3 | 199.2 | - | - | 0 | - |
| - | - | 737.7 | 201.1 | - | - | 0 | - |
| - | - | 1642 | 202.1 | - | - | 0 | - |
| - | - | 3784 | 203.1 | - | - | 0 | - |
| - | - | 1106 | 207.1 | - | - | 0 | - |
| - | - | 8607 | 215.1 | - | - | 0 | - |
| - | - | 710.1 | 219.1 | - | - | 0 | - |
| - | - | 589.8 | 223.2 | - | - | 0 | - |
| - | - | 560.5 | 227.1 | - | - | 0 | - |
| - | - | 499.1 | 228.9 | - | - | 0 | - |
| - | - | 2328 | 229.2 | - | - | 0 | - |
| - | - | 605.6 | 230.2 | - | - | 0 | - |
| - | - | 661.1 | 234.1 | - | - | 0 | - |
| - | - | 1237 | 239.1 | - | - | 0 | - |
| - | - | 4487 | 251.2 | - | - | 0 | - |
| - | - | 481.7 | 269.2 | - | - | 0 | - |
| - | - | 537.4 | 276.3 | - | - | 0 | - |
| - | - | 1927 | 281.1 | - | - | 0 | - |
| - | - | 631.8 | 282 | - | - | 0 | - |
| - | - | 582.4 | 282.2 | - | - | 0 | - |
| - | - | 4265 | 289.1 | - | - | 0 | - |
| - | - | 7562 | 299.1 | - | - | 0 | - |
| - | - | 581.5 | 299.1 | - | - | 0 | - |
| - | - | 1681 | 300.1 | - | - | 0 | - |
| - | - | 1316 | 302.2 | - | - | 0 | - |
| - | - | 3053 | 314.2 | - | - | 0 | - |
| - | - | 579.2 | 318.2 | - | - | 0 | - |
| - | - | 1118 | 326.7 | - | - | 0 | - |
| - | - | 536.4 | 342.2 | - | - | 0 | - |
| - | - | 1902 | 359 | - | - | 0 | - |
| - | - | 839.8 | 360 | - | - | 0 | - |
| 3 | c | 1894 | 379.2 | 0.000761 | 2.007 | +1 | 3 |
| - | - | 1157 | 387.2 | - | - | 0 | - |
| 3 | c | 1925 | 396.2 | 0.0008208 | 2.072 | +1 | 3 |
| - | - | 635.6 | 407.7 | - | - | 0 | - |
| - | - | 654.1 | 414.2 | - | - | 0 | - |
| - | - | 3798 | 415 | - | - | 0 | - |
| - | - | 1257 | 416 | - | - | 0 | - |
| 10 | w | 5510 | 420.2 | 0.004302 | 10.24 | +2 | 6 |
| - | - | 2186 | 420.7 | - | - | 0 | - |
| - | - | 577.5 | 438.2 | - | - | 0 | - |
| - | - | 903.4 | 444.2 | - | - | 0 | - |
| - | - | 841.7 | 444.3 | - | - | 0 | - |
| - | - | 835.1 | 449.2 | - | - | 0 | - |
| 10 | z | 1447 | 449.7 | 0.006197 | 13.78 | +2 | 6 |
| - | - | 602.6 | 450.2 | - | - | 0 | - |
| 10 | y | 620.9 | 457.7 | 0.004234 | 9.249 | +2 | 6 |
| - | - | 3645 | 458.2 | - | - | 0 | - |
| - | - | 1140 | 458.7 | - | - | 0 | - |
| - | - | 1429 | 459.3 | - | - | 0 | - |
| - | - | 716.1 | 460.7 | - | - | 0 | - |
| - | - | 860.1 | 480.7 | - | - | 0 | - |
| - | - | 834.2 | 484.2 | - | - | 0 | - |
| - | - | 876.6 | 485.2 | - | - | 0 | - |
| - | - | 805 | 487.2 | - | - | 0 | - |
| - | - | 1759 | 488.3 | - | - | 0 | - |
| - | - | 2627 | 493.7 | - | - | 0 | - |
| 4 | c | 2206 | 494.2 | 0.007849 | 15.88 | +1 | 4 |
| - | - | 967.8 | 495.3 | - | - | 0 | - |
| - | - | 768.5 | 496.2 | - | - | 0 | - |
| - | - | 2092 | 501.3 | - | - | 0 | - |
| - | - | 805.9 | 501.7 | - | - | 0 | - |
| - | - | 590 | 502.2 | - | - | 0 | - |
| - | - | 2273 | 502.3 | - | - | 0 | - |
| - | - | 2704 | 502.5 | - | - | 0 | - |
| - | - | 5071 | 502.8 | - | - | 0 | - |
| - | - | 2151 | 503 | - | - | 0 | - |
| - | - | 3080 | 503.1 | - | - | 0 | - |
| - | - | 4307 | 503.3 | - | - | 0 | - |
| - | - | 2124 | 503.3 | - | - | 0 | - |
| - | - | 972.5 | 503.8 | - | - | 0 | - |
| - | - | 2030 | 504.1 | - | - | 0 | - |
| - | - | 1252 | 504.3 | - | - | 0 | - |
| 4 | c | 6172 | 511.3 | 0.0006032 | 1.18 | +1 | 4 |
| - | - | 1528 | 512.3 | - | - | 0 | - |
| - | - | 1123 | 515.3 | - | - | 0 | - |
| - | - | 608.8 | 517.3 | - | - | 0 | - |
| 9 | y | 1192 | 521.8 | 0.004779 | 9.159 | +2 | 7 |
| - | - | 888.6 | 522.3 | - | - | 0 | - |
| 8 | z | 875.5 | 542.3 | 0.001693 | 3.123 | +2 | 8 |
| - | - | 664.5 | 545.4 | - | - | 0 | - |
| - | - | 1338 | 547.3 | - | - | 0 | - |
| 8 | y | 1069 | 550.3 | 0.0019 | 3.453 | +2 | 8 |
| - | - | 1514 | 555.3 | - | - | 0 | - |
| 9 | c | 5013 | 555.8 | 0.001012 | 1.821 | +2 | 9 |
| - | - | 1767 | 556.3 | - | - | 0 | - |
| - | - | 906.5 | 556.8 | - | - | 0 | - |
| - | - | 853.9 | 570.3 | - | - | 0 | - |
| - | - | 1726 | 571.3 | - | - | 0 | - |
| 7 | w | 980.9 | 577.3 | 0.002849 | 4.935 | +2 | 9 |
| - | - | 822.2 | 577.8 | - | - | 0 | - |
| - | - | 705.4 | 578.3 | - | - | 0 | - |
| - | - | 784.3 | 596.3 | - | - | 0 | - |
| 7 | y | 2528 | 599.3 | 0.007119 | 11.88 | +2 | 9 |
| 7 | z | 3996 | 599.8 | 0.002592 | 4.321 | +2 | 9 |
| - | - | 864.7 | 600.3 | - | - | 0 | - |
| 12 | z | 4184 | 606.3 | 0.008198 | 13.52 | +1 | 4 |
| 14 | c | 4927 | 607.3 | 0.0008638 | 1.422 | +3 | 14 |
| - | - | 2357 | 607.6 | - | - | 0 | - |
| 7 | y | 4316 | 607.8 | 0.003592 | 5.91 | +2 | 9 |
| - | - | 1790 | 608 | - | - | 0 | - |
| - | - | 2807 | 608.3 | - | - | 0 | - |
| - | - | 1004 | 608.8 | - | - | 0 | - |
| - | - | 1202 | 612.3 | - | - | 0 | - |
| - | - | 641.1 | 612.6 | - | - | 0 | - |
| - | - | 1072 | 613.8 | - | - | 0 | - |
| - | - | 616.5 | 614.1 | - | - | 0 | - |
| - | - | 1748 | 619.8 | - | - | 0 | - |
| 10 | c | 4426 | 620.3 | 0.001139 | 1.836 | +2 | 10 |
| - | - | 3394 | 620.8 | - | - | 0 | - |
| - | - | 1498 | 621.3 | - | - | 0 | - |
| 12 | y | 1882 | 622.3 | 0.008856 | 14.23 | +1 | 4 |
| - | - | 706.4 | 623.3 | - | - | 0 | - |
| 2 | y | 1578 | 626.3 | 0.01186 | 18.94 | +3 | 14 |
| - | - | 865 | 627 | - | - | 0 | - |
| - | - | 1717 | 634.3 | - | - | 0 | - |
| - | - | 922.2 | 634.6 | - | - | 0 | - |
| 6 | w | 2116 | 634.8 | 0.001062 | 1.672 | +2 | 10 |
| - | - | 1184 | 635.3 | - | - | 0 | - |
| - | - | 1641 | 635.8 | - | - | 0 | - |
| - | - | 1167 | 636.7 | - | - | 0 | - |
| - | - | 760.8 | 637.3 | - | - | 0 | - |
| - | - | 2020 | 640 | - | - | 0 | - |
| - | - | 6804 | 640.3 | - | - | 0 | - |
| - | - | 5176 | 640.7 | - | - | 0 | - |
| - | - | 3685 | 641 | - | - | 0 | - |
| - | - | 1493 | 641.3 | - | - | 0 | - |
| - | - | 1181 | 645.6 | - | - | 0 | - |
| - | - | 1026 | 646 | - | - | 0 | - |
| - | - | 1480 | 649.6 | - | - | 0 | - |
| - | - | 4878 | 650 | - | - | 0 | - |
| - | - | 8802 | 650.3 | - | - | 0 | - |
| - | - | 4874 | 650.7 | - | - | 0 | - |
| - | - | 3201 | 651 | - | - | 0 | - |
| - | - | 2207 | 651.3 | - | - | 0 | - |
| - | - | 1161 | 651.6 | - | - | 0 | - |
| - | - | 849.8 | 652 | - | - | 0 | - |
| - | - | 1037 | 653.7 | - | - | 0 | - |
| - | - | 1745 | 654.3 | - | - | 0 | - |
| - | - | 952.3 | 654.7 | - | - | 0 | - |
| - | - | 815.6 | 655 | - | - | 0 | - |
| 6 | z | 1830 | 656.3 | 0.00312 | 4.753 | +2 | 10 |
| - | - | 839.4 | 656.8 | - | - | 0 | - |
| - | - | 1106 | 657 | - | - | 0 | - |
| - | - | 991.6 | 659.7 | - | - | 0 | - |
| - | - | 768.2 | 660 | - | - | 0 | - |
| - | - | 2061 | 664 | - | - | 0 | - |
| 6 | y | 3905 | 664.3 | 0.004425 | 6.661 | +2 | 10 |
| - | - | 2218 | 664.7 | - | - | 0 | - |
| - | - | 1379 | 664.8 | - | - | 0 | - |
| - | - | 798.7 | 665 | - | - | 0 | - |
| - | - | 732.9 | 668 | - | - | 0 | - |
| - | - | 1364 | 669 | - | - | 0 | - |
| - | - | 4908 | 669.3 | - | - | 0 | - |
| - | - | 1.488E+04 | 669.7 | - | - | 0 | - |
| - | - | 1.73E+04 | 670 | - | - | 0 | - |
| - | - | 1.25E+04 | 670.3 | - | - | 0 | - |
| - | - | 5827 | 670.7 | - | - | 0 | - |
| - | - | 4482 | 671 | - | - | 0 | - |
| - | - | 1520 | 672.3 | - | - | 0 | - |
| - | - | 1222 | 674.3 | - | - | 0 | - |
| - | - | 965.4 | 679.8 | - | - | 0 | - |
| 5 | c | 2078 | 680.3 | 0.001487 | 2.186 | +1 | 5 |
| - | - | 839 | 690.3 | - | - | 0 | - |
| 5 | c | 6879 | 697.3 | 0.0006315 | 0.9055 | +1 | 5 |
| - | - | 1930 | 698.3 | - | - | 0 | - |
| - | - | 2583 | 701.3 | - | - | 0 | - |
| 11 | c | 3833 | 701.8 | 0.003349 | 4.772 | +2 | 11 |
| - | - | 2540 | 702.3 | - | - | 0 | - |
| - | - | 1447 | 702.8 | - | - | 0 | - |
| - | - | 3096 | 716.4 | - | - | 0 | - |
| - | - | 1593 | 717.4 | - | - | 0 | - |
| - | - | 785.8 | 726.4 | - | - | 0 | - |
| - | - | 1189 | 744.4 | - | - | 0 | - |
| 5 | z | 1169 | 749.4 | 0.005972 | 7.969 | +2 | 11 |
| - | - | 2057 | 749.9 | - | - | 0 | - |
| - | - | 1232 | 750.4 | - | - | 0 | - |
| - | - | 825.9 | 750.9 | - | - | 0 | - |
| - | - | 1164 | 757.9 | - | - | 0 | - |
| - | - | 918.7 | 765.4 | - | - | 0 | - |
| 12 | c | 850.3 | 765.9 | 0.00439 | 5.733 | +2 | 12 |
| - | - | 1642 | 766.4 | - | - | 0 | - |
| 11 | z | 4944 | 769.4 | 0.01031 | 13.4 | +1 | 5 |
| - | - | 3335 | 770.4 | - | - | 0 | - |
| - | - | 1330 | 771.4 | - | - | 0 | - |
| 11 | y | 2023 | 785.4 | 0.003398 | 4.326 | +1 | 5 |
| - | - | 2043 | 803.4 | - | - | 0 | - |
| 4 | y | 916.3 | 806.4 | 0.003634 | 4.507 | +2 | 12 |
| 4 | z | 2301 | 806.9 | 0.0008278 | 1.026 | +2 | 12 |
| - | - | 1787 | 807.4 | - | - | 0 | - |
| - | - | 987.7 | 807.9 | - | - | 0 | - |
| 6 | c | 1.216E+04 | 810.4 | 0.001016 | 1.254 | +1 | 6 |
| - | - | 5637 | 811.4 | - | - | 0 | - |
| - | - | 1623 | 812.4 | - | - | 0 | - |
| - | - | 2625 | 814.4 | - | - | 0 | - |
| - | - | 1520 | 815.4 | - | - | 0 | - |
| - | - | 748.8 | 817.8 | - | - | 0 | - |
| - | - | 1573 | 831.4 | - | - | 0 | - |
| - | - | 1103 | 832.4 | - | - | 0 | - |
| - | - | 2165 | 837.8 | - | - | 0 | - |
| - | - | 1824 | 838.1 | - | - | 0 | - |
| 10 | w | 1007 | 839.4 | 0.002854 | 3.4 | +1 | 6 |
| 13 | c | 3186 | 846.4 | 0.004198 | 4.959 | +2 | 13 |
| - | - | 2321 | 846.9 | - | - | 0 | - |
| - | - | 1220 | 847.4 | - | - | 0 | - |
| - | - | 910.3 | 847.9 | - | - | 0 | - |
| - | - | 1734 | 867.4 | - | - | 0 | - |
| 3 | y | 1046 | 870.4 | 0.01103 | 12.67 | +2 | 13 |
| 3 | z | 3345 | 870.9 | 0.0007586 | 0.8711 | +2 | 13 |
| - | - | 2102 | 871.4 | - | - | 0 | - |
| - | - | 1331 | 871.9 | - | - | 0 | - |
| - | - | 782.4 | 881.4 | - | - | 0 | - |
| - | - | 840.5 | 887.9 | - | - | 0 | - |
| - | - | 1131 | 888.4 | - | - | 0 | - |
| - | - | 1039 | 889.4 | - | - | 0 | - |
| - | - | 1105 | 890.4 | - | - | 0 | - |
| 10 | z | 6589 | 898.4 | 0.009566 | 10.65 | +1 | 6 |
| - | - | 3668 | 899.4 | - | - | 0 | - |
| - | - | 2344 | 900.4 | - | - | 0 | - |
| 14 | c | 1673 | 910.4 | 0.003164 | 3.475 | +2 | 14 |
| - | - | 2542 | 910.9 | - | - | 0 | - |
| - | - | 814.8 | 915.4 | - | - | 0 | - |
| - | - | 1064 | 916.4 | - | - | 0 | - |
| - | - | 1103 | 924.5 | - | - | 0 | - |
| 7 | c | 1418 | 925.5 | 0.0008493 | 0.9177 | +1 | 7 |
| - | - | 766.5 | 926.5 | - | - | 0 | - |
| - | - | 1496 | 932.5 | - | - | 0 | - |
| - | - | 992.5 | 933.5 | - | - | 0 | - |
| - | - | 1429 | 938.5 | - | - | 0 | - |
| 2 | z | 1902 | 939.4 | 0.009458 | 10.07 | +2 | 14 |
| - | - | 1829 | 939.9 | - | - | 0 | - |
| - | - | 722.4 | 940.3 | - | - | 0 | - |
| - | - | 1691 | 940.4 | - | - | 0 | - |
| - | - | 1017 | 945.5 | - | - | 0 | - |
| - | - | 1334 | 960 | - | - | 0 | - |
| - | - | 2624 | 960.5 | - | - | 0 | - |
| - | - | 1043 | 961 | - | - | 0 | - |
| - | - | 898 | 961.5 | - | - | 0 | - |
| - | - | 688.3 | 965.2 | - | - | 0 | - |
| - | - | 858.7 | 966.5 | - | - | 0 | - |
| - | - | 1018 | 967 | - | - | 0 | - |
| - | - | 863.5 | 967.5 | - | - | 0 | - |
| - | - | 1245 | 968 | - | - | 0 | - |
| - | - | 1620 | 974.5 | - | - | 0 | - |
| - | - | 4281 | 975 | - | - | 0 | - |
| - | - | 3464 | 975.5 | - | - | 0 | - |
| - | - | 2887 | 976 | - | - | 0 | - |
| - | - | 1085 | 976.5 | - | - | 0 | - |
| - | - | 934.5 | 980.5 | - | - | 0 | - |
| - | - | 866.3 | 981 | - | - | 0 | - |
| - | - | 3632 | 981.5 | - | - | 0 | - |
| - | - | 2477 | 982 | - | - | 0 | - |
| 8 | c | 7725 | 982.5 | 0.007279 | 7.408 | +1 | 8 |
| - | - | 1914 | 983 | - | - | 0 | - |
| - | - | 3395 | 983.5 | - | - | 0 | - |
| - | - | 898.6 | 984.5 | - | - | 0 | - |
| - | - | 1019 | 985.5 | - | - | 0 | - |
| - | - | 1769 | 987.5 | - | - | 0 | - |
| - | - | 2786 | 988.5 | - | - | 0 | - |
| - | - | 2671 | 989.5 | - | - | 0 | - |
| - | - | 1070 | 990.5 | - | - | 0 | - |
| - | - | 1108 | 995.5 | - | - | 0 | - |
| - | - | 2746 | 996 | - | - | 0 | - |
| - | - | 7344 | 996.5 | - | - | 0 | - |
| - | - | 6495 | 997 | - | - | 0 | - |
| - | - | 3879 | 997.5 | - | - | 0 | - |
| - | - | 2249 | 998 | - | - | 0 | - |
| - | - | 1663 | 998.5 | - | - | 0 | - |
| - | - | 949.3 | 999.5 | - | - | 0 | - |
| - | - | 2524 | 1004 | - | - | 0 | - |
| - | - | 1728 | 1004 | - | - | 0 | - |
| - | - | 7631 | 1004 | - | - | 0 | - |
| - | - | 7652 | 1005 | - | - | 0 | - |
| - | - | 8400 | 1005 | - | - | 0 | - |
| - | - | 3397 | 1006 | - | - | 0 | - |
| - | - | 3983 | 1006 | - | - | 0 | - |
| - | - | 1026 | 1007 | - | - | 0 | - |
| - | - | 1027 | 1007 | - | - | 0 | - |
| - | - | 1618 | 1008 | - | - | 0 | - |
| 9 | z | 943.7 | 1009 | 0.006194 | 6.142 | +1 | 7 |
| - | - | 2820 | 1028 | - | - | 0 | - |
| - | - | 1120 | 1029 | - | - | 0 | - |
| - | - | 2020 | 1110 | - | - | 0 | - |
| 9 | c | 6697 | 1111 | 0.005516 | 4.967 | +1 | 9 |
| - | - | 3841 | 1112 | - | - | 0 | - |
| - | - | 1725 | 1113 | - | - | 0 | - |
| - | - | 819.2 | 1197 | - | - | 0 | - |
| 7 | z | 827.8 | 1199 | 0.01137 | 9.487 | +1 | 9 |
| - | - | 3061 | 1200 | - | - | 0 | - |
| - | - | 1243 | 1201 | - | - | 0 | - |
| 10 | c | 1829 | 1240 | 0.001986 | 1.602 | +1 | 10 |
| - | - | 1460 | 1241 | - | - | 0 | - |
| - | - | 755.1 | 1242 | - | - | 0 | - |
| - | - | 856.2 | 1257 | - | - | 0 | - |
| 6 | z | 1411 | 1312 | 0.008117 | 6.188 | +1 | 10 |
| - | - | 920.7 | 1313 | - | - | 0 | - |
| - | - | 751.1 | 1326 | - | - | 0 | - |
| - | - | 920.6 | 1404 | - | - | 0 | - |
| 5 | y | 664.8 | 1497 | 0.002669 | 1.783 | +1 | 11 |
| 5 | z | 1133 | 1498 | 0.01682 | 11.23 | +1 | 11 |
| - | - | 954.5 | 1499 | - | - | 0 | - |
| - | - | 1801 | 1614 | - | - | 0 | - |
| 13 | c | 821.4 | 1692 | 0.002 | 1.182 | +1 | 13 |
| 3 | z | 665.2 | 1741 | 0.001931 | 1.109 | +1 | 13 |
| - | - | 833.7 | 1742 | - | - | 0 | - |
| - | - | 881.9 | 1743 | - | - | 0 | - |
| - | - | 1276 | 2009 | - | - | 0 | - |
| - | - | 1277 | 2010 | - | - | 0 | - |

m/z Charge Intensity FragmentType MassShift Position
129.0658721923828 0 601.6983
129.10235595703125 0 989.06775
130.06536865234375 0 1076.3232
131.11788940429688 0 395.4239
132.1020050048828 0 829.98834
132.7487030029297 0 475.7306 c 2
133.0858917236328 0 1551.6216
133.09173583984375 0 422.8467
134.00567626953125 0 432.75745
140.08187866210938 0 458.32172
145.27166748046875 0 444.47092
147.8960723876953 0 368.47137
148.52574157714844 0 390.02893
148.94760131835938 0 669.8233
149.0447540283203 0 1258.3706
155.0928192138672 0 1005.46295
156.07681274414062 0 689.6483
162.34298706054688 0 444.61768
166.06088256835938 0 953.9198
174.0871124267578 0 788.76544
177.1122589111328 0 1358.7135
185.16470336914062 0 2059.2373
187.1077423095703 0 699.452
191.1028289794922 0 1570.7046
194.29266357421875 0 538.3169
199.16888427734375 0 839.2736
201.1233673095703 0 737.69775
202.08200073242188 0 1641.7483
203.1025848388672 0 3784.2676
207.09796142578125 0 1105.8746
215.13894653320312 0 8606.946
219.1337127685547 0 710.14545
223.1553497314453 0 589.7734
227.13919067382812 0 560.46124
228.87734985351562 0 499.13367
229.1545867919922 0 2327.9778
230.15797424316406 0 605.59564
234.12376403808594 0 661.0907
239.09530639648438 0 1237.0802
251.1502685546875 0 4487.046
269.18524169921875 0 481.72452
276.2595520019531 0 537.39014
281.0510559082031 0 1926.8915
282.0493469238281 0 631.82874
282.18023681640625 0 582.37555
289.12164306640625 0 4264.9976
299.0616149902344 0 7561.739
299.13543701171875 0 581.4913
300.0624084472656 0 1680.6085
302.1698913574219 0 1315.538
314.20770263671875 0 3053.2388
318.2001953125 0 579.2011
326.6634216308594 0 1118.092
342.1669921875 0 536.41486
359.02825927734375 0 1901.6151
360.02813720703125 0 839.79047
379.20806884765625 0 1893.7505 c Ammonia loss 2
387.2234802246094 0 1157.1494
396.23455810546875 0 1925.2325 c 2
407.71270751953125 0 635.55676
414.1942138671875 0 654.0914
415.036376953125 0 3798.0483
416.0380859375 0 1256.5502
420.2022399902344 0 5509.6255 w 9
420.7030944824219 0 2185.9722
438.1763916015625 0 577.5285
444.2328796386719 0 903.4463
444.2796936035156 0 841.69476
449.22015380859375 0 835.0916
449.7193908691406 0 1446.9373 z 9
450.2218322753906 0 602.58575
457.71832275390625 0 620.8695 y 9
458.22998046875 0 3644.903
458.72979736328125 0 1140.3862
459.2789306640625 0 1429.1145
460.73486328125 0 716.1023
480.73199462890625 0 860.07404
484.2458801269531 0 834.22943
485.2374572753906 0 876.62665
487.2356872558594 0 805.03937
488.2926940917969 0 1758.8973
493.7494812011719 0 2626.7542
494.2436218261719 0 2206.201 c Ammonia loss 3
495.3029479980469 0 967.8001
496.2464904785156 0 768.4653
501.30474853515625 0 2092.3076
501.7456970214844 0 805.9205
502.2088928222656 0 589.95685
502.2665710449219 0 2272.5166
502.4980773925781 0 2704.03
502.7532043457031 0 5070.8325
502.996826171875 0 2150.5596
503.1086730957031 0 3080.2878
503.2560729980469 0 4306.8145
503.3078918457031 0 2123.8513
503.7582092285156 0 972.49225
504.1095886230469 0 2029.8812
504.2667236328125 0 1251.7391
511.26171875 0 6171.822 c 3
512.2634887695312 0 1528.417
515.3201293945312 0 1123.481
517.2705078125 0 608.78906
521.7652587890625 0 1191.5828 y 8
522.2658081054688 0 888.6042
542.2697143554688 0 875.4735 z 7
545.3758544921875 0 664.45544
547.290283203125 0 1337.9082
550.2788696289062 0 1069.4193 y 7
555.2980346679688 0 1514.1011
555.7891845703125 0 5012.686 c 8
556.2901000976562 0 1767.4902
556.7916259765625 0 906.4842
570.2797241210938 0 853.9468
571.2817993164062 0 1725.7404
577.283203125 0 980.878 w 6
577.784912109375 0 822.15674
578.289306640625 0 705.3684
596.3224487304688 0 784.2806
599.2880859375 0 2527.7595 y Ammonia loss 6
599.7822875976562 0 3996.1396 z 6
600.2783203125 0 864.694
606.3049926757812 0 4184.4165 z 11
607.3004150390625 0 4927.22 c 13
607.6331176757812 0 2357.3499
607.7906494140625 0 4316.3984 y 6
607.9677124023438 0 1789.7391
608.2921752929688 0 2806.9404
608.7905883789062 0 1004.44617
612.3009033203125 0 1202.1016
612.6290283203125 0 641.0784
613.82080078125 0 1071.7653
614.0704345703125 0 616.50507
619.817626953125 0 1747.8398
620.3106079101562 0 4425.799 c 9
620.8113403320312 0 3394.4705
621.3123779296875 0 1498.3953
622.3230590820312 0 1881.592 y 11
623.3264770507812 0 706.4126
626.30419921875 0 1578.341 y Ammonia loss 1
626.9639892578125 0 864.9604
634.3058471679688 0 1716.6261
634.644775390625 0 922.1696
634.7984619140625 0 2116.2698 w 5
635.3035888671875 0 1184.0948
635.7977294921875 0 1641.4829
636.6516723632812 0 1167.2764
637.3008422851562 0 760.82446
639.9921875 0 2020.3215
640.3232421875 0 6803.97
640.6549682617188 0 5175.8857
640.9891967773438 0 3684.5215
641.3226318359375 0 1493.0436
645.6453857421875 0 1181.2451
645.9795532226562 0 1025.957
649.6491088867188 0 1479.6489
649.9775390625 0 4877.92
650.3206787109375 0 8801.629
650.6536865234375 0 4873.947
650.9849853515625 0 3200.5005
651.3153076171875 0 2206.6455
651.6491088867188 0 1161.0944
651.9815063476562 0 849.8252
653.66796875 0 1036.8351
654.341064453125 0 1745.1351
654.6648559570312 0 952.2955
654.9993896484375 0 815.63965
656.3237915039062 0 1829.7134 z 5
656.8250732421875 0 839.3586
656.9736328125 0 1106.4918
659.667724609375 0 991.6012
659.9908447265625 0 768.23517
663.9932250976562 0 2061.065
664.3318481445312 0 3905.1973 y 5
664.6591186523438 0 2217.921
664.8347778320312 0 1378.6979
665.004150390625 0 798.6659
667.9974975585938 0 732.85034
669.0010375976562 0 1363.9745
669.3319702148438 0 4907.5156
669.66259765625 0 14876.789
669.9957275390625 0 17299.746
670.3306884765625 0 12495.343
670.6635131835938 0 5827.401
670.9949340820312 0 4482.3384
672.326904296875 0 1519.9215
674.3214721679688 0 1221.9045
679.8359985351562 0 965.3774
680.3135986328125 0 2077.7593 c Ammonia loss 4
690.3427124023438 0 839.00104
697.3410034179688 0 6878.871 c 4
698.3447875976562 0 1930.4674
701.348876953125 0 2583.0513
701.844482421875 0 3833.1592 c 10
702.34326171875 0 2540.4917
702.8441772460938 0 1446.5967
716.4046630859375 0 3096.1958
717.4059448242188 0 1592.6659
726.3587646484375 0 785.8026
744.3572387695312 0 1189.2992
749.360595703125 0 1169.4916 z 4
749.8656616210938 0 2056.5146
750.3701171875 0 1232.0912
750.872314453125 0 825.88116
757.8758544921875 0 1164.4836
765.3922119140625 0 918.6565
765.8930053710938 0 850.33887 c 11
766.3920288085938 0 1642.2161
769.3662109375 0 4943.8364 z 10
770.3665771484375 0 3335.4048
771.3709716796875 0 1330.0089
785.391845703125 0 2022.974 y 10
803.4292602539062 0 2042.5052
806.3797607421875 0 916.32117 y Ammonia loss 3
806.8792114257812 0 2301.309 z 3
807.3796997070312 0 1786.7345
807.8821411132812 0 987.67786
810.4246826171875 0 12161.127 c 5
811.4276123046875 0 5636.9395
812.4317016601562 0 1622.9559
814.4050903320312 0 2624.7617
815.4013671875 0 1520.382
817.7777709960938 0 748.7588
831.43017578125 0 1572.5427
832.4346313476562 0 1103.4763
837.7918090820312 0 2164.9292
838.1170654296875 0 1824.297
839.4029541015625 0 1006.87244 w 9
846.4024047851562 0 3186.3525 c 12
846.8986206054688 0 2320.7563
847.3995361328125 0 1219.8367
847.8971557617188 0 910.3435
867.4248046875 0 1733.7823
870.4164428710938 0 1046.0765 y Ammonia loss 2
870.9085693359375 0 3344.707 z 2
871.409423828125 0 2102.4312
871.9121704101562 0 1330.5033
881.4457397460938 0 782.3548
887.922119140625 0 840.4966
888.4345703125 0 1130.6271
889.438232421875 0 1039.0474
890.4297485351562 0 1104.8319
898.4095458984375 0 6588.749 z 9
899.415771484375 0 3667.956
900.419677734375 0 2344.4355
910.4488525390625 0 1672.9917 c 13
910.9461669921875 0 2542.13
915.4442138671875 0 814.79706
916.4428100585938 0 1064.2522
924.4569702148438 0 1103.165
925.4534912109375 0 1418.3152 c 6
926.465576171875 0 766.47943
932.4775390625 0 1495.6873
933.4857177734375 0 992.51044
938.467529296875 0 1429.3419
939.4482421875 0 1901.7922 z 1
939.938232421875 0 1828.5067
940.337890625 0 722.36896
940.4485473632812 0 1691.4154
945.48828125 0 1016.6829
959.9816284179688 0 1334.3551
960.4735107421875 0 2624.2446
960.974365234375 0 1043.4235
961.4727783203125 0 897.98694
965.2152709960938 0 688.3231
966.4784545898438 0 858.71466
966.9757080078125 0 1018.042
967.4786987304688 0 863.54803
967.9799194335938 0 1245.4221
974.4862060546875 0 1619.951
974.986083984375 0 4281.159
975.4844970703125 0 3463.8706
975.987548828125 0 2887.4067
976.4862670898438 0 1085.286
980.4979858398438 0 934.457
981.006103515625 0 866.31433
981.4922485351562 0 3631.5723
981.9955444335938 0 2477.2666
982.4813842773438 0 7724.7695 c 7
982.9900512695312 0 1914.4403
983.48046875 0 3394.8027
984.47509765625 0 898.5858
985.4779663085938 0 1018.95074
987.4971313476562 0 1769.1945
988.4865112304688 0 2785.726
989.4803466796875 0 2670.733
990.4723510742188 0 1069.8905
995.4938354492188 0 1107.9344
995.986572265625 0 2746.0813
996.4844360351562 0 7344.463
996.9844360351562 0 6495.1973
997.4871215820312 0 3879.301
997.9815673828125 0 2249.066
998.4794921875 0 1663.4238
999.4832153320312 0 949.27423
1003.5003662109375 0 2523.8398
1003.9873657226562 0 1727.7773
1004.4989624023438 0 7630.8867
1004.9957275390625 0 7651.845
1005.4996337890625 0 8400.192
1005.99755859375 0 3397.4978
1006.4953002929688 0 3982.5354
1006.9963989257812 0 1026.193
1007.4990234375 0 1027.1897
1007.9931640625 0 1617.731
1008.5097045898438 0 943.69604 z Water loss 8
1027.51171875 0 2820.4426
1028.5244140625 0 1119.7917
1109.5802001953125 0 2020.4062
1110.5745849609375 0 6697.132 c 8
1111.5732421875 0 3840.6108
1112.577880859375 0 1724.6877
1196.6019287109375 0 819.15
1198.5738525390625 0 827.7635 z 6
1199.5634765625 0 3060.8123
1200.569091796875 0 1242.6923
1239.6136474609375 0 1829.2101 c 9
1240.6214599609375 0 1459.9824
1241.6275634765625 0 755.0643
1256.697509765625 0 856.244
1311.6546630859375 0 1411.1565 z 5
1312.6455078125 0 920.66876
1325.6558837890625 0 751.1213
1403.6888427734375 0 920.6015
1496.720703125 0 664.7692 y Ammonia loss 4
1497.74267578125 0 1132.8397 z 4
1498.724853515625 0 954.50085
1613.7447509765625 0 1800.5852
1691.7911376953125 0 821.4078 c 12
1740.8094482421875 0 665.2167 z 2
1741.805419921875 0 833.7228
1742.8260498046875 0 881.87555
2008.965576171875 0 1275.627
2009.9595947265625 0 1277.1348

Spectrum Details

|  |  |
| --- | --- |
| Matched peaks? Matched peaks The total absolute number of peaks matched. Additionally in brackets the total fraction of peaks matched and the total number of peaks is shown. | 52 (15.62% of 333) |
| FDR? FDR The false discovery rate estimated for this peptide. It is calculated by matching all theoretical fragments with a non-integer shift with the raw peaks for this spectrum. This is done with 40 different shifts. The resulting percentage is the average number of annotated peaks over the number of annotated peaks with the correct spectrum. | 0.73% |
| Satellite FDR? Satellite FDR See the FDR for details on its calculation. This satellite ion specific FDR only contains the satellite ions (d/w) for I/L/J positions. | 0.00% |
| PSM Score? PSM Score The PSM Score as given by Hecklib to this annotated spectrum. It is shown with three significant figures. | 231 |

#### Spectrum 5013? Spectrum 5013 The raw spectrum of this peptide as annotated by Hecklib. The fragments are coloured according to ion type (see legend). Any peaks with a star '\*' as text can be hovered over to see the full details, first the ion type second the mass shift type. By hovering over the amino acids in the peptide or ions in the legend the corresponding peaks are highlighted. By toggling the 'Unassigned' label you can turn the background (unassigned) peaks on or off in the plot. By updating the slider in the Ion legend you can update the spectrum to only show the top X% of the peaks with labels. The top X% means any peak that is within X% of the highest intensity. By dragging in the spectrum you can zoom in to a specific part of the spectrum and use 'Zoom Out' to get back to the original zoom level. The annotation of the spectrum is based on the given sequence in the peptides file and is done with different software so inconsistencies are likely. The peaks are annotated based on the given sequence, with 20 ppm tolerance.

Copy Data

##### Spectrum 5013 (TSV)

###### Preview

```
Loading example...
```

*Click on the button to copy the data to your clipboard.*

Mz MinMz MaxIntensity Max

WidthHeightPeptide font sizePeptide stroke widthSpectrum font sizeSpectrum stroke widthCompact peptide

Ion legend

wxyz

abcd

OtherUnassignedIon^Charge^_Position_Show for top:%

JHQDWLDGKEYKCKK

07.33e+41.47e+52.20e+52.93e+5

Zoom Out

c^+2^_3_y^+4^_5_y^+1^_1_y^+2^_3_z^+2^_3_c^+2^_4_c^+2^_4_c^+1^_2_z^+2^_4_z^+1^_2_c^+3^_7_z^+2^_4_y^+2^_4_z^+1^_2_y^+4^_10_y^+4^_10_y^+1^_2_c^+5^_13_c^+2^_5_y^+3^_7_y^+3^_7_z^+3^_7_y^+3^_7_z^+3^_8_c^+5^_14_y^+3^_8_z^+3^_8_y^+3^_8_z^+4^_11_y^+4^_11_y^+4^_11_z^+4^_11_c^+1^_3_y^+2^_5_z^+2^_5_w^+4^_12_y^+2^_5_c^+1^_3_c^+2^_6_y^+3^_9_y^+3^_9_z^+3^_9_y^+3^_9_c^+3^_10_w^+2^_6_w^+3^_10_z^+4^_13_z^+4^_13_z^+3^_10_y^+4^_13_y^+4^_13_z^+4^_13_y^+3^_10_y^+3^_10_z^+3^_10_y^+4^_13_z^+2^_6_y^+3^_10_y^+2^_6_y^+2^_6_z^+2^_6_c^+4^_14_c^+2^_7_c^+2^_7_c^+4^_14_y^+2^_6_c^+2^_7_z^+4^_14_z^+4^_14_y^+4^_14_y^+4^_14_z^+4^_14_y^+4^_14_y^+1^_3_z^+1^_3_c^+2^_8_c^+2^_8_w^+2^_7_c^+2^_8_c^+1^_4_z^+3^_11_c^+1^_4_y^+3^_11_y^+3^_11_z^+3^_11_z^+2^_7_c^+3^_12_c^+3^_12_c^+3^_12_c^+1^_4_y^+2^_7_z^+2^_7_y^+2^_7_z^+3^_12_y^+2^_8_y^+2^_8_z^+2^_8_y^+3^_12_w^+1^_4_y^+2^_8_c^+2^_9_w^+3^_13_c^+3^_13_w^+2^_9_y^+3^_13_y^+3^_13_z^+3^_13_y^+3^_13_z^+2^_9_z^+2^_9_y^+2^_9_y^+2^_9_z^+2^_9_c^+3^_14_c^+3^_14_z^+1^_4_c^+3^_14_y^+2^_9_c^+2^_10_c^+2^_10_z^+3^_14_y^+1^_4_z^+3^_14_y^+3^_14_w^+2^_10_z^+2^_10_y^+2^_10_y^+2^_10_z^+2^_10_y^+2^_10_c^+1^_5_c^+1^_5_c^+2^_11_c^+2^_11_c^+1^_5_c^+2^_11_y^+2^_11_y^+2^_11_z^+2^_11_c^+2^_12_c^+2^_12_y^+1^_5_z^+1^_5_w^+2^_12_y^+1^_5_c^+1^_6_z^+2^_12_y^+2^_12_z^+2^_12_c^+1^_6_y^+2^_12_w^+1^_6_w^+2^_13_c^+2^_13_y^+2^_13_z^+2^_13_y^+2^_13_z^+1^_6_y^+1^_6_z^+1^_6_c^+2^_14_c^+1^_7_c^+2^_14_y^+1^_6_c^+1^_7_y^+2^_14_z^+2^_14_c^+1^_8_z^+1^_7_c^+1^_9_y^+1^_8_c^+1^_9_z^+1^_9_c^+1^_10_z^+1^_10_c^+1^_11_

0867173326003466

Fragment Matches Table

Show background peaks

| Position | Ion type | Intensity | mz Theoretical | mz Error (Th) | mz Error (ppm) | Charge | Series Number |
| --- | --- | --- | --- | --- | --- | --- | --- |
| - | - | 389.8 | 120.9 | - | - | 0 | - |
| - | - | 728.7 | 126.1 | - | - | 0 | - |
| - | - | 459.7 | 126.1 | - | - | 0 | - |
| - | - | 728.9 | 127.1 | - | - | 0 | - |
| - | - | 864.4 | 128.1 | - | - | 0 | - |
| - | - | 3774 | 129 | - | - | 0 | - |
| - | - | 694 | 129.1 | - | - | 0 | - |
| - | - | 2.629E+04 | 129.1 | - | - | 0 | - |
| - | - | 9.562E+04 | 130.1 | - | - | 0 | - |
| - | - | 1877 | 130.1 | - | - | 0 | - |
| - | - | 604.7 | 131.1 | - | - | 0 | - |
| - | - | 9297 | 131.1 | - | - | 0 | - |
| - | - | 812.3 | 131.1 | - | - | 0 | - |
| - | - | 1700 | 131.1 | - | - | 0 | - |
| - | - | 563.7 | 132.1 | - | - | 0 | - |
| - | - | 534.3 | 132.1 | - | - | 0 | - |
| - | - | 2877 | 132.1 | - | - | 0 | - |
| - | - | 1524 | 132.1 | - | - | 0 | - |
| - | - | 859 | 133.1 | - | - | 0 | - |
| - | - | 1.392E+04 | 133.1 | - | - | 0 | - |
| - | - | 465.2 | 133.1 | - | - | 0 | - |
| - | - | 1033 | 134.1 | - | - | 0 | - |
| - | - | 499.1 | 136.1 | - | - | 0 | - |
| - | - | 5864 | 136.1 | - | - | 0 | - |
| - | - | 401.3 | 137.1 | - | - | 0 | - |
| - | - | 390.6 | 137.9 | - | - | 0 | - |
| - | - | 1328 | 138.1 | - | - | 0 | - |
| - | - | 2820 | 139 | - | - | 0 | - |
| - | - | 434.9 | 140.3 | - | - | 0 | - |
| - | - | 615.6 | 141.1 | - | - | 0 | - |
| - | - | 585.9 | 143 | - | - | 0 | - |
| - | - | 449.5 | 143.1 | - | - | 0 | - |
| - | - | 2815 | 144.1 | - | - | 0 | - |
| - | - | 2173 | 145.1 | - | - | 0 | - |
| - | - | 1795 | 146.1 | - | - | 0 | - |
| - | - | 414.7 | 146.1 | - | - | 0 | - |
| - | - | 722.8 | 147 | - | - | 0 | - |
| - | - | 820.5 | 147.1 | - | - | 0 | - |
| - | - | 457.8 | 154.1 | - | - | 0 | - |
| - | - | 1.751E+04 | 155.1 | - | - | 0 | - |
| - | - | 759.1 | 156.1 | - | - | 0 | - |
| - | - | 902.7 | 156.1 | - | - | 0 | - |
| - | - | 1590 | 156.1 | - | - | 0 | - |
| - | - | 7593 | 157 | - | - | 0 | - |
| - | - | 669.4 | 157.1 | - | - | 0 | - |
| - | - | 1045 | 158.1 | - | - | 0 | - |
| - | - | 4.701E+04 | 159.1 | - | - | 0 | - |
| - | - | 4371 | 160.1 | - | - | 0 | - |
| - | - | 2189 | 164.1 | - | - | 0 | - |
| - | - | 467.6 | 165.1 | - | - | 0 | - |
| - | - | 1.577E+04 | 166.1 | - | - | 0 | - |
| - | - | 1393 | 167.1 | - | - | 0 | - |
| - | - | 1006 | 168.1 | - | - | 0 | - |
| - | - | 537 | 169.1 | - | - | 0 | - |
| - | - | 1.737E+04 | 170.1 | - | - | 0 | - |
| - | - | 1071 | 171.1 | - | - | 0 | - |
| - | - | 824.9 | 171.1 | - | - | 0 | - |
| - | - | 709.4 | 171.1 | - | - | 0 | - |
| - | - | 3952 | 172.1 | - | - | 0 | - |
| - | - | 1791 | 173.1 | - | - | 0 | - |
| - | - | 389.3 | 173.4 | - | - | 0 | - |
| - | - | 2677 | 173.5 | - | - | 0 | - |
| - | - | 2734 | 174.1 | - | - | 0 | - |
| - | - | 1768 | 176.1 | - | - | 0 | - |
| - | - | 568.9 | 176.6 | - | - | 0 | - |
| - | - | 2673 | 178.1 | - | - | 0 | - |
| - | - | 552.3 | 179.1 | - | - | 0 | - |
| - | - | 534.2 | 181.1 | - | - | 0 | - |
| - | - | 6530 | 183.1 | - | - | 0 | - |
| - | - | 831.7 | 183.1 | - | - | 0 | - |
| - | - | 996.3 | 183.1 | - | - | 0 | - |
| - | - | 859.3 | 183.1 | - | - | 0 | - |
| - | - | 1021 | 185.1 | - | - | 0 | - |
| - | - | 1.008E+04 | 185.1 | - | - | 0 | - |
| - | - | 855.3 | 185.1 | - | - | 0 | - |
| - | - | 1.539E+04 | 185.2 | - | - | 0 | - |
| - | - | 701.4 | 186.1 | - | - | 0 | - |
| - | - | 3193 | 186.1 | - | - | 0 | - |
| - | - | 1189 | 186.2 | - | - | 0 | - |
| - | - | 6943 | 187.1 | - | - | 0 | - |
| - | - | 1404 | 187.1 | - | - | 0 | - |
| - | - | 918.4 | 188.1 | - | - | 0 | - |
| - | - | 802.4 | 189.1 | - | - | 0 | - |
| - | - | 2.91E+04 | 190.1 | - | - | 0 | - |
| 3 | c | 1332 | 190.1 | 0.0006047 | 3.181 | +2 | 3 |
| - | - | 500.1 | 190.6 | - | - | 0 | - |
| - | - | 1091 | 191.1 | - | - | 0 | - |
| - | - | 1891 | 192.1 | - | - | 0 | - |
| - | - | 473.4 | 192.5 | - | - | 0 | - |
| - | - | 752.6 | 195.1 | - | - | 0 | - |
| 11 | y | 521.5 | 197.1 | 0.00355 | 18.01 | +4 | 5 |
| - | - | 8004 | 198.1 | - | - | 0 | - |
| - | - | 515.3 | 199.1 | - | - | 0 | - |
| - | - | 2640 | 199.2 | - | - | 0 | - |
| - | - | 785.2 | 200.1 | - | - | 0 | - |
| - | - | 503.2 | 201.1 | - | - | 0 | - |
| - | - | 958.2 | 202.1 | - | - | 0 | - |
| - | - | 793.7 | 203.1 | - | - | 0 | - |
| - | - | 3139 | 203.1 | - | - | 0 | - |
| - | - | 1069 | 203.1 | - | - | 0 | - |
| - | - | 1286 | 203.2 | - | - | 0 | - |
| - | - | 674.6 | 203.6 | - | - | 0 | - |
| 15 | y | 1.213E+04 | 205.1 | 0.003969 | 19.35 | +1 | 1 |
| - | - | 482.2 | 205.1 | - | - | 0 | - |
| - | - | 1312 | 206.1 | - | - | 0 | - |
| - | - | 1833 | 207.2 | - | - | 0 | - |
| - | - | 1010 | 208.1 | - | - | 0 | - |
| - | - | 4287 | 210.1 | - | - | 0 | - |
| - | - | 848 | 211.1 | - | - | 0 | - |
| - | - | 5958 | 211.6 | - | - | 0 | - |
| - | - | 1735 | 212.1 | - | - | 0 | - |
| - | - | 4945 | 213.1 | - | - | 0 | - |
| - | - | 504.9 | 213.2 | - | - | 0 | - |
| - | - | 967.6 | 215.1 | - | - | 0 | - |
| - | - | 8590 | 217 | - | - | 0 | - |
| - | - | 4113 | 223.1 | - | - | 0 | - |
| - | - | 1.691E+04 | 223.2 | - | - | 0 | - |
| - | - | 604.7 | 224.1 | - | - | 0 | - |
| - | - | 2037 | 224.2 | - | - | 0 | - |
| - | - | 580.4 | 226.1 | - | - | 0 | - |
| - | - | 1510 | 227.1 | - | - | 0 | - |
| - | - | 587.6 | 227.1 | - | - | 0 | - |
| - | - | 746.5 | 228.1 | - | - | 0 | - |
| - | - | 1428 | 228.1 | - | - | 0 | - |
| - | - | 606.8 | 229.1 | - | - | 0 | - |
| - | - | 4485 | 229.1 | - | - | 0 | - |
| - | - | 1152 | 230.2 | - | - | 0 | - |
| - | - | 4667 | 233.1 | - | - | 0 | - |
| - | - | 1227 | 233.6 | - | - | 0 | - |
| - | - | 1.958E+04 | 234.1 | - | - | 0 | - |
| - | - | 2920 | 235.1 | - | - | 0 | - |
| - | - | 2377 | 236.6 | - | - | 0 | - |
| 13 | y | 2289 | 239.1 | 1.441E-06 | 0.006025 | +2 | 3 |
| 13 | z | 1193 | 239.6 | 0.002172 | 9.063 | +2 | 3 |
| - | - | 728.4 | 240.2 | - | - | 0 | - |
| - | - | 556 | 243.1 | - | - | 0 | - |
| - | - | 644.4 | 243.1 | - | - | 0 | - |
| - | - | 2737 | 244.1 | - | - | 0 | - |
| - | - | 507.9 | 245.1 | - | - | 0 | - |
| - | - | 1057 | 245.1 | - | - | 0 | - |
| - | - | 1603 | 246.1 | - | - | 0 | - |
| 4 | c | 7578 | 247.6 | 0.0003015 | 1.218 | +2 | 4 |
| - | - | 1764 | 248.1 | - | - | 0 | - |
| - | - | 2190 | 250.1 | - | - | 0 | - |
| - | - | 2006 | 250.1 | - | - | 0 | - |
| - | - | 2239 | 250.2 | - | - | 0 | - |
| - | - | 9.861E+04 | 251.2 | - | - | 0 | - |
| - | - | 1.325E+04 | 252.2 | - | - | 0 | - |
| - | - | 2908 | 252.6 | - | - | 0 | - |
| - | - | 1124 | 253.2 | - | - | 0 | - |
| - | - | 2924 | 255.1 | - | - | 0 | - |
| - | - | 2025 | 255.8 | - | - | 0 | - |
| 4 | c | 723.3 | 256.1 | 0.004147 | 16.19 | +2 | 4 |
| - | - | 810.2 | 256.6 | - | - | 0 | - |
| - | - | 2051 | 258.2 | - | - | 0 | - |
| - | - | 1124 | 258.7 | - | - | 0 | - |
| - | - | 576.4 | 259.1 | - | - | 0 | - |
| - | - | 1.262E+04 | 259.2 | - | - | 0 | - |
| - | - | 3216 | 261.1 | - | - | 0 | - |
| - | - | 2820 | 261.6 | - | - | 0 | - |
| - | - | 644.6 | 262.1 | - | - | 0 | - |
| - | - | 2331 | 264.2 | - | - | 0 | - |
| - | - | 2481 | 265.1 | - | - | 0 | - |
| - | - | 1370 | 265.6 | - | - | 0 | - |
| - | - | 3460 | 266.1 | - | - | 0 | - |
| - | - | 674.1 | 266.6 | - | - | 0 | - |
| - | - | 577.5 | 267.1 | - | - | 0 | - |
| 2 | c | 1.16E+04 | 268.2 | 0.0006888 | 2.569 | +1 | 2 |
| - | - | 1585 | 269.1 | - | - | 0 | - |
| - | - | 1819 | 269.2 | - | - | 0 | - |
| - | - | 2.02E+04 | 270.1 | - | - | 0 | - |
| - | - | 5388 | 270.6 | - | - | 0 | - |
| - | - | 5176 | 272.1 | - | - | 0 | - |
| - | - | 6374 | 272.2 | - | - | 0 | - |
| - | - | 3824 | 273.1 | - | - | 0 | - |
| - | - | 737.6 | 273.1 | - | - | 0 | - |
| - | - | 958.2 | 273.2 | - | - | 0 | - |
| - | - | 3654 | 274.1 | - | - | 0 | - |
| - | - | 1.743E+04 | 274.1 | - | - | 0 | - |
| - | - | 4706 | 274.7 | - | - | 0 | - |
| - | - | 1764 | 275.1 | - | - | 0 | - |
| - | - | 1027 | 275.2 | - | - | 0 | - |
| - | - | 703.1 | 276.1 | - | - | 0 | - |
| - | - | 729.4 | 277.2 | - | - | 0 | - |
| - | - | 1189 | 282.1 | - | - | 0 | - |
| - | - | 7910 | 282.2 | - | - | 0 | - |
| - | - | 3836 | 283.1 | - | - | 0 | - |
| - | - | 1027 | 283.2 | - | - | 0 | - |
| - | - | 1646 | 284.1 | - | - | 0 | - |
| - | - | 3829 | 284.1 | - | - | 0 | - |
| - | - | 649.5 | 284.1 | - | - | 0 | - |
| - | - | 879.6 | 284.6 | - | - | 0 | - |
| - | - | 1437 | 285.2 | - | - | 0 | - |
| - | - | 1423 | 286.1 | - | - | 0 | - |
| - | - | 6972 | 290.1 | - | - | 0 | - |
| - | - | 1250 | 292.2 | - | - | 0 | - |
| - | - | 1430 | 294.1 | - | - | 0 | - |
| - | - | 798.2 | 294.1 | - | - | 0 | - |
| - | - | 945.1 | 294.2 | - | - | 0 | - |
| - | - | 1033 | 295.1 | - | - | 0 | - |
| 12 | z | 905.6 | 295.1 | 0.0008682 | 2.941 | +2 | 4 |
| - | - | 2160 | 297.1 | - | - | 0 | - |
| - | - | 779 | 297.2 | - | - | 0 | - |
| - | - | 677.6 | 297.6 | - | - | 0 | - |
| 14 | z | 2760 | 300.2 | 0.0009852 | 3.282 | +1 | 2 |
| - | - | 1249 | 301.1 | - | - | 0 | - |
| - | - | 3106 | 301.2 | - | - | 0 | - |
| - | - | 3577 | 302.1 | - | - | 0 | - |
| - | - | 2015 | 302.1 | - | - | 0 | - |
| - | - | 718.6 | 302.5 | - | - | 0 | - |
| - | - | 799.1 | 302.7 | - | - | 0 | - |
| - | - | 611.3 | 303.1 | - | - | 0 | - |
| 7 | c | 1.013E+04 | 303.2 | 0.0002253 | 0.7431 | +3 | 7 |
| - | - | 2968 | 303.2 | - | - | 0 | - |
| 12 | z | 3759 | 303.7 | 0.005387 | 17.74 | +2 | 4 |
| - | - | 2134 | 304.2 | - | - | 0 | - |
| - | - | 882.9 | 305.6 | - | - | 0 | - |
| - | - | 620.3 | 306.1 | - | - | 0 | - |
| - | - | 5467 | 309.1 | - | - | 0 | - |
| - | - | 669.8 | 309.2 | - | - | 0 | - |
| - | - | 1129 | 309.6 | - | - | 0 | - |
| - | - | 838.2 | 309.7 | - | - | 0 | - |
| - | - | 2741 | 311.1 | - | - | 0 | - |
| - | - | 887.5 | 311.6 | - | - | 0 | - |
| 12 | y | 9507 | 311.7 | 0.003641 | 11.68 | +2 | 4 |
| - | - | 1743 | 312.2 | - | - | 0 | - |
| - | - | 558.1 | 312.7 | - | - | 0 | - |
| - | - | 1694 | 315.2 | - | - | 0 | - |
| 14 | z | 4186 | 317.2 | 0.003425 | 10.8 | +1 | 2 |
| - | - | 5005 | 317.7 | - | - | 0 | - |
| - | - | 1.999E+04 | 318.2 | - | - | 0 | - |
| - | - | 1268 | 318.2 | - | - | 0 | - |
| - | - | 9064 | 318.7 | - | - | 0 | - |
| - | - | 719.7 | 318.8 | - | - | 0 | - |
| - | - | 1883 | 319.2 | - | - | 0 | - |
| - | - | 844.1 | 320.2 | - | - | 0 | - |
| - | - | 694.1 | 321.2 | - | - | 0 | - |
| - | - | 1000 | 322.8 | - | - | 0 | - |
| - | - | 3105 | 323.1 | - | - | 0 | - |
| - | - | 1768 | 323.6 | - | - | 0 | - |
| - | - | 870.5 | 324.7 | - | - | 0 | - |
| - | - | 4317 | 326.5 | - | - | 0 | - |
| - | - | 8.255E+04 | 326.7 | - | - | 0 | - |
| - | - | 1754 | 326.8 | - | - | 0 | - |
| - | - | 3.331E+04 | 327.2 | - | - | 0 | - |
| - | - | 1388 | 327.2 | - | - | 0 | - |
| - | - | 8306 | 327.7 | - | - | 0 | - |
| 6 | y | 1219 | 328.2 | 0.0003304 | 1.007 | +4 | 10 |
| - | - | 3849 | 329.2 | - | - | 0 | - |
| - | - | 1225 | 331.5 | - | - | 0 | - |
| - | - | 1.245E+04 | 331.7 | - | - | 0 | - |
| - | - | 774.7 | 331.7 | - | - | 0 | - |
| - | - | 1410 | 331.9 | - | - | 0 | - |
| - | - | 4396 | 332.2 | - | - | 0 | - |
| - | - | 865.5 | 332.7 | - | - | 0 | - |
| 6 | y | 1013 | 332.7 | 0.0006495 | 1.952 | +4 | 10 |
| 14 | y | 5780 | 333.2 | 0.003351 | 10.06 | +1 | 2 |
| - | - | 680.6 | 334.2 | - | - | 0 | - |
| - | - | 639.1 | 335.2 | - | - | 0 | - |
| - | - | 1663 | 337.8 | - | - | 0 | - |
| - | - | 960.6 | 338.1 | - | - | 0 | - |
| - | - | 1387 | 338.2 | - | - | 0 | - |
| - | - | 614.9 | 338.5 | - | - | 0 | - |
| - | - | 1640 | 338.7 | - | - | 0 | - |
| 13 | c | 1443 | 339.2 | 0.00316 | 9.318 | +5 | 13 |
| - | - | 629.1 | 339.7 | - | - | 0 | - |
| - | - | 1075 | 339.7 | - | - | 0 | - |
| - | - | 3441 | 339.7 | - | - | 0 | - |
| - | - | 1785 | 340.2 | - | - | 0 | - |
| 5 | c | 3.36E+04 | 340.7 | 0.00105 | 3.083 | +2 | 5 |
| - | - | 1.843E+04 | 341.2 | - | - | 0 | - |
| - | - | 1079 | 341.2 | - | - | 0 | - |
| - | - | 4248 | 341.7 | - | - | 0 | - |
| - | - | 2977 | 341.8 | - | - | 0 | - |
| 9 | y | 1825 | 342.2 | 0.002812 | 8.219 | +3 | 7 |
| 9 | y | 2401 | 342.5 | 0.000251 | 0.7329 | +3 | 7 |
| 9 | z | 2260 | 342.8 | 0.00111 | 3.237 | +3 | 7 |
| - | - | 1446 | 343.2 | - | - | 0 | - |
| - | - | 858.5 | 346.1 | - | - | 0 | - |
| - | - | 1109 | 347.2 | - | - | 0 | - |
| - | - | 1583 | 347.9 | - | - | 0 | - |
| 9 | y | 1800 | 348.2 | 0.001509 | 4.335 | +3 | 7 |
| - | - | 2181 | 349.2 | - | - | 0 | - |
| - | - | 1112 | 349.7 | - | - | 0 | - |
| - | - | 1919 | 351.2 | - | - | 0 | - |
| - | - | 640.3 | 352.5 | - | - | 0 | - |
| - | - | 640.5 | 352.7 | - | - | 0 | - |
| - | - | 2426 | 353.7 | - | - | 0 | - |
| - | - | 1691 | 354.2 | - | - | 0 | - |
| - | - | 1906 | 355.2 | - | - | 0 | - |
| - | - | 2185 | 355.7 | - | - | 0 | - |
| 8 | z | 801.1 | 355.8 | 0.003217 | 9.04 | +3 | 8 |
| - | - | 933.7 | 356.2 | - | - | 0 | - |
| - | - | 3036 | 356.8 | - | - | 0 | - |
| - | - | 2634 | 357.2 | - | - | 0 | - |
| - | - | 868.1 | 357.7 | - | - | 0 | - |
| - | - | 662.1 | 358.2 | - | - | 0 | - |
| - | - | 2811 | 360.7 | - | - | 0 | - |
| 14 | c | 2409 | 361.2 | 0.000624 | 1.728 | +5 | 14 |
| - | - | 1942 | 361.2 | - | - | 0 | - |
| - | - | 2451 | 361.2 | - | - | 0 | - |
| 8 | y | 1771 | 361.5 | 0.0002645 | 0.7316 | +3 | 8 |
| 8 | z | 7961 | 361.9 | 0.002222 | 6.14 | +3 | 8 |
| - | - | 5243 | 362.2 | - | - | 0 | - |
| - | - | 690.5 | 362.5 | - | - | 0 | - |
| - | - | 806.6 | 362.7 | - | - | 0 | - |
| - | - | 1558 | 363.1 | - | - | 0 | - |
| - | - | 955.4 | 363.2 | - | - | 0 | - |
| - | - | 1564 | 364.1 | - | - | 0 | - |
| - | - | 3435 | 364.2 | - | - | 0 | - |
| - | - | 2038 | 364.7 | - | - | 0 | - |
| - | - | 1071 | 365.7 | - | - | 0 | - |
| - | - | 1687 | 366.2 | - | - | 0 | - |
| 8 | y | 4.785E+04 | 367.2 | 0.002136 | 5.817 | +3 | 8 |
| - | - | 2.843E+04 | 367.5 | - | - | 0 | - |
| - | - | 1440 | 367.7 | - | - | 0 | - |
| - | - | 1152 | 367.7 | - | - | 0 | - |
| - | - | 1.282E+04 | 367.9 | - | - | 0 | - |
| - | - | 4686 | 368.2 | - | - | 0 | - |
| - | - | 1028 | 369.2 | - | - | 0 | - |
| - | - | 1023 | 370.4 | - | - | 0 | - |
| - | - | 654.2 | 370.5 | - | - | 0 | - |
| 5 | z | 1279 | 370.7 | 0.002884 | 7.781 | +4 | 11 |
| - | - | 1613 | 371.2 | - | - | 0 | - |
| - | - | 841.7 | 372.4 | - | - | 0 | - |
| - | - | 4844 | 374.2 | - | - | 0 | - |
| 5 | y | 1.177E+04 | 374.7 | 0.004672 | 12.47 | +4 | 11 |
| 5 | y | 1.062E+04 | 374.9 | 0.001189 | 3.171 | +4 | 11 |
| 5 | z | 9303 | 375.2 | 0.0004864 | 1.296 | +4 | 11 |
| - | - | 4300 | 375.4 | - | - | 0 | - |
| - | - | 1268 | 375.7 | - | - | 0 | - |
| - | - | 619.7 | 375.9 | - | - | 0 | - |
| - | - | 3778 | 376.2 | - | - | 0 | - |
| - | - | 1015 | 376.5 | - | - | 0 | - |
| - | - | 6152 | 378.2 | - | - | 0 | - |
| - | - | 864.6 | 378.2 | - | - | 0 | - |
| 3 | c | 3.222E+04 | 379.2 | 0.001009 | 2.661 | +1 | 3 |
| - | - | 1836 | 379.4 | - | - | 0 | - |
| - | - | 5426 | 380.2 | - | - | 0 | - |
| - | - | 7919 | 381.2 | - | - | 0 | - |
| - | - | 2.179E+04 | 381.2 | - | - | 0 | - |
| - | - | 8988 | 381.5 | - | - | 0 | - |
| - | - | 3423 | 381.9 | - | - | 0 | - |
| - | - | 1327 | 382.2 | - | - | 0 | - |
| - | - | 1110 | 382.2 | - | - | 0 | - |
| - | - | 3.111E+04 | 383.2 | - | - | 0 | - |
| - | - | 1.733E+04 | 383.7 | - | - | 0 | - |
| - | - | 4642 | 384.2 | - | - | 0 | - |
| 11 | y | 2333 | 384.7 | 0.002896 | 7.529 | +2 | 5 |
| - | - | 805.9 | 384.7 | - | - | 0 | - |
| - | - | 2429 | 385.2 | - | - | 0 | - |
| 11 | z | 6248 | 385.2 | 0.001804 | 4.683 | +2 | 5 |
| - | - | 3738 | 385.5 | - | - | 0 | - |
| - | - | 787.3 | 385.7 | - | - | 0 | - |
| - | - | 2606 | 385.9 | - | - | 0 | - |
| - | - | 1919 | 386.2 | - | - | 0 | - |
| - | - | 1166 | 386.5 | - | - | 0 | - |
| - | - | 1394 | 387.2 | - | - | 0 | - |
| - | - | 3.987E+04 | 387.2 | - | - | 0 | - |
| - | - | 1598 | 388.2 | - | - | 0 | - |
| - | - | 8852 | 388.2 | - | - | 0 | - |
| - | - | 3663 | 388.7 | - | - | 0 | - |
| - | - | 792.6 | 388.9 | - | - | 0 | - |
| - | - | 8190 | 389.2 | - | - | 0 | - |
| - | - | 791.9 | 389.2 | - | - | 0 | - |
| - | - | 916.2 | 389.2 | - | - | 0 | - |
| - | - | 1094 | 390 | - | - | 0 | - |
| - | - | 5019 | 390.2 | - | - | 0 | - |
| - | - | 1285 | 390.5 | - | - | 0 | - |
| - | - | 737.3 | 391.1 | - | - | 0 | - |
| - | - | 1708 | 391.2 | - | - | 0 | - |
| - | - | 795.9 | 392.2 | - | - | 0 | - |
| 4 | w | 738.8 | 392.7 | 0.00415 | 10.57 | +4 | 12 |
| 11 | y | 5.024E+04 | 393.2 | 0.003475 | 8.839 | +2 | 5 |
| - | - | 762.9 | 393.5 | - | - | 0 | - |
| - | - | 803.2 | 393.6 | - | - | 0 | - |
| - | - | 2.322E+04 | 393.7 | - | - | 0 | - |
| - | - | 8478 | 394.2 | - | - | 0 | - |
| - | - | 1618 | 394.7 | - | - | 0 | - |
| - | - | 2303 | 395 | - | - | 0 | - |
| - | - | 2153 | 395.1 | - | - | 0 | - |
| - | - | 6039 | 395.2 | - | - | 0 | - |
| - | - | 915.6 | 395.2 | - | - | 0 | - |
| - | - | 6770 | 395.4 | - | - | 0 | - |
| - | - | 3729 | 395.6 | - | - | 0 | - |
| - | - | 2667 | 395.8 | - | - | 0 | - |
| 3 | c | 3.763E+04 | 396.2 | 0.001041 | 2.627 | +1 | 3 |
| 6 | c | 2.52E+04 | 397.2 | 0.001102 | 2.775 | +2 | 6 |
| - | - | 7430 | 397.2 | - | - | 0 | - |
| - | - | 1.187E+04 | 397.7 | - | - | 0 | - |
| - | - | 1801 | 398.2 | - | - | 0 | - |
| - | - | 3108 | 398.2 | - | - | 0 | - |
| - | - | 826.4 | 398.2 | - | - | 0 | - |
| - | - | 1.136E+04 | 398.6 | - | - | 0 | - |
| - | - | 2.814E+04 | 398.8 | - | - | 0 | - |
| - | - | 2.609E+04 | 399 | - | - | 0 | - |
| - | - | 1.702E+04 | 399.2 | - | - | 0 | - |
| - | - | 7520 | 399.4 | - | - | 0 | - |
| 7 | y | 3227 | 399.5 | 0.002153 | 5.389 | +3 | 9 |
| - | - | 3072 | 399.6 | - | - | 0 | - |
| - | - | 812.1 | 399.8 | - | - | 0 | - |
| 7 | y | 3998 | 399.9 | 0.0006904 | 1.727 | +3 | 9 |
| 7 | z | 2799 | 400.2 | 0.0008777 | 2.193 | +3 | 9 |
| - | - | 745.7 | 400.5 | - | - | 0 | - |
| - | - | 764.7 | 400.8 | - | - | 0 | - |
| - | - | 1035 | 401.2 | - | - | 0 | - |
| - | - | 1185 | 401.3 | - | - | 0 | - |
| - | - | 2428 | 402 | - | - | 0 | - |
| - | - | 1.938E+04 | 402.2 | - | - | 0 | - |
| - | - | 2.17E+04 | 402.4 | - | - | 0 | - |
| - | - | 1.622E+04 | 402.6 | - | - | 0 | - |
| - | - | 8306 | 402.8 | - | - | 0 | - |
| - | - | 3323 | 403 | - | - | 0 | - |
| - | - | 2982 | 403.2 | - | - | 0 | - |
| - | - | 2287 | 404.2 | - | - | 0 | - |
| - | - | 1751 | 404.7 | - | - | 0 | - |
| - | - | 1164 | 405.2 | - | - | 0 | - |
| 7 | y | 4.299E+04 | 405.5 | 0.001799 | 4.436 | +3 | 9 |
| - | - | 2.5E+04 | 405.9 | - | - | 0 | - |
| - | - | 1.072E+04 | 406.2 | - | - | 0 | - |
| - | - | 3833 | 406.5 | - | - | 0 | - |
| - | - | 732.3 | 407.2 | - | - | 0 | - |
| 10 | c | 1040 | 408.2 | 0.001161 | 2.845 | +3 | 10 |
| - | - | 2967 | 408.9 | - | - | 0 | - |
| - | - | 1303 | 409.1 | - | - | 0 | - |
| - | - | 709.6 | 409.2 | - | - | 0 | - |
| - | - | 886 | 409.5 | - | - | 0 | - |
| - | - | 1223 | 411.2 | - | - | 0 | - |
| - | - | 996.4 | 411.7 | - | - | 0 | - |
| - | - | 1679 | 412.2 | - | - | 0 | - |
| - | - | 704.2 | 413.1 | - | - | 0 | - |
| - | - | 777.8 | 413.2 | - | - | 0 | - |
| - | - | 1092 | 413.9 | - | - | 0 | - |
| - | - | 976 | 414.2 | - | - | 0 | - |
| - | - | 1354 | 414.6 | - | - | 0 | - |
| - | - | 958.2 | 414.9 | - | - | 0 | - |
| - | - | 779.2 | 415.2 | - | - | 0 | - |
| - | - | 1031 | 415.2 | - | - | 0 | - |
| - | - | 1338 | 415.7 | - | - | 0 | - |
| - | - | 1014 | 416.7 | - | - | 0 | - |
| - | - | 2004 | 418.2 | - | - | 0 | - |
| - | - | 818.3 | 419.7 | - | - | 0 | - |
| 10 | w | 2.028E+05 | 420.2 | 0.003234 | 7.696 | +2 | 6 |
| - | - | 9.086E+04 | 420.7 | - | - | 0 | - |
| - | - | 3.326E+04 | 421.2 | - | - | 0 | - |
| - | - | 1.089E+04 | 421.7 | - | - | 0 | - |
| - | - | 1375 | 422.2 | - | - | 0 | - |
| - | - | 916.6 | 422.7 | - | - | 0 | - |
| 6 | w | 1.64E+04 | 423.5 | 0.001658 | 3.915 | +3 | 10 |
| - | - | 1.133E+04 | 423.9 | - | - | 0 | - |
| - | - | 4455 | 424.2 | - | - | 0 | - |
| - | - | 1327 | 424.5 | - | - | 0 | - |
| - | - | 1228 | 424.7 | - | - | 0 | - |
| - | - | 777 | 424.9 | - | - | 0 | - |
| - | - | 1359 | 425.2 | - | - | 0 | - |
| - | - | 893.9 | 427.2 | - | - | 0 | - |
| - | - | 951.3 | 427.7 | - | - | 0 | - |
| - | - | 7998 | 430.2 | - | - | 0 | - |
| - | - | 3098 | 431.2 | - | - | 0 | - |
| - | - | 1110 | 431.2 | - | - | 0 | - |
| 3 | z | 823 | 431.5 | 0.003116 | 7.222 | +4 | 13 |
| 3 | z | 1707 | 431.7 | 0.004085 | 9.462 | +4 | 13 |
| 6 | z | 986.7 | 432.2 | 0.0001533 | 0.3547 | +3 | 10 |
| - | - | 2453 | 434.2 | - | - | 0 | - |
| - | - | 1933 | 435.2 | - | - | 0 | - |
| - | - | 4375 | 435.2 | - | - | 0 | - |
| 3 | y | 2051 | 435.5 | 0.0007473 | 1.716 | +4 | 13 |
| 3 | y | 1906 | 435.7 | 0.001875 | 4.304 | +4 | 13 |
| 3 | z | 1686 | 436 | 0.0008742 | 2.005 | +4 | 13 |
| - | - | 2169 | 436.2 | - | - | 0 | - |
| 6 | y | 4441 | 437.2 | 0.0007655 | 1.751 | +3 | 10 |
| 6 | y | 6633 | 437.6 | 0.004634 | 10.59 | +3 | 10 |
| 6 | z | 1471 | 437.9 | 0.002371 | 5.415 | +3 | 10 |
| - | - | 1112 | 438.2 | - | - | 0 | - |
| - | - | 1326 | 438.2 | - | - | 0 | - |
| - | - | 692.2 | 438.6 | - | - | 0 | - |
| - | - | 1623 | 438.7 | - | - | 0 | - |
| - | - | 1439 | 439.2 | - | - | 0 | - |
| 3 | y | 1315 | 440 | 0.0009166 | 2.083 | +4 | 13 |
| 10 | z | 892.8 | 440.7 | 0.008671 | 19.68 | +2 | 6 |
| - | - | 883.1 | 441.2 | - | - | 0 | - |
| - | - | 763 | 442.7 | - | - | 0 | - |
| - | - | 3211 | 442.9 | - | - | 0 | - |
| 6 | y | 5.811E+04 | 443.2 | 0.001449 | 3.269 | +3 | 10 |
| - | - | 3.547E+04 | 443.6 | - | - | 0 | - |
| - | - | 1.503E+04 | 443.9 | - | - | 0 | - |
| - | - | 4918 | 444.2 | - | - | 0 | - |
| - | - | 1399 | 444.6 | - | - | 0 | - |
| - | - | 2596 | 445.7 | - | - | 0 | - |
| - | - | 1348 | 446.2 | - | - | 0 | - |
| - | - | 977.7 | 446.6 | - | - | 0 | - |
| - | - | 787.9 | 446.7 | - | - | 0 | - |
| - | - | 737.5 | 446.7 | - | - | 0 | - |
| - | - | 6517 | 447.2 | - | - | 0 | - |
| - | - | 931.4 | 447.7 | - | - | 0 | - |
| - | - | 2756 | 448.2 | - | - | 0 | - |
| 10 | y | 5624 | 448.7 | 0.002919 | 6.504 | +2 | 6 |
| 10 | y | 2640 | 449.2 | 0.005531 | 12.31 | +2 | 6 |
| 10 | z | 3754 | 449.7 | 0.002226 | 4.951 | +2 | 6 |
| - | - | 1611 | 450.2 | - | - | 0 | - |
| - | - | 784.5 | 450.7 | - | - | 0 | - |
| 14 | c | 714.2 | 451.2 | 0.008916 | 19.76 | +4 | 14 |
| - | - | 1431 | 452.2 | - | - | 0 | - |
| - | - | 1040 | 452.5 | - | - | 0 | - |
| - | - | 1132 | 452.6 | - | - | 0 | - |
| - | - | 1234 | 452.7 | - | - | 0 | - |
| - | - | 985 | 453.3 | - | - | 0 | - |
| 7 | c | 1112 | 454.2 | 0.0008609 | 1.895 | +2 | 7 |
| - | - | 747.6 | 454.3 | - | - | 0 | - |
| 7 | c | 3558 | 454.7 | 0.00118 | 2.596 | +2 | 7 |
| - | - | 821.9 | 455 | - | - | 0 | - |
| - | - | 1285 | 455.2 | - | - | 0 | - |
| 14 | c | 1959 | 455.7 | 0.0004083 | 0.896 | +4 | 14 |
| - | - | 1471 | 455.9 | - | - | 0 | - |
| - | - | 1974 | 456 | - | - | 0 | - |
| - | - | 2458 | 456.2 | - | - | 0 | - |
| - | - | 1128 | 457.2 | - | - | 0 | - |
| 10 | y | 5552 | 457.7 | 0.002647 | 5.782 | +2 | 6 |
| - | - | 2166 | 458.2 | - | - | 0 | - |
| - | - | 786.8 | 459.2 | - | - | 0 | - |
| - | - | 774.9 | 459.5 | - | - | 0 | - |
| - | - | 2221 | 460 | - | - | 0 | - |
| - | - | 1748 | 460.2 | - | - | 0 | - |
| - | - | 3.26E+04 | 460.7 | - | - | 0 | - |
| - | - | 1757 | 461 | - | - | 0 | - |
| - | - | 1.666E+04 | 461.2 | - | - | 0 | - |
| - | - | 3390 | 461.5 | - | - | 0 | - |
| - | - | 6834 | 461.7 | - | - | 0 | - |
| - | - | 6304 | 462.2 | - | - | 0 | - |
| - | - | 1711 | 462.2 | - | - | 0 | - |
| - | - | 2601 | 462.7 | - | - | 0 | - |
| 7 | c | 2491 | 463.2 | 0.001913 | 4.131 | +2 | 7 |
| - | - | 1248 | 463.7 | - | - | 0 | - |
| - | - | 1947 | 465.2 | - | - | 0 | - |
| - | - | 7000 | 465.5 | - | - | 0 | - |
| 2 | z | 4567 | 465.7 | 0.0004362 | 0.9366 | +4 | 14 |
| - | - | 1135 | 465.9 | - | - | 0 | - |
| 2 | z | 3408 | 466 | 0.004646 | 9.97 | +4 | 14 |
| - | - | 1898 | 466.2 | - | - | 0 | - |
| - | - | 4225 | 466.2 | - | - | 0 | - |
| - | - | 3351 | 466.6 | - | - | 0 | - |
| - | - | 1898 | 466.9 | - | - | 0 | - |
| - | - | 3662 | 467.2 | - | - | 0 | - |
| - | - | 852.7 | 467.8 | - | - | 0 | - |
| - | - | 1.915E+04 | 468.3 | - | - | 0 | - |
| - | - | 1.168E+04 | 468.8 | - | - | 0 | - |
| - | - | 3611 | 469.2 | - | - | 0 | - |
| - | - | 2691 | 469.3 | - | - | 0 | - |
| - | - | 1423 | 469.5 | - | - | 0 | - |
| 2 | y | 6159 | 469.7 | 0.004178 | 8.895 | +4 | 14 |
| 2 | y | 7716 | 470 | 0.004939 | 10.51 | +4 | 14 |
| 2 | z | 9244 | 470.2 | 0.003654 | 7.772 | +4 | 14 |
| - | - | 5760 | 470.5 | - | - | 0 | - |
| - | - | 2295 | 470.7 | - | - | 0 | - |
| - | - | 708.1 | 471.5 | - | - | 0 | - |
| - | - | 1001 | 471.7 | - | - | 0 | - |
| - | - | 2158 | 472 | - | - | 0 | - |
| - | - | 1785 | 473.2 | - | - | 0 | - |
| - | - | 2143 | 473.7 | - | - | 0 | - |
| - | - | 1976 | 474 | - | - | 0 | - |
| 2 | y | 1.549E+04 | 474.2 | 0.0006604 | 1.393 | +4 | 14 |
| - | - | 1.279E+04 | 474.5 | - | - | 0 | - |
| - | - | 7161 | 474.7 | - | - | 0 | - |
| - | - | 3094 | 475 | - | - | 0 | - |
| - | - | 1779 | 475.2 | - | - | 0 | - |
| - | - | 2800 | 475.7 | - | - | 0 | - |
| - | - | 2157 | 476 | - | - | 0 | - |
| - | - | 2588 | 476.2 | - | - | 0 | - |
| - | - | 2834 | 476.2 | - | - | 0 | - |
| - | - | 3992 | 476.5 | - | - | 0 | - |
| - | - | 2230 | 476.7 | - | - | 0 | - |
| - | - | 2038 | 477 | - | - | 0 | - |
| 13 | y | 5068 | 477.2 | 0.004868 | 10.2 | +1 | 3 |
| - | - | 1581 | 477.7 | - | - | 0 | - |
| - | - | 851.6 | 477.8 | - | - | 0 | - |
| 13 | z | 4719 | 478.2 | 0.001957 | 4.092 | +1 | 3 |
| - | - | 1099 | 478.7 | - | - | 0 | - |
| - | - | 1293 | 479.2 | - | - | 0 | - |
| - | - | 1214 | 479.9 | - | - | 0 | - |
| - | - | 909.6 | 480 | - | - | 0 | - |
| - | - | 3744 | 480.2 | - | - | 0 | - |
| - | - | 5.453E+04 | 480.5 | - | - | 0 | - |
| - | - | 5.195E+04 | 480.7 | - | - | 0 | - |
| - | - | 3.673E+04 | 481 | - | - | 0 | - |
| - | - | 1.368E+04 | 481.2 | - | - | 0 | - |
| - | - | 5081 | 481.5 | - | - | 0 | - |
| - | - | 1814 | 481.7 | - | - | 0 | - |
| - | - | 1388 | 482.2 | - | - | 0 | - |
| 8 | c | 5454 | 482.7 | 0.002659 | 5.508 | +2 | 8 |
| 8 | c | 1.137E+04 | 483.2 | 0.003113 | 6.443 | +2 | 8 |
| - | - | 3061 | 483.5 | - | - | 0 | - |
| - | - | 7353 | 483.7 | - | - | 0 | - |
| - | - | 2007 | 484 | - | - | 0 | - |
| - | - | 1.928E+04 | 484.3 | - | - | 0 | - |
| - | - | 7178 | 484.5 | - | - | 0 | - |
| - | - | 5624 | 484.6 | - | - | 0 | - |
| 9 | w | 8547 | 484.7 | 0.002619 | 5.403 | +2 | 7 |
| - | - | 6199 | 484.8 | - | - | 0 | - |
| - | - | 3773 | 484.9 | - | - | 0 | - |
| - | - | 4038 | 485 | - | - | 0 | - |
| - | - | 4604 | 485.2 | - | - | 0 | - |
| - | - | 3015 | 485.2 | - | - | 0 | - |
| - | - | 975.3 | 485.5 | - | - | 0 | - |
| - | - | 1598 | 485.6 | - | - | 0 | - |
| - | - | 1893 | 485.7 | - | - | 0 | - |
| - | - | 1186 | 485.9 | - | - | 0 | - |
| - | - | 1070 | 486.2 | - | - | 0 | - |
| - | - | 940.4 | 487.2 | - | - | 0 | - |
| - | - | 6594 | 487.5 | - | - | 0 | - |
| - | - | 1.34E+05 | 487.7 | - | - | 0 | - |
| - | - | 1.285E+05 | 488 | - | - | 0 | - |
| - | - | 9.235E+04 | 488.2 | - | - | 0 | - |
| - | - | 3.643E+04 | 488.5 | - | - | 0 | - |
| - | - | 1.131E+04 | 488.7 | - | - | 0 | - |
| - | - | 5237 | 489 | - | - | 0 | - |
| - | - | 5572 | 489.2 | - | - | 0 | - |
| - | - | 1254 | 489.7 | - | - | 0 | - |
| - | - | 936.5 | 490.2 | - | - | 0 | - |
| - | - | 1069 | 490.6 | - | - | 0 | - |
| - | - | 2062 | 491.2 | - | - | 0 | - |
| - | - | 1576 | 491.5 | - | - | 0 | - |
| 8 | c | 2.065E+04 | 491.7 | 0.001619 | 3.291 | +2 | 8 |
| - | - | 2074 | 492 | - | - | 0 | - |
| - | - | 1.385E+04 | 492.2 | - | - | 0 | - |
| - | - | 4306 | 492.7 | - | - | 0 | - |
| 4 | c | 2418 | 493.3 | 0.004138 | 8.388 | +1 | 4 |
| - | - | 1582 | 493.6 | - | - | 0 | - |
| - | - | 1139 | 493.7 | - | - | 0 | - |
| 5 | z | 1673 | 493.9 | 0.005957 | 12.06 | +3 | 11 |
| - | - | 1423 | 494 | - | - | 0 | - |
| 4 | c | 7.547E+04 | 494.2 | 0.001581 | 3.199 | +1 | 4 |
| - | - | 1244 | 494.6 | - | - | 0 | - |
| - | - | 2.004E+04 | 495.2 | - | - | 0 | - |
| - | - | 1998 | 495.8 | - | - | 0 | - |
| - | - | 2388 | 496.2 | - | - | 0 | - |
| - | - | 2398 | 496.2 | - | - | 0 | - |
| - | - | 6371 | 496.8 | - | - | 0 | - |
| - | - | 1889 | 497.2 | - | - | 0 | - |
| - | - | 5456 | 497.3 | - | - | 0 | - |
| - | - | 6683 | 497.8 | - | - | 0 | - |
| - | - | 4967 | 498 | - | - | 0 | - |
| - | - | 1.205E+04 | 498.2 | - | - | 0 | - |
| - | - | 1.1E+04 | 498.5 | - | - | 0 | - |
| - | - | 8669 | 498.7 | - | - | 0 | - |
| - | - | 816.7 | 498.9 | - | - | 0 | - |
| - | - | 4184 | 499 | - | - | 0 | - |
| 5 | y | 1.429E+04 | 499.2 | 0.006238 | 12.49 | +3 | 11 |
| 5 | y | 1.467E+04 | 499.6 | 0.001235 | 2.472 | +3 | 11 |
| - | - | 2024 | 499.7 | - | - | 0 | - |
| 5 | z | 1.826E+04 | 499.9 | 0.00246 | 4.921 | +3 | 11 |
| - | - | 1.031E+04 | 500.2 | - | - | 0 | - |
| - | - | 3883 | 500.6 | - | - | 0 | - |
| - | - | 1049 | 500.9 | - | - | 0 | - |
| - | - | 821.5 | 501.3 | - | - | 0 | - |
| - | - | 2804 | 502 | - | - | 0 | - |
| - | - | 1.227E+04 | 502.3 | - | - | 0 | - |
| - | - | 2.129E+05 | 502.5 | - | - | 0 | - |
| - | - | 2.905E+05 | 502.7 | - | - | 0 | - |
| - | - | 2.2E+05 | 503 | - | - | 0 | - |
| - | - | 1.218E+05 | 503.3 | - | - | 0 | - |
| - | - | 4.07E+04 | 503.5 | - | - | 0 | - |
| - | - | 1.702E+04 | 503.8 | - | - | 0 | - |
| - | - | 5834 | 504 | - | - | 0 | - |
| - | - | 807.1 | 504.2 | - | - | 0 | - |
| - | - | 1191 | 504.2 | - | - | 0 | - |
| - | - | 1335 | 504.6 | - | - | 0 | - |
| 9 | z | 813.7 | 504.8 | 0.0005717 | 1.133 | +2 | 7 |
| 12 | c | 2540 | 504.9 | 0.001765 | 3.496 | +3 | 12 |
| 12 | c | 2.019E+04 | 505.3 | 0.000211 | 0.4175 | +3 | 12 |
| - | - | 1.16E+04 | 505.6 | - | - | 0 | - |
| - | - | 1.061E+04 | 505.7 | - | - | 0 | - |
| - | - | 8157 | 505.9 | - | - | 0 | - |
| - | - | 5305 | 506.2 | - | - | 0 | - |
| - | - | 1103 | 506.6 | - | - | 0 | - |
| - | - | 3993 | 506.7 | - | - | 0 | - |
| - | - | 1433 | 507.2 | - | - | 0 | - |
| - | - | 4343 | 508.6 | - | - | 0 | - |
| - | - | 3355 | 508.9 | - | - | 0 | - |
| - | - | 1725 | 509.2 | - | - | 0 | - |
| - | - | 1405 | 509.6 | - | - | 0 | - |
| - | - | 2490 | 510.6 | - | - | 0 | - |
| 12 | c | 6.716E+04 | 510.9 | 0.00158 | 3.092 | +3 | 12 |
| 4 | c | 1.293E+05 | 511.3 | 0.001533 | 2.999 | +1 | 4 |
| - | - | 2.584E+04 | 511.6 | - | - | 0 | - |
| - | - | 9322 | 511.9 | - | - | 0 | - |
| - | - | 2.297E+04 | 512.3 | - | - | 0 | - |
| 9 | y | 5931 | 512.8 | 0.003098 | 6.041 | +2 | 7 |
| - | - | 9956 | 513.2 | - | - | 0 | - |
| - | - | 4196 | 513.3 | - | - | 0 | - |
| 9 | z | 3.168E+04 | 513.8 | 0.003718 | 7.237 | +2 | 7 |
| - | - | 2.27E+04 | 514.3 | - | - | 0 | - |
| - | - | 4400 | 514.6 | - | - | 0 | - |
| - | - | 8632 | 514.8 | - | - | 0 | - |
| - | - | 2081 | 514.9 | - | - | 0 | - |
| - | - | 2768 | 515.3 | - | - | 0 | - |
| - | - | 9436 | 515.3 | - | - | 0 | - |
| - | - | 1380 | 516.2 | - | - | 0 | - |
| - | - | 3023 | 516.3 | - | - | 0 | - |
| - | - | 2882 | 517.3 | - | - | 0 | - |
| - | - | 1733 | 518.3 | - | - | 0 | - |
| - | - | 2349 | 518.8 | - | - | 0 | - |
| - | - | 2596 | 519.3 | - | - | 0 | - |
| - | - | 918.7 | 519.6 | - | - | 0 | - |
| - | - | 3328 | 519.9 | - | - | 0 | - |
| - | - | 2124 | 520.3 | - | - | 0 | - |
| - | - | 1216 | 520.7 | - | - | 0 | - |
| - | - | 2160 | 521.3 | - | - | 0 | - |
| 9 | y | 3.168E+04 | 521.8 | 0.003131 | 6.001 | +2 | 7 |
| - | - | 1.655E+04 | 522.3 | - | - | 0 | - |
| - | - | 5245 | 522.8 | - | - | 0 | - |
| - | - | 2487 | 523.3 | - | - | 0 | - |
| - | - | 1601 | 523.6 | - | - | 0 | - |
| - | - | 1591 | 525.3 | - | - | 0 | - |
| - | - | 2836 | 526.3 | - | - | 0 | - |
| - | - | 1037 | 526.8 | - | - | 0 | - |
| - | - | 2856 | 527.3 | - | - | 0 | - |
| - | - | 2955 | 527.8 | - | - | 0 | - |
| - | - | 2601 | 532.2 | - | - | 0 | - |
| - | - | 1967 | 532.8 | - | - | 0 | - |
| - | - | 1112 | 533.2 | - | - | 0 | - |
| - | - | 969 | 533.2 | - | - | 0 | - |
| - | - | 1745 | 533.3 | - | - | 0 | - |
| - | - | 2755 | 533.8 | - | - | 0 | - |
| - | - | 2460 | 534.3 | - | - | 0 | - |
| - | - | 1293 | 534.3 | - | - | 0 | - |
| - | - | 2546 | 535.2 | - | - | 0 | - |
| - | - | 1027 | 537.3 | - | - | 0 | - |
| 4 | z | 2449 | 538.3 | 0.001967 | 3.653 | +3 | 12 |
| - | - | 1243 | 538.6 | - | - | 0 | - |
| - | - | 891.2 | 538.8 | - | - | 0 | - |
| - | - | 3820 | 539.2 | - | - | 0 | - |
| - | - | 1105 | 540.2 | - | - | 0 | - |
| 8 | y | 8810 | 541.3 | 0.003149 | 5.817 | +2 | 8 |
| 8 | y | 4744 | 541.8 | 0.006736 | 12.43 | +2 | 8 |
| 8 | z | 3.235E+04 | 542.3 | 0.00206 | 3.798 | +2 | 8 |
| - | - | 1.882E+04 | 542.8 | - | - | 0 | - |
| - | - | 1.083E+04 | 543.3 | - | - | 0 | - |
| 4 | y | 3556 | 543.6 | 0.002586 | 4.758 | +3 | 12 |
| - | - | 1902 | 543.8 | - | - | 0 | - |
| - | - | 2262 | 543.9 | - | - | 0 | - |
| - | - | 2360 | 544.3 | - | - | 0 | - |
| - | - | 1049 | 545.3 | - | - | 0 | - |
| - | - | 782.9 | 545.6 | - | - | 0 | - |
| - | - | 8874 | 546.3 | - | - | 0 | - |
| - | - | 4849 | 546.8 | - | - | 0 | - |
| - | - | 4082 | 547.3 | - | - | 0 | - |
| - | - | 1167 | 547.6 | - | - | 0 | - |
| - | - | 2650 | 547.8 | - | - | 0 | - |
| 12 | w | 2909 | 548.2 | 0.006489 | 11.83 | +1 | 4 |
| - | - | 2074 | 548.3 | - | - | 0 | - |
| - | - | 1246 | 549.2 | - | - | 0 | - |
| - | - | 5840 | 549.6 | - | - | 0 | - |
| - | - | 939.4 | 549.8 | - | - | 0 | - |
| - | - | 5899 | 549.9 | - | - | 0 | - |
| 8 | y | 6.514E+04 | 550.3 | 0.003365 | 6.115 | +2 | 8 |
| - | - | 3.659E+04 | 550.8 | - | - | 0 | - |
| - | - | 1082 | 550.9 | - | - | 0 | - |
| - | - | 1.508E+04 | 551.3 | - | - | 0 | - |
| - | - | 1314 | 551.6 | - | - | 0 | - |
| - | - | 5450 | 551.8 | - | - | 0 | - |
| - | - | 3505 | 552.3 | - | - | 0 | - |
| - | - | 2142 | 553.3 | - | - | 0 | - |
| - | - | 1278 | 553.8 | - | - | 0 | - |
| - | - | 3736 | 554.3 | - | - | 0 | - |
| - | - | 3852 | 554.8 | - | - | 0 | - |
| - | - | 3.074E+04 | 555.3 | - | - | 0 | - |
| 9 | c | 2.098E+05 | 555.8 | 0.0007068 | 1.272 | +2 | 9 |
| - | - | 1.399E+05 | 556.3 | - | - | 0 | - |
| - | - | 4.62E+04 | 556.8 | - | - | 0 | - |
| - | - | 2016 | 556.9 | - | - | 0 | - |
| - | - | 1.214E+04 | 557.3 | - | - | 0 | - |
| - | - | 1260 | 557.6 | - | - | 0 | - |
| - | - | 1951 | 557.8 | - | - | 0 | - |
| - | - | 1570 | 559.3 | - | - | 0 | - |
| - | - | 904.4 | 559.6 | - | - | 0 | - |
| - | - | 975.1 | 559.9 | - | - | 0 | - |
| - | - | 991.3 | 560.9 | - | - | 0 | - |
| - | - | 4204 | 561.3 | - | - | 0 | - |
| 3 | w | 7.43E+04 | 561.6 | 0.001591 | 2.833 | +3 | 13 |
| - | - | 6.046E+04 | 561.9 | - | - | 0 | - |
| - | - | 3.646E+04 | 562.3 | - | - | 0 | - |
| - | - | 1.287E+04 | 562.6 | - | - | 0 | - |
| - | - | 3044 | 562.6 | - | - | 0 | - |
| - | - | 2196 | 562.8 | - | - | 0 | - |
| - | - | 3747 | 562.9 | - | - | 0 | - |
| - | - | 1690 | 563 | - | - | 0 | - |
| - | - | 3690 | 563.3 | - | - | 0 | - |
| - | - | 2769 | 563.3 | - | - | 0 | - |
| - | - | 1297 | 563.6 | - | - | 0 | - |
| - | - | 2.517E+04 | 563.8 | - | - | 0 | - |
| - | - | 872.3 | 563.9 | - | - | 0 | - |
| - | - | 2.131E+04 | 564.3 | - | - | 0 | - |
| 13 | c | 5.036E+04 | 564.6 | 2.71E-05 | 0.04799 | +3 | 13 |
| - | - | 7861 | 564.8 | - | - | 0 | - |
| - | - | 4.745E+04 | 564.9 | - | - | 0 | - |
| - | - | 2.928E+04 | 565.3 | - | - | 0 | - |
| - | - | 9979 | 565.6 | - | - | 0 | - |
| - | - | 4268 | 565.9 | - | - | 0 | - |
| - | - | 6581 | 566.3 | - | - | 0 | - |
| - | - | 3896 | 566.6 | - | - | 0 | - |
| - | - | 3389 | 566.9 | - | - | 0 | - |
| - | - | 1.281E+04 | 567.2 | - | - | 0 | - |
| - | - | 2763 | 567.3 | - | - | 0 | - |
| - | - | 4432 | 568.2 | - | - | 0 | - |
| - | - | 2977 | 568.3 | - | - | 0 | - |
| - | - | 1.517E+04 | 568.8 | - | - | 0 | - |
| - | - | 808.5 | 568.9 | - | - | 0 | - |
| - | - | 1.068E+04 | 569.3 | - | - | 0 | - |
| - | - | 4521 | 569.8 | - | - | 0 | - |
| - | - | 1237 | 569.9 | - | - | 0 | - |
| - | - | 2969 | 570.3 | - | - | 0 | - |
| - | - | 2981 | 570.7 | - | - | 0 | - |
| - | - | 4401 | 570.8 | - | - | 0 | - |
| - | - | 5.411E+04 | 571.3 | - | - | 0 | - |
| - | - | 3.224E+04 | 571.8 | - | - | 0 | - |
| - | - | 1.395E+04 | 572.3 | - | - | 0 | - |
| - | - | 4237 | 572.8 | - | - | 0 | - |
| - | - | 2787 | 573 | - | - | 0 | - |
| - | - | 4586 | 573.3 | - | - | 0 | - |
| - | - | 3400 | 573.6 | - | - | 0 | - |
| - | - | 1703 | 574 | - | - | 0 | - |
| - | - | 1810 | 574.6 | - | - | 0 | - |
| - | - | 4579 | 576.3 | - | - | 0 | - |
| - | - | 3216 | 576.8 | - | - | 0 | - |
| 7 | w | 3664 | 577.3 | 0.005901 | 10.22 | +2 | 9 |
| - | - | 4378 | 577.8 | - | - | 0 | - |
| - | - | 3585 | 578 | - | - | 0 | - |
| - | - | 6274 | 578.3 | - | - | 0 | - |
| - | - | 2998 | 578.6 | - | - | 0 | - |
| - | - | 2.126E+04 | 578.8 | - | - | 0 | - |
| - | - | 1.442E+04 | 579.3 | - | - | 0 | - |
| - | - | 6283 | 579.8 | - | - | 0 | - |
| 3 | y | 3852 | 580.3 | 0.003158 | 5.443 | +3 | 13 |
| 3 | y | 6779 | 580.6 | 0.001691 | 2.913 | +3 | 13 |
| 3 | z | 2.095E+04 | 580.9 | 0.001182 | 2.034 | +3 | 13 |
| - | - | 2.126E+04 | 581.3 | - | - | 0 | - |
| - | - | 1193 | 581.3 | - | - | 0 | - |
| - | - | 1.119E+04 | 581.6 | - | - | 0 | - |
| - | - | 3956 | 581.9 | - | - | 0 | - |
| - | - | 3337 | 582.3 | - | - | 0 | - |
| - | - | 1114 | 583 | - | - | 0 | - |
| - | - | 3261 | 583.3 | - | - | 0 | - |
| - | - | 3297 | 583.6 | - | - | 0 | - |
| - | - | 2567 | 583.9 | - | - | 0 | - |
| - | - | 1743 | 584.8 | - | - | 0 | - |
| - | - | 1715 | 585.3 | - | - | 0 | - |
| - | - | 1329 | 585.8 | - | - | 0 | - |
| 3 | y | 1.164E+04 | 586.3 | 3.021E-06 | 0.005153 | +3 | 13 |
| - | - | 9241 | 586.6 | - | - | 0 | - |
| - | - | 6526 | 586.9 | - | - | 0 | - |
| - | - | 2517 | 587.3 | - | - | 0 | - |
| - | - | 2197 | 587.6 | - | - | 0 | - |
| - | - | 1874 | 588 | - | - | 0 | - |
| - | - | 4575 | 588.3 | - | - | 0 | - |
| - | - | 2949 | 588.6 | - | - | 0 | - |
| - | - | 2230 | 589 | - | - | 0 | - |
| - | - | 2702 | 590.3 | - | - | 0 | - |
| 7 | z | 2683 | 590.8 | 0.001409 | 2.385 | +2 | 9 |
| 7 | z | 4025 | 591.3 | 0.01044 | 17.65 | +2 | 9 |
| - | - | 1029 | 591.8 | - | - | 0 | - |
| - | - | 2217 | 592.3 | - | - | 0 | - |
| - | - | 1.126E+04 | 592.6 | - | - | 0 | - |
| - | - | 1.706E+04 | 593 | - | - | 0 | - |
| - | - | 3352 | 593.3 | - | - | 0 | - |
| - | - | 6918 | 593.3 | - | - | 0 | - |
| - | - | 2861 | 593.6 | - | - | 0 | - |
| - | - | 1172 | 593.8 | - | - | 0 | - |
| - | - | 1305 | 594 | - | - | 0 | - |
| - | - | 2521 | 594.3 | - | - | 0 | - |
| - | - | 1230 | 595.2 | - | - | 0 | - |
| - | - | 2614 | 597 | - | - | 0 | - |
| - | - | 2916 | 597.3 | - | - | 0 | - |
| - | - | 2644 | 597.6 | - | - | 0 | - |
| - | - | 2528 | 597.8 | - | - | 0 | - |
| - | - | 1700 | 598 | - | - | 0 | - |
| - | - | 9861 | 598.3 | - | - | 0 | - |
| 7 | y | 1.26E+04 | 598.8 | 0.0002016 | 0.3367 | +2 | 9 |
| 7 | y | 1.524E+04 | 599.3 | 0.005166 | 8.62 | +2 | 9 |
| 7 | z | 6.612E+04 | 599.8 | 0.002286 | 3.812 | +2 | 9 |
| - | - | 6.295E+04 | 600.3 | - | - | 0 | - |
| - | - | 3.27E+04 | 600.8 | - | - | 0 | - |
| 14 | c | 1.071E+04 | 601.3 | 0.00892 | 14.84 | +3 | 14 |
| 14 | c | 3513 | 601.6 | 0.006153 | 10.23 | +3 | 14 |
| - | - | 3428 | 601.8 | - | - | 0 | - |
| - | - | 2967 | 602 | - | - | 0 | - |
| - | - | 3602 | 602.3 | - | - | 0 | - |
| - | - | 3591 | 602.6 | - | - | 0 | - |
| - | - | 3355 | 602.8 | - | - | 0 | - |
| - | - | 1.079E+04 | 602.9 | - | - | 0 | - |
| - | - | 1.421E+04 | 603.3 | - | - | 0 | - |
| - | - | 9433 | 603.6 | - | - | 0 | - |
| - | - | 1269 | 603.8 | - | - | 0 | - |
| - | - | 2300 | 604 | - | - | 0 | - |
| - | - | 1234 | 604.3 | - | - | 0 | - |
| 12 | z | 3.845E+04 | 606.3 | 0.007404 | 12.21 | +1 | 4 |
| - | - | 2480 | 606.6 | - | - | 0 | - |
| - | - | 1224 | 606.8 | - | - | 0 | - |
| - | - | 1.26E+04 | 607 | - | - | 0 | - |
| 14 | c | 7.565E+04 | 607.3 | 0.001291 | 2.126 | +3 | 14 |
| - | - | 5.927E+04 | 607.6 | - | - | 0 | - |
| 7 | y | 2.229E+05 | 607.8 | 0.00292 | 4.805 | +2 | 9 |
| - | - | 3E+04 | 608 | - | - | 0 | - |
| - | - | 1.613E+05 | 608.3 | - | - | 0 | - |
| - | - | 5755 | 608.6 | - | - | 0 | - |
| - | - | 5.989E+04 | 608.8 | - | - | 0 | - |
| - | - | 1.845E+04 | 609.3 | - | - | 0 | - |
| - | - | 4003 | 609.8 | - | - | 0 | - |
| - | - | 1.298E+04 | 610.3 | - | - | 0 | - |
| - | - | 871.1 | 610.8 | - | - | 0 | - |
| - | - | 4274 | 611.3 | - | - | 0 | - |
| - | - | 3106 | 611.6 | - | - | 0 | - |
| 10 | c | 1.443E+04 | 611.8 | 0.007028 | 11.49 | +2 | 10 |
| - | - | 1.415E+04 | 612 | - | - | 0 | - |
| - | - | 2.665E+04 | 612.3 | - | - | 0 | - |
| - | - | 6476 | 612.6 | - | - | 0 | - |
| - | - | 1.35E+04 | 612.8 | - | - | 0 | - |
| - | - | 2924 | 613 | - | - | 0 | - |
| - | - | 6154 | 613.3 | - | - | 0 | - |
| - | - | 4121 | 613.8 | - | - | 0 | - |
| - | - | 1732 | 614.3 | - | - | 0 | - |
| - | - | 4146 | 615.3 | - | - | 0 | - |
| - | - | 866.1 | 615.6 | - | - | 0 | - |
| - | - | 2112 | 615.8 | - | - | 0 | - |
| - | - | 2316 | 616.3 | - | - | 0 | - |
| - | - | 1028 | 616.6 | - | - | 0 | - |
| - | - | 1610 | 618.3 | - | - | 0 | - |
| - | - | 1121 | 619.3 | - | - | 0 | - |
| - | - | 3112 | 619.8 | - | - | 0 | - |
| 10 | c | 7.665E+04 | 620.3 | 0.002604 | 4.197 | +2 | 10 |
| 2 | z | 7272 | 620.6 | 0.01025 | 16.52 | +3 | 14 |
| - | - | 5.542E+04 | 620.8 | - | - | 0 | - |
| - | - | 8470 | 621 | - | - | 0 | - |
| - | - | 7455 | 621.2 | - | - | 0 | - |
| - | - | 3.327E+04 | 621.3 | - | - | 0 | - |
| - | - | 4186 | 621.6 | - | - | 0 | - |
| - | - | 5495 | 621.8 | - | - | 0 | - |
| - | - | 3415 | 622 | - | - | 0 | - |
| - | - | 2172 | 622.2 | - | - | 0 | - |
| 12 | y | 2.486E+04 | 622.3 | 0.008001 | 12.86 | +1 | 4 |
| - | - | 6355 | 623.3 | - | - | 0 | - |
| - | - | 1938 | 624.3 | - | - | 0 | - |
| - | - | 2784 | 625.3 | - | - | 0 | - |
| - | - | 5889 | 625.7 | - | - | 0 | - |
| - | - | 6271 | 626 | - | - | 0 | - |
| - | - | 8162 | 626.3 | - | - | 0 | - |
| 2 | z | 3.652E+04 | 626.6 | 0.0008079 | 1.289 | +3 | 14 |
| - | - | 1218 | 626.8 | - | - | 0 | - |
| - | - | 3.516E+04 | 627 | - | - | 0 | - |
| - | - | 2.629E+04 | 627.3 | - | - | 0 | - |
| - | - | 1.187E+04 | 627.6 | - | - | 0 | - |
| - | - | 1788 | 627.8 | - | - | 0 | - |
| - | - | 5469 | 628 | - | - | 0 | - |
| - | - | 1203 | 628.3 | - | - | 0 | - |
| - | - | 1217 | 629.3 | - | - | 0 | - |
| - | - | 1799 | 630.3 | - | - | 0 | - |
| - | - | 5563 | 630.6 | - | - | 0 | - |
| - | - | 6091 | 631 | - | - | 0 | - |
| - | - | 2238 | 631.3 | - | - | 0 | - |
| - | - | 1649 | 631.6 | - | - | 0 | - |
| 2 | y | 3114 | 632 | 0.0004056 | 0.6418 | +3 | 14 |
| - | - | 4755 | 632.3 | - | - | 0 | - |
| - | - | 1905 | 632.6 | - | - | 0 | - |
| - | - | 2799 | 633.8 | - | - | 0 | - |
| - | - | 1.62E+04 | 634.3 | - | - | 0 | - |
| - | - | 3210 | 634.6 | - | - | 0 | - |
| 6 | w | 1.744E+04 | 634.8 | 0.002588 | 4.076 | +2 | 10 |
| - | - | 2509 | 634.8 | - | - | 0 | - |
| - | - | 6120 | 635 | - | - | 0 | - |
| - | - | 1.11E+04 | 635.3 | - | - | 0 | - |
| - | - | 5400 | 635.7 | - | - | 0 | - |
| - | - | 4643 | 635.8 | - | - | 0 | - |
| - | - | 1961 | 635.8 | - | - | 0 | - |
| - | - | 3226 | 636 | - | - | 0 | - |
| - | - | 2749 | 636.3 | - | - | 0 | - |
| - | - | 3186 | 636.3 | - | - | 0 | - |
| - | - | 1143 | 636.8 | - | - | 0 | - |
| - | - | 2116 | 637.4 | - | - | 0 | - |
| - | - | 1290 | 638.8 | - | - | 0 | - |
| - | - | 2055 | 639.3 | - | - | 0 | - |
| - | - | 3794 | 640 | - | - | 0 | - |
| - | - | 2.319E+04 | 640.3 | - | - | 0 | - |
| - | - | 2.362E+04 | 640.7 | - | - | 0 | - |
| - | - | 1768 | 640.8 | - | - | 0 | - |
| - | - | 2.324E+04 | 641 | - | - | 0 | - |
| - | - | 1.116E+04 | 641.3 | - | - | 0 | - |
| - | - | 3289 | 641.7 | - | - | 0 | - |
| - | - | 3986 | 641.8 | - | - | 0 | - |
| - | - | 1588 | 642 | - | - | 0 | - |
| - | - | 3852 | 642.3 | - | - | 0 | - |
| - | - | 1275 | 643 | - | - | 0 | - |
| - | - | 1641 | 643.3 | - | - | 0 | - |
| - | - | 5992 | 643.8 | - | - | 0 | - |
| - | - | 8343 | 644.3 | - | - | 0 | - |
| - | - | 6882 | 644.7 | - | - | 0 | - |
| - | - | 2147 | 644.8 | - | - | 0 | - |
| - | - | 9656 | 645 | - | - | 0 | - |
| - | - | 5085 | 645.3 | - | - | 0 | - |
| - | - | 4065 | 645.7 | - | - | 0 | - |
| - | - | 4648 | 646 | - | - | 0 | - |
| - | - | 3966 | 646.3 | - | - | 0 | - |
| - | - | 1638 | 646.7 | - | - | 0 | - |
| 6 | z | 2637 | 647.3 | 0.00211 | 3.259 | +2 | 10 |
| - | - | 2508 | 649.3 | - | - | 0 | - |
| - | - | 7059 | 649.7 | - | - | 0 | - |
| - | - | 1.423E+04 | 650 | - | - | 0 | - |
| - | - | 5.74E+04 | 650.3 | - | - | 0 | - |
| - | - | 5.672E+04 | 650.7 | - | - | 0 | - |
| - | - | 3.985E+04 | 651 | - | - | 0 | - |
| - | - | 1.84E+04 | 651.3 | - | - | 0 | - |
| - | - | 6673 | 651.7 | - | - | 0 | - |
| - | - | 1711 | 651.8 | - | - | 0 | - |
| - | - | 3037 | 652 | - | - | 0 | - |
| - | - | 1.447E+04 | 652.3 | - | - | 0 | - |
| - | - | 6071 | 653.3 | - | - | 0 | - |
| - | - | 2202 | 654.3 | - | - | 0 | - |
| - | - | 4145 | 654.7 | - | - | 0 | - |
| - | - | 7358 | 655 | - | - | 0 | - |
| 6 | y | 1.248E+04 | 655.3 | 0.003415 | 5.211 | +2 | 10 |
| - | - | 9583 | 655.7 | - | - | 0 | - |
| 6 | y | 3977 | 655.8 | 0.002929 | 4.466 | +2 | 10 |
| - | - | 4463 | 656 | - | - | 0 | - |
| 6 | z | 2.96E+04 | 656.3 | 0.002265 | 3.451 | +2 | 10 |
| - | - | 2.021E+04 | 656.8 | - | - | 0 | - |
| - | - | 1.196E+04 | 657.3 | - | - | 0 | - |
| - | - | 7762 | 657.8 | - | - | 0 | - |
| - | - | 4042 | 658.3 | - | - | 0 | - |
| - | - | 6679 | 658.7 | - | - | 0 | - |
| - | - | 2019 | 658.8 | - | - | 0 | - |
| - | - | 5679 | 659 | - | - | 0 | - |
| - | - | 5332 | 659.3 | - | - | 0 | - |
| - | - | 3302 | 659.7 | - | - | 0 | - |
| - | - | 1791 | 660 | - | - | 0 | - |
| - | - | 1236 | 660.3 | - | - | 0 | - |
| - | - | 1170 | 660.7 | - | - | 0 | - |
| - | - | 1253 | 661 | - | - | 0 | - |
| - | - | 2754 | 662.3 | - | - | 0 | - |
| - | - | 7988 | 663.3 | - | - | 0 | - |
| - | - | 3162 | 663.3 | - | - | 0 | - |
| - | - | 1161 | 663.7 | - | - | 0 | - |
| - | - | 6769 | 663.8 | - | - | 0 | - |
| - | - | 2405 | 664 | - | - | 0 | - |
| 6 | y | 6.755E+04 | 664.3 | 0.002106 | 3.17 | +2 | 10 |
| - | - | 9.597E+04 | 664.7 | - | - | 0 | - |
| - | - | 2.795E+04 | 664.8 | - | - | 0 | - |
| - | - | 9.138E+04 | 665 | - | - | 0 | - |
| - | - | 7.157E+04 | 665.3 | - | - | 0 | - |
| - | - | 2.357E+04 | 665.7 | - | - | 0 | - |
| - | - | 5475 | 665.8 | - | - | 0 | - |
| - | - | 1.343E+04 | 666 | - | - | 0 | - |
| - | - | 4790 | 666.3 | - | - | 0 | - |
| - | - | 1702 | 666.7 | - | - | 0 | - |
| - | - | 2086 | 669.3 | - | - | 0 | - |
| - | - | 2.387E+04 | 669.7 | - | - | 0 | - |
| - | - | 4.439E+04 | 670 | - | - | 0 | - |
| - | - | 6.53E+04 | 670.3 | - | - | 0 | - |
| - | - | 4.82E+04 | 670.7 | - | - | 0 | - |
| - | - | 1555 | 670.8 | - | - | 0 | - |
| - | - | 2.705E+04 | 671 | - | - | 0 | - |
| - | - | 1.035E+04 | 671.3 | - | - | 0 | - |
| - | - | 3927 | 671.7 | - | - | 0 | - |
| - | - | 2176 | 671.8 | - | - | 0 | - |
| - | - | 2130 | 672 | - | - | 0 | - |
| - | - | 2652 | 672.3 | - | - | 0 | - |
| - | - | 1402 | 677.8 | - | - | 0 | - |
| 5 | c | 5245 | 679.3 | 0.01183 | 17.41 | +1 | 5 |
| - | - | 3.443E+04 | 679.8 | - | - | 0 | - |
| 5 | c | 1.077E+05 | 680.3 | 0.004738 | 6.965 | +1 | 5 |
| - | - | 1.343E+04 | 680.8 | - | - | 0 | - |
| - | - | 4.486E+04 | 681.3 | - | - | 0 | - |
| - | - | 1367 | 681.8 | - | - | 0 | - |
| - | - | 1.424E+04 | 682.3 | - | - | 0 | - |
| - | - | 1230 | 682.8 | - | - | 0 | - |
| - | - | 3518 | 683.3 | - | - | 0 | - |
| - | - | 1692 | 684.3 | - | - | 0 | - |
| - | - | 1210 | 685.3 | - | - | 0 | - |
| - | - | 2960 | 690.8 | - | - | 0 | - |
| - | - | 3.935E+04 | 691.3 | - | - | 0 | - |
| - | - | 2.64E+04 | 691.8 | - | - | 0 | - |
| - | - | 1.277E+04 | 692.3 | - | - | 0 | - |
| 11 | c | 5761 | 692.8 | 0.002894 | 4.177 | +2 | 11 |
| 11 | c | 2635 | 693.3 | 0.006858 | 9.891 | +2 | 11 |
| - | - | 2303 | 693.8 | - | - | 0 | - |
| - | - | 1887 | 694.3 | - | - | 0 | - |
| - | - | 1037 | 696.3 | - | - | 0 | - |
| - | - | 1013 | 696.8 | - | - | 0 | - |
| 5 | c | 3.234E+04 | 697.3 | 0.001505 | 2.158 | +1 | 5 |
| - | - | 1.429E+04 | 698.3 | - | - | 0 | - |
| - | - | 5279 | 699.3 | - | - | 0 | - |
| - | - | 2951 | 701.3 | - | - | 0 | - |
| 11 | c | 5.521E+04 | 701.8 | 0.00164 | 2.337 | +2 | 11 |
| - | - | 4.57E+04 | 702.3 | - | - | 0 | - |
| - | - | 2.067E+04 | 702.8 | - | - | 0 | - |
| - | - | 6781 | 703.3 | - | - | 0 | - |
| - | - | 1764 | 703.8 | - | - | 0 | - |
| - | - | 1046 | 704.4 | - | - | 0 | - |
| - | - | 1922 | 705.4 | - | - | 0 | - |
| - | - | 1321 | 706.3 | - | - | 0 | - |
| - | - | 3776 | 711.3 | - | - | 0 | - |
| - | - | 1916 | 712.3 | - | - | 0 | - |
| - | - | 1634 | 714.3 | - | - | 0 | - |
| - | - | 1766 | 714.4 | - | - | 0 | - |
| - | - | 1176 | 719.9 | - | - | 0 | - |
| - | - | 5187 | 720.4 | - | - | 0 | - |
| - | - | 3397 | 720.9 | - | - | 0 | - |
| - | - | 5873 | 721.4 | - | - | 0 | - |
| - | - | 3172 | 722.4 | - | - | 0 | - |
| - | - | 2432 | 723.4 | - | - | 0 | - |
| - | - | 1257 | 724.4 | - | - | 0 | - |
| - | - | 2077 | 726.4 | - | - | 0 | - |
| - | - | 3959 | 726.9 | - | - | 0 | - |
| - | - | 2371 | 727.4 | - | - | 0 | - |
| - | - | 1985 | 738.4 | - | - | 0 | - |
| - | - | 8051 | 742.3 | - | - | 0 | - |
| - | - | 5073 | 742.9 | - | - | 0 | - |
| - | - | 3086 | 743.4 | - | - | 0 | - |
| - | - | 2958 | 744.4 | - | - | 0 | - |
| - | - | 2002 | 744.9 | - | - | 0 | - |
| 5 | y | 1249 | 748.4 | 0.009319 | 12.45 | +2 | 11 |
| 5 | y | 4396 | 748.9 | 0.00798 | 10.66 | +2 | 11 |
| 5 | z | 9640 | 749.4 | 0.004324 | 5.77 | +2 | 11 |
| - | - | 1.691E+04 | 749.9 | - | - | 0 | - |
| - | - | 1.149E+04 | 750.4 | - | - | 0 | - |
| - | - | 6734 | 750.9 | - | - | 0 | - |
| - | - | 2529 | 751.4 | - | - | 0 | - |
| - | - | 999.2 | 751.9 | - | - | 0 | - |
| 12 | c | 4470 | 757.4 | 0.002049 | 2.706 | +2 | 12 |
| - | - | 4354 | 757.9 | - | - | 0 | - |
| - | - | 4088 | 758.4 | - | - | 0 | - |
| - | - | 2539 | 762.9 | - | - | 0 | - |
| - | - | 1826 | 763.4 | - | - | 0 | - |
| - | - | 3351 | 765.4 | - | - | 0 | - |
| 12 | c | 6141 | 765.9 | 0.002437 | 3.182 | +2 | 12 |
| - | - | 9337 | 766.4 | - | - | 0 | - |
| - | - | 5245 | 766.9 | - | - | 0 | - |
| - | - | 2061 | 767.4 | - | - | 0 | - |
| 11 | y | 2412 | 768.4 | 0.008343 | 10.86 | +1 | 5 |
| 11 | z | 2.288E+04 | 769.4 | 0.007562 | 9.828 | +1 | 5 |
| - | - | 1.316E+04 | 770.4 | - | - | 0 | - |
| - | - | 2738 | 770.9 | - | - | 0 | - |
| - | - | 6444 | 771.4 | - | - | 0 | - |
| - | - | 2340 | 772.4 | - | - | 0 | - |
| - | - | 966.4 | 773.4 | - | - | 0 | - |
| - | - | 1313 | 775.4 | - | - | 0 | - |
| - | - | 1925 | 776.4 | - | - | 0 | - |
| - | - | 2179 | 777.4 | - | - | 0 | - |
| - | - | 2708 | 777.9 | - | - | 0 | - |
| - | - | 2865 | 778.4 | - | - | 0 | - |
| - | - | 1913 | 778.9 | - | - | 0 | - |
| 4 | w | 2251 | 784.4 | 4.753E-05 | 0.06059 | +2 | 12 |
| - | - | 5143 | 784.9 | - | - | 0 | - |
| 11 | y | 1.599E+04 | 785.4 | 0.008891 | 11.32 | +1 | 5 |
| - | - | 2612 | 785.9 | - | - | 0 | - |
| - | - | 5944 | 786.4 | - | - | 0 | - |
| - | - | 1624 | 787.4 | - | - | 0 | - |
| - | - | 1765 | 791.4 | - | - | 0 | - |
| - | - | 1355 | 792.4 | - | - | 0 | - |
| 6 | c | 5.979E+04 | 793.4 | 0.001607 | 2.026 | +1 | 6 |
| - | - | 3.089E+04 | 794.4 | - | - | 0 | - |
| - | - | 9752 | 795.4 | - | - | 0 | - |
| - | - | 1847 | 796.4 | - | - | 0 | - |
| 4 | z | 1320 | 798.4 | 0.007625 | 9.551 | +2 | 12 |
| 4 | y | 3336 | 806.4 | 0.002474 | 3.069 | +2 | 12 |
| 4 | z | 1.002E+04 | 806.9 | 0.001865 | 2.312 | +2 | 12 |
| - | - | 2.538E+04 | 807.4 | - | - | 0 | - |
| - | - | 1.575E+04 | 807.9 | - | - | 0 | - |
| - | - | 8182 | 808.4 | - | - | 0 | - |
| - | - | 3054 | 808.9 | - | - | 0 | - |
| - | - | 4371 | 809.4 | - | - | 0 | - |
| 6 | c | 9.605E+04 | 810.4 | 0.001364 | 1.683 | +1 | 6 |
| - | - | 4.542E+04 | 811.4 | - | - | 0 | - |
| - | - | 1.323E+04 | 812.4 | - | - | 0 | - |
| - | - | 2443 | 812.9 | - | - | 0 | - |
| - | - | 2099 | 813.4 | - | - | 0 | - |
| - | - | 1181 | 813.9 | - | - | 0 | - |
| 4 | y | 2077 | 814.9 | 0.004025 | 4.94 | +2 | 12 |
| - | - | 1764 | 815.4 | - | - | 0 | - |
| - | - | 2231 | 816.9 | - | - | 0 | - |
| - | - | 3563 | 817.4 | - | - | 0 | - |
| - | - | 2704 | 817.9 | - | - | 0 | - |
| - | - | 1183 | 818.4 | - | - | 0 | - |
| - | - | 1771 | 822.4 | - | - | 0 | - |
| - | - | 3050 | 823.9 | - | - | 0 | - |
| - | - | 3019 | 824.4 | - | - | 0 | - |
| - | - | 1158 | 825.9 | - | - | 0 | - |
| - | - | 2179 | 826.9 | - | - | 0 | - |
| - | - | 1188 | 827.4 | - | - | 0 | - |
| - | - | 1042 | 836.4 | - | - | 0 | - |
| - | - | 2285 | 838.4 | - | - | 0 | - |
| - | - | 1705 | 838.9 | - | - | 0 | - |
| 10 | w | 2795 | 839.4 | 0.007187 | 8.562 | +1 | 6 |
| - | - | 2202 | 840.4 | - | - | 0 | - |
| - | - | 2690 | 841.4 | - | - | 0 | - |
| 3 | w | 2570 | 841.9 | 0.006684 | 7.939 | +2 | 13 |
| - | - | 2079 | 842.4 | - | - | 0 | - |
| - | - | 1248 | 843.4 | - | - | 0 | - |
| - | - | 1053 | 844.4 | - | - | 0 | - |
| 13 | c | 8840 | 846.4 | 0.001173 | 1.386 | +2 | 13 |
| - | - | 9060 | 846.9 | - | - | 0 | - |
| - | - | 5703 | 847.4 | - | - | 0 | - |
| - | - | 2280 | 847.9 | - | - | 0 | - |
| - | - | 2325 | 848.4 | - | - | 0 | - |
| - | - | 1266 | 851.4 | - | - | 0 | - |
| - | - | 1144 | 853.4 | - | - | 0 | - |
| - | - | 1699 | 859.9 | - | - | 0 | - |
| - | - | 2497 | 860.4 | - | - | 0 | - |
| - | - | 3.699E+04 | 867.4 | - | - | 0 | - |
| - | - | 2.012E+04 | 868.4 | - | - | 0 | - |
| - | - | 7309 | 869.4 | - | - | 0 | - |
| 3 | y | 2405 | 870.4 | 0.01261 | 14.49 | +2 | 13 |
| 3 | z | 6003 | 870.9 | 0.002956 | 3.394 | +2 | 13 |
| - | - | 2.193E+04 | 871.4 | - | - | 0 | - |
| - | - | 1.863E+04 | 871.9 | - | - | 0 | - |
| - | - | 1.114E+04 | 872.4 | - | - | 0 | - |
| - | - | 3656 | 872.9 | - | - | 0 | - |
| - | - | 1540 | 873.4 | - | - | 0 | - |
| - | - | 1063 | 873.9 | - | - | 0 | - |
| 3 | y | 2579 | 878.9 | 0.001942 | 2.209 | +2 | 13 |
| - | - | 2206 | 879.4 | - | - | 0 | - |
| - | - | 1086 | 879.9 | - | - | 0 | - |
| 10 | z | 2996 | 880.4 | 0.01705 | 19.37 | +1 | 6 |
| - | - | 2407 | 880.9 | - | - | 0 | - |
| - | - | 4149 | 881.4 | - | - | 0 | - |
| - | - | 3151 | 882.4 | - | - | 0 | - |
| - | - | 2558 | 888.4 | - | - | 0 | - |
| - | - | 8665 | 888.9 | - | - | 0 | - |
| - | - | 7001 | 889.4 | - | - | 0 | - |
| - | - | 5116 | 889.9 | - | - | 0 | - |
| - | - | 1887 | 890.4 | - | - | 0 | - |
| - | - | 1013 | 894.4 | - | - | 0 | - |
| - | - | 1838 | 895.4 | - | - | 0 | - |
| - | - | 1051 | 895.9 | - | - | 0 | - |
| 10 | y | 1392 | 896.4 | 0.0004017 | 0.4481 | +1 | 6 |
| 10 | z | 1.806E+04 | 898.4 | 0.007613 | 8.474 | +1 | 6 |
| - | - | 1.975E+04 | 899.4 | - | - | 0 | - |
| - | - | 8064 | 900.4 | - | - | 0 | - |
| 14 | c | 2258 | 901.4 | 0.01493 | 16.56 | +2 | 14 |
| - | - | 1204 | 902.4 | - | - | 0 | - |
| 7 | c | 8058 | 908.4 | 0.001214 | 1.337 | +1 | 7 |
| - | - | 4600 | 909.4 | - | - | 0 | - |
| 14 | c | 6532 | 910.4 | 0.006052 | 6.648 | +2 | 14 |
| - | - | 4724 | 910.9 | - | - | 0 | - |
| - | - | 2257 | 911.4 | - | - | 0 | - |
| 10 | y | 5020 | 914.4 | 0.008088 | 8.845 | +1 | 6 |
| - | - | 2389 | 915.4 | - | - | 0 | - |
| - | - | 1141 | 916.4 | - | - | 0 | - |
| - | - | 1389 | 922.4 | - | - | 0 | - |
| - | - | 934 | 924.5 | - | - | 0 | - |
| 7 | c | 5854 | 925.5 | 0.001765 | 1.907 | +1 | 7 |
| - | - | 4679 | 926.5 | - | - | 0 | - |
| - | - | 1493 | 927.5 | - | - | 0 | - |
| - | - | 1243 | 931 | - | - | 0 | - |
| - | - | 1096 | 931.5 | - | - | 0 | - |
| 2 | y | 2850 | 938.4 | 0.01868 | 19.91 | +2 | 14 |
| - | - | 1394 | 939 | - | - | 0 | - |
| 2 | z | 5059 | 939.4 | 0.01715 | 18.25 | +2 | 14 |
| - | - | 1.528E+04 | 939.9 | - | - | 0 | - |
| - | - | 1.401E+04 | 940.4 | - | - | 0 | - |
| - | - | 9125 | 940.9 | - | - | 0 | - |
| - | - | 3197 | 941.4 | - | - | 0 | - |
| - | - | 1485 | 941.9 | - | - | 0 | - |
| - | - | 1211 | 952.5 | - | - | 0 | - |
| - | - | 1434 | 953.5 | - | - | 0 | - |
| - | - | 1296 | 961.5 | - | - | 0 | - |
| - | - | 1829 | 966.5 | - | - | 0 | - |
| - | - | 3847 | 967.5 | - | - | 0 | - |
| - | - | 2143 | 968 | - | - | 0 | - |
| - | - | 2104 | 968.5 | - | - | 0 | - |
| - | - | 1549 | 975 | - | - | 0 | - |
| - | - | 1921 | 975.5 | - | - | 0 | - |
| - | - | 2502 | 976 | - | - | 0 | - |
| - | - | 1290 | 976.5 | - | - | 0 | - |
| - | - | 1053 | 977 | - | - | 0 | - |
| - | - | 1196 | 981.5 | - | - | 0 | - |
| 8 | c | 1.348E+04 | 982.5 | 0.002518 | 2.563 | +1 | 8 |
| - | - | 1.187E+04 | 983.5 | - | - | 0 | - |
| - | - | 4140 | 984.5 | - | - | 0 | - |
| - | - | 929 | 985.5 | - | - | 0 | - |
| - | - | 2932 | 996.5 | - | - | 0 | - |
| - | - | 3954 | 997 | - | - | 0 | - |
| - | - | 2543 | 997.5 | - | - | 0 | - |
| - | - | 1063 | 998 | - | - | 0 | - |
| - | - | 996.3 | 1004 | - | - | 0 | - |
| - | - | 2336 | 1005 | - | - | 0 | - |
| - | - | 2386 | 1005 | - | - | 0 | - |
| - | - | 1418 | 1006 | - | - | 0 | - |
| - | - | 2397 | 1010 | - | - | 0 | - |
| - | - | 1845 | 1011 | - | - | 0 | - |
| 9 | z | 1428 | 1027 | 0.008094 | 7.885 | +1 | 7 |
| - | - | 9226 | 1028 | - | - | 0 | - |
| - | - | 4775 | 1029 | - | - | 0 | - |
| - | - | 2278 | 1030 | - | - | 0 | - |
| - | - | 1526 | 1067 | - | - | 0 | - |
| - | - | 1201 | 1068 | - | - | 0 | - |
| - | - | 3688 | 1085 | - | - | 0 | - |
| - | - | 2470 | 1086 | - | - | 0 | - |
| 9 | c | 1017 | 1094 | 0.01969 | 18.01 | +1 | 9 |
| - | - | 2084 | 1095 | - | - | 0 | - |
| - | - | 2200 | 1096 | - | - | 0 | - |
| 8 | y | 1908 | 1100 | 0.009463 | 8.606 | +1 | 8 |
| - | - | 1125 | 1101 | - | - | 0 | - |
| - | - | 991.2 | 1110 | - | - | 0 | - |
| 9 | c | 1.363E+04 | 1111 | 0.0002673 | 0.2407 | +1 | 9 |
| - | - | 1.26E+04 | 1112 | - | - | 0 | - |
| - | - | 4933 | 1113 | - | - | 0 | - |
| - | - | 2259 | 1114 | - | - | 0 | - |
| - | - | 1429 | 1143 | - | - | 0 | - |
| - | - | 1200 | 1197 | - | - | 0 | - |
| 7 | z | 997.4 | 1199 | 0.001727 | 1.441 | +1 | 9 |
| - | - | 4579 | 1200 | - | - | 0 | - |
| - | - | 2403 | 1201 | - | - | 0 | - |
| - | - | 1183 | 1202 | - | - | 0 | - |
| 10 | c | 2310 | 1240 | 0.000944 | 0.7615 | +1 | 10 |
| - | - | 1878 | 1241 | - | - | 0 | - |
| - | - | 1660 | 1242 | - | - | 0 | - |
| - | - | 1095 | 1243 | - | - | 0 | - |
| - | - | 868.9 | 1265 | - | - | 0 | - |
| 6 | z | 847.8 | 1312 | 0.01862 | 14.19 | +1 | 10 |
| - | - | 1941 | 1313 | - | - | 0 | - |
| - | - | 1615 | 1314 | - | - | 0 | - |
| - | - | 1185 | 1315 | - | - | 0 | - |
| - | - | 810.6 | 1359 | - | - | 0 | - |
| - | - | 1894 | 1360 | - | - | 0 | - |
| - | - | 1240 | 1361 | - | - | 0 | - |
| 11 | c | 1730 | 1403 | 0.001162 | 0.8285 | +1 | 11 |
| - | - | 990.4 | 1404 | - | - | 0 | - |
| - | - | 1257 | 1405 | - | - | 0 | - |
| - | - | 700.6 | 1614 | - | - | 0 | - |
| - | - | 739.3 | 1687 | - | - | 0 | - |
| - | - | 726.3 | 2471 | - | - | 0 | - |
| - | - | 1066 | 3046 | - | - | 0 | - |
| - | - | 625.7 | 3289 | - | - | 0 | - |
| - | - | 711.7 | 3432 | - | - | 0 | - |

m/z Charge Intensity FragmentType MassShift Position
120.93453979492188 0 389.80615
126.07940673828125 0 728.698
126.09175872802734 0 459.72244
127.08757781982422 0 728.9094
128.09494018554688 0 864.44885
129.01885986328125 0 3774.081
129.0665283203125 0 693.99603
129.102783203125 0 26292.727
130.06565856933594 0 95623.984
130.10604858398438 0 1877.1478
131.0634002685547 0 604.65094
131.06895446777344 0 9296.7295
131.07424926757812 0 812.3291
131.11843872070312 0 1699.5774
132.05360412597656 0 563.67664
132.066162109375 0 534.27563
132.0813751220703 0 2876.651
132.10247802734375 0 1523.6709
133.0560302734375 0 859.04974
133.06130981445312 0 13924.917
133.08583068847656 0 465.20636
134.06472778320312 0 1032.6588
136.0716552734375 0 499.1095
136.0762176513672 0 5863.5405
137.07931518554688 0 401.326
137.89016723632812 0 390.55136
138.06671142578125 0 1327.7146
139.0032196044922 0 2820.2393
140.3148193359375 0 434.9261
141.1027374267578 0 615.5664
143.04544067382812 0 585.9065
143.11875915527344 0 449.46307
144.0814208984375 0 2814.8604
145.061279296875 0 2173.352
146.09288024902344 0 1795.3385
146.09910583496094 0 414.67255
147.0448760986328 0 722.7545
147.07708740234375 0 820.4801
154.08668518066406 0 457.7947
155.09326171875 0 17514.56
156.06600952148438 0 759.1406
156.09677124023438 0 902.68726
156.13890075683594 0 1590.4714
157.01370239257812 0 7592.603
157.1342315673828 0 669.4096
158.06048583984375 0 1045.4098
159.0922088623047 0 47014.586
160.0956573486328 0 4371.226
164.08245849609375 0 2189.4807
165.10202026367188 0 467.55057
166.06166076660156 0 15766.135
167.06533813476562 0 1393.1162
168.13865661621094 0 1005.9604
169.07659912109375 0 536.9655
170.06057739257812 0 17369.695
171.0631561279297 0 1071.1173
171.0693817138672 0 824.94055
171.11361694335938 0 709.3966
172.0721893310547 0 3951.8257
173.05624389648438 0 1791.3693
173.4222869873047 0 389.32693
173.45095825195312 0 2677.4194
174.07675170898438 0 2734.1028
176.11146545410156 0 1767.9196
176.61293029785156 0 568.8668
178.1343231201172 0 2672.8027
179.13865661621094 0 552.32623
181.0983123779297 0 534.16626
183.11343383789062 0 6529.5693
183.12245178222656 0 831.745
183.12631225585938 0 996.3069
183.14991760253906 0 859.3435
185.07192993164062 0 1020.9185
185.08148193359375 0 10084.902
185.09141540527344 0 855.31665
185.16542053222656 0 15389.882
186.07888793945312 0 701.3816
186.1244659423828 0 3192.6138
186.17376708984375 0 1188.7856
187.08717346191406 0 6942.777
187.10804748535156 0 1404.2396
188.0914306640625 0 918.3896
189.0989990234375 0 802.3699
190.08287048339844 0 29103.648
190.10865783691406 0 1331.8862 c Ammonia loss 2
190.610107421875 0 500.09387
191.08642578125 0 1090.5203
192.07723999023438 0 1890.8328
192.5077362060547 0 473.4322
195.11305236816406 0 752.5992
197.10781860351562 0 521.54755 y 10
198.12820434570312 0 8003.763
199.13198852539062 0 515.30975
199.1698455810547 0 2640.3965
200.14413452148438 0 785.23303
201.1243438720703 0 503.17166
202.1187744140625 0 958.20294
203.10427856445312 0 793.7104
203.11648559570312 0 3139.119
203.13978576660156 0 1069.4877
203.1502227783203 0 1286.0366
203.61753845214844 0 674.58813
205.11883544921875 0 12126.6045 y 14
205.1455841064453 0 482.20346
206.1218719482422 0 1312.0006
207.16091918945312 0 1832.6396
208.14486694335938 0 1009.8839
210.12802124023438 0 4287.253
211.13182067871094 0 848.0402
211.62973022460938 0 5958.4414
212.13148498535156 0 1734.9214
213.07630920410156 0 4944.9126
213.16029357910156 0 504.87198
215.13970947265625 0 967.6332
217.03494262695312 0 8589.877
223.1197052001953 0 4113.045
223.15599060058594 0 16905.068
224.12355041503906 0 604.74066
224.15957641601562 0 2037.376
226.0834197998047 0 580.40674
227.06698608398438 0 1509.998
227.08702087402344 0 587.57806
228.1214599609375 0 746.5193
228.13536071777344 0 1428.1218
229.082763671875 0 606.7668
229.11895751953125 0 4485.1523
230.18707275390625 0 1151.9475
233.14047241210938 0 4666.915
233.62506103515625 0 1227.197
234.1244354248047 0 19577.5
235.12799072265625 0 2920.13
236.63369750976562 0 2377.442
239.1088409423828 0 2289.3489 y Ammonia loss 12
239.61058044433594 0 1192.9451 z 12
240.17198181152344 0 728.4073
243.1102294921875 0 556.0227
243.14996337890625 0 644.4387
244.09359741210938 0 2737.3857
245.09666442871094 0 507.9142
245.1466522216797 0 1057.3656
246.1454315185547 0 1602.6335
247.621826171875 0 7577.788 c Ammonia loss 3
248.12449645996094 0 1764.4604
250.1076202392578 0 2189.959
250.1430206298828 0 2006.262
250.16702270507812 0 2238.7705
251.15101623535156 0 98609.875
252.15420532226562 0 13246.218
252.6034698486328 0 2908.419
253.1568145751953 0 1124.3306
255.14926147460938 0 2924.4507
255.8068084716797 0 2025.3345
256.1389465332031 0 723.30585 c 3
256.63214111328125 0 810.1789
258.16424560546875 0 2051.4612
258.6670837402344 0 1124.4991
259.1096496582031 0 576.418
259.15460205078125 0 12615.326
261.1192932128906 0 3215.9014
261.6087951660156 0 2819.7883
262.11077880859375 0 644.6341
264.1712646484375 0 2331.397
265.1445617675781 0 2481.0864
265.6357727050781 0 1370.4453
266.1251220703125 0 3459.5022
266.6009216308594 0 674.05096
267.1279602050781 0 577.5304
268.177490234375 0 11599.797 c 1
269.138916015625 0 1585.0527
269.1804504394531 0 1819.2725
270.1223449707031 0 20201.574
270.6236267089844 0 5388.3457
272.1070251464844 0 5176.4067
272.1765441894531 0 6374.0107
273.0970153808594 0 3823.8037
273.142822265625 0 737.55646
273.1806640625 0 958.20905
274.1192321777344 0 3654.0396
274.1497497558594 0 17432.543
274.6514587402344 0 4705.758
275.1139221191406 0 1763.658
275.15167236328125 0 1027.3833
276.1319885253906 0 703.1249
277.1558837890625 0 729.35004
282.1335754394531 0 1188.7689
282.1820983886719 0 7909.7686
283.140869140625 0 3835.6255
283.18505859375 0 1027.4244
284.103515625 0 1645.648
284.12030029296875 0 3829.2646
284.1360168457031 0 649.46014
284.62164306640625 0 879.5876
285.15771484375 0 1436.5675
286.14068603515625 0 1423.3779
290.1177978515625 0 6971.9863
292.1665954589844 0 1250.1111
294.1205749511719 0 1430.2327
294.14764404296875 0 798.20575
294.19305419921875 0 945.1269
295.1041259765625 0 1032.8677
295.1478271484375 0 905.6302 z Ammonia loss 11
297.1278991699219 0 2159.9392
297.15655517578125 0 779.04865
297.63134765625 0 677.606
300.1715087890625 0 2759.8445 z Ammonia loss 13
301.1070861816406 0 1248.9329
301.1522521972656 0 3106.224
302.1151123046875 0 3576.7688
302.1451721191406 0 2015.356
302.4929504394531 0 718.57477
302.66119384765625 0 799.1227
303.1186828613281 0 611.2808
303.1524353027344 0 10129.784 c Water loss 6
303.1679992675781 0 2967.7375
303.65484619140625 0 3758.8354 z 11
304.1536865234375 0 2134.0955
305.6466064453125 0 882.8504
306.1448974609375 0 620.3087
309.1457824707031 0 5467.334
309.1939392089844 0 669.8492
309.64825439453125 0 1129.3905
309.6714172363281 0 838.24243
311.1256408691406 0 2740.8499
311.62847900390625 0 887.4616
311.66595458984375 0 9507.014 y 11
312.1671142578125 0 1743.0327
312.6660461425781 0 558.06445
315.1666259765625 0 1693.5399
317.19561767578125 0 4186.1587 z 13
317.6595458984375 0 5005.0977
318.1515808105469 0 19985.607
318.1793518066406 0 1267.6229
318.6533508300781 0 9063.98
318.8331604003906 0 719.71356
319.15496826171875 0 1882.5936
320.19775390625 0 844.0796
321.171142578125 0 694.1377
322.84112548828125 0 1000.49927
323.1440734863281 0 3105.3804
323.6451110839844 0 1767.8784
324.6775817871094 0 870.52545
326.500732421875 0 4317.4326
326.6647644042969 0 82551.16
326.8343505859375 0 1754.1373
327.1663513183594 0 33307.89
327.2023010253906 0 1388.1135
327.6676940917969 0 8306.477
328.1694641113281 0 1219.1381 y Water loss 5
329.1606750488281 0 3849.1394
331.5202331542969 0 1224.6405
331.6570739746094 0 12453.012
331.6749267578125 0 774.70514
331.85504150390625 0 1410.1647
332.1566162109375 0 4395.5347
332.65216064453125 0 865.488
332.67242431640625 0 1013.2631 y 5
333.21441650390625 0 5779.702 y 13
334.21759033203125 0 680.5687
335.16943359375 0 639.11755
337.8477783203125 0 1663.3046
338.1474304199219 0 960.59656
338.18011474609375 0 1387.2062
338.5128479003906 0 614.8901
338.6736145019531 0 1640.233
339.16680908203125 0 1443.1787 c 12
339.6558837890625 0 629.06946
339.6752624511719 0 1074.9313
339.6953125 0 3441.4763
340.1982421875 0 1784.923
340.6622314453125 0 33595.55 c Ammonia loss 4
341.1639709472656 0 18433.887
341.2179870605469 0 1079.1019
341.6659851074219 0 4247.6743
341.8331604003906 0 2976.7314
342.1761169433594 0 1825.0825 y Water loss 8
342.5066833496094 0 2401.341 y Ammonia loss 8
342.8417663574219 0 2259.715 z 8
343.1767272949219 0 1446.423
346.1153564453125 0 858.48376
347.1812438964844 0 1109.277
347.8531188964844 0 1583.3962
348.1839599609375 0 1799.7343 y 8
349.1897277832031 0 2181.347
349.68267822265625 0 1111.6378
351.21514892578125 0 1919.4164
352.51434326171875 0 640.26746
352.68524169921875 0 640.4854
353.6701965332031 0 2425.8142
354.1707763671875 0 1690.922
355.1980285644531 0 1906.4163
355.68133544921875 0 2184.8591
355.8432922363281 0 801.0972 z Water loss 7
356.1824645996094 0 933.6639
356.84027099609375 0 3035.716
357.1768798828125 0 2634.2756
357.693359375 0 868.0828
358.1773986816406 0 662.0936
360.6954040527344 0 2810.9722
361.18115234375 0 2409.258 c Water loss 13
361.1998291015625 0 1941.7992
361.2232971191406 0 2450.7942
361.5138244628906 0 1770.8447 y Ammonia loss 7
361.8478088378906 0 7960.652 z 7
362.183349609375 0 5243.221
362.5140075683594 0 690.5408
362.6882019042969 0 806.6285
363.1427001953125 0 1557.7098
363.2038269042969 0 955.3931
364.12615966796875 0 1564.1145
364.1951904296875 0 3434.768
364.6964111328125 0 2038.0236
365.68804931640625 0 1070.8445
366.18231201171875 0 1686.812
367.1874694824219 0 47847.902 y 7
367.52191162109375 0 28426.693
367.6639404296875 0 1439.5258
367.685791015625 0 1152.3235
367.85565185546875 0 12815.8955
368.18865966796875 0 4686.079
369.21337890625 0 1027.9894
370.4305114746094 0 1022.7008
370.5158386230469 0 654.22736
370.681396484375 0 1279.3247 z Water loss 4
371.1781311035156 0 1612.9973
372.38665771484375 0 841.7087
374.2018737792969 0 4844.2773
374.6936340332031 0 11765.065 y Water loss 4
374.93377685546875 0 10619.104 y Ammonia loss 4
375.1874084472656 0 9303.002 z 4
375.43475341796875 0 4299.81
375.69830322265625 0 1268.356
375.9420471191406 0 619.7111
376.19219970703125 0 3778.0051
376.5223083496094 0 1014.5911
378.2031555175781 0 6151.9897
378.2254943847656 0 864.61725
379.2098388671875 0 32221.996 c Ammonia loss 2
379.4414978027344 0 1836.46
380.2127685546875 0 5425.6484
381.1527404785156 0 7918.8315
381.19110107421875 0 21787.547
381.5253601074219 0 8987.961
381.8594665527344 0 3422.6616
382.15570068359375 0 1327.3546
382.1949157714844 0 1109.696
383.2066955566406 0 31111.379
383.70849609375 0 17333.652
384.210205078125 0 4642.0874
384.6850891113281 0 2333.4246 y Ammonia loss 10
384.7124938964844 0 805.8632
385.15277099609375 0 2429.3137
385.1900939941406 0 6247.5938 z 10
385.5260314941406 0 3737.8965
385.6881408691406 0 787.3313
385.86041259765625 0 2605.7341
386.1940002441406 0 1918.9746
386.5260925292969 0 1165.9254
387.1929626464844 0 1393.8381
387.2248840332031 0 39874
388.1972961425781 0 1598.4288
388.2282409667969 0 8851.569
388.69256591796875 0 3663.4653
388.93646240234375 0 792.6087
389.1874084472656 0 8190.1533
389.2183837890625 0 791.87366
389.2293395996094 0 916.20636
389.9940490722656 0 1094.3662
390.1929626464844 0 5019.2305
390.5298767089844 0 1284.5642
391.1376647949219 0 737.2741
391.199462890625 0 1707.5282
392.2112121582031 0 795.87634
392.6900939941406 0 738.7588 w 3
393.1977844238281 0 50244.473 y 10
393.525634765625 0 762.8733
393.5981140136719 0 803.2104
393.6992492675781 0 23218.621
394.1993713378906 0 8478.203
394.6993103027344 0 1618.2015
394.9955139160156 0 2302.71
395.1363525390625 0 2153.483
395.1949157714844 0 6039.181
395.2230529785156 0 915.6424
395.3935546875 0 6769.7505
395.5942077636719 0 3728.7778
395.79376220703125 0 2666.8372
396.2364196777344 0 37628.867 c 2
397.2043151855469 0 25202.805 c Ammonia loss 5
397.2391662597656 0 7430.295
397.70654296875 0 11869.985
398.1783752441406 0 1801.4454
398.2083435058594 0 3108.0552
398.24005126953125 0 826.4211
398.5976867675781 0 11357.13
398.79638671875 0 28135.318
398.99609375 0 26093.355
399.1958923339844 0 17019.23
399.3970031738281 0 7520.121
399.5262451171875 0 3227.451 y Water loss 6
399.5969543457031 0 3071.7046
399.7966613769531 0 812.1358
399.855712890625 0 3997.914 y Ammonia loss 6
400.19146728515625 0 2798.7473 z 6
400.52557373046875 0 745.709
400.77203369140625 0 764.7458
401.1876525878906 0 1034.5457
401.2619934082031 0 1184.5513
402.0026550292969 0 2428.1184
402.2005615234375 0 19382.348
402.4004821777344 0 21703.781
402.6009826660156 0 16219.506
402.80145263671875 0 8305.992
403.00189208984375 0 3323.4944
403.2013244628906 0 2981.5806
404.2173767089844 0 2287.0342
404.7189636230469 0 1750.8508
405.1974792480469 0 1163.9891
405.5301208496094 0 42985.36 y 6
405.8643493652344 0 24998.105
406.1976318359375 0 10718.732
406.5324401855469 0 3832.6086
407.1987609863281 0 732.29626
408.2010498046875 0 1040.2601 c Ammonia loss 9
408.8705749511719 0 2966.994
409.1468200683594 0 1302.9166
409.20062255859375 0 709.5994
409.5353698730469 0 885.9608
411.19696044921875 0 1223.111
411.6941223144531 0 996.4495
412.1836853027344 0 1678.9135
413.1457214355469 0 704.1711
413.189208984375 0 777.81506
413.8858947753906 0 1091.7173
414.21881103515625 0 975.9884
414.5528869628906 0 1354.16
414.8863830566406 0 958.23663
415.1982116699219 0 779.1799
415.2427673339844 0 1030.9586
415.74578857421875 0 1338.4711
416.73028564453125 0 1014.28955
418.2149658203125 0 2003.7211
419.7059631347656 0 818.2929
420.20330810546875 0 202804.45 w 9
420.7046813964844 0 90859.29
421.2049865722656 0 33264.11
421.7051696777344 0 10891.088
422.20513916015625 0 1375.4781
422.70672607421875 0 916.5526
423.5337829589844 0 16402.033 w 5
423.8677978515625 0 11328.023
424.2019348144531 0 4454.573
424.5338439941406 0 1327.1757
424.72637939453125 0 1227.8384
424.8707580566406 0 776.9719
425.2006530761719 0 1359.0538
427.1897277832031 0 893.85754
427.7142028808594 0 951.3013
430.1945495605469 0 7997.893
431.19940185546875 0 3098.102
431.2299499511719 0 1109.9539
431.4525451660156 0 822.9954 z Water loss 2
431.70574951171875 0 1707.1167 z Ammonia loss 2
432.211669921875 0 986.7326 z Ammonia loss 5
434.2084655761719 0 2453.3047
435.1629943847656 0 1933.2432
435.22491455078125 0 4375.3804
435.4595947265625 0 2051.4 y Water loss 2
435.7082214355469 0 1905.807 y Ammonia loss 2
435.9574279785156 0 1685.8102 z 2
436.227294921875 0 2168.628
437.2223205566406 0 4441.2974 y Water loss 5
437.55572509765625 0 6632.516 y Ammonia loss 5
437.889404296875 0 1470.6302 z 5
438.17340087890625 0 1112.193
438.2219543457031 0 1326.4784
438.5545654296875 0 692.1845
438.7413024902344 0 1622.7041
439.2434387207031 0 1439.3962
439.9620666503906 0 1314.7362 y 2
440.7165832519531 0 892.7954 z Water loss 9
441.2220153808594 0 883.07166
442.7118225097656 0 762.9832
442.8970031738281 0 3211.2568
443.22515869140625 0 58112.695 y 5
443.55938720703125 0 35468.95
443.8930969238281 0 15034.343
444.2261962890625 0 4918.228
444.55999755859375 0 1398.5615
445.71337890625 0 2595.643
446.2064208984375 0 1347.6659
446.5549011230469 0 977.6657
446.7063293457031 0 787.912
446.74505615234375 0 737.484
447.22088623046875 0 6516.9707
447.7445068359375 0 931.3891
448.22174072265625 0 2756.15
448.71435546875 0 5624.297 y Water loss 9
449.2148132324219 0 2640.059 y Ammonia loss 9
449.7109680175781 0 3753.7852 z 9
450.2091369628906 0 1610.6261
450.7104797363281 0 784.5385
451.2327575683594 0 714.23566 c Water loss 13
452.22314453125 0 1430.7544
452.473388671875 0 1040.34
452.5527038574219 0 1132.3043
452.7196044921875 0 1233.8865
453.26202392578125 0 985.0144
454.22381591796875 0 1112.4568 c Water loss 6
454.2587585449219 0 747.6178
454.7178649902344 0 3557.614 c Ammonia loss 6
454.96429443359375 0 821.9146
455.21832275390625 0 1284.6024
455.72607421875 0 1958.9738 c 13
455.8838195800781 0 1470.5914
455.96673583984375 0 1973.83
456.22027587890625 0 2458.0288
457.2209777832031 0 1127.5745
457.71990966796875 0 5552.476 y 9
458.21923828125 0 2165.6006
459.1976623535156 0 786.7616
459.4689025878906 0 774.8642
459.9765625 0 2221.3047
460.2244567871094 0 1747.7001
460.7364196777344 0 32595.262
460.96527099609375 0 1757.2455
461.2381286621094 0 16657.926
461.4697570800781 0 3389.6355
461.7376403808594 0 6833.675
462.2030334472656 0 6303.913
462.2382507324219 0 1710.926
462.7318115234375 0 2600.593
463.23187255859375 0 2490.9683 c 6
463.7346496582031 0 1248.2515
465.225341796875 0 1947.4388
465.4688720703125 0 7000.1597
465.7208251953125 0 4567.1143 z Water loss 1
465.8899230957031 0 1135.3092
465.9710388183594 0 3407.5261 z Ammonia loss 1
466.22222900390625 0 1898.1733
466.2417297363281 0 4224.781
466.55975341796875 0 3350.5566
466.8954772949219 0 1898.3354
467.2477111816406 0 3662.0032
467.75762939453125 0 852.7325
468.2645263671875 0 19151.436
468.7663879394531 0 11683.758
469.2357482910156 0 3610.5793
469.26788330078125 0 2691.4988
469.49139404296875 0 1423.295
469.729248046875 0 6159.3735 y Water loss 1
469.97601318359375 0 7716.1343 y Ammonia loss 1
470.2266845703125 0 9244.325 z 1
470.47711181640625 0 5760.149
470.7268981933594 0 2294.684
471.4855041503906 0 708.08795
471.7320251464844 0 1000.6212
471.983642578125 0 2157.6797
473.23419189453125 0 1784.9901
473.723388671875 0 2142.844
473.9791259765625 0 1976.018
474.22705078125 0 15486.818 y 1
474.47760009765625 0 12790.678
474.728515625 0 7161.1973
474.9786376953125 0 3093.5422
475.22479248046875 0 1778.9471
475.738525390625 0 2800.4607
475.988525390625 0 2157.0813
476.2278747558594 0 2588.2927
476.24530029296875 0 2833.9795
476.4871826171875 0 3991.831
476.73553466796875 0 2230.2363
476.9820251464844 0 2038.4408
477.21527099609375 0 5067.6797 y Ammonia loss 12
477.7242431640625 0 1580.7068
477.75750732421875 0 851.6053
478.2201843261719 0 4719.344 z 12
478.7215881347656 0 1098.7721
479.2384033203125 0 1292.774
479.904296875 0 1214.3218
479.9949951171875 0 909.6267
480.2458801269531 0 3744.0818
480.4940185546875 0 54531.97
480.7445373535156 0 51947.098
480.99517822265625 0 36733.977
481.24517822265625 0 13676.911
481.4955749511719 0 5080.653
481.7453918457031 0 1814.4283
482.2484130859375 0 1387.7867
482.7380676269531 0 5453.9473 c Water loss 7
483.23052978515625 0 11374.277 c Ammonia loss 7
483.49005126953125 0 3060.8076
483.7348937988281 0 7353.1655
483.994140625 0 2006.9674
484.2504577636719 0 19282.713
484.4876403808594 0 7177.9795
484.5774230957031 0 5624.025
484.7252197265625 0 8546.863 w 8
484.75384521484375 0 6199.144
484.91021728515625 0 3772.9946
484.9884948730469 0 4037.851
485.2294616699219 0 4604.089
485.2492370605469 0 3015.4114
485.49151611328125 0 975.3273
485.5845642089844 0 1598.1124
485.7248229980469 0 1893.4199
485.9174499511719 0 1185.8743
486.2490539550781 0 1070.1007
487.2261962890625 0 940.37933
487.4832458496094 0 6593.829
487.73004150390625 0 134034.19
487.98272705078125 0 128529.9
488.23370361328125 0 92350.625
488.48492431640625 0 36427.95
488.7372131347656 0 11308.483
488.9956970214844 0 5237.1436
489.2460632324219 0 5572.0796
489.748779296875 0 1254.2356
490.2446594238281 0 936.47034
490.5777587890625 0 1069.2114
491.2420959472656 0 2061.9902
491.49761962890625 0 1576.3403
491.7423095703125 0 20652.352 c 7
491.99090576171875 0 2073.5317
492.242919921875 0 13851.067
492.7436828613281 0 4306.407
493.24761962890625 0 2418.4326 c Water loss 3
493.5709228515625 0 1581.8765
493.740234375 0 1138.772
493.90399169921875 0 1672.5192 z Water loss 4
493.9893798828125 0 1423.4941
494.23419189453125 0 75471.48 c Ammonia loss 3
494.5723571777344 0 1244.415
495.2380065917969 0 20043.357
495.7669982910156 0 1997.9701
496.2366943359375 0 2387.6282
496.2486877441406 0 2397.9663
496.7757568359375 0 6370.66
497.233154296875 0 1889.4108
497.27789306640625 0 5456.1045
497.7560119628906 0 6683.2915
497.99566650390625 0 4967.08
498.2471618652344 0 12050.22
498.4957275390625 0 11001.563
498.74591064453125 0 8669.181
498.90972900390625 0 816.7019
498.9960632324219 0 4184.4316
499.2432861328125 0 14291.515 y Water loss 4
499.5762939453125 0 14669.115 y Ammonia loss 4
499.74664306640625 0 2023.5946
499.9110107421875 0 18258.93 z 4
500.24609375 0 10313.1455
500.58038330078125 0 3883.4995
500.9137268066406 0 1048.592
501.25128173828125 0 821.5197
501.9936828613281 0 2804.4297
502.2510986328125 0 12267.478
502.4983215332031 0 212894.39
502.7490539550781 0 290462.12
502.9996337890625 0 220023.92
503.25006103515625 0 121829.51
503.5005187988281 0 40702.344
503.7508544921875 0 17016.68
504.0007629394531 0 5833.5283
504.1986999511719 0 807.1139
504.2435302734375 0 1191.2217
504.57806396484375 0 1335.02
504.75482177734375 0 813.67944 z Water loss 8
504.9228820800781 0 2539.7634 c Water loss 11
505.25244140625 0 20194.865 c Ammonia loss 11
505.58587646484375 0 11604.714
505.7371520996094 0 10614.864
505.9200744628906 0 8157.39
506.2409973144531 0 5304.598
506.5867004394531 0 1103.1356
506.73382568359375 0 3993.3774
507.2313232421875 0 1433.1798
508.579345703125 0 4342.9307
508.91436767578125 0 3354.66
509.24603271484375 0 1724.6954
509.5771484375 0 1405.0253
510.5987548828125 0 2490.3948
510.92974853515625 0 67163.01 c 11
511.26385498046875 0 129280.87 c 3
511.5980529785156 0 25842.906
511.9322814941406 0 9321.789
512.2662353515625 0 22970.172
512.7616577148438 0 5931.254 y Water loss 8
513.2349243164062 0 9955.914
513.2682495117188 0 4196.09
513.7569580078125 0 31681.01 z 8
514.256591796875 0 22703.984
514.5814819335938 0 4400.356
514.7596435546875 0 8631.8545
514.9163818359375 0 2080.9148
515.2545776367188 0 2767.7449
515.3199462890625 0 9436.126
516.2211303710938 0 1379.9895
516.3237915039062 0 3023.4814
517.2811279296875 0 2881.794
518.2886352539062 0 1732.5184
518.7587280273438 0 2349.417
519.2627563476562 0 2595.7937
519.6026611328125 0 918.71094
519.9352416992188 0 3327.8152
520.2720336914062 0 2124.3372
520.7378540039062 0 1216.4392
521.2744140625 0 2160.4216
521.7669067382812 0 31676.053 y 8
522.2686767578125 0 16554.854
522.7694091796875 0 5245.2114
523.2671508789062 0 2487.1477
523.5907592773438 0 1601.3057
525.2716674804688 0 1591.4381
526.2622680664062 0 2836.2136
526.7555541992188 0 1036.9865
527.2720947265625 0 2856.3547
527.7717895507812 0 2955.388
532.1942138671875 0 2601.3025
532.763671875 0 1967.492
533.1991577148438 0 1111.5587
533.2469482421875 0 969.01324
533.2781982421875 0 1744.864
533.7711791992188 0 2755.2527
534.2650146484375 0 2459.9653
534.3095703125 0 1292.6915
535.239990234375 0 2546.1265
537.2659301757812 0 1026.9788
538.2577514648438 0 2449.4463 z 3
538.5912475585938 0 1242.6763
538.76708984375 0 891.1713
539.2369384765625 0 3820.087
540.2407836914062 0 1105.3071
541.2723388671875 0 8810.051 y Water loss 7
541.7742309570312 0 4743.741 y Ammonia loss 7
542.2693481445312 0 32345.428 z 7
542.7703247070312 0 18820.934
543.2719116210938 0 10828.74
543.5927734375 0 3555.9114 y 3
543.7698974609375 0 1902.3115
543.929443359375 0 2262.4072
544.2650146484375 0 2359.6165
545.2666015625 0 1048.685
545.5975341796875 0 782.8967
546.2645874023438 0 8873.885
546.7639770507812 0 4848.9736
547.285888671875 0 4082.3936
547.5884399414062 0 1166.813
547.7742919921875 0 2649.6309
548.2410278320312 0 2908.6597 w 11
548.2913818359375 0 2074.371
549.2224731445312 0 1246.0342
549.60888671875 0 5839.555
549.7723999023438 0 939.3529
549.9413452148438 0 5898.684
550.2774047851562 0 65140.16 y 7
550.7789916992188 0 36591.27
550.9443969726562 0 1082.1594
551.2767333984375 0 15076.931
551.6070556640625 0 1313.5206
551.7802124023438 0 5450.3965
552.261962890625 0 3504.694
553.275390625 0 2141.7976
553.7764892578125 0 1278.4779
554.2875366210938 0 3736.1787
554.7825317382812 0 3851.5754
555.2702026367188 0 30738.934
555.7888793945312 0 209843.22 c 8
556.2905883789062 0 139937.34
556.7915649414062 0 46203.656
556.9388427734375 0 2015.5032
557.2919311523438 0 12139.258
557.6127319335938 0 1259.8635
557.7936401367188 0 1951.1001
559.2659912109375 0 1570.4392
559.604736328125 0 904.39
559.9439697265625 0 975.1277
560.9401245117188 0 991.32367
561.2687377929688 0 4203.6284
561.5972900390625 0 74303.875 w 2
561.9317016601562 0 60460.625
562.265625 0 36462.92
562.598388671875 0 12867.823
562.6365356445312 0 3043.6326
562.7765502929688 0 2195.9836
562.9320068359375 0 3747.4893
562.970458984375 0 1689.5929
563.260498046875 0 3689.7935
563.305419921875 0 2768.5505
563.6062622070312 0 1297.2949
563.781982421875 0 25167.361
563.9386596679688 0 872.3395
564.28076171875 0 21313.117
564.6012573242188 0 50357.445 c 12
564.7825927734375 0 7860.8667
564.9351196289062 0 47445.46
565.2701416015625 0 29280.678
565.6031494140625 0 9978.854
565.9378051757812 0 4267.9717
566.2752685546875 0 6580.8047
566.6107177734375 0 3896.1465
566.9451293945312 0 3388.993
567.2323608398438 0 12814.747
567.2781372070312 0 2763.0833
568.2352905273438 0 4431.8667
568.284912109375 0 2977.3013
568.796875 0 15166.528
568.94677734375 0 808.48834
569.2968139648438 0 10679.134
569.7953491210938 0 4520.994
569.94482421875 0 1236.6299
570.2769775390625 0 2968.656
570.7454223632812 0 2980.9338
570.7799682617188 0 4401.448
571.2828979492188 0 54113.81
571.7843017578125 0 32244.02
572.2847290039062 0 13948.982
572.7855224609375 0 4237.254
572.9567260742188 0 2787.2964
573.2882690429688 0 4586.294
573.6162719726562 0 3400.4534
573.9501342773438 0 1703.0522
574.6043701171875 0 1809.5293
576.2750854492188 0 4579.306
576.78076171875 0 3215.8855
577.2801513671875 0 3664.4531 w 6
577.7838134765625 0 4377.6494
577.9614868164062 0 3584.8289
578.289794921875 0 6274.179
578.6283569335938 0 2998.3962
578.7882080078125 0 21262.82
579.28955078125 0 14422.082
579.7890625 0 6282.725
580.2811889648438 0 3852.3457 y Water loss 2
580.6077270507812 0 6779.389 y Ammonia loss 2
580.9407958984375 0 20953.88 z 2
581.2742309570312 0 21256.988
581.3255615234375 0 1192.9672
581.60888671875 0 11191.23
581.941650390625 0 3955.5095
582.267822265625 0 3336.7285
582.9549560546875 0 1114.314
583.2809448242188 0 3261.1921
583.6101684570312 0 3296.9001
583.9426879882812 0 2567.4414
584.7890014648438 0 1743.0614
585.2847290039062 0 1715.1951
585.7941284179688 0 1329.0007
586.2815551757812 0 11638.313 y 2
586.61474609375 0 9241.397
586.9490966796875 0 6526.418
587.2855834960938 0 2516.987
587.6217041015625 0 2197.0598
587.9545288085938 0 1874.3711
588.2864379882812 0 4575.496
588.619140625 0 2948.9827
588.953125 0 2230.2827
590.2843017578125 0 2701.8452
590.781005859375 0 2683.4993 z Water loss 6
591.2820434570312 0 4024.6028 z Ammonia loss 6
591.7822265625 0 1028.6616
592.2976684570312 0 2216.6895
592.6275024414062 0 11263.691
592.961181640625 0 17056.65
593.2548217773438 0 3351.727
593.2958984375 0 6917.9863
593.6258544921875 0 2860.5837
593.77978515625 0 1171.533
593.9609985351562 0 1305.0648
594.2667236328125 0 2521.0437
595.2250366210938 0 1230.0226
596.96240234375 0 2613.711
597.3001708984375 0 2915.6453
597.62744140625 0 2643.584
597.8079833984375 0 2528.0112
597.9599609375 0 1699.9586
598.3056640625 0 9860.854
598.7887573242188 0 12602.799 y Water loss 6
599.2861328125 0 15240.966 y Ammonia loss 6
599.7825927734375 0 66120.695 z 6
600.2843627929688 0 62947.59
600.7854614257812 0 32698.514
601.287109375 0 10707.793 c Water loss 13
601.6301879882812 0 3512.7021 c Ammonia loss 13
601.7864379882812 0 3428.1875
601.95947265625 0 2966.5115
602.294189453125 0 3602.3372
602.6258544921875 0 3591.3845
602.8015747070312 0 3354.9526
602.949951171875 0 10790.73
603.2858276367188 0 14212.682
603.6168823242188 0 9433.395
603.7971801757812 0 1269.2737
603.9502563476562 0 2299.8845
604.2792358398438 0 1233.7625
606.3057861328125 0 38448.637 z 11
606.6334228515625 0 2479.686
606.78955078125 0 1223.6212
606.9695434570312 0 12595.326
607.3008422851562 0 75647.7 c 13
607.6336669921875 0 59268.902
607.7913208007812 0 222935.9 y 6
607.9667358398438 0 30001.635
608.2935180664062 0 161264.64
608.6358642578125 0 5754.918
608.79345703125 0 59892.03
609.29443359375 0 18445.441
609.7949829101562 0 4003.1711
610.2858276367188 0 12983.401
610.8270263671875 0 871.13983
611.2907104492188 0 4273.6714
611.638427734375 0 3105.5894
611.80322265625 0 14431.184 c Ammonia loss 9
611.96435546875 0 14151.324
612.2994995117188 0 26645.98
612.630615234375 0 6475.8447
612.8030395507812 0 13502.614
612.9650268554688 0 2924.1704
613.3018188476562 0 6154.147
613.8015747070312 0 4120.727
614.30517578125 0 1731.8591
615.3041381835938 0 4145.693
615.6390991210938 0 866.1308
615.8036499023438 0 2112.1797
616.3013916015625 0 2316.0479
616.6362915039062 0 1027.7727
618.3335571289062 0 1609.6534
619.3369750976562 0 1121.0483
619.818359375 0 3112.1545
620.3120727539062 0 76652.31 c 9
620.635009765625 0 7271.6 z Water loss 1
620.8135986328125 0 55415.258
620.9728393554688 0 8470.382
621.2432250976562 0 7454.6807
621.314208984375 0 33274.47
621.643310546875 0 4186.352
621.81591796875 0 5494.822
621.9808959960938 0 3414.8838
622.2457885742188 0 2171.6257
622.3239135742188 0 24862.197 y 11
623.3268432617188 0 6355.0264
624.3248901367188 0 1938.435
625.321533203125 0 2784.1653
625.6522827148438 0 5889.3867
625.9857177734375 0 6270.508
626.3131103515625 0 8162.494
626.6290893554688 0 36522.32 z 1
626.82958984375 0 1218.3666
626.9628295898438 0 35156.18
627.296630859375 0 26288.564
627.630126953125 0 11865.003
627.8199462890625 0 1788.031
627.9634399414062 0 5469.4717
628.2933959960938 0 1202.8885
629.307373046875 0 1216.7023
630.3130493164062 0 1798.8328
630.6400146484375 0 5563.106
630.9708251953125 0 6091.2964
631.3071899414062 0 2237.8162
631.6375122070312 0 1649.3191
631.96826171875 0 3113.6216 y 1
632.3027954101562 0 4754.5005
632.6359252929688 0 1904.7767
633.8236083984375 0 2798.9463
634.3274536132812 0 16195.147
634.6494750976562 0 3209.5488
634.7969360351562 0 17442.576 w 5
634.8404541015625 0 2508.5583
634.9837036132812 0 6119.6987
635.2994995117188 0 11096.197
635.6544799804688 0 5399.852
635.7977905273438 0 4643.4697
635.842041015625 0 1960.9668
635.9886474609375 0 3225.571
636.29833984375 0 2748.5024
636.3378295898438 0 3186.0083
636.79541015625 0 1142.889
637.350341796875 0 2115.8442
638.819091796875 0 1290.3752
639.322998046875 0 2054.5833
639.9968872070312 0 3794.371
640.3225708007812 0 23191.824
640.6577758789062 0 23617.207
640.8182373046875 0 1768.0734
640.9911499023438 0 23238.1
641.324951171875 0 11158.033
641.658203125 0 3288.6245
641.8328857421875 0 3986.3923
641.9945678710938 0 1587.6904
642.3351440429688 0 3852.385
642.9799194335938 0 1275.1849
643.3224487304688 0 1640.8776
643.8260498046875 0 5991.557
644.3242797851562 0 8342.893
644.6539306640625 0 6881.7886
644.8256225585938 0 2146.5186
644.9869995117188 0 9656.311
645.3204956054688 0 5085.093
645.650634765625 0 4065.3713
645.9837036132812 0 4648.1943
646.3187255859375 0 3966.379
646.6514282226562 0 1638.3569
647.3195190429688 0 2637.2012 z Water loss 5
649.3245239257812 0 2508.1306
649.6588745117188 0 7059.284
649.9805908203125 0 14228.33
650.324462890625 0 57396.016
650.6593627929688 0 56722.06
650.9938354492188 0 39848.29
651.3277587890625 0 18403.137
651.6615600585938 0 6673.2607
651.805419921875 0 1711.2849
651.9968872070312 0 3037.0566
652.3211669921875 0 14472.981
653.325439453125 0 6071.121
654.3284912109375 0 2202.1404
654.6615600585938 0 4145.176
654.9949340820312 0 7358.248
655.3275756835938 0 12484.252 y Water loss 5
655.6609497070312 0 9582.777
655.825927734375 0 3977.4055 y Ammonia loss 5
655.9945678710938 0 4462.546
656.3246459960938 0 29603.256 z 5
656.8262329101562 0 20206.84
657.3280029296875 0 11955.684
657.8330078125 0 7761.5444
658.3344116210938 0 4042.2927
658.6555786132812 0 6679.1514
658.841064453125 0 2019.2551
658.9864501953125 0 5678.9644
659.3226318359375 0 5331.9355
659.6549682617188 0 3302.0408
659.9841918945312 0 1791.4033
660.318359375 0 1235.7369
660.6611938476562 0 1169.8289
660.9908447265625 0 1252.5365
662.3193359375 0 2754.0586
663.2904663085938 0 7988.104
663.33935546875 0 3162.2764
663.65966796875 0 1160.974
663.831787109375 0 6769.325
663.9891357421875 0 2404.5981
664.3341674804688 0 67547.3 y 5
664.657958984375 0 95967.375
664.8345336914062 0 27949.19
664.9920654296875 0 91378.03
665.3281860351562 0 71568.695
665.6597290039062 0 23570.627
665.8367919921875 0 5474.547
665.9938354492188 0 13428.27
666.3298950195312 0 4789.862
666.659912109375 0 1701.584
669.3270874023438 0 2086.2405
669.661865234375 0 23871.951
669.9966430664062 0 44385.523
670.3316040039062 0 65302.45
670.666259765625 0 48201.246
670.8363037109375 0 1555.2291
671.0000610351562 0 27052.43
671.332763671875 0 10346.431
671.6671142578125 0 3926.8896
671.8260498046875 0 2175.6638
671.9962768554688 0 2130.321
672.3263549804688 0 2652.2898
677.8247680664062 0 1401.8267
679.3428955078125 0 5244.5713 c Water loss 4
679.83642578125 0 34431.586
680.31982421875 0 107667.84 c Ammonia loss 4
680.8397216796875 0 13431.886
681.3217163085938 0 44855.97
681.8427124023438 0 1366.7416
682.3223266601562 0 14241.213
682.8427124023438 0 1229.5848
683.3236694335938 0 3517.9248
684.3260498046875 0 1692.3807
685.3350219726562 0 1210.164
690.842529296875 0 2960.4868
691.3389892578125 0 39354.33
691.83984375 0 26395.291
692.3402099609375 0 12770.346
692.8387451171875 0 5761.176 c Water loss 10
693.334716796875 0 2634.8838 c Ammonia loss 10
693.8342895507812 0 2303.4722
694.3358154296875 0 1886.7849
696.3284912109375 0 1037.3455
696.8460693359375 0 1013.4363
697.3431396484375 0 32337.865 c 4
698.3466796875 0 14288.282
699.3467407226562 0 5279.039
701.3458251953125 0 2951.346
701.8427734375 0 55212.734 c 10
702.3446044921875 0 45696.754
702.84619140625 0 20668.518
703.3488159179688 0 6781.3745
703.8491821289062 0 1764.1549
704.3611450195312 0 1046.0294
705.3576049804688 0 1921.8131
706.345947265625 0 1320.7297
711.3098754882812 0 3776.0784
712.3147583007812 0 1916.2174
714.3162841796875 0 1634.4214
714.3777465820312 0 1766.3708
719.85546875 0 1175.8278
720.3572998046875 0 5187.4087
720.8536987304688 0 3396.7422
721.3526000976562 0 5873.4077
722.359619140625 0 3171.9285
723.3648681640625 0 2432.0513
724.37548828125 0 1257.029
726.3612670898438 0 2077.136
726.8593139648438 0 3959.4985
727.3663330078125 0 2371.2458
738.3759765625 0 1984.9204
742.3492431640625 0 8051.18
742.8504638671875 0 5073.489
743.35107421875 0 3086.4443
744.3864135742188 0 2958.0203
744.8869018554688 0 2002.4441
748.361328125 0 1249.2792 y Water loss 4
748.8546752929688 0 4395.7275 y Ammonia loss 4
749.3622436523438 0 9640.441 z 4
749.8660888671875 0 16905.123
750.3682250976562 0 11485.114
750.8698120117188 0 6733.737
751.3699951171875 0 2528.753
751.8641967773438 0 999.2041
757.373291015625 0 4470.476 c Ammonia loss 11
757.8777465820312 0 4354.4634
758.3773193359375 0 4087.6052
762.868408203125 0 2538.7764
763.3677978515625 0 1826.2294
765.4034423828125 0 3350.9006
765.8910522460938 0 6141.298 c 11
766.398193359375 0 9336.58
766.894775390625 0 5244.8535
767.4043579101562 0 2060.8384
768.3603515625 0 2411.8462 y Ammonia loss 10
769.3689575195312 0 22877.209 z 10
770.3726196289062 0 13163.521
770.8678588867188 0 2737.8804
771.3726806640625 0 6443.5825
772.3694458007812 0 2340.3757
773.3688354492188 0 966.38245
775.3958740234375 0 1313.105
776.3746337890625 0 1924.5847
777.3687133789062 0 2178.7341
777.8748779296875 0 2707.824
778.3735961914062 0 2865.0957
778.8965454101562 0 1913.08
784.3811645507812 0 2250.6633 w 3
784.8814697265625 0 5143.4375
785.3863525390625 0 15990.565 y 10
785.8831787109375 0 2611.609
786.389404296875 0 5944.2603
787.3937377929688 0 1624.169
791.3811645507812 0 1764.6641
792.3851318359375 0 1355.159
793.4007568359375 0 59790.62 c Ammonia loss 5
794.4039306640625 0 30891.502
795.4058837890625 0 9752.434
796.408203125 0 1846.7291
798.3743896484375 0 1319.6047 z Ammonia loss 3
806.3786010742188 0 3335.6033 y Ammonia loss 3
806.878173828125 0 10024.923 z 3
807.3799438476562 0 25383.305
807.882080078125 0 15745.98
808.3832397460938 0 8182.3857
808.8833618164062 0 3053.7075
809.3915405273438 0 4370.992
810.4270629882812 0 96050.78 c 5
811.4301147460938 0 45421.46
812.4321899414062 0 13230.368
812.8815307617188 0 2442.7815
813.4395751953125 0 2099.3823
813.8822631835938 0 1180.9362
814.8853759765625 0 2076.5796 y 3
815.3872680664062 0 1763.9904
816.8836059570312 0 2231.4363
817.394287109375 0 3563.1938
817.893310546875 0 2703.8542
818.39892578125 0 1182.7112
822.414306640625 0 1770.625
823.9072265625 0 3050.4294
824.4048461914062 0 3018.8586
825.8873901367188 0 1157.9596
826.8998413085938 0 2178.5305
827.3995361328125 0 1188.168
836.3810424804688 0 1041.8221
838.39111328125 0 2284.8308
838.8890991210938 0 1704.5216
839.3986206054688 0 2794.7205 w 9
840.4053344726562 0 2202.0022
841.4046630859375 0 2690.1672
841.9013671875 0 2569.9424 w 2
842.4005126953125 0 2079.308
843.4378662109375 0 1248.003
844.4113159179688 0 1053.2563
846.3970336914062 0 8839.62 c 12
846.899658203125 0 9059.541
847.4004516601562 0 5702.937
847.8973388671875 0 2280.0044
848.404296875 0 2325.4153
851.4036865234375 0 1265.8055
853.3636474609375 0 1143.6919
859.9373779296875 0 1699.2249
860.4320068359375 0 2497.1687
867.4246826171875 0 36989.25
868.4285888671875 0 20118.6
869.4308471679688 0 7308.8135
870.4180297851562 0 2405.0083 y Ammonia loss 2
870.9063720703125 0 6003.332 z 2
871.4094848632812 0 21931.41
871.9114379882812 0 18630.598
872.411865234375 0 11140.546
872.913330078125 0 3655.711
873.4144897460938 0 1539.6862
873.9086303710938 0 1063.2598
878.916748046875 0 2579.1091 y 2
879.4188232421875 0 2206.1394
879.9271240234375 0 1086.1604
880.4255981445312 0 2995.5405 z Water loss 9
880.9271850585938 0 2406.6162
881.4381713867188 0 4148.6284
882.4454345703125 0 3151.4868
888.4382934570312 0 2557.6216
888.9418334960938 0 8665.311
889.4422607421875 0 7001.0244
889.9439086914062 0 5116.268
890.4403686523438 0 1886.6396
894.4268798828125 0 1012.8446
895.4327392578125 0 1837.9598
895.934814453125 0 1051.2139
896.4276733398438 0 1391.9202 y Water loss 9
898.4114990234375 0 18064.799 z 9
899.4171142578125 0 19746.293
900.4178466796875 0 8063.882
901.4254760742188 0 2257.8457 c Water loss 13
902.431640625 0 1203.6261
908.4273071289062 0 8057.976 c Ammonia loss 6
909.4323120117188 0 4600.1816
910.4396362304688 0 6531.6543 c 13
910.9415283203125 0 4723.639
911.4450073242188 0 2257.4219
914.4297485351562 0 5020.005 y 9
915.4326782226562 0 2389.3481
916.4295043945312 0 1141.391
922.441162109375 0 1388.9725
924.4711303710938 0 934.0143
925.4544067382812 0 5854.13 c 6
926.4593505859375 0 4678.7197
927.4645385742188 0 1493.0333
930.9719848632812 0 1243.2369
931.46435546875 0 1096.0186
938.4615478515625 0 2849.9775 y Water loss 1
938.967529296875 0 1394.4504
939.4559326171875 0 5059.1084 z 1
939.9391479492188 0 15278.551
940.4418334960938 0 14007.691
940.9414672851562 0 9124.932
941.4435424804688 0 3197.2607
941.9427490234375 0 1485.1324
952.4695434570312 0 1210.5892
953.4761962890625 0 1433.8452
961.485595703125 0 1295.969
966.460693359375 0 1828.5178
967.4766235351562 0 3847.326
967.9804077148438 0 2142.872
968.4849243164062 0 2104.146
974.9860229492188 0 1548.6381
975.4876098632812 0 1920.696
975.9893188476562 0 2502.4058
976.4884033203125 0 1290.2814
976.9965209960938 0 1052.5211
981.4724731445312 0 1196.3749
982.4766235351562 0 13477.216 c 7
983.4801635742188 0 11871.458
984.4821166992188 0 4140.383
985.491455078125 0 928.9919
996.4800415039062 0 2932.2285
996.986083984375 0 3954.0347
997.4849243164062 0 2543.336
997.9850463867188 0 1062.7848
1004.4934692382812 0 996.3333
1004.9924926757812 0 2335.9558
1005.4978637695312 0 2386.2588
1005.99462890625 0 1418.2646
1010.4642944335938 0 2396.7634
1011.4696044921875 0 1844.873
1026.5059814453125 0 1428.152 z 8
1027.513916015625 0 9226.364
1028.5164794921875 0 4775.244
1029.5186767578125 0 2278.1558
1066.55517578125 0 1526.4525
1067.55859375 0 1201.0116
1084.53857421875 0 3687.7864
1085.5361328125 0 2470.1155
1093.5228271484375 0 1016.9858 c Ammonia loss 8
1094.5487060546875 0 2083.9941
1095.55615234375 0 2199.6885
1099.5447998046875 0 1908.4805 y 7
1100.552001953125 0 1125.4843
1109.548828125 0 991.1632
1110.5693359375 0 13632.231 c 8
1111.573486328125 0 12599.508
1112.577880859375 0 4933.0493
1113.577392578125 0 2259.2007
1142.55908203125 0 1429.2269
1196.6185302734375 0 1199.844
1198.564208984375 0 997.43884 z 6
1199.5596923828125 0 4579.458
1200.5645751953125 0 2403.1116
1201.56982421875 0 1182.5435
1239.6107177734375 0 2309.691 c 9
1240.616943359375 0 1878.3679
1241.62158203125 0 1659.829
1242.626953125 0 1094.5087
1264.608642578125 0 868.92206
1311.6279296875 0 847.83594 z 5
1312.642578125 0 1941.4478
1313.6483154296875 0 1614.8304
1314.6429443359375 0 1184.5759
1358.6685791015625 0 810.5722
1359.6639404296875 0 1893.9064
1360.6683349609375 0 1239.8817
1402.673828125 0 1730.037 c 10
1403.6883544921875 0 990.3706
1404.691650390625 0 1256.6218
1613.7657470703125 0 700.5633
1686.7611083984375 0 739.2753
2470.86328125 0 726.3083
3045.662841796875 0 1066.4839
3289.24951171875 0 625.6566
3431.74658203125 0 711.7338

Spectrum Details

|  |  |
| --- | --- |
| Matched peaks? Matched peaks The total absolute number of peaks matched. Additionally in brackets the total fraction of peaks matched and the total number of peaks is shown. | 176 (13.21% of 1332) |
| FDR? FDR The false discovery rate estimated for this peptide. It is calculated by matching all theoretical fragments with a non-integer shift with the raw peaks for this spectrum. This is done with 40 different shifts. The resulting percentage is the average number of annotated peaks over the number of annotated peaks with the correct spectrum. | 1.22% |
| Satellite FDR? Satellite FDR See the FDR for details on its calculation. This satellite ion specific FDR only contains the satellite ions (d/w) for I/L/J positions. | 1.19% |
| PSM Score? PSM Score The PSM Score as given by Hecklib to this annotated spectrum. It is shown with three significant figures. | 452 |

#### Spectrum 5489? Spectrum 5489 The raw spectrum of this peptide as annotated by Hecklib. The fragments are coloured according to ion type (see legend). Any peaks with a star '\*' as text can be hovered over to see the full details, first the ion type second the mass shift type. By hovering over the amino acids in the peptide or ions in the legend the corresponding peaks are highlighted. By toggling the 'Unassigned' label you can turn the background (unassigned) peaks on or off in the plot. By updating the slider in the Ion legend you can update the spectrum to only show the top X% of the peaks with labels. The top X% means any peak that is within X% of the highest intensity. By dragging in the spectrum you can zoom in to a specific part of the spectrum and use 'Zoom Out' to get back to the original zoom level. The annotation of the spectrum is based on the given sequence in the peptides file and is done with different software so inconsistencies are likely. The peaks are annotated based on the given sequence, with 20 ppm tolerance.

Copy Data

##### Spectrum 5489 (TSV)

###### Preview

```
Loading example...
```

*Click on the button to copy the data to your clipboard.*

Mz MinMz MaxIntensity Max

WidthHeightPeptide font sizePeptide stroke widthSpectrum font sizeSpectrum stroke widthCompact peptide

Ion legend

wxyz

abcd

OtherUnassignedIon^Charge^_Position_Show for top:%

JHQDWLDGKEYKCKK

02.61e+45.22e+47.83e+41.04e+5

Zoom Out

c^+1^_2_z^+1^_2_c^+1^_3_c^+1^_4_c^+1^_4_z^+1^_4_c^+3^_14_y^+1^_4_c^+1^_5_z^+1^_5_y^+1^_5_c^+1^_6_z^+2^_13_y^+2^_13_z^+1^_6_c^+1^_7_c^+2^_14_c^+1^_7_z^+2^_14_w^+1^_7_c^+1^_8_z^+1^_7_z^+1^_8_y^+1^_8_c^+1^_9_y^+1^_9_z^+1^_9_c^+1^_10_y^+1^_10_z^+1^_10_c^+1^_11_y^+1^_11_z^+1^_11_c^+1^_12_c^+1^_12_y^+1^_12_z^+1^_12_y^+1^_12_c^+1^_13_z^+1^_13_

0508101715252033

Fragment Matches Table

Show background peaks

| Position | Ion type | Intensity | mz Theoretical | mz Error (Th) | mz Error (ppm) | Charge | Series Number |
| --- | --- | --- | --- | --- | --- | --- | --- |
| - | - | 2116 | 129.1 | - | - | 0 | - |
| - | - | 448.7 | 130.5 | - | - | 0 | - |
| - | - | 882.5 | 134 | - | - | 0 | - |
| - | - | 2601 | 136.1 | - | - | 0 | - |
| - | - | 576.6 | 148.9 | - | - | 0 | - |
| - | - | 808.9 | 148.9 | - | - | 0 | - |
| - | - | 642.4 | 148.9 | - | - | 0 | - |
| - | - | 1109 | 148.9 | - | - | 0 | - |
| - | - | 1302 | 148.9 | - | - | 0 | - |
| - | - | 2659 | 148.9 | - | - | 0 | - |
| - | - | 4380 | 148.9 | - | - | 0 | - |
| - | - | 3406 | 149 | - | - | 0 | - |
| - | - | 1812 | 149 | - | - | 0 | - |
| - | - | 985.6 | 149 | - | - | 0 | - |
| - | - | 901.4 | 149 | - | - | 0 | - |
| - | - | 960.5 | 149 | - | - | 0 | - |
| - | - | 606.1 | 149 | - | - | 0 | - |
| - | - | 534.2 | 149 | - | - | 0 | - |
| - | - | 459.7 | 149 | - | - | 0 | - |
| - | - | 433 | 149 | - | - | 0 | - |
| - | - | 697 | 166.1 | - | - | 0 | - |
| - | - | 857.4 | 173.5 | - | - | 0 | - |
| - | - | 453.8 | 183.9 | - | - | 0 | - |
| - | - | 523.3 | 194.9 | - | - | 0 | - |
| - | - | 533.3 | 212.6 | - | - | 0 | - |
| - | - | 704.9 | 223.2 | - | - | 0 | - |
| - | - | 516.1 | 225 | - | - | 0 | - |
| - | - | 6686 | 232.1 | - | - | 0 | - |
| - | - | 1140 | 239.1 | - | - | 0 | - |
| - | - | 868.5 | 242.1 | - | - | 0 | - |
| - | - | 551.3 | 244.7 | - | - | 0 | - |
| - | - | 644 | 246.1 | - | - | 0 | - |
| - | - | 697.4 | 248.1 | - | - | 0 | - |
| - | - | 7162 | 249.2 | - | - | 0 | - |
| - | - | 924.3 | 250.2 | - | - | 0 | - |
| - | - | 3222 | 251.2 | - | - | 0 | - |
| - | - | 966 | 258.1 | - | - | 0 | - |
| - | - | 600.6 | 263.1 | - | - | 0 | - |
| 2 | c | 1873 | 268.2 | 0.0005362 | 2 | +1 | 2 |
| - | - | 1.456E+04 | 276.1 | - | - | 0 | - |
| - | - | 1559 | 277 | - | - | 0 | - |
| - | - | 1342 | 277.1 | - | - | 0 | - |
| - | - | 1.903E+04 | 277.2 | - | - | 0 | - |
| - | - | 2276 | 278.2 | - | - | 0 | - |
| - | - | 882.4 | 299.1 | - | - | 0 | - |
| - | - | 586.4 | 300.1 | - | - | 0 | - |
| - | - | 1044 | 312.1 | - | - | 0 | - |
| 14 | z | 2142 | 317.2 | 0.005043 | 15.9 | +1 | 2 |
| - | - | 1777 | 318.2 | - | - | 0 | - |
| - | - | 501.2 | 328.5 | - | - | 0 | - |
| - | - | 764 | 330.1 | - | - | 0 | - |
| - | - | 1284 | 331.2 | - | - | 0 | - |
| - | - | 1040 | 338.1 | - | - | 0 | - |
| - | - | 882.2 | 341 | - | - | 0 | - |
| - | - | 790.3 | 342 | - | - | 0 | - |
| - | - | 659.8 | 347.1 | - | - | 0 | - |
| - | - | 1694 | 348.1 | - | - | 0 | - |
| - | - | 3770 | 355.1 | - | - | 0 | - |
| - | - | 2942 | 356.1 | - | - | 0 | - |
| - | - | 1420 | 357.1 | - | - | 0 | - |
| - | - | 755 | 357.1 | - | - | 0 | - |
| - | - | 2607 | 358.1 | - | - | 0 | - |
| - | - | 3520 | 359 | - | - | 0 | - |
| - | - | 895.1 | 359.1 | - | - | 0 | - |
| - | - | 2986 | 360 | - | - | 0 | - |
| - | - | 2665 | 361 | - | - | 0 | - |
| - | - | 689.7 | 362 | - | - | 0 | - |
| - | - | 9573 | 375.1 | - | - | 0 | - |
| - | - | 1.342E+04 | 376.1 | - | - | 0 | - |
| - | - | 1258 | 376.1 | - | - | 0 | - |
| - | - | 603.5 | 376.2 | - | - | 0 | - |
| - | - | 2012 | 377.1 | - | - | 0 | - |
| - | - | 1336 | 378.2 | - | - | 0 | - |
| - | - | 3581 | 387.2 | - | - | 0 | - |
| - | - | 764.7 | 388.2 | - | - | 0 | - |
| - | - | 699.3 | 389.1 | - | - | 0 | - |
| - | - | 1248 | 390.1 | - | - | 0 | - |
| - | - | 9.412E+04 | 393.1 | - | - | 0 | - |
| - | - | 1.489E+04 | 394.1 | - | - | 0 | - |
| 3 | c | 3581 | 396.2 | 0.000363 | 0.9162 | +1 | 3 |
| - | - | 927 | 397.2 | - | - | 0 | - |
| - | - | 1283 | 410.2 | - | - | 0 | - |
| - | - | 557.7 | 417.1 | - | - | 0 | - |
| - | - | 666.9 | 423.2 | - | - | 0 | - |
| - | - | 5565 | 429.1 | - | - | 0 | - |
| - | - | 6911 | 430.1 | - | - | 0 | - |
| - | - | 5906 | 431.1 | - | - | 0 | - |
| - | - | 2352 | 432.1 | - | - | 0 | - |
| - | - | 3.626E+04 | 438.2 | - | - | 0 | - |
| - | - | 7489 | 439.2 | - | - | 0 | - |
| - | - | 789.3 | 451.2 | - | - | 0 | - |
| - | - | 4352 | 460.2 | - | - | 0 | - |
| - | - | 1451 | 461.2 | - | - | 0 | - |
| - | - | 1134 | 467.2 | - | - | 0 | - |
| - | - | 722.1 | 468.3 | - | - | 0 | - |
| - | - | 545.9 | 477.7 | - | - | 0 | - |
| - | - | 570 | 482.7 | - | - | 0 | - |
| - | - | 932.7 | 488.2 | - | - | 0 | - |
| - | - | 2748 | 489.2 | - | - | 0 | - |
| - | - | 975.6 | 490.2 | - | - | 0 | - |
| - | - | 1852 | 491.2 | - | - | 0 | - |
| - | - | 592.4 | 492.3 | - | - | 0 | - |
| 4 | c | 1754 | 494.2 | 0.0039 | 7.892 | +1 | 4 |
| - | - | 614.8 | 496.7 | - | - | 0 | - |
| - | - | 4227 | 506.2 | - | - | 0 | - |
| - | - | 1902 | 507.2 | - | - | 0 | - |
| - | - | 781.6 | 508.2 | - | - | 0 | - |
| - | - | 858.5 | 510.8 | - | - | 0 | - |
| - | - | 775.5 | 511.2 | - | - | 0 | - |
| 4 | c | 6680 | 511.3 | 0.0006642 | 1.299 | +1 | 4 |
| - | - | 1457 | 512.3 | - | - | 0 | - |
| - | - | 2374 | 516.2 | - | - | 0 | - |
| - | - | 790.5 | 516.2 | - | - | 0 | - |
| - | - | 701.9 | 516.3 | - | - | 0 | - |
| - | - | 1417 | 517.2 | - | - | 0 | - |
| - | - | 2958 | 524.2 | - | - | 0 | - |
| - | - | 910.3 | 525.2 | - | - | 0 | - |
| - | - | 2400 | 535.2 | - | - | 0 | - |
| - | - | 792.7 | 536.2 | - | - | 0 | - |
| - | - | 2247 | 538.2 | - | - | 0 | - |
| - | - | 1.357E+04 | 539.2 | - | - | 0 | - |
| - | - | 3790 | 540.2 | - | - | 0 | - |
| - | - | 1.756E+04 | 552.2 | - | - | 0 | - |
| - | - | 1.513E+04 | 553.2 | - | - | 0 | - |
| - | - | 4037 | 554.2 | - | - | 0 | - |
| - | - | 926 | 566.3 | - | - | 0 | - |
| - | - | 2511 | 570.2 | - | - | 0 | - |
| - | - | 1012 | 571.2 | - | - | 0 | - |
| - | - | 3821 | 584.4 | - | - | 0 | - |
| - | - | 700.2 | 585.4 | - | - | 0 | - |
| - | - | 668.6 | 586.2 | - | - | 0 | - |
| - | - | 3053 | 586.3 | - | - | 0 | - |
| - | - | 1095 | 592.3 | - | - | 0 | - |
| - | - | 573.1 | 596.9 | - | - | 0 | - |
| - | - | 2383 | 606.3 | - | - | 0 | - |
| 12 | z | 1930 | 606.3 | 0.007832 | 12.92 | +1 | 4 |
| - | - | 1110 | 607.2 | - | - | 0 | - |
| 14 | c | 1212 | 607.3 | 0.008859 | 14.59 | +3 | 14 |
| - | - | 943.4 | 609.3 | - | - | 0 | - |
| - | - | 2781 | 610.3 | - | - | 0 | - |
| - | - | 1495 | 614.2 | - | - | 0 | - |
| - | - | 1424 | 615.2 | - | - | 0 | - |
| 12 | y | 1035 | 622.3 | 0.01087 | 17.47 | +1 | 4 |
| - | - | 3049 | 623.3 | - | - | 0 | - |
| - | - | 1.509E+04 | 624.3 | - | - | 0 | - |
| - | - | 6516 | 625.3 | - | - | 0 | - |
| - | - | 707.4 | 632.3 | - | - | 0 | - |
| - | - | 791 | 632.6 | - | - | 0 | - |
| - | - | 5105 | 634.3 | - | - | 0 | - |
| - | - | 1110 | 634.6 | - | - | 0 | - |
| - | - | 1995 | 635.3 | - | - | 0 | - |
| - | - | 1190 | 635.3 | - | - | 0 | - |
| - | - | 615.4 | 636.3 | - | - | 0 | - |
| - | - | 654.4 | 648.3 | - | - | 0 | - |
| - | - | 1.067E+04 | 651.3 | - | - | 0 | - |
| - | - | 1156 | 651.3 | - | - | 0 | - |
| - | - | 2.597E+04 | 652.3 | - | - | 0 | - |
| - | - | 1.064E+04 | 653.3 | - | - | 0 | - |
| - | - | 970.5 | 658 | - | - | 0 | - |
| - | - | 763.4 | 663.7 | - | - | 0 | - |
| - | - | 689.4 | 664 | - | - | 0 | - |
| - | - | 1568 | 664.3 | - | - | 0 | - |
| - | - | 763.6 | 664.7 | - | - | 0 | - |
| - | - | 1.033E+05 | 669.3 | - | - | 0 | - |
| - | - | 1.694E+04 | 669.7 | - | - | 0 | - |
| - | - | 1.419E+04 | 670 | - | - | 0 | - |
| - | - | 3.663E+04 | 670.3 | - | - | 0 | - |
| - | - | 5216 | 670.7 | - | - | 0 | - |
| - | - | 2024 | 671 | - | - | 0 | - |
| - | - | 1530 | 671.3 | - | - | 0 | - |
| - | - | 733 | 673.9 | - | - | 0 | - |
| 5 | c | 2862 | 697.3 | 0.002035 | 2.919 | +1 | 5 |
| - | - | 608.1 | 701.6 | - | - | 0 | - |
| - | - | 1217 | 743.5 | - | - | 0 | - |
| 11 | z | 3063 | 769.4 | 0.01214 | 15.78 | +1 | 5 |
| - | - | 1271 | 770.4 | - | - | 0 | - |
| - | - | 1050 | 773.4 | - | - | 0 | - |
| - | - | 1184 | 783.7 | - | - | 0 | - |
| 11 | y | 888.3 | 785.4 | 0.01463 | 18.62 | +1 | 5 |
| - | - | 653.9 | 787.9 | - | - | 0 | - |
| - | - | 595.2 | 789.4 | - | - | 0 | - |
| 6 | c | 2568 | 810.4 | 0.00181 | 2.233 | +1 | 6 |
| - | - | 1168 | 811.4 | - | - | 0 | - |
| - | - | 677.2 | 816.6 | - | - | 0 | - |
| - | - | 898 | 850.4 | - | - | 0 | - |
| 3 | z | 911.6 | 870.9 | 0.003078 | 3.534 | +2 | 13 |
| - | - | 787.6 | 875.5 | - | - | 0 | - |
| 3 | y | 1282 | 878.9 | 0.0004384 | 0.4988 | +2 | 13 |
| - | - | 824.9 | 879.4 | - | - | 0 | - |
| - | - | 809.3 | 879.9 | - | - | 0 | - |
| - | - | 659 | 883.9 | - | - | 0 | - |
| - | - | 865.3 | 887.9 | - | - | 0 | - |
| - | - | 705.6 | 888.4 | - | - | 0 | - |
| - | - | 727.9 | 893.9 | - | - | 0 | - |
| - | - | 969.2 | 894.9 | - | - | 0 | - |
| 10 | z | 2417 | 898.4 | 0.01097 | 12.21 | +1 | 6 |
| - | - | 991.5 | 899.4 | - | - | 0 | - |
| 7 | c | 868 | 908.4 | 0.003961 | 4.36 | +1 | 7 |
| 14 | c | 2295 | 910.4 | 0.003245 | 3.564 | +2 | 14 |
| - | - | 2156 | 910.9 | - | - | 0 | - |
| - | - | 1904 | 911.4 | - | - | 0 | - |
| - | - | 797 | 915.5 | - | - | 0 | - |
| - | - | 729.4 | 921.9 | - | - | 0 | - |
| 7 | c | 1409 | 925.5 | 0.002253 | 2.435 | +1 | 7 |
| - | - | 581.9 | 926.5 | - | - | 0 | - |
| - | - | 1145 | 931.5 | - | - | 0 | - |
| - | - | 866.9 | 939 | - | - | 0 | - |
| 2 | z | 2492 | 939.4 | 0.006162 | 6.56 | +2 | 14 |
| - | - | 917.3 | 939.9 | - | - | 0 | - |
| - | - | 649.1 | 940.4 | - | - | 0 | - |
| - | - | 880.4 | 952.5 | - | - | 0 | - |
| - | - | 651.9 | 952.9 | - | - | 0 | - |
| - | - | 888.5 | 953.5 | - | - | 0 | - |
| - | - | 689.9 | 955.5 | - | - | 0 | - |
| - | - | 744.1 | 956 | - | - | 0 | - |
| - | - | 913.3 | 959 | - | - | 0 | - |
| - | - | 964.6 | 959.5 | - | - | 0 | - |
| - | - | 4906 | 960 | - | - | 0 | - |
| - | - | 5589 | 960.5 | - | - | 0 | - |
| - | - | 2940 | 961 | - | - | 0 | - |
| - | - | 1144 | 961.5 | - | - | 0 | - |
| - | - | 1063 | 962.9 | - | - | 0 | - |
| - | - | 944.9 | 963.5 | - | - | 0 | - |
| - | - | 635.8 | 965 | - | - | 0 | - |
| - | - | 860.7 | 966.5 | - | - | 0 | - |
| - | - | 805 | 967 | - | - | 0 | - |
| - | - | 855.2 | 967.5 | - | - | 0 | - |
| - | - | 2603 | 968 | - | - | 0 | - |
| 9 | w | 2102 | 968.4 | 0.01284 | 13.26 | +1 | 7 |
| - | - | 2543 | 969 | - | - | 0 | - |
| - | - | 1233 | 969.4 | - | - | 0 | - |
| - | - | 751.1 | 972.5 | - | - | 0 | - |
| - | - | 734.6 | 973.5 | - | - | 0 | - |
| - | - | 1437 | 974 | - | - | 0 | - |
| - | - | 4649 | 974.5 | - | - | 0 | - |
| - | - | 7922 | 975 | - | - | 0 | - |
| - | - | 6717 | 975.5 | - | - | 0 | - |
| - | - | 4868 | 976 | - | - | 0 | - |
| - | - | 3291 | 976.5 | - | - | 0 | - |
| - | - | 2123 | 977 | - | - | 0 | - |
| - | - | 1229 | 977.5 | - | - | 0 | - |
| - | - | 672.4 | 979 | - | - | 0 | - |
| - | - | 988 | 979.4 | - | - | 0 | - |
| - | - | 756.3 | 980.5 | - | - | 0 | - |
| - | - | 3914 | 981.5 | - | - | 0 | - |
| - | - | 1900 | 982 | - | - | 0 | - |
| 8 | c | 7282 | 982.5 | 0.002274 | 2.314 | +1 | 8 |
| - | - | 1354 | 983 | - | - | 0 | - |
| - | - | 2313 | 983.5 | - | - | 0 | - |
| - | - | 807.2 | 983.8 | - | - | 0 | - |
| - | - | 1588 | 984.5 | - | - | 0 | - |
| - | - | 1851 | 985 | - | - | 0 | - |
| - | - | 1573 | 985.5 | - | - | 0 | - |
| - | - | 1574 | 985.9 | - | - | 0 | - |
| - | - | 1164 | 986.5 | - | - | 0 | - |
| - | - | 1239 | 987 | - | - | 0 | - |
| - | - | 1915 | 987.5 | - | - | 0 | - |
| - | - | 1473 | 988.4 | - | - | 0 | - |
| - | - | 789.1 | 989 | - | - | 0 | - |
| - | - | 838.9 | 990.5 | - | - | 0 | - |
| - | - | 856.4 | 991 | - | - | 0 | - |
| - | - | 806.5 | 991.4 | - | - | 0 | - |
| - | - | 1169 | 993.5 | - | - | 0 | - |
| - | - | 1862 | 994 | - | - | 0 | - |
| - | - | 1130 | 994.5 | - | - | 0 | - |
| - | - | 1522 | 995.5 | - | - | 0 | - |
| - | - | 3510 | 996 | - | - | 0 | - |
| - | - | 3721 | 996.5 | - | - | 0 | - |
| - | - | 1653 | 997 | - | - | 0 | - |
| - | - | 910.7 | 997.5 | - | - | 0 | - |
| - | - | 967.1 | 998 | - | - | 0 | - |
| - | - | 1227 | 1002 | - | - | 0 | - |
| - | - | 1022 | 1003 | - | - | 0 | - |
| - | - | 3627 | 1003 | - | - | 0 | - |
| - | - | 1.081E+04 | 1004 | - | - | 0 | - |
| - | - | 1.385E+04 | 1004 | - | - | 0 | - |
| - | - | 9371 | 1005 | - | - | 0 | - |
| - | - | 6085 | 1005 | - | - | 0 | - |
| - | - | 2833 | 1006 | - | - | 0 | - |
| - | - | 1084 | 1006 | - | - | 0 | - |
| - | - | 894 | 1022 | - | - | 0 | - |
| 9 | z | 4569 | 1027 | 0.01371 | 13.35 | +1 | 7 |
| - | - | 2868 | 1028 | - | - | 0 | - |
| - | - | 1865 | 1029 | - | - | 0 | - |
| - | - | 803.8 | 1030 | - | - | 0 | - |
| - | - | 820.3 | 1038 | - | - | 0 | - |
| - | - | 1659 | 1039 | - | - | 0 | - |
| - | - | 1072 | 1040 | - | - | 0 | - |
| - | - | 939.8 | 1041 | - | - | 0 | - |
| - | - | 1089 | 1044 | - | - | 0 | - |
| - | - | 1445 | 1044 | - | - | 0 | - |
| - | - | 2616 | 1045 | - | - | 0 | - |
| - | - | 3494 | 1045 | - | - | 0 | - |
| - | - | 2675 | 1045 | - | - | 0 | - |
| - | - | 959.1 | 1046 | - | - | 0 | - |
| - | - | 880.3 | 1050 | - | - | 0 | - |
| - | - | 879.9 | 1055 | - | - | 0 | - |
| - | - | 1197 | 1061 | - | - | 0 | - |
| 8 | z | 1099 | 1084 | 0.0143 | 13.2 | +1 | 8 |
| - | - | 1340 | 1085 | - | - | 0 | - |
| - | - | 962.8 | 1086 | - | - | 0 | - |
| - | - | 760.1 | 1087 | - | - | 0 | - |
| - | - | 825.3 | 1092 | - | - | 0 | - |
| - | - | 729.5 | 1094 | - | - | 0 | - |
| - | - | 1042 | 1095 | - | - | 0 | - |
| - | - | 1141 | 1097 | - | - | 0 | - |
| - | - | 709.4 | 1099 | - | - | 0 | - |
| 8 | y | 1980 | 1100 | 0.01325 | 12.05 | +1 | 8 |
| - | - | 919.8 | 1100 | - | - | 0 | - |
| - | - | 788.4 | 1101 | - | - | 0 | - |
| - | - | 923.6 | 1102 | - | - | 0 | - |
| - | - | 1355 | 1103 | - | - | 0 | - |
| - | - | 1542 | 1104 | - | - | 0 | - |
| - | - | 1409 | 1108 | - | - | 0 | - |
| - | - | 1577 | 1109 | - | - | 0 | - |
| - | - | 946.6 | 1109 | - | - | 0 | - |
| - | - | 848.2 | 1109 | - | - | 0 | - |
| - | - | 1756 | 1110 | - | - | 0 | - |
| 9 | c | 4141 | 1111 | 0.002662 | 2.397 | +1 | 9 |
| - | - | 892 | 1111 | - | - | 0 | - |
| - | - | 2919 | 1112 | - | - | 0 | - |
| - | - | 1153 | 1112 | - | - | 0 | - |
| - | - | 923 | 1113 | - | - | 0 | - |
| - | - | 700.5 | 1115 | - | - | 0 | - |
| - | - | 986.1 | 1115 | - | - | 0 | - |
| - | - | 1613 | 1117 | - | - | 0 | - |
| - | - | 1077 | 1117 | - | - | 0 | - |
| - | - | 1577 | 1117 | - | - | 0 | - |
| - | - | 804.7 | 1118 | - | - | 0 | - |
| - | - | 747.4 | 1118 | - | - | 0 | - |
| - | - | 2255 | 1172 | - | - | 0 | - |
| - | - | 939.3 | 1173 | - | - | 0 | - |
| - | - | 928.6 | 1174 | - | - | 0 | - |
| - | - | 1006 | 1196 | - | - | 0 | - |
| 7 | y | 828.6 | 1198 | 0.002228 | 1.86 | +1 | 9 |
| 7 | z | 2493 | 1199 | 0.01426 | 11.9 | +1 | 9 |
| - | - | 1003 | 1221 | - | - | 0 | - |
| 10 | c | 3404 | 1240 | 0.006925 | 5.587 | +1 | 10 |
| - | - | 2515 | 1241 | - | - | 0 | - |
| - | - | 707.2 | 1274 | - | - | 0 | - |
| - | - | 877.3 | 1275 | - | - | 0 | - |
| - | - | 1914 | 1280 | - | - | 0 | - |
| - | - | 866 | 1281 | - | - | 0 | - |
| - | - | 961 | 1302 | - | - | 0 | - |
| - | - | 1074 | 1303 | - | - | 0 | - |
| 6 | y | 765.3 | 1311 | 0.003101 | 2.366 | +1 | 10 |
| 6 | z | 1741 | 1312 | 0.01447 | 11.03 | +1 | 10 |
| - | - | 1044 | 1313 | - | - | 0 | - |
| - | - | 900 | 1318 | - | - | 0 | - |
| - | - | 1146 | 1337 | - | - | 0 | - |
| - | - | 917.9 | 1340 | - | - | 0 | - |
| - | - | 1550 | 1341 | - | - | 0 | - |
| - | - | 872.5 | 1359 | - | - | 0 | - |
| - | - | 1011 | 1402 | - | - | 0 | - |
| 11 | c | 3136 | 1403 | 0.004092 | 2.917 | +1 | 11 |
| - | - | 2039 | 1404 | - | - | 0 | - |
| - | - | 926.3 | 1407 | - | - | 0 | - |
| - | - | 1350 | 1437 | - | - | 0 | - |
| - | - | 1103 | 1438 | - | - | 0 | - |
| - | - | 697.8 | 1460 | - | - | 0 | - |
| - | - | 1226 | 1465 | - | - | 0 | - |
| - | - | 1813 | 1466 | - | - | 0 | - |
| - | - | 1581 | 1466 | - | - | 0 | - |
| - | - | 1119 | 1474 | - | - | 0 | - |
| - | - | 857.1 | 1494 | - | - | 0 | - |
| - | - | 1211 | 1495 | - | - | 0 | - |
| 5 | y | 1322 | 1497 | 0.01243 | 8.308 | +1 | 11 |
| 5 | z | 3515 | 1498 | 0.01492 | 9.963 | +1 | 11 |
| - | - | 3117 | 1499 | - | - | 0 | - |
| - | - | 1548 | 1500 | - | - | 0 | - |
| 12 | c | 1858 | 1514 | 0.01587 | 10.48 | +1 | 12 |
| - | - | 1197 | 1515 | - | - | 0 | - |
| - | - | 1060 | 1516 | - | - | 0 | - |
| - | - | 1400 | 1530 | - | - | 0 | - |
| 12 | c | 3950 | 1531 | 0.01568 | 10.24 | +1 | 12 |
| - | - | 3063 | 1532 | - | - | 0 | - |
| - | - | 1485 | 1533 | - | - | 0 | - |
| - | - | 973.5 | 1536 | - | - | 0 | - |
| - | - | 942.9 | 1537 | - | - | 0 | - |
| - | - | 1303 | 1544 | - | - | 0 | - |
| - | - | 968.4 | 1544 | - | - | 0 | - |
| - | - | 963.5 | 1545 | - | - | 0 | - |
| - | - | 1123 | 1545 | - | - | 0 | - |
| - | - | 800.7 | 1550 | - | - | 0 | - |
| - | - | 810.4 | 1550 | - | - | 0 | - |
| - | - | 812.2 | 1551 | - | - | 0 | - |
| - | - | 920.2 | 1551 | - | - | 0 | - |
| - | - | 1013 | 1552 | - | - | 0 | - |
| - | - | 896.8 | 1557 | - | - | 0 | - |
| - | - | 2589 | 1558 | - | - | 0 | - |
| - | - | 2968 | 1558 | - | - | 0 | - |
| - | - | 3353 | 1559 | - | - | 0 | - |
| - | - | 1863 | 1559 | - | - | 0 | - |
| - | - | 1124 | 1560 | - | - | 0 | - |
| - | - | 1427 | 1565 | - | - | 0 | - |
| - | - | 1808 | 1566 | - | - | 0 | - |
| - | - | 2445 | 1566 | - | - | 0 | - |
| - | - | 4386 | 1567 | - | - | 0 | - |
| - | - | 2921 | 1567 | - | - | 0 | - |
| - | - | 3191 | 1568 | - | - | 0 | - |
| - | - | 2423 | 1568 | - | - | 0 | - |
| - | - | 654.9 | 1569 | - | - | 0 | - |
| - | - | 987.4 | 1570 | - | - | 0 | - |
| - | - | 782.3 | 1571 | - | - | 0 | - |
| - | - | 1197 | 1575 | - | - | 0 | - |
| - | - | 1230 | 1575 | - | - | 0 | - |
| - | - | 932.3 | 1579 | - | - | 0 | - |
| - | - | 742.4 | 1584 | - | - | 0 | - |
| - | - | 796.3 | 1584 | - | - | 0 | - |
| - | - | 1118 | 1587 | - | - | 0 | - |
| - | - | 931.9 | 1599 | - | - | 0 | - |
| - | - | 903.2 | 1599 | - | - | 0 | - |
| - | - | 1042 | 1603 | - | - | 0 | - |
| - | - | 920.6 | 1604 | - | - | 0 | - |
| - | - | 658.4 | 1608 | - | - | 0 | - |
| - | - | 674.1 | 1609 | - | - | 0 | - |
| - | - | 940.6 | 1610 | - | - | 0 | - |
| - | - | 764.6 | 1611 | - | - | 0 | - |
| 4 | y | 1078 | 1612 | 0.01186 | 7.358 | +1 | 12 |
| 4 | z | 3173 | 1613 | 0.008539 | 5.295 | +1 | 12 |
| - | - | 3412 | 1614 | - | - | 0 | - |
| - | - | 2600 | 1615 | - | - | 0 | - |
| - | - | 810.8 | 1616 | - | - | 0 | - |
| - | - | 790.4 | 1617 | - | - | 0 | - |
| - | - | 1087 | 1619 | - | - | 0 | - |
| - | - | 1114 | 1620 | - | - | 0 | - |
| - | - | 933 | 1624 | - | - | 0 | - |
| - | - | 932.1 | 1625 | - | - | 0 | - |
| - | - | 1264 | 1628 | - | - | 0 | - |
| 4 | y | 1159 | 1629 | 0.006145 | 3.773 | +1 | 12 |
| - | - | 1336 | 1630 | - | - | 0 | - |
| - | - | 1011 | 1632 | - | - | 0 | - |
| - | - | 792.4 | 1632 | - | - | 0 | - |
| - | - | 800.9 | 1633 | - | - | 0 | - |
| - | - | 984.9 | 1633 | - | - | 0 | - |
| - | - | 1378 | 1634 | - | - | 0 | - |
| - | - | 836.5 | 1635 | - | - | 0 | - |
| - | - | 1002 | 1637 | - | - | 0 | - |
| - | - | 897.4 | 1639 | - | - | 0 | - |
| - | - | 1209 | 1640 | - | - | 0 | - |
| - | - | 1126 | 1641 | - | - | 0 | - |
| - | - | 1511 | 1641 | - | - | 0 | - |
| - | - | 1073 | 1642 | - | - | 0 | - |
| - | - | 1047 | 1643 | - | - | 0 | - |
| - | - | 1130 | 1643 | - | - | 0 | - |
| - | - | 930.2 | 1644 | - | - | 0 | - |
| - | - | 792.9 | 1644 | - | - | 0 | - |
| - | - | 1217 | 1646 | - | - | 0 | - |
| - | - | 745 | 1647 | - | - | 0 | - |
| - | - | 1044 | 1647 | - | - | 0 | - |
| - | - | 1144 | 1648 | - | - | 0 | - |
| - | - | 996 | 1650 | - | - | 0 | - |
| - | - | 914.1 | 1651 | - | - | 0 | - |
| - | - | 1641 | 1653 | - | - | 0 | - |
| - | - | 1047 | 1654 | - | - | 0 | - |
| - | - | 1094 | 1655 | - | - | 0 | - |
| - | - | 892.3 | 1655 | - | - | 0 | - |
| - | - | 1206 | 1658 | - | - | 0 | - |
| - | - | 908.3 | 1662 | - | - | 0 | - |
| - | - | 1718 | 1663 | - | - | 0 | - |
| - | - | 1643 | 1663 | - | - | 0 | - |
| - | - | 1614 | 1664 | - | - | 0 | - |
| - | - | 1507 | 1666 | - | - | 0 | - |
| - | - | 1132 | 1667 | - | - | 0 | - |
| - | - | 875 | 1670 | - | - | 0 | - |
| - | - | 1130 | 1672 | - | - | 0 | - |
| - | - | 2010 | 1674 | - | - | 0 | - |
| - | - | 2683 | 1675 | - | - | 0 | - |
| - | - | 1824 | 1675 | - | - | 0 | - |
| - | - | 1507 | 1676 | - | - | 0 | - |
| - | - | 1561 | 1691 | - | - | 0 | - |
| 13 | c | 2814 | 1692 | 0.009108 | 5.384 | +1 | 13 |
| - | - | 3613 | 1693 | - | - | 0 | - |
| - | - | 1448 | 1694 | - | - | 0 | - |
| 3 | z | 1909 | 1741 | 0.02256 | 12.96 | +1 | 13 |
| - | - | 1427 | 1742 | - | - | 0 | - |
| - | - | 1037 | 1743 | - | - | 0 | - |
| - | - | 1107 | 1821 | - | - | 0 | - |
| - | - | 1037 | 1831 | - | - | 0 | - |
| - | - | 935.8 | 1879 | - | - | 0 | - |
| - | - | 1542 | 1918 | - | - | 0 | - |
| - | - | 1558 | 1919 | - | - | 0 | - |
| - | - | 1907 | 1920 | - | - | 0 | - |
| - | - | 1494 | 1921 | - | - | 0 | - |
| - | - | 1549 | 1922 | - | - | 0 | - |
| - | - | 1040 | 1923 | - | - | 0 | - |
| - | - | 1154 | 1933 | - | - | 0 | - |
| - | - | 924.9 | 1938 | - | - | 0 | - |
| - | - | 1072 | 1948 | - | - | 0 | - |
| - | - | 3070 | 1949 | - | - | 0 | - |
| - | - | 3409 | 1950 | - | - | 0 | - |
| - | - | 2277 | 1951 | - | - | 0 | - |
| - | - | 1096 | 1952 | - | - | 0 | - |
| - | - | 1691 | 1953 | - | - | 0 | - |
| - | - | 889.6 | 1954 | - | - | 0 | - |
| - | - | 1200 | 1963 | - | - | 0 | - |
| - | - | 2395 | 1964 | - | - | 0 | - |
| - | - | 1728 | 1965 | - | - | 0 | - |
| - | - | 1359 | 1966 | - | - | 0 | - |
| - | - | 1241 | 1970 | - | - | 0 | - |
| - | - | 986.1 | 1971 | - | - | 0 | - |
| - | - | 1204 | 1980 | - | - | 0 | - |
| - | - | 2167 | 1981 | - | - | 0 | - |
| - | - | 1905 | 1982 | - | - | 0 | - |
| - | - | 1025 | 1983 | - | - | 0 | - |
| - | - | 1030 | 1987 | - | - | 0 | - |
| - | - | 1672 | 1988 | - | - | 0 | - |
| - | - | 1177 | 1990 | - | - | 0 | - |
| - | - | 2821 | 1991 | - | - | 0 | - |
| - | - | 4897 | 1992 | - | - | 0 | - |
| - | - | 4162 | 1993 | - | - | 0 | - |
| - | - | 3637 | 1994 | - | - | 0 | - |
| - | - | 1711 | 1995 | - | - | 0 | - |
| - | - | 1101 | 1996 | - | - | 0 | - |
| - | - | 1831 | 2006 | - | - | 0 | - |
| - | - | 2098 | 2007 | - | - | 0 | - |
| - | - | 9354 | 2008 | - | - | 0 | - |
| - | - | 1.98E+04 | 2009 | - | - | 0 | - |
| - | - | 1.682E+04 | 2010 | - | - | 0 | - |
| - | - | 1.096E+04 | 2011 | - | - | 0 | - |
| - | - | 5460 | 2012 | - | - | 0 | - |
| - | - | 2281 | 2013 | - | - | 0 | - |

m/z Charge Intensity FragmentType MassShift Position
129.10250854492188 0 2115.7969
130.5105438232422 0 448.67108
134.02732849121094 0 882.4501
136.07606506347656 0 2600.8425
148.90086364746094 0 576.58417
148.90768432617188 0 808.941
148.91506958007812 0 642.3968
148.92245483398438 0 1108.9796
148.9295654296875 0 1302.4044
148.93698120117188 0 2659.177
148.94468688964844 0 4380.0693
148.9613800048828 0 3405.505
148.9691162109375 0 1812.3502
148.97625732421875 0 985.5999
148.98365783691406 0 901.4162
148.99069213867188 0 960.5108
148.99794006347656 0 606.07007
149.00579833984375 0 534.1856
149.0125732421875 0 459.7365
149.0352325439453 0 433.0143
166.06137084960938 0 696.9513
173.45018005371094 0 857.42584
183.85836791992188 0 453.80072
194.9304656982422 0 523.30914
212.61233520507812 0 533.2689
223.15541076660156 0 704.9176
225.0435028076172 0 516.1426
232.12945556640625 0 6686.386
239.0948944091797 0 1140.4744
242.1495819091797 0 868.46783
244.7068328857422 0 551.2772
246.10928344726562 0 643.9793
248.0701446533203 0 697.36835
249.16000366210938 0 7161.7563
250.16424560546875 0 924.345
251.150390625 0 3221.871
258.0549621582031 0 965.9577
263.13818359375 0 600.61084
268.1773376464844 0 1873.0077 c 1
276.06500244140625 0 14555.231
277.0487060546875 0 1559.103
277.0666809082031 0 1341.5977
277.1547546386719 0 19029.527
278.1580505371094 0 2276.0042
299.0624084472656 0 882.3915
300.1322021484375 0 586.43195
312.1013488769531 0 1044.4155
317.1940002441406 0 2142.499 z 13
318.20233154296875 0 1776.8568
328.4953918457031 0 501.16504
330.1109619140625 0 763.99475
331.201904296875 0 1284.17
338.1344299316406 0 1040.1041
341.0186767578125 0 882.21747
342.01947021484375 0 790.26587
347.1372375488281 0 659.78827
348.12176513671875 0 1694.2911
355.0700378417969 0 3769.682
356.0706481933594 0 2941.9324
357.0671691894531 0 1419.605
357.1227111816406 0 754.95166
358.1069641113281 0 2607.1816
359.0282287597656 0 3519.5176
359.1102600097656 0 895.05444
360.0292053222656 0 2986.1438
361.02593994140625 0 2665.2705
362.0247497558594 0 689.65344
375.1331481933594 0 9573.055
376.1169738769531 0 13417.25
376.13629150390625 0 1258.3972
376.22418212890625 0 603.5093
377.1196594238281 0 2012.199
378.1653747558594 0 1336.0922
387.223388671875 0 3580.7654
388.22576904296875 0 764.7135
389.1492004394531 0 699.30524
390.1332092285156 0 1248.4688
393.14385986328125 0 94117.31
394.146728515625 0 14894.61
396.2350158691406 0 3581.1125 c 2
397.2379455566406 0 926.95966
410.1737976074219 0 1282.8921
417.14495849609375 0 557.69574
423.22265625 0 666.9162
429.0886535644531 0 5564.6333
430.0893859863281 0 6911.3154
431.0872497558594 0 5905.9863
432.0866394042969 0 2352.1682
438.1694030761719 0 36255.547
439.1726379394531 0 7489.0396
451.2186584472656 0 789.31744
460.2197265625 0 4351.896
461.22088623046875 0 1450.6093
467.2450256347656 0 1133.5314
468.25360107421875 0 722.06226
477.71368408203125 0 545.9359
482.6661071777344 0 569.97284
488.21856689453125 0 932.7205
489.2011413574219 0 2748.177
490.20361328125 0 975.5703
491.24859619140625 0 1851.9718
492.2513732910156 0 592.4252
494.23187255859375 0 1754.2339 c Ammonia loss 3
496.7092590332031 0 614.8148
506.22747802734375 0 4226.6426
507.1903991699219 0 1902.0532
508.1965637207031 0 781.63904
510.78106689453125 0 858.48267
511.18939208984375 0 775.49023
511.26165771484375 0 6679.735 c 3
512.2637939453125 0 1457.2772
516.1656494140625 0 2373.783
516.2072143554688 0 790.50415
516.3247680664062 0 701.8672
517.1677856445312 0 1417.2662
524.2166137695312 0 2957.6404
525.2183837890625 0 910.2634
535.1851196289062 0 2400.4026
536.1856079101562 0 792.72736
538.1964721679688 0 2247.348
539.180419921875 0 13567.765
540.1822509765625 0 3789.8586
552.2113647460938 0 17557.377
553.19775390625 0 15134.793
554.199951171875 0 4037.4387
566.2833251953125 0 926.04645
570.2205200195312 0 2510.5627
571.2197265625 0 1012.2763
584.3614501953125 0 3820.6863
585.366455078125 0 700.19653
586.2040405273438 0 668.63196
586.30810546875 0 3053.4475
592.255615234375 0 1095.4205
596.9476928710938 0 573.09784
606.2577514648438 0 2382.976
606.3053588867188 0 1930.2488 z 11
607.2473754882812 0 1110.2944
607.3084106445312 0 1212.0886 c 13
609.2784423828125 0 943.4429
610.2645874023438 0 2781.3423
614.199951171875 0 1494.5642
615.2034301757812 0 1423.7615
622.321044921875 0 1034.912 y 11
623.28515625 0 3048.9495
624.269775390625 0 15088.8955
625.2733154296875 0 6515.8496
632.2970581054688 0 707.3621
632.63720703125 0 791.0294
634.2529296875 0 5104.81
634.64208984375 0 1109.5828
635.2548828125 0 1994.6782
635.3132934570312 0 1190.0334
636.333251953125 0 615.36584
648.3045654296875 0 654.4218
651.2799682617188 0 10674.941
651.3460693359375 0 1156.1382
652.2654418945312 0 25974.7
653.2674560546875 0 10636.1455
657.9800415039062 0 970.5494
663.650390625 0 763.37646
663.9843139648438 0 689.35016
664.3211059570312 0 1567.626
664.6508178710938 0 763.59534
669.2900390625 0 103336.164
669.6593017578125 0 16937.305
669.9935913085938 0 14186.082
670.2916870117188 0 36628.297
670.6604614257812 0 5216.1353
670.9912719726562 0 2024.4777
671.2872314453125 0 1530.1213
673.8575439453125 0 732.9995
697.339599609375 0 2862.3608 c 4
701.5900268554688 0 608.13666
743.4866943359375 0 1217.2072
769.3643798828125 0 3062.8276 z 10
770.3735961914062 0 1271.0104
773.4281005859375 0 1050.2661
783.651123046875 0 1184.3907
785.380615234375 0 888.3301 y 10
787.9021606445312 0 653.9393
789.4441528320312 0 595.191
810.4238891601562 0 2568.275 c 5
811.4299926757812 0 1168.0137
816.6325073242188 0 677.20856
850.4146728515625 0 897.95435
870.90625 0 911.5647 z 2
875.4743041992188 0 787.5942
878.9191284179688 0 1282.1946 y 2
879.4194946289062 0 824.8701
879.9168090820312 0 809.2834
883.8929443359375 0 658.9884
887.9215698242188 0 865.30005
888.4402465820312 0 705.6491
893.9242553710938 0 727.8869
894.9358520507812 0 969.2274
898.4081420898438 0 2416.7896 z 9
899.4127197265625 0 991.48517
908.4300537109375 0 867.97046 c Ammonia loss 6
910.4424438476562 0 2294.9414 c 13
910.9411010742188 0 2155.811
911.4480590820312 0 1903.8297
915.4559326171875 0 797.03564
921.9378051757812 0 729.4064
925.4548950195312 0 1408.5956 c 6
926.4520263671875 0 581.88226
931.4707641601562 0 1145.3188
938.962890625 0 866.9317
939.4449462890625 0 2492.0747 z 1
939.9262084960938 0 917.27625
940.4241333007812 0 649.1315
952.4843139648438 0 880.4443
952.946533203125 0 651.867
953.4612426757812 0 888.478
955.4667358398438 0 689.9433
955.95556640625 0 744.1026
958.9636840820312 0 913.316
959.48046875 0 964.5917
959.974365234375 0 4905.558
960.4755249023438 0 5588.567
960.9768676757812 0 2939.89
961.4757080078125 0 1143.6376
962.9446411132812 0 1062.62
963.4607543945312 0 944.9355
964.9595336914062 0 635.786
966.4541625976562 0 860.74585
966.966064453125 0 805.009
967.4613647460938 0 855.22577
967.9547119140625 0 2602.7798
968.4612426757812 0 2102.096 w 8
968.9652099609375 0 2542.8293
969.4485473632812 0 1233.4902
972.4564208984375 0 751.09985
973.4616088867188 0 734.5754
973.97412109375 0 1436.699
974.4728393554688 0 4648.9463
974.9766845703125 0 7922.257
975.478271484375 0 6716.9766
975.9696655273438 0 4867.7217
976.4672241210938 0 3290.6516
976.9685668945312 0 2122.7312
977.4614868164062 0 1229.2834
978.9598388671875 0 672.4258
979.4487915039062 0 987.96954
980.4640502929688 0 756.3294
981.4812622070312 0 3914.2021
981.982177734375 0 1900.2589
982.4763793945312 0 7282.399 c 7
982.9732666015625 0 1353.8196
983.4803466796875 0 2313.409
983.8348999023438 0 807.22253
984.467529296875 0 1588.2716
984.9544067382812 0 1850.6196
985.4547729492188 0 1573.2517
985.94677734375 0 1574.2281
986.464599609375 0 1164.0568
986.9634399414062 0 1239.3796
987.4610595703125 0 1915.0475
988.447509765625 0 1473.0613
988.953369140625 0 789.06775
990.4778442382812 0 838.8819
991.0303344726562 0 856.3931
991.4290771484375 0 806.50616
993.4533081054688 0 1169.3564
993.957763671875 0 1862.0853
994.4725952148438 0 1130.3354
995.4758911132812 0 1522.0706
995.9689331054688 0 3509.7886
996.4708862304688 0 3721.4922
996.9639892578125 0 1653.2526
997.4601440429688 0 910.7275
997.9765014648438 0 967.0608
1002.4822387695312 0 1227.4695
1002.9508666992188 0 1022.1983
1003.4856567382812 0 3626.8848
1003.9838256835938 0 10807.415
1004.48486328125 0 13847.507
1004.986083984375 0 9371.202
1005.4866333007812 0 6085.443
1005.9782104492188 0 2833.0518
1006.494140625 0 1084.0475
1021.5146484375 0 894.04266
1026.5003662109375 0 4568.705 z 8
1027.5067138671875 0 2867.5405
1028.50390625 0 1864.5627
1029.86474609375 0 803.7547
1038.193603515625 0 820.27783
1039.1966552734375 0 1659.3405
1039.5367431640625 0 1071.7084
1040.53857421875 0 939.843
1043.8651123046875 0 1089.298
1044.1917724609375 0 1444.528
1044.532958984375 0 2615.691
1044.869873046875 0 3493.8613
1045.204345703125 0 2675.442
1045.5467529296875 0 959.08954
1049.8680419921875 0 880.34656
1054.5244140625 0 879.8979
1060.5128173828125 0 1197.1201
1083.521240234375 0 1098.9457 z 7
1084.53173828125 0 1339.7533
1085.518310546875 0 962.7757
1086.5072021484375 0 760.0704
1091.5445556640625 0 825.34204
1094.180419921875 0 729.4777
1095.189453125 0 1041.79
1097.1890869140625 0 1140.8901
1098.5587158203125 0 709.4022
1099.541015625 0 1979.8983 y 7
1100.2059326171875 0 919.8302
1101.22900390625 0 788.4189
1101.5467529296875 0 923.6162
1102.5400390625 0 1354.7423
1103.53564453125 0 1542.1904
1108.193359375 0 1409.2491
1108.530029296875 0 1577.4681
1108.8575439453125 0 946.59064
1109.188232421875 0 848.2407
1109.5618896484375 0 1756.0907
1110.56640625 0 4140.756 c 8
1111.239013671875 0 891.97565
1111.567138671875 0 2918.8237
1111.888916015625 0 1153.4509
1112.54150390625 0 923.0207
1114.52734375 0 700.5395
1114.86181640625 0 986.1061
1116.5546875 0 1613.4803
1116.899658203125 0 1076.6909
1117.23388671875 0 1576.8093
1117.5653076171875 0 804.6955
1117.893310546875 0 747.4171
1171.5994873046875 0 2255.0193
1172.6048583984375 0 939.27045
1173.60546875 0 928.58417
1195.5777587890625 0 1005.8859
1197.556884765625 0 828.57416 y Ammonia loss 6
1198.5482177734375 0 2493.371 z 6
1220.6187744140625 0 1003.0871
1239.604736328125 0 3404.2708 c 9
1240.6046142578125 0 2515.0693
1274.11328125 0 707.1577
1274.6334228515625 0 877.3041
1279.5911865234375 0 1913.7509
1280.5955810546875 0 865.98645
1301.68359375 0 961.04034
1302.6734619140625 0 1074.3771
1310.6356201171875 0 765.34644 y Ammonia loss 5
1311.632080078125 0 1741.1713 z 5
1312.6357421875 0 1044.3685
1317.6456298828125 0 900.026
1336.6683349609375 0 1146.491
1339.626220703125 0 917.8587
1340.6038818359375 0 1549.5736
1358.6724853515625 0 872.50995
1401.68798828125 0 1011.21747
1402.6708984375 0 3136.1064 c 10
1403.6719970703125 0 2039.2921
1406.7064208984375 0 926.3189
1436.6912841796875 0 1349.8138
1437.7042236328125 0 1103.1399
1460.2159423828125 0 697.83405
1465.230712890625 0 1226.0773
1465.7244873046875 0 1813.3479
1466.2291259765625 0 1580.6355
1473.702392578125 0 1119.3624
1493.731689453125 0 857.1157
1494.7225341796875 0 1210.7723
1496.73046875 0 1321.5947 y Ammonia loss 4
1497.7109375 0 3515.4187 z 4
1498.716796875 0 3116.5078
1499.714111328125 0 1547.8568
1513.7275390625 0 1858.2137 c Ammonia loss 11
1514.7529296875 0 1196.9341
1515.7239990234375 0 1060.3966
1529.7666015625 0 1399.8009
1530.7542724609375 0 3950.1926 c 11
1531.7633056640625 0 3063.059
1532.7552490234375 0 1484.6167
1535.7955322265625 0 973.4555
1537.2537841796875 0 942.9121
1543.7625732421875 0 1302.8503
1544.2625732421875 0 968.44714
1544.7889404296875 0 963.53064
1545.3021240234375 0 1123.2057
1549.768798828125 0 800.72534
1550.2528076171875 0 810.43286
1550.76123046875 0 812.2445
1551.287841796875 0 920.2206
1552.2606201171875 0 1012.55304
1556.761962890625 0 896.83527
1557.781982421875 0 2589.451
1558.2779541015625 0 2968.0935
1558.7869873046875 0 3353.4343
1559.2864990234375 0 1862.9
1560.280029296875 0 1123.9845
1565.27978515625 0 1426.5525
1565.78857421875 0 1808.2485
1566.28369140625 0 2444.8757
1566.80029296875 0 4386.179
1567.298828125 0 2920.9956
1567.801025390625 0 3190.5837
1568.2900390625 0 2422.5269
1568.7608642578125 0 654.91296
1569.7547607421875 0 987.38885
1570.7708740234375 0 782.33594
1574.8125 0 1197.4379
1575.2908935546875 0 1229.6417
1579.2645263671875 0 932.2593
1583.7701416015625 0 742.3635
1584.2752685546875 0 796.25134
1587.288818359375 0 1117.9331
1598.7486572265625 0 931.8623
1599.26318359375 0 903.18414
1603.262451171875 0 1041.556
1603.803955078125 0 920.5533
1607.8028564453125 0 658.4455
1608.78125 0 674.0577
1609.780517578125 0 940.60583
1611.3040771484375 0 764.55316
1611.7568359375 0 1077.5656 y Ammonia loss 3
1612.7442626953125 0 3172.9626 z 3
1613.7349853515625 0 3412.3066
1614.7427978515625 0 2599.8865
1615.7491455078125 0 810.7911
1617.28662109375 0 790.3871
1619.27880859375 0 1086.9902
1620.25341796875 0 1113.549
1624.2542724609375 0 932.97174
1625.2828369140625 0 932.1402
1627.76904296875 0 1263.8119
1628.765380859375 0 1158.5392 y 3
1629.7762451171875 0 1336.1677
1631.7880859375 0 1010.7458
1632.27001953125 0 792.3643
1632.77001953125 0 800.8919
1633.29541015625 0 984.9233
1633.77978515625 0 1378.3108
1634.7535400390625 0 836.4522
1637.2547607421875 0 1002.2166
1638.7911376953125 0 897.3726
1639.7774658203125 0 1209.0632
1640.7779541015625 0 1126.0526
1641.28271484375 0 1511.0878
1642.2852783203125 0 1073.0862
1642.80810546875 0 1046.9059
1643.3109130859375 0 1130.3876
1643.8062744140625 0 930.2493
1644.3297119140625 0 792.93335
1646.302490234375 0 1216.5808
1646.7911376953125 0 745.0034
1647.2791748046875 0 1044.2019
1648.300048828125 0 1143.8121
1649.7894287109375 0 995.95435
1651.30224609375 0 914.0755
1653.3067626953125 0 1641.1797
1653.781005859375 0 1047.429
1654.797607421875 0 1094.0104
1655.27392578125 0 892.2861
1658.314208984375 0 1205.7015
1662.275146484375 0 908.31226
1662.7933349609375 0 1717.5541
1663.2801513671875 0 1643.2876
1663.792724609375 0 1613.9518
1666.3214111328125 0 1506.9131
1666.837646484375 0 1131.6023
1670.3031005859375 0 875.0401
1671.7813720703125 0 1130.2407
1674.3192138671875 0 2009.6798
1674.83056640625 0 2682.9666
1675.3385009765625 0 1824.2601
1675.8341064453125 0 1506.7788
1690.7760009765625 0 1560.9298
1691.780029296875 0 2814.1543 c 12
1692.774658203125 0 3612.9458
1693.780517578125 0 1448.095
1740.788818359375 0 1908.6321 z 2
1741.8099365234375 0 1426.5938
1742.8082275390625 0 1037.254
1820.8812255859375 0 1107.2665
1830.91552734375 0 1037.3376
1878.8662109375 0 935.84296
1917.9541015625 0 1542.2855
1918.957275390625 0 1557.7106
1919.9464111328125 0 1906.8866
1920.9263916015625 0 1493.5261
1921.95166015625 0 1548.729
1922.9432373046875 0 1040.4403
1932.925048828125 0 1154.2112
1937.9281005859375 0 924.9149
1947.9320068359375 0 1071.6046
1948.9356689453125 0 3069.744
1949.9468994140625 0 3409.432
1950.9439697265625 0 2277.1692
1951.9283447265625 0 1096.0714
1952.918212890625 0 1690.6816
1953.935302734375 0 889.573
1962.9605712890625 0 1199.5848
1963.9462890625 0 2394.8193
1964.9515380859375 0 1728.1508
1965.9539794921875 0 1359.2
1969.873779296875 0 1240.648
1970.8836669921875 0 986.09705
1979.9718017578125 0 1204.2223
1980.968994140625 0 2167.17
1981.9736328125 0 1904.8773
1982.966552734375 0 1025.4988
1986.9002685546875 0 1030.3185
1987.9039306640625 0 1672.4594
1989.9193115234375 0 1176.6671
1990.9417724609375 0 2821.1274
1991.93359375 0 4896.647
1992.93017578125 0 4162.002
1993.9364013671875 0 3637.172
1994.9237060546875 0 1710.8544
1995.9327392578125 0 1100.6835
2005.9468994140625 0 1830.811
2006.9354248046875 0 2098.075
2007.9622802734375 0 9354.299
2008.9605712890625 0 19802.256
2009.962646484375 0 16824.111
2010.9630126953125 0 10964.97
2011.959716796875 0 5459.9717
2012.95263671875 0 2281.22

Spectrum Details

|  |  |
| --- | --- |
| Matched peaks? Matched peaks The total absolute number of peaks matched. Additionally in brackets the total fraction of peaks matched and the total number of peaks is shown. | 40 (7.66% of 522) |
| FDR? FDR The false discovery rate estimated for this peptide. It is calculated by matching all theoretical fragments with a non-integer shift with the raw peaks for this spectrum. This is done with 40 different shifts. The resulting percentage is the average number of annotated peaks over the number of annotated peaks with the correct spectrum. | 2.56% |
| Satellite FDR? Satellite FDR See the FDR for details on its calculation. This satellite ion specific FDR only contains the satellite ions (d/w) for I/L/J positions. | - |
| PSM Score? PSM Score The PSM Score as given by Hecklib to this annotated spectrum. It is shown with three significant figures. | 402 |

#### Spectrum 4164? Spectrum 4164 The raw spectrum of this peptide as annotated by Hecklib. The fragments are coloured according to ion type (see legend). Any peaks with a star '\*' as text can be hovered over to see the full details, first the ion type second the mass shift type. By hovering over the amino acids in the peptide or ions in the legend the corresponding peaks are highlighted. By toggling the 'Unassigned' label you can turn the background (unassigned) peaks on or off in the plot. By updating the slider in the Ion legend you can update the spectrum to only show the top X% of the peaks with labels. The top X% means any peak that is within X% of the highest intensity. By dragging in the spectrum you can zoom in to a specific part of the spectrum and use 'Zoom Out' to get back to the original zoom level. The annotation of the spectrum is based on the given sequence in the peptides file and is done with different software so inconsistencies are likely. The peaks are annotated based on the given sequence, with 20 ppm tolerance.

Copy Data

##### Spectrum 4164 (TSV)

###### Preview

```
Loading example...
```

*Click on the button to copy the data to your clipboard.*

Mz MinMz MaxIntensity Max

WidthHeightPeptide font sizePeptide stroke widthSpectrum font sizeSpectrum stroke widthCompact peptide

Ion legend

wxyz

abcd

OtherUnassignedIon^Charge^_Position_Show for top:%

JHQDWLDGKEYKCKK

03.17e+36.34e+39.52e+31.27e+4

Zoom Out

c^+4^_6_z^+2^_3_c^+2^_4_c^+1^_2_z^+2^_4_y^+3^_7_c^+2^_5_y^+3^_8_c^+1^_3_z^+2^_5_y^+2^_5_c^+1^_3_c^+2^_6_c^+2^_6_y^+3^_9_w^+2^_6_w^+3^_10_y^+3^_10_y^+2^_6_c^+4^_14_y^+2^_6_c^+2^_7_y^+4^_14_c^+1^_4_c^+2^_8_y^+3^_11_y^+3^_11_z^+2^_7_c^+1^_4_y^+2^_7_y^+2^_7_z^+2^_7_c^+3^_12_y^+2^_7_z^+2^_8_y^+2^_8_c^+2^_9_w^+3^_13_c^+3^_13_y^+3^_13_z^+2^_9_z^+1^_4_y^+2^_9_c^+3^_14_c^+2^_10_c^+2^_10_z^+3^_14_w^+2^_10_y^+2^_10_z^+2^_10_y^+2^_10_c^+1^_5_c^+2^_11_c^+1^_5_y^+2^_11_y^+2^_11_z^+1^_5_c^+2^_12_y^+1^_5_c^+1^_6_z^+2^_12_c^+1^_6_c^+2^_13_z^+2^_13_z^+1^_6_z^+1^_6_c^+1^_7_c^+1^_7_z^+2^_14_c^+1^_8_z^+1^_7_c^+1^_9_c^+1^_10_

0827165524823310

Fragment Matches Table

Show background peaks

| Position | Ion type | Intensity | mz Theoretical | mz Error (Th) | mz Error (ppm) | Charge | Series Number |
| --- | --- | --- | --- | --- | --- | --- | --- |
| - | - | 393.2 | 127.1 | - | - | 0 | - |
| - | - | 343.5 | 128.2 | - | - | 0 | - |
| - | - | 397.9 | 128.9 | - | - | 0 | - |
| - | - | 2277 | 129 | - | - | 0 | - |
| - | - | 1896 | 129.1 | - | - | 0 | - |
| - | - | 774.8 | 130 | - | - | 0 | - |
| - | - | 785.5 | 130.1 | - | - | 0 | - |
| - | - | 675.5 | 133.1 | - | - | 0 | - |
| - | - | 436.9 | 133.1 | - | - | 0 | - |
| - | - | 1216 | 136.1 | - | - | 0 | - |
| - | - | 1761 | 139 | - | - | 0 | - |
| - | - | 797 | 140 | - | - | 0 | - |
| - | - | 366.8 | 140.6 | - | - | 0 | - |
| - | - | 415.6 | 141.1 | - | - | 0 | - |
| - | - | 574.7 | 146.1 | - | - | 0 | - |
| - | - | 3440 | 149 | - | - | 0 | - |
| - | - | 991.9 | 151.1 | - | - | 0 | - |
| - | - | 1481 | 155.1 | - | - | 0 | - |
| - | - | 4516 | 157 | - | - | 0 | - |
| - | - | 2315 | 158 | - | - | 0 | - |
| - | - | 594.8 | 158.1 | - | - | 0 | - |
| - | - | 739.2 | 159 | - | - | 0 | - |
| - | - | 621.5 | 159.1 | - | - | 0 | - |
| - | - | 1100 | 166.1 | - | - | 0 | - |
| - | - | 496.5 | 167.1 | - | - | 0 | - |
| - | - | 1350 | 173.4 | - | - | 0 | - |
| - | - | 2495 | 175.1 | - | - | 0 | - |
| - | - | 1009 | 175.1 | - | - | 0 | - |
| - | - | 659.6 | 176.1 | - | - | 0 | - |
| - | - | 679.8 | 183.1 | - | - | 0 | - |
| - | - | 779.6 | 185.1 | - | - | 0 | - |
| - | - | 6493 | 185.1 | - | - | 0 | - |
| - | - | 3689 | 185.2 | - | - | 0 | - |
| - | - | 4217 | 186.1 | - | - | 0 | - |
| - | - | 558.9 | 187.1 | - | - | 0 | - |
| - | - | 1738 | 187.1 | - | - | 0 | - |
| - | - | 1046 | 190.1 | - | - | 0 | - |
| - | - | 595.5 | 199.2 | - | - | 0 | - |
| - | - | 489.8 | 199.6 | - | - | 0 | - |
| - | - | 618.3 | 200.2 | - | - | 0 | - |
| 6 | c | 607.6 | 203.1 | 0.0007703 | 3.793 | +4 | 6 |
| - | - | 808.3 | 205.1 | - | - | 0 | - |
| - | - | 2221 | 213.1 | - | - | 0 | - |
| - | - | 600.3 | 213.2 | - | - | 0 | - |
| - | - | 1631 | 214.1 | - | - | 0 | - |
| - | - | 3930 | 217 | - | - | 0 | - |
| - | - | 2922 | 218 | - | - | 0 | - |
| - | - | 653.2 | 218.1 | - | - | 0 | - |
| - | - | 696 | 219.1 | - | - | 0 | - |
| - | - | 2796 | 223.1 | - | - | 0 | - |
| - | - | 938.8 | 223.2 | - | - | 0 | - |
| - | - | 833.8 | 227 | - | - | 0 | - |
| - | - | 648.2 | 230 | - | - | 0 | - |
| 13 | z | 687.6 | 231.1 | 0.002004 | 8.673 | +2 | 3 |
| - | - | 1217 | 234.1 | - | - | 0 | - |
| - | - | 621.7 | 246.4 | - | - | 0 | - |
| 4 | c | 858.8 | 247.6 | 0.0005835 | 2.356 | +2 | 4 |
| - | - | 9572 | 248 | - | - | 0 | - |
| - | - | 728.9 | 248.1 | - | - | 0 | - |
| - | - | 1130 | 249 | - | - | 0 | - |
| - | - | 5486 | 251.2 | - | - | 0 | - |
| - | - | 748.5 | 252.2 | - | - | 0 | - |
| - | - | 4204 | 259.2 | - | - | 0 | - |
| - | - | 7414 | 260.2 | - | - | 0 | - |
| - | - | 1203 | 261.2 | - | - | 0 | - |
| - | - | 797.1 | 263.1 | - | - | 0 | - |
| - | - | 944.5 | 266 | - | - | 0 | - |
| 2 | c | 624.1 | 268.2 | 0.0005973 | 2.227 | +1 | 2 |
| - | - | 626.1 | 269.1 | - | - | 0 | - |
| - | - | 2069 | 270.1 | - | - | 0 | - |
| - | - | 664.1 | 270.2 | - | - | 0 | - |
| - | - | 1623 | 273.1 | - | - | 0 | - |
| - | - | 2315 | 274.1 | - | - | 0 | - |
| - | - | 501.2 | 274.1 | - | - | 0 | - |
| - | - | 2176 | 276 | - | - | 0 | - |
| - | - | 1009 | 281.1 | - | - | 0 | - |
| - | - | 3278 | 297.1 | - | - | 0 | - |
| - | - | 1037 | 297.1 | - | - | 0 | - |
| - | - | 897.6 | 298.1 | - | - | 0 | - |
| - | - | 1478 | 298.2 | - | - | 0 | - |
| - | - | 4063 | 299.1 | - | - | 0 | - |
| - | - | 1297 | 300.1 | - | - | 0 | - |
| - | - | 747.5 | 301.1 | - | - | 0 | - |
| 12 | z | 693.6 | 303.7 | 0.004106 | 13.52 | +2 | 4 |
| - | - | 1476 | 315.1 | - | - | 0 | - |
| - | - | 891.2 | 315.2 | - | - | 0 | - |
| - | - | 647 | 316.1 | - | - | 0 | - |
| - | - | 745 | 317.1 | - | - | 0 | - |
| - | - | 1244 | 330.2 | - | - | 0 | - |
| - | - | 867.6 | 330.2 | - | - | 0 | - |
| - | - | 641.3 | 334.1 | - | - | 0 | - |
| - | - | 2639 | 334.7 | - | - | 0 | - |
| - | - | 626.7 | 335.1 | - | - | 0 | - |
| 9 | y | 713.8 | 342.2 | 0.004338 | 12.68 | +3 | 7 |
| - | - | 592.9 | 343.1 | - | - | 0 | - |
| 5 | c | 2385 | 348.7 | 0.002693 | 7.725 | +2 | 5 |
| - | - | 892.2 | 351.1 | - | - | 0 | - |
| - | - | 3852 | 351.2 | - | - | 0 | - |
| - | - | 673 | 351.9 | - | - | 0 | - |
| - | - | 1186 | 352.2 | - | - | 0 | - |
| - | - | 1155 | 353.2 | - | - | 0 | - |
| - | - | 4153 | 358.1 | - | - | 0 | - |
| - | - | 942.5 | 359.1 | - | - | 0 | - |
| - | - | 2031 | 362.2 | - | - | 0 | - |
| 8 | y | 1959 | 367.2 | 0.003296 | 8.975 | +3 | 8 |
| - | - | 653.5 | 367.5 | - | - | 0 | - |
| 3 | c | 1524 | 379.2 | 0.0002116 | 0.5581 | +1 | 3 |
| - | - | 683.4 | 381.1 | - | - | 0 | - |
| - | - | 819.2 | 381.2 | - | - | 0 | - |
| - | - | 799.3 | 383.2 | - | - | 0 | - |
| - | - | 707.1 | 384.2 | - | - | 0 | - |
| 11 | z | 758.2 | 385.2 | 0.003665 | 9.516 | +2 | 5 |
| - | - | 786.6 | 386.1 | - | - | 0 | - |
| - | - | 719.3 | 387.2 | - | - | 0 | - |
| - | - | 3344 | 387.2 | - | - | 0 | - |
| - | - | 595.8 | 388.2 | - | - | 0 | - |
| - | - | 856.5 | 391.2 | - | - | 0 | - |
| 11 | y | 2434 | 393.2 | 0.00491 | 12.49 | +2 | 5 |
| - | - | 1092 | 393.7 | - | - | 0 | - |
| 3 | c | 3127 | 396.2 | 9.472E-05 | 0.2391 | +1 | 3 |
| - | - | 653.9 | 397.9 | - | - | 0 | - |
| - | - | 634.8 | 398.2 | - | - | 0 | - |
| - | - | 695.9 | 398.4 | - | - | 0 | - |
| - | - | 2943 | 404.1 | - | - | 0 | - |
| - | - | 2097 | 404.2 | - | - | 0 | - |
| - | - | 6619 | 404.2 | - | - | 0 | - |
| 6 | c | 693.9 | 404.7 | 0.003145 | 7.772 | +2 | 6 |
| - | - | 725.2 | 404.9 | - | - | 0 | - |
| - | - | 966.4 | 405.2 | - | - | 0 | - |
| 6 | c | 2181 | 405.2 | 0.00424 | 10.47 | +2 | 6 |
| - | - | 1674 | 405.2 | - | - | 0 | - |
| - | - | 589.7 | 405.4 | - | - | 0 | - |
| 7 | y | 1276 | 405.5 | 0.002806 | 6.919 | +3 | 9 |
| - | - | 810.8 | 405.7 | - | - | 0 | - |
| - | - | 941.5 | 405.9 | - | - | 0 | - |
| - | - | 1426 | 406.2 | - | - | 0 | - |
| - | - | 1075 | 412.2 | - | - | 0 | - |
| 10 | w | 1.256E+04 | 420.2 | 0.004058 | 9.657 | +2 | 6 |
| - | - | 6969 | 420.7 | - | - | 0 | - |
| - | - | 2664 | 421.2 | - | - | 0 | - |
| 6 | w | 714.7 | 423.5 | 7.135E-05 | 0.1685 | +3 | 10 |
| 6 | y | 1074 | 443.2 | 0.002456 | 5.541 | +3 | 10 |
| 10 | y | 942.2 | 448.7 | 0.002522 | 5.62 | +2 | 6 |
| 14 | c | 1220 | 455.2 | 0.005343 | 11.74 | +4 | 14 |
| 10 | y | 616.4 | 457.7 | 0.003959 | 8.649 | +2 | 6 |
| - | - | 2568 | 460.7 | - | - | 0 | - |
| - | - | 2692 | 461.2 | - | - | 0 | - |
| - | - | 1815 | 461.6 | - | - | 0 | - |
| 7 | c | 868 | 462.7 | 0.002579 | 5.574 | +2 | 7 |
| - | - | 799.7 | 466.5 | - | - | 0 | - |
| - | - | 796.4 | 468.2 | - | - | 0 | - |
| 2 | y | 870.8 | 478.2 | 0.00405 | 8.469 | +4 | 14 |
| - | - | 832.4 | 478.5 | - | - | 0 | - |
| - | - | 651.7 | 484.3 | - | - | 0 | - |
| - | - | 2249 | 484.5 | - | - | 0 | - |
| - | - | 1163 | 484.7 | - | - | 0 | - |
| - | - | 1512 | 485 | - | - | 0 | - |
| - | - | 960.8 | 490.2 | - | - | 0 | - |
| - | - | 1140 | 490.7 | - | - | 0 | - |
| - | - | 2962 | 491.7 | - | - | 0 | - |
| - | - | 3590 | 492 | - | - | 0 | - |
| - | - | 835 | 492.2 | - | - | 0 | - |
| - | - | 2244 | 492.5 | - | - | 0 | - |
| - | - | 1131 | 492.7 | - | - | 0 | - |
| 4 | c | 9380 | 494.2 | 0.001001 | 2.026 | +1 | 4 |
| - | - | 2499 | 495.2 | - | - | 0 | - |
| - | - | 875 | 496.2 | - | - | 0 | - |
| - | - | 994.7 | 496.8 | - | - | 0 | - |
| 8 | c | 1504 | 499.7 | 0.003567 | 7.137 | +2 | 8 |
| - | - | 1044 | 500.3 | - | - | 0 | - |
| - | - | 716.3 | 500.7 | - | - | 0 | - |
| - | - | 1044 | 502 | - | - | 0 | - |
| - | - | 1173 | 502.2 | - | - | 0 | - |
| - | - | 868.9 | 502.5 | - | - | 0 | - |
| - | - | 966.3 | 503 | - | - | 0 | - |
| - | - | 1151 | 503.9 | - | - | 0 | - |
| - | - | 4390 | 504.2 | - | - | 0 | - |
| 5 | y | 4025 | 504.6 | 0.000178 | 0.3529 | +3 | 11 |
| 5 | y | 2549 | 504.9 | 0.006228 | 12.34 | +3 | 11 |
| 9 | z | 729.9 | 505.2 | 0.002742 | 5.427 | +2 | 7 |
| - | - | 1901 | 506.2 | - | - | 0 | - |
| - | - | 1.059E+04 | 506.5 | - | - | 0 | - |
| - | - | 1.162E+04 | 506.7 | - | - | 0 | - |
| - | - | 9088 | 507 | - | - | 0 | - |
| - | - | 4787 | 507.2 | - | - | 0 | - |
| - | - | 1444 | 507.5 | - | - | 0 | - |
| 4 | c | 5960 | 511.3 | 0.0001149 | 0.2248 | +1 | 4 |
| - | - | 1135 | 512.3 | - | - | 0 | - |
| 9 | y | 1788 | 512.8 | 0.009201 | 17.94 | +2 | 7 |
| 9 | y | 1493 | 513.3 | 0.003002 | 5.849 | +2 | 7 |
| 9 | z | 1255 | 513.8 | 0.002192 | 4.267 | +2 | 7 |
| - | - | 954.3 | 515.9 | - | - | 0 | - |
| 12 | c | 4957 | 516.3 | 0.002512 | 4.867 | +3 | 12 |
| - | - | 4765 | 516.6 | - | - | 0 | - |
| - | - | 1179 | 516.9 | - | - | 0 | - |
| - | - | 832.3 | 519.3 | - | - | 0 | - |
| 9 | y | 2334 | 521.8 | 0.004718 | 9.042 | +2 | 7 |
| - | - | 785.6 | 522.3 | - | - | 0 | - |
| - | - | 698.8 | 524.8 | - | - | 0 | - |
| - | - | 874.7 | 525.3 | - | - | 0 | - |
| - | - | 928.2 | 535.3 | - | - | 0 | - |
| - | - | 755.2 | 540 | - | - | 0 | - |
| 8 | z | 1881 | 542.3 | 0.002914 | 5.374 | +2 | 8 |
| - | - | 1374 | 542.8 | - | - | 0 | - |
| - | - | 813.2 | 543.3 | - | - | 0 | - |
| 8 | y | 2441 | 550.3 | 0.006478 | 11.77 | +2 | 8 |
| - | - | 1726 | 550.8 | - | - | 0 | - |
| - | - | 813.6 | 553.2 | - | - | 0 | - |
| - | - | 658.9 | 553.3 | - | - | 0 | - |
| - | - | 690.3 | 554.3 | - | - | 0 | - |
| - | - | 1150 | 555.3 | - | - | 0 | - |
| - | - | 1671 | 563.3 | - | - | 0 | - |
| 9 | c | 1.167E+04 | 563.8 | 0.003326 | 5.9 | +2 | 9 |
| - | - | 7757 | 564.3 | - | - | 0 | - |
| - | - | 2734 | 564.8 | - | - | 0 | - |
| 3 | w | 4134 | 566.9 | 0.0007196 | 1.269 | +3 | 13 |
| - | - | 1546 | 567.3 | - | - | 0 | - |
| 13 | c | 3087 | 569.9 | 0.002425 | 4.254 | +3 | 13 |
| - | - | 1825 | 570.3 | - | - | 0 | - |
| - | - | 3805 | 571.3 | - | - | 0 | - |
| - | - | 2194 | 571.8 | - | - | 0 | - |
| - | - | 1398 | 572.3 | - | - | 0 | - |
| - | - | 954.5 | 579.3 | - | - | 0 | - |
| 3 | y | 1056 | 585.9 | 0.002319 | 3.957 | +3 | 13 |
| - | - | 1566 | 596.3 | - | - | 0 | - |
| - | - | 3012 | 596.8 | - | - | 0 | - |
| - | - | 2369 | 597.3 | - | - | 0 | - |
| - | - | 943.7 | 598 | - | - | 0 | - |
| - | - | 917.1 | 598.8 | - | - | 0 | - |
| 7 | z | 4690 | 599.8 | 0.002958 | 4.932 | +2 | 9 |
| - | - | 4680 | 600.3 | - | - | 0 | - |
| - | - | 3003 | 600.8 | - | - | 0 | - |
| - | - | 1323 | 601.3 | - | - | 0 | - |
| 12 | z | 3107 | 606.3 | 0.01143 | 18.86 | +1 | 4 |
| - | - | 1603 | 606.8 | - | - | 0 | - |
| - | - | 2947 | 607.3 | - | - | 0 | - |
| 7 | y | 7632 | 607.8 | 0.00347 | 5.709 | +2 | 9 |
| - | - | 4780 | 608.3 | - | - | 0 | - |
| - | - | 2174 | 608.8 | - | - | 0 | - |
| - | - | 4303 | 609.3 | - | - | 0 | - |
| - | - | 1885 | 609.8 | - | - | 0 | - |
| - | - | 815.3 | 610.3 | - | - | 0 | - |
| 14 | c | 2611 | 612.6 | 0.0009502 | 1.551 | +3 | 14 |
| - | - | 3725 | 613 | - | - | 0 | - |
| - | - | 1704 | 613.3 | - | - | 0 | - |
| - | - | 1090 | 614.3 | - | - | 0 | - |
| 10 | c | 825.2 | 619.8 | 0.004033 | 6.506 | +2 | 10 |
| - | - | 2952 | 622.3 | - | - | 0 | - |
| - | - | 2110 | 626.3 | - | - | 0 | - |
| - | - | 835.8 | 626.6 | - | - | 0 | - |
| - | - | 755.1 | 627.3 | - | - | 0 | - |
| - | - | 1331 | 627.8 | - | - | 0 | - |
| 10 | c | 5104 | 628.3 | 0.003819 | 6.079 | +2 | 10 |
| - | - | 2144 | 628.8 | - | - | 0 | - |
| - | - | 1202 | 629.3 | - | - | 0 | - |
| 2 | z | 1343 | 632 | 0.00113 | 1.788 | +3 | 14 |
| - | - | 1234 | 632.3 | - | - | 0 | - |
| - | - | 1554 | 632.6 | - | - | 0 | - |
| 6 | w | 1703 | 634.8 | 0.004602 | 7.249 | +2 | 10 |
| - | - | 926.3 | 635.3 | - | - | 0 | - |
| - | - | 1082 | 640.3 | - | - | 0 | - |
| - | - | 1719 | 640.4 | - | - | 0 | - |
| - | - | 815.3 | 641.3 | - | - | 0 | - |
| - | - | 2559 | 645.7 | - | - | 0 | - |
| - | - | 3510 | 646 | - | - | 0 | - |
| - | - | 1699 | 646.3 | - | - | 0 | - |
| - | - | 678.6 | 646.7 | - | - | 0 | - |
| - | - | 673.5 | 649.6 | - | - | 0 | - |
| - | - | 1033 | 650.3 | - | - | 0 | - |
| - | - | 1220 | 655 | - | - | 0 | - |
| 6 | y | 971.1 | 655.3 | 0.007016 | 10.71 | +2 | 10 |
| - | - | 6420 | 655.7 | - | - | 0 | - |
| - | - | 5346 | 656 | - | - | 0 | - |
| 6 | z | 5120 | 656.3 | 0.002631 | 4.009 | +2 | 10 |
| - | - | 1417 | 656.7 | - | - | 0 | - |
| - | - | 2045 | 656.8 | - | - | 0 | - |
| - | - | 1075 | 657 | - | - | 0 | - |
| - | - | 725.4 | 660.3 | - | - | 0 | - |
| - | - | 1134 | 661 | - | - | 0 | - |
| 6 | y | 3054 | 664.3 | 0.00589 | 8.866 | +2 | 10 |
| - | - | 820.5 | 669.3 | - | - | 0 | - |
| - | - | 5024 | 669.7 | - | - | 0 | - |
| - | - | 1.064E+04 | 670 | - | - | 0 | - |
| - | - | 8094 | 670.3 | - | - | 0 | - |
| - | - | 3596 | 670.7 | - | - | 0 | - |
| - | - | 1645 | 671 | - | - | 0 | - |
| - | - | 963.4 | 672.3 | - | - | 0 | - |
| - | - | 674.9 | 674.7 | - | - | 0 | - |
| - | - | 2841 | 675 | - | - | 0 | - |
| - | - | 6157 | 675.3 | - | - | 0 | - |
| - | - | 6612 | 675.7 | - | - | 0 | - |
| - | - | 4552 | 676 | - | - | 0 | - |
| - | - | 2941 | 676.3 | - | - | 0 | - |
| - | - | 1264 | 676.7 | - | - | 0 | - |
| - | - | 776.9 | 687.3 | - | - | 0 | - |
| - | - | 1947 | 687.8 | - | - | 0 | - |
| - | - | 2583 | 688.3 | - | - | 0 | - |
| - | - | 860.8 | 688.8 | - | - | 0 | - |
| - | - | 834.4 | 690.8 | - | - | 0 | - |
| - | - | 1141 | 691.3 | - | - | 0 | - |
| - | - | 919.8 | 691.8 | - | - | 0 | - |
| - | - | 1267 | 692.3 | - | - | 0 | - |
| 5 | c | 5036 | 696.3 | 0.005888 | 8.457 | +1 | 5 |
| - | - | 1844 | 697.3 | - | - | 0 | - |
| 11 | c | 3987 | 709.8 | 0.004138 | 5.829 | +2 | 11 |
| - | - | 3068 | 710.3 | - | - | 0 | - |
| - | - | 1922 | 710.8 | - | - | 0 | - |
| - | - | 1035 | 711.5 | - | - | 0 | - |
| 5 | c | 3202 | 713.3 | 0.003082 | 4.32 | +1 | 5 |
| - | - | 981.8 | 714.3 | - | - | 0 | - |
| - | - | 765.1 | 729.3 | - | - | 0 | - |
| - | - | 767.6 | 752.4 | - | - | 0 | - |
| - | - | 1013 | 755.9 | - | - | 0 | - |
| 5 | y | 666.9 | 756.4 | 0.00447 | 5.91 | +2 | 11 |
| 5 | y | 1523 | 756.9 | 0.01374 | 18.16 | +2 | 11 |
| 11 | z | 2495 | 769.4 | 0.008355 | 10.86 | +1 | 5 |
| - | - | 1173 | 770.4 | - | - | 0 | - |
| - | - | 1002 | 771.4 | - | - | 0 | - |
| 12 | c | 1472 | 773.9 | 0.003287 | 4.247 | +2 | 12 |
| - | - | 1604 | 774.4 | - | - | 0 | - |
| 11 | y | 752 | 785.4 | 0.01133 | 14.43 | +1 | 5 |
| - | - | 816.1 | 803.4 | - | - | 0 | - |
| 6 | c | 2818 | 809.4 | 0.01191 | 14.72 | +1 | 6 |
| - | - | 1824 | 810.4 | - | - | 0 | - |
| - | - | 699.8 | 811.4 | - | - | 0 | - |
| - | - | 1316 | 812.4 | - | - | 0 | - |
| 4 | z | 1006 | 814.9 | 0.0007542 | 0.9256 | +2 | 12 |
| - | - | 799 | 815.4 | - | - | 0 | - |
| - | - | 2417 | 815.9 | - | - | 0 | - |
| 6 | c | 8517 | 826.4 | 0.005505 | 6.661 | +1 | 6 |
| - | - | 4230 | 827.4 | - | - | 0 | - |
| - | - | 1224 | 828.4 | - | - | 0 | - |
| 13 | c | 747 | 854.4 | 0.008526 | 9.979 | +2 | 13 |
| - | - | 1735 | 854.9 | - | - | 0 | - |
| 3 | z | 1551 | 878.9 | 0.003143 | 3.576 | +2 | 13 |
| - | - | 3607 | 879.4 | - | - | 0 | - |
| - | - | 2350 | 879.9 | - | - | 0 | - |
| 10 | z | 965.5 | 880.4 | 0.003945 | 4.481 | +1 | 6 |
| - | - | 1119 | 880.9 | - | - | 0 | - |
| - | - | 2738 | 883.4 | - | - | 0 | - |
| - | - | 1648 | 884.4 | - | - | 0 | - |
| - | - | 735.8 | 885.4 | - | - | 0 | - |
| - | - | 1897 | 896.9 | - | - | 0 | - |
| - | - | 1551 | 897.4 | - | - | 0 | - |
| 10 | z | 2869 | 898.4 | 0.006331 | 7.047 | +1 | 6 |
| - | - | 1943 | 899.4 | - | - | 0 | - |
| - | - | 859 | 900.4 | - | - | 0 | - |
| - | - | 1239 | 918.9 | - | - | 0 | - |
| - | - | 719.6 | 919.4 | - | - | 0 | - |
| 7 | c | 953 | 924.4 | 0.004928 | 5.331 | +1 | 7 |
| 7 | c | 819.1 | 941.4 | 0.0003812 | 0.4049 | +1 | 7 |
| - | - | 768 | 946.5 | - | - | 0 | - |
| 2 | z | 833.5 | 947.4 | 0.003289 | 3.471 | +2 | 14 |
| - | - | 1927 | 947.9 | - | - | 0 | - |
| - | - | 1808 | 948.4 | - | - | 0 | - |
| - | - | 1557 | 948.9 | - | - | 0 | - |
| - | - | 787.3 | 949.4 | - | - | 0 | - |
| - | - | 629 | 968.5 | - | - | 0 | - |
| 8 | c | 975.9 | 998.5 | 0.00672 | 6.73 | +1 | 8 |
| - | - | 1434 | 999.5 | - | - | 0 | - |
| - | - | 853.1 | 1000 | - | - | 0 | - |
| - | - | 905.9 | 1004 | - | - | 0 | - |
| - | - | 1819 | 1005 | - | - | 0 | - |
| - | - | 866.6 | 1013 | - | - | 0 | - |
| 9 | z | 871.6 | 1027 | 0.01554 | 15.14 | +1 | 7 |
| - | - | 1528 | 1028 | - | - | 0 | - |
| - | - | 980.5 | 1085 | - | - | 0 | - |
| - | - | 1159 | 1111 | - | - | 0 | - |
| 9 | c | 1681 | 1127 | 0.004652 | 4.129 | +1 | 9 |
| - | - | 1726 | 1128 | - | - | 0 | - |
| 10 | c | 646.7 | 1256 | 0.008934 | 7.115 | +1 | 10 |
| - | - | 1282 | 1420 | - | - | 0 | - |
| - | - | 802.3 | 2254 | - | - | 0 | - |
| - | - | 657.6 | 2466 | - | - | 0 | - |
| - | - | 751.9 | 3277 | - | - | 0 | - |

m/z Charge Intensity FragmentType MassShift Position
127.11176300048828 0 393.2388
128.24969482421875 0 343.49353
128.90475463867188 0 397.93213
129.01846313476562 0 2276.633
129.10243225097656 0 1896.4971
130.0215301513672 0 774.8398
130.06539916992188 0 785.5081
133.0609893798828 0 675.49695
133.0861053466797 0 436.89487
136.07568359375 0 1215.9338
139.0027618408203 0 1760.5504
140.0061798095703 0 797.0342
140.64195251464844 0 366.78732
141.0916748046875 0 415.60425
146.0602569580078 0 574.6753
149.02349853515625 0 3439.5017
151.07554626464844 0 991.8799
155.09286499023438 0 1481.3105
157.01332092285156 0 4515.916
158.01670837402344 0 2314.6462
158.0601806640625 0 594.81354
159.01763916015625 0 739.20874
159.091796875 0 621.52423
166.0610809326172 0 1099.9152
167.05665588378906 0 496.5107
173.43896484375 0 1349.7347
175.0868377685547 0 2494.6992
175.0959930419922 0 1008.61414
176.0904541015625 0 659.6341
183.1134033203125 0 679.8468
185.07180786132812 0 779.588
185.0809326171875 0 6492.532
185.16490173339844 0 3689.223
186.08441162109375 0 4217.255
187.08653259277344 0 558.9043
187.10797119140625 0 1737.7118
190.08238220214844 0 1045.8679
199.16957092285156 0 595.46466
199.5518341064453 0 489.7976
200.17369079589844 0 618.28534
203.1019744873047 0 607.56085 c Ammonia loss 5
205.11825561523438 0 808.345
213.07626342773438 0 2220.9602
213.15951538085938 0 600.2716
214.0794677734375 0 1631.2262
217.03439331054688 0 3930.1646
218.03768920898438 0 2921.7092
218.14918518066406 0 653.19257
219.0801239013672 0 696.036
223.06361389160156 0 2796.2278
223.15576171875 0 938.81635
227.037353515625 0 833.75134
230.0227508544922 0 648.2105
231.09747314453125 0 687.5893 z Ammonia loss 12
234.12457275390625 0 1216.9424
246.36865234375 0 621.7308
247.62094116210938 0 858.8424 c Ammonia loss 3
248.03433227539062 0 9572.209
248.0792999267578 0 728.8915
249.03733825683594 0 1130.0555
251.1505584716797 0 5486.1387
252.15374755859375 0 748.48505
259.1541442871094 0 4203.711
260.1575622558594 0 7413.6836
261.1601867675781 0 1202.8163
263.1493835449219 0 797.1359
266.04473876953125 0 944.46106
268.1773986816406 0 624.0511 c 1
269.1361999511719 0 626.06213
270.09149169921875 0 2068.6328
270.1591491699219 0 664.0857
273.09686279296875 0 1623.1405
274.1004638671875 0 2315.2434
274.14947509765625 0 501.17487
276.02899169921875 0 2175.8267
281.05078125 0 1008.7733
297.0826721191406 0 3278.1643
297.1002502441406 0 1037.0112
298.08367919921875 0 897.5704
298.1545715332031 0 1478.2869
299.0618591308594 0 4063.3687
300.0626525878906 0 1297.0173
301.0599365234375 0 747.5246
303.6561279296875 0 693.5977 z 11
315.0890808105469 0 1476.3312
315.181884765625 0 891.2228
316.0945129394531 0 646.9519
317.1400146484375 0 745.0272
330.1641540527344 0 1244.1372
330.19439697265625 0 867.58746
334.10479736328125 0 641.3292
334.661376953125 0 2638.5862
335.1479187011719 0 626.7191
342.1745910644531 0 713.81366 y Water loss 8
343.0846252441406 0 592.88385
348.65887451171875 0 2384.6763 c Ammonia loss 4
351.1451416015625 0 892.20703
351.2015075683594 0 3851.9185
351.92437744140625 0 672.99396
352.204833984375 0 1186.4955
353.1600646972656 0 1154.6191
358.1441345214844 0 4153.353
359.14599609375 0 942.5471
362.2257080078125 0 2030.5277
367.1863098144531 0 1958.8212 y 7
367.51947021484375 0 653.5033
379.2086181640625 0 1524.496 c Ammonia loss 2
381.14849853515625 0 683.43195
381.1892395019531 0 819.20624
383.20556640625 0 799.2754
384.19342041015625 0 707.072
385.188232421875 0 758.2102 z 10
386.13720703125 0 786.60675
387.1975402832031 0 719.32324
387.22406005859375 0 3344.306
388.22528076171875 0 595.8309
391.2005920410156 0 856.50775
393.19635009765625 0 2434.3523 y 10
393.69537353515625 0 1092.3646
396.2354736328125 0 3126.725 c 2
397.882080078125 0 653.91583
398.2156982421875 0 634.8389
398.39239501953125 0 695.94586
404.1496276855469 0 2943.2598
404.19390869140625 0 2096.8499
404.2286071777344 0 6619.2925
404.7093505859375 0 693.91656 c Water loss 5
404.8885498046875 0 725.1628
405.17523193359375 0 966.43976
405.20245361328125 0 2180.6704 c Ammonia loss 5
405.2307434082031 0 1674.3816
405.4072265625 0 589.65436
405.52911376953125 0 1275.8425 y 6
405.70367431640625 0 810.7831
405.8632507324219 0 941.4794
406.1957702636719 0 1425.6058
412.21917724609375 0 1075.0825
420.2024841308594 0 12561.861 w 9
420.7039794921875 0 6968.838
421.2046813964844 0 2663.9915
423.5353698730469 0 714.7293 w 5
443.2241516113281 0 1073.7239 y 5
448.7147521972656 0 942.21594 y Water loss 9
455.2266845703125 0 1219.8916 c Water loss 13
457.7185974121094 0 616.39655 y 9
460.7359619140625 0 2567.9905
461.2309265136719 0 2692.1472
461.56243896484375 0 1814.5767
462.7142639160156 0 868.04504 c Ammonia loss 6
466.485595703125 0 799.74115
468.2284240722656 0 796.37897
478.2211608886719 0 870.75916 y 1
478.47393798828125 0 832.4012
484.2555847167969 0 651.7073
484.491455078125 0 2249.1267
484.74298095703125 0 1162.5105
484.9916076660156 0 1512.2946
490.24072265625 0 960.83887
490.740966796875 0 1139.8901
491.7284240722656 0 2961.722
491.9797058105469 0 3590.1577
492.2448425292969 0 835.0406
492.4814758300781 0 2244.3638
492.742919921875 0 1131.0675
494.2347717285156 0 9379.725 c Ammonia loss 3
495.2374267578125 0 2498.9753
496.2496032714844 0 874.9603
496.7738952636719 0 994.66296
499.7392578125 0 1503.8735 c 7
500.27508544921875 0 1043.7587
500.7441101074219 0 716.2534
501.99310302734375 0 1044.4939
502.2447814941406 0 1172.9509
502.4951171875 0 868.8937
502.9880065917969 0 966.27826
503.9143371582031 0 1151.4993
504.24664306640625 0 4389.932
504.579345703125 0 4024.6187 y Water loss 4
504.91375732421875 0 2549.0056 y Ammonia loss 4
505.2446594238281 0 729.90814 z Ammonia loss 8
506.2497253417969 0 1901.0568
506.495849609375 0 10594.209
506.74688720703125 0 11617.466
506.9971618652344 0 9087.557
507.247802734375 0 4786.8145
507.4969787597656 0 1444.434
511.26220703125 0 5960.0786 c 3
512.265869140625 0 1135.1366
512.7555541992188 0 1788.3573 y Water loss 8
513.259765625 0 1493.4299 y Ammonia loss 8
513.7584838867188 0 1254.8323 z 8
515.9305419921875 0 954.2707
516.2606811523438 0 4956.842 c 11
516.5958251953125 0 4764.5146
516.9270629882812 0 1179.1837
519.2947387695312 0 832.26324
521.7653198242188 0 2333.6604 y 8
522.2669067382812 0 785.5874
524.7713623046875 0 698.7615
525.2774658203125 0 874.65247
535.2897338867188 0 928.22327
540.0271606445312 0 755.2381
542.2684936523438 0 1880.8572 z 7
542.7687377929688 0 1373.8748
543.268798828125 0 813.2448
550.2742919921875 0 2441.491 y 7
550.7778930664062 0 1725.5078
553.2337646484375 0 813.6321
553.2824096679688 0 658.9005
554.2861328125 0 690.29767
555.3101196289062 0 1150.4434
563.2938232421875 0 1671.2238
563.7864990234375 0 11667.654 c 8
564.2868041992188 0 7757.3433
564.7886962890625 0 2734.0068
566.9281616210938 0 4134.3027 w 2
567.2637329101562 0 1546.3663
569.9336547851562 0 3087.415 c 12
570.2664184570312 0 1825.3245
571.2810668945312 0 3805.311
571.7830200195312 0 2194.4763
572.2858276367188 0 1398.4795
579.2881469726562 0 954.54584
585.9383544921875 0 1055.8859 y Ammonia loss 2
596.3199462890625 0 1566.4377
596.8220825195312 0 3012.1086
597.32275390625 0 2369.3062
597.9610595703125 0 943.738
598.7742309570312 0 917.1337
599.7819213867188 0 4689.9136 z 6
600.2844848632812 0 4679.9688
600.785400390625 0 3002.6858
601.2841796875 0 1323.0667
606.3017578125 0 3107.471 z 11
606.7854614257812 0 1602.7886
607.3026733398438 0 2946.9683
607.790771484375 0 7631.5454 y 6
608.2909545898438 0 4780.1895
608.7935180664062 0 2173.6284
609.3008422851562 0 4302.92
609.8037719726562 0 1884.7026
610.2916870117188 0 815.2848
612.6286010742188 0 2611.3835 c 13
612.9637451171875 0 3724.569
613.2979736328125 0 1703.7322
614.3230590820312 0 1090.2971
619.7952270507812 0 825.2153 c Ammonia loss 9
622.3193969726562 0 2951.5505
626.3054809570312 0 2109.8984
626.6442260742188 0 835.7737
627.3120727539062 0 755.08453
627.8148803710938 0 1331.0227
628.3082885742188 0 5103.9473 c 9
628.8087768554688 0 2143.7812
629.3060302734375 0 1202.4319
631.9594116210938 0 1343.1602 z 1
632.288330078125 0 1233.9493
632.6257934570312 0 1553.6829
634.794921875 0 1702.592 w 5
635.2909545898438 0 926.34076
640.3157958984375 0 1082.2592
640.4133911132812 0 1719.2771
641.31982421875 0 815.28174
645.6532592773438 0 2559.2437
645.9890747070312 0 3509.5503
646.3223876953125 0 1699.0381
646.6603393554688 0 678.5896
649.6461791992188 0 673.49713
650.3187255859375 0 1032.5653
654.9951782226562 0 1219.7711
655.323974609375 0 971.1469 y Water loss 5
655.6538696289062 0 6420.4546
655.9907836914062 0 5346.035
656.3242797851562 0 5119.959 z 5
656.657470703125 0 1416.6187
656.8244018554688 0 2044.8291
656.99560546875 0 1075.4193
660.3229370117188 0 725.3578
660.9902954101562 0 1134.4457
664.3303833007812 0 3053.9705 y 5
669.3234252929688 0 820.4925
669.6583251953125 0 5023.5337
669.98974609375 0 10644.579
670.3244018554688 0 8093.879
670.6563110351562 0 3595.964
670.9910888671875 0 1645.2782
672.278564453125 0 963.3644
674.6658935546875 0 674.9114
674.9927368164062 0 2841.2693
675.3265380859375 0 6157.498
675.661865234375 0 6612.1626
675.9959716796875 0 4552.3276
676.332275390625 0 2940.5298
676.6669921875 0 1264.1786
687.3251953125 0 776.908
687.8328857421875 0 1946.6537
688.335693359375 0 2583.184
688.8362426757812 0 860.8144
690.8471069335938 0 834.40436
691.3381958007812 0 1141.4542
691.8436279296875 0 919.7732
692.3433837890625 0 1267.0953
696.3109741210938 0 5036.2427 c Ammonia loss 4
697.3131713867188 0 1844.3926
709.8402709960938 0 3987.1624 c 10
710.3424072265625 0 3068.0876
710.8414306640625 0 1921.8324
711.4512329101562 0 1034.6636
713.334716796875 0 3202.492 c 4
714.3389282226562 0 981.76874
729.2979125976562 0 765.06824
752.3757934570312 0 767.55115
755.8600463867188 0 1012.9091
756.3701171875 0 666.88214 y Water loss 4
756.8713989257812 0 1523.3801 y Ammonia loss 4
769.3681640625 0 2495.201 z 10
770.3678588867188 0 1173.4824
771.3658447265625 0 1002.4569
773.8869018554688 0 1472.2997 c 11
774.3875732421875 0 1603.8882
785.3839111328125 0 751.95795 y 10
803.430908203125 0 816.1123
809.4010620117188 0 2818.184 c Ammonia loss 5
810.4013061523438 0 1823.7971
811.3966674804688 0 699.8351
812.380615234375 0 1316.4858
814.8757934570312 0 1005.715 z 3
815.3744506835938 0 798.9645
815.8776245117188 0 2417.3909
826.4212036132812 0 8516.823 c 5
827.4240112304688 0 4229.9297
828.4258422851562 0 1223.9288
854.4017333984375 0 747.01416 c 12
854.8984375 0 1735.3947
878.907470703125 0 1551.2468 z 2
879.4083251953125 0 3607.0596
879.907958984375 0 2350.3623
880.4046020507812 0 965.5277 z Water loss 9
880.9082641601562 0 1118.9694
883.4183349609375 0 2737.9065
884.42138671875 0 1647.8483
885.4203491210938 0 735.7612
896.9368286132812 0 1897.2585
897.4430541992188 0 1551.0648
898.4127807617188 0 2869.4297 z 9
899.4127807617188 0 1942.7177
900.4148559570312 0 859.0099
918.9370727539062 0 1238.7395
919.4481201171875 0 719.6405
924.4210205078125 0 952.97943 c Ammonia loss 6
941.4422607421875 0 819.0993 c 6
946.4698486328125 0 767.96814
947.4370727539062 0 833.48315 z 1
947.9354248046875 0 1927.2911
948.4383544921875 0 1807.7753
948.9378051757812 0 1557.2379
949.4358520507812 0 787.3219
968.4904174804688 0 628.96735
998.4708251953125 0 975.8943 c 7
999.4774780273438 0 1434.1556
1000.4682006835938 0 853.1292
1004.4785766601562 0 905.8706
1004.9803466796875 0 1819.13
1012.9937133789062 0 866.57153
1026.49853515625 0 871.6208 z 8
1027.513916015625 0 1527.8638
1084.5322265625 0 980.5249
1110.5426025390625 0 1158.6337
1126.563720703125 0 1681.285 c 8
1127.56298828125 0 1726.0756
1255.610595703125 0 646.68225 c 9
1419.6783447265625 0 1282.3973
2253.73388671875 0 802.25995
2465.8251953125 0 657.5798
3276.918701171875 0 751.89264

Spectrum Details

|  |  |
| --- | --- |
| Matched peaks? Matched peaks The total absolute number of peaks matched. Additionally in brackets the total fraction of peaks matched and the total number of peaks is shown. | 73 (19.47% of 375) |
| FDR? FDR The false discovery rate estimated for this peptide. It is calculated by matching all theoretical fragments with a non-integer shift with the raw peaks for this spectrum. This is done with 40 different shifts. The resulting percentage is the average number of annotated peaks over the number of annotated peaks with the correct spectrum. | 1.63% |
| Satellite FDR? Satellite FDR See the FDR for details on its calculation. This satellite ion specific FDR only contains the satellite ions (d/w) for I/L/J positions. | 0.00% |
| PSM Score? PSM Score The PSM Score as given by Hecklib to this annotated spectrum. It is shown with three significant figures. | 203 |

#### Spectrum 5177? Spectrum 5177 The raw spectrum of this peptide as annotated by Hecklib. The fragments are coloured according to ion type (see legend). Any peaks with a star '\*' as text can be hovered over to see the full details, first the ion type second the mass shift type. By hovering over the amino acids in the peptide or ions in the legend the corresponding peaks are highlighted. By toggling the 'Unassigned' label you can turn the background (unassigned) peaks on or off in the plot. By updating the slider in the Ion legend you can update the spectrum to only show the top X% of the peaks with labels. The top X% means any peak that is within X% of the highest intensity. By dragging in the spectrum you can zoom in to a specific part of the spectrum and use 'Zoom Out' to get back to the original zoom level. The annotation of the spectrum is based on the given sequence in the peptides file and is done with different software so inconsistencies are likely. The peaks are annotated based on the given sequence, with 20 ppm tolerance.

Copy Data

##### Spectrum 5177 (TSV)

###### Preview

```
Loading example...
```

*Click on the button to copy the data to your clipboard.*

Mz MinMz MaxIntensity Max

WidthHeightPeptide font sizePeptide stroke widthSpectrum font sizeSpectrum stroke widthCompact peptide

Ion legend

wxyz

abcd

OtherUnassignedIon^Charge^_Position_Show for top:%

JHQDWLDGKEYKCKK

01.15e+42.31e+43.46e+44.62e+4

Zoom Out

c^+4^_6_c^+1^_2_z^+1^_2_c^+3^_7_z^+2^_4_y^+2^_4_z^+1^_2_y^+1^_2_c^+2^_5_y^+3^_7_c^+3^_9_y^+3^_8_y^+4^_11_y^+4^_11_c^+1^_3_z^+2^_5_y^+2^_5_c^+1^_3_c^+2^_6_y^+3^_9_w^+2^_6_w^+3^_10_y^+3^_10_y^+3^_10_y^+2^_6_z^+2^_6_y^+4^_14_z^+4^_14_y^+4^_14_c^+2^_8_c^+2^_8_w^+2^_7_c^+2^_8_c^+1^_4_y^+3^_11_y^+3^_11_z^+3^_11_c^+3^_12_c^+3^_12_c^+3^_12_c^+1^_4_y^+2^_7_z^+2^_7_y^+2^_7_w^+3^_12_z^+2^_8_c^+2^_9_y^+2^_8_c^+2^_9_w^+3^_13_c^+3^_13_y^+3^_13_z^+3^_13_z^+2^_9_y^+2^_9_y^+2^_9_z^+2^_9_c^+3^_14_c^+3^_14_z^+1^_4_c^+3^_14_y^+2^_9_c^+2^_10_c^+2^_10_z^+3^_14_y^+1^_4_z^+3^_14_y^+3^_14_w^+2^_10_y^+2^_10_z^+2^_10_y^+2^_10_c^+1^_5_c^+1^_5_c^+1^_5_c^+2^_11_z^+2^_11_c^+2^_12_c^+2^_12_z^+1^_5_y^+1^_5_c^+1^_6_y^+2^_12_z^+2^_12_c^+1^_6_w^+2^_13_c^+2^_13_z^+2^_13_z^+1^_6_c^+1^_7_y^+2^_14_z^+2^_14_c^+1^_8_z^+1^_7_c^+1^_9_c^+1^_10_c^+1^_11_

0769153923083077

Fragment Matches Table

Show background peaks

| Position | Ion type | Intensity | mz Theoretical | mz Error (Th) | mz Error (ppm) | Charge | Series Number |
| --- | --- | --- | --- | --- | --- | --- | --- |
| - | - | 588 | 120.1 | - | - | 0 | - |
| - | - | 374.3 | 124.8 | - | - | 0 | - |
| - | - | 841.8 | 127.1 | - | - | 0 | - |
| - | - | 469.9 | 128.1 | - | - | 0 | - |
| - | - | 5982 | 129 | - | - | 0 | - |
| - | - | 730 | 129.1 | - | - | 0 | - |
| - | - | 5067 | 129.1 | - | - | 0 | - |
| - | - | 370.7 | 129.3 | - | - | 0 | - |
| - | - | 1.531E+04 | 130.1 | - | - | 0 | - |
| - | - | 1596 | 131.1 | - | - | 0 | - |
| - | - | 2666 | 133.1 | - | - | 0 | - |
| - | - | 3617 | 133.1 | - | - | 0 | - |
| - | - | 402.5 | 135.2 | - | - | 0 | - |
| - | - | 717.7 | 136.1 | - | - | 0 | - |
| - | - | 4768 | 139 | - | - | 0 | - |
| - | - | 598.1 | 140.1 | - | - | 0 | - |
| - | - | 959.2 | 146.1 | - | - | 0 | - |
| - | - | 4059 | 155.1 | - | - | 0 | - |
| - | - | 642.2 | 155.1 | - | - | 0 | - |
| - | - | 1.537E+04 | 157 | - | - | 0 | - |
| - | - | 566.3 | 157.1 | - | - | 0 | - |
| - | - | 4013 | 159.1 | - | - | 0 | - |
| - | - | 482 | 165.1 | - | - | 0 | - |
| - | - | 3219 | 166.1 | - | - | 0 | - |
| - | - | 634.8 | 168.1 | - | - | 0 | - |
| - | - | 1471 | 170.1 | - | - | 0 | - |
| - | - | 494.2 | 171.1 | - | - | 0 | - |
| - | - | 909 | 172.1 | - | - | 0 | - |
| - | - | 571.9 | 173.1 | - | - | 0 | - |
| - | - | 872.8 | 173.1 | - | - | 0 | - |
| - | - | 3681 | 173.1 | - | - | 0 | - |
| - | - | 2662 | 173.5 | - | - | 0 | - |
| - | - | 1043 | 174.1 | - | - | 0 | - |
| - | - | 4022 | 177.1 | - | - | 0 | - |
| - | - | 704.7 | 181.1 | - | - | 0 | - |
| - | - | 2584 | 183.1 | - | - | 0 | - |
| - | - | 2.116E+04 | 185.1 | - | - | 0 | - |
| - | - | 4019 | 185.2 | - | - | 0 | - |
| - | - | 1300 | 187.1 | - | - | 0 | - |
| - | - | 5076 | 190.1 | - | - | 0 | - |
| - | - | 823.8 | 198.1 | - | - | 0 | - |
| 6 | c | 500.5 | 199.1 | 0.002375 | 11.93 | +4 | 6 |
| - | - | 5812 | 199.2 | - | - | 0 | - |
| - | - | 2.034E+04 | 201.1 | - | - | 0 | - |
| - | - | 2657 | 202.1 | - | - | 0 | - |
| - | - | 5837 | 203.1 | - | - | 0 | - |
| - | - | 1682 | 205.1 | - | - | 0 | - |
| - | - | 524.2 | 208.1 | - | - | 0 | - |
| - | - | 577.3 | 209.1 | - | - | 0 | - |
| - | - | 1.125E+04 | 213.1 | - | - | 0 | - |
| - | - | 765.5 | 215.1 | - | - | 0 | - |
| - | - | 1.776E+04 | 217 | - | - | 0 | - |
| - | - | 1037 | 217.1 | - | - | 0 | - |
| - | - | 843.1 | 218.1 | - | - | 0 | - |
| - | - | 595.8 | 219.1 | - | - | 0 | - |
| - | - | 1368 | 221.1 | - | - | 0 | - |
| - | - | 918.7 | 223.1 | - | - | 0 | - |
| - | - | 2770 | 223.2 | - | - | 0 | - |
| - | - | 652.2 | 225.1 | - | - | 0 | - |
| - | - | 1009 | 225.1 | - | - | 0 | - |
| - | - | 653.6 | 229.1 | - | - | 0 | - |
| - | - | 1601 | 229.2 | - | - | 0 | - |
| - | - | 1333 | 233.1 | - | - | 0 | - |
| - | - | 3957 | 234.1 | - | - | 0 | - |
| - | - | 1031 | 236.6 | - | - | 0 | - |
| - | - | 521.7 | 241.1 | - | - | 0 | - |
| - | - | 809.9 | 243.1 | - | - | 0 | - |
| - | - | 651.6 | 247.1 | - | - | 0 | - |
| - | - | 1.686E+04 | 251.2 | - | - | 0 | - |
| - | - | 2602 | 252.2 | - | - | 0 | - |
| - | - | 739.8 | 256.1 | - | - | 0 | - |
| - | - | 2.429E+04 | 259.2 | - | - | 0 | - |
| - | - | 798.9 | 260.2 | - | - | 0 | - |
| - | - | 783.3 | 261.1 | - | - | 0 | - |
| - | - | 725.4 | 265.1 | - | - | 0 | - |
| - | - | 660.1 | 265.2 | - | - | 0 | - |
| - | - | 690.4 | 266.1 | - | - | 0 | - |
| 2 | c | 2469 | 268.2 | 0.0004142 | 1.544 | +1 | 2 |
| - | - | 2109 | 269.1 | - | - | 0 | - |
| - | - | 1641 | 270.1 | - | - | 0 | - |
| - | - | 7725 | 270.1 | - | - | 0 | - |
| - | - | 867.8 | 271.1 | - | - | 0 | - |
| - | - | 613.4 | 272.1 | - | - | 0 | - |
| - | - | 8765 | 273.1 | - | - | 0 | - |
| - | - | 1958 | 274.1 | - | - | 0 | - |
| - | - | 1164 | 274.7 | - | - | 0 | - |
| - | - | 1647 | 282.2 | - | - | 0 | - |
| - | - | 1046 | 286.1 | - | - | 0 | - |
| - | - | 8763 | 288.2 | - | - | 0 | - |
| - | - | 1726 | 289.2 | - | - | 0 | - |
| - | - | 582.3 | 290.1 | - | - | 0 | - |
| - | - | 1727 | 295.1 | - | - | 0 | - |
| - | - | 1.627E+04 | 296.2 | - | - | 0 | - |
| - | - | 3619 | 297.2 | - | - | 0 | - |
| - | - | 1013 | 298.1 | - | - | 0 | - |
| 14 | z | 681.8 | 300.2 | 0.001565 | 5.214 | +1 | 2 |
| 7 | c | 1221 | 303.2 | 0.0008429 | 2.78 | +3 | 7 |
| 12 | z | 877.6 | 303.7 | 0.004563 | 15.03 | +2 | 4 |
| - | - | 1295 | 309.2 | - | - | 0 | - |
| 12 | y | 1259 | 311.7 | 0.004434 | 14.23 | +2 | 4 |
| - | - | 7519 | 314.2 | - | - | 0 | - |
| - | - | 1685 | 315.2 | - | - | 0 | - |
| - | - | 1001 | 316.2 | - | - | 0 | - |
| 14 | z | 1298 | 317.2 | 0.00367 | 11.57 | +1 | 2 |
| - | - | 1976 | 318.2 | - | - | 0 | - |
| - | - | 711.9 | 318.8 | - | - | 0 | - |
| - | - | 546.4 | 325.2 | - | - | 0 | - |
| - | - | 984.6 | 326.5 | - | - | 0 | - |
| - | - | 9669 | 326.7 | - | - | 0 | - |
| - | - | 3723 | 327.2 | - | - | 0 | - |
| - | - | 2217 | 327.2 | - | - | 0 | - |
| - | - | 838.3 | 327.7 | - | - | 0 | - |
| - | - | 4392 | 329.2 | - | - | 0 | - |
| - | - | 764.4 | 331.7 | - | - | 0 | - |
| 14 | y | 1142 | 333.2 | 0.00451 | 13.54 | +1 | 2 |
| - | - | 2658 | 338.2 | - | - | 0 | - |
| - | - | 1115 | 339.7 | - | - | 0 | - |
| 5 | c | 4103 | 340.7 | 0.000562 | 1.65 | +2 | 5 |
| - | - | 2418 | 341.2 | - | - | 0 | - |
| - | - | 1062 | 341.2 | - | - | 0 | - |
| - | - | 638.3 | 341.7 | - | - | 0 | - |
| - | - | 738.5 | 343.7 | - | - | 0 | - |
| 9 | y | 1013 | 348.2 | 0.006941 | 19.94 | +3 | 7 |
| - | - | 871.3 | 349.2 | - | - | 0 | - |
| - | - | 651.3 | 349.2 | - | - | 0 | - |
| - | - | 756.2 | 355.2 | - | - | 0 | - |
| - | - | 624.8 | 355.7 | - | - | 0 | - |
| - | - | 4786 | 356.2 | - | - | 0 | - |
| - | - | 673.6 | 356.8 | - | - | 0 | - |
| - | - | 845.9 | 357.2 | - | - | 0 | - |
| - | - | 839.5 | 357.2 | - | - | 0 | - |
| - | - | 704.3 | 357.7 | - | - | 0 | - |
| - | - | 849.6 | 359.2 | - | - | 0 | - |
| - | - | 6872 | 361.2 | - | - | 0 | - |
| - | - | 1032 | 362.2 | - | - | 0 | - |
| - | - | 634.4 | 363.2 | - | - | 0 | - |
| 9 | c | 600.2 | 365.2 | 0.007272 | 19.91 | +3 | 9 |
| - | - | 1469 | 365.2 | - | - | 0 | - |
| 8 | y | 1970 | 367.2 | 0.002746 | 7.479 | +3 | 8 |
| - | - | 2664 | 367.2 | - | - | 0 | - |
| - | - | 1754 | 367.5 | - | - | 0 | - |
| - | - | 1068 | 367.9 | - | - | 0 | - |
| - | - | 923.1 | 369.2 | - | - | 0 | - |
| - | - | 5685 | 373.2 | - | - | 0 | - |
| - | - | 1419 | 374.2 | - | - | 0 | - |
| 5 | y | 768.5 | 374.7 | 0.00217 | 5.791 | +4 | 11 |
| 5 | y | 808.6 | 374.9 | 0.003417 | 9.113 | +4 | 11 |
| - | - | 1033 | 376.2 | - | - | 0 | - |
| - | - | 1374 | 378.2 | - | - | 0 | - |
| 3 | c | 5460 | 379.2 | 6.301E-05 | 0.1662 | +1 | 3 |
| - | - | 758.8 | 379.7 | - | - | 0 | - |
| - | - | 4317 | 381.2 | - | - | 0 | - |
| - | - | 501.3 | 381.2 | - | - | 0 | - |
| - | - | 1190 | 381.5 | - | - | 0 | - |
| - | - | 4188 | 383.2 | - | - | 0 | - |
| - | - | 1.499E+04 | 383.2 | - | - | 0 | - |
| - | - | 1304 | 383.7 | - | - | 0 | - |
| - | - | 4887 | 384.2 | - | - | 0 | - |
| 11 | z | 1008 | 385.2 | 0.005374 | 13.95 | +2 | 5 |
| - | - | 2.347E+04 | 385.2 | - | - | 0 | - |
| - | - | 1121 | 385.5 | - | - | 0 | - |
| - | - | 729.6 | 386.2 | - | - | 0 | - |
| - | - | 4620 | 386.2 | - | - | 0 | - |
| - | - | 1.039E+04 | 387.2 | - | - | 0 | - |
| - | - | 2395 | 388.2 | - | - | 0 | - |
| - | - | 1205 | 389.2 | - | - | 0 | - |
| 11 | y | 5614 | 393.2 | 0.00372 | 9.46 | +2 | 5 |
| - | - | 2384 | 393.7 | - | - | 0 | - |
| - | - | 890.9 | 394.2 | - | - | 0 | - |
| 3 | c | 1.011E+04 | 396.2 | 0.0004915 | 1.24 | +1 | 3 |
| 6 | c | 3523 | 397.2 | 2.708E-05 | 0.06817 | +2 | 6 |
| - | - | 2292 | 397.2 | - | - | 0 | - |
| - | - | 1014 | 397.7 | - | - | 0 | - |
| - | - | 1277 | 398.8 | - | - | 0 | - |
| - | - | 1746 | 399 | - | - | 0 | - |
| - | - | 3.607E+04 | 401.2 | - | - | 0 | - |
| - | - | 1.224E+04 | 401.3 | - | - | 0 | - |
| - | - | 8475 | 402.2 | - | - | 0 | - |
| - | - | 8704 | 402.2 | - | - | 0 | - |
| - | - | 2926 | 402.3 | - | - | 0 | - |
| - | - | 2500 | 402.3 | - | - | 0 | - |
| - | - | 816.3 | 402.4 | - | - | 0 | - |
| - | - | 1146 | 402.6 | - | - | 0 | - |
| - | - | 1248 | 403.2 | - | - | 0 | - |
| - | - | 1714 | 403.2 | - | - | 0 | - |
| - | - | 1169 | 403.2 | - | - | 0 | - |
| - | - | 968.3 | 403.3 | - | - | 0 | - |
| 7 | y | 3298 | 405.5 | 0.002806 | 6.919 | +3 | 9 |
| - | - | 2582 | 405.7 | - | - | 0 | - |
| - | - | 1045 | 405.9 | - | - | 0 | - |
| - | - | 1959 | 406.2 | - | - | 0 | - |
| 10 | w | 4.173E+04 | 420.2 | 0.003905 | 9.294 | +2 | 6 |
| - | - | 2.012E+04 | 420.7 | - | - | 0 | - |
| - | - | 6118 | 421.2 | - | - | 0 | - |
| - | - | 992.3 | 421.7 | - | - | 0 | - |
| 6 | w | 2187 | 423.5 | 0.002269 | 5.356 | +3 | 10 |
| - | - | 736 | 424.2 | - | - | 0 | - |
| - | - | 918.5 | 424.7 | - | - | 0 | - |
| - | - | 793.9 | 432.7 | - | - | 0 | - |
| - | - | 3987 | 435.2 | - | - | 0 | - |
| - | - | 827.2 | 436.2 | - | - | 0 | - |
| 6 | y | 781.7 | 437.2 | 0.002108 | 4.822 | +3 | 10 |
| - | - | 853.9 | 438.2 | - | - | 0 | - |
| - | - | 745.8 | 439.7 | - | - | 0 | - |
| 6 | y | 3604 | 443.2 | 0.001388 | 3.131 | +3 | 10 |
| - | - | 2352 | 443.6 | - | - | 0 | - |
| 10 | y | 775.2 | 448.7 | 0.003712 | 8.273 | +2 | 6 |
| 10 | z | 810.1 | 449.7 | 0.003386 | 7.529 | +2 | 6 |
| - | - | 810.2 | 456.2 | - | - | 0 | - |
| - | - | 6659 | 460.7 | - | - | 0 | - |
| - | - | 3921 | 461.2 | - | - | 0 | - |
| - | - | 1115 | 461.7 | - | - | 0 | - |
| - | - | 762.9 | 462.2 | - | - | 0 | - |
| - | - | 2006 | 462.2 | - | - | 0 | - |
| - | - | 813 | 462.7 | - | - | 0 | - |
| - | - | 681.2 | 465.5 | - | - | 0 | - |
| - | - | 1063 | 466.2 | - | - | 0 | - |
| - | - | 952.6 | 467.2 | - | - | 0 | - |
| - | - | 3877 | 468.3 | - | - | 0 | - |
| - | - | 2155 | 468.8 | - | - | 0 | - |
| - | - | 882.5 | 469.3 | - | - | 0 | - |
| 2 | y | 825.5 | 470 | 0.004482 | 9.536 | +4 | 14 |
| 2 | z | 1180 | 470.2 | 0.009148 | 19.45 | +4 | 14 |
| - | - | 1219 | 470.5 | - | - | 0 | - |
| 2 | y | 1517 | 474.2 | 0.0003161 | 0.6667 | +4 | 14 |
| - | - | 1019 | 474.7 | - | - | 0 | - |
| - | - | 1317 | 480.2 | - | - | 0 | - |
| - | - | 6613 | 480.5 | - | - | 0 | - |
| - | - | 6961 | 480.7 | - | - | 0 | - |
| - | - | 3451 | 481 | - | - | 0 | - |
| - | - | 986.3 | 481.5 | - | - | 0 | - |
| 8 | c | 1744 | 482.7 | 0.001377 | 2.853 | +2 | 8 |
| 8 | c | 3763 | 483.2 | 0.001984 | 4.106 | +2 | 8 |
| - | - | 782.8 | 483.5 | - | - | 0 | - |
| - | - | 2002 | 483.7 | - | - | 0 | - |
| - | - | 3760 | 484.2 | - | - | 0 | - |
| - | - | 1282 | 484.5 | - | - | 0 | - |
| - | - | 769.5 | 484.6 | - | - | 0 | - |
| 9 | w | 1799 | 484.7 | 0.001917 | 3.955 | +2 | 7 |
| - | - | 1640 | 485 | - | - | 0 | - |
| - | - | 831.9 | 485.3 | - | - | 0 | - |
| - | - | 1928 | 487.5 | - | - | 0 | - |
| - | - | 1.603E+04 | 487.7 | - | - | 0 | - |
| - | - | 1.86E+04 | 488 | - | - | 0 | - |
| - | - | 7714 | 488.2 | - | - | 0 | - |
| - | - | 2983 | 488.5 | - | - | 0 | - |
| - | - | 894.7 | 488.7 | - | - | 0 | - |
| - | - | 698.2 | 489 | - | - | 0 | - |
| - | - | 942.3 | 491.3 | - | - | 0 | - |
| 8 | c | 3321 | 491.7 | 0.0004283 | 0.8711 | +2 | 8 |
| - | - | 1703 | 492.2 | - | - | 0 | - |
| 4 | c | 1.178E+04 | 494.2 | 0.003076 | 6.224 | +1 | 4 |
| - | - | 2417 | 495.2 | - | - | 0 | - |
| - | - | 1000 | 496.8 | - | - | 0 | - |
| - | - | 756 | 497.8 | - | - | 0 | - |
| - | - | 2178 | 498.2 | - | - | 0 | - |
| - | - | 1385 | 498.5 | - | - | 0 | - |
| 5 | y | 2090 | 499.2 | 0.006848 | 13.72 | +3 | 11 |
| 5 | y | 2410 | 499.6 | 0.002519 | 5.042 | +3 | 11 |
| 5 | z | 677.1 | 499.9 | 0.006061 | 12.12 | +3 | 11 |
| - | - | 1654 | 500.2 | - | - | 0 | - |
| - | - | 2708 | 502.3 | - | - | 0 | - |
| - | - | 2.619E+04 | 502.5 | - | - | 0 | - |
| - | - | 3.498E+04 | 502.7 | - | - | 0 | - |
| - | - | 2.499E+04 | 503 | - | - | 0 | - |
| - | - | 8941 | 503.2 | - | - | 0 | - |
| - | - | 4229 | 503.5 | - | - | 0 | - |
| - | - | 2957 | 503.7 | - | - | 0 | - |
| 12 | c | 907.5 | 504.9 | 0.0004834 | 0.9573 | +3 | 12 |
| 12 | c | 2547 | 505.3 | 0.0004551 | 0.9007 | +3 | 12 |
| - | - | 1182 | 505.6 | - | - | 0 | - |
| - | - | 772.4 | 505.7 | - | - | 0 | - |
| - | - | 1075 | 505.9 | - | - | 0 | - |
| - | - | 741.6 | 510.3 | - | - | 0 | - |
| 12 | c | 1.183E+04 | 510.9 | 0.001214 | 2.375 | +3 | 12 |
| 4 | c | 3.354E+04 | 511.3 | 0.0006785 | 1.327 | +1 | 4 |
| - | - | 7900 | 511.6 | - | - | 0 | - |
| - | - | 976.4 | 511.9 | - | - | 0 | - |
| - | - | 3938 | 512.3 | - | - | 0 | - |
| 9 | y | 830.2 | 512.8 | 0.008225 | 16.04 | +2 | 7 |
| - | - | 1726 | 513.2 | - | - | 0 | - |
| - | - | 1249 | 513.3 | - | - | 0 | - |
| 9 | z | 6756 | 513.8 | 0.004267 | 8.306 | +2 | 7 |
| - | - | 2101 | 514.3 | - | - | 0 | - |
| - | - | 1065 | 514.6 | - | - | 0 | - |
| - | - | 2052 | 514.8 | - | - | 0 | - |
| - | - | 4733 | 515.2 | - | - | 0 | - |
| - | - | 2455 | 515.3 | - | - | 0 | - |
| - | - | 4049 | 515.6 | - | - | 0 | - |
| - | - | 1651 | 515.9 | - | - | 0 | - |
| - | - | 914.8 | 516.2 | - | - | 0 | - |
| - | - | 694.3 | 516.3 | - | - | 0 | - |
| - | - | 718.9 | 518.3 | - | - | 0 | - |
| 9 | y | 6151 | 521.8 | 0.003864 | 7.405 | +2 | 7 |
| - | - | 1554 | 522.3 | - | - | 0 | - |
| - | - | 856.9 | 522.8 | - | - | 0 | - |
| 4 | w | 721.1 | 523.3 | 0.001185 | 2.264 | +3 | 12 |
| - | - | 885.7 | 527.3 | - | - | 0 | - |
| - | - | 907 | 527.8 | - | - | 0 | - |
| - | - | 840.2 | 528.3 | - | - | 0 | - |
| - | - | 687.3 | 528.8 | - | - | 0 | - |
| - | - | 726.6 | 533.8 | - | - | 0 | - |
| - | - | 7308 | 534.9 | - | - | 0 | - |
| - | - | 8838 | 535.3 | - | - | 0 | - |
| - | - | 6012 | 535.6 | - | - | 0 | - |
| - | - | 3190 | 535.9 | - | - | 0 | - |
| - | - | 693 | 537.3 | - | - | 0 | - |
| 8 | z | 4968 | 542.3 | 0.001693 | 3.123 | +2 | 8 |
| - | - | 2351 | 542.8 | - | - | 0 | - |
| - | - | 1079 | 543.3 | - | - | 0 | - |
| - | - | 881.7 | 546.3 | - | - | 0 | - |
| 9 | c | 1841 | 547.3 | 0.005436 | 9.934 | +2 | 9 |
| - | - | 1604 | 549.6 | - | - | 0 | - |
| 8 | y | 9710 | 550.3 | 0.004403 | 8.001 | +2 | 8 |
| - | - | 5626 | 550.8 | - | - | 0 | - |
| - | - | 1980 | 551.3 | - | - | 0 | - |
| - | - | 750.9 | 554.3 | - | - | 0 | - |
| - | - | 882.5 | 554.8 | - | - | 0 | - |
| - | - | 2172 | 555.3 | - | - | 0 | - |
| - | - | 1990 | 555.3 | - | - | 0 | - |
| 9 | c | 4.571E+04 | 555.8 | 0.0002186 | 0.3932 | +2 | 9 |
| - | - | 2.789E+04 | 556.3 | - | - | 0 | - |
| - | - | 8338 | 556.8 | - | - | 0 | - |
| - | - | 2895 | 557.3 | - | - | 0 | - |
| - | - | 1662 | 561.3 | - | - | 0 | - |
| 3 | w | 1.254E+04 | 561.6 | 0.001713 | 3.051 | +3 | 13 |
| - | - | 9521 | 561.9 | - | - | 0 | - |
| - | - | 6225 | 562.3 | - | - | 0 | - |
| - | - | 1299 | 562.6 | - | - | 0 | - |
| - | - | 744 | 563.3 | - | - | 0 | - |
| - | - | 2758 | 563.8 | - | - | 0 | - |
| - | - | 2488 | 564.3 | - | - | 0 | - |
| 13 | c | 7503 | 564.6 | 0.001011 | 1.79 | +3 | 13 |
| - | - | 872 | 564.8 | - | - | 0 | - |
| - | - | 6455 | 564.9 | - | - | 0 | - |
| - | - | 3176 | 565.3 | - | - | 0 | - |
| - | - | 855.2 | 567.2 | - | - | 0 | - |
| - | - | 948.6 | 568.3 | - | - | 0 | - |
| - | - | 1854 | 568.8 | - | - | 0 | - |
| - | - | 764.4 | 569.3 | - | - | 0 | - |
| - | - | 2187 | 570.3 | - | - | 0 | - |
| - | - | 1115 | 570.8 | - | - | 0 | - |
| - | - | 9788 | 571.3 | - | - | 0 | - |
| - | - | 5236 | 571.8 | - | - | 0 | - |
| - | - | 2743 | 572.3 | - | - | 0 | - |
| - | - | 948.8 | 572.8 | - | - | 0 | - |
| - | - | 771.5 | 573.3 | - | - | 0 | - |
| - | - | 820.8 | 577.8 | - | - | 0 | - |
| - | - | 800.6 | 578.3 | - | - | 0 | - |
| - | - | 1863 | 578.8 | - | - | 0 | - |
| - | - | 1184 | 579.3 | - | - | 0 | - |
| 3 | y | 1545 | 580.6 | 0.005048 | 8.695 | +3 | 13 |
| 3 | z | 1610 | 580.9 | 2.19E-05 | 0.03769 | +3 | 13 |
| - | - | 1376 | 581.3 | - | - | 0 | - |
| - | - | 854.2 | 581.6 | - | - | 0 | - |
| - | - | 995.5 | 586.6 | - | - | 0 | - |
| - | - | 774.4 | 586.8 | - | - | 0 | - |
| 7 | z | 948.6 | 590.8 | 0.002986 | 5.054 | +2 | 9 |
| - | - | 1007 | 592.6 | - | - | 0 | - |
| - | - | 2548 | 593 | - | - | 0 | - |
| - | - | 954.7 | 593.3 | - | - | 0 | - |
| - | - | 867.2 | 597.3 | - | - | 0 | - |
| 7 | y | 880.6 | 598.8 | 0.005536 | 9.245 | +2 | 9 |
| 7 | y | 2298 | 599.3 | 0.005532 | 9.232 | +2 | 9 |
| 7 | z | 1.268E+04 | 599.8 | 0.002836 | 4.728 | +2 | 9 |
| - | - | 1.677E+04 | 600.3 | - | - | 0 | - |
| - | - | 8616 | 600.8 | - | - | 0 | - |
| 14 | c | 1924 | 601.3 | 0.0113 | 18.79 | +3 | 14 |
| 14 | c | 766.1 | 601.6 | 0.0003776 | 0.6277 | +3 | 14 |
| - | - | 1250 | 602.3 | - | - | 0 | - |
| - | - | 842.5 | 602.6 | - | - | 0 | - |
| - | - | 2003 | 602.9 | - | - | 0 | - |
| - | - | 2654 | 603.3 | - | - | 0 | - |
| - | - | 1470 | 603.6 | - | - | 0 | - |
| - | - | 704.4 | 604 | - | - | 0 | - |
| 12 | z | 1.043E+04 | 606.3 | 0.008381 | 13.82 | +1 | 4 |
| - | - | 674.5 | 606.6 | - | - | 0 | - |
| - | - | 2107 | 607 | - | - | 0 | - |
| 14 | c | 1.438E+04 | 607.3 | 0.0006807 | 1.121 | +3 | 14 |
| - | - | 1.131E+04 | 607.6 | - | - | 0 | - |
| 7 | y | 1.947E+04 | 607.8 | 0.003836 | 6.311 | +2 | 9 |
| - | - | 4721 | 608 | - | - | 0 | - |
| - | - | 1.489E+04 | 608.3 | - | - | 0 | - |
| - | - | 1361 | 608.6 | - | - | 0 | - |
| - | - | 4390 | 608.8 | - | - | 0 | - |
| - | - | 1882 | 609.3 | - | - | 0 | - |
| - | - | 1641 | 610.3 | - | - | 0 | - |
| 10 | c | 2492 | 611.8 | 0.007883 | 12.88 | +2 | 10 |
| - | - | 2689 | 612 | - | - | 0 | - |
| - | - | 2963 | 612.3 | - | - | 0 | - |
| - | - | 966.7 | 612.6 | - | - | 0 | - |
| - | - | 1747 | 612.8 | - | - | 0 | - |
| - | - | 768 | 613.3 | - | - | 0 | - |
| - | - | 874.6 | 613.3 | - | - | 0 | - |
| - | - | 892.9 | 615.3 | - | - | 0 | - |
| - | - | 1579 | 619.8 | - | - | 0 | - |
| 10 | c | 1.882E+04 | 620.3 | 0.001566 | 2.525 | +2 | 10 |
| 2 | z | 2838 | 620.6 | 0.01062 | 17.11 | +3 | 14 |
| - | - | 1.208E+04 | 620.8 | - | - | 0 | - |
| - | - | 1893 | 621 | - | - | 0 | - |
| - | - | 1135 | 621.2 | - | - | 0 | - |
| - | - | 7928 | 621.3 | - | - | 0 | - |
| - | - | 1476 | 621.6 | - | - | 0 | - |
| - | - | 1348 | 621.8 | - | - | 0 | - |
| - | - | 1079 | 622 | - | - | 0 | - |
| 12 | y | 6444 | 622.3 | 0.007391 | 11.88 | +1 | 4 |
| - | - | 1678 | 623.3 | - | - | 0 | - |
| - | - | 752 | 625.6 | - | - | 0 | - |
| - | - | 976.4 | 626 | - | - | 0 | - |
| - | - | 2325 | 626.3 | - | - | 0 | - |
| 2 | z | 4698 | 626.6 | 0.002822 | 4.504 | +3 | 14 |
| - | - | 7855 | 627 | - | - | 0 | - |
| - | - | 3923 | 627.3 | - | - | 0 | - |
| - | - | 886.6 | 627.6 | - | - | 0 | - |
| - | - | 1187 | 630.6 | - | - | 0 | - |
| - | - | 1086 | 631 | - | - | 0 | - |
| - | - | 720.9 | 631.3 | - | - | 0 | - |
| 2 | y | 891.1 | 632 | 0.001626 | 2.573 | +3 | 14 |
| - | - | 2290 | 634.3 | - | - | 0 | - |
| 6 | w | 2381 | 634.8 | 0.00509 | 8.018 | +2 | 10 |
| - | - | 1701 | 635.3 | - | - | 0 | - |
| - | - | 1147 | 635.7 | - | - | 0 | - |
| - | - | 798 | 636.3 | - | - | 0 | - |
| - | - | 6930 | 640.3 | - | - | 0 | - |
| - | - | 7440 | 640.7 | - | - | 0 | - |
| - | - | 6730 | 641 | - | - | 0 | - |
| - | - | 3501 | 641.3 | - | - | 0 | - |
| - | - | 2042 | 641.7 | - | - | 0 | - |
| - | - | 1363 | 643.8 | - | - | 0 | - |
| - | - | 1085 | 644.3 | - | - | 0 | - |
| - | - | 1231 | 644.7 | - | - | 0 | - |
| - | - | 1766 | 645 | - | - | 0 | - |
| - | - | 2216 | 645.3 | - | - | 0 | - |
| - | - | 1489 | 645.7 | - | - | 0 | - |
| - | - | 2751 | 646.3 | - | - | 0 | - |
| - | - | 942.5 | 649.3 | - | - | 0 | - |
| - | - | 1232 | 649.7 | - | - | 0 | - |
| - | - | 4426 | 650 | - | - | 0 | - |
| - | - | 1.724E+04 | 650.3 | - | - | 0 | - |
| - | - | 1.993E+04 | 650.7 | - | - | 0 | - |
| - | - | 1.091E+04 | 651 | - | - | 0 | - |
| - | - | 6381 | 651.3 | - | - | 0 | - |
| - | - | 1122 | 651.7 | - | - | 0 | - |
| - | - | 2332 | 652.3 | - | - | 0 | - |
| - | - | 1256 | 653.3 | - | - | 0 | - |
| - | - | 948.1 | 654.3 | - | - | 0 | - |
| - | - | 1116 | 654.7 | - | - | 0 | - |
| - | - | 2613 | 655 | - | - | 0 | - |
| 6 | y | 2319 | 655.3 | 0.002927 | 4.466 | +2 | 10 |
| - | - | 2219 | 655.7 | - | - | 0 | - |
| - | - | 2290 | 656 | - | - | 0 | - |
| 6 | z | 3437 | 656.3 | 0.003669 | 5.59 | +2 | 10 |
| - | - | 3359 | 656.8 | - | - | 0 | - |
| - | - | 1454 | 657.3 | - | - | 0 | - |
| - | - | 909.8 | 658.7 | - | - | 0 | - |
| - | - | 1173 | 659 | - | - | 0 | - |
| - | - | 895.4 | 659.3 | - | - | 0 | - |
| - | - | 1261 | 660 | - | - | 0 | - |
| - | - | 902.7 | 662.3 | - | - | 0 | - |
| - | - | 979.5 | 663.8 | - | - | 0 | - |
| - | - | 1275 | 664 | - | - | 0 | - |
| 6 | y | 1.191E+04 | 664.3 | 0.006134 | 9.233 | +2 | 10 |
| - | - | 2.421E+04 | 664.7 | - | - | 0 | - |
| - | - | 3485 | 664.8 | - | - | 0 | - |
| - | - | 2.409E+04 | 665 | - | - | 0 | - |
| - | - | 1.87E+04 | 665.3 | - | - | 0 | - |
| - | - | 7514 | 665.7 | - | - | 0 | - |
| - | - | 3214 | 666 | - | - | 0 | - |
| - | - | 1365 | 666.3 | - | - | 0 | - |
| - | - | 1594 | 668.8 | - | - | 0 | - |
| - | - | 1397 | 669.3 | - | - | 0 | - |
| - | - | 6882 | 669.7 | - | - | 0 | - |
| - | - | 1.22E+04 | 670 | - | - | 0 | - |
| - | - | 1.853E+04 | 670.3 | - | - | 0 | - |
| - | - | 1.5E+04 | 670.7 | - | - | 0 | - |
| - | - | 6875 | 671 | - | - | 0 | - |
| - | - | 3197 | 671.3 | - | - | 0 | - |
| - | - | 893.4 | 671.7 | - | - | 0 | - |
| - | - | 872.1 | 671.8 | - | - | 0 | - |
| 5 | c | 1173 | 679.3 | 0.002975 | 4.38 | +1 | 5 |
| - | - | 5895 | 679.8 | - | - | 0 | - |
| 5 | c | 1.399E+04 | 680.3 | 0.005349 | 7.862 | +1 | 5 |
| - | - | 2726 | 680.8 | - | - | 0 | - |
| - | - | 6278 | 681.3 | - | - | 0 | - |
| - | - | 1247 | 682.3 | - | - | 0 | - |
| - | - | 1238 | 690.8 | - | - | 0 | - |
| - | - | 6705 | 691.3 | - | - | 0 | - |
| - | - | 2625 | 691.8 | - | - | 0 | - |
| - | - | 2635 | 692.3 | - | - | 0 | - |
| - | - | 1407 | 693.8 | - | - | 0 | - |
| 5 | c | 8156 | 697.3 | 0.0004672 | 0.6699 | +1 | 5 |
| - | - | 3733 | 698.3 | - | - | 0 | - |
| - | - | 1194 | 699.3 | - | - | 0 | - |
| - | - | 1481 | 701.3 | - | - | 0 | - |
| 11 | c | 1.611E+04 | 701.8 | 0.0009076 | 1.293 | +2 | 11 |
| - | - | 1.639E+04 | 702.3 | - | - | 0 | - |
| - | - | 5089 | 702.8 | - | - | 0 | - |
| - | - | 1413 | 703.3 | - | - | 0 | - |
| - | - | 758.6 | 711.3 | - | - | 0 | - |
| - | - | 2175 | 714.4 | - | - | 0 | - |
| - | - | 995.2 | 715.4 | - | - | 0 | - |
| - | - | 834.5 | 720.4 | - | - | 0 | - |
| - | - | 833.5 | 720.9 | - | - | 0 | - |
| - | - | 953 | 722.4 | - | - | 0 | - |
| - | - | 1072 | 726.9 | - | - | 0 | - |
| - | - | 1045 | 731.4 | - | - | 0 | - |
| - | - | 1554 | 737.4 | - | - | 0 | - |
| - | - | 1985 | 737.8 | - | - | 0 | - |
| - | - | 1154 | 738.4 | - | - | 0 | - |
| - | - | 1086 | 742.3 | - | - | 0 | - |
| - | - | 1361 | 742.9 | - | - | 0 | - |
| - | - | 1111 | 744.4 | - | - | 0 | - |
| 5 | z | 3650 | 749.4 | 0.001333 | 1.779 | +2 | 11 |
| - | - | 7308 | 749.9 | - | - | 0 | - |
| - | - | 4936 | 750.4 | - | - | 0 | - |
| - | - | 3790 | 750.9 | - | - | 0 | - |
| - | - | 983.7 | 751.4 | - | - | 0 | - |
| 12 | c | 2238 | 757.4 | 0.0008193 | 1.082 | +2 | 12 |
| - | - | 1286 | 757.9 | - | - | 0 | - |
| - | - | 946.8 | 758.4 | - | - | 0 | - |
| - | - | 1049 | 765.4 | - | - | 0 | - |
| 12 | c | 2782 | 765.9 | 0.00378 | 4.936 | +2 | 12 |
| - | - | 4592 | 766.4 | - | - | 0 | - |
| - | - | 2256 | 766.9 | - | - | 0 | - |
| - | - | 1088 | 767.4 | - | - | 0 | - |
| 11 | z | 6017 | 769.4 | 0.008416 | 10.94 | +1 | 5 |
| - | - | 4074 | 770.4 | - | - | 0 | - |
| - | - | 1516 | 771.4 | - | - | 0 | - |
| - | - | 937.6 | 772.4 | - | - | 0 | - |
| - | - | 1213 | 777.9 | - | - | 0 | - |
| - | - | 915 | 784.9 | - | - | 0 | - |
| 11 | y | 2480 | 785.4 | 0.008219 | 10.47 | +1 | 5 |
| - | - | 829.2 | 786.4 | - | - | 0 | - |
| - | - | 1750 | 787.4 | - | - | 0 | - |
| 6 | c | 1.084E+04 | 793.4 | 4.089E-05 | 0.05154 | +1 | 6 |
| - | - | 8189 | 794.4 | - | - | 0 | - |
| - | - | 2081 | 794.9 | - | - | 0 | - |
| - | - | 3752 | 795.4 | - | - | 0 | - |
| - | - | 1240 | 796.4 | - | - | 0 | - |
| - | - | 1968 | 802.9 | - | - | 0 | - |
| - | - | 2236 | 803.4 | - | - | 0 | - |
| - | - | 1151 | 803.9 | - | - | 0 | - |
| 4 | y | 1072 | 806.4 | 0.003202 | 3.971 | +2 | 12 |
| 4 | z | 3724 | 806.9 | 0.003269 | 4.052 | +2 | 12 |
| - | - | 8175 | 807.4 | - | - | 0 | - |
| - | - | 4785 | 807.9 | - | - | 0 | - |
| - | - | 2809 | 808.4 | - | - | 0 | - |
| 6 | c | 3.089E+04 | 810.4 | 0.0002838 | 0.3502 | +1 | 6 |
| - | - | 1.379E+04 | 811.4 | - | - | 0 | - |
| - | - | 4174 | 812.4 | - | - | 0 | - |
| - | - | 1005 | 813.4 | - | - | 0 | - |
| - | - | 1194 | 817.4 | - | - | 0 | - |
| - | - | 1504 | 826.9 | - | - | 0 | - |
| - | - | 1080 | 840.4 | - | - | 0 | - |
| - | - | 1583 | 841.4 | - | - | 0 | - |
| 3 | w | 1169 | 841.9 | 0.006256 | 7.431 | +2 | 13 |
| - | - | 2127 | 842.4 | - | - | 0 | - |
| 13 | c | 2380 | 846.4 | 0.001756 | 2.075 | +2 | 13 |
| - | - | 3507 | 846.9 | - | - | 0 | - |
| - | - | 1694 | 847.4 | - | - | 0 | - |
| - | - | 1006 | 847.9 | - | - | 0 | - |
| - | - | 851.7 | 860.4 | - | - | 0 | - |
| - | - | 1.2E+04 | 867.4 | - | - | 0 | - |
| - | - | 7046 | 868.4 | - | - | 0 | - |
| - | - | 1943 | 869.4 | - | - | 0 | - |
| 3 | z | 3649 | 870.9 | 0.0005841 | 0.6707 | +2 | 13 |
| - | - | 1.124E+04 | 871.4 | - | - | 0 | - |
| - | - | 7575 | 871.9 | - | - | 0 | - |
| - | - | 5171 | 872.4 | - | - | 0 | - |
| - | - | 2079 | 872.9 | - | - | 0 | - |
| - | - | 812.9 | 879.9 | - | - | 0 | - |
| - | - | 1091 | 880.4 | - | - | 0 | - |
| - | - | 1664 | 880.9 | - | - | 0 | - |
| - | - | 2873 | 881.4 | - | - | 0 | - |
| - | - | 1120 | 881.9 | - | - | 0 | - |
| - | - | 912.5 | 888.4 | - | - | 0 | - |
| - | - | 3923 | 888.9 | - | - | 0 | - |
| - | - | 3540 | 889.4 | - | - | 0 | - |
| - | - | 1946 | 889.9 | - | - | 0 | - |
| - | - | 857 | 895.4 | - | - | 0 | - |
| 10 | z | 6719 | 898.4 | 0.007979 | 8.882 | +1 | 6 |
| - | - | 8462 | 899.4 | - | - | 0 | - |
| - | - | 3532 | 900.4 | - | - | 0 | - |
| - | - | 1253 | 901.4 | - | - | 0 | - |
| - | - | 728.6 | 902.9 | - | - | 0 | - |
| - | - | 800.1 | 909.4 | - | - | 0 | - |
| - | - | 1883 | 910.9 | - | - | 0 | - |
| - | - | 719.2 | 923 | - | - | 0 | - |
| 7 | c | 2249 | 925.5 | 0.001338 | 1.445 | +1 | 7 |
| - | - | 1173 | 926.5 | - | - | 0 | - |
| - | - | 1028 | 931 | - | - | 0 | - |
| 2 | y | 1522 | 938.4 | 0.01789 | 19.06 | +2 | 14 |
| - | - | 872.9 | 939 | - | - | 0 | - |
| 2 | z | 3250 | 939.4 | 0.01513 | 16.11 | +2 | 14 |
| - | - | 8294 | 939.9 | - | - | 0 | - |
| - | - | 6288 | 940.4 | - | - | 0 | - |
| - | - | 3657 | 940.9 | - | - | 0 | - |
| - | - | 897.7 | 941.4 | - | - | 0 | - |
| - | - | 844.2 | 941.9 | - | - | 0 | - |
| - | - | 1914 | 952.5 | - | - | 0 | - |
| - | - | 1762 | 953 | - | - | 0 | - |
| - | - | 853.6 | 953.5 | - | - | 0 | - |
| - | - | 1544 | 966.5 | - | - | 0 | - |
| - | - | 775.4 | 967 | - | - | 0 | - |
| - | - | 2999 | 967.5 | - | - | 0 | - |
| - | - | 1991 | 968 | - | - | 0 | - |
| - | - | 1457 | 968.5 | - | - | 0 | - |
| - | - | 1238 | 975 | - | - | 0 | - |
| - | - | 2336 | 975.5 | - | - | 0 | - |
| - | - | 1425 | 976 | - | - | 0 | - |
| - | - | 2147 | 976.5 | - | - | 0 | - |
| - | - | 1377 | 977 | - | - | 0 | - |
| - | - | 830 | 981.5 | - | - | 0 | - |
| 8 | c | 8905 | 982.5 | 0.001297 | 1.32 | +1 | 8 |
| - | - | 6802 | 983.5 | - | - | 0 | - |
| - | - | 2587 | 984.5 | - | - | 0 | - |
| - | - | 1165 | 985.5 | - | - | 0 | - |
| - | - | 3228 | 996.5 | - | - | 0 | - |
| - | - | 3816 | 997 | - | - | 0 | - |
| - | - | 2722 | 997.5 | - | - | 0 | - |
| - | - | 1309 | 998 | - | - | 0 | - |
| - | - | 2033 | 1005 | - | - | 0 | - |
| - | - | 2287 | 1005 | - | - | 0 | - |
| - | - | 1059 | 1006 | - | - | 0 | - |
| - | - | 918.6 | 1006 | - | - | 0 | - |
| 9 | z | 2381 | 1027 | 0.00846 | 8.242 | +1 | 7 |
| - | - | 5646 | 1028 | - | - | 0 | - |
| - | - | 2631 | 1029 | - | - | 0 | - |
| - | - | 965 | 1030 | - | - | 0 | - |
| - | - | 1705 | 1085 | - | - | 0 | - |
| - | - | 1394 | 1095 | - | - | 0 | - |
| - | - | 1756 | 1096 | - | - | 0 | - |
| 9 | c | 6918 | 1111 | 0.001244 | 1.12 | +1 | 9 |
| - | - | 6318 | 1112 | - | - | 0 | - |
| - | - | 4089 | 1113 | - | - | 0 | - |
| - | - | 1052 | 1114 | - | - | 0 | - |
| - | - | 785.4 | 1143 | - | - | 0 | - |
| - | - | 707.6 | 1197 | - | - | 0 | - |
| - | - | 3122 | 1200 | - | - | 0 | - |
| - | - | 1677 | 1201 | - | - | 0 | - |
| - | - | 852.7 | 1225 | - | - | 0 | - |
| 10 | c | 1555 | 1240 | 0.001798 | 1.451 | +1 | 10 |
| - | - | 1961 | 1241 | - | - | 0 | - |
| - | - | 837.1 | 1242 | - | - | 0 | - |
| - | - | 763.9 | 1243 | - | - | 0 | - |
| - | - | 763.2 | 1265 | - | - | 0 | - |
| - | - | 1260 | 1313 | - | - | 0 | - |
| - | - | 893.5 | 1314 | - | - | 0 | - |
| - | - | 816.8 | 1359 | - | - | 0 | - |
| - | - | 1890 | 1360 | - | - | 0 | - |
| - | - | 1449 | 1361 | - | - | 0 | - |
| 11 | c | 1266 | 1403 | 0.001406 | 1.003 | +1 | 11 |
| - | - | 2815 | 1404 | - | - | 0 | - |
| - | - | 1707 | 1405 | - | - | 0 | - |
| - | - | 779.7 | 1500 | - | - | 0 | - |
| - | - | 843.2 | 1615 | - | - | 0 | - |
| - | - | 707.3 | 1712 | - | - | 0 | - |
| - | - | 1161 | 1743 | - | - | 0 | - |
| - | - | 947.5 | 1880 | - | - | 0 | - |
| - | - | 692.7 | 2739 | - | - | 0 | - |
| - | - | 947.7 | 3047 | - | - | 0 | - |

m/z Charge Intensity FragmentType MassShift Position
120.06602478027344 0 588.042
124.79774475097656 0 374.28314
127.11197662353516 0 841.809
128.09524536132812 0 469.85413
129.01846313476562 0 5981.6533
129.05491638183594 0 730.0359
129.10243225097656 0 5067.432
129.32701110839844 0 370.66077
130.06533813476562 0 15308.128
131.06874084472656 0 1596.1394
133.06092834472656 0 2665.8242
133.08615112304688 0 3617.2253
135.1761016845703 0 402.46603
136.07565307617188 0 717.6687
139.00277709960938 0 4768.068
140.0819091796875 0 598.09705
146.09255981445312 0 959.2202
155.0928497314453 0 4059.0806
155.11758422851562 0 642.17065
157.01339721679688 0 15374.153
157.1333465576172 0 566.3215
159.09185791015625 0 4012.957
165.10145568847656 0 482.0146
166.0612335205078 0 3218.5793
168.13829040527344 0 634.7996
170.06031799316406 0 1470.9406
171.1367950439453 0 494.20477
172.07183837890625 0 908.9623
173.05557250976562 0 571.9324
173.09237670898438 0 872.78314
173.12864685058594 0 3680.8535
173.4501495361328 0 2661.9841
174.0767059326172 0 1042.5653
177.11231994628906 0 4021.7207
181.09693908691406 0 704.6912
183.1131134033203 0 2584.2517
185.08103942871094 0 21157.062
185.1651153564453 0 4019.0288
187.10809326171875 0 1299.7086
190.08258056640625 0 5076.312
198.12838745117188 0 823.8052
199.1076202392578 0 500.4838 c Ammonia loss 5
199.16944885253906 0 5811.6045
201.12353515625 0 20336.783
202.12704467773438 0 2656.9578
203.10276794433594 0 5837.1514
205.1184844970703 0 1681.7046
208.14474487304688 0 524.23254
209.09231567382812 0 577.2951
213.0758514404297 0 11251.726
215.13934326171875 0 765.4525
217.03451538085938 0 17756.861
217.08184814453125 0 1037.0968
218.1499481201172 0 843.1001
219.08082580566406 0 595.7763
221.138427734375 0 1368.2026
223.11859130859375 0 918.7225
223.15541076660156 0 2769.922
225.09780883789062 0 652.19635
225.1233367919922 0 1008.6412
229.11874389648438 0 653.6168
229.15476989746094 0 1601.115
233.13909912109375 0 1333.3995
234.12399291992188 0 3956.973
236.63284301757812 0 1030.6786
241.1441650390625 0 521.7381
243.1341094970703 0 809.9168
247.07594299316406 0 651.5527
251.15049743652344 0 16860.643
252.154052734375 0 2601.8494
256.12921142578125 0 739.775
259.1541748046875 0 24289.025
260.1596374511719 0 798.9164
261.1192932128906 0 783.27136
265.1435546875 0 725.4076
265.1636962890625 0 660.0911
266.1254577636719 0 690.4073
268.1772155761719 0 2469.1309 c 1
269.13836669921875 0 2109.0022
270.1214599609375 0 1641.4128
270.1450500488281 0 7725.171
271.1484069824219 0 867.7908
272.1070251464844 0 613.4043
273.0971374511719 0 8764.829
274.1494140625 0 1957.62
274.65069580078125 0 1163.5804
282.18182373046875 0 1647.0133
286.1395568847656 0 1046.2728
288.15582275390625 0 8762.673
289.158935546875 0 1725.8911
290.117431640625 0 582.3484
295.10302734375 0 1727.047
296.1971740722656 0 16273.299
297.2002868652344 0 3618.6047
298.1400451660156 0 1012.5761
300.1709289550781 0 681.83466 z Ammonia loss 13
303.1513671875 0 1220.7332 c Water loss 6
303.6556701660156 0 877.5869 z 11
309.19012451171875 0 1295.1489
311.6651611328125 0 1259.0592 y 11
314.20794677734375 0 7518.672
315.2106018066406 0 1684.6632
316.1505126953125 0 1001.1066
317.19537353515625 0 1297.5452 z 13
318.1517639160156 0 1975.634
318.83441162109375 0 711.8817
325.1872863769531 0 546.3743
326.4990539550781 0 984.636
326.6642150878906 0 9669.161
327.1656799316406 0 3723.3374
327.2017822265625 0 2217.311
327.66619873046875 0 838.3373
329.1602478027344 0 4391.9014
331.65606689453125 0 764.44574
333.2132568359375 0 1142.4082 y 13
338.2073974609375 0 2657.5598
339.6939392089844 0 1114.9828
340.6617431640625 0 4102.944 c Ammonia loss 4
341.16259765625 0 2417.9045
341.183349609375 0 1062.4639
341.664794921875 0 638.2812
343.6947326660156 0 738.4599
348.18939208984375 0 1013.1575 y 8
349.1877136230469 0 871.28143
349.2214660644531 0 651.32294
355.2348937988281 0 756.23596
355.6803283691406 0 624.76733
356.2178039550781 0 4786.0947
356.83917236328125 0 673.5685
357.1794128417969 0 845.8719
357.22064208984375 0 839.499
357.6923828125 0 704.32965
359.1522521972656 0 849.56445
361.2221374511719 0 6871.667
362.1837158203125 0 1031.8481
363.2053527832031 0 634.36664
365.1929626464844 0 600.2326 c Ammonia loss 8
365.21875 0 1469.1833
367.1868591308594 0 1970.491 y 7
367.23297119140625 0 2664.0017
367.5203857421875 0 1753.5474
367.8548583984375 0 1067.9403
369.17327880859375 0 923.14624
373.2447204589844 0 5684.564
374.246826171875 0 1418.9763
374.6911315917969 0 768.49084 y Water loss 4
374.9315490722656 0 808.6231 y Ammonia loss 4
376.19232177734375 0 1033.3368
378.2026672363281 0 1373.7268
379.2088928222656 0 5459.9326 c Ammonia loss 2
379.6909484863281 0 758.795
381.1900939941406 0 4316.8867
381.21038818359375 0 501.33655
381.52520751953125 0 1190.4287
383.2049560546875 0 4187.8755
383.2295227050781 0 14988.15
383.70880126953125 0 1304.3229
384.2323303222656 0 4887.0767
385.1865234375 0 1008.4648 z 10
385.243408203125 0 23468.307
385.5249938964844 0 1121.2407
386.1536865234375 0 729.59015
386.2469482421875 0 4619.6064
387.2241516113281 0 10385.101
388.2275085449219 0 2395.264
389.1866149902344 0 1205.1698
393.1975402832031 0 5613.81 y 10
393.6992492675781 0 2384.073
394.1969299316406 0 890.86145
396.2358703613281 0 10108.291 c 2
397.20318603515625 0 3522.788 c Ammonia loss 5
397.23956298828125 0 2292.4324
397.7060241699219 0 1013.6459
398.7931823730469 0 1276.7668
398.99615478515625 0 1745.5946
401.2398681640625 0 36068.156
401.28759765625 0 12236.833
402.21185302734375 0 8475.3125
402.2425231933594 0 8703.634
402.271240234375 0 2926.1345
402.29180908203125 0 2500.438
402.40130615234375 0 816.3145
402.600830078125 0 1146.0201
403.1899108886719 0 1247.9808
403.2179870605469 0 1714.4899
403.2449951171875 0 1168.8822
403.27325439453125 0 968.27814
405.52911376953125 0 3297.632 y 6
405.7038879394531 0 2581.676
405.8642883300781 0 1044.8591
406.2021179199219 0 1959.1324
420.20263671875 0 41731.6 w 9
420.70416259765625 0 20117.383
421.2044677734375 0 6117.691
421.7056884765625 0 992.2613
423.5331726074219 0 2186.8188 w 5
424.2012939453125 0 735.96497
424.7247009277344 0 918.5124
432.7076721191406 0 793.86334
435.2244873046875 0 3986.7617
436.22686767578125 0 827.1736
437.2209777832031 0 781.6505 y Water loss 5
438.2348937988281 0 853.9291
439.7390441894531 0 745.7583
443.2252197265625 0 3604.1091 y 5
443.55841064453125 0 2351.9355
448.71356201171875 0 775.1684 y Water loss 9
449.7098083496094 0 810.1468 z 9
456.2257385253906 0 810.2467
460.73577880859375 0 6659.273
461.2371826171875 0 3921.3887
461.7388610839844 0 1115.0208
462.201904296875 0 762.86774
462.2441711425781 0 2006.174
462.7471618652344 0 812.9999
465.4708557128906 0 681.20056
466.2411193847656 0 1063.1791
467.24609375 0 952.60406
468.2638854980469 0 3877.3186
468.7668151855469 0 2155.331
469.2689208984375 0 882.50024
469.9755554199219 0 825.45593 y Ammonia loss 1
470.232177734375 0 1180.0208 z 1
470.478271484375 0 1219.2064
474.22802734375 0 1517.3242 y 1
474.7294921875 0 1018.8714
480.2459716796875 0 1317.3418
480.4937438964844 0 6612.9863
480.74420166015625 0 6961.1494
480.9944152832031 0 3450.7915
481.4953308105469 0 986.33246
482.7367858886719 0 1744.489 c Water loss 7
483.2294006347656 0 3763.3154 c Ammonia loss 7
483.48052978515625 0 782.81714
483.73553466796875 0 2001.5812
484.2473449707031 0 3759.9917
484.48681640625 0 1281.8455
484.5732116699219 0 769.48517
484.7259216308594 0 1799.1671 w 8
484.9866638183594 0 1640.3619
485.2512512207031 0 831.92444
487.4842834472656 0 1927.9365
487.7291564941406 0 16032.498
487.9812316894531 0 18595.762
488.2332763671875 0 7714.3364
488.4934997558594 0 2983.4116
488.7423400878906 0 894.6994
488.9955139160156 0 698.15375
491.28826904296875 0 942.28973
491.7411193847656 0 3321.1924 c 7
492.2435607910156 0 1702.728
494.2326965332031 0 11775.318 c Ammonia loss 3
495.23919677734375 0 2416.6936
496.7762451171875 0 1000.4761
497.7550354003906 0 755.96747
498.2455749511719 0 2177.8508
498.49786376953125 0 1385.4745
499.24267578125 0 2089.7769 y Water loss 4
499.5800476074219 0 2410.0981 y Ammonia loss 4
499.90740966796875 0 677.1431 z 4
500.24761962890625 0 1653.8987
502.2511901855469 0 2707.6748
502.49761962890625 0 26186.59
502.7483215332031 0 34979.63
502.9985656738281 0 24986.787
503.2494812011719 0 8940.606
503.5006103515625 0 4229.463
503.7498779296875 0 2957.0562
504.9241638183594 0 907.52716 c Water loss 11
505.252197265625 0 2547.2222 c Ammonia loss 11
505.5865783691406 0 1181.826
505.74310302734375 0 772.4247
505.9196472167969 0 1075.0358
510.2552795410156 0 741.616
510.92938232421875 0 11834.441 c 11
511.26300048828125 0 33539.848 c 3
511.59722900390625 0 7900.3604
511.935302734375 0 976.4117
512.265869140625 0 3938.0203
512.7565307617188 0 830.15533 y Water loss 8
513.234619140625 0 1726.2269
513.2714233398438 0 1248.9072
513.7564086914062 0 6755.7686 z 8
514.2567138671875 0 2101.4832
514.5822143554688 0 1065.0295
514.7586669921875 0 2051.5237
515.2459106445312 0 4733.0474
515.3226318359375 0 2454.5781
515.5789794921875 0 4048.686
515.91357421875 0 1651.1573
516.2462768554688 0 914.81683
516.3247680664062 0 694.31665
518.2886962890625 0 718.86426
521.7661743164062 0 6151.4375 y 8
522.2655639648438 0 1554.4508
522.7698364257812 0 856.86755
523.2577514648438 0 721.1008 w 3
527.2756958007812 0 885.74896
527.7767944335938 0 907.0449
528.2738037109375 0 840.20776
528.7828979492188 0 687.26227
533.7730102539062 0 726.5722
534.9365234375 0 7307.8657
535.2706298828125 0 8838.004
535.6060180664062 0 6012.0215
535.9384765625 0 3189.5835
537.2724609375 0 692.9608
542.2697143554688 0 4968.1123 z 7
542.7682495117188 0 2351.0488
543.2706298828125 0 1079.3387
546.2610473632812 0 881.7114
547.2803344726562 0 1841.2438 c Ammonia loss 8
549.6088256835938 0 1604.4403
550.2763671875 0 9709.603 y 7
550.7783813476562 0 5626.3315
551.2780151367188 0 1979.6075
554.2804565429688 0 750.8604
554.7860107421875 0 882.47107
555.267333984375 0 2172.0386
555.299560546875 0 1990.3512
555.7883911132812 0 45710.047 c 8
556.2900390625 0 27886.666
556.7908325195312 0 8337.6
557.2916870117188 0 2894.9927
561.2721557617188 0 1662.0226
561.59716796875 0 12537.587 w 2
561.9309692382812 0 9520.64
562.2649536132812 0 6224.8237
562.5958862304688 0 1298.8251
563.265869140625 0 743.95386
563.781494140625 0 2757.8281
564.2781982421875 0 2488.4824
564.6002197265625 0 7502.9907 c 12
564.7850952148438 0 872.0299
564.9345703125 0 6455.324
565.2682495117188 0 3175.5908
567.2290649414062 0 855.22705
568.2924194335938 0 948.55817
568.79541015625 0 1853.6019
569.302734375 0 764.3619
570.2735595703125 0 2186.5986
570.7807006835938 0 1115.443
571.2813720703125 0 9787.958
571.783203125 0 5235.5386
572.2817993164062 0 2742.813
572.7826538085938 0 948.759
573.2794799804688 0 771.45404
577.7830810546875 0 820.7702
578.2857666015625 0 800.64935
578.7871704101562 0 1862.514
579.2920532226562 0 1183.6096
580.611083984375 0 1545.2296 y Ammonia loss 2
580.9419555664062 0 1610.1633 z 2
581.2752685546875 0 1375.5504
581.6046752929688 0 854.23035
586.6131591796875 0 995.5482
586.7979736328125 0 774.3978
590.776611328125 0 948.55945 z Water loss 6
592.6298828125 0 1007.38464
592.95947265625 0 2548.3386
593.2969970703125 0 954.66595
597.2861938476562 0 867.2217
598.7944946289062 0 880.5757 y Water loss 6
599.2864990234375 0 2297.9006 y Ammonia loss 6
599.7820434570312 0 12677.38 z 6
600.2835083007812 0 16766.494
600.7852172851562 0 8616.397
601.2847290039062 0 1924.0421 c Water loss 13
601.6236572265625 0 766.1166 c Ammonia loss 13
602.2941284179688 0 1249.8396
602.6282958984375 0 842.50775
602.9490356445312 0 2002.7274
603.2823486328125 0 2653.7114
603.6151123046875 0 1470.352
603.953369140625 0 704.40295
606.3048095703125 0 10426.905 z 11
606.6337280273438 0 674.50836
606.967529296875 0 2107.0342
607.3002319335938 0 14375.996 c 13
607.6322021484375 0 11312.053
607.7904052734375 0 19472.193 y 6
607.9669799804688 0 4721.063
608.29296875 0 14893.146
608.635498046875 0 1361.1761
608.7943115234375 0 4390.1777
609.2982177734375 0 1881.9808
610.283203125 0 1641.2611
611.8040771484375 0 2492.3425 c Ammonia loss 9
611.962158203125 0 2689.3945
612.2987670898438 0 2962.9238
612.6295166015625 0 966.65283
612.8013305664062 0 1746.7614
613.2890625 0 768.01843
613.3345336914062 0 874.5724
615.3114624023438 0 892.8866
619.8156127929688 0 1579.4708
620.31103515625 0 18824.674 c 9
620.6353759765625 0 2837.7766 z Water loss 1
620.8120727539062 0 12077.233
620.9808959960938 0 1893.2003
621.2457275390625 0 1134.783
621.3134765625 0 7928.3145
621.639892578125 0 1475.6262
621.8141479492188 0 1347.872
621.9865112304688 0 1079.1263
622.3245239257812 0 6444.2 y 11
623.328369140625 0 1677.5802
625.6472778320312 0 751.9639
625.9820556640625 0 976.4113
626.3099975585938 0 2324.5803
626.631103515625 0 4698.002 z 1
626.9613037109375 0 7854.5186
627.2962646484375 0 3923.132
627.6267700195312 0 886.6021
630.6372680664062 0 1187.1311
630.9769287109375 0 1086.4446
631.3052368164062 0 720.92017
631.969482421875 0 891.0548 y 1
634.32470703125 0 2290.3235
634.79443359375 0 2380.838 w 5
635.3223876953125 0 1700.9242
635.6544799804688 0 1146.6699
636.301513671875 0 798.0334
640.321533203125 0 6930.016
640.6561279296875 0 7440.291
640.9908447265625 0 6730.3237
641.3237915039062 0 3500.5356
641.6574096679688 0 2042.4835
643.8271484375 0 1362.8423
644.3225708007812 0 1084.6973
644.653564453125 0 1231.4775
644.9895629882812 0 1765.9937
645.3222045898438 0 2215.7454
645.6551513671875 0 1489.0581
646.3154907226562 0 2751.0469
649.3228149414062 0 942.5266
649.6534423828125 0 1232.0773
649.976806640625 0 4426.3857
650.3231811523438 0 17238.53
650.6583251953125 0 19926.744
650.9931030273438 0 10909.149
651.3258056640625 0 6380.548
651.6609497070312 0 1122.1376
652.3193969726562 0 2332.1274
653.32080078125 0 1255.6042
654.3287353515625 0 948.1423
654.6629638671875 0 1116.445
654.9928588867188 0 2612.7104
655.3280639648438 0 2319.122 y Water loss 5
655.6593627929688 0 2219.286
655.9951171875 0 2289.5454
656.3232421875 0 3437.2083 z 5
656.8255004882812 0 3358.9795
657.3262329101562 0 1454.2539
658.6515502929688 0 909.8143
658.9876098632812 0 1173.4961
659.3244018554688 0 895.41614
659.9893798828125 0 1261.2565
662.324462890625 0 902.7194
663.8314208984375 0 979.5298
663.9876098632812 0 1275.1735
664.3301391601562 0 11905.199 y 5
664.6571655273438 0 24208.174
664.8335571289062 0 3484.5964
664.9908447265625 0 24087.482
665.3260498046875 0 18702.17
665.6581420898438 0 7514.34
665.9934692382812 0 3214.3557
666.3231811523438 0 1364.667
668.8212280273438 0 1594.0376
669.3251953125 0 1397.3777
669.661376953125 0 6881.722
669.995849609375 0 12195.069
670.3309936523438 0 18527.47
670.6651000976562 0 15001.013
670.9996948242188 0 6875.4463
671.3325805664062 0 3196.5723
671.6676025390625 0 893.3651
671.8238525390625 0 872.08417
679.3340454101562 0 1172.7839 c Water loss 4
679.8347778320312 0 5895.025
680.3204345703125 0 13986.691 c Ammonia loss 4
680.8388671875 0 2725.5474
681.3208618164062 0 6277.85
682.314208984375 0 1246.5613
690.849365234375 0 1238.0891
691.3372802734375 0 6705.217
691.8394165039062 0 2625.4456
692.3405151367188 0 2635.0227
693.8352661132812 0 1407.2859
697.3421020507812 0 8156.219 c 4
698.345947265625 0 3733.4958
699.3446655273438 0 1193.856
701.3407592773438 0 1481.456
701.842041015625 0 16111.6045 c 10
702.343505859375 0 16386.537
702.8458251953125 0 5088.9756
703.3466186523438 0 1413.4763
711.3076782226562 0 758.5922
714.3795166015625 0 2174.5625
715.3822631835938 0 995.2133
720.3590698242188 0 834.4625
720.8598022460938 0 833.4823
722.3579711914062 0 952.9599
726.8599243164062 0 1071.6449
731.4019165039062 0 1045.3478
737.3548583984375 0 1553.758
737.8499755859375 0 1985.0823
738.3576049804688 0 1153.8936
742.3488159179688 0 1085.9633
742.8535766601562 0 1361.2251
744.38330078125 0 1111
749.365234375 0 3649.604 z 4
749.866455078125 0 7308.274
750.3659057617188 0 4935.6055
750.8692016601562 0 3789.898
751.37646484375 0 983.679
757.3761596679688 0 2237.988 c Ammonia loss 11
757.876953125 0 1286.204
758.3784790039062 0 946.78357
765.4028930664062 0 1048.6302
765.8923950195312 0 2781.6475 c 11
766.3928833007812 0 4591.595
766.8936767578125 0 2256.4272
767.4098510742188 0 1088.2543
769.3681030273438 0 6017.1504 z 10
770.373046875 0 4073.5637
771.3729248046875 0 1516.265
772.375244140625 0 937.6347
777.8677368164062 0 1212.601
784.8826904296875 0 915.0358
785.3870239257812 0 2479.711 y 10
786.3782348632812 0 829.2231
787.3989868164062 0 1750.3733
793.3991088867188 0 10836.151 c Ammonia loss 5
794.3989868164062 0 8189.174
794.8943481445312 0 2080.5457
795.395751953125 0 3752.4497
796.4004516601562 0 1239.6825
802.9087524414062 0 1968.3142
803.4111938476562 0 2236.3389
803.906005859375 0 1151.451
806.3729248046875 0 1071.8767 y Ammonia loss 3
806.8767700195312 0 3723.7805 z 3
807.3801879882812 0 8175.3066
807.8810424804688 0 4785.1465
808.380859375 0 2809.2666
810.4254150390625 0 30889.512 c 5
811.4281616210938 0 13792.908
812.43212890625 0 4174.088
813.4351196289062 0 1005.267
817.3944091796875 0 1193.7975
826.8976440429688 0 1503.9528
840.4044799804688 0 1079.7649
841.4105224609375 0 1582.5792
841.9009399414062 0 1168.6089 w 2
842.4074096679688 0 2127.4785
846.3999633789062 0 2379.7205 c 12
846.8998413085938 0 3507.323
847.3986206054688 0 1694.2311
847.9005737304688 0 1006.0644
860.4310913085938 0 851.6976
867.42431640625 0 12002.634
868.4271240234375 0 7045.874
869.431396484375 0 1943.4989
870.909912109375 0 3649.4402 z 2
871.4099731445312 0 11240.712
871.9096069335938 0 7575.4272
872.41162109375 0 5171.0015
872.9116821289062 0 2079.0378
879.9313354492188 0 812.88617
880.434814453125 0 1090.6976
880.9274291992188 0 1664.1078
881.43896484375 0 2873.0376
881.9429931640625 0 1119.7633
888.4421997070312 0 912.5042
888.939453125 0 3922.9639
889.4432373046875 0 3540.466
889.9428100585938 0 1946.3159
895.4378051757812 0 857.04553
898.4111328125 0 6718.6064 z 9
899.41748046875 0 8462.109
900.4208984375 0 3532.449
901.420166015625 0 1253.0549
902.9451904296875 0 728.608
909.4434814453125 0 800.05884
910.9443969726562 0 1883.4459
922.9652709960938 0 719.248
925.4539794921875 0 2248.5986 c 6
926.4524536132812 0 1172.9601
930.9716796875 0 1028.0638
938.4607543945312 0 1521.578 y Water loss 1
938.96923828125 0 872.8547
939.4539184570312 0 3250.3486 z 1
939.9393920898438 0 8293.737
940.4421997070312 0 6287.8486
940.939453125 0 3657.1057
941.441650390625 0 897.69006
941.9483032226562 0 844.1721
952.4757690429688 0 1913.5507
952.976806640625 0 1762.37
953.4789428710938 0 853.6348
966.4622802734375 0 1543.7987
966.973876953125 0 775.37854
967.4745483398438 0 2998.9006
967.9788818359375 0 1991.3772
968.485595703125 0 1456.6084
974.981689453125 0 1238.4436
975.4848022460938 0 2335.7766
975.9873657226562 0 1424.5099
976.489013671875 0 2147.0706
976.9877319335938 0 1376.7782
981.4874877929688 0 830.0173
982.4754028320312 0 8904.692 c 7
983.479248046875 0 6802.109
984.483642578125 0 2587.36
985.4862060546875 0 1164.893
996.4821166992188 0 3228.1404
996.986328125 0 3815.9517
997.4825439453125 0 2721.7515
997.9844360351562 0 1309.4353
1004.99169921875 0 2033.3369
1005.49609375 0 2286.6436
1005.9982299804688 0 1058.7941
1006.4962158203125 0 918.58154
1026.505615234375 0 2381.3655 z 8
1027.512939453125 0 5645.591
1028.515625 0 2630.5664
1029.5240478515625 0 964.995
1084.5360107421875 0 1704.685
1094.5430908203125 0 1393.666
1095.556396484375 0 1755.8291
1110.5703125 0 6917.7344 c 8
1111.572998046875 0 6318.2627
1112.575927734375 0 4088.5588
1113.5810546875 0 1051.933
1142.5538330078125 0 785.3739
1196.603271484375 0 707.6481
1199.5606689453125 0 3121.6545
1200.56396484375 0 1677.0675
1224.603515625 0 852.74994
1239.60986328125 0 1555.2816 c 9
1240.61767578125 0 1960.5603
1241.616943359375 0 837.0815
1242.6209716796875 0 763.8782
1264.609375 0 763.22424
1312.6317138671875 0 1259.8213
1313.6390380859375 0 893.48645
1358.6510009765625 0 816.8003
1359.665283203125 0 1889.6074
1360.66748046875 0 1448.7689
1402.673583984375 0 1266.1676 c 10
1403.6810302734375 0 2814.7869
1404.6795654296875 0 1707.0471
1499.7381591796875 0 779.666
1614.7569580078125 0 843.18994
1712.006591796875 0 707.2722
1742.814697265625 0 1161.4268
1879.8953857421875 0 947.48114
2739.481201171875 0 692.7038
3046.811279296875 0 947.7079

Spectrum Details

|  |  |
| --- | --- |
| Matched peaks? Matched peaks The total absolute number of peaks matched. Additionally in brackets the total fraction of peaks matched and the total number of peaks is shown. | 97 (14.67% of 661) |
| FDR? FDR The false discovery rate estimated for this peptide. It is calculated by matching all theoretical fragments with a non-integer shift with the raw peaks for this spectrum. This is done with 40 different shifts. The resulting percentage is the average number of annotated peaks over the number of annotated peaks with the correct spectrum. | 0.83% |
| Satellite FDR? Satellite FDR See the FDR for details on its calculation. This satellite ion specific FDR only contains the satellite ions (d/w) for I/L/J positions. | 0.00% |
| PSM Score? PSM Score The PSM Score as given by Hecklib to this annotated spectrum. It is shown with three significant figures. | 245 |

#### Reverse Lookup? Reverse Lookup All places where this read could be placed.

| Group | Segment | Template | Template Part | Read Part | Score | Unique |
| --- | --- | --- | --- | --- | --- | --- |
| Homo sapiens Heavy Chain | IGHC | IGHG1 | [191..206] | [0..15] | 106 | False |
| Homo sapiens Heavy Chain | IGHC | IGHG3 | [238..253] | [0..15] | 106 | False |
| Homo sapiens Heavy Chain | IGHC | IGHG2 | [188..202] | [0..15] | 97 | False |
| Homo sapiens Heavy Chain | IGHC | IGHG4 | [188..203] | [0..15] | 106 | False |

| Recombined | Template Part | Read Part | Score | Unique |
| --- | --- | --- | --- | --- |
| REC-0-1 | [316..331] | [0..15] | 111 | True |

#### Meta Information from Multiple reads

##### Number of combined reads

11

##### Intensity

0.6636

##### TotalArea

1.992E+08

##### Changes to the peptide sequence

JHQDWLDGKEYKCKK

J→LSupport for Leucine based on side chain ions (2 for L 0 for I) (Position: 6)

L→JNo support for either Leucine or Isoleucine based on side chain ions (Position: 6)

J→LSupport for Leucine based on side chain ions (1 for L 0 for I) (Position: 6)

L→JNo support for either Leucine or Isoleucine based on side chain ions (Position: 6)

L→JNo support for either Leucine or Isoleucine based on side chain ions (Position: 1)

#### Positional Score

Copy Data

##### Positional Score (TSV)

###### Preview

```
Loading example...
```

*Click on the button to copy the data to your clipboard.*

1001234567891011121314

Label Value
"0" 0.815
"1" 0.795
"2" 0.753
"3" 0.781
"4" 0.787
"5" 0.807
"6" 0.808
"7" 0.802
"8" 0.805
"9" 0.811
"10" 0.808
"11" 0.769
"12" 0.761
"13" 0.785
"14" 0.761

#### Meta Information from PEAKS

##### Scan Identifier

F1:5173

##### Original sequence

L

H

Q

D

W

L

D

G

K

E

Y

K

C

+58.01

K

K

+58.01

##### Posttranslational Modifications

Carboxymethyl; Carboxymethyl (KW X@N-term)

##### Source File

D:\separate\_stitch\_analyses\xle-disambiguation\raw\20210323\_F1\_UM1\_Peng0013\_SA\_F59\_ingel\_3ug\_ELA.raw

##### Fraction

1

##### Scan Feature

F1:11461

##### De Novo Score

99

##### ConfidenceScore

99

### m/z

669.6603

##### Mass

2005.9568

##### Charge

3

##### Retention Time

26.79

##### Predicted Retention Time

-

##### Area

3.382E+07

##### Parts Per Million

1.1

##### Fragmentation mode

ETHCD

##### Originating file

01 D:\separate\_stitch\_analyses\xle-disambiguation\20210325\_F59\_3ug\_DENOVO\_12.csv

#### Meta Information from PEAKS

##### Scan Identifier

F1:5138

##### Original sequence

L

H

Q

D

W

L

D

G

K

E

Y

K

C

+58.01

K

K

+58.01

##### Posttranslational Modifications

Carboxymethyl; Carboxymethyl (KW X@N-term)

##### Source File

D:\separate\_stitch\_analyses\xle-disambiguation\raw\20210323\_F1\_UM1\_Peng0013\_SA\_F59\_ingel\_3ug\_ELA.raw

##### Fraction

1

##### Scan Feature

F1:4131

##### De Novo Score

98

##### ConfidenceScore

98

### m/z

502.4973

##### Mass

2005.9568

##### Charge

4

##### Retention Time

26.79

##### Predicted Retention Time

-

##### Area

5.82E+07

##### Parts Per Million

1.7

##### Fragmentation mode

ETHCD

##### Originating file

01 D:\separate\_stitch\_analyses\xle-disambiguation\20210325\_F59\_3ug\_DENOVO\_12.csv

#### Meta Information from PEAKS

##### Scan Identifier

F1:4177

##### Original sequence

L

H

Q

D

W

+15.99

L

D

G

K

E

Y

K

C

+58.01

K

K

+58.01

##### Posttranslational Modifications

Oxidation (HW); Carboxymethyl; Carboxymethyl (KW X@N-term)

##### Source File

D:\separate\_stitch\_analyses\xle-disambiguation\raw\20210323\_F1\_UM1\_Peng0013\_SA\_F59\_ingel\_3ug\_ELA.raw

##### Fraction

1

##### Scan Feature

F1:4302

##### De Novo Score

98

##### ConfidenceScore

98

### m/z

506.497

##### Mass

2021.9517

##### Charge

4

##### Retention Time

21.96

##### Predicted Retention Time

-

##### Area

3.62E+06

##### Parts Per Million

3.6

##### Fragmentation mode

ETHCD

##### Originating file

01 D:\separate\_stitch\_analyses\xle-disambiguation\20210325\_F59\_3ug\_DENOVO\_12.csv

#### Meta Information from PEAKS

##### Scan Identifier

F1:5273

##### Original sequence

L

H

Q

D

W

L

D

G

K

E

Y

K

C

+58.01

K

K

+58.01

##### Posttranslational Modifications

Carboxymethyl; Carboxymethyl (KW X@N-term)

##### Source File

D:\separate\_stitch\_analyses\xle-disambiguation\raw\20210323\_F1\_UM1\_Peng0013\_SA\_F59\_ingel\_3ug\_ELA.raw

##### Fraction

1

##### Scan Feature

F1:803

##### De Novo Score

97

##### ConfidenceScore

97

### m/z

402.199

##### Mass

2005.9568

##### Charge

5

##### Retention Time

27.6

##### Predicted Retention Time

-

##### Area

2.295E+06

##### Parts Per Million

0.9

##### Fragmentation mode

ETHCD

##### Originating file

01 D:\separate\_stitch\_analyses\xle-disambiguation\20210325\_F59\_3ug\_DENOVO\_12.csv

#### Meta Information from PEAKS

##### Scan Identifier

F1:5486

##### Original sequence

L

H

Q

D

W

L

D

G

K

E

Y

K

C

+58.01

K

K

+58.01

##### Posttranslational Modifications

Carboxymethyl; Carboxymethyl (KW X@N-term)

##### Source File

D:\separate\_stitch\_analyses\xle-disambiguation\raw\20210323\_F1\_UM1\_Peng0013\_SA\_F59\_ingel\_3ug\_ELA.raw

##### Fraction

1

##### Scan Feature

F1:4129

##### De Novo Score

97

##### ConfidenceScore

97

### m/z

502.4968

##### Mass

2005.9568

##### Charge

4

##### Retention Time

27.6

##### Predicted Retention Time

-

##### Area

4.602E+07

##### Parts Per Million

0.6

##### Fragmentation mode

ETHCD

##### Originating file

01 D:\separate\_stitch\_analyses\xle-disambiguation\20210325\_F59\_3ug\_DENOVO\_12.csv

#### Meta Information from PEAKS

##### Scan Identifier

F1:5469

##### Original sequence

L

H

Q

D

W

L

D

G

K

E

Y

K

C

+58.01

K

K

+58.01

##### Posttranslational Modifications

Carboxymethyl; Carboxymethyl (KW X@N-term)

##### Source File

D:\separate\_stitch\_analyses\xle-disambiguation\raw\20210323\_F1\_UM1\_Peng0013\_SA\_F59\_ingel\_3ug\_ELA.raw

##### Fraction

1

##### Scan Feature

F1:4129

##### De Novo Score

96

##### ConfidenceScore

96

### m/z

502.4968

##### Mass

2005.9568

##### Charge

4

##### Retention Time

27.6

##### Predicted Retention Time

-

##### Area

4.602E+07

##### Parts Per Million

0.6

##### Fragmentation mode

ETHCD

##### Originating file

01 D:\separate\_stitch\_analyses\xle-disambiguation\20210325\_F59\_3ug\_DENOVO\_12.csv

#### Meta Information from PEAKS

##### Scan Identifier

F1:5830

##### Original sequence

L

H

Q

D

W

L

D

G

K

E

Y

K

C

+58.01

K

K

+58.01

##### Posttranslational Modifications

Carboxymethyl; Carboxymethyl (KW X@N-term)

##### Source File

D:\separate\_stitch\_analyses\xle-disambiguation\raw\20210323\_F1\_UM1\_Peng0013\_SA\_F59\_ingel\_3ug\_ELA.raw

##### Fraction

1

##### Scan Feature

-

##### De Novo Score

96

##### ConfidenceScore

96

### m/z

502.4995

##### Mass

2005.9568

##### Charge

4

##### Retention Time

31.23

##### Predicted Retention Time

-

##### Area

0

##### Parts Per Million

6.1

##### Fragmentation mode

ETHCD

##### Originating file

01 D:\separate\_stitch\_analyses\xle-disambiguation\20210325\_F59\_3ug\_DENOVO\_12.csv

#### Meta Information from PEAKS

##### Scan Identifier

F1:5013

##### Original sequence

L

H

Q

D

W

L

D

G

K

E

Y

K

C

+58.01

K

K

+58.01

##### Posttranslational Modifications

Carboxymethyl; Carboxymethyl (KW X@N-term)

##### Source File

D:\separate\_stitch\_analyses\xle-disambiguation\raw\20210323\_F1\_UM1\_Peng0013\_SA\_F59\_ingel\_3ug\_ELA.raw

##### Fraction

1

##### Scan Feature

F1:804

##### De Novo Score

96

##### ConfidenceScore

96

### m/z

402.1993

##### Mass

2005.9568

##### Charge

5

##### Retention Time

26.74

##### Predicted Retention Time

-

##### Area

6.681E+06

##### Parts Per Million

1.6

##### Fragmentation mode

ETHCD

##### Originating file

01 D:\separate\_stitch\_analyses\xle-disambiguation\20210325\_F59\_3ug\_DENOVO\_12.csv

#### Meta Information from PEAKS

##### Scan Identifier

F1:5489

##### Original sequence

L

H

Q

D

W

L

D

G

K

E

Y

K

C

+58.01

K

K

+58.01

##### Posttranslational Modifications

Carboxymethyl; Carboxymethyl (KW X@N-term)

##### Source File

D:\separate\_stitch\_analyses\xle-disambiguation\raw\20210323\_F1\_UM1\_Peng0013\_SA\_F59\_ingel\_3ug\_ELA.raw

##### Fraction

1

##### Scan Feature

-

##### De Novo Score

96

##### ConfidenceScore

96

### m/z

669.6587

##### Mass

2005.9568

##### Charge

3

##### Retention Time

29.36

##### Predicted Retention Time

-

##### Area

0

##### Fragmentation mode

ETHCD

##### Originating file

01 D:\separate\_stitch\_analyses\xle-disambiguation\20210325\_F59\_3ug\_DENOVO\_12.csv

#### Meta Information from PEAKS

##### Scan Identifier

F1:4164

##### Original sequence

L

H

Q

D

W

+15.99

L

D

G

K

E

Y

K

C

+58.01

K

K

+58.01

##### Posttranslational Modifications

Oxidation (HW); Carboxymethyl; Carboxymethyl (KW X@N-term)

##### Source File

D:\separate\_stitch\_analyses\xle-disambiguation\raw\20210323\_F1\_UM1\_Peng0013\_SA\_F59\_ingel\_3ug\_ELA.raw

##### Fraction

1

##### Scan Feature

F1:897

##### De Novo Score

96

##### ConfidenceScore

96

### m/z

405.3983

##### Mass

2021.9517

##### Charge

5

##### Retention Time

21.96

##### Predicted Retention Time

-

##### Area

2.436E+05

##### Parts Per Million

1.8

##### Fragmentation mode

ETHCD

##### Originating file

01 D:\separate\_stitch\_analyses\xle-disambiguation\20210325\_F59\_3ug\_DENOVO\_12.csv

#### Meta Information from PEAKS

##### Scan Identifier

F1:5177

##### Original sequence

L

H

Q

D

W

L

D

G

K

E

Y

K

C

+58.01

K

K

+58.01

##### Posttranslational Modifications

Carboxymethyl; Carboxymethyl (KW X@N-term)

##### Source File

D:\separate\_stitch\_analyses\xle-disambiguation\raw\20210323\_F1\_UM1\_Peng0013\_SA\_F59\_ingel\_3ug\_ELA.raw

##### Fraction

1

##### Scan Feature

F1:803

##### De Novo Score

95

##### ConfidenceScore

95

### m/z

402.199

##### Mass

2005.9568

##### Charge

5

##### Retention Time

27.6

##### Predicted Retention Time

-

##### Area

2.295E+06

##### Parts Per Million

0.9

##### Fragmentation mode

ETHCD

##### Originating file

01 D:\separate\_stitch\_analyses\xle-disambiguation\20210325\_F59\_3ug\_DENOVO\_12.csv
