## Supplementary material for "A handle on mass coincidence errors in *de novo* sequencing of antibodies by bottom-up proteomics": Combined_057.html

Details Combined\_057 | Stitch OverviewUndefined

### Read Combined\_057

#### Sequence (length=11)

JVKDYYPEPVT

#### Spectrum 7672? Spectrum 7672 The raw spectrum of this peptide as annotated by Hecklib. The fragments are coloured according to ion type (see legend). Any peaks with a star '\*' as text can be hovered over to see the full details, first the ion type second the mass shift type. By hovering over the amino acids in the peptide or ions in the legend the corresponding peaks are highlighted. By toggling the 'Unassigned' label you can turn the background (unassigned) peaks on or off in the plot. By updating the slider in the Ion legend you can update the spectrum to only show the top X% of the peaks with labels. The top X% means any peak that is within X% of the highest intensity. By dragging in the spectrum you can zoom in to a specific part of the spectrum and use 'Zoom Out' to get back to the original zoom level. The annotation of the spectrum is based on the given sequence in the peptides file and is done with different software so inconsistencies are likely. The peaks are annotated based on the given sequence, with 20 ppm tolerance.

Copy Data

##### Spectrum 7672 (TSV)

###### Preview

```
Loading example...
```

*Click on the button to copy the data to your clipboard.*

Mz MinMz MaxIntensity Max

WidthHeightPeptide font sizePeptide stroke widthSpectrum font sizeSpectrum stroke widthCompact peptide

Ion legend

wxyz

abcd

OtherUnassignedIon^Charge^_Position_Show for top:%

JVKDYYPEPVT

01.89e+53.78e+55.67e+57.56e+5

Zoom Out

y^+1^_1_y^+1^_2_y^+1^_2_y^+1^_3_y^+1^_3_c^+1^_3_c^+1^_3_y^+1^_4_c^+2^_7_y^+1^_4_c^+1^_4_c^+1^_4_c^+2^_8_y^+1^_5_z^+2^_9_y^+1^_5_y^+2^_9_y^+2^_9_c^+2^_9_y^+2^_9_c^+2^_10_c^+2^_10_c^+1^_5_c^+1^_5_y^+1^_6_c^+1^_6_y^+1^_7_y^+1^_7_c^+1^_7_c^+1^_7_z^+1^_8_y^+1^_8_c^+1^_8_y^+1^_9_y^+1^_9_z^+1^_9_c^+1^_9_c^+1^_9_y^+1^_9_c^+1^_9_y^+1^_10_y^+1^_10_z^+1^_10_c^+1^_10_c^+1^_10_y^+1^_10_c^+1^_10_

0732146421972929

Fragment Matches Table

Show background peaks

| Position | Ion type | Intensity | mz Theoretical | mz Error (Th) | mz Error (ppm) | Charge | Series Number |
| --- | --- | --- | --- | --- | --- | --- | --- |
| 11 | y | 1.908E+04 | 120.1 | 0.0003678 | 3.064 | +1 | 1 |
| - | - | 1912 | 120.1 | - | - | 0 | - |
| - | - | 752.4 | 121.1 | - | - | 0 | - |
| - | - | 536.5 | 128.6 | - | - | 0 | - |
| - | - | 1.196E+04 | 129.1 | - | - | 0 | - |
| - | - | 608.3 | 130.1 | - | - | 0 | - |
| - | - | 389.4 | 138 | - | - | 0 | - |
| - | - | 613.9 | 149 | - | - | 0 | - |
| - | - | 415.8 | 150.8 | - | - | 0 | - |
| - | - | 4065 | 167.1 | - | - | 0 | - |
| - | - | 2.379E+04 | 169.1 | - | - | 0 | - |
| - | - | 1765 | 170.1 | - | - | 0 | - |
| - | - | 2788 | 173.5 | - | - | 0 | - |
| - | - | 742.4 | 181.1 | - | - | 0 | - |
| - | - | 532.4 | 185.1 | - | - | 0 | - |
| - | - | 3.608E+04 | 185.2 | - | - | 0 | - |
| - | - | 4248 | 186.2 | - | - | 0 | - |
| - | - | 6697 | 195.1 | - | - | 0 | - |
| - | - | 991.3 | 196.1 | - | - | 0 | - |
| - | - | 4.83E+04 | 197.1 | - | - | 0 | - |
| - | - | 6207 | 198.1 | - | - | 0 | - |
| - | - | 1207 | 199.1 | - | - | 0 | - |
| 10 | y | 757.2 | 201.1 | 0.0004562 | 2.268 | +1 | 2 |
| - | - | 450.1 | 201.2 | - | - | 0 | - |
| - | - | 485.5 | 205 | - | - | 0 | - |
| - | - | 821.5 | 211.1 | - | - | 0 | - |
| - | - | 2252 | 213.1 | - | - | 0 | - |
| - | - | 2.658E+04 | 213.2 | - | - | 0 | - |
| - | - | 3348 | 214.2 | - | - | 0 | - |
| - | - | 886.8 | 215.1 | - | - | 0 | - |
| 10 | y | 4572 | 219.1 | 0.0004354 | 1.987 | +1 | 2 |
| - | - | 806.9 | 225.1 | - | - | 0 | - |
| - | - | 1.539E+04 | 227.1 | - | - | 0 | - |
| - | - | 1135 | 228.1 | - | - | 0 | - |
| - | - | 923 | 228.2 | - | - | 0 | - |
| - | - | 735.3 | 229.2 | - | - | 0 | - |
| - | - | 1985 | 244.1 | - | - | 0 | - |
| - | - | 505.9 | 258.9 | - | - | 0 | - |
| - | - | 523.8 | 261.3 | - | - | 0 | - |
| - | - | 1363 | 270.2 | - | - | 0 | - |
| - | - | 553.9 | 271.1 | - | - | 0 | - |
| - | - | 819.8 | 279.1 | - | - | 0 | - |
| - | - | 1793 | 280.2 | - | - | 0 | - |
| - | - | 886.1 | 294.1 | - | - | 0 | - |
| - | - | 2389 | 296.2 | - | - | 0 | - |
| 9 | y | 1.225E+04 | 298.2 | 0.0007167 | 2.404 | +1 | 3 |
| - | - | 541.4 | 298.5 | - | - | 0 | - |
| - | - | 1963 | 299.2 | - | - | 0 | - |
| - | - | 1248 | 310.2 | - | - | 0 | - |
| - | - | 7007 | 312.2 | - | - | 0 | - |
| - | - | 763 | 313.2 | - | - | 0 | - |
| - | - | 5683 | 314.2 | - | - | 0 | - |
| - | - | 2916 | 315.3 | - | - | 0 | - |
| 9 | y | 1.559E+05 | 316.2 | 0.0008637 | 2.731 | +1 | 3 |
| - | - | 2.529E+04 | 317.2 | - | - | 0 | - |
| - | - | 3190 | 318.2 | - | - | 0 | - |
| - | - | 1483 | 322.1 | - | - | 0 | - |
| - | - | 660.1 | 323.1 | - | - | 0 | - |
| - | - | 2.205E+04 | 324.2 | - | - | 0 | - |
| - | - | 4175 | 325.2 | - | - | 0 | - |
| - | - | 840.5 | 329.1 | - | - | 0 | - |
| - | - | 4795 | 330.2 | - | - | 0 | - |
| - | - | 4894 | 332.2 | - | - | 0 | - |
| - | - | 980.9 | 333.2 | - | - | 0 | - |
| - | - | 1465 | 337.2 | - | - | 0 | - |
| - | - | 1559 | 338.1 | - | - | 0 | - |
| - | - | 984.1 | 340.2 | - | - | 0 | - |
| 3 | c | 4167 | 341.3 | 0.0006537 | 1.916 | +1 | 3 |
| - | - | 665.9 | 342.3 | - | - | 0 | - |
| - | - | 663.4 | 343.2 | - | - | 0 | - |
| - | - | 1.596E+04 | 357.3 | - | - | 0 | - |
| 3 | c | 8960 | 358.3 | 0.0007489 | 2.09 | +1 | 3 |
| - | - | 797.3 | 359.3 | - | - | 0 | - |
| - | - | 518.1 | 364.2 | - | - | 0 | - |
| - | - | 1979 | 374.2 | - | - | 0 | - |
| - | - | 571.2 | 374.3 | - | - | 0 | - |
| - | - | 692.4 | 375.2 | - | - | 0 | - |
| - | - | 1382 | 391.2 | - | - | 0 | - |
| - | - | 629.8 | 392.2 | - | - | 0 | - |
| - | - | 2518 | 395.2 | - | - | 0 | - |
| - | - | 1238 | 400.2 | - | - | 0 | - |
| - | - | 640.8 | 405.2 | - | - | 0 | - |
| - | - | 1969 | 407.2 | - | - | 0 | - |
| - | - | 2578 | 409.2 | - | - | 0 | - |
| - | - | 603.8 | 409.3 | - | - | 0 | - |
| - | - | 3053 | 421.2 | - | - | 0 | - |
| - | - | 744.3 | 422.2 | - | - | 0 | - |
| - | - | 2279 | 423.2 | - | - | 0 | - |
| - | - | 1.22E+04 | 423.2 | - | - | 0 | - |
| - | - | 735.3 | 424.2 | - | - | 0 | - |
| - | - | 2094 | 424.2 | - | - | 0 | - |
| - | - | 689.9 | 425.2 | - | - | 0 | - |
| - | - | 2051 | 426.2 | - | - | 0 | - |
| - | - | 831.9 | 426.2 | - | - | 0 | - |
| - | - | 942.4 | 426.7 | - | - | 0 | - |
| 8 | y | 1031 | 427.2 | 0.0008482 | 1.985 | +1 | 4 |
| - | - | 2588 | 430.3 | - | - | 0 | - |
| - | - | 1488 | 432.2 | - | - | 0 | - |
| - | - | 2906 | 437.2 | - | - | 0 | - |
| - | - | 750.6 | 438.3 | - | - | 0 | - |
| - | - | 1364 | 439.2 | - | - | 0 | - |
| - | - | 627.4 | 439.3 | - | - | 0 | - |
| 7 | c | 4347 | 440.2 | 0.0002877 | 0.6534 | +2 | 7 |
| - | - | 2016 | 440.7 | - | - | 0 | - |
| - | - | 855.2 | 441.2 | - | - | 0 | - |
| - | - | 2969 | 442.2 | - | - | 0 | - |
| - | - | 815.1 | 443.2 | - | - | 0 | - |
| 8 | y | 4161 | 445.2 | 0.001148 | 2.578 | +1 | 4 |
| - | - | 1348 | 446.2 | - | - | 0 | - |
| - | - | 1103 | 447.2 | - | - | 0 | - |
| - | - | 1030 | 452.3 | - | - | 0 | - |
| 4 | c | 9692 | 456.3 | 0.001238 | 2.712 | +1 | 4 |
| - | - | 2525 | 457.3 | - | - | 0 | - |
| - | - | 919.4 | 459.2 | - | - | 0 | - |
| - | - | 619.3 | 461.2 | - | - | 0 | - |
| - | - | 1331 | 471.2 | - | - | 0 | - |
| - | - | 4349 | 472.3 | - | - | 0 | - |
| 4 | c | 1.285E+04 | 473.3 | 0.001025 | 2.166 | +1 | 4 |
| - | - | 2379 | 474.3 | - | - | 0 | - |
| - | - | 962.1 | 483.3 | - | - | 0 | - |
| - | - | 708.3 | 488.3 | - | - | 0 | - |
| - | - | 684.6 | 489.3 | - | - | 0 | - |
| - | - | 698.8 | 489.8 | - | - | 0 | - |
| - | - | 578.2 | 490.3 | - | - | 0 | - |
| - | - | 1061 | 490.8 | - | - | 0 | - |
| - | - | 1603 | 491.3 | - | - | 0 | - |
| - | - | 815.7 | 491.8 | - | - | 0 | - |
| - | - | 716.3 | 494.7 | - | - | 0 | - |
| - | - | 859.3 | 495.2 | - | - | 0 | - |
| - | - | 1122 | 495.8 | - | - | 0 | - |
| - | - | 1129 | 496.8 | - | - | 0 | - |
| - | - | 1075 | 497.2 | - | - | 0 | - |
| - | - | 4130 | 503.7 | - | - | 0 | - |
| - | - | 3574 | 504.3 | - | - | 0 | - |
| 8 | c | 1.017E+04 | 504.8 | 0.0008722 | 1.728 | +2 | 8 |
| - | - | 6633 | 505.3 | - | - | 0 | - |
| - | - | 1916 | 505.8 | - | - | 0 | - |
| - | - | 1409 | 506.3 | - | - | 0 | - |
| - | - | 576.9 | 509.2 | - | - | 0 | - |
| - | - | 750.1 | 522.3 | - | - | 0 | - |
| 7 | y | 5057 | 524.3 | 0.001826 | 3.483 | +1 | 5 |
| - | - | 1285 | 525.3 | - | - | 0 | - |
| - | - | 625.9 | 529.3 | - | - | 0 | - |
| - | - | 930.6 | 529.8 | - | - | 0 | - |
| - | - | 1132 | 530.3 | - | - | 0 | - |
| - | - | 1825 | 530.8 | - | - | 0 | - |
| - | - | 640.3 | 531.8 | - | - | 0 | - |
| - | - | 1095 | 534.3 | - | - | 0 | - |
| - | - | 1060 | 536.3 | - | - | 0 | - |
| - | - | 2137 | 536.3 | - | - | 0 | - |
| - | - | 3839 | 538.3 | - | - | 0 | - |
| 3 | z | 1578 | 539.3 | 0.003387 | 6.281 | +2 | 9 |
| - | - | 1845 | 539.3 | - | - | 0 | - |
| - | - | 2000 | 539.8 | - | - | 0 | - |
| - | - | 3448 | 540.3 | - | - | 0 | - |
| - | - | 1713 | 541.3 | - | - | 0 | - |
| 7 | y | 6.373E+04 | 542.3 | 0.00121 | 2.231 | +1 | 5 |
| - | - | 1.794E+04 | 543.3 | - | - | 0 | - |
| - | - | 1636 | 543.8 | - | - | 0 | - |
| - | - | 4726 | 544.3 | - | - | 0 | - |
| - | - | 1662 | 544.8 | - | - | 0 | - |
| - | - | 2050 | 545.3 | - | - | 0 | - |
| - | - | 1020 | 545.8 | - | - | 0 | - |
| 3 | y | 1916 | 547.3 | 0.0005566 | 1.017 | +2 | 9 |
| 3 | y | 788.7 | 547.8 | 0.009525 | 17.39 | +2 | 9 |
| - | - | 977.4 | 550.3 | - | - | 0 | - |
| - | - | 1.281E+04 | 552.3 | - | - | 0 | - |
| - | - | 588.3 | 552.7 | - | - | 0 | - |
| - | - | 1.103E+04 | 552.8 | - | - | 0 | - |
| 9 | c | 4.266E+04 | 553.3 | 0.0007964 | 1.439 | +2 | 9 |
| - | - | 2.514E+04 | 553.8 | - | - | 0 | - |
| - | - | 6468 | 554.3 | - | - | 0 | - |
| - | - | 5870 | 554.3 | - | - | 0 | - |
| - | - | 856.5 | 554.8 | - | - | 0 | - |
| - | - | 3966 | 555.3 | - | - | 0 | - |
| 3 | y | 1.767E+04 | 556.3 | 0.006374 | 11.46 | +2 | 9 |
| - | - | 5702 | 557.3 | - | - | 0 | - |
| - | - | 5313 | 558.3 | - | - | 0 | - |
| - | - | 852.8 | 559.3 | - | - | 0 | - |
| - | - | 602.2 | 561 | - | - | 0 | - |
| - | - | 1769 | 567.4 | - | - | 0 | - |
| - | - | 1705 | 568.2 | - | - | 0 | - |
| - | - | 1.498E+04 | 570.3 | - | - | 0 | - |
| - | - | 4794 | 571.3 | - | - | 0 | - |
| - | - | 998.7 | 572.3 | - | - | 0 | - |
| - | - | 1801 | 578.8 | - | - | 0 | - |
| - | - | 881.8 | 579.3 | - | - | 0 | - |
| - | - | 2750 | 579.8 | - | - | 0 | - |
| - | - | 741.4 | 580.3 | - | - | 0 | - |
| - | - | 1735 | 580.3 | - | - | 0 | - |
| - | - | 941.3 | 580.8 | - | - | 0 | - |
| - | - | 839.8 | 584.3 | - | - | 0 | - |
| - | - | 5588 | 587.8 | - | - | 0 | - |
| - | - | 4385 | 588.3 | - | - | 0 | - |
| - | - | 1.362E+04 | 588.8 | - | - | 0 | - |
| - | - | 1.229E+04 | 589.3 | - | - | 0 | - |
| - | - | 4152 | 589.8 | - | - | 0 | - |
| - | - | 3256 | 591.4 | - | - | 0 | - |
| - | - | 1524 | 592.4 | - | - | 0 | - |
| - | - | 719.2 | 593.3 | - | - | 0 | - |
| - | - | 4190 | 593.8 | - | - | 0 | - |
| - | - | 1705 | 594.3 | - | - | 0 | - |
| - | - | 3856 | 601.8 | - | - | 0 | - |
| 10 | c | 3244 | 602.3 | 0.0115 | 19.09 | +2 | 10 |
| 10 | c | 1.284E+04 | 602.8 | 0.001135 | 1.883 | +2 | 10 |
| - | - | 8416 | 603.3 | - | - | 0 | - |
| - | - | 3689 | 603.8 | - | - | 0 | - |
| - | - | 2678 | 607.3 | - | - | 0 | - |
| - | - | 1471 | 615.3 | - | - | 0 | - |
| - | - | 2207 | 617.3 | - | - | 0 | - |
| 5 | c | 1.364E+04 | 619.3 | 0.001569 | 2.533 | +1 | 5 |
| - | - | 4299 | 620.3 | - | - | 0 | - |
| - | - | 1540 | 621.4 | - | - | 0 | - |
| - | - | 713.9 | 622.4 | - | - | 0 | - |
| - | - | 1644 | 631.3 | - | - | 0 | - |
| - | - | 1623 | 632.3 | - | - | 0 | - |
| - | - | 1771 | 633.3 | - | - | 0 | - |
| - | - | 1.131E+04 | 635.4 | - | - | 0 | - |
| - | - | 1.18E+04 | 635.4 | - | - | 0 | - |
| 5 | c | 2.562E+04 | 636.4 | 4.399E-05 | 0.06913 | +1 | 5 |
| - | - | 8151 | 637.4 | - | - | 0 | - |
| - | - | 1939 | 638.4 | - | - | 0 | - |
| - | - | 765 | 638.8 | - | - | 0 | - |
| - | - | 993.1 | 644.3 | - | - | 0 | - |
| - | - | 1018 | 644.8 | - | - | 0 | - |
| - | - | 3466 | 646.4 | - | - | 0 | - |
| - | - | 1647 | 649.3 | - | - | 0 | - |
| - | - | 2244 | 650.3 | - | - | 0 | - |
| - | - | 3.238E+04 | 651.4 | - | - | 0 | - |
| - | - | 1.497E+04 | 652.4 | - | - | 0 | - |
| - | - | 1807 | 652.8 | - | - | 0 | - |
| - | - | 9018 | 653.3 | - | - | 0 | - |
| - | - | 6529 | 653.8 | - | - | 0 | - |
| - | - | 2023 | 654.3 | - | - | 0 | - |
| - | - | 3149 | 661.3 | - | - | 0 | - |
| - | - | 2442 | 661.8 | - | - | 0 | - |
| - | - | 1.384E+04 | 662.3 | - | - | 0 | - |
| - | - | 637.2 | 662.4 | - | - | 0 | - |
| - | - | 9702 | 662.8 | - | - | 0 | - |
| - | - | 4407 | 663.3 | - | - | 0 | - |
| - | - | 867.4 | 663.9 | - | - | 0 | - |
| - | - | 1823 | 667.3 | - | - | 0 | - |
| - | - | 1847 | 669.3 | - | - | 0 | - |
| - | - | 828.9 | 670.3 | - | - | 0 | - |
| - | - | 1065 | 671.3 | - | - | 0 | - |
| - | - | 1048 | 679.3 | - | - | 0 | - |
| - | - | 1198 | 681.3 | - | - | 0 | - |
| - | - | 3428 | 682.4 | - | - | 0 | - |
| - | - | 1209 | 683.4 | - | - | 0 | - |
| - | - | 1110 | 685.3 | - | - | 0 | - |
| - | - | 1304 | 688.3 | - | - | 0 | - |
| - | - | 2.518E+04 | 689.4 | - | - | 0 | - |
| - | - | 1.012E+04 | 690.4 | - | - | 0 | - |
| - | - | 1266 | 691.4 | - | - | 0 | - |
| - | - | 1906 | 697.3 | - | - | 0 | - |
| - | - | 1372 | 698.3 | - | - | 0 | - |
| - | - | 5580 | 703.3 | - | - | 0 | - |
| - | - | 2601 | 704.3 | - | - | 0 | - |
| 6 | y | 9430 | 705.3 | 0.0006865 | 0.9732 | +1 | 6 |
| - | - | 2756 | 706.4 | - | - | 0 | - |
| - | - | 631.3 | 707.4 | - | - | 0 | - |
| - | - | 813.1 | 714.4 | - | - | 0 | - |
| - | - | 724 | 735.4 | - | - | 0 | - |
| - | - | 1384 | 736.4 | - | - | 0 | - |
| - | - | 677.9 | 737.4 | - | - | 0 | - |
| - | - | 3266 | 738.4 | - | - | 0 | - |
| - | - | 7695 | 739.4 | - | - | 0 | - |
| - | - | 6299 | 740.4 | - | - | 0 | - |
| - | - | 3517 | 741.4 | - | - | 0 | - |
| - | - | 1270 | 753.4 | - | - | 0 | - |
| - | - | 6104 | 754.4 | - | - | 0 | - |
| - | - | 1.502E+04 | 755.4 | - | - | 0 | - |
| - | - | 1.452E+04 | 756.4 | - | - | 0 | - |
| - | - | 5682 | 757.4 | - | - | 0 | - |
| - | - | 645.1 | 758.3 | - | - | 0 | - |
| - | - | 6317 | 762.4 | - | - | 0 | - |
| - | - | 2282 | 763.4 | - | - | 0 | - |
| - | - | 6151 | 764.4 | - | - | 0 | - |
| - | - | 1887 | 765.4 | - | - | 0 | - |
| - | - | 3.533E+04 | 766.4 | - | - | 0 | - |
| - | - | 1.727E+04 | 767.4 | - | - | 0 | - |
| - | - | 4956 | 768.4 | - | - | 0 | - |
| - | - | 682.9 | 769.4 | - | - | 0 | - |
| - | - | 1531 | 778.3 | - | - | 0 | - |
| - | - | 4850 | 780.4 | - | - | 0 | - |
| - | - | 2766 | 781.4 | - | - | 0 | - |
| 6 | c | 6.74E+04 | 782.4 | 0.001534 | 1.96 | +1 | 6 |
| - | - | 3.327E+04 | 783.4 | - | - | 0 | - |
| - | - | 8476 | 784.4 | - | - | 0 | - |
| - | - | 1519 | 794.3 | - | - | 0 | - |
| - | - | 963.2 | 795.3 | - | - | 0 | - |
| - | - | 1.646E+04 | 796.4 | - | - | 0 | - |
| - | - | 5808 | 797.4 | - | - | 0 | - |
| - | - | 1678 | 798.4 | - | - | 0 | - |
| - | - | 792.3 | 829.4 | - | - | 0 | - |
| - | - | 993.8 | 839.4 | - | - | 0 | - |
| - | - | 948.7 | 840.4 | - | - | 0 | - |
| 5 | y | 751 | 850.4 | 0.01417 | 16.67 | +1 | 7 |
| - | - | 839.6 | 851.4 | - | - | 0 | - |
| - | - | 2689 | 852.4 | - | - | 0 | - |
| - | - | 1806 | 853.4 | - | - | 0 | - |
| - | - | 733.2 | 859.4 | - | - | 0 | - |
| - | - | 1955 | 863.5 | - | - | 0 | - |
| - | - | 754 | 864.4 | - | - | 0 | - |
| - | - | 757.4 | 864.5 | - | - | 0 | - |
| - | - | 910.7 | 865.4 | - | - | 0 | - |
| - | - | 1875 | 865.5 | - | - | 0 | - |
| - | - | 2630 | 866.4 | - | - | 0 | - |
| - | - | 9380 | 867.4 | - | - | 0 | - |
| 5 | y | 1.322E+04 | 868.4 | 0.0002031 | 0.2339 | +1 | 7 |
| - | - | 5270 | 869.4 | - | - | 0 | - |
| - | - | 1047 | 870.4 | - | - | 0 | - |
| - | - | 2320 | 875.5 | - | - | 0 | - |
| - | - | 2317 | 876.5 | - | - | 0 | - |
| - | - | 1110 | 877.5 | - | - | 0 | - |
| 7 | c | 1457 | 878.5 | 0.001328 | 1.511 | +1 | 7 |
| - | - | 3.073E+04 | 879.5 | - | - | 0 | - |
| - | - | 4.286E+04 | 880.5 | - | - | 0 | - |
| - | - | 2.094E+04 | 881.5 | - | - | 0 | - |
| - | - | 5918 | 882.5 | - | - | 0 | - |
| - | - | 1117 | 883.5 | - | - | 0 | - |
| - | - | 3078 | 891.4 | - | - | 0 | - |
| - | - | 1353 | 892.4 | - | - | 0 | - |
| - | - | 5137 | 893.4 | - | - | 0 | - |
| - | - | 4558 | 893.5 | - | - | 0 | - |
| - | - | 1.723E+04 | 894.5 | - | - | 0 | - |
| - | - | 7.994E+04 | 895.5 | - | - | 0 | - |
| 7 | c | 9.784E+04 | 896.5 | 0.000692 | 0.7719 | +1 | 7 |
| - | - | 4.003E+04 | 897.5 | - | - | 0 | - |
| - | - | 9524 | 898.5 | - | - | 0 | - |
| - | - | 790.4 | 905.4 | - | - | 0 | - |
| - | - | 1354 | 923.4 | - | - | 0 | - |
| - | - | 733.4 | 950.5 | - | - | 0 | - |
| - | - | 926.3 | 960.5 | - | - | 0 | - |
| - | - | 782.5 | 961.5 | - | - | 0 | - |
| - | - | 1576 | 962.5 | - | - | 0 | - |
| - | - | 891.2 | 963.5 | - | - | 0 | - |
| - | - | 1159 | 964.5 | - | - | 0 | - |
| - | - | 1174 | 965.5 | - | - | 0 | - |
| - | - | 932.3 | 966.5 | - | - | 0 | - |
| 4 | z | 2209 | 967.4 | 0.01709 | 17.67 | +1 | 8 |
| - | - | 2938 | 968.4 | - | - | 0 | - |
| - | - | 1191 | 969.4 | - | - | 0 | - |
| - | - | 1042 | 972.5 | - | - | 0 | - |
| - | - | 952.6 | 973.5 | - | - | 0 | - |
| - | - | 1513 | 974.5 | - | - | 0 | - |
| - | - | 725.9 | 975.5 | - | - | 0 | - |
| - | - | 2007 | 978.5 | - | - | 0 | - |
| - | - | 1573 | 979.5 | - | - | 0 | - |
| - | - | 4716 | 980.5 | - | - | 0 | - |
| - | - | 1463 | 981.4 | - | - | 0 | - |
| - | - | 7394 | 981.5 | - | - | 0 | - |
| - | - | 1.011E+04 | 982.4 | - | - | 0 | - |
| - | - | 2119 | 982.5 | - | - | 0 | - |
| 4 | y | 8214 | 983.4 | 0.001328 | 1.351 | +1 | 8 |
| - | - | 887.9 | 983.5 | - | - | 0 | - |
| - | - | 4254 | 984.4 | - | - | 0 | - |
| - | - | 802.8 | 985.4 | - | - | 0 | - |
| - | - | 8793 | 988.5 | - | - | 0 | - |
| - | - | 5579 | 989.5 | - | - | 0 | - |
| - | - | 9510 | 990.5 | - | - | 0 | - |
| - | - | 5388 | 991.5 | - | - | 0 | - |
| - | - | 2.644E+04 | 992.5 | - | - | 0 | - |
| - | - | 1.395E+04 | 993.5 | - | - | 0 | - |
| - | - | 6438 | 994.5 | - | - | 0 | - |
| - | - | 989.8 | 995.5 | - | - | 0 | - |
| - | - | 1.871E+04 | 1006 | - | - | 0 | - |
| - | - | 1.486E+04 | 1007 | - | - | 0 | - |
| 8 | c | 1.047E+05 | 1009 | 0.001636 | 1.622 | +1 | 8 |
| - | - | 6.221E+04 | 1010 | - | - | 0 | - |
| - | - | 2.105E+04 | 1011 | - | - | 0 | - |
| - | - | 2291 | 1012 | - | - | 0 | - |
| - | - | 1475 | 1023 | - | - | 0 | - |
| - | - | 981.2 | 1024 | - | - | 0 | - |
| - | - | 898.4 | 1038 | - | - | 0 | - |
| - | - | 2500 | 1050 | - | - | 0 | - |
| - | - | 2804 | 1051 | - | - | 0 | - |
| - | - | 4736 | 1052 | - | - | 0 | - |
| - | - | 3049 | 1053 | - | - | 0 | - |
| - | - | 1030 | 1054 | - | - | 0 | - |
| - | - | 1028 | 1062 | - | - | 0 | - |
| - | - | 1327 | 1063 | - | - | 0 | - |
| - | - | 911.8 | 1064 | - | - | 0 | - |
| - | - | 1077 | 1076 | - | - | 0 | - |
| - | - | 1196 | 1077 | - | - | 0 | - |
| - | - | 1533 | 1078 | - | - | 0 | - |
| - | - | 1171 | 1079 | - | - | 0 | - |
| - | - | 3334 | 1080 | - | - | 0 | - |
| - | - | 2607 | 1081 | - | - | 0 | - |
| - | - | 1023 | 1082 | - | - | 0 | - |
| - | - | 1003 | 1091 | - | - | 0 | - |
| - | - | 2391 | 1092 | - | - | 0 | - |
| - | - | 1871 | 1093 | - | - | 0 | - |
| 3 | y | 9727 | 1094 | 0.01676 | 15.32 | +1 | 9 |
| 3 | y | 9206 | 1095 | 0.002401 | 2.194 | +1 | 9 |
| 3 | z | 3.775E+04 | 1096 | 0.002389 | 2.181 | +1 | 9 |
| - | - | 2.599E+04 | 1097 | - | - | 0 | - |
| - | - | 1.008E+04 | 1098 | - | - | 0 | - |
| - | - | 1787 | 1099 | - | - | 0 | - |
| - | - | 1129 | 1104 | - | - | 0 | - |
| 9 | c | 1447 | 1105 | 0.01664 | 15.06 | +1 | 9 |
| 9 | c | 2429 | 1106 | 0.01473 | 13.32 | +1 | 9 |
| - | - | 3081 | 1107 | - | - | 0 | - |
| - | - | 1921 | 1108 | - | - | 0 | - |
| - | - | 9006 | 1110 | - | - | 0 | - |
| - | - | 7089 | 1111 | - | - | 0 | - |
| 3 | y | 2.613E+04 | 1112 | 0.001609 | 1.448 | +1 | 9 |
| - | - | 1.704E+04 | 1113 | - | - | 0 | - |
| - | - | 4967 | 1114 | - | - | 0 | - |
| - | - | 5341 | 1121 | - | - | 0 | - |
| - | - | 4009 | 1122 | - | - | 0 | - |
| 9 | c | 1.946E+04 | 1123 | 0.0006308 | 0.5619 | +1 | 9 |
| - | - | 1.193E+04 | 1124 | - | - | 0 | - |
| - | - | 3904 | 1125 | - | - | 0 | - |
| - | - | 717.2 | 1132 | - | - | 0 | - |
| - | - | 888.8 | 1134 | - | - | 0 | - |
| - | - | 1625 | 1141 | - | - | 0 | - |
| - | - | 1042 | 1151 | - | - | 0 | - |
| - | - | 2303 | 1160 | - | - | 0 | - |
| - | - | 5243 | 1161 | - | - | 0 | - |
| - | - | 3536 | 1162 | - | - | 0 | - |
| - | - | 4229 | 1163 | - | - | 0 | - |
| - | - | 4662 | 1164 | - | - | 0 | - |
| - | - | 1819 | 1165 | - | - | 0 | - |
| - | - | 1751 | 1175 | - | - | 0 | - |
| - | - | 8908 | 1176 | - | - | 0 | - |
| - | - | 1.164E+04 | 1177 | - | - | 0 | - |
| - | - | 4.148E+04 | 1178 | - | - | 0 | - |
| - | - | 3.233E+04 | 1179 | - | - | 0 | - |
| - | - | 1.453E+04 | 1180 | - | - | 0 | - |
| - | - | 3147 | 1181 | - | - | 0 | - |
| - | - | 1604 | 1185 | - | - | 0 | - |
| - | - | 2442 | 1187 | - | - | 0 | - |
| - | - | 1844 | 1188 | - | - | 0 | - |
| - | - | 2052 | 1189 | - | - | 0 | - |
| - | - | 1109 | 1190 | - | - | 0 | - |
| 2 | y | 3338 | 1193 | 0.02072 | 17.37 | +1 | 10 |
| 2 | y | 4076 | 1194 | 0.002103 | 1.762 | +1 | 10 |
| 2 | z | 1.695E+04 | 1195 | 0.001602 | 1.341 | +1 | 10 |
| - | - | 1.073E+04 | 1196 | - | - | 0 | - |
| - | - | 4312 | 1197 | - | - | 0 | - |
| - | - | 1589 | 1201 | - | - | 0 | - |
| - | - | 3232 | 1202 | - | - | 0 | - |
| - | - | 8943 | 1203 | - | - | 0 | - |
| 10 | c | 5931 | 1204 | 0.01437 | 11.94 | +1 | 10 |
| 10 | c | 1.629E+04 | 1205 | 0.001613 | 1.339 | +1 | 10 |
| - | - | 1.246E+04 | 1206 | - | - | 0 | - |
| - | - | 1.146E+04 | 1207 | - | - | 0 | - |
| - | - | 7085 | 1208 | - | - | 0 | - |
| - | - | 2131 | 1209 | - | - | 0 | - |
| - | - | 1524 | 1210 | - | - | 0 | - |
| 2 | y | 2587 | 1211 | 0.0006427 | 0.5309 | +1 | 10 |
| - | - | 2023 | 1212 | - | - | 0 | - |
| - | - | 1611 | 1216 | - | - | 0 | - |
| - | - | 1819 | 1217 | - | - | 0 | - |
| - | - | 1601 | 1218 | - | - | 0 | - |
| - | - | 998.5 | 1219 | - | - | 0 | - |
| - | - | 4.303E+04 | 1220 | - | - | 0 | - |
| - | - | 3.274E+04 | 1221 | - | - | 0 | - |
| 10 | c | 1.563E+05 | 1222 | 3.411E-05 | 0.02792 | +1 | 10 |
| - | - | 1.133E+05 | 1223 | - | - | 0 | - |
| - | - | 5.256E+04 | 1224 | - | - | 0 | - |
| - | - | 1.582E+04 | 1225 | - | - | 0 | - |
| - | - | 3016 | 1226 | - | - | 0 | - |
| - | - | 890 | 1231 | - | - | 0 | - |
| - | - | 4635 | 1232 | - | - | 0 | - |
| - | - | 3954 | 1233 | - | - | 0 | - |
| - | - | 3074 | 1234 | - | - | 0 | - |
| - | - | 3653 | 1235 | - | - | 0 | - |
| - | - | 1.105E+04 | 1236 | - | - | 0 | - |
| - | - | 7554 | 1237 | - | - | 0 | - |
| - | - | 4111 | 1238 | - | - | 0 | - |
| - | - | 1539 | 1239 | - | - | 0 | - |
| - | - | 992.9 | 1240 | - | - | 0 | - |
| - | - | 2176 | 1245 | - | - | 0 | - |
| - | - | 2495 | 1246 | - | - | 0 | - |
| - | - | 2970 | 1247 | - | - | 0 | - |
| - | - | 1637 | 1248 | - | - | 0 | - |
| - | - | 6137 | 1249 | - | - | 0 | - |
| - | - | 3823 | 1250 | - | - | 0 | - |
| - | - | 5535 | 1251 | - | - | 0 | - |
| - | - | 4393 | 1252 | - | - | 0 | - |
| - | - | 7614 | 1253 | - | - | 0 | - |
| - | - | 5606 | 1254 | - | - | 0 | - |
| - | - | 2693 | 1255 | - | - | 0 | - |
| - | - | 1172 | 1259 | - | - | 0 | - |
| - | - | 1993 | 1260 | - | - | 0 | - |
| - | - | 1.106E+04 | 1261 | - | - | 0 | - |
| - | - | 1.074E+04 | 1262 | - | - | 0 | - |
| - | - | 1.548E+04 | 1263 | - | - | 0 | - |
| - | - | 1.275E+04 | 1264 | - | - | 0 | - |
| - | - | 5.361E+04 | 1265 | - | - | 0 | - |
| - | - | 4.158E+04 | 1266 | - | - | 0 | - |
| - | - | 2.05E+04 | 1267 | - | - | 0 | - |
| - | - | 5623 | 1268 | - | - | 0 | - |
| - | - | 6153 | 1269 | - | - | 0 | - |
| - | - | 3292 | 1270 | - | - | 0 | - |
| - | - | 1722 | 1271 | - | - | 0 | - |
| - | - | 889.1 | 1272 | - | - | 0 | - |
| - | - | 2856 | 1277 | - | - | 0 | - |
| - | - | 9016 | 1278 | - | - | 0 | - |
| - | - | 2.597E+04 | 1279 | - | - | 0 | - |
| - | - | 2.052E+04 | 1280 | - | - | 0 | - |
| - | - | 2.113E+04 | 1281 | - | - | 0 | - |
| - | - | 1.113E+04 | 1282 | - | - | 0 | - |
| - | - | 4347 | 1283 | - | - | 0 | - |
| - | - | 877.4 | 1284 | - | - | 0 | - |
| - | - | 2064 | 1288 | - | - | 0 | - |
| - | - | 3487 | 1289 | - | - | 0 | - |
| - | - | 4503 | 1290 | - | - | 0 | - |
| - | - | 3015 | 1291 | - | - | 0 | - |
| - | - | 2378 | 1292 | - | - | 0 | - |
| - | - | 1618 | 1293 | - | - | 0 | - |
| - | - | 2003 | 1294 | - | - | 0 | - |
| - | - | 1.487E+04 | 1295 | - | - | 0 | - |
| - | - | 1.043E+04 | 1296 | - | - | 0 | - |
| - | - | 4.103E+04 | 1297 | - | - | 0 | - |
| - | - | 3.41E+04 | 1298 | - | - | 0 | - |
| - | - | 1.3E+04 | 1299 | - | - | 0 | - |
| - | - | 1871 | 1300 | - | - | 0 | - |
| - | - | 4106 | 1304 | - | - | 0 | - |
| - | - | 3.074E+04 | 1305 | - | - | 0 | - |
| - | - | 7.287E+04 | 1306 | - | - | 0 | - |
| - | - | 1.692E+05 | 1307 | - | - | 0 | - |
| - | - | 3.174E+05 | 1308 | - | - | 0 | - |
| - | - | 2.112E+05 | 1309 | - | - | 0 | - |
| - | - | 8.068E+04 | 1310 | - | - | 0 | - |
| - | - | 1.295E+04 | 1311 | - | - | 0 | - |
| - | - | 754.7 | 1321 | - | - | 0 | - |
| - | - | 4.521E+04 | 1322 | - | - | 0 | - |
| - | - | 2.803E+05 | 1323 | - | - | 0 | - |
| - | - | 3.932E+05 | 1324 | - | - | 0 | - |
| - | - | 7.484E+05 | 1325 | - | - | 0 | - |
| - | - | 5.194E+05 | 1326 | - | - | 0 | - |
| - | - | 2.107E+05 | 1327 | - | - | 0 | - |
| - | - | 2.836E+04 | 1328 | - | - | 0 | - |
| - | - | 1357 | 1340 | - | - | 0 | - |
| - | - | 842.8 | 1341 | - | - | 0 | - |
| - | - | 1041 | 1357 | - | - | 0 | - |
| - | - | 1392 | 1358 | - | - | 0 | - |
| - | - | 695.5 | 2900 | - | - | 0 | - |

m/z Charge Intensity FragmentType MassShift Position
120.06588745117188 0 19079.217 y 10
120.0811538696289 0 1912.0985
121.06912994384766 0 752.35065
128.58213806152344 0 536.5045
129.1025848388672 0 11957.271
130.1062469482422 0 608.27496
137.98143005371094 0 389.38403
148.95492553710938 0 613.9414
150.75450134277344 0 415.76797
167.11839294433594 0 4064.6912
169.1339569091797 0 23792.084
170.13734436035156 0 1764.8479
173.4513397216797 0 2788.3364
181.13360595703125 0 742.40656
185.12887573242188 0 532.3705
185.16529846191406 0 36081.18
186.16879272460938 0 4248.209
195.11326599121094 0 6697.359
196.11627197265625 0 991.2816
197.12890625 0 48298.477
198.13230895996094 0 6206.7075
199.10804748535156 0 1207.313
201.1238250732422 0 757.2164 y Water loss 9
201.15853881835938 0 450.13824
205.0348663330078 0 485.5312
211.10791015625 0 821.533
213.12403869628906 0 2252.0286
213.16024780273438 0 26582.396
214.16358947753906 0 3347.941
215.13946533203125 0 886.8144
219.13436889648438 0 4572.363 y 9
225.08734130859375 0 806.8548
227.10316467285156 0 15392.055
228.10618591308594 0 1135.4939
228.17135620117188 0 923.0057
229.15521240234375 0 735.30273
244.1296844482422 0 1984.9105
258.92364501953125 0 505.90176
261.29278564453125 0 523.79297
270.181396484375 0 1363.1912
271.1088562011719 0 553.9249
279.09808349609375 0 819.83374
280.1660461425781 0 1792.9696
294.1455993652344 0 886.1454
296.1616516113281 0 2389.4111
298.1768493652344 0 12248.888 y Water loss 8
298.4950256347656 0 541.4291
299.1799011230469 0 1962.7109
310.21295166015625 0 1248.299
312.15625 0 7007.318
313.16058349609375 0 763.0137
314.1720275878906 0 5683.227
315.2763366699219 0 2915.5984
316.18756103515625 0 155886.05 y 8
317.190673828125 0 25286.615
318.19293212890625 0 3190.166
322.13983154296875 0 1483.2249
323.1432800292969 0 660.12036
324.1563415527344 0 22051.072
325.1597900390625 0 4175.0312
329.1474304199219 0 840.48914
330.16680908203125 0 4795.4287
332.18231201171875 0 4893.7847
333.1856994628906 0 980.8674
337.2243957519531 0 1464.5614
338.135498046875 0 1559.0485
340.15130615234375 0 984.08777
341.25537109375 0 4166.6816 c Ammonia loss 2
342.25787353515625 0 665.9105
343.19921875 0 663.3589
357.274169921875 0 15964.596
358.280517578125 0 8959.514 c 2
359.2842712402344 0 797.3311
364.19573974609375 0 518.0898
374.17169189453125 0 1978.9545
374.2762145996094 0 571.1557
375.1732482910156 0 692.415
391.19873046875 0 1382.0848
392.1821594238281 0 629.8343
395.2301940917969 0 2517.9478
400.1510314941406 0 1238.2882
405.2115173339844 0 640.82324
407.1935729980469 0 1968.9441
409.20941162109375 0 2578.425
409.2820739746094 0 603.77356
421.2095031738281 0 3052.6936
422.2128601074219 0 744.2845
423.1889953613281 0 2279.272
423.2248229980469 0 12200.321
424.1911315917969 0 735.28796
424.2270202636719 0 2094.127
425.2304382324219 0 689.88654
426.1669616699219 0 2051.1362
426.237548828125 0 831.9147
426.74053955078125 0 942.3639
427.2195739746094 0 1030.9783 y Water loss 7
430.3033447265625 0 2588.0588
432.23760986328125 0 1487.8204
437.20428466796875 0 2906.048
438.2714538574219 0 750.5692
439.2218017578125 0 1364.1067
439.2550964355469 0 627.445
440.2344665527344 0 4347.257 c Ammonia loss 6
440.73699951171875 0 2016.2643
441.2359924316406 0 855.24414
442.16217041015625 0 2969.2058
443.1653747558594 0 815.1085
445.2304382324219 0 4161.0728 y 7
446.2333679199219 0 1347.8558
447.2051696777344 0 1103.4886
452.25103759765625 0 1029.6853
456.28289794921875 0 9692.096 c Ammonia loss 3
457.28643798828125 0 2524.5757
459.21002197265625 0 919.4202
461.223388671875 0 619.2537
471.2257995605469 0 1330.6451
472.3011169433594 0 4348.7036
473.3092346191406 0 12851.652 c 3
474.3121337890625 0 2378.964
483.265380859375 0 962.0662
488.3247375488281 0 708.25226
489.3050231933594 0 684.5515
489.75225830078125 0 698.7602
490.2658386230469 0 578.2442
490.7577819824219 0 1060.5801
491.260009765625 0 1603.2266
491.7582702636719 0 815.66223
494.7427673339844 0 716.34314
495.2423400878906 0 859.3198
495.7500305175781 0 1122.212
496.76165771484375 0 1129.3281
497.2057800292969 0 1075.1234
503.74835205078125 0 4129.5557
504.250244140625 0 3574.237
504.75634765625 0 10172.733 c Ammonia loss 7
505.2574768066406 0 6633.484
505.7582092285156 0 1915.5302
506.2623291015625 0 1408.5004
509.2441101074219 0 576.8659
522.2559814453125 0 750.14404
524.2733154296875 0 5056.6235 y Water loss 6
525.2761840820312 0 1284.6418
529.2815551757812 0 625.8852
529.7744140625 0 930.64856
530.2796020507812 0 1131.7422
530.78125 0 1825.022
531.780029296875 0 640.27057
534.2545776367188 0 1095.407
536.2586059570312 0 1060.3818
536.310302734375 0 2136.796
538.2520751953125 0 3839.0483
539.2509155273438 0 1578.438 z Water loss 2
539.2862548828125 0 1844.7148
539.7874755859375 0 2000.4978
540.2671508789062 0 3447.628
541.271728515625 0 1713.0542
542.2832641601562 0 63729.582 y 6
543.2855224609375 0 17940.426
543.7716064453125 0 1636.3834
544.28271484375 0 4726.307
544.7757568359375 0 1661.5211
545.2843017578125 0 2049.531
545.78515625 0 1019.74884
547.2642211914062 0 1915.6587 y Water loss 2
547.7651977539062 0 788.66876 y Ammonia loss 2
550.2509155273438 0 977.3725
552.2753295898438 0 12814.8125
552.6998901367188 0 588.34143
552.7763061523438 0 11034.979
553.2826538085938 0 42661.645 c Ammonia loss 8
553.7844848632812 0 25136.967
554.261474609375 0 6468.049
554.2865600585938 0 5870.238
554.7875366210938 0 856.4978
555.2633666992188 0 3966.4817
556.2625732421875 0 17671.703 y 2
557.265380859375 0 5702.225
558.2755126953125 0 5312.5635
559.28076171875 0 852.7726
560.9935302734375 0 602.1977
567.3521118164062 0 1769.4564
568.2430419921875 0 1705.3389
570.2568359375 0 14984.096
571.2596435546875 0 4793.9653
572.2614135742188 0 998.6722
578.8057250976562 0 1800.5048
579.3038940429688 0 881.8053
579.8146362304688 0 2749.5625
580.2637329101562 0 741.42755
580.3128662109375 0 1735.1339
580.8115234375 0 941.3099
584.2719116210938 0 839.8178
587.812255859375 0 5588.399
588.312744140625 0 4385.164
588.819091796875 0 13616.116
589.3208618164062 0 12286.336
589.8215942382812 0 4151.9673
591.3505859375 0 3256.3704
592.3557739257812 0 1524.0487
593.306396484375 0 719.214
593.8116455078125 0 4189.768
594.313232421875 0 1705.0707
601.8096313476562 0 3855.6228
602.3125610351562 0 3243.851 c Water loss 9
602.8171997070312 0 12842.965 c Ammonia loss 9
603.3184814453125 0 8415.891
603.8197021484375 0 3689.3816
607.3466796875 0 2677.9568
615.31298828125 0 1471.1006
617.3304443359375 0 2206.7607
619.3465576171875 0 13637.569 c Ammonia loss 4
620.34912109375 0 4298.513
621.3515625 0 1540.3743
622.3526611328125 0 713.8524
631.3379516601562 0 1644.1713
632.3424072265625 0 1623.1567
633.3291625976562 0 1770.7743
635.353759765625 0 11305.237
635.3638305664062 0 11802.5
636.37158203125 0 25622.727 c 4
637.3746948242188 0 8150.958
638.3782348632812 0 1938.7302
638.8353881835938 0 765.04364
644.33251953125 0 993.12933
644.8374633789062 0 1017.72205
646.3660888671875 0 3466.2542
649.3485717773438 0 1646.6896
650.3491821289062 0 2243.503
651.3602905273438 0 32382.494
652.364013671875 0 14971.125
652.836669921875 0 1807.323
653.341796875 0 9017.591
653.84326171875 0 6528.9004
654.3389892578125 0 2022.9047
661.3390502929688 0 3149.218
661.8392333984375 0 2442.0417
662.3455810546875 0 13839.691
662.402587890625 0 637.22144
662.8472900390625 0 9702.309
663.3490600585938 0 4406.908
663.8524780273438 0 867.3737
667.3092651367188 0 1823.2383
669.32470703125 0 1847.391
670.3257446289062 0 828.9
671.340576171875 0 1064.8396
679.3117065429688 0 1048.1421
681.326171875 0 1198.3695
682.3788452148438 0 3427.6338
683.3853759765625 0 1209.439
685.32177734375 0 1109.7875
688.3428955078125 0 1303.8593
689.3514404296875 0 25181.512
690.354736328125 0 10115.462
691.3585815429688 0 1266.2483
697.3223266601562 0 1905.6018
698.3214721679688 0 1371.7386
703.3313598632812 0 5580.4194
704.3330078125 0 2601.1143
705.3460693359375 0 9429.54 y 5
706.3507080078125 0 2756.1243
707.3534545898438 0 631.3177
714.41748046875 0 813.0688
735.3984375 0 724.0057
736.3997802734375 0 1383.6088
737.4065551757812 0 677.8608
738.4182739257812 0 3265.9155
739.4263305664062 0 7695.1323
740.4331665039062 0 6298.527
741.438232421875 0 3516.9304
753.4102172851562 0 1269.5586
754.4135131835938 0 6104.331
755.421630859375 0 15019.809
756.42822265625 0 14522.742
757.431640625 0 5682.0474
758.3286743164062 0 645.0701
762.38330078125 0 6316.9463
763.3836059570312 0 2281.9707
764.3975830078125 0 6151.112
765.4049072265625 0 1886.628
766.4146118164062 0 35325.77
767.4183349609375 0 17274.498
768.4207763671875 0 4955.5034
769.426025390625 0 682.8684
778.3377075195312 0 1530.8859
780.3887329101562 0 4849.562
781.3946533203125 0 2765.6492
782.4098510742188 0 67395.35 c Ammonia loss 5
783.4129638671875 0 33267.758
784.4168701171875 0 8475.712
794.3374633789062 0 1518.595
795.3389892578125 0 963.22314
796.3526611328125 0 16457.957
797.3551635742188 0 5807.672
798.3580322265625 0 1678.3562
829.4444580078125 0 792.34766
839.4152221679688 0 993.77026
840.41845703125 0 948.7107
850.3839721679688 0 750.99457 y Water loss 4
851.3919677734375 0 839.5573
852.4083862304688 0 2688.5042
853.4141235351562 0 1806.1085
859.42529296875 0 733.1585
863.4678955078125 0 1955.06
864.3779296875 0 753.966
864.4691162109375 0 757.3717
865.3858642578125 0 910.7434
865.4699096679688 0 1874.6779
866.3955078125 0 2629.818
867.4027709960938 0 9380.378
868.4085083007812 0 13220.059 y 4
869.4130249023438 0 5270.165
870.416015625 0 1047.1876
875.4537353515625 0 2320.3696
876.4585571289062 0 2316.977
877.4557495117188 0 1110.4285
878.4783935546875 0 1457.1067 c Water loss 6
879.4830932617188 0 30730.844
880.4913940429688 0 42857.29
881.4954833984375 0 20935.217
882.4994506835938 0 5918.068
883.5089721679688 0 1117.3035
891.3895874023438 0 3078.0537
892.3917236328125 0 1352.9493
893.402099609375 0 5137.2217
893.4730224609375 0 4557.5425
894.4724731445312 0 17232.719
895.4806518554688 0 79939.58
896.4869384765625 0 97843.195 c 6
897.4906005859375 0 40026.195
898.4945068359375 0 9524.4
905.4033203125 0 790.4404
923.4227905273438 0 1353.8585
950.4629516601562 0 733.39764
960.482666015625 0 926.3165
961.4933471679688 0 782.45435
962.4766845703125 0 1576.3116
963.4846801757812 0 891.2177
964.5006103515625 0 1158.623
965.5161743164062 0 1173.7611
966.5293579101562 0 932.32526
967.4340209960938 0 2209.3591 z 3
968.4295654296875 0 2938.348
969.4334716796875 0 1191.3431
972.4639282226562 0 1042.2095
973.4674682617188 0 952.5585
974.4798583984375 0 1512.5481
975.466064453125 0 725.8844
978.5142211914062 0 2006.8838
979.5083618164062 0 1573.246
980.510498046875 0 4716.271
981.41650390625 0 1462.8737
981.5159912109375 0 7393.8423
982.4277954101562 0 10109.626
982.525390625 0 2118.7717
983.434326171875 0 8214.479 y 3
983.5305786132812 0 887.9488
984.4400024414062 0 4254.453
985.4368286132812 0 802.7925
988.4782104492188 0 8792.74
989.482177734375 0 5578.998
990.48486328125 0 9510.016
991.4894409179688 0 5388.4526
992.5032348632812 0 26435.926
993.5042724609375 0 13945.247
994.5075073242188 0 6437.816
995.5052490234375 0 989.8441
1006.4901733398438 0 18711.23
1007.493896484375 0 14862.828
1008.5053100585938 0 104747.914 c Ammonia loss 7
1009.5087280273438 0 62211.457
1010.5111083984375 0 21046.188
1011.5142211914062 0 2290.6892
1023.4933471679688 0 1474.8934
1024.4945068359375 0 981.2364
1037.518310546875 0 898.42163
1049.5126953125 0 2500.2825
1050.524658203125 0 2803.9556
1051.5260009765625 0 4736.2207
1052.532470703125 0 3049.2253
1053.5380859375 0 1030.159
1061.535400390625 0 1027.6437
1062.544677734375 0 1326.7509
1063.5528564453125 0 911.76447
1075.5225830078125 0 1076.9082
1076.528564453125 0 1196.0922
1077.5457763671875 0 1532.9755
1078.567626953125 0 1171.1167
1079.524658203125 0 3333.5725
1080.5269775390625 0 2607.3003
1081.526123046875 0 1022.96326
1090.55615234375 0 1002.7517
1091.527587890625 0 2390.9187
1092.5115966796875 0 1870.5581
1093.5032958984375 0 9727.266 y Water loss 2
1094.5064697265625 0 9206.407 y Ammonia loss 2
1095.5142822265625 0 37746.906 z 2
1096.5179443359375 0 25985.8
1097.5208740234375 0 10078.821
1098.522216796875 0 1787.1315
1103.5582275390625 0 1129.1979
1104.5557861328125 0 1447.2579 c Water loss 8
1105.5711669921875 0 2428.7166 c Ammonia loss 8
1106.573974609375 0 3081.342
1107.5777587890625 0 1921.141
1109.5169677734375 0 9006.367
1110.5184326171875 0 7089.1636
1111.5322265625 0 26133.863 y 2
1112.5345458984375 0 17044.713
1113.537841796875 0 4967.063
1120.568115234375 0 5340.8633
1121.5697021484375 0 4008.82
1122.5836181640625 0 19464.139 c 8
1123.587646484375 0 11934.047
1124.5902099609375 0 3904.3052
1131.620849609375 0 717.1716
1133.6409912109375 0 888.7829
1140.535888671875 0 1624.9333
1150.5640869140625 0 1042.0989
1159.6209716796875 0 2302.5095
1160.6241455078125 0 5243.1406
1161.6307373046875 0 3535.522
1162.6285400390625 0 4228.896
1163.638427734375 0 4662.497
1164.6409912109375 0 1818.9797
1174.6201171875 0 1751.4299
1175.6229248046875 0 8908.166
1176.628662109375 0 11644.373
1177.638427734375 0 41479.06
1178.639892578125 0 32332.162
1179.6453857421875 0 14533.14
1180.644287109375 0 3146.8198
1184.602783203125 0 1603.6891
1186.6102294921875 0 2442.117
1187.614990234375 0 1843.6952
1188.63037109375 0 2052.2957
1189.6385498046875 0 1108.9489
1192.5677490234375 0 3337.6604 y Water loss 1
1193.5745849609375 0 4075.8545 y Ammonia loss 1
1194.5819091796875 0 16951.752 z 1
1195.5845947265625 0 10729.584
1196.5859375 0 4312.3496
1200.6436767578125 0 1589.4209
1201.6317138671875 0 3232.3757
1202.6185302734375 0 8942.921
1203.62646484375 0 5930.6934 c Water loss 9
1204.62646484375 0 16288.195 c Ammonia loss 9
1205.6326904296875 0 12456.74
1206.6409912109375 0 11459.55
1207.644287109375 0 7084.589
1208.6407470703125 0 2131.3113
1209.5931396484375 0 1524.3993
1210.598388671875 0 2587.4324 y 1
1211.605224609375 0 2022.5764
1215.66455078125 0 1611.2399
1216.656982421875 0 1819.163
1217.6453857421875 0 1600.6042
1218.6199951171875 0 998.5488
1219.6368408203125 0 43032.098
1220.6400146484375 0 32743.52
1221.6513671875 0 156296.72 c 9
1222.6551513671875 0 113307.31
1223.659423828125 0 52564.805
1224.66650390625 0 15818.656
1225.6712646484375 0 3015.5398
1230.65771484375 0 889.95593
1231.6259765625 0 4634.736
1232.6324462890625 0 3953.9263
1233.6287841796875 0 3074.064
1234.655517578125 0 3653.2444
1235.62060546875 0 11053.854
1236.6207275390625 0 7553.7725
1237.62060546875 0 4110.953
1238.624755859375 0 1538.9272
1239.608642578125 0 992.8763
1244.66015625 0 2175.6833
1245.6715087890625 0 2494.6885
1246.657470703125 0 2969.9453
1247.6409912109375 0 1636.7672
1248.6190185546875 0 6136.9053
1249.6187744140625 0 3822.714
1250.64208984375 0 5534.743
1251.648193359375 0 4392.724
1252.6656494140625 0 7613.9326
1253.6685791015625 0 5605.57
1254.67236328125 0 2692.8176
1258.65966796875 0 1171.6667
1259.6583251953125 0 1993.1665
1260.6744384765625 0 11055.684
1261.67529296875 0 10738.65
1262.6627197265625 0 15481.048
1263.659912109375 0 12754.078
1264.647216796875 0 53605.73
1265.6512451171875 0 41582.875
1266.647705078125 0 20495.916
1267.641357421875 0 5622.587
1268.6322021484375 0 6152.671
1269.632568359375 0 3292.4924
1270.6380615234375 0 1722.01
1271.654541015625 0 889.0748
1276.66943359375 0 2856.289
1277.6639404296875 0 9015.744
1278.68212890625 0 25974.568
1279.6856689453125 0 20518.545
1280.695068359375 0 21131.754
1281.6993408203125 0 11130.161
1282.69189453125 0 4346.7217
1283.651123046875 0 877.4388
1287.638916015625 0 2064.3096
1288.667236328125 0 3487.313
1289.6658935546875 0 4503.269
1290.6636962890625 0 3015.0737
1291.6671142578125 0 2377.5476
1292.6690673828125 0 1618.2037
1293.65966796875 0 2003.1274
1294.681884765625 0 14869.597
1295.68359375 0 10429.725
1296.6959228515625 0 41029.04
1297.69970703125 0 34103.434
1298.702392578125 0 12998.411
1299.7108154296875 0 1870.5259
1303.6590576171875 0 4106.379
1304.6636962890625 0 30735.377
1305.6566162109375 0 72873.35
1306.6739501953125 0 169152.69
1307.674072265625 0 317381.53
1308.676513671875 0 211166.1
1309.6768798828125 0 80681.16
1310.677978515625 0 12945.273
1320.6607666015625 0 754.69836
1321.667724609375 0 45207.094
1322.6759033203125 0 280345
1323.6817626953125 0 393217.03
1324.6905517578125 0 748366.94
1325.6934814453125 0 519421.9
1326.6953125 0 210678
1327.6981201171875 0 28364.63
1339.6512451171875 0 1356.9575
1340.6531982421875 0 842.8163
1356.681884765625 0 1040.9194
1357.6859130859375 0 1391.6316
2899.75634765625 0 695.49286

Spectrum Details

|  |  |
| --- | --- |
| Matched peaks? Matched peaks The total absolute number of peaks matched. Additionally in brackets the total fraction of peaks matched and the total number of peaks is shown. | 47 (8.69% of 541) |
| FDR? FDR The false discovery rate estimated for this peptide. It is calculated by matching all theoretical fragments with a non-integer shift with the raw peaks for this spectrum. This is done with 40 different shifts. The resulting percentage is the average number of annotated peaks over the number of annotated peaks with the correct spectrum. | 2.89% |
| Satellite FDR? Satellite FDR See the FDR for details on its calculation. This satellite ion specific FDR only contains the satellite ions (d/w) for I/L/J positions. | - |
| PSM Score? PSM Score The PSM Score as given by Hecklib to this annotated spectrum. It is shown with three significant figures. | 569 |

#### Spectrum 7728? Spectrum 7728 The raw spectrum of this peptide as annotated by Hecklib. The fragments are coloured according to ion type (see legend). Any peaks with a star '\*' as text can be hovered over to see the full details, first the ion type second the mass shift type. By hovering over the amino acids in the peptide or ions in the legend the corresponding peaks are highlighted. By toggling the 'Unassigned' label you can turn the background (unassigned) peaks on or off in the plot. By updating the slider in the Ion legend you can update the spectrum to only show the top X% of the peaks with labels. The top X% means any peak that is within X% of the highest intensity. By dragging in the spectrum you can zoom in to a specific part of the spectrum and use 'Zoom Out' to get back to the original zoom level. The annotation of the spectrum is based on the given sequence in the peptides file and is done with different software so inconsistencies are likely. The peaks are annotated based on the given sequence, with 20 ppm tolerance.

Copy Data

##### Spectrum 7728 (TSV)

###### Preview

```
Loading example...
```

*Click on the button to copy the data to your clipboard.*

Mz MinMz MaxIntensity Max

WidthHeightPeptide font sizePeptide stroke widthSpectrum font sizeSpectrum stroke widthCompact peptide

Ion legend

wxyz

abcd

OtherUnassignedIon^Charge^_Position_Show for top:%

JVKDYYPEPVT

09.06e+41.81e+52.72e+53.63e+5

Zoom Out

y^+1^_1_y^+1^_2_y^+1^_3_y^+1^_3_c^+1^_3_c^+1^_3_c^+2^_7_y^+1^_4_c^+1^_4_c^+1^_4_c^+2^_8_y^+1^_5_z^+2^_9_y^+1^_5_y^+2^_9_c^+2^_9_y^+2^_9_c^+2^_10_c^+1^_5_c^+1^_5_y^+1^_6_c^+1^_6_z^+1^_7_y^+1^_7_c^+1^_7_z^+1^_8_y^+1^_8_c^+1^_8_y^+1^_9_y^+1^_9_z^+1^_9_c^+1^_9_y^+1^_9_c^+1^_9_y^+1^_10_z^+1^_10_c^+1^_10_c^+1^_10_y^+1^_10_c^+1^_10_

0775154923243099

Fragment Matches Table

Show background peaks

| Position | Ion type | Intensity | mz Theoretical | mz Error (Th) | mz Error (ppm) | Charge | Series Number |
| --- | --- | --- | --- | --- | --- | --- | --- |
| 11 | y | 7485 | 120.1 | 0.0002381 | 1.983 | +1 | 1 |
| - | - | 439.1 | 120.1 | - | - | 0 | - |
| - | - | 403.6 | 120.3 | - | - | 0 | - |
| - | - | 469.3 | 122.8 | - | - | 0 | - |
| - | - | 3894 | 129.1 | - | - | 0 | - |
| - | - | 442.2 | 166.5 | - | - | 0 | - |
| - | - | 948.3 | 167.1 | - | - | 0 | - |
| - | - | 8637 | 169.1 | - | - | 0 | - |
| - | - | 727.8 | 170.1 | - | - | 0 | - |
| - | - | 1032 | 173.5 | - | - | 0 | - |
| - | - | 1.18E+04 | 185.2 | - | - | 0 | - |
| - | - | 908.3 | 186.2 | - | - | 0 | - |
| - | - | 496.3 | 193.6 | - | - | 0 | - |
| - | - | 2416 | 195.1 | - | - | 0 | - |
| - | - | 481.9 | 196.6 | - | - | 0 | - |
| - | - | 1.754E+04 | 197.1 | - | - | 0 | - |
| - | - | 1503 | 198.1 | - | - | 0 | - |
| - | - | 749.3 | 199.1 | - | - | 0 | - |
| - | - | 1404 | 213.1 | - | - | 0 | - |
| - | - | 1.04E+04 | 213.2 | - | - | 0 | - |
| - | - | 745 | 214.2 | - | - | 0 | - |
| 10 | y | 1196 | 219.1 | 0.000588 | 2.683 | +1 | 2 |
| - | - | 5377 | 227.1 | - | - | 0 | - |
| - | - | 521.6 | 237.8 | - | - | 0 | - |
| - | - | 898.5 | 244.1 | - | - | 0 | - |
| - | - | 548.8 | 268.2 | - | - | 0 | - |
| - | - | 969.9 | 270.2 | - | - | 0 | - |
| - | - | 681.2 | 280.2 | - | - | 0 | - |
| 9 | y | 3965 | 298.2 | 0.0002589 | 0.8683 | +1 | 3 |
| - | - | 2643 | 312.2 | - | - | 0 | - |
| - | - | 1734 | 314.2 | - | - | 0 | - |
| - | - | 1546 | 315.3 | - | - | 0 | - |
| 9 | y | 5.35E+04 | 316.2 | 0.0002838 | 0.8976 | +1 | 3 |
| - | - | 7451 | 317.2 | - | - | 0 | - |
| - | - | 665 | 318.2 | - | - | 0 | - |
| - | - | 807.6 | 322.1 | - | - | 0 | - |
| - | - | 7857 | 324.2 | - | - | 0 | - |
| - | - | 1444 | 325.2 | - | - | 0 | - |
| - | - | 1363 | 330.2 | - | - | 0 | - |
| - | - | 1725 | 332.2 | - | - | 0 | - |
| 3 | c | 1337 | 341.3 | 0.0002923 | 0.8566 | +1 | 3 |
| - | - | 604.1 | 342.3 | - | - | 0 | - |
| - | - | 8123 | 357.3 | - | - | 0 | - |
| 3 | c | 5001 | 358.3 | 0.001268 | 3.538 | +1 | 3 |
| - | - | 548.9 | 373.3 | - | - | 0 | - |
| - | - | 753.5 | 374.2 | - | - | 0 | - |
| - | - | 1393 | 395.2 | - | - | 0 | - |
| - | - | 733.1 | 407.2 | - | - | 0 | - |
| - | - | 692 | 421.2 | - | - | 0 | - |
| - | - | 4653 | 423.2 | - | - | 0 | - |
| - | - | 972.5 | 424.2 | - | - | 0 | - |
| - | - | 1125 | 426.2 | - | - | 0 | - |
| - | - | 1326 | 430.3 | - | - | 0 | - |
| - | - | 629.9 | 432.2 | - | - | 0 | - |
| - | - | 1288 | 437.2 | - | - | 0 | - |
| 7 | c | 1253 | 440.2 | 0.0001656 | 0.3761 | +2 | 7 |
| - | - | 1000 | 442.2 | - | - | 0 | - |
| 8 | y | 1530 | 445.2 | 0.0009036 | 2.03 | +1 | 4 |
| 4 | c | 3101 | 456.3 | 0.0006272 | 1.375 | +1 | 4 |
| - | - | 611.6 | 457.3 | - | - | 0 | - |
| - | - | 1619 | 472.3 | - | - | 0 | - |
| 4 | c | 6419 | 473.3 | 0.0006839 | 1.445 | +1 | 4 |
| - | - | 1472 | 474.3 | - | - | 0 | - |
| - | - | 1165 | 490.8 | - | - | 0 | - |
| - | - | 633.4 | 491.3 | - | - | 0 | - |
| - | - | 675 | 503.7 | - | - | 0 | - |
| - | - | 792.6 | 504.2 | - | - | 0 | - |
| 8 | c | 2780 | 504.8 | 0.0007147 | 1.416 | +2 | 8 |
| - | - | 2586 | 505.3 | - | - | 0 | - |
| - | - | 838.6 | 505.8 | - | - | 0 | - |
| 7 | y | 1568 | 524.3 | 0.002131 | 4.065 | +1 | 5 |
| - | - | 650.4 | 530.8 | - | - | 0 | - |
| - | - | 1706 | 538.3 | - | - | 0 | - |
| 3 | z | 893.9 | 539.3 | 0.005706 | 10.58 | +2 | 9 |
| - | - | 982.8 | 539.3 | - | - | 0 | - |
| - | - | 1050 | 540.3 | - | - | 0 | - |
| - | - | 1379 | 541.3 | - | - | 0 | - |
| 7 | y | 2.177E+04 | 542.3 | 0.0002943 | 0.5427 | +1 | 5 |
| - | - | 5802 | 543.3 | - | - | 0 | - |
| - | - | 727.3 | 543.8 | - | - | 0 | - |
| - | - | 1840 | 544.3 | - | - | 0 | - |
| - | - | 692 | 544.8 | - | - | 0 | - |
| 3 | y | 657.9 | 547.8 | 0.008366 | 15.27 | +2 | 9 |
| - | - | 4464 | 552.3 | - | - | 0 | - |
| - | - | 4630 | 552.8 | - | - | 0 | - |
| 9 | c | 1.459E+04 | 553.3 | 0.0004243 | 0.7668 | +2 | 9 |
| - | - | 8483 | 553.8 | - | - | 0 | - |
| - | - | 3366 | 554.3 | - | - | 0 | - |
| - | - | 1009 | 554.3 | - | - | 0 | - |
| - | - | 957.2 | 555.3 | - | - | 0 | - |
| 3 | y | 6150 | 556.3 | 0.007106 | 12.77 | +2 | 9 |
| - | - | 1538 | 557.3 | - | - | 0 | - |
| - | - | 1642 | 558.3 | - | - | 0 | - |
| - | - | 5794 | 570.3 | - | - | 0 | - |
| - | - | 1518 | 571.3 | - | - | 0 | - |
| - | - | 726.7 | 579.8 | - | - | 0 | - |
| - | - | 2862 | 587.8 | - | - | 0 | - |
| - | - | 1972 | 588.3 | - | - | 0 | - |
| - | - | 5502 | 588.8 | - | - | 0 | - |
| - | - | 3271 | 589.3 | - | - | 0 | - |
| - | - | 883.7 | 589.8 | - | - | 0 | - |
| - | - | 1199 | 591.3 | - | - | 0 | - |
| - | - | 587.5 | 592.8 | - | - | 0 | - |
| - | - | 970.4 | 593.8 | - | - | 0 | - |
| - | - | 638.9 | 594.3 | - | - | 0 | - |
| - | - | 1342 | 601.8 | - | - | 0 | - |
| - | - | 1672 | 602.3 | - | - | 0 | - |
| 10 | c | 3522 | 602.8 | 0.000403 | 0.6685 | +2 | 10 |
| - | - | 3306 | 603.3 | - | - | 0 | - |
| - | - | 958 | 603.8 | - | - | 0 | - |
| - | - | 801.5 | 615.3 | - | - | 0 | - |
| 5 | c | 4477 | 619.3 | 0.0005676 | 0.9164 | +1 | 5 |
| - | - | 1885 | 620.3 | - | - | 0 | - |
| - | - | 793.9 | 631.3 | - | - | 0 | - |
| - | - | 7406 | 635.4 | - | - | 0 | - |
| 5 | c | 1.269E+04 | 636.4 | 0.001299 | 2.041 | +1 | 5 |
| - | - | 4037 | 637.4 | - | - | 0 | - |
| - | - | 1198 | 650.4 | - | - | 0 | - |
| - | - | 1.47E+04 | 651.4 | - | - | 0 | - |
| - | - | 7045 | 652.4 | - | - | 0 | - |
| - | - | 2470 | 653.3 | - | - | 0 | - |
| - | - | 1946 | 653.8 | - | - | 0 | - |
| - | - | 1041 | 654.3 | - | - | 0 | - |
| - | - | 1127 | 661.3 | - | - | 0 | - |
| - | - | 794 | 661.8 | - | - | 0 | - |
| - | - | 3245 | 662.3 | - | - | 0 | - |
| - | - | 3456 | 662.4 | - | - | 0 | - |
| - | - | 2878 | 662.8 | - | - | 0 | - |
| - | - | 1552 | 663.3 | - | - | 0 | - |
| - | - | 1016 | 663.4 | - | - | 0 | - |
| - | - | 805.3 | 667.3 | - | - | 0 | - |
| - | - | 1648 | 682.4 | - | - | 0 | - |
| - | - | 673.2 | 688.3 | - | - | 0 | - |
| - | - | 1.029E+04 | 689.4 | - | - | 0 | - |
| - | - | 3484 | 690.4 | - | - | 0 | - |
| - | - | 2182 | 703.3 | - | - | 0 | - |
| - | - | 1529 | 704.3 | - | - | 0 | - |
| 6 | y | 2642 | 705.3 | 0.001511 | 2.142 | +1 | 6 |
| - | - | 868.2 | 706.3 | - | - | 0 | - |
| - | - | 1181 | 738.4 | - | - | 0 | - |
| - | - | 3118 | 739.4 | - | - | 0 | - |
| - | - | 3727 | 740.4 | - | - | 0 | - |
| - | - | 1636 | 741.4 | - | - | 0 | - |
| - | - | 2475 | 754.4 | - | - | 0 | - |
| - | - | 7941 | 755.4 | - | - | 0 | - |
| - | - | 7361 | 756.4 | - | - | 0 | - |
| - | - | 2351 | 757.4 | - | - | 0 | - |
| - | - | 1719 | 762.4 | - | - | 0 | - |
| - | - | 2475 | 764.4 | - | - | 0 | - |
| - | - | 1496 | 765.4 | - | - | 0 | - |
| - | - | 1.283E+04 | 766.4 | - | - | 0 | - |
| - | - | 5558 | 767.4 | - | - | 0 | - |
| - | - | 1591 | 768.4 | - | - | 0 | - |
| - | - | 1771 | 780.4 | - | - | 0 | - |
| - | - | 790.6 | 781.4 | - | - | 0 | - |
| 6 | c | 2.426E+04 | 782.4 | 0.0001144 | 0.1462 | +1 | 6 |
| - | - | 1.173E+04 | 783.4 | - | - | 0 | - |
| - | - | 2980 | 784.4 | - | - | 0 | - |
| - | - | 765.8 | 794.3 | - | - | 0 | - |
| - | - | 6129 | 796.4 | - | - | 0 | - |
| - | - | 2347 | 797.4 | - | - | 0 | - |
| 5 | z | 1350 | 852.4 | 0.01559 | 18.29 | +1 | 7 |
| - | - | 953.9 | 866.4 | - | - | 0 | - |
| - | - | 4321 | 867.4 | - | - | 0 | - |
| 5 | y | 5301 | 868.4 | 0.0007524 | 0.8664 | +1 | 7 |
| - | - | 2129 | 869.4 | - | - | 0 | - |
| - | - | 1185 | 875.5 | - | - | 0 | - |
| - | - | 1309 | 876.5 | - | - | 0 | - |
| - | - | 1.509E+04 | 879.5 | - | - | 0 | - |
| - | - | 2.079E+04 | 880.5 | - | - | 0 | - |
| - | - | 8877 | 881.5 | - | - | 0 | - |
| - | - | 2984 | 882.5 | - | - | 0 | - |
| - | - | 1129 | 891.4 | - | - | 0 | - |
| - | - | 1325 | 893.4 | - | - | 0 | - |
| - | - | 1770 | 893.5 | - | - | 0 | - |
| - | - | 9198 | 894.5 | - | - | 0 | - |
| - | - | 3.781E+04 | 895.5 | - | - | 0 | - |
| 7 | c | 4.556E+04 | 896.5 | 0.002645 | 2.95 | +1 | 7 |
| - | - | 1.924E+04 | 897.5 | - | - | 0 | - |
| - | - | 3245 | 898.5 | - | - | 0 | - |
| 4 | z | 1226 | 967.4 | 0.01776 | 18.36 | +1 | 8 |
| - | - | 1376 | 968.4 | - | - | 0 | - |
| - | - | 1073 | 969.4 | - | - | 0 | - |
| - | - | 785.8 | 978.5 | - | - | 0 | - |
| - | - | 907 | 979.5 | - | - | 0 | - |
| - | - | 1823 | 980.5 | - | - | 0 | - |
| - | - | 1237 | 981.4 | - | - | 0 | - |
| - | - | 1689 | 981.5 | - | - | 0 | - |
| - | - | 3460 | 982.4 | - | - | 0 | - |
| - | - | 1226 | 982.5 | - | - | 0 | - |
| 4 | y | 3349 | 983.4 | 0.004685 | 4.764 | +1 | 8 |
| - | - | 1757 | 984.4 | - | - | 0 | - |
| - | - | 3953 | 988.5 | - | - | 0 | - |
| - | - | 1672 | 989.5 | - | - | 0 | - |
| - | - | 3268 | 990.5 | - | - | 0 | - |
| - | - | 1800 | 991.5 | - | - | 0 | - |
| - | - | 8805 | 992.5 | - | - | 0 | - |
| - | - | 4706 | 993.5 | - | - | 0 | - |
| - | - | 2288 | 994.5 | - | - | 0 | - |
| - | - | 6926 | 1006 | - | - | 0 | - |
| - | - | 4898 | 1007 | - | - | 0 | - |
| 8 | c | 3.875E+04 | 1009 | 0.0006837 | 0.6779 | +1 | 8 |
| - | - | 1.859E+04 | 1010 | - | - | 0 | - |
| - | - | 6293 | 1011 | - | - | 0 | - |
| - | - | 1224 | 1050 | - | - | 0 | - |
| - | - | 862.2 | 1051 | - | - | 0 | - |
| - | - | 1992 | 1052 | - | - | 0 | - |
| - | - | 907.7 | 1053 | - | - | 0 | - |
| - | - | 1178 | 1063 | - | - | 0 | - |
| - | - | 1613 | 1079 | - | - | 0 | - |
| - | - | 1966 | 1080 | - | - | 0 | - |
| - | - | 1152 | 1081 | - | - | 0 | - |
| - | - | 935.1 | 1092 | - | - | 0 | - |
| 3 | y | 5726 | 1094 | 0.01895 | 17.33 | +1 | 9 |
| 3 | y | 3969 | 1095 | 0.0007725 | 0.7058 | +1 | 9 |
| 3 | z | 1.916E+04 | 1096 | 0.0006629 | 0.6051 | +1 | 9 |
| - | - | 1.187E+04 | 1097 | - | - | 0 | - |
| - | - | 3930 | 1098 | - | - | 0 | - |
| - | - | 681.2 | 1099 | - | - | 0 | - |
| 9 | c | 709.1 | 1106 | 0.0008963 | 0.8107 | +1 | 9 |
| - | - | 1092 | 1107 | - | - | 0 | - |
| - | - | 831.2 | 1108 | - | - | 0 | - |
| - | - | 3352 | 1110 | - | - | 0 | - |
| - | - | 1928 | 1111 | - | - | 0 | - |
| 3 | y | 8949 | 1112 | 0.001687 | 1.518 | +1 | 9 |
| - | - | 5014 | 1113 | - | - | 0 | - |
| - | - | 1460 | 1114 | - | - | 0 | - |
| - | - | 2662 | 1121 | - | - | 0 | - |
| 9 | c | 9939 | 1123 | 0.002665 | 2.374 | +1 | 9 |
| - | - | 6069 | 1124 | - | - | 0 | - |
| - | - | 2175 | 1125 | - | - | 0 | - |
| - | - | 701.7 | 1141 | - | - | 0 | - |
| - | - | 1355 | 1160 | - | - | 0 | - |
| - | - | 2318 | 1161 | - | - | 0 | - |
| - | - | 1615 | 1162 | - | - | 0 | - |
| - | - | 1740 | 1163 | - | - | 0 | - |
| - | - | 2341 | 1164 | - | - | 0 | - |
| - | - | 891.5 | 1165 | - | - | 0 | - |
| - | - | 5346 | 1176 | - | - | 0 | - |
| - | - | 5108 | 1177 | - | - | 0 | - |
| - | - | 932.9 | 1177 | - | - | 0 | - |
| - | - | 1.681E+04 | 1178 | - | - | 0 | - |
| - | - | 1.312E+04 | 1179 | - | - | 0 | - |
| - | - | 7352 | 1180 | - | - | 0 | - |
| - | - | 1940 | 1181 | - | - | 0 | - |
| - | - | 1147 | 1187 | - | - | 0 | - |
| - | - | 735.2 | 1189 | - | - | 0 | - |
| - | - | 1868 | 1193 | - | - | 0 | - |
| 2 | y | 2320 | 1194 | 0.0007599 | 0.6367 | +1 | 10 |
| 2 | z | 6979 | 1195 | 0.002915 | 2.44 | +1 | 10 |
| - | - | 4909 | 1196 | - | - | 0 | - |
| - | - | 2153 | 1197 | - | - | 0 | - |
| - | - | 1542 | 1202 | - | - | 0 | - |
| - | - | 3995 | 1203 | - | - | 0 | - |
| 10 | c | 2382 | 1204 | 0.02023 | 16.81 | +1 | 10 |
| 10 | c | 6182 | 1205 | 0.001927 | 1.6 | +1 | 10 |
| - | - | 4892 | 1206 | - | - | 0 | - |
| - | - | 4491 | 1207 | - | - | 0 | - |
| - | - | 3255 | 1208 | - | - | 0 | - |
| - | - | 713.5 | 1209 | - | - | 0 | - |
| 2 | y | 1095 | 1211 | 0.004915 | 4.06 | +1 | 10 |
| - | - | 939.3 | 1217 | - | - | 0 | - |
| - | - | 1.894E+04 | 1220 | - | - | 0 | - |
| - | - | 1.468E+04 | 1221 | - | - | 0 | - |
| 10 | c | 7.803E+04 | 1222 | 0.003086 | 2.526 | +1 | 10 |
| - | - | 5.354E+04 | 1223 | - | - | 0 | - |
| - | - | 2.402E+04 | 1224 | - | - | 0 | - |
| - | - | 6997 | 1225 | - | - | 0 | - |
| - | - | 1449 | 1226 | - | - | 0 | - |
| - | - | 1593 | 1232 | - | - | 0 | - |
| - | - | 2381 | 1233 | - | - | 0 | - |
| - | - | 1429 | 1234 | - | - | 0 | - |
| - | - | 1705 | 1235 | - | - | 0 | - |
| - | - | 4648 | 1236 | - | - | 0 | - |
| - | - | 4218 | 1237 | - | - | 0 | - |
| - | - | 1511 | 1238 | - | - | 0 | - |
| - | - | 1135 | 1239 | - | - | 0 | - |
| - | - | 819.5 | 1246 | - | - | 0 | - |
| - | - | 1843 | 1247 | - | - | 0 | - |
| - | - | 848.7 | 1248 | - | - | 0 | - |
| - | - | 2744 | 1249 | - | - | 0 | - |
| - | - | 1986 | 1250 | - | - | 0 | - |
| - | - | 3045 | 1251 | - | - | 0 | - |
| - | - | 1732 | 1252 | - | - | 0 | - |
| - | - | 5136 | 1253 | - | - | 0 | - |
| - | - | 2351 | 1254 | - | - | 0 | - |
| - | - | 1035 | 1255 | - | - | 0 | - |
| - | - | 875.3 | 1259 | - | - | 0 | - |
| - | - | 6022 | 1261 | - | - | 0 | - |
| - | - | 5262 | 1262 | - | - | 0 | - |
| - | - | 7782 | 1263 | - | - | 0 | - |
| - | - | 5665 | 1264 | - | - | 0 | - |
| - | - | 2.693E+04 | 1265 | - | - | 0 | - |
| - | - | 1.756E+04 | 1266 | - | - | 0 | - |
| - | - | 8355 | 1267 | - | - | 0 | - |
| - | - | 2146 | 1268 | - | - | 0 | - |
| - | - | 3060 | 1269 | - | - | 0 | - |
| - | - | 2029 | 1270 | - | - | 0 | - |
| - | - | 1208 | 1271 | - | - | 0 | - |
| - | - | 2036 | 1277 | - | - | 0 | - |
| - | - | 5543 | 1278 | - | - | 0 | - |
| - | - | 1.224E+04 | 1279 | - | - | 0 | - |
| - | - | 8475 | 1280 | - | - | 0 | - |
| - | - | 9183 | 1281 | - | - | 0 | - |
| - | - | 4748 | 1282 | - | - | 0 | - |
| - | - | 1617 | 1283 | - | - | 0 | - |
| - | - | 2145 | 1289 | - | - | 0 | - |
| - | - | 2529 | 1290 | - | - | 0 | - |
| - | - | 1623 | 1291 | - | - | 0 | - |
| - | - | 755.2 | 1292 | - | - | 0 | - |
| - | - | 1091 | 1294 | - | - | 0 | - |
| - | - | 5975 | 1295 | - | - | 0 | - |
| - | - | 5063 | 1296 | - | - | 0 | - |
| - | - | 1.941E+04 | 1297 | - | - | 0 | - |
| - | - | 1.471E+04 | 1298 | - | - | 0 | - |
| - | - | 5158 | 1299 | - | - | 0 | - |
| - | - | 996.5 | 1300 | - | - | 0 | - |
| - | - | 2479 | 1304 | - | - | 0 | - |
| - | - | 1.603E+04 | 1305 | - | - | 0 | - |
| - | - | 3.568E+04 | 1306 | - | - | 0 | - |
| - | - | 8.179E+04 | 1307 | - | - | 0 | - |
| - | - | 1.544E+05 | 1308 | - | - | 0 | - |
| - | - | 1.019E+05 | 1309 | - | - | 0 | - |
| - | - | 3.666E+04 | 1310 | - | - | 0 | - |
| - | - | 5874 | 1311 | - | - | 0 | - |
| - | - | 819.2 | 1321 | - | - | 0 | - |
| - | - | 2.519E+04 | 1322 | - | - | 0 | - |
| - | - | 1.421E+05 | 1323 | - | - | 0 | - |
| - | - | 1.935E+05 | 1324 | - | - | 0 | - |
| - | - | 3.59E+05 | 1325 | - | - | 0 | - |
| - | - | 2.473E+05 | 1326 | - | - | 0 | - |
| - | - | 9.017E+04 | 1327 | - | - | 0 | - |
| - | - | 1.143E+04 | 1328 | - | - | 0 | - |
| - | - | 748.2 | 1340 | - | - | 0 | - |
| - | - | 1064 | 1357 | - | - | 0 | - |
| - | - | 691.6 | 3068 | - | - | 0 | - |

m/z Charge Intensity FragmentType MassShift Position
120.06575775146484 0 7484.8994 y 10
120.08128356933594 0 439.08203
120.27854919433594 0 403.6253
122.77685546875 0 469.34488
129.10247802734375 0 3894.0144
166.48390197753906 0 442.18335
167.11781311035156 0 948.3069
169.1337890625 0 8637.45
170.13682556152344 0 727.76575
173.45225524902344 0 1032.0885
185.16506958007812 0 11800.489
186.1685791015625 0 908.30023
193.63961791992188 0 496.30582
195.11300659179688 0 2415.5684
196.59239196777344 0 481.85974
197.1286163330078 0 17537.52
198.1319580078125 0 1503.1256
199.1075439453125 0 749.29895
213.12347412109375 0 1404.0239
213.15992736816406 0 10395.495
214.16383361816406 0 744.99677
219.134521484375 0 1196.4569 y 9
227.10284423828125 0 5376.78
237.7908477783203 0 521.6481
244.12916564941406 0 898.47614
268.18023681640625 0 548.77625
270.181884765625 0 969.9177
280.16632080078125 0 681.2014
298.1763916015625 0 3964.636 y Water loss 8
312.15570068359375 0 2643.2195
314.1707458496094 0 1733.7745
315.275390625 0 1545.5554
316.1869812011719 0 53504.066 y 8
317.1900634765625 0 7450.976
318.1918029785156 0 664.96826
322.1393127441406 0 807.59064
324.15582275390625 0 7857.3945
325.158935546875 0 1443.8411
330.166259765625 0 1363.4381
332.1815490722656 0 1725.208
341.2544250488281 0 1336.8528 c Ammonia loss 2
342.2586669921875 0 604.07635
357.2736511230469 0 8122.879
358.2799987792969 0 5001.4453 c 2
373.2709045410156 0 548.8798
374.1706237792969 0 753.4926
395.23040771484375 0 1393.0201
407.19049072265625 0 733.0577
421.2080383300781 0 691.9703
423.22406005859375 0 4652.628
424.22650146484375 0 972.466
426.166259765625 0 1125.428
430.3031921386719 0 1325.556
432.23760986328125 0 629.9423
437.2041015625 0 1287.9906
440.2343444824219 0 1252.6895 c Ammonia loss 6
442.1614685058594 0 1000.40326
445.2301940917969 0 1530.4755 y 7
456.28228759765625 0 3101.232 c Ammonia loss 3
457.28277587890625 0 611.6485
472.301025390625 0 1619.3019
473.3075256347656 0 6419.2046 c 3
474.3114318847656 0 1472.242
490.7581787109375 0 1164.5815
491.2578125 0 633.3651
503.74639892578125 0 675.0206
504.2491760253906 0 792.6269
504.7547607421875 0 2780.159 c Ammonia loss 7
505.25726318359375 0 2586.4177
505.7592468261719 0 838.6131
524.2736206054688 0 1568.1765 y Water loss 6
530.7759399414062 0 650.4299
538.2523193359375 0 1706.3975
539.2485961914062 0 893.92554 z Water loss 2
539.2896118164062 0 982.761
540.2640991210938 0 1049.7888
541.2713623046875 0 1378.9924
542.2823486328125 0 21765.057 y 6
543.28466796875 0 5801.797
543.7664184570312 0 727.3031
544.277099609375 0 1840.1631
544.7775268554688 0 691.9514
547.7640380859375 0 657.8509 y Ammonia loss 2
552.273681640625 0 4463.505
552.775390625 0 4629.581
553.2814331054688 0 14588.106 c Ammonia loss 8
553.7838134765625 0 8483.133
554.26123046875 0 3366.183
554.2935180664062 0 1009.42596
555.260498046875 0 957.23224
556.2618408203125 0 6149.951 y 2
557.2645263671875 0 1538.4604
558.2758178710938 0 1642.1393
570.2556762695312 0 5793.718
571.2586669921875 0 1518.0665
579.8124389648438 0 726.7073
587.81005859375 0 2862.0454
588.3116455078125 0 1971.5876
588.8181762695312 0 5501.901
589.3207397460938 0 3270.9514
589.8198852539062 0 883.74677
591.3494262695312 0 1198.5035
592.8040161132812 0 587.5282
593.8120727539062 0 970.39905
594.302978515625 0 638.8851
601.8082275390625 0 1341.6758
602.3101196289062 0 1671.9432
602.8164672851562 0 3522.0923 c Ammonia loss 9
603.3171997070312 0 3305.7393
603.8177490234375 0 957.9566
615.3162841796875 0 801.54266
619.3444213867188 0 4477.369 c Ammonia loss 4
620.3477172851562 0 1885.2734
631.3308715820312 0 793.8709
635.3572387695312 0 7406.384
636.3702392578125 0 12694.027 c 4
637.3740234375 0 4036.9573
650.351806640625 0 1197.5585
651.35888671875 0 14697.593
652.3626708984375 0 7045.352
653.3365478515625 0 2469.7322
653.8418579101562 0 1945.7482
654.3394775390625 0 1040.822
661.3358764648438 0 1127.4093
661.8348388671875 0 794.0192
662.34326171875 0 3244.7822
662.3972778320312 0 3455.532
662.8460083007812 0 2877.9785
663.34521484375 0 1552.136
663.4032592773438 0 1015.6213
667.3046875 0 805.33
682.3760375976562 0 1648.1897
688.341064453125 0 673.1772
689.3500366210938 0 10293.235
690.3544311523438 0 3484.4963
703.3307495117188 0 2181.6628
704.3369140625 0 1529.1884
705.3438720703125 0 2641.954 y 5
706.3468627929688 0 868.19324
738.41748046875 0 1181.0067
739.426513671875 0 3117.9456
740.4323120117188 0 3727.26
741.4336547851562 0 1636.1007
754.4121704101562 0 2474.641
755.420654296875 0 7941.329
756.4273071289062 0 7360.651
757.4315185546875 0 2350.747
762.3802490234375 0 1718.5719
764.3954467773438 0 2475.2688
765.3981323242188 0 1495.9908
766.4127197265625 0 12825.141
767.4168701171875 0 5558.264
768.4201049804688 0 1590.9498
780.3875122070312 0 1770.5919
781.3619995117188 0 790.59766
782.408203125 0 24258.547 c Ammonia loss 5
783.4116821289062 0 11727.303
784.4146118164062 0 2979.6775
794.3346557617188 0 765.8085
796.3507080078125 0 6129.392
797.353515625 0 2347.413
852.4055786132812 0 1349.6455 z 4
866.3863525390625 0 953.94403
867.3997192382812 0 4321.4614
868.407958984375 0 5301.4263 y 4
869.410888671875 0 2129.175
875.4537353515625 0 1185.0366
876.45556640625 0 1308.8094
879.482421875 0 15085.509
880.4901733398438 0 20794.988
881.4945068359375 0 8877.235
882.4968872070312 0 2984.0417
891.3859252929688 0 1128.9122
893.3908081054688 0 1324.5488
893.46875 0 1770.2292
894.47021484375 0 9198.405
895.4783325195312 0 37805.812
896.4849853515625 0 45564.97 c 6
897.4886474609375 0 19236.951
898.4924926757812 0 3244.5994
967.4346923828125 0 1225.5497 z 3
968.4279174804688 0 1376.176
969.4286499023438 0 1072.744
978.5267333984375 0 785.8232
979.5053100585938 0 906.97095
980.5093383789062 0 1823.3748
981.4139404296875 0 1236.7842
981.5174560546875 0 1689.1636
982.4244384765625 0 3459.7266
982.525146484375 0 1226.3054
983.4309692382812 0 3348.6838 y 3
984.4384155273438 0 1757.4558
988.476806640625 0 3952.5413
989.4852905273438 0 1672.3489
990.4862060546875 0 3267.5298
991.4899291992188 0 1800.1759
992.4996337890625 0 8805.483
993.5038452148438 0 4705.609
994.5001220703125 0 2288.3804
1006.4881591796875 0 6925.61
1007.4911499023438 0 4898.3735
1008.5029907226562 0 38752.387 c Ammonia loss 7
1009.505859375 0 18587.373
1010.5089111328125 0 6292.6597
1049.5059814453125 0 1224.1592
1050.5260009765625 0 862.1526
1051.524658203125 0 1991.6763
1052.532958984375 0 907.6788
1062.5584716796875 0 1177.5049
1078.5574951171875 0 1612.5991
1079.5206298828125 0 1965.6477
1080.5177001953125 0 1151.5609
1091.524658203125 0 935.0647
1093.5010986328125 0 5725.6616 y Water loss 2
1094.5032958984375 0 3968.731 y Ammonia loss 2
1095.51123046875 0 19160.361 z 2
1096.5152587890625 0 11865.453
1097.5174560546875 0 3929.9456
1098.5185546875 0 681.23706
1105.5555419921875 0 709.07574 c Ammonia loss 8
1106.5762939453125 0 1092.3579
1107.579345703125 0 831.1947
1109.5166015625 0 3351.577
1110.51513671875 0 1928.2959
1111.5289306640625 0 8948.847 y 2
1112.5330810546875 0 5014.0815
1113.5382080078125 0 1459.6577
1120.5648193359375 0 2661.6233
1122.580322265625 0 9939.066 c 8
1123.585205078125 0 6069.2373
1124.582763671875 0 2174.943
1140.52783203125 0 701.68317
1159.6204833984375 0 1354.5605
1160.6275634765625 0 2318.21
1161.6236572265625 0 1614.6807
1162.6280517578125 0 1739.8501
1163.6365966796875 0 2341.4753
1164.6490478515625 0 891.5052
1175.6207275390625 0 5345.9727
1176.625244140625 0 5108.3955
1177.486328125 0 932.94116
1177.6368408203125 0 16811.422
1178.6361083984375 0 13121.87
1179.6431884765625 0 7351.615
1180.6435546875 0 1940.3467
1186.6024169921875 0 1147.243
1188.6180419921875 0 735.15814
1192.5621337890625 0 1867.8995
1193.5732421875 0 2320.056 y Ammonia loss 1
1194.577392578125 0 6979.4985 z 1
1195.5814208984375 0 4908.698
1196.5855712890625 0 2153.4133
1201.6256103515625 0 1541.6985
1202.6146240234375 0 3995.2654
1203.62060546875 0 2382.437 c Water loss 9
1204.6229248046875 0 6181.6733 c Ammonia loss 9
1205.632080078125 0 4891.813
1206.6397705078125 0 4490.9985
1207.6444091796875 0 3254.8594
1208.6370849609375 0 713.54675
1210.5941162109375 0 1094.7496 y 1
1216.6513671875 0 939.3005
1219.63427734375 0 18936.607
1220.63720703125 0 14678.527
1221.6483154296875 0 78034.44 c 9
1222.6513671875 0 53542.97
1223.6568603515625 0 24023.213
1224.6640625 0 6996.707
1225.667236328125 0 1449.1844
1231.6214599609375 0 1593.0793
1232.6279296875 0 2381.231
1233.6324462890625 0 1429.1207
1234.6541748046875 0 1705.017
1235.61669921875 0 4647.514
1236.6175537109375 0 4217.824
1237.6119384765625 0 1510.5519
1238.6214599609375 0 1134.5784
1245.6610107421875 0 819.54913
1246.642578125 0 1843.183
1247.6502685546875 0 848.72156
1248.6170654296875 0 2743.803
1249.61279296875 0 1985.5824
1250.631103515625 0 3045.2683
1251.6304931640625 0 1731.7213
1252.6624755859375 0 5136.5
1253.6651611328125 0 2350.5244
1254.672607421875 0 1034.9567
1258.6461181640625 0 875.34296
1260.6689453125 0 6021.602
1261.6717529296875 0 5261.7344
1262.65625 0 7782.0166
1263.641357421875 0 5664.978
1264.6439208984375 0 26929.445
1265.64794921875 0 17561.049
1266.646484375 0 8354.55
1267.6370849609375 0 2145.8699
1268.6280517578125 0 3059.947
1269.6339111328125 0 2028.5895
1270.6456298828125 0 1207.997
1276.6590576171875 0 2036.0499
1277.658935546875 0 5542.5493
1278.67822265625 0 12235.539
1279.6820068359375 0 8475.422
1280.6912841796875 0 9182.925
1281.6981201171875 0 4748.3584
1282.6868896484375 0 1616.5875
1288.6558837890625 0 2144.891
1289.661376953125 0 2529.3489
1290.6600341796875 0 1623.4528
1291.6671142578125 0 755.1898
1293.6680908203125 0 1090.7184
1294.674560546875 0 5974.702
1295.6806640625 0 5063.398
1296.692626953125 0 19409.41
1297.696533203125 0 14711.537
1298.69970703125 0 5157.519
1299.6949462890625 0 996.53107
1303.6513671875 0 2479.1714
1304.6605224609375 0 16029.284
1305.6527099609375 0 35680.31
1306.6710205078125 0 81791.45
1307.6712646484375 0 154350.03
1308.6722412109375 0 101906.2
1309.673095703125 0 36659.32
1310.675537109375 0 5873.9634
1320.6552734375 0 819.1999
1321.663330078125 0 25188.166
1322.6719970703125 0 142135.16
1323.677490234375 0 193517.28
1324.6861572265625 0 358986.1
1325.68994140625 0 247339.28
1326.69189453125 0 90172.82
1327.69287109375 0 11433.899
1339.6453857421875 0 748.2084
1356.68701171875 0 1064.0952
3068.18359375 0 691.6067

Spectrum Details

|  |  |
| --- | --- |
| Matched peaks? Matched peaks The total absolute number of peaks matched. Additionally in brackets the total fraction of peaks matched and the total number of peaks is shown. | 40 (11.90% of 336) |
| FDR? FDR The false discovery rate estimated for this peptide. It is calculated by matching all theoretical fragments with a non-integer shift with the raw peaks for this spectrum. This is done with 40 different shifts. The resulting percentage is the average number of annotated peaks over the number of annotated peaks with the correct spectrum. | 1.67% |
| Satellite FDR? Satellite FDR See the FDR for details on its calculation. This satellite ion specific FDR only contains the satellite ions (d/w) for I/L/J positions. | - |
| PSM Score? PSM Score The PSM Score as given by Hecklib to this annotated spectrum. It is shown with three significant figures. | 467 |

#### Reverse Lookup? Reverse Lookup All places where this read could be placed.

| Group | Segment | Template | Template Part | Read Part | Score | Unique |
| --- | --- | --- | --- | --- | --- | --- |
| Homo sapiens Heavy Chain | IGHC | IGHG1 | [27..38] | [0..11] | 79 | False |
| Homo sapiens Heavy Chain | IGHC | IGHG3 | [27..38] | [0..11] | 79 | False |
| Homo sapiens Heavy Chain | IGHC | IGHG2 | [27..38] | [0..11] | 79 | False |
| Homo sapiens Heavy Chain | IGHC | IGHG4 | [27..38] | [0..11] | 79 | False |

| Recombined | Template Part | Read Part | Score | Unique |
| --- | --- | --- | --- | --- |
| REC-0-1 | [149..160] | [0..11] | 79 | True |

#### Meta Information from Multiple reads

##### Number of combined reads

2

##### Intensity

0.8093

##### TotalArea

1.243E+07

##### Changes to the peptide sequence

JVKDYYPEPVT

L→JNo support for either Leucine or Isoleucine based on side chain ions (Position: 1)

#### Positional Score

Copy Data

##### Positional Score (TSV)

###### Preview

```
Loading example...
```

*Click on the button to copy the data to your clipboard.*

00012345678910

Label Value
"0" 0
"1" 0
"2" 0
"3" 0
"4" 0
"5" 0
"6" 0
"7" 0
"8" 0
"9" 0
"10" 0

#### Meta Information from PEAKS

##### Scan Identifier

F2:7672

##### Original sequence

L

V

K

D

Y

Y

P

E

P

V

T

##### Posttranslational Modifications

##### Source File

D:\separate\_stitch\_analyses\xle-disambiguation\raw\20210323\_F1\_UM1\_Peng0013\_SA\_F59\_ingel\_3ug\_TL.raw

##### Fraction

2

##### Scan Feature

F2:10908

##### De Novo Score

98

##### ConfidenceScore

98

### m/z

662.3448

##### Mass

1322.6758

##### Charge

2

##### Retention Time

43.06

##### Predicted Retention Time

-

##### Area

1.243E+07

##### Fragmentation mode

ETHCD

##### Originating file

01 D:\separate\_stitch\_analyses\xle-disambiguation\20210325\_F59\_3ug\_DENOVO\_12.csv

#### Meta Information from PEAKS

##### Scan Identifier

F2:7728

##### Original sequence

L

V

K

D

Y

Y

P

E

P

V

T

##### Posttranslational Modifications

##### Source File

D:\separate\_stitch\_analyses\xle-disambiguation\raw\20210323\_F1\_UM1\_Peng0013\_SA\_F59\_ingel\_3ug\_TL.raw

##### Fraction

2

##### Scan Feature

-

##### De Novo Score

98

##### ConfidenceScore

98

### m/z

662.3445

##### Mass

1322.6758

##### Charge

2

##### Retention Time

43.33

##### Predicted Retention Time

-

##### Area

0

##### Fragmentation mode

ETHCD

##### Originating file

01 D:\separate\_stitch\_analyses\xle-disambiguation\20210325\_F59\_3ug\_DENOVO\_12.csv
