## Supplementary material for "A handle on mass coincidence errors in *de novo* sequencing of antibodies by bottom-up proteomics": Combined_058.html

Details Combined\_058 | Stitch OverviewUndefined

### Read Combined\_058

#### Sequence (length=8)

SLRJDDTA

#### Spectrum 4344? Spectrum 4344 The raw spectrum of this peptide as annotated by Hecklib. The fragments are coloured according to ion type (see legend). Any peaks with a star '\*' as text can be hovered over to see the full details, first the ion type second the mass shift type. By hovering over the amino acids in the peptide or ions in the legend the corresponding peaks are highlighted. By toggling the 'Unassigned' label you can turn the background (unassigned) peaks on or off in the plot. By updating the slider in the Ion legend you can update the spectrum to only show the top X% of the peaks with labels. The top X% means any peak that is within X% of the highest intensity. By dragging in the spectrum you can zoom in to a specific part of the spectrum and use 'Zoom Out' to get back to the original zoom level. The annotation of the spectrum is based on the given sequence in the peptides file and is done with different software so inconsistencies are likely. The peaks are annotated based on the given sequence, with 20 ppm tolerance.

Copy Data

##### Spectrum 4344 (TSV)

###### Preview

```
Loading example...
```

*Click on the button to copy the data to your clipboard.*

Mz MinMz MaxIntensity Max

WidthHeightPeptide font sizePeptide stroke widthSpectrum font sizeSpectrum stroke widthCompact peptide

Ion legend

wxyz

abcd

OtherUnassignedIon^Charge^_Position_Show for top:%

SLRJDDTA

01.20e+62.39e+63.59e+64.78e+6

Zoom Out

y^+1^_2_y^+1^_2_y^+1^_3_c^+2^_5_y^+1^_3_c^+2^_6_c^+1^_3_c^+1^_3_c^+1^_3_y^+2^_7_y^+2^_7_c^+2^_7_y^+2^_7_y^+1^_4_y^+1^_4_c^+1^_4_c^+1^_4_c^+1^_4_y^+1^_5_y^+1^_5_c^+1^_5_c^+1^_5_c^+1^_5_z^+1^_6_y^+1^_6_y^+1^_6_z^+1^_6_y^+1^_6_c^+1^_6_c^+1^_6_w^+1^_7_y^+1^_7_z^+1^_7_c^+1^_7_y^+1^_7_c^+1^_7_

0875175026253500

Fragment Matches Table

Show background peaks

| Position | Ion type | Intensity | mz Theoretical | mz Error (Th) | mz Error (ppm) | Charge | Series Number |
| --- | --- | --- | --- | --- | --- | --- | --- |
| - | - | 1.877E+04 | 120.1 | - | - | 0 | - |
| - | - | 5118 | 124.8 | - | - | 0 | - |
| - | - | 1.34E+04 | 133.1 | - | - | 0 | - |
| - | - | 6289 | 133.1 | - | - | 0 | - |
| - | - | 6276 | 144.9 | - | - | 0 | - |
| - | - | 8949 | 145.1 | - | - | 0 | - |
| - | - | 7652 | 148.9 | - | - | 0 | - |
| - | - | 7116 | 148.9 | - | - | 0 | - |
| - | - | 7458 | 148.9 | - | - | 0 | - |
| - | - | 9156 | 148.9 | - | - | 0 | - |
| - | - | 1.651E+04 | 148.9 | - | - | 0 | - |
| - | - | 1.748E+04 | 148.9 | - | - | 0 | - |
| - | - | 3.687E+04 | 148.9 | - | - | 0 | - |
| - | - | 6.165E+04 | 148.9 | - | - | 0 | - |
| - | - | 5.351E+04 | 149 | - | - | 0 | - |
| - | - | 2.966E+04 | 149 | - | - | 0 | - |
| - | - | 1.889E+04 | 149 | - | - | 0 | - |
| - | - | 1.336E+04 | 149 | - | - | 0 | - |
| - | - | 7677 | 149 | - | - | 0 | - |
| - | - | 6139 | 149 | - | - | 0 | - |
| - | - | 3.268E+04 | 149 | - | - | 0 | - |
| - | - | 6336 | 149.1 | - | - | 0 | - |
| - | - | 6304 | 149.3 | - | - | 0 | - |
| - | - | 2.382E+04 | 157.1 | - | - | 0 | - |
| - | - | 8549 | 158.1 | - | - | 0 | - |
| - | - | 8525 | 167.1 | - | - | 0 | - |
| - | - | 3.248E+04 | 171.1 | - | - | 0 | - |
| 7 | y | 1.058E+05 | 173.1 | 0.0003538 | 2.044 | +1 | 2 |
| - | - | 4.994E+05 | 173.1 | - | - | 0 | - |
| - | - | 1.009E+04 | 174.1 | - | - | 0 | - |
| - | - | 4.592E+04 | 174.1 | - | - | 0 | - |
| - | - | 7167 | 182.3 | - | - | 0 | - |
| - | - | 6.292E+04 | 189.1 | - | - | 0 | - |
| 7 | y | 4.953E+05 | 191.1 | 0.0003787 | 1.982 | +1 | 2 |
| - | - | 3.646E+04 | 192.1 | - | - | 0 | - |
| - | - | 1.9E+04 | 199.1 | - | - | 0 | - |
| - | - | 1.969E+05 | 201.1 | - | - | 0 | - |
| - | - | 1.392E+04 | 202.1 | - | - | 0 | - |
| - | - | 1.357E+04 | 211.1 | - | - | 0 | - |
| - | - | 2.171E+04 | 215.1 | - | - | 0 | - |
| - | - | 1.089E+05 | 217.1 | - | - | 0 | - |
| - | - | 7620 | 218.1 | - | - | 0 | - |
| - | - | 1.322E+04 | 223.1 | - | - | 0 | - |
| - | - | 1.007E+04 | 225 | - | - | 0 | - |
| - | - | 7492 | 225.6 | - | - | 0 | - |
| - | - | 6.92E+04 | 227.2 | - | - | 0 | - |
| - | - | 1.047E+04 | 228.2 | - | - | 0 | - |
| - | - | 4.537E+04 | 229.1 | - | - | 0 | - |
| - | - | 2.089E+04 | 229.2 | - | - | 0 | - |
| - | - | 2.816E+04 | 231.1 | - | - | 0 | - |
| - | - | 7383 | 233.1 | - | - | 0 | - |
| - | - | 6970 | 238.1 | - | - | 0 | - |
| - | - | 1.424E+04 | 239.1 | - | - | 0 | - |
| - | - | 1.223E+04 | 242.1 | - | - | 0 | - |
| - | - | 8116 | 257.1 | - | - | 0 | - |
| - | - | 1.284E+04 | 270.1 | - | - | 0 | - |
| - | - | 1.349E+04 | 270.2 | - | - | 0 | - |
| - | - | 1.834E+04 | 271.2 | - | - | 0 | - |
| - | - | 2.196E+04 | 274.2 | - | - | 0 | - |
| - | - | 8962 | 284.2 | - | - | 0 | - |
| - | - | 1.003E+04 | 286.1 | - | - | 0 | - |
| - | - | 1.707E+04 | 286.2 | - | - | 0 | - |
| - | - | 7379 | 287.4 | - | - | 0 | - |
| 6 | y | 7.256E+04 | 288.1 | 0.0005256 | 1.824 | +1 | 3 |
| - | - | 8930 | 289.1 | - | - | 0 | - |
| 5 | c | 4.884E+04 | 293.2 | 0.0003407 | 1.162 | +2 | 5 |
| - | - | 8794 | 304.1 | - | - | 0 | - |
| 6 | y | 2.314E+04 | 306.1 | 0.0003674 | 1.2 | +1 | 3 |
| - | - | 7907 | 311.6 | - | - | 0 | - |
| - | - | 1.253E+04 | 312.2 | - | - | 0 | - |
| - | - | 1.43E+04 | 314.1 | - | - | 0 | - |
| - | - | 9854 | 314.7 | - | - | 0 | - |
| - | - | 2.019E+04 | 315.2 | - | - | 0 | - |
| - | - | 2.778E+04 | 317.2 | - | - | 0 | - |
| - | - | 3.559E+04 | 329.2 | - | - | 0 | - |
| - | - | 6.078E+05 | 330.2 | - | - | 0 | - |
| - | - | 1.669E+05 | 331.2 | - | - | 0 | - |
| - | - | 2.607E+04 | 332.1 | - | - | 0 | - |
| - | - | 2.203E+04 | 332.2 | - | - | 0 | - |
| - | - | 2.465E+04 | 335.7 | - | - | 0 | - |
| - | - | 1.067E+04 | 336.2 | - | - | 0 | - |
| - | - | 8609 | 338.2 | - | - | 0 | - |
| - | - | 1.042E+04 | 339.2 | - | - | 0 | - |
| - | - | 1.461E+04 | 340.2 | - | - | 0 | - |
| - | - | 9.606E+04 | 341 | - | - | 0 | - |
| - | - | 4.981E+04 | 341.2 | - | - | 0 | - |
| - | - | 2.661E+04 | 341.7 | - | - | 0 | - |
| - | - | 2.809E+04 | 342 | - | - | 0 | - |
| - | - | 1.435E+04 | 342.2 | - | - | 0 | - |
| - | - | 1.203E+04 | 342.2 | - | - | 0 | - |
| - | - | 1.508E+05 | 342.2 | - | - | 0 | - |
| - | - | 1.039E+04 | 342.2 | - | - | 0 | - |
| - | - | 1.279E+04 | 343 | - | - | 0 | - |
| - | - | 2.784E+04 | 343.2 | - | - | 0 | - |
| - | - | 1.474E+04 | 343.7 | - | - | 0 | - |
| - | - | 3.378E+04 | 344.1 | - | - | 0 | - |
| - | - | 8170 | 344.2 | - | - | 0 | - |
| - | - | 2.002E+04 | 344.7 | - | - | 0 | - |
| - | - | 8772 | 345.2 | - | - | 0 | - |
| - | - | 7537 | 348.7 | - | - | 0 | - |
| 6 | c | 2.293E+05 | 350.7 | 0.000358 | 1.021 | +2 | 6 |
| - | - | 5.172E+04 | 351.2 | - | - | 0 | - |
| - | - | 2.403E+04 | 351.7 | - | - | 0 | - |
| - | - | 3.921E+04 | 355.2 | - | - | 0 | - |
| - | - | 1.061E+04 | 356.2 | - | - | 0 | - |
| 3 | c | 2.575E+04 | 356.2 | 0.0005941 | 1.668 | +1 | 3 |
| 3 | c | 2.275E+05 | 357.2 | 0.0004042 | 1.131 | +1 | 3 |
| - | - | 1.269E+04 | 358.2 | - | - | 0 | - |
| - | - | 3.337E+04 | 358.2 | - | - | 0 | - |
| - | - | 3.198E+05 | 359 | - | - | 0 | - |
| - | - | 8311 | 359.3 | - | - | 0 | - |
| - | - | 9.541E+04 | 360 | - | - | 0 | - |
| - | - | 3.413E+04 | 361 | - | - | 0 | - |
| - | - | 1.678E+04 | 362.2 | - | - | 0 | - |
| - | - | 1.389E+04 | 364.7 | - | - | 0 | - |
| - | - | 5.981E+04 | 369.2 | - | - | 0 | - |
| - | - | 9.508E+04 | 369.7 | - | - | 0 | - |
| - | - | 7.446E+04 | 370.2 | - | - | 0 | - |
| - | - | 2.747E+04 | 370.7 | - | - | 0 | - |
| - | - | 1.256E+04 | 372.2 | - | - | 0 | - |
| - | - | 1.253E+06 | 373.2 | - | - | 0 | - |
| 3 | c | 3.234E+06 | 374.3 | 0.0001307 | 0.3492 | +1 | 3 |
| - | - | 5.43E+05 | 375.3 | - | - | 0 | - |
| - | - | 6.152E+04 | 376.3 | - | - | 0 | - |
| - | - | 3.719E+05 | 378.2 | - | - | 0 | - |
| - | - | 2.439E+05 | 378.7 | - | - | 0 | - |
| - | - | 7.022E+04 | 379.2 | - | - | 0 | - |
| - | - | 1.304E+04 | 379.7 | - | - | 0 | - |
| - | - | 9503 | 381.2 | - | - | 0 | - |
| - | - | 9.271E+04 | 383.2 | - | - | 0 | - |
| - | - | 3.434E+04 | 383.7 | - | - | 0 | - |
| - | - | 9961 | 384.2 | - | - | 0 | - |
| - | - | 1.749E+04 | 385.1 | - | - | 0 | - |
| - | - | 2.96E+04 | 385.2 | - | - | 0 | - |
| - | - | 9961 | 386.2 | - | - | 0 | - |
| - | - | 3.686E+05 | 387.2 | - | - | 0 | - |
| - | - | 1.382E+05 | 387.7 | - | - | 0 | - |
| - | - | 3.481E+04 | 388.2 | - | - | 0 | - |
| - | - | 3.611E+04 | 391.7 | - | - | 0 | - |
| - | - | 3.869E+05 | 392.2 | - | - | 0 | - |
| - | - | 1.026E+05 | 392.7 | - | - | 0 | - |
| 2 | y | 4.208E+04 | 393.2 | 0.005211 | 13.25 | +2 | 7 |
| 2 | y | 9028 | 393.7 | 0.005619 | 14.27 | +2 | 7 |
| - | - | 8613 | 395.2 | - | - | 0 | - |
| 7 | c | 7.814E+05 | 401.2 | 0.0007497 | 1.869 | +2 | 7 |
| - | - | 2.966E+05 | 401.7 | - | - | 0 | - |
| - | - | 1.165E+04 | 402.2 | - | - | 0 | - |
| 2 | y | 6.601E+04 | 402.2 | 0.004573 | 11.37 | +2 | 7 |
| 5 | y | 3.238E+04 | 403.1 | 0.0005601 | 1.389 | +1 | 4 |
| - | - | 1.71E+04 | 404.7 | - | - | 0 | - |
| - | - | 1.454E+04 | 405.2 | - | - | 0 | - |
| - | - | 1.195E+04 | 405.7 | - | - | 0 | - |
| - | - | 3.152E+04 | 412.3 | - | - | 0 | - |
| - | - | 2.789E+04 | 413.3 | - | - | 0 | - |
| - | - | 1.083E+05 | 413.7 | - | - | 0 | - |
| - | - | 3.421E+04 | 414.2 | - | - | 0 | - |
| - | - | 1.465E+04 | 414.3 | - | - | 0 | - |
| - | - | 1.547E+04 | 414.7 | - | - | 0 | - |
| - | - | 1.316E+04 | 417.2 | - | - | 0 | - |
| - | - | 7700 | 419.7 | - | - | 0 | - |
| 5 | y | 1.076E+04 | 421.2 | 0.0009103 | 2.162 | +1 | 4 |
| - | - | 2.028E+04 | 425.3 | - | - | 0 | - |
| - | - | 1.363E+04 | 427.2 | - | - | 0 | - |
| - | - | 1.37E+04 | 427.3 | - | - | 0 | - |
| - | - | 1.187E+05 | 427.7 | - | - | 0 | - |
| - | - | 5.408E+04 | 428.2 | - | - | 0 | - |
| - | - | 1.104E+04 | 428.7 | - | - | 0 | - |
| - | - | 8.885E+04 | 429.1 | - | - | 0 | - |
| - | - | 3.421E+04 | 430.1 | - | - | 0 | - |
| - | - | 1.472E+05 | 430.3 | - | - | 0 | - |
| - | - | 1.733E+04 | 431.1 | - | - | 0 | - |
| - | - | 3.106E+04 | 431.3 | - | - | 0 | - |
| - | - | 7.294E+05 | 436.7 | - | - | 0 | - |
| - | - | 3.204E+05 | 437.2 | - | - | 0 | - |
| - | - | 9.824E+04 | 437.7 | - | - | 0 | - |
| - | - | 6.42E+04 | 442.3 | - | - | 0 | - |
| - | - | 2.901E+04 | 443.3 | - | - | 0 | - |
| - | - | 7.288E+04 | 443.3 | - | - | 0 | - |
| - | - | 2.021E+04 | 444.3 | - | - | 0 | - |
| - | - | 2.126E+04 | 445.2 | - | - | 0 | - |
| - | - | 6.371E+04 | 445.7 | - | - | 0 | - |
| - | - | 1.575E+04 | 446.2 | - | - | 0 | - |
| - | - | 2.796E+04 | 446.2 | - | - | 0 | - |
| - | - | 9242 | 446.7 | - | - | 0 | - |
| - | - | 8647 | 447.1 | - | - | 0 | - |
| - | - | 3.626E+04 | 457.2 | - | - | 0 | - |
| - | - | 1.898E+05 | 458.2 | - | - | 0 | - |
| - | - | 3.853E+04 | 459.3 | - | - | 0 | - |
| - | - | 6.552E+04 | 468.3 | - | - | 0 | - |
| 4 | c | 7.029E+04 | 469.3 | 0.0004621 | 0.9846 | +1 | 4 |
| 4 | c | 1.571E+05 | 470.3 | 0.0008739 | 1.858 | +1 | 4 |
| - | - | 4.09E+04 | 471.3 | - | - | 0 | - |
| - | - | 2.32E+04 | 485.3 | - | - | 0 | - |
| - | - | 1.279E+06 | 486.3 | - | - | 0 | - |
| 4 | c | 1.597E+06 | 487.3 | 0.0001015 | 0.2083 | +1 | 4 |
| - | - | 3.664E+05 | 488.3 | - | - | 0 | - |
| - | - | 4.241E+04 | 489.3 | - | - | 0 | - |
| - | - | 2.103E+04 | 498.3 | - | - | 0 | - |
| - | - | 1.563E+04 | 499.3 | - | - | 0 | - |
| - | - | 8.37E+04 | 500.2 | - | - | 0 | - |
| - | - | 2.218E+04 | 501.3 | - | - | 0 | - |
| - | - | 8.379E+04 | 502.3 | - | - | 0 | - |
| - | - | 5.325E+04 | 502.3 | - | - | 0 | - |
| - | - | 1.476E+04 | 503.3 | - | - | 0 | - |
| - | - | 9249 | 513.4 | - | - | 0 | - |
| - | - | 2.93E+04 | 515.3 | - | - | 0 | - |
| 4 | y | 2.686E+04 | 516.2 | 0.00045 | 0.8717 | +1 | 5 |
| - | - | 1.023E+04 | 516.3 | - | - | 0 | - |
| - | - | 4.666E+04 | 517.3 | - | - | 0 | - |
| - | - | 1.054E+04 | 518.3 | - | - | 0 | - |
| 4 | y | 1.998E+04 | 534.2 | 0.001909 | 3.574 | +1 | 5 |
| - | - | 2.537E+04 | 539.3 | - | - | 0 | - |
| - | - | 1.482E+04 | 540.3 | - | - | 0 | - |
| - | - | 2.311E+04 | 541.3 | - | - | 0 | - |
| - | - | 8.175E+04 | 542.3 | - | - | 0 | - |
| - | - | 2.1E+04 | 543.3 | - | - | 0 | - |
| - | - | 1.429E+04 | 545.3 | - | - | 0 | - |
| - | - | 4.677E+04 | 555.3 | - | - | 0 | - |
| - | - | 1.057E+04 | 556.3 | - | - | 0 | - |
| - | - | 1.113E+04 | 557.3 | - | - | 0 | - |
| - | - | 2.365E+04 | 557.3 | - | - | 0 | - |
| - | - | 2.305E+04 | 558.3 | - | - | 0 | - |
| - | - | 2.424E+05 | 558.3 | - | - | 0 | - |
| - | - | 7.902E+04 | 559.4 | - | - | 0 | - |
| - | - | 1.057E+04 | 560.4 | - | - | 0 | - |
| - | - | 1.433E+04 | 567.3 | - | - | 0 | - |
| - | - | 8868 | 568.3 | - | - | 0 | - |
| - | - | 1.003E+05 | 573.3 | - | - | 0 | - |
| - | - | 1.317E+05 | 574.3 | - | - | 0 | - |
| - | - | 4.421E+04 | 575.3 | - | - | 0 | - |
| - | - | 1.468E+04 | 576.3 | - | - | 0 | - |
| - | - | 3.026E+04 | 583.3 | - | - | 0 | - |
| - | - | 3.368E+04 | 583.3 | - | - | 0 | - |
| 5 | c | 2.403E+04 | 584.4 | 0.0002133 | 0.365 | +1 | 5 |
| 5 | c | 4.401E+05 | 585.3 | 0.0008168 | 1.395 | +1 | 5 |
| - | - | 1.528E+05 | 586.3 | - | - | 0 | - |
| - | - | 3.415E+04 | 587.3 | - | - | 0 | - |
| - | - | 2.336E+04 | 587.3 | - | - | 0 | - |
| - | - | 9.71E+04 | 601.3 | - | - | 0 | - |
| - | - | 4.613E+05 | 601.4 | - | - | 0 | - |
| 5 | c | 4.302E+06 | 602.4 | 0.0005128 | 0.8514 | +1 | 5 |
| - | - | 1.256E+06 | 603.4 | - | - | 0 | - |
| - | - | 2.326E+05 | 604.4 | - | - | 0 | - |
| - | - | 1.431E+04 | 605.4 | - | - | 0 | - |
| - | - | 4.739E+04 | 613.3 | - | - | 0 | - |
| - | - | 1.035E+04 | 615.4 | - | - | 0 | - |
| - | - | 4.836E+05 | 618.3 | - | - | 0 | - |
| - | - | 1.22E+05 | 619.3 | - | - | 0 | - |
| - | - | 3.005E+04 | 620.3 | - | - | 0 | - |
| - | - | 9401 | 628.3 | - | - | 0 | - |
| - | - | 3.056E+04 | 628.4 | - | - | 0 | - |
| - | - | 2.311E+04 | 629.3 | - | - | 0 | - |
| - | - | 2.903E+05 | 630.3 | - | - | 0 | - |
| - | - | 8.479E+05 | 631.3 | - | - | 0 | - |
| - | - | 8.89E+04 | 631.3 | - | - | 0 | - |
| - | - | 2.151E+05 | 632.3 | - | - | 0 | - |
| - | - | 1.729E+04 | 632.3 | - | - | 0 | - |
| - | - | 4.364E+04 | 633.3 | - | - | 0 | - |
| - | - | 2.177E+04 | 641.4 | - | - | 0 | - |
| - | - | 1.034E+04 | 646.3 | - | - | 0 | - |
| - | - | 1.385E+04 | 652.3 | - | - | 0 | - |
| - | - | 1.52E+04 | 654.3 | - | - | 0 | - |
| 3 | z | 1.31E+04 | 656.3 | 0.001559 | 2.376 | +1 | 6 |
| - | - | 7.851E+04 | 657.4 | - | - | 0 | - |
| - | - | 1.682E+05 | 658.4 | - | - | 0 | - |
| - | - | 5.982E+04 | 659.4 | - | - | 0 | - |
| - | - | 1.115E+04 | 660.4 | - | - | 0 | - |
| - | - | 4.837E+04 | 670.3 | - | - | 0 | - |
| - | - | 7685 | 670.4 | - | - | 0 | - |
| - | - | 1.459E+04 | 671.3 | - | - | 0 | - |
| 3 | y | 1.165E+05 | 672.3 | 0.000108 | 0.1606 | +1 | 6 |
| - | - | 1.034E+04 | 672.4 | - | - | 0 | - |
| 3 | y | 1.979E+04 | 673.3 | 0.006266 | 9.306 | +1 | 6 |
| - | - | 7.715E+04 | 673.4 | - | - | 0 | - |
| 3 | z | 1.379E+06 | 674.3 | 0.000699 | 1.037 | +1 | 6 |
| - | - | 4.141E+05 | 675.3 | - | - | 0 | - |
| - | - | 2.19E+04 | 675.4 | - | - | 0 | - |
| - | - | 8.652E+04 | 676.3 | - | - | 0 | - |
| - | - | 8.189E+04 | 682.4 | - | - | 0 | - |
| - | - | 3.109E+04 | 683.4 | - | - | 0 | - |
| - | - | 1.366E+04 | 684.4 | - | - | 0 | - |
| - | - | 8.987E+04 | 688.3 | - | - | 0 | - |
| - | - | 2.685E+04 | 689.3 | - | - | 0 | - |
| 3 | y | 8.281E+05 | 690.3 | 0.0004686 | 0.6788 | +1 | 6 |
| - | - | 2.638E+05 | 691.3 | - | - | 0 | - |
| - | - | 6.639E+04 | 692.3 | - | - | 0 | - |
| - | - | 1.709E+04 | 696.4 | - | - | 0 | - |
| 6 | c | 2.073E+06 | 700.4 | 0.0005462 | 0.7798 | +1 | 6 |
| - | - | 7.106E+05 | 701.4 | - | - | 0 | - |
| - | - | 3.161E+05 | 702.4 | - | - | 0 | - |
| - | - | 6.603E+04 | 703.4 | - | - | 0 | - |
| - | - | 1.549E+04 | 714.4 | - | - | 0 | - |
| 6 | c | 2.569E+06 | 717.4 | 0.0004253 | 0.5928 | +1 | 6 |
| - | - | 8.884E+05 | 718.4 | - | - | 0 | - |
| - | - | 1.053E+04 | 719.3 | - | - | 0 | - |
| - | - | 1.962E+05 | 719.4 | - | - | 0 | - |
| - | - | 1.56E+04 | 720.4 | - | - | 0 | - |
| - | - | 9271 | 731.3 | - | - | 0 | - |
| - | - | 1.363E+04 | 743.4 | - | - | 0 | - |
| 2 | w | 6.812E+05 | 744.4 | 0.000463 | 0.622 | +1 | 7 |
| - | - | 2.499E+05 | 745.4 | - | - | 0 | - |
| - | - | 6.321E+04 | 746.4 | - | - | 0 | - |
| - | - | 7.241E+04 | 756.4 | - | - | 0 | - |
| - | - | 1.376E+05 | 757.4 | - | - | 0 | - |
| - | - | 6.055E+04 | 758.4 | - | - | 0 | - |
| - | - | 1.15E+04 | 759.4 | - | - | 0 | - |
| - | - | 1.235E+04 | 765.4 | - | - | 0 | - |
| - | - | 9079 | 773.3 | - | - | 0 | - |
| - | - | 5.038E+04 | 773.4 | - | - | 0 | - |
| - | - | 4.129E+04 | 774.4 | - | - | 0 | - |
| - | - | 6.253E+04 | 775.4 | - | - | 0 | - |
| - | - | 6.788E+04 | 776.4 | - | - | 0 | - |
| - | - | 2.112E+04 | 777.4 | - | - | 0 | - |
| - | - | 1.435E+04 | 783.3 | - | - | 0 | - |
| - | - | 3.524E+05 | 783.4 | - | - | 0 | - |
| - | - | 1.312E+05 | 784.4 | - | - | 0 | - |
| 2 | y | 4.735E+04 | 785.4 | 0.007784 | 9.911 | +1 | 7 |
| 2 | z | 4.626E+05 | 787.4 | 0.0001922 | 0.2441 | +1 | 7 |
| - | - | 1.991E+05 | 788.4 | - | - | 0 | - |
| - | - | 5.891E+04 | 789.4 | - | - | 0 | - |
| - | - | 4.514E+04 | 790.4 | - | - | 0 | - |
| - | - | 1.028E+04 | 791.4 | - | - | 0 | - |
| 7 | c | 1.352E+06 | 801.4 | 0.0007192 | 0.8975 | +1 | 7 |
| - | - | 4.883E+05 | 802.4 | - | - | 0 | - |
| 2 | y | 1.696E+05 | 803.4 | 0.007179 | 8.936 | +1 | 7 |
| - | - | 3.488E+04 | 804.4 | - | - | 0 | - |
| - | - | 5.416E+04 | 813.4 | - | - | 0 | - |
| - | - | 2.124E+04 | 814.4 | - | - | 0 | - |
| - | - | 1.229E+04 | 815.4 | - | - | 0 | - |
| - | - | 2.04E+04 | 817.4 | - | - | 0 | - |
| - | - | 1.899E+06 | 818.4 | - | - | 0 | - |
| 7 | c | 2.32E+06 | 818.4 | 0.002613 | 3.192 | +1 | 7 |
| - | - | 6.873E+05 | 819.4 | - | - | 0 | - |
| - | - | 8.573E+05 | 819.4 | - | - | 0 | - |
| - | - | 1.739E+05 | 820.4 | - | - | 0 | - |
| - | - | 2.322E+05 | 820.4 | - | - | 0 | - |
| - | - | 1.43E+04 | 821.4 | - | - | 0 | - |
| - | - | 1.614E+04 | 821.5 | - | - | 0 | - |
| - | - | 3.406E+04 | 829.5 | - | - | 0 | - |
| - | - | 5.001E+05 | 830.4 | - | - | 0 | - |
| - | - | 4.291E+06 | 831.4 | - | - | 0 | - |
| - | - | 1.704E+06 | 832.4 | - | - | 0 | - |
| - | - | 4.216E+05 | 833.4 | - | - | 0 | - |
| - | - | 4.246E+04 | 834.4 | - | - | 0 | - |
| - | - | 1.602E+05 | 835.4 | - | - | 0 | - |
| - | - | 5.2E+04 | 836.4 | - | - | 0 | - |
| - | - | 1.613E+04 | 844.4 | - | - | 0 | - |
| - | - | 6.554E+04 | 845.5 | - | - | 0 | - |
| - | - | 2.306E+04 | 846.5 | - | - | 0 | - |
| - | - | 2.133E+04 | 847.4 | - | - | 0 | - |
| - | - | 7.913E+05 | 848.4 | - | - | 0 | - |
| - | - | 3.021E+05 | 849.4 | - | - | 0 | - |
| - | - | 1.079E+05 | 850.4 | - | - | 0 | - |
| - | - | 1.112E+04 | 851.4 | - | - | 0 | - |
| - | - | 4.445E+04 | 856.4 | - | - | 0 | - |
| - | - | 6.062E+04 | 857.4 | - | - | 0 | - |
| - | - | 2.391E+04 | 858.4 | - | - | 0 | - |
| - | - | 1.324E+04 | 859.4 | - | - | 0 | - |
| - | - | 1.409E+04 | 863.5 | - | - | 0 | - |
| - | - | 1.119E+05 | 873.5 | - | - | 0 | - |
| - | - | 4.735E+06 | 874.4 | - | - | 0 | - |
| - | - | 1.944E+06 | 875.4 | - | - | 0 | - |
| - | - | 5.485E+05 | 876.4 | - | - | 0 | - |
| - | - | 4.703E+04 | 877.4 | - | - | 0 | - |
| - | - | 9794 | 889.4 | - | - | 0 | - |
| - | - | 2.644E+06 | 890.5 | - | - | 0 | - |
| - | - | 2.091E+06 | 891.5 | - | - | 0 | - |
| - | - | 1.117E+04 | 891.7 | - | - | 0 | - |
| - | - | 6.694E+05 | 892.5 | - | - | 0 | - |
| - | - | 1.197E+05 | 893.5 | - | - | 0 | - |
| - | - | 1.002E+04 | 1289 | - | - | 0 | - |
| - | - | 1.038E+04 | 1360 | - | - | 0 | - |
| - | - | 1.039E+04 | 3465 | - | - | 0 | - |

m/z Charge Intensity FragmentType MassShift Position
120.06573486328125 0 18773.97
124.8177490234375 0 5117.985
133.0611572265625 0 13404.53
133.0865020751953 0 6288.998
144.92153930664062 0 6275.5137
145.0977020263672 0 8948.955
148.89328002929688 0 7651.7993
148.9004364013672 0 7115.7266
148.90737915039062 0 7458.102
148.91522216796875 0 9156.085
148.92257690429688 0 16510.652
148.92965698242188 0 17477.16
148.9368896484375 0 36865.97
148.9447784423828 0 61652.184
148.96156311035156 0 53514.7
148.9694061279297 0 29657.57
148.97671508789062 0 18885.957
148.9839630126953 0 13355.325
148.9915313720703 0 7677.231
149.0056915283203 0 6138.6646
149.0451202392578 0 32680.174
149.07106018066406 0 6335.807
149.25999450683594 0 6303.7124
157.1088104248047 0 23818.316
158.11700439453125 0 8549.281
167.0559844970703 0 8525.337
171.07681274414062 0 32479.45
173.09242248535156 0 105849.86 y Water loss 6
173.1288299560547 0 499432.47
174.0956573486328 0 10088.712
174.13217163085938 0 45923.75
182.28811645507812 0 7166.695
189.08737182617188 0 62924.02
191.10301208496094 0 495257.94 y 6
192.1063232421875 0 36464.44
199.07147216796875 0 18997.451
201.12368774414062 0 196939.62
202.1257781982422 0 13916.772
211.14430236816406 0 13572.998
215.13865661621094 0 21714.488
217.08230590820312 0 108918.87
218.08627319335938 0 7619.561
223.06448364257812 0 13222.163
225.04261779785156 0 10067.491
225.5511932373047 0 7491.852
227.18698120117188 0 69198.42
228.19386291503906 0 10470.632
229.1186981201172 0 45371.395
229.15478515625 0 20886.945
231.06129455566406 0 28157.135
233.14080810546875 0 7383.3486
238.06800842285156 0 6970.1055
239.13937377929688 0 14242.089
242.1134796142578 0 12232.864
257.1490478515625 0 8115.645
270.1094055175781 0 12838.957
270.19293212890625 0 13488.028
271.20062255859375 0 18340.54
274.1751708984375 0 21961.482
284.16607666015625 0 8962.49
286.1036376953125 0 10030.338
286.1507873535156 0 17069.969
287.449462890625 0 7378.525
288.1195373535156 0 72556.66 y Water loss 5
289.1229248046875 0 8929.636
293.1717224121094 0 48835.773 c Ammonia loss 4
304.1143493652344 0 8793.758
306.12994384765625 0 23138.277 y 5
311.59765625 0 7907.4263
312.1784362792969 0 12534.228
314.0991516113281 0 14298.85
314.6929931640625 0 9853.518
315.2267150878906 0 20192.643
317.1811218261719 0 27782.518
329.2308044433594 0 35591.285
330.23797607421875 0 607763.2
331.24334716796875 0 166933.19
332.1101989746094 0 26067.002
332.2471923828125 0 22027.72
335.662109375 0 24646.031
336.16552734375 0 10672.109
338.20556640625 0 8609.166
339.2143859863281 0 10419.766
340.1982116699219 0 14613.615
341.0183410644531 0 96059.01
341.2300720214844 0 49807.44
341.6800537109375 0 26614.14
342.019287109375 0 28091.043
342.18023681640625 0 14346.221
342.1885986328125 0 12032.188
342.21417236328125 0 150753.6
342.2359924316406 0 10388.39
343.0167236328125 0 12785.337
343.21649169921875 0 27835.15
343.69549560546875 0 14739.883
344.1457214355469 0 33781.953
344.2415771484375 0 8170.1064
344.6667785644531 0 20021.254
345.169677734375 0 8771.73
348.6875915527344 0 7536.553
350.6852111816406 0 229277.52 c Ammonia loss 5
351.1868591308594 0 51717.02
351.6869201660156 0 24028.922
355.2330017089844 0 39211.457
356.2177429199219 0 10611.296
356.2410583496094 0 25745.812 c Water loss 2
357.2248840332031 0 227503.03 c Ammonia loss 2
358.1954650878906 0 12691.093
358.22760009765625 0 33371.742
359.0289611816406 0 319768.72
359.28253173828125 0 8311.261
360.0293884277344 0 95413.67
361.026611328125 0 34134.047
362.2276611328125 0 16784.078
364.7004699707031 0 13886.949
369.20135498046875 0 59807.59
369.6952819824219 0 95079.89
370.2079772949219 0 74461.28
370.71063232421875 0 27473.8
372.23516845703125 0 12555.005
373.24359130859375 0 1252671.6
374.25115966796875 0 3233615.8 c 2
375.25421142578125 0 543039.7
376.2567443847656 0 61516.312
378.2063293457031 0 371932.78
378.70111083984375 0 243884.4
379.200927734375 0 70221.24
379.69921875 0 13038.33
381.1777038574219 0 9503.31
383.1982116699219 0 92708.03
383.6988525390625 0 34342.68
384.1990051269531 0 9960.735
385.1355895996094 0 17494.293
385.2198486328125 0 29595.146
386.2270812988281 0 9961.386
387.2117004394531 0 368623.44
387.71282958984375 0 138152.42
388.2142333984375 0 34810.438
391.7128601074219 0 36107.664
392.2040100097656 0 386879
392.70550537109375 0 102618.41
393.2060241699219 0 42077.133 y Water loss 1
393.7088623046875 0 9028.185 y Ammonia loss 1
395.24151611328125 0 8612.817
401.2094421386719 0 781423.75 c Ammonia loss 6
401.7107849121094 0 296583.44
402.18560791015625 0 11651.487
402.2119445800781 0 66006.12 y 1
403.1465148925781 0 32383.773 y Water loss 4
404.7199401855469 0 17099.975
405.2177429199219 0 14536.383
405.70928955078125 0 11945.117
412.2556457519531 0 31515.38
413.25054931640625 0 27889.688
413.7251892089844 0 108345.35
414.2261962890625 0 34206.793
414.2574462890625 0 14646.981
414.7161865234375 0 15473.071
417.1983642578125 0 13161.788
419.71466064453125 0 7700.3184
421.1556091308594 0 10758.509 y 4
425.26300048828125 0 20276.594
427.182861328125 0 13630.224
427.30364990234375 0 13695.064
427.7225341796875 0 118697.74
428.2234802246094 0 54076.188
428.7273254394531 0 11041.879
429.08941650390625 0 88846.52
430.09033203125 0 34208.504
430.26544189453125 0 147222.66
431.08544921875 0 17326.48
431.26812744140625 0 31063.223
436.7279052734375 0 729441.25
437.2296447753906 0 320353.6
437.7301330566406 0 98238.875
442.31463623046875 0 64203.61
443.2735595703125 0 29013.18
443.32177734375 0 72876
444.32843017578125 0 20210.168
445.195556640625 0 21258.062
445.7331237792969 0 63706.11
446.1979064941406 0 15754.274
446.2347412109375 0 27957.377
446.7344970703125 0 9241.731
447.1082763671875 0 8646.977
457.24151611328125 0 36263.07
458.2489013671875 0 189826.45
459.251953125 0 38529.21
468.31719970703125 0 65523.598
469.3240661621094 0 70292.58 c Water loss 3
470.3094177246094 0 157125.72 c Ammonia loss 3
471.311767578125 0 40902.523
485.3206481933594 0 23196.879
486.32794189453125 0 1279119.5
487.3349914550781 0 1597106.4 c 3
488.3378601074219 0 366416.84
489.3411560058594 0 42410.66
498.30572509765625 0 21034.945
499.2615661621094 0 15629.586
500.24737548828125 0 83696.13
501.2508850097656 0 22179.668
502.2605895996094 0 83789.49
502.2903137207031 0 53246.26
503.2636413574219 0 14761.8955
513.3527221679688 0 9249.493
515.2947998046875 0 29297.227
516.23046875 0 26856.781 y Water loss 3
516.2947998046875 0 10231.43
517.2735595703125 0 46663.188
518.2759399414062 0 10535.364
534.2424926757812 0 19980.408 y 3
539.2944946289062 0 25368.4
540.2938232421875 0 14817.83
541.346923828125 0 23105.373
542.330078125 0 81752.266
543.3330078125 0 20999.475
545.2947387695312 0 14285.417
555.2900390625 0 46768.62
556.2904052734375 0 10572.243
557.3001098632812 0 11126.349
557.342041015625 0 23645.43
558.294921875 0 23053.963
558.348876953125 0 242436.64
559.3522338867188 0 79018.16
560.3524169921875 0 10570.346
567.324462890625 0 14328.828
568.3197631835938 0 8867.825
573.2994995117188 0 100343.34
574.2711181640625 0 131697.47
575.2740478515625 0 44207.324
576.2792358398438 0 14679.155
583.2838134765625 0 30256.48
583.3433227539062 0 33680.91
584.3516845703125 0 24031.836 c Water loss 4
585.3363037109375 0 440094.72 c Ammonia loss 4
586.339111328125 0 152786.61
587.279052734375 0 34154.98
587.342529296875 0 23360.172
601.2957153320312 0 97099.34
601.3550415039062 0 461292.7
602.362548828125 0 4302304 c 4
603.3655395507812 0 1256269.4
604.36767578125 0 232611.48
605.3709106445312 0 14311.141
613.3306274414062 0 47387.082
615.3727416992188 0 10345.503
618.2608032226562 0 483646.38
619.2640991210938 0 122026.984
620.2672729492188 0 30048.172
628.3087158203125 0 9400.854
628.3790283203125 0 30562.215
629.3217163085938 0 23107.992
630.333984375 0 290299.6
631.2687377929688 0 847893.25
631.3351440429688 0 88904.28
632.271728515625 0 215075.52
632.3338012695312 0 17293.312
633.2753295898438 0 43640.863
641.3653564453125 0 21769.518
646.3128662109375 0 10342.648
652.3050537109375 0 13846.785
654.3180541992188 0 15196.827
656.31396484375 0 13103.579 z Water loss 2
657.3572387695312 0 78509.31
658.3643798828125 0 168183.05
659.3703002929688 0 59816.875
660.3695068359375 0 11148.042
670.3150634765625 0 48366.055
670.3773193359375 0 7685.1465
671.3222045898438 0 14587.696
672.3312377929688 0 116497.75 y Water loss 2
672.38134765625 0 10340.022
673.3214111328125 0 19785.906 y Ammonia loss 2
673.3764038085938 0 77148.17
674.3236694335938 0 1379327.4 z 2
675.3261108398438 0 414126.7
675.3836059570312 0 21904.043
676.3287963867188 0 86523.41
682.3525390625 0 81888.984
683.3546752929688 0 31092.271
684.3547973632812 0 13662.532
688.3263549804688 0 89871.4
689.3295288085938 0 26853.004
690.3421630859375 0 828121.06 y 2
691.34521484375 0 263793.66
692.3479614257812 0 66391.88
696.3670654296875 0 17093.375
700.3629760742188 0 2072870.4 c Ammonia loss 5
701.3659057617188 0 710555.25
702.3737182617188 0 316102.34
703.3795166015625 0 66032.484
714.3788452148438 0 15491.985
717.389404296875 0 2569269.5 c 5
718.3922729492188 0 888449.75
719.337890625 0 10534.547
719.3951416015625 0 196176.67
720.4020385742188 0 15604.695
731.3388671875 0 9270.598
743.4191284179688 0 13627.314
744.3527221679688 0 681222.5 w 1
745.3558349609375 0 249929.73
746.358154296875 0 63211.91
756.4114990234375 0 72407.62
757.3892822265625 0 137640.61
758.3921508789062 0 60552.438
759.3803100585938 0 11501.76
765.39111328125 0 12346.585
773.3472900390625 0 9079.124
773.4158935546875 0 50381.93
774.4097900390625 0 41287.24
775.3862915039062 0 62533.844
776.3596801757812 0 67882.69
777.3635864257812 0 21122.531
783.3167724609375 0 14347.47
783.3997802734375 0 352356.62
784.403564453125 0 131224.47
785.4074096679688 0 47348.598 y Water loss 1
787.4072265625 0 462636.44 z 1
788.410888671875 0 199128.77
789.41162109375 0 58909.58
790.3709716796875 0 45144.777
791.3714599609375 0 10283.298
801.4108276367188 0 1352317.2 c Ammonia loss 6
802.4136962890625 0 488320.97
803.4185791015625 0 169577.36 y 1
804.4268798828125 0 34876.69
813.4364624023438 0 54158.207
814.428466796875 0 21240.143
815.426025390625 0 12287.282
817.395263671875 0 20397.293
818.3741455078125 0 1898608
818.4392700195312 0 2319731.8 c 6
819.376708984375 0 687339.4
819.4417724609375 0 857255.25
820.3797607421875 0 173892.67
820.44384765625 0 232248.77
821.3768310546875 0 14296.318
821.4536743164062 0 16135.787
829.458251953125 0 34057.855
830.4492797851562 0 500097.06
831.4451293945312 0 4291222
832.4445190429688 0 1704418.5
833.4458618164062 0 421603.06
834.4406127929688 0 42459.31
835.404052734375 0 160234.67
836.406982421875 0 51997.14
844.4270629882812 0 16131.804
845.4600219726562 0 65538.74
846.4603271484375 0 23059.885
847.4323120117188 0 21328
848.4362182617188 0 791326.9
849.4392700195312 0 302052.5
850.4412841796875 0 107885.875
851.4461669921875 0 11118.085
856.4287109375 0 44451.76
857.4251708984375 0 60615.566
858.4247436523438 0 23912.262
859.4234008789062 0 13243.087
863.4676513671875 0 14090.519
873.453857421875 0 111921.62
874.4397583007812 0 4734553
875.4425048828125 0 1943653.5
876.444580078125 0 548497.7
877.4480590820312 0 47026.81
889.4379272460938 0 9794.288
890.4579467773438 0 2644236.2
891.462890625 0 2090916.5
891.6648559570312 0 11171.089
892.4659423828125 0 669380.75
893.469970703125 0 119724.42
1289.2103271484375 0 10021.44
1359.6910400390625 0 10382.89
3465.00634765625 0 10392.781

Spectrum Details

|  |  |
| --- | --- |
| Matched peaks? Matched peaks The total absolute number of peaks matched. Additionally in brackets the total fraction of peaks matched and the total number of peaks is shown. | 36 (9.65% of 373) |
| FDR? FDR The false discovery rate estimated for this peptide. It is calculated by matching all theoretical fragments with a non-integer shift with the raw peaks for this spectrum. This is done with 40 different shifts. The resulting percentage is the average number of annotated peaks over the number of annotated peaks with the correct spectrum. | 2.05% |
| Satellite FDR? Satellite FDR See the FDR for details on its calculation. This satellite ion specific FDR only contains the satellite ions (d/w) for I/L/J positions. | 2.38% |
| PSM Score? PSM Score The PSM Score as given by Hecklib to this annotated spectrum. It is shown with three significant figures. | 454 |

#### Spectrum 4418? Spectrum 4418 The raw spectrum of this peptide as annotated by Hecklib. The fragments are coloured according to ion type (see legend). Any peaks with a star '\*' as text can be hovered over to see the full details, first the ion type second the mass shift type. By hovering over the amino acids in the peptide or ions in the legend the corresponding peaks are highlighted. By toggling the 'Unassigned' label you can turn the background (unassigned) peaks on or off in the plot. By updating the slider in the Ion legend you can update the spectrum to only show the top X% of the peaks with labels. The top X% means any peak that is within X% of the highest intensity. By dragging in the spectrum you can zoom in to a specific part of the spectrum and use 'Zoom Out' to get back to the original zoom level. The annotation of the spectrum is based on the given sequence in the peptides file and is done with different software so inconsistencies are likely. The peaks are annotated based on the given sequence, with 20 ppm tolerance.

Copy Data

##### Spectrum 4418 (TSV)

###### Preview

```
Loading example...
```

*Click on the button to copy the data to your clipboard.*

Mz MinMz MaxIntensity Max

WidthHeightPeptide font sizePeptide stroke widthSpectrum font sizeSpectrum stroke widthCompact peptide

Ion legend

wxyz

abcd

OtherUnassignedIon^Charge^_Position_Show for top:%

SLRJDDTA

07.87e+41.57e+52.36e+53.15e+5

Zoom Out

y^+1^_2_y^+1^_2_y^+1^_3_c^+2^_5_y^+1^_3_c^+2^_6_c^+1^_3_c^+1^_3_c^+1^_3_y^+2^_7_c^+2^_7_c^+2^_7_y^+2^_7_y^+1^_4_c^+1^_4_c^+1^_4_c^+1^_4_y^+1^_5_y^+1^_5_c^+1^_5_c^+1^_5_c^+1^_5_z^+1^_6_y^+1^_6_y^+1^_6_z^+1^_6_y^+1^_6_c^+1^_6_c^+1^_6_w^+1^_7_y^+1^_7_z^+1^_7_c^+1^_7_y^+1^_7_c^+1^_7_

0874174826213495

Fragment Matches Table

Show background peaks

| Position | Ion type | Intensity | mz Theoretical | mz Error (Th) | mz Error (ppm) | Charge | Series Number |
| --- | --- | --- | --- | --- | --- | --- | --- |
| - | - | 870.8 | 120.1 | - | - | 0 | - |
| - | - | 603.5 | 127.1 | - | - | 0 | - |
| - | - | 438.1 | 129.1 | - | - | 0 | - |
| - | - | 457.1 | 130.4 | - | - | 0 | - |
| - | - | 702.9 | 133.1 | - | - | 0 | - |
| - | - | 516.3 | 133.1 | - | - | 0 | - |
| - | - | 399.3 | 135.4 | - | - | 0 | - |
| - | - | 377 | 139.5 | - | - | 0 | - |
| - | - | 407.9 | 142.9 | - | - | 0 | - |
| - | - | 442.5 | 142.9 | - | - | 0 | - |
| - | - | 436.6 | 144.1 | - | - | 0 | - |
| - | - | 1123 | 147.1 | - | - | 0 | - |
| - | - | 4.013E+04 | 149 | - | - | 0 | - |
| - | - | 3264 | 150 | - | - | 0 | - |
| - | - | 1816 | 151 | - | - | 0 | - |
| - | - | 813.1 | 155.1 | - | - | 0 | - |
| - | - | 1612 | 157.1 | - | - | 0 | - |
| - | - | 1.256E+04 | 167.1 | - | - | 0 | - |
| - | - | 733.9 | 167.1 | - | - | 0 | - |
| - | - | 910 | 168.1 | - | - | 0 | - |
| - | - | 883.8 | 169.1 | - | - | 0 | - |
| - | - | 500.3 | 169.4 | - | - | 0 | - |
| - | - | 1491 | 171.1 | - | - | 0 | - |
| 7 | y | 6008 | 173.1 | 0.0002927 | 1.691 | +1 | 2 |
| - | - | 2.838E+04 | 173.1 | - | - | 0 | - |
| - | - | 2029 | 174.1 | - | - | 0 | - |
| - | - | 526.8 | 183.1 | - | - | 0 | - |
| - | - | 4449 | 189.1 | - | - | 0 | - |
| 7 | y | 2.573E+04 | 191.1 | 0.0003482 | 1.822 | +1 | 2 |
| - | - | 2063 | 192.1 | - | - | 0 | - |
| - | - | 1092 | 199.1 | - | - | 0 | - |
| - | - | 1.153E+04 | 201.1 | - | - | 0 | - |
| - | - | 891.7 | 202.1 | - | - | 0 | - |
| - | - | 685.2 | 207 | - | - | 0 | - |
| - | - | 1872 | 209 | - | - | 0 | - |
| - | - | 970.3 | 211.1 | - | - | 0 | - |
| - | - | 3850 | 215.1 | - | - | 0 | - |
| - | - | 777.6 | 216.1 | - | - | 0 | - |
| - | - | 6763 | 217.1 | - | - | 0 | - |
| - | - | 928.1 | 218.1 | - | - | 0 | - |
| - | - | 1.242E+04 | 223.1 | - | - | 0 | - |
| - | - | 1406 | 224.1 | - | - | 0 | - |
| - | - | 1.035E+04 | 225 | - | - | 0 | - |
| - | - | 1652 | 226 | - | - | 0 | - |
| - | - | 721.3 | 227 | - | - | 0 | - |
| - | - | 1054 | 227 | - | - | 0 | - |
| - | - | 4569 | 227.2 | - | - | 0 | - |
| - | - | 3311 | 229.1 | - | - | 0 | - |
| - | - | 1320 | 229.2 | - | - | 0 | - |
| - | - | 1310 | 231.1 | - | - | 0 | - |
| - | - | 2085 | 234.1 | - | - | 0 | - |
| - | - | 849.7 | 239.1 | - | - | 0 | - |
| - | - | 965.1 | 239.1 | - | - | 0 | - |
| - | - | 814.8 | 270.1 | - | - | 0 | - |
| - | - | 634.9 | 271.2 | - | - | 0 | - |
| - | - | 1249 | 274.2 | - | - | 0 | - |
| - | - | 509.9 | 283.1 | - | - | 0 | - |
| - | - | 1120 | 285 | - | - | 0 | - |
| - | - | 933.8 | 286.1 | - | - | 0 | - |
| - | - | 998.9 | 286.2 | - | - | 0 | - |
| 6 | y | 4204 | 288.1 | 0.0004646 | 1.612 | +1 | 3 |
| - | - | 799.1 | 289.1 | - | - | 0 | - |
| 5 | c | 3531 | 293.2 | 0.0004628 | 1.579 | +2 | 5 |
| - | - | 1172 | 293.7 | - | - | 0 | - |
| - | - | 511.8 | 296.4 | - | - | 0 | - |
| - | - | 984.9 | 299.1 | - | - | 0 | - |
| 6 | y | 1089 | 306.1 | 0.0007947 | 2.596 | +1 | 3 |
| - | - | 657.9 | 311 | - | - | 0 | - |
| - | - | 854.6 | 312.2 | - | - | 0 | - |
| - | - | 1165 | 314.1 | - | - | 0 | - |
| - | - | 903.1 | 314.7 | - | - | 0 | - |
| - | - | 920.9 | 315.2 | - | - | 0 | - |
| - | - | 1584 | 317.2 | - | - | 0 | - |
| - | - | 1975 | 318.9 | - | - | 0 | - |
| - | - | 2383 | 325 | - | - | 0 | - |
| - | - | 1312 | 326 | - | - | 0 | - |
| - | - | 1798 | 329.2 | - | - | 0 | - |
| - | - | 3.333E+04 | 330.2 | - | - | 0 | - |
| - | - | 884.8 | 331.2 | - | - | 0 | - |
| - | - | 8129 | 331.2 | - | - | 0 | - |
| - | - | 1479 | 332.1 | - | - | 0 | - |
| - | - | 820.5 | 332.2 | - | - | 0 | - |
| - | - | 902.6 | 332.2 | - | - | 0 | - |
| - | - | 699.9 | 332.7 | - | - | 0 | - |
| - | - | 623.9 | 334.2 | - | - | 0 | - |
| - | - | 594.8 | 339.2 | - | - | 0 | - |
| - | - | 1054 | 340.2 | - | - | 0 | - |
| - | - | 8.229E+04 | 341 | - | - | 0 | - |
| - | - | 1658 | 341.2 | - | - | 0 | - |
| - | - | 1473 | 341.7 | - | - | 0 | - |
| - | - | 2.526E+04 | 342 | - | - | 0 | - |
| - | - | 8123 | 342.2 | - | - | 0 | - |
| - | - | 3199 | 343 | - | - | 0 | - |
| - | - | 1.195E+04 | 343 | - | - | 0 | - |
| - | - | 855.5 | 343.2 | - | - | 0 | - |
| - | - | 1648 | 344 | - | - | 0 | - |
| - | - | 1179 | 344.1 | - | - | 0 | - |
| - | - | 1711 | 344.7 | - | - | 0 | - |
| - | - | 1042 | 345 | - | - | 0 | - |
| 6 | c | 1.14E+04 | 350.7 | 0.0002359 | 0.6728 | +2 | 6 |
| - | - | 3701 | 351.2 | - | - | 0 | - |
| - | - | 6803 | 355.1 | - | - | 0 | - |
| - | - | 2650 | 355.2 | - | - | 0 | - |
| - | - | 1454 | 356.1 | - | - | 0 | - |
| - | - | 669 | 356.2 | - | - | 0 | - |
| 3 | c | 1917 | 356.2 | 0.0002889 | 0.8109 | +1 | 3 |
| - | - | 905.8 | 357.1 | - | - | 0 | - |
| 3 | c | 1.495E+04 | 357.2 | 0.0003736 | 1.046 | +1 | 3 |
| - | - | 2093 | 358.2 | - | - | 0 | - |
| - | - | 3.115E+05 | 359 | - | - | 0 | - |
| - | - | 1.012E+05 | 360 | - | - | 0 | - |
| - | - | 4.739E+04 | 361 | - | - | 0 | - |
| - | - | 1180 | 364.7 | - | - | 0 | - |
| - | - | 734.7 | 367.9 | - | - | 0 | - |
| - | - | 4089 | 369.2 | - | - | 0 | - |
| - | - | 3429 | 369.7 | - | - | 0 | - |
| - | - | 4022 | 370.2 | - | - | 0 | - |
| - | - | 979.9 | 370.7 | - | - | 0 | - |
| - | - | 608.8 | 372.2 | - | - | 0 | - |
| - | - | 6.724E+04 | 373.2 | - | - | 0 | - |
| - | - | 624.5 | 373.9 | - | - | 0 | - |
| 3 | c | 1.829E+05 | 374.3 | 6.966E-05 | 0.1861 | +1 | 3 |
| - | - | 3.234E+04 | 375.3 | - | - | 0 | - |
| - | - | 2797 | 376.3 | - | - | 0 | - |
| - | - | 1.854E+04 | 378.2 | - | - | 0 | - |
| - | - | 1.202E+04 | 378.7 | - | - | 0 | - |
| - | - | 3808 | 379.2 | - | - | 0 | - |
| - | - | 831.2 | 379.7 | - | - | 0 | - |
| - | - | 5568 | 383.2 | - | - | 0 | - |
| - | - | 1548 | 383.7 | - | - | 0 | - |
| - | - | 1124 | 384.2 | - | - | 0 | - |
| - | - | 909.5 | 385.1 | - | - | 0 | - |
| - | - | 1675 | 385.2 | - | - | 0 | - |
| - | - | 414.8 | 387.2 | - | - | 0 | - |
| - | - | 1.869E+04 | 387.2 | - | - | 0 | - |
| - | - | 5681 | 387.7 | - | - | 0 | - |
| - | - | 1690 | 388.2 | - | - | 0 | - |
| - | - | 1405 | 391.7 | - | - | 0 | - |
| - | - | 2.038E+04 | 392.2 | - | - | 0 | - |
| - | - | 6312 | 392.7 | - | - | 0 | - |
| 2 | y | 1829 | 393.2 | 0.00515 | 13.1 | +2 | 7 |
| 7 | c | 789.3 | 400.7 | 0.0006616 | 1.651 | +2 | 7 |
| 7 | c | 3.832E+04 | 401.2 | 0.0005971 | 1.488 | +2 | 7 |
| - | - | 1.204E+04 | 401.7 | - | - | 0 | - |
| 2 | y | 4972 | 402.2 | 0.004268 | 10.61 | +2 | 7 |
| - | - | 1059 | 402.2 | - | - | 0 | - |
| 5 | y | 1286 | 403.1 | 8.076E-05 | 0.2003 | +1 | 4 |
| - | - | 1081 | 403.2 | - | - | 0 | - |
| - | - | 877.3 | 404.7 | - | - | 0 | - |
| - | - | 1073 | 405.2 | - | - | 0 | - |
| - | - | 2003 | 412.3 | - | - | 0 | - |
| - | - | 1667 | 413.2 | - | - | 0 | - |
| - | - | 4479 | 413.7 | - | - | 0 | - |
| - | - | 2096 | 414.2 | - | - | 0 | - |
| - | - | 864.3 | 414.3 | - | - | 0 | - |
| - | - | 1718 | 414.7 | - | - | 0 | - |
| - | - | 1310 | 418.7 | - | - | 0 | - |
| - | - | 617.9 | 424.3 | - | - | 0 | - |
| - | - | 1315 | 425.3 | - | - | 0 | - |
| - | - | 874.9 | 427.3 | - | - | 0 | - |
| - | - | 5600 | 427.7 | - | - | 0 | - |
| - | - | 559.3 | 428.1 | - | - | 0 | - |
| - | - | 3522 | 428.2 | - | - | 0 | - |
| - | - | 783.3 | 428.9 | - | - | 0 | - |
| - | - | 9.643E+04 | 429.1 | - | - | 0 | - |
| - | - | 4.263E+04 | 430.1 | - | - | 0 | - |
| - | - | 666.4 | 430.2 | - | - | 0 | - |
| - | - | 8235 | 430.3 | - | - | 0 | - |
| - | - | 2.101E+04 | 431.1 | - | - | 0 | - |
| - | - | 2144 | 431.3 | - | - | 0 | - |
| - | - | 4.457E+04 | 436.7 | - | - | 0 | - |
| - | - | 1.99E+04 | 437.2 | - | - | 0 | - |
| - | - | 4895 | 437.7 | - | - | 0 | - |
| - | - | 743.1 | 438.2 | - | - | 0 | - |
| - | - | 3720 | 442.3 | - | - | 0 | - |
| - | - | 2021 | 443.3 | - | - | 0 | - |
| - | - | 4247 | 443.3 | - | - | 0 | - |
| - | - | 1455 | 444.3 | - | - | 0 | - |
| - | - | 4650 | 445.1 | - | - | 0 | - |
| - | - | 3745 | 445.7 | - | - | 0 | - |
| - | - | 2625 | 446.1 | - | - | 0 | - |
| - | - | 2372 | 446.2 | - | - | 0 | - |
| - | - | 970.6 | 447.1 | - | - | 0 | - |
| - | - | 2067 | 457.2 | - | - | 0 | - |
| - | - | 9872 | 458.2 | - | - | 0 | - |
| - | - | 2512 | 459.3 | - | - | 0 | - |
| - | - | 3415 | 468.3 | - | - | 0 | - |
| 4 | c | 5049 | 469.3 | 0.0006452 | 1.375 | +1 | 4 |
| 4 | c | 8007 | 470.3 | 0.0007213 | 1.534 | +1 | 4 |
| - | - | 1499 | 471.3 | - | - | 0 | - |
| - | - | 1700 | 485.3 | - | - | 0 | - |
| - | - | 7.397E+04 | 486.3 | - | - | 0 | - |
| 4 | c | 8.661E+04 | 487.3 | 0.0002236 | 0.4588 | +1 | 4 |
| - | - | 2.067E+04 | 488.3 | - | - | 0 | - |
| - | - | 2984 | 489.3 | - | - | 0 | - |
| - | - | 944.5 | 498.3 | - | - | 0 | - |
| - | - | 4314 | 500.2 | - | - | 0 | - |
| - | - | 1365 | 501.3 | - | - | 0 | - |
| - | - | 3505 | 502.3 | - | - | 0 | - |
| - | - | 3339 | 502.3 | - | - | 0 | - |
| - | - | 818.7 | 503.3 | - | - | 0 | - |
| - | - | 1472 | 513.4 | - | - | 0 | - |
| - | - | 1856 | 515.3 | - | - | 0 | - |
| 4 | y | 806.2 | 516.2 | 0.002175 | 4.212 | +1 | 5 |
| - | - | 3692 | 517.3 | - | - | 0 | - |
| - | - | 1842 | 518.3 | - | - | 0 | - |
| 4 | y | 895.3 | 534.2 | 0.00606 | 11.34 | +1 | 5 |
| - | - | 2413 | 539.3 | - | - | 0 | - |
| - | - | 633.4 | 540.3 | - | - | 0 | - |
| - | - | 1315 | 541.3 | - | - | 0 | - |
| - | - | 4939 | 542.3 | - | - | 0 | - |
| - | - | 1312 | 543.3 | - | - | 0 | - |
| - | - | 753.5 | 545.3 | - | - | 0 | - |
| - | - | 1705 | 555.3 | - | - | 0 | - |
| - | - | 1009 | 556.3 | - | - | 0 | - |
| - | - | 1064 | 557.3 | - | - | 0 | - |
| - | - | 2039 | 558.3 | - | - | 0 | - |
| - | - | 1.346E+04 | 558.3 | - | - | 0 | - |
| - | - | 4576 | 559.4 | - | - | 0 | - |
| - | - | 861.4 | 567.3 | - | - | 0 | - |
| - | - | 568.5 | 571.3 | - | - | 0 | - |
| - | - | 6155 | 573.3 | - | - | 0 | - |
| - | - | 8199 | 574.3 | - | - | 0 | - |
| - | - | 2364 | 575.3 | - | - | 0 | - |
| - | - | 2090 | 583.3 | - | - | 0 | - |
| - | - | 1933 | 583.3 | - | - | 0 | - |
| - | - | 961.3 | 584.3 | - | - | 0 | - |
| 5 | c | 2036 | 584.4 | 0.003449 | 5.902 | +1 | 5 |
| 5 | c | 2.58E+04 | 585.3 | 0.0006948 | 1.187 | +1 | 5 |
| - | - | 7543 | 586.3 | - | - | 0 | - |
| - | - | 2069 | 587.3 | - | - | 0 | - |
| - | - | 1800 | 587.3 | - | - | 0 | - |
| - | - | 615.8 | 595.3 | - | - | 0 | - |
| - | - | 4407 | 601.3 | - | - | 0 | - |
| - | - | 2.891E+04 | 601.4 | - | - | 0 | - |
| 5 | c | 2.519E+05 | 602.4 | 0.0003908 | 0.6487 | +1 | 5 |
| - | - | 7.078E+04 | 603.4 | - | - | 0 | - |
| - | - | 723.7 | 603.4 | - | - | 0 | - |
| - | - | 1.419E+04 | 604.4 | - | - | 0 | - |
| - | - | 2796 | 605.3 | - | - | 0 | - |
| - | - | 633.1 | 605.4 | - | - | 0 | - |
| - | - | 698.6 | 606.3 | - | - | 0 | - |
| - | - | 749.1 | 610.3 | - | - | 0 | - |
| - | - | 1986 | 613.3 | - | - | 0 | - |
| - | - | 1201 | 615.4 | - | - | 0 | - |
| - | - | 2.864E+04 | 618.3 | - | - | 0 | - |
| - | - | 6547 | 619.3 | - | - | 0 | - |
| - | - | 1238 | 620.3 | - | - | 0 | - |
| - | - | 1322 | 628.4 | - | - | 0 | - |
| - | - | 1165 | 629.3 | - | - | 0 | - |
| - | - | 1.641E+04 | 630.3 | - | - | 0 | - |
| - | - | 4.697E+04 | 631.3 | - | - | 0 | - |
| - | - | 5004 | 631.3 | - | - | 0 | - |
| - | - | 1.405E+04 | 632.3 | - | - | 0 | - |
| - | - | 939.3 | 632.3 | - | - | 0 | - |
| - | - | 2634 | 633.3 | - | - | 0 | - |
| - | - | 1006 | 641.4 | - | - | 0 | - |
| - | - | 1024 | 654.3 | - | - | 0 | - |
| 3 | z | 593.3 | 657.3 | 0.007839 | 11.93 | +1 | 6 |
| - | - | 3591 | 657.4 | - | - | 0 | - |
| - | - | 9914 | 658.4 | - | - | 0 | - |
| - | - | 3893 | 659.4 | - | - | 0 | - |
| - | - | 847.8 | 660.4 | - | - | 0 | - |
| - | - | 2039 | 670.3 | - | - | 0 | - |
| 3 | y | 5923 | 672.3 | 0.0005023 | 0.7472 | +1 | 6 |
| 3 | y | 1249 | 673.3 | 0.007425 | 11.03 | +1 | 6 |
| - | - | 3132 | 673.4 | - | - | 0 | - |
| 3 | z | 7.706E+04 | 674.3 | 0.0005159 | 0.7651 | +1 | 6 |
| - | - | 2.435E+04 | 675.3 | - | - | 0 | - |
| - | - | 1214 | 675.4 | - | - | 0 | - |
| - | - | 5183 | 676.3 | - | - | 0 | - |
| - | - | 679.5 | 677.3 | - | - | 0 | - |
| - | - | 5120 | 682.4 | - | - | 0 | - |
| - | - | 1190 | 683.4 | - | - | 0 | - |
| - | - | 994.5 | 687.3 | - | - | 0 | - |
| - | - | 5209 | 688.3 | - | - | 0 | - |
| - | - | 1137 | 689.3 | - | - | 0 | - |
| 3 | y | 4.728E+04 | 690.3 | 0.0002245 | 0.3252 | +1 | 6 |
| - | - | 1.554E+04 | 691.3 | - | - | 0 | - |
| - | - | 3431 | 692.3 | - | - | 0 | - |
| - | - | 1340 | 696.4 | - | - | 0 | - |
| 6 | c | 1.157E+05 | 700.4 | 0.000302 | 0.4312 | +1 | 6 |
| - | - | 3.914E+04 | 701.4 | - | - | 0 | - |
| - | - | 1.706E+04 | 702.4 | - | - | 0 | - |
| - | - | 3814 | 703.4 | - | - | 0 | - |
| - | - | 923.5 | 704.4 | - | - | 0 | - |
| - | - | 1143 | 714.4 | - | - | 0 | - |
| 6 | c | 1.507E+05 | 717.4 | 0.0001811 | 0.2525 | +1 | 6 |
| - | - | 4.981E+04 | 718.4 | - | - | 0 | - |
| - | - | 1.184E+04 | 719.4 | - | - | 0 | - |
| - | - | 1076 | 720.4 | - | - | 0 | - |
| - | - | 1470 | 731.3 | - | - | 0 | - |
| 2 | w | 4.021E+04 | 744.4 | 0.000402 | 0.54 | +1 | 7 |
| - | - | 1.207E+04 | 745.4 | - | - | 0 | - |
| - | - | 4084 | 746.4 | - | - | 0 | - |
| - | - | 3836 | 756.4 | - | - | 0 | - |
| - | - | 6850 | 757.4 | - | - | 0 | - |
| - | - | 2872 | 758.4 | - | - | 0 | - |
| - | - | 1568 | 759.4 | - | - | 0 | - |
| - | - | 2490 | 773.4 | - | - | 0 | - |
| - | - | 2743 | 774.4 | - | - | 0 | - |
| - | - | 5105 | 775.4 | - | - | 0 | - |
| - | - | 3837 | 776.4 | - | - | 0 | - |
| - | - | 1018 | 777.4 | - | - | 0 | - |
| - | - | 2.013E+04 | 783.4 | - | - | 0 | - |
| - | - | 958.2 | 783.9 | - | - | 0 | - |
| - | - | 7204 | 784.4 | - | - | 0 | - |
| 2 | y | 2748 | 785.4 | 0.007418 | 9.445 | +1 | 7 |
| 2 | z | 2.961E+04 | 787.4 | 0.0001311 | 0.1665 | +1 | 7 |
| - | - | 9937 | 788.4 | - | - | 0 | - |
| - | - | 3716 | 789.4 | - | - | 0 | - |
| - | - | 2580 | 790.4 | - | - | 0 | - |
| - | - | 949.4 | 791.4 | - | - | 0 | - |
| 7 | c | 7.519E+04 | 801.4 | 0.000353 | 0.4405 | +1 | 7 |
| - | - | 2.9E+04 | 802.4 | - | - | 0 | - |
| 2 | y | 8550 | 803.4 | 0.007485 | 9.316 | +1 | 7 |
| - | - | 1800 | 804.4 | - | - | 0 | - |
| - | - | 2775 | 813.4 | - | - | 0 | - |
| - | - | 1584 | 814.4 | - | - | 0 | - |
| - | - | 1069 | 816.4 | - | - | 0 | - |
| - | - | 728.6 | 817.4 | - | - | 0 | - |
| - | - | 1.041E+05 | 818.4 | - | - | 0 | - |
| 7 | c | 1.346E+05 | 818.4 | 0.002429 | 2.968 | +1 | 7 |
| - | - | 4.037E+04 | 819.4 | - | - | 0 | - |
| - | - | 5.028E+04 | 819.4 | - | - | 0 | - |
| - | - | 1.126E+04 | 820.4 | - | - | 0 | - |
| - | - | 1.329E+04 | 820.4 | - | - | 0 | - |
| - | - | 1819 | 821.4 | - | - | 0 | - |
| - | - | 2018 | 829.5 | - | - | 0 | - |
| - | - | 2.923E+04 | 830.4 | - | - | 0 | - |
| - | - | 2.448E+05 | 831.4 | - | - | 0 | - |
| - | - | 1.008E+05 | 832.4 | - | - | 0 | - |
| - | - | 2.708E+04 | 833.4 | - | - | 0 | - |
| - | - | 2992 | 834.4 | - | - | 0 | - |
| - | - | 9189 | 835.4 | - | - | 0 | - |
| - | - | 3522 | 836.4 | - | - | 0 | - |
| - | - | 784.1 | 837.4 | - | - | 0 | - |
| - | - | 1300 | 844.4 | - | - | 0 | - |
| - | - | 3358 | 845.5 | - | - | 0 | - |
| - | - | 2350 | 846.5 | - | - | 0 | - |
| - | - | 1054 | 847.4 | - | - | 0 | - |
| - | - | 4.333E+04 | 848.4 | - | - | 0 | - |
| - | - | 1.735E+04 | 849.4 | - | - | 0 | - |
| - | - | 4772 | 850.4 | - | - | 0 | - |
| - | - | 1608 | 856.4 | - | - | 0 | - |
| - | - | 3545 | 857.4 | - | - | 0 | - |
| - | - | 2237 | 858.4 | - | - | 0 | - |
| - | - | 756.9 | 859.4 | - | - | 0 | - |
| - | - | 1637 | 863.5 | - | - | 0 | - |
| - | - | 7326 | 873.4 | - | - | 0 | - |
| - | - | 2.726E+05 | 874.4 | - | - | 0 | - |
| - | - | 1.108E+05 | 875.4 | - | - | 0 | - |
| - | - | 3.231E+04 | 876.4 | - | - | 0 | - |
| - | - | 3228 | 877.4 | - | - | 0 | - |
| - | - | 1449 | 883.9 | - | - | 0 | - |
| - | - | 870.7 | 884.4 | - | - | 0 | - |
| - | - | 856.4 | 888.4 | - | - | 0 | - |
| - | - | 3589 | 889.4 | - | - | 0 | - |
| - | - | 1.608E+05 | 890.5 | - | - | 0 | - |
| - | - | 1.142E+05 | 891.5 | - | - | 0 | - |
| - | - | 4.066E+04 | 892.5 | - | - | 0 | - |
| - | - | 5881 | 892.9 | - | - | 0 | - |
| - | - | 8865 | 893.5 | - | - | 0 | - |
| - | - | 1145 | 894.5 | - | - | 0 | - |
| - | - | 745.3 | 922.5 | - | - | 0 | - |
| - | - | 737.8 | 923.5 | - | - | 0 | - |
| - | - | 654.1 | 1175 | - | - | 0 | - |
| - | - | 810.2 | 1768 | - | - | 0 | - |
| - | - | 677.8 | 1769 | - | - | 0 | - |
| - | - | 846.3 | 1770 | - | - | 0 | - |
| - | - | 1346 | 1784 | - | - | 0 | - |
| - | - | 2492 | 1785 | - | - | 0 | - |
| - | - | 1364 | 1786 | - | - | 0 | - |
| - | - | 838.4 | 1787 | - | - | 0 | - |
| - | - | 769.3 | 3043 | - | - | 0 | - |
| - | - | 671.6 | 3112 | - | - | 0 | - |
| - | - | 813 | 3461 | - | - | 0 | - |

m/z Charge Intensity FragmentType MassShift Position
120.06596374511719 0 870.8416
127.08736419677734 0 603.54913
129.10287475585938 0 438.053
130.4119415283203 0 457.14877
133.06106567382812 0 702.90955
133.086181640625 0 516.28906
135.4265594482422 0 399.26096
139.50978088378906 0 376.98068
142.85545349121094 0 407.87842
142.91773986816406 0 442.45728
144.06651306152344 0 436.62167
147.06585693359375 0 1122.7179
149.04518127441406 0 40127.273
150.04473876953125 0 3263.7498
151.0421600341797 0 1815.7352
155.08184814453125 0 813.14954
157.10862731933594 0 1611.8746
167.0557403564453 0 12564.103
167.0639190673828 0 733.9229
168.0553436279297 0 910.0419
169.05345153808594 0 883.76825
169.39089965820312 0 500.33737
171.07699584960938 0 1490.6948
173.0923614501953 0 6007.703 y Water loss 6
173.12876892089844 0 28383.383
174.1324920654297 0 2028.7773
183.1126251220703 0 526.75494
189.0872039794922 0 4449.1235
191.1029815673828 0 25734.309 y 6
192.10646057128906 0 2062.7083
199.07168579101562 0 1092.2407
201.12368774414062 0 11527.874
202.12667846679688 0 891.71423
207.0326385498047 0 685.1797
208.95326232910156 0 1871.5795
211.14378356933594 0 970.27527
215.1390380859375 0 3850.3284
216.0989532470703 0 777.5836
217.0822296142578 0 6762.7695
218.08590698242188 0 928.07056
223.0640411376953 0 12418.083
224.06451416015625 0 1406.4974
225.0433349609375 0 10346.467
226.04302978515625 0 1651.9336
227.02279663085938 0 721.26263
227.04034423828125 0 1054.0947
227.18692016601562 0 4568.9194
229.11866760253906 0 3310.6619
229.15487670898438 0 1320.0427
231.06170654296875 0 1310.196
234.10897827148438 0 2085.1567
239.09487915039062 0 849.6812
239.138427734375 0 965.081
270.10791015625 0 814.83734
271.2010803222656 0 634.9466
274.1751403808594 0 1248.8782
283.0683898925781 0 509.86725
285.00994873046875 0 1120.4125
286.10302734375 0 933.81256
286.15216064453125 0 998.89166
288.1194763183594 0 4203.71 y Water loss 5
289.1221923828125 0 799.1114
293.1718444824219 0 3530.7249 c Ammonia loss 4
293.6728820800781 0 1171.7944
296.4132385253906 0 511.75354
299.0618591308594 0 984.8978
306.13037109375 0 1089.2054 y 5
310.9703369140625 0 657.90295
312.15570068359375 0 854.5969
314.09857177734375 0 1164.641
314.6927490234375 0 903.10547
315.2247009277344 0 920.871
317.1812438964844 0 1584.0714
318.9228210449219 0 1974.6669
324.98699951171875 0 2382.8225
325.98748779296875 0 1311.5922
329.23065185546875 0 1798.092
330.2379150390625 0 33334.902
331.2023010253906 0 884.7755
331.24346923828125 0 8128.588
332.10931396484375 0 1478.8422
332.208251953125 0 820.48694
332.2475891113281 0 902.6141
332.6755065917969 0 699.85754
334.160888671875 0 623.90155
339.2153015136719 0 594.84485
340.199462890625 0 1053.8444
341.01849365234375 0 82289.52
341.22869873046875 0 1658.2307
341.67962646484375 0 1473.3099
342.0188293457031 0 25260.03
342.21380615234375 0 8123.04
342.99658203125 0 3199.4148
343.0163879394531 0 11949.402
343.2152099609375 0 855.5032
343.998779296875 0 1647.8271
344.14404296875 0 1179.0825
344.6670227050781 0 1710.6039
344.99560546875 0 1042.3735
350.6850891113281 0 11404.554 c Ammonia loss 5
351.1866149902344 0 3700.9548
355.07037353515625 0 6802.8647
355.2334289550781 0 2649.9048
356.0712890625 0 1454.1388
356.2175598144531 0 669.00073
356.2407531738281 0 1917.2848 c Water loss 2
357.0677490234375 0 905.7831
357.224853515625 0 14951.16 c Ammonia loss 2
358.2276306152344 0 2093.1538
359.0287780761719 0 311485.94
360.0291442871094 0 101213.414
361.026611328125 0 47391.77
364.7012634277344 0 1180.344
367.8690490722656 0 734.7105
369.2007141113281 0 4088.7534
369.69439697265625 0 3428.9836
370.2080078125 0 4021.6133
370.7118835449219 0 979.8988
372.2370300292969 0 608.82306
373.2435607910156 0 67240.45
373.906494140625 0 624.46277
374.2510986328125 0 182945.61 c 2
375.2539367675781 0 32338.46
376.25567626953125 0 2796.506
378.2060546875 0 18544.844
378.7007141113281 0 12019.65
379.20098876953125 0 3808.1414
379.6999206542969 0 831.16125
383.19818115234375 0 5568.4087
383.69891357421875 0 1547.8273
384.2012939453125 0 1123.6566
385.13623046875 0 909.46643
385.2189636230469 0 1674.7655
387.18994140625 0 414.79352
387.2115783691406 0 18693.273
387.712646484375 0 5680.573
388.21533203125 0 1690.1497
391.71368408203125 0 1404.591
392.2040710449219 0 20384.037
392.7051696777344 0 6311.699
393.2060852050781 0 1828.7195 y Water loss 1
400.71734619140625 0 789.25275 c Water loss 6
401.20928955078125 0 38320.777 c Ammonia loss 6
401.71051025390625 0 12035.361
402.2122497558594 0 4972.291 y 1
402.2394104003906 0 1059.4432
403.1458740234375 0 1286.2539 y Water loss 4
403.2442932128906 0 1081.1566
404.7196044921875 0 877.2692
405.2106628417969 0 1073.3622
412.25506591796875 0 2002.8575
413.2499084472656 0 1666.9728
413.7254638671875 0 4478.648
414.2252197265625 0 2096.1045
414.25726318359375 0 864.3177
414.7176208496094 0 1717.7579
418.7165222167969 0 1309.5023
424.30206298828125 0 617.9196
425.26202392578125 0 1315.188
427.30267333984375 0 874.86774
427.7222900390625 0 5600.098
428.1001281738281 0 559.30273
428.2233581542969 0 3522.2163
428.8908996582031 0 783.3449
429.0894470214844 0 96426.055
430.0899353027344 0 42626.402
430.2338562011719 0 666.4222
430.265380859375 0 8235.424
431.08758544921875 0 21005.268
431.26922607421875 0 2143.6233
436.7278137207031 0 44566.34
437.2293395996094 0 19896.395
437.7307434082031 0 4895.0024
438.229736328125 0 743.0946
442.31427001953125 0 3720.122
443.2731628417969 0 2021.4652
443.3208923339844 0 4246.7944
444.32684326171875 0 1455.1494
445.12060546875 0 4649.8433
445.7332458496094 0 3745.225
446.12176513671875 0 2624.6748
446.2353210449219 0 2371.5596
447.1187438964844 0 970.56525
457.2412414550781 0 2067.4023
458.2485656738281 0 9872.235
459.2520751953125 0 2512.2317
468.31787109375 0 3414.7278
469.3238830566406 0 5048.7134 c Water loss 3
470.30926513671875 0 8007.0703 c Ammonia loss 3
471.3111267089844 0 1498.958
485.3206787109375 0 1700.4198
486.3277587890625 0 73974.586
487.3348693847656 0 86609.586 c 3
488.3375549316406 0 20668.582
489.3418884277344 0 2983.726
498.3030090332031 0 944.5246
500.2469482421875 0 4314.2446
501.2516174316406 0 1364.9302
502.25921630859375 0 3505.2117
502.2889099121094 0 3339.222
503.2917785644531 0 818.7404
513.3516235351562 0 1471.5823
515.2945556640625 0 1856.3777
516.2278442382812 0 806.2459 y Water loss 3
517.271728515625 0 3691.5903
518.273193359375 0 1841.6709
534.2466430664062 0 895.26184 y 3
539.2940063476562 0 2413.4988
540.2936401367188 0 633.4494
541.3472290039062 0 1315.3531
542.3301391601562 0 4939.196
543.3328247070312 0 1312.4694
545.29345703125 0 753.484
555.2890014648438 0 1704.6807
556.2865600585938 0 1009.3111
557.3411865234375 0 1063.6083
558.295166015625 0 2038.7726
558.3489379882812 0 13460.079
559.3526000976562 0 4576.393
567.325439453125 0 861.4027
571.3300170898438 0 568.51355
573.2999877929688 0 6154.776
574.270751953125 0 8198.692
575.274169921875 0 2363.503
583.2833251953125 0 2089.7705
583.3443603515625 0 1933.1019
584.2883911132812 0 961.2637
584.3480224609375 0 2036.3228 c Water loss 4
585.336181640625 0 25795.158 c Ammonia loss 4
586.3387451171875 0 7543.4033
587.2796020507812 0 2069.1318
587.3425903320312 0 1799.7451
595.2921752929688 0 615.82513
601.2948608398438 0 4407.31
601.3546752929688 0 28910.676
602.3624267578125 0 251857.23 c 4
603.3652954101562 0 70783.69
603.4248046875 0 723.7168
604.3673706054688 0 14187.237
605.3040161132812 0 2796.08
605.3585205078125 0 633.0775
606.3098754882812 0 698.60394
610.3018188476562 0 749.0738
613.33056640625 0 1986.1863
615.3726806640625 0 1201.0977
618.2605590820312 0 28643.916
619.263916015625 0 6547.253
620.2693481445312 0 1237.6777
628.3787841796875 0 1322.0997
629.3275756835938 0 1165.3984
630.3338623046875 0 16410.41
631.2684326171875 0 46970.535
631.3341064453125 0 5004.127
632.2715454101562 0 14047.678
632.3314819335938 0 939.3443
633.2731323242188 0 2634.3843
641.36328125 0 1005.7449
654.3207397460938 0 1023.92816
657.3042602539062 0 593.3116 z Ammonia loss 2
657.3569946289062 0 3590.7598
658.3645629882812 0 9913.741
659.3704833984375 0 3893.0469
660.3699340820312 0 847.8374
670.3165893554688 0 2038.5867
672.3306274414062 0 5922.6426 y Water loss 2
673.3225708007812 0 1249.0582 y Ammonia loss 2
673.37744140625 0 3132.2793
674.323486328125 0 77062.516 z 2
675.3255004882812 0 24348.857
675.3875122070312 0 1214.4934
676.3265991210938 0 5183.3945
677.3252563476562 0 679.45245
682.3521118164062 0 5119.895
683.3552856445312 0 1190.4735
687.3426513671875 0 994.50433
688.32763671875 0 5208.517
689.3263549804688 0 1136.836
690.3419189453125 0 47283.793 y 2
691.3448486328125 0 15538.089
692.3468017578125 0 3431.0293
696.3670043945312 0 1340.0704
700.3627319335938 0 115726.27 c Ammonia loss 5
701.3655395507812 0 39141.82
702.3740234375 0 17059.564
703.3792114257812 0 3814.0684
704.3839111328125 0 923.4678
714.3778076171875 0 1143.305
717.38916015625 0 150654.2 c 5
718.3921508789062 0 49806.82
719.3953247070312 0 11837.221
720.3967895507812 0 1075.6206
731.342529296875 0 1469.7067
744.3526611328125 0 40212.715 w 1
745.3555908203125 0 12072.388
746.358642578125 0 4083.816
756.4118041992188 0 3836.2183
757.3893432617188 0 6850.0425
758.3887939453125 0 2871.6736
759.3801879882812 0 1567.9562
773.4149780273438 0 2490.0483
774.387451171875 0 2742.9702
775.3832397460938 0 5105.4487
776.3600463867188 0 3836.9175
777.3609619140625 0 1018.0778
783.3993530273438 0 20131.068
783.8662719726562 0 958.16174
784.4017333984375 0 7204.4893
785.4077758789062 0 2748.463 y Water loss 1
787.4071655273438 0 29605.266 z 1
788.4102783203125 0 9937.43
789.402099609375 0 3716.2996
790.3767700195312 0 2580.218
791.3699951171875 0 949.36523
801.4104614257812 0 75194.15 c Ammonia loss 6
802.4135131835938 0 29001.055
803.4182739257812 0 8549.927 y 1
804.425048828125 0 1800.1758
813.4329223632812 0 2775.3147
814.4335327148438 0 1583.8207
816.4130859375 0 1069.4888
817.3826293945312 0 728.64954
818.3737182617188 0 104084.78
818.4390869140625 0 134644.84 c 6
819.3765869140625 0 40369.26
819.4417724609375 0 50279.535
820.3799438476562 0 11263.519
820.4432983398438 0 13285.045
821.4434814453125 0 1819.4753
829.4584350585938 0 2018.1163
830.4483032226562 0 29230.967
831.4448852539062 0 244790.48
832.4448852539062 0 100791.34
833.44580078125 0 27080.746
834.4364624023438 0 2991.589
835.4037475585938 0 9189.039
836.4052734375 0 3522.469
837.4091796875 0 784.0954
844.427490234375 0 1300.4268
845.4569702148438 0 3357.6108
846.451904296875 0 2349.9268
847.4391479492188 0 1053.5688
848.4357299804688 0 43326.086
849.4384155273438 0 17348.508
850.4420166015625 0 4771.925
856.4287719726562 0 1607.7538
857.4239501953125 0 3545.3188
858.4222412109375 0 2236.7605
859.425048828125 0 756.8539
863.4681396484375 0 1636.698
873.441650390625 0 7326.381
874.439208984375 0 272625.9
875.442138671875 0 110833.195
876.4446411132812 0 32312.672
877.4457397460938 0 3227.8958
883.9171142578125 0 1448.7228
884.4078369140625 0 870.70856
888.4369506835938 0 856.3733
889.4404296875 0 3589.3682
890.4572143554688 0 160829.83
891.4620971679688 0 114182.62
892.4635009765625 0 40658.047
892.9326171875 0 5880.6553
893.4637451171875 0 8865.037
894.4811401367188 0 1145.309
922.4544067382812 0 745.33075
923.4618530273438 0 737.8457
1174.559326171875 0 654.11896
1767.8411865234375 0 810.2164
1768.8299560546875 0 677.822
1769.8441162109375 0 846.32214
1783.857421875 0 1346.0018
1784.8612060546875 0 2491.512
1785.8612060546875 0 1363.9753
1786.8570556640625 0 838.4054
3043.041748046875 0 769.28876
3111.89697265625 0 671.5804
3460.6982421875 0 813.0306

Spectrum Details

|  |  |
| --- | --- |
| Matched peaks? Matched peaks The total absolute number of peaks matched. Additionally in brackets the total fraction of peaks matched and the total number of peaks is shown. | 35 (9.28% of 377) |
| FDR? FDR The false discovery rate estimated for this peptide. It is calculated by matching all theoretical fragments with a non-integer shift with the raw peaks for this spectrum. This is done with 40 different shifts. The resulting percentage is the average number of annotated peaks over the number of annotated peaks with the correct spectrum. | 2.04% |
| Satellite FDR? Satellite FDR See the FDR for details on its calculation. This satellite ion specific FDR only contains the satellite ions (d/w) for I/L/J positions. | 0.00% |
| PSM Score? PSM Score The PSM Score as given by Hecklib to this annotated spectrum. It is shown with three significant figures. | 432 |

#### Reverse Lookup? Reverse Lookup All places where this read could be placed.

| Group | Segment | Template | Template Part | Read Part | Score | Unique |
| --- | --- | --- | --- | --- | --- | --- |
| Homo sapiens Heavy Chain | IGHV | IGHV1-18 | [84..92] | [0..8] | 55 | True |

| Recombined | Template Part | Read Part | Score | Unique |
| --- | --- | --- | --- | --- |
| REC-0-1 | [84..92] | [0..8] | 64 | True |

#### Meta Information from Multiple reads

##### Number of combined reads

2

##### Intensity

0.8056

##### TotalArea

4.053E+08

##### Changes to the peptide sequence

SLRJDDTA

L→JNo support for either Leucine or Isoleucine based on side chain ions (Position: 4)

#### Positional Score

Copy Data

##### Positional Score (TSV)

###### Preview

```
Loading example...
```

*Click on the button to copy the data to your clipboard.*

0001234567

Label Value
"0" 0
"1" 0
"2" 0
"3" 0
"4" 0
"5" 0
"6" 0
"7" 0

#### Meta Information from PEAKS

##### Scan Identifier

F1:4344

##### Original sequence

S

L

R

L

D

D

T

A

##### Posttranslational Modifications

##### Source File

D:\separate\_stitch\_analyses\xle-disambiguation\raw\20210323\_F1\_UM1\_Peng0013\_SA\_F59\_ingel\_3ug\_ELA.raw

##### Fraction

1

##### Scan Feature

F1:2074

##### De Novo Score

98

##### ConfidenceScore

98

### m/z

445.7337

##### Mass

889.4505

##### Charge

2

##### Retention Time

22.86

##### Predicted Retention Time

-

##### Area

2.026E+08

##### Parts Per Million

2.6

##### Fragmentation mode

ETHCD

##### Originating file

01 D:\separate\_stitch\_analyses\xle-disambiguation\20210325\_F59\_3ug\_DENOVO\_12.csv

#### Meta Information from PEAKS

##### Scan Identifier

F1:4418

##### Original sequence

S

L

R

L

D

D

T

A

##### Posttranslational Modifications

##### Source File

D:\separate\_stitch\_analyses\xle-disambiguation\raw\20210323\_F1\_UM1\_Peng0013\_SA\_F59\_ingel\_3ug\_ELA.raw

##### Fraction

1

##### Scan Feature

F1:2074

##### De Novo Score

98

##### ConfidenceScore

98

### m/z

445.7337

##### Mass

889.4505

##### Charge

2

##### Retention Time

22.86

##### Predicted Retention Time

-

##### Area

2.026E+08

##### Parts Per Million

2.6

##### Fragmentation mode

ETHCD

##### Originating file

01 D:\separate\_stitch\_analyses\xle-disambiguation\20210325\_F59\_3ug\_DENOVO\_12.csv
