## Supplementary material for "A handle on mass coincidence errors in *de novo* sequencing of antibodies by bottom-up proteomics": Combined_059.html

Details Combined\_059 | Stitch OverviewUndefined

### Read Combined\_059

#### Sequence (length=8)

JSJTPEQW

#### Spectrum 9728? Spectrum 9728 The raw spectrum of this peptide as annotated by Hecklib. The fragments are coloured according to ion type (see legend). Any peaks with a star '\*' as text can be hovered over to see the full details, first the ion type second the mass shift type. By hovering over the amino acids in the peptide or ions in the legend the corresponding peaks are highlighted. By toggling the 'Unassigned' label you can turn the background (unassigned) peaks on or off in the plot. By updating the slider in the Ion legend you can update the spectrum to only show the top X% of the peaks with labels. The top X% means any peak that is within X% of the highest intensity. By dragging in the spectrum you can zoom in to a specific part of the spectrum and use 'Zoom Out' to get back to the original zoom level. The annotation of the spectrum is based on the given sequence in the peptides file and is done with different software so inconsistencies are likely. The peaks are annotated based on the given sequence, with 20 ppm tolerance.

Copy Data

##### Spectrum 9728 (TSV)

###### Preview

```
Loading example...
```

*Click on the button to copy the data to your clipboard.*

Mz MinMz MaxIntensity Max

WidthHeightPeptide font sizePeptide stroke widthSpectrum font sizeSpectrum stroke widthCompact peptide

Ion legend

wxyz

abcd

OtherUnassignedIon^Charge^_Position_Show for top:%

JSJTPEQW

09.00e+41.80e+52.70e+53.60e+5

Zoom Out

y^+1^_1_y^+1^_2_z^+1^_2_c^+2^_6_c^+1^_3_y^+1^_2_y^+1^_3_y^+1^_3_z^+1^_3_y^+1^_3_c^+1^_5_z^+1^_4_c^+1^_5_y^+1^_4_y^+1^_4_z^+1^_4_y^+1^_4_z^+1^_5_w^+1^_5_c^+1^_6_y^+1^_5_z^+1^_5_c^+1^_6_y^+1^_5_w^+1^_6_y^+1^_6_z^+1^_6_c^+1^_7_c^+1^_7_z^+1^_7_w^+1^_7_z^+1^_7_y^+1^_7_

0827165324803307

Fragment Matches Table

Show background peaks

| Position | Ion type | Intensity | mz Theoretical | mz Error (Th) | mz Error (ppm) | Charge | Series Number |
| --- | --- | --- | --- | --- | --- | --- | --- |
| - | - | 380.4 | 122.5 | - | - | 0 | - |
| - | - | 402.5 | 124.2 | - | - | 0 | - |
| - | - | 954 | 129.1 | - | - | 0 | - |
| - | - | 2862 | 130.1 | - | - | 0 | - |
| - | - | 947.6 | 132.1 | - | - | 0 | - |
| - | - | 919.1 | 133.1 | - | - | 0 | - |
| - | - | 413.4 | 142.1 | - | - | 0 | - |
| - | - | 660 | 142.1 | - | - | 0 | - |
| - | - | 631.5 | 144.1 | - | - | 0 | - |
| - | - | 514.4 | 145.1 | - | - | 0 | - |
| - | - | 1385 | 146.1 | - | - | 0 | - |
| - | - | 686.7 | 146.1 | - | - | 0 | - |
| - | - | 727.8 | 155.1 | - | - | 0 | - |
| - | - | 1.506E+04 | 159.1 | - | - | 0 | - |
| - | - | 1828 | 160.1 | - | - | 0 | - |
| - | - | 511 | 172.6 | - | - | 0 | - |
| - | - | 7846 | 173.1 | - | - | 0 | - |
| - | - | 882.2 | 173.4 | - | - | 0 | - |
| - | - | 1454 | 177.1 | - | - | 0 | - |
| - | - | 1138 | 183.1 | - | - | 0 | - |
| - | - | 1013 | 184.1 | - | - | 0 | - |
| - | - | 515.4 | 185.3 | - | - | 0 | - |
| - | - | 1176 | 187.1 | - | - | 0 | - |
| - | - | 858.4 | 187.1 | - | - | 0 | - |
| - | - | 3748 | 187.1 | - | - | 0 | - |
| - | - | 4.876E+04 | 188.1 | - | - | 0 | - |
| - | - | 5639 | 189.1 | - | - | 0 | - |
| - | - | 495.9 | 190.1 | - | - | 0 | - |
| - | - | 3599 | 199.1 | - | - | 0 | - |
| - | - | 1007 | 201.1 | - | - | 0 | - |
| - | - | 2.02E+04 | 201.1 | - | - | 0 | - |
| - | - | 536.1 | 201.7 | - | - | 0 | - |
| - | - | 1470 | 202.1 | - | - | 0 | - |
| 8 | y | 6.207E+04 | 205.1 | 0.0001359 | 0.6628 | +1 | 1 |
| - | - | 7133 | 206.1 | - | - | 0 | - |
| - | - | 626.7 | 207.1 | - | - | 0 | - |
| - | - | 1.123E+04 | 215.1 | - | - | 0 | - |
| - | - | 524.9 | 218.9 | - | - | 0 | - |
| - | - | 504.5 | 222.3 | - | - | 0 | - |
| - | - | 938.3 | 227.1 | - | - | 0 | - |
| - | - | 1.228E+04 | 227.1 | - | - | 0 | - |
| - | - | 1682 | 228.1 | - | - | 0 | - |
| - | - | 1197 | 240.1 | - | - | 0 | - |
| - | - | 1036 | 240.2 | - | - | 0 | - |
| - | - | 624.6 | 240.4 | - | - | 0 | - |
| - | - | 1374 | 242.7 | - | - | 0 | - |
| - | - | 575.9 | 243.2 | - | - | 0 | - |
| - | - | 3.695E+04 | 244.1 | - | - | 0 | - |
| - | - | 448.5 | 245.1 | - | - | 0 | - |
| - | - | 3731 | 245.1 | - | - | 0 | - |
| - | - | 1337 | 255.1 | - | - | 0 | - |
| - | - | 577.8 | 256.7 | - | - | 0 | - |
| - | - | 1383 | 257.1 | - | - | 0 | - |
| - | - | 2683 | 258.1 | - | - | 0 | - |
| - | - | 1073 | 258.1 | - | - | 0 | - |
| - | - | 642.8 | 269.2 | - | - | 0 | - |
| - | - | 2120 | 270.1 | - | - | 0 | - |
| - | - | 7617 | 284.2 | - | - | 0 | - |
| - | - | 930.6 | 285.2 | - | - | 0 | - |
| - | - | 780.1 | 287.2 | - | - | 0 | - |
| - | - | 1183 | 287.2 | - | - | 0 | - |
| - | - | 2.4E+04 | 296.2 | - | - | 0 | - |
| - | - | 4730 | 297.2 | - | - | 0 | - |
| - | - | 1907 | 298.1 | - | - | 0 | - |
| - | - | 691 | 298.2 | - | - | 0 | - |
| - | - | 659.8 | 300.2 | - | - | 0 | - |
| - | - | 546.2 | 301.2 | - | - | 0 | - |
| - | - | 756.3 | 302.2 | - | - | 0 | - |
| - | - | 1.153E+04 | 302.2 | - | - | 0 | - |
| - | - | 1515 | 303.2 | - | - | 0 | - |
| - | - | 505.4 | 307.2 | - | - | 0 | - |
| - | - | 2044 | 311.1 | - | - | 0 | - |
| - | - | 706.6 | 312.2 | - | - | 0 | - |
| - | - | 1185 | 313.1 | - | - | 0 | - |
| - | - | 9697 | 314.2 | - | - | 0 | - |
| - | - | 1.677E+04 | 315.1 | - | - | 0 | - |
| - | - | 1237 | 315.2 | - | - | 0 | - |
| 7 | y | 2.509E+04 | 316.1 | 0.000151 | 0.4776 | +1 | 2 |
| - | - | 2731 | 316.1 | - | - | 0 | - |
| 7 | z | 3716 | 317.1 | 0.0045 | 14.19 | +1 | 2 |
| - | - | 3830 | 321.2 | - | - | 0 | - |
| - | - | 998.4 | 321.7 | - | - | 0 | - |
| - | - | 670.8 | 327.1 | - | - | 0 | - |
| - | - | 1689 | 327.2 | - | - | 0 | - |
| - | - | 4525 | 328.2 | - | - | 0 | - |
| - | - | 1200 | 329.2 | - | - | 0 | - |
| - | - | 881.5 | 329.2 | - | - | 0 | - |
| 6 | c | 1104 | 329.7 | 0.0001608 | 0.4878 | +2 | 6 |
| - | - | 1461 | 330.2 | - | - | 0 | - |
| - | - | 631.7 | 330.2 | - | - | 0 | - |
| - | - | 578.8 | 330.2 | - | - | 0 | - |
| 3 | c | 5601 | 331.2 | 0.0001489 | 0.4496 | +1 | 3 |
| - | - | 1092 | 332.2 | - | - | 0 | - |
| 7 | y | 2.266E+04 | 333.2 | 0.0002437 | 0.7316 | +1 | 2 |
| - | - | 4229 | 334.2 | - | - | 0 | - |
| - | - | 2190 | 337.2 | - | - | 0 | - |
| - | - | 1438 | 340.2 | - | - | 0 | - |
| - | - | 600.6 | 343.2 | - | - | 0 | - |
| - | - | 3778 | 345.2 | - | - | 0 | - |
| - | - | 1894 | 352.2 | - | - | 0 | - |
| - | - | 668 | 353.2 | - | - | 0 | - |
| - | - | 3085 | 354.2 | - | - | 0 | - |
| - | - | 5.062E+04 | 355.2 | - | - | 0 | - |
| - | - | 7932 | 356.2 | - | - | 0 | - |
| - | - | 1178 | 357.2 | - | - | 0 | - |
| - | - | 674.7 | 365.2 | - | - | 0 | - |
| - | - | 4151 | 369.2 | - | - | 0 | - |
| - | - | 5867 | 370.2 | - | - | 0 | - |
| - | - | 849.8 | 370.3 | - | - | 0 | - |
| - | - | 2260 | 371.2 | - | - | 0 | - |
| - | - | 1.875E+04 | 372.2 | - | - | 0 | - |
| - | - | 837.7 | 373.2 | - | - | 0 | - |
| - | - | 3846 | 373.2 | - | - | 0 | - |
| - | - | 1663 | 379.2 | - | - | 0 | - |
| - | - | 8734 | 387.3 | - | - | 0 | - |
| - | - | 2275 | 388.3 | - | - | 0 | - |
| - | - | 6611 | 396.2 | - | - | 0 | - |
| - | - | 850.1 | 397.2 | - | - | 0 | - |
| - | - | 2.175E+04 | 397.2 | - | - | 0 | - |
| - | - | 1126 | 398.2 | - | - | 0 | - |
| - | - | 4867 | 398.2 | - | - | 0 | - |
| - | - | 679.6 | 400.2 | - | - | 0 | - |
| - | - | 654.6 | 412.2 | - | - | 0 | - |
| - | - | 2642 | 413.2 | - | - | 0 | - |
| - | - | 522 | 414.2 | - | - | 0 | - |
| - | - | 6.154E+04 | 415.3 | - | - | 0 | - |
| - | - | 1.293E+04 | 416.3 | - | - | 0 | - |
| - | - | 666.5 | 417.2 | - | - | 0 | - |
| - | - | 2211 | 417.3 | - | - | 0 | - |
| - | - | 626.6 | 421.2 | - | - | 0 | - |
| - | - | 555.8 | 423.2 | - | - | 0 | - |
| - | - | 1711 | 427.2 | - | - | 0 | - |
| - | - | 677.3 | 428.2 | - | - | 0 | - |
| - | - | 3110 | 429.2 | - | - | 0 | - |
| - | - | 1305 | 430.2 | - | - | 0 | - |
| - | - | 947.9 | 437.7 | - | - | 0 | - |
| - | - | 3307 | 439.2 | - | - | 0 | - |
| - | - | 618.2 | 439.2 | - | - | 0 | - |
| - | - | 1234 | 440.2 | - | - | 0 | - |
| - | - | 929.9 | 440.2 | - | - | 0 | - |
| - | - | 883.5 | 440.3 | - | - | 0 | - |
| - | - | 8928 | 441.2 | - | - | 0 | - |
| - | - | 1124 | 441.2 | - | - | 0 | - |
| - | - | 4679 | 442.2 | - | - | 0 | - |
| - | - | 950.1 | 443.2 | - | - | 0 | - |
| - | - | 1131 | 443.3 | - | - | 0 | - |
| 6 | y | 2366 | 444.2 | 0.0001685 | 0.3793 | +1 | 3 |
| 6 | y | 1628 | 445.2 | 0.0003889 | 0.8736 | +1 | 3 |
| 6 | z | 1892 | 446.2 | 6.572E-05 | 0.1473 | +1 | 3 |
| - | - | 607.2 | 446.7 | - | - | 0 | - |
| - | - | 1499 | 447.2 | - | - | 0 | - |
| - | - | 641 | 447.2 | - | - | 0 | - |
| - | - | 576.8 | 450.3 | - | - | 0 | - |
| - | - | 531.4 | 451.2 | - | - | 0 | - |
| - | - | 1.644E+04 | 456.2 | - | - | 0 | - |
| - | - | 3325 | 457.2 | - | - | 0 | - |
| - | - | 2.199E+04 | 458.2 | - | - | 0 | - |
| - | - | 1121 | 458.3 | - | - | 0 | - |
| - | - | 5018 | 459.2 | - | - | 0 | - |
| - | - | 845.3 | 460.7 | - | - | 0 | - |
| 6 | y | 3181 | 462.2 | 0.0001741 | 0.3766 | +1 | 3 |
| - | - | 4548 | 468.2 | - | - | 0 | - |
| - | - | 1249 | 468.3 | - | - | 0 | - |
| - | - | 1463 | 469.2 | - | - | 0 | - |
| - | - | 1170 | 469.7 | - | - | 0 | - |
| - | - | 779.4 | 470.2 | - | - | 0 | - |
| - | - | 3029 | 471.3 | - | - | 0 | - |
| - | - | 925.7 | 471.3 | - | - | 0 | - |
| - | - | 2963 | 473.2 | - | - | 0 | - |
| - | - | 792.4 | 474.2 | - | - | 0 | - |
| - | - | 789.9 | 478.2 | - | - | 0 | - |
| - | - | 681.8 | 478.7 | - | - | 0 | - |
| - | - | 2285 | 484.2 | - | - | 0 | - |
| - | - | 641.5 | 485.2 | - | - | 0 | - |
| - | - | 2133 | 487.2 | - | - | 0 | - |
| - | - | 580.9 | 487.3 | - | - | 0 | - |
| - | - | 1603 | 488.2 | - | - | 0 | - |
| - | - | 2256 | 494.3 | - | - | 0 | - |
| - | - | 886.9 | 495.3 | - | - | 0 | - |
| - | - | 1124 | 496.2 | - | - | 0 | - |
| - | - | 947.8 | 499.3 | - | - | 0 | - |
| - | - | 889.9 | 500.2 | - | - | 0 | - |
| - | - | 607.8 | 507.2 | - | - | 0 | - |
| - | - | 1098 | 510.3 | - | - | 0 | - |
| 5 | c | 1130 | 511.3 | 9.858E-05 | 0.1928 | +1 | 5 |
| - | - | 3706 | 512.3 | - | - | 0 | - |
| - | - | 951.4 | 513.2 | - | - | 0 | - |
| - | - | 1115 | 513.2 | - | - | 0 | - |
| - | - | 1389 | 513.3 | - | - | 0 | - |
| - | - | 9066 | 514.3 | - | - | 0 | - |
| - | - | 3057 | 515.2 | - | - | 0 | - |
| - | - | 3090 | 515.3 | - | - | 0 | - |
| - | - | 1215 | 516.3 | - | - | 0 | - |
| - | - | 9508 | 524.2 | - | - | 0 | - |
| 5 | z | 3094 | 525.2 | 0.003721 | 7.085 | +1 | 4 |
| - | - | 1.485E+04 | 528.3 | - | - | 0 | - |
| 5 | c | 1.176E+05 | 529.3 | 0.0004399 | 0.831 | +1 | 5 |
| - | - | 1879 | 530.3 | - | - | 0 | - |
| - | - | 3.1E+04 | 530.3 | - | - | 0 | - |
| - | - | 513 | 531.3 | - | - | 0 | - |
| - | - | 4267 | 531.3 | - | - | 0 | - |
| - | - | 782.3 | 534.3 | - | - | 0 | - |
| - | - | 1274 | 537.3 | - | - | 0 | - |
| 5 | y | 5523 | 541.2 | 0.0002285 | 0.4221 | +1 | 4 |
| 5 | y | 2.163E+04 | 542.2 | 8.888E-06 | 0.01639 | +1 | 4 |
| 5 | z | 6080 | 543.2 | 0.004581 | 8.433 | +1 | 4 |
| - | - | 971.3 | 543.3 | - | - | 0 | - |
| - | - | 1066 | 544.2 | - | - | 0 | - |
| - | - | 1212 | 551.3 | - | - | 0 | - |
| - | - | 605.5 | 553.3 | - | - | 0 | - |
| - | - | 1733 | 554.3 | - | - | 0 | - |
| - | - | 3885 | 555.3 | - | - | 0 | - |
| - | - | 3530 | 555.3 | - | - | 0 | - |
| - | - | 1830 | 556.3 | - | - | 0 | - |
| - | - | 1603 | 556.4 | - | - | 0 | - |
| - | - | 8786 | 558.2 | - | - | 0 | - |
| 5 | y | 1.331E+05 | 559.3 | 0.0004782 | 0.8551 | +1 | 4 |
| - | - | 3.772E+04 | 560.3 | - | - | 0 | - |
| - | - | 6907 | 561.3 | - | - | 0 | - |
| - | - | 2577 | 569.3 | - | - | 0 | - |
| - | - | 3977 | 571.3 | - | - | 0 | - |
| - | - | 1066 | 572.3 | - | - | 0 | - |
| - | - | 1379 | 573.2 | - | - | 0 | - |
| - | - | 686.5 | 574.2 | - | - | 0 | - |
| - | - | 904.3 | 586.2 | - | - | 0 | - |
| - | - | 657 | 588.2 | - | - | 0 | - |
| - | - | 3523 | 598.3 | - | - | 0 | - |
| - | - | 1192 | 598.4 | - | - | 0 | - |
| - | - | 7428 | 599.3 | - | - | 0 | - |
| - | - | 5867 | 600.3 | - | - | 0 | - |
| - | - | 4915 | 600.3 | - | - | 0 | - |
| - | - | 1672 | 601.3 | - | - | 0 | - |
| - | - | 1798 | 601.3 | - | - | 0 | - |
| - | - | 831 | 602.3 | - | - | 0 | - |
| - | - | 882.7 | 605.3 | - | - | 0 | - |
| - | - | 1318 | 613.3 | - | - | 0 | - |
| - | - | 1.206E+04 | 614.4 | - | - | 0 | - |
| - | - | 3992 | 615.4 | - | - | 0 | - |
| - | - | 843.1 | 616.4 | - | - | 0 | - |
| - | - | 1147 | 621.3 | - | - | 0 | - |
| - | - | 5571 | 623.3 | - | - | 0 | - |
| - | - | 1913 | 624.3 | - | - | 0 | - |
| - | - | 9603 | 625.3 | - | - | 0 | - |
| 4 | z | 1447 | 626.3 | 0.0009236 | 1.475 | +1 | 5 |
| - | - | 4146 | 626.3 | - | - | 0 | - |
| 4 | w | 723.8 | 627.3 | 0.0005089 | 0.8112 | +1 | 5 |
| - | - | 1005 | 627.3 | - | - | 0 | - |
| - | - | 3193 | 638.3 | - | - | 0 | - |
| - | - | 1069 | 639.3 | - | - | 0 | - |
| - | - | 4571 | 640.3 | - | - | 0 | - |
| 6 | c | 3195 | 640.4 | 0.000979 | 1.529 | +1 | 6 |
| - | - | 7029 | 641.3 | - | - | 0 | - |
| - | - | 1.054E+04 | 641.4 | - | - | 0 | - |
| - | - | 2.691E+04 | 642.3 | - | - | 0 | - |
| - | - | 2102 | 642.3 | - | - | 0 | - |
| - | - | 2980 | 642.4 | - | - | 0 | - |
| 4 | y | 1.228E+04 | 643.3 | 0.004701 | 7.308 | +1 | 5 |
| - | - | 1013 | 643.3 | - | - | 0 | - |
| - | - | 929.8 | 643.4 | - | - | 0 | - |
| 4 | z | 6.85E+04 | 644.3 | 0.0009292 | 1.442 | +1 | 5 |
| - | - | 5.744E+04 | 645.3 | - | - | 0 | - |
| - | - | 465 | 645.3 | - | - | 0 | - |
| - | - | 1.716E+04 | 646.3 | - | - | 0 | - |
| - | - | 3211 | 647.3 | - | - | 0 | - |
| - | - | 2141 | 650.4 | - | - | 0 | - |
| - | - | 9541 | 656.3 | - | - | 0 | - |
| - | - | 6156 | 657.4 | - | - | 0 | - |
| 6 | c | 2.295E+05 | 658.4 | 0.0006135 | 0.9319 | +1 | 6 |
| - | - | 8.103E+04 | 659.4 | - | - | 0 | - |
| 4 | y | 1.509E+04 | 660.3 | 0.001404 | 2.126 | +1 | 5 |
| - | - | 1.699E+04 | 660.4 | - | - | 0 | - |
| - | - | 6443 | 661.3 | - | - | 0 | - |
| - | - | 939.8 | 661.4 | - | - | 0 | - |
| - | - | 954.3 | 662.3 | - | - | 0 | - |
| - | - | 1812 | 668.4 | - | - | 0 | - |
| - | - | 861.9 | 669.4 | - | - | 0 | - |
| - | - | 2023 | 670.4 | - | - | 0 | - |
| - | - | 1092 | 671.4 | - | - | 0 | - |
| - | - | 890.1 | 683.4 | - | - | 0 | - |
| - | - | 3.287E+04 | 684.4 | - | - | 0 | - |
| - | - | 1.102E+04 | 685.4 | - | - | 0 | - |
| - | - | 3176 | 686.4 | - | - | 0 | - |
| - | - | 835.1 | 687.2 | - | - | 0 | - |
| - | - | 2123 | 688.2 | - | - | 0 | - |
| - | - | 780.5 | 688.3 | - | - | 0 | - |
| - | - | 745 | 689.2 | - | - | 0 | - |
| - | - | 724.3 | 697.3 | - | - | 0 | - |
| - | - | 732.2 | 706.4 | - | - | 0 | - |
| - | - | 1341 | 710.3 | - | - | 0 | - |
| - | - | 1554 | 713.4 | - | - | 0 | - |
| 3 | w | 4889 | 714.3 | 0.0006159 | 0.8622 | +1 | 6 |
| - | - | 7336 | 715.3 | - | - | 0 | - |
| - | - | 3101 | 716.3 | - | - | 0 | - |
| - | - | 835.1 | 717.4 | - | - | 0 | - |
| - | - | 1204 | 723.4 | - | - | 0 | - |
| - | - | 684.2 | 724.4 | - | - | 0 | - |
| - | - | 893.2 | 725.4 | - | - | 0 | - |
| - | - | 1.284E+04 | 728.3 | - | - | 0 | - |
| - | - | 4688 | 729.3 | - | - | 0 | - |
| - | - | 1201 | 730.4 | - | - | 0 | - |
| - | - | 2855 | 733.4 | - | - | 0 | - |
| - | - | 2041 | 734.4 | - | - | 0 | - |
| - | - | 908.9 | 735.4 | - | - | 0 | - |
| - | - | 1804 | 741.4 | - | - | 0 | - |
| - | - | 1.229E+04 | 742.4 | - | - | 0 | - |
| - | - | 5264 | 743.4 | - | - | 0 | - |
| - | - | 1579 | 744.4 | - | - | 0 | - |
| - | - | 1.548E+04 | 751.4 | - | - | 0 | - |
| - | - | 8029 | 752.4 | - | - | 0 | - |
| - | - | 7476 | 753.4 | - | - | 0 | - |
| - | - | 4805 | 754.4 | - | - | 0 | - |
| 3 | y | 2461 | 755.4 | 0.0136 | 18 | +1 | 6 |
| 3 | z | 2.393E+04 | 757.4 | 0.00107 | 1.413 | +1 | 6 |
| - | - | 1.563E+04 | 758.4 | - | - | 0 | - |
| - | - | 5405 | 759.4 | - | - | 0 | - |
| - | - | 916.4 | 760.4 | - | - | 0 | - |
| 7 | c | 1.908E+04 | 769.4 | 0.0004154 | 0.5399 | +1 | 7 |
| - | - | 7442 | 770.4 | - | - | 0 | - |
| - | - | 2.08E+04 | 771.4 | - | - | 0 | - |
| - | - | 7636 | 772.4 | - | - | 0 | - |
| - | - | 2.374E+04 | 773.4 | - | - | 0 | - |
| - | - | 9976 | 774.4 | - | - | 0 | - |
| - | - | 3014 | 775.4 | - | - | 0 | - |
| - | - | 1421 | 782.4 | - | - | 0 | - |
| 7 | c | 3.562E+05 | 786.4 | 0.001269 | 1.613 | +1 | 7 |
| - | - | 1.407E+05 | 787.4 | - | - | 0 | - |
| - | - | 1.129E+04 | 788.3 | - | - | 0 | - |
| - | - | 3.485E+04 | 788.4 | - | - | 0 | - |
| - | - | 4207 | 789.3 | - | - | 0 | - |
| - | - | 3011 | 789.4 | - | - | 0 | - |
| - | - | 853.3 | 790.3 | - | - | 0 | - |
| - | - | 1167 | 797.4 | - | - | 0 | - |
| - | - | 2732 | 799.4 | - | - | 0 | - |
| - | - | 1.196E+04 | 800.4 | - | - | 0 | - |
| - | - | 6053 | 801.4 | - | - | 0 | - |
| - | - | 1184 | 802.4 | - | - | 0 | - |
| - | - | 952.3 | 810.4 | - | - | 0 | - |
| - | - | 2180 | 812.4 | - | - | 0 | - |
| - | - | 3184 | 813.4 | - | - | 0 | - |
| - | - | 2534 | 814.5 | - | - | 0 | - |
| - | - | 698.8 | 815.5 | - | - | 0 | - |
| 2 | z | 4712 | 826.4 | 0.001232 | 1.491 | +1 | 7 |
| 2 | w | 1742 | 827.4 | 0.005761 | 6.963 | +1 | 7 |
| - | - | 1553 | 828.4 | - | - | 0 | - |
| - | - | 869.8 | 829.4 | - | - | 0 | - |
| - | - | 1912 | 837.4 | - | - | 0 | - |
| - | - | 1032 | 838.4 | - | - | 0 | - |
| - | - | 4244 | 841.4 | - | - | 0 | - |
| - | - | 6266 | 842.4 | - | - | 0 | - |
| - | - | 1.697E+04 | 843.4 | - | - | 0 | - |
| 2 | z | 7.507E+04 | 844.4 | 0.0007496 | 0.8877 | +1 | 7 |
| - | - | 3.94E+04 | 845.4 | - | - | 0 | - |
| - | - | 9102 | 846.4 | - | - | 0 | - |
| - | - | 831.6 | 847.4 | - | - | 0 | - |
| - | - | 698.8 | 855.4 | - | - | 0 | - |
| - | - | 5.244E+04 | 857.5 | - | - | 0 | - |
| - | - | 2.67E+04 | 858.5 | - | - | 0 | - |
| - | - | 8630 | 859.5 | - | - | 0 | - |
| 2 | y | 1.044E+04 | 860.4 | 0.001896 | 2.203 | +1 | 7 |
| - | - | 6042 | 861.4 | - | - | 0 | - |
| - | - | 1539 | 862.4 | - | - | 0 | - |
| - | - | 2288 | 867.5 | - | - | 0 | - |
| - | - | 955.3 | 868.5 | - | - | 0 | - |
| - | - | 861.7 | 869.5 | - | - | 0 | - |
| - | - | 1353 | 873.4 | - | - | 0 | - |
| - | - | 1.186E+04 | 884.5 | - | - | 0 | - |
| - | - | 2173 | 885.5 | - | - | 0 | - |
| - | - | 5927 | 885.5 | - | - | 0 | - |
| - | - | 3603 | 886.5 | - | - | 0 | - |
| - | - | 1710 | 887.4 | - | - | 0 | - |
| - | - | 656 | 895.5 | - | - | 0 | - |
| - | - | 938.9 | 900.5 | - | - | 0 | - |
| - | - | 8966 | 901.5 | - | - | 0 | - |
| - | - | 5076 | 902.5 | - | - | 0 | - |
| - | - | 3765 | 903.5 | - | - | 0 | - |
| - | - | 1018 | 904.5 | - | - | 0 | - |
| - | - | 1203 | 911.5 | - | - | 0 | - |
| - | - | 1.598E+04 | 912.5 | - | - | 0 | - |
| - | - | 1.064E+04 | 913.5 | - | - | 0 | - |
| - | - | 1.066E+04 | 914.4 | - | - | 0 | - |
| - | - | 2888 | 914.5 | - | - | 0 | - |
| - | - | 3852 | 915.4 | - | - | 0 | - |
| - | - | 1773 | 915.5 | - | - | 0 | - |
| - | - | 1146 | 916.4 | - | - | 0 | - |
| - | - | 1924 | 918.4 | - | - | 0 | - |
| - | - | 732.8 | 920.5 | - | - | 0 | - |
| - | - | 4511 | 928.5 | - | - | 0 | - |
| - | - | 1.647E+04 | 929.5 | - | - | 0 | - |
| - | - | 1.797E+04 | 930.5 | - | - | 0 | - |
| - | - | 9466 | 931.5 | - | - | 0 | - |
| - | - | 3343 | 932.5 | - | - | 0 | - |
| - | - | 4728 | 939.5 | - | - | 0 | - |
| - | - | 2993 | 940.5 | - | - | 0 | - |
| - | - | 1138 | 941.5 | - | - | 0 | - |
| - | - | 2.54E+04 | 956.5 | - | - | 0 | - |
| - | - | 9.126E+04 | 957.5 | - | - | 0 | - |
| - | - | 4.789E+04 | 958.5 | - | - | 0 | - |
| - | - | 1.499E+04 | 959.5 | - | - | 0 | - |
| - | - | 1548 | 960.5 | - | - | 0 | - |
| - | - | 1.275E+05 | 973.5 | - | - | 0 | - |
| - | - | 2.074E+05 | 974.5 | - | - | 0 | - |
| - | - | 1.005E+05 | 975.5 | - | - | 0 | - |
| - | - | 2.703E+04 | 976.5 | - | - | 0 | - |
| - | - | 2513 | 977.5 | - | - | 0 | - |
| - | - | 683.4 | 1232 | - | - | 0 | - |
| - | - | 680.4 | 2762 | - | - | 0 | - |
| - | - | 935.9 | 3082 | - | - | 0 | - |
| - | - | 662.4 | 3089 | - | - | 0 | - |
| - | - | 717 | 3274 | - | - | 0 | - |

m/z Charge Intensity FragmentType MassShift Position
122.49163818359375 0 380.3528
124.22992706298828 0 402.50922
129.0662841796875 0 954.0309
130.06541442871094 0 2861.5977
132.08094787597656 0 947.6269
133.08602905273438 0 919.094
142.05201721191406 0 413.40433
142.12318420410156 0 660.02814
144.08096313476562 0 631.4899
145.1029815673828 0 514.42065
146.0605010986328 0 1385.1306
146.0926513671875 0 686.6775
155.11822509765625 0 727.834
159.0918731689453 0 15062.629
160.0951385498047 0 1828.3809
172.57699584960938 0 511.04697
173.12864685058594 0 7846.2046
173.43751525878906 0 882.1675
177.11233520507812 0 1454.0806
183.11257934570312 0 1137.6749
184.0971221923828 0 1012.8815
185.27792358398438 0 515.42883
187.0628662109375 0 1176.2976
187.08665466308594 0 858.4306
187.14434814453125 0 3747.7825
188.07080078125 0 48756.875
189.07423400878906 0 5639.2183
190.07513427734375 0 495.94754
199.10787963867188 0 3599.0054
201.10206604003906 0 1007.4946
201.12355041503906 0 20197.89
201.65744018554688 0 536.05286
202.12681579589844 0 1470.012
205.0972900390625 0 62070.29 y 7
206.1006317138672 0 7133.408
207.1031494140625 0 626.6533
215.1390838623047 0 11227.579
218.9202880859375 0 524.9292
222.30166625976562 0 504.52097
227.0909881591797 0 938.2739
227.10279846191406 0 12284.06
228.10609436035156 0 1682.177
240.097900390625 0 1196.5686
240.17025756835938 0 1036.2975
240.40521240234375 0 624.6033
242.66055297851562 0 1373.5734
243.16212463378906 0 575.91614
244.12937927246094 0 36950.797
245.11590576171875 0 448.53293
245.132568359375 0 3730.7825
255.1371307373047 0 1336.8872
256.6578063964844 0 577.784
257.1496276855469 0 1383.05
258.1084289550781 0 2682.542
258.1450500488281 0 1073.0697
269.186767578125 0 642.8122
270.12353515625 0 2120.1257
284.1605529785156 0 7617.3213
285.1641540527344 0 930.5824
287.1506042480469 0 780.0953
287.2209167480469 0 1183.0455
296.1970520019531 0 24003.814
297.2003479003906 0 4729.7944
298.1189270019531 0 1906.7668
298.1760559082031 0 690.9602
300.15496826171875 0 659.7668
301.1520080566406 0 546.21954
302.15362548828125 0 756.264
302.1711730957031 0 11531.072
303.1743469238281 0 1515.264
307.18096923828125 0 505.44812
311.12451171875 0 2043.9006
312.15545654296875 0 706.6454
313.1386413574219 0 1185.4321
314.20751953125 0 9697.449
315.1452331542969 0 16770.79
315.21051025390625 0 1236.7817
316.12933349609375 0 25085.512 y Ammonia loss 6
316.1482238769531 0 2730.7434
317.13250732421875 0 3716.407 z 6
321.1789245605469 0 3830.332
321.6808776855469 0 998.42
327.14495849609375 0 670.8345
327.1672058105469 0 1688.7681
328.1504821777344 0 4524.9683
329.1539611816406 0 1200.3643
329.179443359375 0 881.45
329.6919860839844 0 1104.373 c 5
330.1661682128906 0 1460.9436
330.1948547363281 0 631.731
330.2286071777344 0 578.80023
331.234130859375 0 5600.734 c 2
332.2368469238281 0 1091.8193
333.1559753417969 0 22662.564 y 6
334.15899658203125 0 4229.422
337.1510314941406 0 2189.582
340.1879577636719 0 1438.092
343.23345947265625 0 600.59656
345.1770324707031 0 3777.8394
352.2236633300781 0 1894.3976
353.2254943847656 0 667.9641
354.15338134765625 0 3084.8464
355.1611633300781 0 50618.55
356.1642761230469 0 7932.4155
357.1664123535156 0 1178.1118
365.18243408203125 0 674.6969
369.2489929199219 0 4151.1016
370.23333740234375 0 5866.688
370.25531005859375 0 849.7919
371.23077392578125 0 2259.7996
372.1875 0 18750.238
373.1669616699219 0 837.68176
373.1904602050781 0 3845.7441
379.23321533203125 0 1663.0448
387.26025390625 0 8734.265
388.2635803222656 0 2275.149
396.1761779785156 0 6611.4062
397.1798095703125 0 850.1307
397.24456787109375 0 21748.1
398.2039794921875 0 1126.4794
398.24798583984375 0 4866.671
400.18341064453125 0 679.59845
412.1958312988281 0 654.559
413.2029724121094 0 2641.9314
414.24652099609375 0 522.04706
415.255126953125 0 61543.133
416.25830078125 0 12928.84
417.2284851074219 0 666.46967
417.2611083984375 0 2211.0203
421.1728210449219 0 626.57983
423.22552490234375 0 555.778
427.1617736816406 0 1711.0093
428.2175598144531 0 677.29474
429.185302734375 0 3109.8572
430.1879577636719 0 1304.7546
437.7252502441406 0 947.85596
439.1827392578125 0 3307.1497
439.2154235839844 0 618.20557
440.18560791015625 0 1233.7346
440.21807861328125 0 929.8655
440.251953125 0 883.49133
441.1976623535156 0 8927.599
441.2330627441406 0 1124.434
442.1959533691406 0 4678.714
443.19830322265625 0 950.1001
443.2853088378906 0 1131.1202
444.1875915527344 0 2366.2214 y Water loss 5
445.17138671875 0 1627.5254 y Ammonia loss 5
446.1795349121094 0 1891.539 z 5
446.7304992675781 0 607.1975
447.1857604980469 0 1498.7188
447.22930908203125 0 641.032
450.2731018066406 0 576.7838
451.21771240234375 0 531.3884
456.2090148925781 0 16442.879
457.2122497558594 0 3324.674
458.224365234375 0 21988.562
458.26019287109375 0 1121.4125
459.2276611328125 0 5017.5913
460.728515625 0 845.2528
462.1981506347656 0 3180.641 y 5
468.24505615234375 0 4547.857
468.2799072265625 0 1249.1031
469.24749755859375 0 1462.9186
469.73382568359375 0 1170.4141
470.2376403808594 0 779.41144
471.28045654296875 0 3028.993
471.3147277832031 0 925.65375
473.2352294921875 0 2962.6443
474.2397766113281 0 792.383
478.2451171875 0 789.8965
478.74835205078125 0 681.8123
484.23968505859375 0 2285.0835
485.2445983886719 0 641.47864
487.2044677734375 0 2132.832
487.31005859375 0 580.94354
488.2067565917969 0 1603.0739
494.29693603515625 0 2256.4727
495.2996520996094 0 886.93317
496.2182922363281 0 1124.3138
499.27740478515625 0 947.7972
500.2131652832031 0 889.91327
507.20928955078125 0 607.80536
510.25592041015625 0 1097.9357
511.3237609863281 0 1130.0715 c Water loss 4
512.307861328125 0 3706.023
513.207275390625 0 951.3832
513.2466430664062 0 1115.0505
513.3120727539062 0 1389.2655
514.2535400390625 0 9065.88
515.220947265625 0 3056.6199
515.2591552734375 0 3089.7617
516.2646484375 0 1214.6748
524.2138061523438 0 9508.205
525.2180786132812 0 3093.7292 z Water loss 4
528.3262939453125 0 14846.942
529.333984375 0 117551.77 c 4
530.2843627929688 0 1879.1697
530.3367919921875 0 31000.648
531.2938842773438 0 513.0218
531.33935546875 0 4266.532
534.2547607421875 0 782.2751
537.265625 0 1274.2864
541.2402954101562 0 5523.3564 y Water loss 4
542.2245483398438 0 21625.174 y Ammonia loss 4
543.227783203125 0 6080.206 z 4
543.3123779296875 0 971.3013
544.2310180664062 0 1066.0665
551.2831420898438 0 1211.5541
553.2760009765625 0 605.46216
554.2817993164062 0 1733.1918
555.2722778320312 0 3884.7937
555.3497314453125 0 3530.3232
556.2750854492188 0 1829.8102
556.3524169921875 0 1602.9597
558.242431640625 0 8785.737
559.2506103515625 0 133146.53 y 4
560.2536010742188 0 37718.188
561.2560424804688 0 6906.911
569.2916259765625 0 2577.197
571.307861328125 0 3976.5095
572.3080444335938 0 1066.4156
573.244873046875 0 1378.8971
574.2459106445312 0 686.4541
586.239501953125 0 904.3415
588.2482299804688 0 657.0254
598.27392578125 0 3523.1104
598.356201171875 0 1192.355
599.2593383789062 0 7428.1177
600.2660522460938 0 5867.1997
600.285888671875 0 4914.9453
601.2732543945312 0 1672.4756
601.2898559570312 0 1797.8398
602.2752075195312 0 831.015
605.3311767578125 0 882.66455
613.3131103515625 0 1317.9951
614.3626098632812 0 12061.567
615.3676147460938 0 3991.637
616.370361328125 0 843.1092
621.2874755859375 0 1147.2627
623.3392944335938 0 5570.986
624.3438110351562 0 1913.0068
625.331298828125 0 9603.436
626.2685546875 0 1446.6562 z Water loss 3
626.3341674804688 0 4146.051
627.2767944335938 0 723.82996 w 3
627.3394165039062 0 1005.4161
638.3131103515625 0 3193.4392
639.3003540039062 0 1069.2631
640.313232421875 0 4570.968
640.367431640625 0 3194.7742 c Water loss 5
641.3082275390625 0 7028.6807
641.351806640625 0 10544.772
642.2634887695312 0 26914.805
642.3154907226562 0 2102.1257
642.3552856445312 0 2980.2563
643.2675170898438 0 12279.174 y Ammonia loss 3
643.3223876953125 0 1012.67596
643.3648681640625 0 929.83435
644.2791137695312 0 68495.67 z 3
645.2852783203125 0 57441.676
645.3333129882812 0 465.03912
646.2891235351562 0 17164.895
647.2919921875 0 3210.539
650.3504638671875 0 2141.1135
656.32421875 0 9541.34
657.3683471679688 0 6156.059
658.3764038085938 0 229493.53 c 5
659.37890625 0 81025.516
660.29736328125 0 15091.087 y 3
660.3815307617188 0 16985.764
661.301025390625 0 6443.1646
661.3794555664062 0 939.7582
662.2994995117188 0 954.2869
668.3607177734375 0 1812.129
669.360595703125 0 861.94073
670.3974609375 0 2023.45
671.4019775390625 0 1091.545
683.4096069335938 0 890.10535
684.391845703125 0 32871.613
685.3949584960938 0 11024.686
686.3968505859375 0 3176.289
687.1783447265625 0 835.05676
688.180908203125 0 2123.4668
688.2864379882812 0 780.45245
689.1832885742188 0 744.9934
697.3268432617188 0 724.3123
706.3801879882812 0 732.2166
710.3310546875 0 1340.5804
713.3765258789062 0 1554.0391
714.3087158203125 0 4889.245 w 2
715.3358764648438 0 7335.5034
716.3425903320312 0 3101.398
717.3508911132812 0 835.0828
723.4013061523438 0 1204.0521
724.3916015625 0 684.205
725.3907470703125 0 893.156
728.3448486328125 0 12842.683
729.3482666015625 0 4688.0034
730.357421875 0 1201.1659
733.3897705078125 0 2854.9727
734.37939453125 0 2041.0957
735.3841552734375 0 908.8659
741.412109375 0 1804.0327
742.4203491210938 0 12291.596
743.422607421875 0 5264.235
744.4284057617188 0 1578.8279
751.3975219726562 0 15478.158
752.397216796875 0 8029.126
753.3970336914062 0 7476.4746
754.3966674804688 0 4805.12
755.3858642578125 0 2461.1492 y Water loss 2
757.363037109375 0 23933.875 z 2
758.3682861328125 0 15630.105
759.3709106445312 0 5404.9478
760.3760375976562 0 916.403
769.4086303710938 0 19080.104 c Ammonia loss 6
770.4124755859375 0 7441.6855
771.4228515625 0 20801.2
772.4263916015625 0 7635.913
773.3621826171875 0 23739.682
774.3648071289062 0 9975.76
775.3795166015625 0 3013.996
782.3953247070312 0 1420.6268
786.434326171875 0 356238.84 c 6
787.4373168945312 0 140720.16
788.3336181640625 0 11285.533
788.440185546875 0 34846.96
789.3363037109375 0 4206.9624
789.446044921875 0 3011.161
790.3338012695312 0 853.3382
797.4407958984375 0 1167.452
799.376220703125 0 2731.852
800.4039916992188 0 11957.973
801.40673828125 0 6052.736
802.4091796875 0 1183.8729
810.4114379882812 0 952.3468
812.4473876953125 0 2180.1736
813.4389038085938 0 3184.0632
814.458984375 0 2533.6606
815.4636840820312 0 698.78314
826.3843383789062 0 4712.4756 z Water loss 1
827.3876342773438 0 1741.7654 w 1
828.4232177734375 0 1553.1394
829.42724609375 0 869.7585
837.3849487304688 0 1911.5596
838.3931274414062 0 1032.2351
841.4293212890625 0 4243.8257
842.3814086914062 0 6266.312
843.4298095703125 0 16968.316
844.3953857421875 0 75069.305 z 1
845.398193359375 0 39399.36
846.4009399414062 0 9102.487
847.4091796875 0 831.61066
855.4091186523438 0 698.82275
857.4622192382812 0 52436.23
858.4652709960938 0 26703.76
859.4688720703125 0 8629.936
860.4129638671875 0 10443.304 y 1
861.4164428710938 0 6042.305
862.421630859375 0 1539.2906
867.4829711914062 0 2287.6084
868.4891967773438 0 955.28705
869.487060546875 0 861.6892
873.417724609375 0 1353.4901
884.5095825195312 0 11859.104
885.4513549804688 0 2172.9692
885.5107421875 0 5927.4404
886.4612426757812 0 3602.7742
887.4498901367188 0 1709.6714
895.4918212890625 0 655.9924
900.461669921875 0 938.9347
901.4515991210938 0 8966.082
902.4623413085938 0 5076.144
903.4693603515625 0 3764.8105
904.474853515625 0 1018.26605
911.48046875 0 1202.5696
912.5038452148438 0 15979.118
913.5026245117188 0 10642.262
914.4229736328125 0 10655.873
914.5076293945312 0 2888.1724
915.4259643554688 0 3852.23
915.5036010742188 0 1773.0955
916.4203491210938 0 1146.1562
918.4436645507812 0 1923.5508
920.451416015625 0 732.8016
928.4993896484375 0 4511.391
929.4838256835938 0 16474.209
930.5028076171875 0 17974.727
931.5087890625 0 9466.069
932.5140380859375 0 3342.8267
939.4661254882812 0 4728.067
940.4658813476562 0 2992.8716
941.4776000976562 0 1137.6854
956.494140625 0 25401.043
957.4802856445312 0 91256.49
958.4822387695312 0 47887.543
959.4830932617188 0 14987.311
960.4834594726562 0 1547.9337
973.4967651367188 0 127532.86
974.5029296875 0 207402.55
975.5062255859375 0 100492.42
976.5097045898438 0 27025.229
977.5096435546875 0 2512.736
1231.5003662109375 0 683.3846
2762.35888671875 0 680.37537
3081.956787109375 0 935.9252
3088.785888671875 0 662.36896
3273.96142578125 0 716.99347

Spectrum Details

|  |  |
| --- | --- |
| Matched peaks? Matched peaks The total absolute number of peaks matched. Additionally in brackets the total fraction of peaks matched and the total number of peaks is shown. | 33 (8.07% of 409) |
| FDR? FDR The false discovery rate estimated for this peptide. It is calculated by matching all theoretical fragments with a non-integer shift with the raw peaks for this spectrum. This is done with 40 different shifts. The resulting percentage is the average number of annotated peaks over the number of annotated peaks with the correct spectrum. | 5.12% |
| Satellite FDR? Satellite FDR See the FDR for details on its calculation. This satellite ion specific FDR only contains the satellite ions (d/w) for I/L/J positions. | 2.38% |
| PSM Score? PSM Score The PSM Score as given by Hecklib to this annotated spectrum. It is shown with three significant figures. | 446 |

#### Spectrum 9668? Spectrum 9668 The raw spectrum of this peptide as annotated by Hecklib. The fragments are coloured according to ion type (see legend). Any peaks with a star '\*' as text can be hovered over to see the full details, first the ion type second the mass shift type. By hovering over the amino acids in the peptide or ions in the legend the corresponding peaks are highlighted. By toggling the 'Unassigned' label you can turn the background (unassigned) peaks on or off in the plot. By updating the slider in the Ion legend you can update the spectrum to only show the top X% of the peaks with labels. The top X% means any peak that is within X% of the highest intensity. By dragging in the spectrum you can zoom in to a specific part of the spectrum and use 'Zoom Out' to get back to the original zoom level. The annotation of the spectrum is based on the given sequence in the peptides file and is done with different software so inconsistencies are likely. The peaks are annotated based on the given sequence, with 20 ppm tolerance.

Copy Data

##### Spectrum 9668 (TSV)

###### Preview

```
Loading example...
```

*Click on the button to copy the data to your clipboard.*

Mz MinMz MaxIntensity Max

WidthHeightPeptide font sizePeptide stroke widthSpectrum font sizeSpectrum stroke widthCompact peptide

Ion legend

wxyz

abcd

OtherUnassignedIon^Charge^_Position_Show for top:%

JSJTPEQW

02.93e+55.86e+58.80e+51.17e+6

Zoom Out

a^+1^_2_y^+2^_2_a^+1^_2_b^+1^_2_b^+1^_2_y^+1^_1_b^+2^_4_b^+2^_5_b^+2^_5_d^+1^_3_a^+1^_3_y^+2^_4_y^+2^_4_a^+1^_3_b^+1^_3_b^+2^_6_b^+1^_3_y^+1^_2_b^+2^_6_y^+1^_2_b^+2^_7_b^+2^_7_b^+1^_4_b^+1^_4_y^+2^_7_y^+1^_3_y^+1^_3_y^+1^_3_\*\*b^+1^_5_b^+1^_5_y^+1^_4_y^+1^_4_y^+1^_4_b^+1^_6_b^+1^_6_y^+1^_5_y^+1^_5_y^+1^_5_b^+1^_7_y^+1^_6_b^+1^_7_y^+1^_6_y^+1^_7_y^+1^_7_y^+1^_7_

0218436653871

Fragment Matches Table

Show background peaks

| Position | Ion type | Intensity | mz Theoretical | mz Error (Th) | mz Error (ppm) | Charge | Series Number |
| --- | --- | --- | --- | --- | --- | --- | --- |
| - | - | 2151 | 120.1 | - | - | 0 | - |
| - | - | 530.1 | 123.9 | - | - | 0 | - |
| - | - | 4814 | 125.1 | - | - | 0 | - |
| - | - | 1.887E+04 | 125.1 | - | - | 0 | - |
| - | - | 976.9 | 125.1 | - | - | 0 | - |
| - | - | 6069 | 126.1 | - | - | 0 | - |
| - | - | 1426 | 126.1 | - | - | 0 | - |
| - | - | 2341 | 127.1 | - | - | 0 | - |
| - | - | 506.2 | 127.1 | - | - | 0 | - |
| - | - | 2045 | 128.1 | - | - | 0 | - |
| - | - | 7767 | 128.1 | - | - | 0 | - |
| - | - | 4.385E+04 | 129.1 | - | - | 0 | - |
| - | - | 1297 | 129.1 | - | - | 0 | - |
| - | - | 1.511E+04 | 130.1 | - | - | 0 | - |
| - | - | 1.112E+05 | 130.1 | - | - | 0 | - |
| - | - | 9549 | 131.1 | - | - | 0 | - |
| - | - | 979.5 | 131.1 | - | - | 0 | - |
| - | - | 5.494E+04 | 132.1 | - | - | 0 | - |
| - | - | 5949 | 133.1 | - | - | 0 | - |
| - | - | 551.8 | 135.2 | - | - | 0 | - |
| - | - | 934.1 | 136.1 | - | - | 0 | - |
| - | - | 1363 | 137.1 | - | - | 0 | - |
| - | - | 4661 | 138.1 | - | - | 0 | - |
| - | - | 1370 | 138.1 | - | - | 0 | - |
| - | - | 4388 | 139.1 | - | - | 0 | - |
| - | - | 1428 | 141.1 | - | - | 0 | - |
| - | - | 4094 | 142.1 | - | - | 0 | - |
| - | - | 2.767E+04 | 142.1 | - | - | 0 | - |
| - | - | 2306 | 143.1 | - | - | 0 | - |
| - | - | 619.2 | 143.1 | - | - | 0 | - |
| - | - | 1189 | 143.1 | - | - | 0 | - |
| - | - | 2187 | 143.1 | - | - | 0 | - |
| - | - | 943.3 | 144 | - | - | 0 | - |
| - | - | 2.677E+04 | 144.1 | - | - | 0 | - |
| - | - | 2581 | 145.1 | - | - | 0 | - |
| - | - | 1.461E+05 | 146.1 | - | - | 0 | - |
| - | - | 1966 | 146.1 | - | - | 0 | - |
| - | - | 656.5 | 147.1 | - | - | 0 | - |
| - | - | 1.408E+04 | 147.1 | - | - | 0 | - |
| - | - | 1.269E+04 | 147.1 | - | - | 0 | - |
| - | - | 779.4 | 150.4 | - | - | 0 | - |
| - | - | 2.043E+04 | 152.1 | - | - | 0 | - |
| - | - | 1.324E+04 | 153.1 | - | - | 0 | - |
| - | - | 9560 | 153.1 | - | - | 0 | - |
| - | - | 1336 | 153.1 | - | - | 0 | - |
| - | - | 663.7 | 154.1 | - | - | 0 | - |
| - | - | 793 | 154.1 | - | - | 0 | - |
| - | - | 5311 | 155.1 | - | - | 0 | - |
| 2 | a | 2.858E+04 | 155.1 | 0.0005339 | 3.442 | +1 | 2 |
| - | - | 2121 | 156.1 | - | - | 0 | - |
| - | - | 2083 | 156.1 | - | - | 0 | - |
| - | - | 2105 | 158 | - | - | 0 | - |
| - | - | 2.985E+05 | 159.1 | - | - | 0 | - |
| - | - | 5271 | 160.1 | - | - | 0 | - |
| - | - | 1395 | 160.1 | - | - | 0 | - |
| - | - | 3.318E+04 | 160.1 | - | - | 0 | - |
| - | - | 1282 | 161.1 | - | - | 0 | - |
| - | - | 2303 | 163.1 | - | - | 0 | - |
| - | - | 954.3 | 165.1 | - | - | 0 | - |
| 7 | y | 2103 | 167.1 | 0.0007866 | 4.708 | +2 | 2 |
| - | - | 3311 | 169.1 | - | - | 0 | - |
| - | - | 1.208E+04 | 169.1 | - | - | 0 | - |
| - | - | 1.592E+04 | 170.1 | - | - | 0 | - |
| - | - | 2383 | 170.1 | - | - | 0 | - |
| - | - | 1024 | 170.1 | - | - | 0 | - |
| - | - | 2120 | 171.1 | - | - | 0 | - |
| - | - | 2.818E+04 | 171.1 | - | - | 0 | - |
| - | - | 1450 | 171.1 | - | - | 0 | - |
| - | - | 3689 | 171.1 | - | - | 0 | - |
| - | - | 2438 | 172.1 | - | - | 0 | - |
| - | - | 760.5 | 172.2 | - | - | 0 | - |
| 2 | a | 2.173E+05 | 173.1 | 0.0005894 | 3.404 | +1 | 2 |
| - | - | 671.7 | 174.1 | - | - | 0 | - |
| - | - | 1493 | 174.1 | - | - | 0 | - |
| - | - | 1.695E+04 | 174.1 | - | - | 0 | - |
| - | - | 727.3 | 175.1 | - | - | 0 | - |
| - | - | 923.5 | 177.1 | - | - | 0 | - |
| - | - | 1362 | 179.1 | - | - | 0 | - |
| - | - | 3.313E+04 | 181.1 | - | - | 0 | - |
| - | - | 2290 | 182.1 | - | - | 0 | - |
| 2 | b | 5.534E+04 | 183.1 | 0.0006297 | 3.439 | +1 | 2 |
| - | - | 1.687E+04 | 183.1 | - | - | 0 | - |
| - | - | 4628 | 184.1 | - | - | 0 | - |
| - | - | 580.4 | 184.1 | - | - | 0 | - |
| - | - | 2063 | 184.2 | - | - | 0 | - |
| - | - | 1537 | 185.1 | - | - | 0 | - |
| - | - | 716 | 185.1 | - | - | 0 | - |
| - | - | 1804 | 186.1 | - | - | 0 | - |
| - | - | 794.5 | 187.1 | - | - | 0 | - |
| - | - | 3821 | 187.1 | - | - | 0 | - |
| - | - | 2.641E+04 | 187.1 | - | - | 0 | - |
| - | - | 900.7 | 187.1 | - | - | 0 | - |
| - | - | 6.403E+04 | 187.1 | - | - | 0 | - |
| - | - | 1.161E+06 | 188.1 | - | - | 0 | - |
| - | - | 4862 | 188.1 | - | - | 0 | - |
| - | - | 1.329E+05 | 189.1 | - | - | 0 | - |
| - | - | 7462 | 190.1 | - | - | 0 | - |
| - | - | 1660 | 191.1 | - | - | 0 | - |
| - | - | 3460 | 194.1 | - | - | 0 | - |
| - | - | 5578 | 195.1 | - | - | 0 | - |
| - | - | 1793 | 195.1 | - | - | 0 | - |
| - | - | 1972 | 195.1 | - | - | 0 | - |
| - | - | 2879 | 196.1 | - | - | 0 | - |
| - | - | 849.7 | 197.1 | - | - | 0 | - |
| - | - | 3.14E+04 | 197.1 | - | - | 0 | - |
| - | - | 745.8 | 198.1 | - | - | 0 | - |
| - | - | 3562 | 198.1 | - | - | 0 | - |
| - | - | 4877 | 199.1 | - | - | 0 | - |
| - | - | 2.312E+05 | 199.1 | - | - | 0 | - |
| - | - | 1085 | 200.1 | - | - | 0 | - |
| - | - | 2.107E+04 | 200.1 | - | - | 0 | - |
| - | - | 1681 | 200.1 | - | - | 0 | - |
| 2 | b | 2.619E+05 | 201.1 | 0.000563 | 2.799 | +1 | 2 |
| - | - | 2.536E+04 | 202.1 | - | - | 0 | - |
| - | - | 1531 | 203.1 | - | - | 0 | - |
| 8 | y | 4.923E+05 | 205.1 | 0.0006395 | 3.118 | +1 | 1 |
| - | - | 5.35E+04 | 206.1 | - | - | 0 | - |
| - | - | 2477 | 207.1 | - | - | 0 | - |
| - | - | 1168 | 207.1 | - | - | 0 | - |
| - | - | 819.4 | 208.1 | - | - | 0 | - |
| 4 | b | 1038 | 208.1 | 0.001298 | 6.237 | +2 | 4 |
| - | - | 2.327E+04 | 209.1 | - | - | 0 | - |
| - | - | 2278 | 210.1 | - | - | 0 | - |
| - | - | 2566 | 210.1 | - | - | 0 | - |
| - | - | 3288 | 211.1 | - | - | 0 | - |
| - | - | 1.453E+04 | 211.1 | - | - | 0 | - |
| - | - | 6927 | 212.1 | - | - | 0 | - |
| - | - | 988.3 | 212.1 | - | - | 0 | - |
| - | - | 2047 | 213.1 | - | - | 0 | - |
| - | - | 2679 | 213.1 | - | - | 0 | - |
| - | - | 6477 | 214.2 | - | - | 0 | - |
| - | - | 1822 | 215.1 | - | - | 0 | - |
| - | - | 1.582E+05 | 215.1 | - | - | 0 | - |
| - | - | 1.333E+04 | 216.1 | - | - | 0 | - |
| - | - | 1.077E+04 | 217.1 | - | - | 0 | - |
| - | - | 1019 | 217.1 | - | - | 0 | - |
| - | - | 1278 | 217.1 | - | - | 0 | - |
| - | - | 873.6 | 218.2 | - | - | 0 | - |
| - | - | 1546 | 221.1 | - | - | 0 | - |
| - | - | 921.6 | 221.2 | - | - | 0 | - |
| - | - | 1293 | 222.1 | - | - | 0 | - |
| - | - | 9230 | 223.1 | - | - | 0 | - |
| - | - | 2522 | 223.1 | - | - | 0 | - |
| - | - | 999.2 | 223.1 | - | - | 0 | - |
| - | - | 997.2 | 223.2 | - | - | 0 | - |
| - | - | 1966 | 224.1 | - | - | 0 | - |
| - | - | 1.117E+04 | 225.1 | - | - | 0 | - |
| - | - | 6127 | 225.1 | - | - | 0 | - |
| - | - | 1353 | 226.1 | - | - | 0 | - |
| - | - | 1.151E+04 | 226.1 | - | - | 0 | - |
| - | - | 3.76E+05 | 227.1 | - | - | 0 | - |
| - | - | 3.677E+04 | 228.1 | - | - | 0 | - |
| - | - | 939.5 | 228.1 | - | - | 0 | - |
| - | - | 3066 | 229.1 | - | - | 0 | - |
| - | - | 5838 | 229.2 | - | - | 0 | - |
| - | - | 2551 | 230.1 | - | - | 0 | - |
| - | - | 1271 | 231.1 | - | - | 0 | - |
| - | - | 2973 | 233.7 | - | - | 0 | - |
| - | - | 1376 | 237.1 | - | - | 0 | - |
| - | - | 2651 | 237.1 | - | - | 0 | - |
| - | - | 4704 | 238.2 | - | - | 0 | - |
| - | - | 1078 | 238.2 | - | - | 0 | - |
| - | - | 1.754E+04 | 239.1 | - | - | 0 | - |
| - | - | 1313 | 239.2 | - | - | 0 | - |
| - | - | 1221 | 239.2 | - | - | 0 | - |
| - | - | 4338 | 239.6 | - | - | 0 | - |
| - | - | 5.138E+04 | 240.1 | - | - | 0 | - |
| - | - | 2513 | 240.1 | - | - | 0 | - |
| - | - | 1336 | 240.2 | - | - | 0 | - |
| - | - | 1.323E+04 | 241.1 | - | - | 0 | - |
| - | - | 5220 | 241.1 | - | - | 0 | - |
| - | - | 3627 | 241.2 | - | - | 0 | - |
| - | - | 758 | 241.9 | - | - | 0 | - |
| - | - | 1188 | 242.1 | - | - | 0 | - |
| - | - | 9243 | 242.2 | - | - | 0 | - |
| - | - | 1.271E+04 | 242.7 | - | - | 0 | - |
| - | - | 2283 | 243.1 | - | - | 0 | - |
| - | - | 3641 | 243.2 | - | - | 0 | - |
| - | - | 1.109E+05 | 244.1 | - | - | 0 | - |
| - | - | 7517 | 245.1 | - | - | 0 | - |
| - | - | 8950 | 245.1 | - | - | 0 | - |
| - | - | 1520 | 246.1 | - | - | 0 | - |
| 5 | b | 2412 | 247.7 | 0.001195 | 4.824 | +2 | 5 |
| - | - | 1819 | 248.1 | - | - | 0 | - |
| - | - | 912.8 | 248.1 | - | - | 0 | - |
| - | - | 2227 | 249.1 | - | - | 0 | - |
| - | - | 1536 | 249.2 | - | - | 0 | - |
| - | - | 904.9 | 250.2 | - | - | 0 | - |
| - | - | 1101 | 251.1 | - | - | 0 | - |
| - | - | 4778 | 251.1 | - | - | 0 | - |
| - | - | 1.205E+04 | 251.2 | - | - | 0 | - |
| - | - | 5607 | 252.1 | - | - | 0 | - |
| - | - | 1711 | 252.2 | - | - | 0 | - |
| - | - | 8322 | 253.1 | - | - | 0 | - |
| - | - | 1673 | 253.1 | - | - | 0 | - |
| - | - | 1491 | 254.1 | - | - | 0 | - |
| - | - | 1547 | 254.1 | - | - | 0 | - |
| - | - | 2928 | 255.1 | - | - | 0 | - |
| - | - | 1489 | 256.1 | - | - | 0 | - |
| - | - | 1.48E+04 | 256.2 | - | - | 0 | - |
| 5 | b | 3083 | 256.7 | 0.0009935 | 3.871 | +2 | 5 |
| - | - | 1143 | 257.1 | - | - | 0 | - |
| - | - | 2.069E+04 | 257.2 | - | - | 0 | - |
| - | - | 1252 | 257.2 | - | - | 0 | - |
| - | - | 839.4 | 257.2 | - | - | 0 | - |
| - | - | 4.816E+04 | 258.1 | - | - | 0 | - |
| - | - | 9142 | 258.1 | - | - | 0 | - |
| 3 | d | 1272 | 258.2 | 9.625E-05 | 0.3728 | +1 | 3 |
| - | - | 1798 | 259.1 | - | - | 0 | - |
| - | - | 4332 | 259.1 | - | - | 0 | - |
| - | - | 966 | 259.2 | - | - | 0 | - |
| - | - | 1512 | 262.6 | - | - | 0 | - |
| - | - | 4827 | 264.1 | - | - | 0 | - |
| - | - | 1258 | 265.1 | - | - | 0 | - |
| - | - | 4803 | 266.1 | - | - | 0 | - |
| - | - | 9842 | 266.2 | - | - | 0 | - |
| - | - | 3713 | 266.2 | - | - | 0 | - |
| - | - | 8526 | 267.1 | - | - | 0 | - |
| - | - | 1021 | 267.1 | - | - | 0 | - |
| - | - | 1612 | 267.2 | - | - | 0 | - |
| - | - | 1600 | 268.1 | - | - | 0 | - |
| - | - | 1383 | 268.1 | - | - | 0 | - |
| - | - | 1066 | 268.2 | - | - | 0 | - |
| 3 | a | 1.007E+04 | 268.2 | 0.0005306 | 1.978 | +1 | 3 |
| - | - | 1.221E+04 | 269.1 | - | - | 0 | - |
| - | - | 7884 | 269.2 | - | - | 0 | - |
| - | - | 955.5 | 269.2 | - | - | 0 | - |
| - | - | 1.264E+05 | 270.1 | - | - | 0 | - |
| - | - | 3434 | 271.1 | - | - | 0 | - |
| 5 | y | 2.241E+04 | 271.1 | 0.003633 | 13.4 | +2 | 4 |
| 5 | y | 1145 | 271.6 | 0.0008828 | 3.25 | +2 | 4 |
| - | - | 1173 | 272.1 | - | - | 0 | - |
| - | - | 1212 | 272.1 | - | - | 0 | - |
| - | - | 1476 | 272.2 | - | - | 0 | - |
| - | - | 1582 | 274.1 | - | - | 0 | - |
| - | - | 1.023E+04 | 274.2 | - | - | 0 | - |
| - | - | 1252 | 275.2 | - | - | 0 | - |
| - | - | 944.8 | 276.1 | - | - | 0 | - |
| - | - | 1657 | 277.2 | - | - | 0 | - |
| - | - | 1742 | 278.1 | - | - | 0 | - |
| - | - | 2087 | 278.2 | - | - | 0 | - |
| - | - | 1310 | 279.1 | - | - | 0 | - |
| - | - | 1.603E+04 | 280.1 | - | - | 0 | - |
| - | - | 1.045E+04 | 280.2 | - | - | 0 | - |
| - | - | 2790 | 281.1 | - | - | 0 | - |
| - | - | 1283 | 281.2 | - | - | 0 | - |
| - | - | 1.019E+04 | 282.1 | - | - | 0 | - |
| - | - | 5367 | 283.1 | - | - | 0 | - |
| - | - | 1326 | 283.1 | - | - | 0 | - |
| - | - | 5074 | 284.1 | - | - | 0 | - |
| - | - | 1.25E+05 | 284.2 | - | - | 0 | - |
| - | - | 2139 | 285.1 | - | - | 0 | - |
| - | - | 1.798E+04 | 285.2 | - | - | 0 | - |
| - | - | 2255 | 286.1 | - | - | 0 | - |
| - | - | 2078 | 286.1 | - | - | 0 | - |
| - | - | 1690 | 286.2 | - | - | 0 | - |
| 3 | a | 3711 | 286.2 | 0.0007386 | 2.581 | +1 | 3 |
| - | - | 3395 | 287.2 | - | - | 0 | - |
| - | - | 9073 | 292.1 | - | - | 0 | - |
| - | - | 2153 | 293.1 | - | - | 0 | - |
| - | - | 1276 | 293.1 | - | - | 0 | - |
| - | - | 1098 | 293.2 | - | - | 0 | - |
| - | - | 7295 | 294.1 | - | - | 0 | - |
| - | - | 7052 | 294.2 | - | - | 0 | - |
| - | - | 1025 | 295.1 | - | - | 0 | - |
| - | - | 1116 | 295.2 | - | - | 0 | - |
| - | - | 993 | 295.2 | - | - | 0 | - |
| - | - | 3905 | 296.1 | - | - | 0 | - |
| 3 | b | 2.803E+05 | 296.2 | 0.0008553 | 2.887 | +1 | 3 |
| - | - | 4741 | 297.1 | - | - | 0 | - |
| - | - | 4.277E+04 | 297.2 | - | - | 0 | - |
| - | - | 3.977E+04 | 298.1 | - | - | 0 | - |
| - | - | 826.1 | 298.1 | - | - | 0 | - |
| - | - | 1.517E+04 | 298.2 | - | - | 0 | - |
| - | - | 3400 | 298.2 | - | - | 0 | - |
| - | - | 2389 | 299.1 | - | - | 0 | - |
| - | - | 6984 | 299.1 | - | - | 0 | - |
| - | - | 2332 | 299.2 | - | - | 0 | - |
| - | - | 1.574E+04 | 300.2 | - | - | 0 | - |
| - | - | 1411 | 301.2 | - | - | 0 | - |
| - | - | 1.539E+05 | 302.2 | - | - | 0 | - |
| - | - | 2.273E+04 | 303.2 | - | - | 0 | - |
| - | - | 1711 | 304.2 | - | - | 0 | - |
| - | - | 5671 | 307.2 | - | - | 0 | - |
| - | - | 1774 | 307.7 | - | - | 0 | - |
| - | - | 1473 | 308.2 | - | - | 0 | - |
| - | - | 6362 | 309.2 | - | - | 0 | - |
| - | - | 1033 | 309.2 | - | - | 0 | - |
| - | - | 2.534E+04 | 310.1 | - | - | 0 | - |
| - | - | 1322 | 310.2 | - | - | 0 | - |
| - | - | 5314 | 310.2 | - | - | 0 | - |
| - | - | 4046 | 311.1 | - | - | 0 | - |
| - | - | 6659 | 311.2 | - | - | 0 | - |
| - | - | 1513 | 311.2 | - | - | 0 | - |
| - | - | 1.449E+04 | 312.2 | - | - | 0 | - |
| 6 | b | 2666 | 312.2 | 0.001032 | 3.304 | +2 | 6 |
| - | - | 1.04E+04 | 312.2 | - | - | 0 | - |
| - | - | 1022 | 312.7 | - | - | 0 | - |
| - | - | 3300 | 313.2 | - | - | 0 | - |
| - | - | 1762 | 313.2 | - | - | 0 | - |
| - | - | 2104 | 314.1 | - | - | 0 | - |
| 3 | b | 6.358E+04 | 314.2 | 0.0009412 | 2.995 | +1 | 3 |
| - | - | 3.017E+05 | 315.1 | - | - | 0 | - |
| - | - | 9401 | 315.2 | - | - | 0 | - |
| 7 | y | 3.166E+05 | 316.1 | 0.0008529 | 2.698 | +1 | 2 |
| - | - | 4.795E+04 | 316.1 | - | - | 0 | - |
| - | - | 5.369E+04 | 317.1 | - | - | 0 | - |
| - | - | 3123 | 317.2 | - | - | 0 | - |
| - | - | 5616 | 318.1 | - | - | 0 | - |
| - | - | 1567 | 318.2 | - | - | 0 | - |
| - | - | 5809 | 320.1 | - | - | 0 | - |
| - | - | 1448 | 320.2 | - | - | 0 | - |
| - | - | 3238 | 320.7 | - | - | 0 | - |
| - | - | 1369 | 321.1 | - | - | 0 | - |
| 6 | b | 2.206E+04 | 321.2 | 0.001029 | 3.203 | +2 | 6 |
| - | - | 6318 | 321.7 | - | - | 0 | - |
| - | - | 3493 | 322.2 | - | - | 0 | - |
| - | - | 1392 | 324.2 | - | - | 0 | - |
| - | - | 1971 | 325.2 | - | - | 0 | - |
| - | - | 1104 | 325.2 | - | - | 0 | - |
| - | - | 2426 | 326.2 | - | - | 0 | - |
| - | - | 1084 | 326.2 | - | - | 0 | - |
| - | - | 3.18E+04 | 327.2 | - | - | 0 | - |
| - | - | 5.2E+04 | 328.2 | - | - | 0 | - |
| - | - | 3999 | 328.2 | - | - | 0 | - |
| - | - | 1042 | 328.2 | - | - | 0 | - |
| - | - | 7359 | 329.2 | - | - | 0 | - |
| - | - | 3963 | 329.2 | - | - | 0 | - |
| - | - | 4392 | 329.7 | - | - | 0 | - |
| - | - | 7019 | 330.2 | - | - | 0 | - |
| - | - | 1080 | 330.2 | - | - | 0 | - |
| 7 | y | 1.572E+05 | 333.2 | 0.0009456 | 2.838 | +1 | 2 |
| - | - | 2.667E+04 | 334.2 | - | - | 0 | - |
| - | - | 3068 | 334.2 | - | - | 0 | - |
| - | - | 2095 | 335.2 | - | - | 0 | - |
| - | - | 1242 | 335.2 | - | - | 0 | - |
| - | - | 1399 | 336.2 | - | - | 0 | - |
| - | - | 2343 | 336.2 | - | - | 0 | - |
| - | - | 3.331E+04 | 337.2 | - | - | 0 | - |
| - | - | 3963 | 337.2 | - | - | 0 | - |
| - | - | 9821 | 338.1 | - | - | 0 | - |
| - | - | 4921 | 338.2 | - | - | 0 | - |
| - | - | 2415 | 339.1 | - | - | 0 | - |
| - | - | 927.8 | 339.2 | - | - | 0 | - |
| - | - | 1.918E+04 | 340.2 | - | - | 0 | - |
| - | - | 2763 | 341.2 | - | - | 0 | - |
| - | - | 1181 | 342.2 | - | - | 0 | - |
| - | - | 1113 | 343.1 | - | - | 0 | - |
| - | - | 2604 | 343.2 | - | - | 0 | - |
| - | - | 1171 | 344.2 | - | - | 0 | - |
| - | - | 2.086E+04 | 345.2 | - | - | 0 | - |
| - | - | 5270 | 346.2 | - | - | 0 | - |
| - | - | 1419 | 347.2 | - | - | 0 | - |
| - | - | 1956 | 350.2 | - | - | 0 | - |
| - | - | 5463 | 351.2 | - | - | 0 | - |
| - | - | 2.328E+04 | 352.2 | - | - | 0 | - |
| - | - | 1588 | 352.2 | - | - | 0 | - |
| - | - | 4780 | 353.2 | - | - | 0 | - |
| - | - | 6912 | 353.2 | - | - | 0 | - |
| - | - | 1343 | 353.3 | - | - | 0 | - |
| - | - | 943.2 | 354.2 | - | - | 0 | - |
| - | - | 952.2 | 354.2 | - | - | 0 | - |
| - | - | 1375 | 354.2 | - | - | 0 | - |
| - | - | 5.111E+05 | 355.2 | - | - | 0 | - |
| - | - | 7931 | 356.1 | - | - | 0 | - |
| - | - | 8.49E+04 | 356.2 | - | - | 0 | - |
| - | - | 754.4 | 357.1 | - | - | 0 | - |
| - | - | 1.034E+04 | 357.2 | - | - | 0 | - |
| - | - | 975.2 | 357.2 | - | - | 0 | - |
| - | - | 832.1 | 359.2 | - | - | 0 | - |
| - | - | 1674 | 360.7 | - | - | 0 | - |
| - | - | 3060 | 362.2 | - | - | 0 | - |
| - | - | 1058 | 363.1 | - | - | 0 | - |
| - | - | 9669 | 363.2 | - | - | 0 | - |
| - | - | 2442 | 364.2 | - | - | 0 | - |
| - | - | 1201 | 364.2 | - | - | 0 | - |
| - | - | 1.012E+04 | 365.2 | - | - | 0 | - |
| - | - | 2461 | 366.2 | - | - | 0 | - |
| - | - | 1945 | 367.2 | - | - | 0 | - |
| - | - | 892.9 | 367.7 | - | - | 0 | - |
| - | - | 3.519E+04 | 369.3 | - | - | 0 | - |
| - | - | 890.7 | 369.7 | - | - | 0 | - |
| - | - | 4.542E+04 | 370.2 | - | - | 0 | - |
| - | - | 2809 | 371.2 | - | - | 0 | - |
| - | - | 1.717E+04 | 371.2 | - | - | 0 | - |
| - | - | 7231 | 372.2 | - | - | 0 | - |
| - | - | 2831 | 372.2 | - | - | 0 | - |
| - | - | 1.511E+04 | 373.2 | - | - | 0 | - |
| - | - | 1334 | 373.2 | - | - | 0 | - |
| - | - | 2686 | 374.2 | - | - | 0 | - |
| 7 | b | 2687 | 376.2 | 0.000765 | 2.034 | +2 | 7 |
| - | - | 947.7 | 376.7 | - | - | 0 | - |
| - | - | 2.127E+04 | 379.2 | - | - | 0 | - |
| - | - | 1095 | 380.1 | - | - | 0 | - |
| - | - | 3688 | 380.2 | - | - | 0 | - |
| - | - | 2055 | 380.2 | - | - | 0 | - |
| - | - | 1.552E+04 | 381.2 | - | - | 0 | - |
| - | - | 2554 | 381.2 | - | - | 0 | - |
| - | - | 1.167E+04 | 381.2 | - | - | 0 | - |
| - | - | 3071 | 382.2 | - | - | 0 | - |
| - | - | 2160 | 382.2 | - | - | 0 | - |
| - | - | 1176 | 383.2 | - | - | 0 | - |
| - | - | 2013 | 383.3 | - | - | 0 | - |
| - | - | 1071 | 384.3 | - | - | 0 | - |
| 7 | b | 2995 | 385.2 | 0.001159 | 3.009 | +2 | 7 |
| - | - | 1329 | 385.7 | - | - | 0 | - |
| - | - | 5.593E+04 | 387.3 | - | - | 0 | - |
| - | - | 9914 | 388.3 | - | - | 0 | - |
| - | - | 1194 | 391.1 | - | - | 0 | - |
| - | - | 2353 | 391.2 | - | - | 0 | - |
| - | - | 4429 | 393.3 | - | - | 0 | - |
| - | - | 1106 | 394.3 | - | - | 0 | - |
| - | - | 5943 | 395.2 | - | - | 0 | - |
| - | - | 1532 | 395.2 | - | - | 0 | - |
| - | - | 1983 | 395.2 | - | - | 0 | - |
| - | - | 1609 | 396.2 | - | - | 0 | - |
| 4 | b | 1.593E+05 | 397.2 | 0.001028 | 2.589 | +1 | 4 |
| - | - | 3.601E+04 | 398.2 | - | - | 0 | - |
| - | - | 2174 | 399.2 | - | - | 0 | - |
| - | - | 2518 | 399.2 | - | - | 0 | - |
| - | - | 6219 | 399.2 | - | - | 0 | - |
| - | - | 4580 | 399.3 | - | - | 0 | - |
| - | - | 1048 | 404.2 | - | - | 0 | - |
| - | - | 1088 | 405.2 | - | - | 0 | - |
| - | - | 1238 | 405.2 | - | - | 0 | - |
| - | - | 1451 | 405.2 | - | - | 0 | - |
| - | - | 2569 | 407.2 | - | - | 0 | - |
| - | - | 2054 | 407.2 | - | - | 0 | - |
| - | - | 8451 | 409.2 | - | - | 0 | - |
| - | - | 1.398E+04 | 409.2 | - | - | 0 | - |
| - | - | 1228 | 409.2 | - | - | 0 | - |
| - | - | 2100 | 410.2 | - | - | 0 | - |
| - | - | 3758 | 410.2 | - | - | 0 | - |
| - | - | 4143 | 411.2 | - | - | 0 | - |
| - | - | 4521 | 411.3 | - | - | 0 | - |
| - | - | 3429 | 412.2 | - | - | 0 | - |
| - | - | 1714 | 412.2 | - | - | 0 | - |
| - | - | 1297 | 412.3 | - | - | 0 | - |
| - | - | 3762 | 413.2 | - | - | 0 | - |
| 4 | b | 3.238E+05 | 415.3 | 0.001084 | 2.61 | +1 | 4 |
| - | - | 6.766E+04 | 416.3 | - | - | 0 | - |
| - | - | 9322 | 417.3 | - | - | 0 | - |
| - | - | 1817 | 420.2 | - | - | 0 | - |
| - | - | 4346 | 421.2 | - | - | 0 | - |
| 2 | y | 2808 | 422.2 | 0.006766 | 16.03 | +2 | 7 |
| - | - | 1147 | 422.2 | - | - | 0 | - |
| - | - | 1921 | 423.2 | - | - | 0 | - |
| - | - | 9282 | 423.2 | - | - | 0 | - |
| - | - | 1087 | 424.2 | - | - | 0 | - |
| - | - | 3570 | 424.2 | - | - | 0 | - |
| - | - | 2689 | 424.3 | - | - | 0 | - |
| - | - | 1223 | 425.2 | - | - | 0 | - |
| - | - | 2785 | 425.2 | - | - | 0 | - |
| - | - | 1356 | 426.2 | - | - | 0 | - |
| - | - | 2127 | 426.2 | - | - | 0 | - |
| - | - | 4.013E+04 | 427.2 | - | - | 0 | - |
| - | - | 7406 | 427.2 | - | - | 0 | - |
| - | - | 8125 | 428.2 | - | - | 0 | - |
| - | - | 5219 | 428.2 | - | - | 0 | - |
| - | - | 1009 | 429.2 | - | - | 0 | - |
| - | - | 2257 | 430.2 | - | - | 0 | - |
| - | - | 4483 | 431.3 | - | - | 0 | - |
| - | - | 2142 | 437.7 | - | - | 0 | - |
| - | - | 8784 | 438.2 | - | - | 0 | - |
| - | - | 1221 | 438.2 | - | - | 0 | - |
| - | - | 3332 | 439.2 | - | - | 0 | - |
| - | - | 2471 | 440.2 | - | - | 0 | - |
| - | - | 1.389E+04 | 440.3 | - | - | 0 | - |
| - | - | 865.3 | 440.3 | - | - | 0 | - |
| - | - | 8653 | 441.2 | - | - | 0 | - |
| - | - | 3699 | 442.2 | - | - | 0 | - |
| - | - | 2117 | 442.2 | - | - | 0 | - |
| 6 | y | 1.213E+04 | 444.2 | 0.001083 | 2.438 | +1 | 3 |
| 6 | y | 2.951E+04 | 445.2 | 0.001198 | 2.691 | +1 | 3 |
| - | - | 6882 | 446.2 | - | - | 0 | - |
| - | - | 4816 | 446.7 | - | - | 0 | - |
| - | - | 1179 | 447.2 | - | - | 0 | - |
| - | - | 1628 | 447.2 | - | - | 0 | - |
| - | - | 1276 | 449.2 | - | - | 0 | - |
| - | - | 3458 | 450.2 | - | - | 0 | - |
| - | - | 2540 | 450.3 | - | - | 0 | - |
| - | - | 4612 | 451.2 | - | - | 0 | - |
| - | - | 1084 | 451.7 | - | - | 0 | - |
| - | - | 2520 | 452.2 | - | - | 0 | - |
| - | - | 1673 | 455.7 | - | - | 0 | - |
| - | - | 4.013E+04 | 456.2 | - | - | 0 | - |
| - | - | 7825 | 457.2 | - | - | 0 | - |
| - | - | 1156 | 458.2 | - | - | 0 | - |
| - | - | 2136 | 458.3 | - | - | 0 | - |
| - | - | 1730 | 459.3 | - | - | 0 | - |
| - | - | 5094 | 460.7 | - | - | 0 | - |
| - | - | 1281 | 461.2 | - | - | 0 | - |
| 6 | y | 1.446E+04 | 462.2 | 0.001291 | 2.793 | +1 | 3 |
| - | - | 3408 | 463.2 | - | - | 0 | - |
| - | - | 1210 | 463.3 | - | - | 0 | - |
| - | - | 1693 | 464.3 | - | - | 0 | - |
| - | - | 1497 | 465.2 | - | - | 0 | - |
| - | - | 1312 | 466.2 | - | - | 0 | - |
| - | - | 1151 | 466.2 | - | - | 0 | - |
| - | - | 4914 | 466.3 | - | - | 0 | - |
| - | - | 4.171E+04 | 468.2 | - | - | 0 | - |
| - | - | 5831 | 468.3 | - | - | 0 | - |
| - | - | 1.054E+04 | 469.2 | - | - | 0 | - |
| - | - | 4720 | 469.7 | - | - | 0 | - |
| - | - | 2379 | 470.2 | - | - | 0 | - |
| - | - | 1815 | 471.3 | - | - | 0 | - |
| - | - | 1.165E+04 | 472.2 | - | - | 0 | - |
| - | - | 2384 | 473.2 | - | - | 0 | - |
| - | - | 928.6 | 473.2 | - | - | 0 | - |
| - | - | 3052 | 474.2 | - | - | 0 | - |
| - | - | 1964 | 475.2 | - | - | 0 | - |
| - | - | 6602 | 476.3 | - | - | 0 | - |
| - | - | 1750 | 477.3 | - | - | 0 | - |
| - | - | 1.321E+04 | 478.2 | - | - | 0 | - |
| 0 | Precursor | 3065 | 478.2 | 0.0006565 | 1.373 | +2 | -1 |
| - | - | 1578 | 478.8 | - | - | 0 | - |
| - | - | 3250 | 479.2 | - | - | 0 | - |
| - | - | 1271 | 480.2 | - | - | 0 | - |
| - | - | 2796 | 481.3 | - | - | 0 | - |
| - | - | 2824 | 482.3 | - | - | 0 | - |
| - | - | 1210 | 484.2 | - | - | 0 | - |
| - | - | 4625 | 484.3 | - | - | 0 | - |
| - | - | 1512 | 485.3 | - | - | 0 | - |
| - | - | 1464 | 486.3 | - | - | 0 | - |
| 0 | Precursor | 1198 | 487.3 | 0.005903 | 12.11 | +2 | -1 |
| - | - | 1050 | 488.3 | - | - | 0 | - |
| - | - | 5147 | 492.2 | - | - | 0 | - |
| - | - | 1297 | 493.2 | - | - | 0 | - |
| 5 | b | 3.066E+04 | 494.3 | 0.0009683 | 1.959 | +1 | 5 |
| - | - | 1275 | 495.2 | - | - | 0 | - |
| - | - | 7765 | 495.3 | - | - | 0 | - |
| - | - | 1.263E+04 | 496.2 | - | - | 0 | - |
| - | - | 3212 | 497.2 | - | - | 0 | - |
| - | - | 1677 | 498.2 | - | - | 0 | - |
| - | - | 7209 | 499.3 | - | - | 0 | - |
| - | - | 3692 | 500.3 | - | - | 0 | - |
| - | - | 7645 | 506.2 | - | - | 0 | - |
| - | - | 953.8 | 506.3 | - | - | 0 | - |
| - | - | 2749 | 507.2 | - | - | 0 | - |
| - | - | 2443 | 507.3 | - | - | 0 | - |
| - | - | 1647 | 508.2 | - | - | 0 | - |
| - | - | 5576 | 508.3 | - | - | 0 | - |
| - | - | 2993 | 509.3 | - | - | 0 | - |
| - | - | 1.851E+04 | 510.3 | - | - | 0 | - |
| - | - | 4521 | 511.3 | - | - | 0 | - |
| - | - | 2181 | 512.3 | - | - | 0 | - |
| 5 | b | 3.317E+04 | 512.3 | 0.001024 | 1.998 | +1 | 5 |
| - | - | 7463 | 513.2 | - | - | 0 | - |
| - | - | 8871 | 513.3 | - | - | 0 | - |
| - | - | 3320 | 514.2 | - | - | 0 | - |
| - | - | 1390 | 514.3 | - | - | 0 | - |
| - | - | 1449 | 515.2 | - | - | 0 | - |
| - | - | 1117 | 519.3 | - | - | 0 | - |
| - | - | 4679 | 520.2 | - | - | 0 | - |
| - | - | 953.9 | 521.2 | - | - | 0 | - |
| - | - | 4939 | 522.3 | - | - | 0 | - |
| - | - | 2353 | 523.2 | - | - | 0 | - |
| - | - | 1681 | 523.3 | - | - | 0 | - |
| - | - | 9.132E+04 | 524.2 | - | - | 0 | - |
| - | - | 3602 | 524.3 | - | - | 0 | - |
| - | - | 2.659E+04 | 525.2 | - | - | 0 | - |
| - | - | 1243 | 525.3 | - | - | 0 | - |
| - | - | 4323 | 525.3 | - | - | 0 | - |
| - | - | 4161 | 526.2 | - | - | 0 | - |
| - | - | 6027 | 526.3 | - | - | 0 | - |
| - | - | 7234 | 527.3 | - | - | 0 | - |
| - | - | 1.011E+04 | 528.3 | - | - | 0 | - |
| - | - | 2811 | 529.3 | - | - | 0 | - |
| - | - | 1180 | 533.3 | - | - | 0 | - |
| - | - | 3827 | 534.3 | - | - | 0 | - |
| - | - | 1160 | 536.3 | - | - | 0 | - |
| - | - | 1.521E+04 | 537.3 | - | - | 0 | - |
| - | - | 3785 | 538.3 | - | - | 0 | - |
| - | - | 1727 | 539.3 | - | - | 0 | - |
| - | - | 5084 | 540.3 | - | - | 0 | - |
| 5 | y | 4.244E+04 | 541.2 | 0.0008091 | 1.495 | +1 | 4 |
| - | - | 4072 | 541.3 | - | - | 0 | - |
| 5 | y | 1.567E+05 | 542.2 | 0.001474 | 2.718 | +1 | 4 |
| - | - | 4.881E+04 | 543.2 | - | - | 0 | - |
| - | - | 4626 | 543.3 | - | - | 0 | - |
| - | - | 8157 | 544.2 | - | - | 0 | - |
| - | - | 1189 | 544.3 | - | - | 0 | - |
| - | - | 2661 | 545.3 | - | - | 0 | - |
| - | - | 1.336E+04 | 551.3 | - | - | 0 | - |
| - | - | 3343 | 552.3 | - | - | 0 | - |
| - | - | 1201 | 553.3 | - | - | 0 | - |
| - | - | 1.563E+04 | 555.3 | - | - | 0 | - |
| - | - | 4394 | 556.3 | - | - | 0 | - |
| 5 | y | 4.556E+05 | 559.3 | 0.0008035 | 1.437 | +1 | 4 |
| - | - | 1.377E+05 | 560.3 | - | - | 0 | - |
| - | - | 2.535E+04 | 561.3 | - | - | 0 | - |
| - | - | 1321 | 562.3 | - | - | 0 | - |
| - | - | 1.891E+04 | 569.3 | - | - | 0 | - |
| - | - | 1415 | 569.3 | - | - | 0 | - |
| - | - | 5276 | 570.3 | - | - | 0 | - |
| - | - | 1386 | 571.3 | - | - | 0 | - |
| - | - | 1277 | 577.3 | - | - | 0 | - |
| - | - | 2258 | 578.3 | - | - | 0 | - |
| - | - | 1683 | 579.3 | - | - | 0 | - |
| - | - | 1276 | 588.3 | - | - | 0 | - |
| - | - | 3193 | 593.3 | - | - | 0 | - |
| - | - | 2110 | 594.3 | - | - | 0 | - |
| - | - | 878.9 | 595.3 | - | - | 0 | - |
| - | - | 4758 | 595.3 | - | - | 0 | - |
| - | - | 2034 | 596.3 | - | - | 0 | - |
| - | - | 2463 | 597.3 | - | - | 0 | - |
| - | - | 2251 | 597.3 | - | - | 0 | - |
| - | - | 3747 | 603.3 | - | - | 0 | - |
| - | - | 1.04E+04 | 605.3 | - | - | 0 | - |
| - | - | 3678 | 606.3 | - | - | 0 | - |
| - | - | 4351 | 607.3 | - | - | 0 | - |
| - | - | 1327 | 608.3 | - | - | 0 | - |
| - | - | 2513 | 611.3 | - | - | 0 | - |
| - | - | 1744 | 612.3 | - | - | 0 | - |
| - | - | 8578 | 613.4 | - | - | 0 | - |
| - | - | 1212 | 614.3 | - | - | 0 | - |
| - | - | 3764 | 614.4 | - | - | 0 | - |
| - | - | 7602 | 620.3 | - | - | 0 | - |
| - | - | 5006 | 621.3 | - | - | 0 | - |
| - | - | 2119 | 622.3 | - | - | 0 | - |
| - | - | 3485 | 622.4 | - | - | 0 | - |
| 6 | b | 4.672E+04 | 623.3 | 0.001039 | 1.667 | +1 | 6 |
| - | - | 1.736E+04 | 624.3 | - | - | 0 | - |
| - | - | 2880 | 625.3 | - | - | 0 | - |
| - | - | 2965 | 625.3 | - | - | 0 | - |
| - | - | 1294 | 628.3 | - | - | 0 | - |
| - | - | 1394 | 633.3 | - | - | 0 | - |
| - | - | 1790 | 637.4 | - | - | 0 | - |
| - | - | 2.647E+04 | 638.3 | - | - | 0 | - |
| - | - | 9419 | 639.3 | - | - | 0 | - |
| - | - | 1563 | 640.3 | - | - | 0 | - |
| - | - | 1.394E+04 | 640.4 | - | - | 0 | - |
| 6 | b | 4.953E+04 | 641.4 | 0.001827 | 2.848 | +1 | 6 |
| 4 | y | 4718 | 642.3 | 0.0004329 | 0.674 | +1 | 5 |
| - | - | 1.964E+04 | 642.4 | - | - | 0 | - |
| 4 | y | 9138 | 643.3 | 0.002074 | 3.224 | +1 | 5 |
| - | - | 3251 | 643.4 | - | - | 0 | - |
| - | - | 2998 | 644.3 | - | - | 0 | - |
| - | - | 1.587E+04 | 650.4 | - | - | 0 | - |
| - | - | 5084 | 651.4 | - | - | 0 | - |
| - | - | 2046 | 652.4 | - | - | 0 | - |
| - | - | 2.07E+04 | 656.3 | - | - | 0 | - |
| - | - | 6896 | 657.3 | - | - | 0 | - |
| - | - | 1597 | 658.3 | - | - | 0 | - |
| - | - | 2.581E+04 | 658.4 | - | - | 0 | - |
| - | - | 8978 | 659.4 | - | - | 0 | - |
| 4 | y | 5.828E+04 | 660.3 | 0.0009766 | 1.479 | +1 | 5 |
| - | - | 1.996E+04 | 661.3 | - | - | 0 | - |
| - | - | 3628 | 662.3 | - | - | 0 | - |
| - | - | 1145 | 666.3 | - | - | 0 | - |
| - | - | 1.154E+04 | 668.4 | - | - | 0 | - |
| - | - | 3161 | 669.4 | - | - | 0 | - |
| - | - | 1546 | 670.4 | - | - | 0 | - |
| - | - | 1053 | 680.4 | - | - | 0 | - |
| - | - | 1475 | 689.4 | - | - | 0 | - |
| - | - | 1189 | 690.3 | - | - | 0 | - |
| - | - | 1341 | 697.4 | - | - | 0 | - |
| - | - | 4944 | 706.4 | - | - | 0 | - |
| - | - | 5072 | 707.4 | - | - | 0 | - |
| - | - | 1024 | 708.4 | - | - | 0 | - |
| - | - | 2020 | 715.4 | - | - | 0 | - |
| - | - | 3827 | 716.4 | - | - | 0 | - |
| - | - | 1181 | 717.4 | - | - | 0 | - |
| - | - | 8684 | 723.4 | - | - | 0 | - |
| - | - | 6205 | 724.4 | - | - | 0 | - |
| - | - | 7780 | 725.4 | - | - | 0 | - |
| - | - | 2353 | 726.4 | - | - | 0 | - |
| - | - | 1.71E+04 | 733.4 | - | - | 0 | - |
| - | - | 1.551E+04 | 734.4 | - | - | 0 | - |
| - | - | 5407 | 735.4 | - | - | 0 | - |
| - | - | 1150 | 736.4 | - | - | 0 | - |
| - | - | 1516 | 738.3 | - | - | 0 | - |
| - | - | 1.318E+04 | 741.4 | - | - | 0 | - |
| - | - | 7602 | 742.4 | - | - | 0 | - |
| - | - | 1161 | 743.4 | - | - | 0 | - |
| 7 | b | 9.961E+04 | 751.4 | 0.001055 | 1.404 | +1 | 7 |
| - | - | 4.795E+04 | 752.4 | - | - | 0 | - |
| - | - | 1.184E+04 | 753.4 | - | - | 0 | - |
| - | - | 2122 | 754.4 | - | - | 0 | - |
| 3 | y | 5889 | 755.4 | 0.0004754 | 0.6293 | +1 | 6 |
| - | - | 3186 | 756.4 | - | - | 0 | - |
| - | - | 1156 | 757.4 | - | - | 0 | - |
| 7 | b | 7.098E+04 | 769.4 | 0.0008664 | 1.126 | +1 | 7 |
| - | - | 2.989E+04 | 770.4 | - | - | 0 | - |
| - | - | 4815 | 771.4 | - | - | 0 | - |
| 3 | y | 1.06E+04 | 773.4 | 0.000836 | 1.081 | +1 | 6 |
| - | - | 3719 | 774.4 | - | - | 0 | - |
| - | - | 1473 | 807.4 | - | - | 0 | - |
| - | - | 1189 | 824.4 | - | - | 0 | - |
| - | - | 2110 | 825.4 | - | - | 0 | - |
| - | - | 945.9 | 826.4 | - | - | 0 | - |
| 2 | y | 5737 | 842.4 | 0.0009177 | 1.089 | +1 | 7 |
| 2 | y | 3503 | 843.4 | 0.01367 | 16.21 | +1 | 7 |
| 2 | y | 7566 | 860.4 | 0.0001186 | 0.1379 | +1 | 7 |
| - | - | 3208 | 861.4 | - | - | 0 | - |
| - | - | 1085 | 862.4 | - | - | 0 | - |

m/z Charge Intensity FragmentType MassShift Position
120.08124542236328 0 2151.087
123.88644409179688 0 530.13745
125.07144165039062 0 4813.5083
125.10783386230469 0 18865.455
125.11304473876953 0 976.9383
126.0919189453125 0 6068.788
126.11122131347656 0 1426.2091
127.0871810913086 0 2340.9343
127.12349700927734 0 506.19977
128.08224487304688 0 2045.432
128.10748291015625 0 7766.987
129.0663604736328 0 43848.938
129.1028289794922 0 1296.6803
130.05043029785156 0 15108.485
130.06565856933594 0 111154.62
131.06903076171875 0 9548.575
131.07373046875 0 979.48206
132.08131408691406 0 54943.94
133.0852813720703 0 5948.534
135.249755859375 0 551.7589
136.0763397216797 0 934.12244
137.0717315673828 0 1363.4719
138.0918426513672 0 4660.9375
138.12832641601562 0 1370.4729
139.05072021484375 0 4388.045
141.06640625 0 1428.3386
142.06581115722656 0 4093.8645
142.12318420410156 0 27668.229
143.0736541748047 0 2306.1812
143.1068115234375 0 619.1844
143.11854553222656 0 1188.822
143.12664794921875 0 2186.884
144.04510498046875 0 943.2896
144.08135986328125 0 26770.691
145.08457946777344 0 2581.023
146.0606231689453 0 146060.28
146.09304809570312 0 1966.3191
147.05792236328125 0 656.5312
147.06398010253906 0 14083.409
147.07693481445312 0 12690.683
150.4418182373047 0 779.3963
152.1075439453125 0 20431.777
153.06642150878906 0 13238.72
153.10279846191406 0 9560.326
153.11009216308594 0 1336.4055
154.06964111328125 0 663.6698
154.08644104003906 0 793.0074
155.08206176757812 0 5310.57
155.11842346191406 0 28580.818 a Water loss 1
156.10232543945312 0 2120.848
156.12193298339844 0 2083.3125
158.04531860351562 0 2104.8271
159.09225463867188 0 298537.53
160.07611083984375 0 5270.8896
160.0891571044922 0 1395.0361
160.09564208984375 0 33178.21
161.09869384765625 0 1281.7083
163.08714294433594 0 2302.5962
165.10284423828125 0 954.2537
167.08229064941406 0 2103.44 y 6
169.07652282714844 0 3311.0012
169.13412475585938 0 12080.979
170.06060791015625 0 15918.962
170.1183624267578 0 2382.692
170.137451171875 0 1023.8127
171.06410217285156 0 2120.3599
171.11341857910156 0 28179.205
171.12176513671875 0 1449.9489
171.149658203125 0 3688.7734
172.11671447753906 0 2437.5588
172.15310668945312 0 760.5439
173.12904357910156 0 217275.72 a 1
174.0556640625 0 671.7224
174.1250762939453 0 1493.1508
174.13243103027344 0 16954.79
175.134765625 0 727.2502
177.1129608154297 0 923.5085
179.10569763183594 0 1362.1349
181.09779357910156 0 33128.695
182.10113525390625 0 2289.9346
183.11343383789062 0 55338.883 b Water loss 1
183.1497802734375 0 16866.021
184.11685180664062 0 4627.5615
184.13050842285156 0 580.39185
184.15296936035156 0 2063.128
185.09259033203125 0 1537.2499
185.12985229492188 0 715.99445
186.0878448486328 0 1804.169
187.06349182128906 0 794.46826
187.0719757080078 0 3821.024
187.08729553222656 0 26409.912
187.1343536376953 0 900.74225
187.14474487304688 0 64031.973
188.07130432128906 0 1161196.2
188.1481170654297 0 4862.048
189.0745391845703 0 132856.78
190.07711791992188 0 7462.388
191.081787109375 0 1659.6401
194.13412475585938 0 3460.4934
195.07696533203125 0 5578.443
195.113525390625 0 1792.9775
195.1495819091797 0 1972.177
196.14471435546875 0 2879.0195
197.0711669921875 0 849.7221
197.12899780273438 0 31401.51
198.0922393798828 0 745.7835
198.132568359375 0 3561.77
199.07200622558594 0 4876.536
199.1083221435547 0 231167.23
200.07601928710938 0 1085.4968
200.11167907714844 0 21072.102
200.1394805908203 0 1680.8896
201.12393188476562 0 261875.84 b 1
202.12733459472656 0 25358.984
203.12930297851562 0 1531.0632
205.09779357910156 0 492305.66 y 7
206.10110473632812 0 53498.043
207.1036834716797 0 2477.0654
207.1136474609375 0 1167.9921
208.10836791992188 0 819.4299
208.1324920654297 0 1037.8528 b 3
209.09266662597656 0 23273.658
210.09597778320312 0 2277.8647
210.1239013671875 0 2565.6401
211.1080780029297 0 3287.521
211.1447296142578 0 14534.799
212.10360717773438 0 6926.933
212.1482696533203 0 988.29114
213.0878143310547 0 2047.3113
213.1241912841797 0 2679.074
214.1555938720703 0 6477.274
215.0819854736328 0 1821.817
215.13963317871094 0 158194.7
216.14308166503906 0 13332.089
217.0824737548828 0 10768.504
217.11878967285156 0 1018.89886
217.14500427246094 0 1278.3383
218.15065002441406 0 873.62067
221.1283416748047 0 1546.3596
221.16641235351562 0 921.5711
222.1246795654297 0 1293.1495
223.0720672607422 0 9230.215
223.10858154296875 0 2522.3474
223.1449432373047 0 999.22614
223.1811981201172 0 997.24207
224.1397705078125 0 1965.505
225.10299682617188 0 11168.133
225.12405395507812 0 6127.183
226.1070556640625 0 1353.4863
226.11920166015625 0 11510.641
227.10336303710938 0 375969.3
228.106689453125 0 36768.906
228.13417053222656 0 939.4727
229.10829162597656 0 3066.3406
229.1553955078125 0 5838.1772
230.11386108398438 0 2551.313
231.09848022460938 0 1270.9623
233.65550231933594 0 2972.5815
237.10255432128906 0 1376.2427
237.1242218017578 0 2651.3594
238.15553283691406 0 4703.8916
238.19151306152344 0 1077.8986
239.13973999023438 0 17542.996
239.1524200439453 0 1313.4423
239.17626953125 0 1221.2059
239.6087646484375 0 4338.084
240.0985565185547 0 51376.51
240.13540649414062 0 2512.777
240.17025756835938 0 1335.7148
241.0825958251953 0 13229.24
241.10174560546875 0 5219.994
241.19178771972656 0 3627.4756
241.85577392578125 0 758.0163
242.0858154296875 0 1188.4164
242.15061950683594 0 9242.804
242.66079711914062 0 12713.834
243.13455200195312 0 2282.65
243.16256713867188 0 3640.6785
244.1298828125 0 110944.03
245.11386108398438 0 7516.9595
245.1332244873047 0 8950.224
246.12445068359375 0 1519.7214
247.6534881591797 0 2411.94 b Water loss 4
248.10366821289062 0 1819.3817
248.1404571533203 0 912.7606
249.1241455078125 0 2227.1414
249.16053771972656 0 1535.8273
250.1924591064453 0 904.88104
251.10377502441406 0 1101.4886
251.1400146484375 0 4778.484
251.1760711669922 0 12047.426
252.11390686035156 0 5606.7637
252.17977905273438 0 1711.2946
253.0979766845703 0 8322.072
253.11566162109375 0 1672.9694
254.10113525390625 0 1490.6807
254.1136932373047 0 1547.4001
255.13462829589844 0 2927.955
256.1301574707031 0 1488.7933
256.1661376953125 0 14797.057
256.6585693359375 0 3082.9026 b 4
257.1269226074219 0 1143.4006
257.15020751953125 0 20691.037
257.1659851074219 0 1252.0386
257.1720275878906 0 839.4043
258.1090087890625 0 48157.516
258.1455383300781 0 9142.384
258.1811218261719 0 1272.3007 d 2
259.0932312011719 0 1797.5299
259.1120300292969 0 4331.7417
259.15008544921875 0 966.02344
262.6114196777344 0 1511.8529
264.13470458984375 0 4826.6943
265.1182556152344 0 1257.9158
266.1143493652344 0 4803.2
266.1505432128906 0 9841.925
266.18695068359375 0 3712.858
267.1346130371094 0 8526.1045
267.14959716796875 0 1020.74274
267.17108154296875 0 1611.7891
268.09405517578125 0 1600.016
268.12939453125 0 1382.9431
268.1676330566406 0 1065.8735
268.2024841308594 0 10069.925 a Water loss 2
269.14031982421875 0 12212.556
269.1865539550781 0 7884.028
269.20318603515625 0 955.4966
270.12432861328125 0 126383.19
271.1085205078125 0 3433.5408
271.1275329589844 0 22413.723 y Water loss 4
271.6167907714844 0 1145.2778 y Ammonia loss 4
272.1119689941406 0 1172.7178
272.1283264160156 0 1212.1652
272.1608581542969 0 1475.6384
274.118896484375 0 1581.7067
274.1766052246094 0 10230.98
275.17987060546875 0 1251.6962
276.1185607910156 0 944.83124
277.1549987792969 0 1657.1085
278.1145324707031 0 1741.9133
278.1864929199219 0 2087.364
279.09771728515625 0 1310.0636
280.1086730957031 0 16026.486
280.166259765625 0 10450.399
281.112060546875 0 2789.5608
281.1689453125 0 1283.3632
282.1455993652344 0 10194.696
283.1296081542969 0 5367.1255
283.1461181640625 0 1325.8942
284.1249084472656 0 5074.3184
284.1612243652344 0 124980.945
285.145751953125 0 2138.8547
285.16448974609375 0 17976.545
286.1036071777344 0 2255.149
286.1411437988281 0 2077.6292
286.1663513183594 0 1689.9615
286.2132568359375 0 3710.978 a 2
287.1512756347656 0 3394.5852
292.12982177734375 0 9073.333
293.1138610839844 0 2152.5583
293.13226318359375 0 1276.0988
293.16229248046875 0 1097.587
294.1455383300781 0 7294.559
294.182373046875 0 7052.1655
295.1476135253906 0 1024.6768
295.1674499511719 0 1116.4277
295.18719482421875 0 992.96387
296.1250305175781 0 3904.8733
296.1977233886719 0 280337.6 b Water loss 2
297.13543701171875 0 4740.9683
297.20086669921875 0 42774.902
298.1194763183594 0 39768.207
298.1353759765625 0 826.07294
298.177001953125 0 15171.848
298.2033996582031 0 3400.2341
299.10302734375 0 2388.8071
299.12274169921875 0 6984.194
299.1806945800781 0 2332.0828
300.1561584472656 0 15739.674
301.1589050292969 0 1411.0189
302.17193603515625 0 153930.25
303.1751403808594 0 22727.865
304.1775817871094 0 1711.1754
307.18206787109375 0 5670.7476
307.68389892578125 0 1773.5677
308.1614074707031 0 1473.3297
309.156982421875 0 6362.3794
309.20556640625 0 1032.9733
310.1406555175781 0 25339.154
310.1601867675781 0 1322.3654
310.2132263183594 0 5313.7515
311.14410400390625 0 4045.5286
311.172119140625 0 6659.2334
311.2165832519531 0 1512.9834
312.15631103515625 0 14490.039
312.17462158203125 0 2665.7153 b Water loss 5
312.1927795410156 0 10404.448
312.6756591796875 0 1022.3522
313.1591491699219 0 3299.5037
313.1947326660156 0 1762.3734
314.1360778808594 0 2104.3657
314.2083740234375 0 63584.887 b 2
315.1461486816406 0 301695.1
315.21173095703125 0 9401.065
316.1300354003906 0 316574.3 y Ammonia loss 6
316.14923095703125 0 47953.664
317.1332702636719 0 53692.3
317.1506042480469 0 3122.5461
318.13555908203125 0 5615.912
318.1824035644531 0 1567.2498
320.1248779296875 0 5809.093
320.1614074707031 0 1448.0432
320.6880187988281 0 3238.3513
321.127685546875 0 1368.6421
321.1799011230469 0 22060.486 b 5
321.6809387207031 0 6318.106
322.17864990234375 0 3493.4648
324.22845458984375 0 1391.6799
325.1882019042969 0 1970.7003
325.2233581542969 0 1103.6265
326.172119140625 0 2426.1716
326.2085876464844 0 1084.425
327.1672668457031 0 31795.621
328.1512451171875 0 51998.24
328.1695556640625 0 3998.5361
328.2230224609375 0 1042.4664
329.1544189453125 0 7358.8306
329.1828918457031 0 3963.496
329.6938171386719 0 4392.34
330.1664123535156 0 7019.015
330.18829345703125 0 1079.85
333.15667724609375 0 157231.06 y 6
334.1597595214844 0 26667.578
334.2127380371094 0 3067.5347
335.162109375 0 2095.1006
335.2113037109375 0 1242.3687
336.1563415527344 0 1399.1022
336.1929931640625 0 2343.0747
337.1515808105469 0 33310.426
337.1885986328125 0 3963.1565
338.1351623535156 0 9821.47
338.1554870605469 0 4921.4727
339.1390380859375 0 2415.4312
339.2047424316406 0 927.79346
340.18768310546875 0 19180.697
341.19061279296875 0 2762.966
342.2398376464844 0 1180.8164
343.1417236328125 0 1113.132
343.23516845703125 0 2604.2256
344.23858642578125 0 1171.1779
345.1777038574219 0 20862.697
346.1795959472656 0 5269.546
347.17767333984375 0 1419.3582
350.2088928222656 0 1956.1631
351.2402648925781 0 5462.665
352.2237243652344 0 23276.357
352.2444763183594 0 1588.0748
353.1829833984375 0 4779.9893
353.2232360839844 0 6912.06
353.2532958984375 0 1342.8574
354.166259765625 0 943.2107
354.2010498046875 0 952.1896
354.22723388671875 0 1374.6859
355.162109375 0 511083
356.14605712890625 0 7931.1177
356.1652526855469 0 84900.57
357.1445617675781 0 754.3895
357.1671142578125 0 10336.347
357.2127685546875 0 975.1844
359.1589660644531 0 832.0972
360.6720886230469 0 1674.4685
362.20672607421875 0 3059.801
363.14532470703125 0 1057.6583
363.20343017578125 0 9668.972
364.18756103515625 0 2442.2769
364.2101745605469 0 1200.9938
365.1829528808594 0 10124.387
366.18597412109375 0 2461.1304
367.2344970703125 0 1945.3754
367.70257568359375 0 892.87366
369.2503662109375 0 35194.29
369.67889404296875 0 890.6929
370.2344055175781 0 45418.344
371.1920166015625 0 2809.46
371.23284912109375 0 17168.254
372.18890380859375 0 7231.4443
372.2347717285156 0 2831.3418
373.17242431640625 0 15107.7705
373.1929931640625 0 1333.9982
374.1752014160156 0 2685.8767
376.2036437988281 0 2687.115 b Water loss 6
376.7046203613281 0 947.65955
379.2349548339844 0 21267.99
380.1435241699219 0 1094.5507
380.2181091308594 0 3687.794
380.24053955078125 0 2055.4768
381.1564025878906 0 15516.161
381.1824951171875 0 2554.4678
381.21435546875 0 11668.003
382.1600036621094 0 3070.819
382.2174072265625 0 2159.8484
383.1947937011719 0 1175.8671
383.265625 0 2012.9843
384.2677917480469 0 1071.2002
385.2093200683594 0 2995.1138 b 6
385.70989990234375 0 1328.7695
387.2610168457031 0 55934.26
388.2641296386719 0 9914.163
391.1435546875 0 1193.5272
391.1982116699219 0 2353.2996
393.250244140625 0 4428.779
394.25457763671875 0 1106.1058
395.15692138671875 0 5942.5063
395.18389892578125 0 1531.9
395.22998046875 0 1982.5775
396.21356201171875 0 1608.8579
397.2455749511719 0 159300.75 b Water loss 3
398.2487487792969 0 36012.344
399.1659240722656 0 2173.8923
399.1893615722656 0 2517.6345
399.224609375 0 6219.08
399.2518310546875 0 4579.5195
404.1893310546875 0 1047.9227
405.1773986816406 0 1087.9319
405.2142333984375 0 1238.3857
405.2499084472656 0 1451.1079
407.1576232910156 0 2568.9082
407.2301940917969 0 2054.012
409.15191650390625 0 8451.295
409.2091979980469 0 13982.129
409.2366638183594 0 1228.1055
410.1550598144531 0 2099.9092
410.2119445800781 0 3758.473
411.18878173828125 0 4143.17
411.2615051269531 0 4521.192
412.1839294433594 0 3429.474
412.2178039550781 0 1713.7206
412.2643127441406 0 1296.9648
413.240478515625 0 3761.7188
415.2561950683594 0 323791.8 b 3
416.2591857910156 0 67659.766
417.2619323730469 0 9322.327
420.1913757324219 0 1817.1239
421.172607421875 0 4346.015
422.2045593261719 0 2807.7678 y Ammonia loss 1
422.2388610839844 0 1146.9896
423.1898498535156 0 1921.0957
423.2247314453125 0 9281.632
424.18743896484375 0 1086.8235
424.22705078125 0 3569.6895
424.2569885253906 0 2688.698
425.1689147949219 0 1223.4446
425.239501953125 0 2785.128
426.1787109375 0 1356.4629
426.2367858886719 0 2126.6313
427.1622314453125 0 40134.184
427.2196350097656 0 7405.6025
428.16510009765625 0 8125.4077
428.2173156738281 0 5218.8906
429.16802978515625 0 1009.00494
430.1944274902344 0 2257.4492
431.2657775878906 0 4482.686
437.7286071777344 0 2142.4243
438.19940185546875 0 8784.46
438.23052978515625 0 1220.5872
439.184326171875 0 3332.1199
440.2152404785156 0 2470.688
440.25140380859375 0 13889.503
440.28521728515625 0 865.26733
441.2359313964844 0 8653.455
442.19476318359375 0 3698.5566
442.2388916015625 0 2116.993
444.1888427734375 0 12133.034 y Water loss 5
445.1729736328125 0 29511.727 y Ammonia loss 5
446.17559814453125 0 6882.317
446.73236083984375 0 4816.377
447.1769714355469 0 1178.6853
447.2328186035156 0 1628.0162
449.2015686035156 0 1275.8257
450.2351989746094 0 3458.4695
450.27154541015625 0 2540.4087
451.2196044921875 0 4611.5317
451.7271423339844 0 1083.6049
452.2230224609375 0 2520.4302
455.7391052246094 0 1673.4847
456.2099609375 0 40128.766
457.2123107910156 0 7825.3755
458.2156982421875 0 1155.6696
458.2624816894531 0 2136.171
459.26153564453125 0 1729.923
460.72998046875 0 5093.9224
461.2338562011719 0 1280.6958
462.1996154785156 0 14462.749 y 5
463.202880859375 0 3408.3113
463.2574768066406 0 1209.7068
464.2539367675781 0 1693.0208
465.23602294921875 0 1496.793
466.193359375 0 1311.5334
466.2342834472656 0 1150.5824
466.3034973144531 0 4914.1187
468.2463684082031 0 41707.55
468.2829284667969 0 5831.1035
469.24859619140625 0 10539.126
469.73663330078125 0 4720.1953
470.2415771484375 0 2379.2327
471.281494140625 0 1814.663
472.1837158203125 0 11653.36
473.1868591308594 0 2384.3638
473.23504638671875 0 928.5725
474.22216796875 0 3052.4653
475.2200927734375 0 1964.012
476.28851318359375 0 6601.7114
477.2901611328125 0 1750.3671
478.2091064453125 0 13213.152
478.24847412109375 0 3064.9358 Precursor Water loss
478.7501220703125 0 1577.8455
479.21270751953125 0 3250.4402
480.21258544921875 0 1271.2377
481.2671203613281 0 2796.3455
482.263916015625 0 2824.2217
484.2427673339844 0 1209.6558
484.31475830078125 0 4624.9126
485.3150329589844 0 1512.0018
486.2554931640625 0 1463.9652
487.2590026855469 0 1197.713 Precursor
488.25299072265625 0 1050.3851
492.2463684082031 0 5147.4316
493.23199462890625 0 1297.2626
494.29827880859375 0 30657.67 b Water loss 4
495.2373046875 0 1275.2401
495.30145263671875 0 7765.2407
496.2203063964844 0 12631.308
497.2226257324219 0 3212.4446
498.2294006347656 0 1677.1575
499.2779846191406 0 7208.962
500.2774353027344 0 3691.5176
506.20452880859375 0 7645.38
506.26312255859375 0 953.817
507.2095642089844 0 2748.5542
507.29327392578125 0 2443.3254
508.2051086425781 0 1647.0591
508.2787780761719 0 5576.4175
509.27545166015625 0 2993.4253
510.2570495605469 0 18508.43
511.2591552734375 0 4521.205
512.2639770507812 0 2180.8745
512.3088989257812 0 33169.676 b 4
513.2462768554688 0 7462.762
513.3115844726562 0 8871.041
514.2315673828125 0 3319.8433
514.313720703125 0 1390.1265
515.234619140625 0 1449.1532
519.2553100585938 0 1117.0454
520.2413330078125 0 4679.154
521.2489013671875 0 953.85126
522.293701171875 0 4938.822
523.2311401367188 0 2352.777
523.2926025390625 0 1681.1146
524.2149658203125 0 91321.93
524.2684326171875 0 3601.504
525.2180786132812 0 26586.4
525.2666625976562 0 1242.7059
525.304443359375 0 4322.5156
526.2203979492188 0 4160.7046
526.2881469726562 0 6026.645
527.2855834960938 0 7234.307
528.2683715820312 0 10110.334
529.2698974609375 0 2810.8247
533.2719116210938 0 1179.6729
534.2576293945312 0 3826.8384
536.3075561523438 0 1159.674
537.2675170898438 0 15207.217
538.2593994140625 0 3784.925
539.2539672851562 0 1727.0273
540.3038330078125 0 5083.7593
541.2413330078125 0 42439.203 y Water loss 4
541.2979125976562 0 4071.7434
542.2260131835938 0 156698.78 y Ammonia loss 4
543.2286987304688 0 48811.176
543.3135375976562 0 4625.626
544.23095703125 0 8156.7163
544.3128662109375 0 1188.511
545.2928466796875 0 2661.4602
551.2827758789062 0 13361.633
552.2800903320312 0 3343.2256
553.301025390625 0 1201.3014
555.2780151367188 0 15627.56
556.279296875 0 4394.141
559.2518920898438 0 455565.6 y 4
560.2548217773438 0 137661.22
561.2569580078125 0 25345.166
562.2546997070312 0 1321.1747
569.29345703125 0 18909.66
569.339111328125 0 1414.7323
570.2963256835938 0 5276.089
571.2987670898438 0 1385.588
577.33642578125 0 1277.0927
578.3219604492188 0 2258.053
579.3161010742188 0 1683.2074
588.321533203125 0 1276.2189
593.2919311523438 0 3193.3994
594.2854614257812 0 2110.0088
595.2914428710938 0 878.9332
595.3455200195312 0 4758.3784
596.3356323242188 0 2033.5154
597.2691650390625 0 2463.4653
597.3291625976562 0 2250.87
603.2786865234375 0 3746.9683
605.33056640625 0 10396.023
606.3320922851562 0 3678.0835
607.251953125 0 4350.5293
608.2540893554688 0 1326.5834
611.3062133789062 0 2512.5027
612.2999877929688 0 1744.0295
613.3562622070312 0 8578.21
614.2974243164062 0 1211.7516
614.3592529296875 0 3764.3672
620.3049926757812 0 7601.934
621.2962036132812 0 5006.244
622.2941284179688 0 2119.1902
622.3574829101562 0 3484.7725
623.3409423828125 0 46723.31 b Water loss 5
624.3436889648438 0 17357.377
625.264404296875 0 2879.9875
625.3460083007812 0 2964.926
628.3317260742188 0 1294.184
633.3282470703125 0 1394.014
637.3530883789062 0 1789.5808
638.3150634765625 0 26465.895
639.3151245117188 0 9418.883
640.3121337890625 0 1562.8235
640.367431640625 0 13943.718
641.352294921875 0 49534.484 b 5
642.2886352539062 0 4717.7217 y Water loss 3
642.3546752929688 0 19636.79
643.2742919921875 0 9137.867 y Ammonia loss 3
643.3570556640625 0 3250.691
644.2794799804688 0 2997.9834
650.3515625 0 15867.839
651.3521728515625 0 5084.132
652.3566284179688 0 2045.9211
656.325927734375 0 20699.906
657.329345703125 0 6896.196
658.3196411132812 0 1596.9915
658.3779907226562 0 25806.045
659.380859375 0 8977.841
660.2997436523438 0 58276.664 y 3
661.3026733398438 0 19964.674
662.3056030273438 0 3628.0024
666.3091430664062 0 1144.5144
668.362060546875 0 11542.382
669.3658447265625 0 3160.6812
670.3651123046875 0 1546.2124
680.3661499023438 0 1053.073
689.3594360351562 0 1475.2708
690.3472900390625 0 1188.9124
697.3898315429688 0 1341.0625
706.37939453125 0 4943.834
707.3737182617188 0 5072.1265
708.3785400390625 0 1023.80066
715.379150390625 0 2019.5901
716.365478515625 0 3827.2856
717.3618774414062 0 1181.2189
723.4041748046875 0 8683.717
724.3927001953125 0 6205.34
725.3912963867188 0 7779.801
726.3887329101562 0 2353.255
733.3887329101562 0 17100.64
734.3787841796875 0 15512.175
735.3784790039062 0 5406.7285
736.3765258789062 0 1150.1398
738.3497314453125 0 1515.574
741.4149780273438 0 13179.58
742.4176635742188 0 7602.1206
743.4203491210938 0 1160.8597
751.3995361328125 0 99607.44 b Water loss 6
752.3989868164062 0 47950.67
753.3992309570312 0 11840.62
754.39453125 0 2121.8035
755.3727416992188 0 5889.375 y Water loss 2
756.371826171875 0 3186.455
757.3802490234375 0 1156.3284
769.409912109375 0 70977.51 b 6
770.4125366210938 0 29892.34
771.4163818359375 0 4815.3237
773.3836669921875 0 10597.606 y 2
774.3856811523438 0 3718.7085
807.3663330078125 0 1473.3573
824.3907470703125 0 1189.4166
825.3811645507812 0 2110.1443
826.3807373046875 0 945.871
842.4052124023438 0 5736.658 y Water loss 1
843.4019775390625 0 3502.6882 y Ammonia loss 1
860.4149780273438 0 7566.2876 y 1
861.419189453125 0 3207.7307
862.4144287109375 0 1085.041

Spectrum Details

|  |  |
| --- | --- |
| Matched peaks? Matched peaks The total absolute number of peaks matched. Additionally in brackets the total fraction of peaks matched and the total number of peaks is shown. | 47 (6.75% of 696) |
| FDR? FDR The false discovery rate estimated for this peptide. It is calculated by matching all theoretical fragments with a non-integer shift with the raw peaks for this spectrum. This is done with 40 different shifts. The resulting percentage is the average number of annotated peaks over the number of annotated peaks with the correct spectrum. | 1.62% |
| Satellite FDR? Satellite FDR See the FDR for details on its calculation. This satellite ion specific FDR only contains the satellite ions (d/w) for I/L/J positions. | 2.38% |
| PSM Score? PSM Score The PSM Score as given by Hecklib to this annotated spectrum. It is shown with three significant figures. | 603 |

#### Spectrum 9605? Spectrum 9605 The raw spectrum of this peptide as annotated by Hecklib. The fragments are coloured according to ion type (see legend). Any peaks with a star '\*' as text can be hovered over to see the full details, first the ion type second the mass shift type. By hovering over the amino acids in the peptide or ions in the legend the corresponding peaks are highlighted. By toggling the 'Unassigned' label you can turn the background (unassigned) peaks on or off in the plot. By updating the slider in the Ion legend you can update the spectrum to only show the top X% of the peaks with labels. The top X% means any peak that is within X% of the highest intensity. By dragging in the spectrum you can zoom in to a specific part of the spectrum and use 'Zoom Out' to get back to the original zoom level. The annotation of the spectrum is based on the given sequence in the peptides file and is done with different software so inconsistencies are likely. The peaks are annotated based on the given sequence, with 20 ppm tolerance.

Copy Data

##### Spectrum 9605 (TSV)

###### Preview

```
Loading example...
```

*Click on the button to copy the data to your clipboard.*

Mz MinMz MaxIntensity Max

WidthHeightPeptide font sizePeptide stroke widthSpectrum font sizeSpectrum stroke widthCompact peptide

Ion legend

wxyz

abcd

OtherUnassignedIon^Charge^_Position_Show for top:%

JSJTPEQW

02.22e+64.43e+66.65e+68.86e+6

Zoom Out

a^+1^_2_y^+2^_2_a^+1^_2_b^+1^_2_b^+1^_2_y^+1^_1_b^+2^_5_b^+2^_5_d^+1^_3_a^+1^_3_y^+2^_4_a^+1^_3_b^+1^_3_b^+2^_6_b^+1^_3_y^+1^_2_b^+2^_6_y^+1^_2_b^+2^_7_b^+2^_7_b^+1^_4_b^+1^_4_y^+2^_7_y^+1^_3_y^+1^_3_y^+1^_3_\*b^+1^_5_b^+1^_5_y^+1^_4_y^+1^_4_y^+1^_4_b^+1^_6_b^+1^_6_y^+1^_5_y^+1^_5_y^+1^_5_b^+1^_7_y^+1^_6_y^+1^_6_b^+1^_7_y^+1^_6_y^+1^_7_y^+1^_7_

049498914831977

Fragment Matches Table

Show background peaks

| Position | Ion type | Intensity | mz Theoretical | mz Error (Th) | mz Error (ppm) | Charge | Series Number |
| --- | --- | --- | --- | --- | --- | --- | --- |
| - | - | 4.845E+04 | 125.1 | - | - | 0 | - |
| - | - | 1.515E+05 | 125.1 | - | - | 0 | - |
| - | - | 5.187E+04 | 126.1 | - | - | 0 | - |
| - | - | 1.217E+04 | 126.1 | - | - | 0 | - |
| - | - | 1.213E+04 | 127.1 | - | - | 0 | - |
| - | - | 6578 | 127.1 | - | - | 0 | - |
| - | - | 7625 | 127.1 | - | - | 0 | - |
| - | - | 5.928E+04 | 128.1 | - | - | 0 | - |
| - | - | 2.993E+05 | 129.1 | - | - | 0 | - |
| - | - | 4115 | 129.1 | - | - | 0 | - |
| - | - | 1.184E+05 | 130.1 | - | - | 0 | - |
| - | - | 8.168E+05 | 130.1 | - | - | 0 | - |
| - | - | 8524 | 131.1 | - | - | 0 | - |
| - | - | 8.146E+04 | 131.1 | - | - | 0 | - |
| - | - | 9336 | 131.1 | - | - | 0 | - |
| - | - | 4.421E+05 | 132.1 | - | - | 0 | - |
| - | - | 4.302E+04 | 133.1 | - | - | 0 | - |
| - | - | 1.933E+04 | 137.1 | - | - | 0 | - |
| - | - | 2.083E+04 | 138.1 | - | - | 0 | - |
| - | - | 1.35E+04 | 138.1 | - | - | 0 | - |
| - | - | 4.943E+04 | 139.1 | - | - | 0 | - |
| - | - | 5958 | 139.1 | - | - | 0 | - |
| - | - | 6862 | 140.1 | - | - | 0 | - |
| - | - | 7136 | 141.1 | - | - | 0 | - |
| - | - | 5.013E+04 | 142.1 | - | - | 0 | - |
| - | - | 2.139E+05 | 142.1 | - | - | 0 | - |
| - | - | 1.424E+04 | 143.1 | - | - | 0 | - |
| - | - | 8350 | 143.1 | - | - | 0 | - |
| - | - | 1.628E+04 | 143.1 | - | - | 0 | - |
| - | - | 1.864E+05 | 144.1 | - | - | 0 | - |
| - | - | 2.306E+04 | 145.1 | - | - | 0 | - |
| - | - | 1.088E+06 | 146.1 | - | - | 0 | - |
| - | - | 1.15E+04 | 146.1 | - | - | 0 | - |
| - | - | 4866 | 147.1 | - | - | 0 | - |
| - | - | 9.776E+04 | 147.1 | - | - | 0 | - |
| - | - | 9.691E+04 | 147.1 | - | - | 0 | - |
| - | - | 1.334E+05 | 152.1 | - | - | 0 | - |
| - | - | 1.069E+04 | 152.1 | - | - | 0 | - |
| - | - | 1.026E+05 | 153.1 | - | - | 0 | - |
| - | - | 7.235E+04 | 153.1 | - | - | 0 | - |
| - | - | 1.293E+04 | 153.1 | - | - | 0 | - |
| - | - | 6689 | 154.1 | - | - | 0 | - |
| - | - | 8836 | 154.1 | - | - | 0 | - |
| - | - | 8319 | 154.1 | - | - | 0 | - |
| - | - | 3.585E+04 | 155.1 | - | - | 0 | - |
| 2 | a | 2.047E+05 | 155.1 | 0.0005187 | 3.344 | +1 | 2 |
| - | - | 1.379E+04 | 156.1 | - | - | 0 | - |
| - | - | 1.709E+04 | 156.1 | - | - | 0 | - |
| - | - | 1.974E+04 | 158 | - | - | 0 | - |
| - | - | 1.128E+04 | 158.1 | - | - | 0 | - |
| - | - | 2.294E+06 | 159.1 | - | - | 0 | - |
| - | - | 2.644E+04 | 160.1 | - | - | 0 | - |
| - | - | 1.179E+04 | 160.1 | - | - | 0 | - |
| - | - | 2.434E+05 | 160.1 | - | - | 0 | - |
| - | - | 1.068E+04 | 161.1 | - | - | 0 | - |
| - | - | 1.322E+04 | 163.1 | - | - | 0 | - |
| - | - | 8869 | 165.1 | - | - | 0 | - |
| - | - | 5184 | 165.1 | - | - | 0 | - |
| - | - | 5563 | 165.9 | - | - | 0 | - |
| 7 | y | 2.035E+04 | 167.1 | 0.0006646 | 3.977 | +2 | 2 |
| - | - | 2.566E+04 | 169.1 | - | - | 0 | - |
| - | - | 9.47E+04 | 169.1 | - | - | 0 | - |
| - | - | 6096 | 169.2 | - | - | 0 | - |
| - | - | 1.272E+05 | 170.1 | - | - | 0 | - |
| - | - | 2.254E+04 | 170.1 | - | - | 0 | - |
| - | - | 1.892E+04 | 171.1 | - | - | 0 | - |
| - | - | 2.285E+05 | 171.1 | - | - | 0 | - |
| - | - | 3.709E+04 | 171.1 | - | - | 0 | - |
| - | - | 5373 | 172.1 | - | - | 0 | - |
| - | - | 2.269E+04 | 172.1 | - | - | 0 | - |
| 2 | a | 1.653E+06 | 173.1 | 0.0006046 | 3.492 | +1 | 2 |
| - | - | 1.718E+04 | 173.4 | - | - | 0 | - |
| - | - | 1.349E+05 | 174.1 | - | - | 0 | - |
| - | - | 1.272E+04 | 179.1 | - | - | 0 | - |
| - | - | 2.257E+05 | 181.1 | - | - | 0 | - |
| - | - | 7171 | 181.1 | - | - | 0 | - |
| - | - | 1.792E+04 | 182.1 | - | - | 0 | - |
| 2 | b | 3.74E+05 | 183.1 | 0.0006755 | 3.689 | +1 | 2 |
| - | - | 1.271E+05 | 183.1 | - | - | 0 | - |
| - | - | 3.855E+04 | 184.1 | - | - | 0 | - |
| - | - | 7742 | 184.2 | - | - | 0 | - |
| - | - | 1.049E+04 | 185.1 | - | - | 0 | - |
| - | - | 1.017E+04 | 185.1 | - | - | 0 | - |
| - | - | 5874 | 186.1 | - | - | 0 | - |
| - | - | 1.659E+04 | 186.1 | - | - | 0 | - |
| - | - | 2.325E+04 | 187.1 | - | - | 0 | - |
| - | - | 1.782E+05 | 187.1 | - | - | 0 | - |
| - | - | 4.677E+05 | 187.1 | - | - | 0 | - |
| - | - | 8.773E+06 | 188.1 | - | - | 0 | - |
| - | - | 3.763E+04 | 188.1 | - | - | 0 | - |
| - | - | 9.883E+05 | 189.1 | - | - | 0 | - |
| - | - | 5.529E+04 | 190.1 | - | - | 0 | - |
| - | - | 1.01E+04 | 191.1 | - | - | 0 | - |
| - | - | 1.747E+04 | 194.1 | - | - | 0 | - |
| - | - | 4.126E+04 | 195.1 | - | - | 0 | - |
| - | - | 9743 | 195.2 | - | - | 0 | - |
| - | - | 1.512E+04 | 196.1 | - | - | 0 | - |
| - | - | 6644 | 196.2 | - | - | 0 | - |
| - | - | 2.395E+05 | 197.1 | - | - | 0 | - |
| - | - | 8886 | 198.1 | - | - | 0 | - |
| - | - | 7025 | 198.1 | - | - | 0 | - |
| - | - | 2.394E+04 | 198.1 | - | - | 0 | - |
| - | - | 4.067E+04 | 199.1 | - | - | 0 | - |
| - | - | 1.806E+06 | 199.1 | - | - | 0 | - |
| - | - | 1.42E+05 | 200.1 | - | - | 0 | - |
| - | - | 1.823E+04 | 200.1 | - | - | 0 | - |
| 2 | b | 1.96E+06 | 201.1 | 0.000563 | 2.799 | +1 | 2 |
| - | - | 1.694E+05 | 202.1 | - | - | 0 | - |
| - | - | 5902 | 203.1 | - | - | 0 | - |
| 8 | y | 3.726E+06 | 205.1 | 0.0006547 | 3.192 | +1 | 1 |
| - | - | 4.123E+05 | 206.1 | - | - | 0 | - |
| - | - | 2.317E+04 | 207.1 | - | - | 0 | - |
| - | - | 1E+04 | 208.1 | - | - | 0 | - |
| - | - | 1.721E+05 | 209.1 | - | - | 0 | - |
| - | - | 1.847E+04 | 210.1 | - | - | 0 | - |
| - | - | 1.136E+04 | 210.1 | - | - | 0 | - |
| - | - | 2.455E+04 | 211.1 | - | - | 0 | - |
| - | - | 1.022E+05 | 211.1 | - | - | 0 | - |
| - | - | 6.377E+04 | 212.1 | - | - | 0 | - |
| - | - | 1.109E+04 | 212.1 | - | - | 0 | - |
| - | - | 1.625E+04 | 213.1 | - | - | 0 | - |
| - | - | 1.64E+04 | 213.1 | - | - | 0 | - |
| - | - | 4.423E+04 | 214.2 | - | - | 0 | - |
| - | - | 2.132E+04 | 215.1 | - | - | 0 | - |
| - | - | 1.164E+06 | 215.1 | - | - | 0 | - |
| - | - | 1.199E+05 | 216.1 | - | - | 0 | - |
| - | - | 7.98E+04 | 217.1 | - | - | 0 | - |
| - | - | 1.129E+04 | 217.1 | - | - | 0 | - |
| - | - | 6914 | 217.1 | - | - | 0 | - |
| - | - | 7695 | 218.1 | - | - | 0 | - |
| - | - | 1.051E+04 | 218.2 | - | - | 0 | - |
| - | - | 6551 | 219.1 | - | - | 0 | - |
| - | - | 6859 | 220.1 | - | - | 0 | - |
| - | - | 1.154E+04 | 221.1 | - | - | 0 | - |
| - | - | 6453 | 221.2 | - | - | 0 | - |
| - | - | 1.039E+04 | 222.1 | - | - | 0 | - |
| - | - | 8062 | 222.2 | - | - | 0 | - |
| - | - | 5.011E+04 | 223.1 | - | - | 0 | - |
| - | - | 1.8E+04 | 223.1 | - | - | 0 | - |
| - | - | 1.675E+04 | 223.1 | - | - | 0 | - |
| - | - | 1.031E+04 | 223.2 | - | - | 0 | - |
| - | - | 1.1E+04 | 224.1 | - | - | 0 | - |
| - | - | 9.789E+04 | 225.1 | - | - | 0 | - |
| - | - | 4.728E+04 | 225.1 | - | - | 0 | - |
| - | - | 1.127E+04 | 226.1 | - | - | 0 | - |
| - | - | 9.529E+04 | 226.1 | - | - | 0 | - |
| - | - | 2.879E+06 | 227.1 | - | - | 0 | - |
| - | - | 2.834E+05 | 228.1 | - | - | 0 | - |
| - | - | 1.108E+04 | 228.1 | - | - | 0 | - |
| - | - | 2.587E+04 | 229.1 | - | - | 0 | - |
| - | - | 6.541E+04 | 229.2 | - | - | 0 | - |
| - | - | 2.611E+04 | 230.1 | - | - | 0 | - |
| - | - | 7034 | 231.1 | - | - | 0 | - |
| - | - | 3.283E+04 | 233.7 | - | - | 0 | - |
| - | - | 1.574E+04 | 237.1 | - | - | 0 | - |
| - | - | 1.282E+04 | 237.1 | - | - | 0 | - |
| - | - | 2.434E+04 | 238.2 | - | - | 0 | - |
| - | - | 1.418E+05 | 239.1 | - | - | 0 | - |
| - | - | 1.058E+04 | 239.2 | - | - | 0 | - |
| - | - | 1.587E+04 | 239.6 | - | - | 0 | - |
| - | - | 3.697E+05 | 240.1 | - | - | 0 | - |
| - | - | 2.172E+04 | 240.1 | - | - | 0 | - |
| - | - | 6788 | 240.2 | - | - | 0 | - |
| - | - | 9.747E+04 | 241.1 | - | - | 0 | - |
| - | - | 3.624E+04 | 241.1 | - | - | 0 | - |
| - | - | 3.049E+04 | 241.2 | - | - | 0 | - |
| - | - | 7878 | 242.1 | - | - | 0 | - |
| - | - | 6.387E+04 | 242.2 | - | - | 0 | - |
| - | - | 8.158E+04 | 242.7 | - | - | 0 | - |
| - | - | 1.417E+04 | 243.1 | - | - | 0 | - |
| - | - | 1.41E+04 | 243.2 | - | - | 0 | - |
| - | - | 8.32E+05 | 244.1 | - | - | 0 | - |
| - | - | 5.407E+04 | 245.1 | - | - | 0 | - |
| - | - | 8.525E+04 | 245.1 | - | - | 0 | - |
| - | - | 1.257E+04 | 246.1 | - | - | 0 | - |
| 5 | b | 2.166E+04 | 247.7 | 0.0004012 | 1.62 | +2 | 5 |
| - | - | 9406 | 248.1 | - | - | 0 | - |
| - | - | 1.077E+04 | 248.1 | - | - | 0 | - |
| - | - | 8085 | 248.2 | - | - | 0 | - |
| - | - | 1.436E+04 | 248.6 | - | - | 0 | - |
| - | - | 8921 | 249.1 | - | - | 0 | - |
| - | - | 1.003E+04 | 250.1 | - | - | 0 | - |
| - | - | 9037 | 250.2 | - | - | 0 | - |
| - | - | 2.954E+04 | 251.1 | - | - | 0 | - |
| - | - | 7.785E+04 | 251.2 | - | - | 0 | - |
| - | - | 3.624E+04 | 252.1 | - | - | 0 | - |
| - | - | 5.158E+04 | 253.1 | - | - | 0 | - |
| - | - | 8249 | 253.1 | - | - | 0 | - |
| - | - | 1.253E+04 | 254.1 | - | - | 0 | - |
| - | - | 1.034E+04 | 254.1 | - | - | 0 | - |
| - | - | 8420 | 254.2 | - | - | 0 | - |
| - | - | 2.004E+04 | 255.1 | - | - | 0 | - |
| - | - | 1.289E+05 | 256.2 | - | - | 0 | - |
| 5 | b | 2.655E+04 | 256.7 | 0.0004442 | 1.731 | +2 | 5 |
| - | - | 1.103E+04 | 257.1 | - | - | 0 | - |
| - | - | 1.757E+05 | 257.2 | - | - | 0 | - |
| - | - | 3.159E+05 | 258.1 | - | - | 0 | - |
| - | - | 7.28E+04 | 258.1 | - | - | 0 | - |
| 3 | d | 6957 | 258.2 | 2.582E-05 | 0.1 | +1 | 3 |
| - | - | 1.455E+04 | 259.1 | - | - | 0 | - |
| - | - | 3.416E+04 | 259.1 | - | - | 0 | - |
| - | - | 8993 | 262.6 | - | - | 0 | - |
| - | - | 7394 | 263.1 | - | - | 0 | - |
| - | - | 2.916E+04 | 264.1 | - | - | 0 | - |
| - | - | 1.281E+04 | 265.1 | - | - | 0 | - |
| - | - | 3.83E+04 | 266.1 | - | - | 0 | - |
| - | - | 7.135E+04 | 266.2 | - | - | 0 | - |
| - | - | 2.947E+04 | 266.2 | - | - | 0 | - |
| - | - | 7.477E+04 | 267.1 | - | - | 0 | - |
| - | - | 9688 | 267.2 | - | - | 0 | - |
| - | - | 1.25E+04 | 267.2 | - | - | 0 | - |
| - | - | 1.403E+04 | 268.1 | - | - | 0 | - |
| - | - | 1.084E+04 | 268.2 | - | - | 0 | - |
| 3 | a | 8.427E+04 | 268.2 | 0.0006222 | 2.32 | +1 | 3 |
| - | - | 8741 | 269.1 | - | - | 0 | - |
| - | - | 9.356E+04 | 269.1 | - | - | 0 | - |
| - | - | 7.269E+04 | 269.2 | - | - | 0 | - |
| - | - | 9091 | 269.2 | - | - | 0 | - |
| - | - | 9.601E+05 | 270.1 | - | - | 0 | - |
| - | - | 3.038E+04 | 271.1 | - | - | 0 | - |
| 5 | y | 1.445E+05 | 271.1 | 0.003663 | 13.51 | +2 | 4 |
| - | - | 6883 | 272.1 | - | - | 0 | - |
| - | - | 1.286E+04 | 272.2 | - | - | 0 | - |
| - | - | 1.113E+04 | 274.1 | - | - | 0 | - |
| - | - | 6.846E+04 | 274.2 | - | - | 0 | - |
| - | - | 1.192E+04 | 278.2 | - | - | 0 | - |
| - | - | 6656 | 279.1 | - | - | 0 | - |
| - | - | 1.14E+05 | 280.1 | - | - | 0 | - |
| - | - | 8.503E+04 | 280.2 | - | - | 0 | - |
| - | - | 1.932E+04 | 281.1 | - | - | 0 | - |
| - | - | 6874 | 281.1 | - | - | 0 | - |
| - | - | 1.269E+04 | 281.2 | - | - | 0 | - |
| - | - | 7.621E+04 | 282.1 | - | - | 0 | - |
| - | - | 3.859E+04 | 283.1 | - | - | 0 | - |
| - | - | 8615 | 283.1 | - | - | 0 | - |
| - | - | 3.876E+04 | 284.1 | - | - | 0 | - |
| - | - | 9.381E+05 | 284.2 | - | - | 0 | - |
| - | - | 1.99E+04 | 285.1 | - | - | 0 | - |
| - | - | 1.196E+05 | 285.2 | - | - | 0 | - |
| - | - | 1.722E+04 | 286.1 | - | - | 0 | - |
| - | - | 1.18E+04 | 286.1 | - | - | 0 | - |
| - | - | 1.409E+04 | 286.2 | - | - | 0 | - |
| 3 | a | 1.94E+04 | 286.2 | 0.001074 | 3.754 | +1 | 3 |
| - | - | 2.932E+04 | 287.2 | - | - | 0 | - |
| - | - | 6.5E+04 | 292.1 | - | - | 0 | - |
| - | - | 1.643E+04 | 293.1 | - | - | 0 | - |
| - | - | 9232 | 293.2 | - | - | 0 | - |
| - | - | 5.265E+04 | 294.1 | - | - | 0 | - |
| - | - | 8541 | 294.2 | - | - | 0 | - |
| - | - | 5.748E+04 | 294.2 | - | - | 0 | - |
| - | - | 9194 | 295.2 | - | - | 0 | - |
| - | - | 8307 | 295.2 | - | - | 0 | - |
| - | - | 2.695E+04 | 296.1 | - | - | 0 | - |
| 3 | b | 2.054E+06 | 296.2 | 0.0008553 | 2.887 | +1 | 3 |
| - | - | 3.671E+04 | 297.1 | - | - | 0 | - |
| - | - | 3.232E+05 | 297.2 | - | - | 0 | - |
| - | - | 3.027E+05 | 298.1 | - | - | 0 | - |
| - | - | 1.433E+05 | 298.2 | - | - | 0 | - |
| - | - | 2.007E+04 | 298.2 | - | - | 0 | - |
| - | - | 2.362E+04 | 299.1 | - | - | 0 | - |
| - | - | 4.949E+04 | 299.1 | - | - | 0 | - |
| - | - | 1.219E+04 | 299.2 | - | - | 0 | - |
| - | - | 1.1E+05 | 300.2 | - | - | 0 | - |
| - | - | 1.286E+04 | 301.2 | - | - | 0 | - |
| - | - | 1.151E+06 | 302.2 | - | - | 0 | - |
| - | - | 1.589E+05 | 303.2 | - | - | 0 | - |
| - | - | 1.435E+04 | 304.2 | - | - | 0 | - |
| - | - | 3.665E+04 | 307.2 | - | - | 0 | - |
| - | - | 1.603E+04 | 307.7 | - | - | 0 | - |
| - | - | 1.204E+04 | 308.2 | - | - | 0 | - |
| - | - | 1.051E+04 | 308.2 | - | - | 0 | - |
| - | - | 5.548E+04 | 309.2 | - | - | 0 | - |
| - | - | 1.797E+05 | 310.1 | - | - | 0 | - |
| - | - | 1.047E+04 | 310.2 | - | - | 0 | - |
| - | - | 3.741E+04 | 310.2 | - | - | 0 | - |
| - | - | 3.105E+04 | 311.1 | - | - | 0 | - |
| - | - | 5.962E+04 | 311.2 | - | - | 0 | - |
| - | - | 1.16E+05 | 312.2 | - | - | 0 | - |
| 6 | b | 1.567E+04 | 312.2 | 0.0005433 | 1.74 | +2 | 6 |
| - | - | 7.253E+04 | 312.2 | - | - | 0 | - |
| - | - | 9966 | 312.7 | - | - | 0 | - |
| - | - | 1.331E+04 | 313.2 | - | - | 0 | - |
| - | - | 2.059E+04 | 314.1 | - | - | 0 | - |
| 3 | b | 4.557E+05 | 314.2 | 0.0009717 | 3.093 | +1 | 3 |
| - | - | 2.2E+06 | 315.1 | - | - | 0 | - |
| - | - | 6.876E+04 | 315.2 | - | - | 0 | - |
| 7 | y | 2.356E+06 | 316.1 | 0.0008529 | 2.698 | +1 | 2 |
| - | - | 3.513E+05 | 316.1 | - | - | 0 | - |
| - | - | 3.829E+05 | 317.1 | - | - | 0 | - |
| - | - | 2.606E+04 | 317.2 | - | - | 0 | - |
| - | - | 4.049E+04 | 318.1 | - | - | 0 | - |
| - | - | 5.702E+04 | 320.1 | - | - | 0 | - |
| - | - | 1.351E+04 | 320.2 | - | - | 0 | - |
| - | - | 1.471E+04 | 320.7 | - | - | 0 | - |
| 6 | b | 1.835E+05 | 321.2 | 0.0009983 | 3.108 | +2 | 6 |
| - | - | 5.903E+04 | 321.7 | - | - | 0 | - |
| - | - | 2.82E+04 | 322.2 | - | - | 0 | - |
| - | - | 9418 | 324.2 | - | - | 0 | - |
| - | - | 1.395E+04 | 325.2 | - | - | 0 | - |
| - | - | 1.349E+04 | 326.2 | - | - | 0 | - |
| - | - | 1.457E+04 | 326.2 | - | - | 0 | - |
| - | - | 2.437E+05 | 327.2 | - | - | 0 | - |
| - | - | 3.908E+05 | 328.2 | - | - | 0 | - |
| - | - | 2.408E+04 | 328.2 | - | - | 0 | - |
| - | - | 5.506E+04 | 329.2 | - | - | 0 | - |
| - | - | 2.734E+04 | 329.2 | - | - | 0 | - |
| - | - | 4.8E+04 | 329.7 | - | - | 0 | - |
| - | - | 4.674E+04 | 330.2 | - | - | 0 | - |
| - | - | 1.047E+04 | 330.2 | - | - | 0 | - |
| 7 | y | 1.128E+06 | 333.2 | 0.0009151 | 2.747 | +1 | 2 |
| - | - | 1.988E+05 | 334.2 | - | - | 0 | - |
| - | - | 2.23E+04 | 334.2 | - | - | 0 | - |
| - | - | 1.83E+04 | 335.2 | - | - | 0 | - |
| - | - | 1.093E+04 | 336.2 | - | - | 0 | - |
| - | - | 2.403E+05 | 337.2 | - | - | 0 | - |
| - | - | 3.743E+04 | 337.2 | - | - | 0 | - |
| - | - | 8429 | 337.7 | - | - | 0 | - |
| - | - | 8.944E+04 | 338.1 | - | - | 0 | - |
| - | - | 3.262E+04 | 338.2 | - | - | 0 | - |
| - | - | 1.171E+04 | 339.1 | - | - | 0 | - |
| - | - | 1.58E+05 | 340.2 | - | - | 0 | - |
| - | - | 2.178E+04 | 341.2 | - | - | 0 | - |
| - | - | 2.251E+04 | 343.2 | - | - | 0 | - |
| - | - | 1.423E+05 | 345.2 | - | - | 0 | - |
| - | - | 1.204E+04 | 345.2 | - | - | 0 | - |
| - | - | 4.803E+04 | 346.2 | - | - | 0 | - |
| - | - | 1.147E+04 | 347.2 | - | - | 0 | - |
| - | - | 1.201E+04 | 350.2 | - | - | 0 | - |
| - | - | 4.373E+04 | 351.2 | - | - | 0 | - |
| - | - | 1.097E+04 | 352.2 | - | - | 0 | - |
| - | - | 1.758E+05 | 352.2 | - | - | 0 | - |
| - | - | 1.176E+04 | 352.2 | - | - | 0 | - |
| - | - | 2.845E+04 | 353.2 | - | - | 0 | - |
| - | - | 5.362E+04 | 353.2 | - | - | 0 | - |
| - | - | 1.401E+04 | 353.7 | - | - | 0 | - |
| - | - | 1.134E+04 | 354.2 | - | - | 0 | - |
| - | - | 3.668E+06 | 355.2 | - | - | 0 | - |
| - | - | 4.861E+04 | 356.1 | - | - | 0 | - |
| - | - | 5.889E+05 | 356.2 | - | - | 0 | - |
| - | - | 7.059E+04 | 357.2 | - | - | 0 | - |
| - | - | 1.994E+04 | 362.2 | - | - | 0 | - |
| - | - | 8563 | 363.1 | - | - | 0 | - |
| - | - | 6.553E+04 | 363.2 | - | - | 0 | - |
| - | - | 1.743E+04 | 364.2 | - | - | 0 | - |
| - | - | 1.224E+04 | 364.2 | - | - | 0 | - |
| - | - | 6.999E+04 | 365.2 | - | - | 0 | - |
| - | - | 1.961E+04 | 365.3 | - | - | 0 | - |
| - | - | 8770 | 366.2 | - | - | 0 | - |
| - | - | 9830 | 367.2 | - | - | 0 | - |
| - | - | 8159 | 368.2 | - | - | 0 | - |
| - | - | 2.531E+05 | 369.3 | - | - | 0 | - |
| - | - | 3.079E+05 | 370.2 | - | - | 0 | - |
| - | - | 1.796E+04 | 371.2 | - | - | 0 | - |
| - | - | 1.173E+05 | 371.2 | - | - | 0 | - |
| - | - | 5.792E+04 | 372.2 | - | - | 0 | - |
| - | - | 2.022E+04 | 372.2 | - | - | 0 | - |
| - | - | 1.142E+05 | 373.2 | - | - | 0 | - |
| - | - | 9306 | 373.2 | - | - | 0 | - |
| - | - | 2.344E+04 | 374.2 | - | - | 0 | - |
| 7 | b | 1.533E+04 | 376.2 | 0.0002768 | 0.7357 | +2 | 7 |
| - | - | 7552 | 376.7 | - | - | 0 | - |
| - | - | 1.142E+04 | 379.2 | - | - | 0 | - |
| - | - | 1.492E+05 | 379.2 | - | - | 0 | - |
| - | - | 2.336E+04 | 380.2 | - | - | 0 | - |
| - | - | 2.266E+04 | 380.2 | - | - | 0 | - |
| - | - | 1.076E+05 | 381.2 | - | - | 0 | - |
| - | - | 1.263E+04 | 381.2 | - | - | 0 | - |
| - | - | 7.327E+04 | 381.2 | - | - | 0 | - |
| - | - | 1.384E+04 | 382.2 | - | - | 0 | - |
| - | - | 1.388E+04 | 382.2 | - | - | 0 | - |
| - | - | 1.694E+04 | 383.3 | - | - | 0 | - |
| 7 | b | 1.717E+04 | 385.2 | 0.0007012 | 1.82 | +2 | 7 |
| - | - | 6909 | 385.2 | - | - | 0 | - |
| - | - | 3.911E+05 | 387.3 | - | - | 0 | - |
| - | - | 7.599E+04 | 388.3 | - | - | 0 | - |
| - | - | 1.007E+04 | 389.3 | - | - | 0 | - |
| - | - | 3.036E+04 | 391.2 | - | - | 0 | - |
| - | - | 2.519E+04 | 393.3 | - | - | 0 | - |
| - | - | 1.157E+04 | 394.2 | - | - | 0 | - |
| - | - | 2.912E+04 | 395.2 | - | - | 0 | - |
| - | - | 7403 | 395.2 | - | - | 0 | - |
| - | - | 1.647E+04 | 395.2 | - | - | 0 | - |
| 4 | b | 1.2E+06 | 397.2 | 0.001028 | 2.589 | +1 | 4 |
| - | - | 2.503E+05 | 398.2 | - | - | 0 | - |
| - | - | 1.507E+04 | 399.2 | - | - | 0 | - |
| - | - | 2.057E+04 | 399.2 | - | - | 0 | - |
| - | - | 4.322E+04 | 399.2 | - | - | 0 | - |
| - | - | 2.333E+04 | 399.3 | - | - | 0 | - |
| - | - | 1.118E+04 | 400.2 | - | - | 0 | - |
| - | - | 8911 | 405.2 | - | - | 0 | - |
| - | - | 7526 | 405.3 | - | - | 0 | - |
| - | - | 8059 | 406.2 | - | - | 0 | - |
| - | - | 2.099E+04 | 407.2 | - | - | 0 | - |
| - | - | 2.106E+04 | 407.2 | - | - | 0 | - |
| - | - | 5.833E+04 | 409.2 | - | - | 0 | - |
| - | - | 1.092E+05 | 409.2 | - | - | 0 | - |
| - | - | 1.536E+04 | 410.2 | - | - | 0 | - |
| - | - | 2.221E+04 | 410.2 | - | - | 0 | - |
| - | - | 2.915E+04 | 411.2 | - | - | 0 | - |
| - | - | 2.437E+04 | 411.3 | - | - | 0 | - |
| - | - | 2.031E+04 | 412.2 | - | - | 0 | - |
| - | - | 9392 | 412.2 | - | - | 0 | - |
| - | - | 1.083E+04 | 413.2 | - | - | 0 | - |
| - | - | 1.726E+04 | 413.2 | - | - | 0 | - |
| 4 | b | 2.302E+06 | 415.3 | 0.001053 | 2.536 | +1 | 4 |
| - | - | 5.032E+05 | 416.3 | - | - | 0 | - |
| - | - | 6.796E+04 | 417.3 | - | - | 0 | - |
| - | - | 9176 | 420.2 | - | - | 0 | - |
| - | - | 3.295E+04 | 421.2 | - | - | 0 | - |
| 2 | y | 2.402E+04 | 422.2 | 0.006919 | 16.39 | +2 | 7 |
| - | - | 1.058E+04 | 422.2 | - | - | 0 | - |
| - | - | 1.381E+04 | 422.3 | - | - | 0 | - |
| - | - | 1.084E+04 | 423.2 | - | - | 0 | - |
| - | - | 8.181E+04 | 423.2 | - | - | 0 | - |
| - | - | 1.545E+04 | 424.2 | - | - | 0 | - |
| - | - | 2.304E+04 | 424.3 | - | - | 0 | - |
| - | - | 1.642E+04 | 425.2 | - | - | 0 | - |
| - | - | 1.607E+04 | 426.2 | - | - | 0 | - |
| - | - | 1.237E+04 | 426.2 | - | - | 0 | - |
| - | - | 2.943E+05 | 427.2 | - | - | 0 | - |
| - | - | 5.077E+04 | 427.2 | - | - | 0 | - |
| - | - | 6.566E+04 | 428.2 | - | - | 0 | - |
| - | - | 4.197E+04 | 428.2 | - | - | 0 | - |
| - | - | 1.022E+04 | 429.2 | - | - | 0 | - |
| - | - | 3.569E+04 | 431.3 | - | - | 0 | - |
| - | - | 1.951E+04 | 437.7 | - | - | 0 | - |
| - | - | 6.959E+04 | 438.2 | - | - | 0 | - |
| - | - | 1.483E+04 | 438.2 | - | - | 0 | - |
| - | - | 2.729E+04 | 439.2 | - | - | 0 | - |
| - | - | 2.5E+04 | 440.2 | - | - | 0 | - |
| - | - | 1.005E+05 | 440.3 | - | - | 0 | - |
| - | - | 1.292E+04 | 440.3 | - | - | 0 | - |
| - | - | 6.692E+04 | 441.2 | - | - | 0 | - |
| - | - | 3.405E+04 | 442.2 | - | - | 0 | - |
| - | - | 1.334E+04 | 442.2 | - | - | 0 | - |
| - | - | 1.322E+04 | 442.3 | - | - | 0 | - |
| 6 | y | 9.14E+04 | 444.2 | 0.001296 | 2.919 | +1 | 3 |
| - | - | 1.036E+04 | 444.2 | - | - | 0 | - |
| 6 | y | 1.866E+05 | 445.2 | 0.00129 | 2.897 | +1 | 3 |
| - | - | 5.025E+04 | 446.2 | - | - | 0 | - |
| - | - | 1.347E+04 | 446.7 | - | - | 0 | - |
| - | - | 9644 | 447.2 | - | - | 0 | - |
| - | - | 2.301E+04 | 447.2 | - | - | 0 | - |
| - | - | 8279 | 448.3 | - | - | 0 | - |
| - | - | 1.272E+04 | 449.2 | - | - | 0 | - |
| - | - | 1.708E+04 | 449.3 | - | - | 0 | - |
| - | - | 3.272E+04 | 450.2 | - | - | 0 | - |
| - | - | 2.996E+04 | 450.3 | - | - | 0 | - |
| - | - | 2.741E+04 | 451.2 | - | - | 0 | - |
| - | - | 1.708E+04 | 452.2 | - | - | 0 | - |
| - | - | 7596 | 453.3 | - | - | 0 | - |
| - | - | 1.005E+04 | 455.2 | - | - | 0 | - |
| - | - | 3.145E+05 | 456.2 | - | - | 0 | - |
| - | - | 6.37E+04 | 457.2 | - | - | 0 | - |
| - | - | 1.552E+04 | 458.3 | - | - | 0 | - |
| - | - | 3.037E+04 | 460.7 | - | - | 0 | - |
| - | - | 1.318E+04 | 461.2 | - | - | 0 | - |
| 6 | y | 1.023E+05 | 462.2 | 0.000894 | 1.934 | +1 | 3 |
| - | - | 1.986E+04 | 463.2 | - | - | 0 | - |
| - | - | 1.504E+04 | 465.2 | - | - | 0 | - |
| - | - | 1.555E+04 | 466.2 | - | - | 0 | - |
| - | - | 1.066E+04 | 466.2 | - | - | 0 | - |
| - | - | 4.195E+04 | 466.3 | - | - | 0 | - |
| - | - | 9585 | 468.2 | - | - | 0 | - |
| - | - | 2.756E+05 | 468.2 | - | - | 0 | - |
| - | - | 3.105E+04 | 468.3 | - | - | 0 | - |
| - | - | 6.895E+04 | 469.2 | - | - | 0 | - |
| - | - | 1.317E+04 | 469.3 | - | - | 0 | - |
| - | - | 2.599E+04 | 469.7 | - | - | 0 | - |
| - | - | 1.898E+04 | 470.2 | - | - | 0 | - |
| - | - | 8.261E+04 | 472.2 | - | - | 0 | - |
| - | - | 2.577E+04 | 473.2 | - | - | 0 | - |
| - | - | 1.127E+04 | 473.2 | - | - | 0 | - |
| - | - | 9487 | 474.2 | - | - | 0 | - |
| - | - | 1.46E+04 | 475.2 | - | - | 0 | - |
| - | - | 3.864E+04 | 476.3 | - | - | 0 | - |
| - | - | 1.035E+04 | 477.3 | - | - | 0 | - |
| - | - | 9.954E+04 | 478.2 | - | - | 0 | - |
| 0 | Precursor | 1.911E+04 | 478.2 | 0.0009617 | 2.011 | +2 | -1 |
| - | - | 1.36E+04 | 478.8 | - | - | 0 | - |
| - | - | 3.128E+04 | 479.2 | - | - | 0 | - |
| - | - | 2.024E+04 | 481.3 | - | - | 0 | - |
| - | - | 3.43E+04 | 482.3 | - | - | 0 | - |
| - | - | 2.972E+04 | 484.3 | - | - | 0 | - |
| - | - | 1.025E+04 | 485.3 | - | - | 0 | - |
| - | - | 9458 | 485.3 | - | - | 0 | - |
| - | - | 1.903E+04 | 486.3 | - | - | 0 | - |
| - | - | 1.193E+04 | 490.3 | - | - | 0 | - |
| - | - | 4.626E+04 | 492.2 | - | - | 0 | - |
| 5 | b | 2.014E+05 | 494.3 | 0.00106 | 2.144 | +1 | 5 |
| - | - | 7933 | 495.2 | - | - | 0 | - |
| - | - | 6.106E+04 | 495.3 | - | - | 0 | - |
| - | - | 9.244E+04 | 496.2 | - | - | 0 | - |
| - | - | 2.042E+04 | 497.2 | - | - | 0 | - |
| - | - | 1.397E+04 | 498.2 | - | - | 0 | - |
| - | - | 4.403E+04 | 499.3 | - | - | 0 | - |
| - | - | 1.785E+04 | 500.3 | - | - | 0 | - |
| - | - | 6.224E+04 | 506.2 | - | - | 0 | - |
| - | - | 2.098E+04 | 507.2 | - | - | 0 | - |
| - | - | 7757 | 507.3 | - | - | 0 | - |
| - | - | 1.486E+04 | 508.2 | - | - | 0 | - |
| - | - | 2.531E+04 | 508.3 | - | - | 0 | - |
| - | - | 9196 | 509.3 | - | - | 0 | - |
| - | - | 1.3E+05 | 510.3 | - | - | 0 | - |
| - | - | 3.874E+04 | 511.3 | - | - | 0 | - |
| 5 | b | 2.622E+05 | 512.3 | 0.001146 | 2.237 | +1 | 5 |
| - | - | 7.738E+04 | 513.2 | - | - | 0 | - |
| - | - | 6.716E+04 | 513.3 | - | - | 0 | - |
| - | - | 1.321E+04 | 514.2 | - | - | 0 | - |
| - | - | 1.114E+04 | 514.3 | - | - | 0 | - |
| - | - | 1.05E+04 | 519.3 | - | - | 0 | - |
| - | - | 2.793E+04 | 520.2 | - | - | 0 | - |
| - | - | 3.354E+04 | 522.3 | - | - | 0 | - |
| - | - | 1.29E+04 | 523.2 | - | - | 0 | - |
| - | - | 1.419E+04 | 523.3 | - | - | 0 | - |
| - | - | 6.449E+05 | 524.2 | - | - | 0 | - |
| - | - | 3.637E+04 | 524.3 | - | - | 0 | - |
| - | - | 1.849E+05 | 525.2 | - | - | 0 | - |
| - | - | 2.52E+04 | 525.3 | - | - | 0 | - |
| - | - | 2.571E+04 | 526.2 | - | - | 0 | - |
| - | - | 5.524E+04 | 526.3 | - | - | 0 | - |
| - | - | 5.524E+04 | 527.3 | - | - | 0 | - |
| - | - | 5.895E+04 | 528.3 | - | - | 0 | - |
| - | - | 1.848E+04 | 529.3 | - | - | 0 | - |
| - | - | 9704 | 533.3 | - | - | 0 | - |
| - | - | 1.713E+04 | 534.3 | - | - | 0 | - |
| - | - | 1.18E+05 | 537.3 | - | - | 0 | - |
| - | - | 2.864E+04 | 538.3 | - | - | 0 | - |
| - | - | 2.701E+04 | 540.3 | - | - | 0 | - |
| 5 | y | 2.927E+05 | 541.2 | 0.0008091 | 1.495 | +1 | 4 |
| - | - | 3.518E+04 | 541.3 | - | - | 0 | - |
| 5 | y | 1.095E+06 | 542.2 | 0.001474 | 2.718 | +1 | 4 |
| - | - | 3.297E+05 | 543.2 | - | - | 0 | - |
| - | - | 2.686E+04 | 543.3 | - | - | 0 | - |
| - | - | 4.682E+04 | 544.2 | - | - | 0 | - |
| - | - | 8490 | 544.3 | - | - | 0 | - |
| - | - | 1.158E+04 | 545.3 | - | - | 0 | - |
| - | - | 9.154E+04 | 551.3 | - | - | 0 | - |
| - | - | 1.902E+04 | 552.3 | - | - | 0 | - |
| - | - | 1.139E+05 | 555.3 | - | - | 0 | - |
| - | - | 3.286E+04 | 556.3 | - | - | 0 | - |
| - | - | 8634 | 557.3 | - | - | 0 | - |
| 5 | y | 3.275E+06 | 559.3 | 0.0008646 | 1.546 | +1 | 4 |
| - | - | 1.039E+06 | 560.3 | - | - | 0 | - |
| - | - | 1.943E+05 | 561.3 | - | - | 0 | - |
| - | - | 9842 | 562.3 | - | - | 0 | - |
| - | - | 1.566E+05 | 569.3 | - | - | 0 | - |
| - | - | 7866 | 569.3 | - | - | 0 | - |
| - | - | 2.938E+04 | 570.3 | - | - | 0 | - |
| - | - | 1.034E+04 | 577.3 | - | - | 0 | - |
| - | - | 1.131E+04 | 578.3 | - | - | 0 | - |
| - | - | 9775 | 579.3 | - | - | 0 | - |
| - | - | 1.163E+04 | 587.3 | - | - | 0 | - |
| - | - | 2.631E+04 | 593.3 | - | - | 0 | - |
| - | - | 1.438E+04 | 594.3 | - | - | 0 | - |
| - | - | 3.737E+04 | 595.3 | - | - | 0 | - |
| - | - | 2.105E+04 | 596.3 | - | - | 0 | - |
| - | - | 1.124E+04 | 597.3 | - | - | 0 | - |
| - | - | 8438 | 598.3 | - | - | 0 | - |
| - | - | 2.938E+04 | 603.3 | - | - | 0 | - |
| - | - | 7219 | 604.3 | - | - | 0 | - |
| - | - | 5.643E+04 | 605.3 | - | - | 0 | - |
| - | - | 1.779E+04 | 606.3 | - | - | 0 | - |
| - | - | 3.425E+04 | 607.3 | - | - | 0 | - |
| - | - | 1.4E+04 | 608.3 | - | - | 0 | - |
| - | - | 7410 | 610.3 | - | - | 0 | - |
| - | - | 2.053E+04 | 611.3 | - | - | 0 | - |
| - | - | 1.072E+04 | 612.3 | - | - | 0 | - |
| - | - | 5.325E+04 | 613.4 | - | - | 0 | - |
| - | - | 8752 | 614.3 | - | - | 0 | - |
| - | - | 2.105E+04 | 614.4 | - | - | 0 | - |
| - | - | 3.881E+04 | 620.3 | - | - | 0 | - |
| - | - | 3.818E+04 | 621.3 | - | - | 0 | - |
| - | - | 9855 | 622.3 | - | - | 0 | - |
| - | - | 2.58E+04 | 622.4 | - | - | 0 | - |
| 6 | b | 3.288E+05 | 623.3 | 0.001161 | 1.862 | +1 | 6 |
| - | - | 1.181E+05 | 624.3 | - | - | 0 | - |
| - | - | 1.454E+04 | 625.3 | - | - | 0 | - |
| - | - | 1.924E+04 | 625.3 | - | - | 0 | - |
| - | - | 1.022E+04 | 628.3 | - | - | 0 | - |
| - | - | 8447 | 632.3 | - | - | 0 | - |
| - | - | 1.792E+04 | 633.3 | - | - | 0 | - |
| - | - | 1.112E+04 | 637.4 | - | - | 0 | - |
| - | - | 1.973E+05 | 638.3 | - | - | 0 | - |
| - | - | 7.344E+04 | 639.3 | - | - | 0 | - |
| - | - | 9695 | 640.3 | - | - | 0 | - |
| - | - | 9.421E+04 | 640.4 | - | - | 0 | - |
| 6 | b | 3.797E+05 | 641.4 | 0.001705 | 2.658 | +1 | 6 |
| 4 | y | 3.113E+04 | 642.3 | 0.001104 | 1.719 | +1 | 5 |
| - | - | 1.475E+05 | 642.4 | - | - | 0 | - |
| 4 | y | 7.042E+04 | 643.3 | 0.002562 | 3.983 | +1 | 5 |
| - | - | 2.68E+04 | 643.4 | - | - | 0 | - |
| - | - | 2.61E+04 | 644.3 | - | - | 0 | - |
| - | - | 1.233E+05 | 650.4 | - | - | 0 | - |
| - | - | 4.811E+04 | 651.4 | - | - | 0 | - |
| - | - | 9814 | 652.3 | - | - | 0 | - |
| - | - | 1.252E+05 | 656.3 | - | - | 0 | - |
| - | - | 4.764E+04 | 657.3 | - | - | 0 | - |
| - | - | 2.106E+05 | 658.4 | - | - | 0 | - |
| - | - | 6.265E+04 | 659.4 | - | - | 0 | - |
| 4 | y | 4.149E+05 | 660.3 | 0.0009766 | 1.479 | +1 | 5 |
| - | - | 1.609E+05 | 661.3 | - | - | 0 | - |
| - | - | 2.685E+04 | 662.3 | - | - | 0 | - |
| - | - | 9.181E+04 | 668.4 | - | - | 0 | - |
| - | - | 2.837E+04 | 669.4 | - | - | 0 | - |
| - | - | 8488 | 679.4 | - | - | 0 | - |
| - | - | 9975 | 689.4 | - | - | 0 | - |
| - | - | 9365 | 697.4 | - | - | 0 | - |
| - | - | 2.854E+04 | 706.4 | - | - | 0 | - |
| - | - | 3.185E+04 | 707.4 | - | - | 0 | - |
| - | - | 8313 | 715.4 | - | - | 0 | - |
| - | - | 3.242E+04 | 716.4 | - | - | 0 | - |
| - | - | 6.259E+04 | 723.4 | - | - | 0 | - |
| - | - | 4.196E+04 | 724.4 | - | - | 0 | - |
| - | - | 3.307E+04 | 725.4 | - | - | 0 | - |
| - | - | 1.203E+04 | 726.4 | - | - | 0 | - |
| - | - | 9635 | 727.4 | - | - | 0 | - |
| - | - | 1.192E+05 | 733.4 | - | - | 0 | - |
| - | - | 1.183E+05 | 734.4 | - | - | 0 | - |
| - | - | 3.96E+04 | 735.4 | - | - | 0 | - |
| - | - | 9948 | 736.4 | - | - | 0 | - |
| - | - | 1.717E+04 | 738.3 | - | - | 0 | - |
| - | - | 9.502E+04 | 741.4 | - | - | 0 | - |
| - | - | 3.683E+04 | 742.4 | - | - | 0 | - |
| 7 | b | 7.035E+05 | 751.4 | 0.000933 | 1.242 | +1 | 7 |
| - | - | 3.306E+05 | 752.4 | - | - | 0 | - |
| - | - | 6.387E+04 | 753.4 | - | - | 0 | - |
| - | - | 8702 | 754.4 | - | - | 0 | - |
| 3 | y | 4.537E+04 | 755.4 | 0.0006585 | 0.8717 | +1 | 6 |
| 3 | y | 2.098E+04 | 756.4 | 0.01493 | 19.74 | +1 | 6 |
| 7 | b | 4.792E+05 | 769.4 | 0.0006832 | 0.888 | +1 | 7 |
| - | - | 1.919E+05 | 770.4 | - | - | 0 | - |
| - | - | 5.1E+04 | 771.4 | - | - | 0 | - |
| 3 | y | 6.493E+04 | 773.4 | 0.0001646 | 0.2128 | +1 | 6 |
| - | - | 3.245E+04 | 774.4 | - | - | 0 | - |
| - | - | 8333 | 824.4 | - | - | 0 | - |
| - | - | 1.757E+04 | 825.4 | - | - | 0 | - |
| 2 | y | 3.227E+04 | 842.4 | 0.0003641 | 0.4322 | +1 | 7 |
| - | - | 3.331E+04 | 843.4 | - | - | 0 | - |
| - | - | 8383 | 844.4 | - | - | 0 | - |
| 2 | y | 6.135E+04 | 860.4 | 0.0006679 | 0.7763 | +1 | 7 |
| - | - | 1.801E+04 | 861.4 | - | - | 0 | - |
| - | - | 1.286E+04 | 862.4 | - | - | 0 | - |
| - | - | 7679 | 1957 | - | - | 0 | - |

m/z Charge Intensity FragmentType MassShift Position
125.07147216796875 0 48450.1
125.10784149169922 0 151491.42
126.09183502197266 0 51867.547
126.111328125 0 12174.844
127.08697509765625 0 12126.296
127.0952377319336 0 6578.327
127.12333679199219 0 7625.1694
128.1074676513672 0 59276.94
129.06637573242188 0 299290.28
129.1031036376953 0 4114.9873
130.05039978027344 0 118399.29
130.06565856933594 0 816768.44
131.06333923339844 0 8523.669
131.06903076171875 0 81460.68
131.11802673339844 0 9336.3545
132.08131408691406 0 442109.53
133.08468627929688 0 43020.004
137.07151794433594 0 19334.21
138.09197998046875 0 20833.678
138.12841796875 0 13497.019
139.0507354736328 0 49431.848
139.1240997314453 0 5957.9272
140.10702514648438 0 6862.4033
141.0662841796875 0 7136.064
142.06564331054688 0 50130.23
142.12319946289062 0 213886.66
143.07350158691406 0 14236.387
143.1183624267578 0 8350.285
143.12640380859375 0 16279.47
144.08132934570312 0 186440.27
145.0846710205078 0 23062.982
146.0606231689453 0 1087765.5
146.09304809570312 0 11504.075
147.05763244628906 0 4865.5693
147.06396484375 0 97760.766
147.07693481445312 0 96913.48
152.10757446289062 0 133424.33
152.11404418945312 0 10688.869
153.06640625 0 102570.83
153.10275268554688 0 72351.81
153.1101837158203 0 12932.577
154.06991577148438 0 6688.8086
154.08642578125 0 8835.579
154.10604858398438 0 8318.969
155.08189392089844 0 35851.086
155.118408203125 0 204688.72 a Water loss 1
156.10189819335938 0 13788.3955
156.12176513671875 0 17092.121
158.04544067382812 0 19742.518
158.0843048095703 0 11282.274
159.09226989746094 0 2293645.5
160.07615661621094 0 26439.584
160.08946228027344 0 11792.037
160.09559631347656 0 243442.44
161.09889221191406 0 10681.594
163.08729553222656 0 13222.69
165.10275268554688 0 8869.044
165.11074829101562 0 5183.608
165.86500549316406 0 5562.877
167.08216857910156 0 20348.557 y 6
169.0764617919922 0 25656.398
169.1340789794922 0 94697.18
169.21592712402344 0 6096.14
170.0606689453125 0 127209.25
170.1182098388672 0 22536.836
171.0639190673828 0 18915.33
171.1134490966797 0 228512.16
171.14971923828125 0 37093.4
172.1094512939453 0 5373.4844
172.1168212890625 0 22691.557
173.12905883789062 0 1652812.1 a 1
173.43882751464844 0 17179.688
174.13243103027344 0 134899.34
179.10572814941406 0 12720.072
181.0977783203125 0 225687.11
181.13427734375 0 7171.0063
182.10107421875 0 17924.557
183.1134796142578 0 373964.66 b Water loss 1
183.14979553222656 0 127100.95
184.1168975830078 0 38554.305
184.15341186523438 0 7742.227
185.09300231933594 0 10493.757
185.12911987304688 0 10174.793
186.07806396484375 0 5874.438
186.08773803710938 0 16590.69
187.07183837890625 0 23250.246
187.08724975585938 0 178185.61
187.14474487304688 0 467660.28
188.07131958007812 0 8772577
188.14813232421875 0 37627.28
189.07452392578125 0 988298.5
190.07679748535156 0 55287.977
191.0821533203125 0 10098.312
194.13438415527344 0 17471.678
195.0771026611328 0 41262.406
195.15029907226562 0 9742.617
196.1441192626953 0 15120.561
196.19000244140625 0 6643.6123
197.12901306152344 0 239547.83
198.0915985107422 0 8886.316
198.11219787597656 0 7024.575
198.13270568847656 0 23943.791
199.07203674316406 0 40670.836
199.1083221435547 0 1806229.2
200.1116943359375 0 141987.45
200.13987731933594 0 18225.88
201.12393188476562 0 1959738.5 b 1
202.12728881835938 0 169361.36
203.12893676757812 0 5901.895
205.09780883789062 0 3726163 y 7
206.10113525390625 0 412334.34
207.10296630859375 0 23168.547
208.1082000732422 0 10004.802
209.09266662597656 0 172113.97
210.09585571289062 0 18465.598
210.1240997314453 0 11361.198
211.1083221435547 0 24550.504
211.1447296142578 0 102240.39
212.1035919189453 0 63773.695
212.14891052246094 0 11093.502
213.0876922607422 0 16250.71
213.12380981445312 0 16404.578
214.15545654296875 0 44228.547
215.08224487304688 0 21317.166
215.1396484375 0 1163653.1
216.1430206298828 0 119906.92
217.08258056640625 0 79798.5
217.11900329589844 0 11288.228
217.14596557617188 0 6914.026
218.08572387695312 0 7694.6396
218.15048217773438 0 10511.62
219.13421630859375 0 6550.64
220.1085662841797 0 6859.185
221.12791442871094 0 11542.365
221.1668701171875 0 6453.4814
222.12461853027344 0 10394.338
222.1970672607422 0 8062.365
223.07200622558594 0 50105.652
223.10824584960938 0 17998.684
223.1445770263672 0 16749.52
223.18087768554688 0 10309.913
224.1401824951172 0 11000.871
225.1029510498047 0 97887.33
225.12403869628906 0 47283.19
226.10708618164062 0 11265.119
226.1193084716797 0 95285.234
227.1033477783203 0 2879291.2
228.106689453125 0 283430.8
228.1337890625 0 11081.934
229.1083221435547 0 25868.885
229.15528869628906 0 65414.977
230.114013671875 0 26107.846
231.09835815429688 0 7034.1025
233.65548706054688 0 32827.082
237.1033172607422 0 15735.098
237.12307739257812 0 12822.712
238.15602111816406 0 24344.55
239.13966369628906 0 141829.94
239.1760711669922 0 10578.231
239.60855102539062 0 15872.119
240.0985870361328 0 369727.06
240.1351776123047 0 21715.6
240.1701202392578 0 6787.6245
241.0826416015625 0 97471.234
241.10186767578125 0 36240.586
241.19187927246094 0 30490.436
242.08599853515625 0 7878.3643
242.15052795410156 0 63873.707
242.66090393066406 0 81576.55
243.13478088378906 0 14165.701
243.162353515625 0 14100.509
244.12989807128906 0 831974.6
245.1138153076172 0 54070.355
245.1332244873047 0 85252.59
246.12429809570312 0 12565.352
247.65269470214844 0 21655.57 b Water loss 4
248.10406494140625 0 9406.465
248.13975524902344 0 10769.596
248.15419006347656 0 8084.9663
248.61373901367188 0 14356.6455
249.12435913085938 0 8921.162
250.11822509765625 0 10030.774
250.15606689453125 0 9037.089
251.13987731933594 0 29544.56
251.17605590820312 0 77852.445
252.11361694335938 0 36243.94
253.09788513183594 0 51581.773
253.1167449951172 0 8248.925
254.10108947753906 0 12528.184
254.1142120361328 0 10335.758
254.15011596679688 0 8419.738
255.13478088378906 0 20039.791
256.16619873046875 0 128948.945
256.65802001953125 0 26554.518 b 4
257.1274108886719 0 11026.516
257.15020751953125 0 175744.17
258.1090393066406 0 315932.4
258.1455078125 0 72800.49
258.1812438964844 0 6957.2793 d 2
259.0937805175781 0 14548.167
259.1123046875 0 34159.45
262.61114501953125 0 8992.732
263.1131896972656 0 7393.911
264.1346435546875 0 29162.746
265.1181335449219 0 12807.27
266.1142272949219 0 38304.87
266.1505126953125 0 71345.445
266.18634033203125 0 29469.662
267.13446044921875 0 74770.125
267.1523132324219 0 9688.188
267.1701965332031 0 12495.83
268.09375 0 14034.466
268.16534423828125 0 10837.397
268.20257568359375 0 84267.43 a Water loss 2
269.1138916015625 0 8741.402
269.1402587890625 0 93560.38
269.1864929199219 0 72688.76
269.2037353515625 0 9091.391
270.12432861328125 0 960094.06
271.10833740234375 0 30382.252
271.1275634765625 0 144450.75 y Water loss 4
272.13055419921875 0 6883.2983
272.1617736816406 0 12860.456
274.11920166015625 0 11126.85
274.1766357421875 0 68456.26
278.1871337890625 0 11923.706
279.09857177734375 0 6656.3716
280.1087951660156 0 114041.94
280.1663818359375 0 85026.29
281.1119689941406 0 19323.29
281.1274108886719 0 6873.8813
281.1689147949219 0 12692.421
282.14544677734375 0 76205.33
283.1297607421875 0 38585.37
283.1462707519531 0 8614.826
284.1247863769531 0 38757.305
284.1612243652344 0 938096.9
285.14532470703125 0 19895.855
285.1644592285156 0 119553.84
286.103515625 0 17224.91
286.1408996582031 0 11804.299
286.1660461425781 0 14093.186
286.2135925292969 0 19402.135 a 2
287.1510925292969 0 29320.486
292.1300354003906 0 65003.434
293.1133728027344 0 16430.441
293.1619873046875 0 9232.282
294.1457214355469 0 52652.98
294.1651916503906 0 8541.196
294.1819763183594 0 57483.145
295.1667175292969 0 9193.877
295.18572998046875 0 8307.202
296.1248779296875 0 26951.986
296.1977233886719 0 2054384.6 b Water loss 2
297.1350402832031 0 36706.38
297.20086669921875 0 323198.06
298.11944580078125 0 302720.8
298.1769714355469 0 143287.67
298.2038269042969 0 20067.76
299.1039123535156 0 23620.867
299.12261962890625 0 49491.805
299.1798095703125 0 12188.488
300.1562194824219 0 110003.945
301.1577453613281 0 12859.482
302.17193603515625 0 1150804.6
303.17510986328125 0 158919.48
304.1787109375 0 14347.593
307.1820983886719 0 36645.58
307.6839599609375 0 16032.1455
308.1607971191406 0 12041.394
308.1986999511719 0 10507.461
309.1568298339844 0 55476.62
310.1405944824219 0 179663.58
310.1606140136719 0 10474.481
310.21343994140625 0 37409.48
311.14385986328125 0 31051.467
311.1725158691406 0 59624.723
312.1563720703125 0 116038.984
312.17413330078125 0 15667.814 b Water loss 5
312.1926574707031 0 72533.984
312.67864990234375 0 9966.149
313.1590576171875 0 13309.405
314.1362609863281 0 20593.008
314.2084045410156 0 455687.8 b 2
315.1461486816406 0 2200198.2
315.2114562988281 0 68758.33
316.1300354003906 0 2356145.8 y Ammonia loss 6
316.1492614746094 0 351269.38
317.1331787109375 0 382866.84
317.1515808105469 0 26056.346
318.13543701171875 0 40494.016
320.1250915527344 0 57023.57
320.1606750488281 0 13510.309
320.6875 0 14708.707
321.17987060546875 0 183549.4 b 5
321.68145751953125 0 59029.652
322.1787414550781 0 28201.113
324.2290344238281 0 9417.732
325.1882019042969 0 13948.206
326.1720886230469 0 13488.245
326.207763671875 0 14572.881
327.16729736328125 0 243730.9
328.1512451171875 0 390801.25
328.1700439453125 0 24080.365
329.1544189453125 0 55063.266
329.1827392578125 0 27340.047
329.6932373046875 0 48004.137
330.16680908203125 0 46741.805
330.1882019042969 0 10470.216
333.1566467285156 0 1127568.9 y 6
334.1600341796875 0 198836.02
334.2134094238281 0 22301.24
335.16259765625 0 18297.662
336.1930847167969 0 10934.821
337.151611328125 0 240262.98
337.1883239746094 0 37432.348
337.6695861816406 0 8428.9
338.1353759765625 0 89444.3
338.1553955078125 0 32622.143
339.1379699707031 0 11712.525
340.1875915527344 0 158016.44
341.1907653808594 0 21782.525
343.2352294921875 0 22507.883
345.177734375 0 142283.84
345.1998596191406 0 12038.961
346.178955078125 0 48028.56
347.17901611328125 0 11474.673
350.2079772949219 0 12007.654
351.2400817871094 0 43729.383
352.1879577636719 0 10972.1045
352.2239074707031 0 175787.66
352.24639892578125 0 11758.228
353.18304443359375 0 28447.236
353.2220153808594 0 53619.617
353.69281005859375 0 14007.95
354.223388671875 0 11341.167
355.162109375 0 3668229.5
356.14654541015625 0 48605.805
356.1652526855469 0 588906.7
357.1670837402344 0 70589.9
362.20709228515625 0 19935.768
363.1458435058594 0 8562.737
363.2032165527344 0 65532.965
364.1866455078125 0 17430.809
364.2085266113281 0 12242.791
365.1829833984375 0 69991.18
365.2553405761719 0 19612.982
366.1843566894531 0 8770.466
367.2344970703125 0 9830.294
368.1847839355469 0 8158.9004
369.2503356933594 0 253092.7
370.234375 0 307929.56
371.1916809082031 0 17958.453
371.23236083984375 0 117319.06
372.18865966796875 0 57921.92
372.236083984375 0 20218.97
373.1723937988281 0 114216.64
373.1924743652344 0 9305.564
374.17608642578125 0 23438.809
376.2031555175781 0 15332.16 b Water loss 6
376.7052917480469 0 7552.379
379.2027893066406 0 11418.3545
379.2347412109375 0 149150.06
380.2181701660156 0 23361.814
380.23907470703125 0 22655.549
381.1563720703125 0 107607.94
381.181640625 0 12633.443
381.2143859863281 0 73271.66
382.161865234375 0 13843.004
382.2181091308594 0 13883.158
383.2663879394531 0 16940.34
385.2088623046875 0 17165.45 b 6
385.2445068359375 0 6909.0557
387.2611083984375 0 391070.12
388.2640380859375 0 75993.04
389.2651062011719 0 10070.506
391.19830322265625 0 30362.416
393.250244140625 0 25191.066
394.2076721191406 0 11572.976
395.15753173828125 0 29119.49
395.19708251953125 0 7402.8677
395.2301330566406 0 16467.027
397.2455749511719 0 1200107.4 b Water loss 3
398.2488098144531 0 250348.62
399.163330078125 0 15073.351
399.18975830078125 0 20565.531
399.2251281738281 0 43220.848
399.2513122558594 0 23325.346
400.2259216308594 0 11183.867
405.21148681640625 0 8910.616
405.2527770996094 0 7526.2344
406.24517822265625 0 8058.9536
407.1570739746094 0 20991.94
407.23052978515625 0 21061.934
409.15191650390625 0 58333.586
409.2093200683594 0 109188.64
410.1559143066406 0 15363.053
410.2114562988281 0 22207.486
411.18817138671875 0 29149.912
411.260498046875 0 24366.492
412.18536376953125 0 20308.773
412.2189636230469 0 9391.54
413.1664123535156 0 10826.414
413.24090576171875 0 17257.303
415.25616455078125 0 2301956.2 b 3
416.25933837890625 0 503172.53
417.2613220214844 0 67964.39
420.1890563964844 0 9176.303
421.17242431640625 0 32948.73
422.2047119140625 0 24021.775 y Ammonia loss 1
422.23895263671875 0 10583.909
422.27630615234375 0 13809.547
423.19036865234375 0 10838.67
423.2248840332031 0 81807.695
424.2269287109375 0 15449.462
424.2566833496094 0 23036.545
425.2404479980469 0 16415.377
426.17913818359375 0 16066.925
426.23773193359375 0 12369.326
427.162353515625 0 294258.62
427.2195129394531 0 50769.363
428.165283203125 0 65662.39
428.2158508300781 0 41971.887
429.168212890625 0 10222.97
431.2662353515625 0 35687.445
437.7276306152344 0 19513.268
438.1990661621094 0 69592.766
438.2285461425781 0 14825.04
439.18463134765625 0 27289.281
440.21502685546875 0 24997.213
440.2516174316406 0 100493.67
440.2857971191406 0 12918.523
441.23577880859375 0 66915.86
442.194580078125 0 34049.49
442.23541259765625 0 13344.48
442.2684020996094 0 13216.144
444.1890563964844 0 91404.625 y Water loss 5
444.24859619140625 0 10361.952
445.1730651855469 0 186575.9 y Ammonia loss 5
446.1763916015625 0 50254.484
446.7348937988281 0 13471.511
447.1776123046875 0 9643.845
447.23150634765625 0 23009.305
448.2923889160156 0 8278.776
449.20538330078125 0 12722.822
449.2764892578125 0 17076.494
450.23626708984375 0 32719.379
450.27349853515625 0 29957.799
451.2193603515625 0 27405.074
452.2244567871094 0 17076.834
453.27227783203125 0 7596.039
455.24627685546875 0 10047.161
456.21002197265625 0 314505.34
457.2126159667969 0 63701.836
458.26080322265625 0 15520.789
460.7306823730469 0 30373.924
461.2323303222656 0 13184.312
462.19921875 0 102250.29 y 5
463.2025146484375 0 19861.475
465.23614501953125 0 15041.411
466.19439697265625 0 15554.503
466.2304382324219 0 10658.561
466.30322265625 0 41953.87
468.2135925292969 0 9585.29
468.24627685546875 0 275614.44
468.2825622558594 0 31051.002
469.2484436035156 0 68949.9
469.28485107421875 0 13174.108
469.7364501953125 0 25992.633
470.2394714355469 0 18982.338
472.18353271484375 0 82611.305
473.1869201660156 0 25772.928
473.23590087890625 0 11272.311
474.2228698730469 0 9487.433
475.2203674316406 0 14598.043
476.2874450683594 0 38637.656
477.2892150878906 0 10345.164
478.20941162109375 0 99537.46
478.248779296875 0 19110.328 Precursor Water loss
478.7506103515625 0 13595.868
479.2131042480469 0 31282.943
481.26678466796875 0 20239.172
482.2630310058594 0 34299.54
484.31494140625 0 29715.639
485.27618408203125 0 10249.725
485.3154602050781 0 9458.0205
486.2577209472656 0 19028.605
490.2656555175781 0 11926.744
492.24652099609375 0 46256.688
494.2983703613281 0 201400.78 b Water loss 4
495.2382507324219 0 7932.8745
495.3016662597656 0 61058.66
496.2202453613281 0 92442.9
497.22198486328125 0 20415.305
498.2317199707031 0 13972.775
499.27801513671875 0 44028.164
500.2768859863281 0 17851.863
506.2047424316406 0 62240.49
507.2095642089844 0 20980.629
507.2930603027344 0 7756.8574
508.2056579589844 0 14856.468
508.2780456542969 0 25308.697
509.276611328125 0 9196.285
510.2566833496094 0 130049.805
511.26031494140625 0 38740.562
512.3090209960938 0 262212.66 b 4
513.24658203125 0 77376.09
513.3125610351562 0 67162.555
514.22998046875 0 13213.403
514.3134765625 0 11144.949
519.2552490234375 0 10498.102
520.24072265625 0 27927.389
522.2916259765625 0 33536.71
523.2349853515625 0 12902.774
523.29052734375 0 14188.432
524.2150268554688 0 644863.5
524.270263671875 0 36371.023
525.2180786132812 0 184870.89
525.3041381835938 0 25204.078
526.2200927734375 0 25711.256
526.2882690429688 0 55239.26
527.28466796875 0 55243.23
528.2685546875 0 58951.52
529.272216796875 0 18476.965
533.2714233398438 0 9703.534
534.2584228515625 0 17131.773
537.267333984375 0 117950.43
538.2589721679688 0 28637.596
540.3026733398438 0 27006.305
541.2413330078125 0 292673.53 y Water loss 4
541.2991333007812 0 35182.863
542.2260131835938 0 1095161.5 y Ammonia loss 4
543.22900390625 0 329716.16
543.3140869140625 0 26864.797
544.2318725585938 0 46821.66
544.3164672851562 0 8490.258
545.2936401367188 0 11579.655
551.2830810546875 0 91544.64
552.279296875 0 19021.115
555.2780151367188 0 113910.97
556.280517578125 0 32859.062
557.2794189453125 0 8633.741
559.251953125 0 3274775 y 4
560.2548828125 0 1039427.25
561.2571411132812 0 194299.72
562.2573852539062 0 9841.623
569.2935180664062 0 156630.42
569.3330688476562 0 7866.45
570.2957153320312 0 29375.344
577.3353271484375 0 10336.177
578.3241577148438 0 11306.144
579.3175659179688 0 9774.513
587.3169555664062 0 11626.914
593.2945556640625 0 26306.91
594.2899169921875 0 14379.843
595.3452758789062 0 37369.145
596.3411254882812 0 21052.79
597.3257446289062 0 11242.133
598.27294921875 0 8438.01
603.2784423828125 0 29375.549
604.2807006835938 0 7218.853
605.329345703125 0 56425.113
606.3321533203125 0 17789.346
607.2529907226562 0 34252.53
608.2554931640625 0 14003.905
610.31494140625 0 7410.041
611.3067626953125 0 20534.156
612.30029296875 0 10724.29
613.3568725585938 0 53247.523
614.2958374023438 0 8752.009
614.3573608398438 0 21052.32
620.305419921875 0 38813.113
621.2933959960938 0 38182.152
622.29638671875 0 9854.99
622.357666015625 0 25796.654
623.341064453125 0 328814.4 b Water loss 5
624.3439331054688 0 118082.75
625.2628173828125 0 14539.853
625.3475341796875 0 19243.85
628.3308715820312 0 10216.3545
632.3480834960938 0 8447.312
633.3240356445312 0 17922.793
637.3585815429688 0 11120.894
638.3155517578125 0 197286.47
639.3143310546875 0 73441.92
640.3148193359375 0 9694.96
640.3673095703125 0 94213.164
641.3521728515625 0 379725.7 b 5
642.289306640625 0 31127.373 y Water loss 3
642.3546142578125 0 147530.67
643.2747802734375 0 70416.21 y Ammonia loss 3
643.357177734375 0 26797.27
644.275634765625 0 26097.406
650.3519287109375 0 123307.164
651.3511352539062 0 48111.223
652.3428955078125 0 9814.095
656.3259887695312 0 125185.336
657.3289794921875 0 47643.438
658.3780517578125 0 210583.95
659.3812255859375 0 62652.344
660.2997436523438 0 414872.53 y 3
661.302490234375 0 160947.53
662.30419921875 0 26851.605
668.3623657226562 0 91813.56
669.3663940429688 0 28367.436
679.3724365234375 0 8488.206
689.3646850585938 0 9974.89
697.3836059570312 0 9364.935
706.3806762695312 0 28542.094
707.3753051757812 0 31849.473
715.378173828125 0 8313.388
716.3650512695312 0 32420.404
723.4047241210938 0 62592.586
724.3943481445312 0 41956.953
725.387939453125 0 33070.297
726.3839721679688 0 12025.99
727.3795166015625 0 9635.249
733.38818359375 0 119239.97
734.3782348632812 0 118349.74
735.3786010742188 0 39599.484
736.3853759765625 0 9948.186
738.3497314453125 0 17167.104
741.4152221679688 0 95018.37
742.4178466796875 0 36834.176
751.3994140625 0 703548.44 b Water loss 6
752.3987426757812 0 330552.1
753.3994140625 0 63872.812
754.394287109375 0 8701.544
755.3729248046875 0 45365.85 y Water loss 2
756.3712158203125 0 20983.201 y Ammonia loss 2
769.4097290039062 0 479197.2 b 6
770.4127197265625 0 191882.48
771.41552734375 0 50996.61
773.3829956054688 0 64928.895 y 2
774.385009765625 0 32446.164
824.3970947265625 0 8332.572
825.3812866210938 0 17565.129
842.4039306640625 0 32270.02 y Water loss 1
843.4061889648438 0 33305.92
844.4136962890625 0 8382.837
860.41552734375 0 61353.07 y 1
861.41796875 0 18011.14
862.4173583984375 0 12858.142
1957.4564208984375 0 7679.143

Spectrum Details

|  |  |
| --- | --- |
| Matched peaks? Matched peaks The total absolute number of peaks matched. Additionally in brackets the total fraction of peaks matched and the total number of peaks is shown. | 44 (6.83% of 644) |
| FDR? FDR The false discovery rate estimated for this peptide. It is calculated by matching all theoretical fragments with a non-integer shift with the raw peaks for this spectrum. This is done with 40 different shifts. The resulting percentage is the average number of annotated peaks over the number of annotated peaks with the correct spectrum. | 1.52% |
| Satellite FDR? Satellite FDR See the FDR for details on its calculation. This satellite ion specific FDR only contains the satellite ions (d/w) for I/L/J positions. | 2.38% |
| PSM Score? PSM Score The PSM Score as given by Hecklib to this annotated spectrum. It is shown with three significant figures. | 603 |

#### Spectrum 9792? Spectrum 9792 The raw spectrum of this peptide as annotated by Hecklib. The fragments are coloured according to ion type (see legend). Any peaks with a star '\*' as text can be hovered over to see the full details, first the ion type second the mass shift type. By hovering over the amino acids in the peptide or ions in the legend the corresponding peaks are highlighted. By toggling the 'Unassigned' label you can turn the background (unassigned) peaks on or off in the plot. By updating the slider in the Ion legend you can update the spectrum to only show the top X% of the peaks with labels. The top X% means any peak that is within X% of the highest intensity. By dragging in the spectrum you can zoom in to a specific part of the spectrum and use 'Zoom Out' to get back to the original zoom level. The annotation of the spectrum is based on the given sequence in the peptides file and is done with different software so inconsistencies are likely. The peaks are annotated based on the given sequence, with 20 ppm tolerance.

Copy Data

##### Spectrum 9792 (TSV)

###### Preview

```
Loading example...
```

*Click on the button to copy the data to your clipboard.*

Mz MinMz MaxIntensity Max

WidthHeightPeptide font sizePeptide stroke widthSpectrum font sizeSpectrum stroke widthCompact peptide

Ion legend

wxyz

abcd

OtherUnassignedIon^Charge^_Position_Show for top:%

JSJTPEQW

09.44e+41.89e+52.83e+53.78e+5

Zoom Out

a^+1^_2_y^+2^_2_a^+1^_2_b^+1^_2_b^+1^_2_y^+1^_1_b^+2^_5_b^+2^_5_a^+1^_3_y^+2^_4_a^+1^_3_b^+1^_3_b^+2^_6_b^+1^_3_y^+1^_2_b^+2^_6_y^+1^_2_b^+1^_4_b^+1^_4_y^+1^_3_y^+1^_3_y^+1^_3_\*b^+1^_5_b^+1^_5_y^+1^_4_y^+1^_4_y^+1^_4_b^+1^_6_b^+1^_6_y^+1^_5_y^+1^_5_y^+1^_5_b^+1^_7_b^+1^_7_y^+1^_6_b^+1^_7_y^+1^_6_y^+1^_7_y^+1^_7_

0778155723353113

Fragment Matches Table

Show background peaks

| Position | Ion type | Intensity | mz Theoretical | mz Error (Th) | mz Error (ppm) | Charge | Series Number |
| --- | --- | --- | --- | --- | --- | --- | --- |
| - | - | 1583 | 120.1 | - | - | 0 | - |
| - | - | 2162 | 125.1 | - | - | 0 | - |
| - | - | 6243 | 125.1 | - | - | 0 | - |
| - | - | 1568 | 126.1 | - | - | 0 | - |
| - | - | 472.6 | 126.1 | - | - | 0 | - |
| - | - | 1178 | 127.1 | - | - | 0 | - |
| - | - | 485.8 | 127.1 | - | - | 0 | - |
| - | - | 2682 | 128.1 | - | - | 0 | - |
| - | - | 1.411E+04 | 129.1 | - | - | 0 | - |
| - | - | 1387 | 129.1 | - | - | 0 | - |
| - | - | 424.3 | 129.4 | - | - | 0 | - |
| - | - | 4899 | 130.1 | - | - | 0 | - |
| - | - | 3.404E+04 | 130.1 | - | - | 0 | - |
| - | - | 3504 | 131.1 | - | - | 0 | - |
| - | - | 1.633E+04 | 132.1 | - | - | 0 | - |
| - | - | 1141 | 132.1 | - | - | 0 | - |
| - | - | 5471 | 133.1 | - | - | 0 | - |
| - | - | 1011 | 136.1 | - | - | 0 | - |
| - | - | 948.1 | 137.1 | - | - | 0 | - |
| - | - | 1015 | 138.1 | - | - | 0 | - |
| - | - | 1561 | 139.1 | - | - | 0 | - |
| - | - | 376.8 | 139.1 | - | - | 0 | - |
| - | - | 576.8 | 141.1 | - | - | 0 | - |
| - | - | 1504 | 142.1 | - | - | 0 | - |
| - | - | 9189 | 142.1 | - | - | 0 | - |
| - | - | 565.5 | 143.1 | - | - | 0 | - |
| - | - | 529.5 | 144 | - | - | 0 | - |
| - | - | 8055 | 144.1 | - | - | 0 | - |
| - | - | 1116 | 145.1 | - | - | 0 | - |
| - | - | 4.909E+04 | 146.1 | - | - | 0 | - |
| - | - | 4027 | 147.1 | - | - | 0 | - |
| - | - | 4016 | 147.1 | - | - | 0 | - |
| - | - | 770.1 | 149 | - | - | 0 | - |
| - | - | 478.9 | 152 | - | - | 0 | - |
| - | - | 5734 | 152.1 | - | - | 0 | - |
| - | - | 4642 | 153.1 | - | - | 0 | - |
| - | - | 3910 | 153.1 | - | - | 0 | - |
| - | - | 1552 | 155.1 | - | - | 0 | - |
| 2 | a | 7521 | 155.1 | 0.0003203 | 2.065 | +1 | 2 |
| - | - | 645 | 156.1 | - | - | 0 | - |
| - | - | 844 | 156.1 | - | - | 0 | - |
| - | - | 708.1 | 158 | - | - | 0 | - |
| - | - | 9.096E+04 | 159.1 | - | - | 0 | - |
| - | - | 876.3 | 160.1 | - | - | 0 | - |
| - | - | 758.2 | 160.1 | - | - | 0 | - |
| - | - | 9492 | 160.1 | - | - | 0 | - |
| - | - | 1077 | 163.1 | - | - | 0 | - |
| - | - | 545.7 | 165.1 | - | - | 0 | - |
| - | - | 500.5 | 166.1 | - | - | 0 | - |
| 7 | y | 833.9 | 167.1 | 0.0004967 | 2.973 | +2 | 2 |
| - | - | 1061 | 169.1 | - | - | 0 | - |
| - | - | 3504 | 169.1 | - | - | 0 | - |
| - | - | 5104 | 170.1 | - | - | 0 | - |
| - | - | 636 | 170.1 | - | - | 0 | - |
| - | - | 507.7 | 171.1 | - | - | 0 | - |
| - | - | 484.5 | 171.1 | - | - | 0 | - |
| - | - | 9383 | 171.1 | - | - | 0 | - |
| - | - | 1724 | 171.1 | - | - | 0 | - |
| 2 | a | 6.48E+04 | 173.1 | 0.0003452 | 1.994 | +1 | 2 |
| - | - | 2779 | 173.5 | - | - | 0 | - |
| - | - | 580.4 | 174.1 | - | - | 0 | - |
| - | - | 5730 | 174.1 | - | - | 0 | - |
| - | - | 466.3 | 174.3 | - | - | 0 | - |
| - | - | 1458 | 177.1 | - | - | 0 | - |
| - | - | 789.9 | 179.1 | - | - | 0 | - |
| - | - | 568.3 | 180.4 | - | - | 0 | - |
| - | - | 1.145E+04 | 181.1 | - | - | 0 | - |
| - | - | 971.4 | 181.1 | - | - | 0 | - |
| - | - | 463.5 | 182.1 | - | - | 0 | - |
| - | - | 542.7 | 182.1 | - | - | 0 | - |
| 2 | b | 1.659E+04 | 183.1 | 0.0003398 | 1.855 | +1 | 2 |
| - | - | 6022 | 183.1 | - | - | 0 | - |
| - | - | 1828 | 184.1 | - | - | 0 | - |
| - | - | 436.2 | 185.1 | - | - | 0 | - |
| - | - | 920.8 | 186.1 | - | - | 0 | - |
| - | - | 1033 | 187.1 | - | - | 0 | - |
| - | - | 7659 | 187.1 | - | - | 0 | - |
| - | - | 2.108E+04 | 187.1 | - | - | 0 | - |
| - | - | 3.739E+05 | 188.1 | - | - | 0 | - |
| - | - | 1722 | 188.1 | - | - | 0 | - |
| - | - | 4.02E+04 | 189.1 | - | - | 0 | - |
| - | - | 2338 | 190.1 | - | - | 0 | - |
| - | - | 579.9 | 191.1 | - | - | 0 | - |
| - | - | 723.7 | 194.1 | - | - | 0 | - |
| - | - | 609.4 | 194.6 | - | - | 0 | - |
| - | - | 2150 | 195.1 | - | - | 0 | - |
| - | - | 584.2 | 196.1 | - | - | 0 | - |
| - | - | 821.2 | 196.1 | - | - | 0 | - |
| - | - | 1.093E+04 | 197.1 | - | - | 0 | - |
| - | - | 920.4 | 198.1 | - | - | 0 | - |
| - | - | 1660 | 199.1 | - | - | 0 | - |
| - | - | 7.409E+04 | 199.1 | - | - | 0 | - |
| - | - | 6547 | 200.1 | - | - | 0 | - |
| - | - | 671.6 | 200.1 | - | - | 0 | - |
| - | - | 2633 | 201.1 | - | - | 0 | - |
| 2 | b | 7.944E+04 | 201.1 | 0.0002884 | 1.434 | +1 | 2 |
| - | - | 1049 | 202.1 | - | - | 0 | - |
| - | - | 7262 | 202.1 | - | - | 0 | - |
| 8 | y | 1.499E+05 | 205.1 | 0.0003191 | 1.556 | +1 | 1 |
| - | - | 1.644E+04 | 206.1 | - | - | 0 | - |
| - | - | 549.8 | 207 | - | - | 0 | - |
| - | - | 914.4 | 207.1 | - | - | 0 | - |
| - | - | 674.4 | 207.1 | - | - | 0 | - |
| - | - | 7015 | 209.1 | - | - | 0 | - |
| - | - | 1027 | 210.1 | - | - | 0 | - |
| - | - | 743 | 211.1 | - | - | 0 | - |
| - | - | 4483 | 211.1 | - | - | 0 | - |
| - | - | 3286 | 212.1 | - | - | 0 | - |
| - | - | 550.5 | 213.1 | - | - | 0 | - |
| - | - | 1802 | 214.2 | - | - | 0 | - |
| - | - | 535.8 | 215.1 | - | - | 0 | - |
| - | - | 4.997E+04 | 215.1 | - | - | 0 | - |
| - | - | 4782 | 216.1 | - | - | 0 | - |
| - | - | 3919 | 217.1 | - | - | 0 | - |
| - | - | 1125 | 221.1 | - | - | 0 | - |
| - | - | 2290 | 223.1 | - | - | 0 | - |
| - | - | 1337 | 223.1 | - | - | 0 | - |
| - | - | 829 | 223.1 | - | - | 0 | - |
| - | - | 750.3 | 223.2 | - | - | 0 | - |
| - | - | 3443 | 225.1 | - | - | 0 | - |
| - | - | 2483 | 225.1 | - | - | 0 | - |
| - | - | 3803 | 226.1 | - | - | 0 | - |
| - | - | 1.208E+05 | 227.1 | - | - | 0 | - |
| - | - | 1.3E+04 | 228.1 | - | - | 0 | - |
| - | - | 1112 | 229.1 | - | - | 0 | - |
| - | - | 2911 | 229.2 | - | - | 0 | - |
| - | - | 1009 | 230.1 | - | - | 0 | - |
| - | - | 558.5 | 230.8 | - | - | 0 | - |
| - | - | 694.2 | 231.2 | - | - | 0 | - |
| - | - | 1137 | 233.7 | - | - | 0 | - |
| - | - | 971.5 | 237.1 | - | - | 0 | - |
| - | - | 1138 | 238.2 | - | - | 0 | - |
| - | - | 5492 | 239.1 | - | - | 0 | - |
| - | - | 947.7 | 239.6 | - | - | 0 | - |
| - | - | 1.632E+04 | 240.1 | - | - | 0 | - |
| - | - | 902.8 | 240.1 | - | - | 0 | - |
| - | - | 666.9 | 240.1 | - | - | 0 | - |
| - | - | 633.2 | 240.2 | - | - | 0 | - |
| - | - | 3834 | 241.1 | - | - | 0 | - |
| - | - | 1222 | 241.1 | - | - | 0 | - |
| - | - | 1470 | 241.2 | - | - | 0 | - |
| - | - | 2543 | 242.2 | - | - | 0 | - |
| - | - | 3822 | 242.7 | - | - | 0 | - |
| - | - | 736.2 | 243.2 | - | - | 0 | - |
| - | - | 3.303E+04 | 244.1 | - | - | 0 | - |
| - | - | 2305 | 245.1 | - | - | 0 | - |
| - | - | 2978 | 245.1 | - | - | 0 | - |
| - | - | 813.9 | 246.1 | - | - | 0 | - |
| 5 | b | 1913 | 247.7 | 0.0003402 | 1.374 | +2 | 5 |
| - | - | 1808 | 251.1 | - | - | 0 | - |
| - | - | 4385 | 251.2 | - | - | 0 | - |
| - | - | 2170 | 252.1 | - | - | 0 | - |
| - | - | 600.4 | 252.2 | - | - | 0 | - |
| - | - | 2143 | 253.1 | - | - | 0 | - |
| - | - | 909.6 | 255.1 | - | - | 0 | - |
| - | - | 4183 | 256.2 | - | - | 0 | - |
| 5 | b | 1112 | 256.7 | 0.0001085 | 0.4228 | +2 | 5 |
| - | - | 7540 | 257.1 | - | - | 0 | - |
| - | - | 1.496E+04 | 258.1 | - | - | 0 | - |
| - | - | 2502 | 258.1 | - | - | 0 | - |
| - | - | 788.5 | 259.1 | - | - | 0 | - |
| - | - | 1541 | 259.1 | - | - | 0 | - |
| - | - | 571.6 | 259.1 | - | - | 0 | - |
| - | - | 613.4 | 263 | - | - | 0 | - |
| - | - | 984.3 | 263.1 | - | - | 0 | - |
| - | - | 1603 | 264.1 | - | - | 0 | - |
| - | - | 733.1 | 265.1 | - | - | 0 | - |
| - | - | 1774 | 266.1 | - | - | 0 | - |
| - | - | 3826 | 266.1 | - | - | 0 | - |
| - | - | 1710 | 266.2 | - | - | 0 | - |
| - | - | 2883 | 267.1 | - | - | 0 | - |
| 3 | a | 3554 | 268.2 | 0.000317 | 1.182 | +1 | 3 |
| - | - | 3754 | 269.1 | - | - | 0 | - |
| - | - | 2883 | 269.2 | - | - | 0 | - |
| - | - | 3.795E+04 | 270.1 | - | - | 0 | - |
| - | - | 886.8 | 271.1 | - | - | 0 | - |
| 5 | y | 6531 | 271.1 | 0.003236 | 11.94 | +2 | 4 |
| - | - | 668.7 | 274.1 | - | - | 0 | - |
| - | - | 2431 | 274.2 | - | - | 0 | - |
| - | - | 678.8 | 279.1 | - | - | 0 | - |
| - | - | 4766 | 280.1 | - | - | 0 | - |
| - | - | 3785 | 280.2 | - | - | 0 | - |
| - | - | 919.4 | 281.1 | - | - | 0 | - |
| - | - | 3184 | 282.1 | - | - | 0 | - |
| - | - | 1600 | 283.1 | - | - | 0 | - |
| - | - | 4.079E+04 | 284.2 | - | - | 0 | - |
| - | - | 752.9 | 285.1 | - | - | 0 | - |
| - | - | 5241 | 285.2 | - | - | 0 | - |
| 3 | a | 661.9 | 286.2 | 0.001611 | 5.629 | +1 | 3 |
| - | - | 882.5 | 287.2 | - | - | 0 | - |
| - | - | 2760 | 292.1 | - | - | 0 | - |
| - | - | 1080 | 293.1 | - | - | 0 | - |
| - | - | 2522 | 294.1 | - | - | 0 | - |
| - | - | 2865 | 294.2 | - | - | 0 | - |
| - | - | 1734 | 296.1 | - | - | 0 | - |
| 3 | b | 8.428E+04 | 296.2 | 0.0004585 | 1.548 | +1 | 3 |
| - | - | 1475 | 297.1 | - | - | 0 | - |
| - | - | 1.331E+04 | 297.2 | - | - | 0 | - |
| - | - | 851.2 | 298.1 | - | - | 0 | - |
| - | - | 1.122E+04 | 298.1 | - | - | 0 | - |
| - | - | 6138 | 298.2 | - | - | 0 | - |
| - | - | 1435 | 298.2 | - | - | 0 | - |
| - | - | 1158 | 299.1 | - | - | 0 | - |
| - | - | 2122 | 299.1 | - | - | 0 | - |
| - | - | 687.1 | 299.2 | - | - | 0 | - |
| - | - | 4365 | 300.2 | - | - | 0 | - |
| - | - | 4.616E+04 | 302.2 | - | - | 0 | - |
| - | - | 1083 | 303.1 | - | - | 0 | - |
| - | - | 6231 | 303.2 | - | - | 0 | - |
| - | - | 953.5 | 307.2 | - | - | 0 | - |
| - | - | 642 | 308.2 | - | - | 0 | - |
| - | - | 599.4 | 308.2 | - | - | 0 | - |
| - | - | 1923 | 309.2 | - | - | 0 | - |
| - | - | 1461 | 309.2 | - | - | 0 | - |
| - | - | 7659 | 310.1 | - | - | 0 | - |
| - | - | 932.6 | 310.2 | - | - | 0 | - |
| - | - | 1529 | 310.2 | - | - | 0 | - |
| - | - | 753.7 | 311.1 | - | - | 0 | - |
| - | - | 1373 | 311.2 | - | - | 0 | - |
| - | - | 5406 | 312.2 | - | - | 0 | - |
| 6 | b | 973.7 | 312.2 | 0.0001466 | 0.4695 | +2 | 6 |
| - | - | 3311 | 312.2 | - | - | 0 | - |
| - | - | 1164 | 313.2 | - | - | 0 | - |
| - | - | 1226 | 314.1 | - | - | 0 | - |
| 3 | b | 1.893E+04 | 314.2 | 0.0004834 | 1.539 | +1 | 3 |
| - | - | 9.185E+04 | 315.1 | - | - | 0 | - |
| - | - | 3316 | 315.2 | - | - | 0 | - |
| 7 | y | 9.314E+04 | 316.1 | 0.0003341 | 1.057 | +1 | 2 |
| - | - | 1.457E+04 | 316.1 | - | - | 0 | - |
| - | - | 1.595E+04 | 317.1 | - | - | 0 | - |
| - | - | 1281 | 317.2 | - | - | 0 | - |
| - | - | 1540 | 318.1 | - | - | 0 | - |
| - | - | 2053 | 320.1 | - | - | 0 | - |
| - | - | 1006 | 320.2 | - | - | 0 | - |
| - | - | 613 | 321.1 | - | - | 0 | - |
| 6 | b | 5189 | 321.2 | 0.0003879 | 1.208 | +2 | 6 |
| - | - | 2222 | 321.7 | - | - | 0 | - |
| - | - | 1100 | 322.2 | - | - | 0 | - |
| - | - | 1066 | 325.2 | - | - | 0 | - |
| - | - | 906.6 | 326.2 | - | - | 0 | - |
| - | - | 789.9 | 326.2 | - | - | 0 | - |
| - | - | 1.087E+04 | 327.2 | - | - | 0 | - |
| - | - | 1.547E+04 | 328.2 | - | - | 0 | - |
| - | - | 1282 | 328.2 | - | - | 0 | - |
| - | - | 3086 | 329.2 | - | - | 0 | - |
| - | - | 1066 | 329.2 | - | - | 0 | - |
| - | - | 1258 | 329.7 | - | - | 0 | - |
| - | - | 1704 | 330.2 | - | - | 0 | - |
| 7 | y | 4.708E+04 | 333.2 | 0.0004879 | 1.464 | +1 | 2 |
| - | - | 8290 | 334.2 | - | - | 0 | - |
| - | - | 9826 | 337.2 | - | - | 0 | - |
| - | - | 868 | 337.2 | - | - | 0 | - |
| - | - | 741.8 | 337.7 | - | - | 0 | - |
| - | - | 3736 | 338.1 | - | - | 0 | - |
| - | - | 736.9 | 338.2 | - | - | 0 | - |
| - | - | 928.9 | 340.1 | - | - | 0 | - |
| - | - | 6454 | 340.2 | - | - | 0 | - |
| - | - | 964.6 | 341.2 | - | - | 0 | - |
| - | - | 1160 | 343.2 | - | - | 0 | - |
| - | - | 5835 | 345.2 | - | - | 0 | - |
| - | - | 2011 | 346.2 | - | - | 0 | - |
| - | - | 829.1 | 351.2 | - | - | 0 | - |
| - | - | 7382 | 352.2 | - | - | 0 | - |
| - | - | 989.5 | 353.2 | - | - | 0 | - |
| - | - | 1909 | 353.2 | - | - | 0 | - |
| - | - | 897 | 354.1 | - | - | 0 | - |
| - | - | 1.558E+05 | 355.2 | - | - | 0 | - |
| - | - | 2.438E+04 | 356.2 | - | - | 0 | - |
| - | - | 3067 | 357.2 | - | - | 0 | - |
| - | - | 3431 | 359.1 | - | - | 0 | - |
| - | - | 2616 | 363.2 | - | - | 0 | - |
| - | - | 2996 | 365.2 | - | - | 0 | - |
| - | - | 9407 | 369.2 | - | - | 0 | - |
| - | - | 1.185E+04 | 370.2 | - | - | 0 | - |
| - | - | 1224 | 370.3 | - | - | 0 | - |
| - | - | 1453 | 371.2 | - | - | 0 | - |
| - | - | 5392 | 371.2 | - | - | 0 | - |
| - | - | 2413 | 372.2 | - | - | 0 | - |
| - | - | 838 | 372.2 | - | - | 0 | - |
| - | - | 5483 | 373.2 | - | - | 0 | - |
| - | - | 732.5 | 374.2 | - | - | 0 | - |
| - | - | 5739 | 379.2 | - | - | 0 | - |
| - | - | 1043 | 380.2 | - | - | 0 | - |
| - | - | 5191 | 381.2 | - | - | 0 | - |
| - | - | 1260 | 381.2 | - | - | 0 | - |
| - | - | 2575 | 381.2 | - | - | 0 | - |
| - | - | 1185 | 382.2 | - | - | 0 | - |
| - | - | 1.62E+04 | 387.3 | - | - | 0 | - |
| - | - | 3735 | 388.3 | - | - | 0 | - |
| - | - | 895 | 391.2 | - | - | 0 | - |
| - | - | 938.8 | 393.3 | - | - | 0 | - |
| - | - | 658.3 | 394.3 | - | - | 0 | - |
| - | - | 1143 | 395.2 | - | - | 0 | - |
| - | - | 780.9 | 395.2 | - | - | 0 | - |
| 4 | b | 4.612E+04 | 397.2 | 0.0003264 | 0.8217 | +1 | 4 |
| - | - | 9649 | 398.2 | - | - | 0 | - |
| - | - | 877.1 | 399.2 | - | - | 0 | - |
| - | - | 720.2 | 399.2 | - | - | 0 | - |
| - | - | 1477 | 399.2 | - | - | 0 | - |
| - | - | 1360 | 399.3 | - | - | 0 | - |
| - | - | 610.1 | 400.2 | - | - | 0 | - |
| - | - | 619.2 | 405.2 | - | - | 0 | - |
| - | - | 1363 | 407.2 | - | - | 0 | - |
| - | - | 1599 | 409.2 | - | - | 0 | - |
| - | - | 5307 | 409.2 | - | - | 0 | - |
| - | - | 1269 | 411.2 | - | - | 0 | - |
| - | - | 1186 | 411.3 | - | - | 0 | - |
| - | - | 920.2 | 412.2 | - | - | 0 | - |
| - | - | 852.9 | 412.2 | - | - | 0 | - |
| - | - | 659.1 | 413.2 | - | - | 0 | - |
| - | - | 746.1 | 413.2 | - | - | 0 | - |
| 4 | b | 9.315E+04 | 415.3 | 0.0003819 | 0.9196 | +1 | 4 |
| - | - | 2.119E+04 | 416.3 | - | - | 0 | - |
| - | - | 2674 | 417.3 | - | - | 0 | - |
| - | - | 776 | 420.2 | - | - | 0 | - |
| - | - | 1345 | 421.2 | - | - | 0 | - |
| - | - | 3726 | 423.2 | - | - | 0 | - |
| - | - | 896.1 | 424.2 | - | - | 0 | - |
| - | - | 1379 | 424.3 | - | - | 0 | - |
| - | - | 868.3 | 425.2 | - | - | 0 | - |
| - | - | 995.2 | 426.2 | - | - | 0 | - |
| - | - | 1.324E+04 | 427.2 | - | - | 0 | - |
| - | - | 2361 | 427.2 | - | - | 0 | - |
| - | - | 2287 | 428.2 | - | - | 0 | - |
| - | - | 1503 | 428.2 | - | - | 0 | - |
| - | - | 636.3 | 430.2 | - | - | 0 | - |
| - | - | 1057 | 431.3 | - | - | 0 | - |
| - | - | 1023 | 437.7 | - | - | 0 | - |
| - | - | 3054 | 438.2 | - | - | 0 | - |
| - | - | 870.3 | 440.2 | - | - | 0 | - |
| - | - | 4708 | 440.3 | - | - | 0 | - |
| - | - | 3042 | 441.2 | - | - | 0 | - |
| - | - | 579.3 | 442.2 | - | - | 0 | - |
| - | - | 813.1 | 442.2 | - | - | 0 | - |
| - | - | 863.9 | 442.3 | - | - | 0 | - |
| 6 | y | 3945 | 444.2 | 0.0008386 | 1.888 | +1 | 3 |
| 6 | y | 8383 | 445.2 | 0.0006487 | 1.457 | +1 | 3 |
| - | - | 2159 | 446.2 | - | - | 0 | - |
| - | - | 714.3 | 450.2 | - | - | 0 | - |
| - | - | 1082 | 451.2 | - | - | 0 | - |
| - | - | 621.1 | 454.2 | - | - | 0 | - |
| - | - | 665.3 | 455.7 | - | - | 0 | - |
| - | - | 1.21E+04 | 456.2 | - | - | 0 | - |
| - | - | 890.6 | 456.2 | - | - | 0 | - |
| - | - | 2429 | 457.2 | - | - | 0 | - |
| - | - | 1072 | 458.3 | - | - | 0 | - |
| 6 | y | 5038 | 462.2 | 0.0003447 | 0.7459 | +1 | 3 |
| - | - | 994.2 | 463.2 | - | - | 0 | - |
| - | - | 745.7 | 464.2 | - | - | 0 | - |
| - | - | 684.4 | 466.2 | - | - | 0 | - |
| - | - | 1017 | 466.2 | - | - | 0 | - |
| - | - | 1587 | 466.3 | - | - | 0 | - |
| - | - | 1.181E+04 | 468.2 | - | - | 0 | - |
| - | - | 2007 | 468.3 | - | - | 0 | - |
| - | - | 596.8 | 469.2 | - | - | 0 | - |
| - | - | 2404 | 469.2 | - | - | 0 | - |
| - | - | 1580 | 469.7 | - | - | 0 | - |
| - | - | 681.5 | 470.2 | - | - | 0 | - |
| - | - | 1094 | 471.3 | - | - | 0 | - |
| - | - | 3275 | 472.2 | - | - | 0 | - |
| - | - | 752.1 | 473.2 | - | - | 0 | - |
| - | - | 1965 | 476.3 | - | - | 0 | - |
| - | - | 3724 | 478.2 | - | - | 0 | - |
| - | - | 1037 | 479.2 | - | - | 0 | - |
| - | - | 1289 | 482.3 | - | - | 0 | - |
| - | - | 1297 | 484.3 | - | - | 0 | - |
| - | - | 8648 | 486.2 | - | - | 0 | - |
| - | - | 1638 | 487.2 | - | - | 0 | - |
| 0 | Precursor | 631.5 | 487.3 | 0.003309 | 6.791 | +2 | -1 |
| - | - | 1708 | 492.2 | - | - | 0 | - |
| 5 | b | 8767 | 494.3 | 0.0007242 | 1.465 | +1 | 5 |
| - | - | 2321 | 495.3 | - | - | 0 | - |
| - | - | 3376 | 496.2 | - | - | 0 | - |
| - | - | 910.5 | 497.2 | - | - | 0 | - |
| - | - | 2063 | 499.3 | - | - | 0 | - |
| - | - | 874.4 | 500.3 | - | - | 0 | - |
| - | - | 2971 | 506.2 | - | - | 0 | - |
| - | - | 999.5 | 507.2 | - | - | 0 | - |
| - | - | 792.8 | 507.3 | - | - | 0 | - |
| - | - | 1559 | 508.3 | - | - | 0 | - |
| - | - | 657.1 | 509.3 | - | - | 0 | - |
| - | - | 5315 | 510.3 | - | - | 0 | - |
| - | - | 1626 | 511.3 | - | - | 0 | - |
| 5 | b | 1.055E+04 | 512.3 | 1.382E-05 | 0.02698 | +1 | 5 |
| - | - | 2941 | 513.2 | - | - | 0 | - |
| - | - | 2751 | 513.3 | - | - | 0 | - |
| - | - | 1230 | 520.2 | - | - | 0 | - |
| - | - | 2.692E+04 | 524.2 | - | - | 0 | - |
| - | - | 940 | 524.3 | - | - | 0 | - |
| - | - | 8515 | 525.2 | - | - | 0 | - |
| - | - | 1074 | 525.3 | - | - | 0 | - |
| - | - | 1404 | 526.2 | - | - | 0 | - |
| - | - | 1909 | 526.3 | - | - | 0 | - |
| - | - | 2733 | 527.3 | - | - | 0 | - |
| - | - | 3181 | 528.3 | - | - | 0 | - |
| - | - | 1535 | 534.3 | - | - | 0 | - |
| - | - | 3266 | 537.3 | - | - | 0 | - |
| - | - | 761.4 | 538.3 | - | - | 0 | - |
| - | - | 913.1 | 540.3 | - | - | 0 | - |
| 5 | y | 1.292E+04 | 541.2 | 0.0001988 | 0.3673 | +1 | 4 |
| - | - | 1231 | 541.3 | - | - | 0 | - |
| 5 | y | 4.62E+04 | 542.2 | 0.0005582 | 1.029 | +1 | 4 |
| - | - | 1.344E+04 | 543.2 | - | - | 0 | - |
| - | - | 1463 | 543.3 | - | - | 0 | - |
| - | - | 2301 | 544.2 | - | - | 0 | - |
| - | - | 3891 | 551.3 | - | - | 0 | - |
| - | - | 1055 | 552.3 | - | - | 0 | - |
| - | - | 4562 | 555.3 | - | - | 0 | - |
| - | - | 1171 | 556.3 | - | - | 0 | - |
| 5 | y | 1.339E+05 | 559.3 | 0.0001321 | 0.2363 | +1 | 4 |
| - | - | 4.188E+04 | 560.3 | - | - | 0 | - |
| - | - | 7857 | 561.3 | - | - | 0 | - |
| - | - | 6872 | 569.3 | - | - | 0 | - |
| - | - | 1701 | 570.3 | - | - | 0 | - |
| - | - | 670.7 | 579.3 | - | - | 0 | - |
| - | - | 866.4 | 593.3 | - | - | 0 | - |
| - | - | 921.1 | 594.3 | - | - | 0 | - |
| - | - | 1446 | 595.3 | - | - | 0 | - |
| - | - | 1176 | 603.3 | - | - | 0 | - |
| - | - | 2115 | 605.3 | - | - | 0 | - |
| - | - | 1256 | 607.3 | - | - | 0 | - |
| - | - | 609.5 | 612.3 | - | - | 0 | - |
| - | - | 2869 | 613.4 | - | - | 0 | - |
| - | - | 1018 | 614.4 | - | - | 0 | - |
| - | - | 2917 | 620.3 | - | - | 0 | - |
| - | - | 1891 | 621.3 | - | - | 0 | - |
| - | - | 1534 | 622.4 | - | - | 0 | - |
| 6 | b | 1.385E+04 | 623.3 | 0.0004285 | 0.6874 | +1 | 6 |
| - | - | 5337 | 624.3 | - | - | 0 | - |
| - | - | 1129 | 625.3 | - | - | 0 | - |
| - | - | 1423 | 625.3 | - | - | 0 | - |
| - | - | 803 | 628.3 | - | - | 0 | - |
| - | - | 846.2 | 637.4 | - | - | 0 | - |
| - | - | 8454 | 638.3 | - | - | 0 | - |
| - | - | 3360 | 639.3 | - | - | 0 | - |
| - | - | 827.8 | 640.3 | - | - | 0 | - |
| - | - | 4917 | 640.4 | - | - | 0 | - |
| 6 | b | 1.449E+04 | 641.4 | 0.000606 | 0.9448 | +1 | 6 |
| 4 | y | 1794 | 642.3 | 0.0003605 | 0.5613 | +1 | 5 |
| - | - | 4259 | 642.4 | - | - | 0 | - |
| 4 | y | 2739 | 643.3 | 0.00299 | 4.647 | +1 | 5 |
| - | - | 889 | 643.4 | - | - | 0 | - |
| - | - | 4581 | 650.4 | - | - | 0 | - |
| - | - | 2403 | 651.3 | - | - | 0 | - |
| - | - | 675 | 652.4 | - | - | 0 | - |
| - | - | 5222 | 656.3 | - | - | 0 | - |
| - | - | 1991 | 657.3 | - | - | 0 | - |
| - | - | 628.5 | 658.3 | - | - | 0 | - |
| - | - | 7041 | 658.4 | - | - | 0 | - |
| - | - | 2667 | 659.4 | - | - | 0 | - |
| 4 | y | 1.764E+04 | 660.3 | 0.000122 | 0.1848 | +1 | 5 |
| - | - | 4936 | 661.3 | - | - | 0 | - |
| - | - | 1040 | 662.3 | - | - | 0 | - |
| - | - | 3487 | 668.4 | - | - | 0 | - |
| - | - | 1106 | 669.4 | - | - | 0 | - |
| - | - | 1341 | 706.4 | - | - | 0 | - |
| - | - | 741.9 | 707.4 | - | - | 0 | - |
| - | - | 1158 | 716.4 | - | - | 0 | - |
| - | - | 2306 | 723.4 | - | - | 0 | - |
| - | - | 1799 | 724.4 | - | - | 0 | - |
| - | - | 948 | 725.4 | - | - | 0 | - |
| - | - | 5502 | 733.4 | - | - | 0 | - |
| - | - | 4887 | 734.4 | - | - | 0 | - |
| - | - | 1551 | 735.4 | - | - | 0 | - |
| - | - | 4056 | 741.4 | - | - | 0 | - |
| - | - | 1916 | 742.4 | - | - | 0 | - |
| 7 | b | 2.944E+04 | 751.4 | 1.746E-05 | 0.02324 | +1 | 7 |
| 7 | b | 1.248E+04 | 752.4 | 0.01435 | 19.08 | +1 | 7 |
| - | - | 3559 | 753.4 | - | - | 0 | - |
| 3 | y | 1958 | 755.4 | 0.002332 | 3.088 | +1 | 6 |
| 7 | b | 1.911E+04 | 769.4 | 0.0002323 | 0.3019 | +1 | 7 |
| - | - | 8471 | 770.4 | - | - | 0 | - |
| - | - | 2282 | 771.4 | - | - | 0 | - |
| 3 | y | 2103 | 773.4 | 0.0007749 | 1.002 | +1 | 6 |
| - | - | 906.5 | 774.4 | - | - | 0 | - |
| 2 | y | 1191 | 842.4 | 0.0002463 | 0.2924 | +1 | 7 |
| 2 | y | 1779 | 860.4 | 0.0006069 | 0.7053 | +1 | 7 |
| - | - | 968.8 | 861.4 | - | - | 0 | - |
| - | - | 624.5 | 1184 | - | - | 0 | - |
| - | - | 616.4 | 1711 | - | - | 0 | - |
| - | - | 649.7 | 1788 | - | - | 0 | - |
| - | - | 831.1 | 3041 | - | - | 0 | - |
| - | - | 703.1 | 3082 | - | - | 0 | - |

m/z Charge Intensity FragmentType MassShift Position
120.08108520507812 0 1583.047
125.07125091552734 0 2161.5107
125.10765838623047 0 6242.814
126.09162139892578 0 1568.1167
126.11046600341797 0 472.5744
127.08695983886719 0 1177.672
127.12350463867188 0 485.8192
128.10731506347656 0 2682.4268
129.06619262695312 0 14106.758
129.1023712158203 0 1387.3828
129.38717651367188 0 424.29044
130.05020141601562 0 4898.9766
130.0654754638672 0 34044.37
131.0690155029297 0 3503.602
132.0811004638672 0 16327.3
132.10218811035156 0 1140.5109
133.08587646484375 0 5470.9243
136.0761260986328 0 1011.4714
137.0712890625 0 948.145
138.09136962890625 0 1014.7903
139.0507049560547 0 1561.4487
139.123291015625 0 376.7561
141.10269165039062 0 576.7684
142.06541442871094 0 1504.4473
142.12298583984375 0 9188.894
143.0734405517578 0 565.457
144.04476928710938 0 529.4983
144.08116149902344 0 8055.2837
145.08457946777344 0 1115.8036
146.06039428710938 0 49089.4
147.0638427734375 0 4027.0698
147.0767059326172 0 4015.673
149.02359008789062 0 770.1001
151.99134826660156 0 478.9456
152.10733032226562 0 5734.076
153.0662078857422 0 4641.617
153.10250854492188 0 3909.763
155.08213806152344 0 1552.109
155.1182098388672 0 7521.3477 a Water loss 1
156.1022491455078 0 644.97473
156.12196350097656 0 844.01733
158.0452880859375 0 708.1241
159.09202575683594 0 90960.06
160.0759735107422 0 876.2782
160.0886993408203 0 758.1851
160.0954132080078 0 9492.378
163.08714294433594 0 1076.7582
165.10202026367188 0 545.7174
166.08665466308594 0 500.5384
167.08200073242188 0 833.88446 y 6
169.0766143798828 0 1061.1721
169.13394165039062 0 3504.023
170.06036376953125 0 5104.062
170.11807250976562 0 636.01263
171.0638427734375 0 507.72357
171.07659912109375 0 484.47437
171.11309814453125 0 9382.737
171.1494903564453 0 1723.7739
173.12879943847656 0 64803.457 a 1
173.45086669921875 0 2779.3762
174.05589294433594 0 580.43445
174.1322021484375 0 5729.8525
174.29763793945312 0 466.29953
177.11239624023438 0 1457.5282
179.10638427734375 0 789.94995
180.35226440429688 0 568.28
181.09744262695312 0 11452.569
181.10586547851562 0 971.42804
182.0810089111328 0 463.52496
182.10116577148438 0 542.6724
183.11314392089844 0 16589.256 b Water loss 1
183.14956665039062 0 6022.1836
184.1165008544922 0 1827.8689
185.091796875 0 436.2101
186.08746337890625 0 920.8261
187.07229614257812 0 1032.6189
187.0869598388672 0 7659.479
187.1444854736328 0 21084.672
188.07101440429688 0 373852.88
188.14816284179688 0 1722.1279
189.07431030273438 0 40199.22
190.07708740234375 0 2338.226
191.08216857910156 0 579.86426
194.13381958007812 0 723.7018
194.63572692871094 0 609.41644
195.07684326171875 0 2150.0496
196.11988830566406 0 584.19073
196.1443634033203 0 821.19586
197.1287384033203 0 10934.475
198.1323699951172 0 920.375
199.07183837890625 0 1659.6981
199.10800170898438 0 74094.77
200.11148071289062 0 6547.419
200.1404571533203 0 671.63135
201.1022491455078 0 2633.0837
201.1236572265625 0 79442.37 b 1
202.1072998046875 0 1049.0093
202.1270751953125 0 7262.4575
205.09747314453125 0 149895.86 y 7
206.10079956054688 0 16440.984
207.04490661621094 0 549.8117
207.10333251953125 0 914.3674
207.11331176757812 0 674.4451
209.09231567382812 0 7015.282
210.095947265625 0 1026.832
211.10789489746094 0 742.9971
211.14443969726562 0 4483.3315
212.1033477783203 0 3286.423
213.08766174316406 0 550.5402
214.15541076660156 0 1802.3799
215.08181762695312 0 535.79114
215.13934326171875 0 49967.35
216.14288330078125 0 4781.5293
217.08218383789062 0 3919.294
221.12852478027344 0 1124.6929
223.0717010498047 0 2289.5227
223.10836791992188 0 1336.6277
223.14398193359375 0 829.0489
223.18075561523438 0 750.3353
225.1026611328125 0 3443.3684
225.12367248535156 0 2483.0706
226.1188507080078 0 3802.7825
227.10304260253906 0 120796.055
228.10633850097656 0 13004.928
229.1085205078125 0 1111.6542
229.15496826171875 0 2910.9238
230.11329650878906 0 1009.34796
230.76641845703125 0 558.5333
231.1597900390625 0 694.2213
233.655029296875 0 1136.9114
237.123046875 0 971.48663
238.15493774414062 0 1137.5442
239.13929748535156 0 5492.3774
239.6085968017578 0 947.7424
240.0982666015625 0 16316.599
240.11203002929688 0 902.7765
240.13449096679688 0 666.8673
240.1708221435547 0 633.1903
241.08230590820312 0 3834.2148
241.10208129882812 0 1222.2373
241.19175720214844 0 1469.5128
242.15008544921875 0 2542.67
242.66024780273438 0 3821.8813
243.1619110107422 0 736.24896
244.1295166015625 0 33032.086
245.11305236816406 0 2305.0083
245.13299560546875 0 2978.4607
246.12387084960938 0 813.9115
247.6526336669922 0 1912.9858 b Water loss 4
251.13958740234375 0 1807.938
251.17575073242188 0 4384.9717
252.11331176757812 0 2169.6978
252.17974853515625 0 600.4358
253.0974578857422 0 2142.7422
255.13375854492188 0 909.6436
256.165771484375 0 4183.3735
256.6576843261719 0 1112.1547 b 4
257.14990234375 0 7539.6885
258.1087341308594 0 14956.049
258.1450500488281 0 2502.3538
259.093017578125 0 788.4955
259.11236572265625 0 1540.9011
259.1484680175781 0 571.5715
263.0440673828125 0 613.367
263.0713195800781 0 984.2827
264.1344299316406 0 1603.1621
265.1190185546875 0 733.1381
266.1136474609375 0 1774.29
266.1498718261719 0 3826.2205
266.1862487792969 0 1710.4241
267.1343994140625 0 2882.9614
268.2022705078125 0 3554.3442 a Water loss 2
269.1397399902344 0 3753.8271
269.186279296875 0 2882.827
270.1239318847656 0 37947.04
271.10870361328125 0 886.7836
271.12713623046875 0 6531.445 y Water loss 4
274.1184387207031 0 668.72784
274.1761169433594 0 2431.3672
279.0978698730469 0 678.829
280.10845947265625 0 4766.3325
280.1658630371094 0 3784.7915
281.1102294921875 0 919.4114
282.1451416015625 0 3183.5217
283.1300354003906 0 1599.8052
284.1608581542969 0 40785.82
285.1459045410156 0 752.93896
285.1641540527344 0 5240.847
286.2109069824219 0 661.91754 a 2
287.1505432128906 0 882.46106
292.1290588378906 0 2759.9907
293.11395263671875 0 1079.918
294.1454162597656 0 2522.0298
294.1817626953125 0 2865.327
296.1244201660156 0 1734.0975
296.19732666015625 0 84276.914 b Water loss 2
297.1343994140625 0 1475.158
297.200439453125 0 13314.852
298.1007080078125 0 851.2207
298.1190490722656 0 11220.395
298.176513671875 0 6137.9453
298.203125 0 1434.7567
299.1037292480469 0 1158.2787
299.12249755859375 0 2122.004
299.1805725097656 0 687.1273
300.1556701660156 0 4365.073
302.1714782714844 0 46158.855
303.06622314453125 0 1082.6208
303.1745910644531 0 6230.9775
307.18212890625 0 953.4679
308.1629638671875 0 642.0453
308.19970703125 0 599.4009
309.1563720703125 0 1922.8873
309.2045593261719 0 1460.7317
310.14013671875 0 7658.913
310.1585998535156 0 932.60394
310.21246337890625 0 1529.3545
311.14361572265625 0 753.7326
311.1717529296875 0 1373.2915
312.15582275390625 0 5406.3223
312.1737365722656 0 973.6518 b Water loss 5
312.1922912597656 0 3311.1272
313.1594543457031 0 1164.0529
314.13555908203125 0 1225.8483
314.2079162597656 0 18933.473 b 2
315.1455993652344 0 91849.38
315.2110595703125 0 3316.1992
316.1295166015625 0 93140.06 y Ammonia loss 6
316.1487731933594 0 14571.13
317.1327819824219 0 15950.818
317.1531066894531 0 1280.5898
318.1355285644531 0 1539.7279
320.1240539550781 0 2053.417
320.16131591796875 0 1006.3861
321.1300964355469 0 612.96594
321.17926025390625 0 5188.6665 b 5
321.6805419921875 0 2222.4602
322.1766357421875 0 1099.5001
325.1878662109375 0 1065.8407
326.1719055175781 0 906.5739
326.20782470703125 0 789.94867
327.166748046875 0 10874.295
328.1507263183594 0 15467.475
328.1693115234375 0 1281.6884
329.1542053222656 0 3086.0142
329.18060302734375 0 1066.4473
329.69256591796875 0 1258.0924
330.16607666015625 0 1704.045
333.1562194824219 0 47083.664 y 6
334.1593933105469 0 8290.017
337.15118408203125 0 9825.769
337.1864318847656 0 867.9648
337.6692199707031 0 741.8487
338.1348571777344 0 3736.098
338.1539306640625 0 736.936
340.108154296875 0 928.8631
340.18719482421875 0 6453.7705
341.1897888183594 0 964.64435
343.2337951660156 0 1160.4305
345.1771240234375 0 5834.6777
346.1781005859375 0 2011.4738
351.23944091796875 0 829.0596
352.22314453125 0 7381.7715
353.181640625 0 989.5249
353.2216796875 0 1908.6031
354.1246032714844 0 897.03735
355.1615295410156 0 155798.89
356.1645812988281 0 24378.871
357.16748046875 0 3067.0647
359.09234619140625 0 3431.2954
363.2026062011719 0 2615.8247
365.18194580078125 0 2996.418
369.2498779296875 0 9406.957
370.2337341308594 0 11852.526
370.2541198730469 0 1223.6854
371.1923522949219 0 1453.4662
371.2318420410156 0 5392.207
372.1885681152344 0 2413.0098
372.23382568359375 0 838.02765
373.1717834472656 0 5483.3545
374.17572021484375 0 732.46075
379.2342834472656 0 5738.5166
380.21673583984375 0 1043.232
381.1557312011719 0 5190.719
381.1839294433594 0 1260.0249
381.21380615234375 0 2575.0154
382.158203125 0 1184.9685
387.2605895996094 0 16200.122
388.2633972167969 0 3734.9158
391.1975402832031 0 895.00244
393.2500915527344 0 938.8246
394.2514343261719 0 658.3441
395.15673828125 0 1143.3334
395.2305908203125 0 780.90436
397.244873046875 0 46115.152 b Water loss 3
398.2479553222656 0 9648.925
399.1653747558594 0 877.1493
399.1923522949219 0 720.21063
399.223876953125 0 1476.6357
399.2514953613281 0 1360.0588
400.2257080078125 0 610.0643
405.2105712890625 0 619.22034
407.1568908691406 0 1362.9355
409.1513366699219 0 1598.9319
409.20867919921875 0 5306.7544
411.1878356933594 0 1269.1057
411.2611083984375 0 1185.9849
412.185791015625 0 920.1803
412.21759033203125 0 852.88434
413.2055358886719 0 659.06604
413.24102783203125 0 746.14923
415.2554931640625 0 93151.984 b 3
416.2587890625 0 21192.215
417.2610168457031 0 2674.0488
420.1889343261719 0 775.9637
421.17169189453125 0 1345.0131
423.22412109375 0 3725.9092
424.2271423339844 0 896.0641
424.2560119628906 0 1378.7745
425.2386474609375 0 868.34607
426.1766662597656 0 995.15405
427.16162109375 0 13236.829
427.21893310546875 0 2361.4924
428.1653747558594 0 2286.5396
428.21856689453125 0 1502.5269
430.19244384765625 0 636.3378
431.2657165527344 0 1056.9923
437.7252197265625 0 1022.66785
438.1992492675781 0 3054.4517
440.2162170410156 0 870.3016
440.25067138671875 0 4707.5293
441.23492431640625 0 3041.5874
442.1972961425781 0 579.2627
442.23455810546875 0 813.1212
442.2677307128906 0 863.93555
444.1885986328125 0 3945.2373 y Water loss 5
445.17242431640625 0 8382.793 y Ammonia loss 5
446.175537109375 0 2158.786
450.2350158691406 0 714.28015
451.2174377441406 0 1081.7384
454.1705627441406 0 621.0681
455.738525390625 0 665.26373
456.20953369140625 0 12104.359
456.24432373046875 0 890.5653
457.2119445800781 0 2428.7283
458.2608337402344 0 1071.6772
462.19866943359375 0 5037.5103 y 5
463.2008361816406 0 994.2371
464.25 0 745.6999
466.193603515625 0 684.35626
466.2298889160156 0 1016.86456
466.3016662597656 0 1587.1056
468.24554443359375 0 11808.365
468.2823181152344 0 2006.8295
469.1642761230469 0 596.7673
469.24713134765625 0 2404.0806
469.7337341308594 0 1579.8623
470.23944091796875 0 681.5052
471.2816467285156 0 1094.2621
472.1831359863281 0 3274.7285
473.1846923828125 0 752.07916
476.2870178222656 0 1964.9222
478.20843505859375 0 3723.892
479.212158203125 0 1037.0685
482.2640686035156 0 1288.5841
484.3130187988281 0 1297.0029
486.1664123535156 0 8647.903
487.1711730957031 0 1637.8948
487.25640869140625 0 631.4981 Precursor
492.2445068359375 0 1708.0819
494.29803466796875 0 8766.876 b Water loss 4
495.3004455566406 0 2321.1995
496.2190246582031 0 3376.3076
497.2199401855469 0 910.5251
499.2780456542969 0 2063.286
500.27740478515625 0 874.37524
506.2052001953125 0 2970.74
507.2060241699219 0 999.5188
507.2922668457031 0 792.7834
508.27825927734375 0 1558.9036
509.27435302734375 0 657.08673
510.25604248046875 0 5315.082
511.2591857910156 0 1626.1748
512.307861328125 0 10553.952 b 4
513.2452392578125 0 2941.2388
513.3117065429688 0 2751.129
520.2389526367188 0 1230.1361
524.214111328125 0 26915.502
524.2625122070312 0 940.02985
525.21728515625 0 8515.147
525.3074340820312 0 1073.7953
526.220458984375 0 1404.246
526.286865234375 0 1908.7874
527.285400390625 0 2733.4575
528.2673950195312 0 3180.799
534.2556762695312 0 1535.2438
537.2665405273438 0 3266.3518
538.2596435546875 0 761.37714
540.3038330078125 0 913.10547
541.24072265625 0 12915.81 y Water loss 4
541.2933349609375 0 1230.9788
542.22509765625 0 46195.9 y Ammonia loss 4
543.228271484375 0 13438.456
543.3138427734375 0 1462.943
544.230224609375 0 2300.7761
551.282470703125 0 3891.3418
552.2815551757812 0 1054.8307
555.277587890625 0 4561.697
556.2814331054688 0 1171.0094
559.251220703125 0 133878.1 y 4
560.2540893554688 0 41879.9
561.2562866210938 0 7856.99
569.2926635742188 0 6872.204
570.2968139648438 0 1701.0101
579.3164672851562 0 670.6901
593.293701171875 0 866.41034
594.291748046875 0 921.09485
595.3453979492188 0 1446.2919
603.2764892578125 0 1175.591
605.3314819335938 0 2115.4983
607.250732421875 0 1256.3918
612.3009643554688 0 609.4878
613.3555908203125 0 2869.3271
614.3556518554688 0 1018.25903
620.3037109375 0 2916.9006
621.2913818359375 0 1891.0781
622.355712890625 0 1533.6979
623.34033203125 0 13852.379 b Water loss 5
624.3428344726562 0 5337.2817
625.2637329101562 0 1129.4594
625.34375 0 1423.2883
628.3360595703125 0 802.97406
637.35498046875 0 846.2378
638.314453125 0 8454.251
639.3142700195312 0 3359.5144
640.313232421875 0 827.7807
640.3660888671875 0 4916.938
641.35107421875 0 14493.826 b 5
642.287841796875 0 1794.0692 y Water loss 3
642.353759765625 0 4258.6025
643.2752075195312 0 2739.4604 y Ammonia loss 3
643.3558349609375 0 889.01025
650.3513793945312 0 4581.3164
651.3490600585938 0 2402.679
652.357177734375 0 675.0461
656.3258056640625 0 5221.602
657.327880859375 0 1991.1393
658.32958984375 0 628.51935
658.3768310546875 0 7040.6523
659.3793334960938 0 2667.296
660.2986450195312 0 17637.117 y 3
661.302490234375 0 4936.4277
662.3049926757812 0 1040.4597
668.3611450195312 0 3486.8064
669.3619384765625 0 1106.1267
706.3781127929688 0 1340.5913
707.3739624023438 0 741.87756
716.3630981445312 0 1158.4326
723.4032592773438 0 2306.155
724.3968505859375 0 1798.7721
725.3870849609375 0 947.9859
733.3881225585938 0 5501.679
734.3779907226562 0 4887.171
735.380859375 0 1551.2185
741.4140014648438 0 4055.8394
742.4161987304688 0 1915.9486
751.3984985351562 0 29439.275 b Water loss 6
752.3968505859375 0 12477.414 b Ammonia loss 6
753.3991088867188 0 3558.8696
755.3699340820312 0 1958.3435 y Water loss 2
769.4088134765625 0 19111.682 b 6
770.4111328125 0 8471.15
771.4163208007812 0 2282.2903
773.3836059570312 0 2102.6182 y 2
774.3892211914062 0 906.47644
842.404541015625 0 1191.185 y Water loss 1
860.4154663085938 0 1778.9598 y 1
861.4168090820312 0 968.80023
1183.75439453125 0 624.4869
1710.952392578125 0 616.39996
1787.5811767578125 0 649.69824
3041.234619140625 0 831.1255
3082.194091796875 0 703.14374

Spectrum Details

|  |  |
| --- | --- |
| Matched peaks? Matched peaks The total absolute number of peaks matched. Additionally in brackets the total fraction of peaks matched and the total number of peaks is shown. | 40 (8.28% of 483) |
| FDR? FDR The false discovery rate estimated for this peptide. It is calculated by matching all theoretical fragments with a non-integer shift with the raw peaks for this spectrum. This is done with 40 different shifts. The resulting percentage is the average number of annotated peaks over the number of annotated peaks with the correct spectrum. | 1.43% |
| Satellite FDR? Satellite FDR See the FDR for details on its calculation. This satellite ion specific FDR only contains the satellite ions (d/w) for I/L/J positions. | - |
| PSM Score? PSM Score The PSM Score as given by Hecklib to this annotated spectrum. It is shown with three significant figures. | 603 |

#### Spectrum 8241? Spectrum 8241 The raw spectrum of this peptide as annotated by Hecklib. The fragments are coloured according to ion type (see legend). Any peaks with a star '\*' as text can be hovered over to see the full details, first the ion type second the mass shift type. By hovering over the amino acids in the peptide or ions in the legend the corresponding peaks are highlighted. By toggling the 'Unassigned' label you can turn the background (unassigned) peaks on or off in the plot. By updating the slider in the Ion legend you can update the spectrum to only show the top X% of the peaks with labels. The top X% means any peak that is within X% of the highest intensity. By dragging in the spectrum you can zoom in to a specific part of the spectrum and use 'Zoom Out' to get back to the original zoom level. The annotation of the spectrum is based on the given sequence in the peptides file and is done with different software so inconsistencies are likely. The peaks are annotated based on the given sequence, with 20 ppm tolerance.

Copy Data

##### Spectrum 8241 (TSV)

###### Preview

```
Loading example...
```

*Click on the button to copy the data to your clipboard.*

Mz MinMz MaxIntensity Max

WidthHeightPeptide font sizePeptide stroke widthSpectrum font sizeSpectrum stroke widthCompact peptide

Ion legend

wxyz

abcd

OtherUnassignedIon^Charge^_Position_Show for top:%

JSJTPEQW

05.28e+41.06e+51.58e+52.11e+5

Zoom Out

c^+1^_3_y^+1^_2_z^+1^_2_y^+1^_2_y^+2^_7_y^+1^_3_z^+1^_3_y^+1^_3_c^+1^_5_c^+1^_5_z^+1^_4_y^+1^_4_y^+1^_4_z^+1^_4_y^+1^_4_c^+1^_6_z^+1^_5_w^+1^_5_w^+1^_5_c^+1^_6_y^+1^_5_z^+1^_5_y^+1^_5_w^+1^_6_c^+1^_7_c^+1^_7_z^+1^_6_c^+1^_7_y^+1^_6_z^+1^_7_y^+1^_7_z^+1^_7_y^+1^_7_

036572910941458

Fragment Matches Table

Show background peaks

| Position | Ion type | Intensity | mz Theoretical | mz Error (Th) | mz Error (ppm) | Charge | Series Number |
| --- | --- | --- | --- | --- | --- | --- | --- |
| - | - | 376.3 | 121.6 | - | - | 0 | - |
| - | - | 487.3 | 137.3 | - | - | 0 | - |
| - | - | 455.6 | 148.9 | - | - | 0 | - |
| - | - | 414.8 | 148.9 | - | - | 0 | - |
| - | - | 625.6 | 148.9 | - | - | 0 | - |
| - | - | 662.2 | 148.9 | - | - | 0 | - |
| - | - | 660.1 | 148.9 | - | - | 0 | - |
| - | - | 1203 | 148.9 | - | - | 0 | - |
| - | - | 1485 | 148.9 | - | - | 0 | - |
| - | - | 2832 | 148.9 | - | - | 0 | - |
| - | - | 4074 | 148.9 | - | - | 0 | - |
| - | - | 4090 | 149 | - | - | 0 | - |
| - | - | 2254 | 149 | - | - | 0 | - |
| - | - | 1468 | 149 | - | - | 0 | - |
| - | - | 995.7 | 149 | - | - | 0 | - |
| - | - | 672.3 | 149 | - | - | 0 | - |
| - | - | 770.7 | 149 | - | - | 0 | - |
| - | - | 727.5 | 149 | - | - | 0 | - |
| - | - | 511.7 | 149 | - | - | 0 | - |
| - | - | 402.3 | 149.2 | - | - | 0 | - |
| - | - | 615.3 | 155.1 | - | - | 0 | - |
| - | - | 794.7 | 158.1 | - | - | 0 | - |
| - | - | 995.8 | 158.1 | - | - | 0 | - |
| - | - | 4985 | 173.1 | - | - | 0 | - |
| - | - | 7784 | 175.1 | - | - | 0 | - |
| - | - | 614.5 | 176.1 | - | - | 0 | - |
| - | - | 1100 | 183.1 | - | - | 0 | - |
| - | - | 2635 | 187.1 | - | - | 0 | - |
| - | - | 1305 | 197.1 | - | - | 0 | - |
| - | - | 1258 | 199.1 | - | - | 0 | - |
| - | - | 747.6 | 201.1 | - | - | 0 | - |
| - | - | 1.238E+04 | 201.1 | - | - | 0 | - |
| - | - | 2005 | 202.1 | - | - | 0 | - |
| - | - | 1457 | 202.1 | - | - | 0 | - |
| - | - | 3840 | 203.1 | - | - | 0 | - |
| - | - | 7798 | 215.1 | - | - | 0 | - |
| - | - | 3.666E+04 | 221.1 | - | - | 0 | - |
| - | - | 4937 | 222.1 | - | - | 0 | - |
| - | - | 6134 | 227.1 | - | - | 0 | - |
| - | - | 1.039E+04 | 244.1 | - | - | 0 | - |
| - | - | 1140 | 245.1 | - | - | 0 | - |
| - | - | 528.8 | 257 | - | - | 0 | - |
| - | - | 564.4 | 258.1 | - | - | 0 | - |
| - | - | 501.4 | 266.2 | - | - | 0 | - |
| - | - | 4658 | 284.2 | - | - | 0 | - |
| - | - | 572.3 | 289.5 | - | - | 0 | - |
| - | - | 1.622E+04 | 296.2 | - | - | 0 | - |
| - | - | 1942 | 297.2 | - | - | 0 | - |
| - | - | 6542 | 302.2 | - | - | 0 | - |
| - | - | 656.8 | 303.2 | - | - | 0 | - |
| - | - | 605.4 | 311.1 | - | - | 0 | - |
| - | - | 3461 | 314.1 | - | - | 0 | - |
| - | - | 5037 | 314.2 | - | - | 0 | - |
| - | - | 799.3 | 315.1 | - | - | 0 | - |
| - | - | 1443 | 321.2 | - | - | 0 | - |
| - | - | 576.9 | 326.8 | - | - | 0 | - |
| - | - | 1601 | 328.2 | - | - | 0 | - |
| - | - | 630.9 | 330.2 | - | - | 0 | - |
| - | - | 7092 | 331.1 | - | - | 0 | - |
| 3 | c | 1786 | 331.2 | 0.0006372 | 1.924 | +1 | 3 |
| 7 | y | 1.278E+04 | 332.1 | 0.004963 | 14.94 | +1 | 2 |
| - | - | 1203 | 332.1 | - | - | 0 | - |
| 7 | z | 2108 | 333.1 | 0.0004034 | 1.211 | +1 | 2 |
| - | - | 865 | 337.2 | - | - | 0 | - |
| - | - | 618.9 | 340.2 | - | - | 0 | - |
| - | - | 1407 | 345.2 | - | - | 0 | - |
| - | - | 1413 | 348.1 | - | - | 0 | - |
| 7 | y | 1.339E+04 | 349.1 | 0.004598 | 13.17 | +1 | 2 |
| - | - | 2334 | 350.2 | - | - | 0 | - |
| - | - | 952 | 352.2 | - | - | 0 | - |
| - | - | 2.217E+04 | 355.2 | - | - | 0 | - |
| - | - | 3699 | 356.2 | - | - | 0 | - |
| - | - | 714.8 | 357.2 | - | - | 0 | - |
| - | - | 1688 | 369.2 | - | - | 0 | - |
| - | - | 2939 | 370.2 | - | - | 0 | - |
| - | - | 793.7 | 371.2 | - | - | 0 | - |
| - | - | 2784 | 372.2 | - | - | 0 | - |
| - | - | 955.4 | 379.2 | - | - | 0 | - |
| - | - | 4686 | 387.3 | - | - | 0 | - |
| - | - | 988.2 | 388.3 | - | - | 0 | - |
| - | - | 1.085E+04 | 397.2 | - | - | 0 | - |
| - | - | 2261 | 398.2 | - | - | 0 | - |
| - | - | 2.865E+04 | 415.3 | - | - | 0 | - |
| - | - | 6836 | 416.3 | - | - | 0 | - |
| - | - | 1121 | 417.3 | - | - | 0 | - |
| - | - | 1.09E+04 | 429.2 | - | - | 0 | - |
| 2 | y | 1956 | 430.2 | 0.005171 | 12.02 | +2 | 7 |
| - | - | 2285 | 439.2 | - | - | 0 | - |
| - | - | 695.4 | 440.2 | - | - | 0 | - |
| - | - | 2090 | 441.2 | - | - | 0 | - |
| - | - | 688.3 | 441.2 | - | - | 0 | - |
| - | - | 2685 | 442.2 | - | - | 0 | - |
| - | - | 1125 | 443.2 | - | - | 0 | - |
| - | - | 927.2 | 443.2 | - | - | 0 | - |
| - | - | 2591 | 456.2 | - | - | 0 | - |
| - | - | 571.3 | 457.2 | - | - | 0 | - |
| - | - | 1956 | 459.8 | - | - | 0 | - |
| 6 | y | 1151 | 460.2 | 0.004918 | 10.69 | +1 | 3 |
| 6 | z | 3161 | 462.2 | 0.004105 | 8.883 | +1 | 3 |
| - | - | 1432 | 463.2 | - | - | 0 | - |
| - | - | 687.5 | 466.2 | - | - | 0 | - |
| - | - | 1182 | 468.2 | - | - | 0 | - |
| - | - | 704.9 | 468.7 | - | - | 0 | - |
| - | - | 991 | 477.7 | - | - | 0 | - |
| - | - | 2185 | 477.8 | - | - | 0 | - |
| 6 | y | 1760 | 478.2 | 0.005828 | 12.19 | +1 | 3 |
| - | - | 1197 | 486.2 | - | - | 0 | - |
| - | - | 1311 | 494.3 | - | - | 0 | - |
| - | - | 982.6 | 495.8 | - | - | 0 | - |
| - | - | 1165 | 497.2 | - | - | 0 | - |
| - | - | 779.6 | 503.2 | - | - | 0 | - |
| 5 | c | 700.4 | 511.3 | 0.002876 | 5.624 | +1 | 5 |
| - | - | 1313 | 512.3 | - | - | 0 | - |
| - | - | 5499 | 513.2 | - | - | 0 | - |
| - | - | 999.9 | 513.2 | - | - | 0 | - |
| - | - | 1041 | 514.2 | - | - | 0 | - |
| - | - | 3478 | 515.2 | - | - | 0 | - |
| - | - | 872.7 | 516.2 | - | - | 0 | - |
| - | - | 7976 | 528.3 | - | - | 0 | - |
| 5 | c | 5.658E+04 | 529.3 | 0.0003178 | 0.6004 | +1 | 5 |
| - | - | 1963 | 530.2 | - | - | 0 | - |
| - | - | 1.545E+04 | 530.3 | - | - | 0 | - |
| - | - | 2605 | 531.3 | - | - | 0 | - |
| - | - | 3835 | 540.2 | - | - | 0 | - |
| 5 | z | 1096 | 541.2 | 0.0007402 | 1.368 | +1 | 4 |
| - | - | 700.2 | 554.3 | - | - | 0 | - |
| - | - | 764.2 | 555.3 | - | - | 0 | - |
| - | - | 1314 | 555.3 | - | - | 0 | - |
| 5 | y | 3508 | 557.2 | 0.004523 | 8.116 | +1 | 4 |
| 5 | y | 1.184E+04 | 558.2 | 0.004882 | 8.746 | +1 | 4 |
| 5 | z | 4105 | 559.2 | 0.0002307 | 0.4126 | +1 | 4 |
| - | - | 1074 | 569.3 | - | - | 0 | - |
| - | - | 4.078E+04 | 574.2 | - | - | 0 | - |
| 5 | y | 8.923E+04 | 575.2 | 0.003968 | 6.897 | +1 | 4 |
| - | - | 2.301E+04 | 576.2 | - | - | 0 | - |
| - | - | 4567 | 577.3 | - | - | 0 | - |
| - | - | 1117 | 598.3 | - | - | 0 | - |
| - | - | 1652 | 599.3 | - | - | 0 | - |
| - | - | 1556 | 602.2 | - | - | 0 | - |
| - | - | 2100 | 603.2 | - | - | 0 | - |
| - | - | 1194 | 604.2 | - | - | 0 | - |
| - | - | 638.4 | 605.3 | - | - | 0 | - |
| - | - | 3762 | 614.4 | - | - | 0 | - |
| - | - | 3221 | 615.4 | - | - | 0 | - |
| - | - | 1202 | 616.4 | - | - | 0 | - |
| - | - | 1501 | 623.3 | - | - | 0 | - |
| - | - | 2413 | 625.3 | - | - | 0 | - |
| - | - | 1209 | 626.3 | - | - | 0 | - |
| - | - | 1260 | 628.3 | - | - | 0 | - |
| - | - | 796.5 | 640.2 | - | - | 0 | - |
| - | - | 967.2 | 640.3 | - | - | 0 | - |
| 6 | c | 1225 | 640.4 | 0.002683 | 4.19 | +1 | 6 |
| - | - | 7314 | 641.3 | - | - | 0 | - |
| - | - | 2208 | 641.4 | - | - | 0 | - |
| 4 | z | 1137 | 642.3 | 0.003522 | 5.484 | +1 | 5 |
| - | - | 1845 | 642.3 | - | - | 0 | - |
| - | - | 892 | 642.4 | - | - | 0 | - |
| 4 | w | 838.7 | 643.3 | 0.005096 | 7.922 | +1 | 5 |
| 4 | w | 812.1 | 645.2 | 0.00746 | 11.56 | +1 | 5 |
| - | - | 616.7 | 650.4 | - | - | 0 | - |
| - | - | 984.8 | 656.3 | - | - | 0 | - |
| - | - | 1536 | 657.4 | - | - | 0 | - |
| - | - | 1.239E+04 | 658.3 | - | - | 0 | - |
| 6 | c | 7.459E+04 | 658.4 | 0.0005525 | 0.8392 | +1 | 6 |
| 4 | y | 5372 | 659.3 | 0.001515 | 2.298 | +1 | 5 |
| - | - | 2.414E+04 | 659.4 | - | - | 0 | - |
| 4 | z | 3.241E+04 | 660.3 | 0.004066 | 6.158 | +1 | 5 |
| - | - | 4731 | 660.4 | - | - | 0 | - |
| - | - | 2.364E+04 | 661.3 | - | - | 0 | - |
| - | - | 6240 | 662.3 | - | - | 0 | - |
| - | - | 1108 | 663.3 | - | - | 0 | - |
| - | - | 5954 | 675.3 | - | - | 0 | - |
| 4 | y | 1.29E+04 | 676.3 | 0.003652 | 5.401 | +1 | 5 |
| - | - | 3752 | 677.3 | - | - | 0 | - |
| - | - | 997.3 | 678.3 | - | - | 0 | - |
| - | - | 3197 | 684.4 | - | - | 0 | - |
| - | - | 1368 | 685.4 | - | - | 0 | - |
| - | - | 586.7 | 687.4 | - | - | 0 | - |
| - | - | 2238 | 697.3 | - | - | 0 | - |
| - | - | 689.1 | 698.3 | - | - | 0 | - |
| - | - | 1366 | 713.3 | - | - | 0 | - |
| - | - | 1.451E+04 | 715.3 | - | - | 0 | - |
| - | - | 6950 | 716.3 | - | - | 0 | - |
| - | - | 906.7 | 716.4 | - | - | 0 | - |
| - | - | 1156 | 717.3 | - | - | 0 | - |
| - | - | 4698 | 728.3 | - | - | 0 | - |
| - | - | 1698 | 729.3 | - | - | 0 | - |
| 3 | w | 736.8 | 730.3 | 0.002304 | 3.155 | +1 | 6 |
| - | - | 727.7 | 731.3 | - | - | 0 | - |
| - | - | 761.4 | 733.4 | - | - | 0 | - |
| - | - | 694 | 741.4 | - | - | 0 | - |
| - | - | 1399 | 742.4 | - | - | 0 | - |
| - | - | 5009 | 751.4 | - | - | 0 | - |
| - | - | 1612 | 752.4 | - | - | 0 | - |
| - | - | 1.101E+04 | 753.4 | - | - | 0 | - |
| - | - | 5644 | 754.4 | - | - | 0 | - |
| - | - | 2217 | 755.4 | - | - | 0 | - |
| 7 | c | 1302 | 768.4 | 0.003888 | 5.059 | +1 | 7 |
| 7 | c | 6537 | 769.4 | 0.0001712 | 0.2226 | +1 | 7 |
| - | - | 2613 | 770.4 | - | - | 0 | - |
| - | - | 1131 | 771.3 | - | - | 0 | - |
| - | - | 2.289E+04 | 771.4 | - | - | 0 | - |
| - | - | 7878 | 772.4 | - | - | 0 | - |
| 3 | z | 1.721E+04 | 773.4 | 0.003803 | 4.918 | +1 | 6 |
| - | - | 1902 | 773.4 | - | - | 0 | - |
| - | - | 8580 | 774.4 | - | - | 0 | - |
| - | - | 2008 | 775.4 | - | - | 0 | - |
| - | - | 4292 | 783.4 | - | - | 0 | - |
| - | - | 1870 | 784.5 | - | - | 0 | - |
| 7 | c | 2.092E+05 | 786.4 | 0.0009025 | 1.148 | +1 | 7 |
| - | - | 8.982E+04 | 787.4 | - | - | 0 | - |
| - | - | 2087 | 788.4 | - | - | 0 | - |
| - | - | 1.983E+04 | 788.4 | - | - | 0 | - |
| 3 | y | 9416 | 789.4 | 0.01382 | 17.51 | +1 | 6 |
| - | - | 1214 | 789.4 | - | - | 0 | - |
| - | - | 4057 | 790.4 | - | - | 0 | - |
| - | - | 912.5 | 791.4 | - | - | 0 | - |
| - | - | 9413 | 800.4 | - | - | 0 | - |
| - | - | 4341 | 801.4 | - | - | 0 | - |
| - | - | 1180 | 802.4 | - | - | 0 | - |
| - | - | 4425 | 804.3 | - | - | 0 | - |
| - | - | 1798 | 805.3 | - | - | 0 | - |
| - | - | 920.4 | 810.4 | - | - | 0 | - |
| - | - | 937.8 | 812.4 | - | - | 0 | - |
| - | - | 9329 | 812.4 | - | - | 0 | - |
| - | - | 961.3 | 813.4 | - | - | 0 | - |
| - | - | 4693 | 813.5 | - | - | 0 | - |
| - | - | 1054 | 814.4 | - | - | 0 | - |
| - | - | 1.947E+04 | 827.4 | - | - | 0 | - |
| - | - | 922 | 827.5 | - | - | 0 | - |
| - | - | 1.458E+04 | 828.4 | - | - | 0 | - |
| - | - | 5722 | 829.4 | - | - | 0 | - |
| - | - | 955.5 | 830.4 | - | - | 0 | - |
| 2 | z | 4380 | 842.4 | 0.004373 | 5.191 | +1 | 7 |
| - | - | 2.238E+04 | 843.4 | - | - | 0 | - |
| - | - | 1.072E+04 | 844.4 | - | - | 0 | - |
| - | - | 1.52E+04 | 845.4 | - | - | 0 | - |
| - | - | 5776 | 846.5 | - | - | 0 | - |
| - | - | 1443 | 847.5 | - | - | 0 | - |
| - | - | 7508 | 857.5 | - | - | 0 | - |
| - | - | 2048 | 858.4 | - | - | 0 | - |
| - | - | 4095 | 858.5 | - | - | 0 | - |
| 2 | y | 1603 | 859.4 | 0.003037 | 3.534 | +1 | 7 |
| - | - | 1157 | 859.5 | - | - | 0 | - |
| 2 | z | 4.639E+04 | 860.4 | 0.003696 | 4.296 | +1 | 7 |
| - | - | 2.183E+04 | 861.4 | - | - | 0 | - |
| - | - | 5408 | 862.4 | - | - | 0 | - |
| - | - | 782.6 | 863.4 | - | - | 0 | - |
| - | - | 2150 | 873.5 | - | - | 0 | - |
| - | - | 2046 | 875.5 | - | - | 0 | - |
| 2 | y | 3688 | 876.4 | 0.003893 | 4.442 | +1 | 7 |
| - | - | 2595 | 877.4 | - | - | 0 | - |
| - | - | 1879 | 910.5 | - | - | 0 | - |
| - | - | 1737 | 911.5 | - | - | 0 | - |
| - | - | 872.8 | 918.5 | - | - | 0 | - |
| - | - | 3.994E+04 | 928.5 | - | - | 0 | - |
| - | - | 2.195E+04 | 929.5 | - | - | 0 | - |
| - | - | 2443 | 930.4 | - | - | 0 | - |
| - | - | 5640 | 930.5 | - | - | 0 | - |
| - | - | 2400 | 931.5 | - | - | 0 | - |
| - | - | 840 | 932.5 | - | - | 0 | - |
| - | - | 2.716E+04 | 945.5 | - | - | 0 | - |
| - | - | 1.306E+04 | 946.5 | - | - | 0 | - |
| - | - | 4986 | 947.5 | - | - | 0 | - |
| - | - | 4676 | 955.5 | - | - | 0 | - |
| - | - | 2862 | 956.5 | - | - | 0 | - |
| - | - | 1580 | 957.5 | - | - | 0 | - |
| - | - | 1210 | 962.5 | - | - | 0 | - |
| - | - | 3.315E+04 | 972.5 | - | - | 0 | - |
| - | - | 5.727E+04 | 973.5 | - | - | 0 | - |
| - | - | 2.662E+04 | 974.5 | - | - | 0 | - |
| - | - | 2050 | 974.6 | - | - | 0 | - |
| - | - | 8446 | 975.5 | - | - | 0 | - |
| - | - | 823.9 | 988.6 | - | - | 0 | - |
| - | - | 6.219E+04 | 989.5 | - | - | 0 | - |
| - | - | 1.354E+05 | 990.5 | - | - | 0 | - |
| - | - | 6.659E+04 | 991.5 | - | - | 0 | - |
| - | - | 2.018E+04 | 992.5 | - | - | 0 | - |
| - | - | 1283 | 993.5 | - | - | 0 | - |
| - | - | 980 | 1028 | - | - | 0 | - |
| - | - | 803.7 | 1444 | - | - | 0 | - |

m/z Charge Intensity FragmentType MassShift Position
121.57476806640625 0 376.2849
137.2867431640625 0 487.3151
148.87136840820312 0 455.64694
148.8782196044922 0 414.81488
148.900146484375 0 625.6081
148.90695190429688 0 662.189
148.9140625 0 660.1237
148.9217987060547 0 1202.6763
148.92889404296875 0 1484.6636
148.93626403808594 0 2832.2712
148.9440155029297 0 4073.771
148.96067810058594 0 4089.858
148.96849060058594 0 2253.8796
148.9757080078125 0 1467.5283
148.98265075683594 0 995.67163
148.99014282226562 0 672.2567
148.9976043701172 0 770.6843
149.00457763671875 0 727.46124
149.01939392089844 0 511.67892
149.20858764648438 0 402.3235
155.1177978515625 0 615.3144
158.0602264404297 0 794.6937
158.08424377441406 0 995.7786
173.12857055664062 0 4984.686
175.0867156982422 0 7784.466
176.09005737304688 0 614.47046
183.1128387451172 0 1099.6088
187.14419555664062 0 2634.8528
197.12820434570312 0 1305.4412
199.10772705078125 0 1258.1918
201.1129150390625 0 747.5541
201.1234130859375 0 12383.3545
202.0739288330078 0 2004.8256
202.12646484375 0 1456.6744
203.08143615722656 0 3839.6682
215.13905334472656 0 7798.3047
221.09219360351562 0 36664.715
222.09559631347656 0 4936.6807
227.10263061523438 0 6134.236
244.12921142578125 0 10392.888
245.1328887939453 0 1140.0554
257.01715087890625 0 528.75916
258.1089782714844 0 564.38855
266.1508483886719 0 501.3616
284.1604919433594 0 4658.1143
289.5162353515625 0 572.3471
296.1968994140625 0 16218.17
297.2002258300781 0 1941.7856
302.1713562011719 0 6541.921
303.17437744140625 0 656.83014
311.1242980957031 0 605.3975
314.1134338378906 0 3461.3652
314.2074890136719 0 5036.999
315.11541748046875 0 799.33417
321.1790466308594 0 1442.7875
326.82501220703125 0 576.8995
328.15032958984375 0 1601.401
330.1676025390625 0 630.8788
331.1401062011719 0 7092.111
331.234619140625 0 1786.3607 c 2
332.1241455078125 0 12776.758 y Ammonia loss 6
332.1432800292969 0 1203.0447
333.1274108886719 0 2108.2712 z 6
337.15142822265625 0 865.0072
340.185302734375 0 618.85016
345.1766052246094 0 1407.3322
348.1429748535156 0 1413.3875
349.15032958984375 0 13385.87 y 6
350.154052734375 0 2334.021
352.2226257324219 0 952.0352
355.16107177734375 0 22167.686
356.1637268066406 0 3699.1572
357.1759338378906 0 714.76904
369.24969482421875 0 1688.0978
370.233642578125 0 2939.007
371.23101806640625 0 793.6834
372.1873779296875 0 2783.7776
379.23358154296875 0 955.4371
387.26031494140625 0 4686.207
388.26434326171875 0 988.24927
397.2444763183594 0 10850.082
398.2474670410156 0 2261.0789
415.255126953125 0 28654.205
416.25836181640625 0 6836.267
417.2596130371094 0 1120.677
429.1854553222656 0 10900.439
430.1876220703125 0 1956.4476 y Ammonia loss 1
439.18341064453125 0 2285.1782
440.18756103515625 0 695.3808
441.1973571777344 0 2089.9216
441.2303771972656 0 688.2942
442.19439697265625 0 2684.941
443.157470703125 0 1124.8627
443.1977233886719 0 927.21265
456.2099914550781 0 2590.9307
457.21551513671875 0 571.29877
459.8128356933594 0 1956.1451
460.18267822265625 0 1151.4064 y Water loss 5
462.1737060546875 0 3161.0117 z 5
463.1809387207031 0 1431.8368
466.1549987792969 0 687.5411
468.2442626953125 0 1182.4202
468.7270812988281 0 704.9216
477.7319030761719 0 990.9654
477.8223876953125 0 2184.6777
478.19415283203125 0 1759.8783 y 5
486.24639892578125 0 1197.2742
494.2979736328125 0 1310.9822
495.8352966308594 0 982.57574
497.2119445800781 0 1164.9049
503.20111083984375 0 779.6077
511.32098388671875 0 700.40466 c Water loss 4
512.30859375 0 1313.0186
513.2062377929688 0 5499.403
513.2457275390625 0 999.9343
514.208984375 0 1040.5789
515.2214965820312 0 3478.156
516.22314453125 0 872.68805
528.3262939453125 0 7976.1074
529.3341064453125 0 56583.14 c 4
530.2356567382812 0 1963.0703
530.3368530273438 0 15451.363
531.33935546875 0 2604.816
540.208984375 0 3835.2656
541.2110595703125 0 1096.2842 z Water loss 4
554.2789916992188 0 700.23553
555.2785034179688 0 764.2053
555.349853515625 0 1313.9791
557.2350463867188 0 3507.5554 y Water loss 4
558.2194213867188 0 11839.492 y Ammonia loss 4
559.2225952148438 0 4104.712 z 4
569.2899169921875 0 1074.3103
574.2378540039062 0 40782.016
575.2450561523438 0 89227.25 y 4
576.248046875 0 23005.242
577.2504272460938 0 4567.289
598.2760009765625 0 1116.5538
599.2817993164062 0 1651.7048
602.2345581054688 0 1556.1814
603.2390747070312 0 2099.5786
604.2423706054688 0 1194.3209
605.3392333984375 0 638.40686
614.3633422851562 0 3762.024
615.3685302734375 0 3221.271
616.3724975585938 0 1201.5924
623.33984375 0 1501.0997
625.3314208984375 0 2413.226
626.3343505859375 0 1208.6624
628.3063354492188 0 1260.2917
640.2483520507812 0 796.5129
640.304931640625 0 967.20575
640.36376953125 0 1225.3179 c Water loss 5
641.3133544921875 0 7313.767
641.3605346679688 0 2208.1204
642.2630004882812 0 1137.1407 z Water loss 3
642.316650390625 0 1845.4143
642.364990234375 0 892.0281
643.26220703125 0 838.7414 w 3
645.2540283203125 0 812.0552 w 3
650.3556518554688 0 616.6658
656.328369140625 0 984.78595
657.3702392578125 0 1535.5164
658.2593383789062 0 12393.49
658.37646484375 0 74587.58 c 5
659.2637329101562 0 5371.6987 y Ammonia loss 3
659.3792724609375 0 24142.873
660.2741088867188 0 32412.445 z 3
660.38232421875 0 4731.46
661.2796020507812 0 23639.818
662.2835693359375 0 6239.648
663.2867431640625 0 1107.697
675.2849731445312 0 5954.357
676.2924194335938 0 12896.109 y 3
677.2944946289062 0 3752.434
678.299560546875 0 997.2582
684.3923950195312 0 3197.3582
685.3984985351562 0 1368.4365
687.3544311523438 0 586.72144
697.3286743164062 0 2237.9568
698.3248901367188 0 689.0961
713.3212280273438 0 1366.4695
715.337158203125 0 14511.619
716.3404541015625 0 6949.946
716.4061889648438 0 906.6975
717.3458251953125 0 1155.9489
728.3461303710938 0 4698.4517
729.3465576171875 0 1698.0728
730.3016357421875 0 736.79425 w 2
731.306640625 0 727.7174
733.389892578125 0 761.35455
741.411376953125 0 693.95026
742.419921875 0 1398.6416
751.3963623046875 0 5009.308
752.39404296875 0 1612.0372
753.4120483398438 0 11012.373
754.4067993164062 0 5644.4585
755.4099731445312 0 2217.197
768.421142578125 0 1301.5602 c Water loss 6
769.4088745117188 0 6537.145 c Ammonia loss 6
770.4118041992188 0 2612.777
771.3446655273438 0 1131.4856
771.4236450195312 0 22890.154
772.4264526367188 0 7878.132
773.35791015625 0 17206.83 z 2
773.4300537109375 0 1901.7562
774.3617553710938 0 8579.869
775.3671875 0 2007.5237
783.4473876953125 0 4291.517
784.4518432617188 0 1869.6182
786.4346923828125 0 209178.02 c 6
787.437744140625 0 89820.35
788.3701782226562 0 2087.0947
788.440673828125 0 19834.299
789.3590087890625 0 9415.968 y 2
789.4442138671875 0 1213.784
790.3630981445312 0 4056.5908
791.3751831054688 0 912.5177
800.4244995117188 0 9412.631
801.4273681640625 0 4340.545
802.43603515625 0 1180.3851
804.3267822265625 0 4424.6284
805.3304443359375 0 1797.5209
810.41357421875 0 920.3824
812.3692016601562 0 937.82556
812.44970703125 0 9329.058
813.37548828125 0 961.3131
813.4527587890625 0 4692.7856
814.447265625 0 1053.6356
827.4373779296875 0 19472.076
827.526611328125 0 922.0292
828.4323120117188 0 14579.31
829.4330444335938 0 5722.0913
830.4342651367188 0 955.5165
842.3799438476562 0 4379.7437 z Water loss 1
843.4295043945312 0 22375.557
844.4329223632812 0 10721.884
845.4451904296875 0 15197.819
846.4501342773438 0 5776.059
847.4537963867188 0 1443.3634
857.4616088867188 0 7508.0107
858.3758544921875 0 2047.7644
858.4644165039062 0 4094.7087
859.38134765625 0 1602.5464 y Ammonia loss 1
859.466552734375 0 1156.9258
860.3898315429688 0 46389.9 z 1
861.3931274414062 0 21829.385
862.3952026367188 0 5408.171
863.4012451171875 0 782.6395
873.4556274414062 0 2150.4932
875.4703979492188 0 2045.5496
876.4087524414062 0 3687.858 y 1
877.4141235351562 0 2594.5845
910.4928588867188 0 1878.6979
911.4866333007812 0 1737.1693
918.4962768554688 0 872.8432
928.4998168945312 0 39939.59
929.5031127929688 0 21947.17
930.415283203125 0 2443.309
930.5064697265625 0 5640.123
931.4913330078125 0 2399.516
932.49755859375 0 840.00916
945.4791259765625 0 27160.03
946.4832153320312 0 13056.49
947.486083984375 0 4985.849
955.4635620117188 0 4676.433
956.4662475585938 0 2861.5264
957.4660034179688 0 1579.9225
962.5081176757812 0 1209.5302
972.4893798828125 0 33145.836
973.4788818359375 0 57267.547
974.4790649414062 0 26618.254
974.5712890625 0 2049.6062
975.4814453125 0 8445.527
988.551025390625 0 823.8824
989.4925537109375 0 62190.465
990.498779296875 0 135405.16
991.502197265625 0 66588.65
992.5060424804688 0 20183.824
993.5121459960938 0 1283.4542
1027.5814208984375 0 980.04016
1443.6092529296875 0 803.70306

Spectrum Details

|  |  |
| --- | --- |
| Matched peaks? Matched peaks The total absolute number of peaks matched. Additionally in brackets the total fraction of peaks matched and the total number of peaks is shown. | 33 (11.74% of 281) |
| FDR? FDR The false discovery rate estimated for this peptide. It is calculated by matching all theoretical fragments with a non-integer shift with the raw peaks for this spectrum. This is done with 40 different shifts. The resulting percentage is the average number of annotated peaks over the number of annotated peaks with the correct spectrum. | 4.26% |
| Satellite FDR? Satellite FDR See the FDR for details on its calculation. This satellite ion specific FDR only contains the satellite ions (d/w) for I/L/J positions. | 14.29% |
| PSM Score? PSM Score The PSM Score as given by Hecklib to this annotated spectrum. It is shown with three significant figures. | 446 |

#### Spectrum 10159? Spectrum 10159 The raw spectrum of this peptide as annotated by Hecklib. The fragments are coloured according to ion type (see legend). Any peaks with a star '\*' as text can be hovered over to see the full details, first the ion type second the mass shift type. By hovering over the amino acids in the peptide or ions in the legend the corresponding peaks are highlighted. By toggling the 'Unassigned' label you can turn the background (unassigned) peaks on or off in the plot. By updating the slider in the Ion legend you can update the spectrum to only show the top X% of the peaks with labels. The top X% means any peak that is within X% of the highest intensity. By dragging in the spectrum you can zoom in to a specific part of the spectrum and use 'Zoom Out' to get back to the original zoom level. The annotation of the spectrum is based on the given sequence in the peptides file and is done with different software so inconsistencies are likely. The peaks are annotated based on the given sequence, with 20 ppm tolerance.

Copy Data

##### Spectrum 10159 (TSV)

###### Preview

```
Loading example...
```

*Click on the button to copy the data to your clipboard.*

Mz MinMz MaxIntensity Max

WidthHeightPeptide font sizePeptide stroke widthSpectrum font sizeSpectrum stroke widthCompact peptide

Ion legend

wxyz

abcd

OtherUnassignedIon^Charge^_Position_Show for top:%

JSJTPEQW

02.77e+45.54e+48.30e+41.11e+5

Zoom Out

a^+1^_2_a^+1^_2_b^+1^_2_b^+1^_2_y^+1^_1_a^+1^_3_y^+2^_4_b^+1^_3_b^+1^_3_y^+1^_2_b^+2^_6_y^+1^_2_b^+1^_4_b^+1^_4_y^+1^_3_y^+1^_3_y^+1^_3_\*b^+1^_5_b^+1^_5_y^+1^_4_y^+1^_4_y^+1^_4_b^+1^_6_b^+1^_6_y^+1^_5_b^+1^_7_b^+1^_7_y^+1^_6_b^+1^_7_y^+1^_6_y^+1^_7_y^+1^_7_

02515017521002

Fragment Matches Table

Show background peaks

| Position | Ion type | Intensity | mz Theoretical | mz Error (Th) | mz Error (ppm) | Charge | Series Number |
| --- | --- | --- | --- | --- | --- | --- | --- |
| - | - | 2503 | 120.1 | - | - | 0 | - |
| - | - | 358.4 | 121.1 | - | - | 0 | - |
| - | - | 890.9 | 125.1 | - | - | 0 | - |
| - | - | 1826 | 125.1 | - | - | 0 | - |
| - | - | 860 | 126.1 | - | - | 0 | - |
| - | - | 686.5 | 127.1 | - | - | 0 | - |
| - | - | 446.4 | 127.1 | - | - | 0 | - |
| - | - | 418.4 | 128.1 | - | - | 0 | - |
| - | - | 1220 | 128.1 | - | - | 0 | - |
| - | - | 3844 | 129.1 | - | - | 0 | - |
| - | - | 349 | 129.1 | - | - | 0 | - |
| - | - | 2088 | 129.1 | - | - | 0 | - |
| - | - | 2236 | 130.1 | - | - | 0 | - |
| - | - | 1.127E+04 | 130.1 | - | - | 0 | - |
| - | - | 1038 | 131.1 | - | - | 0 | - |
| - | - | 4523 | 132.1 | - | - | 0 | - |
| - | - | 1.592E+04 | 133.1 | - | - | 0 | - |
| - | - | 477.6 | 134.4 | - | - | 0 | - |
| - | - | 1263 | 136.1 | - | - | 0 | - |
| - | - | 556.8 | 137.1 | - | - | 0 | - |
| - | - | 424.1 | 141.1 | - | - | 0 | - |
| - | - | 475.8 | 141.9 | - | - | 0 | - |
| - | - | 2304 | 142.1 | - | - | 0 | - |
| - | - | 410.1 | 143.1 | - | - | 0 | - |
| - | - | 2247 | 144.1 | - | - | 0 | - |
| - | - | 407.7 | 145.3 | - | - | 0 | - |
| - | - | 1.461E+04 | 146.1 | - | - | 0 | - |
| - | - | 1636 | 147.1 | - | - | 0 | - |
| - | - | 1337 | 147.1 | - | - | 0 | - |
| - | - | 809.2 | 149 | - | - | 0 | - |
| - | - | 1794 | 152.1 | - | - | 0 | - |
| - | - | 869 | 153.1 | - | - | 0 | - |
| - | - | 1472 | 153.1 | - | - | 0 | - |
| 2 | a | 2288 | 155.1 | 0.0002898 | 1.868 | +1 | 2 |
| - | - | 572.2 | 157.1 | - | - | 0 | - |
| - | - | 535.7 | 157.7 | - | - | 0 | - |
| - | - | 2.957E+04 | 159.1 | - | - | 0 | - |
| - | - | 830.1 | 160.1 | - | - | 0 | - |
| - | - | 1860 | 160.1 | - | - | 0 | - |
| - | - | 412.7 | 162.1 | - | - | 0 | - |
| - | - | 548.6 | 163.1 | - | - | 0 | - |
| - | - | 665.3 | 169.1 | - | - | 0 | - |
| - | - | 1371 | 169.1 | - | - | 0 | - |
| - | - | 1473 | 170.1 | - | - | 0 | - |
| - | - | 533.1 | 170.1 | - | - | 0 | - |
| - | - | 2940 | 171.1 | - | - | 0 | - |
| - | - | 460.6 | 173.1 | - | - | 0 | - |
| 2 | a | 2.208E+04 | 173.1 | 0.00033 | 1.906 | +1 | 2 |
| - | - | 2605 | 173.4 | - | - | 0 | - |
| - | - | 830.9 | 174.1 | - | - | 0 | - |
| - | - | 1404 | 174.1 | - | - | 0 | - |
| - | - | 539.8 | 175.1 | - | - | 0 | - |
| - | - | 1333 | 175.1 | - | - | 0 | - |
| - | - | 5334 | 177.1 | - | - | 0 | - |
| - | - | 2575 | 181.1 | - | - | 0 | - |
| 2 | b | 5051 | 183.1 | 0.0003092 | 1.689 | +1 | 2 |
| - | - | 1890 | 183.1 | - | - | 0 | - |
| - | - | 576.7 | 187.1 | - | - | 0 | - |
| - | - | 3028 | 187.1 | - | - | 0 | - |
| - | - | 6119 | 187.1 | - | - | 0 | - |
| - | - | 1.096E+05 | 188.1 | - | - | 0 | - |
| - | - | 738.5 | 188.1 | - | - | 0 | - |
| - | - | 1.223E+04 | 189.1 | - | - | 0 | - |
| - | - | 535.3 | 190.1 | - | - | 0 | - |
| - | - | 444.2 | 190.8 | - | - | 0 | - |
| - | - | 609.2 | 193.7 | - | - | 0 | - |
| - | - | 535.3 | 195.1 | - | - | 0 | - |
| - | - | 3288 | 197.1 | - | - | 0 | - |
| - | - | 2.377E+04 | 199.1 | - | - | 0 | - |
| - | - | 1757 | 200.1 | - | - | 0 | - |
| - | - | 3311 | 201.1 | - | - | 0 | - |
| 2 | b | 2.376E+04 | 201.1 | 0.0002426 | 1.206 | +1 | 2 |
| - | - | 1669 | 202.1 | - | - | 0 | - |
| 8 | y | 4.594E+04 | 205.1 | 0.0002428 | 1.184 | +1 | 1 |
| - | - | 4448 | 206.1 | - | - | 0 | - |
| - | - | 2416 | 209.1 | - | - | 0 | - |
| - | - | 1921 | 211.1 | - | - | 0 | - |
| - | - | 754.7 | 212.1 | - | - | 0 | - |
| - | - | 484.2 | 213.1 | - | - | 0 | - |
| - | - | 600.6 | 213.1 | - | - | 0 | - |
| - | - | 1.491E+04 | 215.1 | - | - | 0 | - |
| - | - | 1402 | 216.1 | - | - | 0 | - |
| - | - | 1004 | 217.1 | - | - | 0 | - |
| - | - | 1584 | 221.1 | - | - | 0 | - |
| - | - | 676.6 | 223.1 | - | - | 0 | - |
| - | - | 875.7 | 225.1 | - | - | 0 | - |
| - | - | 922.2 | 225.1 | - | - | 0 | - |
| - | - | 1473 | 226.1 | - | - | 0 | - |
| - | - | 3.435E+04 | 227.1 | - | - | 0 | - |
| - | - | 3879 | 228.1 | - | - | 0 | - |
| - | - | 936.2 | 229.2 | - | - | 0 | - |
| - | - | 1396 | 239.1 | - | - | 0 | - |
| - | - | 3836 | 240.1 | - | - | 0 | - |
| - | - | 1222 | 241.1 | - | - | 0 | - |
| - | - | 491 | 241.1 | - | - | 0 | - |
| - | - | 863.4 | 242.1 | - | - | 0 | - |
| - | - | 645.3 | 243.2 | - | - | 0 | - |
| - | - | 1.073E+04 | 244.1 | - | - | 0 | - |
| - | - | 1359 | 245.1 | - | - | 0 | - |
| - | - | 1192 | 245.1 | - | - | 0 | - |
| - | - | 590.4 | 248.1 | - | - | 0 | - |
| - | - | 969.9 | 251.1 | - | - | 0 | - |
| - | - | 942.2 | 251.2 | - | - | 0 | - |
| - | - | 737.2 | 252.1 | - | - | 0 | - |
| - | - | 707.4 | 253.1 | - | - | 0 | - |
| - | - | 1399 | 256.2 | - | - | 0 | - |
| - | - | 2834 | 257.1 | - | - | 0 | - |
| - | - | 4271 | 258.1 | - | - | 0 | - |
| - | - | 541.1 | 258.1 | - | - | 0 | - |
| - | - | 1596 | 258.1 | - | - | 0 | - |
| - | - | 504.7 | 259.1 | - | - | 0 | - |
| - | - | 677.9 | 266.1 | - | - | 0 | - |
| - | - | 662.3 | 266.1 | - | - | 0 | - |
| - | - | 701 | 267.1 | - | - | 0 | - |
| 3 | a | 919 | 268.2 | 0.00166 | 6.188 | +1 | 3 |
| - | - | 909.9 | 269.1 | - | - | 0 | - |
| - | - | 1125 | 269.2 | - | - | 0 | - |
| - | - | 1.192E+04 | 270.1 | - | - | 0 | - |
| 5 | y | 1731 | 271.1 | 0.00348 | 12.84 | +2 | 4 |
| - | - | 561.1 | 277 | - | - | 0 | - |
| - | - | 1357 | 280.1 | - | - | 0 | - |
| - | - | 894.9 | 280.2 | - | - | 0 | - |
| - | - | 1088 | 282.1 | - | - | 0 | - |
| - | - | 714.3 | 283.1 | - | - | 0 | - |
| - | - | 1180 | 283.2 | - | - | 0 | - |
| - | - | 1.173E+04 | 284.2 | - | - | 0 | - |
| - | - | 2172 | 285.2 | - | - | 0 | - |
| - | - | 907.2 | 294.2 | - | - | 0 | - |
| - | - | 793.7 | 296.1 | - | - | 0 | - |
| 3 | b | 2.561E+04 | 296.2 | 0.000367 | 1.239 | +1 | 3 |
| - | - | 3422 | 297.2 | - | - | 0 | - |
| - | - | 3641 | 298.1 | - | - | 0 | - |
| - | - | 1029 | 298.2 | - | - | 0 | - |
| - | - | 666.6 | 299.1 | - | - | 0 | - |
| - | - | 1067 | 300.2 | - | - | 0 | - |
| - | - | 1.428E+04 | 302.2 | - | - | 0 | - |
| - | - | 1641 | 303.2 | - | - | 0 | - |
| - | - | 1718 | 309.2 | - | - | 0 | - |
| - | - | 1924 | 310.1 | - | - | 0 | - |
| - | - | 823.4 | 310.2 | - | - | 0 | - |
| - | - | 1332 | 312.2 | - | - | 0 | - |
| - | - | 1270 | 312.2 | - | - | 0 | - |
| 3 | b | 5506 | 314.2 | 0.0003614 | 1.15 | +1 | 3 |
| - | - | 509 | 314.2 | - | - | 0 | - |
| - | - | 2.826E+04 | 315.1 | - | - | 0 | - |
| - | - | 651.7 | 315.2 | - | - | 0 | - |
| 7 | y | 2.759E+04 | 316.1 | 0.000212 | 0.6707 | +1 | 2 |
| - | - | 4190 | 316.1 | - | - | 0 | - |
| - | - | 4936 | 317.1 | - | - | 0 | - |
| - | - | 650.3 | 318.1 | - | - | 0 | - |
| - | - | 613.3 | 318.2 | - | - | 0 | - |
| - | - | 622.8 | 320.1 | - | - | 0 | - |
| 6 | b | 1360 | 321.2 | 0.000314 | 0.9776 | +2 | 6 |
| - | - | 866.7 | 321.7 | - | - | 0 | - |
| - | - | 590.8 | 326.2 | - | - | 0 | - |
| - | - | 3742 | 327.2 | - | - | 0 | - |
| - | - | 8658 | 327.2 | - | - | 0 | - |
| - | - | 4232 | 328.2 | - | - | 0 | - |
| - | - | 480.8 | 328.2 | - | - | 0 | - |
| 7 | y | 1.238E+04 | 333.2 | 0.0004574 | 1.373 | +1 | 2 |
| - | - | 2686 | 334.2 | - | - | 0 | - |
| - | - | 2327 | 337.2 | - | - | 0 | - |
| - | - | 1363 | 338.1 | - | - | 0 | - |
| - | - | 1617 | 340.2 | - | - | 0 | - |
| - | - | 1497 | 345.2 | - | - | 0 | - |
| - | - | 649.6 | 346.2 | - | - | 0 | - |
| - | - | 2162 | 352.2 | - | - | 0 | - |
| - | - | 1135 | 353.2 | - | - | 0 | - |
| - | - | 4.507E+04 | 355.2 | - | - | 0 | - |
| - | - | 879 | 356.1 | - | - | 0 | - |
| - | - | 7577 | 356.2 | - | - | 0 | - |
| - | - | 896.3 | 357.2 | - | - | 0 | - |
| - | - | 1160 | 365.2 | - | - | 0 | - |
| - | - | 2731 | 369.2 | - | - | 0 | - |
| - | - | 3039 | 370.2 | - | - | 0 | - |
| - | - | 1014 | 371.2 | - | - | 0 | - |
| - | - | 819.2 | 372.2 | - | - | 0 | - |
| - | - | 2066 | 373.2 | - | - | 0 | - |
| - | - | 1347 | 379.2 | - | - | 0 | - |
| - | - | 1187 | 381.2 | - | - | 0 | - |
| - | - | 934 | 381.2 | - | - | 0 | - |
| - | - | 2058 | 381.2 | - | - | 0 | - |
| - | - | 5429 | 387.3 | - | - | 0 | - |
| - | - | 683.9 | 388.3 | - | - | 0 | - |
| 4 | b | 1.326E+04 | 397.2 | 5.178E-05 | 0.1303 | +1 | 4 |
| - | - | 2535 | 398.2 | - | - | 0 | - |
| - | - | 3058 | 399.3 | - | - | 0 | - |
| - | - | 975.4 | 409.2 | - | - | 0 | - |
| - | - | 740.5 | 411.2 | - | - | 0 | - |
| - | - | 571.5 | 414.2 | - | - | 0 | - |
| 4 | b | 2.457E+04 | 415.3 | 0.0002293 | 0.5521 | +1 | 4 |
| - | - | 5354 | 416.3 | - | - | 0 | - |
| - | - | 1192 | 423.2 | - | - | 0 | - |
| - | - | 3613 | 427.2 | - | - | 0 | - |
| - | - | 740.6 | 438.2 | - | - | 0 | - |
| - | - | 1072 | 440.3 | - | - | 0 | - |
| - | - | 863.1 | 441.2 | - | - | 0 | - |
| 6 | y | 1825 | 444.2 | 0.0008386 | 1.888 | +1 | 3 |
| 6 | y | 2626 | 445.2 | 0.0004351 | 0.9774 | +1 | 3 |
| - | - | 4428 | 456.2 | - | - | 0 | - |
| - | - | 681.3 | 457.2 | - | - | 0 | - |
| 6 | y | 1128 | 462.2 | 0.0001616 | 0.3497 | +1 | 3 |
| - | - | 4085 | 468.2 | - | - | 0 | - |
| - | - | 889.4 | 469.2 | - | - | 0 | - |
| - | - | 689.8 | 469.7 | - | - | 0 | - |
| - | - | 4881 | 471.3 | - | - | 0 | - |
| - | - | 743.6 | 472.2 | - | - | 0 | - |
| - | - | 1000 | 478.2 | - | - | 0 | - |
| 0 | Precursor | 692 | 487.3 | 0.003614 | 7.417 | +2 | -1 |
| - | - | 1274 | 488.3 | - | - | 0 | - |
| 5 | b | 2303 | 494.3 | 0.0001913 | 0.3871 | +1 | 5 |
| - | - | 950.1 | 495.3 | - | - | 0 | - |
| - | - | 1362 | 496.2 | - | - | 0 | - |
| - | - | 1352 | 510.3 | - | - | 0 | - |
| 5 | b | 2819 | 512.3 | 0.0004411 | 0.8609 | +1 | 5 |
| - | - | 1086 | 513.3 | - | - | 0 | - |
| - | - | 8012 | 524.2 | - | - | 0 | - |
| - | - | 1834 | 525.2 | - | - | 0 | - |
| - | - | 889.7 | 526.3 | - | - | 0 | - |
| - | - | 647.4 | 528.3 | - | - | 0 | - |
| - | - | 1140 | 537.3 | - | - | 0 | - |
| 5 | y | 2870 | 541.2 | 0.001083 | 2.001 | +1 | 4 |
| 5 | y | 1.173E+04 | 542.2 | 0.0006192 | 1.142 | +1 | 4 |
| - | - | 3037 | 543.2 | - | - | 0 | - |
| - | - | 1037 | 551.3 | - | - | 0 | - |
| - | - | 765.3 | 555.3 | - | - | 0 | - |
| 5 | y | 3.572E+04 | 559.3 | 0.000173 | 0.3094 | +1 | 4 |
| - | - | 1.145E+04 | 560.3 | - | - | 0 | - |
| - | - | 2215 | 561.3 | - | - | 0 | - |
| - | - | 1752 | 569.3 | - | - | 0 | - |
| - | - | 742.3 | 605.3 | - | - | 0 | - |
| - | - | 653.1 | 615.3 | - | - | 0 | - |
| 6 | b | 3086 | 623.3 | 0.0001233 | 0.1978 | +1 | 6 |
| - | - | 1484 | 624.3 | - | - | 0 | - |
| - | - | 2212 | 638.3 | - | - | 0 | - |
| - | - | 945.5 | 640.4 | - | - | 0 | - |
| 6 | b | 3584 | 641.4 | 0.0003096 | 0.4827 | +1 | 6 |
| - | - | 1025 | 642.4 | - | - | 0 | - |
| - | - | 981.7 | 650.3 | - | - | 0 | - |
| - | - | 2010 | 656.3 | - | - | 0 | - |
| - | - | 2076 | 658.4 | - | - | 0 | - |
| - | - | 866.9 | 659.4 | - | - | 0 | - |
| 4 | y | 4807 | 660.3 | 6.097E-05 | 0.09234 | +1 | 5 |
| - | - | 1759 | 661.3 | - | - | 0 | - |
| - | - | 692.1 | 668.4 | - | - | 0 | - |
| - | - | 749 | 733.4 | - | - | 0 | - |
| - | - | 1115 | 734.4 | - | - | 0 | - |
| - | - | 716.2 | 735.4 | - | - | 0 | - |
| - | - | 1091 | 741.4 | - | - | 0 | - |
| 7 | b | 6928 | 751.4 | 0.0009591 | 1.276 | +1 | 7 |
| 7 | b | 4416 | 752.4 | 0.01289 | 17.13 | +1 | 7 |
| - | - | 1148 | 753.4 | - | - | 0 | - |
| 3 | y | 894.6 | 755.4 | 0.001112 | 1.472 | +1 | 6 |
| 7 | b | 5921 | 769.4 | 0.0002933 | 0.3812 | +1 | 7 |
| - | - | 2190 | 770.4 | - | - | 0 | - |
| 3 | y | 1100 | 773.4 | 0.001202 | 1.554 | +1 | 6 |
| 2 | y | 742.2 | 842.4 | 0.002317 | 2.751 | +1 | 7 |
| 2 | y | 721.2 | 860.4 | 0.000919 | 1.068 | +1 | 7 |
| - | - | 647.7 | 992.5 | - | - | 0 | - |

m/z Charge Intensity FragmentType MassShift Position
120.08108520507812 0 2503.3318
121.08417510986328 0 358.43674
125.07150268554688 0 890.9232
125.1076889038086 0 1826.1333
126.09158325195312 0 859.9709
127.07585906982422 0 686.4717
127.08671569824219 0 446.3657
128.07064819335938 0 418.3929
128.1072998046875 0 1220.329
129.0662384033203 0 3843.6133
129.070068359375 0 348.955
129.10263061523438 0 2087.5464
130.0502471923828 0 2236.1687
130.0654296875 0 11270.873
131.07003784179688 0 1038.2582
132.08103942871094 0 4522.549
133.0862274169922 0 15918.916
134.3995361328125 0 477.56003
136.07614135742188 0 1263.3556
137.0712432861328 0 556.8351
141.10250854492188 0 424.13092
141.94317626953125 0 475.83087
142.12306213378906 0 2304.0293
143.0699462890625 0 410.07797
144.08120727539062 0 2246.6956
145.27886962890625 0 407.6794
146.06036376953125 0 14610.366
147.0636444091797 0 1635.5818
147.07669067382812 0 1336.519
148.95387268066406 0 809.2048
152.10739135742188 0 1794.3308
153.0661163330078 0 868.9915
153.1026611328125 0 1472.022
155.11817932128906 0 2287.6377 a Water loss 1
157.08567810058594 0 572.21356
157.7144012451172 0 535.69476
159.0919952392578 0 29573.572
160.07586669921875 0 830.0975
160.09523010253906 0 1860.3375
162.05377197265625 0 412.68585
163.08677673339844 0 548.5802
169.09674072265625 0 665.316
169.13369750976562 0 1370.802
170.06028747558594 0 1473.4995
170.1183624267578 0 533.09125
171.11318969726562 0 2940.0022
173.0919647216797 0 460.55286
173.1287841796875 0 22079.668 a 1
173.4402618408203 0 2605.0852
174.0551300048828 0 830.9273
174.13243103027344 0 1404.1531
175.08627319335938 0 539.7639
175.09658813476562 0 1332.6372
177.11236572265625 0 5334.0415
181.09742736816406 0 2575.0334
183.1131134033203 0 5051.417 b Water loss 1
183.1494140625 0 1889.9263
187.07180786132812 0 576.74164
187.0869598388672 0 3027.7744
187.1444549560547 0 6119.2563
188.0709228515625 0 109606.3
188.1476593017578 0 738.4617
189.0742950439453 0 12229.682
190.07582092285156 0 535.2989
190.7716064453125 0 444.23138
193.6612548828125 0 609.22394
195.07662963867188 0 535.2814
197.12852478027344 0 3288.3562
199.1079864501953 0 23769.736
200.1116180419922 0 1756.8254
201.1023712158203 0 3310.9814
201.1236114501953 0 23764.416 b 1
202.1273193359375 0 1668.667
205.09739685058594 0 45936.38 y 7
206.1007537841797 0 4447.5767
209.09231567382812 0 2416.495
211.14434814453125 0 1921.4532
212.10325622558594 0 754.7178
213.08729553222656 0 484.17178
213.12359619140625 0 600.565
215.1392822265625 0 14907.414
216.14324951171875 0 1401.824
217.08248901367188 0 1004.2925
221.1387481689453 0 1584.1821
223.0724639892578 0 676.62
225.1024169921875 0 875.7316
225.12355041503906 0 922.1965
226.11865234375 0 1473.3031
227.1029510498047 0 34347.35
228.10630798339844 0 3878.8164
229.15545654296875 0 936.1632
239.13951110839844 0 1396.1089
240.0980987548828 0 3835.6333
241.08197021484375 0 1221.6425
241.10009765625 0 491.00522
242.14964294433594 0 863.3559
243.1612548828125 0 645.3231
244.12948608398438 0 10725.6045
245.11329650878906 0 1358.6953
245.13282775878906 0 1192.4806
248.1042938232422 0 590.4422
251.1394500732422 0 969.87115
251.17567443847656 0 942.1689
252.11253356933594 0 737.2075
253.09732055664062 0 707.43384
256.1658020019531 0 1398.814
257.1495666503906 0 2833.7605
258.1085510253906 0 4271.3677
258.1217346191406 0 541.09436
258.1451721191406 0 1595.7832
259.11187744140625 0 504.68307
266.1125183105469 0 677.8829
266.14990234375 0 662.2852
267.1334533691406 0 701.0317
268.20361328125 0 918.9556 a Water loss 2
269.13916015625 0 909.8979
269.1864318847656 0 1125.354
270.1239318847656 0 11918.839
271.12738037109375 0 1730.7959 y Water loss 4
277.01727294921875 0 561.1267
280.1085510253906 0 1357.186
280.1663513183594 0 894.86774
282.1451721191406 0 1088.3274
283.1309814453125 0 714.2893
283.17425537109375 0 1179.514
284.16070556640625 0 11732.655
285.1641540527344 0 2171.96
294.1806945800781 0 907.19293
296.1244812011719 0 793.66705
296.1972351074219 0 25614.54 b Water loss 2
297.20037841796875 0 3421.717
298.1188659667969 0 3640.9578
298.176513671875 0 1029.3973
299.1227722167969 0 666.57324
300.1544189453125 0 1066.7677
302.1713562011719 0 14284.108
303.1744689941406 0 1640.8472
309.2038269042969 0 1717.7449
310.1399230957031 0 1924.1375
310.21136474609375 0 823.4206
312.1556091308594 0 1331.6161
312.1921081542969 0 1269.942
314.2077941894531 0 5506.3306 b 2
314.22357177734375 0 508.99643
315.1455078125 0 28264.898
315.2111511230469 0 651.658
316.12939453125 0 27585.074 y Ammonia loss 6
316.1488037109375 0 4189.7515
317.13262939453125 0 4935.618
318.13641357421875 0 650.254
318.18115234375 0 613.2779
320.1239013671875 0 622.83417
321.1785583496094 0 1359.7123 b 5
321.68121337890625 0 866.7037
326.1720275878906 0 590.8059
327.16656494140625 0 3741.827
327.2016296386719 0 8658.227
328.15069580078125 0 4231.812
328.1676330566406 0 480.7624
333.15618896484375 0 12380.295 y 6
334.1595153808594 0 2686.327
337.15142822265625 0 2327.2827
338.1341247558594 0 1362.8281
340.18670654296875 0 1616.8512
345.17724609375 0 1496.9785
346.1771240234375 0 649.5999
352.2227783203125 0 2162.4302
353.2194519042969 0 1135.0635
355.16131591796875 0 45065.24
356.14617919921875 0 879.0145
356.1640930175781 0 7576.789
357.1684265136719 0 896.32294
365.1817932128906 0 1159.6718
369.24908447265625 0 2730.991
370.2331237792969 0 3038.7214
371.2315673828125 0 1013.99194
372.1858825683594 0 819.21625
373.172119140625 0 2066.1526
379.232421875 0 1346.6267
381.1553649902344 0 1187.2578
381.215576171875 0 934.0138
381.24761962890625 0 2057.9421
387.26043701171875 0 5428.954
388.2643737792969 0 683.868
397.2445983886719 0 13263.276 b Water loss 3
398.2473449707031 0 2535.4517
399.25921630859375 0 3058.3455
409.2080078125 0 975.3909
411.1871643066406 0 740.53815
414.20654296875 0 571.4947
415.2553405761719 0 24568.408 b 3
416.2586669921875 0 5354.052
423.2231140136719 0 1191.7556
427.1614990234375 0 3613.0493
438.198974609375 0 740.55115
440.25048828125 0 1072.4028
441.2340087890625 0 863.0934
444.1885986328125 0 1825.0984 y Water loss 5
445.1722106933594 0 2626.0046 y Ammonia loss 5
456.20916748046875 0 4427.6343
457.21221923828125 0 681.3299
462.198486328125 0 1128.2792 y 5
468.24542236328125 0 4084.5215
469.248046875 0 889.4436
469.7331237792969 0 689.7887
471.3165588378906 0 4880.5103
472.18267822265625 0 743.55444
478.2087097167969 0 1000.4607
487.2567138671875 0 692.0171 Precursor
488.344482421875 0 1274.2992
494.297119140625 0 2302.6125 b Water loss 4
495.3006286621094 0 950.14526
496.2185974121094 0 1361.6925
510.25634765625 0 1351.7751
512.3074340820312 0 2818.923 b 4
513.3109130859375 0 1085.9791
524.2136840820312 0 8011.558
525.2164306640625 0 1833.7363
526.2877807617188 0 889.68506
528.2685546875 0 647.3762
537.266845703125 0 1140.0713
541.2394409179688 0 2870.4949 y Water loss 4
542.2251586914062 0 11733.98 y Ammonia loss 4
543.2274169921875 0 3037.0796
551.2811889648438 0 1037.4122
555.2760620117188 0 765.27344
559.2509155273438 0 35721.06 y 4
560.2539672851562 0 11449.751
561.2553100585938 0 2214.8193
569.293701171875 0 1751.8684
605.3304443359375 0 742.31006
615.2752685546875 0 653.0695
623.3400268554688 0 3086.2095 b Water loss 5
624.3430786132812 0 1484.2085
638.3125 0 2212.0645
640.3681030273438 0 945.53766
641.3501586914062 0 3583.6938 b 5
642.3529052734375 0 1024.9862
650.3457641601562 0 981.7092
656.3261108398438 0 2010.274
658.3783569335938 0 2075.9695
659.3822631835938 0 866.90405
660.2987060546875 0 4807.351 y 3
661.3025512695312 0 1759.3325
668.363037109375 0 692.05493
733.3864135742188 0 748.9525
734.3809204101562 0 1114.667
735.373291015625 0 716.20166
741.4141845703125 0 1091.2288
751.3975219726562 0 6927.74 b Water loss 6
752.3953857421875 0 4416.246 b Ammonia loss 6
753.398193359375 0 1147.6655
755.3711547851562 0 894.55884 y Water loss 2
769.4087524414062 0 5920.579 b 6
770.411376953125 0 2190.301
773.384033203125 0 1099.5294 y 2
842.4019775390625 0 742.1504 y Water loss 1
860.4139404296875 0 721.1905 y 1
992.5420532226562 0 647.6828

Spectrum Details

|  |  |
| --- | --- |
| Matched peaks? Matched peaks The total absolute number of peaks matched. Additionally in brackets the total fraction of peaks matched and the total number of peaks is shown. | 33 (12.74% of 259) |
| FDR? FDR The false discovery rate estimated for this peptide. It is calculated by matching all theoretical fragments with a non-integer shift with the raw peaks for this spectrum. This is done with 40 different shifts. The resulting percentage is the average number of annotated peaks over the number of annotated peaks with the correct spectrum. | 0.65% |
| Satellite FDR? Satellite FDR See the FDR for details on its calculation. This satellite ion specific FDR only contains the satellite ions (d/w) for I/L/J positions. | - |
| PSM Score? PSM Score The PSM Score as given by Hecklib to this annotated spectrum. It is shown with three significant figures. | 523 |

#### Spectrum 9852? Spectrum 9852 The raw spectrum of this peptide as annotated by Hecklib. The fragments are coloured according to ion type (see legend). Any peaks with a star '\*' as text can be hovered over to see the full details, first the ion type second the mass shift type. By hovering over the amino acids in the peptide or ions in the legend the corresponding peaks are highlighted. By toggling the 'Unassigned' label you can turn the background (unassigned) peaks on or off in the plot. By updating the slider in the Ion legend you can update the spectrum to only show the top X% of the peaks with labels. The top X% means any peak that is within X% of the highest intensity. By dragging in the spectrum you can zoom in to a specific part of the spectrum and use 'Zoom Out' to get back to the original zoom level. The annotation of the spectrum is based on the given sequence in the peptides file and is done with different software so inconsistencies are likely. The peaks are annotated based on the given sequence, with 20 ppm tolerance.

Copy Data

##### Spectrum 9852 (TSV)

###### Preview

```
Loading example...
```

*Click on the button to copy the data to your clipboard.*

Mz MinMz MaxIntensity Max

WidthHeightPeptide font sizePeptide stroke widthSpectrum font sizeSpectrum stroke widthCompact peptide

Ion legend

wxyz

abcd

OtherUnassignedIon^Charge^_Position_Show for top:%

JSJTPEQW

07.16e+41.43e+52.15e+52.87e+5

Zoom Out

a^+1^_2_a^+1^_2_b^+1^_2_b^+1^_2_y^+1^_1_b^+2^_5_a^+1^_3_y^+2^_4_a^+1^_3_b^+1^_3_b^+2^_6_b^+1^_3_y^+1^_2_b^+2^_6_y^+1^_2_b^+1^_4_b^+1^_4_y^+1^_3_y^+1^_3_y^+1^_3_\*b^+1^_5_b^+1^_5_y^+1^_4_y^+1^_4_y^+1^_4_b^+1^_6_b^+1^_6_y^+1^_5_y^+1^_5_y^+1^_5_b^+1^_7_b^+1^_7_y^+1^_6_y^+1^_6_b^+1^_7_y^+1^_6_y^+1^_7_y^+1^_7_

041382712401653

Fragment Matches Table

Show background peaks

| Position | Ion type | Intensity | mz Theoretical | mz Error (Th) | mz Error (ppm) | Charge | Series Number |
| --- | --- | --- | --- | --- | --- | --- | --- |
| - | - | 2120 | 120.1 | - | - | 0 | - |
| - | - | 333.9 | 120.8 | - | - | 0 | - |
| - | - | 1337 | 125.1 | - | - | 0 | - |
| - | - | 4458 | 125.1 | - | - | 0 | - |
| - | - | 1463 | 126.1 | - | - | 0 | - |
| - | - | 388.3 | 126.4 | - | - | 0 | - |
| - | - | 500.2 | 127.1 | - | - | 0 | - |
| - | - | 420 | 127.1 | - | - | 0 | - |
| - | - | 512.2 | 128.1 | - | - | 0 | - |
| - | - | 2090 | 128.1 | - | - | 0 | - |
| - | - | 1.093E+04 | 129.1 | - | - | 0 | - |
| - | - | 1423 | 129.1 | - | - | 0 | - |
| - | - | 3974 | 130.1 | - | - | 0 | - |
| - | - | 2.569E+04 | 130.1 | - | - | 0 | - |
| - | - | 2510 | 131.1 | - | - | 0 | - |
| - | - | 1.335E+04 | 132.1 | - | - | 0 | - |
| - | - | 1911 | 132.1 | - | - | 0 | - |
| - | - | 5961 | 133.1 | - | - | 0 | - |
| - | - | 425.9 | 135.7 | - | - | 0 | - |
| - | - | 676.1 | 136.1 | - | - | 0 | - |
| - | - | 472.3 | 137.1 | - | - | 0 | - |
| - | - | 471.8 | 137.1 | - | - | 0 | - |
| - | - | 1109 | 138.1 | - | - | 0 | - |
| - | - | 1280 | 139.1 | - | - | 0 | - |
| - | - | 552.6 | 139.7 | - | - | 0 | - |
| - | - | 434.9 | 141.1 | - | - | 0 | - |
| - | - | 847.5 | 142.1 | - | - | 0 | - |
| - | - | 6313 | 142.1 | - | - | 0 | - |
| - | - | 1046 | 143.1 | - | - | 0 | - |
| - | - | 5738 | 144.1 | - | - | 0 | - |
| - | - | 366.2 | 144.5 | - | - | 0 | - |
| - | - | 422.8 | 145.1 | - | - | 0 | - |
| - | - | 3.735E+04 | 146.1 | - | - | 0 | - |
| - | - | 3786 | 147.1 | - | - | 0 | - |
| - | - | 2537 | 147.1 | - | - | 0 | - |
| - | - | 417.7 | 148 | - | - | 0 | - |
| - | - | 4190 | 152.1 | - | - | 0 | - |
| - | - | 3054 | 153.1 | - | - | 0 | - |
| - | - | 3036 | 153.1 | - | - | 0 | - |
| - | - | 532.8 | 154.1 | - | - | 0 | - |
| - | - | 1480 | 155.1 | - | - | 0 | - |
| - | - | 525.9 | 155.1 | - | - | 0 | - |
| 2 | a | 5950 | 155.1 | 0.000305 | 1.967 | +1 | 2 |
| - | - | 835.4 | 156.1 | - | - | 0 | - |
| - | - | 569.5 | 156.1 | - | - | 0 | - |
| - | - | 878.9 | 158 | - | - | 0 | - |
| - | - | 7.453E+04 | 159.1 | - | - | 0 | - |
| - | - | 1129 | 160.1 | - | - | 0 | - |
| - | - | 7500 | 160.1 | - | - | 0 | - |
| - | - | 521 | 161.1 | - | - | 0 | - |
| - | - | 430.6 | 161.3 | - | - | 0 | - |
| - | - | 1069 | 163.1 | - | - | 0 | - |
| - | - | 573.6 | 166.1 | - | - | 0 | - |
| - | - | 943.4 | 169.1 | - | - | 0 | - |
| - | - | 2519 | 169.1 | - | - | 0 | - |
| - | - | 4233 | 170.1 | - | - | 0 | - |
| - | - | 8206 | 171.1 | - | - | 0 | - |
| - | - | 941.5 | 171.1 | - | - | 0 | - |
| 2 | a | 5.34E+04 | 173.1 | 0.00033 | 1.906 | +1 | 2 |
| - | - | 811.7 | 173.5 | - | - | 0 | - |
| - | - | 862.8 | 174.1 | - | - | 0 | - |
| - | - | 4807 | 174.1 | - | - | 0 | - |
| - | - | 1976 | 177.1 | - | - | 0 | - |
| - | - | 510.3 | 180.2 | - | - | 0 | - |
| - | - | 6545 | 181.1 | - | - | 0 | - |
| - | - | 631.6 | 182.1 | - | - | 0 | - |
| 2 | b | 1.215E+04 | 183.1 | 0.0002787 | 1.522 | +1 | 2 |
| - | - | 4109 | 183.1 | - | - | 0 | - |
| - | - | 1687 | 184.1 | - | - | 0 | - |
| - | - | 1058 | 187.1 | - | - | 0 | - |
| - | - | 6248 | 187.1 | - | - | 0 | - |
| - | - | 1.574E+04 | 187.1 | - | - | 0 | - |
| - | - | 2.837E+05 | 188.1 | - | - | 0 | - |
| - | - | 1469 | 188.1 | - | - | 0 | - |
| - | - | 3.119E+04 | 189.1 | - | - | 0 | - |
| - | - | 1868 | 190.1 | - | - | 0 | - |
| - | - | 1338 | 195.1 | - | - | 0 | - |
| - | - | 589.8 | 195.1 | - | - | 0 | - |
| - | - | 676.3 | 196.1 | - | - | 0 | - |
| - | - | 8417 | 197.1 | - | - | 0 | - |
| - | - | 865.5 | 198.1 | - | - | 0 | - |
| - | - | 1448 | 199.1 | - | - | 0 | - |
| - | - | 5.565E+04 | 199.1 | - | - | 0 | - |
| - | - | 5627 | 200.1 | - | - | 0 | - |
| - | - | 584.2 | 200.1 | - | - | 0 | - |
| - | - | 2558 | 201.1 | - | - | 0 | - |
| 2 | b | 6.209E+04 | 201.1 | 0.0002579 | 1.282 | +1 | 2 |
| - | - | 5917 | 202.1 | - | - | 0 | - |
| 8 | y | 1.195E+05 | 205.1 | 0.0003191 | 1.556 | +1 | 1 |
| - | - | 1.366E+04 | 206.1 | - | - | 0 | - |
| - | - | 735.7 | 207.1 | - | - | 0 | - |
| - | - | 6835 | 209.1 | - | - | 0 | - |
| - | - | 751.1 | 210.1 | - | - | 0 | - |
| - | - | 1136 | 211.1 | - | - | 0 | - |
| - | - | 3426 | 211.1 | - | - | 0 | - |
| - | - | 2210 | 212.1 | - | - | 0 | - |
| - | - | 844.9 | 213.1 | - | - | 0 | - |
| - | - | 728.8 | 213.1 | - | - | 0 | - |
| - | - | 1149 | 214.2 | - | - | 0 | - |
| - | - | 545.6 | 215.1 | - | - | 0 | - |
| - | - | 3.781E+04 | 215.1 | - | - | 0 | - |
| - | - | 3385 | 216.1 | - | - | 0 | - |
| - | - | 2559 | 217.1 | - | - | 0 | - |
| - | - | 512.8 | 222.1 | - | - | 0 | - |
| - | - | 1204 | 223.1 | - | - | 0 | - |
| - | - | 618.6 | 223.1 | - | - | 0 | - |
| - | - | 3604 | 225.1 | - | - | 0 | - |
| - | - | 1455 | 225.1 | - | - | 0 | - |
| - | - | 1009 | 226.1 | - | - | 0 | - |
| - | - | 4023 | 226.1 | - | - | 0 | - |
| - | - | 8.512E+04 | 227.1 | - | - | 0 | - |
| - | - | 8681 | 228.1 | - | - | 0 | - |
| - | - | 1089 | 229.1 | - | - | 0 | - |
| - | - | 1627 | 229.2 | - | - | 0 | - |
| - | - | 949.9 | 230.1 | - | - | 0 | - |
| - | - | 726.5 | 231.1 | - | - | 0 | - |
| - | - | 484.2 | 231.9 | - | - | 0 | - |
| - | - | 804.9 | 233.7 | - | - | 0 | - |
| - | - | 620.7 | 238.2 | - | - | 0 | - |
| - | - | 4624 | 239.1 | - | - | 0 | - |
| - | - | 1.185E+04 | 240.1 | - | - | 0 | - |
| - | - | 924.7 | 240.1 | - | - | 0 | - |
| - | - | 700.1 | 240.2 | - | - | 0 | - |
| - | - | 3156 | 241.1 | - | - | 0 | - |
| - | - | 1220 | 241.1 | - | - | 0 | - |
| - | - | 604.6 | 241.1 | - | - | 0 | - |
| - | - | 1412 | 242.2 | - | - | 0 | - |
| - | - | 2894 | 242.7 | - | - | 0 | - |
| - | - | 1145 | 243.1 | - | - | 0 | - |
| - | - | 2.423E+04 | 244.1 | - | - | 0 | - |
| - | - | 1961 | 245.1 | - | - | 0 | - |
| - | - | 2933 | 245.1 | - | - | 0 | - |
| - | - | 544.4 | 248.2 | - | - | 0 | - |
| - | - | 800.7 | 251.1 | - | - | 0 | - |
| - | - | 2742 | 251.2 | - | - | 0 | - |
| - | - | 1690 | 252.1 | - | - | 0 | - |
| - | - | 540.4 | 252.2 | - | - | 0 | - |
| - | - | 1899 | 253.1 | - | - | 0 | - |
| - | - | 3776 | 256.2 | - | - | 0 | - |
| 5 | b | 1331 | 256.7 | 0.0001356 | 0.5284 | +2 | 5 |
| - | - | 5942 | 257.1 | - | - | 0 | - |
| - | - | 9272 | 258.1 | - | - | 0 | - |
| - | - | 1869 | 258.1 | - | - | 0 | - |
| - | - | 595.8 | 259.1 | - | - | 0 | - |
| - | - | 1380 | 264.1 | - | - | 0 | - |
| - | - | 541.5 | 265.1 | - | - | 0 | - |
| - | - | 978.5 | 266.1 | - | - | 0 | - |
| - | - | 2252 | 266.2 | - | - | 0 | - |
| - | - | 3100 | 267.1 | - | - | 0 | - |
| - | - | 543.2 | 267.1 | - | - | 0 | - |
| 3 | a | 2434 | 268.2 | 0.0002254 | 0.8406 | +1 | 3 |
| - | - | 2700 | 269.1 | - | - | 0 | - |
| - | - | 2443 | 269.2 | - | - | 0 | - |
| - | - | 2.947E+04 | 270.1 | - | - | 0 | - |
| - | - | 877.7 | 271.1 | - | - | 0 | - |
| 5 | y | 4022 | 271.1 | 0.003358 | 12.39 | +2 | 4 |
| - | - | 656.3 | 272.2 | - | - | 0 | - |
| - | - | 1952 | 274.2 | - | - | 0 | - |
| - | - | 709.4 | 278.2 | - | - | 0 | - |
| - | - | 3779 | 280.1 | - | - | 0 | - |
| - | - | 2455 | 280.2 | - | - | 0 | - |
| - | - | 785.2 | 281.1 | - | - | 0 | - |
| - | - | 1764 | 282.1 | - | - | 0 | - |
| - | - | 917.1 | 283.1 | - | - | 0 | - |
| - | - | 3.414E+04 | 284.2 | - | - | 0 | - |
| - | - | 958.8 | 285.1 | - | - | 0 | - |
| - | - | 3586 | 285.2 | - | - | 0 | - |
| - | - | 729.6 | 286.1 | - | - | 0 | - |
| - | - | 830.2 | 286.1 | - | - | 0 | - |
| 3 | a | 620.7 | 286.2 | 0.0002198 | 0.7681 | +1 | 3 |
| - | - | 994.4 | 287.2 | - | - | 0 | - |
| - | - | 1918 | 292.1 | - | - | 0 | - |
| - | - | 1259 | 294.1 | - | - | 0 | - |
| - | - | 1537 | 294.2 | - | - | 0 | - |
| - | - | 832.9 | 296.1 | - | - | 0 | - |
| 3 | b | 6.274E+04 | 296.2 | 0.0003975 | 1.342 | +1 | 3 |
| - | - | 1106 | 297.1 | - | - | 0 | - |
| - | - | 1.148E+04 | 297.2 | - | - | 0 | - |
| - | - | 9146 | 298.1 | - | - | 0 | - |
| - | - | 3992 | 298.2 | - | - | 0 | - |
| - | - | 918.8 | 298.2 | - | - | 0 | - |
| - | - | 1095 | 299.1 | - | - | 0 | - |
| - | - | 1157 | 299.1 | - | - | 0 | - |
| - | - | 546.7 | 299.2 | - | - | 0 | - |
| - | - | 2904 | 300.2 | - | - | 0 | - |
| - | - | 3.601E+04 | 302.2 | - | - | 0 | - |
| - | - | 7001 | 303.2 | - | - | 0 | - |
| - | - | 1230 | 307.2 | - | - | 0 | - |
| - | - | 1255 | 309.2 | - | - | 0 | - |
| - | - | 958.7 | 309.2 | - | - | 0 | - |
| - | - | 5545 | 310.1 | - | - | 0 | - |
| - | - | 1598 | 310.2 | - | - | 0 | - |
| - | - | 1130 | 311.1 | - | - | 0 | - |
| - | - | 1639 | 311.2 | - | - | 0 | - |
| - | - | 3944 | 312.2 | - | - | 0 | - |
| 6 | b | 1266 | 312.2 | 0.0005248 | 1.681 | +2 | 6 |
| - | - | 2458 | 312.2 | - | - | 0 | - |
| - | - | 555.6 | 313.2 | - | - | 0 | - |
| - | - | 834.4 | 314.1 | - | - | 0 | - |
| 3 | b | 1.4E+04 | 314.2 | 0.0004529 | 1.441 | +1 | 3 |
| - | - | 6.811E+04 | 315.1 | - | - | 0 | - |
| - | - | 2686 | 315.2 | - | - | 0 | - |
| 7 | y | 7.445E+04 | 316.1 | 0.0003036 | 0.9603 | +1 | 2 |
| - | - | 9773 | 316.1 | - | - | 0 | - |
| - | - | 1.174E+04 | 317.1 | - | - | 0 | - |
| - | - | 1228 | 317.2 | - | - | 0 | - |
| - | - | 1494 | 318.1 | - | - | 0 | - |
| - | - | 1147 | 320.1 | - | - | 0 | - |
| 6 | b | 5083 | 321.2 | 0.0002964 | 0.9227 | +2 | 6 |
| - | - | 1378 | 321.7 | - | - | 0 | - |
| - | - | 637.1 | 326.2 | - | - | 0 | - |
| - | - | 7735 | 327.2 | - | - | 0 | - |
| - | - | 1.233E+04 | 328.2 | - | - | 0 | - |
| - | - | 823.2 | 328.2 | - | - | 0 | - |
| - | - | 1735 | 329.2 | - | - | 0 | - |
| - | - | 655.5 | 329.2 | - | - | 0 | - |
| - | - | 778.3 | 329.7 | - | - | 0 | - |
| - | - | 1690 | 330.2 | - | - | 0 | - |
| 7 | y | 3.592E+04 | 333.2 | 0.0004574 | 1.373 | +1 | 2 |
| - | - | 7064 | 334.2 | - | - | 0 | - |
| - | - | 8427 | 337.2 | - | - | 0 | - |
| - | - | 985.5 | 337.2 | - | - | 0 | - |
| - | - | 2702 | 338.1 | - | - | 0 | - |
| - | - | 648.4 | 340.1 | - | - | 0 | - |
| - | - | 4413 | 340.2 | - | - | 0 | - |
| - | - | 770.3 | 341.2 | - | - | 0 | - |
| - | - | 666.5 | 343.2 | - | - | 0 | - |
| - | - | 743.1 | 344.1 | - | - | 0 | - |
| - | - | 4025 | 345.2 | - | - | 0 | - |
| - | - | 1731 | 346.2 | - | - | 0 | - |
| - | - | 1205 | 351.2 | - | - | 0 | - |
| - | - | 5406 | 352.2 | - | - | 0 | - |
| - | - | 1242 | 353.2 | - | - | 0 | - |
| - | - | 1979 | 353.2 | - | - | 0 | - |
| - | - | 1.174E+05 | 355.2 | - | - | 0 | - |
| - | - | 1.965E+04 | 356.2 | - | - | 0 | - |
| - | - | 2579 | 357.2 | - | - | 0 | - |
| - | - | 753.1 | 359.1 | - | - | 0 | - |
| - | - | 590.4 | 361.2 | - | - | 0 | - |
| - | - | 2534 | 363.2 | - | - | 0 | - |
| - | - | 2548 | 365.2 | - | - | 0 | - |
| - | - | 7235 | 369.2 | - | - | 0 | - |
| - | - | 1.045E+04 | 370.2 | - | - | 0 | - |
| - | - | 2773 | 371.2 | - | - | 0 | - |
| - | - | 2163 | 372.2 | - | - | 0 | - |
| - | - | 557.8 | 372.2 | - | - | 0 | - |
| - | - | 4617 | 373.2 | - | - | 0 | - |
| - | - | 778 | 373.2 | - | - | 0 | - |
| - | - | 865 | 374.2 | - | - | 0 | - |
| - | - | 5277 | 379.2 | - | - | 0 | - |
| - | - | 1394 | 380.2 | - | - | 0 | - |
| - | - | 3109 | 381.2 | - | - | 0 | - |
| - | - | 693.6 | 381.2 | - | - | 0 | - |
| - | - | 2106 | 381.2 | - | - | 0 | - |
| - | - | 638.4 | 383.3 | - | - | 0 | - |
| - | - | 1.32E+04 | 387.3 | - | - | 0 | - |
| - | - | 2441 | 388.3 | - | - | 0 | - |
| - | - | 664.7 | 389.3 | - | - | 0 | - |
| - | - | 810.6 | 391.2 | - | - | 0 | - |
| - | - | 948.9 | 395.2 | - | - | 0 | - |
| - | - | 659.1 | 395.2 | - | - | 0 | - |
| 4 | b | 3.697E+04 | 397.2 | 0.0002654 | 0.6681 | +1 | 4 |
| - | - | 8375 | 398.2 | - | - | 0 | - |
| - | - | 582.8 | 399.2 | - | - | 0 | - |
| - | - | 1519 | 399.2 | - | - | 0 | - |
| - | - | 1079 | 399.3 | - | - | 0 | - |
| - | - | 554.5 | 405.2 | - | - | 0 | - |
| - | - | 605 | 407.2 | - | - | 0 | - |
| - | - | 1080 | 409.2 | - | - | 0 | - |
| - | - | 3650 | 409.2 | - | - | 0 | - |
| - | - | 847.9 | 410.2 | - | - | 0 | - |
| - | - | 883.5 | 411.2 | - | - | 0 | - |
| - | - | 1113 | 411.3 | - | - | 0 | - |
| - | - | 801 | 412.2 | - | - | 0 | - |
| 4 | b | 7.461E+04 | 415.3 | 0.0003208 | 0.7726 | +1 | 4 |
| - | - | 1.616E+04 | 416.3 | - | - | 0 | - |
| - | - | 2096 | 417.3 | - | - | 0 | - |
| - | - | 2286 | 423.2 | - | - | 0 | - |
| - | - | 1232 | 424.3 | - | - | 0 | - |
| - | - | 1.022E+04 | 427.2 | - | - | 0 | - |
| - | - | 2564 | 427.2 | - | - | 0 | - |
| - | - | 2006 | 428.2 | - | - | 0 | - |
| - | - | 1462 | 428.2 | - | - | 0 | - |
| - | - | 836.6 | 431.3 | - | - | 0 | - |
| - | - | 629.4 | 433.9 | - | - | 0 | - |
| - | - | 805.7 | 437.7 | - | - | 0 | - |
| - | - | 1940 | 438.2 | - | - | 0 | - |
| - | - | 3231 | 440.3 | - | - | 0 | - |
| - | - | 1815 | 441.2 | - | - | 0 | - |
| - | - | 1427 | 442.2 | - | - | 0 | - |
| - | - | 634.4 | 442.3 | - | - | 0 | - |
| 6 | y | 2808 | 444.2 | 0.0008691 | 1.957 | +1 | 3 |
| 6 | y | 5222 | 445.2 | 0.0005877 | 1.32 | +1 | 3 |
| - | - | 929.4 | 446.2 | - | - | 0 | - |
| - | - | 1161 | 446.7 | - | - | 0 | - |
| - | - | 1256 | 450.2 | - | - | 0 | - |
| - | - | 734.5 | 450.3 | - | - | 0 | - |
| - | - | 1007 | 451.2 | - | - | 0 | - |
| - | - | 9112 | 456.2 | - | - | 0 | - |
| - | - | 2231 | 457.2 | - | - | 0 | - |
| - | - | 690.6 | 458.3 | - | - | 0 | - |
| 6 | y | 3171 | 462.2 | 0.0005278 | 1.142 | +1 | 3 |
| - | - | 1427 | 466.3 | - | - | 0 | - |
| - | - | 8767 | 468.2 | - | - | 0 | - |
| - | - | 1290 | 468.3 | - | - | 0 | - |
| - | - | 1957 | 469.2 | - | - | 0 | - |
| - | - | 1166 | 469.7 | - | - | 0 | - |
| - | - | 1119 | 470.2 | - | - | 0 | - |
| - | - | 2521 | 472.2 | - | - | 0 | - |
| - | - | 904.4 | 473.2 | - | - | 0 | - |
| - | - | 605.8 | 473.2 | - | - | 0 | - |
| - | - | 1015 | 476.3 | - | - | 0 | - |
| - | - | 3122 | 478.2 | - | - | 0 | - |
| - | - | 627.1 | 481.2 | - | - | 0 | - |
| - | - | 2171 | 486.1 | - | - | 0 | - |
| - | - | 1362 | 486.3 | - | - | 0 | - |
| 0 | Precursor | 698.4 | 487.3 | 0.009016 | 18.5 | +2 | -1 |
| - | - | 1055 | 492.2 | - | - | 0 | - |
| 5 | b | 6018 | 494.3 | 0.0005411 | 1.095 | +1 | 5 |
| - | - | 1515 | 495.3 | - | - | 0 | - |
| - | - | 3082 | 496.2 | - | - | 0 | - |
| - | - | 995.7 | 499.3 | - | - | 0 | - |
| - | - | 669.2 | 500.3 | - | - | 0 | - |
| - | - | 2338 | 506.2 | - | - | 0 | - |
| - | - | 981.7 | 508.3 | - | - | 0 | - |
| - | - | 4633 | 510.3 | - | - | 0 | - |
| - | - | 1158 | 511.3 | - | - | 0 | - |
| 5 | b | 6819 | 512.3 | 7.486E-05 | 0.1461 | +1 | 5 |
| - | - | 1746 | 513.2 | - | - | 0 | - |
| - | - | 2329 | 513.3 | - | - | 0 | - |
| - | - | 808.3 | 514.2 | - | - | 0 | - |
| - | - | 1298 | 520.2 | - | - | 0 | - |
| - | - | 2.1E+04 | 524.2 | - | - | 0 | - |
| - | - | 6132 | 525.2 | - | - | 0 | - |
| - | - | 1001 | 525.3 | - | - | 0 | - |
| - | - | 761.8 | 526.2 | - | - | 0 | - |
| - | - | 1174 | 526.3 | - | - | 0 | - |
| - | - | 2142 | 527.3 | - | - | 0 | - |
| - | - | 1782 | 528.3 | - | - | 0 | - |
| - | - | 869.3 | 529.3 | - | - | 0 | - |
| - | - | 600.3 | 533.3 | - | - | 0 | - |
| - | - | 3260 | 537.3 | - | - | 0 | - |
| - | - | 676.9 | 540.3 | - | - | 0 | - |
| 5 | y | 9807 | 541.2 | 0.0001064 | 0.1966 | +1 | 4 |
| 5 | y | 3.579E+04 | 542.2 | 0.0004972 | 0.9169 | +1 | 4 |
| - | - | 1.028E+04 | 543.2 | - | - | 0 | - |
| - | - | 1071 | 543.3 | - | - | 0 | - |
| - | - | 1627 | 544.2 | - | - | 0 | - |
| - | - | 705.5 | 545.3 | - | - | 0 | - |
| - | - | 2967 | 551.3 | - | - | 0 | - |
| - | - | 704.1 | 552.3 | - | - | 0 | - |
| - | - | 3202 | 555.3 | - | - | 0 | - |
| 5 | y | 1.018E+05 | 559.3 | 1.008E-05 | 0.01802 | +1 | 4 |
| - | - | 3.19E+04 | 560.3 | - | - | 0 | - |
| - | - | 5775 | 561.3 | - | - | 0 | - |
| - | - | 1031 | 562.3 | - | - | 0 | - |
| - | - | 4348 | 569.3 | - | - | 0 | - |
| - | - | 1396 | 570.3 | - | - | 0 | - |
| - | - | 920.4 | 593.3 | - | - | 0 | - |
| - | - | 994.9 | 595.3 | - | - | 0 | - |
| - | - | 2623 | 605.3 | - | - | 0 | - |
| - | - | 589.9 | 606.3 | - | - | 0 | - |
| - | - | 949.8 | 607.2 | - | - | 0 | - |
| - | - | 783.8 | 608.3 | - | - | 0 | - |
| - | - | 1511 | 613.4 | - | - | 0 | - |
| - | - | 664.3 | 614.4 | - | - | 0 | - |
| - | - | 1723 | 620.3 | - | - | 0 | - |
| - | - | 1583 | 621.3 | - | - | 0 | - |
| - | - | 1172 | 622.4 | - | - | 0 | - |
| 6 | b | 9137 | 623.3 | 0.0003064 | 0.4915 | +1 | 6 |
| - | - | 3441 | 624.3 | - | - | 0 | - |
| - | - | 878.8 | 625.3 | - | - | 0 | - |
| - | - | 5335 | 638.3 | - | - | 0 | - |
| - | - | 2248 | 639.3 | - | - | 0 | - |
| - | - | 2719 | 640.4 | - | - | 0 | - |
| 6 | b | 1.025E+04 | 641.4 | 0.0005449 | 0.8497 | +1 | 6 |
| 4 | y | 1631 | 642.3 | 5.537E-05 | 0.08621 | +1 | 5 |
| - | - | 3680 | 642.4 | - | - | 0 | - |
| 4 | y | 1893 | 643.3 | 0.001403 | 2.181 | +1 | 5 |
| - | - | 3084 | 650.4 | - | - | 0 | - |
| - | - | 1437 | 651.3 | - | - | 0 | - |
| - | - | 4694 | 656.3 | - | - | 0 | - |
| - | - | 1302 | 657.3 | - | - | 0 | - |
| - | - | 5650 | 658.4 | - | - | 0 | - |
| - | - | 2635 | 659.4 | - | - | 0 | - |
| 4 | y | 1.14E+04 | 660.3 | 0.000183 | 0.2772 | +1 | 5 |
| - | - | 4061 | 661.3 | - | - | 0 | - |
| - | - | 1057 | 662.3 | - | - | 0 | - |
| - | - | 3023 | 668.4 | - | - | 0 | - |
| - | - | 1069 | 669.4 | - | - | 0 | - |
| - | - | 895.5 | 707.4 | - | - | 0 | - |
| - | - | 1247 | 716.4 | - | - | 0 | - |
| - | - | 1204 | 723.4 | - | - | 0 | - |
| - | - | 1490 | 724.4 | - | - | 0 | - |
| - | - | 1557 | 725.4 | - | - | 0 | - |
| - | - | 4187 | 733.4 | - | - | 0 | - |
| - | - | 3178 | 734.4 | - | - | 0 | - |
| - | - | 1412 | 735.4 | - | - | 0 | - |
| - | - | 702.6 | 736.4 | - | - | 0 | - |
| - | - | 2238 | 741.4 | - | - | 0 | - |
| - | - | 1029 | 742.4 | - | - | 0 | - |
| 7 | b | 2.151E+04 | 751.4 | 0.0003487 | 0.4641 | +1 | 7 |
| 7 | b | 1.081E+04 | 752.4 | 0.01417 | 18.83 | +1 | 7 |
| - | - | 3225 | 753.4 | - | - | 0 | - |
| 3 | y | 1205 | 755.4 | 0.002515 | 3.33 | +1 | 6 |
| 3 | y | 1023 | 756.4 | 0.005168 | 6.833 | +1 | 6 |
| 7 | b | 1.473E+04 | 769.4 | 0.0004764 | 0.6192 | +1 | 7 |
| - | - | 6205 | 770.4 | - | - | 0 | - |
| - | - | 1371 | 771.4 | - | - | 0 | - |
| 3 | y | 1640 | 773.4 | 7.954E-05 | 0.1029 | +1 | 6 |
| - | - | 891.4 | 774.4 | - | - | 0 | - |
| - | - | 734.8 | 797.4 | - | - | 0 | - |
| 2 | y | 983.5 | 842.4 | 0.002439 | 2.896 | +1 | 7 |
| 2 | y | 1352 | 860.4 | 0.00214 | 2.487 | +1 | 7 |
| - | - | 618.1 | 1625 | - | - | 0 | - |
| - | - | 790 | 1637 | - | - | 0 | - |

m/z Charge Intensity FragmentType MassShift Position
120.08114624023438 0 2120.0974
120.84408569335938 0 333.9008
125.07119750976562 0 1336.799
125.10763549804688 0 4457.865
126.09172821044922 0 1462.7856
126.35767364501953 0 388.31223
127.07575225830078 0 500.17847
127.08708190917969 0 419.97076
128.08262634277344 0 512.17456
128.10728454589844 0 2089.742
129.06617736816406 0 10925.476
129.10264587402344 0 1422.8115
130.05026245117188 0 3973.5942
130.06544494628906 0 25689.95
131.069091796875 0 2509.677
132.0811004638672 0 13351.558
132.10227966308594 0 1911.3588
133.08602905273438 0 5961.499
135.74765014648438 0 425.89548
136.07606506347656 0 676.13214
137.0586395263672 0 472.29648
137.07142639160156 0 471.76855
138.0917510986328 0 1108.8457
139.0506134033203 0 1280.4656
139.70379638671875 0 552.552
141.10226440429688 0 434.85245
142.06568908691406 0 847.54535
142.12298583984375 0 6312.503
143.1181640625 0 1046.1049
144.08111572265625 0 5737.6797
144.5458526611328 0 366.18335
145.08477783203125 0 422.82245
146.0603790283203 0 37350.504
147.06393432617188 0 3785.5571
147.07667541503906 0 2536.5762
148.0161895751953 0 417.66763
152.10733032226562 0 4190.2134
153.0660858154297 0 3054.1533
153.1025390625 0 3035.5688
154.0863037109375 0 532.80493
155.08193969726562 0 1479.8328
155.0989990234375 0 525.90295
155.11819458007812 0 5949.829 a Water loss 1
156.10208129882812 0 835.43134
156.12142944335938 0 569.482
158.0451202392578 0 878.9091
159.0919952392578 0 74533.695
160.07589721679688 0 1129.3195
160.09536743164062 0 7499.736
161.09767150878906 0 521.0364
161.33963012695312 0 430.63553
163.0868682861328 0 1068.7461
166.08633422851562 0 573.62244
169.076171875 0 943.39404
169.1338653564453 0 2518.5522
170.0603790283203 0 4232.902
171.11312866210938 0 8206.485
171.14988708496094 0 941.47424
173.1287841796875 0 53399.402 a 1
173.4522247314453 0 811.72266
174.0554656982422 0 862.7716
174.1321563720703 0 4806.6943
177.11236572265625 0 1976.1588
180.23751831054688 0 510.305
181.0974884033203 0 6545.119
182.10134887695312 0 631.6266
183.1130828857422 0 12149.124 b Water loss 1
183.14944458007812 0 4108.805
184.11659240722656 0 1686.972
187.07177734375 0 1058.3896
187.0869598388672 0 6248.248
187.1444854736328 0 15736.135
188.0709991455078 0 283704.7
188.1476593017578 0 1469.114
189.0742950439453 0 31192.611
190.0769500732422 0 1867.882
195.07669067382812 0 1338.1235
195.11343383789062 0 589.8129
196.1438446044922 0 676.3496
197.1287841796875 0 8417.041
198.1317138671875 0 865.5349
199.07228088378906 0 1448.2947
199.1079864501953 0 55645.54
200.11143493652344 0 5627.1733
200.1389923095703 0 584.1676
201.1022186279297 0 2557.6028
201.12362670898438 0 62091.31 b 1
202.12704467773438 0 5916.698
205.09747314453125 0 119468.13 y 7
206.10076904296875 0 13663.867
207.10198974609375 0 735.69666
209.09237670898438 0 6835.4805
210.09616088867188 0 751.0642
211.1079559326172 0 1136.2179
211.14450073242188 0 3425.7917
212.10324096679688 0 2210.1306
213.08700561523438 0 844.911
213.1230926513672 0 728.81006
214.1554412841797 0 1149.2039
215.0814208984375 0 545.61304
215.13934326171875 0 37806.91
216.14254760742188 0 3384.6375
217.08229064941406 0 2559.3865
222.12335205078125 0 512.84937
223.0718231201172 0 1203.6681
223.1085662841797 0 618.6334
225.10232543945312 0 3603.6028
225.12371826171875 0 1454.6934
226.10693359375 0 1008.7962
226.11892700195312 0 4022.732
227.10301208496094 0 85118.96
228.10623168945312 0 8681.3
229.10794067382812 0 1089.0665
229.15521240234375 0 1627.2527
230.1136474609375 0 949.88007
231.08676147460938 0 726.4524
231.9134979248047 0 484.19373
233.6561279296875 0 804.887
238.15443420410156 0 620.7122
239.13937377929688 0 4623.7993
240.09820556640625 0 11851.338
240.13401794433594 0 924.6625
240.1712646484375 0 700.0798
241.082275390625 0 3156.331
241.1015625 0 1220.3154
241.11807250976562 0 604.5808
242.1504669189453 0 1411.8108
242.66018676757812 0 2894.0745
243.13467407226562 0 1145.245
244.1295166015625 0 24232.227
245.11326599121094 0 1960.6638
245.13282775878906 0 2932.9126
248.1532745361328 0 544.3964
251.13906860351562 0 800.7373
251.1756134033203 0 2742.4485
252.11297607421875 0 1690.1057
252.1791229248047 0 540.4348
253.09735107421875 0 1898.5986
256.16583251953125 0 3775.6228
256.6574401855469 0 1330.8427 b 4
257.1497802734375 0 5941.69
258.1085510253906 0 9271.563
258.1456604003906 0 1868.5654
259.1112976074219 0 595.8308
264.134765625 0 1379.5964
265.1206970214844 0 541.45184
266.1134948730469 0 978.51886
266.1501770019531 0 2252.386
267.1339111328125 0 3100.0156
267.1482238769531 0 543.2048
268.2021789550781 0 2434.327 a Water loss 2
269.1401062011719 0 2699.971
269.18603515625 0 2442.9038
270.1239318847656 0 29472.197
271.1107177734375 0 877.7253
271.12725830078125 0 4022.053 y Water loss 4
272.1611328125 0 656.32416
274.176025390625 0 1951.9445
278.18731689453125 0 709.36255
280.1084899902344 0 3779.2305
280.1661682128906 0 2455.2083
281.1128234863281 0 785.1852
282.1446533203125 0 1764.0769
283.1292724609375 0 917.0923
284.1607971191406 0 34136.223
285.1475830078125 0 958.8303
285.1639709472656 0 3586.3054
286.103271484375 0 729.6488
286.1395263671875 0 830.16345
286.2127380371094 0 620.6746 a 2
287.1503601074219 0 994.396
292.1300964355469 0 1917.9694
294.1446228027344 0 1259.4517
294.18145751953125 0 1537.284
296.124755859375 0 832.945
296.197265625 0 62735.94 b Water loss 2
297.1342468261719 0 1106.0592
297.2005310058594 0 11476.107
298.1189270019531 0 9146.232
298.1764221191406 0 3992.1301
298.20220947265625 0 918.7721
299.1034240722656 0 1095.2816
299.12213134765625 0 1157.46
299.1773376464844 0 546.6842
300.155517578125 0 2904.427
302.17144775390625 0 36006.38
303.1746520996094 0 7000.753
307.18096923828125 0 1230.2635
309.15594482421875 0 1255.091
309.2041320800781 0 958.6801
310.1400451660156 0 5545.305
310.2115173339844 0 1597.7059
311.14422607421875 0 1130.0656
311.1710510253906 0 1638.8896
312.15576171875 0 3943.5166
312.1730651855469 0 1266.4208 b Water loss 5
312.1921691894531 0 2457.984
313.1593933105469 0 555.5556
314.1348571777344 0 834.35333
314.2078857421875 0 13995.114 b 2
315.1455383300781 0 68108.055
315.2111511230469 0 2685.5522
316.1294860839844 0 74450.12 y Ammonia loss 6
316.1485595703125 0 9773.484
317.13275146484375 0 11736.839
317.1527099609375 0 1227.6765
318.1349792480469 0 1493.8492
320.12506103515625 0 1147.1915
321.1791687011719 0 5083.2446 b 5
321.68084716796875 0 1377.6327
326.1715393066406 0 637.1266
327.16668701171875 0 7734.96
328.15057373046875 0 12326.509
328.16888427734375 0 823.2243
329.1541442871094 0 1734.8418
329.1800842285156 0 655.46625
329.6927490234375 0 778.293
330.166015625 0 1689.9773
333.15618896484375 0 35919.832 y 6
334.1593322753906 0 7064.0977
337.1508483886719 0 8427.129
337.18707275390625 0 985.5032
338.1351623535156 0 2701.9048
340.10888671875 0 648.39984
340.1870422363281 0 4412.747
341.1891174316406 0 770.25006
343.2333984375 0 666.4971
344.1197204589844 0 743.0918
345.1769714355469 0 4025.039
346.1784362792969 0 1731.3428
351.240234375 0 1204.7661
352.22344970703125 0 5405.7085
353.18212890625 0 1241.5029
353.2201843261719 0 1979.1024
355.1614074707031 0 117383.77
356.1644592285156 0 19650.207
357.1671447753906 0 2578.9888
359.0926208496094 0 753.08136
361.1728515625 0 590.38513
363.20263671875 0 2533.5527
365.1820068359375 0 2547.8298
369.2496643066406 0 7234.7344
370.2336730957031 0 10451.984
371.2317810058594 0 2772.505
372.1878967285156 0 2163.1206
372.2328796386719 0 557.7748
373.1719665527344 0 4616.6665
373.19793701171875 0 778.0015
374.1756896972656 0 864.9984
379.2342529296875 0 5277.4023
380.2378845214844 0 1394.3402
381.1554870605469 0 3108.7754
381.18157958984375 0 693.64526
381.21417236328125 0 2105.959
383.2662048339844 0 638.401
387.2603759765625 0 13199.995
388.2632141113281 0 2441.0647
389.2669982910156 0 664.70483
391.195068359375 0 810.5909
395.1566162109375 0 948.9393
395.1942138671875 0 659.1399
397.24481201171875 0 36974.05 b Water loss 3
398.2476501464844 0 8375.422
399.1904296875 0 582.8479
399.2237854003906 0 1518.6598
399.2510681152344 0 1078.6573
405.21435546875 0 554.5253
407.1571350097656 0 604.98517
409.15093994140625 0 1080.3987
409.2083435058594 0 3649.6758
410.2113342285156 0 847.8838
411.18817138671875 0 883.45966
411.2602233886719 0 1113.47
412.1846923828125 0 801.0408
415.25543212890625 0 74609.99 b 3
416.25872802734375 0 16159.485
417.2598571777344 0 2095.5803
423.2243347167969 0 2285.9448
424.2548828125 0 1231.9133
427.1616516113281 0 10218.65
427.2193298339844 0 2563.7202
428.16363525390625 0 2005.8662
428.2186279296875 0 1461.7388
431.2629699707031 0 836.5544
433.9101867675781 0 629.43866
437.727294921875 0 805.708
438.1991271972656 0 1940.4664
440.2510070800781 0 3230.8762
441.23431396484375 0 1814.5032
442.1930236816406 0 1427.4653
442.2671203613281 0 634.42444
444.1886291503906 0 2807.9768 y Water loss 5
445.17236328125 0 5222.004 y Ammonia loss 5
446.1761474609375 0 929.3535
446.73095703125 0 1161.3502
450.2344970703125 0 1256.2628
450.2720642089844 0 734.48944
451.21856689453125 0 1007.4158
456.2092590332031 0 9112.206
457.2123718261719 0 2230.7007
458.2616271972656 0 690.56415
462.1988525390625 0 3171.441 y 5
466.30120849609375 0 1426.917
468.2455139160156 0 8767.483
468.28076171875 0 1289.6654
469.24798583984375 0 1956.8668
469.73272705078125 0 1165.7053
470.23724365234375 0 1119.4503
472.18316650390625 0 2521.2698
473.188720703125 0 904.4138
473.2336120605469 0 605.8169
476.2858581542969 0 1015.2511
478.2087097167969 0 3122.4912
481.1974182128906 0 627.1473
486.1496276855469 0 2171.4478
486.2568359375 0 1361.5483
487.2621154785156 0 698.4 Precursor
492.2453308105469 0 1054.7839
494.2978515625 0 6018.092 b Water loss 4
495.3000183105469 0 1515.1766
496.22003173828125 0 3081.6636
499.2773132324219 0 995.70526
500.2716369628906 0 669.2022
506.20306396484375 0 2338.1584
508.2782287597656 0 981.65656
510.2560729980469 0 4632.687
511.2606201171875 0 1157.5284
512.3078002929688 0 6819.2544 b 4
513.2446899414062 0 1745.8694
513.3111572265625 0 2329.387
514.2327270507812 0 808.3206
520.2396850585938 0 1298.4674
524.2139282226562 0 20998.264
525.2174682617188 0 6132.4497
525.3043823242188 0 1001.418
526.2188720703125 0 761.8077
526.2864990234375 0 1174.2305
527.2827758789062 0 2141.8564
528.2684936523438 0 1782.336
529.2700805664062 0 869.299
533.2708129882812 0 600.2591
537.2667236328125 0 3260.3052
540.3011474609375 0 676.91205
541.2404174804688 0 9806.602 y Water loss 4
542.2250366210938 0 35788.957 y Ammonia loss 4
543.2277221679688 0 10280.448
543.3109130859375 0 1071.486
544.229736328125 0 1626.5024
545.2907104492188 0 705.52295
551.2818603515625 0 2966.708
552.2871704101562 0 704.12115
555.2774047851562 0 3202.2065
559.2510986328125 0 101758.35 y 4
560.2540283203125 0 31901.273
561.2557373046875 0 5774.6807
562.2991943359375 0 1031.2437
569.2927856445312 0 4347.528
570.2953491210938 0 1395.9904
593.2939453125 0 920.43634
595.3462524414062 0 994.9164
605.328857421875 0 2623.3875
606.3272094726562 0 589.92865
607.2461547851562 0 949.7783
608.2553100585938 0 783.81256
613.3550415039062 0 1511.1554
614.35302734375 0 664.2701
620.3021240234375 0 1722.5298
621.2918701171875 0 1583.3413
622.3547973632812 0 1171.5433
623.3402099609375 0 9136.644 b Water loss 5
624.3421630859375 0 3440.6868
625.2579956054688 0 878.81177
638.3139038085938 0 5334.6167
639.3123779296875 0 2247.755
640.3659057617188 0 2719.1394
641.3510131835938 0 10245.1875 b 5
642.2881469726562 0 1631.2773 y Water loss 3
642.3541259765625 0 3680.1804
643.2736206054688 0 1893.0092 y Ammonia loss 3
650.3507690429688 0 3083.67
651.3497924804688 0 1436.8352
656.3253173828125 0 4693.905
657.32763671875 0 1301.8783
658.3768310546875 0 5650.203
659.378662109375 0 2634.5447
660.298583984375 0 11403.897 y 3
661.302490234375 0 4060.7644
662.303955078125 0 1057.3336
668.361572265625 0 3023.2866
669.3619995117188 0 1069.4362
707.3739624023438 0 895.5248
716.3609008789062 0 1247.389
723.4049682617188 0 1203.7506
724.3964233398438 0 1490.461
725.386474609375 0 1556.8802
733.3871459960938 0 4186.9326
734.3770141601562 0 3177.9563
735.3777465820312 0 1411.5374
736.3821411132812 0 702.6433
741.4134521484375 0 2238.288
742.4202880859375 0 1029.4988
751.3981323242188 0 21510.8 b Water loss 6
752.3966674804688 0 10814.156 b Ammonia loss 6
753.396728515625 0 3224.7092
755.3697509765625 0 1204.5444 y Water loss 2
756.3614501953125 0 1023.48834 y Ammonia loss 2
769.4085693359375 0 14726.074 b 6
770.4114990234375 0 6205.0464
771.41357421875 0 1370.6628
773.3827514648438 0 1640.3938 y 2
774.3850708007812 0 891.449
797.4212036132812 0 734.83124
842.40185546875 0 983.4722 y Water loss 1
860.4127197265625 0 1351.635 y 1
1625.107177734375 0 618.14343
1636.98828125 0 789.99396

Spectrum Details

|  |  |
| --- | --- |
| Matched peaks? Matched peaks The total absolute number of peaks matched. Additionally in brackets the total fraction of peaks matched and the total number of peaks is shown. | 39 (9.38% of 416) |
| FDR? FDR The false discovery rate estimated for this peptide. It is calculated by matching all theoretical fragments with a non-integer shift with the raw peaks for this spectrum. This is done with 40 different shifts. The resulting percentage is the average number of annotated peaks over the number of annotated peaks with the correct spectrum. | 1.34% |
| Satellite FDR? Satellite FDR See the FDR for details on its calculation. This satellite ion specific FDR only contains the satellite ions (d/w) for I/L/J positions. | - |
| PSM Score? PSM Score The PSM Score as given by Hecklib to this annotated spectrum. It is shown with three significant figures. | 631 |

#### Spectrum 9975? Spectrum 9975 The raw spectrum of this peptide as annotated by Hecklib. The fragments are coloured according to ion type (see legend). Any peaks with a star '\*' as text can be hovered over to see the full details, first the ion type second the mass shift type. By hovering over the amino acids in the peptide or ions in the legend the corresponding peaks are highlighted. By toggling the 'Unassigned' label you can turn the background (unassigned) peaks on or off in the plot. By updating the slider in the Ion legend you can update the spectrum to only show the top X% of the peaks with labels. The top X% means any peak that is within X% of the highest intensity. By dragging in the spectrum you can zoom in to a specific part of the spectrum and use 'Zoom Out' to get back to the original zoom level. The annotation of the spectrum is based on the given sequence in the peptides file and is done with different software so inconsistencies are likely. The peaks are annotated based on the given sequence, with 20 ppm tolerance.

Copy Data

##### Spectrum 9975 (TSV)

###### Preview

```
Loading example...
```

*Click on the button to copy the data to your clipboard.*

Mz MinMz MaxIntensity Max

WidthHeightPeptide font sizePeptide stroke widthSpectrum font sizeSpectrum stroke widthCompact peptide

Ion legend

wxyz

abcd

OtherUnassignedIon^Charge^_Position_Show for top:%

JSJTPEQW

04.55e+49.10e+41.37e+51.82e+5

Zoom Out

a^+1^_2_y^+2^_2_a^+1^_2_b^+1^_2_b^+1^_2_y^+1^_1_b^+2^_5_b^+2^_5_a^+1^_3_y^+2^_4_b^+1^_3_b^+1^_3_y^+1^_2_b^+2^_6_y^+1^_2_b^+1^_4_b^+1^_4_y^+1^_3_y^+1^_3_y^+1^_3_b^+1^_5_b^+1^_5_y^+1^_4_y^+1^_4_y^+1^_4_b^+1^_6_b^+1^_6_y^+1^_5_y^+1^_5_b^+1^_7_b^+1^_7_b^+1^_7_y^+1^_6_y^+1^_7_

0796159323893186

Fragment Matches Table

Show background peaks

| Position | Ion type | Intensity | mz Theoretical | mz Error (Th) | mz Error (ppm) | Charge | Series Number |
| --- | --- | --- | --- | --- | --- | --- | --- |
| - | - | 2857 | 120.1 | - | - | 0 | - |
| - | - | 321.9 | 122 | - | - | 0 | - |
| - | - | 1393 | 125.1 | - | - | 0 | - |
| - | - | 3179 | 125.1 | - | - | 0 | - |
| - | - | 1350 | 126.1 | - | - | 0 | - |
| - | - | 535.5 | 127.1 | - | - | 0 | - |
| - | - | 1410 | 128.1 | - | - | 0 | - |
| - | - | 7573 | 129.1 | - | - | 0 | - |
| - | - | 1776 | 129.1 | - | - | 0 | - |
| - | - | 1583 | 130.1 | - | - | 0 | - |
| - | - | 1.655E+04 | 130.1 | - | - | 0 | - |
| - | - | 1725 | 131.1 | - | - | 0 | - |
| - | - | 887.5 | 132.1 | - | - | 0 | - |
| - | - | 8956 | 132.1 | - | - | 0 | - |
| - | - | 1397 | 132.1 | - | - | 0 | - |
| - | - | 1.048E+04 | 133.1 | - | - | 0 | - |
| - | - | 490.2 | 135.1 | - | - | 0 | - |
| - | - | 1314 | 136.1 | - | - | 0 | - |
| - | - | 1203 | 139.1 | - | - | 0 | - |
| - | - | 432.2 | 139.1 | - | - | 0 | - |
| - | - | 506.7 | 142.1 | - | - | 0 | - |
| - | - | 3823 | 142.1 | - | - | 0 | - |
| - | - | 457.9 | 143.1 | - | - | 0 | - |
| - | - | 520.2 | 143.1 | - | - | 0 | - |
| - | - | 454.2 | 144 | - | - | 0 | - |
| - | - | 4118 | 144.1 | - | - | 0 | - |
| - | - | 1025 | 145.1 | - | - | 0 | - |
| - | - | 2.369E+04 | 146.1 | - | - | 0 | - |
| - | - | 1125 | 146.1 | - | - | 0 | - |
| - | - | 2119 | 147.1 | - | - | 0 | - |
| - | - | 1640 | 147.1 | - | - | 0 | - |
| - | - | 536.1 | 147.1 | - | - | 0 | - |
| - | - | 462.7 | 149.3 | - | - | 0 | - |
| - | - | 2521 | 152.1 | - | - | 0 | - |
| - | - | 420.9 | 152.5 | - | - | 0 | - |
| - | - | 2325 | 153.1 | - | - | 0 | - |
| - | - | 1531 | 153.1 | - | - | 0 | - |
| - | - | 1292 | 155.1 | - | - | 0 | - |
| - | - | 442.2 | 155.1 | - | - | 0 | - |
| 2 | a | 4299 | 155.1 | 0.0003203 | 2.065 | +1 | 2 |
| - | - | 4.874E+04 | 159.1 | - | - | 0 | - |
| - | - | 5420 | 160.1 | - | - | 0 | - |
| - | - | 435.9 | 167.1 | - | - | 0 | - |
| 7 | y | 954.1 | 167.1 | 0.000161 | 0.9637 | +2 | 2 |
| - | - | 1277 | 169.1 | - | - | 0 | - |
| - | - | 1813 | 169.1 | - | - | 0 | - |
| - | - | 3506 | 170.1 | - | - | 0 | - |
| - | - | 451.7 | 170.7 | - | - | 0 | - |
| - | - | 4743 | 171.1 | - | - | 0 | - |
| - | - | 1062 | 171.1 | - | - | 0 | - |
| 2 | a | 3.341E+04 | 173.1 | 0.0003452 | 1.994 | +1 | 2 |
| - | - | 763.8 | 174.1 | - | - | 0 | - |
| - | - | 2584 | 174.1 | - | - | 0 | - |
| - | - | 446.2 | 177.1 | - | - | 0 | - |
| - | - | 2741 | 177.1 | - | - | 0 | - |
| - | - | 518.5 | 178.1 | - | - | 0 | - |
| - | - | 4713 | 181.1 | - | - | 0 | - |
| - | - | 783 | 183.1 | - | - | 0 | - |
| 2 | b | 8833 | 183.1 | 0.0002787 | 1.522 | +1 | 2 |
| - | - | 2125 | 183.1 | - | - | 0 | - |
| - | - | 1009 | 184.1 | - | - | 0 | - |
| - | - | 1108 | 185.1 | - | - | 0 | - |
| - | - | 654.4 | 187.1 | - | - | 0 | - |
| - | - | 3766 | 187.1 | - | - | 0 | - |
| - | - | 9564 | 187.1 | - | - | 0 | - |
| - | - | 1.802E+05 | 188.1 | - | - | 0 | - |
| - | - | 743.7 | 188.1 | - | - | 0 | - |
| - | - | 2.109E+04 | 189.1 | - | - | 0 | - |
| - | - | 629.9 | 189.1 | - | - | 0 | - |
| - | - | 912.3 | 190.1 | - | - | 0 | - |
| - | - | 565.4 | 191.1 | - | - | 0 | - |
| - | - | 678.6 | 194.1 | - | - | 0 | - |
| - | - | 570.2 | 195.1 | - | - | 0 | - |
| - | - | 1178 | 195.1 | - | - | 0 | - |
| - | - | 4971 | 197.1 | - | - | 0 | - |
| - | - | 1312 | 199.1 | - | - | 0 | - |
| - | - | 3.765E+04 | 199.1 | - | - | 0 | - |
| - | - | 2872 | 200.1 | - | - | 0 | - |
| - | - | 3344 | 201.1 | - | - | 0 | - |
| 2 | b | 4.025E+04 | 201.1 | 0.0002731 | 1.358 | +1 | 2 |
| - | - | 724 | 202.1 | - | - | 0 | - |
| - | - | 3776 | 202.1 | - | - | 0 | - |
| 8 | y | 7.547E+04 | 205.1 | 0.0002885 | 1.407 | +1 | 1 |
| - | - | 7759 | 206.1 | - | - | 0 | - |
| - | - | 509.5 | 207.1 | - | - | 0 | - |
| - | - | 3509 | 209.1 | - | - | 0 | - |
| - | - | 770.4 | 211.1 | - | - | 0 | - |
| - | - | 2303 | 211.1 | - | - | 0 | - |
| - | - | 888.8 | 212.1 | - | - | 0 | - |
| - | - | 800.2 | 213.1 | - | - | 0 | - |
| - | - | 651.6 | 213.1 | - | - | 0 | - |
| - | - | 890.5 | 214.2 | - | - | 0 | - |
| - | - | 2.501E+04 | 215.1 | - | - | 0 | - |
| - | - | 506.7 | 216.1 | - | - | 0 | - |
| - | - | 1985 | 216.1 | - | - | 0 | - |
| - | - | 1528 | 217.1 | - | - | 0 | - |
| - | - | 646.7 | 221.1 | - | - | 0 | - |
| - | - | 1019 | 223.1 | - | - | 0 | - |
| - | - | 1245 | 224.1 | - | - | 0 | - |
| - | - | 1859 | 225.1 | - | - | 0 | - |
| - | - | 1201 | 225.1 | - | - | 0 | - |
| - | - | 1678 | 226.1 | - | - | 0 | - |
| - | - | 5.606E+04 | 227.1 | - | - | 0 | - |
| - | - | 5506 | 228.1 | - | - | 0 | - |
| - | - | 1384 | 229.2 | - | - | 0 | - |
| - | - | 647.1 | 230.1 | - | - | 0 | - |
| - | - | 572.7 | 235.7 | - | - | 0 | - |
| - | - | 638.5 | 238.2 | - | - | 0 | - |
| - | - | 2432 | 239.1 | - | - | 0 | - |
| - | - | 6683 | 240.1 | - | - | 0 | - |
| - | - | 2251 | 241.1 | - | - | 0 | - |
| - | - | 560.5 | 241.1 | - | - | 0 | - |
| - | - | 568 | 242.1 | - | - | 0 | - |
| - | - | 1002 | 242.2 | - | - | 0 | - |
| - | - | 1995 | 242.7 | - | - | 0 | - |
| - | - | 1.709E+04 | 244.1 | - | - | 0 | - |
| - | - | 1138 | 245.1 | - | - | 0 | - |
| - | - | 1900 | 245.1 | - | - | 0 | - |
| 5 | b | 547.9 | 247.7 | 0.0003159 | 1.276 | +2 | 5 |
| - | - | 593.7 | 251.1 | - | - | 0 | - |
| - | - | 1900 | 251.2 | - | - | 0 | - |
| - | - | 878 | 252.1 | - | - | 0 | - |
| - | - | 939.7 | 253.1 | - | - | 0 | - |
| - | - | 628.9 | 254.1 | - | - | 0 | - |
| - | - | 671.8 | 255.1 | - | - | 0 | - |
| - | - | 1685 | 256.2 | - | - | 0 | - |
| 5 | b | 585.1 | 256.7 | 0.0008104 | 3.158 | +2 | 5 |
| - | - | 3391 | 257.1 | - | - | 0 | - |
| - | - | 7036 | 258.1 | - | - | 0 | - |
| - | - | 1534 | 258.1 | - | - | 0 | - |
| - | - | 631.4 | 264.1 | - | - | 0 | - |
| - | - | 586.5 | 265.1 | - | - | 0 | - |
| - | - | 1097 | 266.1 | - | - | 0 | - |
| - | - | 1716 | 266.2 | - | - | 0 | - |
| - | - | 1166 | 267.1 | - | - | 0 | - |
| 3 | a | 807.8 | 268.2 | 1.182E-05 | 0.04406 | +1 | 3 |
| - | - | 1986 | 269.1 | - | - | 0 | - |
| - | - | 1260 | 269.2 | - | - | 0 | - |
| - | - | 2.014E+04 | 270.1 | - | - | 0 | - |
| 5 | y | 2422 | 271.1 | 0.00287 | 10.59 | +2 | 4 |
| - | - | 884.8 | 274.2 | - | - | 0 | - |
| - | - | 3068 | 280.1 | - | - | 0 | - |
| - | - | 2201 | 280.2 | - | - | 0 | - |
| - | - | 683.2 | 281.1 | - | - | 0 | - |
| - | - | 1568 | 282.1 | - | - | 0 | - |
| - | - | 806.8 | 283.2 | - | - | 0 | - |
| - | - | 742.5 | 284.1 | - | - | 0 | - |
| - | - | 974.6 | 284.1 | - | - | 0 | - |
| - | - | 1.924E+04 | 284.2 | - | - | 0 | - |
| - | - | 846.8 | 285.1 | - | - | 0 | - |
| - | - | 2396 | 285.2 | - | - | 0 | - |
| - | - | 849.8 | 287.2 | - | - | 0 | - |
| - | - | 1585 | 292.1 | - | - | 0 | - |
| - | - | 1254 | 294.1 | - | - | 0 | - |
| - | - | 1002 | 294.2 | - | - | 0 | - |
| - | - | 772.7 | 296.1 | - | - | 0 | - |
| 3 | b | 4.009E+04 | 296.2 | 0.0002754 | 0.9298 | +1 | 3 |
| - | - | 1184 | 297.1 | - | - | 0 | - |
| - | - | 6767 | 297.2 | - | - | 0 | - |
| - | - | 5330 | 298.1 | - | - | 0 | - |
| - | - | 2937 | 298.2 | - | - | 0 | - |
| - | - | 749.8 | 298.2 | - | - | 0 | - |
| - | - | 549.8 | 298.6 | - | - | 0 | - |
| - | - | 1037 | 299.1 | - | - | 0 | - |
| - | - | 2221 | 300.2 | - | - | 0 | - |
| - | - | 586.6 | 301.1 | - | - | 0 | - |
| - | - | 2.087E+04 | 302.2 | - | - | 0 | - |
| - | - | 3373 | 303.2 | - | - | 0 | - |
| - | - | 920 | 307.2 | - | - | 0 | - |
| - | - | 1603 | 309.2 | - | - | 0 | - |
| - | - | 1222 | 309.2 | - | - | 0 | - |
| - | - | 3243 | 310.1 | - | - | 0 | - |
| - | - | 705.1 | 310.2 | - | - | 0 | - |
| - | - | 949.3 | 311.2 | - | - | 0 | - |
| - | - | 2553 | 312.2 | - | - | 0 | - |
| - | - | 1551 | 312.2 | - | - | 0 | - |
| - | - | 594.4 | 314.1 | - | - | 0 | - |
| 3 | b | 8469 | 314.2 | 0.0002088 | 0.6645 | +1 | 3 |
| - | - | 4.299E+04 | 315.1 | - | - | 0 | - |
| - | - | 1017 | 315.2 | - | - | 0 | - |
| 7 | y | 4.472E+04 | 316.1 | 0.0002425 | 0.7672 | +1 | 2 |
| - | - | 6985 | 316.1 | - | - | 0 | - |
| - | - | 7841 | 317.1 | - | - | 0 | - |
| - | - | 1035 | 318.1 | - | - | 0 | - |
| - | - | 1129 | 320.1 | - | - | 0 | - |
| 6 | b | 2600 | 321.2 | 0.0001309 | 0.4075 | +2 | 6 |
| - | - | 787.2 | 321.7 | - | - | 0 | - |
| - | - | 581.3 | 323.2 | - | - | 0 | - |
| - | - | 877.8 | 326.2 | - | - | 0 | - |
| - | - | 4260 | 327.2 | - | - | 0 | - |
| - | - | 6919 | 328.2 | - | - | 0 | - |
| - | - | 684.3 | 328.2 | - | - | 0 | - |
| - | - | 1420 | 329.2 | - | - | 0 | - |
| - | - | 708.2 | 329.2 | - | - | 0 | - |
| - | - | 1344 | 329.7 | - | - | 0 | - |
| 7 | y | 2.16E+04 | 333.2 | 0.0002743 | 0.8232 | +1 | 2 |
| - | - | 3643 | 334.2 | - | - | 0 | - |
| - | - | 594.7 | 334.2 | - | - | 0 | - |
| - | - | 4134 | 337.2 | - | - | 0 | - |
| - | - | 1256 | 338.1 | - | - | 0 | - |
| - | - | 689 | 338.2 | - | - | 0 | - |
| - | - | 2500 | 340.2 | - | - | 0 | - |
| - | - | 2657 | 345.2 | - | - | 0 | - |
| - | - | 621.4 | 346.2 | - | - | 0 | - |
| - | - | 872.4 | 351.2 | - | - | 0 | - |
| - | - | 3363 | 352.2 | - | - | 0 | - |
| - | - | 785.3 | 353.2 | - | - | 0 | - |
| - | - | 701.5 | 353.2 | - | - | 0 | - |
| - | - | 7.487E+04 | 355.2 | - | - | 0 | - |
| - | - | 1227 | 356.1 | - | - | 0 | - |
| - | - | 1.155E+04 | 356.2 | - | - | 0 | - |
| - | - | 1041 | 357.2 | - | - | 0 | - |
| - | - | 567.4 | 359.4 | - | - | 0 | - |
| - | - | 559.3 | 360.9 | - | - | 0 | - |
| - | - | 1366 | 363.2 | - | - | 0 | - |
| - | - | 1458 | 365.2 | - | - | 0 | - |
| - | - | 4501 | 369.2 | - | - | 0 | - |
| - | - | 5849 | 370.2 | - | - | 0 | - |
| - | - | 625.3 | 370.3 | - | - | 0 | - |
| - | - | 2198 | 371.2 | - | - | 0 | - |
| - | - | 1162 | 372.2 | - | - | 0 | - |
| - | - | 2939 | 373.2 | - | - | 0 | - |
| - | - | 2474 | 379.2 | - | - | 0 | - |
| - | - | 1547 | 381.2 | - | - | 0 | - |
| - | - | 1249 | 381.2 | - | - | 0 | - |
| - | - | 7188 | 387.3 | - | - | 0 | - |
| - | - | 1992 | 388.3 | - | - | 0 | - |
| 4 | b | 2.318E+04 | 397.2 | 8.229E-05 | 0.2072 | +1 | 4 |
| - | - | 1106 | 397.3 | - | - | 0 | - |
| - | - | 4560 | 398.2 | - | - | 0 | - |
| - | - | 1189 | 399.2 | - | - | 0 | - |
| - | - | 800.5 | 399.3 | - | - | 0 | - |
| - | - | 946.9 | 409.2 | - | - | 0 | - |
| - | - | 1947 | 409.2 | - | - | 0 | - |
| - | - | 731.7 | 411.2 | - | - | 0 | - |
| 4 | b | 4.486E+04 | 415.3 | 4.617E-05 | 0.1112 | +1 | 4 |
| - | - | 9749 | 416.3 | - | - | 0 | - |
| - | - | 945.3 | 417.2 | - | - | 0 | - |
| - | - | 1556 | 417.3 | - | - | 0 | - |
| - | - | 1875 | 423.2 | - | - | 0 | - |
| - | - | 5687 | 427.2 | - | - | 0 | - |
| - | - | 993.5 | 427.2 | - | - | 0 | - |
| - | - | 1358 | 428.2 | - | - | 0 | - |
| - | - | 754.3 | 431.3 | - | - | 0 | - |
| - | - | 1301 | 438.2 | - | - | 0 | - |
| - | - | 1403 | 440.3 | - | - | 0 | - |
| - | - | 965.2 | 441.2 | - | - | 0 | - |
| 6 | y | 1461 | 444.2 | 0.0002893 | 0.6513 | +1 | 3 |
| 6 | y | 4039 | 445.2 | 0.0001142 | 0.2566 | +1 | 3 |
| - | - | 1147 | 446.2 | - | - | 0 | - |
| - | - | 793.1 | 450.2 | - | - | 0 | - |
| - | - | 795.8 | 451.2 | - | - | 0 | - |
| - | - | 6345 | 456.2 | - | - | 0 | - |
| - | - | 1348 | 457.2 | - | - | 0 | - |
| - | - | 1148 | 460.3 | - | - | 0 | - |
| 6 | y | 1200 | 462.2 | 0.001108 | 2.397 | +1 | 3 |
| - | - | 4556 | 468.2 | - | - | 0 | - |
| - | - | 1039 | 468.3 | - | - | 0 | - |
| - | - | 1522 | 469.2 | - | - | 0 | - |
| - | - | 625 | 469.7 | - | - | 0 | - |
| - | - | 2392 | 470.2 | - | - | 0 | - |
| - | - | 690.9 | 471.3 | - | - | 0 | - |
| - | - | 1169 | 472.2 | - | - | 0 | - |
| - | - | 1178 | 476.3 | - | - | 0 | - |
| - | - | 2120 | 478.2 | - | - | 0 | - |
| - | - | 731.4 | 481.3 | - | - | 0 | - |
| - | - | 633.6 | 484.3 | - | - | 0 | - |
| - | - | 748.5 | 488.2 | - | - | 0 | - |
| - | - | 864.6 | 492.2 | - | - | 0 | - |
| 5 | b | 3774 | 494.3 | 9.977E-05 | 0.2018 | +1 | 5 |
| - | - | 1743 | 495.3 | - | - | 0 | - |
| - | - | 1185 | 496.2 | - | - | 0 | - |
| - | - | 1164 | 499.3 | - | - | 0 | - |
| - | - | 687.7 | 500.3 | - | - | 0 | - |
| - | - | 1341 | 506.2 | - | - | 0 | - |
| - | - | 1097 | 508.3 | - | - | 0 | - |
| - | - | 2241 | 510.3 | - | - | 0 | - |
| 5 | b | 4817 | 512.3 | 0.0004745 | 0.9261 | +1 | 5 |
| - | - | 859.5 | 513.2 | - | - | 0 | - |
| - | - | 933.1 | 513.3 | - | - | 0 | - |
| - | - | 608.2 | 523.3 | - | - | 0 | - |
| - | - | 1.307E+04 | 524.2 | - | - | 0 | - |
| - | - | 3351 | 525.2 | - | - | 0 | - |
| - | - | 683.4 | 525.3 | - | - | 0 | - |
| - | - | 857.3 | 526.2 | - | - | 0 | - |
| - | - | 1475 | 526.3 | - | - | 0 | - |
| - | - | 974.8 | 528.3 | - | - | 0 | - |
| - | - | 1743 | 537.3 | - | - | 0 | - |
| 5 | y | 5566 | 541.2 | 0.0005947 | 1.099 | +1 | 4 |
| - | - | 1001 | 541.3 | - | - | 0 | - |
| 5 | y | 2.043E+04 | 542.2 | 0.000253 | 0.4666 | +1 | 4 |
| - | - | 7170 | 543.2 | - | - | 0 | - |
| - | - | 1249 | 544.2 | - | - | 0 | - |
| - | - | 2239 | 551.3 | - | - | 0 | - |
| - | - | 1948 | 555.3 | - | - | 0 | - |
| 5 | y | 6.103E+04 | 559.3 | 0.0004782 | 0.8551 | +1 | 4 |
| - | - | 2.001E+04 | 560.3 | - | - | 0 | - |
| - | - | 4204 | 561.3 | - | - | 0 | - |
| - | - | 982.3 | 562.3 | - | - | 0 | - |
| - | - | 1945 | 569.3 | - | - | 0 | - |
| - | - | 929.2 | 570.3 | - | - | 0 | - |
| - | - | 1552 | 605.3 | - | - | 0 | - |
| - | - | 1011 | 607.2 | - | - | 0 | - |
| - | - | 1435 | 613.4 | - | - | 0 | - |
| - | - | 1006 | 620.3 | - | - | 0 | - |
| - | - | 724.7 | 622.3 | - | - | 0 | - |
| - | - | 1131 | 622.4 | - | - | 0 | - |
| 6 | b | 6025 | 623.3 | 0.0006091 | 0.9772 | +1 | 6 |
| - | - | 1435 | 624.3 | - | - | 0 | - |
| - | - | 649.7 | 625.3 | - | - | 0 | - |
| - | - | 4023 | 638.3 | - | - | 0 | - |
| - | - | 735.6 | 639.3 | - | - | 0 | - |
| - | - | 2666 | 640.4 | - | - | 0 | - |
| 6 | b | 6622 | 641.4 | 0.0004316 | 0.673 | +1 | 6 |
| - | - | 1895 | 642.4 | - | - | 0 | - |
| 4 | y | 1108 | 643.3 | 0.0003063 | 0.4762 | +1 | 5 |
| - | - | 765.5 | 643.4 | - | - | 0 | - |
| - | - | 1916 | 650.3 | - | - | 0 | - |
| - | - | 2465 | 656.3 | - | - | 0 | - |
| - | - | 954.8 | 657.3 | - | - | 0 | - |
| - | - | 3837 | 658.4 | - | - | 0 | - |
| - | - | 1260 | 659.4 | - | - | 0 | - |
| 4 | y | 6169 | 660.3 | 0.0004882 | 0.7394 | +1 | 5 |
| - | - | 2969 | 661.3 | - | - | 0 | - |
| - | - | 1780 | 668.4 | - | - | 0 | - |
| - | - | 1110 | 723.4 | - | - | 0 | - |
| - | - | 1060 | 724.4 | - | - | 0 | - |
| - | - | 775.9 | 725.4 | - | - | 0 | - |
| - | - | 1925 | 733.4 | - | - | 0 | - |
| - | - | 1831 | 734.4 | - | - | 0 | - |
| - | - | 2496 | 741.4 | - | - | 0 | - |
| 7 | b | 1.42E+04 | 751.4 | 0.001692 | 2.251 | +1 | 7 |
| 7 | b | 6705 | 752.4 | 0.01344 | 17.86 | +1 | 7 |
| - | - | 959 | 753.4 | - | - | 0 | - |
| 7 | b | 9321 | 769.4 | 0.001087 | 1.412 | +1 | 7 |
| - | - | 3729 | 770.4 | - | - | 0 | - |
| - | - | 1077 | 771.4 | - | - | 0 | - |
| 3 | y | 1780 | 773.4 | 0.003253 | 4.207 | +1 | 6 |
| - | - | 652.2 | 774.4 | - | - | 0 | - |
| 2 | y | 968.1 | 860.4 | 0.0004848 | 0.5635 | +1 | 7 |
| - | - | 611.6 | 970.3 | - | - | 0 | - |
| - | - | 626.5 | 1132 | - | - | 0 | - |
| - | - | 649.5 | 1226 | - | - | 0 | - |
| - | - | 577.9 | 1239 | - | - | 0 | - |
| - | - | 650.4 | 1559 | - | - | 0 | - |
| - | - | 607.5 | 2297 | - | - | 0 | - |
| - | - | 639.1 | 2405 | - | - | 0 | - |
| - | - | 747.2 | 2730 | - | - | 0 | - |
| - | - | 747.8 | 2804 | - | - | 0 | - |
| - | - | 706.2 | 3118 | - | - | 0 | - |
| - | - | 819.5 | 3154 | - | - | 0 | - |

m/z Charge Intensity FragmentType MassShift Position
120.08113861083984 0 2857.3645
122.04698181152344 0 321.93588
125.07122802734375 0 1393.1466
125.10772705078125 0 3178.6692
126.0916976928711 0 1350.064
127.08706665039062 0 535.5425
128.10731506347656 0 1409.6824
129.0662384033203 0 7573.193
129.1025390625 0 1775.649
130.05029296875 0 1582.5024
130.0654754638672 0 16549.984
131.06930541992188 0 1725.3723
132.07591247558594 0 887.5038
132.0811309814453 0 8956.179
132.10227966308594 0 1397.3136
133.08619689941406 0 10478.71
135.08084106445312 0 490.24167
136.07606506347656 0 1314.464
139.05067443847656 0 1203.2177
139.11111450195312 0 432.1763
142.0651092529297 0 506.6868
142.12303161621094 0 3823.0024
143.10691833496094 0 457.91043
143.12623596191406 0 520.21313
144.0443878173828 0 454.17743
144.08116149902344 0 4118.307
145.08570861816406 0 1024.745
146.0603790283203 0 23689.963
146.06704711914062 0 1125.0725
147.0638427734375 0 2118.8503
147.07681274414062 0 1639.7755
147.11294555664062 0 536.13837
149.29637145996094 0 462.74222
152.10733032226562 0 2520.7634
152.4986114501953 0 420.9122
153.06613159179688 0 2324.9016
153.10263061523438 0 1530.8108
155.08172607421875 0 1291.6378
155.1002197265625 0 442.20737
155.1182098388672 0 4299.393 a Water loss 1
159.09202575683594 0 48735.344
160.0953826904297 0 5420.2505
167.0703582763672 0 435.9072
167.0816650390625 0 954.08997 y 6
169.09730529785156 0 1277.1393
169.13375854492188 0 1813.4988
170.06033325195312 0 3506.3413
170.72146606445312 0 451.6758
171.11305236816406 0 4742.581
171.1493682861328 0 1061.7562
173.12879943847656 0 33408.445 a 1
174.0554656982422 0 763.8142
174.13201904296875 0 2583.5332
177.10379028320312 0 446.20316
177.11248779296875 0 2741.0425
178.11585998535156 0 518.4807
181.09751892089844 0 4713.292
183.10348510742188 0 783.04285
183.1130828857422 0 8832.647 b Water loss 1
183.1494140625 0 2124.9402
184.11639404296875 0 1008.9007
185.0924835205078 0 1108.2362
187.07183837890625 0 654.4059
187.08697509765625 0 3766.1218
187.14447021484375 0 9564.018
188.0709686279297 0 180224.28
188.14794921875 0 743.6653
189.07432556152344 0 21094.51
189.1117401123047 0 629.88354
190.077392578125 0 912.27545
191.08206176757812 0 565.411
194.1338653564453 0 678.55493
195.07595825195312 0 570.2135
195.1230010986328 0 1178.4446
197.12864685058594 0 4971.1694
199.07159423828125 0 1311.559
199.10797119140625 0 37654.33
200.11135864257812 0 2871.531
201.10231018066406 0 3344.4458
201.12364196777344 0 40245.402 b 1
202.10826110839844 0 724.0253
202.12705993652344 0 3775.745
205.09744262695312 0 75467.57 y 7
206.1007843017578 0 7759.4077
207.10281372070312 0 509.49622
209.09237670898438 0 3509.0947
211.10794067382812 0 770.3618
211.1441650390625 0 2303.373
212.10250854492188 0 888.8179
213.0870819091797 0 800.1689
213.12319946289062 0 651.5725
214.15545654296875 0 890.52875
215.13934326171875 0 25005.29
216.09768676757812 0 506.68695
216.14276123046875 0 1985.3054
217.0821075439453 0 1528.1404
221.1388702392578 0 646.71985
223.07186889648438 0 1018.8935
224.1029815673828 0 1245.4565
225.10255432128906 0 1858.7393
225.12356567382812 0 1200.8185
226.11907958984375 0 1678.3516
227.1029510498047 0 56060.26
228.10630798339844 0 5506.405
229.15478515625 0 1384.4729
230.11309814453125 0 647.12897
235.69366455078125 0 572.7097
238.15597534179688 0 638.54236
239.1390838623047 0 2432.2798
240.09817504882812 0 6682.912
241.08251953125 0 2251.06
241.10205078125 0 560.51935
242.11279296875 0 568.0149
242.1502685546875 0 1001.8512
242.6605224609375 0 1995.0988
244.1294708251953 0 17091.686
245.11337280273438 0 1138.0452
245.13253784179688 0 1900.0697
247.6519775390625 0 547.8681 b Water loss 4
251.1392364501953 0 593.71844
251.17593383789062 0 1899.6309
252.1134490966797 0 877.98883
253.09718322753906 0 939.6922
254.11422729492188 0 628.8856
255.13343811035156 0 671.76154
256.1656494140625 0 1684.5288
256.65838623046875 0 585.0734 b 4
257.14984130859375 0 3390.6719
258.1085510253906 0 7036.0693
258.14501953125 0 1533.5581
264.1336669921875 0 631.3654
265.118896484375 0 586.4608
266.113525390625 0 1096.7747
266.1500244140625 0 1715.5494
267.13433837890625 0 1165.5765
268.20196533203125 0 807.7859 a Water loss 2
269.1391296386719 0 1986.0024
269.18634033203125 0 1259.5585
270.1238708496094 0 20140.012
271.12677001953125 0 2422.4875 y Water loss 4
274.1766052246094 0 884.77606
280.10845947265625 0 3068.3604
280.16607666015625 0 2200.847
281.1126708984375 0 683.2279
282.1452331542969 0 1567.8977
283.1751403808594 0 806.82117
284.1242370605469 0 742.53424
284.1430358886719 0 974.63806
284.16070556640625 0 19242.426
285.1469421386719 0 846.8096
285.1639709472656 0 2395.523
287.15032958984375 0 849.80597
292.1293640136719 0 1584.8127
294.14556884765625 0 1253.6473
294.18145751953125 0 1002.1223
296.1247863769531 0 772.7064
296.1971435546875 0 40088.477 b Water loss 2
297.1348571777344 0 1183.848
297.20025634765625 0 6766.7197
298.1188049316406 0 5330.147
298.1766662597656 0 2936.5913
298.2025451660156 0 749.7536
298.5511779785156 0 549.8213
299.1224060058594 0 1036.5168
300.1557922363281 0 2220.5427
301.1407470703125 0 586.60144
302.17132568359375 0 20870.385
303.1741027832031 0 3372.9116
307.18206787109375 0 920.03375
309.1549987792969 0 1603.1143
309.2045593261719 0 1221.8539
310.14013671875 0 3242.579
310.2127685546875 0 705.09235
311.1716613769531 0 949.3027
312.155517578125 0 2552.7136
312.1915283203125 0 1550.6633
314.1347351074219 0 594.37683
314.2076416015625 0 8469.303 b 2
315.1454162597656 0 42988.82
315.21173095703125 0 1016.7846
316.1294250488281 0 44719.15 y Ammonia loss 6
316.14874267578125 0 6984.5864
317.13262939453125 0 7841.1533
318.1343688964844 0 1034.9427
320.12396240234375 0 1129.4794
321.1787414550781 0 2600.458 b 5
321.6806640625 0 787.24255
323.1693420410156 0 581.31757
326.1716613769531 0 877.8446
327.1663513183594 0 4259.899
328.1505432128906 0 6918.695
328.1688537597656 0 684.33307
329.1533203125 0 1420.1577
329.1828918457031 0 708.2147
329.69244384765625 0 1344.2124
333.156005859375 0 21596.688 y 6
334.1595153808594 0 3642.5212
334.21490478515625 0 594.7497
337.15093994140625 0 4134.2856
338.1343078613281 0 1255.8474
338.15435791015625 0 688.9838
340.1869812011719 0 2500.3906
345.1766357421875 0 2657.2986
346.17681884765625 0 621.36865
351.240966796875 0 872.40594
352.2229919433594 0 3362.905
353.182861328125 0 785.31805
353.2198791503906 0 701.5246
355.1612854003906 0 74874.02
356.1459655761719 0 1226.8596
356.16424560546875 0 11551.79
357.16815185546875 0 1040.9253
359.4363098144531 0 567.38416
360.8933410644531 0 559.34064
363.2036437988281 0 1366.4552
365.18212890625 0 1458.4294
369.2494812011719 0 4501.0444
370.233154296875 0 5849.365
370.25531005859375 0 625.2639
371.2308654785156 0 2198.3013
372.18951416015625 0 1162.156
373.17138671875 0 2938.7783
379.23394775390625 0 2474.27
381.1555480957031 0 1546.7717
381.2125549316406 0 1249.3273
387.2605285644531 0 7188.484
388.2633972167969 0 1992.034
397.24462890625 0 23176.84 b Water loss 3
397.2743225097656 0 1105.9481
398.2479553222656 0 4559.5435
399.2222595214844 0 1188.5409
399.25164794921875 0 800.46857
409.15338134765625 0 946.91675
409.207763671875 0 1947.0361
411.1882019042969 0 731.6906
415.2551574707031 0 44856.86 b 3
416.25823974609375 0 9749.146
417.2290344238281 0 945.347
417.2611999511719 0 1555.9724
423.22412109375 0 1875.2468
427.1614074707031 0 5686.622
427.2176818847656 0 993.4635
428.16485595703125 0 1358.2148
431.26422119140625 0 754.3492
438.1983642578125 0 1300.699
440.2501525878906 0 1403.2465
441.2345275878906 0 965.22986
444.18804931640625 0 1461.4369 y Water loss 5
445.1716613769531 0 4038.8218 y Ammonia loss 5
446.176025390625 0 1147.279
450.2377014160156 0 793.1136
451.22406005859375 0 795.8489
456.2090148925781 0 6345.4478
457.2100830078125 0 1347.7164
460.2561950683594 0 1148.3397
462.1994323730469 0 1199.9672 y 5
468.2450866699219 0 4555.9165
468.2810974121094 0 1039.2494
469.2442321777344 0 1522.1484
469.7373352050781 0 625.00275
470.2385559082031 0 2392.276
471.2787170410156 0 690.90717
472.183349609375 0 1169.0881
476.2885437011719 0 1178.1676
478.20831298828125 0 2119.9136
481.264404296875 0 731.40405
484.3162536621094 0 633.5845
488.24951171875 0 748.46985
492.2438659667969 0 864.6052
494.2972106933594 0 3773.963 b Water loss 4
495.2997741699219 0 1743.4631
496.219482421875 0 1185.1088
499.2765197753906 0 1163.6237
500.2779235839844 0 687.68384
506.2048034667969 0 1340.5662
508.27508544921875 0 1096.5986
510.2561340332031 0 2240.585
512.308349609375 0 4816.5386 b 4
513.244384765625 0 859.53394
513.3118896484375 0 933.07837
523.2908325195312 0 608.2168
524.2136840820312 0 13065.373
525.2169799804688 0 3350.8096
525.3034057617188 0 683.36487
526.2213134765625 0 857.2994
526.2863159179688 0 1474.558
528.2673950195312 0 974.7865
537.2655639648438 0 1742.6501
541.2399291992188 0 5566.3857 y Water loss 4
541.3008422851562 0 1001.36914
542.2247924804688 0 20426.38 y Ammonia loss 4
543.2273559570312 0 7169.825
544.2302856445312 0 1249.2399
551.2818603515625 0 2238.7869
555.2776489257812 0 1947.9695
559.2506103515625 0 61027.215 y 4
560.2532958984375 0 20011.059
561.255615234375 0 4203.8867
562.2984008789062 0 982.31586
569.2925415039062 0 1944.7875
570.2972412109375 0 929.173
605.3278198242188 0 1552.2367
607.2495727539062 0 1011.3359
613.3538208007812 0 1435.1887
620.3047485351562 0 1006.15814
622.298828125 0 724.70966
622.3553466796875 0 1131.1667
623.3392944335938 0 6025.1567 b Water loss 5
624.3422241210938 0 1434.8486
625.3453979492188 0 649.67725
638.3146362304688 0 4022.794
639.3067016601562 0 735.6336
640.3656005859375 0 2666.3257
641.3500366210938 0 6622.1323 b 5
642.3516235351562 0 1895.0836
643.2719116210938 0 1107.7228 y Ammonia loss 3
643.3557739257812 0 765.49695
650.3496704101562 0 1915.8876
656.3237915039062 0 2464.8342
657.3267822265625 0 954.7768
658.3757934570312 0 3837.0833
659.37890625 0 1260.4343
660.2982788085938 0 6168.936 y 3
661.3007202148438 0 2969.4614
668.3585815429688 0 1779.7499
723.4005126953125 0 1110.0743
724.396240234375 0 1060.0834
725.3825073242188 0 775.89154
733.3869018554688 0 1924.5767
734.3773193359375 0 1831.0177
741.4119873046875 0 2496.2983
751.3967895507812 0 14198.027 b Water loss 6
752.3959350585938 0 6705.2617 b Ammonia loss 6
753.3954467773438 0 959.0128
769.407958984375 0 9320.516 b 6
770.4105224609375 0 3728.9858
771.4090576171875 0 1077.3036
773.3795776367188 0 1780.2299 y 2
774.38623046875 0 652.1716
860.4153442382812 0 968.14594 y 1
970.3051147460938 0 611.60516
1132.15966796875 0 626.5205
1225.9010009765625 0 649.46246
1238.82763671875 0 577.87665
1558.63037109375 0 650.3866
2296.691650390625 0 607.48303
2405.159423828125 0 639.07764
2730.171630859375 0 747.16455
2804.2529296875 0 747.8225
3117.526611328125 0 706.1665
3154.1171875 0 819.4533

Spectrum Details

|  |  |
| --- | --- |
| Matched peaks? Matched peaks The total absolute number of peaks matched. Additionally in brackets the total fraction of peaks matched and the total number of peaks is shown. | 34 (9.69% of 351) |
| FDR? FDR The false discovery rate estimated for this peptide. It is calculated by matching all theoretical fragments with a non-integer shift with the raw peaks for this spectrum. This is done with 40 different shifts. The resulting percentage is the average number of annotated peaks over the number of annotated peaks with the correct spectrum. | 0.63% |
| Satellite FDR? Satellite FDR See the FDR for details on its calculation. This satellite ion specific FDR only contains the satellite ions (d/w) for I/L/J positions. | - |
| PSM Score? PSM Score The PSM Score as given by Hecklib to this annotated spectrum. It is shown with three significant figures. | 498 |

#### Spectrum 7901? Spectrum 7901 The raw spectrum of this peptide as annotated by Hecklib. The fragments are coloured according to ion type (see legend). Any peaks with a star '\*' as text can be hovered over to see the full details, first the ion type second the mass shift type. By hovering over the amino acids in the peptide or ions in the legend the corresponding peaks are highlighted. By toggling the 'Unassigned' label you can turn the background (unassigned) peaks on or off in the plot. By updating the slider in the Ion legend you can update the spectrum to only show the top X% of the peaks with labels. The top X% means any peak that is within X% of the highest intensity. By dragging in the spectrum you can zoom in to a specific part of the spectrum and use 'Zoom Out' to get back to the original zoom level. The annotation of the spectrum is based on the given sequence in the peptides file and is done with different software so inconsistencies are likely. The peaks are annotated based on the given sequence, with 20 ppm tolerance.

Copy Data

##### Spectrum 7901 (TSV)

###### Preview

```
Loading example...
```

*Click on the button to copy the data to your clipboard.*

Mz MinMz MaxIntensity Max

WidthHeightPeptide font sizePeptide stroke widthSpectrum font sizeSpectrum stroke widthCompact peptide

Ion legend

wxyz

abcd

OtherUnassignedIon^Charge^_Position_Show for top:%

JSJTPEQW

02.33e+44.67e+47.00e+49.33e+4

Zoom Out

y^+1^_2_z^+1^_2_y^+1^_2_y^+2^_7_y^+1^_3_c^+1^_5_z^+1^_4_y^+1^_4_y^+1^_4_z^+1^_4_y^+1^_4_c^+1^_6_c^+1^_6_y^+1^_5_z^+1^_5_y^+1^_5_w^+1^_6_c^+1^_7_z^+1^_6_c^+1^_7_y^+1^_6_z^+1^_7_y^+1^_7_z^+1^_7_y^+1^_7_

0777155523323110

Fragment Matches Table

Show background peaks

| Position | Ion type | Intensity | mz Theoretical | mz Error (Th) | mz Error (ppm) | Charge | Series Number |
| --- | --- | --- | --- | --- | --- | --- | --- |
| - | - | 348.2 | 135 | - | - | 0 | - |
| - | - | 436.3 | 143.4 | - | - | 0 | - |
| - | - | 428.1 | 148.9 | - | - | 0 | - |
| - | - | 545.1 | 148.9 | - | - | 0 | - |
| - | - | 565.8 | 148.9 | - | - | 0 | - |
| - | - | 544.2 | 148.9 | - | - | 0 | - |
| - | - | 463.4 | 148.9 | - | - | 0 | - |
| - | - | 1025 | 148.9 | - | - | 0 | - |
| - | - | 994.5 | 148.9 | - | - | 0 | - |
| - | - | 2470 | 148.9 | - | - | 0 | - |
| - | - | 3980 | 148.9 | - | - | 0 | - |
| - | - | 4352 | 149 | - | - | 0 | - |
| - | - | 2354 | 149 | - | - | 0 | - |
| - | - | 1329 | 149 | - | - | 0 | - |
| - | - | 1126 | 149 | - | - | 0 | - |
| - | - | 847 | 149 | - | - | 0 | - |
| - | - | 820.5 | 149 | - | - | 0 | - |
| - | - | 756 | 149 | - | - | 0 | - |
| - | - | 573.2 | 149 | - | - | 0 | - |
| - | - | 582 | 149 | - | - | 0 | - |
| - | - | 566.7 | 149 | - | - | 0 | - |
| - | - | 504.6 | 149 | - | - | 0 | - |
| - | - | 473.1 | 149.1 | - | - | 0 | - |
| - | - | 404 | 152.3 | - | - | 0 | - |
| - | - | 405.3 | 155.4 | - | - | 0 | - |
| - | - | 411.1 | 157.1 | - | - | 0 | - |
| - | - | 2741 | 173.1 | - | - | 0 | - |
| - | - | 1051 | 173.4 | - | - | 0 | - |
| - | - | 4264 | 175.1 | - | - | 0 | - |
| - | - | 487.4 | 186.3 | - | - | 0 | - |
| - | - | 1211 | 187.1 | - | - | 0 | - |
| - | - | 504.6 | 194.6 | - | - | 0 | - |
| - | - | 610.4 | 197.1 | - | - | 0 | - |
| - | - | 5860 | 201.1 | - | - | 0 | - |
| - | - | 1315 | 203.1 | - | - | 0 | - |
| - | - | 761.6 | 211.1 | - | - | 0 | - |
| - | - | 4018 | 215.1 | - | - | 0 | - |
| - | - | 1.966E+04 | 221.1 | - | - | 0 | - |
| - | - | 1847 | 222.1 | - | - | 0 | - |
| - | - | 2304 | 227.1 | - | - | 0 | - |
| - | - | 540.1 | 231 | - | - | 0 | - |
| - | - | 5048 | 244.1 | - | - | 0 | - |
| - | - | 614.3 | 245.1 | - | - | 0 | - |
| - | - | 2057 | 284.2 | - | - | 0 | - |
| - | - | 822.6 | 285.2 | - | - | 0 | - |
| - | - | 540.5 | 295.3 | - | - | 0 | - |
| - | - | 7884 | 296.2 | - | - | 0 | - |
| - | - | 956.9 | 297.2 | - | - | 0 | - |
| - | - | 3110 | 302.2 | - | - | 0 | - |
| - | - | 1703 | 314.1 | - | - | 0 | - |
| - | - | 2698 | 314.2 | - | - | 0 | - |
| - | - | 1210 | 328.2 | - | - | 0 | - |
| - | - | 4279 | 331.1 | - | - | 0 | - |
| 7 | y | 5681 | 332.1 | 0.005329 | 16.05 | +1 | 2 |
| - | - | 810.7 | 332.1 | - | - | 0 | - |
| 7 | z | 1056 | 333.1 | 0.0002508 | 0.7528 | +1 | 2 |
| - | - | 1083 | 348.1 | - | - | 0 | - |
| 7 | y | 6441 | 349.1 | 0.004903 | 14.04 | +1 | 2 |
| - | - | 845.5 | 350.2 | - | - | 0 | - |
| - | - | 1.151E+04 | 355.2 | - | - | 0 | - |
| - | - | 1832 | 356.2 | - | - | 0 | - |
| - | - | 867.7 | 369.3 | - | - | 0 | - |
| - | - | 1105 | 370.2 | - | - | 0 | - |
| - | - | 1363 | 372.2 | - | - | 0 | - |
| - | - | 743.8 | 379.2 | - | - | 0 | - |
| - | - | 2265 | 387.3 | - | - | 0 | - |
| - | - | 4954 | 397.2 | - | - | 0 | - |
| - | - | 747.7 | 398.2 | - | - | 0 | - |
| - | - | 592.1 | 405.2 | - | - | 0 | - |
| - | - | 1.606E+04 | 415.3 | - | - | 0 | - |
| - | - | 2007 | 416.3 | - | - | 0 | - |
| - | - | 6710 | 429.2 | - | - | 0 | - |
| 2 | y | 1521 | 430.2 | 0.003584 | 8.332 | +2 | 7 |
| - | - | 1719 | 439.2 | - | - | 0 | - |
| - | - | 713.6 | 441.2 | - | - | 0 | - |
| - | - | 2162 | 442.2 | - | - | 0 | - |
| - | - | 797.8 | 443.2 | - | - | 0 | - |
| - | - | 1037 | 456.2 | - | - | 0 | - |
| - | - | 2076 | 459.8 | - | - | 0 | - |
| - | - | 1128 | 463.2 | - | - | 0 | - |
| - | - | 2901 | 477.8 | - | - | 0 | - |
| 6 | y | 750 | 478.2 | 0.005706 | 11.93 | +1 | 3 |
| - | - | 966.8 | 495.8 | - | - | 0 | - |
| - | - | 731.8 | 503.2 | - | - | 0 | - |
| - | - | 727.5 | 512.3 | - | - | 0 | - |
| - | - | 4374 | 513.2 | - | - | 0 | - |
| - | - | 1596 | 514.2 | - | - | 0 | - |
| - | - | 2219 | 515.2 | - | - | 0 | - |
| - | - | 4287 | 528.3 | - | - | 0 | - |
| 5 | c | 3.143E+04 | 529.3 | 0.0001347 | 0.2545 | +1 | 5 |
| - | - | 1116 | 530.2 | - | - | 0 | - |
| - | - | 9465 | 530.3 | - | - | 0 | - |
| - | - | 1648 | 531.3 | - | - | 0 | - |
| - | - | 1774 | 540.2 | - | - | 0 | - |
| 5 | z | 703.6 | 541.2 | 0.003609 | 6.668 | +1 | 4 |
| 5 | y | 1526 | 557.2 | 0.003363 | 6.035 | +1 | 4 |
| 5 | y | 4711 | 558.2 | 0.005675 | 10.17 | +1 | 4 |
| 5 | z | 1742 | 559.2 | 4.763E-05 | 0.08516 | +1 | 4 |
| - | - | 1.842E+04 | 574.2 | - | - | 0 | - |
| 5 | y | 4.75E+04 | 575.2 | 0.004334 | 7.534 | +1 | 4 |
| - | - | 1.384E+04 | 576.2 | - | - | 0 | - |
| - | - | 2530 | 577.3 | - | - | 0 | - |
| - | - | 1007 | 603.2 | - | - | 0 | - |
| - | - | 1365 | 614.4 | - | - | 0 | - |
| - | - | 2148 | 615.4 | - | - | 0 | - |
| - | - | 1003 | 623.3 | - | - | 0 | - |
| - | - | 2491 | 625.3 | - | - | 0 | - |
| - | - | 1424 | 626.3 | - | - | 0 | - |
| - | - | 924.3 | 628.3 | - | - | 0 | - |
| - | - | 937.4 | 640.3 | - | - | 0 | - |
| 6 | c | 886.8 | 640.4 | 0.002195 | 3.427 | +1 | 6 |
| - | - | 3099 | 641.3 | - | - | 0 | - |
| - | - | 1402 | 641.4 | - | - | 0 | - |
| - | - | 1005 | 642.3 | - | - | 0 | - |
| - | - | 1.068E+04 | 658.3 | - | - | 0 | - |
| 6 | c | 3.518E+04 | 658.4 | 6.422E-05 | 0.09755 | +1 | 6 |
| 4 | y | 5462 | 659.3 | 0.002186 | 3.316 | +1 | 5 |
| - | - | 1.142E+04 | 659.4 | - | - | 0 | - |
| 4 | z | 2.363E+04 | 660.3 | 0.00431 | 6.528 | +1 | 5 |
| - | - | 2985 | 660.4 | - | - | 0 | - |
| - | - | 1.426E+04 | 661.3 | - | - | 0 | - |
| - | - | 4001 | 662.3 | - | - | 0 | - |
| - | - | 767.5 | 663.3 | - | - | 0 | - |
| - | - | 2356 | 675.3 | - | - | 0 | - |
| 4 | y | 4846 | 676.3 | 0.004141 | 6.123 | +1 | 5 |
| - | - | 856.9 | 677.3 | - | - | 0 | - |
| - | - | 639.4 | 678.3 | - | - | 0 | - |
| - | - | 703.6 | 684.4 | - | - | 0 | - |
| - | - | 791.7 | 697.3 | - | - | 0 | - |
| - | - | 6157 | 715.3 | - | - | 0 | - |
| - | - | 1982 | 716.3 | - | - | 0 | - |
| - | - | 2882 | 728.3 | - | - | 0 | - |
| - | - | 1010 | 729.3 | - | - | 0 | - |
| 3 | w | 996.2 | 730.3 | 0.000534 | 0.7312 | +1 | 6 |
| - | - | 643.6 | 742.4 | - | - | 0 | - |
| - | - | 1021 | 749.4 | - | - | 0 | - |
| - | - | 3025 | 751.4 | - | - | 0 | - |
| - | - | 1473 | 752.4 | - | - | 0 | - |
| - | - | 9704 | 753.4 | - | - | 0 | - |
| - | - | 4073 | 754.4 | - | - | 0 | - |
| - | - | 1315 | 755.4 | - | - | 0 | - |
| 7 | c | 4275 | 769.4 | 0.0008053 | 1.047 | +1 | 7 |
| - | - | 1126 | 770.4 | - | - | 0 | - |
| - | - | 1.718E+04 | 771.4 | - | - | 0 | - |
| - | - | 7302 | 772.4 | - | - | 0 | - |
| 3 | z | 9271 | 773.4 | 0.004902 | 6.338 | +1 | 6 |
| - | - | 1297 | 773.4 | - | - | 0 | - |
| - | - | 5829 | 774.4 | - | - | 0 | - |
| - | - | 2251 | 775.4 | - | - | 0 | - |
| - | - | 4314 | 783.4 | - | - | 0 | - |
| - | - | 1551 | 784.4 | - | - | 0 | - |
| 7 | c | 9.239E+04 | 786.4 | 0.0001701 | 0.2162 | +1 | 7 |
| - | - | 3.583E+04 | 787.4 | - | - | 0 | - |
| - | - | 8659 | 788.4 | - | - | 0 | - |
| 3 | y | 8242 | 789.4 | 0.01553 | 19.68 | +1 | 6 |
| - | - | 2913 | 790.4 | - | - | 0 | - |
| - | - | 2926 | 800.4 | - | - | 0 | - |
| - | - | 1113 | 801.4 | - | - | 0 | - |
| - | - | 2264 | 804.3 | - | - | 0 | - |
| - | - | 3966 | 812.5 | - | - | 0 | - |
| - | - | 2702 | 813.5 | - | - | 0 | - |
| - | - | 747.7 | 814.4 | - | - | 0 | - |
| - | - | 6583 | 827.4 | - | - | 0 | - |
| - | - | 6550 | 828.4 | - | - | 0 | - |
| - | - | 2356 | 829.4 | - | - | 0 | - |
| 2 | z | 1737 | 842.4 | 0.004678 | 5.554 | +1 | 7 |
| - | - | 1.397E+04 | 843.4 | - | - | 0 | - |
| - | - | 7249 | 844.4 | - | - | 0 | - |
| - | - | 8975 | 845.4 | - | - | 0 | - |
| - | - | 3130 | 846.4 | - | - | 0 | - |
| - | - | 3336 | 857.5 | - | - | 0 | - |
| 2 | y | 890.3 | 858.4 | 0.0163 | 18.99 | +1 | 7 |
| - | - | 2172 | 858.5 | - | - | 0 | - |
| 2 | z | 1.898E+04 | 860.4 | 0.004551 | 5.289 | +1 | 7 |
| - | - | 9671 | 861.4 | - | - | 0 | - |
| - | - | 2629 | 862.4 | - | - | 0 | - |
| - | - | 2215 | 873.5 | - | - | 0 | - |
| - | - | 851.8 | 874.5 | - | - | 0 | - |
| - | - | 3083 | 875.5 | - | - | 0 | - |
| 2 | y | 1524 | 876.4 | 0.001295 | 1.478 | +1 | 7 |
| - | - | 1488 | 876.5 | - | - | 0 | - |
| - | - | 1130 | 877.4 | - | - | 0 | - |
| - | - | 1.136E+04 | 928.5 | - | - | 0 | - |
| - | - | 5426 | 929.5 | - | - | 0 | - |
| - | - | 1986 | 930.5 | - | - | 0 | - |
| - | - | 1089 | 931.5 | - | - | 0 | - |
| - | - | 929.8 | 932.5 | - | - | 0 | - |
| - | - | 7968 | 945.5 | - | - | 0 | - |
| - | - | 4741 | 946.5 | - | - | 0 | - |
| - | - | 1878 | 947.5 | - | - | 0 | - |
| - | - | 2423 | 955.5 | - | - | 0 | - |
| - | - | 905.5 | 956.5 | - | - | 0 | - |
| - | - | 765.6 | 962.5 | - | - | 0 | - |
| - | - | 1.18E+04 | 972.5 | - | - | 0 | - |
| - | - | 2.505E+04 | 973.5 | - | - | 0 | - |
| - | - | 1.27E+04 | 974.5 | - | - | 0 | - |
| - | - | 3755 | 975.5 | - | - | 0 | - |
| - | - | 855.7 | 976.5 | - | - | 0 | - |
| - | - | 2.788E+04 | 989.5 | - | - | 0 | - |
| - | - | 7.271E+04 | 990.5 | - | - | 0 | - |
| - | - | 3.756E+04 | 991.5 | - | - | 0 | - |
| - | - | 1.14E+04 | 992.5 | - | - | 0 | - |
| - | - | 1145 | 993.5 | - | - | 0 | - |
| - | - | 687.7 | 1349 | - | - | 0 | - |
| - | - | 660.9 | 2520 | - | - | 0 | - |
| - | - | 833.4 | 3079 | - | - | 0 | - |

m/z Charge Intensity FragmentType MassShift Position
135.04049682617188 0 348.23804
143.37783813476562 0 436.2669
148.8779296875 0 428.08984
148.884765625 0 545.0832
148.89915466308594 0 565.82996
148.90625 0 544.2376
148.91383361816406 0 463.3665
148.92132568359375 0 1024.5603
148.9287567138672 0 994.48236
148.93548583984375 0 2470.1616
148.9432373046875 0 3979.7964
148.9598846435547 0 4352.263
148.96771240234375 0 2353.5352
148.97463989257812 0 1328.831
148.9821014404297 0 1125.8593
148.98948669433594 0 847.003
148.99652099609375 0 820.48395
149.0037078857422 0 755.95087
149.01133728027344 0 573.23755
149.0188751220703 0 582.0313
149.0260009765625 0 566.69086
149.04736328125 0 504.60324
149.06207275390625 0 473.13416
152.3017120361328 0 403.97128
155.40249633789062 0 405.27972
157.0757598876953 0 411.1054
173.1286163330078 0 2741.2354
173.44163513183594 0 1050.6716
175.08677673339844 0 4263.904
186.2598876953125 0 487.42908
187.14410400390625 0 1210.6539
194.61566162109375 0 504.59122
197.12892150878906 0 610.4366
201.12351989746094 0 5860.2124
203.0817413330078 0 1314.8285
211.0945587158203 0 761.6414
215.1390380859375 0 4017.6458
221.09219360351562 0 19657.652
222.09552001953125 0 1847.2379
227.1026153564453 0 2303.795
230.9820556640625 0 540.0645
244.1294403076172 0 5047.5254
245.13211059570312 0 614.30615
284.1608581542969 0 2056.913
285.1646728515625 0 822.6482
295.29840087890625 0 540.49866
296.1970520019531 0 7883.7827
297.2002258300781 0 956.9251
302.1715393066406 0 3109.8142
314.1138000488281 0 1703.1996
314.2075500488281 0 2697.918
328.1502380371094 0 1209.9268
331.14013671875 0 4279.0225
332.12451171875 0 5681.4224 y Ammonia loss 6
332.1440734863281 0 810.6882
333.12725830078125 0 1055.94 z 6
348.1426696777344 0 1083.1711
349.150634765625 0 6441.0703 y 6
350.1534729003906 0 845.5364
355.16119384765625 0 11511.152
356.164794921875 0 1831.5835
369.2505798339844 0 867.67084
370.2344055175781 0 1104.5665
372.1869812011719 0 1362.7489
379.23431396484375 0 743.75867
387.26068115234375 0 2264.8645
397.24456787109375 0 4954.364
398.2482604980469 0 747.6955
405.1781921386719 0 592.14233
415.2554016113281 0 16061.059
416.2585144042969 0 2007.3008
429.185546875 0 6710.2627
430.189208984375 0 1520.5591 y Ammonia loss 1
439.1823425292969 0 1718.9053
441.1988830566406 0 713.5909
442.1940002441406 0 2161.6987
443.1553955078125 0 797.79895
456.20941162109375 0 1036.897
459.81298828125 0 2075.7415
463.1819152832031 0 1128.2515
477.82342529296875 0 2900.822
478.19403076171875 0 749.954 y 5
495.8342590332031 0 966.79614
503.2035217285156 0 731.7827
512.310546875 0 727.46545
513.2066040039062 0 4373.5396
514.2108764648438 0 1595.9714
515.2205200195312 0 2219.487
528.326416015625 0 4287.431
529.3342895507812 0 31431.021 c 4
530.234619140625 0 1116.245
530.3372802734375 0 9464.898
531.341552734375 0 1648.2688
540.2089233398438 0 1773.6742
541.2081909179688 0 703.57275 z Water loss 4
557.23388671875 0 1525.786 y Water loss 4
558.22021484375 0 4711.2944 y Ammonia loss 4
559.222412109375 0 1742.1526 z 4
574.2381591796875 0 18422.342
575.2454223632812 0 47497.027 y 4
576.2489624023438 0 13840.476
577.2511596679688 0 2530.1187
603.240234375 0 1006.7752
614.3627319335938 0 1364.5352
615.3712158203125 0 2148.1968
623.34033203125 0 1002.8635
625.3314208984375 0 2491.2183
626.3367919921875 0 1424.4817
628.3056030273438 0 924.25507
640.305419921875 0 937.44934
640.3642578125 0 886.79956 c Water loss 5
641.3121337890625 0 3099.31
641.3607177734375 0 1402.2705
642.3146362304688 0 1005.4221
658.2595825195312 0 10676.415
658.376953125 0 35177.055 c 5
659.264404296875 0 5462.4546 y Ammonia loss 3
659.3804321289062 0 11418.596
660.2743530273438 0 23625.566 z 3
660.3816528320312 0 2985.3108
661.2801513671875 0 14257.089
662.282470703125 0 4000.8474
663.2899169921875 0 767.5341
675.28564453125 0 2355.5195
676.2929077148438 0 4846.119 y 3
677.2955932617188 0 856.87036
678.3037109375 0 639.37
684.3936157226562 0 703.5527
697.3245239257812 0 791.67535
715.3379516601562 0 6156.9927
716.342529296875 0 1981.7212
728.3460693359375 0 2881.703
729.3496704101562 0 1010.4059
730.2998657226562 0 996.2423 w 2
742.4254760742188 0 643.5757
749.394775390625 0 1020.56244
751.3954467773438 0 3024.994
752.3966064453125 0 1473.1174
753.41259765625 0 9704.096
754.4105224609375 0 4073.4685
755.4158325195312 0 1315.0244
769.4098510742188 0 4274.575 c Ammonia loss 6
770.4088134765625 0 1125.6151
771.4244995117188 0 17182.312
772.4275512695312 0 7301.9937
773.3590087890625 0 9270.607 z 2
773.4302368164062 0 1296.557
774.3628540039062 0 5829.2188
775.365234375 0 2250.9316
783.44775390625 0 4314.2075
784.44970703125 0 1551.0857
786.4354248046875 0 92385.71 c 6
787.4384155273438 0 35834.934
788.44140625 0 8658.594
789.3572998046875 0 8242.436 y 2
790.3580932617188 0 2912.6099
800.4268188476562 0 2925.8906
801.42919921875 0 1112.8423
804.328857421875 0 2263.5317
812.4508666992188 0 3965.986
813.4527587890625 0 2702.2551
814.4483032226562 0 747.7293
827.4376220703125 0 6583.38
828.4317016601562 0 6549.7295
829.4323120117188 0 2356.313
842.3802490234375 0 1736.9594 z Water loss 1
843.4321899414062 0 13972.955
844.4353637695312 0 7249.4155
845.4454956054688 0 8974.642
846.4495239257812 0 3129.7458
857.45849609375 0 3336.08
858.3779907226562 0 890.2971 y Water loss 1
858.4661254882812 0 2172.117
860.3906860351562 0 18976.383 z 1
861.3943481445312 0 9671.45
862.3981323242188 0 2628.5818
873.4588623046875 0 2214.7542
874.4653930664062 0 851.7779
875.471435546875 0 3083.1064
876.403564453125 0 1523.95 y 1
876.4810791015625 0 1487.8661
877.4203491210938 0 1129.8564
928.5006103515625 0 11357.248
929.50244140625 0 5425.5435
930.5062866210938 0 1985.8483
931.48974609375 0 1088.7083
932.4935302734375 0 929.8281
945.4794311523438 0 7967.5044
946.485107421875 0 4740.9663
947.4976196289062 0 1878.1779
955.4653930664062 0 2423.402
956.4679565429688 0 905.52136
962.50634765625 0 765.6226
972.4901123046875 0 11802.928
973.4780883789062 0 25052.105
974.4801025390625 0 12704.727
975.4818725585938 0 3755.3057
976.491943359375 0 855.7179
989.4931030273438 0 27878.223
990.4996337890625 0 72712.81
991.503173828125 0 37561.01
992.5072631835938 0 11396.295
993.5111083984375 0 1145.1455
1349.2197265625 0 687.7286
2519.6630859375 0 660.9115
3078.791015625 0 833.43726

Spectrum Details

|  |  |
| --- | --- |
| Matched peaks? Matched peaks The total absolute number of peaks matched. Additionally in brackets the total fraction of peaks matched and the total number of peaks is shown. | 25 (12.14% of 206) |
| FDR? FDR The false discovery rate estimated for this peptide. It is calculated by matching all theoretical fragments with a non-integer shift with the raw peaks for this spectrum. This is done with 40 different shifts. The resulting percentage is the average number of annotated peaks over the number of annotated peaks with the correct spectrum. | 3.52% |
| Satellite FDR? Satellite FDR See the FDR for details on its calculation. This satellite ion specific FDR only contains the satellite ions (d/w) for I/L/J positions. | 14.29% |
| PSM Score? PSM Score The PSM Score as given by Hecklib to this annotated spectrum. It is shown with three significant figures. | 323 |

#### Spectrum 9915? Spectrum 9915 The raw spectrum of this peptide as annotated by Hecklib. The fragments are coloured according to ion type (see legend). Any peaks with a star '\*' as text can be hovered over to see the full details, first the ion type second the mass shift type. By hovering over the amino acids in the peptide or ions in the legend the corresponding peaks are highlighted. By toggling the 'Unassigned' label you can turn the background (unassigned) peaks on or off in the plot. By updating the slider in the Ion legend you can update the spectrum to only show the top X% of the peaks with labels. The top X% means any peak that is within X% of the highest intensity. By dragging in the spectrum you can zoom in to a specific part of the spectrum and use 'Zoom Out' to get back to the original zoom level. The annotation of the spectrum is based on the given sequence in the peptides file and is done with different software so inconsistencies are likely. The peaks are annotated based on the given sequence, with 20 ppm tolerance.

Copy Data

##### Spectrum 9915 (TSV)

###### Preview

```
Loading example...
```

*Click on the button to copy the data to your clipboard.*

Mz MinMz MaxIntensity Max

WidthHeightPeptide font sizePeptide stroke widthSpectrum font sizeSpectrum stroke widthCompact peptide

Ion legend

wxyz

abcd

OtherUnassignedIon^Charge^_Position_Show for top:%

JSJTPEQW

06.22e+41.24e+51.87e+52.49e+5

Zoom Out

a^+1^_2_y^+2^_2_a^+1^_2_b^+1^_2_b^+1^_2_y^+1^_1_b^+2^_5_b^+2^_5_d^+1^_3_a^+1^_3_y^+2^_4_a^+1^_3_b^+1^_3_b^+1^_3_y^+1^_2_b^+2^_6_y^+1^_2_b^+1^_4_b^+1^_4_y^+1^_3_y^+1^_3_y^+1^_3_b^+1^_5_b^+1^_5_y^+1^_4_y^+1^_4_y^+1^_4_b^+1^_6_b^+1^_6_y^+1^_5_y^+1^_5_b^+1^_7_b^+1^_7_y^+1^_6_b^+1^_7_y^+1^_6_y^+1^_7_y^+1^_7_

0778155623353113

Fragment Matches Table

Show background peaks

| Position | Ion type | Intensity | mz Theoretical | mz Error (Th) | mz Error (ppm) | Charge | Series Number |
| --- | --- | --- | --- | --- | --- | --- | --- |
| - | - | 2000 | 120.1 | - | - | 0 | - |
| - | - | 614.7 | 121 | - | - | 0 | - |
| - | - | 1659 | 125.1 | - | - | 0 | - |
| - | - | 3667 | 125.1 | - | - | 0 | - |
| - | - | 881.9 | 126.1 | - | - | 0 | - |
| - | - | 393.4 | 127.1 | - | - | 0 | - |
| - | - | 662.8 | 127.1 | - | - | 0 | - |
| - | - | 1600 | 128.1 | - | - | 0 | - |
| - | - | 9531 | 129.1 | - | - | 0 | - |
| - | - | 2289 | 129.1 | - | - | 0 | - |
| - | - | 3249 | 130.1 | - | - | 0 | - |
| - | - | 2.371E+04 | 130.1 | - | - | 0 | - |
| - | - | 2091 | 131.1 | - | - | 0 | - |
| - | - | 1.151E+04 | 132.1 | - | - | 0 | - |
| - | - | 1355 | 132.1 | - | - | 0 | - |
| - | - | 3135 | 133.1 | - | - | 0 | - |
| - | - | 1349 | 136.1 | - | - | 0 | - |
| - | - | 771.8 | 138.1 | - | - | 0 | - |
| - | - | 1040 | 139.1 | - | - | 0 | - |
| - | - | 394.9 | 141.1 | - | - | 0 | - |
| - | - | 553.4 | 141.1 | - | - | 0 | - |
| - | - | 997.6 | 142.1 | - | - | 0 | - |
| - | - | 5667 | 142.1 | - | - | 0 | - |
| - | - | 492.1 | 142.4 | - | - | 0 | - |
| - | - | 575.5 | 143.1 | - | - | 0 | - |
| - | - | 425.1 | 143.1 | - | - | 0 | - |
| - | - | 778.9 | 143.1 | - | - | 0 | - |
| - | - | 5804 | 144.1 | - | - | 0 | - |
| - | - | 3.306E+04 | 146.1 | - | - | 0 | - |
| - | - | 2846 | 147.1 | - | - | 0 | - |
| - | - | 3173 | 147.1 | - | - | 0 | - |
| - | - | 496.2 | 149.1 | - | - | 0 | - |
| - | - | 3895 | 152.1 | - | - | 0 | - |
| - | - | 2722 | 153.1 | - | - | 0 | - |
| - | - | 1805 | 153.1 | - | - | 0 | - |
| - | - | 1120 | 155.1 | - | - | 0 | - |
| 2 | a | 6114 | 155.1 | 0.0003508 | 2.262 | +1 | 2 |
| - | - | 574.2 | 156.9 | - | - | 0 | - |
| - | - | 6.644E+04 | 159.1 | - | - | 0 | - |
| - | - | 1200 | 160.1 | - | - | 0 | - |
| - | - | 6174 | 160.1 | - | - | 0 | - |
| - | - | 966.5 | 163.1 | - | - | 0 | - |
| 7 | y | 563.6 | 167.1 | 0.0004814 | 2.881 | +2 | 2 |
| - | - | 1267 | 169.1 | - | - | 0 | - |
| - | - | 1150 | 169.1 | - | - | 0 | - |
| - | - | 2432 | 169.1 | - | - | 0 | - |
| - | - | 4138 | 170.1 | - | - | 0 | - |
| - | - | 5715 | 171.1 | - | - | 0 | - |
| - | - | 674.5 | 172.1 | - | - | 0 | - |
| 2 | a | 4.337E+04 | 173.1 | 0.0003452 | 1.994 | +1 | 2 |
| - | - | 537.9 | 174.1 | - | - | 0 | - |
| - | - | 672.5 | 174.1 | - | - | 0 | - |
| - | - | 3679 | 174.1 | - | - | 0 | - |
| - | - | 769.2 | 177.1 | - | - | 0 | - |
| - | - | 7238 | 181.1 | - | - | 0 | - |
| - | - | 572.3 | 182.1 | - | - | 0 | - |
| 2 | b | 1.185E+04 | 183.1 | 0.0003245 | 1.772 | +1 | 2 |
| - | - | 3960 | 183.1 | - | - | 0 | - |
| - | - | 989.2 | 184.1 | - | - | 0 | - |
| - | - | 1127 | 187.1 | - | - | 0 | - |
| - | - | 5921 | 187.1 | - | - | 0 | - |
| - | - | 1.249E+04 | 187.1 | - | - | 0 | - |
| - | - | 2.462E+05 | 188.1 | - | - | 0 | - |
| - | - | 1090 | 188.1 | - | - | 0 | - |
| - | - | 2.497E+04 | 189.1 | - | - | 0 | - |
| - | - | 1236 | 190.1 | - | - | 0 | - |
| - | - | 750.2 | 191.1 | - | - | 0 | - |
| - | - | 848.8 | 195.1 | - | - | 0 | - |
| - | - | 677.2 | 195.1 | - | - | 0 | - |
| - | - | 548.2 | 196.1 | - | - | 0 | - |
| - | - | 7581 | 197.1 | - | - | 0 | - |
| - | - | 1081 | 199.1 | - | - | 0 | - |
| - | - | 4.9E+04 | 199.1 | - | - | 0 | - |
| - | - | 4197 | 200.1 | - | - | 0 | - |
| - | - | 957.1 | 200.1 | - | - | 0 | - |
| - | - | 3304 | 201.1 | - | - | 0 | - |
| 2 | b | 4.98E+04 | 201.1 | 0.0002579 | 1.282 | +1 | 2 |
| - | - | 487.5 | 202.1 | - | - | 0 | - |
| - | - | 3828 | 202.1 | - | - | 0 | - |
| 8 | y | 1.008E+05 | 205.1 | 0.0003191 | 1.556 | +1 | 1 |
| - | - | 9799 | 206.1 | - | - | 0 | - |
| - | - | 830.4 | 207.1 | - | - | 0 | - |
| - | - | 5159 | 209.1 | - | - | 0 | - |
| - | - | 692.7 | 210.1 | - | - | 0 | - |
| - | - | 620 | 211.1 | - | - | 0 | - |
| - | - | 2949 | 211.1 | - | - | 0 | - |
| - | - | 1504 | 212.1 | - | - | 0 | - |
| - | - | 602.1 | 213.1 | - | - | 0 | - |
| - | - | 890.5 | 214.2 | - | - | 0 | - |
| - | - | 3.201E+04 | 215.1 | - | - | 0 | - |
| - | - | 2258 | 216.1 | - | - | 0 | - |
| - | - | 2623 | 217.1 | - | - | 0 | - |
| - | - | 528.2 | 217.1 | - | - | 0 | - |
| - | - | 499.6 | 221.1 | - | - | 0 | - |
| - | - | 2098 | 221.1 | - | - | 0 | - |
| - | - | 1116 | 223.1 | - | - | 0 | - |
| - | - | 646.5 | 223.1 | - | - | 0 | - |
| - | - | 1898 | 225.1 | - | - | 0 | - |
| - | - | 1331 | 225.1 | - | - | 0 | - |
| - | - | 896.4 | 226.1 | - | - | 0 | - |
| - | - | 2409 | 226.1 | - | - | 0 | - |
| - | - | 7.336E+04 | 227.1 | - | - | 0 | - |
| - | - | 7966 | 228.1 | - | - | 0 | - |
| - | - | 1205 | 229.2 | - | - | 0 | - |
| - | - | 717.1 | 231.1 | - | - | 0 | - |
| - | - | 597.3 | 237.1 | - | - | 0 | - |
| - | - | 992.3 | 238.2 | - | - | 0 | - |
| - | - | 3572 | 239.1 | - | - | 0 | - |
| - | - | 580.9 | 239.2 | - | - | 0 | - |
| - | - | 8772 | 240.1 | - | - | 0 | - |
| - | - | 723.4 | 240.1 | - | - | 0 | - |
| - | - | 516.8 | 240.1 | - | - | 0 | - |
| - | - | 523.6 | 240.2 | - | - | 0 | - |
| - | - | 3202 | 241.1 | - | - | 0 | - |
| - | - | 535.5 | 241.1 | - | - | 0 | - |
| - | - | 740.8 | 241.2 | - | - | 0 | - |
| - | - | 1268 | 242.2 | - | - | 0 | - |
| - | - | 2938 | 242.7 | - | - | 0 | - |
| - | - | 707.5 | 243.1 | - | - | 0 | - |
| - | - | 863.1 | 244.1 | - | - | 0 | - |
| - | - | 1.728E+04 | 244.1 | - | - | 0 | - |
| - | - | 1098 | 245.1 | - | - | 0 | - |
| - | - | 1637 | 245.1 | - | - | 0 | - |
| 5 | b | 593.5 | 247.7 | 1.074E-05 | 0.04338 | +2 | 5 |
| - | - | 725.9 | 249.1 | - | - | 0 | - |
| - | - | 569.2 | 249.9 | - | - | 0 | - |
| - | - | 485 | 251.1 | - | - | 0 | - |
| - | - | 2946 | 251.2 | - | - | 0 | - |
| - | - | 916.2 | 252.1 | - | - | 0 | - |
| - | - | 526 | 252.3 | - | - | 0 | - |
| - | - | 1778 | 253.1 | - | - | 0 | - |
| - | - | 724.2 | 254.1 | - | - | 0 | - |
| - | - | 2590 | 256.2 | - | - | 0 | - |
| 5 | b | 1012 | 256.7 | 0.0004442 | 1.731 | +2 | 5 |
| - | - | 4069 | 257.1 | - | - | 0 | - |
| - | - | 9343 | 258.1 | - | - | 0 | - |
| - | - | 2126 | 258.1 | - | - | 0 | - |
| 3 | d | 678.6 | 258.2 | 0.0006667 | 2.582 | +1 | 3 |
| - | - | 609.1 | 259.1 | - | - | 0 | - |
| - | - | 1351 | 259.1 | - | - | 0 | - |
| - | - | 839.8 | 266.1 | - | - | 0 | - |
| - | - | 2299 | 266.2 | - | - | 0 | - |
| - | - | 1682 | 267.1 | - | - | 0 | - |
| 3 | a | 2167 | 268.2 | 7.974E-05 | 0.2973 | +1 | 3 |
| - | - | 2304 | 269.1 | - | - | 0 | - |
| - | - | 1406 | 269.2 | - | - | 0 | - |
| - | - | 2.375E+04 | 270.1 | - | - | 0 | - |
| - | - | 881.2 | 270.1 | - | - | 0 | - |
| - | - | 796 | 271.1 | - | - | 0 | - |
| 5 | y | 4012 | 271.1 | 0.003389 | 12.5 | +2 | 4 |
| - | - | 1421 | 274.2 | - | - | 0 | - |
| - | - | 3092 | 280.1 | - | - | 0 | - |
| - | - | 1778 | 280.2 | - | - | 0 | - |
| - | - | 2040 | 282.1 | - | - | 0 | - |
| - | - | 990 | 283.1 | - | - | 0 | - |
| - | - | 967.1 | 284.1 | - | - | 0 | - |
| - | - | 2.381E+04 | 284.2 | - | - | 0 | - |
| - | - | 1035 | 285.1 | - | - | 0 | - |
| - | - | 2906 | 285.2 | - | - | 0 | - |
| 3 | a | 836.4 | 286.2 | 0.001257 | 4.393 | +1 | 3 |
| - | - | 1608 | 292.1 | - | - | 0 | - |
| - | - | 1400 | 294.1 | - | - | 0 | - |
| - | - | 1412 | 294.2 | - | - | 0 | - |
| - | - | 938.3 | 296.1 | - | - | 0 | - |
| 3 | b | 5.064E+04 | 296.2 | 0.000428 | 1.445 | +1 | 3 |
| - | - | 634.7 | 297.1 | - | - | 0 | - |
| - | - | 7596 | 297.2 | - | - | 0 | - |
| - | - | 7745 | 298.1 | - | - | 0 | - |
| - | - | 3901 | 298.2 | - | - | 0 | - |
| - | - | 1058 | 298.2 | - | - | 0 | - |
| - | - | 1175 | 299.1 | - | - | 0 | - |
| - | - | 2924 | 300.2 | - | - | 0 | - |
| - | - | 618.2 | 302.1 | - | - | 0 | - |
| - | - | 3.01E+04 | 302.2 | - | - | 0 | - |
| - | - | 4556 | 303.2 | - | - | 0 | - |
| - | - | 1088 | 307.2 | - | - | 0 | - |
| - | - | 628.3 | 308.2 | - | - | 0 | - |
| - | - | 1248 | 309.2 | - | - | 0 | - |
| - | - | 1306 | 309.2 | - | - | 0 | - |
| - | - | 5931 | 310.1 | - | - | 0 | - |
| - | - | 954.9 | 310.2 | - | - | 0 | - |
| - | - | 1144 | 311.1 | - | - | 0 | - |
| - | - | 1254 | 311.2 | - | - | 0 | - |
| - | - | 2628 | 312.2 | - | - | 0 | - |
| - | - | 1017 | 312.2 | - | - | 0 | - |
| - | - | 694.2 | 313.2 | - | - | 0 | - |
| - | - | 633.6 | 313.2 | - | - | 0 | - |
| - | - | 611.4 | 314.1 | - | - | 0 | - |
| 3 | b | 1.114E+04 | 314.2 | 0.0004529 | 1.441 | +1 | 3 |
| - | - | 5.435E+04 | 315.1 | - | - | 0 | - |
| - | - | 2308 | 315.2 | - | - | 0 | - |
| 7 | y | 5.891E+04 | 316.1 | 0.0003341 | 1.057 | +1 | 2 |
| - | - | 9016 | 316.1 | - | - | 0 | - |
| - | - | 1.045E+04 | 317.1 | - | - | 0 | - |
| - | - | 787.8 | 317.2 | - | - | 0 | - |
| - | - | 812.9 | 318.1 | - | - | 0 | - |
| - | - | 1201 | 320.1 | - | - | 0 | - |
| - | - | 622.8 | 320.2 | - | - | 0 | - |
| 6 | b | 4521 | 321.2 | 0.0003269 | 1.018 | +2 | 6 |
| - | - | 892 | 321.7 | - | - | 0 | - |
| - | - | 847.5 | 326.2 | - | - | 0 | - |
| - | - | 6415 | 327.2 | - | - | 0 | - |
| - | - | 1.117E+04 | 328.2 | - | - | 0 | - |
| - | - | 877.4 | 328.2 | - | - | 0 | - |
| - | - | 1537 | 329.2 | - | - | 0 | - |
| - | - | 1813 | 330.2 | - | - | 0 | - |
| 7 | y | 2.687E+04 | 333.2 | 0.0004879 | 1.464 | +1 | 2 |
| - | - | 1098 | 334.1 | - | - | 0 | - |
| - | - | 4799 | 334.2 | - | - | 0 | - |
| - | - | 839.4 | 334.2 | - | - | 0 | - |
| - | - | 5695 | 337.2 | - | - | 0 | - |
| - | - | 2760 | 338.1 | - | - | 0 | - |
| - | - | 828.3 | 338.2 | - | - | 0 | - |
| - | - | 621 | 339.1 | - | - | 0 | - |
| - | - | 604.6 | 339.1 | - | - | 0 | - |
| - | - | 3986 | 340.2 | - | - | 0 | - |
| - | - | 940.6 | 341.2 | - | - | 0 | - |
| - | - | 922.9 | 343.2 | - | - | 0 | - |
| - | - | 3218 | 345.2 | - | - | 0 | - |
| - | - | 1372 | 346.2 | - | - | 0 | - |
| - | - | 920.9 | 351.2 | - | - | 0 | - |
| - | - | 3868 | 352.2 | - | - | 0 | - |
| - | - | 1048 | 353.2 | - | - | 0 | - |
| - | - | 9.01E+04 | 355.2 | - | - | 0 | - |
| - | - | 7478 | 356.1 | - | - | 0 | - |
| - | - | 1.234E+04 | 356.2 | - | - | 0 | - |
| - | - | 1148 | 357.1 | - | - | 0 | - |
| - | - | 1630 | 357.2 | - | - | 0 | - |
| - | - | 2099 | 363.2 | - | - | 0 | - |
| - | - | 2465 | 364.2 | - | - | 0 | - |
| - | - | 1519 | 365.2 | - | - | 0 | - |
| - | - | 7386 | 369.2 | - | - | 0 | - |
| - | - | 9003 | 370.2 | - | - | 0 | - |
| - | - | 807.9 | 370.3 | - | - | 0 | - |
| - | - | 648.3 | 371.2 | - | - | 0 | - |
| - | - | 3613 | 371.2 | - | - | 0 | - |
| - | - | 1633 | 372.2 | - | - | 0 | - |
| - | - | 2693 | 373.2 | - | - | 0 | - |
| - | - | 4711 | 379.2 | - | - | 0 | - |
| - | - | 3234 | 381.2 | - | - | 0 | - |
| - | - | 2064 | 381.2 | - | - | 0 | - |
| - | - | 635.9 | 381.7 | - | - | 0 | - |
| - | - | 9642 | 387.3 | - | - | 0 | - |
| - | - | 2076 | 388.3 | - | - | 0 | - |
| - | - | 852.7 | 391.1 | - | - | 0 | - |
| - | - | 583.9 | 391.2 | - | - | 0 | - |
| - | - | 999.3 | 393.2 | - | - | 0 | - |
| - | - | 965.4 | 395.2 | - | - | 0 | - |
| - | - | 742.4 | 395.2 | - | - | 0 | - |
| - | - | 832.8 | 396.2 | - | - | 0 | - |
| 4 | b | 2.938E+04 | 397.2 | 0.0002654 | 0.6681 | +1 | 4 |
| - | - | 6691 | 398.2 | - | - | 0 | - |
| - | - | 1273 | 399.2 | - | - | 0 | - |
| - | - | 1150 | 399.3 | - | - | 0 | - |
| - | - | 1580 | 409.2 | - | - | 0 | - |
| - | - | 2482 | 409.2 | - | - | 0 | - |
| - | - | 686.5 | 410.2 | - | - | 0 | - |
| - | - | 792.7 | 411.2 | - | - | 0 | - |
| - | - | 819.4 | 411.3 | - | - | 0 | - |
| - | - | 861.5 | 413.2 | - | - | 0 | - |
| 4 | b | 5.888E+04 | 415.3 | 0.0003513 | 0.8461 | +1 | 4 |
| - | - | 1.322E+04 | 416.3 | - | - | 0 | - |
| - | - | 1311 | 417.3 | - | - | 0 | - |
| - | - | 778.9 | 421.2 | - | - | 0 | - |
| - | - | 2105 | 423.2 | - | - | 0 | - |
| - | - | 874.8 | 424.3 | - | - | 0 | - |
| - | - | 760.4 | 426.2 | - | - | 0 | - |
| - | - | 6405 | 427.2 | - | - | 0 | - |
| - | - | 1705 | 427.2 | - | - | 0 | - |
| - | - | 1463 | 428.2 | - | - | 0 | - |
| - | - | 1120 | 428.2 | - | - | 0 | - |
| - | - | 882.7 | 431.3 | - | - | 0 | - |
| - | - | 747.6 | 434.9 | - | - | 0 | - |
| - | - | 914.5 | 438.2 | - | - | 0 | - |
| - | - | 902.8 | 440.2 | - | - | 0 | - |
| - | - | 2686 | 440.3 | - | - | 0 | - |
| - | - | 1642 | 441.2 | - | - | 0 | - |
| 6 | y | 2065 | 444.2 | 0.0003516 | 0.7915 | +1 | 3 |
| 6 | y | 5250 | 445.2 | 0.0005266 | 1.183 | +1 | 3 |
| - | - | 957.1 | 446.2 | - | - | 0 | - |
| - | - | 1051 | 451.2 | - | - | 0 | - |
| - | - | 1075 | 451.7 | - | - | 0 | - |
| - | - | 6293 | 456.2 | - | - | 0 | - |
| - | - | 1504 | 457.2 | - | - | 0 | - |
| 6 | y | 2346 | 462.2 | 9.039E-06 | 0.01956 | +1 | 3 |
| - | - | 1004 | 466.3 | - | - | 0 | - |
| - | - | 7610 | 468.2 | - | - | 0 | - |
| - | - | 735.8 | 468.3 | - | - | 0 | - |
| - | - | 955.5 | 469.3 | - | - | 0 | - |
| - | - | 1499 | 469.7 | - | - | 0 | - |
| - | - | 766.2 | 470.2 | - | - | 0 | - |
| - | - | 1956 | 472.2 | - | - | 0 | - |
| - | - | 604.1 | 474.2 | - | - | 0 | - |
| - | - | 932.7 | 476.3 | - | - | 0 | - |
| - | - | 2481 | 478.2 | - | - | 0 | - |
| - | - | 922.3 | 486.2 | - | - | 0 | - |
| - | - | 730.1 | 487.3 | - | - | 0 | - |
| - | - | 1866 | 488.3 | - | - | 0 | - |
| 5 | b | 5665 | 494.3 | 0.0002359 | 0.4773 | +1 | 5 |
| - | - | 1523 | 495.3 | - | - | 0 | - |
| - | - | 2307 | 496.2 | - | - | 0 | - |
| - | - | 801.4 | 499.3 | - | - | 0 | - |
| - | - | 2253 | 506.2 | - | - | 0 | - |
| - | - | 758.2 | 508.3 | - | - | 0 | - |
| - | - | 3223 | 510.3 | - | - | 0 | - |
| 5 | b | 5373 | 512.3 | 0.0001693 | 0.3304 | +1 | 5 |
| - | - | 1309 | 513.2 | - | - | 0 | - |
| - | - | 2205 | 513.3 | - | - | 0 | - |
| - | - | 676.9 | 514.2 | - | - | 0 | - |
| - | - | 1.619E+04 | 524.2 | - | - | 0 | - |
| - | - | 5563 | 525.2 | - | - | 0 | - |
| - | - | 1180 | 526.3 | - | - | 0 | - |
| - | - | 1459 | 527.3 | - | - | 0 | - |
| - | - | 2235 | 528.3 | - | - | 0 | - |
| - | - | 740.7 | 529.3 | - | - | 0 | - |
| - | - | 653.6 | 535.3 | - | - | 0 | - |
| - | - | 1630 | 537.3 | - | - | 0 | - |
| - | - | 1037 | 538.3 | - | - | 0 | - |
| 5 | y | 6775 | 541.2 | 0.0003819 | 0.7056 | +1 | 4 |
| - | - | 816.5 | 541.3 | - | - | 0 | - |
| 5 | y | 2.719E+04 | 542.2 | 0.0006192 | 1.142 | +1 | 4 |
| - | - | 8559 | 543.2 | - | - | 0 | - |
| - | - | 796 | 543.3 | - | - | 0 | - |
| - | - | 1153 | 544.2 | - | - | 0 | - |
| - | - | 838.8 | 545.3 | - | - | 0 | - |
| - | - | 1980 | 551.3 | - | - | 0 | - |
| - | - | 655.5 | 552.3 | - | - | 0 | - |
| - | - | 2921 | 555.3 | - | - | 0 | - |
| 5 | y | 7.287E+04 | 559.3 | 7.111E-05 | 0.1272 | +1 | 4 |
| - | - | 2.248E+04 | 560.3 | - | - | 0 | - |
| - | - | 4720 | 561.3 | - | - | 0 | - |
| - | - | 1118 | 562.3 | - | - | 0 | - |
| - | - | 2842 | 569.3 | - | - | 0 | - |
| - | - | 1051 | 570.3 | - | - | 0 | - |
| - | - | 920.1 | 577.3 | - | - | 0 | - |
| - | - | 1056 | 593.3 | - | - | 0 | - |
| - | - | 1227 | 595.3 | - | - | 0 | - |
| - | - | 1384 | 605.3 | - | - | 0 | - |
| - | - | 955.6 | 607.3 | - | - | 0 | - |
| - | - | 1205 | 613.4 | - | - | 0 | - |
| - | - | 1442 | 620.3 | - | - | 0 | - |
| 6 | b | 8623 | 623.3 | 0.0003674 | 0.5895 | +1 | 6 |
| - | - | 1944 | 624.3 | - | - | 0 | - |
| - | - | 641.1 | 625.3 | - | - | 0 | - |
| - | - | 786.7 | 625.3 | - | - | 0 | - |
| - | - | 5120 | 638.3 | - | - | 0 | - |
| - | - | 1938 | 639.3 | - | - | 0 | - |
| - | - | 2460 | 640.4 | - | - | 0 | - |
| 6 | b | 8369 | 641.4 | 0.0007891 | 1.23 | +1 | 6 |
| - | - | 2897 | 642.4 | - | - | 0 | - |
| 4 | y | 1846 | 643.3 | 0.005065 | 7.873 | +1 | 5 |
| - | - | 3380 | 650.3 | - | - | 0 | - |
| - | - | 1008 | 651.3 | - | - | 0 | - |
| - | - | 2793 | 656.3 | - | - | 0 | - |
| - | - | 875.3 | 657.3 | - | - | 0 | - |
| - | - | 3504 | 658.4 | - | - | 0 | - |
| - | - | 1652 | 659.4 | - | - | 0 | - |
| 4 | y | 9060 | 660.3 | 0.000122 | 0.1848 | +1 | 5 |
| - | - | 3036 | 661.3 | - | - | 0 | - |
| - | - | 681.9 | 662.3 | - | - | 0 | - |
| - | - | 1454 | 668.4 | - | - | 0 | - |
| - | - | 704.4 | 716.4 | - | - | 0 | - |
| - | - | 1300 | 723.4 | - | - | 0 | - |
| - | - | 1113 | 724.4 | - | - | 0 | - |
| - | - | 938.4 | 725.4 | - | - | 0 | - |
| - | - | 1313 | 727.5 | - | - | 0 | - |
| - | - | 3505 | 733.4 | - | - | 0 | - |
| - | - | 2981 | 734.4 | - | - | 0 | - |
| - | - | 1308 | 735.4 | - | - | 0 | - |
| - | - | 2056 | 741.4 | - | - | 0 | - |
| - | - | 1380 | 742.4 | - | - | 0 | - |
| 7 | b | 1.821E+04 | 751.4 | 0.0004708 | 0.6266 | +1 | 7 |
| 7 | b | 8100 | 752.4 | 0.01374 | 18.27 | +1 | 7 |
| - | - | 2707 | 753.4 | - | - | 0 | - |
| 3 | y | 908.8 | 755.4 | 0.002245 | 2.973 | +1 | 6 |
| - | - | 824.4 | 769.3 | - | - | 0 | - |
| 7 | b | 1.212E+04 | 769.4 | 0.0003544 | 0.4606 | +1 | 7 |
| - | - | 5660 | 770.4 | - | - | 0 | - |
| - | - | 982.1 | 771.4 | - | - | 0 | - |
| 3 | y | 1235 | 773.4 | 0.001972 | 2.549 | +1 | 6 |
| 2 | y | 679.2 | 843.4 | 0.006099 | 7.231 | +1 | 7 |
| 2 | y | 694.3 | 860.4 | 0.0003086 | 0.3587 | +1 | 7 |
| - | - | 667.5 | 1056 | - | - | 0 | - |
| - | - | 557.1 | 1082 | - | - | 0 | - |
| - | - | 742 | 1223 | - | - | 0 | - |
| - | - | 532.9 | 1259 | - | - | 0 | - |
| - | - | 682.6 | 1751 | - | - | 0 | - |
| - | - | 650.7 | 2348 | - | - | 0 | - |
| - | - | 681.6 | 2551 | - | - | 0 | - |
| - | - | 739.1 | 3082 | - | - | 0 | - |

m/z Charge Intensity FragmentType MassShift Position
120.0810546875 0 2000.0172
121.0282974243164 0 614.666
125.0714111328125 0 1658.5043
125.10773468017578 0 3666.7698
126.09163665771484 0 881.8894
127.07573699951172 0 393.4173
127.08684539794922 0 662.76044
128.107421875 0 1600.1261
129.06617736816406 0 9531.094
129.10256958007812 0 2289.0933
130.05014038085938 0 3248.8467
130.06546020507812 0 23708.129
131.06895446777344 0 2091.0137
132.0811004638672 0 11514.895
132.1022491455078 0 1354.7025
133.08607482910156 0 3134.5332
136.0760040283203 0 1348.7124
138.09164428710938 0 771.7782
139.05038452148438 0 1039.9569
141.09149169921875 0 394.92856
141.10276794433594 0 553.3834
142.0653533935547 0 997.5948
142.12301635742188 0 5667.1807
142.4151611328125 0 492.0949
143.0725555419922 0 575.51166
143.1065673828125 0 425.1446
143.11807250976562 0 778.92584
144.08116149902344 0 5804.2266
146.06039428710938 0 33062.195
147.06370544433594 0 2846.3
147.07676696777344 0 3173.21
149.09725952148438 0 496.21622
152.10733032226562 0 3895.3635
153.066162109375 0 2721.9668
153.10256958007812 0 1805.1411
155.08189392089844 0 1119.69
155.1182403564453 0 6113.897 a Water loss 1
156.92111206054688 0 574.2456
159.09201049804688 0 66435.516
160.07595825195312 0 1199.8094
160.0954132080078 0 6173.9575
163.0867919921875 0 966.5189
167.0819854736328 0 563.6294 y 6
169.076416015625 0 1266.8964
169.0977020263672 0 1149.7665
169.13381958007812 0 2431.9329
170.060302734375 0 4137.6914
171.11312866210938 0 5714.9736
172.11622619628906 0 674.50085
173.12879943847656 0 43366.688 a 1
174.05511474609375 0 537.91064
174.12496948242188 0 672.546
174.1321563720703 0 3678.8662
177.11260986328125 0 769.1942
181.0974884033203 0 7237.7886
182.10081481933594 0 572.31006
183.11312866210938 0 11854.451 b Water loss 1
183.1495819091797 0 3959.6184
184.11669921875 0 989.2315
187.0720672607422 0 1127.4806
187.08697509765625 0 5920.84
187.14447021484375 0 12487.884
188.07101440429688 0 246208.47
188.14817810058594 0 1089.7273
189.07432556152344 0 24971.627
190.07640075683594 0 1236.348
191.08169555664062 0 750.2242
195.07664489746094 0 848.81683
195.1491241455078 0 677.20483
196.1449432373047 0 548.1901
197.1287841796875 0 7581.3623
199.07139587402344 0 1081.3914
199.10801696777344 0 48995.97
200.11141967773438 0 4196.996
200.13983154296875 0 957.08655
201.1023712158203 0 3303.7554
201.12362670898438 0 49797.445 b 1
202.1072540283203 0 487.50223
202.12716674804688 0 3827.647
205.09747314453125 0 100795.99 y 7
206.1007537841797 0 9799.173
207.10252380371094 0 830.39557
209.09234619140625 0 5158.6406
210.12451171875 0 692.6796
211.10797119140625 0 619.9718
211.1444091796875 0 2948.9822
212.10330200195312 0 1504.2233
213.08724975585938 0 602.0508
214.1551055908203 0 890.4734
215.1393280029297 0 32005.88
216.14273071289062 0 2257.5486
217.0819854736328 0 2622.6653
217.0922088623047 0 528.2303
221.0799102783203 0 499.56287
221.12864685058594 0 2098.4175
223.0718994140625 0 1116.3322
223.14439392089844 0 646.4555
225.102783203125 0 1897.5477
225.12364196777344 0 1331.004
226.1069793701172 0 896.391
226.11915588378906 0 2408.734
227.10302734375 0 73358.52
228.1062774658203 0 7966.1836
229.15521240234375 0 1204.6389
231.09934997558594 0 717.127
237.12289428710938 0 597.34033
238.1556396484375 0 992.31006
239.13922119140625 0 3572.155
239.15081787109375 0 580.90424
240.0982208251953 0 8772.496
240.1117706298828 0 723.41626
240.13458251953125 0 516.8216
240.1702117919922 0 523.57513
241.0825653076172 0 3201.7488
241.10284423828125 0 535.5289
241.1912078857422 0 740.7831
242.15052795410156 0 1268.0488
242.6602783203125 0 2937.515
243.134521484375 0 707.53674
244.1153106689453 0 863.0819
244.12954711914062 0 17282.57
245.11363220214844 0 1098.0646
245.13294982910156 0 1636.6781
247.65228271484375 0 593.5443 b Water loss 4
249.12425231933594 0 725.91876
249.94471740722656 0 569.1795
251.140869140625 0 484.98462
251.1757049560547 0 2946.0374
252.11326599121094 0 916.22345
252.3122100830078 0 526.0228
253.09762573242188 0 1777.9849
254.11373901367188 0 724.1863
256.16595458984375 0 2589.631
256.65802001953125 0 1011.6892 b 4
257.1497802734375 0 4069.284
258.10870361328125 0 9342.853
258.1456604003906 0 2125.893
258.181884765625 0 678.5823 d 2
259.0921630859375 0 609.14886
259.11163330078125 0 1350.5925
266.1136779785156 0 839.7749
266.15032958984375 0 2299.1013
267.13433837890625 0 1681.9163
268.2018737792969 0 2167.445 a Water loss 2
269.1398010253906 0 2303.8655
269.1860656738281 0 1405.5836
270.1239318847656 0 23753.635
270.14056396484375 0 881.1942
271.1083984375 0 796.035
271.1272888183594 0 4011.6086 y Water loss 4
274.1766052246094 0 1421.0703
280.10870361328125 0 3091.5127
280.1661376953125 0 1778.3528
282.1448059082031 0 2040.1182
283.12939453125 0 989.9918
284.124267578125 0 967.1173
284.1607971191406 0 23808.54
285.1458740234375 0 1035.2322
285.1639709472656 0 2906.244
286.2137756347656 0 836.4164 a 2
292.1297912597656 0 1607.9697
294.14569091796875 0 1400.4003
294.1815185546875 0 1411.7407
296.1239013671875 0 938.3408
296.1972961425781 0 50637.25 b Water loss 2
297.13446044921875 0 634.6908
297.2005615234375 0 7595.891
298.11895751953125 0 7745.474
298.17645263671875 0 3900.8857
298.203125 0 1057.7964
299.1224365234375 0 1175.3342
300.1559143066406 0 2923.8906
302.1156311035156 0 618.1606
302.1714782714844 0 30099.633
303.1746520996094 0 4556.1934
307.1810607910156 0 1087.5841
308.1966247558594 0 628.2658
309.1576232910156 0 1248.0819
309.2037048339844 0 1306.1381
310.1402282714844 0 5931.42
310.212646484375 0 954.9345
311.1451416015625 0 1144.008
311.17169189453125 0 1254.4598
312.1555480957031 0 2628.0469
312.1925048828125 0 1017.0712
313.15936279296875 0 694.23645
313.19537353515625 0 633.6228
314.13427734375 0 611.4185
314.2078857421875 0 11139.408 b 2
315.14556884765625 0 54345.902
315.2112121582031 0 2307.653
316.1295166015625 0 58905.28 y Ammonia loss 6
316.1488342285156 0 9016.415
317.1328125 0 10449.091
317.1527404785156 0 787.7657
318.13555908203125 0 812.85876
320.12506103515625 0 1200.881
320.1610107421875 0 622.8025
321.17919921875 0 4520.7725 b 5
321.68072509765625 0 892.02325
326.1720886230469 0 847.4598
327.1668395996094 0 6414.9575
328.15069580078125 0 11165.281
328.16961669921875 0 877.4185
329.1531677246094 0 1537.3805
330.166015625 0 1812.7064
333.1562194824219 0 26868.531 y 6
334.1391906738281 0 1098.4733
334.1593017578125 0 4798.7
334.21234130859375 0 839.43195
337.1513671875 0 5695.1763
338.13531494140625 0 2759.677
338.15509033203125 0 828.2779
339.1062927246094 0 620.9693
339.1384582519531 0 604.63336
340.1870422363281 0 3986.0906
341.18988037109375 0 940.55096
343.2343444824219 0 922.90936
345.1775207519531 0 3218.3794
346.1784362792969 0 1372.1106
351.2408447265625 0 920.85004
352.22296142578125 0 3868.266
353.2222595214844 0 1047.6888
355.16143798828125 0 90096.03
356.1445007324219 0 7478.383
356.1649169921875 0 12341.612
357.1463928222656 0 1148.4252
357.16790771484375 0 1630.0042
363.20318603515625 0 2098.791
364.2392578125 0 2465.2417
365.1820373535156 0 1518.5265
369.2499084472656 0 7386.299
370.23370361328125 0 9002.628
370.25311279296875 0 807.941
371.1719970703125 0 648.3454
371.23004150390625 0 3613.435
372.1876220703125 0 1632.6995
373.17193603515625 0 2692.5083
379.2344055175781 0 4711.3906
381.1561584472656 0 3233.6636
381.21429443359375 0 2064.4866
381.68499755859375 0 635.8836
387.2605285644531 0 9642.39
388.2629699707031 0 2076.1375
391.1392822265625 0 852.68616
391.19873046875 0 583.89105
393.2488708496094 0 999.3252
395.1573486328125 0 965.39307
395.2291564941406 0 742.4313
396.215087890625 0 832.82764
397.24481201171875 0 29383.762 b Water loss 3
398.2478942871094 0 6690.9517
399.2220764160156 0 1273.424
399.2504577636719 0 1149.7412
409.1506652832031 0 1580.1718
409.2086181640625 0 2481.8518
410.2110900878906 0 686.4674
411.1882629394531 0 792.74164
411.2607116699219 0 819.3667
413.239501953125 0 861.50854
415.2554626464844 0 58876.633 b 3
416.25848388671875 0 13220.959
417.2610168457031 0 1310.7593
421.1691589355469 0 778.90796
423.2231750488281 0 2104.9092
424.25482177734375 0 874.84595
426.238037109375 0 760.38336
427.16168212890625 0 6404.7075
427.2188415527344 0 1705.4746
428.1643371582031 0 1462.8625
428.2145080566406 0 1119.8975
431.26763916015625 0 882.6977
434.8704833984375 0 747.60156
438.1991882324219 0 914.5072
440.2188720703125 0 902.84796
440.2517395019531 0 2686.3682
441.2344970703125 0 1642.061
444.1874084472656 0 2065.0469 y Water loss 5
445.17230224609375 0 5249.8774 y Ammonia loss 5
446.174560546875 0 957.1455
451.2183532714844 0 1050.9471
451.72314453125 0 1074.6794
456.20904541015625 0 6293.411
457.2119140625 0 1504.2495
462.1983337402344 0 2345.778 y 5
466.3047790527344 0 1004.4299
468.2455749511719 0 7610.0024
468.2806091308594 0 735.8429
469.2506408691406 0 955.4835
469.7345275878906 0 1499.4078
470.2344665527344 0 766.2259
472.1833801269531 0 1955.961
474.2178039550781 0 604.068
476.2859191894531 0 932.6649
478.2087707519531 0 2480.732
486.1501770019531 0 922.2827
487.3050537109375 0 730.098
488.3056335449219 0 1865.9636
494.29754638671875 0 5664.511 b Water loss 4
495.2987976074219 0 1522.9603
496.22052001953125 0 2307.1328
499.2762145996094 0 801.3761
506.2039489746094 0 2253.1455
508.2770690917969 0 758.2231
510.2559814453125 0 3222.795
512.3080444335938 0 5373.342 b 4
513.2465209960938 0 1308.7546
513.3118286132812 0 2204.906
514.2313232421875 0 676.8595
524.214111328125 0 16193.146
525.217529296875 0 5562.849
526.2857055664062 0 1180.2836
527.2841796875 0 1458.9714
528.2667236328125 0 2235.1936
529.2720947265625 0 740.7466
535.2802734375 0 653.643
537.2666015625 0 1630.4456
538.2529296875 0 1036.616
541.2409057617188 0 6775.159 y Water loss 4
541.2999267578125 0 816.5419
542.2251586914062 0 27194.074 y Ammonia loss 4
543.2274169921875 0 8558.864
543.3096923828125 0 795.9533
544.2300415039062 0 1152.8223
545.292724609375 0 838.77673
551.2816162109375 0 1980.0012
552.2802124023438 0 655.5022
555.2781982421875 0 2920.8408
559.2511596679688 0 72872.02 y 4
560.251953125 0 22479.717
561.2523193359375 0 4719.8335
562.2978515625 0 1117.6682
569.2922973632812 0 2842.4692
570.2928466796875 0 1050.7847
577.3345336914062 0 920.14813
593.2946166992188 0 1056.4536
595.3458862304688 0 1227.1575
605.3327026367188 0 1384.0823
607.25146484375 0 955.5665
613.3540649414062 0 1205.0897
620.3034057617188 0 1442.4635
623.3402709960938 0 8623.439 b Water loss 5
624.3425903320312 0 1943.8665
625.264404296875 0 641.1111
625.34375 0 786.7212
638.3131713867188 0 5120.4956
639.3118286132812 0 1937.5264
640.3665771484375 0 2459.8125
641.3512573242188 0 8368.62 b 5
642.35400390625 0 2897.1335
643.2772827148438 0 1846.2965 y Ammonia loss 3
650.3497924804688 0 3379.8528
651.3432006835938 0 1008.44147
656.32470703125 0 2793.4946
657.324462890625 0 875.2623
658.376220703125 0 3503.892
659.3795166015625 0 1651.5677
660.2986450195312 0 9059.763 y 3
661.3004760742188 0 3036.484
662.3032836914062 0 681.8672
668.36181640625 0 1454.3922
716.3646850585938 0 704.36646
723.4053344726562 0 1299.54
724.3936157226562 0 1113.0665
725.3836669921875 0 938.3935
727.4708862304688 0 1312.7828
733.3869018554688 0 3504.971
734.3749389648438 0 2981.4543
735.3723754882812 0 1307.8186
741.4130859375 0 2055.6897
742.41748046875 0 1380.4498
751.3980102539062 0 18206.21 b Water loss 6
752.396240234375 0 8100.0234 b Ammonia loss 6
753.3927001953125 0 2706.8733
755.37451171875 0 908.7873 y Water loss 2
769.3355712890625 0 824.4344
769.40869140625 0 12121.379 b 6
770.4090576171875 0 5660.1978
771.4116821289062 0 982.0745
773.380859375 0 1235.0856 y 2
843.3944091796875 0 679.2007 y Ammonia loss 1
860.41455078125 0 694.3495 y 1
1055.633544921875 0 667.46594
1081.59814453125 0 557.13226
1222.5565185546875 0 741.97754
1259.0589599609375 0 532.8799
1751.141845703125 0 682.56415
2347.755859375 0 650.6809
2551.229248046875 0 681.57385
3082.17333984375 0 739.07275

Spectrum Details

|  |  |
| --- | --- |
| Matched peaks? Matched peaks The total absolute number of peaks matched. Additionally in brackets the total fraction of peaks matched and the total number of peaks is shown. | 38 (9.74% of 390) |
| FDR? FDR The false discovery rate estimated for this peptide. It is calculated by matching all theoretical fragments with a non-integer shift with the raw peaks for this spectrum. This is done with 40 different shifts. The resulting percentage is the average number of annotated peaks over the number of annotated peaks with the correct spectrum. | 1.00% |
| Satellite FDR? Satellite FDR See the FDR for details on its calculation. This satellite ion specific FDR only contains the satellite ions (d/w) for I/L/J positions. | 4.76% |
| PSM Score? PSM Score The PSM Score as given by Hecklib to this annotated spectrum. It is shown with three significant figures. | 576 |

#### Spectrum 10510? Spectrum 10510 The raw spectrum of this peptide as annotated by Hecklib. The fragments are coloured according to ion type (see legend). Any peaks with a star '\*' as text can be hovered over to see the full details, first the ion type second the mass shift type. By hovering over the amino acids in the peptide or ions in the legend the corresponding peaks are highlighted. By toggling the 'Unassigned' label you can turn the background (unassigned) peaks on or off in the plot. By updating the slider in the Ion legend you can update the spectrum to only show the top X% of the peaks with labels. The top X% means any peak that is within X% of the highest intensity. By dragging in the spectrum you can zoom in to a specific part of the spectrum and use 'Zoom Out' to get back to the original zoom level. The annotation of the spectrum is based on the given sequence in the peptides file and is done with different software so inconsistencies are likely. The peaks are annotated based on the given sequence, with 20 ppm tolerance.

Copy Data

##### Spectrum 10510 (TSV)

###### Preview

```
Loading example...
```

*Click on the button to copy the data to your clipboard.*

Mz MinMz MaxIntensity Max

WidthHeightPeptide font sizePeptide stroke widthSpectrum font sizeSpectrum stroke widthCompact peptide

Ion legend

wxyz

abcd

OtherUnassignedIon^Charge^_Position_Show for top:%

JSJTPEQW

07.41e+31.48e+42.22e+42.97e+4

Zoom Out

y^+1^_1_y^+1^_2_y^+1^_2_y^+1^_3_c^+1^_5_y^+1^_4_y^+1^_4_z^+1^_5_c^+1^_6_y^+1^_5_z^+1^_6_c^+1^_7_y^+1^_7_z^+1^_7_y^+1^_7_

0649129819472595

Fragment Matches Table

Show background peaks

| Position | Ion type | Intensity | mz Theoretical | mz Error (Th) | mz Error (ppm) | Charge | Series Number |
| --- | --- | --- | --- | --- | --- | --- | --- |
| - | - | 376.5 | 120.7 | - | - | 0 | - |
| - | - | 373.2 | 122 | - | - | 0 | - |
| - | - | 976.3 | 133.1 | - | - | 0 | - |
| - | - | 410.6 | 136.9 | - | - | 0 | - |
| - | - | 422.7 | 137.1 | - | - | 0 | - |
| - | - | 463.7 | 149 | - | - | 0 | - |
| - | - | 930.3 | 159.1 | - | - | 0 | - |
| - | - | 427.9 | 162.8 | - | - | 0 | - |
| - | - | 452.5 | 167.5 | - | - | 0 | - |
| - | - | 564 | 177.1 | - | - | 0 | - |
| - | - | 4138 | 188.1 | - | - | 0 | - |
| - | - | 2106 | 201.1 | - | - | 0 | - |
| - | - | 2109 | 201.1 | - | - | 0 | - |
| 8 | y | 5122 | 205.1 | 2.913E-05 | 0.1421 | +1 | 1 |
| - | - | 743.7 | 206.1 | - | - | 0 | - |
| - | - | 1355 | 215.1 | - | - | 0 | - |
| - | - | 1253 | 227.1 | - | - | 0 | - |
| - | - | 2875 | 244.1 | - | - | 0 | - |
| - | - | 577.2 | 258.1 | - | - | 0 | - |
| - | - | 2263 | 296.2 | - | - | 0 | - |
| - | - | 577.5 | 297.2 | - | - | 0 | - |
| - | - | 648.9 | 302.2 | - | - | 0 | - |
| - | - | 556.9 | 309.1 | - | - | 0 | - |
| - | - | 1154 | 309.2 | - | - | 0 | - |
| - | - | 645.4 | 310.2 | - | - | 0 | - |
| - | - | 494.6 | 313.2 | - | - | 0 | - |
| - | - | 544.3 | 314.2 | - | - | 0 | - |
| - | - | 918.3 | 315.1 | - | - | 0 | - |
| 7 | y | 1897 | 316.1 | 0.0001815 | 0.5742 | +1 | 2 |
| 7 | y | 1949 | 333.2 | 0.0001827 | 0.5484 | +1 | 2 |
| - | - | 4353 | 355.2 | - | - | 0 | - |
| - | - | 791.2 | 356.2 | - | - | 0 | - |
| - | - | 558 | 366.1 | - | - | 0 | - |
| - | - | 666.6 | 370.2 | - | - | 0 | - |
| - | - | 2520 | 372.2 | - | - | 0 | - |
| - | - | 1889 | 397.2 | - | - | 0 | - |
| - | - | 5658 | 415.3 | - | - | 0 | - |
| - | - | 974.5 | 416.3 | - | - | 0 | - |
| - | - | 677.8 | 416.9 | - | - | 0 | - |
| - | - | 843.1 | 434.9 | - | - | 0 | - |
| - | - | 747.4 | 441.2 | - | - | 0 | - |
| - | - | 735.6 | 441.2 | - | - | 0 | - |
| 6 | y | 574.3 | 444.2 | 0.00212 | 4.774 | +1 | 3 |
| - | - | 595.5 | 451.2 | - | - | 0 | - |
| - | - | 2104 | 456.2 | - | - | 0 | - |
| - | - | 2291 | 458.2 | - | - | 0 | - |
| - | - | 600.5 | 470.2 | - | - | 0 | - |
| - | - | 620.1 | 485.7 | - | - | 0 | - |
| - | - | 793.2 | 486.2 | - | - | 0 | - |
| - | - | 651.5 | 488.3 | - | - | 0 | - |
| - | - | 1059 | 514.3 | - | - | 0 | - |
| - | - | 1381 | 528.3 | - | - | 0 | - |
| 5 | c | 9214 | 529.3 | 1.263E-05 | 0.02386 | +1 | 5 |
| - | - | 3081 | 530.3 | - | - | 0 | - |
| 5 | y | 1617 | 542.2 | 0.0009066 | 1.672 | +1 | 4 |
| 5 | y | 1.105E+04 | 559.3 | 5.096E-05 | 0.09111 | +1 | 4 |
| - | - | 2986 | 560.3 | - | - | 0 | - |
| - | - | 627.5 | 599.3 | - | - | 0 | - |
| - | - | 868.6 | 614.4 | - | - | 0 | - |
| - | - | 816.5 | 641.4 | - | - | 0 | - |
| - | - | 2875 | 642.3 | - | - | 0 | - |
| 4 | z | 6334 | 644.3 | 0.0002578 | 0.4001 | +1 | 5 |
| - | - | 5556 | 645.3 | - | - | 0 | - |
| - | - | 1694 | 646.3 | - | - | 0 | - |
| - | - | 676.9 | 647.3 | - | - | 0 | - |
| 6 | c | 1.909E+04 | 658.4 | 5.785E-05 | 0.08786 | +1 | 6 |
| - | - | 8706 | 659.4 | - | - | 0 | - |
| 4 | y | 1328 | 660.3 | 0.001465 | 2.218 | +1 | 5 |
| - | - | 1576 | 660.4 | - | - | 0 | - |
| - | - | 617.5 | 661.3 | - | - | 0 | - |
| - | - | 3152 | 684.4 | - | - | 0 | - |
| - | - | 905.7 | 685.4 | - | - | 0 | - |
| - | - | 695.5 | 728.3 | - | - | 0 | - |
| - | - | 731.4 | 741.4 | - | - | 0 | - |
| - | - | 728 | 751.4 | - | - | 0 | - |
| 3 | z | 1356 | 757.4 | 0.001066 | 1.408 | +1 | 6 |
| - | - | 1536 | 758.4 | - | - | 0 | - |
| - | - | 785.5 | 770.4 | - | - | 0 | - |
| - | - | 1866 | 771.4 | - | - | 0 | - |
| - | - | 701.7 | 772.4 | - | - | 0 | - |
| - | - | 2033 | 773.4 | - | - | 0 | - |
| - | - | 1151 | 774.4 | - | - | 0 | - |
| 7 | c | 2.936E+04 | 786.4 | 1.305E-05 | 0.01659 | +1 | 7 |
| - | - | 1.125E+04 | 787.4 | - | - | 0 | - |
| - | - | 533.9 | 788.3 | - | - | 0 | - |
| - | - | 2949 | 788.4 | - | - | 0 | - |
| - | - | 723.4 | 789.4 | - | - | 0 | - |
| - | - | 609.1 | 800.4 | - | - | 0 | - |
| - | - | 736.8 | 828.4 | - | - | 0 | - |
| 2 | y | 694.3 | 842.4 | 0.00281 | 3.335 | +1 | 7 |
| - | - | 1153 | 843.4 | - | - | 0 | - |
| 2 | z | 5591 | 844.4 | 0.0002613 | 0.3095 | +1 | 7 |
| - | - | 2993 | 845.4 | - | - | 0 | - |
| - | - | 828.2 | 846.4 | - | - | 0 | - |
| - | - | 3661 | 857.5 | - | - | 0 | - |
| - | - | 1658 | 858.5 | - | - | 0 | - |
| - | - | 735.2 | 859.5 | - | - | 0 | - |
| 2 | y | 1125 | 860.4 | 0.0006748 | 0.7843 | +1 | 7 |
| - | - | 627.1 | 875 | - | - | 0 | - |
| - | - | 1316 | 884.5 | - | - | 0 | - |
| - | - | 937.7 | 901.5 | - | - | 0 | - |
| - | - | 604.3 | 903.5 | - | - | 0 | - |
| - | - | 1602 | 912.5 | - | - | 0 | - |
| - | - | 993.2 | 914.4 | - | - | 0 | - |
| - | - | 1100 | 929.5 | - | - | 0 | - |
| - | - | 841.4 | 930.5 | - | - | 0 | - |
| - | - | 783 | 931.5 | - | - | 0 | - |
| - | - | 1558 | 956.5 | - | - | 0 | - |
| - | - | 6819 | 957.5 | - | - | 0 | - |
| - | - | 3122 | 958.5 | - | - | 0 | - |
| - | - | 643.2 | 959.5 | - | - | 0 | - |
| - | - | 9839 | 973.5 | - | - | 0 | - |
| - | - | 1.581E+04 | 974.5 | - | - | 0 | - |
| - | - | 7638 | 975.5 | - | - | 0 | - |
| - | - | 523.6 | 975.6 | - | - | 0 | - |
| - | - | 3036 | 976.5 | - | - | 0 | - |
| - | - | 2085 | 1459 | - | - | 0 | - |
| - | - | 1441 | 1460 | - | - | 0 | - |
| - | - | 773.1 | 2570 | - | - | 0 | - |

m/z Charge Intensity FragmentType MassShift Position
120.6772689819336 0 376.5369
122.00000762939453 0 373.22238
133.08627319335938 0 976.298
136.8639373779297 0 410.5956
137.0963592529297 0 422.6673
148.9550018310547 0 463.74496
159.09170532226562 0 930.27026
162.75784301757812 0 427.8847
167.4859619140625 0 452.54242
177.11172485351562 0 564.03546
188.07070922851562 0 4137.9443
201.1019744873047 0 2106.2595
201.12364196777344 0 2108.7673
205.09718322753906 0 5122.0806 y 7
206.1002655029297 0 743.65814
215.13937377929688 0 1355.2206
227.10243225097656 0 1253.1512
244.12930297851562 0 2875.4297
258.1448059082031 0 577.1966
296.1968078613281 0 2263.4087
297.19915771484375 0 577.47327
302.171630859375 0 648.8834
309.1371765136719 0 556.9025
309.2033996582031 0 1153.6376
310.2066650390625 0 645.41455
313.2369079589844 0 494.56064
314.20660400390625 0 544.3188
315.1457214355469 0 918.3088
316.1293640136719 0 1896.7732 y Ammonia loss 6
333.1559143066406 0 1949.3948 y 6
355.16131591796875 0 4353.0757
356.1650390625 0 791.2124
366.09039306640625 0 557.98505
370.2332458496094 0 666.5802
372.1882629394531 0 2520.36
397.24468994140625 0 1889.0774
415.2551574707031 0 5657.64
416.2592468261719 0 974.4979
416.8586730957031 0 677.82635
434.8694763183594 0 843.0982
441.20068359375 0 747.37085
441.23468017578125 0 735.5531
444.18988037109375 0 574.26794 y Water loss 5
451.19647216796875 0 595.5051
456.2084655761719 0 2103.6187
458.2242431640625 0 2290.8484
470.1959228515625 0 600.49146
485.7112121582031 0 620.1191
486.1873474121094 0 793.21814
488.2960205078125 0 651.54785
514.2522583007812 0 1059.3889
528.3276977539062 0 1380.6375
529.3344116210938 0 9213.723 c 4
530.3380737304688 0 3080.8853
542.2236328125 0 1617.4934 y Ammonia loss 4
559.2510375976562 0 11047.026 y 4
560.2536010742188 0 2985.7808
599.2596435546875 0 627.4983
614.3639526367188 0 868.5529
641.3565673828125 0 816.4888
642.2637939453125 0 2875.3906
644.27978515625 0 6334.104 z 3
645.2854614257812 0 5556.279
646.2919921875 0 1694.3597
647.292236328125 0 676.8977
658.3770751953125 0 19089.473 c 5
659.3796997070312 0 8706.3
660.2973022460938 0 1328.0537 y 3
660.381591796875 0 1576.0951
661.2913208007812 0 617.5211
684.392578125 0 3151.7546
685.3958740234375 0 905.69324
728.342041015625 0 695.4775
741.4135131835938 0 731.42365
751.3966064453125 0 727.9782
757.3651733398438 0 1356.4739 z 2
758.3695068359375 0 1535.5706
770.4078979492188 0 785.49506
771.4235229492188 0 1866.3027
772.4268798828125 0 701.70746
773.3622436523438 0 2033.4889
774.3643798828125 0 1151.4117
786.4356079101562 0 29358.055 c 6
787.4384765625 0 11253.5625
788.343505859375 0 533.9278
788.4407348632812 0 2948.9167
789.4467163085938 0 723.3631
800.4077758789062 0 609.0895
828.4207153320312 0 736.7748
842.4071044921875 0 694.3193 y Water loss 1
843.4285278320312 0 1153.0167
844.3958740234375 0 5590.598 z 1
845.4014892578125 0 2993.1602
846.3992919921875 0 828.2163
857.4663696289062 0 3660.529
858.4652709960938 0 1657.55
859.4707641601562 0 735.23474
860.4141845703125 0 1125.3756 y 1
875.0042114257812 0 627.1378
884.51171875 0 1315.5768
901.4560546875 0 937.71954
903.4851684570312 0 604.28
912.5077514648438 0 1601.7798
914.4219360351562 0 993.1591
929.4910278320312 0 1100.3729
930.508544921875 0 841.3656
931.51416015625 0 782.9834
956.4910278320312 0 1557.64
957.4822387695312 0 6818.7754
958.4835815429688 0 3121.6477
959.4818115234375 0 643.2354
973.4981079101562 0 9839.437
974.5050659179688 0 15813.424
975.507568359375 0 7637.9673
975.603271484375 0 523.5737
976.5098876953125 0 3036.3684
1458.8565673828125 0 2084.7642
1459.8480224609375 0 1440.9182
2569.680908203125 0 773.09015

Spectrum Details

|  |  |
| --- | --- |
| Matched peaks? Matched peaks The total absolute number of peaks matched. Additionally in brackets the total fraction of peaks matched and the total number of peaks is shown. | 15 (12.61% of 119) |
| FDR? FDR The false discovery rate estimated for this peptide. It is calculated by matching all theoretical fragments with a non-integer shift with the raw peaks for this spectrum. This is done with 40 different shifts. The resulting percentage is the average number of annotated peaks over the number of annotated peaks with the correct spectrum. | 3.17% |
| Satellite FDR? Satellite FDR See the FDR for details on its calculation. This satellite ion specific FDR only contains the satellite ions (d/w) for I/L/J positions. | - |
| PSM Score? PSM Score The PSM Score as given by Hecklib to this annotated spectrum. It is shown with three significant figures. | 179 |

#### Spectrum 7735? Spectrum 7735 The raw spectrum of this peptide as annotated by Hecklib. The fragments are coloured according to ion type (see legend). Any peaks with a star '\*' as text can be hovered over to see the full details, first the ion type second the mass shift type. By hovering over the amino acids in the peptide or ions in the legend the corresponding peaks are highlighted. By toggling the 'Unassigned' label you can turn the background (unassigned) peaks on or off in the plot. By updating the slider in the Ion legend you can update the spectrum to only show the top X% of the peaks with labels. The top X% means any peak that is within X% of the highest intensity. By dragging in the spectrum you can zoom in to a specific part of the spectrum and use 'Zoom Out' to get back to the original zoom level. The annotation of the spectrum is based on the given sequence in the peptides file and is done with different software so inconsistencies are likely. The peaks are annotated based on the given sequence, with 20 ppm tolerance.

Copy Data

##### Spectrum 7735 (TSV)

###### Preview

```
Loading example...
```

*Click on the button to copy the data to your clipboard.*

Mz MinMz MaxIntensity Max

WidthHeightPeptide font sizePeptide stroke widthSpectrum font sizeSpectrum stroke widthCompact peptide

Ion legend

wxyz

abcd

OtherUnassignedIon^Charge^_Position_Show for top:%

JSJTPEQW

01.68e+43.35e+45.03e+46.70e+4

Zoom Out

y^+1^_2_z^+1^_2_y^+1^_2_y^+1^_3_z^+1^_3_c^+1^_5_y^+1^_4_y^+1^_4_z^+1^_4_y^+1^_4_c^+1^_6_y^+1^_5_z^+1^_5_y^+1^_5_c^+1^_7_z^+1^_6_c^+1^_7_z^+1^_7_z^+1^_7_y^+1^_7_

0868173626043472

Fragment Matches Table

Show background peaks

| Position | Ion type | Intensity | mz Theoretical | mz Error (Th) | mz Error (ppm) | Charge | Series Number |
| --- | --- | --- | --- | --- | --- | --- | --- |
| - | - | 370.2 | 121.6 | - | - | 0 | - |
| - | - | 449.8 | 169.7 | - | - | 0 | - |
| - | - | 503.9 | 169.8 | - | - | 0 | - |
| - | - | 1482 | 173.1 | - | - | 0 | - |
| - | - | 982.2 | 173.4 | - | - | 0 | - |
| - | - | 3185 | 175.1 | - | - | 0 | - |
| - | - | 905 | 187.1 | - | - | 0 | - |
| - | - | 4837 | 201.1 | - | - | 0 | - |
| - | - | 1429 | 203.1 | - | - | 0 | - |
| - | - | 2580 | 215.1 | - | - | 0 | - |
| - | - | 1.57E+04 | 221.1 | - | - | 0 | - |
| - | - | 840.2 | 222.1 | - | - | 0 | - |
| - | - | 538.8 | 223.9 | - | - | 0 | - |
| - | - | 2848 | 227.1 | - | - | 0 | - |
| - | - | 3830 | 244.1 | - | - | 0 | - |
| - | - | 503.5 | 244.9 | - | - | 0 | - |
| - | - | 561.5 | 255.5 | - | - | 0 | - |
| - | - | 656.8 | 258.1 | - | - | 0 | - |
| - | - | 557 | 282.9 | - | - | 0 | - |
| - | - | 1819 | 284.2 | - | - | 0 | - |
| - | - | 7666 | 296.2 | - | - | 0 | - |
| - | - | 1326 | 297.2 | - | - | 0 | - |
| - | - | 468.8 | 300.2 | - | - | 0 | - |
| - | - | 2844 | 302.2 | - | - | 0 | - |
| - | - | 668.7 | 309.2 | - | - | 0 | - |
| - | - | 1380 | 314.1 | - | - | 0 | - |
| - | - | 1778 | 314.2 | - | - | 0 | - |
| - | - | 2870 | 331.1 | - | - | 0 | - |
| 7 | y | 4469 | 332.1 | 0.004688 | 14.12 | +1 | 2 |
| 7 | z | 913.1 | 333.1 | 0.000739 | 2.219 | +1 | 2 |
| - | - | 1577 | 348.1 | - | - | 0 | - |
| 7 | y | 5878 | 349.1 | 0.004262 | 12.21 | +1 | 2 |
| - | - | 1712 | 350.2 | - | - | 0 | - |
| - | - | 7952 | 355.2 | - | - | 0 | - |
| - | - | 1053 | 356.2 | - | - | 0 | - |
| - | - | 1074 | 369.3 | - | - | 0 | - |
| - | - | 801.3 | 370.2 | - | - | 0 | - |
| - | - | 788.5 | 372.2 | - | - | 0 | - |
| - | - | 1665 | 387.3 | - | - | 0 | - |
| - | - | 559.9 | 388.3 | - | - | 0 | - |
| - | - | 4970 | 397.2 | - | - | 0 | - |
| - | - | 937.7 | 398.2 | - | - | 0 | - |
| - | - | 575.2 | 411.6 | - | - | 0 | - |
| - | - | 1.042E+04 | 415.3 | - | - | 0 | - |
| - | - | 2156 | 416.3 | - | - | 0 | - |
| - | - | 738 | 417.3 | - | - | 0 | - |
| - | - | 4170 | 429.2 | - | - | 0 | - |
| - | - | 1388 | 439.2 | - | - | 0 | - |
| - | - | 894.7 | 441.2 | - | - | 0 | - |
| - | - | 1637 | 442.2 | - | - | 0 | - |
| - | - | 623.1 | 443.2 | - | - | 0 | - |
| - | - | 1217 | 456.2 | - | - | 0 | - |
| - | - | 1657 | 459.8 | - | - | 0 | - |
| 6 | y | 943.2 | 460.2 | 0.006139 | 13.34 | +1 | 3 |
| 6 | z | 763.2 | 462.2 | 0.004655 | 10.07 | +1 | 3 |
| - | - | 605 | 474.3 | - | - | 0 | - |
| - | - | 624.5 | 477.7 | - | - | 0 | - |
| - | - | 2069 | 477.8 | - | - | 0 | - |
| - | - | 834.5 | 495.8 | - | - | 0 | - |
| - | - | 898.6 | 503.2 | - | - | 0 | - |
| - | - | 663.9 | 504.2 | - | - | 0 | - |
| - | - | 4751 | 513.2 | - | - | 0 | - |
| - | - | 860.1 | 514.2 | - | - | 0 | - |
| - | - | 1811 | 515.2 | - | - | 0 | - |
| - | - | 566.4 | 520.3 | - | - | 0 | - |
| - | - | 566.9 | 526.7 | - | - | 0 | - |
| - | - | 3821 | 528.3 | - | - | 0 | - |
| 5 | c | 2.296E+04 | 529.3 | 0.000684 | 1.292 | +1 | 5 |
| - | - | 6453 | 530.3 | - | - | 0 | - |
| - | - | 1353 | 540.2 | - | - | 0 | - |
| 5 | y | 1583 | 557.2 | 0.003546 | 6.364 | +1 | 4 |
| 5 | y | 3974 | 558.2 | 0.004699 | 8.418 | +1 | 4 |
| 5 | z | 1433 | 559.2 | 0.0009289 | 1.661 | +1 | 4 |
| - | - | 1.292E+04 | 574.2 | - | - | 0 | - |
| 5 | y | 4.01E+04 | 575.2 | 0.003723 | 6.473 | +1 | 4 |
| - | - | 1.084E+04 | 576.2 | - | - | 0 | - |
| - | - | 2216 | 577.3 | - | - | 0 | - |
| - | - | 556.3 | 598.3 | - | - | 0 | - |
| - | - | 770.2 | 602.2 | - | - | 0 | - |
| - | - | 901.7 | 614.4 | - | - | 0 | - |
| - | - | 1411 | 615.4 | - | - | 0 | - |
| - | - | 2386 | 625.3 | - | - | 0 | - |
| - | - | 766.8 | 628.3 | - | - | 0 | - |
| - | - | 2069 | 641.3 | - | - | 0 | - |
| - | - | 786.8 | 641.4 | - | - | 0 | - |
| - | - | 9956 | 658.3 | - | - | 0 | - |
| 6 | c | 2.77E+04 | 658.4 | 0.0008577 | 1.303 | +1 | 6 |
| 4 | y | 3671 | 659.3 | 0.0008436 | 1.28 | +1 | 5 |
| - | - | 1.018E+04 | 659.4 | - | - | 0 | - |
| 4 | z | 1.819E+04 | 660.3 | 0.0037 | 5.603 | +1 | 5 |
| - | - | 2679 | 660.4 | - | - | 0 | - |
| - | - | 1.097E+04 | 661.3 | - | - | 0 | - |
| - | - | 2617 | 662.3 | - | - | 0 | - |
| - | - | 1349 | 675.3 | - | - | 0 | - |
| 4 | y | 4016 | 676.3 | 0.00408 | 6.032 | +1 | 5 |
| - | - | 620.1 | 677.3 | - | - | 0 | - |
| - | - | 1112 | 684.4 | - | - | 0 | - |
| - | - | 3008 | 715.3 | - | - | 0 | - |
| - | - | 1558 | 716.3 | - | - | 0 | - |
| - | - | 2051 | 728.3 | - | - | 0 | - |
| - | - | 1125 | 729.3 | - | - | 0 | - |
| - | - | 775.1 | 742.4 | - | - | 0 | - |
| - | - | 1722 | 751.4 | - | - | 0 | - |
| - | - | 755.9 | 752.4 | - | - | 0 | - |
| - | - | 8228 | 753.4 | - | - | 0 | - |
| - | - | 3737 | 754.4 | - | - | 0 | - |
| - | - | 974.7 | 755.4 | - | - | 0 | - |
| 7 | c | 2661 | 769.4 | 0.0005001 | 0.65 | +1 | 7 |
| - | - | 1.567E+04 | 771.4 | - | - | 0 | - |
| - | - | 6576 | 772.4 | - | - | 0 | - |
| 3 | z | 1.007E+04 | 773.4 | 0.004414 | 5.707 | +1 | 6 |
| - | - | 1207 | 773.4 | - | - | 0 | - |
| - | - | 5281 | 774.4 | - | - | 0 | - |
| - | - | 1900 | 775.4 | - | - | 0 | - |
| - | - | 3499 | 783.4 | - | - | 0 | - |
| - | - | 826.6 | 784.5 | - | - | 0 | - |
| 7 | c | 6.637E+04 | 786.4 | 0.001208 | 1.536 | +1 | 7 |
| - | - | 2.623E+04 | 787.4 | - | - | 0 | - |
| - | - | 994.9 | 788.4 | - | - | 0 | - |
| - | - | 6743 | 788.4 | - | - | 0 | - |
| - | - | 6621 | 789.4 | - | - | 0 | - |
| - | - | 2624 | 790.4 | - | - | 0 | - |
| - | - | 1412 | 800.4 | - | - | 0 | - |
| - | - | 1859 | 804.3 | - | - | 0 | - |
| - | - | 777.1 | 810.4 | - | - | 0 | - |
| - | - | 4580 | 812.5 | - | - | 0 | - |
| - | - | 1776 | 813.5 | - | - | 0 | - |
| - | - | 3925 | 827.4 | - | - | 0 | - |
| - | - | 3919 | 828.4 | - | - | 0 | - |
| - | - | 1184 | 829.4 | - | - | 0 | - |
| 2 | z | 1262 | 842.4 | 0.005289 | 6.278 | +1 | 7 |
| - | - | 1.547E+04 | 843.4 | - | - | 0 | - |
| - | - | 8270 | 844.4 | - | - | 0 | - |
| - | - | 6538 | 845.4 | - | - | 0 | - |
| - | - | 2843 | 846.5 | - | - | 0 | - |
| - | - | 7104 | 857.5 | - | - | 0 | - |
| - | - | 703.8 | 858.4 | - | - | 0 | - |
| - | - | 1809 | 858.5 | - | - | 0 | - |
| 2 | z | 1.544E+04 | 860.4 | 0.003574 | 4.154 | +1 | 7 |
| - | - | 6641 | 861.4 | - | - | 0 | - |
| - | - | 2081 | 862.4 | - | - | 0 | - |
| - | - | 644.1 | 863.6 | - | - | 0 | - |
| - | - | 1131 | 873.5 | - | - | 0 | - |
| - | - | 3891 | 875.5 | - | - | 0 | - |
| 2 | y | 1327 | 876.4 | 0.0016 | 1.826 | +1 | 7 |
| - | - | 1021 | 876.5 | - | - | 0 | - |
| - | - | 782.9 | 877.5 | - | - | 0 | - |
| - | - | 1.245E+04 | 928.5 | - | - | 0 | - |
| - | - | 7898 | 929.5 | - | - | 0 | - |
| - | - | 1887 | 930.5 | - | - | 0 | - |
| - | - | 1013 | 931.5 | - | - | 0 | - |
| - | - | 715 | 932.5 | - | - | 0 | - |
| - | - | 3896 | 945.5 | - | - | 0 | - |
| - | - | 3262 | 946.5 | - | - | 0 | - |
| - | - | 1026 | 947.5 | - | - | 0 | - |
| - | - | 2178 | 955.5 | - | - | 0 | - |
| - | - | 1497 | 956.5 | - | - | 0 | - |
| - | - | 6128 | 972.5 | - | - | 0 | - |
| - | - | 1.858E+04 | 973.5 | - | - | 0 | - |
| - | - | 8068 | 974.5 | - | - | 0 | - |
| - | - | 667.3 | 974.6 | - | - | 0 | - |
| - | - | 3551 | 975.5 | - | - | 0 | - |
| - | - | 2151 | 987.5 | - | - | 0 | - |
| - | - | 1232 | 987.6 | - | - | 0 | - |
| - | - | 1387 | 988.5 | - | - | 0 | - |
| - | - | 2.182E+04 | 989.5 | - | - | 0 | - |
| - | - | 5.564E+04 | 990.5 | - | - | 0 | - |
| - | - | 3.106E+04 | 991.5 | - | - | 0 | - |
| - | - | 8658 | 992.5 | - | - | 0 | - |
| - | - | 1080 | 1486 | - | - | 0 | - |
| - | - | 679.8 | 1550 | - | - | 0 | - |
| - | - | 657.2 | 2691 | - | - | 0 | - |
| - | - | 693.6 | 3081 | - | - | 0 | - |
| - | - | 722.8 | 3347 | - | - | 0 | - |
| - | - | 784.5 | 3438 | - | - | 0 | - |

m/z Charge Intensity FragmentType MassShift Position
121.5859603881836 0 370.23282
169.72113037109375 0 449.79715
169.808349609375 0 503.85556
173.1282196044922 0 1482.1841
173.4394989013672 0 982.23
175.08639526367188 0 3185.2517
187.14413452148438 0 905.04376
201.12319946289062 0 4837.1045
203.08119201660156 0 1429.2317
215.13885498046875 0 2579.9973
221.0919189453125 0 15704.851
222.09548950195312 0 840.21295
223.87904357910156 0 538.7978
227.10244750976562 0 2847.936
244.1289825439453 0 3829.9648
244.8595428466797 0 503.53485
255.546142578125 0 561.5449
258.1084289550781 0 656.83606
282.8887634277344 0 556.9917
284.16015625 0 1818.7935
296.19659423828125 0 7665.9546
297.1994323730469 0 1325.6368
300.15838623046875 0 468.79907
302.17059326171875 0 2843.5237
309.20367431640625 0 668.69745
314.11370849609375 0 1379.9935
314.2073059082031 0 1777.6608
331.1400146484375 0 2870.1523
332.1238708496094 0 4468.8887 y Ammonia loss 6
333.12774658203125 0 913.0576 z 6
348.1427001953125 0 1577.0964
349.1499938964844 0 5877.5747 y 6
350.1545104980469 0 1712.2197
355.1607666015625 0 7951.726
356.1636657714844 0 1053.4675
369.2500305175781 0 1074.2943
370.2326965332031 0 801.2667
372.1866455078125 0 788.4763
387.2601318359375 0 1665.3066
388.262451171875 0 559.8748
397.244140625 0 4970.0415
398.2476806640625 0 937.68713
411.599853515625 0 575.163
415.2547912597656 0 10424.911
416.25714111328125 0 2156.447
417.2599182128906 0 738.00555
429.184814453125 0 4169.503
439.1830749511719 0 1388.2725
441.1963195800781 0 894.7137
442.1928405761719 0 1636.688
443.15289306640625 0 623.0765
456.2081298828125 0 1217.47
459.8126220703125 0 1657.3033
460.18389892578125 0 943.18494 y Water loss 5
462.17425537109375 0 763.24316 z 5
474.3102722167969 0 605.0417
477.7296447753906 0 624.46545
477.8226318359375 0 2069.3398
495.8313293457031 0 834.5012
503.2001953125 0 898.5742
504.208251953125 0 663.90674
513.2061157226562 0 4751.0303
514.2098999023438 0 860.0959
515.2208862304688 0 1810.9048
520.3282470703125 0 566.35187
526.747314453125 0 566.91516
528.326904296875 0 3820.628
529.333740234375 0 22962.902 c 4
530.3365478515625 0 6452.8965
540.2084350585938 0 1352.68
557.2340698242188 0 1582.9679 y Water loss 4
558.21923828125 0 3974.234 y Ammonia loss 4
559.221435546875 0 1433.305 z 4
574.2376708984375 0 12924.155
575.2448120117188 0 40103.742 y 4
576.2479248046875 0 10842.864
577.250732421875 0 2216.3994
598.27490234375 0 556.25397
602.2369384765625 0 770.1678
614.361083984375 0 901.6794
615.36962890625 0 1410.6968
625.3298950195312 0 2386.3
628.3056030273438 0 766.82794
641.3123779296875 0 2069.2761
641.3599243164062 0 786.7553
658.259033203125 0 9955.949
658.3761596679688 0 27702.818 c 5
659.2630615234375 0 3671.1926 y Ammonia loss 3
659.37890625 0 10180.289
660.2737426757812 0 18190.998 z 3
660.3821411132812 0 2678.82
661.2789916992188 0 10965.448
662.283203125 0 2616.8813
675.2840576171875 0 1348.9242
676.2928466796875 0 4015.9473 y 3
677.298095703125 0 620.1358
684.3961791992188 0 1112.2484
715.337158203125 0 3008.0588
716.341064453125 0 1557.8629
728.346923828125 0 2050.8245
729.3472900390625 0 1125.2751
742.4141235351562 0 775.0512
751.3990478515625 0 1721.6935
752.3919677734375 0 755.8599
753.411865234375 0 8227.667
754.40966796875 0 3736.6074
755.4130859375 0 974.71655
769.4095458984375 0 2661.2788 c Ammonia loss 6
771.4236450195312 0 15667.42
772.4258422851562 0 6575.994
773.3585205078125 0 10074.595 z 2
773.42919921875 0 1206.5236
774.3623657226562 0 5280.555
775.364990234375 0 1899.9895
783.447265625 0 3498.7178
784.4501342773438 0 826.6184
786.4343872070312 0 66367.234 c 6
787.4374389648438 0 26234.322
788.3681640625 0 994.8924
788.4404296875 0 6742.999
789.3541870117188 0 6621.1377
790.3587036132812 0 2624.4211
800.4198608398438 0 1411.9923
804.3292846679688 0 1859.0007
810.4118041992188 0 777.14264
812.450927734375 0 4579.7627
813.4542236328125 0 1775.7982
827.43798828125 0 3925.1743
828.4288330078125 0 3919.3413
829.4301147460938 0 1183.7474
842.380859375 0 1262.3179 z Water loss 1
843.4313354492188 0 15465.036
844.4345703125 0 8269.514
845.4450073242188 0 6537.6655
846.4501953125 0 2843.2192
857.4600830078125 0 7104.073
858.3735961914062 0 703.8233
858.46240234375 0 1809.2805
860.3897094726562 0 15438.484 z 1
861.3933715820312 0 6641.2583
862.396484375 0 2080.6064
863.6065063476562 0 644.1079
873.4599609375 0 1131.2476
875.471435546875 0 3890.8247
876.4032592773438 0 1326.7411 y 1
876.4822387695312 0 1021.304
877.4564819335938 0 782.9016
928.5001831054688 0 12452.77
929.5018310546875 0 7898.1777
930.5061645507812 0 1886.8275
931.500244140625 0 1012.57367
932.4913330078125 0 714.9969
945.4779052734375 0 3895.5657
946.4930419921875 0 3261.6985
947.4996948242188 0 1025.8812
955.4594116210938 0 2178.1147
956.4661254882812 0 1497.1494
972.489013671875 0 6127.9277
973.47705078125 0 18579.5
974.4788818359375 0 8068.199
974.57568359375 0 667.2583
975.4837646484375 0 3551.424
987.4942626953125 0 2150.8804
987.6154174804688 0 1232.363
988.4948120117188 0 1386.8721
989.4921875 0 21822.027
990.4990234375 0 55635.277
991.50244140625 0 31055.871
992.50634765625 0 8658.048
1485.8448486328125 0 1080.2614
1550.01513671875 0 679.8194
2690.730712890625 0 657.2445
3081.37451171875 0 693.5957
3346.786865234375 0 722.8052
3437.990478515625 0 784.54333

Spectrum Details

|  |  |
| --- | --- |
| Matched peaks? Matched peaks The total absolute number of peaks matched. Additionally in brackets the total fraction of peaks matched and the total number of peaks is shown. | 20 (11.43% of 175) |
| FDR? FDR The false discovery rate estimated for this peptide. It is calculated by matching all theoretical fragments with a non-integer shift with the raw peaks for this spectrum. This is done with 40 different shifts. The resulting percentage is the average number of annotated peaks over the number of annotated peaks with the correct spectrum. | 4.29% |
| Satellite FDR? Satellite FDR See the FDR for details on its calculation. This satellite ion specific FDR only contains the satellite ions (d/w) for I/L/J positions. | ∞ |
| PSM Score? PSM Score The PSM Score as given by Hecklib to this annotated spectrum. It is shown with three significant figures. | 267 |

#### Spectrum 10395? Spectrum 10395 The raw spectrum of this peptide as annotated by Hecklib. The fragments are coloured according to ion type (see legend). Any peaks with a star '\*' as text can be hovered over to see the full details, first the ion type second the mass shift type. By hovering over the amino acids in the peptide or ions in the legend the corresponding peaks are highlighted. By toggling the 'Unassigned' label you can turn the background (unassigned) peaks on or off in the plot. By updating the slider in the Ion legend you can update the spectrum to only show the top X% of the peaks with labels. The top X% means any peak that is within X% of the highest intensity. By dragging in the spectrum you can zoom in to a specific part of the spectrum and use 'Zoom Out' to get back to the original zoom level. The annotation of the spectrum is based on the given sequence in the peptides file and is done with different software so inconsistencies are likely. The peaks are annotated based on the given sequence, with 20 ppm tolerance.

Copy Data

##### Spectrum 10395 (TSV)

###### Preview

```
Loading example...
```

*Click on the button to copy the data to your clipboard.*

Mz MinMz MaxIntensity Max

WidthHeightPeptide font sizePeptide stroke widthSpectrum font sizeSpectrum stroke widthCompact peptide

Ion legend

wxyz

abcd

OtherUnassignedIon^Charge^_Position_Show for top:%

JSJTPEQW

08.38e+31.68e+42.51e+43.35e+4

Zoom Out

z^+1^_1_y^+1^_1_y^+1^_2_c^+1^_3_y^+1^_2_c^+1^_5_y^+1^_4_y^+1^_4_y^+1^_5_z^+1^_5_c^+1^_6_y^+1^_5_z^+1^_6_c^+1^_7_c^+1^_7_z^+1^_7_y^+1^_7_

0527105415812108

Fragment Matches Table

Show background peaks

| Position | Ion type | Intensity | mz Theoretical | mz Error (Th) | mz Error (ppm) | Charge | Series Number |
| --- | --- | --- | --- | --- | --- | --- | --- |
| - | - | 417.6 | 127.1 | - | - | 0 | - |
| - | - | 2252 | 133.1 | - | - | 0 | - |
| - | - | 486.4 | 152.8 | - | - | 0 | - |
| - | - | 440.3 | 155.8 | - | - | 0 | - |
| - | - | 1528 | 159.1 | - | - | 0 | - |
| - | - | 465.4 | 164 | - | - | 0 | - |
| - | - | 627.1 | 173.1 | - | - | 0 | - |
| - | - | 2036 | 173.5 | - | - | 0 | - |
| - | - | 458 | 176.3 | - | - | 0 | - |
| - | - | 1075 | 177.1 | - | - | 0 | - |
| - | - | 4601 | 188.1 | - | - | 0 | - |
| 8 | z | 557.1 | 189.1 | 0.003754 | 19.85 | +1 | 1 |
| - | - | 1016 | 201.1 | - | - | 0 | - |
| - | - | 1531 | 201.1 | - | - | 0 | - |
| - | - | 2143 | 201.1 | - | - | 0 | - |
| 8 | y | 6268 | 205.1 | 0.0001359 | 0.6628 | +1 | 1 |
| - | - | 1115 | 215.1 | - | - | 0 | - |
| - | - | 478 | 215.3 | - | - | 0 | - |
| - | - | 1230 | 227.1 | - | - | 0 | - |
| - | - | 2998 | 244.1 | - | - | 0 | - |
| - | - | 541.9 | 250.7 | - | - | 0 | - |
| - | - | 698.2 | 255.1 | - | - | 0 | - |
| - | - | 496.2 | 264 | - | - | 0 | - |
| - | - | 501 | 267.6 | - | - | 0 | - |
| - | - | 737.2 | 284.2 | - | - | 0 | - |
| - | - | 2770 | 296.2 | - | - | 0 | - |
| - | - | 1096 | 302.2 | - | - | 0 | - |
| - | - | 877.9 | 314.2 | - | - | 0 | - |
| - | - | 1720 | 315.1 | - | - | 0 | - |
| 7 | y | 2175 | 316.1 | 0.0002425 | 0.7672 | +1 | 2 |
| - | - | 965.6 | 327.2 | - | - | 0 | - |
| 3 | c | 648.5 | 331.2 | 0.001522 | 4.596 | +1 | 3 |
| 7 | y | 1486 | 333.2 | 0.0003056 | 0.9172 | +1 | 2 |
| - | - | 4651 | 355.2 | - | - | 0 | - |
| - | - | 795.1 | 356.2 | - | - | 0 | - |
| - | - | 599.6 | 367.5 | - | - | 0 | - |
| - | - | 786.2 | 370.2 | - | - | 0 | - |
| - | - | 2502 | 372.2 | - | - | 0 | - |
| - | - | 780.7 | 381.2 | - | - | 0 | - |
| - | - | 609.8 | 396.2 | - | - | 0 | - |
| - | - | 1547 | 397.2 | - | - | 0 | - |
| - | - | 615.8 | 400.4 | - | - | 0 | - |
| - | - | 6335 | 415.3 | - | - | 0 | - |
| - | - | 945.7 | 416.3 | - | - | 0 | - |
| - | - | 524.8 | 424.9 | - | - | 0 | - |
| - | - | 878 | 434.9 | - | - | 0 | - |
| - | - | 565 | 450 | - | - | 0 | - |
| - | - | 538.7 | 453.2 | - | - | 0 | - |
| - | - | 1523 | 456.2 | - | - | 0 | - |
| - | - | 2348 | 458.2 | - | - | 0 | - |
| - | - | 667.4 | 458.3 | - | - | 0 | - |
| - | - | 782 | 468.2 | - | - | 0 | - |
| - | - | 961.3 | 469.2 | - | - | 0 | - |
| - | - | 1179 | 471.3 | - | - | 0 | - |
| - | - | 742.9 | 486.3 | - | - | 0 | - |
| - | - | 793.1 | 487.1 | - | - | 0 | - |
| - | - | 732 | 487.3 | - | - | 0 | - |
| - | - | 878.1 | 488.2 | - | - | 0 | - |
| - | - | 744.7 | 514.3 | - | - | 0 | - |
| - | - | 792.3 | 515.3 | - | - | 0 | - |
| - | - | 823.4 | 524.2 | - | - | 0 | - |
| - | - | 1923 | 528.3 | - | - | 0 | - |
| 5 | c | 1.187E+04 | 529.3 | 7.367E-05 | 0.1392 | +1 | 5 |
| - | - | 3091 | 530.3 | - | - | 0 | - |
| - | - | 634.3 | 531.3 | - | - | 0 | - |
| 5 | y | 1452 | 542.2 | 0.0004972 | 0.9169 | +1 | 4 |
| - | - | 653 | 555.3 | - | - | 0 | - |
| 5 | y | 1.333E+04 | 559.3 | 0.0001932 | 0.3454 | +1 | 4 |
| - | - | 2998 | 560.3 | - | - | 0 | - |
| - | - | 679.9 | 571.3 | - | - | 0 | - |
| - | - | 781.6 | 599.3 | - | - | 0 | - |
| - | - | 603.1 | 600.3 | - | - | 0 | - |
| - | - | 803 | 614.4 | - | - | 0 | - |
| - | - | 1086 | 625.3 | - | - | 0 | - |
| - | - | 1856 | 642.3 | - | - | 0 | - |
| 4 | y | 1372 | 643.3 | 0.004213 | 6.549 | +1 | 5 |
| 4 | z | 7367 | 644.3 | 0.0003188 | 0.4949 | +1 | 5 |
| - | - | 6528 | 645.3 | - | - | 0 | - |
| - | - | 1739 | 646.3 | - | - | 0 | - |
| - | - | 641.8 | 650.3 | - | - | 0 | - |
| 6 | c | 2.466E+04 | 658.4 | 5.785E-05 | 0.08786 | +1 | 6 |
| - | - | 8168 | 659.4 | - | - | 0 | - |
| 4 | y | 1365 | 660.3 | 0.002441 | 3.697 | +1 | 5 |
| - | - | 2738 | 660.4 | - | - | 0 | - |
| - | - | 3311 | 684.4 | - | - | 0 | - |
| - | - | 1575 | 685.4 | - | - | 0 | - |
| - | - | 1428 | 728.3 | - | - | 0 | - |
| - | - | 862.1 | 729.9 | - | - | 0 | - |
| - | - | 1004 | 742.4 | - | - | 0 | - |
| - | - | 898.2 | 743.4 | - | - | 0 | - |
| - | - | 1087 | 751.4 | - | - | 0 | - |
| - | - | 759.4 | 753.4 | - | - | 0 | - |
| 3 | z | 2524 | 757.4 | 0.002165 | 2.859 | +1 | 6 |
| - | - | 1612 | 758.4 | - | - | 0 | - |
| 7 | c | 2406 | 769.4 | 0.0007206 | 0.9365 | +1 | 7 |
| - | - | 1654 | 771.4 | - | - | 0 | - |
| - | - | 670.7 | 772.4 | - | - | 0 | - |
| - | - | 2901 | 773.4 | - | - | 0 | - |
| - | - | 738.5 | 775.4 | - | - | 0 | - |
| 7 | c | 3.319E+04 | 786.4 | 1.305E-05 | 0.01659 | +1 | 7 |
| - | - | 1.591E+04 | 787.4 | - | - | 0 | - |
| - | - | 933.1 | 788.3 | - | - | 0 | - |
| - | - | 3494 | 788.4 | - | - | 0 | - |
| - | - | 1530 | 800.4 | - | - | 0 | - |
| - | - | 600 | 817.3 | - | - | 0 | - |
| - | - | 831.1 | 841.4 | - | - | 0 | - |
| - | - | 2169 | 843.4 | - | - | 0 | - |
| 2 | z | 7963 | 844.4 | 0.001387 | 1.642 | +1 | 7 |
| - | - | 3210 | 845.4 | - | - | 0 | - |
| - | - | 1083 | 846.4 | - | - | 0 | - |
| - | - | 4383 | 857.5 | - | - | 0 | - |
| - | - | 2465 | 858.5 | - | - | 0 | - |
| - | - | 630.8 | 859.5 | - | - | 0 | - |
| 2 | y | 1076 | 860.4 | 0.0005459 | 0.6344 | +1 | 7 |
| - | - | 1100 | 884.5 | - | - | 0 | - |
| - | - | 937.7 | 901.5 | - | - | 0 | - |
| - | - | 1752 | 912.5 | - | - | 0 | - |
| - | - | 945.1 | 913.5 | - | - | 0 | - |
| - | - | 1497 | 914.4 | - | - | 0 | - |
| - | - | 1138 | 929.5 | - | - | 0 | - |
| - | - | 2065 | 930.5 | - | - | 0 | - |
| - | - | 987.3 | 931.5 | - | - | 0 | - |
| - | - | 2002 | 956.5 | - | - | 0 | - |
| - | - | 7875 | 957.5 | - | - | 0 | - |
| - | - | 4913 | 958.5 | - | - | 0 | - |
| - | - | 810.2 | 958.6 | - | - | 0 | - |
| - | - | 1457 | 959.5 | - | - | 0 | - |
| - | - | 740.7 | 972.6 | - | - | 0 | - |
| - | - | 1.34E+04 | 973.5 | - | - | 0 | - |
| - | - | 1.846E+04 | 974.5 | - | - | 0 | - |
| - | - | 1548 | 974.6 | - | - | 0 | - |
| - | - | 1.136E+04 | 975.5 | - | - | 0 | - |
| - | - | 935.8 | 975.6 | - | - | 0 | - |
| - | - | 2203 | 976.5 | - | - | 0 | - |
| - | - | 1893 | 1442 | - | - | 0 | - |
| - | - | 1480 | 1443 | - | - | 0 | - |
| - | - | 5160 | 1459 | - | - | 0 | - |
| - | - | 4218 | 1460 | - | - | 0 | - |
| - | - | 1392 | 1461 | - | - | 0 | - |
| - | - | 723.5 | 1462 | - | - | 0 | - |
| - | - | 610.6 | 1713 | - | - | 0 | - |
| - | - | 637.2 | 1910 | - | - | 0 | - |
| - | - | 745.9 | 2087 | - | - | 0 | - |

m/z Charge Intensity FragmentType MassShift Position
127.07572937011719 0 417.55185
133.0860595703125 0 2251.6145
152.77540588378906 0 486.4468
155.8026580810547 0 440.2971
159.09185791015625 0 1528.0776
164.03843688964844 0 465.43942
173.12884521484375 0 627.1257
173.45147705078125 0 2035.5771
176.31234741210938 0 457.9922
177.1123046875 0 1075.258
188.0707244873047 0 4600.774
189.07467651367188 0 557.1073 z 7
201.1024932861328 0 1016.2879
201.11224365234375 0 1530.785
201.12347412109375 0 2143.0713
205.0972900390625 0 6268.3975 y 7
215.139404296875 0 1115.1011
215.33056640625 0 478.04025
227.1019744873047 0 1229.5001
244.12924194335938 0 2998.2175
250.6910858154297 0 541.87866
255.1363983154297 0 698.1874
263.95458984375 0 496.17905
267.64654541015625 0 500.98462
284.1613464355469 0 737.2008
296.1971130371094 0 2770.0251
302.1715087890625 0 1096.3535
314.20782470703125 0 877.92523
315.14495849609375 0 1719.7941
316.1294250488281 0 2174.582 y Ammonia loss 6
327.20086669921875 0 965.6023
331.2355041503906 0 648.4984 c 2
333.1554260253906 0 1485.9869 y 6
355.1611022949219 0 4650.5747
356.16497802734375 0 795.14996
367.50567626953125 0 599.61426
370.23358154296875 0 786.2469
372.187744140625 0 2502.4893
381.24749755859375 0 780.7473
396.17718505859375 0 609.8054
397.2444763183594 0 1546.9692
400.3539123535156 0 615.79486
415.2551574707031 0 6334.6855
416.2573547363281 0 945.673
424.8943176269531 0 524.76624
434.868896484375 0 877.9683
449.959716796875 0 564.98926
453.20928955078125 0 538.6722
456.2098083496094 0 1523.2686
458.2242736816406 0 2347.8982
458.258056640625 0 667.44464
468.24481201171875 0 782.0401
469.1689453125 0 961.28516
471.31683349609375 0 1179.2864
486.2906799316406 0 742.8769
487.0914611816406 0 793.1325
487.2978820800781 0 732.0008
488.1641845703125 0 878.12366
514.2521362304688 0 744.6758
515.2589721679688 0 792.3157
524.2146606445312 0 823.38214
528.3273315429688 0 1923.0757
529.3343505859375 0 11868.217 c 4
530.33740234375 0 3091.374
531.334716796875 0 634.2729
542.2250366210938 0 1451.9343 y Ammonia loss 4
555.3482055664062 0 652.981
559.2512817382812 0 13331.198 y 4
560.25439453125 0 2997.5613
571.3062133789062 0 679.911
599.2630004882812 0 781.5657
600.2619018554688 0 603.0739
614.3615112304688 0 803.0303
625.3309936523438 0 1085.7598
642.2635498046875 0 1855.542
643.2680053710938 0 1371.6157 y Ammonia loss 3
644.2797241210938 0 7366.636 z 3
645.2857055664062 0 6527.7866
646.289794921875 0 1738.9742
650.3477783203125 0 641.7953
658.3770751953125 0 24663.959 c 5
659.3804931640625 0 8168.282
660.2963256835938 0 1364.6451 y 3
660.3838500976562 0 2738.4304
684.3929443359375 0 3310.7742
685.3950805664062 0 1575.4093
728.3430786132812 0 1427.9452
729.9338989257812 0 862.1088
742.4207153320312 0 1004.2478
743.4234008789062 0 898.23834
751.3934936523438 0 1087.2745
753.393798828125 0 759.4094
757.3662719726562 0 2524.1328 z 2
758.367919921875 0 1611.8674
769.4083251953125 0 2406.0125 c Ammonia loss 6
771.4249877929688 0 1653.9553
772.4261474609375 0 670.66583
773.3621215820312 0 2901.482
775.389404296875 0 738.5001
786.4356079101562 0 33192.21 c 6
787.4385375976562 0 15906.269
788.3308715820312 0 933.12415
788.4412841796875 0 3494.306
800.406005859375 0 1530.2058
817.2981567382812 0 599.96625
841.4219970703125 0 831.1251
843.4328002929688 0 2169.1204
844.3975219726562 0 7962.5483 z 1
845.4000244140625 0 3210.0244
846.4028930664062 0 1082.9817
857.4645385742188 0 4382.9307
858.4680786132812 0 2465.107
859.4649047851562 0 630.8365
860.4154052734375 0 1076.3167 y 1
884.50732421875 0 1100.349
901.4554443359375 0 937.69214
912.5054321289062 0 1751.6561
913.4957885742188 0 945.1238
914.4230346679688 0 1496.8378
929.4940795898438 0 1137.7042
930.5107421875 0 2064.7766
931.5194091796875 0 987.2791
956.4940795898438 0 2001.5013
957.4829711914062 0 7874.598
958.4840698242188 0 4912.6973
958.5751953125 0 810.16394
959.4804077148438 0 1457.2649
972.5874633789062 0 740.66833
973.49853515625 0 13401.538
974.5045166015625 0 18461.598
974.6049194335938 0 1548.2385
975.50732421875 0 11355.59
975.6065673828125 0 935.79407
976.5093383789062 0 2202.6145
1441.8260498046875 0 1892.6487
1442.83447265625 0 1479.752
1458.8592529296875 0 5159.8076
1459.8612060546875 0 4218.2876
1460.856689453125 0 1391.7684
1461.866455078125 0 723.524
1712.96630859375 0 610.5827
1910.482177734375 0 637.24713
2087.046630859375 0 745.9056

Spectrum Details

|  |  |
| --- | --- |
| Matched peaks? Matched peaks The total absolute number of peaks matched. Additionally in brackets the total fraction of peaks matched and the total number of peaks is shown. | 17 (11.89% of 143) |
| FDR? FDR The false discovery rate estimated for this peptide. It is calculated by matching all theoretical fragments with a non-integer shift with the raw peaks for this spectrum. This is done with 40 different shifts. The resulting percentage is the average number of annotated peaks over the number of annotated peaks with the correct spectrum. | 3.78% |
| Satellite FDR? Satellite FDR See the FDR for details on its calculation. This satellite ion specific FDR only contains the satellite ions (d/w) for I/L/J positions. | - |
| PSM Score? PSM Score The PSM Score as given by Hecklib to this annotated spectrum. It is shown with three significant figures. | 213 |

#### Spectrum 8179? Spectrum 8179 The raw spectrum of this peptide as annotated by Hecklib. The fragments are coloured according to ion type (see legend). Any peaks with a star '\*' as text can be hovered over to see the full details, first the ion type second the mass shift type. By hovering over the amino acids in the peptide or ions in the legend the corresponding peaks are highlighted. By toggling the 'Unassigned' label you can turn the background (unassigned) peaks on or off in the plot. By updating the slider in the Ion legend you can update the spectrum to only show the top X% of the peaks with labels. The top X% means any peak that is within X% of the highest intensity. By dragging in the spectrum you can zoom in to a specific part of the spectrum and use 'Zoom Out' to get back to the original zoom level. The annotation of the spectrum is based on the given sequence in the peptides file and is done with different software so inconsistencies are likely. The peaks are annotated based on the given sequence, with 20 ppm tolerance.

Copy Data

##### Spectrum 8179 (TSV)

###### Preview

```
Loading example...
```

*Click on the button to copy the data to your clipboard.*

Mz MinMz MaxIntensity Max

WidthHeightPeptide font sizePeptide stroke widthSpectrum font sizeSpectrum stroke widthCompact peptide

Ion legend

wxyz

abcd

OtherUnassignedIon^Charge^_Position_Show for top:%

JSJTPEQW

05.51e+41.10e+51.65e+52.20e+5

Zoom Out

a^+1^_2_a^+1^_2_b^+1^_2_b^+1^_2_b^+2^_5_a^+1^_3_y^+2^_4_y^+2^_4_b^+1^_3_b^+2^_6_b^+1^_3_b^+2^_6_y^+1^_2_y^+1^_2_b^+1^_4_b^+1^_4_y^+1^_3_y^+1^_3_y^+1^_3_\*b^+1^_5_b^+1^_5_y^+1^_4_y^+1^_4_y^+1^_4_b^+1^_6_b^+1^_6_y^+1^_5_y^+1^_5_y^+1^_5_b^+1^_7_b^+1^_7_b^+1^_7_y^+1^_6_y^+1^_6_y^+1^_6_y^+1^_7_y^+1^_7_y^+1^_7_

0840167925193359

Fragment Matches Table

Show background peaks

| Position | Ion type | Intensity | mz Theoretical | mz Error (Th) | mz Error (ppm) | Charge | Series Number |
| --- | --- | --- | --- | --- | --- | --- | --- |
| - | - | 4784 | 120.1 | - | - | 0 | - |
| - | - | 639.2 | 121.1 | - | - | 0 | - |
| - | - | 1267 | 125.1 | - | - | 0 | - |
| - | - | 6053 | 125.1 | - | - | 0 | - |
| - | - | 1752 | 126.1 | - | - | 0 | - |
| - | - | 798.6 | 126.1 | - | - | 0 | - |
| - | - | 1105 | 127.1 | - | - | 0 | - |
| - | - | 407.3 | 128.1 | - | - | 0 | - |
| - | - | 3092 | 128.1 | - | - | 0 | - |
| - | - | 1.125E+04 | 129.1 | - | - | 0 | - |
| - | - | 3250 | 129.1 | - | - | 0 | - |
| - | - | 4863 | 130.1 | - | - | 0 | - |
| - | - | 2.726E+04 | 130.1 | - | - | 0 | - |
| - | - | 2474 | 131.1 | - | - | 0 | - |
| - | - | 6427 | 132 | - | - | 0 | - |
| - | - | 2929 | 132.1 | - | - | 0 | - |
| - | - | 806.3 | 132.1 | - | - | 0 | - |
| - | - | 1434 | 133.1 | - | - | 0 | - |
| - | - | 2587 | 134.1 | - | - | 0 | - |
| - | - | 3139 | 136.1 | - | - | 0 | - |
| - | - | 530.3 | 137.1 | - | - | 0 | - |
| - | - | 488.1 | 137.1 | - | - | 0 | - |
| - | - | 1150 | 138.1 | - | - | 0 | - |
| - | - | 872.7 | 139.1 | - | - | 0 | - |
| - | - | 454.3 | 139.1 | - | - | 0 | - |
| - | - | 1133 | 141.1 | - | - | 0 | - |
| - | - | 9141 | 142.1 | - | - | 0 | - |
| - | - | 592.9 | 143.1 | - | - | 0 | - |
| - | - | 1.102E+04 | 146.1 | - | - | 0 | - |
| - | - | 1029 | 147.1 | - | - | 0 | - |
| - | - | 3566 | 147.1 | - | - | 0 | - |
| - | - | 2322 | 148.1 | - | - | 0 | - |
| - | - | 888.6 | 148.9 | - | - | 0 | - |
| - | - | 483 | 151 | - | - | 0 | - |
| - | - | 5624 | 152.1 | - | - | 0 | - |
| - | - | 474 | 153.1 | - | - | 0 | - |
| - | - | 2691 | 153.1 | - | - | 0 | - |
| - | - | 2805 | 153.1 | - | - | 0 | - |
| - | - | 675.7 | 153.1 | - | - | 0 | - |
| - | - | 453.6 | 154.1 | - | - | 0 | - |
| - | - | 2068 | 155.1 | - | - | 0 | - |
| 2 | a | 8286 | 155.1 | 0.0003508 | 2.262 | +1 | 2 |
| - | - | 437.6 | 156 | - | - | 0 | - |
| - | - | 943.5 | 156.1 | - | - | 0 | - |
| - | - | 526 | 157.1 | - | - | 0 | - |
| - | - | 1.569E+04 | 157.1 | - | - | 0 | - |
| - | - | 3.324E+04 | 158.1 | - | - | 0 | - |
| - | - | 1599 | 158.1 | - | - | 0 | - |
| - | - | 3569 | 159.1 | - | - | 0 | - |
| - | - | 6425 | 159.1 | - | - | 0 | - |
| - | - | 683 | 160.1 | - | - | 0 | - |
| - | - | 1048 | 160.1 | - | - | 0 | - |
| - | - | 1045 | 163.1 | - | - | 0 | - |
| - | - | 998.7 | 166.1 | - | - | 0 | - |
| - | - | 558.1 | 167.1 | - | - | 0 | - |
| - | - | 1393 | 168.1 | - | - | 0 | - |
| - | - | 3809 | 169.1 | - | - | 0 | - |
| - | - | 4051 | 169.1 | - | - | 0 | - |
| - | - | 720.8 | 170.1 | - | - | 0 | - |
| - | - | 8529 | 171.1 | - | - | 0 | - |
| - | - | 802.3 | 171.1 | - | - | 0 | - |
| - | - | 1137 | 172.1 | - | - | 0 | - |
| 2 | a | 7.367E+04 | 173.1 | 0.000391 | 2.258 | +1 | 2 |
| - | - | 2998 | 173.5 | - | - | 0 | - |
| - | - | 480.1 | 174.1 | - | - | 0 | - |
| - | - | 476 | 174.1 | - | - | 0 | - |
| - | - | 5564 | 174.1 | - | - | 0 | - |
| - | - | 1.437E+05 | 175.1 | - | - | 0 | - |
| - | - | 629.5 | 176.1 | - | - | 0 | - |
| - | - | 1.363E+04 | 176.1 | - | - | 0 | - |
| - | - | 665 | 177.1 | - | - | 0 | - |
| - | - | 545 | 177.1 | - | - | 0 | - |
| - | - | 1.477E+04 | 181.1 | - | - | 0 | - |
| - | - | 8967 | 181.1 | - | - | 0 | - |
| - | - | 1154 | 182.1 | - | - | 0 | - |
| - | - | 965.3 | 182.1 | - | - | 0 | - |
| 2 | b | 1.773E+04 | 183.1 | 0.0003245 | 1.772 | +1 | 2 |
| - | - | 3269 | 183.1 | - | - | 0 | - |
| - | - | 1537 | 184.1 | - | - | 0 | - |
| - | - | 3786 | 185.1 | - | - | 0 | - |
| - | - | 562.7 | 185.1 | - | - | 0 | - |
| - | - | 3698 | 186.1 | - | - | 0 | - |
| - | - | 643.9 | 187.1 | - | - | 0 | - |
| - | - | 1.899E+04 | 187.1 | - | - | 0 | - |
| - | - | 4035 | 188.1 | - | - | 0 | - |
| - | - | 1328 | 188.1 | - | - | 0 | - |
| - | - | 1222 | 195.1 | - | - | 0 | - |
| - | - | 9025 | 197.1 | - | - | 0 | - |
| - | - | 968.4 | 198.1 | - | - | 0 | - |
| - | - | 1472 | 199.1 | - | - | 0 | - |
| - | - | 6.554E+04 | 199.1 | - | - | 0 | - |
| - | - | 538.8 | 200.1 | - | - | 0 | - |
| - | - | 6364 | 200.1 | - | - | 0 | - |
| - | - | 610.2 | 200.1 | - | - | 0 | - |
| 2 | b | 7.993E+04 | 201.1 | 0.0003036 | 1.51 | +1 | 2 |
| - | - | 8444 | 202.1 | - | - | 0 | - |
| - | - | 7.228E+04 | 203.1 | - | - | 0 | - |
| - | - | 9148 | 204.1 | - | - | 0 | - |
| - | - | 8498 | 204.1 | - | - | 0 | - |
| - | - | 833.6 | 205.1 | - | - | 0 | - |
| - | - | 1149 | 205.1 | - | - | 0 | - |
| - | - | 6028 | 209.1 | - | - | 0 | - |
| - | - | 1153 | 210.1 | - | - | 0 | - |
| - | - | 883.1 | 210.1 | - | - | 0 | - |
| - | - | 777 | 211.1 | - | - | 0 | - |
| - | - | 3148 | 211.1 | - | - | 0 | - |
| - | - | 1676 | 212.1 | - | - | 0 | - |
| - | - | 1207 | 213.1 | - | - | 0 | - |
| - | - | 704 | 213.1 | - | - | 0 | - |
| - | - | 623.6 | 214.1 | - | - | 0 | - |
| - | - | 758.2 | 214.2 | - | - | 0 | - |
| - | - | 4.719E+04 | 215.1 | - | - | 0 | - |
| - | - | 4401 | 216.1 | - | - | 0 | - |
| - | - | 3396 | 217.1 | - | - | 0 | - |
| - | - | 672.1 | 217.1 | - | - | 0 | - |
| - | - | 2.181E+05 | 221.1 | - | - | 0 | - |
| - | - | 2.502E+04 | 222.1 | - | - | 0 | - |
| - | - | 2507 | 223.1 | - | - | 0 | - |
| - | - | 1190 | 223.1 | - | - | 0 | - |
| - | - | 647.7 | 223.1 | - | - | 0 | - |
| - | - | 613.7 | 223.1 | - | - | 0 | - |
| - | - | 629.4 | 225.1 | - | - | 0 | - |
| - | - | 1234 | 225.1 | - | - | 0 | - |
| - | - | 2877 | 226.1 | - | - | 0 | - |
| - | - | 1210 | 226.2 | - | - | 0 | - |
| - | - | 9.618E+04 | 227.1 | - | - | 0 | - |
| - | - | 8355 | 228.1 | - | - | 0 | - |
| - | - | 780.2 | 229.1 | - | - | 0 | - |
| - | - | 2008 | 229.2 | - | - | 0 | - |
| - | - | 828.4 | 230.1 | - | - | 0 | - |
| - | - | 957.2 | 231.1 | - | - | 0 | - |
| - | - | 707.3 | 233.1 | - | - | 0 | - |
| - | - | 1351 | 233.2 | - | - | 0 | - |
| - | - | 1374 | 233.7 | - | - | 0 | - |
| - | - | 855.1 | 237.1 | - | - | 0 | - |
| - | - | 1057 | 238.2 | - | - | 0 | - |
| - | - | 4265 | 239.1 | - | - | 0 | - |
| - | - | 1.164E+04 | 240.1 | - | - | 0 | - |
| - | - | 821.1 | 240.1 | - | - | 0 | - |
| - | - | 3711 | 241.1 | - | - | 0 | - |
| - | - | 2622 | 241.1 | - | - | 0 | - |
| - | - | 1827 | 242.2 | - | - | 0 | - |
| - | - | 1793 | 242.7 | - | - | 0 | - |
| - | - | 2.437E+04 | 244.1 | - | - | 0 | - |
| - | - | 1668 | 245.1 | - | - | 0 | - |
| - | - | 2308 | 245.1 | - | - | 0 | - |
| - | - | 743.8 | 246.1 | - | - | 0 | - |
| - | - | 789.1 | 250.1 | - | - | 0 | - |
| - | - | 711.7 | 251.1 | - | - | 0 | - |
| - | - | 1138 | 251.1 | - | - | 0 | - |
| - | - | 3543 | 251.2 | - | - | 0 | - |
| - | - | 1493 | 252.1 | - | - | 0 | - |
| - | - | 697.1 | 253.1 | - | - | 0 | - |
| - | - | 561 | 255.1 | - | - | 0 | - |
| - | - | 4523 | 256.2 | - | - | 0 | - |
| 5 | b | 758.7 | 256.7 | 0.0007494 | 2.92 | +2 | 5 |
| - | - | 857.5 | 257.1 | - | - | 0 | - |
| - | - | 6174 | 257.1 | - | - | 0 | - |
| - | - | 1.039E+04 | 258.1 | - | - | 0 | - |
| - | - | 3521 | 258.1 | - | - | 0 | - |
| - | - | 1062 | 259.1 | - | - | 0 | - |
| - | - | 1260 | 264.1 | - | - | 0 | - |
| - | - | 686.5 | 265.1 | - | - | 0 | - |
| - | - | 1592 | 266.1 | - | - | 0 | - |
| - | - | 2971 | 266.1 | - | - | 0 | - |
| - | - | 1004 | 266.2 | - | - | 0 | - |
| - | - | 1995 | 267.1 | - | - | 0 | - |
| - | - | 587.6 | 267.2 | - | - | 0 | - |
| - | - | 841.8 | 267.2 | - | - | 0 | - |
| - | - | 875.5 | 268.1 | - | - | 0 | - |
| - | - | 3846 | 268.1 | - | - | 0 | - |
| 3 | a | 2610 | 268.2 | 0.0002865 | 1.068 | +1 | 3 |
| - | - | 859.4 | 269.1 | - | - | 0 | - |
| - | - | 732.5 | 269.1 | - | - | 0 | - |
| - | - | 985.9 | 269.1 | - | - | 0 | - |
| - | - | 2358 | 269.2 | - | - | 0 | - |
| - | - | 4761 | 270.1 | - | - | 0 | - |
| - | - | 987.3 | 271.1 | - | - | 0 | - |
| - | - | 763.7 | 274.1 | - | - | 0 | - |
| - | - | 2142 | 274.2 | - | - | 0 | - |
| - | - | 554.2 | 276.2 | - | - | 0 | - |
| - | - | 585.5 | 279.1 | - | - | 0 | - |
| 5 | y | 654 | 279.6 | 0.002373 | 8.488 | +2 | 4 |
| - | - | 2137 | 280.2 | - | - | 0 | - |
| - | - | 2955 | 282.1 | - | - | 0 | - |
| - | - | 971.4 | 283.1 | - | - | 0 | - |
| - | - | 3.465E+04 | 284.2 | - | - | 0 | - |
| - | - | 1738 | 285.1 | - | - | 0 | - |
| - | - | 4878 | 285.2 | - | - | 0 | - |
| - | - | 1.713E+04 | 286.1 | - | - | 0 | - |
| - | - | 1017 | 286.1 | - | - | 0 | - |
| - | - | 3201 | 287.1 | - | - | 0 | - |
| - | - | 1067 | 287.2 | - | - | 0 | - |
| 5 | y | 784.1 | 288.1 | 0.002618 | 9.086 | +2 | 4 |
| - | - | 3719 | 292.1 | - | - | 0 | - |
| - | - | 783.7 | 293.1 | - | - | 0 | - |
| - | - | 640.9 | 293.1 | - | - | 0 | - |
| - | - | 1352 | 294.1 | - | - | 0 | - |
| - | - | 2360 | 294.2 | - | - | 0 | - |
| - | - | 7180 | 296.1 | - | - | 0 | - |
| - | - | 1118 | 296.1 | - | - | 0 | - |
| 3 | b | 7.851E+04 | 296.2 | 0.000367 | 1.239 | +1 | 3 |
| - | - | 2376 | 297.1 | - | - | 0 | - |
| - | - | 707.3 | 297.1 | - | - | 0 | - |
| - | - | 1.326E+04 | 297.2 | - | - | 0 | - |
| - | - | 2747 | 298.2 | - | - | 0 | - |
| - | - | 1160 | 298.2 | - | - | 0 | - |
| - | - | 2874 | 300.2 | - | - | 0 | - |
| - | - | 3.948E+04 | 302.2 | - | - | 0 | - |
| - | - | 5801 | 303.2 | - | - | 0 | - |
| - | - | 547 | 304.2 | - | - | 0 | - |
| - | - | 730.3 | 307.2 | - | - | 0 | - |
| - | - | 2214 | 309.2 | - | - | 0 | - |
| - | - | 743.7 | 309.2 | - | - | 0 | - |
| - | - | 7121 | 310.1 | - | - | 0 | - |
| - | - | 1472 | 310.2 | - | - | 0 | - |
| - | - | 1176 | 311.1 | - | - | 0 | - |
| - | - | 882.4 | 311.2 | - | - | 0 | - |
| - | - | 4462 | 312.2 | - | - | 0 | - |
| 6 | b | 808.4 | 312.2 | 0.0007264 | 2.327 | +2 | 6 |
| - | - | 1925 | 312.2 | - | - | 0 | - |
| - | - | 4759 | 313.1 | - | - | 0 | - |
| - | - | 947.2 | 313.2 | - | - | 0 | - |
| - | - | 4.205E+04 | 314.1 | - | - | 0 | - |
| 3 | b | 1.672E+04 | 314.2 | 0.0004224 | 1.344 | +1 | 3 |
| - | - | 555.2 | 314.9 | - | - | 0 | - |
| - | - | 6941 | 315.1 | - | - | 0 | - |
| - | - | 2643 | 315.2 | - | - | 0 | - |
| - | - | 2186 | 320.1 | - | - | 0 | - |
| 6 | b | 2349 | 321.2 | 0.0004184 | 1.303 | +2 | 6 |
| - | - | 971.2 | 321.7 | - | - | 0 | - |
| - | - | 742.2 | 322.2 | - | - | 0 | - |
| - | - | 560 | 325.2 | - | - | 0 | - |
| - | - | 801.2 | 326.2 | - | - | 0 | - |
| - | - | 612.9 | 326.4 | - | - | 0 | - |
| - | - | 7682 | 327.2 | - | - | 0 | - |
| - | - | 1.144E+04 | 328.2 | - | - | 0 | - |
| - | - | 1118 | 328.2 | - | - | 0 | - |
| - | - | 1822 | 329.2 | - | - | 0 | - |
| - | - | 1690 | 330.2 | - | - | 0 | - |
| - | - | 7.995E+04 | 331.1 | - | - | 0 | - |
| 7 | y | 7.213E+04 | 332.1 | 0.005177 | 15.59 | +1 | 2 |
| - | - | 1.167E+04 | 332.1 | - | - | 0 | - |
| - | - | 1.147E+04 | 333.1 | - | - | 0 | - |
| - | - | 996.1 | 333.1 | - | - | 0 | - |
| - | - | 1575 | 334.1 | - | - | 0 | - |
| - | - | 680.2 | 334.2 | - | - | 0 | - |
| - | - | 6650 | 337.2 | - | - | 0 | - |
| - | - | 642.8 | 337.2 | - | - | 0 | - |
| - | - | 566.2 | 337.5 | - | - | 0 | - |
| - | - | 3447 | 338.1 | - | - | 0 | - |
| - | - | 1005 | 338.2 | - | - | 0 | - |
| - | - | 3517 | 340.2 | - | - | 0 | - |
| - | - | 896 | 341.2 | - | - | 0 | - |
| - | - | 1024 | 343.2 | - | - | 0 | - |
| - | - | 4678 | 345.2 | - | - | 0 | - |
| - | - | 935.9 | 346.2 | - | - | 0 | - |
| 7 | y | 4.139E+04 | 349.1 | 0.005178 | 14.83 | +1 | 2 |
| - | - | 6057 | 350.2 | - | - | 0 | - |
| - | - | 686.8 | 351.2 | - | - | 0 | - |
| - | - | 1342 | 351.2 | - | - | 0 | - |
| - | - | 6601 | 352.2 | - | - | 0 | - |
| - | - | 774.6 | 353.2 | - | - | 0 | - |
| - | - | 1525 | 353.2 | - | - | 0 | - |
| - | - | 1.196E+05 | 355.2 | - | - | 0 | - |
| - | - | 1733 | 356.1 | - | - | 0 | - |
| - | - | 1.915E+04 | 356.2 | - | - | 0 | - |
| - | - | 1177 | 357.2 | - | - | 0 | - |
| - | - | 2641 | 363.2 | - | - | 0 | - |
| - | - | 702.3 | 364.2 | - | - | 0 | - |
| - | - | 1975 | 365.2 | - | - | 0 | - |
| - | - | 1.039E+04 | 369.2 | - | - | 0 | - |
| - | - | 1.17E+04 | 370.2 | - | - | 0 | - |
| - | - | 3890 | 371.2 | - | - | 0 | - |
| - | - | 1486 | 372.2 | - | - | 0 | - |
| - | - | 1231 | 372.2 | - | - | 0 | - |
| - | - | 4727 | 373.2 | - | - | 0 | - |
| - | - | 602.7 | 374.2 | - | - | 0 | - |
| - | - | 605.8 | 379.1 | - | - | 0 | - |
| - | - | 5157 | 379.2 | - | - | 0 | - |
| - | - | 1035 | 380.2 | - | - | 0 | - |
| - | - | 1384 | 381.2 | - | - | 0 | - |
| - | - | 1980 | 381.2 | - | - | 0 | - |
| - | - | 1.278E+04 | 387.3 | - | - | 0 | - |
| - | - | 2863 | 388.3 | - | - | 0 | - |
| - | - | 645.9 | 391.2 | - | - | 0 | - |
| - | - | 2685 | 397.2 | - | - | 0 | - |
| 4 | b | 3.73E+04 | 397.2 | 0.0001738 | 0.4376 | +1 | 4 |
| - | - | 846.3 | 398.2 | - | - | 0 | - |
| - | - | 7370 | 398.2 | - | - | 0 | - |
| - | - | 683.2 | 399.2 | - | - | 0 | - |
| - | - | 1055 | 399.2 | - | - | 0 | - |
| - | - | 1161 | 399.3 | - | - | 0 | - |
| - | - | 2555 | 409.2 | - | - | 0 | - |
| - | - | 932.9 | 412.2 | - | - | 0 | - |
| - | - | 743.7 | 413.2 | - | - | 0 | - |
| 4 | b | 7.842E+04 | 415.3 | 0.0001072 | 0.2582 | +1 | 4 |
| - | - | 1.747E+04 | 416.3 | - | - | 0 | - |
| - | - | 2218 | 417.3 | - | - | 0 | - |
| - | - | 974.9 | 421.2 | - | - | 0 | - |
| - | - | 2562 | 423.2 | - | - | 0 | - |
| - | - | 657.8 | 424.2 | - | - | 0 | - |
| - | - | 2388 | 425.1 | - | - | 0 | - |
| - | - | 2122 | 427.2 | - | - | 0 | - |
| - | - | 1519 | 428.2 | - | - | 0 | - |
| - | - | 2393 | 438.2 | - | - | 0 | - |
| - | - | 1374 | 439.8 | - | - | 0 | - |
| - | - | 947.4 | 440.2 | - | - | 0 | - |
| - | - | 3033 | 440.3 | - | - | 0 | - |
| - | - | 1528 | 441.2 | - | - | 0 | - |
| - | - | 1447 | 442.2 | - | - | 0 | - |
| - | - | 1.109E+04 | 443.2 | - | - | 0 | - |
| - | - | 2600 | 444.2 | - | - | 0 | - |
| - | - | 1657 | 448.3 | - | - | 0 | - |
| - | - | 582 | 450.3 | - | - | 0 | - |
| - | - | 1.011E+04 | 456.2 | - | - | 0 | - |
| - | - | 1550 | 457.2 | - | - | 0 | - |
| 6 | y | 4394 | 460.2 | 0.00504 | 10.95 | +1 | 3 |
| 6 | y | 6116 | 461.2 | 0.00543 | 11.78 | +1 | 3 |
| - | - | 1736 | 462.2 | - | - | 0 | - |
| - | - | 1181 | 466.3 | - | - | 0 | - |
| - | - | 7245 | 468.2 | - | - | 0 | - |
| - | - | 980.4 | 468.3 | - | - | 0 | - |
| - | - | 1065 | 468.7 | - | - | 0 | - |
| - | - | 836 | 469.3 | - | - | 0 | - |
| - | - | 641.7 | 475.2 | - | - | 0 | - |
| - | - | 1374 | 476.3 | - | - | 0 | - |
| - | - | 1787 | 477.7 | - | - | 0 | - |
| - | - | 1679 | 477.8 | - | - | 0 | - |
| 6 | y | 5318 | 478.2 | 0.005004 | 10.46 | +1 | 3 |
| - | - | 1655 | 478.2 | - | - | 0 | - |
| - | - | 1026 | 479.2 | - | - | 0 | - |
| - | - | 644.8 | 481.3 | - | - | 0 | - |
| - | - | 651.2 | 481.3 | - | - | 0 | - |
| - | - | 884 | 483.2 | - | - | 0 | - |
| - | - | 822.5 | 484.3 | - | - | 0 | - |
| 0 | Precursor | 1745 | 486.2 | 0.002696 | 5.545 | +2 | -1 |
| - | - | 1530 | 486.7 | - | - | 0 | - |
| - | - | 1052 | 487.1 | - | - | 0 | - |
| - | - | 3433 | 488.2 | - | - | 0 | - |
| - | - | 787.4 | 489.2 | - | - | 0 | - |
| - | - | 1470 | 492.2 | - | - | 0 | - |
| - | - | 2004 | 494.2 | - | - | 0 | - |
| - | - | 731.5 | 494.3 | - | - | 0 | - |
| 5 | b | 7388 | 494.3 | 8.333E-05 | 0.1686 | +1 | 5 |
| - | - | 722.7 | 495.3 | - | - | 0 | - |
| - | - | 1483 | 495.3 | - | - | 0 | - |
| - | - | 9356 | 495.8 | - | - | 0 | - |
| - | - | 1212 | 496.2 | - | - | 0 | - |
| - | - | 1033 | 499.3 | - | - | 0 | - |
| - | - | 975.2 | 508.3 | - | - | 0 | - |
| - | - | 4618 | 510.3 | - | - | 0 | - |
| - | - | 1924 | 512.2 | - | - | 0 | - |
| 5 | b | 4606 | 512.3 | 0.0001359 | 0.2653 | +1 | 5 |
| - | - | 2351 | 513.2 | - | - | 0 | - |
| - | - | 1024 | 513.3 | - | - | 0 | - |
| - | - | 2386 | 522.2 | - | - | 0 | - |
| - | - | 869.2 | 522.3 | - | - | 0 | - |
| - | - | 833.4 | 523.2 | - | - | 0 | - |
| - | - | 651.2 | 524.3 | - | - | 0 | - |
| - | - | 967.4 | 525.3 | - | - | 0 | - |
| - | - | 1166 | 526.3 | - | - | 0 | - |
| - | - | 1464 | 527.3 | - | - | 0 | - |
| - | - | 2005 | 528.3 | - | - | 0 | - |
| - | - | 1296 | 529.2 | - | - | 0 | - |
| - | - | 885.3 | 534.3 | - | - | 0 | - |
| - | - | 2262 | 537.3 | - | - | 0 | - |
| - | - | 597.3 | 537.4 | - | - | 0 | - |
| - | - | 768.2 | 538.3 | - | - | 0 | - |
| - | - | 2637 | 539.2 | - | - | 0 | - |
| - | - | 2.235E+04 | 540.2 | - | - | 0 | - |
| - | - | 905.1 | 540.3 | - | - | 0 | - |
| - | - | 5692 | 541.2 | - | - | 0 | - |
| - | - | 1101 | 541.3 | - | - | 0 | - |
| - | - | 831.2 | 542.2 | - | - | 0 | - |
| - | - | 700.4 | 543.3 | - | - | 0 | - |
| - | - | 2611 | 551.3 | - | - | 0 | - |
| - | - | 2982 | 555.3 | - | - | 0 | - |
| - | - | 897.7 | 556.3 | - | - | 0 | - |
| 5 | y | 1.586E+04 | 557.2 | 0.004461 | 8.007 | +1 | 4 |
| 5 | y | 4.558E+04 | 558.2 | 0.004943 | 8.855 | +1 | 4 |
| - | - | 1.318E+04 | 559.2 | - | - | 0 | - |
| - | - | 2217 | 560.2 | - | - | 0 | - |
| - | - | 3109 | 569.3 | - | - | 0 | - |
| - | - | 1725 | 570.3 | - | - | 0 | - |
| - | - | 1116 | 573.3 | - | - | 0 | - |
| 5 | y | 1.72E+05 | 575.2 | 0.004334 | 7.534 | +1 | 4 |
| - | - | 4.965E+04 | 576.2 | - | - | 0 | - |
| - | - | 9929 | 577.3 | - | - | 0 | - |
| - | - | 944.8 | 578.3 | - | - | 0 | - |
| - | - | 664.5 | 578.3 | - | - | 0 | - |
| - | - | 797.9 | 595.3 | - | - | 0 | - |
| - | - | 701.5 | 603.3 | - | - | 0 | - |
| - | - | 1893 | 605.3 | - | - | 0 | - |
| - | - | 1797 | 613.4 | - | - | 0 | - |
| - | - | 1220 | 620.3 | - | - | 0 | - |
| - | - | 1033 | 621.3 | - | - | 0 | - |
| - | - | 905.3 | 622.4 | - | - | 0 | - |
| - | - | 1520 | 623.2 | - | - | 0 | - |
| 6 | b | 9082 | 623.3 | 0.000304 | 0.4876 | +1 | 6 |
| - | - | 3163 | 624.3 | - | - | 0 | - |
| - | - | 901.7 | 634.9 | - | - | 0 | - |
| - | - | 4915 | 638.3 | - | - | 0 | - |
| - | - | 1653 | 639.3 | - | - | 0 | - |
| - | - | 847.6 | 639.9 | - | - | 0 | - |
| - | - | 3182 | 640.4 | - | - | 0 | - |
| - | - | 1395 | 641.3 | - | - | 0 | - |
| 6 | b | 8052 | 641.4 | 0.0001787 | 0.2787 | +1 | 6 |
| - | - | 3112 | 642.4 | - | - | 0 | - |
| - | - | 766.4 | 643.9 | - | - | 0 | - |
| - | - | 1526 | 650.4 | - | - | 0 | - |
| - | - | 1444 | 651.3 | - | - | 0 | - |
| - | - | 5050 | 656.3 | - | - | 0 | - |
| - | - | 1457 | 657.3 | - | - | 0 | - |
| 4 | y | 2098 | 658.3 | 0.002987 | 4.537 | +1 | 5 |
| - | - | 3692 | 658.4 | - | - | 0 | - |
| 4 | y | 4740 | 659.3 | 0.006276 | 9.519 | +1 | 5 |
| - | - | 1633 | 659.4 | - | - | 0 | - |
| - | - | 1773 | 660.3 | - | - | 0 | - |
| - | - | 765.5 | 660.4 | - | - | 0 | - |
| - | - | 1933 | 668.4 | - | - | 0 | - |
| 4 | y | 2.239E+04 | 676.3 | 0.004141 | 6.123 | +1 | 5 |
| - | - | 8103 | 677.3 | - | - | 0 | - |
| - | - | 2312 | 678.3 | - | - | 0 | - |
| - | - | 1377 | 723.4 | - | - | 0 | - |
| - | - | 1020 | 724.4 | - | - | 0 | - |
| - | - | 2904 | 733.4 | - | - | 0 | - |
| - | - | 3437 | 734.4 | - | - | 0 | - |
| - | - | 1115 | 735.4 | - | - | 0 | - |
| - | - | 1854 | 741.4 | - | - | 0 | - |
| 7 | b | 1.669E+04 | 751.4 | 0.001447 | 1.926 | +1 | 7 |
| 7 | b | 7603 | 752.4 | 0.01326 | 17.62 | +1 | 7 |
| - | - | 1636 | 753.4 | - | - | 0 | - |
| - | - | 745 | 754.3 | - | - | 0 | - |
| 7 | b | 8999 | 769.4 | 0.001331 | 1.73 | +1 | 7 |
| - | - | 4100 | 770.4 | - | - | 0 | - |
| 3 | y | 1829 | 771.4 | 0.001503 | 1.949 | +1 | 6 |
| 3 | y | 907 | 772.3 | 0.01187 | 15.37 | +1 | 6 |
| 3 | y | 3984 | 789.4 | 0.002474 | 3.134 | +1 | 6 |
| - | - | 1937 | 790.4 | - | - | 0 | - |
| 2 | y | 1254 | 858.4 | 0.007683 | 8.95 | +1 | 7 |
| 2 | y | 980.4 | 859.4 | 0.01048 | 12.2 | +1 | 7 |
| 2 | y | 3277 | 876.4 | 0.00432 | 4.93 | +1 | 7 |
| - | - | 1402 | 877.4 | - | - | 0 | - |
| - | - | 1121 | 895.5 | - | - | 0 | - |
| - | - | 1001 | 3080 | - | - | 0 | - |
| - | - | 712.3 | 3325 | - | - | 0 | - |

m/z Charge Intensity FragmentType MassShift Position
120.08114624023438 0 4783.722
121.08455657958984 0 639.2263
125.07120513916016 0 1266.9747
125.10771179199219 0 6052.504
126.0916976928711 0 1752.0114
126.11109161376953 0 798.5642
127.08686065673828 0 1104.9669
128.08236694335938 0 407.32114
128.10736083984375 0 3091.5999
129.0662384033203 0 11246.194
129.10269165039062 0 3249.6553
130.05026245117188 0 4863.14
130.0655059814453 0 27258.771
131.06890869140625 0 2474.1433
132.04478454589844 0 6426.7725
132.0811004638672 0 2928.6274
132.1023712158203 0 806.26465
133.0860595703125 0 1434.324
134.0603790283203 0 2586.5413
136.0760955810547 0 3139.4368
137.07177734375 0 530.3353
137.07875061035156 0 488.0761
138.0916748046875 0 1149.9543
139.05055236816406 0 872.6985
139.08685302734375 0 454.30463
141.06622314453125 0 1132.7633
142.123046875 0 9141.025
143.12698364257812 0 592.8549
146.06040954589844 0 11018.455
147.0637969970703 0 1029.3312
147.07679748535156 0 3566.1257
148.07598876953125 0 2322.3372
148.947265625 0 888.55914
151.0499725341797 0 483.00458
152.10731506347656 0 5624.205
153.054931640625 0 473.9933
153.06614685058594 0 2690.915
153.10267639160156 0 2804.7966
153.1099853515625 0 675.6936
154.08682250976562 0 453.56277
155.08192443847656 0 2068.444
155.1182403564453 0 8285.743 a Water loss 1
155.97926330566406 0 437.62778
156.12152099609375 0 943.46277
157.06068420410156 0 526.0444
157.07638549804688 0 15692.947
158.0603790283203 0 33240.81
158.07986450195312 0 1599.3672
159.063720703125 0 3569.2148
159.0920867919922 0 6424.663
160.07626342773438 0 682.9925
160.0953369140625 0 1047.9335
163.08685302734375 0 1045.0712
166.08645629882812 0 998.73706
167.081787109375 0 558.1045
168.0771942138672 0 1393.229
169.06112670898438 0 3808.9775
169.13389587402344 0 4050.803
170.11795043945312 0 720.7566
171.1131591796875 0 8529.18
171.14968872070312 0 802.29065
172.1165008544922 0 1137.2228
173.12884521484375 0 73670.78 a 1
173.4503631591797 0 2998.3875
174.05560302734375 0 480.1035
174.12509155273438 0 475.96033
174.1322021484375 0 5564.2563
175.0869903564453 0 143690.81
176.0712432861328 0 629.4563
176.09033203125 0 13627.082
177.09280395507812 0 665.0277
177.11231994628906 0 545.0457
181.06112670898438 0 14765.387
181.0974578857422 0 8967.429
182.06431579589844 0 1153.552
182.10145568847656 0 965.31573
183.11312866210938 0 17729.838 b Water loss 1
183.14956665039062 0 3268.7998
184.11666870117188 0 1536.8582
185.07138061523438 0 3785.9973
185.1284942626953 0 562.70905
186.0553741455078 0 3697.8242
187.07154846191406 0 643.8515
187.1444549560547 0 18985.959
188.07102966308594 0 4035.3254
188.14761352539062 0 1328.0583
195.07667541503906 0 1221.8364
197.12881469726562 0 9025.2705
198.13247680664062 0 968.3818
199.0717315673828 0 1472.3453
199.10801696777344 0 65541.195
200.10324096679688 0 538.83655
200.1114501953125 0 6364.4863
200.13966369628906 0 610.17914
201.12367248535156 0 79933.555 b 1
202.1270751953125 0 8444.092
203.08181762695312 0 72280.46
204.06581115722656 0 9147.7
204.08517456054688 0 8497.918
205.06887817382812 0 833.5838
205.09762573242188 0 1148.5525
209.09242248535156 0 6028.2666
210.09573364257812 0 1153.1826
210.12393188476562 0 883.05963
211.108154296875 0 776.99713
211.14453125 0 3148.1953
212.1033477783203 0 1675.6335
213.08758544921875 0 1206.7317
213.12400817871094 0 704.0143
214.11888122558594 0 623.63
214.15467834472656 0 758.19855
215.13934326171875 0 47192.074
216.14276123046875 0 4401.062
217.08221435546875 0 3396.1836
217.11907958984375 0 672.0567
221.0924530029297 0 218136.61
222.0957489013672 0 25023.422
223.0716552734375 0 2506.7798
223.09783935546875 0 1189.7128
223.1093292236328 0 647.66565
223.14422607421875 0 613.7404
225.10281372070312 0 629.3924
225.1228485107422 0 1234.3872
226.11911010742188 0 2876.7664
226.15528869628906 0 1209.6656
227.10301208496094 0 96175.055
228.1062774658203 0 8355.439
229.10804748535156 0 780.1913
229.155029296875 0 2007.7546
230.1137237548828 0 828.3981
231.07643127441406 0 957.22485
233.14987182617188 0 707.31635
233.16493225097656 0 1350.8832
233.6552276611328 0 1374.0029
237.12342834472656 0 855.05896
238.15538024902344 0 1056.687
239.1393585205078 0 4265.4976
240.0982666015625 0 11637.564
240.13534545898438 0 821.1479
241.0821533203125 0 3711.4495
241.09817504882812 0 2621.7156
242.15025329589844 0 1827.1697
242.66058349609375 0 1792.9856
244.12954711914062 0 24369.643
245.1134033203125 0 1668.2745
245.13259887695312 0 2308.1392
246.12411499023438 0 743.8463
250.11915588378906 0 789.0794
251.08050537109375 0 711.71466
251.14019775390625 0 1138.4489
251.1757354736328 0 3543.4985
252.11312866210938 0 1492.8595
253.0960693359375 0 697.0706
255.13449096679688 0 561.0421
256.16583251953125 0 4523.0996
256.6583251953125 0 758.69 b 4
257.1256103515625 0 857.4934
257.14971923828125 0 6173.9097
258.108642578125 0 10394.903
258.1451416015625 0 3520.5317
259.111083984375 0 1062.1647
264.134033203125 0 1260.3107
265.1204833984375 0 686.5294
266.1136169433594 0 1592.0107
266.1499328613281 0 2970.5557
266.18658447265625 0 1003.62726
267.1344299316406 0 1995.0133
267.1528625488281 0 587.61945
267.1703796386719 0 841.7717
268.0940246582031 0 875.4976
268.1083068847656 0 3845.6174
268.2022399902344 0 2609.6912 a Water loss 2
269.0921630859375 0 859.3973
269.11248779296875 0 732.4872
269.1391296386719 0 985.8933
269.1861877441406 0 2357.8342
270.12396240234375 0 4761.343
271.1281433105469 0 987.2836
274.1186828613281 0 763.69965
274.1758728027344 0 2142.2908
276.1707458496094 0 554.21356
279.0984802246094 0 585.483
279.61328125 0 653.9722 y Ammonia loss 4
280.1654052734375 0 2136.7185
282.14483642578125 0 2954.7002
283.1297912597656 0 971.4335
284.1607360839844 0 34653.363
285.1346740722656 0 1737.8561
285.1640319824219 0 4878.189
286.11895751953125 0 17125.121
286.1374816894531 0 1016.87537
287.1221923828125 0 3201.282
287.15008544921875 0 1066.6444
288.1268005371094 0 784.12225 y 4
292.1295471191406 0 3719.0964
293.1134338378906 0 783.7149
293.13262939453125 0 640.9361
294.1451721191406 0 1351.7405
294.1817626953125 0 2360.1948
296.1032409667969 0 7180.427
296.1229553222656 0 1117.9244
296.1972351074219 0 78511.66 b Water loss 2
297.0870666503906 0 2375.8306
297.1051940917969 0 707.29645
297.200439453125 0 13261.5625
298.1763000488281 0 2747.376
298.20233154296875 0 1159.884
300.15545654296875 0 2873.5603
302.1714172363281 0 39476.008
303.17486572265625 0 5800.9346
304.179443359375 0 547.0232
307.1807556152344 0 730.2876
309.1558837890625 0 2213.7751
309.20391845703125 0 743.7203
310.1402587890625 0 7120.76
310.2126159667969 0 1472.1318
311.1425476074219 0 1175.8319
311.17218017578125 0 882.42487
312.155517578125 0 4461.545
312.17431640625 0 808.384 b Water loss 5
312.1922607421875 0 1925.1077
313.1298522949219 0 4758.645
313.1591491699219 0 947.2156
314.1138916015625 0 42047.484
314.2078552246094 0 16719.88 b 2
314.8863525390625 0 555.2079
315.1170959472656 0 6940.534
315.21075439453125 0 2643.1748
320.12451171875 0 2185.61
321.1792907714844 0 2348.9014 b 5
321.681640625 0 971.228
322.177001953125 0 742.19525
325.225341796875 0 560.0417
326.1734313964844 0 801.1558
326.35443115234375 0 612.9387
327.16668701171875 0 7681.8667
328.1505432128906 0 11441.827
328.1706848144531 0 1117.5396
329.1529235839844 0 1822.0044
330.1663818359375 0 1689.6636
331.140380859375 0 79945.51
332.1243591308594 0 72130.17 y Ammonia loss 6
332.14398193359375 0 11670.781
333.12725830078125 0 11469.488
333.14642333984375 0 996.12823
334.1304016113281 0 1574.8936
334.21240234375 0 680.18335
337.1507263183594 0 6649.622
337.1863708496094 0 642.7979
337.4805908203125 0 566.1913
338.1348571777344 0 3446.6604
338.1550598144531 0 1005.34375
340.18658447265625 0 3516.8447
341.1898193359375 0 896.00183
343.2344055175781 0 1024.1276
345.1768798828125 0 4678.1016
346.177978515625 0 935.9167
349.1509094238281 0 41394.574 y 6
350.1539306640625 0 6056.695
351.1567687988281 0 686.7848
351.2398681640625 0 1342.1344
352.223388671875 0 6600.8086
353.1825256347656 0 774.6466
353.22125244140625 0 1525.0005
355.16131591796875 0 119605.82
356.1455993652344 0 1732.7478
356.1644592285156 0 19152.986
357.1674499511719 0 1176.8125
363.2021179199219 0 2640.8008
364.2072448730469 0 702.29895
365.1819152832031 0 1975.0919
369.2496032714844 0 10391.704
370.23358154296875 0 11696.02
371.2330017089844 0 3889.5662
372.1881103515625 0 1485.8328
372.23419189453125 0 1230.541
373.1716003417969 0 4727.3545
374.1764221191406 0 602.6882
379.13970947265625 0 605.7589
379.2340393066406 0 5156.725
380.2391662597656 0 1034.773
381.1560363769531 0 1383.938
381.21380615234375 0 1979.5675
387.2604064941406 0 12780.436
388.2627868652344 0 2862.9426
391.1975402832031 0 645.9342
397.1508483886719 0 2684.5608
397.2447204589844 0 37303.855 b Water loss 3
398.1538391113281 0 846.29315
398.24755859375 0 7370.4375
399.1873779296875 0 683.1559
399.2239685058594 0 1054.5126
399.2502746582031 0 1161.1497
409.2088317871094 0 2554.832
412.18341064453125 0 932.8875
413.2390441894531 0 743.66925
415.2552185058594 0 78415.54 b 3
416.25836181640625 0 17468.13
417.26116943359375 0 2218.3547
421.1723327636719 0 974.91864
423.2247314453125 0 2562.088
424.2271423339844 0 657.8231
425.1457824707031 0 2387.671
427.2191162109375 0 2122.1055
428.2156066894531 0 1518.7612
438.19952392578125 0 2392.5288
439.84375 0 1373.6832
440.21917724609375 0 947.38104
440.2505798339844 0 3033.4585
441.2348327636719 0 1527.8857
442.1715393066406 0 1446.5249
443.1562194824219 0 11090.297
444.1585693359375 0 2600.233
448.2655944824219 0 1657.3314
450.27374267578125 0 582.04517
456.20880126953125 0 10114.33
457.2118225097656 0 1549.9843
460.18280029296875 0 4393.846 y Water loss 5
461.1672058105469 0 6115.7075 y Ammonia loss 5
462.1713562011719 0 1736.0431
466.30438232421875 0 1180.7732
468.24493408203125 0 7244.928
468.28118896484375 0 980.38306
468.727294921875 0 1064.8586
469.2510986328125 0 835.9942
475.2185974121094 0 641.6601
476.28851318359375 0 1374.2822
477.7322082519531 0 1786.6238
477.82403564453125 0 1678.5822
478.1933288574219 0 5318.2866 y 5
478.2329406738281 0 1655.1771
479.19512939453125 0 1025.6182
481.26947021484375 0 644.8485
481.3141174316406 0 651.21027
483.2451477050781 0 883.9549
484.31512451171875 0 822.46655
486.2455139160156 0 1745.0006 Precursor Water loss
486.7447204589844 0 1530.3259
487.1064147949219 0 1051.6559
488.1781005859375 0 3432.5654
489.1788635253906 0 787.4243
492.2467346191406 0 1469.8887
494.2041015625 0 2004.3685
494.2532043457031 0 731.51044
494.2973937988281 0 7388.233 b Water loss 4
495.26336669921875 0 722.73706
495.3011169433594 0 1482.705
495.83349609375 0 9356.327
496.21710205078125 0 1211.7748
499.2767028808594 0 1032.8755
508.2738037109375 0 975.2385
510.2552185058594 0 4618.4653
512.215576171875 0 1923.7926
512.3077392578125 0 4605.92 b 4
513.2459716796875 0 2350.921
513.3115844726562 0 1023.70667
522.1978759765625 0 2385.8818
522.2965698242188 0 869.2155
523.2025756835938 0 833.375
524.2711181640625 0 651.21735
525.302978515625 0 967.35425
526.2857666015625 0 1165.6058
527.2835083007812 0 1464.0409
528.2662353515625 0 2005.0891
529.2384643554688 0 1295.7249
534.2573852539062 0 885.3153
537.2666625976562 0 2261.7727
537.3662109375 0 597.3491
538.2587280273438 0 768.15924
539.2239379882812 0 2637.427
540.2086181640625 0 22346.586
540.3036499023438 0 905.0643
541.2115478515625 0 5691.932
541.2984008789062 0 1100.9828
542.2179565429688 0 831.2496
543.3136596679688 0 700.41016
551.2819213867188 0 2610.8325
555.2767944335938 0 2982.3948
556.2794189453125 0 897.7225
557.2349853515625 0 15860.23 y Water loss 4
558.219482421875 0 45575.227 y Ammonia loss 4
559.2223510742188 0 13175.707
560.224365234375 0 2217.0635
569.292724609375 0 3109.023
570.2924194335938 0 1724.7985
573.3241577148438 0 1116.1976
575.2454223632812 0 171950.61 y 4
576.2485961914062 0 49646.6
577.25048828125 0 9928.654
578.2636108398438 0 944.7832
578.3333740234375 0 664.4891
595.345947265625 0 797.9335
603.2774658203125 0 701.4577
605.3301391601562 0 1893.4261
613.3547973632812 0 1796.9207
620.3026733398438 0 1220.1475
621.2866821289062 0 1033.3558
622.3561401367188 0 905.3301
623.2459716796875 0 1520.3997
623.339599609375 0 9082.114 b Water loss 5
624.3428955078125 0 3162.9912
634.8850708007812 0 901.67194
638.3138427734375 0 4915.0596
639.3143920898438 0 1652.9364
639.8731689453125 0 847.6024
640.3660888671875 0 3182.0679
641.25732421875 0 1394.7952
641.3506469726562 0 8051.757 b 5
642.3549194335938 0 3111.5535
643.8533935546875 0 766.36456
650.3504028320312 0 1525.5953
651.3497924804688 0 1444.292
656.3239135742188 0 5050.157
657.3264770507812 0 1457.4001
658.2811889648438 0 2097.5476 y Water loss 3
658.3766479492188 0 3691.8376
659.2684936523438 0 4740.275 y Ammonia loss 3
659.3782348632812 0 1632.6794
660.270751953125 0 1773.3652
660.3815307617188 0 765.4671
668.36083984375 0 1932.507
676.2929077148438 0 22389.518 y 3
677.2957153320312 0 8102.7075
678.2988891601562 0 2312.2969
723.4047241210938 0 1377.4698
724.4063720703125 0 1020.0531
733.3876342773438 0 2904.089
734.3782348632812 0 3436.625
735.3760986328125 0 1115.1891
741.41259765625 0 1853.614
751.3970336914062 0 16688.842 b Water loss 6
752.395751953125 0 7603.3677 b Ammonia loss 6
753.3976440429688 0 1635.5278
754.3399658203125 0 745.0468
769.40771484375 0 8999.075 b 6
770.4100952148438 0 4100.463
771.36376953125 0 1829.316 y Water loss 2
772.358154296875 0 906.9848 y Ammonia loss 2
789.3753051757812 0 3983.8113 y 2
790.3767700195312 0 1937.0092
858.4019775390625 0 1254.1608 y Water loss 1
859.3887939453125 0 980.3663 y Ammonia loss 1
876.4091796875 0 3277.4075 y 1
877.4138793945312 0 1402.4275
895.5248413085938 0 1121.4741
3080.126220703125 0 1001.19025
3325.35888671875 0 712.3397

Spectrum Details

|  |  |
| --- | --- |
| Matched peaks? Matched peaks The total absolute number of peaks matched. Additionally in brackets the total fraction of peaks matched and the total number of peaks is shown. | 39 (8.72% of 447) |
| FDR? FDR The false discovery rate estimated for this peptide. It is calculated by matching all theoretical fragments with a non-integer shift with the raw peaks for this spectrum. This is done with 40 different shifts. The resulting percentage is the average number of annotated peaks over the number of annotated peaks with the correct spectrum. | 1.83% |
| Satellite FDR? Satellite FDR See the FDR for details on its calculation. This satellite ion specific FDR only contains the satellite ions (d/w) for I/L/J positions. | - |
| PSM Score? PSM Score The PSM Score as given by Hecklib to this annotated spectrum. It is shown with three significant figures. | 603 |

#### Spectrum 10331? Spectrum 10331 The raw spectrum of this peptide as annotated by Hecklib. The fragments are coloured according to ion type (see legend). Any peaks with a star '\*' as text can be hovered over to see the full details, first the ion type second the mass shift type. By hovering over the amino acids in the peptide or ions in the legend the corresponding peaks are highlighted. By toggling the 'Unassigned' label you can turn the background (unassigned) peaks on or off in the plot. By updating the slider in the Ion legend you can update the spectrum to only show the top X% of the peaks with labels. The top X% means any peak that is within X% of the highest intensity. By dragging in the spectrum you can zoom in to a specific part of the spectrum and use 'Zoom Out' to get back to the original zoom level. The annotation of the spectrum is based on the given sequence in the peptides file and is done with different software so inconsistencies are likely. The peaks are annotated based on the given sequence, with 20 ppm tolerance.

Copy Data

##### Spectrum 10331 (TSV)

###### Preview

```
Loading example...
```

*Click on the button to copy the data to your clipboard.*

Mz MinMz MaxIntensity Max

WidthHeightPeptide font sizePeptide stroke widthSpectrum font sizeSpectrum stroke widthCompact peptide

Ion legend

wxyz

abcd

OtherUnassignedIon^Charge^_Position_Show for top:%

JSJTPEQW

09.86e+31.97e+42.96e+43.95e+4

Zoom Out

y^+1^_1_y^+1^_2_y^+1^_2_c^+1^_5_y^+1^_4_z^+1^_4_y^+1^_4_y^+1^_5_z^+1^_5_c^+1^_6_y^+1^_5_z^+1^_6_c^+1^_7_c^+1^_7_z^+1^_7_y^+1^_7_

0778155523333111

Fragment Matches Table

Show background peaks

| Position | Ion type | Intensity | mz Theoretical | mz Error (Th) | mz Error (ppm) | Charge | Series Number |
| --- | --- | --- | --- | --- | --- | --- | --- |
| - | - | 397.6 | 120.4 | - | - | 0 | - |
| - | - | 1241 | 131.1 | - | - | 0 | - |
| - | - | 537.2 | 132.1 | - | - | 0 | - |
| - | - | 1.787E+04 | 133.1 | - | - | 0 | - |
| - | - | 415 | 137.3 | - | - | 0 | - |
| - | - | 431.9 | 147.3 | - | - | 0 | - |
| - | - | 841.3 | 149 | - | - | 0 | - |
| - | - | 511.2 | 155.1 | - | - | 0 | - |
| - | - | 1561 | 159.1 | - | - | 0 | - |
| - | - | 543 | 160.1 | - | - | 0 | - |
| - | - | 833.8 | 173.1 | - | - | 0 | - |
| - | - | 2142 | 173.5 | - | - | 0 | - |
| - | - | 1611 | 175.1 | - | - | 0 | - |
| - | - | 9529 | 177.1 | - | - | 0 | - |
| - | - | 499.9 | 182.7 | - | - | 0 | - |
| - | - | 779 | 187.1 | - | - | 0 | - |
| - | - | 6947 | 188.1 | - | - | 0 | - |
| - | - | 543.7 | 189.1 | - | - | 0 | - |
| - | - | 681.1 | 195.1 | - | - | 0 | - |
| - | - | 1269 | 201.1 | - | - | 0 | - |
| - | - | 760.1 | 201.1 | - | - | 0 | - |
| - | - | 1944 | 201.1 | - | - | 0 | - |
| 8 | y | 6135 | 205.1 | 4.716E-05 | 0.2299 | +1 | 1 |
| - | - | 707.8 | 206.1 | - | - | 0 | - |
| - | - | 1396 | 215.1 | - | - | 0 | - |
| - | - | 4656 | 221.1 | - | - | 0 | - |
| - | - | 1284 | 227.1 | - | - | 0 | - |
| - | - | 513.3 | 234.6 | - | - | 0 | - |
| - | - | 5444 | 244.1 | - | - | 0 | - |
| - | - | 1874 | 265.2 | - | - | 0 | - |
| - | - | 913.5 | 283.2 | - | - | 0 | - |
| - | - | 663.1 | 284.2 | - | - | 0 | - |
| - | - | 2551 | 296.2 | - | - | 0 | - |
| - | - | 1537 | 302.2 | - | - | 0 | - |
| - | - | 3946 | 309.2 | - | - | 0 | - |
| - | - | 809.3 | 314.2 | - | - | 0 | - |
| - | - | 1630 | 315.1 | - | - | 0 | - |
| 7 | y | 2462 | 316.1 | 0.0002731 | 0.8638 | +1 | 2 |
| - | - | 638.8 | 321.2 | - | - | 0 | - |
| - | - | 1.846E+04 | 327.2 | - | - | 0 | - |
| 7 | y | 2832 | 333.2 | 0.0004574 | 1.373 | +1 | 2 |
| - | - | 5852 | 355.2 | - | - | 0 | - |
| - | - | 1398 | 355.2 | - | - | 0 | - |
| - | - | 1005 | 356.2 | - | - | 0 | - |
| - | - | 2339 | 372.2 | - | - | 0 | - |
| - | - | 912.9 | 373.2 | - | - | 0 | - |
| - | - | 1.053E+04 | 381.2 | - | - | 0 | - |
| - | - | 1779 | 397.2 | - | - | 0 | - |
| - | - | 6761 | 399.3 | - | - | 0 | - |
| - | - | 585.4 | 415.2 | - | - | 0 | - |
| - | - | 6022 | 415.3 | - | - | 0 | - |
| - | - | 1504 | 416.3 | - | - | 0 | - |
| - | - | 515.6 | 433.2 | - | - | 0 | - |
| - | - | 874.3 | 442.3 | - | - | 0 | - |
| - | - | 1650 | 453.3 | - | - | 0 | - |
| - | - | 1954 | 456.2 | - | - | 0 | - |
| - | - | 3245 | 458.2 | - | - | 0 | - |
| - | - | 1072 | 459.2 | - | - | 0 | - |
| - | - | 565.7 | 461.3 | - | - | 0 | - |
| - | - | 737.7 | 468.3 | - | - | 0 | - |
| - | - | 773.1 | 469.2 | - | - | 0 | - |
| - | - | 1.738E+04 | 471.3 | - | - | 0 | - |
| - | - | 676.6 | 486.2 | - | - | 0 | - |
| - | - | 1206 | 486.3 | - | - | 0 | - |
| - | - | 854.2 | 487.2 | - | - | 0 | - |
| - | - | 643.4 | 488.3 | - | - | 0 | - |
| - | - | 3763 | 488.3 | - | - | 0 | - |
| - | - | 601.6 | 510.5 | - | - | 0 | - |
| - | - | 523 | 511.3 | - | - | 0 | - |
| - | - | 1153 | 514.3 | - | - | 0 | - |
| - | - | 923.8 | 524.2 | - | - | 0 | - |
| - | - | 1450 | 528.3 | - | - | 0 | - |
| 5 | c | 1.318E+04 | 529.3 | 0.0005367 | 1.014 | +1 | 5 |
| - | - | 2583 | 530.3 | - | - | 0 | - |
| 5 | y | 2117 | 542.2 | 0.000253 | 0.4666 | +1 | 4 |
| 5 | z | 793.1 | 543.2 | 0.003788 | 6.973 | +1 | 4 |
| - | - | 1011 | 558.2 | - | - | 0 | - |
| 5 | y | 1.519E+04 | 559.3 | 0.0005594 | 1 | +1 | 4 |
| - | - | 3694 | 560.3 | - | - | 0 | - |
| - | - | 959.3 | 561.3 | - | - | 0 | - |
| - | - | 835.5 | 571.3 | - | - | 0 | - |
| - | - | 834 | 599.3 | - | - | 0 | - |
| - | - | 703.5 | 600.3 | - | - | 0 | - |
| - | - | 1286 | 614.4 | - | - | 0 | - |
| - | - | 942.4 | 625.3 | - | - | 0 | - |
| - | - | 652.5 | 638.3 | - | - | 0 | - |
| - | - | 664.3 | 640.3 | - | - | 0 | - |
| - | - | 1558 | 641.3 | - | - | 0 | - |
| - | - | 1157 | 641.4 | - | - | 0 | - |
| - | - | 2477 | 642.3 | - | - | 0 | - |
| 4 | y | 2365 | 643.3 | 0.001405 | 2.184 | +1 | 5 |
| 4 | z | 8700 | 644.3 | 0.0002915 | 0.4525 | +1 | 5 |
| - | - | 6988 | 645.3 | - | - | 0 | - |
| - | - | 2247 | 646.3 | - | - | 0 | - |
| - | - | 769.2 | 650.9 | - | - | 0 | - |
| - | - | 1135 | 656.4 | - | - | 0 | - |
| - | - | 1749 | 657.4 | - | - | 0 | - |
| 6 | c | 2.538E+04 | 658.4 | 0.001034 | 1.571 | +1 | 6 |
| - | - | 1.125E+04 | 659.4 | - | - | 0 | - |
| 4 | y | 1712 | 660.3 | 0.001465 | 2.219 | +1 | 5 |
| - | - | 2112 | 660.4 | - | - | 0 | - |
| - | - | 796.7 | 661.3 | - | - | 0 | - |
| - | - | 3461 | 684.4 | - | - | 0 | - |
| - | - | 1855 | 685.4 | - | - | 0 | - |
| - | - | 1106 | 715.3 | - | - | 0 | - |
| - | - | 1116 | 728.3 | - | - | 0 | - |
| - | - | 1973 | 742.4 | - | - | 0 | - |
| - | - | 1110 | 743.4 | - | - | 0 | - |
| - | - | 1282 | 751.4 | - | - | 0 | - |
| - | - | 1008 | 753.4 | - | - | 0 | - |
| - | - | 726.7 | 754.4 | - | - | 0 | - |
| 3 | z | 3076 | 757.4 | 0.002043 | 2.697 | +1 | 6 |
| - | - | 1331 | 758.4 | - | - | 0 | - |
| - | - | 737.9 | 759.4 | - | - | 0 | - |
| 7 | c | 1952 | 769.4 | 0.0009884 | 1.285 | +1 | 7 |
| - | - | 795.3 | 770.4 | - | - | 0 | - |
| - | - | 2008 | 771.4 | - | - | 0 | - |
| - | - | 788.7 | 772.4 | - | - | 0 | - |
| - | - | 2100 | 773.4 | - | - | 0 | - |
| - | - | 1232 | 774.4 | - | - | 0 | - |
| - | - | 1665 | 784.4 | - | - | 0 | - |
| - | - | 727.6 | 785.4 | - | - | 0 | - |
| 7 | c | 3.906E+04 | 786.4 | 0.001661 | 2.112 | +1 | 7 |
| - | - | 1.656E+04 | 787.4 | - | - | 0 | - |
| - | - | 1253 | 788.3 | - | - | 0 | - |
| - | - | 3749 | 788.4 | - | - | 0 | - |
| - | - | 1033 | 800.4 | - | - | 0 | - |
| - | - | 1973 | 808.4 | - | - | 0 | - |
| - | - | 993.8 | 809.4 | - | - | 0 | - |
| - | - | 721.6 | 836.4 | - | - | 0 | - |
| - | - | 716.3 | 837.4 | - | - | 0 | - |
| - | - | 655.6 | 841.4 | - | - | 0 | - |
| - | - | 832.5 | 842.4 | - | - | 0 | - |
| - | - | 2016 | 843.4 | - | - | 0 | - |
| 2 | z | 7972 | 844.4 | 0.002241 | 2.654 | +1 | 7 |
| - | - | 3236 | 845.4 | - | - | 0 | - |
| - | - | 691.6 | 846.4 | - | - | 0 | - |
| - | - | 5914 | 857.5 | - | - | 0 | - |
| - | - | 2803 | 858.5 | - | - | 0 | - |
| - | - | 831.7 | 859.5 | - | - | 0 | - |
| 2 | y | 1332 | 860.4 | 0.002255 | 2.621 | +1 | 7 |
| - | - | 700.4 | 861.4 | - | - | 0 | - |
| - | - | 1793 | 884.5 | - | - | 0 | - |
| - | - | 1181 | 901.5 | - | - | 0 | - |
| - | - | 661.9 | 909.4 | - | - | 0 | - |
| - | - | 813.7 | 910.4 | - | - | 0 | - |
| - | - | 2027 | 912.5 | - | - | 0 | - |
| - | - | 779.9 | 913.5 | - | - | 0 | - |
| - | - | 1079 | 914.4 | - | - | 0 | - |
| - | - | 817.4 | 915.4 | - | - | 0 | - |
| - | - | 2100 | 929.5 | - | - | 0 | - |
| - | - | 2335 | 930.5 | - | - | 0 | - |
| - | - | 820.4 | 939.5 | - | - | 0 | - |
| - | - | 2181 | 954.4 | - | - | 0 | - |
| - | - | 1283 | 955.4 | - | - | 0 | - |
| - | - | 2188 | 956.5 | - | - | 0 | - |
| - | - | 8136 | 957.5 | - | - | 0 | - |
| - | - | 5086 | 958.5 | - | - | 0 | - |
| - | - | 1587 | 959.5 | - | - | 0 | - |
| - | - | 665.8 | 965.4 | - | - | 0 | - |
| - | - | 3155 | 971.4 | - | - | 0 | - |
| - | - | 860.6 | 972.4 | - | - | 0 | - |
| - | - | 1.563E+04 | 973.5 | - | - | 0 | - |
| - | - | 2.28E+04 | 974.5 | - | - | 0 | - |
| - | - | 1117 | 974.6 | - | - | 0 | - |
| - | - | 8256 | 975.5 | - | - | 0 | - |
| - | - | 3273 | 976.5 | - | - | 0 | - |
| - | - | 679.5 | 1295 | - | - | 0 | - |
| - | - | 634.3 | 1451 | - | - | 0 | - |
| - | - | 1353 | 1459 | - | - | 0 | - |
| - | - | 1045 | 1460 | - | - | 0 | - |
| - | - | 887.7 | 1461 | - | - | 0 | - |
| - | - | 703.7 | 1992 | - | - | 0 | - |
| - | - | 707.7 | 2593 | - | - | 0 | - |
| - | - | 917.8 | 3080 | - | - | 0 | - |

m/z Charge Intensity FragmentType MassShift Position
120.35992431640625 0 397.60742
131.07046508789062 0 1240.5065
132.1022186279297 0 537.17993
133.08595275878906 0 17871.957
137.31494140625 0 414.95834
147.25225830078125 0 431.93872
148.954833984375 0 841.26984
155.07090759277344 0 511.17368
159.09169006347656 0 1561.2247
160.08753967285156 0 543.03735
173.128662109375 0 833.76526
173.451171875 0 2142.2861
175.0962371826172 0 1611.3546
177.11215209960938 0 9529.323
182.66383361816406 0 499.87933
187.1441192626953 0 779.0034
188.07064819335938 0 6946.866
189.07440185546875 0 543.66437
195.1230926513672 0 681.069
201.10240173339844 0 1269.4729
201.1122283935547 0 760.0915
201.1231689453125 0 1943.8724
205.09710693359375 0 6135.3433 y 7
206.10079956054688 0 707.8131
215.1392364501953 0 1395.8422
221.1384735107422 0 4655.758
227.10272216796875 0 1283.6586
234.64369201660156 0 513.3337
244.1293487548828 0 5443.9824
265.16436767578125 0 1873.636
283.17474365234375 0 913.51337
284.1606750488281 0 663.0695
296.197265625 0 2551.251
302.17095947265625 0 1536.6381
309.1912536621094 0 3945.6074
314.2077941894531 0 809.2978
315.1458740234375 0 1630.4436
316.12945556640625 0 2462.2979 y Ammonia loss 6
321.18182373046875 0 638.8442
327.20172119140625 0 18456.875
333.15618896484375 0 2831.8196 y 6
355.16131591796875 0 5851.5957
355.2235107421875 0 1398.1738
356.164306640625 0 1004.5994
372.1873779296875 0 2338.9822
373.23443603515625 0 912.8923
381.2483825683594 0 10528.91
397.2445373535156 0 1779.1808
399.25946044921875 0 6760.6733
415.1687316894531 0 585.38385
415.2557678222656 0 6021.87
416.25921630859375 0 1504.2362
433.21905517578125 0 515.5526
442.29534912109375 0 874.3382
453.3061828613281 0 1649.7029
456.2101135253906 0 1953.7335
458.2256774902344 0 3245.2537
459.2286376953125 0 1072.1648
461.34295654296875 0 565.70807
468.30706787109375 0 737.6976
469.16595458984375 0 773.1283
471.31707763671875 0 17382.318
486.22674560546875 0 676.562
486.3191223144531 0 1206.1208
487.1745910644531 0 854.171
488.2932434082031 0 643.36816
488.3432312011719 0 3763.367
510.485595703125 0 601.5666
511.254638671875 0 523.0413
514.2547607421875 0 1153.3901
524.2161865234375 0 923.8012
528.3267211914062 0 1450.2341
529.3349609375 0 13183.713 c 4
530.339111328125 0 2582.6733
542.2247924804688 0 2117.2217 y Ammonia loss 4
543.2285766601562 0 793.0528 z 4
558.2457275390625 0 1010.82605
559.2516479492188 0 15191.706 y 4
560.2548828125 0 3693.5664
561.2576293945312 0 959.32904
571.3095703125 0 835.4875
599.2567138671875 0 834.0233
600.2760620117188 0 703.4506
614.3641357421875 0 1286.2887
625.3364868164062 0 942.43945
638.3132934570312 0 652.50964
640.32080078125 0 664.3397
641.3123779296875 0 1558.2
641.3607788085938 0 1157.0393
642.2659301757812 0 2476.6572
643.2708129882812 0 2364.581 y Ammonia loss 3
644.2803344726562 0 8699.536 z 3
645.287109375 0 6987.686
646.2896118164062 0 2247.135
650.8590698242188 0 769.18085
656.36474609375 0 1135.4011
657.370849609375 0 1749.2278
658.3780517578125 0 25376.031 c 5
659.38134765625 0 11248.977
660.3002319335938 0 1711.5204 y 3
660.3838500976562 0 2112.4084
661.3052368164062 0 796.65796
684.3942260742188 0 3461.3806
685.3928833007812 0 1855.168
715.3392333984375 0 1106.2552
728.3458862304688 0 1115.8712
742.4234008789062 0 1972.8788
743.419677734375 0 1110.3167
751.3998413085938 0 1282.3994
753.3922729492188 0 1008.0095
754.39697265625 0 726.728
757.3661499023438 0 3075.7505 z 2
758.367919921875 0 1330.859
759.37353515625 0 737.8836
769.4100341796875 0 1952.4501 c Ammonia loss 6
770.4114379882812 0 795.3048
771.4283447265625 0 2008.3141
772.4292602539062 0 788.69434
773.3648071289062 0 2099.8433
774.3638916015625 0 1231.8601
784.421875 0 1665.0056
785.4227905273438 0 727.5677
786.437255859375 0 39061.45 c 6
787.4396362304688 0 16562.457
788.335205078125 0 1253.4269
788.4429931640625 0 3749.3228
800.4093627929688 0 1032.9778
808.3937377929688 0 1973.1346
809.4078979492188 0 993.8443
836.3987426757812 0 721.584
837.3895874023438 0 716.27954
841.4317626953125 0 655.5969
842.3817138671875 0 832.5415
843.4332275390625 0 2016.0035
844.3983764648438 0 7971.8022 z 1
845.4025268554688 0 3236.1384
846.3987426757812 0 691.6342
857.4661254882812 0 5913.944
858.4686889648438 0 2802.7139
859.4730224609375 0 831.68854
860.4171142578125 0 1331.7426 y 1
861.4194946289062 0 700.3826
884.511474609375 0 1792.9312
901.4567260742188 0 1180.9933
909.3731689453125 0 661.9177
910.3784790039062 0 813.70544
912.505859375 0 2026.5687
913.5020751953125 0 779.8525
914.43212890625 0 1078.5668
915.4270629882812 0 817.42896
929.4873657226562 0 2100.144
930.5039672851562 0 2334.725
939.4739379882812 0 820.41406
954.416748046875 0 2181.3296
955.421142578125 0 1283.0679
956.497314453125 0 2188.419
957.485107421875 0 8135.939
958.4869995117188 0 5086.0137
959.4872436523438 0 1587.1433
965.4252319335938 0 665.8035
971.4229125976562 0 3155.103
972.4172973632812 0 860.6412
973.5004272460938 0 15630.286
974.5068969726562 0 22803.684
974.6168212890625 0 1116.9905
975.5093383789062 0 8256.441
976.5111083984375 0 3272.5684
1295.47216796875 0 679.5397
1450.6224365234375 0 634.33
1458.865966796875 0 1352.7677
1459.8553466796875 0 1045.1765
1460.8001708984375 0 887.68176
1992.266845703125 0 703.7407
2592.6875 0 707.69916
3080.154541015625 0 917.8106

Spectrum Details

|  |  |
| --- | --- |
| Matched peaks? Matched peaks The total absolute number of peaks matched. Additionally in brackets the total fraction of peaks matched and the total number of peaks is shown. | 16 (9.14% of 175) |
| FDR? FDR The false discovery rate estimated for this peptide. It is calculated by matching all theoretical fragments with a non-integer shift with the raw peaks for this spectrum. This is done with 40 different shifts. The resulting percentage is the average number of annotated peaks over the number of annotated peaks with the correct spectrum. | 3.42% |
| Satellite FDR? Satellite FDR See the FDR for details on its calculation. This satellite ion specific FDR only contains the satellite ions (d/w) for I/L/J positions. | - |
| PSM Score? PSM Score The PSM Score as given by Hecklib to this annotated spectrum. It is shown with three significant figures. | 196 |

#### Spectrum 7205? Spectrum 7205 The raw spectrum of this peptide as annotated by Hecklib. The fragments are coloured according to ion type (see legend). Any peaks with a star '\*' as text can be hovered over to see the full details, first the ion type second the mass shift type. By hovering over the amino acids in the peptide or ions in the legend the corresponding peaks are highlighted. By toggling the 'Unassigned' label you can turn the background (unassigned) peaks on or off in the plot. By updating the slider in the Ion legend you can update the spectrum to only show the top X% of the peaks with labels. The top X% means any peak that is within X% of the highest intensity. By dragging in the spectrum you can zoom in to a specific part of the spectrum and use 'Zoom Out' to get back to the original zoom level. The annotation of the spectrum is based on the given sequence in the peptides file and is done with different software so inconsistencies are likely. The peaks are annotated based on the given sequence, with 20 ppm tolerance.

Copy Data

##### Spectrum 7205 (TSV)

###### Preview

```
Loading example...
```

*Click on the button to copy the data to your clipboard.*

Mz MinMz MaxIntensity Max

WidthHeightPeptide font sizePeptide stroke widthSpectrum font sizeSpectrum stroke widthCompact peptide

Ion legend

wxyz

abcd

OtherUnassignedIon^Charge^_Position_Show for top:%

JSJTPEQW

01.47e+42.94e+44.41e+45.88e+4

Zoom Out

y^+1^_2_y^+1^_2_c^+1^_5_y^+1^_4_y^+1^_4_z^+1^_4_y^+1^_4_c^+1^_6_y^+1^_5_z^+1^_5_y^+1^_5_c^+1^_7_c^+1^_7_z^+1^_6_c^+1^_7_z^+1^_7_y^+1^_7_y^+1^_7_z^+1^_7_

0501100215042005

Fragment Matches Table

Show background peaks

| Position | Ion type | Intensity | mz Theoretical | mz Error (Th) | mz Error (ppm) | Charge | Series Number |
| --- | --- | --- | --- | --- | --- | --- | --- |
| - | - | 345 | 121.3 | - | - | 0 | - |
| - | - | 574.2 | 130.1 | - | - | 0 | - |
| - | - | 443.1 | 132.9 | - | - | 0 | - |
| - | - | 428.4 | 136.1 | - | - | 0 | - |
| - | - | 864.4 | 149 | - | - | 0 | - |
| - | - | 465.9 | 149.4 | - | - | 0 | - |
| - | - | 473.8 | 150.1 | - | - | 0 | - |
| - | - | 548.6 | 157.1 | - | - | 0 | - |
| - | - | 1017 | 158.1 | - | - | 0 | - |
| - | - | 487.7 | 173.1 | - | - | 0 | - |
| - | - | 1832 | 173.1 | - | - | 0 | - |
| - | - | 1546 | 175.1 | - | - | 0 | - |
| - | - | 445.1 | 182.6 | - | - | 0 | - |
| - | - | 532.9 | 183.1 | - | - | 0 | - |
| - | - | 690.9 | 187.1 | - | - | 0 | - |
| - | - | 4007 | 201.1 | - | - | 0 | - |
| - | - | 3561 | 202.1 | - | - | 0 | - |
| - | - | 570.4 | 202.1 | - | - | 0 | - |
| - | - | 1.093E+04 | 203.1 | - | - | 0 | - |
| - | - | 1606 | 204.1 | - | - | 0 | - |
| - | - | 455.2 | 214.4 | - | - | 0 | - |
| - | - | 1640 | 215.1 | - | - | 0 | - |
| - | - | 8148 | 221.1 | - | - | 0 | - |
| - | - | 1207 | 222.1 | - | - | 0 | - |
| - | - | 1588 | 227.1 | - | - | 0 | - |
| - | - | 652 | 229.2 | - | - | 0 | - |
| - | - | 500.9 | 233.9 | - | - | 0 | - |
| - | - | 4343 | 244.1 | - | - | 0 | - |
| - | - | 696.2 | 256.4 | - | - | 0 | - |
| - | - | 457.7 | 276.6 | - | - | 0 | - |
| - | - | 1324 | 284.2 | - | - | 0 | - |
| - | - | 4631 | 296.2 | - | - | 0 | - |
| - | - | 1699 | 302.2 | - | - | 0 | - |
| - | - | 1397 | 314.1 | - | - | 0 | - |
| - | - | 1613 | 314.2 | - | - | 0 | - |
| - | - | 470.8 | 321.2 | - | - | 0 | - |
| - | - | 486.4 | 324.1 | - | - | 0 | - |
| - | - | 554.5 | 328.2 | - | - | 0 | - |
| - | - | 1520 | 331.1 | - | - | 0 | - |
| 7 | y | 1928 | 332.1 | 0.004566 | 13.75 | +1 | 2 |
| - | - | 537.3 | 344.4 | - | - | 0 | - |
| 7 | y | 2832 | 349.1 | 0.005147 | 14.74 | +1 | 2 |
| - | - | 1.275E+04 | 355.2 | - | - | 0 | - |
| - | - | 746.4 | 355.2 | - | - | 0 | - |
| - | - | 1539 | 356.2 | - | - | 0 | - |
| - | - | 964.4 | 370.2 | - | - | 0 | - |
| - | - | 1774 | 372.2 | - | - | 0 | - |
| - | - | 1117 | 387.3 | - | - | 0 | - |
| - | - | 3115 | 397.2 | - | - | 0 | - |
| - | - | 1045 | 398.2 | - | - | 0 | - |
| - | - | 481.9 | 399 | - | - | 0 | - |
| - | - | 585.2 | 400.2 | - | - | 0 | - |
| - | - | 6733 | 415.3 | - | - | 0 | - |
| - | - | 1541 | 416.3 | - | - | 0 | - |
| - | - | 551.3 | 423.7 | - | - | 0 | - |
| - | - | 600.4 | 439.2 | - | - | 0 | - |
| - | - | 2732 | 441.2 | - | - | 0 | - |
| - | - | 563 | 441.9 | - | - | 0 | - |
| - | - | 579.2 | 444.2 | - | - | 0 | - |
| - | - | 1757 | 456.2 | - | - | 0 | - |
| - | - | 1528 | 459.8 | - | - | 0 | - |
| - | - | 1103 | 471.3 | - | - | 0 | - |
| - | - | 2529 | 477.8 | - | - | 0 | - |
| - | - | 526.7 | 485.7 | - | - | 0 | - |
| - | - | 810.5 | 495.8 | - | - | 0 | - |
| - | - | 586.9 | 524.3 | - | - | 0 | - |
| - | - | 747.5 | 528.3 | - | - | 0 | - |
| 5 | c | 8237 | 529.3 | 4.84E-05 | 0.09144 | +1 | 5 |
| - | - | 1396 | 530.2 | - | - | 0 | - |
| - | - | 1900 | 530.3 | - | - | 0 | - |
| - | - | 1168 | 540.2 | - | - | 0 | - |
| - | - | 1319 | 554.3 | - | - | 0 | - |
| 5 | y | 1412 | 557.2 | 0.005743 | 10.31 | +1 | 4 |
| 5 | y | 2233 | 558.2 | 0.006713 | 12.03 | +1 | 4 |
| 5 | z | 1350 | 559.2 | 0.001268 | 2.268 | +1 | 4 |
| - | - | 604.2 | 569.3 | - | - | 0 | - |
| - | - | 1839 | 574.2 | - | - | 0 | - |
| 5 | y | 1.094E+04 | 575.2 | 0.004578 | 7.958 | +1 | 4 |
| - | - | 3180 | 576.2 | - | - | 0 | - |
| - | - | 949.2 | 599.3 | - | - | 0 | - |
| - | - | 3943 | 602.2 | - | - | 0 | - |
| - | - | 1146 | 603.2 | - | - | 0 | - |
| - | - | 594.2 | 603.7 | - | - | 0 | - |
| - | - | 683.6 | 614.4 | - | - | 0 | - |
| - | - | 1099 | 623.3 | - | - | 0 | - |
| - | - | 628.3 | 624.3 | - | - | 0 | - |
| - | - | 903.3 | 638.3 | - | - | 0 | - |
| - | - | 6488 | 641.3 | - | - | 0 | - |
| - | - | 2606 | 642.3 | - | - | 0 | - |
| - | - | 3078 | 658.3 | - | - | 0 | - |
| 6 | c | 1.912E+04 | 658.4 | 0.0001189 | 0.1806 | +1 | 6 |
| 4 | y | 2157 | 659.3 | 0.001271 | 1.928 | +1 | 5 |
| - | - | 7496 | 659.4 | - | - | 0 | - |
| 4 | z | 5739 | 660.3 | 0.004066 | 6.158 | +1 | 5 |
| - | - | 1781 | 660.4 | - | - | 0 | - |
| - | - | 3671 | 661.3 | - | - | 0 | - |
| 4 | y | 2243 | 676.3 | 0.004507 | 6.664 | +1 | 5 |
| - | - | 732.3 | 704.3 | - | - | 0 | - |
| - | - | 618.7 | 734.4 | - | - | 0 | - |
| - | - | 609.4 | 741.9 | - | - | 0 | - |
| - | - | 804.2 | 742.9 | - | - | 0 | - |
| - | - | 2155 | 751.4 | - | - | 0 | - |
| - | - | 932.7 | 752.4 | - | - | 0 | - |
| - | - | 878.9 | 753.4 | - | - | 0 | - |
| - | - | 3222 | 754.4 | - | - | 0 | - |
| - | - | 1394 | 755.4 | - | - | 0 | - |
| 7 | c | 861.6 | 768.4 | 0.01341 | 17.45 | +1 | 7 |
| 7 | c | 4621 | 769.4 | 0.0008426 | 1.095 | +1 | 7 |
| - | - | 1413 | 770.4 | - | - | 0 | - |
| - | - | 852.5 | 771.4 | - | - | 0 | - |
| 3 | z | 1568 | 773.4 | 0.005146 | 6.654 | +1 | 6 |
| - | - | 760.1 | 780.4 | - | - | 0 | - |
| - | - | 1020 | 781.4 | - | - | 0 | - |
| - | - | 779.1 | 782.4 | - | - | 0 | - |
| 7 | c | 5.824E+04 | 786.4 | 0.0001701 | 0.2162 | +1 | 7 |
| - | - | 2.299E+04 | 787.4 | - | - | 0 | - |
| - | - | 5707 | 788.4 | - | - | 0 | - |
| - | - | 1010 | 789.4 | - | - | 0 | - |
| - | - | 817.5 | 798.4 | - | - | 0 | - |
| - | - | 1426 | 800.4 | - | - | 0 | - |
| - | - | 1079 | 801.4 | - | - | 0 | - |
| - | - | 1177 | 804.3 | - | - | 0 | - |
| - | - | 3490 | 812.4 | - | - | 0 | - |
| - | - | 3215 | 812.5 | - | - | 0 | - |
| - | - | 955.8 | 813.4 | - | - | 0 | - |
| - | - | 1187 | 813.5 | - | - | 0 | - |
| - | - | 1616 | 824.4 | - | - | 0 | - |
| - | - | 653.6 | 825.5 | - | - | 0 | - |
| - | - | 2394 | 827.5 | - | - | 0 | - |
| - | - | 900.3 | 828.5 | - | - | 0 | - |
| 2 | z | 2996 | 842.4 | 0.005838 | 6.93 | +1 | 7 |
| - | - | 6653 | 843.4 | - | - | 0 | - |
| - | - | 3253 | 844.4 | - | - | 0 | - |
| - | - | 4095 | 857.5 | - | - | 0 | - |
| 2 | y | 844.7 | 858.4 | 0.01429 | 16.65 | +1 | 7 |
| - | - | 1176 | 858.5 | - | - | 0 | - |
| 2 | y | 811.1 | 859.4 | 0.01537 | 17.88 | +1 | 7 |
| 2 | z | 1.52E+04 | 860.4 | 0.004612 | 5.36 | +1 | 7 |
| - | - | 982.5 | 860.5 | - | - | 0 | - |
| - | - | 7870 | 861.4 | - | - | 0 | - |
| - | - | 2237 | 862.4 | - | - | 0 | - |
| - | - | 2.613E+04 | 875.5 | - | - | 0 | - |
| - | - | 1.212E+04 | 876.5 | - | - | 0 | - |
| - | - | 3719 | 877.5 | - | - | 0 | - |
| - | - | 1939 | 900.5 | - | - | 0 | - |
| - | - | 2342 | 910.5 | - | - | 0 | - |
| - | - | 1880 | 911.5 | - | - | 0 | - |
| - | - | 718.6 | 917.5 | - | - | 0 | - |
| - | - | 859.3 | 918.5 | - | - | 0 | - |
| - | - | 625.3 | 920.5 | - | - | 0 | - |
| - | - | 4.842E+04 | 928.5 | - | - | 0 | - |
| - | - | 2.523E+04 | 929.5 | - | - | 0 | - |
| - | - | 2855 | 930.4 | - | - | 0 | - |
| - | - | 6987 | 930.5 | - | - | 0 | - |
| - | - | 946.2 | 931.4 | - | - | 0 | - |
| - | - | 640.3 | 937.4 | - | - | 0 | - |
| - | - | 851.6 | 945.5 | - | - | 0 | - |
| - | - | 1.063E+04 | 946.5 | - | - | 0 | - |
| - | - | 5734 | 947.5 | - | - | 0 | - |
| - | - | 1952 | 948.5 | - | - | 0 | - |
| - | - | 7432 | 955.5 | - | - | 0 | - |
| - | - | 4527 | 956.5 | - | - | 0 | - |
| - | - | 1608 | 957.5 | - | - | 0 | - |
| - | - | 902.1 | 970.4 | - | - | 0 | - |
| - | - | 4623 | 972.5 | - | - | 0 | - |
| - | - | 3.06E+04 | 973.5 | - | - | 0 | - |
| - | - | 1.479E+04 | 974.5 | - | - | 0 | - |
| - | - | 4365 | 975.5 | - | - | 0 | - |
| - | - | 862 | 978 | - | - | 0 | - |
| - | - | 3134 | 978.5 | - | - | 0 | - |
| - | - | 1763 | 979 | - | - | 0 | - |
| - | - | 894 | 979.5 | - | - | 0 | - |
| - | - | 778.3 | 984 | - | - | 0 | - |
| - | - | 1090 | 987.1 | - | - | 0 | - |
| - | - | 708.4 | 987.6 | - | - | 0 | - |
| - | - | 1.863E+04 | 989.5 | - | - | 0 | - |
| - | - | 1.912E+04 | 990.5 | - | - | 0 | - |
| - | - | 1.007E+04 | 991.5 | - | - | 0 | - |
| - | - | 1067 | 992 | - | - | 0 | - |
| - | - | 1560 | 992.5 | - | - | 0 | - |
| - | - | 2040 | 993 | - | - | 0 | - |
| - | - | 1487 | 993.1 | - | - | 0 | - |
| - | - | 1024 | 993.6 | - | - | 0 | - |
| - | - | 698.9 | 1051 | - | - | 0 | - |
| - | - | 740.1 | 1467 | - | - | 0 | - |
| - | - | 1644 | 1468 | - | - | 0 | - |
| - | - | 1872 | 1469 | - | - | 0 | - |
| - | - | 1434 | 1484 | - | - | 0 | - |
| - | - | 1699 | 1485 | - | - | 0 | - |
| - | - | 2552 | 1486 | - | - | 0 | - |
| - | - | 1120 | 1487 | - | - | 0 | - |
| - | - | 683.9 | 1958 | - | - | 0 | - |
| - | - | 838.8 | 1962 | - | - | 0 | - |
| - | - | 750.7 | 1964 | - | - | 0 | - |
| - | - | 800 | 1967 | - | - | 0 | - |
| - | - | 1143 | 1968 | - | - | 0 | - |
| - | - | 709.7 | 1976 | - | - | 0 | - |
| - | - | 1179 | 1983 | - | - | 0 | - |
| - | - | 1802 | 1985 | - | - | 0 | - |

m/z Charge Intensity FragmentType MassShift Position
121.25521850585938 0 344.9957
130.0653076171875 0 574.1941
132.90077209472656 0 443.10867
136.07623291015625 0 428.39484
148.95558166503906 0 864.39874
149.36590576171875 0 465.94394
150.08535766601562 0 473.79028
157.07595825195312 0 548.61096
158.08392333984375 0 1017.2967
173.10812377929688 0 487.72028
173.12838745117188 0 1831.9592
175.08688354492188 0 1546.3584
182.58599853515625 0 445.14197
183.11257934570312 0 532.90564
187.14462280273438 0 690.8958
201.12347412109375 0 4007.398
202.0736541748047 0 3560.841
202.1271209716797 0 570.359
203.0814971923828 0 10928.155
204.0846710205078 0 1606.2483
214.3704833984375 0 455.1934
215.13946533203125 0 1640.3844
221.09213256835938 0 8148.107
222.09542846679688 0 1206.556
227.1022186279297 0 1587.8215
229.1549072265625 0 651.95337
233.890380859375 0 500.88855
244.12936401367188 0 4342.7217
256.4054870605469 0 696.23615
276.5655517578125 0 457.71323
284.1606750488281 0 1324.4952
296.1968994140625 0 4631.359
302.1706237792969 0 1699.1831
314.1139831542969 0 1396.9169
314.2068176269531 0 1613.3784
321.1787414550781 0 470.7556
324.05987548828125 0 486.39508
328.1514892578125 0 554.45044
331.14105224609375 0 1519.6469
332.1237487792969 0 1928.2651 y Ammonia loss 6
344.3645935058594 0 537.2725
349.15087890625 0 2832.3845 y 6
355.1611633300781 0 12751.297
355.18621826171875 0 746.4312
356.1639404296875 0 1539.2184
370.2334289550781 0 964.3789
372.18780517578125 0 1774.1366
387.2593688964844 0 1116.5667
397.2445983886719 0 3114.7244
398.2043151855469 0 1044.9999
398.9595642089844 0 481.88013
400.17376708984375 0 585.238
415.2552795410156 0 6732.86
416.2585144042969 0 1541.3486
423.6654052734375 0 551.29834
439.1826171875 0 600.38654
441.198486328125 0 2731.7307
441.8515319824219 0 562.9985
444.2088928222656 0 579.2061
456.20953369140625 0 1756.7352
459.81243896484375 0 1528.428
471.3046569824219 0 1103.3794
477.82281494140625 0 2529.0557
485.71124267578125 0 526.67883
495.8320617675781 0 810.5034
524.327392578125 0 586.85815
528.3262329101562 0 747.4668
529.33447265625 0 8236.877 c 4
530.2478637695312 0 1395.6355
530.3367309570312 0 1899.854
540.2094116210938 0 1167.6283
554.28466796875 0 1319.446
557.2362670898438 0 1412.4282 y Water loss 4
558.2212524414062 0 2232.9504 y Ammonia loss 4
559.2236328125 0 1349.6543 z 4
569.2925415039062 0 604.2194
574.2383422851562 0 1839.0327
575.2456665039062 0 10941.676 y 4
576.2493286132812 0 3180.4155
599.2813110351562 0 949.1632
602.2328491210938 0 3943.345
603.23583984375 0 1145.629
603.667724609375 0 594.22174
614.3673095703125 0 683.56995
623.3001098632812 0 1099.3132
624.3040771484375 0 628.2967
638.3189086914062 0 903.2894
641.3141479492188 0 6488.1836
642.3171997070312 0 2605.5234
658.259033203125 0 3077.6277
658.3771362304688 0 19123.1 c 5
659.2634887695312 0 2157.1924 y Ammonia loss 3
659.3804321289062 0 7496.3257
660.2741088867188 0 5738.937 z 3
660.3810424804688 0 1781.3477
661.2789916992188 0 3670.5007
676.2932739257812 0 2242.5935 y 3
704.2860717773438 0 732.3311
734.369384765625 0 618.6904
741.9185180664062 0 609.39166
742.9232177734375 0 804.2322
751.3973999023438 0 2155.3293
752.4030151367188 0 932.70807
753.4014282226562 0 878.9313
754.3988037109375 0 3222.4744
755.398193359375 0 1394.0955
768.41162109375 0 861.64954 c Water loss 6
769.408203125 0 4620.906 c Ammonia loss 6
770.4103393554688 0 1413.4108
771.41845703125 0 852.5033
773.3592529296875 0 1567.7012 z 2
780.377197265625 0 760.1308
781.3875732421875 0 1019.69714
782.3992919921875 0 779.1138
786.4354248046875 0 58239.207 c 6
787.4384765625 0 22986.562
788.4408569335938 0 5706.881
789.437744140625 0 1010.2126
798.4411010742188 0 817.46844
800.4042358398438 0 1425.9147
801.417236328125 0 1078.9886
804.3301391601562 0 1177.1812
812.369873046875 0 3490.1028
812.451904296875 0 3214.9102
813.3717651367188 0 955.7949
813.4518432617188 0 1187.217
824.3710327148438 0 1616.435
825.5275268554688 0 653.561
827.4529418945312 0 2393.9575
828.4522094726562 0 900.26556
842.3814086914062 0 2995.865 z Water loss 1
843.40966796875 0 6652.9976
844.4134521484375 0 3253.1414
857.4627685546875 0 4094.863
858.3800048828125 0 844.7228 y Water loss 1
858.4663696289062 0 1176.4595
859.3936767578125 0 811.1058 y Ammonia loss 1
860.3907470703125 0 15197.293 z 1
860.4804077148438 0 982.5076
861.3947143554688 0 7869.9526
862.3948974609375 0 2236.6716
875.4747924804688 0 26130.521
876.4769287109375 0 12123.753
877.4789428710938 0 3718.6345
900.4711303710938 0 1939.4038
910.4876708984375 0 2341.8281
911.4825439453125 0 1880.4365
917.4625244140625 0 718.5916
918.4830322265625 0 859.2956
920.4678955078125 0 625.3419
928.5006103515625 0 48421.277
929.5032348632812 0 25226.045
930.416748046875 0 2855.366
930.508056640625 0 6987.3193
931.424072265625 0 946.1565
937.4454345703125 0 640.2674
945.4902954101562 0 851.607
946.5106201171875 0 10625.974
947.5147094726562 0 5734.2983
948.5137329101562 0 1952.3479
955.4649658203125 0 7432.2856
956.4682006835938 0 4526.99
957.4737548828125 0 1607.6909
970.4466552734375 0 902.11084
972.4890747070312 0 4622.919
973.4766235351562 0 30600.77
974.4791259765625 0 14790.194
975.4813232421875 0 4364.822
977.9747924804688 0 862.0459
978.4757690429688 0 3134.3003
978.9780883789062 0 1763.4341
979.4862670898438 0 894.01215
983.9600219726562 0 778.2872
987.0657958984375 0 1089.9246
987.5606689453125 0 708.4297
989.4924926757812 0 18630.404
990.5011596679688 0 19117.107
991.5044555664062 0 10070.651
991.96142578125 0 1066.8651
992.4951782226562 0 1560.123
992.9555053710938 0 2040.3826
993.06201171875 0 1486.6475
993.5601806640625 0 1023.52625
1051.1505126953125 0 698.9164
1466.8419189453125 0 740.14166
1467.7991943359375 0 1644.1818
1468.81201171875 0 1871.6349
1483.82958984375 0 1433.7303
1484.84033203125 0 1698.5271
1485.845947265625 0 2552.303
1486.8487548828125 0 1120.3923
1957.9686279296875 0 683.91895
1961.9063720703125 0 838.79663
1963.8985595703125 0 750.6688
1966.907958984375 0 799.9531
1967.8944091796875 0 1143.3865
1975.9522705078125 0 709.6602
1982.92138671875 0 1178.936
1984.9053955078125 0 1802.1898

Spectrum Details

|  |  |
| --- | --- |
| Matched peaks? Matched peaks The total absolute number of peaks matched. Additionally in brackets the total fraction of peaks matched and the total number of peaks is shown. | 19 (9.55% of 199) |
| FDR? FDR The false discovery rate estimated for this peptide. It is calculated by matching all theoretical fragments with a non-integer shift with the raw peaks for this spectrum. This is done with 40 different shifts. The resulting percentage is the average number of annotated peaks over the number of annotated peaks with the correct spectrum. | 3.76% |
| Satellite FDR? Satellite FDR See the FDR for details on its calculation. This satellite ion specific FDR only contains the satellite ions (d/w) for I/L/J positions. | ∞ |
| PSM Score? PSM Score The PSM Score as given by Hecklib to this annotated spectrum. It is shown with three significant figures. | 248 |

#### Spectrum 7837? Spectrum 7837 The raw spectrum of this peptide as annotated by Hecklib. The fragments are coloured according to ion type (see legend). Any peaks with a star '\*' as text can be hovered over to see the full details, first the ion type second the mass shift type. By hovering over the amino acids in the peptide or ions in the legend the corresponding peaks are highlighted. By toggling the 'Unassigned' label you can turn the background (unassigned) peaks on or off in the plot. By updating the slider in the Ion legend you can update the spectrum to only show the top X% of the peaks with labels. The top X% means any peak that is within X% of the highest intensity. By dragging in the spectrum you can zoom in to a specific part of the spectrum and use 'Zoom Out' to get back to the original zoom level. The annotation of the spectrum is based on the given sequence in the peptides file and is done with different software so inconsistencies are likely. The peaks are annotated based on the given sequence, with 20 ppm tolerance.

Copy Data

##### Spectrum 7837 (TSV)

###### Preview

```
Loading example...
```

*Click on the button to copy the data to your clipboard.*

Mz MinMz MaxIntensity Max

WidthHeightPeptide font sizePeptide stroke widthSpectrum font sizeSpectrum stroke widthCompact peptide

Ion legend

wxyz

abcd

OtherUnassignedIon^Charge^_Position_Show for top:%

JSJTPEQW

03.85e+47.70e+41.16e+51.54e+5

Zoom Out

a^+1^_2_d^+1^_2_a^+1^_2_b^+1^_2_b^+1^_2_a^+1^_3_y^+2^_4_b^+1^_3_b^+1^_3_b^+2^_6_y^+1^_2_y^+1^_2_b^+1^_4_b^+1^_4_y^+2^_7_y^+1^_3_y^+1^_3_y^+1^_3_\*b^+1^_5_\*b^+1^_5_y^+1^_4_y^+1^_4_y^+1^_4_b^+1^_6_b^+1^_6_y^+1^_5_y^+1^_5_y^+1^_5_b^+1^_7_b^+1^_7_y^+1^_6_y^+1^_7_

0863172625903453

Fragment Matches Table

Show background peaks

| Position | Ion type | Intensity | mz Theoretical | mz Error (Th) | mz Error (ppm) | Charge | Series Number |
| --- | --- | --- | --- | --- | --- | --- | --- |
| - | - | 3877 | 120.1 | - | - | 0 | - |
| - | - | 3545 | 125.1 | - | - | 0 | - |
| - | - | 1429 | 126.1 | - | - | 0 | - |
| - | - | 697.6 | 127.1 | - | - | 0 | - |
| - | - | 1650 | 128.1 | - | - | 0 | - |
| - | - | 6947 | 129.1 | - | - | 0 | - |
| - | - | 2731 | 129.1 | - | - | 0 | - |
| - | - | 1847 | 130.1 | - | - | 0 | - |
| - | - | 1.646E+04 | 130.1 | - | - | 0 | - |
| - | - | 985.8 | 130.1 | - | - | 0 | - |
| - | - | 1732 | 131.1 | - | - | 0 | - |
| - | - | 427.5 | 131.2 | - | - | 0 | - |
| - | - | 3624 | 132 | - | - | 0 | - |
| - | - | 487.3 | 132.1 | - | - | 0 | - |
| - | - | 1283 | 132.1 | - | - | 0 | - |
| - | - | 604.4 | 133 | - | - | 0 | - |
| - | - | 1696 | 134.1 | - | - | 0 | - |
| - | - | 392 | 135.1 | - | - | 0 | - |
| - | - | 1530 | 136.1 | - | - | 0 | - |
| - | - | 551.8 | 141.1 | - | - | 0 | - |
| - | - | 822.5 | 141.1 | - | - | 0 | - |
| - | - | 5239 | 142.1 | - | - | 0 | - |
| - | - | 522 | 143.1 | - | - | 0 | - |
| - | - | 427.4 | 143.1 | - | - | 0 | - |
| - | - | 6780 | 146.1 | - | - | 0 | - |
| - | - | 749.5 | 147.1 | - | - | 0 | - |
| - | - | 2171 | 147.1 | - | - | 0 | - |
| - | - | 579.2 | 148.8 | - | - | 0 | - |
| - | - | 424.7 | 148.9 | - | - | 0 | - |
| - | - | 441.3 | 148.9 | - | - | 0 | - |
| - | - | 571.6 | 148.9 | - | - | 0 | - |
| - | - | 475.6 | 148.9 | - | - | 0 | - |
| - | - | 559.7 | 148.9 | - | - | 0 | - |
| - | - | 906.3 | 148.9 | - | - | 0 | - |
| - | - | 907.1 | 148.9 | - | - | 0 | - |
| - | - | 1173 | 148.9 | - | - | 0 | - |
| - | - | 1459 | 148.9 | - | - | 0 | - |
| - | - | 3530 | 148.9 | - | - | 0 | - |
| - | - | 5435 | 149 | - | - | 0 | - |
| - | - | 3559 | 149 | - | - | 0 | - |
| - | - | 1479 | 149 | - | - | 0 | - |
| - | - | 933.2 | 149 | - | - | 0 | - |
| - | - | 956.2 | 149 | - | - | 0 | - |
| - | - | 1050 | 149 | - | - | 0 | - |
| - | - | 682.8 | 149 | - | - | 0 | - |
| - | - | 755.8 | 149 | - | - | 0 | - |
| - | - | 551.3 | 149 | - | - | 0 | - |
| - | - | 405.4 | 149 | - | - | 0 | - |
| - | - | 455.6 | 149 | - | - | 0 | - |
| - | - | 2451 | 152.1 | - | - | 0 | - |
| - | - | 1628 | 153.1 | - | - | 0 | - |
| - | - | 650.1 | 153.1 | - | - | 0 | - |
| - | - | 527.4 | 155.1 | - | - | 0 | - |
| 2 | a | 5195 | 155.1 | 0.0002593 | 1.671 | +1 | 2 |
| - | - | 459.4 | 156.1 | - | - | 0 | - |
| - | - | 7560 | 157.1 | - | - | 0 | - |
| 2 | d | 478.2 | 157.1 | 0.0006004 | 3.821 | +1 | 2 |
| - | - | 1.955E+04 | 158.1 | - | - | 0 | - |
| - | - | 933 | 158.1 | - | - | 0 | - |
| - | - | 2351 | 159.1 | - | - | 0 | - |
| - | - | 1105 | 159.1 | - | - | 0 | - |
| - | - | 442.7 | 160.1 | - | - | 0 | - |
| - | - | 698.8 | 166.1 | - | - | 0 | - |
| - | - | 2746 | 169.1 | - | - | 0 | - |
| - | - | 534.3 | 169.1 | - | - | 0 | - |
| - | - | 2543 | 169.1 | - | - | 0 | - |
| - | - | 4652 | 171.1 | - | - | 0 | - |
| - | - | 1485 | 171.1 | - | - | 0 | - |
| 2 | a | 3.835E+04 | 173.1 | 0.0003605 | 2.082 | +1 | 2 |
| - | - | 633.1 | 173.5 | - | - | 0 | - |
| - | - | 2824 | 174.1 | - | - | 0 | - |
| - | - | 468.7 | 174.1 | - | - | 0 | - |
| - | - | 1.041E+05 | 175.1 | - | - | 0 | - |
| - | - | 593.1 | 176.1 | - | - | 0 | - |
| - | - | 594 | 176.1 | - | - | 0 | - |
| - | - | 9374 | 176.1 | - | - | 0 | - |
| - | - | 536.7 | 177.1 | - | - | 0 | - |
| - | - | 9910 | 181.1 | - | - | 0 | - |
| - | - | 4687 | 181.1 | - | - | 0 | - |
| 2 | b | 1.028E+04 | 183.1 | 0.0003245 | 1.772 | +1 | 2 |
| - | - | 819.8 | 183.1 | - | - | 0 | - |
| - | - | 1990 | 183.1 | - | - | 0 | - |
| - | - | 926.1 | 184.1 | - | - | 0 | - |
| - | - | 631.2 | 185.1 | - | - | 0 | - |
| - | - | 1500 | 186.1 | - | - | 0 | - |
| - | - | 1.25E+04 | 187.1 | - | - | 0 | - |
| - | - | 1701 | 188.1 | - | - | 0 | - |
| - | - | 995.5 | 188.1 | - | - | 0 | - |
| - | - | 481.6 | 189.7 | - | - | 0 | - |
| - | - | 5390 | 197.1 | - | - | 0 | - |
| - | - | 707.2 | 199.1 | - | - | 0 | - |
| - | - | 3.637E+04 | 199.1 | - | - | 0 | - |
| - | - | 3254 | 200.1 | - | - | 0 | - |
| - | - | 1202 | 201.1 | - | - | 0 | - |
| 2 | b | 4.582E+04 | 201.1 | 0.0003036 | 1.51 | +1 | 2 |
| - | - | 4400 | 202.1 | - | - | 0 | - |
| - | - | 1.548E+04 | 203.1 | - | - | 0 | - |
| - | - | 3087 | 204.1 | - | - | 0 | - |
| - | - | 1672 | 204.1 | - | - | 0 | - |
| - | - | 896.9 | 205.1 | - | - | 0 | - |
| - | - | 562 | 209.1 | - | - | 0 | - |
| - | - | 3129 | 209.1 | - | - | 0 | - |
| - | - | 1278 | 211.1 | - | - | 0 | - |
| - | - | 771 | 211.1 | - | - | 0 | - |
| - | - | 2151 | 211.1 | - | - | 0 | - |
| - | - | 996.3 | 212.1 | - | - | 0 | - |
| - | - | 678 | 213.1 | - | - | 0 | - |
| - | - | 2.711E+04 | 215.1 | - | - | 0 | - |
| - | - | 3052 | 216.1 | - | - | 0 | - |
| - | - | 2485 | 217.1 | - | - | 0 | - |
| - | - | 1.526E+05 | 221.1 | - | - | 0 | - |
| - | - | 1.63E+04 | 222.1 | - | - | 0 | - |
| - | - | 916.9 | 223.1 | - | - | 0 | - |
| - | - | 1669 | 223.1 | - | - | 0 | - |
| - | - | 688.4 | 225.1 | - | - | 0 | - |
| - | - | 1739 | 226.1 | - | - | 0 | - |
| - | - | 779.7 | 226.2 | - | - | 0 | - |
| - | - | 5.331E+04 | 227.1 | - | - | 0 | - |
| - | - | 6132 | 228.1 | - | - | 0 | - |
| - | - | 550.7 | 228.1 | - | - | 0 | - |
| - | - | 567.6 | 229.1 | - | - | 0 | - |
| - | - | 1412 | 229.2 | - | - | 0 | - |
| - | - | 965.3 | 233.2 | - | - | 0 | - |
| - | - | 886.4 | 233.2 | - | - | 0 | - |
| - | - | 908.5 | 233.7 | - | - | 0 | - |
| - | - | 542.8 | 237.1 | - | - | 0 | - |
| - | - | 551.8 | 238.2 | - | - | 0 | - |
| - | - | 3281 | 239.1 | - | - | 0 | - |
| - | - | 5936 | 240.1 | - | - | 0 | - |
| - | - | 1666 | 241.1 | - | - | 0 | - |
| - | - | 1882 | 241.1 | - | - | 0 | - |
| - | - | 736.8 | 241.2 | - | - | 0 | - |
| - | - | 921.3 | 242.2 | - | - | 0 | - |
| - | - | 1066 | 242.7 | - | - | 0 | - |
| - | - | 1.207E+04 | 244.1 | - | - | 0 | - |
| - | - | 1044 | 245.1 | - | - | 0 | - |
| - | - | 1389 | 245.1 | - | - | 0 | - |
| - | - | 723.8 | 249.2 | - | - | 0 | - |
| - | - | 884.4 | 251.1 | - | - | 0 | - |
| - | - | 2130 | 251.2 | - | - | 0 | - |
| - | - | 1836 | 256.2 | - | - | 0 | - |
| - | - | 3544 | 257.1 | - | - | 0 | - |
| - | - | 6505 | 258.1 | - | - | 0 | - |
| - | - | 1794 | 258.1 | - | - | 0 | - |
| - | - | 486.5 | 259.1 | - | - | 0 | - |
| - | - | 953.9 | 264.1 | - | - | 0 | - |
| - | - | 504.9 | 266.1 | - | - | 0 | - |
| - | - | 1845 | 266.1 | - | - | 0 | - |
| - | - | 1106 | 267.1 | - | - | 0 | - |
| - | - | 1205 | 268.1 | - | - | 0 | - |
| 3 | a | 1795 | 268.2 | 7.974E-05 | 0.2973 | +1 | 3 |
| - | - | 1315 | 269.2 | - | - | 0 | - |
| - | - | 1372 | 274.2 | - | - | 0 | - |
| - | - | 1054 | 280.2 | - | - | 0 | - |
| - | - | 1049 | 282.1 | - | - | 0 | - |
| - | - | 1.942E+04 | 284.2 | - | - | 0 | - |
| - | - | 1509 | 285.1 | - | - | 0 | - |
| - | - | 2683 | 285.2 | - | - | 0 | - |
| - | - | 1.268E+04 | 286.1 | - | - | 0 | - |
| - | - | 651.8 | 286.1 | - | - | 0 | - |
| - | - | 1842 | 287.1 | - | - | 0 | - |
| 5 | y | 628.8 | 288.1 | 0.002801 | 9.722 | +2 | 4 |
| - | - | 1642 | 292.1 | - | - | 0 | - |
| - | - | 1171 | 294.2 | - | - | 0 | - |
| - | - | 1907 | 296.1 | - | - | 0 | - |
| - | - | 705.9 | 296.1 | - | - | 0 | - |
| 3 | b | 4.384E+04 | 296.2 | 0.0003975 | 1.342 | +1 | 3 |
| - | - | 5916 | 297.2 | - | - | 0 | - |
| - | - | 1734 | 298.2 | - | - | 0 | - |
| - | - | 1559 | 300.2 | - | - | 0 | - |
| - | - | 2.511E+04 | 302.2 | - | - | 0 | - |
| - | - | 3386 | 303.2 | - | - | 0 | - |
| - | - | 868.4 | 309.2 | - | - | 0 | - |
| - | - | 2899 | 310.1 | - | - | 0 | - |
| - | - | 942.9 | 310.2 | - | - | 0 | - |
| - | - | 3084 | 312.2 | - | - | 0 | - |
| - | - | 1116 | 312.2 | - | - | 0 | - |
| - | - | 1494 | 313.1 | - | - | 0 | - |
| - | - | 2.268E+04 | 314.1 | - | - | 0 | - |
| 3 | b | 1.06E+04 | 314.2 | 0.0004529 | 1.441 | +1 | 3 |
| - | - | 3781 | 315.1 | - | - | 0 | - |
| - | - | 1520 | 315.2 | - | - | 0 | - |
| - | - | 1217 | 320.1 | - | - | 0 | - |
| 6 | b | 1391 | 321.2 | 0.0005405 | 1.683 | +2 | 6 |
| - | - | 714.2 | 323.2 | - | - | 0 | - |
| - | - | 3922 | 327.2 | - | - | 0 | - |
| - | - | 5622 | 328.2 | - | - | 0 | - |
| - | - | 1281 | 330.2 | - | - | 0 | - |
| - | - | 4.989E+04 | 331.1 | - | - | 0 | - |
| 7 | y | 4.942E+04 | 332.1 | 0.005238 | 15.77 | +1 | 2 |
| - | - | 6486 | 332.1 | - | - | 0 | - |
| - | - | 9116 | 333.1 | - | - | 0 | - |
| - | - | 864.6 | 333.1 | - | - | 0 | - |
| - | - | 636.9 | 334.2 | - | - | 0 | - |
| - | - | 4243 | 337.2 | - | - | 0 | - |
| - | - | 2744 | 338.1 | - | - | 0 | - |
| - | - | 583.7 | 338.2 | - | - | 0 | - |
| - | - | 705.1 | 339.1 | - | - | 0 | - |
| - | - | 2415 | 340.2 | - | - | 0 | - |
| - | - | 1156 | 343.2 | - | - | 0 | - |
| - | - | 3729 | 345.2 | - | - | 0 | - |
| - | - | 761 | 346.2 | - | - | 0 | - |
| 7 | y | 2.567E+04 | 349.1 | 0.005239 | 15 | +1 | 2 |
| - | - | 4498 | 350.2 | - | - | 0 | - |
| - | - | 891.2 | 351.2 | - | - | 0 | - |
| - | - | 3544 | 352.2 | - | - | 0 | - |
| - | - | 1427 | 353.2 | - | - | 0 | - |
| - | - | 6.551E+04 | 355.2 | - | - | 0 | - |
| - | - | 1291 | 356.1 | - | - | 0 | - |
| - | - | 1.054E+04 | 356.2 | - | - | 0 | - |
| - | - | 1569 | 357.2 | - | - | 0 | - |
| - | - | 1137 | 363.2 | - | - | 0 | - |
| - | - | 913.5 | 365.2 | - | - | 0 | - |
| - | - | 5133 | 369.3 | - | - | 0 | - |
| - | - | 7523 | 370.2 | - | - | 0 | - |
| - | - | 598.6 | 370.3 | - | - | 0 | - |
| - | - | 2299 | 371.2 | - | - | 0 | - |
| - | - | 1078 | 372.2 | - | - | 0 | - |
| - | - | 638.3 | 372.2 | - | - | 0 | - |
| - | - | 1685 | 373.2 | - | - | 0 | - |
| - | - | 2782 | 379.2 | - | - | 0 | - |
| - | - | 1463 | 381.2 | - | - | 0 | - |
| - | - | 8028 | 387.3 | - | - | 0 | - |
| - | - | 1129 | 388.3 | - | - | 0 | - |
| - | - | 1871 | 397.2 | - | - | 0 | - |
| 4 | b | 2.516E+04 | 397.2 | 0.0002959 | 0.7449 | +1 | 4 |
| - | - | 5721 | 398.2 | - | - | 0 | - |
| - | - | 738.4 | 399.2 | - | - | 0 | - |
| - | - | 772.9 | 399.3 | - | - | 0 | - |
| - | - | 2122 | 409.2 | - | - | 0 | - |
| 4 | b | 4.877E+04 | 415.3 | 0.0003208 | 0.7726 | +1 | 4 |
| - | - | 1.004E+04 | 416.3 | - | - | 0 | - |
| - | - | 1458 | 417.3 | - | - | 0 | - |
| - | - | 1066 | 423.2 | - | - | 0 | - |
| - | - | 1756 | 425.1 | - | - | 0 | - |
| 2 | y | 982.6 | 429.7 | 0.003896 | 9.066 | +2 | 7 |
| - | - | 689.2 | 434.9 | - | - | 0 | - |
| - | - | 1286 | 438.2 | - | - | 0 | - |
| - | - | 1632 | 439.8 | - | - | 0 | - |
| - | - | 1634 | 440.3 | - | - | 0 | - |
| - | - | 846.9 | 441.2 | - | - | 0 | - |
| - | - | 9700 | 443.2 | - | - | 0 | - |
| - | - | 1908 | 444.2 | - | - | 0 | - |
| - | - | 696.1 | 445.7 | - | - | 0 | - |
| - | - | 1123 | 448.3 | - | - | 0 | - |
| - | - | 644.8 | 450.3 | - | - | 0 | - |
| - | - | 616.4 | 451.3 | - | - | 0 | - |
| - | - | 5339 | 456.2 | - | - | 0 | - |
| - | - | 1322 | 457.2 | - | - | 0 | - |
| 6 | y | 2902 | 460.2 | 0.004888 | 10.62 | +1 | 3 |
| 6 | y | 3808 | 461.2 | 0.00543 | 11.78 | +1 | 3 |
| - | - | 955.6 | 462.2 | - | - | 0 | - |
| - | - | 1060 | 466.3 | - | - | 0 | - |
| - | - | 4870 | 468.2 | - | - | 0 | - |
| - | - | 899.1 | 468.3 | - | - | 0 | - |
| - | - | 955.2 | 476.3 | - | - | 0 | - |
| - | - | 1691 | 477.7 | - | - | 0 | - |
| - | - | 1067 | 477.8 | - | - | 0 | - |
| 6 | y | 2655 | 478.2 | 0.004546 | 9.508 | +1 | 3 |
| 0 | Precursor | 925.2 | 486.2 | 0.0009658 | 1.986 | +2 | -1 |
| - | - | 1221 | 486.9 | - | - | 0 | - |
| - | - | 1229 | 488.2 | - | - | 0 | - |
| - | - | 825.4 | 492.2 | - | - | 0 | - |
| - | - | 1546 | 494.2 | - | - | 0 | - |
| 5 | b | 3101 | 494.3 | 0.001992 | 4.03 | +1 | 5 |
| - | - | 851.6 | 494.6 | - | - | 0 | - |
| 0 | Precursor | 669.4 | 495.2 | 0.005379 | 10.86 | +2 | -1 |
| - | - | 1187 | 495.3 | - | - | 0 | - |
| - | - | 9255 | 495.8 | - | - | 0 | - |
| - | - | 840.1 | 496.2 | - | - | 0 | - |
| - | - | 2018 | 510.3 | - | - | 0 | - |
| - | - | 750.4 | 511.3 | - | - | 0 | - |
| - | - | 983.6 | 512.2 | - | - | 0 | - |
| 5 | b | 3103 | 512.3 | 0.0002914 | 0.5687 | +1 | 5 |
| - | - | 1103 | 513.3 | - | - | 0 | - |
| - | - | 586.2 | 522.3 | - | - | 0 | - |
| - | - | 595.7 | 525.3 | - | - | 0 | - |
| - | - | 1539 | 537.3 | - | - | 0 | - |
| - | - | 1.22E+04 | 540.2 | - | - | 0 | - |
| - | - | 3058 | 541.2 | - | - | 0 | - |
| - | - | 629.6 | 541.3 | - | - | 0 | - |
| - | - | 839.3 | 543.3 | - | - | 0 | - |
| - | - | 1597 | 551.3 | - | - | 0 | - |
| - | - | 841.7 | 552.3 | - | - | 0 | - |
| - | - | 1403 | 555.3 | - | - | 0 | - |
| - | - | 781.3 | 556.3 | - | - | 0 | - |
| 5 | y | 7905 | 557.2 | 0.003912 | 7.021 | +1 | 4 |
| 5 | y | 2.606E+04 | 558.2 | 0.005431 | 9.73 | +1 | 4 |
| - | - | 8970 | 559.2 | - | - | 0 | - |
| - | - | 1497 | 560.2 | - | - | 0 | - |
| - | - | 1665 | 569.3 | - | - | 0 | - |
| - | - | 833.3 | 570.3 | - | - | 0 | - |
| 5 | y | 9.764E+04 | 575.2 | 0.004761 | 8.277 | +1 | 4 |
| - | - | 3.109E+04 | 576.2 | - | - | 0 | - |
| - | - | 969.2 | 576.3 | - | - | 0 | - |
| - | - | 6105 | 577.3 | - | - | 0 | - |
| - | - | 596.1 | 603.9 | - | - | 0 | - |
| - | - | 1102 | 605.3 | - | - | 0 | - |
| - | - | 837.8 | 613.4 | - | - | 0 | - |
| - | - | 738.4 | 620.3 | - | - | 0 | - |
| - | - | 741 | 622.4 | - | - | 0 | - |
| - | - | 631.2 | 623.2 | - | - | 0 | - |
| 6 | b | 3695 | 623.3 | 0.0001233 | 0.1978 | +1 | 6 |
| - | - | 1144 | 624.3 | - | - | 0 | - |
| - | - | 2283 | 638.3 | - | - | 0 | - |
| - | - | 837.1 | 639.3 | - | - | 0 | - |
| - | - | 698.2 | 639.9 | - | - | 0 | - |
| - | - | 797.8 | 640.4 | - | - | 0 | - |
| - | - | 1076 | 641.3 | - | - | 0 | - |
| 6 | b | 3911 | 641.4 | 0.001705 | 2.658 | +1 | 6 |
| - | - | 1755 | 642.4 | - | - | 0 | - |
| - | - | 1709 | 650.3 | - | - | 0 | - |
| - | - | 1438 | 656.3 | - | - | 0 | - |
| - | - | 858.3 | 657.3 | - | - | 0 | - |
| 4 | y | 1034 | 658.3 | 0.007442 | 11.31 | +1 | 5 |
| - | - | 2124 | 658.4 | - | - | 0 | - |
| 4 | y | 2797 | 659.3 | 0.006764 | 10.26 | +1 | 5 |
| - | - | 1126 | 668.4 | - | - | 0 | - |
| 4 | y | 1.126E+04 | 676.3 | 0.00469 | 6.935 | +1 | 5 |
| - | - | 3894 | 677.3 | - | - | 0 | - |
| - | - | 599 | 678.3 | - | - | 0 | - |
| - | - | 1012 | 724.4 | - | - | 0 | - |
| - | - | 2211 | 733.4 | - | - | 0 | - |
| - | - | 1426 | 734.4 | - | - | 0 | - |
| - | - | 696.3 | 735.4 | - | - | 0 | - |
| - | - | 1693 | 741.4 | - | - | 0 | - |
| 7 | b | 9224 | 751.4 | 0.0004708 | 0.6266 | +1 | 7 |
| - | - | 3572 | 752.4 | - | - | 0 | - |
| 7 | b | 4549 | 769.4 | 0.0009037 | 1.174 | +1 | 7 |
| - | - | 2536 | 770.4 | - | - | 0 | - |
| 3 | y | 2138 | 789.4 | 0.003329 | 4.217 | +1 | 6 |
| - | - | 951.8 | 790.4 | - | - | 0 | - |
| 2 | y | 1001 | 876.4 | 0.00371 | 4.233 | +1 | 7 |
| - | - | 735.2 | 877.4 | - | - | 0 | - |
| - | - | 608.4 | 885.5 | - | - | 0 | - |
| - | - | 667 | 2007 | - | - | 0 | - |
| - | - | 761.8 | 3081 | - | - | 0 | - |
| - | - | 698 | 3419 | - | - | 0 | - |

m/z Charge Intensity FragmentType MassShift Position
120.08108520507812 0 3877.2837
125.10762786865234 0 3545.3994
126.091552734375 0 1428.8878
127.08673095703125 0 697.59863
128.1072998046875 0 1649.574
129.06619262695312 0 6946.74
129.10255432128906 0 2731.138
130.05027770996094 0 1846.5591
130.06546020507812 0 16462.592
130.07066345214844 0 985.80066
131.06883239746094 0 1732.3628
131.22311401367188 0 427.53952
132.04483032226562 0 3624.496
132.0813751220703 0 487.31204
132.1020965576172 0 1282.6278
133.0479736328125 0 604.3857
134.06045532226562 0 1696.0739
135.0799560546875 0 391.95755
136.07606506347656 0 1530.3551
141.06655883789062 0 551.78406
141.10264587402344 0 822.50964
142.12295532226562 0 5239.282
143.11764526367188 0 522.0471
143.12570190429688 0 427.3525
146.06040954589844 0 6779.6143
147.0640106201172 0 749.4808
147.0768280029297 0 2171.4966
148.83082580566406 0 579.17596
148.86776733398438 0 424.70367
148.88233947753906 0 441.29776
148.88951110839844 0 571.62787
148.8965301513672 0 475.63824
148.90371704101562 0 559.6585
148.9105224609375 0 906.26166
148.918212890625 0 907.1028
148.92543029785156 0 1173.1704
148.93260192871094 0 1458.6023
148.94041442871094 0 3529.5342
148.9569854736328 0 5434.6167
148.9648895263672 0 3558.9136
148.97239685058594 0 1479.2898
148.9790496826172 0 933.1682
148.98707580566406 0 956.1755
148.9942169189453 0 1050.3679
149.00181579589844 0 682.82367
149.0090789794922 0 755.8072
149.023193359375 0 551.32544
149.03045654296875 0 405.39352
149.0384979248047 0 455.60327
152.10723876953125 0 2451.152
153.10231018066406 0 1628.3032
153.109619140625 0 650.1479
155.08140563964844 0 527.4078
155.11814880371094 0 5195.0537 a Water loss 1
156.12158203125 0 459.42758
157.07638549804688 0 7560.4785
157.13414001464844 0 478.20078 d 1
158.0603790283203 0 19554.457
158.07980346679688 0 932.95734
159.06373596191406 0 2351.2566
159.09210205078125 0 1105.0815
160.1242218017578 0 442.65384
166.0868682861328 0 698.7565
169.06109619140625 0 2746.4023
169.0977325439453 0 534.2918
169.1337890625 0 2542.775
171.1131134033203 0 4651.5566
171.14967346191406 0 1485.4795
173.12881469726562 0 38346.934 a 1
173.45050048828125 0 633.1444
174.1321563720703 0 2824.3271
174.13961791992188 0 468.65683
175.0869598388672 0 104103.195
176.07077026367188 0 593.0911
176.0819549560547 0 594.009
176.0903778076172 0 9373.511
177.09327697753906 0 536.68365
181.0611114501953 0 9909.571
181.097412109375 0 4687.206
183.11312866210938 0 10277.719 b Water loss 1
183.12179565429688 0 819.844
183.14947509765625 0 1989.8789
184.1171112060547 0 926.1348
185.09228515625 0 631.2172
186.05531311035156 0 1499.6626
187.14447021484375 0 12496.823
188.07093811035156 0 1700.6389
188.14779663085938 0 995.45715
189.7252197265625 0 481.63892
197.12875366210938 0 5389.8193
199.0713653564453 0 707.2494
199.10800170898438 0 36372.29
200.11131286621094 0 3254.0989
201.06634521484375 0 1201.7769
201.12367248535156 0 45818.062 b 1
202.12710571289062 0 4400.204
203.08181762695312 0 15480.811
204.06588745117188 0 3086.7605
204.08517456054688 0 1672.0139
205.0975341796875 0 896.9022
209.08090209960938 0 562.00555
209.0924072265625 0 3128.6643
211.09439086914062 0 1277.9456
211.1064453125 0 771.04
211.1441650390625 0 2150.843
212.1038055419922 0 996.3213
213.12376403808594 0 677.9989
215.13934326171875 0 27109.379
216.14279174804688 0 3052.1987
217.0823211669922 0 2485.4067
221.0924530029297 0 152559.25
222.09571838378906 0 16298.708
223.0720977783203 0 916.92957
223.09718322753906 0 1668.7678
225.1233673095703 0 688.3928
226.1189422607422 0 1739.1898
226.15467834472656 0 779.6816
227.10299682617188 0 53305.336
228.1063232421875 0 6132.2827
228.13221740722656 0 550.71484
229.1092071533203 0 567.5744
229.1549835205078 0 1411.9685
233.15016174316406 0 965.2913
233.1646728515625 0 886.3688
233.65492248535156 0 908.47766
237.1248016357422 0 542.79456
238.15460205078125 0 551.82996
239.13925170898438 0 3280.7607
240.09829711914062 0 5935.9883
241.0826416015625 0 1665.9668
241.09828186035156 0 1882.3644
241.19119262695312 0 736.7674
242.15037536621094 0 921.2655
242.6612548828125 0 1065.7434
244.12957763671875 0 12065.722
245.1132354736328 0 1043.7399
245.13233947753906 0 1388.7761
249.1602325439453 0 723.78986
251.13954162597656 0 884.3878
251.1754150390625 0 2129.9204
256.1658935546875 0 1836.3829
257.1497497558594 0 3544.4707
258.1087341308594 0 6505.2163
258.14508056640625 0 1794.2544
259.0916748046875 0 486.522
264.1338195800781 0 953.9497
266.11297607421875 0 504.92688
266.1498107910156 0 1845.4407
267.1338195800781 0 1105.6774
268.1076354980469 0 1205.0671
268.2018737792969 0 1795.3625 a Water loss 2
269.1859130859375 0 1314.6052
274.17657470703125 0 1372.2518
280.1663513183594 0 1054.4908
282.14501953125 0 1049.0989
284.16082763671875 0 19421.635
285.1351623535156 0 1509.4052
285.16412353515625 0 2683.0728
286.1189270019531 0 12683.03
286.13525390625 0 651.7954
287.12213134765625 0 1841.8617
288.1269836425781 0 628.7939 y 4
292.12982177734375 0 1642.083
294.18206787109375 0 1170.5536
296.1030578613281 0 1907.2742
296.1236877441406 0 705.87866
296.197265625 0 43835.82 b Water loss 2
297.2005615234375 0 5916.3613
298.17584228515625 0 1734.381
300.1560363769531 0 1559.2198
302.1714782714844 0 25109.373
303.1745300292969 0 3385.7927
309.15557861328125 0 868.4001
310.1399841308594 0 2899.1582
310.2127380371094 0 942.8513
312.15594482421875 0 3083.5938
312.1930236816406 0 1116.2565
313.1297607421875 0 1494.0768
314.1138916015625 0 22677.12
314.2078857421875 0 10599.354 b 2
315.1169128417969 0 3781.1665
315.21099853515625 0 1519.7928
320.12420654296875 0 1216.8031
321.1794128417969 0 1391.0531 b 5
323.2069091796875 0 714.20856
327.16656494140625 0 3922.3865
328.1507263183594 0 5622.1704
330.1671447753906 0 1280.6079
331.14056396484375 0 49890.496
332.1244201660156 0 49421.727 y Ammonia loss 6
332.14398193359375 0 6485.999
333.1276550292969 0 9116.36
333.1468811035156 0 864.6005
334.2129821777344 0 636.8548
337.1512145996094 0 4243.2593
338.1349792480469 0 2744.0327
338.1557312011719 0 583.6794
339.1371154785156 0 705.0861
340.1871032714844 0 2415.3562
343.2352600097656 0 1155.5914
345.1772766113281 0 3729.001
346.1777038574219 0 760.9803
349.1509704589844 0 25672.771 y 6
350.15362548828125 0 4498.3022
351.2388610839844 0 891.15845
352.2235107421875 0 3544.3318
353.2210388183594 0 1426.9625
355.1614074707031 0 65505.633
356.1451110839844 0 1290.5293
356.1645202636719 0 10544.538
357.1664733886719 0 1568.8384
363.2029113769531 0 1137.1805
365.18304443359375 0 913.49414
369.2500305175781 0 5132.5054
370.2336730957031 0 7523.2026
370.25390625 0 598.5587
371.23468017578125 0 2298.6292
372.18841552734375 0 1078.3103
372.2346496582031 0 638.3332
373.1714782714844 0 1685.135
379.2345275878906 0 2781.5576
381.2142333984375 0 1462.9114
387.2601318359375 0 8027.625
388.264892578125 0 1129.267
397.1506652832031 0 1870.5879
397.2448425292969 0 25155.879 b Water loss 3
398.24774169921875 0 5720.997
399.2243347167969 0 738.39984
399.251708984375 0 772.91364
409.208740234375 0 2121.5383
415.25543212890625 0 48769.367 b 3
416.2586975097656 0 10036.779
417.26214599609375 0 1457.6335
423.22412109375 0 1066.0363
425.1466369628906 0 1756.011
429.7046813964844 0 982.6156 y Water loss 1
434.8529357910156 0 689.22076
438.1978759765625 0 1285.8064
439.8440246582031 0 1632.4288
440.25091552734375 0 1634.272
441.2349853515625 0 846.8714
443.156494140625 0 9699.662
444.1597595214844 0 1908.4067
445.7260437011719 0 696.14703
448.2643737792969 0 1123.2739
450.2721252441406 0 644.75433
451.2577819824219 0 616.43085
456.20965576171875 0 5339.2485
457.21142578125 0 1322.1212
460.1826477050781 0 2901.6877 y Water loss 5
461.1672058105469 0 3807.6616 y Ammonia loss 5
462.1702880859375 0 955.6414
466.3011169433594 0 1060.2693
468.24542236328125 0 4870.194
468.2799072265625 0 899.077
476.29010009765625 0 955.2163
477.7341003417969 0 1691.005
477.8210754394531 0 1066.9513
478.19287109375 0 2655.452 y 5
486.2418518066406 0 925.23895 Precursor Water loss
486.9422302246094 0 1221.2651
488.1783752441406 0 1229.3263
492.2462463378906 0 825.4376
494.20391845703125 0 1545.8142
494.2953186035156 0 3100.5532 b Water loss 4
494.6205139160156 0 851.63696
495.25347900390625 0 669.4259 Precursor
495.29925537109375 0 1186.9794
495.8337097167969 0 9254.554
496.2240295410156 0 840.1118
510.2558288574219 0 2017.7538
511.25811767578125 0 750.4185
512.2139282226562 0 983.627
512.3081665039062 0 3102.626 b 4
513.3098754882812 0 1103.3292
522.292236328125 0 586.1684
525.3076171875 0 595.7263
537.2680053710938 0 1539.0522
540.208984375 0 12204.147
541.2116088867188 0 3057.5845
541.2984008789062 0 629.58704
543.3137817382812 0 839.31537
551.2826538085938 0 1596.897
552.2763671875 0 841.6915
555.275146484375 0 1403.4735
556.281982421875 0 781.32806
557.2344360351562 0 7905.2896 y Water loss 4
558.219970703125 0 26062.984 y Ammonia loss 4
559.22265625 0 8970.162
560.2266845703125 0 1497.1008
569.2942504882812 0 1665.4678
570.2943725585938 0 833.25745
575.245849609375 0 97641.46 y 4
576.2490234375 0 31086.506
576.3018188476562 0 969.15
577.251220703125 0 6104.662
603.8521118164062 0 596.135
605.330078125 0 1102.3514
613.356201171875 0 837.8261
620.3029174804688 0 738.364
622.3552856445312 0 740.9887
623.2481689453125 0 631.1672
623.3400268554688 0 3694.7307 b Water loss 5
624.3427124023438 0 1143.6857
638.3143920898438 0 2283.2764
639.3087768554688 0 837.0904
639.8674926757812 0 698.22156
640.3692626953125 0 797.82324
641.2566528320312 0 1076.3363
641.3521728515625 0 3910.5615 b 5
642.3525390625 0 1755.3168
650.3491821289062 0 1708.5565
656.32275390625 0 1437.9734
657.327880859375 0 858.34625
658.28564453125 0 1033.5077 y Water loss 3
658.377685546875 0 2123.7341
659.2689819335938 0 2796.7407 y Ammonia loss 3
668.363037109375 0 1125.8582
676.29345703125 0 11257.3545 y 3
677.2965087890625 0 3894.103
678.3017578125 0 599.04
724.3963012695312 0 1012.19745
733.38720703125 0 2210.6138
734.3745727539062 0 1426.0979
735.3782958984375 0 696.2536
741.4116821289062 0 1693.328
751.3980102539062 0 9224.122 b Water loss 6
752.3986206054688 0 3572.2163
769.4081420898438 0 4548.9116 b 6
770.4074096679688 0 2536.4556
789.3761596679688 0 2138.4124 y 2
790.3826293945312 0 951.7883
876.4085693359375 0 1001.1102 y 1
877.4091796875 0 735.24426
885.517822265625 0 608.4146
2007.0191650390625 0 666.9581
3081.18310546875 0 761.8466
3418.599853515625 0 697.9925

Spectrum Details

|  |  |
| --- | --- |
| Matched peaks? Matched peaks The total absolute number of peaks matched. Additionally in brackets the total fraction of peaks matched and the total number of peaks is shown. | 34 (10.06% of 338) |
| FDR? FDR The false discovery rate estimated for this peptide. It is calculated by matching all theoretical fragments with a non-integer shift with the raw peaks for this spectrum. This is done with 40 different shifts. The resulting percentage is the average number of annotated peaks over the number of annotated peaks with the correct spectrum. | 1.68% |
| Satellite FDR? Satellite FDR See the FDR for details on its calculation. This satellite ion specific FDR only contains the satellite ions (d/w) for I/L/J positions. | - |
| PSM Score? PSM Score The PSM Score as given by Hecklib to this annotated spectrum. It is shown with three significant figures. | 474 |

#### Reverse Lookup? Reverse Lookup All places where this read could be placed.

| Group | Segment | Template | Template Part | Read Part | Score | Unique |
| --- | --- | --- | --- | --- | --- | --- |
| Homo sapiens Light Chain | IGLC | IGLC2 | [71..79] | [0..8] | 64 | False |
| Homo sapiens Light Chain | IGLC | IGLC3 | [69..77] | [0..8] | 64 | False |
| Homo sapiens Light Chain | IGLC | IGLC6 | [71..79] | [0..8] | 64 | False |
| Homo sapiens Light Chain | IGLC | IGLC7 | [71..79] | [0..8] | 64 | False |

| Recombined | Template Part | Read Part | Score | Unique |
| --- | --- | --- | --- | --- |
| REC-0-1\_002 | [182..190] | [0..8] | 64 | True |

#### Meta Information from Multiple reads

##### Number of combined reads

17

##### Intensity

0.8532

##### TotalArea

2.257E+09

##### Changes to the peptide sequence

JSJTPEQW

I→JNo support for either Leucine or Isoleucine based on side chain ions (Position: 3)

L→ISupport for Isoleucine based on side chain ions (1 for I 0 for L) (Position: 3)

J→LSupport for Leucine based on side chain ions (1 for L 0 for I) (Position: 3)

L→JNo support for either Leucine or Isoleucine based on side chain ions (Position: 3)

J→LSupport for Leucine based on side chain ions (1 for L 0 for I) (Position: 3)

I→JNo support for either Leucine or Isoleucine based on side chain ions (Position: 3)

L→ISupport for Isoleucine based on side chain ions (1 for I 0 for L) (Position: 3)

L→JNo support for either Leucine or Isoleucine based on side chain ions (Position: 1)

#### Positional Score

Copy Data

##### Positional Score (TSV)

###### Preview

```
Loading example...
```

*Click on the button to copy the data to your clipboard.*

1001234567

Label Value
"0" 0.836
"1" 0.814
"2" 0.861
"3" 0.868
"4" 0.864
"5" 0.867
"6" 0.866
"7" 0.882

#### Meta Information from PEAKS

##### Scan Identifier

F3:9728

##### Original sequence

L

S

L

T

P

E

Q

W

##### Posttranslational Modifications

##### Source File

D:\separate\_stitch\_analyses\xle-disambiguation\raw\20210323\_F1\_UM1\_Peng0013\_SA\_F59\_ingel\_3ug\_chymo.raw

##### Fraction

3

##### Scan Feature

F3:3577

##### De Novo Score

98

##### ConfidenceScore

98

### m/z

487.2539

##### Mass

972.4916

##### Charge

2

##### Retention Time

53.66

##### Predicted Retention Time

-

##### Area

7.064E+08

##### Parts Per Million

1.7

##### Fragmentation mode

ETHCD

##### Originating file

01 D:\separate\_stitch\_analyses\xle-disambiguation\20210325\_F59\_3ug\_DENOVO\_12.csv

#### Meta Information from PEAKS

##### Scan Identifier

F3:9668

##### Original sequence

L

S

L

T

P

E

Q

W

##### Posttranslational Modifications

##### Source File

D:\separate\_stitch\_analyses\xle-disambiguation\raw\20210323\_F1\_UM1\_Peng0013\_SA\_F59\_ingel\_3ug\_chymo.raw

##### Fraction

3

##### Scan Feature

F3:3577

##### De Novo Score

98

##### ConfidenceScore

98

### m/z

487.2539

##### Mass

972.4916

##### Charge

2

##### Retention Time

53.66

##### Predicted Retention Time

-

##### Area

7.064E+08

##### Parts Per Million

1.7

##### Fragmentation mode

HCD

##### Originating file

01 D:\separate\_stitch\_analyses\xle-disambiguation\20210325\_F59\_3ug\_DENOVO\_12.csv

#### Meta Information from PEAKS

##### Scan Identifier

F3:9605

##### Original sequence

L

S

L

T

P

E

Q

W

##### Posttranslational Modifications

##### Source File

D:\separate\_stitch\_analyses\xle-disambiguation\raw\20210323\_F1\_UM1\_Peng0013\_SA\_F59\_ingel\_3ug\_chymo.raw

##### Fraction

3

##### Scan Feature

F3:3577

##### De Novo Score

98

##### ConfidenceScore

98

### m/z

487.2539

##### Mass

972.4916

##### Charge

2

##### Retention Time

53.66

##### Predicted Retention Time

-

##### Area

7.064E+08

##### Parts Per Million

1.7

##### Fragmentation mode

HCD

##### Originating file

01 D:\separate\_stitch\_analyses\xle-disambiguation\20210325\_F59\_3ug\_DENOVO\_12.csv

#### Meta Information from PEAKS

##### Scan Identifier

F3:9792

##### Original sequence

L

S

L

T

P

E

Q

W

##### Posttranslational Modifications

##### Source File

D:\separate\_stitch\_analyses\xle-disambiguation\raw\20210323\_F1\_UM1\_Peng0013\_SA\_F59\_ingel\_3ug\_chymo.raw

##### Fraction

3

##### Scan Feature

-

##### De Novo Score

98

##### ConfidenceScore

98

### m/z

487.2539

##### Mass

972.4916

##### Charge

2

##### Retention Time

54.95

##### Predicted Retention Time

-

##### Area

0

##### Parts Per Million

1.7

##### Fragmentation mode

HCD

##### Originating file

01 D:\separate\_stitch\_analyses\xle-disambiguation\20210325\_F59\_3ug\_DENOVO\_12.csv

#### Meta Information from PEAKS

##### Scan Identifier

F3:8241

##### Original sequence

L

S

L

T

P

E

Q

W

+15.99

##### Posttranslational Modifications

Oxidation (HW)

##### Source File

D:\separate\_stitch\_analyses\xle-disambiguation\raw\20210323\_F1\_UM1\_Peng0013\_SA\_F59\_ingel\_3ug\_chymo.raw

##### Fraction

3

##### Scan Feature

F3:3855

##### De Novo Score

98

##### ConfidenceScore

98

### m/z

495.2515

##### Mass

988.4866

##### Charge

2

##### Retention Time

46.14

##### Predicted Retention Time

-

##### Area

2.715E+07

##### Parts Per Million

2

##### Fragmentation mode

ETHCD

##### Originating file

01 D:\separate\_stitch\_analyses\xle-disambiguation\20210325\_F59\_3ug\_DENOVO\_12.csv

#### Meta Information from PEAKS

##### Scan Identifier

F3:10159

##### Original sequence

L

S

L

T

P

E

Q

W

##### Posttranslational Modifications

##### Source File

D:\separate\_stitch\_analyses\xle-disambiguation\raw\20210323\_F1\_UM1\_Peng0013\_SA\_F59\_ingel\_3ug\_chymo.raw

##### Fraction

3

##### Scan Feature

-

##### De Novo Score

98

##### ConfidenceScore

98

### m/z

487.2538

##### Mass

972.4916

##### Charge

2

##### Retention Time

57.17

##### Predicted Retention Time

-

##### Area

0

##### Parts Per Million

1.5

##### Fragmentation mode

HCD

##### Originating file

01 D:\separate\_stitch\_analyses\xle-disambiguation\20210325\_F59\_3ug\_DENOVO\_12.csv

#### Meta Information from PEAKS

##### Scan Identifier

F3:9852

##### Original sequence

L

S

L

T

P

E

Q

W

##### Posttranslational Modifications

##### Source File

D:\separate\_stitch\_analyses\xle-disambiguation\raw\20210323\_F1\_UM1\_Peng0013\_SA\_F59\_ingel\_3ug\_chymo.raw

##### Fraction

3

##### Scan Feature

-

##### De Novo Score

97

##### ConfidenceScore

97

### m/z

487.2539

##### Mass

972.4916

##### Charge

2

##### Retention Time

55.31

##### Predicted Retention Time

-

##### Area

0

##### Parts Per Million

1.7

##### Fragmentation mode

HCD

##### Originating file

01 D:\separate\_stitch\_analyses\xle-disambiguation\20210325\_F59\_3ug\_DENOVO\_12.csv

#### Meta Information from PEAKS

##### Scan Identifier

F3:9975

##### Original sequence

L

S

L

T

P

E

Q

W

##### Posttranslational Modifications

##### Source File

D:\separate\_stitch\_analyses\xle-disambiguation\raw\20210323\_F1\_UM1\_Peng0013\_SA\_F59\_ingel\_3ug\_chymo.raw

##### Fraction

3

##### Scan Feature

-

##### De Novo Score

97

##### ConfidenceScore

97

### m/z

487.254

##### Mass

972.4916

##### Charge

2

##### Retention Time

56.05

##### Predicted Retention Time

-

##### Area

0

##### Parts Per Million

1.9

##### Fragmentation mode

HCD

##### Originating file

01 D:\separate\_stitch\_analyses\xle-disambiguation\20210325\_F59\_3ug\_DENOVO\_12.csv

#### Meta Information from PEAKS

##### Scan Identifier

F3:7901

##### Original sequence

L

S

L

T

P

E

Q

W

+15.99

##### Posttranslational Modifications

Oxidation (HW)

##### Source File

D:\separate\_stitch\_analyses\xle-disambiguation\raw\20210323\_F1\_UM1\_Peng0013\_SA\_F59\_ingel\_3ug\_chymo.raw

##### Fraction

3

##### Scan Feature

F3:3855

##### De Novo Score

97

##### ConfidenceScore

97

### m/z

495.2515

##### Mass

988.4866

##### Charge

2

##### Retention Time

46.14

##### Predicted Retention Time

-

##### Area

2.715E+07

##### Parts Per Million

2

##### Fragmentation mode

ETHCD

##### Originating file

01 D:\separate\_stitch\_analyses\xle-disambiguation\20210325\_F59\_3ug\_DENOVO\_12.csv

#### Meta Information from PEAKS

##### Scan Identifier

F3:9915

##### Original sequence

L

S

L

T

P

E

Q

W

##### Posttranslational Modifications

##### Source File

D:\separate\_stitch\_analyses\xle-disambiguation\raw\20210323\_F1\_UM1\_Peng0013\_SA\_F59\_ingel\_3ug\_chymo.raw

##### Fraction

3

##### Scan Feature

-

##### De Novo Score

97

##### ConfidenceScore

97

### m/z

487.254

##### Mass

972.4916

##### Charge

2

##### Retention Time

55.68

##### Predicted Retention Time

-

##### Area

0

##### Parts Per Million

1.9

##### Fragmentation mode

HCD

##### Originating file

01 D:\separate\_stitch\_analyses\xle-disambiguation\20210325\_F59\_3ug\_DENOVO\_12.csv

#### Meta Information from PEAKS

##### Scan Identifier

F3:10510

##### Original sequence

L

S

L

T

P

E

Q

W

##### Posttranslational Modifications

##### Source File

D:\separate\_stitch\_analyses\xle-disambiguation\raw\20210323\_F1\_UM1\_Peng0013\_SA\_F59\_ingel\_3ug\_chymo.raw

##### Fraction

3

##### Scan Feature

-

##### De Novo Score

97

##### ConfidenceScore

97

### m/z

487.2539

##### Mass

972.4916

##### Charge

2

##### Retention Time

59.34

##### Predicted Retention Time

-

##### Area

0

##### Parts Per Million

1.7

##### Fragmentation mode

ETHCD

##### Originating file

01 D:\separate\_stitch\_analyses\xle-disambiguation\20210325\_F59\_3ug\_DENOVO\_12.csv

#### Meta Information from PEAKS

##### Scan Identifier

F3:7735

##### Original sequence

L

S

L

T

P

E

Q

W

+15.99

##### Posttranslational Modifications

Oxidation (HW)

##### Source File

D:\separate\_stitch\_analyses\xle-disambiguation\raw\20210323\_F1\_UM1\_Peng0013\_SA\_F59\_ingel\_3ug\_chymo.raw

##### Fraction

3

##### Scan Feature

F3:3855

##### De Novo Score

97

##### ConfidenceScore

97

### m/z

495.2515

##### Mass

988.4866

##### Charge

2

##### Retention Time

46.14

##### Predicted Retention Time

-

##### Area

2.715E+07

##### Parts Per Million

2

##### Fragmentation mode

ETHCD

##### Originating file

01 D:\separate\_stitch\_analyses\xle-disambiguation\20210325\_F59\_3ug\_DENOVO\_12.csv

#### Meta Information from PEAKS

##### Scan Identifier

F3:10395

##### Original sequence

L

S

L

T

P

E

Q

W

##### Posttranslational Modifications

##### Source File

D:\separate\_stitch\_analyses\xle-disambiguation\raw\20210323\_F1\_UM1\_Peng0013\_SA\_F59\_ingel\_3ug\_chymo.raw

##### Fraction

3

##### Scan Feature

-

##### De Novo Score

96

##### ConfidenceScore

96

### m/z

487.2539

##### Mass

972.4916

##### Charge

2

##### Retention Time

58.61

##### Predicted Retention Time

-

##### Area

0

##### Parts Per Million

1.6

##### Fragmentation mode

ETHCD

##### Originating file

01 D:\separate\_stitch\_analyses\xle-disambiguation\20210325\_F59\_3ug\_DENOVO\_12.csv

#### Meta Information from PEAKS

##### Scan Identifier

F3:8179

##### Original sequence

L

S

L

T

P

E

Q

W

+15.99

##### Posttranslational Modifications

Oxidation (HW)

##### Source File

D:\separate\_stitch\_analyses\xle-disambiguation\raw\20210323\_F1\_UM1\_Peng0013\_SA\_F59\_ingel\_3ug\_chymo.raw

##### Fraction

3

##### Scan Feature

F3:3855

##### De Novo Score

96

##### ConfidenceScore

96

### m/z

495.2515

##### Mass

988.4866

##### Charge

2

##### Retention Time

46.14

##### Predicted Retention Time

-

##### Area

2.715E+07

##### Parts Per Million

2

##### Fragmentation mode

HCD

##### Originating file

01 D:\separate\_stitch\_analyses\xle-disambiguation\20210325\_F59\_3ug\_DENOVO\_12.csv

#### Meta Information from PEAKS

##### Scan Identifier

F3:10331

##### Original sequence

L

S

L

T

P

E

Q

W

##### Posttranslational Modifications

##### Source File

D:\separate\_stitch\_analyses\xle-disambiguation\raw\20210323\_F1\_UM1\_Peng0013\_SA\_F59\_ingel\_3ug\_chymo.raw

##### Fraction

3

##### Scan Feature

-

##### De Novo Score

95

##### ConfidenceScore

95

### m/z

487.2539

##### Mass

972.4916

##### Charge

2

##### Retention Time

58.24

##### Predicted Retention Time

-

##### Area

0

##### Parts Per Million

1.7

##### Fragmentation mode

ETHCD

##### Originating file

01 D:\separate\_stitch\_analyses\xle-disambiguation\20210325\_F59\_3ug\_DENOVO\_12.csv

#### Meta Information from PEAKS

##### Scan Identifier

F3:7205

##### Original sequence

L

S

L

T

P

E

Q

W

+15.99

##### Posttranslational Modifications

Oxidation (HW)

##### Source File

D:\separate\_stitch\_analyses\xle-disambiguation\raw\20210323\_F1\_UM1\_Peng0013\_SA\_F59\_ingel\_3ug\_chymo.raw

##### Fraction

3

##### Scan Feature

F3:3852

##### De Novo Score

95

##### ConfidenceScore

95

### m/z

495.2509

##### Mass

988.4866

##### Charge

2

##### Retention Time

40.15

##### Predicted Retention Time

-

##### Area

1.623E+06

##### Parts Per Million

0.7

##### Fragmentation mode

ETHCD

##### Originating file

01 D:\separate\_stitch\_analyses\xle-disambiguation\20210325\_F59\_3ug\_DENOVO\_12.csv

#### Meta Information from PEAKS

##### Scan Identifier

F3:7837

##### Original sequence

L

S

L

T

P

E

Q

W

+15.99

##### Posttranslational Modifications

Oxidation (HW)

##### Source File

D:\separate\_stitch\_analyses\xle-disambiguation\raw\20210323\_F1\_UM1\_Peng0013\_SA\_F59\_ingel\_3ug\_chymo.raw

##### Fraction

3

##### Scan Feature

F3:3855

##### De Novo Score

95

##### ConfidenceScore

95

### m/z

495.2515

##### Mass

988.4866

##### Charge

2

##### Retention Time

46.14

##### Predicted Retention Time

-

##### Area

2.715E+07

##### Parts Per Million

2

##### Fragmentation mode

HCD

##### Originating file

01 D:\separate\_stitch\_analyses\xle-disambiguation\20210325\_F59\_3ug\_DENOVO\_12.csv
