## Supplementary material for "A handle on mass coincidence errors in *de novo* sequencing of antibodies by bottom-up proteomics": Combined_060.html

Details Combined\_060 | Stitch OverviewUndefined

### Read Combined\_060

#### Sequence (length=7)

FDAVGVK

#### Spectrum 3941? Spectrum 3941 The raw spectrum of this peptide as annotated by Hecklib. The fragments are coloured according to ion type (see legend). Any peaks with a star '\*' as text can be hovered over to see the full details, first the ion type second the mass shift type. By hovering over the amino acids in the peptide or ions in the legend the corresponding peaks are highlighted. By toggling the 'Unassigned' label you can turn the background (unassigned) peaks on or off in the plot. By updating the slider in the Ion legend you can update the spectrum to only show the top X% of the peaks with labels. The top X% means any peak that is within X% of the highest intensity. By dragging in the spectrum you can zoom in to a specific part of the spectrum and use 'Zoom Out' to get back to the original zoom level. The annotation of the spectrum is based on the given sequence in the peptides file and is done with different software so inconsistencies are likely. The peaks are annotated based on the given sequence, with 20 ppm tolerance.

Copy Data

##### Spectrum 3941 (TSV)

###### Preview

```
Loading example...
```

*Click on the button to copy the data to your clipboard.*

Mz MinMz MaxIntensity Max

WidthHeightPeptide font sizePeptide stroke widthSpectrum font sizeSpectrum stroke widthCompact peptide

Ion legend

wxyz

abcd

OtherUnassignedIon^Charge^_Position_Show for top:%

FDAVGVK

03.98e+47.95e+41.19e+51.59e+5

Zoom Out

y^+1^_1_z^+1^_1_y^+1^_1_w^+1^_2_y^+1^_2_z^+1^_2_y^+1^_2_z^+1^_3_y^+1^_3_w^+1^_4_y^+1^_4_z^+1^_4_y^+1^_4_c^+1^_4_z^+1^_5_y^+1^_5_c^+1^_5_w^+1^_6_z^+1^_6_y^+1^_6_z^+1^_6_y^+1^_6_c^+1^_6_

034769310401387

Fragment Matches Table

Show background peaks

| Position | Ion type | Intensity | mz Theoretical | mz Error (Th) | mz Error (ppm) | Charge | Series Number |
| --- | --- | --- | --- | --- | --- | --- | --- |
| - | - | 7.084E+04 | 120.1 | - | - | 0 | - |
| - | - | 5215 | 121.1 | - | - | 0 | - |
| - | - | 8039 | 129.1 | - | - | 0 | - |
| 7 | y | 1.066E+04 | 130.1 | 0.0002928 | 2.251 | +1 | 1 |
| - | - | 505.4 | 130.1 | - | - | 0 | - |
| 7 | z | 5298 | 131.1 | 0.0003108 | 2.371 | +1 | 1 |
| - | - | 381.7 | 131.8 | - | - | 0 | - |
| - | - | 393.3 | 132.1 | - | - | 0 | - |
| - | - | 410.3 | 133.1 | - | - | 0 | - |
| - | - | 362.5 | 135.9 | - | - | 0 | - |
| - | - | 446.1 | 141.6 | - | - | 0 | - |
| - | - | 417.7 | 142.1 | - | - | 0 | - |
| - | - | 2797 | 142.1 | - | - | 0 | - |
| - | - | 1071 | 143.1 | - | - | 0 | - |
| 7 | y | 2.503E+04 | 147.1 | 0.000294 | 1.998 | +1 | 1 |
| - | - | 1501 | 148.1 | - | - | 0 | - |
| - | - | 5125 | 149 | - | - | 0 | - |
| - | - | 400.8 | 150.7 | - | - | 0 | - |
| - | - | 407.6 | 153 | - | - | 0 | - |
| - | - | 518.3 | 156.1 | - | - | 0 | - |
| - | - | 935.8 | 157.1 | - | - | 0 | - |
| - | - | 402 | 160 | - | - | 0 | - |
| - | - | 417.4 | 161.4 | - | - | 0 | - |
| - | - | 2326 | 167.1 | - | - | 0 | - |
| - | - | 1419 | 171.1 | - | - | 0 | - |
| - | - | 1007 | 173.5 | - | - | 0 | - |
| - | - | 464.2 | 174.1 | - | - | 0 | - |
| - | - | 882.3 | 174.1 | - | - | 0 | - |
| - | - | 512 | 175 | - | - | 0 | - |
| - | - | 573.3 | 175.1 | - | - | 0 | - |
| - | - | 658 | 187.1 | - | - | 0 | - |
| - | - | 2657 | 191.1 | - | - | 0 | - |
| - | - | 497.8 | 193.7 | - | - | 0 | - |
| - | - | 771.7 | 199.2 | - | - | 0 | - |
| - | - | 650.1 | 212.1 | - | - | 0 | - |
| - | - | 614 | 213.2 | - | - | 0 | - |
| - | - | 521.2 | 214 | - | - | 0 | - |
| 6 | w | 1121 | 215.1 | 0.0004922 | 2.288 | +1 | 2 |
| - | - | 662.6 | 216.1 | - | - | 0 | - |
| - | - | 1.007E+04 | 217.1 | - | - | 0 | - |
| - | - | 1035 | 218.1 | - | - | 0 | - |
| - | - | 538.1 | 219.2 | - | - | 0 | - |
| - | - | 3253 | 228.1 | - | - | 0 | - |
| - | - | 818.9 | 228.2 | - | - | 0 | - |
| 6 | y | 888 | 229.2 | 0.0003603 | 1.572 | +1 | 2 |
| - | - | 511.2 | 230 | - | - | 0 | - |
| 6 | z | 3197 | 230.2 | 0.0002105 | 0.9144 | +1 | 2 |
| - | - | 845.8 | 231.1 | - | - | 0 | - |
| - | - | 1448 | 231.2 | - | - | 0 | - |
| - | - | 1.724E+04 | 235.1 | - | - | 0 | - |
| - | - | 1707 | 236.1 | - | - | 0 | - |
| - | - | 1021 | 245.2 | - | - | 0 | - |
| - | - | 5225 | 245.2 | - | - | 0 | - |
| 6 | y | 8652 | 246.2 | 0.0001632 | 0.6627 | +1 | 2 |
| - | - | 753.2 | 247.2 | - | - | 0 | - |
| - | - | 914.2 | 253.2 | - | - | 0 | - |
| - | - | 597.3 | 255.1 | - | - | 0 | - |
| - | - | 3690 | 262.1 | - | - | 0 | - |
| - | - | 974.5 | 263.1 | - | - | 0 | - |
| - | - | 1.941E+04 | 263.1 | - | - | 0 | - |
| - | - | 2085 | 264.1 | - | - | 0 | - |
| - | - | 760.6 | 270.2 | - | - | 0 | - |
| - | - | 1605 | 273.2 | - | - | 0 | - |
| - | - | 1548 | 273.2 | - | - | 0 | - |
| - | - | 1173 | 281.1 | - | - | 0 | - |
| - | - | 736.3 | 283.2 | - | - | 0 | - |
| - | - | 549.4 | 283.2 | - | - | 0 | - |
| - | - | 1066 | 284.2 | - | - | 0 | - |
| - | - | 2889 | 284.2 | - | - | 0 | - |
| - | - | 713.9 | 285.2 | - | - | 0 | - |
| - | - | 2010 | 286.1 | - | - | 0 | - |
| 5 | z | 3.611E+04 | 287.2 | 0.0003379 | 1.177 | +1 | 3 |
| - | - | 8.927E+04 | 288.2 | - | - | 0 | - |
| - | - | 9158 | 289.2 | - | - | 0 | - |
| - | - | 1197 | 290.2 | - | - | 0 | - |
| - | - | 510.1 | 298.3 | - | - | 0 | - |
| - | - | 5193 | 299.1 | - | - | 0 | - |
| - | - | 4688 | 299.2 | - | - | 0 | - |
| - | - | 1.491E+04 | 302.2 | - | - | 0 | - |
| 5 | y | 1.96E+04 | 303.2 | 1.456E-05 | 0.04802 | +1 | 3 |
| - | - | 3183 | 304.2 | - | - | 0 | - |
| - | - | 562.4 | 305.2 | - | - | 0 | - |
| - | - | 1598 | 311.2 | - | - | 0 | - |
| - | - | 1643 | 312.2 | - | - | 0 | - |
| - | - | 531.8 | 323.2 | - | - | 0 | - |
| - | - | 532.3 | 325.2 | - | - | 0 | - |
| - | - | 4251 | 327.1 | - | - | 0 | - |
| - | - | 3944 | 327.2 | - | - | 0 | - |
| - | - | 6788 | 330.2 | - | - | 0 | - |
| - | - | 2146 | 331.2 | - | - | 0 | - |
| - | - | 1003 | 333.1 | - | - | 0 | - |
| - | - | 1.072E+04 | 334.1 | - | - | 0 | - |
| - | - | 1588 | 335.1 | - | - | 0 | - |
| - | - | 7471 | 343.2 | - | - | 0 | - |
| - | - | 1528 | 344.2 | - | - | 0 | - |
| - | - | 812.1 | 344.2 | - | - | 0 | - |
| - | - | 1122 | 354.2 | - | - | 0 | - |
| - | - | 2707 | 355.2 | - | - | 0 | - |
| - | - | 1203 | 359.2 | - | - | 0 | - |
| - | - | 983.1 | 367.1 | - | - | 0 | - |
| - | - | 769.8 | 367.2 | - | - | 0 | - |
| - | - | 3904 | 368.2 | - | - | 0 | - |
| - | - | 1782 | 368.7 | - | - | 0 | - |
| - | - | 3018 | 369.1 | - | - | 0 | - |
| - | - | 2212 | 369.3 | - | - | 0 | - |
| 4 | w | 2.652E+04 | 371.2 | 7.684E-05 | 0.207 | +1 | 4 |
| - | - | 5624 | 372.2 | - | - | 0 | - |
| - | - | 1295 | 373.2 | - | - | 0 | - |
| 4 | y | 1956 | 385.2 | 3.978E-05 | 0.1032 | +1 | 4 |
| - | - | 1503 | 386.2 | - | - | 0 | - |
| 4 | z | 3092 | 386.3 | 0.000388 | 1.005 | +1 | 4 |
| - | - | 1980 | 387.2 | - | - | 0 | - |
| - | - | 1110 | 391.2 | - | - | 0 | - |
| - | - | 737.2 | 398.2 | - | - | 0 | - |
| - | - | 2517 | 399.2 | - | - | 0 | - |
| - | - | 4254 | 400.3 | - | - | 0 | - |
| - | - | 830.5 | 401.3 | - | - | 0 | - |
| 4 | y | 1.089E+04 | 402.3 | 0.0002971 | 0.7386 | +1 | 4 |
| - | - | 2007 | 403.3 | - | - | 0 | - |
| - | - | 654.6 | 404.3 | - | - | 0 | - |
| - | - | 6868 | 405.2 | - | - | 0 | - |
| - | - | 1615 | 406.2 | - | - | 0 | - |
| - | - | 1061 | 414.2 | - | - | 0 | - |
| - | - | 1071 | 414.3 | - | - | 0 | - |
| - | - | 1001 | 415.2 | - | - | 0 | - |
| - | - | 5714 | 415.2 | - | - | 0 | - |
| - | - | 1008 | 416.2 | - | - | 0 | - |
| - | - | 1092 | 419.2 | - | - | 0 | - |
| - | - | 632.3 | 427.8 | - | - | 0 | - |
| - | - | 2267 | 432.2 | - | - | 0 | - |
| - | - | 2.333E+04 | 433.2 | - | - | 0 | - |
| - | - | 5360 | 434.2 | - | - | 0 | - |
| - | - | 932.8 | 435.2 | - | - | 0 | - |
| - | - | 1386 | 438.3 | - | - | 0 | - |
| - | - | 6960 | 439.3 | - | - | 0 | - |
| - | - | 1335 | 440.3 | - | - | 0 | - |
| - | - | 3992 | 442.2 | - | - | 0 | - |
| - | - | 1.192E+04 | 442.3 | - | - | 0 | - |
| - | - | 1038 | 443.2 | - | - | 0 | - |
| - | - | 2672 | 443.3 | - | - | 0 | - |
| - | - | 544.8 | 444.2 | - | - | 0 | - |
| - | - | 547.9 | 444.3 | - | - | 0 | - |
| - | - | 2753 | 449.2 | - | - | 0 | - |
| 4 | c | 552.1 | 450.2 | 0.00299 | 6.642 | +1 | 4 |
| 3 | z | 4.273E+04 | 457.3 | 0.000401 | 0.877 | +1 | 5 |
| - | - | 8744 | 458.3 | - | - | 0 | - |
| - | - | 1986 | 459.3 | - | - | 0 | - |
| - | - | 973.2 | 462.2 | - | - | 0 | - |
| - | - | 8919 | 463.2 | - | - | 0 | - |
| - | - | 3670 | 464.2 | - | - | 0 | - |
| - | - | 1326 | 471.3 | - | - | 0 | - |
| - | - | 704.3 | 472.2 | - | - | 0 | - |
| - | - | 809.8 | 472.3 | - | - | 0 | - |
| 3 | y | 3.897E+04 | 473.3 | 0.0005063 | 1.07 | +1 | 5 |
| - | - | 9556 | 474.3 | - | - | 0 | - |
| - | - | 1320 | 475.3 | - | - | 0 | - |
| - | - | 1138 | 485.3 | - | - | 0 | - |
| - | - | 1235 | 486.3 | - | - | 0 | - |
| - | - | 1252 | 489.2 | - | - | 0 | - |
| - | - | 1.029E+04 | 490.2 | - | - | 0 | - |
| - | - | 2303 | 491.2 | - | - | 0 | - |
| - | - | 2603 | 500.3 | - | - | 0 | - |
| - | - | 849.8 | 501.2 | - | - | 0 | - |
| - | - | 1.357E+04 | 506.2 | - | - | 0 | - |
| 5 | c | 2.792E+04 | 507.3 | 0.0003146 | 0.6201 | +1 | 5 |
| - | - | 6994 | 508.3 | - | - | 0 | - |
| - | - | 1541 | 509.3 | - | - | 0 | - |
| 2 | w | 915.6 | 527.3 | 0.002246 | 4.259 | +1 | 6 |
| - | - | 2.757E+04 | 528.3 | - | - | 0 | - |
| - | - | 6110 | 529.3 | - | - | 0 | - |
| - | - | 997 | 530.3 | - | - | 0 | - |
| - | - | 523.8 | 535.3 | - | - | 0 | - |
| - | - | 882.8 | 544.3 | - | - | 0 | - |
| - | - | 898.8 | 545.3 | - | - | 0 | - |
| - | - | 1766 | 546.3 | - | - | 0 | - |
| - | - | 1443 | 553.8 | - | - | 0 | - |
| 2 | z | 685.1 | 554.3 | 0.003279 | 5.916 | +1 | 6 |
| - | - | 6141 | 561.3 | - | - | 0 | - |
| - | - | 4725 | 562.3 | - | - | 0 | - |
| - | - | 3304 | 563.3 | - | - | 0 | - |
| - | - | 667.2 | 565.4 | - | - | 0 | - |
| 2 | y | 2985 | 570.3 | 0.0007906 | 1.386 | +1 | 6 |
| - | - | 5322 | 571.3 | - | - | 0 | - |
| 2 | z | 2.397E+04 | 572.3 | 2.221E-05 | 0.03882 | +1 | 6 |
| - | - | 6455 | 573.3 | - | - | 0 | - |
| - | - | 1181 | 574.3 | - | - | 0 | - |
| 2 | y | 5.21E+04 | 588.3 | 0.0001306 | 0.2219 | +1 | 6 |
| - | - | 6.258E+04 | 589.3 | - | - | 0 | - |
| - | - | 1.011E+04 | 589.3 | - | - | 0 | - |
| - | - | 1.625E+04 | 590.3 | - | - | 0 | - |
| - | - | 1367 | 590.3 | - | - | 0 | - |
| - | - | 4326 | 591.3 | - | - | 0 | - |
| - | - | 685.8 | 598.3 | - | - | 0 | - |
| - | - | 1178 | 605.3 | - | - | 0 | - |
| 6 | c | 7.818E+04 | 606.3 | 0.0003023 | 0.4986 | +1 | 6 |
| - | - | 2.393E+04 | 607.3 | - | - | 0 | - |
| - | - | 5465 | 608.3 | - | - | 0 | - |
| - | - | 835.5 | 644.3 | - | - | 0 | - |
| - | - | 1167 | 658.4 | - | - | 0 | - |
| - | - | 710.1 | 674.3 | - | - | 0 | - |
| - | - | 838.8 | 675.3 | - | - | 0 | - |
| - | - | 1.142E+04 | 675.4 | - | - | 0 | - |
| - | - | 1.03E+04 | 676.4 | - | - | 0 | - |
| - | - | 896.9 | 677.3 | - | - | 0 | - |
| - | - | 4756 | 677.4 | - | - | 0 | - |
| - | - | 768.9 | 678.4 | - | - | 0 | - |
| - | - | 1072 | 690.4 | - | - | 0 | - |
| - | - | 641.9 | 691.4 | - | - | 0 | - |
| - | - | 935.8 | 697.4 | - | - | 0 | - |
| - | - | 1804 | 708.4 | - | - | 0 | - |
| - | - | 590.8 | 709.4 | - | - | 0 | - |
| - | - | 1.575E+05 | 719.4 | - | - | 0 | - |
| - | - | 5.677E+04 | 720.4 | - | - | 0 | - |
| - | - | 1.608E+04 | 721.4 | - | - | 0 | - |
| - | - | 881.2 | 733.4 | - | - | 0 | - |
| - | - | 2441 | 734.4 | - | - | 0 | - |
| - | - | 7.471E+04 | 735.4 | - | - | 0 | - |
| - | - | 4.081E+04 | 736.4 | - | - | 0 | - |
| - | - | 882.4 | 737.3 | - | - | 0 | - |
| - | - | 1.053E+04 | 737.4 | - | - | 0 | - |
| - | - | 712 | 738.4 | - | - | 0 | - |
| - | - | 614 | 787.6 | - | - | 0 | - |
| - | - | 729.1 | 869.5 | - | - | 0 | - |
| - | - | 575.4 | 902 | - | - | 0 | - |
| - | - | 649.2 | 929.5 | - | - | 0 | - |
| - | - | 1037 | 1028 | - | - | 0 | - |
| - | - | 894.1 | 1029 | - | - | 0 | - |
| - | - | 640.4 | 1059 | - | - | 0 | - |
| - | - | 2432 | 1069 | - | - | 0 | - |
| - | - | 1803 | 1070 | - | - | 0 | - |
| - | - | 1093 | 1071 | - | - | 0 | - |
| - | - | 816 | 1086 | - | - | 0 | - |
| - | - | 821.8 | 1087 | - | - | 0 | - |
| - | - | 629.1 | 1091 | - | - | 0 | - |
| - | - | 973.9 | 1107 | - | - | 0 | - |
| - | - | 3166 | 1108 | - | - | 0 | - |
| - | - | 928.6 | 1109 | - | - | 0 | - |
| - | - | 788.9 | 1373 | - | - | 0 | - |

m/z Charge Intensity FragmentType MassShift Position
120.08108520507812 0 70835.57
121.08440399169922 0 5214.888
129.10250854492188 0 8038.6616
130.0865478515625 0 10663.464 y Ammonia loss 6
130.10565185546875 0 505.40698
131.09439086914062 0 5298.011 z 6
131.7864532470703 0 381.6503
132.09719848632812 0 393.28192
133.08665466308594 0 410.2739
135.85452270507812 0 362.51584
141.6382598876953 0 446.13953
142.08114624023438 0 417.65155
142.0865936279297 0 2796.6997
143.11827087402344 0 1071.2047
147.11309814453125 0 25026.174 y 6
148.1165008544922 0 1501.081
149.0452117919922 0 5125.161
150.7186737060547 0 400.76724
153.01258850097656 0 407.63272
156.08970642089844 0 518.32776
157.097412109375 0 935.8161
159.99253845214844 0 401.97748
161.37350463867188 0 417.37073
167.0557403564453 0 2325.6958
171.11309814453125 0 1419.076
173.4517364501953 0 1007.15955
174.1009979248047 0 464.24203
174.12413024902344 0 882.3022
175.02975463867188 0 512.012
175.08694458007812 0 573.25323
187.07167053222656 0 658.0459
191.1181182861328 0 2657.079
193.72264099121094 0 497.80438
199.16957092285156 0 771.7217
212.13973999023438 0 650.0859
213.16064453125 0 614.03955
214.0404510498047 0 521.1675
215.13951110839844 0 1120.5847 w 5
216.0572967529297 0 662.59644
217.09744262695312 0 10072.928
218.10008239746094 0 1035.1946
219.23727416992188 0 538.13727
228.13450622558594 0 3253.4702
228.17103576660156 0 818.9094
229.155029296875 0 887.9705 y Ammonia loss 5
230.03436279296875 0 511.16486
230.16270446777344 0 3197.4878 z 5
231.12181091308594 0 845.76324
231.1700439453125 0 1448.1299
235.10806274414062 0 17237.281
236.11106872558594 0 1706.719
245.16091918945312 0 1020.76666
245.1737518310547 0 5224.703
246.18138122558594 0 8651.9375 y 5
247.18505859375 0 753.21985
253.16598510742188 0 914.1864
255.0656280517578 0 597.32214
262.094970703125 0 3690.142
263.08685302734375 0 974.53217
263.102783203125 0 19410.082
264.10614013671875 0 2084.8457
270.1809387207031 0 760.5746
273.1681823730469 0 1605.0215
273.1919250488281 0 1548.0385
281.0516357421875 0 1173.2861
283.1893005371094 0 736.2953
283.23443603515625 0 549.4048
284.1606750488281 0 1065.901
284.197021484375 0 2888.6067
285.2042541503906 0 713.86426
286.14019775390625 0 2010.4519
287.1842956542969 0 36108.574 z 4
288.1919860839844 0 89268.85
289.1951599121094 0 9157.883
290.1973876953125 0 1196.7488
298.2909851074219 0 510.07303
299.06201171875 0 5193.499
299.20806884765625 0 4687.7417
302.1952819824219 0 14913.388
303.2026672363281 0 19598.629 y 4
304.20623779296875 0 3183.3047
305.20623779296875 0 562.3551
311.1846008300781 0 1598.487
312.191650390625 0 1643.1901
323.1687316894531 0 531.81683
325.150634765625 0 532.27155
327.0783996582031 0 4251.158
327.20306396484375 0 3943.5242
330.1902160644531 0 6787.835
331.19647216796875 0 2146.1472
333.1322326660156 0 1003.0933
334.1400146484375 0 10718.639
335.1439514160156 0 1588.0273
343.16143798828125 0 7471.344
344.166259765625 0 1528.057
344.2296142578125 0 812.05035
354.2276916503906 0 1122.2396
355.23406982421875 0 2707.342
359.20166015625 0 1202.881
367.07086181640625 0 983.1015
367.2433776855469 0 769.828
368.20513916015625 0 3903.729
368.7066650390625 0 1781.6317
369.12451171875 0 3017.534
369.2613220214844 0 2211.9614
371.2289733886719 0 26520.395 w 3
372.2320251464844 0 5623.9473
373.2346496582031 0 1295.4696
385.2445068359375 0 1956.3767 y Ammonia loss 3
386.216552734375 0 1502.6698
386.2519836425781 0 3091.9202 z 3
387.203369140625 0 1980.2653
391.1617126464844 0 1109.7908
398.2163391113281 0 737.20105
399.22357177734375 0 2516.6543
400.2556457519531 0 4253.6772
401.2598571777344 0 830.52606
402.2713928222656 0 10892.776 y 3
403.27496337890625 0 2006.6771
404.2765197753906 0 654.6022
405.21356201171875 0 6868.2085
406.2172546386719 0 1615.128
414.2347106933594 0 1060.6204
414.28302001953125 0 1071.205
415.1973571777344 0 1000.99976
415.24298095703125 0 5714.038
416.244873046875 0 1007.89185
419.1933288574219 0 1092.3771
427.7994384765625 0 632.26373
432.2005310058594 0 2267.263
433.2086181640625 0 23333.145
434.2116394042969 0 5360.2163
435.215576171875 0 932.83496
438.2715759277344 0 1385.5127
439.2790222167969 0 6959.816
440.2828369140625 0 1334.9114
442.2302551269531 0 3992.0298
442.26666259765625 0 11916.466
443.2340087890625 0 1037.9781
443.2693786621094 0 2671.7837
444.2301025390625 0 544.80884
444.2718200683594 0 547.9404
449.2279052734375 0 2753.1394
450.2317199707031 0 552.1385 c 3
457.2898864746094 0 42725.03 z 2
458.2930908203125 0 8743.919
459.2960510253906 0 1985.6145
462.23406982421875 0 973.1846
463.24298095703125 0 8918.883
464.2485656738281 0 3669.8987
471.2924499511719 0 1326.3475
472.21954345703125 0 704.2853
472.29766845703125 0 809.77637
473.3087158203125 0 38969.06 y 2
474.3112487792969 0 9555.672
475.3146667480469 0 1319.9277
485.3214111328125 0 1138.323
486.2811279296875 0 1235.0848
489.22149658203125 0 1252.1635
490.2301025390625 0 10285.368
491.2341003417969 0 2302.6133
500.2500305175781 0 2602.7573
501.24993896484375 0 849.8312
506.24847412109375 0 13568.555
507.255859375 0 27919.154 c 4
508.2586669921875 0 6994.1924
509.2617492675781 0 1541.1736
527.3165283203125 0 915.6212 w 1
528.3268432617188 0 27572.268
529.3282470703125 0 6109.911
530.3336181640625 0 996.962
535.3063354492188 0 523.80304
544.2770385742188 0 882.8091
545.28173828125 0 898.80804
546.2915649414062 0 1766.4745
553.8074340820312 0 1443.46
554.3091430664062 0 685.1458 z Water loss 1
561.3036499023438 0 6141.189
562.3097534179688 0 4724.645
563.3169555664062 0 3304.3113
565.3795776367188 0 667.20593
570.3253784179688 0 2984.5608 y Water loss 1
571.28857421875 0 5321.631
572.31640625 0 23968.053 z 1
573.3195190429688 0 6455.285
574.3221435546875 0 1180.5018
588.3350219726562 0 52096.406 y 1
589.2977905273438 0 62577.855
589.3397216796875 0 10110.147
590.3006591796875 0 16250.781
590.3368530273438 0 1366.7478
591.3037109375 0 4326.013
598.3206176757812 0 685.7791
605.3150634765625 0 1177.5021
606.3248901367188 0 78177.96 c 5
607.3279418945312 0 23927.996
608.3306274414062 0 5465.3423
644.3486938476562 0 835.5377
658.3648681640625 0 1166.7639
674.3271484375 0 710.1012
675.3363037109375 0 838.78674
675.3950805664062 0 11424.253
676.3919677734375 0 10300.818
677.339111328125 0 896.9235
677.3951416015625 0 4756.3496
678.3966674804688 0 768.9332
690.4070434570312 0 1071.8118
691.3987426757812 0 641.9109
697.4404296875 0 935.8098
708.4158325195312 0 1804.3322
709.4183349609375 0 590.812
719.3846435546875 0 157470.05
720.3875732421875 0 56774.008
721.3900756835938 0 16076.837
733.3601684570312 0 881.2016
734.3904418945312 0 2441.3318
735.4033203125 0 74710.91
736.4073486328125 0 40809.215
737.3384399414062 0 882.41406
737.4105224609375 0 10528.165
738.4108276367188 0 712.02875
787.6439819335938 0 614.04004
869.5498046875 0 729.06464
901.95263671875 0 575.40106
929.4888305664062 0 649.1972
1027.586669921875 0 1036.7751
1028.5828857421875 0 894.1213
1058.5908203125 0 640.43176
1068.6064453125 0 2432.316
1069.600341796875 0 1802.5198
1070.5965576171875 0 1093.3741
1085.6087646484375 0 815.959
1086.6092529296875 0 821.7691
1090.5765380859375 0 629.05585
1106.5948486328125 0 973.91003
1107.61474609375 0 3166.0537
1108.60986328125 0 928.6264
1372.9390869140625 0 788.8624

Spectrum Details

|  |  |
| --- | --- |
| Matched peaks? Matched peaks The total absolute number of peaks matched. Additionally in brackets the total fraction of peaks matched and the total number of peaks is shown. | 23 (9.66% of 238) |
| FDR? FDR The false discovery rate estimated for this peptide. It is calculated by matching all theoretical fragments with a non-integer shift with the raw peaks for this spectrum. This is done with 40 different shifts. The resulting percentage is the average number of annotated peaks over the number of annotated peaks with the correct spectrum. | 0.72% |
| Satellite FDR? Satellite FDR See the FDR for details on its calculation. This satellite ion specific FDR only contains the satellite ions (d/w) for I/L/J positions. | - |
| PSM Score? PSM Score The PSM Score as given by Hecklib to this annotated spectrum. It is shown with three significant figures. | 302 |

#### Spectrum 4107? Spectrum 4107 The raw spectrum of this peptide as annotated by Hecklib. The fragments are coloured according to ion type (see legend). Any peaks with a star '\*' as text can be hovered over to see the full details, first the ion type second the mass shift type. By hovering over the amino acids in the peptide or ions in the legend the corresponding peaks are highlighted. By toggling the 'Unassigned' label you can turn the background (unassigned) peaks on or off in the plot. By updating the slider in the Ion legend you can update the spectrum to only show the top X% of the peaks with labels. The top X% means any peak that is within X% of the highest intensity. By dragging in the spectrum you can zoom in to a specific part of the spectrum and use 'Zoom Out' to get back to the original zoom level. The annotation of the spectrum is based on the given sequence in the peptides file and is done with different software so inconsistencies are likely. The peaks are annotated based on the given sequence, with 20 ppm tolerance.

Copy Data

##### Spectrum 4107 (TSV)

###### Preview

```
Loading example...
```

*Click on the button to copy the data to your clipboard.*

Mz MinMz MaxIntensity Max

WidthHeightPeptide font sizePeptide stroke widthSpectrum font sizeSpectrum stroke widthCompact peptide

Ion legend

wxyz

abcd

OtherUnassignedIon^Charge^_Position_Show for top:%

FDAVGVK

02.97e+45.94e+48.91e+41.19e+5

Zoom Out

y^+1^_1_y^+1^_1_d^+1^_2_a^+1^_2_a^+1^_2_y^+1^_2_b^+1^_2_a^+1^_3_y^+1^_3_b^+1^_3_\*\*y^+1^_4_y^+1^_5_y^+1^_6_y^+1^_6_y^+1^_6_

0781156223433124

Fragment Matches Table

Show background peaks

| Position | Ion type | Intensity | mz Theoretical | mz Error (Th) | mz Error (ppm) | Charge | Series Number |
| --- | --- | --- | --- | --- | --- | --- | --- |
| - | - | 514.9 | 120 | - | - | 0 | - |
| - | - | 1.176E+05 | 120.1 | - | - | 0 | - |
| - | - | 8895 | 121.1 | - | - | 0 | - |
| - | - | 350.7 | 121.1 | - | - | 0 | - |
| - | - | 452.6 | 122.1 | - | - | 0 | - |
| - | - | 1283 | 123.1 | - | - | 0 | - |
| - | - | 559.7 | 124 | - | - | 0 | - |
| - | - | 377.8 | 126.5 | - | - | 0 | - |
| - | - | 2074 | 128.1 | - | - | 0 | - |
| - | - | 1.971E+04 | 129.1 | - | - | 0 | - |
| - | - | 645.7 | 130.1 | - | - | 0 | - |
| 7 | y | 1.066E+04 | 130.1 | 0.0003386 | 2.603 | +1 | 1 |
| - | - | 1360 | 130.1 | - | - | 0 | - |
| - | - | 548.6 | 132 | - | - | 0 | - |
| - | - | 941 | 136 | - | - | 0 | - |
| - | - | 1206 | 136.1 | - | - | 0 | - |
| - | - | 552 | 137.1 | - | - | 0 | - |
| - | - | 746.6 | 138.1 | - | - | 0 | - |
| - | - | 602.2 | 139.1 | - | - | 0 | - |
| - | - | 836.8 | 140.1 | - | - | 0 | - |
| - | - | 695.4 | 141.1 | - | - | 0 | - |
| - | - | 532.8 | 142.1 | - | - | 0 | - |
| - | - | 621.4 | 143.1 | - | - | 0 | - |
| - | - | 428.1 | 143.1 | - | - | 0 | - |
| - | - | 4218 | 143.1 | - | - | 0 | - |
| - | - | 459.2 | 145 | - | - | 0 | - |
| - | - | 727.8 | 145.1 | - | - | 0 | - |
| - | - | 910.1 | 146.1 | - | - | 0 | - |
| - | - | 445.6 | 147 | - | - | 0 | - |
| 7 | y | 1.349E+04 | 147.1 | 0.0003398 | 2.31 | +1 | 1 |
| - | - | 700.5 | 148.1 | - | - | 0 | - |
| - | - | 476 | 149 | - | - | 0 | - |
| - | - | 1039 | 149 | - | - | 0 | - |
| - | - | 6695 | 149 | - | - | 0 | - |
| - | - | 749.6 | 152.1 | - | - | 0 | - |
| - | - | 559.3 | 153.1 | - | - | 0 | - |
| - | - | 501.8 | 156 | - | - | 0 | - |
| - | - | 749.3 | 157.1 | - | - | 0 | - |
| - | - | 1826 | 157.1 | - | - | 0 | - |
| - | - | 508.6 | 159 | - | - | 0 | - |
| - | - | 5081 | 159.1 | - | - | 0 | - |
| - | - | 825.1 | 161 | - | - | 0 | - |
| - | - | 904.7 | 163 | - | - | 0 | - |
| - | - | 517.1 | 163.1 | - | - | 0 | - |
| - | - | 462.4 | 164.1 | - | - | 0 | - |
| - | - | 461.3 | 165.1 | - | - | 0 | - |
| - | - | 492.5 | 166.1 | - | - | 0 | - |
| - | - | 1.023E+04 | 167.1 | - | - | 0 | - |
| - | - | 415.8 | 167.1 | - | - | 0 | - |
| - | - | 699.6 | 169.1 | - | - | 0 | - |
| - | - | 520.6 | 170 | - | - | 0 | - |
| - | - | 919.1 | 171 | - | - | 0 | - |
| - | - | 4739 | 171.1 | - | - | 0 | - |
| - | - | 443.6 | 172.7 | - | - | 0 | - |
| - | - | 548.9 | 173 | - | - | 0 | - |
| - | - | 448.1 | 173 | - | - | 0 | - |
| - | - | 1215 | 173.5 | - | - | 0 | - |
| - | - | 490 | 174 | - | - | 0 | - |
| - | - | 1970 | 175 | - | - | 0 | - |
| - | - | 1360 | 175.1 | - | - | 0 | - |
| - | - | 514 | 177 | - | - | 0 | - |
| - | - | 472.2 | 180.1 | - | - | 0 | - |
| - | - | 430.7 | 181.3 | - | - | 0 | - |
| - | - | 518.8 | 183.1 | - | - | 0 | - |
| - | - | 490.4 | 185.6 | - | - | 0 | - |
| - | - | 485.1 | 187 | - | - | 0 | - |
| - | - | 5602 | 187.1 | - | - | 0 | - |
| - | - | 468.4 | 187.1 | - | - | 0 | - |
| - | - | 1227 | 187.1 | - | - | 0 | - |
| - | - | 1083 | 191 | - | - | 0 | - |
| 2 | d | 3282 | 191.1 | 0.0003813 | 1.995 | +1 | 2 |
| - | - | 1365 | 197.1 | - | - | 0 | - |
| - | - | 2315 | 199.2 | - | - | 0 | - |
| - | - | 1197 | 202 | - | - | 0 | - |
| - | - | 2018 | 202.1 | - | - | 0 | - |
| - | - | 614.1 | 204 | - | - | 0 | - |
| - | - | 651.1 | 213 | - | - | 0 | - |
| - | - | 712.3 | 214 | - | - | 0 | - |
| - | - | 811 | 215.1 | - | - | 0 | - |
| - | - | 728 | 216.1 | - | - | 0 | - |
| - | - | 976.8 | 217.1 | - | - | 0 | - |
| 2 | a | 1.149E+04 | 217.1 | 0.0002733 | 1.259 | +1 | 2 |
| - | - | 1155 | 218.1 | - | - | 0 | - |
| - | - | 1357 | 221.1 | - | - | 0 | - |
| - | - | 512 | 224.1 | - | - | 0 | - |
| - | - | 881.3 | 225 | - | - | 0 | - |
| - | - | 888.4 | 226.1 | - | - | 0 | - |
| - | - | 3139 | 226.2 | - | - | 0 | - |
| - | - | 5648 | 228.1 | - | - | 0 | - |
| - | - | 826.1 | 232.1 | - | - | 0 | - |
| - | - | 640.8 | 233 | - | - | 0 | - |
| - | - | 2113 | 233.2 | - | - | 0 | - |
| 2 | a | 1.291E+04 | 235.1 | 0.0002829 | 1.203 | +1 | 2 |
| - | - | 1407 | 236.1 | - | - | 0 | - |
| - | - | 882.3 | 240.1 | - | - | 0 | - |
| - | - | 1822 | 241.1 | - | - | 0 | - |
| - | - | 557.6 | 245.9 | - | - | 0 | - |
| 6 | y | 3406 | 246.2 | 0.0002852 | 1.159 | +1 | 2 |
| - | - | 812.3 | 255.1 | - | - | 0 | - |
| - | - | 2422 | 256.1 | - | - | 0 | - |
| 2 | b | 1.176E+04 | 263.1 | 0.0001193 | 0.4534 | +1 | 2 |
| - | - | 1377 | 264.1 | - | - | 0 | - |
| - | - | 1395 | 268.1 | - | - | 0 | - |
| - | - | 843.3 | 271.1 | - | - | 0 | - |
| - | - | 864.6 | 272.1 | - | - | 0 | - |
| - | - | 960.1 | 281.1 | - | - | 0 | - |
| - | - | 1452 | 283 | - | - | 0 | - |
| - | - | 624.2 | 285 | - | - | 0 | - |
| - | - | 2136 | 286.1 | - | - | 0 | - |
| - | - | 886.5 | 287.1 | - | - | 0 | - |
| 3 | a | 634.5 | 288.1 | 0.001444 | 5.011 | +1 | 3 |
| - | - | 9186 | 299.1 | - | - | 0 | - |
| - | - | 3404 | 299.2 | - | - | 0 | - |
| - | - | 628.9 | 300.2 | - | - | 0 | - |
| 5 | y | 1.375E+04 | 303.2 | 0.0003517 | 1.16 | +1 | 3 |
| - | - | 1787 | 304.2 | - | - | 0 | - |
| - | - | 717.5 | 315.2 | - | - | 0 | - |
| - | - | 600.5 | 323.2 | - | - | 0 | - |
| - | - | 531.9 | 323.2 | - | - | 0 | - |
| - | - | 705.6 | 325.1 | - | - | 0 | - |
| - | - | 8967 | 327.1 | - | - | 0 | - |
| - | - | 1182 | 327.2 | - | - | 0 | - |
| 3 | b | 2433 | 334.1 | 9.875E-05 | 0.2955 | +1 | 3 |
| - | - | 8268 | 343.2 | - | - | 0 | - |
| - | - | 1547 | 344.2 | - | - | 0 | - |
| - | - | 505.4 | 355.4 | - | - | 0 | - |
| 0 | Precursor | 748.7 | 359.2 | 0.004247 | 11.82 | +2 | -1 |
| - | - | 607.2 | 367.1 | - | - | 0 | - |
| 0 | Precursor | 1541 | 368.2 | 3.815E-05 | 0.1036 | +2 | -1 |
| - | - | 2658 | 369.1 | - | - | 0 | - |
| - | - | 1150 | 387.2 | - | - | 0 | - |
| - | - | 849.2 | 391.7 | - | - | 0 | - |
| 4 | y | 6883 | 402.3 | 3.858E-05 | 0.09592 | +1 | 4 |
| - | - | 1283 | 403.3 | - | - | 0 | - |
| - | - | 2019 | 403.7 | - | - | 0 | - |
| - | - | 594.5 | 413.3 | - | - | 0 | - |
| - | - | 1619 | 414.2 | - | - | 0 | - |
| - | - | 1037 | 438.3 | - | - | 0 | - |
| - | - | 1494 | 442.2 | - | - | 0 | - |
| - | - | 1981 | 460.2 | - | - | 0 | - |
| 3 | y | 2.126E+04 | 473.3 | 0.0001401 | 0.296 | +1 | 5 |
| - | - | 4675 | 474.3 | - | - | 0 | - |
| - | - | 856.4 | 475.3 | - | - | 0 | - |
| - | - | 673.7 | 494.2 | - | - | 0 | - |
| 2 | y | 1978 | 570.3 | 0.000247 | 0.4331 | +1 | 6 |
| 2 | y | 621.4 | 571.3 | 0.005789 | 10.13 | +1 | 6 |
| 2 | y | 2.638E+04 | 588.3 | 0.0004357 | 0.7406 | +1 | 6 |
| - | - | 9306 | 589.3 | - | - | 0 | - |
| - | - | 2057 | 590.3 | - | - | 0 | - |
| - | - | 555.1 | 650.3 | - | - | 0 | - |
| - | - | 550.2 | 685.4 | - | - | 0 | - |
| - | - | 1139 | 709.4 | - | - | 0 | - |
| - | - | 585.9 | 773.3 | - | - | 0 | - |
| - | - | 605.2 | 933.7 | - | - | 0 | - |
| - | - | 616.8 | 1232 | - | - | 0 | - |
| - | - | 723.1 | 1316 | - | - | 0 | - |
| - | - | 613.5 | 1341 | - | - | 0 | - |
| - | - | 637.2 | 1563 | - | - | 0 | - |
| - | - | 821.2 | 1871 | - | - | 0 | - |
| - | - | 731.1 | 2326 | - | - | 0 | - |
| - | - | 700.8 | 3074 | - | - | 0 | - |
| - | - | 750.1 | 3075 | - | - | 0 | - |
| - | - | 727.5 | 3093 | - | - | 0 | - |

m/z Charge Intensity FragmentType MassShift Position
120.02408599853516 0 514.8555
120.0811767578125 0 117646.82
121.08446502685547 0 8895.439
121.10124969482422 0 350.67035
122.08732604980469 0 452.5551
123.1170425415039 0 1283.2377
124.01891326904297 0 559.6638
126.45226287841797 0 377.75143
128.1073455810547 0 2074.17
129.10255432128906 0 19705.164
130.06541442871094 0 645.65955
130.0865936279297 0 10660.764 y Ammonia loss 6
130.10597229003906 0 1360.4633
131.97439575195312 0 548.6294
136.018798828125 0 940.9888
136.07603454589844 0 1206.0408
137.13258361816406 0 551.9832
138.06642150878906 0 746.61774
139.08705139160156 0 602.20465
140.0821990966797 0 836.81287
141.06610107421875 0 695.39935
142.06103515625 0 532.7921
143.0941619873047 0 621.4493
143.10858154296875 0 428.09183
143.11825561523438 0 4218.3125
145.0499725341797 0 459.1931
145.12229919433594 0 727.7561
146.0968475341797 0 910.1277
147.0349884033203 0 445.64313
147.11314392089844 0 13493.966 y 6
148.1165771484375 0 700.5072
148.95469665527344 0 475.9809
149.02403259277344 0 1039.3302
149.0452117919922 0 6694.534
152.07162475585938 0 749.6437
153.1026611328125 0 559.29913
155.97732543945312 0 501.8461
157.0648651123047 0 749.2821
157.09732055664062 0 1825.8215
159.0345001220703 0 508.60666
159.07675170898438 0 5081.4106
161.0137481689453 0 825.06775
163.02928161621094 0 904.68646
163.14878845214844 0 517.0686
164.08265686035156 0 462.37564
165.10145568847656 0 461.27292
166.07882690429688 0 492.4811
167.05575561523438 0 10230.61
167.06996154785156 0 415.81055
169.09750366210938 0 699.55914
169.9776153564453 0 520.59534
171.0079345703125 0 919.0946
171.1132049560547 0 4738.5464
172.66368103027344 0 443.647
173.0133056640625 0 548.8664
173.02345275878906 0 448.14804
173.45147705078125 0 1214.8029
174.04562377929688 0 489.99396
175.0296173095703 0 1970.0544
175.0867919921875 0 1360.1442
177.04534912109375 0 514.01025
180.1029815673828 0 472.21738
181.32862854003906 0 430.72516
183.1129150390625 0 518.7943
185.6392364501953 0 490.38742
187.0294952392578 0 485.11752
187.07167053222656 0 5602.088
187.1077880859375 0 468.43243
187.1444549560547 0 1226.6914
191.0247802734375 0 1082.5244
191.11827087402344 0 3281.8892 d 1
197.11813354492188 0 1365.3062
199.16934204101562 0 2314.787
202.04095458984375 0 1196.5876
202.10763549804688 0 2018.109
203.98345947265625 0 614.06177
212.9977264404297 0 651.1055
214.0403594970703 0 712.3356
215.1393585205078 0 810.97174
216.05636596679688 0 727.98505
217.08485412597656 0 976.80615
217.09742736816406 0 11492.692 a Water loss 1
218.1009521484375 0 1155.2804
221.09255981445312 0 1357.1544
224.1397247314453 0 512.0058
225.04278564453125 0 881.25977
226.11880493164062 0 888.3788
226.15521240234375 0 3138.6824
228.134521484375 0 5647.8003
232.05101013183594 0 826.1065
233.0361785888672 0 640.7595
233.1649932861328 0 2113.4814
235.10800170898438 0 12910.432 a 1
236.111328125 0 1406.9966
240.1339111328125 0 882.30884
241.1190948486328 0 1822.4664
245.9239959716797 0 557.62946
246.18150329589844 0 3405.9744 y 5
255.0652618408203 0 812.33075
256.1290283203125 0 2421.621
263.1027526855469 0 11756.205 b 1
264.10693359375 0 1376.6418
268.1295471191406 0 1395.464
271.0619812011719 0 843.29285
272.1241760253906 0 864.5691
281.0518798828125 0 960.1154
283.0307312011719 0 1452.3262
285.0097961425781 0 624.2083
286.1396179199219 0 2136.026
287.0566101074219 0 886.4767
288.1357116699219 0 634.5068 a Water loss 2
299.0619812011719 0 9185.554
299.2079772949219 0 3404.3628
300.2112121582031 0 628.8526
303.2030334472656 0 13745.916 y 4
304.206787109375 0 1786.91
315.1654052734375 0 717.493
323.1697082519531 0 600.4511
323.20745849609375 0 531.8908
325.1499938964844 0 705.61237
327.07843017578125 0 8966.753
327.20367431640625 0 1181.9589
334.1396484375 0 2433.0657 b 2
343.16131591796875 0 8267.613
344.1639099121094 0 1547.0695
355.40264892578125 0 505.43866
359.1958923339844 0 748.71985 Precursor Water loss
367.07061767578125 0 607.19
368.20538330078125 0 1541.4019 Precursor
369.1242980957031 0 2657.5747
387.20245361328125 0 1149.8749
391.7225341796875 0 849.16785
402.27105712890625 0 6882.6167 y 3
403.2735290527344 0 1283.2852
403.72314453125 0 2018.6753
413.2718505859375 0 594.548
414.2342529296875 0 1619.4652
438.27166748046875 0 1037.1382
442.2297058105469 0 1493.8433
460.2414855957031 0 1981.1338
473.308349609375 0 21259.639 y 2
474.3113708496094 0 4675.193
475.31390380859375 0 856.38245
494.2335205078125 0 673.6822
570.3243408203125 0 1978.2944 y Water loss 1
571.3143920898438 0 621.40247 y Ammonia loss 1
588.334716796875 0 26376.71 y 1
589.3375854492188 0 9306.404
590.3397827148438 0 2056.9143
650.2935180664062 0 555.134
685.3848266601562 0 550.24585
709.3869018554688 0 1138.6605
773.2640991210938 0 585.9106
933.700927734375 0 605.24963
1232.434326171875 0 616.764
1315.9326171875 0 723.10236
1340.6942138671875 0 613.5041
1563.2635498046875 0 637.1568
1871.235595703125 0 821.2387
2326.09619140625 0 731.0984
3074.279296875 0 700.7864
3075.000732421875 0 750.13464
3092.819091796875 0 727.54333

Spectrum Details

|  |  |
| --- | --- |
| Matched peaks? Matched peaks The total absolute number of peaks matched. Additionally in brackets the total fraction of peaks matched and the total number of peaks is shown. | 17 (10.43% of 163) |
| FDR? FDR The false discovery rate estimated for this peptide. It is calculated by matching all theoretical fragments with a non-integer shift with the raw peaks for this spectrum. This is done with 40 different shifts. The resulting percentage is the average number of annotated peaks over the number of annotated peaks with the correct spectrum. | 0.98% |
| Satellite FDR? Satellite FDR See the FDR for details on its calculation. This satellite ion specific FDR only contains the satellite ions (d/w) for I/L/J positions. | - |
| PSM Score? PSM Score The PSM Score as given by Hecklib to this annotated spectrum. It is shown with three significant figures. | 199 |

#### Spectrum 4009? Spectrum 4009 The raw spectrum of this peptide as annotated by Hecklib. The fragments are coloured according to ion type (see legend). Any peaks with a star '\*' as text can be hovered over to see the full details, first the ion type second the mass shift type. By hovering over the amino acids in the peptide or ions in the legend the corresponding peaks are highlighted. By toggling the 'Unassigned' label you can turn the background (unassigned) peaks on or off in the plot. By updating the slider in the Ion legend you can update the spectrum to only show the top X% of the peaks with labels. The top X% means any peak that is within X% of the highest intensity. By dragging in the spectrum you can zoom in to a specific part of the spectrum and use 'Zoom Out' to get back to the original zoom level. The annotation of the spectrum is based on the given sequence in the peptides file and is done with different software so inconsistencies are likely. The peaks are annotated based on the given sequence, with 20 ppm tolerance.

Copy Data

##### Spectrum 4009 (TSV)

###### Preview

```
Loading example...
```

*Click on the button to copy the data to your clipboard.*

Mz MinMz MaxIntensity Max

WidthHeightPeptide font sizePeptide stroke widthSpectrum font sizeSpectrum stroke widthCompact peptide

Ion legend

wxyz

abcd

OtherUnassignedIon^Charge^_Position_Show for top:%

FDAVGVK

01.48e+42.96e+44.44e+45.92e+4

Zoom Out

y^+1^_1_z^+1^_1_y^+1^_1_w^+1^_2_y^+1^_2_z^+1^_2_y^+1^_2_z^+1^_3_y^+1^_3_w^+1^_4_y^+1^_4_z^+1^_4_y^+1^_4_z^+1^_5_y^+1^_5_c^+1^_5_z^+1^_6_y^+1^_6_z^+1^_6_y^+1^_6_c^+1^_6_

0775155123263102

Fragment Matches Table

Show background peaks

| Position | Ion type | Intensity | mz Theoretical | mz Error (Th) | mz Error (ppm) | Charge | Series Number |
| --- | --- | --- | --- | --- | --- | --- | --- |
| - | - | 2.329E+04 | 120.1 | - | - | 0 | - |
| - | - | 1671 | 121.1 | - | - | 0 | - |
| - | - | 480.1 | 127.1 | - | - | 0 | - |
| - | - | 429.5 | 128.1 | - | - | 0 | - |
| - | - | 3612 | 129.1 | - | - | 0 | - |
| 7 | y | 3995 | 130.1 | 0.0002165 | 1.664 | +1 | 1 |
| 7 | z | 2078 | 131.1 | 0.000326 | 2.487 | +1 | 1 |
| - | - | 369.5 | 133.1 | - | - | 0 | - |
| - | - | 469.8 | 140.1 | - | - | 0 | - |
| - | - | 903.8 | 142.1 | - | - | 0 | - |
| 7 | y | 8619 | 147.1 | 0.000233 | 1.583 | +1 | 1 |
| - | - | 466.2 | 147.8 | - | - | 0 | - |
| - | - | 665.7 | 148.9 | - | - | 0 | - |
| - | - | 543.1 | 148.9 | - | - | 0 | - |
| - | - | 635 | 148.9 | - | - | 0 | - |
| - | - | 567.4 | 148.9 | - | - | 0 | - |
| - | - | 497 | 148.9 | - | - | 0 | - |
| - | - | 990.9 | 148.9 | - | - | 0 | - |
| - | - | 1306 | 148.9 | - | - | 0 | - |
| - | - | 1322 | 148.9 | - | - | 0 | - |
| - | - | 3258 | 148.9 | - | - | 0 | - |
| - | - | 5716 | 148.9 | - | - | 0 | - |
| - | - | 3958 | 149 | - | - | 0 | - |
| - | - | 1640 | 149 | - | - | 0 | - |
| - | - | 1341 | 149 | - | - | 0 | - |
| - | - | 1104 | 149 | - | - | 0 | - |
| - | - | 517.5 | 149 | - | - | 0 | - |
| - | - | 824.9 | 149 | - | - | 0 | - |
| - | - | 617.1 | 149 | - | - | 0 | - |
| - | - | 485.6 | 149 | - | - | 0 | - |
| - | - | 428.8 | 149 | - | - | 0 | - |
| - | - | 516.4 | 149 | - | - | 0 | - |
| - | - | 515.9 | 149 | - | - | 0 | - |
| - | - | 6916 | 149 | - | - | 0 | - |
| - | - | 491.2 | 154.8 | - | - | 0 | - |
| - | - | 2504 | 167.1 | - | - | 0 | - |
| - | - | 461 | 169.4 | - | - | 0 | - |
| - | - | 455.5 | 171.1 | - | - | 0 | - |
| - | - | 745.3 | 177.1 | - | - | 0 | - |
| - | - | 478.8 | 177.9 | - | - | 0 | - |
| - | - | 887 | 191.1 | - | - | 0 | - |
| - | - | 863.8 | 199.2 | - | - | 0 | - |
| - | - | 560 | 203 | - | - | 0 | - |
| - | - | 1172 | 212.1 | - | - | 0 | - |
| - | - | 590.8 | 214 | - | - | 0 | - |
| 6 | w | 707.7 | 215.1 | 0.00095 | 4.416 | +1 | 2 |
| - | - | 2861 | 217.1 | - | - | 0 | - |
| - | - | 779.3 | 225.2 | - | - | 0 | - |
| - | - | 1023 | 228.1 | - | - | 0 | - |
| 6 | y | 622.7 | 229.2 | 0.0002383 | 1.04 | +1 | 2 |
| 6 | z | 2015 | 230.2 | 2.736E-05 | 0.1189 | +1 | 2 |
| - | - | 715.3 | 231.2 | - | - | 0 | - |
| - | - | 845.6 | 232.1 | - | - | 0 | - |
| - | - | 704.1 | 235.1 | - | - | 0 | - |
| - | - | 4779 | 235.1 | - | - | 0 | - |
| - | - | 627.7 | 236.1 | - | - | 0 | - |
| - | - | 1789 | 245.2 | - | - | 0 | - |
| 6 | y | 3015 | 246.2 | 0.0001937 | 0.7867 | +1 | 2 |
| - | - | 850.1 | 255.1 | - | - | 0 | - |
| - | - | 480.2 | 261.2 | - | - | 0 | - |
| - | - | 1258 | 262.1 | - | - | 0 | - |
| - | - | 6085 | 263.1 | - | - | 0 | - |
| - | - | 610.2 | 269 | - | - | 0 | - |
| - | - | 530.1 | 271.1 | - | - | 0 | - |
| - | - | 584.2 | 273.2 | - | - | 0 | - |
| - | - | 606.5 | 273.2 | - | - | 0 | - |
| - | - | 554.5 | 278.2 | - | - | 0 | - |
| - | - | 1576 | 281.1 | - | - | 0 | - |
| - | - | 1488 | 283.2 | - | - | 0 | - |
| - | - | 857.2 | 284.2 | - | - | 0 | - |
| 5 | z | 1.363E+04 | 287.2 | 0.0001548 | 0.5391 | +1 | 3 |
| - | - | 3.036E+04 | 288.2 | - | - | 0 | - |
| - | - | 2884 | 289.2 | - | - | 0 | - |
| - | - | 610 | 295.2 | - | - | 0 | - |
| - | - | 6247 | 299.1 | - | - | 0 | - |
| - | - | 1125 | 299.2 | - | - | 0 | - |
| - | - | 4102 | 302.2 | - | - | 0 | - |
| 5 | y | 7159 | 303.2 | 0.0001671 | 0.5513 | +1 | 3 |
| - | - | 1182 | 304.2 | - | - | 0 | - |
| - | - | 525.1 | 314.2 | - | - | 0 | - |
| - | - | 5263 | 327.1 | - | - | 0 | - |
| - | - | 967.1 | 327.2 | - | - | 0 | - |
| - | - | 2195 | 330.2 | - | - | 0 | - |
| - | - | 4210 | 334.1 | - | - | 0 | - |
| - | - | 739.6 | 335.1 | - | - | 0 | - |
| - | - | 3155 | 343.2 | - | - | 0 | - |
| - | - | 922.6 | 355.2 | - | - | 0 | - |
| - | - | 831.4 | 367.1 | - | - | 0 | - |
| - | - | 553.7 | 368.2 | - | - | 0 | - |
| - | - | 1443 | 368.2 | - | - | 0 | - |
| - | - | 2770 | 369.1 | - | - | 0 | - |
| 4 | w | 9097 | 371.2 | 0.0001063 | 0.2863 | +1 | 4 |
| - | - | 638.1 | 371.6 | - | - | 0 | - |
| - | - | 1636 | 372.2 | - | - | 0 | - |
| - | - | 606.7 | 380.3 | - | - | 0 | - |
| 4 | y | 1545 | 385.2 | 8.229E-05 | 0.2136 | +1 | 4 |
| 4 | z | 1153 | 386.3 | 0.0009853 | 2.551 | +1 | 4 |
| - | - | 821.1 | 391.7 | - | - | 0 | - |
| - | - | 690.1 | 398.2 | - | - | 0 | - |
| - | - | 923.9 | 399.2 | - | - | 0 | - |
| - | - | 1851 | 400.3 | - | - | 0 | - |
| 4 | y | 3306 | 402.3 | 0.0004802 | 1.194 | +1 | 4 |
| - | - | 753.9 | 404.2 | - | - | 0 | - |
| - | - | 1846 | 405.2 | - | - | 0 | - |
| - | - | 1921 | 415.2 | - | - | 0 | - |
| - | - | 582.9 | 416.2 | - | - | 0 | - |
| - | - | 546.2 | 426.9 | - | - | 0 | - |
| - | - | 786.6 | 432.2 | - | - | 0 | - |
| - | - | 7150 | 433.2 | - | - | 0 | - |
| - | - | 955.5 | 434.2 | - | - | 0 | - |
| - | - | 1271 | 439.3 | - | - | 0 | - |
| - | - | 589.3 | 439.9 | - | - | 0 | - |
| - | - | 2072 | 442.2 | - | - | 0 | - |
| - | - | 4895 | 442.3 | - | - | 0 | - |
| - | - | 1032 | 443.3 | - | - | 0 | - |
| 3 | z | 1.713E+04 | 457.3 | 9.587E-05 | 0.2096 | +1 | 5 |
| - | - | 2752 | 458.3 | - | - | 0 | - |
| - | - | 2324 | 463.2 | - | - | 0 | - |
| - | - | 949.9 | 464.2 | - | - | 0 | - |
| - | - | 890.6 | 471.3 | - | - | 0 | - |
| 3 | y | 1.218E+04 | 473.3 | 0.0002927 | 0.6184 | +1 | 5 |
| - | - | 2796 | 474.3 | - | - | 0 | - |
| - | - | 3642 | 490.2 | - | - | 0 | - |
| - | - | 1139 | 491.2 | - | - | 0 | - |
| - | - | 1333 | 492.3 | - | - | 0 | - |
| - | - | 949.5 | 500.3 | - | - | 0 | - |
| - | - | 4616 | 506.2 | - | - | 0 | - |
| 5 | c | 9944 | 507.3 | 0.0007113 | 1.402 | +1 | 5 |
| - | - | 2519 | 508.3 | - | - | 0 | - |
| - | - | 765.7 | 513.3 | - | - | 0 | - |
| - | - | 776.6 | 513.8 | - | - | 0 | - |
| - | - | 9987 | 528.3 | - | - | 0 | - |
| - | - | 2560 | 529.3 | - | - | 0 | - |
| - | - | 814.1 | 545.3 | - | - | 0 | - |
| - | - | 761.6 | 546.3 | - | - | 0 | - |
| - | - | 701.3 | 553.3 | - | - | 0 | - |
| - | - | 5420 | 553.8 | - | - | 0 | - |
| 2 | z | 1861 | 554.3 | 0.005232 | 9.44 | +1 | 6 |
| - | - | 1503 | 561.3 | - | - | 0 | - |
| - | - | 2085 | 562.3 | - | - | 0 | - |
| - | - | 1139 | 563.3 | - | - | 0 | - |
| 2 | y | 809.6 | 570.3 | 0.003293 | 5.774 | +1 | 6 |
| - | - | 2265 | 571.3 | - | - | 0 | - |
| 2 | z | 7287 | 572.3 | 0.0007546 | 1.319 | +1 | 6 |
| - | - | 2222 | 573.3 | - | - | 0 | - |
| 2 | y | 1.826E+04 | 588.3 | 0.0002526 | 0.4294 | +1 | 6 |
| - | - | 2.008E+04 | 589.3 | - | - | 0 | - |
| - | - | 3126 | 589.3 | - | - | 0 | - |
| - | - | 6096 | 590.3 | - | - | 0 | - |
| - | - | 812.1 | 591.3 | - | - | 0 | - |
| - | - | 663.6 | 603.3 | - | - | 0 | - |
| 6 | c | 2.41E+04 | 606.3 | 6.392E-05 | 0.1054 | +1 | 6 |
| - | - | 8900 | 607.3 | - | - | 0 | - |
| - | - | 1101 | 608.3 | - | - | 0 | - |
| - | - | 622.8 | 614.3 | - | - | 0 | - |
| - | - | 2995 | 675.4 | - | - | 0 | - |
| - | - | 3704 | 676.4 | - | - | 0 | - |
| - | - | 1146 | 677.4 | - | - | 0 | - |
| - | - | 559.2 | 709.4 | - | - | 0 | - |
| - | - | 597.8 | 713.9 | - | - | 0 | - |
| - | - | 5.858E+04 | 719.4 | - | - | 0 | - |
| - | - | 2E+04 | 720.4 | - | - | 0 | - |
| - | - | 6059 | 721.4 | - | - | 0 | - |
| - | - | 553.5 | 731.2 | - | - | 0 | - |
| - | - | 636.2 | 734.4 | - | - | 0 | - |
| - | - | 2.62E+04 | 735.4 | - | - | 0 | - |
| - | - | 1.239E+04 | 736.4 | - | - | 0 | - |
| - | - | 4492 | 737.4 | - | - | 0 | - |
| - | - | 1174 | 760.4 | - | - | 0 | - |
| - | - | 943.6 | 809.5 | - | - | 0 | - |
| - | - | 600.8 | 825.5 | - | - | 0 | - |
| - | - | 728.9 | 849.5 | - | - | 0 | - |
| - | - | 742.6 | 889.6 | - | - | 0 | - |
| - | - | 579.7 | 965.6 | - | - | 0 | - |
| - | - | 878.8 | 1005 | - | - | 0 | - |
| - | - | 772.8 | 1006 | - | - | 0 | - |
| - | - | 760.8 | 1009 | - | - | 0 | - |
| - | - | 1194 | 1027 | - | - | 0 | - |
| - | - | 972.6 | 1028 | - | - | 0 | - |
| - | - | 1037 | 1029 | - | - | 0 | - |
| - | - | 676.4 | 1045 | - | - | 0 | - |
| - | - | 1841 | 1069 | - | - | 0 | - |
| - | - | 1851 | 1070 | - | - | 0 | - |
| - | - | 796.6 | 1071 | - | - | 0 | - |
| - | - | 970.7 | 1086 | - | - | 0 | - |
| - | - | 955.6 | 1087 | - | - | 0 | - |
| - | - | 586.5 | 1091 | - | - | 0 | - |
| - | - | 2436 | 1107 | - | - | 0 | - |
| - | - | 1.327E+04 | 1108 | - | - | 0 | - |
| - | - | 6007 | 1109 | - | - | 0 | - |
| - | - | 613 | 1266 | - | - | 0 | - |
| - | - | 636 | 2535 | - | - | 0 | - |
| - | - | 717.9 | 2726 | - | - | 0 | - |
| - | - | 745.7 | 3071 | - | - | 0 | - |

m/z Charge Intensity FragmentType MassShift Position
120.08100128173828 0 23292.146
121.08428192138672 0 1670.7656
127.11044311523438 0 480.11572
128.10716247558594 0 429.46588
129.10247802734375 0 3612.202
130.0864715576172 0 3994.896 y Ammonia loss 6
131.0944061279297 0 2078.0864 z 6
133.08648681640625 0 369.47913
140.08250427246094 0 469.83087
142.08656311035156 0 903.8414
147.113037109375 0 8619.192 y 6
147.80096435546875 0 466.19897
148.88040161132812 0 665.714
148.88731384277344 0 543.05554
148.8946990966797 0 634.97174
148.90182495117188 0 567.44226
148.90939331054688 0 497.03595
148.91639709472656 0 990.89905
148.92359924316406 0 1306.1449
148.93077087402344 0 1322.089
148.93832397460938 0 3258.2766
148.94610595703125 0 5715.7285
148.96273803710938 0 3957.8352
148.97044372558594 0 1640.1003
148.97750854492188 0 1341.1603
148.9849853515625 0 1104.2444
148.99209594726562 0 517.4776
148.9991912841797 0 824.8627
149.0070037841797 0 617.05237
149.01400756835938 0 485.64084
149.02207946777344 0 428.82367
149.03562927246094 0 516.40466
149.03785705566406 0 515.9213
149.0449676513672 0 6915.954
154.78463745117188 0 491.23355
167.0554962158203 0 2504.2734
169.37142944335938 0 461.0226
171.11383056640625 0 455.4603
177.11277770996094 0 745.2918
177.885498046875 0 478.77997
191.1180877685547 0 887.02216
199.16966247558594 0 863.7625
203.0242919921875 0 559.95844
212.13958740234375 0 1172.1808
214.04043579101562 0 590.85
215.1399688720703 0 707.65704 w 5
217.0971221923828 0 2861.0056
225.159912109375 0 779.3449
228.13441467285156 0 1022.6324
229.1549072265625 0 622.7074 y Ammonia loss 5
230.1625213623047 0 2015.1204 z 5
231.17103576660156 0 715.3083
232.0509033203125 0 845.6217
235.0952911376953 0 704.13367
235.10781860351562 0 4778.959
236.11109924316406 0 627.72363
245.1736602783203 0 1789.0378
246.18141174316406 0 3015.1206 y 5
255.06640625 0 850.07263
261.1592712402344 0 480.206
262.0945739746094 0 1257.7292
263.1025085449219 0 6084.6045
269.0456237792969 0 610.2069
271.0618591308594 0 530.1069
273.1691589355469 0 584.19446
273.1922302246094 0 606.54205
278.15692138671875 0 554.51385
281.0515441894531 0 1575.9329
283.1748352050781 0 1488.4248
284.19708251953125 0 857.21326
287.1841125488281 0 13628.617 z 4
288.19183349609375 0 30362.994
289.1946716308594 0 2884.0586
295.1946716308594 0 609.97736
299.0616149902344 0 6246.8315
299.2079772949219 0 1125.3627
302.1952819824219 0 4101.9385
303.2025146484375 0 7159.1367 y 4
304.2067565917969 0 1182.276
314.1581726074219 0 525.1247
327.078369140625 0 5263.4033
327.2030944824219 0 967.06805
330.190673828125 0 2195.0618
334.1402893066406 0 4209.5127
335.1407165527344 0 739.64026
343.1612243652344 0 3155.0356
355.23297119140625 0 922.64703
367.0682373046875 0 831.433
368.1795959472656 0 553.6555
368.204833984375 0 1443.464
369.1246032714844 0 2769.6558
371.2287902832031 0 9097.04 w 3
371.5964660644531 0 638.14246
372.23089599609375 0 1636.4011
380.2833251953125 0 606.67365
385.24462890625 0 1545.1863 y Ammonia loss 3
386.25335693359375 0 1152.827 z 3
391.7257080078125 0 821.071
398.2147216796875 0 690.1261
399.2239685058594 0 923.89856
400.255859375 0 1851.3354
402.2715759277344 0 3305.9966 y 3
404.2259826660156 0 753.90845
405.21343994140625 0 1845.9045
415.2430114746094 0 1920.8778
416.2464599609375 0 582.8767
426.93365478515625 0 546.16406
432.2018737792969 0 786.5983
433.2080383300781 0 7150.1074
434.2110290527344 0 955.47156
439.2803649902344 0 1270.9956
439.9383239746094 0 589.2821
442.2298889160156 0 2071.5835
442.2662048339844 0 4894.8853
443.2695007324219 0 1032.4379
457.2895812988281 0 17134.676 z 2
458.2929382324219 0 2751.9453
463.2427673339844 0 2323.9106
464.247802734375 0 949.85187
471.2934875488281 0 890.6485
473.3085021972656 0 12178.404 y 2
474.31085205078125 0 2796.042
490.2301330566406 0 3641.8845
491.2330322265625 0 1139.3037
492.2645568847656 0 1332.5078
500.252197265625 0 949.52454
506.2478942871094 0 4615.689
507.2554626464844 0 9944.432 c 4
508.2600402832031 0 2519.011
513.264404296875 0 765.73456
513.7699584960938 0 776.6031
528.3263549804688 0 9986.542
529.3302001953125 0 2560.065
545.29150390625 0 814.14075
546.29443359375 0 761.55914
553.3047485351562 0 701.2602
553.80908203125 0 5419.6543
554.3110961914062 0 1860.7308 z Water loss 1
561.3030395507812 0 1503.3848
562.3092651367188 0 2084.798
563.3162841796875 0 1138.6135
570.327880859375 0 809.60095 y Water loss 1
571.2872314453125 0 2265.37
572.315673828125 0 7287.308 z 1
573.3201293945312 0 2222.1653
588.3348999023438 0 18263.623 y 1
589.29736328125 0 20084.518
589.339599609375 0 3125.574
590.3003540039062 0 6096.4443
591.3056030273438 0 812.0776
603.3269653320312 0 663.6347
606.3245239257812 0 24097.248 c 5
607.3268432617188 0 8899.935
608.3311157226562 0 1101.3141
614.319091796875 0 622.79913
675.3930053710938 0 2995.0063
676.392578125 0 3703.751
677.3943481445312 0 1145.9543
709.4168090820312 0 559.20154
713.9011840820312 0 597.77234
719.3840942382812 0 58579.727
720.3870849609375 0 20004.375
721.389404296875 0 6058.785
731.2385864257812 0 553.51544
734.3853759765625 0 636.16925
735.4027099609375 0 26196.857
736.4066772460938 0 12385.253
737.41015625 0 4491.573
760.4491577148438 0 1174.466
809.5181884765625 0 943.588
825.5372314453125 0 600.77045
849.5345458984375 0 728.9294
889.5602416992188 0 742.5503
965.5554809570312 0 579.681
1004.5733642578125 0 878.75916
1005.578857421875 0 772.80963
1008.58935546875 0 760.8467
1026.5718994140625 0 1194.0676
1027.575927734375 0 972.64264
1028.5823974609375 0 1036.697
1044.5911865234375 0 676.40436
1068.603515625 0 1840.6249
1069.600341796875 0 1851.0732
1070.59814453125 0 796.6479
1085.60791015625 0 970.7401
1086.6121826171875 0 955.61975
1090.5634765625 0 586.4861
1106.6090087890625 0 2435.634
1107.6168212890625 0 13274.926
1108.620849609375 0 6006.913
1265.5010986328125 0 612.9879
2535.419189453125 0 636.0434
2726.1572265625 0 717.89343
3071.2548828125 0 745.7124

Spectrum Details

|  |  |
| --- | --- |
| Matched peaks? Matched peaks The total absolute number of peaks matched. Additionally in brackets the total fraction of peaks matched and the total number of peaks is shown. | 21 (10.82% of 194) |
| FDR? FDR The false discovery rate estimated for this peptide. It is calculated by matching all theoretical fragments with a non-integer shift with the raw peaks for this spectrum. This is done with 40 different shifts. The resulting percentage is the average number of annotated peaks over the number of annotated peaks with the correct spectrum. | 0.57% |
| Satellite FDR? Satellite FDR See the FDR for details on its calculation. This satellite ion specific FDR only contains the satellite ions (d/w) for I/L/J positions. | - |
| PSM Score? PSM Score The PSM Score as given by Hecklib to this annotated spectrum. It is shown with three significant figures. | 281 |

#### Spectrum 4278? Spectrum 4278 The raw spectrum of this peptide as annotated by Hecklib. The fragments are coloured according to ion type (see legend). Any peaks with a star '\*' as text can be hovered over to see the full details, first the ion type second the mass shift type. By hovering over the amino acids in the peptide or ions in the legend the corresponding peaks are highlighted. By toggling the 'Unassigned' label you can turn the background (unassigned) peaks on or off in the plot. By updating the slider in the Ion legend you can update the spectrum to only show the top X% of the peaks with labels. The top X% means any peak that is within X% of the highest intensity. By dragging in the spectrum you can zoom in to a specific part of the spectrum and use 'Zoom Out' to get back to the original zoom level. The annotation of the spectrum is based on the given sequence in the peptides file and is done with different software so inconsistencies are likely. The peaks are annotated based on the given sequence, with 20 ppm tolerance.

Copy Data

##### Spectrum 4278 (TSV)

###### Preview

```
Loading example...
```

*Click on the button to copy the data to your clipboard.*

Mz MinMz MaxIntensity Max

WidthHeightPeptide font sizePeptide stroke widthSpectrum font sizeSpectrum stroke widthCompact peptide

Ion legend

wxyz

abcd

OtherUnassignedIon^Charge^_Position_Show for top:%

FDAVGVK

03.05e+36.10e+39.15e+31.22e+4

Zoom Out

y^+1^_1_y^+1^_1_w^+1^_2_z^+1^_2_y^+1^_2_z^+1^_3_y^+1^_3_w^+1^_4_y^+1^_4_z^+1^_5_y^+1^_5_c^+1^_5_z^+1^_6_y^+1^_6_c^+1^_6_

02805598391119

Fragment Matches Table

Show background peaks

| Position | Ion type | Intensity | mz Theoretical | mz Error (Th) | mz Error (ppm) | Charge | Series Number |
| --- | --- | --- | --- | --- | --- | --- | --- |
| - | - | 6168 | 120.1 | - | - | 0 | - |
| - | - | 459.4 | 121.1 | - | - | 0 | - |
| - | - | 359.2 | 128.8 | - | - | 0 | - |
| - | - | 814.9 | 129.1 | - | - | 0 | - |
| 7 | y | 975.6 | 130.1 | 0.0002012 | 1.547 | +1 | 1 |
| - | - | 400.3 | 131.7 | - | - | 0 | - |
| - | - | 1115 | 133.1 | - | - | 0 | - |
| - | - | 591.8 | 136 | - | - | 0 | - |
| - | - | 613.7 | 140.1 | - | - | 0 | - |
| - | - | 473.3 | 140.6 | - | - | 0 | - |
| 7 | y | 1508 | 147.1 | 9.562E-05 | 0.65 | +1 | 1 |
| - | - | 871.5 | 148.9 | - | - | 0 | - |
| - | - | 1161 | 149 | - | - | 0 | - |
| - | - | 6799 | 149 | - | - | 0 | - |
| - | - | 445.7 | 151.2 | - | - | 0 | - |
| - | - | 430.7 | 158.8 | - | - | 0 | - |
| - | - | 2478 | 167.1 | - | - | 0 | - |
| - | - | 866.1 | 175.1 | - | - | 0 | - |
| - | - | 713.8 | 180.9 | - | - | 0 | - |
| - | - | 1440 | 199.2 | - | - | 0 | - |
| - | - | 694 | 201 | - | - | 0 | - |
| - | - | 519.9 | 208.9 | - | - | 0 | - |
| - | - | 468.9 | 212 | - | - | 0 | - |
| - | - | 1011 | 212.1 | - | - | 0 | - |
| - | - | 637.5 | 214 | - | - | 0 | - |
| 6 | w | 652.1 | 215.1 | 0.0002328 | 1.082 | +1 | 2 |
| - | - | 639.5 | 216.1 | - | - | 0 | - |
| - | - | 819.6 | 217.1 | - | - | 0 | - |
| - | - | 562.1 | 217.2 | - | - | 0 | - |
| - | - | 537.1 | 230 | - | - | 0 | - |
| 6 | z | 549.1 | 230.2 | 0.000644 | 2.798 | +1 | 2 |
| - | - | 1428 | 235.1 | - | - | 0 | - |
| - | - | 561.5 | 245.2 | - | - | 0 | - |
| 6 | y | 653.9 | 246.2 | 0.0005388 | 2.188 | +1 | 2 |
| - | - | 868.4 | 249.1 | - | - | 0 | - |
| - | - | 779.1 | 255.1 | - | - | 0 | - |
| - | - | 770.4 | 261.2 | - | - | 0 | - |
| - | - | 1128 | 263.1 | - | - | 0 | - |
| - | - | 695.4 | 269 | - | - | 0 | - |
| - | - | 965.7 | 271.1 | - | - | 0 | - |
| - | - | 2129 | 281.1 | - | - | 0 | - |
| - | - | 1320 | 283.2 | - | - | 0 | - |
| - | - | 761.6 | 285 | - | - | 0 | - |
| 5 | z | 4132 | 287.2 | 0.0001243 | 0.4328 | +1 | 3 |
| - | - | 7820 | 288.2 | - | - | 0 | - |
| - | - | 944.4 | 289.2 | - | - | 0 | - |
| - | - | 6291 | 299.1 | - | - | 0 | - |
| - | - | 1348 | 302.2 | - | - | 0 | - |
| 5 | y | 2110 | 303.2 | 0.0002892 | 0.9539 | +1 | 3 |
| - | - | 570.4 | 311.9 | - | - | 0 | - |
| - | - | 654.9 | 322.2 | - | - | 0 | - |
| - | - | 4976 | 327.1 | - | - | 0 | - |
| - | - | 832 | 334.1 | - | - | 0 | - |
| - | - | 569.4 | 338.8 | - | - | 0 | - |
| - | - | 640.8 | 352.2 | - | - | 0 | - |
| - | - | 750.7 | 367.1 | - | - | 0 | - |
| - | - | 3112 | 369.1 | - | - | 0 | - |
| - | - | 574.2 | 369.1 | - | - | 0 | - |
| 4 | w | 2010 | 371.2 | 0.0006567 | 1.769 | +1 | 4 |
| - | - | 692.9 | 372.2 | - | - | 0 | - |
| 4 | y | 1473 | 402.3 | 0.0002666 | 0.6627 | +1 | 4 |
| - | - | 615.6 | 405.2 | - | - | 0 | - |
| - | - | 560.3 | 415.2 | - | - | 0 | - |
| - | - | 1640 | 433.2 | - | - | 0 | - |
| - | - | 1090 | 442.3 | - | - | 0 | - |
| 3 | z | 2927 | 457.3 | 0.0006147 | 1.344 | +1 | 5 |
| - | - | 809.7 | 458.3 | - | - | 0 | - |
| 3 | y | 2848 | 473.3 | 0.0003232 | 0.6829 | +1 | 5 |
| - | - | 1035 | 490.2 | - | - | 0 | - |
| 5 | c | 2580 | 507.3 | 0.0003873 | 0.7636 | +1 | 5 |
| - | - | 2373 | 528.3 | - | - | 0 | - |
| 2 | z | 2150 | 572.3 | 0.001198 | 2.094 | +1 | 6 |
| 2 | y | 4780 | 588.3 | 0.0001136 | 0.1931 | +1 | 6 |
| - | - | 4963 | 589.3 | - | - | 0 | - |
| - | - | 793.4 | 589.3 | - | - | 0 | - |
| - | - | 1568 | 590.3 | - | - | 0 | - |
| 6 | c | 7932 | 606.3 | 0.0002413 | 0.3979 | +1 | 6 |
| - | - | 1538 | 607.3 | - | - | 0 | - |
| - | - | 1413 | 675.4 | - | - | 0 | - |
| - | - | 739.3 | 676.4 | - | - | 0 | - |
| - | - | 1043 | 697.4 | - | - | 0 | - |
| - | - | 1.208E+04 | 719.4 | - | - | 0 | - |
| - | - | 4832 | 720.4 | - | - | 0 | - |
| - | - | 985.4 | 721.4 | - | - | 0 | - |
| - | - | 5858 | 735.4 | - | - | 0 | - |
| - | - | 3242 | 736.4 | - | - | 0 | - |
| - | - | 1005 | 737.4 | - | - | 0 | - |
| - | - | 620.3 | 884.5 | - | - | 0 | - |
| - | - | 628.1 | 999.6 | - | - | 0 | - |
| - | - | 629.8 | 1027 | - | - | 0 | - |
| - | - | 692.8 | 1028 | - | - | 0 | - |
| - | - | 979.1 | 1043 | - | - | 0 | - |
| - | - | 906.4 | 1044 | - | - | 0 | - |
| - | - | 743.1 | 1068 | - | - | 0 | - |
| - | - | 833.1 | 1069 | - | - | 0 | - |
| - | - | 920.8 | 1071 | - | - | 0 | - |
| - | - | 1172 | 1103 | - | - | 0 | - |
| - | - | 1035 | 1104 | - | - | 0 | - |
| - | - | 626.2 | 1105 | - | - | 0 | - |
| - | - | 929.9 | 1108 | - | - | 0 | - |

m/z Charge Intensity FragmentType MassShift Position
120.08094787597656 0 6167.7646
121.08464050292969 0 459.39627
128.78306579589844 0 359.22583
129.1024627685547 0 814.85284
130.08645629882812 0 975.5556 y Ammonia loss 6
131.72213745117188 0 400.26572
133.08615112304688 0 1114.8875
136.0183868408203 0 591.83746
140.08233642578125 0 613.74335
140.64608764648438 0 473.32938
147.11289978027344 0 1508.1552 y 6
148.9471893310547 0 871.54944
149.02359008789062 0 1161.4889
149.04505920410156 0 6798.5454
151.15509033203125 0 445.665
158.82887268066406 0 430.65125
167.0554962158203 0 2477.8901
175.13279724121094 0 866.0783
180.85943603515625 0 713.78204
199.16937255859375 0 1440.4352
200.9965057373047 0 693.981
208.9211883544922 0 519.8619
211.9764862060547 0 468.91434
212.13992309570312 0 1010.5317
214.04006958007812 0 637.5184
215.13925170898438 0 652.1482 w 5
216.0565643310547 0 639.5209
217.09690856933594 0 819.5925
217.1937713623047 0 562.0956
230.03475952148438 0 537.09283
230.16184997558594 0 549.0975 z 5
235.10792541503906 0 1428.2354
245.17320251464844 0 561.52124
246.18067932128906 0 653.89105 y 5
249.1121368408203 0 868.3531
255.06552124023438 0 779.12134
261.16009521484375 0 770.40765
263.1027526855469 0 1128.283
269.0462951660156 0 695.35693
271.0623779296875 0 965.70245
281.05145263671875 0 2128.8918
283.1749572753906 0 1320.1876
285.0100402832031 0 761.6003
287.18408203125 0 4132.2373 z 4
288.1918029785156 0 7820.1377
289.1954040527344 0 944.40454
299.0619201660156 0 6290.509
302.1942138671875 0 1348.4062
303.202392578125 0 2110.335 y 4
311.94317626953125 0 570.4086
322.1862487792969 0 654.88617
327.0785217285156 0 4975.541
334.13922119140625 0 832.0449
338.8282165527344 0 569.3513
352.1965637207031 0 640.844
367.0682678222656 0 750.6772
369.1248779296875 0 3111.914
369.14910888671875 0 574.22
371.22955322265625 0 2010.4675 w 3
372.2315673828125 0 692.9016
402.2713623046875 0 1473.068 y 3
405.21343994140625 0 615.5814
415.2441711425781 0 560.31537
433.2080383300781 0 1640.2167
442.26702880859375 0 1090.31
457.29010009765625 0 2927.0679 z 2
458.2908935546875 0 809.72504
473.30853271484375 0 2848.2021 y 2
490.2299499511719 0 1035.4342
507.2565612792969 0 2580.0083 c 4
528.3280029296875 0 2372.658
572.317626953125 0 2149.7498 z 1
588.3352661132812 0 4779.9854 y 1
589.2974853515625 0 4963.371
589.341552734375 0 793.37976
590.30224609375 0 1567.7664
606.3248291015625 0 7932.4097 c 5
607.3292236328125 0 1537.7653
675.3975830078125 0 1413.2076
676.39013671875 0 739.27826
697.4420166015625 0 1042.6198
719.3846435546875 0 12084.222
720.3882446289062 0 4831.521
721.3932495117188 0 985.37067
735.403564453125 0 5857.8403
736.4071655273438 0 3242.195
737.4172973632812 0 1004.53656
884.5406494140625 0 620.27216
999.5758666992188 0 628.1183
1026.5670166015625 0 629.76886
1027.57373046875 0 692.78986
1042.59765625 0 979.11554
1043.5927734375 0 906.4297
1067.589599609375 0 743.1234
1068.60400390625 0 833.09216
1070.591796875 0 920.8172
1102.6197509765625 0 1171.9526
1103.6097412109375 0 1035.4368
1104.599609375 0 626.1765
1107.6112060546875 0 929.9184

Spectrum Details

|  |  |
| --- | --- |
| Matched peaks? Matched peaks The total absolute number of peaks matched. Additionally in brackets the total fraction of peaks matched and the total number of peaks is shown. | 15 (15.00% of 100) |
| FDR? FDR The false discovery rate estimated for this peptide. It is calculated by matching all theoretical fragments with a non-integer shift with the raw peaks for this spectrum. This is done with 40 different shifts. The resulting percentage is the average number of annotated peaks over the number of annotated peaks with the correct spectrum. | 0.95% |
| Satellite FDR? Satellite FDR See the FDR for details on its calculation. This satellite ion specific FDR only contains the satellite ions (d/w) for I/L/J positions. | - |
| PSM Score? PSM Score The PSM Score as given by Hecklib to this annotated spectrum. It is shown with three significant figures. | 167 |

#### Spectrum 4671? Spectrum 4671 The raw spectrum of this peptide as annotated by Hecklib. The fragments are coloured according to ion type (see legend). Any peaks with a star '\*' as text can be hovered over to see the full details, first the ion type second the mass shift type. By hovering over the amino acids in the peptide or ions in the legend the corresponding peaks are highlighted. By toggling the 'Unassigned' label you can turn the background (unassigned) peaks on or off in the plot. By updating the slider in the Ion legend you can update the spectrum to only show the top X% of the peaks with labels. The top X% means any peak that is within X% of the highest intensity. By dragging in the spectrum you can zoom in to a specific part of the spectrum and use 'Zoom Out' to get back to the original zoom level. The annotation of the spectrum is based on the given sequence in the peptides file and is done with different software so inconsistencies are likely. The peaks are annotated based on the given sequence, with 20 ppm tolerance.

Copy Data

##### Spectrum 4671 (TSV)

###### Preview

```
Loading example...
```

*Click on the button to copy the data to your clipboard.*

Mz MinMz MaxIntensity Max

WidthHeightPeptide font sizePeptide stroke widthSpectrum font sizeSpectrum stroke widthCompact peptide

Ion legend

wxyz

abcd

OtherUnassignedIon^Charge^_Position_Show for top:%

FDAVGVK

07.26e+31.45e+42.18e+42.90e+4

Zoom Out

y^+1^_1_y^+1^_1_d^+1^_2_a^+1^_2_a^+1^_2_y^+1^_2_b^+1^_2_y^+1^_3_y^+1^_4_y^+1^_5_y^+1^_6_

0559111816772236

Fragment Matches Table

Show background peaks

| Position | Ion type | Intensity | mz Theoretical | mz Error (Th) | mz Error (ppm) | Charge | Series Number |
| --- | --- | --- | --- | --- | --- | --- | --- |
| - | - | 542.6 | 120 | - | - | 0 | - |
| - | - | 2.875E+04 | 120.1 | - | - | 0 | - |
| - | - | 378.5 | 121 | - | - | 0 | - |
| - | - | 479.8 | 121.1 | - | - | 0 | - |
| - | - | 2495 | 121.1 | - | - | 0 | - |
| - | - | 318.6 | 121.3 | - | - | 0 | - |
| - | - | 511.5 | 123.1 | - | - | 0 | - |
| - | - | 1400 | 123.1 | - | - | 0 | - |
| - | - | 541.4 | 124.1 | - | - | 0 | - |
| - | - | 419 | 125.1 | - | - | 0 | - |
| - | - | 465.9 | 127.1 | - | - | 0 | - |
| - | - | 486.9 | 127.1 | - | - | 0 | - |
| - | - | 483.5 | 128.1 | - | - | 0 | - |
| - | - | 988.4 | 128.1 | - | - | 0 | - |
| - | - | 606.1 | 129.1 | - | - | 0 | - |
| - | - | 7232 | 129.1 | - | - | 0 | - |
| - | - | 2372 | 130 | - | - | 0 | - |
| - | - | 1150 | 130.1 | - | - | 0 | - |
| 7 | y | 1918 | 130.1 | 7.917E-05 | 0.6086 | +1 | 1 |
| - | - | 579.9 | 130.1 | - | - | 0 | - |
| - | - | 492.7 | 132 | - | - | 0 | - |
| - | - | 415.8 | 132 | - | - | 0 | - |
| - | - | 585.5 | 132.1 | - | - | 0 | - |
| - | - | 496.8 | 133.1 | - | - | 0 | - |
| - | - | 488.3 | 133.1 | - | - | 0 | - |
| - | - | 369.5 | 135.4 | - | - | 0 | - |
| - | - | 1231 | 136 | - | - | 0 | - |
| - | - | 971.5 | 136.1 | - | - | 0 | - |
| - | - | 472.3 | 137.1 | - | - | 0 | - |
| - | - | 553.8 | 138 | - | - | 0 | - |
| - | - | 634.6 | 138.1 | - | - | 0 | - |
| - | - | 1081 | 140.1 | - | - | 0 | - |
| - | - | 611.4 | 142.1 | - | - | 0 | - |
| - | - | 491.4 | 142.1 | - | - | 0 | - |
| - | - | 1303 | 143.1 | - | - | 0 | - |
| - | - | 1186 | 143.1 | - | - | 0 | - |
| - | - | 852.2 | 145 | - | - | 0 | - |
| - | - | 896.7 | 145.1 | - | - | 0 | - |
| - | - | 1027 | 146.1 | - | - | 0 | - |
| - | - | 1918 | 147.1 | - | - | 0 | - |
| 7 | y | 2765 | 147.1 | 0.0002177 | 1.48 | +1 | 1 |
| - | - | 659.1 | 148 | - | - | 0 | - |
| - | - | 1586 | 149 | - | - | 0 | - |
| - | - | 7731 | 149 | - | - | 0 | - |
| - | - | 532.1 | 152 | - | - | 0 | - |
| - | - | 822.5 | 152.1 | - | - | 0 | - |
| - | - | 612.3 | 155 | - | - | 0 | - |
| - | - | 746.8 | 156.1 | - | - | 0 | - |
| - | - | 735.6 | 157.1 | - | - | 0 | - |
| - | - | 652.1 | 157.1 | - | - | 0 | - |
| - | - | 566 | 159 | - | - | 0 | - |
| - | - | 1082 | 159.1 | - | - | 0 | - |
| - | - | 1669 | 159.1 | - | - | 0 | - |
| - | - | 514.3 | 159.1 | - | - | 0 | - |
| - | - | 585.4 | 163 | - | - | 0 | - |
| - | - | 1.347E+04 | 167.1 | - | - | 0 | - |
| - | - | 673.3 | 167.1 | - | - | 0 | - |
| - | - | 1148 | 171.1 | - | - | 0 | - |
| - | - | 944 | 171.1 | - | - | 0 | - |
| - | - | 1086 | 175 | - | - | 0 | - |
| - | - | 3113 | 175.1 | - | - | 0 | - |
| - | - | 711.9 | 177 | - | - | 0 | - |
| - | - | 1264 | 177 | - | - | 0 | - |
| - | - | 489 | 178 | - | - | 0 | - |
| - | - | 556.1 | 179.1 | - | - | 0 | - |
| - | - | 600.3 | 184.1 | - | - | 0 | - |
| - | - | 985.1 | 185.2 | - | - | 0 | - |
| - | - | 1350 | 187.1 | - | - | 0 | - |
| - | - | 752.3 | 187.1 | - | - | 0 | - |
| - | - | 1787 | 187.1 | - | - | 0 | - |
| - | - | 710.7 | 189 | - | - | 0 | - |
| - | - | 1342 | 191 | - | - | 0 | - |
| 2 | d | 466.2 | 191.1 | 0.0009612 | 5.029 | +1 | 2 |
| - | - | 617.2 | 195.1 | - | - | 0 | - |
| - | - | 3062 | 199.2 | - | - | 0 | - |
| - | - | 935.3 | 201.1 | - | - | 0 | - |
| - | - | 876.2 | 202 | - | - | 0 | - |
| - | - | 2510 | 202.1 | - | - | 0 | - |
| - | - | 1183 | 214 | - | - | 0 | - |
| - | - | 1604 | 215.1 | - | - | 0 | - |
| - | - | 807.2 | 216.1 | - | - | 0 | - |
| - | - | 598 | 216.1 | - | - | 0 | - |
| 2 | a | 3372 | 217.1 | 7.491E-05 | 0.3451 | +1 | 2 |
| - | - | 1294 | 225 | - | - | 0 | - |
| - | - | 1459 | 226.2 | - | - | 0 | - |
| - | - | 717.3 | 227 | - | - | 0 | - |
| - | - | 1147 | 228.1 | - | - | 0 | - |
| - | - | 726 | 230 | - | - | 0 | - |
| - | - | 1083 | 232.1 | - | - | 0 | - |
| - | - | 745.9 | 233 | - | - | 0 | - |
| - | - | 552.7 | 233.2 | - | - | 0 | - |
| 2 | a | 1848 | 235.1 | 9.854E-05 | 0.4191 | +1 | 2 |
| 6 | y | 1359 | 246.2 | 0.0001268 | 0.5149 | +1 | 2 |
| - | - | 983.8 | 249.1 | - | - | 0 | - |
| - | - | 902.6 | 251.2 | - | - | 0 | - |
| - | - | 1696 | 255.1 | - | - | 0 | - |
| - | - | 572.3 | 259 | - | - | 0 | - |
| 2 | b | 2483 | 263.1 | 3.33E-05 | 0.1266 | +1 | 2 |
| - | - | 555 | 264.6 | - | - | 0 | - |
| - | - | 862.3 | 269 | - | - | 0 | - |
| - | - | 977.4 | 271.1 | - | - | 0 | - |
| - | - | 1357 | 281.1 | - | - | 0 | - |
| - | - | 1192 | 283 | - | - | 0 | - |
| - | - | 606.9 | 283.1 | - | - | 0 | - |
| - | - | 1.113E+04 | 299.1 | - | - | 0 | - |
| - | - | 702.7 | 299.2 | - | - | 0 | - |
| 5 | y | 2936 | 303.2 | 0.0001061 | 0.35 | +1 | 3 |
| - | - | 583.2 | 306.9 | - | - | 0 | - |
| - | - | 1.272E+04 | 327.1 | - | - | 0 | - |
| - | - | 2055 | 343.2 | - | - | 0 | - |
| - | - | 545.6 | 349.1 | - | - | 0 | - |
| - | - | 705.9 | 367.1 | - | - | 0 | - |
| - | - | 2971 | 369.1 | - | - | 0 | - |
| 4 | y | 764.5 | 402.3 | 8.349E-05 | 0.2075 | +1 | 4 |
| - | - | 683.5 | 403.3 | - | - | 0 | - |
| - | - | 668.3 | 460.2 | - | - | 0 | - |
| 3 | y | 5229 | 473.3 | 0.0005063 | 1.07 | +1 | 5 |
| - | - | 792.3 | 474.3 | - | - | 0 | - |
| - | - | 1026 | 494.2 | - | - | 0 | - |
| - | - | 662.7 | 541.6 | - | - | 0 | - |
| - | - | 551.4 | 561.3 | - | - | 0 | - |
| 2 | y | 6398 | 588.3 | 0.0001306 | 0.2219 | +1 | 6 |
| - | - | 1582 | 589.3 | - | - | 0 | - |
| - | - | 571.9 | 600.2 | - | - | 0 | - |
| - | - | 558.5 | 809.9 | - | - | 0 | - |
| - | - | 561.1 | 885.7 | - | - | 0 | - |
| - | - | 676.4 | 1067 | - | - | 0 | - |
| - | - | 608.1 | 1407 | - | - | 0 | - |
| - | - | 680.9 | 2214 | - | - | 0 | - |

m/z Charge Intensity FragmentType MassShift Position
120.0238037109375 0 542.6389
120.08092498779297 0 28747.834
121.03961181640625 0 378.45462
121.06491088867188 0 479.75693
121.08425903320312 0 2494.6355
121.3470687866211 0 318.6099
123.0802001953125 0 511.52518
123.1170425415039 0 1400.2275
124.08700561523438 0 541.3893
125.05966186523438 0 418.99667
127.05059051513672 0 465.88483
127.07549285888672 0 486.89478
128.07057189941406 0 483.4533
128.1071014404297 0 988.36096
129.06576538085938 0 606.0918
129.1023712158203 0 7232.215
130.0499725341797 0 2372.385
130.06536865234375 0 1150.3318
130.08633422851562 0 1918.2494 y Ammonia loss 6
130.10562133789062 0 579.9305
131.97438049316406 0 492.68134
132.02468872070312 0 415.793
132.08096313476562 0 585.51843
133.0606689453125 0 496.79224
133.10142517089844 0 488.30634
135.3824005126953 0 369.4718
136.01869201660156 0 1231.3206
136.07595825195312 0 971.46216
137.08306884765625 0 472.30917
138.0343475341797 0 553.7601
138.0664825439453 0 634.6242
140.08197021484375 0 1081.0465
142.09796142578125 0 611.4155
142.1223602294922 0 491.43973
143.0941619873047 0 1302.6042
143.1181640625 0 1185.5515
145.04954528808594 0 852.1594
145.12242126464844 0 896.7391
146.060302734375 0 1027.0951
147.07664489746094 0 1917.5752
147.11302185058594 0 2764.8018 y 6
148.0184326171875 0 659.082
149.02352905273438 0 1585.6666
149.04495239257812 0 7731.384
152.0136260986328 0 532.1323
152.07015991210938 0 822.4822
155.01300048828125 0 612.34985
156.07681274414062 0 746.8151
157.0653533935547 0 735.6327
157.0976104736328 0 652.0622
159.03477478027344 0 565.9943
159.0764617919922 0 1082.457
159.091796875 0 1669.023
159.1129150390625 0 514.26514
163.0293731689453 0 585.395
167.05543518066406 0 13467.284
167.06808471679688 0 673.2627
171.11297607421875 0 1147.9852
171.1492919921875 0 943.96
175.0294647216797 0 1085.744
175.0867156982422 0 3112.9492
177.00863647460938 0 711.8848
177.045166015625 0 1263.7382
178.02890014648438 0 488.9526
179.1184844970703 0 556.14404
184.13409423828125 0 600.3139
185.16514587402344 0 985.1384
187.0715789794922 0 1350.4075
187.10818481445312 0 752.3491
187.1443328857422 0 1786.9309
189.04544067382812 0 710.6552
191.0244140625 0 1342.3467
191.1188507080078 0 466.15137 d 1
195.0879364013672 0 617.2281
199.16944885253906 0 3062.2986
201.06582641601562 0 935.3147
202.040283203125 0 876.1972
202.1074676513672 0 2509.8962
214.0402374267578 0 1182.7458
215.13877868652344 0 1603.8611
216.05580139160156 0 807.155
216.12326049804688 0 598.00134
217.09722900390625 0 3371.6155 a Water loss 1
225.04269409179688 0 1294.0569
226.1548309326172 0 1458.9956
227.0220184326172 0 717.3129
228.13479614257812 0 1146.834
230.0352020263672 0 726.00793
232.05104064941406 0 1083.2632
233.03628540039062 0 745.92773
233.16464233398438 0 552.67664
235.1076202392578 0 1848.3237 a 1
246.18109130859375 0 1358.8279 y 5
249.1123809814453 0 983.7673
251.15029907226562 0 902.59106
255.0649871826172 0 1695.7308
259.03607177734375 0 572.2761
263.10260009765625 0 2483.4763 b 1
264.6226806640625 0 555.0479
268.97796630859375 0 862.31177
271.06219482421875 0 977.3982
281.05157470703125 0 1356.5748
283.030517578125 0 1191.6053
283.0609436035156 0 606.9195
299.0617980957031 0 11129.356
299.2084655761719 0 702.7372
303.20257568359375 0 2935.9956 y 4
306.9031677246094 0 583.2393
327.0782470703125 0 12721.811
343.1614685058594 0 2054.8682
349.1485900878906 0 545.6153
367.06988525390625 0 705.9241
369.1233825683594 0 2970.5146
402.27117919921875 0 764.4713 y 3
403.2734375 0 683.5238
460.2380065917969 0 668.29156
473.3087158203125 0 5228.5757 y 2
474.3111877441406 0 792.27594
494.2357482910156 0 1026.1948
541.62451171875 0 662.70795
561.2674560546875 0 551.4133
588.3350219726562 0 6398.1143 y 1
589.337158203125 0 1582.2684
600.24169921875 0 571.89197
809.8988647460938 0 558.4701
885.7167358398438 0 561.0795
1067.27392578125 0 676.4313
1407.0760498046875 0 608.06305
2213.64306640625 0 680.8942

Spectrum Details

|  |  |
| --- | --- |
| Matched peaks? Matched peaks The total absolute number of peaks matched. Additionally in brackets the total fraction of peaks matched and the total number of peaks is shown. | 11 (8.53% of 129) |
| FDR? FDR The false discovery rate estimated for this peptide. It is calculated by matching all theoretical fragments with a non-integer shift with the raw peaks for this spectrum. This is done with 40 different shifts. The resulting percentage is the average number of annotated peaks over the number of annotated peaks with the correct spectrum. | 1.08% |
| Satellite FDR? Satellite FDR See the FDR for details on its calculation. This satellite ion specific FDR only contains the satellite ions (d/w) for I/L/J positions. | - |
| PSM Score? PSM Score The PSM Score as given by Hecklib to this annotated spectrum. It is shown with three significant figures. | 126 |

#### Reverse Lookup? Reverse Lookup All places where this read could be placed.

| Group | Segment | Template | Template Part | Read Part | Score | Unique |
| --- | --- | --- | --- | --- | --- | --- |
| Decoy | Decoy | THER | [541..548] | [0..7] | 56 | True |

| Recombined | Template Part | Read Part | Score | Unique |
| --- | --- | --- | --- | --- |
| THER | [541..548] | [0..7] | 56 | True |

#### Meta Information from Multiple reads

##### Number of combined reads

5

##### Intensity

0.6498

##### TotalArea

3.989E+07

#### Positional Score

Copy Data

##### Positional Score (TSV)

###### Preview

```
Loading example...
```

*Click on the button to copy the data to your clipboard.*

100123456

Label Value
"0" 0.582
"1" 0.582
"2" 0.564
"3" 0.562
"4" 0.58
"5" 0.592
"6" 0.596

#### Meta Information from PEAKS

##### Scan Identifier

F2:3941

##### Original sequence

F

D

A

V

G

V

K

##### Posttranslational Modifications

##### Source File

D:\separate\_stitch\_analyses\xle-disambiguation\raw\20210323\_F1\_UM1\_Peng0013\_SA\_F59\_ingel\_3ug\_TL.raw

##### Fraction

2

##### Scan Feature

F2:305

##### De Novo Score

98

##### ConfidenceScore

98

### m/z

368.2054

##### Mass

734.3962

##### Charge

2

##### Retention Time

20.66

##### Predicted Retention Time

-

##### Area

1.318E+07

##### Parts Per Million

0

##### Fragmentation mode

ETHCD

##### Originating file

01 D:\separate\_stitch\_analyses\xle-disambiguation\20210325\_F59\_3ug\_DENOVO\_12.csv

#### Meta Information from PEAKS

##### Scan Identifier

F2:4107

##### Original sequence

F

D

A

V

G

V

K

##### Posttranslational Modifications

##### Source File

D:\separate\_stitch\_analyses\xle-disambiguation\raw\20210323\_F1\_UM1\_Peng0013\_SA\_F59\_ingel\_3ug\_TL.raw

##### Fraction

2

##### Scan Feature

F2:305

##### De Novo Score

98

##### ConfidenceScore

98

### m/z

368.2054

##### Mass

734.3962

##### Charge

2

##### Retention Time

20.66

##### Predicted Retention Time

-

##### Area

1.318E+07

##### Parts Per Million

0

##### Fragmentation mode

HCD

##### Originating file

01 D:\separate\_stitch\_analyses\xle-disambiguation\20210325\_F59\_3ug\_DENOVO\_12.csv

#### Meta Information from PEAKS

##### Scan Identifier

F2:4009

##### Original sequence

F

D

A

V

G

V

K

##### Posttranslational Modifications

##### Source File

D:\separate\_stitch\_analyses\xle-disambiguation\raw\20210323\_F1\_UM1\_Peng0013\_SA\_F59\_ingel\_3ug\_TL.raw

##### Fraction

2

##### Scan Feature

F2:305

##### De Novo Score

97

##### ConfidenceScore

97

### m/z

368.2054

##### Mass

734.3962

##### Charge

2

##### Retention Time

20.66

##### Predicted Retention Time

-

##### Area

1.318E+07

##### Parts Per Million

0

##### Fragmentation mode

ETHCD

##### Originating file

01 D:\separate\_stitch\_analyses\xle-disambiguation\20210325\_F59\_3ug\_DENOVO\_12.csv

#### Meta Information from PEAKS

##### Scan Identifier

F2:4278

##### Original sequence

F

D

A

V

G

V

K

##### Posttranslational Modifications

##### Source File

D:\separate\_stitch\_analyses\xle-disambiguation\raw\20210323\_F1\_UM1\_Peng0013\_SA\_F59\_ingel\_3ug\_TL.raw

##### Fraction

2

##### Scan Feature

-

##### De Novo Score

97

##### ConfidenceScore

97

### m/z

368.2061

##### Mass

734.3962

##### Charge

2

##### Retention Time

22.8

##### Predicted Retention Time

-

##### Area

0

##### Parts Per Million

2

##### Fragmentation mode

ETHCD

##### Originating file

01 D:\separate\_stitch\_analyses\xle-disambiguation\20210325\_F59\_3ug\_DENOVO\_12.csv

#### Meta Information from PEAKS

##### Scan Identifier

F2:4671

##### Original sequence

F

D

A

V

G

V

K

##### Posttranslational Modifications

##### Source File

D:\separate\_stitch\_analyses\xle-disambiguation\raw\20210323\_F1\_UM1\_Peng0013\_SA\_F59\_ingel\_3ug\_TL.raw

##### Fraction

2

##### Scan Feature

F2:304

##### De Novo Score

95

##### ConfidenceScore

95

### m/z

368.2052

##### Mass

734.3962

##### Charge

2

##### Retention Time

23.65

##### Predicted Retention Time

-

##### Area

3.594E+05

##### Fragmentation mode

HCD

##### Originating file

01 D:\separate\_stitch\_analyses\xle-disambiguation\20210325\_F59\_3ug\_DENOVO\_12.csv
