## Supplementary material for "A handle on mass coincidence errors in *de novo* sequencing of antibodies by bottom-up proteomics": Combined_061.html

Details Combined\_061 | Stitch OverviewUndefined

### Read Combined\_061

#### Sequence (length=9)

JFPPSSEEJ

#### Spectrum 8530? Spectrum 8530 The raw spectrum of this peptide as annotated by Hecklib. The fragments are coloured according to ion type (see legend). Any peaks with a star '\*' as text can be hovered over to see the full details, first the ion type second the mass shift type. By hovering over the amino acids in the peptide or ions in the legend the corresponding peaks are highlighted. By toggling the 'Unassigned' label you can turn the background (unassigned) peaks on or off in the plot. By updating the slider in the Ion legend you can update the spectrum to only show the top X% of the peaks with labels. The top X% means any peak that is within X% of the highest intensity. By dragging in the spectrum you can zoom in to a specific part of the spectrum and use 'Zoom Out' to get back to the original zoom level. The annotation of the spectrum is based on the given sequence in the peptides file and is done with different software so inconsistencies are likely. The peaks are annotated based on the given sequence, with 20 ppm tolerance.

Copy Data

##### Spectrum 8530 (TSV)

###### Preview

```
Loading example...
```

*Click on the button to copy the data to your clipboard.*

Mz MinMz MaxIntensity Max

WidthHeightPeptide font sizePeptide stroke widthSpectrum font sizeSpectrum stroke widthCompact peptide

Ion legend

wxyz

abcd

OtherUnassignedIon^Charge^_Position_Show for top:%

JFPPSSEEJ

06.77e+41.35e+52.03e+52.71e+5

Zoom Out

y^+1^_1_y^+1^_2_y^+1^_2_c^+1^_2_z^+2^_6_c^+1^_3_y^+2^_7_y^+1^_3_c^+2^_7_c^+2^_7_y^+1^_3_c^+1^_4_c^+1^_4_y^+1^_4_c^+1^_5_c^+1^_5_y^+1^_5_c^+1^_6_y^+1^_6_z^+1^_6_c^+1^_6_y^+1^_6_y^+1^_7_c^+1^_7_c^+1^_7_z^+1^_8_c^+1^_8_z^+1^_8_c^+1^_8_y^+1^_8_

038777311601546

Fragment Matches Table

Show background peaks

| Position | Ion type | Intensity | mz Theoretical | mz Error (Th) | mz Error (ppm) | Charge | Series Number |
| --- | --- | --- | --- | --- | --- | --- | --- |
| - | - | 5812 | 120.1 | - | - | 0 | - |
| - | - | 1192 | 130.1 | - | - | 0 | - |
| 9 | y | 3.72E+04 | 132.1 | 0.0002067 | 1.565 | +1 | 1 |
| - | - | 2581 | 133.1 | - | - | 0 | - |
| - | - | 437.8 | 144.6 | - | - | 0 | - |
| - | - | 465.4 | 154.3 | - | - | 0 | - |
| - | - | 460.5 | 164.1 | - | - | 0 | - |
| - | - | 419.4 | 175.8 | - | - | 0 | - |
| - | - | 527.8 | 195.1 | - | - | 0 | - |
| - | - | 1992 | 197.1 | - | - | 0 | - |
| - | - | 1634 | 199.1 | - | - | 0 | - |
| - | - | 466.2 | 199.1 | - | - | 0 | - |
| - | - | 3718 | 212.1 | - | - | 0 | - |
| - | - | 830.7 | 217.1 | - | - | 0 | - |
| - | - | 535.6 | 221.1 | - | - | 0 | - |
| - | - | 547.1 | 225 | - | - | 0 | - |
| - | - | 3.088E+04 | 233.2 | - | - | 0 | - |
| - | - | 4276 | 234.2 | - | - | 0 | - |
| - | - | 1514 | 241.1 | - | - | 0 | - |
| 8 | y | 2.007E+04 | 243.1 | 0.0001912 | 0.7865 | +1 | 2 |
| - | - | 2156 | 244.1 | - | - | 0 | - |
| - | - | 1581 | 259.1 | - | - | 0 | - |
| 8 | y | 5886 | 261.1 | 0.0001196 | 0.4578 | +1 | 2 |
| 2 | c | 2.179E+04 | 261.2 | 5.172E-06 | 0.0198 | +1 | 2 |
| - | - | 733.2 | 262.1 | - | - | 0 | - |
| - | - | 3216 | 262.2 | - | - | 0 | - |
| - | - | 1293 | 264.1 | - | - | 0 | - |
| - | - | 6811 | 282.1 | - | - | 0 | - |
| - | - | 738.6 | 283.1 | - | - | 0 | - |
| - | - | 1043 | 296.1 | - | - | 0 | - |
| - | - | 1.6E+04 | 299.2 | - | - | 0 | - |
| - | - | 1766 | 300.2 | - | - | 0 | - |
| - | - | 547.3 | 309.2 | - | - | 0 | - |
| 4 | z | 1600 | 314.1 | 0.005818 | 18.52 | +2 | 6 |
| - | - | 647.1 | 314.6 | - | - | 0 | - |
| - | - | 611.4 | 328.1 | - | - | 0 | - |
| - | - | 1205 | 338.7 | - | - | 0 | - |
| - | - | 1527 | 346.1 | - | - | 0 | - |
| - | - | 1983 | 351.2 | - | - | 0 | - |
| - | - | 551.4 | 352.2 | - | - | 0 | - |
| - | - | 796.3 | 354.2 | - | - | 0 | - |
| - | - | 509.4 | 357.2 | - | - | 0 | - |
| 3 | c | 2507 | 358.2 | 0.0005736 | 1.601 | +1 | 3 |
| - | - | 498.6 | 359.2 | - | - | 0 | - |
| - | - | 3917 | 369.2 | - | - | 0 | - |
| 3 | y | 905.9 | 370.7 | 0.005801 | 15.65 | +2 | 7 |
| 7 | y | 1033 | 372.2 | 0.0003486 | 0.9368 | +1 | 3 |
| 7 | c | 732.9 | 379.7 | 0.006957 | 18.32 | +2 | 7 |
| - | - | 1074 | 380.2 | - | - | 0 | - |
| - | - | 3.718E+04 | 386.2 | - | - | 0 | - |
| - | - | 6308 | 387.2 | - | - | 0 | - |
| 7 | c | 1006 | 388.2 | 0.004458 | 11.48 | +2 | 7 |
| 7 | y | 1075 | 390.2 | 0.0002932 | 0.7515 | +1 | 3 |
| - | - | 789.6 | 401.2 | - | - | 0 | - |
| - | - | 763.7 | 412.2 | - | - | 0 | - |
| - | - | 846.5 | 427.3 | - | - | 0 | - |
| - | - | 6086 | 428.3 | - | - | 0 | - |
| - | - | 2942 | 429.3 | - | - | 0 | - |
| - | - | 680.4 | 430.3 | - | - | 0 | - |
| - | - | 666 | 433.2 | - | - | 0 | - |
| - | - | 639 | 453.9 | - | - | 0 | - |
| 4 | c | 1954 | 455.3 | 0.0002064 | 0.4533 | +1 | 4 |
| - | - | 1549 | 456.3 | - | - | 0 | - |
| - | - | 1042 | 462.2 | - | - | 0 | - |
| - | - | 1.483E+04 | 471.3 | - | - | 0 | - |
| 4 | c | 1.513E+04 | 472.3 | 0.0008155 | 1.727 | +1 | 4 |
| - | - | 4861 | 473.3 | - | - | 0 | - |
| - | - | 1844 | 474.3 | - | - | 0 | - |
| 6 | y | 787 | 477.2 | 0.001919 | 4.021 | +1 | 4 |
| - | - | 3219 | 480.2 | - | - | 0 | - |
| - | - | 721 | 481.2 | - | - | 0 | - |
| - | - | 1086 | 497.3 | - | - | 0 | - |
| - | - | 1.665E+04 | 498.2 | - | - | 0 | - |
| - | - | 1657 | 498.3 | - | - | 0 | - |
| - | - | 3884 | 499.2 | - | - | 0 | - |
| - | - | 756 | 500.2 | - | - | 0 | - |
| - | - | 588.5 | 509.5 | - | - | 0 | - |
| - | - | 828.5 | 512.2 | - | - | 0 | - |
| - | - | 1.741E+04 | 515.2 | - | - | 0 | - |
| - | - | 3721 | 516.2 | - | - | 0 | - |
| - | - | 3234 | 516.3 | - | - | 0 | - |
| - | - | 568.8 | 516.4 | - | - | 0 | - |
| - | - | 1636 | 530.2 | - | - | 0 | - |
| - | - | 677.7 | 531.2 | - | - | 0 | - |
| 5 | c | 2932 | 542.3 | 0.0003134 | 0.5779 | +1 | 5 |
| - | - | 949.2 | 543.3 | - | - | 0 | - |
| - | - | 3707 | 544.3 | - | - | 0 | - |
| - | - | 1363 | 545.3 | - | - | 0 | - |
| - | - | 4.101E+04 | 558.3 | - | - | 0 | - |
| 5 | c | 3.159E+04 | 559.3 | 0.002021 | 3.614 | +1 | 5 |
| - | - | 9192 | 560.3 | - | - | 0 | - |
| - | - | 2261 | 561.3 | - | - | 0 | - |
| 5 | y | 1231 | 564.3 | 0.002308 | 4.09 | +1 | 5 |
| - | - | 721.1 | 575.2 | - | - | 0 | - |
| - | - | 1008 | 591.2 | - | - | 0 | - |
| - | - | 751 | 593.3 | - | - | 0 | - |
| - | - | 1883 | 599.3 | - | - | 0 | - |
| - | - | 894.3 | 601.3 | - | - | 0 | - |
| - | - | 831.4 | 601.3 | - | - | 0 | - |
| - | - | 654.7 | 602.3 | - | - | 0 | - |
| - | - | 1463 | 603.4 | - | - | 0 | - |
| - | - | 4370 | 609.3 | - | - | 0 | - |
| - | - | 1076 | 610.3 | - | - | 0 | - |
| - | - | 2754 | 611.3 | - | - | 0 | - |
| - | - | 691.8 | 612.3 | - | - | 0 | - |
| - | - | 2315 | 626.3 | - | - | 0 | - |
| - | - | 4.473E+04 | 627.3 | - | - | 0 | - |
| - | - | 1.378E+04 | 628.3 | - | - | 0 | - |
| - | - | 2490 | 629.3 | - | - | 0 | - |
| 6 | c | 5825 | 629.3 | 6.828E-06 | 0.01085 | +1 | 6 |
| - | - | 698.3 | 630.3 | - | - | 0 | - |
| - | - | 1789 | 630.3 | - | - | 0 | - |
| 4 | y | 1077 | 643.3 | 0.002368 | 3.681 | +1 | 6 |
| - | - | 1790 | 644.3 | - | - | 0 | - |
| 4 | z | 977.5 | 645.3 | 0.006316 | 9.788 | +1 | 6 |
| - | - | 1197 | 645.3 | - | - | 0 | - |
| 6 | c | 6.845E+04 | 646.4 | 5.302E-05 | 0.08202 | +1 | 6 |
| - | - | 2.37E+04 | 647.4 | - | - | 0 | - |
| - | - | 4496 | 648.4 | - | - | 0 | - |
| - | - | 1729 | 660.3 | - | - | 0 | - |
| 4 | y | 1.667E+04 | 661.3 | 0.0003231 | 0.4886 | +1 | 6 |
| - | - | 6341 | 662.3 | - | - | 0 | - |
| - | - | 1106 | 663.3 | - | - | 0 | - |
| - | - | 3698 | 672.4 | - | - | 0 | - |
| - | - | 1595 | 673.4 | - | - | 0 | - |
| - | - | 1181 | 685.3 | - | - | 0 | - |
| - | - | 824.4 | 686.4 | - | - | 0 | - |
| - | - | 1030 | 689.3 | - | - | 0 | - |
| - | - | 663.8 | 699.3 | - | - | 0 | - |
| - | - | 1749 | 701.3 | - | - | 0 | - |
| - | - | 772.2 | 712.4 | - | - | 0 | - |
| - | - | 1206 | 714.3 | - | - | 0 | - |
| - | - | 1548 | 722.3 | - | - | 0 | - |
| - | - | 776.3 | 723.3 | - | - | 0 | - |
| - | - | 2220 | 730.4 | - | - | 0 | - |
| - | - | 1051 | 731.4 | - | - | 0 | - |
| - | - | 1895 | 732.4 | - | - | 0 | - |
| - | - | 687.8 | 733.4 | - | - | 0 | - |
| - | - | 8145 | 739.3 | - | - | 0 | - |
| 3 | y | 1.067E+04 | 740.3 | 0.0008737 | 1.18 | +1 | 7 |
| - | - | 3593 | 741.4 | - | - | 0 | - |
| - | - | 1971 | 742.4 | - | - | 0 | - |
| - | - | 1214 | 743.4 | - | - | 0 | - |
| - | - | 5.264E+04 | 757.3 | - | - | 0 | - |
| 7 | c | 1.934E+05 | 758.4 | 0.01482 | 19.54 | +1 | 7 |
| - | - | 6.778E+04 | 759.4 | - | - | 0 | - |
| - | - | 1.762E+04 | 760.4 | - | - | 0 | - |
| - | - | 2946 | 761.4 | - | - | 0 | - |
| - | - | 709.9 | 774.3 | - | - | 0 | - |
| 7 | c | 2.021E+05 | 775.4 | 0.0004708 | 0.6072 | +1 | 7 |
| - | - | 8.891E+04 | 776.4 | - | - | 0 | - |
| - | - | 2.065E+04 | 777.4 | - | - | 0 | - |
| - | - | 2297 | 778.4 | - | - | 0 | - |
| - | - | 1307 | 785.3 | - | - | 0 | - |
| - | - | 630.1 | 787.4 | - | - | 0 | - |
| - | - | 763.7 | 788.4 | - | - | 0 | - |
| - | - | 682.5 | 799.4 | - | - | 0 | - |
| - | - | 1.541E+04 | 801.4 | - | - | 0 | - |
| - | - | 7419 | 802.4 | - | - | 0 | - |
| - | - | 1084 | 803.4 | - | - | 0 | - |
| - | - | 1060 | 814.4 | - | - | 0 | - |
| - | - | 695.4 | 815.4 | - | - | 0 | - |
| - | - | 615.1 | 834.4 | - | - | 0 | - |
| - | - | 818.1 | 843.4 | - | - | 0 | - |
| - | - | 885.7 | 851.4 | - | - | 0 | - |
| - | - | 751.6 | 857.5 | - | - | 0 | - |
| - | - | 3679 | 859.4 | - | - | 0 | - |
| - | - | 4931 | 860.4 | - | - | 0 | - |
| - | - | 3219 | 861.4 | - | - | 0 | - |
| - | - | 963.5 | 862.4 | - | - | 0 | - |
| - | - | 1673 | 869.4 | - | - | 0 | - |
| - | - | 1341 | 870.4 | - | - | 0 | - |
| 2 | z | 4708 | 871.4 | 0.0004392 | 0.5041 | +1 | 8 |
| - | - | 2408 | 872.4 | - | - | 0 | - |
| - | - | 781.6 | 873.4 | - | - | 0 | - |
| 8 | c | 2.818E+04 | 887.4 | 0.001073 | 1.209 | +1 | 8 |
| - | - | 1.213E+04 | 888.4 | - | - | 0 | - |
| 2 | z | 1.677E+04 | 889.4 | 0.00196 | 2.203 | +1 | 8 |
| - | - | 7626 | 890.4 | - | - | 0 | - |
| - | - | 2069 | 891.4 | - | - | 0 | - |
| - | - | 696.5 | 901.5 | - | - | 0 | - |
| 8 | c | 2.682E+05 | 904.4 | 0.0004003 | 0.4426 | +1 | 8 |
| 2 | y | 1.297E+05 | 905.4 | 0.01803 | 19.91 | +1 | 8 |
| - | - | 3.895E+04 | 906.4 | - | - | 0 | - |
| - | - | 3320 | 907.4 | - | - | 0 | - |
| - | - | 685.6 | 917.4 | - | - | 0 | - |
| - | - | 731.1 | 933.5 | - | - | 0 | - |
| - | - | 1.391E+04 | 947.5 | - | - | 0 | - |
| - | - | 8010 | 948.5 | - | - | 0 | - |
| - | - | 2517 | 949.5 | - | - | 0 | - |
| - | - | 999.2 | 957.5 | - | - | 0 | - |
| - | - | 944.6 | 958.5 | - | - | 0 | - |
| - | - | 947.1 | 959.4 | - | - | 0 | - |
| - | - | 4183 | 963.5 | - | - | 0 | - |
| - | - | 2095 | 964.5 | - | - | 0 | - |
| - | - | 702.7 | 970.8 | - | - | 0 | - |
| - | - | 873.9 | 971.5 | - | - | 0 | - |
| - | - | 988.4 | 972.5 | - | - | 0 | - |
| - | - | 1616 | 973.5 | - | - | 0 | - |
| - | - | 1730 | 974.5 | - | - | 0 | - |
| - | - | 936.2 | 975.5 | - | - | 0 | - |
| - | - | 4790 | 984.5 | - | - | 0 | - |
| - | - | 2910 | 985.5 | - | - | 0 | - |
| - | - | 3856 | 991.5 | - | - | 0 | - |
| - | - | 1783 | 992.5 | - | - | 0 | - |
| - | - | 757.4 | 1000 | - | - | 0 | - |
| - | - | 2486 | 1001 | - | - | 0 | - |
| - | - | 2.676E+04 | 1002 | - | - | 0 | - |
| - | - | 1.638E+04 | 1003 | - | - | 0 | - |
| - | - | 5645 | 1004 | - | - | 0 | - |
| - | - | 1575 | 1005 | - | - | 0 | - |
| - | - | 6.121E+04 | 1019 | - | - | 0 | - |
| - | - | 2.482E+05 | 1020 | - | - | 0 | - |
| - | - | 1.322E+05 | 1021 | - | - | 0 | - |
| - | - | 4.133E+04 | 1022 | - | - | 0 | - |
| - | - | 4033 | 1023 | - | - | 0 | - |
| - | - | 1722 | 1052 | - | - | 0 | - |
| - | - | 861.2 | 1053 | - | - | 0 | - |
| - | - | 666.9 | 1056 | - | - | 0 | - |
| - | - | 1367 | 1374 | - | - | 0 | - |
| - | - | 1119 | 1513 | - | - | 0 | - |
| - | - | 1025 | 1514 | - | - | 0 | - |
| - | - | 764.6 | 1516 | - | - | 0 | - |
| - | - | 755.9 | 1531 | - | - | 0 | - |

m/z Charge Intensity FragmentType MassShift Position
120.08098602294922 0 5812.241
130.0501251220703 0 1191.516
132.10211181640625 0 37195.902 y 8
133.10543823242188 0 2581.2874
144.6355438232422 0 437.82538
154.27322387695312 0 465.35422
164.07693481445312 0 460.5362
175.81573486328125 0 419.4051
195.1126251220703 0 527.80774
197.12867736816406 0 1991.888
199.07174682617188 0 1634.2302
199.12661743164062 0 466.18842
212.13946533203125 0 3717.6057
217.1459503173828 0 830.663
221.0924530029297 0 535.5746
224.9877166748047 0 547.1299
233.1649627685547 0 30876.648
234.16854858398438 0 4275.929
241.08201599121094 0 1513.8442
243.13412475585938 0 20074.357 y Water loss 7
244.1371612548828 0 2155.764
259.0921325683594 0 1581.0604
261.1443786621094 0 5886.4966 y 7
261.1597595214844 0 21792.463 c Ammonia loss 1
262.1485595703125 0 733.2487
262.1632080078125 0 3215.6453
264.1343688964844 0 1292.7379
282.14501953125 0 6810.663
283.1484375 0 738.5602
296.12481689453125 0 1043.1506
299.171630859375 0 16003.622
300.174560546875 0 1765.6257
309.2038269042969 0 547.25073
314.1351318359375 0 1600.2368 z Water loss 3
314.63763427734375 0 647.1492
328.1163330078125 0 611.357
338.6687927246094 0 1204.6493
346.1244201660156 0 1527.2124
351.1661376953125 0 1983.4843
352.16998291015625 0 551.4065
354.1659851074219 0 796.3177
357.2054748535156 0 509.41742
358.2119445800781 0 2507.2974 c Ammonia loss 2
359.2134704589844 0 498.60873
369.177001953125 0 3916.9382
370.6824951171875 0 905.8741 y Water loss 2
372.1761779785156 0 1032.8839 y Water loss 6
379.6826477050781 0 732.87805 c Ammonia loss 6
380.1852722167969 0 1073.675
386.2035827636719 0 37184.918
387.2066650390625 0 6307.6436
388.20733642578125 0 1005.7667 c 6
390.1867980957031 0 1074.6477 y 6
401.1669006347656 0 789.6476
412.2087097167969 0 763.6839
427.2734069824219 0 846.47687
428.27813720703125 0 6086.4194
429.2833251953125 0 2941.908
430.2859191894531 0 680.3564
433.15948486328125 0 666.04175
453.94464111328125 0 638.99164
455.26507568359375 0 1953.7411 c Ammonia loss 3
456.2722473144531 0 1549.338
462.1978759765625 0 1041.8674
471.28436279296875 0 14827.894
472.291015625 0 15131.643 c 3
473.2951354980469 0 4861.175
474.30474853515625 0 1843.6755
477.2210388183594 0 786.99725 y 5
480.2099609375 0 3218.6921
481.20977783203125 0 721.0475
497.3001708984375 0 1086.2233
498.2197570800781 0 16654.416
498.3067321777344 0 1657.4333
499.2233581542969 0 3884.2869
500.22137451171875 0 756.0122
509.5024108886719 0 588.52106
512.196533203125 0 828.5005
515.2459106445312 0 17414.334
516.2483520507812 0 3720.5332
516.3168334960938 0 3234.263
516.3552856445312 0 568.78357
530.20849609375 0 1635.6987
531.2124633789062 0 677.66833
542.2969970703125 0 2932.0195 c Ammonia loss 4
543.3021240234375 0 949.15497
544.3125610351562 0 3707.1917
545.3153076171875 0 1363.0559
558.3160400390625 0 41008.26
559.3218383789062 0 31585.535 c 4
560.3256225585938 0 9192.332
561.33544921875 0 2261.0078
564.2488403320312 0 1230.9379 y 4
575.24072265625 0 721.0776
591.2421875 0 1007.93335
593.3027954101562 0 751.04895
599.2672119140625 0 1883.065
601.2767333984375 0 894.3133
601.3314208984375 0 831.38916
602.3414916992188 0 654.72577
603.3501586914062 0 1462.6897
609.2518310546875 0 4370.0166
610.2553100585938 0 1075.8391
611.3146362304688 0 2753.6824
612.316650390625 0 691.80176
626.2556762695312 0 2314.6904
627.261962890625 0 44733.25
628.2647094726562 0 13779.129
629.2685546875 0 2489.9966
629.329345703125 0 5824.576 c Ammonia loss 5
630.2750244140625 0 698.28546
630.3323364257812 0 1789.279
643.2957153320312 0 1077.1058 y Water loss 3
644.28955078125 0 1790.3623
645.29150390625 0 977.5098 z 3
645.3474731445312 0 1197.157
646.3558349609375 0 68447.234 c 5
647.3585815429688 0 23702.06
648.36181640625 0 4496.277
660.2955932617188 0 1729.3671
661.3035888671875 0 16673.635 y 3
662.3065795898438 0 6340.916
663.3118286132812 0 1105.9686
672.3703002929688 0 3697.8447
673.3734130859375 0 1594.9023
685.3268432617188 0 1180.8733
686.3754272460938 0 824.3664
689.2984008789062 0 1030.448
699.3385620117188 0 663.82263
701.287841796875 0 1748.8596
712.352783203125 0 772.179
714.2946166992188 0 1206.3228
722.3455200195312 0 1547.8057
723.338623046875 0 776.2769
730.377197265625 0 2219.6516
731.3838500976562 0 1050.9309
732.3895874023438 0 1894.8285
733.3976440429688 0 687.84454
739.3372802734375 0 8144.978
740.3469848632812 0 10669.173 y Water loss 2
741.3513793945312 0 3593.2239
742.3532104492188 0 1971.2303
743.3534545898438 0 1213.5212
757.3483276367188 0 52639.492
758.3571166992188 0 193436.67 c Ammonia loss 6
759.3601684570312 0 67775.055
760.3651123046875 0 17622.65
761.370849609375 0 2946.1707
774.3280639648438 0 709.8839
775.3980102539062 0 202117.08 c 6
776.4012451171875 0 88909.63
777.404052734375 0 20647.256
778.4075317382812 0 2297.2996
785.3439331054688 0 1306.5071
787.3551635742188 0 630.0539
788.4077758789062 0 763.68787
799.4208984375 0 682.53925
801.4139404296875 0 15407.817
802.416748046875 0 7418.727
803.417236328125 0 1084.0585
814.4302978515625 0 1060.1881
815.43310546875 0 695.3776
834.357421875 0 615.07874
843.4285888671875 0 818.0737
851.3932495117188 0 885.7004
857.4652709960938 0 751.57117
859.419189453125 0 3679.4573
860.4259643554688 0 4931.4995
861.4304809570312 0 3218.584
862.4395141601562 0 963.5244
869.39990234375 0 1673.4482
870.409423828125 0 1340.7355
871.396240234375 0 4708.1157 z Water loss 1
872.39892578125 0 2408.2444
873.402587890625 0 781.6411
887.4134521484375 0 28175.096 c Ammonia loss 7
888.4168701171875 0 12131.856
889.4083251953125 0 16769.672 z 1
890.4094848632812 0 7626.411
891.4110717773438 0 2068.996
901.4859008789062 0 696.5223
904.440673828125 0 268160.12 c 7
905.443115234375 0 129685.74 y 1
906.4459838867188 0 38952.53
907.4470825195312 0 3319.5024
917.4468994140625 0 685.61926
933.4888916015625 0 731.0578
947.4949951171875 0 13907.221
948.4973754882812 0 8009.616
949.49755859375 0 2516.8171
957.520263671875 0 999.1861
958.5279541015625 0 944.6206
959.4324340820312 0 947.13995
963.4523315429688 0 4182.744
964.4549560546875 0 2094.677
970.8251342773438 0 702.68744
971.50390625 0 873.8685
972.4962158203125 0 988.3503
973.5060424804688 0 1616.3973
974.5125732421875 0 1729.9186
975.512939453125 0 936.23846
984.4780883789062 0 4790.1323
985.4820556640625 0 2910.423
991.523681640625 0 3856.4907
992.5262451171875 0 1783.1378
1000.4778442382812 0 757.3945
1001.4993896484375 0 2485.5928
1002.4902954101562 0 26758.875
1003.492919921875 0 16382.581
1004.494873046875 0 5644.8716
1005.4906005859375 0 1574.9635
1018.507568359375 0 61211.426
1019.51513671875 0 248215.52
1020.5184936523438 0 132234.58
1021.5215454101562 0 41329.617
1022.5235595703125 0 4033.4185
1051.508056640625 0 1722.4053
1052.515625 0 861.19275
1056.3970947265625 0 666.9147
1373.7034912109375 0 1367.1674
1512.8717041015625 0 1118.7377
1513.8690185546875 0 1024.5803
1515.77197265625 0 764.57263
1530.860107421875 0 755.92664

Spectrum Details

|  |  |
| --- | --- |
| Matched peaks? Matched peaks The total absolute number of peaks matched. Additionally in brackets the total fraction of peaks matched and the total number of peaks is shown. | 30 (13.39% of 224) |
| FDR? FDR The false discovery rate estimated for this peptide. It is calculated by matching all theoretical fragments with a non-integer shift with the raw peaks for this spectrum. This is done with 40 different shifts. The resulting percentage is the average number of annotated peaks over the number of annotated peaks with the correct spectrum. | 1.03% |
| Satellite FDR? Satellite FDR See the FDR for details on its calculation. This satellite ion specific FDR only contains the satellite ions (d/w) for I/L/J positions. | - |
| PSM Score? PSM Score The PSM Score as given by Hecklib to this annotated spectrum. It is shown with three significant figures. | 387 |

#### Spectrum 8467? Spectrum 8467 The raw spectrum of this peptide as annotated by Hecklib. The fragments are coloured according to ion type (see legend). Any peaks with a star '\*' as text can be hovered over to see the full details, first the ion type second the mass shift type. By hovering over the amino acids in the peptide or ions in the legend the corresponding peaks are highlighted. By toggling the 'Unassigned' label you can turn the background (unassigned) peaks on or off in the plot. By updating the slider in the Ion legend you can update the spectrum to only show the top X% of the peaks with labels. The top X% means any peak that is within X% of the highest intensity. By dragging in the spectrum you can zoom in to a specific part of the spectrum and use 'Zoom Out' to get back to the original zoom level. The annotation of the spectrum is based on the given sequence in the peptides file and is done with different software so inconsistencies are likely. The peaks are annotated based on the given sequence, with 20 ppm tolerance.

Copy Data

##### Spectrum 8467 (TSV)

###### Preview

```
Loading example...
```

*Click on the button to copy the data to your clipboard.*

Mz MinMz MaxIntensity Max

WidthHeightPeptide font sizePeptide stroke widthSpectrum font sizeSpectrum stroke widthCompact peptide

Ion legend

wxyz

abcd

OtherUnassignedIon^Charge^_Position_Show for top:%

JFPPSSEEJ

01.16e+52.33e+53.49e+54.65e+5

Zoom Out

y^+1^_1_y^+1^_2_y^+1^_2_c^+1^_2_z^+2^_6_c^+1^_3_y^+2^_7_y^+1^_3_c^+2^_7_c^+2^_7_y^+1^_3_c^+2^_8_c^+1^_4_y^+1^_4_c^+1^_4_y^+1^_4_c^+1^_5_c^+1^_5_y^+1^_5_z^+1^_6_c^+1^_6_c^+1^_6_y^+1^_6_z^+1^_6_c^+1^_6_y^+1^_6_y^+1^_7_c^+1^_7_c^+1^_7_z^+1^_8_c^+1^_8_z^+1^_8_c^+1^_8_

040180312041605

Fragment Matches Table

Show background peaks

| Position | Ion type | Intensity | mz Theoretical | mz Error (Th) | mz Error (ppm) | Charge | Series Number |
| --- | --- | --- | --- | --- | --- | --- | --- |
| - | - | 1.014E+04 | 120.1 | - | - | 0 | - |
| - | - | 891.2 | 121.1 | - | - | 0 | - |
| - | - | 401.9 | 128.3 | - | - | 0 | - |
| - | - | 447.7 | 129.1 | - | - | 0 | - |
| - | - | 430.8 | 129.1 | - | - | 0 | - |
| - | - | 2209 | 130.1 | - | - | 0 | - |
| 9 | y | 7.486E+04 | 132.1 | 0.000283 | 2.142 | +1 | 1 |
| - | - | 582.1 | 133.1 | - | - | 0 | - |
| - | - | 518.9 | 133.1 | - | - | 0 | - |
| - | - | 3457 | 133.1 | - | - | 0 | - |
| - | - | 411.4 | 166.3 | - | - | 0 | - |
| - | - | 448.2 | 167.1 | - | - | 0 | - |
| - | - | 600.4 | 168.1 | - | - | 0 | - |
| - | - | 471.9 | 171.8 | - | - | 0 | - |
| - | - | 473.7 | 184.6 | - | - | 0 | - |
| - | - | 488.2 | 189.9 | - | - | 0 | - |
| - | - | 3911 | 197.1 | - | - | 0 | - |
| - | - | 2186 | 199.1 | - | - | 0 | - |
| - | - | 506.6 | 209.4 | - | - | 0 | - |
| - | - | 4859 | 212.1 | - | - | 0 | - |
| - | - | 603.5 | 215.1 | - | - | 0 | - |
| - | - | 2282 | 217.1 | - | - | 0 | - |
| - | - | 954.2 | 217.1 | - | - | 0 | - |
| - | - | 902.9 | 221.1 | - | - | 0 | - |
| - | - | 474.5 | 229.1 | - | - | 0 | - |
| - | - | 5.955E+04 | 233.2 | - | - | 0 | - |
| - | - | 9639 | 234.2 | - | - | 0 | - |
| - | - | 3256 | 241.1 | - | - | 0 | - |
| 8 | y | 3.857E+04 | 243.1 | 0.0003286 | 1.351 | +1 | 2 |
| - | - | 1157 | 243.1 | - | - | 0 | - |
| - | - | 4306 | 244.1 | - | - | 0 | - |
| - | - | 3503 | 259.1 | - | - | 0 | - |
| 8 | y | 1.141E+04 | 261.1 | 2.516E-06 | 0.009636 | +1 | 2 |
| 2 | c | 3.816E+04 | 261.2 | 0.0001272 | 0.4872 | +1 | 2 |
| - | - | 848.2 | 262.1 | - | - | 0 | - |
| - | - | 5923 | 262.2 | - | - | 0 | - |
| - | - | 645.3 | 263.2 | - | - | 0 | - |
| - | - | 1702 | 264.1 | - | - | 0 | - |
| - | - | 495 | 276.2 | - | - | 0 | - |
| - | - | 826 | 282.1 | - | - | 0 | - |
| - | - | 1.296E+04 | 282.1 | - | - | 0 | - |
| - | - | 1624 | 283.1 | - | - | 0 | - |
| - | - | 752.5 | 286.1 | - | - | 0 | - |
| - | - | 752.8 | 287.1 | - | - | 0 | - |
| - | - | 1826 | 296.1 | - | - | 0 | - |
| - | - | 969.1 | 296.2 | - | - | 0 | - |
| - | - | 859.5 | 297.2 | - | - | 0 | - |
| - | - | 746.7 | 298.2 | - | - | 0 | - |
| - | - | 2.354E+04 | 299.2 | - | - | 0 | - |
| - | - | 3430 | 300.2 | - | - | 0 | - |
| - | - | 578.3 | 304.1 | - | - | 0 | - |
| - | - | 1290 | 305.1 | - | - | 0 | - |
| - | - | 566.3 | 305.6 | - | - | 0 | - |
| - | - | 464.4 | 306.8 | - | - | 0 | - |
| - | - | 537.5 | 308.1 | - | - | 0 | - |
| 4 | z | 2198 | 314.1 | 0.005543 | 17.65 | +2 | 6 |
| - | - | 1051 | 314.6 | - | - | 0 | - |
| - | - | 522 | 322.9 | - | - | 0 | - |
| - | - | 727 | 326.2 | - | - | 0 | - |
| - | - | 907.9 | 329.1 | - | - | 0 | - |
| - | - | 1038 | 333.2 | - | - | 0 | - |
| - | - | 3268 | 338.7 | - | - | 0 | - |
| - | - | 940.8 | 339.2 | - | - | 0 | - |
| - | - | 2720 | 346.1 | - | - | 0 | - |
| - | - | 764.8 | 348.2 | - | - | 0 | - |
| - | - | 3964 | 351.2 | - | - | 0 | - |
| - | - | 1134 | 352.2 | - | - | 0 | - |
| - | - | 1150 | 354.2 | - | - | 0 | - |
| - | - | 1161 | 355.2 | - | - | 0 | - |
| - | - | 654.4 | 357.2 | - | - | 0 | - |
| 3 | c | 6005 | 358.2 | 0.0001588 | 0.4433 | +1 | 3 |
| - | - | 1433 | 359.2 | - | - | 0 | - |
| - | - | 774.4 | 361.7 | - | - | 0 | - |
| - | - | 6828 | 369.2 | - | - | 0 | - |
| - | - | 937.5 | 370.2 | - | - | 0 | - |
| 3 | y | 2350 | 370.7 | 0.006442 | 17.38 | +2 | 7 |
| - | - | 1568 | 371.2 | - | - | 0 | - |
| 7 | y | 2031 | 372.2 | 0.001055 | 2.835 | +1 | 3 |
| 7 | c | 2238 | 379.7 | 0.005461 | 14.38 | +2 | 7 |
| - | - | 2256 | 380.2 | - | - | 0 | - |
| - | - | 1219 | 380.2 | - | - | 0 | - |
| - | - | 5.972E+04 | 386.2 | - | - | 0 | - |
| - | - | 1.077E+04 | 387.2 | - | - | 0 | - |
| 7 | c | 964.9 | 388.2 | 0.003634 | 9.36 | +2 | 7 |
| 7 | y | 1405 | 390.2 | 0.0004697 | 1.204 | +1 | 3 |
| - | - | 733.6 | 397.2 | - | - | 0 | - |
| - | - | 837 | 398.2 | - | - | 0 | - |
| - | - | 981.1 | 401.2 | - | - | 0 | - |
| - | - | 1152 | 412.2 | - | - | 0 | - |
| - | - | 1055 | 415.3 | - | - | 0 | - |
| - | - | 796.9 | 417.2 | - | - | 0 | - |
| - | - | 604.8 | 426.7 | - | - | 0 | - |
| - | - | 1205 | 427.3 | - | - | 0 | - |
| - | - | 1.277E+04 | 428.3 | - | - | 0 | - |
| - | - | 5662 | 429.3 | - | - | 0 | - |
| - | - | 684.8 | 431.2 | - | - | 0 | - |
| - | - | 1984 | 433.2 | - | - | 0 | - |
| 8 | c | 1490 | 444.2 | 0.0003724 | 0.8384 | +2 | 8 |
| - | - | 688.9 | 455.2 | - | - | 0 | - |
| 4 | c | 3228 | 455.3 | 0.0001148 | 0.2522 | +1 | 4 |
| - | - | 2849 | 456.3 | - | - | 0 | - |
| - | - | 992.6 | 457.3 | - | - | 0 | - |
| 6 | y | 702.1 | 459.2 | 0.000704 | 1.533 | +1 | 4 |
| - | - | 1312 | 462.2 | - | - | 0 | - |
| - | - | 813.4 | 471.2 | - | - | 0 | - |
| - | - | 2.402E+04 | 471.3 | - | - | 0 | - |
| 4 | c | 2.489E+04 | 472.3 | 0.0006935 | 1.468 | +1 | 4 |
| - | - | 6379 | 473.3 | - | - | 0 | - |
| - | - | 4884 | 474.3 | - | - | 0 | - |
| - | - | 675.1 | 475.3 | - | - | 0 | - |
| 6 | y | 1882 | 477.2 | 0.00256 | 5.364 | +1 | 4 |
| - | - | 596.8 | 478.2 | - | - | 0 | - |
| - | - | 6048 | 480.2 | - | - | 0 | - |
| - | - | 1479 | 481.2 | - | - | 0 | - |
| - | - | 2059 | 497.3 | - | - | 0 | - |
| - | - | 3.382E+04 | 498.2 | - | - | 0 | - |
| - | - | 2561 | 498.3 | - | - | 0 | - |
| - | - | 8223 | 499.2 | - | - | 0 | - |
| - | - | 689.9 | 499.3 | - | - | 0 | - |
| - | - | 1130 | 500.2 | - | - | 0 | - |
| - | - | 1140 | 502.3 | - | - | 0 | - |
| - | - | 681 | 507.2 | - | - | 0 | - |
| - | - | 2.798E+04 | 515.2 | - | - | 0 | - |
| - | - | 7788 | 516.2 | - | - | 0 | - |
| - | - | 6447 | 516.3 | - | - | 0 | - |
| - | - | 1471 | 517.3 | - | - | 0 | - |
| - | - | 1670 | 517.3 | - | - | 0 | - |
| - | - | 872.4 | 524.3 | - | - | 0 | - |
| - | - | 972.6 | 529.3 | - | - | 0 | - |
| - | - | 3552 | 530.2 | - | - | 0 | - |
| 5 | c | 4337 | 542.3 | 0.0001139 | 0.2099 | +1 | 5 |
| - | - | 974.7 | 543.3 | - | - | 0 | - |
| - | - | 5443 | 544.3 | - | - | 0 | - |
| - | - | 2267 | 545.3 | - | - | 0 | - |
| - | - | 579.8 | 556.7 | - | - | 0 | - |
| - | - | 1338 | 557.3 | - | - | 0 | - |
| - | - | 6.411E+04 | 558.3 | - | - | 0 | - |
| 5 | c | 5.369E+04 | 559.3 | 0.001655 | 2.959 | +1 | 5 |
| - | - | 1.399E+04 | 560.3 | - | - | 0 | - |
| - | - | 3593 | 561.3 | - | - | 0 | - |
| - | - | 1164 | 562.3 | - | - | 0 | - |
| 5 | y | 2291 | 564.3 | 0.001049 | 1.859 | +1 | 5 |
| - | - | 722.8 | 565.3 | - | - | 0 | - |
| - | - | 652 | 566.8 | - | - | 0 | - |
| - | - | 861 | 574.2 | - | - | 0 | - |
| - | - | 915.1 | 575.2 | - | - | 0 | - |
| - | - | 1917 | 591.2 | - | - | 0 | - |
| - | - | 1053 | 593.3 | - | - | 0 | - |
| - | - | 3439 | 599.3 | - | - | 0 | - |
| - | - | 803.4 | 600.3 | - | - | 0 | - |
| - | - | 1172 | 601.3 | - | - | 0 | - |
| - | - | 898.2 | 601.3 | - | - | 0 | - |
| - | - | 991.5 | 602.2 | - | - | 0 | - |
| - | - | 4846 | 603.2 | - | - | 0 | - |
| - | - | 1560 | 603.4 | - | - | 0 | - |
| - | - | 932 | 604.2 | - | - | 0 | - |
| - | - | 816.2 | 604.4 | - | - | 0 | - |
| - | - | 7288 | 609.3 | - | - | 0 | - |
| - | - | 2628 | 610.3 | - | - | 0 | - |
| - | - | 895 | 611.3 | - | - | 0 | - |
| - | - | 5212 | 611.3 | - | - | 0 | - |
| - | - | 1555 | 612.3 | - | - | 0 | - |
| - | - | 711.7 | 613.3 | - | - | 0 | - |
| - | - | 1813 | 626.3 | - | - | 0 | - |
| 4 | z | 8.357E+04 | 627.3 | 0.01236 | 19.7 | +1 | 6 |
| - | - | 2.657E+04 | 628.3 | - | - | 0 | - |
| 6 | c | 895.9 | 628.3 | 0.007616 | 12.12 | +1 | 6 |
| - | - | 817.8 | 628.3 | - | - | 0 | - |
| - | - | 7222 | 629.3 | - | - | 0 | - |
| 6 | c | 9661 | 629.3 | 0.001044 | 1.66 | +1 | 6 |
| - | - | 1020 | 630.3 | - | - | 0 | - |
| - | - | 3833 | 630.3 | - | - | 0 | - |
| 4 | y | 2220 | 643.3 | 0.0009642 | 1.499 | +1 | 6 |
| - | - | 4753 | 644.3 | - | - | 0 | - |
| 4 | z | 1987 | 645.3 | 0.002959 | 4.586 | +1 | 6 |
| - | - | 2948 | 645.3 | - | - | 0 | - |
| 6 | c | 1.147E+05 | 646.4 | 0.0003132 | 0.4846 | +1 | 6 |
| - | - | 4.3E+04 | 647.4 | - | - | 0 | - |
| - | - | 8655 | 648.4 | - | - | 0 | - |
| - | - | 2028 | 658.4 | - | - | 0 | - |
| - | - | 2281 | 660.3 | - | - | 0 | - |
| 4 | y | 3.216E+04 | 661.3 | 4.309E-05 | 0.06516 | +1 | 6 |
| - | - | 1.068E+04 | 662.3 | - | - | 0 | - |
| - | - | 2299 | 663.3 | - | - | 0 | - |
| - | - | 8201 | 672.4 | - | - | 0 | - |
| - | - | 3178 | 673.4 | - | - | 0 | - |
| - | - | 1351 | 685.3 | - | - | 0 | - |
| - | - | 1036 | 686.4 | - | - | 0 | - |
| - | - | 680.7 | 687.3 | - | - | 0 | - |
| - | - | 1053 | 688.3 | - | - | 0 | - |
| - | - | 1525 | 689.3 | - | - | 0 | - |
| - | - | 764.5 | 690.3 | - | - | 0 | - |
| - | - | 902.7 | 696.4 | - | - | 0 | - |
| - | - | 1060 | 698.3 | - | - | 0 | - |
| - | - | 2786 | 701.3 | - | - | 0 | - |
| - | - | 880 | 702.3 | - | - | 0 | - |
| - | - | 985.2 | 704.3 | - | - | 0 | - |
| - | - | 1327 | 712.4 | - | - | 0 | - |
| - | - | 1862 | 713.4 | - | - | 0 | - |
| - | - | 1697 | 714.3 | - | - | 0 | - |
| - | - | 827.5 | 714.4 | - | - | 0 | - |
| - | - | 3768 | 722.3 | - | - | 0 | - |
| - | - | 1450 | 723.3 | - | - | 0 | - |
| - | - | 4942 | 730.4 | - | - | 0 | - |
| - | - | 3329 | 731.4 | - | - | 0 | - |
| - | - | 2057 | 732.4 | - | - | 0 | - |
| - | - | 1.254E+04 | 739.3 | - | - | 0 | - |
| 3 | y | 2.183E+04 | 740.3 | 0.001057 | 1.427 | +1 | 7 |
| - | - | 7354 | 741.4 | - | - | 0 | - |
| - | - | 3954 | 742.4 | - | - | 0 | - |
| - | - | 1358 | 743.4 | - | - | 0 | - |
| - | - | 644.9 | 756.3 | - | - | 0 | - |
| - | - | 8.536E+04 | 757.3 | - | - | 0 | - |
| 7 | c | 3.279E+05 | 758.4 | 0.0142 | 18.73 | +1 | 7 |
| - | - | 1.204E+05 | 759.4 | - | - | 0 | - |
| - | - | 3.329E+04 | 760.4 | - | - | 0 | - |
| - | - | 3733 | 761.4 | - | - | 0 | - |
| - | - | 973.8 | 764.4 | - | - | 0 | - |
| - | - | 690.4 | 764.9 | - | - | 0 | - |
| - | - | 1332 | 765.4 | - | - | 0 | - |
| - | - | 930.4 | 767.3 | - | - | 0 | - |
| - | - | 1646 | 774.3 | - | - | 0 | - |
| 7 | c | 3.349E+05 | 775.4 | 4.357E-05 | 0.05619 | +1 | 7 |
| - | - | 1.515E+05 | 776.4 | - | - | 0 | - |
| - | - | 3.539E+04 | 777.4 | - | - | 0 | - |
| - | - | 2605 | 778.4 | - | - | 0 | - |
| - | - | 1079 | 785.3 | - | - | 0 | - |
| - | - | 1807 | 786.4 | - | - | 0 | - |
| - | - | 1735 | 786.4 | - | - | 0 | - |
| - | - | 798.6 | 787.4 | - | - | 0 | - |
| - | - | 940.7 | 787.4 | - | - | 0 | - |
| - | - | 1227 | 788.4 | - | - | 0 | - |
| - | - | 2.219E+04 | 801.4 | - | - | 0 | - |
| - | - | 1.124E+04 | 802.4 | - | - | 0 | - |
| - | - | 2534 | 803.4 | - | - | 0 | - |
| - | - | 2962 | 814.4 | - | - | 0 | - |
| - | - | 920.7 | 815.4 | - | - | 0 | - |
| - | - | 1119 | 817.4 | - | - | 0 | - |
| - | - | 1134 | 843.4 | - | - | 0 | - |
| - | - | 1657 | 851.4 | - | - | 0 | - |
| - | - | 6228 | 859.4 | - | - | 0 | - |
| - | - | 1.026E+04 | 860.4 | - | - | 0 | - |
| - | - | 5948 | 861.4 | - | - | 0 | - |
| - | - | 1176 | 862.4 | - | - | 0 | - |
| - | - | 4096 | 869.4 | - | - | 0 | - |
| - | - | 1935 | 870.4 | - | - | 0 | - |
| 2 | z | 8354 | 871.4 | 0.0003172 | 0.364 | +1 | 8 |
| - | - | 4168 | 872.4 | - | - | 0 | - |
| - | - | 1810 | 873.4 | - | - | 0 | - |
| 8 | c | 5.339E+04 | 887.4 | 0.0007677 | 0.8651 | +1 | 8 |
| - | - | 2.562E+04 | 888.4 | - | - | 0 | - |
| 2 | z | 2.896E+04 | 889.4 | 0.002631 | 2.958 | +1 | 8 |
| - | - | 1.24E+04 | 890.4 | - | - | 0 | - |
| - | - | 4307 | 891.4 | - | - | 0 | - |
| - | - | 1160 | 892.4 | - | - | 0 | - |
| - | - | 807.4 | 901.5 | - | - | 0 | - |
| 8 | c | 4.608E+05 | 904.4 | 2.69E-05 | 0.02975 | +1 | 8 |
| - | - | 2.281E+05 | 905.4 | - | - | 0 | - |
| - | - | 5.932E+04 | 906.4 | - | - | 0 | - |
| - | - | 6163 | 907.4 | - | - | 0 | - |
| - | - | 750 | 929.5 | - | - | 0 | - |
| - | - | 1058 | 930.5 | - | - | 0 | - |
| - | - | 1032 | 933.5 | - | - | 0 | - |
| - | - | 2.66E+04 | 947.5 | - | - | 0 | - |
| - | - | 1.29E+04 | 948.5 | - | - | 0 | - |
| - | - | 4191 | 949.5 | - | - | 0 | - |
| - | - | 1079 | 955.5 | - | - | 0 | - |
| - | - | 3237 | 956.5 | - | - | 0 | - |
| - | - | 2563 | 957.5 | - | - | 0 | - |
| - | - | 1545 | 958.5 | - | - | 0 | - |
| - | - | 2690 | 959.4 | - | - | 0 | - |
| - | - | 1118 | 960.5 | - | - | 0 | - |
| - | - | 5795 | 963.5 | - | - | 0 | - |
| - | - | 3252 | 964.5 | - | - | 0 | - |
| - | - | 1041 | 965.5 | - | - | 0 | - |
| - | - | 870.5 | 965.6 | - | - | 0 | - |
| - | - | 1430 | 971.5 | - | - | 0 | - |
| - | - | 914.9 | 972.5 | - | - | 0 | - |
| - | - | 3952 | 973.5 | - | - | 0 | - |
| - | - | 1765 | 974.5 | - | - | 0 | - |
| - | - | 1022 | 975.5 | - | - | 0 | - |
| - | - | 883.7 | 976.5 | - | - | 0 | - |
| - | - | 7456 | 984.5 | - | - | 0 | - |
| - | - | 3690 | 985.5 | - | - | 0 | - |
| - | - | 1478 | 986.5 | - | - | 0 | - |
| - | - | 1476 | 989.5 | - | - | 0 | - |
| - | - | 999.3 | 990.5 | - | - | 0 | - |
| - | - | 5958 | 991.5 | - | - | 0 | - |
| - | - | 3149 | 992.5 | - | - | 0 | - |
| - | - | 1316 | 993.5 | - | - | 0 | - |
| - | - | 1877 | 1000 | - | - | 0 | - |
| - | - | 5025 | 1001 | - | - | 0 | - |
| - | - | 4.346E+04 | 1002 | - | - | 0 | - |
| - | - | 2.522E+04 | 1003 | - | - | 0 | - |
| - | - | 9568 | 1004 | - | - | 0 | - |
| - | - | 1287 | 1005 | - | - | 0 | - |
| - | - | 2246 | 1017 | - | - | 0 | - |
| - | - | 1.091E+05 | 1019 | - | - | 0 | - |
| - | - | 4.209E+05 | 1020 | - | - | 0 | - |
| - | - | 2.261E+05 | 1021 | - | - | 0 | - |
| - | - | 6.861E+04 | 1022 | - | - | 0 | - |
| - | - | 6781 | 1023 | - | - | 0 | - |
| - | - | 3507 | 1052 | - | - | 0 | - |
| - | - | 1934 | 1053 | - | - | 0 | - |
| - | - | 891.7 | 1513 | - | - | 0 | - |
| - | - | 1362 | 1514 | - | - | 0 | - |
| - | - | 1153 | 1530 | - | - | 0 | - |
| - | - | 1173 | 1531 | - | - | 0 | - |
| - | - | 663.1 | 1590 | - | - | 0 | - |

m/z Charge Intensity FragmentType MassShift Position
120.08106994628906 0 10144.281
121.08444213867188 0 891.1854
128.2966766357422 0 401.9329
129.06610107421875 0 447.6858
129.10243225097656 0 430.7561
130.0501251220703 0 2209.0474
132.10218811035156 0 74860.01 y 8
133.08621215820312 0 582.0779
133.10057067871094 0 518.94293
133.10557556152344 0 3457.2708
166.2949981689453 0 411.4221
167.1177520751953 0 448.20605
168.1261444091797 0 600.4321
171.83006286621094 0 471.93738
184.5930633544922 0 473.6617
189.9167938232422 0 488.23926
197.1288604736328 0 3911.1455
199.0714874267578 0 2186.3838
209.42147827148438 0 506.6409
212.1396026611328 0 4858.5435
215.14022827148438 0 603.52075
217.08216857910156 0 2281.6296
217.14617919921875 0 954.2405
221.0930633544922 0 902.92365
229.13101196289062 0 474.5498
233.16510009765625 0 59554.258
234.16859436035156 0 9639.152
241.08241271972656 0 3255.655
243.13426208496094 0 38573.5 y Water loss 7
243.14881896972656 0 1156.5782
244.13723754882812 0 4305.994
259.09271240234375 0 3503.4692
261.1445007324219 0 11408.667 y 7
261.1598815917969 0 38161.266 c Ammonia loss 1
262.14892578125 0 848.16174
262.1631774902344 0 5923.4585
263.16595458984375 0 645.3172
264.1343078613281 0 1701.8823
276.1829528808594 0 494.98654
282.1277160644531 0 826.0366
282.14508056640625 0 12958.89
283.1480712890625 0 1623.9583
286.10394287109375 0 752.54663
287.1026306152344 0 752.8442
296.1243896484375 0 1825.7129
296.1966247558594 0 969.1085
297.1588134765625 0 859.45526
298.1641540527344 0 746.6733
299.1717529296875 0 23544.969
300.17486572265625 0 3430.0994
304.1133117675781 0 578.3297
305.1304016113281 0 1290.0638
305.6318664550781 0 566.2507
306.8336486816406 0 464.35785
308.1442565917969 0 537.4989
314.1354064941406 0 2197.6445 z Water loss 3
314.63726806640625 0 1050.5677
322.9450988769531 0 521.9906
326.1595153808594 0 726.97217
329.1219787597656 0 907.93774
333.1560363769531 0 1038.3771
338.6689147949219 0 3268.499
339.1697998046875 0 940.83057
346.1246337890625 0 2720.2332
348.1755065917969 0 764.821
351.16668701171875 0 3963.7986
352.1695251464844 0 1133.809
354.16650390625 0 1150.2155
355.16229248046875 0 1161.1246
357.2051086425781 0 654.39734
358.2126770019531 0 6004.7026 c Ammonia loss 2
359.2154541015625 0 1433.1174
361.67657470703125 0 774.4035
369.1768798828125 0 6828.0264
370.17926025390625 0 937.4944
370.6831359863281 0 2349.799 y Water loss 2
371.1839294433594 0 1567.9717
372.1775817871094 0 2030.9518 y Water loss 6
379.68414306640625 0 2238.1748 c Ammonia loss 6
380.1561584472656 0 2255.9138
380.18341064453125 0 1218.9824
386.2036437988281 0 59715.617
387.20672607421875 0 10770.784
388.2065124511719 0 964.90344 c 6
390.18756103515625 0 1405.0677 y 6
397.2436828613281 0 733.59247
398.16851806640625 0 836.97614
401.16705322265625 0 981.12646
412.2105712890625 0 1152.414
415.25701904296875 0 1054.6416
417.19354248046875 0 796.90894
426.7033386230469 0 604.8099
427.2708435058594 0 1205.4678
428.2784729003906 0 12770.756
429.2843933105469 0 5662.1895
431.2304992675781 0 684.81244
433.1568908691406 0 1983.7076
444.2112731933594 0 1489.679 c Ammonia loss 7
455.2298889160156 0 688.93115
455.2651672363281 0 3227.9795 c Ammonia loss 3
456.2722473144531 0 2849.3967
457.27740478515625 0 992.61926
459.2092590332031 0 702.1475 y Water loss 5
462.198486328125 0 1311.9098
471.2259521484375 0 813.3995
471.284423828125 0 24020.266
472.2911376953125 0 24888.021 c 3
473.29559326171875 0 6379.458
474.307373046875 0 4883.895
475.31146240234375 0 675.0589
477.2216796875 0 1882.496 y 5
478.2231140136719 0 596.8181
480.20953369140625 0 6047.5127
481.2117004394531 0 1478.8258
497.29925537109375 0 2058.6375
498.22003173828125 0 33819.523
498.3063659667969 0 2560.788
499.2230224609375 0 8223.326
499.3130798339844 0 689.87537
500.2223815917969 0 1129.5547
502.25457763671875 0 1140.2837
507.2120666503906 0 680.9646
515.2462768554688 0 27981.768
516.2491455078125 0 7788.211
516.3185424804688 0 6447.1587
517.2530517578125 0 1471.1088
517.3225708007812 0 1670.4944
524.2860107421875 0 872.35144
529.3399047851562 0 972.55975
530.2106323242188 0 3552.42
542.2974243164062 0 4336.525 c Ammonia loss 4
543.3026123046875 0 974.6915
544.3128662109375 0 5443.434
545.3165893554688 0 2267.4714
556.676513671875 0 579.7747
557.328857421875 0 1338.3483
558.3161010742188 0 64110.37
559.3222045898438 0 53692.973 c 4
560.3256225585938 0 13988.925
561.3353881835938 0 3593.1255
562.3411254882812 0 1164.4944
564.252197265625 0 2290.743 y 4
565.2534790039062 0 722.8493
566.7679443359375 0 652.04486
574.2413940429688 0 860.9634
575.2421875 0 915.0514
591.2423706054688 0 1917.2253
593.30224609375 0 1052.7308
599.267822265625 0 3439.2258
600.2689819335938 0 803.4359
601.280029296875 0 1171.5137
601.332763671875 0 898.177
602.2321166992188 0 991.5307
603.2406005859375 0 4845.5854
603.3504638671875 0 1560.0488
604.2422485351562 0 931.95764
604.35400390625 0 816.17017
609.2523193359375 0 7287.717
610.2555541992188 0 2627.507
611.2593383789062 0 895.0325
611.3151245117188 0 5212.139
612.3200073242188 0 1555.4402
613.3179931640625 0 711.7292
626.2556762695312 0 1812.5809
627.2622680664062 0 83573.82 z Water loss 3
628.26513671875 0 26565.73
628.3377075195312 0 895.89557 c Water loss 5
628.3472290039062 0 817.7884
629.2678833007812 0 7221.537
629.3303833007812 0 9661.138 c Ammonia loss 5
630.2759399414062 0 1020.3479
630.332275390625 0 3832.7366
643.2943115234375 0 2219.616 y Water loss 3
644.28955078125 0 4752.987
645.2881469726562 0 1987.0343 z 3
645.348388671875 0 2948.295
646.356201171875 0 114722.17 c 5
647.3590087890625 0 42997.816
648.3613891601562 0 8654.967
658.376953125 0 2027.8219
660.2929077148438 0 2280.9956
661.303955078125 0 32162.893 y 3
662.306884765625 0 10680.494
663.3084716796875 0 2299.2217
672.37158203125 0 8200.928
673.375 0 3178.4355
685.3296508789062 0 1350.8025
686.3770141601562 0 1035.6758
687.2617797851562 0 680.6553
688.2799682617188 0 1052.8182
689.2946166992188 0 1525.243
690.3024291992188 0 764.5299
696.3572387695312 0 902.66364
698.335693359375 0 1060.3381
701.2881469726562 0 2785.775
702.296630859375 0 879.96234
704.32958984375 0 985.2481
712.3556518554688 0 1327.0259
713.3541259765625 0 1862.2483
714.2953491210938 0 1696.8243
714.3641357421875 0 827.527
722.34228515625 0 3768.2085
723.3438720703125 0 1449.8542
730.3776245117188 0 4942.0386
731.3820190429688 0 3328.8354
732.3909912109375 0 2057.2798
739.3382568359375 0 12536.693
740.34716796875 0 21825.678 y Water loss 2
741.3519897460938 0 7353.5547
742.3536376953125 0 3954.187
743.3555908203125 0 1358.1606
756.3139038085938 0 644.93713
757.348876953125 0 85359.19
758.3577270507812 0 327887.56 c Ammonia loss 6
759.360595703125 0 120407.15
760.3655395507812 0 33288.215
761.3720092773438 0 3733.4304
764.4380493164062 0 973.8218
764.9481201171875 0 690.4357
765.447265625 0 1332.0177
767.3330688476562 0 930.437
774.3294067382812 0 1646.348
775.3984375 0 334922.16 c 6
776.4014282226562 0 151535.38
777.404052734375 0 35387.094
778.4067993164062 0 2604.8496
785.348388671875 0 1078.8196
786.3505249023438 0 1807.4089
786.4380493164062 0 1734.6937
787.3531494140625 0 798.57196
787.4359130859375 0 940.7273
788.40478515625 0 1226.6571
801.4141235351562 0 22188.152
802.4168701171875 0 11244.951
803.4213256835938 0 2533.7803
814.4307861328125 0 2961.6758
815.4347534179688 0 920.7477
817.3836669921875 0 1118.9746
843.4259033203125 0 1134.4636
851.3949584960938 0 1657.4244
859.4194946289062 0 6227.6646
860.4258422851562 0 10264.445
861.4314575195312 0 5948.225
862.439208984375 0 1175.5044
869.4035034179688 0 4096.0923
870.4034423828125 0 1935.3744
871.3961181640625 0 8354.238 z Water loss 1
872.3994140625 0 4167.5947
873.4100341796875 0 1809.8438
887.4137573242188 0 53392.71 c Ammonia loss 7
888.4160766601562 0 25621.637
889.4089965820312 0 28955.533 z 1
890.4107666015625 0 12402.411
891.41455078125 0 4307.3047
892.4207153320312 0 1159.7811
901.4859008789062 0 807.38727
904.4411010742188 0 460769.6 c 7
905.443359375 0 228092.23
906.446044921875 0 59316.22
907.4481201171875 0 6163.2295
929.5023193359375 0 749.9526
930.4697265625 0 1058.0673
933.4949951171875 0 1031.7274
947.4951782226562 0 26596.795
948.4976806640625 0 12899.915
949.5003662109375 0 4191.271
955.5064086914062 0 1078.7423
956.4932861328125 0 3237.4897
957.5084228515625 0 2562.8188
958.5023193359375 0 1544.976
959.4357299804688 0 2690.2383
960.514892578125 0 1117.6654
963.4533081054688 0 5795.451
964.457275390625 0 3252.1997
965.4619140625 0 1040.813
965.6105346679688 0 870.5208
971.4909057617188 0 1430.4497
972.4916381835938 0 914.8977
973.508056640625 0 3952.1362
974.5069580078125 0 1764.9028
975.5030517578125 0 1022.407
976.4747924804688 0 883.65704
984.4788208007812 0 7455.588
985.4842529296875 0 3689.9888
986.4880981445312 0 1477.8417
989.512451171875 0 1476.1366
990.5136108398438 0 999.26996
991.5198364257812 0 5957.5938
992.524169921875 0 3148.561
993.5318603515625 0 1315.5657
1000.485107421875 0 1876.741
1001.4954833984375 0 5025.269
1002.4901123046875 0 43457.79
1003.4927368164062 0 25218.266
1004.4951171875 0 9567.539
1005.4949340820312 0 1287.1161
1017.4851684570312 0 2245.618
1018.5072631835938 0 109050.695
1019.515625 0 420870.06
1020.518798828125 0 226140.94
1021.5220947265625 0 68609.63
1022.524658203125 0 6781.3643
1051.508056640625 0 3507.0532
1052.50830078125 0 1933.5677
1512.8695068359375 0 891.6576
1513.8717041015625 0 1361.561
1529.89794921875 0 1153.1567
1530.897705078125 0 1172.7842
1589.5821533203125 0 663.06323

Spectrum Details

|  |  |
| --- | --- |
| Matched peaks? Matched peaks The total absolute number of peaks matched. Additionally in brackets the total fraction of peaks matched and the total number of peaks is shown. | 33 (10.68% of 309) |
| FDR? FDR The false discovery rate estimated for this peptide. It is calculated by matching all theoretical fragments with a non-integer shift with the raw peaks for this spectrum. This is done with 40 different shifts. The resulting percentage is the average number of annotated peaks over the number of annotated peaks with the correct spectrum. | 1.88% |
| Satellite FDR? Satellite FDR See the FDR for details on its calculation. This satellite ion specific FDR only contains the satellite ions (d/w) for I/L/J positions. | - |
| PSM Score? PSM Score The PSM Score as given by Hecklib to this annotated spectrum. It is shown with three significant figures. | 429 |

#### Spectrum 8589? Spectrum 8589 The raw spectrum of this peptide as annotated by Hecklib. The fragments are coloured according to ion type (see legend). Any peaks with a star '\*' as text can be hovered over to see the full details, first the ion type second the mass shift type. By hovering over the amino acids in the peptide or ions in the legend the corresponding peaks are highlighted. By toggling the 'Unassigned' label you can turn the background (unassigned) peaks on or off in the plot. By updating the slider in the Ion legend you can update the spectrum to only show the top X% of the peaks with labels. The top X% means any peak that is within X% of the highest intensity. By dragging in the spectrum you can zoom in to a specific part of the spectrum and use 'Zoom Out' to get back to the original zoom level. The annotation of the spectrum is based on the given sequence in the peptides file and is done with different software so inconsistencies are likely. The peaks are annotated based on the given sequence, with 20 ppm tolerance.

Copy Data

##### Spectrum 8589 (TSV)

###### Preview

```
Loading example...
```

*Click on the button to copy the data to your clipboard.*

Mz MinMz MaxIntensity Max

WidthHeightPeptide font sizePeptide stroke widthSpectrum font sizeSpectrum stroke widthCompact peptide

Ion legend

wxyz

abcd

OtherUnassignedIon^Charge^_Position_Show for top:%

JFPPSSEEJ

04.05e+48.11e+41.22e+51.62e+5

Zoom Out

y^+1^_1_y^+1^_2_y^+1^_2_c^+1^_2_c^+1^_3_y^+2^_7_c^+1^_4_c^+1^_4_c^+1^_5_c^+1^_5_y^+1^_5_c^+1^_6_y^+1^_6_z^+1^_6_c^+1^_6_y^+1^_6_y^+1^_7_y^+1^_7_c^+1^_7_z^+1^_8_c^+1^_8_z^+1^_8_c^+1^_8_y^+1^_8_

0781156223423123

Fragment Matches Table

Show background peaks

| Position | Ion type | Intensity | mz Theoretical | mz Error (Th) | mz Error (ppm) | Charge | Series Number |
| --- | --- | --- | --- | --- | --- | --- | --- |
| - | - | 3749 | 120.1 | - | - | 0 | - |
| - | - | 387 | 120.4 | - | - | 0 | - |
| - | - | 456.9 | 121.1 | - | - | 0 | - |
| - | - | 353.4 | 128.5 | - | - | 0 | - |
| - | - | 414 | 129.6 | - | - | 0 | - |
| - | - | 547.1 | 130.1 | - | - | 0 | - |
| - | - | 371.7 | 131.1 | - | - | 0 | - |
| 9 | y | 2.218E+04 | 132.1 | 0.0001914 | 1.449 | +1 | 1 |
| - | - | 1297 | 133.1 | - | - | 0 | - |
| - | - | 423.7 | 138.3 | - | - | 0 | - |
| - | - | 414.6 | 140.4 | - | - | 0 | - |
| - | - | 977.9 | 148.9 | - | - | 0 | - |
| - | - | 1701 | 173.4 | - | - | 0 | - |
| - | - | 451.3 | 183 | - | - | 0 | - |
| - | - | 1582 | 197.1 | - | - | 0 | - |
| - | - | 940.6 | 199.1 | - | - | 0 | - |
| - | - | 504.3 | 208.8 | - | - | 0 | - |
| - | - | 1493 | 212.1 | - | - | 0 | - |
| - | - | 459.2 | 217.1 | - | - | 0 | - |
| - | - | 556.6 | 229.7 | - | - | 0 | - |
| - | - | 1.81E+04 | 233.2 | - | - | 0 | - |
| - | - | 2709 | 234.2 | - | - | 0 | - |
| - | - | 827.4 | 241.1 | - | - | 0 | - |
| 8 | y | 1.318E+04 | 243.1 | 2.338E-05 | 0.09616 | +1 | 2 |
| - | - | 1149 | 244.1 | - | - | 0 | - |
| - | - | 830.2 | 259.1 | - | - | 0 | - |
| 8 | y | 3338 | 261.1 | 0.0003027 | 1.159 | +1 | 2 |
| 2 | c | 1.172E+04 | 261.2 | 8.638E-05 | 0.3308 | +1 | 2 |
| - | - | 1801 | 262.2 | - | - | 0 | - |
| - | - | 4648 | 282.1 | - | - | 0 | - |
| - | - | 518.5 | 282.2 | - | - | 0 | - |
| - | - | 857.2 | 296.1 | - | - | 0 | - |
| - | - | 8315 | 299.2 | - | - | 0 | - |
| - | - | 1291 | 300.2 | - | - | 0 | - |
| - | - | 704 | 309.2 | - | - | 0 | - |
| - | - | 614.4 | 312.7 | - | - | 0 | - |
| - | - | 605.6 | 316.1 | - | - | 0 | - |
| - | - | 822.9 | 327.1 | - | - | 0 | - |
| - | - | 863.5 | 338.7 | - | - | 0 | - |
| - | - | 684.3 | 346.1 | - | - | 0 | - |
| - | - | 1430 | 351.2 | - | - | 0 | - |
| 3 | c | 1639 | 358.2 | 0.0002074 | 0.579 | +1 | 3 |
| - | - | 1570 | 369.2 | - | - | 0 | - |
| 3 | y | 1038 | 370.7 | 0.006747 | 18.2 | +2 | 7 |
| - | - | 2.196E+04 | 386.2 | - | - | 0 | - |
| - | - | 4368 | 387.2 | - | - | 0 | - |
| - | - | 519.3 | 396.9 | - | - | 0 | - |
| - | - | 542.1 | 412.9 | - | - | 0 | - |
| - | - | 538.2 | 415.5 | - | - | 0 | - |
| - | - | 617.5 | 419 | - | - | 0 | - |
| - | - | 4851 | 428.3 | - | - | 0 | - |
| - | - | 684.2 | 429.2 | - | - | 0 | - |
| - | - | 1998 | 429.3 | - | - | 0 | - |
| - | - | 626.5 | 431.2 | - | - | 0 | - |
| 4 | c | 1296 | 455.3 | 0.0008312 | 1.826 | +1 | 4 |
| - | - | 1394 | 456.3 | - | - | 0 | - |
| - | - | 7585 | 471.3 | - | - | 0 | - |
| 4 | c | 9348 | 472.3 | 0.0009986 | 2.114 | +1 | 4 |
| - | - | 2359 | 473.3 | - | - | 0 | - |
| - | - | 2117 | 474.3 | - | - | 0 | - |
| - | - | 681.9 | 475.3 | - | - | 0 | - |
| - | - | 1802 | 480.2 | - | - | 0 | - |
| - | - | 1051 | 481.2 | - | - | 0 | - |
| - | - | 1.081E+04 | 498.2 | - | - | 0 | - |
| - | - | 650.9 | 498.3 | - | - | 0 | - |
| - | - | 2474 | 499.2 | - | - | 0 | - |
| - | - | 822.4 | 509.3 | - | - | 0 | - |
| - | - | 1.018E+04 | 515.2 | - | - | 0 | - |
| - | - | 2914 | 516.2 | - | - | 0 | - |
| - | - | 1774 | 516.3 | - | - | 0 | - |
| - | - | 842 | 530.2 | - | - | 0 | - |
| 5 | c | 1268 | 542.3 | 6.925E-05 | 0.1277 | +1 | 5 |
| - | - | 803.9 | 543.3 | - | - | 0 | - |
| - | - | 2291 | 544.3 | - | - | 0 | - |
| - | - | 2.347E+04 | 558.3 | - | - | 0 | - |
| 5 | c | 1.783E+04 | 559.3 | 0.002082 | 3.723 | +1 | 5 |
| - | - | 4333 | 560.3 | - | - | 0 | - |
| - | - | 794.5 | 561.3 | - | - | 0 | - |
| 5 | y | 767.5 | 564.3 | 0.002674 | 4.739 | +1 | 5 |
| - | - | 1193 | 599.3 | - | - | 0 | - |
| - | - | 625.5 | 604.4 | - | - | 0 | - |
| - | - | 2886 | 609.3 | - | - | 0 | - |
| - | - | 1218 | 611.3 | - | - | 0 | - |
| - | - | 562.9 | 612.3 | - | - | 0 | - |
| - | - | 572.4 | 618.6 | - | - | 0 | - |
| - | - | 839.9 | 626.3 | - | - | 0 | - |
| - | - | 2.524E+04 | 627.3 | - | - | 0 | - |
| - | - | 6935 | 628.3 | - | - | 0 | - |
| - | - | 1616 | 629.3 | - | - | 0 | - |
| 6 | c | 3338 | 629.3 | 6.786E-05 | 0.1078 | +1 | 6 |
| - | - | 1086 | 630.3 | - | - | 0 | - |
| 4 | y | 1091 | 643.3 | 0.003894 | 6.053 | +1 | 6 |
| - | - | 1937 | 644.3 | - | - | 0 | - |
| 4 | z | 846.8 | 645.3 | 0.003203 | 4.964 | +1 | 6 |
| - | - | 715.2 | 645.3 | - | - | 0 | - |
| 6 | c | 4.052E+04 | 646.4 | 0.0004192 | 0.6486 | +1 | 6 |
| - | - | 1.498E+04 | 647.4 | - | - | 0 | - |
| - | - | 3046 | 648.4 | - | - | 0 | - |
| 4 | y | 1.088E+04 | 661.3 | 0.0006893 | 1.042 | +1 | 6 |
| - | - | 2777 | 662.3 | - | - | 0 | - |
| - | - | 2237 | 672.4 | - | - | 0 | - |
| - | - | 1595 | 673.4 | - | - | 0 | - |
| - | - | 617.4 | 698.3 | - | - | 0 | - |
| - | - | 1083 | 701.3 | - | - | 0 | - |
| - | - | 928.4 | 714.3 | - | - | 0 | - |
| - | - | 793.5 | 722.3 | - | - | 0 | - |
| - | - | 910.8 | 723.3 | - | - | 0 | - |
| - | - | 1632 | 730.4 | - | - | 0 | - |
| - | - | 942.8 | 731.4 | - | - | 0 | - |
| - | - | 1186 | 732.4 | - | - | 0 | - |
| - | - | 4424 | 739.3 | - | - | 0 | - |
| 3 | y | 7191 | 740.3 | 0.002644 | 3.571 | +1 | 7 |
| - | - | 2060 | 741.4 | - | - | 0 | - |
| - | - | 1485 | 742.4 | - | - | 0 | - |
| - | - | 3.002E+04 | 757.3 | - | - | 0 | - |
| - | - | 682.1 | 757.9 | - | - | 0 | - |
| 3 | y | 1.12E+05 | 758.4 | 0.0003526 | 0.4649 | +1 | 7 |
| - | - | 4.359E+04 | 759.4 | - | - | 0 | - |
| - | - | 1.048E+04 | 760.4 | - | - | 0 | - |
| - | - | 1495 | 761.4 | - | - | 0 | - |
| - | - | 1172 | 764.4 | - | - | 0 | - |
| - | - | 1407 | 764.9 | - | - | 0 | - |
| - | - | 951.6 | 765.4 | - | - | 0 | - |
| - | - | 4802 | 766.4 | - | - | 0 | - |
| - | - | 3612 | 766.9 | - | - | 0 | - |
| - | - | 662.9 | 774.3 | - | - | 0 | - |
| 7 | c | 1.223E+05 | 775.4 | 0.001081 | 1.394 | +1 | 7 |
| - | - | 5.041E+04 | 776.4 | - | - | 0 | - |
| - | - | 1.31E+04 | 777.4 | - | - | 0 | - |
| - | - | 916.8 | 778.4 | - | - | 0 | - |
| - | - | 694.5 | 786.4 | - | - | 0 | - |
| - | - | 764.5 | 788.4 | - | - | 0 | - |
| - | - | 8180 | 801.4 | - | - | 0 | - |
| - | - | 3602 | 802.4 | - | - | 0 | - |
| - | - | 1574 | 803.4 | - | - | 0 | - |
| - | - | 1964 | 814.4 | - | - | 0 | - |
| - | - | 1084 | 815.4 | - | - | 0 | - |
| - | - | 661.9 | 843.4 | - | - | 0 | - |
| - | - | 967.9 | 851.4 | - | - | 0 | - |
| - | - | 708.8 | 857.5 | - | - | 0 | - |
| - | - | 1708 | 859.4 | - | - | 0 | - |
| - | - | 2924 | 860.4 | - | - | 0 | - |
| - | - | 1654 | 861.4 | - | - | 0 | - |
| - | - | 1117 | 869.4 | - | - | 0 | - |
| - | - | 1376 | 870.4 | - | - | 0 | - |
| 2 | z | 3284 | 871.4 | 0.0006834 | 0.7842 | +1 | 8 |
| - | - | 2219 | 872.4 | - | - | 0 | - |
| 8 | c | 1.683E+04 | 887.4 | 0.001683 | 1.897 | +1 | 8 |
| - | - | 7889 | 888.4 | - | - | 0 | - |
| 2 | z | 1.12E+04 | 889.4 | 0.001349 | 1.517 | +1 | 8 |
| - | - | 5395 | 890.4 | - | - | 0 | - |
| - | - | 1366 | 891.4 | - | - | 0 | - |
| 8 | c | 1.605E+05 | 904.4 | 0.001316 | 1.455 | +1 | 8 |
| 2 | y | 8.536E+04 | 905.4 | 0.01723 | 19.03 | +1 | 8 |
| - | - | 2.465E+04 | 906.4 | - | - | 0 | - |
| - | - | 1993 | 907.4 | - | - | 0 | - |
| - | - | 820.2 | 939.5 | - | - | 0 | - |
| - | - | 728.7 | 940.5 | - | - | 0 | - |
| - | - | 8805 | 947.5 | - | - | 0 | - |
| - | - | 4810 | 948.5 | - | - | 0 | - |
| - | - | 934.2 | 949.5 | - | - | 0 | - |
| - | - | 634.7 | 954.5 | - | - | 0 | - |
| - | - | 844.7 | 957.5 | - | - | 0 | - |
| - | - | 1172 | 959.4 | - | - | 0 | - |
| - | - | 2184 | 963.5 | - | - | 0 | - |
| - | - | 1077 | 964.5 | - | - | 0 | - |
| - | - | 566.3 | 965.5 | - | - | 0 | - |
| - | - | 685.9 | 973.5 | - | - | 0 | - |
| - | - | 1184 | 974.5 | - | - | 0 | - |
| - | - | 2125 | 984.5 | - | - | 0 | - |
| - | - | 1464 | 985.5 | - | - | 0 | - |
| - | - | 1712 | 991.5 | - | - | 0 | - |
| - | - | 1102 | 1001 | - | - | 0 | - |
| - | - | 1.474E+04 | 1002 | - | - | 0 | - |
| - | - | 1084 | 1003 | - | - | 0 | - |
| - | - | 1.064E+04 | 1003 | - | - | 0 | - |
| - | - | 3799 | 1004 | - | - | 0 | - |
| - | - | 1516 | 1005 | - | - | 0 | - |
| - | - | 3889 | 1017 | - | - | 0 | - |
| - | - | 2378 | 1018 | - | - | 0 | - |
| - | - | 3.771E+04 | 1019 | - | - | 0 | - |
| - | - | 1.453E+05 | 1020 | - | - | 0 | - |
| - | - | 8.064E+04 | 1021 | - | - | 0 | - |
| - | - | 809.4 | 1021 | - | - | 0 | - |
| - | - | 2.465E+04 | 1022 | - | - | 0 | - |
| - | - | 2464 | 1023 | - | - | 0 | - |
| - | - | 1058 | 1052 | - | - | 0 | - |
| - | - | 773.9 | 1315 | - | - | 0 | - |
| - | - | 1154 | 1374 | - | - | 0 | - |
| - | - | 970.7 | 1488 | - | - | 0 | - |
| - | - | 707.8 | 1490 | - | - | 0 | - |
| - | - | 1071 | 1498 | - | - | 0 | - |
| - | - | 1338 | 1505 | - | - | 0 | - |
| - | - | 925.1 | 1509 | - | - | 0 | - |
| - | - | 1392 | 1512 | - | - | 0 | - |
| - | - | 984.5 | 1514 | - | - | 0 | - |
| - | - | 679.8 | 1529 | - | - | 0 | - |
| - | - | 660 | 1531 | - | - | 0 | - |
| - | - | 745.5 | 1531 | - | - | 0 | - |
| - | - | 1777 | 1532 | - | - | 0 | - |
| - | - | 2890 | 1533 | - | - | 0 | - |
| - | - | 1373 | 1534 | - | - | 0 | - |
| - | - | 697.6 | 3092 | - | - | 0 | - |

m/z Charge Intensity FragmentType MassShift Position
120.08098602294922 0 3749.3667
120.40402221679688 0 386.96237
121.08438873291016 0 456.9282
128.53782653808594 0 353.37292
129.6173095703125 0 413.9584
130.05023193359375 0 547.06067
131.07040405273438 0 371.7296
132.1020965576172 0 22181.021 y 8
133.10556030273438 0 1297.3163
138.2909698486328 0 423.68826
140.4077606201172 0 414.6488
148.9474639892578 0 977.852
173.439208984375 0 1701.0101
183.0051727294922 0 451.28033
197.12843322753906 0 1582.1101
199.0713653564453 0 940.6487
208.7689971923828 0 504.32684
212.13931274414062 0 1493.199
217.0818328857422 0 459.15503
229.73936462402344 0 556.6133
233.16490173339844 0 18099.906
234.16802978515625 0 2709.3828
241.08238220214844 0 827.41815
243.1339569091797 0 13177.624 y Water loss 7
244.13795471191406 0 1148.6105
259.09259033203125 0 830.19434
261.1441955566406 0 3338.3179 y 7
261.15966796875 0 11722.515 c Ammonia loss 1
262.163330078125 0 1800.584
282.14501953125 0 4647.8804
282.2492980957031 0 518.5176
296.1234130859375 0 857.2404
299.1716003417969 0 8314.838
300.1745910644531 0 1290.5615
309.20343017578125 0 703.953
312.6753845214844 0 614.4064
316.14910888671875 0 605.5894
327.1427001953125 0 822.9103
338.6683654785156 0 863.4715
346.1257629394531 0 684.2684
351.1668701171875 0 1429.7633
358.2123107910156 0 1639.4897 c Ammonia loss 2
369.1764831542969 0 1569.5159
370.6834411621094 0 1037.5876 y Water loss 2
386.203369140625 0 21955.281
387.20709228515625 0 4367.803
396.86688232421875 0 519.30963
412.8753662109375 0 542.081
415.467041015625 0 538.2014
418.9822082519531 0 617.45874
428.2779846191406 0 4850.8345
429.2142333984375 0 684.1538
429.2846374511719 0 1998.2437
431.2269592285156 0 626.47314
455.26611328125 0 1295.9944 c Ammonia loss 3
456.272216796875 0 1394.1322
471.2840576171875 0 7584.614
472.29083251953125 0 9348.187 c 3
473.295654296875 0 2358.8567
474.30511474609375 0 2117.367
475.3099060058594 0 681.9492
480.20819091796875 0 1802.3243
481.2106628417969 0 1051.3597
498.2196044921875 0 10811.777
498.306640625 0 650.8653
499.22265625 0 2473.7644
509.25433349609375 0 822.3511
515.2457275390625 0 10179.213
516.2484741210938 0 2913.6204
516.3184204101562 0 1774.0199
530.207763671875 0 842.0198
542.2972412109375 0 1267.8225 c Ammonia loss 4
543.3038330078125 0 803.8985
544.3125 0 2290.87
558.3157348632812 0 23474.625
559.32177734375 0 17834.379 c 4
560.3255615234375 0 4332.841
561.3341064453125 0 794.47345
564.2484741210938 0 767.54144 y 4
599.2647094726562 0 1193.1398
604.3531494140625 0 625.5461
609.2510375976562 0 2886.4702
611.3163452148438 0 1217.7738
612.3095703125 0 562.8677
618.5508422851562 0 572.42633
626.2539672851562 0 839.91907
627.2617797851562 0 25240.332
628.2645874023438 0 6934.5815
629.2664794921875 0 1616.2931
629.3294067382812 0 3338.0735 c Ammonia loss 5
630.3363037109375 0 1086.2037
643.2972412109375 0 1091.0702 y Water loss 3
644.28857421875 0 1936.6068
645.2883911132812 0 846.75037 z 3
645.3472900390625 0 715.1798
646.35546875 0 40524.11 c 5
647.3583984375 0 14977.791
648.36083984375 0 3045.615
661.30322265625 0 10882.087 y 3
662.3080444335938 0 2777.177
672.3701782226562 0 2237.1125
673.3754272460938 0 1594.8702
698.3384399414062 0 617.3899
701.2864990234375 0 1082.5116
714.2962036132812 0 928.4185
722.3418579101562 0 793.5443
723.3480834960938 0 910.77515
730.3754272460938 0 1632.4486
731.3815307617188 0 942.7564
732.3899536132812 0 1185.842
739.3373413085938 0 4423.8823
740.3487548828125 0 7190.8296 y Water loss 2
741.3519897460938 0 2059.6924
742.3563232421875 0 1485.3907
757.3479614257812 0 30024.035
757.9000244140625 0 682.1213
758.3563232421875 0 111983.664 y 2
759.359619140625 0 43587.043
760.3648071289062 0 10479.497
761.3657836914062 0 1495.2941
764.3615112304688 0 1171.7866
764.86279296875 0 1407.46
765.3693237304688 0 951.6412
766.3571166992188 0 4801.9375
766.859130859375 0 3611.8367
774.3262939453125 0 662.939
775.3973999023438 0 122259.38 c 6
776.4006958007812 0 50407.82
777.4029541015625 0 13102.91
778.4041137695312 0 916.83856
786.43701171875 0 694.51996
788.4025268554688 0 764.5496
801.4136352539062 0 8179.5376
802.4143676757812 0 3601.6814
803.4203491210938 0 1573.8478
814.427978515625 0 1963.9624
815.4375610351562 0 1084.208
843.43115234375 0 661.935
851.3983154296875 0 967.8682
857.4678344726562 0 708.85
859.4202270507812 0 1707.6917
860.426513671875 0 2924.168
861.43408203125 0 1654.3486
869.402587890625 0 1116.7767
870.4031982421875 0 1375.5465
871.396484375 0 3284.0315 z Water loss 1
872.39892578125 0 2218.8423
887.412841796875 0 16829.145 c Ammonia loss 7
888.414794921875 0 7889.2065
889.40771484375 0 11203.038 z 1
890.4099731445312 0 5394.791
891.4194946289062 0 1365.9733
904.4397583007812 0 160497.88 c 7
905.4423217773438 0 85355.15 y 1
906.445556640625 0 24654.12
907.448486328125 0 1993.05
939.4638671875 0 820.2185
940.4634399414062 0 728.68085
947.4942626953125 0 8805.209
948.49658203125 0 4810.036
949.5015258789062 0 934.212
954.4931030273438 0 634.6811
957.50390625 0 844.67773
959.43408203125 0 1172.4023
963.4509887695312 0 2183.7542
964.4573364257812 0 1077.2705
965.4569091796875 0 566.3162
973.520263671875 0 685.92676
974.5121459960938 0 1183.8457
984.4773559570312 0 2125.393
985.4815673828125 0 1463.8018
991.5182495117188 0 1711.8479
1001.481689453125 0 1101.7783
1002.4884033203125 0 14739.267
1002.5972900390625 0 1083.6382
1003.4937744140625 0 10639.586
1004.494140625 0 3798.734
1005.4883422851562 0 1515.5767
1016.5828247070312 0 3888.5413
1017.5813598632812 0 2377.992
1018.5067138671875 0 37710.05
1019.5143432617188 0 145280.61
1020.5173950195312 0 80638.01
1020.6447143554688 0 809.4445
1021.5206909179688 0 24653.158
1022.5199584960938 0 2464.1924
1051.5052490234375 0 1058.4874
1314.816650390625 0 773.86664
1373.695068359375 0 1154.3864
1487.7401123046875 0 970.72955
1489.7415771484375 0 707.7876
1497.666748046875 0 1071.3439
1504.7347412109375 0 1338.3256
1508.7635498046875 0 925.1305
1511.70849609375 0 1392.1067
1513.8790283203125 0 984.47565
1528.7064208984375 0 679.75555
1530.6815185546875 0 659.97314
1530.9085693359375 0 745.52136
1531.708984375 0 1776.7155
1532.7099609375 0 2890.117
1533.698486328125 0 1372.9967
3092.087890625 0 697.6051

Spectrum Details

|  |  |
| --- | --- |
| Matched peaks? Matched peaks The total absolute number of peaks matched. Additionally in brackets the total fraction of peaks matched and the total number of peaks is shown. | 24 (11.82% of 203) |
| FDR? FDR The false discovery rate estimated for this peptide. It is calculated by matching all theoretical fragments with a non-integer shift with the raw peaks for this spectrum. This is done with 40 different shifts. The resulting percentage is the average number of annotated peaks over the number of annotated peaks with the correct spectrum. | 1.29% |
| Satellite FDR? Satellite FDR See the FDR for details on its calculation. This satellite ion specific FDR only contains the satellite ions (d/w) for I/L/J positions. | - |
| PSM Score? PSM Score The PSM Score as given by Hecklib to this annotated spectrum. It is shown with three significant figures. | 326 |

#### Spectrum 8125? Spectrum 8125 The raw spectrum of this peptide as annotated by Hecklib. The fragments are coloured according to ion type (see legend). Any peaks with a star '\*' as text can be hovered over to see the full details, first the ion type second the mass shift type. By hovering over the amino acids in the peptide or ions in the legend the corresponding peaks are highlighted. By toggling the 'Unassigned' label you can turn the background (unassigned) peaks on or off in the plot. By updating the slider in the Ion legend you can update the spectrum to only show the top X% of the peaks with labels. The top X% means any peak that is within X% of the highest intensity. By dragging in the spectrum you can zoom in to a specific part of the spectrum and use 'Zoom Out' to get back to the original zoom level. The annotation of the spectrum is based on the given sequence in the peptides file and is done with different software so inconsistencies are likely. The peaks are annotated based on the given sequence, with 20 ppm tolerance.

Copy Data

##### Spectrum 8125 (TSV)

###### Preview

```
Loading example...
```

*Click on the button to copy the data to your clipboard.*

Mz MinMz MaxIntensity Max

WidthHeightPeptide font sizePeptide stroke widthSpectrum font sizeSpectrum stroke widthCompact peptide

Ion legend

wxyz

abcd

OtherUnassignedIon^Charge^_Position_Show for top:%

JFPPSSEEJ

01.48e+52.97e+54.45e+55.94e+5

Zoom Out

y^+1^_1_y^+1^_2_z^+1^_2_y^+1^_2_c^+1^_2_z^+2^_6_c^+1^_3_y^+2^_7_y^+1^_3_c^+2^_7_c^+2^_7_y^+1^_3_c^+2^_8_c^+1^_4_y^+1^_4_c^+1^_4_y^+1^_4_c^+1^_5_y^+1^_5_c^+1^_5_y^+1^_5_z^+1^_6_c^+1^_6_c^+1^_6_y^+1^_6_z^+1^_6_c^+1^_6_y^+1^_6_w^+1^_7_y^+1^_7_c^+1^_7_c^+1^_7_z^+1^_8_c^+1^_8_z^+1^_8_c^+1^_8_

0659131719762634

Fragment Matches Table

Show background peaks

| Position | Ion type | Intensity | mz Theoretical | mz Error (Th) | mz Error (ppm) | Charge | Series Number |
| --- | --- | --- | --- | --- | --- | --- | --- |
| - | - | 1.297E+04 | 120.1 | - | - | 0 | - |
| - | - | 370.6 | 120.9 | - | - | 0 | - |
| - | - | 1033 | 121.1 | - | - | 0 | - |
| - | - | 442.5 | 129.9 | - | - | 0 | - |
| - | - | 3325 | 130.1 | - | - | 0 | - |
| 9 | y | 1.019E+05 | 132.1 | 0.0003288 | 2.489 | +1 | 1 |
| - | - | 6392 | 133.1 | - | - | 0 | - |
| - | - | 517.2 | 137.5 | - | - | 0 | - |
| - | - | 397.6 | 139.9 | - | - | 0 | - |
| - | - | 505.7 | 148.8 | - | - | 0 | - |
| - | - | 479.9 | 148.9 | - | - | 0 | - |
| - | - | 548.2 | 148.9 | - | - | 0 | - |
| - | - | 524.7 | 148.9 | - | - | 0 | - |
| - | - | 754.4 | 148.9 | - | - | 0 | - |
| - | - | 782 | 148.9 | - | - | 0 | - |
| - | - | 1088 | 148.9 | - | - | 0 | - |
| - | - | 1256 | 148.9 | - | - | 0 | - |
| - | - | 1883 | 148.9 | - | - | 0 | - |
| - | - | 3816 | 148.9 | - | - | 0 | - |
| - | - | 4567 | 149 | - | - | 0 | - |
| - | - | 2701 | 149 | - | - | 0 | - |
| - | - | 1455 | 149 | - | - | 0 | - |
| - | - | 1121 | 149 | - | - | 0 | - |
| - | - | 663.1 | 149 | - | - | 0 | - |
| - | - | 858.2 | 149 | - | - | 0 | - |
| - | - | 565.7 | 149 | - | - | 0 | - |
| - | - | 403.3 | 149 | - | - | 0 | - |
| - | - | 404.3 | 152.4 | - | - | 0 | - |
| - | - | 506.5 | 157.9 | - | - | 0 | - |
| - | - | 849.7 | 168.1 | - | - | 0 | - |
| - | - | 432.2 | 170.5 | - | - | 0 | - |
| - | - | 541.8 | 171.2 | - | - | 0 | - |
| - | - | 452.6 | 182.6 | - | - | 0 | - |
| - | - | 541.6 | 189.1 | - | - | 0 | - |
| - | - | 3879 | 197.1 | - | - | 0 | - |
| - | - | 711.9 | 198.1 | - | - | 0 | - |
| - | - | 2592 | 199.1 | - | - | 0 | - |
| - | - | 593.4 | 202.1 | - | - | 0 | - |
| - | - | 496.9 | 205.1 | - | - | 0 | - |
| - | - | 5752 | 212.1 | - | - | 0 | - |
| - | - | 689.8 | 213.1 | - | - | 0 | - |
| - | - | 3175 | 217.1 | - | - | 0 | - |
| - | - | 1645 | 217.1 | - | - | 0 | - |
| - | - | 1697 | 225.1 | - | - | 0 | - |
| - | - | 572.9 | 233 | - | - | 0 | - |
| - | - | 8.384E+04 | 233.2 | - | - | 0 | - |
| - | - | 1.088E+04 | 234.2 | - | - | 0 | - |
| - | - | 725.4 | 235.2 | - | - | 0 | - |
| - | - | 505.5 | 238.8 | - | - | 0 | - |
| - | - | 4160 | 241.1 | - | - | 0 | - |
| 8 | y | 5.522E+04 | 243.1 | 0.0003896 | 1.602 | +1 | 2 |
| - | - | 5835 | 244.1 | - | - | 0 | - |
| 8 | z | 1025 | 245.1 | 0.002201 | 8.98 | +1 | 2 |
| - | - | 4841 | 259.1 | - | - | 0 | - |
| 8 | y | 1.519E+04 | 261.1 | 0.0001856 | 0.7108 | +1 | 2 |
| 2 | c | 5.919E+04 | 261.2 | 0.0003103 | 1.188 | +1 | 2 |
| - | - | 1895 | 262.1 | - | - | 0 | - |
| - | - | 9402 | 262.2 | - | - | 0 | - |
| - | - | 1004 | 263.2 | - | - | 0 | - |
| - | - | 2908 | 264.1 | - | - | 0 | - |
| - | - | 675.3 | 282.1 | - | - | 0 | - |
| - | - | 1.494E+04 | 282.1 | - | - | 0 | - |
| - | - | 2428 | 283.1 | - | - | 0 | - |
| - | - | 525 | 284.1 | - | - | 0 | - |
| - | - | 1144 | 286.1 | - | - | 0 | - |
| - | - | 2405 | 296.1 | - | - | 0 | - |
| - | - | 968.4 | 297.2 | - | - | 0 | - |
| - | - | 569.4 | 298.2 | - | - | 0 | - |
| - | - | 3.269E+04 | 299.2 | - | - | 0 | - |
| - | - | 3621 | 300.2 | - | - | 0 | - |
| - | - | 972.3 | 305.1 | - | - | 0 | - |
| - | - | 628.2 | 305.6 | - | - | 0 | - |
| - | - | 1430 | 306.2 | - | - | 0 | - |
| - | - | 760.7 | 308.2 | - | - | 0 | - |
| - | - | 554.4 | 308.2 | - | - | 0 | - |
| - | - | 556.6 | 310.1 | - | - | 0 | - |
| 4 | z | 4575 | 314.1 | 0.005787 | 18.42 | +2 | 6 |
| - | - | 874.9 | 314.6 | - | - | 0 | - |
| - | - | 1181 | 328.1 | - | - | 0 | - |
| - | - | 846.1 | 329.7 | - | - | 0 | - |
| - | - | 942.3 | 331.2 | - | - | 0 | - |
| - | - | 1360 | 333.2 | - | - | 0 | - |
| - | - | 5339 | 338.7 | - | - | 0 | - |
| - | - | 2094 | 339.2 | - | - | 0 | - |
| - | - | 600.4 | 341.2 | - | - | 0 | - |
| - | - | 3267 | 346.1 | - | - | 0 | - |
| - | - | 724.3 | 347.7 | - | - | 0 | - |
| - | - | 6059 | 351.2 | - | - | 0 | - |
| - | - | 950.6 | 352.2 | - | - | 0 | - |
| - | - | 818.5 | 352.7 | - | - | 0 | - |
| - | - | 758.2 | 353.2 | - | - | 0 | - |
| - | - | 2444 | 354.2 | - | - | 0 | - |
| 3 | c | 7348 | 358.2 | 0.0002809 | 0.7841 | +1 | 3 |
| - | - | 1649 | 359.2 | - | - | 0 | - |
| - | - | 561.5 | 360.5 | - | - | 0 | - |
| - | - | 1962 | 361.7 | - | - | 0 | - |
| - | - | 1168 | 362.2 | - | - | 0 | - |
| - | - | 8718 | 369.2 | - | - | 0 | - |
| - | - | 1811 | 370.2 | - | - | 0 | - |
| 3 | y | 4674 | 370.7 | 0.004977 | 13.43 | +2 | 7 |
| - | - | 2024 | 371.2 | - | - | 0 | - |
| - | - | 819.3 | 371.7 | - | - | 0 | - |
| 7 | y | 2936 | 372.2 | 0.0003227 | 0.8672 | +1 | 3 |
| - | - | 865.8 | 373.2 | - | - | 0 | - |
| 7 | c | 2411 | 379.7 | 0.004942 | 13.02 | +2 | 7 |
| - | - | 1131 | 380.2 | - | - | 0 | - |
| - | - | 7.775E+04 | 386.2 | - | - | 0 | - |
| - | - | 1.428E+04 | 387.2 | - | - | 0 | - |
| - | - | 626.1 | 387.7 | - | - | 0 | - |
| 7 | c | 1995 | 388.2 | 0.006045 | 15.57 | +2 | 7 |
| 7 | y | 2395 | 390.2 | 0.0004087 | 1.047 | +1 | 3 |
| - | - | 2214 | 401.2 | - | - | 0 | - |
| - | - | 1320 | 412.2 | - | - | 0 | - |
| - | - | 1799 | 427.3 | - | - | 0 | - |
| - | - | 1.466E+04 | 428.3 | - | - | 0 | - |
| - | - | 876.9 | 429.2 | - | - | 0 | - |
| - | - | 6070 | 429.3 | - | - | 0 | - |
| - | - | 875.2 | 430.3 | - | - | 0 | - |
| - | - | 2693 | 433.2 | - | - | 0 | - |
| - | - | 547.9 | 434.2 | - | - | 0 | - |
| - | - | 989.1 | 435.2 | - | - | 0 | - |
| - | - | 1219 | 441.2 | - | - | 0 | - |
| 8 | c | 1527 | 444.2 | 0.0004945 | 1.113 | +2 | 8 |
| - | - | 872.2 | 452.2 | - | - | 0 | - |
| 4 | c | 6141 | 455.3 | 0.0003429 | 0.7533 | +1 | 4 |
| - | - | 4631 | 456.3 | - | - | 0 | - |
| - | - | 1278 | 457.3 | - | - | 0 | - |
| - | - | 557.6 | 458.3 | - | - | 0 | - |
| 6 | y | 1084 | 459.2 | 0.001772 | 3.859 | +1 | 4 |
| - | - | 2294 | 462.2 | - | - | 0 | - |
| - | - | 2172 | 470.2 | - | - | 0 | - |
| - | - | 1052 | 471.2 | - | - | 0 | - |
| - | - | 3.114E+04 | 471.3 | - | - | 0 | - |
| 4 | c | 3.132E+04 | 472.3 | 0.0005714 | 1.21 | +1 | 4 |
| - | - | 9859 | 473.3 | - | - | 0 | - |
| - | - | 4857 | 474.3 | - | - | 0 | - |
| - | - | 975.3 | 475.3 | - | - | 0 | - |
| 6 | y | 3014 | 477.2 | 0.0009731 | 2.039 | +1 | 4 |
| - | - | 874 | 478.2 | - | - | 0 | - |
| - | - | 1.049E+04 | 480.2 | - | - | 0 | - |
| - | - | 1720 | 481.2 | - | - | 0 | - |
| - | - | 597.1 | 484.2 | - | - | 0 | - |
| - | - | 635.7 | 491.4 | - | - | 0 | - |
| - | - | 3067 | 497.3 | - | - | 0 | - |
| - | - | 4.337E+04 | 498.2 | - | - | 0 | - |
| - | - | 3071 | 498.3 | - | - | 0 | - |
| - | - | 1.103E+04 | 499.2 | - | - | 0 | - |
| - | - | 1309 | 499.3 | - | - | 0 | - |
| - | - | 1869 | 500.2 | - | - | 0 | - |
| - | - | 1094 | 502.3 | - | - | 0 | - |
| - | - | 774.6 | 509.2 | - | - | 0 | - |
| - | - | 1561 | 512.2 | - | - | 0 | - |
| - | - | 3.856E+04 | 515.2 | - | - | 0 | - |
| - | - | 8824 | 516.2 | - | - | 0 | - |
| - | - | 8580 | 516.3 | - | - | 0 | - |
| - | - | 1903 | 517.3 | - | - | 0 | - |
| - | - | 2214 | 517.3 | - | - | 0 | - |
| - | - | 1141 | 524.3 | - | - | 0 | - |
| - | - | 4747 | 530.2 | - | - | 0 | - |
| - | - | 1135 | 531.2 | - | - | 0 | - |
| - | - | 612 | 538.4 | - | - | 0 | - |
| 5 | c | 7224 | 542.3 | 0.0005411 | 0.9978 | +1 | 5 |
| - | - | 2398 | 543.3 | - | - | 0 | - |
| - | - | 7939 | 544.3 | - | - | 0 | - |
| - | - | 3152 | 545.3 | - | - | 0 | - |
| 5 | y | 1094 | 546.2 | 0.0003223 | 0.5901 | +1 | 5 |
| - | - | 8.542E+04 | 558.3 | - | - | 0 | - |
| 5 | c | 6.587E+04 | 559.3 | 0.00135 | 2.413 | +1 | 5 |
| - | - | 1.883E+04 | 560.3 | - | - | 0 | - |
| - | - | 4651 | 561.3 | - | - | 0 | - |
| - | - | 1524 | 562.3 | - | - | 0 | - |
| 5 | y | 4622 | 564.3 | 4.95E-05 | 0.08773 | +1 | 5 |
| - | - | 927.7 | 565.3 | - | - | 0 | - |
| - | - | 1026 | 573.2 | - | - | 0 | - |
| - | - | 902 | 583.3 | - | - | 0 | - |
| - | - | 820.4 | 584.3 | - | - | 0 | - |
| - | - | 591.1 | 585.3 | - | - | 0 | - |
| - | - | 2402 | 591.2 | - | - | 0 | - |
| - | - | 1212 | 593.3 | - | - | 0 | - |
| - | - | 4068 | 599.3 | - | - | 0 | - |
| - | - | 1431 | 600.3 | - | - | 0 | - |
| - | - | 1356 | 601.3 | - | - | 0 | - |
| - | - | 1479 | 601.3 | - | - | 0 | - |
| - | - | 1983 | 603.4 | - | - | 0 | - |
| - | - | 1043 | 604.4 | - | - | 0 | - |
| - | - | 1.109E+04 | 609.3 | - | - | 0 | - |
| - | - | 4286 | 610.3 | - | - | 0 | - |
| - | - | 1349 | 611.3 | - | - | 0 | - |
| - | - | 6484 | 611.3 | - | - | 0 | - |
| - | - | 2149 | 612.3 | - | - | 0 | - |
| - | - | 2689 | 626.3 | - | - | 0 | - |
| 4 | z | 1.156E+05 | 627.3 | 0.01217 | 19.4 | +1 | 6 |
| - | - | 3.488E+04 | 628.3 | - | - | 0 | - |
| - | - | 817 | 628.3 | - | - | 0 | - |
| 6 | c | 1049 | 628.3 | 0.002821 | 4.49 | +1 | 6 |
| - | - | 7626 | 629.3 | - | - | 0 | - |
| 6 | c | 1.236E+04 | 629.3 | 0.001105 | 1.757 | +1 | 6 |
| - | - | 1503 | 630.3 | - | - | 0 | - |
| - | - | 4994 | 630.3 | - | - | 0 | - |
| 4 | y | 3070 | 643.3 | 0.001453 | 2.258 | +1 | 6 |
| - | - | 5004 | 644.3 | - | - | 0 | - |
| 4 | z | 3840 | 645.3 | 0.002898 | 4.491 | +1 | 6 |
| - | - | 3716 | 645.3 | - | - | 0 | - |
| 6 | c | 1.477E+05 | 646.4 | 0.0005573 | 0.8623 | +1 | 6 |
| - | - | 5.552E+04 | 647.4 | - | - | 0 | - |
| - | - | 1.081E+04 | 648.4 | - | - | 0 | - |
| - | - | 620.7 | 649.4 | - | - | 0 | - |
| - | - | 2033 | 660.3 | - | - | 0 | - |
| 4 | y | 4.032E+04 | 661.3 | 0.0003483 | 0.5266 | +1 | 6 |
| - | - | 1.346E+04 | 662.3 | - | - | 0 | - |
| - | - | 2571 | 663.3 | - | - | 0 | - |
| - | - | 1.067E+04 | 672.4 | - | - | 0 | - |
| - | - | 4673 | 673.4 | - | - | 0 | - |
| - | - | 1201 | 685.3 | - | - | 0 | - |
| - | - | 755.3 | 686.3 | - | - | 0 | - |
| - | - | 2028 | 689.3 | - | - | 0 | - |
| - | - | 807.7 | 696.4 | - | - | 0 | - |
| - | - | 1798 | 698.3 | - | - | 0 | - |
| - | - | 4540 | 701.3 | - | - | 0 | - |
| - | - | 1978 | 702.3 | - | - | 0 | - |
| - | - | 2016 | 704.3 | - | - | 0 | - |
| - | - | 2098 | 712.4 | - | - | 0 | - |
| - | - | 1689 | 713.4 | - | - | 0 | - |
| - | - | 2484 | 714.3 | - | - | 0 | - |
| 3 | w | 979.4 | 715.3 | 0.004479 | 6.262 | +1 | 7 |
| - | - | 4461 | 722.3 | - | - | 0 | - |
| - | - | 1589 | 723.3 | - | - | 0 | - |
| - | - | 937.2 | 724.3 | - | - | 0 | - |
| - | - | 1040 | 729.4 | - | - | 0 | - |
| - | - | 7101 | 730.4 | - | - | 0 | - |
| - | - | 3694 | 731.4 | - | - | 0 | - |
| - | - | 3005 | 732.4 | - | - | 0 | - |
| - | - | 1313 | 733.4 | - | - | 0 | - |
| - | - | 1.8E+04 | 739.3 | - | - | 0 | - |
| 3 | y | 3.045E+04 | 740.3 | 0.002094 | 2.829 | +1 | 7 |
| - | - | 1.118E+04 | 741.4 | - | - | 0 | - |
| - | - | 4798 | 742.4 | - | - | 0 | - |
| - | - | 2309 | 743.4 | - | - | 0 | - |
| - | - | 1.136E+05 | 757.3 | - | - | 0 | - |
| 7 | c | 4.264E+05 | 758.4 | 0.01359 | 17.93 | +1 | 7 |
| - | - | 1.654E+05 | 759.4 | - | - | 0 | - |
| - | - | 4.147E+04 | 760.4 | - | - | 0 | - |
| - | - | 5818 | 761.4 | - | - | 0 | - |
| - | - | 1163 | 762.4 | - | - | 0 | - |
| - | - | 868.8 | 766.3 | - | - | 0 | - |
| - | - | 2128 | 774.3 | - | - | 0 | - |
| 7 | c | 4.392E+05 | 775.4 | 0.0003837 | 0.4948 | +1 | 7 |
| - | - | 1.879E+05 | 776.4 | - | - | 0 | - |
| - | - | 4.623E+04 | 777.4 | - | - | 0 | - |
| - | - | 3937 | 778.4 | - | - | 0 | - |
| - | - | 1042 | 784.3 | - | - | 0 | - |
| - | - | 1057 | 785.3 | - | - | 0 | - |
| - | - | 1864 | 786.4 | - | - | 0 | - |
| - | - | 666.3 | 786.4 | - | - | 0 | - |
| - | - | 1212 | 787.4 | - | - | 0 | - |
| - | - | 1484 | 788.4 | - | - | 0 | - |
| - | - | 935.7 | 789.4 | - | - | 0 | - |
| - | - | 855.2 | 790.4 | - | - | 0 | - |
| - | - | 699.5 | 799.4 | - | - | 0 | - |
| - | - | 2.969E+04 | 801.4 | - | - | 0 | - |
| - | - | 1.383E+04 | 802.4 | - | - | 0 | - |
| - | - | 4112 | 803.4 | - | - | 0 | - |
| - | - | 968.9 | 814.4 | - | - | 0 | - |
| - | - | 1854 | 817.4 | - | - | 0 | - |
| - | - | 743.1 | 818.4 | - | - | 0 | - |
| - | - | 689.7 | 841.4 | - | - | 0 | - |
| - | - | 961.6 | 842.4 | - | - | 0 | - |
| - | - | 925.2 | 843.4 | - | - | 0 | - |
| - | - | 1309 | 851.4 | - | - | 0 | - |
| - | - | 9651 | 859.4 | - | - | 0 | - |
| - | - | 1.245E+04 | 860.4 | - | - | 0 | - |
| - | - | 7665 | 861.4 | - | - | 0 | - |
| - | - | 1689 | 862.4 | - | - | 0 | - |
| - | - | 6666 | 869.4 | - | - | 0 | - |
| - | - | 3870 | 870.4 | - | - | 0 | - |
| 2 | z | 1.132E+04 | 871.4 | 0.001111 | 1.275 | +1 | 8 |
| - | - | 5980 | 872.4 | - | - | 0 | - |
| - | - | 2520 | 873.4 | - | - | 0 | - |
| 8 | c | 7.803E+04 | 887.4 | 0.0002795 | 0.3149 | +1 | 8 |
| - | - | 3.664E+04 | 888.4 | - | - | 0 | - |
| 2 | z | 3.927E+04 | 889.4 | 0.003302 | 3.713 | +1 | 8 |
| - | - | 1.792E+04 | 890.4 | - | - | 0 | - |
| - | - | 6126 | 891.4 | - | - | 0 | - |
| - | - | 960.8 | 892.4 | - | - | 0 | - |
| - | - | 1289 | 901.5 | - | - | 0 | - |
| - | - | 1057 | 902.5 | - | - | 0 | - |
| - | - | 754.4 | 903.5 | - | - | 0 | - |
| 8 | c | 5.881E+05 | 904.4 | 0.0006373 | 0.7046 | +1 | 8 |
| - | - | 2.993E+05 | 905.4 | - | - | 0 | - |
| - | - | 8.478E+04 | 906.4 | - | - | 0 | - |
| - | - | 7120 | 907.4 | - | - | 0 | - |
| - | - | 1103 | 917.4 | - | - | 0 | - |
| - | - | 828.4 | 918.4 | - | - | 0 | - |
| - | - | 612.8 | 919.5 | - | - | 0 | - |
| - | - | 1664 | 929.5 | - | - | 0 | - |
| - | - | 1573 | 930.5 | - | - | 0 | - |
| - | - | 791.2 | 931.5 | - | - | 0 | - |
| - | - | 694.5 | 932.5 | - | - | 0 | - |
| - | - | 1086 | 933.5 | - | - | 0 | - |
| - | - | 819.8 | 934.4 | - | - | 0 | - |
| - | - | 842.9 | 946.4 | - | - | 0 | - |
| - | - | 3.213E+04 | 947.5 | - | - | 0 | - |
| - | - | 1.833E+04 | 948.5 | - | - | 0 | - |
| - | - | 6007 | 949.5 | - | - | 0 | - |
| - | - | 712.9 | 956.5 | - | - | 0 | - |
| - | - | 1211 | 957.5 | - | - | 0 | - |
| - | - | 4215 | 959.4 | - | - | 0 | - |
| - | - | 903.9 | 960.4 | - | - | 0 | - |
| - | - | 955.4 | 960.5 | - | - | 0 | - |
| - | - | 815.5 | 961.5 | - | - | 0 | - |
| - | - | 7761 | 963.5 | - | - | 0 | - |
| - | - | 5146 | 964.5 | - | - | 0 | - |
| - | - | 1860 | 965.5 | - | - | 0 | - |
| - | - | 950.4 | 971.5 | - | - | 0 | - |
| - | - | 1461 | 972.5 | - | - | 0 | - |
| - | - | 4641 | 973.5 | - | - | 0 | - |
| - | - | 3045 | 974.5 | - | - | 0 | - |
| - | - | 1118 | 975.5 | - | - | 0 | - |
| - | - | 723.8 | 976.5 | - | - | 0 | - |
| - | - | 851 | 977.5 | - | - | 0 | - |
| - | - | 661 | 979.4 | - | - | 0 | - |
| - | - | 1.03E+04 | 984.5 | - | - | 0 | - |
| - | - | 6504 | 985.5 | - | - | 0 | - |
| - | - | 1885 | 986.5 | - | - | 0 | - |
| - | - | 1816 | 989.5 | - | - | 0 | - |
| - | - | 8017 | 991.5 | - | - | 0 | - |
| - | - | 4371 | 992.5 | - | - | 0 | - |
| - | - | 1476 | 993.5 | - | - | 0 | - |
| - | - | 7099 | 1002 | - | - | 0 | - |
| - | - | 5.951E+04 | 1002 | - | - | 0 | - |
| - | - | 3.442E+04 | 1003 | - | - | 0 | - |
| - | - | 1.038E+04 | 1004 | - | - | 0 | - |
| - | - | 2206 | 1005 | - | - | 0 | - |
| - | - | 1.349E+05 | 1019 | - | - | 0 | - |
| - | - | 5.412E+05 | 1020 | - | - | 0 | - |
| - | - | 2.893E+05 | 1021 | - | - | 0 | - |
| - | - | 9.22E+04 | 1022 | - | - | 0 | - |
| - | - | 1.024E+04 | 1023 | - | - | 0 | - |
| - | - | 4278 | 1052 | - | - | 0 | - |
| - | - | 2287 | 1053 | - | - | 0 | - |
| - | - | 705 | 1582 | - | - | 0 | - |
| - | - | 713.8 | 2608 | - | - | 0 | - |

m/z Charge Intensity FragmentType MassShift Position
120.08109283447266 0 12973.089
120.91679382324219 0 370.63055
121.08446502685547 0 1032.5051
129.8535614013672 0 442.47815
130.0501251220703 0 3324.7427
132.10223388671875 0 101936.76 y 8
133.1055450439453 0 6391.7183
137.504638671875 0 517.23553
139.8654327392578 0 397.6295
148.76048278808594 0 505.656
148.88436889648438 0 479.92435
148.89089965820312 0 548.1526
148.89825439453125 0 524.66833
148.90533447265625 0 754.42535
148.91259765625 0 782.0114
148.9199676513672 0 1088.2339
148.92727661132812 0 1255.5017
148.9342803955078 0 1883.2097
148.94200134277344 0 3816.2
148.95867919921875 0 4567.339
148.96652221679688 0 2701.22
148.97377014160156 0 1455.469
148.9806671142578 0 1120.8071
148.98825073242188 0 663.1144
148.99530029296875 0 858.23663
149.0021514892578 0 565.71124
149.0103759765625 0 403.27362
152.44688415527344 0 404.30737
157.87449645996094 0 506.51245
168.12588500976562 0 849.7057
170.5107879638672 0 432.194
171.23089599609375 0 541.8495
182.607177734375 0 452.56766
189.08839416503906 0 541.6026
197.1287841796875 0 3878.625
198.13233947753906 0 711.8561
199.0716094970703 0 2591.7898
202.1185760498047 0 593.3975
205.12368774414062 0 496.85184
212.13966369628906 0 5751.9614
213.1433563232422 0 689.8372
217.082275390625 0 3175.4292
217.14637756347656 0 1644.7268
225.12353515625 0 1697.3376
233.0255584716797 0 572.85364
233.16522216796875 0 83837.37
234.1685791015625 0 10880.097
235.17141723632812 0 725.3617
238.82131958007812 0 505.5394
241.082275390625 0 4160.1875
243.1343231201172 0 55222.16 y Water loss 7
244.13731384277344 0 5835.226
245.1279754638672 0 1025.3651 z 7
259.09271240234375 0 4840.8965
261.1446838378906 0 15188.666 y 7
261.1600646972656 0 59187.277 c Ammonia loss 1
262.14813232421875 0 1894.7122
262.163330078125 0 9402.131
263.16693115234375 0 1003.5769
264.13458251953125 0 2908.1074
282.1279296875 0 675.31384
282.1451721191406 0 14942.908
283.1481018066406 0 2427.5781
284.1474304199219 0 525.0302
286.1036682128906 0 1143.6038
296.1246643066406 0 2405.2031
297.1582336425781 0 968.3533
298.1632385253906 0 569.3811
299.1718444824219 0 32686.984
300.1750183105469 0 3620.9548
305.1286926269531 0 972.3284
305.6314697265625 0 628.18445
306.16400146484375 0 1430.2936
308.15936279296875 0 760.7233
308.1952209472656 0 554.4046
310.10498046875 0 556.6427
314.1351623535156 0 4575.2197 z Water loss 3
314.6353759765625 0 874.93604
328.1143798828125 0 1180.758
329.66339111328125 0 846.1106
331.226806640625 0 942.2527
333.1560363769531 0 1359.6285
338.6690673828125 0 5338.7373
339.1706848144531 0 2094.1943
341.1837158203125 0 600.4211
346.1247253417969 0 3266.9028
347.6741638183594 0 724.3453
351.16680908203125 0 6059.276
352.1697082519531 0 950.64465
352.66619873046875 0 818.5449
353.169189453125 0 758.1589
354.1659851074219 0 2444.0117
358.2127990722656 0 7347.55 c Ammonia loss 2
359.2158203125 0 1648.7498
360.5106506347656 0 561.48126
361.6763000488281 0 1962.1693
362.1776428222656 0 1168.2003
369.1769104003906 0 8718.166
370.1793518066406 0 1811.1263
370.6816711425781 0 4674.485 y Water loss 2
371.18548583984375 0 2024.2196
371.68609619140625 0 819.3404
372.1768493652344 0 2936.4597 y Water loss 6
373.1813659667969 0 865.7605
379.6846618652344 0 2411.481 c Ammonia loss 6
380.18939208984375 0 1130.9879
386.20379638671875 0 77752.41
387.206787109375 0 14277.94
387.6673278808594 0 626.1373
388.20892333984375 0 1994.7944 c 6
390.1875 0 2394.8787 y 6
401.1671142578125 0 2214.1538
412.20892333984375 0 1319.7156
427.272216796875 0 1799.2424
428.2787170410156 0 14664.3545
429.2141418457031 0 876.9239
429.2839050292969 0 6070.0024
430.2909851074219 0 875.1971
433.15643310546875 0 2692.687
434.204833984375 0 547.8901
435.2056884765625 0 989.08344
441.1984558105469 0 1219.3789
444.2113952636719 0 1526.8193 c Ammonia loss 7
452.2143249511719 0 872.1544
455.265625 0 6141.033 c Ammonia loss 3
456.2721252441406 0 4631.42
457.2771301269531 0 1278.0085
458.282958984375 0 557.62823
459.2103271484375 0 1084.2329 y Water loss 5
462.1990661621094 0 2293.8906
470.22552490234375 0 2172.3455
471.2454528808594 0 1052.2205
471.2845153808594 0 31138.5
472.291259765625 0 31318.768 c 3
473.29571533203125 0 9858.52
474.3062438964844 0 4856.731
475.3098449707031 0 975.3388
477.2200927734375 0 3013.6174 y 5
478.22186279296875 0 874.0015
480.20989990234375 0 10485.969
481.21343994140625 0 1719.6376
484.203857421875 0 597.1286
491.4392395019531 0 635.70624
497.2999267578125 0 3066.807
498.22021484375 0 43368.285
498.30694580078125 0 3070.6064
499.2230224609375 0 11027.244
499.3133544921875 0 1308.5109
500.2264404296875 0 1869.0051
502.252685546875 0 1093.605
509.2494201660156 0 774.64215
512.2005004882812 0 1560.5216
515.2467041015625 0 38564.25
516.2492065429688 0 8824.361
516.3186645507812 0 8580.234
517.251953125 0 1903.3086
517.322021484375 0 2214.4438
524.2857666015625 0 1140.5492
530.2095947265625 0 4746.7563
531.2110595703125 0 1134.6743
538.3515014648438 0 611.99316
542.2978515625 0 7223.6626 c Ammonia loss 4
543.3014526367188 0 2398.3403
544.3131713867188 0 7939.1855
545.3167114257812 0 3152.2092
546.2409057617188 0 1093.9204 y Water loss 4
558.3165283203125 0 85417.48
559.322509765625 0 65866.49 c 4
560.3258666992188 0 18832.762
561.3356323242188 0 4650.981
562.3428955078125 0 1524.022
564.2510986328125 0 4621.8247 y 4
565.2574462890625 0 927.6595
573.2307739257812 0 1025.8763
583.3128051757812 0 902.04175
584.3319702148438 0 820.354
585.337158203125 0 591.11316
591.2418823242188 0 2402.4563
593.3046875 0 1211.6912
599.2677001953125 0 4068.197
600.2708129882812 0 1430.9025
601.2811889648438 0 1355.5343
601.3350830078125 0 1478.8685
603.3504028320312 0 1982.6375
604.3521728515625 0 1042.7731
609.251708984375 0 11088.978
610.255859375 0 4286.2817
611.2578125 0 1348.8562
611.3151245117188 0 6484.03
612.3197631835938 0 2148.6138
626.2557373046875 0 2688.841
627.262451171875 0 115557.8 z Water loss 3
628.2655029296875 0 34875.566
628.3368530273438 0 816.9774
628.34814453125 0 1048.7739 c Water loss 5
629.26806640625 0 7625.795
629.3304443359375 0 12356.706 c Ammonia loss 5
630.2705688476562 0 1503.0221
630.3328247070312 0 4994.47
643.2947998046875 0 3069.8254 y Water loss 3
644.2897338867188 0 5004.2085
645.2880859375 0 3840.3877 z 3
645.3491821289062 0 3715.5144
646.3564453125 0 147720.34 c 5
647.3592529296875 0 55516.418
648.36181640625 0 10806.251
649.3689575195312 0 620.74255
660.2945556640625 0 2033.2167
661.3042602539062 0 40324.504 y 3
662.3071899414062 0 13457.497
663.3107299804688 0 2571.4622
672.3720092773438 0 10667.774
673.375244140625 0 4672.7285
685.3270874023438 0 1200.7289
686.31884765625 0 755.3008
689.2987060546875 0 2028.3827
696.3587036132812 0 807.7166
698.3353271484375 0 1798.4355
701.2872314453125 0 4540.2295
702.2891845703125 0 1978.1964
704.3270263671875 0 2016.1947
712.3555908203125 0 2098.273
713.3584594726562 0 1688.8308
714.2942504882812 0 2484.1646
715.3099975585938 0 979.39703 w 2
722.3404541015625 0 4461.2734
723.34423828125 0 1588.8904
724.3479614257812 0 937.16223
729.3687744140625 0 1039.7521
730.3770141601562 0 7101.3984
731.3821411132812 0 3694.349
732.3926391601562 0 3004.9963
733.3964233398438 0 1312.6453
739.3387451171875 0 17999.166
740.3482055664062 0 30451.186 y Water loss 2
741.3519287109375 0 11177.482
742.35400390625 0 4797.6562
743.3607177734375 0 2309.032
757.349365234375 0 113572.98
758.3583374023438 0 426377.8 c Ammonia loss 6
759.3612670898438 0 165370.6
760.36572265625 0 41474.54
761.372802734375 0 5818.153
762.373291015625 0 1163.1682
766.3275146484375 0 868.8379
774.331298828125 0 2128.0115
775.3988647460938 0 439199.28 c 6
776.40185546875 0 187873.55
777.4044189453125 0 46227.867
778.4080200195312 0 3937.0369
784.328857421875 0 1042.2106
785.342041015625 0 1057.2565
786.3502197265625 0 1863.5667
786.4305419921875 0 666.3215
787.3568115234375 0 1211.7205
788.40625 0 1483.9929
789.412109375 0 935.70337
790.3590087890625 0 855.1992
799.3881225585938 0 699.50134
801.4144897460938 0 29688.254
802.4176025390625 0 13828.29
803.42041015625 0 4112.386
814.4342041015625 0 968.92596
817.3884887695312 0 1854.3947
818.3965454101562 0 743.11
841.4165649414062 0 689.7301
842.40283203125 0 961.6246
843.428466796875 0 925.1558
851.3989868164062 0 1308.9515
859.4196166992188 0 9651.459
860.4263916015625 0 12445.187
861.431640625 0 7664.7427
862.4348754882812 0 1688.8429
869.4039306640625 0 6665.8667
870.4086303710938 0 3870.3022
871.3969116210938 0 11324.207 z Water loss 1
872.399658203125 0 5979.7183
873.412353515625 0 2519.7175
887.4142456054688 0 78026.805 c Ammonia loss 7
888.4179077148438 0 36641.57
889.40966796875 0 39271.844 z 1
890.4107055664062 0 17921.332
891.4149780273438 0 6126.2334
892.4237060546875 0 960.8281
901.4863891601562 0 1289.3588
902.49853515625 0 1056.6178
903.5099487304688 0 754.4482
904.4417114257812 0 588104.9 c 7
905.4440307617188 0 299274.3
906.4466552734375 0 84776.67
907.448486328125 0 7120.478
917.4403076171875 0 1102.7429
918.44921875 0 828.37427
919.495849609375 0 612.84576
929.4988403320312 0 1664.235
930.47021484375 0 1572.8185
931.466064453125 0 791.1836
932.4751586914062 0 694.46344
933.4880981445312 0 1085.9346
934.41748046875 0 819.7799
946.4356079101562 0 842.89606
947.49560546875 0 32126.953
948.49853515625 0 18326.922
949.5029907226562 0 6006.6626
956.493896484375 0 712.8628
957.52001953125 0 1210.925
959.4360961914062 0 4214.8896
960.4227905273438 0 903.927
960.5130004882812 0 955.37994
961.5205078125 0 815.5042
963.4537353515625 0 7761.495
964.4574584960938 0 5145.585
965.4620971679688 0 1859.9249
971.5023193359375 0 950.3622
972.49462890625 0 1460.9604
973.5106201171875 0 4640.9175
974.5107421875 0 3044.681
975.513916015625 0 1118.1057
976.4733276367188 0 723.8145
977.4700317382812 0 851.0214
979.4421997070312 0 660.9503
984.4801635742188 0 10300.479
985.4822998046875 0 6503.764
986.4847412109375 0 1884.6495
989.5054321289062 0 1816.0815
991.521728515625 0 8016.7188
992.5256958007812 0 4371.266
993.530517578125 0 1476.2769
1001.5072021484375 0 7098.8325
1002.4910278320312 0 59507.527
1003.494140625 0 34419.8
1004.4962768554688 0 10379.867
1005.4844360351562 0 2205.9836
1018.5089111328125 0 134934.55
1019.516845703125 0 541174.5
1020.51953125 0 289307.8
1021.5228271484375 0 92198.63
1022.5220947265625 0 10240.791
1051.5076904296875 0 4277.9727
1052.5091552734375 0 2286.9211
1582.0927734375 0 705.02136
2608.064453125 0 713.8106

Spectrum Details

|  |  |
| --- | --- |
| Matched peaks? Matched peaks The total absolute number of peaks matched. Additionally in brackets the total fraction of peaks matched and the total number of peaks is shown. | 36 (10.53% of 342) |
| FDR? FDR The false discovery rate estimated for this peptide. It is calculated by matching all theoretical fragments with a non-integer shift with the raw peaks for this spectrum. This is done with 40 different shifts. The resulting percentage is the average number of annotated peaks over the number of annotated peaks with the correct spectrum. | 1.59% |
| Satellite FDR? Satellite FDR See the FDR for details on its calculation. This satellite ion specific FDR only contains the satellite ions (d/w) for I/L/J positions. | - |
| PSM Score? PSM Score The PSM Score as given by Hecklib to this annotated spectrum. It is shown with three significant figures. | 472 |

#### Spectrum 8215? Spectrum 8215 The raw spectrum of this peptide as annotated by Hecklib. The fragments are coloured according to ion type (see legend). Any peaks with a star '\*' as text can be hovered over to see the full details, first the ion type second the mass shift type. By hovering over the amino acids in the peptide or ions in the legend the corresponding peaks are highlighted. By toggling the 'Unassigned' label you can turn the background (unassigned) peaks on or off in the plot. By updating the slider in the Ion legend you can update the spectrum to only show the top X% of the peaks with labels. The top X% means any peak that is within X% of the highest intensity. By dragging in the spectrum you can zoom in to a specific part of the spectrum and use 'Zoom Out' to get back to the original zoom level. The annotation of the spectrum is based on the given sequence in the peptides file and is done with different software so inconsistencies are likely. The peaks are annotated based on the given sequence, with 20 ppm tolerance.

Copy Data

##### Spectrum 8215 (TSV)

###### Preview

```
Loading example...
```

*Click on the button to copy the data to your clipboard.*

Mz MinMz MaxIntensity Max

WidthHeightPeptide font sizePeptide stroke widthSpectrum font sizeSpectrum stroke widthCompact peptide

Ion legend

wxyz

abcd

OtherUnassignedIon^Charge^_Position_Show for top:%

JFPPSSEEJ

02.66e+55.32e+57.98e+51.06e+6

Zoom Out

y^+1^_1_y^+1^_2_z^+1^_2_y^+1^_2_c^+1^_2_z^+2^_6_c^+1^_3_y^+2^_7_y^+1^_3_c^+2^_7_c^+2^_7_y^+1^_3_c^+2^_8_y^+2^_8_c^+1^_4_y^+1^_4_c^+1^_4_y^+1^_4_c^+1^_5_y^+1^_5_c^+1^_5_y^+1^_5_z^+1^_6_c^+1^_6_y^+1^_6_z^+1^_6_c^+1^_6_y^+1^_6_w^+1^_7_y^+1^_7_c^+1^_7_c^+1^_7_z^+1^_8_c^+1^_8_c^+1^_8_z^+1^_8_c^+1^_8_

0777155523323110

Fragment Matches Table

Show background peaks

| Position | Ion type | Intensity | mz Theoretical | mz Error (Th) | mz Error (ppm) | Charge | Series Number |
| --- | --- | --- | --- | --- | --- | --- | --- |
| - | - | 3.891E+04 | 120.1 | - | - | 0 | - |
| - | - | 2744 | 121.1 | - | - | 0 | - |
| - | - | 525.9 | 121.1 | - | - | 0 | - |
| - | - | 557.6 | 121.8 | - | - | 0 | - |
| - | - | 679 | 126.1 | - | - | 0 | - |
| - | - | 8931 | 130.1 | - | - | 0 | - |
| 9 | y | 2.589E+05 | 132.1 | 0.0003593 | 2.72 | +1 | 1 |
| - | - | 1118 | 133.1 | - | - | 0 | - |
| - | - | 1.77E+04 | 133.1 | - | - | 0 | - |
| - | - | 704 | 134.1 | - | - | 0 | - |
| - | - | 1184 | 136.1 | - | - | 0 | - |
| - | - | 608.1 | 145 | - | - | 0 | - |
| - | - | 1721 | 147.1 | - | - | 0 | - |
| - | - | 805.9 | 148.1 | - | - | 0 | - |
| - | - | 563.8 | 148.9 | - | - | 0 | - |
| - | - | 632 | 148.9 | - | - | 0 | - |
| - | - | 832.5 | 148.9 | - | - | 0 | - |
| - | - | 794.2 | 148.9 | - | - | 0 | - |
| - | - | 1878 | 148.9 | - | - | 0 | - |
| - | - | 1598 | 148.9 | - | - | 0 | - |
| - | - | 2587 | 148.9 | - | - | 0 | - |
| - | - | 4751 | 148.9 | - | - | 0 | - |
| - | - | 6543 | 149 | - | - | 0 | - |
| - | - | 3762 | 149 | - | - | 0 | - |
| - | - | 1896 | 149 | - | - | 0 | - |
| - | - | 1414 | 149 | - | - | 0 | - |
| - | - | 1231 | 149 | - | - | 0 | - |
| - | - | 760.2 | 149 | - | - | 0 | - |
| - | - | 707.3 | 149 | - | - | 0 | - |
| - | - | 661.4 | 149 | - | - | 0 | - |
| - | - | 593.8 | 163.1 | - | - | 0 | - |
| - | - | 8804 | 166.1 | - | - | 0 | - |
| - | - | 1029 | 167.1 | - | - | 0 | - |
| - | - | 1180 | 168.1 | - | - | 0 | - |
| - | - | 987.4 | 169.1 | - | - | 0 | - |
| - | - | 1299 | 173.5 | - | - | 0 | - |
| - | - | 1286 | 175.1 | - | - | 0 | - |
| - | - | 573.9 | 177 | - | - | 0 | - |
| - | - | 1877 | 189.1 | - | - | 0 | - |
| - | - | 2503 | 195.1 | - | - | 0 | - |
| - | - | 1.237E+04 | 197.1 | - | - | 0 | - |
| - | - | 940.1 | 198.1 | - | - | 0 | - |
| - | - | 7311 | 199.1 | - | - | 0 | - |
| - | - | 651.1 | 211.1 | - | - | 0 | - |
| - | - | 1.174E+04 | 212.1 | - | - | 0 | - |
| - | - | 707.6 | 213.1 | - | - | 0 | - |
| - | - | 1155 | 213.1 | - | - | 0 | - |
| - | - | 913.6 | 215.1 | - | - | 0 | - |
| - | - | 7196 | 217.1 | - | - | 0 | - |
| - | - | 2349 | 217.1 | - | - | 0 | - |
| - | - | 2943 | 225.1 | - | - | 0 | - |
| - | - | 1264 | 231.1 | - | - | 0 | - |
| - | - | 2.169E+05 | 233.2 | - | - | 0 | - |
| - | - | 3.285E+04 | 234.2 | - | - | 0 | - |
| - | - | 774.3 | 235.1 | - | - | 0 | - |
| - | - | 1829 | 235.2 | - | - | 0 | - |
| - | - | 917.7 | 240.6 | - | - | 0 | - |
| - | - | 9178 | 241.1 | - | - | 0 | - |
| - | - | 1000 | 242.1 | - | - | 0 | - |
| 8 | y | 1.399E+05 | 243.1 | 0.0005269 | 2.167 | +1 | 2 |
| - | - | 1.697E+04 | 244.1 | - | - | 0 | - |
| 8 | z | 2531 | 245.1 | 0.003224 | 13.15 | +1 | 2 |
| - | - | 829.6 | 245.1 | - | - | 0 | - |
| - | - | 903.3 | 249.6 | - | - | 0 | - |
| - | - | 1.066E+04 | 253.1 | - | - | 0 | - |
| - | - | 836.6 | 254.2 | - | - | 0 | - |
| - | - | 1.235E+04 | 259.1 | - | - | 0 | - |
| - | - | 519.6 | 259.1 | - | - | 0 | - |
| - | - | 1518 | 260.1 | - | - | 0 | - |
| 8 | y | 4.126E+04 | 261.1 | 0.0002772 | 1.061 | +1 | 2 |
| 2 | c | 1.57E+05 | 261.2 | 0.0004629 | 1.773 | +1 | 2 |
| - | - | 4971 | 262.1 | - | - | 0 | - |
| - | - | 2.269E+04 | 262.2 | - | - | 0 | - |
| - | - | 685.7 | 263.2 | - | - | 0 | - |
| - | - | 723.8 | 263.2 | - | - | 0 | - |
| - | - | 7218 | 264.1 | - | - | 0 | - |
| - | - | 1160 | 265.1 | - | - | 0 | - |
| - | - | 1000 | 272.1 | - | - | 0 | - |
| - | - | 1161 | 273.1 | - | - | 0 | - |
| - | - | 706.6 | 274.7 | - | - | 0 | - |
| - | - | 4.304E+04 | 282.1 | - | - | 0 | - |
| - | - | 5046 | 283.1 | - | - | 0 | - |
| - | - | 1842 | 286.1 | - | - | 0 | - |
| - | - | 3795 | 296.1 | - | - | 0 | - |
| - | - | 1610 | 296.6 | - | - | 0 | - |
| - | - | 1940 | 297.2 | - | - | 0 | - |
| - | - | 1656 | 297.7 | - | - | 0 | - |
| - | - | 1387 | 298.2 | - | - | 0 | - |
| - | - | 5.952E+04 | 299.2 | - | - | 0 | - |
| - | - | 806.7 | 300.1 | - | - | 0 | - |
| - | - | 1238 | 300.2 | - | - | 0 | - |
| - | - | 9074 | 300.2 | - | - | 0 | - |
| - | - | 922 | 301.2 | - | - | 0 | - |
| - | - | 1747 | 304.1 | - | - | 0 | - |
| - | - | 3845 | 305.1 | - | - | 0 | - |
| - | - | 1475 | 305.6 | - | - | 0 | - |
| - | - | 1982 | 306.2 | - | - | 0 | - |
| - | - | 1039 | 308.2 | - | - | 0 | - |
| 4 | z | 1.238E+04 | 314.1 | 0.005635 | 17.94 | +2 | 6 |
| - | - | 2567 | 314.6 | - | - | 0 | - |
| - | - | 979.8 | 315.1 | - | - | 0 | - |
| - | - | 1048 | 320.1 | - | - | 0 | - |
| - | - | 760.1 | 323.2 | - | - | 0 | - |
| - | - | 776.6 | 325.2 | - | - | 0 | - |
| - | - | 727.5 | 326.2 | - | - | 0 | - |
| - | - | 4237 | 328.1 | - | - | 0 | - |
| - | - | 967.9 | 329.1 | - | - | 0 | - |
| - | - | 1430 | 329.7 | - | - | 0 | - |
| - | - | 1879 | 330.2 | - | - | 0 | - |
| - | - | 3252 | 331.2 | - | - | 0 | - |
| - | - | 788.3 | 332.2 | - | - | 0 | - |
| - | - | 2836 | 333.2 | - | - | 0 | - |
| - | - | 1018 | 334.2 | - | - | 0 | - |
| - | - | 1171 | 338.1 | - | - | 0 | - |
| - | - | 1.098E+04 | 338.7 | - | - | 0 | - |
| - | - | 3457 | 339.2 | - | - | 0 | - |
| - | - | 1207 | 340.2 | - | - | 0 | - |
| - | - | 2007 | 341.2 | - | - | 0 | - |
| - | - | 2416 | 342.2 | - | - | 0 | - |
| - | - | 7970 | 346.1 | - | - | 0 | - |
| - | - | 1521 | 347.1 | - | - | 0 | - |
| - | - | 3458 | 347.7 | - | - | 0 | - |
| - | - | 1.387E+04 | 351.2 | - | - | 0 | - |
| - | - | 2755 | 352.2 | - | - | 0 | - |
| - | - | 2554 | 352.7 | - | - | 0 | - |
| - | - | 1850 | 353.2 | - | - | 0 | - |
| - | - | 8501 | 354.2 | - | - | 0 | - |
| - | - | 2274 | 355.2 | - | - | 0 | - |
| - | - | 808.4 | 356.7 | - | - | 0 | - |
| - | - | 1512 | 357.2 | - | - | 0 | - |
| 3 | c | 1.704E+04 | 358.2 | 0.000464 | 1.295 | +1 | 3 |
| - | - | 2798 | 359.2 | - | - | 0 | - |
| - | - | 4471 | 361.7 | - | - | 0 | - |
| - | - | 2463 | 362.2 | - | - | 0 | - |
| - | - | 1103 | 362.7 | - | - | 0 | - |
| - | - | 761.5 | 365.1 | - | - | 0 | - |
| - | - | 939.8 | 365.7 | - | - | 0 | - |
| - | - | 722.7 | 368.2 | - | - | 0 | - |
| - | - | 2.371E+04 | 369.2 | - | - | 0 | - |
| - | - | 4243 | 370.2 | - | - | 0 | - |
| 3 | y | 1.135E+04 | 370.7 | 0.006168 | 16.64 | +2 | 7 |
| - | - | 4921 | 371.2 | - | - | 0 | - |
| 7 | y | 7683 | 372.2 | 0.0004448 | 1.195 | +1 | 3 |
| - | - | 1130 | 373.2 | - | - | 0 | - |
| - | - | 664.3 | 374.2 | - | - | 0 | - |
| - | - | 712.2 | 375.7 | - | - | 0 | - |
| 7 | c | 8916 | 379.7 | 0.004912 | 12.94 | +2 | 7 |
| - | - | 2620 | 380.2 | - | - | 0 | - |
| - | - | 1609 | 383.2 | - | - | 0 | - |
| - | - | 1.378E+05 | 386.2 | - | - | 0 | - |
| - | - | 2.674E+04 | 387.2 | - | - | 0 | - |
| 7 | c | 3626 | 388.2 | 0.006014 | 15.49 | +2 | 7 |
| 7 | y | 4901 | 390.2 | 7.299E-05 | 0.1871 | +1 | 3 |
| - | - | 1670 | 391.2 | - | - | 0 | - |
| - | - | 1177 | 393.2 | - | - | 0 | - |
| - | - | 6266 | 401.2 | - | - | 0 | - |
| - | - | 1115 | 402.2 | - | - | 0 | - |
| - | - | 995.8 | 405.2 | - | - | 0 | - |
| - | - | 1999 | 412.2 | - | - | 0 | - |
| - | - | 1395 | 413.2 | - | - | 0 | - |
| - | - | 1431 | 415.1 | - | - | 0 | - |
| - | - | 1197 | 415.2 | - | - | 0 | - |
| - | - | 1995 | 417.2 | - | - | 0 | - |
| - | - | 972.2 | 417.7 | - | - | 0 | - |
| - | - | 926.3 | 419.2 | - | - | 0 | - |
| - | - | 1506 | 423.2 | - | - | 0 | - |
| - | - | 1891 | 426.2 | - | - | 0 | - |
| - | - | 1824 | 426.7 | - | - | 0 | - |
| - | - | 4431 | 427.3 | - | - | 0 | - |
| - | - | 2.808E+04 | 428.3 | - | - | 0 | - |
| - | - | 2810 | 429.2 | - | - | 0 | - |
| - | - | 1.228E+04 | 429.3 | - | - | 0 | - |
| - | - | 1772 | 430.2 | - | - | 0 | - |
| - | - | 2515 | 430.3 | - | - | 0 | - |
| - | - | 1546 | 431.2 | - | - | 0 | - |
| - | - | 5936 | 433.2 | - | - | 0 | - |
| - | - | 1354 | 434.2 | - | - | 0 | - |
| - | - | 2105 | 434.2 | - | - | 0 | - |
| - | - | 1721 | 435.2 | - | - | 0 | - |
| - | - | 1058 | 439.2 | - | - | 0 | - |
| - | - | 2554 | 441.2 | - | - | 0 | - |
| 8 | c | 4753 | 444.2 | 0.0004945 | 1.113 | +2 | 8 |
| - | - | 1669 | 444.7 | - | - | 0 | - |
| - | - | 913.8 | 446.2 | - | - | 0 | - |
| - | - | 2242 | 452.2 | - | - | 0 | - |
| 2 | y | 831.3 | 453.2 | 0.002445 | 5.394 | +2 | 8 |
| - | - | 956.3 | 455.2 | - | - | 0 | - |
| 4 | c | 1.471E+04 | 455.3 | 0.0009228 | 2.027 | +1 | 4 |
| - | - | 1333 | 455.3 | - | - | 0 | - |
| - | - | 4530 | 456.2 | - | - | 0 | - |
| - | - | 7772 | 456.3 | - | - | 0 | - |
| - | - | 3849 | 457.3 | - | - | 0 | - |
| 6 | y | 3832 | 459.2 | 0.001284 | 2.796 | +1 | 4 |
| - | - | 5938 | 462.2 | - | - | 0 | - |
| - | - | 1200 | 463.2 | - | - | 0 | - |
| - | - | 1321 | 468.2 | - | - | 0 | - |
| - | - | 5326 | 470.2 | - | - | 0 | - |
| - | - | 5.392E+04 | 471.3 | - | - | 0 | - |
| 4 | c | 5.748E+04 | 472.3 | 0.0001441 | 0.3052 | +1 | 4 |
| - | - | 1.596E+04 | 473.3 | - | - | 0 | - |
| - | - | 1148 | 473.7 | - | - | 0 | - |
| - | - | 9907 | 474.3 | - | - | 0 | - |
| - | - | 2403 | 475.3 | - | - | 0 | - |
| 6 | y | 4172 | 477.2 | 0.0014 | 2.934 | +1 | 4 |
| - | - | 1608 | 478.2 | - | - | 0 | - |
| - | - | 2.386E+04 | 480.2 | - | - | 0 | - |
| - | - | 5790 | 481.2 | - | - | 0 | - |
| - | - | 1438 | 482.2 | - | - | 0 | - |
| - | - | 1127 | 482.3 | - | - | 0 | - |
| - | - | 1596 | 484.2 | - | - | 0 | - |
| - | - | 1219 | 491.7 | - | - | 0 | - |
| - | - | 4648 | 497.3 | - | - | 0 | - |
| - | - | 1.127E+05 | 498.2 | - | - | 0 | - |
| - | - | 5643 | 498.3 | - | - | 0 | - |
| - | - | 3.106E+04 | 499.2 | - | - | 0 | - |
| - | - | 2417 | 499.3 | - | - | 0 | - |
| - | - | 5080 | 500.2 | - | - | 0 | - |
| - | - | 947.3 | 500.3 | - | - | 0 | - |
| - | - | 1209 | 501.2 | - | - | 0 | - |
| - | - | 2048 | 502.2 | - | - | 0 | - |
| - | - | 1939 | 502.3 | - | - | 0 | - |
| - | - | 1149 | 510.3 | - | - | 0 | - |
| - | - | 3076 | 512.2 | - | - | 0 | - |
| - | - | 1108 | 514.3 | - | - | 0 | - |
| - | - | 6.554E+04 | 515.2 | - | - | 0 | - |
| - | - | 1771 | 515.3 | - | - | 0 | - |
| - | - | 1.911E+04 | 516.2 | - | - | 0 | - |
| - | - | 1.442E+04 | 516.3 | - | - | 0 | - |
| - | - | 3311 | 517.3 | - | - | 0 | - |
| - | - | 3542 | 517.3 | - | - | 0 | - |
| - | - | 2604 | 524.3 | - | - | 0 | - |
| - | - | 1218 | 525.3 | - | - | 0 | - |
| - | - | 1020 | 528.2 | - | - | 0 | - |
| - | - | 1.169E+04 | 530.2 | - | - | 0 | - |
| - | - | 2594 | 531.2 | - | - | 0 | - |
| - | - | 2168 | 536.2 | - | - | 0 | - |
| - | - | 2909 | 540.3 | - | - | 0 | - |
| 5 | c | 1.374E+04 | 542.3 | 0.0006021 | 1.11 | +1 | 5 |
| - | - | 7000 | 543.3 | - | - | 0 | - |
| - | - | 1.305E+04 | 544.3 | - | - | 0 | - |
| - | - | 5267 | 545.3 | - | - | 0 | - |
| 5 | y | 2561 | 546.2 | 0.001604 | 2.937 | +1 | 5 |
| - | - | 802.3 | 547.2 | - | - | 0 | - |
| - | - | 1010 | 554.2 | - | - | 0 | - |
| - | - | 1.479E+05 | 558.3 | - | - | 0 | - |
| 5 | c | 1.264E+05 | 559.3 | 0.001106 | 1.977 | +1 | 5 |
| - | - | 3.428E+04 | 560.3 | - | - | 0 | - |
| - | - | 8609 | 561.3 | - | - | 0 | - |
| - | - | 2067 | 562.3 | - | - | 0 | - |
| 5 | y | 8133 | 564.3 | 0.000744 | 1.318 | +1 | 5 |
| - | - | 1620 | 565.3 | - | - | 0 | - |
| - | - | 2483 | 567.3 | - | - | 0 | - |
| - | - | 7981 | 571.2 | - | - | 0 | - |
| - | - | 2185 | 573.2 | - | - | 0 | - |
| - | - | 824.1 | 574.2 | - | - | 0 | - |
| - | - | 884.1 | 578.3 | - | - | 0 | - |
| - | - | 3397 | 581.3 | - | - | 0 | - |
| - | - | 1812 | 582.3 | - | - | 0 | - |
| - | - | 1852 | 583.3 | - | - | 0 | - |
| - | - | 6021 | 591.2 | - | - | 0 | - |
| - | - | 2006 | 592.2 | - | - | 0 | - |
| - | - | 2632 | 593.3 | - | - | 0 | - |
| - | - | 1.017E+04 | 599.3 | - | - | 0 | - |
| - | - | 3335 | 600.3 | - | - | 0 | - |
| - | - | 1798 | 601.3 | - | - | 0 | - |
| - | - | 3282 | 601.3 | - | - | 0 | - |
| - | - | 2085 | 602.3 | - | - | 0 | - |
| - | - | 4092 | 603.4 | - | - | 0 | - |
| - | - | 1264 | 604.4 | - | - | 0 | - |
| - | - | 1373 | 606.3 | - | - | 0 | - |
| - | - | 842.8 | 607.3 | - | - | 0 | - |
| - | - | 2.959E+04 | 609.3 | - | - | 0 | - |
| - | - | 8416 | 610.3 | - | - | 0 | - |
| - | - | 1333 | 611.3 | - | - | 0 | - |
| - | - | 1.681E+04 | 611.3 | - | - | 0 | - |
| - | - | 5560 | 612.3 | - | - | 0 | - |
| - | - | 855.3 | 613.3 | - | - | 0 | - |
| - | - | 1708 | 625.3 | - | - | 0 | - |
| - | - | 5603 | 626.3 | - | - | 0 | - |
| 4 | z | 2.845E+05 | 627.3 | 0.01144 | 18.24 | +1 | 6 |
| - | - | 9.041E+04 | 628.3 | - | - | 0 | - |
| - | - | 2.391E+04 | 629.3 | - | - | 0 | - |
| 6 | c | 3.239E+04 | 629.3 | 0.001411 | 2.241 | +1 | 6 |
| - | - | 2400 | 630.3 | - | - | 0 | - |
| - | - | 1.055E+04 | 630.3 | - | - | 0 | - |
| - | - | 1493 | 631.3 | - | - | 0 | - |
| - | - | 1078 | 639.3 | - | - | 0 | - |
| 4 | y | 6386 | 643.3 | 0.003406 | 5.294 | +1 | 6 |
| - | - | 1.073E+04 | 644.3 | - | - | 0 | - |
| 4 | z | 8896 | 645.3 | 9.04E-05 | 0.1401 | +1 | 6 |
| - | - | 7474 | 645.3 | - | - | 0 | - |
| 6 | c | 2.589E+05 | 646.4 | 0.00129 | 1.995 | +1 | 6 |
| - | - | 9.959E+04 | 647.4 | - | - | 0 | - |
| - | - | 2.015E+04 | 648.4 | - | - | 0 | - |
| - | - | 3664 | 649.3 | - | - | 0 | - |
| - | - | 1236 | 649.4 | - | - | 0 | - |
| - | - | 1135 | 650.3 | - | - | 0 | - |
| - | - | 1228 | 651.3 | - | - | 0 | - |
| - | - | 1533 | 652.3 | - | - | 0 | - |
| - | - | 4865 | 660.3 | - | - | 0 | - |
| 4 | y | 8.15E+04 | 661.3 | 0.00102 | 1.542 | +1 | 6 |
| - | - | 2.666E+04 | 662.3 | - | - | 0 | - |
| - | - | 5186 | 663.3 | - | - | 0 | - |
| - | - | 7694 | 667.3 | - | - | 0 | - |
| - | - | 2152 | 668.3 | - | - | 0 | - |
| - | - | 1.642E+04 | 672.4 | - | - | 0 | - |
| - | - | 7323 | 673.4 | - | - | 0 | - |
| - | - | 1024 | 674.4 | - | - | 0 | - |
| - | - | 2524 | 683.3 | - | - | 0 | - |
| - | - | 2996 | 684.3 | - | - | 0 | - |
| - | - | 3991 | 685.3 | - | - | 0 | - |
| - | - | 2440 | 686.3 | - | - | 0 | - |
| - | - | 1018 | 687.3 | - | - | 0 | - |
| - | - | 3689 | 689.3 | - | - | 0 | - |
| - | - | 1414 | 696.4 | - | - | 0 | - |
| - | - | 4292 | 698.3 | - | - | 0 | - |
| - | - | 1967 | 699.3 | - | - | 0 | - |
| - | - | 8241 | 701.3 | - | - | 0 | - |
| - | - | 3141 | 702.3 | - | - | 0 | - |
| - | - | 4005 | 704.3 | - | - | 0 | - |
| - | - | 1488 | 705.3 | - | - | 0 | - |
| - | - | 4883 | 712.4 | - | - | 0 | - |
| - | - | 2608 | 713.4 | - | - | 0 | - |
| - | - | 4795 | 714.3 | - | - | 0 | - |
| - | - | 1026 | 714.4 | - | - | 0 | - |
| 3 | w | 2411 | 715.3 | 0.01125 | 15.73 | +1 | 7 |
| - | - | 1.312E+04 | 722.3 | - | - | 0 | - |
| - | - | 4983 | 723.3 | - | - | 0 | - |
| - | - | 2513 | 724.3 | - | - | 0 | - |
| - | - | 1.387E+04 | 730.4 | - | - | 0 | - |
| - | - | 6553 | 731.4 | - | - | 0 | - |
| - | - | 1490 | 732.3 | - | - | 0 | - |
| - | - | 6500 | 732.4 | - | - | 0 | - |
| - | - | 1617 | 733.4 | - | - | 0 | - |
| - | - | 2.978E+04 | 739.3 | - | - | 0 | - |
| 3 | y | 7.377E+04 | 740.3 | 0.003498 | 4.725 | +1 | 7 |
| - | - | 2.43E+04 | 741.4 | - | - | 0 | - |
| - | - | 9463 | 742.4 | - | - | 0 | - |
| - | - | 3512 | 743.4 | - | - | 0 | - |
| - | - | 1057 | 744.4 | - | - | 0 | - |
| - | - | 2670 | 750.3 | - | - | 0 | - |
| - | - | 1434 | 752.3 | - | - | 0 | - |
| - | - | 1879 | 753.3 | - | - | 0 | - |
| - | - | 1246 | 756.3 | - | - | 0 | - |
| - | - | 2.055E+05 | 757.4 | - | - | 0 | - |
| 7 | c | 8.753E+05 | 758.4 | 0.01219 | 16.07 | +1 | 7 |
| - | - | 3.368E+05 | 759.4 | - | - | 0 | - |
| - | - | 8.729E+04 | 760.4 | - | - | 0 | - |
| - | - | 1.218E+04 | 761.4 | - | - | 0 | - |
| - | - | 1463 | 762.4 | - | - | 0 | - |
| - | - | 1458 | 766.3 | - | - | 0 | - |
| - | - | 1249 | 767.3 | - | - | 0 | - |
| - | - | 4975 | 768.3 | - | - | 0 | - |
| - | - | 1292 | 769.3 | - | - | 0 | - |
| - | - | 1345 | 770.4 | - | - | 0 | - |
| - | - | 6024 | 774.3 | - | - | 0 | - |
| 7 | c | 7.825E+05 | 775.4 | 0.001665 | 2.148 | +1 | 7 |
| - | - | 3.391E+05 | 776.4 | - | - | 0 | - |
| - | - | 8.13E+04 | 777.4 | - | - | 0 | - |
| - | - | 6575 | 778.4 | - | - | 0 | - |
| - | - | 2030 | 784.3 | - | - | 0 | - |
| - | - | 9862 | 785.4 | - | - | 0 | - |
| - | - | 3725 | 786.4 | - | - | 0 | - |
| - | - | 2223 | 787.4 | - | - | 0 | - |
| - | - | 2985 | 788.4 | - | - | 0 | - |
| - | - | 1216 | 789.4 | - | - | 0 | - |
| - | - | 1873 | 790.4 | - | - | 0 | - |
| - | - | 1210 | 791.4 | - | - | 0 | - |
| - | - | 5.081E+04 | 801.4 | - | - | 0 | - |
| - | - | 2.241E+04 | 802.4 | - | - | 0 | - |
| - | - | 1227 | 803.3 | - | - | 0 | - |
| - | - | 5080 | 803.4 | - | - | 0 | - |
| - | - | 2265 | 812.4 | - | - | 0 | - |
| - | - | 1031 | 813.4 | - | - | 0 | - |
| - | - | 3179 | 817.4 | - | - | 0 | - |
| - | - | 1603 | 818.4 | - | - | 0 | - |
| - | - | 1034 | 819.4 | - | - | 0 | - |
| - | - | 1475 | 821.3 | - | - | 0 | - |
| - | - | 7323 | 830.4 | - | - | 0 | - |
| - | - | 2634 | 831.4 | - | - | 0 | - |
| - | - | 1458 | 837.4 | - | - | 0 | - |
| - | - | 1828 | 842.4 | - | - | 0 | - |
| - | - | 1238 | 843.4 | - | - | 0 | - |
| - | - | 4156 | 851.4 | - | - | 0 | - |
| - | - | 1700 | 852.4 | - | - | 0 | - |
| - | - | 1878 | 855.4 | - | - | 0 | - |
| - | - | 2.091E+04 | 859.4 | - | - | 0 | - |
| - | - | 2.541E+04 | 860.4 | - | - | 0 | - |
| - | - | 1.329E+04 | 861.4 | - | - | 0 | - |
| - | - | 4069 | 862.4 | - | - | 0 | - |
| - | - | 1.659E+04 | 869.4 | - | - | 0 | - |
| - | - | 7356 | 870.4 | - | - | 0 | - |
| 2 | z | 1.918E+04 | 871.4 | 0.002637 | 3.026 | +1 | 8 |
| - | - | 2.51E+04 | 872.4 | - | - | 0 | - |
| - | - | 6352 | 873.4 | - | - | 0 | - |
| 8 | c | 1066 | 886.4 | 0.001925 | 2.171 | +1 | 8 |
| 8 | c | 1.876E+05 | 887.4 | 0.001185 | 1.336 | +1 | 8 |
| - | - | 8.637E+04 | 888.4 | - | - | 0 | - |
| 2 | z | 7.96E+04 | 889.4 | 0.005194 | 5.84 | +1 | 8 |
| - | - | 3.088E+04 | 890.4 | - | - | 0 | - |
| - | - | 1.071E+04 | 891.4 | - | - | 0 | - |
| - | - | 1384 | 892.4 | - | - | 0 | - |
| - | - | 2008 | 901.5 | - | - | 0 | - |
| - | - | 1435 | 902.5 | - | - | 0 | - |
| 8 | c | 1.053E+06 | 904.4 | 0.001858 | 2.054 | +1 | 8 |
| - | - | 5.342E+05 | 905.4 | - | - | 0 | - |
| - | - | 1.512E+05 | 906.4 | - | - | 0 | - |
| - | - | 1.274E+04 | 907.4 | - | - | 0 | - |
| - | - | 1909 | 917.4 | - | - | 0 | - |
| - | - | 1146 | 929.5 | - | - | 0 | - |
| - | - | 3345 | 930.5 | - | - | 0 | - |
| - | - | 1549 | 931.5 | - | - | 0 | - |
| - | - | 2123 | 933.5 | - | - | 0 | - |
| - | - | 975.9 | 945.5 | - | - | 0 | - |
| - | - | 1306 | 946.4 | - | - | 0 | - |
| - | - | 5.851E+04 | 947.5 | - | - | 0 | - |
| - | - | 3.36E+04 | 948.5 | - | - | 0 | - |
| - | - | 9402 | 949.5 | - | - | 0 | - |
| - | - | 981.5 | 950.5 | - | - | 0 | - |
| - | - | 2862 | 957.5 | - | - | 0 | - |
| - | - | 2089 | 958.5 | - | - | 0 | - |
| - | - | 7280 | 959.4 | - | - | 0 | - |
| - | - | 2424 | 960.5 | - | - | 0 | - |
| - | - | 7068 | 961.4 | - | - | 0 | - |
| - | - | 1362 | 962.4 | - | - | 0 | - |
| - | - | 1.532E+04 | 963.5 | - | - | 0 | - |
| - | - | 6678 | 964.5 | - | - | 0 | - |
| - | - | 2590 | 965.5 | - | - | 0 | - |
| - | - | 2234 | 971.5 | - | - | 0 | - |
| - | - | 1483 | 972.5 | - | - | 0 | - |
| - | - | 7366 | 973.5 | - | - | 0 | - |
| - | - | 4320 | 974.5 | - | - | 0 | - |
| - | - | 1938 | 975.5 | - | - | 0 | - |
| - | - | 2395 | 977.5 | - | - | 0 | - |
| - | - | 1137 | 983.5 | - | - | 0 | - |
| - | - | 1.566E+04 | 984.5 | - | - | 0 | - |
| - | - | 9307 | 985.5 | - | - | 0 | - |
| - | - | 2083 | 986.5 | - | - | 0 | - |
| - | - | 928.7 | 987.4 | - | - | 0 | - |
| - | - | 3548 | 989.5 | - | - | 0 | - |
| - | - | 1065 | 990.5 | - | - | 0 | - |
| - | - | 1.306E+04 | 991.5 | - | - | 0 | - |
| - | - | 6671 | 992.5 | - | - | 0 | - |
| - | - | 2542 | 993.5 | - | - | 0 | - |
| - | - | 1.102E+04 | 1002 | - | - | 0 | - |
| - | - | 9.823E+04 | 1002 | - | - | 0 | - |
| - | - | 5.867E+04 | 1003 | - | - | 0 | - |
| - | - | 1.766E+04 | 1004 | - | - | 0 | - |
| - | - | 2306 | 1005 | - | - | 0 | - |
| - | - | 959.2 | 1016 | - | - | 0 | - |
| - | - | 2.361E+05 | 1019 | - | - | 0 | - |
| - | - | 9.713E+05 | 1020 | - | - | 0 | - |
| - | - | 5.185E+05 | 1021 | - | - | 0 | - |
| - | - | 1.546E+05 | 1022 | - | - | 0 | - |
| - | - | 1.474E+04 | 1023 | - | - | 0 | - |
| - | - | 5775 | 1052 | - | - | 0 | - |
| - | - | 3557 | 1053 | - | - | 0 | - |
| - | - | 1582 | 1054 | - | - | 0 | - |
| - | - | 902.7 | 1658 | - | - | 0 | - |
| - | - | 987.3 | 2630 | - | - | 0 | - |
| - | - | 879.9 | 3030 | - | - | 0 | - |
| - | - | 874.2 | 3079 | - | - | 0 | - |

m/z Charge Intensity FragmentType MassShift Position
120.08110809326172 0 38908.234
121.08451080322266 0 2743.8484
121.08879852294922 0 525.8692
121.80667877197266 0 557.5793
126.0555648803711 0 678.96387
130.0502471923828 0 8930.53
132.10226440429688 0 258942.22 y 8
133.09986877441406 0 1117.5209
133.1055908203125 0 17701.094
134.1063232421875 0 703.95514
136.07606506347656 0 1184.1338
145.0198211669922 0 608.059
147.07667541503906 0 1721.028
148.0609893798828 0 805.91
148.86912536621094 0 563.7733
148.89047241210938 0 632.04755
148.8981475830078 0 832.54956
148.9049072265625 0 794.1789
148.91958618164062 0 1878.1088
148.92698669433594 0 1598.0234
148.93397521972656 0 2587.1226
148.94183349609375 0 4751.171
148.9584503173828 0 6543.15
148.96620178222656 0 3762.0356
148.97360229492188 0 1896.4489
148.9807891845703 0 1414.4869
148.98814392089844 0 1231.0145
148.99554443359375 0 760.15485
149.0172119140625 0 707.27966
149.02423095703125 0 661.37225
163.08627319335938 0 593.7741
166.0865478515625 0 8803.7295
167.11817932128906 0 1028.6078
168.1264190673828 0 1180.2003
169.13401794433594 0 987.3539
173.4510040283203 0 1299.0627
175.0718231201172 0 1286.3665
177.04742431640625 0 573.9383
189.08737182617188 0 1876.773
195.1132049560547 0 2502.7595
197.1288299560547 0 12371.594
198.1321258544922 0 940.07874
199.0717010498047 0 7311.11
211.13194274902344 0 651.0755
212.13970947265625 0 11738.226
213.0877685546875 0 707.63617
213.14373779296875 0 1155.0197
215.14064025878906 0 913.564
217.08218383789062 0 7195.8145
217.14678955078125 0 2349.2634
225.1238250732422 0 2942.81
231.09812927246094 0 1263.7759
233.1653594970703 0 216946.94
234.16867065429688 0 32851.066
235.1080322265625 0 774.29755
235.1716766357422 0 1829.0726
240.60848999023438 0 917.72064
241.0824432373047 0 9177.808
242.086669921875 0 1000.2642
243.13446044921875 0 139875.2 y Water loss 7
244.13755798339844 0 16973.05
245.12899780273438 0 2531.1707 z 7
245.1414337158203 0 829.6213
249.6136474609375 0 903.31555
253.1187286376953 0 10662.1455
254.15052795410156 0 836.5611
259.0928955078125 0 12346.433
259.1076965332031 0 519.6075
260.09649658203125 0 1517.6731
261.144775390625 0 41257.84 y 7
261.16021728515625 0 157012.73 c Ammonia loss 1
262.1481628417969 0 4970.553
262.1634826660156 0 22690.074
263.1502380371094 0 685.7005
263.1658935546875 0 723.7594
264.134765625 0 7217.611
265.137451171875 0 1159.7434
272.1239013671875 0 1000.1528
273.12408447265625 0 1160.7745
274.6832275390625 0 706.5826
282.1453552246094 0 43040.1
283.1485595703125 0 5046.4897
286.10382080078125 0 1841.9003
296.1251220703125 0 3794.7686
296.62579345703125 0 1609.9742
297.1579895019531 0 1940.0098
297.6602783203125 0 1656.091
298.1640319824219 0 1387.2054
299.1719665527344 0 59517.37
300.1375732421875 0 806.6695
300.1559753417969 0 1237.5502
300.1753845214844 0 9074.376
301.177490234375 0 922.039
304.1147155761719 0 1747.344
305.129638671875 0 3845.1025
305.6312255859375 0 1475.267
306.1642150878906 0 1981.8082
308.1976623535156 0 1038.7744
314.13531494140625 0 12381.441 z Water loss 3
314.6371765136719 0 2566.847
315.1373596191406 0 979.8149
320.123779296875 0 1047.966
323.1728820800781 0 760.1176
325.1865539550781 0 776.5821
326.1626892089844 0 727.5189
328.1147155761719 0 4236.692
329.11834716796875 0 967.91296
329.6644287109375 0 1430.0259
330.16748046875 0 1879.1326
331.2259216308594 0 3251.9597
332.2336120605469 0 788.28094
333.15643310546875 0 2836.3428
334.16021728515625 0 1018.07666
338.1340026855469 0 1170.6288
338.6693115234375 0 10983.571
339.1701965332031 0 3457.239
340.178466796875 0 1206.9244
341.18359375 0 2007.3757
342.18243408203125 0 2415.8345
346.1250915527344 0 7970.109
347.1280517578125 0 1520.8495
347.6748962402344 0 3457.7117
351.1669921875 0 13874.758
352.1700134277344 0 2754.769
352.66705322265625 0 2554.4077
353.16876220703125 0 1849.9186
354.1663818359375 0 8501.104
355.1639404296875 0 2273.577
356.6855773925781 0 808.35925
357.2059020996094 0 1511.9154
358.2129821777344 0 17040.674 c Ammonia loss 2
359.2164001464844 0 2797.791
361.67620849609375 0 4470.6445
362.1770324707031 0 2462.8755
362.68218994140625 0 1103.4651
365.1459655761719 0 761.4912
365.6939697265625 0 939.7691
368.1933898925781 0 722.73584
369.1773681640625 0 23705.367
370.18017578125 0 4242.7456
370.682861328125 0 11351.644 y Water loss 2
371.18524169921875 0 4921.2114
372.1769714355469 0 7682.5347 y Water loss 6
373.1778564453125 0 1129.9746
374.176025390625 0 664.29474
375.6672668457031 0 712.214
379.6846923828125 0 8915.763 c Ammonia loss 6
380.1866760253906 0 2619.9036
383.1548767089844 0 1609.1473
386.2040710449219 0 137844.47
387.2073669433594 0 26743.865
388.2088928222656 0 3626.2207 c 6
390.1871643066406 0 4900.8896 y 6
391.19232177734375 0 1670.4843
393.177734375 0 1177.131
401.167724609375 0 6265.54
402.1697998046875 0 1114.514
405.16363525390625 0 995.7605
412.210693359375 0 1998.616
413.2210388183594 0 1395.3748
415.14599609375 0 1431.002
415.2326354980469 0 1197.2635
417.1965637207031 0 1995.3568
417.6965026855469 0 972.1998
419.184814453125 0 926.2964
423.1876220703125 0 1506.0303
426.20159912109375 0 1891.3763
426.704345703125 0 1823.9551
427.27099609375 0 4431.1396
428.2789611816406 0 28079.201
429.21380615234375 0 2810.3274
429.2842102050781 0 12276.113
430.21527099609375 0 1772.3242
430.28948974609375 0 2515.1006
431.2301330566406 0 1546.3546
433.15771484375 0 5935.9814
434.1604919433594 0 1353.7202
434.2041320800781 0 2105.4749
435.2078857421875 0 1720.6094
439.18115234375 0 1057.5834
441.1981201171875 0 2554.2305
444.2113952636719 0 4752.8823 c Ammonia loss 7
444.71234130859375 0 1668.7031
446.2420349121094 0 913.7697
452.2146301269531 0 2242.2598
453.2186279296875 0 831.2607 y 1
455.2073059082031 0 956.276
455.2662048339844 0 14711.981 c Ammonia loss 3
455.2995910644531 0 1332.7701
456.21002197265625 0 4529.9717
456.2722473144531 0 7771.566
457.2784118652344 0 3848.5364
459.2098388671875 0 3831.8752 y Water loss 5
462.1997375488281 0 5938.3647
463.20123291015625 0 1199.9319
468.2082214355469 0 1321.4532
470.22607421875 0 5326.01
471.28497314453125 0 53921.07
472.29168701171875 0 57477.805 c 3
473.29571533203125 0 15955.039
473.7381286621094 0 1147.9302
474.3067321777344 0 9907.384
475.3118591308594 0 2403.119
477.22052001953125 0 4171.636 y 5
478.22296142578125 0 1607.9973
480.2100830078125 0 23858.953
481.21295166015625 0 5789.814
482.2163391113281 0 1437.89
482.26007080078125 0 1127.4188
484.203369140625 0 1595.7966
491.7490234375 0 1219.4004
497.300048828125 0 4647.6313
498.2206115722656 0 112737.27
498.3069763183594 0 5643.1494
499.22357177734375 0 31058.648
499.3129577636719 0 2416.6213
500.2264709472656 0 5080.003
500.32452392578125 0 947.289
501.2272644042969 0 1209.0759
502.21490478515625 0 2047.6766
502.2545471191406 0 1939.4865
510.2931823730469 0 1148.6871
512.1986083984375 0 3076.4656
514.3311767578125 0 1107.9387
515.2470092773438 0 65535.023
515.3154296875 0 1770.8812
516.2495727539062 0 19107.832
516.3192749023438 0 14415.129
517.2503051757812 0 3310.9546
517.3214721679688 0 3542.2158
524.2886352539062 0 2604.3445
525.288818359375 0 1218.1283
528.2314453125 0 1019.7473
530.2098999023438 0 11691.674
531.2125854492188 0 2593.6396
536.2008056640625 0 2167.6787
540.3057861328125 0 2908.6719
542.2979125976562 0 13735.548 c Ammonia loss 4
543.3014526367188 0 7000.343
544.3135986328125 0 13050.55
545.3168334960938 0 5266.9155
546.2421875 0 2560.554 y Water loss 4
547.243896484375 0 802.2677
554.2084350585938 0 1009.60565
558.3170776367188 0 147909.02
559.32275390625 0 126434.086 c 4
560.3267211914062 0 34281.973
561.3357543945312 0 8609.328
562.3408203125 0 2067.465
564.2518920898438 0 8133.1553 y 4
565.2586059570312 0 1620.377
567.2658081054688 0 2483.1428
571.2364501953125 0 7980.521
573.232421875 0 2184.5652
574.2366333007812 0 824.07855
578.261474609375 0 884.09894
581.2582397460938 0 3397.1633
582.278564453125 0 1811.5068
583.3147583007812 0 1851.717
591.2424926757812 0 6020.884
592.2437133789062 0 2006.417
593.3058471679688 0 2632.105
599.2685546875 0 10168.137
600.272216796875 0 3335.0789
601.2800903320312 0 1797.7736
601.33544921875 0 3281.8843
602.3414916992188 0 2084.6663
603.3515014648438 0 4092.3315
604.3523559570312 0 1264.1031
606.277587890625 0 1373.3031
607.2809448242188 0 842.84064
609.2528076171875 0 29589.95
610.2550048828125 0 8415.826
611.2574462890625 0 1333.1481
611.314697265625 0 16811.441
612.3189697265625 0 5559.653
613.3203735351562 0 855.28827
625.2847900390625 0 1707.8416
626.2549438476562 0 5603.247
627.26318359375 0 284458.25 z Water loss 3
628.2661743164062 0 90413.836
629.2686767578125 0 23914.596
629.3307495117188 0 32386.473 c Ammonia loss 5
630.2722778320312 0 2399.9185
630.3343505859375 0 10552.591
631.3299560546875 0 1492.7653
639.2999267578125 0 1078.1624
643.2967529296875 0 6386.203 y Water loss 3
644.2908935546875 0 10734.919
645.2852783203125 0 8895.594 z 3
645.3492431640625 0 7474.009
646.357177734375 0 258878.5 c 5
647.35986328125 0 99594.7
648.362548828125 0 20147.916
649.2857666015625 0 3663.723
649.3666381835938 0 1235.8953
650.2952880859375 0 1135.3157
651.2757568359375 0 1227.6592
652.282958984375 0 1533.1107
660.2969360351562 0 4864.963
661.304931640625 0 81497.65 y 3
662.308349609375 0 26664.246
663.3092041015625 0 5186.1465
667.2952270507812 0 7694.467
668.3064575195312 0 2151.9412
672.3726196289062 0 16417.885
673.3756713867188 0 7322.5254
674.3821411132812 0 1024.3563
683.3180541992188 0 2524.231
684.3211669921875 0 2996.2603
685.3289184570312 0 3991.3127
686.3220825195312 0 2439.53
687.3259887695312 0 1018.0861
689.2998657226562 0 3688.771
696.3563232421875 0 1414.0897
698.3375244140625 0 4291.9663
699.3431396484375 0 1967.4915
701.28759765625 0 8240.797
702.2936401367188 0 3141.244
704.3286743164062 0 4004.879
705.32958984375 0 1488.0803
712.3568115234375 0 4883.138
713.3563842773438 0 2608.3186
714.2960205078125 0 4795.1987
714.3555297851562 0 1025.6702
715.30322265625 0 2411.063 w 2
722.3433837890625 0 13124.993
723.3466186523438 0 4982.695
724.34814453125 0 2513.2605
730.3787231445312 0 13867.683
731.3822021484375 0 6552.5967
732.328369140625 0 1490.0154
732.3914794921875 0 6499.9717
733.3973388671875 0 1616.5408
739.3397216796875 0 29776.75
740.349609375 0 73766.766 y Water loss 2
741.3538208007812 0 24296.19
742.3555908203125 0 9462.914
743.3598022460938 0 3512.3762
744.3557739257812 0 1056.72
750.3309326171875 0 2669.724
752.3216552734375 0 1433.8816
753.3276977539062 0 1878.7172
756.3202514648438 0 1245.9365
757.3504638671875 0 205451.16
758.3597412109375 0 875332.06 c Ammonia loss 6
759.3628540039062 0 336831.06
760.366455078125 0 87292.07
761.373291015625 0 12181.335
762.3756713867188 0 1462.7646
766.3262329101562 0 1458.1656
767.3341674804688 0 1249.042
768.341796875 0 4974.9766
769.3486328125 0 1291.5823
770.357177734375 0 1344.6234
774.3313598632812 0 6023.806
775.400146484375 0 782463.06 c 6
776.4027709960938 0 339118.8
777.4053955078125 0 81303.28
778.4072265625 0 6574.7563
784.3311157226562 0 2030.2101
785.3651123046875 0 9862.157
786.3571166992188 0 3724.6755
787.3598022460938 0 2223.0422
788.4067993164062 0 2985.2563
789.4072265625 0 1215.6995
790.3583984375 0 1873.2001
791.363037109375 0 1209.5958
801.4155883789062 0 50812.613
802.41845703125 0 22411.79
803.3424072265625 0 1227.268
803.4234619140625 0 5079.711
812.3594360351562 0 2264.9656
813.366943359375 0 1031.2255
817.3861694335938 0 3179.2593
818.3926391601562 0 1603.266
819.3604125976562 0 1033.9995
821.340576171875 0 1475.0879
830.3698120117188 0 7323.3716
831.3754272460938 0 2633.5012
837.3676147460938 0 1458.2242
842.4025268554688 0 1828.2958
843.4060668945312 0 1238.3401
851.392822265625 0 4155.543
852.3955688476562 0 1700.4813
855.3751831054688 0 1878.1611
859.4210815429688 0 20914.389
860.427001953125 0 25406.43
861.4320068359375 0 13293.728
862.4373779296875 0 4069.0872
869.4047241210938 0 16589.346
870.4072265625 0 7356.47
871.3984375 0 19179.451 z Water loss 1
872.4013671875 0 25103.81
873.4049072265625 0 6352.475
886.4324340820312 0 1066.4626 c Water loss 7
887.4157104492188 0 187599.94 c Ammonia loss 7
888.4188842773438 0 86367.13
889.4115600585938 0 79604.89 z 1
890.4129638671875 0 30878.816
891.4151000976562 0 10713.007
892.4229125976562 0 1384.3782
901.49072265625 0 2008.3895
902.4963989257812 0 1434.6935
904.4429321289062 0 1053247.8 c 7
905.4451904296875 0 534150.3
906.4478759765625 0 151165.34
907.449951171875 0 12744.865
917.4420776367188 0 1909.0992
929.4934692382812 0 1146.4358
930.473388671875 0 3344.7153
931.4786987304688 0 1548.7709
933.4872436523438 0 2122.512
945.4807739257812 0 975.9364
946.426513671875 0 1305.7579
947.4971923828125 0 58510.57
948.4998779296875 0 33601.516
949.5037841796875 0 9401.553
950.5077514648438 0 981.5045
957.5177612304688 0 2862.2332
958.5189208984375 0 2089.0076
959.4358520507812 0 7280.2446
960.5128784179688 0 2423.705
961.440185546875 0 7067.7046
962.448486328125 0 1362.3142
963.4561767578125 0 15320.411
964.4603271484375 0 6678.3867
965.4588012695312 0 2590.2256
971.5010375976562 0 2234.0146
972.4956665039062 0 1483.3401
973.5113525390625 0 7365.676
974.5150756835938 0 4320.02
975.5218505859375 0 1937.65
977.4740600585938 0 2395.2122
983.5060424804688 0 1137.2781
984.4815063476562 0 15658.419
985.4835815429688 0 9306.886
986.4944458007812 0 2082.8164
987.422119140625 0 928.7092
989.5073852539062 0 3548.383
990.5133056640625 0 1065.3741
991.5232543945312 0 13056.047
992.525390625 0 6670.9917
993.5288696289062 0 2541.9126
1001.507568359375 0 11018.305
1002.4931030273438 0 98234.33
1003.4948120117188 0 58667.184
1004.4976196289062 0 17658.365
1005.4957275390625 0 2305.836
1016.4834594726562 0 959.23596
1018.5105590820312 0 236143.62
1019.5184936523438 0 971313.94
1020.5213012695312 0 518504.16
1021.5237426757812 0 154631.17
1022.5247192382812 0 14740.595
1051.5091552734375 0 5774.8813
1052.510986328125 0 3556.992
1053.5067138671875 0 1582.4308
1658.2830810546875 0 902.6768
2630.32275390625 0 987.3192
3029.703125 0 879.89325
3079.101806640625 0 874.248

Spectrum Details

|  |  |
| --- | --- |
| Matched peaks? Matched peaks The total absolute number of peaks matched. Additionally in brackets the total fraction of peaks matched and the total number of peaks is shown. | 37 (8.01% of 462) |
| FDR? FDR The false discovery rate estimated for this peptide. It is calculated by matching all theoretical fragments with a non-integer shift with the raw peaks for this spectrum. This is done with 40 different shifts. The resulting percentage is the average number of annotated peaks over the number of annotated peaks with the correct spectrum. | 1.74% |
| Satellite FDR? Satellite FDR See the FDR for details on its calculation. This satellite ion specific FDR only contains the satellite ions (d/w) for I/L/J positions. | - |
| PSM Score? PSM Score The PSM Score as given by Hecklib to this annotated spectrum. It is shown with three significant figures. | 472 |

#### Spectrum 8292? Spectrum 8292 The raw spectrum of this peptide as annotated by Hecklib. The fragments are coloured according to ion type (see legend). Any peaks with a star '\*' as text can be hovered over to see the full details, first the ion type second the mass shift type. By hovering over the amino acids in the peptide or ions in the legend the corresponding peaks are highlighted. By toggling the 'Unassigned' label you can turn the background (unassigned) peaks on or off in the plot. By updating the slider in the Ion legend you can update the spectrum to only show the top X% of the peaks with labels. The top X% means any peak that is within X% of the highest intensity. By dragging in the spectrum you can zoom in to a specific part of the spectrum and use 'Zoom Out' to get back to the original zoom level. The annotation of the spectrum is based on the given sequence in the peptides file and is done with different software so inconsistencies are likely. The peaks are annotated based on the given sequence, with 20 ppm tolerance.

Copy Data

##### Spectrum 8292 (TSV)

###### Preview

```
Loading example...
```

*Click on the button to copy the data to your clipboard.*

Mz MinMz MaxIntensity Max

WidthHeightPeptide font sizePeptide stroke widthSpectrum font sizeSpectrum stroke widthCompact peptide

Ion legend

wxyz

abcd

OtherUnassignedIon^Charge^_Position_Show for top:%

JFPPSSEEJ

02.32e+54.63e+56.95e+59.26e+5

Zoom Out

y^+1^_1_a^+2^_3_a^+2^_3_b^+2^_3_a^+1^_2_a^+1^_2_y^+1^_2_b^+1^_2_y^+1^_2_b^+1^_2_y^+2^_5_b^+2^_6_a^+1^_3_b^+1^_3_b^+1^_3_b^+2^_7_b^+2^_7_y^+1^_3_b^+2^_7_y^+1^_3_b^+2^_8_b^+2^_8_b^+2^_8_y^+2^_8_b^+1^_4_y^+1^_4_y^+1^_4_b^+1^_5_b^+1^_5_y^+1^_5_y^+1^_5_b^+1^_6_b^+1^_6_b^+1^_6_y^+1^_6_y^+1^_6_b^+1^_7_b^+1^_7_b^+1^_7_b^+1^_8_b^+1^_8_y^+1^_8_

0778155623343112

Fragment Matches Table

Show background peaks

| Position | Ion type | Intensity | mz Theoretical | mz Error (Th) | mz Error (ppm) | Charge | Series Number |
| --- | --- | --- | --- | --- | --- | --- | --- |
| - | - | 5.585E+05 | 120.1 | - | - | 0 | - |
| - | - | 4.552E+04 | 121.1 | - | - | 0 | - |
| - | - | 711.5 | 122.1 | - | - | 0 | - |
| - | - | 5.904E+04 | 126.1 | - | - | 0 | - |
| - | - | 701.3 | 126.3 | - | - | 0 | - |
| - | - | 3135 | 127.1 | - | - | 0 | - |
| - | - | 1.77E+04 | 129.1 | - | - | 0 | - |
| - | - | 2480 | 129.1 | - | - | 0 | - |
| - | - | 5.96E+04 | 130.1 | - | - | 0 | - |
| - | - | 3284 | 131.1 | - | - | 0 | - |
| 9 | y | 8.938E+05 | 132.1 | 0.000344 | 2.604 | +1 | 1 |
| - | - | 2091 | 133.1 | - | - | 0 | - |
| - | - | 2214 | 133.1 | - | - | 0 | - |
| - | - | 5.519E+04 | 133.1 | - | - | 0 | - |
| - | - | 2565 | 134.1 | - | - | 0 | - |
| - | - | 2300 | 136.1 | - | - | 0 | - |
| - | - | 1.304E+04 | 138.1 | - | - | 0 | - |
| - | - | 2507 | 139.1 | - | - | 0 | - |
| - | - | 1.083E+04 | 139.1 | - | - | 0 | - |
| - | - | 1004 | 140.1 | - | - | 0 | - |
| - | - | 1039 | 140.1 | - | - | 0 | - |
| - | - | 8004 | 141.1 | - | - | 0 | - |
| - | - | 694.7 | 143.4 | - | - | 0 | - |
| - | - | 994.7 | 144.1 | - | - | 0 | - |
| - | - | 1.342E+04 | 147.1 | - | - | 0 | - |
| - | - | 4505 | 148.1 | - | - | 0 | - |
| - | - | 1677 | 148.1 | - | - | 0 | - |
| - | - | 945.8 | 149 | - | - | 0 | - |
| - | - | 884.3 | 151.1 | - | - | 0 | - |
| - | - | 1975 | 151.1 | - | - | 0 | - |
| - | - | 991.1 | 152.1 | - | - | 0 | - |
| - | - | 2463 | 152.1 | - | - | 0 | - |
| - | - | 2942 | 153.1 | - | - | 0 | - |
| - | - | 4739 | 154.1 | - | - | 0 | - |
| - | - | 3.388E+04 | 154.1 | - | - | 0 | - |
| - | - | 2327 | 155.1 | - | - | 0 | - |
| - | - | 2724 | 155.1 | - | - | 0 | - |
| - | - | 1356 | 155.1 | - | - | 0 | - |
| - | - | 2.011E+04 | 157.1 | - | - | 0 | - |
| 3 | a | 5.042E+04 | 157.1 | 0.001738 | 11.06 | +2 | 3 |
| - | - | 1074 | 158.1 | - | - | 0 | - |
| - | - | 3348 | 158.1 | - | - | 0 | - |
| - | - | 1167 | 159.1 | - | - | 0 | - |
| - | - | 1124 | 162.1 | - | - | 0 | - |
| - | - | 844.7 | 165.1 | - | - | 0 | - |
| - | - | 2021 | 165.1 | - | - | 0 | - |
| 3 | a | 1218 | 165.6 | 0.0001546 | 0.9334 | +2 | 3 |
| - | - | 5225 | 166.1 | - | - | 0 | - |
| - | - | 1523 | 167 | - | - | 0 | - |
| - | - | 4.74E+04 | 167.1 | - | - | 0 | - |
| - | - | 4.82E+04 | 167.1 | - | - | 0 | - |
| - | - | 2080 | 168.1 | - | - | 0 | - |
| - | - | 3990 | 168.1 | - | - | 0 | - |
| - | - | 3947 | 168.1 | - | - | 0 | - |
| - | - | 908 | 169.1 | - | - | 0 | - |
| - | - | 894.4 | 169.1 | - | - | 0 | - |
| - | - | 2051 | 170 | - | - | 0 | - |
| - | - | 1169 | 170.1 | - | - | 0 | - |
| - | - | 1.108E+04 | 171.1 | - | - | 0 | - |
| 3 | b | 937.4 | 171.1 | 0.002339 | 13.67 | +2 | 3 |
| - | - | 3.114E+04 | 172.1 | - | - | 0 | - |
| - | - | 3338 | 173.1 | - | - | 0 | - |
| - | - | 2779 | 173.1 | - | - | 0 | - |
| - | - | 1447 | 174.1 | - | - | 0 | - |
| - | - | 2.334E+04 | 175.1 | - | - | 0 | - |
| - | - | 1014 | 175.1 | - | - | 0 | - |
| - | - | 4361 | 175.1 | - | - | 0 | - |
| - | - | 727 | 175.8 | - | - | 0 | - |
| - | - | 1306 | 176.1 | - | - | 0 | - |
| - | - | 3017 | 177.1 | - | - | 0 | - |
| - | - | 1183 | 177.1 | - | - | 0 | - |
| - | - | 1164 | 179.1 | - | - | 0 | - |
| - | - | 1459 | 180.1 | - | - | 0 | - |
| - | - | 805 | 180.2 | - | - | 0 | - |
| - | - | 2653 | 181.1 | - | - | 0 | - |
| - | - | 5434 | 181.1 | - | - | 0 | - |
| - | - | 1743 | 182.1 | - | - | 0 | - |
| - | - | 829.3 | 183.1 | - | - | 0 | - |
| - | - | 969.1 | 183.1 | - | - | 0 | - |
| - | - | 5107 | 183.1 | - | - | 0 | - |
| - | - | 8.966E+04 | 185.1 | - | - | 0 | - |
| - | - | 760.6 | 185.4 | - | - | 0 | - |
| - | - | 1.049E+04 | 186.1 | - | - | 0 | - |
| - | - | 884 | 187.1 | - | - | 0 | - |
| - | - | 968.6 | 188.1 | - | - | 0 | - |
| - | - | 4701 | 188.1 | - | - | 0 | - |
| - | - | 3.407E+04 | 189.1 | - | - | 0 | - |
| - | - | 1118 | 193.1 | - | - | 0 | - |
| - | - | 1.219E+04 | 195.1 | - | - | 0 | - |
| - | - | 9.713E+04 | 195.1 | - | - | 0 | - |
| - | - | 930.8 | 195.7 | - | - | 0 | - |
| - | - | 1030 | 196.1 | - | - | 0 | - |
| - | - | 9657 | 196.1 | - | - | 0 | - |
| - | - | 2.2E+05 | 197.1 | - | - | 0 | - |
| - | - | 1175 | 198.1 | - | - | 0 | - |
| - | - | 2.032E+04 | 198.1 | - | - | 0 | - |
| - | - | 5.279E+04 | 199.1 | - | - | 0 | - |
| - | - | 7483 | 199.1 | - | - | 0 | - |
| - | - | 3724 | 200.1 | - | - | 0 | - |
| - | - | 2010 | 201.1 | - | - | 0 | - |
| - | - | 887.5 | 201.1 | - | - | 0 | - |
| - | - | 2760 | 202.1 | - | - | 0 | - |
| - | - | 6502 | 203.1 | - | - | 0 | - |
| - | - | 1760 | 205.2 | - | - | 0 | - |
| - | - | 1239 | 208.1 | - | - | 0 | - |
| - | - | 1464 | 208.1 | - | - | 0 | - |
| - | - | 1.308E+04 | 209.1 | - | - | 0 | - |
| - | - | 1033 | 209.1 | - | - | 0 | - |
| - | - | 6718 | 211.1 | - | - | 0 | - |
| - | - | 1102 | 212.1 | - | - | 0 | - |
| - | - | 1.263E+04 | 212.1 | - | - | 0 | - |
| - | - | 1.579E+04 | 213.1 | - | - | 0 | - |
| - | - | 1.038E+04 | 213.1 | - | - | 0 | - |
| - | - | 1153 | 214.1 | - | - | 0 | - |
| - | - | 1047 | 214.1 | - | - | 0 | - |
| - | - | 3439 | 214.1 | - | - | 0 | - |
| - | - | 6918 | 215.1 | - | - | 0 | - |
| - | - | 971.1 | 216.1 | - | - | 0 | - |
| 2 | a | 1132 | 216.1 | 0.00267 | 12.35 | +1 | 2 |
| - | - | 9.729E+04 | 217.1 | - | - | 0 | - |
| - | - | 2.069E+04 | 217.1 | - | - | 0 | - |
| - | - | 5806 | 218.1 | - | - | 0 | - |
| - | - | 2190 | 218.1 | - | - | 0 | - |
| - | - | 1279 | 221.1 | - | - | 0 | - |
| - | - | 1033 | 221.1 | - | - | 0 | - |
| - | - | 2712 | 223.1 | - | - | 0 | - |
| - | - | 4426 | 224.1 | - | - | 0 | - |
| - | - | 8098 | 225.1 | - | - | 0 | - |
| - | - | 3.016E+04 | 225.1 | - | - | 0 | - |
| - | - | 5058 | 226.1 | - | - | 0 | - |
| - | - | 1389 | 227.1 | - | - | 0 | - |
| - | - | 1.62E+04 | 227.1 | - | - | 0 | - |
| - | - | 1650 | 227.1 | - | - | 0 | - |
| - | - | 1816 | 228.1 | - | - | 0 | - |
| - | - | 1931 | 229.2 | - | - | 0 | - |
| - | - | 1.538E+04 | 231.1 | - | - | 0 | - |
| - | - | 951.2 | 231.1 | - | - | 0 | - |
| - | - | 1510 | 231.6 | - | - | 0 | - |
| - | - | 1107 | 232.1 | - | - | 0 | - |
| 2 | a | 8.013E+05 | 233.2 | 0.0003824 | 1.64 | +1 | 2 |
| - | - | 1.205E+05 | 234.2 | - | - | 0 | - |
| - | - | 1681 | 235.1 | - | - | 0 | - |
| - | - | 8557 | 235.2 | - | - | 0 | - |
| - | - | 2359 | 235.6 | - | - | 0 | - |
| - | - | 4.113E+04 | 236.1 | - | - | 0 | - |
| - | - | 3921 | 236.1 | - | - | 0 | - |
| - | - | 3228 | 237.1 | - | - | 0 | - |
| - | - | 2617 | 237.1 | - | - | 0 | - |
| - | - | 1075 | 239.1 | - | - | 0 | - |
| - | - | 4532 | 240.1 | - | - | 0 | - |
| - | - | 3448 | 240.6 | - | - | 0 | - |
| - | - | 7.87E+04 | 241.1 | - | - | 0 | - |
| - | - | 1.065E+04 | 242.1 | - | - | 0 | - |
| - | - | 7316 | 242.1 | - | - | 0 | - |
| 8 | y | 5.631E+05 | 243.1 | 0.0004048 | 1.665 | +1 | 2 |
| 2 | b | 5.702E+04 | 244.1 | 0.003925 | 16.08 | +1 | 2 |
| - | - | 2.207E+04 | 245.1 | - | - | 0 | - |
| - | - | 3549 | 245.1 | - | - | 0 | - |
| - | - | 2578 | 246.1 | - | - | 0 | - |
| - | - | 2667 | 249.6 | - | - | 0 | - |
| - | - | 2718 | 250.1 | - | - | 0 | - |
| - | - | 1258 | 250.1 | - | - | 0 | - |
| - | - | 1908 | 251.1 | - | - | 0 | - |
| - | - | 1.142E+04 | 252.1 | - | - | 0 | - |
| - | - | 1891 | 253.1 | - | - | 0 | - |
| - | - | 1623 | 253.1 | - | - | 0 | - |
| - | - | 2020 | 253.2 | - | - | 0 | - |
| - | - | 1.315E+05 | 254.1 | - | - | 0 | - |
| - | - | 1.455E+04 | 254.2 | - | - | 0 | - |
| - | - | 1.556E+04 | 255.1 | - | - | 0 | - |
| - | - | 1914 | 255.2 | - | - | 0 | - |
| - | - | 1652 | 256.1 | - | - | 0 | - |
| - | - | 1782 | 256.1 | - | - | 0 | - |
| - | - | 5280 | 258.1 | - | - | 0 | - |
| - | - | 6.747E+04 | 259.1 | - | - | 0 | - |
| - | - | 7047 | 260.1 | - | - | 0 | - |
| 8 | y | 7.579E+04 | 261.1 | 0.0001551 | 0.5939 | +1 | 2 |
| 2 | b | 3.362E+05 | 261.2 | 0.0002188 | 0.8378 | +1 | 2 |
| - | - | 5976 | 262.1 | - | - | 0 | - |
| - | - | 5.256E+04 | 262.2 | - | - | 0 | - |
| - | - | 4209 | 263.2 | - | - | 0 | - |
| - | - | 4852 | 264.1 | - | - | 0 | - |
| - | - | 6.306E+04 | 264.1 | - | - | 0 | - |
| - | - | 8782 | 265.1 | - | - | 0 | - |
| - | - | 1263 | 267.1 | - | - | 0 | - |
| - | - | 1.022E+04 | 268.1 | - | - | 0 | - |
| - | - | 3504 | 268.1 | - | - | 0 | - |
| - | - | 1537 | 269.1 | - | - | 0 | - |
| - | - | 1756 | 269.1 | - | - | 0 | - |
| - | - | 1838 | 269.1 | - | - | 0 | - |
| - | - | 1427 | 271.2 | - | - | 0 | - |
| - | - | 7.082E+04 | 272.1 | - | - | 0 | - |
| - | - | 1.027E+04 | 273.1 | - | - | 0 | - |
| - | - | 1074 | 274.1 | - | - | 0 | - |
| - | - | 3372 | 276.1 | - | - | 0 | - |
| - | - | 1507 | 277.1 | - | - | 0 | - |
| - | - | 2373 | 278.1 | - | - | 0 | - |
| - | - | 1165 | 278.1 | - | - | 0 | - |
| - | - | 1341 | 279.1 | - | - | 0 | - |
| - | - | 1083 | 280.2 | - | - | 0 | - |
| - | - | 4063 | 281.1 | - | - | 0 | - |
| - | - | 3104 | 281.2 | - | - | 0 | - |
| - | - | 6347 | 282.1 | - | - | 0 | - |
| - | - | 2.549E+05 | 282.1 | - | - | 0 | - |
| 5 | y | 2051 | 282.6 | 0.001344 | 4.754 | +2 | 5 |
| - | - | 1009 | 283.1 | - | - | 0 | - |
| - | - | 3.335E+04 | 283.1 | - | - | 0 | - |
| - | - | 1611 | 284.1 | - | - | 0 | - |
| - | - | 7784 | 285.2 | - | - | 0 | - |
| - | - | 4.789E+04 | 286.1 | - | - | 0 | - |
| - | - | 1373 | 286.2 | - | - | 0 | - |
| - | - | 3667 | 287.1 | - | - | 0 | - |
| - | - | 2484 | 287.1 | - | - | 0 | - |
| - | - | 1404 | 287.1 | - | - | 0 | - |
| - | - | 1074 | 288.2 | - | - | 0 | - |
| - | - | 2794 | 288.2 | - | - | 0 | - |
| - | - | 2580 | 289.2 | - | - | 0 | - |
| - | - | 1486 | 290.1 | - | - | 0 | - |
| - | - | 3401 | 291.1 | - | - | 0 | - |
| - | - | 3204 | 292.1 | - | - | 0 | - |
| - | - | 1705 | 292.2 | - | - | 0 | - |
| - | - | 2019 | 294.1 | - | - | 0 | - |
| - | - | 3415 | 296.1 | - | - | 0 | - |
| - | - | 2.568E+04 | 296.1 | - | - | 0 | - |
| - | - | 4144 | 296.6 | - | - | 0 | - |
| - | - | 2091 | 297.1 | - | - | 0 | - |
| - | - | 6482 | 297.2 | - | - | 0 | - |
| - | - | 2183 | 297.7 | - | - | 0 | - |
| - | - | 3729 | 299.1 | - | - | 0 | - |
| - | - | 1.228E+04 | 299.2 | - | - | 0 | - |
| - | - | 8311 | 300.1 | - | - | 0 | - |
| - | - | 1190 | 300.1 | - | - | 0 | - |
| - | - | 1.011E+04 | 300.2 | - | - | 0 | - |
| - | - | 1695 | 300.2 | - | - | 0 | - |
| - | - | 2612 | 301.2 | - | - | 0 | - |
| - | - | 1031 | 302.2 | - | - | 0 | - |
| - | - | 2.167E+04 | 304.1 | - | - | 0 | - |
| - | - | 1447 | 304.1 | - | - | 0 | - |
| - | - | 6891 | 305.1 | - | - | 0 | - |
| - | - | 2026 | 305.2 | - | - | 0 | - |
| - | - | 2289 | 305.6 | - | - | 0 | - |
| - | - | 8292 | 306.1 | - | - | 0 | - |
| 6 | b | 4013 | 306.2 | 0.0006405 | 2.092 | +2 | 6 |
| - | - | 1460 | 306.7 | - | - | 0 | - |
| - | - | 1262 | 307.1 | - | - | 0 | - |
| - | - | 2060 | 307.2 | - | - | 0 | - |
| - | - | 1.304E+04 | 308.2 | - | - | 0 | - |
| - | - | 7411 | 308.2 | - | - | 0 | - |
| - | - | 1707 | 309.2 | - | - | 0 | - |
| - | - | 2032 | 309.2 | - | - | 0 | - |
| - | - | 1.494E+04 | 310.1 | - | - | 0 | - |
| - | - | 1361 | 310.1 | - | - | 0 | - |
| - | - | 1971 | 311.1 | - | - | 0 | - |
| - | - | 3752 | 311.2 | - | - | 0 | - |
| - | - | 3586 | 312.1 | - | - | 0 | - |
| - | - | 4081 | 312.2 | - | - | 0 | - |
| 3 | a | 4643 | 313.2 | 7.547E-05 | 0.241 | +1 | 3 |
| - | - | 4436 | 314.1 | - | - | 0 | - |
| - | - | 2.754E+04 | 314.1 | - | - | 0 | - |
| - | - | 5070 | 314.6 | - | - | 0 | - |
| - | - | 2557 | 315.1 | - | - | 0 | - |
| - | - | 1763 | 317.2 | - | - | 0 | - |
| - | - | 2960 | 318.1 | - | - | 0 | - |
| - | - | 2529 | 319.1 | - | - | 0 | - |
| - | - | 2111 | 320.1 | - | - | 0 | - |
| - | - | 2342 | 320.7 | - | - | 0 | - |
| - | - | 1.035E+04 | 321.2 | - | - | 0 | - |
| - | - | 1617 | 322.2 | - | - | 0 | - |
| - | - | 1.174E+04 | 323.2 | - | - | 0 | - |
| - | - | 1.118E+04 | 324.2 | - | - | 0 | - |
| - | - | 1842 | 324.2 | - | - | 0 | - |
| - | - | 1855 | 325.2 | - | - | 0 | - |
| - | - | 3112 | 326.2 | - | - | 0 | - |
| - | - | 1455 | 327.2 | - | - | 0 | - |
| - | - | 3.232E+04 | 328.1 | - | - | 0 | - |
| - | - | 4900 | 329.1 | - | - | 0 | - |
| - | - | 6368 | 329.7 | - | - | 0 | - |
| - | - | 5966 | 330.2 | - | - | 0 | - |
| - | - | 3.916E+04 | 333.2 | - | - | 0 | - |
| - | - | 7763 | 334.2 | - | - | 0 | - |
| - | - | 1092 | 335.2 | - | - | 0 | - |
| - | - | 3242 | 336.2 | - | - | 0 | - |
| - | - | 1.265E+04 | 337.2 | - | - | 0 | - |
| - | - | 1297 | 338.1 | - | - | 0 | - |
| - | - | 6965 | 338.1 | - | - | 0 | - |
| - | - | 2095 | 338.2 | - | - | 0 | - |
| - | - | 2.7E+04 | 338.7 | - | - | 0 | - |
| - | - | 3.302E+04 | 339.2 | - | - | 0 | - |
| - | - | 1272 | 339.7 | - | - | 0 | - |
| - | - | 3423 | 340.2 | - | - | 0 | - |
| 3 | b | 4.75E+04 | 341.2 | 0.003291 | 9.645 | +1 | 3 |
| - | - | 1.115E+04 | 342.2 | - | - | 0 | - |
| - | - | 4254 | 343.2 | - | - | 0 | - |
| - | - | 1248 | 343.2 | - | - | 0 | - |
| - | - | 2455 | 343.7 | - | - | 0 | - |
| - | - | 2517 | 344.2 | - | - | 0 | - |
| - | - | 5.697E+04 | 346.1 | - | - | 0 | - |
| - | - | 1.064E+04 | 347.1 | - | - | 0 | - |
| - | - | 5403 | 347.7 | - | - | 0 | - |
| - | - | 1364 | 348.1 | - | - | 0 | - |
| - | - | 1228 | 348.2 | - | - | 0 | - |
| - | - | 1436 | 348.2 | - | - | 0 | - |
| - | - | 1.848E+05 | 351.2 | - | - | 0 | - |
| - | - | 3.22E+04 | 352.2 | - | - | 0 | - |
| - | - | 3628 | 352.7 | - | - | 0 | - |
| - | - | 1694 | 353.1 | - | - | 0 | - |
| - | - | 5734 | 353.2 | - | - | 0 | - |
| - | - | 2.745E+04 | 354.2 | - | - | 0 | - |
| - | - | 3.092E+04 | 355.2 | - | - | 0 | - |
| - | - | 6851 | 356.1 | - | - | 0 | - |
| - | - | 3359 | 356.2 | - | - | 0 | - |
| - | - | 1277 | 357.1 | - | - | 0 | - |
| - | - | 1460 | 357.2 | - | - | 0 | - |
| 3 | b | 3.099E+04 | 358.2 | 0.0002809 | 0.7841 | +1 | 3 |
| - | - | 6109 | 359.2 | - | - | 0 | - |
| - | - | 1.01E+04 | 361.7 | - | - | 0 | - |
| - | - | 4749 | 362.2 | - | - | 0 | - |
| - | - | 3.057E+04 | 365.1 | - | - | 0 | - |
| - | - | 2419 | 365.2 | - | - | 0 | - |
| - | - | 1812 | 365.7 | - | - | 0 | - |
| - | - | 3339 | 366.1 | - | - | 0 | - |
| - | - | 2062 | 366.2 | - | - | 0 | - |
| - | - | 2660 | 367.2 | - | - | 0 | - |
| - | - | 1206 | 367.2 | - | - | 0 | - |
| - | - | 4653 | 368.2 | - | - | 0 | - |
| - | - | 2.139E+05 | 369.2 | - | - | 0 | - |
| - | - | 1413 | 370.1 | - | - | 0 | - |
| - | - | 3.429E+04 | 370.2 | - | - | 0 | - |
| 7 | b | 1.516E+04 | 370.7 | 0.002681 | 7.233 | +2 | 7 |
| - | - | 6358 | 371.2 | - | - | 0 | - |
| 7 | b | 1.026E+04 | 371.2 | 0.006959 | 18.75 | +2 | 7 |
| - | - | 1340 | 371.7 | - | - | 0 | - |
| 7 | y | 2.483E+04 | 372.2 | 0.0002922 | 0.7852 | +1 | 3 |
| - | - | 5.237E+04 | 373.2 | - | - | 0 | - |
| - | - | 9174 | 374.2 | - | - | 0 | - |
| - | - | 3030 | 375.2 | - | - | 0 | - |
| - | - | 1582 | 377.2 | - | - | 0 | - |
| - | - | 1609 | 379.1 | - | - | 0 | - |
| 7 | b | 8597 | 379.7 | 0.004637 | 12.21 | +2 | 7 |
| - | - | 2872 | 380.2 | - | - | 0 | - |
| - | - | 981.8 | 380.7 | - | - | 0 | - |
| - | - | 5.736E+04 | 383.2 | - | - | 0 | - |
| - | - | 1.046E+04 | 384.2 | - | - | 0 | - |
| - | - | 1223 | 384.2 | - | - | 0 | - |
| - | - | 1547 | 385.2 | - | - | 0 | - |
| - | - | 2.279E+04 | 386.2 | - | - | 0 | - |
| - | - | 3177 | 387.2 | - | - | 0 | - |
| - | - | 3500 | 387.2 | - | - | 0 | - |
| - | - | 1757 | 387.2 | - | - | 0 | - |
| - | - | 1041 | 388.2 | - | - | 0 | - |
| 7 | y | 7185 | 390.2 | 0.000134 | 0.3435 | +1 | 3 |
| - | - | 1915 | 391.2 | - | - | 0 | - |
| - | - | 1776 | 393.1 | - | - | 0 | - |
| - | - | 6044 | 393.2 | - | - | 0 | - |
| - | - | 1114 | 393.2 | - | - | 0 | - |
| - | - | 2090 | 394.2 | - | - | 0 | - |
| - | - | 1796 | 395.2 | - | - | 0 | - |
| - | - | 2284 | 395.7 | - | - | 0 | - |
| - | - | 1909 | 396.2 | - | - | 0 | - |
| - | - | 8316 | 397.1 | - | - | 0 | - |
| - | - | 1348 | 398.1 | - | - | 0 | - |
| - | - | 1468 | 399.2 | - | - | 0 | - |
| - | - | 1.034E+05 | 401.2 | - | - | 0 | - |
| - | - | 3005 | 401.3 | - | - | 0 | - |
| - | - | 1.836E+04 | 402.2 | - | - | 0 | - |
| - | - | 2182 | 403.2 | - | - | 0 | - |
| - | - | 4673 | 405.2 | - | - | 0 | - |
| - | - | 1361 | 405.2 | - | - | 0 | - |
| - | - | 1493 | 406.2 | - | - | 0 | - |
| - | - | 1021 | 409.2 | - | - | 0 | - |
| - | - | 2734 | 411.2 | - | - | 0 | - |
| - | - | 1975 | 411.2 | - | - | 0 | - |
| - | - | 1281 | 412.2 | - | - | 0 | - |
| - | - | 1.601E+04 | 415.1 | - | - | 0 | - |
| - | - | 2641 | 416.2 | - | - | 0 | - |
| - | - | 3537 | 416.2 | - | - | 0 | - |
| - | - | 3834 | 417.2 | - | - | 0 | - |
| - | - | 1378 | 417.7 | - | - | 0 | - |
| - | - | 1578 | 418.2 | - | - | 0 | - |
| - | - | 5260 | 423.2 | - | - | 0 | - |
| - | - | 1343 | 424.2 | - | - | 0 | - |
| - | - | 3060 | 425.1 | - | - | 0 | - |
| - | - | 3219 | 425.2 | - | - | 0 | - |
| - | - | 1683 | 425.2 | - | - | 0 | - |
| - | - | 3125 | 426.2 | - | - | 0 | - |
| - | - | 1.282E+04 | 427.3 | - | - | 0 | - |
| - | - | 3727 | 428.3 | - | - | 0 | - |
| - | - | 1295 | 429.2 | - | - | 0 | - |
| - | - | 7771 | 429.2 | - | - | 0 | - |
| - | - | 2918 | 430.2 | - | - | 0 | - |
| - | - | 3.25E+04 | 433.2 | - | - | 0 | - |
| - | - | 5069 | 434.2 | - | - | 0 | - |
| - | - | 2.056E+04 | 434.2 | - | - | 0 | - |
| 8 | b | 5703 | 435.2 | 0.0007414 | 1.704 | +2 | 8 |
| 8 | b | 1877 | 435.7 | 0.007696 | 17.66 | +2 | 8 |
| - | - | 2894 | 436.3 | - | - | 0 | - |
| - | - | 1.127E+04 | 440.2 | - | - | 0 | - |
| - | - | 7714 | 441.2 | - | - | 0 | - |
| - | - | 1771 | 442.2 | - | - | 0 | - |
| - | - | 3071 | 443.1 | - | - | 0 | - |
| - | - | 2448 | 443.2 | - | - | 0 | - |
| - | - | 1239 | 444.1 | - | - | 0 | - |
| - | - | 6788 | 444.2 | - | - | 0 | - |
| 8 | b | 2045 | 444.2 | 0.006476 | 14.58 | +2 | 8 |
| - | - | 1706 | 444.7 | - | - | 0 | - |
| - | - | 1565 | 446.2 | - | - | 0 | - |
| - | - | 2558 | 448.2 | - | - | 0 | - |
| - | - | 1.372E+04 | 450.2 | - | - | 0 | - |
| - | - | 3443 | 451.2 | - | - | 0 | - |
| - | - | 4.677E+04 | 452.2 | - | - | 0 | - |
| - | - | 1428 | 453.2 | - | - | 0 | - |
| 2 | y | 9210 | 453.2 | 0.0006138 | 1.354 | +2 | 8 |
| - | - | 1793 | 454.2 | - | - | 0 | - |
| - | - | 7750 | 454.3 | - | - | 0 | - |
| 4 | b | 3.263E+04 | 455.3 | 0.0004345 | 0.9544 | +1 | 4 |
| - | - | 8528 | 456.3 | - | - | 0 | - |
| - | - | 1633 | 457.3 | - | - | 0 | - |
| 6 | y | 1.437E+04 | 459.2 | 0.001375 | 2.995 | +1 | 4 |
| - | - | 2801 | 460.2 | - | - | 0 | - |
| - | - | 4.02E+04 | 462.2 | - | - | 0 | - |
| - | - | 1.228E+04 | 463.2 | - | - | 0 | - |
| - | - | 1173 | 464.2 | - | - | 0 | - |
| - | - | 4437 | 464.3 | - | - | 0 | - |
| - | - | 1195 | 465.3 | - | - | 0 | - |
| - | - | 5904 | 466.2 | - | - | 0 | - |
| - | - | 2.522E+04 | 468.2 | - | - | 0 | - |
| - | - | 1279 | 468.2 | - | - | 0 | - |
| - | - | 4974 | 469.2 | - | - | 0 | - |
| - | - | 2158 | 469.2 | - | - | 0 | - |
| - | - | 1.801E+05 | 470.2 | - | - | 0 | - |
| - | - | 4.186E+04 | 471.2 | - | - | 0 | - |
| - | - | 7188 | 472.2 | - | - | 0 | - |
| - | - | 5507 | 476.2 | - | - | 0 | - |
| 6 | y | 1.056E+04 | 477.2 | 0.001004 | 2.103 | +1 | 4 |
| - | - | 1388 | 477.9 | - | - | 0 | - |
| - | - | 3238 | 478.2 | - | - | 0 | - |
| - | - | 1.602E+05 | 480.2 | - | - | 0 | - |
| - | - | 3.935E+04 | 481.2 | - | - | 0 | - |
| - | - | 6738 | 482.2 | - | - | 0 | - |
| - | - | 8127 | 482.3 | - | - | 0 | - |
| - | - | 1823 | 483.3 | - | - | 0 | - |
| - | - | 1239 | 483.9 | - | - | 0 | - |
| - | - | 1.749E+04 | 484.2 | - | - | 0 | - |
| - | - | 3119 | 485.2 | - | - | 0 | - |
| - | - | 5819 | 486.3 | - | - | 0 | - |
| - | - | 2556 | 487.3 | - | - | 0 | - |
| - | - | 1301 | 488.3 | - | - | 0 | - |
| - | - | 1360 | 491.7 | - | - | 0 | - |
| - | - | 1577 | 492.2 | - | - | 0 | - |
| - | - | 1.294E+04 | 494.2 | - | - | 0 | - |
| - | - | 4114 | 495.2 | - | - | 0 | - |
| - | - | 2548 | 496.2 | - | - | 0 | - |
| - | - | 5482 | 497.2 | - | - | 0 | - |
| - | - | 5.545E+05 | 498.2 | - | - | 0 | - |
| - | - | 1.412E+05 | 499.2 | - | - | 0 | - |
| - | - | 2.117E+04 | 500.2 | - | - | 0 | - |
| - | - | 3241 | 501.2 | - | - | 0 | - |
| - | - | 2.746E+04 | 502.2 | - | - | 0 | - |
| - | - | 7890 | 503.2 | - | - | 0 | - |
| - | - | 1396 | 504.2 | - | - | 0 | - |
| - | - | 1254 | 504.3 | - | - | 0 | - |
| - | - | 1880 | 508.3 | - | - | 0 | - |
| - | - | 2136 | 510.2 | - | - | 0 | - |
| - | - | 1670 | 511.2 | - | - | 0 | - |
| - | - | 2.772E+04 | 512.2 | - | - | 0 | - |
| - | - | 1182 | 512.3 | - | - | 0 | - |
| - | - | 6429 | 513.2 | - | - | 0 | - |
| - | - | 1715 | 514.2 | - | - | 0 | - |
| - | - | 6097 | 514.3 | - | - | 0 | - |
| - | - | 4817 | 514.3 | - | - | 0 | - |
| - | - | 1.288E+04 | 515.2 | - | - | 0 | - |
| - | - | 1177 | 515.3 | - | - | 0 | - |
| - | - | 1332 | 515.3 | - | - | 0 | - |
| - | - | 7229 | 516.2 | - | - | 0 | - |
| - | - | 1224 | 517.2 | - | - | 0 | - |
| - | - | 1397 | 518.2 | - | - | 0 | - |
| - | - | 1565 | 519.2 | - | - | 0 | - |
| 5 | b | 7633 | 524.3 | 2.614E-06 | 0.004986 | +1 | 5 |
| - | - | 2739 | 525.3 | - | - | 0 | - |
| - | - | 2336 | 526.3 | - | - | 0 | - |
| - | - | 4295 | 528.2 | - | - | 0 | - |
| - | - | 2005 | 529.2 | - | - | 0 | - |
| - | - | 8.903E+04 | 530.2 | - | - | 0 | - |
| - | - | 1.093E+04 | 530.3 | - | - | 0 | - |
| - | - | 2.231E+04 | 531.2 | - | - | 0 | - |
| - | - | 1752 | 531.3 | - | - | 0 | - |
| - | - | 3570 | 532.2 | - | - | 0 | - |
| - | - | 1249 | 532.3 | - | - | 0 | - |
| - | - | 1266 | 534.2 | - | - | 0 | - |
| 5 | b | 2.51E+04 | 542.3 | 0.0001139 | 0.2099 | +1 | 5 |
| - | - | 8788 | 543.3 | - | - | 0 | - |
| - | - | 1213 | 544.3 | - | - | 0 | - |
| - | - | 2737 | 545.2 | - | - | 0 | - |
| 5 | y | 8202 | 546.2 | 0.0003491 | 0.639 | +1 | 5 |
| - | - | 1983 | 547.2 | - | - | 0 | - |
| - | - | 1062 | 548.1 | - | - | 0 | - |
| - | - | 7973 | 548.2 | - | - | 0 | - |
| - | - | 1654 | 549.2 | - | - | 0 | - |
| - | - | 1943 | 553.3 | - | - | 0 | - |
| - | - | 1899 | 554.2 | - | - | 0 | - |
| - | - | 1942 | 556.2 | - | - | 0 | - |
| - | - | 8814 | 563.2 | - | - | 0 | - |
| 5 | y | 1.387E+04 | 564.3 | 0.001148 | 2.035 | +1 | 5 |
| - | - | 1810 | 565.3 | - | - | 0 | - |
| - | - | 2980 | 565.3 | - | - | 0 | - |
| - | - | 2113 | 566.3 | - | - | 0 | - |
| - | - | 1647 | 572.3 | - | - | 0 | - |
| - | - | 1.001E+04 | 573.2 | - | - | 0 | - |
| - | - | 848.5 | 573.3 | - | - | 0 | - |
| - | - | 3199 | 574.2 | - | - | 0 | - |
| - | - | 1248 | 574.3 | - | - | 0 | - |
| - | - | 2538 | 575.3 | - | - | 0 | - |
| - | - | 2235 | 579.2 | - | - | 0 | - |
| - | - | 2017 | 579.3 | - | - | 0 | - |
| - | - | 3.098E+04 | 581.3 | - | - | 0 | - |
| - | - | 9427 | 582.3 | - | - | 0 | - |
| - | - | 1314 | 583.3 | - | - | 0 | - |
| - | - | 3.07E+04 | 583.3 | - | - | 0 | - |
| - | - | 1.299E+04 | 584.3 | - | - | 0 | - |
| - | - | 1761 | 585.3 | - | - | 0 | - |
| - | - | 2148 | 588.3 | - | - | 0 | - |
| - | - | 1547 | 590.3 | - | - | 0 | - |
| - | - | 2.965E+04 | 591.2 | - | - | 0 | - |
| - | - | 1885 | 591.3 | - | - | 0 | - |
| - | - | 9933 | 592.2 | - | - | 0 | - |
| - | - | 1435 | 593.2 | - | - | 0 | - |
| - | - | 1.176E+04 | 593.3 | - | - | 0 | - |
| - | - | 4546 | 594.3 | - | - | 0 | - |
| - | - | 2036 | 597.2 | - | - | 0 | - |
| - | - | 1866 | 597.3 | - | - | 0 | - |
| - | - | 1.788E+05 | 599.3 | - | - | 0 | - |
| - | - | 5.901E+04 | 600.3 | - | - | 0 | - |
| - | - | 1341 | 600.3 | - | - | 0 | - |
| - | - | 1.125E+04 | 601.3 | - | - | 0 | - |
| - | - | 1.938E+04 | 601.3 | - | - | 0 | - |
| - | - | 6774 | 602.3 | - | - | 0 | - |
| - | - | 1122 | 603.3 | - | - | 0 | - |
| - | - | 3841 | 606.3 | - | - | 0 | - |
| - | - | 4431 | 607.3 | - | - | 0 | - |
| - | - | 3075 | 608.3 | - | - | 0 | - |
| - | - | 1.079E+05 | 609.3 | - | - | 0 | - |
| - | - | 3.377E+04 | 610.3 | - | - | 0 | - |
| - | - | 1638 | 610.3 | - | - | 0 | - |
| - | - | 3249 | 611.3 | - | - | 0 | - |
| 6 | b | 5.637E+04 | 611.3 | 0.007983 | 13.06 | +1 | 6 |
| 6 | b | 2.139E+04 | 612.3 | 0.01081 | 17.65 | +1 | 6 |
| - | - | 3528 | 613.3 | - | - | 0 | - |
| - | - | 6792 | 613.4 | - | - | 0 | - |
| - | - | 7055 | 615.3 | - | - | 0 | - |
| - | - | 4081 | 616.3 | - | - | 0 | - |
| - | - | 3772 | 617.3 | - | - | 0 | - |
| - | - | 9722 | 625.3 | - | - | 0 | - |
| - | - | 4166 | 626.3 | - | - | 0 | - |
| - | - | 9.032E+05 | 627.3 | - | - | 0 | - |
| - | - | 2.774E+05 | 628.3 | - | - | 0 | - |
| - | - | 5.556E+04 | 629.3 | - | - | 0 | - |
| 6 | b | 5.789E+04 | 629.3 | 0.000312 | 0.4958 | +1 | 6 |
| - | - | 4239 | 630.3 | - | - | 0 | - |
| - | - | 2.138E+04 | 630.3 | - | - | 0 | - |
| - | - | 4979 | 631.3 | - | - | 0 | - |
| - | - | 2442 | 633.3 | - | - | 0 | - |
| - | - | 1690 | 634.3 | - | - | 0 | - |
| 4 | y | 2.63E+04 | 643.3 | 0.0009642 | 1.499 | +1 | 6 |
| - | - | 8906 | 644.3 | - | - | 0 | - |
| - | - | 4.137E+04 | 645.3 | - | - | 0 | - |
| - | - | 1.474E+04 | 646.3 | - | - | 0 | - |
| - | - | 2059 | 647.3 | - | - | 0 | - |
| - | - | 1935 | 653.3 | - | - | 0 | - |
| - | - | 995.3 | 658.3 | - | - | 0 | - |
| 4 | y | 1.045E+05 | 661.3 | 0.0003483 | 0.5266 | +1 | 6 |
| - | - | 3.445E+04 | 662.3 | - | - | 0 | - |
| - | - | 6133 | 663.3 | - | - | 0 | - |
| - | - | 1238 | 676.3 | - | - | 0 | - |
| - | - | 8466 | 677.4 | - | - | 0 | - |
| - | - | 4588 | 686.3 | - | - | 0 | - |
| - | - | 1376 | 692.3 | - | - | 0 | - |
| - | - | 7730 | 694.3 | - | - | 0 | - |
| - | - | 3225 | 695.3 | - | - | 0 | - |
| - | - | 1791 | 696.4 | - | - | 0 | - |
| - | - | 1.258E+04 | 704.3 | - | - | 0 | - |
| - | - | 6167 | 705.3 | - | - | 0 | - |
| - | - | 1966 | 706.3 | - | - | 0 | - |
| - | - | 2985 | 710.3 | - | - | 0 | - |
| - | - | 1094 | 711.3 | - | - | 0 | - |
| - | - | 5.343E+04 | 712.4 | - | - | 0 | - |
| - | - | 2.497E+04 | 713.4 | - | - | 0 | - |
| - | - | 7796 | 714.4 | - | - | 0 | - |
| - | - | 3.985E+04 | 722.3 | - | - | 0 | - |
| - | - | 2.016E+04 | 723.3 | - | - | 0 | - |
| - | - | 4778 | 724.3 | - | - | 0 | - |
| - | - | 2283 | 728.3 | - | - | 0 | - |
| - | - | 3721 | 729.3 | - | - | 0 | - |
| - | - | 7.123E+04 | 730.4 | - | - | 0 | - |
| - | - | 2.874E+04 | 731.4 | - | - | 0 | - |
| - | - | 4661 | 732.4 | - | - | 0 | - |
| - | - | 1479 | 738.3 | - | - | 0 | - |
| 7 | b | 1.684E+05 | 740.4 | 0.01231 | 16.62 | +1 | 7 |
| 7 | b | 7.317E+04 | 741.3 | 0.006729 | 9.077 | +1 | 7 |
| - | - | 1.806E+04 | 742.4 | - | - | 0 | - |
| - | - | 1457 | 743.4 | - | - | 0 | - |
| - | - | 3627 | 746.3 | - | - | 0 | - |
| - | - | 1884 | 747.3 | - | - | 0 | - |
| - | - | 1740 | 754.3 | - | - | 0 | - |
| - | - | 4640 | 756.3 | - | - | 0 | - |
| - | - | 1946 | 757.3 | - | - | 0 | - |
| 7 | b | 9.173E+05 | 758.4 | 0.01219 | 16.07 | +1 | 7 |
| - | - | 3.826E+05 | 759.4 | - | - | 0 | - |
| - | - | 9.221E+04 | 760.4 | - | - | 0 | - |
| - | - | 6890 | 761.4 | - | - | 0 | - |
| - | - | 2733 | 762.4 | - | - | 0 | - |
| - | - | 2893 | 772.4 | - | - | 0 | - |
| - | - | 1152 | 773.4 | - | - | 0 | - |
| - | - | 1.858E+04 | 774.3 | - | - | 0 | - |
| - | - | 7584 | 775.3 | - | - | 0 | - |
| - | - | 2042 | 776.3 | - | - | 0 | - |
| - | - | 4958 | 790.4 | - | - | 0 | - |
| - | - | 2286 | 791.4 | - | - | 0 | - |
| - | - | 1547 | 833.4 | - | - | 0 | - |
| - | - | 6466 | 841.4 | - | - | 0 | - |
| - | - | 8260 | 842.4 | - | - | 0 | - |
| - | - | 3719 | 843.4 | - | - | 0 | - |
| - | - | 1.307E+04 | 851.4 | - | - | 0 | - |
| - | - | 5700 | 852.4 | - | - | 0 | - |
| - | - | 3125 | 854.4 | - | - | 0 | - |
| - | - | 6.572E+04 | 859.4 | - | - | 0 | - |
| - | - | 2.739E+04 | 860.4 | - | - | 0 | - |
| - | - | 8669 | 861.4 | - | - | 0 | - |
| - | - | 1672 | 862.4 | - | - | 0 | - |
| 8 | b | 3.636E+04 | 869.4 | 0.0007621 | 0.8766 | +1 | 8 |
| - | - | 1.784E+04 | 870.4 | - | - | 0 | - |
| - | - | 5847 | 871.4 | - | - | 0 | - |
| 8 | b | 2.947E+05 | 887.4 | 0.001134 | 1.278 | +1 | 8 |
| - | - | 1.599E+05 | 888.4 | - | - | 0 | - |
| - | - | 4.238E+04 | 889.4 | - | - | 0 | - |
| - | - | 4182 | 890.4 | - | - | 0 | - |
| 2 | y | 4257 | 905.4 | 0.001506 | 1.663 | +1 | 8 |
| - | - | 2801 | 906.4 | - | - | 0 | - |
| - | - | 1154 | 954 | - | - | 0 | - |
| - | - | 1093 | 1028 | - | - | 0 | - |
| - | - | 1164 | 1349 | - | - | 0 | - |
| - | - | 1235 | 1773 | - | - | 0 | - |
| - | - | 1108 | 1938 | - | - | 0 | - |
| - | - | 1187 | 2054 | - | - | 0 | - |
| - | - | 1218 | 3081 | - | - | 0 | - |

m/z Charge Intensity FragmentType MassShift Position
120.08114624023438 0 558457.44
121.08443450927734 0 45516.63
122.08776092529297 0 711.4928
126.05526733398438 0 59043.363
126.2887954711914 0 701.2555
127.05862426757812 0 3135.4312
129.06617736816406 0 17698.148
129.10252380371094 0 2479.782
130.05020141601562 0 59601.652
131.0535430908203 0 3283.597
132.1022491455078 0 893784.3 y 8
133.08642578125 0 2090.738
133.09988403320312 0 2213.8691
133.10556030273438 0 55191.586
134.10662841796875 0 2564.9412
136.07626342773438 0 2300.1375
138.12802124023438 0 13039.5625
139.05038452148438 0 2507.4707
139.0869598388672 0 10827.477
140.07054138183594 0 1004.24414
140.09030151367188 0 1038.5781
141.06613159179688 0 8004.1587
143.4122772216797 0 694.66565
144.06591796875 0 994.7465
147.07672119140625 0 13415.69
148.06080627441406 0 4504.939
148.0754852294922 0 1676.5271
148.95420837402344 0 945.84644
151.08766174316406 0 884.2798
151.1232147216797 0 1975.0913
152.08204650878906 0 991.0957
152.10739135742188 0 2463.3562
153.0660858154297 0 2941.8455
154.05023193359375 0 4738.828
154.08656311035156 0 33875.113
155.08187866210938 0 2326.5908
155.08998107910156 0 2724.4749
155.11776733398438 0 1355.9663
157.06103515625 0 20113.041
157.09742736816406 0 50419.69 a Ammonia loss 2
158.09292602539062 0 1073.7017
158.10079956054688 0 3348.2031
159.09181213378906 0 1166.6521
162.0914764404297 0 1123.6708
165.06700134277344 0 844.7165
165.10256958007812 0 2021.2517
165.6125946044922 0 1217.9921 a 2
166.0865478515625 0 5224.8086
167.04559326171875 0 1522.759
167.08181762695312 0 47397.754
167.1181640625 0 48197.156
168.06570434570312 0 2080.0015
168.08522033691406 0 3990.4905
168.12171936035156 0 3947.2427
169.09767150878906 0 908.0446
169.13427734375 0 894.3639
170.04539489746094 0 2051.0676
170.11756896972656 0 1168.7979
171.07676696777344 0 11080.665
171.0942840576172 0 937.39496 b Ammonia loss 2
172.11233520507812 0 31141.334
173.11561584472656 0 3337.8406
173.12867736816406 0 2779.0842
174.0550994873047 0 1447.398
175.07162475585938 0 23335.832
175.08511352539062 0 1013.6666
175.1192626953125 0 4360.641
175.809326171875 0 727.01764
176.075439453125 0 1305.8448
177.1027374267578 0 3017.303
177.11170959472656 0 1182.6871
179.1179962158203 0 1164.0521
180.10203552246094 0 1459.0294
180.16331481933594 0 804.98846
181.06089782714844 0 2652.978
181.09730529785156 0 5433.87
182.08135986328125 0 1742.8763
183.08584594726562 0 829.2873
183.11326599121094 0 969.1393
183.14947509765625 0 5106.549
185.0923614501953 0 89657.92
185.4304962158203 0 760.5818
186.09573364257812 0 10494.859
187.0982208251953 0 884.02936
188.06983947753906 0 968.5663
188.14373779296875 0 4700.911
189.0872344970703 0 34068.832
193.121337890625 0 1118.2244
195.07666015625 0 12185.684
195.11306762695312 0 97125.984
195.67031860351562 0 930.7962
196.06021118164062 0 1029.7292
196.11634826660156 0 9656.622
197.12872314453125 0 219955.16
198.11326599121094 0 1175.2849
198.1321563720703 0 20316.81
199.0715789794922 0 52791.863
199.10784912109375 0 7482.8623
200.07473754882812 0 3723.9365
201.12384033203125 0 2009.6455
201.1382598876953 0 887.5327
202.11891174316406 0 2759.8845
203.10281372070312 0 6501.9644
205.17047119140625 0 1760.4644
208.07171630859375 0 1238.9149
208.10862731933594 0 1463.9648
209.09239196777344 0 13080.274
209.1282196044922 0 1032.5844
211.1442108154297 0 6718.1855
212.10293579101562 0 1102.3656
212.13970947265625 0 12626.254
213.0872039794922 0 15790.8955
213.1237030029297 0 10384.77
214.09132385253906 0 1152.5355
214.1284637451172 0 1046.6791
214.13917541503906 0 3438.657
215.13919067382812 0 6917.7427
216.09800720214844 0 971.06256
216.14096069335938 0 1132.2943 a Ammonia loss 1
217.08216857910156 0 97290.66
217.1337432861328 0 20688.393
218.0854949951172 0 5806.4043
218.13816833496094 0 2189.959
221.0919952392578 0 1278.5531
221.12875366210938 0 1032.5693
223.07142639160156 0 2711.5908
224.10328674316406 0 4425.8926
225.09835815429688 0 8098.3945
225.12359619140625 0 30158.176
226.11903381347656 0 5057.9683
227.06639099121094 0 1389.0251
227.10301208496094 0 16198.752
227.11691284179688 0 1649.5098
228.10496520996094 0 1816.2466
229.155517578125 0 1930.9392
231.097900390625 0 15377.811
231.11050415039062 0 951.22986
231.60281372070312 0 1509.8263
232.10086059570312 0 1107.1342
233.16522216796875 0 801332.44 a 1
234.16848754882812 0 120540.29
235.14462280273438 0 1681.3839
235.17149353027344 0 8557.436
235.61607360839844 0 2359.4895
236.10324096679688 0 41132.754
236.13937377929688 0 3920.5776
237.10679626464844 0 3227.8267
237.1234893798828 0 2616.5981
239.13807678222656 0 1074.7045
240.09799194335938 0 4531.9194
240.60870361328125 0 3447.5398
241.08221435546875 0 78698.38
242.08541870117188 0 10647.585
242.11402893066406 0 7315.979
243.13433837890625 0 563144.7 y Water loss 7
244.1371307373047 0 57017.656 b Ammonia loss 1
245.12887573242188 0 22068.447
245.1391143798828 0 3548.9355
246.1322479248047 0 2578.0479
249.6136932373047 0 2666.902
250.08204650878906 0 2717.9373
250.1162109375 0 1257.9387
251.1397247314453 0 1908.4447
252.1346893310547 0 11423.323
253.0825958251953 0 1891.4963
253.1181182861328 0 1623.1577
253.16973876953125 0 2019.9482
254.11380004882812 0 131511.5
254.1500701904297 0 14546.743
255.116943359375 0 15555.772
255.15377807617188 0 1913.9058
256.0924987792969 0 1651.5568
256.11883544921875 0 1781.9558
258.1087646484375 0 5279.517
259.0926513671875 0 67473.64
260.0961608886719 0 7047.1416
261.1446533203125 0 75788.49 y 7
261.15997314453125 0 336160.75 b 1
262.1479187011719 0 5975.552
262.16326904296875 0 52564.08
263.16595458984375 0 4208.6064
264.0982971191406 0 4852.305
264.1344909667969 0 63064.04
265.1376647949219 0 8782.27
267.09735107421875 0 1263.4235
268.09307861328125 0 10222.468
268.1292419433594 0 3503.8237
269.0766906738281 0 1536.9552
269.0970764160156 0 1756.0957
269.1131591796875 0 1837.5349
271.1771240234375 0 1426.7225
272.1242370605469 0 70817.836
273.1251220703125 0 10274.315
274.103515625 0 1073.6847
276.1190185546875 0 3372.0808
277.1189270019531 0 1507.2709
278.113525390625 0 2372.8694
278.14990234375 0 1165.2935
279.1349182128906 0 1340.7164
280.1653137207031 0 1083.4677
281.0772705078125 0 4062.5474
281.1611633300781 0 3103.9336
282.1083984375 0 6347.466
282.1451721191406 0 254870.69
282.62786865234375 0 2051.088 y 4
283.09417724609375 0 1008.8359
283.1482849121094 0 33350.094
284.1487121582031 0 1610.6427
285.19659423828125 0 7784.377
286.10369873046875 0 47888.99
286.1993103027344 0 1373.0488
287.1062927246094 0 3667.0264
287.1198425292969 0 2483.5027
287.136474609375 0 1403.729
288.1528625488281 0 1073.8961
288.2037048339844 0 2793.5264
289.1514892578125 0 2579.729
290.134521484375 0 1486.1512
291.1322326660156 0 3401.3125
292.1297912597656 0 3204.3933
292.1664123535156 0 1704.5403
294.1201171875 0 2018.512
296.0878601074219 0 3415.0044
296.1243591308594 0 25677.105
296.6261291503906 0 4143.9175
297.12713623046875 0 2090.954
297.1582946777344 0 6482.126
297.659423828125 0 2182.8423
299.08758544921875 0 3729.0337
299.17169189453125 0 12276.01
300.11944580078125 0 8311.393
300.1376647949219 0 1189.7538
300.1557312011719 0 10111.263
300.1741638183594 0 1695.3579
301.1575622558594 0 2611.8657
302.17181396484375 0 1031.3007
304.1142578125 0 21670.225
304.1329650878906 0 1447.1105
305.12957763671875 0 6890.702
305.1602478027344 0 2026.4196
305.631591796875 0 2289.1902
306.14495849609375 0 8292.218
306.1636657714844 0 4012.5796 b Water loss 5
306.6666564941406 0 1459.6514
307.1470642089844 0 1261.6313
307.1875305175781 0 2059.7844
308.16082763671875 0 13039.13
308.1973876953125 0 7410.6157
309.158447265625 0 1706.7292
309.2010803222656 0 2032.0516
310.10369873046875 0 14935.2705
310.1382141113281 0 1360.9397
311.1057434082031 0 1971.32
311.171875 0 3752.4138
312.1302490234375 0 3585.707
312.1555480957031 0 4080.8923
313.19097900390625 0 4643.029 a Ammonia loss 2
314.0990905761719 0 4435.727
314.1351013183594 0 27538.129
314.6365661621094 0 5069.7773
315.1392517089844 0 2557.3938
317.1864013671875 0 1762.5717
318.12982177734375 0 2959.9692
319.1394348144531 0 2528.9507
320.1243896484375 0 2111.1892
320.6586608886719 0 2341.989
321.15673828125 0 10349.852
322.1591796875 0 1617.4263
323.1716613769531 0 11735.252
324.15545654296875 0 11179.685
324.174560546875 0 1842.318
325.1575012207031 0 1854.712
326.17071533203125 0 3112.4385
327.1697998046875 0 1454.6943
328.1143798828125 0 32316.434
329.11846923828125 0 4899.961
329.66351318359375 0 6368.205
330.1656494140625 0 5965.9604
333.15618896484375 0 39164.805
334.15924072265625 0 7763.194
335.17047119140625 0 1092.0859
336.1552734375 0 3242.3254
337.1510925292969 0 12654.806
338.09881591796875 0 1297.1354
338.1351623535156 0 6964.642
338.1545715332031 0 2095.3396
338.6690368652344 0 26999.785
339.16766357421875 0 33016.71
339.6703796386719 0 1271.6779
340.1692810058594 0 3422.5745
341.18267822265625 0 47497.496 b Ammonia loss 2
342.1842346191406 0 11146.765
343.16156005859375 0 4254.1562
343.185791015625 0 1248.0585
343.6605224609375 0 2455.3572
344.16314697265625 0 2516.7065
346.1247863769531 0 56974.723
347.1300964355469 0 10640.736
347.6749267578125 0 5402.5884
348.1317138671875 0 1363.622
348.1553039550781 0 1227.5608
348.17645263671875 0 1436.1942
351.1666259765625 0 184788.94
352.169677734375 0 32198.63
352.66571044921875 0 3627.9497
353.1465148925781 0 1693.7463
353.1710205078125 0 5734.354
354.1661682128906 0 27453.857
355.162109375 0 30923.186
356.1453552246094 0 6851.483
356.16583251953125 0 3359.0999
357.1483459472656 0 1277.4215
357.2154235839844 0 1459.5283
358.2127990722656 0 30988.38 b 2
359.2159118652344 0 6108.8364
361.6762390136719 0 10096.211
362.1788024902344 0 4748.8003
365.14581298828125 0 30574.639
365.18341064453125 0 2418.6846
365.6896667480469 0 1811.6173
366.1486511230469 0 3338.5164
366.1692199707031 0 2062.386
367.1963806152344 0 2660.0981
367.2352600097656 0 1206.201
368.1942138671875 0 4652.529
369.1771240234375 0 213854.16
370.1233825683594 0 1412.5541
370.1799621582031 0 34287.83
370.681640625 0 15161.447 b Water loss 6
371.1560363769531 0 6358.233
371.18328857421875 0 10262.075 b Ammonia loss 6
371.68426513671875 0 1339.6342
372.17681884765625 0 24832.805 y Water loss 6
373.1723327636719 0 52373.32
374.17523193359375 0 9174.352
375.167724609375 0 3029.7393
377.18267822265625 0 1582.3011
379.12640380859375 0 1609.1954
379.6849670410156 0 8596.556 b 6
380.1897888183594 0 2871.5317
380.6896057128906 0 981.8098
383.1564636230469 0 57357.117
384.1592102050781 0 10459.679
384.18865966796875 0 1222.8049
385.20794677734375 0 1547.0587
386.2039489746094 0 22786.688
387.1505432128906 0 3177.0857
387.1874694824219 0 3500.4475
387.2102966308594 0 1757.2965
388.19219970703125 0 1041.4492
390.1872253417969 0 7185.175 y 6
391.1910095214844 0 1914.7805
393.1429443359375 0 1776.2152
393.1772766113281 0 6043.723
393.2033386230469 0 1114.4998
394.16058349609375 0 2090.314
395.19219970703125 0 1796.4873
395.7294616699219 0 2283.9973
396.2326965332031 0 1908.931
397.1353454589844 0 8315.638
398.1191101074219 0 1348.4479
399.18634033203125 0 1468.1615
401.16705322265625 0 103359.516
401.2877502441406 0 3004.7822
402.1698913574219 0 18357.1
403.1722717285156 0 2181.8877
405.1619873046875 0 4673.3086
405.2115783691406 0 1361.175
406.1640319824219 0 1493.0819
409.2065124511719 0 1021.3241
411.1507873535156 0 2734.3875
411.2039489746094 0 1974.6045
412.1724548339844 0 1280.7596
415.14642333984375 0 16012.968
416.1505432128906 0 2640.5044
416.1927795410156 0 3536.7522
417.19635009765625 0 3834.1855
417.6969299316406 0 1377.8284
418.1954345703125 0 1578.3088
423.1878662109375 0 5259.517
424.1907043457031 0 1342.7504
425.130615234375 0 3059.7104
425.1674499511719 0 3218.5938
425.2048034667969 0 1683.0366
426.2018737792969 0 3124.8828
427.27081298828125 0 12815.76
428.2735595703125 0 3727.158
429.1626892089844 0 1295.1179
429.2135925292969 0 7770.6416
430.21551513671875 0 2917.5078
433.1568298339844 0 32499.076
434.1606140136719 0 5068.859
434.20379638671875 0 20558.074
435.20635986328125 0 5703.256 b Water loss 7
435.705322265625 0 1876.742 b Ammonia loss 7
436.2554931640625 0 2893.814
440.21417236328125 0 11272.637
441.1991271972656 0 7713.8564
442.20050048828125 0 1770.5258
443.1409606933594 0 3070.6277
443.1781311035156 0 2448.1145
444.1481628417969 0 1239.1057
444.1868896484375 0 6788.0347
444.2173767089844 0 2044.6381 b 7
444.7131042480469 0 1705.7576
446.2408752441406 0 1565.1365
448.18316650390625 0 2558.3733
450.1987609863281 0 13720.235
451.2001953125 0 3443.3613
452.21429443359375 0 46772.54
453.1634826660156 0 1427.9316
453.216796875 0 9209.563 y 1
454.22747802734375 0 1793.2662
454.2660217285156 0 7749.9487
455.2657165527344 0 32629.303 b 3
456.26861572265625 0 8527.581
457.2718811035156 0 1633.4316
459.2099304199219 0 14374.967 y Water loss 5
460.213623046875 0 2800.9692
462.1988830566406 0 40198.926
463.20172119140625 0 12275.456
464.2149353027344 0 1173.0417
464.2512512207031 0 4437.268
465.2546081542969 0 1194.8917
466.1939697265625 0 5903.835
468.20928955078125 0 25223.896
468.2435302734375 0 1278.5736
469.21319580078125 0 4974.3696
469.2459411621094 0 2158.2087
470.2250061035156 0 180107.02
471.227783203125 0 41864.41
472.2295837402344 0 7187.5615
476.177978515625 0 5506.8867
477.2201232910156 0 10562.449 y 5
477.9244384765625 0 1388.4116
478.22625732421875 0 3238.4053
480.20928955078125 0 160174.77
481.21246337890625 0 39346.117
482.217041015625 0 6737.641
482.2612609863281 0 8126.814
483.26556396484375 0 1822.7538
483.9281921386719 0 1239.3472
484.20416259765625 0 17485.766
485.2058410644531 0 3118.8955
486.2563781738281 0 5818.6562
487.25921630859375 0 2556.2205
488.2547912597656 0 1300.5146
491.7469787597656 0 1359.9313
492.2082824707031 0 1577.0781
494.18890380859375 0 12944.247
495.19232177734375 0 4113.913
496.2417297363281 0 2548.3052
497.236572265625 0 5482.01
498.22003173828125 0 554517.4
499.2227478027344 0 141192.97
500.22552490234375 0 21166.55
501.2274475097656 0 3240.6658
502.2145690917969 0 27455.084
503.21746826171875 0 7890.22
504.21588134765625 0 1396.3595
504.2821044921875 0 1253.6976
508.25225830078125 0 1880.4196
510.2193298339844 0 2135.7944
511.220947265625 0 1670.03
512.1992797851562 0 27720.53
512.3187866210938 0 1182.3535
513.2027587890625 0 6429.1577
514.2073974609375 0 1715.2501
514.254150390625 0 6097.3623
514.3030395507812 0 4816.658
515.2470092773438 0 12881.2295
515.2902221679688 0 1177.0046
515.3060913085938 0 1332.209
516.243408203125 0 7229.108
517.2442016601562 0 1224.213
518.2459106445312 0 1396.9167
519.2457275390625 0 1565.174
524.2867431640625 0 7632.7124 b Water loss 4
525.2887573242188 0 2739.365
526.2614135742188 0 2335.6746
528.2313232421875 0 4294.9287
529.2344970703125 0 2004.8794
530.2094116210938 0 89031.51
530.3300170898438 0 10930.428
531.2125854492188 0 22309.752
531.33349609375 0 1751.9851
532.215087890625 0 3570.0847
532.2794799804688 0 1249.3607
534.2366333007812 0 1266.0406
542.2974243164062 0 25103.346 b 4
543.30029296875 0 8788.191
544.3025512695312 0 1212.7677
545.2366333007812 0 2736.9148
546.240234375 0 8201.614 y Water loss 4
547.2440795898438 0 1983.2148
548.0692138671875 0 1062.4437
548.220947265625 0 7973.023
549.2256469726562 0 1653.9314
553.26318359375 0 1942.6302
554.2454833984375 0 1898.9812
556.2235717773438 0 1942.0997
563.24609375 0 8813.871
564.25 0 13871.729 y 4
565.2565307617188 0 1810.3994
565.30029296875 0 2980.341
566.297119140625 0 2112.5764
572.2718505859375 0 1647.0728
573.2301635742188 0 10011.768
573.2720947265625 0 848.50494
574.2344360351562 0 3199.4429
574.2857055664062 0 1247.9218
575.2840576171875 0 2537.8904
579.2395629882812 0 2234.9172
579.2828369140625 0 2017.3588
581.2567138671875 0 30981.072
582.25537109375 0 9427.39
583.2691040039062 0 1313.8914
583.3097534179688 0 30699.086
584.3137817382812 0 12986.119
585.3162231445312 0 1760.894
588.2666625976562 0 2147.6323
590.2632446289062 0 1547.2926
591.2409057617188 0 29647.611
591.2924194335938 0 1885.1129
592.2442016601562 0 9933.184
593.244873046875 0 1434.7241
593.30029296875 0 11755.588
594.3035888671875 0 4545.794
597.2462158203125 0 2036.1799
597.29443359375 0 1865.716
599.2671508789062 0 178803
600.2698364257812 0 59014.18
600.3255615234375 0 1341.0604
601.2723999023438 0 11248.813
601.3345336914062 0 19381.115
602.3375244140625 0 6773.7236
603.3404541015625 0 1122.0358
606.2777099609375 0 3841.3772
607.274169921875 0 4431.227
608.2691040039062 0 3074.992
609.2518310546875 0 107905.72
610.254638671875 0 33771.645
610.3118896484375 0 1638.4058
611.2545776367188 0 3249.2988
611.310791015625 0 56366.62 b Water loss 5
612.3135986328125 0 21394.104 b Ammonia loss 5
613.3153076171875 0 3528.2532
613.3674926757812 0 6792.2744
615.2997436523438 0 7054.6387
616.301025390625 0 4081.4138
617.29443359375 0 3771.977
625.285400390625 0 9722.046
626.2860717773438 0 4166.287
627.2623291015625 0 903215.06
628.2649536132812 0 277405.3
629.2673950195312 0 55556.49
629.3296508789062 0 57890.4 b 5
630.27001953125 0 4238.724
630.332275390625 0 21376.469
631.3353271484375 0 4979.115
633.3233032226562 0 2441.666
634.3285522460938 0 1689.9624
643.2943115234375 0 26298.688 y Water loss 3
644.296630859375 0 8906.273
645.2767944335938 0 41374.605
646.2783813476562 0 14743.7705
647.2854614257812 0 2059.369
653.3306884765625 0 1934.8024
658.318603515625 0 995.33813
661.3042602539062 0 104465.805 y 3
662.3074951171875 0 34446.164
663.3111572265625 0 6133.365
676.3308715820312 0 1237.7229
677.3650512695312 0 8465.977
686.3146362304688 0 4587.5806
692.322998046875 0 1376.0842
694.3425903320312 0 7729.6064
695.3430786132812 0 3224.9575
696.3551635742188 0 1791.2052
704.3284912109375 0 12576.467
705.3299560546875 0 6167.332
706.3297729492188 0 1965.7858
710.3321533203125 0 2985.003
711.3492431640625 0 1094.2181
712.3532104492188 0 53433.16
713.35400390625 0 24974.342
714.35986328125 0 7795.5576
722.3406372070312 0 39848.227
723.342041015625 0 20158.588
724.3458862304688 0 4778.1
728.3439331054688 0 2282.8884
729.3104248046875 0 3720.9617
730.3770141601562 0 71228.79
731.3796997070312 0 28736.807
732.3818969726562 0 4660.735
738.3084716796875 0 1478.6372
740.3490600585938 0 168390.25 b Water loss 6
741.3521118164062 0 73166.68 b Ammonia loss 6
742.3546142578125 0 18064.32
743.3580932617188 0 1456.7267
746.3350830078125 0 3627.2383
747.3388061523438 0 1883.9712
754.3438110351562 0 1739.7422
756.3182373046875 0 4640.1943
757.322265625 0 1946.4552
758.3597412109375 0 917252.25 b 6
759.3627319335938 0 382615.56
760.3653564453125 0 92213.96
761.3671264648438 0 6890.1025
762.36572265625 0 2733.0884
772.3516235351562 0 2893.4265
773.35791015625 0 1151.9954
774.330322265625 0 18577.207
775.3331298828125 0 7584.369
776.3365478515625 0 2041.6123
790.3602905273438 0 4957.7095
791.3657836914062 0 2286.3716
833.3821411132812 0 1547.3015
841.4095458984375 0 6465.8364
842.396240234375 0 8260.168
843.4013061523438 0 3718.548
851.3919677734375 0 13071.458
852.39599609375 0 5699.868
854.4366455078125 0 3125.472
859.41845703125 0 65721.98
860.4209594726562 0 27392.438
861.42431640625 0 8668.95
862.427734375 0 1672.113
869.4031982421875 0 36356.91 b Water loss 7
870.4058837890625 0 17839.113
871.40966796875 0 5846.654
887.4133911132812 0 294657.1 b 7
888.4163208007812 0 159856.67
889.4188842773438 0 42375.906
890.4207153320312 0 4181.9727
905.423583984375 0 4257.122 y 1
906.42822265625 0 2801.4048
954.0159301757812 0 1153.8391
1027.6248779296875 0 1093.4493
1349.4219970703125 0 1164.4617
1772.8955078125 0 1234.9764
1937.812744140625 0 1108.0608
2054.398193359375 0 1186.7819
3081.450439453125 0 1218.058

Spectrum Details

|  |  |
| --- | --- |
| Matched peaks? Matched peaks The total absolute number of peaks matched. Additionally in brackets the total fraction of peaks matched and the total number of peaks is shown. | 42 (6.52% of 644) |
| FDR? FDR The false discovery rate estimated for this peptide. It is calculated by matching all theoretical fragments with a non-integer shift with the raw peaks for this spectrum. This is done with 40 different shifts. The resulting percentage is the average number of annotated peaks over the number of annotated peaks with the correct spectrum. | 0.45% |
| Satellite FDR? Satellite FDR See the FDR for details on its calculation. This satellite ion specific FDR only contains the satellite ions (d/w) for I/L/J positions. | - |
| PSM Score? PSM Score The PSM Score as given by Hecklib to this annotated spectrum. It is shown with three significant figures. | 517 |

#### Spectrum 8371? Spectrum 8371 The raw spectrum of this peptide as annotated by Hecklib. The fragments are coloured according to ion type (see legend). Any peaks with a star '\*' as text can be hovered over to see the full details, first the ion type second the mass shift type. By hovering over the amino acids in the peptide or ions in the legend the corresponding peaks are highlighted. By toggling the 'Unassigned' label you can turn the background (unassigned) peaks on or off in the plot. By updating the slider in the Ion legend you can update the spectrum to only show the top X% of the peaks with labels. The top X% means any peak that is within X% of the highest intensity. By dragging in the spectrum you can zoom in to a specific part of the spectrum and use 'Zoom Out' to get back to the original zoom level. The annotation of the spectrum is based on the given sequence in the peptides file and is done with different software so inconsistencies are likely. The peaks are annotated based on the given sequence, with 20 ppm tolerance.

Copy Data

##### Spectrum 8371 (TSV)

###### Preview

```
Loading example...
```

*Click on the button to copy the data to your clipboard.*

Mz MinMz MaxIntensity Max

WidthHeightPeptide font sizePeptide stroke widthSpectrum font sizeSpectrum stroke widthCompact peptide

Ion legend

wxyz

abcd

OtherUnassignedIon^Charge^_Position_Show for top:%

JFPPSSEEJ

01.42e+52.83e+54.25e+55.67e+5

Zoom Out

y^+1^_1_a^+2^_3_a^+1^_2_y^+1^_2_b^+1^_2_y^+1^_2_b^+1^_2_y^+2^_5_y^+2^_5_b^+2^_6_a^+1^_3_b^+1^_3_b^+1^_3_b^+2^_7_b^+2^_7_y^+1^_3_b^+2^_7_y^+1^_3_b^+2^_8_y^+2^_8_b^+1^_4_y^+1^_4_y^+1^_4_b^+1^_5_b^+1^_5_y^+1^_5_y^+1^_5_b^+1^_6_b^+1^_6_b^+1^_6_y^+1^_6_y^+1^_6_b^+1^_7_b^+1^_7_b^+1^_7_b^+1^_8_b^+1^_8_y^+1^_8_

0688137520632751

Fragment Matches Table

Show background peaks

| Position | Ion type | Intensity | mz Theoretical | mz Error (Th) | mz Error (ppm) | Charge | Series Number |
| --- | --- | --- | --- | --- | --- | --- | --- |
| - | - | 3.745E+05 | 120.1 | - | - | 0 | - |
| - | - | 3.015E+04 | 121.1 | - | - | 0 | - |
| - | - | 4.003E+04 | 126.1 | - | - | 0 | - |
| - | - | 2648 | 127.1 | - | - | 0 | - |
| - | - | 508.9 | 128.1 | - | - | 0 | - |
| - | - | 487.7 | 128.1 | - | - | 0 | - |
| - | - | 685.7 | 129.1 | - | - | 0 | - |
| - | - | 1.167E+04 | 129.1 | - | - | 0 | - |
| - | - | 2475 | 129.1 | - | - | 0 | - |
| - | - | 3.687E+04 | 130.1 | - | - | 0 | - |
| - | - | 892.9 | 130.1 | - | - | 0 | - |
| - | - | 1914 | 131.1 | - | - | 0 | - |
| 9 | y | 5.483E+05 | 132.1 | 0.0004203 | 3.182 | +1 | 1 |
| - | - | 2356 | 133.1 | - | - | 0 | - |
| - | - | 1467 | 133.1 | - | - | 0 | - |
| - | - | 3.484E+04 | 133.1 | - | - | 0 | - |
| - | - | 1159 | 134.1 | - | - | 0 | - |
| - | - | 503.9 | 135.1 | - | - | 0 | - |
| - | - | 3225 | 136.1 | - | - | 0 | - |
| - | - | 6621 | 138.1 | - | - | 0 | - |
| - | - | 1207 | 139.1 | - | - | 0 | - |
| - | - | 6455 | 139.1 | - | - | 0 | - |
| - | - | 909.4 | 139.1 | - | - | 0 | - |
| - | - | 526.3 | 140.1 | - | - | 0 | - |
| - | - | 661.2 | 140.1 | - | - | 0 | - |
| - | - | 5847 | 141.1 | - | - | 0 | - |
| - | - | 2463 | 143.1 | - | - | 0 | - |
| - | - | 819.1 | 144.1 | - | - | 0 | - |
| - | - | 520.2 | 144.1 | - | - | 0 | - |
| - | - | 805.4 | 145.1 | - | - | 0 | - |
| - | - | 686.2 | 145.5 | - | - | 0 | - |
| - | - | 787.3 | 146.1 | - | - | 0 | - |
| - | - | 8924 | 147.1 | - | - | 0 | - |
| - | - | 2715 | 148.1 | - | - | 0 | - |
| - | - | 1424 | 149 | - | - | 0 | - |
| - | - | 730.2 | 151.1 | - | - | 0 | - |
| - | - | 1720 | 151.1 | - | - | 0 | - |
| - | - | 1744 | 152.1 | - | - | 0 | - |
| - | - | 1595 | 153.1 | - | - | 0 | - |
| - | - | 2469 | 154.1 | - | - | 0 | - |
| - | - | 2.121E+04 | 154.1 | - | - | 0 | - |
| - | - | 2539 | 155.1 | - | - | 0 | - |
| - | - | 1669 | 155.1 | - | - | 0 | - |
| - | - | 782.4 | 156.1 | - | - | 0 | - |
| - | - | 1.403E+04 | 157.1 | - | - | 0 | - |
| 3 | a | 3.267E+04 | 157.1 | 0.001662 | 10.58 | +2 | 3 |
| - | - | 689.6 | 158.1 | - | - | 0 | - |
| - | - | 1639 | 158.1 | - | - | 0 | - |
| - | - | 601.4 | 158.2 | - | - | 0 | - |
| - | - | 574.2 | 163.6 | - | - | 0 | - |
| - | - | 1437 | 165.1 | - | - | 0 | - |
| - | - | 2318 | 166.1 | - | - | 0 | - |
| - | - | 1401 | 167 | - | - | 0 | - |
| - | - | 3.114E+04 | 167.1 | - | - | 0 | - |
| - | - | 2.93E+04 | 167.1 | - | - | 0 | - |
| - | - | 3388 | 168.1 | - | - | 0 | - |
| - | - | 2675 | 168.1 | - | - | 0 | - |
| - | - | 979.5 | 170 | - | - | 0 | - |
| - | - | 876 | 170.1 | - | - | 0 | - |
| - | - | 7459 | 171.1 | - | - | 0 | - |
| - | - | 954.7 | 171.1 | - | - | 0 | - |
| - | - | 1040 | 172.1 | - | - | 0 | - |
| - | - | 2.073E+04 | 172.1 | - | - | 0 | - |
| - | - | 2944 | 173.1 | - | - | 0 | - |
| - | - | 3174 | 173.1 | - | - | 0 | - |
| - | - | 601.6 | 174.1 | - | - | 0 | - |
| - | - | 1.59E+04 | 175.1 | - | - | 0 | - |
| - | - | 1098 | 175.1 | - | - | 0 | - |
| - | - | 1110 | 175.1 | - | - | 0 | - |
| - | - | 666.9 | 176.1 | - | - | 0 | - |
| - | - | 2256 | 177.1 | - | - | 0 | - |
| - | - | 923.9 | 177.1 | - | - | 0 | - |
| - | - | 737 | 179.1 | - | - | 0 | - |
| - | - | 1502 | 181.1 | - | - | 0 | - |
| - | - | 3242 | 181.1 | - | - | 0 | - |
| - | - | 1966 | 182.1 | - | - | 0 | - |
| - | - | 1464 | 183.1 | - | - | 0 | - |
| - | - | 3217 | 183.1 | - | - | 0 | - |
| - | - | 1125 | 185.1 | - | - | 0 | - |
| - | - | 5.709E+04 | 185.1 | - | - | 0 | - |
| - | - | 1765 | 185.2 | - | - | 0 | - |
| - | - | 5180 | 186.1 | - | - | 0 | - |
| - | - | 645.5 | 186.2 | - | - | 0 | - |
| - | - | 746.7 | 188.1 | - | - | 0 | - |
| - | - | 1783 | 188.1 | - | - | 0 | - |
| - | - | 2.205E+04 | 189.1 | - | - | 0 | - |
| - | - | 3014 | 189.1 | - | - | 0 | - |
| - | - | 1438 | 190.1 | - | - | 0 | - |
| - | - | 877.4 | 191.1 | - | - | 0 | - |
| - | - | 776.8 | 193.1 | - | - | 0 | - |
| - | - | 572.4 | 193.5 | - | - | 0 | - |
| - | - | 8951 | 195.1 | - | - | 0 | - |
| - | - | 6.026E+04 | 195.1 | - | - | 0 | - |
| - | - | 6071 | 196.1 | - | - | 0 | - |
| - | - | 1.356E+05 | 197.1 | - | - | 0 | - |
| - | - | 806.5 | 198.1 | - | - | 0 | - |
| - | - | 861.7 | 198.1 | - | - | 0 | - |
| - | - | 1.39E+04 | 198.1 | - | - | 0 | - |
| - | - | 3.238E+04 | 199.1 | - | - | 0 | - |
| - | - | 988.7 | 199.1 | - | - | 0 | - |
| - | - | 4725 | 199.1 | - | - | 0 | - |
| - | - | 788.1 | 199.1 | - | - | 0 | - |
| - | - | 598.1 | 199.1 | - | - | 0 | - |
| - | - | 3317 | 200.1 | - | - | 0 | - |
| - | - | 2223 | 201.1 | - | - | 0 | - |
| - | - | 2314 | 202.1 | - | - | 0 | - |
| - | - | 616.7 | 203.1 | - | - | 0 | - |
| - | - | 3598 | 203.1 | - | - | 0 | - |
| - | - | 845.1 | 205.2 | - | - | 0 | - |
| - | - | 1203 | 208.1 | - | - | 0 | - |
| - | - | 8534 | 209.1 | - | - | 0 | - |
| - | - | 1015 | 210.1 | - | - | 0 | - |
| - | - | 3972 | 211.1 | - | - | 0 | - |
| - | - | 8751 | 212.1 | - | - | 0 | - |
| - | - | 1.006E+04 | 213.1 | - | - | 0 | - |
| - | - | 5604 | 213.1 | - | - | 0 | - |
| - | - | 1049 | 213.1 | - | - | 0 | - |
| - | - | 1086 | 213.2 | - | - | 0 | - |
| - | - | 1079 | 214.1 | - | - | 0 | - |
| - | - | 1042 | 214.1 | - | - | 0 | - |
| - | - | 4257 | 215.1 | - | - | 0 | - |
| - | - | 701 | 216.1 | - | - | 0 | - |
| - | - | 6.13E+04 | 217.1 | - | - | 0 | - |
| - | - | 1.404E+04 | 217.1 | - | - | 0 | - |
| - | - | 793.5 | 217.6 | - | - | 0 | - |
| - | - | 4875 | 218.1 | - | - | 0 | - |
| - | - | 1409 | 218.1 | - | - | 0 | - |
| - | - | 1932 | 221.1 | - | - | 0 | - |
| - | - | 2039 | 223.1 | - | - | 0 | - |
| - | - | 2635 | 224.1 | - | - | 0 | - |
| - | - | 1.721E+04 | 225.1 | - | - | 0 | - |
| - | - | 2632 | 226.1 | - | - | 0 | - |
| - | - | 1258 | 226.1 | - | - | 0 | - |
| - | - | 1002 | 227.1 | - | - | 0 | - |
| - | - | 9287 | 227.1 | - | - | 0 | - |
| - | - | 1308 | 229.1 | - | - | 0 | - |
| - | - | 1.016E+04 | 231.1 | - | - | 0 | - |
| - | - | 624.8 | 231.1 | - | - | 0 | - |
| - | - | 1517 | 231.6 | - | - | 0 | - |
| - | - | 1258 | 232.1 | - | - | 0 | - |
| 2 | a | 5.023E+05 | 233.2 | 0.0005045 | 2.164 | +1 | 2 |
| - | - | 7.773E+04 | 234.2 | - | - | 0 | - |
| - | - | 592 | 234.2 | - | - | 0 | - |
| - | - | 1160 | 235.1 | - | - | 0 | - |
| - | - | 3999 | 235.2 | - | - | 0 | - |
| - | - | 2.319E+04 | 236.1 | - | - | 0 | - |
| - | - | 1653 | 236.1 | - | - | 0 | - |
| - | - | 1023 | 237.1 | - | - | 0 | - |
| - | - | 2917 | 237.1 | - | - | 0 | - |
| - | - | 1501 | 237.1 | - | - | 0 | - |
| - | - | 1075 | 238.1 | - | - | 0 | - |
| - | - | 730.4 | 238.2 | - | - | 0 | - |
| - | - | 1035 | 239.1 | - | - | 0 | - |
| - | - | 3164 | 240.1 | - | - | 0 | - |
| - | - | 2554 | 240.6 | - | - | 0 | - |
| - | - | 4.767E+04 | 241.1 | - | - | 0 | - |
| - | - | 5252 | 242.1 | - | - | 0 | - |
| - | - | 4450 | 242.1 | - | - | 0 | - |
| 8 | y | 3.439E+05 | 243.1 | 0.0004964 | 2.042 | +1 | 2 |
| 2 | b | 3.523E+04 | 244.1 | 0.004017 | 16.45 | +1 | 2 |
| - | - | 696 | 245.1 | - | - | 0 | - |
| - | - | 1.244E+04 | 245.1 | - | - | 0 | - |
| - | - | 2276 | 246.1 | - | - | 0 | - |
| - | - | 1700 | 249.6 | - | - | 0 | - |
| - | - | 1988 | 250.1 | - | - | 0 | - |
| - | - | 1424 | 250.1 | - | - | 0 | - |
| - | - | 838.7 | 252.1 | - | - | 0 | - |
| - | - | 5822 | 252.1 | - | - | 0 | - |
| - | - | 929.4 | 253.1 | - | - | 0 | - |
| - | - | 916.7 | 253.1 | - | - | 0 | - |
| - | - | 1674 | 253.2 | - | - | 0 | - |
| - | - | 8.188E+04 | 254.1 | - | - | 0 | - |
| - | - | 9586 | 254.2 | - | - | 0 | - |
| - | - | 1.085E+04 | 255.1 | - | - | 0 | - |
| - | - | 1378 | 255.2 | - | - | 0 | - |
| - | - | 1149 | 256.1 | - | - | 0 | - |
| - | - | 3073 | 258.1 | - | - | 0 | - |
| - | - | 705.3 | 258.1 | - | - | 0 | - |
| - | - | 4.174E+04 | 259.1 | - | - | 0 | - |
| - | - | 5360 | 260.1 | - | - | 0 | - |
| 8 | y | 4.85E+04 | 261.1 | 0.0002772 | 1.061 | +1 | 2 |
| 2 | b | 2.077E+05 | 261.2 | 0.0003409 | 1.305 | +1 | 2 |
| - | - | 5588 | 262.1 | - | - | 0 | - |
| - | - | 3.062E+04 | 262.2 | - | - | 0 | - |
| - | - | 2283 | 263.2 | - | - | 0 | - |
| - | - | 3234 | 264.1 | - | - | 0 | - |
| - | - | 3.864E+04 | 264.1 | - | - | 0 | - |
| - | - | 5243 | 265.1 | - | - | 0 | - |
| - | - | 901 | 267.1 | - | - | 0 | - |
| - | - | 6700 | 268.1 | - | - | 0 | - |
| - | - | 2176 | 268.1 | - | - | 0 | - |
| - | - | 1355 | 269.1 | - | - | 0 | - |
| - | - | 844.8 | 269.1 | - | - | 0 | - |
| - | - | 1192 | 269.1 | - | - | 0 | - |
| - | - | 888.9 | 270.1 | - | - | 0 | - |
| - | - | 852.8 | 270.1 | - | - | 0 | - |
| - | - | 3.984E+04 | 272.1 | - | - | 0 | - |
| - | - | 7782 | 273.1 | - | - | 0 | - |
| 5 | y | 1683 | 273.6 | 2.858E-05 | 0.1045 | +2 | 5 |
| - | - | 881.1 | 274.1 | - | - | 0 | - |
| - | - | 1953 | 276.1 | - | - | 0 | - |
| - | - | 978.1 | 277.1 | - | - | 0 | - |
| - | - | 1602 | 278.1 | - | - | 0 | - |
| - | - | 1570 | 278.1 | - | - | 0 | - |
| - | - | 877 | 280.2 | - | - | 0 | - |
| - | - | 2368 | 281.1 | - | - | 0 | - |
| - | - | 2047 | 281.2 | - | - | 0 | - |
| - | - | 4549 | 282.1 | - | - | 0 | - |
| - | - | 5581 | 282.1 | - | - | 0 | - |
| - | - | 1.635E+05 | 282.1 | - | - | 0 | - |
| 5 | y | 1139 | 282.6 | 0.0009469 | 3.35 | +2 | 5 |
| - | - | 769.8 | 283.1 | - | - | 0 | - |
| - | - | 804.8 | 283.1 | - | - | 0 | - |
| - | - | 2.328E+04 | 283.1 | - | - | 0 | - |
| - | - | 1478 | 284.2 | - | - | 0 | - |
| - | - | 5802 | 285.2 | - | - | 0 | - |
| - | - | 2.857E+04 | 286.1 | - | - | 0 | - |
| - | - | 3451 | 287.1 | - | - | 0 | - |
| - | - | 958.9 | 287.1 | - | - | 0 | - |
| - | - | 1023 | 287.2 | - | - | 0 | - |
| - | - | 1237 | 287.6 | - | - | 0 | - |
| - | - | 880.6 | 288.2 | - | - | 0 | - |
| - | - | 2845 | 289.2 | - | - | 0 | - |
| - | - | 1347 | 291.1 | - | - | 0 | - |
| - | - | 930 | 292.1 | - | - | 0 | - |
| - | - | 831.6 | 294.1 | - | - | 0 | - |
| - | - | 2671 | 296.1 | - | - | 0 | - |
| - | - | 1.527E+04 | 296.1 | - | - | 0 | - |
| - | - | 1626 | 296.6 | - | - | 0 | - |
| - | - | 1215 | 297.1 | - | - | 0 | - |
| - | - | 5008 | 297.2 | - | - | 0 | - |
| - | - | 1935 | 297.7 | - | - | 0 | - |
| - | - | 863.3 | 298.2 | - | - | 0 | - |
| - | - | 3994 | 299.1 | - | - | 0 | - |
| - | - | 7983 | 299.2 | - | - | 0 | - |
| - | - | 4356 | 300.1 | - | - | 0 | - |
| - | - | 1383 | 300.1 | - | - | 0 | - |
| - | - | 6859 | 300.2 | - | - | 0 | - |
| - | - | 1568 | 300.2 | - | - | 0 | - |
| - | - | 755.6 | 300.6 | - | - | 0 | - |
| - | - | 1332 | 301.2 | - | - | 0 | - |
| - | - | 1.572E+04 | 304.1 | - | - | 0 | - |
| - | - | 5485 | 305.1 | - | - | 0 | - |
| - | - | 1531 | 305.2 | - | - | 0 | - |
| - | - | 1647 | 305.6 | - | - | 0 | - |
| - | - | 4705 | 306.1 | - | - | 0 | - |
| 6 | b | 3320 | 306.2 | 0.001098 | 3.587 | +2 | 6 |
| - | - | 883.5 | 306.7 | - | - | 0 | - |
| - | - | 1253 | 307.1 | - | - | 0 | - |
| - | - | 7988 | 308.2 | - | - | 0 | - |
| - | - | 4895 | 308.2 | - | - | 0 | - |
| - | - | 1283 | 309.2 | - | - | 0 | - |
| - | - | 8304 | 310.1 | - | - | 0 | - |
| - | - | 943.5 | 310.1 | - | - | 0 | - |
| - | - | 1225 | 311.1 | - | - | 0 | - |
| - | - | 943.7 | 311.1 | - | - | 0 | - |
| - | - | 2285 | 311.2 | - | - | 0 | - |
| - | - | 3022 | 312.2 | - | - | 0 | - |
| 3 | a | 1808 | 313.2 | 0.001115 | 3.559 | +1 | 3 |
| - | - | 2572 | 314.1 | - | - | 0 | - |
| - | - | 1.452E+04 | 314.1 | - | - | 0 | - |
| - | - | 2725 | 314.6 | - | - | 0 | - |
| - | - | 1442 | 318.1 | - | - | 0 | - |
| - | - | 2833 | 319.1 | - | - | 0 | - |
| - | - | 1212 | 320.1 | - | - | 0 | - |
| - | - | 1716 | 320.7 | - | - | 0 | - |
| - | - | 5430 | 321.2 | - | - | 0 | - |
| - | - | 1030 | 322.2 | - | - | 0 | - |
| - | - | 7119 | 323.2 | - | - | 0 | - |
| - | - | 6815 | 324.2 | - | - | 0 | - |
| - | - | 1305 | 324.2 | - | - | 0 | - |
| - | - | 861.1 | 325.2 | - | - | 0 | - |
| - | - | 1473 | 326.2 | - | - | 0 | - |
| - | - | 1.966E+04 | 328.1 | - | - | 0 | - |
| - | - | 2593 | 329.1 | - | - | 0 | - |
| - | - | 858.8 | 329.2 | - | - | 0 | - |
| - | - | 5614 | 329.7 | - | - | 0 | - |
| - | - | 3526 | 330.2 | - | - | 0 | - |
| - | - | 2.725E+04 | 333.2 | - | - | 0 | - |
| - | - | 3838 | 334.2 | - | - | 0 | - |
| - | - | 725.7 | 335.1 | - | - | 0 | - |
| - | - | 1783 | 336.2 | - | - | 0 | - |
| - | - | 7869 | 337.2 | - | - | 0 | - |
| - | - | 5022 | 338.1 | - | - | 0 | - |
| - | - | 1277 | 338.2 | - | - | 0 | - |
| - | - | 1.413E+04 | 338.7 | - | - | 0 | - |
| - | - | 2.078E+04 | 339.2 | - | - | 0 | - |
| - | - | 933.1 | 339.7 | - | - | 0 | - |
| - | - | 2958 | 340.2 | - | - | 0 | - |
| 3 | b | 2.966E+04 | 341.2 | 0.003199 | 9.377 | +1 | 3 |
| - | - | 8944 | 342.2 | - | - | 0 | - |
| - | - | 1298 | 343.2 | - | - | 0 | - |
| - | - | 1225 | 343.2 | - | - | 0 | - |
| - | - | 3.472E+04 | 346.1 | - | - | 0 | - |
| - | - | 5636 | 347.1 | - | - | 0 | - |
| - | - | 3428 | 347.7 | - | - | 0 | - |
| - | - | 1965 | 348.2 | - | - | 0 | - |
| - | - | 1041 | 350.2 | - | - | 0 | - |
| - | - | 1.167E+05 | 351.2 | - | - | 0 | - |
| - | - | 1110 | 352.1 | - | - | 0 | - |
| - | - | 2.061E+04 | 352.2 | - | - | 0 | - |
| - | - | 1876 | 352.7 | - | - | 0 | - |
| - | - | 2000 | 353.1 | - | - | 0 | - |
| - | - | 2377 | 353.2 | - | - | 0 | - |
| - | - | 1.747E+04 | 354.2 | - | - | 0 | - |
| - | - | 1.801E+04 | 355.2 | - | - | 0 | - |
| - | - | 946.8 | 356.1 | - | - | 0 | - |
| - | - | 5696 | 356.1 | - | - | 0 | - |
| - | - | 2213 | 356.2 | - | - | 0 | - |
| - | - | 1278 | 356.7 | - | - | 0 | - |
| - | - | 800.2 | 357.1 | - | - | 0 | - |
| - | - | 776.3 | 357.2 | - | - | 0 | - |
| 3 | b | 1.728E+04 | 358.2 | 0.000464 | 1.295 | +1 | 3 |
| - | - | 3420 | 359.2 | - | - | 0 | - |
| - | - | 4626 | 361.7 | - | - | 0 | - |
| - | - | 2849 | 362.2 | - | - | 0 | - |
| - | - | 1.604E+04 | 365.1 | - | - | 0 | - |
| - | - | 769.5 | 365.7 | - | - | 0 | - |
| - | - | 1554 | 366.1 | - | - | 0 | - |
| - | - | 1084 | 366.2 | - | - | 0 | - |
| - | - | 1134 | 367.2 | - | - | 0 | - |
| - | - | 3499 | 368.2 | - | - | 0 | - |
| - | - | 1.272E+05 | 369.2 | - | - | 0 | - |
| - | - | 2.528E+04 | 370.2 | - | - | 0 | - |
| 7 | b | 9735 | 370.7 | 0.002101 | 5.669 | +2 | 7 |
| - | - | 2896 | 371.2 | - | - | 0 | - |
| 7 | b | 6127 | 371.2 | 0.00644 | 17.35 | +2 | 7 |
| - | - | 994.5 | 371.2 | - | - | 0 | - |
| 7 | y | 1.506E+04 | 372.2 | 0.0003227 | 0.8672 | +1 | 3 |
| - | - | 3.148E+04 | 373.2 | - | - | 0 | - |
| - | - | 6325 | 374.2 | - | - | 0 | - |
| - | - | 1125 | 375.2 | - | - | 0 | - |
| - | - | 1870 | 379.1 | - | - | 0 | - |
| 7 | b | 4181 | 379.7 | 0.004668 | 12.29 | +2 | 7 |
| - | - | 1848 | 380.2 | - | - | 0 | - |
| - | - | 1332 | 381.2 | - | - | 0 | - |
| - | - | 3.712E+04 | 383.2 | - | - | 0 | - |
| - | - | 6786 | 384.2 | - | - | 0 | - |
| - | - | 1309 | 384.2 | - | - | 0 | - |
| - | - | 1.343E+04 | 386.2 | - | - | 0 | - |
| - | - | 1738 | 387.2 | - | - | 0 | - |
| - | - | 2547 | 387.2 | - | - | 0 | - |
| - | - | 1161 | 388.1 | - | - | 0 | - |
| 7 | y | 5319 | 390.2 | 0.0008359 | 2.142 | +1 | 3 |
| - | - | 1175 | 391.2 | - | - | 0 | - |
| - | - | 3603 | 393.2 | - | - | 0 | - |
| - | - | 1558 | 394.2 | - | - | 0 | - |
| - | - | 1518 | 395.2 | - | - | 0 | - |
| - | - | 5105 | 397.1 | - | - | 0 | - |
| - | - | 852.8 | 398.1 | - | - | 0 | - |
| - | - | 1045 | 399.2 | - | - | 0 | - |
| - | - | 6.436E+04 | 401.2 | - | - | 0 | - |
| - | - | 1761 | 401.2 | - | - | 0 | - |
| - | - | 1.287E+04 | 402.2 | - | - | 0 | - |
| - | - | 2367 | 403.2 | - | - | 0 | - |
| - | - | 2870 | 405.2 | - | - | 0 | - |
| - | - | 776 | 406.2 | - | - | 0 | - |
| - | - | 897.2 | 407.2 | - | - | 0 | - |
| - | - | 2942 | 411.2 | - | - | 0 | - |
| - | - | 1054 | 412.2 | - | - | 0 | - |
| - | - | 1154 | 413.5 | - | - | 0 | - |
| - | - | 981.7 | 413.9 | - | - | 0 | - |
| - | - | 9126 | 415.1 | - | - | 0 | - |
| - | - | 2378 | 416.1 | - | - | 0 | - |
| - | - | 3829 | 416.2 | - | - | 0 | - |
| - | - | 1814 | 417.2 | - | - | 0 | - |
| - | - | 955.8 | 418.2 | - | - | 0 | - |
| - | - | 1127 | 422.2 | - | - | 0 | - |
| - | - | 3394 | 423.2 | - | - | 0 | - |
| - | - | 1210 | 424.2 | - | - | 0 | - |
| - | - | 1304 | 424.2 | - | - | 0 | - |
| - | - | 1889 | 425.1 | - | - | 0 | - |
| - | - | 2354 | 425.2 | - | - | 0 | - |
| - | - | 995.4 | 425.2 | - | - | 0 | - |
| - | - | 2834 | 426.2 | - | - | 0 | - |
| - | - | 899.5 | 426.7 | - | - | 0 | - |
| - | - | 9172 | 427.3 | - | - | 0 | - |
| - | - | 3503 | 428.3 | - | - | 0 | - |
| - | - | 5491 | 429.2 | - | - | 0 | - |
| - | - | 2316 | 430.2 | - | - | 0 | - |
| - | - | 862.1 | 431.2 | - | - | 0 | - |
| - | - | 1055 | 432.2 | - | - | 0 | - |
| - | - | 1.806E+04 | 433.2 | - | - | 0 | - |
| - | - | 3415 | 434.2 | - | - | 0 | - |
| - | - | 1.305E+04 | 434.2 | - | - | 0 | - |
| 8 | b | 1807 | 435.2 | 0.002206 | 5.07 | +2 | 8 |
| - | - | 978.9 | 435.7 | - | - | 0 | - |
| - | - | 1237 | 436.3 | - | - | 0 | - |
| - | - | 5317 | 440.2 | - | - | 0 | - |
| - | - | 5663 | 441.2 | - | - | 0 | - |
| - | - | 1693 | 442.2 | - | - | 0 | - |
| - | - | 1247 | 443.1 | - | - | 0 | - |
| - | - | 2644 | 443.2 | - | - | 0 | - |
| - | - | 4299 | 444.2 | - | - | 0 | - |
| - | - | 1163 | 444.7 | - | - | 0 | - |
| - | - | 1748 | 448.2 | - | - | 0 | - |
| - | - | 9742 | 450.2 | - | - | 0 | - |
| - | - | 2221 | 451.2 | - | - | 0 | - |
| - | - | 2.722E+04 | 452.2 | - | - | 0 | - |
| 2 | y | 6533 | 453.2 | 0.00098 | 2.162 | +2 | 8 |
| - | - | 892.6 | 454.2 | - | - | 0 | - |
| - | - | 3084 | 454.3 | - | - | 0 | - |
| 4 | b | 1.997E+04 | 455.3 | 0.0005261 | 1.155 | +1 | 4 |
| - | - | 5877 | 456.3 | - | - | 0 | - |
| - | - | 931.1 | 457.3 | - | - | 0 | - |
| 6 | y | 7511 | 459.2 | 0.001192 | 2.596 | +1 | 4 |
| - | - | 1596 | 460.2 | - | - | 0 | - |
| - | - | 867.5 | 461.2 | - | - | 0 | - |
| - | - | 2.661E+04 | 462.2 | - | - | 0 | - |
| - | - | 6797 | 463.2 | - | - | 0 | - |
| - | - | 962.6 | 464.2 | - | - | 0 | - |
| - | - | 2695 | 464.3 | - | - | 0 | - |
| - | - | 3280 | 466.2 | - | - | 0 | - |
| - | - | 1033 | 467.2 | - | - | 0 | - |
| - | - | 1.401E+04 | 468.2 | - | - | 0 | - |
| - | - | 2750 | 469.2 | - | - | 0 | - |
| - | - | 1.088E+05 | 470.2 | - | - | 0 | - |
| - | - | 2.542E+04 | 471.2 | - | - | 0 | - |
| - | - | 4311 | 472.2 | - | - | 0 | - |
| - | - | 2876 | 476.2 | - | - | 0 | - |
| 6 | y | 4114 | 477.2 | 0.002224 | 4.661 | +1 | 4 |
| - | - | 1468 | 478.2 | - | - | 0 | - |
| - | - | 975.8 | 479.2 | - | - | 0 | - |
| - | - | 9.696E+04 | 480.2 | - | - | 0 | - |
| - | - | 2.584E+04 | 481.2 | - | - | 0 | - |
| - | - | 3562 | 482.2 | - | - | 0 | - |
| - | - | 5124 | 482.3 | - | - | 0 | - |
| - | - | 8684 | 484.2 | - | - | 0 | - |
| - | - | 1318 | 485.2 | - | - | 0 | - |
| - | - | 3499 | 486.3 | - | - | 0 | - |
| - | - | 1428 | 487.3 | - | - | 0 | - |
| - | - | 8806 | 494.2 | - | - | 0 | - |
| - | - | 1351 | 495.2 | - | - | 0 | - |
| - | - | 1831 | 496.2 | - | - | 0 | - |
| - | - | 3724 | 497.2 | - | - | 0 | - |
| - | - | 942.8 | 497.3 | - | - | 0 | - |
| - | - | 3.449E+05 | 498.2 | - | - | 0 | - |
| - | - | 8.775E+04 | 499.2 | - | - | 0 | - |
| - | - | 1.201E+04 | 500.2 | - | - | 0 | - |
| - | - | 1.714E+04 | 502.2 | - | - | 0 | - |
| - | - | 3937 | 503.2 | - | - | 0 | - |
| - | - | 862.8 | 504.2 | - | - | 0 | - |
| - | - | 874.6 | 504.3 | - | - | 0 | - |
| - | - | 2091 | 510.2 | - | - | 0 | - |
| - | - | 1.456E+04 | 512.2 | - | - | 0 | - |
| - | - | 4105 | 513.2 | - | - | 0 | - |
| - | - | 3435 | 514.3 | - | - | 0 | - |
| - | - | 2948 | 514.3 | - | - | 0 | - |
| - | - | 8977 | 515.2 | - | - | 0 | - |
| - | - | 1516 | 515.3 | - | - | 0 | - |
| - | - | 4078 | 516.2 | - | - | 0 | - |
| - | - | 1172 | 517.2 | - | - | 0 | - |
| 5 | b | 5676 | 524.3 | 0.0001195 | 0.2278 | +1 | 5 |
| - | - | 1719 | 525.3 | - | - | 0 | - |
| - | - | 3234 | 528.2 | - | - | 0 | - |
| - | - | 5.078E+04 | 530.2 | - | - | 0 | - |
| - | - | 1.272E+04 | 531.2 | - | - | 0 | - |
| - | - | 2141 | 532.2 | - | - | 0 | - |
| - | - | 1166 | 533.3 | - | - | 0 | - |
| 5 | b | 1.55E+04 | 542.3 | 0.000297 | 0.5476 | +1 | 5 |
| - | - | 5153 | 543.3 | - | - | 0 | - |
| - | - | 1357 | 544.3 | - | - | 0 | - |
| 5 | y | 5452 | 546.2 | 0.0001392 | 0.2549 | +1 | 5 |
| - | - | 1246 | 547.2 | - | - | 0 | - |
| - | - | 4539 | 548.2 | - | - | 0 | - |
| - | - | 1380 | 554.2 | - | - | 0 | - |
| - | - | 3754 | 563.2 | - | - | 0 | - |
| 5 | y | 7601 | 564.3 | 0.001209 | 2.143 | +1 | 5 |
| - | - | 2594 | 565.3 | - | - | 0 | - |
| - | - | 1187 | 565.3 | - | - | 0 | - |
| - | - | 1254 | 566.3 | - | - | 0 | - |
| - | - | 1457 | 566.3 | - | - | 0 | - |
| - | - | 6406 | 573.2 | - | - | 0 | - |
| - | - | 1441 | 574.2 | - | - | 0 | - |
| - | - | 1378 | 575.2 | - | - | 0 | - |
| - | - | 1073 | 575.3 | - | - | 0 | - |
| - | - | 1.804E+04 | 581.3 | - | - | 0 | - |
| - | - | 6510 | 582.3 | - | - | 0 | - |
| - | - | 1048 | 583.3 | - | - | 0 | - |
| - | - | 1.695E+04 | 583.3 | - | - | 0 | - |
| - | - | 5553 | 584.3 | - | - | 0 | - |
| - | - | 2417 | 585.3 | - | - | 0 | - |
| - | - | 1163 | 588.3 | - | - | 0 | - |
| - | - | 831.7 | 589.3 | - | - | 0 | - |
| - | - | 2.009E+04 | 591.2 | - | - | 0 | - |
| - | - | 5180 | 592.2 | - | - | 0 | - |
| - | - | 1156 | 593.2 | - | - | 0 | - |
| - | - | 6176 | 593.3 | - | - | 0 | - |
| - | - | 2238 | 594.3 | - | - | 0 | - |
| - | - | 2395 | 597.3 | - | - | 0 | - |
| - | - | 1.089E+05 | 599.3 | - | - | 0 | - |
| - | - | 3.808E+04 | 600.3 | - | - | 0 | - |
| - | - | 1349 | 600.3 | - | - | 0 | - |
| - | - | 6952 | 601.3 | - | - | 0 | - |
| - | - | 1.025E+04 | 601.3 | - | - | 0 | - |
| - | - | 4244 | 602.3 | - | - | 0 | - |
| - | - | 2292 | 606.3 | - | - | 0 | - |
| - | - | 2013 | 607.3 | - | - | 0 | - |
| - | - | 1813 | 608.3 | - | - | 0 | - |
| - | - | 6.676E+04 | 609.3 | - | - | 0 | - |
| - | - | 2.331E+04 | 610.3 | - | - | 0 | - |
| - | - | 2191 | 611.3 | - | - | 0 | - |
| 6 | b | 3.711E+04 | 611.3 | 0.007739 | 12.66 | +1 | 6 |
| 6 | b | 1.419E+04 | 612.3 | 0.009771 | 15.96 | +1 | 6 |
| - | - | 1209 | 613.3 | - | - | 0 | - |
| - | - | 3763 | 615.3 | - | - | 0 | - |
| - | - | 2384 | 616.3 | - | - | 0 | - |
| - | - | 1792 | 617.3 | - | - | 0 | - |
| - | - | 4502 | 625.3 | - | - | 0 | - |
| - | - | 2334 | 626.3 | - | - | 0 | - |
| - | - | 5.613E+05 | 627.3 | - | - | 0 | - |
| - | - | 1.727E+05 | 628.3 | - | - | 0 | - |
| - | - | 1076 | 628.4 | - | - | 0 | - |
| - | - | 949.7 | 628.4 | - | - | 0 | - |
| - | - | 3.227E+04 | 629.3 | - | - | 0 | - |
| 6 | b | 3.618E+04 | 629.3 | 0.0004951 | 0.7867 | +1 | 6 |
| - | - | 2491 | 630.3 | - | - | 0 | - |
| - | - | 1.156E+04 | 630.3 | - | - | 0 | - |
| - | - | 2412 | 631.3 | - | - | 0 | - |
| - | - | 1990 | 633.3 | - | - | 0 | - |
| 4 | y | 1.627E+04 | 643.3 | 0.001025 | 1.594 | +1 | 6 |
| - | - | 1445 | 643.4 | - | - | 0 | - |
| - | - | 6820 | 644.3 | - | - | 0 | - |
| - | - | 2.651E+04 | 645.3 | - | - | 0 | - |
| - | - | 8443 | 646.3 | - | - | 0 | - |
| - | - | 1223 | 647.3 | - | - | 0 | - |
| 4 | y | 6.592E+04 | 661.3 | 0.0005924 | 0.8958 | +1 | 6 |
| - | - | 2.188E+04 | 662.3 | - | - | 0 | - |
| - | - | 4250 | 663.3 | - | - | 0 | - |
| - | - | 1077 | 676.3 | - | - | 0 | - |
| - | - | 4024 | 686.3 | - | - | 0 | - |
| - | - | 1497 | 687.3 | - | - | 0 | - |
| - | - | 4996 | 694.3 | - | - | 0 | - |
| - | - | 2522 | 695.3 | - | - | 0 | - |
| - | - | 4686 | 700.4 | - | - | 0 | - |
| - | - | 1796 | 701.4 | - | - | 0 | - |
| - | - | 5633 | 704.3 | - | - | 0 | - |
| - | - | 3032 | 705.3 | - | - | 0 | - |
| - | - | 1731 | 710.3 | - | - | 0 | - |
| - | - | 3.299E+04 | 712.4 | - | - | 0 | - |
| - | - | 1.464E+04 | 713.4 | - | - | 0 | - |
| - | - | 3495 | 714.4 | - | - | 0 | - |
| - | - | 2.489E+04 | 722.3 | - | - | 0 | - |
| - | - | 9266 | 723.3 | - | - | 0 | - |
| - | - | 1803 | 724.3 | - | - | 0 | - |
| - | - | 1740 | 729.3 | - | - | 0 | - |
| - | - | 4.095E+04 | 730.4 | - | - | 0 | - |
| - | - | 1.821E+04 | 731.4 | - | - | 0 | - |
| - | - | 4420 | 732.4 | - | - | 0 | - |
| - | - | 1388 | 738.3 | - | - | 0 | - |
| 7 | b | 1.05E+05 | 740.4 | 0.01219 | 16.46 | +1 | 7 |
| 7 | b | 4.437E+04 | 741.3 | 0.007461 | 10.06 | +1 | 7 |
| - | - | 1.052E+04 | 742.4 | - | - | 0 | - |
| - | - | 947.7 | 744.4 | - | - | 0 | - |
| - | - | 1893 | 746.3 | - | - | 0 | - |
| - | - | 3382 | 756.3 | - | - | 0 | - |
| - | - | 986.9 | 757.3 | - | - | 0 | - |
| 7 | b | 5.576E+05 | 758.4 | 0.01213 | 15.99 | +1 | 7 |
| - | - | 2.225E+05 | 759.4 | - | - | 0 | - |
| - | - | 5.463E+04 | 760.4 | - | - | 0 | - |
| - | - | 3711 | 761.4 | - | - | 0 | - |
| - | - | 898.5 | 762.4 | - | - | 0 | - |
| - | - | 1875 | 772.4 | - | - | 0 | - |
| - | - | 1118 | 773.4 | - | - | 0 | - |
| - | - | 1.157E+04 | 774.3 | - | - | 0 | - |
| - | - | 4524 | 775.3 | - | - | 0 | - |
| - | - | 2272 | 783.4 | - | - | 0 | - |
| - | - | 4803 | 790.4 | - | - | 0 | - |
| - | - | 1978 | 791.4 | - | - | 0 | - |
| - | - | 1935 | 801.4 | - | - | 0 | - |
| - | - | 1436 | 833.4 | - | - | 0 | - |
| - | - | 2727 | 841.4 | - | - | 0 | - |
| - | - | 5880 | 842.4 | - | - | 0 | - |
| - | - | 3237 | 843.4 | - | - | 0 | - |
| - | - | 3879 | 844.4 | - | - | 0 | - |
| - | - | 1487 | 845.5 | - | - | 0 | - |
| - | - | 1964 | 847.4 | - | - | 0 | - |
| - | - | 7171 | 851.4 | - | - | 0 | - |
| - | - | 2894 | 852.4 | - | - | 0 | - |
| - | - | 1784 | 854.4 | - | - | 0 | - |
| - | - | 4.051E+04 | 859.4 | - | - | 0 | - |
| - | - | 1.919E+04 | 860.4 | - | - | 0 | - |
| - | - | 4891 | 861.4 | - | - | 0 | - |
| - | - | 907.4 | 862.4 | - | - | 0 | - |
| 8 | b | 2.342E+04 | 869.4 | 0.000518 | 0.5958 | +1 | 8 |
| - | - | 1.273E+04 | 870.4 | - | - | 0 | - |
| - | - | 3725 | 871.4 | - | - | 0 | - |
| - | - | 9799 | 872.4 | - | - | 0 | - |
| - | - | 3622 | 873.4 | - | - | 0 | - |
| 8 | b | 1.851E+05 | 887.4 | 0.0008288 | 0.9339 | +1 | 8 |
| - | - | 8.954E+04 | 888.4 | - | - | 0 | - |
| - | - | 2.626E+04 | 889.4 | - | - | 0 | - |
| - | - | 2788 | 890.4 | - | - | 0 | - |
| 2 | y | 2853 | 905.4 | 0.000102 | 0.1126 | +1 | 8 |
| - | - | 1973 | 906.4 | - | - | 0 | - |
| - | - | 1180 | 907.4 | - | - | 0 | - |
| - | - | 854.7 | 1145 | - | - | 0 | - |
| - | - | 763 | 1249 | - | - | 0 | - |
| - | - | 865.8 | 1359 | - | - | 0 | - |
| - | - | 800.8 | 1538 | - | - | 0 | - |
| - | - | 1005 | 2724 | - | - | 0 | - |

m/z Charge Intensity FragmentType MassShift Position
120.08120727539062 0 374481.12
121.08448028564453 0 30154.889
126.0553207397461 0 40031.688
127.05870056152344 0 2647.5122
128.10702514648438 0 508.93384
128.1126708984375 0 487.73798
129.0608367919922 0 685.7324
129.0662078857422 0 11665.248
129.10247802734375 0 2475.1816
130.0502471923828 0 36871.785
130.0654754638672 0 892.9386
131.05369567871094 0 1914.49
132.10232543945312 0 548335.2 y 8
133.08628845214844 0 2355.7922
133.0998992919922 0 1466.9615
133.10562133789062 0 34840.77
134.10653686523438 0 1158.8641
135.12342834472656 0 503.87598
136.07601928710938 0 3225.2327
138.1280975341797 0 6620.707
139.05030822753906 0 1207.4006
139.08702087402344 0 6455.015
139.1314697265625 0 909.38916
140.07081604003906 0 526.2649
140.0900421142578 0 661.1501
141.0662841796875 0 5846.74
143.11839294433594 0 2463.2346
144.06564331054688 0 819.0715
144.12123107910156 0 520.16693
145.09738159179688 0 805.3861
145.49781799316406 0 686.2291
146.06051635742188 0 787.26654
147.07676696777344 0 8923.847
148.060791015625 0 2714.7395
148.9538116455078 0 1423.5344
151.0966339111328 0 730.1861
151.12353515625 0 1720.2714
152.10740661621094 0 1744.3674
153.06631469726562 0 1594.6217
154.05015563964844 0 2469.1853
154.0865936279297 0 21209.814
155.08209228515625 0 2539.423
155.09048461914062 0 1669.4791
156.0847930908203 0 782.37225
157.06114196777344 0 14029.162
157.09750366210938 0 32672.115 a Ammonia loss 2
158.0653533935547 0 689.614
158.10055541992188 0 1638.8544
158.2281036376953 0 601.4353
163.59475708007812 0 574.24066
165.1028594970703 0 1436.5311
166.08670043945312 0 2318.1003
167.04534912109375 0 1400.8185
167.08187866210938 0 31141.37
167.1182403564453 0 29300.486
168.0855255126953 0 3388.2087
168.12156677246094 0 2674.7617
170.044921875 0 979.5359
170.09642028808594 0 876.0031
171.07679748535156 0 7459.0537
171.11317443847656 0 954.6617
172.08004760742188 0 1040.3213
172.1124267578125 0 20725.648
173.11578369140625 0 2944.2766
173.1288299560547 0 3174.0178
174.0553436279297 0 601.56415
175.07176208496094 0 15902.038
175.0867462158203 0 1098.1249
175.11932373046875 0 1109.725
176.07525634765625 0 666.8779
177.1026153564453 0 2256.216
177.11138916015625 0 923.87384
179.11795043945312 0 736.9969
181.06146240234375 0 1501.7357
181.09744262695312 0 3241.8235
182.0816650390625 0 1965.6792
183.1128387451172 0 1464.1263
183.14947509765625 0 3217.183
185.0561065673828 0 1124.6581
185.09243774414062 0 57088.453
185.16506958007812 0 1765.4634
186.09588623046875 0 5179.9995
186.16842651367188 0 645.4658
188.07144165039062 0 746.71436
188.14369201660156 0 1782.7567
189.08737182617188 0 22046.244
189.12368774414062 0 3014.1008
190.127197265625 0 1437.7512
191.1183624267578 0 877.41986
193.12155151367188 0 776.8173
193.52391052246094 0 572.4182
195.07675170898438 0 8950.834
195.1131591796875 0 60257.023
196.1166534423828 0 6070.8926
197.12879943847656 0 135592.98
198.08741760253906 0 806.46277
198.11293029785156 0 861.66046
198.13209533691406 0 13899.801
199.07164001464844 0 32378.188
199.0985107421875 0 988.68854
199.10806274414062 0 4725.173
199.13389587402344 0 788.1422
199.1440887451172 0 598.1069
200.0752410888672 0 3316.7734
201.12367248535156 0 2222.9695
202.11904907226562 0 2314.259
203.08218383789062 0 616.68536
203.1028594970703 0 3598.069
205.17051696777344 0 845.06665
208.10836791992188 0 1202.6929
209.09237670898438 0 8533.698
210.0963134765625 0 1014.5496
211.14462280273438 0 3971.652
212.13975524902344 0 8751.019
213.08726501464844 0 10064.483
213.123779296875 0 5603.767
213.14254760742188 0 1049.377
213.15977478027344 0 1085.511
214.0907440185547 0 1078.946
214.11878967285156 0 1041.6515
215.13941955566406 0 4257.136
216.09812927246094 0 700.9669
217.08226013183594 0 61304.516
217.13388061523438 0 14044.849
217.60586547851562 0 793.5316
218.08558654785156 0 4875.0317
218.1379852294922 0 1409.0815
221.09246826171875 0 1931.61
223.0719757080078 0 2039.1622
224.10353088378906 0 2634.785
225.12374877929688 0 17205.738
226.11863708496094 0 2632.2915
226.12892150878906 0 1258.4255
227.06639099121094 0 1002.4969
227.1030731201172 0 9286.91
229.11900329589844 0 1307.6528
231.09800720214844 0 10161.797
231.1103973388672 0 624.8267
231.60263061523438 0 1517.2959
232.10235595703125 0 1257.97
233.16534423828125 0 502275.2 a 1
234.1685791015625 0 77733.67
234.17962646484375 0 592.047
235.1443328857422 0 1159.9613
235.17108154296875 0 3999.008
236.10336303710938 0 23193.285
236.1396026611328 0 1653.0372
237.08692932128906 0 1023.4695
237.1066436767578 0 2916.688
237.12261962890625 0 1501.1958
238.08201599121094 0 1074.6807
238.2166290283203 0 730.36414
239.14976501464844 0 1035.355
240.09815979003906 0 3164.424
240.6084442138672 0 2553.5308
241.0823516845703 0 47666.95
242.08566284179688 0 5251.885
242.114013671875 0 4450.1978
243.13442993164062 0 343892.97 y Water loss 7
244.13722229003906 0 35232.133 b Ammonia loss 1
245.0766143798828 0 696.0227
245.12913513183594 0 12437.114
246.1321258544922 0 2276.2021
249.61412048339844 0 1699.5637
250.08290100097656 0 1988.0305
250.1169891357422 0 1423.5221
252.0990447998047 0 838.6523
252.13482666015625 0 5821.62
253.08187866210938 0 929.37573
253.13735961914062 0 916.67596
253.16998291015625 0 1674.404
254.1139373779297 0 81878.03
254.1502685546875 0 9586.215
255.11712646484375 0 10846.395
255.15403747558594 0 1377.8588
256.11846923828125 0 1148.7
258.10894775390625 0 3073.415
258.14471435546875 0 705.2863
259.0928039550781 0 41743.516
260.09600830078125 0 5360.2393
261.144775390625 0 48496.42 y 7
261.16009521484375 0 207728.22 b 1
262.1480407714844 0 5587.862
262.16339111328125 0 30616.303
263.166015625 0 2283.0044
264.0981750488281 0 3234.068
264.1345520019531 0 38641.477
265.1376647949219 0 5242.6753
267.09881591796875 0 900.95636
268.09307861328125 0 6700.096
268.1294860839844 0 2175.7385
269.0761413574219 0 1354.5582
269.09454345703125 0 844.83105
269.1129150390625 0 1192.316
270.1092529296875 0 888.91736
270.1447448730469 0 852.8483
272.1243591308594 0 39836.61
273.1247253417969 0 7781.657
273.6239013671875 0 1682.9159 y Water loss 4
274.1031799316406 0 881.0519
276.1192626953125 0 1953.4255
277.10357666015625 0 978.0837
278.1138000488281 0 1602.4243
278.14996337890625 0 1569.6857
280.16619873046875 0 876.95905
281.0767517089844 0 2367.684
281.16156005859375 0 2046.9716
282.1083679199219 0 4549.1245
282.1291809082031 0 5580.5854
282.1452941894531 0 163483.48
282.6282653808594 0 1138.6295 y 4
283.0935974121094 0 769.81445
283.1131591796875 0 804.76624
283.14837646484375 0 23276.975
284.15045166015625 0 1478.405
285.19671630859375 0 5802.241
286.10382080078125 0 28573.484
287.1070251464844 0 3451.4558
287.1349182128906 0 958.85706
287.1509704589844 0 1023.2283
287.6214599609375 0 1237.2739
288.2042236328125 0 880.57745
289.1517028808594 0 2844.8542
291.1328125 0 1347.3567
292.1295471191406 0 930.0357
294.1460876464844 0 831.59375
296.0885925292969 0 2671.328
296.1244812011719 0 15271.012
296.626953125 0 1625.7657
297.1283264160156 0 1214.9021
297.158203125 0 5008.326
297.6598815917969 0 1935.0034
298.1769714355469 0 863.25287
299.087890625 0 3993.937
299.1719970703125 0 7982.6807
300.1192626953125 0 4356.0557
300.1365051269531 0 1383.1614
300.15570068359375 0 6859.033
300.1750183105469 0 1568.2284
300.63995361328125 0 755.6174
301.1554870605469 0 1332.0995
304.11431884765625 0 15720.354
305.12994384765625 0 5484.704
305.1611633300781 0 1531.3732
305.6317443847656 0 1647.2472
306.145263671875 0 4705.3286
306.16412353515625 0 3319.841 b Water loss 5
306.6648254394531 0 883.47156
307.1492004394531 0 1252.7797
308.1611022949219 0 7988.4424
308.1974792480469 0 4895.045
309.2010192871094 0 1283.1672
310.10406494140625 0 8303.972
310.1378173828125 0 943.5013
311.08740234375 0 1225.0892
311.10650634765625 0 943.73865
311.17138671875 0 2285.1155
312.1558532714844 0 3022.0427
313.1921691894531 0 1807.7168 a Ammonia loss 2
314.0995788574219 0 2572.3237
314.1352233886719 0 14524.667
314.6361083984375 0 2725.2354
318.1295471191406 0 1442.1185
319.1397705078125 0 2833.34
320.1252746582031 0 1212.0562
320.65911865234375 0 1716.4751
321.15631103515625 0 5430.2754
322.1592712402344 0 1029.6027
323.17181396484375 0 7119.0527
324.15570068359375 0 6815.0674
324.1748352050781 0 1304.841
325.1585998535156 0 861.08685
326.17138671875 0 1473.4163
328.1144714355469 0 19661.406
329.11785888671875 0 2593.068
329.1512451171875 0 858.84406
329.6640625 0 5614.4087
330.16546630859375 0 3526.4111
333.15631103515625 0 27253.41
334.1589050292969 0 3838.147
335.1347351074219 0 725.747
336.15576171875 0 1783.4111
337.1512756347656 0 7868.8604
338.13507080078125 0 5021.5137
338.1541442871094 0 1277.1578
338.66925048828125 0 14133.388
339.1676025390625 0 20783.887
339.6721496582031 0 933.0785
340.1694641113281 0 2958.2437
341.1827697753906 0 29655.748 b Ammonia loss 2
342.1842346191406 0 8944.091
343.1627502441406 0 1297.9542
343.1856384277344 0 1224.5748
346.1249084472656 0 34722.74
347.1296691894531 0 5635.647
347.6748962402344 0 3427.7136
348.17578125 0 1964.8081
350.1845703125 0 1041.4901
351.166748046875 0 116735.3
352.1147766113281 0 1109.556
352.1697082519531 0 20608.63
352.66729736328125 0 1875.6044
353.14544677734375 0 1999.5116
353.1712646484375 0 2376.6514
354.16619873046875 0 17468.846
355.1622619628906 0 18013.543
356.1087951660156 0 946.814
356.1455078125 0 5696.3755
356.1653747558594 0 2212.8467
356.6841735839844 0 1277.5864
357.1471252441406 0 800.23663
357.2116394042969 0 776.34955
358.2129821777344 0 17283.527 b 2
359.21624755859375 0 3420.44
361.6761474609375 0 4625.677
362.1800231933594 0 2849.1
365.1457824707031 0 16043.684
365.6919860839844 0 769.5446
366.1478271484375 0 1554.0916
366.16925048828125 0 1084.0403
367.1964416503906 0 1133.9364
368.1929931640625 0 3498.9924
369.17724609375 0 127179.28
370.1802673339844 0 25275.438
370.6822204589844 0 9734.538 b Water loss 6
371.1560363769531 0 2895.9138
371.1827697753906 0 6126.7876 b Ammonia loss 6
371.2257995605469 0 994.4967
372.1768493652344 0 15064.175 y Water loss 6
373.17242431640625 0 31475.6
374.1751708984375 0 6324.5547
375.1684265136719 0 1124.834
379.1252746582031 0 1869.6146
379.6849365234375 0 4180.5244 b 6
380.18756103515625 0 1848.1741
381.17822265625 0 1331.8605
383.1565246582031 0 37116.99
384.1592102050781 0 6785.6885
384.1878967285156 0 1309.1509
386.20416259765625 0 13429.811
387.1510314941406 0 1738.4069
387.20751953125 0 2547.3572
388.1355285644531 0 1161.3466
390.18792724609375 0 5318.6313 y 6
391.19195556640625 0 1174.6732
393.17694091796875 0 3603.2375
394.162841796875 0 1558.2802
395.22821044921875 0 1517.6252
397.1357727050781 0 5105.369
398.1186218261719 0 852.7606
399.18896484375 0 1045.4164
401.1671447753906 0 64361.055
401.2158203125 0 1760.7458
402.17022705078125 0 12865.886
403.1724853515625 0 2367.4746
405.16204833984375 0 2870.0398
406.1659851074219 0 775.9674
407.1571044921875 0 897.19916
411.15167236328125 0 2942.2024
412.17315673828125 0 1054.2491
413.51690673828125 0 1153.5356
413.8527526855469 0 981.7087
415.1465148925781 0 9125.973
416.14923095703125 0 2377.9104
416.1929931640625 0 3829.4163
417.19720458984375 0 1814.3989
418.1936950683594 0 955.8489
422.20263671875 0 1127.1552
423.18780517578125 0 3393.8281
424.1922302246094 0 1209.7163
424.21942138671875 0 1304.3235
425.1322326660156 0 1888.9374
425.1673583984375 0 2353.8745
425.1982727050781 0 995.4341
426.2020568847656 0 2833.689
426.7017517089844 0 899.51324
427.2708740234375 0 9171.72
428.2740478515625 0 3503.0337
429.2130126953125 0 5490.6113
430.21466064453125 0 2316.262
431.2148132324219 0 862.079
432.18890380859375 0 1054.8933
433.1568603515625 0 18058.688
434.16094970703125 0 3414.9004
434.2037353515625 0 13048.713
435.20782470703125 0 1807.3641 b Water loss 7
435.70855712890625 0 978.8876
436.2550048828125 0 1237.2078
440.21429443359375 0 5316.7437
441.19879150390625 0 5663.068
442.20050048828125 0 1693.1233
443.1433410644531 0 1247.0593
443.17852783203125 0 2644.3337
444.18731689453125 0 4299.402
444.7142333984375 0 1163.4889
448.18341064453125 0 1747.7222
450.19903564453125 0 9741.992
451.2027893066406 0 2221.2615
452.21441650390625 0 27224.98
453.2171630859375 0 6532.774 y 1
454.22418212890625 0 892.64667
454.265625 0 3084.3052
455.26580810546875 0 19972.926 b 3
456.2687072753906 0 5876.7476
457.2737731933594 0 931.1389
459.2097473144531 0 7510.6714 y Water loss 5
460.2134094238281 0 1596.273
461.2138977050781 0 867.498
462.1988830566406 0 26606.053
463.20184326171875 0 6796.6875
464.20965576171875 0 962.5742
464.25067138671875 0 2694.619
466.1932373046875 0 3279.8499
467.1991882324219 0 1032.7137
468.2091979980469 0 14006.705
469.2125549316406 0 2749.7002
470.2251281738281 0 108796.46
471.2278747558594 0 25417.746
472.23028564453125 0 4310.9473
476.17864990234375 0 2875.737
477.2213439941406 0 4114.159 y 5
478.22808837890625 0 1467.6024
479.2231140136719 0 975.8168
480.20953369140625 0 96960.2
481.21240234375 0 25844.34
482.216796875 0 3561.5996
482.2624206542969 0 5123.897
484.2043762207031 0 8683.7295
485.20440673828125 0 1318.2223
486.25616455078125 0 3498.6924
487.2585754394531 0 1428.1829
494.18902587890625 0 8805.731
495.1931457519531 0 1350.7911
496.2410888671875 0 1830.624
497.2367858886719 0 3724.3293
497.27349853515625 0 942.768
498.22015380859375 0 344935.75
499.2229919433594 0 87752.64
500.22589111328125 0 12009.559
502.2149353027344 0 17141.45
503.2176208496094 0 3936.5781
504.2173767089844 0 862.8308
504.28350830078125 0 874.5584
510.221435546875 0 2091.1938
512.19921875 0 14564.546
513.2019653320312 0 4105.075
514.2531127929688 0 3434.5493
514.30224609375 0 2947.6655
515.2466430664062 0 8977.228
515.3067016601562 0 1516.321
516.2443237304688 0 4077.7969
517.2462158203125 0 1171.9954
524.286865234375 0 5676.2993 b Water loss 4
525.288330078125 0 1718.5284
528.23046875 0 3233.867
530.2095336914062 0 50782.83
531.2127685546875 0 12716.922
532.2144165039062 0 2140.5818
533.273681640625 0 1166.1111
542.297607421875 0 15504.518 b 4
543.3008422851562 0 5152.6387
544.30224609375 0 1357.4772
546.24072265625 0 5451.625 y Water loss 4
547.2418212890625 0 1246.0555
548.2206420898438 0 4538.6445
554.2481689453125 0 1380.0764
563.2459106445312 0 3754.3862
564.2499389648438 0 7600.833 y 4
565.25390625 0 2594.0798
565.3013305664062 0 1186.9569
566.2537231445312 0 1253.9161
566.3018798828125 0 1456.571
573.2301635742188 0 6405.859
574.2330322265625 0 1441.3507
575.2415771484375 0 1378.2609
575.2861328125 0 1073.4072
581.2567138671875 0 18041.291
582.255859375 0 6510.449
583.2625732421875 0 1048.3485
583.31103515625 0 16948.855
584.3123779296875 0 5552.607
585.331298828125 0 2416.632
588.2655029296875 0 1163.0674
589.2637939453125 0 831.7212
591.2412109375 0 20093.684
592.244140625 0 5179.9326
593.2477416992188 0 1155.7247
593.3008422851562 0 6175.7334
594.3036499023438 0 2238.4822
597.2503051757812 0 2395.4048
599.267333984375 0 108902.38
600.2702026367188 0 38079.016
600.3273315429688 0 1349.3046
601.2720947265625 0 6952.0005
601.3345947265625 0 10249.666
602.337646484375 0 4243.559
606.279052734375 0 2291.964
607.2741088867188 0 2013.0789
608.2689819335938 0 1813.27
609.2520751953125 0 66764.02
610.2549438476562 0 23307.62
611.2550659179688 0 2190.6714
611.31103515625 0 37110.973 b Water loss 5
612.3125610351562 0 14193.12 b Ammonia loss 5
613.3204956054688 0 1208.5204
615.2996215820312 0 3763.4636
616.299560546875 0 2384.2869
617.2937622070312 0 1791.9379
625.2859497070312 0 4502.486
626.2868041992188 0 2333.8152
627.262451171875 0 561301.9
628.2652587890625 0 172682.53
628.3626708984375 0 1076.0485
628.3827514648438 0 949.6882
629.267578125 0 32273.262
629.329833984375 0 36180.61 b 5
630.2681884765625 0 2490.8206
630.3320922851562 0 11562.534
631.33642578125 0 2411.5164
633.326171875 0 1990.1036
643.2943725585938 0 16270.2295 y Water loss 3
643.3619384765625 0 1444.7953
644.2970581054688 0 6820.1226
645.2761840820312 0 26513.238
646.2808837890625 0 8443.179
647.2813720703125 0 1222.9985
661.3045043945312 0 65924.27 y 3
662.307861328125 0 21883.164
663.3099365234375 0 4250.454
676.327880859375 0 1077.0227
686.3137817382812 0 4024.4631
687.31640625 0 1497.3339
694.3445434570312 0 4996.2876
695.3429565429688 0 2522.0815
700.3626708984375 0 4686.119
701.3624877929688 0 1796.2732
704.3271484375 0 5633.2573
705.331787109375 0 3032.036
710.337158203125 0 1731.0138
712.3529663085938 0 32988.383
713.35498046875 0 14638.365
714.3592529296875 0 3494.6062
722.3399658203125 0 24885.285
723.34375 0 9266.452
724.34521484375 0 1802.7291
729.3089599609375 0 1740.435
730.376953125 0 40953.703
731.3800048828125 0 18206.562
732.3850708007812 0 4419.541
738.3052368164062 0 1388.47
740.3491821289062 0 105037.19 b Water loss 6
741.3528442382812 0 44372.926 b Ammonia loss 6
742.354248046875 0 10516.299
744.3517456054688 0 947.7035
746.3343505859375 0 1893.2134
756.3226928710938 0 3382.204
757.3193359375 0 986.89166
758.3598022460938 0 557557.6 b 6
759.3629150390625 0 222452.6
760.3652954101562 0 54626
761.36865234375 0 3710.977
762.369873046875 0 898.5301
772.3526000976562 0 1875.1016
773.4260864257812 0 1118.1279
774.3297119140625 0 11568.825
775.3330688476562 0 4523.5864
783.4028930664062 0 2271.7078
790.360107421875 0 4803.346
791.3643188476562 0 1978.3304
801.4109497070312 0 1934.5393
833.3828125 0 1435.9048
841.4093017578125 0 2727.2412
842.39697265625 0 5879.911
843.4076538085938 0 3237.018
844.4478759765625 0 3879.0847
845.4581298828125 0 1487.3823
847.4327392578125 0 1963.5312
851.392578125 0 7170.572
852.396240234375 0 2894.2654
854.43603515625 0 1783.9498
859.4187622070312 0 40505.17
860.422119140625 0 19192.44
861.425537109375 0 4890.903
862.4352416992188 0 907.3817
869.4034423828125 0 23421.654 b Water loss 7
870.4058227539062 0 12726.801
871.4091186523438 0 3725.33
872.4461059570312 0 9798.709
873.4494018554688 0 3621.536
887.4136962890625 0 185121.77 b 7
888.4166870117188 0 89539.9
889.4193725585938 0 26262.785
890.4207153320312 0 2787.909
905.4249877929688 0 2853.0242 y 1
906.4296875 0 1973.4996
907.4285888671875 0 1180.0138
1145.3795166015625 0 854.6521
1249.0174560546875 0 762.96155
1359.1812744140625 0 865.8395
1537.73876953125 0 800.7647
2723.601318359375 0 1004.8044

Spectrum Details

|  |  |
| --- | --- |
| Matched peaks? Matched peaks The total absolute number of peaks matched. Additionally in brackets the total fraction of peaks matched and the total number of peaks is shown. | 38 (6.32% of 601) |
| FDR? FDR The false discovery rate estimated for this peptide. It is calculated by matching all theoretical fragments with a non-integer shift with the raw peaks for this spectrum. This is done with 40 different shifts. The resulting percentage is the average number of annotated peaks over the number of annotated peaks with the correct spectrum. | 0.56% |
| Satellite FDR? Satellite FDR See the FDR for details on its calculation. This satellite ion specific FDR only contains the satellite ions (d/w) for I/L/J positions. | - |
| PSM Score? PSM Score The PSM Score as given by Hecklib to this annotated spectrum. It is shown with three significant figures. | 492 |

#### Reverse Lookup? Reverse Lookup All places where this read could be placed.

| Group | Segment | Template | Template Part | Read Part | Score | Unique |
| --- | --- | --- | --- | --- | --- | --- |
| Homo sapiens Light Chain | IGLC | IGLC2 | [10..19] | [0..9] | 72 | False |
| Homo sapiens Light Chain | IGLC | IGLC3 | [8..17] | [0..9] | 72 | False |
| Homo sapiens Light Chain | IGLC | IGLC6 | [10..19] | [0..9] | 72 | False |
| Homo sapiens Light Chain | IGLC | IGLC7 | [10..19] | [0..9] | 72 | False |

| Recombined | Template Part | Read Part | Score | Unique |
| --- | --- | --- | --- | --- |
| REC-0-1\_002 | [121..130] | [0..9] | 72 | True |

#### Meta Information from Multiple reads

##### Number of combined reads

7

##### Intensity

0.7954

##### TotalArea

1.218E+09

##### Changes to the peptide sequence

JFPPSSEEJ

L→JNo support for either Leucine or Isoleucine based on side chain ions (Position: 9)

L→JNo support for either Leucine or Isoleucine based on side chain ions (Position: 1)

#### Positional Score

Copy Data

##### Positional Score (TSV)

###### Preview

```
Loading example...
```

*Click on the button to copy the data to your clipboard.*

10012345678

Label Value
"0" 0.683
"1" 0.661
"2" 0.699
"3" 0.706
"4" 0.7
"5" 0.687
"6" 0.701
"7" 0.711
"8" 0.714

#### Meta Information from PEAKS

##### Scan Identifier

F3:8530

##### Original sequence

L

F

P

P

S

S

E

E

L

##### Posttranslational Modifications

##### Source File

D:\separate\_stitch\_analyses\xle-disambiguation\raw\20210323\_F1\_UM1\_Peng0013\_SA\_F59\_ingel\_3ug\_chymo.raw

##### Fraction

3

##### Scan Feature

F3:4445

##### De Novo Score

98

##### ConfidenceScore

98

### m/z

509.7585

##### Mass

1017.5018

##### Charge

2

##### Retention Time

46.24

##### Predicted Retention Time

-

##### Area

1.74E+08

##### Parts Per Million

0.6

##### Fragmentation mode

ETHCD

##### Originating file

01 D:\separate\_stitch\_analyses\xle-disambiguation\20210325\_F59\_3ug\_DENOVO\_12.csv

#### Meta Information from PEAKS

##### Scan Identifier

F3:8467

##### Original sequence

L

F

P

P

S

S

E

E

L

##### Posttranslational Modifications

##### Source File

D:\separate\_stitch\_analyses\xle-disambiguation\raw\20210323\_F1\_UM1\_Peng0013\_SA\_F59\_ingel\_3ug\_chymo.raw

##### Fraction

3

##### Scan Feature

F3:4445

##### De Novo Score

98

##### ConfidenceScore

98

### m/z

509.7585

##### Mass

1017.5018

##### Charge

2

##### Retention Time

46.24

##### Predicted Retention Time

-

##### Area

1.74E+08

##### Parts Per Million

0.6

##### Fragmentation mode

ETHCD

##### Originating file

01 D:\separate\_stitch\_analyses\xle-disambiguation\20210325\_F59\_3ug\_DENOVO\_12.csv

#### Meta Information from PEAKS

##### Scan Identifier

F3:8589

##### Original sequence

L

F

P

P

S

S

E

E

L

##### Posttranslational Modifications

##### Source File

D:\separate\_stitch\_analyses\xle-disambiguation\raw\20210323\_F1\_UM1\_Peng0013\_SA\_F59\_ingel\_3ug\_chymo.raw

##### Fraction

3

##### Scan Feature

F3:4445

##### De Novo Score

98

##### ConfidenceScore

98

### m/z

509.7585

##### Mass

1017.5018

##### Charge

2

##### Retention Time

46.24

##### Predicted Retention Time

-

##### Area

1.74E+08

##### Parts Per Million

0.6

##### Fragmentation mode

ETHCD

##### Originating file

01 D:\separate\_stitch\_analyses\xle-disambiguation\20210325\_F59\_3ug\_DENOVO\_12.csv

#### Meta Information from PEAKS

##### Scan Identifier

F3:8125

##### Original sequence

L

F

P

P

S

S

E

E

L

##### Posttranslational Modifications

##### Source File

D:\separate\_stitch\_analyses\xle-disambiguation\raw\20210323\_F1\_UM1\_Peng0013\_SA\_F59\_ingel\_3ug\_chymo.raw

##### Fraction

3

##### Scan Feature

F3:4445

##### De Novo Score

97

##### ConfidenceScore

97

### m/z

509.7585

##### Mass

1017.5018

##### Charge

2

##### Retention Time

46.24

##### Predicted Retention Time

-

##### Area

1.74E+08

##### Parts Per Million

0.6

##### Fragmentation mode

ETHCD

##### Originating file

01 D:\separate\_stitch\_analyses\xle-disambiguation\20210325\_F59\_3ug\_DENOVO\_12.csv

#### Meta Information from PEAKS

##### Scan Identifier

F3:8215

##### Original sequence

L

F

P

P

S

S

E

E

L

##### Posttranslational Modifications

##### Source File

D:\separate\_stitch\_analyses\xle-disambiguation\raw\20210323\_F1\_UM1\_Peng0013\_SA\_F59\_ingel\_3ug\_chymo.raw

##### Fraction

3

##### Scan Feature

F3:4445

##### De Novo Score

97

##### ConfidenceScore

97

### m/z

509.7585

##### Mass

1017.5018

##### Charge

2

##### Retention Time

46.24

##### Predicted Retention Time

-

##### Area

1.74E+08

##### Parts Per Million

0.6

##### Fragmentation mode

ETHCD

##### Originating file

01 D:\separate\_stitch\_analyses\xle-disambiguation\20210325\_F59\_3ug\_DENOVO\_12.csv

#### Meta Information from PEAKS

##### Scan Identifier

F3:8292

##### Original sequence

L

F

P

P

S

S

E

E

L

##### Posttranslational Modifications

##### Source File

D:\separate\_stitch\_analyses\xle-disambiguation\raw\20210323\_F1\_UM1\_Peng0013\_SA\_F59\_ingel\_3ug\_chymo.raw

##### Fraction

3

##### Scan Feature

F3:4445

##### De Novo Score

97

##### ConfidenceScore

97

### m/z

509.7585

##### Mass

1017.5018

##### Charge

2

##### Retention Time

46.24

##### Predicted Retention Time

-

##### Area

1.74E+08

##### Parts Per Million

0.6

##### Fragmentation mode

HCD

##### Originating file

01 D:\separate\_stitch\_analyses\xle-disambiguation\20210325\_F59\_3ug\_DENOVO\_12.csv

#### Meta Information from PEAKS

##### Scan Identifier

F3:8371

##### Original sequence

L

F

P

P

S

S

E

E

L

##### Posttranslational Modifications

##### Source File

D:\separate\_stitch\_analyses\xle-disambiguation\raw\20210323\_F1\_UM1\_Peng0013\_SA\_F59\_ingel\_3ug\_chymo.raw

##### Fraction

3

##### Scan Feature

F3:4445

##### De Novo Score

96

##### ConfidenceScore

96

### m/z

509.7585

##### Mass

1017.5018

##### Charge

2

##### Retention Time

46.24

##### Predicted Retention Time

-

##### Area

1.74E+08

##### Parts Per Million

0.6

##### Fragmentation mode

HCD

##### Originating file

01 D:\separate\_stitch\_analyses\xle-disambiguation\20210325\_F59\_3ug\_DENOVO\_12.csv
