## Supplementary material for "A handle on mass coincidence errors in *de novo* sequencing of antibodies by bottom-up proteomics": Combined_062.html

Details Combined\_062 | Stitch OverviewUndefined

### Read Combined\_062

#### Sequence (length=18)

VKAGVETTTPSKQSNNKY

#### Spectrum 2479? Spectrum 2479 The raw spectrum of this peptide as annotated by Hecklib. The fragments are coloured according to ion type (see legend). Any peaks with a star '\*' as text can be hovered over to see the full details, first the ion type second the mass shift type. By hovering over the amino acids in the peptide or ions in the legend the corresponding peaks are highlighted. By toggling the 'Unassigned' label you can turn the background (unassigned) peaks on or off in the plot. By updating the slider in the Ion legend you can update the spectrum to only show the top X% of the peaks with labels. The top X% means any peak that is within X% of the highest intensity. By dragging in the spectrum you can zoom in to a specific part of the spectrum and use 'Zoom Out' to get back to the original zoom level. The annotation of the spectrum is based on the given sequence in the peptides file and is done with different software so inconsistencies are likely. The peaks are annotated based on the given sequence, with 20 ppm tolerance.

Copy Data

##### Spectrum 2479 (TSV)

###### Preview

```
Loading example...
```

*Click on the button to copy the data to your clipboard.*

Mz MinMz MaxIntensity Max

WidthHeightPeptide font sizePeptide stroke widthSpectrum font sizeSpectrum stroke widthCompact peptide

Ion legend

wxyz

abcd

OtherUnassignedIon^Charge^_Position_Show for top:%

VKAGVETTTPSKQSNNKY

01.46e+42.92e+44.38e+45.83e+4

Zoom Out

c^+1^_2_z^+1^_2_c^+1^_3_c^+1^_3_c^+1^_4_c^+1^_4_z^+1^_3_c^+1^_5_c^+1^_5_w^+1^_4_z^+1^_4_z^+2^_10_c^+1^_6_y^+1^_5_z^+1^_5_y^+3^_17_y^+1^_5_z^+2^_11_y^+2^_12_y^+2^_12_z^+2^_12_w^+1^_6_c^+1^_7_z^+1^_6_y^+2^_13_y^+1^_6_c^+1^_8_z^+2^_15_c^+2^_16_y^+2^_15_c^+2^_16_z^+2^_16_z^+2^_16_y^+2^_16_y^+1^_7_y^+1^_7_z^+1^_7_y^+1^_7_c^+1^_9_c^+2^_17_z^+2^_17_z^+1^_8_y^+1^_8_c^+1^_10_c^+1^_11_y^+1^_10_y^+1^_10_z^+1^_10_y^+1^_10_c^+1^_12_z^+1^_11_y^+1^_11_c^+1^_13_z^+1^_12_y^+1^_12_c^+1^_14_z^+1^_13_y^+1^_13_c^+1^_15_z^+1^_14_y^+1^_14_z^+1^_15_y^+1^_15_c^+1^_16_z^+1^_16_y^+1^_16_c^+1^_17_z^+1^_17_

0824164824723296

Fragment Matches Table

Show background peaks

| Position | Ion type | Intensity | mz Theoretical | mz Error (Th) | mz Error (ppm) | Charge | Series Number |
| --- | --- | --- | --- | --- | --- | --- | --- |
| - | - | 421.7 | 125.4 | - | - | 0 | - |
| - | - | 549.5 | 129.1 | - | - | 0 | - |
| - | - | 7171 | 129.1 | - | - | 0 | - |
| - | - | 436.3 | 133 | - | - | 0 | - |
| - | - | 481.4 | 148.5 | - | - | 0 | - |
| - | - | 495.5 | 148.8 | - | - | 0 | - |
| - | - | 755.7 | 148.9 | - | - | 0 | - |
| - | - | 501 | 148.9 | - | - | 0 | - |
| - | - | 634.8 | 148.9 | - | - | 0 | - |
| - | - | 630 | 148.9 | - | - | 0 | - |
| - | - | 710.1 | 148.9 | - | - | 0 | - |
| - | - | 1165 | 148.9 | - | - | 0 | - |
| - | - | 1178 | 148.9 | - | - | 0 | - |
| - | - | 2427 | 148.9 | - | - | 0 | - |
| - | - | 3937 | 148.9 | - | - | 0 | - |
| - | - | 4314 | 149 | - | - | 0 | - |
| - | - | 2556 | 149 | - | - | 0 | - |
| - | - | 1419 | 149 | - | - | 0 | - |
| - | - | 1241 | 149 | - | - | 0 | - |
| - | - | 935.1 | 149 | - | - | 0 | - |
| - | - | 697.7 | 149 | - | - | 0 | - |
| - | - | 608.9 | 149 | - | - | 0 | - |
| - | - | 676.1 | 149 | - | - | 0 | - |
| - | - | 514 | 149 | - | - | 0 | - |
| - | - | 468.5 | 149.2 | - | - | 0 | - |
| - | - | 521 | 187.1 | - | - | 0 | - |
| - | - | 603.3 | 188.1 | - | - | 0 | - |
| - | - | 501.6 | 210.3 | - | - | 0 | - |
| - | - | 582.8 | 224.7 | - | - | 0 | - |
| - | - | 579.2 | 227.2 | - | - | 0 | - |
| - | - | 821.6 | 228.1 | - | - | 0 | - |
| 2 | c | 3674 | 228.2 | 0.0001212 | 0.531 | +1 | 2 |
| - | - | 608.9 | 253.2 | - | - | 0 | - |
| - | - | 825.1 | 273.2 | - | - | 0 | - |
| 17 | z | 4528 | 294.2 | 9.051E-05 | 0.3077 | +1 | 2 |
| - | - | 1136 | 295.2 | - | - | 0 | - |
| 3 | c | 918.4 | 299.2 | 0.0002476 | 0.8277 | +1 | 3 |
| - | - | 553.2 | 306.2 | - | - | 0 | - |
| - | - | 2741 | 315.2 | - | - | 0 | - |
| 3 | c | 1885 | 316.2 | 0.0004296 | 1.358 | +1 | 3 |
| - | - | 675.3 | 329.2 | - | - | 0 | - |
| - | - | 780.3 | 330.2 | - | - | 0 | - |
| - | - | 602.3 | 337.1 | - | - | 0 | - |
| - | - | 1360 | 355.1 | - | - | 0 | - |
| 4 | c | 1285 | 356.2 | 1.715E-05 | 0.04814 | +1 | 4 |
| - | - | 1171 | 364.2 | - | - | 0 | - |
| 4 | c | 1.341E+04 | 373.3 | 0.0002563 | 0.6867 | +1 | 4 |
| - | - | 2032 | 374.3 | - | - | 0 | - |
| - | - | 869.9 | 384.2 | - | - | 0 | - |
| 16 | z | 5503 | 408.2 | 4.921E-05 | 0.1205 | +1 | 3 |
| - | - | 7643 | 409.2 | - | - | 0 | - |
| - | - | 1918 | 410.2 | - | - | 0 | - |
| - | - | 1146 | 427.3 | - | - | 0 | - |
| - | - | 2227 | 429.1 | - | - | 0 | - |
| - | - | 967.2 | 429.3 | - | - | 0 | - |
| - | - | 847.7 | 430.1 | - | - | 0 | - |
| 5 | c | 1805 | 455.3 | 0.0003593 | 0.7892 | +1 | 5 |
| 5 | c | 1.216E+04 | 472.3 | 0.0001278 | 0.2705 | +1 | 5 |
| - | - | 2706 | 473.3 | - | - | 0 | - |
| 15 | w | 1571 | 478.2 | 0.0004073 | 0.8517 | +1 | 4 |
| - | - | 668.4 | 485.6 | - | - | 0 | - |
| - | - | 1323 | 498.3 | - | - | 0 | - |
| - | - | 905.2 | 506.9 | - | - | 0 | - |
| - | - | 674.6 | 507.2 | - | - | 0 | - |
| 15 | z | 7644 | 522.2 | 8.365E-05 | 0.1602 | +1 | 4 |
| - | - | 7440 | 523.2 | - | - | 0 | - |
| - | - | 2124 | 524.3 | - | - | 0 | - |
| - | - | 744 | 533.8 | - | - | 0 | - |
| - | - | 1625 | 546.8 | - | - | 0 | - |
| - | - | 594.3 | 556.9 | - | - | 0 | - |
| - | - | 2052 | 557.4 | - | - | 0 | - |
| - | - | 738.6 | 558.3 | - | - | 0 | - |
| - | - | 763.1 | 558.4 | - | - | 0 | - |
| 9 | z | 813.4 | 575.8 | 0.0009646 | 1.675 | +2 | 10 |
| - | - | 737.9 | 585 | - | - | 0 | - |
| - | - | 1446 | 600.4 | - | - | 0 | - |
| 6 | c | 1.21E+04 | 601.4 | 0.0005761 | 0.9579 | +1 | 6 |
| - | - | 4019 | 602.4 | - | - | 0 | - |
| - | - | 732.8 | 603.4 | - | - | 0 | - |
| - | - | 1012 | 607.3 | - | - | 0 | - |
| 14 | y | 809.5 | 608.3 | 0.0008436 | 1.387 | +1 | 5 |
| 14 | z | 2346 | 609.3 | 0.001488 | 2.443 | +1 | 5 |
| - | - | 1.149E+04 | 610.3 | - | - | 0 | - |
| - | - | 3722 | 611.3 | - | - | 0 | - |
| 2 | y | 982.3 | 618.3 | 0.002549 | 4.122 | +3 | 17 |
| - | - | 1350 | 618.7 | - | - | 0 | - |
| - | - | 1720 | 619 | - | - | 0 | - |
| 14 | y | 1720 | 625.3 | 0.001577 | 2.522 | +1 | 5 |
| 8 | z | 853.4 | 626.3 | 0.001 | 1.597 | +2 | 11 |
| - | - | 864.7 | 637.3 | - | - | 0 | - |
| - | - | 888.2 | 638.8 | - | - | 0 | - |
| - | - | 1032 | 640.4 | - | - | 0 | - |
| - | - | 1577 | 645.3 | - | - | 0 | - |
| - | - | 3466 | 645.7 | - | - | 0 | - |
| - | - | 1314 | 646 | - | - | 0 | - |
| - | - | 1129 | 646.3 | - | - | 0 | - |
| - | - | 1168 | 651.3 | - | - | 0 | - |
| - | - | 4.589E+04 | 651.3 | - | - | 0 | - |
| - | - | 4.742E+04 | 651.7 | - | - | 0 | - |
| - | - | 2.629E+04 | 652 | - | - | 0 | - |
| - | - | 9459 | 652.3 | - | - | 0 | - |
| - | - | 793.8 | 652.6 | - | - | 0 | - |
| - | - | 2942 | 652.7 | - | - | 0 | - |
| 7 | y | 711.1 | 675.8 | 0.009022 | 13.35 | +2 | 12 |
| 7 | y | 5369 | 676.3 | 0.001274 | 1.883 | +2 | 12 |
| 7 | z | 2932 | 676.8 | 0.003599 | 5.318 | +2 | 12 |
| - | - | 1166 | 677.3 | - | - | 0 | - |
| 13 | w | 4486 | 679.3 | 1.526E-05 | 0.02246 | +1 | 6 |
| - | - | 1117 | 680.3 | - | - | 0 | - |
| - | - | 679.3 | 693.3 | - | - | 0 | - |
| - | - | 1791 | 701.4 | - | - | 0 | - |
| - | - | 673.6 | 702.3 | - | - | 0 | - |
| 7 | c | 1.12E+04 | 702.4 | 0.000342 | 0.4868 | +1 | 7 |
| - | - | 3819 | 703.4 | - | - | 0 | - |
| 13 | z | 5048 | 737.3 | 0.0003733 | 0.5064 | +1 | 6 |
| - | - | 2271 | 738.3 | - | - | 0 | - |
| - | - | 3356 | 741.4 | - | - | 0 | - |
| - | - | 841.5 | 742.4 | - | - | 0 | - |
| 6 | y | 1470 | 749.4 | 0.001204 | 1.606 | +2 | 13 |
| - | - | 704.6 | 749.9 | - | - | 0 | - |
| 13 | y | 939.4 | 753.4 | 0.002984 | 3.961 | +1 | 6 |
| - | - | 668.2 | 759.4 | - | - | 0 | - |
| - | - | 1069 | 799.4 | - | - | 0 | - |
| - | - | 1010 | 799.9 | - | - | 0 | - |
| - | - | 1935 | 802.5 | - | - | 0 | - |
| 8 | c | 1.767E+04 | 803.5 | 0.000413 | 0.514 | +1 | 8 |
| - | - | 7368 | 804.5 | - | - | 0 | - |
| - | - | 1760 | 805.5 | - | - | 0 | - |
| - | - | 616.3 | 808.9 | - | - | 0 | - |
| 4 | z | 1406 | 819.4 | 0.002513 | 3.067 | +2 | 15 |
| - | - | 1908 | 819.9 | - | - | 0 | - |
| - | - | 1758 | 820.4 | - | - | 0 | - |
| 16 | c | 903.2 | 821.4 | 0.002176 | 2.649 | +2 | 16 |
| - | - | 973.7 | 822.4 | - | - | 0 | - |
| - | - | 3096 | 826.9 | - | - | 0 | - |
| 4 | y | 6118 | 827.4 | 0.00217 | 2.623 | +2 | 15 |
| - | - | 4987 | 827.9 | - | - | 0 | - |
| - | - | 2980 | 828.4 | - | - | 0 | - |
| - | - | 2189 | 829.9 | - | - | 0 | - |
| 16 | c | 2.236E+04 | 830.4 | 0.0003172 | 0.3819 | +2 | 16 |
| - | - | 1.987E+04 | 830.9 | - | - | 0 | - |
| - | - | 9388 | 831.4 | - | - | 0 | - |
| - | - | 3489 | 831.9 | - | - | 0 | - |
| 3 | z | 1043 | 846.4 | 0.01253 | 14.8 | +2 | 16 |
| 3 | z | 1.458E+04 | 854.9 | 0.0003177 | 0.3716 | +2 | 16 |
| - | - | 9726 | 855.4 | - | - | 0 | - |
| - | - | 8111 | 855.9 | - | - | 0 | - |
| - | - | 2183 | 856.4 | - | - | 0 | - |
| - | - | 2807 | 862.4 | - | - | 0 | - |
| 3 | y | 1.876E+04 | 862.9 | 0.0007076 | 0.82 | +2 | 16 |
| 12 | y | 1.782E+04 | 863.4 | 0.009685 | 11.22 | +1 | 7 |
| - | - | 8391 | 863.9 | - | - | 0 | - |
| 12 | y | 3511 | 864.4 | 0.009046 | 10.47 | +1 | 7 |
| - | - | 751.9 | 864.9 | - | - | 0 | - |
| 12 | z | 6608 | 865.4 | 0.001281 | 1.48 | +1 | 7 |
| - | - | 1137 | 865.9 | - | - | 0 | - |
| - | - | 2838 | 866.4 | - | - | 0 | - |
| - | - | 2088 | 867.4 | - | - | 0 | - |
| - | - | 826.3 | 868.4 | - | - | 0 | - |
| - | - | 2702 | 872.5 | - | - | 0 | - |
| - | - | 3020 | 873 | - | - | 0 | - |
| - | - | 2300 | 873.5 | - | - | 0 | - |
| 12 | y | 2141 | 881.4 | 0.001878 | 2.13 | +1 | 7 |
| - | - | 1518 | 883.4 | - | - | 0 | - |
| - | - | 696.5 | 886.5 | - | - | 0 | - |
| 9 | c | 1172 | 887.5 | 0.003049 | 3.435 | +1 | 9 |
| 17 | c | 2.773E+04 | 894.5 | 0.0004354 | 0.4867 | +2 | 17 |
| - | - | 2.757E+04 | 895 | - | - | 0 | - |
| - | - | 1.46E+04 | 895.5 | - | - | 0 | - |
| - | - | 4589 | 896 | - | - | 0 | - |
| - | - | 1086 | 896.5 | - | - | 0 | - |
| - | - | 1391 | 897 | - | - | 0 | - |
| - | - | 1171 | 897.5 | - | - | 0 | - |
| - | - | 1153 | 898 | - | - | 0 | - |
| - | - | 730.8 | 908.5 | - | - | 0 | - |
| 2 | z | 4835 | 919 | 0.0003138 | 0.3415 | +2 | 17 |
| - | - | 4184 | 919.5 | - | - | 0 | - |
| - | - | 2463 | 920 | - | - | 0 | - |
| - | - | 1294 | 920.5 | - | - | 0 | - |
| - | - | 703.5 | 923.5 | - | - | 0 | - |
| - | - | 817.4 | 934.5 | - | - | 0 | - |
| - | - | 1124 | 946.5 | - | - | 0 | - |
| - | - | 1606 | 947.5 | - | - | 0 | - |
| - | - | 1790 | 948 | - | - | 0 | - |
| - | - | 1023 | 948.5 | - | - | 0 | - |
| 11 | z | 7801 | 952.5 | 0.0003208 | 0.3368 | +1 | 8 |
| - | - | 7911 | 953.5 | - | - | 0 | - |
| - | - | 982.8 | 954 | - | - | 0 | - |
| - | - | 6889 | 954.5 | - | - | 0 | - |
| - | - | 6917 | 955 | - | - | 0 | - |
| - | - | 4941 | 955.5 | - | - | 0 | - |
| - | - | 1855 | 956 | - | - | 0 | - |
| - | - | 1513 | 968 | - | - | 0 | - |
| 11 | y | 4082 | 968.5 | 0.01236 | 12.76 | +1 | 8 |
| - | - | 5157 | 969 | - | - | 0 | - |
| - | - | 3503 | 969.5 | - | - | 0 | - |
| - | - | 1272 | 970 | - | - | 0 | - |
| - | - | 2.189E+04 | 976.5 | - | - | 0 | - |
| - | - | 2.84E+04 | 977 | - | - | 0 | - |
| - | - | 2.052E+04 | 977.5 | - | - | 0 | - |
| - | - | 9322 | 978 | - | - | 0 | - |
| - | - | 2897 | 978.5 | - | - | 0 | - |
| - | - | 2818 | 1001 | - | - | 0 | - |
| 10 | c | 1.184E+04 | 1002 | 0.0004525 | 0.4518 | +1 | 10 |
| - | - | 6020 | 1003 | - | - | 0 | - |
| - | - | 2295 | 1004 | - | - | 0 | - |
| - | - | 1420 | 1045 | - | - | 0 | - |
| - | - | 812.7 | 1046 | - | - | 0 | - |
| 11 | c | 1.18E+04 | 1089 | 0.0003764 | 0.3458 | +1 | 11 |
| - | - | 7026 | 1090 | - | - | 0 | - |
| - | - | 2231 | 1091 | - | - | 0 | - |
| - | - | 976.3 | 1107 | - | - | 0 | - |
| - | - | 821.7 | 1108 | - | - | 0 | - |
| 9 | y | 854.1 | 1149 | 0.01978 | 17.22 | +1 | 10 |
| 9 | y | 2781 | 1150 | 0.0007438 | 0.647 | +1 | 10 |
| 9 | z | 1.286E+04 | 1151 | 0.001367 | 1.188 | +1 | 10 |
| - | - | 1.052E+04 | 1152 | - | - | 0 | - |
| - | - | 5111 | 1153 | - | - | 0 | - |
| - | - | 1355 | 1154 | - | - | 0 | - |
| 9 | y | 816.7 | 1167 | 0.008107 | 6.95 | +1 | 10 |
| - | - | 717 | 1168 | - | - | 0 | - |
| - | - | 726.1 | 1188 | - | - | 0 | - |
| 12 | c | 1.233E+04 | 1217 | 0.0003687 | 0.3031 | +1 | 12 |
| - | - | 6857 | 1218 | - | - | 0 | - |
| - | - | 3081 | 1219 | - | - | 0 | - |
| 8 | z | 6929 | 1252 | 0.001072 | 0.8561 | +1 | 11 |
| - | - | 6442 | 1253 | - | - | 0 | - |
| - | - | 2386 | 1254 | - | - | 0 | - |
| 8 | y | 887.8 | 1268 | 0.005147 | 4.06 | +1 | 11 |
| - | - | 676.3 | 1296 | - | - | 0 | - |
| - | - | 778.7 | 1301 | - | - | 0 | - |
| - | - | 7241 | 1344 | - | - | 0 | - |
| 13 | c | 9558 | 1345 | 0.004991 | 3.712 | +1 | 13 |
| - | - | 4915 | 1346 | - | - | 0 | - |
| - | - | 2009 | 1347 | - | - | 0 | - |
| 7 | z | 1.046E+04 | 1353 | 0.001753 | 1.296 | +1 | 12 |
| - | - | 1.042E+04 | 1354 | - | - | 0 | - |
| - | - | 4951 | 1355 | - | - | 0 | - |
| - | - | 1240 | 1356 | - | - | 0 | - |
| 7 | y | 3499 | 1369 | 0.0009457 | 0.691 | +1 | 12 |
| - | - | 2536 | 1370 | - | - | 0 | - |
| - | - | 699.2 | 1419 | - | - | 0 | - |
| - | - | 3479 | 1431 | - | - | 0 | - |
| 14 | c | 1.255E+04 | 1432 | 0.003328 | 2.324 | +1 | 14 |
| - | - | 8439 | 1433 | - | - | 0 | - |
| - | - | 4614 | 1434 | - | - | 0 | - |
| - | - | 1106 | 1475 | - | - | 0 | - |
| 6 | z | 1.298E+04 | 1482 | 0.002842 | 1.918 | +1 | 13 |
| - | - | 1.185E+04 | 1483 | - | - | 0 | - |
| - | - | 5529 | 1484 | - | - | 0 | - |
| - | - | 1307 | 1485 | - | - | 0 | - |
| 6 | y | 972 | 1498 | 0.004923 | 3.287 | +1 | 13 |
| - | - | 1055 | 1510 | - | - | 0 | - |
| - | - | 7284 | 1545 | - | - | 0 | - |
| 15 | c | 1.537E+04 | 1546 | 0.005972 | 3.864 | +1 | 15 |
| - | - | 1.197E+04 | 1547 | - | - | 0 | - |
| - | - | 4385 | 1548 | - | - | 0 | - |
| - | - | 1407 | 1549 | - | - | 0 | - |
| 5 | z | 7768 | 1581 | 0.002775 | 1.755 | +1 | 14 |
| - | - | 1.14E+04 | 1582 | - | - | 0 | - |
| - | - | 6705 | 1583 | - | - | 0 | - |
| - | - | 1581 | 1584 | - | - | 0 | - |
| 5 | y | 1318 | 1597 | 0.0002299 | 0.144 | +1 | 14 |
| - | - | 971.4 | 1598 | - | - | 0 | - |
| - | - | 758.8 | 1618 | - | - | 0 | - |
| 4 | z | 2103 | 1638 | 0.001777 | 1.085 | +1 | 15 |
| - | - | 9088 | 1639 | - | - | 0 | - |
| - | - | 6208 | 1640 | - | - | 0 | - |
| - | - | 3802 | 1641 | - | - | 0 | - |
| - | - | 1589 | 1642 | - | - | 0 | - |
| - | - | 855.2 | 1653 | - | - | 0 | - |
| 4 | y | 2993 | 1654 | 0.002557 | 1.546 | +1 | 15 |
| - | - | 2441 | 1655 | - | - | 0 | - |
| - | - | 1370 | 1656 | - | - | 0 | - |
| - | - | 846.9 | 1659 | - | - | 0 | - |
| 16 | c | 1.672E+04 | 1660 | 0.0004379 | 0.2638 | +1 | 16 |
| - | - | 1.363E+04 | 1661 | - | - | 0 | - |
| - | - | 6736 | 1662 | - | - | 0 | - |
| - | - | 2392 | 1663 | - | - | 0 | - |
| 3 | z | 2193 | 1709 | 0.0004155 | 0.2431 | +1 | 16 |
| - | - | 1.432E+04 | 1710 | - | - | 0 | - |
| - | - | 9751 | 1711 | - | - | 0 | - |
| - | - | 5112 | 1712 | - | - | 0 | - |
| - | - | 1776 | 1713 | - | - | 0 | - |
| 3 | y | 2030 | 1725 | 0.003786 | 2.195 | +1 | 16 |
| - | - | 2210 | 1726 | - | - | 0 | - |
| - | - | 1076 | 1727 | - | - | 0 | - |
| - | - | 797.4 | 1737 | - | - | 0 | - |
| - | - | 2223 | 1745 | - | - | 0 | - |
| - | - | 988.7 | 1746 | - | - | 0 | - |
| - | - | 1153 | 1747 | - | - | 0 | - |
| 17 | c | 2907 | 1788 | 0.004337 | 2.425 | +1 | 17 |
| - | - | 5392 | 1789 | - | - | 0 | - |
| - | - | 4761 | 1790 | - | - | 0 | - |
| - | - | 2162 | 1791 | - | - | 0 | - |
| 2 | z | 996.9 | 1837 | 0.00848 | 4.616 | +1 | 17 |
| - | - | 6016 | 1838 | - | - | 0 | - |
| - | - | 5201 | 1839 | - | - | 0 | - |
| - | - | 2883 | 1840 | - | - | 0 | - |
| - | - | 810.8 | 1841 | - | - | 0 | - |
| - | - | 942.4 | 1895 | - | - | 0 | - |
| - | - | 1641 | 1908 | - | - | 0 | - |
| - | - | 6157 | 1909 | - | - | 0 | - |
| - | - | 5639 | 1910 | - | - | 0 | - |
| - | - | 3819 | 1911 | - | - | 0 | - |
| - | - | 880.3 | 1912 | - | - | 0 | - |
| - | - | 1103 | 1925 | - | - | 0 | - |
| - | - | 5656 | 1926 | - | - | 0 | - |
| - | - | 4111 | 1927 | - | - | 0 | - |
| - | - | 3009 | 1928 | - | - | 0 | - |
| - | - | 2939 | 1936 | - | - | 0 | - |
| - | - | 7370 | 1937 | - | - | 0 | - |
| - | - | 5086 | 1938 | - | - | 0 | - |
| - | - | 2892 | 1939 | - | - | 0 | - |
| - | - | 1138 | 1940 | - | - | 0 | - |
| - | - | 3271 | 1952 | - | - | 0 | - |
| - | - | 1.579E+04 | 1953 | - | - | 0 | - |
| - | - | 5.776E+04 | 1954 | - | - | 0 | - |
| - | - | 5.27E+04 | 1955 | - | - | 0 | - |
| - | - | 2.947E+04 | 1956 | - | - | 0 | - |
| - | - | 1.278E+04 | 1957 | - | - | 0 | - |
| - | - | 2649 | 1958 | - | - | 0 | - |
| - | - | 692.1 | 3264 | - | - | 0 | - |

m/z Charge Intensity FragmentType MassShift Position
125.42489624023438 0 421.67044
129.09786987304688 0 549.54474
129.1021728515625 0 7171.3584
133.00648498535156 0 436.29825
148.45616149902344 0 481.36313
148.8418426513672 0 495.4685
148.88514709472656 0 755.71564
148.892822265625 0 500.9976
148.8995819091797 0 634.84326
148.90676879882812 0 629.95087
148.91378784179688 0 710.09924
148.92115783691406 0 1165.1993
148.92832946777344 0 1178.3838
148.9354705810547 0 2427.1025
148.94317626953125 0 3937.2927
148.95970153808594 0 4313.845
148.96742248535156 0 2556.2178
148.97451782226562 0 1419.2957
148.98159790039062 0 1240.6931
148.98922729492188 0 935.0716
148.996337890625 0 697.68384
149.0108642578125 0 608.9481
149.01780700683594 0 676.0919
149.0325164794922 0 514.0444
149.19688415527344 0 468.51328
187.14395141601562 0 520.976
188.11517333984375 0 603.30597
210.3145294189453 0 501.62024
224.74156188964844 0 582.82306
227.16354370117188 0 579.1815
228.1340789794922 0 821.64294
228.1705322265625 0 3674.0066 c Ammonia loss 1
253.21580505371094 0 608.90314
273.22723388671875 0 825.0527
294.1573181152344 0 4527.9062 z 16
295.1630859375 0 1136.2195
299.20751953125 0 918.4413 c Ammonia loss 2
306.2005310058594 0 553.1504
315.2266540527344 0 2740.6802
316.23388671875 0 1884.7183 c 2
329.2416076660156 0 675.3122
330.2490539550781 0 780.33386
337.127685546875 0 602.27277
355.0696105957031 0 1360.0092
356.229248046875 0 1285.093 c Ammonia loss 3
364.2106628417969 0 1171.2795
373.2555236816406 0 13414.918 c 3
374.258544921875 0 2031.8467
384.2255859375 0 869.92035
408.2002868652344 0 5502.919 z 15
409.2076721191406 0 7643.122
410.21124267578125 0 1918.038
427.3031311035156 0 1146.2104
429.0885009765625 0 2226.9065
429.3193664550781 0 967.1791
430.0885314941406 0 847.677
455.2980041503906 0 1804.8768 c Ammonia loss 4
472.3240661621094 0 12156.052 c 4
473.3268737792969 0 2705.5486
478.2292175292969 0 1571.2189 w 14
485.5995788574219 0 668.39777
498.3398132324219 0 1322.6566
506.91650390625 0 905.18884
507.24591064453125 0 674.5755
522.2433471679688 0 7643.7 z 14
523.2496337890625 0 7440.47
524.2518920898438 0 2123.673
533.7730712890625 0 743.95197
546.7616577148438 0 1624.7037
556.9356689453125 0 594.33234
557.3536987304688 0 2052.27
558.2704467773438 0 738.6258
558.35595703125 0 763.12683
575.7833251953125 0 813.38104 z 8
584.97998046875 0 737.88275
600.3582763671875 0 1445.5804
601.3662109375 0 12100.481 c 5
602.369873046875 0 4018.6924
603.3743896484375 0 732.8349
607.2568359375 0 1012.34033
608.268310546875 0 809.5043 y Ammonia loss 13
609.2738037109375 0 2346.4124 z 13
610.2827758789062 0 11492.224
611.285400390625 0 3721.9624
618.3156127929688 0 982.25146 y 1
618.6521606445312 0 1350.2719
618.9868774414062 0 1720.109
625.2955932617188 0 1719.9562 y 13
626.30712890625 0 853.39557 z 7
637.3290405273438 0 864.7044
638.8197631835938 0 888.15546
640.386474609375 0 1031.5905
645.3389892578125 0 1577.3931
645.6683959960938 0 3466.2798
646.0009765625 0 1313.672
646.3319702148438 0 1129.4368
651.2785034179688 0 1168.3119
651.3406372070312 0 45892.23
651.6749267578125 0 47415.71
652.0089111328125 0 26290.21
652.343017578125 0 9458.89
652.6195068359375 0 793.7944
652.6788330078125 0 2942.315
675.8270263671875 0 711.0643 y Water loss 6
676.3267822265625 0 5368.678 y Ammonia loss 6
676.828369140625 0 2931.6648 z 6
677.3294067382812 0 1165.7366
679.3045654296875 0 4485.7393 w 12
680.3093872070312 0 1116.7509
693.3229370117188 0 679.27997
701.4071655273438 0 1790.5243
702.34375 0 673.62555
702.4141235351562 0 11196.336 c 6
703.4169921875 0 3818.6423
737.33349609375 0 5048.4336 z 12
738.3380126953125 0 2271.3684
741.4381713867188 0 3356.1174
742.4467163085938 0 841.4555
749.3638305664062 0 1470.0125 y 5
749.8715209960938 0 704.5939
753.349609375 0 939.4473 y 12
759.3826904296875 0 668.2495
799.3968505859375 0 1069.4636
799.9057006835938 0 1009.80035
802.45458984375 0 1935.4646
803.4617309570312 0 17669.002 c 7
804.4656982421875 0 7367.869
805.46630859375 0 1759.6947
808.9217529296875 0 616.3253
819.3956909179688 0 1406.4658 z 3
819.9043579101562 0 1908.3236
820.3992309570312 0 1758.361
821.42919921875 0 903.1877 c Water loss 15
822.4338989257812 0 973.7422
826.9031982421875 0 3095.9946
827.4053955078125 0 6118.154 y 3
827.908203125 0 4986.8164
828.4108276367188 0 2979.6165
829.9343872070312 0 2188.9658
830.4363403320312 0 22355.285 c 15
830.937255859375 0 19869.396
831.4386596679688 0 9387.713
831.940185546875 0 3488.7454
846.416015625 0 1042.5334 z Ammonia loss 2
854.9164428710938 0 14580.005 z 2
855.4178466796875 0 9726.102
855.9189453125 0 8110.737
856.4214477539062 0 2183.2253
862.4200439453125 0 2807.2373
862.9254150390625 0 18764.844 y 2
863.4273071289062 0 17822.27 y Water loss 11
863.9280395507812 0 8391.379
864.4300537109375 0 3511.2053 y Ammonia loss 11
864.9339599609375 0 751.9225
865.4275512695312 0 6607.697 z 11
865.9442749023438 0 1137.0765
866.432373046875 0 2838.104
867.4319458007812 0 2087.993
868.430419921875 0 826.2763
872.4757080078125 0 2702.075
872.9764404296875 0 3020.426
873.4775390625 0 2299.8333
881.4456787109375 0 2140.523 y 11
883.4483642578125 0 1517.9609
886.4773559570312 0 696.4962
887.480224609375 0 1172.3118 c Ammonia loss 8
894.4837036132812 0 27734.092 c 16
894.9846801757812 0 27570.012
895.4856567382812 0 14603.431
895.9862060546875 0 4589.2305
896.4892578125 0 1085.7028
896.9696044921875 0 1390.8094
897.4703369140625 0 1171.0023
897.9669799804688 0 1153.2512
908.4744873046875 0 730.7873
918.9639282226562 0 4835.0605 z 1
919.46484375 0 4183.8423
919.9677734375 0 2463.2048
920.4667358398438 0 1293.7568
923.4901733398438 0 703.46857
934.4890747070312 0 817.41046
946.50390625 0 1123.5306
947.4925537109375 0 1606.1208
947.99658203125 0 1790.2965
948.4976196289062 0 1022.8856
952.461181640625 0 7800.6475 z 10
953.4667358398438 0 7910.715
954.0136108398438 0 982.83276
954.4927368164062 0 6888.552
955.0035400390625 0 6916.7417
955.5015258789062 0 4941.4478
956.0086669921875 0 1854.5845
968.0027465820312 0 1513.1051
968.491943359375 0 4081.6223 y 10
969.0013427734375 0 5157.0493
969.5008544921875 0 3502.9316
970.00439453125 0 1271.867
976.5071411132812 0 21893.357
977.009521484375 0 28395.441
977.5110473632812 0 20515.162
978.0123291015625 0 9321.917
978.5155029296875 0 2897.2249
1000.55615234375 0 2817.7998
1001.5621337890625 0 11838.778 c 9
1002.5656127929688 0 6020.404
1003.5672607421875 0 2295.2668
1044.58056640625 0 1420.2977
1045.5880126953125 0 812.6858
1088.59423828125 0 11801.826 c 10
1089.5975341796875 0 7026.3896
1090.5987548828125 0 2231.4114
1106.556640625 0 976.2546
1107.5516357421875 0 821.7115
1148.5496826171875 0 854.11597 y Water loss 8
1149.552734375 0 2780.7073 y Ammonia loss 8
1150.5599365234375 0 12855.005 z 8
1151.56591796875 0 10524.849
1152.56787109375 0 5111.1377
1153.5684814453125 0 1354.5166
1166.588134765625 0 816.7042 y 8
1167.5904541015625 0 717.03375
1187.6121826171875 0 726.13916
1216.689208984375 0 12334.062 c 11
1217.6912841796875 0 6856.899
1218.6968994140625 0 3081.1846
1251.60791015625 0 6929.191 z 7
1252.612060546875 0 6442.4346
1253.6129150390625 0 2386.2075
1267.62255859375 0 887.8387 y 7
1295.5936279296875 0 676.3401
1300.6700439453125 0 778.65894
1343.73876953125 0 7241.3823
1344.7431640625 0 9558.331 c 12
1345.7476806640625 0 4915.1133
1346.7545166015625 0 2008.5579
1352.6549072265625 0 10455.348 z 6
1353.6588134765625 0 10419.56
1354.6627197265625 0 4951.0645
1355.664794921875 0 1239.5481
1368.6744384765625 0 3499.4038 y 6
1369.6761474609375 0 2535.8286
1418.7147216796875 0 699.23346
1430.7705078125 0 3478.5137
1431.77685546875 0 12547.591 c 13
1432.781005859375 0 8438.693
1433.7822265625 0 4613.588
1474.7435302734375 0 1105.7892
1481.6964111328125 0 12975.231 z 5
1482.7012939453125 0 11845.8125
1483.70361328125 0 5528.675
1484.704833984375 0 1307.3492
1497.722900390625 0 971.9729 y 5
1509.72119140625 0 1055.4388
1544.808837890625 0 7284.156
1545.817138671875 0 15367.374 c 14
1546.8187255859375 0 11968.445
1547.8271484375 0 4384.8076
1548.8228759765625 0 1407.1989
1580.764892578125 0 7768.4067 z 4
1581.7672119140625 0 11397.38
1582.7725830078125 0 6705.427
1583.7728271484375 0 1581.4261
1596.78662109375 0 1318.1844 y 4
1597.7900390625 0 971.4325
1617.8433837890625 0 758.82355
1637.787353515625 0 2102.8296 z 3
1638.794921875 0 9087.967
1639.799072265625 0 6207.5024
1640.7987060546875 0 3801.6257
1641.8006591796875 0 1588.9121
1652.81201171875 0 855.19006
1653.8052978515625 0 2992.605 y 3
1654.8123779296875 0 2440.5906
1655.8067626953125 0 1369.7146
1658.85595703125 0 846.9172
1659.8656005859375 0 16721.297 c 15
1660.866455078125 0 13633.997
1661.8702392578125 0 6736.214
1662.8720703125 0 2392.4949
1708.82666015625 0 2193.1663 z 2
1709.831787109375 0 14322.971
1710.8345947265625 0 9750.617
1711.838134765625 0 5111.905
1712.84521484375 0 1775.8683
1724.8487548828125 0 2029.5032 y 2
1725.843505859375 0 2210.199
1726.856689453125 0 1076.285
1736.8739013671875 0 797.37286
1744.962646484375 0 2222.9968
1745.949462890625 0 988.71466
1746.9471435546875 0 1152.7976
1787.9566650390625 0 2906.8403 c 16
1788.961669921875 0 5391.9575
1789.9658203125 0 4760.8525
1790.9674072265625 0 2162.4192
1836.9296875 0 996.86194 z 1
1837.9251708984375 0 6015.858
1838.9298095703125 0 5200.7764
1839.93017578125 0 2882.5178
1840.929443359375 0 810.7842
1894.9732666015625 0 942.4224
1908.0113525390625 0 1641.4553
1909.002685546875 0 6157.406
1910.0050048828125 0 5638.6895
1911.0098876953125 0 3819.1833
1912.0133056640625 0 880.25397
1925.02001953125 0 1103.1328
1926.0228271484375 0 5656.1704
1927.03173828125 0 4110.8145
1928.03466796875 0 3009.134
1936.0054931640625 0 2938.6045
1936.9969482421875 0 7369.536
1938.001220703125 0 5086.0503
1939.0006103515625 0 2892.3308
1940.010498046875 0 1138.1235
1951.9937744140625 0 3271.4404
1953.0107421875 0 15790.651
1954.0179443359375 0 57758.22
1955.02197265625 0 52698.18
1956.0245361328125 0 29466.34
1957.0269775390625 0 12781.426
1958.0247802734375 0 2648.6343
3263.7900390625 0 692.06683

Spectrum Details

|  |  |
| --- | --- |
| Matched peaks? Matched peaks The total absolute number of peaks matched. Additionally in brackets the total fraction of peaks matched and the total number of peaks is shown. | 68 (21.05% of 323) |
| FDR? FDR The false discovery rate estimated for this peptide. It is calculated by matching all theoretical fragments with a non-integer shift with the raw peaks for this spectrum. This is done with 40 different shifts. The resulting percentage is the average number of annotated peaks over the number of annotated peaks with the correct spectrum. | 3.36% |
| Satellite FDR? Satellite FDR See the FDR for details on its calculation. This satellite ion specific FDR only contains the satellite ions (d/w) for I/L/J positions. | - |
| PSM Score? PSM Score The PSM Score as given by Hecklib to this annotated spectrum. It is shown with three significant figures. | 627 |

#### Spectrum 2473? Spectrum 2473 The raw spectrum of this peptide as annotated by Hecklib. The fragments are coloured according to ion type (see legend). Any peaks with a star '\*' as text can be hovered over to see the full details, first the ion type second the mass shift type. By hovering over the amino acids in the peptide or ions in the legend the corresponding peaks are highlighted. By toggling the 'Unassigned' label you can turn the background (unassigned) peaks on or off in the plot. By updating the slider in the Ion legend you can update the spectrum to only show the top X% of the peaks with labels. The top X% means any peak that is within X% of the highest intensity. By dragging in the spectrum you can zoom in to a specific part of the spectrum and use 'Zoom Out' to get back to the original zoom level. The annotation of the spectrum is based on the given sequence in the peptides file and is done with different software so inconsistencies are likely. The peaks are annotated based on the given sequence, with 20 ppm tolerance.

Copy Data

##### Spectrum 2473 (TSV)

###### Preview

```
Loading example...
```

*Click on the button to copy the data to your clipboard.*

Mz MinMz MaxIntensity Max

WidthHeightPeptide font sizePeptide stroke widthSpectrum font sizeSpectrum stroke widthCompact peptide

Ion legend

wxyz

abcd

OtherUnassignedIon^Charge^_Position_Show for top:%

VKAGVETTTPSKQSNNKY

01.38e+42.76e+44.13e+45.51e+4

Zoom Out

y^+1^_1_c^+1^_2_c^+1^_2_c^+2^_6_z^+1^_2_c^+1^_3_y^+1^_2_c^+1^_3_y^+3^_9_c^+1^_4_w^+1^_3_c^+1^_4_y^+3^_10_z^+1^_3_y^+3^_11_y^+1^_3_z^+2^_7_y^+2^_7_c^+1^_5_w^+2^_8_c^+1^_5_y^+2^_8_z^+2^_8_w^+1^_4_y^+2^_8_w^+2^_9_z^+1^_4_y^+2^_9_y^+1^_4_c^+2^_11_w^+2^_10_w^+2^_10_y^+2^_10_z^+2^_10_y^+2^_10_c^+1^_6_w^+1^_5_c^+3^_17_c^+1^_6_y^+1^_5_c^+2^_12_z^+1^_5_z^+3^_17_y^+3^_17_w^+2^_11_y^+1^_5_y^+2^_11_z^+2^_11_y^+2^_11_z^+2^_12_z^+2^_12_w^+2^_12_c^+2^_13_y^+2^_12_y^+2^_12_z^+2^_12_w^+1^_6_y^+2^_12_c^+1^_7_w^+2^_13_c^+2^_14_z^+2^_13_z^+1^_6_y^+2^_13_z^+2^_13_y^+2^_13_y^+1^_6_c^+2^_15_c^+1^_8_y^+2^_14_z^+2^_14_y^+2^_14_c^+1^_8_w^+1^_7_z^+2^_15_y^+2^_15_c^+2^_16_z^+2^_16_y^+2^_16_y^+1^_7_z^+1^_7_y^+1^_7_c^+1^_9_w^+2^_17_c^+2^_17_z^+2^_17_y^+2^_17_w^+1^_8_y^+1^_8_z^+1^_8_c^+1^_10_c^+1^_11_z^+1^_10_y^+1^_10_z^+1^_10_y^+1^_10_c^+1^_12_z^+1^_11_z^+1^_12_c^+1^_13_z^+1^_12_c^+1^_14_z^+1^_13_c^+1^_15_z^+1^_14_z^+1^_15_y^+1^_15_c^+1^_16_

0628125518832510

Fragment Matches Table

Show background peaks

| Position | Ion type | Intensity | mz Theoretical | mz Error (Th) | mz Error (ppm) | Charge | Series Number |
| --- | --- | --- | --- | --- | --- | --- | --- |
| - | - | 1212 | 120.1 | - | - | 0 | - |
| - | - | 371.6 | 121.9 | - | - | 0 | - |
| - | - | 530 | 127.1 | - | - | 0 | - |
| - | - | 670.5 | 128.1 | - | - | 0 | - |
| - | - | 1.849E+04 | 129.1 | - | - | 0 | - |
| - | - | 855.2 | 130.1 | - | - | 0 | - |
| - | - | 2416 | 131.1 | - | - | 0 | - |
| - | - | 501.4 | 131.1 | - | - | 0 | - |
| - | - | 1752 | 136.1 | - | - | 0 | - |
| - | - | 397.7 | 138.5 | - | - | 0 | - |
| - | - | 446.3 | 140.6 | - | - | 0 | - |
| - | - | 489.8 | 141.4 | - | - | 0 | - |
| - | - | 385.6 | 141.5 | - | - | 0 | - |
| - | - | 436.6 | 143.8 | - | - | 0 | - |
| - | - | 1467 | 146.1 | - | - | 0 | - |
| - | - | 925.5 | 149 | - | - | 0 | - |
| - | - | 1786 | 155.1 | - | - | 0 | - |
| - | - | 1485 | 164 | - | - | 0 | - |
| - | - | 640.4 | 171.1 | - | - | 0 | - |
| - | - | 3316 | 173.4 | - | - | 0 | - |
| 18 | y | 1283 | 182.1 | 0.0003286 | 1.805 | +1 | 1 |
| - | - | 436 | 183.1 | - | - | 0 | - |
| - | - | 894.5 | 183.1 | - | - | 0 | - |
| - | - | 3373 | 188.1 | - | - | 0 | - |
| - | - | 716.4 | 189.1 | - | - | 0 | - |
| - | - | 472.6 | 204.6 | - | - | 0 | - |
| - | - | 488.8 | 207.3 | - | - | 0 | - |
| - | - | 885.9 | 210.1 | - | - | 0 | - |
| - | - | 1089 | 211.1 | - | - | 0 | - |
| - | - | 778.6 | 217.2 | - | - | 0 | - |
| - | - | 536.9 | 217.4 | - | - | 0 | - |
| - | - | 1228 | 227.1 | - | - | 0 | - |
| - | - | 1644 | 228.1 | - | - | 0 | - |
| 2 | c | 7288 | 228.2 | 1.435E-05 | 0.06289 | +1 | 2 |
| - | - | 899.9 | 229.2 | - | - | 0 | - |
| - | - | 521.5 | 231 | - | - | 0 | - |
| - | - | 1668 | 245.2 | - | - | 0 | - |
| 2 | c | 581.3 | 245.2 | 5.893E-05 | 0.2404 | +1 | 2 |
| - | - | 1504 | 249.2 | - | - | 0 | - |
| - | - | 831.1 | 250.2 | - | - | 0 | - |
| - | - | 530.7 | 265.2 | - | - | 0 | - |
| - | - | 597.1 | 266.2 | - | - | 0 | - |
| 6 | c | 781.3 | 292.7 | 0.0004066 | 1.389 | +2 | 6 |
| - | - | 653 | 293.1 | - | - | 0 | - |
| 17 | z | 1561 | 294.2 | 0.0001536 | 0.5223 | +1 | 2 |
| 3 | c | 1138 | 299.2 | 0.0003392 | 1.134 | +1 | 3 |
| - | - | 2734 | 302.2 | - | - | 0 | - |
| 17 | y | 1221 | 310.2 | 1.575E-05 | 0.05078 | +1 | 2 |
| - | - | 491.4 | 311.4 | - | - | 0 | - |
| 3 | c | 1666 | 316.2 | 0.0004249 | 1.344 | +1 | 3 |
| - | - | 904.4 | 337.1 | - | - | 0 | - |
| 10 | y | 1353 | 355.8 | 0.001311 | 3.684 | +3 | 9 |
| - | - | 757.9 | 356.2 | - | - | 0 | - |
| 4 | c | 881.3 | 356.2 | 0.001326 | 3.721 | +1 | 4 |
| - | - | 908.7 | 359.2 | - | - | 0 | - |
| - | - | 715.7 | 363.2 | - | - | 0 | - |
| 16 | w | 680.8 | 364.2 | 0.001443 | 3.964 | +1 | 3 |
| - | - | 672.9 | 370.7 | - | - | 0 | - |
| 4 | c | 8048 | 373.3 | 0.0002258 | 0.605 | +1 | 4 |
| - | - | 1114 | 374.3 | - | - | 0 | - |
| - | - | 1437 | 384.2 | - | - | 0 | - |
| - | - | 874.2 | 384.7 | - | - | 0 | - |
| - | - | 740.6 | 385.2 | - | - | 0 | - |
| - | - | 585.5 | 388.8 | - | - | 0 | - |
| 9 | y | 905 | 389.5 | 0.001314 | 3.374 | +3 | 10 |
| - | - | 647.3 | 404.2 | - | - | 0 | - |
| - | - | 1263 | 404.7 | - | - | 0 | - |
| 16 | z | 4602 | 408.2 | 0.0002865 | 0.7018 | +1 | 3 |
| - | - | 2210 | 409.2 | - | - | 0 | - |
| - | - | 1439 | 416.2 | - | - | 0 | - |
| 8 | y | 775.1 | 423.2 | 0.003393 | 8.017 | +3 | 11 |
| 16 | y | 1798 | 424.2 | 9.652E-05 | 0.2275 | +1 | 3 |
| - | - | 612.2 | 427.2 | - | - | 0 | - |
| - | - | 1241 | 427.3 | - | - | 0 | - |
| - | - | 739.8 | 428.3 | - | - | 0 | - |
| - | - | 600.6 | 428.9 | - | - | 0 | - |
| - | - | 558 | 429.1 | - | - | 0 | - |
| - | - | 835.1 | 430.9 | - | - | 0 | - |
| 12 | z | 633 | 433.2 | 0.0006955 | 1.606 | +2 | 7 |
| - | - | 681.2 | 435.2 | - | - | 0 | - |
| 12 | y | 2077 | 441.2 | 0.0004277 | 0.9694 | +2 | 7 |
| - | - | 1191 | 441.7 | - | - | 0 | - |
| - | - | 988.6 | 442.7 | - | - | 0 | - |
| - | - | 674 | 446.2 | - | - | 0 | - |
| - | - | 834 | 448.2 | - | - | 0 | - |
| - | - | 958.1 | 454.7 | - | - | 0 | - |
| - | - | 744 | 455.2 | - | - | 0 | - |
| 5 | c | 3415 | 455.3 | 0.0001457 | 0.32 | +1 | 5 |
| 11 | w | 4030 | 468.2 | 3.282E-05 | 0.07009 | +2 | 8 |
| - | - | 2439 | 468.7 | - | - | 0 | - |
| - | - | 822.7 | 469 | - | - | 0 | - |
| - | - | 610.3 | 470.7 | - | - | 0 | - |
| 5 | c | 3.145E+04 | 472.3 | 2.483E-05 | 0.05258 | +1 | 5 |
| - | - | 581.9 | 473.2 | - | - | 0 | - |
| - | - | 8678 | 473.3 | - | - | 0 | - |
| - | - | 596.6 | 474.3 | - | - | 0 | - |
| - | - | 612 | 475.8 | - | - | 0 | - |
| 11 | y | 1043 | 476.2 | 0.0002977 | 0.6252 | +2 | 8 |
| 11 | z | 801.8 | 476.7 | 0.001344 | 2.819 | +2 | 8 |
| - | - | 698.6 | 477.2 | - | - | 0 | - |
| 15 | w | 1.022E+04 | 478.2 | 0.0003556 | 0.7437 | +1 | 4 |
| - | - | 3445 | 479.2 | - | - | 0 | - |
| - | - | 3601 | 479.8 | - | - | 0 | - |
| - | - | 3148 | 480 | - | - | 0 | - |
| - | - | 1144 | 480.3 | - | - | 0 | - |
| - | - | 595.5 | 480.5 | - | - | 0 | - |
| - | - | 1265 | 484.3 | - | - | 0 | - |
| 11 | y | 1078 | 484.7 | 0.004128 | 8.516 | +2 | 8 |
| - | - | 747.4 | 488.2 | - | - | 0 | - |
| - | - | 1161 | 489.1 | - | - | 0 | - |
| - | - | 839.4 | 493.3 | - | - | 0 | - |
| - | - | 1937 | 498.3 | - | - | 0 | - |
| - | - | 965.1 | 499.3 | - | - | 0 | - |
| - | - | 2651 | 503.2 | - | - | 0 | - |
| - | - | 923.6 | 504.2 | - | - | 0 | - |
| 10 | w | 1784 | 511.7 | 0.0003409 | 0.6662 | +2 | 9 |
| - | - | 993.5 | 512.3 | - | - | 0 | - |
| - | - | 683.5 | 515.3 | - | - | 0 | - |
| - | - | 1020 | 521.8 | - | - | 0 | - |
| 15 | z | 4258 | 522.2 | 0.000694 | 1.329 | +1 | 4 |
| - | - | 618.4 | 522.8 | - | - | 0 | - |
| - | - | 2543 | 523.2 | - | - | 0 | - |
| - | - | 1086 | 532.3 | - | - | 0 | - |
| - | - | 580.5 | 532.8 | - | - | 0 | - |
| 10 | y | 1.201E+04 | 533.3 | 0.0001593 | 0.2987 | +2 | 9 |
| - | - | 6763 | 533.8 | - | - | 0 | - |
| - | - | 2929 | 534.3 | - | - | 0 | - |
| 15 | y | 1697 | 538.3 | 0.0008908 | 1.655 | +1 | 4 |
| 11 | c | 893.8 | 544.8 | 0.0002254 | 0.4137 | +2 | 11 |
| - | - | 1033 | 546.8 | - | - | 0 | - |
| - | - | 1267 | 547.3 | - | - | 0 | - |
| - | - | 680 | 552.8 | - | - | 0 | - |
| - | - | 976.9 | 553.3 | - | - | 0 | - |
| - | - | 3037 | 553.8 | - | - | 0 | - |
| - | - | 1970 | 554.3 | - | - | 0 | - |
| - | - | 637.4 | 554.8 | - | - | 0 | - |
| - | - | 1191 | 557.4 | - | - | 0 | - |
| - | - | 892.9 | 558.4 | - | - | 0 | - |
| - | - | 931.6 | 559.3 | - | - | 0 | - |
| - | - | 5638 | 565.3 | - | - | 0 | - |
| - | - | 1542 | 566.3 | - | - | 0 | - |
| 9 | w | 2788 | 567.3 | 8.308E-05 | 0.1465 | +2 | 10 |
| - | - | 2348 | 567.8 | - | - | 0 | - |
| 9 | w | 6383 | 568.3 | 0.0009463 | 1.665 | +2 | 10 |
| - | - | 2125 | 568.8 | - | - | 0 | - |
| - | - | 1778 | 569.3 | - | - | 0 | - |
| - | - | 3974 | 574.8 | - | - | 0 | - |
| 9 | y | 2.157E+04 | 575.3 | 0.000287 | 0.4988 | +2 | 10 |
| 9 | z | 1.552E+04 | 575.8 | 0.002246 | 3.901 | +2 | 10 |
| - | - | 7739 | 576.3 | - | - | 0 | - |
| - | - | 2752 | 576.8 | - | - | 0 | - |
| - | - | 1096 | 577.3 | - | - | 0 | - |
| - | - | 1442 | 582 | - | - | 0 | - |
| - | - | 1323 | 582.3 | - | - | 0 | - |
| 9 | y | 6901 | 583.8 | 4.931E-05 | 0.08447 | +2 | 10 |
| - | - | 5678 | 584.3 | - | - | 0 | - |
| 6 | c | 760.5 | 584.3 | 0.0008214 | 1.406 | +1 | 6 |
| - | - | 2412 | 584.8 | - | - | 0 | - |
| 14 | w | 2453 | 592.3 | 0.001373 | 2.318 | +1 | 5 |
| - | - | 1158 | 593.3 | - | - | 0 | - |
| 17 | c | 3.256E+04 | 596.7 | 0.000254 | 0.4257 | +3 | 17 |
| - | - | 2.537E+04 | 597 | - | - | 0 | - |
| - | - | 1.429E+04 | 597.3 | - | - | 0 | - |
| - | - | 4974 | 597.7 | - | - | 0 | - |
| - | - | 2800 | 598 | - | - | 0 | - |
| 6 | c | 4.146E+04 | 601.4 | 0.0001488 | 0.2475 | +1 | 6 |
| - | - | 1.422E+04 | 602.4 | - | - | 0 | - |
| - | - | 606.8 | 603.3 | - | - | 0 | - |
| - | - | 2339 | 603.4 | - | - | 0 | - |
| - | - | 1591 | 607.3 | - | - | 0 | - |
| - | - | 787.2 | 607.6 | - | - | 0 | - |
| 14 | y | 1630 | 608.3 | 0.0007433 | 1.222 | +1 | 5 |
| 12 | c | 2.032E+04 | 608.8 | 0.0001605 | 0.2636 | +2 | 12 |
| 14 | z | 1.258E+04 | 609.3 | 0.0005117 | 0.8398 | +1 | 5 |
| - | - | 1.273E+04 | 609.3 | - | - | 0 | - |
| - | - | 5021 | 609.9 | - | - | 0 | - |
| - | - | 5794 | 610.3 | - | - | 0 | - |
| - | - | 1456 | 610.4 | - | - | 0 | - |
| - | - | 1475 | 611.3 | - | - | 0 | - |
| 2 | z | 724.8 | 613 | 0.00469 | 7.65 | +3 | 17 |
| - | - | 1487 | 613.3 | - | - | 0 | - |
| - | - | 1023 | 613.6 | - | - | 0 | - |
| - | - | 2304 | 615.7 | - | - | 0 | - |
| - | - | 1183 | 616 | - | - | 0 | - |
| - | - | 3014 | 616.3 | - | - | 0 | - |
| 2 | y | 743.4 | 618.3 | 0.002122 | 3.431 | +3 | 17 |
| - | - | 934.5 | 618.7 | - | - | 0 | - |
| 8 | w | 711.6 | 618.8 | 0.008357 | 13.51 | +2 | 11 |
| - | - | 2868 | 621.9 | - | - | 0 | - |
| - | - | 771.9 | 622.4 | - | - | 0 | - |
| - | - | 702 | 623.8 | - | - | 0 | - |
| - | - | 1204 | 624.3 | - | - | 0 | - |
| 14 | y | 5016 | 625.3 | 0.001943 | 3.108 | +1 | 5 |
| 8 | y | 5543 | 625.8 | 7.836E-05 | 0.1252 | +2 | 11 |
| 8 | z | 6801 | 626.3 | 0.003503 | 5.592 | +2 | 11 |
| - | - | 2109 | 626.8 | - | - | 0 | - |
| - | - | 770.5 | 627.3 | - | - | 0 | - |
| - | - | 1549 | 631.7 | - | - | 0 | - |
| - | - | 813.7 | 632 | - | - | 0 | - |
| - | - | 1500 | 632.3 | - | - | 0 | - |
| - | - | 967.5 | 632.7 | - | - | 0 | - |
| - | - | 846.5 | 633 | - | - | 0 | - |
| - | - | 952.3 | 633.3 | - | - | 0 | - |
| 8 | y | 4516 | 634.3 | 0.0001359 | 0.2142 | +2 | 11 |
| - | - | 2474 | 634.8 | - | - | 0 | - |
| - | - | 802.9 | 635.3 | - | - | 0 | - |
| - | - | 1160 | 636.3 | - | - | 0 | - |
| - | - | 5352 | 636.7 | - | - | 0 | - |
| - | - | 5262 | 637 | - | - | 0 | - |
| - | - | 2594 | 637.3 | - | - | 0 | - |
| - | - | 797.3 | 637.7 | - | - | 0 | - |
| - | - | 764.4 | 640 | - | - | 0 | - |
| - | - | 3136 | 640.4 | - | - | 0 | - |
| - | - | 7139 | 641 | - | - | 0 | - |
| - | - | 9483 | 641.3 | - | - | 0 | - |
| - | - | 646.9 | 641.4 | - | - | 0 | - |
| - | - | 5408 | 641.7 | - | - | 0 | - |
| - | - | 3047 | 642 | - | - | 0 | - |
| - | - | 2469 | 645.7 | - | - | 0 | - |
| - | - | 1.69E+04 | 646 | - | - | 0 | - |
| - | - | 1.91E+04 | 646.3 | - | - | 0 | - |
| - | - | 9051 | 646.7 | - | - | 0 | - |
| - | - | 3524 | 647 | - | - | 0 | - |
| - | - | 2115 | 647.3 | - | - | 0 | - |
| - | - | 997.2 | 647.7 | - | - | 0 | - |
| - | - | 1162 | 650.7 | - | - | 0 | - |
| - | - | 2664 | 651 | - | - | 0 | - |
| - | - | 4.808E+04 | 651.3 | - | - | 0 | - |
| - | - | 5.458E+04 | 651.7 | - | - | 0 | - |
| - | - | 2.603E+04 | 652 | - | - | 0 | - |
| - | - | 1.097E+04 | 652.3 | - | - | 0 | - |
| - | - | 4440 | 652.7 | - | - | 0 | - |
| - | - | 1483 | 653 | - | - | 0 | - |
| - | - | 973.2 | 653.3 | - | - | 0 | - |
| - | - | 1441 | 654.8 | - | - | 0 | - |
| - | - | 691.8 | 655.3 | - | - | 0 | - |
| - | - | 790.9 | 658.4 | - | - | 0 | - |
| 7 | z | 2678 | 667.8 | 8.684E-05 | 0.13 | +2 | 12 |
| 7 | z | 3126 | 668.3 | 0.01114 | 16.67 | +2 | 12 |
| - | - | 1921 | 668.8 | - | - | 0 | - |
| 7 | w | 1828 | 669.3 | 0.004842 | 7.235 | +2 | 12 |
| - | - | 1293 | 669.8 | - | - | 0 | - |
| - | - | 1232 | 672.4 | - | - | 0 | - |
| 13 | c | 3.402E+04 | 672.9 | 0.0001528 | 0.2271 | +2 | 13 |
| - | - | 2.53E+04 | 673.4 | - | - | 0 | - |
| - | - | 1.054E+04 | 673.9 | - | - | 0 | - |
| - | - | 4519 | 674.4 | - | - | 0 | - |
| 7 | y | 1986 | 675.8 | 0.01152 | 17.05 | +2 | 12 |
| 7 | y | 8330 | 676.3 | 0.0004191 | 0.6196 | +2 | 12 |
| 7 | z | 1.33E+04 | 676.8 | 0.001219 | 1.801 | +2 | 12 |
| - | - | 6931 | 677.3 | - | - | 0 | - |
| - | - | 3902 | 677.8 | - | - | 0 | - |
| - | - | 1473 | 678.3 | - | - | 0 | - |
| 13 | w | 5076 | 679.3 | 0.000351 | 0.5166 | +1 | 6 |
| - | - | 1699 | 680.3 | - | - | 0 | - |
| 7 | y | 3939 | 684.8 | 0.0001614 | 0.2356 | +2 | 12 |
| - | - | 2607 | 685.3 | - | - | 0 | - |
| - | - | 1513 | 685.8 | - | - | 0 | - |
| - | - | 1221 | 701.4 | - | - | 0 | - |
| 7 | c | 1.031E+04 | 702.4 | 0.0005251 | 0.7475 | +1 | 7 |
| - | - | 4499 | 703.4 | - | - | 0 | - |
| 6 | w | 9077 | 711.8 | 0.0002382 | 0.3346 | +2 | 13 |
| - | - | 8723 | 712.3 | - | - | 0 | - |
| - | - | 4188 | 712.8 | - | - | 0 | - |
| - | - | 1096 | 713.3 | - | - | 0 | - |
| - | - | 1454 | 715.4 | - | - | 0 | - |
| 14 | c | 2.429E+04 | 716.4 | 0.0001754 | 0.2448 | +2 | 14 |
| - | - | 1.811E+04 | 716.9 | - | - | 0 | - |
| - | - | 8287 | 717.4 | - | - | 0 | - |
| - | - | 2252 | 717.9 | - | - | 0 | - |
| - | - | 586.7 | 719.3 | - | - | 0 | - |
| 6 | z | 692.5 | 732.8 | 0.008887 | 12.13 | +2 | 13 |
| - | - | 610.3 | 733.3 | - | - | 0 | - |
| 13 | z | 9864 | 737.3 | 0.0001292 | 0.1752 | +1 | 6 |
| - | - | 3524 | 738.3 | - | - | 0 | - |
| - | - | 1232 | 739.3 | - | - | 0 | - |
| 6 | y | 692.9 | 740.4 | 0.0114 | 15.39 | +2 | 13 |
| 6 | z | 1.171E+04 | 741.4 | 0.0004816 | 0.6497 | +2 | 13 |
| - | - | 3508 | 741.4 | - | - | 0 | - |
| - | - | 7818 | 741.9 | - | - | 0 | - |
| - | - | 3686 | 742.4 | - | - | 0 | - |
| - | - | 2053 | 742.4 | - | - | 0 | - |
| - | - | 1653 | 742.9 | - | - | 0 | - |
| - | - | 673.5 | 745.4 | - | - | 0 | - |
| 6 | y | 1258 | 749.4 | 0.002302 | 3.072 | +2 | 13 |
| - | - | 2465 | 749.9 | - | - | 0 | - |
| - | - | 768.7 | 751.4 | - | - | 0 | - |
| 13 | y | 1678 | 753.4 | 0.00196 | 2.601 | +1 | 6 |
| - | - | 784.8 | 755.9 | - | - | 0 | - |
| - | - | 863.9 | 768.4 | - | - | 0 | - |
| - | - | 1095 | 772.4 | - | - | 0 | - |
| - | - | 1505 | 772.9 | - | - | 0 | - |
| 15 | c | 1.924E+04 | 773.4 | 0.001131 | 1.463 | +2 | 15 |
| - | - | 1.583E+04 | 773.9 | - | - | 0 | - |
| - | - | 7946 | 774.4 | - | - | 0 | - |
| - | - | 2239 | 774.9 | - | - | 0 | - |
| - | - | 1110 | 775.4 | - | - | 0 | - |
| - | - | 708.1 | 780.9 | - | - | 0 | - |
| - | - | 1110 | 781.4 | - | - | 0 | - |
| - | - | 1051 | 783.9 | - | - | 0 | - |
| 8 | c | 869 | 786.4 | 0.0003793 | 0.4823 | +1 | 8 |
| - | - | 985.5 | 788.4 | - | - | 0 | - |
| 5 | y | 1090 | 789.9 | 0.0121 | 15.31 | +2 | 14 |
| 5 | z | 9163 | 790.9 | 0.0001427 | 0.1804 | +2 | 14 |
| - | - | 8827 | 791.4 | - | - | 0 | - |
| - | - | 3762 | 791.9 | - | - | 0 | - |
| - | - | 1461 | 792.4 | - | - | 0 | - |
| 5 | y | 1.161E+04 | 798.9 | 0.0002609 | 0.3266 | +2 | 14 |
| - | - | 8827 | 799.4 | - | - | 0 | - |
| - | - | 3419 | 799.9 | - | - | 0 | - |
| - | - | 1624 | 800.4 | - | - | 0 | - |
| - | - | 1314 | 802.5 | - | - | 0 | - |
| 8 | c | 2.486E+04 | 803.5 | 1.424E-05 | 0.01772 | +1 | 8 |
| - | - | 9202 | 804.5 | - | - | 0 | - |
| - | - | 2612 | 805.5 | - | - | 0 | - |
| 12 | w | 1353 | 807.4 | 0.0007314 | 0.906 | +1 | 7 |
| - | - | 721.2 | 811.4 | - | - | 0 | - |
| 4 | z | 7696 | 819.4 | 0.001048 | 1.279 | +2 | 15 |
| - | - | 4709 | 819.9 | - | - | 0 | - |
| - | - | 2725 | 820.4 | - | - | 0 | - |
| - | - | 1010 | 820.9 | - | - | 0 | - |
| 4 | y | 1.188E+04 | 827.4 | 0.0007054 | 0.8525 | +2 | 15 |
| - | - | 1.135E+04 | 827.9 | - | - | 0 | - |
| - | - | 6028 | 828.4 | - | - | 0 | - |
| - | - | 2230 | 828.9 | - | - | 0 | - |
| - | - | 1528 | 829.4 | - | - | 0 | - |
| - | - | 1339 | 829.9 | - | - | 0 | - |
| 16 | c | 3.531E+04 | 830.4 | 0.0005003 | 0.6024 | +2 | 16 |
| - | - | 2.834E+04 | 830.9 | - | - | 0 | - |
| - | - | 1.23E+04 | 831.4 | - | - | 0 | - |
| - | - | 5430 | 831.9 | - | - | 0 | - |
| - | - | 1545 | 832.4 | - | - | 0 | - |
| - | - | 771.4 | 832.9 | - | - | 0 | - |
| - | - | 683.8 | 833.9 | - | - | 0 | - |
| 3 | z | 1.217E+04 | 854.9 | 0.0002567 | 0.3002 | +2 | 16 |
| - | - | 1.131E+04 | 855.4 | - | - | 0 | - |
| - | - | 5936 | 855.9 | - | - | 0 | - |
| - | - | 2103 | 856.4 | - | - | 0 | - |
| 3 | y | 3577 | 862.9 | 0.0005855 | 0.6785 | +2 | 16 |
| 12 | y | 2579 | 863.4 | 0.00999 | 11.57 | +1 | 7 |
| - | - | 1181 | 863.9 | - | - | 0 | - |
| - | - | 696.4 | 864.5 | - | - | 0 | - |
| 12 | z | 4790 | 865.4 | 0.000733 | 0.8469 | +1 | 7 |
| - | - | 5325 | 866.4 | - | - | 0 | - |
| - | - | 2819 | 867.4 | - | - | 0 | - |
| - | - | 990.1 | 868.4 | - | - | 0 | - |
| - | - | 761.1 | 869.5 | - | - | 0 | - |
| - | - | 3605 | 872.5 | - | - | 0 | - |
| - | - | 6687 | 873 | - | - | 0 | - |
| - | - | 4855 | 873.5 | - | - | 0 | - |
| - | - | 1327 | 874 | - | - | 0 | - |
| 12 | y | 2174 | 881.4 | 0.002944 | 3.34 | +1 | 7 |
| - | - | 1145 | 882.5 | - | - | 0 | - |
| - | - | 1290 | 886.5 | - | - | 0 | - |
| 9 | c | 1575 | 887.5 | 0.002011 | 2.266 | +1 | 9 |
| 2 | w | 1627 | 889.9 | 0.0001746 | 0.1963 | +2 | 17 |
| - | - | 2501 | 894.4 | - | - | 0 | - |
| 17 | c | 5161 | 894.5 | 0.0003581 | 0.4003 | +2 | 17 |
| - | - | 4971 | 895 | - | - | 0 | - |
| - | - | 1730 | 895.4 | - | - | 0 | - |
| - | - | 4119 | 895.5 | - | - | 0 | - |
| - | - | 776.2 | 896 | - | - | 0 | - |
| 2 | z | 2809 | 919 | 0.0002965 | 0.3226 | +2 | 17 |
| - | - | 7802 | 919.5 | - | - | 0 | - |
| - | - | 5634 | 920 | - | - | 0 | - |
| - | - | 3399 | 920.5 | - | - | 0 | - |
| - | - | 716.7 | 921 | - | - | 0 | - |
| - | - | 870.6 | 925.5 | - | - | 0 | - |
| - | - | 775.7 | 926.5 | - | - | 0 | - |
| 2 | y | 978.4 | 927 | 0.005827 | 6.286 | +2 | 17 |
| 11 | w | 860.2 | 935.5 | 0.0007411 | 0.7922 | +1 | 8 |
| - | - | 2127 | 939.5 | - | - | 0 | - |
| - | - | 2202 | 940 | - | - | 0 | - |
| - | - | 1736 | 940.5 | - | - | 0 | - |
| - | - | 976.6 | 941 | - | - | 0 | - |
| - | - | 1483 | 946.5 | - | - | 0 | - |
| - | - | 1337 | 947 | - | - | 0 | - |
| - | - | 822.9 | 947.5 | - | - | 0 | - |
| - | - | 1648 | 948 | - | - | 0 | - |
| - | - | 1505 | 948.5 | - | - | 0 | - |
| - | - | 1131 | 949 | - | - | 0 | - |
| 11 | y | 1200 | 951.5 | 0.008822 | 9.272 | +1 | 8 |
| 11 | z | 3130 | 952.5 | 0.0005948 | 0.6244 | +1 | 8 |
| - | - | 8912 | 953.5 | - | - | 0 | - |
| - | - | 4238 | 954.5 | - | - | 0 | - |
| - | - | 9586 | 955 | - | - | 0 | - |
| - | - | 9705 | 955.5 | - | - | 0 | - |
| - | - | 5033 | 956 | - | - | 0 | - |
| - | - | 2759 | 956.5 | - | - | 0 | - |
| - | - | 845.6 | 960 | - | - | 0 | - |
| - | - | 1020 | 960.5 | - | - | 0 | - |
| - | - | 774 | 961.5 | - | - | 0 | - |
| - | - | 1480 | 968 | - | - | 0 | - |
| - | - | 5780 | 968.5 | - | - | 0 | - |
| - | - | 3.785E+04 | 969 | - | - | 0 | - |
| - | - | 2.963E+04 | 969.5 | - | - | 0 | - |
| - | - | 1.58E+04 | 970 | - | - | 0 | - |
| - | - | 7981 | 970.5 | - | - | 0 | - |
| - | - | 2800 | 971 | - | - | 0 | - |
| - | - | 899.2 | 973.5 | - | - | 0 | - |
| - | - | 733.2 | 974.5 | - | - | 0 | - |
| - | - | 5367 | 976.5 | - | - | 0 | - |
| - | - | 6757 | 977 | - | - | 0 | - |
| - | - | 2.287E+04 | 977.5 | - | - | 0 | - |
| - | - | 1.787E+04 | 978 | - | - | 0 | - |
| - | - | 1.098E+04 | 978.5 | - | - | 0 | - |
| - | - | 5628 | 979 | - | - | 0 | - |
| - | - | 1818 | 979.5 | - | - | 0 | - |
| - | - | 1079 | 986.5 | - | - | 0 | - |
| - | - | 1083 | 999.6 | - | - | 0 | - |
| - | - | 1174 | 1001 | - | - | 0 | - |
| 10 | c | 2.046E+04 | 1002 | 0.0001473 | 0.1471 | +1 | 10 |
| - | - | 1.028E+04 | 1003 | - | - | 0 | - |
| - | - | 3588 | 1004 | - | - | 0 | - |
| - | - | 1413 | 1045 | - | - | 0 | - |
| - | - | 1347 | 1046 | - | - | 0 | - |
| - | - | 1228 | 1088 | - | - | 0 | - |
| 11 | c | 1.374E+04 | 1089 | 0.0006206 | 0.5701 | +1 | 11 |
| - | - | 8767 | 1090 | - | - | 0 | - |
| - | - | 2954 | 1091 | - | - | 0 | - |
| - | - | 888.5 | 1102 | - | - | 0 | - |
| 9 | z | 925.6 | 1134 | 0.02042 | 18.02 | +1 | 10 |
| 9 | y | 1491 | 1150 | 0.002941 | 2.558 | +1 | 10 |
| 9 | z | 3617 | 1151 | 0.00332 | 2.885 | +1 | 10 |
| - | - | 7156 | 1152 | - | - | 0 | - |
| - | - | 3849 | 1153 | - | - | 0 | - |
| - | - | 1002 | 1154 | - | - | 0 | - |
| 9 | y | 865.1 | 1167 | 0.006663 | 5.712 | +1 | 10 |
| - | - | 1075 | 1174 | - | - | 0 | - |
| - | - | 2066 | 1189 | - | - | 0 | - |
| - | - | 1358 | 1190 | - | - | 0 | - |
| - | - | 712.3 | 1201 | - | - | 0 | - |
| 12 | c | 3947 | 1217 | 0.000857 | 0.7044 | +1 | 12 |
| - | - | 4370 | 1218 | - | - | 0 | - |
| - | - | 1626 | 1219 | - | - | 0 | - |
| 8 | z | 1663 | 1252 | 0.002102 | 1.68 | +1 | 11 |
| - | - | 3228 | 1253 | - | - | 0 | - |
| - | - | 2005 | 1254 | - | - | 0 | - |
| - | - | 1383 | 1255 | - | - | 0 | - |
| - | - | 1655 | 1301 | - | - | 0 | - |
| - | - | 1249 | 1302 | - | - | 0 | - |
| - | - | 1569 | 1318 | - | - | 0 | - |
| - | - | 1204 | 1319 | - | - | 0 | - |
| 7 | z | 799.6 | 1336 | 0.01466 | 10.98 | +1 | 12 |
| - | - | 917.1 | 1337 | - | - | 0 | - |
| - | - | 1480 | 1344 | - | - | 0 | - |
| 13 | c | 4532 | 1345 | 0.0004746 | 0.3529 | +1 | 13 |
| - | - | 6383 | 1346 | - | - | 0 | - |
| - | - | 2841 | 1347 | - | - | 0 | - |
| - | - | 1246 | 1348 | - | - | 0 | - |
| 7 | z | 3301 | 1353 | 0.00627 | 4.635 | +1 | 12 |
| - | - | 1.381E+04 | 1354 | - | - | 0 | - |
| - | - | 7128 | 1355 | - | - | 0 | - |
| - | - | 3073 | 1356 | - | - | 0 | - |
| - | - | 830.6 | 1357 | - | - | 0 | - |
| - | - | 1107 | 1431 | - | - | 0 | - |
| 14 | c | 5716 | 1432 | 0.006258 | 4.371 | +1 | 14 |
| - | - | 6266 | 1433 | - | - | 0 | - |
| - | - | 2719 | 1434 | - | - | 0 | - |
| - | - | 1239 | 1435 | - | - | 0 | - |
| 6 | z | 2398 | 1482 | 8.758E-05 | 0.05911 | +1 | 13 |
| - | - | 9502 | 1483 | - | - | 0 | - |
| - | - | 7518 | 1484 | - | - | 0 | - |
| - | - | 2383 | 1485 | - | - | 0 | - |
| - | - | 1618 | 1486 | - | - | 0 | - |
| - | - | 929.2 | 1545 | - | - | 0 | - |
| 15 | c | 4075 | 1546 | 0.007193 | 4.653 | +1 | 15 |
| - | - | 4290 | 1547 | - | - | 0 | - |
| - | - | 2477 | 1548 | - | - | 0 | - |
| - | - | 749.6 | 1549 | - | - | 0 | - |
| 5 | z | 1746 | 1581 | 0.0006994 | 0.4424 | +1 | 14 |
| - | - | 4743 | 1582 | - | - | 0 | - |
| - | - | 3618 | 1583 | - | - | 0 | - |
| - | - | 2061 | 1584 | - | - | 0 | - |
| - | - | 828.8 | 1585 | - | - | 0 | - |
| - | - | 1873 | 1598 | - | - | 0 | - |
| - | - | 858.4 | 1599 | - | - | 0 | - |
| 4 | z | 780.3 | 1638 | 0.00373 | 2.278 | +1 | 15 |
| - | - | 3403 | 1639 | - | - | 0 | - |
| - | - | 1830 | 1640 | - | - | 0 | - |
| - | - | 1909 | 1641 | - | - | 0 | - |
| 4 | y | 1081 | 1654 | 0.005866 | 3.547 | +1 | 15 |
| - | - | 2134 | 1655 | - | - | 0 | - |
| - | - | 1156 | 1657 | - | - | 0 | - |
| 16 | c | 3115 | 1660 | 0.001027 | 0.6187 | +1 | 16 |
| - | - | 5022 | 1661 | - | - | 0 | - |
| - | - | 3786 | 1662 | - | - | 0 | - |
| - | - | 1260 | 1663 | - | - | 0 | - |
| - | - | 4553 | 1710 | - | - | 0 | - |
| - | - | 4087 | 1711 | - | - | 0 | - |
| - | - | 2183 | 1712 | - | - | 0 | - |
| - | - | 1082 | 1745 | - | - | 0 | - |
| - | - | 1164 | 1790 | - | - | 0 | - |
| - | - | 1019 | 1791 | - | - | 0 | - |
| - | - | 1533 | 1839 | - | - | 0 | - |
| - | - | 753.1 | 1840 | - | - | 0 | - |
| - | - | 1711 | 1910 | - | - | 0 | - |
| - | - | 900.7 | 1911 | - | - | 0 | - |
| - | - | 1969 | 1937 | - | - | 0 | - |
| - | - | 3189 | 1938 | - | - | 0 | - |
| - | - | 2115 | 1939 | - | - | 0 | - |
| - | - | 1424 | 1940 | - | - | 0 | - |
| - | - | 1155 | 1953 | - | - | 0 | - |
| - | - | 3282 | 1954 | - | - | 0 | - |
| - | - | 4300 | 1955 | - | - | 0 | - |
| - | - | 3167 | 1956 | - | - | 0 | - |
| - | - | 1191 | 1957 | - | - | 0 | - |
| - | - | 644.2 | 2486 | - | - | 0 | - |

m/z Charge Intensity FragmentType MassShift Position
120.05716705322266 0 1211.5443
121.92160034179688 0 371.55017
127.08700561523438 0 529.9626
128.0943603515625 0 670.50903
129.1022186279297 0 18486.61
130.10556030273438 0 855.2196
131.1179656982422 0 2416.3677
131.12281799316406 0 501.39664
136.07566833496094 0 1751.9415
138.5199432373047 0 397.65274
140.57362365722656 0 446.33157
141.44189453125 0 489.8149
141.50994873046875 0 385.6087
143.84205627441406 0 436.579
146.12869262695312 0 1467.1086
148.9556121826172 0 925.4581
155.11781311035156 0 1786.0167
164.04676818847656 0 1484.9807
171.0884552001953 0 640.35565
173.44032287597656 0 3315.7087
182.08084106445312 0 1283.387 y 17
183.1126251220703 0 436.00775
183.14927673339844 0 894.4723
188.11541748046875 0 3372.8093
189.1195831298828 0 716.41394
204.56951904296875 0 472.55948
207.26402282714844 0 488.8267
210.09967041015625 0 885.88916
211.1077880859375 0 1089.4192
217.1656951904297 0 778.56287
217.4361572265625 0 536.8603
227.1261444091797 0 1228.0859
228.1340789794922 0 1643.5703
228.17063903808594 0 7288.002 c Ammonia loss 1
229.17311096191406 0 899.91693
231.04135131835938 0 521.50104
245.16033935546875 0 1667.7498
245.1971435546875 0 581.29694 c 1
249.15992736816406 0 1504.2896
250.16688537597656 0 831.1401
265.1516418457031 0 530.6605
266.15142822265625 0 597.0715
292.6741638183594 0 781.27527 c Ammonia loss 5
293.1350402832031 0 653.03217
294.1575622558594 0 1561.125 z 16
299.2074279785156 0 1138.1525 c Ammonia loss 2
302.15863037109375 0 2733.9805
310.1761169433594 0 1221.0227 y 16
311.361083984375 0 491.43466
316.2347412109375 0 1665.5258 c 2
337.1266784667969 0 904.4249
355.84765625 0 1353.145 y 9
356.1833190917969 0 757.94104
356.2279052734375 0 881.3006 c Ammonia loss 3
359.20330810546875 0 908.72394
363.2033386230469 0 715.68164
364.1881408691406 0 680.77606 w 15
370.7159729003906 0 672.9068
373.25555419921875 0 8047.5605 c 3
374.2583923339844 0 1113.9558
384.2235107421875 0 1437.207
384.7144775390625 0 874.20154
385.2274475097656 0 740.6482
388.77056884765625 0 585.4686
389.53021240234375 0 905.0122 y 8
404.20269775390625 0 647.28424
404.7041015625 0 1262.5557
408.20062255859375 0 4601.9473 z 15
409.2062683105469 0 2209.9238
416.2008361816406 0 1438.6559
423.210693359375 0 775.12646 y 7
424.2189636230469 0 1797.6016 y 15
427.19927978515625 0 612.2165
427.30322265625 0 1240.6748
428.3104248046875 0 739.8274
428.89044189453125 0 600.59467
429.08709716796875 0 558.0315
430.8877868652344 0 835.06244
433.21875 0 633.0164 z 11
435.2401123046875 0 681.15424
441.22784423828125 0 2076.5918 y 11
441.7304382324219 0 1191.4786
442.732666015625 0 988.603
446.2335510253906 0 674.00726
448.2181396484375 0 834.04456
454.728759765625 0 958.14246
455.2335205078125 0 743.9616
455.29779052734375 0 3414.5864 c Ammonia loss 4
468.232666015625 0 4029.5354 w 10
468.7352294921875 0 2438.5125
468.9952697753906 0 822.67566
470.7440185546875 0 610.3373
472.32421875 0 31447.492 c 4
473.2499084472656 0 581.94415
473.3272399902344 0 8678.325
474.3300476074219 0 596.5908
475.75299072265625 0 612.04144
476.2298583984375 0 1043.1283 y Ammonia loss 10
476.73541259765625 0 801.75226 z 10
477.23504638671875 0 698.6454
478.22998046875 0 10220.727 w 14
479.2335510253906 0 3444.6172
479.7510986328125 0 3601.1033
480.0032653808594 0 3148.22
480.2532043457031 0 1144.3995
480.50567626953125 0 595.4807
484.2544250488281 0 1265.0502
484.74755859375 0 1077.8539 y 10
488.23638916015625 0 747.44293
489.0555419921875 0 1160.7014
493.2590026855469 0 839.40857
498.34027099609375 0 1936.7422
499.3424072265625 0 965.1456
503.2337951660156 0 2651.1748
504.2367248535156 0 923.60657
511.7490539550781 0 1784.4796 w 9
512.2521362304688 0 993.4564
515.2554321289062 0 683.46643
521.7554321289062 0 1020.1203
522.2439575195312 0 4258.325 z 14
522.7638549804688 0 618.3868
523.2487182617188 0 2542.5615
532.2601928710938 0 1085.5594
532.7684326171875 0 580.4669
533.2696533203125 0 12007.439 y 9
533.7713012695312 0 6762.6904
534.271728515625 0 2928.644
538.2628784179688 0 1696.5336 y 14
544.8007202148438 0 893.7586 c 10
546.7655029296875 0 1033.0765
547.2689819335938 0 1266.8646
552.7784423828125 0 680.00665
553.280517578125 0 976.9272
553.7803344726562 0 3036.5798
554.2806396484375 0 1970.4451
554.7828369140625 0 637.38544
557.3544311523438 0 1190.6362
558.3563232421875 0 892.9123
559.2639770507812 0 931.5654
565.2617797851562 0 5637.515
566.2658081054688 0 1542.1699
567.2828369140625 0 2788.275 w 8
567.7855834960938 0 2347.8296
568.2734985351562 0 6383.269 w 8
568.7755737304688 0 2125.295
569.27783203125 0 1777.7623
574.7761840820312 0 3973.783
575.2800903320312 0 21573.92 y Ammonia loss 8
575.7820434570312 0 15519.824 z 8
576.2855224609375 0 7738.9375
576.7879028320312 0 2751.6975
577.2874755859375 0 1095.833
581.9873657226562 0 1442.2916
582.3218994140625 0 1322.9396
583.793701171875 0 6900.5435 y 8
584.2947387695312 0 5677.8857
584.3394165039062 0 760.4861 c Ammonia loss 5
584.7953491210938 0 2412.2612
592.2711791992188 0 2453.4836 w 13
593.2735595703125 0 1158.4663
596.6582641601562 0 32562.295 c 16
596.9925537109375 0 25370.846
597.3267211914062 0 14293.734
597.6604614257812 0 4974.2017
597.9949951171875 0 2800.1182
601.3666381835938 0 41464.574 c 5
602.36962890625 0 14216.6455
603.2953491210938 0 606.77466
603.372314453125 0 2339.1296
607.2596435546875 0 1591.4834
607.6387329101562 0 787.15027
608.2667236328125 0 1629.5775 y Ammonia loss 13
608.8482666015625 0 20317.693 c 11
609.2747802734375 0 12583.715 z 13
609.349853515625 0 12728.172
609.8507690429688 0 5020.781
610.27880859375 0 5793.568
610.35009765625 0 1455.8099
611.285400390625 0 1475.1383
612.9832763671875 0 724.8496 z 1
613.3150634765625 0 1486.897
613.6448974609375 0 1023.49493
615.656982421875 0 2303.9475
615.9884033203125 0 1182.5939
616.3281860351562 0 3013.9253
618.3160400390625 0 743.4429 y 1
618.6548461914062 0 934.4623
618.8047485351562 0 711.61536 w 7
621.8541259765625 0 2867.868
622.3588256835938 0 771.8567
623.8090209960938 0 701.98456
624.3106079101562 0 1203.8738
625.2959594726562 0 5016.271 y 13
625.8041381835938 0 5543.4663 y Ammonia loss 7
626.3046264648438 0 6800.7065 z 7
626.808349609375 0 2109.2449
627.3050537109375 0 770.50195
631.6622924804688 0 1549.3013
631.9990844726562 0 813.6873
632.3358154296875 0 1500.4159
632.6700439453125 0 967.54236
633.0026245117188 0 846.4555
633.26220703125 0 952.3077
634.317626953125 0 4516.2314 y 7
634.8177490234375 0 2474.4402
635.3209228515625 0 802.9275
636.3381958007812 0 1159.823
636.6697387695312 0 5351.7207
637.0050659179688 0 5261.8467
637.340576171875 0 2593.613
637.673583984375 0 797.3001
639.9976196289062 0 764.371
640.3908081054688 0 3136.31
640.9932861328125 0 7139.1426
641.32666015625 0 9483.313
641.387939453125 0 646.90283
641.6616821289062 0 5407.598
641.9959106445312 0 3046.5264
645.6696166992188 0 2468.5806
646.0010375976562 0 16896.506
646.33544921875 0 19101.107
646.6693115234375 0 9050.816
647.0051879882812 0 3523.5325
647.3366088867188 0 2115.4634
647.6676635742188 0 997.19977
650.6694946289062 0 1161.6031
651.0033569335938 0 2663.9038
651.3406372070312 0 48083.29
651.6751098632812 0 54576.355
652.00927734375 0 26034.852
652.3438720703125 0 10968.446
652.6781616210938 0 4440.4795
653.0110473632812 0 1482.9161
653.3438110351562 0 973.21155
654.83154296875 0 1440.83
655.33642578125 0 691.7835
658.400146484375 0 790.8521
667.8265991210938 0 2678.4746 z Water loss 6
668.329833984375 0 3125.967 z Ammonia loss 6
668.8309326171875 0 1921.4872
669.3250732421875 0 1827.5529 w 6
669.8277587890625 0 1293.4747
672.3710327148438 0 1232.2543
672.8778686523438 0 34024.84 c 12
673.3790283203125 0 25299.781
673.8806762695312 0 10540.52
674.3822021484375 0 4519.472
675.8245239257812 0 1985.7097 y Water loss 6
676.32763671875 0 8330.043 y Ammonia loss 6
676.8307495117188 0 13303.75 z 6
677.3326416015625 0 6931.1377
677.8333129882812 0 3901.9712
678.33544921875 0 1473.3547
679.304931640625 0 5075.631 w 12
680.3082885742188 0 1699.0477
684.8414916992188 0 3939.172 y 6
685.3418579101562 0 2607.162
685.8431396484375 0 1512.8904
701.3787841796875 0 1220.5828
702.4139404296875 0 10313.088 c 6
703.416748046875 0 4499.2627
711.8463745117188 0 9076.714 w 5
712.34765625 0 8723.492
712.848388671875 0 4187.887
713.349853515625 0 1095.6289
715.3850708007812 0 1453.7239
716.3935546875 0 24290.3 c 13
716.894775390625 0 18105.826
717.396728515625 0 8286.849
717.8974609375 0 2252.2788
719.3380737304688 0 586.71265
732.848876953125 0 692.45435 z Ammonia loss 5
733.3472900390625 0 610.3384
737.333740234375 0 9863.727 z 12
738.3369140625 0 3523.5652
739.3391723632812 0 1231.9124
740.345947265625 0 692.8857 y Water loss 5
741.352783203125 0 11711.04 z 5
741.4373168945312 0 3507.502
741.8541870117188 0 7818.131
742.3555908203125 0 3686.2927
742.4414672851562 0 2053.2551
742.8562622070312 0 1652.6055
745.3814697265625 0 673.4643
749.3649291992188 0 1257.6902 y 5
749.8643798828125 0 2464.8582
751.4038696289062 0 768.74976
753.3545532226562 0 1677.8195 y 12
755.882568359375 0 784.81116
768.4256591796875 0 863.8668
772.403076171875 0 1094.649
772.8966674804688 0 1504.7776
773.4140625 0 19235.684 c 14
773.916015625 0 15834.006
774.417724609375 0 7946.312
774.916259765625 0 2239.4302
775.4196166992188 0 1110.0411
780.8959350585938 0 708.1471
781.4035034179688 0 1109.516
783.8732299804688 0 1050.9432
786.4359741210938 0 868.9525 c Ammonia loss 7
788.4130249023438 0 985.4818
789.8794555664062 0 1089.8287 y Water loss 4
790.8873291015625 0 9162.861 z 4
791.3889770507812 0 8826.55
791.889404296875 0 3762.0952
792.3905029296875 0 1460.6824
798.8970947265625 0 11608.48 y 4
799.3980102539062 0 8827.363
799.8995971679688 0 3419.1736
800.4002685546875 0 1623.5393
802.4542236328125 0 1313.8328
803.462158203125 0 24859.736 c 7
804.4652099609375 0 9201.657
805.4678344726562 0 2612.3848
807.3624267578125 0 1353.2034 w 11
811.4004516601562 0 721.20154
819.3971557617188 0 7696.4536 z 3
819.8997192382812 0 4708.807
820.3992919921875 0 2725.1506
820.9052734375 0 1009.90356
827.4068603515625 0 11881.305 y 3
827.9091186523438 0 11353.497
828.410888671875 0 6027.7485
828.91064453125 0 2229.8384
829.420166015625 0 1528.0955
829.9290161132812 0 1338.8422
830.4361572265625 0 35305.562 c 15
830.9381713867188 0 28339.459
831.4396362304688 0 12303.216
831.9403686523438 0 5429.709
832.4461669921875 0 1545.4816
832.9251098632812 0 771.4217
833.9043579101562 0 683.783
854.91650390625 0 12168.185 z 2
855.4183959960938 0 11305.399
855.9204711914062 0 5935.647
856.4205322265625 0 2103.4634
862.925537109375 0 3576.8284 y 2
863.427001953125 0 2578.9487 y Water loss 11
863.9238891601562 0 1180.7719
864.4660034179688 0 696.40857
865.4295654296875 0 4789.9507 z 11
866.4354858398438 0 5325.0317
867.4353637695312 0 2819.1345
868.439208984375 0 990.14636
869.472412109375 0 761.1474
872.4771118164062 0 3604.8494
872.9794921875 0 6687.228
873.4801635742188 0 4854.609
873.9842529296875 0 1327.4718
881.4505004882812 0 2174.3455 y 11
882.4547729492188 0 1145.0123
886.4810791015625 0 1289.786
887.4812622070312 0 1574.9879 c Ammonia loss 8
889.9315795898438 0 1627.0656 w 1
894.3925170898438 0 2500.8423
894.4844970703125 0 5161.477 c 16
894.9849853515625 0 4971.241
895.3963623046875 0 1730.0863
895.4882202148438 0 4119.393
895.9886474609375 0 776.2
918.9645385742188 0 2809.3638 z 1
919.4672241210938 0 7801.883
919.9678955078125 0 5633.5005
920.470458984375 0 3398.9368
920.9677124023438 0 716.6859
925.4773559570312 0 870.64905
926.4712524414062 0 775.67456
926.9794311523438 0 978.3605 y 1
935.4588623046875 0 860.1585 w 10
939.4677124023438 0 2126.9526
939.9673461914062 0 2201.9373
940.4666137695312 0 1736.4425
940.9662475585938 0 976.628
946.4917602539062 0 1483.3442
947.00146484375 0 1336.7397
947.4971923828125 0 822.90826
947.9966430664062 0 1648.1151
948.5013427734375 0 1505.1072
948.9949340820312 0 1130.5927
951.4442138671875 0 1200.2013 y Ammonia loss 10
952.4602661132812 0 3129.645 z 10
953.4664916992188 0 8912.413
954.4783935546875 0 4238.329
955.004638671875 0 9585.972
955.5042114257812 0 9705.429
956.0071411132812 0 5032.802
956.50634765625 0 2759.3518
960.0045776367188 0 845.5843
960.490966796875 0 1019.58923
961.5101928710938 0 773.9925
968.0010375976562 0 1480.1505
968.4992065429688 0 5779.5854
969.0018920898438 0 37846.07
969.503173828125 0 29630.479
970.0045166015625 0 15803.594
970.5057373046875 0 7981.338
971.0075073242188 0 2800.2466
973.5087890625 0 899.2416
974.5100708007812 0 733.1877
976.5068359375 0 5366.9253
977.009033203125 0 6757.081
977.5147705078125 0 22867.508
978.01611328125 0 17869.885
978.5173950195312 0 10976.121
979.0186157226562 0 5627.704
979.5224609375 0 1818.1991
986.5210571289062 0 1079.2797
999.5542602539062 0 1083.2908
1000.551513671875 0 1173.8868
1001.5624389648438 0 20462.355 c 9
1002.564697265625 0 10275.592
1003.5682373046875 0 3587.5518
1044.57568359375 0 1413.2964
1045.5811767578125 0 1346.9177
1087.5711669921875 0 1228.4464
1088.593994140625 0 13742.116 c 10
1089.5966796875 0 8766.983
1090.5989990234375 0 2953.9214
1101.57861328125 0 888.4815
1133.55517578125 0 925.5807 z Ammonia loss 8
1149.550537109375 0 1491.1036 y Ammonia loss 8
1150.5579833984375 0 3617.4746 z 8
1151.5672607421875 0 7156.4795
1152.570556640625 0 3848.5027
1153.584228515625 0 1002.0317
1166.5733642578125 0 865.1005 y 8
1173.67041015625 0 1074.8542
1188.597412109375 0 2065.642
1189.60546875 0 1358.2782
1200.6876220703125 0 712.3222
1216.688720703125 0 3947.4365 c 11
1217.6929931640625 0 4370.253
1218.6966552734375 0 1626.1177
1251.611083984375 0 1662.7117 z 7
1252.614013671875 0 3227.6226
1253.61376953125 0 2005.1654
1254.619873046875 0 1382.642
1300.73486328125 0 1655.0146
1301.72900390625 0 1248.7767
1317.6497802734375 0 1568.8667
1318.65380859375 0 1203.9961
1335.644775390625 0 799.5814 z Ammonia loss 6
1336.6495361328125 0 917.1291
1343.7396240234375 0 1480.2859
1344.7476806640625 0 4531.9414 c 12
1345.751953125 0 6383.2124
1346.7569580078125 0 2840.9482
1347.7672119140625 0 1245.7894
1352.650390625 0 3301.4646 z 6
1353.662353515625 0 13806.913
1354.664794921875 0 7127.923
1355.6724853515625 0 3072.9265
1356.6680908203125 0 830.55475
1430.7569580078125 0 1106.5531
1431.77392578125 0 5716.175 c 13
1432.7830810546875 0 6265.8623
1433.78369140625 0 2718.5115
1434.7861328125 0 1238.8651
1481.6993408203125 0 2397.6658 z 5
1482.7025146484375 0 9501.947
1483.7080078125 0 7517.7676
1484.71337890625 0 2382.9062
1485.711181640625 0 1618.3137
1544.7978515625 0 929.2281
1545.81591796875 0 4074.9724 c 14
1546.82080078125 0 4289.5186
1547.8271484375 0 2477.0369
1548.8306884765625 0 749.57324
1580.7669677734375 0 1745.6865 z 4
1581.769775390625 0 4743.366
1582.7763671875 0 3618.0278
1583.7818603515625 0 2061.088
1584.781494140625 0 828.79803
1597.7921142578125 0 1872.5969
1598.7886962890625 0 858.4224
1637.785400390625 0 780.32184 z 3
1638.79833984375 0 3403.0017
1639.80029296875 0 1829.572
1640.8023681640625 0 1908.6887
1653.813720703125 0 1081.4081 y 3
1654.81005859375 0 2134.4207
1656.8138427734375 0 1156.2831
1659.8670654296875 0 3114.9846 c 15
1660.8709716796875 0 5022.2314
1661.8724365234375 0 3786.0312
1662.8828125 0 1259.6542
1709.83447265625 0 4553.343
1710.8365478515625 0 4086.8708
1711.8394775390625 0 2182.8005
1744.9547119140625 0 1081.6915
1789.9747314453125 0 1164.2438
1790.9793701171875 0 1018.6651
1838.9188232421875 0 1532.6406
1839.9251708984375 0 753.1285
1910.0069580078125 0 1711.4615
1911.03369140625 0 900.74384
1936.9954833984375 0 1968.8997
1937.99560546875 0 3189.1375
1938.999267578125 0 2115.1082
1940.001220703125 0 1423.6975
1953.00537109375 0 1155.1465
1954.0184326171875 0 3281.8687
1955.025634765625 0 4299.8335
1956.0281982421875 0 3167.2527
1957.02783203125 0 1190.708
2485.512451171875 0 644.24225

Spectrum Details

|  |  |
| --- | --- |
| Matched peaks? Matched peaks The total absolute number of peaks matched. Additionally in brackets the total fraction of peaks matched and the total number of peaks is shown. | 108 (21.26% of 508) |
| FDR? FDR The false discovery rate estimated for this peptide. It is calculated by matching all theoretical fragments with a non-integer shift with the raw peaks for this spectrum. This is done with 40 different shifts. The resulting percentage is the average number of annotated peaks over the number of annotated peaks with the correct spectrum. | 2.54% |
| Satellite FDR? Satellite FDR See the FDR for details on its calculation. This satellite ion specific FDR only contains the satellite ions (d/w) for I/L/J positions. | - |
| PSM Score? PSM Score The PSM Score as given by Hecklib to this annotated spectrum. It is shown with three significant figures. | 609 |

#### Reverse Lookup? Reverse Lookup All places where this read could be placed.

| Group | Segment | Template | Template Part | Read Part | Score | Unique |
| --- | --- | --- | --- | --- | --- | --- |
| Homo sapiens Light Chain | IGLC | IGLC2 | [48..66] | [0..18] | 144 | False |
| Homo sapiens Light Chain | IGLC | IGLC3 | [46..64] | [0..18] | 144 | False |

| Recombined | Template Part | Read Part | Score | Unique |
| --- | --- | --- | --- | --- |
| REC-0-1\_002 | [159..177] | [0..18] | 144 | True |

#### Meta Information from Multiple reads

##### Number of combined reads

2

##### Intensity

0.4687

##### TotalArea

2.653E+06

#### Positional Score

Copy Data

##### Positional Score (TSV)

###### Preview

```
Loading example...
```

*Click on the button to copy the data to your clipboard.*

0001234567891011121314151617

Label Value
"0" 0
"1" 0
"2" 0
"3" 0
"4" 0
"5" 0
"6" 0
"7" 0
"8" 0
"9" 0
"10" 0
"11" 0
"12" 0
"13" 0
"14" 0
"15" 0
"16" 0
"17" 0

#### Meta Information from PEAKS

##### Scan Identifier

F2:2479

##### Original sequence

V

K

A

G

V

E

T

T

T

P

S

K

Q

S

N

N

K

Y

##### Posttranslational Modifications

##### Source File

D:\separate\_stitch\_analyses\xle-disambiguation\raw\20210323\_F1\_UM1\_Peng0013\_SA\_F59\_ingel\_3ug\_TL.raw

##### Fraction

2

##### Scan Feature

F2:10431

##### De Novo Score

99

##### ConfidenceScore

99

### m/z

651.3417

##### Mass

1951.001

##### Charge

3

##### Retention Time

12.59

##### Predicted Retention Time

-

##### Area

1.256E+06

##### Parts Per Million

1.1

##### Fragmentation mode

ETHCD

##### Originating file

01 D:\separate\_stitch\_analyses\xle-disambiguation\20210325\_F59\_3ug\_DENOVO\_12.csv

#### Meta Information from PEAKS

##### Scan Identifier

F2:2473

##### Original sequence

V

K

A

G

V

E

T

T

T

P

S

K

Q

S

N

N

K

Y

##### Posttranslational Modifications

##### Source File

D:\separate\_stitch\_analyses\xle-disambiguation\raw\20210323\_F1\_UM1\_Peng0013\_SA\_F59\_ingel\_3ug\_TL.raw

##### Fraction

2

##### Scan Feature

F2:3663

##### De Novo Score

98

##### ConfidenceScore

98

### m/z

488.7578

##### Mass

1951.001

##### Charge

4

##### Retention Time

12.64

##### Predicted Retention Time

-

##### Area

1.397E+06

##### Parts Per Million

0.6

##### Fragmentation mode

ETHCD

##### Originating file

01 D:\separate\_stitch\_analyses\xle-disambiguation\20210325\_F59\_3ug\_DENOVO\_12.csv
