## Supplementary material for "A handle on mass coincidence errors in *de novo* sequencing of antibodies by bottom-up proteomics": Combined_063.html

Details Combined\_063 | Stitch OverviewUndefined

### Read Combined\_063

#### Sequence (length=10)

JYSKJTVDKS

#### Spectrum 3715? Spectrum 3715 The raw spectrum of this peptide as annotated by Hecklib. The fragments are coloured according to ion type (see legend). Any peaks with a star '\*' as text can be hovered over to see the full details, first the ion type second the mass shift type. By hovering over the amino acids in the peptide or ions in the legend the corresponding peaks are highlighted. By toggling the 'Unassigned' label you can turn the background (unassigned) peaks on or off in the plot. By updating the slider in the Ion legend you can update the spectrum to only show the top X% of the peaks with labels. The top X% means any peak that is within X% of the highest intensity. By dragging in the spectrum you can zoom in to a specific part of the spectrum and use 'Zoom Out' to get back to the original zoom level. The annotation of the spectrum is based on the given sequence in the peptides file and is done with different software so inconsistencies are likely. The peaks are annotated based on the given sequence, with 20 ppm tolerance.

Copy Data

##### Spectrum 3715 (TSV)

###### Preview

```
Loading example...
```

*Click on the button to copy the data to your clipboard.*

Mz MinMz MaxIntensity Max

WidthHeightPeptide font sizePeptide stroke widthSpectrum font sizeSpectrum stroke widthCompact peptide

Ion legend

wxyz

abcd

OtherUnassignedIon^Charge^_Position_Show for top:%

JYSKJTVDKS

02.77e+45.53e+48.30e+41.11e+5

Zoom Out

w^+1^_2_y^+1^_2_z^+1^_2_y^+1^_2_y^+2^_5_w^+1^_3_z^+1^_3_y^+1^_3_c^+1^_3_c^+1^_3_y^+2^_7_w^+1^_4_y^+2^_8_z^+2^_8_z^+1^_4_y^+2^_8_y^+1^_4_c^+1^_4_z^+2^_9_c^+1^_4_y^+2^_9_y^+2^_9_z^+2^_9_w^+1^_5_w^+1^_5_y^+2^_9_c^+2^_9_c^+2^_9_y^+1^_5_z^+1^_5_c^+2^_9_y^+1^_5_w^+1^_6_c^+1^_5_c^+1^_5_y^+1^_6_z^+1^_6_y^+1^_6_w^+1^_7_c^+1^_6_y^+1^_7_z^+1^_7_y^+1^_7_c^+1^_7_z^+1^_8_y^+1^_8_c^+1^_8_c^+1^_8_z^+1^_9_y^+1^_9_c^+1^_9_

0838167625153353

Fragment Matches Table

Show background peaks

| Position | Ion type | Intensity | mz Theoretical | mz Error (Th) | mz Error (ppm) | Charge | Series Number |
| --- | --- | --- | --- | --- | --- | --- | --- |
| - | - | 453.6 | 121 | - | - | 0 | - |
| - | - | 4013 | 127.1 | - | - | 0 | - |
| - | - | 592.9 | 128.1 | - | - | 0 | - |
| - | - | 4840 | 128.1 | - | - | 0 | - |
| - | - | 1.286E+04 | 129.1 | - | - | 0 | - |
| - | - | 367.3 | 129.7 | - | - | 0 | - |
| - | - | 696.6 | 130.1 | - | - | 0 | - |
| - | - | 3536 | 131.1 | - | - | 0 | - |
| - | - | 2.159E+04 | 136.1 | - | - | 0 | - |
| - | - | 2491 | 142.1 | - | - | 0 | - |
| - | - | 739.5 | 143.1 | - | - | 0 | - |
| - | - | 616.2 | 146.1 | - | - | 0 | - |
| - | - | 903.2 | 149 | - | - | 0 | - |
| - | - | 1362 | 149 | - | - | 0 | - |
| - | - | 583.8 | 154.7 | - | - | 0 | - |
| - | - | 613.9 | 156.1 | - | - | 0 | - |
| - | - | 449.5 | 158.1 | - | - | 0 | - |
| 9 | w | 1.332E+04 | 160.1 | 0.0001432 | 0.8944 | +1 | 2 |
| - | - | 874.5 | 161.1 | - | - | 0 | - |
| - | - | 1853 | 173.1 | - | - | 0 | - |
| - | - | 1369 | 173.5 | - | - | 0 | - |
| - | - | 851.3 | 174.1 | - | - | 0 | - |
| - | - | 937.3 | 175.1 | - | - | 0 | - |
| - | - | 3990 | 181.1 | - | - | 0 | - |
| - | - | 1837 | 184.1 | - | - | 0 | - |
| - | - | 638.2 | 187.1 | - | - | 0 | - |
| - | - | 532.7 | 189 | - | - | 0 | - |
| - | - | 1016 | 189.1 | - | - | 0 | - |
| - | - | 823.1 | 198.1 | - | - | 0 | - |
| - | - | 574 | 201.1 | - | - | 0 | - |
| - | - | 1909 | 215.1 | - | - | 0 | - |
| - | - | 804.9 | 215.1 | - | - | 0 | - |
| 9 | y | 4982 | 216.1 | 0.0002689 | 1.244 | +1 | 2 |
| 9 | z | 4107 | 218.1 | 8.169E-05 | 0.3745 | +1 | 2 |
| - | - | 608.5 | 219.1 | - | - | 0 | - |
| - | - | 1348 | 221.1 | - | - | 0 | - |
| - | - | 802.5 | 223.1 | - | - | 0 | - |
| - | - | 531.2 | 224.4 | - | - | 0 | - |
| - | - | 2783 | 225 | - | - | 0 | - |
| - | - | 2516 | 226.1 | - | - | 0 | - |
| - | - | 907.4 | 228.1 | - | - | 0 | - |
| - | - | 1163 | 233.2 | - | - | 0 | - |
| 9 | y | 4551 | 234.1 | 0.0001564 | 0.6682 | +1 | 2 |
| - | - | 2465 | 239.1 | - | - | 0 | - |
| - | - | 600.7 | 239.1 | - | - | 0 | - |
| - | - | 3253 | 244.1 | - | - | 0 | - |
| - | - | 3.581E+04 | 249.2 | - | - | 0 | - |
| - | - | 1149 | 250.1 | - | - | 0 | - |
| - | - | 5887 | 250.2 | - | - | 0 | - |
| - | - | 1711 | 251.1 | - | - | 0 | - |
| - | - | 1655 | 268.1 | - | - | 0 | - |
| - | - | 776 | 269.1 | - | - | 0 | - |
| - | - | 774.9 | 275 | - | - | 0 | - |
| 6 | y | 592.4 | 275.1 | 0.0009353 | 3.399 | +2 | 5 |
| - | - | 1.878E+04 | 277.2 | - | - | 0 | - |
| - | - | 3080 | 278.2 | - | - | 0 | - |
| - | - | 536.6 | 278.2 | - | - | 0 | - |
| 8 | w | 673.1 | 288.2 | 0.0002154 | 0.7474 | +1 | 3 |
| - | - | 4843 | 289.2 | - | - | 0 | - |
| - | - | 3607 | 295.1 | - | - | 0 | - |
| - | - | 954.4 | 296.1 | - | - | 0 | - |
| - | - | 2005 | 299.1 | - | - | 0 | - |
| - | - | 547.4 | 307.7 | - | - | 0 | - |
| - | - | 937.9 | 311.2 | - | - | 0 | - |
| - | - | 665.8 | 319.2 | - | - | 0 | - |
| - | - | 1783 | 324.1 | - | - | 0 | - |
| 8 | z | 4450 | 333.2 | 5.516E-05 | 0.1656 | +1 | 3 |
| - | - | 718.2 | 334.2 | - | - | 0 | - |
| - | - | 649.2 | 338.2 | - | - | 0 | - |
| - | - | 565.4 | 342.2 | - | - | 0 | - |
| 8 | y | 2651 | 349.2 | 0.0001604 | 0.4595 | +1 | 3 |
| 3 | c | 2933 | 363.2 | 0.0004113 | 1.132 | +1 | 3 |
| - | - | 3213 | 364.2 | - | - | 0 | - |
| - | - | 1095 | 365.2 | - | - | 0 | - |
| - | - | 2186 | 367.1 | - | - | 0 | - |
| - | - | 3403 | 367.2 | - | - | 0 | - |
| - | - | 1820 | 367.3 | - | - | 0 | - |
| - | - | 1405 | 368.2 | - | - | 0 | - |
| - | - | 752.1 | 369.1 | - | - | 0 | - |
| - | - | 978.4 | 379.2 | - | - | 0 | - |
| 3 | c | 1248 | 381.2 | 0.000285 | 0.7476 | +1 | 3 |
| - | - | 1089 | 385.1 | - | - | 0 | - |
| - | - | 971.8 | 385.2 | - | - | 0 | - |
| - | - | 867.7 | 385.6 | - | - | 0 | - |
| - | - | 728.4 | 386.1 | - | - | 0 | - |
| - | - | 612.4 | 386.9 | - | - | 0 | - |
| - | - | 900.4 | 388.2 | - | - | 0 | - |
| - | - | 627.8 | 394.4 | - | - | 0 | - |
| - | - | 824.6 | 395.2 | - | - | 0 | - |
| 4 | y | 1920 | 395.7 | 0.0007072 | 1.787 | +2 | 7 |
| - | - | 864.7 | 401.2 | - | - | 0 | - |
| - | - | 1460 | 402.2 | - | - | 0 | - |
| 7 | w | 1.432E+04 | 417.2 | 0.0002213 | 0.5305 | +1 | 4 |
| - | - | 2400 | 418.2 | - | - | 0 | - |
| - | - | 829.6 | 429.2 | - | - | 0 | - |
| 3 | y | 1558 | 430.7 | 0.004926 | 11.44 | +2 | 8 |
| 3 | z | 698.3 | 431.2 | 0.001581 | 3.666 | +2 | 8 |
| - | - | 1101 | 431.7 | - | - | 0 | - |
| 7 | z | 7865 | 432.2 | 0.0003668 | 0.8487 | +1 | 4 |
| - | - | 1291 | 433.2 | - | - | 0 | - |
| 3 | y | 8695 | 439.3 | 0.000196 | 0.4461 | +2 | 8 |
| - | - | 2982 | 439.8 | - | - | 0 | - |
| - | - | 1414 | 440.3 | - | - | 0 | - |
| - | - | 885.1 | 444.2 | - | - | 0 | - |
| - | - | 1681 | 446.3 | - | - | 0 | - |
| - | - | 985.2 | 447.3 | - | - | 0 | - |
| 7 | y | 2594 | 448.2 | 0.0002908 | 0.6488 | +1 | 4 |
| - | - | 1321 | 462.2 | - | - | 0 | - |
| - | - | 677.8 | 463.2 | - | - | 0 | - |
| - | - | 754.3 | 465.3 | - | - | 0 | - |
| - | - | 747.5 | 471.3 | - | - | 0 | - |
| - | - | 581.9 | 472.3 | - | - | 0 | - |
| - | - | 1497 | 474.3 | - | - | 0 | - |
| - | - | 1191 | 477.3 | - | - | 0 | - |
| - | - | 882.7 | 484.7 | - | - | 0 | - |
| - | - | 1686 | 489.3 | - | - | 0 | - |
| 4 | c | 1741 | 492.3 | 0.0009324 | 1.894 | +1 | 4 |
| - | - | 732.4 | 493.3 | - | - | 0 | - |
| - | - | 740 | 494.3 | - | - | 0 | - |
| - | - | 1514 | 501.2 | - | - | 0 | - |
| - | - | 670.6 | 503.3 | - | - | 0 | - |
| 2 | z | 1163 | 503.8 | 0.00129 | 2.562 | +2 | 9 |
| - | - | 691.5 | 508.3 | - | - | 0 | - |
| 4 | c | 7.343E+04 | 509.3 | 0.0001706 | 0.335 | +1 | 4 |
| - | - | 2.129E+04 | 510.3 | - | - | 0 | - |
| - | - | 3757 | 511.3 | - | - | 0 | - |
| 2 | y | 2074 | 511.8 | 0.001219 | 2.382 | +2 | 9 |
| 2 | y | 1116 | 512.3 | 0.00799 | 15.6 | +2 | 9 |
| 2 | z | 1977 | 512.8 | 0.000965 | 1.882 | +2 | 9 |
| - | - | 1050 | 513.3 | - | - | 0 | - |
| - | - | 840.5 | 513.8 | - | - | 0 | - |
| - | - | 1123 | 514.3 | - | - | 0 | - |
| - | - | 761 | 515.3 | - | - | 0 | - |
| - | - | 641.1 | 515.8 | - | - | 0 | - |
| 6 | w | 1.297E+04 | 516.3 | 7.523E-05 | 0.1457 | +1 | 5 |
| - | - | 3038 | 517.3 | - | - | 0 | - |
| 6 | w | 9536 | 518.2 | 0.0003029 | 0.5844 | +1 | 5 |
| - | - | 2672 | 519.2 | - | - | 0 | - |
| 2 | y | 1.1E+04 | 520.8 | 0.0001573 | 0.302 | +2 | 9 |
| - | - | 4795 | 521.3 | - | - | 0 | - |
| - | - | 2399 | 521.8 | - | - | 0 | - |
| - | - | 1221 | 523.8 | - | - | 0 | - |
| 9 | c | 1132 | 524.3 | 0.002457 | 4.686 | +2 | 9 |
| 9 | c | 2021 | 524.8 | 0.0004696 | 0.8948 | +2 | 9 |
| - | - | 884.1 | 525.3 | - | - | 0 | - |
| - | - | 942.6 | 525.8 | - | - | 0 | - |
| - | - | 5740 | 531.3 | - | - | 0 | - |
| 6 | y | 6438 | 532.3 | 0.0006377 | 1.198 | +1 | 5 |
| 6 | z | 1.356E+04 | 533.3 | 0.0004484 | 0.8408 | +1 | 5 |
| 9 | c | 8.314E+04 | 533.3 | 1.243E-05 | 0.02331 | +2 | 9 |
| - | - | 4.946E+04 | 533.8 | - | - | 0 | - |
| - | - | 4838 | 534.3 | - | - | 0 | - |
| - | - | 1.85E+04 | 534.3 | - | - | 0 | - |
| - | - | 3363 | 534.8 | - | - | 0 | - |
| - | - | 888 | 535.3 | - | - | 0 | - |
| - | - | 2365 | 541.3 | - | - | 0 | - |
| - | - | 1904 | 541.8 | - | - | 0 | - |
| - | - | 1.095E+05 | 547.8 | - | - | 0 | - |
| - | - | 6.519E+04 | 548.3 | - | - | 0 | - |
| - | - | 2.387E+04 | 548.8 | - | - | 0 | - |
| 6 | y | 9760 | 549.3 | 0.003026 | 5.508 | +1 | 5 |
| - | - | 920.4 | 549.8 | - | - | 0 | - |
| - | - | 1218 | 550.3 | - | - | 0 | - |
| - | - | 1092 | 554.8 | - | - | 0 | - |
| - | - | 1790 | 558.3 | - | - | 0 | - |
| - | - | 1645 | 559.3 | - | - | 0 | - |
| - | - | 713.2 | 560.8 | - | - | 0 | - |
| - | - | 3287 | 569.3 | - | - | 0 | - |
| - | - | 2376 | 569.8 | - | - | 0 | - |
| - | - | 688 | 574.4 | - | - | 0 | - |
| - | - | 952.9 | 575.3 | - | - | 0 | - |
| - | - | 850.4 | 576.4 | - | - | 0 | - |
| - | - | 1.052E+05 | 577.3 | - | - | 0 | - |
| - | - | 6.915E+04 | 577.8 | - | - | 0 | - |
| - | - | 2.124E+04 | 578.3 | - | - | 0 | - |
| - | - | 761 | 578.4 | - | - | 0 | - |
| - | - | 7124 | 578.8 | - | - | 0 | - |
| - | - | 2296 | 579.3 | - | - | 0 | - |
| - | - | 603.6 | 592.3 | - | - | 0 | - |
| - | - | 693.2 | 593.3 | - | - | 0 | - |
| - | - | 1212 | 601.4 | - | - | 0 | - |
| - | - | 953.9 | 602.4 | - | - | 0 | - |
| 5 | w | 4.615E+04 | 603.3 | 0.0002123 | 0.352 | +1 | 6 |
| - | - | 1.505E+04 | 604.3 | - | - | 0 | - |
| - | - | 2721 | 605.3 | - | - | 0 | - |
| 5 | c | 1886 | 605.4 | 0.0011 | 1.818 | +1 | 5 |
| - | - | 784.5 | 606.3 | - | - | 0 | - |
| - | - | 899.8 | 606.4 | - | - | 0 | - |
| - | - | 2923 | 621.4 | - | - | 0 | - |
| 5 | c | 6.637E+04 | 622.4 | 0.0002447 | 0.3931 | +1 | 5 |
| - | - | 2.549E+04 | 623.4 | - | - | 0 | - |
| - | - | 5046 | 624.4 | - | - | 0 | - |
| - | - | 872.1 | 625.4 | - | - | 0 | - |
| 5 | y | 1281 | 644.4 | 0.0002223 | 0.3449 | +1 | 6 |
| 5 | z | 1.778E+04 | 646.4 | 2.548E-06 | 0.003941 | +1 | 6 |
| - | - | 7457 | 647.4 | - | - | 0 | - |
| - | - | 902.4 | 648.3 | - | - | 0 | - |
| - | - | 1827 | 648.4 | - | - | 0 | - |
| 5 | y | 1.287E+04 | 662.4 | 0.0002889 | 0.4362 | +1 | 6 |
| - | - | 4518 | 663.4 | - | - | 0 | - |
| - | - | 1359 | 664.3 | - | - | 0 | - |
| - | - | 1672 | 664.4 | - | - | 0 | - |
| - | - | 991.3 | 670.4 | - | - | 0 | - |
| - | - | 966.5 | 681.3 | - | - | 0 | - |
| - | - | 5131 | 686.4 | - | - | 0 | - |
| - | - | 2025 | 687.4 | - | - | 0 | - |
| - | - | 650.3 | 692.4 | - | - | 0 | - |
| - | - | 1294 | 702.4 | - | - | 0 | - |
| - | - | 1788 | 712.4 | - | - | 0 | - |
| - | - | 866.1 | 713.5 | - | - | 0 | - |
| 4 | w | 882.4 | 716.4 | 0.006908 | 9.642 | +1 | 7 |
| - | - | 805.9 | 718.4 | - | - | 0 | - |
| - | - | 777.9 | 722.4 | - | - | 0 | - |
| 6 | c | 2.681E+04 | 723.4 | 0.0003768 | 0.5208 | +1 | 6 |
| - | - | 1.207E+04 | 724.4 | - | - | 0 | - |
| - | - | 2795 | 725.4 | - | - | 0 | - |
| - | - | 1497 | 730.5 | - | - | 0 | - |
| - | - | 1418 | 767.9 | - | - | 0 | - |
| - | - | 800.5 | 768.9 | - | - | 0 | - |
| - | - | 901.1 | 769.3 | - | - | 0 | - |
| - | - | 876.5 | 772.4 | - | - | 0 | - |
| 4 | y | 1125 | 772.5 | 0.002227 | 2.883 | +1 | 7 |
| - | - | 3307 | 773.5 | - | - | 0 | - |
| 4 | z | 5.096E+04 | 774.4 | 0.0003154 | 0.4073 | +1 | 7 |
| - | - | 2.091E+04 | 775.5 | - | - | 0 | - |
| - | - | 5783 | 776.5 | - | - | 0 | - |
| - | - | 858.4 | 777.5 | - | - | 0 | - |
| - | - | 821.3 | 778.5 | - | - | 0 | - |
| - | - | 2150 | 789.5 | - | - | 0 | - |
| 4 | y | 3.158E+04 | 790.5 | 0.0002681 | 0.3392 | +1 | 7 |
| - | - | 1.29E+04 | 791.5 | - | - | 0 | - |
| - | - | 3518 | 792.5 | - | - | 0 | - |
| - | - | 1730 | 807.4 | - | - | 0 | - |
| - | - | 771.5 | 808.4 | - | - | 0 | - |
| - | - | 1646 | 817.5 | - | - | 0 | - |
| - | - | 4194 | 821.5 | - | - | 0 | - |
| 7 | c | 6.066E+04 | 822.5 | 0.0005534 | 0.6728 | +1 | 7 |
| - | - | 2.779E+04 | 823.5 | - | - | 0 | - |
| - | - | 8214 | 824.5 | - | - | 0 | - |
| - | - | 1274 | 825.5 | - | - | 0 | - |
| 3 | z | 3.446E+04 | 861.5 | 0.0006461 | 0.75 | +1 | 8 |
| - | - | 1.657E+04 | 862.5 | - | - | 0 | - |
| - | - | 376.5 | 862.6 | - | - | 0 | - |
| - | - | 3709 | 863.5 | - | - | 0 | - |
| - | - | 797.3 | 864.5 | - | - | 0 | - |
| 3 | y | 3547 | 877.5 | 0.0003882 | 0.4424 | +1 | 8 |
| - | - | 2016 | 878.5 | - | - | 0 | - |
| - | - | 2439 | 893.5 | - | - | 0 | - |
| - | - | 1015 | 894.5 | - | - | 0 | - |
| 8 | c | 1439 | 920.5 | 0.001191 | 1.294 | +1 | 8 |
| - | - | 1234 | 921.5 | - | - | 0 | - |
| - | - | 2009 | 936.5 | - | - | 0 | - |
| 8 | c | 1.006E+05 | 937.5 | 0.0005799 | 0.6185 | +1 | 8 |
| - | - | 5.704E+04 | 938.5 | - | - | 0 | - |
| - | - | 1.569E+04 | 939.5 | - | - | 0 | - |
| - | - | 2774 | 940.5 | - | - | 0 | - |
| - | - | 940.1 | 941.5 | - | - | 0 | - |
| - | - | 714.3 | 968.9 | - | - | 0 | - |
| - | - | 770.2 | 1007 | - | - | 0 | - |
| - | - | 1336 | 1022 | - | - | 0 | - |
| - | - | 1666 | 1023 | - | - | 0 | - |
| - | - | 747.4 | 1024 | - | - | 0 | - |
| 2 | z | 2044 | 1025 | 0.005442 | 5.312 | +1 | 9 |
| - | - | 6553 | 1026 | - | - | 0 | - |
| - | - | 2625 | 1027 | - | - | 0 | - |
| 2 | y | 771.2 | 1041 | 0.008541 | 8.208 | +1 | 9 |
| - | - | 2922 | 1050 | - | - | 0 | - |
| - | - | 2351 | 1051 | - | - | 0 | - |
| - | - | 1192 | 1052 | - | - | 0 | - |
| 9 | c | 1.302E+04 | 1066 | 0.001732 | 1.625 | +1 | 9 |
| - | - | 1.722E+04 | 1067 | - | - | 0 | - |
| - | - | 7543 | 1068 | - | - | 0 | - |
| - | - | 2243 | 1069 | - | - | 0 | - |
| - | - | 686.7 | 1070 | - | - | 0 | - |
| - | - | 1627 | 1083 | - | - | 0 | - |
| - | - | 625.5 | 1084 | - | - | 0 | - |
| - | - | 876.2 | 1095 | - | - | 0 | - |
| - | - | 1703 | 1096 | - | - | 0 | - |
| - | - | 1010 | 1097 | - | - | 0 | - |
| - | - | 853.7 | 1109 | - | - | 0 | - |
| - | - | 2407 | 1110 | - | - | 0 | - |
| - | - | 1431 | 1111 | - | - | 0 | - |
| - | - | 659.9 | 1112 | - | - | 0 | - |
| - | - | 727.8 | 1121 | - | - | 0 | - |
| - | - | 1.111E+04 | 1138 | - | - | 0 | - |
| - | - | 6.071E+04 | 1139 | - | - | 0 | - |
| - | - | 4.107E+04 | 1140 | - | - | 0 | - |
| - | - | 1.389E+04 | 1141 | - | - | 0 | - |
| - | - | 4526 | 1142 | - | - | 0 | - |
| - | - | 7900 | 1154 | - | - | 0 | - |
| - | - | 1.595E+04 | 1155 | - | - | 0 | - |
| - | - | 832.4 | 1156 | - | - | 0 | - |
| - | - | 1.076E+04 | 1156 | - | - | 0 | - |
| - | - | 3589 | 1157 | - | - | 0 | - |
| - | - | 854.2 | 1158 | - | - | 0 | - |
| - | - | 610.6 | 1730 | - | - | 0 | - |
| - | - | 798 | 3040 | - | - | 0 | - |
| - | - | 669.1 | 3320 | - | - | 0 | - |

m/z Charge Intensity FragmentType MassShift Position
121.02889251708984 0 453.63623
127.08671569824219 0 4012.5088
128.090087890625 0 592.8875
128.09457397460938 0 4840.1987
129.1024169921875 0 12859.934
129.71929931640625 0 367.25522
130.1057891845703 0 696.6238
131.11805725097656 0 3536.0496
136.0758514404297 0 21589.342
142.05001831054688 0 2490.729
143.0943603515625 0 739.48755
146.12823486328125 0 616.2476
148.95472717285156 0 903.16284
149.04495239257812 0 1361.6665
154.69656372070312 0 583.7875
156.10223388671875 0 613.8956
158.14100646972656 0 449.52686
160.06057739257812 0 13324.916 w 8
161.06399536132812 0 874.53375
173.1286163330078 0 1852.878
173.45138549804688 0 1369.2659
174.13658142089844 0 851.2724
175.13328552246094 0 937.3336
181.097412109375 0 3989.6807
184.12063598632812 0 1836.9452
187.1444091796875 0 638.24384
188.9955596923828 0 532.65576
189.10043334960938 0 1015.9005
198.12400817871094 0 823.1266
201.1234893798828 0 573.9985
215.10269165039062 0 1908.9596
215.13912963867188 0 804.9307
216.13453674316406 0 4982.113 y Water loss 8
218.12619018554688 0 4107.0444 z 8
219.13034057617188 0 608.4851
221.0843505859375 0 1347.7407
223.1077423095703 0 802.5419
224.3513641357422 0 531.19336
225.0430908203125 0 2783.1548
226.1188201904297 0 2516.0933
228.1342010498047 0 907.4195
233.1607666015625 0 1162.9384
234.14498901367188 0 4551.4136 y 8
239.09510803222656 0 2464.7043
239.11276245117188 0 600.732
244.12939453125 0 3252.565
249.15988159179688 0 35805.523
250.11895751953125 0 1149.4635
250.16319274902344 0 5887.3643
251.10281372070312 0 1710.8917
268.128662109375 0 1655.4695
269.1133728027344 0 775.96063
275.03289794921875 0 774.86475
275.1466369628906 0 592.4312 y 5
277.1546630859375 0 18777.854
278.15802001953125 0 3079.663
278.1745300292969 0 536.57605
288.1551818847656 0 673.1392 w 7
289.1633605957031 0 4842.5107
295.1032409667969 0 3607.1023
296.105224609375 0 954.4011
299.0616455078125 0 2004.5371
307.6866760253906 0 547.4491
311.2072448730469 0 937.93604
319.1881408691406 0 665.81134
324.1462097167969 0 1783.1198
333.1531066894531 0 4450.4917 z 7
334.1561279296875 0 718.19574
338.2043762207031 0 649.235
342.1557922363281 0 565.3808
349.17193603515625 0 2651.075 y 7
363.2022705078125 0 2933.4744 c Water loss 2
364.1864013671875 0 3212.7793
365.189453125 0 1095.3665
367.0694885253906 0 2186.0918
367.19573974609375 0 3403.3057
367.26959228515625 0 1819.9814
368.1990051269531 0 1404.8876
369.1241760253906 0 752.13416
379.19708251953125 0 978.4262
381.2135314941406 0 1247.723 c 2
385.1462097167969 0 1088.6115
385.2186279296875 0 971.7657
385.5550842285156 0 867.6801
386.1291809082031 0 728.39575
386.8822937011719 0 612.38916
388.232177734375 0 900.3926
394.4405212402344 0 627.7718
395.24481201171875 0 824.56244
395.73779296875 0 1919.5558 y 3
401.23907470703125 0 864.71954
402.2467956542969 0 1459.6318
417.1982116699219 0 14322.429 w 6
418.20208740234375 0 2399.8667
429.2326965332031 0 829.563
430.7447509765625 0 1558.0333 y Ammonia loss 2
431.2421569824219 0 698.2546 z 2
431.74395751953125 0 1100.7454
432.2218322753906 0 7864.7026 z 6
433.2237854003906 0 1290.5985
439.2532958984375 0 8695.116 y 2
439.7552795410156 0 2981.7827
440.2559814453125 0 1413.7101
444.2439880371094 0 885.05066
446.2610168457031 0 1680.7811
447.26495361328125 0 985.187
448.2398986816406 0 2594.4102 y 6
462.1965026855469 0 1320.9813
463.20013427734375 0 677.83264
465.2945556640625 0 754.34186
471.2668762207031 0 747.5422
472.2786865234375 0 581.87573
474.2720031738281 0 1496.8456
477.2705383300781 0 1191.487
484.743408203125 0 882.6604
489.279296875 0 1685.8074
492.2825927734375 0 1741.1609 c Ammonia loss 3
493.2886047363281 0 732.39764
494.29766845703125 0 739.97833
501.2290954589844 0 1513.7454
503.2734680175781 0 670.62976
503.7688293457031 0 1163.2883 z Water loss 1
508.30352783203125 0 691.5216
509.3083801269531 0 73433 c 3
510.3115539550781 0 21290.668
511.3139343261719 0 3756.9246
511.78070068359375 0 2074.058 y Water loss 1
512.2794799804688 0 1116.255 y Ammonia loss 1
512.7763671875 0 1977.3585 z 1
513.2817993164062 0 1050.4365
513.7782592773438 0 840.4703
514.3226928710938 0 1123.4762
515.325927734375 0 760.9825
515.7955932617188 0 641.09186
516.2664794921875 0 12974.172 w 5
517.2701416015625 0 3037.7302
518.2459716796875 0 9535.66 w 5
519.2486572265625 0 2671.8005
520.7846069335938 0 11004.917 y 1
521.2870483398438 0 4795.145
521.7866821289062 0 2399.077
523.8016357421875 0 1220.8088
524.31103515625 0 1131.9537 c Water loss 8
524.8059692382812 0 2021.0768 c Ammonia loss 8
525.3074340820312 0 884.07733
525.8103637695312 0 942.5983
531.2540893554688 0 5739.512
532.2606811523438 0 6438.2407 y Ammonia loss 5
533.2695922851562 0 13557.079 z 5
533.3187866210938 0 83139.42 c 8
533.8204345703125 0 49461.76
534.27197265625 0 4837.997
534.3215942382812 0 18496.867
534.8233032226562 0 3363.0867
535.2760620117188 0 888.04034
541.2877807617188 0 2364.6084
541.7885131835938 0 1903.7577
547.7899780273438 0 109506.38
548.29150390625 0 65194.76
548.7928466796875 0 23867.521
549.2908935546875 0 9760.364 y 5
549.794677734375 0 920.43555
550.2924194335938 0 1218.0215
554.827392578125 0 1091.7555
558.3358154296875 0 1789.9336
559.3438720703125 0 1645.1373
560.8321533203125 0 713.1973
569.3199462890625 0 3286.509
569.8196411132812 0 2375.8787
574.35400390625 0 688.0096
575.2820434570312 0 952.8726
576.3690185546875 0 850.3655
577.326904296875 0 105151.914
577.8283081054688 0 69147.87
578.3295288085938 0 21240.193
578.3746948242188 0 761.0205
578.8311767578125 0 7124.4463
579.3331298828125 0 2295.6428
592.3013916015625 0 603.60864
593.3316650390625 0 693.2322
601.356689453125 0 1211.5853
602.3597412109375 0 953.9472
603.2986450195312 0 46147.918 w 4
604.3016967773438 0 15051.85
605.3045043945312 0 2720.8645
605.3646240234375 0 1885.8323 c Ammonia loss 4
606.310546875 0 784.5217
606.3685302734375 0 899.7654
621.38427734375 0 2923.1548
622.3920288085938 0 66367.32 c 4
623.3953857421875 0 25493.535
624.3980712890625 0 5045.929
625.3992919921875 0 872.12146
644.3611450195312 0 1280.7124 y Water loss 4
646.3532104492188 0 17775.05 z 4
647.3562622070312 0 7457.459
648.27294921875 0 902.4045
648.3583984375 0 1826.9224
662.3716430664062 0 12873.581 y 4
663.3751220703125 0 4518.252
664.2930297851562 0 1358.5026
664.3759765625 0 1671.9817
670.4151611328125 0 991.2506
681.3203735351562 0 966.5393
686.4315795898438 0 5131.312
687.4365234375 0 2024.5537
692.3939819335938 0 650.32526
702.447509765625 0 1293.5225
712.449951171875 0 1788.0511
713.4505004882812 0 866.1302
716.389404296875 0 882.42 w 3
718.3819580078125 0 805.8866
722.43017578125 0 777.8523
723.4395751953125 0 26807.53 c 5
724.4425048828125 0 12074.601
725.444580078125 0 2794.7996
730.4570922851562 0 1496.8035
767.897705078125 0 1417.9136
768.9012451171875 0 800.48267
769.3341064453125 0 901.11633
772.384033203125 0 876.50616
772.4585571289062 0 1125.3998 y Water loss 3
773.4635620117188 0 3306.6282
774.448486328125 0 50955.79 z 3
775.4518432617188 0 20908.387
776.454833984375 0 5783.1274
777.47021484375 0 858.39923
778.488037109375 0 821.2739
789.471923828125 0 2150.4514
790.4671630859375 0 31584.908 y 3
791.4698486328125 0 12900.347
792.4715576171875 0 3518.2769
807.4241943359375 0 1730.1426
808.4255981445312 0 771.4646
817.4878540039062 0 1645.592
821.500732421875 0 4194.191
822.5078125 0 60655.945 c 6
823.5110473632812 0 27790.814
824.5137329101562 0 8213.821
825.5216674804688 0 1274.4443
861.4795532226562 0 34457.05 z 2
862.4829711914062 0 16572.45
862.5562744140625 0 376.51337
863.4837036132812 0 3709.2327
864.4907836914062 0 797.31055
877.49853515625 0 3546.916 y 2
878.49951171875 0 2016.3317
893.5223388671875 0 2439.421
894.52490234375 0 1014.9979
920.507568359375 0 1438.9342 c Ammonia loss 7
921.511962890625 0 1233.5209
936.5235595703125 0 2009.0262
937.5347290039062 0 100644.41 c 7
938.5377197265625 0 57035.473
939.5400390625 0 15690.128
940.54296875 0 2774.3494
941.5428466796875 0 940.1462
968.91259765625 0 714.2628
1006.6084594726562 0 770.1928
1021.6206665039062 0 1335.9962
1022.6260375976562 0 1665.5271
1023.62841796875 0 747.375
1024.5380859375 0 2044.1688 z 1
1025.549072265625 0 6552.8457
1026.5523681640625 0 2625.3213
1040.5537109375 0 771.2024 y 1
1049.6124267578125 0 2922.104
1050.612548828125 0 2351.0674
1051.617919921875 0 1191.8248
1065.6285400390625 0 13020.58 c 8
1066.6343994140625 0 17219.955
1067.639404296875 0 7543.201
1068.6427001953125 0 2242.5046
1069.6488037109375 0 686.6579
1082.571044921875 0 1626.6494
1083.5845947265625 0 625.5464
1094.5792236328125 0 876.19476
1095.6292724609375 0 1703.3832
1096.629638671875 0 1010.2743
1108.6468505859375 0 853.715
1109.655029296875 0 2407.1936
1110.6546630859375 0 1431.259
1111.656494140625 0 659.89636
1120.6376953125 0 727.83514
1137.6270751953125 0 11109.639
1138.6341552734375 0 60706.824
1139.6363525390625 0 41066.918
1140.64013671875 0 13891.74
1141.6422119140625 0 4525.9736
1153.64453125 0 7899.999
1154.6510009765625 0 15951.153
1155.520751953125 0 832.38727
1155.6558837890625 0 10755.498
1156.6591796875 0 3589.395
1157.6767578125 0 854.24384
1730.4549560546875 0 610.6102
3039.9208984375 0 797.97015
3319.749267578125 0 669.06445

Spectrum Details

|  |  |
| --- | --- |
| Matched peaks? Matched peaks The total absolute number of peaks matched. Additionally in brackets the total fraction of peaks matched and the total number of peaks is shown. | 51 (17.11% of 298) |
| FDR? FDR The false discovery rate estimated for this peptide. It is calculated by matching all theoretical fragments with a non-integer shift with the raw peaks for this spectrum. This is done with 40 different shifts. The resulting percentage is the average number of annotated peaks over the number of annotated peaks with the correct spectrum. | 1.21% |
| Satellite FDR? Satellite FDR See the FDR for details on its calculation. This satellite ion specific FDR only contains the satellite ions (d/w) for I/L/J positions. | 9.52% |
| PSM Score? PSM Score The PSM Score as given by Hecklib to this annotated spectrum. It is shown with three significant figures. | 416 |

#### Spectrum 3725? Spectrum 3725 The raw spectrum of this peptide as annotated by Hecklib. The fragments are coloured according to ion type (see legend). Any peaks with a star '\*' as text can be hovered over to see the full details, first the ion type second the mass shift type. By hovering over the amino acids in the peptide or ions in the legend the corresponding peaks are highlighted. By toggling the 'Unassigned' label you can turn the background (unassigned) peaks on or off in the plot. By updating the slider in the Ion legend you can update the spectrum to only show the top X% of the peaks with labels. The top X% means any peak that is within X% of the highest intensity. By dragging in the spectrum you can zoom in to a specific part of the spectrum and use 'Zoom Out' to get back to the original zoom level. The annotation of the spectrum is based on the given sequence in the peptides file and is done with different software so inconsistencies are likely. The peaks are annotated based on the given sequence, with 20 ppm tolerance.

Copy Data

##### Spectrum 3725 (TSV)

###### Preview

```
Loading example...
```

*Click on the button to copy the data to your clipboard.*

Mz MinMz MaxIntensity Max

WidthHeightPeptide font sizePeptide stroke widthSpectrum font sizeSpectrum stroke widthCompact peptide

Ion legend

wxyz

abcd

OtherUnassignedIon^Charge^_Position_Show for top:%

JYSKJTVDKS

01.37e+42.73e+44.10e+45.47e+4

Zoom Out

y^+1^_2_y^+1^_2_y^+1^_2_a^+1^_2_b^+1^_2_y^+1^_3_y^+1^_3_b^+1^_3_y^+2^_8_y^+1^_4_b^+1^_4_b^+1^_4_b^+2^_9_y^+2^_9_b^+2^_9_y^+1^_5_y^+1^_5_\*\*\*b^+1^_5_y^+1^_6_y^+1^_6_b^+1^_6_b^+1^_6_y^+1^_7_b^+1^_7_y^+1^_7_b^+1^_7_y^+1^_8_y^+1^_8_b^+1^_8_y^+1^_9_

0585117017552340

Fragment Matches Table

Show background peaks

| Position | Ion type | Intensity | mz Theoretical | mz Error (Th) | mz Error (ppm) | Charge | Series Number |
| --- | --- | --- | --- | --- | --- | --- | --- |
| - | - | 647.9 | 120.1 | - | - | 0 | - |
| - | - | 1508 | 120.1 | - | - | 0 | - |
| - | - | 397.2 | 120.8 | - | - | 0 | - |
| - | - | 402.9 | 121.7 | - | - | 0 | - |
| - | - | 371.9 | 124 | - | - | 0 | - |
| - | - | 484.5 | 129.1 | - | - | 0 | - |
| - | - | 3.812E+04 | 129.1 | - | - | 0 | - |
| - | - | 490.9 | 130 | - | - | 0 | - |
| - | - | 2598 | 130.1 | - | - | 0 | - |
| - | - | 698.6 | 134 | - | - | 0 | - |
| - | - | 3.418E+04 | 136.1 | - | - | 0 | - |
| - | - | 399.5 | 136.2 | - | - | 0 | - |
| - | - | 412.3 | 136.9 | - | - | 0 | - |
| - | - | 2506 | 137.1 | - | - | 0 | - |
| - | - | 534.5 | 139.1 | - | - | 0 | - |
| - | - | 550.2 | 148.9 | - | - | 0 | - |
| - | - | 534.3 | 153.1 | - | - | 0 | - |
| - | - | 479.1 | 154.6 | - | - | 0 | - |
| - | - | 1314 | 157.1 | - | - | 0 | - |
| - | - | 421.3 | 167.1 | - | - | 0 | - |
| - | - | 536.8 | 169.1 | - | - | 0 | - |
| - | - | 708.8 | 169.1 | - | - | 0 | - |
| - | - | 944.7 | 171.1 | - | - | 0 | - |
| - | - | 490.4 | 173.1 | - | - | 0 | - |
| - | - | 3508 | 173.1 | - | - | 0 | - |
| - | - | 1482 | 173.5 | - | - | 0 | - |
| - | - | 745.9 | 178.1 | - | - | 0 | - |
| - | - | 453.9 | 181 | - | - | 0 | - |
| - | - | 673 | 181.1 | - | - | 0 | - |
| - | - | 1229 | 183.1 | - | - | 0 | - |
| - | - | 765.1 | 183.1 | - | - | 0 | - |
| - | - | 761.3 | 184.1 | - | - | 0 | - |
| - | - | 441.8 | 184.5 | - | - | 0 | - |
| - | - | 808.4 | 185.1 | - | - | 0 | - |
| - | - | 480.1 | 186.1 | - | - | 0 | - |
| - | - | 979.4 | 187.1 | - | - | 0 | - |
| - | - | 1623 | 187.1 | - | - | 0 | - |
| - | - | 1164 | 189.1 | - | - | 0 | - |
| - | - | 941.8 | 197.1 | - | - | 0 | - |
| - | - | 657.7 | 197.2 | - | - | 0 | - |
| - | - | 1988 | 198.1 | - | - | 0 | - |
| - | - | 1883 | 199.1 | - | - | 0 | - |
| - | - | 1443 | 201.1 | - | - | 0 | - |
| - | - | 906.1 | 202.1 | - | - | 0 | - |
| - | - | 779.7 | 203.1 | - | - | 0 | - |
| - | - | 701.5 | 204.1 | - | - | 0 | - |
| - | - | 1931 | 209.1 | - | - | 0 | - |
| - | - | 610.8 | 215.1 | - | - | 0 | - |
| - | - | 5386 | 215.1 | - | - | 0 | - |
| 9 | y | 7883 | 216.1 | 6.683E-05 | 0.3092 | +1 | 2 |
| - | - | 704.3 | 217.1 | - | - | 0 | - |
| 9 | y | 1223 | 217.1 | 0.0004704 | 2.166 | +1 | 2 |
| - | - | 690.9 | 217.1 | - | - | 0 | - |
| - | - | 2095 | 223.1 | - | - | 0 | - |
| - | - | 677.2 | 224.2 | - | - | 0 | - |
| - | - | 6977 | 226.1 | - | - | 0 | - |
| - | - | 722.7 | 227.1 | - | - | 0 | - |
| - | - | 697.5 | 229.1 | - | - | 0 | - |
| 9 | y | 5587 | 234.1 | 8.015E-05 | 0.3423 | +1 | 2 |
| - | - | 1243 | 235.1 | - | - | 0 | - |
| - | - | 1425 | 239.1 | - | - | 0 | - |
| - | - | 1744 | 240.1 | - | - | 0 | - |
| - | - | 647.1 | 242.2 | - | - | 0 | - |
| - | - | 5102 | 244.1 | - | - | 0 | - |
| - | - | 1115 | 248.2 | - | - | 0 | - |
| 2 | a | 2.125E+04 | 249.2 | 1.009E-05 | 0.04048 | +1 | 2 |
| - | - | 2897 | 250.2 | - | - | 0 | - |
| - | - | 1688 | 251.1 | - | - | 0 | - |
| - | - | 616.2 | 251.2 | - | - | 0 | - |
| - | - | 571.2 | 265 | - | - | 0 | - |
| - | - | 592.6 | 267.1 | - | - | 0 | - |
| - | - | 1130 | 270.2 | - | - | 0 | - |
| 2 | b | 6131 | 277.2 | 9.744E-05 | 0.3516 | +1 | 2 |
| - | - | 788.7 | 278.2 | - | - | 0 | - |
| - | - | 579.7 | 282.1 | - | - | 0 | - |
| - | - | 3127 | 285 | - | - | 0 | - |
| - | - | 667.7 | 290.1 | - | - | 0 | - |
| - | - | 1120 | 296.2 | - | - | 0 | - |
| - | - | 2547 | 299.1 | - | - | 0 | - |
| - | - | 579 | 303.1 | - | - | 0 | - |
| - | - | 1984 | 311.2 | - | - | 0 | - |
| - | - | 721.3 | 312.2 | - | - | 0 | - |
| - | - | 1122 | 316.2 | - | - | 0 | - |
| - | - | 1283 | 329.2 | - | - | 0 | - |
| 8 | y | 536.4 | 331.2 | 0.000105 | 0.3171 | +1 | 3 |
| 8 | y | 2358 | 349.2 | 0.0008161 | 2.337 | +1 | 3 |
| - | - | 8276 | 355.1 | - | - | 0 | - |
| - | - | 2187 | 356.1 | - | - | 0 | - |
| - | - | 555.4 | 361 | - | - | 0 | - |
| 3 | b | 1190 | 364.2 | 0.0003143 | 0.8631 | +1 | 3 |
| - | - | 625.5 | 379.2 | - | - | 0 | - |
| - | - | 737.8 | 415.8 | - | - | 0 | - |
| - | - | 619.7 | 415.8 | - | - | 0 | - |
| - | - | 679.9 | 420 | - | - | 0 | - |
| - | - | 757.8 | 427.2 | - | - | 0 | - |
| - | - | 1047 | 429.2 | - | - | 0 | - |
| - | - | 935.4 | 430.3 | - | - | 0 | - |
| 3 | y | 3326 | 439.3 | 0.0005975 | 1.36 | +2 | 8 |
| - | - | 1714 | 439.8 | - | - | 0 | - |
| 7 | y | 2015 | 448.2 | 0.0001688 | 0.3765 | +1 | 4 |
| 4 | b | 1111 | 474.3 | 2.245E-05 | 0.04734 | +1 | 4 |
| 4 | b | 1806 | 492.3 | 0.0006272 | 1.274 | +1 | 4 |
| - | - | 630.4 | 493.3 | - | - | 0 | - |
| - | - | 601.8 | 497.2 | - | - | 0 | - |
| - | - | 1246 | 511.3 | - | - | 0 | - |
| 9 | b | 654.3 | 515.8 | 0.00322 | 6.243 | +2 | 9 |
| 2 | y | 3075 | 520.8 | 0.0005235 | 1.005 | +2 | 9 |
| - | - | 1705 | 521.3 | - | - | 0 | - |
| 9 | b | 916.2 | 524.8 | 0.004986 | 9.501 | +2 | 9 |
| - | - | 1216 | 525.3 | - | - | 0 | - |
| - | - | 2214 | 529.3 | - | - | 0 | - |
| 6 | y | 794.4 | 531.3 | 0.004903 | 9.229 | +1 | 5 |
| - | - | 1140 | 533.8 | - | - | 0 | - |
| - | - | 719.2 | 534.3 | - | - | 0 | - |
| 6 | y | 3392 | 549.3 | 0.0005231 | 0.9524 | +1 | 5 |
| - | - | 834.3 | 550.3 | - | - | 0 | - |
| - | - | 2131 | 557.3 | - | - | 0 | - |
| - | - | 658.3 | 559.8 | - | - | 0 | - |
| - | - | 762.3 | 560.3 | - | - | 0 | - |
| 0 | Precursor | 4056 | 568.3 | 0.0001414 | 0.2489 | +2 | -1 |
| 0 | Precursor | 2867 | 568.8 | 0.008622 | 15.16 | +2 | -1 |
| - | - | 821.9 | 575.6 | - | - | 0 | - |
| - | - | 711.5 | 576.3 | - | - | 0 | - |
| 0 | Precursor | 5.413E+04 | 577.3 | 0.0001971 | 0.3413 | +2 | -1 |
| - | - | 3.755E+04 | 577.8 | - | - | 0 | - |
| - | - | 1.427E+04 | 578.3 | - | - | 0 | - |
| - | - | 1884 | 578.8 | - | - | 0 | - |
| - | - | 594.4 | 584 | - | - | 0 | - |
| - | - | 2006 | 602.8 | - | - | 0 | - |
| - | - | 649.4 | 603.3 | - | - | 0 | - |
| 5 | b | 1592 | 605.4 | 0.0009748 | 1.61 | +1 | 5 |
| - | - | 1067 | 643.3 | - | - | 0 | - |
| 5 | y | 2544 | 644.4 | 0.0005274 | 0.8185 | +1 | 6 |
| - | - | 1096 | 645.4 | - | - | 0 | - |
| - | - | 929.1 | 653.9 | - | - | 0 | - |
| - | - | 779.6 | 654.4 | - | - | 0 | - |
| - | - | 829.9 | 659.8 | - | - | 0 | - |
| 5 | y | 3505 | 662.4 | 0.0008993 | 1.358 | +1 | 6 |
| - | - | 880.7 | 663.4 | - | - | 0 | - |
| - | - | 705.6 | 685.4 | - | - | 0 | - |
| 6 | b | 700.8 | 688.4 | 0.0006165 | 0.8955 | +1 | 6 |
| - | - | 944.1 | 692.4 | - | - | 0 | - |
| 6 | b | 701.3 | 706.4 | 0.003674 | 5.201 | +1 | 6 |
| - | - | 610.2 | 709.9 | - | - | 0 | - |
| - | - | 990.9 | 720.4 | - | - | 0 | - |
| - | - | 755 | 721.4 | - | - | 0 | - |
| - | - | 1060 | 722.4 | - | - | 0 | - |
| - | - | 835.8 | 728.4 | - | - | 0 | - |
| - | - | 975.8 | 739.3 | - | - | 0 | - |
| - | - | 878.8 | 754.4 | - | - | 0 | - |
| 4 | y | 4562 | 772.5 | 0.0007639 | 0.9889 | +1 | 7 |
| - | - | 2042 | 773.5 | - | - | 0 | - |
| - | - | 1558 | 782.4 | - | - | 0 | - |
| - | - | 1664 | 782.9 | - | - | 0 | - |
| - | - | 799.3 | 785.4 | - | - | 0 | - |
| 7 | b | 795.6 | 787.5 | 0.0002438 | 0.3096 | +1 | 7 |
| 4 | y | 2015 | 790.5 | 0.001258 | 1.591 | +1 | 7 |
| 7 | b | 1378 | 805.5 | 0.002935 | 3.644 | +1 | 7 |
| - | - | 706.3 | 806.5 | - | - | 0 | - |
| - | - | 768.1 | 820.4 | - | - | 0 | - |
| - | - | 684.7 | 842.5 | - | - | 0 | - |
| 3 | y | 1616 | 859.5 | 0.003862 | 4.493 | +1 | 8 |
| 3 | y | 1.095E+04 | 877.5 | 0.0006934 | 0.7902 | +1 | 8 |
| - | - | 5225 | 878.5 | - | - | 0 | - |
| - | - | 1293 | 879.5 | - | - | 0 | - |
| - | - | 728.4 | 903.5 | - | - | 0 | - |
| 8 | b | 3745 | 920.5 | 0.001741 | 1.891 | +1 | 8 |
| - | - | 1910 | 921.5 | - | - | 0 | - |
| - | - | 663.7 | 922.5 | - | - | 0 | - |
| - | - | 1677 | 935.5 | - | - | 0 | - |
| - | - | 689.3 | 936.3 | - | - | 0 | - |
| - | - | 600.2 | 938.4 | - | - | 0 | - |
| - | - | 1276 | 950.5 | - | - | 0 | - |
| - | - | 2071 | 1038 | - | - | 0 | - |
| 2 | y | 1693 | 1041 | 0.0001181 | 0.1135 | +1 | 9 |
| - | - | 1651 | 1042 | - | - | 0 | - |
| - | - | 658.7 | 1195 | - | - | 0 | - |
| - | - | 631.3 | 1393 | - | - | 0 | - |
| - | - | 698.2 | 1538 | - | - | 0 | - |
| - | - | 705.6 | 1550 | - | - | 0 | - |
| - | - | 663.9 | 2316 | - | - | 0 | - |

m/z Charge Intensity FragmentType MassShift Position
120.06562805175781 0 647.8776
120.0809555053711 0 1508.2219
120.79678344726562 0 397.19012
121.71919250488281 0 402.85495
123.97521209716797 0 371.86868
129.05267333984375 0 484.48038
129.1023406982422 0 38116.59
130.0496368408203 0 490.88284
130.10565185546875 0 2598.3845
134.0272674560547 0 698.58624
136.0757598876953 0 34184.46
136.15757751464844 0 399.50552
136.88490295410156 0 412.25677
137.07919311523438 0 2506.2908
139.08677673339844 0 534.5006
148.94741821289062 0 550.2175
153.1024932861328 0 534.32434
154.58226013183594 0 479.1046
157.09718322753906 0 1313.8407
167.08177185058594 0 421.31708
169.0609588623047 0 536.77454
169.13351440429688 0 708.8327
171.1126708984375 0 944.6563
173.09185791015625 0 490.44183
173.1286163330078 0 3507.7605
173.45167541503906 0 1481.6578
178.0864715576172 0 745.86456
181.046875 0 453.9231
181.09725952148438 0 673.00555
183.11297607421875 0 1229.2814
183.1488800048828 0 765.0746
184.10838317871094 0 761.2742
184.50546264648438 0 441.77515
185.0919647216797 0 808.365
186.08712768554688 0 480.09628
187.10752868652344 0 979.3587
187.14405822753906 0 1623.191
189.0872344970703 0 1164.4934
197.12892150878906 0 941.8397
197.164306640625 0 657.6941
198.12344360351562 0 1987.9039
199.1079559326172 0 1882.5846
201.12326049804688 0 1442.8375
202.1269989013672 0 906.1234
203.10243225097656 0 779.6649
204.13406372070312 0 701.51605
209.0921173095703 0 1930.8966
215.10317993164062 0 610.7862
215.13894653320312 0 5385.502
216.1342010498047 0 7882.97 y Water loss 8
217.08282470703125 0 704.2904
217.11781311035156 0 1223.3467 y Ammonia loss 8
217.14622497558594 0 690.93713
223.10775756835938 0 2094.7927
224.1751251220703 0 677.1721
226.1186065673828 0 6977.3423
227.1226348876953 0 722.70667
229.11868286132812 0 697.4728
234.14491271972656 0 5587.006 y 8
235.14788818359375 0 1243.1163
239.09449768066406 0 1424.5382
240.13400268554688 0 1744.2369
242.18670654296875 0 647.0967
244.12913513183594 0 5101.548
248.1609649658203 0 1114.7762
249.1597442626953 0 21253.125 a 1
250.16273498535156 0 2896.8445
251.10287475585938 0 1687.5691
251.16583251953125 0 616.1521
264.95654296875 0 571.1704
267.10955810546875 0 592.63055
270.18096923828125 0 1129.9238
277.1545715332031 0 6130.745 b 1
278.1567687988281 0 788.6639
282.14422607421875 0 579.7221
285.0095520019531 0 3127.2454
290.0807189941406 0 667.69214
296.1966247558594 0 1120.2874
299.0613098144531 0 2546.872
303.1300354003906 0 579.0065
311.2072448730469 0 1984.1642
312.20989990234375 0 721.3325
316.15032958984375 0 1121.9077
329.2182922363281 0 1283.3042
331.16131591796875 0 536.37036 y Water loss 7
349.17095947265625 0 2357.8264 y 7
355.0697021484375 0 8276.077
356.06976318359375 0 2187.2412
361.00848388671875 0 555.35565
364.18701171875 0 1190.3329 b 2
379.1978454589844 0 625.52295
415.78192138671875 0 737.83295
415.8349304199219 0 619.71405
419.9953918457031 0 679.9264
427.1932373046875 0 757.7515
429.23193359375 0 1046.5721
430.2679138183594 0 935.40717
439.25250244140625 0 3326.1326 y 2
439.7544860839844 0 1713.7311
448.2400207519531 0 2015.1705 y 6
474.2711181640625 0 1110.6254 b Water loss 3
492.28228759765625 0 1806.4912 b 3
493.2850036621094 0 630.393
497.24224853515625 0 601.75055
511.32391357421875 0 1245.7987
515.7969970703125 0 654.2586 b Water loss 8
520.7842407226562 0 3075.0374 y 1
521.2855224609375 0 1705.4056
524.8104858398438 0 916.22235 b 8
525.30810546875 0 1215.6733
529.3339233398438 0 2213.5598
531.2723999023438 0 794.4407 y Water loss 5
533.8108520507812 0 1140.3195
534.3123168945312 0 719.20465
549.2883911132812 0 3392.4395 y 5
550.2891235351562 0 834.2531
557.328125 0 2130.6765
559.8126220703125 0 658.31036
560.3008422851562 0 762.33575
568.3216552734375 0 4056.346 Precursor Water loss
568.8221435546875 0 2866.6538 Precursor Ammonia loss
575.6211547851562 0 821.85706
576.28564453125 0 711.4814
577.3265991210938 0 54128.867 Precursor
577.8280029296875 0 37554.074
578.3294067382812 0 14273.957
578.8291015625 0 1884.0444
584.0226440429688 0 594.3865
602.7847290039062 0 2005.983
603.2861938476562 0 649.3951
605.36669921875 0 1592.3469 b 4
643.2907104492188 0 1067.2828
644.36083984375 0 2544.2112 y Water loss 4
645.365478515625 0 1096.4827
653.8651123046875 0 929.0603
654.364990234375 0 779.5941
659.8275146484375 0 829.8825
662.3710327148438 0 3504.9832 y 4
663.3765869140625 0 880.7099
685.4228515625 0 705.63586
688.4022216796875 0 700.7628 b Water loss 5
692.3973388671875 0 944.09015
706.4097290039062 0 701.28595 b 5
709.8533935546875 0 610.2018
720.3602905273438 0 990.8984
721.3650512695312 0 754.993
722.3645629882812 0 1059.8424
728.3787841796875 0 835.7987
739.3375244140625 0 975.79553
754.4443359375 0 878.8185
772.45556640625 0 4562.3145 y Water loss 3
773.4581298828125 0 2042.1239
782.3534545898438 0 1557.9556
782.8516235351562 0 1664.2473
785.37890625 0 799.27515
787.4710083007812 0 795.55383 b Water loss 6
790.4656372070312 0 2014.7635 y 3
805.4788818359375 0 1378.3313 b 6
806.4857788085938 0 706.3253
820.4381713867188 0 768.095
842.4557495117188 0 684.6787
859.4844970703125 0 1615.7988 y Water loss 2
877.4982299804688 0 10951.757 y 2
878.5010375976562 0 5224.644
879.5054931640625 0 1292.7355
903.5062866210938 0 728.4019
920.5070190429688 0 3745.2698 b 7
921.508056640625 0 1909.9908
922.517333984375 0 663.70746
935.5169677734375 0 1677.3267
936.3252563476562 0 689.3196
938.43408203125 0 600.17957
950.4762573242188 0 1275.5562
1037.5074462890625 0 2071.081
1040.5621337890625 0 1693.2212 y 1
1041.5634765625 0 1650.855
1195.2677001953125 0 658.67615
1392.7391357421875 0 631.32086
1538.0579833984375 0 698.16034
1550.3582763671875 0 705.5717
2316.422119140625 0 663.9247

Spectrum Details

|  |  |
| --- | --- |
| Matched peaks? Matched peaks The total absolute number of peaks matched. Additionally in brackets the total fraction of peaks matched and the total number of peaks is shown. | 33 (18.23% of 181) |
| FDR? FDR The false discovery rate estimated for this peptide. It is calculated by matching all theoretical fragments with a non-integer shift with the raw peaks for this spectrum. This is done with 40 different shifts. The resulting percentage is the average number of annotated peaks over the number of annotated peaks with the correct spectrum. | 0.65% |
| Satellite FDR? Satellite FDR See the FDR for details on its calculation. This satellite ion specific FDR only contains the satellite ions (d/w) for I/L/J positions. | - |
| PSM Score? PSM Score The PSM Score as given by Hecklib to this annotated spectrum. It is shown with three significant figures. | 371 |

#### Reverse Lookup? Reverse Lookup All places where this read could be placed.

| Group | Segment | Template | Template Part | Read Part | Score | Unique |
| --- | --- | --- | --- | --- | --- | --- |
| Homo sapiens Heavy Chain | IGHC | IGHG1 | [288..298] | [0..10] | 80 | False |
| Homo sapiens Heavy Chain | IGHC | IGHG3 | [335..345] | [0..10] | 80 | False |
| Homo sapiens Heavy Chain | IGHC | IGHG2 | [284..294] | [0..10] | 80 | False |

| Recombined | Template Part | Read Part | Score | Unique |
| --- | --- | --- | --- | --- |
| REC-0-1 | [413..423] | [0..10] | 80 | True |

#### Meta Information from Multiple reads

##### Number of combined reads

2

##### Intensity

0.38

##### TotalArea

8.496E+05

##### Changes to the peptide sequence

JYSKJTVDKS

L→JNo support for either Leucine or Isoleucine based on side chain ions (Position: 5)

L→JNo support for either Leucine or Isoleucine based on side chain ions (Position: 1)

#### Positional Score

Copy Data

##### Positional Score (TSV)

###### Preview

```
Loading example...
```

*Click on the button to copy the data to your clipboard.*

000123456789

Label Value
"0" 0
"1" 0
"2" 0
"3" 0
"4" 0
"5" 0
"6" 0
"7" 0
"8" 0
"9" 0

#### Meta Information from PEAKS

##### Scan Identifier

F1:3715

##### Original sequence

L

Y

S

K

L

T

V

D

K

S

##### Posttranslational Modifications

##### Source File

D:\separate\_stitch\_analyses\xle-disambiguation\raw\20210323\_F1\_UM1\_Peng0013\_SA\_F59\_ingel\_3ug\_ELA.raw

##### Fraction

1

##### Scan Feature

F1:524

##### De Novo Score

99

##### ConfidenceScore

99

### m/z

385.2202

##### Mass

1152.6389

##### Charge

3

##### Retention Time

19.35

##### Predicted Retention Time

-

##### Area

6.626E+05

##### Fragmentation mode

ETHCD

##### Originating file

01 D:\separate\_stitch\_analyses\xle-disambiguation\20210325\_F59\_3ug\_DENOVO\_12.csv

#### Meta Information from PEAKS

##### Scan Identifier

F1:3725

##### Original sequence

L

Y

S

K

L

T

V

D

K

S

##### Posttranslational Modifications

##### Source File

D:\separate\_stitch\_analyses\xle-disambiguation\raw\20210323\_F1\_UM1\_Peng0013\_SA\_F59\_ingel\_3ug\_ELA.raw

##### Fraction

1

##### Scan Feature

F1:7466

##### De Novo Score

98

##### ConfidenceScore

98

### m/z

577.3265

##### Mass

1152.6389

##### Charge

2

##### Retention Time

19.35

##### Predicted Retention Time

-

##### Area

1.87E+05

##### Fragmentation mode

HCD

##### Originating file

01 D:\separate\_stitch\_analyses\xle-disambiguation\20210325\_F59\_3ug\_DENOVO\_12.csv
