## Supplementary material for "A handle on mass coincidence errors in *de novo* sequencing of antibodies by bottom-up proteomics": Combined_064.html

Details Combined\_064 | Stitch OverviewUndefined

### Read Combined\_064

#### Sequence (length=10)

JSSPATJDSR

#### Spectrum 3844? Spectrum 3844 The raw spectrum of this peptide as annotated by Hecklib. The fragments are coloured according to ion type (see legend). Any peaks with a star '\*' as text can be hovered over to see the full details, first the ion type second the mass shift type. By hovering over the amino acids in the peptide or ions in the legend the corresponding peaks are highlighted. By toggling the 'Unassigned' label you can turn the background (unassigned) peaks on or off in the plot. By updating the slider in the Ion legend you can update the spectrum to only show the top X% of the peaks with labels. The top X% means any peak that is within X% of the highest intensity. By dragging in the spectrum you can zoom in to a specific part of the spectrum and use 'Zoom Out' to get back to the original zoom level. The annotation of the spectrum is based on the given sequence in the peptides file and is done with different software so inconsistencies are likely. The peaks are annotated based on the given sequence, with 20 ppm tolerance.

Copy Data

##### Spectrum 3844 (TSV)

###### Preview

```
Loading example...
```

*Click on the button to copy the data to your clipboard.*

Mz MinMz MaxIntensity Max

WidthHeightPeptide font sizePeptide stroke widthSpectrum font sizeSpectrum stroke widthCompact peptide

Ion legend

wxyz

abcd

OtherUnassignedIon^Charge^_Position_Show for top:%

JSSPATJDSR

02.59e+45.19e+47.78e+41.04e+5

Zoom Out

a^+1^_2_y^+1^_1_a^+1^_2_y^+1^_1_b^+1^_2_b^+1^_2_y^+2^_4_a^+1^_3_y^+1^_2_y^+1^_2_y^+1^_2_b^+1^_3_b^+1^_3_y^+2^_6_y^+1^_3_y^+1^_3_y^+2^_7_y^+1^_3_y^+2^_7_y^+2^_8_y^+2^_8_b^+1^_5_b^+1^_5_y^+2^_9_y^+1^_4_y^+1^_4_\*\*b^+1^_6_y^+1^_5_y^+1^_5_y^+1^_6_y^+1^_6_y^+1^_7_y^+1^_7_y^+1^_7_y^+1^_8_y^+1^_8_y^+1^_8_y^+1^_9_y^+1^_9_y^+1^_9_

0840168025203360

Fragment Matches Table

Show background peaks

| Position | Ion type | Intensity | mz Theoretical | mz Error (Th) | mz Error (ppm) | Charge | Series Number |
| --- | --- | --- | --- | --- | --- | --- | --- |
| - | - | 431.1 | 123.1 | - | - | 0 | - |
| - | - | 441.2 | 124.1 | - | - | 0 | - |
| - | - | 1885 | 125.1 | - | - | 0 | - |
| - | - | 368.2 | 125.4 | - | - | 0 | - |
| - | - | 392.3 | 127.7 | - | - | 0 | - |
| - | - | 1086 | 128.1 | - | - | 0 | - |
| - | - | 1091 | 129.1 | - | - | 0 | - |
| - | - | 1515 | 129.1 | - | - | 0 | - |
| - | - | 4315 | 130.1 | - | - | 0 | - |
| - | - | 621.1 | 130.1 | - | - | 0 | - |
| - | - | 625.5 | 130.1 | - | - | 0 | - |
| - | - | 423.4 | 130.8 | - | - | 0 | - |
| - | - | 477.4 | 131.5 | - | - | 0 | - |
| - | - | 393.6 | 131.7 | - | - | 0 | - |
| - | - | 7134 | 136.1 | - | - | 0 | - |
| - | - | 693.3 | 137.1 | - | - | 0 | - |
| - | - | 471.9 | 138.1 | - | - | 0 | - |
| - | - | 3207 | 139.1 | - | - | 0 | - |
| - | - | 718.4 | 141.1 | - | - | 0 | - |
| - | - | 1.214E+04 | 141.1 | - | - | 0 | - |
| - | - | 474.6 | 143 | - | - | 0 | - |
| - | - | 466 | 143.1 | - | - | 0 | - |
| - | - | 928.7 | 145.1 | - | - | 0 | - |
| - | - | 423.1 | 146.2 | - | - | 0 | - |
| - | - | 1012 | 147.1 | - | - | 0 | - |
| - | - | 491.1 | 147.1 | - | - | 0 | - |
| - | - | 428.7 | 148.9 | - | - | 0 | - |
| - | - | 522 | 148.9 | - | - | 0 | - |
| - | - | 898.7 | 148.9 | - | - | 0 | - |
| - | - | 967.8 | 148.9 | - | - | 0 | - |
| - | - | 977.2 | 148.9 | - | - | 0 | - |
| - | - | 1279 | 148.9 | - | - | 0 | - |
| - | - | 1552 | 148.9 | - | - | 0 | - |
| - | - | 3385 | 148.9 | - | - | 0 | - |
| - | - | 6067 | 149 | - | - | 0 | - |
| - | - | 3743 | 149 | - | - | 0 | - |
| - | - | 1499 | 149 | - | - | 0 | - |
| - | - | 1147 | 149 | - | - | 0 | - |
| - | - | 1015 | 149 | - | - | 0 | - |
| - | - | 629.7 | 149 | - | - | 0 | - |
| - | - | 546.6 | 149 | - | - | 0 | - |
| - | - | 787.4 | 149 | - | - | 0 | - |
| - | - | 553.5 | 149 | - | - | 0 | - |
| - | - | 563.9 | 149 | - | - | 0 | - |
| - | - | 1022 | 149 | - | - | 0 | - |
| - | - | 414.7 | 149.1 | - | - | 0 | - |
| - | - | 541.4 | 151 | - | - | 0 | - |
| - | - | 523.4 | 151.4 | - | - | 0 | - |
| - | - | 664.8 | 155.1 | - | - | 0 | - |
| 2 | a | 4061 | 155.1 | 0.0001982 | 1.278 | +1 | 2 |
| - | - | 2701 | 157.1 | - | - | 0 | - |
| - | - | 1.189E+04 | 157.1 | - | - | 0 | - |
| - | - | 1145 | 157.1 | - | - | 0 | - |
| 10 | y | 8928 | 158.1 | 0.000172 | 1.088 | +1 | 1 |
| - | - | 1078 | 158.1 | - | - | 0 | - |
| - | - | 619.4 | 159.1 | - | - | 0 | - |
| - | - | 941.6 | 167.1 | - | - | 0 | - |
| - | - | 3350 | 167.1 | - | - | 0 | - |
| - | - | 1.264E+04 | 169.1 | - | - | 0 | - |
| - | - | 686.5 | 169.1 | - | - | 0 | - |
| - | - | 8513 | 169.1 | - | - | 0 | - |
| - | - | 972.9 | 170.1 | - | - | 0 | - |
| - | - | 511.6 | 170.1 | - | - | 0 | - |
| - | - | 1606 | 173.1 | - | - | 0 | - |
| 2 | a | 1.711E+04 | 173.1 | 0.0001926 | 1.113 | +1 | 2 |
| - | - | 1520 | 174.1 | - | - | 0 | - |
| - | - | 1637 | 175.1 | - | - | 0 | - |
| 10 | y | 1.965E+04 | 175.1 | 0.0001885 | 1.076 | +1 | 1 |
| - | - | 1560 | 176.1 | - | - | 0 | - |
| - | - | 510.6 | 182.1 | - | - | 0 | - |
| 2 | b | 2.516E+04 | 183.1 | 0.0001414 | 0.7722 | +1 | 2 |
| - | - | 797.5 | 184.1 | - | - | 0 | - |
| - | - | 2300 | 184.1 | - | - | 0 | - |
| - | - | 1.018E+04 | 185.1 | - | - | 0 | - |
| - | - | 817.9 | 185.1 | - | - | 0 | - |
| - | - | 453.3 | 186.1 | - | - | 0 | - |
| - | - | 6090 | 187.1 | - | - | 0 | - |
| - | - | 581.4 | 188.1 | - | - | 0 | - |
| - | - | 662.6 | 189.1 | - | - | 0 | - |
| - | - | 3543 | 197.1 | - | - | 0 | - |
| - | - | 746.2 | 201.1 | - | - | 0 | - |
| 2 | b | 1.321E+04 | 201.1 | 4.424E-05 | 0.22 | +1 | 2 |
| - | - | 541.1 | 201.1 | - | - | 0 | - |
| - | - | 719.1 | 202.1 | - | - | 0 | - |
| - | - | 2414 | 202.1 | - | - | 0 | - |
| - | - | 969.3 | 202.1 | - | - | 0 | - |
| - | - | 671.8 | 203.1 | - | - | 0 | - |
| - | - | 858.6 | 203.1 | - | - | 0 | - |
| - | - | 601.5 | 208.1 | - | - | 0 | - |
| - | - | 563.5 | 209.1 | - | - | 0 | - |
| - | - | 917.4 | 210.1 | - | - | 0 | - |
| - | - | 2060 | 210.1 | - | - | 0 | - |
| - | - | 1050 | 212.1 | - | - | 0 | - |
| - | - | 2141 | 215.1 | - | - | 0 | - |
| - | - | 647.6 | 216.7 | - | - | 0 | - |
| - | - | 1155 | 220.1 | - | - | 0 | - |
| - | - | 1096 | 221.1 | - | - | 0 | - |
| - | - | 713 | 222.1 | - | - | 0 | - |
| - | - | 649 | 224.1 | - | - | 0 | - |
| - | - | 1429 | 224.1 | - | - | 0 | - |
| - | - | 690.6 | 225 | - | - | 0 | - |
| - | - | 2840 | 225.1 | - | - | 0 | - |
| - | - | 1397 | 226.1 | - | - | 0 | - |
| - | - | 1530 | 227.1 | - | - | 0 | - |
| - | - | 1130 | 228.1 | - | - | 0 | - |
| - | - | 674.5 | 229.1 | - | - | 0 | - |
| - | - | 872.5 | 230.1 | - | - | 0 | - |
| - | - | 589.5 | 232.1 | - | - | 0 | - |
| - | - | 3714 | 233.1 | - | - | 0 | - |
| - | - | 3139 | 235.1 | - | - | 0 | - |
| 7 | y | 702.3 | 237.1 | 0.00201 | 8.476 | +2 | 4 |
| - | - | 2.207E+04 | 238.1 | - | - | 0 | - |
| - | - | 676.7 | 239.1 | - | - | 0 | - |
| - | - | 2076 | 239.1 | - | - | 0 | - |
| - | - | 526 | 240.2 | - | - | 0 | - |
| 3 | a | 1372 | 242.1 | 0.0001217 | 0.5027 | +1 | 3 |
| 9 | y | 2506 | 244.1 | 0.0002091 | 0.8565 | +1 | 2 |
| 9 | y | 1.516E+04 | 245.1 | 0.0001871 | 0.7631 | +1 | 2 |
| - | - | 2349 | 246.1 | - | - | 0 | - |
| - | - | 700.5 | 247.1 | - | - | 0 | - |
| - | - | 5403 | 252.1 | - | - | 0 | - |
| - | - | 1119 | 253.1 | - | - | 0 | - |
| - | - | 809.4 | 254.1 | - | - | 0 | - |
| - | - | 5610 | 256.1 | - | - | 0 | - |
| - | - | 650.1 | 257.1 | - | - | 0 | - |
| - | - | 1256 | 261.1 | - | - | 0 | - |
| 9 | y | 5.377E+04 | 262.2 | 2.04E-05 | 0.07781 | +1 | 2 |
| - | - | 4946 | 263.2 | - | - | 0 | - |
| - | - | 934.8 | 268.1 | - | - | 0 | - |
| - | - | 1631 | 268.2 | - | - | 0 | - |
| 3 | b | 2.908E+04 | 270.1 | 6.489E-05 | 0.2402 | +1 | 3 |
| - | - | 3907 | 271.1 | - | - | 0 | - |
| - | - | 885.2 | 284.2 | - | - | 0 | - |
| 3 | b | 2449 | 288.2 | 2.877E-05 | 0.09985 | +1 | 3 |
| - | - | 2537 | 299.1 | - | - | 0 | - |
| - | - | 583.4 | 299.1 | - | - | 0 | - |
| - | - | 2349 | 300.1 | - | - | 0 | - |
| - | - | 4014 | 301.1 | - | - | 0 | - |
| - | - | 680.4 | 301.4 | - | - | 0 | - |
| - | - | 2099 | 302.1 | - | - | 0 | - |
| - | - | 1013 | 303.1 | - | - | 0 | - |
| - | - | 1028 | 312.2 | - | - | 0 | - |
| - | - | 597.4 | 313.2 | - | - | 0 | - |
| - | - | 883.8 | 322.2 | - | - | 0 | - |
| 5 | y | 823.4 | 322.7 | 0.001502 | 4.656 | +2 | 6 |
| - | - | 986.4 | 324.1 | - | - | 0 | - |
| - | - | 1074 | 330.2 | - | - | 0 | - |
| - | - | 1006 | 337.2 | - | - | 0 | - |
| - | - | 3061 | 339.2 | - | - | 0 | - |
| - | - | 1005 | 341.2 | - | - | 0 | - |
| - | - | 2609 | 342.1 | - | - | 0 | - |
| - | - | 746.5 | 343.2 | - | - | 0 | - |
| - | - | 599.3 | 346.2 | - | - | 0 | - |
| - | - | 783.2 | 347 | - | - | 0 | - |
| - | - | 708.8 | 355.2 | - | - | 0 | - |
| - | - | 620.2 | 356.6 | - | - | 0 | - |
| - | - | 3133 | 357.2 | - | - | 0 | - |
| - | - | 603.9 | 358.2 | - | - | 0 | - |
| - | - | 1332 | 358.2 | - | - | 0 | - |
| 8 | y | 3260 | 359.2 | 9.208E-05 | 0.2564 | +1 | 3 |
| 8 | y | 2295 | 360.2 | 0.0001904 | 0.5287 | +1 | 3 |
| - | - | 1068 | 361 | - | - | 0 | - |
| - | - | 1622 | 365.2 | - | - | 0 | - |
| - | - | 632.9 | 368.2 | - | - | 0 | - |
| - | - | 667.6 | 369.7 | - | - | 0 | - |
| - | - | 969 | 370.2 | - | - | 0 | - |
| - | - | 4944 | 370.7 | - | - | 0 | - |
| 4 | y | 4469 | 371.2 | 0.00164 | 4.419 | +2 | 7 |
| - | - | 2386 | 371.7 | - | - | 0 | - |
| - | - | 769.4 | 372.2 | - | - | 0 | - |
| - | - | 3230 | 375.2 | - | - | 0 | - |
| - | - | 6005 | 376.2 | - | - | 0 | - |
| - | - | 1051 | 376.2 | - | - | 0 | - |
| 8 | y | 6835 | 377.2 | 6.129E-06 | 0.01625 | +1 | 3 |
| - | - | 727.3 | 378.2 | - | - | 0 | - |
| - | - | 728.5 | 378.7 | - | - | 0 | - |
| - | - | 2.902E+04 | 379.7 | - | - | 0 | - |
| 4 | y | 3.034E+04 | 380.2 | 0.00176 | 4.628 | +2 | 7 |
| - | - | 9924 | 380.7 | - | - | 0 | - |
| - | - | 2671 | 381.2 | - | - | 0 | - |
| - | - | 663.4 | 383.1 | - | - | 0 | - |
| - | - | 1562 | 383.2 | - | - | 0 | - |
| - | - | 6173 | 402.2 | - | - | 0 | - |
| - | - | 759.8 | 403.2 | - | - | 0 | - |
| - | - | 1677 | 403.2 | - | - | 0 | - |
| - | - | 1391 | 405.2 | - | - | 0 | - |
| - | - | 2324 | 405.7 | - | - | 0 | - |
| - | - | 1490 | 406.2 | - | - | 0 | - |
| - | - | 2.212E+04 | 414.2 | - | - | 0 | - |
| 3 | y | 2.803E+04 | 414.7 | 0.002197 | 5.298 | +2 | 8 |
| - | - | 1.234E+04 | 415.2 | - | - | 0 | - |
| - | - | 2450 | 415.7 | - | - | 0 | - |
| - | - | 941.2 | 416.2 | - | - | 0 | - |
| - | - | 921.2 | 419 | - | - | 0 | - |
| - | - | 732.2 | 421 | - | - | 0 | - |
| - | - | 1.799E+04 | 423.2 | - | - | 0 | - |
| 3 | y | 2.097E+04 | 423.7 | 0.002286 | 5.395 | +2 | 8 |
| - | - | 1.131E+04 | 424.2 | - | - | 0 | - |
| - | - | 2627 | 424.7 | - | - | 0 | - |
| - | - | 1.441E+04 | 431.2 | - | - | 0 | - |
| - | - | 5501 | 432.3 | - | - | 0 | - |
| - | - | 869.3 | 433.3 | - | - | 0 | - |
| - | - | 1002 | 435.6 | - | - | 0 | - |
| - | - | 1343 | 438.2 | - | - | 0 | - |
| 5 | b | 695.6 | 438.2 | 0.003509 | 8.007 | +1 | 5 |
| - | - | 2071 | 440.3 | - | - | 0 | - |
| - | - | 2574 | 441.2 | - | - | 0 | - |
| - | - | 1343 | 442.6 | - | - | 0 | - |
| - | - | 747.8 | 444.6 | - | - | 0 | - |
| - | - | 633.2 | 448.7 | - | - | 0 | - |
| - | - | 1046 | 452.3 | - | - | 0 | - |
| 5 | b | 818 | 456.2 | 0.0008494 | 1.862 | +1 | 5 |
| - | - | 8.366E+04 | 458.3 | - | - | 0 | - |
| - | - | 1091 | 459.1 | - | - | 0 | - |
| - | - | 2.978E+04 | 459.3 | - | - | 0 | - |
| - | - | 3910 | 460.3 | - | - | 0 | - |
| - | - | 1132 | 461.6 | - | - | 0 | - |
| - | - | 609.6 | 466.6 | - | - | 0 | - |
| 2 | y | 623.4 | 467.2 | 0.003383 | 7.24 | +2 | 9 |
| - | - | 763.2 | 470.3 | - | - | 0 | - |
| 7 | y | 938.6 | 473.2 | 0.001164 | 2.46 | +1 | 4 |
| - | - | 689.9 | 473.6 | - | - | 0 | - |
| - | - | 1103 | 474.1 | - | - | 0 | - |
| - | - | 798.4 | 481.1 | - | - | 0 | - |
| - | - | 800.9 | 483.3 | - | - | 0 | - |
| - | - | 815.4 | 483.7 | - | - | 0 | - |
| - | - | 756.3 | 489.1 | - | - | 0 | - |
| - | - | 8318 | 489.3 | - | - | 0 | - |
| 7 | y | 6548 | 490.3 | 0.001104 | 2.253 | +1 | 4 |
| - | - | 1171 | 491.3 | - | - | 0 | - |
| - | - | 648.5 | 497.7 | - | - | 0 | - |
| - | - | 3214 | 498.3 | - | - | 0 | - |
| - | - | 779.3 | 504.3 | - | - | 0 | - |
| - | - | 786.7 | 505.3 | - | - | 0 | - |
| - | - | 2358 | 505.8 | - | - | 0 | - |
| - | - | 740.5 | 506.3 | - | - | 0 | - |
| - | - | 999.9 | 507.2 | - | - | 0 | - |
| - | - | 1534 | 507.2 | - | - | 0 | - |
| - | - | 784 | 513.8 | - | - | 0 | - |
| - | - | 1632 | 514.3 | - | - | 0 | - |
| 0 | Precursor | 2159 | 514.8 | 0.001555 | 3.02 | +2 | -1 |
| - | - | 1080 | 521.3 | - | - | 0 | - |
| - | - | 1.028E+05 | 522.3 | - | - | 0 | - |
| - | - | 4.587E+04 | 523.3 | - | - | 0 | - |
| 0 | Precursor | 3303 | 523.8 | 0.004146 | 7.915 | +2 | -1 |
| - | - | 1.329E+04 | 524.3 | - | - | 0 | - |
| - | - | 1939 | 525.3 | - | - | 0 | - |
| 6 | b | 803.1 | 539.3 | 0.002726 | 5.054 | +1 | 6 |
| - | - | 3602 | 545.8 | - | - | 0 | - |
| - | - | 2376 | 546.3 | - | - | 0 | - |
| - | - | 2370 | 567.3 | - | - | 0 | - |
| - | - | 834 | 572.3 | - | - | 0 | - |
| 6 | y | 786.1 | 573.3 | 0.005342 | 9.318 | +1 | 5 |
| - | - | 1473 | 585.3 | - | - | 0 | - |
| - | - | 2.668E+04 | 590.3 | - | - | 0 | - |
| 6 | y | 2.413E+04 | 591.3 | 0.002407 | 4.07 | +1 | 5 |
| - | - | 6490 | 592.3 | - | - | 0 | - |
| - | - | 1425 | 593.3 | - | - | 0 | - |
| - | - | 928.6 | 595.8 | - | - | 0 | - |
| - | - | 957.9 | 627.3 | - | - | 0 | - |
| - | - | 2822 | 643.4 | - | - | 0 | - |
| 5 | y | 3129 | 644.3 | 0.003079 | 4.779 | +1 | 6 |
| - | - | 1042 | 645.3 | - | - | 0 | - |
| - | - | 780 | 646.3 | - | - | 0 | - |
| - | - | 2360 | 652.8 | - | - | 0 | - |
| - | - | 1950 | 653.3 | - | - | 0 | - |
| - | - | 2.28E+04 | 661.4 | - | - | 0 | - |
| 5 | y | 2.014E+04 | 662.3 | 0.003318 | 5.009 | +1 | 6 |
| - | - | 6535 | 663.4 | - | - | 0 | - |
| - | - | 1068 | 664.3 | - | - | 0 | - |
| - | - | 774.6 | 665.7 | - | - | 0 | - |
| - | - | 1200 | 671.3 | - | - | 0 | - |
| - | - | 760.9 | 672.3 | - | - | 0 | - |
| - | - | 693.7 | 673.3 | - | - | 0 | - |
| - | - | 750.1 | 734.4 | - | - | 0 | - |
| - | - | 1153 | 734.9 | - | - | 0 | - |
| 4 | y | 1673 | 741.4 | 0.003782 | 5.102 | +1 | 7 |
| 4 | y | 1417 | 742.4 | 0.008414 | 11.33 | +1 | 7 |
| - | - | 3.548E+04 | 758.4 | - | - | 0 | - |
| 4 | y | 4.29E+04 | 759.4 | 0.003899 | 5.134 | +1 | 7 |
| - | - | 1.517E+04 | 760.4 | - | - | 0 | - |
| - | - | 3330 | 761.4 | - | - | 0 | - |
| - | - | 630.2 | 826.5 | - | - | 0 | - |
| - | - | 860.9 | 827.4 | - | - | 0 | - |
| 3 | y | 1464 | 828.4 | 0.0008676 | 1.047 | +1 | 8 |
| 3 | y | 665.9 | 829.4 | 0.01453 | 17.52 | +1 | 8 |
| - | - | 1.517E+04 | 845.4 | - | - | 0 | - |
| 3 | y | 2.371E+04 | 846.4 | 0.002998 | 3.542 | +1 | 8 |
| - | - | 9558 | 847.4 | - | - | 0 | - |
| - | - | 2758 | 848.4 | - | - | 0 | - |
| - | - | 1570 | 914.5 | - | - | 0 | - |
| 2 | y | 2186 | 915.5 | 0.001493 | 1.631 | +1 | 9 |
| 2 | y | 1213 | 916.4 | 0.01656 | 18.07 | +1 | 9 |
| - | - | 1.251E+04 | 932.5 | - | - | 0 | - |
| 2 | y | 1.868E+04 | 933.5 | 0.003013 | 3.228 | +1 | 9 |
| - | - | 7058 | 934.5 | - | - | 0 | - |
| - | - | 1862 | 935.5 | - | - | 0 | - |
| - | - | 1442 | 942.5 | - | - | 0 | - |
| - | - | 1178 | 943.5 | - | - | 0 | - |
| - | - | 766.1 | 1027 | - | - | 0 | - |
| - | - | 634.8 | 2299 | - | - | 0 | - |
| - | - | 684.7 | 3326 | - | - | 0 | - |

m/z Charge Intensity FragmentType MassShift Position
123.09173583984375 0 431.1155
124.11256408691406 0 441.2188
125.10767364501953 0 1885.4725
125.44654083251953 0 368.21997
127.6700668334961 0 392.26224
128.107177734375 0 1085.9297
129.06610107421875 0 1090.8453
129.10255432128906 0 1515.0131
130.061279296875 0 4315.311
130.0863800048828 0 621.0625
130.09800720214844 0 625.5065
130.78628540039062 0 423.41562
131.54966735839844 0 477.42963
131.73336791992188 0 393.58603
136.07591247558594 0 7133.5723
137.07933044433594 0 693.29913
138.09210205078125 0 471.8724
139.0868377685547 0 3207.0566
141.06600952148438 0 718.43945
141.10244750976562 0 12143.397
143.0338134765625 0 474.59824
143.11827087402344 0 466.0491
145.09754943847656 0 928.667
146.185791015625 0 423.11795
147.07667541503906 0 1011.8642
147.11309814453125 0 491.1491
148.88189697265625 0 428.68753
148.88905334472656 0 522.0269
148.90316772460938 0 898.664
148.910400390625 0 967.83374
148.91770935058594 0 977.1661
148.9250946044922 0 1279.0503
148.93223571777344 0 1551.8008
148.9398956298828 0 3385.3792
148.95645141601562 0 6066.891
148.9642333984375 0 3743.1924
148.97190856933594 0 1499.2482
148.9790802001953 0 1146.7528
148.98648071289062 0 1014.98157
148.99354553222656 0 629.6683
149.00079345703125 0 546.5912
149.0076446533203 0 787.39325
149.0152130126953 0 553.52795
149.03025817871094 0 563.91205
149.04457092285156 0 1022.1193
149.05870056152344 0 414.7026
151.04254150390625 0 541.3693
151.3609619140625 0 523.4069
155.08168029785156 0 664.7761
155.1180877685547 0 4061.2644 a Water loss 1
157.06092834472656 0 2700.705
157.09732055664062 0 11889.897
157.10848999023438 0 1145.0497
158.0925750732422 0 8928.005 y Ammonia loss 9
158.10018920898438 0 1078.3943
159.07691955566406 0 619.3976
167.05543518066406 0 941.6206
167.08169555664062 0 3350.068
169.09732055664062 0 12635.568
169.12594604492188 0 686.5473
169.1337127685547 0 8513.125
170.1004180908203 0 972.90375
170.1371307373047 0 511.61017
173.09222412109375 0 1605.7808
173.12864685058594 0 17114.4 a 1
174.13198852539062 0 1519.8448
175.07139587402344 0 1636.7498
175.119140625 0 19653.252 y 9
176.12232971191406 0 1560.4453
182.09214782714844 0 510.61493
183.11294555664062 0 25162.36 b Water loss 1
184.10824584960938 0 797.52203
184.11647033691406 0 2299.5498
185.0922088623047 0 10180.441
185.13973999023438 0 817.9289
186.1234588623047 0 453.28854
187.1442413330078 0 6090.116
188.14779663085938 0 581.37396
189.08705139160156 0 662.5641
197.12864685058594 0 3543.3313
201.09884643554688 0 746.2282
201.1234130859375 0 13214.974 b 1
201.13348388671875 0 541.0912
202.08265686035156 0 719.063
202.11878967285156 0 2414.3032
202.1280975341797 0 969.26154
203.0655975341797 0 671.8031
203.10279846191406 0 858.6373
208.10829162597656 0 601.462
209.10256958007812 0 563.54004
210.08731079101562 0 917.44305
210.12380981445312 0 2059.9973
212.10272216796875 0 1050.1448
215.1393280029297 0 2140.6736
216.72198486328125 0 647.5662
220.1295623779297 0 1155.1162
221.09242248535156 0 1096.0787
222.1244354248047 0 713.0286
224.10281372070312 0 648.99115
224.13934326171875 0 1429.027
225.0438995361328 0 690.5781
225.12353515625 0 2840.357
226.1189422607422 0 1396.5232
227.11378479003906 0 1529.9318
228.13461303710938 0 1129.795
229.1178436279297 0 674.5101
230.12574768066406 0 872.53296
232.13955688476562 0 589.5296
233.1318359375 0 3713.749
235.14395141601562 0 3138.5723
237.1233673095703 0 702.3425 y Ammonia loss 6
238.1188201904297 0 22072.309
239.0896759033203 0 676.6866
239.12210083007812 0 2075.6353
240.17047119140625 0 525.967
242.15003967285156 0 1371.9438 a Water loss 2
244.140625 0 2505.7341 y Water loss 8
245.12461853027344 0 15155.548 y Ammonia loss 8
246.12852478027344 0 2348.7922
247.1088104248047 0 700.45
252.13449096679688 0 5403.4995
253.13734436035156 0 1119.45
254.1144256591797 0 809.3534
256.12908935546875 0 5610.08
257.13360595703125 0 650.1058
261.1258544921875 0 1256.4958
262.1510009765625 0 53770.855 y 8
263.1540222167969 0 4946.0415
268.129638671875 0 934.78204
268.166015625 0 1630.5399
270.1448974609375 0 29076.414 b Water loss 2
271.1479187011719 0 3907.2822
284.1607666015625 0 885.1942
288.1554260253906 0 2449.4922 b 2
299.06170654296875 0 2537.4753
299.1355895996094 0 583.39575
300.0619812011719 0 2348.959
301.05987548828125 0 4013.8008
301.43017578125 0 680.36835
302.0605773925781 0 2099.1584
303.0567626953125 0 1013.1491
312.1558837890625 0 1028.1501
313.15020751953125 0 597.39325
322.1802062988281 0 883.84534
322.6732482910156 0 823.41034 y Water loss 4
324.1307678222656 0 986.38153
330.1652526855469 0 1074.2799
337.22528076171875 0 1005.7134
339.1664733886719 0 3060.906
341.1573181152344 0 1005.36487
342.140625 0 2609.3267
343.16265869140625 0 746.4704
346.1774597167969 0 599.26154
346.9742126464844 0 783.1513
355.1964111328125 0 708.80945
356.6369323730469 0 620.15485
357.17626953125 0 3132.9543
358.1827392578125 0 603.93756
358.2079772949219 0 1332.2307
359.1672668457031 0 3260.103 y Water loss 7
360.15118408203125 0 2294.8477 y Ammonia loss 7
361.0241394042969 0 1067.7805
365.21844482421875 0 1622.2137
368.1920166015625 0 632.8791
369.697509765625 0 667.60895
370.1920166015625 0 968.9934
370.7052917480469 0 4944.297
371.19976806640625 0 4468.5327 y Water loss 3
371.70111083984375 0 2386.253
372.2010803222656 0 769.43884
375.2349548339844 0 3229.6462
376.1938781738281 0 6004.8823
376.2353820800781 0 1051.4529
377.17791748046875 0 6835.3853 y 7
378.1785888671875 0 727.30365
378.7033386230469 0 728.5301
379.7112121582031 0 29020.896
380.2051696777344 0 30344.652 y 3
380.7060852050781 0 9923.834
381.205810546875 0 2671.395
383.14251708984375 0 663.3548
383.2292785644531 0 1561.6128
402.2095642089844 0 6172.626
403.1641540527344 0 759.7901
403.2115173339844 0 1676.5415
405.2173156738281 0 1391.3674
405.7110900878906 0 2323.6853
406.209716796875 0 1490.3406
414.22210693359375 0 22116.209
414.7163391113281 0 28025.219 y Water loss 2
415.21697998046875 0 12340.564
415.7174377441406 0 2450.4668
416.1834716796875 0 941.23645
419.0305480957031 0 921.24554
420.9945068359375 0 732.15594
423.22772216796875 0 17991.77
423.7217102050781 0 20973.85 y 2
424.2220153808594 0 11314.589
424.7226257324219 0 2626.596
431.2487487792969 0 14411.61
432.25201416015625 0 5501.235
433.251953125 0 869.254
435.6373596191406 0 1002.1722
438.19708251953125 0 1343.296
438.231201171875 0 695.5809 b Water loss 4
440.2622375488281 0 2071.294
441.2458801269531 0 2573.5576
442.6405029296875 0 1342.743
444.6428527832031 0 747.8204
448.7364501953125 0 633.24005
452.25091552734375 0 1045.9485
456.2461242675781 0 818.0451 b 4
458.2723693847656 0 83660.766
459.1333312988281 0 1090.6826
459.27508544921875 0 29783.963
460.27789306640625 0 3909.6545
461.63897705078125 0 1132.2119
466.626953125 0 609.5866
467.2320556640625 0 623.3813 y 1
470.2606506347656 0 763.1818
473.2366027832031 0 938.6485 y Ammonia loss 6
473.6086120605469 0 689.91174
474.1117858886719 0 1102.8063
481.14495849609375 0 798.3679
483.2547607421875 0 800.8669
483.6506652832031 0 815.4463
489.1435241699219 0 756.3455
489.2778015136719 0 8317.729
490.2630920410156 0 6548.2153 y 6
491.2643127441406 0 1170.5106
497.73809814453125 0 648.51184
498.2563781738281 0 3214.2507
504.2572937011719 0 779.33594
505.2746276855469 0 786.6681
505.7682800292969 0 2357.5798
506.2669372558594 0 740.45636
507.175537109375 0 999.91473
507.2463684082031 0 1534.2693
513.7657470703125 0 784.0035
514.2747192382812 0 1631.5085
514.7737426757812 0 2159.1257 Precursor Water loss
521.2711181640625 0 1080.0875
522.2703247070312 0 102754.07
523.2733154296875 0 45873.76
523.7816162109375 0 3302.843 Precursor
524.2713012695312 0 13289.65
525.26904296875 0 1938.7375
539.2796630859375 0 803.14594 b Water loss 5
545.7626342773438 0 3601.8157
546.2655029296875 0 2375.8113
567.2765502929688 0 2370.4385
572.31591796875 0 834.0065
573.304443359375 0 786.0934 y Water loss 5
585.2926025390625 0 1473.0155
590.3253784179688 0 26680.293
591.3120727539062 0 24131.875 y 5
592.3134155273438 0 6490.002
593.3140869140625 0 1424.5498
595.80224609375 0 928.56836
627.305908203125 0 957.8772
643.3519287109375 0 2821.9956
644.3392944335938 0 3129.2861 y Water loss 4
645.3341674804688 0 1041.8578
646.324462890625 0 779.9506
652.8286743164062 0 2360.1917
653.3305053710938 0 1950.4086
661.3624267578125 0 22799.355
662.35009765625 0 20140.588 y 4
663.3522338867188 0 6534.626
664.349609375 0 1067.7228
665.7069091796875 0 774.6463
671.3450317382812 0 1199.5635
672.3300170898438 0 760.93994
673.333251953125 0 693.7017
734.3663940429688 0 750.1403
734.8597412109375 0 1152.6124
741.3927612304688 0 1673.486 y Water loss 3
742.3814086914062 0 1416.8008 y Ammonia loss 3
758.4148559570312 0 35481.23
759.4034423828125 0 42895.9 y 3
760.4046630859375 0 15169.914
761.4077758789062 0 3330.2544
826.5211791992188 0 630.21484
827.4361572265625 0 860.8685
828.421875 0 1463.6173 y Water loss 2
829.4195556640625 0 665.94824 y Ammonia loss 2
845.4462890625 0 15174.487
846.4345703125 0 23714.658 y 2
847.4371948242188 0 9558.027
848.43603515625 0 2757.8186
914.465087890625 0 1569.87
915.4545288085938 0 2185.9148 y Water loss 1
916.45361328125 0 1213.4857 y Ammonia loss 1
932.4775390625 0 12508.435
933.4666137695312 0 18675.77 y 1
934.4700927734375 0 7057.6875
935.4654541015625 0 1861.9299
942.4642333984375 0 1442.0117
943.4586181640625 0 1177.9419
1026.5069580078125 0 766.10504
2298.646728515625 0 634.76025
3326.2919921875 0 684.7221

Spectrum Details

|  |  |
| --- | --- |
| Matched peaks? Matched peaks The total absolute number of peaks matched. Additionally in brackets the total fraction of peaks matched and the total number of peaks is shown. | 42 (13.91% of 302) |
| FDR? FDR The false discovery rate estimated for this peptide. It is calculated by matching all theoretical fragments with a non-integer shift with the raw peaks for this spectrum. This is done with 40 different shifts. The resulting percentage is the average number of annotated peaks over the number of annotated peaks with the correct spectrum. | 0.62% |
| Satellite FDR? Satellite FDR See the FDR for details on its calculation. This satellite ion specific FDR only contains the satellite ions (d/w) for I/L/J positions. | - |
| PSM Score? PSM Score The PSM Score as given by Hecklib to this annotated spectrum. It is shown with three significant figures. | 535 |

#### Spectrum 3642? Spectrum 3642 The raw spectrum of this peptide as annotated by Hecklib. The fragments are coloured according to ion type (see legend). Any peaks with a star '\*' as text can be hovered over to see the full details, first the ion type second the mass shift type. By hovering over the amino acids in the peptide or ions in the legend the corresponding peaks are highlighted. By toggling the 'Unassigned' label you can turn the background (unassigned) peaks on or off in the plot. By updating the slider in the Ion legend you can update the spectrum to only show the top X% of the peaks with labels. The top X% means any peak that is within X% of the highest intensity. By dragging in the spectrum you can zoom in to a specific part of the spectrum and use 'Zoom Out' to get back to the original zoom level. The annotation of the spectrum is based on the given sequence in the peptides file and is done with different software so inconsistencies are likely. The peaks are annotated based on the given sequence, with 20 ppm tolerance.

Copy Data

##### Spectrum 3642 (TSV)

###### Preview

```
Loading example...
```

*Click on the button to copy the data to your clipboard.*

Mz MinMz MaxIntensity Max

WidthHeightPeptide font sizePeptide stroke widthSpectrum font sizeSpectrum stroke widthCompact peptide

Ion legend

wxyz

abcd

OtherUnassignedIon^Charge^_Position_Show for top:%

JSSPATJDSR

01.67e+53.34e+55.01e+56.68e+5

Zoom Out

a^+1^_2_y^+1^_1_a^+1^_2_y^+1^_1_b^+1^_2_b^+1^_2_a^+1^_3_y^+1^_2_y^+1^_2_y^+1^_2_b^+1^_3_y^+2^_5_b^+1^_3_y^+2^_5_y^+2^_6_y^+2^_6_y^+1^_3_y^+1^_3_b^+1^_4_y^+2^_7_y^+1^_3_y^+2^_7_b^+1^_4_y^+2^_8_y^+2^_8_b^+1^_5_b^+1^_5_y^+2^_9_y^+2^_9_y^+1^_4_y^+1^_4_y^+1^_4_\*\*\*b^+1^_6_b^+1^_6_y^+1^_5_y^+1^_5_y^+1^_6_b^+1^_7_y^+1^_6_b^+1^_7_y^+1^_7_y^+1^_7_y^+1^_7_b^+1^_8_b^+1^_8_y^+1^_8_y^+1^_8_y^+1^_8_y^+1^_9_y^+1^_9_y^+1^_9_

0840167925193359

Fragment Matches Table

Show background peaks

| Position | Ion type | Intensity | mz Theoretical | mz Error (Th) | mz Error (ppm) | Charge | Series Number |
| --- | --- | --- | --- | --- | --- | --- | --- |
| - | - | 329 | 122.6 | - | - | 0 | - |
| - | - | 1631 | 123.1 | - | - | 0 | - |
| - | - | 1.258E+04 | 125.1 | - | - | 0 | - |
| - | - | 879.7 | 126.1 | - | - | 0 | - |
| - | - | 1899 | 127.1 | - | - | 0 | - |
| - | - | 367.5 | 127.1 | - | - | 0 | - |
| - | - | 956.1 | 128.1 | - | - | 0 | - |
| - | - | 4697 | 128.1 | - | - | 0 | - |
| - | - | 3871 | 129.1 | - | - | 0 | - |
| - | - | 898.6 | 129.1 | - | - | 0 | - |
| - | - | 1044 | 129.1 | - | - | 0 | - |
| - | - | 1239 | 130.1 | - | - | 0 | - |
| - | - | 1.094E+04 | 130.1 | - | - | 0 | - |
| - | - | 467 | 130.1 | - | - | 0 | - |
| - | - | 1917 | 130.1 | - | - | 0 | - |
| - | - | 740.5 | 131.1 | - | - | 0 | - |
| - | - | 804.6 | 133.1 | - | - | 0 | - |
| - | - | 474.3 | 133.1 | - | - | 0 | - |
| - | - | 395.1 | 135.9 | - | - | 0 | - |
| - | - | 5526 | 136.1 | - | - | 0 | - |
| - | - | 405.1 | 136.6 | - | - | 0 | - |
| - | - | 970.8 | 137.1 | - | - | 0 | - |
| - | - | 513.5 | 137.1 | - | - | 0 | - |
| - | - | 2501 | 138.1 | - | - | 0 | - |
| - | - | 361.8 | 139 | - | - | 0 | - |
| - | - | 706.2 | 139.1 | - | - | 0 | - |
| - | - | 1.913E+04 | 139.1 | - | - | 0 | - |
| - | - | 893.3 | 140.1 | - | - | 0 | - |
| - | - | 1444 | 140.1 | - | - | 0 | - |
| - | - | 2410 | 141.1 | - | - | 0 | - |
| - | - | 7.165E+04 | 141.1 | - | - | 0 | - |
| - | - | 5108 | 142.1 | - | - | 0 | - |
| - | - | 900.7 | 142.1 | - | - | 0 | - |
| - | - | 529 | 143.1 | - | - | 0 | - |
| - | - | 976 | 143.1 | - | - | 0 | - |
| - | - | 5142 | 145.1 | - | - | 0 | - |
| - | - | 6900 | 147.1 | - | - | 0 | - |
| - | - | 728.6 | 147.1 | - | - | 0 | - |
| - | - | 567.3 | 148.9 | - | - | 0 | - |
| - | - | 561.5 | 148.9 | - | - | 0 | - |
| - | - | 741.7 | 148.9 | - | - | 0 | - |
| - | - | 936.2 | 148.9 | - | - | 0 | - |
| - | - | 1368 | 148.9 | - | - | 0 | - |
| - | - | 1210 | 148.9 | - | - | 0 | - |
| - | - | 2968 | 148.9 | - | - | 0 | - |
| - | - | 4546 | 148.9 | - | - | 0 | - |
| - | - | 3878 | 149 | - | - | 0 | - |
| - | - | 2365 | 149 | - | - | 0 | - |
| - | - | 1319 | 149 | - | - | 0 | - |
| - | - | 1393 | 149 | - | - | 0 | - |
| - | - | 1039 | 149 | - | - | 0 | - |
| - | - | 804.4 | 149 | - | - | 0 | - |
| - | - | 565.1 | 149 | - | - | 0 | - |
| - | - | 661 | 149 | - | - | 0 | - |
| - | - | 698.7 | 149 | - | - | 0 | - |
| - | - | 1173 | 149 | - | - | 0 | - |
| - | - | 491.3 | 150.1 | - | - | 0 | - |
| - | - | 869.4 | 151.1 | - | - | 0 | - |
| - | - | 1122 | 153.1 | - | - | 0 | - |
| - | - | 801.4 | 153.1 | - | - | 0 | - |
| - | - | 730 | 155 | - | - | 0 | - |
| - | - | 3736 | 155.1 | - | - | 0 | - |
| 2 | a | 2.117E+04 | 155.1 | 0.0002288 | 1.475 | +1 | 2 |
| - | - | 581.5 | 156.1 | - | - | 0 | - |
| - | - | 1075 | 156.1 | - | - | 0 | - |
| - | - | 1733 | 156.1 | - | - | 0 | - |
| - | - | 488.6 | 156.6 | - | - | 0 | - |
| - | - | 1.812E+04 | 157.1 | - | - | 0 | - |
| - | - | 6.366E+04 | 157.1 | - | - | 0 | - |
| - | - | 4267 | 157.1 | - | - | 0 | - |
| - | - | 1124 | 158.1 | - | - | 0 | - |
| 10 | y | 3.084E+04 | 158.1 | 0.000233 | 1.474 | +1 | 1 |
| - | - | 4327 | 158.1 | - | - | 0 | - |
| - | - | 625.5 | 159.1 | - | - | 0 | - |
| - | - | 2131 | 159.1 | - | - | 0 | - |
| - | - | 464.9 | 160.5 | - | - | 0 | - |
| - | - | 755.9 | 166.1 | - | - | 0 | - |
| - | - | 1134 | 167.1 | - | - | 0 | - |
| - | - | 1.899E+04 | 167.1 | - | - | 0 | - |
| - | - | 753.2 | 167.1 | - | - | 0 | - |
| - | - | 1711 | 168.1 | - | - | 0 | - |
| - | - | 644.7 | 169.1 | - | - | 0 | - |
| - | - | 902.5 | 169.1 | - | - | 0 | - |
| - | - | 8.106E+04 | 169.1 | - | - | 0 | - |
| - | - | 2805 | 169.1 | - | - | 0 | - |
| - | - | 734.6 | 170.1 | - | - | 0 | - |
| - | - | 7012 | 170.1 | - | - | 0 | - |
| - | - | 549.9 | 171.1 | - | - | 0 | - |
| - | - | 828.2 | 171.1 | - | - | 0 | - |
| - | - | 1.147E+04 | 173.1 | - | - | 0 | - |
| 2 | a | 8.921E+04 | 173.1 | 0.0002384 | 1.377 | +1 | 2 |
| - | - | 1324 | 174.1 | - | - | 0 | - |
| - | - | 1110 | 174.1 | - | - | 0 | - |
| - | - | 829.7 | 174.1 | - | - | 0 | - |
| - | - | 7616 | 174.1 | - | - | 0 | - |
| - | - | 9794 | 175.1 | - | - | 0 | - |
| 10 | y | 6.546E+04 | 175.1 | 0.000219 | 1.25 | +1 | 1 |
| - | - | 835.3 | 176.1 | - | - | 0 | - |
| - | - | 455 | 176.1 | - | - | 0 | - |
| - | - | 3887 | 176.1 | - | - | 0 | - |
| - | - | 1065 | 179.1 | - | - | 0 | - |
| - | - | 514.6 | 179.1 | - | - | 0 | - |
| - | - | 1139 | 181.1 | - | - | 0 | - |
| - | - | 1411 | 182.1 | - | - | 0 | - |
| 2 | b | 1.447E+05 | 183.1 | 0.000233 | 1.272 | +1 | 2 |
| - | - | 1767 | 184.1 | - | - | 0 | - |
| - | - | 1.271E+04 | 184.1 | - | - | 0 | - |
| - | - | 2100 | 185.1 | - | - | 0 | - |
| - | - | 5.991E+04 | 185.1 | - | - | 0 | - |
| - | - | 4382 | 186.1 | - | - | 0 | - |
| - | - | 2074 | 186.1 | - | - | 0 | - |
| - | - | 1111 | 187.1 | - | - | 0 | - |
| - | - | 2255 | 187.1 | - | - | 0 | - |
| - | - | 1.891E+04 | 187.1 | - | - | 0 | - |
| - | - | 2730 | 188.1 | - | - | 0 | - |
| - | - | 555 | 192.1 | - | - | 0 | - |
| - | - | 1418 | 193.1 | - | - | 0 | - |
| - | - | 1899 | 195.1 | - | - | 0 | - |
| - | - | 829.6 | 196.1 | - | - | 0 | - |
| - | - | 548.2 | 196.1 | - | - | 0 | - |
| - | - | 522.4 | 197.1 | - | - | 0 | - |
| - | - | 465.5 | 197.1 | - | - | 0 | - |
| - | - | 4897 | 197.1 | - | - | 0 | - |
| - | - | 1227 | 199.1 | - | - | 0 | - |
| - | - | 1073 | 200.1 | - | - | 0 | - |
| 2 | b | 8.932E+04 | 201.1 | 0.0001511 | 0.7511 | +1 | 2 |
| - | - | 4140 | 202.1 | - | - | 0 | - |
| - | - | 3963 | 202.1 | - | - | 0 | - |
| - | - | 8337 | 202.1 | - | - | 0 | - |
| - | - | 4433 | 203.1 | - | - | 0 | - |
| - | - | 4936 | 203.1 | - | - | 0 | - |
| - | - | 6101 | 208.1 | - | - | 0 | - |
| - | - | 567.8 | 208.1 | - | - | 0 | - |
| - | - | 499.3 | 208.3 | - | - | 0 | - |
| - | - | 1521 | 209.1 | - | - | 0 | - |
| - | - | 985.4 | 209.1 | - | - | 0 | - |
| - | - | 2132 | 210.1 | - | - | 0 | - |
| - | - | 9725 | 210.1 | - | - | 0 | - |
| - | - | 2216 | 211.1 | - | - | 0 | - |
| - | - | 864.4 | 211.1 | - | - | 0 | - |
| - | - | 1236 | 211.1 | - | - | 0 | - |
| - | - | 1203 | 215.1 | - | - | 0 | - |
| - | - | 9703 | 215.1 | - | - | 0 | - |
| - | - | 569.6 | 216.1 | - | - | 0 | - |
| - | - | 678.8 | 218.2 | - | - | 0 | - |
| - | - | 571.9 | 220.1 | - | - | 0 | - |
| - | - | 2900 | 220.1 | - | - | 0 | - |
| - | - | 3137 | 221.1 | - | - | 0 | - |
| - | - | 825.9 | 222.1 | - | - | 0 | - |
| - | - | 2024 | 222.1 | - | - | 0 | - |
| - | - | 1579 | 223.1 | - | - | 0 | - |
| - | - | 592.5 | 223.1 | - | - | 0 | - |
| - | - | 1902 | 224.1 | - | - | 0 | - |
| - | - | 5971 | 224.1 | - | - | 0 | - |
| - | - | 597.6 | 225 | - | - | 0 | - |
| - | - | 563.9 | 225 | - | - | 0 | - |
| - | - | 1318 | 225.1 | - | - | 0 | - |
| - | - | 1.97E+04 | 225.1 | - | - | 0 | - |
| - | - | 8450 | 226.1 | - | - | 0 | - |
| - | - | 983 | 226.1 | - | - | 0 | - |
| - | - | 6570 | 227.1 | - | - | 0 | - |
| - | - | 2874 | 228.1 | - | - | 0 | - |
| - | - | 3107 | 228.1 | - | - | 0 | - |
| - | - | 1587 | 229.1 | - | - | 0 | - |
| - | - | 5750 | 232.1 | - | - | 0 | - |
| - | - | 2662 | 235.1 | - | - | 0 | - |
| - | - | 2659 | 236.1 | - | - | 0 | - |
| - | - | 1029 | 237.1 | - | - | 0 | - |
| - | - | 1.369E+05 | 238.1 | - | - | 0 | - |
| - | - | 1.495E+04 | 239.1 | - | - | 0 | - |
| - | - | 945.1 | 240.1 | - | - | 0 | - |
| - | - | 755 | 240.1 | - | - | 0 | - |
| - | - | 2350 | 240.2 | - | - | 0 | - |
| - | - | 1179 | 242.1 | - | - | 0 | - |
| 3 | a | 9903 | 242.1 | 0.000198 | 0.8178 | +1 | 3 |
| - | - | 1635 | 243.1 | - | - | 0 | - |
| - | - | 859.4 | 243.2 | - | - | 0 | - |
| 9 | y | 1.33E+04 | 244.1 | 0.0001176 | 0.4815 | +1 | 2 |
| 9 | y | 3.642E+04 | 245.1 | 0.0002328 | 0.9498 | +1 | 2 |
| - | - | 1850 | 246.1 | - | - | 0 | - |
| - | - | 3310 | 246.1 | - | - | 0 | - |
| - | - | 2163 | 250.1 | - | - | 0 | - |
| - | - | 1530 | 252.1 | - | - | 0 | - |
| - | - | 3.914E+04 | 252.1 | - | - | 0 | - |
| - | - | 591.7 | 253.1 | - | - | 0 | - |
| - | - | 2283 | 253.1 | - | - | 0 | - |
| - | - | 5651 | 253.1 | - | - | 0 | - |
| - | - | 5985 | 254.1 | - | - | 0 | - |
| - | - | 618.2 | 254.1 | - | - | 0 | - |
| - | - | 3.655E+04 | 256.1 | - | - | 0 | - |
| - | - | 3766 | 257.1 | - | - | 0 | - |
| 9 | y | 6.614E+05 | 262.2 | 0.000173 | 0.6599 | +1 | 2 |
| - | - | 6.416E+04 | 263.2 | - | - | 0 | - |
| - | - | 2869 | 264.1 | - | - | 0 | - |
| - | - | 6593 | 264.2 | - | - | 0 | - |
| - | - | 2360 | 268.1 | - | - | 0 | - |
| - | - | 6547 | 268.2 | - | - | 0 | - |
| - | - | 1383 | 269.1 | - | - | 0 | - |
| - | - | 790.9 | 269.2 | - | - | 0 | - |
| - | - | 5624 | 270.1 | - | - | 0 | - |
| 3 | b | 1.951E+05 | 270.1 | 9.541E-05 | 0.3532 | +1 | 3 |
| - | - | 2.438E+04 | 271.1 | - | - | 0 | - |
| - | - | 878 | 271.2 | - | - | 0 | - |
| - | - | 1912 | 272.1 | - | - | 0 | - |
| - | - | 2547 | 272.2 | - | - | 0 | - |
| - | - | 669.5 | 278.1 | - | - | 0 | - |
| - | - | 959 | 280.1 | - | - | 0 | - |
| - | - | 674.5 | 280.2 | - | - | 0 | - |
| - | - | 1578 | 281.1 | - | - | 0 | - |
| - | - | 2673 | 282.1 | - | - | 0 | - |
| - | - | 1572 | 282.2 | - | - | 0 | - |
| - | - | 886.2 | 284.2 | - | - | 0 | - |
| - | - | 1882 | 286.2 | - | - | 0 | - |
| - | - | 1048 | 286.2 | - | - | 0 | - |
| - | - | 804.5 | 286.7 | - | - | 0 | - |
| 6 | y | 1949 | 287.2 | 0.000345 | 1.202 | +2 | 5 |
| - | - | 573.9 | 287.2 | - | - | 0 | - |
| - | - | 5680 | 288.1 | - | - | 0 | - |
| 3 | b | 1.388E+04 | 288.2 | 0.0001814 | 0.6294 | +1 | 3 |
| - | - | 1243 | 289.2 | - | - | 0 | - |
| - | - | 1622 | 294.1 | - | - | 0 | - |
| - | - | 969 | 295.1 | - | - | 0 | - |
| 6 | y | 3031 | 296.2 | 0.001227 | 4.144 | +2 | 5 |
| - | - | 1000 | 297.1 | - | - | 0 | - |
| - | - | 780.3 | 297.2 | - | - | 0 | - |
| - | - | 626.7 | 298.1 | - | - | 0 | - |
| - | - | 1046 | 298.2 | - | - | 0 | - |
| - | - | 1561 | 299.1 | - | - | 0 | - |
| - | - | 2368 | 299.1 | - | - | 0 | - |
| - | - | 1371 | 300.1 | - | - | 0 | - |
| - | - | 1843 | 300.1 | - | - | 0 | - |
| - | - | 3104 | 301.1 | - | - | 0 | - |
| - | - | 725.3 | 301.1 | - | - | 0 | - |
| - | - | 2162 | 302.1 | - | - | 0 | - |
| - | - | 1052 | 303.1 | - | - | 0 | - |
| - | - | 1408 | 304.2 | - | - | 0 | - |
| - | - | 907.7 | 307.1 | - | - | 0 | - |
| - | - | 1409 | 307.1 | - | - | 0 | - |
| - | - | 1074 | 308.1 | - | - | 0 | - |
| - | - | 1401 | 311.2 | - | - | 0 | - |
| - | - | 2347 | 311.2 | - | - | 0 | - |
| - | - | 2094 | 312.1 | - | - | 0 | - |
| - | - | 3576 | 312.2 | - | - | 0 | - |
| - | - | 1681 | 313.2 | - | - | 0 | - |
| - | - | 1977 | 314.1 | - | - | 0 | - |
| - | - | 688.2 | 314.2 | - | - | 0 | - |
| - | - | 2283 | 315.2 | - | - | 0 | - |
| - | - | 1221 | 316.2 | - | - | 0 | - |
| - | - | 1969 | 317.1 | - | - | 0 | - |
| - | - | 5205 | 321.2 | - | - | 0 | - |
| - | - | 1167 | 322.1 | - | - | 0 | - |
| - | - | 1032 | 322.2 | - | - | 0 | - |
| - | - | 1935 | 322.2 | - | - | 0 | - |
| 5 | y | 4419 | 322.7 | 0.0002818 | 0.8733 | +2 | 6 |
| - | - | 2735 | 323.2 | - | - | 0 | - |
| - | - | 546.3 | 323.7 | - | - | 0 | - |
| - | - | 2739 | 324.1 | - | - | 0 | - |
| - | - | 644.4 | 324.2 | - | - | 0 | - |
| - | - | 2271 | 325.1 | - | - | 0 | - |
| - | - | 3642 | 325.2 | - | - | 0 | - |
| - | - | 944.7 | 326.2 | - | - | 0 | - |
| - | - | 707.5 | 328.2 | - | - | 0 | - |
| - | - | 3999 | 329.2 | - | - | 0 | - |
| - | - | 2126 | 330.2 | - | - | 0 | - |
| - | - | 1.113E+04 | 331.2 | - | - | 0 | - |
| 5 | y | 2634 | 331.7 | 0.0006147 | 1.853 | +2 | 6 |
| - | - | 4060 | 332.2 | - | - | 0 | - |
| - | - | 1968 | 332.2 | - | - | 0 | - |
| - | - | 722.6 | 333.2 | - | - | 0 | - |
| - | - | 1047 | 335.1 | - | - | 0 | - |
| - | - | 1028 | 335.2 | - | - | 0 | - |
| - | - | 1755 | 335.2 | - | - | 0 | - |
| - | - | 6116 | 337.2 | - | - | 0 | - |
| - | - | 1308 | 338.2 | - | - | 0 | - |
| - | - | 1.874E+04 | 339.2 | - | - | 0 | - |
| - | - | 1998 | 339.2 | - | - | 0 | - |
| - | - | 3119 | 340.2 | - | - | 0 | - |
| - | - | 2224 | 341.2 | - | - | 0 | - |
| - | - | 1.653E+04 | 342.1 | - | - | 0 | - |
| - | - | 734 | 342.2 | - | - | 0 | - |
| - | - | 969.8 | 343.1 | - | - | 0 | - |
| - | - | 835.9 | 343.1 | - | - | 0 | - |
| - | - | 3652 | 343.2 | - | - | 0 | - |
| - | - | 637.8 | 347 | - | - | 0 | - |
| - | - | 1230 | 349.2 | - | - | 0 | - |
| - | - | 963.5 | 349.2 | - | - | 0 | - |
| - | - | 1088 | 353.1 | - | - | 0 | - |
| - | - | 2537 | 355.2 | - | - | 0 | - |
| - | - | 7123 | 355.2 | - | - | 0 | - |
| - | - | 1581 | 356.2 | - | - | 0 | - |
| - | - | 1.575E+04 | 357.2 | - | - | 0 | - |
| - | - | 3442 | 358.2 | - | - | 0 | - |
| - | - | 1848 | 358.2 | - | - | 0 | - |
| 8 | y | 2.253E+04 | 359.2 | 9.208E-05 | 0.2564 | +1 | 3 |
| 8 | y | 2.629E+04 | 360.2 | 6.836E-05 | 0.1898 | +1 | 3 |
| - | - | 4006 | 361.2 | - | - | 0 | - |
| - | - | 804.1 | 361.2 | - | - | 0 | - |
| - | - | 1152 | 362.2 | - | - | 0 | - |
| - | - | 1054 | 363.2 | - | - | 0 | - |
| - | - | 8275 | 365.2 | - | - | 0 | - |
| - | - | 1720 | 366.2 | - | - | 0 | - |
| - | - | 1210 | 367.2 | - | - | 0 | - |
| 4 | b | 2280 | 367.2 | 0.000249 | 0.6782 | +1 | 4 |
| - | - | 1737 | 369.7 | - | - | 0 | - |
| - | - | 4373 | 370.2 | - | - | 0 | - |
| - | - | 1.663E+04 | 370.7 | - | - | 0 | - |
| 4 | y | 5.016E+04 | 371.2 | 0.0005722 | 1.542 | +2 | 7 |
| - | - | 1.981E+04 | 371.7 | - | - | 0 | - |
| - | - | 5435 | 372.2 | - | - | 0 | - |
| - | - | 1.997E+04 | 376.2 | - | - | 0 | - |
| - | - | 583.8 | 376.9 | - | - | 0 | - |
| 8 | y | 1.29E+05 | 377.2 | 5.491E-05 | 0.1456 | +1 | 3 |
| - | - | 1.986E+04 | 378.2 | - | - | 0 | - |
| - | - | 3946 | 378.7 | - | - | 0 | - |
| - | - | 8072 | 379.2 | - | - | 0 | - |
| - | - | 1.078E+05 | 379.7 | - | - | 0 | - |
| 4 | y | 2.737E+05 | 380.2 | 0.000722 | 1.899 | +2 | 7 |
| - | - | 1.114E+05 | 380.7 | - | - | 0 | - |
| - | - | 1158 | 381.2 | - | - | 0 | - |
| - | - | 2.327E+04 | 381.2 | - | - | 0 | - |
| - | - | 1780 | 381.7 | - | - | 0 | - |
| - | - | 691 | 382.2 | - | - | 0 | - |
| - | - | 893.2 | 383.2 | - | - | 0 | - |
| - | - | 7671 | 383.2 | - | - | 0 | - |
| - | - | 1322 | 384.2 | - | - | 0 | - |
| 4 | b | 2056 | 385.2 | 0.006776 | 17.59 | +1 | 4 |
| - | - | 917.3 | 385.7 | - | - | 0 | - |
| - | - | 1459 | 387.2 | - | - | 0 | - |
| - | - | 1200 | 390.2 | - | - | 0 | - |
| - | - | 2167 | 391.2 | - | - | 0 | - |
| - | - | 934.2 | 392.2 | - | - | 0 | - |
| - | - | 815.7 | 392.2 | - | - | 0 | - |
| - | - | 1703 | 396.7 | - | - | 0 | - |
| - | - | 1462 | 397.2 | - | - | 0 | - |
| - | - | 904.9 | 397.7 | - | - | 0 | - |
| - | - | 924.6 | 398.2 | - | - | 0 | - |
| - | - | 759.6 | 399.2 | - | - | 0 | - |
| - | - | 1017 | 400.2 | - | - | 0 | - |
| - | - | 1076 | 401.2 | - | - | 0 | - |
| - | - | 1210 | 401.2 | - | - | 0 | - |
| - | - | 4217 | 405.2 | - | - | 0 | - |
| - | - | 1.874E+04 | 405.7 | - | - | 0 | - |
| - | - | 6709 | 406.2 | - | - | 0 | - |
| - | - | 992 | 406.2 | - | - | 0 | - |
| - | - | 2061 | 406.7 | - | - | 0 | - |
| - | - | 995.2 | 407.2 | - | - | 0 | - |
| - | - | 4384 | 408.2 | - | - | 0 | - |
| - | - | 974.7 | 409.2 | - | - | 0 | - |
| - | - | 1011 | 410.2 | - | - | 0 | - |
| - | - | 6.702E+04 | 414.2 | - | - | 0 | - |
| 3 | y | 2.301E+05 | 414.7 | 0.0009154 | 2.207 | +2 | 8 |
| - | - | 9.472E+04 | 415.2 | - | - | 0 | - |
| - | - | 2.306E+04 | 415.7 | - | - | 0 | - |
| - | - | 3196 | 416.2 | - | - | 0 | - |
| - | - | 708.7 | 419 | - | - | 0 | - |
| - | - | 804.9 | 419 | - | - | 0 | - |
| - | - | 1208 | 420 | - | - | 0 | - |
| - | - | 1843 | 420.2 | - | - | 0 | - |
| - | - | 6.446E+04 | 423.2 | - | - | 0 | - |
| 3 | y | 1.973E+05 | 423.7 | 0.001004 | 2.37 | +2 | 8 |
| - | - | 8.216E+04 | 424.2 | - | - | 0 | - |
| - | - | 2.368E+04 | 424.7 | - | - | 0 | - |
| - | - | 1086 | 425.2 | - | - | 0 | - |
| - | - | 3667 | 426.2 | - | - | 0 | - |
| - | - | 1698 | 428.3 | - | - | 0 | - |
| - | - | 941.6 | 430.2 | - | - | 0 | - |
| - | - | 3696 | 434.2 | - | - | 0 | - |
| - | - | 1443 | 436.2 | - | - | 0 | - |
| - | - | 1141 | 438.2 | - | - | 0 | - |
| 5 | b | 2334 | 438.2 | 0.0009465 | 2.16 | +1 | 5 |
| - | - | 660.7 | 439.2 | - | - | 0 | - |
| - | - | 719 | 440.2 | - | - | 0 | - |
| - | - | 2012 | 442.3 | - | - | 0 | - |
| - | - | 807.6 | 443.2 | - | - | 0 | - |
| - | - | 865.8 | 443.3 | - | - | 0 | - |
| - | - | 2325 | 444.2 | - | - | 0 | - |
| - | - | 946.9 | 448.7 | - | - | 0 | - |
| - | - | 3083 | 449.2 | - | - | 0 | - |
| - | - | 1607 | 449.7 | - | - | 0 | - |
| - | - | 1.04E+04 | 452.3 | - | - | 0 | - |
| - | - | 1863 | 453.3 | - | - | 0 | - |
| - | - | 2296 | 455.2 | - | - | 0 | - |
| - | - | 1396 | 456.2 | - | - | 0 | - |
| 5 | b | 4082 | 456.2 | 8.643E-05 | 0.1894 | +1 | 5 |
| - | - | 663.4 | 457.2 | - | - | 0 | - |
| - | - | 1174 | 457.2 | - | - | 0 | - |
| - | - | 1791 | 457.7 | - | - | 0 | - |
| 2 | y | 5389 | 458.2 | 0.001289 | 2.813 | +2 | 9 |
| - | - | 2401 | 458.7 | - | - | 0 | - |
| - | - | 663.3 | 459.2 | - | - | 0 | - |
| - | - | 1873 | 462.7 | - | - | 0 | - |
| - | - | 2758 | 463.2 | - | - | 0 | - |
| - | - | 1036 | 463.7 | - | - | 0 | - |
| - | - | 1425 | 466.7 | - | - | 0 | - |
| 2 | y | 3905 | 467.2 | 0.002782 | 5.954 | +2 | 9 |
| - | - | 1496 | 467.7 | - | - | 0 | - |
| - | - | 613.9 | 470.2 | - | - | 0 | - |
| - | - | 4110 | 470.3 | - | - | 0 | - |
| - | - | 2656 | 471.3 | - | - | 0 | - |
| - | - | 1725 | 471.7 | - | - | 0 | - |
| - | - | 2237 | 472.2 | - | - | 0 | - |
| 7 | y | 2934 | 472.3 | 0.001659 | 3.514 | +1 | 4 |
| - | - | 2834 | 472.7 | - | - | 0 | - |
| 7 | y | 7237 | 473.2 | 0.0003005 | 0.6351 | +1 | 4 |
| - | - | 910 | 474.2 | - | - | 0 | - |
| - | - | 930.4 | 476.3 | - | - | 0 | - |
| - | - | 764.6 | 479.3 | - | - | 0 | - |
| - | - | 4922 | 480.2 | - | - | 0 | - |
| - | - | 1405 | 481.2 | - | - | 0 | - |
| - | - | 851.1 | 489.2 | - | - | 0 | - |
| - | - | 3.123E+04 | 489.3 | - | - | 0 | - |
| 7 | y | 7.295E+04 | 490.3 | 0.0005551 | 1.132 | +1 | 4 |
| - | - | 1.776E+04 | 491.3 | - | - | 0 | - |
| - | - | 2658 | 492.3 | - | - | 0 | - |
| - | - | 678 | 495.3 | - | - | 0 | - |
| - | - | 1219 | 497.3 | - | - | 0 | - |
| - | - | 1.711E+04 | 498.3 | - | - | 0 | - |
| - | - | 4273 | 499.3 | - | - | 0 | - |
| - | - | 651.6 | 499.8 | - | - | 0 | - |
| - | - | 646.5 | 500.3 | - | - | 0 | - |
| - | - | 2245 | 503.3 | - | - | 0 | - |
| - | - | 857 | 504.2 | - | - | 0 | - |
| - | - | 1882 | 505.3 | - | - | 0 | - |
| - | - | 9615 | 505.8 | - | - | 0 | - |
| - | - | 4853 | 506.3 | - | - | 0 | - |
| - | - | 2419 | 506.8 | - | - | 0 | - |
| - | - | 656.1 | 508.8 | - | - | 0 | - |
| - | - | 1246 | 513.8 | - | - | 0 | - |
| - | - | 5441 | 514.3 | - | - | 0 | - |
| 0 | Precursor | 1.826E+04 | 514.8 | 0.001494 | 2.901 | +2 | -1 |
| 0 | Precursor | 8517 | 515.3 | 0.009181 | 17.82 | +2 | -1 |
| - | - | 2945 | 515.8 | - | - | 0 | - |
| - | - | 6689 | 521.3 | - | - | 0 | - |
| - | - | 1608 | 522.3 | - | - | 0 | - |
| - | - | 1.115E+04 | 523.3 | - | - | 0 | - |
| 0 | Precursor | 3.966E+04 | 523.8 | 0.0009719 | 1.856 | +2 | -1 |
| - | - | 1.863E+04 | 524.3 | - | - | 0 | - |
| - | - | 6334 | 524.8 | - | - | 0 | - |
| - | - | 968.4 | 525.3 | - | - | 0 | - |
| - | - | 1172 | 538.3 | - | - | 0 | - |
| 6 | b | 8102 | 539.3 | 0.0003872 | 0.718 | +1 | 6 |
| - | - | 1515 | 540.3 | - | - | 0 | - |
| - | - | 5007 | 545.8 | - | - | 0 | - |
| - | - | 2922 | 546.3 | - | - | 0 | - |
| - | - | 2773 | 546.3 | - | - | 0 | - |
| - | - | 1139 | 546.8 | - | - | 0 | - |
| - | - | 2301 | 547.3 | - | - | 0 | - |
| - | - | 2269 | 549.3 | - | - | 0 | - |
| - | - | 1380 | 550.3 | - | - | 0 | - |
| - | - | 1715 | 555.3 | - | - | 0 | - |
| - | - | 2436 | 556.3 | - | - | 0 | - |
| 6 | b | 3889 | 557.3 | 0.001327 | 2.382 | +1 | 6 |
| - | - | 1101 | 558.3 | - | - | 0 | - |
| - | - | 8887 | 567.3 | - | - | 0 | - |
| - | - | 3307 | 568.3 | - | - | 0 | - |
| - | - | 3517 | 572.3 | - | - | 0 | - |
| 6 | y | 9462 | 573.3 | 0.000154 | 0.2686 | +1 | 5 |
| - | - | 3069 | 574.3 | - | - | 0 | - |
| - | - | 849.1 | 575.3 | - | - | 0 | - |
| - | - | 933.4 | 583.3 | - | - | 0 | - |
| - | - | 1728 | 584.3 | - | - | 0 | - |
| - | - | 7732 | 585.3 | - | - | 0 | - |
| - | - | 2444 | 586.3 | - | - | 0 | - |
| - | - | 9.161E+04 | 590.3 | - | - | 0 | - |
| 6 | y | 1.84E+05 | 591.3 | 0.0008808 | 1.49 | +1 | 5 |
| - | - | 5.326E+04 | 592.3 | - | - | 0 | - |
| - | - | 1.051E+04 | 593.3 | - | - | 0 | - |
| - | - | 690.3 | 594.3 | - | - | 0 | - |
| - | - | 1821 | 600.3 | - | - | 0 | - |
| - | - | 2148 | 601.3 | - | - | 0 | - |
| - | - | 1888 | 602.3 | - | - | 0 | - |
| - | - | 622.8 | 603.3 | - | - | 0 | - |
| - | - | 961.2 | 616.3 | - | - | 0 | - |
| - | - | 1609 | 617.3 | - | - | 0 | - |
| - | - | 762.2 | 618.3 | - | - | 0 | - |
| - | - | 1254 | 626.3 | - | - | 0 | - |
| - | - | 1926 | 627.3 | - | - | 0 | - |
| - | - | 1830 | 634.4 | - | - | 0 | - |
| - | - | 836.1 | 635.4 | - | - | 0 | - |
| - | - | 863.8 | 636.3 | - | - | 0 | - |
| - | - | 672.3 | 642.4 | - | - | 0 | - |
| - | - | 8498 | 643.4 | - | - | 0 | - |
| 5 | y | 1.709E+04 | 644.3 | 0.000882 | 1.369 | +1 | 6 |
| - | - | 6592 | 645.3 | - | - | 0 | - |
| - | - | 1551 | 646.3 | - | - | 0 | - |
| - | - | 679 | 647.3 | - | - | 0 | - |
| 7 | b | 4007 | 652.4 | 0.0008521 | 1.306 | +1 | 7 |
| - | - | 2988 | 652.8 | - | - | 0 | - |
| - | - | 1825 | 653.3 | - | - | 0 | - |
| - | - | 1201 | 653.8 | - | - | 0 | - |
| - | - | 1614 | 654.3 | - | - | 0 | - |
| - | - | 1287 | 661.3 | - | - | 0 | - |
| - | - | 7.381E+04 | 661.4 | - | - | 0 | - |
| 5 | y | 1.402E+05 | 662.3 | 0.001426 | 2.153 | +1 | 6 |
| - | - | 4.437E+04 | 663.4 | - | - | 0 | - |
| - | - | 8480 | 664.4 | - | - | 0 | - |
| 7 | b | 1712 | 670.4 | 3.189E-06 | 0.004757 | +1 | 7 |
| - | - | 2098 | 671.3 | - | - | 0 | - |
| - | - | 6175 | 672.3 | - | - | 0 | - |
| - | - | 1928 | 673.3 | - | - | 0 | - |
| - | - | 825.6 | 674.3 | - | - | 0 | - |
| - | - | 840.5 | 698.4 | - | - | 0 | - |
| - | - | 989.4 | 699.4 | - | - | 0 | - |
| - | - | 1711 | 715.4 | - | - | 0 | - |
| - | - | 756 | 716.4 | - | - | 0 | - |
| - | - | 756.2 | 719 | - | - | 0 | - |
| - | - | 811.1 | 723.4 | - | - | 0 | - |
| - | - | 1133 | 724.4 | - | - | 0 | - |
| - | - | 929.7 | 725.4 | - | - | 0 | - |
| - | - | 945.2 | 734.4 | - | - | 0 | - |
| - | - | 3150 | 740.4 | - | - | 0 | - |
| 4 | y | 1.243E+04 | 741.4 | 0.0002422 | 0.3266 | +1 | 7 |
| 4 | y | 6641 | 742.4 | 0.01409 | 18.98 | +1 | 7 |
| - | - | 780.9 | 742.5 | - | - | 0 | - |
| - | - | 2029 | 743.4 | - | - | 0 | - |
| - | - | 1123 | 749.4 | - | - | 0 | - |
| - | - | 635.3 | 756.4 | - | - | 0 | - |
| - | - | 1621 | 757.4 | - | - | 0 | - |
| - | - | 1036 | 758.3 | - | - | 0 | - |
| - | - | 1.181E+05 | 758.4 | - | - | 0 | - |
| 4 | y | 2.813E+05 | 759.4 | 0.001579 | 2.08 | +1 | 7 |
| - | - | 1.076E+05 | 760.4 | - | - | 0 | - |
| - | - | 2.71E+04 | 761.4 | - | - | 0 | - |
| - | - | 1852 | 762.4 | - | - | 0 | - |
| 8 | b | 1149 | 767.4 | 0.003028 | 3.945 | +1 | 8 |
| 8 | b | 2727 | 785.4 | 0.001006 | 1.281 | +1 | 8 |
| - | - | 1688 | 786.4 | - | - | 0 | - |
| - | - | 827.3 | 810.4 | - | - | 0 | - |
| - | - | 1000 | 815.4 | - | - | 0 | - |
| - | - | 1089 | 816.4 | - | - | 0 | - |
| - | - | 800.4 | 817.4 | - | - | 0 | - |
| - | - | 2666 | 827.4 | - | - | 0 | - |
| 3 | y | 8481 | 828.4 | 0.0009896 | 1.195 | +1 | 8 |
| 3 | y | 4039 | 829.4 | 0.01551 | 18.7 | +1 | 8 |
| - | - | 1097 | 830.4 | - | - | 0 | - |
| - | - | 1121 | 845.4 | - | - | 0 | - |
| - | - | 5.6E+04 | 845.4 | - | - | 0 | - |
| 3 | y | 1.403E+05 | 846.4 | 0.001106 | 1.307 | +1 | 8 |
| - | - | 5.867E+04 | 847.4 | - | - | 0 | - |
| - | - | 1.496E+04 | 848.4 | - | - | 0 | - |
| - | - | 1556 | 849.4 | - | - | 0 | - |
| - | - | 1726 | 856.4 | - | - | 0 | - |
| - | - | 1896 | 897.4 | - | - | 0 | - |
| - | - | 1432 | 898.4 | - | - | 0 | - |
| - | - | 800.5 | 902.5 | - | - | 0 | - |
| - | - | 1139 | 903.5 | - | - | 0 | - |
| - | - | 3670 | 914.5 | - | - | 0 | - |
| 2 | y | 1.298E+04 | 915.5 | 0.001127 | 1.231 | +1 | 9 |
| 2 | y | 6682 | 916.4 | 0.01827 | 19.94 | +1 | 9 |
| - | - | 908.1 | 917.5 | - | - | 0 | - |
| - | - | 1332 | 925.4 | - | - | 0 | - |
| - | - | 4.184E+04 | 932.5 | - | - | 0 | - |
| 2 | y | 1.074E+05 | 933.5 | 0.001487 | 1.593 | +1 | 9 |
| - | - | 4.894E+04 | 934.5 | - | - | 0 | - |
| - | - | 1.461E+04 | 935.5 | - | - | 0 | - |
| - | - | 1406 | 936.5 | - | - | 0 | - |
| - | - | 4230 | 942.5 | - | - | 0 | - |
| - | - | 1.015E+04 | 943.5 | - | - | 0 | - |
| - | - | 5476 | 944.5 | - | - | 0 | - |
| - | - | 1412 | 945.5 | - | - | 0 | - |
| - | - | 1157 | 1091 | - | - | 0 | - |
| - | - | 672.2 | 1707 | - | - | 0 | - |
| - | - | 609.8 | 1829 | - | - | 0 | - |
| - | - | 721.3 | 3080 | - | - | 0 | - |
| - | - | 678.4 | 3325 | - | - | 0 | - |

m/z Charge Intensity FragmentType MassShift Position
122.58350372314453 0 329.02994
123.09193420410156 0 1631.3757
125.10758972167969 0 12581.451
126.11083221435547 0 879.6909
127.08690643310547 0 1899.0605
127.11433410644531 0 367.5348
128.07081604003906 0 956.0945
128.10723876953125 0 4696.589
129.06610107421875 0 3870.585
129.10252380371094 0 898.6259
129.11375427246094 0 1044.2917
130.05003356933594 0 1239.1927
130.0613555908203 0 10941.167
130.06947326660156 0 466.99835
130.09779357910156 0 1917.4619
131.11814880371094 0 740.5313
133.08595275878906 0 804.58466
133.09764099121094 0 474.288
135.87771606445312 0 395.07233
136.0759735107422 0 5525.6655
136.5862274169922 0 405.06534
137.07125854492188 0 970.82043
137.0792694091797 0 513.51685
138.09149169921875 0 2501.215
138.97019958496094 0 361.76138
139.05047607421875 0 706.18384
139.0868377685547 0 19129.691
140.08189392089844 0 893.26556
140.09010314941406 0 1444.1057
141.0662384033203 0 2409.6643
141.1024932861328 0 71654.09
142.1057891845703 0 5108.284
142.1227264404297 0 900.65125
143.08123779296875 0 528.97363
143.11810302734375 0 975.9944
145.097412109375 0 5142.339
147.07669067382812 0 6900.2886
147.11282348632812 0 728.6107
148.8934783935547 0 567.3355
148.9009552001953 0 561.4873
148.90792846679688 0 741.69055
148.91519165039062 0 936.23804
148.92221069335938 0 1367.9958
148.92933654785156 0 1209.9734
148.93658447265625 0 2968.16
148.94422912597656 0 4545.9473
148.9606170654297 0 3877.9387
148.96823120117188 0 2365.397
148.9753875732422 0 1318.5347
148.9825439453125 0 1393.1445
148.9895782470703 0 1039.4897
148.99705505371094 0 804.42145
149.00418090820312 0 565.10986
149.01119995117188 0 660.96643
149.01809692382812 0 698.7246
149.0455322265625 0 1172.6105
150.05503845214844 0 491.2896
151.08673095703125 0 869.4324
153.0663299560547 0 1121.9692
153.1024932861328 0 801.377
155.0449981689453 0 729.9531
155.0817413330078 0 3735.794
155.1181182861328 0 21170.498 a Water loss 1
156.0846405029297 0 581.5342
156.1021270751953 0 1075.384
156.12168884277344 0 1733.1786
156.57638549804688 0 488.64178
157.0609893798828 0 18123.713
157.0973663330078 0 63657.652
157.1087188720703 0 4266.6274
158.0644989013672 0 1124.1165
158.09263610839844 0 30837.22 y Ammonia loss 9
158.1005096435547 0 4326.6963
159.0770263671875 0 625.5322
159.09605407714844 0 2131.0369
160.47613525390625 0 464.89008
166.09779357910156 0 755.932
167.0556182861328 0 1134.3875
167.0817413330078 0 18989.59
167.1182861328125 0 753.1898
168.0849151611328 0 1710.5784
169.0531005859375 0 644.6647
169.06097412109375 0 902.53064
169.0973663330078 0 81056.84
169.13377380371094 0 2804.881
170.09373474121094 0 734.6023
170.1007537841797 0 7011.8193
171.07687377929688 0 549.90076
171.11277770996094 0 828.15796
173.09225463867188 0 11471.641
173.12869262695312 0 89206.31 a 1
174.0878448486328 0 1324.4355
174.09547424316406 0 1109.9827
174.1248321533203 0 829.74243
174.13198852539062 0 7616.265
175.07156372070312 0 9794.213
175.11917114257812 0 65464.68 y 9
176.074951171875 0 835.3305
176.11572265625 0 454.97815
176.1226043701172 0 3887.4014
179.0817413330078 0 1065.191
179.11767578125 0 514.6238
181.09701538085938 0 1139.1719
182.0924835205078 0 1411.4236
183.113037109375 0 144685.81 b Water loss 1
184.10861206054688 0 1766.5173
184.11642456054688 0 12711.251
185.05584716796875 0 2099.6282
185.09226989746094 0 59910.43
186.09576416015625 0 4381.776
186.12367248535156 0 2073.7837
187.07081604003906 0 1110.5057
187.1078338623047 0 2255.0063
187.14431762695312 0 18907.514
188.14779663085938 0 2729.6008
192.1136474609375 0 554.96
193.09750366210938 0 1418.393
195.07652282714844 0 1899.4902
196.108154296875 0 829.553
196.14443969726562 0 548.2289
197.0933074951172 0 522.3602
197.1037139892578 0 465.53024
197.128662109375 0 4897.1904
199.1189727783203 0 1226.612
200.10311889648438 0 1073.3353
201.12351989746094 0 89322.69 b 1
202.08250427246094 0 4139.838
202.11851501464844 0 3962.864
202.1270294189453 0 8337.091
203.06639099121094 0 4432.9194
203.10275268554688 0 4935.746
208.10818481445312 0 6101.215
208.1450653076172 0 567.78406
208.25650024414062 0 499.31485
209.09222412109375 0 1521.3794
209.1030731201172 0 985.4001
210.08726501464844 0 2131.882
210.12387084960938 0 9725.152
211.10781860351562 0 2215.9265
211.12745666503906 0 864.36304
211.14447021484375 0 1236.247
215.11410522460938 0 1203.1821
215.13922119140625 0 9703.037
216.14334106445312 0 569.58765
218.16119384765625 0 678.76776
220.10940551757812 0 571.9066
220.1293182373047 0 2900.313
221.09219360351562 0 3137.3733
222.09567260742188 0 825.89935
222.12368774414062 0 2023.6255
223.10777282714844 0 1578.9712
223.14373779296875 0 592.4884
224.103515625 0 1902.1948
224.13961791992188 0 5971.4756
225.0066680908203 0 597.63385
225.04345703125 0 563.87695
225.08717346191406 0 1318.4783
225.12356567382812 0 19703.922
226.11880493164062 0 8449.781
226.13018798828125 0 982.96136
227.11399841308594 0 6569.8877
228.09815979003906 0 2874.0632
228.13442993164062 0 3106.6902
229.11842346191406 0 1587.3167
232.14064025878906 0 5749.637
235.14425659179688 0 2662.1099
236.1031036376953 0 2658.7083
237.0872802734375 0 1029.1837
238.11888122558594 0 136929.39
239.12217712402344 0 14949.074
240.12310791015625 0 945.06946
240.13494873046875 0 754.9824
240.17062377929688 0 2349.7493
242.11354064941406 0 1178.65
242.15011596679688 0 9902.92 a Water loss 2
243.13375854492188 0 1635.057
243.15274047851562 0 859.42004
244.14053344726562 0 13304.825 y Water loss 8
245.12466430664062 0 36419.723 y Ammonia loss 8
246.1086883544922 0 1850.3577
246.12802124023438 0 3310.0842
250.11863708496094 0 2162.517
252.0976104736328 0 1529.9153
252.13449096679688 0 39143.65
253.1046905517578 0 591.6816
253.11866760253906 0 2282.9788
253.13768005371094 0 5651.1406
254.1136932373047 0 5985.1777
254.13760375976562 0 618.19116
256.1293029785156 0 36551.074
257.1326599121094 0 3766.083
262.1511535644531 0 661397.9 y 8
263.1541442871094 0 64164.95
264.0978088378906 0 2868.8801
264.1559753417969 0 6592.586
268.1291198730469 0 2360.3552
268.1657409667969 0 6547.1206
269.1236267089844 0 1382.8754
269.1619567871094 0 790.878
270.1092834472656 0 5624.0176
270.1449279785156 0 195116.25 b Water loss 2
271.14813232421875 0 24379.936
271.17486572265625 0 878.0197
272.1239929199219 0 1912.2268
272.1502990722656 0 2547.101
278.1239929199219 0 669.4635
280.1282653808594 0 959.0401
280.16668701171875 0 674.52435
281.125 0 1578.4474
282.1084289550781 0 2673.4233
282.1812744140625 0 1572.232
284.1593322753906 0 886.1616
286.15118408203125 0 1882.3965
286.1763916015625 0 1048.4933
286.6615295410156 0 804.53876
287.1535339355469 0 1948.87 y Water loss 5
287.1701354980469 0 573.90326
288.1304931640625 0 5679.6904
288.15557861328125 0 13875.101 b 2
289.1590576171875 0 1243.365
294.1451110839844 0 1621.6685
295.14019775390625 0 968.9797
296.1596984863281 0 3031.3457 y 5
297.1192626953125 0 1000.28265
297.15667724609375 0 780.28845
298.13824462890625 0 626.6513
298.1761474609375 0 1045.8527
299.06158447265625 0 1561.3705
299.1349792480469 0 2367.7988
300.0625305175781 0 1370.7523
300.119140625 0 1843.1488
301.0597229003906 0 3104.4316
301.1495056152344 0 725.31757
302.0588073730469 0 2161.5693
303.05731201171875 0 1052.479
304.16290283203125 0 1407.6543
307.1036682128906 0 907.7241
307.1402282714844 0 1408.922
308.1239318847656 0 1074.2
311.17193603515625 0 1401.313
311.2082824707031 0 2346.9602
312.13043212890625 0 2093.915
312.1555480957031 0 3576.1948
313.1534118652344 0 1680.538
314.1463623046875 0 1976.6289
314.16973876953125 0 688.2
315.1669006347656 0 2282.754
316.1646423339844 0 1220.7386
317.1458435058594 0 1968.7761
321.1558532714844 0 5205.3867
322.13995361328125 0 1167.4001
322.1592102050781 0 1032.1381
322.1785583496094 0 1934.7054
322.6720275878906 0 4418.7534 y Water loss 4
323.17327880859375 0 2734.7732
323.6748046875 0 546.3274
324.1303405761719 0 2739.4895
324.1929626464844 0 644.37286
325.1148681640625 0 2270.943
325.1514587402344 0 3641.5283
326.1546630859375 0 944.6764
328.16192626953125 0 707.4867
329.18267822265625 0 3998.8433
330.16632080078125 0 2126.1284
331.1725769042969 0 11125.979
331.6776428222656 0 2634.3052 y 4
332.1565856933594 0 4060.274
332.17706298828125 0 1967.644
333.177734375 0 722.5634
335.13555908203125 0 1046.5068
335.15582275390625 0 1028.0404
335.2077331542969 0 1755.2083
337.22369384765625 0 6115.8823
338.22625732421875 0 1307.9027
339.1664123535156 0 18736.102
339.202880859375 0 1997.7014
340.16937255859375 0 3119.2756
341.1567077636719 0 2224.309
342.1410217285156 0 16534.852
342.1588134765625 0 733.95123
343.14093017578125 0 969.8354
343.146728515625 0 835.91833
343.1612854003906 0 3651.8604
346.97454833984375 0 637.78125
349.150146484375 0 1230.2794
349.1861267089844 0 963.54205
353.14508056640625 0 1088.4009
355.1971435546875 0 2536.8352
355.2341003417969 0 7123.2417
356.2376403808594 0 1581.3752
357.1767272949219 0 15749.592
358.18048095703125 0 3442.1301
358.20892333984375 0 1847.5575
359.1672668457031 0 22533.25 y Water loss 7
360.15130615234375 0 26289.4 y Ammonia loss 7
361.1543273925781 0 4006.3328
361.17926025390625 0 804.05707
362.1940002441406 0 1152.4949
363.2037353515625 0 1053.8138
365.2183837890625 0 8275.097
366.2216491699219 0 1719.9216
367.16241455078125 0 1210.4904
367.1978454589844 0 2279.83 b Water loss 3
369.6980285644531 0 1736.5427
370.19122314453125 0 4373.286
370.7060852050781 0 16630.514
371.1986999511719 0 50164.58 y Water loss 3
371.699951171875 0 19806.215
372.2013244628906 0 5435.475
376.19384765625 0 19969.207
376.8617858886719 0 583.7719
377.177978515625 0 128971.57 y 7
378.1807861328125 0 19859.781
378.7033386230469 0 3946.1597
379.1958923339844 0 8071.7695
379.7113952636719 0 107759.01
380.2041320800781 0 273706.25 y 3
380.7051696777344 0 111380.805
381.17791748046875 0 1157.5927
381.20660400390625 0 23273.7
381.7073974609375 0 1780.33
382.20733642578125 0 691.0449
383.197265625 0 893.1657
383.22906494140625 0 7670.6924
384.2312927246094 0 1321.7428
385.2013854980469 0 2056.3252 b 3
385.6965026855469 0 917.25476
387.1622314453125 0 1458.6913
390.1784973144531 0 1199.763
391.1616516113281 0 2167.1604
392.1644287109375 0 934.15674
392.2303771972656 0 815.7188
396.70526123046875 0 1703.2423
397.1969299316406 0 1461.7191
397.6986389160156 0 904.9162
398.2389831542969 0 924.60895
399.1891174316406 0 759.6093
400.2184143066406 0 1016.93866
401.2026672363281 0 1076.1971
401.2400817871094 0 1210.3058
405.2176513671875 0 4217.2544
405.7096252441406 0 18743.006
406.2099304199219 0 6709.179
406.2444152832031 0 991.9908
406.7107849121094 0 2060.984
407.22869873046875 0 995.2489
408.18756103515625 0 4383.678
409.17083740234375 0 974.683
410.23974609375 0 1010.96375
414.2223205566406 0 67023.195
414.7150573730469 0 230107.53 y Water loss 2
415.2162170410156 0 94720.31
415.7171325683594 0 23057.082
416.21875 0 3196.33
418.9950866699219 0 708.6695
419.03448486328125 0 804.8703
419.99688720703125 0 1207.5215
420.2236022949219 0 1843.2109
423.2275695800781 0 64462.145
423.7204284667969 0 197274.31 y 2
424.2215576171875 0 82156.64
424.7224426269531 0 23678.492
425.2192687988281 0 1085.9747
426.19879150390625 0 3667.1172
428.2501525878906 0 1697.9762
430.2292175292969 0 941.6407
434.2399597167969 0 3695.507
436.18212890625 0 1442.8955
438.2021179199219 0 1140.5599
438.23565673828125 0 2333.722 b Water loss 4
439.237060546875 0 660.72345
440.214599609375 0 719.00256
442.26654052734375 0 2011.9381
443.226806640625 0 807.6085
443.270263671875 0 865.80145
444.2091979980469 0 2325.3794
448.7333984375 0 946.902
449.2259826660156 0 3082.6575
449.72430419921875 0 1607.1958
452.25048828125 0 10398.223
453.2532653808594 0 1862.9196
455.2256774902344 0 2295.5378
456.20867919921875 0 1396.1868
456.245361328125 0 4081.6829 b 4
457.2154846191406 0 663.4343
457.2491149902344 0 1174.371
457.73797607421875 0 1790.6906
458.2314453125 0 5388.5674 y Water loss 1
458.73187255859375 0 2400.6233
459.232421875 0 663.25024
462.73004150390625 0 1872.6927
463.2238464355469 0 2757.6428
463.7259826660156 0 1035.6014
466.7444152832031 0 1425.0656
467.23822021484375 0 3904.5676 y 1
467.7375183105469 0 1496.1781
470.2271423339844 0 613.91266
470.2614440917969 0 4109.6187
471.26458740234375 0 2656.2556
471.7367858886719 0 1724.6715
472.22540283203125 0 2236.574
472.2530822753906 0 2934.193 y Water loss 6
472.729736328125 0 2834.3696
473.2351379394531 0 7237.1704 y Ammonia loss 6
474.2360534667969 0 909.9717
476.2520751953125 0 930.35455
479.26171875 0 764.6463
480.2456359863281 0 4921.69
481.2486267089844 0 1405.2817
489.23651123046875 0 851.0687
489.2779541015625 0 31234.436
490.2625427246094 0 72949.21 y 6
491.26495361328125 0 17762.824
492.2670593261719 0 2658.0547
495.2568664550781 0 678.0156
497.2654113769531 0 1219.2369
498.2561340332031 0 17109.117
499.2586669921875 0 4273.2295
499.7662353515625 0 651.63043
500.2524108886719 0 646.4686
503.2610168457031 0 2244.825
504.2478332519531 0 856.9812
505.273193359375 0 1882.1594
505.7676696777344 0 9615.495
506.2677917480469 0 4852.636
506.7693176269531 0 2419.089
508.7751770019531 0 656.109
513.7644653320312 0 1246.2251
514.2788696289062 0 5441.0728
514.773681640625 0 18255.357 Precursor Water loss
515.2733764648438 0 8516.529 Precursor Ammonia loss
515.7760620117188 0 2944.9844
521.2724609375 0 6688.9585
522.2721557617188 0 1607.6614
523.2850952148438 0 11149.566
523.7784423828125 0 39662.48 Precursor
524.2799072265625 0 18634.262
524.781494140625 0 6333.736
525.27978515625 0 968.3664
538.2617797851562 0 1171.7567
539.2827758789062 0 8102.3945 b Water loss 5
540.283447265625 0 1515.1227
545.7632446289062 0 5007.113
546.2630004882812 0 2922.189
546.30126953125 0 2772.5942
546.7654418945312 0 1138.9991
547.2841186523438 0 2301.1382
549.2678833007812 0 2268.9353
550.2694091796875 0 1380.3134
555.289306640625 0 1715.1669
556.2738647460938 0 2435.6707
557.2916259765625 0 3889.0425 b 5
558.2933349609375 0 1100.7169
567.2769165039062 0 8887.192
568.2799682617188 0 3307.3418
572.314453125 0 3516.7373
573.2992553710938 0 9461.662 y Water loss 5
574.2952270507812 0 3069.1565
575.2864379882812 0 849.1068
583.2822265625 0 933.35364
584.3009033203125 0 1728.3688
585.2883911132812 0 7731.516
586.2905883789062 0 2443.6506
590.325439453125 0 91612.68
591.310546875 0 184017.83 y 5
592.3131103515625 0 53255.13
593.3150024414062 0 10508.759
594.317138671875 0 690.33875
600.309814453125 0 1821.451
601.292236328125 0 2147.583
602.305908203125 0 1888.3413
603.3093872070312 0 622.7973
616.343505859375 0 961.22217
617.3388061523438 0 1609.0377
618.32275390625 0 762.21564
626.3264770507812 0 1253.5608
627.312744140625 0 1925.8737
634.3546752929688 0 1830.3953
635.356201171875 0 836.11456
636.2965087890625 0 863.8254
642.38232421875 0 672.3284
643.3518676757812 0 8498.5
644.3370971679688 0 17085.959 y Water loss 4
645.3339233398438 0 6591.7207
646.3281860351562 0 1550.5951
647.3272705078125 0 678.9688
652.3656005859375 0 4007.0142 b Water loss 6
652.8282470703125 0 2988.246
653.3284301757812 0 1824.7256
653.8297119140625 0 1201.4185
654.3189086914062 0 1613.624
661.2963256835938 0 1287.4093
661.362548828125 0 73810.23
662.3482055664062 0 140156.75 y 4
663.350341796875 0 44369.25
664.3521118164062 0 8479.875
670.3770141601562 0 1712.4375 b 6
671.3466186523438 0 2097.9438
672.3307495117188 0 6175.4673
673.33251953125 0 1927.9722
674.341064453125 0 825.5993
698.3755493164062 0 840.48474
699.36865234375 0 989.44464
715.37646484375 0 1711.3157
716.3818969726562 0 756.0054
719.0211181640625 0 756.2176
723.3772583007812 0 811.08777
724.3688354492188 0 1132.5792
725.3580322265625 0 929.7341
734.3641357421875 0 945.1705
740.4032592773438 0 3149.6047
741.3892211914062 0 12432.249 y Water loss 3
742.3870849609375 0 6640.9473 y Ammonia loss 3
742.4654541015625 0 780.92737
743.3836059570312 0 2029.0813
749.381103515625 0 1123.4856
756.3928833007812 0 635.3413
757.3856811523438 0 1621.3381
758.3328857421875 0 1035.769
758.4148559570312 0 118088.266
759.401123046875 0 281335.22 y 3
760.4029541015625 0 107585.18
761.405029296875 0 27101.555
762.4087524414062 0 1851.9519
767.3964233398438 0 1148.9172 b Water loss 7
785.4029541015625 0 2727.2402 b 7
786.406494140625 0 1688.0117
810.4024047851562 0 827.3292
815.4365234375 0 1000.0482
816.4248046875 0 1088.7241
817.4259643554688 0 800.3577
827.4352416992188 0 2666.2888
828.4219970703125 0 8480.68 y Water loss 2
829.4205322265625 0 4038.978 y Ammonia loss 2
830.4195556640625 0 1096.546
845.3516845703125 0 1120.5509
845.4463500976562 0 55999.523
846.4326782226562 0 140335.17 y 2
847.4347534179688 0 58672.523
848.4374389648438 0 14962.666
849.4389038085938 0 1556.146
856.416748046875 0 1725.5613
897.4468383789062 0 1895.6167
898.4360961914062 0 1431.832
902.4680786132812 0 800.5131
903.4613037109375 0 1138.7712
914.4661865234375 0 3670.4734
915.4541625976562 0 12975.621 y Water loss 1
916.455322265625 0 6681.6523 y Ammonia loss 1
917.4556884765625 0 908.1101
925.4378051757812 0 1331.8989
932.477783203125 0 41839.125
933.465087890625 0 107371.055 y 1
934.466552734375 0 48938.41
935.468017578125 0 14612.416
936.4683227539062 0 1405.9949
942.4622802734375 0 4229.5664
943.4500732421875 0 10150.144
944.4508056640625 0 5475.79
945.45849609375 0 1411.5742
1090.511474609375 0 1156.7704
1707.0126953125 0 672.1559
1828.7396240234375 0 609.84686
3079.732421875 0 721.3099
3325.493408203125 0 678.43353

Spectrum Details

|  |  |
| --- | --- |
| Matched peaks? Matched peaks The total absolute number of peaks matched. Additionally in brackets the total fraction of peaks matched and the total number of peaks is shown. | 54 (9.56% of 565) |
| FDR? FDR The false discovery rate estimated for this peptide. It is calculated by matching all theoretical fragments with a non-integer shift with the raw peaks for this spectrum. This is done with 40 different shifts. The resulting percentage is the average number of annotated peaks over the number of annotated peaks with the correct spectrum. | 0.40% |
| Satellite FDR? Satellite FDR See the FDR for details on its calculation. This satellite ion specific FDR only contains the satellite ions (d/w) for I/L/J positions. | - |
| PSM Score? PSM Score The PSM Score as given by Hecklib to this annotated spectrum. It is shown with three significant figures. | 734 |

#### Reverse Lookup? Reverse Lookup All places where this read could be placed.

| Group | Segment | Template | Template Part | Read Part | Score | Unique |
| --- | --- | --- | --- | --- | --- | --- |
| Decoy | Decoy | TRYP | [97..107] | [0..10] | 75 | True |

| Recombined | Template Part | Read Part | Score | Unique |
| --- | --- | --- | --- | --- |
| TRYP | [97..107] | [0..10] | 75 | True |

#### Meta Information from Multiple reads

##### Number of combined reads

2

##### Intensity

0.5791

##### TotalArea

1.613E+07

##### Changes to the peptide sequence

JSSPATJDSR

L→JNo support for either Leucine or Isoleucine based on side chain ions (Position: 7)

L→JNo support for either Leucine or Isoleucine based on side chain ions (Position: 1)

#### Positional Score

Copy Data

##### Positional Score (TSV)

###### Preview

```
Loading example...
```

*Click on the button to copy the data to your clipboard.*

000123456789

Label Value
"0" 0
"1" 0
"2" 0
"3" 0
"4" 0
"5" 0
"6" 0
"7" 0
"8" 0
"9" 0

#### Meta Information from PEAKS

##### Scan Identifier

F4:3844

##### Original sequence

L

S

S

P

A

T

L

D

S

R

##### Posttranslational Modifications

##### Source File

D:\separate\_stitch\_analyses\xle-disambiguation\raw\20210323\_F1\_UM1\_Peng0013\_SA\_F59\_ingel\_3ug\_tryp.raw

##### Fraction

4

##### Scan Feature

F4:3759

##### De Novo Score

98

##### ConfidenceScore

98

### m/z

523.7781

##### Mass

1045.5403

##### Charge

2

##### Retention Time

20.42

##### Predicted Retention Time

22.61

##### Area

3.856E+06

##### Parts Per Million

1.3

##### Fragmentation mode

HCD

##### Originating file

01 D:\separate\_stitch\_analyses\xle-disambiguation\20210325\_F59\_3ug\_DENOVO\_12.csv

#### Meta Information from PEAKS

##### Scan Identifier

F4:3642

##### Original sequence

L

S

S

P

A

T

L

D

S

R

##### Posttranslational Modifications

##### Source File

D:\separate\_stitch\_analyses\xle-disambiguation\raw\20210323\_F1\_UM1\_Peng0013\_SA\_F59\_ingel\_3ug\_tryp.raw

##### Fraction

4

##### Scan Feature

F4:3760

##### De Novo Score

98

##### ConfidenceScore

99

### m/z

523.7786

##### Mass

1045.5403

##### Charge

2

##### Retention Time

19.39

##### Predicted Retention Time

22.61

##### Area

1.228E+07

##### Parts Per Million

2.2

##### Fragmentation mode

HCD

##### Originating file

01 D:\separate\_stitch\_analyses\xle-disambiguation\20210325\_F59\_3ug\_DENOVO\_12.csv
