## Supplementary material for "A handle on mass coincidence errors in *de novo* sequencing of antibodies by bottom-up proteomics": Combined_065.html

Details Combined\_065 | Stitch OverviewUndefined

### Read Combined\_065

#### Sequence (length=14)

VVTVPSSSJGTQTY

#### Spectrum 7044? Spectrum 7044 The raw spectrum of this peptide as annotated by Hecklib. The fragments are coloured according to ion type (see legend). Any peaks with a star '\*' as text can be hovered over to see the full details, first the ion type second the mass shift type. By hovering over the amino acids in the peptide or ions in the legend the corresponding peaks are highlighted. By toggling the 'Unassigned' label you can turn the background (unassigned) peaks on or off in the plot. By updating the slider in the Ion legend you can update the spectrum to only show the top X% of the peaks with labels. The top X% means any peak that is within X% of the highest intensity. By dragging in the spectrum you can zoom in to a specific part of the spectrum and use 'Zoom Out' to get back to the original zoom level. The annotation of the spectrum is based on the given sequence in the peptides file and is done with different software so inconsistencies are likely. The peaks are annotated based on the given sequence, with 20 ppm tolerance.

Copy Data

##### Spectrum 7044 (TSV)

###### Preview

```
Loading example...
```

*Click on the button to copy the data to your clipboard.*

Mz MinMz MaxIntensity Max

WidthHeightPeptide font sizePeptide stroke widthSpectrum font sizeSpectrum stroke widthCompact peptide

Ion legend

wxyz

abcd

OtherUnassignedIon^Charge^_Position_Show for top:%

VVTVPSSSJGTQTY

01.94e+43.88e+45.82e+47.76e+4

Zoom Out

y^+1^_1_y^+1^_2_y^+1^_3_y^+1^_3_y^+2^_10_c^+1^_5_y^+1^_5_y^+1^_5_c^+1^_6_y^+2^_12_y^+1^_6_c^+1^_7_y^+1^_7_c^+1^_8_y^+1^_8_c^+1^_9_c^+1^_10_y^+1^_10_y^+1^_10_z^+1^_10_y^+1^_10_c^+1^_11_z^+1^_11_y^+1^_11_c^+1^_12_c^+1^_12_z^+1^_12_z^+1^_12_z^+1^_12_y^+1^_12_c^+1^_13_z^+1^_13_z^+1^_13_z^+1^_13_

0816163324493265

Fragment Matches Table

Show background peaks

| Position | Ion type | Intensity | mz Theoretical | mz Error (Th) | mz Error (ppm) | Charge | Series Number |
| --- | --- | --- | --- | --- | --- | --- | --- |
| - | - | 335 | 121.8 | - | - | 0 | - |
| - | - | 823.6 | 133.1 | - | - | 0 | - |
| - | - | 474.4 | 137.7 | - | - | 0 | - |
| - | - | 467.6 | 145.3 | - | - | 0 | - |
| - | - | 459.9 | 147.3 | - | - | 0 | - |
| - | - | 477.9 | 153.1 | - | - | 0 | - |
| - | - | 562.4 | 155.1 | - | - | 0 | - |
| - | - | 2971 | 171.1 | - | - | 0 | - |
| - | - | 790.9 | 173.1 | - | - | 0 | - |
| - | - | 1306 | 173.4 | - | - | 0 | - |
| - | - | 566.5 | 177 | - | - | 0 | - |
| - | - | 444.3 | 178 | - | - | 0 | - |
| - | - | 446 | 181.6 | - | - | 0 | - |
| 14 | y | 3236 | 182.1 | 0.0002054 | 1.128 | +1 | 1 |
| - | - | 463.4 | 196.6 | - | - | 0 | - |
| - | - | 4780 | 199.1 | - | - | 0 | - |
| - | - | 586.8 | 200.1 | - | - | 0 | - |
| - | - | 585.4 | 201.1 | - | - | 0 | - |
| - | - | 949.8 | 230.1 | - | - | 0 | - |
| - | - | 506.8 | 263.2 | - | - | 0 | - |
| - | - | 7865 | 282.2 | - | - | 0 | - |
| 13 | y | 9910 | 283.1 | 5.81E-05 | 0.2052 | +1 | 2 |
| - | - | 710.6 | 283.2 | - | - | 0 | - |
| - | - | 1185 | 284.1 | - | - | 0 | - |
| - | - | 7326 | 300.2 | - | - | 0 | - |
| - | - | 1556 | 301.2 | - | - | 0 | - |
| - | - | 932.3 | 353.3 | - | - | 0 | - |
| - | - | 668.4 | 355.2 | - | - | 0 | - |
| - | - | 2120 | 371.3 | - | - | 0 | - |
| - | - | 700.1 | 372.3 | - | - | 0 | - |
| - | - | 671.1 | 374.3 | - | - | 0 | - |
| - | - | 1.418E+04 | 381.2 | - | - | 0 | - |
| - | - | 2091 | 382.3 | - | - | 0 | - |
| - | - | 1091 | 388.2 | - | - | 0 | - |
| 12 | y | 965.4 | 393.2 | 1.161E-05 | 0.02953 | +1 | 3 |
| - | - | 4899 | 399.3 | - | - | 0 | - |
| - | - | 1169 | 400.3 | - | - | 0 | - |
| 12 | y | 1435 | 411.2 | 0.0004405 | 1.071 | +1 | 3 |
| - | - | 600.2 | 466.1 | - | - | 0 | - |
| - | - | 806.2 | 472.2 | - | - | 0 | - |
| 5 | y | 1909 | 511.7 | 0.0008307 | 1.623 | +2 | 10 |
| - | - | 868.4 | 512.2 | - | - | 0 | - |
| 5 | c | 1150 | 513.3 | 8.998E-05 | 0.1753 | +1 | 5 |
| - | - | 972.2 | 534.2 | - | - | 0 | - |
| 10 | y | 1004 | 551.2 | 0.0007009 | 1.272 | +1 | 5 |
| 10 | y | 1177 | 569.3 | 0.0004033 | 0.7085 | +1 | 5 |
| - | - | 1046 | 574.3 | - | - | 0 | - |
| - | - | 1793 | 599.4 | - | - | 0 | - |
| 6 | c | 7718 | 600.4 | 0.0001391 | 0.2317 | +1 | 6 |
| - | - | 2694 | 601.4 | - | - | 0 | - |
| 3 | y | 1130 | 612.3 | 0.00464 | 7.578 | +2 | 12 |
| - | - | 968.2 | 630.3 | - | - | 0 | - |
| - | - | 1287 | 644.4 | - | - | 0 | - |
| - | - | 577.1 | 651.4 | - | - | 0 | - |
| - | - | 661.6 | 652.4 | - | - | 0 | - |
| - | - | 797.5 | 668.4 | - | - | 0 | - |
| - | - | 1791 | 675.4 | - | - | 0 | - |
| - | - | 702.3 | 676.4 | - | - | 0 | - |
| - | - | 581.9 | 681.3 | - | - | 0 | - |
| 9 | y | 1060 | 682.3 | 0.001154 | 1.692 | +1 | 6 |
| - | - | 1.47E+04 | 686.4 | - | - | 0 | - |
| 7 | c | 1.187E+04 | 687.4 | 0.002627 | 3.821 | +1 | 7 |
| - | - | 2150 | 688.4 | - | - | 0 | - |
| - | - | 760.6 | 702.3 | - | - | 0 | - |
| - | - | 1340 | 703.3 | - | - | 0 | - |
| - | - | 6406 | 720.3 | - | - | 0 | - |
| - | - | 2386 | 720.4 | - | - | 0 | - |
| - | - | 1472 | 721.3 | - | - | 0 | - |
| - | - | 1171 | 740.4 | - | - | 0 | - |
| - | - | 609.5 | 748.7 | - | - | 0 | - |
| - | - | 2380 | 758.4 | - | - | 0 | - |
| - | - | 2042 | 768.4 | - | - | 0 | - |
| 8 | y | 4000 | 769.4 | 0.002604 | 3.385 | +1 | 7 |
| - | - | 1120 | 770.4 | - | - | 0 | - |
| - | - | 662.7 | 771.1 | - | - | 0 | - |
| - | - | 3402 | 773.4 | - | - | 0 | - |
| 8 | c | 1.436E+04 | 774.4 | 0.0007194 | 0.9289 | +1 | 8 |
| - | - | 4605 | 775.4 | - | - | 0 | - |
| - | - | 1365 | 776.4 | - | - | 0 | - |
| - | - | 1431 | 841.4 | - | - | 0 | - |
| - | - | 1201 | 842.4 | - | - | 0 | - |
| - | - | 1280 | 843.5 | - | - | 0 | - |
| - | - | 9363 | 844.5 | - | - | 0 | - |
| - | - | 3533 | 845.5 | - | - | 0 | - |
| - | - | 978.9 | 846.5 | - | - | 0 | - |
| - | - | 1651 | 852.5 | - | - | 0 | - |
| 7 | y | 1113 | 856.4 | 0.001439 | 1.681 | +1 | 8 |
| - | - | 3783 | 859.4 | - | - | 0 | - |
| - | - | 1547 | 860.4 | - | - | 0 | - |
| - | - | 4769 | 886.5 | - | - | 0 | - |
| 9 | c | 4611 | 887.5 | 0.002813 | 3.17 | +1 | 9 |
| - | - | 2074 | 888.5 | - | - | 0 | - |
| - | - | 824.6 | 889.5 | - | - | 0 | - |
| - | - | 859.6 | 900.5 | - | - | 0 | - |
| - | - | 1201 | 901.5 | - | - | 0 | - |
| - | - | 1547 | 902.5 | - | - | 0 | - |
| - | - | 1007 | 903.5 | - | - | 0 | - |
| - | - | 666 | 927.5 | - | - | 0 | - |
| - | - | 708 | 942.5 | - | - | 0 | - |
| - | - | 1.117E+04 | 943.5 | - | - | 0 | - |
| 10 | c | 1.013E+04 | 944.5 | 0.003464 | 3.667 | +1 | 10 |
| - | - | 3240 | 945.5 | - | - | 0 | - |
| - | - | 2242 | 946.6 | - | - | 0 | - |
| - | - | 773 | 947.6 | - | - | 0 | - |
| - | - | 628.4 | 959.9 | - | - | 0 | - |
| - | - | 781.3 | 1005 | - | - | 0 | - |
| - | - | 800.5 | 1011 | - | - | 0 | - |
| - | - | 647.4 | 1021 | - | - | 0 | - |
| 5 | y | 5102 | 1022 | 0.001011 | 0.9887 | +1 | 10 |
| 5 | y | 4957 | 1023 | 0.0165 | 16.12 | +1 | 10 |
| 5 | z | 1302 | 1024 | 0.005317 | 5.19 | +1 | 10 |
| - | - | 1022 | 1029 | - | - | 0 | - |
| - | - | 1312 | 1039 | - | - | 0 | - |
| 5 | y | 1.901E+04 | 1040 | 0.001444 | 1.388 | +1 | 10 |
| - | - | 1.101E+04 | 1041 | - | - | 0 | - |
| - | - | 3164 | 1042 | - | - | 0 | - |
| - | - | 3352 | 1045 | - | - | 0 | - |
| 11 | c | 2.267E+04 | 1046 | 0.0007883 | 0.754 | +1 | 11 |
| - | - | 1.328E+04 | 1047 | - | - | 0 | - |
| - | - | 2663 | 1048 | - | - | 0 | - |
| - | - | 645.8 | 1058 | - | - | 0 | - |
| - | - | 842.7 | 1060 | - | - | 0 | - |
| 4 | z | 1.274E+04 | 1124 | 0.0005966 | 0.531 | +1 | 11 |
| - | - | 8838 | 1125 | - | - | 0 | - |
| - | - | 2247 | 1126 | - | - | 0 | - |
| - | - | 883.8 | 1131 | - | - | 0 | - |
| - | - | 855.2 | 1132 | - | - | 0 | - |
| - | - | 850.9 | 1139 | - | - | 0 | - |
| 4 | y | 1011 | 1140 | 0.007724 | 6.778 | +1 | 11 |
| - | - | 1547 | 1141 | - | - | 0 | - |
| - | - | 876.7 | 1142 | - | - | 0 | - |
| - | - | 1402 | 1143 | - | - | 0 | - |
| 12 | c | 2426 | 1157 | 0.000224 | 0.1937 | +1 | 12 |
| - | - | 1413 | 1158 | - | - | 0 | - |
| - | - | 1458 | 1159 | - | - | 0 | - |
| - | - | 1427 | 1160 | - | - | 0 | - |
| - | - | 1116 | 1172 | - | - | 0 | - |
| - | - | 2938 | 1173 | - | - | 0 | - |
| 12 | c | 6.23E+04 | 1174 | 0.001138 | 0.9699 | +1 | 12 |
| - | - | 3.928E+04 | 1175 | - | - | 0 | - |
| - | - | 1.25E+04 | 1176 | - | - | 0 | - |
| - | - | 2068 | 1177 | - | - | 0 | - |
| 3 | z | 700 | 1207 | 0.00416 | 3.448 | +1 | 12 |
| 3 | z | 856.5 | 1208 | 0.0127 | 10.52 | +1 | 12 |
| 3 | z | 4946 | 1225 | 0.0009193 | 0.7507 | +1 | 12 |
| - | - | 4764 | 1226 | - | - | 0 | - |
| - | - | 1105 | 1227 | - | - | 0 | - |
| - | - | 608.9 | 1227 | - | - | 0 | - |
| - | - | 618.1 | 1240 | - | - | 0 | - |
| 3 | y | 2978 | 1241 | 0.004412 | 3.556 | +1 | 12 |
| - | - | 2011 | 1242 | - | - | 0 | - |
| - | - | 779.2 | 1244 | - | - | 0 | - |
| - | - | 864.9 | 1260 | - | - | 0 | - |
| 13 | c | 2.152E+04 | 1275 | 0.002796 | 2.194 | +1 | 13 |
| - | - | 1.608E+04 | 1276 | - | - | 0 | - |
| - | - | 5566 | 1277 | - | - | 0 | - |
| - | - | 756.7 | 1278 | - | - | 0 | - |
| 2 | z | 2168 | 1306 | 0.007035 | 5.388 | +1 | 13 |
| 2 | z | 1187 | 1307 | 0.01985 | 15.19 | +1 | 13 |
| 2 | z | 1.024E+04 | 1324 | 0.003041 | 2.298 | +1 | 13 |
| - | - | 8970 | 1325 | - | - | 0 | - |
| - | - | 2389 | 1326 | - | - | 0 | - |
| - | - | 928.2 | 1369 | - | - | 0 | - |
| - | - | 895.4 | 1370 | - | - | 0 | - |
| - | - | 905.4 | 1378 | - | - | 0 | - |
| - | - | 1002 | 1384 | - | - | 0 | - |
| - | - | 862.2 | 1385 | - | - | 0 | - |
| - | - | 1968 | 1394 | - | - | 0 | - |
| - | - | 4800 | 1395 | - | - | 0 | - |
| - | - | 3024 | 1396 | - | - | 0 | - |
| - | - | 1473 | 1397 | - | - | 0 | - |
| - | - | 868.1 | 1398 | - | - | 0 | - |
| - | - | 4384 | 1405 | - | - | 0 | - |
| - | - | 2793 | 1406 | - | - | 0 | - |
| - | - | 1846 | 1407 | - | - | 0 | - |
| - | - | 3472 | 1412 | - | - | 0 | - |
| - | - | 2305 | 1413 | - | - | 0 | - |
| - | - | 4906 | 1422 | - | - | 0 | - |
| - | - | 1.613E+04 | 1423 | - | - | 0 | - |
| - | - | 1.297E+04 | 1424 | - | - | 0 | - |
| - | - | 4639 | 1425 | - | - | 0 | - |
| - | - | 953.6 | 1426 | - | - | 0 | - |
| - | - | 2691 | 1438 | - | - | 0 | - |
| - | - | 2.211E+04 | 1439 | - | - | 0 | - |
| - | - | 7.681E+04 | 1440 | - | - | 0 | - |
| - | - | 5.698E+04 | 1441 | - | - | 0 | - |
| - | - | 1.935E+04 | 1442 | - | - | 0 | - |
| - | - | 2492 | 1443 | - | - | 0 | - |
| - | - | 837.4 | 2141 | - | - | 0 | - |
| - | - | 802.2 | 2142 | - | - | 0 | - |
| - | - | 795.4 | 2157 | - | - | 0 | - |
| - | - | 1738 | 2158 | - | - | 0 | - |
| - | - | 1194 | 2159 | - | - | 0 | - |
| - | - | 979.5 | 2160 | - | - | 0 | - |
| - | - | 676.8 | 3003 | - | - | 0 | - |
| - | - | 637.5 | 3233 | - | - | 0 | - |

m/z Charge Intensity FragmentType MassShift Position
121.82434844970703 0 334.98105
133.08619689941406 0 823.55365
137.69515991210938 0 474.42004
145.2974395751953 0 467.62344
147.27198791503906 0 459.86505
153.11715698242188 0 477.9387
155.07064819335938 0 562.417
171.14932250976562 0 2970.5305
173.12835693359375 0 790.86096
173.43836975097656 0 1306.4436
177.00209045410156 0 566.51935
178.02511596679688 0 444.27585
181.58453369140625 0 446.00653
182.0813751220703 0 3235.7605 y 13
196.55813598632812 0 463.3734
199.14430236816406 0 4779.9917
200.1475830078125 0 586.8247
201.12307739257812 0 585.35596
230.11370849609375 0 949.8085
263.1695251464844 0 506.7956
282.1812744140625 0 7864.8535
283.12890625 0 9910.097 y 12
283.18536376953125 0 710.5984
284.13226318359375 0 1184.7571
300.1919250488281 0 7326.3975
301.195556640625 0 1556.0265
353.2539367675781 0 932.25696
355.2353210449219 0 668.36707
371.2644348144531 0 2120.2527
372.2685852050781 0 700.09894
374.258056640625 0 671.0569
381.2497253417969 0 14181.756
382.2525939941406 0 2091.1082
388.1829528808594 0 1090.565
393.1768493652344 0 965.35754 y Water loss 11
399.26055908203125 0 4898.5854
400.26397705078125 0 1169.1151
411.1878662109375 0 1435.1344 y 11
466.0982666015625 0 600.1977
472.23828125 0 806.21436
511.7439270019531 0 1909.1378 y Water loss 4
512.2462768554688 0 868.3924
513.339599609375 0 1150.0651 c 4
534.2297973632812 0 972.2376
551.2467041015625 0 1004.4852 y Water loss 9
569.2561645507812 0 1177.4908 y 9
574.2628784179688 0 1046.4163
599.3645629882812 0 1793.1735
600.3713989257812 0 7717.695 c 5
601.3746948242188 0 2694.1895
612.2977905273438 0 1130.3583 y Ammonia loss 2
630.3107299804688 0 968.1599
644.3960571289062 0 1287.3508
651.3560791015625 0 577.1445
652.3765258789062 0 661.61743
668.3847045898438 0 797.4927
675.3792114257812 0 1791.2885
676.3848876953125 0 702.2623
681.3269653320312 0 581.8527
682.3394775390625 0 1059.5007 y 8
686.3953857421875 0 14695.139
687.4009399414062 0 11872.106 c 6
688.40283203125 0 2149.9626
702.3171997070312 0 760.5755
703.3060302734375 0 1339.6421
720.3314819335938 0 6405.6323
720.4370727539062 0 2386.3608
721.330810546875 0 1472.4574
740.3529663085938 0 1171.2106
748.71435546875 0 609.5305
758.36669921875 0 2379.7588
768.3646850585938 0 2041.5076
769.3700561523438 0 3999.7498 y 7
770.372314453125 0 1120.1215
771.0919799804688 0 662.6715
773.426513671875 0 3402.405
774.4348754882812 0 14362.801 c 7
775.4379272460938 0 4604.5547
776.4412231445312 0 1364.6412
841.4057006835938 0 1430.659
842.4037475585938 0 1201.1349
843.50244140625 0 1279.5022
844.5135498046875 0 9362.645
845.5170288085938 0 3533.1199
846.5147705078125 0 978.93964
852.4824829101562 0 1650.9579
856.4061279296875 0 1112.9314 y 6
859.4152221679688 0 3783.3794
860.416015625 0 1547.3306
886.510986328125 0 4768.8125
887.516845703125 0 4611.088 c 8
888.517822265625 0 2074.4937
889.5161743164062 0 824.62714
900.5311889648438 0 859.6145
901.5316162109375 0 1201.4111
902.53662109375 0 1547.0746
903.5493774414062 0 1006.5711
927.5161743164062 0 666.0045
942.468017578125 0 708.0214
943.53271484375 0 11174.129
944.5376586914062 0 10130.76 c 9
945.5438842773438 0 3240.193
946.551025390625 0 2241.7986
947.5612182617188 0 773.0363
959.8963012695312 0 628.3954
1005.4713134765625 0 781.30634
1010.562744140625 0 800.49475
1021.478271484375 0 647.36566
1022.4779052734375 0 5102.4365 y Water loss 4
1023.4794311523438 0 4957.2085 y Ammonia loss 4
1024.47607421875 0 1302.4742 z 4
1028.5684814453125 0 1022.1761
1039.48046875 0 1312.3973
1040.488037109375 0 19005.31 y 4
1041.492431640625 0 11006.108
1042.4967041015625 0 3163.8513
1044.579833984375 0 3351.9102
1045.5880126953125 0 22672.26 c 10
1046.5911865234375 0 13283.024
1047.59375 0 2663.22
1057.5296630859375 0 645.8482
1059.5389404296875 0 842.6549
1123.53857421875 0 12744.466 z 3
1124.5413818359375 0 8837.809
1125.5418701171875 0 2246.712
1130.632080078125 0 883.8168
1131.639404296875 0 855.2348
1138.6024169921875 0 850.8719
1139.5501708984375 0 1010.8746 y 3
1140.5906982421875 0 1546.7152
1141.602294921875 0 876.6764
1142.593505859375 0 1401.6218
1156.62060546875 0 2426.375 c Ammonia loss 11
1157.618896484375 0 1413.3751
1158.629638671875 0 1457.7113
1159.641845703125 0 1426.5007
1171.6192626953125 0 1116.3315
1172.6328125 0 2937.7847
1173.646240234375 0 62297.39 c 11
1174.64892578125 0 39283.277
1175.6522216796875 0 12502.218
1176.656005859375 0 2067.7202
1206.5804443359375 0 699.9634 z Water loss 2
1207.572998046875 0 856.49854 z Ammonia loss 2
1224.5877685546875 0 4946.0083 z 2
1225.589111328125 0 4763.5977
1226.5850830078125 0 1104.5405
1227.084228515625 0 608.86304
1239.6103515625 0 618.1143
1240.6099853515625 0 2978.4336 y 2
1241.6226806640625 0 2010.5731
1243.655517578125 0 779.1531
1259.68017578125 0 864.9403
1274.6922607421875 0 21519.145 c 12
1275.6949462890625 0 16078.752
1276.7003173828125 0 5566.281
1277.7027587890625 0 756.673
1305.6517333984375 0 2167.5388 z Water loss 1
1306.6485595703125 0 1186.9492 z Ammonia loss 1
1323.6522216796875 0 10238.883 z 1
1324.65576171875 0 8969.58
1325.660400390625 0 2388.862
1368.7083740234375 0 928.2261
1369.709716796875 0 895.4129
1377.725830078125 0 905.3852
1383.689208984375 0 1001.6471
1384.692138671875 0 862.22095
1393.7391357421875 0 1968.2734
1394.728515625 0 4800.33
1395.7337646484375 0 3024.313
1396.7396240234375 0 1473.4886
1397.71240234375 0 868.0599
1404.7098388671875 0 4383.746
1405.7108154296875 0 2793.198
1406.7093505859375 0 1845.7737
1411.75146484375 0 3472.0854
1412.75341796875 0 2305.0564
1421.7330322265625 0 4906.2827
1422.72265625 0 16125.045
1423.7220458984375 0 12968.503
1424.721435546875 0 4639.4287
1425.7166748046875 0 953.5521
1437.71875 0 2690.7402
1438.73779296875 0 22114.6
1439.7454833984375 0 76813.414
1440.7481689453125 0 56980.934
1441.750732421875 0 19350.285
1442.752197265625 0 2491.7063
2141.0126953125 0 837.39655
2142.017333984375 0 802.1917
2157.05517578125 0 795.4023
2158.03076171875 0 1738.1353
2159.0458984375 0 1194.2473
2160.049072265625 0 979.50586
3002.504150390625 0 676.83624
3232.9599609375 0 637.47687

Spectrum Details

|  |  |
| --- | --- |
| Matched peaks? Matched peaks The total absolute number of peaks matched. Additionally in brackets the total fraction of peaks matched and the total number of peaks is shown. | 34 (17.35% of 196) |
| FDR? FDR The false discovery rate estimated for this peptide. It is calculated by matching all theoretical fragments with a non-integer shift with the raw peaks for this spectrum. This is done with 40 different shifts. The resulting percentage is the average number of annotated peaks over the number of annotated peaks with the correct spectrum. | 4.97% |
| Satellite FDR? Satellite FDR See the FDR for details on its calculation. This satellite ion specific FDR only contains the satellite ions (d/w) for I/L/J positions. | - |
| PSM Score? PSM Score The PSM Score as given by Hecklib to this annotated spectrum. It is shown with three significant figures. | 386 |

#### Spectrum 7102? Spectrum 7102 The raw spectrum of this peptide as annotated by Hecklib. The fragments are coloured according to ion type (see legend). Any peaks with a star '\*' as text can be hovered over to see the full details, first the ion type second the mass shift type. By hovering over the amino acids in the peptide or ions in the legend the corresponding peaks are highlighted. By toggling the 'Unassigned' label you can turn the background (unassigned) peaks on or off in the plot. By updating the slider in the Ion legend you can update the spectrum to only show the top X% of the peaks with labels. The top X% means any peak that is within X% of the highest intensity. By dragging in the spectrum you can zoom in to a specific part of the spectrum and use 'Zoom Out' to get back to the original zoom level. The annotation of the spectrum is based on the given sequence in the peptides file and is done with different software so inconsistencies are likely. The peaks are annotated based on the given sequence, with 20 ppm tolerance.

Copy Data

##### Spectrum 7102 (TSV)

###### Preview

```
Loading example...
```

*Click on the button to copy the data to your clipboard.*

Mz MinMz MaxIntensity Max

WidthHeightPeptide font sizePeptide stroke widthSpectrum font sizeSpectrum stroke widthCompact peptide

Ion legend

wxyz

abcd

OtherUnassignedIon^Charge^_Position_Show for top:%

VVTVPSSSJGTQTY

01.15e+42.29e+43.44e+44.58e+4

Zoom Out

y^+1^_1_y^+1^_2_y^+1^_4_c^+1^_5_y^+1^_5_c^+1^_6_y^+2^_12_y^+1^_6_c^+1^_7_y^+1^_7_c^+1^_8_y^+1^_8_c^+1^_9_c^+1^_10_z^+1^_10_y^+1^_10_y^+1^_10_z^+1^_10_y^+1^_10_c^+1^_11_z^+1^_11_y^+1^_11_c^+1^_12_c^+1^_12_z^+1^_12_z^+1^_12_y^+1^_12_c^+1^_13_z^+1^_13_z^+1^_13_

0813162624383251

Fragment Matches Table

Show background peaks

| Position | Ion type | Intensity | mz Theoretical | mz Error (Th) | mz Error (ppm) | Charge | Series Number |
| --- | --- | --- | --- | --- | --- | --- | --- |
| - | - | 469.6 | 127.5 | - | - | 0 | - |
| - | - | 717.8 | 136.1 | - | - | 0 | - |
| - | - | 476.7 | 140.5 | - | - | 0 | - |
| - | - | 457.7 | 152.2 | - | - | 0 | - |
| - | - | 442.3 | 159.5 | - | - | 0 | - |
| - | - | 2001 | 171.1 | - | - | 0 | - |
| - | - | 651.5 | 173.1 | - | - | 0 | - |
| - | - | 1855 | 173.5 | - | - | 0 | - |
| 14 | y | 1904 | 182.1 | 0.0003275 | 1.799 | +1 | 1 |
| - | - | 2334 | 199.1 | - | - | 0 | - |
| - | - | 596.6 | 201.1 | - | - | 0 | - |
| - | - | 448.8 | 222.8 | - | - | 0 | - |
| - | - | 485.4 | 230 | - | - | 0 | - |
| - | - | 491.5 | 250.1 | - | - | 0 | - |
| - | - | 531.7 | 277.5 | - | - | 0 | - |
| - | - | 4816 | 282.2 | - | - | 0 | - |
| 13 | y | 5198 | 283.1 | 3.345E-05 | 0.1182 | +1 | 2 |
| - | - | 804.1 | 283.2 | - | - | 0 | - |
| - | - | 1021 | 284.1 | - | - | 0 | - |
| - | - | 501 | 287.5 | - | - | 0 | - |
| - | - | 4881 | 300.2 | - | - | 0 | - |
| - | - | 686.9 | 301.2 | - | - | 0 | - |
| - | - | 870.3 | 348.2 | - | - | 0 | - |
| - | - | 963.4 | 371.3 | - | - | 0 | - |
| - | - | 7947 | 381.2 | - | - | 0 | - |
| - | - | 1794 | 382.3 | - | - | 0 | - |
| - | - | 791.3 | 388.2 | - | - | 0 | - |
| - | - | 2464 | 399.3 | - | - | 0 | - |
| - | - | 931.7 | 487.2 | - | - | 0 | - |
| 11 | y | 712.5 | 512.2 | 0.003238 | 6.322 | +1 | 4 |
| 5 | c | 683.8 | 513.3 | 0.001253 | 2.44 | +1 | 5 |
| - | - | 1029 | 534.2 | - | - | 0 | - |
| 10 | y | 1001 | 551.2 | 0.000825 | 1.497 | +1 | 5 |
| - | - | 1222 | 556.3 | - | - | 0 | - |
| - | - | 970.4 | 574.3 | - | - | 0 | - |
| - | - | 676.7 | 599.4 | - | - | 0 | - |
| 6 | c | 4574 | 600.4 | 0.0005933 | 0.9882 | +1 | 6 |
| - | - | 1604 | 601.4 | - | - | 0 | - |
| 3 | y | 566.7 | 612.3 | 0.008607 | 14.06 | +2 | 12 |
| - | - | 593.1 | 629.8 | - | - | 0 | - |
| - | - | 843.5 | 644.4 | - | - | 0 | - |
| - | - | 574.7 | 652.4 | - | - | 0 | - |
| 9 | y | 1224 | 682.3 | 0.0004936 | 0.7235 | +1 | 6 |
| - | - | 7831 | 686.4 | - | - | 0 | - |
| 7 | c | 5937 | 687.4 | 0.003176 | 4.62 | +1 | 7 |
| - | - | 1184 | 688.4 | - | - | 0 | - |
| - | - | 1277 | 703.3 | - | - | 0 | - |
| - | - | 786.1 | 718.3 | - | - | 0 | - |
| - | - | 6786 | 720.3 | - | - | 0 | - |
| - | - | 585.6 | 720.4 | - | - | 0 | - |
| - | - | 1155 | 721.3 | - | - | 0 | - |
| - | - | 569.8 | 731.4 | - | - | 0 | - |
| - | - | 1175 | 758.4 | - | - | 0 | - |
| - | - | 880.2 | 768.4 | - | - | 0 | - |
| 8 | y | 1841 | 769.4 | 0.004618 | 6.003 | +1 | 7 |
| - | - | 649.5 | 770.4 | - | - | 0 | - |
| - | - | 1989 | 773.4 | - | - | 0 | - |
| 8 | c | 7514 | 774.4 | 0.0005973 | 0.7713 | +1 | 8 |
| - | - | 2106 | 775.4 | - | - | 0 | - |
| - | - | 652.6 | 779.4 | - | - | 0 | - |
| - | - | 541.9 | 802.2 | - | - | 0 | - |
| - | - | 784.5 | 841.4 | - | - | 0 | - |
| - | - | 4782 | 844.5 | - | - | 0 | - |
| - | - | 1519 | 845.5 | - | - | 0 | - |
| 7 | y | 766.1 | 856.4 | 9.649E-05 | 0.1127 | +1 | 8 |
| - | - | 2266 | 859.4 | - | - | 0 | - |
| - | - | 771.6 | 860.4 | - | - | 0 | - |
| - | - | 3668 | 886.5 | - | - | 0 | - |
| 9 | c | 3134 | 887.5 | 0.002569 | 2.895 | +1 | 9 |
| - | - | 1282 | 888.5 | - | - | 0 | - |
| - | - | 921.1 | 900.5 | - | - | 0 | - |
| - | - | 821.1 | 901.5 | - | - | 0 | - |
| - | - | 917.4 | 902.5 | - | - | 0 | - |
| - | - | 6918 | 943.5 | - | - | 0 | - |
| 10 | c | 7142 | 944.5 | 0.002243 | 2.375 | +1 | 10 |
| - | - | 2244 | 945.5 | - | - | 0 | - |
| - | - | 877.4 | 946.5 | - | - | 0 | - |
| - | - | 793.1 | 959.5 | - | - | 0 | - |
| - | - | 1023 | 1004 | - | - | 0 | - |
| 5 | z | 716.9 | 1006 | 0.004076 | 4.05 | +1 | 10 |
| - | - | 877.8 | 1011 | - | - | 0 | - |
| 5 | y | 4054 | 1022 | 0.0004616 | 0.4515 | +1 | 10 |
| 5 | y | 1867 | 1023 | 0.01626 | 15.88 | +1 | 10 |
| 5 | z | 796.5 | 1024 | 0.01948 | 19.01 | +1 | 10 |
| 5 | y | 1.214E+04 | 1040 | 0.0004673 | 0.4491 | +1 | 10 |
| - | - | 6138 | 1041 | - | - | 0 | - |
| - | - | 1151 | 1043 | - | - | 0 | - |
| - | - | 1891 | 1045 | - | - | 0 | - |
| 11 | c | 1.235E+04 | 1046 | 0.001155 | 1.104 | +1 | 11 |
| - | - | 7769 | 1047 | - | - | 0 | - |
| - | - | 2289 | 1048 | - | - | 0 | - |
| - | - | 741.6 | 1071 | - | - | 0 | - |
| 4 | z | 6769 | 1124 | 0.001939 | 1.726 | +1 | 11 |
| - | - | 4575 | 1125 | - | - | 0 | - |
| - | - | 1336 | 1126 | - | - | 0 | - |
| 4 | y | 730.3 | 1140 | 0.009244 | 8.112 | +1 | 11 |
| - | - | 823.6 | 1141 | - | - | 0 | - |
| - | - | 1046 | 1143 | - | - | 0 | - |
| 12 | c | 1356 | 1157 | 0.0008343 | 0.7214 | +1 | 12 |
| - | - | 1149 | 1159 | - | - | 0 | - |
| - | - | 2061 | 1173 | - | - | 0 | - |
| 12 | c | 3.843E+04 | 1174 | 0.001138 | 0.9699 | +1 | 12 |
| - | - | 2.133E+04 | 1175 | - | - | 0 | - |
| - | - | 7621 | 1176 | - | - | 0 | - |
| - | - | 1187 | 1177 | - | - | 0 | - |
| 3 | z | 713.7 | 1207 | 0.004526 | 3.751 | +1 | 12 |
| 3 | z | 3132 | 1225 | 0.0006752 | 0.5513 | +1 | 12 |
| - | - | 1703 | 1226 | - | - | 0 | - |
| 3 | y | 2204 | 1241 | 0.002459 | 1.982 | +1 | 12 |
| - | - | 807.8 | 1242 | - | - | 0 | - |
| - | - | 762.8 | 1243 | - | - | 0 | - |
| 13 | c | 1.27E+04 | 1275 | 0.002186 | 1.715 | +1 | 13 |
| - | - | 9357 | 1276 | - | - | 0 | - |
| - | - | 3323 | 1277 | - | - | 0 | - |
| 2 | z | 824 | 1306 | 0.00203 | 1.555 | +1 | 13 |
| 2 | z | 6023 | 1324 | 0.0009663 | 0.73 | +1 | 13 |
| - | - | 3946 | 1325 | - | - | 0 | - |
| - | - | 2074 | 1326 | - | - | 0 | - |
| - | - | 897.3 | 1394 | - | - | 0 | - |
| - | - | 2439 | 1395 | - | - | 0 | - |
| - | - | 2070 | 1396 | - | - | 0 | - |
| - | - | 2021 | 1405 | - | - | 0 | - |
| - | - | 1655 | 1406 | - | - | 0 | - |
| - | - | 1951 | 1412 | - | - | 0 | - |
| - | - | 1446 | 1413 | - | - | 0 | - |
| - | - | 3208 | 1422 | - | - | 0 | - |
| - | - | 9187 | 1423 | - | - | 0 | - |
| - | - | 6048 | 1424 | - | - | 0 | - |
| - | - | 3524 | 1425 | - | - | 0 | - |
| - | - | 948.5 | 1437 | - | - | 0 | - |
| - | - | 952.9 | 1438 | - | - | 0 | - |
| - | - | 1.377E+04 | 1439 | - | - | 0 | - |
| - | - | 4.537E+04 | 1440 | - | - | 0 | - |
| - | - | 3.406E+04 | 1441 | - | - | 0 | - |
| - | - | 1.186E+04 | 1442 | - | - | 0 | - |
| - | - | 1535 | 1443 | - | - | 0 | - |
| - | - | 644.3 | 2141 | - | - | 0 | - |
| - | - | 1958 | 2158 | - | - | 0 | - |
| - | - | 1448 | 2159 | - | - | 0 | - |
| - | - | 785.8 | 2160 | - | - | 0 | - |
| - | - | 677.5 | 2161 | - | - | 0 | - |
| - | - | 635.2 | 2268 | - | - | 0 | - |
| - | - | 712.9 | 2612 | - | - | 0 | - |
| - | - | 756 | 3070 | - | - | 0 | - |
| - | - | 780.6 | 3072 | - | - | 0 | - |
| - | - | 683.5 | 3219 | - | - | 0 | - |

m/z Charge Intensity FragmentType MassShift Position
127.53412628173828 0 469.5851
136.07601928710938 0 717.7775
140.4606475830078 0 476.71213
152.19631958007812 0 457.6805
159.4684295654297 0 442.33197
171.1492919921875 0 2001.3616
173.12869262695312 0 651.5083
173.45169067382812 0 1855.4818
182.0814971923828 0 1903.8604 y 13
199.1441650390625 0 2334.4292
201.12393188476562 0 596.582
222.7985382080078 0 448.80954
230.0423126220703 0 485.39377
250.12025451660156 0 491.48035
277.5387878417969 0 531.6718
282.1813659667969 0 4816.1406
283.1288146972656 0 5198.361 y 12
283.1854248046875 0 804.12354
284.1328125 0 1020.98987
287.4754333496094 0 500.9824
300.1921691894531 0 4881.09
301.1959533691406 0 686.8531
348.1507873535156 0 870.3362
371.2646179199219 0 963.3544
381.2493896484375 0 7947.4976
382.2527770996094 0 1794.4685
388.18280029296875 0 791.34174
399.2605285644531 0 2463.982
487.2296142578125 0 931.7113
512.2383422851562 0 712.4942 y 10
513.3382568359375 0 683.814 c 4
534.2290649414062 0 1028.8594
551.2451782226562 0 1001.2699 y Water loss 9
556.25390625 0 1221.7422
574.2637329101562 0 970.38074
599.3654174804688 0 676.67676
600.3721313476562 0 4573.6377 c 5
601.3749389648438 0 1604.1494
612.3017578125 0 566.7051 y Ammonia loss 2
629.8301391601562 0 593.0666
644.4002075195312 0 843.5026
652.3668823242188 0 574.69763
682.3411254882812 0 1223.9858 y 8
686.39599609375 0 7830.591
687.400390625 0 5937.373 c 6
688.4032592773438 0 1184.1505
703.3073120117188 0 1276.6707
718.3378295898438 0 786.126
720.33203125 0 6786.1436
720.3916015625 0 585.5849
721.3260498046875 0 1155.0382
731.4317626953125 0 569.80975
758.3673706054688 0 1175.4167
768.3671875 0 880.16974
769.3680419921875 0 1840.9485 y 7
770.3782958984375 0 649.5304
773.427490234375 0 1988.9803
774.4349975585938 0 7513.621 c 7
775.4384155273438 0 2105.9092
779.4266357421875 0 652.6486
802.21875 0 541.85504
841.4074096679688 0 784.4813
844.5143432617188 0 4781.5054
845.5171508789062 0 1519.1229
856.40478515625 0 766.1264 y 6
859.4163818359375 0 2266.2454
860.4171142578125 0 771.5855
886.5111083984375 0 3667.7524
887.51708984375 0 3133.8762 c 8
888.5176391601562 0 1281.7219
900.5293579101562 0 921.0783
901.5283203125 0 821.07837
902.5457763671875 0 917.40686
943.5324096679688 0 6918.352
944.5388793945312 0 7141.5386 c 9
945.5418701171875 0 2244.2075
946.5458374023438 0 877.3656
959.4834594726562 0 793.1012
1004.4711303710938 0 1022.5751
1006.4561157226562 0 716.8865 z Water loss 4
1010.56884765625 0 877.81903
1022.4784545898438 0 4053.5532 y Water loss 4
1023.4791870117188 0 1867.3008 y Ammonia loss 4
1024.490234375 0 796.51373 z 4
1040.489013671875 0 12144.258 y 4
1041.4915771484375 0 6137.7026
1042.5003662109375 0 1151.3593
1044.5784912109375 0 1890.7722
1045.587646484375 0 12353.512 c 10
1046.591552734375 0 7768.581
1047.59619140625 0 2288.6106
1071.0126953125 0 741.6135
1123.5372314453125 0 6769.478 z 3
1124.5400390625 0 4574.827
1125.5416259765625 0 1336.3572
1139.567138671875 0 730.2994 y 3
1140.58447265625 0 823.6107
1142.6002197265625 0 1046.4012
1156.6199951171875 0 1356.4287 c Ammonia loss 11
1158.6361083984375 0 1149.112
1172.6365966796875 0 2061.0562
1173.646240234375 0 38426.734 c 11
1174.6490478515625 0 21328.957
1175.65380859375 0 7621.066
1176.6614990234375 0 1187.2582
1206.580810546875 0 713.7078 z Water loss 2
1224.5875244140625 0 3132.1868 z 2
1225.5897216796875 0 1703.0099
1240.6080322265625 0 2204.094 y 2
1241.6280517578125 0 807.7876
1242.6490478515625 0 762.7945
1274.69287109375 0 12698.055 c 12
1275.6949462890625 0 9356.536
1276.699951171875 0 3322.5613
1305.646728515625 0 824.0264 z Water loss 1
1323.654296875 0 6023.071 z 1
1324.6572265625 0 3946.359
1325.6591796875 0 2073.9797
1393.7398681640625 0 897.2576
1394.731201171875 0 2439.346
1395.72998046875 0 2069.9329
1404.7083740234375 0 2021.2986
1405.720703125 0 1655.3672
1411.750244140625 0 1950.9932
1412.7598876953125 0 1445.9648
1421.727294921875 0 3207.5378
1422.7227783203125 0 9187.258
1423.7242431640625 0 6048.386
1424.7218017578125 0 3523.825
1436.701416015625 0 948.45496
1437.7430419921875 0 952.8569
1438.73779296875 0 13772.75
1439.7462158203125 0 45374.75
1440.748291015625 0 34061.086
1441.750244140625 0 11860.141
1442.74951171875 0 1534.852
2140.9931640625 0 644.27985
2158.02685546875 0 1957.9897
2159.042724609375 0 1447.699
2160.049560546875 0 785.81555
2161.03515625 0 677.54315
2267.84765625 0 635.19476
2611.95947265625 0 712.9209
3070.238525390625 0 756.00336
3071.509765625 0 780.5911
3219.085693359375 0 683.5168

Spectrum Details

|  |  |
| --- | --- |
| Matched peaks? Matched peaks The total absolute number of peaks matched. Additionally in brackets the total fraction of peaks matched and the total number of peaks is shown. | 30 (20.55% of 146) |
| FDR? FDR The false discovery rate estimated for this peptide. It is calculated by matching all theoretical fragments with a non-integer shift with the raw peaks for this spectrum. This is done with 40 different shifts. The resulting percentage is the average number of annotated peaks over the number of annotated peaks with the correct spectrum. | 3.17% |
| Satellite FDR? Satellite FDR See the FDR for details on its calculation. This satellite ion specific FDR only contains the satellite ions (d/w) for I/L/J positions. | - |
| PSM Score? PSM Score The PSM Score as given by Hecklib to this annotated spectrum. It is shown with three significant figures. | 336 |

#### Reverse Lookup? Reverse Lookup All places where this read could be placed.

| Group | Segment | Template | Template Part | Read Part | Score | Unique |
| --- | --- | --- | --- | --- | --- | --- |
| Homo sapiens Heavy Chain | IGHC | IGHG1 | [67..81] | [0..14] | 112 | False |
| Homo sapiens Heavy Chain | IGHC | IGHG3 | [67..81] | [0..14] | 112 | False |
| Homo sapiens Heavy Chain | IGHC | IGHG4 | [67..81] | [0..14] | 103 | False |

| Recombined | Template Part | Read Part | Score | Unique |
| --- | --- | --- | --- | --- |
| REC-0-1 | [189..203] | [0..14] | 112 | True |

#### Meta Information from Multiple reads

##### Number of combined reads

2

##### Intensity

0.8946

##### TotalArea

1.586E+08

##### Changes to the peptide sequence

VVTVPSSSJGTQTY

L→JNo support for either Leucine or Isoleucine based on side chain ions (Position: 9)

#### Positional Score

Copy Data

##### Positional Score (TSV)

###### Preview

```
Loading example...
```

*Click on the button to copy the data to your clipboard.*

00012345678910111213

Label Value
"0" 0
"1" 0
"2" 0
"3" 0
"4" 0
"5" 0
"6" 0
"7" 0
"8" 0
"9" 0
"10" 0
"11" 0
"12" 0
"13" 0

#### Meta Information from PEAKS

##### Scan Identifier

F2:7044

##### Original sequence

V

V

T

V

P

S

S

S

L

G

T

Q

T

Y

##### Posttranslational Modifications

##### Source File

D:\separate\_stitch\_analyses\xle-disambiguation\raw\20210323\_F1\_UM1\_Peng0013\_SA\_F59\_ingel\_3ug\_TL.raw

##### Fraction

2

##### Scan Feature

F2:13097

##### De Novo Score

98

##### ConfidenceScore

98

### m/z

719.8757

##### Mass

1437.7351

##### Charge

2

##### Retention Time

38.43

##### Predicted Retention Time

-

##### Area

1.586E+08

##### Parts Per Million

1.2

##### Fragmentation mode

ETHCD

##### Originating file

01 D:\separate\_stitch\_analyses\xle-disambiguation\20210325\_F59\_3ug\_DENOVO\_12.csv

#### Meta Information from PEAKS

##### Scan Identifier

F2:7102

##### Original sequence

V

V

T

V

P

S

S

S

L

G

T

Q

T

Y

##### Posttranslational Modifications

##### Source File

D:\separate\_stitch\_analyses\xle-disambiguation\raw\20210323\_F1\_UM1\_Peng0013\_SA\_F59\_ingel\_3ug\_TL.raw

##### Fraction

2

##### Scan Feature

-

##### De Novo Score

98

##### ConfidenceScore

98

### m/z

719.8759

##### Mass

1437.7351

##### Charge

2

##### Retention Time

39.55

##### Predicted Retention Time

-

##### Area

0

##### Parts Per Million

1.4

##### Fragmentation mode

ETHCD

##### Originating file

01 D:\separate\_stitch\_analyses\xle-disambiguation\20210325\_F59\_3ug\_DENOVO\_12.csv
