## Supplementary material for "A handle on mass coincidence errors in *de novo* sequencing of antibodies by bottom-up proteomics": Combined_066.html

Details Combined\_066 | Stitch OverviewUndefined

### Read Combined\_066

#### Sequence (length=17)

APEJVESGGGLAQPGTS

#### Spectrum 6376? Spectrum 6376 The raw spectrum of this peptide as annotated by Hecklib. The fragments are coloured according to ion type (see legend). Any peaks with a star '\*' as text can be hovered over to see the full details, first the ion type second the mass shift type. By hovering over the amino acids in the peptide or ions in the legend the corresponding peaks are highlighted. By toggling the 'Unassigned' label you can turn the background (unassigned) peaks on or off in the plot. By updating the slider in the Ion legend you can update the spectrum to only show the top X% of the peaks with labels. The top X% means any peak that is within X% of the highest intensity. By dragging in the spectrum you can zoom in to a specific part of the spectrum and use 'Zoom Out' to get back to the original zoom level. The annotation of the spectrum is based on the given sequence in the peptides file and is done with different software so inconsistencies are likely. The peaks are annotated based on the given sequence, with 20 ppm tolerance.

Copy Data

##### Spectrum 6376 (TSV)

###### Preview

```
Loading example...
```

*Click on the button to copy the data to your clipboard.*

Mz MinMz MaxIntensity Max

WidthHeightPeptide font sizePeptide stroke widthSpectrum font sizeSpectrum stroke widthCompact peptide

Ion legend

wxyz

abcd

OtherUnassignedIon^Charge^_Position_Show for top:%

APEJVESGGGLAQPGTS

01.40e+52.80e+54.20e+55.60e+5

Zoom Out

y^+1^_2_y^+1^_2_y^+1^_3_y^+1^_3_y^+1^_4_y^+1^_4_y^+1^_5_y^+1^_5_y^+1^_5_y^+1^_6_y^+1^_6_y^+2^_13_z^+2^_13_c^+1^_5_c^+2^_13_y^+1^_7_c^+2^_14_y^+1^_8_c^+1^_6_y^+1^_8_c^+2^_16_y^+1^_9_y^+1^_9_c^+1^_7_z^+1^_10_y^+1^_10_y^+1^_10_c^+1^_8_z^+1^_11_y^+1^_11_y^+1^_11_c^+1^_9_y^+1^_11_c^+1^_10_z^+1^_12_y^+1^_12_c^+1^_11_c^+1^_12_y^+1^_13_z^+1^_13_c^+1^_12_y^+1^_13_z^+1^_14_z^+1^_14_z^+1^_14_c^+1^_13_c^+1^_13_y^+1^_14_z^+1^_15_c^+1^_14_y^+1^_15_z^+1^_15_y^+1^_15_c^+1^_15_c^+1^_15_c^+1^_16_

0743148622292972

Fragment Matches Table

Show background peaks

| Position | Ion type | Intensity | mz Theoretical | mz Error (Th) | mz Error (ppm) | Charge | Series Number |
| --- | --- | --- | --- | --- | --- | --- | --- |
| - | - | 534.9 | 122.5 | - | - | 0 | - |
| - | - | 611.1 | 126.1 | - | - | 0 | - |
| - | - | 621.3 | 127.1 | - | - | 0 | - |
| - | - | 629.1 | 129 | - | - | 0 | - |
| - | - | 690.2 | 131.1 | - | - | 0 | - |
| - | - | 631.5 | 135.7 | - | - | 0 | - |
| - | - | 618.2 | 138.2 | - | - | 0 | - |
| - | - | 586.4 | 140 | - | - | 0 | - |
| - | - | 576.1 | 148.8 | - | - | 0 | - |
| - | - | 600.9 | 148.9 | - | - | 0 | - |
| - | - | 671.6 | 148.9 | - | - | 0 | - |
| - | - | 663.4 | 148.9 | - | - | 0 | - |
| - | - | 863.5 | 148.9 | - | - | 0 | - |
| - | - | 1088 | 148.9 | - | - | 0 | - |
| - | - | 1074 | 148.9 | - | - | 0 | - |
| - | - | 879.5 | 148.9 | - | - | 0 | - |
| - | - | 1369 | 148.9 | - | - | 0 | - |
| - | - | 2282 | 148.9 | - | - | 0 | - |
| - | - | 3903 | 148.9 | - | - | 0 | - |
| - | - | 5922 | 148.9 | - | - | 0 | - |
| - | - | 5651 | 149 | - | - | 0 | - |
| - | - | 3261 | 149 | - | - | 0 | - |
| - | - | 1831 | 149 | - | - | 0 | - |
| - | - | 1604 | 149 | - | - | 0 | - |
| - | - | 1051 | 149 | - | - | 0 | - |
| - | - | 671.9 | 149 | - | - | 0 | - |
| - | - | 892.5 | 149 | - | - | 0 | - |
| - | - | 981.7 | 149 | - | - | 0 | - |
| - | - | 590.2 | 149 | - | - | 0 | - |
| - | - | 6847 | 155.1 | - | - | 0 | - |
| - | - | 899.1 | 156.1 | - | - | 0 | - |
| - | - | 607.6 | 159.1 | - | - | 0 | - |
| 16 | y | 4051 | 189.1 | 0.0002512 | 1.328 | +1 | 2 |
| - | - | 4852 | 200.1 | - | - | 0 | - |
| 16 | y | 1.085E+04 | 207.1 | 0.000154 | 0.7436 | +1 | 2 |
| - | - | 839.4 | 209.1 | - | - | 0 | - |
| - | - | 1473 | 210.1 | - | - | 0 | - |
| - | - | 1074 | 227.1 | - | - | 0 | - |
| - | - | 6729 | 228.1 | - | - | 0 | - |
| - | - | 2724 | 238.1 | - | - | 0 | - |
| 15 | y | 1495 | 246.1 | 8.871E-05 | 0.3604 | +1 | 3 |
| - | - | 3.368E+04 | 256.1 | - | - | 0 | - |
| - | - | 2676 | 257.1 | - | - | 0 | - |
| 15 | y | 3094 | 264.1 | 0.000312 | 1.181 | +1 | 3 |
| - | - | 752.2 | 283.1 | - | - | 0 | - |
| - | - | 755.2 | 291.1 | - | - | 0 | - |
| - | - | 1020 | 295.1 | - | - | 0 | - |
| - | - | 1643 | 299.1 | - | - | 0 | - |
| - | - | 881.4 | 299.1 | - | - | 0 | - |
| - | - | 3117 | 325.2 | - | - | 0 | - |
| - | - | 4848 | 338.1 | - | - | 0 | - |
| - | - | 1353 | 340.2 | - | - | 0 | - |
| 14 | y | 5.392E+04 | 343.2 | 0.0004102 | 1.195 | +1 | 4 |
| - | - | 7390 | 344.2 | - | - | 0 | - |
| - | - | 917.4 | 345.2 | - | - | 0 | - |
| - | - | 789.2 | 349.2 | - | - | 0 | - |
| - | - | 922.4 | 354.2 | - | - | 0 | - |
| - | - | 4139 | 355.1 | - | - | 0 | - |
| - | - | 1270 | 356.1 | - | - | 0 | - |
| - | - | 2110 | 359.2 | - | - | 0 | - |
| 14 | y | 2.971E+05 | 361.2 | 0.0002825 | 0.7822 | +1 | 4 |
| - | - | 4.509E+04 | 362.2 | - | - | 0 | - |
| - | - | 4238 | 363.2 | - | - | 0 | - |
| - | - | 926.9 | 366.2 | - | - | 0 | - |
| - | - | 3010 | 367.2 | - | - | 0 | - |
| - | - | 1522 | 384.2 | - | - | 0 | - |
| - | - | 4356 | 423.2 | - | - | 0 | - |
| - | - | 1219 | 424.2 | - | - | 0 | - |
| - | - | 903.1 | 435.7 | - | - | 0 | - |
| - | - | 1243 | 439.3 | - | - | 0 | - |
| - | - | 1391 | 443.2 | - | - | 0 | - |
| - | - | 1.108E+04 | 451.2 | - | - | 0 | - |
| - | - | 2640 | 452.2 | - | - | 0 | - |
| - | - | 1848 | 454.2 | - | - | 0 | - |
| - | - | 1552 | 455.2 | - | - | 0 | - |
| - | - | 4542 | 469.2 | - | - | 0 | - |
| 13 | y | 7571 | 471.2 | 0.0007621 | 1.617 | +1 | 5 |
| 13 | y | 1206 | 472.2 | 0.003258 | 6.899 | +1 | 5 |
| - | - | 840.1 | 478.7 | - | - | 0 | - |
| - | - | 1717 | 484.3 | - | - | 0 | - |
| - | - | 946.9 | 487.2 | - | - | 0 | - |
| 13 | y | 1.574E+04 | 489.2 | 0.0006955 | 1.422 | +1 | 5 |
| - | - | 4289 | 490.2 | - | - | 0 | - |
| - | - | 2447 | 501.2 | - | - | 0 | - |
| - | - | 4555 | 522.3 | - | - | 0 | - |
| - | - | 1266 | 523.3 | - | - | 0 | - |
| - | - | 1251 | 529.2 | - | - | 0 | - |
| - | - | 802.4 | 532.3 | - | - | 0 | - |
| - | - | 5390 | 534.8 | - | - | 0 | - |
| - | - | 2634 | 535.3 | - | - | 0 | - |
| 12 | y | 1435 | 542.3 | 0.002375 | 4.38 | +1 | 6 |
| - | - | 2.032E+04 | 550.3 | - | - | 0 | - |
| - | - | 7016 | 551.3 | - | - | 0 | - |
| - | - | 1253 | 552.3 | - | - | 0 | - |
| - | - | 1015 | 553.3 | - | - | 0 | - |
| - | - | 2611 | 554.3 | - | - | 0 | - |
| 12 | y | 3281 | 560.3 | 0.0004164 | 0.7432 | +1 | 6 |
| - | - | 1263 | 561.3 | - | - | 0 | - |
| - | - | 1.35E+04 | 568.3 | - | - | 0 | - |
| - | - | 3193 | 569.3 | - | - | 0 | - |
| - | - | 8999 | 570.3 | - | - | 0 | - |
| - | - | 5103 | 570.8 | - | - | 0 | - |
| 5 | y | 7586 | 571.3 | 0.006894 | 12.07 | +2 | 13 |
| 5 | z | 4536 | 572.3 | 0.001477 | 2.581 | +2 | 13 |
| 5 | c | 6027 | 585.3 | 0.003396 | 5.801 | +1 | 5 |
| - | - | 2341 | 586.3 | - | - | 0 | - |
| - | - | 1203 | 589.3 | - | - | 0 | - |
| - | - | 1294 | 600.3 | - | - | 0 | - |
| - | - | 814.5 | 601.3 | - | - | 0 | - |
| - | - | 961.9 | 625.3 | - | - | 0 | - |
| 13 | c | 1025 | 634.3 | 0.0007803 | 1.23 | +2 | 13 |
| - | - | 1001 | 634.8 | - | - | 0 | - |
| - | - | 2026 | 642.3 | - | - | 0 | - |
| - | - | 884.3 | 643.3 | - | - | 0 | - |
| - | - | 1521 | 655.3 | - | - | 0 | - |
| - | - | 1284 | 663.3 | - | - | 0 | - |
| - | - | 1391 | 670.4 | - | - | 0 | - |
| - | - | 1591 | 671.3 | - | - | 0 | - |
| - | - | 1362 | 672.3 | - | - | 0 | - |
| 11 | y | 1116 | 673.4 | 0.003755 | 5.576 | +1 | 7 |
| - | - | 9305 | 679.3 | - | - | 0 | - |
| - | - | 3436 | 680.3 | - | - | 0 | - |
| - | - | 1370 | 682.3 | - | - | 0 | - |
| 14 | c | 1320 | 682.8 | 0.001405 | 2.058 | +2 | 14 |
| - | - | 1.048E+04 | 697.3 | - | - | 0 | - |
| - | - | 2857 | 698.3 | - | - | 0 | - |
| - | - | 1.243E+04 | 700.3 | - | - | 0 | - |
| - | - | 4589 | 701.3 | - | - | 0 | - |
| 10 | y | 7019 | 713.3 | 0.01287 | 18.04 | +1 | 8 |
| 6 | c | 3.656E+04 | 714.4 | 0.003936 | 5.509 | +1 | 6 |
| - | - | 1.33E+04 | 715.4 | - | - | 0 | - |
| - | - | 2846 | 716.4 | - | - | 0 | - |
| 10 | y | 2700 | 730.4 | 0.00477 | 6.53 | +1 | 8 |
| - | - | 959.8 | 731.4 | - | - | 0 | - |
| - | - | 1161 | 739.4 | - | - | 0 | - |
| - | - | 1236 | 752.9 | - | - | 0 | - |
| - | - | 3357 | 758.4 | - | - | 0 | - |
| - | - | 1052 | 759.4 | - | - | 0 | - |
| 16 | c | 1958 | 761.9 | 8.777E-05 | 0.1152 | +2 | 16 |
| - | - | 1232 | 762.4 | - | - | 0 | - |
| - | - | 3684 | 766.4 | - | - | 0 | - |
| - | - | 2698 | 767.4 | - | - | 0 | - |
| 9 | y | 1727 | 769.4 | 0.001447 | 1.881 | +1 | 9 |
| - | - | 1045 | 775.4 | - | - | 0 | - |
| - | - | 1220 | 775.9 | - | - | 0 | - |
| - | - | 4594 | 784.4 | - | - | 0 | - |
| - | - | 1406 | 785.4 | - | - | 0 | - |
| - | - | 7506 | 786.4 | - | - | 0 | - |
| 9 | y | 6702 | 787.4 | 0.002613 | 3.318 | +1 | 9 |
| - | - | 1880 | 788.4 | - | - | 0 | - |
| - | - | 1431 | 796.4 | - | - | 0 | - |
| - | - | 2541 | 796.9 | - | - | 0 | - |
| - | - | 3778 | 799.4 | - | - | 0 | - |
| - | - | 8526 | 800.4 | - | - | 0 | - |
| 7 | c | 2.126E+04 | 801.4 | 0.004226 | 5.273 | +1 | 7 |
| - | - | 8951 | 802.4 | - | - | 0 | - |
| - | - | 1446 | 803.4 | - | - | 0 | - |
| - | - | 9126 | 805.4 | - | - | 0 | - |
| - | - | 6231 | 805.9 | - | - | 0 | - |
| - | - | 3296 | 806.4 | - | - | 0 | - |
| 8 | z | 1155 | 811.4 | 0.01213 | 14.95 | +1 | 10 |
| - | - | 1206 | 812.4 | - | - | 0 | - |
| - | - | 1675 | 814.4 | - | - | 0 | - |
| - | - | 1319 | 814.9 | - | - | 0 | - |
| - | - | 1.233E+04 | 815.4 | - | - | 0 | - |
| - | - | 2007 | 815.9 | - | - | 0 | - |
| - | - | 4449 | 816.4 | - | - | 0 | - |
| - | - | 1279 | 823.4 | - | - | 0 | - |
| - | - | 1308 | 825.4 | - | - | 0 | - |
| 8 | y | 4340 | 826.4 | 2.289E-05 | 0.0277 | +1 | 10 |
| - | - | 1638 | 827.4 | - | - | 0 | - |
| - | - | 3719 | 829.4 | - | - | 0 | - |
| - | - | 3337 | 841.4 | - | - | 0 | - |
| - | - | 5733 | 843.4 | - | - | 0 | - |
| 8 | y | 9330 | 844.4 | 0.0003337 | 0.3952 | +1 | 10 |
| - | - | 3420 | 845.4 | - | - | 0 | - |
| - | - | 1.088E+04 | 857.4 | - | - | 0 | - |
| 8 | c | 2.28E+04 | 858.4 | 0.00451 | 5.254 | +1 | 8 |
| - | - | 8245 | 859.4 | - | - | 0 | - |
| - | - | 1872 | 860.4 | - | - | 0 | - |
| - | - | 937.2 | 862.4 | - | - | 0 | - |
| - | - | 3185 | 871.4 | - | - | 0 | - |
| - | - | 1.145E+04 | 872.4 | - | - | 0 | - |
| - | - | 4477 | 873.4 | - | - | 0 | - |
| - | - | 2266 | 880.4 | - | - | 0 | - |
| - | - | 1212 | 881.4 | - | - | 0 | - |
| - | - | 1012 | 895.4 | - | - | 0 | - |
| - | - | 1023 | 896.4 | - | - | 0 | - |
| 7 | z | 4159 | 898.4 | 0.01181 | 13.15 | +1 | 11 |
| - | - | 2568 | 899.4 | - | - | 0 | - |
| - | - | 906.6 | 911.4 | - | - | 0 | - |
| - | - | 1169 | 912.5 | - | - | 0 | - |
| - | - | 810.7 | 913.2 | - | - | 0 | - |
| 7 | y | 4033 | 913.4 | 0.002205 | 2.414 | +1 | 11 |
| 7 | y | 1.119E+04 | 914.4 | 0.01347 | 14.74 | +1 | 11 |
| 9 | c | 1.014E+04 | 915.4 | 0.00626 | 6.838 | +1 | 9 |
| - | - | 4042 | 916.4 | - | - | 0 | - |
| - | - | 1510 | 917.4 | - | - | 0 | - |
| - | - | 1.132E+04 | 928.4 | - | - | 0 | - |
| - | - | 2.615E+04 | 929.5 | - | - | 0 | - |
| - | - | 1.466E+04 | 930.5 | - | - | 0 | - |
| 7 | y | 2.644E+04 | 931.4 | 0.001512 | 1.624 | +1 | 11 |
| - | - | 1.053E+04 | 932.5 | - | - | 0 | - |
| - | - | 2031 | 933.5 | - | - | 0 | - |
| - | - | 8201 | 937.4 | - | - | 0 | - |
| - | - | 3501 | 938.4 | - | - | 0 | - |
| - | - | 2312 | 939.5 | - | - | 0 | - |
| - | - | 1.331E+04 | 955.4 | - | - | 0 | - |
| - | - | 6226 | 956.4 | - | - | 0 | - |
| - | - | 1835 | 957.4 | - | - | 0 | - |
| - | - | 2.702E+04 | 971.5 | - | - | 0 | - |
| 10 | c | 9.096E+04 | 972.5 | 0.004042 | 4.156 | +1 | 10 |
| - | - | 3.895E+04 | 973.5 | - | - | 0 | - |
| - | - | 9143 | 974.5 | - | - | 0 | - |
| - | - | 2294 | 1011 | - | - | 0 | - |
| - | - | 1362 | 1022 | - | - | 0 | - |
| - | - | 1175 | 1025 | - | - | 0 | - |
| 6 | z | 1556 | 1026 | 0.01494 | 14.56 | +1 | 12 |
| - | - | 1252 | 1028 | - | - | 0 | - |
| - | - | 1136 | 1032 | - | - | 0 | - |
| - | - | 4640 | 1041 | - | - | 0 | - |
| - | - | 1.391E+04 | 1042 | - | - | 0 | - |
| - | - | 8154 | 1043 | - | - | 0 | - |
| - | - | 2656 | 1044 | - | - | 0 | - |
| - | - | 1048 | 1045 | - | - | 0 | - |
| - | - | 7051 | 1051 | - | - | 0 | - |
| - | - | 4743 | 1052 | - | - | 0 | - |
| - | - | 2941 | 1053 | - | - | 0 | - |
| - | - | 1097 | 1055 | - | - | 0 | - |
| - | - | 898.9 | 1058 | - | - | 0 | - |
| - | - | 8955 | 1059 | - | - | 0 | - |
| 6 | y | 1.695E+04 | 1060 | 0.0004313 | 0.4067 | +1 | 12 |
| - | - | 7246 | 1061 | - | - | 0 | - |
| - | - | 2000 | 1062 | - | - | 0 | - |
| - | - | 2.62E+04 | 1069 | - | - | 0 | - |
| - | - | 1.349E+04 | 1070 | - | - | 0 | - |
| - | - | 4760 | 1071 | - | - | 0 | - |
| - | - | 2.965E+04 | 1085 | - | - | 0 | - |
| 11 | c | 4.376E+04 | 1086 | 0.005464 | 5.034 | +1 | 11 |
| - | - | 2.25E+04 | 1087 | - | - | 0 | - |
| - | - | 6220 | 1088 | - | - | 0 | - |
| - | - | 1417 | 1112 | - | - | 0 | - |
| - | - | 1.327E+04 | 1113 | - | - | 0 | - |
| - | - | 1.56E+04 | 1114 | - | - | 0 | - |
| - | - | 6356 | 1115 | - | - | 0 | - |
| - | - | 1952 | 1116 | - | - | 0 | - |
| - | - | 8043 | 1122 | - | - | 0 | - |
| - | - | 4468 | 1123 | - | - | 0 | - |
| - | - | 1889 | 1124 | - | - | 0 | - |
| - | - | 1919 | 1125 | - | - | 0 | - |
| 12 | c | 2601 | 1139 | 0.004181 | 3.672 | +1 | 12 |
| - | - | 3.259E+04 | 1140 | - | - | 0 | - |
| - | - | 2.008E+04 | 1141 | - | - | 0 | - |
| 5 | y | 7401 | 1142 | 0.01093 | 9.578 | +1 | 13 |
| - | - | 2246 | 1143 | - | - | 0 | - |
| 5 | z | 1517 | 1144 | 0.002491 | 2.179 | +1 | 13 |
| - | - | 1551 | 1145 | - | - | 0 | - |
| - | - | 1325 | 1155 | - | - | 0 | - |
| - | - | 2.783E+04 | 1156 | - | - | 0 | - |
| 12 | c | 1.318E+05 | 1157 | 0.00376 | 3.251 | +1 | 12 |
| - | - | 7.392E+04 | 1158 | - | - | 0 | - |
| - | - | 2.234E+04 | 1159 | - | - | 0 | - |
| 5 | y | 1.023E+04 | 1160 | 0.004885 | 4.213 | +1 | 13 |
| - | - | 5820 | 1161 | - | - | 0 | - |
| - | - | 1190 | 1162 | - | - | 0 | - |
| - | - | 1408 | 1169 | - | - | 0 | - |
| - | - | 2100 | 1183 | - | - | 0 | - |
| - | - | 942.7 | 1196 | - | - | 0 | - |
| - | - | 1399 | 1223 | - | - | 0 | - |
| - | - | 3723 | 1232 | - | - | 0 | - |
| - | - | 2240 | 1233 | - | - | 0 | - |
| 4 | z | 2477 | 1239 | 0.0008177 | 0.6601 | +1 | 14 |
| 4 | z | 6988 | 1240 | 0.02286 | 18.44 | +1 | 14 |
| - | - | 6.068E+04 | 1241 | - | - | 0 | - |
| - | - | 4.503E+04 | 1242 | - | - | 0 | - |
| - | - | 1.666E+04 | 1243 | - | - | 0 | - |
| - | - | 3923 | 1244 | - | - | 0 | - |
| - | - | 1.73E+04 | 1250 | - | - | 0 | - |
| - | - | 1.351E+04 | 1251 | - | - | 0 | - |
| - | - | 1.807E+04 | 1252 | - | - | 0 | - |
| - | - | 9589 | 1253 | - | - | 0 | - |
| - | - | 2625 | 1254 | - | - | 0 | - |
| 4 | z | 2.489E+04 | 1257 | 0.0007622 | 0.6066 | +1 | 14 |
| - | - | 1.502E+04 | 1258 | - | - | 0 | - |
| - | - | 5386 | 1259 | - | - | 0 | - |
| 13 | c | 3278 | 1267 | 0.005874 | 4.637 | +1 | 13 |
| 13 | c | 1.3E+05 | 1268 | 0.003806 | 3.002 | +1 | 13 |
| - | - | 8.474E+04 | 1269 | - | - | 0 | - |
| - | - | 2.933E+04 | 1270 | - | - | 0 | - |
| - | - | 3275 | 1271 | - | - | 0 | - |
| - | - | 7366 | 1272 | - | - | 0 | - |
| 4 | y | 1.29E+04 | 1273 | 0.003739 | 2.938 | +1 | 14 |
| - | - | 1.052E+04 | 1274 | - | - | 0 | - |
| - | - | 3037 | 1275 | - | - | 0 | - |
| - | - | 2185 | 1323 | - | - | 0 | - |
| - | - | 1347 | 1324 | - | - | 0 | - |
| - | - | 2580 | 1325 | - | - | 0 | - |
| - | - | 971.2 | 1326 | - | - | 0 | - |
| - | - | 858.4 | 1330 | - | - | 0 | - |
| - | - | 5945 | 1338 | - | - | 0 | - |
| - | - | 4615 | 1339 | - | - | 0 | - |
| - | - | 2698 | 1340 | - | - | 0 | - |
| - | - | 1158 | 1352 | - | - | 0 | - |
| 3 | z | 1897 | 1368 | 0.001052 | 0.7695 | +1 | 15 |
| - | - | 1400 | 1369 | - | - | 0 | - |
| - | - | 4351 | 1381 | - | - | 0 | - |
| 14 | c | 1.264E+04 | 1382 | 0.005604 | 4.056 | +1 | 14 |
| - | - | 9149 | 1383 | - | - | 0 | - |
| 3 | y | 2928 | 1384 | 0.01892 | 13.67 | +1 | 15 |
| 3 | z | 1.878E+04 | 1386 | 0.0002645 | 0.1909 | +1 | 15 |
| - | - | 1.403E+04 | 1387 | - | - | 0 | - |
| - | - | 5407 | 1388 | - | - | 0 | - |
| - | - | 1279 | 1389 | - | - | 0 | - |
| - | - | 2.033E+04 | 1395 | - | - | 0 | - |
| - | - | 1.883E+04 | 1396 | - | - | 0 | - |
| - | - | 1.35E+04 | 1397 | - | - | 0 | - |
| - | - | 4465 | 1398 | - | - | 0 | - |
| - | - | 1226 | 1399 | - | - | 0 | - |
| 3 | y | 7677 | 1402 | 0.001655 | 1.18 | +1 | 15 |
| - | - | 6185 | 1403 | - | - | 0 | - |
| - | - | 2781 | 1404 | - | - | 0 | - |
| 15 | c | 1585 | 1422 | 0.009858 | 6.934 | +1 | 15 |
| - | - | 2093 | 1423 | - | - | 0 | - |
| - | - | 4096 | 1424 | - | - | 0 | - |
| - | - | 3431 | 1425 | - | - | 0 | - |
| - | - | 1359 | 1426 | - | - | 0 | - |
| - | - | 1421 | 1438 | - | - | 0 | - |
| 15 | c | 5.861E+04 | 1439 | 0.005461 | 3.796 | +1 | 15 |
| - | - | 4.905E+04 | 1440 | - | - | 0 | - |
| - | - | 2.184E+04 | 1441 | - | - | 0 | - |
| - | - | 4208 | 1442 | - | - | 0 | - |
| - | - | 7124 | 1478 | - | - | 0 | - |
| - | - | 5380 | 1479 | - | - | 0 | - |
| - | - | 1983 | 1480 | - | - | 0 | - |
| - | - | 8659 | 1496 | - | - | 0 | - |
| - | - | 5564 | 1497 | - | - | 0 | - |
| - | - | 3378 | 1498 | - | - | 0 | - |
| - | - | 2022 | 1508 | - | - | 0 | - |
| - | - | 2024 | 1512 | - | - | 0 | - |
| - | - | 1536 | 1514 | - | - | 0 | - |
| - | - | 1721 | 1525 | - | - | 0 | - |
| - | - | 1168 | 1526 | - | - | 0 | - |
| - | - | 1593 | 1527 | - | - | 0 | - |
| - | - | 1423 | 1528 | - | - | 0 | - |
| - | - | 3919 | 1529 | - | - | 0 | - |
| - | - | 5903 | 1530 | - | - | 0 | - |
| - | - | 3318 | 1531 | - | - | 0 | - |
| - | - | 1134 | 1532 | - | - | 0 | - |
| - | - | 1120 | 1539 | - | - | 0 | - |
| 16 | c | 7.403E+04 | 1540 | 0.006387 | 4.148 | +1 | 16 |
| - | - | 6.218E+04 | 1541 | - | - | 0 | - |
| - | - | 2.984E+04 | 1542 | - | - | 0 | - |
| - | - | 5810 | 1543 | - | - | 0 | - |
| - | - | 1318 | 1550 | - | - | 0 | - |
| - | - | 1108 | 1551 | - | - | 0 | - |
| - | - | 6079 | 1553 | - | - | 0 | - |
| - | - | 4448 | 1554 | - | - | 0 | - |
| - | - | 2256 | 1555 | - | - | 0 | - |
| - | - | 3510 | 1556 | - | - | 0 | - |
| - | - | 7863 | 1557 | - | - | 0 | - |
| - | - | 1.589E+04 | 1558 | - | - | 0 | - |
| - | - | 1.368E+04 | 1559 | - | - | 0 | - |
| - | - | 3906 | 1560 | - | - | 0 | - |
| - | - | 1117 | 1566 | - | - | 0 | - |
| - | - | 3004 | 1567 | - | - | 0 | - |
| - | - | 1909 | 1568 | - | - | 0 | - |
| - | - | 1134 | 1569 | - | - | 0 | - |
| - | - | 2381 | 1570 | - | - | 0 | - |
| - | - | 1090 | 1571 | - | - | 0 | - |
| - | - | 1177 | 1572 | - | - | 0 | - |
| - | - | 9881 | 1573 | - | - | 0 | - |
| - | - | 7238 | 1574 | - | - | 0 | - |
| - | - | 4662 | 1575 | - | - | 0 | - |
| - | - | 1562 | 1582 | - | - | 0 | - |
| - | - | 9922 | 1583 | - | - | 0 | - |
| - | - | 8.287E+04 | 1584 | - | - | 0 | - |
| - | - | 7.097E+04 | 1585 | - | - | 0 | - |
| - | - | 3.376E+04 | 1586 | - | - | 0 | - |
| - | - | 7639 | 1587 | - | - | 0 | - |
| - | - | 1059 | 1588 | - | - | 0 | - |
| - | - | 4588 | 1594 | - | - | 0 | - |
| - | - | 3998 | 1595 | - | - | 0 | - |
| - | - | 2398 | 1596 | - | - | 0 | - |
| - | - | 1292 | 1597 | - | - | 0 | - |
| - | - | 1415 | 1599 | - | - | 0 | - |
| - | - | 1517 | 1600 | - | - | 0 | - |
| - | - | 5.153E+04 | 1601 | - | - | 0 | - |
| - | - | 5.358E+04 | 1602 | - | - | 0 | - |
| - | - | 2.394E+04 | 1603 | - | - | 0 | - |
| - | - | 2886 | 1604 | - | - | 0 | - |
| - | - | 2068 | 1609 | - | - | 0 | - |
| - | - | 2834 | 1610 | - | - | 0 | - |
| - | - | 2.239E+04 | 1611 | - | - | 0 | - |
| - | - | 1.628E+05 | 1612 | - | - | 0 | - |
| - | - | 1.451E+05 | 1613 | - | - | 0 | - |
| - | - | 6.548E+04 | 1614 | - | - | 0 | - |
| - | - | 1.012E+04 | 1615 | - | - | 0 | - |
| - | - | 2133 | 1626 | - | - | 0 | - |
| - | - | 1.153E+04 | 1627 | - | - | 0 | - |
| - | - | 1.349E+05 | 1628 | - | - | 0 | - |
| - | - | 5.545E+05 | 1629 | - | - | 0 | - |
| - | - | 4.48E+05 | 1630 | - | - | 0 | - |
| - | - | 1.862E+05 | 1631 | - | - | 0 | - |
| - | - | 2.485E+04 | 1632 | - | - | 0 | - |
| - | - | 1356 | 1644 | - | - | 0 | - |
| - | - | 1659 | 1645 | - | - | 0 | - |
| - | - | 1099 | 2263 | - | - | 0 | - |
| - | - | 1031 | 2512 | - | - | 0 | - |
| - | - | 973.5 | 2942 | - | - | 0 | - |

m/z Charge Intensity FragmentType MassShift Position
122.46458435058594 0 534.86383
126.10679626464844 0 611.1293
127.08657836914062 0 621.3365
128.98988342285156 0 629.094
131.0816192626953 0 690.2073
135.65731811523438 0 631.4755
138.21603393554688 0 618.17126
140.00204467773438 0 586.36176
148.82839965820312 0 576.05865
148.85650634765625 0 600.9036
148.86422729492188 0 671.596
148.88534545898438 0 663.40106
148.8927764892578 0 863.54865
148.89984130859375 0 1088.081
148.9072265625 0 1074.3541
148.91436767578125 0 879.5124
148.92193603515625 0 1369.3004
148.92897033691406 0 2282.123
148.93612670898438 0 3902.9006
148.94387817382812 0 5922.408
148.96051025390625 0 5650.6826
148.96832275390625 0 3260.6606
148.9755096435547 0 1830.5416
148.9826202392578 0 1604.3159
148.9899444580078 0 1051.1542
148.99710083007812 0 671.92206
149.00453186035156 0 892.5455
149.01177978515625 0 981.655
149.018798828125 0 590.2095
155.08172607421875 0 6847.2666
156.0853729248047 0 899.07544
159.07643127441406 0 607.5868
189.0872344970703 0 4051.342 y Water loss 15
200.10311889648438 0 4852.129
207.0977020263672 0 10852.585 y 15
209.09201049804688 0 839.3822
210.1233367919922 0 1472.5555
227.1027069091797 0 1074.2267
228.1343994140625 0 6729.092
238.11907958984375 0 2723.6394
246.10853576660156 0 1495.3604 y Water loss 14
256.12933349609375 0 33682.258
257.1320495605469 0 2676.3594
264.11932373046875 0 3094.4355 y 14
283.1400146484375 0 752.2053
291.07373046875 0 755.22516
295.1043395996094 0 1020.01416
299.0619812011719 0 1642.9578
299.135009765625 0 881.44684
325.15087890625 0 3117.2595
338.13507080078125 0 4847.8716
340.18731689453125 0 1352.9343
343.16162109375 0 53921.832 y Water loss 13
344.1644287109375 0 7389.8735
345.1654052734375 0 917.4193
349.15167236328125 0 789.1834
354.17669677734375 0 922.42316
355.0700988769531 0 4139.0425
356.144287109375 0 1269.899
359.1557312011719 0 2109.9214
361.17205810546875 0 297117.66 y 13
362.175048828125 0 45094.492
363.1773986816406 0 4237.565
366.1758728027344 0 926.85596
367.1617736816406 0 3009.611
384.18792724609375 0 1521.6417
423.22418212890625 0 4356.0674
424.2288513183594 0 1219.0477
435.7076721191406 0 903.0771
439.2569274902344 0 1242.767
443.2257385253906 0 1390.5782
451.2190246582031 0 11076.661
452.222900390625 0 2639.7842
454.1944274902344 0 1848.3636
455.22503662109375 0 1551.9703
469.2303771972656 0 4542.376
471.2205505371094 0 7570.88 y Water loss 12
472.2070617675781 0 1206.4207 y Ammonia loss 12
478.72149658203125 0 840.1491
484.2527770996094 0 1717.0276
487.2149658203125 0 946.938
489.2310485839844 0 15742.032 y 12
490.2342834472656 0 4288.5864
501.2319641113281 0 2446.5508
522.2926635742188 0 4555.435
523.2957153320312 0 1265.7318
529.192626953125 0 1250.8297
532.2822265625 0 802.41675
534.764892578125 0 5390.1567
535.2659301757812 0 2634.3564
542.25927734375 0 1435.0125 y Water loss 11
550.2877807617188 0 20322.824
551.2911376953125 0 7015.8784
552.2919311523438 0 1253.4137
553.2732543945312 0 1015.1052
554.2566528320312 0 2611.291
560.2678833007812 0 3281.1077 y 11
561.2732543945312 0 1262.6832
568.29833984375 0 13501.925
569.3014526367188 0 3193.0159
570.2835693359375 0 8998.923
570.7853393554688 0 5103.281
571.2847290039062 0 7586.203 y Water loss 4
572.2722778320312 0 4536.0684 z 4
585.3253784179688 0 6027.2354 c 4
586.3278198242188 0 2340.9265
589.2943115234375 0 1202.905
600.298828125 0 1294.1635
601.3007202148438 0 814.5374
625.303955078125 0 961.9303
634.3133544921875 0 1024.9032 c Ammonia loss 12
634.8119506835938 0 1000.5848
642.2781372070312 0 2026.3829
643.2808227539062 0 884.32007
655.3255004882812 0 1520.7367
663.33837890625 0 1284.4828
670.3524780273438 0 1391.3408
671.3466186523438 0 1591.1418
672.3405151367188 0 1362.3965
673.3552856445312 0 1115.847 y 10
679.3307495117188 0 9305.368
680.3322143554688 0 3436.1543
682.3163452148438 0 1369.7113
682.839111328125 0 1320.2753 c Ammonia loss 13
697.3427124023438 0 10484.396
698.3472290039062 0 2856.9385
700.3269653320312 0 12429.859
701.328125 0 4588.6035
713.3593139648438 0 7019.126 y Ammonia loss 9
714.367431640625 0 36557.555 c 5
715.3699951171875 0 13302.0205
716.3740844726562 0 2845.7046
730.3682250976562 0 2699.9045 y 9
731.3719482421875 0 959.81244
739.3718872070312 0 1160.8926
752.8780517578125 0 1235.9652
758.39306640625 0 3357.1602
759.3932495117188 0 1052.0836
761.875 0 1957.5328 c Ammonia loss 15
762.3762817382812 0 1232.4568
766.364990234375 0 3684.3608
767.3692626953125 0 2698.4966
769.3824462890625 0 1726.9984 y Water loss 8
775.39697265625 0 1045.4115
775.90283203125 0 1220.135
784.3733520507812 0 4593.973
785.3681030273438 0 1406.3646
786.3873901367188 0 7506.149
787.391845703125 0 6702.1934 y 8
788.3858642578125 0 1880.3843
796.3831176757812 0 1430.5137
796.8797607421875 0 2540.7605
799.3953247070312 0 3777.5825
800.3923950195312 0 8525.682
801.399169921875 0 21257.834 c 6
802.4025268554688 0 8950.9375
803.4072875976562 0 1445.5992
805.3870239257812 0 9125.588
805.8909912109375 0 6231.023
806.3907470703125 0 3296.253
811.3585205078125 0 1155.3414 z Ammonia loss 7
812.3623046875 0 1206.0208
814.4039916992188 0 1674.6733
814.9006958007812 0 1318.612
815.4163208007812 0 12332.703
815.9201049804688 0 2006.7015
816.4178466796875 0 4448.604
823.3822021484375 0 1279.1776
825.4000854492188 0 1308.13
826.4053344726562 0 4339.73 y Water loss 7
827.4078369140625 0 1637.6392
829.3699951171875 0 3719.235
841.3970336914062 0 3337.4978
843.4095458984375 0 5732.6597
844.4155883789062 0 9329.79 y 7
845.4165649414062 0 3420.3647
857.4130249023438 0 10880.259
858.4203491210938 0 22801.355 c 7
859.4234008789062 0 8245.259
860.4267578125 0 1871.6353
862.3931274414062 0 937.15533
871.4285278320312 0 3185.37
872.4349975585938 0 11446.987
873.439208984375 0 4477.3545
880.4033813476562 0 2266.1572
881.408935546875 0 1212.0072
895.43505859375 0 1012.01196
896.4307250976562 0 1023.4593
898.4144897460938 0 4158.792 z Ammonia loss 6
899.418701171875 0 2568.0544
911.4343872070312 0 906.602
912.4810791015625 0 1168.9187
913.2276000976562 0 810.69073
913.4351806640625 0 4033.0364 y Water loss 6
914.4348754882812 0 11191.266 y Ammonia loss 6
915.4400634765625 0 10136.58 c 8
916.4439697265625 0 4042.2822
917.4484252929688 0 1509.5719
928.4481811523438 0 11324.598
929.4569702148438 0 26147.908
930.4526977539062 0 14663.345
931.449462890625 0 26435.982 y 6
932.4501342773438 0 10527.958
933.4526977539062 0 2030.8081
937.4271240234375 0 8201.359
938.4271240234375 0 3500.526
939.4747924804688 0 2312.4304
955.4393310546875 0 13312.573
956.4412231445312 0 6225.8604
957.4425659179688 0 1834.7977
971.4561767578125 0 27023.865
972.4637451171875 0 90955.3 c 9
973.4666137695312 0 38951.953
974.4696044921875 0 9142.689
1010.517822265625 0 2293.7212
1022.4993286132812 0 1361.8619
1025.4779052734375 0 1175.2001
1026.4761962890625 0 1555.7606 z Water loss 5
1028.4749755859375 0 1251.5293
1032.5 0 1135.5834
1040.5264892578125 0 4639.938
1041.5316162109375 0 13914.516
1042.5325927734375 0 8154.042
1043.5396728515625 0 2656.337
1045.470703125 0 1047.9683
1050.5098876953125 0 7051.454
1051.5181884765625 0 4743.3994
1052.5120849609375 0 2941.1018
1054.52392578125 0 1096.6224
1057.5042724609375 0 898.91644
1059.483154296875 0 8954.718
1060.4901123046875 0 16952.713 y 5
1061.4930419921875 0 7245.748
1062.4952392578125 0 2000.3136
1068.52197265625 0 26203.326
1069.5240478515625 0 13489.212
1070.5301513671875 0 4760.054
1084.5406494140625 0 29653.234
1085.54638671875 0 43764.74 c 10
1086.550048828125 0 22501.283
1087.55322265625 0 6219.663
1111.565673828125 0 1416.535
1112.5714111328125 0 13265.495
1113.576416015625 0 15601.843
1114.580810546875 0 6355.9834
1115.5836181640625 0 1951.9573
1121.548828125 0 8043.0674
1122.553466796875 0 4467.7935
1123.543701171875 0 1889.3218
1124.5391845703125 0 1918.6636
1138.57421875 0 2601.103 c Water loss 11
1139.5595703125 0 32589.467
1140.5621337890625 0 20082.883
1141.559326171875 0 7400.9556 y Water loss 4
1142.5589599609375 0 2246.2124
1143.542724609375 0 1517.0927 z 4
1144.5526123046875 0 1550.6263
1154.541015625 0 1325.1687
1155.5760498046875 0 27830.08
1156.585205078125 0 131841.78 c 11
1157.5875244140625 0 73915.06
1158.5880126953125 0 22340.555
1159.5638427734375 0 10233.157 y 4
1160.5628662109375 0 5819.9805
1161.568359375 0 1189.8071
1168.558349609375 0 1408.4707
1182.6070556640625 0 2100.1274
1195.5770263671875 0 942.7461
1222.6085205078125 0 1398.722
1231.5948486328125 0 3722.9438
1232.5977783203125 0 2240.4358
1238.6129150390625 0 2476.6646 z Water loss 3
1239.62060546875 0 6987.8193 z Ammonia loss 3
1240.6287841796875 0 60676.727
1241.633056640625 0 45027.22
1242.63671875 0 16659.41
1243.64306640625 0 3923.3586
1249.6058349609375 0 17299.385
1250.60302734375 0 13512.709
1251.5999755859375 0 18072.207
1252.6007080078125 0 9588.844
1253.6043701171875 0 2625.3206
1256.62353515625 0 24888.23 z 3
1257.6263427734375 0 15017.667
1258.6297607421875 0 5386.256
1266.631103515625 0 3278.3179 c Water loss 12
1267.6171875 0 130040.36 c Ammonia loss 12
1268.6197509765625 0 84743.86
1269.62158203125 0 29326.824
1270.6197509765625 0 3275.0923
1271.63525390625 0 7366.481
1272.6392822265625 0 12902.188 y 3
1273.6429443359375 0 10517.657
1274.646728515625 0 3036.8555
1322.6795654296875 0 2185.133
1323.6741943359375 0 1347.3044
1324.6732177734375 0 2579.5293
1325.6453857421875 0 971.21484
1329.5941162109375 0 858.3848
1337.6820068359375 0 5944.825
1338.67822265625 0 4614.5547
1339.684326171875 0 2698.334
1351.6962890625 0 1158.2711
1367.6552734375 0 1897.1705 z Water loss 2
1368.6820068359375 0 1399.7347
1380.6898193359375 0 4351.432
1381.6947021484375 0 12635.083 c 13
1382.6968994140625 0 9148.668
1383.6939697265625 0 2927.609 y Water loss 2
1385.6666259765625 0 18779.352 z 2
1386.6697998046875 0 14029.437
1387.6727294921875 0 5407.032
1388.6846923828125 0 1279.1404
1394.702392578125 0 20328.531
1395.7073974609375 0 18826.535
1396.715087890625 0 13500.484
1397.7171630859375 0 4465.161
1398.7203369140625 0 1226.2109
1401.6839599609375 0 7677.0664 y 2
1402.6865234375 0 6184.79
1403.689453125 0 2781.453
1421.705078125 0 1585.0126 c Ammonia loss 14
1422.6939697265625 0 2092.622
1423.708740234375 0 4095.5984
1424.7152099609375 0 3430.9492
1425.7177734375 0 1359.1536
1437.6845703125 0 1421.4218
1438.71630859375 0 58605.066 c 14
1439.7193603515625 0 49054.227
1440.723876953125 0 21837.885
1441.7325439453125 0 4207.905
1477.7403564453125 0 7123.562
1478.741455078125 0 5379.8706
1479.74560546875 0 1982.983
1495.7498779296875 0 8659.218
1496.749755859375 0 5563.881
1497.759521484375 0 3378.4006
1507.7308349609375 0 2021.6973
1511.7454833984375 0 2024.4883
1513.752685546875 0 1535.5822
1524.7530517578125 0 1720.7362
1525.7452392578125 0 1168.216
1526.73876953125 0 1592.8063
1527.729736328125 0 1422.726
1528.7608642578125 0 3919.1182
1529.7626953125 0 5902.961
1530.7625732421875 0 3318.2153
1531.7786865234375 0 1134.0403
1538.771728515625 0 1119.6508
1539.7630615234375 0 74033.11 c 15
1540.7657470703125 0 62177.895
1541.7694091796875 0 29839.072
1542.7674560546875 0 5810.414
1549.7705078125 0 1317.6654
1550.734130859375 0 1108.1193
1552.7471923828125 0 6079.168
1553.7496337890625 0 4448.449
1554.752197265625 0 2255.8381
1555.7239990234375 0 3509.5618
1556.76025390625 0 7862.813
1557.7554931640625 0 15891.854
1558.759521484375 0 13676.036
1559.7646484375 0 3906.209
1565.763671875 0 1116.8129
1566.7860107421875 0 3003.5098
1567.7786865234375 0 1909.454
1568.802978515625 0 1134.0342
1569.7662353515625 0 2380.5425
1570.7777099609375 0 1090.3302
1571.755615234375 0 1176.5085
1572.7301025390625 0 9881.397
1573.7315673828125 0 7237.8936
1574.742919921875 0 4662.447
1581.748291015625 0 1562.4729
1582.779541015625 0 9922.163
1583.7689208984375 0 82869.94
1584.77001953125 0 70970.2
1585.7696533203125 0 33755.582
1586.7611083984375 0 7638.59
1587.736572265625 0 1059.0813
1593.756103515625 0 4587.7144
1594.7578125 0 3998.1782
1595.748779296875 0 2397.7913
1596.739501953125 0 1292.4912
1598.7701416015625 0 1415.4554
1599.780517578125 0 1516.6194
1600.792724609375 0 51529.484
1601.79541015625 0 53578.293
1602.7987060546875 0 23939.934
1603.80224609375 0 2886.3943
1608.78173828125 0 2068.1104
1609.7698974609375 0 2834.333
1610.7752685546875 0 22391.414
1611.7628173828125 0 162810.27
1612.7650146484375 0 145149.97
1613.7667236328125 0 65483.81
1614.7686767578125 0 10115.217
1625.7799072265625 0 2133.205
1626.774169921875 0 11532.414
1627.7794189453125 0 134910.8
1628.7867431640625 0 554469.5
1629.7904052734375 0 448021.62
1630.7930908203125 0 186222.11
1631.7965087890625 0 24853.611
1643.7406005859375 0 1355.9814
1644.741943359375 0 1659.4995
2262.533203125 0 1099.4099
2511.644775390625 0 1030.7845
2942.199462890625 0 973.5155

Spectrum Details

|  |  |
| --- | --- |
| Matched peaks? Matched peaks The total absolute number of peaks matched. Additionally in brackets the total fraction of peaks matched and the total number of peaks is shown. | 56 (13.69% of 409) |
| FDR? FDR The false discovery rate estimated for this peptide. It is calculated by matching all theoretical fragments with a non-integer shift with the raw peaks for this spectrum. This is done with 40 different shifts. The resulting percentage is the average number of annotated peaks over the number of annotated peaks with the correct spectrum. | 0.43% |
| Satellite FDR? Satellite FDR See the FDR for details on its calculation. This satellite ion specific FDR only contains the satellite ions (d/w) for I/L/J positions. | - |
| PSM Score? PSM Score The PSM Score as given by Hecklib to this annotated spectrum. It is shown with three significant figures. | 677 |

#### Spectrum 6271? Spectrum 6271 The raw spectrum of this peptide as annotated by Hecklib. The fragments are coloured according to ion type (see legend). Any peaks with a star '\*' as text can be hovered over to see the full details, first the ion type second the mass shift type. By hovering over the amino acids in the peptide or ions in the legend the corresponding peaks are highlighted. By toggling the 'Unassigned' label you can turn the background (unassigned) peaks on or off in the plot. By updating the slider in the Ion legend you can update the spectrum to only show the top X% of the peaks with labels. The top X% means any peak that is within X% of the highest intensity. By dragging in the spectrum you can zoom in to a specific part of the spectrum and use 'Zoom Out' to get back to the original zoom level. The annotation of the spectrum is based on the given sequence in the peptides file and is done with different software so inconsistencies are likely. The peaks are annotated based on the given sequence, with 20 ppm tolerance.

Copy Data

##### Spectrum 6271 (TSV)

###### Preview

```
Loading example...
```

*Click on the button to copy the data to your clipboard.*

Mz MinMz MaxIntensity Max

WidthHeightPeptide font sizePeptide stroke widthSpectrum font sizeSpectrum stroke widthCompact peptide

Ion legend

wxyz

abcd

OtherUnassignedIon^Charge^_Position_Show for top:%

APEJVESGGGLAQPGTS

01.86e+53.72e+55.58e+57.45e+5

Zoom Out

y^+1^_2_y^+1^_2_y^+3^_7_z^+2^_5_c^+1^_2_y^+1^_3_y^+3^_9_y^+1^_3_y^+2^_6_y^+2^_6_y^+3^_10_c^+2^_5_y^+2^_7_w^+3^_12_y^+1^_4_z^+2^_8_c^+1^_3_y^+2^_8_y^+2^_8_c^+2^_6_y^+1^_4_c^+1^_3_y^+2^_9_c^+3^_12_y^+2^_9_y^+2^_10_w^+1^_5_z^+2^_11_z^+1^_5_y^+2^_11_c^+2^_9_y^+1^_5_z^+1^_5_c^+1^_4_y^+1^_5_y^+2^_12_z^+2^_12_c^+2^_11_y^+1^_6_y^+1^_6_y^+2^_13_y^+2^_13_z^+2^_13_c^+2^_12_c^+1^_5_w^+1^_7_y^+2^_14_y^+2^_14_c^+2^_13_c^+2^_13_y^+2^_14_z^+1^_7_y^+1^_7_z^+1^_7_y^+1^_7_c^+2^_14_c^+2^_14_y^+2^_15_y^+2^_15_z^+1^_8_y^+2^_15_c^+2^_15_c^+2^_15_y^+1^_8_c^+1^_6_c^+2^_15_y^+1^_8_y^+2^_16_z^+2^_16_y^+2^_16_z^+1^_9_c^+2^_16_c^+2^_16_y^+1^_9_c^+2^_16_z^+1^_9_c^+1^_7_y^+1^_9_c^+1^_7_z^+1^_10_y^+1^_10_y^+1^_10_z^+1^_10_c^+1^_8_y^+1^_10_c^+1^_8_c^+1^_9_z^+1^_11_y^+1^_11_y^+1^_11_c^+1^_9_y^+1^_11_c^+1^_10_c^+1^_10_w^+1^_12_z^+1^_12_z^+1^_12_y^+1^_12_c^+1^_11_z^+1^_13_y^+1^_13_z^+1^_13_c^+1^_12_y^+1^_13_w^+1^_14_z^+1^_14_z^+1^_14_y^+1^_14_y^+1^_14_z^+1^_14_c^+1^_13_y^+1^_14_c^+1^_14_z^+1^_15_z^+1^_15_c^+1^_14_y^+1^_15_z^+1^_15_y^+1^_15_c^+1^_15_c^+1^_15_y^+1^_16_y^+1^_16_y^+1^_16_c^+1^_16_

041382512381650

Fragment Matches Table

Show background peaks

| Position | Ion type | Intensity | mz Theoretical | mz Error (Th) | mz Error (ppm) | Charge | Series Number |
| --- | --- | --- | --- | --- | --- | --- | --- |
| - | - | 1343 | 120.1 | - | - | 0 | - |
| - | - | 464.6 | 120.1 | - | - | 0 | - |
| - | - | 450.1 | 122.9 | - | - | 0 | - |
| - | - | 1152 | 124.1 | - | - | 0 | - |
| - | - | 508.1 | 126.1 | - | - | 0 | - |
| - | - | 1065 | 127.1 | - | - | 0 | - |
| - | - | 1181 | 128 | - | - | 0 | - |
| - | - | 935.5 | 128.1 | - | - | 0 | - |
| - | - | 779.7 | 128.1 | - | - | 0 | - |
| - | - | 3.63E+04 | 129.1 | - | - | 0 | - |
| - | - | 1269 | 129.1 | - | - | 0 | - |
| - | - | 2484 | 130.1 | - | - | 0 | - |
| - | - | 473.8 | 130.1 | - | - | 0 | - |
| - | - | 2223 | 130.1 | - | - | 0 | - |
| - | - | 1659 | 131.1 | - | - | 0 | - |
| - | - | 486.9 | 131.1 | - | - | 0 | - |
| - | - | 3804 | 133.1 | - | - | 0 | - |
| - | - | 785.7 | 133.1 | - | - | 0 | - |
| - | - | 6748 | 143.1 | - | - | 0 | - |
| - | - | 2620 | 145.1 | - | - | 0 | - |
| - | - | 1.406E+04 | 147.1 | - | - | 0 | - |
| - | - | 1299 | 148.1 | - | - | 0 | - |
| - | - | 969 | 149 | - | - | 0 | - |
| - | - | 4835 | 152.1 | - | - | 0 | - |
| - | - | 859.2 | 153.1 | - | - | 0 | - |
| - | - | 581.4 | 153.1 | - | - | 0 | - |
| - | - | 1.469E+04 | 155.1 | - | - | 0 | - |
| - | - | 1421 | 156.1 | - | - | 0 | - |
| - | - | 592.2 | 156.1 | - | - | 0 | - |
| - | - | 710 | 157.1 | - | - | 0 | - |
| - | - | 673.9 | 157.1 | - | - | 0 | - |
| - | - | 1679 | 157.1 | - | - | 0 | - |
| - | - | 484.1 | 158.4 | - | - | 0 | - |
| - | - | 2502 | 159.1 | - | - | 0 | - |
| - | - | 627.1 | 165.1 | - | - | 0 | - |
| - | - | 2941 | 166.1 | - | - | 0 | - |
| - | - | 2427 | 171.1 | - | - | 0 | - |
| - | - | 4467 | 171.1 | - | - | 0 | - |
| - | - | 928.9 | 172.1 | - | - | 0 | - |
| - | - | 8.535E+04 | 172.1 | - | - | 0 | - |
| - | - | 838.9 | 173.1 | - | - | 0 | - |
| - | - | 738.5 | 173.1 | - | - | 0 | - |
| - | - | 6510 | 173.1 | - | - | 0 | - |
| - | - | 781.9 | 181.1 | - | - | 0 | - |
| - | - | 1578 | 181.1 | - | - | 0 | - |
| - | - | 5.19E+04 | 181.1 | - | - | 0 | - |
| - | - | 4632 | 182.1 | - | - | 0 | - |
| - | - | 3525 | 183.1 | - | - | 0 | - |
| - | - | 1619 | 183.1 | - | - | 0 | - |
| - | - | 471.5 | 183.1 | - | - | 0 | - |
| - | - | 5663 | 183.1 | - | - | 0 | - |
| - | - | 3534 | 184.1 | - | - | 0 | - |
| - | - | 1625 | 185.1 | - | - | 0 | - |
| - | - | 2953 | 185.1 | - | - | 0 | - |
| - | - | 1018 | 185.2 | - | - | 0 | - |
| - | - | 1014 | 187.1 | - | - | 0 | - |
| 16 | y | 1.232E+04 | 189.1 | 0.0005258 | 2.781 | +1 | 2 |
| - | - | 730.7 | 190.1 | - | - | 0 | - |
| - | - | 608.1 | 190.1 | - | - | 0 | - |
| - | - | 1617 | 193.1 | - | - | 0 | - |
| - | - | 2173 | 197.1 | - | - | 0 | - |
| - | - | 2229 | 199.1 | - | - | 0 | - |
| - | - | 8381 | 199.1 | - | - | 0 | - |
| - | - | 3.596E+04 | 200.1 | - | - | 0 | - |
| - | - | 710.7 | 200.1 | - | - | 0 | - |
| - | - | 3311 | 201.1 | - | - | 0 | - |
| - | - | 1295 | 201.1 | - | - | 0 | - |
| - | - | 1986 | 202.1 | - | - | 0 | - |
| - | - | 694.9 | 203.1 | - | - | 0 | - |
| 16 | y | 3.97E+04 | 207.1 | 0.0004897 | 2.365 | +1 | 2 |
| - | - | 2682 | 208.1 | - | - | 0 | - |
| - | - | 1.051E+05 | 209.1 | - | - | 0 | - |
| - | - | 1.424E+04 | 210.1 | - | - | 0 | - |
| - | - | 4017 | 210.1 | - | - | 0 | - |
| - | - | 648.5 | 211.1 | - | - | 0 | - |
| - | - | 922.6 | 211.1 | - | - | 0 | - |
| - | - | 3403 | 211.1 | - | - | 0 | - |
| - | - | 1.157E+04 | 211.1 | - | - | 0 | - |
| - | - | 776.4 | 212.1 | - | - | 0 | - |
| - | - | 2771 | 212.1 | - | - | 0 | - |
| - | - | 2053 | 212.1 | - | - | 0 | - |
| - | - | 3251 | 213.1 | - | - | 0 | - |
| - | - | 1598 | 213.2 | - | - | 0 | - |
| - | - | 491.4 | 214.1 | - | - | 0 | - |
| - | - | 664.5 | 214.1 | - | - | 0 | - |
| - | - | 2711 | 215.1 | - | - | 0 | - |
| - | - | 1328 | 217.1 | - | - | 0 | - |
| - | - | 514.4 | 225.1 | - | - | 0 | - |
| - | - | 2870 | 225.1 | - | - | 0 | - |
| 11 | y | 2143 | 225.1 | 0.001965 | 8.729 | +3 | 7 |
| - | - | 6234 | 226.1 | - | - | 0 | - |
| - | - | 7419 | 227.1 | - | - | 0 | - |
| - | - | 474.1 | 227.1 | - | - | 0 | - |
| - | - | 792.9 | 227.1 | - | - | 0 | - |
| - | - | 7643 | 227.1 | - | - | 0 | - |
| - | - | 1815 | 228.1 | - | - | 0 | - |
| - | - | 1.887E+04 | 228.1 | - | - | 0 | - |
| - | - | 4938 | 229.1 | - | - | 0 | - |
| - | - | 677.1 | 230.1 | - | - | 0 | - |
| - | - | 3609 | 233.2 | - | - | 0 | - |
| 13 | z | 2228 | 237.1 | 0.001723 | 7.266 | +2 | 5 |
| - | - | 691.1 | 238.1 | - | - | 0 | - |
| - | - | 9866 | 238.1 | - | - | 0 | - |
| - | - | 803.5 | 241.1 | - | - | 0 | - |
| - | - | 536.2 | 242.1 | - | - | 0 | - |
| - | - | 4524 | 242.2 | - | - | 0 | - |
| - | - | 635.2 | 243.1 | - | - | 0 | - |
| - | - | 7952 | 243.1 | - | - | 0 | - |
| - | - | 847.6 | 243.1 | - | - | 0 | - |
| - | - | 922.1 | 244.1 | - | - | 0 | - |
| 2 | c | 1.838E+04 | 244.1 | 0.003881 | 15.9 | +1 | 2 |
| - | - | 4474 | 244.2 | - | - | 0 | - |
| - | - | 484.6 | 245.1 | - | - | 0 | - |
| - | - | 1738 | 245.1 | - | - | 0 | - |
| - | - | 576.7 | 245.2 | - | - | 0 | - |
| 15 | y | 6166 | 246.1 | 0.0007754 | 3.15 | +1 | 3 |
| - | - | 793.4 | 250.1 | - | - | 0 | - |
| - | - | 587.8 | 251.1 | - | - | 0 | - |
| - | - | 649.5 | 251.2 | - | - | 0 | - |
| - | - | 2712 | 252.1 | - | - | 0 | - |
| - | - | 1674 | 254.1 | - | - | 0 | - |
| - | - | 638.2 | 254.1 | - | - | 0 | - |
| - | - | 1563 | 255.1 | - | - | 0 | - |
| - | - | 3453 | 255.1 | - | - | 0 | - |
| - | - | 2745 | 256.1 | - | - | 0 | - |
| - | - | 6.854E+04 | 256.1 | - | - | 0 | - |
| 9 | y | 8269 | 257.1 | 0.0001191 | 0.463 | +3 | 9 |
| - | - | 3809 | 257.2 | - | - | 0 | - |
| - | - | 692.5 | 258.1 | - | - | 0 | - |
| - | - | 2597 | 259.1 | - | - | 0 | - |
| - | - | 1136 | 259.2 | - | - | 0 | - |
| - | - | 2237 | 260.2 | - | - | 0 | - |
| - | - | 2221 | 261.2 | - | - | 0 | - |
| - | - | 2438 | 262.6 | - | - | 0 | - |
| - | - | 1026 | 262.6 | - | - | 0 | - |
| 15 | y | 4434 | 264.1 | 0.0004951 | 1.874 | +1 | 3 |
| - | - | 1087 | 264.1 | - | - | 0 | - |
| - | - | 2256 | 266.1 | - | - | 0 | - |
| - | - | 4123 | 267.1 | - | - | 0 | - |
| - | - | 5417 | 268.1 | - | - | 0 | - |
| - | - | 5203 | 270.1 | - | - | 0 | - |
| - | - | 676.3 | 270.7 | - | - | 0 | - |
| - | - | 1277 | 271.2 | - | - | 0 | - |
| 12 | y | 7473 | 271.6 | 0.000479 | 1.764 | +2 | 6 |
| - | - | 2147 | 272.1 | - | - | 0 | - |
| - | - | 8.27E+04 | 273.2 | - | - | 0 | - |
| - | - | 832.4 | 274.1 | - | - | 0 | - |
| - | - | 1060 | 274.1 | - | - | 0 | - |
| - | - | 1.061E+04 | 274.2 | - | - | 0 | - |
| - | - | 939.2 | 275.2 | - | - | 0 | - |
| - | - | 811.3 | 275.2 | - | - | 0 | - |
| - | - | 1584 | 280.2 | - | - | 0 | - |
| 12 | y | 723.8 | 280.6 | 0.001117 | 3.981 | +2 | 6 |
| 8 | y | 4589 | 282.1 | 0.001986 | 7.038 | +3 | 10 |
| - | - | 4404 | 283.1 | - | - | 0 | - |
| - | - | 3570 | 284.1 | - | - | 0 | - |
| 5 | c | 1.734E+04 | 284.2 | 0.00161 | 5.666 | +2 | 5 |
| - | - | 5516 | 285.2 | - | - | 0 | - |
| - | - | 793.3 | 291.2 | - | - | 0 | - |
| - | - | 1673 | 292.1 | - | - | 0 | - |
| - | - | 1062 | 296.2 | - | - | 0 | - |
| - | - | 1358 | 297.2 | - | - | 0 | - |
| - | - | 2473 | 297.2 | - | - | 0 | - |
| - | - | 8725 | 298.1 | - | - | 0 | - |
| - | - | 693.9 | 298.1 | - | - | 0 | - |
| - | - | 4567 | 298.2 | - | - | 0 | - |
| - | - | 1651 | 299.1 | - | - | 0 | - |
| - | - | 8899 | 299.2 | - | - | 0 | - |
| - | - | 1119 | 300.1 | - | - | 0 | - |
| - | - | 7605 | 300.2 | - | - | 0 | - |
| - | - | 866.7 | 301.2 | - | - | 0 | - |
| - | - | 1186 | 301.2 | - | - | 0 | - |
| - | - | 1484 | 301.2 | - | - | 0 | - |
| - | - | 1290 | 307.1 | - | - | 0 | - |
| - | - | 4.983E+04 | 310.1 | - | - | 0 | - |
| - | - | 6639 | 311.1 | - | - | 0 | - |
| - | - | 2307 | 312.1 | - | - | 0 | - |
| - | - | 1905 | 313.1 | - | - | 0 | - |
| - | - | 5899 | 313.2 | - | - | 0 | - |
| - | - | 1034 | 315.2 | - | - | 0 | - |
| - | - | 1763 | 316.2 | - | - | 0 | - |
| - | - | 749.1 | 316.2 | - | - | 0 | - |
| - | - | 1268 | 319.2 | - | - | 0 | - |
| - | - | 842.2 | 319.7 | - | - | 0 | - |
| - | - | 647.3 | 320.2 | - | - | 0 | - |
| - | - | 615.6 | 322.2 | - | - | 0 | - |
| - | - | 3389 | 324.1 | - | - | 0 | - |
| - | - | 645.9 | 324.2 | - | - | 0 | - |
| - | - | 1.101E+04 | 325.2 | - | - | 0 | - |
| - | - | 1208 | 326.1 | - | - | 0 | - |
| - | - | 1343 | 326.2 | - | - | 0 | - |
| - | - | 555.3 | 327.2 | - | - | 0 | - |
| - | - | 644.2 | 328.2 | - | - | 0 | - |
| 11 | y | 1.105E+04 | 328.2 | 0.000958 | 2.919 | +2 | 7 |
| - | - | 1624 | 328.2 | - | - | 0 | - |
| - | - | 4024 | 328.7 | - | - | 0 | - |
| 6 | w | 1543 | 329.2 | 0.000819 | 2.488 | +3 | 12 |
| - | - | 815.8 | 329.2 | - | - | 0 | - |
| - | - | 1071 | 330.2 | - | - | 0 | - |
| - | - | 3666 | 331.1 | - | - | 0 | - |
| - | - | 520.6 | 333.1 | - | - | 0 | - |
| - | - | 3042 | 337.2 | - | - | 0 | - |
| - | - | 7.034E+04 | 338.1 | - | - | 0 | - |
| - | - | 1.129E+04 | 339.1 | - | - | 0 | - |
| - | - | 2843 | 339.2 | - | - | 0 | - |
| - | - | 1090 | 339.2 | - | - | 0 | - |
| - | - | 2171 | 340.1 | - | - | 0 | - |
| - | - | 1776 | 340.1 | - | - | 0 | - |
| - | - | 1323 | 340.2 | - | - | 0 | - |
| - | - | 3155 | 340.2 | - | - | 0 | - |
| - | - | 818.5 | 340.7 | - | - | 0 | - |
| - | - | 1583 | 341.1 | - | - | 0 | - |
| - | - | 3279 | 341.2 | - | - | 0 | - |
| - | - | 664.3 | 341.7 | - | - | 0 | - |
| - | - | 2.341E+04 | 342.2 | - | - | 0 | - |
| - | - | 3324 | 342.2 | - | - | 0 | - |
| 14 | y | 1.223E+05 | 343.2 | 0.0007154 | 2.085 | +1 | 4 |
| - | - | 2.144E+04 | 344.2 | - | - | 0 | - |
| - | - | 2536 | 344.2 | - | - | 0 | - |
| - | - | 653.5 | 345.1 | - | - | 0 | - |
| - | - | 2739 | 345.2 | - | - | 0 | - |
| - | - | 1721 | 347.7 | - | - | 0 | - |
| - | - | 1059 | 349.2 | - | - | 0 | - |
| 10 | z | 2698 | 349.2 | 0.006909 | 19.79 | +2 | 8 |
| - | - | 1146 | 349.7 | - | - | 0 | - |
| - | - | 826 | 350.2 | - | - | 0 | - |
| - | - | 1260 | 351.1 | - | - | 0 | - |
| - | - | 1279 | 351.2 | - | - | 0 | - |
| - | - | 831.3 | 352.1 | - | - | 0 | - |
| - | - | 4401 | 353.1 | - | - | 0 | - |
| - | - | 7937 | 354.2 | - | - | 0 | - |
| 3 | c | 7946 | 355.2 | 0.004019 | 11.32 | +1 | 3 |
| - | - | 1.979E+04 | 355.2 | - | - | 0 | - |
| - | - | 6569 | 356.1 | - | - | 0 | - |
| - | - | 939.9 | 356.2 | - | - | 0 | - |
| - | - | 6846 | 356.2 | - | - | 0 | - |
| 10 | y | 4168 | 356.7 | 0.0009989 | 2.8 | +2 | 8 |
| - | - | 2099 | 357.2 | - | - | 0 | - |
| 10 | y | 1061 | 357.2 | 0.003834 | 10.73 | +2 | 8 |
| 6 | c | 660.4 | 357.7 | 0.001486 | 4.155 | +2 | 6 |
| - | - | 6006 | 359.2 | - | - | 0 | - |
| - | - | 9464 | 360.2 | - | - | 0 | - |
| 14 | y | 7.372E+05 | 361.2 | 0.0006792 | 1.881 | +1 | 4 |
| - | - | 1.236E+05 | 362.2 | - | - | 0 | - |
| - | - | 1.838E+04 | 363.2 | - | - | 0 | - |
| - | - | 1747 | 364.2 | - | - | 0 | - |
| - | - | 1060 | 365.2 | - | - | 0 | - |
| - | - | 2267 | 366.2 | - | - | 0 | - |
| - | - | 5431 | 367.2 | - | - | 0 | - |
| - | - | 720.3 | 367.2 | - | - | 0 | - |
| - | - | 6748 | 368.2 | - | - | 0 | - |
| - | - | 1312 | 368.2 | - | - | 0 | - |
| - | - | 3956 | 369.2 | - | - | 0 | - |
| - | - | 1198 | 369.7 | - | - | 0 | - |
| - | - | 4891 | 370.1 | - | - | 0 | - |
| - | - | 4076 | 370.2 | - | - | 0 | - |
| - | - | 711.7 | 371.1 | - | - | 0 | - |
| - | - | 7435 | 371.2 | - | - | 0 | - |
| - | - | 4246 | 371.2 | - | - | 0 | - |
| - | - | 1081 | 372.2 | - | - | 0 | - |
| - | - | 6074 | 372.2 | - | - | 0 | - |
| 3 | c | 1.036E+04 | 373.2 | 0.004025 | 10.78 | +1 | 3 |
| - | - | 1257 | 373.2 | - | - | 0 | - |
| - | - | 1792 | 374.2 | - | - | 0 | - |
| - | - | 5486 | 374.7 | - | - | 0 | - |
| - | - | 2157 | 375.2 | - | - | 0 | - |
| - | - | 621 | 375.7 | - | - | 0 | - |
| - | - | 1143 | 376.2 | - | - | 0 | - |
| - | - | 9361 | 379.2 | - | - | 0 | - |
| - | - | 1690 | 380.2 | - | - | 0 | - |
| - | - | 5364 | 383.7 | - | - | 0 | - |
| - | - | 591.3 | 384.2 | - | - | 0 | - |
| - | - | 1.367E+04 | 384.2 | - | - | 0 | - |
| - | - | 1433 | 384.7 | - | - | 0 | - |
| 9 | y | 6349 | 385.2 | 0.0005167 | 1.341 | +2 | 9 |
| - | - | 1859 | 385.7 | - | - | 0 | - |
| 12 | c | 2222 | 386.2 | 0.001617 | 4.186 | +3 | 12 |
| - | - | 4777 | 388.1 | - | - | 0 | - |
| - | - | 2060 | 389.2 | - | - | 0 | - |
| - | - | 651.1 | 391.2 | - | - | 0 | - |
| - | - | 1097 | 391.7 | - | - | 0 | - |
| - | - | 935.8 | 391.7 | - | - | 0 | - |
| - | - | 1893 | 392.7 | - | - | 0 | - |
| - | - | 610.3 | 393.2 | - | - | 0 | - |
| - | - | 1129 | 394.2 | - | - | 0 | - |
| 9 | y | 1246 | 394.2 | 0.001343 | 3.408 | +2 | 9 |
| - | - | 1002 | 396.2 | - | - | 0 | - |
| - | - | 5.792E+04 | 397.2 | - | - | 0 | - |
| - | - | 1.275E+04 | 398.2 | - | - | 0 | - |
| - | - | 2665 | 399.2 | - | - | 0 | - |
| - | - | 829.9 | 399.2 | - | - | 0 | - |
| - | - | 7997 | 401.2 | - | - | 0 | - |
| - | - | 1298 | 402.2 | - | - | 0 | - |
| - | - | 7116 | 403.2 | - | - | 0 | - |
| - | - | 2916 | 403.7 | - | - | 0 | - |
| - | - | 873.6 | 404.2 | - | - | 0 | - |
| - | - | 1002 | 404.7 | - | - | 0 | - |
| - | - | 1554 | 405.2 | - | - | 0 | - |
| - | - | 1890 | 405.2 | - | - | 0 | - |
| - | - | 1133 | 408.2 | - | - | 0 | - |
| - | - | 1778 | 409.2 | - | - | 0 | - |
| - | - | 912.1 | 410.2 | - | - | 0 | - |
| - | - | 4890 | 410.2 | - | - | 0 | - |
| - | - | 1571 | 411.2 | - | - | 0 | - |
| - | - | 9993 | 412.2 | - | - | 0 | - |
| - | - | 1.941E+04 | 412.2 | - | - | 0 | - |
| - | - | 7650 | 412.7 | - | - | 0 | - |
| - | - | 1109 | 413.2 | - | - | 0 | - |
| - | - | 4420 | 413.2 | - | - | 0 | - |
| 8 | y | 5351 | 413.7 | 0.0005007 | 1.21 | +2 | 10 |
| - | - | 943.5 | 414.2 | - | - | 0 | - |
| 13 | w | 1.441E+04 | 415.2 | 0.00107 | 2.578 | +1 | 5 |
| - | - | 2919 | 416.2 | - | - | 0 | - |
| - | - | 716.6 | 420.2 | - | - | 0 | - |
| - | - | 2638 | 421.2 | - | - | 0 | - |
| - | - | 1410 | 421.7 | - | - | 0 | - |
| - | - | 3.29E+04 | 423.2 | - | - | 0 | - |
| - | - | 9212 | 424.2 | - | - | 0 | - |
| - | - | 2214 | 425.2 | - | - | 0 | - |
| - | - | 600 | 425.2 | - | - | 0 | - |
| - | - | 1162 | 426.2 | - | - | 0 | - |
| - | - | 945.8 | 426.2 | - | - | 0 | - |
| - | - | 8428 | 427.2 | - | - | 0 | - |
| - | - | 1184 | 428.2 | - | - | 0 | - |
| - | - | 1919 | 430.2 | - | - | 0 | - |
| - | - | 1709 | 431.7 | - | - | 0 | - |
| - | - | 612.6 | 432.2 | - | - | 0 | - |
| - | - | 2681 | 433.2 | - | - | 0 | - |
| - | - | 695.4 | 433.2 | - | - | 0 | - |
| - | - | 4022 | 436.2 | - | - | 0 | - |
| - | - | 656.5 | 436.2 | - | - | 0 | - |
| - | - | 972.7 | 437.2 | - | - | 0 | - |
| - | - | 1495 | 437.2 | - | - | 0 | - |
| - | - | 931.7 | 438.2 | - | - | 0 | - |
| - | - | 3681 | 439.3 | - | - | 0 | - |
| - | - | 623.4 | 440.2 | - | - | 0 | - |
| - | - | 6542 | 440.7 | - | - | 0 | - |
| - | - | 2266 | 441.2 | - | - | 0 | - |
| - | - | 3199 | 441.2 | - | - | 0 | - |
| - | - | 2189 | 442.2 | - | - | 0 | - |
| - | - | 3206 | 443.2 | - | - | 0 | - |
| - | - | 766.7 | 443.3 | - | - | 0 | - |
| - | - | 717.2 | 446.2 | - | - | 0 | - |
| - | - | 2641 | 448.2 | - | - | 0 | - |
| 7 | z | 7606 | 449.7 | 0.007517 | 16.72 | +2 | 11 |
| - | - | 5668 | 450.2 | - | - | 0 | - |
| - | - | 1520 | 450.7 | - | - | 0 | - |
| - | - | 5.548E+04 | 451.2 | - | - | 0 | - |
| - | - | 1485 | 451.9 | - | - | 0 | - |
| - | - | 1.311E+04 | 452.2 | - | - | 0 | - |
| - | - | 2570 | 452.3 | - | - | 0 | - |
| - | - | 1.791E+04 | 453.2 | - | - | 0 | - |
| - | - | 3809 | 454.2 | - | - | 0 | - |
| - | - | 5640 | 454.2 | - | - | 0 | - |
| 13 | z | 2713 | 455.2 | 0.003707 | 8.144 | +1 | 5 |
| - | - | 1.27E+04 | 455.2 | - | - | 0 | - |
| - | - | 676 | 455.7 | - | - | 0 | - |
| - | - | 1.22E+04 | 456.2 | - | - | 0 | - |
| 7 | y | 5319 | 457.2 | 0.002971 | 6.497 | +2 | 11 |
| - | - | 1657 | 457.7 | - | - | 0 | - |
| 9 | c | 1127 | 458.2 | 0.004174 | 9.109 | +2 | 9 |
| - | - | 1952 | 460.2 | - | - | 0 | - |
| - | - | 937.2 | 460.3 | - | - | 0 | - |
| - | - | 2699 | 467.2 | - | - | 0 | - |
| - | - | 5140 | 468.2 | - | - | 0 | - |
| - | - | 2.022E+04 | 469.2 | - | - | 0 | - |
| - | - | 1328 | 469.3 | - | - | 0 | - |
| - | - | 2324 | 469.7 | - | - | 0 | - |
| - | - | 938.7 | 470.2 | - | - | 0 | - |
| - | - | 4016 | 470.2 | - | - | 0 | - |
| 13 | y | 3.213E+04 | 471.2 | 0.00125 | 2.654 | +1 | 5 |
| - | - | 5591 | 472.2 | - | - | 0 | - |
| - | - | 1123 | 472.3 | - | - | 0 | - |
| 13 | z | 2426 | 473.2 | 0.001088 | 2.3 | +1 | 5 |
| - | - | 1527 | 473.2 | - | - | 0 | - |
| - | - | 1097 | 474.2 | - | - | 0 | - |
| - | - | 760.6 | 476.3 | - | - | 0 | - |
| - | - | 9359 | 478.2 | - | - | 0 | - |
| - | - | 4648 | 478.7 | - | - | 0 | - |
| - | - | 2097 | 479.2 | - | - | 0 | - |
| - | - | 6659 | 483.2 | - | - | 0 | - |
| - | - | 1429 | 484.2 | - | - | 0 | - |
| - | - | 4918 | 484.3 | - | - | 0 | - |
| - | - | 4929 | 485.3 | - | - | 0 | - |
| 4 | c | 1.168E+04 | 486.3 | 0.003219 | 6.62 | +1 | 4 |
| - | - | 3676 | 487.2 | - | - | 0 | - |
| - | - | 2438 | 487.3 | - | - | 0 | - |
| - | - | 5044 | 488.2 | - | - | 0 | - |
| - | - | 898.3 | 488.3 | - | - | 0 | - |
| 13 | y | 9.435E+04 | 489.2 | 0.001092 | 2.232 | +1 | 5 |
| - | - | 2.21E+04 | 490.2 | - | - | 0 | - |
| - | - | 1.137E+04 | 491.2 | - | - | 0 | - |
| - | - | 2717 | 492.2 | - | - | 0 | - |
| - | - | 4586 | 493.3 | - | - | 0 | - |
| - | - | 1244 | 494.3 | - | - | 0 | - |
| - | - | 1083 | 498.3 | - | - | 0 | - |
| - | - | 6843 | 501.2 | - | - | 0 | - |
| - | - | 1419 | 502.2 | - | - | 0 | - |
| - | - | 1061 | 502.8 | - | - | 0 | - |
| - | - | 1080 | 503.7 | - | - | 0 | - |
| - | - | 1071 | 504.2 | - | - | 0 | - |
| - | - | 2967 | 504.3 | - | - | 0 | - |
| - | - | 1127 | 507.2 | - | - | 0 | - |
| - | - | 2733 | 508.3 | - | - | 0 | - |
| - | - | 886.5 | 508.3 | - | - | 0 | - |
| - | - | 4482 | 509.2 | - | - | 0 | - |
| - | - | 2885 | 509.3 | - | - | 0 | - |
| - | - | 1058 | 510.2 | - | - | 0 | - |
| - | - | 899.4 | 510.3 | - | - | 0 | - |
| - | - | 1085 | 511.8 | - | - | 0 | - |
| - | - | 742.2 | 512.3 | - | - | 0 | - |
| - | - | 2615 | 512.7 | - | - | 0 | - |
| - | - | 1703 | 513.2 | - | - | 0 | - |
| - | - | 741.8 | 517.2 | - | - | 0 | - |
| - | - | 685.2 | 518.8 | - | - | 0 | - |
| - | - | 1178 | 520.8 | - | - | 0 | - |
| 6 | y | 3287 | 521.7 | 0.001917 | 3.674 | +2 | 12 |
| - | - | 1403 | 522.3 | - | - | 0 | - |
| - | - | 1.712E+04 | 522.3 | - | - | 0 | - |
| 6 | z | 883.2 | 522.7 | 0.006851 | 13.11 | +2 | 12 |
| - | - | 5893 | 523.3 | - | - | 0 | - |
| - | - | 8271 | 524.2 | - | - | 0 | - |
| - | - | 1985 | 524.3 | - | - | 0 | - |
| - | - | 2590 | 525.3 | - | - | 0 | - |
| - | - | 1531 | 525.3 | - | - | 0 | - |
| - | - | 4259 | 525.8 | - | - | 0 | - |
| - | - | 2657 | 526.3 | - | - | 0 | - |
| - | - | 2417 | 526.3 | - | - | 0 | - |
| - | - | 832.5 | 527.2 | - | - | 0 | - |
| - | - | 771.5 | 527.3 | - | - | 0 | - |
| - | - | 670.4 | 527.3 | - | - | 0 | - |
| - | - | 678 | 528.3 | - | - | 0 | - |
| - | - | 3873 | 532.3 | - | - | 0 | - |
| - | - | 1031 | 533.3 | - | - | 0 | - |
| - | - | 676.3 | 533.8 | - | - | 0 | - |
| 11 | c | 5809 | 534.3 | 0.006175 | 11.56 | +2 | 11 |
| - | - | 1.239E+04 | 534.8 | - | - | 0 | - |
| - | - | 1002 | 534.8 | - | - | 0 | - |
| - | - | 6519 | 535.3 | - | - | 0 | - |
| - | - | 2887 | 535.8 | - | - | 0 | - |
| - | - | 1879 | 538.3 | - | - | 0 | - |
| - | - | 1.144E+04 | 540.3 | - | - | 0 | - |
| - | - | 760.7 | 541.3 | - | - | 0 | - |
| - | - | 4114 | 541.3 | - | - | 0 | - |
| 12 | y | 4.161E+04 | 542.3 | 0.001093 | 2.016 | +1 | 6 |
| - | - | 2109 | 542.8 | - | - | 0 | - |
| - | - | 9780 | 543.3 | - | - | 0 | - |
| - | - | 3392 | 543.3 | - | - | 0 | - |
| - | - | 1088 | 543.8 | - | - | 0 | - |
| - | - | 1846 | 544.3 | - | - | 0 | - |
| - | - | 8.277E+04 | 550.3 | - | - | 0 | - |
| - | - | 2.497E+04 | 551.3 | - | - | 0 | - |
| - | - | 2.174E+04 | 552.3 | - | - | 0 | - |
| - | - | 1.344E+04 | 553.3 | - | - | 0 | - |
| - | - | 4387 | 554.3 | - | - | 0 | - |
| - | - | 2087 | 554.3 | - | - | 0 | - |
| - | - | 1.732E+04 | 555.3 | - | - | 0 | - |
| - | - | 1052 | 555.3 | - | - | 0 | - |
| - | - | 1.44E+04 | 556.3 | - | - | 0 | - |
| - | - | 3161 | 557.3 | - | - | 0 | - |
| - | - | 2356 | 558.3 | - | - | 0 | - |
| - | - | 1682 | 559.3 | - | - | 0 | - |
| - | - | 903.5 | 559.3 | - | - | 0 | - |
| 12 | y | 3.746E+04 | 560.3 | 0.0009657 | 1.724 | +1 | 6 |
| - | - | 1.606E+04 | 561.3 | - | - | 0 | - |
| - | - | 3480 | 561.8 | - | - | 0 | - |
| - | - | 1.274E+04 | 562.3 | - | - | 0 | - |
| - | - | 1710 | 562.8 | - | - | 0 | - |
| - | - | 3608 | 563.3 | - | - | 0 | - |
| - | - | 3819 | 565.3 | - | - | 0 | - |
| - | - | 2330 | 566.3 | - | - | 0 | - |
| - | - | 2133 | 567.3 | - | - | 0 | - |
| - | - | 5.953E+04 | 568.3 | - | - | 0 | - |
| - | - | 1.924E+04 | 569.3 | - | - | 0 | - |
| - | - | 1.433E+04 | 570.3 | - | - | 0 | - |
| - | - | 8918 | 570.8 | - | - | 0 | - |
| 5 | y | 1.739E+04 | 571.3 | 0.006528 | 11.43 | +2 | 13 |
| 5 | y | 3374 | 571.8 | 0.01135 | 19.84 | +2 | 13 |
| 5 | z | 7993 | 572.3 | 0.001111 | 1.941 | +2 | 13 |
| - | - | 1706 | 573.3 | - | - | 0 | - |
| - | - | 634.5 | 578.3 | - | - | 0 | - |
| 12 | c | 2045 | 578.8 | 0.001734 | 2.996 | +2 | 12 |
| - | - | 972.3 | 579.3 | - | - | 0 | - |
| - | - | 714.4 | 580.3 | - | - | 0 | - |
| - | - | 1788 | 581.3 | - | - | 0 | - |
| - | - | 3726 | 583.3 | - | - | 0 | - |
| - | - | 1511 | 584.3 | - | - | 0 | - |
| 5 | c | 5.22E+04 | 585.3 | 0.00303 | 5.176 | +1 | 5 |
| - | - | 1.784E+04 | 586.3 | - | - | 0 | - |
| - | - | 3335 | 587.3 | - | - | 0 | - |
| - | - | 720.1 | 588.3 | - | - | 0 | - |
| - | - | 743.9 | 589.3 | - | - | 0 | - |
| - | - | 8751 | 591.3 | - | - | 0 | - |
| - | - | 3052 | 592.3 | - | - | 0 | - |
| - | - | 3179 | 595.3 | - | - | 0 | - |
| - | - | 7617 | 596.3 | - | - | 0 | - |
| - | - | 1618 | 597.3 | - | - | 0 | - |
| - | - | 804 | 598.3 | - | - | 0 | - |
| - | - | 810.6 | 598.3 | - | - | 0 | - |
| - | - | 1754 | 599.8 | - | - | 0 | - |
| - | - | 4258 | 600.3 | - | - | 0 | - |
| - | - | 827.9 | 601.3 | - | - | 0 | - |
| - | - | 2116 | 609.3 | - | - | 0 | - |
| - | - | 2029 | 610.3 | - | - | 0 | - |
| - | - | 1.851E+04 | 611.3 | - | - | 0 | - |
| - | - | 6851 | 612.3 | - | - | 0 | - |
| - | - | 2019 | 613.4 | - | - | 0 | - |
| 11 | w | 2.12E+04 | 614.3 | 0.001204 | 1.96 | +1 | 7 |
| - | - | 7173 | 615.3 | - | - | 0 | - |
| - | - | 684.7 | 616.3 | - | - | 0 | - |
| - | - | 1071 | 618.8 | - | - | 0 | - |
| - | - | 1070 | 619.3 | - | - | 0 | - |
| - | - | 1464 | 621.3 | - | - | 0 | - |
| - | - | 6016 | 622.3 | - | - | 0 | - |
| - | - | 1819 | 623.3 | - | - | 0 | - |
| - | - | 3690 | 625.3 | - | - | 0 | - |
| - | - | 2307 | 625.8 | - | - | 0 | - |
| - | - | 1989 | 626.3 | - | - | 0 | - |
| - | - | 2376 | 626.8 | - | - | 0 | - |
| - | - | 1594 | 627.3 | - | - | 0 | - |
| 4 | y | 2334 | 627.8 | 0.0002017 | 0.3213 | +2 | 14 |
| 4 | y | 2035 | 628.3 | 0.01082 | 17.22 | +2 | 14 |
| - | - | 897.5 | 630.8 | - | - | 0 | - |
| - | - | 1149 | 632.3 | - | - | 0 | - |
| 13 | c | 814.3 | 633.8 | 0.01134 | 17.89 | +2 | 13 |
| 13 | c | 1.041E+04 | 634.3 | 0.0009634 | 1.519 | +2 | 13 |
| - | - | 5543 | 634.8 | - | - | 0 | - |
| - | - | 3285 | 635.3 | - | - | 0 | - |
| - | - | 1738 | 635.8 | - | - | 0 | - |
| 4 | y | 791 | 636.8 | 0.003098 | 4.865 | +2 | 14 |
| - | - | 1789 | 637.3 | - | - | 0 | - |
| - | - | 3376 | 638.3 | - | - | 0 | - |
| 11 | z | 3919 | 639.3 | 0.003319 | 5.192 | +1 | 7 |
| - | - | 2183 | 640.3 | - | - | 0 | - |
| - | - | 940 | 641.3 | - | - | 0 | - |
| - | - | 806.1 | 642.4 | - | - | 0 | - |
| - | - | 4327 | 643.3 | - | - | 0 | - |
| - | - | 1556 | 644.3 | - | - | 0 | - |
| - | - | 2543 | 646.4 | - | - | 0 | - |
| - | - | 759.2 | 646.8 | - | - | 0 | - |
| - | - | 886.9 | 648.3 | - | - | 0 | - |
| - | - | 2294 | 652.3 | - | - | 0 | - |
| - | - | 1681 | 653.3 | - | - | 0 | - |
| - | - | 1137 | 654.3 | - | - | 0 | - |
| 11 | y | 9024 | 655.3 | 0.0007696 | 1.174 | +1 | 7 |
| - | - | 3818 | 656.3 | - | - | 0 | - |
| - | - | 739.7 | 656.8 | - | - | 0 | - |
| 11 | z | 1953 | 657.3 | 0.000201 | 0.3058 | +1 | 7 |
| - | - | 691.7 | 658.3 | - | - | 0 | - |
| - | - | 812.1 | 660.3 | - | - | 0 | - |
| - | - | 1038 | 661.3 | - | - | 0 | - |
| - | - | 1037 | 664.3 | - | - | 0 | - |
| - | - | 3234 | 666.3 | - | - | 0 | - |
| - | - | 1493 | 667.3 | - | - | 0 | - |
| - | - | 721.2 | 668.3 | - | - | 0 | - |
| - | - | 1665 | 669.3 | - | - | 0 | - |
| - | - | 1.116E+04 | 670.4 | - | - | 0 | - |
| - | - | 4905 | 671.4 | - | - | 0 | - |
| - | - | 1975 | 672.4 | - | - | 0 | - |
| 11 | y | 2.62E+04 | 673.4 | 0.001496 | 2.222 | +1 | 7 |
| - | - | 8402 | 674.4 | - | - | 0 | - |
| - | - | 761.4 | 674.8 | - | - | 0 | - |
| - | - | 1878 | 675.4 | - | - | 0 | - |
| - | - | 2664 | 677.3 | - | - | 0 | - |
| - | - | 1114 | 678.3 | - | - | 0 | - |
| - | - | 2.262E+04 | 679.3 | - | - | 0 | - |
| - | - | 9573 | 680.3 | - | - | 0 | - |
| - | - | 1.415E+04 | 681.3 | - | - | 0 | - |
| - | - | 3197 | 681.8 | - | - | 0 | - |
| - | - | 1.341E+04 | 682.3 | - | - | 0 | - |
| 14 | c | 2802 | 682.8 | 0.0003036 | 0.4447 | +2 | 14 |
| - | - | 6631 | 683.3 | - | - | 0 | - |
| - | - | 1578 | 683.8 | - | - | 0 | - |
| - | - | 2006 | 684.3 | - | - | 0 | - |
| - | - | 2510 | 685.3 | - | - | 0 | - |
| - | - | 1.641E+04 | 686.3 | - | - | 0 | - |
| - | - | 1428 | 689.3 | - | - | 0 | - |
| 14 | c | 896.6 | 691.4 | 0.002351 | 3.4 | +2 | 14 |
| 3 | y | 1776 | 692.3 | 0.0003428 | 0.4952 | +2 | 15 |
| 3 | y | 845.7 | 692.8 | 0.007894 | 11.39 | +2 | 15 |
| - | - | 2478 | 694.3 | - | - | 0 | - |
| - | - | 1923 | 695.4 | - | - | 0 | - |
| 10 | z | 7331 | 696.3 | 0.002913 | 4.183 | +1 | 8 |
| - | - | 4.189E+04 | 697.3 | - | - | 0 | - |
| - | - | 830.6 | 697.9 | - | - | 0 | - |
| - | - | 1.517E+04 | 698.3 | - | - | 0 | - |
| - | - | 5606 | 699.4 | - | - | 0 | - |
| - | - | 1.648E+04 | 700.3 | - | - | 0 | - |
| 3 | y | 6293 | 701.3 | 0.01039 | 14.81 | +2 | 15 |
| - | - | 2397 | 702.3 | - | - | 0 | - |
| - | - | 1255 | 702.8 | - | - | 0 | - |
| - | - | 1826 | 703.3 | - | - | 0 | - |
| - | - | 1239 | 703.8 | - | - | 0 | - |
| - | - | 807.4 | 704.3 | - | - | 0 | - |
| - | - | 1320 | 705.3 | - | - | 0 | - |
| - | - | 842.9 | 707.9 | - | - | 0 | - |
| - | - | 4951 | 709.3 | - | - | 0 | - |
| - | - | 2945 | 710.3 | - | - | 0 | - |
| 15 | c | 1157 | 710.9 | 0.00426 | 5.993 | +2 | 15 |
| 15 | c | 7959 | 711.4 | 0.001944 | 2.733 | +2 | 15 |
| - | - | 1.234E+04 | 711.9 | - | - | 0 | - |
| 10 | y | 2.127E+04 | 712.4 | 0.003177 | 4.46 | +1 | 8 |
| - | - | 5191 | 712.9 | - | - | 0 | - |
| - | - | 6879 | 713.4 | - | - | 0 | - |
| 6 | c | 1.275E+05 | 714.4 | 0.003081 | 4.313 | +1 | 6 |
| - | - | 4.806E+04 | 715.4 | - | - | 0 | - |
| - | - | 1.388E+04 | 716.4 | - | - | 0 | - |
| - | - | 3087 | 717.4 | - | - | 0 | - |
| 15 | c | 8011 | 719.9 | 0.001242 | 1.725 | +2 | 15 |
| - | - | 7057 | 720.4 | - | - | 0 | - |
| - | - | 8889 | 720.9 | - | - | 0 | - |
| - | - | 6124 | 721.4 | - | - | 0 | - |
| - | - | 2152 | 721.9 | - | - | 0 | - |
| - | - | 1601 | 722.4 | - | - | 0 | - |
| - | - | 2400 | 723.4 | - | - | 0 | - |
| - | - | 1.021E+04 | 724.4 | - | - | 0 | - |
| - | - | 8241 | 724.9 | - | - | 0 | - |
| - | - | 7370 | 725.4 | - | - | 0 | - |
| - | - | 1453 | 725.9 | - | - | 0 | - |
| - | - | 2428 | 726.4 | - | - | 0 | - |
| - | - | 1189 | 726.9 | - | - | 0 | - |
| - | - | 3441 | 727.4 | - | - | 0 | - |
| - | - | 1129 | 728.4 | - | - | 0 | - |
| - | - | 1953 | 729.3 | - | - | 0 | - |
| - | - | 1912 | 729.4 | - | - | 0 | - |
| - | - | 4815 | 729.9 | - | - | 0 | - |
| 10 | y | 3.82E+04 | 730.4 | 0.0008457 | 1.158 | +1 | 8 |
| - | - | 1127 | 730.9 | - | - | 0 | - |
| - | - | 1.397E+04 | 731.4 | - | - | 0 | - |
| - | - | 1220 | 731.9 | - | - | 0 | - |
| - | - | 3865 | 732.4 | - | - | 0 | - |
| - | - | 1100 | 733.4 | - | - | 0 | - |
| - | - | 899.5 | 735.4 | - | - | 0 | - |
| - | - | 1103 | 738.4 | - | - | 0 | - |
| - | - | 1.289E+04 | 739.4 | - | - | 0 | - |
| - | - | 9768 | 739.9 | - | - | 0 | - |
| - | - | 9518 | 740.4 | - | - | 0 | - |
| 2 | y | 4653 | 740.9 | 0.006784 | 9.156 | +2 | 16 |
| - | - | 3107 | 741.4 | - | - | 0 | - |
| 2 | z | 1879 | 741.9 | 0.00952 | 12.83 | +2 | 16 |
| - | - | 2816 | 742.4 | - | - | 0 | - |
| - | - | 1192 | 743.4 | - | - | 0 | - |
| - | - | 3069 | 744.9 | - | - | 0 | - |
| - | - | 2820 | 745.4 | - | - | 0 | - |
| - | - | 1282 | 745.9 | - | - | 0 | - |
| - | - | 961.3 | 746.4 | - | - | 0 | - |
| - | - | 720.6 | 747.4 | - | - | 0 | - |
| - | - | 1477 | 747.9 | - | - | 0 | - |
| - | - | 4682 | 748.4 | - | - | 0 | - |
| - | - | 2193 | 748.9 | - | - | 0 | - |
| - | - | 1765 | 749.4 | - | - | 0 | - |
| 2 | y | 869.9 | 749.9 | 0.002051 | 2.734 | +2 | 16 |
| - | - | 1236 | 750.3 | - | - | 0 | - |
| - | - | 1730 | 751.4 | - | - | 0 | - |
| - | - | 2754 | 751.9 | - | - | 0 | - |
| - | - | 1576 | 752.4 | - | - | 0 | - |
| - | - | 4497 | 752.9 | - | - | 0 | - |
| 9 | z | 1.48E+04 | 753.4 | 0.002872 | 3.813 | +1 | 9 |
| - | - | 2.872E+04 | 753.9 | - | - | 0 | - |
| - | - | 2.836E+04 | 754.4 | - | - | 0 | - |
| - | - | 1.38E+04 | 754.9 | - | - | 0 | - |
| - | - | 4814 | 755.4 | - | - | 0 | - |
| - | - | 1997 | 755.9 | - | - | 0 | - |
| - | - | 3088 | 756.4 | - | - | 0 | - |
| - | - | 918.4 | 756.9 | - | - | 0 | - |
| - | - | 4419 | 757.4 | - | - | 0 | - |
| - | - | 1188 | 757.9 | - | - | 0 | - |
| - | - | 3032 | 758.4 | - | - | 0 | - |
| - | - | 1043 | 758.9 | - | - | 0 | - |
| - | - | 2576 | 759.4 | - | - | 0 | - |
| 16 | c | 1.621E+04 | 761.4 | 0.00216 | 2.836 | +2 | 16 |
| 16 | c | 4.457E+04 | 761.9 | 0.001133 | 1.487 | +2 | 16 |
| - | - | 4.598E+04 | 762.4 | - | - | 0 | - |
| - | - | 2.573E+04 | 762.9 | - | - | 0 | - |
| - | - | 1.235E+04 | 763.4 | - | - | 0 | - |
| - | - | 5077 | 763.9 | - | - | 0 | - |
| - | - | 3935 | 764.4 | - | - | 0 | - |
| - | - | 1995 | 764.9 | - | - | 0 | - |
| - | - | 8109 | 765.4 | - | - | 0 | - |
| - | - | 1.415E+04 | 766.4 | - | - | 0 | - |
| - | - | 7124 | 767.4 | - | - | 0 | - |
| - | - | 8822 | 767.9 | - | - | 0 | - |
| - | - | 1.788E+04 | 768.4 | - | - | 0 | - |
| - | - | 5572 | 768.9 | - | - | 0 | - |
| 9 | y | 1.245E+04 | 769.4 | 0.01146 | 14.89 | +1 | 9 |
| - | - | 3194 | 769.9 | - | - | 0 | - |
| 16 | c | 3.01E+04 | 770.4 | 0.003597 | 4.669 | +2 | 16 |
| - | - | 1.889E+04 | 770.9 | - | - | 0 | - |
| 9 | z | 2.442E+04 | 771.4 | 0.008177 | 10.6 | +1 | 9 |
| - | - | 4523 | 771.9 | - | - | 0 | - |
| - | - | 5738 | 772.4 | - | - | 0 | - |
| - | - | 1810 | 773.4 | - | - | 0 | - |
| - | - | 1083 | 774.9 | - | - | 0 | - |
| - | - | 6365 | 775.4 | - | - | 0 | - |
| - | - | 1350 | 776.4 | - | - | 0 | - |
| - | - | 2.925E+04 | 776.9 | - | - | 0 | - |
| - | - | 2.372E+04 | 777.4 | - | - | 0 | - |
| - | - | 1.378E+04 | 777.9 | - | - | 0 | - |
| - | - | 6125 | 778.4 | - | - | 0 | - |
| - | - | 7305 | 778.9 | - | - | 0 | - |
| - | - | 1.382E+04 | 779.4 | - | - | 0 | - |
| - | - | 9069 | 779.9 | - | - | 0 | - |
| - | - | 3545 | 780.4 | - | - | 0 | - |
| - | - | 1008 | 780.9 | - | - | 0 | - |
| - | - | 2829 | 781.4 | - | - | 0 | - |
| - | - | 862.5 | 782.4 | - | - | 0 | - |
| - | - | 1225 | 782.4 | - | - | 0 | - |
| - | - | 2849 | 782.9 | - | - | 0 | - |
| 7 | c | 5674 | 783.4 | 0.007516 | 9.594 | +1 | 7 |
| - | - | 5106 | 783.9 | - | - | 0 | - |
| - | - | 1.651E+04 | 784.4 | - | - | 0 | - |
| - | - | 2139 | 784.9 | - | - | 0 | - |
| - | - | 8528 | 785.4 | - | - | 0 | - |
| - | - | 2051 | 785.9 | - | - | 0 | - |
| - | - | 2946 | 786.4 | - | - | 0 | - |
| - | - | 4822 | 786.9 | - | - | 0 | - |
| 9 | y | 3.219E+04 | 787.4 | 0.0006595 | 0.8376 | +1 | 9 |
| - | - | 2977 | 787.9 | - | - | 0 | - |
| - | - | 1.403E+04 | 788.4 | - | - | 0 | - |
| - | - | 3022 | 788.9 | - | - | 0 | - |
| - | - | 5114 | 789.4 | - | - | 0 | - |
| - | - | 6005 | 791.9 | - | - | 0 | - |
| - | - | 7280 | 792.4 | - | - | 0 | - |
| - | - | 3998 | 792.9 | - | - | 0 | - |
| - | - | 3177 | 793.4 | - | - | 0 | - |
| - | - | 1224 | 793.9 | - | - | 0 | - |
| - | - | 3167 | 794.4 | - | - | 0 | - |
| - | - | 921 | 795.4 | - | - | 0 | - |
| - | - | 1997 | 796.4 | - | - | 0 | - |
| - | - | 6783 | 796.9 | - | - | 0 | - |
| - | - | 1.999E+04 | 797.4 | - | - | 0 | - |
| - | - | 1.575E+04 | 797.9 | - | - | 0 | - |
| - | - | 9160 | 798.4 | - | - | 0 | - |
| - | - | 2751 | 798.9 | - | - | 0 | - |
| - | - | 6019 | 799.4 | - | - | 0 | - |
| - | - | 2973 | 800.4 | - | - | 0 | - |
| 7 | c | 6.435E+04 | 801.4 | 0.0027 | 3.369 | +1 | 7 |
| - | - | 2.513E+04 | 802.4 | - | - | 0 | - |
| - | - | 6884 | 803.4 | - | - | 0 | - |
| - | - | 1307 | 804.9 | - | - | 0 | - |
| - | - | 2.699E+04 | 805.4 | - | - | 0 | - |
| - | - | 8.192E+04 | 805.9 | - | - | 0 | - |
| - | - | 1.035E+05 | 806.4 | - | - | 0 | - |
| - | - | 7.457E+04 | 806.9 | - | - | 0 | - |
| - | - | 3.382E+04 | 807.4 | - | - | 0 | - |
| - | - | 9479 | 807.9 | - | - | 0 | - |
| - | - | 7936 | 808.4 | - | - | 0 | - |
| - | - | 2546 | 809.4 | - | - | 0 | - |
| 8 | z | 5264 | 810.4 | 0.001489 | 1.838 | +1 | 10 |
| - | - | 2889 | 811.4 | - | - | 0 | - |
| - | - | 1660 | 812.4 | - | - | 0 | - |
| - | - | 1629 | 813.4 | - | - | 0 | - |
| - | - | 3237 | 813.9 | - | - | 0 | - |
| - | - | 6.744E+04 | 814.4 | - | - | 0 | - |
| - | - | 1.954E+05 | 814.9 | - | - | 0 | - |
| - | - | 1.533E+05 | 815.4 | - | - | 0 | - |
| - | - | 7.116E+04 | 815.9 | - | - | 0 | - |
| - | - | 3.253E+04 | 816.4 | - | - | 0 | - |
| - | - | 1.281E+04 | 816.9 | - | - | 0 | - |
| - | - | 878 | 817.4 | - | - | 0 | - |
| - | - | 1.024E+04 | 823.4 | - | - | 0 | - |
| - | - | 5556 | 824.4 | - | - | 0 | - |
| - | - | 4198 | 825.4 | - | - | 0 | - |
| 8 | y | 1.166E+04 | 826.4 | 0.003441 | 4.164 | +1 | 10 |
| 8 | y | 5210 | 827.4 | 0.01639 | 19.81 | +1 | 10 |
| 8 | z | 1.889E+04 | 828.4 | 0.00301 | 3.633 | +1 | 10 |
| - | - | 8178 | 829.4 | - | - | 0 | - |
| - | - | 2534 | 830.4 | - | - | 0 | - |
| 8 | c | 902.2 | 840.4 | 0.002002 | 2.382 | +1 | 8 |
| - | - | 9767 | 841.4 | - | - | 0 | - |
| - | - | 4930 | 842.4 | - | - | 0 | - |
| - | - | 3716 | 843.4 | - | - | 0 | - |
| 8 | y | 3.779E+04 | 844.4 | 0.00131 | 1.552 | +1 | 10 |
| - | - | 1.749E+04 | 845.4 | - | - | 0 | - |
| - | - | 5463 | 846.4 | - | - | 0 | - |
| - | - | 1070 | 847.4 | - | - | 0 | - |
| - | - | 5749 | 853.4 | - | - | 0 | - |
| - | - | 4691 | 854.4 | - | - | 0 | - |
| - | - | 2180 | 855.4 | - | - | 0 | - |
| - | - | 5398 | 856.4 | - | - | 0 | - |
| - | - | 3985 | 857.4 | - | - | 0 | - |
| 8 | c | 1.291E+05 | 858.4 | 0.002496 | 2.908 | +1 | 8 |
| - | - | 5.679E+04 | 859.4 | - | - | 0 | - |
| - | - | 1.758E+04 | 860.4 | - | - | 0 | - |
| - | - | 2544 | 861.4 | - | - | 0 | - |
| - | - | 1607 | 862.4 | - | - | 0 | - |
| - | - | 898.9 | 863.4 | - | - | 0 | - |
| - | - | 809.9 | 868.4 | - | - | 0 | - |
| - | - | 1038 | 869.4 | - | - | 0 | - |
| - | - | 2.226E+04 | 871.4 | - | - | 0 | - |
| - | - | 9623 | 872.4 | - | - | 0 | - |
| - | - | 3897 | 873.4 | - | - | 0 | - |
| - | - | 862.2 | 874.4 | - | - | 0 | - |
| - | - | 848.3 | 879.4 | - | - | 0 | - |
| - | - | 9704 | 880.4 | - | - | 0 | - |
| - | - | 4150 | 881.4 | - | - | 0 | - |
| - | - | 8463 | 882.4 | - | - | 0 | - |
| - | - | 5609 | 883.4 | - | - | 0 | - |
| - | - | 2128 | 884.4 | - | - | 0 | - |
| - | - | 912.8 | 885.4 | - | - | 0 | - |
| - | - | 1689 | 887.4 | - | - | 0 | - |
| - | - | 1173 | 894.5 | - | - | 0 | - |
| - | - | 1708 | 895.4 | - | - | 0 | - |
| - | - | 1774 | 896.4 | - | - | 0 | - |
| 9 | c | 1.319E+04 | 897.4 | 0.01394 | 15.54 | +1 | 9 |
| 7 | z | 1.826E+04 | 898.4 | 0.01779 | 19.81 | +1 | 11 |
| - | - | 7696 | 899.4 | - | - | 0 | - |
| - | - | 1775 | 900.4 | - | - | 0 | - |
| - | - | 1138 | 901.4 | - | - | 0 | - |
| - | - | 5644 | 904.4 | - | - | 0 | - |
| - | - | 815.5 | 909.4 | - | - | 0 | - |
| - | - | 1290 | 910.4 | - | - | 0 | - |
| - | - | 1027 | 911.4 | - | - | 0 | - |
| - | - | 6082 | 912.5 | - | - | 0 | - |
| 7 | y | 8832 | 913.4 | 0.01032 | 11.3 | +1 | 11 |
| 7 | y | 6980 | 914.4 | 0.01073 | 11.73 | +1 | 11 |
| 9 | c | 2.026E+05 | 915.4 | 0.005588 | 6.104 | +1 | 9 |
| - | - | 9.739E+04 | 916.4 | - | - | 0 | - |
| - | - | 3.214E+04 | 917.4 | - | - | 0 | - |
| - | - | 5562 | 918.4 | - | - | 0 | - |
| - | - | 5265 | 919.4 | - | - | 0 | - |
| - | - | 3442 | 920.4 | - | - | 0 | - |
| - | - | 871.8 | 921.4 | - | - | 0 | - |
| - | - | 1953 | 923.4 | - | - | 0 | - |
| - | - | 805 | 924.4 | - | - | 0 | - |
| - | - | 1646 | 927.4 | - | - | 0 | - |
| - | - | 1.899E+04 | 928.5 | - | - | 0 | - |
| - | - | 9159 | 929.5 | - | - | 0 | - |
| - | - | 3849 | 930.5 | - | - | 0 | - |
| 7 | y | 3.943E+04 | 931.4 | 0.001329 | 1.427 | +1 | 11 |
| - | - | 2.233E+04 | 932.5 | - | - | 0 | - |
| - | - | 8004 | 933.5 | - | - | 0 | - |
| - | - | 1587 | 934.5 | - | - | 0 | - |
| - | - | 2.214E+04 | 937.4 | - | - | 0 | - |
| - | - | 1.123E+04 | 938.4 | - | - | 0 | - |
| - | - | 1.532E+04 | 939.4 | - | - | 0 | - |
| - | - | 7971 | 940.4 | - | - | 0 | - |
| - | - | 7324 | 941.4 | - | - | 0 | - |
| - | - | 2371 | 942.4 | - | - | 0 | - |
| - | - | 824.9 | 953.4 | - | - | 0 | - |
| 10 | c | 2180 | 954.5 | 0.01283 | 13.44 | +1 | 10 |
| - | - | 9.028E+04 | 955.4 | - | - | 0 | - |
| - | - | 4.599E+04 | 956.4 | - | - | 0 | - |
| - | - | 1.644E+04 | 957.4 | - | - | 0 | - |
| - | - | 3796 | 958.5 | - | - | 0 | - |
| - | - | 4859 | 959.4 | - | - | 0 | - |
| - | - | 1761 | 960.4 | - | - | 0 | - |
| - | - | 1074 | 961.4 | - | - | 0 | - |
| 10 | c | 1.687E+05 | 972.5 | 0.002821 | 2.901 | +1 | 10 |
| - | - | 9.013E+04 | 973.5 | - | - | 0 | - |
| - | - | 3.003E+04 | 974.5 | - | - | 0 | - |
| - | - | 5934 | 975.5 | - | - | 0 | - |
| - | - | 1270 | 976.5 | - | - | 0 | - |
| - | - | 1662 | 982.4 | - | - | 0 | - |
| 6 | w | 1619 | 985.5 | 0.0008354 | 0.8478 | +1 | 12 |
| - | - | 4223 | 987.5 | - | - | 0 | - |
| - | - | 1870 | 988.4 | - | - | 0 | - |
| - | - | 932.8 | 989.4 | - | - | 0 | - |
| - | - | 1.071E+04 | 993.5 | - | - | 0 | - |
| - | - | 7253 | 994.5 | - | - | 0 | - |
| - | - | 2599 | 995.5 | - | - | 0 | - |
| - | - | 946.6 | 1016 | - | - | 0 | - |
| - | - | 3544 | 1018 | - | - | 0 | - |
| - | - | 1044 | 1024 | - | - | 0 | - |
| - | - | 1631 | 1025 | - | - | 0 | - |
| - | - | 2556 | 1025 | - | - | 0 | - |
| 6 | z | 9747 | 1026 | 0.006152 | 5.994 | +1 | 12 |
| - | - | 6095 | 1027 | - | - | 0 | - |
| - | - | 2859 | 1028 | - | - | 0 | - |
| - | - | 1234 | 1029 | - | - | 0 | - |
| - | - | 1676 | 1033 | - | - | 0 | - |
| - | - | 1070 | 1034 | - | - | 0 | - |
| - | - | 1116 | 1034 | - | - | 0 | - |
| - | - | 1296 | 1039 | - | - | 0 | - |
| - | - | 842.2 | 1040 | - | - | 0 | - |
| - | - | 2927 | 1041 | - | - | 0 | - |
| - | - | 8539 | 1042 | - | - | 0 | - |
| - | - | 4717 | 1043 | - | - | 0 | - |
| - | - | 1820 | 1043 | - | - | 0 | - |
| 6 | z | 5.955E+04 | 1044 | 0.001569 | 1.502 | +1 | 12 |
| - | - | 3.067E+04 | 1045 | - | - | 0 | - |
| - | - | 1.192E+04 | 1046 | - | - | 0 | - |
| - | - | 1670 | 1047 | - | - | 0 | - |
| - | - | 8494 | 1051 | - | - | 0 | - |
| - | - | 7411 | 1052 | - | - | 0 | - |
| - | - | 3.824E+04 | 1053 | - | - | 0 | - |
| - | - | 2.663E+04 | 1054 | - | - | 0 | - |
| - | - | 1.072E+04 | 1055 | - | - | 0 | - |
| - | - | 3503 | 1056 | - | - | 0 | - |
| - | - | 1778 | 1057 | - | - | 0 | - |
| - | - | 1553 | 1059 | - | - | 0 | - |
| 6 | y | 3.845E+04 | 1060 | 0.001278 | 1.205 | +1 | 12 |
| - | - | 940.4 | 1061 | - | - | 0 | - |
| - | - | 2.578E+04 | 1061 | - | - | 0 | - |
| - | - | 9501 | 1062 | - | - | 0 | - |
| - | - | 2871 | 1064 | - | - | 0 | - |
| - | - | 2.261E+04 | 1065 | - | - | 0 | - |
| - | - | 1.282E+04 | 1066 | - | - | 0 | - |
| - | - | 6206 | 1067 | - | - | 0 | - |
| - | - | 8370 | 1068 | - | - | 0 | - |
| - | - | 4.074E+04 | 1069 | - | - | 0 | - |
| - | - | 2.383E+04 | 1070 | - | - | 0 | - |
| - | - | 1.035E+04 | 1071 | - | - | 0 | - |
| - | - | 3456 | 1072 | - | - | 0 | - |
| - | - | 2047 | 1073 | - | - | 0 | - |
| - | - | 1311 | 1074 | - | - | 0 | - |
| - | - | 1718 | 1075 | - | - | 0 | - |
| - | - | 988.1 | 1084 | - | - | 0 | - |
| - | - | 2212 | 1085 | - | - | 0 | - |
| 11 | c | 1.392E+05 | 1086 | 0.003511 | 3.234 | +1 | 11 |
| - | - | 8.289E+04 | 1087 | - | - | 0 | - |
| - | - | 3.282E+04 | 1088 | - | - | 0 | - |
| - | - | 7690 | 1089 | - | - | 0 | - |
| - | - | 1126 | 1090 | - | - | 0 | - |
| - | - | 1295 | 1091 | - | - | 0 | - |
| - | - | 1777 | 1095 | - | - | 0 | - |
| - | - | 1228 | 1096 | - | - | 0 | - |
| - | - | 813.4 | 1097 | - | - | 0 | - |
| - | - | 1634 | 1104 | - | - | 0 | - |
| - | - | 5048 | 1113 | - | - | 0 | - |
| - | - | 3330 | 1114 | - | - | 0 | - |
| - | - | 1057 | 1115 | - | - | 0 | - |
| - | - | 7043 | 1122 | - | - | 0 | - |
| - | - | 6572 | 1123 | - | - | 0 | - |
| - | - | 5.71E+04 | 1124 | - | - | 0 | - |
| - | - | 3.97E+04 | 1125 | - | - | 0 | - |
| 5 | z | 1.742E+04 | 1126 | 0.01489 | 13.23 | +1 | 13 |
| - | - | 5924 | 1127 | - | - | 0 | - |
| - | - | 1207 | 1128 | - | - | 0 | - |
| - | - | 1.612E+04 | 1139 | - | - | 0 | - |
| - | - | 5.018E+04 | 1140 | - | - | 0 | - |
| - | - | 3.322E+04 | 1141 | - | - | 0 | - |
| 5 | y | 1.405E+04 | 1142 | 0.01508 | 13.21 | +1 | 13 |
| - | - | 5212 | 1143 | - | - | 0 | - |
| 5 | z | 5.267E+04 | 1144 | 0.002125 | 1.858 | +1 | 13 |
| - | - | 3.405E+04 | 1145 | - | - | 0 | - |
| - | - | 1.554E+04 | 1146 | - | - | 0 | - |
| - | - | 3117 | 1147 | - | - | 0 | - |
| - | - | 1042 | 1151 | - | - | 0 | - |
| - | - | 1896 | 1152 | - | - | 0 | - |
| - | - | 1212 | 1153 | - | - | 0 | - |
| - | - | 1878 | 1155 | - | - | 0 | - |
| - | - | 3503 | 1156 | - | - | 0 | - |
| 12 | c | 2.31E+05 | 1157 | 0.002905 | 2.512 | +1 | 12 |
| - | - | 1.531E+05 | 1158 | - | - | 0 | - |
| - | - | 5.704E+04 | 1159 | - | - | 0 | - |
| 5 | y | 2.776E+04 | 1160 | 0.01477 | 12.74 | +1 | 13 |
| - | - | 1.393E+04 | 1161 | - | - | 0 | - |
| - | - | 6645 | 1162 | - | - | 0 | - |
| - | - | 1915 | 1163 | - | - | 0 | - |
| - | - | 3649 | 1167 | - | - | 0 | - |
| - | - | 2376 | 1168 | - | - | 0 | - |
| - | - | 2061 | 1169 | - | - | 0 | - |
| - | - | 1635 | 1170 | - | - | 0 | - |
| - | - | 811.3 | 1175 | - | - | 0 | - |
| - | - | 899.6 | 1176 | - | - | 0 | - |
| - | - | 1200 | 1180 | - | - | 0 | - |
| - | - | 3174 | 1183 | - | - | 0 | - |
| - | - | 1616 | 1184 | - | - | 0 | - |
| - | - | 792.4 | 1185 | - | - | 0 | - |
| - | - | 3268 | 1186 | - | - | 0 | - |
| - | - | 2005 | 1187 | - | - | 0 | - |
| - | - | 2383 | 1188 | - | - | 0 | - |
| - | - | 2065 | 1189 | - | - | 0 | - |
| - | - | 1.285E+04 | 1193 | - | - | 0 | - |
| - | - | 9908 | 1194 | - | - | 0 | - |
| - | - | 8075 | 1195 | - | - | 0 | - |
| - | - | 4137 | 1196 | - | - | 0 | - |
| - | - | 2503 | 1197 | - | - | 0 | - |
| - | - | 2092 | 1198 | - | - | 0 | - |
| - | - | 1839 | 1199 | - | - | 0 | - |
| 4 | w | 2373 | 1214 | 0.005429 | 4.474 | +1 | 14 |
| - | - | 1948 | 1215 | - | - | 0 | - |
| - | - | 1103 | 1216 | - | - | 0 | - |
| - | - | 3575 | 1217 | - | - | 0 | - |
| - | - | 7912 | 1223 | - | - | 0 | - |
| - | - | 8513 | 1224 | - | - | 0 | - |
| - | - | 7997 | 1225 | - | - | 0 | - |
| - | - | 3058 | 1226 | - | - | 0 | - |
| - | - | 1659 | 1227 | - | - | 0 | - |
| - | - | 906.9 | 1232 | - | - | 0 | - |
| - | - | 851.8 | 1233 | - | - | 0 | - |
| - | - | 3673 | 1234 | - | - | 0 | - |
| - | - | 1842 | 1235 | - | - | 0 | - |
| - | - | 1229 | 1236 | - | - | 0 | - |
| 4 | z | 7262 | 1239 | 0.0005251 | 0.424 | +1 | 14 |
| 4 | z | 6959 | 1240 | 0.02274 | 18.34 | +1 | 14 |
| - | - | 4.062E+04 | 1241 | - | - | 0 | - |
| - | - | 2.994E+04 | 1242 | - | - | 0 | - |
| - | - | 1.111E+04 | 1243 | - | - | 0 | - |
| - | - | 2505 | 1244 | - | - | 0 | - |
| - | - | 5773 | 1250 | - | - | 0 | - |
| - | - | 8639 | 1251 | - | - | 0 | - |
| - | - | 3.925E+04 | 1252 | - | - | 0 | - |
| - | - | 2.739E+04 | 1253 | - | - | 0 | - |
| - | - | 1.158E+04 | 1254 | - | - | 0 | - |
| 4 | y | 5829 | 1255 | 0.0204 | 16.26 | +1 | 14 |
| 4 | y | 2817 | 1256 | 0.003523 | 2.806 | +1 | 14 |
| 4 | z | 1.899E+05 | 1257 | 0.0005805 | 0.462 | +1 | 14 |
| - | - | 1.325E+05 | 1258 | - | - | 0 | - |
| - | - | 5.574E+04 | 1259 | - | - | 0 | - |
| - | - | 1.391E+04 | 1260 | - | - | 0 | - |
| - | - | 2799 | 1261 | - | - | 0 | - |
| - | - | 955.2 | 1266 | - | - | 0 | - |
| - | - | 1.058E+04 | 1267 | - | - | 0 | - |
| 13 | c | 1.911E+05 | 1268 | 0.00405 | 3.195 | +1 | 13 |
| - | - | 1.505E+05 | 1269 | - | - | 0 | - |
| - | - | 7.02E+04 | 1270 | - | - | 0 | - |
| - | - | 2.124E+04 | 1271 | - | - | 0 | - |
| - | - | 6722 | 1272 | - | - | 0 | - |
| 4 | y | 5.786E+04 | 1273 | 4.495E-05 | 0.03532 | +1 | 14 |
| - | - | 4.453E+04 | 1274 | - | - | 0 | - |
| - | - | 1.863E+04 | 1275 | - | - | 0 | - |
| - | - | 6478 | 1276 | - | - | 0 | - |
| - | - | 1188 | 1277 | - | - | 0 | - |
| - | - | 960.4 | 1301 | - | - | 0 | - |
| - | - | 1226 | 1312 | - | - | 0 | - |
| - | - | 1118 | 1313 | - | - | 0 | - |
| - | - | 782.7 | 1315 | - | - | 0 | - |
| - | - | 2626 | 1330 | - | - | 0 | - |
| - | - | 4276 | 1331 | - | - | 0 | - |
| 14 | c | 949.6 | 1365 | 7.123E-05 | 0.05219 | +1 | 14 |
| - | - | 1478 | 1366 | - | - | 0 | - |
| - | - | 2290 | 1367 | - | - | 0 | - |
| 3 | z | 2580 | 1368 | 0.01677 | 12.26 | +1 | 15 |
| 3 | z | 3129 | 1369 | 0.02592 | 18.94 | +1 | 15 |
| - | - | 3358 | 1381 | - | - | 0 | - |
| 14 | c | 1.347E+04 | 1382 | 0.005116 | 3.702 | +1 | 14 |
| - | - | 9653 | 1383 | - | - | 0 | - |
| 3 | y | 6075 | 1384 | 0.02515 | 18.17 | +1 | 15 |
| - | - | 1504 | 1385 | - | - | 0 | - |
| 3 | z | 4.4E+04 | 1386 | 0.00059 | 0.4258 | +1 | 15 |
| - | - | 3.383E+04 | 1387 | - | - | 0 | - |
| - | - | 1.542E+04 | 1388 | - | - | 0 | - |
| - | - | 4818 | 1389 | - | - | 0 | - |
| - | - | 2688 | 1395 | - | - | 0 | - |
| - | - | 3235 | 1396 | - | - | 0 | - |
| - | - | 1812 | 1397 | - | - | 0 | - |
| 3 | y | 6964 | 1402 | 0.000678 | 0.4837 | +1 | 15 |
| - | - | 5114 | 1403 | - | - | 0 | - |
| - | - | 3098 | 1404 | - | - | 0 | - |
| - | - | 1445 | 1412 | - | - | 0 | - |
| 15 | c | 3596 | 1422 | 0.0001523 | 0.1071 | +1 | 15 |
| - | - | 1.09E+04 | 1423 | - | - | 0 | - |
| - | - | 8485 | 1424 | - | - | 0 | - |
| - | - | 3908 | 1425 | - | - | 0 | - |
| - | - | 1637 | 1426 | - | - | 0 | - |
| 15 | c | 4.224E+04 | 1439 | 0.005827 | 4.05 | +1 | 15 |
| - | - | 4.073E+04 | 1440 | - | - | 0 | - |
| - | - | 2.114E+04 | 1441 | - | - | 0 | - |
| - | - | 7070 | 1442 | - | - | 0 | - |
| - | - | 1673 | 1443 | - | - | 0 | - |
| - | - | 1657 | 1462 | - | - | 0 | - |
| - | - | 2278 | 1478 | - | - | 0 | - |
| - | - | 6158 | 1479 | - | - | 0 | - |
| - | - | 5493 | 1480 | - | - | 0 | - |
| 2 | y | 2252 | 1481 | 0.01694 | 11.44 | +1 | 16 |
| 2 | y | 1068 | 1482 | 0.02767 | 18.68 | +1 | 16 |
| - | - | 3128 | 1496 | - | - | 0 | - |
| - | - | 2635 | 1497 | - | - | 0 | - |
| - | - | 1056 | 1498 | - | - | 0 | - |
| 2 | y | 1172 | 1499 | 0.000147 | 0.09807 | +1 | 16 |
| - | - | 1106 | 1507 | - | - | 0 | - |
| - | - | 1109 | 1508 | - | - | 0 | - |
| - | - | 1099 | 1509 | - | - | 0 | - |
| - | - | 2316 | 1513 | - | - | 0 | - |
| - | - | 2264 | 1514 | - | - | 0 | - |
| - | - | 1.235E+04 | 1524 | - | - | 0 | - |
| - | - | 1.015E+04 | 1525 | - | - | 0 | - |
| - | - | 6157 | 1526 | - | - | 0 | - |
| - | - | 1516 | 1527 | - | - | 0 | - |
| - | - | 909.9 | 1528 | - | - | 0 | - |
| 16 | c | 5265 | 1540 | 0.005044 | 3.276 | +1 | 16 |
| - | - | 1.717E+04 | 1541 | - | - | 0 | - |
| - | - | 1.449E+04 | 1542 | - | - | 0 | - |
| - | - | 6422 | 1543 | - | - | 0 | - |
| - | - | 1170 | 1544 | - | - | 0 | - |
| - | - | 1083 | 1545 | - | - | 0 | - |
| - | - | 1298 | 1558 | - | - | 0 | - |
| - | - | 2397 | 1567 | - | - | 0 | - |
| - | - | 2670 | 1568 | - | - | 0 | - |
| - | - | 1858 | 1569 | - | - | 0 | - |
| - | - | 9809 | 1584 | - | - | 0 | - |
| - | - | 1.115E+04 | 1585 | - | - | 0 | - |
| - | - | 6628 | 1586 | - | - | 0 | - |
| - | - | 2956 | 1587 | - | - | 0 | - |
| - | - | 6020 | 1595 | - | - | 0 | - |
| - | - | 5062 | 1596 | - | - | 0 | - |
| - | - | 3932 | 1597 | - | - | 0 | - |
| - | - | 2117 | 1598 | - | - | 0 | - |
| - | - | 1007 | 1599 | - | - | 0 | - |
| - | - | 4213 | 1601 | - | - | 0 | - |
| - | - | 5092 | 1602 | - | - | 0 | - |
| - | - | 3374 | 1603 | - | - | 0 | - |
| - | - | 1609 | 1604 | - | - | 0 | - |
| - | - | 1036 | 1605 | - | - | 0 | - |
| - | - | 2223 | 1611 | - | - | 0 | - |
| - | - | 1.23E+04 | 1612 | - | - | 0 | - |
| - | - | 9.59E+04 | 1613 | - | - | 0 | - |
| - | - | 8.86E+04 | 1614 | - | - | 0 | - |
| - | - | 4.198E+04 | 1615 | - | - | 0 | - |
| - | - | 1.697E+04 | 1616 | - | - | 0 | - |
| - | - | 5374 | 1617 | - | - | 0 | - |
| - | - | 5164 | 1628 | - | - | 0 | - |
| - | - | 2.401E+04 | 1629 | - | - | 0 | - |
| - | - | 7.534E+04 | 1630 | - | - | 0 | - |
| - | - | 6.047E+04 | 1631 | - | - | 0 | - |
| - | - | 2.915E+04 | 1632 | - | - | 0 | - |
| - | - | 1.173E+04 | 1633 | - | - | 0 | - |
| - | - | 4181 | 1634 | - | - | 0 | - |

m/z Charge Intensity FragmentType MassShift Position
120.06594848632812 0 1342.6655
120.08126831054688 0 464.62985
122.85167694091797 0 450.10287
124.0761947631836 0 1151.6576
126.05555725097656 0 508.099
127.08686828613281 0 1064.6996
128.03443908691406 0 1180.7009
128.09478759765625 0 935.5066
128.1075897216797 0 779.7184
129.06625366210938 0 36295.434
129.10296630859375 0 1269.271
130.05026245117188 0 2484.421
130.0641326904297 0 473.84604
130.06967163085938 0 2223.0479
131.0819854736328 0 1658.7119
131.11849975585938 0 486.86102
133.0612335205078 0 3803.8765
133.08665466308594 0 785.70245
143.11831665039062 0 6747.6216
145.0612335205078 0 2620.2703
147.07687377929688 0 14062.9
148.08053588867188 0 1298.8544
148.95469665527344 0 968.9643
152.071044921875 0 4834.6553
153.06671142578125 0 859.21136
153.07421875 0 581.44073
155.08192443847656 0 14694.629
156.08531188964844 0 1420.685
156.1022491455078 0 592.1821
157.0739288330078 0 709.9645
157.09793090820312 0 673.9263
157.1337890625 0 1679.2953
158.3658905029297 0 484.11737
159.076904296875 0 2501.6948
165.0786590576172 0 627.10815
166.08665466308594 0 2940.9062
171.07687377929688 0 2426.631
171.11328125 0 4466.7886
172.07225036621094 0 928.8755
172.10853576660156 0 85352.62
173.092529296875 0 838.9173
173.10598754882812 0 738.51324
173.11190795898438 0 6510.132
181.06124877929688 0 781.9278
181.08351135253906 0 1578.225
181.09764099121094 0 51899.605
182.10121154785156 0 4632.4956
183.07684326171875 0 3525.0974
183.08932495117188 0 1618.7567
183.10638427734375 0 471.51056
183.1131591796875 0 5662.643
184.08477783203125 0 3534.034
185.09251403808594 0 1624.8511
185.12879943847656 0 2953.16
185.16464233398438 0 1017.6965
187.1082305908203 0 1014.2507
189.08750915527344 0 12323.931 y Water loss 15
190.0825958251953 0 730.6811
190.09144592285156 0 608.1024
193.07386779785156 0 1617.0447
197.1288604736328 0 2172.901
199.07188415527344 0 2229.0916
199.10813903808594 0 8380.952
200.1034393310547 0 35961.11
200.13955688476562 0 710.7315
201.10687255859375 0 3310.7097
201.12310791015625 0 1295.0964
202.0826416015625 0 1985.6686
203.10311889648438 0 694.9277
207.09803771972656 0 39696.93 y 15
208.10140991210938 0 2681.5425
209.09259033203125 0 105094.35
210.09837341308594 0 14237.684
210.12399291992188 0 4017.3828
211.0850067138672 0 648.50323
211.09649658203125 0 922.64575
211.10789489746094 0 3402.6934
211.1319122314453 0 11567.0625
212.09295654296875 0 776.42145
212.10353088378906 0 2770.858
212.13905334472656 0 2052.9102
213.12384033203125 0 3250.636
213.1604461669922 0 1597.9734
214.0826416015625 0 491.39136
214.11927795410156 0 664.4662
215.13963317871094 0 2710.7034
217.0826416015625 0 1328.1599
225.05191040039062 0 514.40564
225.11134338378906 0 2870.0195
225.12399291992188 0 2143.2412 y 10
226.11895751953125 0 6234.3765
227.10324096679688 0 7418.5137
227.12161254882812 0 474.1434
227.127197265625 0 792.91174
227.13958740234375 0 7643.473
228.0984344482422 0 1815.0901
228.13482666015625 0 18874.074
229.118896484375 0 4938.3823
230.1220245361328 0 677.05005
233.1656036376953 0 3609.3408
237.11117553710938 0 2228.0405 z 12
238.0828399658203 0 691.0993
238.11912536621094 0 9866.192
241.09397888183594 0 803.49506
242.12709045410156 0 536.1527
242.150390625 0 4523.7593
243.11102294921875 0 635.18463
243.13449096679688 0 7952.1646
243.1473846435547 0 847.56146
244.11537170410156 0 922.1033
244.12982177734375 0 18379.787 c 1
244.1663360595703 0 4474.011
245.10055541992188 0 484.6399
245.13323974609375 0 1737.8562
245.16851806640625 0 576.6537
246.10922241210938 0 6165.6743 y Water loss 14
250.0952911376953 0 793.4389
251.1019287109375 0 587.8355
251.1770782470703 0 649.4744
252.1348419189453 0 2711.5886
254.11416625976562 0 1674.2157
254.13784790039062 0 638.1925
255.12159729003906 0 1563.4844
255.14549255371094 0 3452.6936
256.09381103515625 0 2744.8108
256.1297607421875 0 68544.07
257.1329345703125 0 8269.474 y Water loss 8
257.16143798828125 0 3809.4055
258.13397216796875 0 692.5013
259.1046142578125 0 2596.9358
259.1777038574219 0 1136.2529
260.1607971191406 0 2237.3499
261.1602783203125 0 2221.489
262.6269836425781 0 2437.562
262.64410400390625 0 1025.6356
264.1195068359375 0 4434.2734 y 14
264.13397216796875 0 1086.6455
266.11419677734375 0 2256.3953
267.121826171875 0 4122.6333
268.1295471191406 0 5417.109
270.1454772949219 0 5202.904
270.659912109375 0 676.2944
271.17681884765625 0 1276.5872
271.632568359375 0 7472.9272 y Water loss 11
272.1334533691406 0 2146.9172
273.1562805175781 0 82696.02
274.1034240722656 0 832.43616
274.1416015625 0 1059.7903
274.1593933105469 0 10612.53
275.1607666015625 0 939.2355
275.17694091796875 0 811.30615
280.1539306640625 0 1583.8221
280.63848876953125 0 723.8172 y 11
282.1454772949219 0 4589.1694 y 7
283.1409912109375 0 4404.0933
284.1247863769531 0 3569.9878
284.1611328125 0 17341.428 c Water loss 4
285.1572265625 0 5515.607
291.1569519042969 0 793.31067
292.12969970703125 0 1673.4512
296.1983947753906 0 1061.6914
297.1537780761719 0 1358.3616
297.169189453125 0 2473.023
298.1278381347656 0 8725.352
298.14404296875 0 693.9155
298.1766052246094 0 4567.2104
299.1335754394531 0 1651.4458
299.17218017578125 0 8899.228
300.1185302734375 0 1118.6533
300.1675109863281 0 7605.007
301.15234375 0 866.7425
301.1702575683594 0 1186.0461
301.18780517578125 0 1483.6494
307.11712646484375 0 1289.6072
310.1405334472656 0 49829.35
311.1435546875 0 6638.5938
312.1449890136719 0 2307.455
313.1146240234375 0 1904.5065
313.18817138671875 0 5898.9287
315.1679992675781 0 1033.9111
316.1510314941406 0 1763.4573
316.1986083984375 0 749.0655
319.1697692871094 0 1267.5687
319.66595458984375 0 842.24634
320.1722412109375 0 647.2948
322.16339111328125 0 615.551
324.14434814453125 0 3389.088
324.1949768066406 0 645.9252
325.1513977050781 0 11008.205
326.1349182128906 0 1208.4426
326.1549072265625 0 1342.6495
327.1662292480469 0 555.2619
328.15313720703125 0 644.23755
328.1750793457031 0 11045.662 y Water loss 10
328.1976013183594 0 1624.2397
328.6766662597656 0 4023.898
329.15850830078125 0 1542.7676 w 5
329.1786193847656 0 815.8389
330.1668701171875 0 1070.5148
331.1257629394531 0 3665.6553
333.134521484375 0 520.61194
337.1517028808594 0 3042.431
338.1355895996094 0 70343.77
339.1386413574219 0 11285.629
339.1663818359375 0 2842.8477
339.1889343261719 0 1089.9642
340.1260681152344 0 2171.1375
340.143798828125 0 1776.1085
340.1697082519531 0 1322.6615
340.1878662109375 0 3154.582
340.6708984375 0 818.49493
341.1463317871094 0 1582.907
341.183349609375 0 3279.1455
341.6786193847656 0 664.2936
342.1541442871094 0 23411.477
342.2032775878906 0 3324.259
343.16192626953125 0 122305.81 y Water loss 13
344.1651611328125 0 21436.264
344.1935119628906 0 2536.0557
345.1441345214844 0 653.50793
345.16851806640625 0 2739.072
347.6799621582031 0 1720.7386
349.1527404785156 0 1059.1315
349.1744079589844 0 2698.4688 z Ammonia loss 9
349.6758117675781 0 1146.3091
350.15704345703125 0 825.9599
351.14373779296875 0 1260.4624
351.1658630371094 0 1279.3627
352.1480407714844 0 831.33636
353.1464538574219 0 4401.0005
354.1783447265625 0 7937.1675
355.1617126464844 0 7945.512 c Water loss 2
355.1983642578125 0 19790.66
356.1458435058594 0 6568.8516
356.1665954589844 0 939.8964
356.1962890625 0 6845.7236
356.68585205078125 0 4168.1816 y Water loss 9
357.150390625 0 2098.9543
357.1806945800781 0 1060.9845 y Ammonia loss 9
357.6878356933594 0 660.43365 c 5
359.1566467285156 0 6005.946
360.16455078125 0 9463.599
361.1724548339844 0 737194.75 y 13
362.17547607421875 0 123623.74
363.1799011230469 0 18375.105
364.1820983886719 0 1747.2207
365.16986083984375 0 1059.8484
366.1776428222656 0 2267.2495
367.16180419921875 0 5431.0845
367.1834411621094 0 720.2536
368.16900634765625 0 6748.4507
368.1930847167969 0 1312.2303
369.1757507324219 0 3955.7666
369.68896484375 0 1198.3154
370.1365661621094 0 4891.4243
370.2096862792969 0 4075.9475
371.1370544433594 0 711.69434
371.15704345703125 0 7435.436
371.2055969238281 0 4246.065
372.162841796875 0 1080.9688
372.1884765625 0 6073.851
373.1722717285156 0 10361.706 c 2
373.21502685546875 0 1256.8589
374.176513671875 0 1792.0566
374.67987060546875 0 5486.379
375.1809387207031 0 2157.473
375.68292236328125 0 620.9916
376.190185546875 0 1142.6321
379.1621398925781 0 9361.078
380.1644287109375 0 1690.0708
383.6853332519531 0 5363.7183
384.1642150878906 0 591.3356
384.1883239746094 0 13668.106
384.6868896484375 0 1433.2324
385.195068359375 0 6349.338 y Water loss 8
385.6977233886719 0 1858.7026
386.2027893066406 0 2221.7742 c 11
388.14703369140625 0 4776.8403
389.1676025390625 0 2059.616
391.19720458984375 0 651.1155
391.69671630859375 0 1097.3485
391.7255859375 0 935.8138
392.69061279296875 0 1893.1254
393.1912536621094 0 610.2714
394.150390625 0 1129.228
394.19952392578125 0 1245.671 y 8
396.188720703125 0 1002.065
397.1727294921875 0 57915.36
398.17596435546875 0 12751.291
399.182861328125 0 2664.671
399.2351989746094 0 829.89844
401.2154235839844 0 7996.9263
402.218017578125 0 1297.9673
403.1905212402344 0 7115.8496
403.6924133300781 0 2915.5217
404.1922912597656 0 873.6046
404.7021179199219 0 1002.4345
405.1630554199219 0 1554.2352
405.2120361328125 0 1890.1072
408.1882019042969 0 1132.5966
409.2206726074219 0 1777.9431
410.1775207519531 0 912.1266
410.20416259765625 0 4889.697
411.2074890136719 0 1571.4175
412.1947326660156 0 9993.108
412.2206726074219 0 19411.65
412.6976013183594 0 7650.1875
413.1983947753906 0 1108.5856
413.2231750488281 0 4420.3213
413.7068176269531 0 5351.076 y Water loss 7
414.2273254394531 0 943.48315
415.18341064453125 0 14407.863 w 12
416.1865539550781 0 2918.663
420.1898498535156 0 716.55365
421.2015380859375 0 2638.0378
421.7022399902344 0 1410.2128
423.22491455078125 0 32896.133
424.22845458984375 0 9211.511
425.2160339355469 0 2214.1228
425.2366943359375 0 600.0404
426.1885986328125 0 1161.5862
426.2205810546875 0 945.82916
427.23126220703125 0 8427.607
428.2110900878906 0 1184.078
430.1942443847656 0 1918.5715
431.70172119140625 0 1708.966
432.20135498046875 0 612.64667
433.2089538574219 0 2681.465
433.23907470703125 0 695.42975
436.18353271484375 0 4022.025
436.2119445800781 0 656.4712
437.1888427734375 0 972.703
437.21783447265625 0 1495.0935
438.1741943359375 0 931.6787
439.2568054199219 0 3680.651
440.2174377441406 0 623.4236
440.7070007324219 0 6542.4536
441.2079772949219 0 2265.8362
441.2361145019531 0 3199.0771
442.24176025390625 0 2189.4346
443.22601318359375 0 3206.341
443.2557373046875 0 766.67255
446.2213439941406 0 717.23773
448.21844482421875 0 2640.8145
449.7124938964844 0 7605.692 z Ammonia loss 6
450.2141418457031 0 5667.7866
450.7156066894531 0 1519.8737
451.2198486328125 0 55481.383
451.8931884765625 0 1484.8917
452.2228088378906 0 13111.378
452.2525634765625 0 2569.7278
453.2118225097656 0 17906.586
454.1918640136719 0 3808.8032
454.2179260253906 0 5640.4307
455.1973571777344 0 2713.335 z Water loss 12
455.2254333496094 0 12700.634
455.7238464355469 0 676.03784
456.2103271484375 0 12199.325
457.2193603515625 0 5319.4927 y Water loss 6
457.72528076171875 0 1657.2216
458.2226257324219 0 1127.1776 c 8
460.21392822265625 0 1951.7638
460.25128173828125 0 937.24304
467.2353210449219 0 2699.41
468.2368469238281 0 5140.4443
469.2320861816406 0 20215.994
469.2793884277344 0 1328.0723
469.71978759765625 0 2323.6946
470.209228515625 0 938.69385
470.2363586425781 0 4016.2488
471.2210388183594 0 32129.732 y Water loss 12
472.2227783203125 0 5590.9766
472.25439453125 0 1123.4746
473.2105407714844 0 2425.9358 z 12
473.2391662597656 0 1526.5269
474.2170104980469 0 1096.8468
476.25469970703125 0 760.59296
478.2236633300781 0 9359.102
478.7251281738281 0 4647.7764
479.2262268066406 0 2097.303
483.2206726074219 0 6658.821
484.221435546875 0 1429.2198
484.25335693359375 0 4918.1626
485.2521667480469 0 4928.6006
486.25714111328125 0 11675.6875 c 3
487.21575927734375 0 3675.9028
487.2619934082031 0 2438.4568
488.22296142578125 0 5044.4644
488.2587585449219 0 898.28815
489.2314453125 0 94354.92 y 12
490.23468017578125 0 22097.82
491.2443542480469 0 11374.429
492.2493591308594 0 2716.698
493.26641845703125 0 4586.0566
494.26953125 0 1244.3795
498.2569580078125 0 1083.1772
501.231201171875 0 6843.2295
502.235107421875 0 1419.4661
502.75640869140625 0 1060.9707
503.7351989746094 0 1080.326
504.240234375 0 1070.8336
504.27947998046875 0 2967.2961
507.22332763671875 0 1126.9557
508.26507568359375 0 2733.4016
508.3023681640625 0 886.45435
509.23626708984375 0 4481.8223
509.2725524902344 0 2885.3538
510.2385559082031 0 1058.1509
510.2763671875 0 899.38135
511.76287841796875 0 1085.3307
512.265625 0 742.1823
512.7388916015625 0 2614.5981
513.240234375 0 1703.2023
517.1904907226562 0 741.83203
518.7578125 0 685.21356
520.7675170898438 0 1178.2001
521.7455444335938 0 3287.2744 y Water loss 5
522.2503051757812 0 1403.3054
522.29345703125 0 17115.002
522.7463989257812 0 883.2096 z 5
523.2960815429688 0 5892.791
524.2476196289062 0 8270.853
524.2907104492188 0 1985.1256
525.2518310546875 0 2589.5593
525.2952270507812 0 1531.3243
525.7598266601562 0 4259.4
526.2590942382812 0 2657.3274
526.3003540039062 0 2416.8284
527.2086181640625 0 832.471
527.2544555664062 0 771.46857
527.305908203125 0 670.3683
528.2650756835938 0 677.9829
532.2783203125 0 3873.1938
533.279052734375 0 1031.3899
533.7966918945312 0 676.28687
534.2804565429688 0 5808.8955 c Water loss 10
534.765380859375 0 12391.607
534.8064575195312 0 1002.49097
535.2689208984375 0 6519.44
535.768798828125 0 2886.622
538.2760620117188 0 1878.6082
540.3042602539062 0 11439.831
541.259033203125 0 760.7473
541.30859375 0 4114.0723
542.2579956054688 0 41613.637 y Water loss 11
542.8057861328125 0 2109.0325
543.25927734375 0 9780.423
543.3119506835938 0 3391.872
543.8091430664062 0 1087.6335
544.2683715820312 0 1846.2947
550.2884521484375 0 82767.695
551.29150390625 0 24969.113
552.2825927734375 0 21743.838
553.2846069335938 0 13440.274
554.2579956054688 0 4386.9697
554.2951049804688 0 2087.3467
555.2654418945312 0 17315.555
555.3101196289062 0 1052.0292
556.2725830078125 0 14402.565
557.275146484375 0 3161.4197
558.2830200195312 0 2355.8574
559.261474609375 0 1682.0271
559.3290405273438 0 903.4888
560.2684326171875 0 37458.8 y 11
561.2739868164062 0 16058.101
561.7803955078125 0 3480.1562
562.2799072265625 0 12740.895
562.7764892578125 0 1709.7439
563.28515625 0 3608.0693
565.287109375 0 3818.676
566.2922973632812 0 2329.794
567.2911987304688 0 2133.2776
568.2991333007812 0 59533.816
569.3020629882812 0 19236.113
570.2849731445312 0 14332.655
570.7855834960938 0 8918.112
571.2843627929688 0 17393.898 y Water loss 4
571.7811889648438 0 3374.323 y Ammonia loss 4
572.2726440429688 0 7993.281 z 4
573.2723999023438 0 1706.4789
578.2621459960938 0 634.5446
578.79638671875 0 2044.8102 c 11
579.2974853515625 0 972.32996
580.30712890625 0 714.38727
581.306640625 0 1787.7462
583.3206787109375 0 3725.566
584.318359375 0 1510.704
585.3257446289062 0 52198.22 c 4
586.3284912109375 0 17844.967
587.3319091796875 0 3334.8352
588.2672729492188 0 720.0575
589.2958984375 0 743.88666
591.3021240234375 0 8750.506
592.3056640625 0 3052.3254
595.2889404296875 0 3178.532
596.2703247070312 0 7617.3413
597.2736206054688 0 1617.5333
598.2764892578125 0 804.0213
598.3340454101562 0 810.6272
599.8074340820312 0 1753.6335
600.2985229492188 0 4258.001
601.3059692382812 0 827.90375
609.31494140625 0 2116.4863
610.3173828125 0 2028.9265
611.341064453125 0 18511.607
612.3439331054688 0 6850.7246
613.351318359375 0 2019.4458
614.2792358398438 0 21202.309 w 10
615.283203125 0 7173.4624
616.2722778320312 0 684.67615
618.8169555664062 0 1071.3398
619.3162231445312 0 1069.9281
621.3115844726562 0 1463.5159
622.3097534179688 0 6016.038
623.31396484375 0 1819.4663
625.3225708007812 0 3689.9736
625.80810546875 0 2307.0247
626.3154907226562 0 1989.3357
626.8124389648438 0 2376.4795
627.263671875 0 1593.7938
627.820068359375 0 2334.4336 y Water loss 3
628.3226928710938 0 2034.6343 y Ammonia loss 3
630.8142700195312 0 897.5476
632.3380126953125 0 1149.351
633.810791015625 0 814.3419 c Water loss 12
634.3131713867188 0 10405.549 c Ammonia loss 12
634.8151245117188 0 5542.9
635.3175659179688 0 3284.5784
635.8172607421875 0 1738.2463
636.8282470703125 0 791.045 y 3
637.3290405273438 0 1789.27
638.3279418945312 0 3375.6504
639.3255615234375 0 3919.2253 z Water loss 10
640.3318481445312 0 2182.5105
641.3369140625 0 939.98474
642.3541870117188 0 806.11383
643.3196411132812 0 4327.115
644.3206176757812 0 1556.3588
646.3570556640625 0 2543.1506
646.8267822265625 0 759.24927
648.3248901367188 0 886.8731
652.32275390625 0 2294.3252
653.3236694335938 0 1680.7579
654.3145751953125 0 1136.5289
655.3417358398438 0 9023.501 y Water loss 10
656.3427124023438 0 3818.496
656.8346557617188 0 739.73224
657.3330078125 0 1953.3752 z 10
658.344482421875 0 691.74274
660.3291625976562 0 812.11926
661.3229370117188 0 1037.9108
664.3404541015625 0 1037.1542
666.3341064453125 0 3234.2695
667.34130859375 0 1492.9724
668.3345947265625 0 721.19556
669.3472290039062 0 1664.565
670.3534545898438 0 11160.233
671.353271484375 0 4905.1357
672.3657836914062 0 1975.0698
673.35302734375 0 26195.379 y 10
674.3540649414062 0 8402.48
674.8364868164062 0 761.4296
675.35498046875 0 1878.4542
677.3355102539062 0 2664.0264
678.3281860351562 0 1114.061
679.3309936523438 0 22621.652
680.3336791992188 0 9572.825
681.3245849609375 0 14146.465
681.8345947265625 0 3197.0225
682.330322265625 0 13407.65
682.8408203125 0 2801.8892 c Ammonia loss 13
683.3355102539062 0 6631.4976
683.8414306640625 0 1577.9982
684.313720703125 0 2005.8516
685.3157958984375 0 2510.3215
686.3428344726562 0 16411.355
689.3473510742188 0 1428.2733
691.3514404296875 0 896.55035 c 13
692.3408203125 0 1776.2856 y Water loss 2
692.841064453125 0 845.6606 y Ammonia loss 2
694.3458862304688 0 2478.4941
695.3513793945312 0 1922.5298
696.3466186523438 0 7331.1997 z Water loss 9
697.3423461914062 0 41894.027
697.85546875 0 830.56146
698.3458251953125 0 15172.43
699.3514404296875 0 5605.931
700.3323364257812 0 16481.506
701.3360595703125 0 6292.931 y 2
702.3387451171875 0 2396.8062
702.8453979492188 0 1255.4795
703.345703125 0 1826.357
703.8482666015625 0 1238.6686
704.3477172851562 0 807.42804
705.3429565429688 0 1319.7452
707.876708984375 0 842.92365
709.34130859375 0 4950.922
710.3468017578125 0 2944.6853
710.85498046875 0 1157.3179 c Water loss 14
711.3493041992188 0 7959.2324 c Ammonia loss 14
711.8534545898438 0 12342.246
712.3592529296875 0 21268.426 y Water loss 9
712.8580322265625 0 5190.7744
713.3651123046875 0 6878.674
714.3682861328125 0 127525.63 c 5
715.3707275390625 0 48061.168
716.3741455078125 0 13875.429
717.378173828125 0 3086.823
719.86328125 0 8010.9043 c 14
720.3643188476562 0 7057.405
720.8689575195312 0 8889.244
721.3704833984375 0 6124.0117
721.877197265625 0 2152.0515
722.3720703125 0 1600.7151
723.3565673828125 0 2400.4348
724.3590087890625 0 10211.073
724.8601684570312 0 8240.967
725.3600463867188 0 7369.557
725.8589477539062 0 1453.4329
726.3609008789062 0 2427.901
726.85791015625 0 1188.7286
727.3724365234375 0 3440.5938
728.3760986328125 0 1129.12
729.341064453125 0 1952.5894
729.4025268554688 0 1911.618
729.8838500976562 0 4815.4707
730.3738403320312 0 38202.508 y 9
730.8715209960938 0 1127.2175
731.3766479492188 0 13970.921
731.8646850585938 0 1220.1252
732.3815307617188 0 3864.8936
733.3742065429688 0 1100.2406
735.3516235351562 0 899.4642
738.3694458007812 0 1103.3586
739.3729858398438 0 12888.465
739.8742065429688 0 9767.511
740.371337890625 0 9517.906
740.8743286132812 0 4652.9844 y Water loss 1
741.3748168945312 0 3107.1433
741.8729858398438 0 1879.1616 z 1
742.384521484375 0 2815.5679
743.3865966796875 0 1191.7635
744.8612670898438 0 3069.3806
745.3606567382812 0 2819.7776
745.8610229492188 0 1281.6249
746.355712890625 0 961.25055
747.3735961914062 0 720.58496
747.8797607421875 0 1477.2634
748.3563842773438 0 4682.0557
748.883056640625 0 2193.2053
749.3578491210938 0 1765.2024
749.8748779296875 0 869.9019 y 1
750.3468017578125 0 1236.4176
751.3634643554688 0 1730.414
751.8665161132812 0 2754.1504
752.3744506835938 0 1576.1442
752.8707275390625 0 4496.8306
753.3680419921875 0 14798.22 z Water loss 8
753.8660888671875 0 28716.582
754.3671264648438 0 28359.912
754.866943359375 0 13802.468
755.3697509765625 0 4814.4766
755.8743896484375 0 1997.0444
756.3823852539062 0 3088.0952
756.8823852539062 0 918.3671
757.3659057617188 0 4418.54
757.898193359375 0 1188.1721
758.3659057617188 0 3031.5872
758.8692016601562 0 1042.5543
759.3648681640625 0 2576.4841
761.3809204101562 0 16209.3 c Water loss 15
761.876220703125 0 44570.85 c Ammonia loss 15
762.3775634765625 0 45980.414
762.8793334960938 0 25731.602
763.380859375 0 12348.148
763.8734741210938 0 5076.806
764.3697509765625 0 3934.759
764.8689575195312 0 1995.0432
765.4025268554688 0 8108.9644
766.3642578125 0 14147.33
767.3670654296875 0 7124.1294
767.8739013671875 0 8821.876
768.36279296875 0 17877.83
768.8765869140625 0 5571.525
769.3724365234375 0 12450.156 y Water loss 8
769.8787231445312 0 3193.9673
770.384765625 0 30096.912 c 15
770.8876342773438 0 18891.432
771.3839111328125 0 24424.938 z 8
771.8931274414062 0 4522.9956
772.3834838867188 0 5738.0405
773.3842163085938 0 1809.5931
774.8841552734375 0 1082.8002
775.3985595703125 0 6364.5757
776.3857421875 0 1349.7905
776.8795166015625 0 29251.777
777.3810424804688 0 23718.807
777.88037109375 0 13778.808
778.3776245117188 0 6125.237
778.885498046875 0 7304.595
779.3851318359375 0 13818.69
779.8843383789062 0 9069.058
780.385986328125 0 3544.8435
780.9055786132812 0 1008.40326
781.3854370117188 0 2829.221
782.3748168945312 0 862.4679
782.44775390625 0 1225.144
782.8906860351562 0 2849.204
783.3853149414062 0 5673.9106 c Water loss 6
783.887939453125 0 5105.9053
784.3760986328125 0 16508.408
784.8824462890625 0 2138.5608
785.379638671875 0 8528.119
785.8862915039062 0 2051.2651
786.387939453125 0 2946.245
786.8707885742188 0 4822.4316
787.393798828125 0 32193.791 y 8
787.8739013671875 0 2977.033
788.3944702148438 0 14031.373
788.876220703125 0 3021.5676
789.3941650390625 0 5114.4126
791.89599609375 0 6005.0015
792.3915405273438 0 7279.6206
792.8961791992188 0 3997.793
793.3822631835938 0 3176.554
793.8711547851562 0 1223.6282
794.4075317382812 0 3166.8923
795.3825073242188 0 920.9538
796.3831787109375 0 1996.8049
796.887939453125 0 6782.7603
797.3834228515625 0 19991.482
797.8841552734375 0 15746.538
798.385986328125 0 9160.154
798.8863525390625 0 2750.84
799.3990478515625 0 6019.336
800.3998413085938 0 2972.5347
801.4006958007812 0 64347.848 c 6
802.4032592773438 0 25125.195
803.4054565429688 0 6884.055
804.8873291015625 0 1306.7023
805.3873291015625 0 26994.277
805.893798828125 0 81922.04
806.3914184570312 0 103509.66
806.891357421875 0 74571.55
807.3873901367188 0 33821.137
807.8955078125 0 9478.936
808.3821411132812 0 7935.92
809.381103515625 0 2545.5164
810.3881225585938 0 5264.052 z Water loss 7
811.38916015625 0 2888.544
812.4107055664062 0 1659.7784
813.3911743164062 0 1628.8923
813.8908081054688 0 3237.096
814.3967895507812 0 67437.8
814.8987426757812 0 195367.9
815.4002685546875 0 153304.56
815.9019165039062 0 71159.11
816.407470703125 0 32534.941
816.9146118164062 0 12808.482
817.403076171875 0 877.99475
823.3840942382812 0 10240.943
824.387451171875 0 5556.0806
825.3814697265625 0 4198.4326
826.4019165039062 0 11658.348 y Water loss 7
827.40576171875 0 5210.189 y Ammonia loss 7
828.4002075195312 0 18892.64 z 7
829.4024047851562 0 8177.8276
830.40380859375 0 2533.5774
840.4122924804688 0 902.1982 c Water loss 7
841.3941040039062 0 9766.553
842.4014892578125 0 4929.9517
843.41455078125 0 3715.561
844.4146118164062 0 37789.17 y 7
845.4180908203125 0 17494.266
846.4223022460938 0 5463.4644
847.4210815429688 0 1069.7203
853.4015502929688 0 5749.1133
854.4011840820312 0 4691.3203
855.4031982421875 0 2179.5637
856.419921875 0 5397.7397
857.41943359375 0 3984.6575
858.42236328125 0 129106.95 c 7
859.4248046875 0 56787.973
860.4273681640625 0 17582.43
861.4303588867188 0 2543.8394
862.39306640625 0 1606.7253
863.4006958007812 0 898.8914
868.4008178710938 0 809.9355
869.4249877929688 0 1038.1372
871.4290161132812 0 22262.621
872.4300537109375 0 9622.851
873.43017578125 0 3896.789
874.4332885742188 0 862.2049
879.412109375 0 848.3394
880.405029296875 0 9703.785
881.4083251953125 0 4149.89
882.4005737304688 0 8463.1875
883.4033203125 0 5608.923
884.4052734375 0 2128.297
885.4053344726562 0 912.7594
887.4190673828125 0 1688.7498
894.4725341796875 0 1173.1201
895.4249267578125 0 1708.2161
896.4486083984375 0 1774.4866
897.4218139648438 0 13185.753 c Water loss 8
898.4204711914062 0 18258.688 z Ammonia loss 6
899.4234619140625 0 7695.9985
900.427001953125 0 1775.3914
901.4239501953125 0 1138.3134
904.4417114257812 0 5643.8896
909.433837890625 0 815.48303
910.433349609375 0 1289.5881
911.4459228515625 0 1027.0103
912.4793090820312 0 6081.6777
913.4270629882812 0 8831.867 y Water loss 6
914.43212890625 0 6980.0254 y Ammonia loss 6
915.4407348632812 0 202563.5 c 8
916.4437866210938 0 97387.89
917.44580078125 0 32136.527
918.447509765625 0 5562.232
919.419189453125 0 5264.935
920.4189453125 0 3441.6519
921.4135131835938 0 871.7601
923.4213256835938 0 1953.4062
924.4249267578125 0 804.9624
927.444091796875 0 1645.7109
928.4509887695312 0 18991.293
929.4537353515625 0 9158.93
930.4533081054688 0 3849.4634
931.4492797851562 0 39428.105 y 6
932.4522705078125 0 22333.678
933.4559936523438 0 8003.625
934.4613647460938 0 1586.8823
937.4279174804688 0 22138.982
938.4314575195312 0 11225.08
939.4244995117188 0 15323.245
940.4247436523438 0 7970.5986
941.4324951171875 0 7324.384
942.4386596679688 0 2370.884
953.427001953125 0 824.93585
954.4443969726562 0 2179.7734 c Water loss 9
955.4389038085938 0 90282.734
956.44189453125 0 45987.887
957.4461059570312 0 16438.66
958.4515991210938 0 3795.959
959.4459228515625 0 4858.847
960.4483642578125 0 1761.3643
961.4478149414062 0 1074.1476
972.4649658203125 0 168679.52 c 9
973.4671630859375 0 90130.8
974.4696655273438 0 30034.29
975.4701538085938 0 5933.6943
976.4742431640625 0 1269.5066
982.43896484375 0 1661.5027
985.4593505859375 0 1619.3574 w 5
987.4894409179688 0 4223.2046
988.4173583984375 0 1869.9592
989.4181518554688 0 932.81165
993.4901733398438 0 10708.545
994.4930419921875 0 7252.9937
995.4971313476562 0 2598.8076
1015.504638671875 0 946.6012
1017.5255126953125 0 3543.6248
1023.5247192382812 0 1043.5918
1024.521484375 0 1631.1672
1025.4979248046875 0 2555.985
1026.4674072265625 0 9746.756 z Water loss 5
1027.4697265625 0 6095.0938
1028.4703369140625 0 2859.0312
1029.479248046875 0 1234.0037
1032.5050048828125 0 1676.238
1033.5025634765625 0 1070.3983
1034.490478515625 0 1116.3472
1038.5054931640625 0 1296.0262
1039.518310546875 0 842.20654
1040.5284423828125 0 2927.1768
1041.5322265625 0 8538.708
1042.5262451171875 0 4717.426
1043.4923095703125 0 1819.7899
1044.473388671875 0 59551.492 z 5
1045.4764404296875 0 30668.133
1046.478759765625 0 11917.507
1047.4827880859375 0 1669.9495
1050.5133056640625 0 8494.457
1051.5106201171875 0 7411.271
1052.5047607421875 0 38235.71
1053.5074462890625 0 26633.807
1054.5106201171875 0 10717.938
1055.512939453125 0 3502.9543
1056.5225830078125 0 1777.9342
1059.4827880859375 0 1553.0801
1060.4918212890625 0 38450.066 y 5
1061.364990234375 0 940.3581
1061.49560546875 0 25777.22
1062.498779296875 0 9501.166
1063.5025634765625 0 2870.693
1064.5267333984375 0 22610.547
1065.5311279296875 0 12822.6455
1066.5325927734375 0 6205.572
1067.516845703125 0 8370.216
1068.5238037109375 0 40743.402
1069.5274658203125 0 23832.5
1070.529541015625 0 10352.602
1071.53515625 0 3455.7917
1072.5201416015625 0 2046.7357
1073.517578125 0 1311.4136
1074.523681640625 0 1718.2054
1083.5516357421875 0 988.10376
1084.5328369140625 0 2212.148
1085.54833984375 0 139180.89 c 10
1086.551513671875 0 82889.88
1087.5555419921875 0 32819.23
1088.556396484375 0 7690.091
1089.56396484375 0 1126.4163
1090.531982421875 0 1295.4381
1094.5604248046875 0 1776.6877
1095.560546875 0 1228.313
1096.5682373046875 0 813.36444
1103.5303955078125 0 1634.3652
1112.57373046875 0 5048.2944
1113.5743408203125 0 3330.4216
1114.584228515625 0 1057.0455
1121.54638671875 0 7042.652
1122.548583984375 0 6572.2524
1123.540771484375 0 57097.69
1124.544189453125 0 39701.254
1125.5445556640625 0 17421.727 z Water loss 4
1126.5455322265625 0 5923.541
1127.5408935546875 0 1207.1863
1138.554443359375 0 16120.191
1139.558349609375 0 50175.785
1140.5615234375 0 33217.32
1141.5634765625 0 14052.486 y Water loss 4
1142.5606689453125 0 5212.007
1143.5423583984375 0 52669.613 z 4
1144.5447998046875 0 34048.402
1145.55712890625 0 15544.6
1146.5518798828125 0 3116.9639
1150.5972900390625 0 1042.4045
1151.5899658203125 0 1896.0706
1152.5880126953125 0 1212.2261
1154.579345703125 0 1878.4948
1155.5712890625 0 3503.0046
1156.5860595703125 0 230971.36 c 11
1157.588623046875 0 153069.36
1158.5908203125 0 57042.72
1159.57373046875 0 27760.725 y 4
1160.567626953125 0 13925.474
1161.56640625 0 6645.2573
1162.5701904296875 0 1914.8195
1166.5594482421875 0 3648.826
1167.5635986328125 0 2376.0002
1168.58984375 0 2060.9558
1169.6134033203125 0 1635.0981
1174.58251953125 0 811.2923
1175.5823974609375 0 899.5565
1179.5863037109375 0 1199.5768
1182.599609375 0 3173.9617
1183.6031494140625 0 1615.8517
1184.5869140625 0 792.37646
1185.595703125 0 3268.4255
1186.5831298828125 0 2005.127
1187.5556640625 0 2382.5073
1188.5574951171875 0 2064.7473
1192.5860595703125 0 12848.043
1193.589111328125 0 9908.434
1194.5902099609375 0 8075.294
1195.593017578125 0 4136.9634
1196.5980224609375 0 2503.2576
1197.60546875 0 2091.6733
1198.6119384765625 0 1838.6254
1213.574951171875 0 2373.0876 w 3
1214.5738525390625 0 1947.5469
1215.58740234375 0 1102.6342
1216.6214599609375 0 3575.0437
1222.6187744140625 0 7911.7646
1223.61865234375 0 8512.946
1224.61962890625 0 7997.4634
1225.6202392578125 0 3057.7932
1226.6234130859375 0 1658.6898
1231.5986328125 0 906.91925
1232.619384765625 0 851.7945
1233.59130859375 0 3672.6216
1234.58984375 0 1841.5454
1235.595703125 0 1229.1704
1238.6142578125 0 7262.072 z Water loss 3
1239.6204833984375 0 6958.7075 z Ammonia loss 3
1240.62890625 0 40624.55
1241.6329345703125 0 29936.934
1242.6370849609375 0 11114.308
1243.63525390625 0 2505.1497
1249.6051025390625 0 5773.4995
1250.606689453125 0 8639.153
1251.60009765625 0 39247.54
1252.6019287109375 0 27386.113
1253.605224609375 0 11576.703
1254.612060546875 0 5829.262 y Water loss 3
1255.6199951171875 0 2816.6497 y Ammonia loss 3
1256.6248779296875 0 189876.27 z 3
1257.62744140625 0 132511.73
1258.630126953125 0 55743.41
1259.6329345703125 0 13909.477
1260.6361083984375 0 2799.002
1265.602783203125 0 955.2064
1266.6085205078125 0 10582.922
1267.616943359375 0 191101.6 c Ammonia loss 12
1268.6204833984375 0 150518.5
1269.6234130859375 0 70201.195
1270.6273193359375 0 21235.2
1271.631591796875 0 6721.5615
1272.64306640625 0 57862.445 y 3
1273.646484375 0 44526.67
1274.64794921875 0 18627.03
1275.651123046875 0 6477.877
1276.650390625 0 1187.8213
1300.6468505859375 0 960.41144
1311.652587890625 0 1225.513
1312.6636962890625 0 1118.1538
1314.6505126953125 0 782.70233
1329.6051025390625 0 2625.6938
1330.654052734375 0 4276.249
1364.673828125 0 949.5944 c Ammonia loss 13
1365.673828125 0 1478.2716
1366.6749267578125 0 2290.3896
1367.673095703125 0 2580.0144 z Water loss 2
1368.666259765625 0 3129.0144 z Ammonia loss 2
1380.685546875 0 3357.5847
1381.6951904296875 0 13467.136 c 13
1382.69873046875 0 9652.86
1383.7001953125 0 6075.2847 y Water loss 2
1384.6875 0 1503.6456
1385.66748046875 0 43997.47 z 2
1386.67041015625 0 33834.516
1387.672607421875 0 15422.385
1388.6741943359375 0 4818.402
1394.7022705078125 0 2687.5886
1395.7069091796875 0 3234.888
1396.7049560546875 0 1812.2291
1401.6849365234375 0 6964.0015 y 2
1402.6898193359375 0 5114.0303
1403.690673828125 0 3097.645
1411.729248046875 0 1444.8636
1421.695068359375 0 3595.781 c Ammonia loss 14
1422.6971435546875 0 10899.613
1423.7008056640625 0 8485.002
1424.705078125 0 3908.3494
1425.7034912109375 0 1637.2955
1438.7159423828125 0 42239.53 c 14
1439.7198486328125 0 40730.03
1440.72314453125 0 21136.521
1441.7296142578125 0 7069.9336
1442.7291259765625 0 1673.251
1461.7335205078125 0 1657.3296
1477.74462890625 0 2278.272
1478.745361328125 0 6158.048
1479.7459716796875 0 5493.211
1480.7447509765625 0 2252.26 y Water loss 1
1481.739501953125 0 1068.1884 y Ammonia loss 1
1495.753173828125 0 3127.6423
1496.75146484375 0 2634.7341
1497.76513671875 0 1055.8396
1498.738525390625 0 1171.8569 y 1
1506.7099609375 0 1105.9873
1507.710693359375 0 1108.7966
1508.721923828125 0 1098.885
1512.7763671875 0 2315.9028
1513.780029296875 0 2264.448
1523.74609375 0 12345.2705
1524.7486572265625 0 10147.303
1525.748291015625 0 6157.0513
1526.7503662109375 0 1515.9498
1527.7589111328125 0 909.9418
1539.764404296875 0 5264.798 c 15
1540.77001953125 0 17173.67
1541.7735595703125 0 14488.013
1542.7767333984375 0 6422.261
1543.771728515625 0 1169.5697
1544.7696533203125 0 1082.754
1557.767822265625 0 1298.1469
1566.762451171875 0 2396.9807
1567.76416015625 0 2669.6216
1568.7554931640625 0 1857.6478
1583.7860107421875 0 9809.192
1584.782470703125 0 11151.526
1585.7862548828125 0 6627.67
1586.79296875 0 2956.1914
1594.756103515625 0 6019.942
1595.7586669921875 0 5061.6064
1596.756591796875 0 3931.5317
1597.7625732421875 0 2117.2673
1598.755126953125 0 1007.3841
1600.7926025390625 0 4213.28
1601.7994384765625 0 5092.1567
1602.7989501953125 0 3374.3005
1603.8028564453125 0 1608.6978
1604.810546875 0 1036.1572
1610.760009765625 0 2223.3418
1611.76806640625 0 12297.764
1612.768798828125 0 95903.555
1613.771240234375 0 88602.35
1614.7747802734375 0 41975.15
1615.7791748046875 0 16973.486
1616.7904052734375 0 5374.157
1627.77734375 0 5164.391
1628.7861328125 0 24012.734
1629.7928466796875 0 75339.1
1630.796142578125 0 60474.57
1631.8004150390625 0 29151.674
1632.8062744140625 0 11725.697
1633.8245849609375 0 4180.8633

Spectrum Details

|  |  |
| --- | --- |
| Matched peaks? Matched peaks The total absolute number of peaks matched. Additionally in brackets the total fraction of peaks matched and the total number of peaks is shown. | 125 (11.26% of 1110) |
| FDR? FDR The false discovery rate estimated for this peptide. It is calculated by matching all theoretical fragments with a non-integer shift with the raw peaks for this spectrum. This is done with 40 different shifts. The resulting percentage is the average number of annotated peaks over the number of annotated peaks with the correct spectrum. | 0.04% |
| Satellite FDR? Satellite FDR See the FDR for details on its calculation. This satellite ion specific FDR only contains the satellite ions (d/w) for I/L/J positions. | 0.00% |
| PSM Score? PSM Score The PSM Score as given by Hecklib to this annotated spectrum. It is shown with three significant figures. | 1.16E+03 |

#### Reverse Lookup? Reverse Lookup All places where this read could be placed.

| Group | Segment | Template | Template Part | Read Part | Score | Unique |
| --- | --- | --- | --- | --- | --- | --- |
| Homo sapiens Heavy Chain | IGHV | IGHV3-9 | [2..17] | [0..17] | 95 | False |
| Homo sapiens Heavy Chain | IGHV | IGHV3-48 | [2..17] | [0..17] | 95 | False |
| Homo sapiens Heavy Chain | IGHV | IGHV3-21 | [2..17] | [0..17] | 86 | False |
| Homo sapiens Heavy Chain | IGHV | IGHV3-7 | [2..17] | [0..17] | 95 | False |
| Homo sapiens Heavy Chain | IGHV | IGHV3-13 | [2..17] | [0..17] | 95 | False |
| Homo sapiens Heavy Chain | IGHV | IGHV3-11 | [2..17] | [0..17] | 86 | False |
| Homo sapiens Heavy Chain | IGHV | IGHV3-74 | [2..17] | [0..17] | 95 | False |
| Homo sapiens Heavy Chain | IGHV | IGHV3-23 | [2..17] | [0..17] | 86 | False |
| Homo sapiens Heavy Chain | IGHV | IGHV3-53 | [2..17] | [0..17] | 95 | False |
| Homo sapiens Heavy Chain | IGHV | IGHV3-66 | [2..17] | [0..17] | 95 | False |
| Homo sapiens Heavy Chain | IGHV | IGHV3-NL1 | [2..17] | [0..17] | 86 | False |
| Homo sapiens Heavy Chain | IGHV | IGHV3-30-5 | [2..17] | [0..17] | 86 | False |
| Homo sapiens Heavy Chain | IGHV | IGHV3-72 | [2..17] | [0..17] | 95 | False |
| Homo sapiens Heavy Chain | IGHV | IGHV3-30 | [2..17] | [0..17] | 86 | False |
| Homo sapiens Heavy Chain | IGHV | IGHV3-33 | [2..17] | [0..17] | 86 | False |
| Homo sapiens Heavy Chain | IGHV | IGHV3-64 | [2..17] | [0..17] | 95 | False |
| Homo sapiens Heavy Chain | IGHV | IGHV3-15 | [2..17] | [0..17] | 86 | False |
| Homo sapiens Heavy Chain | IGHV | IGHV3-49 | [2..17] | [0..17] | 95 | False |
| Homo sapiens Heavy Chain | IGHV | IGHV3-73 | [2..17] | [0..17] | 95 | False |

| Recombined | Template Part | Read Part | Score | Unique |
| --- | --- | --- | --- | --- |
| REC-0-1 | [1..17] | [0..17] | 127 | True |

#### Meta Information from Multiple reads

##### Number of combined reads

2

##### Intensity

1

##### TotalArea

0

##### Changes to the peptide sequence

APEJVESGGGLAQPGTS

J→LSupport for Leucine based on side chain ions (1 for L 0 for I) (Position: 11)

L→JNo support for either Leucine or Isoleucine based on side chain ions (Position: 11)

L→JNo support for either Leucine or Isoleucine based on side chain ions (Position: 4)

#### Positional Score

Copy Data

##### Positional Score (TSV)

###### Preview

```
Loading example...
```

*Click on the button to copy the data to your clipboard.*

00012345678910111213141516

Label Value
"0" 0
"1" 0
"2" 0
"3" 0
"4" 0
"5" 0
"6" 0
"7" 0
"8" 0
"9" 0
"10" 0
"11" 0
"12" 0
"13" 0
"14" 0
"15" 0
"16" 0

#### Meta Information from PEAKS

##### Scan Identifier

F2:6376

##### Original sequence

A

+58.01

P

E

L

V

E

S

G

G

G

L

A

Q

P

G

T

S

##### Posttranslational Modifications

Carboxymethyl (KW X@N-term)

##### Source File

D:\separate\_stitch\_analyses\xle-disambiguation\raw\20210323\_F1\_UM1\_Peng0013\_SA\_F59\_ingel\_3ug\_TL.raw

##### Fraction

2

##### Scan Feature

-

##### De Novo Score

98

##### ConfidenceScore

98

### m/z

814.3962

##### Mass

1626.7737

##### Charge

2

##### Retention Time

35.1

##### Predicted Retention Time

-

##### Area

0

##### Parts Per Million

2.5

##### Fragmentation mode

ETHCD

##### Originating file

01 D:\separate\_stitch\_analyses\xle-disambiguation\20210325\_F59\_3ug\_DENOVO\_12.csv

#### Meta Information from PEAKS

##### Scan Identifier

F2:6271

##### Original sequence

A

+58.01

P

E

L

V

E

S

G

G

G

L

A

Q

P

G

T

S

##### Posttranslational Modifications

Carboxymethyl (KW X@N-term)

##### Source File

D:\separate\_stitch\_analyses\xle-disambiguation\raw\20210323\_F1\_UM1\_Peng0013\_SA\_F59\_ingel\_3ug\_TL.raw

##### Fraction

2

##### Scan Feature

-

##### De Novo Score

98

##### ConfidenceScore

98

### m/z

543.2664

##### Mass

1626.7737

##### Charge

3

##### Retention Time

34.5

##### Predicted Retention Time

-

##### Area

0

##### Parts Per Million

2.2

##### Fragmentation mode

ETHCD

##### Originating file

01 D:\separate\_stitch\_analyses\xle-disambiguation\20210325\_F59\_3ug\_DENOVO\_12.csv
