## Supplementary material for "A handle on mass coincidence errors in *de novo* sequencing of antibodies by bottom-up proteomics": Combined_067.html

Details Combined\_067 | Stitch OverviewUndefined

### Read Combined\_067

#### Sequence (length=8)

SDSVEGRF

#### Spectrum 3834? Spectrum 3834 The raw spectrum of this peptide as annotated by Hecklib. The fragments are coloured according to ion type (see legend). Any peaks with a star '\*' as text can be hovered over to see the full details, first the ion type second the mass shift type. By hovering over the amino acids in the peptide or ions in the legend the corresponding peaks are highlighted. By toggling the 'Unassigned' label you can turn the background (unassigned) peaks on or off in the plot. By updating the slider in the Ion legend you can update the spectrum to only show the top X% of the peaks with labels. The top X% means any peak that is within X% of the highest intensity. By dragging in the spectrum you can zoom in to a specific part of the spectrum and use 'Zoom Out' to get back to the original zoom level. The annotation of the spectrum is based on the given sequence in the peptides file and is done with different software so inconsistencies are likely. The peaks are annotated based on the given sequence, with 20 ppm tolerance.

Copy Data

##### Spectrum 3834 (TSV)

###### Preview

```
Loading example...
```

*Click on the button to copy the data to your clipboard.*

Mz MinMz MaxIntensity Max

WidthHeightPeptide font sizePeptide stroke widthSpectrum font sizeSpectrum stroke widthCompact peptide

Ion legend

wxyz

abcd

OtherUnassignedIon^Charge^_Position_Show for top:%

SDSVEGRF

07.44e+61.49e+72.23e+72.97e+7

Zoom Out

d^+1^_2_y^+2^_2_a^+1^_2_y^+1^_1_a^+1^_2_b^+1^_2_b^+1^_2_a^+1^_3_y^+2^_4_y^+2^_4_y^+2^_4_a^+1^_3_b^+1^_3_b^+1^_3_y^+2^_5_y^+2^_5_y^+1^_2_y^+1^_2_y^+2^_6_y^+2^_6_b^+2^_7_y^+1^_3_b^+2^_7_b^+1^_4_y^+1^_3_b^+1^_4_y^+2^_7_y^+2^_7_\*\*y^+1^_4_y^+1^_4_b^+1^_5_y^+1^_4_b^+1^_5_b^+1^_6_y^+1^_5_y^+1^_5_y^+1^_5_y^+1^_6_y^+1^_6_y^+1^_6_b^+1^_7_b^+1^_7_y^+1^_7_y^+1^_7_

040180312041605

Fragment Matches Table

Show background peaks

| Position | Ion type | Intensity | mz Theoretical | mz Error (Th) | mz Error (ppm) | Charge | Series Number |
| --- | --- | --- | --- | --- | --- | --- | --- |
| - | - | 1.292E+06 | 120.1 | - | - | 0 | - |
| - | - | 1.119E+05 | 121.1 | - | - | 0 | - |
| - | - | 2.69E+04 | 122 | - | - | 0 | - |
| - | - | 1.142E+04 | 122.1 | - | - | 0 | - |
| - | - | 1.309E+04 | 123.1 | - | - | 0 | - |
| - | - | 9257 | 123.1 | - | - | 0 | - |
| - | - | 1.786E+04 | 124 | - | - | 0 | - |
| - | - | 1.702E+04 | 124.1 | - | - | 0 | - |
| - | - | 3.365E+05 | 124.1 | - | - | 0 | - |
| - | - | 3.036E+04 | 124.1 | - | - | 0 | - |
| - | - | 1.345E+04 | 125 | - | - | 0 | - |
| - | - | 2.949E+04 | 125.1 | - | - | 0 | - |
| - | - | 2.661E+04 | 125.1 | - | - | 0 | - |
| - | - | 1.463E+04 | 126.1 | - | - | 0 | - |
| - | - | 3.774E+04 | 127.1 | - | - | 0 | - |
| - | - | 1.239E+05 | 127.1 | - | - | 0 | - |
| - | - | 1.188E+05 | 129.1 | - | - | 0 | - |
| - | - | 5.902E+04 | 129.1 | - | - | 0 | - |
| - | - | 1.501E+04 | 129.1 | - | - | 0 | - |
| - | - | 6.381E+05 | 130.1 | - | - | 0 | - |
| - | - | 3.473E+04 | 130.1 | - | - | 0 | - |
| - | - | 1.396E+04 | 131 | - | - | 0 | - |
| - | - | 2.736E+04 | 131.1 | - | - | 0 | - |
| 2 | d | 2.479E+06 | 131.1 | 0.0004662 | 3.556 | +1 | 2 |
| - | - | 2.382E+04 | 132.1 | - | - | 0 | - |
| - | - | 1.343E+05 | 132.1 | - | - | 0 | - |
| - | - | 1.633E+04 | 133.1 | - | - | 0 | - |
| - | - | 9793 | 133.7 | - | - | 0 | - |
| - | - | 1.678E+04 | 134 | - | - | 0 | - |
| - | - | 8.081E+04 | 136.1 | - | - | 0 | - |
| - | - | 1.712E+04 | 138.1 | - | - | 0 | - |
| - | - | 5.529E+04 | 138.1 | - | - | 0 | - |
| - | - | 1.392E+04 | 138.1 | - | - | 0 | - |
| - | - | 2.275E+05 | 139.1 | - | - | 0 | - |
| - | - | 8762 | 139.1 | - | - | 0 | - |
| - | - | 7.045E+04 | 140 | - | - | 0 | - |
| - | - | 1.022E+06 | 140.1 | - | - | 0 | - |
| - | - | 1.966E+05 | 141.1 | - | - | 0 | - |
| - | - | 6.568E+04 | 141.1 | - | - | 0 | - |
| - | - | 1.634E+06 | 141.1 | - | - | 0 | - |
| - | - | 1.955E+04 | 142.1 | - | - | 0 | - |
| - | - | 2.145E+04 | 142.1 | - | - | 0 | - |
| - | - | 1.641E+04 | 142.1 | - | - | 0 | - |
| - | - | 1.092E+05 | 142.1 | - | - | 0 | - |
| - | - | 7.525E+04 | 143 | - | - | 0 | - |
| - | - | 1.232E+04 | 145.1 | - | - | 0 | - |
| - | - | 1.577E+04 | 148.1 | - | - | 0 | - |
| - | - | 1.221E+04 | 149 | - | - | 0 | - |
| - | - | 3.599E+04 | 149.1 | - | - | 0 | - |
| - | - | 1.314E+04 | 150 | - | - | 0 | - |
| - | - | 1.433E+04 | 151.1 | - | - | 0 | - |
| - | - | 2.096E+04 | 151.1 | - | - | 0 | - |
| - | - | 1.44E+04 | 152.1 | - | - | 0 | - |
| - | - | 1.971E+05 | 152.1 | - | - | 0 | - |
| - | - | 4.65E+04 | 153.1 | - | - | 0 | - |
| 7 | y | 1.59E+04 | 153.1 | 0.001586 | 10.36 | +2 | 2 |
| - | - | 2.514E+04 | 153.1 | - | - | 0 | - |
| - | - | 3.444E+04 | 154.1 | - | - | 0 | - |
| - | - | 5.604E+04 | 155 | - | - | 0 | - |
| - | - | 1.669E+05 | 155.1 | - | - | 0 | - |
| - | - | 1.607E+05 | 156.1 | - | - | 0 | - |
| - | - | 1.818E+04 | 156.1 | - | - | 0 | - |
| 2 | a | 1.3E+07 | 157.1 | 0.000526 | 3.349 | +1 | 2 |
| - | - | 1.405E+05 | 157.1 | - | - | 0 | - |
| - | - | 1.021E+05 | 158 | - | - | 0 | - |
| - | - | 8.268E+04 | 158.1 | - | - | 0 | - |
| - | - | 8.425E+05 | 158.1 | - | - | 0 | - |
| - | - | 9.357E+05 | 158.1 | - | - | 0 | - |
| - | - | 1.331E+04 | 158.1 | - | - | 0 | - |
| - | - | 6.339E+04 | 159.1 | - | - | 0 | - |
| - | - | 1.105E+05 | 159.1 | - | - | 0 | - |
| - | - | 8.721E+06 | 159.1 | - | - | 0 | - |
| - | - | 4.548E+04 | 160.1 | - | - | 0 | - |
| - | - | 6.455E+05 | 160.1 | - | - | 0 | - |
| - | - | 2.426E+04 | 161.1 | - | - | 0 | - |
| - | - | 1.215E+04 | 165.1 | - | - | 0 | - |
| - | - | 2.026E+04 | 166.1 | - | - | 0 | - |
| 8 | y | 1.076E+06 | 166.1 | 0.0005064 | 3.049 | +1 | 1 |
| - | - | 2.037E+05 | 167 | - | - | 0 | - |
| - | - | 2.937E+04 | 167.1 | - | - | 0 | - |
| - | - | 9.647E+04 | 167.1 | - | - | 0 | - |
| - | - | 1.528E+04 | 167.1 | - | - | 0 | - |
| - | - | 1.893E+04 | 168 | - | - | 0 | - |
| - | - | 6.099E+04 | 168.1 | - | - | 0 | - |
| - | - | 9.916E+05 | 169.1 | - | - | 0 | - |
| - | - | 7.62E+05 | 169.1 | - | - | 0 | - |
| - | - | 1.148E+05 | 169.1 | - | - | 0 | - |
| - | - | 9867 | 170 | - | - | 0 | - |
| - | - | 6.041E+04 | 170.1 | - | - | 0 | - |
| - | - | 5.111E+04 | 170.1 | - | - | 0 | - |
| - | - | 2.666E+04 | 171.1 | - | - | 0 | - |
| - | - | 1.876E+04 | 171.1 | - | - | 0 | - |
| - | - | 7.785E+05 | 172.1 | - | - | 0 | - |
| - | - | 4.021E+04 | 173.1 | - | - | 0 | - |
| - | - | 2.565E+05 | 173.1 | - | - | 0 | - |
| - | - | 1.323E+04 | 173.1 | - | - | 0 | - |
| - | - | 9144 | 173.1 | - | - | 0 | - |
| - | - | 4.12E+04 | 173.1 | - | - | 0 | - |
| 2 | a | 9.836E+06 | 175.1 | 0.0005814 | 3.321 | +1 | 2 |
| - | - | 9.444E+05 | 175.1 | - | - | 0 | - |
| - | - | 6.11E+05 | 176.1 | - | - | 0 | - |
| - | - | 6.628E+04 | 176.1 | - | - | 0 | - |
| - | - | 7.713E+04 | 177.1 | - | - | 0 | - |
| - | - | 2.017E+04 | 179 | - | - | 0 | - |
| - | - | 1.114E+04 | 179.6 | - | - | 0 | - |
| - | - | 1.983E+05 | 180.1 | - | - | 0 | - |
| - | - | 4.334E+04 | 181.1 | - | - | 0 | - |
| - | - | 1.635E+04 | 181.1 | - | - | 0 | - |
| - | - | 2.551E+04 | 182.1 | - | - | 0 | - |
| - | - | 4.161E+04 | 182.1 | - | - | 0 | - |
| - | - | 1.259E+04 | 183.1 | - | - | 0 | - |
| - | - | 8.793E+04 | 183.1 | - | - | 0 | - |
| - | - | 6.991E+04 | 184.1 | - | - | 0 | - |
| - | - | 1.86E+04 | 184.1 | - | - | 0 | - |
| 2 | b | 1.042E+06 | 185.1 | 0.0006065 | 3.277 | +1 | 2 |
| - | - | 1.268E+04 | 185.1 | - | - | 0 | - |
| - | - | 1.223E+04 | 185.1 | - | - | 0 | - |
| - | - | 3.718E+04 | 186 | - | - | 0 | - |
| - | - | 7.42E+04 | 186.1 | - | - | 0 | - |
| - | - | 3.094E+05 | 186.1 | - | - | 0 | - |
| - | - | 1.859E+05 | 187.1 | - | - | 0 | - |
| - | - | 2.686E+06 | 187.1 | - | - | 0 | - |
| - | - | 2.344E+05 | 188.1 | - | - | 0 | - |
| - | - | 4.315E+04 | 190.1 | - | - | 0 | - |
| - | - | 1.214E+04 | 191 | - | - | 0 | - |
| - | - | 1.78E+04 | 193.1 | - | - | 0 | - |
| - | - | 5.524E+04 | 194.1 | - | - | 0 | - |
| - | - | 1.501E+04 | 194.1 | - | - | 0 | - |
| - | - | 3.18E+04 | 196.1 | - | - | 0 | - |
| - | - | 2.022E+04 | 196.1 | - | - | 0 | - |
| - | - | 1.726E+04 | 196.1 | - | - | 0 | - |
| - | - | 4.604E+04 | 197.1 | - | - | 0 | - |
| - | - | 8.02E+04 | 197.1 | - | - | 0 | - |
| - | - | 2.316E+06 | 197.1 | - | - | 0 | - |
| - | - | 1.419E+05 | 198.1 | - | - | 0 | - |
| - | - | 1.753E+05 | 198.1 | - | - | 0 | - |
| - | - | 1.345E+04 | 198.1 | - | - | 0 | - |
| - | - | 7.849E+04 | 199.1 | - | - | 0 | - |
| - | - | 3.099E+04 | 199.1 | - | - | 0 | - |
| - | - | 2.155E+04 | 200.1 | - | - | 0 | - |
| - | - | 6.091E+05 | 201.1 | - | - | 0 | - |
| - | - | 4.721E+04 | 202.1 | - | - | 0 | - |
| - | - | 6.303E+04 | 202.1 | - | - | 0 | - |
| 2 | b | 1.482E+07 | 203.1 | 0.0005398 | 2.658 | +1 | 2 |
| - | - | 9.992E+05 | 204.1 | - | - | 0 | - |
| - | - | 6.225E+04 | 204.1 | - | - | 0 | - |
| - | - | 1.144E+05 | 205.1 | - | - | 0 | - |
| - | - | 2.344E+04 | 206.1 | - | - | 0 | - |
| - | - | 6.137E+04 | 207.1 | - | - | 0 | - |
| - | - | 7.26E+04 | 208.1 | - | - | 0 | - |
| - | - | 1.989E+04 | 208.1 | - | - | 0 | - |
| - | - | 1.091E+04 | 208.5 | - | - | 0 | - |
| - | - | 2.897E+04 | 209.1 | - | - | 0 | - |
| - | - | 1.417E+04 | 209.1 | - | - | 0 | - |
| - | - | 1.172E+05 | 210.1 | - | - | 0 | - |
| - | - | 5.684E+04 | 211.1 | - | - | 0 | - |
| - | - | 1.205E+04 | 211.1 | - | - | 0 | - |
| - | - | 4.71E+04 | 212.1 | - | - | 0 | - |
| - | - | 1.021E+05 | 213.1 | - | - | 0 | - |
| - | - | 1.448E+04 | 214.1 | - | - | 0 | - |
| - | - | 1.624E+04 | 214.1 | - | - | 0 | - |
| - | - | 5.967E+05 | 214.1 | - | - | 0 | - |
| - | - | 6.33E+05 | 215.1 | - | - | 0 | - |
| - | - | 4.09E+04 | 215.1 | - | - | 0 | - |
| - | - | 1.457E+04 | 216.1 | - | - | 0 | - |
| - | - | 3.278E+04 | 216.1 | - | - | 0 | - |
| - | - | 8.976E+04 | 217.1 | - | - | 0 | - |
| - | - | 4.489E+04 | 217.1 | - | - | 0 | - |
| - | - | 4.521E+04 | 220.1 | - | - | 0 | - |
| - | - | 4.482E+04 | 221.1 | - | - | 0 | - |
| - | - | 7.176E+04 | 221.1 | - | - | 0 | - |
| - | - | 1.605E+04 | 222.1 | - | - | 0 | - |
| - | - | 3.163E+04 | 222.6 | - | - | 0 | - |
| - | - | 1.945E+04 | 223.1 | - | - | 0 | - |
| - | - | 1.155E+04 | 223.1 | - | - | 0 | - |
| - | - | 5.154E+04 | 224.1 | - | - | 0 | - |
| - | - | 4.039E+04 | 224.1 | - | - | 0 | - |
| - | - | 1.682E+04 | 224.1 | - | - | 0 | - |
| - | - | 1.06E+05 | 225.1 | - | - | 0 | - |
| - | - | 1.318E+04 | 225.1 | - | - | 0 | - |
| - | - | 1.882E+04 | 226.1 | - | - | 0 | - |
| - | - | 3.349E+05 | 226.1 | - | - | 0 | - |
| - | - | 3.799E+04 | 226.1 | - | - | 0 | - |
| - | - | 5.742E+04 | 227.1 | - | - | 0 | - |
| - | - | 3.819E+04 | 227.1 | - | - | 0 | - |
| - | - | 2.008E+04 | 227.1 | - | - | 0 | - |
| - | - | 1.935E+04 | 228.1 | - | - | 0 | - |
| - | - | 1.24E+04 | 228.2 | - | - | 0 | - |
| - | - | 6.943E+05 | 229.1 | - | - | 0 | - |
| - | - | 7.561E+04 | 230.1 | - | - | 0 | - |
| - | - | 4.624E+06 | 232.1 | - | - | 0 | - |
| - | - | 4.004E+05 | 233.1 | - | - | 0 | - |
| - | - | 1.779E+04 | 234.1 | - | - | 0 | - |
| - | - | 1.671E+04 | 234.1 | - | - | 0 | - |
| - | - | 2.036E+05 | 236.1 | - | - | 0 | - |
| - | - | 3.728E+04 | 237.1 | - | - | 0 | - |
| - | - | 2.456E+04 | 237.1 | - | - | 0 | - |
| - | - | 1.34E+04 | 237.1 | - | - | 0 | - |
| - | - | 2.998E+04 | 237.1 | - | - | 0 | - |
| - | - | 2.151E+05 | 238.1 | - | - | 0 | - |
| - | - | 1.034E+05 | 239.1 | - | - | 0 | - |
| - | - | 2.415E+04 | 239.1 | - | - | 0 | - |
| - | - | 1.554E+04 | 240.1 | - | - | 0 | - |
| - | - | 1.37E+04 | 241.1 | - | - | 0 | - |
| - | - | 2.662E+04 | 241.1 | - | - | 0 | - |
| - | - | 1.893E+05 | 242.1 | - | - | 0 | - |
| - | - | 8.496E+04 | 242.1 | - | - | 0 | - |
| - | - | 2.801E+04 | 243.1 | - | - | 0 | - |
| - | - | 3.348E+04 | 243.1 | - | - | 0 | - |
| 3 | a | 6.604E+05 | 244.1 | 0.0007241 | 2.967 | +1 | 3 |
| - | - | 3.013E+04 | 244.1 | - | - | 0 | - |
| - | - | 3E+05 | 245.1 | - | - | 0 | - |
| - | - | 6.352E+04 | 245.1 | - | - | 0 | - |
| - | - | 1.795E+04 | 245.1 | - | - | 0 | - |
| 5 | y | 1.798E+05 | 245.6 | 0.0007038 | 2.865 | +2 | 4 |
| - | - | 3.192E+04 | 246.1 | - | - | 0 | - |
| - | - | 1.099E+04 | 246.1 | - | - | 0 | - |
| 5 | y | 4.554E+04 | 246.1 | 0.003191 | 12.96 | +2 | 4 |
| - | - | 3.035E+04 | 246.1 | - | - | 0 | - |
| - | - | 3.405E+04 | 248.1 | - | - | 0 | - |
| - | - | 1.302E+04 | 251.1 | - | - | 0 | - |
| - | - | 1.567E+04 | 251.1 | - | - | 0 | - |
| - | - | 1.074E+05 | 252.1 | - | - | 0 | - |
| - | - | 4.126E+04 | 253.1 | - | - | 0 | - |
| - | - | 8.575E+05 | 254.1 | - | - | 0 | - |
| 5 | y | 6.334E+04 | 254.6 | 0.0005331 | 2.094 | +2 | 4 |
| - | - | 1.581E+04 | 255.1 | - | - | 0 | - |
| - | - | 6.405E+04 | 255.1 | - | - | 0 | - |
| - | - | 1.914E+04 | 255.1 | - | - | 0 | - |
| - | - | 3.475E+04 | 255.1 | - | - | 0 | - |
| - | - | 9.599E+05 | 256.1 | - | - | 0 | - |
| - | - | 1.284E+05 | 257.1 | - | - | 0 | - |
| - | - | 1.074E+05 | 257.1 | - | - | 0 | - |
| - | - | 1.317E+04 | 257.6 | - | - | 0 | - |
| - | - | 3.095E+05 | 259.2 | - | - | 0 | - |
| - | - | 3.627E+04 | 260.1 | - | - | 0 | - |
| - | - | 5.253E+04 | 260.2 | - | - | 0 | - |
| 3 | a | 5.538E+05 | 262.1 | 0.0005812 | 2.217 | +1 | 3 |
| - | - | 2.194E+04 | 262.1 | - | - | 0 | - |
| - | - | 3.229E+05 | 262.2 | - | - | 0 | - |
| - | - | 4.097E+04 | 263.1 | - | - | 0 | - |
| - | - | 4.637E+05 | 263.1 | - | - | 0 | - |
| - | - | 3.83E+04 | 263.2 | - | - | 0 | - |
| - | - | 2.266E+04 | 264.1 | - | - | 0 | - |
| - | - | 5.645E+04 | 264.1 | - | - | 0 | - |
| - | - | 3.479E+05 | 265.1 | - | - | 0 | - |
| - | - | 1.305E+05 | 266.1 | - | - | 0 | - |
| - | - | 8.137E+04 | 266.1 | - | - | 0 | - |
| - | - | 1.551E+04 | 266.6 | - | - | 0 | - |
| - | - | 2.71E+04 | 267.1 | - | - | 0 | - |
| - | - | 1.907E+04 | 267.1 | - | - | 0 | - |
| - | - | 1.915E+04 | 267.1 | - | - | 0 | - |
| - | - | 2.91E+04 | 268.1 | - | - | 0 | - |
| - | - | 1.658E+04 | 269.1 | - | - | 0 | - |
| - | - | 2.314E+04 | 270.1 | - | - | 0 | - |
| - | - | 7.404E+05 | 271.1 | - | - | 0 | - |
| - | - | 3.024E+04 | 271.1 | - | - | 0 | - |
| 3 | b | 7.015E+06 | 272.1 | 0.0005757 | 2.116 | +1 | 3 |
| - | - | 7.434E+05 | 273.1 | - | - | 0 | - |
| - | - | 8.941E+04 | 274.1 | - | - | 0 | - |
| - | - | 7.175E+04 | 274.1 | - | - | 0 | - |
| - | - | 3.311E+04 | 274.2 | - | - | 0 | - |
| - | - | 2.965E+04 | 276.2 | - | - | 0 | - |
| - | - | 1.745E+04 | 278.1 | - | - | 0 | - |
| - | - | 1.423E+05 | 280.1 | - | - | 0 | - |
| - | - | 2.069E+04 | 280.2 | - | - | 0 | - |
| - | - | 1.816E+04 | 281.1 | - | - | 0 | - |
| - | - | 3.543E+04 | 281.1 | - | - | 0 | - |
| - | - | 1.912E+04 | 281.1 | - | - | 0 | - |
| - | - | 3.153E+05 | 283.1 | - | - | 0 | - |
| - | - | 1.013E+06 | 284.1 | - | - | 0 | - |
| - | - | 3.846E+04 | 284.1 | - | - | 0 | - |
| - | - | 2.198E+05 | 285.1 | - | - | 0 | - |
| - | - | 1.321E+04 | 286.1 | - | - | 0 | - |
| - | - | 8.536E+04 | 286.1 | - | - | 0 | - |
| - | - | 1.7E+04 | 287.1 | - | - | 0 | - |
| - | - | 1.563E+05 | 287.2 | - | - | 0 | - |
| - | - | 1.304E+05 | 288.1 | - | - | 0 | - |
| - | - | 5.148E+04 | 288.2 | - | - | 0 | - |
| - | - | 2.656E+04 | 289.1 | - | - | 0 | - |
| - | - | 2.632E+04 | 289.1 | - | - | 0 | - |
| 3 | b | 4.749E+06 | 290.1 | 0.0008143 | 2.807 | +1 | 3 |
| - | - | 5.19E+05 | 291.1 | - | - | 0 | - |
| - | - | 3.392E+04 | 291.1 | - | - | 0 | - |
| - | - | 5.853E+04 | 292.1 | - | - | 0 | - |
| - | - | 1.747E+04 | 294.1 | - | - | 0 | - |
| - | - | 2.578E+04 | 295.1 | - | - | 0 | - |
| 4 | y | 2.813E+04 | 295.2 | 0.001211 | 4.101 | +2 | 5 |
| - | - | 2.197E+04 | 296.1 | - | - | 0 | - |
| - | - | 4.592E+04 | 296.1 | - | - | 0 | - |
| - | - | 4.245E+04 | 297.2 | - | - | 0 | - |
| - | - | 1.652E+05 | 298.1 | - | - | 0 | - |
| - | - | 4.697E+04 | 298.2 | - | - | 0 | - |
| - | - | 1.86E+04 | 299.1 | - | - | 0 | - |
| - | - | 6.764E+04 | 299.7 | - | - | 0 | - |
| - | - | 2.51E+04 | 300.2 | - | - | 0 | - |
| - | - | 1.853E+05 | 301.2 | - | - | 0 | - |
| - | - | 1.999E+05 | 302.1 | - | - | 0 | - |
| - | - | 3.285E+04 | 302.2 | - | - | 0 | - |
| - | - | 3.315E+04 | 303.1 | - | - | 0 | - |
| 4 | y | 4.484E+05 | 304.2 | 0.00075 | 2.466 | +2 | 5 |
| - | - | 1.363E+05 | 304.7 | - | - | 0 | - |
| 7 | y | 2.8E+06 | 305.2 | 0.0008346 | 2.735 | +1 | 2 |
| - | - | 4.795E+05 | 306.2 | - | - | 0 | - |
| - | - | 1.903E+05 | 307.1 | - | - | 0 | - |
| - | - | 4.531E+04 | 307.2 | - | - | 0 | - |
| - | - | 2.044E+06 | 308.1 | - | - | 0 | - |
| - | - | 8.427E+04 | 309.1 | - | - | 0 | - |
| - | - | 2.986E+05 | 309.1 | - | - | 0 | - |
| - | - | 1.688E+04 | 309.2 | - | - | 0 | - |
| - | - | 4.784E+04 | 310.1 | - | - | 0 | - |
| - | - | 3.005E+04 | 311.1 | - | - | 0 | - |
| - | - | 7.706E+04 | 312.1 | - | - | 0 | - |
| - | - | 3.812E+04 | 313.2 | - | - | 0 | - |
| - | - | 1.571E+04 | 313.6 | - | - | 0 | - |
| - | - | 4.747E+04 | 314.1 | - | - | 0 | - |
| - | - | 7.377E+04 | 314.1 | - | - | 0 | - |
| - | - | 2.105E+05 | 315.2 | - | - | 0 | - |
| - | - | 4.3E+04 | 315.7 | - | - | 0 | - |
| - | - | 3.944E+05 | 316.2 | - | - | 0 | - |
| - | - | 2.557E+04 | 316.2 | - | - | 0 | - |
| - | - | 6.587E+04 | 317.2 | - | - | 0 | - |
| - | - | 2.513E+04 | 319.2 | - | - | 0 | - |
| - | - | 1.142E+05 | 319.2 | - | - | 0 | - |
| - | - | 2.325E+05 | 320.2 | - | - | 0 | - |
| - | - | 1.514E+04 | 320.2 | - | - | 0 | - |
| - | - | 4.137E+04 | 321.2 | - | - | 0 | - |
| 7 | y | 5.043E+05 | 322.2 | 0.0008969 | 2.784 | +1 | 2 |
| - | - | 4.524E+04 | 322.7 | - | - | 0 | - |
| - | - | 6.919E+04 | 323.2 | - | - | 0 | - |
| - | - | 6.288E+04 | 324.7 | - | - | 0 | - |
| - | - | 1.289E+04 | 324.7 | - | - | 0 | - |
| - | - | 1.836E+06 | 325.2 | - | - | 0 | - |
| - | - | 1.296E+06 | 326.1 | - | - | 0 | - |
| - | - | 2.173E+05 | 327.1 | - | - | 0 | - |
| - | - | 1.652E+04 | 328.1 | - | - | 0 | - |
| - | - | 4.993E+04 | 328.2 | - | - | 0 | - |
| - | - | 8.313E+04 | 329.7 | - | - | 0 | - |
| - | - | 3.182E+04 | 330.1 | - | - | 0 | - |
| - | - | 2.153E+04 | 330.2 | - | - | 0 | - |
| - | - | 7.62E+04 | 332.1 | - | - | 0 | - |
| - | - | 2.401E+04 | 332.2 | - | - | 0 | - |
| - | - | 3.071E+04 | 333.2 | - | - | 0 | - |
| - | - | 1.385E+04 | 334.2 | - | - | 0 | - |
| - | - | 3.134E+04 | 335.1 | - | - | 0 | - |
| - | - | 2.28E+04 | 336.7 | - | - | 0 | - |
| - | - | 2.078E+04 | 337.1 | - | - | 0 | - |
| - | - | 1.708E+04 | 337.2 | - | - | 0 | - |
| - | - | 5.263E+04 | 337.2 | - | - | 0 | - |
| - | - | 2.082E+04 | 338.1 | - | - | 0 | - |
| - | - | 1.518E+04 | 338.2 | - | - | 0 | - |
| 3 | y | 8.53E+05 | 338.7 | 0.000974 | 2.876 | +2 | 6 |
| - | - | 2.992E+05 | 339.2 | - | - | 0 | - |
| - | - | 6.167E+04 | 339.7 | - | - | 0 | - |
| - | - | 3.889E+06 | 343.2 | - | - | 0 | - |
| - | - | 5.954E+04 | 343.7 | - | - | 0 | - |
| - | - | 5.18E+05 | 344.2 | - | - | 0 | - |
| - | - | 2.764E+05 | 345.2 | - | - | 0 | - |
| - | - | 4.921E+04 | 345.2 | - | - | 0 | - |
| - | - | 3.654E+04 | 346.2 | - | - | 0 | - |
| 3 | y | 4.864E+05 | 347.7 | 0.0008186 | 2.354 | +2 | 6 |
| - | - | 1.976E+04 | 348.2 | - | - | 0 | - |
| - | - | 1.748E+05 | 348.2 | - | - | 0 | - |
| - | - | 1.488E+04 | 348.7 | - | - | 0 | - |
| - | - | 3.038E+04 | 348.7 | - | - | 0 | - |
| - | - | 1.101E+05 | 352.7 | - | - | 0 | - |
| - | - | 2.472E+05 | 353.1 | - | - | 0 | - |
| - | - | 5.788E+04 | 354.1 | - | - | 0 | - |
| - | - | 1.859E+04 | 355.1 | - | - | 0 | - |
| - | - | 2.126E+04 | 355.2 | - | - | 0 | - |
| - | - | 6.62E+04 | 356.1 | - | - | 0 | - |
| - | - | 1.294E+04 | 357.1 | - | - | 0 | - |
| 7 | b | 9.109E+04 | 357.2 | 0.00051 | 1.428 | +2 | 7 |
| - | - | 4.249E+04 | 357.7 | - | - | 0 | - |
| - | - | 5.594E+06 | 361.2 | - | - | 0 | - |
| 6 | y | 1.411E+06 | 362.2 | 0.002412 | 6.659 | +1 | 3 |
| - | - | 2.066E+05 | 363.2 | - | - | 0 | - |
| - | - | 1.638E+04 | 364.2 | - | - | 0 | - |
| 7 | b | 1.223E+05 | 366.2 | 0.0008124 | 2.219 | +2 | 7 |
| - | - | 3.929E+04 | 366.7 | - | - | 0 | - |
| - | - | 2.198E+04 | 367.2 | - | - | 0 | - |
| - | - | 1.742E+04 | 368.1 | - | - | 0 | - |
| 4 | b | 1.907E+06 | 371.2 | 0.0008874 | 2.391 | +1 | 4 |
| - | - | 3.19E+05 | 372.2 | - | - | 0 | - |
| - | - | 1.628E+05 | 373.2 | - | - | 0 | - |
| - | - | 5.903E+04 | 373.7 | - | - | 0 | - |
| - | - | 2.734E+04 | 374.1 | - | - | 0 | - |
| - | - | 1.351E+04 | 374.2 | - | - | 0 | - |
| - | - | 2.43E+04 | 377.2 | - | - | 0 | - |
| - | - | 1.944E+04 | 378.7 | - | - | 0 | - |
| 6 | y | 1.178E+07 | 379.2 | 0.0009175 | 2.42 | +1 | 3 |
| - | - | 2.227E+06 | 380.2 | - | - | 0 | - |
| - | - | 2.986E+04 | 381.2 | - | - | 0 | - |
| - | - | 2.622E+05 | 381.2 | - | - | 0 | - |
| - | - | 2.284E+04 | 382.2 | - | - | 0 | - |
| - | - | 8.399E+04 | 383.1 | - | - | 0 | - |
| - | - | 1.847E+04 | 383.2 | - | - | 0 | - |
| - | - | 2.093E+04 | 384.2 | - | - | 0 | - |
| - | - | 7.289E+04 | 385.2 | - | - | 0 | - |
| - | - | 1.707E+04 | 386.2 | - | - | 0 | - |
| - | - | 6.02E+04 | 387.2 | - | - | 0 | - |
| - | - | 3.268E+04 | 387.7 | - | - | 0 | - |
| 4 | b | 3.676E+05 | 389.2 | 0.0008513 | 2.187 | +1 | 4 |
| - | - | 2.652E+04 | 389.2 | - | - | 0 | - |
| - | - | 6.956E+04 | 390.2 | - | - | 0 | - |
| - | - | 1.686E+04 | 390.2 | - | - | 0 | - |
| - | - | 1.895E+04 | 390.7 | - | - | 0 | - |
| - | - | 1.79E+04 | 395.2 | - | - | 0 | - |
| 2 | y | 7.709E+05 | 396.2 | 0.0008386 | 2.117 | +2 | 7 |
| - | - | 3.048E+05 | 396.7 | - | - | 0 | - |
| - | - | 7.487E+04 | 397.2 | - | - | 0 | - |
| - | - | 2.05E+04 | 397.2 | - | - | 0 | - |
| - | - | 2.129E+04 | 400.2 | - | - | 0 | - |
| - | - | 1.325E+05 | 401.1 | - | - | 0 | - |
| - | - | 1.91E+04 | 401.2 | - | - | 0 | - |
| - | - | 1.876E+04 | 402.1 | - | - | 0 | - |
| - | - | 1.785E+04 | 403.2 | - | - | 0 | - |
| 2 | y | 8.997E+04 | 405.2 | 0.0008969 | 2.213 | +2 | 7 |
| - | - | 2.181E+04 | 405.7 | - | - | 0 | - |
| - | - | 1.747E+04 | 407.2 | - | - | 0 | - |
| - | - | 3.51E+04 | 408.2 | - | - | 0 | - |
| - | - | 3.506E+04 | 409.2 | - | - | 0 | - |
| - | - | 3.233E+05 | 410.2 | - | - | 0 | - |
| - | - | 1.304E+05 | 410.7 | - | - | 0 | - |
| - | - | 4.03E+04 | 411.2 | - | - | 0 | - |
| - | - | 8.34E+04 | 413.2 | - | - | 0 | - |
| - | - | 2.375E+04 | 414.2 | - | - | 0 | - |
| - | - | 7.767E+04 | 414.2 | - | - | 0 | - |
| - | - | 1.66E+04 | 415.2 | - | - | 0 | - |
| - | - | 2.009E+04 | 415.7 | - | - | 0 | - |
| - | - | 2.61E+04 | 416.2 | - | - | 0 | - |
| - | - | 6.963E+04 | 416.7 | - | - | 0 | - |
| - | - | 1.663E+04 | 417.2 | - | - | 0 | - |
| - | - | 4.678E+04 | 419.1 | - | - | 0 | - |
| - | - | 5.537E+04 | 421.7 | - | - | 0 | - |
| - | - | 1.018E+05 | 422.2 | - | - | 0 | - |
| - | - | 6.112E+04 | 422.7 | - | - | 0 | - |
| - | - | 3.26E+04 | 424.2 | - | - | 0 | - |
| - | - | 3.699E+04 | 424.7 | - | - | 0 | - |
| - | - | 2.382E+05 | 425.2 | - | - | 0 | - |
| - | - | 3.48E+04 | 426.2 | - | - | 0 | - |
| - | - | 2.148E+04 | 427.2 | - | - | 0 | - |
| - | - | 8.915E+04 | 427.2 | - | - | 0 | - |
| - | - | 1.801E+05 | 430.2 | - | - | 0 | - |
| - | - | 3.709E+05 | 430.7 | - | - | 0 | - |
| - | - | 3.982E+05 | 431.2 | - | - | 0 | - |
| - | - | 3.306E+04 | 431.7 | - | - | 0 | - |
| - | - | 6.875E+04 | 432.2 | - | - | 0 | - |
| - | - | 4.766E+04 | 436.2 | - | - | 0 | - |
| - | - | 2.342E+04 | 437.2 | - | - | 0 | - |
| 0 | Precursor | 1.289E+06 | 439.7 | 0.001121 | 2.549 | +2 | -1 |
| - | - | 5.649E+05 | 440.2 | - | - | 0 | - |
| - | - | 1.522E+05 | 440.7 | - | - | 0 | - |
| - | - | 2.033E+04 | 441.2 | - | - | 0 | - |
| - | - | 3.159E+05 | 442.2 | - | - | 0 | - |
| - | - | 3.85E+04 | 443.2 | - | - | 0 | - |
| - | - | 2.18E+04 | 443.3 | - | - | 0 | - |
| - | - | 1.725E+04 | 444.2 | - | - | 0 | - |
| - | - | 1.456E+04 | 445.2 | - | - | 0 | - |
| - | - | 8.788E+04 | 448.2 | - | - | 0 | - |
| 0 | Precursor | 1.504E+05 | 448.7 | 0.001362 | 3.036 | +2 | -1 |
| - | - | 1.418E+05 | 449.2 | - | - | 0 | - |
| - | - | 1.967E+04 | 449.7 | - | - | 0 | - |
| - | - | 3.446E+04 | 450.2 | - | - | 0 | - |
| - | - | 9.55E+04 | 454.2 | - | - | 0 | - |
| - | - | 4.839E+04 | 455.2 | - | - | 0 | - |
| - | - | 8.286E+04 | 455.2 | - | - | 0 | - |
| - | - | 2.909E+04 | 456.2 | - | - | 0 | - |
| - | - | 1.526E+06 | 460.3 | - | - | 0 | - |
| - | - | 3.219E+05 | 461.3 | - | - | 0 | - |
| - | - | 7.325E+04 | 462.3 | - | - | 0 | - |
| - | - | 1.621E+04 | 463.3 | - | - | 0 | - |
| - | - | 4.668E+04 | 466.2 | - | - | 0 | - |
| - | - | 1.826E+04 | 467.2 | - | - | 0 | - |
| - | - | 2.464E+04 | 470.2 | - | - | 0 | - |
| - | - | 7.893E+04 | 470.2 | - | - | 0 | - |
| - | - | 1.403E+05 | 472.2 | - | - | 0 | - |
| - | - | 1.774E+04 | 472.2 | - | - | 0 | - |
| - | - | 2.798E+05 | 473.2 | - | - | 0 | - |
| - | - | 8.279E+04 | 474.2 | - | - | 0 | - |
| - | - | 8.904E+04 | 482.2 | - | - | 0 | - |
| - | - | 2.202E+04 | 484.3 | - | - | 0 | - |
| - | - | 3.975E+04 | 487.3 | - | - | 0 | - |
| 5 | y | 2.43E+06 | 490.2 | 0.001238 | 2.525 | +1 | 4 |
| 5 | y | 3.558E+05 | 491.2 | 0.00529 | 10.77 | +1 | 4 |
| - | - | 4.287E+05 | 491.2 | - | - | 0 | - |
| - | - | 1.181E+05 | 492.2 | - | - | 0 | - |
| - | - | 2.124E+04 | 493.2 | - | - | 0 | - |
| - | - | 3.41E+04 | 494.2 | - | - | 0 | - |
| 5 | b | 2.383E+05 | 500.2 | 0.0009274 | 1.854 | +1 | 5 |
| - | - | 6.542E+04 | 501.2 | - | - | 0 | - |
| - | - | 1.767E+05 | 501.3 | - | - | 0 | - |
| - | - | 1.728E+04 | 502.2 | - | - | 0 | - |
| - | - | 4.024E+04 | 502.3 | - | - | 0 | - |
| - | - | 1.651E+04 | 505.3 | - | - | 0 | - |
| 5 | y | 2.945E+07 | 508.3 | 0.001202 | 2.364 | +1 | 4 |
| - | - | 7.44E+06 | 509.3 | - | - | 0 | - |
| - | - | 1.906E+04 | 509.4 | - | - | 0 | - |
| - | - | 1.183E+06 | 510.3 | - | - | 0 | - |
| - | - | 1.261E+05 | 511.3 | - | - | 0 | - |
| - | - | 1.631E+05 | 512.2 | - | - | 0 | - |
| - | - | 1.816E+04 | 513.2 | - | - | 0 | - |
| - | - | 8.087E+04 | 517.3 | - | - | 0 | - |
| 5 | b | 5.092E+04 | 518.2 | 0.004938 | 9.528 | +1 | 5 |
| - | - | 1.652E+05 | 518.2 | - | - | 0 | - |
| - | - | 1.982E+04 | 518.3 | - | - | 0 | - |
| - | - | 2.112E+04 | 519.2 | - | - | 0 | - |
| - | - | 3.997E+04 | 519.2 | - | - | 0 | - |
| - | - | 8.953E+05 | 529.3 | - | - | 0 | - |
| - | - | 1.961E+05 | 530.3 | - | - | 0 | - |
| - | - | 2.311E+04 | 531.3 | - | - | 0 | - |
| - | - | 2.241E+04 | 539.3 | - | - | 0 | - |
| - | - | 3.501E+06 | 547.3 | - | - | 0 | - |
| - | - | 8.408E+05 | 548.3 | - | - | 0 | - |
| - | - | 1.461E+05 | 549.3 | - | - | 0 | - |
| 6 | b | 1.603E+04 | 557.2 | 0.003939 | 7.069 | +1 | 6 |
| - | - | 7.168E+04 | 557.3 | - | - | 0 | - |
| - | - | 1.744E+04 | 561.3 | - | - | 0 | - |
| - | - | 1.041E+05 | 571.3 | - | - | 0 | - |
| - | - | 1.335E+05 | 572.3 | - | - | 0 | - |
| - | - | 2.118E+04 | 573.3 | - | - | 0 | - |
| 4 | y | 6.426E+04 | 589.3 | 0.0003592 | 0.6095 | +1 | 5 |
| 4 | y | 1.372E+05 | 590.3 | 0.002672 | 4.526 | +1 | 5 |
| - | - | 3.527E+04 | 591.3 | - | - | 0 | - |
| 4 | y | 8.635E+06 | 607.3 | 0.001208 | 1.989 | +1 | 5 |
| - | - | 2.736E+06 | 608.3 | - | - | 0 | - |
| - | - | 5.243E+05 | 609.3 | - | - | 0 | - |
| - | - | 1.666E+04 | 610.3 | - | - | 0 | - |
| - | - | 2.383E+04 | 616.3 | - | - | 0 | - |
| - | - | 2.288E+05 | 617.3 | - | - | 0 | - |
| - | - | 7.887E+04 | 618.3 | - | - | 0 | - |
| - | - | 2.099E+04 | 619.3 | - | - | 0 | - |
| - | - | 5.189E+04 | 626.3 | - | - | 0 | - |
| - | - | 3.587E+04 | 627.3 | - | - | 0 | - |
| - | - | 2.633E+04 | 629.3 | - | - | 0 | - |
| - | - | 2.532E+04 | 631.3 | - | - | 0 | - |
| - | - | 1.907E+04 | 632.3 | - | - | 0 | - |
| - | - | 3.275E+04 | 634.3 | - | - | 0 | - |
| - | - | 2.068E+05 | 644.3 | - | - | 0 | - |
| - | - | 6.657E+04 | 645.3 | - | - | 0 | - |
| - | - | 3.862E+04 | 648.3 | - | - | 0 | - |
| - | - | 1.951E+04 | 652.3 | - | - | 0 | - |
| - | - | 2.857E+04 | 658.3 | - | - | 0 | - |
| - | - | 8.84E+04 | 659.3 | - | - | 0 | - |
| - | - | 3.895E+04 | 660.3 | - | - | 0 | - |
| - | - | 2.884E+05 | 662.3 | - | - | 0 | - |
| - | - | 1.025E+05 | 663.3 | - | - | 0 | - |
| - | - | 1.295E+05 | 664.3 | - | - | 0 | - |
| - | - | 3.596E+04 | 665.3 | - | - | 0 | - |
| - | - | 2.396E+04 | 672.3 | - | - | 0 | - |
| - | - | 1.606E+04 | 674.3 | - | - | 0 | - |
| 3 | y | 9.614E+05 | 676.3 | 0.0009236 | 1.366 | +1 | 6 |
| 3 | y | 4.68E+05 | 677.3 | 0.01245 | 18.38 | +1 | 6 |
| - | - | 1.25E+05 | 678.3 | - | - | 0 | - |
| - | - | 1.918E+04 | 679.3 | - | - | 0 | - |
| - | - | 2.852E+04 | 686.3 | - | - | 0 | - |
| - | - | 5.569E+04 | 692.3 | - | - | 0 | - |
| - | - | 1.84E+04 | 693.3 | - | - | 0 | - |
| 3 | y | 2.092E+07 | 694.4 | 0.001162 | 1.674 | +1 | 6 |
| - | - | 7.424E+06 | 695.4 | - | - | 0 | - |
| - | - | 1.659E+06 | 696.4 | - | - | 0 | - |
| - | - | 7.879E+04 | 697.4 | - | - | 0 | - |
| - | - | 3.599E+04 | 703.3 | - | - | 0 | - |
| - | - | 2.758E+05 | 704.3 | - | - | 0 | - |
| - | - | 8.796E+04 | 705.3 | - | - | 0 | - |
| - | - | 3.769E+04 | 706.3 | - | - | 0 | - |
| 7 | b | 8.099E+04 | 713.3 | 0.0007891 | 1.106 | +1 | 7 |
| - | - | 3.249E+04 | 714.3 | - | - | 0 | - |
| 7 | b | 3.511E+05 | 731.3 | 0.0009666 | 1.322 | +1 | 7 |
| - | - | 1.228E+05 | 732.3 | - | - | 0 | - |
| - | - | 2.29E+04 | 733.3 | - | - | 0 | - |
| - | - | 2.312E+04 | 749.3 | - | - | 0 | - |
| - | - | 3.21E+04 | 761.3 | - | - | 0 | - |
| - | - | 1.861E+04 | 762.3 | - | - | 0 | - |
| - | - | 3.859E+04 | 773.4 | - | - | 0 | - |
| - | - | 5.294E+04 | 774.3 | - | - | 0 | - |
| - | - | 1.814E+04 | 775.4 | - | - | 0 | - |
| - | - | 4.375E+04 | 779.3 | - | - | 0 | - |
| - | - | 2.65E+04 | 780.3 | - | - | 0 | - |
| 2 | y | 6.183E+05 | 791.4 | 0.0007139 | 0.9022 | +1 | 7 |
| - | - | 2.575E+05 | 792.4 | - | - | 0 | - |
| - | - | 6.393E+04 | 793.4 | - | - | 0 | - |
| - | - | 2.202E+04 | 801.4 | - | - | 0 | - |
| 2 | y | 1.958E+06 | 809.4 | 0.0005252 | 0.6489 | +1 | 7 |
| - | - | 8.704E+05 | 810.4 | - | - | 0 | - |
| - | - | 2.06E+05 | 811.4 | - | - | 0 | - |
| - | - | 1.805E+05 | 819.4 | - | - | 0 | - |
| - | - | 7.957E+04 | 820.4 | - | - | 0 | - |
| - | - | 1.252E+04 | 1589 | - | - | 0 | - |

m/z Charge Intensity FragmentType MassShift Position
120.08125305175781 0 1291746.6
121.08464050292969 0 111922.64
122.02436828613281 0 26895.832
122.0711669921875 0 11415
123.0555419921875 0 13091.872
123.09241485595703 0 9257.091
124.0400390625 0 17857.66
124.07107543945312 0 17024.71
124.0761947631836 0 336492.44
124.08731842041016 0 30358.514
125.03494262695312 0 13446.174
125.0714340209961 0 29492.37
125.0795669555664 0 26605.016
126.05525970458984 0 14628.966
127.05062103271484 0 37738.14
127.08708190917969 0 123912.586
129.0663604736328 0 118803.836
129.10281372070312 0 59023.74
129.11387634277344 0 15007.668
130.05035400390625 0 638124.94
130.09799194335938 0 34728.832
131.04934692382812 0 13960.034
131.05398559570312 0 27362.68
131.08197021484375 0 2478769.5 d 1
132.07997131347656 0 23824.336
132.0853729248047 0 134340.17
133.09764099121094 0 16325.834
133.67628479003906 0 9793.225
134.0450897216797 0 16783.361
136.08741760253906 0 80805.625
138.05557250976562 0 17121.13
138.06668090820312 0 55289.31
138.09217834472656 0 13920.68
139.05067443847656 0 227479.03
139.08726501464844 0 8761.711
140.0347137451172 0 70447.65
140.08233642578125 0 1022192.2
141.0663299560547 0 196611.97
141.0858154296875 0 65678.47
141.10272216796875 0 1633723.5
142.07003784179688 0 19547.56
142.08705139160156 0 21451.965
142.10009765625 0 16406.557
142.1061553955078 0 109156.67
143.04566955566406 0 75251.01
145.06163024902344 0 12319.329
148.06124877929688 0 15772.389
149.0452880859375 0 12205.346
149.0601348876953 0 35987.883
150.01942443847656 0 13137.663
151.0867919921875 0 14331.594
151.09829711914062 0 20955.143
152.0718994140625 0 14402.929
152.08245849609375 0 197076.94
153.06642150878906 0 46498.496
153.08563232421875 0 15902.53 y Ammonia loss 6
153.1138458251953 0 25139.102
154.09791564941406 0 34440.684
155.0456085205078 0 56041.816
155.08203125 0 166856.78
156.07730102539062 0 160698.84
156.10215759277344 0 18175.97
157.06129455566406 0 13004620 a Water loss 1
157.10879516601562 0 140513.8
158.0452423095703 0 102137.21
158.05821228027344 0 82679.87
158.0646514892578 0 842485.9
158.09288024902344 0 935694.5
158.11265563964844 0 13306.344
159.0657958984375 0 63392.516
159.0769805908203 0 110491.305
159.11328125 0 8721291
160.1103973388672 0 45477.945
160.11666870117188 0 645502.3
161.11865234375 0 24259.5
165.1029052734375 0 12151.563
166.06187438964844 0 20262.182
166.08676147460938 0 1075609.2 y 7
167.04562377929688 0 203718.33
167.0823211669922 0 29374.1
167.09014892578125 0 96468.12
167.1183319091797 0 15276.968
168.04925537109375 0 18934.238
168.0771942138672 0 60985.324
169.06126403808594 0 991595.5
169.09765625 0 761974.2
169.10882568359375 0 114783.46
170.04588317871094 0 9867.139
170.06466674804688 0 60411.29
170.1008758544922 0 51108.52
171.07696533203125 0 26664.84
171.11297607421875 0 18759.82
172.1085968017578 0 778453.44
173.05628967285156 0 40207.49
173.0926055908203 0 256521.02
173.1020965576172 0 13227.55
173.10423278808594 0 9144.372
173.11212158203125 0 41198.695
175.07191467285156 0 9836232 a 1
175.11941528320312 0 944437.3
176.07522583007812 0 611046.4
176.12298583984375 0 66278.555
177.07626342773438 0 77127.86
179.0456085205078 0 20172.201
179.61422729492188 0 11135.633
180.07725524902344 0 198294.27
181.06109619140625 0 43339.598
181.08078002929688 0 16347.832
182.0567626953125 0 25514.596
182.0929412841797 0 41607.9
183.0762481689453 0 12585.006
183.11329650878906 0 87933.86
184.07217407226562 0 69907.21
184.09701538085938 0 18597.822
185.05628967285156 0 1041596.9 b Water loss 1
185.10313415527344 0 12684.994
185.1282196044922 0 12230.817
186.04025268554688 0 37184.754
186.0595245361328 0 74200.664
186.087890625 0 309407.03
187.0718994140625 0 185918.08
187.10826110839844 0 2685704.5
188.1117401123047 0 234362.83
190.11927795410156 0 43152.805
191.04498291015625 0 12139.4
193.0980224609375 0 17800.416
194.05661010742188 0 55236.938
194.12948608398438 0 15005.258
196.07186889648438 0 31804.74
196.1090545654297 0 20215.252
196.11883544921875 0 17264.322
197.0561065673828 0 46042.668
197.0933074951172 0 80200.92
197.10382080078125 0 2316171.8
198.0876922607422 0 141868.05
198.10719299316406 0 175277.98
198.12400817871094 0 13447.138
199.07176208496094 0 78493.445
199.1083526611328 0 30987.07
200.10687255859375 0 21552.324
201.12387084960938 0 609139
202.08274841308594 0 47210.117
202.12730407714844 0 63031.477
203.06678771972656 0 14822571 b 1
204.07015991210938 0 999233.4
204.09841918945312 0 62246.07
205.07131958007812 0 114426.05
206.0930938720703 0 23439.77
207.1134033203125 0 61367.42
208.07241821289062 0 72600.76
208.10865783691406 0 19885.477
208.45030212402344 0 10908.69
209.05624389648438 0 28967.367
209.09329223632812 0 14165.708
210.12429809570312 0 117238.73
211.10841369628906 0 56844.582
211.1272430419922 0 12048.289
212.06698608398438 0 47098.25
213.05128479003906 0 102113.48
214.0545654296875 0 14482.959
214.0831298828125 0 16235.467
214.13040161132812 0 596711.56
215.1144561767578 0 633005.9
215.13388061523438 0 40902.91
216.0989532470703 0 14567.916
216.11798095703125 0 32775.883
217.08238220214844 0 89756.414
217.1343231201172 0 44890.43
220.10862731933594 0 45210.07
221.0779266357422 0 44818.05
221.0928192138672 0 71760.05
222.09576416015625 0 16046.257
222.62208557128906 0 31627.998
223.10787963867188 0 19445.885
223.12257385253906 0 11548.224
224.0673370361328 0 51537.89
224.10350036621094 0 40393.418
224.14027404785156 0 16823.244
225.05120849609375 0 105994.54
225.0994873046875 0 13175.67
226.0550079345703 0 18819.688
226.0829315185547 0 334924.44
226.11941528320312 0 37985.508
227.06674194335938 0 57420.652
227.08670043945312 0 38191.27
227.11758422851562 0 20079.045
228.13499450683594 0 19349.066
228.19351196289062 0 12398.067
229.1189422607422 0 694340.6
230.1223602294922 0 75609.72
232.1410675048828 0 4623500
233.14427185058594 0 400408.3
234.1251983642578 0 17787.469
234.1458740234375 0 16713.385
236.06723022460938 0 203625.81
237.05130004882812 0 37283.19
237.07032775878906 0 24564.691
237.08653259277344 0 13404.139
237.1351318359375 0 29977.707
238.11932373046875 0 215102.8
239.10299682617188 0 103413.43
239.12277221679688 0 24148.994
240.10728454589844 0 15541.516
241.08106994628906 0 13701.785
241.0938262939453 0 26622.928
242.07774353027344 0 189302.23
242.12786865234375 0 84959.125
243.0614013671875 0 28008.992
243.13363647460938 0 33482.105
244.09352111816406 0 660447 a Water loss 2
244.145263671875 0 30127
245.0775146484375 0 299988.53
245.0965576171875 0 63521.992
245.1298370361328 0 17948.87
245.62477111816406 0 179824.84 y Water loss 4
246.0811004638672 0 31916.17
246.09849548339844 0 10985.617
246.11288452148438 0 45538.688 y Ammonia loss 4
246.12652587890625 0 30347.81
248.1033477783203 0 34045.54
251.06639099121094 0 13023.935
251.10333251953125 0 15672.775
252.13497924804688 0 107407.38
253.09356689453125 0 41262.316
254.07781982421875 0 857453.44
254.6298828125 0 63341.2 y 4
255.05999755859375 0 15805.806
255.08087158203125 0 64047.57
255.12933349609375 0 19140.158
255.14569091796875 0 34750.58
256.1298522949219 0 959887.06
257.1136779785156 0 128412.31
257.1326904296875 0 107372.91
257.6240234375 0 13171.951
259.1558837890625 0 309487.38
260.0884704589844 0 36265.383
260.1590270996094 0 52526.355
262.10394287109375 0 553823.25 a 2
262.1300964355469 0 21940.219
262.155517578125 0 322934.78
263.10736083984375 0 40966.824
263.1395263671875 0 463711.9
263.1578369140625 0 38297.465
264.0984802246094 0 22655.152
264.1429443359375 0 56452.633
265.13006591796875 0 347904.78
266.1141052246094 0 130526.13
266.1355285644531 0 81367.09
266.6298828125 0 15510.246
267.0975646972656 0 27096.895
267.1162414550781 0 19069.441
267.13201904296875 0 19146.19
268.129638671875 0 29098.846
269.0771179199219 0 16576.73
270.146484375 0 23136.568
271.1042785644531 0 740373
271.130615234375 0 30238.084
272.0882873535156 0 7014615.5 b Water loss 2
273.0915832519531 0 743420.44
274.09326171875 0 89412.125
274.1402282714844 0 71752.83
274.1556091308594 0 33109.625
276.1824035644531 0 29652.023
278.1140441894531 0 17453.094
280.1410217285156 0 142298.84
280.1659240722656 0 20688.754
281.1116638183594 0 18156.87
281.1257629394531 0 35428.754
281.14349365234375 0 19124.352
283.1407470703125 0 315264.88
284.12481689453125 0 1012611.56
284.1410827636719 0 38458.758
285.1261901855469 0 219827.56
286.1255798339844 0 13210.884
286.1404113769531 0 85356.27
287.0877380371094 0 17002.078
287.1507568359375 0 156299.84
288.1351623535156 0 130426.96
288.1550598144531 0 51477.23
289.1151123046875 0 26562.697
289.1388244628906 0 26323.703
290.0990905761719 0 4748863 b 2
291.1023254394531 0 518990.66
291.1342468261719 0 33919.375
292.10382080078125 0 58529.67
294.1086120605469 0 17465.285
295.1415100097656 0 25782.465
295.15948486328125 0 28134.775 y Water loss 3
296.0885009765625 0 21968.986
296.12432861328125 0 45921.586
297.16766357421875 0 42447.562
298.1405944824219 0 165200.69
298.1552429199219 0 46971.676
299.14447021484375 0 18597.912
299.6517028808594 0 67637.48
300.15374755859375 0 25101.045
301.1512756347656 0 185276.06
302.13531494140625 0 199929.6
302.15277099609375 0 32848.406
303.1374206542969 0 33153.098
304.164306640625 0 448385.22 y 3
304.666015625 0 136259.9
305.1616516113281 0 2800127 y Ammonia loss 6
306.16473388671875 0 479533.6
307.14117431640625 0 190292.28
307.16656494140625 0 45314.367
308.13616943359375 0 2043719
309.1201171875 0 84272.18
309.13934326171875 0 298566.2
309.1556396484375 0 16884.504
310.1405029296875 0 47840.375
311.1360168457031 0 30050.975
312.1198425292969 0 77058.47
313.15130615234375 0 38121.652
313.64886474609375 0 15708.371
314.0996398925781 0 47466.49
314.1353759765625 0 73766.9
315.1783142089844 0 210529.94
315.6727600097656 0 42999.812
316.1510314941406 0 394403.22
316.1819763183594 0 25566.465
317.1546630859375 0 65869.29
319.1597595214844 0 25129.18
319.17742919921875 0 114173.32
320.1614074707031 0 232523.7
320.18072509765625 0 15139.315
321.1632080078125 0 41372.586
322.1882629394531 0 504321.94 y 6
322.65447998046875 0 45237.027
323.1912841796875 0 69191.95
324.6781311035156 0 62881.63
324.6968078613281 0 12891.598
325.1517028808594 0 1836285.9
326.1473083496094 0 1295903.4
327.1493835449219 0 217322.34
328.129638671875 0 16519.334
328.15093994140625 0 49930.742
329.6700134277344 0 83133.29
330.1291809082031 0 31824.988
330.1716613769531 0 21531.02
332.1098327636719 0 76201.766
332.17047119140625 0 24013.781
333.204345703125 0 30708.385
334.16058349609375 0 13853.77
335.1353454589844 0 31340.395
336.6513671875 0 22802.184
337.1159362792969 0 20777.273
337.1526794433594 0 17076.031
337.1883544921875 0 52629.867
338.0996398925781 0 20821.459
338.1924743652344 0 15176.584
338.6752624511719 0 852963.75 y Water loss 2
339.17657470703125 0 299227.22
339.6783447265625 0 61671.617
343.1632080078125 0 3888516.5
343.66717529296875 0 59542
344.1618957519531 0 518043.12
345.15753173828125 0 276378.88
345.1758728027344 0 49206.973
346.15985107421875 0 36540.508
347.6803894042969 0 486419.97 y 2
348.16009521484375 0 19759.992
348.18170166015625 0 174798.5
348.6598815917969 0 14883.288
348.6824645996094 0 30378.008
352.6723327636719 0 110105.484
353.1464538574219 0 247183.3
354.1492919921875 0 57876.984
355.1258544921875 0 18585.268
355.16192626953125 0 21260.824
356.1097717285156 0 66202.93
357.1145935058594 0 12940.035
357.164794921875 0 91092.23 b Water loss 6
357.6659240722656 0 42488.344
361.1835021972656 0 5594468.5
362.1846923828125 0 1411360.4 y Ammonia loss 5
363.18682861328125 0 206620.81
364.19732666015625 0 16378.4
366.1703796386719 0 122325.88 b 6
366.67156982421875 0 39294.945
367.16510009765625 0 21975.895
368.14703369140625 0 17417.406
371.1570129394531 0 1907253.8 b Water loss 3
372.1602783203125 0 318963.78
373.1853332519531 0 162837.45
373.6876220703125 0 59029.316
374.12030029296875 0 27343.775
374.189208984375 0 13509.305
377.15667724609375 0 24304.492
378.66961669921875 0 19441.658
379.2097473144531 0 11775358 y 5
380.2125549316406 0 2226622.2
381.1837463378906 0 29862.613
381.21527099609375 0 262191.84
382.19091796875 0 22839.625
383.12066650390625 0 83991.266
383.1577453613281 0 18473.764
384.2041320800781 0 20926.139
385.1871643066406 0 72893.22
386.1905212402344 0 17071.54
387.1830749511719 0 60199.742
387.67962646484375 0 32676.076
389.16754150390625 0 367600.2 b 3
389.1947021484375 0 26522.764
390.17242431640625 0 69559.02
390.1972351074219 0 16860.973
390.6721496582031 0 18950.746
395.16778564453125 0 17899.303
396.1885986328125 0 770935.2 y Water loss 1
396.6899108886719 0 304840.97
397.1914367675781 0 74867.36
397.22021484375 0 20501.479
400.2208251953125 0 21292.488
401.1314697265625 0 132517.66
401.1795959472656 0 19100.127
402.1320495605469 0 18764.611
403.1832580566406 0 17848.27
405.1939392089844 0 89974.45 y 1
405.694580078125 0 21809.758
407.2033386230469 0 17470.02
408.1888122558594 0 35100.93
409.17266845703125 0 35057.797
410.1858215332031 0 323272.94
410.6879577636719 0 130443.72
411.1883544921875 0 40300.16
413.1693115234375 0 83402.32
414.1708068847656 0 23748.742
414.24688720703125 0 77673.46
415.2480163574219 0 16600.287
415.69366455078125 0 20093.496
416.1876220703125 0 26104.998
416.7013244628906 0 69628.59
417.2059020996094 0 16634.91
419.1414489746094 0 46778.727
421.6939697265625 0 55370.8
422.18695068359375 0 101839.27
422.68792724609375 0 61120.094
424.2314453125 0 32598.58
424.7003479003906 0 36986.99
425.215087890625 0 238209.84
426.2154846191406 0 34800.297
427.1831970214844 0 21476.072
427.21044921875 0 89150.88
430.2094421386719 0 180136.22
430.6996154785156 0 370907.56
431.1956481933594 0 398168.72
431.70184326171875 0 33064.99
432.1963195800781 0 68747.3
436.18450927734375 0 47661.83
437.168212890625 0 23416.508
439.70489501953125 0 1289085.6 Precursor Water loss
440.2061767578125 0 564914.1
440.7073059082031 0 152240.8
441.20947265625 0 20333.328
442.2419128417969 0 315945.75
443.2248229980469 0 38503.79
443.25042724609375 0 21801.775
444.2276916503906 0 17253.436
445.22308349609375 0 14556.91
448.22021484375 0 87875.91
448.7104187011719 0 150387.66 Precursor
449.20782470703125 0 141824.38
449.712646484375 0 19673.62
450.20849609375 0 34459.44
454.19482421875 0 95496.516
455.1768493652344 0 48388.734
455.20452880859375 0 82858.92
456.1830139160156 0 29088.773
460.25250244140625 0 1526314.4
461.25518798828125 0 321918.5
462.2510070800781 0 73248.71
463.2516174316406 0 16213.804
466.23504638671875 0 46679.797
467.2304382324219 0 18258.957
470.1939697265625 0 24636.838
470.2373352050781 0 78926.71
472.204833984375 0 140349.53
472.239013671875 0 17741.076
473.21551513671875 0 279782.84
474.2168273925781 0 82792.555
482.1888122558594 0 89037.91
484.25262451171875 0 22016.943
487.2510681152344 0 39747.75
490.2420959472656 0 2429551.8 y Water loss 4
491.23016357421875 0 355818.28 y Ammonia loss 4
491.243896484375 0 428686.06
492.2302551269531 0 118124.83
493.2341003417969 0 21244.266
494.2345275878906 0 34096.39
500.19964599609375 0 238268.17 b Water loss 4
501.20458984375 0 65423.344
501.279541015625 0 176669.64
502.2049865722656 0 17281.28
502.28216552734375 0 40244.23
505.26385498046875 0 16514.348
508.25262451171875 0 29453226 y 4
509.25537109375 0 7440349.5
509.36712646484375 0 19063.11
510.2576599121094 0 1182641.8
511.2621154785156 0 126137.34
512.24755859375 0 163094.2
513.2437744140625 0 18159.158
517.2745361328125 0 80872.4
518.204345703125 0 50916.113 b 4
518.2369995117188 0 165167.88
518.278076171875 0 19815.488
519.205322265625 0 21122.484
519.24169921875 0 39970.938
529.2738647460938 0 895258.56
530.2742919921875 0 196086.55
531.2781982421875 0 23105.498
539.2557983398438 0 22410.086
547.2844848632812 0 3501309.2
548.2869262695312 0 840780
549.2889404296875 0 146135.61
557.22412109375 0 16033.67 b Water loss 5
557.2686767578125 0 71684.47
561.31591796875 0 17438.861
571.2989501953125 0 104069.89
572.2847900390625 0 133476.64
573.2955932617188 0 21177.47
589.3096313476562 0 64262.72 y Water loss 3
590.2959594726562 0 137225.1 y Ammonia loss 3
591.2988891601562 0 35274.164
607.321044921875 0 8634630 y 3
608.32373046875 0 2735781.8
609.3262329101562 0 524260.16
610.3267211914062 0 16660.02
616.3077392578125 0 23829.865
617.3050537109375 0 228827.83
618.3077392578125 0 78872.75
619.31884765625 0 20986.256
626.2906494140625 0 51890.395
627.2766723632812 0 35866.35
629.3048706054688 0 26333.621
631.3135986328125 0 25321.33
632.2610473632812 0 19071.41
634.3199462890625 0 32746.252
644.3006591796875 0 206819.89
645.30029296875 0 66569.586
648.34619140625 0 38620.676
652.32958984375 0 19505.285
658.334716796875 0 28565.275
659.3182373046875 0 88396.66
660.3138427734375 0 38949.156
662.3114013671875 0 288397.28
663.3138427734375 0 102526.36
664.3405151367188 0 129522.74
665.3446655273438 0 35961.973
672.298583984375 0 23959.135
674.3301391601562 0 16061.051
676.3422241210938 0 961371.75 y Water loss 2
677.3377685546875 0 468022.06 y Ammonia loss 2
678.3391723632812 0 125049.04
679.331787109375 0 19179.994
686.3314819335938 0 28521.045
692.3004150390625 0 55691.027
693.3058471679688 0 18397.3
694.35302734375 0 20919032 y 2
695.355712890625 0 7423522.5
696.3583984375 0 1659184.9
697.3603515625 0 78789.586
703.3368530273438 0 35985.504
704.3370361328125 0 275806.16
705.3408813476562 0 87964.08
706.3416748046875 0 37693.78
713.3220825195312 0 80989.89 b Water loss 6
714.3224487304688 0 32485.283
731.3328247070312 0 351081.53 b 6
732.335205078125 0 122787.734
733.3388671875 0 22895.299
749.343017578125 0 23124.973
761.3253173828125 0 32097.62
762.32470703125 0 18609.783
773.3606567382812 0 38589.26
774.3486328125 0 52936.004
775.3689575195312 0 18137.639
779.3325805664062 0 43747.902
780.3405151367188 0 26495.309
791.3689575195312 0 618300.75 y Water loss 1
792.3699951171875 0 257522.08
793.37255859375 0 63931.516
801.3636474609375 0 22015.508
809.3793334960938 0 1957733 y 1
810.3821411132812 0 870436.7
811.384521484375 0 206017.11
819.3631591796875 0 180505.34
820.3662109375 0 79574.734
1589.4481201171875 0 12522.372

Spectrum Details

|  |  |
| --- | --- |
| Matched peaks? Matched peaks The total absolute number of peaks matched. Additionally in brackets the total fraction of peaks matched and the total number of peaks is shown. | 46 (7.81% of 589) |
| FDR? FDR The false discovery rate estimated for this peptide. It is calculated by matching all theoretical fragments with a non-integer shift with the raw peaks for this spectrum. This is done with 40 different shifts. The resulting percentage is the average number of annotated peaks over the number of annotated peaks with the correct spectrum. | 0.05% |
| Satellite FDR? Satellite FDR See the FDR for details on its calculation. This satellite ion specific FDR only contains the satellite ions (d/w) for I/L/J positions. | - |
| PSM Score? PSM Score The PSM Score as given by Hecklib to this annotated spectrum. It is shown with three significant figures. | 557 |

#### Spectrum 4171? Spectrum 4171 The raw spectrum of this peptide as annotated by Hecklib. The fragments are coloured according to ion type (see legend). Any peaks with a star '\*' as text can be hovered over to see the full details, first the ion type second the mass shift type. By hovering over the amino acids in the peptide or ions in the legend the corresponding peaks are highlighted. By toggling the 'Unassigned' label you can turn the background (unassigned) peaks on or off in the plot. By updating the slider in the Ion legend you can update the spectrum to only show the top X% of the peaks with labels. The top X% means any peak that is within X% of the highest intensity. By dragging in the spectrum you can zoom in to a specific part of the spectrum and use 'Zoom Out' to get back to the original zoom level. The annotation of the spectrum is based on the given sequence in the peptides file and is done with different software so inconsistencies are likely. The peaks are annotated based on the given sequence, with 20 ppm tolerance.

Copy Data

##### Spectrum 4171 (TSV)

###### Preview

```
Loading example...
```

*Click on the button to copy the data to your clipboard.*

Mz MinMz MaxIntensity Max

WidthHeightPeptide font sizePeptide stroke widthSpectrum font sizeSpectrum stroke widthCompact peptide

Ion legend

wxyz

abcd

OtherUnassignedIon^Charge^_Position_Show for top:%

SDSVEGRF

04.07e+48.14e+41.22e+51.63e+5

Zoom Out

d^+1^_2_a^+1^_2_y^+1^_1_a^+1^_2_b^+1^_2_b^+1^_2_a^+1^_3_y^+2^_4_a^+1^_3_b^+1^_3_b^+1^_3_y^+2^_5_y^+1^_2_y^+1^_2_y^+2^_6_y^+2^_6_b^+2^_7_y^+1^_3_b^+1^_4_y^+1^_3_b^+1^_4_y^+2^_7_\*\*y^+1^_4_y^+1^_4_b^+1^_5_y^+1^_4_b^+1^_5_y^+1^_5_y^+1^_6_y^+1^_6_y^+1^_6_b^+1^_7_y^+1^_7_y^+1^_7_

0796159123873183

Fragment Matches Table

Show background peaks

| Position | Ion type | Intensity | mz Theoretical | mz Error (Th) | mz Error (ppm) | Charge | Series Number |
| --- | --- | --- | --- | --- | --- | --- | --- |
| - | - | 9287 | 120.1 | - | - | 0 | - |
| - | - | 1009 | 121.1 | - | - | 0 | - |
| - | - | 420.4 | 122.7 | - | - | 0 | - |
| - | - | 2211 | 124.1 | - | - | 0 | - |
| - | - | 452.7 | 126.1 | - | - | 0 | - |
| - | - | 1195 | 127.1 | - | - | 0 | - |
| - | - | 1059 | 129.1 | - | - | 0 | - |
| - | - | 7463 | 129.1 | - | - | 0 | - |
| - | - | 3963 | 130.1 | - | - | 0 | - |
| - | - | 523.3 | 130.1 | - | - | 0 | - |
| - | - | 549.3 | 130.1 | - | - | 0 | - |
| - | - | 1.125E+04 | 131.1 | - | - | 0 | - |
| 2 | d | 1.522E+04 | 131.1 | 0.0003441 | 2.625 | +1 | 2 |
| - | - | 565.2 | 132.1 | - | - | 0 | - |
| - | - | 834.8 | 132.1 | - | - | 0 | - |
| - | - | 8301 | 133.1 | - | - | 0 | - |
| - | - | 526.8 | 134.1 | - | - | 0 | - |
| - | - | 3950 | 136.1 | - | - | 0 | - |
| - | - | 1839 | 139.1 | - | - | 0 | - |
| - | - | 900.4 | 139.1 | - | - | 0 | - |
| - | - | 6635 | 140.1 | - | - | 0 | - |
| - | - | 1386 | 141.1 | - | - | 0 | - |
| - | - | 1.029E+04 | 141.1 | - | - | 0 | - |
| - | - | 505.2 | 142.1 | - | - | 0 | - |
| - | - | 421 | 144.4 | - | - | 0 | - |
| - | - | 933.6 | 146.1 | - | - | 0 | - |
| - | - | 520.3 | 147 | - | - | 0 | - |
| - | - | 428.9 | 148.6 | - | - | 0 | - |
| - | - | 890.6 | 148.9 | - | - | 0 | - |
| - | - | 1.102E+04 | 149 | - | - | 0 | - |
| - | - | 7974 | 149 | - | - | 0 | - |
| - | - | 1369 | 150 | - | - | 0 | - |
| - | - | 1345 | 150 | - | - | 0 | - |
| - | - | 3154 | 151 | - | - | 0 | - |
| - | - | 468.3 | 151 | - | - | 0 | - |
| - | - | 1138 | 152.1 | - | - | 0 | - |
| - | - | 826 | 155.1 | - | - | 0 | - |
| - | - | 529 | 155.1 | - | - | 0 | - |
| - | - | 1843 | 156.1 | - | - | 0 | - |
| 2 | a | 7.766E+04 | 157.1 | 0.0003581 | 2.28 | +1 | 2 |
| - | - | 3115 | 157.1 | - | - | 0 | - |
| - | - | 841 | 157.1 | - | - | 0 | - |
| - | - | 622.7 | 158 | - | - | 0 | - |
| - | - | 5187 | 158.1 | - | - | 0 | - |
| - | - | 5791 | 158.1 | - | - | 0 | - |
| - | - | 730.3 | 159.1 | - | - | 0 | - |
| - | - | 696.6 | 159.1 | - | - | 0 | - |
| - | - | 1430 | 159.1 | - | - | 0 | - |
| - | - | 5.374E+04 | 159.1 | - | - | 0 | - |
| - | - | 4157 | 160.1 | - | - | 0 | - |
| - | - | 576.7 | 161.1 | - | - | 0 | - |
| - | - | 428.2 | 162.2 | - | - | 0 | - |
| - | - | 748.2 | 165.1 | - | - | 0 | - |
| 8 | y | 6776 | 166.1 | 0.0003691 | 2.222 | +1 | 1 |
| - | - | 1148 | 167 | - | - | 0 | - |
| - | - | 1304 | 167 | - | - | 0 | - |
| - | - | 1.156E+04 | 167.1 | - | - | 0 | - |
| - | - | 763 | 167.1 | - | - | 0 | - |
| - | - | 2942 | 168.1 | - | - | 0 | - |
| - | - | 4925 | 169.1 | - | - | 0 | - |
| - | - | 6005 | 169.1 | - | - | 0 | - |
| - | - | 5144 | 169.1 | - | - | 0 | - |
| - | - | 852.4 | 169.1 | - | - | 0 | - |
| - | - | 500.6 | 171.1 | - | - | 0 | - |
| - | - | 464.3 | 171.1 | - | - | 0 | - |
| - | - | 4651 | 172.1 | - | - | 0 | - |
| - | - | 504.1 | 173.1 | - | - | 0 | - |
| - | - | 1307 | 173.1 | - | - | 0 | - |
| - | - | 6867 | 173.1 | - | - | 0 | - |
| - | - | 2165 | 173.4 | - | - | 0 | - |
| 2 | a | 6.194E+04 | 175.1 | 0.0003525 | 2.014 | +1 | 2 |
| - | - | 4711 | 175.1 | - | - | 0 | - |
| - | - | 5690 | 175.1 | - | - | 0 | - |
| - | - | 4151 | 176.1 | - | - | 0 | - |
| - | - | 4060 | 177.1 | - | - | 0 | - |
| - | - | 1685 | 180.1 | - | - | 0 | - |
| - | - | 1122 | 182.1 | - | - | 0 | - |
| - | - | 880.2 | 183.1 | - | - | 0 | - |
| - | - | 968.8 | 183.1 | - | - | 0 | - |
| - | - | 907.9 | 184.1 | - | - | 0 | - |
| 2 | b | 6169 | 185.1 | 0.0002403 | 1.298 | +1 | 2 |
| - | - | 657.4 | 186.1 | - | - | 0 | - |
| - | - | 1837 | 186.1 | - | - | 0 | - |
| - | - | 1165 | 187.1 | - | - | 0 | - |
| - | - | 797 | 187.1 | - | - | 0 | - |
| - | - | 1.611E+04 | 187.1 | - | - | 0 | - |
| - | - | 2244 | 188.1 | - | - | 0 | - |
| - | - | 1135 | 188.1 | - | - | 0 | - |
| - | - | 715.4 | 195.1 | - | - | 0 | - |
| - | - | 746.3 | 197.1 | - | - | 0 | - |
| - | - | 1.351E+04 | 197.1 | - | - | 0 | - |
| - | - | 597.9 | 198.1 | - | - | 0 | - |
| - | - | 1129 | 198.1 | - | - | 0 | - |
| - | - | 532.2 | 199.1 | - | - | 0 | - |
| - | - | 709.4 | 199.1 | - | - | 0 | - |
| - | - | 3007 | 201.1 | - | - | 0 | - |
| - | - | 4286 | 201.1 | - | - | 0 | - |
| - | - | 806 | 202.1 | - | - | 0 | - |
| 2 | b | 8.854E+04 | 203.1 | 0.0002804 | 1.381 | +1 | 2 |
| - | - | 6710 | 204.1 | - | - | 0 | - |
| - | - | 616.9 | 205.1 | - | - | 0 | - |
| - | - | 1073 | 205.1 | - | - | 0 | - |
| - | - | 509.7 | 207.1 | - | - | 0 | - |
| - | - | 503.8 | 208.1 | - | - | 0 | - |
| - | - | 1971 | 209 | - | - | 0 | - |
| - | - | 586.8 | 209.1 | - | - | 0 | - |
| - | - | 1744 | 211 | - | - | 0 | - |
| - | - | 3460 | 214.1 | - | - | 0 | - |
| - | - | 2688 | 215.1 | - | - | 0 | - |
| - | - | 660.4 | 217.1 | - | - | 0 | - |
| - | - | 7943 | 219.1 | - | - | 0 | - |
| - | - | 1171 | 220.1 | - | - | 0 | - |
| - | - | 809.4 | 221.1 | - | - | 0 | - |
| - | - | 1626 | 221.1 | - | - | 0 | - |
| - | - | 3874 | 223.1 | - | - | 0 | - |
| - | - | 2417 | 224.1 | - | - | 0 | - |
| - | - | 6589 | 225 | - | - | 0 | - |
| - | - | 1343 | 225.1 | - | - | 0 | - |
| - | - | 2895 | 226 | - | - | 0 | - |
| - | - | 1979 | 226.1 | - | - | 0 | - |
| - | - | 750.5 | 226.2 | - | - | 0 | - |
| - | - | 1217 | 227 | - | - | 0 | - |
| - | - | 4313 | 227 | - | - | 0 | - |
| - | - | 843 | 228 | - | - | 0 | - |
| - | - | 3407 | 229.1 | - | - | 0 | - |
| - | - | 2.616E+04 | 232.1 | - | - | 0 | - |
| - | - | 1484 | 233.1 | - | - | 0 | - |
| - | - | 1073 | 236.1 | - | - | 0 | - |
| - | - | 2313 | 238.1 | - | - | 0 | - |
| - | - | 611.2 | 239.2 | - | - | 0 | - |
| - | - | 1132 | 240.1 | - | - | 0 | - |
| - | - | 743.2 | 242.1 | - | - | 0 | - |
| 3 | a | 5639 | 244.1 | 0.0004647 | 1.904 | +1 | 3 |
| - | - | 1781 | 245.1 | - | - | 0 | - |
| - | - | 681.8 | 245.1 | - | - | 0 | - |
| 5 | y | 1047 | 245.6 | 0.0008869 | 3.611 | +2 | 4 |
| - | - | 4636 | 254.1 | - | - | 0 | - |
| - | - | 588.4 | 255.1 | - | - | 0 | - |
| - | - | 5587 | 256.1 | - | - | 0 | - |
| - | - | 1855 | 259.2 | - | - | 0 | - |
| 3 | a | 2601 | 262.1 | 0.0003065 | 1.17 | +1 | 3 |
| - | - | 1539 | 262.2 | - | - | 0 | - |
| - | - | 1782 | 263.1 | - | - | 0 | - |
| - | - | 1312 | 265.1 | - | - | 0 | - |
| - | - | 644 | 266.1 | - | - | 0 | - |
| - | - | 5100 | 271.1 | - | - | 0 | - |
| 3 | b | 4.176E+04 | 272.1 | 0.000179 | 0.6579 | +1 | 3 |
| - | - | 4143 | 273.1 | - | - | 0 | - |
| - | - | 846.9 | 280.1 | - | - | 0 | - |
| - | - | 2032 | 283.1 | - | - | 0 | - |
| - | - | 1123 | 283.2 | - | - | 0 | - |
| - | - | 6333 | 284.1 | - | - | 0 | - |
| - | - | 777.6 | 284.1 | - | - | 0 | - |
| - | - | 798.2 | 286.1 | - | - | 0 | - |
| - | - | 810 | 287 | - | - | 0 | - |
| - | - | 826.9 | 288.1 | - | - | 0 | - |
| - | - | 1884 | 289.2 | - | - | 0 | - |
| 3 | b | 2.854E+04 | 290.1 | 0.0003565 | 1.229 | +1 | 3 |
| - | - | 2273 | 291.1 | - | - | 0 | - |
| - | - | 1442 | 298.1 | - | - | 0 | - |
| - | - | 953.7 | 301.2 | - | - | 0 | - |
| - | - | 961 | 302.1 | - | - | 0 | - |
| 4 | y | 1630 | 304.2 | 0.0002617 | 0.8605 | +2 | 5 |
| 7 | y | 1.524E+04 | 305.2 | 0.0004379 | 1.435 | +1 | 2 |
| - | - | 2333 | 306.2 | - | - | 0 | - |
| - | - | 1146 | 307.1 | - | - | 0 | - |
| - | - | 1563 | 307.2 | - | - | 0 | - |
| - | - | 1.11E+04 | 308.1 | - | - | 0 | - |
| - | - | 1908 | 309.1 | - | - | 0 | - |
| - | - | 638.9 | 313.2 | - | - | 0 | - |
| - | - | 843.6 | 315.2 | - | - | 0 | - |
| - | - | 771.2 | 315.7 | - | - | 0 | - |
| - | - | 1739 | 316.2 | - | - | 0 | - |
| - | - | 699.8 | 318.9 | - | - | 0 | - |
| - | - | 692.5 | 319.2 | - | - | 0 | - |
| - | - | 938.5 | 320.2 | - | - | 0 | - |
| - | - | 1543 | 320.9 | - | - | 0 | - |
| 7 | y | 3193 | 322.2 | 0.0006222 | 1.931 | +1 | 2 |
| - | - | 838.8 | 325 | - | - | 0 | - |
| - | - | 1.074E+04 | 325.2 | - | - | 0 | - |
| - | - | 866.3 | 326 | - | - | 0 | - |
| - | - | 6836 | 326.1 | - | - | 0 | - |
| - | - | 3104 | 327 | - | - | 0 | - |
| - | - | 988.3 | 327.1 | - | - | 0 | - |
| - | - | 1042 | 328 | - | - | 0 | - |
| - | - | 636.7 | 332.1 | - | - | 0 | - |
| - | - | 578 | 336.7 | - | - | 0 | - |
| 3 | y | 4983 | 338.7 | 0.0003636 | 1.074 | +2 | 6 |
| - | - | 1019 | 339.2 | - | - | 0 | - |
| - | - | 552.7 | 340.9 | - | - | 0 | - |
| - | - | 700.2 | 341 | - | - | 0 | - |
| - | - | 859.8 | 342 | - | - | 0 | - |
| - | - | 3753 | 343 | - | - | 0 | - |
| - | - | 2.228E+04 | 343.2 | - | - | 0 | - |
| - | - | 1294 | 344 | - | - | 0 | - |
| - | - | 1464 | 344 | - | - | 0 | - |
| - | - | 3033 | 344.2 | - | - | 0 | - |
| - | - | 3894 | 345 | - | - | 0 | - |
| - | - | 1898 | 345.2 | - | - | 0 | - |
| - | - | 1539 | 346 | - | - | 0 | - |
| 3 | y | 2649 | 347.7 | 0.0001275 | 0.3667 | +2 | 6 |
| - | - | 1722 | 353.1 | - | - | 0 | - |
| - | - | 682.3 | 356.1 | - | - | 0 | - |
| - | - | 931.7 | 357.1 | - | - | 0 | - |
| 7 | b | 1021 | 357.2 | 0.000375 | 1.05 | +2 | 7 |
| - | - | 9802 | 359 | - | - | 0 | - |
| - | - | 1.263E+04 | 360 | - | - | 0 | - |
| - | - | 4.787E+04 | 361 | - | - | 0 | - |
| - | - | 3.067E+04 | 361.2 | - | - | 0 | - |
| - | - | 2.016E+04 | 362 | - | - | 0 | - |
| 6 | y | 7122 | 362.2 | 0.002442 | 6.743 | +1 | 3 |
| - | - | 8086 | 363 | - | - | 0 | - |
| - | - | 1010 | 363.2 | - | - | 0 | - |
| - | - | 577.7 | 364 | - | - | 0 | - |
| - | - | 542.9 | 370.2 | - | - | 0 | - |
| 4 | b | 1.182E+04 | 371.2 | 0.000216 | 0.582 | +1 | 4 |
| - | - | 1743 | 372.2 | - | - | 0 | - |
| - | - | 1281 | 373.7 | - | - | 0 | - |
| - | - | 819.8 | 377.2 | - | - | 0 | - |
| 6 | y | 6.61E+04 | 379.2 | 0.000124 | 0.3271 | +1 | 3 |
| - | - | 1.247E+04 | 380.2 | - | - | 0 | - |
| - | - | 632.9 | 381.1 | - | - | 0 | - |
| - | - | 1719 | 381.2 | - | - | 0 | - |
| - | - | 631.8 | 387.2 | - | - | 0 | - |
| 4 | b | 2160 | 389.2 | 0.0006987 | 1.795 | +1 | 4 |
| - | - | 1.195E+04 | 395.2 | - | - | 0 | - |
| 2 | y | 3095 | 396.2 | 0.0001685 | 0.4252 | +2 | 7 |
| - | - | 2970 | 396.2 | - | - | 0 | - |
| - | - | 1697 | 396.7 | - | - | 0 | - |
| - | - | 658.5 | 398.1 | - | - | 0 | - |
| - | - | 731.2 | 409.1 | - | - | 0 | - |
| - | - | 1879 | 410.2 | - | - | 0 | - |
| - | - | 645.6 | 421.6 | - | - | 0 | - |
| - | - | 1118 | 422.2 | - | - | 0 | - |
| - | - | 1199 | 425.2 | - | - | 0 | - |
| - | - | 1416 | 429.1 | - | - | 0 | - |
| - | - | 1279 | 430.1 | - | - | 0 | - |
| - | - | 743.7 | 430.3 | - | - | 0 | - |
| - | - | 2323 | 430.7 | - | - | 0 | - |
| - | - | 757 | 430.9 | - | - | 0 | - |
| - | - | 6599 | 431.1 | - | - | 0 | - |
| - | - | 2401 | 431.2 | - | - | 0 | - |
| - | - | 647.2 | 431.8 | - | - | 0 | - |
| - | - | 4723 | 432.1 | - | - | 0 | - |
| - | - | 919 | 432.9 | - | - | 0 | - |
| - | - | 1430 | 433.1 | - | - | 0 | - |
| 0 | Precursor | 7519 | 439.7 | 0.000541 | 1.23 | +2 | -1 |
| - | - | 3251 | 440.2 | - | - | 0 | - |
| - | - | 2005 | 442.2 | - | - | 0 | - |
| - | - | 3131 | 448.3 | - | - | 0 | - |
| 0 | Precursor | 759.8 | 448.7 | 0.003154 | 7.03 | +2 | -1 |
| - | - | 1039 | 448.8 | - | - | 0 | - |
| - | - | 698.7 | 449.3 | - | - | 0 | - |
| - | - | 650.4 | 454.2 | - | - | 0 | - |
| - | - | 7221 | 460.3 | - | - | 0 | - |
| - | - | 2197 | 461.3 | - | - | 0 | - |
| - | - | 2076 | 473.2 | - | - | 0 | - |
| 5 | y | 1.389E+04 | 490.2 | 0.0002001 | 0.4082 | +1 | 4 |
| 5 | y | 2337 | 491.2 | 0.005046 | 10.27 | +1 | 4 |
| - | - | 2257 | 491.2 | - | - | 0 | - |
| 5 | b | 1933 | 500.2 | 0.0007443 | 1.488 | +1 | 5 |
| 5 | y | 1.611E+05 | 508.3 | 0.0001945 | 0.3827 | +1 | 4 |
| - | - | 4.285E+04 | 509.3 | - | - | 0 | - |
| - | - | 7374 | 510.3 | - | - | 0 | - |
| - | - | 772.7 | 512.2 | - | - | 0 | - |
| 5 | b | 821.1 | 518.2 | 0.002801 | 5.406 | +1 | 5 |
| - | - | 4347 | 529.3 | - | - | 0 | - |
| - | - | 991.8 | 530.3 | - | - | 0 | - |
| - | - | 580.5 | 536.6 | - | - | 0 | - |
| - | - | 1.838E+04 | 547.3 | - | - | 0 | - |
| - | - | 5484 | 548.3 | - | - | 0 | - |
| - | - | 1049 | 549.3 | - | - | 0 | - |
| - | - | 1196 | 555.3 | - | - | 0 | - |
| - | - | 1657 | 571.3 | - | - | 0 | - |
| - | - | 1083 | 571.8 | - | - | 0 | - |
| 4 | y | 4.742E+04 | 607.3 | 0.0001095 | 0.1802 | +1 | 5 |
| - | - | 1.578E+04 | 608.3 | - | - | 0 | - |
| - | - | 3902 | 609.3 | - | - | 0 | - |
| - | - | 1277 | 617.3 | - | - | 0 | - |
| - | - | 846.4 | 618.3 | - | - | 0 | - |
| - | - | 1163 | 644.3 | - | - | 0 | - |
| - | - | 631.3 | 645.3 | - | - | 0 | - |
| - | - | 2010 | 662.3 | - | - | 0 | - |
| 3 | y | 5193 | 676.3 | 0.0005574 | 0.8241 | +1 | 6 |
| 3 | y | 2551 | 677.3 | 0.008302 | 12.26 | +1 | 6 |
| 3 | y | 1.218E+05 | 694.4 | 0.0002417 | 0.3481 | +1 | 6 |
| - | - | 4.309E+04 | 695.4 | - | - | 0 | - |
| - | - | 8792 | 696.4 | - | - | 0 | - |
| - | - | 2090 | 704.3 | - | - | 0 | - |
| 7 | b | 2232 | 731.3 | 0.001231 | 1.683 | +1 | 7 |
| - | - | 754.8 | 732.3 | - | - | 0 | - |
| 2 | y | 3127 | 791.4 | 0.000995 | 1.257 | +1 | 7 |
| - | - | 1001 | 792.4 | - | - | 0 | - |
| 2 | y | 1.076E+04 | 809.4 | 0.0002682 | 0.3314 | +1 | 7 |
| - | - | 3648 | 810.4 | - | - | 0 | - |
| - | - | 1518 | 811.4 | - | - | 0 | - |
| - | - | 657.4 | 819.4 | - | - | 0 | - |
| - | - | 636.8 | 1012 | - | - | 0 | - |
| - | - | 752.8 | 1265 | - | - | 0 | - |
| - | - | 705.1 | 1468 | - | - | 0 | - |
| - | - | 598.6 | 1838 | - | - | 0 | - |
| - | - | 640.3 | 1878 | - | - | 0 | - |
| - | - | 593.8 | 1922 | - | - | 0 | - |
| - | - | 663 | 2063 | - | - | 0 | - |
| - | - | 810.9 | 2623 | - | - | 0 | - |
| - | - | 694.8 | 2646 | - | - | 0 | - |
| - | - | 758.6 | 3151 | - | - | 0 | - |

m/z Charge Intensity FragmentType MassShift Position
120.08110046386719 0 9286.614
121.0843505859375 0 1008.9209
122.66899108886719 0 420.3691
124.0760498046875 0 2211.4902
126.05512237548828 0 452.7452
127.08698272705078 0 1195.0175
129.06622314453125 0 1058.9948
129.10255432128906 0 7463.4785
130.05020141601562 0 3963.3545
130.06544494628906 0 523.313
130.1060791015625 0 549.29834
131.07058715820312 0 11246.514
131.08184814453125 0 15215.734 d 1
132.0738525390625 0 565.1569
132.08535766601562 0 834.83014
133.08627319335938 0 8300.984
134.089599609375 0 526.81506
136.07601928710938 0 3950.198
139.05055236816406 0 1839.0562
139.07546997070312 0 900.3802
140.08216857910156 0 6635.4106
141.06614685058594 0 1386.2003
141.10256958007812 0 10294.818
142.1063690185547 0 505.1894
144.41082763671875 0 420.98538
146.0602569580078 0 933.60455
147.04461669921875 0 520.32214
148.55091857910156 0 428.88666
148.947021484375 0 890.5742
149.02369689941406 0 11023.0205
149.04522705078125 0 7974.1367
150.02708435058594 0 1368.9572
150.0443878173828 0 1345.1912
151.04214477539062 0 3153.7578
151.04786682128906 0 468.29724
152.08230590820312 0 1138.0745
155.0817413330078 0 826.025
155.11810302734375 0 528.9915
156.07723999023438 0 1842.7429
157.06112670898438 0 77660.984 a Water loss 1
157.08631896972656 0 3114.927
157.10858154296875 0 841.00854
158.0450439453125 0 622.73615
158.0644989013672 0 5186.6084
158.09263610839844 0 5790.8457
159.06515502929688 0 730.30457
159.0770263671875 0 696.5581
159.0919952392578 0 1429.7916
159.11314392089844 0 53744.387
160.11642456054688 0 4157.3213
161.11793518066406 0 576.6716
162.1879425048828 0 428.24008
165.05455017089844 0 748.1832
166.0866241455078 0 6776.4307 y 7
167.03433227539062 0 1147.5112
167.0457000732422 0 1303.8317
167.0557403564453 0 11560.9
167.0902557373047 0 762.9755
168.0557861328125 0 2941.5986
169.052490234375 0 4925.0796
169.06112670898438 0 6005.3394
169.09744262695312 0 5143.638
169.10874938964844 0 852.38666
171.07656860351562 0 500.58835
171.1377410888672 0 464.31546
172.10848999023438 0 4651.071
173.0563507080078 0 504.13126
173.09228515625 0 1307.0327
173.12876892089844 0 6866.661
173.4398193359375 0 2165.324
175.07168579101562 0 61937.367 a 1
175.0967559814453 0 4710.5107
175.11935424804688 0 5689.7197
176.07505798339844 0 4150.6504
177.11236572265625 0 4059.866
180.07704162597656 0 1685.0143
182.08192443847656 0 1122.1447
183.1132049560547 0 880.24725
183.14903259277344 0 968.7678
184.07154846191406 0 907.8619
185.05592346191406 0 6169.0244 b Water loss 1
186.05897521972656 0 657.3797
186.087646484375 0 1836.8568
187.0714111328125 0 1165.4749
187.0984344482422 0 797.0425
187.1080322265625 0 16109.749
188.07106018066406 0 2243.9727
188.11184692382812 0 1135.4908
195.12405395507812 0 715.3832
197.09327697753906 0 746.32324
197.1036376953125 0 13507.139
198.087646484375 0 597.8595
198.10728454589844 0 1128.7449
199.07289123535156 0 532.2303
199.10916137695312 0 709.3807
201.1124725341797 0 3007.2156
201.12368774414062 0 4286.407
202.08251953125 0 806.0048
203.0665283203125 0 88535.984 b 1
204.07008361816406 0 6709.91
205.07147216796875 0 616.9188
205.09742736816406 0 1073.0986
207.11380004882812 0 509.6672
208.0713653564453 0 503.76144
208.9536895751953 0 1970.8341
209.09259033203125 0 586.76215
210.95021057128906 0 1743.9836
214.13015747070312 0 3459.7314
215.11416625976562 0 2687.633
217.08197021484375 0 660.4181
219.123046875 0 7943.3257
220.12681579589844 0 1170.7814
221.09230041503906 0 809.44867
221.13880920410156 0 1626.4973
223.06382751464844 0 3874.3335
224.06507873535156 0 2416.7375
225.04318237304688 0 6588.9155
225.0613250732422 0 1342.9258
226.0435028076172 0 2894.8289
226.08251953125 0 1978.7593
226.1553192138672 0 750.5493
227.0221710205078 0 1217.1969
227.04039001464844 0 4313.022
228.03994750976562 0 843.0191
229.1186065673828 0 3406.8103
232.14077758789062 0 26155.729
233.1442413330078 0 1483.6544
236.06689453125 0 1072.7662
238.11900329589844 0 2313.2346
239.1504669189453 0 611.2447
240.1350860595703 0 1131.8733
242.0775909423828 0 743.2491
244.09326171875 0 5639.4307 a Water loss 2
245.0769500732422 0 1781.3561
245.09535217285156 0 681.78033
245.6249542236328 0 1046.8182 y Water loss 4
254.07752990722656 0 4636.0137
255.0800323486328 0 588.426
256.1295471191406 0 5586.7065
259.1557922363281 0 1854.5272
262.1036682128906 0 2600.7122 a 2
262.15533447265625 0 1539.3286
263.1396179199219 0 1781.8993
265.1292724609375 0 1311.9324
266.1347351074219 0 644.0212
271.1039123535156 0 5100.2266
272.087890625 0 41762.523 b Water loss 2
273.0909118652344 0 4143.0005
280.14190673828125 0 846.91376
283.14031982421875 0 2031.6039
283.1748046875 0 1123.1609
284.1244201660156 0 6333.3467
284.1424560546875 0 777.56274
286.1396789550781 0 798.2011
287.0063171386719 0 810.00507
288.1340637207031 0 826.94183
289.16424560546875 0 1884.2516
290.0986328125 0 28543.39 b 2
291.1021728515625 0 2272.859
298.14007568359375 0 1442.2019
301.1507873535156 0 953.7487
302.1351318359375 0 960.95435
304.163818359375 0 1629.7069 y 3
305.1612548828125 0 15239.19 y Ammonia loss 6
306.1644592285156 0 2332.592
307.14019775390625 0 1145.9232
307.1755676269531 0 1562.6207
308.1357116699219 0 11099.632
309.1385803222656 0 1908.3156
313.1502685546875 0 638.9335
315.1786193847656 0 843.58014
315.6719665527344 0 771.2125
316.15045166015625 0 1739.1687
318.92254638671875 0 699.7779
319.1777038574219 0 692.503
320.1607360839844 0 938.4726
320.9197998046875 0 1543.2555
322.18798828125 0 3193.109 y 6
324.9872741699219 0 838.8398
325.1512145996094 0 10735.986
325.98577880859375 0 866.27924
326.1471252441406 0 6836.262
326.9844970703125 0 3103.5215
327.14996337890625 0 988.31116
327.9842834472656 0 1042.3457
332.10809326171875 0 636.67706
336.652099609375 0 578.0065
338.6746520996094 0 4983.346 y Water loss 2
339.1763000488281 0 1018.69366
340.9091491699219 0 552.6795
341.0193176269531 0 700.2018
342.0175476074219 0 859.7891
343.01654052734375 0 3752.5503
343.162353515625 0 22284.709
343.9964294433594 0 1293.8079
344.01788330078125 0 1464.2102
344.16845703125 0 3033.2012
344.9951477050781 0 3894.0557
345.1567077636719 0 1898.4567
345.99652099609375 0 1538.5363
347.679443359375 0 2648.9297 y 2
353.1456604003906 0 1721.5731
356.10699462890625 0 682.3247
357.0699462890625 0 931.7439
357.1639099121094 0 1020.7473 b Water loss 6
359.02850341796875 0 9801.922
360.0290222167969 0 12633.959
361.0265808105469 0 47873.78
361.1828918457031 0 30672.432
362.0265808105469 0 20161.223
362.1847229003906 0 7121.5176 y Ammonia loss 5
363.02435302734375 0 8085.61
363.1861572265625 0 1010.12213
364.0242004394531 0 577.68695
370.2399597167969 0 542.9203
371.1563415527344 0 11815.338 b Water loss 3
372.1592712402344 0 1742.7678
373.68511962890625 0 1280.6091
377.2164001464844 0 819.7971
379.2089538574219 0 66104.44 y 5
380.2118225097656 0 12474.286
381.0833435058594 0 632.8831
381.21453857421875 0 1718.8579
387.1795349121094 0 631.83673
389.1673889160156 0 2160.3218 b 3
395.22784423828125 0 11945.75
396.1875915527344 0 3095.0022 y Water loss 1
396.2311706542969 0 2970.0112
396.6896057128906 0 1697.0745
398.09381103515625 0 658.4725
409.131103515625 0 731.1955
410.1852722167969 0 1878.7435
421.6117248535156 0 645.5502
422.1873474121094 0 1117.8263
425.2151794433594 0 1198.7031
429.0899658203125 0 1416.2167
430.08978271484375 0 1279.1975
430.2656555175781 0 743.7458
430.69903564453125 0 2322.9844
430.8873596191406 0 757.0081
431.0876770019531 0 6599.401
431.19427490234375 0 2400.8772
431.84356689453125 0 647.2172
432.0881042480469 0 4722.648
432.88616943359375 0 918.9878
433.0872802734375 0 1429.8059
439.7043151855469 0 7518.7266 Precursor Water loss
440.2058410644531 0 3251.122
442.2405090332031 0 2004.6044
448.27496337890625 0 3131.2646
448.7059020996094 0 759.81464 Precursor
448.76727294921875 0 1038.9132
449.27606201171875 0 698.69653
454.195556640625 0 650.37836
460.2521667480469 0 7220.751
461.2541198730469 0 2196.581
473.2139587402344 0 2075.68
490.2410583496094 0 13890.466 y Water loss 4
491.22991943359375 0 2336.9067 y Ammonia loss 4
491.2392272949219 0 2257.2437
500.199462890625 0 1932.7109 b Water loss 4
508.2516174316406 0 161104.42 y 4
509.254638671875 0 42848.004
510.2567138671875 0 7374.3164
512.243408203125 0 772.6707
518.2064819335938 0 821.05396 b 4
529.2731323242188 0 4346.633
530.2745361328125 0 991.76575
536.5983276367188 0 580.48883
547.2836303710938 0 18382.922
548.286376953125 0 5484.035
549.2857055664062 0 1048.8448
555.2727661132812 0 1195.7175
571.2589721679688 0 1656.9359
571.760986328125 0 1082.9907
607.3199462890625 0 47419.38 y 3
608.3223876953125 0 15776.17
609.3259887695312 0 3901.8728
617.3040161132812 0 1277.1093
618.3093872070312 0 846.42316
644.300537109375 0 1162.6982
645.3118286132812 0 631.3036
662.3098754882812 0 2009.8398
676.3418579101562 0 5193.076 y Water loss 2
677.3336181640625 0 2550.786 y Ammonia loss 2
694.3516235351562 0 121839.02 y 2
695.354248046875 0 43085.305
696.35693359375 0 8792.433
704.3358764648438 0 2089.7095
731.3306274414062 0 2232.1616 b 6
732.331787109375 0 754.77026
791.3672485351562 0 3127.0676 y Water loss 1
792.36865234375 0 1000.9448
809.3785400390625 0 10758.836 y 1
810.379150390625 0 3647.5852
811.3798828125 0 1518.325
819.3649291992188 0 657.3857
1012.4805297851562 0 636.81244
1265.1324462890625 0 752.83093
1467.5440673828125 0 705.0792
1837.67333984375 0 598.563
1878.4034423828125 0 640.275
1922.3199462890625 0 593.7685
2062.640625 0 663.029
2623.234130859375 0 810.8582
2645.72900390625 0 694.7615
3151.0615234375 0 758.5904

Spectrum Details

|  |  |
| --- | --- |
| Matched peaks? Matched peaks The total absolute number of peaks matched. Additionally in brackets the total fraction of peaks matched and the total number of peaks is shown. | 36 (11.73% of 307) |
| FDR? FDR The false discovery rate estimated for this peptide. It is calculated by matching all theoretical fragments with a non-integer shift with the raw peaks for this spectrum. This is done with 40 different shifts. The resulting percentage is the average number of annotated peaks over the number of annotated peaks with the correct spectrum. | 0.73% |
| Satellite FDR? Satellite FDR See the FDR for details on its calculation. This satellite ion specific FDR only contains the satellite ions (d/w) for I/L/J positions. | - |
| PSM Score? PSM Score The PSM Score as given by Hecklib to this annotated spectrum. It is shown with three significant figures. | 455 |

#### Reverse Lookup? Reverse Lookup All places where this read could be placed.

| Group | Segment | Template | Template Part | Read Part | Score | Unique |
| --- | --- | --- | --- | --- | --- | --- |
| Homo sapiens Heavy Chain | IGHV | IGHV3-9 | [61..68] | [0..8] | 46 | False |
| Homo sapiens Heavy Chain | IGHV | IGHV3-48 | [61..68] | [0..8] | 46 | False |
| Homo sapiens Heavy Chain | IGHV | IGHV3-21 | [61..68] | [0..8] | 46 | False |
| Homo sapiens Heavy Chain | IGHV | IGHV3-7 | [61..68] | [0..8] | 46 | False |
| Homo sapiens Heavy Chain | IGHV | IGHV3-43 | [61..68] | [0..8] | 46 | False |
| Homo sapiens Heavy Chain | IGHV | IGHV3-11 | [61..68] | [0..8] | 46 | False |
| Homo sapiens Heavy Chain | IGHV | IGHV3-74 | [61..68] | [0..8] | 46 | False |
| Homo sapiens Heavy Chain | IGHV | IGHV3-20 | [61..68] | [0..8] | 46 | False |
| Homo sapiens Heavy Chain | IGHV | IGHV3-23 | [61..68] | [0..8] | 46 | False |
| Homo sapiens Heavy Chain | IGHV | IGHV3-53 | [60..67] | [0..8] | 46 | False |
| Homo sapiens Heavy Chain | IGHV | IGHV3-66 | [60..67] | [0..8] | 46 | False |
| Homo sapiens Heavy Chain | IGHV | IGHV3-NL1 | [61..68] | [0..8] | 46 | False |
| Homo sapiens Heavy Chain | IGHV | IGHV3-30-5 | [61..68] | [0..8] | 46 | False |
| Homo sapiens Heavy Chain | IGHV | IGHV3-30 | [61..68] | [0..8] | 46 | False |
| Homo sapiens Heavy Chain | IGHV | IGHV3-33 | [61..68] | [0..8] | 46 | False |

| Recombined | Template Part | Read Part | Score | Unique |
| --- | --- | --- | --- | --- |
| REC-0-1 | [60..68] | [0..8] | 64 | True |

#### Meta Information from Multiple reads

##### Number of combined reads

2

##### Intensity

0.937

##### TotalArea

5.632E+08

#### Positional Score

Copy Data

##### Positional Score (TSV)

###### Preview

```
Loading example...
```

*Click on the button to copy the data to your clipboard.*

0001234567

Label Value
"0" 0
"1" 0
"2" 0
"3" 0
"4" 0
"5" 0
"6" 0
"7" 0

#### Meta Information from PEAKS

##### Scan Identifier

F3:3834

##### Original sequence

S

D

S

V

E

G

R

F

##### Posttranslational Modifications

##### Source File

D:\separate\_stitch\_analyses\xle-disambiguation\raw\20210323\_F1\_UM1\_Peng0013\_SA\_F59\_ingel\_3ug\_chymo.raw

##### Fraction

3

##### Scan Feature

F3:2271

##### De Novo Score

98

##### ConfidenceScore

98

### m/z

448.7096

##### Mass

895.4035

##### Charge

2

##### Retention Time

21

##### Predicted Retention Time

-

##### Area

5.632E+08

##### Parts Per Million

1.2

##### Fragmentation mode

HCD

##### Originating file

01 D:\separate\_stitch\_analyses\xle-disambiguation\20210325\_F59\_3ug\_DENOVO\_12.csv

#### Meta Information from PEAKS

##### Scan Identifier

F3:4171

##### Original sequence

S

D

S

V

E

G

R

F

##### Posttranslational Modifications

##### Source File

D:\separate\_stitch\_analyses\xle-disambiguation\raw\20210323\_F1\_UM1\_Peng0013\_SA\_F59\_ingel\_3ug\_chymo.raw

##### Fraction

3

##### Scan Feature

-

##### De Novo Score

98

##### ConfidenceScore

98

### m/z

448.7098

##### Mass

895.4035

##### Charge

2

##### Retention Time

22.92

##### Predicted Retention Time

-

##### Area

0

##### Parts Per Million

1.7

##### Fragmentation mode

HCD

##### Originating file

01 D:\separate\_stitch\_analyses\xle-disambiguation\20210325\_F59\_3ug\_DENOVO\_12.csv
