## Supplementary material for "A handle on mass coincidence errors in *de novo* sequencing of antibodies by bottom-up proteomics": Combined_068.html

Details Combined\_068 | Stitch OverviewUndefined

### Read Combined\_068

#### Sequence (length=7)

JSLSPGK

#### Spectrum 3938? Spectrum 3938 The raw spectrum of this peptide as annotated by Hecklib. The fragments are coloured according to ion type (see legend). Any peaks with a star '\*' as text can be hovered over to see the full details, first the ion type second the mass shift type. By hovering over the amino acids in the peptide or ions in the legend the corresponding peaks are highlighted. By toggling the 'Unassigned' label you can turn the background (unassigned) peaks on or off in the plot. By updating the slider in the Ion legend you can update the spectrum to only show the top X% of the peaks with labels. The top X% means any peak that is within X% of the highest intensity. By dragging in the spectrum you can zoom in to a specific part of the spectrum and use 'Zoom Out' to get back to the original zoom level. The annotation of the spectrum is based on the given sequence in the peptides file and is done with different software so inconsistencies are likely. The peaks are annotated based on the given sequence, with 20 ppm tolerance.

Copy Data

##### Spectrum 3938 (TSV)

###### Preview

```
Loading example...
```

*Click on the button to copy the data to your clipboard.*

Mz MinMz MaxIntensity Max

WidthHeightPeptide font sizePeptide stroke widthSpectrum font sizeSpectrum stroke widthCompact peptide

Ion legend

wxyz

abcd

OtherUnassignedIon^Charge^_Position_Show for top:%

JSLSPGK

05.17e+41.03e+51.55e+52.07e+5

Zoom Out

y^+1^_1_y^+1^_1_y^+2^_3_a^+1^_2_a^+1^_2_b^+1^_2_y^+1^_2_b^+1^_2_y^+1^_2_y^+2^_5_y^+1^_3_b^+1^_3_y^+1^_3_\*\*y^+1^_4_b^+1^_4_y^+1^_4_y^+1^_5_y^+1^_5_y^+1^_6_y^+1^_6_

0565112916942258

Fragment Matches Table

Show background peaks

| Position | Ion type | Intensity | mz Theoretical | mz Error (Th) | mz Error (ppm) | Charge | Series Number |
| --- | --- | --- | --- | --- | --- | --- | --- |
| - | - | 1163 | 120.1 | - | - | 0 | - |
| - | - | 1372 | 120.1 | - | - | 0 | - |
| - | - | 522.7 | 121 | - | - | 0 | - |
| - | - | 534.7 | 121 | - | - | 0 | - |
| - | - | 648 | 121.1 | - | - | 0 | - |
| - | - | 1872 | 123.1 | - | - | 0 | - |
| - | - | 675.7 | 124.1 | - | - | 0 | - |
| - | - | 458 | 125.1 | - | - | 0 | - |
| - | - | 9125 | 125.1 | - | - | 0 | - |
| - | - | 885.5 | 126.1 | - | - | 0 | - |
| - | - | 456.1 | 126.1 | - | - | 0 | - |
| - | - | 469.3 | 126.1 | - | - | 0 | - |
| - | - | 470.1 | 127.1 | - | - | 0 | - |
| - | - | 5411 | 127.1 | - | - | 0 | - |
| - | - | 468.8 | 127.1 | - | - | 0 | - |
| - | - | 5022 | 128.1 | - | - | 0 | - |
| - | - | 437.2 | 128.5 | - | - | 0 | - |
| - | - | 3.774E+04 | 129.1 | - | - | 0 | - |
| - | - | 945.2 | 130.1 | - | - | 0 | - |
| 7 | y | 1.8E+04 | 130.1 | 0.0003386 | 2.603 | +1 | 1 |
| - | - | 2124 | 130.1 | - | - | 0 | - |
| - | - | 808.9 | 131.1 | - | - | 0 | - |
| - | - | 890.3 | 131.1 | - | - | 0 | - |
| - | - | 629.8 | 132.1 | - | - | 0 | - |
| - | - | 747.2 | 133.1 | - | - | 0 | - |
| - | - | 369.1 | 134.8 | - | - | 0 | - |
| - | - | 3731 | 136.1 | - | - | 0 | - |
| - | - | 570.5 | 137.1 | - | - | 0 | - |
| - | - | 582.2 | 137.1 | - | - | 0 | - |
| - | - | 1927 | 138.1 | - | - | 0 | - |
| - | - | 1177 | 139.1 | - | - | 0 | - |
| - | - | 488.3 | 140.1 | - | - | 0 | - |
| - | - | 696 | 141.1 | - | - | 0 | - |
| - | - | 4002 | 141.1 | - | - | 0 | - |
| - | - | 1229 | 143.1 | - | - | 0 | - |
| - | - | 427.8 | 144.1 | - | - | 0 | - |
| - | - | 1103 | 145 | - | - | 0 | - |
| 7 | y | 3.224E+04 | 147.1 | 0.0003398 | 2.31 | +1 | 1 |
| - | - | 2038 | 148.1 | - | - | 0 | - |
| - | - | 499.1 | 148.9 | - | - | 0 | - |
| - | - | 1238 | 149 | - | - | 0 | - |
| - | - | 455 | 149.1 | - | - | 0 | - |
| - | - | 774.4 | 150 | - | - | 0 | - |
| - | - | 491.5 | 151.1 | - | - | 0 | - |
| - | - | 521.4 | 151.1 | - | - | 0 | - |
| 5 | y | 1116 | 151.1 | 0.0002303 | 1.524 | +2 | 3 |
| - | - | 1007 | 153.1 | - | - | 0 | - |
| - | - | 681.3 | 153.1 | - | - | 0 | - |
| - | - | 1.202E+04 | 155.1 | - | - | 0 | - |
| 2 | a | 9376 | 155.1 | 0.0002593 | 1.671 | +1 | 2 |
| - | - | 422.9 | 155.8 | - | - | 0 | - |
| - | - | 614.8 | 156.1 | - | - | 0 | - |
| - | - | 1664 | 156.1 | - | - | 0 | - |
| - | - | 1160 | 156.1 | - | - | 0 | - |
| - | - | 1.779E+04 | 157.1 | - | - | 0 | - |
| - | - | 870.9 | 158.1 | - | - | 0 | - |
| - | - | 523.3 | 159.1 | - | - | 0 | - |
| - | - | 1324 | 159.1 | - | - | 0 | - |
| - | - | 2778 | 159.1 | - | - | 0 | - |
| - | - | 478.6 | 161.1 | - | - | 0 | - |
| - | - | 468.9 | 165.1 | - | - | 0 | - |
| - | - | 494 | 166.1 | - | - | 0 | - |
| - | - | 641.9 | 167.1 | - | - | 0 | - |
| - | - | 2332 | 167.1 | - | - | 0 | - |
| - | - | 529.5 | 168 | - | - | 0 | - |
| - | - | 413.7 | 169.1 | - | - | 0 | - |
| - | - | 2313 | 169.1 | - | - | 0 | - |
| - | - | 642.6 | 170.1 | - | - | 0 | - |
| - | - | 1417 | 172.1 | - | - | 0 | - |
| - | - | 699.5 | 173.1 | - | - | 0 | - |
| - | - | 1.087E+04 | 173.1 | - | - | 0 | - |
| 2 | a | 8.876E+04 | 173.1 | 0.00033 | 1.906 | +1 | 2 |
| - | - | 1443 | 173.4 | - | - | 0 | - |
| - | - | 814.4 | 174.1 | - | - | 0 | - |
| - | - | 829.7 | 174.1 | - | - | 0 | - |
| - | - | 8193 | 174.1 | - | - | 0 | - |
| - | - | 651.3 | 175.1 | - | - | 0 | - |
| - | - | 826.6 | 179.1 | - | - | 0 | - |
| - | - | 499.5 | 181.1 | - | - | 0 | - |
| - | - | 594.4 | 181.1 | - | - | 0 | - |
| 2 | b | 7907 | 183.1 | 0.0002635 | 1.439 | +1 | 2 |
| - | - | 1942 | 183.1 | - | - | 0 | - |
| - | - | 649.5 | 183.1 | - | - | 0 | - |
| - | - | 595.6 | 184.1 | - | - | 0 | - |
| - | - | 580.1 | 184.1 | - | - | 0 | - |
| - | - | 1.095E+04 | 185.1 | - | - | 0 | - |
| - | - | 655.2 | 186.1 | - | - | 0 | - |
| - | - | 1.683E+04 | 186.1 | - | - | 0 | - |
| - | - | 569 | 187.1 | - | - | 0 | - |
| 6 | y | 6985 | 187.1 | 0.000405 | 2.165 | +1 | 2 |
| - | - | 1019 | 187.1 | - | - | 0 | - |
| - | - | 569.5 | 193.1 | - | - | 0 | - |
| - | - | 513.6 | 193.3 | - | - | 0 | - |
| - | - | 578 | 194.1 | - | - | 0 | - |
| - | - | 1084 | 196.1 | - | - | 0 | - |
| 2 | b | 4.576E+04 | 201.1 | 0.0002426 | 1.206 | +1 | 2 |
| - | - | 3234 | 202.1 | - | - | 0 | - |
| - | - | 4384 | 202.1 | - | - | 0 | - |
| - | - | 1114 | 203.1 | - | - | 0 | - |
| - | - | 562.5 | 203.1 | - | - | 0 | - |
| 6 | y | 7.578E+04 | 204.1 | 0.0002994 | 1.467 | +1 | 2 |
| - | - | 906.4 | 205.1 | - | - | 0 | - |
| - | - | 6228 | 205.1 | - | - | 0 | - |
| - | - | 1376 | 206.1 | - | - | 0 | - |
| - | - | 1103 | 207.1 | - | - | 0 | - |
| - | - | 815.9 | 209.2 | - | - | 0 | - |
| - | - | 1072 | 211.1 | - | - | 0 | - |
| - | - | 667.5 | 214.1 | - | - | 0 | - |
| - | - | 576.2 | 215.1 | - | - | 0 | - |
| - | - | 536.6 | 216.1 | - | - | 0 | - |
| - | - | 630.8 | 217.1 | - | - | 0 | - |
| - | - | 765.5 | 219.1 | - | - | 0 | - |
| - | - | 623.5 | 223.1 | - | - | 0 | - |
| - | - | 6141 | 224.1 | - | - | 0 | - |
| - | - | 773.9 | 225.1 | - | - | 0 | - |
| - | - | 584.1 | 225.1 | - | - | 0 | - |
| - | - | 767.1 | 226.1 | - | - | 0 | - |
| - | - | 466.1 | 227.5 | - | - | 0 | - |
| - | - | 843.5 | 228.2 | - | - | 0 | - |
| - | - | 664.5 | 236.1 | - | - | 0 | - |
| - | - | 649.1 | 240.1 | - | - | 0 | - |
| - | - | 7247 | 242.1 | - | - | 0 | - |
| 3 | y | 798.8 | 242.1 | 0.0007321 | 3.023 | +2 | 5 |
| - | - | 839.4 | 243.1 | - | - | 0 | - |
| - | - | 1285 | 243.1 | - | - | 0 | - |
| - | - | 1205 | 251.2 | - | - | 0 | - |
| - | - | 657.4 | 252.1 | - | - | 0 | - |
| - | - | 554.6 | 254.1 | - | - | 0 | - |
| - | - | 3103 | 260.1 | - | - | 0 | - |
| - | - | 633.8 | 268.2 | - | - | 0 | - |
| - | - | 4058 | 270.1 | - | - | 0 | - |
| - | - | 998.2 | 279.2 | - | - | 0 | - |
| - | - | 4144 | 282.2 | - | - | 0 | - |
| - | - | 1580 | 283.1 | - | - | 0 | - |
| - | - | 1.544E+04 | 283.2 | - | - | 0 | - |
| 5 | y | 704.9 | 284.2 | 0.002085 | 7.336 | +1 | 3 |
| - | - | 2219 | 284.2 | - | - | 0 | - |
| - | - | 4914 | 288.2 | - | - | 0 | - |
| 3 | b | 2541 | 296.2 | 0.0001839 | 0.6207 | +1 | 3 |
| 5 | y | 2.049E+05 | 301.2 | 0.0004072 | 1.352 | +1 | 3 |
| - | - | 2.976E+04 | 302.2 | - | - | 0 | - |
| - | - | 1581 | 303.2 | - | - | 0 | - |
| - | - | 854 | 303.2 | - | - | 0 | - |
| - | - | 683.8 | 308.2 | - | - | 0 | - |
| - | - | 2113 | 309.2 | - | - | 0 | - |
| - | - | 552.1 | 315.9 | - | - | 0 | - |
| - | - | 622.7 | 316.9 | - | - | 0 | - |
| - | - | 505 | 322.2 | - | - | 0 | - |
| - | - | 720.1 | 325.2 | - | - | 0 | - |
| - | - | 1386 | 334.9 | - | - | 0 | - |
| - | - | 1813 | 335.2 | - | - | 0 | - |
| - | - | 1967 | 339.3 | - | - | 0 | - |
| 0 | Precursor | 722.8 | 342.2 | 0.001085 | 3.172 | +2 | -1 |
| - | - | 927.2 | 350.9 | - | - | 0 | - |
| - | - | 5251 | 351.1 | - | - | 0 | - |
| 0 | Precursor | 1279 | 351.2 | 0.0008441 | 2.403 | +2 | -1 |
| - | - | 624.5 | 351.4 | - | - | 0 | - |
| - | - | 2301 | 352.2 | - | - | 0 | - |
| - | - | 1.293E+04 | 352.2 | - | - | 0 | - |
| - | - | 644.3 | 352.9 | - | - | 0 | - |
| - | - | 2251 | 353.2 | - | - | 0 | - |
| - | - | 1126 | 355.2 | - | - | 0 | - |
| - | - | 1370 | 358.2 | - | - | 0 | - |
| - | - | 742.7 | 367.2 | - | - | 0 | - |
| - | - | 2919 | 367.2 | - | - | 0 | - |
| - | - | 588.1 | 368.2 | - | - | 0 | - |
| - | - | 723.7 | 369.1 | - | - | 0 | - |
| 4 | y | 1.021E+04 | 370.2 | 9.092E-05 | 0.2456 | +1 | 4 |
| - | - | 2165 | 371.2 | - | - | 0 | - |
| 4 | b | 726.5 | 383.2 | 0.0009924 | 2.589 | +1 | 4 |
| - | - | 1864 | 384.3 | - | - | 0 | - |
| - | - | 750.5 | 385.2 | - | - | 0 | - |
| 4 | y | 6.517E+04 | 388.2 | 0.0002087 | 0.5375 | +1 | 4 |
| - | - | 1.151E+04 | 389.2 | - | - | 0 | - |
| - | - | 1603 | 390.2 | - | - | 0 | - |
| - | - | 3376 | 398.2 | - | - | 0 | - |
| - | - | 774.1 | 406.2 | - | - | 0 | - |
| - | - | 944.5 | 424.2 | - | - | 0 | - |
| - | - | 1046 | 442.2 | - | - | 0 | - |
| - | - | 1049 | 456.3 | - | - | 0 | - |
| - | - | 1175 | 465.3 | - | - | 0 | - |
| - | - | 2141 | 467.3 | - | - | 0 | - |
| - | - | 617 | 468.3 | - | - | 0 | - |
| 3 | y | 1.111E+04 | 483.3 | 0.0001957 | 0.4049 | +1 | 5 |
| - | - | 2172 | 484.3 | - | - | 0 | - |
| - | - | 2191 | 484.3 | - | - | 0 | - |
| - | - | 816.6 | 485.3 | - | - | 0 | - |
| - | - | 3969 | 495.3 | - | - | 0 | - |
| 3 | y | 4.238E+04 | 501.3 | 0.000129 | 0.2574 | +1 | 5 |
| - | - | 1.01E+04 | 502.3 | - | - | 0 | - |
| - | - | 2158 | 503.3 | - | - | 0 | - |
| - | - | 781 | 535.3 | - | - | 0 | - |
| - | - | 1065 | 537.3 | - | - | 0 | - |
| - | - | 1111 | 552.3 | - | - | 0 | - |
| - | - | 686.4 | 553.3 | - | - | 0 | - |
| - | - | 1919 | 554.3 | - | - | 0 | - |
| - | - | 603.2 | 555.3 | - | - | 0 | - |
| - | - | 636.8 | 564.3 | - | - | 0 | - |
| - | - | 1071 | 565.3 | - | - | 0 | - |
| 2 | y | 8397 | 570.3 | 2.887E-06 | 0.005062 | +1 | 6 |
| - | - | 2773 | 571.3 | - | - | 0 | - |
| - | - | 692.8 | 572.3 | - | - | 0 | - |
| - | - | 908.5 | 580.3 | - | - | 0 | - |
| - | - | 1456 | 582.3 | - | - | 0 | - |
| - | - | 649.2 | 583.3 | - | - | 0 | - |
| 2 | y | 1.097E+05 | 588.3 | 0.0002526 | 0.4294 | +1 | 6 |
| - | - | 2.968E+04 | 589.3 | - | - | 0 | - |
| - | - | 1111 | 589.4 | - | - | 0 | - |
| - | - | 6620 | 590.3 | - | - | 0 | - |
| - | - | 7298 | 598.3 | - | - | 0 | - |
| - | - | 3167 | 599.3 | - | - | 0 | - |
| - | - | 716 | 839.7 | - | - | 0 | - |
| - | - | 543.3 | 865.7 | - | - | 0 | - |
| - | - | 522.7 | 869.4 | - | - | 0 | - |
| - | - | 725.3 | 2236 | - | - | 0 | - |

m/z Charge Intensity FragmentType MassShift Position
120.06584167480469 0 1162.7181
120.08112335205078 0 1372.3967
120.99555969238281 0 522.6697
121.02876281738281 0 534.6724
121.06514739990234 0 647.978
123.1172103881836 0 1872.2239
124.07595825195312 0 675.69604
125.060302734375 0 457.9959
125.10767364501953 0 9124.602
126.05532836914062 0 885.5139
126.09156036376953 0 456.0571
126.11090850830078 0 469.34833
127.07585906982422 0 470.1294
127.08692169189453 0 5411.175
127.12300109863281 0 468.77145
128.10726928710938 0 5021.684
128.53359985351562 0 437.2203
129.10256958007812 0 37735.348
130.06539916992188 0 945.23254
130.0865936279297 0 17998.873 y Ammonia loss 6
130.10595703125 0 2124.3855
131.07061767578125 0 808.9026
131.09007263183594 0 890.33215
132.10231018066406 0 629.81335
133.08604431152344 0 747.245
134.8173065185547 0 369.1264
136.07603454589844 0 3731.2158
137.0598602294922 0 570.4541
137.0712127685547 0 582.22205
138.09153747558594 0 1927.4811
139.08677673339844 0 1176.5569
140.0828857421875 0 488.30435
141.0704345703125 0 695.9895
141.10256958007812 0 4002.1306
143.09425354003906 0 1229.326
144.0803680419922 0 427.7608
145.04989624023438 0 1103.1564
147.11314392089844 0 32242.48 y 6
148.11651611328125 0 2038.193
148.94715881347656 0 499.09378
149.02362060546875 0 1237.8745
149.07118225097656 0 455.04144
150.0269317626953 0 774.3758
151.0757598876953 0 491.55
151.08639526367188 0 521.3806
151.096923828125 0 1116.2307 y 4
153.05496215820312 0 1006.8058
153.06634521484375 0 681.30396
155.0818328857422 0 12015.485
155.11814880371094 0 9376.39 a Water loss 1
155.82525634765625 0 422.94714
156.08547973632812 0 614.82495
156.10214233398438 0 1663.7423
156.12168884277344 0 1160.4731
157.0974884033203 0 17787.473
158.10096740722656 0 870.87054
159.0661163330078 0 523.26373
159.09176635742188 0 1323.8743
159.1133270263672 0 2777.7153
161.05955505371094 0 478.5558
165.06622314453125 0 468.94
166.09764099121094 0 493.9945
167.07040405273438 0 641.9254
167.08160400390625 0 2331.9568
168.02261352539062 0 529.5016
169.07595825195312 0 413.66525
169.0974884033203 0 2312.7998
170.060302734375 0 642.5816
172.10836791992188 0 1417.3397
173.05593872070312 0 699.50525
173.09234619140625 0 10866.302
173.1287841796875 0 88761.28 a 1
173.4398956298828 0 1442.6892
174.09619140625 0 814.40753
174.1246795654297 0 829.6664
174.1322479248047 0 8193.064
175.13397216796875 0 651.3299
179.08172607421875 0 826.58716
181.09718322753906 0 499.54535
181.13363647460938 0 594.41174
183.11306762695312 0 7907.3022 b Water loss 1
183.1243133544922 0 1941.9612
183.14938354492188 0 649.5337
184.11672973632812 0 595.57086
184.12721252441406 0 580.0772
185.09242248535156 0 10952.646
186.0954132080078 0 655.2314
186.1240234375 0 16834.664
187.08917236328125 0 568.9972
187.10812377929688 0 6985.2583 y Ammonia loss 5
187.1280975341797 0 1018.54816
193.13900756835938 0 569.49884
193.34510803222656 0 513.63983
194.09213256835938 0 577.9904
196.10877990722656 0 1084.4136
201.1236114501953 0 45763.74 b 1
202.1079559326172 0 3233.811
202.12705993652344 0 4384.1343
203.066162109375 0 1113.7177
203.1289825439453 0 562.4591
204.1345672607422 0 75782.734 y 5
205.11932373046875 0 906.35205
205.13787841796875 0 6227.7563
206.09254455566406 0 1375.8843
207.0767364501953 0 1103.3901
209.1651611328125 0 815.8546
211.1084442138672 0 1072.1003
214.1195068359375 0 667.5319
215.13990783691406 0 576.2129
216.07687377929688 0 536.55414
217.0829315185547 0 630.7568
219.0982666015625 0 765.4866
223.10711669921875 0 623.4628
224.10324096679688 0 6140.6133
225.10728454589844 0 773.87036
225.12327575683594 0 584.1346
226.11859130859375 0 767.12573
227.54147338867188 0 466.12302
228.17063903808594 0 843.4528
236.140625 0 664.4565
240.13491821289062 0 649.0869
242.11390686035156 0 7246.595
242.15065002441406 0 798.7988 y Water loss 2
243.11871337890625 0 839.4204
243.14678955078125 0 1285.3439
251.18685913085938 0 1204.9595
252.13453674316406 0 657.3592
254.11378479003906 0 554.5542
260.12445068359375 0 3102.505
268.21343994140625 0 633.8463
270.1450500488281 0 4057.6443
279.1817626953125 0 998.21924
282.160400390625 0 4143.972
283.1399841308594 0 1580.0447
283.1767883300781 0 15444.509
284.1625671386719 0 704.8905 y Ammonia loss 4
284.18115234375 0 2219.0193
288.1558532714844 0 4913.728
296.1970520019531 0 2540.9026 b Water loss 2
301.18743896484375 0 204891.47 y 4
302.1905212402344 0 29755.486
303.1898193359375 0 1581.4531
303.2049560546875 0 853.9831
308.1619567871094 0 683.8028
309.2035217285156 0 2112.8406
315.89349365234375 0 552.10876
316.89959716796875 0 622.65717
322.18914794921875 0 505.01065
325.18780517578125 0 720.0949
334.9120178222656 0 1385.836
335.171142578125 0 1813.2594
339.2508850097656 0 1966.9453
342.2068786621094 0 722.7506 Precursor Water loss
350.94720458984375 0 927.17664
351.1411437988281 0 5251.3833
351.21240234375 0 1278.5295 Precursor
351.36297607421875 0 624.4621
352.1981506347656 0 2301.4814
352.2383728027344 0 12929.099
352.9215087890625 0 644.334
353.1817321777344 0 2251.2588
355.1976013183594 0 1125.9663
358.20867919921875 0 1370.2699
367.19879150390625 0 742.74316
367.24505615234375 0 2918.5508
368.2460021972656 0 588.0768
369.06109619140625 0 723.7479
370.2084045410156 0 10211.928 y Water loss 3
371.2121276855469 0 2165.2634
383.2298889160156 0 726.5476 b Water loss 3
384.2715759277344 0 1863.877
385.20892333984375 0 750.5077
388.2192687988281 0 65173.438 y 3
389.221923828125 0 11505.082
390.2247619628906 0 1602.6624
398.2029113769531 0 3376.357
406.20880126953125 0 774.1139
424.21954345703125 0 944.5428
442.2293701171875 0 1045.8046
456.2576599121094 0 1048.9822
465.2825927734375 0 1175.4939
467.30914306640625 0 2141.3093
468.31341552734375 0 617.0363
483.2927551269531 0 11107.029 y Water loss 2
484.25213623046875 0 2172.4487
484.2960510253906 0 2190.6733
485.2525329589844 0 816.5543
495.3039245605469 0 3968.7212
501.3032531738281 0 42378.902 y 2
502.30615234375 0 10101.607
503.30816650390625 0 2157.5608
535.2886352539062 0 781.0093
537.3158569335938 0 1064.8977
552.313232421875 0 1111.0873
553.2971801757812 0 686.44073
554.3405151367188 0 1918.7006
555.3276977539062 0 603.18634
564.3255004882812 0 636.7948
565.3109741210938 0 1071.4153
570.3245849609375 0 8397.29 y Water loss 1
571.3275756835938 0 2773.1208
572.3294067382812 0 692.8447
580.307373046875 0 908.4624
582.3358154296875 0 1456.0864
583.3351440429688 0 649.2113
588.3348999023438 0 109738.02 y 1
589.3380126953125 0 29677.387
589.3923950195312 0 1110.7448
590.340576171875 0 6620.1797
598.3191528320312 0 7297.993
599.3222045898438 0 3166.8342
839.7376708984375 0 716.00525
865.6665649414062 0 543.25195
869.4278564453125 0 522.7427
2236.04443359375 0 725.283

Spectrum Details

|  |  |
| --- | --- |
| Matched peaks? Matched peaks The total absolute number of peaks matched. Additionally in brackets the total fraction of peaks matched and the total number of peaks is shown. | 22 (10.23% of 215) |
| FDR? FDR The false discovery rate estimated for this peptide. It is calculated by matching all theoretical fragments with a non-integer shift with the raw peaks for this spectrum. This is done with 40 different shifts. The resulting percentage is the average number of annotated peaks over the number of annotated peaks with the correct spectrum. | 0.76% |
| Satellite FDR? Satellite FDR See the FDR for details on its calculation. This satellite ion specific FDR only contains the satellite ions (d/w) for I/L/J positions. | - |
| PSM Score? PSM Score The PSM Score as given by Hecklib to this annotated spectrum. It is shown with three significant figures. | 274 |

#### Spectrum 3875? Spectrum 3875 The raw spectrum of this peptide as annotated by Hecklib. The fragments are coloured according to ion type (see legend). Any peaks with a star '\*' as text can be hovered over to see the full details, first the ion type second the mass shift type. By hovering over the amino acids in the peptide or ions in the legend the corresponding peaks are highlighted. By toggling the 'Unassigned' label you can turn the background (unassigned) peaks on or off in the plot. By updating the slider in the Ion legend you can update the spectrum to only show the top X% of the peaks with labels. The top X% means any peak that is within X% of the highest intensity. By dragging in the spectrum you can zoom in to a specific part of the spectrum and use 'Zoom Out' to get back to the original zoom level. The annotation of the spectrum is based on the given sequence in the peptides file and is done with different software so inconsistencies are likely. The peaks are annotated based on the given sequence, with 20 ppm tolerance.

Copy Data

##### Spectrum 3875 (TSV)

###### Preview

```
Loading example...
```

*Click on the button to copy the data to your clipboard.*

Mz MinMz MaxIntensity Max

WidthHeightPeptide font sizePeptide stroke widthSpectrum font sizeSpectrum stroke widthCompact peptide

Ion legend

wxyz

abcd

OtherUnassignedIon^Charge^_Position_Show for top:%

JSLSPGK

01.38e+52.77e+54.15e+55.54e+5

Zoom Out

y^+1^_1_y^+1^_1_y^+2^_3_z^+1^_2_y^+1^_2_y^+2^_5_w^+1^_3_y^+1^_3_y^+2^_6_y^+2^_6_y^+1^_3_c^+1^_3_z^+1^_4_w^+1^_4_y^+1^_4_y^+1^_4_z^+1^_4_y^+1^_4_w^+1^_5_z^+1^_5_y^+1^_5_z^+1^_5_y^+1^_5_c^+1^_5_c^+1^_6_w^+1^_6_y^+1^_6_c^+1^_6_y^+1^_6_

0828165624843313

Fragment Matches Table

Show background peaks

| Position | Ion type | Intensity | mz Theoretical | mz Error (Th) | mz Error (ppm) | Charge | Series Number |
| --- | --- | --- | --- | --- | --- | --- | --- |
| - | - | 3700 | 125.1 | - | - | 0 | - |
| - | - | 677.5 | 128.1 | - | - | 0 | - |
| - | - | 1437 | 128.1 | - | - | 0 | - |
| - | - | 5020 | 129.1 | - | - | 0 | - |
| - | - | 493.5 | 130.1 | - | - | 0 | - |
| 7 | y | 1756 | 130.1 | 0.0004606 | 3.541 | +1 | 1 |
| - | - | 1717 | 131.1 | - | - | 0 | - |
| - | - | 525.2 | 131.1 | - | - | 0 | - |
| - | - | 380.6 | 133 | - | - | 0 | - |
| - | - | 1176 | 133.1 | - | - | 0 | - |
| - | - | 994 | 142.1 | - | - | 0 | - |
| 7 | y | 6542 | 147.1 | 0.0003855 | 2.621 | +1 | 1 |
| - | - | 587.8 | 148.1 | - | - | 0 | - |
| - | - | 403.7 | 150.2 | - | - | 0 | - |
| 5 | y | 952.4 | 151.1 | 1.383E-06 | 0.009151 | +2 | 3 |
| - | - | 1403 | 155.1 | - | - | 0 | - |
| - | - | 5357 | 155.1 | - | - | 0 | - |
| - | - | 548.9 | 157.1 | - | - | 0 | - |
| - | - | 1529 | 157.1 | - | - | 0 | - |
| - | - | 5351 | 169.1 | - | - | 0 | - |
| - | - | 1542 | 170.1 | - | - | 0 | - |
| - | - | 3.978E+04 | 172.1 | - | - | 0 | - |
| - | - | 2414 | 173.1 | - | - | 0 | - |
| - | - | 2289 | 173.1 | - | - | 0 | - |
| - | - | 4.79E+04 | 173.1 | - | - | 0 | - |
| - | - | 3176 | 174.1 | - | - | 0 | - |
| - | - | 3509 | 183.1 | - | - | 0 | - |
| - | - | 1632 | 185.1 | - | - | 0 | - |
| - | - | 1930 | 186.1 | - | - | 0 | - |
| 6 | z | 4375 | 188.1 | 0.0005603 | 2.979 | +1 | 2 |
| - | - | 1785 | 189.1 | - | - | 0 | - |
| - | - | 6389 | 200.1 | - | - | 0 | - |
| - | - | 4.227E+04 | 201.1 | - | - | 0 | - |
| - | - | 3089 | 202.1 | - | - | 0 | - |
| - | - | 3725 | 202.1 | - | - | 0 | - |
| 6 | y | 1.214E+04 | 204.1 | 0.0003604 | 1.766 | +1 | 2 |
| - | - | 599.6 | 204.1 | - | - | 0 | - |
| - | - | 1028 | 205.1 | - | - | 0 | - |
| - | - | 624.7 | 211.1 | - | - | 0 | - |
| - | - | 793.9 | 224.1 | - | - | 0 | - |
| - | - | 1462 | 226.1 | - | - | 0 | - |
| - | - | 592.8 | 227.1 | - | - | 0 | - |
| - | - | 8119 | 229.1 | - | - | 0 | - |
| - | - | 1054 | 230.1 | - | - | 0 | - |
| - | - | 688 | 231.1 | - | - | 0 | - |
| - | - | 1115 | 232.1 | - | - | 0 | - |
| - | - | 2833 | 242.1 | - | - | 0 | - |
| - | - | 2121 | 245.1 | - | - | 0 | - |
| - | - | 531.2 | 247 | - | - | 0 | - |
| 3 | y | 1979 | 251.2 | 0.0002868 | 1.142 | +2 | 5 |
| 5 | w | 9764 | 258.1 | 0.0003396 | 1.315 | +1 | 3 |
| - | - | 8890 | 259.1 | - | - | 0 | - |
| - | - | 1083 | 260.1 | - | - | 0 | - |
| - | - | 500.3 | 260.1 | - | - | 0 | - |
| - | - | 1821 | 270.1 | - | - | 0 | - |
| - | - | 777.8 | 271.2 | - | - | 0 | - |
| - | - | 2239 | 272.2 | - | - | 0 | - |
| - | - | 9717 | 273.2 | - | - | 0 | - |
| - | - | 820.8 | 273.2 | - | - | 0 | - |
| - | - | 987.5 | 274.2 | - | - | 0 | - |
| - | - | 911.2 | 283.1 | - | - | 0 | - |
| - | - | 1122 | 283.2 | - | - | 0 | - |
| - | - | 9280 | 283.2 | - | - | 0 | - |
| 5 | y | 527.8 | 284.2 | 0.0002653 | 0.9338 | +1 | 3 |
| - | - | 673.3 | 284.2 | - | - | 0 | - |
| 2 | y | 512.6 | 285.7 | 0.0004353 | 1.524 | +2 | 6 |
| - | - | 735.6 | 287.2 | - | - | 0 | - |
| - | - | 2291 | 288.2 | - | - | 0 | - |
| 2 | y | 563.1 | 294.7 | 0.001607 | 5.452 | +2 | 6 |
| - | - | 732.9 | 295.2 | - | - | 0 | - |
| - | - | 7322 | 296.2 | - | - | 0 | - |
| - | - | 1476 | 297.2 | - | - | 0 | - |
| - | - | 1.944E+04 | 299.2 | - | - | 0 | - |
| - | - | 1.202E+04 | 300.2 | - | - | 0 | - |
| - | - | 2706 | 301.1 | - | - | 0 | - |
| 5 | y | 1.417E+05 | 301.2 | 0.0006209 | 2.061 | +1 | 3 |
| - | - | 2.146E+04 | 302.2 | - | - | 0 | - |
| - | - | 2172 | 303.2 | - | - | 0 | - |
| - | - | 590 | 313.2 | - | - | 0 | - |
| - | - | 1657 | 314.1 | - | - | 0 | - |
| - | - | 2034 | 314.2 | - | - | 0 | - |
| - | - | 5056 | 315.2 | - | - | 0 | - |
| - | - | 835.7 | 316.2 | - | - | 0 | - |
| - | - | 1202 | 320.2 | - | - | 0 | - |
| - | - | 1.357E+04 | 328.2 | - | - | 0 | - |
| - | - | 4894 | 329.2 | - | - | 0 | - |
| - | - | 979.1 | 330.2 | - | - | 0 | - |
| 3 | c | 7377 | 331.2 | 0.0008203 | 2.477 | +1 | 3 |
| - | - | 1494 | 332.2 | - | - | 0 | - |
| - | - | 872.1 | 333.7 | - | - | 0 | - |
| - | - | 904.8 | 334.9 | - | - | 0 | - |
| - | - | 642 | 335.2 | - | - | 0 | - |
| - | - | 734.4 | 336.2 | - | - | 0 | - |
| - | - | 1075 | 337.2 | - | - | 0 | - |
| - | - | 5332 | 342.2 | - | - | 0 | - |
| - | - | 2598 | 342.2 | - | - | 0 | - |
| - | - | 948.3 | 343.2 | - | - | 0 | - |
| - | - | 1291 | 344.2 | - | - | 0 | - |
| - | - | 1811 | 350.2 | - | - | 0 | - |
| - | - | 746.5 | 351 | - | - | 0 | - |
| - | - | 2932 | 351.1 | - | - | 0 | - |
| - | - | 4004 | 351.2 | - | - | 0 | - |
| - | - | 1709 | 351.7 | - | - | 0 | - |
| - | - | 945.2 | 352.1 | - | - | 0 | - |
| - | - | 3093 | 352.2 | - | - | 0 | - |
| - | - | 1160 | 352.2 | - | - | 0 | - |
| - | - | 6832 | 352.2 | - | - | 0 | - |
| - | - | 4152 | 353.2 | - | - | 0 | - |
| 4 | z | 1485 | 354.2 | 0.001539 | 4.345 | +1 | 4 |
| 4 | w | 4.025E+04 | 355.2 | 0.0005847 | 1.646 | +1 | 4 |
| - | - | 7034 | 356.2 | - | - | 0 | - |
| - | - | 790.5 | 357.2 | - | - | 0 | - |
| - | - | 608.1 | 367.2 | - | - | 0 | - |
| - | - | 701.1 | 369.2 | - | - | 0 | - |
| - | - | 1.97E+04 | 370.2 | - | - | 0 | - |
| 4 | y | 5334 | 370.2 | 0.001313 | 3.546 | +1 | 4 |
| 4 | y | 1.728E+04 | 371.2 | 0.0004944 | 1.332 | +1 | 4 |
| 4 | z | 1.987E+04 | 372.2 | 0.0001408 | 0.3782 | +1 | 4 |
| - | - | 2153 | 372.2 | - | - | 0 | - |
| - | - | 2.075E+04 | 373.2 | - | - | 0 | - |
| - | - | 1551 | 373.2 | - | - | 0 | - |
| - | - | 2677 | 374.2 | - | - | 0 | - |
| - | - | 5506 | 383.2 | - | - | 0 | - |
| - | - | 854.8 | 384.2 | - | - | 0 | - |
| - | - | 648.7 | 385.2 | - | - | 0 | - |
| 4 | y | 5.806E+04 | 388.2 | 0.0006664 | 1.717 | +1 | 4 |
| - | - | 1.012E+04 | 389.2 | - | - | 0 | - |
| - | - | 1839 | 390.2 | - | - | 0 | - |
| - | - | 2935 | 396.2 | - | - | 0 | - |
| - | - | 2984 | 398.2 | - | - | 0 | - |
| - | - | 915.9 | 399.2 | - | - | 0 | - |
| - | - | 8520 | 400.2 | - | - | 0 | - |
| - | - | 4.873E+04 | 401.2 | - | - | 0 | - |
| - | - | 9133 | 402.2 | - | - | 0 | - |
| - | - | 1019 | 403.2 | - | - | 0 | - |
| - | - | 1323 | 411.2 | - | - | 0 | - |
| - | - | 5150 | 414.2 | - | - | 0 | - |
| - | - | 1375 | 415.2 | - | - | 0 | - |
| - | - | 798.3 | 416.2 | - | - | 0 | - |
| - | - | 1141 | 424.2 | - | - | 0 | - |
| - | - | 711.7 | 425.2 | - | - | 0 | - |
| - | - | 824 | 427.2 | - | - | 0 | - |
| - | - | 1724 | 429.2 | - | - | 0 | - |
| 3 | w | 1.508E+05 | 442.2 | 0.000966 | 2.184 | +1 | 5 |
| - | - | 3.33E+04 | 443.2 | - | - | 0 | - |
| - | - | 5424 | 444.2 | - | - | 0 | - |
| - | - | 3019 | 455.3 | - | - | 0 | - |
| - | - | 6864 | 458.2 | - | - | 0 | - |
| - | - | 2164 | 459.3 | - | - | 0 | - |
| - | - | 1297 | 465.3 | - | - | 0 | - |
| - | - | 626.4 | 466.3 | - | - | 0 | - |
| 3 | z | 1879 | 467.3 | 0.001403 | 3.002 | +1 | 5 |
| - | - | 1249 | 470.3 | - | - | 0 | - |
| - | - | 1057 | 471.3 | - | - | 0 | - |
| - | - | 913.1 | 472.3 | - | - | 0 | - |
| - | - | 1908 | 479.3 | - | - | 0 | - |
| - | - | 1462 | 480.3 | - | - | 0 | - |
| 3 | y | 1.202E+04 | 483.3 | 0.0008366 | 1.731 | +1 | 5 |
| - | - | 3810 | 484.3 | - | - | 0 | - |
| 3 | z | 2.463E+04 | 485.3 | 0.001031 | 2.124 | +1 | 5 |
| - | - | 1205 | 486.2 | - | - | 0 | - |
| - | - | 6795 | 486.3 | - | - | 0 | - |
| - | - | 1623 | 487.3 | - | - | 0 | - |
| - | - | 2139 | 496.3 | - | - | 0 | - |
| - | - | 2246 | 497.3 | - | - | 0 | - |
| - | - | 1275 | 498.2 | - | - | 0 | - |
| - | - | 3891 | 498.3 | - | - | 0 | - |
| - | - | 1082 | 499.3 | - | - | 0 | - |
| - | - | 1793 | 501.2 | - | - | 0 | - |
| 3 | y | 6.834E+04 | 501.3 | 0.0008615 | 1.718 | +1 | 5 |
| - | - | 680.2 | 502.2 | - | - | 0 | - |
| - | - | 1.652E+04 | 502.3 | - | - | 0 | - |
| - | - | 2709 | 503.3 | - | - | 0 | - |
| - | - | 1930 | 514.2 | - | - | 0 | - |
| - | - | 2.262E+04 | 514.3 | - | - | 0 | - |
| 5 | c | 2.251E+04 | 515.3 | 0.0001096 | 0.2128 | +1 | 5 |
| - | - | 1.753E+04 | 516.3 | - | - | 0 | - |
| - | - | 6293 | 516.3 | - | - | 0 | - |
| - | - | 4906 | 517.3 | - | - | 0 | - |
| - | - | 881.4 | 519.3 | - | - | 0 | - |
| - | - | 2116 | 527.2 | - | - | 0 | - |
| - | - | 4428 | 527.7 | - | - | 0 | - |
| - | - | 3216 | 528.2 | - | - | 0 | - |
| - | - | 3940 | 528.3 | - | - | 0 | - |
| - | - | 1.661E+04 | 529.3 | - | - | 0 | - |
| - | - | 5405 | 529.3 | - | - | 0 | - |
| - | - | 4237 | 530.3 | - | - | 0 | - |
| - | - | 1288 | 530.3 | - | - | 0 | - |
| - | - | 942.4 | 531.3 | - | - | 0 | - |
| - | - | 1443 | 535.3 | - | - | 0 | - |
| - | - | 2710 | 542.3 | - | - | 0 | - |
| - | - | 1202 | 553.3 | - | - | 0 | - |
| 6 | c | 4882 | 554.3 | 0.001243 | 2.242 | +1 | 6 |
| 2 | w | 9947 | 555.3 | 0.001253 | 2.256 | +1 | 6 |
| - | - | 2649 | 556.3 | - | - | 0 | - |
| - | - | 1126 | 558.3 | - | - | 0 | - |
| 2 | y | 9637 | 570.3 | 0.0003691 | 0.6472 | +1 | 6 |
| - | - | 4629 | 571.3 | - | - | 0 | - |
| 6 | c | 4.112E+05 | 572.3 | 0.0007655 | 1.338 | +1 | 6 |
| - | - | 1.206E+05 | 573.3 | - | - | 0 | - |
| - | - | 2.669E+04 | 574.3 | - | - | 0 | - |
| - | - | 725.6 | 575.3 | - | - | 0 | - |
| - | - | 1274 | 580.3 | - | - | 0 | - |
| 2 | y | 1.476E+05 | 588.3 | 0.0009681 | 1.645 | +1 | 6 |
| - | - | 4.633E+04 | 589.3 | - | - | 0 | - |
| - | - | 8500 | 590.3 | - | - | 0 | - |
| - | - | 2.55E+04 | 598.3 | - | - | 0 | - |
| - | - | 8716 | 599.3 | - | - | 0 | - |
| - | - | 1811 | 600.3 | - | - | 0 | - |
| - | - | 815.5 | 607.3 | - | - | 0 | - |
| - | - | 1241 | 614.3 | - | - | 0 | - |
| - | - | 870.1 | 615.3 | - | - | 0 | - |
| - | - | 1027 | 617.3 | - | - | 0 | - |
| - | - | 863.4 | 624.3 | - | - | 0 | - |
| - | - | 692.5 | 628.4 | - | - | 0 | - |
| - | - | 5688 | 629.3 | - | - | 0 | - |
| - | - | 2056 | 630.3 | - | - | 0 | - |
| - | - | 1175 | 631.4 | - | - | 0 | - |
| - | - | 3.393E+05 | 642.3 | - | - | 0 | - |
| - | - | 1.149E+05 | 643.3 | - | - | 0 | - |
| - | - | 2.661E+04 | 644.4 | - | - | 0 | - |
| - | - | 1093 | 645.4 | - | - | 0 | - |
| - | - | 1.84E+04 | 646.4 | - | - | 0 | - |
| - | - | 5072 | 647.4 | - | - | 0 | - |
| - | - | 1431 | 648.4 | - | - | 0 | - |
| - | - | 5354 | 655.4 | - | - | 0 | - |
| - | - | 1492 | 656.4 | - | - | 0 | - |
| - | - | 794.9 | 657.4 | - | - | 0 | - |
| - | - | 3397 | 659.4 | - | - | 0 | - |
| - | - | 1537 | 660.4 | - | - | 0 | - |
| - | - | 4603 | 667.4 | - | - | 0 | - |
| - | - | 2231 | 668.4 | - | - | 0 | - |
| - | - | 853.1 | 672.4 | - | - | 0 | - |
| - | - | 2757 | 684.4 | - | - | 0 | - |
| - | - | 2.27E+05 | 685.4 | - | - | 0 | - |
| - | - | 8.21E+04 | 686.4 | - | - | 0 | - |
| - | - | 2.268E+04 | 687.4 | - | - | 0 | - |
| - | - | 1733 | 688.4 | - | - | 0 | - |
| - | - | 2.409E+05 | 701.4 | - | - | 0 | - |
| - | - | 5.483E+05 | 702.4 | - | - | 0 | - |
| - | - | 1.919E+05 | 703.4 | - | - | 0 | - |
| - | - | 1109 | 703.5 | - | - | 0 | - |
| - | - | 4.477E+04 | 704.4 | - | - | 0 | - |
| - | - | 2983 | 705.4 | - | - | 0 | - |
| - | - | 1854 | 717.4 | - | - | 0 | - |
| - | - | 1928 | 761.3 | - | - | 0 | - |
| - | - | 643.9 | 924 | - | - | 0 | - |
| - | - | 1728 | 996.4 | - | - | 0 | - |
| - | - | 1014 | 997.4 | - | - | 0 | - |
| - | - | 2615 | 1007 | - | - | 0 | - |
| - | - | 1983 | 1008 | - | - | 0 | - |
| - | - | 940.1 | 1009 | - | - | 0 | - |
| - | - | 927.4 | 1019 | - | - | 0 | - |
| - | - | 959.6 | 1020 | - | - | 0 | - |
| - | - | 1348 | 1037 | - | - | 0 | - |
| - | - | 1069 | 1038 | - | - | 0 | - |
| - | - | 1849 | 1054 | - | - | 0 | - |
| - | - | 8566 | 1055 | - | - | 0 | - |
| - | - | 5347 | 1056 | - | - | 0 | - |
| - | - | 1091 | 1057 | - | - | 0 | - |
| - | - | 658.9 | 1361 | - | - | 0 | - |
| - | - | 712.8 | 1864 | - | - | 0 | - |
| - | - | 883.1 | 2964 | - | - | 0 | - |
| - | - | 771.8 | 3041 | - | - | 0 | - |
| - | - | 839.7 | 3043 | - | - | 0 | - |
| - | - | 720.5 | 3161 | - | - | 0 | - |
| - | - | 677.7 | 3280 | - | - | 0 | - |

m/z Charge Intensity FragmentType MassShift Position
125.10769653320312 0 3700.3462
128.0711212158203 0 677.50354
128.10733032226562 0 1436.5289
129.10264587402344 0 5020.435
130.06556701660156 0 493.46204
130.0867156982422 0 1755.8076 y Ammonia loss 6
131.0707244873047 0 1717.4108
131.11813354492188 0 525.1682
132.98095703125 0 380.60638
133.0862274169922 0 1175.5598
142.08694458007812 0 994.0482
147.11318969726562 0 6541.8853 y 6
148.1166534423828 0 587.8418
150.18954467773438 0 403.69528
151.09715270996094 0 952.36035 y 4
155.08180236816406 0 1402.9336
155.1183319091797 0 5356.8315
157.0866241455078 0 548.85297
157.0974884033203 0 1529.0914
169.07371520996094 0 5350.501
170.0816192626953 0 1541.5349
172.10848999023438 0 39780.55
173.0925750732422 0 2413.5354
173.11175537109375 0 2288.7463
173.1288604736328 0 47901.72
174.13229370117188 0 3176.418
183.1131591796875 0 3508.5222
185.0924530029297 0 1631.9883
186.12435913085938 0 1930.3955
188.11610412597656 0 4374.695 z 5
189.1234893798828 0 1785.0486
200.11605834960938 0 6388.5815
201.12374877929688 0 42265.695
202.11871337890625 0 3088.954
202.1273651123047 0 3725.2266
204.13462829589844 0 12135.749 y 5
204.14524841308594 0 599.61096
205.13821411132812 0 1027.7329
211.1084747314453 0 624.6819
224.1027069091797 0 793.8641
226.0949249267578 0 1462.2307
227.10299682617188 0 592.8479
229.1062469482422 0 8119.4834
230.1129913330078 0 1053.9833
231.12118530273438 0 688.0157
232.1193389892578 0 1115.3582
242.11404418945312 0 2832.5085
245.1374969482422 0 2121.3855
247.0321502685547 0 531.18365
251.15548706054688 0 1978.8513 y 2
258.1451721191406 0 9763.997 w 4
259.140625 0 8890.435
260.1249694824219 0 1083.4413
260.1426086425781 0 500.29276
270.1446533203125 0 1821.0966
271.1531066894531 0 777.75653
272.1604919433594 0 2238.9346
273.168701171875 0 9717.164
273.18402099609375 0 820.8308
274.1724548339844 0 987.46875
283.11724853515625 0 911.2075
283.15362548828125 0 1121.9436
283.1769714355469 0 9279.913
284.16021728515625 0 527.77936 y Ammonia loss 4
284.17919921875 0 673.32587
285.6654968261719 0 512.56805 y Water loss 1
287.21978759765625 0 735.6025
288.156005859375 0 2290.5564
294.6728210449219 0 563.068 y 1
295.19000244140625 0 732.9282
296.1974792480469 0 7322.063
297.1999816894531 0 1476.1251
299.17205810546875 0 19444.744
300.1791687011719 0 12015.25
301.1272888183594 0 2705.8057
301.1876525878906 0 141703.64 y 4
302.1908874511719 0 21460.285
303.1934509277344 0 2172.1875
313.20166015625 0 590.04486
314.1349792480469 0 1657.2084
314.2078857421875 0 2034.199
315.20355224609375 0 5055.88
316.20654296875 0 835.7186
320.2217102050781 0 1202.0154
328.1748962402344 0 13573.304
329.1807861328125 0 4894.2227
330.18487548828125 0 979.1393
331.23480224609375 0 7377.111 c 2
332.2377014160156 0 1494.1012
333.6957702636719 0 872.1484
334.91082763671875 0 904.81915
335.173583984375 0 642.00586
336.1789855957031 0 734.4054
337.1872253417969 0 1074.7482
342.1898193359375 0 5332.2593
342.2094421386719 0 2598.2898
343.1925964355469 0 948.2943
344.1697692871094 0 1291.4603
350.1871643066406 0 1811.4602
350.9866638183594 0 746.4735
351.14190673828125 0 2931.7441
351.2140808105469 0 4004.4246
351.71527099609375 0 1709.0894
352.1493225097656 0 945.2331
352.1741943359375 0 3092.5938
352.1969299316406 0 1159.546
352.23895263671875 0 6831.8613
353.182373046875 0 4152.3936
354.188232421875 0 1484.7726 z Water loss 3
355.19818115234375 0 40252.113 w 3
356.20098876953125 0 7034.0933
357.2004699707031 0 790.4669
367.1980285644531 0 608.1204
369.20440673828125 0 701.1243
370.1850891113281 0 19698.443
370.2098083496094 0 5334.2583 y Water loss 3
371.1920166015625 0 17277.121 y Ammonia loss 3
372.2001953125 0 19868.32 z 3
372.22467041015625 0 2153.153
373.20806884765625 0 20749.848
373.2450256347656 0 1551.0828
374.211181640625 0 2677.4907
383.2287902832031 0 5506.2295
384.2304382324219 0 854.7995
385.20501708984375 0 648.72015
388.2197265625 0 58058.15 y 3
389.2227783203125 0 10122.547
390.2260437011719 0 1839.447
396.1893005371094 0 2934.8354
398.2038269042969 0 2984.3462
399.2096252441406 0 915.88525
400.232421875 0 8520.432
401.2400817871094 0 48733.734
402.2429504394531 0 9132.973
403.24371337890625 0 1018.7356
411.2112121582031 0 1322.5939
414.2113952636719 0 5150.447
415.2113037109375 0 1375.1935
416.21337890625 0 798.32794
424.2208557128906 0 1141.0095
425.2200622558594 0 711.69934
427.21990966796875 0 823.9788
429.2233581542969 0 1723.8196
442.2305908203125 0 150802.88 w 2
443.2334289550781 0 33302.598
444.2361145019531 0 5423.6406
455.2746887207031 0 3018.57
458.249267578125 0 6864.246
459.2531433105469 0 2163.6394
465.28192138671875 0 1297.1149
466.26788330078125 0 626.3799
467.2752380371094 0 1878.5234 z Water loss 2
470.2976989746094 0 1248.9259
471.30316162109375 0 1056.5975
472.30218505859375 0 913.0861
479.27484130859375 0 1907.6802
480.28173828125 0 1462.0732
483.29339599609375 0 12018.419 y Water loss 2
484.29736328125 0 3810.2224
485.2854309082031 0 24627.838 z 2
486.2469482421875 0 1204.552
486.28875732421875 0 6795.0645
487.2918701171875 0 1623.2823
496.30133056640625 0 2139.0054
497.28570556640625 0 2245.8977
498.24658203125 0 1275.2028
498.2919921875 0 3890.6892
499.2927551269531 0 1082.18
501.2380065917969 0 1792.6213
501.3039855957031 0 68342.78 y 2
502.24542236328125 0 680.20667
502.3071594238281 0 16517.51
503.30999755859375 0 2708.539
514.24072265625 0 1929.8016
514.3118286132812 0 22620.307
515.3186645507812 0 22509.639 c 4
516.2545166015625 0 17532.162
516.3221435546875 0 6293.0034
517.2584838867188 0 4905.6807
519.2949829101562 0 881.43945
527.2315673828125 0 2116.2617
527.7349853515625 0 4428.0903
528.2373046875 0 3215.7576
528.3269653320312 0 3940.2493
529.2623291015625 0 16610.986
529.3342895507812 0 5404.632
530.2649536132812 0 4236.961
530.3375244140625 0 1288.0979
531.2664184570312 0 942.3848
535.2882690429688 0 1442.5145
542.3084716796875 0 2710.4204
553.3023071289062 0 1201.9781
554.3284301757812 0 4881.7427 c Water loss 5
555.31494140625 0 9946.824 w 1
556.318115234375 0 2649.1575
558.3226318359375 0 1125.9486
570.32421875 0 9636.83 y Water loss 1
571.3273315429688 0 4629.039
572.3410034179688 0 411196.03 c 5
573.343505859375 0 120600.58
574.34619140625 0 26688.236
575.3466796875 0 725.5585
580.3086547851562 0 1274.454
588.3361206054688 0 147627.3 y 1
589.3389282226562 0 46325.984
590.3416137695312 0 8499.801
598.3204345703125 0 25503.998
599.3233642578125 0 8716.431
600.3305053710938 0 1811.2592
607.2538452148438 0 815.5472
614.3317260742188 0 1241.2178
615.3267822265625 0 870.07324
617.3402099609375 0 1026.7716
624.2802124023438 0 863.39557
628.3534545898438 0 692.5245
629.3392944335938 0 5688.144
630.34033203125 0 2055.653
631.3538818359375 0 1174.672
642.3468627929688 0 339284.16
643.3497314453125 0 114946.01
644.35205078125 0 26610.582
645.3538208007812 0 1093.1713
646.365234375 0 18400.654
647.367919921875 0 5071.5464
648.3701171875 0 1430.9626
655.3908081054688 0 5353.5483
656.4072265625 0 1492.2645
657.4216918945312 0 794.89075
659.3751831054688 0 3397.4001
660.3754272460938 0 1537.3552
667.3909912109375 0 4603.492
668.3958129882812 0 2231.2703
672.411865234375 0 853.0888
684.4175415039062 0 2756.9412
685.4016723632812 0 227046.9
686.4046630859375 0 82100.16
687.406982421875 0 22675.441
688.4075927734375 0 1732.8832
701.4202270507812 0 240874.7
702.4273071289062 0 548301.7
703.4302978515625 0 191859.52
703.505126953125 0 1108.5063
704.43310546875 0 44774.7
705.434814453125 0 2983.4375
717.3900146484375 0 1853.5092
761.3364868164062 0 1927.6176
924.0321044921875 0 643.85425
996.4351806640625 0 1727.5853
997.4396362304688 0 1013.9192
1007.4490356445312 0 2615.319
1008.4512939453125 0 1982.5483
1009.46044921875 0 940.1121
1019.4567260742188 0 927.4102
1020.44775390625 0 959.62213
1037.4609375 0 1348.3141
1038.4583740234375 0 1068.6862
1054.462890625 0 1848.7003
1055.46826171875 0 8565.753
1056.47265625 0 5347.2905
1057.477783203125 0 1090.5659
1361.3798828125 0 658.94556
1864.1885986328125 0 712.76855
2964.04931640625 0 883.0788
3040.889404296875 0 771.80365
3042.56884765625 0 839.6936
3160.67236328125 0 720.5088
3279.720947265625 0 677.65717

Spectrum Details

|  |  |
| --- | --- |
| Matched peaks? Matched peaks The total absolute number of peaks matched. Additionally in brackets the total fraction of peaks matched and the total number of peaks is shown. | 29 (10.86% of 267) |
| FDR? FDR The false discovery rate estimated for this peptide. It is calculated by matching all theoretical fragments with a non-integer shift with the raw peaks for this spectrum. This is done with 40 different shifts. The resulting percentage is the average number of annotated peaks over the number of annotated peaks with the correct spectrum. | 0.25% |
| Satellite FDR? Satellite FDR See the FDR for details on its calculation. This satellite ion specific FDR only contains the satellite ions (d/w) for I/L/J positions. | 0.00% |
| PSM Score? PSM Score The PSM Score as given by Hecklib to this annotated spectrum. It is shown with three significant figures. | 310 |

#### Reverse Lookup? Reverse Lookup All places where this read could be placed.

| Group | Segment | Template | Template Part | Read Part | Score | Unique |
| --- | --- | --- | --- | --- | --- | --- |
| Homo sapiens Heavy Chain | IGHC | IGHG1 | [323..330] | [0..7] | 56 | False |
| Homo sapiens Heavy Chain | IGHC | IGHG3 | [370..377] | [0..7] | 56 | False |
| Homo sapiens Heavy Chain | IGHC | IGHG2 | [319..326] | [0..7] | 56 | False |

| Recombined | Template Part | Read Part | Score | Unique |
| --- | --- | --- | --- | --- |
| REC-0-1 | [448..455] | [0..7] | 47 | True |

#### Meta Information from Multiple reads

##### Number of combined reads

2

##### Intensity

1

##### TotalArea

0

##### Changes to the peptide sequence

JSLSPGK

J→LSupport for Leucine based on side chain ions (1 for L 0 for I) (Position: 3)

L→JNo support for either Leucine or Isoleucine based on side chain ions (Position: 3)

L→JNo support for either Leucine or Isoleucine based on side chain ions (Position: 1)

#### Positional Score

Copy Data

##### Positional Score (TSV)

###### Preview

```
Loading example...
```

*Click on the button to copy the data to your clipboard.*

000123456

Label Value
"0" 0
"1" 0
"2" 0
"3" 0
"4" 0
"5" 0
"6" 0

#### Meta Information from PEAKS

##### Scan Identifier

F1:3938

##### Original sequence

L

S

L

S

P

G

K

##### Posttranslational Modifications

##### Source File

D:\separate\_stitch\_analyses\xle-disambiguation\raw\20210323\_F1\_UM1\_Peng0013\_SA\_F59\_ingel\_3ug\_ELA.raw

##### Fraction

1

##### Scan Feature

-

##### De Novo Score

99

##### ConfidenceScore

99

### m/z

351.2136

##### Mass

700.4119

##### Charge

2

##### Retention Time

20.64

##### Predicted Retention Time

-

##### Area

0

##### Parts Per Million

1

##### Fragmentation mode

HCD

##### Originating file

01 D:\separate\_stitch\_analyses\xle-disambiguation\20210325\_F59\_3ug\_DENOVO\_12.csv

#### Meta Information from PEAKS

##### Scan Identifier

F1:3875

##### Original sequence

L

S

L

S

P

G

K

##### Posttranslational Modifications

##### Source File

D:\separate\_stitch\_analyses\xle-disambiguation\raw\20210323\_F1\_UM1\_Peng0013\_SA\_F59\_ingel\_3ug\_ELA.raw

##### Fraction

1

##### Scan Feature

-

##### De Novo Score

98

##### ConfidenceScore

98

### m/z

351.2139

##### Mass

700.4119

##### Charge

2

##### Retention Time

20.28

##### Predicted Retention Time

-

##### Area

0

##### Parts Per Million

2

##### Fragmentation mode

ETHCD

##### Originating file

01 D:\separate\_stitch\_analyses\xle-disambiguation\20210325\_F59\_3ug\_DENOVO\_12.csv
