## Supplementary material for "A handle on mass coincidence errors in *de novo* sequencing of antibodies by bottom-up proteomics": Combined_069.html

Details Combined\_069 | Stitch OverviewUndefined

### Read Combined\_069

#### Sequence (length=16)

VYTLPPSREEMTKNQK

#### Spectrum 4198? Spectrum 4198 The raw spectrum of this peptide as annotated by Hecklib. The fragments are coloured according to ion type (see legend). Any peaks with a star '\*' as text can be hovered over to see the full details, first the ion type second the mass shift type. By hovering over the amino acids in the peptide or ions in the legend the corresponding peaks are highlighted. By toggling the 'Unassigned' label you can turn the background (unassigned) peaks on or off in the plot. By updating the slider in the Ion legend you can update the spectrum to only show the top X% of the peaks with labels. The top X% means any peak that is within X% of the highest intensity. By dragging in the spectrum you can zoom in to a specific part of the spectrum and use 'Zoom Out' to get back to the original zoom level. The annotation of the spectrum is based on the given sequence in the peptides file and is done with different software so inconsistencies are likely. The peaks are annotated based on the given sequence, with 20 ppm tolerance.

Copy Data

##### Spectrum 4198 (TSV)

###### Preview

```
Loading example...
```

*Click on the button to copy the data to your clipboard.*

Mz MinMz MaxIntensity Max

WidthHeightPeptide font sizePeptide stroke widthSpectrum font sizeSpectrum stroke widthCompact peptide

Ion legend

wxyz

abcd

OtherUnassignedIon^Charge^_Position_Show for top:%

VYTLPPSREEMTKNQK

02.05e+44.09e+46.14e+48.19e+4

Zoom Out

y^+1^_2_y^+1^_3_y^+3^_12_z^+1^_4_y^+3^_14_y^+3^_14_y^+1^_4_y^+3^_15_z^+3^_15_w^+1^_5_w^+1^_5_y^+2^_10_y^+1^_5_z^+1^_5_y^+2^_10_y^+1^_5_c^+1^_6_z^+2^_11_y^+2^_11_y^+2^_12_z^+2^_12_y^+2^_12_c^+1^_7_c^+2^_13_w^+2^_13_z^+1^_6_z^+1^_6_z^+2^_13_y^+2^_13_y^+1^_6_c^+2^_14_y^+2^_14_y^+2^_14_z^+2^_14_y^+2^_14_w^+1^_7_c^+2^_15_c^+2^_15_c^+1^_8_z^+1^_7_y^+2^_15_z^+2^_15_y^+2^_15_y^+1^_7_w^+1^_8_c^+1^_9_z^+1^_8_y^+1^_8_c^+1^_10_z^+1^_9_y^+1^_9_z^+1^_10_y^+1^_10_c^+1^_11_y^+1^_11_c^+1^_12_c^+1^_13_z^+1^_13_y^+1^_13_c^+1^_14_z^+1^_14_y^+1^_14_c^+1^_15_z^+1^_15_

0506101115172022

Fragment Matches Table

Show background peaks

| Position | Ion type | Intensity | mz Theoretical | mz Error (Th) | mz Error (ppm) | Charge | Series Number |
| --- | --- | --- | --- | --- | --- | --- | --- |
| - | - | 369.6 | 121.7 | - | - | 0 | - |
| - | - | 349.6 | 124.2 | - | - | 0 | - |
| - | - | 360.6 | 127.5 | - | - | 0 | - |
| - | - | 871.2 | 129.1 | - | - | 0 | - |
| - | - | 2618 | 136.1 | - | - | 0 | - |
| - | - | 452.8 | 145.1 | - | - | 0 | - |
| - | - | 448.6 | 147.1 | - | - | 0 | - |
| - | - | 928.1 | 148.9 | - | - | 0 | - |
| - | - | 419.9 | 151.6 | - | - | 0 | - |
| - | - | 880.5 | 169.1 | - | - | 0 | - |
| - | - | 1775 | 187.1 | - | - | 0 | - |
| - | - | 459.7 | 192.2 | - | - | 0 | - |
| - | - | 512.1 | 194.3 | - | - | 0 | - |
| - | - | 1043 | 197.1 | - | - | 0 | - |
| - | - | 450.1 | 203.6 | - | - | 0 | - |
| - | - | 536.1 | 205.7 | - | - | 0 | - |
| - | - | 1314 | 215.1 | - | - | 0 | - |
| - | - | 782.6 | 225 | - | - | 0 | - |
| - | - | 1.426E+04 | 235.1 | - | - | 0 | - |
| - | - | 1971 | 236.1 | - | - | 0 | - |
| - | - | 1074 | 239.1 | - | - | 0 | - |
| - | - | 492.2 | 240.4 | - | - | 0 | - |
| - | - | 695.2 | 263.1 | - | - | 0 | - |
| - | - | 1.12E+04 | 263.1 | - | - | 0 | - |
| - | - | 1473 | 264.1 | - | - | 0 | - |
| - | - | 1079 | 283.1 | - | - | 0 | - |
| - | - | 956 | 295.1 | - | - | 0 | - |
| - | - | 498.9 | 299.1 | - | - | 0 | - |
| - | - | 2994 | 299.2 | - | - | 0 | - |
| - | - | 714.6 | 320.2 | - | - | 0 | - |
| 15 | y | 1054 | 333.2 | 0.003678 | 11.04 | +1 | 2 |
| - | - | 1298 | 341 | - | - | 0 | - |
| - | - | 568.5 | 342.9 | - | - | 0 | - |
| - | - | 8157 | 346.2 | - | - | 0 | - |
| - | - | 1789 | 347.2 | - | - | 0 | - |
| - | - | 4361 | 355.1 | - | - | 0 | - |
| - | - | 4995 | 359 | - | - | 0 | - |
| - | - | 803.9 | 360.2 | - | - | 0 | - |
| - | - | 1.086E+04 | 364.2 | - | - | 0 | - |
| - | - | 2422 | 365.2 | - | - | 0 | - |
| - | - | 592.2 | 387.7 | - | - | 0 | - |
| - | - | 4059 | 388.2 | - | - | 0 | - |
| - | - | 1.32E+04 | 429.1 | - | - | 0 | - |
| - | - | 3470 | 431.3 | - | - | 0 | - |
| 14 | y | 1536 | 447.2 | 0.002935 | 6.562 | +1 | 3 |
| - | - | 2300 | 449.3 | - | - | 0 | - |
| - | - | 594.2 | 450.3 | - | - | 0 | - |
| - | - | 684.4 | 458.3 | - | - | 0 | - |
| - | - | 7817 | 459.3 | - | - | 0 | - |
| - | - | 2725 | 460.3 | - | - | 0 | - |
| - | - | 566.2 | 473.1 | - | - | 0 | - |
| - | - | 592.7 | 476.3 | - | - | 0 | - |
| - | - | 3432 | 477.3 | - | - | 0 | - |
| - | - | 1085 | 478.3 | - | - | 0 | - |
| 5 | y | 5156 | 506.9 | 0.0003148 | 0.621 | +3 | 12 |
| - | - | 653.2 | 507.1 | - | - | 0 | - |
| - | - | 3854 | 507.2 | - | - | 0 | - |
| - | - | 1310 | 507.6 | - | - | 0 | - |
| - | - | 693.1 | 507.9 | - | - | 0 | - |
| - | - | 621.2 | 514.2 | - | - | 0 | - |
| - | - | 2879 | 515.3 | - | - | 0 | - |
| - | - | 974.7 | 516.3 | - | - | 0 | - |
| - | - | 2059 | 545.4 | - | - | 0 | - |
| 13 | z | 8164 | 559.3 | 0.004467 | 7.986 | +1 | 4 |
| - | - | 3678 | 560.3 | - | - | 0 | - |
| - | - | 1091 | 561.3 | - | - | 0 | - |
| 3 | y | 1152 | 572.3 | 0.00186 | 3.249 | +3 | 14 |
| 3 | y | 1430 | 572.6 | 0.006134 | 10.71 | +3 | 14 |
| 13 | y | 7849 | 575.3 | 0.003781 | 6.573 | +1 | 4 |
| - | - | 1068 | 576.3 | - | - | 0 | - |
| - | - | 614 | 586.3 | - | - | 0 | - |
| - | - | 1197 | 592.9 | - | - | 0 | - |
| - | - | 2010 | 593.2 | - | - | 0 | - |
| - | - | 741.5 | 601.2 | - | - | 0 | - |
| - | - | 1608 | 602.4 | - | - | 0 | - |
| - | - | 1022 | 603.4 | - | - | 0 | - |
| - | - | 1222 | 615.3 | - | - | 0 | - |
| - | - | 2120 | 616.3 | - | - | 0 | - |
| - | - | 639.7 | 617.3 | - | - | 0 | - |
| - | - | 3170 | 619.3 | - | - | 0 | - |
| - | - | 1141 | 624.6 | - | - | 0 | - |
| - | - | 631.1 | 625.7 | - | - | 0 | - |
| 2 | y | 1504 | 627 | 0.005756 | 9.18 | +3 | 15 |
| 2 | z | 1153 | 627.3 | 0.003615 | 5.763 | +3 | 15 |
| 12 | w | 2491 | 643.3 | 0.004667 | 7.254 | +1 | 5 |
| - | - | 763.2 | 644.3 | - | - | 0 | - |
| 12 | w | 2136 | 645.3 | 0.003401 | 5.271 | +1 | 5 |
| 7 | y | 895.6 | 654.3 | 0.0009419 | 1.439 | +2 | 10 |
| - | - | 675.9 | 656.8 | - | - | 0 | - |
| - | - | 815 | 657.8 | - | - | 0 | - |
| - | - | 1508 | 658.3 | - | - | 0 | - |
| 12 | y | 3267 | 659.3 | 0.005746 | 8.714 | +1 | 5 |
| - | - | 6771 | 659.7 | - | - | 0 | - |
| - | - | 9083 | 660 | - | - | 0 | - |
| 12 | z | 1.248E+04 | 660.3 | 0.009787 | 14.82 | +1 | 5 |
| - | - | 2672 | 660.7 | - | - | 0 | - |
| - | - | 4217 | 661.3 | - | - | 0 | - |
| 7 | y | 707.9 | 662.8 | 0.0008593 | 1.296 | +2 | 10 |
| - | - | 1145 | 666.3 | - | - | 0 | - |
| - | - | 1686 | 666.3 | - | - | 0 | - |
| - | - | 768.5 | 666.9 | - | - | 0 | - |
| - | - | 845.5 | 667.3 | - | - | 0 | - |
| - | - | 1.039E+04 | 667.4 | - | - | 0 | - |
| - | - | 649.7 | 674.4 | - | - | 0 | - |
| - | - | 802.5 | 675.4 | - | - | 0 | - |
| 12 | y | 3467 | 676.4 | 0.005134 | 7.591 | +1 | 5 |
| - | - | 1534 | 677.4 | - | - | 0 | - |
| - | - | 744.2 | 678.4 | - | - | 0 | - |
| - | - | 706.7 | 679.4 | - | - | 0 | - |
| - | - | 1027 | 687.4 | - | - | 0 | - |
| 6 | c | 3111 | 688.4 | 0.002081 | 3.023 | +1 | 6 |
| - | - | 836.6 | 689.4 | - | - | 0 | - |
| - | - | 823.7 | 693.3 | - | - | 0 | - |
| 6 | z | 657.3 | 694.3 | 0.002916 | 4.199 | +2 | 11 |
| - | - | 1445 | 697.4 | - | - | 0 | - |
| - | - | 652.9 | 698.4 | - | - | 0 | - |
| - | - | 887.3 | 701.4 | - | - | 0 | - |
| - | - | 4327 | 707.3 | - | - | 0 | - |
| - | - | 3781 | 707.8 | - | - | 0 | - |
| - | - | 2429 | 708.3 | - | - | 0 | - |
| 6 | y | 3509 | 711.3 | 0.0001732 | 0.2434 | +2 | 11 |
| - | - | 2092 | 711.8 | - | - | 0 | - |
| - | - | 1238 | 712.3 | - | - | 0 | - |
| - | - | 2954 | 713.4 | - | - | 0 | - |
| - | - | 2355 | 714.4 | - | - | 0 | - |
| - | - | 714.5 | 717.4 | - | - | 0 | - |
| - | - | 927.9 | 718.3 | - | - | 0 | - |
| - | - | 1010 | 727.4 | - | - | 0 | - |
| - | - | 1981 | 727.9 | - | - | 0 | - |
| - | - | 2679 | 728.4 | - | - | 0 | - |
| - | - | 708.1 | 728.9 | - | - | 0 | - |
| - | - | 722.7 | 729.4 | - | - | 0 | - |
| - | - | 3517 | 744.4 | - | - | 0 | - |
| - | - | 1.197E+04 | 745.4 | - | - | 0 | - |
| - | - | 5358 | 746.4 | - | - | 0 | - |
| - | - | 1205 | 747.4 | - | - | 0 | - |
| 5 | y | 2514 | 751.4 | 0.006231 | 8.293 | +2 | 12 |
| 5 | z | 1064 | 751.9 | 0.001647 | 2.19 | +2 | 12 |
| 5 | y | 5.809E+04 | 759.9 | 0.0003415 | 0.4495 | +2 | 12 |
| - | - | 4.444E+04 | 760.4 | - | - | 0 | - |
| - | - | 2.716E+04 | 760.9 | - | - | 0 | - |
| - | - | 8015 | 761.4 | - | - | 0 | - |
| - | - | 2437 | 761.9 | - | - | 0 | - |
| - | - | 1222 | 764.9 | - | - | 0 | - |
| - | - | 1433 | 766.4 | - | - | 0 | - |
| - | - | 1201 | 772.4 | - | - | 0 | - |
| - | - | 846.9 | 774.9 | - | - | 0 | - |
| 7 | c | 1.466E+04 | 775.4 | 0.0006193 | 0.7986 | +1 | 7 |
| - | - | 7025 | 776.4 | - | - | 0 | - |
| - | - | 1908 | 777.4 | - | - | 0 | - |
| - | - | 1145 | 780.9 | - | - | 0 | - |
| - | - | 1.039E+04 | 782.9 | - | - | 0 | - |
| 13 | c | 1.829E+04 | 783.4 | 0.001622 | 2.07 | +2 | 13 |
| - | - | 1.374E+04 | 783.9 | - | - | 0 | - |
| - | - | 7640 | 784.4 | - | - | 0 | - |
| - | - | 2610 | 784.9 | - | - | 0 | - |
| - | - | 1112 | 785.4 | - | - | 0 | - |
| 4 | w | 1.441E+04 | 786.9 | 0.0006744 | 0.8571 | +2 | 13 |
| - | - | 1.052E+04 | 787.4 | - | - | 0 | - |
| - | - | 8094 | 787.9 | - | - | 0 | - |
| - | - | 3641 | 788.4 | - | - | 0 | - |
| - | - | 1049 | 788.9 | - | - | 0 | - |
| 11 | z | 806.5 | 789.4 | 0.001788 | 2.265 | +1 | 6 |
| - | - | 783.1 | 794.8 | - | - | 0 | - |
| - | - | 862.3 | 800.4 | - | - | 0 | - |
| 11 | z | 3250 | 807.4 | 0.003613 | 4.475 | +1 | 6 |
| 4 | z | 2.262E+04 | 808.4 | 0.003398 | 4.203 | +2 | 13 |
| - | - | 1.42E+04 | 808.9 | - | - | 0 | - |
| - | - | 1.257E+04 | 809.4 | - | - | 0 | - |
| - | - | 4802 | 809.9 | - | - | 0 | - |
| - | - | 3530 | 810.4 | - | - | 0 | - |
| - | - | 2278 | 814.4 | - | - | 0 | - |
| - | - | 969.1 | 814.9 | - | - | 0 | - |
| - | - | 693.2 | 815.4 | - | - | 0 | - |
| - | - | 923.9 | 815.9 | - | - | 0 | - |
| 4 | y | 9927 | 816.4 | 0.0002407 | 0.2948 | +2 | 13 |
| - | - | 7838 | 816.9 | - | - | 0 | - |
| - | - | 4924 | 817.4 | - | - | 0 | - |
| - | - | 2195 | 817.9 | - | - | 0 | - |
| - | - | 1084 | 818.4 | - | - | 0 | - |
| - | - | 2159 | 818.9 | - | - | 0 | - |
| 11 | y | 1283 | 823.4 | 0.005763 | 6.999 | +1 | 6 |
| - | - | 685.3 | 830.4 | - | - | 0 | - |
| - | - | 2502 | 830.9 | - | - | 0 | - |
| - | - | 2798 | 831.4 | - | - | 0 | - |
| - | - | 779 | 831.9 | - | - | 0 | - |
| - | - | 893 | 834.4 | - | - | 0 | - |
| - | - | 1070 | 835.4 | - | - | 0 | - |
| - | - | 2271 | 839.9 | - | - | 0 | - |
| 14 | c | 3.425E+04 | 840.4 | 0.002436 | 2.898 | +2 | 14 |
| - | - | 3.023E+04 | 840.9 | - | - | 0 | - |
| - | - | 1.768E+04 | 841.4 | - | - | 0 | - |
| - | - | 8158 | 841.9 | - | - | 0 | - |
| - | - | 2463 | 842.4 | - | - | 0 | - |
| - | - | 745.2 | 842.9 | - | - | 0 | - |
| - | - | 816.3 | 853.9 | - | - | 0 | - |
| - | - | 1689 | 857.5 | - | - | 0 | - |
| 3 | y | 1242 | 857.9 | 0.007818 | 9.113 | +2 | 14 |
| 3 | y | 2659 | 858.4 | 0.002066 | 2.407 | +2 | 14 |
| 3 | z | 1.305E+04 | 858.9 | 0.0006257 | 0.7284 | +2 | 14 |
| - | - | 968 | 859.3 | - | - | 0 | - |
| - | - | 1.286E+04 | 859.4 | - | - | 0 | - |
| - | - | 5751 | 859.9 | - | - | 0 | - |
| - | - | 3630 | 860.4 | - | - | 0 | - |
| 3 | y | 2.439E+04 | 866.9 | 0.0002052 | 0.2366 | +2 | 14 |
| - | - | 2.564E+04 | 867.4 | - | - | 0 | - |
| - | - | 1.471E+04 | 867.9 | - | - | 0 | - |
| - | - | 7332 | 868.4 | - | - | 0 | - |
| - | - | 2460 | 868.9 | - | - | 0 | - |
| - | - | 985.9 | 870.9 | - | - | 0 | - |
| - | - | 900.1 | 871.4 | - | - | 0 | - |
| - | - | 1765 | 873.4 | - | - | 0 | - |
| - | - | 1765 | 874.4 | - | - | 0 | - |
| 10 | w | 8058 | 877.4 | 0.0003863 | 0.4403 | +1 | 7 |
| - | - | 4126 | 878.4 | - | - | 0 | - |
| - | - | 1199 | 879.4 | - | - | 0 | - |
| - | - | 1628 | 880.4 | - | - | 0 | - |
| - | - | 1736 | 880.9 | - | - | 0 | - |
| - | - | 1525 | 881.4 | - | - | 0 | - |
| - | - | 951.3 | 881.9 | - | - | 0 | - |
| - | - | 1685 | 882.4 | - | - | 0 | - |
| - | - | 718.5 | 887.1 | - | - | 0 | - |
| - | - | 872.7 | 887.4 | - | - | 0 | - |
| - | - | 919.3 | 887.5 | - | - | 0 | - |
| - | - | 1076 | 888.5 | - | - | 0 | - |
| - | - | 2750 | 888.9 | - | - | 0 | - |
| - | - | 2789 | 889.3 | - | - | 0 | - |
| - | - | 840.1 | 889.9 | - | - | 0 | - |
| - | - | 726.7 | 890.3 | - | - | 0 | - |
| 15 | c | 997.4 | 895.9 | 0.007296 | 8.143 | +2 | 15 |
| - | - | 1955 | 896.5 | - | - | 0 | - |
| - | - | 1911 | 897 | - | - | 0 | - |
| - | - | 972.3 | 897.5 | - | - | 0 | - |
| - | - | 951.3 | 901.5 | - | - | 0 | - |
| - | - | 1064 | 902.5 | - | - | 0 | - |
| - | - | 1042 | 903.9 | - | - | 0 | - |
| 15 | c | 1.06E+04 | 904.5 | 0.002017 | 2.23 | +2 | 15 |
| - | - | 1.098E+04 | 905 | - | - | 0 | - |
| - | - | 6562 | 905.5 | - | - | 0 | - |
| - | - | 2743 | 906 | - | - | 0 | - |
| - | - | 918 | 909.4 | - | - | 0 | - |
| - | - | 1327 | 909.9 | - | - | 0 | - |
| - | - | 1725 | 910.5 | - | - | 0 | - |
| - | - | 827 | 911.5 | - | - | 0 | - |
| - | - | 2426 | 912.4 | - | - | 0 | - |
| - | - | 1633 | 912.9 | - | - | 0 | - |
| - | - | 920.2 | 916.4 | - | - | 0 | - |
| - | - | 1059 | 917.4 | - | - | 0 | - |
| - | - | 935.1 | 923 | - | - | 0 | - |
| - | - | 968.1 | 924.5 | - | - | 0 | - |
| - | - | 3425 | 925.4 | - | - | 0 | - |
| - | - | 3296 | 925.9 | - | - | 0 | - |
| - | - | 2675 | 926.4 | - | - | 0 | - |
| - | - | 1536 | 928.5 | - | - | 0 | - |
| - | - | 813.6 | 930.5 | - | - | 0 | - |
| - | - | 955.8 | 931 | - | - | 0 | - |
| - | - | 1165 | 931.4 | - | - | 0 | - |
| 8 | c | 2.418E+04 | 931.5 | 0.0003993 | 0.4287 | +1 | 8 |
| - | - | 1405 | 931.9 | - | - | 0 | - |
| - | - | 3379 | 932 | - | - | 0 | - |
| - | - | 1.495E+04 | 932.5 | - | - | 0 | - |
| - | - | 2624 | 933 | - | - | 0 | - |
| - | - | 4046 | 933.5 | - | - | 0 | - |
| - | - | 938.1 | 934.5 | - | - | 0 | - |
| 10 | z | 1.842E+04 | 936.4 | 0.0003266 | 0.3488 | +1 | 7 |
| - | - | 1.101E+04 | 937.4 | - | - | 0 | - |
| - | - | 4279 | 938.4 | - | - | 0 | - |
| 2 | y | 1013 | 939.5 | 0.01507 | 16.04 | +2 | 15 |
| - | - | 2180 | 940 | - | - | 0 | - |
| 2 | z | 8303 | 940.5 | 0.001462 | 1.555 | +2 | 15 |
| - | - | 9445 | 941 | - | - | 0 | - |
| - | - | 7803 | 941.5 | - | - | 0 | - |
| - | - | 3488 | 942 | - | - | 0 | - |
| - | - | 1601 | 942.4 | - | - | 0 | - |
| - | - | 917.6 | 944.5 | - | - | 0 | - |
| - | - | 1185 | 945.5 | - | - | 0 | - |
| - | - | 1623 | 946 | - | - | 0 | - |
| - | - | 3123 | 946.5 | - | - | 0 | - |
| - | - | 2932 | 947 | - | - | 0 | - |
| - | - | 1380 | 947.5 | - | - | 0 | - |
| - | - | 2054 | 948 | - | - | 0 | - |
| 2 | y | 4442 | 948.5 | 0.001073 | 1.131 | +2 | 15 |
| - | - | 3560 | 949 | - | - | 0 | - |
| - | - | 1084 | 949.5 | - | - | 0 | - |
| - | - | 1230 | 950 | - | - | 0 | - |
| 10 | y | 1239 | 952.4 | 0.001918 | 2.014 | +1 | 7 |
| - | - | 1381 | 953 | - | - | 0 | - |
| - | - | 732.8 | 953.5 | - | - | 0 | - |
| - | - | 3.533E+04 | 954 | - | - | 0 | - |
| - | - | 3.928E+04 | 954.5 | - | - | 0 | - |
| - | - | 2.736E+04 | 955 | - | - | 0 | - |
| - | - | 1.379E+04 | 955.5 | - | - | 0 | - |
| - | - | 4814 | 956 | - | - | 0 | - |
| - | - | 809.9 | 959.5 | - | - | 0 | - |
| - | - | 964.9 | 960 | - | - | 0 | - |
| - | - | 778.6 | 960.5 | - | - | 0 | - |
| - | - | 1106 | 961 | - | - | 0 | - |
| - | - | 2220 | 961.5 | - | - | 0 | - |
| - | - | 4959 | 962 | - | - | 0 | - |
| - | - | 7099 | 962.5 | - | - | 0 | - |
| - | - | 4861 | 963 | - | - | 0 | - |
| - | - | 3232 | 963.5 | - | - | 0 | - |
| - | - | 1368 | 964 | - | - | 0 | - |
| - | - | 1513 | 966.5 | - | - | 0 | - |
| - | - | 1129 | 966.9 | - | - | 0 | - |
| - | - | 2117 | 967 | - | - | 0 | - |
| - | - | 1154 | 967.4 | - | - | 0 | - |
| - | - | 3718 | 967.5 | - | - | 0 | - |
| - | - | 2532 | 968 | - | - | 0 | - |
| - | - | 1158 | 968.5 | - | - | 0 | - |
| - | - | 7289 | 969 | - | - | 0 | - |
| - | - | 4342 | 969.5 | - | - | 0 | - |
| - | - | 3363 | 970 | - | - | 0 | - |
| - | - | 3592 | 970.5 | - | - | 0 | - |
| - | - | 3835 | 971 | - | - | 0 | - |
| - | - | 2647 | 971.5 | - | - | 0 | - |
| - | - | 822.7 | 972 | - | - | 0 | - |
| - | - | 1096 | 973.5 | - | - | 0 | - |
| - | - | 2077 | 975.5 | - | - | 0 | - |
| - | - | 1.275E+04 | 976 | - | - | 0 | - |
| - | - | 1.585E+04 | 976.5 | - | - | 0 | - |
| - | - | 9373 | 977 | - | - | 0 | - |
| - | - | 6915 | 977.5 | - | - | 0 | - |
| - | - | 2146 | 978 | - | - | 0 | - |
| - | - | 1089 | 978.5 | - | - | 0 | - |
| - | - | 1573 | 981 | - | - | 0 | - |
| - | - | 2242 | 981.5 | - | - | 0 | - |
| - | - | 1549 | 982 | - | - | 0 | - |
| - | - | 2796 | 982.5 | - | - | 0 | - |
| - | - | 2585 | 983 | - | - | 0 | - |
| - | - | 1042 | 984.6 | - | - | 0 | - |
| - | - | 1251 | 988.4 | - | - | 0 | - |
| - | - | 6144 | 989.5 | - | - | 0 | - |
| - | - | 4.445E+04 | 990 | - | - | 0 | - |
| - | - | 4.994E+04 | 990.5 | - | - | 0 | - |
| - | - | 3.045E+04 | 991 | - | - | 0 | - |
| - | - | 1.824E+04 | 991.5 | - | - | 0 | - |
| - | - | 7024 | 992 | - | - | 0 | - |
| - | - | 2331 | 992.5 | - | - | 0 | - |
| - | - | 997.4 | 993.4 | - | - | 0 | - |
| - | - | 1702 | 995.9 | - | - | 0 | - |
| - | - | 5474 | 996.4 | - | - | 0 | - |
| - | - | 4952 | 996.9 | - | - | 0 | - |
| - | - | 3490 | 997.4 | - | - | 0 | - |
| - | - | 3.185E+04 | 998 | - | - | 0 | - |
| - | - | 7.06E+04 | 998.5 | - | - | 0 | - |
| - | - | 6.185E+04 | 999 | - | - | 0 | - |
| - | - | 3.606E+04 | 999.5 | - | - | 0 | - |
| - | - | 1.69E+04 | 1000 | - | - | 0 | - |
| - | - | 6140 | 1001 | - | - | 0 | - |
| - | - | 1244 | 1001 | - | - | 0 | - |
| 9 | w | 5124 | 1006 | 0.0003958 | 0.3932 | +1 | 8 |
| - | - | 2194 | 1007 | - | - | 0 | - |
| - | - | 1037 | 1035 | - | - | 0 | - |
| - | - | 975.7 | 1058 | - | - | 0 | - |
| 9 | c | 3.367E+04 | 1061 | 0.0005309 | 0.5005 | +1 | 9 |
| - | - | 1.974E+04 | 1062 | - | - | 0 | - |
| - | - | 7705 | 1063 | - | - | 0 | - |
| - | - | 1574 | 1064 | - | - | 0 | - |
| 9 | z | 1.564E+04 | 1065 | 0.001313 | 1.232 | +1 | 8 |
| - | - | 8443 | 1066 | - | - | 0 | - |
| - | - | 4101 | 1067 | - | - | 0 | - |
| - | - | 1410 | 1068 | - | - | 0 | - |
| 9 | y | 2470 | 1081 | 0.002519 | 2.329 | +1 | 8 |
| - | - | 1669 | 1082 | - | - | 0 | - |
| - | - | 974.5 | 1086 | - | - | 0 | - |
| - | - | 980.6 | 1093 | - | - | 0 | - |
| - | - | 1801 | 1100 | - | - | 0 | - |
| - | - | 1228 | 1101 | - | - | 0 | - |
| - | - | 1146 | 1102 | - | - | 0 | - |
| - | - | 1021 | 1102 | - | - | 0 | - |
| - | - | 832.6 | 1102 | - | - | 0 | - |
| - | - | 1071 | 1106 | - | - | 0 | - |
| - | - | 1276 | 1107 | - | - | 0 | - |
| - | - | 1413 | 1107 | - | - | 0 | - |
| - | - | 1480 | 1108 | - | - | 0 | - |
| - | - | 1072 | 1111 | - | - | 0 | - |
| - | - | 1602 | 1112 | - | - | 0 | - |
| - | - | 1306 | 1112 | - | - | 0 | - |
| - | - | 3927 | 1189 | - | - | 0 | - |
| 10 | c | 3.029E+04 | 1190 | 0.0003142 | 0.2641 | +1 | 10 |
| - | - | 1.926E+04 | 1191 | - | - | 0 | - |
| - | - | 5315 | 1192 | - | - | 0 | - |
| - | - | 1917 | 1193 | - | - | 0 | - |
| 8 | z | 1.143E+04 | 1222 | 0.0009096 | 0.7446 | +1 | 9 |
| - | - | 1.069E+04 | 1223 | - | - | 0 | - |
| - | - | 5616 | 1224 | - | - | 0 | - |
| - | - | 2474 | 1225 | - | - | 0 | - |
| - | - | 2361 | 1228 | - | - | 0 | - |
| - | - | 1478 | 1229 | - | - | 0 | - |
| - | - | 877.6 | 1231 | - | - | 0 | - |
| - | - | 1245 | 1237 | - | - | 0 | - |
| 8 | y | 5778 | 1238 | 0.001213 | 0.9801 | +1 | 9 |
| - | - | 4591 | 1239 | - | - | 0 | - |
| - | - | 2119 | 1240 | - | - | 0 | - |
| - | - | 905.7 | 1246 | - | - | 0 | - |
| - | - | 834.1 | 1283 | - | - | 0 | - |
| - | - | 740.8 | 1288 | - | - | 0 | - |
| - | - | 2774 | 1293 | - | - | 0 | - |
| - | - | 2416 | 1294 | - | - | 0 | - |
| - | - | 1956 | 1295 | - | - | 0 | - |
| - | - | 896.3 | 1299 | - | - | 0 | - |
| - | - | 2052 | 1300 | - | - | 0 | - |
| - | - | 1191 | 1301 | - | - | 0 | - |
| 7 | z | 2401 | 1309 | 0.005746 | 4.391 | +1 | 10 |
| - | - | 4448 | 1310 | - | - | 0 | - |
| - | - | 2566 | 1311 | - | - | 0 | - |
| - | - | 1337 | 1312 | - | - | 0 | - |
| - | - | 1166 | 1313 | - | - | 0 | - |
| - | - | 3309 | 1316 | - | - | 0 | - |
| - | - | 811.7 | 1316 | - | - | 0 | - |
| - | - | 1.028E+04 | 1317 | - | - | 0 | - |
| - | - | 6392 | 1318 | - | - | 0 | - |
| - | - | 997.8 | 1323 | - | - | 0 | - |
| 7 | y | 1111 | 1325 | 0.0118 | 8.91 | +1 | 10 |
| - | - | 1060 | 1326 | - | - | 0 | - |
| - | - | 1028 | 1329 | - | - | 0 | - |
| - | - | 975.9 | 1330 | - | - | 0 | - |
| - | - | 956.7 | 1330 | - | - | 0 | - |
| - | - | 1778 | 1331 | - | - | 0 | - |
| - | - | 1489 | 1332 | - | - | 0 | - |
| - | - | 1060 | 1333 | - | - | 0 | - |
| - | - | 1557 | 1334 | - | - | 0 | - |
| - | - | 754.3 | 1335 | - | - | 0 | - |
| - | - | 1344 | 1336 | - | - | 0 | - |
| 11 | c | 1.978E+04 | 1337 | 0.003259 | 2.438 | +1 | 11 |
| - | - | 1.423E+04 | 1338 | - | - | 0 | - |
| - | - | 6980 | 1339 | - | - | 0 | - |
| - | - | 2601 | 1340 | - | - | 0 | - |
| - | - | 914.2 | 1341 | - | - | 0 | - |
| - | - | 865.8 | 1359 | - | - | 0 | - |
| - | - | 1011 | 1361 | - | - | 0 | - |
| - | - | 904.8 | 1366 | - | - | 0 | - |
| - | - | 1043 | 1376 | - | - | 0 | - |
| - | - | 914.5 | 1378 | - | - | 0 | - |
| - | - | 987.2 | 1388 | - | - | 0 | - |
| - | - | 1056 | 1388 | - | - | 0 | - |
| - | - | 2118 | 1389 | - | - | 0 | - |
| - | - | 2087 | 1394 | - | - | 0 | - |
| - | - | 2521 | 1395 | - | - | 0 | - |
| - | - | 1232 | 1396 | - | - | 0 | - |
| 6 | y | 2888 | 1422 | 0.0005427 | 0.3817 | +1 | 11 |
| - | - | 1581 | 1423 | - | - | 0 | - |
| - | - | 802.6 | 1424 | - | - | 0 | - |
| - | - | 759.5 | 1425 | - | - | 0 | - |
| - | - | 1349 | 1437 | - | - | 0 | - |
| 12 | c | 1.365E+04 | 1438 | 0.00331 | 2.302 | +1 | 12 |
| - | - | 1.026E+04 | 1439 | - | - | 0 | - |
| - | - | 5447 | 1440 | - | - | 0 | - |
| - | - | 1804 | 1441 | - | - | 0 | - |
| - | - | 893.4 | 1453 | - | - | 0 | - |
| - | - | 996.3 | 1488 | - | - | 0 | - |
| - | - | 992.3 | 1489 | - | - | 0 | - |
| - | - | 847.1 | 1494 | - | - | 0 | - |
| - | - | 765.2 | 1496 | - | - | 0 | - |
| - | - | 750.4 | 1521 | - | - | 0 | - |
| - | - | 838.6 | 1523 | - | - | 0 | - |
| - | - | 1135 | 1529 | - | - | 0 | - |
| - | - | 838.2 | 1531 | - | - | 0 | - |
| - | - | 926.2 | 1552 | - | - | 0 | - |
| - | - | 1191 | 1552 | - | - | 0 | - |
| - | - | 1423 | 1553 | - | - | 0 | - |
| - | - | 1002 | 1553 | - | - | 0 | - |
| - | - | 1043 | 1565 | - | - | 0 | - |
| 13 | c | 4082 | 1566 | 0.006247 | 3.99 | +1 | 13 |
| - | - | 3191 | 1567 | - | - | 0 | - |
| - | - | 1701 | 1568 | - | - | 0 | - |
| - | - | 924.7 | 1574 | - | - | 0 | - |
| 4 | z | 3947 | 1616 | 0.001716 | 1.062 | +1 | 13 |
| - | - | 1.887E+04 | 1617 | - | - | 0 | - |
| - | - | 1.807E+04 | 1618 | - | - | 0 | - |
| - | - | 8077 | 1619 | - | - | 0 | - |
| - | - | 4054 | 1620 | - | - | 0 | - |
| - | - | 1740 | 1621 | - | - | 0 | - |
| - | - | 822.1 | 1623 | - | - | 0 | - |
| - | - | 966.3 | 1631 | - | - | 0 | - |
| 4 | y | 1586 | 1632 | 0.006049 | 3.707 | +1 | 13 |
| - | - | 1420 | 1633 | - | - | 0 | - |
| - | - | 1028 | 1634 | - | - | 0 | - |
| - | - | 747.7 | 1638 | - | - | 0 | - |
| - | - | 1617 | 1639 | - | - | 0 | - |
| - | - | 1593 | 1652 | - | - | 0 | - |
| - | - | 944 | 1652 | - | - | 0 | - |
| - | - | 2187 | 1653 | - | - | 0 | - |
| - | - | 1207 | 1658 | - | - | 0 | - |
| - | - | 1255 | 1659 | - | - | 0 | - |
| - | - | 1408 | 1660 | - | - | 0 | - |
| - | - | 2302 | 1660 | - | - | 0 | - |
| - | - | 2609 | 1661 | - | - | 0 | - |
| - | - | 2681 | 1661 | - | - | 0 | - |
| - | - | 1907 | 1662 | - | - | 0 | - |
| - | - | 1527 | 1667 | - | - | 0 | - |
| - | - | 1181 | 1667 | - | - | 0 | - |
| - | - | 2075 | 1668 | - | - | 0 | - |
| - | - | 1152 | 1668 | - | - | 0 | - |
| - | - | 1189 | 1679 | - | - | 0 | - |
| 14 | c | 1.406E+04 | 1680 | 0.005556 | 3.308 | +1 | 14 |
| - | - | 1.289E+04 | 1681 | - | - | 0 | - |
| - | - | 8356 | 1682 | - | - | 0 | - |
| - | - | 3328 | 1683 | - | - | 0 | - |
| - | - | 1370 | 1684 | - | - | 0 | - |
| 3 | z | 1323 | 1717 | 0.006304 | 3.672 | +1 | 14 |
| - | - | 5696 | 1718 | - | - | 0 | - |
| - | - | 4932 | 1719 | - | - | 0 | - |
| - | - | 2361 | 1720 | - | - | 0 | - |
| - | - | 1375 | 1721 | - | - | 0 | - |
| 3 | y | 1332 | 1733 | 0.003536 | 2.041 | +1 | 14 |
| - | - | 1626 | 1734 | - | - | 0 | - |
| 15 | c | 1647 | 1808 | 0.01631 | 9.024 | +1 | 15 |
| - | - | 3314 | 1809 | - | - | 0 | - |
| - | - | 2691 | 1810 | - | - | 0 | - |
| - | - | 1935 | 1811 | - | - | 0 | - |
| - | - | 3419 | 1864 | - | - | 0 | - |
| - | - | 3642 | 1865 | - | - | 0 | - |
| - | - | 2377 | 1866 | - | - | 0 | - |
| - | - | 1082 | 1867 | - | - | 0 | - |
| 2 | z | 1343 | 1880 | 0.0218 | 11.6 | +1 | 15 |
| - | - | 8471 | 1881 | - | - | 0 | - |
| - | - | 9121 | 1882 | - | - | 0 | - |
| - | - | 6839 | 1883 | - | - | 0 | - |
| - | - | 2801 | 1884 | - | - | 0 | - |
| - | - | 1041 | 1891 | - | - | 0 | - |
| - | - | 1148 | 1893 | - | - | 0 | - |
| - | - | 4760 | 1907 | - | - | 0 | - |
| - | - | 1.247E+04 | 1908 | - | - | 0 | - |
| - | - | 9904 | 1909 | - | - | 0 | - |
| - | - | 6735 | 1910 | - | - | 0 | - |
| - | - | 2950 | 1911 | - | - | 0 | - |
| - | - | 1680 | 1934 | - | - | 0 | - |
| - | - | 2433 | 1935 | - | - | 0 | - |
| - | - | 1142 | 1936 | - | - | 0 | - |
| - | - | 1645 | 1937 | - | - | 0 | - |
| - | - | 3476 | 1938 | - | - | 0 | - |
| - | - | 2114 | 1939 | - | - | 0 | - |
| - | - | 1734 | 1940 | - | - | 0 | - |
| - | - | 1120 | 1941 | - | - | 0 | - |
| - | - | 3001 | 1951 | - | - | 0 | - |
| - | - | 1.106E+04 | 1952 | - | - | 0 | - |
| - | - | 1.047E+04 | 1953 | - | - | 0 | - |
| - | - | 6604 | 1954 | - | - | 0 | - |
| - | - | 4084 | 1955 | - | - | 0 | - |
| - | - | 991.1 | 1956 | - | - | 0 | - |
| - | - | 1389 | 1963 | - | - | 0 | - |
| - | - | 2087 | 1964 | - | - | 0 | - |
| - | - | 972.8 | 1966 | - | - | 0 | - |
| - | - | 2194 | 1968 | - | - | 0 | - |
| - | - | 6718 | 1969 | - | - | 0 | - |
| - | - | 5585 | 1970 | - | - | 0 | - |
| - | - | 4053 | 1971 | - | - | 0 | - |
| - | - | 2432 | 1972 | - | - | 0 | - |
| - | - | 1097 | 1976 | - | - | 0 | - |
| - | - | 9117 | 1979 | - | - | 0 | - |
| - | - | 3.303E+04 | 1980 | - | - | 0 | - |
| - | - | 3.327E+04 | 1981 | - | - | 0 | - |
| - | - | 2.054E+04 | 1982 | - | - | 0 | - |
| - | - | 1.11E+04 | 1983 | - | - | 0 | - |
| - | - | 4079 | 1984 | - | - | 0 | - |
| - | - | 1491 | 1992 | - | - | 0 | - |
| - | - | 4626 | 1993 | - | - | 0 | - |
| - | - | 6343 | 1994 | - | - | 0 | - |
| - | - | 4308 | 1995 | - | - | 0 | - |
| - | - | 1.968E+04 | 1996 | - | - | 0 | - |
| - | - | 8.109E+04 | 1997 | - | - | 0 | - |
| - | - | 7.798E+04 | 1998 | - | - | 0 | - |
| - | - | 5.465E+04 | 1999 | - | - | 0 | - |
| - | - | 2.346E+04 | 2000 | - | - | 0 | - |
| - | - | 8529 | 2001 | - | - | 0 | - |
| - | - | 2907 | 2002 | - | - | 0 | - |

m/z Charge Intensity FragmentType MassShift Position
121.72193908691406 0 369.61768
124.19652557373047 0 349.6162
127.46900177001953 0 360.57532
129.10240173339844 0 871.1896
136.07569885253906 0 2618.2397
145.12887573242188 0 452.78098
147.0762176513672 0 448.5614
148.9471435546875 0 928.07654
151.61671447753906 0 419.91418
169.13345336914062 0 880.48486
187.14393615722656 0 1775.3486
192.1538848876953 0 459.6936
194.28594970703125 0 512.0753
197.1283721923828 0 1043.1641
203.58641052246094 0 450.08417
205.6564178466797 0 536.08966
215.13870239257812 0 1314.443
225.04356384277344 0 782.5691
235.14398193359375 0 14256.707
236.1473388671875 0 1970.9958
239.09474182128906 0 1074.4009
240.39463806152344 0 492.20844
263.12359619140625 0 695.23083
263.13885498046875 0 11201.307
264.1423034667969 0 1473.2064
283.14068603515625 0 1078.5923
295.1022644042969 0 956.0088
299.09698486328125 0 498.93225
299.17108154296875 0 2993.6987
320.1610412597656 0 714.59015
333.1777038574219 0 1053.8517 y 14
341.01800537109375 0 1297.8811
342.9356384277344 0 568.4995
346.17608642578125 0 8157.0303
347.1787414550781 0 1788.8167
355.06964111328125 0 4361.1636
359.0283203125 0 4995.216
360.1914367675781 0 803.87866
364.1865539550781 0 10856.518
365.18963623046875 0 2421.6992
387.7482604980469 0 592.1598
388.1832275390625 0 4058.654
429.08880615234375 0 13195.044
431.26531982421875 0 3469.87
447.22137451171875 0 1535.6953 y 13
449.2763671875 0 2299.512
450.27886962890625 0 594.1844
458.2594909667969 0 684.3883
459.26055908203125 0 7817.1973
460.26373291015625 0 2724.8728
473.0938720703125 0 566.2297
476.2654724121094 0 592.71497
477.2707824707031 0 3432.403
478.2757263183594 0 1084.6156
506.9122619628906 0 5155.8877 y 4
507.12774658203125 0 653.20544
507.24560546875 0 3854.0383
507.5793762207031 0 1309.6628
507.91552734375 0 693.1169
514.228759765625 0 621.1783
515.282470703125 0 2879.1987
516.2822875976562 0 974.7109
545.3770141601562 0 2058.9673
559.2960815429688 0 8163.823 z 12
560.3003540039062 0 3678.009
561.3029174804688 0 1090.5979
572.2838134765625 0 1151.545 y Water loss 2
572.6198120117188 0 1429.7928 y Ammonia loss 2
575.3154907226562 0 7849.1045 y 12
576.3209838867188 0 1067.7922
586.2794799804688 0 613.99304
592.9044189453125 0 1196.6204
593.2373046875 0 2010.3809
601.2432250976562 0 741.5281
602.3982543945312 0 1607.934
603.4031372070312 0 1021.78625
615.346923828125 0 1222.0435
616.3302001953125 0 2119.5405
617.3272705078125 0 639.66644
619.2679443359375 0 3170.1484
624.647216796875 0 1141.4935
625.653076171875 0 631.0545
626.973876953125 0 1504.4128 y Ammonia loss 1
627.3076782226562 0 1152.6641 z 1
643.3408203125 0 2490.9531 w 11
644.33740234375 0 763.2176
645.3213500976562 0 2136.1409 w 11
654.2973022460938 0 895.62494 y Ammonia loss 6
656.7891235351562 0 675.89624
657.8062133789062 0 815.0109
658.3284301757812 0 1508.4695
659.3346557617188 0 3267.2563 y Ammonia loss 11
659.6630859375 0 6770.9697
659.9969482421875 0 9083.482
660.3384399414062 0 12475.274 z 11
660.6644897460938 0 2672.254
661.3474731445312 0 4217.276
662.8123779296875 0 707.9272 y 6
666.2774047851562 0 1145.3256
666.3460693359375 0 1685.6123
666.861328125 0 768.4724
667.2711791992188 0 845.5337
667.4144897460938 0 10385.53
674.357421875 0 649.6728
675.3543090820312 0 802.5354
676.36181640625 0 3466.7534 y 11
677.3634643554688 0 1534.2567
678.3689575195312 0 744.1736
679.3812255859375 0 706.72235
687.3942260742188 0 1026.8839
688.4007568359375 0 3111.2075 c 5
689.4076538085938 0 836.5816
693.34814453125 0 823.6893
694.326171875 0 657.2887 z Water loss 5
697.3899536132812 0 1445.3887
698.3901977539062 0 652.86163
701.3703002929688 0 887.2797
707.343994140625 0 4326.7744
707.843017578125 0 3780.641
708.3439331054688 0 2428.8782
711.3380737304688 0 3508.5803 y 5
711.8391723632812 0 2091.8445
712.33935546875 0 1237.604
713.4100952148438 0 2953.9475
714.4141845703125 0 2355.2532
717.359619140625 0 714.5086
718.3359985351562 0 927.92267
727.3864135742188 0 1009.85034
727.864990234375 0 1980.8495
728.3668823242188 0 2678.7173
728.8606567382812 0 708.0731
729.3734130859375 0 722.74
744.3881225585938 0 3516.689
745.3954467773438 0 11974.577
746.3978271484375 0 5358.1636
747.4019775390625 0 1205.0093
751.3572387695312 0 2514.3218 y Ammonia loss 4
751.8565673828125 0 1063.7129 z 4
759.8646240234375 0 58089.125 y 4
760.3651733398438 0 44444.906
760.8662719726562 0 27163.453
761.36767578125 0 8015.0283
761.8695068359375 0 2437.3767
764.88037109375 0 1222.3239
766.3989868164062 0 1432.9436
772.388916015625 0 1201.2908
774.8626098632812 0 846.88666
775.4354858398438 0 14664.565 c 6
776.4379272460938 0 7025.3765
777.4395141601562 0 1908.228
780.8644409179688 0 1145.2339
782.8995971679688 0 10389.138
783.4024047851562 0 18294.992 c 12
783.90380859375 0 13738.58
784.4056396484375 0 7640.4683
784.90673828125 0 2610.0366
785.40771484375 0 1112.4094
786.8702392578125 0 14413.355 w 3
787.37060546875 0 10521.387
787.871337890625 0 8094.3506
788.3717651367188 0 3641.3706
788.8778076171875 0 1048.6637
789.3699340820312 0 806.5444 z Water loss 10
794.7750854492188 0 783.1109
800.412109375 0 862.3266
807.38232421875 0 3250.3242 z 10
808.3935546875 0 22616.146 z 3
808.8978271484375 0 14202.345
809.3960571289062 0 12568.537
809.89892578125 0 4802.1943
810.3985595703125 0 3530.1514
814.4105834960938 0 2278.4238
814.9104614257812 0 969.10333
815.4071044921875 0 693.21173
815.9024658203125 0 923.9046
816.4065551757812 0 9927.311 y 3
816.9083862304688 0 7838.269
817.4071044921875 0 4923.6436
817.9107666015625 0 2194.9202
818.4196166992188 0 1083.506
818.9166870117188 0 2158.9731
823.4031982421875 0 1283.2758 y 10
830.382568359375 0 685.295
830.8934326171875 0 2501.5225
831.3880615234375 0 2798.3105
831.892333984375 0 779.0266
834.3784790039062 0 892.97736
835.4307861328125 0 1069.6633
839.91943359375 0 2270.99
840.4246826171875 0 34248.594 c 13
840.9259643554688 0 30226.393
841.426513671875 0 17681.29
841.9273681640625 0 8158.158
842.426025390625 0 2462.705
842.9303588867188 0 745.2458
853.9368896484375 0 816.33624
857.46728515625 0 1689.2925
857.9170532226562 0 1241.9915 y Water loss 2
858.4189453125 0 2658.6743 y Ammonia loss 2
858.920166015625 0 13052.513 z 2
859.3258056640625 0 967.99884
859.4219360351562 0 12857.586
859.923828125 0 5751.048
860.4214477539062 0 3630.2715
866.9303588867188 0 24392.004 y 2
867.431640625 0 25637.719
867.9320068359375 0 14708.955
868.43115234375 0 7332.4062
868.9321899414062 0 2459.9905
870.91455078125 0 985.90076
871.4124145507812 0 900.08246
873.4324340820312 0 1765.3243
874.4267578125 0 1764.9053
877.4083862304688 0 8057.9946 w 9
878.4107666015625 0 4125.6655
879.40576171875 0 1198.578
880.4151611328125 0 1628.1528
880.9273681640625 0 1735.7107
881.4252319335938 0 1524.5737
881.9320068359375 0 951.28613
882.4418334960938 0 1684.5828
887.1242065429688 0 718.47705
887.4326782226562 0 872.6653
887.52294921875 0 919.2577
888.5279541015625 0 1075.6675
888.8521728515625 0 2749.5671
889.3489379882812 0 2788.6843
889.8564453125 0 840.137
890.3446655273438 0 726.69586
895.945556640625 0 997.3985 c Ammonia loss 14
896.4537353515625 0 1954.5463
896.9558715820312 0 1911.3489
897.4564208984375 0 972.33344
901.4942626953125 0 951.3181
902.49609375 0 1064.4644
903.9473876953125 0 1042.4645
904.4535522460938 0 10599.825 c 14
904.9542236328125 0 10983.2
905.4526977539062 0 6562.176
905.9556274414062 0 2742.647
909.4410400390625 0 917.9724
909.9459838867188 0 1327.3652
910.4629516601562 0 1725.251
911.4547119140625 0 826.9927
912.4210205078125 0 2426.2932
912.927490234375 0 1632.8157
916.4409790039062 0 920.1848
917.4296264648438 0 1058.5779
922.9658813476562 0 935.12213
924.4740600585938 0 968.1286
925.4405517578125 0 3424.8872
925.9362182617188 0 3295.967
926.4384765625 0 2674.5923
928.4716186523438 0 1536.0168
930.5326538085938 0 813.6467
930.9830322265625 0 955.81055
931.3792724609375 0 1165.4193
931.536376953125 0 24176.477 c 7
931.89208984375 0 1405.2734
931.9822998046875 0 3378.9038
932.537841796875 0 14947.139
932.9776611328125 0 2624.477
933.5416259765625 0 4046.272
934.5464477539062 0 938.1275
936.421630859375 0 18423.232 z 9
937.424072265625 0 11006.549
938.4246826171875 0 4279.2876
939.4414672851562 0 1012.96 y Water loss 1
939.971435546875 0 2180.1548
940.4539184570312 0 8303.316 z 1
940.9535522460938 0 9444.907
941.4548950195312 0 7803.3716
941.9525146484375 0 3488.3718
942.4454345703125 0 1601.4095
944.4820556640625 0 917.6365
945.4824829101562 0 1185.0969
945.9738159179688 0 1623.3263
946.4798583984375 0 3122.69
946.980224609375 0 2932.0908
947.4730834960938 0 1379.9268
947.9666748046875 0 2054.3264
948.462890625 0 4441.6353 y 1
948.9608764648438 0 3559.7053
949.46240234375 0 1083.6561
949.9633178710938 0 1230.3358
952.4381103515625 0 1238.6008 y 9
952.9804077148438 0 1380.6539
953.4614868164062 0 732.8434
953.987548828125 0 35325.59
954.4887084960938 0 39282.04
954.98974609375 0 27363.45
955.4898071289062 0 13785.232
955.9909057617188 0 4813.8667
959.4811401367188 0 809.9013
959.9943237304688 0 964.9488
960.4796752929688 0 778.58795
960.9833374023438 0 1105.6864
961.4689331054688 0 2220.0076
961.95947265625 0 4958.943
962.4642333984375 0 7099.4683
962.9696044921875 0 4861.1865
963.4721069335938 0 3232.4597
963.9764404296875 0 1368.3917
966.496337890625 0 1513.4403
966.8860473632812 0 1128.6711
967.0046997070312 0 2117.4814
967.389892578125 0 1153.6226
967.5004272460938 0 3718.0056
967.9909057617188 0 2532.27
968.4951782226562 0 1158.2175
968.9832153320312 0 7289.019
969.4796142578125 0 4342.285
969.9833984375 0 3363.2368
970.4761352539062 0 3592.2307
970.9733276367188 0 3834.77
971.4727172851562 0 2646.7644
971.9740600585938 0 822.6764
973.4588623046875 0 1096.1334
975.4902954101562 0 2077.2524
975.990966796875 0 12753.669
976.4912719726562 0 15850.739
976.9899291992188 0 9372.959
977.4880981445312 0 6914.7334
977.9902954101562 0 2146.1829
978.4847412109375 0 1089.1671
980.984130859375 0 1572.6002
981.487060546875 0 2242.2512
981.9839477539062 0 1548.6824
982.4794921875 0 2795.5386
982.97705078125 0 2584.627
984.6095581054688 0 1042.4436
988.413818359375 0 1250.6533
989.4944458007812 0 6143.919
989.9895629882812 0 44448.406
990.489990234375 0 49942.51
990.9908447265625 0 30446.162
991.4921875 0 18238.977
991.992919921875 0 7023.84
992.4830322265625 0 2331.0498
993.4371337890625 0 997.43506
995.915771484375 0 1702.1691
996.4219970703125 0 5474.267
996.92041015625 0 4951.6724
997.4210205078125 0 3490.454
997.9967041015625 0 31846.031
998.4986572265625 0 70604.44
999.0001220703125 0 61852.066
999.5011596679688 0 36056.496
1000.0016479492188 0 16900.238
1000.5012817382812 0 6140.4927
1001.000732421875 0 1243.9181
1006.4509887695312 0 5123.775 w 8
1007.4520263671875 0 2193.5369
1034.864501953125 0 1037.1327
1058.223876953125 0 975.74066
1060.5791015625 0 33670.555 c 8
1061.5826416015625 0 19736.94
1062.5849609375 0 7704.788
1063.584716796875 0 1573.6559
1065.4652099609375 0 15643.986 z 8
1066.465576171875 0 8443.351
1067.4686279296875 0 4100.7637
1068.47314453125 0 1410.3729
1081.4801025390625 0 2470.3103 y 8
1082.4869384765625 0 1668.5588
1085.5006103515625 0 974.47974
1092.572998046875 0 980.6099
1099.576904296875 0 1800.9458
1100.568359375 0 1228.1554
1101.5733642578125 0 1146.1362
1101.909912109375 0 1020.7144
1102.2437744140625 0 832.56775
1105.575927734375 0 1070.8281
1106.56005859375 0 1276.3916
1106.91064453125 0 1412.8583
1107.5841064453125 0 1480.0889
1111.24267578125 0 1072.258
1111.5789794921875 0 1602.4669
1111.9119873046875 0 1305.7776
1188.611572265625 0 3926.5022
1189.620849609375 0 30288.434 c 9
1190.6234130859375 0 19264.357
1191.62890625 0 5315.472
1192.6329345703125 0 1916.8971
1221.56591796875 0 11425.059 z 7
1222.5706787109375 0 10689.208
1223.5682373046875 0 5615.5728
1224.568359375 0 2473.95
1227.6334228515625 0 2361.2117
1228.6241455078125 0 1477.6167
1230.6632080078125 0 877.5869
1236.5758056640625 0 1245.2866
1237.58251953125 0 5777.6104 y 7
1238.5845947265625 0 4590.502
1239.58154296875 0 2118.8154
1245.6046142578125 0 905.6961
1282.60498046875 0 834.0826
1287.62353515625 0 740.7988
1292.637939453125 0 2774.3052
1293.6444091796875 0 2416.0332
1294.6431884765625 0 1955.657
1298.6395263671875 0 896.3109
1299.63232421875 0 2051.9177
1300.635498046875 0 1191.4844
1308.602783203125 0 2400.7458 z 6
1309.6046142578125 0 4448.039
1310.6087646484375 0 2566.4736
1311.6104736328125 0 1337.3309
1312.6082763671875 0 1166.244
1315.6478271484375 0 3308.5122
1316.1650390625 0 811.6708
1316.653076171875 0 10276.646
1317.6572265625 0 6391.9956
1323.1566162109375 0 997.80444
1324.6275634765625 0 1110.7748 y 6
1325.615966796875 0 1059.9341
1328.6429443359375 0 1027.9408
1329.6875 0 975.8907
1330.19091796875 0 956.66473
1330.6517333984375 0 1777.7931
1331.684814453125 0 1489.4144
1332.6781005859375 0 1059.6082
1333.6439208984375 0 1556.7904
1334.63232421875 0 754.32404
1335.656982421875 0 1344.3804
1336.6549072265625 0 19779.912 c 10
1337.6580810546875 0 14225.206
1338.66064453125 0 6980.3765
1339.66015625 0 2600.9807
1340.6590576171875 0 914.216
1359.1727294921875 0 865.8078
1360.685791015625 0 1010.9193
1366.1929931640625 0 904.8352
1375.6761474609375 0 1043.4177
1377.6810302734375 0 914.5192
1387.6944580078125 0 987.2071
1388.192626953125 0 1056.181
1388.68896484375 0 2117.707
1393.6837158203125 0 2087.0044
1394.6885986328125 0 2521.481
1395.6990966796875 0 1232.3306
1421.6690673828125 0 2888.464 y 5
1422.67041015625 0 1580.5991
1423.6590576171875 0 802.6085
1424.665771484375 0 759.47784
1436.701171875 0 1348.9332
1437.70263671875 0 13651.704 c 11
1438.70703125 0 10261.518
1439.70849609375 0 5447.146
1440.71044921875 0 1804.4869
1452.746826171875 0 893.44794
1488.2591552734375 0 996.2574
1488.759765625 0 992.3321
1494.247802734375 0 847.0684
1495.754150390625 0 765.1857
1520.7227783203125 0 750.35693
1522.7850341796875 0 838.6293
1528.7694091796875 0 1135.1201
1530.7757568359375 0 838.2407
1551.8087158203125 0 926.18933
1552.294677734375 0 1190.5481
1552.8009033203125 0 1422.9092
1553.3079833984375 0 1002.1932
1564.790771484375 0 1043.461
1565.800537109375 0 4082.0903 c 12
1566.7994384765625 0 3190.6838
1567.8070068359375 0 1701.3207
1573.78125 0 924.7491
1615.784912109375 0 3946.772 z 3
1616.79443359375 0 18873.166
1617.7950439453125 0 18074.701
1618.794921875 0 8077.244
1619.7989501953125 0 4054.1113
1620.80322265625 0 1739.6902
1623.3448486328125 0 822.08905
1630.8184814453125 0 966.2696
1631.8114013671875 0 1586.3806 y 3
1632.7921142578125 0 1420.4836
1633.8123779296875 0 1027.8933
1637.8416748046875 0 747.71094
1638.852294921875 0 1617.4991
1651.8721923828125 0 1593.4364
1652.361572265625 0 943.9885
1652.85888671875 0 2187.4375
1658.354248046875 0 1207.371
1658.864501953125 0 1254.8822
1659.8583984375 0 1407.5607
1660.3702392578125 0 2301.8833
1660.8636474609375 0 2609.4841
1661.3721923828125 0 2680.6785
1661.8687744140625 0 1906.8245
1666.863037109375 0 1527.4443
1667.362548828125 0 1180.702
1667.8658447265625 0 2075.4548
1668.377685546875 0 1152.3674
1678.8262939453125 0 1188.7329
1679.8427734375 0 14063.523 c 13
1680.843994140625 0 12894.787
1681.845458984375 0 8356.128
1682.8497314453125 0 3328.1812
1683.8582763671875 0 1369.6611
1716.8280029296875 0 1322.5381 z 2
1717.8411865234375 0 5695.863
1718.84619140625 0 4931.805
1719.8431396484375 0 2360.7427
1720.8458251953125 0 1374.8582
1732.8565673828125 0 1331.9127 y 2
1733.8538818359375 0 1626.0076
1807.912109375 0 1646.8208 c 14
1808.909912109375 0 3314.4092
1809.906494140625 0 2690.8281
1810.90673828125 0 1934.726
1863.95458984375 0 3419.4414
1864.9573974609375 0 3642.374
1865.9591064453125 0 2376.8262
1866.9525146484375 0 1082.0795
1879.91943359375 0 1342.8267 z 1
1880.9053955078125 0 8471.39
1881.905517578125 0 9120.912
1882.90771484375 0 6838.7603
1883.9090576171875 0 2801.1235
1890.945556640625 0 1040.7136
1892.9517822265625 0 1148.339
1906.9671630859375 0 4759.8994
1907.9757080078125 0 12473.786
1908.9775390625 0 9904.107
1909.976806640625 0 6734.885
1910.981689453125 0 2949.6196
1933.991455078125 0 1680.2043
1934.9881591796875 0 2433.1353
1935.961181640625 0 1142.2572
1936.95849609375 0 1645.1964
1937.957275390625 0 3476.0142
1938.96240234375 0 2114.3242
1939.9583740234375 0 1733.6746
1940.962158203125 0 1120.049
1950.9747314453125 0 3001.1343
1951.978759765625 0 11061.144
1952.9813232421875 0 10471.368
1953.9832763671875 0 6603.761
1954.98046875 0 4084.0557
1955.9991455078125 0 991.1124
1962.9627685546875 0 1389.1045
1963.9599609375 0 2086.9272
1965.95703125 0 972.7888
1968.00341796875 0 2193.6338
1969.0042724609375 0 6717.597
1970.001708984375 0 5585.4243
1971.0072021484375 0 4053.2246
1972.0028076171875 0 2431.8608
1975.8074951171875 0 1097.2853
1978.969482421875 0 9116.675
1979.973876953125 0 33026.727
1980.975341796875 0 33268.46
1981.975830078125 0 20535.549
1982.9783935546875 0 11099.623
1983.9754638671875 0 4078.5054
1991.841552734375 0 1490.8942
1992.845458984375 0 4626.13
1993.8463134765625 0 6343.429
1994.9302978515625 0 4308.2866
1995.9876708984375 0 19680.277
1996.998046875 0 81086.56
1998.0001220703125 0 77978.09
1999.001708984375 0 54649.285
2000.0025634765625 0 23464.002
2001.004150390625 0 8529.335
2002.0045166015625 0 2906.7202

Spectrum Details

|  |  |
| --- | --- |
| Matched peaks? Matched peaks The total absolute number of peaks matched. Additionally in brackets the total fraction of peaks matched and the total number of peaks is shown. | 64 (11.27% of 568) |
| FDR? FDR The false discovery rate estimated for this peptide. It is calculated by matching all theoretical fragments with a non-integer shift with the raw peaks for this spectrum. This is done with 40 different shifts. The resulting percentage is the average number of annotated peaks over the number of annotated peaks with the correct spectrum. | 2.46% |
| Satellite FDR? Satellite FDR See the FDR for details on its calculation. This satellite ion specific FDR only contains the satellite ions (d/w) for I/L/J positions. | 0.00% |
| PSM Score? PSM Score The PSM Score as given by Hecklib to this annotated spectrum. It is shown with three significant figures. | 391 |

#### Spectrum 4132? Spectrum 4132 The raw spectrum of this peptide as annotated by Hecklib. The fragments are coloured according to ion type (see legend). Any peaks with a star '\*' as text can be hovered over to see the full details, first the ion type second the mass shift type. By hovering over the amino acids in the peptide or ions in the legend the corresponding peaks are highlighted. By toggling the 'Unassigned' label you can turn the background (unassigned) peaks on or off in the plot. By updating the slider in the Ion legend you can update the spectrum to only show the top X% of the peaks with labels. The top X% means any peak that is within X% of the highest intensity. By dragging in the spectrum you can zoom in to a specific part of the spectrum and use 'Zoom Out' to get back to the original zoom level. The annotation of the spectrum is based on the given sequence in the peptides file and is done with different software so inconsistencies are likely. The peaks are annotated based on the given sequence, with 20 ppm tolerance.

Copy Data

##### Spectrum 4132 (TSV)

###### Preview

```
Loading example...
```

*Click on the button to copy the data to your clipboard.*

Mz MinMz MaxIntensity Max

WidthHeightPeptide font sizePeptide stroke widthSpectrum font sizeSpectrum stroke widthCompact peptide

Ion legend

wxyz

abcd

OtherUnassignedIon^Charge^_Position_Show for top:%

VYTLPPSREEMTKNQK

02.35e+44.69e+47.04e+49.39e+4

Zoom Out

y^+1^_2_y^+1^_3_y^+3^_12_y^+3^_13_z^+1^_4_y^+3^_14_y^+3^_14_z^+3^_14_y^+1^_4_y^+3^_15_y^+3^_15_w^+1^_5_w^+1^_5_y^+1^_5_z^+1^_5_y^+1^_5_c^+1^_6_y^+2^_11_w^+2^_12_y^+2^_12_y^+2^_12_z^+2^_12_y^+2^_12_c^+2^_13_c^+1^_7_c^+2^_13_w^+2^_13_z^+1^_6_z^+1^_6_y^+2^_13_z^+2^_13_y^+2^_13_y^+1^_6_c^+2^_14_c^+2^_14_y^+2^_14_y^+2^_14_z^+2^_14_y^+2^_14_w^+1^_7_c^+2^_15_c^+2^_15_c^+1^_8_z^+1^_7_y^+2^_15_z^+2^_15_y^+2^_15_y^+1^_7_w^+1^_8_c^+1^_9_z^+1^_8_y^+1^_8_c^+1^_10_z^+1^_9_y^+1^_9_y^+1^_10_z^+1^_10_y^+1^_10_c^+1^_11_y^+1^_11_c^+1^_12_c^+1^_13_y^+1^_13_z^+1^_13_y^+1^_13_c^+1^_14_z^+1^_14_y^+1^_14_c^+1^_15_z^+1^_15_

0506101115172022

Fragment Matches Table

Show background peaks

| Position | Ion type | Intensity | mz Theoretical | mz Error (Th) | mz Error (ppm) | Charge | Series Number |
| --- | --- | --- | --- | --- | --- | --- | --- |
| - | - | 398 | 123.2 | - | - | 0 | - |
| - | - | 1964 | 129.1 | - | - | 0 | - |
| - | - | 3848 | 136.1 | - | - | 0 | - |
| - | - | 684.8 | 148.9 | - | - | 0 | - |
| - | - | 468.3 | 155.1 | - | - | 0 | - |
| - | - | 470.7 | 160.9 | - | - | 0 | - |
| - | - | 447.7 | 173.1 | - | - | 0 | - |
| - | - | 2954 | 173.5 | - | - | 0 | - |
| - | - | 1242 | 187.1 | - | - | 0 | - |
| - | - | 670.7 | 197.1 | - | - | 0 | - |
| - | - | 528.8 | 211.5 | - | - | 0 | - |
| - | - | 1597 | 215.1 | - | - | 0 | - |
| - | - | 656.1 | 216.1 | - | - | 0 | - |
| - | - | 574.5 | 219.8 | - | - | 0 | - |
| - | - | 670.5 | 225 | - | - | 0 | - |
| - | - | 561.9 | 230.3 | - | - | 0 | - |
| - | - | 1.657E+04 | 235.1 | - | - | 0 | - |
| - | - | 2460 | 236.1 | - | - | 0 | - |
| - | - | 1455 | 239.1 | - | - | 0 | - |
| - | - | 1.405E+04 | 263.1 | - | - | 0 | - |
| - | - | 1185 | 264.1 | - | - | 0 | - |
| - | - | 928.5 | 283.1 | - | - | 0 | - |
| - | - | 1073 | 295.1 | - | - | 0 | - |
| - | - | 963.1 | 299.1 | - | - | 0 | - |
| - | - | 3591 | 299.2 | - | - | 0 | - |
| - | - | 597 | 300.2 | - | - | 0 | - |
| - | - | 579.5 | 318.2 | - | - | 0 | - |
| - | - | 1159 | 320.2 | - | - | 0 | - |
| - | - | 1384 | 325.2 | - | - | 0 | - |
| 15 | y | 653.4 | 333.2 | 0.004776 | 14.34 | +1 | 2 |
| - | - | 579 | 339.2 | - | - | 0 | - |
| - | - | 715.5 | 341 | - | - | 0 | - |
| - | - | 643.2 | 342.2 | - | - | 0 | - |
| - | - | 588.5 | 344.2 | - | - | 0 | - |
| - | - | 9763 | 346.2 | - | - | 0 | - |
| - | - | 1731 | 347.2 | - | - | 0 | - |
| - | - | 4427 | 355.1 | - | - | 0 | - |
| - | - | 671.2 | 356.1 | - | - | 0 | - |
| - | - | 5078 | 359 | - | - | 0 | - |
| - | - | 1534 | 360.2 | - | - | 0 | - |
| - | - | 1.421E+04 | 364.2 | - | - | 0 | - |
| - | - | 2460 | 365.2 | - | - | 0 | - |
| - | - | 798.8 | 382.2 | - | - | 0 | - |
| - | - | 4424 | 388.2 | - | - | 0 | - |
| - | - | 791 | 389.2 | - | - | 0 | - |
| - | - | 723.3 | 396.2 | - | - | 0 | - |
| - | - | 1076 | 400.3 | - | - | 0 | - |
| - | - | 507.2 | 405.2 | - | - | 0 | - |
| - | - | 1.15E+04 | 429.1 | - | - | 0 | - |
| - | - | 2724 | 431.3 | - | - | 0 | - |
| - | - | 790.3 | 432.2 | - | - | 0 | - |
| - | - | 642.6 | 433.2 | - | - | 0 | - |
| 14 | y | 1602 | 447.2 | 0.004552 | 10.18 | +1 | 3 |
| - | - | 531 | 448.2 | - | - | 0 | - |
| - | - | 2244 | 449.3 | - | - | 0 | - |
| - | - | 898 | 450.3 | - | - | 0 | - |
| - | - | 1.24E+04 | 459.3 | - | - | 0 | - |
| - | - | 3183 | 460.3 | - | - | 0 | - |
| - | - | 3908 | 477.3 | - | - | 0 | - |
| - | - | 771.2 | 497.2 | - | - | 0 | - |
| 5 | y | 5751 | 506.9 | 0.0003453 | 0.6812 | +3 | 12 |
| - | - | 3655 | 507.2 | - | - | 0 | - |
| - | - | 2265 | 507.6 | - | - | 0 | - |
| - | - | 641.3 | 507.9 | - | - | 0 | - |
| - | - | 958.8 | 511.3 | - | - | 0 | - |
| - | - | 2871 | 515.3 | - | - | 0 | - |
| - | - | 785.8 | 519 | - | - | 0 | - |
| 4 | y | 738.3 | 544.6 | 0.0001898 | 0.3486 | +3 | 13 |
| - | - | 2984 | 545.4 | - | - | 0 | - |
| - | - | 1044 | 546.4 | - | - | 0 | - |
| - | - | 574 | 556.3 | - | - | 0 | - |
| 13 | z | 1.034E+04 | 559.3 | 0.004528 | 8.095 | +1 | 4 |
| - | - | 4131 | 560.3 | - | - | 0 | - |
| - | - | 705.4 | 561.3 | - | - | 0 | - |
| 3 | y | 2171 | 572.3 | 0.0005168 | 0.9031 | +3 | 14 |
| 3 | y | 1780 | 572.6 | 0.006744 | 11.78 | +3 | 14 |
| 3 | z | 1165 | 572.9 | 0.005153 | 8.994 | +3 | 14 |
| - | - | 908.7 | 574.3 | - | - | 0 | - |
| 13 | y | 7606 | 575.3 | 0.003476 | 6.042 | +1 | 4 |
| - | - | 2573 | 576.3 | - | - | 0 | - |
| - | - | 3782 | 602.4 | - | - | 0 | - |
| - | - | 1159 | 603.4 | - | - | 0 | - |
| - | - | 608.9 | 611.4 | - | - | 0 | - |
| - | - | 1633 | 616.3 | - | - | 0 | - |
| - | - | 669.4 | 617.3 | - | - | 0 | - |
| - | - | 3380 | 619.3 | - | - | 0 | - |
| - | - | 741.3 | 620.3 | - | - | 0 | - |
| - | - | 701.1 | 625 | - | - | 0 | - |
| 2 | y | 1705 | 626.6 | 0.0007727 | 1.233 | +3 | 15 |
| 2 | y | 898.1 | 627 | 0.005573 | 8.888 | +3 | 15 |
| 12 | w | 4039 | 643.3 | 0.003873 | 6.02 | +1 | 5 |
| - | - | 1116 | 644.3 | - | - | 0 | - |
| - | - | 1081 | 644.4 | - | - | 0 | - |
| 12 | w | 3147 | 645.3 | 0.004439 | 6.879 | +1 | 5 |
| - | - | 1003 | 646.3 | - | - | 0 | - |
| - | - | 603.1 | 648.3 | - | - | 0 | - |
| - | - | 663.6 | 656.3 | - | - | 0 | - |
| - | - | 1502 | 658.3 | - | - | 0 | - |
| 12 | y | 3718 | 659.3 | 0.005441 | 8.252 | +1 | 5 |
| - | - | 7623 | 659.7 | - | - | 0 | - |
| - | - | 8564 | 660 | - | - | 0 | - |
| 12 | z | 1.456E+04 | 660.3 | 0.01021 | 15.47 | +1 | 5 |
| - | - | 4724 | 660.7 | - | - | 0 | - |
| - | - | 2871 | 661 | - | - | 0 | - |
| - | - | 3754 | 661.3 | - | - | 0 | - |
| - | - | 1482 | 662.4 | - | - | 0 | - |
| - | - | 801.8 | 664.8 | - | - | 0 | - |
| - | - | 827.6 | 665.3 | - | - | 0 | - |
| - | - | 845 | 666.4 | - | - | 0 | - |
| - | - | 1011 | 666.8 | - | - | 0 | - |
| - | - | 1820 | 667.3 | - | - | 0 | - |
| - | - | 1010 | 674.4 | - | - | 0 | - |
| - | - | 671 | 675.4 | - | - | 0 | - |
| 12 | y | 3688 | 676.4 | 0.005012 | 7.41 | +1 | 5 |
| - | - | 1659 | 677.4 | - | - | 0 | - |
| - | - | 697.3 | 679.4 | - | - | 0 | - |
| - | - | 667 | 684.3 | - | - | 0 | - |
| - | - | 1538 | 687.4 | - | - | 0 | - |
| 6 | c | 4290 | 688.4 | 0.0002503 | 0.3636 | +1 | 6 |
| - | - | 671.4 | 689.4 | - | - | 0 | - |
| - | - | 2314 | 697.4 | - | - | 0 | - |
| - | - | 782 | 698.4 | - | - | 0 | - |
| - | - | 769.7 | 701.4 | - | - | 0 | - |
| - | - | 5943 | 707.3 | - | - | 0 | - |
| - | - | 3664 | 707.8 | - | - | 0 | - |
| - | - | 2297 | 708.3 | - | - | 0 | - |
| - | - | 898.4 | 708.8 | - | - | 0 | - |
| - | - | 831.6 | 710.3 | - | - | 0 | - |
| 6 | y | 4918 | 711.3 | 0.0005593 | 0.7862 | +2 | 11 |
| - | - | 2424 | 711.8 | - | - | 0 | - |
| - | - | 1458 | 712.3 | - | - | 0 | - |
| - | - | 2690 | 713.4 | - | - | 0 | - |
| - | - | 1020 | 714.4 | - | - | 0 | - |
| - | - | 2418 | 714.4 | - | - | 0 | - |
| - | - | 683.1 | 717.4 | - | - | 0 | - |
| - | - | 5086 | 727.9 | - | - | 0 | - |
| - | - | 2283 | 728.4 | - | - | 0 | - |
| - | - | 1122 | 728.9 | - | - | 0 | - |
| - | - | 1539 | 732.4 | - | - | 0 | - |
| 5 | w | 704 | 738.3 | 1.776E-05 | 0.02406 | +2 | 12 |
| - | - | 830 | 743.4 | - | - | 0 | - |
| - | - | 4536 | 744.4 | - | - | 0 | - |
| - | - | 1.591E+04 | 745.4 | - | - | 0 | - |
| - | - | 6689 | 746.4 | - | - | 0 | - |
| - | - | 1554 | 747.4 | - | - | 0 | - |
| 5 | y | 1696 | 750.9 | 0.004752 | 6.329 | +2 | 12 |
| 5 | y | 2663 | 751.4 | 0.006353 | 8.455 | +2 | 12 |
| 5 | z | 1530 | 751.9 | 0.0011 | 1.463 | +2 | 12 |
| - | - | 1273 | 752.4 | - | - | 0 | - |
| - | - | 961.9 | 752.9 | - | - | 0 | - |
| - | - | 1223 | 758.9 | - | - | 0 | - |
| - | - | 905.8 | 759.4 | - | - | 0 | - |
| 5 | y | 6.876E+04 | 759.9 | 0.0004026 | 0.5298 | +2 | 12 |
| - | - | 5.996E+04 | 760.4 | - | - | 0 | - |
| - | - | 3.263E+04 | 760.9 | - | - | 0 | - |
| - | - | 1.116E+04 | 761.4 | - | - | 0 | - |
| - | - | 3730 | 761.9 | - | - | 0 | - |
| - | - | 1142 | 762.4 | - | - | 0 | - |
| - | - | 1015 | 763.9 | - | - | 0 | - |
| - | - | 847.6 | 765.4 | - | - | 0 | - |
| - | - | 1012 | 772.4 | - | - | 0 | - |
| - | - | 964.6 | 774.4 | - | - | 0 | - |
| 13 | c | 700.1 | 774.9 | 0.01086 | 14.02 | +2 | 13 |
| 7 | c | 1.948E+04 | 775.4 | 0.0003751 | 0.4838 | +1 | 7 |
| - | - | 7900 | 776.4 | - | - | 0 | - |
| - | - | 1382 | 777.4 | - | - | 0 | - |
| - | - | 8523 | 782.9 | - | - | 0 | - |
| 13 | c | 2.052E+04 | 783.4 | 0.002049 | 2.615 | +2 | 13 |
| - | - | 1.546E+04 | 783.9 | - | - | 0 | - |
| - | - | 8433 | 784.4 | - | - | 0 | - |
| - | - | 3093 | 784.9 | - | - | 0 | - |
| - | - | 969.8 | 785.4 | - | - | 0 | - |
| 4 | w | 1.578E+04 | 786.9 | 0.0006744 | 0.8571 | +2 | 13 |
| - | - | 1.519E+04 | 787.4 | - | - | 0 | - |
| - | - | 8194 | 787.9 | - | - | 0 | - |
| - | - | 3551 | 788.4 | - | - | 0 | - |
| 11 | z | 831.5 | 789.4 | 0.0004096 | 0.519 | +1 | 6 |
| 11 | z | 3877 | 807.4 | 0.001843 | 2.283 | +1 | 6 |
| 4 | y | 885.9 | 807.9 | 0.003208 | 3.971 | +2 | 13 |
| 4 | z | 2.588E+04 | 808.4 | 0.002726 | 3.373 | +2 | 13 |
| - | - | 1.845E+04 | 808.9 | - | - | 0 | - |
| - | - | 1.484E+04 | 809.4 | - | - | 0 | - |
| - | - | 4785 | 809.9 | - | - | 0 | - |
| - | - | 1787 | 810.4 | - | - | 0 | - |
| - | - | 1254 | 814.4 | - | - | 0 | - |
| - | - | 1535 | 814.9 | - | - | 0 | - |
| - | - | 1082 | 815.4 | - | - | 0 | - |
| - | - | 984.5 | 815.9 | - | - | 0 | - |
| 4 | y | 1.396E+04 | 816.4 | 0.0003017 | 0.3696 | +2 | 13 |
| - | - | 9514 | 816.9 | - | - | 0 | - |
| - | - | 5147 | 817.4 | - | - | 0 | - |
| - | - | 3859 | 817.9 | - | - | 0 | - |
| - | - | 2752 | 818.4 | - | - | 0 | - |
| - | - | 1722 | 818.9 | - | - | 0 | - |
| - | - | 806.7 | 819.4 | - | - | 0 | - |
| 11 | y | 2476 | 823.4 | 0.003627 | 4.405 | +1 | 6 |
| - | - | 1040 | 824.4 | - | - | 0 | - |
| - | - | 953.1 | 829.9 | - | - | 0 | - |
| - | - | 849.2 | 830.2 | - | - | 0 | - |
| - | - | 3670 | 830.9 | - | - | 0 | - |
| - | - | 3488 | 831.4 | - | - | 0 | - |
| 14 | c | 1570 | 831.9 | 0.01401 | 16.85 | +2 | 14 |
| - | - | 624.5 | 833.4 | - | - | 0 | - |
| - | - | 1950 | 834.9 | - | - | 0 | - |
| - | - | 1010 | 835.4 | - | - | 0 | - |
| - | - | 2159 | 839.9 | - | - | 0 | - |
| 14 | c | 3.988E+04 | 840.4 | 0.002375 | 2.826 | +2 | 14 |
| - | - | 3.622E+04 | 840.9 | - | - | 0 | - |
| - | - | 1.976E+04 | 841.4 | - | - | 0 | - |
| - | - | 7535 | 841.9 | - | - | 0 | - |
| - | - | 2181 | 842.4 | - | - | 0 | - |
| - | - | 1009 | 849.4 | - | - | 0 | - |
| - | - | 1002 | 851.5 | - | - | 0 | - |
| - | - | 842 | 853.9 | - | - | 0 | - |
| - | - | 1115 | 857.5 | - | - | 0 | - |
| 3 | y | 1086 | 857.9 | 0.0001164 | 0.1357 | +2 | 14 |
| 3 | y | 5656 | 858.4 | 0.001456 | 1.696 | +2 | 14 |
| 3 | z | 1.347E+04 | 858.9 | 0.0006257 | 0.7284 | +2 | 14 |
| - | - | 1.334E+04 | 859.4 | - | - | 0 | - |
| - | - | 7576 | 859.9 | - | - | 0 | - |
| - | - | 4470 | 860.4 | - | - | 0 | - |
| - | - | 1421 | 860.9 | - | - | 0 | - |
| - | - | 786.3 | 861.4 | - | - | 0 | - |
| - | - | 987.6 | 863.4 | - | - | 0 | - |
| 3 | y | 3.236E+04 | 866.9 | 0.0002662 | 0.3071 | +2 | 14 |
| - | - | 2.927E+04 | 867.4 | - | - | 0 | - |
| - | - | 1.624E+04 | 867.9 | - | - | 0 | - |
| - | - | 8361 | 868.4 | - | - | 0 | - |
| - | - | 2925 | 868.9 | - | - | 0 | - |
| - | - | 1175 | 869.4 | - | - | 0 | - |
| - | - | 1077 | 870.9 | - | - | 0 | - |
| - | - | 1954 | 871.4 | - | - | 0 | - |
| - | - | 1318 | 871.9 | - | - | 0 | - |
| - | - | 1530 | 873.4 | - | - | 0 | - |
| - | - | 1659 | 874.4 | - | - | 0 | - |
| 10 | w | 8537 | 877.4 | 0.0005084 | 0.5794 | +1 | 7 |
| - | - | 4682 | 878.4 | - | - | 0 | - |
| - | - | 1525 | 879.4 | - | - | 0 | - |
| - | - | 1539 | 880.4 | - | - | 0 | - |
| - | - | 951.7 | 880.9 | - | - | 0 | - |
| - | - | 1030 | 881.4 | - | - | 0 | - |
| - | - | 1509 | 881.9 | - | - | 0 | - |
| - | - | 1198 | 882.4 | - | - | 0 | - |
| - | - | 853.6 | 883 | - | - | 0 | - |
| 15 | c | 1572 | 895.9 | 0.01431 | 15.98 | +2 | 15 |
| - | - | 1878 | 896.5 | - | - | 0 | - |
| - | - | 2066 | 897 | - | - | 0 | - |
| - | - | 1756 | 897.5 | - | - | 0 | - |
| - | - | 1078 | 904 | - | - | 0 | - |
| 15 | c | 1.1E+04 | 904.5 | 0.001833 | 2.027 | +2 | 15 |
| - | - | 1.328E+04 | 905 | - | - | 0 | - |
| - | - | 8248 | 905.5 | - | - | 0 | - |
| - | - | 4195 | 906 | - | - | 0 | - |
| - | - | 2307 | 906.5 | - | - | 0 | - |
| - | - | 832.4 | 909.5 | - | - | 0 | - |
| - | - | 1253 | 910.5 | - | - | 0 | - |
| - | - | 1549 | 911.5 | - | - | 0 | - |
| - | - | 2992 | 912.4 | - | - | 0 | - |
| - | - | 2023 | 912.9 | - | - | 0 | - |
| - | - | 1244 | 925.5 | - | - | 0 | - |
| - | - | 1713 | 928.5 | - | - | 0 | - |
| - | - | 1084 | 930.5 | - | - | 0 | - |
| - | - | 800.2 | 931 | - | - | 0 | - |
| 8 | c | 3.246E+04 | 931.5 | 0.0001552 | 0.1666 | +1 | 8 |
| - | - | 4666 | 932 | - | - | 0 | - |
| - | - | 1.615E+04 | 932.5 | - | - | 0 | - |
| - | - | 3520 | 933 | - | - | 0 | - |
| - | - | 4503 | 933.5 | - | - | 0 | - |
| 10 | z | 2.073E+04 | 936.4 | 0.0007539 | 0.805 | +1 | 7 |
| - | - | 1.27E+04 | 937.4 | - | - | 0 | - |
| - | - | 5278 | 938.4 | - | - | 0 | - |
| 2 | y | 1442 | 939.5 | 0.01739 | 18.51 | +2 | 15 |
| - | - | 1628 | 940 | - | - | 0 | - |
| 2 | z | 1.006E+04 | 940.5 | 0.0025 | 2.658 | +2 | 15 |
| - | - | 1.291E+04 | 941 | - | - | 0 | - |
| - | - | 8171 | 941.5 | - | - | 0 | - |
| - | - | 4546 | 942 | - | - | 0 | - |
| - | - | 1502 | 942.4 | - | - | 0 | - |
| - | - | 946.4 | 944.5 | - | - | 0 | - |
| - | - | 963.2 | 945 | - | - | 0 | - |
| - | - | 1157 | 945.5 | - | - | 0 | - |
| - | - | 2107 | 946 | - | - | 0 | - |
| - | - | 2978 | 946.5 | - | - | 0 | - |
| - | - | 4081 | 947 | - | - | 0 | - |
| - | - | 1770 | 947.5 | - | - | 0 | - |
| - | - | 1930 | 948 | - | - | 0 | - |
| 2 | y | 5273 | 948.5 | 0.0008285 | 0.8735 | +2 | 15 |
| - | - | 4933 | 949 | - | - | 0 | - |
| - | - | 2195 | 949.5 | - | - | 0 | - |
| - | - | 980.4 | 950 | - | - | 0 | - |
| - | - | 678.5 | 950.5 | - | - | 0 | - |
| 10 | y | 2855 | 952.4 | 0.002965 | 3.113 | +1 | 7 |
| - | - | 1133 | 953 | - | - | 0 | - |
| - | - | 1112 | 953.5 | - | - | 0 | - |
| - | - | 4.238E+04 | 954 | - | - | 0 | - |
| - | - | 4.842E+04 | 954.5 | - | - | 0 | - |
| - | - | 3.194E+04 | 955 | - | - | 0 | - |
| - | - | 1.722E+04 | 955.5 | - | - | 0 | - |
| - | - | 5498 | 956 | - | - | 0 | - |
| - | - | 1045 | 956.5 | - | - | 0 | - |
| - | - | 798.5 | 957.5 | - | - | 0 | - |
| - | - | 1075 | 959 | - | - | 0 | - |
| - | - | 1101 | 960.5 | - | - | 0 | - |
| - | - | 1188 | 961 | - | - | 0 | - |
| - | - | 1083 | 961.5 | - | - | 0 | - |
| - | - | 6966 | 962 | - | - | 0 | - |
| - | - | 7758 | 962.5 | - | - | 0 | - |
| - | - | 6150 | 963 | - | - | 0 | - |
| - | - | 3036 | 963.5 | - | - | 0 | - |
| - | - | 1645 | 964 | - | - | 0 | - |
| - | - | 1260 | 966.5 | - | - | 0 | - |
| - | - | 2043 | 967 | - | - | 0 | - |
| - | - | 3165 | 967.5 | - | - | 0 | - |
| - | - | 2513 | 968 | - | - | 0 | - |
| - | - | 2672 | 968.5 | - | - | 0 | - |
| - | - | 7159 | 969 | - | - | 0 | - |
| - | - | 4573 | 969.5 | - | - | 0 | - |
| - | - | 2938 | 970 | - | - | 0 | - |
| - | - | 3945 | 970.5 | - | - | 0 | - |
| - | - | 3442 | 971 | - | - | 0 | - |
| - | - | 3694 | 971.5 | - | - | 0 | - |
| - | - | 1121 | 972 | - | - | 0 | - |
| - | - | 1060 | 972.5 | - | - | 0 | - |
| - | - | 990.5 | 973.5 | - | - | 0 | - |
| - | - | 2118 | 975.5 | - | - | 0 | - |
| - | - | 1.53E+04 | 976 | - | - | 0 | - |
| - | - | 1.754E+04 | 976.5 | - | - | 0 | - |
| - | - | 1.267E+04 | 977 | - | - | 0 | - |
| - | - | 7811 | 977.5 | - | - | 0 | - |
| - | - | 3284 | 978 | - | - | 0 | - |
| - | - | 1613 | 978.5 | - | - | 0 | - |
| - | - | 1928 | 981 | - | - | 0 | - |
| - | - | 2271 | 981.5 | - | - | 0 | - |
| - | - | 3066 | 982 | - | - | 0 | - |
| - | - | 2522 | 982.5 | - | - | 0 | - |
| - | - | 1739 | 983 | - | - | 0 | - |
| - | - | 1142 | 983.5 | - | - | 0 | - |
| - | - | 858 | 983.6 | - | - | 0 | - |
| - | - | 1117 | 989 | - | - | 0 | - |
| - | - | 8441 | 989.5 | - | - | 0 | - |
| - | - | 5.187E+04 | 990 | - | - | 0 | - |
| - | - | 6.061E+04 | 990.5 | - | - | 0 | - |
| - | - | 3.9E+04 | 991 | - | - | 0 | - |
| - | - | 1.863E+04 | 991.5 | - | - | 0 | - |
| - | - | 6482 | 992 | - | - | 0 | - |
| - | - | 2898 | 992.5 | - | - | 0 | - |
| - | - | 1464 | 997 | - | - | 0 | - |
| - | - | 2220 | 997.5 | - | - | 0 | - |
| - | - | 3.598E+04 | 998 | - | - | 0 | - |
| - | - | 8.009E+04 | 998.5 | - | - | 0 | - |
| - | - | 6.648E+04 | 999 | - | - | 0 | - |
| - | - | 4.235E+04 | 999.5 | - | - | 0 | - |
| - | - | 1.845E+04 | 1000 | - | - | 0 | - |
| - | - | 5030 | 1001 | - | - | 0 | - |
| - | - | 1548 | 1001 | - | - | 0 | - |
| 9 | w | 4660 | 1006 | 0.001983 | 1.97 | +1 | 8 |
| - | - | 2482 | 1007 | - | - | 0 | - |
| - | - | 1010 | 1009 | - | - | 0 | - |
| - | - | 1351 | 1035 | - | - | 0 | - |
| 9 | c | 3.543E+04 | 1061 | 0.0001646 | 0.1552 | +1 | 9 |
| - | - | 2.24E+04 | 1062 | - | - | 0 | - |
| - | - | 7986 | 1063 | - | - | 0 | - |
| - | - | 1806 | 1064 | - | - | 0 | - |
| 9 | z | 1.873E+04 | 1065 | 0.001435 | 1.347 | +1 | 8 |
| - | - | 1.174E+04 | 1066 | - | - | 0 | - |
| - | - | 4766 | 1067 | - | - | 0 | - |
| - | - | 1885 | 1068 | - | - | 0 | - |
| 9 | y | 1525 | 1081 | 0.001298 | 1.2 | +1 | 8 |
| - | - | 2173 | 1082 | - | - | 0 | - |
| - | - | 1035 | 1085 | - | - | 0 | - |
| - | - | 1111 | 1092 | - | - | 0 | - |
| - | - | 1204 | 1092 | - | - | 0 | - |
| - | - | 980.3 | 1096 | - | - | 0 | - |
| - | - | 1975 | 1100 | - | - | 0 | - |
| - | - | 1253 | 1101 | - | - | 0 | - |
| - | - | 1447 | 1102 | - | - | 0 | - |
| - | - | 1416 | 1102 | - | - | 0 | - |
| - | - | 1975 | 1102 | - | - | 0 | - |
| - | - | 854.9 | 1105 | - | - | 0 | - |
| - | - | 1019 | 1105 | - | - | 0 | - |
| - | - | 1144 | 1106 | - | - | 0 | - |
| - | - | 2257 | 1106 | - | - | 0 | - |
| - | - | 1008 | 1106 | - | - | 0 | - |
| - | - | 1467 | 1107 | - | - | 0 | - |
| - | - | 1246 | 1107 | - | - | 0 | - |
| - | - | 2296 | 1107 | - | - | 0 | - |
| - | - | 3048 | 1108 | - | - | 0 | - |
| - | - | 1611 | 1108 | - | - | 0 | - |
| - | - | 1445 | 1111 | - | - | 0 | - |
| - | - | 2372 | 1111 | - | - | 0 | - |
| - | - | 2548 | 1112 | - | - | 0 | - |
| - | - | 1955 | 1112 | - | - | 0 | - |
| - | - | 1810 | 1112 | - | - | 0 | - |
| - | - | 879.6 | 1160 | - | - | 0 | - |
| - | - | 1072 | 1169 | - | - | 0 | - |
| - | - | 5815 | 1189 | - | - | 0 | - |
| 10 | c | 3.131E+04 | 1190 | 0.0004363 | 0.3667 | +1 | 10 |
| - | - | 1.998E+04 | 1191 | - | - | 0 | - |
| - | - | 7682 | 1192 | - | - | 0 | - |
| - | - | 2036 | 1193 | - | - | 0 | - |
| 8 | z | 1.555E+04 | 1222 | 0.0005552 | 0.4545 | +1 | 9 |
| - | - | 1.271E+04 | 1223 | - | - | 0 | - |
| - | - | 7710 | 1224 | - | - | 0 | - |
| - | - | 2430 | 1225 | - | - | 0 | - |
| - | - | 2608 | 1228 | - | - | 0 | - |
| - | - | 1277 | 1229 | - | - | 0 | - |
| - | - | 827.3 | 1231 | - | - | 0 | - |
| - | - | 991.2 | 1237 | - | - | 0 | - |
| 8 | y | 6559 | 1238 | 7.797E-06 | 0.0063 | +1 | 9 |
| - | - | 5299 | 1239 | - | - | 0 | - |
| - | - | 2234 | 1240 | - | - | 0 | - |
| - | - | 1028 | 1241 | - | - | 0 | - |
| - | - | 3705 | 1293 | - | - | 0 | - |
| - | - | 3429 | 1294 | - | - | 0 | - |
| - | - | 1903 | 1295 | - | - | 0 | - |
| - | - | 1719 | 1300 | - | - | 0 | - |
| - | - | 865.7 | 1301 | - | - | 0 | - |
| 7 | y | 861.6 | 1308 | 0.003274 | 2.504 | +1 | 10 |
| 7 | z | 2189 | 1309 | 0.00123 | 0.9398 | +1 | 10 |
| - | - | 5046 | 1310 | - | - | 0 | - |
| - | - | 3128 | 1311 | - | - | 0 | - |
| - | - | 1803 | 1312 | - | - | 0 | - |
| - | - | 1360 | 1315 | - | - | 0 | - |
| - | - | 2771 | 1316 | - | - | 0 | - |
| - | - | 1.01E+04 | 1317 | - | - | 0 | - |
| - | - | 5511 | 1318 | - | - | 0 | - |
| 7 | y | 1028 | 1325 | 0.01803 | 13.61 | +1 | 10 |
| - | - | 2455 | 1331 | - | - | 0 | - |
| - | - | 2046 | 1332 | - | - | 0 | - |
| - | - | 2032 | 1333 | - | - | 0 | - |
| - | - | 2101 | 1334 | - | - | 0 | - |
| 11 | c | 2.018E+04 | 1337 | 0.004357 | 3.26 | +1 | 11 |
| - | - | 1.568E+04 | 1338 | - | - | 0 | - |
| - | - | 8154 | 1339 | - | - | 0 | - |
| - | - | 3210 | 1340 | - | - | 0 | - |
| - | - | 1789 | 1359 | - | - | 0 | - |
| - | - | 1530 | 1360 | - | - | 0 | - |
| - | - | 1911 | 1360 | - | - | 0 | - |
| - | - | 1228 | 1361 | - | - | 0 | - |
| - | - | 1268 | 1366 | - | - | 0 | - |
| - | - | 1058 | 1367 | - | - | 0 | - |
| - | - | 1150 | 1367 | - | - | 0 | - |
| - | - | 1735 | 1376 | - | - | 0 | - |
| - | - | 1567 | 1377 | - | - | 0 | - |
| - | - | 1090 | 1381 | - | - | 0 | - |
| - | - | 1135 | 1387 | - | - | 0 | - |
| - | - | 2261 | 1388 | - | - | 0 | - |
| - | - | 2826 | 1388 | - | - | 0 | - |
| - | - | 1604 | 1389 | - | - | 0 | - |
| - | - | 3579 | 1394 | - | - | 0 | - |
| - | - | 898.8 | 1394 | - | - | 0 | - |
| - | - | 2735 | 1395 | - | - | 0 | - |
| - | - | 2472 | 1396 | - | - | 0 | - |
| 6 | y | 2974 | 1422 | 0.0008001 | 0.5628 | +1 | 11 |
| - | - | 2177 | 1423 | - | - | 0 | - |
| - | - | 979.2 | 1424 | - | - | 0 | - |
| - | - | 1925 | 1437 | - | - | 0 | - |
| 12 | c | 1.747E+04 | 1438 | 0.003798 | 2.642 | +1 | 12 |
| - | - | 1.329E+04 | 1439 | - | - | 0 | - |
| - | - | 6133 | 1440 | - | - | 0 | - |
| - | - | 2934 | 1441 | - | - | 0 | - |
| - | - | 1093 | 1452 | - | - | 0 | - |
| - | - | 1413 | 1453 | - | - | 0 | - |
| - | - | 1191 | 1453 | - | - | 0 | - |
| - | - | 822.9 | 1459 | - | - | 0 | - |
| - | - | 1085 | 1460 | - | - | 0 | - |
| - | - | 1023 | 1487 | - | - | 0 | - |
| - | - | 1132 | 1488 | - | - | 0 | - |
| - | - | 775.3 | 1502 | - | - | 0 | - |
| - | - | 1022 | 1509 | - | - | 0 | - |
| - | - | 1072 | 1520 | - | - | 0 | - |
| - | - | 821.9 | 1521 | - | - | 0 | - |
| - | - | 1072 | 1528 | - | - | 0 | - |
| - | - | 1927 | 1529 | - | - | 0 | - |
| - | - | 1706 | 1530 | - | - | 0 | - |
| - | - | 2194 | 1552 | - | - | 0 | - |
| - | - | 2492 | 1552 | - | - | 0 | - |
| - | - | 1216 | 1553 | - | - | 0 | - |
| - | - | 1014 | 1558 | - | - | 0 | - |
| - | - | 1811 | 1559 | - | - | 0 | - |
| - | - | 1422 | 1565 | - | - | 0 | - |
| 13 | c | 5045 | 1566 | 0.001975 | 1.261 | +1 | 13 |
| - | - | 3462 | 1567 | - | - | 0 | - |
| - | - | 2133 | 1568 | - | - | 0 | - |
| - | - | 816.5 | 1569 | - | - | 0 | - |
| - | - | 909.3 | 1573 | - | - | 0 | - |
| 4 | y | 1868 | 1615 | 0.01221 | 7.563 | +1 | 13 |
| 4 | z | 4867 | 1616 | 0.001228 | 0.76 | +1 | 13 |
| - | - | 2.088E+04 | 1617 | - | - | 0 | - |
| - | - | 1.853E+04 | 1618 | - | - | 0 | - |
| - | - | 1.007E+04 | 1619 | - | - | 0 | - |
| - | - | 5146 | 1620 | - | - | 0 | - |
| - | - | 1667 | 1621 | - | - | 0 | - |
| - | - | 1491 | 1630 | - | - | 0 | - |
| - | - | 917.9 | 1631 | - | - | 0 | - |
| - | - | 867.1 | 1631 | - | - | 0 | - |
| 4 | y | 3205 | 1632 | 0.01069 | 6.549 | +1 | 13 |
| - | - | 1441 | 1633 | - | - | 0 | - |
| - | - | 1017 | 1636 | - | - | 0 | - |
| - | - | 1036 | 1637 | - | - | 0 | - |
| - | - | 2159 | 1638 | - | - | 0 | - |
| - | - | 1226 | 1638 | - | - | 0 | - |
| - | - | 1485 | 1639 | - | - | 0 | - |
| - | - | 1125 | 1639 | - | - | 0 | - |
| - | - | 803.5 | 1644 | - | - | 0 | - |
| - | - | 1078 | 1645 | - | - | 0 | - |
| - | - | 859.8 | 1645 | - | - | 0 | - |
| - | - | 1562 | 1652 | - | - | 0 | - |
| - | - | 2927 | 1653 | - | - | 0 | - |
| - | - | 2209 | 1653 | - | - | 0 | - |
| - | - | 1030 | 1658 | - | - | 0 | - |
| - | - | 1655 | 1658 | - | - | 0 | - |
| - | - | 2108 | 1659 | - | - | 0 | - |
| - | - | 1216 | 1659 | - | - | 0 | - |
| - | - | 2145 | 1660 | - | - | 0 | - |
| - | - | 1979 | 1660 | - | - | 0 | - |
| - | - | 4086 | 1661 | - | - | 0 | - |
| - | - | 4808 | 1661 | - | - | 0 | - |
| - | - | 3275 | 1662 | - | - | 0 | - |
| - | - | 1278 | 1662 | - | - | 0 | - |
| - | - | 972 | 1666 | - | - | 0 | - |
| - | - | 2327 | 1666 | - | - | 0 | - |
| - | - | 1934 | 1667 | - | - | 0 | - |
| - | - | 2579 | 1667 | - | - | 0 | - |
| - | - | 2827 | 1668 | - | - | 0 | - |
| - | - | 1909 | 1668 | - | - | 0 | - |
| 14 | c | 1.384E+04 | 1680 | 0.00458 | 2.726 | +1 | 14 |
| - | - | 1.448E+04 | 1681 | - | - | 0 | - |
| - | - | 7345 | 1682 | - | - | 0 | - |
| - | - | 3718 | 1683 | - | - | 0 | - |
| - | - | 1651 | 1684 | - | - | 0 | - |
| 3 | z | 1135 | 1717 | 0.002119 | 1.234 | +1 | 14 |
| - | - | 6962 | 1718 | - | - | 0 | - |
| - | - | 5932 | 1719 | - | - | 0 | - |
| - | - | 3815 | 1720 | - | - | 0 | - |
| - | - | 1206 | 1721 | - | - | 0 | - |
| 3 | y | 1438 | 1733 | 0.002079 | 1.2 | +1 | 14 |
| - | - | 1486 | 1734 | - | - | 0 | - |
| - | - | 1237 | 1735 | - | - | 0 | - |
| - | - | 920.6 | 1793 | - | - | 0 | - |
| 15 | c | 1570 | 1808 | 0.002887 | 1.597 | +1 | 15 |
| - | - | 3570 | 1809 | - | - | 0 | - |
| - | - | 3076 | 1810 | - | - | 0 | - |
| - | - | 1547 | 1811 | - | - | 0 | - |
| - | - | 1228 | 1812 | - | - | 0 | - |
| - | - | 1426 | 1863 | - | - | 0 | - |
| - | - | 4150 | 1864 | - | - | 0 | - |
| - | - | 4297 | 1865 | - | - | 0 | - |
| - | - | 1512 | 1866 | - | - | 0 | - |
| - | - | 1256 | 1867 | - | - | 0 | - |
| 2 | z | 1723 | 1880 | 0.008492 | 4.518 | +1 | 15 |
| - | - | 1.081E+04 | 1881 | - | - | 0 | - |
| - | - | 1.13E+04 | 1882 | - | - | 0 | - |
| - | - | 6417 | 1883 | - | - | 0 | - |
| - | - | 2189 | 1884 | - | - | 0 | - |
| - | - | 1238 | 1885 | - | - | 0 | - |
| - | - | 1007 | 1891 | - | - | 0 | - |
| - | - | 1812 | 1892 | - | - | 0 | - |
| - | - | 1560 | 1893 | - | - | 0 | - |
| - | - | 1234 | 1894 | - | - | 0 | - |
| - | - | 1387 | 1897 | - | - | 0 | - |
| - | - | 4385 | 1907 | - | - | 0 | - |
| - | - | 1.178E+04 | 1908 | - | - | 0 | - |
| - | - | 1.217E+04 | 1909 | - | - | 0 | - |
| - | - | 7314 | 1910 | - | - | 0 | - |
| - | - | 3413 | 1911 | - | - | 0 | - |
| - | - | 1622 | 1912 | - | - | 0 | - |
| - | - | 1523 | 1933 | - | - | 0 | - |
| - | - | 1730 | 1934 | - | - | 0 | - |
| - | - | 1718 | 1935 | - | - | 0 | - |
| - | - | 1809 | 1936 | - | - | 0 | - |
| - | - | 2807 | 1937 | - | - | 0 | - |
| - | - | 4160 | 1938 | - | - | 0 | - |
| - | - | 3335 | 1939 | - | - | 0 | - |
| - | - | 2172 | 1940 | - | - | 0 | - |
| - | - | 3814 | 1951 | - | - | 0 | - |
| - | - | 1.101E+04 | 1952 | - | - | 0 | - |
| - | - | 1.293E+04 | 1953 | - | - | 0 | - |
| - | - | 7240 | 1954 | - | - | 0 | - |
| - | - | 3550 | 1955 | - | - | 0 | - |
| - | - | 1428 | 1956 | - | - | 0 | - |
| - | - | 1641 | 1963 | - | - | 0 | - |
| - | - | 1401 | 1964 | - | - | 0 | - |
| - | - | 1683 | 1965 | - | - | 0 | - |
| - | - | 2531 | 1968 | - | - | 0 | - |
| - | - | 8396 | 1969 | - | - | 0 | - |
| - | - | 7761 | 1970 | - | - | 0 | - |
| - | - | 5364 | 1971 | - | - | 0 | - |
| - | - | 2228 | 1972 | - | - | 0 | - |
| - | - | 1649 | 1978 | - | - | 0 | - |
| - | - | 1.013E+04 | 1979 | - | - | 0 | - |
| - | - | 3.642E+04 | 1980 | - | - | 0 | - |
| - | - | 3.559E+04 | 1981 | - | - | 0 | - |
| - | - | 2.502E+04 | 1982 | - | - | 0 | - |
| - | - | 1.344E+04 | 1983 | - | - | 0 | - |
| - | - | 5046 | 1984 | - | - | 0 | - |
| - | - | 1011 | 1985 | - | - | 0 | - |
| - | - | 5347 | 1995 | - | - | 0 | - |
| - | - | 2.27E+04 | 1996 | - | - | 0 | - |
| - | - | 9.293E+04 | 1997 | - | - | 0 | - |
| - | - | 9.093E+04 | 1998 | - | - | 0 | - |
| - | - | 6.033E+04 | 1999 | - | - | 0 | - |
| - | - | 2.766E+04 | 2000 | - | - | 0 | - |
| - | - | 9066 | 2001 | - | - | 0 | - |
| - | - | 2688 | 2002 | - | - | 0 | - |

m/z Charge Intensity FragmentType MassShift Position
123.2157974243164 0 397.9948
129.10227966308594 0 1964.3021
136.07571411132812 0 3848.1877
148.9477996826172 0 684.8036
155.11790466308594 0 468.31882
160.9010772705078 0 470.69846
173.1297607421875 0 447.65912
173.45095825195312 0 2953.8857
187.14471435546875 0 1242.1549
197.1280059814453 0 670.7186
211.54884338378906 0 528.77313
215.13865661621094 0 1596.7544
216.1343994140625 0 656.1019
219.81387329101562 0 574.468
225.0438995361328 0 670.4903
230.33151245117188 0 561.93506
235.14405822753906 0 16574.85
236.1475830078125 0 2459.745
239.09474182128906 0 1455.104
263.13885498046875 0 14051.693
264.141845703125 0 1184.5001
283.1405029296875 0 928.48987
295.1031188964844 0 1072.7908
299.06134033203125 0 963.08203
299.171630859375 0 3590.7588
300.17437744140625 0 597.009
318.181396484375 0 579.49023
320.16094970703125 0 1158.7588
325.1868591308594 0 1384.3402
333.1766052246094 0 653.37225 y 14
339.1647644042969 0 578.96295
341.0184631347656 0 715.52905
342.17742919921875 0 643.201
344.1912536621094 0 588.5365
346.17608642578125 0 9763.077
347.17852783203125 0 1730.7793
355.06964111328125 0 4426.6274
356.1446228027344 0 671.1974
359.0284118652344 0 5077.933
360.1916198730469 0 1533.5555
364.1865234375 0 14211.188
365.1900634765625 0 2460.2205
382.246337890625 0 798.7788
388.1829833984375 0 4423.6025
389.186767578125 0 791.0371
396.187255859375 0 723.3205
400.25567626953125 0 1076.1918
405.249755859375 0 507.2
429.0888977050781 0 11500.435
431.2649841308594 0 2724.4724
432.20867919921875 0 790.31665
433.24505615234375 0 642.5853
447.2197570800781 0 1601.7072 y 13
448.2216796875 0 530.9978
449.2768859863281 0 2244.3975
450.28082275390625 0 898.0011
459.26031494140625 0 12399.824
460.26385498046875 0 3182.9583
477.2712097167969 0 3908.2117
497.2364807128906 0 771.1948
506.91229248046875 0 5750.5293 y 4
507.2459411621094 0 3654.8452
507.5802307128906 0 2264.6108
507.9111022949219 0 641.265
511.2878112792969 0 958.82715
515.2828979492188 0 2871.0793
519.0394897460938 0 785.80524
544.6064453125 0 738.3084 y 3
545.3760986328125 0 2984.007
546.376953125 0 1044.017
556.3170776367188 0 573.99634
559.2960205078125 0 10344.424 z 12
560.3012084960938 0 4131.3003
561.3055419921875 0 705.3918
572.28515625 0 2170.6064 y Water loss 2
572.6204223632812 0 1779.5038 y Ammonia loss 2
572.9547729492188 0 1165.4064 z 2
574.3204956054688 0 908.7414
575.3157958984375 0 7605.774 y 12
576.3175048828125 0 2573.2786
602.398193359375 0 3782.321
603.4028930664062 0 1159.1915
611.3524780273438 0 608.9296
616.3294067382812 0 1633.45
617.3316650390625 0 669.4187
619.2686767578125 0 3379.6035
620.2728271484375 0 741.3442
624.9810791015625 0 701.1133
626.6393432617188 0 1705.4927 y Water loss 1
626.9736938476562 0 898.13214 y Ammonia loss 1
643.3416137695312 0 4038.6487 w 11
644.3379516601562 0 1115.549
644.3916625976562 0 1080.6185
645.3203125 0 3147.1895 w 11
646.3232421875 0 1002.85236
648.3287353515625 0 603.1351
656.3153076171875 0 663.6045
658.3228149414062 0 1501.5026
659.3349609375 0 3717.7825 y Ammonia loss 11
659.6624145507812 0 7622.7305
659.9973754882812 0 8564.008
660.3380126953125 0 14562.135 z 11
660.6656494140625 0 4724.048
660.999267578125 0 2871.0537
661.3472290039062 0 3754.0276
662.3533325195312 0 1481.7518
664.8341064453125 0 801.761
665.3408203125 0 827.5576
666.3562622070312 0 844.982
666.8494262695312 0 1011.32965
667.3107299804688 0 1820.2747
674.3583374023438 0 1009.6937
675.3600463867188 0 671.04614
676.3619384765625 0 3688.4597 y 11
677.3629150390625 0 1659.1997
679.3761596679688 0 697.3139
684.3290405273438 0 667.00964
687.3948974609375 0 1537.792
688.402587890625 0 4289.574 c 5
689.4012451171875 0 671.4257
697.3890380859375 0 2314.3533
698.3953857421875 0 782.0142
701.3765258789062 0 769.67163
707.3432006835938 0 5943.4062
707.8440551757812 0 3663.664
708.3450317382812 0 2297.0747
708.8467407226562 0 898.44446
710.3322143554688 0 831.60474
711.3373413085938 0 4917.878 y 5
711.8385009765625 0 2423.837
712.3383178710938 0 1458.2758
713.41015625 0 2689.5889
714.35009765625 0 1020.2379
714.414794921875 0 2417.7393
717.3628540039062 0 683.1404
727.864501953125 0 5085.627
728.3656005859375 0 2282.7886
728.8654174804688 0 1122.0675
732.4287109375 0 1539.0087
738.3432006835938 0 703.9996 w 4
743.380615234375 0 830.04095
744.3880615234375 0 4535.8154
745.3954467773438 0 15907.789
746.3976440429688 0 6688.798
747.398681640625 0 1553.8728
750.854248046875 0 1696.0012 y Water loss 4
751.3573608398438 0 2663.255 y Ammonia loss 4
751.8538208007812 0 1529.5034 z 4
752.3538208007812 0 1273.026
752.8507080078125 0 961.8523
758.856689453125 0 1223.0576
759.36474609375 0 905.76715
759.8646850585938 0 68764.83 y 4
760.3654174804688 0 59955.047
760.866455078125 0 32625.297
761.3678588867188 0 11156.934
761.872802734375 0 3729.526
762.3912353515625 0 1142.46
763.8800659179688 0 1015.07855
765.3748168945312 0 847.57635
772.3877563476562 0 1011.9318
774.35595703125 0 964.59784
774.8766479492188 0 700.10657 c Ammonia loss 12
775.4352416992188 0 19478.701 c 6
776.4376220703125 0 7899.9756
777.443115234375 0 1381.8132
782.8992309570312 0 8522.501
783.40283203125 0 20515.652 c 12
783.9041748046875 0 15462.931
784.4054565429688 0 8433.322
784.905517578125 0 3093.2317
785.4140625 0 969.80743
786.8702392578125 0 15776.255 w 3
787.3707275390625 0 15185.996
787.87060546875 0 8193.705
788.3717041015625 0 3550.5913
789.3677368164062 0 831.5058 z Water loss 10
807.3805541992188 0 3877.0994 z 10
807.8898315429688 0 885.9113 y Ammonia loss 3
808.3942260742188 0 25882.562 z 3
808.8981323242188 0 18450.494
809.396240234375 0 14835.809
809.8984985351562 0 4784.613
810.394287109375 0 1787.411
814.4080810546875 0 1254.114
814.9152221679688 0 1535.4738
815.409912109375 0 1082.2606
815.9046020507812 0 984.47107
816.4066162109375 0 13964.51 y 3
816.9080200195312 0 9514.195
817.407958984375 0 5146.8896
817.9102172851562 0 3858.92
818.41552734375 0 2751.9604
818.91455078125 0 1721.9913
819.4187622070312 0 806.6792
823.4010620117188 0 2476.356 y 10
824.3966064453125 0 1039.9137
829.8822631835938 0 953.1417
830.1885375976562 0 849.1884
830.8920288085938 0 3670.302
831.3944091796875 0 3487.6548
831.8949584960938 0 1569.966 c Ammonia loss 13
833.4028930664062 0 624.4635
834.9314575195312 0 1950.4128
835.4290161132812 0 1009.9968
839.92041015625 0 2158.5706
840.4246215820312 0 39880.137 c 13
840.9259033203125 0 36215.656
841.4266967773438 0 19759.521
841.926025390625 0 7534.663
842.4257202148438 0 2180.6636
849.4144287109375 0 1008.7843
851.4838256835938 0 1002.02985
853.9317626953125 0 842.04193
857.4600219726562 0 1114.8287
857.9249877929688 0 1085.5109 y Water loss 2
858.4183349609375 0 5655.9 y Ammonia loss 2
858.920166015625 0 13467.171 z 2
859.4222412109375 0 13342.333
859.922607421875 0 7575.7466
860.4246215820312 0 4470.266
860.9253540039062 0 1421.3354
861.429931640625 0 786.2706
863.3983764648438 0 987.59424
866.930419921875 0 32360.275 y 2
867.4314575195312 0 29273.709
867.9320678710938 0 16236.701
868.4324951171875 0 8360.994
868.93408203125 0 2925.036
869.43310546875 0 1174.9443
870.9124755859375 0 1076.7628
871.4161376953125 0 1954.0367
871.9176025390625 0 1318.4409
873.4305419921875 0 1530.305
874.4328002929688 0 1658.5358
877.4085083007812 0 8536.537 w 9
878.4060668945312 0 4682.387
879.4134521484375 0 1524.94
880.4190673828125 0 1538.769
880.9244384765625 0 951.7246
881.4260864257812 0 1030.4065
881.9322509765625 0 1509.3032
882.4385375976562 0 1197.7584
882.9510498046875 0 853.5523
895.9525756835938 0 1571.6152 c Ammonia loss 14
896.462646484375 0 1878.0747
896.9623413085938 0 2066.3608
897.4566040039062 0 1755.9739
903.9542236328125 0 1077.6351
904.453369140625 0 10997.129 c 14
904.9542236328125 0 13277.099
905.4542236328125 0 8247.603
905.9552612304688 0 4194.7705
906.4601440429688 0 2307.3574
909.4580688476562 0 832.3952
910.4631958007812 0 1253.4127
911.4547729492188 0 1548.8611
912.4237670898438 0 2992.3926
912.922119140625 0 2023.2184
925.4611206054688 0 1244.1057
928.4745483398438 0 1713.1411
930.5238647460938 0 1084.3363
930.9818725585938 0 800.1798
931.5361328125 0 32460.078 c 7
931.9796752929688 0 4666.184
932.5379028320312 0 16151.347
932.9768676757812 0 3520.245
933.5409545898438 0 4502.837
936.4220581054688 0 20734.426 z 9
937.4239501953125 0 12695.699
938.4248046875 0 5277.6753
939.4391479492188 0 1442.4452 y Water loss 1
939.97119140625 0 1628.1467
940.4549560546875 0 10062.19 z 1
940.9542846679688 0 12914.819
941.4544067382812 0 8171.3057
941.9537963867188 0 4545.8447
942.4490966796875 0 1502.1724
944.4853515625 0 946.3686
944.9852905273438 0 963.1502
945.4890747070312 0 1156.535
945.9785766601562 0 2107.2246
946.4825439453125 0 2977.5525
946.9803466796875 0 4080.9185
947.4757080078125 0 1770.0892
947.9674072265625 0 1930.3385
948.462646484375 0 5273.0264 y 1
948.9617919921875 0 4933.2397
949.4613647460938 0 2194.7654
949.9640502929688 0 980.3673
950.4646606445312 0 678.4957
952.4429931640625 0 2855.278 y 9
952.982421875 0 1133.1603
953.4755249023438 0 1111.6016
953.9881591796875 0 42383.492
954.4890747070312 0 48417.164
954.9898071289062 0 31944.639
955.488525390625 0 17215.703
955.9906616210938 0 5498.1157
956.4854125976562 0 1045.1755
957.5470581054688 0 798.5203
958.9840087890625 0 1074.9792
960.4908447265625 0 1100.522
960.9736938476562 0 1188.4636
961.4744873046875 0 1082.8834
961.9593505859375 0 6966.359
962.4623413085938 0 7757.564
962.9729614257812 0 6150.3022
963.4835205078125 0 3036.2578
963.9708251953125 0 1644.6259
966.4862670898438 0 1259.5538
966.9985961914062 0 2043.1617
967.4952392578125 0 3165.4094
967.9906616210938 0 2513.2224
968.4908447265625 0 2671.9045
968.9823608398438 0 7159.0444
969.48095703125 0 4572.9375
969.9845581054688 0 2938.4253
970.4735717773438 0 3945.2468
970.9698486328125 0 3441.6077
971.4730834960938 0 3694.1848
971.9719848632812 0 1120.7911
972.4801025390625 0 1060.2255
973.4732055664062 0 990.50323
975.4921264648438 0 2118.3308
975.9901123046875 0 15304.131
976.4906005859375 0 17543.018
976.990234375 0 12672.935
977.4883422851562 0 7810.7773
977.9893798828125 0 3284.099
978.4892578125 0 1612.7838
980.9879150390625 0 1927.8866
981.4819946289062 0 2270.904
981.9851684570312 0 3066.4158
982.4834594726562 0 2522.2805
982.9791259765625 0 1738.9144
983.4876708984375 0 1141.6382
983.6060791015625 0 857.9717
988.983154296875 0 1116.5481
989.4920654296875 0 8441.381
989.9895629882812 0 51868.77
990.4901123046875 0 60612.49
990.9908447265625 0 38996.793
991.491943359375 0 18627.684
991.9915161132812 0 6481.7773
992.48876953125 0 2898.0098
996.9908447265625 0 1464.0837
997.4891967773438 0 2219.6128
997.9963989257812 0 35981.5
998.4984741210938 0 80090.06
999.0001831054688 0 66478.39
999.501220703125 0 42345.16
1000.0011596679688 0 18447.686
1000.5038452148438 0 5030.208
1001.0048217773438 0 1548.3896
1006.4525756835938 0 4659.539 w 8
1007.4559326171875 0 2481.703
1009.4629516601562 0 1009.7494
1035.197998046875 0 1351.2285
1060.5787353515625 0 35426.34 c 8
1061.581787109375 0 22403.863
1062.585693359375 0 7985.711
1063.58203125 0 1805.8679
1065.46533203125 0 18727.158 z 8
1066.46630859375 0 11736.559
1067.46923828125 0 4766.178
1068.468994140625 0 1885.2712
1081.4813232421875 0 1524.9973 y 8
1082.484130859375 0 2172.7478
1085.493408203125 0 1035.11
1091.89892578125 0 1111.2896
1092.242919921875 0 1203.7018
1096.2431640625 0 980.27075
1099.5750732421875 0 1975.2246
1100.57568359375 0 1252.6621
1101.584228515625 0 1446.8132
1101.9166259765625 0 1416.3662
1102.2462158203125 0 1974.7917
1104.9012451171875 0 854.9407
1105.22265625 0 1019.3427
1105.567626953125 0 1144.469
1105.901611328125 0 2256.9402
1106.236572265625 0 1008.37317
1106.575927734375 0 1466.9397
1106.914794921875 0 1246.4606
1107.2510986328125 0 2296.3396
1107.5830078125 0 3048.0208
1107.9141845703125 0 1610.8032
1110.9029541015625 0 1444.7585
1111.2396240234375 0 2372.0183
1111.5758056640625 0 2547.884
1111.90771484375 0 1955.0874
1112.248046875 0 1810.4536
1159.5694580078125 0 879.60455
1168.5531005859375 0 1072.2809
1188.6109619140625 0 5814.59
1189.6207275390625 0 31311.426 c 9
1190.6243896484375 0 19982.611
1191.62744140625 0 7682.167
1192.6302490234375 0 2036.4508
1221.564453125 0 15553.218 z 7
1222.5673828125 0 12710.489
1223.5689697265625 0 7710.106
1224.5679931640625 0 2429.864
1227.635009765625 0 2608.0317
1228.6328125 0 1276.7743
1230.6507568359375 0 827.26294
1236.5782470703125 0 991.2088
1237.583740234375 0 6559.042 y 7
1238.5850830078125 0 5298.639
1239.5865478515625 0 2233.9558
1240.5819091796875 0 1028.1119
1292.64208984375 0 3705.4087
1293.6480712890625 0 3429.2556
1294.64794921875 0 1902.5792
1299.63720703125 0 1719.1973
1300.622802734375 0 865.74994
1307.5859375 0 861.624 y Ammonia loss 6
1308.5982666015625 0 2188.9614 z 6
1309.605712890625 0 5046.4805
1310.60400390625 0 3128.3123
1311.6055908203125 0 1802.8379
1314.6492919921875 0 1359.8239
1315.6436767578125 0 2770.929
1316.6541748046875 0 10099.614
1317.6552734375 0 5511.2383
1324.6337890625 0 1027.8939 y 6
1330.6744384765625 0 2454.7327
1331.652099609375 0 2046.151
1332.656494140625 0 2032.4137
1333.6510009765625 0 2101.2307
1336.656005859375 0 20183.312 c 10
1337.659423828125 0 15676.798
1338.6583251953125 0 8153.5034
1339.66162109375 0 3209.5227
1359.1746826171875 0 1788.5585
1359.6761474609375 0 1530.2162
1360.1708984375 0 1910.9332
1360.6773681640625 0 1227.8492
1366.1663818359375 0 1267.5264
1366.6737060546875 0 1057.77
1367.168701171875 0 1149.6802
1375.6748046875 0 1735.2743
1376.679443359375 0 1567.1293
1380.72216796875 0 1089.6029
1387.1837158203125 0 1134.7219
1387.6788330078125 0 2261.2188
1388.193115234375 0 2826.2615
1388.6942138671875 0 1604.2596
1393.685546875 0 3578.696
1394.1705322265625 0 898.75195
1394.6898193359375 0 2735.4866
1395.6866455078125 0 2472.0886
1421.667724609375 0 2973.8193 y 5
1422.6644287109375 0 2177.0515
1423.68701171875 0 979.1596
1436.6943359375 0 1925.0231
1437.703125 0 17467.467 c 11
1438.7064208984375 0 13288.935
1439.7081298828125 0 6133.0654
1440.7098388671875 0 2933.5847
1452.2412109375 0 1092.7126
1452.743896484375 0 1412.9718
1453.233642578125 0 1191.2299
1459.2296142578125 0 822.8736
1459.7381591796875 0 1085.1156
1487.265625 0 1022.81396
1487.76171875 0 1131.8254
1502.302734375 0 775.3287
1508.781982421875 0 1021.7144
1519.7291259765625 0 1071.764
1520.7138671875 0 821.93805
1527.7681884765625 0 1071.899
1528.7689208984375 0 1927.3185
1529.7738037109375 0 1705.6666
1551.80517578125 0 2193.6758
1552.2996826171875 0 2491.562
1552.79736328125 0 1215.926
1558.2958984375 0 1013.7828
1558.7930908203125 0 1811.1709
1564.78466796875 0 1422.1208
1565.7962646484375 0 5045.41 c 12
1566.798583984375 0 3462.1113
1567.8052978515625 0 2132.6365
1568.7984619140625 0 816.5401
1572.79541015625 0 909.2798
1614.791015625 0 1867.8427 y Ammonia loss 3
1615.785400390625 0 4867.4604 z 3
1616.79296875 0 20876.818
1617.7930908203125 0 18530.629
1618.794189453125 0 10066.256
1619.799560546875 0 5145.9404
1620.7939453125 0 1667.2029
1629.8525390625 0 1491.0043
1630.843017578125 0 917.8879
1631.382568359375 0 867.1241
1631.8160400390625 0 3204.9404 y 3
1632.804931640625 0 1440.9596
1635.85546875 0 1017.31476
1636.827880859375 0 1036.2029
1637.8394775390625 0 2158.6958
1638.355712890625 0 1225.5536
1638.8546142578125 0 1485.0281
1639.3680419921875 0 1125.2682
1643.8472900390625 0 803.5436
1644.8636474609375 0 1078.433
1645.3487548828125 0 859.84546
1652.3734130859375 0 1562.3313
1652.856201171875 0 2927.126
1653.3675537109375 0 2209.3184
1657.8375244140625 0 1029.8512
1658.3468017578125 0 1655.3134
1658.858154296875 0 2108.2917
1659.354736328125 0 1215.5021
1659.85498046875 0 2145.2417
1660.373046875 0 1979.1553
1660.8709716796875 0 4085.9597
1661.37109375 0 4807.6963
1661.8658447265625 0 3274.601
1662.3726806640625 0 1278.2169
1665.866943359375 0 972.0353
1666.3614501953125 0 2326.5393
1666.866455078125 0 1933.7172
1667.3670654296875 0 2578.9458
1667.866455078125 0 2827.1724
1668.365234375 0 1909.4125
1679.841796875 0 13844.761 c 13
1680.843994140625 0 14484.487
1681.8460693359375 0 7344.7856
1682.8507080078125 0 3718.2554
1683.856201171875 0 1651.49
1716.83642578125 0 1134.9877 z 2
1717.8414306640625 0 6962.346
1718.8431396484375 0 5932.2134
1719.843994140625 0 3815.3838
1720.8349609375 0 1205.5142
1732.8509521484375 0 1437.5807 y 2
1733.8604736328125 0 1486.1132
1734.870361328125 0 1236.508
1792.888916015625 0 920.63556
1807.898681640625 0 1569.6658 c 14
1808.9058837890625 0 3570.4905
1809.90283203125 0 3075.5342
1810.906005859375 0 1546.7739
1811.9130859375 0 1228.3966
1862.9442138671875 0 1426.1227
1863.9559326171875 0 4150.043
1864.96142578125 0 4296.7295
1865.9598388671875 0 1512.3813
1866.978759765625 0 1256.3818
1879.9061279296875 0 1722.6077 z 1
1880.9029541015625 0 10809.222
1881.905029296875 0 11299.871
1882.9078369140625 0 6417.134
1883.9119873046875 0 2189.3682
1884.907958984375 0 1238.3253
1890.95556640625 0 1007.3632
1891.9583740234375 0 1812.3029
1892.958251953125 0 1559.9872
1893.9486083984375 0 1233.8853
1896.911865234375 0 1386.9341
1906.9693603515625 0 4384.8545
1907.9737548828125 0 11782.829
1908.97607421875 0 12172.316
1909.980224609375 0 7313.664
1910.969970703125 0 3413.4968
1911.9710693359375 0 1621.8086
1932.9913330078125 0 1523.108
1934.003173828125 0 1730.3414
1934.9808349609375 0 1718.1852
1935.9697265625 0 1808.7585
1936.9603271484375 0 2807.3804
1937.95166015625 0 4160.0454
1938.9566650390625 0 3335.005
1939.9508056640625 0 2172.0945
1950.9779052734375 0 3814.0674
1951.978271484375 0 11005.125
1952.9791259765625 0 12930.803
1953.9827880859375 0 7240.437
1954.9781494140625 0 3550.0557
1955.97509765625 0 1428.0616
1962.9505615234375 0 1641.264
1963.956298828125 0 1401.0447
1964.956787109375 0 1683.386
1967.990478515625 0 2530.839
1969.0028076171875 0 8396.034
1970.0035400390625 0 7760.8633
1971.0059814453125 0 5364.495
1972.0062255859375 0 2228.311
1977.953369140625 0 1649.2045
1978.971435546875 0 10134.43
1979.9737548828125 0 36424.055
1980.9755859375 0 35587.098
1981.9744873046875 0 25021.814
1982.976806640625 0 13439.725
1983.972900390625 0 5046.493
1984.9727783203125 0 1010.73157
1994.9765625 0 5347.001
1995.988525390625 0 22703.348
1996.9967041015625 0 92926.89
1997.9996337890625 0 90930.234
1999.00048828125 0 60332.598
2000.000732421875 0 27655.924
2001.00244140625 0 9065.681
2001.99609375 0 2688.206

Spectrum Details

|  |  |
| --- | --- |
| Matched peaks? Matched peaks The total absolute number of peaks matched. Additionally in brackets the total fraction of peaks matched and the total number of peaks is shown. | 70 (11.57% of 605) |
| FDR? FDR The false discovery rate estimated for this peptide. It is calculated by matching all theoretical fragments with a non-integer shift with the raw peaks for this spectrum. This is done with 40 different shifts. The resulting percentage is the average number of annotated peaks over the number of annotated peaks with the correct spectrum. | 1.77% |
| Satellite FDR? Satellite FDR See the FDR for details on its calculation. This satellite ion specific FDR only contains the satellite ions (d/w) for I/L/J positions. | 0.00% |
| PSM Score? PSM Score The PSM Score as given by Hecklib to this annotated spectrum. It is shown with three significant figures. | 424 |

#### Spectrum 3932? Spectrum 3932 The raw spectrum of this peptide as annotated by Hecklib. The fragments are coloured according to ion type (see legend). Any peaks with a star '\*' as text can be hovered over to see the full details, first the ion type second the mass shift type. By hovering over the amino acids in the peptide or ions in the legend the corresponding peaks are highlighted. By toggling the 'Unassigned' label you can turn the background (unassigned) peaks on or off in the plot. By updating the slider in the Ion legend you can update the spectrum to only show the top X% of the peaks with labels. The top X% means any peak that is within X% of the highest intensity. By dragging in the spectrum you can zoom in to a specific part of the spectrum and use 'Zoom Out' to get back to the original zoom level. The annotation of the spectrum is based on the given sequence in the peptides file and is done with different software so inconsistencies are likely. The peaks are annotated based on the given sequence, with 20 ppm tolerance.

Copy Data

##### Spectrum 3932 (TSV)

###### Preview

```
Loading example...
```

*Click on the button to copy the data to your clipboard.*

Mz MinMz MaxIntensity Max

WidthHeightPeptide font sizePeptide stroke widthSpectrum font sizeSpectrum stroke widthCompact peptide

Ion legend

wxyz

abcd

OtherUnassignedIon^Charge^_Position_Show for top:%

VYTLPPSREEMTKNQK

03.20e+46.41e+49.61e+41.28e+5

Zoom Out

y^+1^_2_y^+1^_3_c^+2^_8_y^+3^_12_y^+3^_13_z^+1^_4_y^+3^_14_y^+3^_14_z^+3^_14_c^+1^_5_y^+1^_4_c^+3^_15_y^+3^_15_y^+3^_15_z^+3^_15_w^+1^_5_w^+1^_5_y^+2^_10_z^+2^_10_y^+1^_5_z^+1^_5_y^+2^_10_c^+1^_6_y^+1^_5_c^+1^_6_y^+2^_11_y^+2^_12_y^+2^_12_z^+2^_12_y^+2^_12_c^+1^_7_c^+2^_13_w^+2^_13_z^+1^_6_y^+1^_6_y^+2^_13_z^+2^_13_y^+2^_13_y^+1^_6_c^+2^_14_c^+2^_14_y^+2^_14_y^+2^_14_z^+2^_14_y^+2^_14_w^+1^_7_c^+2^_15_c^+2^_15_c^+1^_8_z^+1^_7_y^+2^_15_z^+2^_15_y^+2^_15_y^+1^_7_w^+1^_8_c^+1^_9_c^+1^_9_z^+1^_8_y^+1^_8_c^+1^_10_z^+1^_9_y^+1^_9_z^+1^_10_y^+1^_10_c^+1^_11_y^+1^_11_c^+1^_12_c^+1^_13_c^+1^_14_z^+1^_14_z^+1^_15_

0839167725163355

Fragment Matches Table

Show background peaks

| Position | Ion type | Intensity | mz Theoretical | mz Error (Th) | mz Error (ppm) | Charge | Series Number |
| --- | --- | --- | --- | --- | --- | --- | --- |
| - | - | 502.5 | 128.1 | - | - | 0 | - |
| - | - | 1.646E+04 | 129.1 | - | - | 0 | - |
| - | - | 370.3 | 130.1 | - | - | 0 | - |
| - | - | 1489 | 130.1 | - | - | 0 | - |
| - | - | 6377 | 136.1 | - | - | 0 | - |
| - | - | 402.8 | 143.2 | - | - | 0 | - |
| - | - | 2211 | 146.1 | - | - | 0 | - |
| - | - | 1806 | 155.1 | - | - | 0 | - |
| - | - | 726.6 | 169.1 | - | - | 0 | - |
| - | - | 512.9 | 171.1 | - | - | 0 | - |
| - | - | 1887 | 173.1 | - | - | 0 | - |
| - | - | 3567 | 173.4 | - | - | 0 | - |
| - | - | 866.9 | 183.1 | - | - | 0 | - |
| - | - | 2447 | 187.1 | - | - | 0 | - |
| - | - | 1503 | 197.1 | - | - | 0 | - |
| - | - | 1261 | 198.1 | - | - | 0 | - |
| - | - | 2408 | 200.1 | - | - | 0 | - |
| - | - | 2878 | 201.1 | - | - | 0 | - |
| - | - | 758.3 | 203.2 | - | - | 0 | - |
| - | - | 619.6 | 205.1 | - | - | 0 | - |
| - | - | 505.9 | 210.7 | - | - | 0 | - |
| - | - | 1640 | 212.1 | - | - | 0 | - |
| - | - | 1965 | 215.1 | - | - | 0 | - |
| - | - | 1.111E+04 | 216.1 | - | - | 0 | - |
| - | - | 1061 | 216.1 | - | - | 0 | - |
| - | - | 1327 | 217.1 | - | - | 0 | - |
| - | - | 2903 | 217.2 | - | - | 0 | - |
| - | - | 743.3 | 218.2 | - | - | 0 | - |
| - | - | 575.2 | 228.1 | - | - | 0 | - |
| - | - | 657.1 | 230.2 | - | - | 0 | - |
| - | - | 2266 | 233.2 | - | - | 0 | - |
| - | - | 2.776E+04 | 235.1 | - | - | 0 | - |
| - | - | 4506 | 236.1 | - | - | 0 | - |
| - | - | 829.6 | 243.1 | - | - | 0 | - |
| - | - | 812.5 | 243.1 | - | - | 0 | - |
| - | - | 472.2 | 252.1 | - | - | 0 | - |
| - | - | 2.206E+04 | 263.1 | - | - | 0 | - |
| - | - | 3880 | 264.1 | - | - | 0 | - |
| - | - | 1034 | 269.2 | - | - | 0 | - |
| - | - | 606.6 | 271.2 | - | - | 0 | - |
| - | - | 831.8 | 274.2 | - | - | 0 | - |
| - | - | 521.6 | 297.1 | - | - | 0 | - |
| - | - | 504.6 | 297.2 | - | - | 0 | - |
| - | - | 613.4 | 298.1 | - | - | 0 | - |
| - | - | 3923 | 299.2 | - | - | 0 | - |
| - | - | 668.8 | 300.2 | - | - | 0 | - |
| - | - | 1988 | 304.2 | - | - | 0 | - |
| - | - | 933.4 | 311.2 | - | - | 0 | - |
| - | - | 1050 | 315.2 | - | - | 0 | - |
| - | - | 1338 | 320.2 | - | - | 0 | - |
| - | - | 1404 | 326.2 | - | - | 0 | - |
| - | - | 691 | 327.2 | - | - | 0 | - |
| - | - | 1209 | 328.2 | - | - | 0 | - |
| - | - | 4948 | 329.2 | - | - | 0 | - |
| - | - | 1273 | 330.2 | - | - | 0 | - |
| 15 | y | 1770 | 333.2 | 0.004075 | 12.23 | +1 | 2 |
| - | - | 607.7 | 341 | - | - | 0 | - |
| - | - | 1351 | 342.2 | - | - | 0 | - |
| - | - | 651.7 | 344.2 | - | - | 0 | - |
| - | - | 1209 | 345.3 | - | - | 0 | - |
| - | - | 1.542E+04 | 346.2 | - | - | 0 | - |
| - | - | 3876 | 346.2 | - | - | 0 | - |
| - | - | 3775 | 347.2 | - | - | 0 | - |
| - | - | 1477 | 347.2 | - | - | 0 | - |
| - | - | 2276 | 355.1 | - | - | 0 | - |
| - | - | 1878 | 359 | - | - | 0 | - |
| - | - | 1768 | 360.2 | - | - | 0 | - |
| - | - | 2.644E+04 | 364.2 | - | - | 0 | - |
| - | - | 4215 | 365.2 | - | - | 0 | - |
| - | - | 1162 | 366.2 | - | - | 0 | - |
| - | - | 5352 | 382.2 | - | - | 0 | - |
| - | - | 3489 | 383.2 | - | - | 0 | - |
| - | - | 717.3 | 383.3 | - | - | 0 | - |
| - | - | 5329 | 384.2 | - | - | 0 | - |
| - | - | 1140 | 385.2 | - | - | 0 | - |
| - | - | 780.3 | 386.2 | - | - | 0 | - |
| - | - | 3816 | 388.2 | - | - | 0 | - |
| - | - | 3036 | 392.3 | - | - | 0 | - |
| - | - | 1018 | 392.8 | - | - | 0 | - |
| - | - | 1025 | 396.2 | - | - | 0 | - |
| - | - | 1.208E+04 | 400.3 | - | - | 0 | - |
| - | - | 3058 | 401.3 | - | - | 0 | - |
| - | - | 4155 | 417.3 | - | - | 0 | - |
| - | - | 1221 | 418.3 | - | - | 0 | - |
| - | - | 1336 | 425.2 | - | - | 0 | - |
| - | - | 829.5 | 427.3 | - | - | 0 | - |
| - | - | 5012 | 429.1 | - | - | 0 | - |
| - | - | 6098 | 431.3 | - | - | 0 | - |
| - | - | 1716 | 432.2 | - | - | 0 | - |
| - | - | 1343 | 432.3 | - | - | 0 | - |
| - | - | 674 | 433.2 | - | - | 0 | - |
| - | - | 754.2 | 433.2 | - | - | 0 | - |
| - | - | 3242 | 435.8 | - | - | 0 | - |
| - | - | 2170 | 436.3 | - | - | 0 | - |
| - | - | 1389 | 441.3 | - | - | 0 | - |
| - | - | 692.9 | 442.3 | - | - | 0 | - |
| - | - | 539.7 | 442.7 | - | - | 0 | - |
| - | - | 978.3 | 443.3 | - | - | 0 | - |
| 14 | y | 4149 | 447.2 | 0.003209 | 7.176 | +1 | 3 |
| - | - | 658.5 | 448.2 | - | - | 0 | - |
| - | - | 4779 | 449.3 | - | - | 0 | - |
| - | - | 1267 | 450.3 | - | - | 0 | - |
| - | - | 3221 | 456.3 | - | - | 0 | - |
| 8 | c | 749.3 | 457.3 | 0.005536 | 12.11 | +2 | 8 |
| - | - | 2.045E+04 | 459.3 | - | - | 0 | - |
| - | - | 6083 | 460.3 | - | - | 0 | - |
| - | - | 781.3 | 461.3 | - | - | 0 | - |
| - | - | 787.3 | 472.3 | - | - | 0 | - |
| - | - | 6393 | 477.3 | - | - | 0 | - |
| - | - | 2008 | 478.3 | - | - | 0 | - |
| - | - | 1052 | 483.3 | - | - | 0 | - |
| - | - | 1680 | 483.8 | - | - | 0 | - |
| - | - | 8727 | 492.3 | - | - | 0 | - |
| - | - | 8912 | 492.8 | - | - | 0 | - |
| - | - | 1710 | 493.3 | - | - | 0 | - |
| - | - | 1678 | 497.2 | - | - | 0 | - |
| - | - | 1513 | 501.3 | - | - | 0 | - |
| - | - | 878.6 | 502.4 | - | - | 0 | - |
| - | - | 1152 | 504.2 | - | - | 0 | - |
| - | - | 1002 | 505.3 | - | - | 0 | - |
| - | - | 1270 | 505.8 | - | - | 0 | - |
| 5 | y | 1.102E+04 | 506.9 | 0.001108 | 2.186 | +3 | 12 |
| - | - | 7080 | 507.2 | - | - | 0 | - |
| - | - | 4856 | 507.6 | - | - | 0 | - |
| - | - | 709.9 | 507.9 | - | - | 0 | - |
| - | - | 955.1 | 514.2 | - | - | 0 | - |
| - | - | 8211 | 515.3 | - | - | 0 | - |
| - | - | 2457 | 516.3 | - | - | 0 | - |
| - | - | 8754 | 528.4 | - | - | 0 | - |
| - | - | 2985 | 529.4 | - | - | 0 | - |
| - | - | 1509 | 544.4 | - | - | 0 | - |
| 4 | y | 757 | 544.6 | 0.0009088 | 1.669 | +3 | 13 |
| - | - | 714.7 | 544.9 | - | - | 0 | - |
| - | - | 4.113E+04 | 545.4 | - | - | 0 | - |
| - | - | 918.2 | 546.3 | - | - | 0 | - |
| - | - | 1.663E+04 | 546.4 | - | - | 0 | - |
| - | - | 3579 | 547.4 | - | - | 0 | - |
| - | - | 793.3 | 556.3 | - | - | 0 | - |
| 13 | z | 1.559E+04 | 559.3 | 0.003368 | 6.022 | +1 | 4 |
| - | - | 7677 | 560.3 | - | - | 0 | - |
| - | - | 1623 | 561.3 | - | - | 0 | - |
| - | - | 744.6 | 562.3 | - | - | 0 | - |
| - | - | 855.5 | 564.2 | - | - | 0 | - |
| - | - | 843.1 | 568.3 | - | - | 0 | - |
| 3 | y | 3962 | 572.3 | 0.0005818 | 1.017 | +3 | 14 |
| 3 | y | 1635 | 572.6 | 0.005462 | 9.539 | +3 | 14 |
| 3 | z | 890.2 | 572.9 | 0.006801 | 11.87 | +3 | 14 |
| 5 | c | 892.5 | 573.3 | 0.008089 | 14.11 | +1 | 5 |
| - | - | 604.5 | 574.3 | - | - | 0 | - |
| 13 | y | 1.305E+04 | 575.3 | 0.002805 | 4.875 | +1 | 4 |
| - | - | 3828 | 576.3 | - | - | 0 | - |
| - | - | 704.2 | 577.3 | - | - | 0 | - |
| - | - | 3931 | 581.8 | - | - | 0 | - |
| - | - | 4385 | 582.3 | - | - | 0 | - |
| - | - | 1065 | 582.8 | - | - | 0 | - |
| - | - | 590 | 583.3 | - | - | 0 | - |
| - | - | 2705 | 584.4 | - | - | 0 | - |
| - | - | 1787 | 585.4 | - | - | 0 | - |
| 15 | c | 745.5 | 597.6 | 0.004819 | 8.063 | +3 | 15 |
| - | - | 893 | 601.4 | - | - | 0 | - |
| - | - | 4.443E+04 | 602.4 | - | - | 0 | - |
| - | - | 2.128E+04 | 603.4 | - | - | 0 | - |
| - | - | 3786 | 604.4 | - | - | 0 | - |
| - | - | 1879 | 605.4 | - | - | 0 | - |
| - | - | 867 | 605.4 | - | - | 0 | - |
| - | - | 1883 | 605.9 | - | - | 0 | - |
| - | - | 1014 | 606.4 | - | - | 0 | - |
| - | - | 877.6 | 615.3 | - | - | 0 | - |
| - | - | 814 | 615.6 | - | - | 0 | - |
| - | - | 2829 | 616.3 | - | - | 0 | - |
| - | - | 1291 | 617.3 | - | - | 0 | - |
| - | - | 921.3 | 619 | - | - | 0 | - |
| - | - | 1540 | 619.3 | - | - | 0 | - |
| - | - | 1513 | 619.8 | - | - | 0 | - |
| - | - | 1397 | 625 | - | - | 0 | - |
| - | - | 871.3 | 625.3 | - | - | 0 | - |
| 2 | y | 2303 | 626.6 | 0.002096 | 3.345 | +3 | 15 |
| 2 | y | 1801 | 627 | 0.01003 | 15.99 | +3 | 15 |
| 2 | z | 1369 | 627.3 | 0.007522 | 11.99 | +3 | 15 |
| - | - | 1184 | 628.4 | - | - | 0 | - |
| - | - | 954.7 | 631.3 | - | - | 0 | - |
| - | - | 1314 | 633.3 | - | - | 0 | - |
| - | - | 810.6 | 639.4 | - | - | 0 | - |
| - | - | 795.7 | 640.4 | - | - | 0 | - |
| 12 | w | 5545 | 643.3 | 0.002652 | 4.123 | +1 | 5 |
| - | - | 1649 | 644.3 | - | - | 0 | - |
| - | - | 1140 | 644.4 | - | - | 0 | - |
| 12 | w | 5524 | 645.3 | 0.002242 | 3.474 | +1 | 5 |
| - | - | 1974 | 646.3 | - | - | 0 | - |
| - | - | 786.8 | 646.8 | - | - | 0 | - |
| - | - | 706.9 | 647.3 | - | - | 0 | - |
| 7 | y | 887.2 | 654.3 | 0.004551 | 6.956 | +2 | 10 |
| 7 | z | 1208 | 654.8 | 0.002653 | 4.052 | +2 | 10 |
| - | - | 1165 | 655.3 | - | - | 0 | - |
| - | - | 3946 | 655.4 | - | - | 0 | - |
| - | - | 2113 | 656.4 | - | - | 0 | - |
| - | - | 2777 | 657.8 | - | - | 0 | - |
| - | - | 3253 | 658.3 | - | - | 0 | - |
| 12 | y | 9042 | 659.3 | 0.004464 | 6.77 | +1 | 5 |
| - | - | 1.555E+04 | 659.7 | - | - | 0 | - |
| - | - | 1.848E+04 | 660 | - | - | 0 | - |
| 12 | z | 2.606E+04 | 660.3 | 0.00942 | 14.27 | +1 | 5 |
| - | - | 4101 | 660.7 | - | - | 0 | - |
| - | - | 1256 | 660.9 | - | - | 0 | - |
| - | - | 1308 | 661 | - | - | 0 | - |
| - | - | 6791 | 661.3 | - | - | 0 | - |
| - | - | 2064 | 662.3 | - | - | 0 | - |
| 7 | y | 1300 | 662.8 | 0.002629 | 3.967 | +2 | 10 |
| - | - | 780.9 | 663.3 | - | - | 0 | - |
| - | - | 1169 | 665.3 | - | - | 0 | - |
| - | - | 976.4 | 666.2 | - | - | 0 | - |
| - | - | 2635 | 666.3 | - | - | 0 | - |
| - | - | 2006 | 666.3 | - | - | 0 | - |
| - | - | 4612 | 669.4 | - | - | 0 | - |
| - | - | 3895 | 669.9 | - | - | 0 | - |
| 6 | c | 2505 | 670.4 | 0.00116 | 1.731 | +1 | 6 |
| - | - | 1404 | 670.9 | - | - | 0 | - |
| - | - | 1244 | 674.4 | - | - | 0 | - |
| - | - | 1374 | 675.4 | - | - | 0 | - |
| - | - | 761.2 | 675.8 | - | - | 0 | - |
| 12 | y | 7506 | 676.4 | 0.002632 | 3.891 | +1 | 5 |
| - | - | 3128 | 677.4 | - | - | 0 | - |
| - | - | 974.1 | 678.4 | - | - | 0 | - |
| - | - | 1212 | 679.4 | - | - | 0 | - |
| - | - | 1284 | 684.3 | - | - | 0 | - |
| - | - | 923.2 | 685.3 | - | - | 0 | - |
| - | - | 2560 | 687.4 | - | - | 0 | - |
| 6 | c | 3961 | 688.4 | 0.0003113 | 0.4522 | +1 | 6 |
| - | - | 2019 | 689.4 | - | - | 0 | - |
| - | - | 1104 | 693.3 | - | - | 0 | - |
| - | - | 689.8 | 696.3 | - | - | 0 | - |
| - | - | 2005 | 696.4 | - | - | 0 | - |
| - | - | 1785 | 696.9 | - | - | 0 | - |
| - | - | 2015 | 697.4 | - | - | 0 | - |
| - | - | 2281 | 701.4 | - | - | 0 | - |
| - | - | 817.3 | 702.4 | - | - | 0 | - |
| - | - | 8770 | 707.3 | - | - | 0 | - |
| - | - | 7051 | 707.8 | - | - | 0 | - |
| - | - | 2104 | 708.3 | - | - | 0 | - |
| - | - | 1547 | 710.4 | - | - | 0 | - |
| - | - | 1962 | 710.9 | - | - | 0 | - |
| 6 | y | 8808 | 711.3 | 0.0007225 | 1.016 | +2 | 11 |
| - | - | 1270 | 711.4 | - | - | 0 | - |
| - | - | 5226 | 711.8 | - | - | 0 | - |
| - | - | 2576 | 712.3 | - | - | 0 | - |
| - | - | 731.1 | 712.8 | - | - | 0 | - |
| - | - | 4481 | 713.4 | - | - | 0 | - |
| - | - | 1227 | 714.4 | - | - | 0 | - |
| - | - | 620 | 715.4 | - | - | 0 | - |
| - | - | 1034 | 717.4 | - | - | 0 | - |
| - | - | 752.7 | 718.4 | - | - | 0 | - |
| - | - | 4539 | 718.9 | - | - | 0 | - |
| - | - | 3336 | 719.4 | - | - | 0 | - |
| - | - | 1772 | 719.9 | - | - | 0 | - |
| - | - | 1813 | 727.4 | - | - | 0 | - |
| - | - | 5798 | 727.9 | - | - | 0 | - |
| - | - | 4436 | 728.4 | - | - | 0 | - |
| - | - | 2458 | 728.9 | - | - | 0 | - |
| - | - | 779.5 | 729.9 | - | - | 0 | - |
| - | - | 1921 | 732.4 | - | - | 0 | - |
| - | - | 914.5 | 739.3 | - | - | 0 | - |
| - | - | 810.3 | 739.9 | - | - | 0 | - |
| - | - | 773.6 | 740.1 | - | - | 0 | - |
| - | - | 648.5 | 741.9 | - | - | 0 | - |
| - | - | 1921 | 743.4 | - | - | 0 | - |
| - | - | 1.122E+04 | 744.4 | - | - | 0 | - |
| - | - | 2.629E+04 | 745.4 | - | - | 0 | - |
| - | - | 1.083E+04 | 746.4 | - | - | 0 | - |
| - | - | 2656 | 747.4 | - | - | 0 | - |
| 5 | y | 2077 | 750.9 | 0.00422 | 5.62 | +2 | 12 |
| - | - | 2821 | 751.1 | - | - | 0 | - |
| - | - | 2192 | 751.2 | - | - | 0 | - |
| 5 | y | 3439 | 751.4 | 0.005376 | 7.156 | +2 | 12 |
| - | - | 7952 | 751.4 | - | - | 0 | - |
| - | - | 1121 | 751.6 | - | - | 0 | - |
| - | - | 8126 | 751.8 | - | - | 0 | - |
| 5 | z | 1610 | 751.9 | 0.002809 | 3.736 | +2 | 12 |
| - | - | 5033 | 752.1 | - | - | 0 | - |
| - | - | 1024 | 752.4 | - | - | 0 | - |
| - | - | 3220 | 752.4 | - | - | 0 | - |
| - | - | 810.7 | 752.9 | - | - | 0 | - |
| - | - | 767.1 | 756.1 | - | - | 0 | - |
| - | - | 972.2 | 756.4 | - | - | 0 | - |
| - | - | 908.5 | 758.9 | - | - | 0 | - |
| - | - | 1510 | 759.4 | - | - | 0 | - |
| 5 | y | 1.269E+05 | 759.9 | 0.001806 | 2.377 | +2 | 12 |
| - | - | 9.512E+04 | 760.4 | - | - | 0 | - |
| - | - | 4.909E+04 | 760.9 | - | - | 0 | - |
| - | - | 1.843E+04 | 761.4 | - | - | 0 | - |
| - | - | 2553 | 761.9 | - | - | 0 | - |
| - | - | 1405 | 763.9 | - | - | 0 | - |
| - | - | 1110 | 764.4 | - | - | 0 | - |
| - | - | 1727 | 764.9 | - | - | 0 | - |
| - | - | 1469 | 767.5 | - | - | 0 | - |
| - | - | 1449 | 768.5 | - | - | 0 | - |
| - | - | 1229 | 772.9 | - | - | 0 | - |
| - | - | 859.8 | 773.9 | - | - | 0 | - |
| - | - | 1688 | 774.4 | - | - | 0 | - |
| 7 | c | 2.054E+04 | 775.4 | 0.001596 | 2.058 | +1 | 7 |
| - | - | 1476 | 775.6 | - | - | 0 | - |
| - | - | 2759 | 775.9 | - | - | 0 | - |
| - | - | 1244 | 776.2 | - | - | 0 | - |
| - | - | 8375 | 776.4 | - | - | 0 | - |
| - | - | 2096 | 777.4 | - | - | 0 | - |
| - | - | 1279 | 778.4 | - | - | 0 | - |
| - | - | 1159 | 779.7 | - | - | 0 | - |
| - | - | 940.9 | 780.2 | - | - | 0 | - |
| - | - | 1235 | 781.4 | - | - | 0 | - |
| - | - | 1.248E+04 | 782.9 | - | - | 0 | - |
| - | - | 2972 | 783.2 | - | - | 0 | - |
| 13 | c | 2.833E+04 | 783.4 | 0.004307 | 5.498 | +2 | 13 |
| - | - | 1055 | 783.5 | - | - | 0 | - |
| - | - | 1764 | 783.7 | - | - | 0 | - |
| - | - | 1.619E+04 | 783.9 | - | - | 0 | - |
| - | - | 9301 | 784.4 | - | - | 0 | - |
| - | - | 4714 | 784.9 | - | - | 0 | - |
| - | - | 1555 | 785.4 | - | - | 0 | - |
| - | - | 836.9 | 786.4 | - | - | 0 | - |
| 4 | w | 2.412E+04 | 786.9 | 0.001773 | 2.253 | +2 | 13 |
| - | - | 2.24E+04 | 787.4 | - | - | 0 | - |
| - | - | 1.529E+04 | 787.9 | - | - | 0 | - |
| - | - | 5577 | 788.4 | - | - | 0 | - |
| - | - | 2016 | 788.9 | - | - | 0 | - |
| 11 | z | 1133 | 789.4 | 0.002302 | 2.916 | +1 | 6 |
| - | - | 1215 | 792.7 | - | - | 0 | - |
| - | - | 908.7 | 792.9 | - | - | 0 | - |
| - | - | 5644 | 794.1 | - | - | 0 | - |
| - | - | 1.506E+04 | 794.4 | - | - | 0 | - |
| - | - | 1.189E+04 | 794.8 | - | - | 0 | - |
| - | - | 7122 | 795.1 | - | - | 0 | - |
| - | - | 4114 | 795.4 | - | - | 0 | - |
| - | - | 905.7 | 795.8 | - | - | 0 | - |
| - | - | 964.8 | 797.4 | - | - | 0 | - |
| - | - | 899.5 | 797.7 | - | - | 0 | - |
| - | - | 1081 | 799.1 | - | - | 0 | - |
| - | - | 2223 | 799.4 | - | - | 0 | - |
| - | - | 1191 | 799.7 | - | - | 0 | - |
| - | - | 3274 | 800.4 | - | - | 0 | - |
| - | - | 5232 | 801 | - | - | 0 | - |
| - | - | 1467 | 801.2 | - | - | 0 | - |
| - | - | 2434 | 801.5 | - | - | 0 | - |
| - | - | 780.6 | 802 | - | - | 0 | - |
| - | - | 839.5 | 802.7 | - | - | 0 | - |
| - | - | 3172 | 803.4 | - | - | 0 | - |
| - | - | 2118 | 803.7 | - | - | 0 | - |
| - | - | 2765 | 803.9 | - | - | 0 | - |
| - | - | 2212 | 804.2 | - | - | 0 | - |
| - | - | 2232 | 804.4 | - | - | 0 | - |
| - | - | 1778 | 804.9 | - | - | 0 | - |
| 11 | y | 2181 | 805.4 | 0.01425 | 17.7 | +1 | 6 |
| - | - | 781.3 | 806.9 | - | - | 0 | - |
| 4 | y | 6062 | 807.4 | 0.01535 | 19.01 | +2 | 13 |
| - | - | 3670 | 807.7 | - | - | 0 | - |
| - | - | 4822 | 807.9 | - | - | 0 | - |
| - | - | 3788 | 808.2 | - | - | 0 | - |
| 4 | z | 3.693E+04 | 808.4 | 0.0001019 | 0.126 | +2 | 13 |
| - | - | 2309 | 808.7 | - | - | 0 | - |
| - | - | 2.52E+04 | 808.9 | - | - | 0 | - |
| - | - | 1.607E+04 | 809.4 | - | - | 0 | - |
| - | - | 5289 | 809.9 | - | - | 0 | - |
| - | - | 3901 | 810.4 | - | - | 0 | - |
| - | - | 863.5 | 810.9 | - | - | 0 | - |
| - | - | 2541 | 811.4 | - | - | 0 | - |
| - | - | 1610 | 811.7 | - | - | 0 | - |
| - | - | 2118 | 811.9 | - | - | 0 | - |
| - | - | 2042 | 812.2 | - | - | 0 | - |
| - | - | 3778 | 812.4 | - | - | 0 | - |
| - | - | 2750 | 812.7 | - | - | 0 | - |
| - | - | 1721 | 812.9 | - | - | 0 | - |
| - | - | 1089 | 813.2 | - | - | 0 | - |
| - | - | 1191 | 813.4 | - | - | 0 | - |
| - | - | 1843 | 814.4 | - | - | 0 | - |
| - | - | 2172 | 814.9 | - | - | 0 | - |
| - | - | 738.8 | 815.2 | - | - | 0 | - |
| - | - | 2234 | 815.4 | - | - | 0 | - |
| - | - | 3282 | 815.7 | - | - | 0 | - |
| - | - | 3677 | 815.9 | - | - | 0 | - |
| - | - | 2884 | 816.2 | - | - | 0 | - |
| 4 | y | 1.857E+04 | 816.4 | 0.002987 | 3.659 | +2 | 13 |
| - | - | 1342 | 816.7 | - | - | 0 | - |
| - | - | 1.868E+04 | 816.9 | - | - | 0 | - |
| - | - | 7529 | 817.4 | - | - | 0 | - |
| - | - | 4169 | 817.9 | - | - | 0 | - |
| - | - | 1107 | 818.2 | - | - | 0 | - |
| - | - | 4645 | 818.4 | - | - | 0 | - |
| - | - | 4585 | 818.7 | - | - | 0 | - |
| - | - | 7269 | 818.9 | - | - | 0 | - |
| - | - | 5639 | 819.2 | - | - | 0 | - |
| - | - | 6114 | 819.4 | - | - | 0 | - |
| - | - | 2213 | 819.7 | - | - | 0 | - |
| - | - | 1742 | 819.9 | - | - | 0 | - |
| - | - | 1043 | 820.2 | - | - | 0 | - |
| - | - | 1185 | 821.7 | - | - | 0 | - |
| - | - | 1398 | 821.9 | - | - | 0 | - |
| - | - | 991.9 | 822.4 | - | - | 0 | - |
| 11 | y | 3883 | 823.4 | 0.00204 | 2.477 | +1 | 6 |
| - | - | 1648 | 824.4 | - | - | 0 | - |
| - | - | 872.3 | 825.2 | - | - | 0 | - |
| - | - | 947.7 | 825.4 | - | - | 0 | - |
| - | - | 4514 | 825.7 | - | - | 0 | - |
| - | - | 1.094E+04 | 825.9 | - | - | 0 | - |
| - | - | 4959 | 826.2 | - | - | 0 | - |
| - | - | 5559 | 826.4 | - | - | 0 | - |
| - | - | 2341 | 826.7 | - | - | 0 | - |
| - | - | 893.1 | 827.2 | - | - | 0 | - |
| - | - | 3255 | 827.5 | - | - | 0 | - |
| - | - | 1396 | 828.5 | - | - | 0 | - |
| - | - | 1808 | 829.4 | - | - | 0 | - |
| - | - | 1.147E+04 | 829.7 | - | - | 0 | - |
| - | - | 3.138E+04 | 829.9 | - | - | 0 | - |
| - | - | 2.969E+04 | 830.2 | - | - | 0 | - |
| - | - | 2.19E+04 | 830.4 | - | - | 0 | - |
| - | - | 1.04E+04 | 830.7 | - | - | 0 | - |
| - | - | 2962 | 830.9 | - | - | 0 | - |
| - | - | 3393 | 830.9 | - | - | 0 | - |
| - | - | 1579 | 831.2 | - | - | 0 | - |
| - | - | 4101 | 831.4 | - | - | 0 | - |
| 14 | c | 2985 | 831.9 | 0.01444 | 17.36 | +2 | 14 |
| - | - | 749.8 | 832.2 | - | - | 0 | - |
| - | - | 1306 | 832.9 | - | - | 0 | - |
| - | - | 3582 | 833.2 | - | - | 0 | - |
| - | - | 4738 | 833.4 | - | - | 0 | - |
| - | - | 2691 | 833.7 | - | - | 0 | - |
| - | - | 2738 | 833.9 | - | - | 0 | - |
| - | - | 1410 | 834.5 | - | - | 0 | - |
| - | - | 2127 | 834.9 | - | - | 0 | - |
| - | - | 2081 | 835.4 | - | - | 0 | - |
| - | - | 5679 | 837.1 | - | - | 0 | - |
| - | - | 1.234E+04 | 837.5 | - | - | 0 | - |
| - | - | 1.357E+04 | 837.8 | - | - | 0 | - |
| - | - | 8767 | 838.1 | - | - | 0 | - |
| - | - | 1922 | 838.5 | - | - | 0 | - |
| - | - | 768.5 | 838.8 | - | - | 0 | - |
| - | - | 2880 | 839.9 | - | - | 0 | - |
| 14 | c | 4.545E+04 | 840.4 | 0.004023 | 4.786 | +2 | 14 |
| - | - | 4.144E+04 | 840.9 | - | - | 0 | - |
| - | - | 2.43E+04 | 841.4 | - | - | 0 | - |
| - | - | 8695 | 841.9 | - | - | 0 | - |
| - | - | 2353 | 842.4 | - | - | 0 | - |
| - | - | 906.2 | 842.8 | - | - | 0 | - |
| - | - | 1269 | 843 | - | - | 0 | - |
| - | - | 1059 | 843.5 | - | - | 0 | - |
| - | - | 687.9 | 844.9 | - | - | 0 | - |
| - | - | 646.7 | 845.9 | - | - | 0 | - |
| - | - | 1659 | 849.4 | - | - | 0 | - |
| - | - | 9653 | 851 | - | - | 0 | - |
| - | - | 1.688E+04 | 851.5 | - | - | 0 | - |
| - | - | 9049 | 852 | - | - | 0 | - |
| - | - | 4823 | 852.5 | - | - | 0 | - |
| - | - | 2451 | 853 | - | - | 0 | - |
| - | - | 924.8 | 853.4 | - | - | 0 | - |
| - | - | 1347 | 855.5 | - | - | 0 | - |
| - | - | 851.6 | 857.5 | - | - | 0 | - |
| 3 | y | 1201 | 857.9 | 0.01166 | 13.59 | +2 | 14 |
| 3 | y | 7724 | 858.4 | 0.003958 | 4.611 | +2 | 14 |
| 3 | z | 2.221E+04 | 858.9 | 0.001388 | 1.617 | +2 | 14 |
| - | - | 2.037E+04 | 859.4 | - | - | 0 | - |
| - | - | 1.026E+04 | 859.9 | - | - | 0 | - |
| - | - | 5351 | 860.4 | - | - | 0 | - |
| - | - | 1184 | 860.9 | - | - | 0 | - |
| - | - | 1200 | 865.9 | - | - | 0 | - |
| - | - | 1360 | 866.4 | - | - | 0 | - |
| 3 | y | 5.59E+04 | 866.9 | 0.00228 | 2.63 | +2 | 14 |
| - | - | 4.719E+04 | 867.4 | - | - | 0 | - |
| - | - | 2.579E+04 | 867.9 | - | - | 0 | - |
| - | - | 1257 | 868.1 | - | - | 0 | - |
| - | - | 1.291E+04 | 868.4 | - | - | 0 | - |
| - | - | 2339 | 868.9 | - | - | 0 | - |
| - | - | 1863 | 870.9 | - | - | 0 | - |
| - | - | 2452 | 871.4 | - | - | 0 | - |
| - | - | 3154 | 871.9 | - | - | 0 | - |
| - | - | 1197 | 872.1 | - | - | 0 | - |
| - | - | 2202 | 872.5 | - | - | 0 | - |
| - | - | 3531 | 873.4 | - | - | 0 | - |
| - | - | 2923 | 874.4 | - | - | 0 | - |
| - | - | 1216 | 875.4 | - | - | 0 | - |
| - | - | 802.5 | 876.4 | - | - | 0 | - |
| - | - | 841.1 | 877.1 | - | - | 0 | - |
| 10 | w | 1.556E+04 | 877.4 | 0.003255 | 3.71 | +1 | 7 |
| - | - | 9908 | 878.4 | - | - | 0 | - |
| - | - | 3926 | 879.4 | - | - | 0 | - |
| - | - | 2706 | 880.4 | - | - | 0 | - |
| - | - | 1062 | 880.8 | - | - | 0 | - |
| - | - | 1321 | 880.9 | - | - | 0 | - |
| - | - | 1039 | 881.1 | - | - | 0 | - |
| - | - | 2795 | 881.4 | - | - | 0 | - |
| - | - | 1346 | 881.8 | - | - | 0 | - |
| - | - | 1273 | 881.9 | - | - | 0 | - |
| - | - | 739.6 | 882.1 | - | - | 0 | - |
| - | - | 847.2 | 882.4 | - | - | 0 | - |
| - | - | 789.6 | 885.8 | - | - | 0 | - |
| - | - | 892.1 | 886 | - | - | 0 | - |
| - | - | 3654 | 886.1 | - | - | 0 | - |
| - | - | 1.386E+04 | 886.5 | - | - | 0 | - |
| - | - | 2.03E+04 | 886.8 | - | - | 0 | - |
| - | - | 1.432E+04 | 887.1 | - | - | 0 | - |
| - | - | 5074 | 887.5 | - | - | 0 | - |
| - | - | 3019 | 887.8 | - | - | 0 | - |
| - | - | 1218 | 888.5 | - | - | 0 | - |
| - | - | 784.7 | 889.4 | - | - | 0 | - |
| - | - | 998.9 | 890.8 | - | - | 0 | - |
| - | - | 1348 | 891.5 | - | - | 0 | - |
| - | - | 923.9 | 891.8 | - | - | 0 | - |
| 15 | c | 2177 | 895.9 | 0.003694 | 4.124 | +2 | 15 |
| - | - | 1102 | 896.1 | - | - | 0 | - |
| - | - | 5018 | 896.5 | - | - | 0 | - |
| - | - | 739.1 | 896.8 | - | - | 0 | - |
| - | - | 3202 | 897 | - | - | 0 | - |
| - | - | 1668 | 897.5 | - | - | 0 | - |
| - | - | 1375 | 899 | - | - | 0 | - |
| - | - | 3126 | 899.5 | - | - | 0 | - |
| - | - | 1078 | 899.8 | - | - | 0 | - |
| - | - | 3367 | 900 | - | - | 0 | - |
| - | - | 1233 | 900.5 | - | - | 0 | - |
| - | - | 1374 | 900.8 | - | - | 0 | - |
| - | - | 3981 | 901.1 | - | - | 0 | - |
| - | - | 4095 | 901.4 | - | - | 0 | - |
| - | - | 3746 | 901.8 | - | - | 0 | - |
| - | - | 1513 | 902.1 | - | - | 0 | - |
| - | - | 1254 | 903.8 | - | - | 0 | - |
| - | - | 888.4 | 904 | - | - | 0 | - |
| 15 | c | 1.887E+04 | 904.5 | 0.005129 | 5.671 | +2 | 15 |
| - | - | 1205 | 904.8 | - | - | 0 | - |
| - | - | 1.519E+04 | 905 | - | - | 0 | - |
| - | - | 1908 | 905.1 | - | - | 0 | - |
| - | - | 1.341E+04 | 905.5 | - | - | 0 | - |
| - | - | 6317 | 905.8 | - | - | 0 | - |
| - | - | 4426 | 906 | - | - | 0 | - |
| - | - | 7162 | 906.1 | - | - | 0 | - |
| - | - | 6817 | 906.5 | - | - | 0 | - |
| - | - | 3169 | 906.8 | - | - | 0 | - |
| - | - | 1055 | 907.1 | - | - | 0 | - |
| - | - | 923.6 | 910 | - | - | 0 | - |
| - | - | 3424 | 910.5 | - | - | 0 | - |
| - | - | 831.1 | 910.8 | - | - | 0 | - |
| - | - | 1210 | 911 | - | - | 0 | - |
| - | - | 993.2 | 911.1 | - | - | 0 | - |
| - | - | 2168 | 911.5 | - | - | 0 | - |
| - | - | 2265 | 912.4 | - | - | 0 | - |
| - | - | 2496 | 912.9 | - | - | 0 | - |
| - | - | 1483 | 913.4 | - | - | 0 | - |
| - | - | 840.2 | 918.8 | - | - | 0 | - |
| - | - | 1217 | 919.4 | - | - | 0 | - |
| - | - | 835.4 | 919.8 | - | - | 0 | - |
| - | - | 6006 | 920.2 | - | - | 0 | - |
| - | - | 9912 | 920.5 | - | - | 0 | - |
| - | - | 5761 | 920.8 | - | - | 0 | - |
| - | - | 3621 | 921.2 | - | - | 0 | - |
| - | - | 938.9 | 921.5 | - | - | 0 | - |
| - | - | 986.4 | 923 | - | - | 0 | - |
| - | - | 3307 | 924.5 | - | - | 0 | - |
| - | - | 1.157E+04 | 924.8 | - | - | 0 | - |
| - | - | 1.731E+04 | 925.1 | - | - | 0 | - |
| - | - | 1.122E+04 | 925.5 | - | - | 0 | - |
| - | - | 5589 | 925.8 | - | - | 0 | - |
| - | - | 1021 | 926.1 | - | - | 0 | - |
| - | - | 1763 | 926.5 | - | - | 0 | - |
| - | - | 1096 | 929.1 | - | - | 0 | - |
| - | - | 1554 | 929.8 | - | - | 0 | - |
| - | - | 4902 | 930.1 | - | - | 0 | - |
| - | - | 5408 | 930.5 | - | - | 0 | - |
| - | - | 3062 | 930.8 | - | - | 0 | - |
| - | - | 900.5 | 931 | - | - | 0 | - |
| - | - | 1565 | 931.1 | - | - | 0 | - |
| 8 | c | 3.028E+04 | 931.5 | 0.002169 | 2.329 | +1 | 8 |
| - | - | 4868 | 932 | - | - | 0 | - |
| - | - | 1.388E+04 | 932.5 | - | - | 0 | - |
| - | - | 3953 | 933 | - | - | 0 | - |
| - | - | 3112 | 933.5 | - | - | 0 | - |
| - | - | 807.6 | 934 | - | - | 0 | - |
| - | - | 1471 | 934.5 | - | - | 0 | - |
| 10 | z | 3.408E+04 | 936.4 | 0.002768 | 2.956 | +1 | 7 |
| - | - | 2.004E+04 | 937.4 | - | - | 0 | - |
| - | - | 9477 | 938.4 | - | - | 0 | - |
| - | - | 887.4 | 939.2 | - | - | 0 | - |
| - | - | 2668 | 939.4 | - | - | 0 | - |
| 2 | y | 2173 | 939.9 | 0.01728 | 18.38 | +2 | 15 |
| 2 | z | 1.232E+04 | 940.5 | 0.003721 | 3.956 | +2 | 15 |
| - | - | 1.638E+04 | 941 | - | - | 0 | - |
| - | - | 1.235E+04 | 941.5 | - | - | 0 | - |
| - | - | 6102 | 942 | - | - | 0 | - |
| - | - | 2288 | 942.5 | - | - | 0 | - |
| - | - | 2190 | 943.8 | - | - | 0 | - |
| - | - | 3099 | 944.1 | - | - | 0 | - |
| - | - | 1071 | 944.5 | - | - | 0 | - |
| - | - | 2796 | 945.5 | - | - | 0 | - |
| - | - | 1729 | 946 | - | - | 0 | - |
| - | - | 5093 | 946.5 | - | - | 0 | - |
| - | - | 3778 | 947 | - | - | 0 | - |
| - | - | 3587 | 947.5 | - | - | 0 | - |
| - | - | 2714 | 948 | - | - | 0 | - |
| 2 | y | 8428 | 948.5 | 0.00266 | 2.804 | +2 | 15 |
| - | - | 851.6 | 948.8 | - | - | 0 | - |
| - | - | 6506 | 949 | - | - | 0 | - |
| - | - | 835.5 | 949.2 | - | - | 0 | - |
| - | - | 3679 | 949.5 | - | - | 0 | - |
| - | - | 2145 | 950 | - | - | 0 | - |
| 10 | y | 3049 | 952.4 | 0.005162 | 5.42 | +1 | 7 |
| - | - | 1905 | 953 | - | - | 0 | - |
| - | - | 873.6 | 953.2 | - | - | 0 | - |
| - | - | 3332 | 953.5 | - | - | 0 | - |
| - | - | 4.66E+04 | 954 | - | - | 0 | - |
| - | - | 5.19E+04 | 954.5 | - | - | 0 | - |
| - | - | 3.279E+04 | 955 | - | - | 0 | - |
| - | - | 1.542E+04 | 955.5 | - | - | 0 | - |
| - | - | 6621 | 956 | - | - | 0 | - |
| - | - | 1024 | 958.5 | - | - | 0 | - |
| - | - | 1217 | 960.5 | - | - | 0 | - |
| - | - | 1477 | 961 | - | - | 0 | - |
| - | - | 2042 | 961.5 | - | - | 0 | - |
| - | - | 9768 | 962 | - | - | 0 | - |
| - | - | 915 | 962.1 | - | - | 0 | - |
| - | - | 9636 | 962.5 | - | - | 0 | - |
| - | - | 9611 | 962.9 | - | - | 0 | - |
| - | - | 7249 | 963 | - | - | 0 | - |
| - | - | 1.461E+04 | 963.2 | - | - | 0 | - |
| - | - | 1.148E+04 | 963.5 | - | - | 0 | - |
| - | - | 4153 | 963.9 | - | - | 0 | - |
| - | - | 1737 | 964 | - | - | 0 | - |
| - | - | 3243 | 964.2 | - | - | 0 | - |
| - | - | 1374 | 964.5 | - | - | 0 | - |
| - | - | 929.9 | 964.9 | - | - | 0 | - |
| - | - | 2316 | 966.5 | - | - | 0 | - |
| - | - | 1867 | 967 | - | - | 0 | - |
| - | - | 1773 | 967.2 | - | - | 0 | - |
| - | - | 9607 | 967.5 | - | - | 0 | - |
| - | - | 883.7 | 967.6 | - | - | 0 | - |
| - | - | 7370 | 967.8 | - | - | 0 | - |
| - | - | 1233 | 968 | - | - | 0 | - |
| - | - | 8586 | 968.2 | - | - | 0 | - |
| - | - | 5338 | 968.5 | - | - | 0 | - |
| - | - | 1187 | 968.6 | - | - | 0 | - |
| - | - | 1457 | 968.8 | - | - | 0 | - |
| - | - | 7182 | 969 | - | - | 0 | - |
| - | - | 753.8 | 969.2 | - | - | 0 | - |
| - | - | 7658 | 969.5 | - | - | 0 | - |
| - | - | 963 | 969.6 | - | - | 0 | - |
| - | - | 4391 | 970 | - | - | 0 | - |
| - | - | 4961 | 970.5 | - | - | 0 | - |
| - | - | 3067 | 971 | - | - | 0 | - |
| - | - | 2459 | 971.5 | - | - | 0 | - |
| - | - | 1086 | 971.8 | - | - | 0 | - |
| - | - | 1290 | 972 | - | - | 0 | - |
| - | - | 1553 | 972.2 | - | - | 0 | - |
| - | - | 3503 | 972.5 | - | - | 0 | - |
| - | - | 4160 | 972.8 | - | - | 0 | - |
| - | - | 7551 | 973.2 | - | - | 0 | - |
| - | - | 3561 | 973.5 | - | - | 0 | - |
| - | - | 1995 | 973.8 | - | - | 0 | - |
| - | - | 1323 | 975 | - | - | 0 | - |
| - | - | 2564 | 975.5 | - | - | 0 | - |
| - | - | 1.873E+04 | 976 | - | - | 0 | - |
| - | - | 1.599E+04 | 976.5 | - | - | 0 | - |
| - | - | 1.51E+04 | 977 | - | - | 0 | - |
| - | - | 978.7 | 977.2 | - | - | 0 | - |
| - | - | 7609 | 977.5 | - | - | 0 | - |
| - | - | 1571 | 978 | - | - | 0 | - |
| - | - | 962.5 | 978.5 | - | - | 0 | - |
| - | - | 773.4 | 980.5 | - | - | 0 | - |
| - | - | 3190 | 981 | - | - | 0 | - |
| - | - | 2915 | 981.5 | - | - | 0 | - |
| - | - | 2747 | 982 | - | - | 0 | - |
| - | - | 2622 | 982.5 | - | - | 0 | - |
| - | - | 1997 | 983 | - | - | 0 | - |
| - | - | 852 | 983.5 | - | - | 0 | - |
| - | - | 1.511E+04 | 983.6 | - | - | 0 | - |
| - | - | 1.363E+04 | 984.6 | - | - | 0 | - |
| - | - | 4636 | 985.6 | - | - | 0 | - |
| - | - | 1005 | 986.5 | - | - | 0 | - |
| - | - | 2007 | 986.6 | - | - | 0 | - |
| - | - | 1905 | 989 | - | - | 0 | - |
| - | - | 1.129E+04 | 989.5 | - | - | 0 | - |
| - | - | 6.298E+04 | 990 | - | - | 0 | - |
| - | - | 6.835E+04 | 990.5 | - | - | 0 | - |
| - | - | 5.173E+04 | 991 | - | - | 0 | - |
| - | - | 3885 | 991.2 | - | - | 0 | - |
| - | - | 2.784E+04 | 991.5 | - | - | 0 | - |
| - | - | 6709 | 991.8 | - | - | 0 | - |
| - | - | 8331 | 992 | - | - | 0 | - |
| - | - | 2216 | 992.2 | - | - | 0 | - |
| - | - | 2573 | 992.5 | - | - | 0 | - |
| - | - | 1108 | 993.4 | - | - | 0 | - |
| - | - | 1119 | 994.4 | - | - | 0 | - |
| - | - | 2490 | 996.5 | - | - | 0 | - |
| - | - | 1638 | 996.8 | - | - | 0 | - |
| - | - | 1358 | 997.2 | - | - | 0 | - |
| - | - | 1916 | 997.5 | - | - | 0 | - |
| - | - | 4.772E+04 | 998 | - | - | 0 | - |
| - | - | 9.448E+04 | 998.5 | - | - | 0 | - |
| - | - | 8.224E+04 | 999 | - | - | 0 | - |
| - | - | 4.433E+04 | 999.5 | - | - | 0 | - |
| - | - | 2.012E+04 | 1000 | - | - | 0 | - |
| - | - | 6503 | 1001 | - | - | 0 | - |
| - | - | 1.45E+04 | 1001 | - | - | 0 | - |
| - | - | 1.855E+04 | 1001 | - | - | 0 | - |
| - | - | 1.33E+04 | 1002 | - | - | 0 | - |
| - | - | 8337 | 1002 | - | - | 0 | - |
| - | - | 3854 | 1002 | - | - | 0 | - |
| - | - | 2118 | 1003 | - | - | 0 | - |
| - | - | 2382 | 1005 | - | - | 0 | - |
| - | - | 7351 | 1005 | - | - | 0 | - |
| - | - | 1869 | 1005 | - | - | 0 | - |
| - | - | 5761 | 1006 | - | - | 0 | - |
| - | - | 2297 | 1006 | - | - | 0 | - |
| - | - | 2901 | 1006 | - | - | 0 | - |
| 9 | w | 7444 | 1006 | 0.003203 | 3.183 | +1 | 8 |
| - | - | 1143 | 1007 | - | - | 0 | - |
| - | - | 4268 | 1007 | - | - | 0 | - |
| - | - | 2402 | 1008 | - | - | 0 | - |
| - | - | 2584 | 1010 | - | - | 0 | - |
| - | - | 3965 | 1011 | - | - | 0 | - |
| - | - | 1624 | 1011 | - | - | 0 | - |
| - | - | 1621 | 1011 | - | - | 0 | - |
| - | - | 1350 | 1012 | - | - | 0 | - |
| - | - | 983.1 | 1013 | - | - | 0 | - |
| - | - | 2021 | 1014 | - | - | 0 | - |
| - | - | 1196 | 1015 | - | - | 0 | - |
| - | - | 952.4 | 1016 | - | - | 0 | - |
| - | - | 1386 | 1018 | - | - | 0 | - |
| - | - | 1097 | 1018 | - | - | 0 | - |
| - | - | 1904 | 1020 | - | - | 0 | - |
| - | - | 1533 | 1020 | - | - | 0 | - |
| - | - | 1278 | 1020 | - | - | 0 | - |
| - | - | 2955 | 1027 | - | - | 0 | - |
| - | - | 2106 | 1027 | - | - | 0 | - |
| - | - | 1174 | 1028 | - | - | 0 | - |
| - | - | 957.4 | 1029 | - | - | 0 | - |
| - | - | 3187 | 1029 | - | - | 0 | - |
| - | - | 1947 | 1029 | - | - | 0 | - |
| - | - | 1681 | 1030 | - | - | 0 | - |
| - | - | 1135 | 1034 | - | - | 0 | - |
| - | - | 1.379E+04 | 1034 | - | - | 0 | - |
| - | - | 3.697E+04 | 1034 | - | - | 0 | - |
| - | - | 4.624E+04 | 1035 | - | - | 0 | - |
| - | - | 2.951E+04 | 1035 | - | - | 0 | - |
| - | - | 1.478E+04 | 1035 | - | - | 0 | - |
| - | - | 7310 | 1036 | - | - | 0 | - |
| - | - | 3295 | 1036 | - | - | 0 | - |
| - | - | 2683 | 1039 | - | - | 0 | - |
| - | - | 2267 | 1039 | - | - | 0 | - |
| - | - | 3918 | 1039 | - | - | 0 | - |
| - | - | 5487 | 1040 | - | - | 0 | - |
| - | - | 2530 | 1040 | - | - | 0 | - |
| 9 | c | 3404 | 1044 | 0.001484 | 1.423 | +1 | 9 |
| - | - | 5851 | 1044 | - | - | 0 | - |
| - | - | 6472 | 1044 | - | - | 0 | - |
| - | - | 3757 | 1045 | - | - | 0 | - |
| - | - | 2200 | 1045 | - | - | 0 | - |
| - | - | 1358 | 1046 | - | - | 0 | - |
| - | - | 6883 | 1048 | - | - | 0 | - |
| - | - | 1.908E+04 | 1049 | - | - | 0 | - |
| - | - | 1.684E+04 | 1049 | - | - | 0 | - |
| - | - | 6375 | 1050 | - | - | 0 | - |
| - | - | 3689 | 1050 | - | - | 0 | - |
| - | - | 1216 | 1051 | - | - | 0 | - |
| - | - | 974 | 1053 | - | - | 0 | - |
| - | - | 3010 | 1055 | - | - | 0 | - |
| - | - | 1789 | 1055 | - | - | 0 | - |
| - | - | 1912 | 1056 | - | - | 0 | - |
| - | - | 1281 | 1057 | - | - | 0 | - |
| - | - | 5335 | 1058 | - | - | 0 | - |
| - | - | 2674 | 1058 | - | - | 0 | - |
| - | - | 2209 | 1059 | - | - | 0 | - |
| 9 | c | 4.204E+04 | 1061 | 0.00285 | 2.687 | +1 | 9 |
| - | - | 2.478E+04 | 1062 | - | - | 0 | - |
| - | - | 1130 | 1062 | - | - | 0 | - |
| - | - | 3354 | 1062 | - | - | 0 | - |
| - | - | 1.105E+04 | 1063 | - | - | 0 | - |
| - | - | 7067 | 1063 | - | - | 0 | - |
| - | - | 1.284E+04 | 1063 | - | - | 0 | - |
| - | - | 1.671E+04 | 1064 | - | - | 0 | - |
| - | - | 1.356E+04 | 1064 | - | - | 0 | - |
| - | - | 6473 | 1064 | - | - | 0 | - |
| - | - | 1791 | 1065 | - | - | 0 | - |
| - | - | 2172 | 1065 | - | - | 0 | - |
| 9 | z | 2.471E+04 | 1065 | 0.003754 | 3.523 | +1 | 8 |
| - | - | 2.02E+04 | 1066 | - | - | 0 | - |
| - | - | 1703 | 1067 | - | - | 0 | - |
| - | - | 8443 | 1067 | - | - | 0 | - |
| - | - | 1865 | 1068 | - | - | 0 | - |
| - | - | 3750 | 1068 | - | - | 0 | - |
| - | - | 1990 | 1068 | - | - | 0 | - |
| - | - | 2243 | 1069 | - | - | 0 | - |
| - | - | 1201 | 1069 | - | - | 0 | - |
| - | - | 1669 | 1072 | - | - | 0 | - |
| - | - | 1838 | 1072 | - | - | 0 | - |
| - | - | 2072 | 1072 | - | - | 0 | - |
| - | - | 2222 | 1077 | - | - | 0 | - |
| - | - | 5853 | 1077 | - | - | 0 | - |
| - | - | 7122 | 1077 | - | - | 0 | - |
| - | - | 5291 | 1078 | - | - | 0 | - |
| - | - | 2433 | 1078 | - | - | 0 | - |
| - | - | 1694 | 1078 | - | - | 0 | - |
| - | - | 985.2 | 1081 | - | - | 0 | - |
| 9 | y | 3354 | 1081 | 0.005049 | 4.669 | +1 | 8 |
| - | - | 1817 | 1082 | - | - | 0 | - |
| - | - | 1850 | 1082 | - | - | 0 | - |
| - | - | 2117 | 1082 | - | - | 0 | - |
| - | - | 2063 | 1083 | - | - | 0 | - |
| - | - | 2492 | 1083 | - | - | 0 | - |
| - | - | 1289 | 1083 | - | - | 0 | - |
| - | - | 1502 | 1083 | - | - | 0 | - |
| - | - | 1600 | 1085 | - | - | 0 | - |
| - | - | 2469 | 1086 | - | - | 0 | - |
| - | - | 4068 | 1086 | - | - | 0 | - |
| - | - | 4541 | 1086 | - | - | 0 | - |
| - | - | 7161 | 1087 | - | - | 0 | - |
| - | - | 9585 | 1087 | - | - | 0 | - |
| - | - | 1.028E+04 | 1087 | - | - | 0 | - |
| - | - | 1.074E+04 | 1088 | - | - | 0 | - |
| - | - | 3583 | 1088 | - | - | 0 | - |
| - | - | 2996 | 1088 | - | - | 0 | - |
| - | - | 1463 | 1089 | - | - | 0 | - |
| - | - | 1238 | 1089 | - | - | 0 | - |
| - | - | 1177 | 1090 | - | - | 0 | - |
| - | - | 1174 | 1090 | - | - | 0 | - |
| - | - | 2303 | 1091 | - | - | 0 | - |
| - | - | 1691 | 1091 | - | - | 0 | - |
| - | - | 4171 | 1091 | - | - | 0 | - |
| - | - | 1.62E+04 | 1092 | - | - | 0 | - |
| - | - | 2.39E+04 | 1092 | - | - | 0 | - |
| - | - | 1.828E+04 | 1092 | - | - | 0 | - |
| - | - | 1.057E+04 | 1093 | - | - | 0 | - |
| - | - | 5437 | 1093 | - | - | 0 | - |
| - | - | 2288 | 1093 | - | - | 0 | - |
| - | - | 2152 | 1095 | - | - | 0 | - |
| - | - | 4880 | 1095 | - | - | 0 | - |
| - | - | 4935 | 1095 | - | - | 0 | - |
| - | - | 5101 | 1096 | - | - | 0 | - |
| - | - | 5941 | 1096 | - | - | 0 | - |
| - | - | 4114 | 1096 | - | - | 0 | - |
| - | - | 5106 | 1097 | - | - | 0 | - |
| - | - | 2839 | 1097 | - | - | 0 | - |
| - | - | 1107 | 1098 | - | - | 0 | - |
| - | - | 2404 | 1100 | - | - | 0 | - |
| - | - | 1716 | 1100 | - | - | 0 | - |
| - | - | 1559 | 1100 | - | - | 0 | - |
| - | - | 8850 | 1101 | - | - | 0 | - |
| - | - | 3.452E+04 | 1101 | - | - | 0 | - |
| - | - | 6.051E+04 | 1101 | - | - | 0 | - |
| - | - | 4.715E+04 | 1102 | - | - | 0 | - |
| - | - | 3.35E+04 | 1102 | - | - | 0 | - |
| - | - | 1.28E+04 | 1102 | - | - | 0 | - |
| - | - | 8037 | 1103 | - | - | 0 | - |
| - | - | 1486 | 1103 | - | - | 0 | - |
| - | - | 1653 | 1103 | - | - | 0 | - |
| - | - | 1511 | 1105 | - | - | 0 | - |
| - | - | 2006 | 1105 | - | - | 0 | - |
| - | - | 6411 | 1106 | - | - | 0 | - |
| - | - | 1.246E+04 | 1106 | - | - | 0 | - |
| - | - | 1.64E+04 | 1106 | - | - | 0 | - |
| - | - | 5.781E+04 | 1107 | - | - | 0 | - |
| - | - | 9.054E+04 | 1107 | - | - | 0 | - |
| - | - | 8.137E+04 | 1107 | - | - | 0 | - |
| - | - | 4.871E+04 | 1108 | - | - | 0 | - |
| - | - | 1.83E+04 | 1108 | - | - | 0 | - |
| - | - | 1.058E+04 | 1108 | - | - | 0 | - |
| - | - | 2389 | 1109 | - | - | 0 | - |
| - | - | 1108 | 1109 | - | - | 0 | - |
| - | - | 1653 | 1110 | - | - | 0 | - |
| - | - | 1963 | 1111 | - | - | 0 | - |
| - | - | 5925 | 1111 | - | - | 0 | - |
| - | - | 1.011E+04 | 1111 | - | - | 0 | - |
| - | - | 1.245E+04 | 1112 | - | - | 0 | - |
| - | - | 5638 | 1112 | - | - | 0 | - |
| - | - | 1420 | 1112 | - | - | 0 | - |
| - | - | 3904 | 1126 | - | - | 0 | - |
| - | - | 9339 | 1127 | - | - | 0 | - |
| - | - | 1.176E+04 | 1127 | - | - | 0 | - |
| - | - | 8506 | 1128 | - | - | 0 | - |
| - | - | 2721 | 1128 | - | - | 0 | - |
| - | - | 1134 | 1135 | - | - | 0 | - |
| - | - | 5924 | 1138 | - | - | 0 | - |
| - | - | 4710 | 1139 | - | - | 0 | - |
| - | - | 3514 | 1139 | - | - | 0 | - |
| - | - | 1306 | 1140 | - | - | 0 | - |
| - | - | 1161 | 1148 | - | - | 0 | - |
| - | - | 7420 | 1167 | - | - | 0 | - |
| - | - | 2.521E+04 | 1168 | - | - | 0 | - |
| - | - | 2.977E+04 | 1168 | - | - | 0 | - |
| - | - | 2.082E+04 | 1169 | - | - | 0 | - |
| - | - | 8462 | 1169 | - | - | 0 | - |
| - | - | 2035 | 1170 | - | - | 0 | - |
| - | - | 2690 | 1175 | - | - | 0 | - |
| - | - | 2324 | 1175 | - | - | 0 | - |
| - | - | 2068 | 1176 | - | - | 0 | - |
| - | - | 1069 | 1180 | - | - | 0 | - |
| - | - | 6876 | 1189 | - | - | 0 | - |
| 10 | c | 3.414E+04 | 1190 | 0.00286 | 2.404 | +1 | 10 |
| - | - | 2.578E+04 | 1191 | - | - | 0 | - |
| - | - | 9475 | 1191 | - | - | 0 | - |
| - | - | 1.847E+04 | 1192 | - | - | 0 | - |
| - | - | 9029 | 1192 | - | - | 0 | - |
| - | - | 5048 | 1193 | - | - | 0 | - |
| - | - | 1308 | 1193 | - | - | 0 | - |
| - | - | 976 | 1199 | - | - | 0 | - |
| - | - | 1051 | 1210 | - | - | 0 | - |
| - | - | 1813 | 1211 | - | - | 0 | - |
| - | - | 1543 | 1221 | - | - | 0 | - |
| 8 | z | 1.876E+04 | 1222 | 0.003595 | 2.943 | +1 | 9 |
| - | - | 2.068E+04 | 1223 | - | - | 0 | - |
| - | - | 1.426E+04 | 1224 | - | - | 0 | - |
| - | - | 5865 | 1225 | - | - | 0 | - |
| - | - | 2402 | 1226 | - | - | 0 | - |
| - | - | 1616 | 1228 | - | - | 0 | - |
| - | - | 1169 | 1231 | - | - | 0 | - |
| - | - | 1370 | 1234 | - | - | 0 | - |
| - | - | 1047 | 1234 | - | - | 0 | - |
| - | - | 1261 | 1235 | - | - | 0 | - |
| - | - | 2283 | 1237 | - | - | 0 | - |
| 8 | y | 8348 | 1238 | 0.002083 | 1.683 | +1 | 9 |
| - | - | 6360 | 1239 | - | - | 0 | - |
| - | - | 2922 | 1240 | - | - | 0 | - |
| - | - | 1948 | 1246 | - | - | 0 | - |
| - | - | 2456 | 1246 | - | - | 0 | - |
| - | - | 2279 | 1247 | - | - | 0 | - |
| - | - | 1730 | 1247 | - | - | 0 | - |
| - | - | 1049 | 1248 | - | - | 0 | - |
| - | - | 1041 | 1248 | - | - | 0 | - |
| - | - | 3003 | 1255 | - | - | 0 | - |
| - | - | 8806 | 1256 | - | - | 0 | - |
| - | - | 1.104E+04 | 1256 | - | - | 0 | - |
| - | - | 7905 | 1257 | - | - | 0 | - |
| - | - | 5620 | 1257 | - | - | 0 | - |
| - | - | 1616 | 1258 | - | - | 0 | - |
| - | - | 1197 | 1272 | - | - | 0 | - |
| - | - | 1113 | 1273 | - | - | 0 | - |
| - | - | 4074 | 1293 | - | - | 0 | - |
| - | - | 4419 | 1294 | - | - | 0 | - |
| - | - | 2849 | 1295 | - | - | 0 | - |
| - | - | 1124 | 1302 | - | - | 0 | - |
| - | - | 2467 | 1302 | - | - | 0 | - |
| - | - | 2566 | 1303 | - | - | 0 | - |
| - | - | 1139 | 1307 | - | - | 0 | - |
| - | - | 3027 | 1308 | - | - | 0 | - |
| - | - | 2540 | 1308 | - | - | 0 | - |
| 7 | z | 1298 | 1309 | 0.01049 | 8.015 | +1 | 10 |
| - | - | 8315 | 1310 | - | - | 0 | - |
| - | - | 6856 | 1311 | - | - | 0 | - |
| - | - | 3063 | 1312 | - | - | 0 | - |
| - | - | 1188 | 1313 | - | - | 0 | - |
| - | - | 1269 | 1315 | - | - | 0 | - |
| - | - | 1185 | 1315 | - | - | 0 | - |
| - | - | 2764 | 1316 | - | - | 0 | - |
| - | - | 5862 | 1317 | - | - | 0 | - |
| - | - | 926.3 | 1322 | - | - | 0 | - |
| - | - | 1315 | 1322 | - | - | 0 | - |
| 7 | y | 2084 | 1325 | 0.001671 | 1.261 | +1 | 10 |
| - | - | 1939 | 1326 | - | - | 0 | - |
| - | - | 1286 | 1327 | - | - | 0 | - |
| - | - | 1295 | 1329 | - | - | 0 | - |
| - | - | 5309 | 1329 | - | - | 0 | - |
| - | - | 7722 | 1330 | - | - | 0 | - |
| - | - | 4711 | 1330 | - | - | 0 | - |
| - | - | 5109 | 1331 | - | - | 0 | - |
| - | - | 2619 | 1331 | - | - | 0 | - |
| - | - | 1646 | 1336 | - | - | 0 | - |
| - | - | 1726 | 1336 | - | - | 0 | - |
| 11 | c | 2.213E+04 | 1337 | 0.007531 | 5.634 | +1 | 11 |
| - | - | 1061 | 1337 | - | - | 0 | - |
| - | - | 1.355E+04 | 1338 | - | - | 0 | - |
| - | - | 6932 | 1339 | - | - | 0 | - |
| - | - | 1469 | 1340 | - | - | 0 | - |
| - | - | 1881 | 1340 | - | - | 0 | - |
| - | - | 2128 | 1351 | - | - | 0 | - |
| - | - | 2132 | 1351 | - | - | 0 | - |
| - | - | 1313 | 1352 | - | - | 0 | - |
| - | - | 1038 | 1357 | - | - | 0 | - |
| - | - | 1913 | 1358 | - | - | 0 | - |
| - | - | 1.005E+04 | 1358 | - | - | 0 | - |
| - | - | 3.136E+04 | 1359 | - | - | 0 | - |
| - | - | 3.534E+04 | 1359 | - | - | 0 | - |
| - | - | 2.692E+04 | 1360 | - | - | 0 | - |
| - | - | 1.14E+04 | 1360 | - | - | 0 | - |
| - | - | 4685 | 1361 | - | - | 0 | - |
| - | - | 1154 | 1361 | - | - | 0 | - |
| - | - | 1249 | 1365 | - | - | 0 | - |
| - | - | 1854 | 1365 | - | - | 0 | - |
| - | - | 4134 | 1366 | - | - | 0 | - |
| - | - | 4041 | 1366 | - | - | 0 | - |
| - | - | 3194 | 1367 | - | - | 0 | - |
| - | - | 1221 | 1367 | - | - | 0 | - |
| - | - | 1419 | 1372 | - | - | 0 | - |
| - | - | 1309 | 1373 | - | - | 0 | - |
| - | - | 1644 | 1376 | - | - | 0 | - |
| - | - | 1522 | 1378 | - | - | 0 | - |
| - | - | 2716 | 1379 | - | - | 0 | - |
| - | - | 3778 | 1379 | - | - | 0 | - |
| - | - | 5515 | 1380 | - | - | 0 | - |
| - | - | 7926 | 1380 | - | - | 0 | - |
| - | - | 5998 | 1381 | - | - | 0 | - |
| - | - | 3353 | 1381 | - | - | 0 | - |
| - | - | 1524 | 1382 | - | - | 0 | - |
| - | - | 1167 | 1386 | - | - | 0 | - |
| - | - | 8165 | 1387 | - | - | 0 | - |
| - | - | 3.081E+04 | 1387 | - | - | 0 | - |
| - | - | 4.308E+04 | 1388 | - | - | 0 | - |
| - | - | 3.295E+04 | 1388 | - | - | 0 | - |
| - | - | 1.629E+04 | 1389 | - | - | 0 | - |
| - | - | 6323 | 1389 | - | - | 0 | - |
| - | - | 2486 | 1390 | - | - | 0 | - |
| - | - | 1399 | 1393 | - | - | 0 | - |
| - | - | 4621 | 1394 | - | - | 0 | - |
| - | - | 2169 | 1394 | - | - | 0 | - |
| - | - | 5214 | 1395 | - | - | 0 | - |
| - | - | 6712 | 1395 | - | - | 0 | - |
| - | - | 2452 | 1396 | - | - | 0 | - |
| - | - | 3288 | 1396 | - | - | 0 | - |
| - | - | 1278 | 1397 | - | - | 0 | - |
| 6 | y | 1810 | 1422 | 0.0147 | 10.34 | +1 | 11 |
| - | - | 2271 | 1423 | - | - | 0 | - |
| - | - | 1148 | 1430 | - | - | 0 | - |
| - | - | 1144 | 1436 | - | - | 0 | - |
| - | - | 1661 | 1437 | - | - | 0 | - |
| 12 | c | 1.536E+04 | 1438 | 0.006484 | 4.51 | +1 | 12 |
| - | - | 1.198E+04 | 1439 | - | - | 0 | - |
| - | - | 6150 | 1440 | - | - | 0 | - |
| - | - | 1825 | 1441 | - | - | 0 | - |
| - | - | 2489 | 1443 | - | - | 0 | - |
| - | - | 2752 | 1443 | - | - | 0 | - |
| - | - | 3909 | 1444 | - | - | 0 | - |
| - | - | 3977 | 1444 | - | - | 0 | - |
| - | - | 3523 | 1445 | - | - | 0 | - |
| - | - | 1568 | 1445 | - | - | 0 | - |
| - | - | 1676 | 1450 | - | - | 0 | - |
| - | - | 5618 | 1451 | - | - | 0 | - |
| - | - | 1.558E+04 | 1451 | - | - | 0 | - |
| - | - | 2.599E+04 | 1452 | - | - | 0 | - |
| - | - | 1.82E+04 | 1452 | - | - | 0 | - |
| - | - | 1.078E+04 | 1453 | - | - | 0 | - |
| - | - | 3727 | 1453 | - | - | 0 | - |
| - | - | 1413 | 1454 | - | - | 0 | - |
| - | - | 1625 | 1458 | - | - | 0 | - |
| - | - | 2247 | 1458 | - | - | 0 | - |
| - | - | 2509 | 1459 | - | - | 0 | - |
| - | - | 4474 | 1459 | - | - | 0 | - |
| - | - | 3054 | 1460 | - | - | 0 | - |
| - | - | 2337 | 1460 | - | - | 0 | - |
| - | - | 1548 | 1465 | - | - | 0 | - |
| - | - | 1762 | 1466 | - | - | 0 | - |
| - | - | 1154 | 1473 | - | - | 0 | - |
| - | - | 2222 | 1479 | - | - | 0 | - |
| - | - | 2117 | 1479 | - | - | 0 | - |
| - | - | 2879 | 1480 | - | - | 0 | - |
| - | - | 4010 | 1486 | - | - | 0 | - |
| - | - | 1.532E+04 | 1487 | - | - | 0 | - |
| - | - | 1.928E+04 | 1487 | - | - | 0 | - |
| - | - | 1.821E+04 | 1488 | - | - | 0 | - |
| - | - | 9507 | 1488 | - | - | 0 | - |
| - | - | 5190 | 1489 | - | - | 0 | - |
| - | - | 2873 | 1489 | - | - | 0 | - |
| - | - | 1050 | 1493 | - | - | 0 | - |
| - | - | 2319 | 1493 | - | - | 0 | - |
| - | - | 3222 | 1494 | - | - | 0 | - |
| - | - | 2967 | 1494 | - | - | 0 | - |
| - | - | 2906 | 1495 | - | - | 0 | - |
| - | - | 2509 | 1495 | - | - | 0 | - |
| - | - | 1639 | 1496 | - | - | 0 | - |
| - | - | 3791 | 1501 | - | - | 0 | - |
| - | - | 8049 | 1501 | - | - | 0 | - |
| - | - | 8031 | 1502 | - | - | 0 | - |
| - | - | 8198 | 1502 | - | - | 0 | - |
| - | - | 2591 | 1503 | - | - | 0 | - |
| - | - | 1102 | 1503 | - | - | 0 | - |
| - | - | 1495 | 1507 | - | - | 0 | - |
| - | - | 2380 | 1520 | - | - | 0 | - |
| - | - | 1848 | 1522 | - | - | 0 | - |
| - | - | 1086 | 1523 | - | - | 0 | - |
| - | - | 1634 | 1528 | - | - | 0 | - |
| - | - | 3990 | 1529 | - | - | 0 | - |
| - | - | 1798 | 1529 | - | - | 0 | - |
| - | - | 3591 | 1530 | - | - | 0 | - |
| - | - | 1195 | 1531 | - | - | 0 | - |
| - | - | 1268 | 1538 | - | - | 0 | - |
| - | - | 1089 | 1542 | - | - | 0 | - |
| - | - | 2040 | 1542 | - | - | 0 | - |
| - | - | 3556 | 1543 | - | - | 0 | - |
| - | - | 3833 | 1543 | - | - | 0 | - |
| - | - | 2523 | 1544 | - | - | 0 | - |
| - | - | 2108 | 1544 | - | - | 0 | - |
| - | - | 1091 | 1545 | - | - | 0 | - |
| - | - | 1371 | 1550 | - | - | 0 | - |
| - | - | 4405 | 1550 | - | - | 0 | - |
| - | - | 2.015E+04 | 1551 | - | - | 0 | - |
| - | - | 3.944E+04 | 1551 | - | - | 0 | - |
| - | - | 3.779E+04 | 1552 | - | - | 0 | - |
| - | - | 2.421E+04 | 1552 | - | - | 0 | - |
| - | - | 1.26E+04 | 1553 | - | - | 0 | - |
| - | - | 4297 | 1553 | - | - | 0 | - |
| - | - | 2689 | 1554 | - | - | 0 | - |
| - | - | 1705 | 1557 | - | - | 0 | - |
| - | - | 2628 | 1558 | - | - | 0 | - |
| - | - | 3532 | 1558 | - | - | 0 | - |
| - | - | 4295 | 1559 | - | - | 0 | - |
| - | - | 2166 | 1559 | - | - | 0 | - |
| - | - | 1905 | 1565 | - | - | 0 | - |
| - | - | 2456 | 1565 | - | - | 0 | - |
| 13 | c | 5668 | 1566 | 0.02004 | 12.8 | +1 | 13 |
| - | - | 3295 | 1566 | - | - | 0 | - |
| - | - | 4723 | 1567 | - | - | 0 | - |
| - | - | 1818 | 1568 | - | - | 0 | - |
| - | - | 1669 | 1572 | - | - | 0 | - |
| - | - | 2308 | 1573 | - | - | 0 | - |
| - | - | 1234 | 1574 | - | - | 0 | - |
| - | - | 1919 | 1585 | - | - | 0 | - |
| - | - | 2528 | 1586 | - | - | 0 | - |
| - | - | 2750 | 1586 | - | - | 0 | - |
| - | - | 3471 | 1587 | - | - | 0 | - |
| - | - | 2828 | 1587 | - | - | 0 | - |
| - | - | 2948 | 1593 | - | - | 0 | - |
| - | - | 4344 | 1594 | - | - | 0 | - |
| - | - | 4924 | 1594 | - | - | 0 | - |
| - | - | 1.334E+04 | 1595 | - | - | 0 | - |
| - | - | 1.309E+04 | 1595 | - | - | 0 | - |
| - | - | 9311 | 1596 | - | - | 0 | - |
| - | - | 3456 | 1596 | - | - | 0 | - |
| - | - | 2680 | 1597 | - | - | 0 | - |
| - | - | 1078 | 1599 | - | - | 0 | - |
| - | - | 1828 | 1600 | - | - | 0 | - |
| - | - | 1654 | 1601 | - | - | 0 | - |
| - | - | 1826 | 1602 | - | - | 0 | - |
| - | - | 1349 | 1602 | - | - | 0 | - |
| - | - | 2235 | 1607 | - | - | 0 | - |
| - | - | 1888 | 1608 | - | - | 0 | - |
| - | - | 1514 | 1608 | - | - | 0 | - |
| - | - | 1544 | 1609 | - | - | 0 | - |
| - | - | 1576 | 1614 | - | - | 0 | - |
| - | - | 1285 | 1615 | - | - | 0 | - |
| - | - | 4619 | 1615 | - | - | 0 | - |
| - | - | 9327 | 1616 | - | - | 0 | - |
| - | - | 5511 | 1616 | - | - | 0 | - |
| - | - | 1.838E+04 | 1617 | - | - | 0 | - |
| - | - | 1640 | 1617 | - | - | 0 | - |
| - | - | 1.742E+04 | 1618 | - | - | 0 | - |
| - | - | 1.145E+04 | 1619 | - | - | 0 | - |
| - | - | 3913 | 1620 | - | - | 0 | - |
| - | - | 2729 | 1621 | - | - | 0 | - |
| - | - | 1878 | 1621 | - | - | 0 | - |
| - | - | 1888 | 1622 | - | - | 0 | - |
| - | - | 1633 | 1622 | - | - | 0 | - |
| - | - | 2735 | 1623 | - | - | 0 | - |
| - | - | 2447 | 1623 | - | - | 0 | - |
| - | - | 1370 | 1628 | - | - | 0 | - |
| - | - | 2532 | 1628 | - | - | 0 | - |
| - | - | 2950 | 1629 | - | - | 0 | - |
| - | - | 4641 | 1629 | - | - | 0 | - |
| - | - | 7928 | 1630 | - | - | 0 | - |
| - | - | 1.188E+04 | 1630 | - | - | 0 | - |
| - | - | 9800 | 1631 | - | - | 0 | - |
| - | - | 6558 | 1631 | - | - | 0 | - |
| - | - | 4218 | 1632 | - | - | 0 | - |
| - | - | 1686 | 1632 | - | - | 0 | - |
| - | - | 1032 | 1634 | - | - | 0 | - |
| - | - | 1515 | 1636 | - | - | 0 | - |
| - | - | 1944 | 1636 | - | - | 0 | - |
| - | - | 5220 | 1637 | - | - | 0 | - |
| - | - | 1.024E+04 | 1637 | - | - | 0 | - |
| - | - | 1.723E+04 | 1638 | - | - | 0 | - |
| - | - | 1.212E+04 | 1638 | - | - | 0 | - |
| - | - | 8171 | 1639 | - | - | 0 | - |
| - | - | 4723 | 1639 | - | - | 0 | - |
| - | - | 2119 | 1640 | - | - | 0 | - |
| - | - | 2555 | 1642 | - | - | 0 | - |
| - | - | 5135 | 1643 | - | - | 0 | - |
| - | - | 6685 | 1643 | - | - | 0 | - |
| - | - | 8623 | 1644 | - | - | 0 | - |
| - | - | 3789 | 1644 | - | - | 0 | - |
| - | - | 3189 | 1645 | - | - | 0 | - |
| - | - | 3455 | 1645 | - | - | 0 | - |
| - | - | 4353 | 1646 | - | - | 0 | - |
| - | - | 4201 | 1646 | - | - | 0 | - |
| - | - | 4287 | 1647 | - | - | 0 | - |
| - | - | 1636 | 1647 | - | - | 0 | - |
| - | - | 1666 | 1650 | - | - | 0 | - |
| - | - | 9428 | 1651 | - | - | 0 | - |
| - | - | 2.418E+04 | 1651 | - | - | 0 | - |
| - | - | 3.135E+04 | 1652 | - | - | 0 | - |
| - | - | 2.473E+04 | 1652 | - | - | 0 | - |
| - | - | 1.563E+04 | 1653 | - | - | 0 | - |
| - | - | 7167 | 1653 | - | - | 0 | - |
| - | - | 3966 | 1654 | - | - | 0 | - |
| - | - | 1536 | 1654 | - | - | 0 | - |
| - | - | 2814 | 1658 | - | - | 0 | - |
| - | - | 4722 | 1658 | - | - | 0 | - |
| - | - | 6864 | 1659 | - | - | 0 | - |
| - | - | 2.424E+04 | 1659 | - | - | 0 | - |
| - | - | 5.919E+04 | 1660 | - | - | 0 | - |
| - | - | 6.942E+04 | 1660 | - | - | 0 | - |
| - | - | 5.064E+04 | 1661 | - | - | 0 | - |
| - | - | 2.849E+04 | 1661 | - | - | 0 | - |
| - | - | 1.187E+04 | 1662 | - | - | 0 | - |
| - | - | 4629 | 1662 | - | - | 0 | - |
| - | - | 1737 | 1663 | - | - | 0 | - |
| - | - | 3377 | 1666 | - | - | 0 | - |
| - | - | 6669 | 1666 | - | - | 0 | - |
| - | - | 7150 | 1667 | - | - | 0 | - |
| - | - | 7455 | 1667 | - | - | 0 | - |
| - | - | 5212 | 1668 | - | - | 0 | - |
| 14 | c | 7250 | 1680 | 0.01117 | 6.65 | +1 | 14 |
| - | - | 5877 | 1681 | - | - | 0 | - |
| - | - | 3865 | 1682 | - | - | 0 | - |
| - | - | 1280 | 1683 | - | - | 0 | - |
| - | - | 2351 | 1701 | - | - | 0 | - |
| - | - | 3898 | 1702 | - | - | 0 | - |
| - | - | 2935 | 1703 | - | - | 0 | - |
| 3 | z | 1590 | 1717 | 0.008711 | 5.074 | +1 | 14 |
| - | - | 5730 | 1718 | - | - | 0 | - |
| - | - | 5549 | 1719 | - | - | 0 | - |
| - | - | 2971 | 1720 | - | - | 0 | - |
| - | - | 1284 | 1761 | - | - | 0 | - |
| - | - | 1646 | 1809 | - | - | 0 | - |
| - | - | 1699 | 1810 | - | - | 0 | - |
| - | - | 1713 | 1864 | - | - | 0 | - |
| - | - | 2171 | 1865 | - | - | 0 | - |
| 2 | z | 1681 | 1880 | 0.009103 | 4.842 | +1 | 15 |
| - | - | 6910 | 1881 | - | - | 0 | - |
| - | - | 6536 | 1882 | - | - | 0 | - |
| - | - | 4622 | 1883 | - | - | 0 | - |
| - | - | 2139 | 1884 | - | - | 0 | - |
| - | - | 2482 | 1907 | - | - | 0 | - |
| - | - | 5867 | 1908 | - | - | 0 | - |
| - | - | 6263 | 1909 | - | - | 0 | - |
| - | - | 4095 | 1910 | - | - | 0 | - |
| - | - | 2309 | 1911 | - | - | 0 | - |
| - | - | 2256 | 1938 | - | - | 0 | - |
| - | - | 1907 | 1939 | - | - | 0 | - |
| - | - | 1986 | 1951 | - | - | 0 | - |
| - | - | 4903 | 1952 | - | - | 0 | - |
| - | - | 6000 | 1953 | - | - | 0 | - |
| - | - | 3747 | 1954 | - | - | 0 | - |
| - | - | 2266 | 1955 | - | - | 0 | - |
| - | - | 1693 | 1968 | - | - | 0 | - |
| - | - | 2861 | 1969 | - | - | 0 | - |
| - | - | 2832 | 1970 | - | - | 0 | - |
| - | - | 1588 | 1971 | - | - | 0 | - |
| - | - | 1261 | 1972 | - | - | 0 | - |
| - | - | 6223 | 1979 | - | - | 0 | - |
| - | - | 1.947E+04 | 1980 | - | - | 0 | - |
| - | - | 2.013E+04 | 1981 | - | - | 0 | - |
| - | - | 1.269E+04 | 1982 | - | - | 0 | - |
| - | - | 6442 | 1983 | - | - | 0 | - |
| - | - | 2580 | 1995 | - | - | 0 | - |
| - | - | 1.113E+04 | 1996 | - | - | 0 | - |
| - | - | 4.552E+04 | 1997 | - | - | 0 | - |
| - | - | 3.982E+04 | 1998 | - | - | 0 | - |
| - | - | 2.617E+04 | 1999 | - | - | 0 | - |
| - | - | 1.094E+04 | 2000 | - | - | 0 | - |
| - | - | 3627 | 2001 | - | - | 0 | - |
| - | - | 1333 | 2053 | - | - | 0 | - |
| - | - | 1210 | 2095 | - | - | 0 | - |
| - | - | 2538 | 2096 | - | - | 0 | - |
| - | - | 2790 | 2097 | - | - | 0 | - |
| - | - | 2771 | 2098 | - | - | 0 | - |
| - | - | 1680 | 2110 | - | - | 0 | - |
| - | - | 1117 | 2254 | - | - | 0 | - |
| - | - | 1288 | 2333 | - | - | 0 | - |
| - | - | 4548 | 2334 | - | - | 0 | - |
| - | - | 6542 | 2335 | - | - | 0 | - |
| - | - | 7733 | 2336 | - | - | 0 | - |
| - | - | 3702 | 2337 | - | - | 0 | - |
| - | - | 1689 | 2338 | - | - | 0 | - |
| - | - | 1252 | 2383 | - | - | 0 | - |
| - | - | 1383 | 2384 | - | - | 0 | - |
| - | - | 1368 | 2512 | - | - | 0 | - |
| - | - | 1270 | 2716 | - | - | 0 | - |
| - | - | 1736 | 2717 | - | - | 0 | - |
| - | - | 1187 | 2718 | - | - | 0 | - |
| - | - | 2502 | 2774 | - | - | 0 | - |
| - | - | 2864 | 2775 | - | - | 0 | - |
| - | - | 1794 | 2776 | - | - | 0 | - |
| - | - | 1087 | 3102 | - | - | 0 | - |
| - | - | 981.5 | 3322 | - | - | 0 | - |

m/z Charge Intensity FragmentType MassShift Position
128.09486389160156 0 502.53568
129.1025390625 0 16461.412
130.09869384765625 0 370.3455
130.1058807373047 0 1489.3458
136.0760498046875 0 6377.176
143.22535705566406 0 402.79132
146.12890625 0 2211.2124
155.11801147460938 0 1805.5616
169.1339874267578 0 726.59985
171.135498046875 0 512.9111
173.12884521484375 0 1887.0486
173.44015502929688 0 3567.2773
183.1134796142578 0 866.94666
187.14466857910156 0 2446.5244
197.12921142578125 0 1502.9731
198.12399291992188 0 1260.54
200.13999938964844 0 2408.4912
201.1236572265625 0 2877.6436
203.15072631835938 0 758.3203
205.1177978515625 0 619.63434
210.68634033203125 0 505.93808
212.1395721435547 0 1639.6277
215.1392822265625 0 1964.7227
216.13470458984375 0 11114.297
216.14537048339844 0 1061.2385
217.13816833496094 0 1327.483
217.16612243652344 0 2903.1057
218.15060424804688 0 743.3211
228.13412475585938 0 575.244
230.15057373046875 0 657.0776
233.16114807128906 0 2266.3293
235.14459228515625 0 27763.916
236.14810180664062 0 4506.1445
243.1095733642578 0 829.57434
243.14614868164062 0 812.4519
252.13284301757812 0 472.22006
263.139404296875 0 22062.994
264.142822265625 0 3880.3418
269.1612243652344 0 1034.065
271.18994140625 0 606.643
274.188720703125 0 831.7872
297.1197509765625 0 521.56647
297.1571350097656 0 504.5511
298.12786865234375 0 613.4181
299.17193603515625 0 3923.4482
300.17584228515625 0 668.8
304.1986083984375 0 1987.6991
311.2088317871094 0 933.44147
315.16668701171875 0 1050.4384
320.1609191894531 0 1338.0884
326.1596374511719 0 1404.2737
327.164794921875 0 691.03937
328.2353820800781 0 1208.634
329.2189636230469 0 4947.815
330.22161865234375 0 1272.573
333.17730712890625 0 1769.9606 y 14
341.0186767578125 0 607.71136
342.1769714355469 0 1350.5927
344.19439697265625 0 651.74786
345.26153564453125 0 1208.6643
346.17694091796875 0 15417.523
346.2453308105469 0 3876.3962
347.1803894042969 0 3775.2175
347.2490539550781 0 1476.5579
355.0708312988281 0 2275.5435
359.02899169921875 0 1878.1588
360.1922607421875 0 1767.5256
364.1872863769531 0 26440.264
365.1904296875 0 4215.2124
366.1936950683594 0 1161.9581
382.2454833984375 0 5352.154
383.227783203125 0 3489.3813
383.2517395019531 0 717.3139
384.235595703125 0 5328.858
385.238525390625 0 1139.7637
386.2420959472656 0 780.3216
388.18377685546875 0 3815.9377
392.2521667480469 0 3036.3906
392.7520751953125 0 1018.138
396.18792724609375 0 1025.3016
400.25634765625 0 12078.889
401.2590026855469 0 3057.9988
417.28289794921875 0 4155.015
418.2854309082031 0 1221.2589
425.2270202636719 0 1335.5726
427.2646789550781 0 829.5359
429.0894775390625 0 5012.053
431.2667236328125 0 6098.4263
432.20953369140625 0 1716.2883
432.26983642578125 0 1342.8894
433.21209716796875 0 674.0135
433.2441711425781 0 754.2023
435.7683410644531 0 3242.3145
436.27069091796875 0 2169.95
441.2570495605469 0 1388.9117
442.26226806640625 0 692.92456
442.7106628417969 0 539.664
443.2984619140625 0 978.33954
447.2210998535156 0 4149.185 y 13
448.2225036621094 0 658.48785
449.2772216796875 0 4779.3276
450.2807922363281 0 1267.0444
456.268798828125 0 3221.0608
457.2718811035156 0 749.34485 c Water loss 7
459.2615966796875 0 20452.111
460.2647399902344 0 6083.1924
461.2671813964844 0 781.2873
472.2906494140625 0 787.26917
477.2720031738281 0 6393.1997
478.2752990722656 0 2007.699
483.30621337890625 0 1051.8292
483.80194091796875 0 1679.5436
492.31011962890625 0 8727.261
492.8118896484375 0 8912.456
493.3131103515625 0 1709.7965
497.2360534667969 0 1677.7502
501.2684020996094 0 1512.7872
502.3707580566406 0 878.6059
504.2142639160156 0 1152.0864
505.31646728515625 0 1002.2156
505.81976318359375 0 1270.0262
506.9130554199219 0 11023.213 y 4
507.24786376953125 0 7079.8228
507.5806579589844 0 4855.5024
507.9115905761719 0 709.9067
514.2300415039062 0 955.0991
515.2837524414062 0 8211.335
516.2868041992188 0 2457.1802
528.3517456054688 0 8754.024
529.353515625 0 2985.337
544.3701171875 0 1509.4379
544.6075439453125 0 756.97186 y 3
544.9462890625 0 714.7441
545.3782348632812 0 41125.625
546.2694702148438 0 918.2166
546.3811645507812 0 16629.262
547.3833618164062 0 3578.6602
556.3140258789062 0 793.30066
559.2971801757812 0 15588.799 z 12
560.3025512695312 0 7677.224
561.3056030273438 0 1622.8416
562.311767578125 0 744.6249
564.246337890625 0 855.50006
568.3477783203125 0 843.0589
572.2862548828125 0 3961.5945 y Water loss 2
572.619140625 0 1634.588 y Ammonia loss 2
572.9564208984375 0 890.202 z 2
573.3314208984375 0 892.45654 c Water loss 4
574.3203125 0 604.5302
575.3164672851562 0 13049.559 y 12
576.31884765625 0 3827.6462
577.318603515625 0 704.1535
581.7816772460938 0 3931.0115
582.2818603515625 0 4385.1577
582.7778930664062 0 1065.2659
583.27978515625 0 590.0281
584.3648681640625 0 2705.275
585.3704223632812 0 1786.9019
597.6327514648438 0 745.51294 c Ammonia loss 14
601.3927612304688 0 892.9851
602.3997802734375 0 44433.152
603.402587890625 0 21277.346
604.4052124023438 0 3785.5083
605.3583984375 0 1879.3586
605.406494140625 0 867.0345
605.8587036132812 0 1882.6715
606.3609008789062 0 1013.8631
615.31787109375 0 877.61444
615.6480712890625 0 814.02045
616.3313598632812 0 2828.9668
617.3341064453125 0 1290.9231
618.9796752929688 0 921.33765
619.2669067382812 0 1539.6133
619.7971801757812 0 1512.7345
624.9841918945312 0 1396.7065
625.3108520507812 0 871.34863
626.6422119140625 0 2303.0588 y Water loss 1
626.9781494140625 0 1800.6355 y Ammonia loss 1
627.3115844726562 0 1368.6986 z 1
628.416259765625 0 1183.6888
631.330810546875 0 954.69116
633.3048095703125 0 1313.6537
639.3807373046875 0 810.5724
640.38916015625 0 795.68066
643.3428344726562 0 5544.8364 w 11
644.3423461914062 0 1648.7023
644.3941040039062 0 1140.4192
645.322509765625 0 5523.5786 w 11
646.3215942382812 0 1973.6167
646.8028564453125 0 786.81006
647.3281860351562 0 706.91547
654.3027954101562 0 887.23505 y Ammonia loss 6
654.8048095703125 0 1208.0087 z 6
655.3060302734375 0 1165.4583
655.4009399414062 0 3945.7617
656.403564453125 0 2113.175
657.81005859375 0 2777.2913
658.325927734375 0 3252.7932
659.3359375 0 9042.027 y Ammonia loss 11
659.6646118164062 0 15550.471
659.9987182617188 0 18483.572
660.3388061523438 0 26058.62 z 11
660.6666259765625 0 4101.45
660.8750610351562 0 1255.6407
661.0023193359375 0 1307.8949
661.3494262695312 0 6791.4077
662.3494873046875 0 2064.05
662.8141479492188 0 1300.4788 y 6
663.3148193359375 0 780.91785
665.3422241210938 0 1169.0571
666.2130737304688 0 976.3954
666.2774047851562 0 2635.2966
666.3382568359375 0 2005.6172
669.3882446289062 0 4611.9917
669.8884887695312 0 3895.3845
670.39111328125 0 2504.983 c Water loss 5
670.889404296875 0 1404.2892
674.357666015625 0 1243.5662
675.3563232421875 0 1374.4337
675.8461303710938 0 761.2041
676.3643188476562 0 7505.752 y 11
677.3643798828125 0 3127.531
678.3712768554688 0 974.06256
679.3789672851562 0 1212.3961
684.331298828125 0 1283.5111
685.3336791992188 0 923.24225
687.3951416015625 0 2560.0703
688.4025268554688 0 3960.829 c 5
689.40478515625 0 2019.325
693.345703125 0 1103.8735
696.34375 0 689.8329
696.41357421875 0 2005.1235
696.9138793945312 0 1785.1211
697.3927612304688 0 2015.4912
701.3754272460938 0 2281.2095
702.3792724609375 0 817.3342
707.3443603515625 0 8770.084
707.845947265625 0 7050.544
708.348388671875 0 2104.388
710.4085693359375 0 1546.6648
710.9096069335938 0 1962.1235
711.338623046875 0 8807.793 y 5
711.4077758789062 0 1269.991
711.841064453125 0 5225.5513
712.3424072265625 0 2575.7297
712.835693359375 0 731.1036
713.4130859375 0 4481.0347
714.4212036132812 0 1226.5776
715.4207763671875 0 620.03406
717.365234375 0 1034.4663
718.352783203125 0 752.7198
718.921142578125 0 4539.3115
719.4224853515625 0 3336.4624
719.925537109375 0 1771.8796
727.38720703125 0 1813.3389
727.8660278320312 0 5797.883
728.3726806640625 0 4435.5723
728.8670654296875 0 2457.6877
729.87060546875 0 779.48486
732.4307861328125 0 1921.1892
739.3463134765625 0 914.4681
739.8745727539062 0 810.32074
740.1406860351562 0 773.624
741.8741455078125 0 648.52185
743.3831176757812 0 1920.8452
744.3894653320312 0 11216.648
745.3970336914062 0 26287.783
746.3999633789062 0 10829.088
747.40380859375 0 2655.72
750.8632202148438 0 2077.0496 y Water loss 4
751.09423828125 0 2821.0752
751.1527099609375 0 2191.5437
751.3563842773438 0 3439.1013 y Ammonia loss 4
751.4296264648438 0 7952.407
751.6488037109375 0 1121.2604
751.7644653320312 0 8125.7485
751.8521118164062 0 1610.0635 z 4
752.0980224609375 0 5032.557
752.3626708984375 0 1024.2351
752.432861328125 0 3220.4646
752.8543090820312 0 810.74677
756.0901489257812 0 767.1395
756.4247436523438 0 972.1524
758.858642578125 0 908.4952
759.3616943359375 0 1509.5295
759.8660888671875 0 126908.086 y 4
760.3676147460938 0 95121.984
760.8681030273438 0 49085.953
761.3712158203125 0 18428.49
761.8742065429688 0 2552.5366
763.886474609375 0 1404.8055
764.3856811523438 0 1109.9553
764.8832397460938 0 1727.3684
767.4767456054688 0 1469.0769
768.4828491210938 0 1448.5259
772.8864135742188 0 1229.1772
773.8704223632812 0 859.77045
774.365966796875 0 1688.0342
775.4364624023438 0 20542.277 c 6
775.648681640625 0 1475.7529
775.9010009765625 0 2759.1743
776.1537475585938 0 1244.2014
776.4393920898438 0 8375.28
777.4419555664062 0 2096.2026
778.4487915039062 0 1279.2173
779.6563720703125 0 1158.8397
780.1531982421875 0 940.8789
781.3675537109375 0 1234.6707
782.9011840820312 0 12475.637
783.1621704101562 0 2972.4175
783.4050903320312 0 28328.73 c 12
783.4888916015625 0 1055.0083
783.6710205078125 0 1764.2551
783.9057006835938 0 16191.986
784.407470703125 0 9300.529
784.9083251953125 0 4713.83
785.40625 0 1554.8096
786.40673828125 0 836.9283
786.871337890625 0 24115.53 w 3
787.3729858398438 0 22395.527
787.8734741210938 0 15294.126
788.3728637695312 0 5577.205
788.8754272460938 0 2016.2739
789.3658447265625 0 1133.2715 z Water loss 10
792.67333984375 0 1214.8966
792.9271850585938 0 908.7193
794.109375 0 5644.24
794.44384765625 0 15063.725
794.7786254882812 0 11892.7705
795.1113891601562 0 7122.407
795.4434814453125 0 4114.167
795.778076171875 0 905.7429
797.4098510742188 0 964.77686
797.6727905273438 0 899.49963
799.1004638671875 0 1080.8643
799.4306030273438 0 2223.201
799.6736450195312 0 1191.4265
800.4490966796875 0 3273.776
800.9520263671875 0 5231.828
801.17529296875 0 1466.7668
801.451904296875 0 2433.532
801.9557495117188 0 780.6336
802.6686401367188 0 839.46936
803.4281616210938 0 3172.0576
803.6799926757812 0 2117.908
803.9275512695312 0 2765.295
804.177734375 0 2212.4802
804.428955078125 0 2231.891
804.91650390625 0 1778.3035
805.401123046875 0 2181.09 y Water loss 10
806.9168090820312 0 781.2817
807.3856811523438 0 6062.416 y Water loss 3
807.6820068359375 0 3670.0835
807.927001953125 0 4822.1636
808.1806640625 0 3787.627
808.3968505859375 0 36931.094 z 3
808.679931640625 0 2309.2058
808.9007568359375 0 25195.844
809.3986206054688 0 16074.718
809.9017944335938 0 5288.918
810.4054565429688 0 3900.642
810.9293212890625 0 863.4591
811.4195556640625 0 2541.2136
811.6739501953125 0 1610.4172
811.9241943359375 0 2118.0127
812.1787719726562 0 2042.3962
812.4286499023438 0 3777.5183
812.67724609375 0 2750.2893
812.9241333007812 0 1720.75
813.1810302734375 0 1089.1953
813.4278564453125 0 1190.9021
814.4126586914062 0 1842.5332
814.9127807617188 0 2171.5032
815.1806640625 0 738.7677
815.4218139648438 0 2234.176
815.680908203125 0 3281.643
815.9219360351562 0 3677.125
816.173828125 0 2883.9229
816.4093017578125 0 18570.803 y 3
816.6787109375 0 1341.537
816.9096069335938 0 18679.656
817.410400390625 0 7528.7593
817.9136352539062 0 4169.3384
818.1831665039062 0 1107.0233
818.4223022460938 0 4644.658
818.6754150390625 0 4584.992
818.9242553710938 0 7268.657
819.1764526367188 0 5638.8076
819.4276123046875 0 6114.2285
819.6813354492188 0 2212.9163
819.9273681640625 0 1741.7776
820.1766967773438 0 1042.8582
821.676513671875 0 1184.8567
821.9295043945312 0 1398.4988
822.4364013671875 0 991.8953
823.3994750976562 0 3882.8386 y 10
824.4002685546875 0 1647.8132
825.181884765625 0 872.3329
825.4322509765625 0 947.6902
825.6843872070312 0 4514.344
825.9344482421875 0 10937.478
826.1834106445312 0 4958.843
826.433837890625 0 5558.859
826.6863403320312 0 2341.1802
827.1807250976562 0 893.08203
827.5133056640625 0 3255.0413
828.5149536132812 0 1395.7168
829.431884765625 0 1807.853
829.685791015625 0 11473.918
829.9375 0 31381.31
830.1876220703125 0 29694.992
830.4381713867188 0 21895.014
830.6883544921875 0 10402.209
830.887451171875 0 2961.877
830.9420776367188 0 3392.859
831.1939086914062 0 1578.604
831.3928833007812 0 4100.8853
831.89453125 0 2984.9268 c Ammonia loss 13
832.1997680664062 0 749.84564
832.9286499023438 0 1305.7954
833.182373046875 0 3581.7434
833.4352416992188 0 4738.3315
833.684814453125 0 2690.5051
833.946044921875 0 2738.3115
834.4608764648438 0 1409.7964
834.9376220703125 0 2127.158
835.4310913085938 0 2080.9048
837.1240234375 0 5678.7993
837.4572143554688 0 12344.3125
837.7932739257812 0 13570.143
838.1264038085938 0 8766.514
838.4601440429688 0 1922.2954
838.793212890625 0 768.51245
839.9219970703125 0 2879.5647
840.42626953125 0 45445.133 c 13
840.927978515625 0 41442.37
841.4290771484375 0 24299.873
841.92919921875 0 8694.745
842.4397583007812 0 2353.3533
842.7869873046875 0 906.1994
842.97119140625 0 1269.4261
843.4679565429688 0 1059.1469
844.92578125 0 687.8803
845.9151611328125 0 646.72534
849.4140014648438 0 1658.9529
850.9774780273438 0 9653.295
851.4783935546875 0 16884.955
851.9798583984375 0 9049.493
852.4818725585938 0 4822.5938
852.9822998046875 0 2450.6787
853.4315185546875 0 924.8462
855.5158081054688 0 1346.7473
857.4515380859375 0 851.58545
857.9132080078125 0 1201.4557 y Water loss 2
858.4208374023438 0 7723.744 y Ammonia loss 2
858.9221801757812 0 22214.684 z 2
859.4241333007812 0 20370.326
859.9260864257812 0 10257.444
860.4271240234375 0 5351.252
860.924560546875 0 1184.0396
865.9181518554688 0 1199.9476
866.4232788085938 0 1359.8728
866.9324340820312 0 55900.11 y 2
867.43359375 0 47194.72
867.93408203125 0 25791.861
868.0875244140625 0 1257.0482
868.43505859375 0 12908.616
868.933837890625 0 2338.705
870.9160766601562 0 1863.2242
871.4196166992188 0 2452.2163
871.9222412109375 0 3154.132
872.1332397460938 0 1196.9644
872.458740234375 0 2201.8528
873.4327392578125 0 3530.616
874.4324951171875 0 2922.6863
875.434326171875 0 1216.4114
876.4229736328125 0 802.4944
877.1082763671875 0 841.1215
877.4112548828125 0 15563.318 w 9
878.4119262695312 0 9908.392
879.411376953125 0 3926.37
880.4205322265625 0 2705.9082
880.776611328125 0 1061.6991
880.9302978515625 0 1320.8265
881.1172485351562 0 1038.5247
881.4496459960938 0 2795.0579
881.791259765625 0 1345.799
881.9295654296875 0 1273.2345
882.1058349609375 0 739.64044
882.4305419921875 0 847.2071
885.7822265625 0 789.6407
886.0143432617188 0 892.0545
886.119873046875 0 3654.3105
886.4569702148438 0 13863.186
886.7860107421875 0 20300.473
887.1213989257812 0 14316.82
887.4555053710938 0 5074.292
887.7849731445312 0 3018.878
888.5289306640625 0 1218.063
889.447265625 0 784.7378
890.7833251953125 0 998.9029
891.451171875 0 1347.9733
891.772705078125 0 923.9048
895.9419555664062 0 2176.7407 c Ammonia loss 14
896.1126098632812 0 1102.4364
896.4567260742188 0 5017.9746
896.7884521484375 0 739.0992
896.9564819335938 0 3202.091
897.4639892578125 0 1667.8324
899.0075073242188 0 1375.4767
899.506591796875 0 3126.4675
899.8026123046875 0 1078.4423
900.0093994140625 0 3367.4468
900.5028076171875 0 1233.1437
900.7802124023438 0 1373.6533
901.1138305664062 0 3981.422
901.4478149414062 0 4094.658
901.7799072265625 0 3745.7583
902.1160278320312 0 1513.1375
903.7962646484375 0 1254.2843
903.9564819335938 0 888.4007
904.4566650390625 0 18868.525 c 14
904.7962036132812 0 1204.8514
904.956298828125 0 15191.041
905.1217041015625 0 1907.8685
905.455810546875 0 13411.847
905.7879638671875 0 6316.632
905.9566650390625 0 4426.177
906.118408203125 0 7162.2056
906.4524536132812 0 6817.392
906.787353515625 0 3168.701
907.1151123046875 0 1054.8507
909.959228515625 0 923.5573
910.455322265625 0 3423.8174
910.7845458984375 0 831.0509
910.9502563476562 0 1209.7955
911.1133422851562 0 993.18585
911.4610595703125 0 2167.803
912.431396484375 0 2265.171
912.9244995117188 0 2495.7402
913.42626953125 0 1482.5603
918.7885131835938 0 840.16705
919.447265625 0 1217.325
919.8087158203125 0 835.3515
920.1524047851562 0 6006.46
920.4864501953125 0 9911.647
920.8204956054688 0 5760.7866
921.1528930664062 0 3620.7031
921.4857788085938 0 938.9041
922.973388671875 0 986.363
924.4644165039062 0 3307.183
924.7942504882812 0 11567.591
925.1275634765625 0 17307.518
925.4619750976562 0 11220.729
925.7936401367188 0 5589.23
926.1317138671875 0 1021.4024
926.4610595703125 0 1762.7703
929.1221923828125 0 1096.1033
929.791748046875 0 1554.041
930.1298828125 0 4902.2114
930.4715576171875 0 5408.4683
930.798583984375 0 3061.514
930.9804077148438 0 900.4663
931.1349487304688 0 1564.7408
931.5381469726562 0 30278.53 c 7
931.9815063476562 0 4867.605
932.539306640625 0 13881.373
932.9786376953125 0 3953.002
933.5414428710938 0 3112.0933
933.98291015625 0 807.63446
934.543701171875 0 1471.2997
936.424072265625 0 34082.652 z 9
937.427001953125 0 20042.06
938.4274291992188 0 9476.553
939.1613159179688 0 887.41003
939.4325561523438 0 2668.154
939.9658203125 0 2173.2546 y Ammonia loss 1
940.4561767578125 0 12321.675 z 1
940.957275390625 0 16381.567
941.455322265625 0 12354.985
941.9542236328125 0 6101.8267
942.4555053710938 0 2287.5024
943.8129272460938 0 2189.5251
944.1427612304688 0 3099.4873
944.4891357421875 0 1071.2129
945.4814453125 0 2795.9346
945.9848022460938 0 1729.2712
946.4815673828125 0 5092.6514
946.98388671875 0 3777.9612
947.4874267578125 0 3587.1997
947.9627075195312 0 2713.8386
948.4644775390625 0 8427.63 y 1
948.8424682617188 0 851.58453
948.962158203125 0 6505.664
949.1687622070312 0 835.456
949.466796875 0 3678.6587
949.9618530273438 0 2144.732
952.4451904296875 0 3049.257 y 9
952.98095703125 0 1904.6589
953.1522216796875 0 873.609
953.4667358398438 0 3331.9758
953.990234375 0 46598.7
954.4912109375 0 51904.81
954.9924926757812 0 32790.938
955.4918823242188 0 15423.306
955.995361328125 0 6621.096
958.487060546875 0 1023.8899
960.4820556640625 0 1216.8816
960.9769287109375 0 1477.4755
961.47021484375 0 2042.204
961.9624633789062 0 9768.138
962.1411743164062 0 915.0187
962.47216796875 0 9635.855
962.850341796875 0 9611.26
962.9752807617188 0 7248.604
963.185546875 0 14605.16
963.516357421875 0 11480.566
963.85302734375 0 4152.9785
963.9730834960938 0 1737.4851
964.1870727539062 0 3243.2317
964.5172729492188 0 1374.1409
964.8641357421875 0 929.9099
966.4990844726562 0 2315.8235
966.990966796875 0 1866.7974
967.1661376953125 0 1773.4797
967.4943237304688 0 9606.923
967.593505859375 0 883.6559
967.8291015625 0 7370.046
968.0010375976562 0 1232.9736
968.1612548828125 0 8586.069
968.4918212890625 0 5337.8105
968.598388671875 0 1186.7958
968.8201904296875 0 1457.0333
968.9819946289062 0 7182.086
969.1543579101562 0 753.8228
969.486328125 0 7658.286
969.5934448242188 0 963.04553
969.9815673828125 0 4390.5913
970.4793701171875 0 4960.5625
970.971923828125 0 3067.315
971.4797973632812 0 2459.117
971.8195190429688 0 1085.6539
971.97412109375 0 1289.9335
972.15673828125 0 1553.419
972.4950561523438 0 3503.288
972.8291015625 0 4159.505
973.1631469726562 0 7550.7383
973.493408203125 0 3560.7595
973.8320922851562 0 1995.3427
974.9850463867188 0 1323.0549
975.4973754882812 0 2564.1538
975.992919921875 0 18726.734
976.492919921875 0 15985.981
976.9931030273438 0 15103.095
977.1578369140625 0 978.7254
977.490966796875 0 7608.5146
977.9940795898438 0 1571.1113
978.46875 0 962.4571
980.47509765625 0 773.4452
980.9871215820312 0 3190.2793
981.4837646484375 0 2915.167
981.9892578125 0 2747.1655
982.488525390625 0 2621.657
982.99169921875 0 1996.5707
983.5083618164062 0 852.03955
983.6123046875 0 15107.73
984.6161499023438 0 13626.904
985.6199340820312 0 4636.0444
986.5114135742188 0 1005.09985
986.6256713867188 0 2006.7365
988.9789428710938 0 1904.607
989.4942016601562 0 11287.039
989.991943359375 0 62978.156
990.49267578125 0 68348.664
990.993408203125 0 51725.74
991.1771850585938 0 3884.9
991.4967041015625 0 27840.219
991.8380126953125 0 6708.8125
991.9940185546875 0 8330.504
992.1734008789062 0 2216.0193
992.5010986328125 0 2573.2832
993.4464111328125 0 1108.2969
994.4392700195312 0 1118.9218
996.5104370117188 0 2490.1167
996.8421630859375 0 1637.9257
997.1783447265625 0 1358.2506
997.49072265625 0 1916.1813
997.9984130859375 0 47716.387
998.5012817382812 0 94481.055
999.0023803710938 0 82235.766
999.504150390625 0 44333.555
1000.00439453125 0 20123.97
1000.5111694335938 0 6503.3223
1000.8649291992188 0 14503.7
1001.1986694335938 0 18546.629
1001.53369140625 0 13298.944
1001.8666381835938 0 8337.444
1002.202392578125 0 3853.754
1002.5377807617188 0 2117.933
1004.5701904296875 0 2381.501
1005.0736083984375 0 7351.187
1005.1915283203125 0 1869.087
1005.57080078125 0 5760.754
1005.8636474609375 0 2296.581
1006.0751342773438 0 2901.3672
1006.4537963867188 0 7444.094 w 8
1006.5677490234375 0 1143.4946
1007.4571533203125 0 4267.9453
1008.4608154296875 0 2402.3179
1010.1900024414062 0 2584.375
1010.5221557617188 0 3965.2026
1010.8612060546875 0 1624.433
1011.19140625 0 1621.0416
1011.5216674804688 0 1350.3363
1013.2040405273438 0 983.0558
1013.5286254882812 0 2021.2125
1015.1895751953125 0 1195.8328
1015.8571166992188 0 952.3549
1017.57958984375 0 1386.3329
1018.0787353515625 0 1097.2255
1019.537353515625 0 1903.6819
1019.8612060546875 0 1532.9459
1020.1909790039062 0 1277.9753
1026.581787109375 0 2955.438
1027.0784912109375 0 2105.8745
1027.85302734375 0 1173.8654
1028.541748046875 0 957.3546
1028.864501953125 0 3187.183
1029.19970703125 0 1947.4767
1029.53662109375 0 1680.667
1033.533935546875 0 1134.5298
1033.8685302734375 0 13788.099
1034.2032470703125 0 36965.33
1034.5369873046875 0 46239.543
1034.8717041015625 0 29513.838
1035.2054443359375 0 14778.338
1035.537841796875 0 7310.3203
1035.8734130859375 0 3295.372
1038.5352783203125 0 2683.1362
1038.8701171875 0 2266.6855
1039.1986083984375 0 3917.9312
1039.535400390625 0 5486.6475
1039.8763427734375 0 2530.0332
1043.550537109375 0 3403.9526 c Ammonia loss 8
1043.8839111328125 0 5851.4688
1044.21826171875 0 6472.1216
1044.552490234375 0 3757.2974
1044.8851318359375 0 2199.7642
1045.5535888671875 0 1357.8032
1048.087646484375 0 6882.6494
1048.5888671875 0 19075.736
1049.0911865234375 0 16841.975
1049.59130859375 0 6374.8433
1050.0950927734375 0 3688.998
1050.5887451171875 0 1215.8257
1053.2171630859375 0 973.97894
1054.5216064453125 0 3010.2505
1055.0291748046875 0 1788.7725
1055.5198974609375 0 1912.1809
1057.22998046875 0 1281.0408
1057.547119140625 0 5335.1616
1057.8797607421875 0 2674.1099
1058.537109375 0 2209.1333
1060.5814208984375 0 42041.832 c 8
1061.5841064453125 0 24783.947
1061.9014892578125 0 1130.2571
1062.2354736328125 0 3353.982
1062.5819091796875 0 11052.863
1062.89794921875 0 7066.7925
1063.2156982421875 0 12841.556
1063.5535888671875 0 16714.865
1063.883544921875 0 13557.63
1064.2169189453125 0 6472.5444
1064.55908203125 0 1791.3021
1064.889404296875 0 2172.016
1065.4676513671875 0 24706.549 z 8
1066.4691162109375 0 20197.432
1067.236083984375 0 1703.1971
1067.470947265625 0 8442.813
1067.8834228515625 0 1864.8975
1068.215576171875 0 3749.956
1068.462158203125 0 1989.7037
1068.5552978515625 0 2242.6982
1068.8848876953125 0 1201.3595
1071.5743408203125 0 1669.2131
1071.9022216796875 0 1837.8214
1072.2342529296875 0 2072.1152
1076.5723876953125 0 2222.0347
1076.908447265625 0 5853.1284
1077.23779296875 0 7121.601
1077.5709228515625 0 5290.785
1077.9073486328125 0 2433.4897
1078.2359619140625 0 1693.7382
1081.237548828125 0 985.18884
1081.4876708984375 0 3353.7876 y 8
1081.8922119140625 0 1817.167
1082.23486328125 0 1849.5328
1082.482666015625 0 2116.8325
1082.5816650390625 0 2062.9775
1082.90478515625 0 2492.3179
1083.086669921875 0 1289.2312
1083.237548828125 0 1502.3197
1085.247314453125 0 1600.4163
1085.580078125 0 2468.6177
1085.9090576171875 0 4067.5864
1086.2462158203125 0 4540.802
1086.5743408203125 0 7160.788
1086.9036865234375 0 9584.557
1087.2362060546875 0 10281.561
1087.57177734375 0 10740.775
1087.9063720703125 0 3582.6714
1088.2381591796875 0 2995.5742
1088.57666015625 0 1462.9216
1088.9078369140625 0 1237.9044
1089.5660400390625 0 1176.5847
1089.89892578125 0 1173.5739
1090.5714111328125 0 2303.3394
1090.904541015625 0 1690.6995
1091.2432861328125 0 4170.652
1091.5770263671875 0 16196.307
1091.91162109375 0 23903.357
1092.2447509765625 0 18281.715
1092.5782470703125 0 10570.862
1092.912841796875 0 5437.148
1093.242431640625 0 2287.8096
1094.5643310546875 0 2152.085
1094.9072265625 0 4880.122
1095.2420654296875 0 4934.5166
1095.5726318359375 0 5101.4507
1095.906005859375 0 5940.992
1096.2379150390625 0 4114.2417
1096.573486328125 0 5106.2114
1096.9100341796875 0 2839.2217
1097.5806884765625 0 1107.1633
1099.565185546875 0 2403.9019
1099.9031982421875 0 1716.4412
1100.23291015625 0 1558.9548
1100.5770263671875 0 8849.634
1100.91015625 0 34519.023
1101.243408203125 0 60512.855
1101.5775146484375 0 47151.336
1101.911376953125 0 33499.504
1102.2462158203125 0 12803.786
1102.578369140625 0 8036.6675
1102.9100341796875 0 1485.7651
1103.2496337890625 0 1653.1218
1104.625 0 1511.0299
1105.2353515625 0 2005.6783
1105.575927734375 0 6411.363
1105.909423828125 0 12464.295
1106.2474365234375 0 16397.463
1106.5845947265625 0 57813.656
1106.919189453125 0 90539.49
1107.252685546875 0 81371.5
1107.5869140625 0 48713.32
1107.920654296875 0 18298.176
1108.254150390625 0 10584.6045
1108.591064453125 0 2389.1685
1109.2508544921875 0 1107.9557
1110.2369384765625 0 1652.9252
1110.5712890625 0 1963.4442
1110.9097900390625 0 5924.698
1111.24365234375 0 10108.421
1111.57861328125 0 12446.44
1111.9139404296875 0 5637.7324
1112.051513671875 0 1420.2551
1126.140625 0 3904.495
1126.638427734375 0 9339.402
1127.141357421875 0 11758.88
1127.6429443359375 0 8506.262
1128.1422119140625 0 2721.399
1134.558349609375 0 1134.2905
1138.0599365234375 0 5924.3804
1138.5633544921875 0 4709.788
1139.0611572265625 0 3514.4736
1139.554443359375 0 1306.2617
1147.618896484375 0 1161.2601
1167.067138671875 0 7419.6426
1167.5675048828125 0 25211.707
1168.0684814453125 0 29773.938
1168.569580078125 0 20820.832
1169.0689697265625 0 8462.44
1169.562744140625 0 2035.4705
1174.60400390625 0 2689.73
1175.071533203125 0 2324.0134
1175.5679931640625 0 2068.2964
1179.58154296875 0 1069.0432
1188.615234375 0 6876.326
1189.6240234375 0 34143.855 c 9
1190.6322021484375 0 25776.104
1191.159912109375 0 9474.647
1191.6529541015625 0 18471.146
1192.16552734375 0 9028.6045
1192.65576171875 0 5048.1636
1193.172119140625 0 1308.4816
1198.6529541015625 0 975.9962
1209.7071533203125 0 1050.8575
1210.705322265625 0 1813.35
1221.1387939453125 0 1543.3298
1221.568603515625 0 18761.723 z 7
1222.572509765625 0 20683.6
1223.5740966796875 0 14259.058
1224.57470703125 0 5864.727
1225.5753173828125 0 2402.3303
1227.6343994140625 0 1616.1652
1230.6556396484375 0 1169.0953
1233.665771484375 0 1369.5192
1234.1734619140625 0 1046.9348
1234.684326171875 0 1261.1917
1236.5816650390625 0 2282.6846
1237.5858154296875 0 8348.431 y 7
1238.587158203125 0 6359.785
1239.5882568359375 0 2922.0251
1245.6226806640625 0 1948.114
1246.1177978515625 0 2456.33
1246.6221923828125 0 2279.3103
1247.12646484375 0 1730.2833
1247.667236328125 0 1049.3557
1248.1639404296875 0 1041.0746
1255.1824951171875 0 3002.5496
1255.6832275390625 0 8805.991
1256.1851806640625 0 11036.135
1256.68603515625 0 7905.3906
1257.1890869140625 0 5619.9355
1258.18212890625 0 1616.1587
1271.665771484375 0 1197.2471
1272.6612548828125 0 1112.5485
1292.6473388671875 0 4074.072
1293.6494140625 0 4418.9863
1294.6495361328125 0 2849.334
1301.6566162109375 0 1123.5123
1302.1541748046875 0 2467.1394
1302.6539306640625 0 2566.3406
1307.1939697265625 0 1138.9551
1307.6961669921875 0 3026.6099
1308.19384765625 0 2539.5095
1308.5865478515625 0 1298.2144 z 6
1309.607177734375 0 8315.279
1310.6077880859375 0 6855.6577
1311.6112060546875 0 3062.6487
1312.6080322265625 0 1188.0913
1314.65966796875 0 1269.2891
1315.1658935546875 0 1184.7207
1315.6573486328125 0 2764.202
1316.6600341796875 0 5861.6104
1321.6680908203125 0 926.26465
1322.18798828125 0 1314.8496
1324.617431640625 0 2084.0588 y 6
1325.6214599609375 0 1938.8916
1326.6248779296875 0 1286.4019
1328.6722412109375 0 1295.1826
1329.1953125 0 5309.2627
1329.697998046875 0 7722.4053
1330.19921875 0 4711.245
1330.697509765625 0 5109.4077
1331.1976318359375 0 2619.4253
1335.6702880859375 0 1645.9512
1336.18017578125 0 1725.6085
1336.6591796875 0 22127.447 c 10
1337.1661376953125 0 1060.64
1337.6640625 0 13548.64
1338.660400390625 0 6932.0957
1339.643310546875 0 1468.8119
1339.7838134765625 0 1880.7958
1350.682373046875 0 2128.1135
1351.1695556640625 0 2131.8975
1351.686279296875 0 1313.1547
1357.16015625 0 1038.0471
1357.69091796875 0 1912.9963
1358.1785888671875 0 10049.857
1358.678466796875 0 31355.992
1359.178955078125 0 35344.957
1359.68017578125 0 26922.39
1360.179931640625 0 11404.866
1360.68115234375 0 4685.0894
1361.1768798828125 0 1154.3052
1364.689453125 0 1249.0615
1365.1724853515625 0 1854.2466
1365.6695556640625 0 4134.1973
1366.17578125 0 4040.8918
1366.6771240234375 0 3194.4253
1367.181640625 0 1221.0948
1372.211669921875 0 1418.931
1372.720947265625 0 1308.77
1375.6837158203125 0 1643.8312
1378.1873779296875 0 1521.9137
1378.6856689453125 0 2715.8772
1379.1944580078125 0 3778.2275
1379.7073974609375 0 5514.9683
1380.2191162109375 0 7926.435
1380.72900390625 0 5997.848
1381.2247314453125 0 3352.9067
1381.7242431640625 0 1524.3234
1385.724853515625 0 1167.2675
1386.6845703125 0 8165.2593
1387.18701171875 0 30814.646
1387.6884765625 0 43084.71
1388.189208984375 0 32949.52
1388.6910400390625 0 16289.773
1389.1893310546875 0 6323.196
1389.6912841796875 0 2486.255
1393.211669921875 0 1398.7657
1393.695068359375 0 4621.3047
1394.189453125 0 2168.5603
1394.6910400390625 0 5214.0903
1395.1898193359375 0 6711.6655
1395.699462890625 0 2452.4863
1396.198486328125 0 3287.945
1396.6932373046875 0 1277.9197
1421.6832275390625 0 1810.3809 y 5
1422.6915283203125 0 2271.153
1429.7296142578125 0 1147.6019
1435.7589111328125 0 1143.5139
1436.6962890625 0 1661.4294
1437.705810546875 0 15355.364 c 11
1438.7088623046875 0 11979.041
1439.70703125 0 6149.694
1440.7119140625 0 1825.3672
1442.7352294921875 0 2489.2556
1443.249267578125 0 2752.2485
1443.7584228515625 0 3909.2507
1444.2623291015625 0 3977.4006
1444.762451171875 0 3522.6838
1445.2708740234375 0 1568.1136
1450.23291015625 0 1676.2377
1450.7332763671875 0 5617.8726
1451.233642578125 0 15580.367
1451.7357177734375 0 25992.152
1452.23583984375 0 18195.81
1452.7354736328125 0 10775.316
1453.2381591796875 0 3726.9062
1453.743408203125 0 1413.0599
1457.73388671875 0 1625.4432
1458.229736328125 0 2247.0842
1458.736083984375 0 2508.9893
1459.2451171875 0 4474.088
1459.741943359375 0 3053.9146
1460.240234375 0 2336.957
1465.236083984375 0 1547.7373
1465.7431640625 0 1762.4907
1472.767578125 0 1154.4862
1478.75048828125 0 2222.233
1479.253173828125 0 2116.8416
1479.7674560546875 0 2879.3455
1486.255615234375 0 4010.3376
1486.754150390625 0 15320.514
1487.2542724609375 0 19275.188
1487.755859375 0 18209.12
1488.25537109375 0 9506.874
1488.75537109375 0 5190.19
1489.2552490234375 0 2872.6343
1492.78076171875 0 1050.2947
1493.2684326171875 0 2319.333
1493.765869140625 0 3222.1123
1494.2589111328125 0 2967.1897
1494.7537841796875 0 2905.9004
1495.2587890625 0 2508.5852
1495.7689208984375 0 1638.7151
1500.788818359375 0 3790.9397
1501.29345703125 0 8048.5615
1501.7947998046875 0 8031.2686
1502.2965087890625 0 8197.512
1502.794677734375 0 2590.809
1503.306640625 0 1102.1744
1507.2879638671875 0 1495.2599
1519.736328125 0 2380.2803
1521.7861328125 0 1847.6929
1522.7718505859375 0 1085.875
1527.7677001953125 0 1634.3368
1528.7847900390625 0 3990.236
1529.2947998046875 0 1798.0685
1529.79052734375 0 3590.6628
1530.773193359375 0 1194.8466
1537.8004150390625 0 1267.6895
1541.7950439453125 0 1088.601
1542.2972412109375 0 2039.728
1542.792724609375 0 3556.0288
1543.2926025390625 0 3833.4575
1543.796875 0 2523.106
1544.2996826171875 0 2108.324
1544.7950439453125 0 1091.1691
1549.8050537109375 0 1371.1971
1550.2967529296875 0 4404.7676
1550.8001708984375 0 20153.262
1551.302001953125 0 39441.547
1551.8023681640625 0 37786.113
1552.3045654296875 0 24213.191
1552.8037109375 0 12595.238
1553.3040771484375 0 4297.084
1553.8031005859375 0 2689.4768
1557.3089599609375 0 1705.044
1557.796875 0 2628.4492
1558.301513671875 0 3531.5159
1558.7987060546875 0 4295.4814
1559.3004150390625 0 2165.6274
1564.798828125 0 1905.1166
1565.3184814453125 0 2455.9084
1565.8143310546875 0 5668.205 c 12
1566.324951171875 0 3294.803
1566.81494140625 0 4723.3105
1567.8065185546875 0 1818.3303
1572.326171875 0 1669.1862
1572.8095703125 0 2308.467
1573.825439453125 0 1234.2343
1585.3323974609375 0 1919.45
1585.81982421875 0 2527.9778
1586.323974609375 0 2750.4182
1586.8204345703125 0 3470.7935
1587.3228759765625 0 2828.186
1593.343994140625 0 2948.128
1593.846923828125 0 4343.8516
1594.3328857421875 0 4924.2847
1594.8232421875 0 13344.331
1595.322265625 0 13092.62
1595.8231201171875 0 9310.834
1596.3282470703125 0 3455.9668
1596.8170166015625 0 2680.4731
1599.3272705078125 0 1078.4994
1599.8681640625 0 1827.7751
1600.8612060546875 0 1653.6906
1601.8846435546875 0 1826.3312
1602.3309326171875 0 1348.5841
1607.357666015625 0 2235.431
1607.863525390625 0 1887.7493
1608.3441162109375 0 1513.8232
1609.358154296875 0 1544.0734
1614.3724365234375 0 1575.5073
1614.836669921875 0 1285.3184
1615.35693359375 0 4619.4775
1615.83740234375 0 9327.168
1616.3612060546875 0 5510.831
1616.802490234375 0 18384.514
1617.360107421875 0 1640.0956
1617.7996826171875 0 17419.58
1618.7996826171875 0 11454.646
1619.802978515625 0 3913.0737
1620.820556640625 0 2729.4714
1621.357421875 0 1877.866
1621.837646484375 0 1887.8578
1622.350341796875 0 1633.4054
1622.846923828125 0 2735.158
1623.3480224609375 0 2446.535
1627.85693359375 0 1370.3882
1628.3612060546875 0 2532.0405
1628.85888671875 0 2949.6958
1629.3544921875 0 4640.756
1629.8514404296875 0 7928.447
1630.3502197265625 0 11876.116
1630.8505859375 0 9800.127
1631.3515625 0 6558.0527
1631.8394775390625 0 4217.643
1632.354736328125 0 1686.0148
1633.8319091796875 0 1031.7477
1635.857421875 0 1515.3397
1636.35400390625 0 1943.563
1636.8565673828125 0 5220.1963
1637.364501953125 0 10243.051
1637.86181640625 0 17230.75
1638.36376953125 0 12117.719
1638.86669921875 0 8170.983
1639.365966796875 0 4722.6753
1639.870361328125 0 2118.8342
1642.3514404296875 0 2554.5603
1642.8553466796875 0 5134.667
1643.357666015625 0 6684.5645
1643.853271484375 0 8623.345
1644.3533935546875 0 3789.028
1644.859619140625 0 3189.0886
1645.3634033203125 0 3454.7195
1645.8734130859375 0 4353.0293
1646.3797607421875 0 4201.484
1646.875 0 4286.9614
1647.3773193359375 0 1636.3662
1650.3531494140625 0 1666.4221
1650.8648681640625 0 9427.849
1651.3643798828125 0 24183.992
1651.86572265625 0 31348.709
1652.3660888671875 0 24734.77
1652.8653564453125 0 15626.609
1653.3651123046875 0 7166.912
1653.8638916015625 0 3965.736
1654.3681640625 0 1536.1223
1657.85791015625 0 2814.099
1658.3607177734375 0 4721.778
1658.8621826171875 0 6864.1006
1659.3707275390625 0 24244.8
1659.8740234375 0 59191.176
1660.376220703125 0 69423.14
1660.877197265625 0 50638.72
1661.3785400390625 0 28494.293
1661.879150390625 0 11871.844
1662.375732421875 0 4628.5596
1662.8798828125 0 1737.4391
1665.86279296875 0 3376.9292
1666.3623046875 0 6669.186
1666.86669921875 0 7149.652
1667.367919921875 0 7454.626
1667.8740234375 0 5211.9116
1679.848388671875 0 7250.0713 c 13
1680.85107421875 0 5876.717
1681.8519287109375 0 3864.682
1682.84619140625 0 1279.5867
1700.9381103515625 0 2351.1929
1701.9453125 0 3898.1868
1702.952392578125 0 2935.3503
1716.843017578125 0 1589.6423 z 2
1717.846435546875 0 5729.5864
1718.8450927734375 0 5549.261
1719.850341796875 0 2970.611
1760.9202880859375 0 1283.5798
1808.9029541015625 0 1646.109
1809.91162109375 0 1698.9274
1863.9608154296875 0 1712.5985
1864.9622802734375 0 2171.1084
1879.90673828125 0 1680.6742 z 1
1880.9071044921875 0 6910.317
1881.9102783203125 0 6535.6865
1882.9093017578125 0 4621.813
1883.9031982421875 0 2138.874
1906.978515625 0 2482.2612
1907.9759521484375 0 5867.24
1908.97509765625 0 6262.8315
1909.9866943359375 0 4095.4973
1910.9833984375 0 2308.9966
1937.9652099609375 0 2255.821
1938.963134765625 0 1907.4314
1950.9888916015625 0 1986.1895
1951.982421875 0 4902.945
1952.983154296875 0 5999.6084
1953.9974365234375 0 3746.6301
1954.9912109375 0 2266.0947
1968.0169677734375 0 1693.3821
1969.0025634765625 0 2861.0635
1970.01025390625 0 2831.7368
1971.0120849609375 0 1587.6874
1972.0076904296875 0 1261.1626
1978.9735107421875 0 6222.87
1979.9765625 0 19469.934
1980.97802734375 0 20129.365
1981.9771728515625 0 12693.412
1982.9818115234375 0 6442.491
1994.984375 0 2580.348
1995.993408203125 0 11128.105
1997.000244140625 0 45522.83
1998.0037841796875 0 39818.918
1999.0057373046875 0 26165.727
2000.0057373046875 0 10944.723
2001.0050048828125 0 3626.5596
2053.159423828125 0 1332.9637
2095.163818359375 0 1209.7522
2096.16796875 0 2537.8455
2097.17529296875 0 2789.9246
2098.187255859375 0 2770.9316
2110.05322265625 0 1679.9017
2254.286376953125 0 1117.1708
2333.13037109375 0 1288.459
2334.130126953125 0 4547.657
2335.12939453125 0 6542.0796
2336.1318359375 0 7733.4478
2337.131591796875 0 3701.9116
2338.12890625 0 1689.0001
2383.3056640625 0 1252.3868
2384.32958984375 0 1382.5372
2512.33837890625 0 1367.8759
2716.322021484375 0 1269.5013
2717.37744140625 0 1736.443
2718.34716796875 0 1187.1024
2774.3798828125 0 2502.3662
2775.387939453125 0 2864.186
2776.370361328125 0 1794.1283
3101.59423828125 0 1087.0681
3321.77490234375 0 981.49835

Spectrum Details

|  |  |
| --- | --- |
| Matched peaks? Matched peaks The total absolute number of peaks matched. Additionally in brackets the total fraction of peaks matched and the total number of peaks is shown. | 71 (5.58% of 1272) |
| FDR? FDR The false discovery rate estimated for this peptide. It is calculated by matching all theoretical fragments with a non-integer shift with the raw peaks for this spectrum. This is done with 40 different shifts. The resulting percentage is the average number of annotated peaks over the number of annotated peaks with the correct spectrum. | 8.89% |
| Satellite FDR? Satellite FDR See the FDR for details on its calculation. This satellite ion specific FDR only contains the satellite ions (d/w) for I/L/J positions. | 16.67% |
| PSM Score? PSM Score The PSM Score as given by Hecklib to this annotated spectrum. It is shown with three significant figures. | 391 |

#### Reverse Lookup? Reverse Lookup All places where this read could be placed.

| Group | Segment | Template | Template Part | Read Part | Score | Unique |
| --- | --- | --- | --- | --- | --- | --- |
| Homo sapiens Heavy Chain | IGHC | IGHG3 | [277..293] | [0..16] | 119 | False |
| Homo sapiens Heavy Chain | IGHC | IGHG2 | [226..242] | [0..16] | 119 | False |
| Homo sapiens Heavy Chain | IGHC | IGHG4 | [227..243] | [0..16] | 110 | False |

| Recombined | Template Part | Read Part | Score | Unique |
| --- | --- | --- | --- | --- |
| REC-0-1 | [355..371] | [0..16] | 105 | True |

#### Meta Information from Multiple reads

##### Number of combined reads

3

##### Intensity

0.7242

##### TotalArea

1.803E+08

#### Positional Score

Copy Data

##### Positional Score (TSV)

###### Preview

```
Loading example...
```

*Click on the button to copy the data to your clipboard.*

100123456789101112131415

Label Value
"0" 0.333
"1" 0.333
"2" 0.33
"3" 0.31
"4" 0.283
"5" 0.31
"6" 0.317
"7" 0.323
"8" 0.327
"9" 0.327
"10" 0.327
"11" 0.327
"12" 0.327
"13" 0.31
"14" 0.303
"15" 0.283

#### Meta Information from PEAKS

##### Scan Identifier

F2:4198

##### Original sequence

V

Y

T

L

P

P

S

R

E

E

M

+15.99

T

K

N

Q

K

+58.01

##### Posttranslational Modifications

Oxidation (M); Carboxymethyl (KW X@N-term)

##### Source File

D:\separate\_stitch\_analyses\xle-disambiguation\raw\20210323\_F1\_UM1\_Peng0013\_SA\_F59\_ingel\_3ug\_TL.raw

##### Fraction

2

##### Scan Feature

F2:11026

##### De Novo Score

98

##### ConfidenceScore

98

### m/z

665.6665

##### Mass

1993.9778

##### Charge

3

##### Retention Time

20.97

##### Predicted Retention Time

-

##### Area

6.009E+07

##### Parts Per Million

0

##### Fragmentation mode

ETHCD

##### Originating file

01 D:\separate\_stitch\_analyses\xle-disambiguation\20210325\_F59\_3ug\_DENOVO\_12.csv

#### Meta Information from PEAKS

##### Scan Identifier

F2:4132

##### Original sequence

V

Y

T

L

P

P

S

R

E

E

M

+15.99

T

K

N

Q

K

+58.01

##### Posttranslational Modifications

Oxidation (M); Carboxymethyl (KW X@N-term)

##### Source File

D:\separate\_stitch\_analyses\xle-disambiguation\raw\20210323\_F1\_UM1\_Peng0013\_SA\_F59\_ingel\_3ug\_TL.raw

##### Fraction

2

##### Scan Feature

F2:11026

##### De Novo Score

98

##### ConfidenceScore

98

### m/z

665.6665

##### Mass

1993.9778

##### Charge

3

##### Retention Time

20.97

##### Predicted Retention Time

-

##### Area

6.009E+07

##### Parts Per Million

0

##### Fragmentation mode

ETHCD

##### Originating file

01 D:\separate\_stitch\_analyses\xle-disambiguation\20210325\_F59\_3ug\_DENOVO\_12.csv

#### Meta Information from PEAKS

##### Scan Identifier

F2:3932

##### Original sequence

V

Y

T

L

P

P

S

R

E

E

M

+15.99

T

K

N

Q

K

+58.01

##### Posttranslational Modifications

Oxidation (M); Carboxymethyl (KW X@N-term)

##### Source File

D:\separate\_stitch\_analyses\xle-disambiguation\raw\20210323\_F1\_UM1\_Peng0013\_SA\_F59\_ingel\_3ug\_TL.raw

##### Fraction

2

##### Scan Feature

F2:11026

##### De Novo Score

95

##### ConfidenceScore

95

### m/z

665.6665

##### Mass

1993.9778

##### Charge

3

##### Retention Time

20.97

##### Predicted Retention Time

-

##### Area

6.009E+07

##### Parts Per Million

0

##### Fragmentation mode

ETHCD

##### Originating file

01 D:\separate\_stitch\_analyses\xle-disambiguation\20210325\_F59\_3ug\_DENOVO\_12.csv
