## Supplementary material for "A handle on mass coincidence errors in *de novo* sequencing of antibodies by bottom-up proteomics": Combined_070.html

Details Combined\_070 | Stitch OverviewUndefined

### Read Combined\_070

#### Sequence (length=8)

AEFAEVSK

#### Spectrum 3462? Spectrum 3462 The raw spectrum of this peptide as annotated by Hecklib. The fragments are coloured according to ion type (see legend). Any peaks with a star '\*' as text can be hovered over to see the full details, first the ion type second the mass shift type. By hovering over the amino acids in the peptide or ions in the legend the corresponding peaks are highlighted. By toggling the 'Unassigned' label you can turn the background (unassigned) peaks on or off in the plot. By updating the slider in the Ion legend you can update the spectrum to only show the top X% of the peaks with labels. The top X% means any peak that is within X% of the highest intensity. By dragging in the spectrum you can zoom in to a specific part of the spectrum and use 'Zoom Out' to get back to the original zoom level. The annotation of the spectrum is based on the given sequence in the peptides file and is done with different software so inconsistencies are likely. The peaks are annotated based on the given sequence, with 20 ppm tolerance.

Copy Data

##### Spectrum 3462 (TSV)

###### Preview

```
Loading example...
```

*Click on the button to copy the data to your clipboard.*

Mz MinMz MaxIntensity Max

WidthHeightPeptide font sizePeptide stroke widthSpectrum font sizeSpectrum stroke widthCompact peptide

Ion legend

wxyz

abcd

OtherUnassignedIon^Charge^_Position_Show for top:%

AEFAEVSK

02.40e+34.80e+37.19e+39.59e+3

Zoom Out

y^+1^_2_y^+1^_2_y^+1^_3_w^+1^_4_z^+1^_4_y^+1^_4_z^+1^_5_y^+1^_5_c^+1^_5_z^+1^_6_y^+1^_6_c^+1^_6_y^+1^_6_c^+1^_7_y^+1^_7_z^+1^_7_

0776155323293106

Fragment Matches Table

Show background peaks

| Position | Ion type | Intensity | mz Theoretical | mz Error (Th) | mz Error (ppm) | Charge | Series Number |
| --- | --- | --- | --- | --- | --- | --- | --- |
| - | - | 337.9 | 121.5 | - | - | 0 | - |
| - | - | 404.5 | 151.2 | - | - | 0 | - |
| - | - | 1082 | 153.1 | - | - | 0 | - |
| - | - | 407.9 | 154.7 | - | - | 0 | - |
| - | - | 1022 | 155.1 | - | - | 0 | - |
| - | - | 880.3 | 156.1 | - | - | 0 | - |
| - | - | 3839 | 173.1 | - | - | 0 | - |
| - | - | 566.8 | 183.6 | - | - | 0 | - |
| - | - | 496.3 | 198.1 | - | - | 0 | - |
| - | - | 3528 | 201.1 | - | - | 0 | - |
| - | - | 536.2 | 201.9 | - | - | 0 | - |
| - | - | 513.3 | 202.1 | - | - | 0 | - |
| 7 | y | 1387 | 217.1 | 0.0002567 | 1.182 | +1 | 2 |
| - | - | 493.4 | 220.5 | - | - | 0 | - |
| 7 | y | 1013 | 234.1 | 0.0004769 | 2.037 | +1 | 2 |
| - | - | 499.1 | 237.7 | - | - | 0 | - |
| - | - | 520.7 | 240.1 | - | - | 0 | - |
| - | - | 2216 | 318.2 | - | - | 0 | - |
| 6 | y | 1241 | 333.2 | 0.0006 | 1.801 | +1 | 3 |
| - | - | 712.5 | 341.2 | - | - | 0 | - |
| 5 | w | 3666 | 387.2 | 0.0001573 | 0.4063 | +1 | 4 |
| - | - | 763.6 | 388.2 | - | - | 0 | - |
| - | - | 1797 | 405.2 | - | - | 0 | - |
| - | - | 1262 | 421.8 | - | - | 0 | - |
| - | - | 859.8 | 422.8 | - | - | 0 | - |
| - | - | 3319 | 439.8 | - | - | 0 | - |
| - | - | 1185 | 440.2 | - | - | 0 | - |
| - | - | 1170 | 440.8 | - | - | 0 | - |
| - | - | 917.6 | 442.3 | - | - | 0 | - |
| 5 | z | 2744 | 446.2 | 0.0001587 | 0.3555 | +1 | 4 |
| - | - | 1975 | 447.2 | - | - | 0 | - |
| - | - | 759.9 | 458.3 | - | - | 0 | - |
| 5 | y | 862.9 | 462.3 | 0.001393 | 3.014 | +1 | 4 |
| 4 | z | 1573 | 517.3 | 0.0003035 | 0.5868 | +1 | 5 |
| - | - | 673.5 | 518.3 | - | - | 0 | - |
| - | - | 533.1 | 521.8 | - | - | 0 | - |
| 4 | y | 1548 | 533.3 | 0.0008699 | 1.631 | +1 | 5 |
| - | - | 1088 | 564.3 | - | - | 0 | - |
| 5 | c | 1563 | 565.3 | 0.001521 | 2.692 | +1 | 5 |
| - | - | 639.2 | 566.3 | - | - | 0 | - |
| - | - | 2104 | 620.3 | - | - | 0 | - |
| - | - | 1498 | 621.3 | - | - | 0 | - |
| - | - | 720.2 | 634.3 | - | - | 0 | - |
| - | - | 842.4 | 641.8 | - | - | 0 | - |
| - | - | 622.8 | 642.3 | - | - | 0 | - |
| - | - | 1043 | 642.8 | - | - | 0 | - |
| 3 | z | 657.2 | 647.3 | 0.01263 | 19.51 | +1 | 6 |
| - | - | 689.3 | 651.3 | - | - | 0 | - |
| - | - | 584.1 | 660.3 | - | - | 0 | - |
| 3 | y | 2836 | 663.3 | 0.01261 | 19.02 | +1 | 6 |
| 6 | c | 8475 | 664.3 | 0.0113 | 17.01 | +1 | 6 |
| - | - | 2505 | 665.3 | - | - | 0 | - |
| 3 | y | 7344 | 680.4 | 3.916E-05 | 0.05756 | +1 | 6 |
| - | - | 2520 | 681.4 | - | - | 0 | - |
| - | - | 780.9 | 682.4 | - | - | 0 | - |
| 7 | c | 6005 | 751.4 | 3.502E-05 | 0.04661 | +1 | 7 |
| - | - | 2301 | 752.4 | - | - | 0 | - |
| - | - | 571.1 | 772.6 | - | - | 0 | - |
| 2 | y | 611.8 | 791.4 | 0.004052 | 5.121 | +1 | 7 |
| 2 | z | 3998 | 793.4 | 0.0008721 | 1.099 | +1 | 7 |
| - | - | 1180 | 794.4 | - | - | 0 | - |
| - | - | 9495 | 864.4 | - | - | 0 | - |
| - | - | 3890 | 865.4 | - | - | 0 | - |
| - | - | 7512 | 880.4 | - | - | 0 | - |
| - | - | 4454 | 881.4 | - | - | 0 | - |
| - | - | 2011 | 882.4 | - | - | 0 | - |
| - | - | 720.6 | 882.6 | - | - | 0 | - |
| - | - | 753.6 | 883.5 | - | - | 0 | - |
| - | - | 864.5 | 1236 | - | - | 0 | - |
| - | - | 1462 | 1284 | - | - | 0 | - |
| - | - | 1824 | 1285 | - | - | 0 | - |
| - | - | 1214 | 1286 | - | - | 0 | - |
| - | - | 736.1 | 1301 | - | - | 0 | - |
| - | - | 1177 | 1302 | - | - | 0 | - |
| - | - | 1376 | 1303 | - | - | 0 | - |
| - | - | 676.3 | 1304 | - | - | 0 | - |
| - | - | 678.4 | 1305 | - | - | 0 | - |
| - | - | 747.1 | 1320 | - | - | 0 | - |
| - | - | 713.2 | 1325 | - | - | 0 | - |
| - | - | 762.7 | 2038 | - | - | 0 | - |
| - | - | 580.2 | 2640 | - | - | 0 | - |
| - | - | 853.5 | 3075 | - | - | 0 | - |

m/z Charge Intensity FragmentType MassShift Position
121.50997161865234 0 337.85458
151.19342041015625 0 404.54456
153.0772247314453 0 1081.9952
154.69459533691406 0 407.94125
155.0818328857422 0 1021.53076
156.07713317871094 0 880.30634
173.09219360351562 0 3838.797
183.6058807373047 0 566.77795
198.0715789794922 0 496.28647
201.08717346191406 0 3527.9495
201.9192352294922 0 536.16895
202.0902099609375 0 513.28735
217.11802673339844 0 1387.0057 y Ammonia loss 6
220.49371337890625 0 493.44254
234.1453094482422 0 1013.0932 y 6
237.6512908935547 0 499.1055
240.09486389160156 0 520.7079
318.20233154296875 0 2215.8877
333.212646484375 0 1241.4542 y 5
341.185791015625 0 712.49475
387.2239685058594 0 3665.7554 w 4
388.2276306152344 0 763.6057
405.22613525390625 0 1796.8481
421.83306884765625 0 1261.7825
422.8351135253906 0 859.78674
439.84368896484375 0 3318.5269
440.1901550292969 0 1185.3297
440.8446350097656 0 1170.1312
442.2647399902344 0 917.5714
446.2372741699219 0 2744.2207 z 4
447.2449645996094 0 1975.4843
458.2615966796875 0 759.9093
462.2572326660156 0 862.8736 y 4
517.27392578125 0 1573.436 z 3
518.284423828125 0 673.5096
521.7965087890625 0 533.06494
533.2938232421875 0 1547.908 y 3
564.2538452148438 0 1088.0543
565.2601318359375 0 1563.3748 c 4
566.2647094726562 0 639.17236
620.3156127929688 0 2103.615
621.3291625976562 0 1497.9668
634.3154907226562 0 720.1552
641.8009643554688 0 842.3769
642.3043823242188 0 622.819
642.8037719726562 0 1043.1501
647.303466796875 0 657.2468 z Ammonia loss 2
651.3123168945312 0 689.2532
660.3363037109375 0 584.14215
663.3222045898438 0 2836.0066 y Ammonia loss 2
664.3413696289062 0 8474.924 c 5
665.34423828125 0 2504.7983
680.361328125 0 7344.045 y 2
681.3651123046875 0 2520.1018
682.3654174804688 0 780.90826
751.362060546875 0 6004.569 c 6
752.3638916015625 0 2300.737
772.5518188476562 0 571.0742
791.3893432617188 0 611.77814 y Water loss 1
793.3861083984375 0 3998.2239 z 1
794.3851928710938 0 1179.7589
864.4223022460938 0 9495.126
865.425048828125 0 3890.3318
880.4403076171875 0 7512.411
881.4468994140625 0 4453.9062
882.4456176757812 0 2011.4768
882.56640625 0 720.5606
883.4635620117188 0 753.6432
1235.565185546875 0 864.53217
1283.5966796875 0 1462.2863
1284.5980224609375 0 1823.8955
1285.6083984375 0 1214.0874
1300.68603515625 0 736.102
1301.6202392578125 0 1177.261
1302.611083984375 0 1375.5864
1303.623291015625 0 676.3224
1304.6346435546875 0 678.41425
1319.6722412109375 0 747.09717
1324.67822265625 0 713.1709
2037.8857421875 0 762.66144
2640.118408203125 0 580.20526
3075.123046875 0 853.4613

Spectrum Details

|  |  |
| --- | --- |
| Matched peaks? Matched peaks The total absolute number of peaks matched. Additionally in brackets the total fraction of peaks matched and the total number of peaks is shown. | 16 (19.51% of 82) |
| FDR? FDR The false discovery rate estimated for this peptide. It is calculated by matching all theoretical fragments with a non-integer shift with the raw peaks for this spectrum. This is done with 40 different shifts. The resulting percentage is the average number of annotated peaks over the number of annotated peaks with the correct spectrum. | 0.00% |
| Satellite FDR? Satellite FDR See the FDR for details on its calculation. This satellite ion specific FDR only contains the satellite ions (d/w) for I/L/J positions. | - |
| PSM Score? PSM Score The PSM Score as given by Hecklib to this annotated spectrum. It is shown with three significant figures. | 175 |

#### Spectrum 3397? Spectrum 3397 The raw spectrum of this peptide as annotated by Hecklib. The fragments are coloured according to ion type (see legend). Any peaks with a star '\*' as text can be hovered over to see the full details, first the ion type second the mass shift type. By hovering over the amino acids in the peptide or ions in the legend the corresponding peaks are highlighted. By toggling the 'Unassigned' label you can turn the background (unassigned) peaks on or off in the plot. By updating the slider in the Ion legend you can update the spectrum to only show the top X% of the peaks with labels. The top X% means any peak that is within X% of the highest intensity. By dragging in the spectrum you can zoom in to a specific part of the spectrum and use 'Zoom Out' to get back to the original zoom level. The annotation of the spectrum is based on the given sequence in the peptides file and is done with different software so inconsistencies are likely. The peaks are annotated based on the given sequence, with 20 ppm tolerance.

Copy Data

##### Spectrum 3397 (TSV)

###### Preview

```
Loading example...
```

*Click on the button to copy the data to your clipboard.*

Mz MinMz MaxIntensity Max

WidthHeightPeptide font sizePeptide stroke widthSpectrum font sizeSpectrum stroke widthCompact peptide

Ion legend

wxyz

abcd

OtherUnassignedIon^Charge^_Position_Show for top:%

AEFAEVSK

01.84e+43.67e+45.51e+47.34e+4

Zoom Out

y^+1^_1_y^+1^_1_a^+1^_2_a^+1^_2_b^+1^_2_y^+1^_2_y^+1^_2_y^+1^_2_y^+2^_5_a^+1^_3_y^+1^_3_a^+1^_3_b^+1^_3_y^+1^_3_y^+2^_6_b^+1^_3_\*y^+1^_4_y^+1^_4_y^+1^_5_y^+1^_5_b^+1^_5_y^+1^_6_y^+1^_6_y^+1^_7_y^+1^_7_

0561112216832244

Fragment Matches Table

Show background peaks

| Position | Ion type | Intensity | mz Theoretical | mz Error (Th) | mz Error (ppm) | Charge | Series Number |
| --- | --- | --- | --- | --- | --- | --- | --- |
| - | - | 3.915E+04 | 120.1 | - | - | 0 | - |
| - | - | 3100 | 121.1 | - | - | 0 | - |
| - | - | 1074 | 127.1 | - | - | 0 | - |
| - | - | 2267 | 128.1 | - | - | 0 | - |
| - | - | 338.7 | 128.9 | - | - | 0 | - |
| - | - | 1.16E+04 | 129.1 | - | - | 0 | - |
| - | - | 517.4 | 130 | - | - | 0 | - |
| 8 | y | 5547 | 130.1 | 0.0002165 | 1.664 | +1 | 1 |
| - | - | 572 | 130.1 | - | - | 0 | - |
| - | - | 1045 | 136.1 | - | - | 0 | - |
| - | - | 2334 | 138.1 | - | - | 0 | - |
| - | - | 468.9 | 144.6 | - | - | 0 | - |
| 8 | y | 8594 | 147.1 | 0.0001872 | 1.272 | +1 | 1 |
| - | - | 505.9 | 148.1 | - | - | 0 | - |
| 2 | a | 2.069E+04 | 155.1 | 0.000161 | 1.038 | +1 | 2 |
| - | - | 1944 | 156.1 | - | - | 0 | - |
| - | - | 1729 | 156.1 | - | - | 0 | - |
| - | - | 855.3 | 159.1 | - | - | 0 | - |
| - | - | 1172 | 171.1 | - | - | 0 | - |
| 2 | a | 7.2E+04 | 173.1 | 0.0001401 | 0.8097 | +1 | 2 |
| - | - | 973.6 | 173.4 | - | - | 0 | - |
| - | - | 4273 | 174.1 | - | - | 0 | - |
| - | - | 3264 | 183.1 | - | - | 0 | - |
| - | - | 1699 | 184.1 | - | - | 0 | - |
| - | - | 506.8 | 184.6 | - | - | 0 | - |
| - | - | 1006 | 186.1 | - | - | 0 | - |
| - | - | 1532 | 187.1 | - | - | 0 | - |
| - | - | 1.412E+04 | 191.1 | - | - | 0 | - |
| - | - | 1218 | 192.1 | - | - | 0 | - |
| - | - | 935.2 | 198.1 | - | - | 0 | - |
| - | - | 1688 | 199.1 | - | - | 0 | - |
| 2 | b | 5.052E+04 | 201.1 | 3.754E-05 | 0.1867 | +1 | 2 |
| - | - | 2751 | 201.1 | - | - | 0 | - |
| - | - | 4062 | 202.1 | - | - | 0 | - |
| - | - | 733.4 | 207.1 | - | - | 0 | - |
| - | - | 660.3 | 211.1 | - | - | 0 | - |
| 7 | y | 6728 | 216.1 | 5.796E-06 | 0.02682 | +1 | 2 |
| - | - | 528.4 | 217.1 | - | - | 0 | - |
| 7 | y | 788.7 | 217.1 | 0.000552 | 2.542 | +1 | 2 |
| - | - | 2.237E+04 | 219.1 | - | - | 0 | - |
| - | - | 2911 | 220.1 | - | - | 0 | - |
| - | - | 1695 | 229.1 | - | - | 0 | - |
| - | - | 5184 | 231.1 | - | - | 0 | - |
| 7 | y | 2.896E+04 | 234.1 | 6.489E-05 | 0.2772 | +1 | 2 |
| - | - | 2677 | 235.1 | - | - | 0 | - |
| - | - | 588 | 240.4 | - | - | 0 | - |
| - | - | 2489 | 249.1 | - | - | 0 | - |
| - | - | 1397 | 259.1 | - | - | 0 | - |
| 4 | y | 736.4 | 267.2 | 0.0003041 | 1.138 | +2 | 5 |
| - | - | 1079 | 277.1 | - | - | 0 | - |
| - | - | 2373 | 298.1 | - | - | 0 | - |
| - | - | 675 | 298.9 | - | - | 0 | - |
| - | - | 2414 | 300.2 | - | - | 0 | - |
| 3 | a | 1029 | 302.1 | 0.0001682 | 0.5566 | +1 | 3 |
| - | - | 1693 | 309.2 | - | - | 0 | - |
| 6 | y | 1457 | 315.2 | 0.0004737 | 1.503 | +1 | 3 |
| - | - | 760.1 | 316.1 | - | - | 0 | - |
| 3 | a | 1442 | 320.2 | 0.0004485 | 1.401 | +1 | 3 |
| 3 | b | 3008 | 330.1 | 0.0001487 | 0.4505 | +1 | 3 |
| 6 | y | 1.569E+04 | 333.2 | 5.068E-05 | 0.1521 | +1 | 3 |
| - | - | 1945 | 334.2 | - | - | 0 | - |
| 3 | y | 6063 | 340.7 | 0.0003009 | 0.8831 | +2 | 6 |
| - | - | 2295 | 341.2 | - | - | 0 | - |
| - | - | 588.3 | 344.9 | - | - | 0 | - |
| - | - | 554.4 | 347.9 | - | - | 0 | - |
| 3 | b | 2.101E+04 | 348.2 | 0.0001543 | 0.4433 | +1 | 3 |
| - | - | 3370 | 349.2 | - | - | 0 | - |
| - | - | 714.8 | 350.2 | - | - | 0 | - |
| - | - | 1311 | 387.2 | - | - | 0 | - |
| - | - | 671.1 | 390.2 | - | - | 0 | - |
| - | - | 615.2 | 404.1 | - | - | 0 | - |
| - | - | 1110 | 410.2 | - | - | 0 | - |
| - | - | 2314 | 419.2 | - | - | 0 | - |
| - | - | 654.8 | 420.2 | - | - | 0 | - |
| 0 | Precursor | 1069 | 431.7 | 0.00176 | 4.078 | +2 | -1 |
| - | - | 1128 | 435.2 | - | - | 0 | - |
| - | - | 6932 | 439.8 | - | - | 0 | - |
| - | - | 1109 | 440.2 | - | - | 0 | - |
| - | - | 2741 | 440.8 | - | - | 0 | - |
| - | - | 1343 | 441.3 | - | - | 0 | - |
| - | - | 733 | 441.8 | - | - | 0 | - |
| 5 | y | 4654 | 444.2 | 0.0003103 | 0.6985 | +1 | 4 |
| - | - | 2706 | 447.2 | - | - | 0 | - |
| - | - | 1917 | 459.2 | - | - | 0 | - |
| 5 | y | 1E+04 | 462.3 | 1.072E-05 | 0.0232 | +1 | 4 |
| - | - | 2843 | 463.3 | - | - | 0 | - |
| - | - | 656 | 487.9 | - | - | 0 | - |
| - | - | 585.2 | 502.8 | - | - | 0 | - |
| 4 | y | 992.8 | 515.3 | 0.0009556 | 1.854 | +1 | 5 |
| - | - | 736.3 | 516.2 | - | - | 0 | - |
| 4 | y | 2.308E+04 | 533.3 | 0.000595 | 1.116 | +1 | 5 |
| - | - | 1145 | 534.3 | - | - | 0 | - |
| - | - | 6570 | 534.3 | - | - | 0 | - |
| - | - | 829.7 | 535.3 | - | - | 0 | - |
| - | - | 636.5 | 547.9 | - | - | 0 | - |
| 5 | b | 612.8 | 548.2 | 0.0009798 | 1.787 | +1 | 5 |
| - | - | 1042 | 558.3 | - | - | 0 | - |
| 3 | y | 1783 | 662.4 | 0.001804 | 2.723 | +1 | 6 |
| 3 | y | 7.27E+04 | 680.4 | 0.001199 | 1.762 | +1 | 6 |
| - | - | 2.705E+04 | 681.4 | - | - | 0 | - |
| - | - | 6072 | 682.4 | - | - | 0 | - |
| 2 | y | 3208 | 791.4 | 0.001306 | 1.65 | +1 | 7 |
| - | - | 1517 | 792.4 | - | - | 0 | - |
| 2 | y | 5353 | 809.4 | 0.001189 | 1.469 | +1 | 7 |
| - | - | 1814 | 810.4 | - | - | 0 | - |
| - | - | 1112 | 811.4 | - | - | 0 | - |
| - | - | 722.3 | 1655 | - | - | 0 | - |
| - | - | 724.3 | 1665 | - | - | 0 | - |
| - | - | 641.1 | 2135 | - | - | 0 | - |
| - | - | 627 | 2222 | - | - | 0 | - |

m/z Charge Intensity FragmentType MassShift Position
120.08099365234375 0 39148.996
121.0843505859375 0 3100.3992
127.08686828613281 0 1074.0603
128.0707244873047 0 2267.3054
128.92758178710938 0 338.65286
129.1024169921875 0 11596.417
130.0497283935547 0 517.3956
130.0864715576172 0 5547.34 y Ammonia loss 7
130.10568237304688 0 571.96204
136.07586669921875 0 1044.7123
138.05514526367188 0 2333.6523
144.60792541503906 0 468.86118
147.1129913330078 0 8593.935 y 7
148.11659240722656 0 505.94608
155.0816650390625 0 20693.123 a Water loss 1
156.0657958984375 0 1944.2441
156.08506774902344 0 1728.5262
159.1128387451172 0 855.2532
171.11270141601562 0 1172.3132
173.0922088623047 0 71997.92 a 1
173.4386749267578 0 973.6194
174.0955047607422 0 4273.085
183.11293029785156 0 3263.979
184.06040954589844 0 1698.7745
184.57704162597656 0 506.84677
186.09129333496094 0 1006.4052
187.10784912109375 0 1532.4723
191.1180419921875 0 14116.444
192.1211395263672 0 1218.2944
198.12393188476562 0 935.18695
199.10751342773438 0 1688.2323
201.08702087402344 0 50519.16 b 1
201.12338256835938 0 2750.6868
202.09046936035156 0 4062.0876
207.11285400390625 0 733.40356
211.10699462890625 0 660.32074
216.13426208496094 0 6728.341 y Water loss 6
217.0821533203125 0 528.4394
217.11883544921875 0 788.65656 y Ammonia loss 6
219.1128692626953 0 22367.059
220.11611938476562 0 2910.9387
229.11802673339844 0 1695.1249
231.11294555664062 0 5184.4526
234.1448974609375 0 28963.879 y 6
235.14816284179688 0 2676.7822
240.42835998535156 0 588.0464
249.12330627441406 0 2488.9946
259.1075439453125 0 1397.1431
267.1498107910156 0 736.4387 y 3
277.1188659667969 0 1078.8583
298.13958740234375 0 2372.5486
298.9220275878906 0 674.99115
300.1553039550781 0 2413.5332
302.1497497558594 0 1029.2981 a Water loss 2
309.2032165527344 0 1693.3016
315.2031555175781 0 1457.4325 y Water loss 5
316.148681640625 0 760.1117
320.1600341796875 0 1441.7008 a 2
330.1446838378906 0 3007.7966 b Water loss 2
333.21319580078125 0 15691.9795 y 5
334.21563720703125 0 1944.5511
340.68402099609375 0 6063.2964 y 2
341.18585205078125 0 2294.845
344.91839599609375 0 588.34906
347.9393310546875 0 554.3611
348.1552429199219 0 21009.242 b 2
349.1580505371094 0 3370.238
350.1603088378906 0 714.76636
387.1867980957031 0 1310.5022
390.1676940917969 0 671.06683
404.0699768066406 0 615.2008
410.1976013183594 0 1109.5985
419.2286682128906 0 2313.625
420.1927490234375 0 654.7771
431.7171325683594 0 1069.1029 Precursor Water loss
435.1877136230469 0 1127.9431
439.8439025878906 0 6932.2476
440.1916809082031 0 1109.3594
440.84490966796875 0 2740.9885
441.2976379394531 0 1342.5166
441.8426513671875 0 733.03754
444.2449645996094 0 4653.844 y Water loss 4
447.2227783203125 0 2706.2517
459.1858825683594 0 1916.5149
462.2558288574219 0 10002.543 y 4
463.2588806152344 0 2843.226
487.8512878417969 0 656.0426
502.7541809082031 0 585.2034
515.2814331054688 0 992.84595 y Water loss 3
516.24365234375 0 736.2893
533.2923583984375 0 23075.645 y 3
534.2526245117188 0 1145.1749
534.2951049804688 0 6570.123
535.29931640625 0 829.66345
547.8883666992188 0 636.51025
548.236083984375 0 612.836 b 4
558.2540893554688 0 1041.524
662.3489990234375 0 1783.0383 y Water loss 2
680.3601684570312 0 72699.36 y 2
681.36328125 0 27051.268
682.3665771484375 0 6072.2266
791.39208984375 0 3207.561 y Water loss 1
792.3938598632812 0 1517.0205
809.4027709960938 0 5353.2104 y 1
810.4072875976562 0 1814.4462
811.4085083007812 0 1112.3984
1655.1158447265625 0 722.25494
1665.45458984375 0 724.30115
2135.1015625 0 641.1185
2221.77294921875 0 627.02576

Spectrum Details

|  |  |
| --- | --- |
| Matched peaks? Matched peaks The total absolute number of peaks matched. Additionally in brackets the total fraction of peaks matched and the total number of peaks is shown. | 26 (23.64% of 110) |
| FDR? FDR The false discovery rate estimated for this peptide. It is calculated by matching all theoretical fragments with a non-integer shift with the raw peaks for this spectrum. This is done with 40 different shifts. The resulting percentage is the average number of annotated peaks over the number of annotated peaks with the correct spectrum. | 0.09% |
| Satellite FDR? Satellite FDR See the FDR for details on its calculation. This satellite ion specific FDR only contains the satellite ions (d/w) for I/L/J positions. | - |
| PSM Score? PSM Score The PSM Score as given by Hecklib to this annotated spectrum. It is shown with three significant figures. | 355 |

#### Spectrum 3535? Spectrum 3535 The raw spectrum of this peptide as annotated by Hecklib. The fragments are coloured according to ion type (see legend). Any peaks with a star '\*' as text can be hovered over to see the full details, first the ion type second the mass shift type. By hovering over the amino acids in the peptide or ions in the legend the corresponding peaks are highlighted. By toggling the 'Unassigned' label you can turn the background (unassigned) peaks on or off in the plot. By updating the slider in the Ion legend you can update the spectrum to only show the top X% of the peaks with labels. The top X% means any peak that is within X% of the highest intensity. By dragging in the spectrum you can zoom in to a specific part of the spectrum and use 'Zoom Out' to get back to the original zoom level. The annotation of the spectrum is based on the given sequence in the peptides file and is done with different software so inconsistencies are likely. The peaks are annotated based on the given sequence, with 20 ppm tolerance.

Copy Data

##### Spectrum 3535 (TSV)

###### Preview

```
Loading example...
```

*Click on the button to copy the data to your clipboard.*

Mz MinMz MaxIntensity Max

WidthHeightPeptide font sizePeptide stroke widthSpectrum font sizeSpectrum stroke widthCompact peptide

Ion legend

wxyz

abcd

OtherUnassignedIon^Charge^_Position_Show for top:%

AEFAEVSK

03.95e+47.89e+41.18e+51.58e+5

Zoom Out

y^+1^_1_y^+1^_1_a^+1^_2_a^+1^_2_b^+1^_2_b^+1^_2_y^+1^_2_y^+1^_2_y^+2^_4_y^+1^_2_y^+2^_5_a^+1^_3_y^+1^_3_a^+1^_3_b^+1^_3_y^+1^_3_y^+2^_6_b^+1^_3_y^+2^_7_y^+2^_7_\*\*\*y^+1^_4_y^+1^_4_y^+1^_5_y^+1^_5_b^+1^_5_y^+1^_6_y^+1^_6_y^+1^_7_y^+1^_7_

034268410261368

Fragment Matches Table

Show background peaks

| Position | Ion type | Intensity | mz Theoretical | mz Error (Th) | mz Error (ppm) | Charge | Series Number |
| --- | --- | --- | --- | --- | --- | --- | --- |
| - | - | 1559 | 120.1 | - | - | 0 | - |
| - | - | 7.951E+04 | 120.1 | - | - | 0 | - |
| - | - | 6275 | 121.1 | - | - | 0 | - |
| - | - | 389.7 | 122 | - | - | 0 | - |
| - | - | 1420 | 127.1 | - | - | 0 | - |
| - | - | 366.8 | 127.1 | - | - | 0 | - |
| - | - | 4895 | 128.1 | - | - | 0 | - |
| - | - | 2.724E+04 | 129.1 | - | - | 0 | - |
| - | - | 934.1 | 130.1 | - | - | 0 | - |
| 8 | y | 1.115E+04 | 130.1 | 0.0003691 | 2.837 | +1 | 1 |
| - | - | 1868 | 130.1 | - | - | 0 | - |
| - | - | 631.4 | 131.1 | - | - | 0 | - |
| - | - | 420.7 | 134.1 | - | - | 0 | - |
| - | - | 4228 | 136.1 | - | - | 0 | - |
| - | - | 390.1 | 136.8 | - | - | 0 | - |
| - | - | 590 | 137.1 | - | - | 0 | - |
| - | - | 4385 | 138.1 | - | - | 0 | - |
| - | - | 631 | 138.1 | - | - | 0 | - |
| - | - | 478.3 | 143.1 | - | - | 0 | - |
| - | - | 434.8 | 145.1 | - | - | 0 | - |
| 8 | y | 1.94E+04 | 147.1 | 0.0003703 | 2.517 | +1 | 1 |
| - | - | 1154 | 148.1 | - | - | 0 | - |
| - | - | 417.9 | 149.4 | - | - | 0 | - |
| - | - | 400.7 | 152.4 | - | - | 0 | - |
| - | - | 498.6 | 153.1 | - | - | 0 | - |
| 2 | a | 4.292E+04 | 155.1 | 0.0003899 | 2.514 | +1 | 2 |
| - | - | 3023 | 156.1 | - | - | 0 | - |
| - | - | 3148 | 156.1 | - | - | 0 | - |
| - | - | 489.5 | 156.1 | - | - | 0 | - |
| - | - | 467.3 | 159.1 | - | - | 0 | - |
| - | - | 2306 | 159.1 | - | - | 0 | - |
| - | - | 1475 | 171.1 | - | - | 0 | - |
| 2 | a | 1.548E+05 | 173.1 | 0.0004606 | 2.661 | +1 | 2 |
| - | - | 913.8 | 174.1 | - | - | 0 | - |
| - | - | 1.148E+04 | 174.1 | - | - | 0 | - |
| - | - | 838.7 | 175.1 | - | - | 0 | - |
| - | - | 976.7 | 181.1 | - | - | 0 | - |
| - | - | 889.4 | 182.1 | - | - | 0 | - |
| 2 | b | 869 | 183.1 | 0.0004246 | 2.319 | +1 | 2 |
| - | - | 6083 | 183.1 | - | - | 0 | - |
| - | - | 2451 | 184.1 | - | - | 0 | - |
| - | - | 466 | 184.1 | - | - | 0 | - |
| - | - | 2185 | 186.1 | - | - | 0 | - |
| - | - | 3994 | 187.1 | - | - | 0 | - |
| - | - | 2.904E+04 | 191.1 | - | - | 0 | - |
| - | - | 3334 | 192.1 | - | - | 0 | - |
| - | - | 553.5 | 193.1 | - | - | 0 | - |
| - | - | 1847 | 198.1 | - | - | 0 | - |
| - | - | 3243 | 199.1 | - | - | 0 | - |
| 2 | b | 1.071E+05 | 201.1 | 0.0004343 | 2.16 | +1 | 2 |
| - | - | 7534 | 201.1 | - | - | 0 | - |
| - | - | 7523 | 202.1 | - | - | 0 | - |
| - | - | 651.1 | 203.1 | - | - | 0 | - |
| - | - | 552.3 | 204.1 | - | - | 0 | - |
| - | - | 633.7 | 207.1 | - | - | 0 | - |
| - | - | 1223 | 211.1 | - | - | 0 | - |
| - | - | 649.2 | 213.1 | - | - | 0 | - |
| - | - | 648.5 | 215.1 | - | - | 0 | - |
| 7 | y | 1.395E+04 | 216.1 | 0.0004367 | 2.021 | +1 | 2 |
| 7 | y | 2010 | 217.1 | 0.000552 | 2.542 | +1 | 2 |
| - | - | 1196 | 217.1 | - | - | 0 | - |
| - | - | 4.19E+04 | 219.1 | - | - | 0 | - |
| - | - | 4558 | 220.1 | - | - | 0 | - |
| - | - | 636.4 | 223.1 | - | - | 0 | - |
| 5 | y | 549.7 | 223.1 | 0.004046 | 18.13 | +2 | 4 |
| - | - | 895.2 | 226.1 | - | - | 0 | - |
| - | - | 677.2 | 226.2 | - | - | 0 | - |
| - | - | 640.6 | 227.1 | - | - | 0 | - |
| - | - | 3430 | 229.1 | - | - | 0 | - |
| - | - | 1.076E+04 | 231.1 | - | - | 0 | - |
| - | - | 1360 | 232.1 | - | - | 0 | - |
| - | - | 676.3 | 233.1 | - | - | 0 | - |
| 7 | y | 5.802E+04 | 234.1 | 0.0004158 | 1.776 | +1 | 2 |
| - | - | 4725 | 235.1 | - | - | 0 | - |
| - | - | 1201 | 240.1 | - | - | 0 | - |
| - | - | 1001 | 244.1 | - | - | 0 | - |
| - | - | 6808 | 249.1 | - | - | 0 | - |
| - | - | 623.3 | 250.1 | - | - | 0 | - |
| - | - | 937.4 | 255.1 | - | - | 0 | - |
| - | - | 3673 | 259.1 | - | - | 0 | - |
| - | - | 626.1 | 260.1 | - | - | 0 | - |
| 4 | y | 1150 | 267.2 | 0.001067 | 3.994 | +2 | 5 |
| - | - | 577.8 | 270.2 | - | - | 0 | - |
| - | - | 772.3 | 272.2 | - | - | 0 | - |
| - | - | 597 | 275.1 | - | - | 0 | - |
| - | - | 2653 | 277.1 | - | - | 0 | - |
| - | - | 1180 | 280.1 | - | - | 0 | - |
| - | - | 998.7 | 282.1 | - | - | 0 | - |
| - | - | 696.2 | 283.1 | - | - | 0 | - |
| - | - | 672.7 | 283.1 | - | - | 0 | - |
| - | - | 794.6 | 285.1 | - | - | 0 | - |
| - | - | 560.7 | 288.1 | - | - | 0 | - |
| - | - | 3311 | 298.1 | - | - | 0 | - |
| - | - | 725.4 | 299.1 | - | - | 0 | - |
| - | - | 5736 | 300.2 | - | - | 0 | - |
| - | - | 1215 | 301.2 | - | - | 0 | - |
| 3 | a | 3841 | 302.1 | 0.0003201 | 1.059 | +1 | 3 |
| - | - | 2944 | 303.1 | - | - | 0 | - |
| - | - | 797.1 | 303.2 | - | - | 0 | - |
| - | - | 1115 | 306.1 | - | - | 0 | - |
| - | - | 1840 | 309.2 | - | - | 0 | - |
| 6 | y | 2118 | 315.2 | 0.0004432 | 1.406 | +1 | 3 |
| - | - | 832.8 | 316.2 | - | - | 0 | - |
| 3 | a | 2913 | 320.2 | 0.0006197 | 1.935 | +1 | 3 |
| 3 | b | 7422 | 330.1 | 0.0004006 | 1.213 | +1 | 3 |
| - | - | 1319 | 331.1 | - | - | 0 | - |
| 6 | y | 3.302E+04 | 333.2 | 0.0004681 | 1.405 | +1 | 3 |
| - | - | 4499 | 334.2 | - | - | 0 | - |
| 3 | y | 1.241E+04 | 340.7 | 0.0005231 | 1.535 | +2 | 6 |
| - | - | 5340 | 341.2 | - | - | 0 | - |
| - | - | 641 | 341.7 | - | - | 0 | - |
| 3 | b | 4.501E+04 | 348.2 | 0.0003339 | 0.9592 | +1 | 3 |
| - | - | 7199 | 349.2 | - | - | 0 | - |
| - | - | 696.2 | 350.2 | - | - | 0 | - |
| - | - | 1603 | 369.2 | - | - | 0 | - |
| - | - | 679.2 | 375.9 | - | - | 0 | - |
| - | - | 1161 | 376.2 | - | - | 0 | - |
| - | - | 3623 | 387.2 | - | - | 0 | - |
| - | - | 1609 | 390.2 | - | - | 0 | - |
| 2 | y | 649.4 | 396.2 | 0.0004696 | 1.185 | +2 | 7 |
| - | - | 619.9 | 402.2 | - | - | 0 | - |
| 2 | y | 919.4 | 405.2 | 0.002969 | 7.328 | +2 | 7 |
| - | - | 701.5 | 405.8 | - | - | 0 | - |
| - | - | 1338 | 410.2 | - | - | 0 | - |
| - | - | 1461 | 417.2 | - | - | 0 | - |
| - | - | 6516 | 419.2 | - | - | 0 | - |
| - | - | 1656 | 420.2 | - | - | 0 | - |
| - | - | 889.7 | 421.8 | - | - | 0 | - |
| - | - | 657.6 | 422.8 | - | - | 0 | - |
| - | - | 2354 | 423.2 | - | - | 0 | - |
| - | - | 841 | 423.7 | - | - | 0 | - |
| - | - | 704.6 | 426.2 | - | - | 0 | - |
| 0 | Precursor | 952.6 | 431.7 | 0.0009557 | 2.214 | +2 | -1 |
| 0 | Precursor | 692 | 432.2 | 0.007666 | 17.74 | +2 | -1 |
| - | - | 2816 | 435.2 | - | - | 0 | - |
| - | - | 8412 | 439.8 | - | - | 0 | - |
| - | - | 2707 | 440.2 | - | - | 0 | - |
| 0 | Precursor | 1103 | 440.7 | 0.0008003 | 1.816 | +2 | -1 |
| - | - | 4737 | 440.8 | - | - | 0 | - |
| - | - | 690.6 | 441.2 | - | - | 0 | - |
| - | - | 5667 | 441.3 | - | - | 0 | - |
| - | - | 1789 | 441.8 | - | - | 0 | - |
| 5 | y | 8546 | 444.2 | 0.0004832 | 1.088 | +1 | 4 |
| - | - | 1972 | 445.2 | - | - | 0 | - |
| - | - | 7055 | 447.2 | - | - | 0 | - |
| - | - | 1772 | 448.2 | - | - | 0 | - |
| - | - | 624.5 | 458.4 | - | - | 0 | - |
| - | - | 4830 | 459.2 | - | - | 0 | - |
| - | - | 1305 | 460.2 | - | - | 0 | - |
| 5 | y | 2.442E+04 | 462.3 | 0.0005691 | 1.231 | +1 | 4 |
| - | - | 4184 | 463.3 | - | - | 0 | - |
| - | - | 636.3 | 472.2 | - | - | 0 | - |
| - | - | 1911 | 477.2 | - | - | 0 | - |
| - | - | 780.8 | 488.3 | - | - | 0 | - |
| - | - | 835.7 | 489.2 | - | - | 0 | - |
| - | - | 714.8 | 497.3 | - | - | 0 | - |
| - | - | 1132 | 506.3 | - | - | 0 | - |
| 4 | y | 2623 | 515.3 | 0.0005093 | 0.9883 | +1 | 5 |
| - | - | 1013 | 516.2 | - | - | 0 | - |
| - | - | 656.3 | 516.3 | - | - | 0 | - |
| 4 | y | 5.355E+04 | 533.3 | 0.0001375 | 0.2577 | +1 | 5 |
| - | - | 2138 | 534.3 | - | - | 0 | - |
| - | - | 1.366E+04 | 534.3 | - | - | 0 | - |
| - | - | 1228 | 535.3 | - | - | 0 | - |
| - | - | 3162 | 535.3 | - | - | 0 | - |
| 5 | b | 795.5 | 548.2 | 0.002865 | 5.227 | +1 | 5 |
| - | - | 1709 | 558.3 | - | - | 0 | - |
| - | - | 682.7 | 662.3 | - | - | 0 | - |
| 3 | y | 2921 | 662.4 | 0.0004547 | 0.6865 | +1 | 6 |
| - | - | 1461 | 663.4 | - | - | 0 | - |
| - | - | 2489 | 664.3 | - | - | 0 | - |
| 3 | y | 1.563E+05 | 680.4 | 2.188E-05 | 0.03215 | +1 | 6 |
| - | - | 5.55E+04 | 681.4 | - | - | 0 | - |
| - | - | 1.401E+04 | 682.4 | - | - | 0 | - |
| - | - | 1144 | 683.4 | - | - | 0 | - |
| 2 | y | 8243 | 791.4 | 0.001123 | 1.419 | +1 | 7 |
| - | - | 3382 | 792.4 | - | - | 0 | - |
| 2 | y | 1.288E+04 | 809.4 | 0.0009452 | 1.168 | +1 | 7 |
| - | - | 4366 | 810.4 | - | - | 0 | - |
| - | - | 1435 | 811.4 | - | - | 0 | - |
| - | - | 581.4 | 1354 | - | - | 0 | - |

m/z Charge Intensity FragmentType MassShift Position
120.06597137451172 0 1559.2363
120.08119201660156 0 79510.76
121.08456420898438 0 6274.5635
122.01673126220703 0 389.7498
127.08695220947266 0 1419.909
127.1133041381836 0 366.84433
128.07098388671875 0 4894.726
129.10263061523438 0 27240.248
130.05038452148438 0 934.0692
130.0866241455078 0 11151.465 y Ammonia loss 7
130.1060028076172 0 1867.7594
131.0900115966797 0 631.4359
134.11880493164062 0 420.69043
136.07608032226562 0 4228.349
136.84413146972656 0 390.0574
137.07960510253906 0 589.9657
138.05535888671875 0 4385.2207
138.09149169921875 0 630.98865
143.1183319091797 0 478.32596
145.06106567382812 0 434.75528
147.11317443847656 0 19397.209 y 7
148.116455078125 0 1153.8446
149.40133666992188 0 417.94324
152.40658569335938 0 400.7089
153.10301208496094 0 498.63376
155.08189392089844 0 42919.39 a Water loss 1
156.06591796875 0 3022.8467
156.08523559570312 0 3147.719
156.1298065185547 0 489.5328
159.07650756835938 0 467.286
159.11293029785156 0 2305.779
171.1130828857422 0 1475.2057
173.092529296875 0 154811.28 a 1
174.0879669189453 0 913.7781
174.09588623046875 0 11476.149
175.09689331054688 0 838.68225
181.09764099121094 0 976.69037
182.0814971923828 0 889.4315
183.07684326171875 0 868.9693 b Water loss 1
183.1131591796875 0 6083.002
184.0607452392578 0 2450.646
184.06964111328125 0 465.9959
186.09165954589844 0 2185.2017
187.108154296875 0 3993.7693
191.1182861328125 0 29040.684
192.12161254882812 0 3334.198
193.09671020507812 0 553.50824
198.12403869628906 0 1846.9172
199.10797119140625 0 3242.7488
201.08741760253906 0 107132.305 b 1
201.12367248535156 0 7534.4316
202.09078979492188 0 7523.312
203.09246826171875 0 651.0538
204.1024169921875 0 552.265
207.11251831054688 0 633.67786
211.10841369628906 0 1223.3759
213.08773803710938 0 649.22034
215.13938903808594 0 648.523
216.13470458984375 0 13945.867 y Water loss 6
217.11883544921875 0 2009.647 y Ammonia loss 6
217.13780212402344 0 1195.7356
219.11317443847656 0 41900.203
220.1163787841797 0 4557.8433
223.10931396484375 0 636.3758
223.12232971191406 0 549.7443 y Ammonia loss 4
226.11915588378906 0 895.1536
226.15538024902344 0 677.2039
227.1033172607422 0 640.60864
229.11874389648438 0 3429.859
231.11317443847656 0 10759.518
232.11647033691406 0 1359.8085
233.12838745117188 0 676.3485
234.14524841308594 0 58018.887 y 6
235.14859008789062 0 4725.0474
240.13436889648438 0 1200.7537
244.12962341308594 0 1001.0026
249.1238555908203 0 6807.6104
250.1268768310547 0 623.30524
255.13426208496094 0 937.4439
259.1081848144531 0 3672.8362
260.111328125 0 626.0651
267.1490478515625 0 1150.1768 y 3
270.1814880371094 0 577.8114
272.1605529785156 0 772.282
275.1395568847656 0 597.0221
277.1187744140625 0 2653.2515
280.1290588378906 0 1179.9935
282.14532470703125 0 998.70184
283.1297607421875 0 696.2028
283.14697265625 0 672.72754
285.12176513671875 0 794.6249
288.13525390625 0 560.701
298.1402282714844 0 3311.455
299.1446533203125 0 725.4495
300.15570068359375 0 5736.1587
301.15826416015625 0 1215.2705
302.1502380371094 0 3841.4592 a Water loss 2
303.134033203125 0 2944.2925
303.15252685546875 0 797.10254
306.1446533203125 0 1114.646
309.2041015625 0 1840.2057
315.203125 0 2118.2668 y Water loss 5
316.150146484375 0 832.7555
320.1611022949219 0 2913.438 a 2
330.1452331542969 0 7421.704 b Water loss 2
331.1484069824219 0 1319.3706
333.2137145996094 0 33022.49 y 5
334.21673583984375 0 4498.6226
340.6848449707031 0 12412.682 y 2
341.185791015625 0 5340.104
341.6872253417969 0 640.9552
348.1557312011719 0 45009.8 b 2
349.1589050292969 0 7199.1475
350.1611328125 0 696.2112
369.176513671875 0 1602.5193
375.87384033203125 0 679.2025
376.15179443359375 0 1160.5063
387.1883850097656 0 3622.7695
390.1666564941406 0 1609.2793
396.2008056640625 0 649.35425 y Water loss 1
402.2041931152344 0 619.92584
405.2085876464844 0 919.42456 y 1
405.81182861328125 0 701.51544
410.1989440917969 0 1337.5117
417.1781005859375 0 1461.4476
419.2295837402344 0 6516.2573
420.2323913574219 0 1656.1075
421.8321228027344 0 889.7431
422.83319091796875 0 657.5825
423.20574951171875 0 2354.4268
423.7095947265625 0 841.0441
426.23590087890625 0 704.62463
431.7198486328125 0 952.5574 Precursor Water loss
432.21856689453125 0 692.0229 Precursor Ammonia loss
435.18804931640625 0 2816.2551
439.8441162109375 0 8411.574
440.1923828125 0 2706.7842
440.7249755859375 0 1102.675 Precursor
440.8449401855469 0 4736.9683
441.19378662109375 0 690.6153
441.2978515625 0 5666.9336
441.8445739746094 0 1788.5436
444.2457580566406 0 8545.7295 y Water loss 4
445.24981689453125 0 1972.1958
447.2242736816406 0 7054.8306
448.2279357910156 0 1772.1315
458.40728759765625 0 624.4751
459.1880798339844 0 4830.496
460.1916809082031 0 1304.785
462.25640869140625 0 24420.275 y 4
463.259765625 0 4184.199
472.2398681640625 0 636.30536
477.19830322265625 0 1910.982
488.25323486328125 0 780.8282
489.2360534667969 0 835.73785
497.2705383300781 0 714.7993
506.2602844238281 0 1131.8574
515.2828979492188 0 2622.64 y Water loss 3
516.2449340820312 0 1012.64465
516.2861938476562 0 656.34515
533.2930908203125 0 53547.51 y 3
534.2551879882812 0 2137.6484
534.2962036132812 0 13662.115
535.2583618164062 0 1228.3683
535.2984008789062 0 3162.2063
548.2322387695312 0 795.47656 b 4
558.25537109375 0 1709.3195
662.2954711914062 0 682.6697
662.3512573242188 0 2920.7668 y Water loss 2
663.3566284179688 0 1461.0997
664.303955078125 0 2489.0938
680.3613891601562 0 156270.61 y 2
681.3642578125 0 55500.805
682.3671264648438 0 14005.957
683.3744506835938 0 1143.8384
791.3922729492188 0 8242.8545 y Water loss 1
792.3964233398438 0 3382.261
809.4030151367188 0 12883.645 y 1
810.4058227539062 0 4365.508
811.4089965820312 0 1435.0072
1354.3936767578125 0 581.40466

Spectrum Details

|  |  |
| --- | --- |
| Matched peaks? Matched peaks The total absolute number of peaks matched. Additionally in brackets the total fraction of peaks matched and the total number of peaks is shown. | 32 (17.68% of 181) |
| FDR? FDR The false discovery rate estimated for this peptide. It is calculated by matching all theoretical fragments with a non-integer shift with the raw peaks for this spectrum. This is done with 40 different shifts. The resulting percentage is the average number of annotated peaks over the number of annotated peaks with the correct spectrum. | 0.07% |
| Satellite FDR? Satellite FDR See the FDR for details on its calculation. This satellite ion specific FDR only contains the satellite ions (d/w) for I/L/J positions. | - |
| PSM Score? PSM Score The PSM Score as given by Hecklib to this annotated spectrum. It is shown with three significant figures. | 377 |

#### Reverse Lookup? Reverse Lookup All places where this read could be placed.

| Group | Segment | Template | Template Part | Read Part | Score | Unique |
| --- | --- | --- | --- | --- | --- | --- |
| Homo sapiens Heavy Chain | IGHV | IGHV1-18 | [6..13] | [0..8] | 32 | False |
| Homo sapiens Heavy Chain | IGHV | IGHV1-45 | [6..13] | [0..8] | 32 | False |
| Homo sapiens Heavy Chain | IGHV | IGHV5-51 | [6..13] | [0..8] | 32 | False |
| Homo sapiens Heavy Chain | IGHV | IGHV5-10-1 | [6..13] | [0..8] | 32 | False |
| Homo sapiens Heavy Chain | IGHV | IGHV1-24 | [6..13] | [0..8] | 32 | False |
| Homo sapiens Heavy Chain | IGHV | IGHV1-69 | [6..13] | [0..8] | 32 | False |
| Homo sapiens Heavy Chain | IGHV | IGHV1-2 | [6..13] | [0..8] | 32 | False |
| Homo sapiens Heavy Chain | IGHV | IGHV1-69-2 | [6..13] | [0..8] | 32 | False |
| Homo sapiens Heavy Chain | IGHV | IGHV1-3 | [6..13] | [0..8] | 32 | False |
| Homo sapiens Heavy Chain | IGHV | IGHV1-46 | [6..13] | [0..8] | 32 | False |
| Homo sapiens Heavy Chain | IGHV | IGHV1-8 | [6..13] | [0..8] | 32 | False |

#### Meta Information from Multiple reads

##### Number of combined reads

3

##### Intensity

0.5832

##### TotalArea

7.343E+05

#### Positional Score

Copy Data

##### Positional Score (TSV)

###### Preview

```
Loading example...
```

*Click on the button to copy the data to your clipboard.*

1001234567

Label Value
"0" 0.323
"1" 0.33
"2" 0.33
"3" 0.33
"4" 0.33
"5" 0.33
"6" 0.333
"7" 0.333

#### Meta Information from PEAKS

##### Scan Identifier

F1:3535

##### Original sequence

A

E

F

A

E

V

S

K

##### Posttranslational Modifications

##### Source File

D:\separate\_stitch\_analyses\xle-disambiguation\raw\20210323\_F1\_UM1\_Peng0013\_SA\_F59\_ingel\_3ug\_ELA.raw

##### Fraction

1

##### Scan Feature

F1:1913

##### De Novo Score

99

##### ConfidenceScore

99

### m/z

440.725

##### Mass

879.4338

##### Charge

2

##### Retention Time

18.34

##### Predicted Retention Time

-

##### Area

5.373E+05

##### Parts Per Million

1.8

##### Fragmentation mode

HCD

##### Originating file

01 D:\separate\_stitch\_analyses\xle-disambiguation\20210325\_F59\_3ug\_DENOVO\_12.csv

#### Meta Information from PEAKS

##### Scan Identifier

F2:3462

##### Original sequence

A

E

F

A

E

V

S

K

##### Posttranslational Modifications

##### Source File

D:\separate\_stitch\_analyses\xle-disambiguation\raw\20210323\_F1\_UM1\_Peng0013\_SA\_F59\_ingel\_3ug\_TL.raw

##### Fraction

2

##### Scan Feature

F2:1903

##### De Novo Score

98

##### ConfidenceScore

98

### m/z

440.7246

##### Mass

879.4338

##### Charge

2

##### Retention Time

18.18

##### Predicted Retention Time

-

##### Area

1.97E+05

##### Parts Per Million

1

##### Fragmentation mode

ETHCD

##### Originating file

01 D:\separate\_stitch\_analyses\xle-disambiguation\20210325\_F59\_3ug\_DENOVO\_12.csv

#### Meta Information from PEAKS

##### Scan Identifier

F4:3397

##### Original sequence

A

E

F

A

E

V

S

K

##### Posttranslational Modifications

##### Source File

D:\separate\_stitch\_analyses\xle-disambiguation\raw\20210323\_F1\_UM1\_Peng0013\_SA\_F59\_ingel\_3ug\_tryp.raw

##### Fraction

4

##### Scan Feature

-

##### De Novo Score

96

##### ConfidenceScore

99

### m/z

440.7247

##### Mass

879.4338

##### Charge

2

##### Retention Time

18.12

##### Predicted Retention Time

13.07

##### Area

0

##### Parts Per Million

1.3

##### Fragmentation mode

HCD

##### Originating file

01 D:\separate\_stitch\_analyses\xle-disambiguation\20210325\_F59\_3ug\_DENOVO\_12.csv
