## Supplementary material for "A handle on mass coincidence errors in *de novo* sequencing of antibodies by bottom-up proteomics": Combined_071.html

Details Combined\_071 | Stitch OverviewUndefined

### Read Combined\_071

#### Sequence (length=8)

KJSTAASF

#### Spectrum 4426? Spectrum 4426 The raw spectrum of this peptide as annotated by Hecklib. The fragments are coloured according to ion type (see legend). Any peaks with a star '\*' as text can be hovered over to see the full details, first the ion type second the mass shift type. By hovering over the amino acids in the peptide or ions in the legend the corresponding peaks are highlighted. By toggling the 'Unassigned' label you can turn the background (unassigned) peaks on or off in the plot. By updating the slider in the Ion legend you can update the spectrum to only show the top X% of the peaks with labels. The top X% means any peak that is within X% of the highest intensity. By dragging in the spectrum you can zoom in to a specific part of the spectrum and use 'Zoom Out' to get back to the original zoom level. The annotation of the spectrum is based on the given sequence in the peptides file and is done with different software so inconsistencies are likely. The peaks are annotated based on the given sequence, with 20 ppm tolerance.

Copy Data

##### Spectrum 4426 (TSV)

###### Preview

```
Loading example...
```

*Click on the button to copy the data to your clipboard.*

Mz MinMz MaxIntensity Max

WidthHeightPeptide font sizePeptide stroke widthSpectrum font sizeSpectrum stroke widthCompact peptide

Ion legend

wxyz

abcd

OtherUnassignedIon^Charge^_Position_Show for top:%

KJSTAASF

01.92e+63.83e+65.75e+67.66e+6

Zoom Out

y^+1^_1_c^+2^_4_y^+1^_2_c^+1^_2_c^+2^_5_y^+1^_2_c^+1^_2_c^+2^_6_y^+1^_3_y^+1^_3_c^+1^_3_c^+1^_3_c^+2^_7_c^+2^_7_c^+1^_3_y^+1^_4_c^+1^_4_c^+1^_4_c^+1^_4_y^+1^_5_y^+1^_5_c^+1^_5_c^+1^_5_c^+1^_5_y^+1^_6_c^+1^_6_y^+1^_6_c^+1^_6_w^+1^_7_c^+1^_7_z^+1^_7_c^+1^_7_y^+1^_7_z^+1^_7_y^+1^_7_

0779155823383117

Fragment Matches Table

Show background peaks

| Position | Ion type | Intensity | mz Theoretical | mz Error (Th) | mz Error (ppm) | Charge | Series Number |
| --- | --- | --- | --- | --- | --- | --- | --- |
| - | - | 1.094E+06 | 120.1 | - | - | 0 | - |
| - | - | 8.643E+04 | 121.1 | - | - | 0 | - |
| - | - | 6172 | 126.1 | - | - | 0 | - |
| - | - | 3.261E+04 | 128.1 | - | - | 0 | - |
| - | - | 4.324E+05 | 129.1 | - | - | 0 | - |
| - | - | 1.646E+04 | 130.1 | - | - | 0 | - |
| - | - | 3918 | 130.3 | - | - | 0 | - |
| - | - | 1.418E+04 | 131 | - | - | 0 | - |
| - | - | 2.192E+04 | 131.1 | - | - | 0 | - |
| - | - | 4330 | 134.8 | - | - | 0 | - |
| - | - | 4169 | 135.2 | - | - | 0 | - |
| - | - | 4663 | 139.1 | - | - | 0 | - |
| - | - | 1.372E+04 | 141.1 | - | - | 0 | - |
| - | - | 2.262E+04 | 145.1 | - | - | 0 | - |
| - | - | 8923 | 146.1 | - | - | 0 | - |
| - | - | 1.334E+05 | 146.1 | - | - | 0 | - |
| - | - | 6306 | 147.1 | - | - | 0 | - |
| - | - | 1.05E+04 | 149 | - | - | 0 | - |
| - | - | 2.307E+04 | 149.1 | - | - | 0 | - |
| - | - | 1.172E+04 | 153.1 | - | - | 0 | - |
| - | - | 9.117E+04 | 155.1 | - | - | 0 | - |
| - | - | 7.913E+04 | 159.1 | - | - | 0 | - |
| 8 | y | 2.756E+06 | 166.1 | 0.0005369 | 3.233 | +1 | 1 |
| - | - | 2.399E+05 | 167.1 | - | - | 0 | - |
| - | - | 8511 | 168.1 | - | - | 0 | - |
| - | - | 6.351E+04 | 171.1 | - | - | 0 | - |
| - | - | 9037 | 172.1 | - | - | 0 | - |
| - | - | 7713 | 172.1 | - | - | 0 | - |
| - | - | 7737 | 172.1 | - | - | 0 | - |
| - | - | 1.917E+04 | 173.1 | - | - | 0 | - |
| - | - | 8193 | 173.5 | - | - | 0 | - |
| - | - | 5218 | 180.1 | - | - | 0 | - |
| - | - | 4537 | 181.1 | - | - | 0 | - |
| - | - | 4401 | 184.9 | - | - | 0 | - |
| - | - | 4859 | 185.1 | - | - | 0 | - |
| - | - | 9488 | 189.1 | - | - | 0 | - |
| - | - | 6.245E+04 | 189.1 | - | - | 0 | - |
| - | - | 6406 | 190.1 | - | - | 0 | - |
| - | - | 1.174E+04 | 196.2 | - | - | 0 | - |
| - | - | 5.937E+04 | 197.2 | - | - | 0 | - |
| - | - | 5116 | 198.1 | - | - | 0 | - |
| - | - | 5316 | 198.1 | - | - | 0 | - |
| - | - | 6631 | 198.2 | - | - | 0 | - |
| - | - | 1.714E+04 | 200.1 | - | - | 0 | - |
| - | - | 1.819E+04 | 201.1 | - | - | 0 | - |
| - | - | 5749 | 202.1 | - | - | 0 | - |
| - | - | 1.691E+05 | 207.1 | - | - | 0 | - |
| - | - | 1.544E+04 | 208.1 | - | - | 0 | - |
| - | - | 4611 | 210.4 | - | - | 0 | - |
| - | - | 6128 | 212.1 | - | - | 0 | - |
| - | - | 6013 | 213.1 | - | - | 0 | - |
| - | - | 5.126E+04 | 214.2 | - | - | 0 | - |
| - | - | 1.513E+05 | 215.2 | - | - | 0 | - |
| - | - | 8034 | 216.1 | - | - | 0 | - |
| - | - | 6.702E+05 | 216.2 | - | - | 0 | - |
| - | - | 8254 | 217.1 | - | - | 0 | - |
| - | - | 7.112E+04 | 217.2 | - | - | 0 | - |
| - | - | 2.041E+04 | 218.2 | - | - | 0 | - |
| 4 | c | 9293 | 224.1 | 0.002411 | 10.75 | +2 | 4 |
| - | - | 3.989E+04 | 224.2 | - | - | 0 | - |
| - | - | 3.104E+04 | 225.2 | - | - | 0 | - |
| - | - | 2.653E+04 | 228.2 | - | - | 0 | - |
| - | - | 1.89E+04 | 230.1 | - | - | 0 | - |
| 7 | y | 2.253E+05 | 235.1 | 0.0005423 | 2.307 | +1 | 2 |
| - | - | 2.484E+04 | 236.1 | - | - | 0 | - |
| - | - | 8.477E+04 | 241.2 | - | - | 0 | - |
| - | - | 3.676E+04 | 242.1 | - | - | 0 | - |
| - | - | 2.914E+04 | 242.2 | - | - | 0 | - |
| 2 | c | 2.378E+05 | 242.2 | 0.0004489 | 1.853 | +1 | 2 |
| - | - | 2.642E+04 | 243.2 | - | - | 0 | - |
| - | - | 6190 | 243.5 | - | - | 0 | - |
| - | - | 2.541E+04 | 244.1 | - | - | 0 | - |
| - | - | 6473 | 244.2 | - | - | 0 | - |
| 5 | c | 1.716E+04 | 251.2 | 0.0005157 | 2.053 | +2 | 5 |
| 7 | y | 1.628E+06 | 253.1 | 0.000552 | 2.181 | +1 | 2 |
| - | - | 1.891E+05 | 254.1 | - | - | 0 | - |
| - | - | 1.545E+04 | 255.1 | - | - | 0 | - |
| - | - | 2.04E+05 | 258.2 | - | - | 0 | - |
| 2 | c | 6.439E+05 | 259.2 | 0.0003127 | 1.206 | +1 | 2 |
| - | - | 5.811E+04 | 260.1 | - | - | 0 | - |
| - | - | 7795 | 260.1 | - | - | 0 | - |
| - | - | 6.08E+04 | 260.2 | - | - | 0 | - |
| - | - | 2.597E+04 | 263.1 | - | - | 0 | - |
| - | - | 6327 | 266.2 | - | - | 0 | - |
| - | - | 6512 | 268.2 | - | - | 0 | - |
| - | - | 6248 | 269.2 | - | - | 0 | - |
| - | - | 1.683E+04 | 269.2 | - | - | 0 | - |
| - | - | 5760 | 272.2 | - | - | 0 | - |
| - | - | 1.468E+04 | 277.2 | - | - | 0 | - |
| - | - | 4903 | 279.8 | - | - | 0 | - |
| - | - | 1.079E+04 | 283.2 | - | - | 0 | - |
| - | - | 8943 | 283.2 | - | - | 0 | - |
| - | - | 1.951E+04 | 284.2 | - | - | 0 | - |
| - | - | 2.651E+04 | 284.2 | - | - | 0 | - |
| - | - | 6.102E+04 | 285.2 | - | - | 0 | - |
| - | - | 7596 | 286.1 | - | - | 0 | - |
| - | - | 6850 | 286.2 | - | - | 0 | - |
| 6 | c | 4.18E+04 | 286.7 | 0.0004677 | 1.631 | +2 | 6 |
| - | - | 9879 | 287.2 | - | - | 0 | - |
| - | - | 8999 | 287.2 | - | - | 0 | - |
| - | - | 7722 | 288.1 | - | - | 0 | - |
| - | - | 1.931E+04 | 289.2 | - | - | 0 | - |
| - | - | 1.406E+04 | 293.2 | - | - | 0 | - |
| - | - | 7896 | 294.2 | - | - | 0 | - |
| - | - | 7821 | 295.1 | - | - | 0 | - |
| - | - | 9403 | 296.2 | - | - | 0 | - |
| - | - | 1.361E+04 | 298.7 | - | - | 0 | - |
| - | - | 1.138E+04 | 301.2 | - | - | 0 | - |
| - | - | 3.572E+04 | 301.2 | - | - | 0 | - |
| - | - | 7974 | 302.2 | - | - | 0 | - |
| - | - | 8389 | 302.2 | - | - | 0 | - |
| - | - | 2.284E+04 | 302.2 | - | - | 0 | - |
| - | - | 2.043E+04 | 303.2 | - | - | 0 | - |
| - | - | 2.7E+05 | 303.2 | - | - | 0 | - |
| - | - | 1.064E+04 | 304.2 | - | - | 0 | - |
| - | - | 3.462E+04 | 304.2 | - | - | 0 | - |
| 6 | y | 1.667E+05 | 306.1 | 0.0006752 | 2.206 | +1 | 3 |
| - | - | 1.098E+04 | 306.2 | - | - | 0 | - |
| - | - | 2.5E+04 | 307.1 | - | - | 0 | - |
| - | - | 4.608E+04 | 307.2 | - | - | 0 | - |
| - | - | 3.86E+04 | 307.7 | - | - | 0 | - |
| - | - | 7716 | 308.2 | - | - | 0 | - |
| - | - | 3.233E+04 | 311.2 | - | - | 0 | - |
| - | - | 2.057E+04 | 312.2 | - | - | 0 | - |
| - | - | 6209 | 312.7 | - | - | 0 | - |
| - | - | 4.986E+04 | 313.2 | - | - | 0 | - |
| - | - | 1.83E+04 | 313.2 | - | - | 0 | - |
| - | - | 4.506E+04 | 313.2 | - | - | 0 | - |
| - | - | 5363 | 313.4 | - | - | 0 | - |
| - | - | 5408 | 314.2 | - | - | 0 | - |
| - | - | 7972 | 314.2 | - | - | 0 | - |
| - | - | 9840 | 316.2 | - | - | 0 | - |
| - | - | 2.922E+04 | 319.2 | - | - | 0 | - |
| - | - | 3.921E+04 | 321.2 | - | - | 0 | - |
| - | - | 9213 | 321.7 | - | - | 0 | - |
| 6 | y | 6.073E+04 | 324.2 | 0.0007002 | 2.16 | +1 | 3 |
| - | - | 1.091E+04 | 325.2 | - | - | 0 | - |
| - | - | 8409 | 327.2 | - | - | 0 | - |
| - | - | 7840 | 327.2 | - | - | 0 | - |
| 3 | c | 2.332E+04 | 328.2 | 0.0004555 | 1.388 | +1 | 3 |
| 3 | c | 1.621E+05 | 329.2 | 0.0007233 | 2.197 | +1 | 3 |
| 7 | c | 1.868E+04 | 329.7 | 0.0006617 | 2.007 | +2 | 7 |
| 7 | c | 7.413E+04 | 330.2 | 0.0007499 | 2.271 | +2 | 7 |
| - | - | 2.073E+04 | 330.2 | - | - | 0 | - |
| - | - | 1.971E+04 | 330.7 | - | - | 0 | - |
| - | - | 2.731E+05 | 331.2 | - | - | 0 | - |
| - | - | 2.436E+04 | 331.2 | - | - | 0 | - |
| - | - | 3.069E+04 | 332.2 | - | - | 0 | - |
| - | - | 1.074E+04 | 339.2 | - | - | 0 | - |
| - | - | 1.806E+04 | 343.2 | - | - | 0 | - |
| - | - | 1.121E+04 | 345.2 | - | - | 0 | - |
| - | - | 5.235E+05 | 345.2 | - | - | 0 | - |
| 3 | c | 7.71E+05 | 346.2 | 0.0002667 | 0.7704 | +1 | 3 |
| - | - | 1.299E+05 | 347.2 | - | - | 0 | - |
| - | - | 1.55E+04 | 348.3 | - | - | 0 | - |
| - | - | 9262 | 354.2 | - | - | 0 | - |
| - | - | 9968 | 354.3 | - | - | 0 | - |
| - | - | 4.598E+04 | 355.2 | - | - | 0 | - |
| - | - | 7812 | 355.2 | - | - | 0 | - |
| - | - | 8561 | 356.2 | - | - | 0 | - |
| - | - | 1.708E+04 | 356.3 | - | - | 0 | - |
| - | - | 7326 | 360.2 | - | - | 0 | - |
| - | - | 1.093E+04 | 367.2 | - | - | 0 | - |
| - | - | 1.417E+04 | 368.2 | - | - | 0 | - |
| - | - | 2.763E+04 | 372.2 | - | - | 0 | - |
| - | - | 1.131E+04 | 372.3 | - | - | 0 | - |
| - | - | 2.123E+04 | 372.7 | - | - | 0 | - |
| - | - | 6467 | 373.2 | - | - | 0 | - |
| - | - | 4.798E+04 | 373.2 | - | - | 0 | - |
| - | - | 8534 | 374.2 | - | - | 0 | - |
| 5 | y | 4.402E+04 | 377.2 | 0.0003657 | 0.9694 | +1 | 4 |
| - | - | 1.174E+04 | 378.2 | - | - | 0 | - |
| - | - | 9632 | 378.2 | - | - | 0 | - |
| - | - | 9431 | 380.2 | - | - | 0 | - |
| - | - | 9.793E+04 | 380.7 | - | - | 0 | - |
| - | - | 4.794E+04 | 381.2 | - | - | 0 | - |
| - | - | 2.013E+04 | 382.2 | - | - | 0 | - |
| - | - | 5365 | 382.2 | - | - | 0 | - |
| - | - | 2.001E+04 | 382.2 | - | - | 0 | - |
| - | - | 4.332E+04 | 384.3 | - | - | 0 | - |
| - | - | 9.756E+05 | 385.3 | - | - | 0 | - |
| - | - | 2.992E+05 | 386.3 | - | - | 0 | - |
| - | - | 4.671E+04 | 387.3 | - | - | 0 | - |
| - | - | 1.115E+04 | 389.7 | - | - | 0 | - |
| - | - | 1.586E+04 | 390.2 | - | - | 0 | - |
| - | - | 3.522E+04 | 394.2 | - | - | 0 | - |
| - | - | 2.153E+04 | 394.7 | - | - | 0 | - |
| - | - | 1.1E+05 | 395.2 | - | - | 0 | - |
| - | - | 4.71E+04 | 395.7 | - | - | 0 | - |
| - | - | 1.578E+04 | 396.2 | - | - | 0 | - |
| - | - | 5946 | 398.2 | - | - | 0 | - |
| - | - | 1.892E+04 | 398.2 | - | - | 0 | - |
| - | - | 6.265E+04 | 400.2 | - | - | 0 | - |
| - | - | 5.314E+04 | 400.3 | - | - | 0 | - |
| - | - | 1.468E+04 | 401.2 | - | - | 0 | - |
| - | - | 9347 | 401.3 | - | - | 0 | - |
| - | - | 1.771E+04 | 402.3 | - | - | 0 | - |
| - | - | 6159 | 403.2 | - | - | 0 | - |
| - | - | 6526 | 403.3 | - | - | 0 | - |
| - | - | 4.392E+05 | 403.7 | - | - | 0 | - |
| - | - | 1.74E+05 | 404.2 | - | - | 0 | - |
| - | - | 2.768E+05 | 404.3 | - | - | 0 | - |
| - | - | 5.826E+04 | 404.7 | - | - | 0 | - |
| - | - | 6.35E+04 | 405.3 | - | - | 0 | - |
| - | - | 1.221E+04 | 408.2 | - | - | 0 | - |
| - | - | 2.233E+04 | 411.3 | - | - | 0 | - |
| - | - | 6.102E+04 | 412.3 | - | - | 0 | - |
| - | - | 1.701E+04 | 413.3 | - | - | 0 | - |
| - | - | 2.769E+04 | 414.2 | - | - | 0 | - |
| - | - | 7044 | 414.3 | - | - | 0 | - |
| - | - | 3.696E+04 | 416.3 | - | - | 0 | - |
| - | - | 1.073E+04 | 418.2 | - | - | 0 | - |
| - | - | 1.432E+05 | 418.2 | - | - | 0 | - |
| - | - | 2.943E+04 | 419.2 | - | - | 0 | - |
| - | - | 1.766E+04 | 424.2 | - | - | 0 | - |
| - | - | 5839 | 424.3 | - | - | 0 | - |
| - | - | 2.712E+04 | 426.2 | - | - | 0 | - |
| - | - | 1.342E+04 | 426.3 | - | - | 0 | - |
| - | - | 1.036E+04 | 428.2 | - | - | 0 | - |
| - | - | 9469 | 429.2 | - | - | 0 | - |
| 4 | c | 6.738E+04 | 429.3 | 0.0006591 | 1.535 | +1 | 4 |
| 4 | c | 1.187E+05 | 430.3 | 0.001049 | 2.438 | +1 | 4 |
| - | - | 2.994E+04 | 431.3 | - | - | 0 | - |
| - | - | 6818 | 432.3 | - | - | 0 | - |
| - | - | 6826 | 436.3 | - | - | 0 | - |
| - | - | 8997 | 438.3 | - | - | 0 | - |
| - | - | 7584 | 439.3 | - | - | 0 | - |
| - | - | 1.018E+04 | 441.2 | - | - | 0 | - |
| - | - | 1.181E+04 | 442.2 | - | - | 0 | - |
| - | - | 1.194E+04 | 443.3 | - | - | 0 | - |
| - | - | 4.821E+04 | 444.2 | - | - | 0 | - |
| - | - | 1.175E+04 | 445.2 | - | - | 0 | - |
| - | - | 1.85E+04 | 446.3 | - | - | 0 | - |
| 4 | c | 3.814E+06 | 447.3 | 0.0009281 | 2.075 | +1 | 4 |
| - | - | 8.708E+05 | 448.3 | - | - | 0 | - |
| - | - | 1.138E+05 | 449.3 | - | - | 0 | - |
| - | - | 1.358E+04 | 453.3 | - | - | 0 | - |
| - | - | 9445 | 454.2 | - | - | 0 | - |
| - | - | 9512 | 455.3 | - | - | 0 | - |
| - | - | 8883 | 457.3 | - | - | 0 | - |
| - | - | 2.126E+04 | 459.3 | - | - | 0 | - |
| - | - | 9840 | 460.2 | - | - | 0 | - |
| - | - | 9111 | 461.3 | - | - | 0 | - |
| - | - | 4.783E+04 | 465.3 | - | - | 0 | - |
| - | - | 5916 | 466.3 | - | - | 0 | - |
| - | - | 3.982E+04 | 471.3 | - | - | 0 | - |
| - | - | 8573 | 472.2 | - | - | 0 | - |
| - | - | 6745 | 472.3 | - | - | 0 | - |
| - | - | 7.27E+04 | 473.3 | - | - | 0 | - |
| - | - | 3.052E+04 | 474.3 | - | - | 0 | - |
| - | - | 7013 | 475.2 | - | - | 0 | - |
| - | - | 9727 | 475.3 | - | - | 0 | - |
| 4 | y | 2.518E+04 | 478.2 | 0.001088 | 2.275 | +1 | 5 |
| - | - | 6182 | 481.3 | - | - | 0 | - |
| - | - | 1.162E+04 | 482.3 | - | - | 0 | - |
| - | - | 1.223E+04 | 483.3 | - | - | 0 | - |
| - | - | 8.991E+04 | 483.3 | - | - | 0 | - |
| - | - | 8784 | 484.3 | - | - | 0 | - |
| - | - | 1.448E+04 | 484.3 | - | - | 0 | - |
| - | - | 1.897E+04 | 485.3 | - | - | 0 | - |
| - | - | 1.447E+04 | 487.3 | - | - | 0 | - |
| - | - | 8289 | 492.3 | - | - | 0 | - |
| - | - | 1.179E+04 | 493.2 | - | - | 0 | - |
| - | - | 8458 | 495.3 | - | - | 0 | - |
| 4 | y | 3.783E+04 | 496.2 | 0.0008994 | 1.812 | +1 | 5 |
| - | - | 2.217E+04 | 497.2 | - | - | 0 | - |
| - | - | 1.617E+04 | 499.3 | - | - | 0 | - |
| 5 | c | 3.812E+04 | 500.3 | 0.0008377 | 1.674 | +1 | 5 |
| 5 | c | 5.251E+05 | 501.3 | 0.000892 | 1.779 | +1 | 5 |
| - | - | 1.295E+05 | 502.3 | - | - | 0 | - |
| - | - | 2.198E+04 | 503.3 | - | - | 0 | - |
| - | - | 1.154E+04 | 505.2 | - | - | 0 | - |
| - | - | 6200 | 508.3 | - | - | 0 | - |
| - | - | 1.42E+04 | 509.3 | - | - | 0 | - |
| - | - | 8135 | 510.3 | - | - | 0 | - |
| - | - | 2.043E+04 | 510.3 | - | - | 0 | - |
| - | - | 1.246E+04 | 511.3 | - | - | 0 | - |
| - | - | 1.55E+04 | 513.3 | - | - | 0 | - |
| - | - | 1.921E+04 | 515.3 | - | - | 0 | - |
| - | - | 1.201E+04 | 516.3 | - | - | 0 | - |
| 5 | c | 5.668E+06 | 518.3 | 0.0008932 | 1.723 | +1 | 5 |
| - | - | 1.403E+06 | 519.3 | - | - | 0 | - |
| - | - | 6128 | 520.2 | - | - | 0 | - |
| - | - | 2.468E+05 | 520.3 | - | - | 0 | - |
| - | - | 7678 | 521.3 | - | - | 0 | - |
| - | - | 3.796E+04 | 526.3 | - | - | 0 | - |
| - | - | 7945 | 527.3 | - | - | 0 | - |
| - | - | 1.171E+04 | 528.3 | - | - | 0 | - |
| - | - | 2.838E+04 | 528.3 | - | - | 0 | - |
| - | - | 2.792E+04 | 530.3 | - | - | 0 | - |
| - | - | 1.86E+04 | 531.3 | - | - | 0 | - |
| - | - | 8382 | 531.3 | - | - | 0 | - |
| - | - | 9743 | 535.3 | - | - | 0 | - |
| - | - | 8.18E+04 | 536.3 | - | - | 0 | - |
| - | - | 2.79E+04 | 537.3 | - | - | 0 | - |
| - | - | 7418 | 541.3 | - | - | 0 | - |
| - | - | 8777 | 543.3 | - | - | 0 | - |
| - | - | 2.602E+05 | 544.3 | - | - | 0 | - |
| - | - | 1.553E+05 | 545.4 | - | - | 0 | - |
| - | - | 1.273E+04 | 546.2 | - | - | 0 | - |
| - | - | 2.306E+04 | 546.3 | - | - | 0 | - |
| - | - | 3.402E+04 | 546.4 | - | - | 0 | - |
| - | - | 7967 | 547.3 | - | - | 0 | - |
| - | - | 8375 | 552.3 | - | - | 0 | - |
| - | - | 6506 | 553.3 | - | - | 0 | - |
| - | - | 2.67E+05 | 554.3 | - | - | 0 | - |
| - | - | 6.56E+04 | 555.3 | - | - | 0 | - |
| - | - | 2.039E+04 | 556.3 | - | - | 0 | - |
| - | - | 1.468E+04 | 558.3 | - | - | 0 | - |
| - | - | 1.662E+05 | 564.3 | - | - | 0 | - |
| 3 | y | 1.146E+05 | 565.3 | 0.00146 | 2.584 | +1 | 6 |
| - | - | 3.252E+04 | 566.3 | - | - | 0 | - |
| - | - | 1.372E+04 | 568.3 | - | - | 0 | - |
| - | - | 2.57E+04 | 570.3 | - | - | 0 | - |
| - | - | 1.804E+04 | 571.3 | - | - | 0 | - |
| 6 | c | 1.381E+06 | 572.3 | 0.0007655 | 1.338 | +1 | 6 |
| - | - | 3.731E+05 | 573.3 | - | - | 0 | - |
| - | - | 8.626E+04 | 574.3 | - | - | 0 | - |
| - | - | 1.51E+04 | 575.4 | - | - | 0 | - |
| - | - | 7674 | 578.3 | - | - | 0 | - |
| - | - | 6.664E+05 | 582.3 | - | - | 0 | - |
| 3 | y | 6.575E+05 | 583.3 | 0.0004284 | 0.7344 | +1 | 6 |
| - | - | 1.722E+05 | 584.3 | - | - | 0 | - |
| - | - | 2.696E+04 | 585.3 | - | - | 0 | - |
| - | - | 3.061E+04 | 588.3 | - | - | 0 | - |
| 6 | c | 5.568E+06 | 589.4 | 0.001011 | 1.715 | +1 | 6 |
| - | - | 1.614E+06 | 590.4 | - | - | 0 | - |
| - | - | 3.16E+05 | 591.4 | - | - | 0 | - |
| - | - | 1.679E+04 | 592.4 | - | - | 0 | - |
| - | - | 8267 | 595.4 | - | - | 0 | - |
| - | - | 1.936E+04 | 596.3 | - | - | 0 | - |
| - | - | 1.948E+04 | 597.3 | - | - | 0 | - |
| - | - | 1.805E+04 | 605.3 | - | - | 0 | - |
| - | - | 2.257E+04 | 606.3 | - | - | 0 | - |
| - | - | 8098 | 611.4 | - | - | 0 | - |
| - | - | 4.413E+04 | 613.4 | - | - | 0 | - |
| - | - | 1.196E+05 | 614.4 | - | - | 0 | - |
| - | - | 4.296E+04 | 615.4 | - | - | 0 | - |
| - | - | 1.408E+04 | 616.4 | - | - | 0 | - |
| - | - | 1.246E+05 | 623.4 | - | - | 0 | - |
| - | - | 2.974E+04 | 624.3 | - | - | 0 | - |
| - | - | 1.48E+04 | 625.3 | - | - | 0 | - |
| - | - | 2.226E+05 | 631.4 | - | - | 0 | - |
| - | - | 1.313E+05 | 632.4 | - | - | 0 | - |
| - | - | 3.533E+04 | 633.4 | - | - | 0 | - |
| 2 | w | 5.266E+04 | 637.3 | 0.0006647 | 1.043 | +1 | 7 |
| - | - | 8162 | 638.3 | - | - | 0 | - |
| - | - | 5.446E+05 | 641.4 | - | - | 0 | - |
| - | - | 1.794E+05 | 642.4 | - | - | 0 | - |
| - | - | 3.876E+04 | 643.4 | - | - | 0 | - |
| 7 | c | 1.284E+06 | 659.4 | 0.0005364 | 0.8135 | +1 | 7 |
| - | - | 3.876E+05 | 660.4 | - | - | 0 | - |
| - | - | 1.118E+04 | 660.4 | - | - | 0 | - |
| - | - | 9.887E+04 | 661.4 | - | - | 0 | - |
| 2 | z | 6.451E+04 | 662.3 | 9.425E-05 | 0.1423 | +1 | 7 |
| - | - | 1.864E+04 | 662.4 | - | - | 0 | - |
| - | - | 2.651E+04 | 663.3 | - | - | 0 | - |
| 7 | c | 7.584E+06 | 676.4 | 0.0008428 | 1.246 | +1 | 7 |
| - | - | 2.517E+06 | 677.4 | - | - | 0 | - |
| 2 | y | 3.676E+04 | 678.3 | 0.002671 | 3.938 | +1 | 7 |
| - | - | 5.446E+05 | 678.4 | - | - | 0 | - |
| - | - | 1.6E+04 | 679.4 | - | - | 0 | - |
| - | - | 3.098E+04 | 679.4 | - | - | 0 | - |
| 2 | z | 2.101E+05 | 680.3 | 0.0008211 | 1.207 | +1 | 7 |
| - | - | 1.087E+05 | 681.3 | - | - | 0 | - |
| - | - | 2.795E+04 | 682.3 | - | - | 0 | - |
| 2 | y | 8141 | 696.4 | 0.006328 | 9.087 | +1 | 7 |
| - | - | 1.264E+05 | 733.4 | - | - | 0 | - |
| - | - | 5.088E+04 | 734.4 | - | - | 0 | - |
| - | - | 2.091E+04 | 735.4 | - | - | 0 | - |
| - | - | 1.388E+04 | 735.5 | - | - | 0 | - |
| - | - | 1.234E+04 | 747.4 | - | - | 0 | - |
| - | - | 1.505E+04 | 749.4 | - | - | 0 | - |
| - | - | 1.04E+04 | 750.4 | - | - | 0 | - |
| - | - | 1.802E+05 | 752.4 | - | - | 0 | - |
| - | - | 6.238E+04 | 753.4 | - | - | 0 | - |
| - | - | 1.948E+04 | 754.4 | - | - | 0 | - |
| - | - | 1.335E+04 | 761.4 | - | - | 0 | - |
| - | - | 7003 | 762.4 | - | - | 0 | - |
| - | - | 6.645E+04 | 763.5 | - | - | 0 | - |
| - | - | 2.444E+04 | 764.5 | - | - | 0 | - |
| - | - | 1.235E+05 | 765.4 | - | - | 0 | - |
| - | - | 8140 | 765.4 | - | - | 0 | - |
| - | - | 3.96E+04 | 766.4 | - | - | 0 | - |
| - | - | 1.335E+04 | 767.4 | - | - | 0 | - |
| - | - | 1.041E+04 | 769.4 | - | - | 0 | - |
| - | - | 2.42E+05 | 779.5 | - | - | 0 | - |
| - | - | 9.048E+04 | 780.5 | - | - | 0 | - |
| - | - | 2.433E+04 | 781.5 | - | - | 0 | - |
| - | - | 2.66E+05 | 782.4 | - | - | 0 | - |
| - | - | 9.25E+04 | 783.4 | - | - | 0 | - |
| - | - | 2.317E+04 | 784.4 | - | - | 0 | - |
| - | - | 4.23E+04 | 790.4 | - | - | 0 | - |
| - | - | 2.393E+04 | 791.4 | - | - | 0 | - |
| - | - | 4.12E+05 | 797.5 | - | - | 0 | - |
| - | - | 1.725E+05 | 798.5 | - | - | 0 | - |
| - | - | 4.479E+04 | 799.5 | - | - | 0 | - |
| - | - | 5.438E+04 | 807.4 | - | - | 0 | - |
| - | - | 2.874E+06 | 808.4 | - | - | 0 | - |
| - | - | 1.174E+06 | 809.4 | - | - | 0 | - |
| - | - | 3.65E+05 | 810.4 | - | - | 0 | - |
| - | - | 2.176E+04 | 811.4 | - | - | 0 | - |
| - | - | 3.375E+06 | 824.5 | - | - | 0 | - |
| - | - | 3.275E+06 | 825.5 | - | - | 0 | - |
| - | - | 1.23E+06 | 826.5 | - | - | 0 | - |
| - | - | 2.612E+05 | 827.5 | - | - | 0 | - |
| - | - | 1.924E+04 | 828.5 | - | - | 0 | - |
| - | - | 8809 | 840.4 | - | - | 0 | - |
| - | - | 6119 | 878.5 | - | - | 0 | - |
| - | - | 5924 | 927.7 | - | - | 0 | - |
| - | - | 5823 | 1958 | - | - | 0 | - |
| - | - | 7491 | 2446 | - | - | 0 | - |
| - | - | 6509 | 2470 | - | - | 0 | - |
| - | - | 9859 | 3085 | - | - | 0 | - |
| - | - | 8398 | 3086 | - | - | 0 | - |

m/z Charge Intensity FragmentType MassShift Position
120.08124542236328 0 1093538.2
121.0845718383789 0 86428.414
126.10303497314453 0 6172.159
128.09487915039062 0 32610.934
129.10269165039062 0 432393.66
130.1060791015625 0 16455.479
130.2517852783203 0 3917.643
131.04959106445312 0 14175.022
131.08193969726562 0 21915.365
134.83804321289062 0 4329.6304
135.21824645996094 0 4169.3735
139.0868377685547 0 4662.948
141.0664520263672 0 13716.144
145.1214141845703 0 22617.812
146.12327575683594 0 8923.236
146.1292266845703 0 133408.55
147.1327362060547 0 6306.3135
148.9547576904297 0 10498.97
149.06007385253906 0 23074.135
153.1028289794922 0 11717.3955
155.11831665039062 0 91165.26
159.0768585205078 0 79131
166.0867919921875 0 2756044.5 y 7
167.09007263183594 0 239883.78
168.0913543701172 0 8510.753
171.11329650878906 0 63508.387
172.07640075683594 0 9037.387
172.11691284179688 0 7712.6465
172.14511108398438 0 7737.1187
173.09237670898438 0 19170.965
173.45314025878906 0 8192.72
180.1141815185547 0 5218.004
181.13323974609375 0 4537.351
184.9217529296875 0 4400.988
185.1281280517578 0 4858.564
189.08766174316406 0 9487.776
189.1026611328125 0 62452.324
190.1060333251953 0 6406.1255
196.156982421875 0 11738.656
197.16531372070312 0 59373.93
198.1242218017578 0 5116.2915
198.14830017089844 0 5316.2974
198.16940307617188 0 6631.0063
200.139892578125 0 17139.176
201.1238250732422 0 18186.512
202.14279174804688 0 5748.539
207.11331176757812 0 169068.52
208.11680603027344 0 15439.441
210.37979125976562 0 4611.2007
212.10313415527344 0 6128.424
213.14698791503906 0 6013.225
214.19187927246094 0 51257.434
215.19964599609375 0 151289.45
216.1348419189453 0 8034.163
216.20748901367188 0 670184.5
217.0975341796875 0 8254.216
217.2108917236328 0 71118.4
218.15057373046875 0 20410.008
224.15232849121094 0 9293.105 c 3
224.1763153076172 0 39886.246
225.1602020263672 0 31040.31
228.1586151123047 0 26526.096
230.1143035888672 0 18903.271
235.10826110839844 0 225260.33 y Water loss 6
236.11190795898438 0 24838.744
241.17909240722656 0 84766.8
242.1140899658203 0 36764.703
242.15037536621094 0 29137.43
242.18675231933594 0 237798.98 c Ammonia loss 1
243.1899871826172 0 26417.365
243.53150939941406 0 6190.3193
244.12954711914062 0 25411.73
244.20236206054688 0 6473.4067
251.1557159423828 0 17156.055 c Ammonia loss 4
253.11883544921875 0 1628225.9 y 6
254.12208557128906 0 189089
255.1241455078125 0 15445.234
258.2054443359375 0 204001.86
259.2131652832031 0 643894.5 c 1
260.1244812011719 0 58106.09
260.1391906738281 0 7794.598
260.2164306640625 0 60798.367
263.1029968261719 0 25967.459
266.185302734375 0 6326.526
268.2032165527344 0 6512.205
269.1613464355469 0 6248.0693
269.2102966308594 0 16828.99
272.16058349609375 0 5759.723
277.15106201171875 0 14681.676
279.7665710449219 0 4902.8877
283.1769714355469 0 10792.275
283.2136535644531 0 8943.21
284.16107177734375 0 19506.828
284.22113037109375 0 26508.75
285.2289123535156 0 61019.395
286.1415100097656 0 7595.958
286.2320556640625 0 6849.6763
286.6742248535156 0 41796.953 c Ammonia loss 5
287.17340087890625 0 9879.204
287.22088623046875 0 8999.135
288.1341857910156 0 7721.5996
289.1748352050781 0 19313.482
293.1974792480469 0 14062.628
294.1829528808594 0 7896.3867
295.1413879394531 0 7821.1494
296.1980285644531 0 9402.548
298.67462158203125 0 13613.385
301.1886291503906 0 11382.983
301.22406005859375 0 35723.375
302.1690979003906 0 7974.186
302.18572998046875 0 8388.845
302.2314453125 0 22843.21
303.167236328125 0 20434.418
303.2396545410156 0 269974.84
304.17474365234375 0 10636.503
304.24267578125 0 34619.984
306.1455078125 0 166735.11 y Water loss 5
306.1648254394531 0 10978.31
307.1485900878906 0 25004.201
307.18768310546875 0 46081
307.6799621582031 0 38599.945
308.18084716796875 0 7716.169
311.2084045410156 0 32331.885
312.1925354003906 0 20568.96
312.67333984375 0 6209.4243
313.1513671875 0 49859.617
313.1874084472656 0 18296.014
313.22412109375 0 45055.59
313.3619384765625 0 5362.681
314.15484619140625 0 5407.716
314.2275695800781 0 7971.544
316.19256591796875 0 9840.175
319.197998046875 0 29217.578
321.1853332519531 0 39214.46
321.6796569824219 0 9213.457
324.1560974121094 0 60726.027 y 5
325.1593322753906 0 10905.201
327.2027893066406 0 8409.256
327.225830078125 0 7840.0654
328.2347717285156 0 23320.809 c Water loss 2
329.21905517578125 0 162093.94 c Ammonia loss 2
329.69842529296875 0 18676.838 c Water loss 6
330.1905212402344 0 74126.64 c Ammonia loss 6
330.2227783203125 0 20727.346
330.6924133300781 0 19712.566
331.16192626953125 0 273083.22
331.2343444824219 0 24359.533
332.16473388671875 0 30687.229
339.23974609375 0 10741.271
343.1985168457031 0 18060.473
345.2182922363281 0 11212.769
345.23773193359375 0 523471.34
346.2451477050781 0 770978.06 c 2
347.2484130859375 0 129850.07
348.2510070800781 0 15496.34
354.2142028808594 0 9261.612
354.250244140625 0 9967.631
355.1982421875 0 45983.285
355.2347106933594 0 7811.807
356.20111083984375 0 8561.365
356.2651672363281 0 17075.645
360.22235107421875 0 7325.631
367.2341003417969 0 10934.487
368.24053955078125 0 14168.449
372.2093505859375 0 27627.035
372.26116943359375 0 11306.246
372.7100524902344 0 21226.645
373.17303466796875 0 6467.308
373.20880126953125 0 47975.023
374.21270751953125 0 8533.736
377.18231201171875 0 44023.742 y Water loss 4
378.1869812011719 0 11740.099
378.21337890625 0 9632.416
380.2317810058594 0 9431.089
380.7215881347656 0 97926.3
381.2226257324219 0 47938.42
382.1723937988281 0 20134.53
382.2148132324219 0 5365.3887
382.2457275390625 0 20005.516
384.2611083984375 0 43316.984
385.26898193359375 0 975603.9
386.27398681640625 0 299191.38
387.2789001464844 0 46713.26
389.7269592285156 0 11151.92
390.2328186035156 0 15860.686
394.2459411621094 0 35224.65
394.7192687988281 0 21525.762
395.2117614746094 0 109956.83
395.7132263183594 0 47097.305
396.2166748046875 0 15783.9795
398.20648193359375 0 5945.512
398.24066162109375 0 18915.375
400.1835021972656 0 62646.684
400.2561340332031 0 53141.168
401.1876525878906 0 14681.993
401.260498046875 0 9347.323
402.27130126953125 0 17705.596
403.2334289550781 0 6158.5864
403.277099609375 0 6526.072
403.7246398925781 0 439152.16
404.2261657714844 0 173991.34
404.2874755859375 0 276827.7
404.72735595703125 0 58256.008
405.2908630371094 0 63504.39
408.22198486328125 0 12208.803
411.2720947265625 0 22331.172
412.2559814453125 0 61019.86
413.2605285644531 0 17013.807
414.23553466796875 0 27689.613
414.2687072753906 0 7043.635
416.2508850097656 0 36959.188
418.16339111328125 0 10732.481
418.194091796875 0 143164.3
419.19659423828125 0 29426.64
424.2199401855469 0 17660.707
424.2543640136719 0 5838.8887
426.2354431152344 0 27123.043
426.2710266113281 0 13415.291
428.2261047363281 0 10362.328
429.2332458496094 0 9469.184
429.28265380859375 0 67377.44 c Water loss 3
430.2670593261719 0 118688.69 c Ammonia loss 3
431.26971435546875 0 29943.48
432.2685852050781 0 6818.3813
436.2586975097656 0 6825.8774
438.270751953125 0 8996.798
439.2690124511719 0 7584.119
441.2467041015625 0 10183.813
442.2314147949219 0 11805.924
443.2955627441406 0 11941.683
444.24591064453125 0 48213.66
445.24749755859375 0 11746.172
446.2852783203125 0 18497.408
447.2934875488281 0 3813641.8 c 3
448.2962646484375 0 870785.8
449.2986755371094 0 113844.305
453.2813415527344 0 13581.415
454.230224609375 0 9444.519
455.29803466796875 0 9512.187
457.2805480957031 0 8883.094
459.2576599121094 0 21262.707
460.2178649902344 0 9840.129
461.2722473144531 0 9111.396
465.2832946777344 0 47826.49
466.27618408203125 0 5915.553
471.29351806640625 0 39817.355
472.20458984375 0 8573.255
472.2997131347656 0 6745.218
473.3089599609375 0 72704.02
474.3123779296875 0 30524.498
475.22796630859375 0 7013.227
475.3213806152344 0 9727.298
478.230712890625 0 25184.084 y Water loss 3
481.275390625 0 6182.22
482.310302734375 0 11621.098
483.25616455078125 0 12233.784
483.29388427734375 0 89910.5
484.27947998046875 0 8784.347
484.2959289550781 0 14476.928
485.3087158203125 0 18971.84
487.2866516113281 0 14467.436
492.29339599609375 0 8289.214
493.24090576171875 0 11794.186
495.259521484375 0 8458.484
496.2410888671875 0 37826.926 y 3
497.24505615234375 0 22168.98
499.2881164550781 0 16171.074
500.3199462890625 0 38115.52 c Water loss 4
501.30401611328125 0 525061.5 c Ammonia loss 4
502.3070373535156 0 129528.71
503.30999755859375 0 21977.139
505.21453857421875 0 11541.093
508.32421875 0 6199.8486
509.30780029296875 0 14196.781
510.26776123046875 0 8135.1167
510.30584716796875 0 20427.775
511.2510986328125 0 12460.338
513.26806640625 0 15497.634
515.2598266601562 0 19205.105
516.2651977539062 0 12012.963
518.33056640625 0 5668014 c 4
519.3333129882812 0 1403135
520.2276000976562 0 6128.1084
520.3357543945312 0 246808.12
521.3369750976562 0 7678.118
526.3356323242188 0 37963.684
527.3203735351562 0 7945.1094
528.2778930664062 0 11710.178
528.316162109375 0 28384.287
530.3422241210938 0 27923.652
531.27880859375 0 18599.488
531.3462524414062 0 8381.673
535.2890014648438 0 9742.94
536.31982421875 0 81802.63
537.318359375 0 27901.455
541.263916015625 0 7418.4443
543.312744140625 0 8776.615
544.3461303710938 0 260189.55
545.351806640625 0 155270.75
546.2451782226562 0 12730.071
546.2886962890625 0 23064.812
546.3556518554688 0 34022.21
547.2936401367188 0 7966.8076
552.3130493164062 0 8375.358
553.2994995117188 0 6505.7056
554.3302001953125 0 267039
555.3318481445312 0 65604.77
556.3330078125 0 20387.754
558.323974609375 0 14682.051
564.2547607421875 0 166248.17
565.2601928710938 0 114634.61 y Water loss 2
566.2633056640625 0 32519.686
568.2621459960938 0 13715.7
570.3245849609375 0 25697.168
571.3272705078125 0 18036.28
572.3410034179688 0 1380513 c Ammonia loss 5
573.3436279296875 0 373066.28
574.3490600585938 0 86262.19
575.3545532226562 0 15097.14
578.3298950195312 0 7674.1777
582.2650146484375 0 666365.1
583.2717895507812 0 657482.4 y 2
584.2747192382812 0 172185.61
585.2774658203125 0 26962.375
588.3356323242188 0 30607.105
589.3677978515625 0 5568425 c 5
590.3701171875 0 1614218.9
591.372802734375 0 316029.9
592.3756713867188 0 16785.213
595.3560791015625 0 8266.77
596.3435668945312 0 19362.006
597.3401489257812 0 19482.334
605.3412475585938 0 18049.66
606.3287353515625 0 22569.828
611.3504028320312 0 8097.8857
613.3673706054688 0 44132.094
614.3731079101562 0 119600.21
615.3775024414062 0 42964.195
616.3748779296875 0 14080.097
623.3515014648438 0 124649.73
624.3466796875 0 29742.406
625.3472290039062 0 14799.429
631.3778686523438 0 222550.39
632.3831176757812 0 131309.72
633.3864135742188 0 35329.426
637.283447265625 0 52660.594 w 1
638.2860717773438 0 8162.3086
641.3621826171875 0 544587.9
642.3646240234375 0 179440.17
643.364990234375 0 38756.727
659.372802734375 0 1283626.6 c Ammonia loss 6
660.3758544921875 0 387639.97
660.4428100585938 0 11178.403
661.3788452148438 0 98868.266
662.3270874023438 0 64513.023 z Water loss 1
662.392333984375 0 18642.932
663.3303833007812 0 26506.834
676.399658203125 0 7584339.5 c 6
677.4021606445312 0 2517495
678.348388671875 0 36755.97 y Water loss 1
678.4046630859375 0 544570.94
679.3507080078125 0 15999.608
679.4075317382812 0 30980.695
680.33837890625 0 210060.61 z 1
681.3429565429688 0 108737.45
682.3441772460938 0 27951.611
696.3626098632812 0 8140.834 y 1
733.39697265625 0 126383.66
734.3998413085938 0 50880.69
735.4020385742188 0 20911.691
735.4671630859375 0 13880.135
747.4161987304688 0 12343.676
749.3601684570312 0 15046.764
750.364013671875 0 10398.309
752.3699340820312 0 180198.6
753.3731689453125 0 62380.508
754.375 0 19478.54
761.44580078125 0 13353.468
762.4473266601562 0 7003.2915
763.4569702148438 0 66453.12
764.4547729492188 0 24444.354
765.3778686523438 0 123508.766
765.437255859375 0 8139.706
766.3821411132812 0 39598.336
767.3831787109375 0 13345.325
769.3980712890625 0 10405.716
779.4534301757812 0 241975.33
780.4567260742188 0 90479.48
781.4591064453125 0 24327.666
782.404541015625 0 266023.38
783.40771484375 0 92496.555
784.4097900390625 0 23173.258
790.4227294921875 0 42296.586
791.4263305664062 0 23934.271
797.4642333984375 0 412037.94
798.4671630859375 0 172451.44
799.4703369140625 0 44790.35
807.4481201171875 0 54380.656
808.4328002929688 0 2873722.8
809.4359130859375 0 1174245.1
810.4383544921875 0 364998.78
811.4395141601562 0 21758.113
824.4515380859375 0 3374779.5
825.4573974609375 0 3275005
826.4603271484375 0 1230262.6
827.4641723632812 0 261217.05
828.4682006835938 0 19241.316
840.4322509765625 0 8809.146
878.5269165039062 0 6119.073
927.7279663085938 0 5924.2515
1957.5030517578125 0 5822.792
2446.34423828125 0 7491.224
2470.0927734375 0 6509.36
3085.445556640625 0 9859.484
3086.048828125 0 8398.267

Spectrum Details

|  |  |
| --- | --- |
| Matched peaks? Matched peaks The total absolute number of peaks matched. Additionally in brackets the total fraction of peaks matched and the total number of peaks is shown. | 35 (8.43% of 415) |
| FDR? FDR The false discovery rate estimated for this peptide. It is calculated by matching all theoretical fragments with a non-integer shift with the raw peaks for this spectrum. This is done with 40 different shifts. The resulting percentage is the average number of annotated peaks over the number of annotated peaks with the correct spectrum. | 5.65% |
| Satellite FDR? Satellite FDR See the FDR for details on its calculation. This satellite ion specific FDR only contains the satellite ions (d/w) for I/L/J positions. | 7.14% |
| PSM Score? PSM Score The PSM Score as given by Hecklib to this annotated spectrum. It is shown with three significant figures. | 472 |

#### Spectrum 5039? Spectrum 5039 The raw spectrum of this peptide as annotated by Hecklib. The fragments are coloured according to ion type (see legend). Any peaks with a star '\*' as text can be hovered over to see the full details, first the ion type second the mass shift type. By hovering over the amino acids in the peptide or ions in the legend the corresponding peaks are highlighted. By toggling the 'Unassigned' label you can turn the background (unassigned) peaks on or off in the plot. By updating the slider in the Ion legend you can update the spectrum to only show the top X% of the peaks with labels. The top X% means any peak that is within X% of the highest intensity. By dragging in the spectrum you can zoom in to a specific part of the spectrum and use 'Zoom Out' to get back to the original zoom level. The annotation of the spectrum is based on the given sequence in the peptides file and is done with different software so inconsistencies are likely. The peaks are annotated based on the given sequence, with 20 ppm tolerance.

Copy Data

##### Spectrum 5039 (TSV)

###### Preview

```
Loading example...
```

*Click on the button to copy the data to your clipboard.*

Mz MinMz MaxIntensity Max

WidthHeightPeptide font sizePeptide stroke widthSpectrum font sizeSpectrum stroke widthCompact peptide

Ion legend

wxyz

abcd

OtherUnassignedIon^Charge^_Position_Show for top:%

KJSTAASF

01.55e+43.10e+44.65e+46.19e+4

Zoom Out

y^+1^_1_y^+1^_2_c^+1^_2_y^+1^_2_c^+1^_2_y^+1^_3_c^+1^_3_c^+1^_3_c^+1^_4_c^+1^_4_c^+1^_5_c^+1^_5_c^+1^_6_y^+1^_6_c^+1^_6_c^+1^_7_c^+1^_7_z^+1^_7_

043286412961727

Fragment Matches Table

Show background peaks

| Position | Ion type | Intensity | mz Theoretical | mz Error (Th) | mz Error (ppm) | Charge | Series Number |
| --- | --- | --- | --- | --- | --- | --- | --- |
| - | - | 453.2 | 120.1 | - | - | 0 | - |
| - | - | 5910 | 120.1 | - | - | 0 | - |
| - | - | 568.5 | 121.1 | - | - | 0 | - |
| - | - | 440.7 | 124.7 | - | - | 0 | - |
| - | - | 902.2 | 129.1 | - | - | 0 | - |
| - | - | 3791 | 129.1 | - | - | 0 | - |
| - | - | 396.3 | 129.3 | - | - | 0 | - |
| - | - | 349.3 | 129.4 | - | - | 0 | - |
| - | - | 401.1 | 140.9 | - | - | 0 | - |
| - | - | 417.3 | 144.9 | - | - | 0 | - |
| - | - | 649.7 | 146.1 | - | - | 0 | - |
| - | - | 408 | 150.7 | - | - | 0 | - |
| - | - | 851.2 | 166.1 | - | - | 0 | - |
| 8 | y | 1.438E+04 | 166.1 | 7.341E-05 | 0.442 | +1 | 1 |
| - | - | 1642 | 167.1 | - | - | 0 | - |
| - | - | 519 | 171.1 | - | - | 0 | - |
| - | - | 2346 | 173.5 | - | - | 0 | - |
| - | - | 512.5 | 179.5 | - | - | 0 | - |
| - | - | 796.2 | 184.1 | - | - | 0 | - |
| - | - | 490.4 | 196.1 | - | - | 0 | - |
| - | - | 476.8 | 197.2 | - | - | 0 | - |
| - | - | 939.7 | 207.1 | - | - | 0 | - |
| - | - | 792.9 | 215.2 | - | - | 0 | - |
| - | - | 5499 | 216.2 | - | - | 0 | - |
| 7 | y | 656.1 | 235.1 | 0.0007865 | 3.345 | +1 | 2 |
| 2 | c | 1601 | 242.2 | 0.0004056 | 1.675 | +1 | 2 |
| - | - | 525.2 | 247.4 | - | - | 0 | - |
| 7 | y | 8938 | 253.1 | 0.0001804 | 0.7129 | +1 | 2 |
| - | - | 1109 | 254.1 | - | - | 0 | - |
| - | - | 1209 | 258.2 | - | - | 0 | - |
| 2 | c | 4688 | 259.2 | 0.0001756 | 0.6772 | +1 | 2 |
| - | - | 843.9 | 260.2 | - | - | 0 | - |
| - | - | 686.7 | 269.2 | - | - | 0 | - |
| - | - | 2635 | 303.2 | - | - | 0 | - |
| 6 | y | 587.9 | 306.1 | 0.0005455 | 1.782 | +1 | 3 |
| - | - | 744.4 | 313.1 | - | - | 0 | - |
| 3 | c | 824.3 | 329.2 | 0.0006195 | 1.882 | +1 | 3 |
| - | - | 1217 | 331.2 | - | - | 0 | - |
| - | - | 4409 | 345.2 | - | - | 0 | - |
| 3 | c | 6196 | 346.2 | 0.0006488 | 1.874 | +1 | 3 |
| - | - | 1285 | 347.2 | - | - | 0 | - |
| - | - | 900.8 | 375.9 | - | - | 0 | - |
| - | - | 847.4 | 376.9 | - | - | 0 | - |
| - | - | 567.2 | 380.7 | - | - | 0 | - |
| - | - | 8184 | 385.3 | - | - | 0 | - |
| - | - | 2924 | 386.3 | - | - | 0 | - |
| - | - | 711.6 | 393.8 | - | - | 0 | - |
| - | - | 1644 | 395.2 | - | - | 0 | - |
| - | - | 1968 | 403.7 | - | - | 0 | - |
| - | - | 2179 | 404.3 | - | - | 0 | - |
| - | - | 1686 | 411.8 | - | - | 0 | - |
| - | - | 1053 | 412.8 | - | - | 0 | - |
| - | - | 6245 | 413.3 | - | - | 0 | - |
| 4 | c | 1129 | 430.3 | 0.0005379 | 1.25 | +1 | 4 |
| 4 | c | 3.088E+04 | 447.3 | 7.897E-05 | 0.1765 | +1 | 4 |
| - | - | 7029 | 448.3 | - | - | 0 | - |
| - | - | 990.4 | 449.3 | - | - | 0 | - |
| - | - | 569.4 | 456.2 | - | - | 0 | - |
| - | - | 576.7 | 477.7 | - | - | 0 | - |
| 5 | c | 2869 | 501.3 | 0.0002372 | 0.4731 | +1 | 5 |
| - | - | 1189 | 502.3 | - | - | 0 | - |
| 5 | c | 4.419E+04 | 518.3 | 0.0004496 | 0.8674 | +1 | 5 |
| - | - | 1.243E+04 | 519.3 | - | - | 0 | - |
| - | - | 1853 | 520.3 | - | - | 0 | - |
| - | - | 1323 | 544.3 | - | - | 0 | - |
| - | - | 1001 | 545.4 | - | - | 0 | - |
| - | - | 738.5 | 554.3 | - | - | 0 | - |
| - | - | 1768 | 564.3 | - | - | 0 | - |
| 6 | c | 6554 | 572.3 | 2.795E-05 | 0.04884 | +1 | 6 |
| - | - | 2334 | 573.3 | - | - | 0 | - |
| - | - | 649 | 574.3 | - | - | 0 | - |
| - | - | 4236 | 582.3 | - | - | 0 | - |
| 3 | y | 4719 | 583.3 | 0.001405 | 2.409 | +1 | 6 |
| - | - | 1774 | 584.3 | - | - | 0 | - |
| 6 | c | 4.614E+04 | 589.4 | 0.000515 | 0.8739 | +1 | 6 |
| - | - | 1.158E+04 | 590.4 | - | - | 0 | - |
| - | - | 2462 | 591.4 | - | - | 0 | - |
| - | - | 1120 | 611.8 | - | - | 0 | - |
| - | - | 831.4 | 614.4 | - | - | 0 | - |
| - | - | 2647 | 619.8 | - | - | 0 | - |
| - | - | 1999 | 620.3 | - | - | 0 | - |
| - | - | 1609 | 631.4 | - | - | 0 | - |
| - | - | 731 | 632.4 | - | - | 0 | - |
| - | - | 2041 | 641.4 | - | - | 0 | - |
| - | - | 1075 | 642.4 | - | - | 0 | - |
| 7 | c | 5252 | 659.4 | 0.000196 | 0.2973 | +1 | 7 |
| - | - | 1295 | 660.4 | - | - | 0 | - |
| - | - | 774 | 674.4 | - | - | 0 | - |
| 7 | c | 6.133E+04 | 676.4 | 0.0005 | 0.7392 | +1 | 7 |
| - | - | 1.935E+04 | 677.4 | - | - | 0 | - |
| - | - | 4086 | 678.4 | - | - | 0 | - |
| 2 | z | 1637 | 680.3 | 0.0003996 | 0.5874 | +1 | 7 |
| - | - | 982.6 | 681.3 | - | - | 0 | - |
| - | - | 1239 | 733.4 | - | - | 0 | - |
| - | - | 1339 | 752.4 | - | - | 0 | - |
| - | - | 1155 | 765.4 | - | - | 0 | - |
| - | - | 1226 | 779.5 | - | - | 0 | - |
| - | - | 653.9 | 780.5 | - | - | 0 | - |
| - | - | 2436 | 782.4 | - | - | 0 | - |
| - | - | 1419 | 783.4 | - | - | 0 | - |
| - | - | 2410 | 797.5 | - | - | 0 | - |
| - | - | 969.9 | 798.5 | - | - | 0 | - |
| - | - | 2.19E+04 | 808.4 | - | - | 0 | - |
| - | - | 8956 | 809.4 | - | - | 0 | - |
| - | - | 2447 | 810.4 | - | - | 0 | - |
| - | - | 2.613E+04 | 824.5 | - | - | 0 | - |
| - | - | 2.635E+04 | 825.5 | - | - | 0 | - |
| - | - | 8906 | 826.5 | - | - | 0 | - |
| - | - | 2187 | 827.5 | - | - | 0 | - |
| - | - | 576.2 | 971.8 | - | - | 0 | - |
| - | - | 571.6 | 1064 | - | - | 0 | - |
| - | - | 593.3 | 1111 | - | - | 0 | - |
| - | - | 1603 | 1223 | - | - | 0 | - |
| - | - | 1371 | 1224 | - | - | 0 | - |
| - | - | 921.1 | 1225 | - | - | 0 | - |
| - | - | 1067 | 1238 | - | - | 0 | - |
| - | - | 3208 | 1239 | - | - | 0 | - |
| - | - | 7345 | 1240 | - | - | 0 | - |
| - | - | 5740 | 1241 | - | - | 0 | - |
| - | - | 2777 | 1242 | - | - | 0 | - |
| - | - | 672.9 | 1490 | - | - | 0 | - |
| - | - | 662.6 | 1710 | - | - | 0 | - |

m/z Charge Intensity FragmentType MassShift Position
120.07707214355469 0 453.20068
120.08077239990234 0 5910.264
121.0842514038086 0 568.4647
124.66239166259766 0 440.69946
129.0547332763672 0 902.2384
129.10218811035156 0 3790.8135
129.2718505859375 0 396.32544
129.40855407714844 0 349.26468
140.87864685058594 0 401.0843
144.88043212890625 0 417.33813
146.12918090820312 0 649.667
150.66015625 0 408.04105
166.07894897460938 0 851.1693
166.086181640625 0 14375.618 y 7
167.08970642089844 0 1642.0391
171.11251831054688 0 519.0194
173.45050048828125 0 2346.289
179.54737854003906 0 512.5398
184.0728302001953 0 796.1954
196.12571716308594 0 490.36462
197.1648406982422 0 476.82913
207.1125030517578 0 939.6604
215.19915771484375 0 792.9205
216.20692443847656 0 5499.0117
235.10850524902344 0 656.09326 y Water loss 6
242.18589782714844 0 1600.7913 c Ammonia loss 1
247.4104766845703 0 525.1597
253.11810302734375 0 8937.979 y 6
254.12152099609375 0 1109.09
258.2052001953125 0 1208.7778
259.2126770019531 0 4687.6465 c 1
260.21636962890625 0 843.8993
269.20843505859375 0 686.71716
303.2388916015625 0 2634.8193
306.144287109375 0 587.93805 y Water loss 5
313.149658203125 0 744.40533
329.21771240234375 0 824.25476 c Ammonia loss 2
331.1607971191406 0 1216.7118
345.2370910644531 0 4408.863
346.2442321777344 0 6195.676 c 2
347.24676513671875 0 1285.2432
375.8602294921875 0 900.8143
376.8657531738281 0 847.38617
380.72137451171875 0 567.19257
385.2681884765625 0 8183.7715
386.2725524902344 0 2923.7126
393.83685302734375 0 711.5646
395.2091979980469 0 1643.8794
403.72369384765625 0 1968.3114
404.2865295410156 0 2178.8367
411.84912109375 0 1685.798
412.8493957519531 0 1052.6338
413.2660217285156 0 6244.8994
430.2654724121094 0 1129.2721 c Ammonia loss 3
447.29248046875 0 30879.428 c 3
448.29522705078125 0 7028.7954
449.3001403808594 0 990.3807
456.2132568359375 0 569.4302
477.736572265625 0 576.6799
501.3028869628906 0 2868.8394 c Ammonia loss 4
502.3041687011719 0 1188.5933
518.3292236328125 0 44193.555 c 4
519.3323974609375 0 12434.855
520.3329467773438 0 1853.3903
544.3455200195312 0 1323.1677
545.350341796875 0 1000.9512
554.3313598632812 0 738.5188
564.2523803710938 0 1768.3713
572.3402099609375 0 6554.1724 c Ammonia loss 5
573.3428344726562 0 2334.2344
574.3439331054688 0 648.99664
582.263671875 0 4236.1943
583.2708129882812 0 4718.624 y 2
584.2723388671875 0 1774.4608
589.3662719726562 0 46135.9 c 5
590.3692626953125 0 11583.404
591.3712158203125 0 2461.583
611.8375244140625 0 1119.6907
614.3766479492188 0 831.44867
619.8467407226562 0 2646.976
620.3496704101562 0 1999.1084
631.3775634765625 0 1608.6106
632.3821411132812 0 731.00745
641.3616333007812 0 2041.4805
642.3633422851562 0 1075.1782
659.3720703125 0 5252.4033 c Ammonia loss 6
660.3753051757812 0 1295.4293
674.385009765625 0 774.03033
676.3983154296875 0 61330.4 c 6
677.4014282226562 0 19345.297
678.4035034179688 0 4086.1536
680.337158203125 0 1637.2855 z 1
681.3417358398438 0 982.62964
733.3988647460938 0 1239.1772
752.3693237304688 0 1339.4707
765.3778076171875 0 1155.4795
779.454833984375 0 1225.5833
780.4623413085938 0 653.94434
782.40380859375 0 2436.2102
783.4054565429688 0 1419.1554
797.4622192382812 0 2409.9695
798.463134765625 0 969.9295
808.4317626953125 0 21895.719
809.4346923828125 0 8955.986
810.4370727539062 0 2446.878
824.4501953125 0 26132.459
825.4562377929688 0 26351.643
826.4595947265625 0 8906.191
827.4645385742188 0 2187.0823
971.7732543945312 0 576.18854
1064.39013671875 0 571.5587
1110.6209716796875 0 593.31555
1222.6715087890625 0 1603.2843
1223.67822265625 0 1370.8694
1224.6822509765625 0 921.11926
1237.69140625 0 1067.27
1238.686767578125 0 3208.2766
1239.6929931640625 0 7345.2197
1240.6995849609375 0 5740.2075
1241.70263671875 0 2776.5454
1489.5294189453125 0 672.90405
1710.3326416015625 0 662.555

Spectrum Details

|  |  |
| --- | --- |
| Matched peaks? Matched peaks The total absolute number of peaks matched. Additionally in brackets the total fraction of peaks matched and the total number of peaks is shown. | 18 (14.75% of 122) |
| FDR? FDR The false discovery rate estimated for this peptide. It is calculated by matching all theoretical fragments with a non-integer shift with the raw peaks for this spectrum. This is done with 40 different shifts. The resulting percentage is the average number of annotated peaks over the number of annotated peaks with the correct spectrum. | 3.84% |
| Satellite FDR? Satellite FDR See the FDR for details on its calculation. This satellite ion specific FDR only contains the satellite ions (d/w) for I/L/J positions. | ∞ |
| PSM Score? PSM Score The PSM Score as given by Hecklib to this annotated spectrum. It is shown with three significant figures. | 243 |

#### Spectrum 4759? Spectrum 4759 The raw spectrum of this peptide as annotated by Hecklib. The fragments are coloured according to ion type (see legend). Any peaks with a star '\*' as text can be hovered over to see the full details, first the ion type second the mass shift type. By hovering over the amino acids in the peptide or ions in the legend the corresponding peaks are highlighted. By toggling the 'Unassigned' label you can turn the background (unassigned) peaks on or off in the plot. By updating the slider in the Ion legend you can update the spectrum to only show the top X% of the peaks with labels. The top X% means any peak that is within X% of the highest intensity. By dragging in the spectrum you can zoom in to a specific part of the spectrum and use 'Zoom Out' to get back to the original zoom level. The annotation of the spectrum is based on the given sequence in the peptides file and is done with different software so inconsistencies are likely. The peaks are annotated based on the given sequence, with 20 ppm tolerance.

Copy Data

##### Spectrum 4759 (TSV)

###### Preview

```
Loading example...
```

*Click on the button to copy the data to your clipboard.*

Mz MinMz MaxIntensity Max

WidthHeightPeptide font sizePeptide stroke widthSpectrum font sizeSpectrum stroke widthCompact peptide

Ion legend

wxyz

abcd

OtherUnassignedIon^Charge^_Position_Show for top:%

KJSTAASF

03.55e+47.11e+41.07e+51.42e+5

Zoom Out

y^+1^_1_y^+1^_2_c^+1^_2_y^+1^_2_c^+1^_2_y^+1^_3_y^+1^_3_c^+1^_3_c^+2^_7_c^+1^_3_y^+1^_4_c^+1^_4_c^+1^_4_c^+1^_4_c^+1^_5_c^+1^_5_c^+1^_5_y^+1^_6_c^+1^_6_y^+1^_6_c^+1^_6_w^+1^_7_c^+1^_7_z^+1^_7_c^+1^_7_y^+1^_7_z^+1^_7_

0725145121762901

Fragment Matches Table

Show background peaks

| Position | Ion type | Intensity | mz Theoretical | mz Error (Th) | mz Error (ppm) | Charge | Series Number |
| --- | --- | --- | --- | --- | --- | --- | --- |
| - | - | 1.293E+04 | 120.1 | - | - | 0 | - |
| - | - | 788.3 | 121.1 | - | - | 0 | - |
| - | - | 797.9 | 128.1 | - | - | 0 | - |
| - | - | 468.9 | 129.1 | - | - | 0 | - |
| - | - | 8520 | 129.1 | - | - | 0 | - |
| - | - | 473.1 | 130.1 | - | - | 0 | - |
| - | - | 325.6 | 131.1 | - | - | 0 | - |
| - | - | 390.4 | 136.9 | - | - | 0 | - |
| - | - | 3349 | 146.1 | - | - | 0 | - |
| - | - | 739.4 | 155.1 | - | - | 0 | - |
| - | - | 1007 | 159.1 | - | - | 0 | - |
| 8 | y | 3.284E+04 | 166.1 | 0.0002318 | 1.395 | +1 | 1 |
| - | - | 3296 | 167.1 | - | - | 0 | - |
| - | - | 876.4 | 171.1 | - | - | 0 | - |
| - | - | 1025 | 173.5 | - | - | 0 | - |
| - | - | 474.7 | 189.1 | - | - | 0 | - |
| - | - | 587.4 | 189.1 | - | - | 0 | - |
| - | - | 507.7 | 192.1 | - | - | 0 | - |
| - | - | 421.9 | 193.6 | - | - | 0 | - |
| - | - | 693.7 | 197.2 | - | - | 0 | - |
| - | - | 499.7 | 197.7 | - | - | 0 | - |
| - | - | 2316 | 207.1 | - | - | 0 | - |
| - | - | 915.9 | 212.1 | - | - | 0 | - |
| - | - | 3385 | 215.2 | - | - | 0 | - |
| - | - | 1.388E+04 | 216.2 | - | - | 0 | - |
| - | - | 1080 | 217.2 | - | - | 0 | - |
| - | - | 574.1 | 225.2 | - | - | 0 | - |
| - | - | 762.4 | 228.2 | - | - | 0 | - |
| 7 | y | 2675 | 235.1 | 0.0006491 | 2.761 | +1 | 2 |
| - | - | 561.1 | 239.3 | - | - | 0 | - |
| - | - | 2100 | 241.2 | - | - | 0 | - |
| - | - | 734.5 | 242.1 | - | - | 0 | - |
| 2 | c | 4045 | 242.2 | 9.792E-05 | 0.4043 | +1 | 2 |
| - | - | 500.8 | 247.5 | - | - | 0 | - |
| - | - | 1074 | 253.1 | - | - | 0 | - |
| 7 | y | 1.721E+04 | 253.1 | 0.0002621 | 1.035 | +1 | 2 |
| - | - | 1804 | 254.1 | - | - | 0 | - |
| - | - | 4178 | 258.2 | - | - | 0 | - |
| 2 | c | 1.187E+04 | 259.2 | 2.296E-05 | 0.08859 | +1 | 2 |
| - | - | 1423 | 260.2 | - | - | 0 | - |
| - | - | 5545 | 303.2 | - | - | 0 | - |
| - | - | 989.5 | 304.2 | - | - | 0 | - |
| 6 | y | 2958 | 306.1 | 0.0003013 | 0.9842 | +1 | 3 |
| - | - | 807.8 | 307.2 | - | - | 0 | - |
| 6 | y | 609.7 | 324.2 | 0.0005816 | 1.794 | +1 | 3 |
| 3 | c | 2944 | 329.2 | 0.0003266 | 0.992 | +1 | 3 |
| 7 | c | 1232 | 330.2 | 0.0004403 | 1.334 | +2 | 7 |
| - | - | 2913 | 331.2 | - | - | 0 | - |
| - | - | 625.6 | 332.2 | - | - | 0 | - |
| - | - | 669.2 | 333.2 | - | - | 0 | - |
| - | - | 9722 | 345.2 | - | - | 0 | - |
| 3 | c | 1.358E+04 | 346.2 | 9.947E-05 | 0.2873 | +1 | 3 |
| - | - | 2948 | 347.2 | - | - | 0 | - |
| - | - | 613.3 | 358.2 | - | - | 0 | - |
| - | - | 583.8 | 365.1 | - | - | 0 | - |
| - | - | 603.8 | 373.2 | - | - | 0 | - |
| - | - | 710 | 375.9 | - | - | 0 | - |
| - | - | 1325 | 376.9 | - | - | 0 | - |
| 5 | y | 687.6 | 377.2 | 0.001251 | 3.316 | +1 | 4 |
| - | - | 1529 | 380.7 | - | - | 0 | - |
| - | - | 623.8 | 381.2 | - | - | 0 | - |
| - | - | 1.82E+04 | 385.3 | - | - | 0 | - |
| - | - | 5451 | 386.3 | - | - | 0 | - |
| - | - | 615.9 | 387.3 | - | - | 0 | - |
| - | - | 1046 | 394.8 | - | - | 0 | - |
| - | - | 1527 | 394.9 | - | - | 0 | - |
| - | - | 1554 | 395.2 | - | - | 0 | - |
| - | - | 602.6 | 401.2 | - | - | 0 | - |
| - | - | 4638 | 403.7 | - | - | 0 | - |
| - | - | 2539 | 404.2 | - | - | 0 | - |
| - | - | 4686 | 404.3 | - | - | 0 | - |
| - | - | 1559 | 405.3 | - | - | 0 | - |
| - | - | 1576 | 411.8 | - | - | 0 | - |
| - | - | 6656 | 413.3 | - | - | 0 | - |
| - | - | 1472 | 418.2 | - | - | 0 | - |
| 4 | c | 1297 | 429.3 | 0.0001344 | 0.3131 | +1 | 4 |
| 4 | c | 2339 | 430.3 | 0.0004692 | 1.09 | +1 | 4 |
| 4 | c | 7.404E+04 | 447.3 | 0.0003788 | 0.8469 | +1 | 4 |
| - | - | 1.472E+04 | 448.3 | - | - | 0 | - |
| - | - | 2625 | 449.3 | - | - | 0 | - |
| - | - | 1008 | 483.3 | - | - | 0 | - |
| - | - | 2002 | 488.3 | - | - | 0 | - |
| 5 | c | 860 | 500.3 | 0.0001664 | 0.3325 | +1 | 5 |
| 5 | c | 7156 | 501.3 | 0.0005258 | 1.049 | +1 | 5 |
| - | - | 1960 | 502.3 | - | - | 0 | - |
| 5 | c | 1.041E+05 | 518.3 | 3.869E-05 | 0.07464 | +1 | 5 |
| - | - | 2.547E+04 | 519.3 | - | - | 0 | - |
| - | - | 4803 | 520.3 | - | - | 0 | - |
| - | - | 988.7 | 525.8 | - | - | 0 | - |
| - | - | 606.1 | 530.3 | - | - | 0 | - |
| - | - | 922.1 | 536.3 | - | - | 0 | - |
| - | - | 2433 | 544.3 | - | - | 0 | - |
| - | - | 2672 | 545.4 | - | - | 0 | - |
| - | - | 2519 | 554.3 | - | - | 0 | - |
| - | - | 681.6 | 555.3 | - | - | 0 | - |
| - | - | 3558 | 564.3 | - | - | 0 | - |
| 3 | y | 1692 | 565.3 | 0.001766 | 3.123 | +1 | 6 |
| 6 | c | 1.559E+04 | 572.3 | 9.412E-05 | 0.1644 | +1 | 6 |
| - | - | 4632 | 573.3 | - | - | 0 | - |
| - | - | 1592 | 574.3 | - | - | 0 | - |
| - | - | 9088 | 582.3 | - | - | 0 | - |
| 3 | y | 1.311E+04 | 583.3 | 0.001222 | 2.095 | +1 | 6 |
| - | - | 1039 | 583.3 | - | - | 0 | - |
| - | - | 3244 | 584.3 | - | - | 0 | - |
| - | - | 880 | 585.3 | - | - | 0 | - |
| 6 | c | 9.898E+04 | 589.4 | 8.779E-05 | 0.149 | +1 | 6 |
| - | - | 2.818E+04 | 590.4 | - | - | 0 | - |
| - | - | 5720 | 591.4 | - | - | 0 | - |
| - | - | 663.9 | 613.4 | - | - | 0 | - |
| - | - | 2089 | 614.4 | - | - | 0 | - |
| - | - | 994.7 | 618.3 | - | - | 0 | - |
| - | - | 1045 | 619.4 | - | - | 0 | - |
| - | - | 892.9 | 619.8 | - | - | 0 | - |
| - | - | 1450 | 620.4 | - | - | 0 | - |
| - | - | 861.6 | 620.9 | - | - | 0 | - |
| - | - | 1160 | 623.3 | - | - | 0 | - |
| - | - | 534.1 | 624.3 | - | - | 0 | - |
| - | - | 2420 | 631.4 | - | - | 0 | - |
| - | - | 1612 | 632.4 | - | - | 0 | - |
| - | - | 593.2 | 633.4 | - | - | 0 | - |
| - | - | 791.3 | 635.3 | - | - | 0 | - |
| 2 | w | 1388 | 637.3 | 0.003669 | 5.757 | +1 | 7 |
| - | - | 5278 | 641.4 | - | - | 0 | - |
| - | - | 2070 | 642.4 | - | - | 0 | - |
| 7 | c | 1.265E+04 | 659.4 | 0.000135 | 0.2047 | +1 | 7 |
| - | - | 4081 | 660.4 | - | - | 0 | - |
| - | - | 715.3 | 661.2 | - | - | 0 | - |
| - | - | 1661 | 661.4 | - | - | 0 | - |
| 2 | z | 815.6 | 662.3 | 0.0001553 | 0.2345 | +1 | 7 |
| - | - | 5228 | 674.4 | - | - | 0 | - |
| - | - | 1736 | 675.4 | - | - | 0 | - |
| 7 | c | 1.407E+05 | 676.4 | 7.275E-05 | 0.1076 | +1 | 7 |
| - | - | 4.467E+04 | 677.4 | - | - | 0 | - |
| 2 | y | 910.5 | 678.3 | 0.001268 | 1.869 | +1 | 7 |
| - | - | 1.028E+04 | 678.4 | - | - | 0 | - |
| 2 | z | 4104 | 680.3 | 0.000699 | 1.027 | +1 | 7 |
| - | - | 2373 | 681.3 | - | - | 0 | - |
| - | - | 876.5 | 682.3 | - | - | 0 | - |
| - | - | 2942 | 733.4 | - | - | 0 | - |
| - | - | 899.1 | 734.4 | - | - | 0 | - |
| - | - | 3484 | 752.4 | - | - | 0 | - |
| - | - | 1095 | 753.4 | - | - | 0 | - |
| - | - | 1134 | 763.5 | - | - | 0 | - |
| - | - | 789 | 764.5 | - | - | 0 | - |
| - | - | 1748 | 765.4 | - | - | 0 | - |
| - | - | 939.2 | 768.4 | - | - | 0 | - |
| - | - | 4281 | 779.5 | - | - | 0 | - |
| - | - | 1605 | 780.5 | - | - | 0 | - |
| - | - | 654 | 781.5 | - | - | 0 | - |
| - | - | 4045 | 782.4 | - | - | 0 | - |
| - | - | 2166 | 783.4 | - | - | 0 | - |
| - | - | 692.1 | 784.4 | - | - | 0 | - |
| - | - | 675.6 | 790.4 | - | - | 0 | - |
| - | - | 6889 | 797.5 | - | - | 0 | - |
| - | - | 2939 | 798.5 | - | - | 0 | - |
| - | - | 1988 | 806.4 | - | - | 0 | - |
| - | - | 1873 | 807.4 | - | - | 0 | - |
| - | - | 5.14E+04 | 808.4 | - | - | 0 | - |
| - | - | 2.089E+04 | 809.4 | - | - | 0 | - |
| - | - | 8007 | 810.4 | - | - | 0 | - |
| - | - | 2548 | 822.4 | - | - | 0 | - |
| - | - | 777 | 823.3 | - | - | 0 | - |
| - | - | 3093 | 823.4 | - | - | 0 | - |
| - | - | 6.343E+04 | 824.5 | - | - | 0 | - |
| - | - | 5.685E+04 | 825.5 | - | - | 0 | - |
| - | - | 2.049E+04 | 826.5 | - | - | 0 | - |
| - | - | 4836 | 827.5 | - | - | 0 | - |
| - | - | 625.6 | 888.4 | - | - | 0 | - |
| - | - | 678.1 | 895.2 | - | - | 0 | - |
| - | - | 1384 | 905.4 | - | - | 0 | - |
| - | - | 3539 | 906.4 | - | - | 0 | - |
| - | - | 1825 | 907.4 | - | - | 0 | - |
| - | - | 808.8 | 1042 | - | - | 0 | - |
| - | - | 578.1 | 1070 | - | - | 0 | - |
| - | - | 1199 | 1111 | - | - | 0 | - |
| - | - | 972.5 | 1112 | - | - | 0 | - |
| - | - | 1738 | 1225 | - | - | 0 | - |
| - | - | 1078 | 1226 | - | - | 0 | - |
| - | - | 1445 | 1238 | - | - | 0 | - |
| - | - | 1899 | 1239 | - | - | 0 | - |
| - | - | 703.6 | 1240 | - | - | 0 | - |
| - | - | 3698 | 1241 | - | - | 0 | - |
| - | - | 3601 | 1242 | - | - | 0 | - |
| - | - | 782.2 | 1342 | - | - | 0 | - |
| - | - | 644 | 2015 | - | - | 0 | - |
| - | - | 760.5 | 2873 | - | - | 0 | - |

m/z Charge Intensity FragmentType MassShift Position
120.0810317993164 0 12932.251
121.08446502685547 0 788.3314
128.09463500976562 0 797.94086
129.05490112304688 0 468.87106
129.1024627685547 0 8519.694
130.10598754882812 0 473.1045
131.08175659179688 0 325.59427
136.9339141845703 0 390.3827
146.1290283203125 0 3348.831
155.11831665039062 0 739.41174
159.0766143798828 0 1006.7917
166.08648681640625 0 32838 y 7
167.08990478515625 0 3295.7024
171.1131134033203 0 876.44977
173.45175170898438 0 1025.4592
189.08676147460938 0 474.73126
189.1032257080078 0 587.38257
192.0912322998047 0 507.73938
193.5643768310547 0 421.85596
197.16485595703125 0 693.7375
197.70716857910156 0 499.6627
207.1127471923828 0 2316.291
212.13929748535156 0 915.8859
215.19924926757812 0 3385.1104
216.2072296142578 0 13876.3545
217.21102905273438 0 1079.8318
225.15972900390625 0 574.06586
228.15774536132812 0 762.41064
235.10836791992188 0 2674.968 y Water loss 6
239.3134765625 0 561.1339
241.17849731445312 0 2100.3906
242.11378479003906 0 734.54315
242.1864013671875 0 4044.5017 c Ammonia loss 1
247.4697265625 0 500.83942
253.10365295410156 0 1074.0355
253.11854553222656 0 17205.252 y 6
254.12173461914062 0 1804.3031
258.20526123046875 0 4178.1113
259.21282958984375 0 11869.018 c 1
260.2162170410156 0 1423.3591
303.23931884765625 0 5544.9697
304.2423095703125 0 989.51544
306.14453125 0 2958.414 y Water loss 5
307.1874084472656 0 807.7833
324.1548156738281 0 609.6979 y 5
329.2186584472656 0 2943.822 c Ammonia loss 2
330.1893310546875 0 1232.1157 c Ammonia loss 6
331.1612548828125 0 2913.3228
332.1642761230469 0 625.647
333.1995544433594 0 669.2443
345.2374572753906 0 9722.454
346.2447814941406 0 13578.161 c 2
347.2483825683594 0 2947.762
358.19549560546875 0 613.31665
365.06524658203125 0 583.7819
373.20843505859375 0 603.83606
375.85980224609375 0 709.98096
376.8670959472656 0 1325.438
377.1831970214844 0 687.5675 y Water loss 4
380.720703125 0 1529.1224
381.2222595214844 0 623.8488
385.2684326171875 0 18195.607
386.27337646484375 0 5451.059
387.279296875 0 615.9198
394.8352966308594 0 1045.8066
394.8777770996094 0 1526.5353
395.21124267578125 0 1553.8431
401.1901550292969 0 602.5733
403.7239685058594 0 4637.541
404.22576904296875 0 2539.2646
404.28668212890625 0 4685.5186
405.2909240722656 0 1558.6848
411.84912109375 0 1575.9296
413.2663879394531 0 6655.6553
418.1930847167969 0 1471.775
429.2818603515625 0 1297.2938 c Water loss 3
430.2664794921875 0 2338.721 c Ammonia loss 3
447.2929382324219 0 74042.05 c 3
448.29583740234375 0 14715.016
449.2981872558594 0 2625.3027
483.2926025390625 0 1007.64996
488.2832336425781 0 2001.9758
500.31927490234375 0 860.0236 c Water loss 4
501.30364990234375 0 7155.968 c Ammonia loss 4
502.30645751953125 0 1960.164
518.3297119140625 0 104070.3 c 4
519.3325805664062 0 25466.656
520.3351440429688 0 4802.8184
525.77197265625 0 988.6597
530.3402099609375 0 606.1069
536.3206176757812 0 922.1035
544.345458984375 0 2432.8152
545.3507690429688 0 2671.592
554.3309326171875 0 2518.876
555.326904296875 0 681.5785
564.254150390625 0 3557.7617
565.2598876953125 0 1691.6425 y Water loss 2
572.34033203125 0 15593.74 c Ammonia loss 5
573.34423828125 0 4631.894
574.3485717773438 0 1592.2162
582.2640991210938 0 9088.021
583.27099609375 0 13109.649 y 2
583.3241577148438 0 1039.3947
584.2745361328125 0 3244.047
585.2788696289062 0 880.04205
589.36669921875 0 98975.81 c 5
590.3695678710938 0 28183.084
591.3728637695312 0 5720.443
613.367919921875 0 663.86816
614.373291015625 0 2088.7156
618.3377685546875 0 994.6956
619.3535766601562 0 1044.7985
619.8056030273438 0 892.9194
620.358154296875 0 1450.2827
620.8572387695312 0 861.56396
623.34912109375 0 1160.0555
624.342041015625 0 534.0923
631.3777465820312 0 2419.8125
632.3827514648438 0 1612.01
633.3865966796875 0 593.1614
635.2630615234375 0 791.2613
637.2791137695312 0 1387.9849 w 1
641.361328125 0 5278.334
642.3643798828125 0 2070.2937
659.3721313476562 0 12651.353 c Ammonia loss 6
660.3751220703125 0 4080.7888
661.2261962890625 0 715.266
661.37744140625 0 1661.3032
662.3271484375 0 815.6409 z Water loss 1
674.3828125 0 5228.0312
675.3848876953125 0 1735.8
676.3987426757812 0 140726.33 c 6
677.4014282226562 0 44666.38
678.3469848632812 0 910.51245 y Water loss 1
678.404296875 0 10276.021
680.3382568359375 0 4104.327 z 1
681.3416748046875 0 2372.6648
682.3494262695312 0 876.51056
733.3966674804688 0 2942.0996
734.3970336914062 0 899.1244
752.36962890625 0 3483.5073
753.3717651367188 0 1095.0957
763.4600830078125 0 1134.3846
764.45166015625 0 788.9872
765.3793334960938 0 1747.9712
768.4370727539062 0 939.2106
779.4537353515625 0 4280.692
780.4577026367188 0 1604.9137
781.45947265625 0 653.97687
782.4042358398438 0 4045.1265
783.4078369140625 0 2166.337
784.4096069335938 0 692.1406
790.420654296875 0 675.6243
797.46337890625 0 6889.084
798.4678344726562 0 2938.797
806.4209594726562 0 1987.6243
807.4369506835938 0 1872.7384
808.4320068359375 0 51399.168
809.4349975585938 0 20890.1
810.4371948242188 0 8006.9453
822.4349975585938 0 2547.86
823.2518310546875 0 776.9549
823.4404296875 0 3093.0295
824.4505615234375 0 63428.38
825.4563598632812 0 56848.914
826.4598388671875 0 20487.648
827.4633178710938 0 4836.0024
888.4401245117188 0 625.5862
895.2493896484375 0 678.14606
905.4420166015625 0 1384.1134
906.4495849609375 0 3539.0754
907.4491577148438 0 1825.0466
1041.6043701171875 0 808.8417
1069.663818359375 0 578.1272
1110.6314697265625 0 1198.6561
1111.63232421875 0 972.47656
1224.68505859375 0 1737.5607
1225.68408203125 0 1078.0311
1237.69921875 0 1444.9011
1238.6981201171875 0 1898.826
1239.704833984375 0 703.5923
1240.712646484375 0 3698.2788
1241.7181396484375 0 3600.7688
1342.175048828125 0 782.173
2015.10546875 0 644.0314
2872.642333984375 0 760.50867

Spectrum Details

|  |  |
| --- | --- |
| Matched peaks? Matched peaks The total absolute number of peaks matched. Additionally in brackets the total fraction of peaks matched and the total number of peaks is shown. | 27 (14.52% of 186) |
| FDR? FDR The false discovery rate estimated for this peptide. It is calculated by matching all theoretical fragments with a non-integer shift with the raw peaks for this spectrum. This is done with 40 different shifts. The resulting percentage is the average number of annotated peaks over the number of annotated peaks with the correct spectrum. | 3.97% |
| Satellite FDR? Satellite FDR See the FDR for details on its calculation. This satellite ion specific FDR only contains the satellite ions (d/w) for I/L/J positions. | 4.76% |
| PSM Score? PSM Score The PSM Score as given by Hecklib to this annotated spectrum. It is shown with three significant figures. | 383 |

#### Spectrum 5518? Spectrum 5518 The raw spectrum of this peptide as annotated by Hecklib. The fragments are coloured according to ion type (see legend). Any peaks with a star '\*' as text can be hovered over to see the full details, first the ion type second the mass shift type. By hovering over the amino acids in the peptide or ions in the legend the corresponding peaks are highlighted. By toggling the 'Unassigned' label you can turn the background (unassigned) peaks on or off in the plot. By updating the slider in the Ion legend you can update the spectrum to only show the top X% of the peaks with labels. The top X% means any peak that is within X% of the highest intensity. By dragging in the spectrum you can zoom in to a specific part of the spectrum and use 'Zoom Out' to get back to the original zoom level. The annotation of the spectrum is based on the given sequence in the peptides file and is done with different software so inconsistencies are likely. The peaks are annotated based on the given sequence, with 20 ppm tolerance.

Copy Data

##### Spectrum 5518 (TSV)

###### Preview

```
Loading example...
```

*Click on the button to copy the data to your clipboard.*

Mz MinMz MaxIntensity Max

WidthHeightPeptide font sizePeptide stroke widthSpectrum font sizeSpectrum stroke widthCompact peptide

Ion legend

wxyz

abcd

OtherUnassignedIon^Charge^_Position_Show for top:%

KJSTAASF

01.08e+42.15e+43.23e+44.31e+4

Zoom Out

y^+1^_1_y^+1^_2_c^+1^_2_y^+1^_2_c^+1^_2_y^+1^_3_c^+1^_3_c^+1^_3_c^+1^_4_c^+1^_4_c^+1^_4_c^+1^_5_c^+1^_5_c^+1^_6_y^+1^_6_c^+1^_6_c^+1^_7_c^+1^_7_z^+1^_7_

0688137620642752

Fragment Matches Table

Show background peaks

| Position | Ion type | Intensity | mz Theoretical | mz Error (Th) | mz Error (ppm) | Charge | Series Number |
| --- | --- | --- | --- | --- | --- | --- | --- |
| - | - | 359.1 | 120.1 | - | - | 0 | - |
| - | - | 4379 | 120.1 | - | - | 0 | - |
| - | - | 561.5 | 121.1 | - | - | 0 | - |
| - | - | 454.2 | 129.1 | - | - | 0 | - |
| - | - | 2889 | 129.1 | - | - | 0 | - |
| - | - | 363.9 | 129.4 | - | - | 0 | - |
| - | - | 717.4 | 146.1 | - | - | 0 | - |
| - | - | 672.6 | 166.1 | - | - | 0 | - |
| 8 | y | 9821 | 166.1 | 2.764E-05 | 0.1664 | +1 | 1 |
| - | - | 958.4 | 167.1 | - | - | 0 | - |
| - | - | 430 | 172.1 | - | - | 0 | - |
| - | - | 471.3 | 175.1 | - | - | 0 | - |
| - | - | 444.5 | 185.3 | - | - | 0 | - |
| - | - | 445 | 213.2 | - | - | 0 | - |
| - | - | 1128 | 215.2 | - | - | 0 | - |
| - | - | 3185 | 216.2 | - | - | 0 | - |
| - | - | 533.5 | 217.1 | - | - | 0 | - |
| - | - | 571.7 | 232.1 | - | - | 0 | - |
| - | - | 550.3 | 233.2 | - | - | 0 | - |
| 7 | y | 639.7 | 235.1 | 0.001366 | 5.811 | +1 | 2 |
| 2 | c | 1082 | 242.2 | 9.792E-05 | 0.4043 | +1 | 2 |
| - | - | 538.1 | 242.8 | - | - | 0 | - |
| 7 | y | 5886 | 253.1 | 4.844E-05 | 0.1914 | +1 | 2 |
| - | - | 502.9 | 253.4 | - | - | 0 | - |
| - | - | 625.6 | 254.1 | - | - | 0 | - |
| - | - | 1714 | 258.2 | - | - | 0 | - |
| 2 | c | 4088 | 259.2 | 0.000145 | 0.5595 | +1 | 2 |
| - | - | 506.3 | 267.7 | - | - | 0 | - |
| - | - | 555.4 | 283.2 | - | - | 0 | - |
| - | - | 525 | 288.7 | - | - | 0 | - |
| - | - | 507.6 | 297.8 | - | - | 0 | - |
| - | - | 758.7 | 301.1 | - | - | 0 | - |
| - | - | 1181 | 301.1 | - | - | 0 | - |
| - | - | 1663 | 303.2 | - | - | 0 | - |
| 6 | y | 797.7 | 306.1 | 0.0002785 | 0.9098 | +1 | 3 |
| 3 | c | 1082 | 329.2 | 0.001517 | 4.607 | +1 | 3 |
| - | - | 1084 | 331.2 | - | - | 0 | - |
| - | - | 697.9 | 334.9 | - | - | 0 | - |
| - | - | 2393 | 345.2 | - | - | 0 | - |
| 3 | c | 5225 | 346.2 | 0.0002826 | 0.8161 | +1 | 3 |
| - | - | 801.8 | 375.9 | - | - | 0 | - |
| - | - | 3352 | 376.9 | - | - | 0 | - |
| - | - | 673.9 | 377.9 | - | - | 0 | - |
| - | - | 563 | 383.7 | - | - | 0 | - |
| - | - | 5308 | 385.3 | - | - | 0 | - |
| - | - | 941.5 | 386.3 | - | - | 0 | - |
| - | - | 495 | 394.4 | - | - | 0 | - |
| - | - | 632.1 | 394.8 | - | - | 0 | - |
| - | - | 3055 | 394.9 | - | - | 0 | - |
| - | - | 1749 | 403.7 | - | - | 0 | - |
| - | - | 2233 | 404.3 | - | - | 0 | - |
| - | - | 1057 | 411.8 | - | - | 0 | - |
| - | - | 611 | 413.2 | - | - | 0 | - |
| - | - | 1.269E+04 | 413.3 | - | - | 0 | - |
| - | - | 696.9 | 414.3 | - | - | 0 | - |
| 4 | c | 648.6 | 429.3 | 4.871E-05 | 0.1135 | +1 | 4 |
| 4 | c | 814.6 | 430.3 | 0.0001412 | 0.3282 | +1 | 4 |
| 4 | c | 2.203E+04 | 447.3 | 1.258E-05 | 0.02813 | +1 | 4 |
| - | - | 4646 | 448.3 | - | - | 0 | - |
| 5 | c | 1812 | 501.3 | 9.853E-05 | 0.1966 | +1 | 5 |
| 5 | c | 3.193E+04 | 518.3 | 0.0002665 | 0.5141 | +1 | 5 |
| - | - | 7207 | 519.3 | - | - | 0 | - |
| - | - | 759.9 | 520.3 | - | - | 0 | - |
| - | - | 913.6 | 544.3 | - | - | 0 | - |
| 6 | c | 5911 | 572.3 | 8.899E-05 | 0.1555 | +1 | 6 |
| - | - | 1172 | 573.3 | - | - | 0 | - |
| - | - | 776.2 | 574.3 | - | - | 0 | - |
| - | - | 3458 | 582.3 | - | - | 0 | - |
| 3 | y | 3746 | 583.3 | 0.0009777 | 1.676 | +1 | 6 |
| - | - | 1271 | 584.3 | - | - | 0 | - |
| 6 | c | 3.184E+04 | 589.4 | 0.000454 | 0.7703 | +1 | 6 |
| - | - | 8901 | 590.4 | - | - | 0 | - |
| - | - | 1732 | 591.4 | - | - | 0 | - |
| - | - | 1394 | 610.8 | - | - | 0 | - |
| - | - | 1736 | 611.3 | - | - | 0 | - |
| - | - | 663.2 | 631.4 | - | - | 0 | - |
| - | - | 1712 | 641.4 | - | - | 0 | - |
| 7 | c | 3789 | 659.4 | 0.000196 | 0.2973 | +1 | 7 |
| - | - | 2160 | 660.4 | - | - | 0 | - |
| - | - | 661.4 | 674.4 | - | - | 0 | - |
| 7 | c | 4.263E+04 | 676.4 | 0.0003169 | 0.4685 | +1 | 7 |
| - | - | 1.361E+04 | 677.4 | - | - | 0 | - |
| - | - | 3147 | 678.4 | - | - | 0 | - |
| 2 | z | 847.1 | 680.3 | 0.001309 | 1.925 | +1 | 7 |
| - | - | 811.7 | 681.3 | - | - | 0 | - |
| - | - | 560.3 | 733.4 | - | - | 0 | - |
| - | - | 839.6 | 752.4 | - | - | 0 | - |
| - | - | 1714 | 779.5 | - | - | 0 | - |
| - | - | 1652 | 782.4 | - | - | 0 | - |
| - | - | 695.1 | 789.6 | - | - | 0 | - |
| - | - | 1878 | 797.5 | - | - | 0 | - |
| - | - | 608.9 | 798.5 | - | - | 0 | - |
| - | - | 1.428E+04 | 808.4 | - | - | 0 | - |
| - | - | 1020 | 808.5 | - | - | 0 | - |
| - | - | 7736 | 809.4 | - | - | 0 | - |
| - | - | 2277 | 810.4 | - | - | 0 | - |
| - | - | 566.2 | 821.3 | - | - | 0 | - |
| - | - | 1.785E+04 | 824.5 | - | - | 0 | - |
| - | - | 653.8 | 825.3 | - | - | 0 | - |
| - | - | 1.819E+04 | 825.5 | - | - | 0 | - |
| - | - | 5631 | 826.5 | - | - | 0 | - |
| - | - | 1332 | 827.5 | - | - | 0 | - |
| - | - | 3865 | 1204 | - | - | 0 | - |
| - | - | 2613 | 1205 | - | - | 0 | - |
| - | - | 1059 | 1222 | - | - | 0 | - |
| - | - | 826.6 | 1238 | - | - | 0 | - |
| - | - | 1993 | 1239 | - | - | 0 | - |
| - | - | 1018 | 1240 | - | - | 0 | - |
| - | - | 755.8 | 1241 | - | - | 0 | - |
| - | - | 687.5 | 1243 | - | - | 0 | - |
| - | - | 585.8 | 1775 | - | - | 0 | - |
| - | - | 712 | 2031 | - | - | 0 | - |
| - | - | 670.2 | 2725 | - | - | 0 | - |

m/z Charge Intensity FragmentType MassShift Position
120.0771484375 0 359.08536
120.08082580566406 0 4378.9355
121.08428192138672 0 561.51697
129.05459594726562 0 454.17313
129.10223388671875 0 2888.9563
129.3690948486328 0 363.88513
146.12913513183594 0 717.39624
166.07894897460938 0 672.6429
166.0862274169922 0 9821.11 y 7
167.08995056152344 0 958.4016
172.11929321289062 0 429.9688
175.13206481933594 0 471.32706
185.324951171875 0 444.50986
213.1589813232422 0 445.0061
215.19927978515625 0 1128.2064
216.2068328857422 0 3185.367
217.09268188476562 0 533.4776
232.07899475097656 0 571.7417
233.18846130371094 0 550.33014
235.1090850830078 0 639.6721 y Water loss 6
242.1864013671875 0 1082.3896 c Ammonia loss 1
242.84640502929688 0 538.0732
253.1183319091797 0 5886.1914 y 6
253.38232421875 0 502.8648
254.12234497070312 0 625.6499
258.2046813964844 0 1713.7639
259.21270751953125 0 4087.8943 c 1
267.74017333984375 0 506.32843
283.1749267578125 0 555.3647
288.7388610839844 0 525.04333
297.76287841796875 0 507.56543
301.09619140625 0 758.71625
301.1409606933594 0 1181.0975
303.23883056640625 0 1662.6177
306.1451110839844 0 797.7047 y Water loss 5
329.2198486328125 0 1081.8401 c Ammonia loss 2
331.16131591796875 0 1083.7617
334.91058349609375 0 697.94556
345.23699951171875 0 2392.6145
346.2445983886719 0 5224.98 c 2
375.8589782714844 0 801.8337
376.8668212890625 0 3352.123
377.8653564453125 0 673.9142
383.7356872558594 0 562.98364
385.2682800292969 0 5307.541
386.2733459472656 0 941.54486
394.4253234863281 0 494.977
394.8425598144531 0 632.1243
394.877197265625 0 3054.9639
403.72406005859375 0 1749.3639
404.2866516113281 0 2232.6313
411.8489074707031 0 1057.1138
413.1817626953125 0 611.005
413.26629638671875 0 12694.685
414.26373291015625 0 696.87146
429.28204345703125 0 648.59467 c Water loss 3
430.265869140625 0 814.6038 c Ammonia loss 3
447.2925720214844 0 22029.217 c 3
448.2956848144531 0 4645.7695
501.30322265625 0 1812.1166 c Ammonia loss 4
518.3294067382812 0 31934.943 c 4
519.3319702148438 0 7206.527
520.3360595703125 0 759.94147
544.3469848632812 0 913.6232
572.3401489257812 0 5911.123 c Ammonia loss 5
573.3450927734375 0 1172.2288
574.3499145507812 0 776.2388
582.26416015625 0 3458.3716
583.271240234375 0 3745.7717 y 2
584.275390625 0 1271.4713
589.3663330078125 0 31836.139 c 5
590.36962890625 0 8901.228
591.3716430664062 0 1732.3064
610.8035888671875 0 1394.4607
611.3049926757812 0 1736.4836
631.3788452148438 0 663.15247
641.3621215820312 0 1711.598
659.3720703125 0 3789.3848 c Ammonia loss 6
660.3782958984375 0 2159.9324
674.3839111328125 0 661.41034
676.3984985351562 0 42634.613 c 6
677.4012451171875 0 13608.095
678.4041137695312 0 3146.9272
680.3388671875 0 847.05164 z 1
681.3431396484375 0 811.7065
733.3984375 0 560.2722
752.3660888671875 0 839.6368
779.451904296875 0 1714.2415
782.4053344726562 0 1652.1991
789.5661010742188 0 695.1369
797.4647827148438 0 1878.2114
798.4755249023438 0 608.85706
808.4318237304688 0 14276.169
808.513671875 0 1019.9117
809.4348754882812 0 7736.2104
810.4400634765625 0 2276.8333
821.263427734375 0 566.2106
824.4503784179688 0 17848.672
825.2719116210938 0 653.76337
825.4554443359375 0 18187.83
826.459716796875 0 5630.5396
827.4617919921875 0 1331.8835
1203.5936279296875 0 3864.5798
1204.6002197265625 0 2613.4634
1221.60205078125 0 1058.9854
1237.72314453125 0 826.5597
1238.71240234375 0 1993.2717
1239.7117919921875 0 1017.81146
1240.7103271484375 0 755.7564
1242.6876220703125 0 687.5104
1775.32763671875 0 585.7779
2030.940673828125 0 712.02325
2724.8486328125 0 670.16046

Spectrum Details

|  |  |
| --- | --- |
| Matched peaks? Matched peaks The total absolute number of peaks matched. Additionally in brackets the total fraction of peaks matched and the total number of peaks is shown. | 19 (16.81% of 113) |
| FDR? FDR The false discovery rate estimated for this peptide. It is calculated by matching all theoretical fragments with a non-integer shift with the raw peaks for this spectrum. This is done with 40 different shifts. The resulting percentage is the average number of annotated peaks over the number of annotated peaks with the correct spectrum. | 3.88% |
| Satellite FDR? Satellite FDR See the FDR for details on its calculation. This satellite ion specific FDR only contains the satellite ions (d/w) for I/L/J positions. | ∞ |
| PSM Score? PSM Score The PSM Score as given by Hecklib to this annotated spectrum. It is shown with three significant figures. | 262 |

#### Reverse Lookup? Reverse Lookup All places where this read could be placed.

| Group | Segment | Template | Template Part | Read Part | Score | Unique |
| --- | --- | --- | --- | --- | --- | --- |
| Homo sapiens Heavy Chain | IGHV | IGHV3-73 | [18..26] | [0..8] | 46 | True |

| Recombined | Template Part | Read Part | Score | Unique |
| --- | --- | --- | --- | --- |
| TRYP | [96..104] | [0..8] | 32 | True |

#### Meta Information from Multiple reads

##### Number of combined reads

4

##### Intensity

0.9508

##### TotalArea

1.95E+08

##### Changes to the peptide sequence

KJSTAASF

L→JNo support for either Leucine or Isoleucine based on side chain ions (Position: 2)

J→LSupport for Leucine based on side chain ions (1 for L 0 for I) (Position: 2)

L→JNo support for either Leucine or Isoleucine based on side chain ions (Position: 2)

#### Positional Score

Copy Data

##### Positional Score (TSV)

###### Preview

```
Loading example...
```

*Click on the button to copy the data to your clipboard.*

1001234567

Label Value
"0" 0.468
"1" 0.46
"2" 0.465
"3" 0.475
"4" 0.487
"5" 0.497
"6" 0.5
"7" 0.5

#### Meta Information from PEAKS

##### Scan Identifier

F3:4426

##### Original sequence

K

L

S

T

A

A

S

F

##### Posttranslational Modifications

##### Source File

D:\separate\_stitch\_analyses\xle-disambiguation\raw\20210323\_F1\_UM1\_Peng0013\_SA\_F59\_ingel\_3ug\_chymo.raw

##### Fraction

3

##### Scan Feature

F3:1096

##### De Novo Score

98

##### ConfidenceScore

98

### m/z

412.7301

##### Mass

823.4439

##### Charge

2

##### Retention Time

24.3

##### Predicted Retention Time

-

##### Area

1.95E+08

##### Parts Per Million

2.1

##### Fragmentation mode

ETHCD

##### Originating file

01 D:\separate\_stitch\_analyses\xle-disambiguation\20210325\_F59\_3ug\_DENOVO\_12.csv

#### Meta Information from PEAKS

##### Scan Identifier

F3:5039

##### Original sequence

K

L

S

T

A

A

S

F

##### Posttranslational Modifications

##### Source File

D:\separate\_stitch\_analyses\xle-disambiguation\raw\20210323\_F1\_UM1\_Peng0013\_SA\_F59\_ingel\_3ug\_chymo.raw

##### Fraction

3

##### Scan Feature

-

##### De Novo Score

98

##### ConfidenceScore

98

### m/z

412.7296

##### Mass

823.4439

##### Charge

2

##### Retention Time

27.84

##### Predicted Retention Time

-

##### Area

0

##### Parts Per Million

0.9

##### Fragmentation mode

ETHCD

##### Originating file

01 D:\separate\_stitch\_analyses\xle-disambiguation\20210325\_F59\_3ug\_DENOVO\_12.csv

#### Meta Information from PEAKS

##### Scan Identifier

F3:4759

##### Original sequence

K

L

S

T

A

A

S

F

##### Posttranslational Modifications

##### Source File

D:\separate\_stitch\_analyses\xle-disambiguation\raw\20210323\_F1\_UM1\_Peng0013\_SA\_F59\_ingel\_3ug\_chymo.raw

##### Fraction

3

##### Scan Feature

-

##### De Novo Score

96

##### ConfidenceScore

96

### m/z

412.7298

##### Mass

823.4439

##### Charge

2

##### Retention Time

26.24

##### Predicted Retention Time

-

##### Area

0

##### Parts Per Million

1.3

##### Fragmentation mode

ETHCD

##### Originating file

01 D:\separate\_stitch\_analyses\xle-disambiguation\20210325\_F59\_3ug\_DENOVO\_12.csv

#### Meta Information from PEAKS

##### Scan Identifier

F3:5518

##### Original sequence

K

L

S

T

A

A

S

F

##### Posttranslational Modifications

##### Source File

D:\separate\_stitch\_analyses\xle-disambiguation\raw\20210323\_F1\_UM1\_Peng0013\_SA\_F59\_ingel\_3ug\_chymo.raw

##### Fraction

3

##### Scan Feature

-

##### De Novo Score

96

##### ConfidenceScore

96

### m/z

412.7297

##### Mass

823.4439

##### Charge

2

##### Retention Time

30.53

##### Predicted Retention Time

-

##### Area

0

##### Parts Per Million

1.2

##### Fragmentation mode

ETHCD

##### Originating file

01 D:\separate\_stitch\_analyses\xle-disambiguation\20210325\_F59\_3ug\_DENOVO\_12.csv
