## Supplementary material for "A handle on mass coincidence errors in *de novo* sequencing of antibodies by bottom-up proteomics": Combined_072.html

Details Combined\_072 | Stitch OverviewUndefined

### Read Combined\_072

#### Sequence (length=11)

JSEVSDRPSGV

#### Spectrum 3712? Spectrum 3712 The raw spectrum of this peptide as annotated by Hecklib. The fragments are coloured according to ion type (see legend). Any peaks with a star '\*' as text can be hovered over to see the full details, first the ion type second the mass shift type. By hovering over the amino acids in the peptide or ions in the legend the corresponding peaks are highlighted. By toggling the 'Unassigned' label you can turn the background (unassigned) peaks on or off in the plot. By updating the slider in the Ion legend you can update the spectrum to only show the top X% of the peaks with labels. The top X% means any peak that is within X% of the highest intensity. By dragging in the spectrum you can zoom in to a specific part of the spectrum and use 'Zoom Out' to get back to the original zoom level. The annotation of the spectrum is based on the given sequence in the peptides file and is done with different software so inconsistencies are likely. The peaks are annotated based on the given sequence, with 20 ppm tolerance.

Copy Data

##### Spectrum 3712 (TSV)

###### Preview

```
Loading example...
```

*Click on the button to copy the data to your clipboard.*

Mz MinMz MaxIntensity Max

WidthHeightPeptide font sizePeptide stroke widthSpectrum font sizeSpectrum stroke widthCompact peptide

Ion legend

wxyz

abcd

OtherUnassignedIon^Charge^_Position_Show for top:%

JSEVSDRPSGV

03.91e+67.83e+61.17e+71.57e+7

Zoom Out

a^+1^_2_a^+1^_2_y^+1^_2_b^+1^_2_b^+1^_2_y^+1^_3_y^+1^_3_a^+1^_3_a^+1^_3_b^+1^_3_b^+2^_6_b^+1^_3_y^+1^_4_y^+2^_7_y^+1^_4_y^+2^_8_y^+2^_8_y^+2^_8_b^+1^_4_b^+1^_4_y^+2^_9_y^+2^_9_b^+2^_9_y^+1^_5_y^+1^_5_b^+2^_10_b^+2^_10_y^+2^_10_b^+2^_10_y^+1^_5_b^+1^_5_y^+2^_10_\*\*\*y^+1^_6_b^+1^_6_y^+1^_6_y^+1^_7_y^+1^_7_b^+1^_7_b^+1^_7_y^+1^_8_y^+1^_8_y^+1^_8_b^+1^_8_y^+1^_9_y^+1^_9_y^+1^_9_b^+1^_9_y^+1^_10_y^+1^_10_b^+1^_10_y^+1^_10_

02645297931058

Fragment Matches Table

Show background peaks

| Position | Ion type | Intensity | mz Theoretical | mz Error (Th) | mz Error (ppm) | Charge | Series Number |
| --- | --- | --- | --- | --- | --- | --- | --- |
| - | - | 1.3E+05 | 125.1 | - | - | 0 | - |
| - | - | 2.392E+05 | 126.1 | - | - | 0 | - |
| - | - | 1.775E+04 | 126.1 | - | - | 0 | - |
| - | - | 1.275E+04 | 128 | - | - | 0 | - |
| - | - | 1.224E+05 | 128.1 | - | - | 0 | - |
| - | - | 2.005E+05 | 129.1 | - | - | 0 | - |
| - | - | 6.51E+04 | 130.1 | - | - | 0 | - |
| - | - | 2.81E+04 | 131.1 | - | - | 0 | - |
| - | - | 2.258E+04 | 132.1 | - | - | 0 | - |
| - | - | 1.635E+04 | 132.7 | - | - | 0 | - |
| - | - | 2.05E+04 | 133.1 | - | - | 0 | - |
| - | - | 5.754E+04 | 138.1 | - | - | 0 | - |
| - | - | 1.144E+04 | 138.1 | - | - | 0 | - |
| - | - | 3.078E+04 | 139.1 | - | - | 0 | - |
| - | - | 5.724E+04 | 140.1 | - | - | 0 | - |
| - | - | 1.956E+04 | 140.1 | - | - | 0 | - |
| - | - | 1.956E+04 | 141.1 | - | - | 0 | - |
| - | - | 1.476E+04 | 141.2 | - | - | 0 | - |
| - | - | 5.483E+04 | 144.1 | - | - | 0 | - |
| - | - | 9.877E+04 | 145.1 | - | - | 0 | - |
| - | - | 1.402E+04 | 152.3 | - | - | 0 | - |
| - | - | 3.729E+04 | 153.1 | - | - | 0 | - |
| - | - | 1.981E+05 | 154.1 | - | - | 0 | - |
| - | - | 1.826E+04 | 155.1 | - | - | 0 | - |
| 2 | a | 2.853E+05 | 155.1 | 0.0003661 | 2.36 | +1 | 2 |
| - | - | 1.006E+05 | 156.1 | - | - | 0 | - |
| - | - | 1.542E+05 | 157.1 | - | - | 0 | - |
| - | - | 4.914E+05 | 157.1 | - | - | 0 | - |
| - | - | 1.741E+05 | 157.1 | - | - | 0 | - |
| - | - | 1.845E+04 | 158.1 | - | - | 0 | - |
| - | - | 2.479E+04 | 158.1 | - | - | 0 | - |
| - | - | 1.203E+05 | 159.1 | - | - | 0 | - |
| - | - | 2.638E+04 | 166.1 | - | - | 0 | - |
| - | - | 8.545E+04 | 167.1 | - | - | 0 | - |
| - | - | 1.972E+04 | 169.1 | - | - | 0 | - |
| - | - | 2.239E+04 | 169.1 | - | - | 0 | - |
| - | - | 1.588E+04 | 170 | - | - | 0 | - |
| - | - | 2.596E+05 | 171.1 | - | - | 0 | - |
| - | - | 4.647E+04 | 172.1 | - | - | 0 | - |
| - | - | 2.398E+04 | 172.1 | - | - | 0 | - |
| 2 | a | 3.709E+06 | 173.1 | 0.000452 | 2.611 | +1 | 2 |
| - | - | 8.702E+04 | 173.5 | - | - | 0 | - |
| - | - | 2.607E+05 | 174.1 | - | - | 0 | - |
| - | - | 1.598E+05 | 175.1 | - | - | 0 | - |
| 10 | y | 6.181E+05 | 175.1 | 0.0004203 | 2.4 | +1 | 2 |
| - | - | 2.902E+04 | 176.1 | - | - | 0 | - |
| - | - | 2.276E+04 | 182.1 | - | - | 0 | - |
| 2 | b | 3.881E+06 | 183.1 | 0.0004924 | 2.689 | +1 | 2 |
| - | - | 2.086E+04 | 184.1 | - | - | 0 | - |
| - | - | 2.723E+04 | 184.1 | - | - | 0 | - |
| - | - | 3.875E+05 | 184.1 | - | - | 0 | - |
| - | - | 1.881E+04 | 185.1 | - | - | 0 | - |
| - | - | 9.452E+05 | 185.1 | - | - | 0 | - |
| - | - | 2.068E+04 | 185.1 | - | - | 0 | - |
| - | - | 7.499E+04 | 186.1 | - | - | 0 | - |
| - | - | 2.561E+05 | 187.1 | - | - | 0 | - |
| - | - | 2.45E+04 | 188.1 | - | - | 0 | - |
| - | - | 1.416E+06 | 189.1 | - | - | 0 | - |
| - | - | 9.071E+04 | 190.1 | - | - | 0 | - |
| - | - | 6.022E+04 | 192.1 | - | - | 0 | - |
| - | - | 6.999E+04 | 194.1 | - | - | 0 | - |
| - | - | 1.733E+04 | 194.1 | - | - | 0 | - |
| - | - | 3.617E+04 | 195.1 | - | - | 0 | - |
| - | - | 3.697E+04 | 195.1 | - | - | 0 | - |
| - | - | 9.347E+05 | 199.1 | - | - | 0 | - |
| - | - | 5.516E+04 | 200.1 | - | - | 0 | - |
| 2 | b | 5.303E+06 | 201.1 | 0.0004562 | 2.268 | +1 | 2 |
| - | - | 4.492E+05 | 202.1 | - | - | 0 | - |
| - | - | 5.426E+05 | 203.1 | - | - | 0 | - |
| - | - | 3.462E+04 | 203.1 | - | - | 0 | - |
| - | - | 3.014E+04 | 204.1 | - | - | 0 | - |
| - | - | 6.129E+04 | 209.1 | - | - | 0 | - |
| - | - | 3.594E+05 | 211.1 | - | - | 0 | - |
| - | - | 1.663E+04 | 211.1 | - | - | 0 | - |
| - | - | 2.091E+05 | 212.1 | - | - | 0 | - |
| - | - | 1.655E+04 | 212.1 | - | - | 0 | - |
| - | - | 1.995E+04 | 213.1 | - | - | 0 | - |
| - | - | 2.746E+04 | 213.2 | - | - | 0 | - |
| - | - | 2.537E+04 | 215.1 | - | - | 0 | - |
| - | - | 2.199E+04 | 216.1 | - | - | 0 | - |
| - | - | 2.745E+06 | 217.1 | - | - | 0 | - |
| - | - | 2.317E+05 | 218.1 | - | - | 0 | - |
| - | - | 2.588E+04 | 219.1 | - | - | 0 | - |
| - | - | 5.073E+04 | 220.1 | - | - | 0 | - |
| - | - | 3.232E+04 | 220.1 | - | - | 0 | - |
| - | - | 2.729E+04 | 223.1 | - | - | 0 | - |
| - | - | 1.79E+05 | 224.1 | - | - | 0 | - |
| - | - | 1.599E+04 | 224.1 | - | - | 0 | - |
| - | - | 2.023E+04 | 225.1 | - | - | 0 | - |
| - | - | 6.155E+04 | 227.1 | - | - | 0 | - |
| - | - | 3.456E+04 | 227.2 | - | - | 0 | - |
| - | - | 5.194E+05 | 229.1 | - | - | 0 | - |
| - | - | 1.644E+05 | 230.1 | - | - | 0 | - |
| - | - | 7.142E+04 | 237.1 | - | - | 0 | - |
| - | - | 2.06E+05 | 237.1 | - | - | 0 | - |
| - | - | 2.125E+04 | 240.1 | - | - | 0 | - |
| - | - | 3.1E+05 | 242.1 | - | - | 0 | - |
| - | - | 2.837E+04 | 243.1 | - | - | 0 | - |
| - | - | 1.877E+04 | 243.1 | - | - | 0 | - |
| - | - | 1.76E+04 | 243.6 | - | - | 0 | - |
| 9 | y | 3.347E+04 | 244.1 | 0.0003036 | 1.244 | +1 | 3 |
| - | - | 1.782E+04 | 246.8 | - | - | 0 | - |
| - | - | 3.824E+04 | 252.1 | - | - | 0 | - |
| - | - | 2.577E+04 | 253.1 | - | - | 0 | - |
| - | - | 6.696E+04 | 254.2 | - | - | 0 | - |
| - | - | 7.989E+05 | 255.1 | - | - | 0 | - |
| - | - | 4.014E+04 | 255.2 | - | - | 0 | - |
| - | - | 7.961E+04 | 256.1 | - | - | 0 | - |
| - | - | 1.823E+04 | 259.9 | - | - | 0 | - |
| - | - | 4.174E+05 | 260.1 | - | - | 0 | - |
| - | - | 3.146E+04 | 261.1 | - | - | 0 | - |
| 9 | y | 4.821E+05 | 262.1 | 0.0003895 | 1.486 | +1 | 3 |
| - | - | 2.734E+04 | 263.1 | - | - | 0 | - |
| - | - | 5.846E+04 | 267.1 | - | - | 0 | - |
| - | - | 3.218E+04 | 269.1 | - | - | 0 | - |
| - | - | 1.23E+05 | 270.1 | - | - | 0 | - |
| - | - | 2.524E+04 | 271.1 | - | - | 0 | - |
| - | - | 2.365E+04 | 271.1 | - | - | 0 | - |
| - | - | 1.111E+05 | 272.1 | - | - | 0 | - |
| - | - | 1.23E+05 | 272.1 | - | - | 0 | - |
| - | - | 2.719E+04 | 273.1 | - | - | 0 | - |
| - | - | 7.128E+04 | 280.1 | - | - | 0 | - |
| - | - | 2.886E+04 | 281.1 | - | - | 0 | - |
| - | - | 3.154E+04 | 282.2 | - | - | 0 | - |
| - | - | 7.89E+04 | 284.1 | - | - | 0 | - |
| 3 | a | 3.205E+05 | 284.2 | 0.0005891 | 2.073 | +1 | 3 |
| - | - | 3.667E+04 | 285.2 | - | - | 0 | - |
| - | - | 2.17E+04 | 286.1 | - | - | 0 | - |
| - | - | 1.415E+05 | 288.2 | - | - | 0 | - |
| - | - | 2.378E+04 | 289.2 | - | - | 0 | - |
| - | - | 5.393E+04 | 294.1 | - | - | 0 | - |
| - | - | 3.744E+04 | 296.1 | - | - | 0 | - |
| - | - | 4.004E+04 | 296.2 | - | - | 0 | - |
| - | - | 4.144E+05 | 298.1 | - | - | 0 | - |
| - | - | 1.549E+05 | 299.1 | - | - | 0 | - |
| - | - | 3.127E+04 | 300.1 | - | - | 0 | - |
| - | - | 1.776E+04 | 301.1 | - | - | 0 | - |
| - | - | 7.346E+04 | 302.1 | - | - | 0 | - |
| 3 | a | 8.072E+04 | 302.2 | 0.0007666 | 2.537 | +1 | 3 |
| - | - | 2.825E+04 | 306.2 | - | - | 0 | - |
| - | - | 1.76E+04 | 307.1 | - | - | 0 | - |
| 3 | b | 4.861E+06 | 312.2 | 0.0008222 | 2.634 | +1 | 3 |
| - | - | 7.626E+05 | 313.2 | - | - | 0 | - |
| - | - | 9.5E+04 | 313.2 | - | - | 0 | - |
| - | - | 1.76E+04 | 314 | - | - | 0 | - |
| - | - | 3.892E+04 | 314.1 | - | - | 0 | - |
| - | - | 5.113E+04 | 314.2 | - | - | 0 | - |
| - | - | 2.611E+04 | 314.2 | - | - | 0 | - |
| 6 | b | 1.198E+06 | 316.2 | 0.0006586 | 2.083 | +2 | 6 |
| - | - | 1.754E+05 | 317.2 | - | - | 0 | - |
| - | - | 3.168E+04 | 318.2 | - | - | 0 | - |
| - | - | 2.343E+04 | 319.2 | - | - | 0 | - |
| - | - | 1.709E+04 | 321.3 | - | - | 0 | - |
| - | - | 1.566E+05 | 324.1 | - | - | 0 | - |
| - | - | 4.402E+04 | 324.2 | - | - | 0 | - |
| - | - | 3.773E+04 | 325.1 | - | - | 0 | - |
| - | - | 8.396E+04 | 326.1 | - | - | 0 | - |
| - | - | 2.344E+04 | 329.2 | - | - | 0 | - |
| 3 | b | 2.682E+06 | 330.2 | 0.0007861 | 2.381 | +1 | 3 |
| - | - | 6.731E+05 | 331.2 | - | - | 0 | - |
| - | - | 6.002E+04 | 332.2 | - | - | 0 | - |
| - | - | 7.011E+04 | 335.1 | - | - | 0 | - |
| - | - | 5.026E+04 | 336.2 | - | - | 0 | - |
| - | - | 1.8E+04 | 338.1 | - | - | 0 | - |
| - | - | 4.931E+04 | 341.2 | - | - | 0 | - |
| 8 | y | 2.138E+05 | 341.2 | 0.0008845 | 2.592 | +1 | 4 |
| - | - | 9.874E+05 | 342.1 | - | - | 0 | - |
| - | - | 3.822E+04 | 342.2 | - | - | 0 | - |
| - | - | 1.264E+05 | 343.1 | - | - | 0 | - |
| - | - | 3.896E+04 | 344.1 | - | - | 0 | - |
| - | - | 2.851E+04 | 349.2 | - | - | 0 | - |
| - | - | 2.087E+04 | 350.1 | - | - | 0 | - |
| 5 | y | 3.65E+04 | 350.2 | 0.0002741 | 0.7828 | +2 | 7 |
| - | - | 3.762E+04 | 352.1 | - | - | 0 | - |
| - | - | 2.398E+04 | 352.2 | - | - | 0 | - |
| - | - | 5.467E+04 | 357.2 | - | - | 0 | - |
| - | - | 8.359E+05 | 359.2 | - | - | 0 | - |
| 8 | y | 2.747E+06 | 359.2 | 0.0005126 | 1.427 | +1 | 4 |
| - | - | 9.833E+04 | 359.7 | - | - | 0 | - |
| - | - | 1.059E+05 | 360.2 | - | - | 0 | - |
| - | - | 4.365E+05 | 360.2 | - | - | 0 | - |
| - | - | 6.454E+04 | 361.2 | - | - | 0 | - |
| - | - | 9.745E+04 | 367.2 | - | - | 0 | - |
| - | - | 2.758E+04 | 370.2 | - | - | 0 | - |
| - | - | 2.06E+04 | 372.2 | - | - | 0 | - |
| - | - | 2.596E+04 | 377.2 | - | - | 0 | - |
| - | - | 3.108E+04 | 380.2 | - | - | 0 | - |
| - | - | 1.879E+04 | 380.2 | - | - | 0 | - |
| - | - | 2.856E+04 | 381.2 | - | - | 0 | - |
| - | - | 7.912E+04 | 383.2 | - | - | 0 | - |
| - | - | 3.864E+04 | 384.2 | - | - | 0 | - |
| - | - | 1.875E+05 | 385.2 | - | - | 0 | - |
| - | - | 2.294E+04 | 386.2 | - | - | 0 | - |
| - | - | 2.559E+04 | 393.2 | - | - | 0 | - |
| - | - | 2.948E+04 | 393.2 | - | - | 0 | - |
| - | - | 6.028E+04 | 395.2 | - | - | 0 | - |
| - | - | 4.225E+04 | 396.2 | - | - | 0 | - |
| - | - | 3.396E+04 | 396.7 | - | - | 0 | - |
| - | - | 8.176E+04 | 398.2 | - | - | 0 | - |
| - | - | 3.689E+05 | 399.2 | - | - | 0 | - |
| 4 | y | 3.777E+04 | 399.7 | 0.001864 | 4.664 | +2 | 8 |
| 4 | y | 6.656E+04 | 400.2 | 0.002319 | 5.794 | +2 | 8 |
| - | - | 9.604E+05 | 401.2 | - | - | 0 | - |
| - | - | 1.777E+05 | 402.2 | - | - | 0 | - |
| - | - | 1.654E+05 | 403.2 | - | - | 0 | - |
| - | - | 3.655E+04 | 403.2 | - | - | 0 | - |
| - | - | 2.875E+04 | 404.2 | - | - | 0 | - |
| 4 | y | 2.663E+05 | 408.7 | 0.0006713 | 1.642 | +2 | 8 |
| - | - | 2.381E+04 | 408.7 | - | - | 0 | - |
| - | - | 8.11E+04 | 409.2 | - | - | 0 | - |
| 4 | b | 1.768E+05 | 411.2 | 0.0009508 | 2.312 | +1 | 4 |
| - | - | 2.017E+04 | 412.2 | - | - | 0 | - |
| - | - | 4.278E+04 | 412.2 | - | - | 0 | - |
| - | - | 8.941E+04 | 413.2 | - | - | 0 | - |
| - | - | 4.322E+05 | 416.2 | - | - | 0 | - |
| - | - | 5.463E+04 | 417.2 | - | - | 0 | - |
| - | - | 1.107E+05 | 417.2 | - | - | 0 | - |
| - | - | 2.52E+04 | 423.2 | - | - | 0 | - |
| - | - | 2.374E+04 | 424.2 | - | - | 0 | - |
| 4 | b | 1.148E+06 | 429.2 | 0.0009147 | 2.131 | +1 | 4 |
| - | - | 4.166E+05 | 430.2 | - | - | 0 | - |
| - | - | 2.3E+04 | 431.2 | - | - | 0 | - |
| - | - | 7.837E+04 | 431.2 | - | - | 0 | - |
| - | - | 9.23E+04 | 439.2 | - | - | 0 | - |
| - | - | 2.376E+04 | 440.2 | - | - | 0 | - |
| - | - | 1.811E+05 | 441.2 | - | - | 0 | - |
| - | - | 2.551E+04 | 441.7 | - | - | 0 | - |
| - | - | 5.135E+04 | 442.2 | - | - | 0 | - |
| - | - | 2.36E+04 | 446.2 | - | - | 0 | - |
| - | - | 3.872E+04 | 450.2 | - | - | 0 | - |
| - | - | 1.885E+04 | 450.3 | - | - | 0 | - |
| - | - | 2.19E+04 | 453.2 | - | - | 0 | - |
| - | - | 1.365E+05 | 455.2 | - | - | 0 | - |
| - | - | 7.676E+04 | 455.7 | - | - | 0 | - |
| - | - | 2.575E+04 | 456.2 | - | - | 0 | - |
| - | - | 6.478E+04 | 456.2 | - | - | 0 | - |
| - | - | 3.063E+04 | 457.2 | - | - | 0 | - |
| - | - | 5.153E+05 | 458.2 | - | - | 0 | - |
| - | - | 7.946E+04 | 459.2 | - | - | 0 | - |
| 3 | y | 1.739E+05 | 464.2 | 0.0009535 | 2.054 | +2 | 9 |
| - | - | 8.526E+04 | 464.7 | - | - | 0 | - |
| - | - | 2.262E+04 | 465.2 | - | - | 0 | - |
| - | - | 2.76E+04 | 469.3 | - | - | 0 | - |
| - | - | 2.595E+04 | 470.3 | - | - | 0 | - |
| - | - | 2.804E+04 | 471.2 | - | - | 0 | - |
| 3 | y | 4.893E+05 | 473.2 | 0.001073 | 2.267 | +2 | 9 |
| - | - | 3.003E+04 | 473.3 | - | - | 0 | - |
| - | - | 2.525E+05 | 473.7 | - | - | 0 | - |
| - | - | 4.037E+04 | 474.2 | - | - | 0 | - |
| - | - | 2.64E+04 | 479.3 | - | - | 0 | - |
| - | - | 9.179E+04 | 480.2 | - | - | 0 | - |
| - | - | 1.953E+04 | 481.2 | - | - | 0 | - |
| - | - | 1.885E+04 | 482.2 | - | - | 0 | - |
| - | - | 3.913E+04 | 484.2 | - | - | 0 | - |
| - | - | 2.445E+04 | 484.7 | - | - | 0 | - |
| - | - | 1.816E+04 | 485.2 | - | - | 0 | - |
| 9 | b | 7.66E+04 | 486.2 | 0.0009381 | 1.929 | +2 | 9 |
| - | - | 7.974E+04 | 486.7 | - | - | 0 | - |
| - | - | 5.622E+04 | 488.3 | - | - | 0 | - |
| - | - | 2.296E+04 | 495.3 | - | - | 0 | - |
| - | - | 3.715E+04 | 496.2 | - | - | 0 | - |
| 7 | y | 1.32E+05 | 497.3 | 0.0006951 | 1.398 | +1 | 5 |
| 7 | y | 6.699E+05 | 498.3 | 0.0001967 | 0.3949 | +1 | 5 |
| - | - | 9.202E+04 | 498.7 | - | - | 0 | - |
| - | - | 1.238E+05 | 499.3 | - | - | 0 | - |
| - | - | 4.923E+04 | 500.2 | - | - | 0 | - |
| - | - | 1.835E+04 | 500.3 | - | - | 0 | - |
| 10 | b | 2.573E+04 | 505.7 | 8.633E-05 | 0.1707 | +2 | 10 |
| 10 | b | 4.675E+04 | 506.2 | 0.009371 | 18.51 | +2 | 10 |
| 2 | y | 2.291E+05 | 507.7 | 0.0006863 | 1.352 | +2 | 10 |
| - | - | 2.264E+04 | 508.2 | - | - | 0 | - |
| - | - | 1.179E+05 | 508.2 | - | - | 0 | - |
| - | - | 3.566E+04 | 513.2 | - | - | 0 | - |
| 10 | b | 1.677E+05 | 514.8 | 0.001071 | 2.08 | +2 | 10 |
| - | - | 9.97E+04 | 515.3 | - | - | 0 | - |
| 7 | y | 2.697E+06 | 515.3 | 0.0009946 | 1.93 | +1 | 5 |
| 5 | b | 9.506E+04 | 516.3 | 0.002244 | 4.347 | +1 | 5 |
| - | - | 6.877E+05 | 516.3 | - | - | 0 | - |
| 2 | y | 6.36E+05 | 516.8 | 0.0008666 | 1.677 | +2 | 10 |
| - | - | 3.106E+05 | 517.3 | - | - | 0 | - |
| - | - | 9.362E+04 | 517.3 | - | - | 0 | - |
| - | - | 1.044E+05 | 517.8 | - | - | 0 | - |
| - | - | 3.652E+04 | 518.2 | - | - | 0 | - |
| - | - | 2.366E+04 | 519.2 | - | - | 0 | - |
| - | - | 3.863E+04 | 521.7 | - | - | 0 | - |
| - | - | 2.593E+04 | 522.7 | - | - | 0 | - |
| - | - | 5.157E+04 | 526.2 | - | - | 0 | - |
| - | - | 1.553E+05 | 531.3 | - | - | 0 | - |
| - | - | 3.757E+04 | 538.3 | - | - | 0 | - |
| - | - | 4.415E+04 | 541.3 | - | - | 0 | - |
| - | - | 1.19E+05 | 543.3 | - | - | 0 | - |
| - | - | 7.858E+04 | 550.3 | - | - | 0 | - |
| - | - | 5.617E+04 | 550.8 | - | - | 0 | - |
| - | - | 1.254E+05 | 555.3 | - | - | 0 | - |
| - | - | 6.625E+04 | 555.8 | - | - | 0 | - |
| - | - | 2.372E+04 | 556.3 | - | - | 0 | - |
| - | - | 7.595E+04 | 559.3 | - | - | 0 | - |
| - | - | 4.636E+04 | 560.3 | - | - | 0 | - |
| 0 | Precursor | 6.819E+05 | 564.3 | 0.0007991 | 1.416 | +2 | -1 |
| 0 | Precursor | 4.049E+05 | 564.8 | 0.009768 | 17.3 | +2 | -1 |
| - | - | 2.251E+04 | 565.2 | - | - | 0 | - |
| - | - | 1.573E+05 | 565.3 | - | - | 0 | - |
| - | - | 4.516E+04 | 565.8 | - | - | 0 | - |
| - | - | 3.908E+05 | 569.3 | - | - | 0 | - |
| - | - | 1.062E+05 | 570.3 | - | - | 0 | - |
| - | - | 3.219E+04 | 572.3 | - | - | 0 | - |
| 0 | Precursor | 2.504E+06 | 573.3 | 0.0008268 | 1.442 | +2 | -1 |
| - | - | 1.633E+06 | 573.8 | - | - | 0 | - |
| - | - | 5.984E+05 | 574.3 | - | - | 0 | - |
| - | - | 7.86E+04 | 574.8 | - | - | 0 | - |
| - | - | 2.843E+04 | 582.3 | - | - | 0 | - |
| - | - | 5.291E+04 | 583.2 | - | - | 0 | - |
| - | - | 3.035E+05 | 587.3 | - | - | 0 | - |
| - | - | 1.104E+05 | 588.3 | - | - | 0 | - |
| - | - | 3.152E+04 | 595.3 | - | - | 0 | - |
| - | - | 2.367E+05 | 600.3 | - | - | 0 | - |
| - | - | 5.866E+04 | 601.3 | - | - | 0 | - |
| 6 | y | 5.49E+04 | 612.3 | 0.0004301 | 0.7024 | +1 | 6 |
| 6 | b | 1.024E+05 | 613.3 | 0.01025 | 16.71 | +1 | 6 |
| - | - | 7.824E+05 | 618.3 | - | - | 0 | - |
| - | - | 2.334E+05 | 619.3 | - | - | 0 | - |
| - | - | 3.751E+04 | 620.3 | - | - | 0 | - |
| - | - | 2.827E+04 | 624.3 | - | - | 0 | - |
| 6 | y | 2.623E+06 | 630.3 | 0.00109 | 1.73 | +1 | 6 |
| - | - | 7.977E+05 | 631.3 | - | - | 0 | - |
| - | - | 1.561E+05 | 632.3 | - | - | 0 | - |
| - | - | 2.459E+04 | 640.3 | - | - | 0 | - |
| - | - | 6.649E+04 | 642.3 | - | - | 0 | - |
| - | - | 1.241E+05 | 646.3 | - | - | 0 | - |
| - | - | 4.321E+04 | 647.3 | - | - | 0 | - |
| - | - | 8.027E+04 | 656.3 | - | - | 0 | - |
| - | - | 3.903E+04 | 657.3 | - | - | 0 | - |
| - | - | 5.806E+04 | 658.3 | - | - | 0 | - |
| - | - | 2.374E+04 | 664.3 | - | - | 0 | - |
| - | - | 7.382E+04 | 671.3 | - | - | 0 | - |
| - | - | 2.729E+04 | 672.3 | - | - | 0 | - |
| - | - | 4.815E+05 | 674.3 | - | - | 0 | - |
| - | - | 1.605E+05 | 675.3 | - | - | 0 | - |
| - | - | 3.104E+04 | 676.3 | - | - | 0 | - |
| - | - | 3.007E+04 | 682.3 | - | - | 0 | - |
| - | - | 1.225E+05 | 687.3 | - | - | 0 | - |
| - | - | 2.24E+04 | 688.3 | - | - | 0 | - |
| 5 | y | 4.921E+05 | 699.3 | 0.0006836 | 0.9775 | +1 | 7 |
| - | - | 1.958E+05 | 700.3 | - | - | 0 | - |
| - | - | 4.955E+04 | 701.3 | - | - | 0 | - |
| 5 | y | 1.55E+07 | 717.4 | 0.001166 | 1.626 | +1 | 7 |
| - | - | 5.453E+06 | 718.4 | - | - | 0 | - |
| - | - | 1.128E+06 | 719.4 | - | - | 0 | - |
| - | - | 1.018E+05 | 720.4 | - | - | 0 | - |
| - | - | 4.819E+04 | 725.4 | - | - | 0 | - |
| - | - | 2.997E+04 | 726.4 | - | - | 0 | - |
| - | - | 4.855E+04 | 727.3 | - | - | 0 | - |
| - | - | 4.798E+04 | 735.3 | - | - | 0 | - |
| - | - | 2.524E+05 | 753.4 | - | - | 0 | - |
| - | - | 1.146E+05 | 754.4 | - | - | 0 | - |
| - | - | 4.14E+04 | 755.4 | - | - | 0 | - |
| - | - | 2.803E+04 | 757.4 | - | - | 0 | - |
| - | - | 7.699E+04 | 759.4 | - | - | 0 | - |
| - | - | 1.961E+04 | 760.4 | - | - | 0 | - |
| 7 | b | 6.438E+04 | 769.4 | 1.749E-05 | 0.02273 | +1 | 7 |
| - | - | 3.325E+04 | 770.4 | - | - | 0 | - |
| - | - | 8.566E+04 | 771.4 | - | - | 0 | - |
| - | - | 3.018E+04 | 772.4 | - | - | 0 | - |
| - | - | 7.888E+04 | 782.4 | - | - | 0 | - |
| - | - | 4.718E+04 | 786.4 | - | - | 0 | - |
| 7 | b | 1.014E+06 | 787.4 | 0.001233 | 1.565 | +1 | 7 |
| - | - | 4.02E+05 | 788.4 | - | - | 0 | - |
| - | - | 7.647E+04 | 789.4 | - | - | 0 | - |
| - | - | 4.609E+04 | 792.4 | - | - | 0 | - |
| - | - | 2.626E+04 | 793.4 | - | - | 0 | - |
| 4 | y | 2.151E+05 | 798.4 | 0.000507 | 0.635 | +1 | 8 |
| 4 | y | 1.141E+05 | 799.4 | 0.01564 | 19.56 | +1 | 8 |
| - | - | 4.827E+04 | 800.4 | - | - | 0 | - |
| - | - | 3.833E+05 | 810.4 | - | - | 0 | - |
| - | - | 1.951E+05 | 811.4 | - | - | 0 | - |
| - | - | 3.567E+04 | 812.4 | - | - | 0 | - |
| 4 | y | 8.586E+06 | 816.4 | 0.001112 | 1.362 | +1 | 8 |
| - | - | 3.696E+06 | 817.4 | - | - | 0 | - |
| - | - | 8.318E+05 | 818.4 | - | - | 0 | - |
| - | - | 8.336E+04 | 819.4 | - | - | 0 | - |
| - | - | 2.625E+04 | 827.4 | - | - | 0 | - |
| - | - | 1.717E+05 | 828.4 | - | - | 0 | - |
| - | - | 8.698E+04 | 829.4 | - | - | 0 | - |
| - | - | 5.536E+04 | 830.4 | - | - | 0 | - |
| - | - | 3.103E+04 | 840.4 | - | - | 0 | - |
| - | - | 3.12E+05 | 846.4 | - | - | 0 | - |
| - | - | 1.036E+05 | 847.4 | - | - | 0 | - |
| - | - | 3.728E+04 | 848.4 | - | - | 0 | - |
| - | - | 1.123E+05 | 858.4 | - | - | 0 | - |
| - | - | 4.27E+04 | 859.4 | - | - | 0 | - |
| 8 | b | 4.431E+04 | 884.4 | 0.002119 | 2.395 | +1 | 8 |
| - | - | 2.091E+04 | 886.4 | - | - | 0 | - |
| - | - | 3.147E+04 | 887.4 | - | - | 0 | - |
| - | - | 2.505E+04 | 897.4 | - | - | 0 | - |
| - | - | 3.422E+04 | 899.5 | - | - | 0 | - |
| - | - | 2.937E+04 | 903.4 | - | - | 0 | - |
| - | - | 2.39E+04 | 909.4 | - | - | 0 | - |
| - | - | 2.27E+05 | 915.4 | - | - | 0 | - |
| - | - | 9.048E+04 | 916.4 | - | - | 0 | - |
| - | - | 2.43E+04 | 917.4 | - | - | 0 | - |
| 3 | y | 9.65E+05 | 927.5 | 0.000155 | 0.1671 | +1 | 9 |
| 3 | y | 4.941E+05 | 928.4 | 0.01815 | 19.55 | +1 | 9 |
| - | - | 1.397E+05 | 929.5 | - | - | 0 | - |
| - | - | 2.822E+04 | 930.4 | - | - | 0 | - |
| - | - | 4.686E+05 | 933.4 | - | - | 0 | - |
| - | - | 2.198E+05 | 934.4 | - | - | 0 | - |
| - | - | 3.806E+04 | 935.4 | - | - | 0 | - |
| - | - | 4.942E+04 | 943.5 | - | - | 0 | - |
| 3 | y | 7.446E+06 | 945.5 | 0.0006939 | 0.7339 | +1 | 9 |
| - | - | 3.59E+06 | 946.5 | - | - | 0 | - |
| - | - | 9.198E+05 | 947.5 | - | - | 0 | - |
| - | - | 9.741E+04 | 948.5 | - | - | 0 | - |
| 9 | b | 2.362E+05 | 971.5 | 0.0006078 | 0.6257 | +1 | 9 |
| - | - | 1.179E+05 | 972.5 | - | - | 0 | - |
| - | - | 3.675E+04 | 973.5 | - | - | 0 | - |
| - | - | 4.584E+04 | 979.5 | - | - | 0 | - |
| - | - | 3.774E+04 | 984.5 | - | - | 0 | - |
| - | - | 2.525E+04 | 985.5 | - | - | 0 | - |
| - | - | 2.15E+04 | 986.5 | - | - | 0 | - |
| - | - | 1.084E+05 | 996.5 | - | - | 0 | - |
| - | - | 5.996E+04 | 997.5 | - | - | 0 | - |
| - | - | 2.332E+04 | 1001 | - | - | 0 | - |
| - | - | 8.181E+04 | 1002 | - | - | 0 | - |
| - | - | 3.719E+04 | 1003 | - | - | 0 | - |
| - | - | 2.882E+04 | 1012 | - | - | 0 | - |
| 2 | y | 1.096E+06 | 1014 | 0.00102 | 1.005 | +1 | 10 |
| 2 | y | 5.873E+05 | 1015 | 0.01829 | 18.01 | +1 | 10 |
| - | - | 1.94E+05 | 1016 | - | - | 0 | - |
| - | - | 3.334E+04 | 1017 | - | - | 0 | - |
| - | - | 5.444E+04 | 1024 | - | - | 0 | - |
| - | - | 3.215E+04 | 1025 | - | - | 0 | - |
| 10 | b | 2.005E+05 | 1029 | 0.001361 | 1.323 | +1 | 10 |
| - | - | 1.104E+05 | 1030 | - | - | 0 | - |
| - | - | 3.918E+04 | 1031 | - | - | 0 | - |
| 2 | y | 1.056E+07 | 1032 | 0.001197 | 1.16 | +1 | 10 |
| - | - | 5.471E+06 | 1033 | - | - | 0 | - |
| - | - | 1.618E+06 | 1035 | - | - | 0 | - |
| - | - | 1.719E+05 | 1036 | - | - | 0 | - |
| - | - | 4.674E+05 | 1042 | - | - | 0 | - |
| - | - | 2.958E+05 | 1043 | - | - | 0 | - |
| - | - | 8.051E+04 | 1044 | - | - | 0 | - |
| - | - | 3.624E+04 | 1047 | - | - | 0 | - |
| - | - | 3.069E+04 | 1048 | - | - | 0 | - |

m/z Charge Intensity FragmentType MassShift Position
125.10773468017578 0 130025.86
126.05535888671875 0 239214.27
126.11088562011719 0 17752.373
127.96387481689453 0 12747.324
128.10731506347656 0 122449.36
129.10263061523438 0 200465.28
130.05029296875 0 65102.812
131.08200073242188 0 28098.326
132.1023406982422 0 22579.998
132.7283477783203 0 16346.705
133.06106567382812 0 20497.865
138.09181213378906 0 57539.098
138.1348114013672 0 11439.289
139.08702087402344 0 30783.71
140.08238220214844 0 57238.023
140.14389038085938 0 19558.682
141.10272216796875 0 19557.555
141.15451049804688 0 14763.513
144.0661163330078 0 54830.004
145.06117248535156 0 98770.82
152.28712463378906 0 14021.83
153.0663604736328 0 37286.645
154.05029296875 0 198063.22
155.1111602783203 0 18261.525
155.11825561523438 0 285333 a Water loss 1
156.10231018066406 0 100566.3
157.06121826171875 0 154173.08
157.09756469726562 0 491355.44
157.1088104248047 0 174106.9
158.0929718017578 0 18447.588
158.10084533691406 0 24793.867
159.1132049560547 0 120275.72
166.08636474609375 0 26383.658
167.08197021484375 0 85453.95
169.09768676757812 0 19715.346
169.13426208496094 0 22388.559
170.04554748535156 0 15882.991
171.0768585205078 0 259638.5
172.06105041503906 0 46470.996
172.08018493652344 0 23976.588
173.12890625 0 3708797 a 1
173.450927734375 0 87022.6
174.1322784423828 0 260681.33
175.07179260253906 0 159845.03
175.10813903808594 0 618142.75 y 9
176.11183166503906 0 29021.55
182.0934600830078 0 22764.145
183.11329650878906 0 3880924.8 b Water loss 1
184.097412109375 0 20862.18
184.10882568359375 0 27228.791
184.11671447753906 0 387534.47
185.05593872070312 0 18813.45
185.0924835205078 0 945224.7
185.11795043945312 0 20679.486
186.09596252441406 0 74992.39
187.10812377929688 0 256076.38
188.111572265625 0 24500.65
189.08746337890625 0 1416419
190.09091186523438 0 90710.59
192.11351013183594 0 60216.555
194.09271240234375 0 69992.01
194.12925720214844 0 17329.857
195.0770263671875 0 36168.695
195.1127471923828 0 36974.41
199.0717315673828 0 934705.7
200.07542419433594 0 55162.523
201.1238250732422 0 5303161.5 b 1
202.12722778320312 0 449214.53
203.06668090820312 0 542595.1
203.12901306152344 0 34623.336
204.06971740722656 0 30143.19
209.14028930664062 0 61294.35
211.10821533203125 0 359403.1
211.11856079101562 0 16630.197
212.1033477783203 0 209068.66
212.13865661621094 0 16550.389
213.08738708496094 0 19953.646
213.16058349609375 0 27456.824
215.13941955566406 0 25368.799
216.13487243652344 0 21987.105
217.08238220214844 0 2745358
218.0858154296875 0 231691.95
219.08778381347656 0 25881.86
220.09332275390625 0 50733.863
220.10784912109375 0 32324.53
223.1082305908203 0 27291.326
224.1034393310547 0 179016.92
224.11227416992188 0 15993.689
225.1074676513672 0 20229.117
227.06678771972656 0 61547.367
227.1759796142578 0 34563.434
229.11883544921875 0 519351.72
230.1139373779297 0 164440.78
237.0988311767578 0 71416.01
237.13497924804688 0 206011.73
240.09783935546875 0 21247.688
242.11402893066406 0 310003.28
243.11819458007812 0 28366.33
243.13270568847656 0 18770.602
243.60556030273438 0 17595.914
244.12948608398438 0 33466.33 y Water loss 8
246.81361389160156 0 17816.938
252.13522338867188 0 38235.77
253.1186981201172 0 25771.586
254.16184997558594 0 66960.49
255.10934448242188 0 798945.8
255.1712188720703 0 40139.926
256.11248779296875 0 79613.914
259.93115234375 0 18230.348
260.12451171875 0 417438
261.1278076171875 0 31458.14
262.14013671875 0 482063.44 y 8
263.14398193359375 0 27341.523
267.1341857910156 0 58457.934
269.1131896972656 0 32178.326
270.14532470703125 0 122961.94
271.1299743652344 0 25242.463
271.1468505859375 0 23646.375
272.1239929199219 0 111140.84
272.13604736328125 0 123029.234
273.127197265625 0 27194.775
280.1298522949219 0 71277.484
281.12530517578125 0 28861.754
282.18133544921875 0 31535.506
284.1244201660156 0 78899.61
284.16107177734375 0 320511.28 a Water loss 2
285.16400146484375 0 36670.336
286.10430908203125 0 21701.953
288.1560363769531 0 141482.4
289.15960693359375 0 23784.1
294.1459045410156 0 53933.625
296.1366271972656 0 37444.855
296.172119140625 0 40044.926
298.1403503417969 0 414437.84
299.1365661621094 0 154857.58
300.1312561035156 0 31274.422
301.1487121582031 0 17756.057
302.1354675292969 0 73458.836
302.17181396484375 0 80724.04 a 2
306.15606689453125 0 28245.36
307.104736328125 0 17595.582
312.1562194824219 0 4860621.5 b Water loss 2
313.1593322753906 0 762554.3
313.1882629394531 0 94998.22
314.03851318359375 0 17601.738
314.14501953125 0 38922.6
314.16217041015625 0 51127.76
314.1907958984375 0 26110.79
316.1509704589844 0 1197926.8 b 5
317.1507263183594 0 175421.23
318.1536560058594 0 31678.316
319.1614074707031 0 23429.15
321.2785339355469 0 17091.217
324.1310119628906 0 156623.28
324.167236328125 0 44019.99
325.1144714355469 0 37728.742
326.1356201171875 0 83962.07
329.1827697753906 0 23438.262
330.166748046875 0 2681678.8 b 2
331.1710510253906 0 673117.56
332.17364501953125 0 60018
335.1360778808594 0 70110.67
336.17828369140625 0 50256.176
338.1357116699219 0 18003.715
341.1569519042969 0 49306.527
341.1828308105469 0 213776.98 y Water loss 7
342.14166259765625 0 987388.5
342.1828308105469 0 38218.406
343.1441345214844 0 126382.87
344.1465148925781 0 38957.76
349.1513671875 0 28509.291
350.134765625 0 20870.002
350.1749267578125 0 36496.434 y Water loss 4
352.1257019042969 0 37620.18
352.1618347167969 0 23982.89
357.1770935058594 0 54669.566
359.1679382324219 0 835893.4
359.1930236816406 0 2746708.2 y 7
359.68157958984375 0 98329.305
360.1708679199219 0 105939.66
360.1960754394531 0 436518.9
361.1980285644531 0 64541.88
367.1619873046875 0 97446.96
370.2200622558594 0 27576.332
372.17596435546875 0 20599.68
377.18048095703125 0 25955.635
380.1569519042969 0 31082.232
380.20086669921875 0 18793.463
381.18695068359375 0 28555.572
383.2297668457031 0 79120.12
384.21148681640625 0 38637.137
385.17236328125 0 187494.95
386.174072265625 0 22941.87
393.1880798339844 0 25589.02
393.21441650390625 0 29478.982
395.1565856933594 0 60279.133
396.18792724609375 0 42254.086
396.6861267089844 0 33960.56
398.2152099609375 0 81764.3
399.1986999511719 0 368931.4
399.7107238769531 0 37772.633 y Water loss 3
400.20318603515625 0 66563.26 y Ammonia loss 3
401.240234375 0 960375.4
402.243408203125 0 177716.14
403.18341064453125 0 165436.66
403.2455139160156 0 36551.637
404.18426513671875 0 28754.26
408.7148132324219 0 266275.72 y 3
408.7425231933594 0 23806.1
409.2165222167969 0 81099.945
411.2247619628906 0 176766.83 b Water loss 3
412.1824951171875 0 20174.924
412.22833251953125 0 42777.426
413.16790771484375 0 89406.52
416.225830078125 0 432192.88
417.1988525390625 0 54631.336
417.2292175292969 0 110675.88
423.1985778808594 0 25198.871
424.1866760253906 0 23735.645
429.23529052734375 0 1147754.4 b 3
430.23968505859375 0 416606.97
431.1743469238281 0 23001.893
431.2407531738281 0 78371.47
439.1947021484375 0 92301.69
440.2257385253906 0 23758.6
441.21063232421875 0 181111.9
441.727783203125 0 25511.588
442.2137451171875 0 51347.438
446.21966552734375 0 23598.596
450.23406982421875 0 38718.305
450.29754638671875 0 18847.4
453.23773193359375 0 21895.564
455.2261047363281 0 136469.72
455.7274169921875 0 76762.33
456.2153625488281 0 25748.082
456.2471008300781 0 64781.42
457.2490539550781 0 30631.643
458.2366943359375 0 515334.4
459.2392883300781 0 79462.62
464.2311096191406 0 173921.05 y Water loss 2
464.7335205078125 0 85263.86
465.2361755371094 0 22623.283
469.2908020019531 0 27603.033
470.2633056640625 0 25951.057
471.24395751953125 0 28037.357
473.23651123046875 0 489338.16 y 2
473.2707214355469 0 30031.885
473.73773193359375 0 252535.94
474.23712158203125 0 40374.566
479.2737731933594 0 26400.73
480.249755859375 0 91785.09
481.2477111816406 0 19525.627
482.18865966796875 0 18851.336
484.2314758300781 0 39131.31
484.7303161621094 0 24447.588
485.2103271484375 0 18160.041
486.24420166015625 0 76595.14 b 8
486.7465515136719 0 79741.414
488.27337646484375 0 56221.1
495.2547912597656 0 22958.537
496.21392822265625 0 37153.8
497.28375244140625 0 132016.53 y Water loss 6
498.2668762207031 0 669881.6 y Ammonia loss 6
498.74114990234375 0 92019.484
499.2696838378906 0 123754.66
500.1985168457031 0 49230.816
500.275390625 0 18345.348
505.7486267089844 0 25729.22 b Water loss 9
506.2500915527344 0 46746.94 b Ammonia loss 9
507.7468566894531 0 229085.05 y Water loss 1
508.212158203125 0 22636.639
508.2493896484375 0 117938.73
513.241455078125 0 35655.06
514.7550659179688 0 167723.94 b 9
515.2576293945312 0 99695.46
515.2946166992188 0 2697125.2 y 6
516.26416015625 0 95055.49 b 4
516.2978515625 0 687717.5
516.7523193359375 0 635972.56 y 1
517.25390625 0 310622.4
517.3003540039062 0 93617.91
517.7551879882812 0 104378.57
518.2103271484375 0 36520.97
519.21240234375 0 23661.746
521.7451171875 0 38626.77
522.7481079101562 0 25929.51
526.2265625 0 51573.562
531.2529907226562 0 155311.58
538.2618408203125 0 37568.69
541.2753295898438 0 44145.445
543.2522583007812 0 119026.914
550.2921142578125 0 78580.48
550.7918090820312 0 56170.273
555.2840576171875 0 125384.79
555.78173828125 0 66252.22
556.2808227539062 0 23720.748
559.2848510742188 0 75946.516
560.2854614257812 0 46360.332
564.2890014648438 0 681875.1 Precursor Water loss
564.7899780273438 0 404919.38 Precursor Ammonia loss
565.2393188476562 0 22512.15
565.2921752929688 0 157338.12
565.791259765625 0 45161.215
569.2686767578125 0 390810.84
570.2684936523438 0 106224.48
572.2771606445312 0 32193.008
573.2943115234375 0 2503662 Precursor
573.7957763671875 0 1633026.1
574.296875 0 598370.25
574.7988891601562 0 78604.82
582.2630615234375 0 28433.432
583.2483520507812 0 52909.97
587.279052734375 0 303508.44
588.2818603515625 0 110405.9
595.2850341796875 0 31517.447
600.2743530273438 0 236744.45
601.2761840820312 0 58664.188
612.3095703125 0 54900.23 y Water loss 5
613.2930297851562 0 102419.04 b Water loss 5
618.2850341796875 0 782414.5
619.2880859375 0 233443.03
620.2908325195312 0 37507.848
624.3048706054688 0 28268.459
630.3216552734375 0 2622809 y 5
631.3240966796875 0 797683.5
632.3262329101562 0 156149.61
640.3047485351562 0 24591.404
642.3189086914062 0 66494.8
646.31640625 0 124060.4
647.3189086914062 0 43208.195
656.3003540039062 0 80268.64
657.2905883789062 0 39026.645
658.3057250976562 0 58058.1
664.3020629882812 0 23735.018
671.3467407226562 0 73815.27
672.3491821289062 0 27285.625
674.3109741210938 0 481505.47
675.3162231445312 0 160471.56
676.3218994140625 0 31044.678
682.314453125 0 30070.408
687.3436279296875 0 122510.19
688.3439331054688 0 22397.186
699.3427124023438 0 492064.97 y Water loss 4
700.3427124023438 0 195759.81
701.3424682617188 0 49553
717.353759765625 0 15503751 y 4
718.3564453125 0 5452586
719.3585815429688 0 1127958
720.3603515625 0 101768.33
725.3604125976562 0 48186.37
726.3585205078125 0 29972.111
727.3390502929688 0 48551.887
735.3402709960938 0 47983.355
753.353759765625 0 252369.3
754.35546875 0 114645.195
755.3552856445312 0 41402.625
757.371337890625 0 28027.89
759.4002685546875 0 76994.32
760.4080810546875 0 19608.877
769.3839111328125 0 64376.99 b Water loss 6
770.3946533203125 0 33251.75
771.3633422851562 0 85660.17
772.3628540039062 0 30177.012
782.378662109375 0 78877.34
786.4093017578125 0 47180.77
787.3956909179688 0 1014462.1 b 6
788.39794921875 0 402033.62
789.3964233398438 0 76470.9
792.3621215820312 0 46086.957
793.3572387695312 0 26263.145
798.4109497070312 0 215090.6 y Water loss 3
799.4100952148438 0 114090.65 y Ammonia loss 3
800.3999633789062 0 48266.984
810.375244140625 0 383333.75
811.3770141601562 0 195061.27
812.3773193359375 0 35672.7
816.422119140625 0 8585914 y 3
817.4248657226562 0 3696455.2
818.4271850585938 0 831757.9
819.4283447265625 0 83364.14
827.4105224609375 0 26253.484
828.3863525390625 0 171720.34
829.385986328125 0 86976.93
830.3984375 0 55362.934
840.3856201171875 0 31028.89
846.3963012695312 0 312026.72
847.401123046875 0 103551.04
848.4013671875 0 37277.996
858.3963012695312 0 112279.11
859.3980102539062 0 42695.92
884.4493408203125 0 44313.117 b 7
886.4312744140625 0 20913.748
887.4207153320312 0 31474.164
897.4078979492188 0 25045.33
899.45849609375 0 34224.875
903.421875 0 29372.562
909.4339599609375 0 23899.87
915.4195556640625 0 227043.55
916.4207153320312 0 90481.19
917.416259765625 0 24297.742
927.452880859375 0 964977.75 y Water loss 2
928.4552001953125 0 494127.53 y Ammonia loss 2
929.4542846679688 0 139695.08
930.44873046875 0 28216.23
933.4277954101562 0 468590.34
934.431396484375 0 219822.28
935.4376220703125 0 38058.34
943.486083984375 0 49423.74
945.4642944335938 0 7446226 y 2
946.46728515625 0 3590004.8
947.469482421875 0 919812
948.472412109375 0 97406.016
971.4798583984375 0 236247.03 b 8
972.4828491210938 0 117914.266
973.466552734375 0 36746.848
979.4611206054688 0 45838.258
984.4827880859375 0 37737.67
985.4805297851562 0 25252.482
986.4876098632812 0 21498.516
996.4762573242188 0 108367.79
997.471923828125 0 59962.918
1000.501220703125 0 23318.309
1002.4878540039062 0 81812.37
1003.4918212890625 0 37189.57
1012.4684448242188 0 28824.285
1014.486083984375 0 1095996.4 y Water loss 1
1015.4873657226562 0 587325.8 y Ammonia loss 1
1016.4879760742188 0 194039.02
1017.4959716796875 0 33336.246
1024.46923828125 0 54443.637
1025.4705810546875 0 32153.977
1028.5020751953125 0 200488.33 b 9
1029.502685546875 0 110441.74
1030.5072021484375 0 39176.86
1032.496826171875 0 10562329 y 1
1033.4993896484375 0 5470522.5
1034.501708984375 0 1618316.4
1035.50244140625 0 171886.94
1042.48046875 0 467366.25
1043.48388671875 0 295821.9
1044.4854736328125 0 80508.11
1046.5130615234375 0 36236.97
1047.5115966796875 0 30693.309

Spectrum Details

|  |  |
| --- | --- |
| Matched peaks? Matched peaks The total absolute number of peaks matched. Additionally in brackets the total fraction of peaks matched and the total number of peaks is shown. | 54 (12.19% of 443) |
| FDR? FDR The false discovery rate estimated for this peptide. It is calculated by matching all theoretical fragments with a non-integer shift with the raw peaks for this spectrum. This is done with 40 different shifts. The resulting percentage is the average number of annotated peaks over the number of annotated peaks with the correct spectrum. | 0.09% |
| Satellite FDR? Satellite FDR See the FDR for details on its calculation. This satellite ion specific FDR only contains the satellite ions (d/w) for I/L/J positions. | - |
| PSM Score? PSM Score The PSM Score as given by Hecklib to this annotated spectrum. It is shown with three significant figures. | 633 |

#### Spectrum 3800? Spectrum 3800 The raw spectrum of this peptide as annotated by Hecklib. The fragments are coloured according to ion type (see legend). Any peaks with a star '\*' as text can be hovered over to see the full details, first the ion type second the mass shift type. By hovering over the amino acids in the peptide or ions in the legend the corresponding peaks are highlighted. By toggling the 'Unassigned' label you can turn the background (unassigned) peaks on or off in the plot. By updating the slider in the Ion legend you can update the spectrum to only show the top X% of the peaks with labels. The top X% means any peak that is within X% of the highest intensity. By dragging in the spectrum you can zoom in to a specific part of the spectrum and use 'Zoom Out' to get back to the original zoom level. The annotation of the spectrum is based on the given sequence in the peptides file and is done with different software so inconsistencies are likely. The peaks are annotated based on the given sequence, with 20 ppm tolerance.

Copy Data

##### Spectrum 3800 (TSV)

###### Preview

```
Loading example...
```

*Click on the button to copy the data to your clipboard.*

Mz MinMz MaxIntensity Max

WidthHeightPeptide font sizePeptide stroke widthSpectrum font sizeSpectrum stroke widthCompact peptide

Ion legend

wxyz

abcd

OtherUnassignedIon^Charge^_Position_Show for top:%

JSEVSDRPSGV

05.21e+41.04e+51.56e+52.08e+5

Zoom Out

y^+1^_2_w^+1^_3_w^+1^_4_y^+1^_4_y^+2^_8_y^+2^_9_y^+2^_9_c^+2^_9_z^+1^_5_y^+2^_10_c^+2^_10_y^+1^_5_y^+2^_10_z^+1^_6_c^+1^_6_y^+1^_7_z^+1^_7_y^+1^_7_z^+1^_8_w^+1^_8_c^+1^_7_y^+1^_8_z^+1^_8_y^+1^_8_w^+1^_9_c^+1^_8_z^+1^_9_y^+1^_9_z^+1^_9_y^+1^_9_c^+1^_9_c^+1^_9_y^+1^_10_y^+1^_10_z^+1^_10_y^+1^_10_c^+1^_10_

0699139720962794

Fragment Matches Table

Show background peaks

| Position | Ion type | Intensity | mz Theoretical | mz Error (Th) | mz Error (ppm) | Charge | Series Number |
| --- | --- | --- | --- | --- | --- | --- | --- |
| - | - | 406.1 | 122.3 | - | - | 0 | - |
| - | - | 513.8 | 129.1 | - | - | 0 | - |
| - | - | 394.6 | 132.1 | - | - | 0 | - |
| - | - | 340.7 | 133.8 | - | - | 0 | - |
| - | - | 418.6 | 137.8 | - | - | 0 | - |
| - | - | 474.9 | 143.7 | - | - | 0 | - |
| - | - | 385.9 | 145.6 | - | - | 0 | - |
| - | - | 471.1 | 145.6 | - | - | 0 | - |
| - | - | 423.7 | 148.6 | - | - | 0 | - |
| - | - | 553.6 | 148.9 | - | - | 0 | - |
| - | - | 625.7 | 148.9 | - | - | 0 | - |
| - | - | 772.6 | 148.9 | - | - | 0 | - |
| - | - | 970.5 | 148.9 | - | - | 0 | - |
| - | - | 1089 | 148.9 | - | - | 0 | - |
| - | - | 1200 | 148.9 | - | - | 0 | - |
| - | - | 3050 | 148.9 | - | - | 0 | - |
| - | - | 4677 | 148.9 | - | - | 0 | - |
| - | - | 3389 | 149 | - | - | 0 | - |
| - | - | 1649 | 149 | - | - | 0 | - |
| - | - | 1155 | 149 | - | - | 0 | - |
| - | - | 1229 | 149 | - | - | 0 | - |
| - | - | 1043 | 149 | - | - | 0 | - |
| - | - | 839.3 | 149 | - | - | 0 | - |
| - | - | 564.5 | 149 | - | - | 0 | - |
| - | - | 423.3 | 149.1 | - | - | 0 | - |
| - | - | 535.2 | 156.3 | - | - | 0 | - |
| - | - | 416.9 | 159.1 | - | - | 0 | - |
| - | - | 444.7 | 162.3 | - | - | 0 | - |
| - | - | 1.046E+04 | 173.1 | - | - | 0 | - |
| - | - | 1032 | 173.4 | - | - | 0 | - |
| - | - | 1136 | 174.1 | - | - | 0 | - |
| 10 | y | 2584 | 175.1 | 0.0002982 | 1.703 | +1 | 2 |
| - | - | 6425 | 183.1 | - | - | 0 | - |
| - | - | 459 | 185.6 | - | - | 0 | - |
| - | - | 574.8 | 189.1 | - | - | 0 | - |
| - | - | 1563 | 199.1 | - | - | 0 | - |
| - | - | 2.744E+04 | 201.1 | - | - | 0 | - |
| - | - | 2609 | 202.1 | - | - | 0 | - |
| - | - | 1034 | 203.1 | - | - | 0 | - |
| - | - | 1159 | 211.1 | - | - | 0 | - |
| - | - | 5.307E+04 | 215.1 | - | - | 0 | - |
| - | - | 4775 | 217.1 | - | - | 0 | - |
| 9 | w | 2141 | 229.1 | 0.0003689 | 1.61 | +1 | 3 |
| - | - | 527.4 | 231.1 | - | - | 0 | - |
| - | - | 547.1 | 246.3 | - | - | 0 | - |
| - | - | 465.6 | 255.1 | - | - | 0 | - |
| - | - | 477.7 | 255.4 | - | - | 0 | - |
| - | - | 558.6 | 280.1 | - | - | 0 | - |
| - | - | 1303 | 284.2 | - | - | 0 | - |
| - | - | 644.7 | 289.5 | - | - | 0 | - |
| - | - | 766.9 | 298.1 | - | - | 0 | - |
| - | - | 2744 | 302.2 | - | - | 0 | - |
| - | - | 2.133E+04 | 312.2 | - | - | 0 | - |
| - | - | 3545 | 313.2 | - | - | 0 | - |
| - | - | 1140 | 314.1 | - | - | 0 | - |
| - | - | 754.8 | 314.2 | - | - | 0 | - |
| 8 | w | 4770 | 316.2 | 0.0003839 | 1.214 | +1 | 4 |
| - | - | 571.8 | 317.2 | - | - | 0 | - |
| - | - | 2.027E+04 | 330.2 | - | - | 0 | - |
| - | - | 2975 | 331.2 | - | - | 0 | - |
| - | - | 2773 | 332.1 | - | - | 0 | - |
| - | - | 1026 | 335.1 | - | - | 0 | - |
| - | - | 1197 | 342.1 | - | - | 0 | - |
| 8 | y | 7382 | 359.2 | 5.488E-05 | 0.1528 | +1 | 4 |
| - | - | 740.7 | 359.7 | - | - | 0 | - |
| - | - | 1896 | 360.1 | - | - | 0 | - |
| - | - | 779.7 | 360.2 | - | - | 0 | - |
| - | - | 638.3 | 383.2 | - | - | 0 | - |
| - | - | 665.2 | 385.2 | - | - | 0 | - |
| - | - | 595.8 | 397.2 | - | - | 0 | - |
| - | - | 752.4 | 399.2 | - | - | 0 | - |
| - | - | 9476 | 401.2 | - | - | 0 | - |
| - | - | 1644 | 402.2 | - | - | 0 | - |
| 4 | y | 1647 | 408.7 | 0.0001832 | 0.4482 | +2 | 8 |
| - | - | 877.1 | 409.2 | - | - | 0 | - |
| - | - | 1120 | 411.2 | - | - | 0 | - |
| - | - | 1.01E+04 | 429.2 | - | - | 0 | - |
| - | - | 1993 | 430.2 | - | - | 0 | - |
| - | - | 746.9 | 439.1 | - | - | 0 | - |
| 3 | y | 1993 | 464.2 | 0.0005873 | 1.265 | +2 | 9 |
| - | - | 972.8 | 464.7 | - | - | 0 | - |
| 3 | y | 5058 | 473.2 | 0.001134 | 2.396 | +2 | 9 |
| - | - | 2641 | 473.7 | - | - | 0 | - |
| 9 | c | 1029 | 486.2 | 0.001274 | 2.62 | +2 | 9 |
| - | - | 1129 | 498.3 | - | - | 0 | - |
| 7 | z | 4.83E+04 | 499.3 | 0.0005842 | 1.17 | +1 | 5 |
| - | - | 2.57E+04 | 500.3 | - | - | 0 | - |
| - | - | 5782 | 501.3 | - | - | 0 | - |
| - | - | 749.4 | 502.3 | - | - | 0 | - |
| 2 | y | 1267 | 507.7 | 7.599E-05 | 0.1497 | +2 | 10 |
| - | - | 625.1 | 508.3 | - | - | 0 | - |
| - | - | 758.9 | 513.3 | - | - | 0 | - |
| - | - | 1293 | 514.3 | - | - | 0 | - |
| 10 | c | 1209 | 514.8 | 0.0008216 | 1.596 | +2 | 10 |
| - | - | 1078 | 515.3 | - | - | 0 | - |
| 7 | y | 1.289E+04 | 515.3 | 0.0004453 | 0.8642 | +1 | 5 |
| - | - | 3053 | 516.3 | - | - | 0 | - |
| 2 | y | 6229 | 516.8 | 0.0006225 | 1.205 | +2 | 10 |
| - | - | 2737 | 517.3 | - | - | 0 | - |
| - | - | 1192 | 517.8 | - | - | 0 | - |
| - | - | 748.9 | 521.7 | - | - | 0 | - |
| - | - | 1387 | 538.2 | - | - | 0 | - |
| - | - | 644.6 | 542.3 | - | - | 0 | - |
| - | - | 715.7 | 556.3 | - | - | 0 | - |
| - | - | 6944 | 564.3 | - | - | 0 | - |
| - | - | 4539 | 564.8 | - | - | 0 | - |
| - | - | 875.2 | 565.3 | - | - | 0 | - |
| - | - | 1098 | 569.3 | - | - | 0 | - |
| - | - | 2512 | 570.3 | - | - | 0 | - |
| - | - | 1376 | 571.3 | - | - | 0 | - |
| - | - | 2446 | 572.3 | - | - | 0 | - |
| - | - | 3.113E+04 | 573.3 | - | - | 0 | - |
| - | - | 2.078E+04 | 573.8 | - | - | 0 | - |
| - | - | 5747 | 574.3 | - | - | 0 | - |
| - | - | 585.8 | 585.8 | - | - | 0 | - |
| 6 | z | 1.964E+04 | 614.3 | 0.000283 | 0.4607 | +1 | 6 |
| - | - | 3.596E+04 | 615.3 | - | - | 0 | - |
| - | - | 8988 | 616.3 | - | - | 0 | - |
| - | - | 1685 | 617.3 | - | - | 0 | - |
| - | - | 3854 | 629.3 | - | - | 0 | - |
| 6 | c | 3.926E+04 | 630.3 | 0.01153 | 18.29 | +1 | 6 |
| - | - | 1.244E+04 | 631.3 | - | - | 0 | - |
| - | - | 1750 | 632.3 | - | - | 0 | - |
| - | - | 572 | 655.3 | - | - | 0 | - |
| - | - | 729.6 | 657.3 | - | - | 0 | - |
| - | - | 1827 | 658.3 | - | - | 0 | - |
| - | - | 605.1 | 659.3 | - | - | 0 | - |
| - | - | 733.4 | 677.7 | - | - | 0 | - |
| - | - | 6951 | 699.3 | - | - | 0 | - |
| 5 | y | 7346 | 700.3 | 0.0007881 | 1.125 | +1 | 7 |
| 5 | z | 1.593E+04 | 701.3 | 0.0004344 | 0.6194 | +1 | 7 |
| - | - | 4.007E+04 | 702.3 | - | - | 0 | - |
| - | - | 1.236E+04 | 703.3 | - | - | 0 | - |
| - | - | 2811 | 704.3 | - | - | 0 | - |
| - | - | 4011 | 716.3 | - | - | 0 | - |
| 5 | y | 9.15E+04 | 717.4 | 0.0002507 | 0.3495 | +1 | 7 |
| - | - | 3.03E+04 | 718.4 | - | - | 0 | - |
| - | - | 6637 | 719.4 | - | - | 0 | - |
| - | - | 1705 | 725.3 | - | - | 0 | - |
| - | - | 681.9 | 730.4 | - | - | 0 | - |
| - | - | 998.5 | 744.3 | - | - | 0 | - |
| - | - | 840.5 | 753.4 | - | - | 0 | - |
| - | - | 782.4 | 755.4 | - | - | 0 | - |
| - | - | 1972 | 756.4 | - | - | 0 | - |
| - | - | 1054 | 759.4 | - | - | 0 | - |
| - | - | 4.094E+04 | 760.4 | - | - | 0 | - |
| - | - | 1.594E+04 | 761.4 | - | - | 0 | - |
| - | - | 4360 | 762.4 | - | - | 0 | - |
| - | - | 574.4 | 770.4 | - | - | 0 | - |
| - | - | 684.7 | 773.4 | - | - | 0 | - |
| 4 | z | 1632 | 782.4 | 0.0001781 | 0.2277 | +1 | 8 |
| - | - | 1201 | 783.4 | - | - | 0 | - |
| 4 | w | 2914 | 785.4 | 0.001685 | 2.145 | +1 | 8 |
| - | - | 1248 | 786.4 | - | - | 0 | - |
| 7 | c | 7391 | 787.4 | 0.0004154 | 0.5275 | +1 | 7 |
| - | - | 2663 | 788.4 | - | - | 0 | - |
| - | - | 772.2 | 789.4 | - | - | 0 | - |
| 4 | y | 738.3 | 798.4 | 0.004437 | 5.557 | +1 | 8 |
| 4 | z | 3.948E+04 | 800.4 | 0.0006707 | 0.838 | +1 | 8 |
| - | - | 2.133E+04 | 801.4 | - | - | 0 | - |
| - | - | 5140 | 802.4 | - | - | 0 | - |
| - | - | 972.5 | 803.4 | - | - | 0 | - |
| - | - | 947.9 | 810.4 | - | - | 0 | - |
| - | - | 687 | 815.4 | - | - | 0 | - |
| 4 | y | 4.372E+04 | 816.4 | 0.0002572 | 0.3151 | +1 | 8 |
| - | - | 1.917E+04 | 817.4 | - | - | 0 | - |
| - | - | 3532 | 818.4 | - | - | 0 | - |
| - | - | 733.7 | 829.4 | - | - | 0 | - |
| - | - | 9958 | 857.5 | - | - | 0 | - |
| - | - | 586.5 | 858.4 | - | - | 0 | - |
| - | - | 4192 | 858.5 | - | - | 0 | - |
| - | - | 1306 | 859.5 | - | - | 0 | - |
| - | - | 1372 | 861.4 | - | - | 0 | - |
| - | - | 1329 | 861.9 | - | - | 0 | - |
| 3 | w | 1.856E+04 | 870.4 | 0.0001295 | 0.1488 | +1 | 9 |
| - | - | 7605 | 871.4 | - | - | 0 | - |
| - | - | 2077 | 872.4 | - | - | 0 | - |
| - | - | 669 | 879.8 | - | - | 0 | - |
| - | - | 5975 | 885.5 | - | - | 0 | - |
| - | - | 3231 | 886.5 | - | - | 0 | - |
| - | - | 750.1 | 887.5 | - | - | 0 | - |
| - | - | 7108 | 900.5 | - | - | 0 | - |
| 8 | c | 3.686E+04 | 901.5 | 0.0003498 | 0.388 | +1 | 8 |
| - | - | 1.66E+04 | 902.5 | - | - | 0 | - |
| - | - | 4725 | 903.5 | - | - | 0 | - |
| - | - | 891.5 | 904.5 | - | - | 0 | - |
| 3 | z | 1458 | 911.4 | 0.001573 | 1.725 | +1 | 9 |
| - | - | 1048 | 926.5 | - | - | 0 | - |
| 3 | y | 2605 | 927.5 | 0.01492 | 16.09 | +1 | 9 |
| 3 | z | 5.717E+04 | 929.4 | 0.0001919 | 0.2065 | +1 | 9 |
| - | - | 2.872E+04 | 930.4 | - | - | 0 | - |
| - | - | 8021 | 931.4 | - | - | 0 | - |
| - | - | 820 | 932.4 | - | - | 0 | - |
| - | - | 1.627E+04 | 944.5 | - | - | 0 | - |
| 3 | y | 4.779E+04 | 945.5 | 0.003196 | 3.381 | +1 | 9 |
| - | - | 2.22E+04 | 946.5 | - | - | 0 | - |
| - | - | 5964 | 947.5 | - | - | 0 | - |
| - | - | 711.1 | 948.5 | - | - | 0 | - |
| - | - | 1870 | 957.5 | - | - | 0 | - |
| - | - | 1269 | 958.5 | - | - | 0 | - |
| 9 | c | 1539 | 971.5 | 0.002012 | 2.071 | +1 | 9 |
| - | - | 1072 | 972.5 | - | - | 0 | - |
| - | - | 1128 | 973.5 | - | - | 0 | - |
| 9 | c | 2.559E+04 | 988.5 | 0.0002428 | 0.2456 | +1 | 9 |
| - | - | 1.226E+04 | 989.5 | - | - | 0 | - |
| - | - | 3350 | 990.5 | - | - | 0 | - |
| - | - | 2.151E+04 | 1002 | - | - | 0 | - |
| - | - | 1.02E+04 | 1003 | - | - | 0 | - |
| - | - | 3686 | 1004 | - | - | 0 | - |
| - | - | 705.7 | 1013 | - | - | 0 | - |
| 2 | y | 4260 | 1014 | 0.001727 | 1.702 | +1 | 10 |
| 2 | y | 2121 | 1015 | 0.01816 | 17.89 | +1 | 10 |
| 2 | z | 3.972E+04 | 1016 | 0.0005121 | 0.5038 | +1 | 10 |
| - | - | 2.003E+04 | 1017 | - | - | 0 | - |
| - | - | 5456 | 1018 | - | - | 0 | - |
| - | - | 1944 | 1031 | - | - | 0 | - |
| - | - | 1075 | 1032 | - | - | 0 | - |
| 2 | y | 3.635E+04 | 1032 | 0.0005869 | 0.5684 | +1 | 10 |
| - | - | 1.771E+04 | 1033 | - | - | 0 | - |
| - | - | 6243 | 1035 | - | - | 0 | - |
| - | - | 853.1 | 1036 | - | - | 0 | - |
| - | - | 1543 | 1042 | - | - | 0 | - |
| 10 | c | 8.743E+04 | 1046 | 0.0003245 | 0.3103 | +1 | 10 |
| - | - | 4.972E+04 | 1047 | - | - | 0 | - |
| - | - | 1.426E+04 | 1048 | - | - | 0 | - |
| - | - | 2139 | 1049 | - | - | 0 | - |
| - | - | 722.9 | 1071 | - | - | 0 | - |
| - | - | 1221 | 1085 | - | - | 0 | - |
| - | - | 2879 | 1086 | - | - | 0 | - |
| - | - | 8281 | 1087 | - | - | 0 | - |
| - | - | 8115 | 1088 | - | - | 0 | - |
| - | - | 4977 | 1089 | - | - | 0 | - |
| - | - | 1609 | 1090 | - | - | 0 | - |
| - | - | 2451 | 1098 | - | - | 0 | - |
| - | - | 1077 | 1099 | - | - | 0 | - |
| - | - | 1254 | 1100 | - | - | 0 | - |
| - | - | 1.168E+04 | 1101 | - | - | 0 | - |
| - | - | 6154 | 1102 | - | - | 0 | - |
| - | - | 3564 | 1103 | - | - | 0 | - |
| - | - | 6929 | 1104 | - | - | 0 | - |
| - | - | 4521 | 1105 | - | - | 0 | - |
| - | - | 1778 | 1106 | - | - | 0 | - |
| - | - | 817.1 | 1112 | - | - | 0 | - |
| - | - | 679.5 | 1113 | - | - | 0 | - |
| - | - | 835.4 | 1114 | - | - | 0 | - |
| - | - | 1199 | 1115 | - | - | 0 | - |
| - | - | 1276 | 1116 | - | - | 0 | - |
| - | - | 3287 | 1119 | - | - | 0 | - |
| - | - | 1410 | 1120 | - | - | 0 | - |
| - | - | 1.203E+04 | 1129 | - | - | 0 | - |
| - | - | 2.017E+05 | 1130 | - | - | 0 | - |
| - | - | 1.162E+05 | 1131 | - | - | 0 | - |
| - | - | 4.032E+04 | 1132 | - | - | 0 | - |
| - | - | 5681 | 1133 | - | - | 0 | - |
| - | - | 997.7 | 1145 | - | - | 0 | - |
| - | - | 614.7 | 1145 | - | - | 0 | - |
| - | - | 8.395E+04 | 1146 | - | - | 0 | - |
| - | - | 2.062E+05 | 1147 | - | - | 0 | - |
| - | - | 1.152E+05 | 1148 | - | - | 0 | - |
| - | - | 1193 | 1148 | - | - | 0 | - |
| - | - | 3.533E+04 | 1149 | - | - | 0 | - |
| - | - | 5099 | 1150 | - | - | 0 | - |
| - | - | 1415 | 1179 | - | - | 0 | - |
| - | - | 846 | 1720 | - | - | 0 | - |
| - | - | 1486 | 1722 | - | - | 0 | - |
| - | - | 2007 | 1723 | - | - | 0 | - |
| - | - | 1484 | 1724 | - | - | 0 | - |
| - | - | 697.5 | 1751 | - | - | 0 | - |
| - | - | 688.1 | 2767 | - | - | 0 | - |

m/z Charge Intensity FragmentType MassShift Position
122.26419067382812 0 406.12653
129.10231018066406 0 513.7508
132.12730407714844 0 394.62332
133.7930450439453 0 340.74643
137.83200073242188 0 418.5958
143.66552734375 0 474.876
145.5753173828125 0 385.8782
145.6005401611328 0 471.1319
148.55751037597656 0 423.68463
148.89468383789062 0 553.59283
148.90174865722656 0 625.6912
148.9090118408203 0 772.60815
148.91622924804688 0 970.46295
148.92323303222656 0 1088.9961
148.930419921875 0 1199.5713
148.9379119873047 0 3050.105
148.9456024169922 0 4676.7246
148.96202087402344 0 3389.3745
148.9698028564453 0 1648.626
148.97706604003906 0 1155.2079
148.98435974121094 0 1229.1536
148.9913330078125 0 1043.3121
148.99887084960938 0 839.2742
149.00619506835938 0 564.4582
149.08592224121094 0 423.34882
156.26376342773438 0 535.2146
159.08497619628906 0 416.8805
162.34754943847656 0 444.72897
173.12872314453125 0 10460.199
173.4412841796875 0 1032.1261
174.1317138671875 0 1136.3279
175.10801696777344 0 2584.3145 y 9
183.11306762695312 0 6424.5537
185.60626220703125 0 458.98367
189.08773803710938 0 574.7505
199.07139587402344 0 1563.3688
201.12356567382812 0 27435.201
202.12689208984375 0 2608.7183
203.10302734375 0 1033.5781
211.1084747314453 0 1158.8402
215.13926696777344 0 53070.566
217.0821075439453 0 4775.3237
229.11865234375 0 2140.7852 w 8
231.06105041503906 0 527.38074
246.34478759765625 0 547.0632
255.0618896484375 0 465.63794
255.4033660888672 0 477.71518
280.14361572265625 0 558.56616
284.16156005859375 0 1303.4874
289.4521484375 0 644.7331
298.1383972167969 0 766.91077
302.1717224121094 0 2743.9756
312.15576171875 0 21329.459
313.158935546875 0 3544.7524
314.0988464355469 0 1139.848
314.1593933105469 0 754.8064
316.15069580078125 0 4769.8247 w 7
317.15350341796875 0 571.75684
330.1664733886719 0 20271.873
331.1694641113281 0 2975.1753
332.10968017578125 0 2773.2537
335.1401062011719 0 1026.1875
342.0918884277344 0 1196.5002
359.19256591796875 0 7381.8057 y 7
359.6817932128906 0 740.69604
360.1036376953125 0 1895.5364
360.1954650878906 0 779.7426
383.22998046875 0 638.2787
385.17047119140625 0 665.18646
397.19293212890625 0 595.80646
399.1879577636719 0 752.3641
401.2400207519531 0 9475.975
402.2431945800781 0 1644.4995
408.7139587402344 0 1646.8887 y 3
409.21600341796875 0 877.05164
411.22406005859375 0 1119.5403
429.2350158691406 0 10100.296
430.237548828125 0 1993.2797
439.14581298828125 0 746.8996
464.2307434082031 0 1993.1698 y Water loss 2
464.73211669921875 0 972.80475
473.236572265625 0 5057.8164 y 2
473.7373352050781 0 2640.8271
486.2445373535156 0 1028.9047 c Ammonia loss 8
498.2563171386719 0 1129.4426
499.2754821777344 0 48297.777 z 6
500.2810363769531 0 25697.176
501.2849426269531 0 5781.817
502.287109375 0 749.40106
507.7462463378906 0 1266.6317 y Water loss 1
508.2516784667969 0 625.0567
513.2919311523438 0 758.89685
514.288330078125 0 1292.6096
514.753173828125 0 1209.3967 c Ammonia loss 9
515.2511596679688 0 1077.886
515.2940673828125 0 12887.804 y 6
516.2970581054688 0 3052.9663
516.7520751953125 0 6228.744 y 1
517.2547607421875 0 2736.5725
517.7547607421875 0 1191.9093
521.74169921875 0 748.9474
538.2144775390625 0 1387.0813
542.2828979492188 0 644.6349
556.2720336914062 0 715.6687
564.2888793945312 0 6943.7866
564.7900390625 0 4538.8706
565.2926635742188 0 875.17883
569.2666625976562 0 1097.8103
570.3126831054688 0 2512.2556
571.317626953125 0 1376.0785
572.2831420898438 0 2445.7666
573.293701171875 0 31134.998
573.7952270507812 0 20776.002
574.2954711914062 0 5746.703
585.8313598632812 0 585.77094
614.3021240234375 0 19644.3 z 5
615.309326171875 0 35961.76
616.3125 0 8988.204
617.31494140625 0 1685.1595
629.3130493164062 0 3853.975
630.3208618164062 0 39257.418 c Water loss 5
631.3235473632812 0 12438.895
632.3253173828125 0 1750.124
655.3263549804688 0 572.0221
657.3029174804688 0 729.5601
658.31640625 0 1827.0398
659.3250732421875 0 605.1412
677.6848754882812 0 733.42706
699.3201904296875 0 6950.5977
700.3252563476562 0 7345.8984 y Ammonia loss 4
701.3334350585938 0 15932.8545 z 4
702.3412475585938 0 40068.3
703.3448486328125 0 12361.743
704.3453979492188 0 2811.474
716.3451538085938 0 4010.5007
717.3528442382812 0 91502.4 y 4
718.35595703125 0 30295.469
719.35791015625 0 6636.588
725.321533203125 0 1705.3643
730.368408203125 0 681.86304
744.3394775390625 0 998.4522
753.35498046875 0 840.5005
755.4050903320312 0 782.44366
756.4139404296875 0 1972.4052
759.4035034179688 0 1054.2611
760.4078369140625 0 40939.008
761.4111938476562 0 15942.565
762.4124145507812 0 4359.97
770.3889770507812 0 574.37164
773.3779296875 0 684.6698
782.3915405273438 0 1632.3666 z Water loss 3
783.396240234375 0 1201.2402
785.3804931640625 0 2914.379 w 3
786.3821411132812 0 1248.3677
787.39404296875 0 7391.493 c Ammonia loss 6
788.3988647460938 0 2663.0576
789.3977661132812 0 772.1649
798.406005859375 0 738.3265 y Water loss 3
800.4029541015625 0 39479.42 z 3
801.4065551757812 0 21328.166
802.40966796875 0 5139.74
803.4151611328125 0 972.466
810.36962890625 0 947.8947
815.4151611328125 0 686.9909
816.4212646484375 0 43717.41 y 3
817.42431640625 0 19166.715
818.4237670898438 0 3532.068
829.3958740234375 0 733.7219
857.4607543945312 0 9957.584
858.3922119140625 0 586.5323
858.4651489257812 0 4191.524
859.4702758789062 0 1305.8827
861.4197387695312 0 1371.5802
861.926025390625 0 1328.8558
870.4317016601562 0 18556.354 w 2
871.4342651367188 0 7604.606
872.4373779296875 0 2077.3425
879.7722778320312 0 668.952
885.454345703125 0 5975.389
886.458740234375 0 3230.6028
887.4635620117188 0 750.1362
900.4710693359375 0 7107.97
901.47412109375 0 36856.586 c 7
902.4768676757812 0 16596.715
903.4798583984375 0 4725.165
904.4845581054688 0 891.4702
911.4327392578125 0 1457.8873 z Water loss 2
926.4739379882812 0 1048.4995
927.4679565429688 0 2605.4749 y Water loss 2
929.445068359375 0 57171.832 z 2
930.4479370117188 0 28718.732
931.4497680664062 0 8021.278
932.449951171875 0 819.96594
944.4911499023438 0 16271.175
945.466796875 0 47791.918 y 2
946.4685668945312 0 22196.67
947.4703979492188 0 5963.9937
948.4631958007812 0 711.11786
957.5226440429688 0 1869.7229
958.5259399414062 0 1269.3949
971.4812622070312 0 1539.4124 c Ammonia loss 8
972.4825439453125 0 1072.2473
973.4918212890625 0 1128.026
988.5060424804688 0 25593.271 c 8
989.5084228515625 0 12262.676
990.5086059570312 0 3349.5283
1001.5138549804688 0 21512.447
1002.516357421875 0 10199.805
1003.5184326171875 0 3686.285
1012.5040893554688 0 705.7165
1014.4833374023438 0 4259.613 y Water loss 1
1015.4872436523438 0 2121.1555 y Ammonia loss 1
1016.4774169921875 0 39722.34 z 1
1017.4801635742188 0 20030.5
1018.4806518554688 0 5456.4893
1030.5098876953125 0 1944.0507
1031.520263671875 0 1075.2762
1032.4962158203125 0 36346.06 y 1
1033.49853515625 0 17712.768
1034.5013427734375 0 6242.618
1035.504150390625 0 853.0604
1042.480712890625 0 1543.3215
1045.527587890625 0 87429.3 c 9
1046.5303955078125 0 49720.574
1047.532470703125 0 14263.949
1048.5347900390625 0 2138.8198
1070.54833984375 0 722.8751
1084.59375 0 1221.1692
1085.57861328125 0 2878.56
1086.569580078125 0 8280.974
1087.547607421875 0 8114.961
1088.54638671875 0 4976.501
1089.544921875 0 1609.0543
1097.55712890625 0 2450.902
1098.563232421875 0 1077.466
1099.557861328125 0 1254.3265
1100.5726318359375 0 11680.055
1101.570556640625 0 6154.188
1102.584228515625 0 3563.8357
1103.5584716796875 0 6928.7295
1104.558837890625 0 4521.003
1105.5611572265625 0 1777.8317
1111.5423583984375 0 817.1491
1112.54833984375 0 679.47705
1113.55517578125 0 835.40314
1114.547119140625 0 1199.1079
1115.5413818359375 0 1276.2028
1118.5916748046875 0 3287.4817
1119.5997314453125 0 1409.5018
1128.577392578125 0 12026.612
1129.5618896484375 0 201701.33
1130.564208984375 0 116153.79
1131.5662841796875 0 40318.24
1132.5689697265625 0 5681.2593
1144.53466796875 0 997.6789
1145.107177734375 0 614.739
1145.57958984375 0 83953.88
1146.5865478515625 0 206192.58
1147.58935546875 0 115170.77
1147.74609375 0 1192.5557
1148.5919189453125 0 35329.97
1149.5869140625 0 5099.2715
1178.576416015625 0 1414.5942
1719.831298828125 0 846.0139
1721.8609619140625 0 1486.2319
1722.8406982421875 0 2007.2831
1723.8125 0 1483.5735
1751.42822265625 0 697.47015
2766.547607421875 0 688.1305

Spectrum Details

|  |  |
| --- | --- |
| Matched peaks? Matched peaks The total absolute number of peaks matched. Additionally in brackets the total fraction of peaks matched and the total number of peaks is shown. | 37 (13.75% of 269) |
| FDR? FDR The false discovery rate estimated for this peptide. It is calculated by matching all theoretical fragments with a non-integer shift with the raw peaks for this spectrum. This is done with 40 different shifts. The resulting percentage is the average number of annotated peaks over the number of annotated peaks with the correct spectrum. | 0.45% |
| Satellite FDR? Satellite FDR See the FDR for details on its calculation. This satellite ion specific FDR only contains the satellite ions (d/w) for I/L/J positions. | - |
| PSM Score? PSM Score The PSM Score as given by Hecklib to this annotated spectrum. It is shown with three significant figures. | 331 |

#### Spectrum 3858? Spectrum 3858 The raw spectrum of this peptide as annotated by Hecklib. The fragments are coloured according to ion type (see legend). Any peaks with a star '\*' as text can be hovered over to see the full details, first the ion type second the mass shift type. By hovering over the amino acids in the peptide or ions in the legend the corresponding peaks are highlighted. By toggling the 'Unassigned' label you can turn the background (unassigned) peaks on or off in the plot. By updating the slider in the Ion legend you can update the spectrum to only show the top X% of the peaks with labels. The top X% means any peak that is within X% of the highest intensity. By dragging in the spectrum you can zoom in to a specific part of the spectrum and use 'Zoom Out' to get back to the original zoom level. The annotation of the spectrum is based on the given sequence in the peptides file and is done with different software so inconsistencies are likely. The peaks are annotated based on the given sequence, with 20 ppm tolerance.

Copy Data

##### Spectrum 3858 (TSV)

###### Preview

```
Loading example...
```

*Click on the button to copy the data to your clipboard.*

Mz MinMz MaxIntensity Max

WidthHeightPeptide font sizePeptide stroke widthSpectrum font sizeSpectrum stroke widthCompact peptide

Ion legend

wxyz

abcd

OtherUnassignedIon^Charge^_Position_Show for top:%

JSEVSDRPSGV

03.03e+46.06e+49.09e+41.21e+5

Zoom Out

y^+1^_2_w^+1^_3_w^+1^_4_y^+1^_4_y^+2^_8_c^+1^_4_y^+2^_9_y^+2^_9_c^+2^_9_z^+1^_5_y^+2^_10_y^+1^_5_y^+2^_10_c^+1^_5_z^+1^_6_c^+1^_6_y^+1^_7_z^+1^_7_y^+1^_7_z^+1^_8_w^+1^_8_c^+1^_7_z^+1^_8_y^+1^_8_w^+1^_9_c^+1^_8_y^+1^_9_z^+1^_9_y^+1^_9_c^+1^_9_c^+1^_9_y^+1^_10_y^+1^_10_z^+1^_10_c^+1^_10_y^+1^_10_c^+1^_10_

043586913041738

Fragment Matches Table

Show background peaks

| Position | Ion type | Intensity | mz Theoretical | mz Error (Th) | mz Error (ppm) | Charge | Series Number |
| --- | --- | --- | --- | --- | --- | --- | --- |
| - | - | 385.8 | 126 | - | - | 0 | - |
| - | - | 407.8 | 127.5 | - | - | 0 | - |
| - | - | 457.2 | 129.1 | - | - | 0 | - |
| - | - | 1060 | 136.1 | - | - | 0 | - |
| - | - | 505.2 | 149 | - | - | 0 | - |
| - | - | 461.2 | 165.4 | - | - | 0 | - |
| - | - | 6177 | 173.1 | - | - | 0 | - |
| - | - | 1183 | 173.4 | - | - | 0 | - |
| - | - | 537.2 | 175.1 | - | - | 0 | - |
| 10 | y | 1426 | 175.1 | 0.0002982 | 1.703 | +1 | 2 |
| - | - | 3622 | 183.1 | - | - | 0 | - |
| - | - | 496 | 184.1 | - | - | 0 | - |
| - | - | 652.5 | 189.1 | - | - | 0 | - |
| - | - | 646 | 199.1 | - | - | 0 | - |
| - | - | 1.445E+04 | 201.1 | - | - | 0 | - |
| - | - | 1055 | 201.1 | - | - | 0 | - |
| - | - | 1959 | 202.1 | - | - | 0 | - |
| - | - | 601.1 | 211.1 | - | - | 0 | - |
| - | - | 2.511E+04 | 215.1 | - | - | 0 | - |
| - | - | 2590 | 217.1 | - | - | 0 | - |
| 9 | w | 1483 | 229.1 | 0.0003078 | 1.344 | +1 | 3 |
| - | - | 776.7 | 261.2 | - | - | 0 | - |
| - | - | 530.7 | 266.7 | - | - | 0 | - |
| - | - | 1.178E+04 | 312.2 | - | - | 0 | - |
| - | - | 2006 | 313.2 | - | - | 0 | - |
| - | - | 1099 | 314.1 | - | - | 0 | - |
| 8 | w | 2719 | 316.2 | 0.0006891 | 2.18 | +1 | 4 |
| - | - | 9956 | 330.2 | - | - | 0 | - |
| - | - | 1273 | 331.2 | - | - | 0 | - |
| - | - | 1186 | 332.1 | - | - | 0 | - |
| - | - | 515.3 | 339.1 | - | - | 0 | - |
| - | - | 935.6 | 342.1 | - | - | 0 | - |
| 8 | y | 4709 | 359.2 | 2.436E-05 | 0.06782 | +1 | 4 |
| - | - | 970.6 | 360.1 | - | - | 0 | - |
| - | - | 723.6 | 360.2 | - | - | 0 | - |
| - | - | 606.3 | 396.1 | - | - | 0 | - |
| - | - | 5477 | 401.2 | - | - | 0 | - |
| - | - | 1083 | 402.2 | - | - | 0 | - |
| 4 | y | 1805 | 408.7 | 0.0003358 | 0.8216 | +2 | 8 |
| - | - | 772.7 | 411.2 | - | - | 0 | - |
| - | - | 583 | 416.9 | - | - | 0 | - |
| - | - | 6583 | 429.2 | - | - | 0 | - |
| - | - | 807 | 430.2 | - | - | 0 | - |
| - | - | 1188 | 439.1 | - | - | 0 | - |
| 4 | c | 1372 | 446.3 | 0.001252 | 2.805 | +1 | 4 |
| 3 | y | 1323 | 464.2 | 0.0006788 | 1.462 | +2 | 9 |
| 3 | y | 2732 | 473.2 | 0.001134 | 2.396 | +2 | 9 |
| - | - | 1202 | 473.7 | - | - | 0 | - |
| - | - | 994 | 485.7 | - | - | 0 | - |
| 9 | c | 918.1 | 486.2 | 0.0008929 | 1.836 | +2 | 9 |
| - | - | 530.1 | 493.4 | - | - | 0 | - |
| - | - | 577.2 | 498.5 | - | - | 0 | - |
| 7 | z | 2.998E+04 | 499.3 | 0.0007978 | 1.598 | +1 | 5 |
| - | - | 1.497E+04 | 500.3 | - | - | 0 | - |
| - | - | 3537 | 501.3 | - | - | 0 | - |
| - | - | 629.3 | 502.3 | - | - | 0 | - |
| 2 | y | 1425 | 507.7 | 0.001815 | 3.576 | +2 | 10 |
| - | - | 583.4 | 508.8 | - | - | 0 | - |
| - | - | 966.1 | 514.3 | - | - | 0 | - |
| 7 | y | 6351 | 515.3 | 0.0004453 | 0.8642 | +1 | 5 |
| - | - | 1637 | 516.3 | - | - | 0 | - |
| 2 | y | 3990 | 516.8 | 0.0006835 | 1.323 | +2 | 10 |
| - | - | 1474 | 517.3 | - | - | 0 | - |
| - | - | 661.5 | 518.2 | - | - | 0 | - |
| 5 | c | 1879 | 533.3 | 0.001663 | 3.119 | +1 | 5 |
| - | - | 877.6 | 554.8 | - | - | 0 | - |
| - | - | 4570 | 564.3 | - | - | 0 | - |
| - | - | 1630 | 564.8 | - | - | 0 | - |
| - | - | 780.9 | 565.3 | - | - | 0 | - |
| - | - | 850.9 | 569.3 | - | - | 0 | - |
| - | - | 1338 | 570.3 | - | - | 0 | - |
| - | - | 855.9 | 571.3 | - | - | 0 | - |
| - | - | 2555 | 572.3 | - | - | 0 | - |
| - | - | 1.741E+04 | 573.3 | - | - | 0 | - |
| - | - | 1.118E+04 | 573.8 | - | - | 0 | - |
| - | - | 3874 | 574.3 | - | - | 0 | - |
| - | - | 820.7 | 574.8 | - | - | 0 | - |
| - | - | 1085 | 590.3 | - | - | 0 | - |
| - | - | 4401 | 605.3 | - | - | 0 | - |
| - | - | 921.6 | 606.3 | - | - | 0 | - |
| 6 | z | 1.171E+04 | 614.3 | 0.0009544 | 1.554 | +1 | 6 |
| - | - | 2.044E+04 | 615.3 | - | - | 0 | - |
| - | - | 6658 | 616.3 | - | - | 0 | - |
| - | - | 1418 | 617.3 | - | - | 0 | - |
| - | - | 2500 | 629.3 | - | - | 0 | - |
| 6 | c | 2.285E+04 | 630.3 | 0.01196 | 18.97 | +1 | 6 |
| - | - | 6085 | 631.3 | - | - | 0 | - |
| - | - | 1707 | 632.3 | - | - | 0 | - |
| - | - | 571 | 642.3 | - | - | 0 | - |
| - | - | 1476 | 658.3 | - | - | 0 | - |
| - | - | 2291 | 659.3 | - | - | 0 | - |
| - | - | 559 | 660.3 | - | - | 0 | - |
| - | - | 3690 | 699.3 | - | - | 0 | - |
| 5 | y | 3876 | 700.3 | 0.0001167 | 0.1666 | +1 | 7 |
| 5 | z | 9182 | 701.3 | 0.0004811 | 0.686 | +1 | 7 |
| - | - | 2.066E+04 | 702.3 | - | - | 0 | - |
| - | - | 6901 | 703.3 | - | - | 0 | - |
| - | - | 1976 | 704.3 | - | - | 0 | - |
| - | - | 2520 | 716.3 | - | - | 0 | - |
| 5 | y | 5.186E+04 | 717.4 | 0.0004338 | 0.6048 | +1 | 7 |
| - | - | 1.847E+04 | 718.4 | - | - | 0 | - |
| - | - | 4507 | 719.4 | - | - | 0 | - |
| - | - | 1134 | 723.4 | - | - | 0 | - |
| - | - | 1144 | 725.3 | - | - | 0 | - |
| - | - | 1265 | 756.4 | - | - | 0 | - |
| - | - | 882.2 | 757.4 | - | - | 0 | - |
| - | - | 2.438E+04 | 760.4 | - | - | 0 | - |
| - | - | 9200 | 761.4 | - | - | 0 | - |
| - | - | 2152 | 762.4 | - | - | 0 | - |
| - | - | 537.4 | 768.8 | - | - | 0 | - |
| - | - | 757.1 | 770.4 | - | - | 0 | - |
| - | - | 581.1 | 775.4 | - | - | 0 | - |
| 4 | z | 732.6 | 782.4 | 0.002263 | 2.893 | +1 | 8 |
| 4 | w | 1822 | 785.4 | 0.0026 | 3.311 | +1 | 8 |
| 7 | c | 3509 | 787.4 | 0.001111 | 1.41 | +1 | 7 |
| - | - | 1666 | 788.4 | - | - | 0 | - |
| - | - | 707.3 | 792.4 | - | - | 0 | - |
| 4 | z | 2.394E+04 | 800.4 | 0.0008538 | 1.067 | +1 | 8 |
| - | - | 1.298E+04 | 801.4 | - | - | 0 | - |
| - | - | 3520 | 802.4 | - | - | 0 | - |
| - | - | 604.2 | 806.4 | - | - | 0 | - |
| - | - | 731.4 | 807.4 | - | - | 0 | - |
| - | - | 581.2 | 809.4 | - | - | 0 | - |
| - | - | 1275 | 810.4 | - | - | 0 | - |
| 4 | y | 2.562E+04 | 816.4 | 0.0005624 | 0.6889 | +1 | 8 |
| - | - | 1.082E+04 | 817.4 | - | - | 0 | - |
| - | - | 2977 | 818.4 | - | - | 0 | - |
| - | - | 4839 | 857.5 | - | - | 0 | - |
| - | - | 2753 | 858.5 | - | - | 0 | - |
| 3 | w | 9460 | 870.4 | 0.001106 | 1.271 | +1 | 9 |
| - | - | 3471 | 871.4 | - | - | 0 | - |
| - | - | 1686 | 872.4 | - | - | 0 | - |
| - | - | 3639 | 885.5 | - | - | 0 | - |
| - | - | 1831 | 886.5 | - | - | 0 | - |
| - | - | 3864 | 900.5 | - | - | 0 | - |
| 8 | c | 1.932E+04 | 901.5 | 0.001143 | 1.268 | +1 | 8 |
| - | - | 864.9 | 902.4 | - | - | 0 | - |
| - | - | 1.095E+04 | 902.5 | - | - | 0 | - |
| - | - | 2380 | 903.5 | - | - | 0 | - |
| - | - | 862.4 | 904.5 | - | - | 0 | - |
| - | - | 829.7 | 912.4 | - | - | 0 | - |
| - | - | 666.8 | 926.5 | - | - | 0 | - |
| 3 | y | 1455 | 927.5 | 0.00723 | 7.796 | +1 | 9 |
| 3 | z | 3.38E+04 | 929.4 | 0.000314 | 0.3378 | +1 | 9 |
| - | - | 1.788E+04 | 930.4 | - | - | 0 | - |
| - | - | 3745 | 931.4 | - | - | 0 | - |
| - | - | 9674 | 944.5 | - | - | 0 | - |
| 3 | y | 2.56E+04 | 945.5 | 0.004356 | 4.607 | +1 | 9 |
| - | - | 1.301E+04 | 946.5 | - | - | 0 | - |
| - | - | 3699 | 947.5 | - | - | 0 | - |
| - | - | 1049 | 954.4 | - | - | 0 | - |
| - | - | 951.8 | 957.5 | - | - | 0 | - |
| 9 | c | 1245 | 971.5 | 0.006411 | 6.599 | +1 | 9 |
| - | - | 663.1 | 972.2 | - | - | 0 | - |
| - | - | 1184 | 972.5 | - | - | 0 | - |
| 9 | c | 1.593E+04 | 988.5 | 0.00067 | 0.6778 | +1 | 9 |
| - | - | 7297 | 989.5 | - | - | 0 | - |
| - | - | 2940 | 990.5 | - | - | 0 | - |
| - | - | 1.289E+04 | 1002 | - | - | 0 | - |
| - | - | 6316 | 1003 | - | - | 0 | - |
| - | - | 2061 | 1004 | - | - | 0 | - |
| - | - | 704 | 1005 | - | - | 0 | - |
| 2 | y | 1700 | 1014 | 0.00252 | 2.484 | +1 | 10 |
| 2 | y | 1040 | 1015 | 0.01658 | 16.32 | +1 | 10 |
| 2 | z | 1.993E+04 | 1016 | 0.001428 | 1.405 | +1 | 10 |
| - | - | 1.019E+04 | 1017 | - | - | 0 | - |
| - | - | 3847 | 1018 | - | - | 0 | - |
| 10 | c | 750.1 | 1029 | 0.008405 | 8.172 | +1 | 10 |
| - | - | 4670 | 1030 | - | - | 0 | - |
| - | - | 2995 | 1031 | - | - | 0 | - |
| - | - | 1506 | 1032 | - | - | 0 | - |
| 2 | y | 2.317E+04 | 1032 | 0.001197 | 1.16 | +1 | 10 |
| - | - | 1.11E+04 | 1033 | - | - | 0 | - |
| - | - | 3148 | 1035 | - | - | 0 | - |
| - | - | 1537 | 1041 | - | - | 0 | - |
| - | - | 1787 | 1042 | - | - | 0 | - |
| - | - | 805.9 | 1043 | - | - | 0 | - |
| 10 | c | 4.862E+04 | 1046 | 0.0005686 | 0.5439 | +1 | 10 |
| - | - | 2.818E+04 | 1047 | - | - | 0 | - |
| - | - | 9597 | 1048 | - | - | 0 | - |
| - | - | 1143 | 1085 | - | - | 0 | - |
| - | - | 1812 | 1086 | - | - | 0 | - |
| - | - | 4844 | 1087 | - | - | 0 | - |
| - | - | 4055 | 1088 | - | - | 0 | - |
| - | - | 2561 | 1089 | - | - | 0 | - |
| - | - | 888 | 1090 | - | - | 0 | - |
| - | - | 1378 | 1098 | - | - | 0 | - |
| - | - | 1532 | 1100 | - | - | 0 | - |
| - | - | 7115 | 1101 | - | - | 0 | - |
| - | - | 4167 | 1102 | - | - | 0 | - |
| - | - | 1123 | 1103 | - | - | 0 | - |
| - | - | 5621 | 1104 | - | - | 0 | - |
| - | - | 3248 | 1105 | - | - | 0 | - |
| - | - | 1028 | 1106 | - | - | 0 | - |
| - | - | 617.3 | 1116 | - | - | 0 | - |
| - | - | 1547 | 1119 | - | - | 0 | - |
| - | - | 774.3 | 1120 | - | - | 0 | - |
| - | - | 1132 | 1128 | - | - | 0 | - |
| - | - | 8190 | 1129 | - | - | 0 | - |
| - | - | 1.114E+05 | 1130 | - | - | 0 | - |
| - | - | 6.841E+04 | 1131 | - | - | 0 | - |
| - | - | 2.172E+04 | 1132 | - | - | 0 | - |
| - | - | 3538 | 1133 | - | - | 0 | - |
| - | - | 2004 | 1145 | - | - | 0 | - |
| - | - | 5.009E+04 | 1146 | - | - | 0 | - |
| - | - | 1.2E+05 | 1147 | - | - | 0 | - |
| - | - | 6.469E+04 | 1148 | - | - | 0 | - |
| - | - | 2.217E+04 | 1149 | - | - | 0 | - |
| - | - | 3242 | 1150 | - | - | 0 | - |
| - | - | 752.4 | 1179 | - | - | 0 | - |
| - | - | 826.1 | 1721 | - | - | 0 | - |

m/z Charge Intensity FragmentType MassShift Position
126.0135726928711 0 385.75742
127.53914642333984 0 407.80594
129.10250854492188 0 457.19144
136.0760498046875 0 1059.8573
148.95440673828125 0 505.24985
165.35595703125 0 461.1532
173.12876892089844 0 6176.7983
173.43829345703125 0 1182.8494
175.07180786132812 0 537.24286
175.10801696777344 0 1426.0076 y 9
183.11325073242188 0 3621.9958
184.11680603027344 0 496.02972
189.08729553222656 0 652.46875
199.0719451904297 0 646.0448
201.1236572265625 0 14449.53
201.1337127685547 0 1054.6764
202.12716674804688 0 1959.1649
211.10829162597656 0 601.14105
215.1393585205078 0 25112.086
217.08229064941406 0 2589.8516
229.11859130859375 0 1482.6182 w 8
261.1565856933594 0 776.6659
266.7084045410156 0 530.69763
312.1560363769531 0 11775.231
313.158935546875 0 2006.346
314.0992736816406 0 1099.2185
316.1510009765625 0 2719.4128 w 7
330.1665344238281 0 9956.111
331.17022705078125 0 1273.0173
332.1094970703125 0 1185.6257
339.0919494628906 0 515.3469
342.0937194824219 0 935.5864
359.1925354003906 0 4709.1436 y 7
360.1044616699219 0 970.55475
360.1968688964844 0 723.6409
396.0744323730469 0 606.28406
401.239990234375 0 5476.947
402.2422790527344 0 1082.5358
408.71380615234375 0 1804.6257 y 3
411.2251281738281 0 772.7178
416.9217224121094 0 582.9599
429.23516845703125 0 6583.0317
430.23748779296875 0 806.9676
439.1476135253906 0 1187.5173
446.2621765136719 0 1371.5337 c 3
464.2308349609375 0 1323.0452 y Water loss 2
473.236572265625 0 2731.7456 y 2
473.7368469238281 0 1202.3088
485.7354736328125 0 993.9613
486.24237060546875 0 918.1242 c Ammonia loss 8
493.3859558105469 0 530.12933
498.544921875 0 577.2208
499.27569580078125 0 29982.645 z 6
500.2812194824219 0 14969.492
501.2855529785156 0 3536.713
502.2902526855469 0 629.29877
507.74798583984375 0 1424.8737 y Water loss 1
508.8418884277344 0 583.3848
514.2870483398438 0 966.11224
515.2940673828125 0 6351.4937 y 6
516.2984619140625 0 1637.3953
516.7521362304688 0 3989.6025 y 1
517.2554931640625 0 1473.7227
518.2083740234375 0 661.456
533.2946166992188 0 1878.895 c 4
554.8153686523438 0 877.57086
564.2890014648438 0 4569.692
564.791259765625 0 1630.001
565.2903442382812 0 780.876
569.2662963867188 0 850.94916
570.3133544921875 0 1337.6863
571.31884765625 0 855.9281
572.2852783203125 0 2555.2988
573.2939453125 0 17409.46
573.7955322265625 0 11179.182
574.2975463867188 0 3873.5442
574.800048828125 0 820.66315
590.314453125 0 1084.838
605.3256225585938 0 4400.6987
606.3287353515625 0 921.5715
614.3027954101562 0 11712.824 z 5
615.309814453125 0 20438.982
616.312744140625 0 6658.388
617.3169555664062 0 1417.5137
629.3134765625 0 2499.5173
630.3212890625 0 22851.229 c Water loss 5
631.3235473632812 0 6085.0522
632.3255004882812 0 1707.4573
642.3317260742188 0 571.0351
658.3189697265625 0 1475.6117
659.3356323242188 0 2290.9705
660.3423461914062 0 558.97327
699.3207397460938 0 3689.784
700.325927734375 0 3875.589 y Ammonia loss 4
701.3343505859375 0 9182.237 z 4
702.3413696289062 0 20655.814
703.3440551757812 0 6901.378
704.34716796875 0 1976.0256
716.3460083007812 0 2519.9797
717.35302734375 0 51855.87 y 4
718.3560180664062 0 18474.654
719.3582763671875 0 4506.8965
723.3623046875 0 1133.8616
725.3215942382812 0 1144.2015
756.4119262695312 0 1264.7673
757.4147338867188 0 882.2254
760.4082641601562 0 24381.648
761.4114990234375 0 9200.05
762.4125366210938 0 2152.0645
768.7955932617188 0 537.3867
770.3880615234375 0 757.0581
775.3839721679688 0 581.14844
782.3939819335938 0 732.6092 z Water loss 3
785.3814086914062 0 1822.1423 w 3
787.3955688476562 0 3508.658 c Ammonia loss 6
788.3975830078125 0 1666.1561
792.400146484375 0 707.33026
800.4031372070312 0 23942.14 z 3
801.4071044921875 0 12984.472
802.4091796875 0 3520.3108
806.3951416015625 0 604.20245
807.4048461914062 0 731.3948
809.4071044921875 0 581.2452
810.3701782226562 0 1274.797
816.4215698242188 0 25615.121 y 3
817.4244995117188 0 10815.994
818.4280395507812 0 2976.822
857.4599609375 0 4838.6396
858.4641723632812 0 2753.214
870.4326782226562 0 9460.226 w 2
871.4349365234375 0 3471.4976
872.4384155273438 0 1685.9475
885.455810546875 0 3638.8684
886.4548950195312 0 1830.5107
900.4732055664062 0 3863.825
901.4749145507812 0 19316.27 c 7
902.3931274414062 0 864.86145
902.4782104492188 0 10953.467
903.4856567382812 0 2380.385
904.4854736328125 0 862.3912
912.4397583007812 0 829.66016
926.48388671875 0 666.79944
927.4602661132812 0 1454.6655 y Water loss 2
929.4451904296875 0 33801.12 z 2
930.448486328125 0 17881.451
931.4495239257812 0 3745.0085
944.489501953125 0 9673.897
945.4679565429688 0 25599.398 y 2
946.46826171875 0 13009.013
947.4705200195312 0 3698.917
954.443603515625 0 1049.4236
957.5245361328125 0 951.80554
971.4728393554688 0 1244.582 c Ammonia loss 8
972.2448120117188 0 663.0629
972.4847412109375 0 1183.6819
988.5064697265625 0 15930.7 c 8
989.5086669921875 0 7296.931
990.513427734375 0 2939.52
1001.5138549804688 0 12891.16
1002.5164794921875 0 6315.697
1003.518798828125 0 2060.8323
1004.5194091796875 0 704.0001
1014.4825439453125 0 1700.129 y Water loss 1
1015.4856567382812 0 1039.5453 y Ammonia loss 1
1016.4783325195312 0 19932.357 z 1
1017.4805297851562 0 10190.962
1018.4832153320312 0 3846.5913
1028.4923095703125 0 750.1163 c Ammonia loss 9
1029.5020751953125 0 4669.7686
1030.50537109375 0 2994.9534
1031.512451171875 0 1505.8154
1032.496826171875 0 23174.502 y 1
1033.4993896484375 0 11097.007
1034.5015869140625 0 3147.5596
1041.4757080078125 0 1536.7471
1042.47900390625 0 1786.6691
1043.4786376953125 0 805.8736
1045.52783203125 0 48622.94 c 9
1046.531005859375 0 28183.502
1047.532958984375 0 9597.042
1084.585693359375 0 1143.1886
1085.57568359375 0 1812.2246
1086.567626953125 0 4843.9155
1087.549072265625 0 4054.755
1088.549560546875 0 2561.2476
1089.5399169921875 0 888.0168
1097.5594482421875 0 1378.4048
1099.5579833984375 0 1531.8037
1100.572265625 0 7115.176
1101.5760498046875 0 4167.134
1102.5947265625 0 1122.5281
1103.5611572265625 0 5621.208
1104.558349609375 0 3248.1038
1105.552734375 0 1027.7576
1115.525146484375 0 617.2654
1118.596435546875 0 1547.2292
1119.5960693359375 0 774.2817
1127.51611328125 0 1132.1243
1128.57080078125 0 8189.701
1129.561767578125 0 111426.57
1130.5640869140625 0 68408.56
1131.5665283203125 0 21717.725
1132.56640625 0 3538.1675
1144.5321044921875 0 2004.4497
1145.578125 0 50090.207
1146.5863037109375 0 119971.02
1147.589599609375 0 64685.625
1148.5927734375 0 22165.418
1149.5941162109375 0 3242.4097
1178.5733642578125 0 752.37506
1720.81787109375 0 826.1296

Spectrum Details

|  |  |
| --- | --- |
| Matched peaks? Matched peaks The total absolute number of peaks matched. Additionally in brackets the total fraction of peaks matched and the total number of peaks is shown. | 37 (17.54% of 211) |
| FDR? FDR The false discovery rate estimated for this peptide. It is calculated by matching all theoretical fragments with a non-integer shift with the raw peaks for this spectrum. This is done with 40 different shifts. The resulting percentage is the average number of annotated peaks over the number of annotated peaks with the correct spectrum. | 0.32% |
| Satellite FDR? Satellite FDR See the FDR for details on its calculation. This satellite ion specific FDR only contains the satellite ions (d/w) for I/L/J positions. | - |
| PSM Score? PSM Score The PSM Score as given by Hecklib to this annotated spectrum. It is shown with three significant figures. | 349 |

#### Spectrum 4054? Spectrum 4054 The raw spectrum of this peptide as annotated by Hecklib. The fragments are coloured according to ion type (see legend). Any peaks with a star '\*' as text can be hovered over to see the full details, first the ion type second the mass shift type. By hovering over the amino acids in the peptide or ions in the legend the corresponding peaks are highlighted. By toggling the 'Unassigned' label you can turn the background (unassigned) peaks on or off in the plot. By updating the slider in the Ion legend you can update the spectrum to only show the top X% of the peaks with labels. The top X% means any peak that is within X% of the highest intensity. By dragging in the spectrum you can zoom in to a specific part of the spectrum and use 'Zoom Out' to get back to the original zoom level. The annotation of the spectrum is based on the given sequence in the peptides file and is done with different software so inconsistencies are likely. The peaks are annotated based on the given sequence, with 20 ppm tolerance.

Copy Data

##### Spectrum 4054 (TSV)

###### Preview

```
Loading example...
```

*Click on the button to copy the data to your clipboard.*

Mz MinMz MaxIntensity Max

WidthHeightPeptide font sizePeptide stroke widthSpectrum font sizeSpectrum stroke widthCompact peptide

Ion legend

wxyz

abcd

OtherUnassignedIon^Charge^_Position_Show for top:%

JSEVSDRPSGV

01.30e+42.59e+43.89e+45.19e+4

Zoom Out

y^+1^_2_w^+1^_3_y^+1^_4_y^+2^_9_z^+1^_5_c^+2^_10_y^+1^_5_y^+2^_10_z^+1^_6_c^+1^_6_y^+1^_7_z^+1^_7_y^+1^_7_w^+1^_8_c^+1^_7_z^+1^_8_y^+1^_8_w^+1^_9_c^+1^_8_z^+1^_9_y^+1^_9_c^+1^_9_c^+1^_9_y^+1^_10_y^+1^_10_z^+1^_10_y^+1^_10_c^+1^_10_

0767153523023070

Fragment Matches Table

Show background peaks

| Position | Ion type | Intensity | mz Theoretical | mz Error (Th) | mz Error (ppm) | Charge | Series Number |
| --- | --- | --- | --- | --- | --- | --- | --- |
| - | - | 387.9 | 129.1 | - | - | 0 | - |
| - | - | 493.2 | 149.3 | - | - | 0 | - |
| - | - | 1923 | 173.1 | - | - | 0 | - |
| - | - | 2110 | 173.4 | - | - | 0 | - |
| 10 | y | 1043 | 175.1 | 0.0001138 | 0.6499 | +1 | 2 |
| - | - | 1642 | 183.1 | - | - | 0 | - |
| - | - | 520.3 | 193.1 | - | - | 0 | - |
| - | - | 6667 | 201.1 | - | - | 0 | - |
| - | - | 2378 | 210 | - | - | 0 | - |
| - | - | 3130 | 215.1 | - | - | 0 | - |
| - | - | 1198 | 217.1 | - | - | 0 | - |
| 9 | w | 501 | 229.1 | 0.0008213 | 3.585 | +1 | 3 |
| - | - | 537.2 | 230.1 | - | - | 0 | - |
| - | - | 568 | 231 | - | - | 0 | - |
| - | - | 780.1 | 239.1 | - | - | 0 | - |
| - | - | 599.4 | 247 | - | - | 0 | - |
| - | - | 5765 | 312.2 | - | - | 0 | - |
| - | - | 1002 | 313.2 | - | - | 0 | - |
| - | - | 3801 | 330.2 | - | - | 0 | - |
| - | - | 565.7 | 334.2 | - | - | 0 | - |
| - | - | 541.5 | 344.3 | - | - | 0 | - |
| 8 | y | 1610 | 359.2 | 0.001715 | 4.775 | +1 | 4 |
| - | - | 595.5 | 382.2 | - | - | 0 | - |
| - | - | 714.3 | 386.2 | - | - | 0 | - |
| - | - | 2448 | 401.2 | - | - | 0 | - |
| - | - | 704.4 | 410.2 | - | - | 0 | - |
| - | - | 1988 | 429.2 | - | - | 0 | - |
| - | - | 556 | 430.2 | - | - | 0 | - |
| - | - | 611.9 | 463.3 | - | - | 0 | - |
| 3 | y | 679.9 | 473.2 | 8.693E-05 | 0.1837 | +2 | 9 |
| - | - | 811 | 473.7 | - | - | 0 | - |
| 7 | z | 1.237E+04 | 499.3 | 0.000606 | 1.214 | +1 | 5 |
| - | - | 5875 | 500.3 | - | - | 0 | - |
| - | - | 1305 | 501.3 | - | - | 0 | - |
| 10 | c | 597.1 | 514.8 | 0.001193 | 2.317 | +2 | 10 |
| 7 | y | 3333 | 515.3 | 0.0008364 | 1.623 | +1 | 5 |
| 2 | y | 1151 | 516.8 | 0.001111 | 2.15 | +2 | 10 |
| - | - | 719 | 520.2 | - | - | 0 | - |
| - | - | 650.2 | 521.2 | - | - | 0 | - |
| - | - | 654.7 | 536.8 | - | - | 0 | - |
| - | - | 2053 | 538.3 | - | - | 0 | - |
| - | - | 1089 | 539.2 | - | - | 0 | - |
| - | - | 633 | 545.6 | - | - | 0 | - |
| - | - | 561.1 | 554.3 | - | - | 0 | - |
| - | - | 5805 | 556.3 | - | - | 0 | - |
| - | - | 765.5 | 557.3 | - | - | 0 | - |
| - | - | 904 | 564.3 | - | - | 0 | - |
| - | - | 833.6 | 564.8 | - | - | 0 | - |
| - | - | 822.1 | 570.3 | - | - | 0 | - |
| - | - | 842.4 | 571.3 | - | - | 0 | - |
| - | - | 5087 | 572.3 | - | - | 0 | - |
| - | - | 7775 | 573.3 | - | - | 0 | - |
| - | - | 5541 | 573.8 | - | - | 0 | - |
| - | - | 4058 | 574.3 | - | - | 0 | - |
| 6 | z | 4250 | 614.3 | 8.322E-05 | 0.1355 | +1 | 6 |
| - | - | 9047 | 615.3 | - | - | 0 | - |
| - | - | 2083 | 616.3 | - | - | 0 | - |
| 6 | c | 8351 | 630.3 | 0.01025 | 16.26 | +1 | 6 |
| - | - | 2677 | 631.3 | - | - | 0 | - |
| - | - | 1734 | 699.3 | - | - | 0 | - |
| 5 | y | 1363 | 700.3 | 0.0003608 | 0.5152 | +1 | 7 |
| 5 | z | 4230 | 701.3 | 0.00251 | 3.578 | +1 | 7 |
| - | - | 9507 | 702.3 | - | - | 0 | - |
| - | - | 2925 | 703.3 | - | - | 0 | - |
| - | - | 922.6 | 716.3 | - | - | 0 | - |
| 5 | y | 2.256E+04 | 717.4 | 0.0006038 | 0.8417 | +1 | 7 |
| - | - | 7510 | 718.4 | - | - | 0 | - |
| - | - | 2184 | 719.4 | - | - | 0 | - |
| - | - | 538.8 | 737.3 | - | - | 0 | - |
| - | - | 703.5 | 744.3 | - | - | 0 | - |
| - | - | 9215 | 760.4 | - | - | 0 | - |
| - | - | 4675 | 761.4 | - | - | 0 | - |
| - | - | 1010 | 762.4 | - | - | 0 | - |
| 4 | w | 934 | 785.4 | 0.002173 | 2.767 | +1 | 8 |
| 7 | c | 1085 | 787.4 | 4.915E-05 | 0.06242 | +1 | 7 |
| 4 | z | 8852 | 800.4 | 0.0001825 | 0.228 | +1 | 8 |
| - | - | 6128 | 801.4 | - | - | 0 | - |
| - | - | 667.6 | 801.8 | - | - | 0 | - |
| - | - | 1466 | 802.4 | - | - | 0 | - |
| 4 | y | 1.053E+04 | 816.4 | 0.0008414 | 1.031 | +1 | 8 |
| - | - | 4598 | 817.4 | - | - | 0 | - |
| - | - | 1059 | 818.4 | - | - | 0 | - |
| - | - | 790.7 | 852 | - | - | 0 | - |
| - | - | 2896 | 857.5 | - | - | 0 | - |
| 3 | w | 3921 | 870.4 | 0.001289 | 1.481 | +1 | 9 |
| - | - | 2319 | 871.4 | - | - | 0 | - |
| - | - | 1625 | 885.5 | - | - | 0 | - |
| - | - | 1491 | 900.5 | - | - | 0 | - |
| 8 | c | 8895 | 901.5 | 0.0003498 | 0.388 | +1 | 8 |
| - | - | 3935 | 902.5 | - | - | 0 | - |
| - | - | 1517 | 903.5 | - | - | 0 | - |
| 3 | z | 1.461E+04 | 929.4 | 0.0001133 | 0.1219 | +1 | 9 |
| - | - | 7371 | 930.4 | - | - | 0 | - |
| - | - | 1505 | 931.4 | - | - | 0 | - |
| - | - | 4595 | 944.5 | - | - | 0 | - |
| 3 | y | 9505 | 945.5 | 0.004173 | 4.414 | +1 | 9 |
| - | - | 4441 | 946.5 | - | - | 0 | - |
| - | - | 852.8 | 947.5 | - | - | 0 | - |
| - | - | 756.5 | 957.5 | - | - | 0 | - |
| 9 | c | 657.9 | 971.5 | 0.006777 | 6.976 | +1 | 9 |
| - | - | 690.6 | 972.5 | - | - | 0 | - |
| 9 | c | 6451 | 988.5 | 0.0002455 | 0.2483 | +1 | 9 |
| - | - | 3342 | 989.5 | - | - | 0 | - |
| - | - | 5807 | 1002 | - | - | 0 | - |
| - | - | 2610 | 1003 | - | - | 0 | - |
| 2 | y | 1202 | 1014 | 0.006121 | 6.034 | +1 | 10 |
| 2 | y | 749.4 | 1015 | 0.01017 | 10.01 | +1 | 10 |
| 2 | z | 9727 | 1016 | 0.0003901 | 0.3837 | +1 | 10 |
| - | - | 4550 | 1017 | - | - | 0 | - |
| - | - | 1574 | 1018 | - | - | 0 | - |
| - | - | 782.4 | 1031 | - | - | 0 | - |
| 2 | y | 8506 | 1032 | 0.0001455 | 0.141 | +1 | 10 |
| - | - | 5320 | 1034 | - | - | 0 | - |
| - | - | 989.5 | 1034 | - | - | 0 | - |
| 10 | c | 2.131E+04 | 1046 | 0.001629 | 1.558 | +1 | 10 |
| - | - | 1.287E+04 | 1047 | - | - | 0 | - |
| - | - | 2780 | 1048 | - | - | 0 | - |
| - | - | 790.8 | 1086 | - | - | 0 | - |
| - | - | 1710 | 1087 | - | - | 0 | - |
| - | - | 1653 | 1088 | - | - | 0 | - |
| - | - | 1192 | 1089 | - | - | 0 | - |
| - | - | 3468 | 1101 | - | - | 0 | - |
| - | - | 1648 | 1102 | - | - | 0 | - |
| - | - | 696.8 | 1103 | - | - | 0 | - |
| - | - | 2084 | 1104 | - | - | 0 | - |
| - | - | 1374 | 1105 | - | - | 0 | - |
| - | - | 874.9 | 1117 | - | - | 0 | - |
| - | - | 1026 | 1119 | - | - | 0 | - |
| - | - | 3845 | 1129 | - | - | 0 | - |
| - | - | 4.832E+04 | 1130 | - | - | 0 | - |
| - | - | 3.028E+04 | 1131 | - | - | 0 | - |
| - | - | 8873 | 1132 | - | - | 0 | - |
| - | - | 1565 | 1133 | - | - | 0 | - |
| - | - | 1.918E+04 | 1146 | - | - | 0 | - |
| - | - | 5.134E+04 | 1147 | - | - | 0 | - |
| - | - | 2.736E+04 | 1148 | - | - | 0 | - |
| - | - | 9291 | 1149 | - | - | 0 | - |
| - | - | 1648 | 1150 | - | - | 0 | - |
| - | - | 1961 | 1716 | - | - | 0 | - |
| - | - | 1735 | 1717 | - | - | 0 | - |
| - | - | 1844 | 1718 | - | - | 0 | - |
| - | - | 751.1 | 1720 | - | - | 0 | - |
| - | - | 1830 | 1721 | - | - | 0 | - |
| - | - | 1467 | 1722 | - | - | 0 | - |
| - | - | 780.8 | 3039 | - | - | 0 | - |

m/z Charge Intensity FragmentType MassShift Position
129.10215759277344 0 387.90546
149.30552673339844 0 493.1586
173.12811279296875 0 1922.7815
173.43943786621094 0 2110.3716
175.10760498046875 0 1043.4922 y 9
183.1123504638672 0 1642.1217
193.0819091796875 0 520.30786
201.12289428710938 0 6666.7974
210.02474975585938 0 2377.9368
215.13861083984375 0 3130.477
217.08108520507812 0 1197.9387
229.11746215820312 0 501.0127 w 8
230.11276245117188 0 537.20996
231.04598999023438 0 567.96893
239.11367797851562 0 780.0556
246.96775817871094 0 599.3854
312.1549377441406 0 5764.8594
313.1581726074219 0 1002.0751
330.1657409667969 0 3800.7085
334.175048828125 0 565.72784
344.2682800292969 0 541.4544
359.1907958984375 0 1610.3905 y 7
382.2089538574219 0 595.5012
386.1842956542969 0 714.3489
401.2389831542969 0 2448.436
410.2044677734375 0 704.42615
429.2341003417969 0 1987.539
430.23931884765625 0 555.9825
463.25274658203125 0 611.9181
473.2353515625 0 679.9071 y 2
473.7362976074219 0 811.01337
499.2742919921875 0 12366.853 z 6
500.280029296875 0 5875.295
501.2837829589844 0 1304.7399
514.7551879882812 0 597.1192 c Ammonia loss 9
515.2927856445312 0 3332.734 y 6
516.7525634765625 0 1150.824 y 1
520.2498779296875 0 718.97534
521.2340087890625 0 650.24677
536.8032836914062 0 654.7262
538.260986328125 0 2053.161
539.2462158203125 0 1088.991
545.625 0 633.00635
554.2720947265625 0 561.1458
556.2723388671875 0 5805.257
557.2537231445312 0 765.5183
564.2893676757812 0 903.97156
564.7858276367188 0 833.58624
570.3098754882812 0 822.1077
571.317138671875 0 842.4426
572.28271484375 0 5087.0723
573.2914428710938 0 7774.6963
573.7943115234375 0 5540.8667
574.286865234375 0 4057.966
614.3017578125 0 4250.028 z 5
615.308349609375 0 9046.834
616.31201171875 0 2082.5723
630.319580078125 0 8351.011 c Water loss 5
631.3228149414062 0 2676.957
699.3203125 0 1733.8351
700.32568359375 0 1363.0636 y Ammonia loss 4
701.3313598632812 0 4229.9023 z 4
702.3406372070312 0 9507.454
703.3426513671875 0 2925.4995
716.3389282226562 0 922.6458
717.3519897460938 0 22558.625 y 4
718.3539428710938 0 7509.52
719.3572387695312 0 2184.1733
737.3170166015625 0 538.8248
744.3440551757812 0 703.539
760.40625 0 9215.053
761.4096069335938 0 4674.9224
762.4122314453125 0 1010.1434
785.3809814453125 0 934.0258 w 3
787.3944091796875 0 1085.4738 c Ammonia loss 6
800.4024658203125 0 8851.898 z 3
801.4056396484375 0 6128.37
801.80126953125 0 667.64496
802.4108276367188 0 1465.6592
816.420166015625 0 10532.84 y 3
817.4239501953125 0 4597.6577
818.4212036132812 0 1058.5331
851.95068359375 0 790.6749
857.460693359375 0 2896.4678
870.432861328125 0 3920.9946 w 2
871.4337158203125 0 2319.1
885.4526977539062 0 1624.8489
900.4651489257812 0 1490.5144
901.47412109375 0 8895.157 c 7
902.4765014648438 0 3935.195
903.4817504882812 0 1516.7108
929.4447631835938 0 14607.067 z 2
930.4473266601562 0 7371.1445
931.4474487304688 0 1505.4425
944.4902954101562 0 4595.08
945.4677734375 0 9504.577 y 2
946.4671630859375 0 4440.558
947.4749145507812 0 852.8362
957.5203857421875 0 756.4626
971.4724731445312 0 657.9059 c Ammonia loss 8
972.4780883789062 0 690.5884
988.5055541992188 0 6450.8687 c 8
989.5065307617188 0 3341.8416
1001.5120849609375 0 5807.492
1002.515380859375 0 2609.9304
1014.4789428710938 0 1201.7108 y Water loss 1
1015.479248046875 0 749.3598 y Ammonia loss 1
1016.477294921875 0 9727.19 z 1
1017.4791259765625 0 4550.216
1018.4789428710938 0 1574.1946
1030.50390625 0 782.4206
1032.4954833984375 0 8506.445 y 1
1033.5 0 5319.9565
1034.4937744140625 0 989.4871
1045.525634765625 0 21307.387 c 9
1046.5299072265625 0 12870.471
1047.5308837890625 0 2780.1606
1085.572021484375 0 790.79407
1086.5687255859375 0 1709.751
1087.546875 0 1653.3651
1088.551513671875 0 1192.1061
1100.5736083984375 0 3467.9668
1101.5721435546875 0 1647.6245
1102.556640625 0 696.7561
1103.5545654296875 0 2083.7024
1104.560546875 0 1373.9686
1116.5531005859375 0 874.8775
1118.5943603515625 0 1026.4526
1128.572998046875 0 3844.8767
1129.561279296875 0 48323.707
1130.563720703125 0 30275.572
1131.5654296875 0 8872.9
1132.5667724609375 0 1564.6534
1145.5791015625 0 19178.113
1146.586181640625 0 51337.05
1147.588623046875 0 27363.43
1148.5902099609375 0 9290.936
1149.589599609375 0 1647.893
1715.77880859375 0 1961.4039
1716.8006591796875 0 1735.0305
1717.7926025390625 0 1844.1007
1719.7940673828125 0 751.0599
1720.8436279296875 0 1829.6243
1721.843994140625 0 1467.2593
3039.19873046875 0 780.7716

Spectrum Details

|  |  |
| --- | --- |
| Matched peaks? Matched peaks The total absolute number of peaks matched. Additionally in brackets the total fraction of peaks matched and the total number of peaks is shown. | 28 (19.31% of 145) |
| FDR? FDR The false discovery rate estimated for this peptide. It is calculated by matching all theoretical fragments with a non-integer shift with the raw peaks for this spectrum. This is done with 40 different shifts. The resulting percentage is the average number of annotated peaks over the number of annotated peaks with the correct spectrum. | 0.09% |
| Satellite FDR? Satellite FDR See the FDR for details on its calculation. This satellite ion specific FDR only contains the satellite ions (d/w) for I/L/J positions. | - |
| PSM Score? PSM Score The PSM Score as given by Hecklib to this annotated spectrum. It is shown with three significant figures. | 263 |

#### Spectrum 3922? Spectrum 3922 The raw spectrum of this peptide as annotated by Hecklib. The fragments are coloured according to ion type (see legend). Any peaks with a star '\*' as text can be hovered over to see the full details, first the ion type second the mass shift type. By hovering over the amino acids in the peptide or ions in the legend the corresponding peaks are highlighted. By toggling the 'Unassigned' label you can turn the background (unassigned) peaks on or off in the plot. By updating the slider in the Ion legend you can update the spectrum to only show the top X% of the peaks with labels. The top X% means any peak that is within X% of the highest intensity. By dragging in the spectrum you can zoom in to a specific part of the spectrum and use 'Zoom Out' to get back to the original zoom level. The annotation of the spectrum is based on the given sequence in the peptides file and is done with different software so inconsistencies are likely. The peaks are annotated based on the given sequence, with 20 ppm tolerance.

Copy Data

##### Spectrum 3922 (TSV)

###### Preview

```
Loading example...
```

*Click on the button to copy the data to your clipboard.*

Mz MinMz MaxIntensity Max

WidthHeightPeptide font sizePeptide stroke widthSpectrum font sizeSpectrum stroke widthCompact peptide

Ion legend

wxyz

abcd

OtherUnassignedIon^Charge^_Position_Show for top:%

JSEVSDRPSGV

03.30e+46.60e+49.90e+41.32e+5

Zoom Out

a^+1^_2_a^+1^_2_y^+1^_2_b^+1^_2_b^+1^_2_y^+1^_3_a^+1^_3_a^+1^_3_b^+1^_3_b^+2^_6_b^+1^_3_y^+1^_4_y^+1^_4_y^+2^_8_y^+2^_8_b^+1^_4_b^+1^_4_y^+2^_9_y^+2^_9_y^+1^_5_y^+1^_5_y^+2^_10_b^+2^_10_y^+1^_5_y^+2^_10_\*\*\*y^+1^_6_b^+1^_6_y^+1^_6_y^+1^_7_y^+1^_7_y^+1^_7_b^+1^_7_y^+1^_8_y^+1^_8_y^+1^_8_y^+1^_9_y^+1^_9_b^+1^_9_y^+1^_10_y^+1^_10_b^+1^_10_y^+1^_10_

0769153823073076

Fragment Matches Table

Show background peaks

| Position | Ion type | Intensity | mz Theoretical | mz Error (Th) | mz Error (ppm) | Charge | Series Number |
| --- | --- | --- | --- | --- | --- | --- | --- |
| - | - | 613.6 | 120.1 | - | - | 0 | - |
| - | - | 1242 | 120.1 | - | - | 0 | - |
| - | - | 716.1 | 125.1 | - | - | 0 | - |
| - | - | 2147 | 126.1 | - | - | 0 | - |
| - | - | 739.9 | 127.1 | - | - | 0 | - |
| - | - | 1098 | 128.1 | - | - | 0 | - |
| - | - | 699 | 129.1 | - | - | 0 | - |
| - | - | 1.066E+04 | 129.1 | - | - | 0 | - |
| - | - | 1008 | 130.1 | - | - | 0 | - |
| - | - | 433 | 130.1 | - | - | 0 | - |
| - | - | 482.9 | 130.1 | - | - | 0 | - |
| - | - | 465.7 | 130.1 | - | - | 0 | - |
| - | - | 496.5 | 131.1 | - | - | 0 | - |
| - | - | 504.3 | 133.1 | - | - | 0 | - |
| - | - | 4362 | 136.1 | - | - | 0 | - |
| - | - | 433.7 | 137 | - | - | 0 | - |
| - | - | 739 | 138.1 | - | - | 0 | - |
| - | - | 427.3 | 139.1 | - | - | 0 | - |
| - | - | 589.9 | 140.1 | - | - | 0 | - |
| - | - | 435.6 | 141.1 | - | - | 0 | - |
| - | - | 571.3 | 143.1 | - | - | 0 | - |
| - | - | 1135 | 145.1 | - | - | 0 | - |
| - | - | 451.9 | 145.1 | - | - | 0 | - |
| - | - | 552 | 147.1 | - | - | 0 | - |
| - | - | 535.7 | 148.9 | - | - | 0 | - |
| - | - | 501.3 | 148.9 | - | - | 0 | - |
| - | - | 590.3 | 148.9 | - | - | 0 | - |
| - | - | 490 | 148.9 | - | - | 0 | - |
| - | - | 924.1 | 148.9 | - | - | 0 | - |
| - | - | 643.6 | 148.9 | - | - | 0 | - |
| - | - | 955.4 | 148.9 | - | - | 0 | - |
| - | - | 1158 | 148.9 | - | - | 0 | - |
| - | - | 2720 | 148.9 | - | - | 0 | - |
| - | - | 4246 | 148.9 | - | - | 0 | - |
| - | - | 3500 | 149 | - | - | 0 | - |
| - | - | 1632 | 149 | - | - | 0 | - |
| - | - | 887.2 | 149 | - | - | 0 | - |
| - | - | 929 | 149 | - | - | 0 | - |
| - | - | 935.4 | 149 | - | - | 0 | - |
| - | - | 513.6 | 149 | - | - | 0 | - |
| - | - | 688.6 | 153.1 | - | - | 0 | - |
| - | - | 2185 | 154.1 | - | - | 0 | - |
| - | - | 528.4 | 155 | - | - | 0 | - |
| - | - | 609.8 | 155.1 | - | - | 0 | - |
| 2 | a | 2963 | 155.1 | 0.0002745 | 1.77 | +1 | 2 |
| - | - | 1113 | 156.1 | - | - | 0 | - |
| - | - | 1339 | 157.1 | - | - | 0 | - |
| - | - | 4131 | 157.1 | - | - | 0 | - |
| - | - | 1624 | 157.1 | - | - | 0 | - |
| - | - | 521 | 158.1 | - | - | 0 | - |
| - | - | 636.1 | 159.1 | - | - | 0 | - |
| - | - | 690.3 | 159.1 | - | - | 0 | - |
| - | - | 1332 | 159.1 | - | - | 0 | - |
| - | - | 553.1 | 166.1 | - | - | 0 | - |
| - | - | 473.6 | 166.6 | - | - | 0 | - |
| - | - | 1562 | 167.1 | - | - | 0 | - |
| - | - | 533.2 | 167.1 | - | - | 0 | - |
| - | - | 461.9 | 168.1 | - | - | 0 | - |
| - | - | 652.4 | 169.1 | - | - | 0 | - |
| - | - | 416.9 | 170.8 | - | - | 0 | - |
| - | - | 3240 | 171.1 | - | - | 0 | - |
| - | - | 536.8 | 171.1 | - | - | 0 | - |
| - | - | 516.1 | 172.1 | - | - | 0 | - |
| - | - | 1950 | 173.1 | - | - | 0 | - |
| 2 | a | 3.656E+04 | 173.1 | 0.0003452 | 1.994 | +1 | 2 |
| - | - | 2731 | 174.1 | - | - | 0 | - |
| - | - | 1691 | 175.1 | - | - | 0 | - |
| - | - | 1173 | 175.1 | - | - | 0 | - |
| 10 | y | 5403 | 175.1 | 0.0002982 | 1.703 | +1 | 2 |
| - | - | 732.5 | 181.1 | - | - | 0 | - |
| 2 | b | 3.797E+04 | 183.1 | 0.0003245 | 1.772 | +1 | 2 |
| - | - | 3093 | 184.1 | - | - | 0 | - |
| - | - | 604.7 | 185.1 | - | - | 0 | - |
| - | - | 8101 | 185.1 | - | - | 0 | - |
| - | - | 598.6 | 185.1 | - | - | 0 | - |
| - | - | 582 | 186.1 | - | - | 0 | - |
| - | - | 896.1 | 186.1 | - | - | 0 | - |
| - | - | 3817 | 187.1 | - | - | 0 | - |
| - | - | 824.7 | 187.1 | - | - | 0 | - |
| - | - | 1.341E+04 | 189.1 | - | - | 0 | - |
| - | - | 984 | 190.1 | - | - | 0 | - |
| - | - | 810.7 | 195.1 | - | - | 0 | - |
| - | - | 9166 | 199.1 | - | - | 0 | - |
| - | - | 1177 | 199.1 | - | - | 0 | - |
| - | - | 646.6 | 200.1 | - | - | 0 | - |
| 2 | b | 5.083E+04 | 201.1 | 0.0002884 | 1.434 | +1 | 2 |
| - | - | 4042 | 202.1 | - | - | 0 | - |
| - | - | 4490 | 203.1 | - | - | 0 | - |
| - | - | 1423 | 203.1 | - | - | 0 | - |
| - | - | 498.7 | 204.1 | - | - | 0 | - |
| - | - | 602.8 | 207.1 | - | - | 0 | - |
| - | - | 624 | 209.1 | - | - | 0 | - |
| - | - | 2561 | 210 | - | - | 0 | - |
| - | - | 3254 | 211.1 | - | - | 0 | - |
| - | - | 521.7 | 211.1 | - | - | 0 | - |
| - | - | 1836 | 212.1 | - | - | 0 | - |
| - | - | 1038 | 213.1 | - | - | 0 | - |
| - | - | 593.7 | 213.2 | - | - | 0 | - |
| - | - | 1.683E+04 | 215.1 | - | - | 0 | - |
| - | - | 2.613E+04 | 217.1 | - | - | 0 | - |
| - | - | 1967 | 218.1 | - | - | 0 | - |
| - | - | 821 | 223.1 | - | - | 0 | - |
| - | - | 2489 | 224.1 | - | - | 0 | - |
| - | - | 1026 | 227.1 | - | - | 0 | - |
| - | - | 644.7 | 227.1 | - | - | 0 | - |
| - | - | 665.9 | 227.2 | - | - | 0 | - |
| - | - | 4801 | 229.1 | - | - | 0 | - |
| - | - | 1710 | 230.1 | - | - | 0 | - |
| - | - | 496.7 | 231.1 | - | - | 0 | - |
| - | - | 604.4 | 235.1 | - | - | 0 | - |
| - | - | 1845 | 237.1 | - | - | 0 | - |
| - | - | 1446 | 240.1 | - | - | 0 | - |
| - | - | 2488 | 242.1 | - | - | 0 | - |
| - | - | 962.4 | 251.2 | - | - | 0 | - |
| - | - | 857.4 | 254.2 | - | - | 0 | - |
| - | - | 925.5 | 255.1 | - | - | 0 | - |
| - | - | 6215 | 255.1 | - | - | 0 | - |
| - | - | 3909 | 260.1 | - | - | 0 | - |
| - | - | 645.8 | 261.2 | - | - | 0 | - |
| 9 | y | 3678 | 262.1 | 0.0002369 | 0.9039 | +1 | 3 |
| - | - | 599.2 | 269.1 | - | - | 0 | - |
| - | - | 1350 | 270.1 | - | - | 0 | - |
| - | - | 1151 | 272.1 | - | - | 0 | - |
| - | - | 1143 | 272.1 | - | - | 0 | - |
| - | - | 695.8 | 280.1 | - | - | 0 | - |
| - | - | 920.1 | 284.1 | - | - | 0 | - |
| 3 | a | 3021 | 284.2 | 0.0004976 | 1.751 | +1 | 3 |
| - | - | 1288 | 288.2 | - | - | 0 | - |
| - | - | 3783 | 298.1 | - | - | 0 | - |
| - | - | 1651 | 299.1 | - | - | 0 | - |
| 3 | a | 1375 | 302.2 | 6.474E-05 | 0.2143 | +1 | 3 |
| 3 | b | 4.484E+04 | 312.2 | 0.0005171 | 1.656 | +1 | 3 |
| - | - | 7217 | 313.2 | - | - | 0 | - |
| - | - | 1075 | 313.2 | - | - | 0 | - |
| - | - | 867.3 | 314.1 | - | - | 0 | - |
| 6 | b | 1.008E+04 | 316.2 | 0.0004144 | 1.311 | +2 | 6 |
| - | - | 1665 | 317.2 | - | - | 0 | - |
| - | - | 688.5 | 323.2 | - | - | 0 | - |
| - | - | 1650 | 324.1 | - | - | 0 | - |
| 3 | b | 2.358E+04 | 330.2 | 0.000542 | 1.641 | +1 | 3 |
| - | - | 6127 | 331.2 | - | - | 0 | - |
| - | - | 676.4 | 332.2 | - | - | 0 | - |
| - | - | 591.6 | 337.1 | - | - | 0 | - |
| 8 | y | 1869 | 341.2 | 0.001068 | 3.129 | +1 | 4 |
| - | - | 8846 | 342.1 | - | - | 0 | - |
| - | - | 1456 | 343.1 | - | - | 0 | - |
| - | - | 580.9 | 355.2 | - | - | 0 | - |
| - | - | 8352 | 359.2 | - | - | 0 | - |
| 8 | y | 2.568E+04 | 359.2 | 0.0003601 | 1.002 | +1 | 4 |
| - | - | 643.9 | 359.7 | - | - | 0 | - |
| - | - | 1208 | 360.2 | - | - | 0 | - |
| - | - | 4011 | 360.2 | - | - | 0 | - |
| - | - | 565.6 | 366.1 | - | - | 0 | - |
| - | - | 1031 | 383.2 | - | - | 0 | - |
| - | - | 2160 | 385.2 | - | - | 0 | - |
| - | - | 586 | 387.2 | - | - | 0 | - |
| - | - | 2928 | 399.2 | - | - | 0 | - |
| 4 | y | 637.2 | 400.2 | 0.00113 | 2.823 | +2 | 8 |
| - | - | 1.041E+04 | 401.2 | - | - | 0 | - |
| - | - | 1710 | 402.2 | - | - | 0 | - |
| - | - | 1395 | 403.2 | - | - | 0 | - |
| - | - | 669.5 | 404.2 | - | - | 0 | - |
| 4 | y | 2022 | 408.7 | 0.0006408 | 1.568 | +2 | 8 |
| 4 | b | 984.7 | 411.2 | 0.0005235 | 1.273 | +1 | 4 |
| - | - | 974.7 | 413.2 | - | - | 0 | - |
| - | - | 3970 | 416.2 | - | - | 0 | - |
| - | - | 774.3 | 422.2 | - | - | 0 | - |
| 4 | b | 9957 | 429.2 | 0.0007926 | 1.847 | +1 | 4 |
| - | - | 2438 | 430.2 | - | - | 0 | - |
| - | - | 2038 | 441.2 | - | - | 0 | - |
| - | - | 1061 | 455.2 | - | - | 0 | - |
| - | - | 866.7 | 455.7 | - | - | 0 | - |
| - | - | 1022 | 456.2 | - | - | 0 | - |
| - | - | 3385 | 458.2 | - | - | 0 | - |
| - | - | 1274 | 459.2 | - | - | 0 | - |
| 3 | y | 1935 | 464.2 | 0.000923 | 1.988 | +2 | 9 |
| - | - | 1026 | 464.7 | - | - | 0 | - |
| 3 | y | 3304 | 473.2 | 0.001073 | 2.267 | +2 | 9 |
| - | - | 2089 | 473.7 | - | - | 0 | - |
| - | - | 1152 | 486.7 | - | - | 0 | - |
| - | - | 672.5 | 494.7 | - | - | 0 | - |
| 7 | y | 1198 | 497.3 | 0.001472 | 2.959 | +1 | 5 |
| 7 | y | 5086 | 498.3 | 0.0008681 | 1.742 | +1 | 5 |
| - | - | 1164 | 499.3 | - | - | 0 | - |
| 2 | y | 1786 | 507.7 | 0.001327 | 2.614 | +2 | 10 |
| - | - | 1297 | 508.2 | - | - | 0 | - |
| - | - | 696.5 | 508.7 | - | - | 0 | - |
| - | - | 697.2 | 514.2 | - | - | 0 | - |
| 10 | b | 932.3 | 514.8 | 0.0001502 | 0.2917 | +2 | 10 |
| - | - | 999.5 | 515.3 | - | - | 0 | - |
| 7 | y | 2.377E+04 | 515.3 | 0.0006284 | 1.22 | +1 | 5 |
| - | - | 6213 | 516.3 | - | - | 0 | - |
| 2 | y | 5656 | 516.8 | 0.0003784 | 0.7322 | +2 | 10 |
| - | - | 2285 | 517.3 | - | - | 0 | - |
| - | - | 1071 | 517.3 | - | - | 0 | - |
| - | - | 1073 | 517.8 | - | - | 0 | - |
| - | - | 1462 | 531.3 | - | - | 0 | - |
| - | - | 769.3 | 538.3 | - | - | 0 | - |
| - | - | 996.8 | 543.3 | - | - | 0 | - |
| - | - | 772.6 | 555.3 | - | - | 0 | - |
| - | - | 697.8 | 560.3 | - | - | 0 | - |
| 0 | Precursor | 7312 | 564.3 | 0.000616 | 1.092 | +2 | -1 |
| - | - | 785.9 | 564.7 | - | - | 0 | - |
| 0 | Precursor | 4333 | 564.8 | 0.00989 | 17.51 | +2 | -1 |
| - | - | 1188 | 565.3 | - | - | 0 | - |
| - | - | 3496 | 569.3 | - | - | 0 | - |
| - | - | 706.7 | 570.3 | - | - | 0 | - |
| - | - | 2053 | 572.3 | - | - | 0 | - |
| 0 | Precursor | 2.262E+04 | 573.3 | 0.0002165 | 0.3776 | +2 | -1 |
| - | - | 1.435E+04 | 573.8 | - | - | 0 | - |
| - | - | 5084 | 574.3 | - | - | 0 | - |
| - | - | 1096 | 574.5 | - | - | 0 | - |
| - | - | 706.1 | 574.6 | - | - | 0 | - |
| - | - | 639.2 | 574.8 | - | - | 0 | - |
| - | - | 3223 | 587.3 | - | - | 0 | - |
| - | - | 1086 | 588.3 | - | - | 0 | - |
| - | - | 1771 | 600.3 | - | - | 0 | - |
| 6 | y | 901.8 | 612.3 | 0.0002413 | 0.3941 | +1 | 6 |
| 6 | b | 652.1 | 613.3 | 0.008538 | 13.92 | +1 | 6 |
| - | - | 5855 | 618.3 | - | - | 0 | - |
| - | - | 1604 | 619.3 | - | - | 0 | - |
| - | - | 888 | 626.3 | - | - | 0 | - |
| 6 | y | 2.367E+04 | 630.3 | 0.000724 | 1.149 | +1 | 6 |
| - | - | 5941 | 631.3 | - | - | 0 | - |
| - | - | 1121 | 632.3 | - | - | 0 | - |
| - | - | 900.4 | 646.3 | - | - | 0 | - |
| - | - | 3932 | 674.3 | - | - | 0 | - |
| - | - | 1060 | 675.3 | - | - | 0 | - |
| - | - | 1013 | 687.3 | - | - | 0 | - |
| - | - | 612.3 | 695.8 | - | - | 0 | - |
| 5 | y | 4337 | 699.3 | 7.322E-05 | 0.1047 | +1 | 7 |
| 5 | y | 1851 | 700.3 | 0.01221 | 17.44 | +1 | 7 |
| 5 | y | 1.308E+05 | 717.4 | 0.0003118 | 0.4346 | +1 | 7 |
| - | - | 4.876E+04 | 718.4 | - | - | 0 | - |
| - | - | 1.027E+04 | 719.4 | - | - | 0 | - |
| - | - | 758.3 | 727.3 | - | - | 0 | - |
| - | - | 637.3 | 728.4 | - | - | 0 | - |
| - | - | 2683 | 753.4 | - | - | 0 | - |
| - | - | 1131 | 754.4 | - | - | 0 | - |
| 7 | b | 8294 | 787.4 | 0.0003171 | 0.4027 | +1 | 7 |
| - | - | 3926 | 788.4 | - | - | 0 | - |
| - | - | 1422 | 789.4 | - | - | 0 | - |
| 4 | y | 2857 | 798.4 | 0.000629 | 0.7879 | +1 | 8 |
| 4 | y | 965.2 | 799.4 | 0.01436 | 17.96 | +1 | 8 |
| - | - | 685.5 | 802.9 | - | - | 0 | - |
| - | - | 3139 | 810.4 | - | - | 0 | - |
| - | - | 1513 | 811.4 | - | - | 0 | - |
| 4 | y | 7.514E+04 | 816.4 | 7.411E-05 | 0.09078 | +1 | 8 |
| - | - | 3.315E+04 | 817.4 | - | - | 0 | - |
| - | - | 7493 | 818.4 | - | - | 0 | - |
| - | - | 1316 | 828.4 | - | - | 0 | - |
| - | - | 3169 | 846.4 | - | - | 0 | - |
| - | - | 942.9 | 847.4 | - | - | 0 | - |
| - | - | 888.7 | 858.4 | - | - | 0 | - |
| - | - | 1941 | 915.4 | - | - | 0 | - |
| - | - | 745.9 | 916.4 | - | - | 0 | - |
| 3 | y | 7457 | 927.5 | 0.0004554 | 0.491 | +1 | 9 |
| - | - | 3585 | 928.5 | - | - | 0 | - |
| - | - | 1697 | 929.5 | - | - | 0 | - |
| - | - | 4852 | 933.4 | - | - | 0 | - |
| - | - | 1880 | 934.4 | - | - | 0 | - |
| 3 | y | 6.403E+04 | 945.5 | 3.851E-05 | 0.04074 | +1 | 9 |
| - | - | 2.869E+04 | 946.5 | - | - | 0 | - |
| - | - | 8971 | 947.5 | - | - | 0 | - |
| 9 | b | 1781 | 971.5 | 0.000735 | 0.7565 | +1 | 9 |
| - | - | 1109 | 996.5 | - | - | 0 | - |
| - | - | 867.3 | 997.5 | - | - | 0 | - |
| - | - | 906.7 | 1002 | - | - | 0 | - |
| 2 | y | 1.016E+04 | 1014 | 0.0001652 | 0.1629 | +1 | 10 |
| 2 | y | 4608 | 1015 | 0.01694 | 16.69 | +1 | 10 |
| - | - | 811 | 1016 | - | - | 0 | - |
| 10 | b | 1778 | 1029 | 0.0006285 | 0.611 | +1 | 10 |
| - | - | 1014 | 1030 | - | - | 0 | - |
| 2 | y | 8.77E+04 | 1032 | 9.86E-05 | 0.0955 | +1 | 10 |
| - | - | 4.813E+04 | 1033 | - | - | 0 | - |
| - | - | 1.46E+04 | 1035 | - | - | 0 | - |
| - | - | 2761 | 1036 | - | - | 0 | - |
| - | - | 3565 | 1042 | - | - | 0 | - |
| - | - | 2426 | 1043 | - | - | 0 | - |
| - | - | 643.6 | 1683 | - | - | 0 | - |
| - | - | 702 | 1969 | - | - | 0 | - |
| - | - | 714.4 | 3045 | - | - | 0 | - |

m/z Charge Intensity FragmentType MassShift Position
120.06550598144531 0 613.6348
120.0811538696289 0 1241.9207
125.10759735107422 0 716.0977
126.05525970458984 0 2146.916
127.0505599975586 0 739.93964
128.1072998046875 0 1098.3143
129.06614685058594 0 698.95325
129.10255432128906 0 10662.485
130.05038452148438 0 1007.91486
130.06509399414062 0 432.99536
130.08653259277344 0 482.9092
130.10594177246094 0 465.73404
131.08189392089844 0 496.46844
133.06076049804688 0 504.31003
136.0760040283203 0 4362.049
136.99374389648438 0 433.71603
138.0918426513672 0 739.0264
139.08721923828125 0 427.25137
140.08209228515625 0 589.9088
141.10235595703125 0 435.62405
143.08135986328125 0 571.2545
145.06118774414062 0 1135.157
145.0678253173828 0 451.9499
147.07701110839844 0 552.0173
148.86521911621094 0 535.693
148.87905883789062 0 501.311
148.88638305664062 0 590.2646
148.893798828125 0 489.98398
148.90142822265625 0 924.0948
148.90797424316406 0 643.6093
148.9228515625 0 955.35986
148.93026733398438 0 1158.2612
148.93751525878906 0 2719.5784
148.9452362060547 0 4245.8247
148.96189880371094 0 3500.114
148.96963500976562 0 1631.8761
148.97665405273438 0 887.23724
148.98406982421875 0 929.03125
148.9912872314453 0 935.4382
148.99832153320312 0 513.5628
153.06600952148438 0 688.5822
154.0501708984375 0 2185.1875
155.04595947265625 0 528.3635
155.08212280273438 0 609.808
155.1181640625 0 2962.652 a Water loss 1
156.10177612304688 0 1112.8247
157.0610809326172 0 1339.4386
157.0974884033203 0 4130.67
157.1088104248047 0 1624.2291
158.0932159423828 0 521.0208
159.07684326171875 0 636.1118
159.09156799316406 0 690.33405
159.1131591796875 0 1331.7858
166.061279296875 0 553.0682
166.57632446289062 0 473.61392
167.08193969726562 0 1561.6697
167.11854553222656 0 533.2214
168.1473846435547 0 461.8904
169.096923828125 0 652.44794
170.84860229492188 0 416.85077
171.0767822265625 0 3240.1602
171.1129150390625 0 536.75073
172.07984924316406 0 516.14154
173.09254455566406 0 1949.75
173.12879943847656 0 36560.363 a 1
174.1321563720703 0 2731.199
175.07176208496094 0 1691.1777
175.0870361328125 0 1172.5272
175.10801696777344 0 5403.4624 y 9
181.06153869628906 0 732.5437
183.11312866210938 0 37966.434 b Water loss 1
184.11648559570312 0 3093.144
185.05642700195312 0 604.693
185.0924072265625 0 8101.075
185.12734985351562 0 598.64923
186.0963592529297 0 582.00653
186.1241455078125 0 896.05774
187.10789489746094 0 3817.032
187.14430236816406 0 824.6755
189.08731079101562 0 13410.507
190.09095764160156 0 983.9542
195.11363220214844 0 810.68695
199.0717315673828 0 9166.263
199.1083221435547 0 1176.6598
200.0756378173828 0 646.61115
201.1236572265625 0 50830.844 b 1
202.12701416015625 0 4042.3481
203.06661987304688 0 4489.7456
203.1027374267578 0 1423.2733
204.1345977783203 0 498.69193
207.1126251220703 0 602.8397
209.10243225097656 0 623.96295
210.02574157714844 0 2561.4517
211.10801696777344 0 3253.8743
211.11814880371094 0 521.72235
212.10308837890625 0 1835.9071
213.08766174316406 0 1037.9542
213.1602783203125 0 593.7467
215.13934326171875 0 16825.518
217.08221435546875 0 26129.607
218.08575439453125 0 1967.2686
223.10787963867188 0 820.9603
224.10317993164062 0 2489.392
227.0669708251953 0 1025.6124
227.10267639160156 0 644.68036
227.1761474609375 0 665.9264
229.1187744140625 0 4800.8784
230.1139678955078 0 1709.9182
231.09872436523438 0 496.71985
235.10806274414062 0 604.3981
237.1352081298828 0 1845.1903
240.13433837890625 0 1446.178
242.11399841308594 0 2488.3157
251.15028381347656 0 962.4291
254.16180419921875 0 857.4468
255.0611114501953 0 925.4895
255.10923767089844 0 6215.319
260.124267578125 0 3909.3337
261.1556091308594 0 645.77997
262.1399841308594 0 3677.9976 y 8
269.1284484863281 0 599.221
270.1452331542969 0 1350.3347
272.1224670410156 0 1150.8728
272.137451171875 0 1143.3491
280.13037109375 0 695.8198
284.12420654296875 0 920.1025
284.1609802246094 0 3021.3208 a Water loss 2
288.1558837890625 0 1288.1317
298.1401672363281 0 3783.3506
299.1362609863281 0 1651.3079
302.1711120605469 0 1374.9948 a 2
312.1559143066406 0 44837.418 b Water loss 2
313.158935546875 0 7217.456
313.18695068359375 0 1074.7098
314.09820556640625 0 867.33124
316.1507263183594 0 10076.342 b 5
317.15185546875 0 1664.7119
323.1707458496094 0 688.4508
324.1307678222656 0 1650.2937
330.16650390625 0 23578.172 b 2
331.1706848144531 0 6126.772
332.1729736328125 0 676.42303
337.1499938964844 0 591.5916
341.1830139160156 0 1868.7501 y Water loss 7
342.14129638671875 0 8846.459
343.1448669433594 0 1455.75
355.1610107421875 0 580.93317
359.1676330566406 0 8352.492
359.19287109375 0 25684.367 y 7
359.679931640625 0 643.89453
360.17132568359375 0 1207.9519
360.1961975097656 0 4011.0889
366.1426086425781 0 565.6475
383.2292175292969 0 1031.0211
385.1717834472656 0 2159.5488
387.1959228515625 0 585.9504
399.1982116699219 0 2927.8455
400.1997375488281 0 637.16943 y Ammonia loss 3
401.2401123046875 0 10406.259
402.243408203125 0 1709.5526
403.1839294433594 0 1395.034
404.18511962890625 0 669.4757
408.71478271484375 0 2022.1813 y 3
411.2243347167969 0 984.7253 b Water loss 3
413.1656799316406 0 974.73804
416.2259216308594 0 3970.373
422.1669616699219 0 774.2533
429.23516845703125 0 9956.805 b 3
430.23919677734375 0 2438.1313
441.2098083496094 0 2038.3326
455.2246398925781 0 1061.3022
455.7284851074219 0 866.7182
456.2464599609375 0 1021.7114
458.23681640625 0 3385.101
459.2394104003906 0 1274.2775
464.2310791015625 0 1935.108 y Water loss 2
464.73431396484375 0 1025.8805
473.23651123046875 0 3303.8696 y 2
473.73809814453125 0 2089.0005
486.74530029296875 0 1151.7753
494.7364196777344 0 672.4974
497.2815856933594 0 1198.0869 y Water loss 6
498.2662048339844 0 5086.2925 y Ammonia loss 6
499.26739501953125 0 1163.5264
507.74749755859375 0 1786.4663 y Water loss 1
508.2499084472656 0 1296.8777
508.74920654296875 0 696.50116
514.2235107421875 0 697.2331
514.7538452148438 0 932.2791 b 9
515.2573852539062 0 999.524
515.2942504882812 0 23769.674 y 6
516.2975463867188 0 6212.591
516.7518310546875 0 5656.2812 y 1
517.2562255859375 0 2284.5435
517.3003540039062 0 1070.6605
517.7540283203125 0 1073.2891
531.2549438476562 0 1462.2406
538.2648315429688 0 769.3111
543.2554931640625 0 996.7732
555.28271484375 0 772.6043
560.2835083007812 0 697.75885
564.288818359375 0 7311.5 Precursor Water loss
564.7464599609375 0 785.9055
564.7901000976562 0 4333.1294 Precursor Ammonia loss
565.296875 0 1188.4694
569.267578125 0 3496.0085
570.2708740234375 0 706.7471
572.2825317382812 0 2052.5984
573.293701171875 0 22621.205 Precursor
573.7950439453125 0 14353.979
574.295166015625 0 5083.8716
574.54150390625 0 1095.8914
574.6067504882812 0 706.1137
574.7857055664062 0 639.15765
587.2786254882812 0 3222.5845
588.3143920898438 0 1086.3914
600.2752685546875 0 1770.566
612.3102416992188 0 901.7986 y Water loss 5
613.2913208007812 0 652.11096 b Water loss 5
618.2848510742188 0 5855.186
619.2891235351562 0 1603.7366
626.3220825195312 0 888.00494
630.3212890625 0 23668.363 y 5
631.32275390625 0 5941.1504
632.3256225585938 0 1120.8467
646.3162231445312 0 900.36835
674.31005859375 0 3931.9604
675.3130493164062 0 1060.2263
687.3439331054688 0 1013.48865
695.8355102539062 0 612.3027
699.3421020507812 0 4336.5957 y Water loss 4
700.3382568359375 0 1850.6678 y Ammonia loss 4
717.3529052734375 0 130757.58 y 4
718.3556518554688 0 48756.004
719.3579711914062 0 10274.858
727.3382568359375 0 758.30756
728.3528442382812 0 637.28687
753.3525390625 0 2682.9019
754.3515014648438 0 1131.4135
787.394775390625 0 8293.949 b 6
788.396728515625 0 3926.453
789.3939819335938 0 1422.0786
798.4110717773438 0 2857.1028 y Water loss 3
799.4088134765625 0 965.1711 y Ammonia loss 3
802.8687744140625 0 685.4998
810.3734130859375 0 3138.5786
811.3762817382812 0 1512.9102
816.4210815429688 0 75144.09 y 3
817.4239501953125 0 33149.508
818.4258422851562 0 7493.1865
828.3846435546875 0 1315.5242
846.3947143554688 0 3169.2275
847.4013061523438 0 942.9406
858.3942260742188 0 888.7335
915.42138671875 0 1940.9856
916.4120483398438 0 745.9049
927.4534912109375 0 7457.2314 y Water loss 2
928.4562377929688 0 3585.0203
929.4583740234375 0 1696.7344
933.4275512695312 0 4852.386
934.4312744140625 0 1880.3326
945.4635620117188 0 64033.68 y 2
946.466796875 0 28688.637
947.4679565429688 0 8971.191
971.478515625 0 1781.1952 b 8
996.477294921875 0 1109.3569
997.47314453125 0 867.2578
1002.4891357421875 0 906.69995
1014.4852294921875 0 10164.1875 y Water loss 1
1015.4860229492188 0 4608.298 y Ammonia loss 1
1016.4876708984375 0 811.01385
1028.5013427734375 0 1777.7922 b 9
1029.5 0 1014.4065
1032.4957275390625 0 87702.84 y 1
1033.4984130859375 0 48134.707
1034.5008544921875 0 14599.772
1035.5032958984375 0 2761.055
1042.4801025390625 0 3564.8826
1043.484375 0 2425.6978
1682.733154296875 0 643.6436
1969.0552978515625 0 702.01825
3045.30322265625 0 714.44366

Spectrum Details

|  |  |
| --- | --- |
| Matched peaks? Matched peaks The total absolute number of peaks matched. Additionally in brackets the total fraction of peaks matched and the total number of peaks is shown. | 45 (15.96% of 282) |
| FDR? FDR The false discovery rate estimated for this peptide. It is calculated by matching all theoretical fragments with a non-integer shift with the raw peaks for this spectrum. This is done with 40 different shifts. The resulting percentage is the average number of annotated peaks over the number of annotated peaks with the correct spectrum. | 0.05% |
| Satellite FDR? Satellite FDR See the FDR for details on its calculation. This satellite ion specific FDR only contains the satellite ions (d/w) for I/L/J positions. | - |
| PSM Score? PSM Score The PSM Score as given by Hecklib to this annotated spectrum. It is shown with three significant figures. | 540 |

#### Spectrum 4123? Spectrum 4123 The raw spectrum of this peptide as annotated by Hecklib. The fragments are coloured according to ion type (see legend). Any peaks with a star '\*' as text can be hovered over to see the full details, first the ion type second the mass shift type. By hovering over the amino acids in the peptide or ions in the legend the corresponding peaks are highlighted. By toggling the 'Unassigned' label you can turn the background (unassigned) peaks on or off in the plot. By updating the slider in the Ion legend you can update the spectrum to only show the top X% of the peaks with labels. The top X% means any peak that is within X% of the highest intensity. By dragging in the spectrum you can zoom in to a specific part of the spectrum and use 'Zoom Out' to get back to the original zoom level. The annotation of the spectrum is based on the given sequence in the peptides file and is done with different software so inconsistencies are likely. The peaks are annotated based on the given sequence, with 20 ppm tolerance.

Copy Data

##### Spectrum 4123 (TSV)

###### Preview

```
Loading example...
```

*Click on the button to copy the data to your clipboard.*

Mz MinMz MaxIntensity Max

WidthHeightPeptide font sizePeptide stroke widthSpectrum font sizeSpectrum stroke widthCompact peptide

Ion legend

wxyz

abcd

OtherUnassignedIon^Charge^_Position_Show for top:%

JSEVSDRPSGV

01.42e+42.84e+44.25e+45.67e+4

Zoom Out

y^+1^_2_w^+1^_3_w^+1^_4_y^+1^_4_y^+2^_9_z^+1^_5_y^+2^_10_c^+2^_10_y^+1^_5_y^+2^_10_z^+1^_6_c^+1^_6_y^+1^_7_y^+1^_7_z^+1^_7_y^+1^_7_c^+1^_7_y^+1^_8_z^+1^_8_y^+1^_8_w^+1^_9_c^+1^_8_z^+1^_9_y^+1^_9_c^+1^_9_c^+1^_9_z^+1^_10_y^+1^_10_c^+1^_10_

0580115917392318

Fragment Matches Table

Show background peaks

| Position | Ion type | Intensity | mz Theoretical | mz Error (Th) | mz Error (ppm) | Charge | Series Number |
| --- | --- | --- | --- | --- | --- | --- | --- |
| - | - | 464.2 | 123.1 | - | - | 0 | - |
| - | - | 537.4 | 129.1 | - | - | 0 | - |
| - | - | 436.1 | 134 | - | - | 0 | - |
| - | - | 395.2 | 144.6 | - | - | 0 | - |
| - | - | 409 | 145.6 | - | - | 0 | - |
| - | - | 432.2 | 150.5 | - | - | 0 | - |
| - | - | 466 | 152.7 | - | - | 0 | - |
| - | - | 488 | 160.1 | - | - | 0 | - |
| - | - | 1.818E+04 | 173.1 | - | - | 0 | - |
| - | - | 2157 | 173.1 | - | - | 0 | - |
| - | - | 1243 | 174.1 | - | - | 0 | - |
| 10 | y | 1011 | 175.1 | 0.0004648 | 2.654 | +1 | 2 |
| - | - | 447.4 | 182.8 | - | - | 0 | - |
| - | - | 1355 | 183.1 | - | - | 0 | - |
| - | - | 1794 | 187.1 | - | - | 0 | - |
| - | - | 6657 | 201.1 | - | - | 0 | - |
| - | - | 692.9 | 202.1 | - | - | 0 | - |
| - | - | 456.3 | 203.9 | - | - | 0 | - |
| - | - | 487.3 | 212 | - | - | 0 | - |
| - | - | 2283 | 215.1 | - | - | 0 | - |
| - | - | 493.3 | 216.1 | - | - | 0 | - |
| - | - | 929.1 | 217.1 | - | - | 0 | - |
| - | - | 561.6 | 217.4 | - | - | 0 | - |
| - | - | 546.2 | 224.3 | - | - | 0 | - |
| 9 | w | 569.2 | 229.1 | 0.0001499 | 0.6543 | +1 | 3 |
| - | - | 6353 | 239.1 | - | - | 0 | - |
| - | - | 1.24E+04 | 260.1 | - | - | 0 | - |
| - | - | 940.8 | 261.1 | - | - | 0 | - |
| - | - | 975.3 | 270.1 | - | - | 0 | - |
| - | - | 1377 | 288.2 | - | - | 0 | - |
| - | - | 903.2 | 296.2 | - | - | 0 | - |
| - | - | 4397 | 312.2 | - | - | 0 | - |
| - | - | 1265 | 314.2 | - | - | 0 | - |
| 8 | w | 1162 | 316.2 | 0.0006586 | 2.083 | +1 | 4 |
| - | - | 552.3 | 322 | - | - | 0 | - |
| - | - | 2066 | 324.1 | - | - | 0 | - |
| - | - | 3817 | 330.2 | - | - | 0 | - |
| - | - | 634 | 331.2 | - | - | 0 | - |
| - | - | 3924 | 334.2 | - | - | 0 | - |
| - | - | 1062 | 335.2 | - | - | 0 | - |
| - | - | 627 | 338.2 | - | - | 0 | - |
| - | - | 1565 | 342.2 | - | - | 0 | - |
| - | - | 528.5 | 346.9 | - | - | 0 | - |
| 8 | y | 2029 | 359.2 | 0.0004029 | 1.122 | +1 | 4 |
| - | - | 1172 | 368.2 | - | - | 0 | - |
| - | - | 2746 | 373.2 | - | - | 0 | - |
| - | - | 1.183E+04 | 383.2 | - | - | 0 | - |
| - | - | 2161 | 384.2 | - | - | 0 | - |
| - | - | 8839 | 386.2 | - | - | 0 | - |
| - | - | 1597 | 387.2 | - | - | 0 | - |
| - | - | 2.629E+04 | 401.2 | - | - | 0 | - |
| - | - | 4870 | 402.2 | - | - | 0 | - |
| - | - | 2487 | 429.2 | - | - | 0 | - |
| - | - | 669.8 | 430.2 | - | - | 0 | - |
| - | - | 629.1 | 460.2 | - | - | 0 | - |
| 3 | y | 1057 | 473.2 | 0.001714 | 3.621 | +2 | 9 |
| 7 | z | 9453 | 499.3 | 0.0006147 | 1.231 | +1 | 5 |
| - | - | 5157 | 500.3 | - | - | 0 | - |
| - | - | 689 | 501.3 | - | - | 0 | - |
| 2 | y | 739.1 | 507.7 | 0.00206 | 4.058 | +2 | 10 |
| 10 | c | 641.8 | 514.8 | 0.0008216 | 1.596 | +2 | 10 |
| 7 | y | 2519 | 515.3 | 0.0002622 | 0.5089 | +1 | 5 |
| 2 | y | 1712 | 516.8 | 1.215E-05 | 0.02351 | +2 | 10 |
| - | - | 819.1 | 537.3 | - | - | 0 | - |
| - | - | 4403 | 554.3 | - | - | 0 | - |
| - | - | 701.4 | 554.8 | - | - | 0 | - |
| - | - | 1734 | 555.3 | - | - | 0 | - |
| - | - | 2153 | 555.3 | - | - | 0 | - |
| - | - | 1019 | 556.3 | - | - | 0 | - |
| - | - | 1033 | 564.3 | - | - | 0 | - |
| - | - | 1018 | 564.8 | - | - | 0 | - |
| - | - | 610.4 | 571.3 | - | - | 0 | - |
| - | - | 3.129E+04 | 572.3 | - | - | 0 | - |
| - | - | 1.038E+04 | 573.3 | - | - | 0 | - |
| - | - | 5.614E+04 | 573.3 | - | - | 0 | - |
| - | - | 2527 | 573.8 | - | - | 0 | - |
| - | - | 2674 | 574.3 | - | - | 0 | - |
| - | - | 1.766E+04 | 574.3 | - | - | 0 | - |
| 6 | z | 4540 | 614.3 | 0.0005271 | 0.8581 | +1 | 6 |
| - | - | 6958 | 615.3 | - | - | 0 | - |
| - | - | 2504 | 616.3 | - | - | 0 | - |
| - | - | 1079 | 617.3 | - | - | 0 | - |
| 6 | c | 7354 | 630.3 | 0.01068 | 16.94 | +1 | 6 |
| - | - | 2403 | 631.3 | - | - | 0 | - |
| - | - | 645.3 | 676.4 | - | - | 0 | - |
| 5 | y | 1050 | 699.3 | 0.01396 | 19.97 | +1 | 7 |
| 5 | y | 1639 | 700.3 | 0.001165 | 1.664 | +1 | 7 |
| 5 | z | 3024 | 701.3 | 0.001594 | 2.273 | +1 | 7 |
| - | - | 7308 | 702.3 | - | - | 0 | - |
| - | - | 2245 | 703.3 | - | - | 0 | - |
| - | - | 1591 | 716.4 | - | - | 0 | - |
| 5 | y | 1.66E+04 | 717.4 | 0.0004817 | 0.6715 | +1 | 7 |
| - | - | 6765 | 718.4 | - | - | 0 | - |
| - | - | 926.8 | 719.4 | - | - | 0 | - |
| - | - | 611.9 | 749.8 | - | - | 0 | - |
| - | - | 634.4 | 756.4 | - | - | 0 | - |
| - | - | 8021 | 760.4 | - | - | 0 | - |
| - | - | 4032 | 761.4 | - | - | 0 | - |
| - | - | 894.5 | 777.3 | - | - | 0 | - |
| 7 | c | 1462 | 787.4 | 0.001941 | 2.465 | +1 | 7 |
| 4 | y | 1895 | 798.4 | 0.003193 | 3.999 | +1 | 8 |
| 4 | z | 9248 | 800.4 | 0.0001825 | 0.228 | +1 | 8 |
| - | - | 5303 | 801.4 | - | - | 0 | - |
| - | - | 1607 | 802.4 | - | - | 0 | - |
| - | - | 2109 | 815.4 | - | - | 0 | - |
| 4 | y | 9263 | 816.4 | 0.0005624 | 0.6889 | +1 | 8 |
| - | - | 3081 | 817.4 | - | - | 0 | - |
| - | - | 816.6 | 818.4 | - | - | 0 | - |
| - | - | 1832 | 857.5 | - | - | 0 | - |
| - | - | 973.5 | 858.4 | - | - | 0 | - |
| - | - | 1118 | 858.5 | - | - | 0 | - |
| - | - | 670.8 | 859.4 | - | - | 0 | - |
| 3 | w | 3547 | 870.4 | 0.0001295 | 0.1488 | +1 | 9 |
| - | - | 1982 | 871.4 | - | - | 0 | - |
| - | - | 844.9 | 885.5 | - | - | 0 | - |
| - | - | 828.5 | 886.5 | - | - | 0 | - |
| - | - | 1116 | 900.5 | - | - | 0 | - |
| 8 | c | 7554 | 901.5 | 0.001115 | 1.237 | +1 | 8 |
| - | - | 3070 | 902.5 | - | - | 0 | - |
| - | - | 1054 | 903.5 | - | - | 0 | - |
| - | - | 1147 | 911.5 | - | - | 0 | - |
| - | - | 1804 | 926.5 | - | - | 0 | - |
| - | - | 860.1 | 927.5 | - | - | 0 | - |
| 3 | z | 1.155E+04 | 929.4 | 0.0008457 | 0.9099 | +1 | 9 |
| - | - | 4864 | 930.4 | - | - | 0 | - |
| - | - | 2373 | 931.4 | - | - | 0 | - |
| - | - | 3169 | 944.5 | - | - | 0 | - |
| 3 | y | 9800 | 945.5 | 0.002708 | 2.864 | +1 | 9 |
| - | - | 4480 | 946.5 | - | - | 0 | - |
| - | - | 629.4 | 947.5 | - | - | 0 | - |
| 9 | c | 737.8 | 971.5 | 0.004453 | 4.584 | +1 | 9 |
| 9 | c | 5559 | 988.5 | 0.002016 | 2.039 | +1 | 9 |
| - | - | 3218 | 989.5 | - | - | 0 | - |
| - | - | 959.6 | 990.5 | - | - | 0 | - |
| - | - | 4077 | 1002 | - | - | 0 | - |
| - | - | 1459 | 1003 | - | - | 0 | - |
| - | - | 823.1 | 1004 | - | - | 0 | - |
| 2 | z | 7478 | 1016 | 0.001502 | 1.478 | +1 | 10 |
| - | - | 4093 | 1017 | - | - | 0 | - |
| - | - | 1512 | 1018 | - | - | 0 | - |
| 2 | y | 7276 | 1032 | 0.0001455 | 0.141 | +1 | 10 |
| - | - | 2516 | 1033 | - | - | 0 | - |
| - | - | 942.9 | 1034 | - | - | 0 | - |
| 10 | c | 1.607E+04 | 1046 | 0.001995 | 1.908 | +1 | 10 |
| - | - | 1.104E+04 | 1047 | - | - | 0 | - |
| - | - | 1959 | 1048 | - | - | 0 | - |
| - | - | 1973 | 1057 | - | - | 0 | - |
| - | - | 1631 | 1058 | - | - | 0 | - |
| - | - | 620.7 | 1064 | - | - | 0 | - |
| - | - | 1531 | 1087 | - | - | 0 | - |
| - | - | 1724 | 1088 | - | - | 0 | - |
| - | - | 1159 | 1089 | - | - | 0 | - |
| - | - | 2840 | 1101 | - | - | 0 | - |
| - | - | 1053 | 1102 | - | - | 0 | - |
| - | - | 1265 | 1103 | - | - | 0 | - |
| - | - | 2071 | 1104 | - | - | 0 | - |
| - | - | 888.8 | 1105 | - | - | 0 | - |
| - | - | 718 | 1118 | - | - | 0 | - |
| - | - | 823.2 | 1126 | - | - | 0 | - |
| - | - | 6096 | 1129 | - | - | 0 | - |
| - | - | 4.207E+04 | 1130 | - | - | 0 | - |
| - | - | 2.481E+04 | 1131 | - | - | 0 | - |
| - | - | 7156 | 1132 | - | - | 0 | - |
| - | - | 1287 | 1133 | - | - | 0 | - |
| - | - | 737.2 | 1144 | - | - | 0 | - |
| - | - | 1577 | 1145 | - | - | 0 | - |
| - | - | 1.882E+04 | 1146 | - | - | 0 | - |
| - | - | 4.27E+04 | 1147 | - | - | 0 | - |
| - | - | 1361 | 1147 | - | - | 0 | - |
| - | - | 2.513E+04 | 1148 | - | - | 0 | - |
| - | - | 1810 | 1148 | - | - | 0 | - |
| - | - | 7141 | 1149 | - | - | 0 | - |
| - | - | 1776 | 1150 | - | - | 0 | - |
| - | - | 876.8 | 1659 | - | - | 0 | - |
| - | - | 892.9 | 1701 | - | - | 0 | - |
| - | - | 791.3 | 1705 | - | - | 0 | - |
| - | - | 784.3 | 1716 | - | - | 0 | - |
| - | - | 2360 | 1717 | - | - | 0 | - |
| - | - | 1793 | 1718 | - | - | 0 | - |
| - | - | 1489 | 1719 | - | - | 0 | - |
| - | - | 815.8 | 1720 | - | - | 0 | - |
| - | - | 1063 | 1721 | - | - | 0 | - |
| - | - | 1821 | 1722 | - | - | 0 | - |
| - | - | 692.2 | 2004 | - | - | 0 | - |
| - | - | 782.7 | 2294 | - | - | 0 | - |
| - | - | 961.3 | 2295 | - | - | 0 | - |

m/z Charge Intensity FragmentType MassShift Position
123.13774871826172 0 464.20135
129.1021728515625 0 537.427
133.95748901367188 0 436.0565
144.56954956054688 0 395.17844
145.6404571533203 0 409.03616
150.50936889648438 0 432.1879
152.6637725830078 0 465.95572
160.07582092285156 0 488.0275
173.09230041503906 0 18178.656
173.1287078857422 0 2157.3267
174.0956268310547 0 1243.3392
175.1072540283203 0 1011.42084 y 9
182.8031463623047 0 447.41196
183.11312866210938 0 1354.7303
187.1078338623047 0 1794.4526
201.1235809326172 0 6657.0996
202.12667846679688 0 692.8853
203.8563232421875 0 456.33383
211.98875427246094 0 487.34708
215.1392059326172 0 2282.8916
216.0992431640625 0 493.2789
217.08206176757812 0 929.0522
217.3632049560547 0 561.6366
224.33462524414062 0 546.1921
229.11813354492188 0 569.15375 w 8
239.11407470703125 0 6352.5054
260.1242370605469 0 12404.065
261.1280517578125 0 940.8388
270.1447448730469 0 975.2935
288.155029296875 0 1377.258
296.19683837890625 0 903.20215
312.1556396484375 0 4396.6157
314.20721435546875 0 1264.576
316.1509704589844 0 1162.123 w 7
322.019287109375 0 552.33417
324.1459655761719 0 2065.929
330.16656494140625 0 3816.7437
331.1700134277344 0 633.95435
334.1763610839844 0 3923.6377
335.179931640625 0 1062.3009
338.2070617675781 0 626.95874
342.15655517578125 0 1564.5841
346.87921142578125 0 528.4905
359.1921081542969 0 2029.4883 y 7
368.1726379394531 0 1171.8959
373.244384765625 0 2745.8757
383.2291259765625 0 11827.3125
384.23211669921875 0 2161.113
386.1824951171875 0 8838.922
387.1859436035156 0 1596.6455
401.2397155761719 0 26285.361
402.2429504394531 0 4869.7866
429.23486328125 0 2486.5715
430.2352600097656 0 669.8489
460.23736572265625 0 629.1261
473.2371520996094 0 1057.1661 y 2
499.2755126953125 0 9453.04 z 6
500.28106689453125 0 5157.323
501.28338623046875 0 688.98956
507.7441101074219 0 739.0998 y Water loss 1
514.753173828125 0 641.81744 c Ammonia loss 9
515.2938842773438 0 2518.7273 y 6
516.75146484375 0 1711.5417 y 1
537.3046875 0 819.0717
554.2726440429688 0 4402.907
554.8192138671875 0 701.4135
555.2750854492188 0 1733.5688
555.31494140625 0 2152.6445
556.317138671875 0 1019.4404
564.2894287109375 0 1033.0896
564.7881469726562 0 1017.93964
571.3140869140625 0 610.3838
572.282958984375 0 31294.293
573.2861938476562 0 10378.868
573.324462890625 0 56143.695
573.794921875 0 2527.0684
574.2905883789062 0 2673.833
574.3271484375 0 17661.246
614.3023681640625 0 4539.7793 z 5
615.308837890625 0 6957.6855
616.30517578125 0 2504.0044
617.312255859375 0 1078.5199
630.3200073242188 0 7354.1006 c Water loss 5
631.3231201171875 0 2402.8345
676.4107055664062 0 645.33215
699.3280639648438 0 1049.7905 y Water loss 4
700.3272094726562 0 1639.4893 y Ammonia loss 4
701.332275390625 0 3024.1987 z 4
702.3406372070312 0 7307.9062
703.344970703125 0 2244.8032
716.35791015625 0 1591.0385
717.3521118164062 0 16596.11 y 4
718.3556518554688 0 6764.6646
719.3560791015625 0 926.818
749.8251342773438 0 611.86646
756.4108276367188 0 634.39825
760.4074096679688 0 8020.807
761.4100952148438 0 4032.1084
777.346923828125 0 894.4623
787.3925170898438 0 1462.0107 c Ammonia loss 6
798.4136352539062 0 1894.7095 y Water loss 3
800.4024658203125 0 9248.398 z 3
801.4069213867188 0 5303.0864
802.4110717773438 0 1607.1229
815.4329223632812 0 2108.502
816.4215698242188 0 9262.505 y 3
817.4234619140625 0 3080.5068
818.4257202148438 0 816.63995
857.4602661132812 0 1831.5886
858.38232421875 0 973.5276
858.4690551757812 0 1117.6992
859.377197265625 0 670.8304
870.4317016601562 0 3546.9927 w 2
871.4328002929688 0 1981.8141
885.4547729492188 0 844.90045
886.4578857421875 0 828.50543
900.4788208007812 0 1115.9955
901.47265625 0 7554.013 c 7
902.4767456054688 0 3069.935
903.4820556640625 0 1054.2638
911.4930419921875 0 1147.2467
926.5153198242188 0 1804.0323
927.5200805664062 0 860.1429
929.4440307617188 0 11546.476 z 2
930.4478149414062 0 4863.5615
931.4481201171875 0 2372.817
944.4866333007812 0 3169.28
945.46630859375 0 9800.474 y 2
946.4683837890625 0 4480.3623
947.4644165039062 0 629.4476
971.4837036132812 0 737.75134 c Ammonia loss 8
988.5037841796875 0 5559.0176 c 8
989.5074462890625 0 3218.3118
990.505859375 0 959.6402
1001.5118408203125 0 4077.2812
1002.5149536132812 0 1459.3105
1003.5202026367188 0 823.11774
1016.4754028320312 0 7478.179 z 1
1017.4789428710938 0 4093.3857
1018.4848022460938 0 1512.3506
1032.4954833984375 0 7276.3457 y 1
1033.498046875 0 2515.6504
1034.492919921875 0 942.8739
1045.5252685546875 0 16074.735 c 9
1046.5277099609375 0 11035.079
1047.531005859375 0 1958.7018
1056.5782470703125 0 1973.0695
1057.5811767578125 0 1630.9188
1064.484619140625 0 620.7325
1086.561767578125 0 1530.8127
1087.5482177734375 0 1724.0941
1088.5491943359375 0 1158.7797
1100.5723876953125 0 2839.7705
1101.5623779296875 0 1053.0515
1102.54052734375 0 1265.44
1103.558837890625 0 2070.6934
1104.5582275390625 0 888.78125
1117.6070556640625 0 718.0172
1126.49755859375 0 823.20404
1128.5751953125 0 6095.5215
1129.560546875 0 42072.844
1130.563232421875 0 24806.152
1131.5643310546875 0 7156.157
1132.5650634765625 0 1286.6952
1143.54833984375 0 737.15607
1144.5950927734375 0 1576.6512
1145.5810546875 0 18821.246
1146.5848388671875 0 42697.703
1147.06982421875 0 1360.998
1147.5863037109375 0 25133.459
1148.06298828125 0 1810.1881
1148.58837890625 0 7140.617
1149.5794677734375 0 1776.2513
1658.759765625 0 876.7554
1700.740966796875 0 892.8875
1704.8057861328125 0 791.2693
1715.751953125 0 784.31635
1716.7685546875 0 2359.7612
1717.7706298828125 0 1793.1292
1718.766357421875 0 1489.025
1719.820068359375 0 815.84814
1720.8138427734375 0 1062.8693
1721.8004150390625 0 1820.7345
2003.9832763671875 0 692.2133
2294.11083984375 0 782.7292
2295.116455078125 0 961.26917

Spectrum Details

|  |  |
| --- | --- |
| Matched peaks? Matched peaks The total absolute number of peaks matched. Additionally in brackets the total fraction of peaks matched and the total number of peaks is shown. | 29 (15.59% of 186) |
| FDR? FDR The false discovery rate estimated for this peptide. It is calculated by matching all theoretical fragments with a non-integer shift with the raw peaks for this spectrum. This is done with 40 different shifts. The resulting percentage is the average number of annotated peaks over the number of annotated peaks with the correct spectrum. | 0.33% |
| Satellite FDR? Satellite FDR See the FDR for details on its calculation. This satellite ion specific FDR only contains the satellite ions (d/w) for I/L/J positions. | - |
| PSM Score? PSM Score The PSM Score as given by Hecklib to this annotated spectrum. It is shown with three significant figures. | 263 |

#### Reverse Lookup? Reverse Lookup All places where this read could be placed.

| Group | Segment | Template | Template Part | Read Part | Score | Unique |
| --- | --- | --- | --- | --- | --- | --- |
| Homo sapiens Light Chain | IGLV | IGLV2-14 | [49..60] | [0..11] | 74 | False |
| Homo sapiens Light Chain | IGLV | IGLV2-18 | [49..60] | [0..11] | 74 | False |
| Homo sapiens Light Chain | IGLV | IGLV2-8 | [49..60] | [0..11] | 70 | False |

| Recombined | Template Part | Read Part | Score | Unique |
| --- | --- | --- | --- | --- |
| REC-0-1\_002 | [49..60] | [0..11] | 88 | True |

#### Meta Information from Multiple reads

##### Number of combined reads

6

##### Intensity

0.8876

##### TotalArea

3.077E+08

##### Changes to the peptide sequence

JSEVSDRPSGV

L→JNo support for either Leucine or Isoleucine based on side chain ions (Position: 1)

#### Positional Score

Copy Data

##### Positional Score (TSV)

###### Preview

```
Loading example...
```

*Click on the button to copy the data to your clipboard.*

10012345678910

Label Value
"0" 0.65
"1" 0.637
"2" 0.66
"3" 0.658
"4" 0.66
"5" 0.658
"6" 0.642
"7" 0.632
"8" 0.642
"9" 0.62
"10" 0.637

#### Meta Information from PEAKS

##### Scan Identifier

F1:3712

##### Original sequence

L

S

E

V

S

D

R

P

S

G

V

##### Posttranslational Modifications

##### Source File

D:\separate\_stitch\_analyses\xle-disambiguation\raw\20210323\_F1\_UM1\_Peng0013\_SA\_F59\_ingel\_3ug\_ELA.raw

##### Fraction

1

##### Scan Feature

F1:7258

##### De Novo Score

98

##### ConfidenceScore

98

### m/z

573.2948

##### Mass

1144.5723

##### Charge

2

##### Retention Time

19.27

##### Predicted Retention Time

-

##### Area

3.058E+08

##### Parts Per Million

2.4

##### Fragmentation mode

HCD

##### Originating file

01 D:\separate\_stitch\_analyses\xle-disambiguation\20210325\_F59\_3ug\_DENOVO\_12.csv

#### Meta Information from PEAKS

##### Scan Identifier

F1:3800

##### Original sequence

L

S

E

V

S

D

R

P

S

G

V

##### Posttranslational Modifications

##### Source File

D:\separate\_stitch\_analyses\xle-disambiguation\raw\20210323\_F1\_UM1\_Peng0013\_SA\_F59\_ingel\_3ug\_ELA.raw

##### Fraction

1

##### Scan Feature

-

##### De Novo Score

98

##### ConfidenceScore

98

### m/z

573.2946

##### Mass

1144.5723

##### Charge

2

##### Retention Time

19.84

##### Predicted Retention Time

-

##### Area

0

##### Parts Per Million

2

##### Fragmentation mode

ETHCD

##### Originating file

01 D:\separate\_stitch\_analyses\xle-disambiguation\20210325\_F59\_3ug\_DENOVO\_12.csv

#### Meta Information from PEAKS

##### Scan Identifier

F1:3858

##### Original sequence

L

S

E

V

S

D

R

P

S

G

V

##### Posttranslational Modifications

##### Source File

D:\separate\_stitch\_analyses\xle-disambiguation\raw\20210323\_F1\_UM1\_Peng0013\_SA\_F59\_ingel\_3ug\_ELA.raw

##### Fraction

1

##### Scan Feature

-

##### De Novo Score

98

##### ConfidenceScore

98

### m/z

573.2943

##### Mass

1144.5723

##### Charge

2

##### Retention Time

20.18

##### Predicted Retention Time

-

##### Area

0

##### Parts Per Million

1.6

##### Fragmentation mode

ETHCD

##### Originating file

01 D:\separate\_stitch\_analyses\xle-disambiguation\20210325\_F59\_3ug\_DENOVO\_12.csv

#### Meta Information from PEAKS

##### Scan Identifier

F1:4054

##### Original sequence

L

S

E

V

S

D

R

P

S

G

V

##### Posttranslational Modifications

##### Source File

D:\separate\_stitch\_analyses\xle-disambiguation\raw\20210323\_F1\_UM1\_Peng0013\_SA\_F59\_ingel\_3ug\_ELA.raw

##### Fraction

1

##### Scan Feature

-

##### De Novo Score

97

##### ConfidenceScore

97

### m/z

573.2946

##### Mass

1144.5723

##### Charge

2

##### Retention Time

21.3

##### Predicted Retention Time

-

##### Area

0

##### Parts Per Million

2.1

##### Fragmentation mode

ETHCD

##### Originating file

01 D:\separate\_stitch\_analyses\xle-disambiguation\20210325\_F59\_3ug\_DENOVO\_12.csv

#### Meta Information from PEAKS

##### Scan Identifier

F1:3922

##### Original sequence

L

S

E

V

S

D

R

P

S

G

V

##### Posttranslational Modifications

##### Source File

D:\separate\_stitch\_analyses\xle-disambiguation\raw\20210323\_F1\_UM1\_Peng0013\_SA\_F59\_ingel\_3ug\_ELA.raw

##### Fraction

1

##### Scan Feature

-

##### De Novo Score

96

##### ConfidenceScore

96

### m/z

573.2944

##### Mass

1144.5723

##### Charge

2

##### Retention Time

20.56

##### Predicted Retention Time

-

##### Area

0

##### Parts Per Million

1.8

##### Fragmentation mode

HCD

##### Originating file

01 D:\separate\_stitch\_analyses\xle-disambiguation\20210325\_F59\_3ug\_DENOVO\_12.csv

#### Meta Information from PEAKS

##### Scan Identifier

F1:4123

##### Original sequence

L

S

E

V

S

D

R

P

S

G

V

##### Posttranslational Modifications

##### Source File

D:\separate\_stitch\_analyses\xle-disambiguation\raw\20210323\_F1\_UM1\_Peng0013\_SA\_F59\_ingel\_3ug\_ELA.raw

##### Fraction

1

##### Scan Feature

F1:7254

##### De Novo Score

95

##### ConfidenceScore

95

### m/z

573.2928

##### Mass

1144.5723

##### Charge

2

##### Retention Time

21.79

##### Predicted Retention Time

-

##### Area

1.892E+06

##### Fragmentation mode

ETHCD

##### Originating file

01 D:\separate\_stitch\_analyses\xle-disambiguation\20210325\_F59\_3ug\_DENOVO\_12.csv
