## Supplementary material for "A handle on mass coincidence errors in *de novo* sequencing of antibodies by bottom-up proteomics": Combined_073.html

Details Combined\_073 | Stitch OverviewUndefined

### Read Combined\_073

#### Sequence (length=10)

JTSGVHTFPA

#### Spectrum 5220? Spectrum 5220 The raw spectrum of this peptide as annotated by Hecklib. The fragments are coloured according to ion type (see legend). Any peaks with a star '\*' as text can be hovered over to see the full details, first the ion type second the mass shift type. By hovering over the amino acids in the peptide or ions in the legend the corresponding peaks are highlighted. By toggling the 'Unassigned' label you can turn the background (unassigned) peaks on or off in the plot. By updating the slider in the Ion legend you can update the spectrum to only show the top X% of the peaks with labels. The top X% means any peak that is within X% of the highest intensity. By dragging in the spectrum you can zoom in to a specific part of the spectrum and use 'Zoom Out' to get back to the original zoom level. The annotation of the spectrum is based on the given sequence in the peptides file and is done with different software so inconsistencies are likely. The peaks are annotated based on the given sequence, with 20 ppm tolerance.

Copy Data

##### Spectrum 5220 (TSV)

###### Preview

```
Loading example...
```

*Click on the button to copy the data to your clipboard.*

Mz MinMz MaxIntensity Max

WidthHeightPeptide font sizePeptide stroke widthSpectrum font sizeSpectrum stroke widthCompact peptide

Ion legend

wxyz

abcd

OtherUnassignedIon^Charge^_Position_Show for top:%

JTSGVHTFPA

01.83e+63.66e+65.49e+67.32e+6

Zoom Out

y^+1^_2_c^+1^_2_y^+1^_3_c^+1^_4_y^+2^_8_y^+2^_8_c^+2^_8_y^+1^_4_y^+2^_9_c^+1^_5_y^+2^_9_c^+2^_9_c^+1^_5_z^+1^_5_y^+1^_5_z^+1^_5_y^+1^_5_c^+1^_6_c^+1^_6_c^+1^_6_z^+1^_6_w^+1^_6_z^+1^_6_y^+1^_6_z^+1^_7_c^+1^_7_c^+1^_7_y^+1^_7_z^+1^_7_c^+1^_7_y^+1^_7_z^+1^_8_w^+1^_8_y^+1^_8_z^+1^_8_y^+1^_8_c^+1^_8_z^+1^_9_w^+1^_9_z^+1^_9_y^+1^_9_c^+1^_9_c^+1^_9_

0606121218182424

Fragment Matches Table

Show background peaks

| Position | Ion type | Intensity | mz Theoretical | mz Error (Th) | mz Error (ppm) | Charge | Series Number |
| --- | --- | --- | --- | --- | --- | --- | --- |
| - | - | 7613 | 120.1 | - | - | 0 | - |
| - | - | 3761 | 122.4 | - | - | 0 | - |
| - | - | 1.639E+04 | 125.1 | - | - | 0 | - |
| - | - | 3428 | 127.6 | - | - | 0 | - |
| - | - | 4587 | 130.3 | - | - | 0 | - |
| - | - | 3994 | 131.1 | - | - | 0 | - |
| - | - | 7.871E+04 | 140.1 | - | - | 0 | - |
| - | - | 5112 | 141.1 | - | - | 0 | - |
| - | - | 6.521E+04 | 142.1 | - | - | 0 | - |
| - | - | 4285 | 148.6 | - | - | 0 | - |
| - | - | 3.475E+04 | 152.1 | - | - | 0 | - |
| - | - | 5302 | 153.1 | - | - | 0 | - |
| - | - | 2.564E+04 | 155.1 | - | - | 0 | - |
| - | - | 4.827E+04 | 169.1 | - | - | 0 | - |
| - | - | 4231 | 171.2 | - | - | 0 | - |
| - | - | 1.066E+04 | 173.4 | - | - | 0 | - |
| - | - | 4584 | 178.1 | - | - | 0 | - |
| - | - | 5296 | 186.3 | - | - | 0 | - |
| 9 | y | 1.163E+06 | 187.1 | 0.0003134 | 1.675 | +1 | 2 |
| - | - | 4.899E+05 | 187.1 | - | - | 0 | - |
| - | - | 8.431E+04 | 188.1 | - | - | 0 | - |
| - | - | 3.822E+04 | 188.1 | - | - | 0 | - |
| - | - | 5824 | 189.1 | - | - | 0 | - |
| - | - | 6021 | 189.1 | - | - | 0 | - |
| - | - | 7919 | 194.1 | - | - | 0 | - |
| - | - | 1.892E+04 | 195.1 | - | - | 0 | - |
| - | - | 6.091E+04 | 197.1 | - | - | 0 | - |
| 2 | c | 5.207E+05 | 215.1 | 0.0002938 | 1.366 | +1 | 2 |
| - | - | 5.277E+04 | 216.1 | - | - | 0 | - |
| - | - | 7601 | 224.1 | - | - | 0 | - |
| - | - | 9423 | 239.1 | - | - | 0 | - |
| - | - | 5941 | 239.2 | - | - | 0 | - |
| - | - | 5232 | 249.8 | - | - | 0 | - |
| - | - | 9645 | 251.2 | - | - | 0 | - |
| - | - | 4.542E+04 | 252.2 | - | - | 0 | - |
| - | - | 4.842E+04 | 284.2 | - | - | 0 | - |
| - | - | 7002 | 285.2 | - | - | 0 | - |
| - | - | 1.205E+04 | 295.2 | - | - | 0 | - |
| - | - | 1.128E+05 | 296.2 | - | - | 0 | - |
| - | - | 1.549E+04 | 297.2 | - | - | 0 | - |
| - | - | 8256 | 308.2 | - | - | 0 | - |
| - | - | 5418 | 323.6 | - | - | 0 | - |
| - | - | 5545 | 323.7 | - | - | 0 | - |
| - | - | 6787 | 325.2 | - | - | 0 | - |
| - | - | 5244 | 325.9 | - | - | 0 | - |
| 8 | y | 6.063E+04 | 334.2 | 0.0003199 | 0.9574 | +1 | 3 |
| - | - | 1.408E+04 | 335.2 | - | - | 0 | - |
| - | - | 1.269E+05 | 339.2 | - | - | 0 | - |
| - | - | 1.937E+04 | 340.2 | - | - | 0 | - |
| - | - | 6918 | 341.2 | - | - | 0 | - |
| - | - | 9805 | 343.2 | - | - | 0 | - |
| - | - | 8158 | 352.2 | - | - | 0 | - |
| - | - | 6286 | 352.2 | - | - | 0 | - |
| - | - | 7089 | 353.2 | - | - | 0 | - |
| - | - | 1.081E+04 | 354.7 | - | - | 0 | - |
| 4 | c | 5795 | 359.2 | 0.001074 | 2.991 | +1 | 4 |
| - | - | 7023 | 363.7 | - | - | 0 | - |
| - | - | 7361 | 365.7 | - | - | 0 | - |
| - | - | 6145 | 368.2 | - | - | 0 | - |
| - | - | 1.715E+04 | 371.2 | - | - | 0 | - |
| - | - | 6026 | 380.2 | - | - | 0 | - |
| - | - | 6.96E+04 | 381.2 | - | - | 0 | - |
| - | - | 2.063E+04 | 382.2 | - | - | 0 | - |
| - | - | 8.633E+04 | 383.2 | - | - | 0 | - |
| - | - | 1.703E+04 | 384.2 | - | - | 0 | - |
| - | - | 2.089E+04 | 386.2 | - | - | 0 | - |
| - | - | 8261 | 388.2 | - | - | 0 | - |
| - | - | 8058 | 395.2 | - | - | 0 | - |
| - | - | 1.196E+04 | 397.2 | - | - | 0 | - |
| - | - | 1.33E+04 | 398.2 | - | - | 0 | - |
| 3 | y | 1.907E+04 | 399.2 | 0.0002615 | 0.655 | +2 | 8 |
| - | - | 6472 | 399.2 | - | - | 0 | - |
| - | - | 1.503E+04 | 399.7 | - | - | 0 | - |
| - | - | 8067 | 400.2 | - | - | 0 | - |
| - | - | 1.067E+04 | 401.3 | - | - | 0 | - |
| - | - | 2.169E+04 | 404.2 | - | - | 0 | - |
| - | - | 1.223E+04 | 404.7 | - | - | 0 | - |
| - | - | 2.145E+04 | 406.2 | - | - | 0 | - |
| 3 | y | 1.739E+04 | 408.2 | 0.002645 | 6.479 | +2 | 8 |
| - | - | 1.847E+04 | 408.2 | - | - | 0 | - |
| - | - | 1.375E+04 | 408.7 | - | - | 0 | - |
| - | - | 6172 | 409.2 | - | - | 0 | - |
| - | - | 1.781E+04 | 413.2 | - | - | 0 | - |
| - | - | 9178 | 413.7 | - | - | 0 | - |
| - | - | 9345 | 414.2 | - | - | 0 | - |
| 8 | c | 3.693E+04 | 422.2 | 0.0002905 | 0.688 | +2 | 8 |
| - | - | 1.253E+04 | 422.7 | - | - | 0 | - |
| - | - | 1.903E+04 | 430.3 | - | - | 0 | - |
| 7 | y | 7894 | 435.2 | 0.003362 | 7.724 | +1 | 4 |
| - | - | 3.101E+04 | 440.2 | - | - | 0 | - |
| - | - | 2.243E+04 | 440.7 | - | - | 0 | - |
| - | - | 9413 | 441.2 | - | - | 0 | - |
| 2 | y | 7275 | 449.7 | 0.001931 | 4.293 | +2 | 9 |
| - | - | 1.537E+04 | 452.7 | - | - | 0 | - |
| - | - | 8679 | 453.2 | - | - | 0 | - |
| - | - | 6641 | 454.7 | - | - | 0 | - |
| - | - | 6765 | 455.2 | - | - | 0 | - |
| - | - | 1.692E+04 | 457.3 | - | - | 0 | - |
| 5 | c | 2.008E+04 | 458.3 | 0.0009152 | 1.997 | +1 | 5 |
| 2 | y | 2.422E+04 | 458.7 | 0.0003105 | 0.6769 | +2 | 9 |
| - | - | 2.158E+04 | 459.2 | - | - | 0 | - |
| - | - | 1.742E+04 | 461.7 | - | - | 0 | - |
| - | - | 1.24E+04 | 462.2 | - | - | 0 | - |
| - | - | 1.674E+04 | 464.2 | - | - | 0 | - |
| - | - | 6158 | 469.2 | - | - | 0 | - |
| 9 | c | 2.307E+04 | 470.7 | 5.995E-05 | 0.1274 | +2 | 9 |
| - | - | 2.045E+04 | 471.2 | - | - | 0 | - |
| - | - | 1.067E+04 | 471.8 | - | - | 0 | - |
| 5 | c | 6212 | 475.3 | 0.001955 | 4.113 | +1 | 5 |
| - | - | 9686 | 481.2 | - | - | 0 | - |
| - | - | 1.293E+05 | 482.2 | - | - | 0 | - |
| - | - | 3.392E+04 | 483.2 | - | - | 0 | - |
| - | - | 2.579E+04 | 484.3 | - | - | 0 | - |
| - | - | 7548 | 485.3 | - | - | 0 | - |
| - | - | 7532 | 488.3 | - | - | 0 | - |
| - | - | 5899 | 488.8 | - | - | 0 | - |
| - | - | 6092 | 489.3 | - | - | 0 | - |
| - | - | 8.272E+04 | 497.3 | - | - | 0 | - |
| - | - | 5.938E+04 | 497.8 | - | - | 0 | - |
| - | - | 1.764E+04 | 498.3 | - | - | 0 | - |
| - | - | 1.048E+05 | 499.3 | - | - | 0 | - |
| - | - | 3.315E+04 | 500.3 | - | - | 0 | - |
| - | - | 9.216E+04 | 506.3 | - | - | 0 | - |
| - | - | 4.153E+04 | 506.8 | - | - | 0 | - |
| - | - | 2.155E+04 | 507.2 | - | - | 0 | - |
| - | - | 8926 | 507.3 | - | - | 0 | - |
| - | - | 1.438E+04 | 508.2 | - | - | 0 | - |
| - | - | 5515 | 509.2 | - | - | 0 | - |
| - | - | 1.088E+05 | 515.3 | - | - | 0 | - |
| - | - | 6.768E+04 | 515.8 | - | - | 0 | - |
| - | - | 2.987E+04 | 516.3 | - | - | 0 | - |
| - | - | 1.702E+04 | 524.3 | - | - | 0 | - |
| - | - | 8339 | 525.3 | - | - | 0 | - |
| - | - | 2.348E+04 | 526.3 | - | - | 0 | - |
| - | - | 3.276E+04 | 527.3 | - | - | 0 | - |
| - | - | 9137 | 528.3 | - | - | 0 | - |
| 6 | z | 2.487E+04 | 538.3 | 0.001143 | 2.124 | +1 | 5 |
| - | - | 2.909E+04 | 539.3 | - | - | 0 | - |
| - | - | 4.354E+04 | 542.3 | - | - | 0 | - |
| - | - | 1.412E+04 | 543.3 | - | - | 0 | - |
| - | - | 5987 | 544 | - | - | 0 | - |
| - | - | 9035 | 548.3 | - | - | 0 | - |
| - | - | 1.068E+04 | 548.8 | - | - | 0 | - |
| - | - | 3.154E+04 | 553.3 | - | - | 0 | - |
| 6 | y | 1.32E+04 | 554.3 | 0.001345 | 2.427 | +1 | 5 |
| - | - | 1.01E+04 | 554.3 | - | - | 0 | - |
| - | - | 1.148E+04 | 555.3 | - | - | 0 | - |
| 6 | z | 1.946E+05 | 556.3 | 2.191E-05 | 0.03939 | +1 | 5 |
| - | - | 1.037E+06 | 557.3 | - | - | 0 | - |
| - | - | 3.546E+05 | 558.3 | - | - | 0 | - |
| - | - | 5.2E+04 | 559.3 | - | - | 0 | - |
| - | - | 2.162E+04 | 567.3 | - | - | 0 | - |
| - | - | 4.343E+04 | 568.3 | - | - | 0 | - |
| - | - | 1.095E+05 | 568.3 | - | - | 0 | - |
| - | - | 1.663E+04 | 569.3 | - | - | 0 | - |
| - | - | 5.702E+04 | 569.3 | - | - | 0 | - |
| - | - | 1.027E+04 | 570.3 | - | - | 0 | - |
| - | - | 4.013E+04 | 571.3 | - | - | 0 | - |
| 6 | y | 1.568E+05 | 572.3 | 0.0002523 | 0.4409 | +1 | 5 |
| - | - | 5.31E+04 | 573.3 | - | - | 0 | - |
| - | - | 7870 | 577.3 | - | - | 0 | - |
| - | - | 2.169E+04 | 583.3 | - | - | 0 | - |
| - | - | 6727 | 584.3 | - | - | 0 | - |
| - | - | 1.717E+04 | 585.3 | - | - | 0 | - |
| - | - | 3.269E+04 | 586.3 | - | - | 0 | - |
| - | - | 1.652E+04 | 587.3 | - | - | 0 | - |
| - | - | 9660 | 593.3 | - | - | 0 | - |
| 6 | c | 1.956E+04 | 594.3 | 0.002752 | 4.631 | +1 | 6 |
| 6 | c | 6.745E+04 | 595.3 | 0.001208 | 2.029 | +1 | 6 |
| - | - | 3.374E+04 | 596.3 | - | - | 0 | - |
| - | - | 2.893E+04 | 597.3 | - | - | 0 | - |
| - | - | 1.028E+04 | 598.3 | - | - | 0 | - |
| - | - | 1.168E+04 | 600.3 | - | - | 0 | - |
| - | - | 2.941E+04 | 601.3 | - | - | 0 | - |
| - | - | 1.541E+04 | 610.3 | - | - | 0 | - |
| - | - | 1.016E+05 | 611.3 | - | - | 0 | - |
| - | - | 5.282E+05 | 611.3 | - | - | 0 | - |
| 6 | c | 1.612E+06 | 612.3 | 0.0003166 | 0.517 | +1 | 6 |
| - | - | 4.926E+05 | 613.3 | - | - | 0 | - |
| - | - | 8.516E+04 | 614.4 | - | - | 0 | - |
| - | - | 1.812E+04 | 626.4 | - | - | 0 | - |
| - | - | 6401 | 627.4 | - | - | 0 | - |
| - | - | 3.046E+05 | 629.3 | - | - | 0 | - |
| - | - | 9.46E+04 | 630.3 | - | - | 0 | - |
| - | - | 2.333E+04 | 631.3 | - | - | 0 | - |
| 5 | z | 9510 | 637.3 | 5.122E-05 | 0.08037 | +1 | 6 |
| 5 | w | 3.798E+04 | 640.3 | 0.0004494 | 0.7019 | +1 | 6 |
| - | - | 8699 | 641.3 | - | - | 0 | - |
| - | - | 4.441E+04 | 651.4 | - | - | 0 | - |
| - | - | 2.003E+04 | 652.4 | - | - | 0 | - |
| 5 | z | 2.507E+05 | 655.3 | 0.0002898 | 0.4422 | +1 | 6 |
| - | - | 3.927E+05 | 656.3 | - | - | 0 | - |
| - | - | 1.154E+05 | 657.3 | - | - | 0 | - |
| - | - | 1.916E+04 | 658.3 | - | - | 0 | - |
| - | - | 1.018E+04 | 669.3 | - | - | 0 | - |
| - | - | 3.929E+04 | 669.4 | - | - | 0 | - |
| - | - | 2.069E+04 | 670.3 | - | - | 0 | - |
| - | - | 9151 | 670.4 | - | - | 0 | - |
| 5 | y | 6.52E+04 | 671.4 | 6.273E-05 | 0.09344 | +1 | 6 |
| - | - | 1.232E+04 | 672.4 | - | - | 0 | - |
| - | - | 3.714E+04 | 678.4 | - | - | 0 | - |
| - | - | 1.473E+04 | 679.4 | - | - | 0 | - |
| - | - | 1.125E+04 | 687.3 | - | - | 0 | - |
| 4 | z | 2.703E+04 | 694.3 | 0.002675 | 3.852 | +1 | 7 |
| - | - | 1.368E+04 | 695.3 | - | - | 0 | - |
| 7 | c | 8929 | 695.4 | 0.01109 | 15.95 | +1 | 7 |
| 7 | c | 7.528E+04 | 696.4 | 0.0003888 | 0.5584 | +1 | 7 |
| - | - | 2.404E+04 | 697.4 | - | - | 0 | - |
| - | - | 1.227E+04 | 698.4 | - | - | 0 | - |
| - | - | 1.37E+04 | 698.9 | - | - | 0 | - |
| - | - | 6649 | 699.4 | - | - | 0 | - |
| - | - | 9869 | 700.9 | - | - | 0 | - |
| 4 | y | 7595 | 710.4 | 0.002832 | 3.987 | +1 | 7 |
| - | - | 1.541E+04 | 711.4 | - | - | 0 | - |
| 4 | z | 1.113E+05 | 712.4 | 0.003291 | 4.619 | +1 | 7 |
| - | - | 1.205E+05 | 712.4 | - | - | 0 | - |
| 7 | c | 1.013E+06 | 713.4 | 0.00173 | 2.426 | +1 | 7 |
| - | - | 4.055E+05 | 714.4 | - | - | 0 | - |
| - | - | 8.35E+04 | 715.4 | - | - | 0 | - |
| - | - | 9295 | 716.4 | - | - | 0 | - |
| - | - | 1.195E+04 | 719.9 | - | - | 0 | - |
| - | - | 7396 | 722.3 | - | - | 0 | - |
| - | - | 1.134E+04 | 726.4 | - | - | 0 | - |
| - | - | 3.382E+04 | 727.4 | - | - | 0 | - |
| 4 | y | 1.663E+05 | 728.4 | 0.0001031 | 0.1416 | +1 | 7 |
| - | - | 6.999E+04 | 729.4 | - | - | 0 | - |
| - | - | 9.489E+04 | 730.4 | - | - | 0 | - |
| - | - | 4.161E+04 | 731.4 | - | - | 0 | - |
| - | - | 8170 | 732.4 | - | - | 0 | - |
| - | - | 9571 | 740.3 | - | - | 0 | - |
| - | - | 1.143E+04 | 765.4 | - | - | 0 | - |
| - | - | 6833 | 765.9 | - | - | 0 | - |
| - | - | 6206 | 771.4 | - | - | 0 | - |
| - | - | 1.492E+04 | 771.9 | - | - | 0 | - |
| - | - | 1.161E+04 | 772.4 | - | - | 0 | - |
| - | - | 8380 | 773.9 | - | - | 0 | - |
| 3 | z | 5.259E+04 | 781.4 | 0.002162 | 2.767 | +1 | 8 |
| 3 | w | 3.882E+04 | 782.4 | 0.00194 | 2.479 | +1 | 8 |
| - | - | 1.026E+04 | 783.4 | - | - | 0 | - |
| - | - | 1.849E+04 | 789.4 | - | - | 0 | - |
| - | - | 8033 | 790.4 | - | - | 0 | - |
| 3 | y | 6.731E+04 | 797.4 | 0.003561 | 4.466 | +1 | 8 |
| - | - | 5.016E+04 | 798.4 | - | - | 0 | - |
| 3 | z | 8.772E+05 | 799.4 | 0.0005696 | 0.7125 | +1 | 8 |
| - | - | 4.414E+05 | 800.4 | - | - | 0 | - |
| - | - | 1.011E+05 | 801.4 | - | - | 0 | - |
| - | - | 1.612E+04 | 802.4 | - | - | 0 | - |
| - | - | 8.035E+04 | 807.4 | - | - | 0 | - |
| - | - | 4.391E+04 | 808.4 | - | - | 0 | - |
| - | - | 7673 | 813.4 | - | - | 0 | - |
| 3 | y | 8.429E+05 | 815.4 | 0.0007664 | 0.9399 | +1 | 8 |
| - | - | 3.841E+05 | 816.4 | - | - | 0 | - |
| - | - | 9.36E+04 | 817.4 | - | - | 0 | - |
| - | - | 2.762E+05 | 825.4 | - | - | 0 | - |
| - | - | 1.103E+05 | 826.4 | - | - | 0 | - |
| - | - | 3.241E+04 | 827.4 | - | - | 0 | - |
| - | - | 8169 | 839.4 | - | - | 0 | - |
| - | - | 9369 | 841.4 | - | - | 0 | - |
| 8 | c | 4.962E+05 | 843.4 | 0.0003213 | 0.3809 | +1 | 8 |
| - | - | 2.243E+05 | 844.4 | - | - | 0 | - |
| - | - | 5.495E+04 | 845.4 | - | - | 0 | - |
| - | - | 2.626E+04 | 880.4 | - | - | 0 | - |
| - | - | 1.707E+04 | 881.4 | - | - | 0 | - |
| 2 | z | 4.896E+04 | 882.4 | 0.001847 | 2.093 | +1 | 9 |
| 2 | w | 3.515E+04 | 883.4 | 0.005917 | 6.698 | +1 | 9 |
| - | - | 1.043E+05 | 898.4 | - | - | 0 | - |
| - | - | 1.738E+05 | 899.4 | - | - | 0 | - |
| 2 | z | 4.222E+05 | 900.4 | 0.001088 | 1.209 | +1 | 9 |
| - | - | 2.002E+05 | 901.4 | - | - | 0 | - |
| - | - | 6.808E+04 | 902.4 | - | - | 0 | - |
| - | - | 1.536E+04 | 913.5 | - | - | 0 | - |
| - | - | 1.076E+04 | 914.5 | - | - | 0 | - |
| 2 | y | 2.948E+05 | 916.5 | 0.0005864 | 0.6398 | +1 | 9 |
| - | - | 1.49E+05 | 917.5 | - | - | 0 | - |
| - | - | 4.707E+04 | 918.5 | - | - | 0 | - |
| - | - | 9941 | 921.5 | - | - | 0 | - |
| - | - | 4.261E+04 | 926.4 | - | - | 0 | - |
| - | - | 2.795E+04 | 927.4 | - | - | 0 | - |
| - | - | 8272 | 938.5 | - | - | 0 | - |
| 9 | c | 1.023E+04 | 939.5 | 0.005898 | 6.278 | +1 | 9 |
| - | - | 1.951E+04 | 955.5 | - | - | 0 | - |
| - | - | 9924 | 956.5 | - | - | 0 | - |
| 9 | c | 1.763E+06 | 957.5 | 0.0003496 | 0.3651 | +1 | 9 |
| - | - | 9.593E+05 | 958.5 | - | - | 0 | - |
| - | - | 2.872E+05 | 959.5 | - | - | 0 | - |
| - | - | 3.834E+04 | 960.5 | - | - | 0 | - |
| - | - | 5.624E+05 | 970.5 | - | - | 0 | - |
| - | - | 3.016E+05 | 971.5 | - | - | 0 | - |
| - | - | 1.061E+05 | 972.5 | - | - | 0 | - |
| - | - | 1.379E+04 | 973.5 | - | - | 0 | - |
| - | - | 2.65E+04 | 974.5 | - | - | 0 | - |
| - | - | 2.305E+04 | 984.5 | - | - | 0 | - |
| - | - | 1.538E+04 | 985.5 | - | - | 0 | - |
| - | - | 9.081E+04 | 986.6 | - | - | 0 | - |
| - | - | 5.443E+04 | 987.6 | - | - | 0 | - |
| - | - | 2.082E+04 | 988.6 | - | - | 0 | - |
| - | - | 5.576E+04 | 995.5 | - | - | 0 | - |
| - | - | 2.349E+04 | 996.5 | - | - | 0 | - |
| - | - | 8936 | 997.5 | - | - | 0 | - |
| - | - | 3.562E+04 | 1012 | - | - | 0 | - |
| - | - | 5.117E+04 | 1013 | - | - | 0 | - |
| - | - | 2.264E+06 | 1014 | - | - | 0 | - |
| - | - | 1.331E+06 | 1015 | - | - | 0 | - |
| - | - | 4.037E+05 | 1016 | - | - | 0 | - |
| - | - | 3.682E+04 | 1017 | - | - | 0 | - |
| - | - | 3.592E+04 | 1028 | - | - | 0 | - |
| - | - | 1.148E+05 | 1029 | - | - | 0 | - |
| - | - | 1.771E+06 | 1030 | - | - | 0 | - |
| - | - | 7.252E+06 | 1031 | - | - | 0 | - |
| - | - | 3.907E+06 | 1032 | - | - | 0 | - |
| - | - | 1.18E+06 | 1033 | - | - | 0 | - |
| - | - | 1.397E+05 | 1034 | - | - | 0 | - |
| - | - | 1.221E+04 | 1046 | - | - | 0 | - |
| - | - | 7851 | 1047 | - | - | 0 | - |
| - | - | 1.036E+04 | 1096 | - | - | 0 | - |
| - | - | 1.17E+04 | 1143 | - | - | 0 | - |
| - | - | 1.304E+04 | 1147 | - | - | 0 | - |
| - | - | 8628 | 1439 | - | - | 0 | - |
| - | - | 2.25E+04 | 1527 | - | - | 0 | - |
| - | - | 1.897E+04 | 1528 | - | - | 0 | - |
| - | - | 1.117E+04 | 1530 | - | - | 0 | - |
| - | - | 2.068E+04 | 1531 | - | - | 0 | - |
| - | - | 1.126E+04 | 1532 | - | - | 0 | - |
| - | - | 9023 | 1542 | - | - | 0 | - |
| - | - | 1.922E+04 | 1544 | - | - | 0 | - |
| - | - | 8852 | 1545 | - | - | 0 | - |
| - | - | 1.85E+04 | 1548 | - | - | 0 | - |
| - | - | 8546 | 1549 | - | - | 0 | - |
| - | - | 7300 | 1582 | - | - | 0 | - |
| - | - | 7132 | 1947 | - | - | 0 | - |
| - | - | 6972 | 2400 | - | - | 0 | - |

m/z Charge Intensity FragmentType MassShift Position
120.08112335205078 0 7613.084
122.36892700195312 0 3761.0837
125.10765838623047 0 16386.44
127.56163024902344 0 3427.5503
130.26683044433594 0 4586.9067
131.1141357421875 0 3994.1182
140.08213806152344 0 78707.26
141.0853271484375 0 5112.078
142.1229248046875 0 65214.773
148.559326171875 0 4284.5283
152.1072998046875 0 34746.44
153.1108856201172 0 5302.409
155.09292602539062 0 25643.62
169.13385009765625 0 48272.938
171.24021911621094 0 4230.9233
173.4388427734375 0 10658.565
178.14915466308594 0 4583.7637
186.33358764648438 0 5295.777
187.1080322265625 0 1162887 y 8
187.1444091796875 0 489852.28
188.11141967773438 0 84308.445
188.1477508544922 0 38221.184
189.08726501464844 0 5824.193
189.11276245117188 0 6021.265
194.12925720214844 0 7919.1655
195.13671875 0 18915.055
197.1287384033203 0 60906.68
215.13931274414062 0 520670.12 c Ammonia loss 1
216.1427764892578 0 52767.105
224.1033477783203 0 7600.825
239.1142120361328 0 9422.503
239.15029907226562 0 5940.9307
249.75308227539062 0 5232.2705
251.15028381347656 0 9645.395
252.1583709716797 0 45422.69
284.16064453125 0 48418.855
285.1632995605469 0 7002.369
295.16357421875 0 12051.15
296.1720886230469 0 112759.08
297.1754455566406 0 15491.464
308.1637268066406 0 8256.326
323.5780334472656 0 5417.526
323.6531677246094 0 5545.267
325.1661682128906 0 6786.971
325.871826171875 0 5244.396
334.17645263671875 0 60633.535 y 7
335.17919921875 0 14076.559
339.1904296875 0 126901.51
340.1935729980469 0 19371.324
341.1816101074219 0 6917.5225
343.17767333984375 0 9804.876
352.15240478515625 0 8157.5557
352.1972961425781 0 6286.2554
353.1603088378906 0 7089.324
354.6775207519531 0 10813.184
359.1914367675781 0 5795.369 c Ammonia loss 3
363.68206787109375 0 7022.7065
365.6788024902344 0 7360.9688
368.17333984375 0 6145.3027
371.1719665527344 0 17145.887
380.1813049316406 0 6025.614
381.1885070800781 0 69602.16
382.1944580078125 0 20628.363
383.20391845703125 0 86325.95
384.2075500488281 0 17029.967
386.181884765625 0 20888.732
388.20855712890625 0 8260.545
395.20416259765625 0 8058.4146
397.2142333984375 0 11963.793
398.2150573730469 0 13296.646
399.2004089355469 0 19067.08 y Water loss 2
399.22430419921875 0 6472.184
399.70172119140625 0 15026.103
400.206298828125 0 8067.007
401.2619323730469 0 10668.87
404.21185302734375 0 21687.7
404.7147216796875 0 12232.904
406.18377685546875 0 21446.908
408.20330810546875 0 17393.783 y 2
408.2271728515625 0 18474.35
408.7276916503906 0 13751.245
409.2292175292969 0 6171.8564
413.2164306640625 0 17811.287
413.716552734375 0 9177.615
414.2060546875 0 9344.555
422.2218933105469 0 36932.11 c Ammonia loss 7
422.7242736816406 0 12527.934
430.2658996582031 0 19029.668
435.2271728515625 0 7893.9937 y 6
440.2387390136719 0 31012.895
440.7198181152344 0 22427.94
441.2435302734375 0 9412.555
449.7264404296875 0 7275.264 y Water loss 1
452.7372741699219 0 15368.508
453.2375793457031 0 8679.268
454.7171325683594 0 6641.331
455.24169921875 0 6764.609
457.2529296875 0 16924.977
458.260009765625 0 20082.736 c Ammonia loss 4
458.7301025390625 0 24223.926 y 1
459.2322082519531 0 21581.309
461.7438049316406 0 17423.254
462.2444763183594 0 12396.789
464.22528076171875 0 16740.424
469.2300720214844 0 6158.184
470.7479248046875 0 23069.004 c Ammonia loss 8
471.2491149902344 0 20453.717
471.7514953613281 0 10671.247
475.2894287109375 0 6212.418 c 4
481.2273254394531 0 9686.329
482.2362365722656 0 129343.28
483.2397766113281 0 33922.594
484.2520751953125 0 25787.424
485.2525939941406 0 7547.9243
488.2568054199219 0 7531.691
488.75347900390625 0 5899.061
489.2510681152344 0 6091.8013
497.2615051269531 0 82721.99
497.7628479003906 0 59375.668
498.26239013671875 0 17643.629
499.2669677734375 0 104807.266
500.2705078125 0 33152.88
506.2669372558594 0 92157.336
506.76861572265625 0 41526.164
507.2324523925781 0 21548.234
507.265380859375 0 8926.143
508.23956298828125 0 14378.799
509.2419128417969 0 5515.169
515.272216796875 0 108835.98
515.773193359375 0 67678.16
516.2743530273438 0 29867.732
524.2611083984375 0 17019.543
525.2735595703125 0 8338.534
526.2537231445312 0 23477.191
527.2603149414062 0 32757.107
528.2633666992188 0 9136.695
538.2545776367188 0 24872.984 z Water loss 5
539.2611083984375 0 29086.11
542.2726440429688 0 43541.65
543.2754516601562 0 14118.06
544.024169921875 0 5986.9717
548.3201293945312 0 9035.222
548.8240966796875 0 10675.385
553.3212890625 0 31542.43
554.2708129882812 0 13201.706 y Water loss 5
554.321533203125 0 10095.448
555.272705078125 0 11484.25
556.2639770507812 0 194642.62 z 5
557.271728515625 0 1036535
558.2750244140625 0 354563.38
559.278564453125 0 51995.027
567.3236694335938 0 21619.918
568.2833862304688 0 43425.094
568.33251953125 0 109544.93
569.2862548828125 0 16634.379
569.3385009765625 0 57017.05
570.342041015625 0 10267.787
571.2749633789062 0 40131.566
572.282470703125 0 156824.4 y 5
573.2854614257812 0 53098.508
577.3106079101562 0 7870.289
583.2855834960938 0 21688.139
584.28955078125 0 6727.42
585.2786865234375 0 17173.24
586.296630859375 0 32691.436
587.3021240234375 0 16522.035
593.32373046875 0 9660.3
594.3330688476562 0 19558.854 c Water loss 5
595.321044921875 0 67448.56 c Ammonia loss 5
596.3226928710938 0 33740.45
597.334716796875 0 28932.387
598.3369750976562 0 10277.833
600.2777709960938 0 11684.691
601.3106689453125 0 29409.129
610.3314819335938 0 15406.324
611.2921142578125 0 101644.45
611.3389282226562 0 528198.2
612.3460693359375 0 1611549 c 5
613.3494262695312 0 492644.28
614.351806640625 0 85164.77
626.37451171875 0 18124.516
627.3760375976562 0 6400.566
629.304443359375 0 304648.22
630.307373046875 0 94599.086
631.3095703125 0 23333.973
637.3218994140625 0 9509.848 z Water loss 4
640.3093872070312 0 37979.074 w 4
641.31396484375 0 8699.3545
651.368896484375 0 44408.715
652.3712158203125 0 20031.988
655.3327026367188 0 250735.16 z 4
656.339111328125 0 392653.22
657.3425903320312 0 115384.51
658.343017578125 0 19164.26
669.3244018554688 0 10181.022
669.3812255859375 0 39291.855
670.3434448242188 0 20688.758
670.3955078125 0 9151.474
671.35107421875 0 65203.023 y 4
672.358642578125 0 12323.29
678.3571166992188 0 37141.855
679.3604125976562 0 14725.087
687.339599609375 0 11245.157
694.3406372070312 0 27029.611 z Water loss 3
695.33544921875 0 13681.109
695.3945922851562 0 8928.555 c Water loss 6
696.3671264648438 0 75275.69 c Ammonia loss 6
697.3729858398438 0 24039.14
698.3792724609375 0 12274.22
698.8901977539062 0 13703.209
699.3917846679688 0 6649.2363
700.8882446289062 0 9869.213
710.3648681640625 0 7594.8257 y Water loss 3
711.3759765625 0 15414.902
712.3505859375 0 111258.49 z 3
712.38525390625 0 120491.84
713.392333984375 0 1013376.25 c 6
714.3952026367188 0 405477.1
715.3977661132812 0 83495.16
716.4014282226562 0 9294.749
719.89404296875 0 11946.005
722.3218383789062 0 7396.3965
726.3514404296875 0 11340.222
727.3648681640625 0 33817.996
728.3724975585938 0 166260.39 y 3
729.3748168945312 0 69988.38
730.3525390625 0 94894.945
731.3551635742188 0 41612.637
732.3583374023438 0 8170.4883
740.3314208984375 0 9570.547
765.3911743164062 0 11431.127
765.8983764648438 0 6833.3657
771.4017944335938 0 6206.2295
771.917724609375 0 14918.796
772.4180908203125 0 11605.514
773.912841796875 0 8380.071
781.3775024414062 0 52590.336 z Water loss 2
782.3812255859375 0 38816.766 w 2
783.3876953125 0 10261.702
789.40771484375 0 18487.646
790.3989868164062 0 8033.3716
797.3905029296875 0 67305.94 y Water loss 2
798.3850708007812 0 50164.062
799.386474609375 0 877209.5 z 2
800.3903198242188 0 441413.34
801.3938598632812 0 101093.33
802.3994140625 0 16122.668
807.4148559570312 0 80351.914
808.4180297851562 0 43910.473
813.3792724609375 0 7673.3257
815.4053955078125 0 842892.8 y 2
816.4083862304688 0 384130.44
817.41064453125 0 93595.33
825.4248046875 0 276154.7
826.4285888671875 0 110334.89
827.4273071289062 0 32410.43
839.4432373046875 0 8169.044
841.425048828125 0 9369.164
843.4356079101562 0 496189 c Ammonia loss 7
844.4383544921875 0 224289.55
845.4405517578125 0 54954.133
880.4232177734375 0 26259.488
881.429931640625 0 17069.451
882.4248657226562 0 48958.668 z Water loss 1
883.4249267578125 0 35147.695 w 1
898.4213256835938 0 104318.79
899.42578125 0 173820.62
900.4324951171875 0 422245.1 z 1
901.4361572265625 0 200238.88
902.4384765625 0 68082.195
913.498779296875 0 15355.282
914.507568359375 0 10762.571
916.4517211914062 0 294776.5 y 1
917.4547119140625 0 148959.2
918.4572143554688 0 47066.69
921.4615478515625 0 9940.792
926.4359130859375 0 42605.09
927.4407958984375 0 27950.86
938.4800415039062 0 8271.855
939.498779296875 0 10230.837 c Water loss 8
955.4982299804688 0 19505.688
956.501708984375 0 9923.538
957.514892578125 0 1762710.5 c 8
958.5179443359375 0 959264.8
959.5205688476562 0 287202.5
960.5211791992188 0 38344.402
970.462890625 0 562366.4
971.4652709960938 0 301612.53
972.4676513671875 0 106118.414
973.4661865234375 0 13794.419
974.4783935546875 0 26496.129
984.537353515625 0 23045.248
985.5369262695312 0 15380.154
986.5532836914062 0 90808.07
987.5562744140625 0 54432.535
988.5563354492188 0 20817.305
995.5047607421875 0 55763.8
996.5073852539062 0 23493.988
997.518798828125 0 8936.185
1011.5034790039062 0 35618.82
1012.5244140625 0 51165.797
1013.51708984375 0 2263977
1014.5204467773438 0 1331290.5
1015.5225219726562 0 403693.7
1016.5238647460938 0 36819.69
1027.5159912109375 0 35917.203
1028.5264892578125 0 114779.95
1029.5357666015625 0 1770902.1
1030.5433349609375 0 7251617
1031.546142578125 0 3907042.8
1032.5491943359375 0 1179755.1
1033.550537109375 0 139705.97
1045.502685546875 0 12207.2
1046.506103515625 0 7851.1543
1095.6383056640625 0 10356.137
1142.570556640625 0 11697.186
1146.568115234375 0 13038.883
1438.78076171875 0 8628.137
1526.8079833984375 0 22503.826
1527.80908203125 0 18968.021
1529.8284912109375 0 11173.451
1530.80615234375 0 20675.215
1531.791748046875 0 11264.946
1541.8338623046875 0 9023.453
1543.8272705078125 0 19216.395
1544.827880859375 0 8851.992
1547.8272705078125 0 18503.805
1548.8448486328125 0 8545.785
1582.2978515625 0 7300.239
1947.337158203125 0 7132.0977
2400.173583984375 0 6972.0215

Spectrum Details

|  |  |
| --- | --- |
| Matched peaks? Matched peaks The total absolute number of peaks matched. Additionally in brackets the total fraction of peaks matched and the total number of peaks is shown. | 43 (12.99% of 331) |
| FDR? FDR The false discovery rate estimated for this peptide. It is calculated by matching all theoretical fragments with a non-integer shift with the raw peaks for this spectrum. This is done with 40 different shifts. The resulting percentage is the average number of annotated peaks over the number of annotated peaks with the correct spectrum. | 0.06% |
| Satellite FDR? Satellite FDR See the FDR for details on its calculation. This satellite ion specific FDR only contains the satellite ions (d/w) for I/L/J positions. | - |
| PSM Score? PSM Score The PSM Score as given by Hecklib to this annotated spectrum. It is shown with three significant figures. | 536 |

#### Spectrum 5282? Spectrum 5282 The raw spectrum of this peptide as annotated by Hecklib. The fragments are coloured according to ion type (see legend). Any peaks with a star '\*' as text can be hovered over to see the full details, first the ion type second the mass shift type. By hovering over the amino acids in the peptide or ions in the legend the corresponding peaks are highlighted. By toggling the 'Unassigned' label you can turn the background (unassigned) peaks on or off in the plot. By updating the slider in the Ion legend you can update the spectrum to only show the top X% of the peaks with labels. The top X% means any peak that is within X% of the highest intensity. By dragging in the spectrum you can zoom in to a specific part of the spectrum and use 'Zoom Out' to get back to the original zoom level. The annotation of the spectrum is based on the given sequence in the peptides file and is done with different software so inconsistencies are likely. The peaks are annotated based on the given sequence, with 20 ppm tolerance.

Copy Data

##### Spectrum 5282 (TSV)

###### Preview

```
Loading example...
```

*Click on the button to copy the data to your clipboard.*

Mz MinMz MaxIntensity Max

WidthHeightPeptide font sizePeptide stroke widthSpectrum font sizeSpectrum stroke widthCompact peptide

Ion legend

wxyz

abcd

OtherUnassignedIon^Charge^_Position_Show for top:%

JTSGVHTFPA

02.67e+55.35e+58.02e+51.07e+6

Zoom Out

y^+1^_2_c^+1^_2_c^+1^_3_y^+1^_3_c^+2^_7_c^+1^_4_y^+2^_8_y^+2^_8_c^+2^_8_y^+1^_4_y^+2^_9_c^+1^_5_y^+2^_9_c^+2^_9_z^+1^_5_y^+1^_5_z^+1^_5_y^+1^_5_c^+1^_6_c^+1^_6_c^+1^_6_w^+1^_6_z^+1^_6_y^+1^_6_z^+1^_7_c^+1^_7_y^+1^_7_z^+1^_7_c^+1^_7_y^+1^_7_z^+1^_8_w^+1^_8_y^+1^_8_z^+1^_8_y^+1^_8_c^+1^_8_z^+1^_9_w^+1^_9_w^+1^_9_y^+1^_9_z^+1^_9_y^+1^_9_c^+1^_9_

039178211721563

Fragment Matches Table

Show background peaks

| Position | Ion type | Intensity | mz Theoretical | mz Error (Th) | mz Error (ppm) | Charge | Series Number |
| --- | --- | --- | --- | --- | --- | --- | --- |
| - | - | 1878 | 120.1 | - | - | 0 | - |
| - | - | 425.4 | 121 | - | - | 0 | - |
| - | - | 593.2 | 122.4 | - | - | 0 | - |
| - | - | 536.9 | 123.1 | - | - | 0 | - |
| - | - | 3607 | 125.1 | - | - | 0 | - |
| - | - | 903.5 | 129.1 | - | - | 0 | - |
| - | - | 1423 | 131.1 | - | - | 0 | - |
| - | - | 1042 | 133.1 | - | - | 0 | - |
| - | - | 1.794E+04 | 133.1 | - | - | 0 | - |
| - | - | 619.3 | 139.5 | - | - | 0 | - |
| - | - | 1.146E+04 | 140.1 | - | - | 0 | - |
| - | - | 600.1 | 141.3 | - | - | 0 | - |
| - | - | 1.275E+04 | 142.1 | - | - | 0 | - |
| - | - | 1169 | 143.1 | - | - | 0 | - |
| - | - | 777.7 | 149 | - | - | 0 | - |
| - | - | 6052 | 152.1 | - | - | 0 | - |
| - | - | 752.2 | 153.1 | - | - | 0 | - |
| - | - | 4969 | 155.1 | - | - | 0 | - |
| - | - | 889.3 | 157.1 | - | - | 0 | - |
| - | - | 2025 | 163.1 | - | - | 0 | - |
| - | - | 9680 | 169.1 | - | - | 0 | - |
| - | - | 583.4 | 170.1 | - | - | 0 | - |
| - | - | 839.7 | 171.1 | - | - | 0 | - |
| - | - | 533.5 | 172.7 | - | - | 0 | - |
| - | - | 624.2 | 173.8 | - | - | 0 | - |
| - | - | 1817 | 175.1 | - | - | 0 | - |
| - | - | 1.394E+04 | 177.1 | - | - | 0 | - |
| - | - | 1320 | 178.1 | - | - | 0 | - |
| - | - | 676.3 | 183.1 | - | - | 0 | - |
| 9 | y | 1.973E+05 | 187.1 | 0.000466 | 2.491 | +1 | 2 |
| - | - | 8.595E+04 | 187.1 | - | - | 0 | - |
| - | - | 1.642E+04 | 188.1 | - | - | 0 | - |
| - | - | 589.8 | 188.1 | - | - | 0 | - |
| - | - | 8225 | 188.1 | - | - | 0 | - |
| - | - | 1129 | 189.1 | - | - | 0 | - |
| - | - | 1310 | 194.1 | - | - | 0 | - |
| - | - | 3746 | 195.1 | - | - | 0 | - |
| - | - | 1.13E+04 | 197.1 | - | - | 0 | - |
| - | - | 898.6 | 198.1 | - | - | 0 | - |
| - | - | 3424 | 201.1 | - | - | 0 | - |
| - | - | 6916 | 213.2 | - | - | 0 | - |
| 2 | c | 8.894E+04 | 215.1 | 0.0004617 | 2.146 | +1 | 2 |
| - | - | 9197 | 216.1 | - | - | 0 | - |
| - | - | 1148 | 221.1 | - | - | 0 | - |
| - | - | 4903 | 221.1 | - | - | 0 | - |
| - | - | 1.967E+04 | 223.1 | - | - | 0 | - |
| - | - | 1107 | 224.1 | - | - | 0 | - |
| - | - | 738.6 | 234.1 | - | - | 0 | - |
| - | - | 2017 | 239.1 | - | - | 0 | - |
| - | - | 3754 | 239.2 | - | - | 0 | - |
| - | - | 3.798E+04 | 241.2 | - | - | 0 | - |
| - | - | 961.5 | 245.1 | - | - | 0 | - |
| - | - | 1394 | 251.2 | - | - | 0 | - |
| - | - | 8174 | 252.2 | - | - | 0 | - |
| - | - | 1059 | 256.2 | - | - | 0 | - |
| - | - | 718.5 | 260.2 | - | - | 0 | - |
| - | - | 2326 | 265.2 | - | - | 0 | - |
| - | - | 2048 | 273.2 | - | - | 0 | - |
| - | - | 1.663E+04 | 276.2 | - | - | 0 | - |
| - | - | 920 | 283.2 | - | - | 0 | - |
| - | - | 1.159E+04 | 284.2 | - | - | 0 | - |
| - | - | 1493 | 285.2 | - | - | 0 | - |
| - | - | 748.4 | 294.2 | - | - | 0 | - |
| - | - | 1271 | 295.2 | - | - | 0 | - |
| - | - | 1.621E+04 | 296.2 | - | - | 0 | - |
| - | - | 2409 | 297.2 | - | - | 0 | - |
| - | - | 995.7 | 299.2 | - | - | 0 | - |
| 3 | c | 1009 | 302.2 | 0.0007361 | 2.436 | +1 | 3 |
| - | - | 670.4 | 304.2 | - | - | 0 | - |
| - | - | 934.4 | 308.2 | - | - | 0 | - |
| - | - | 2277 | 309.2 | - | - | 0 | - |
| - | - | 731.5 | 310.2 | - | - | 0 | - |
| - | - | 1101 | 310.2 | - | - | 0 | - |
| - | - | 1037 | 322.2 | - | - | 0 | - |
| - | - | 1007 | 323.2 | - | - | 0 | - |
| - | - | 1136 | 325.2 | - | - | 0 | - |
| - | - | 1043 | 326.1 | - | - | 0 | - |
| - | - | 1318 | 326.2 | - | - | 0 | - |
| - | - | 1800 | 327.2 | - | - | 0 | - |
| 8 | y | 9826 | 334.2 | 0.0006556 | 1.962 | +1 | 3 |
| - | - | 1949 | 335.2 | - | - | 0 | - |
| - | - | 3.982E+04 | 336.2 | - | - | 0 | - |
| - | - | 1.558E+04 | 339.2 | - | - | 0 | - |
| - | - | 912.9 | 339.7 | - | - | 0 | - |
| - | - | 3107 | 340.2 | - | - | 0 | - |
| - | - | 1046 | 341.2 | - | - | 0 | - |
| - | - | 1381 | 343.2 | - | - | 0 | - |
| 7 | c | 766.9 | 348.7 | 1.794E-05 | 0.05146 | +2 | 7 |
| - | - | 827.3 | 349.2 | - | - | 0 | - |
| - | - | 862.3 | 352.2 | - | - | 0 | - |
| - | - | 1360 | 353.2 | - | - | 0 | - |
| - | - | 1325 | 353.2 | - | - | 0 | - |
| - | - | 923.3 | 353.2 | - | - | 0 | - |
| - | - | 834.7 | 354.2 | - | - | 0 | - |
| - | - | 2.208E+04 | 354.2 | - | - | 0 | - |
| 4 | c | 1729 | 359.2 | 0.001184 | 3.296 | +1 | 4 |
| - | - | 2851 | 363.7 | - | - | 0 | - |
| - | - | 938.8 | 366.2 | - | - | 0 | - |
| - | - | 2975 | 371.2 | - | - | 0 | - |
| - | - | 2246 | 371.2 | - | - | 0 | - |
| - | - | 867.1 | 375.2 | - | - | 0 | - |
| - | - | 2366 | 380.2 | - | - | 0 | - |
| - | - | 1.029E+04 | 381.2 | - | - | 0 | - |
| - | - | 3262 | 382.2 | - | - | 0 | - |
| - | - | 1.236E+04 | 383.2 | - | - | 0 | - |
| - | - | 2367 | 384.2 | - | - | 0 | - |
| - | - | 3161 | 386.2 | - | - | 0 | - |
| - | - | 1139 | 387.2 | - | - | 0 | - |
| - | - | 1587 | 388.2 | - | - | 0 | - |
| - | - | 957 | 390.2 | - | - | 0 | - |
| - | - | 2660 | 391.7 | - | - | 0 | - |
| - | - | 878.8 | 392.2 | - | - | 0 | - |
| - | - | 1793 | 395.2 | - | - | 0 | - |
| - | - | 812.9 | 396.2 | - | - | 0 | - |
| - | - | 1592 | 397.2 | - | - | 0 | - |
| - | - | 2401 | 398.2 | - | - | 0 | - |
| 3 | y | 4326 | 399.2 | 0.0003794 | 0.9503 | +2 | 8 |
| - | - | 2262 | 399.7 | - | - | 0 | - |
| - | - | 841.4 | 400.2 | - | - | 0 | - |
| - | - | 1.093E+04 | 401.3 | - | - | 0 | - |
| - | - | 2606 | 402.3 | - | - | 0 | - |
| - | - | 862.6 | 403.2 | - | - | 0 | - |
| - | - | 4203 | 404.2 | - | - | 0 | - |
| - | - | 1432 | 405.2 | - | - | 0 | - |
| - | - | 1751 | 406.2 | - | - | 0 | - |
| 3 | y | 2658 | 408.2 | 0.002706 | 6.628 | +2 | 8 |
| - | - | 4077 | 408.2 | - | - | 0 | - |
| - | - | 2060 | 408.7 | - | - | 0 | - |
| - | - | 3487 | 413.2 | - | - | 0 | - |
| - | - | 1591 | 413.3 | - | - | 0 | - |
| - | - | 2275 | 413.7 | - | - | 0 | - |
| - | - | 2528 | 414.2 | - | - | 0 | - |
| - | - | 2266 | 414.7 | - | - | 0 | - |
| - | - | 3177 | 415.3 | - | - | 0 | - |
| 8 | c | 7557 | 422.2 | 0.001114 | 2.639 | +2 | 8 |
| - | - | 1912 | 422.7 | - | - | 0 | - |
| - | - | 4237 | 423.3 | - | - | 0 | - |
| - | - | 1301 | 427.3 | - | - | 0 | - |
| - | - | 757.1 | 427.8 | - | - | 0 | - |
| - | - | 2488 | 430.3 | - | - | 0 | - |
| 7 | y | 2070 | 435.2 | 0.002568 | 5.901 | +1 | 4 |
| - | - | 885.4 | 437.2 | - | - | 0 | - |
| - | - | 1157 | 438.2 | - | - | 0 | - |
| - | - | 827.3 | 438.7 | - | - | 0 | - |
| - | - | 5037 | 440.2 | - | - | 0 | - |
| - | - | 3369 | 440.7 | - | - | 0 | - |
| - | - | 1531 | 441.2 | - | - | 0 | - |
| - | - | 2.451E+04 | 441.3 | - | - | 0 | - |
| - | - | 2507 | 445.2 | - | - | 0 | - |
| - | - | 860 | 446.2 | - | - | 0 | - |
| - | - | 1028 | 447.2 | - | - | 0 | - |
| - | - | 891.2 | 448.7 | - | - | 0 | - |
| - | - | 1079 | 449.2 | - | - | 0 | - |
| 2 | y | 1422 | 449.7 | 0.001565 | 3.479 | +2 | 9 |
| - | - | 880.5 | 450.2 | - | - | 0 | - |
| - | - | 1373 | 451.2 | - | - | 0 | - |
| - | - | 1851 | 452.7 | - | - | 0 | - |
| - | - | 1912 | 456.8 | - | - | 0 | - |
| - | - | 1300 | 457.3 | - | - | 0 | - |
| 5 | c | 2230 | 458.3 | 0.001618 | 3.53 | +1 | 5 |
| 2 | y | 6767 | 458.7 | 0.001257 | 2.739 | +2 | 9 |
| - | - | 1915 | 459.2 | - | - | 0 | - |
| - | - | 4.808E+04 | 459.3 | - | - | 0 | - |
| - | - | 1671 | 459.7 | - | - | 0 | - |
| - | - | 1.041E+04 | 460.3 | - | - | 0 | - |
| - | - | 2935 | 461.7 | - | - | 0 | - |
| - | - | 1878 | 462.2 | - | - | 0 | - |
| - | - | 3503 | 464.2 | - | - | 0 | - |
| - | - | 1136 | 465.2 | - | - | 0 | - |
| - | - | 1112 | 470.2 | - | - | 0 | - |
| 9 | c | 4574 | 470.7 | 0.001466 | 3.114 | +2 | 9 |
| - | - | 3886 | 471.2 | - | - | 0 | - |
| - | - | 872.6 | 471.8 | - | - | 0 | - |
| - | - | 857 | 473.3 | - | - | 0 | - |
| - | - | 782.9 | 475.7 | - | - | 0 | - |
| - | - | 2190 | 481.2 | - | - | 0 | - |
| - | - | 2.108E+04 | 482.2 | - | - | 0 | - |
| - | - | 5332 | 483.2 | - | - | 0 | - |
| - | - | 4770 | 483.8 | - | - | 0 | - |
| - | - | 3407 | 484.3 | - | - | 0 | - |
| - | - | 2384 | 484.3 | - | - | 0 | - |
| - | - | 1354 | 485.3 | - | - | 0 | - |
| - | - | 987.2 | 488.3 | - | - | 0 | - |
| - | - | 1327 | 488.7 | - | - | 0 | - |
| - | - | 1.373E+04 | 497.3 | - | - | 0 | - |
| - | - | 8631 | 497.8 | - | - | 0 | - |
| - | - | 1.372E+04 | 498.3 | - | - | 0 | - |
| - | - | 904.3 | 498.8 | - | - | 0 | - |
| - | - | 1.427E+04 | 499.3 | - | - | 0 | - |
| - | - | 4895 | 500.3 | - | - | 0 | - |
| - | - | 1003 | 501.3 | - | - | 0 | - |
| - | - | 1.884E+04 | 506.3 | - | - | 0 | - |
| - | - | 9406 | 506.8 | - | - | 0 | - |
| - | - | 3798 | 507.2 | - | - | 0 | - |
| - | - | 3222 | 507.3 | - | - | 0 | - |
| - | - | 811.1 | 508.2 | - | - | 0 | - |
| - | - | 1365 | 509.3 | - | - | 0 | - |
| - | - | 3318 | 514.3 | - | - | 0 | - |
| - | - | 1.939E+04 | 515.3 | - | - | 0 | - |
| - | - | 1361 | 515.3 | - | - | 0 | - |
| - | - | 1.349E+04 | 515.8 | - | - | 0 | - |
| - | - | 1485 | 516.2 | - | - | 0 | - |
| - | - | 3.869E+04 | 516.3 | - | - | 0 | - |
| - | - | 1485 | 521.8 | - | - | 0 | - |
| - | - | 2680 | 524.3 | - | - | 0 | - |
| - | - | 1472 | 525.3 | - | - | 0 | - |
| - | - | 3783 | 526.3 | - | - | 0 | - |
| - | - | 6018 | 527.3 | - | - | 0 | - |
| - | - | 1606 | 528.3 | - | - | 0 | - |
| - | - | 914.4 | 532.8 | - | - | 0 | - |
| 6 | z | 4361 | 538.3 | 0.001204 | 2.238 | +1 | 5 |
| - | - | 3652 | 539.3 | - | - | 0 | - |
| - | - | 4888 | 539.8 | - | - | 0 | - |
| - | - | 2784 | 540.3 | - | - | 0 | - |
| - | - | 1662 | 540.8 | - | - | 0 | - |
| - | - | 7603 | 542.3 | - | - | 0 | - |
| - | - | 2281 | 543.3 | - | - | 0 | - |
| - | - | 9867 | 548.3 | - | - | 0 | - |
| - | - | 5645 | 548.8 | - | - | 0 | - |
| - | - | 1326 | 549.3 | - | - | 0 | - |
| - | - | 5896 | 553.3 | - | - | 0 | - |
| 6 | y | 1104 | 554.3 | 0.004153 | 7.493 | +1 | 5 |
| - | - | 884.1 | 554.3 | - | - | 0 | - |
| - | - | 1893 | 555.3 | - | - | 0 | - |
| 6 | z | 2.832E+04 | 556.3 | 0.001077 | 1.936 | +1 | 5 |
| - | - | 1.508E+05 | 557.3 | - | - | 0 | - |
| - | - | 4.828E+04 | 558.3 | - | - | 0 | - |
| - | - | 7458 | 559.3 | - | - | 0 | - |
| - | - | 2764 | 567.3 | - | - | 0 | - |
| - | - | 6910 | 568.3 | - | - | 0 | - |
| - | - | 1.837E+04 | 568.3 | - | - | 0 | - |
| - | - | 1665 | 569.3 | - | - | 0 | - |
| - | - | 9438 | 569.3 | - | - | 0 | - |
| - | - | 853.2 | 570.3 | - | - | 0 | - |
| - | - | 1786 | 570.3 | - | - | 0 | - |
| - | - | 8099 | 571.3 | - | - | 0 | - |
| 6 | y | 2.487E+04 | 572.3 | 0.0005411 | 0.9456 | +1 | 5 |
| - | - | 9086 | 573.3 | - | - | 0 | - |
| - | - | 1318 | 574.3 | - | - | 0 | - |
| - | - | 1273 | 577.3 | - | - | 0 | - |
| - | - | 4192 | 583.3 | - | - | 0 | - |
| - | - | 1693 | 584.3 | - | - | 0 | - |
| - | - | 2289 | 585.3 | - | - | 0 | - |
| - | - | 5705 | 586.3 | - | - | 0 | - |
| - | - | 2765 | 587.3 | - | - | 0 | - |
| - | - | 1168 | 593.3 | - | - | 0 | - |
| - | - | 1350 | 593.3 | - | - | 0 | - |
| 6 | c | 2822 | 594.3 | 0.004583 | 7.712 | +1 | 6 |
| 6 | c | 1.157E+04 | 595.3 | 0.001757 | 2.952 | +1 | 6 |
| - | - | 4879 | 596.3 | - | - | 0 | - |
| - | - | 4649 | 597.3 | - | - | 0 | - |
| - | - | 879.1 | 598.3 | - | - | 0 | - |
| - | - | 933.3 | 600.3 | - | - | 0 | - |
| - | - | 4598 | 601.3 | - | - | 0 | - |
| - | - | 1720 | 602.3 | - | - | 0 | - |
| - | - | 900.8 | 605.3 | - | - | 0 | - |
| - | - | 871.2 | 606.3 | - | - | 0 | - |
| - | - | 2084 | 610.3 | - | - | 0 | - |
| - | - | 1.876E+04 | 611.3 | - | - | 0 | - |
| - | - | 7.527E+04 | 611.3 | - | - | 0 | - |
| 6 | c | 2.372E+05 | 612.3 | 0.0002938 | 0.4797 | +1 | 6 |
| - | - | 7.15E+04 | 613.3 | - | - | 0 | - |
| - | - | 1769 | 614.3 | - | - | 0 | - |
| - | - | 1.27E+04 | 614.4 | - | - | 0 | - |
| - | - | 927.9 | 615.4 | - | - | 0 | - |
| - | - | 1005 | 619.4 | - | - | 0 | - |
| - | - | 880.6 | 624.3 | - | - | 0 | - |
| - | - | 1.218E+04 | 626.4 | - | - | 0 | - |
| - | - | 2341 | 627.4 | - | - | 0 | - |
| - | - | 5.134E+04 | 629.3 | - | - | 0 | - |
| - | - | 1.803E+04 | 630.3 | - | - | 0 | - |
| - | - | 4781 | 631.3 | - | - | 0 | - |
| 5 | w | 5179 | 640.3 | 0.001548 | 2.418 | +1 | 6 |
| - | - | 1059 | 640.8 | - | - | 0 | - |
| - | - | 1057 | 641.3 | - | - | 0 | - |
| - | - | 730.7 | 643.9 | - | - | 0 | - |
| - | - | 1247 | 649.9 | - | - | 0 | - |
| - | - | 936.8 | 650.4 | - | - | 0 | - |
| - | - | 6168 | 651.4 | - | - | 0 | - |
| - | - | 1871 | 651.9 | - | - | 0 | - |
| - | - | 2383 | 652.4 | - | - | 0 | - |
| - | - | 1225 | 654.3 | - | - | 0 | - |
| 5 | z | 3.712E+04 | 655.3 | 0.0008391 | 1.28 | +1 | 6 |
| - | - | 5.854E+04 | 656.3 | - | - | 0 | - |
| - | - | 1.984E+04 | 657.3 | - | - | 0 | - |
| - | - | 3784 | 658.3 | - | - | 0 | - |
| - | - | 1519 | 660.4 | - | - | 0 | - |
| - | - | 1107 | 668.4 | - | - | 0 | - |
| - | - | 1367 | 669.3 | - | - | 0 | - |
| - | - | 7076 | 669.4 | - | - | 0 | - |
| - | - | 3137 | 670.3 | - | - | 0 | - |
| - | - | 2514 | 670.4 | - | - | 0 | - |
| 5 | y | 1.008E+04 | 671.4 | 6.273E-05 | 0.09344 | +1 | 6 |
| - | - | 3429 | 672.4 | - | - | 0 | - |
| - | - | 1335 | 676.4 | - | - | 0 | - |
| - | - | 1300 | 677.4 | - | - | 0 | - |
| - | - | 6531 | 678.4 | - | - | 0 | - |
| - | - | 2757 | 679.4 | - | - | 0 | - |
| - | - | 792.4 | 680.4 | - | - | 0 | - |
| - | - | 1264 | 683.4 | - | - | 0 | - |
| - | - | 1382 | 684.4 | - | - | 0 | - |
| - | - | 868.6 | 686.3 | - | - | 0 | - |
| - | - | 1043 | 688.4 | - | - | 0 | - |
| - | - | 1994 | 690.9 | - | - | 0 | - |
| 4 | z | 4935 | 694.3 | 0.002919 | 4.204 | +1 | 7 |
| - | - | 1136 | 695.3 | - | - | 0 | - |
| 7 | c | 1.255E+04 | 696.4 | 0.00132 | 1.896 | +1 | 7 |
| - | - | 4942 | 697.4 | - | - | 0 | - |
| - | - | 860.4 | 697.9 | - | - | 0 | - |
| - | - | 1.134E+04 | 698.4 | - | - | 0 | - |
| - | - | 2.31E+04 | 698.9 | - | - | 0 | - |
| - | - | 1.145E+04 | 699.4 | - | - | 0 | - |
| - | - | 5500 | 699.9 | - | - | 0 | - |
| - | - | 1212 | 700.4 | - | - | 0 | - |
| - | - | 1598 | 705.9 | - | - | 0 | - |
| 4 | y | 1220 | 710.4 | 0.000952 | 1.34 | +1 | 7 |
| - | - | 2866 | 711.4 | - | - | 0 | - |
| 4 | z | 1.795E+04 | 712.4 | 0.002741 | 3.848 | +1 | 7 |
| - | - | 1.989E+04 | 712.4 | - | - | 0 | - |
| 7 | c | 1.62E+05 | 713.4 | 0.0007539 | 1.057 | +1 | 7 |
| - | - | 5.881E+04 | 714.4 | - | - | 0 | - |
| - | - | 1.249E+04 | 715.4 | - | - | 0 | - |
| - | - | 1301 | 716.4 | - | - | 0 | - |
| - | - | 2103 | 726.4 | - | - | 0 | - |
| - | - | 4554 | 727.4 | - | - | 0 | - |
| 4 | y | 2.852E+04 | 728.4 | 0.0005683 | 0.7802 | +1 | 7 |
| - | - | 9941 | 729.4 | - | - | 0 | - |
| - | - | 1.931E+04 | 730.4 | - | - | 0 | - |
| - | - | 5838 | 731.4 | - | - | 0 | - |
| - | - | 1397 | 732.4 | - | - | 0 | - |
| - | - | 1618 | 740.3 | - | - | 0 | - |
| - | - | 3963 | 742.4 | - | - | 0 | - |
| - | - | 3512 | 742.9 | - | - | 0 | - |
| - | - | 2559 | 743.4 | - | - | 0 | - |
| - | - | 1155 | 744.4 | - | - | 0 | - |
| - | - | 1696 | 749.4 | - | - | 0 | - |
| - | - | 2340 | 750.4 | - | - | 0 | - |
| - | - | 1627 | 750.9 | - | - | 0 | - |
| - | - | 1226 | 751.4 | - | - | 0 | - |
| - | - | 1083 | 751.9 | - | - | 0 | - |
| - | - | 1107 | 754.9 | - | - | 0 | - |
| - | - | 1697 | 756.4 | - | - | 0 | - |
| - | - | 1330 | 762.4 | - | - | 0 | - |
| - | - | 3928 | 762.9 | - | - | 0 | - |
| - | - | 1.172E+04 | 763.4 | - | - | 0 | - |
| - | - | 7591 | 763.9 | - | - | 0 | - |
| - | - | 4306 | 764.4 | - | - | 0 | - |
| - | - | 1677 | 764.9 | - | - | 0 | - |
| - | - | 993.1 | 765.4 | - | - | 0 | - |
| - | - | 2642 | 770.4 | - | - | 0 | - |
| - | - | 8569 | 770.9 | - | - | 0 | - |
| - | - | 1.114E+04 | 771.4 | - | - | 0 | - |
| - | - | 1.328E+04 | 771.9 | - | - | 0 | - |
| - | - | 8117 | 772.4 | - | - | 0 | - |
| - | - | 2103 | 772.9 | - | - | 0 | - |
| - | - | 1954 | 773.4 | - | - | 0 | - |
| - | - | 992.5 | 774.4 | - | - | 0 | - |
| 3 | z | 9959 | 781.4 | 0.003505 | 4.486 | +1 | 8 |
| 3 | w | 5307 | 782.4 | 0.000658 | 0.8411 | +1 | 8 |
| - | - | 1261 | 782.5 | - | - | 0 | - |
| - | - | 1582 | 783.4 | - | - | 0 | - |
| - | - | 874.6 | 783.5 | - | - | 0 | - |
| - | - | 2084 | 789.4 | - | - | 0 | - |
| - | - | 1107 | 790.4 | - | - | 0 | - |
| 3 | y | 1.123E+04 | 797.4 | 0.001242 | 1.558 | +1 | 8 |
| - | - | 9027 | 798.4 | - | - | 0 | - |
| 3 | z | 1.343E+05 | 799.4 | 0.001668 | 2.087 | +1 | 8 |
| - | - | 6.254E+04 | 800.4 | - | - | 0 | - |
| - | - | 1.687E+04 | 801.4 | - | - | 0 | - |
| - | - | 1772 | 802.4 | - | - | 0 | - |
| - | - | 1.323E+04 | 807.4 | - | - | 0 | - |
| - | - | 5627 | 808.4 | - | - | 0 | - |
| - | - | 2095 | 809.4 | - | - | 0 | - |
| - | - | 1045 | 813.4 | - | - | 0 | - |
| - | - | 1305 | 814.4 | - | - | 0 | - |
| 3 | y | 1.469E+05 | 815.4 | 0.001499 | 1.838 | +1 | 8 |
| - | - | 6.466E+04 | 816.4 | - | - | 0 | - |
| - | - | 1.685E+04 | 817.4 | - | - | 0 | - |
| - | - | 1936 | 818.4 | - | - | 0 | - |
| - | - | 4.441E+04 | 825.4 | - | - | 0 | - |
| - | - | 2.208E+04 | 826.4 | - | - | 0 | - |
| - | - | 5546 | 827.4 | - | - | 0 | - |
| - | - | 2017 | 839.4 | - | - | 0 | - |
| - | - | 1957 | 841.4 | - | - | 0 | - |
| 8 | c | 8.985E+04 | 843.4 | 0.0005942 | 0.7045 | +1 | 8 |
| - | - | 3.881E+04 | 844.4 | - | - | 0 | - |
| - | - | 1.113E+04 | 845.4 | - | - | 0 | - |
| - | - | 998.8 | 846.4 | - | - | 0 | - |
| - | - | 3462 | 880.4 | - | - | 0 | - |
| - | - | 2141 | 881.4 | - | - | 0 | - |
| 2 | z | 7131 | 882.4 | 0.001603 | 1.816 | +1 | 9 |
| 2 | w | 4532 | 883.4 | 0.002926 | 3.312 | +1 | 9 |
| 2 | w | 1238 | 885.4 | 0.01097 | 12.39 | +1 | 9 |
| - | - | 909.2 | 887.5 | - | - | 0 | - |
| 2 | y | 1.388E+04 | 898.4 | 0.01651 | 18.38 | +1 | 9 |
| - | - | 2.108E+04 | 899.4 | - | - | 0 | - |
| 2 | z | 6.5E+04 | 900.4 | 0.0002949 | 0.3275 | +1 | 9 |
| - | - | 2.987E+04 | 901.4 | - | - | 0 | - |
| - | - | 9408 | 902.4 | - | - | 0 | - |
| - | - | 1021 | 903.4 | - | - | 0 | - |
| - | - | 1712 | 908.4 | - | - | 0 | - |
| - | - | 977.3 | 909.4 | - | - | 0 | - |
| - | - | 2855 | 913.5 | - | - | 0 | - |
| 2 | y | 5.339E+04 | 916.5 | 0.0005733 | 0.6256 | +1 | 9 |
| - | - | 3.13E+04 | 917.5 | - | - | 0 | - |
| - | - | 1.093E+04 | 918.5 | - | - | 0 | - |
| - | - | 2046 | 919.5 | - | - | 0 | - |
| - | - | 6970 | 926.4 | - | - | 0 | - |
| - | - | 3156 | 927.4 | - | - | 0 | - |
| - | - | 4415 | 951.4 | - | - | 0 | - |
| - | - | 2188 | 952.4 | - | - | 0 | - |
| - | - | 954.5 | 953.4 | - | - | 0 | - |
| - | - | 2228 | 955.5 | - | - | 0 | - |
| - | - | 1063 | 956.5 | - | - | 0 | - |
| 9 | c | 2.629E+05 | 957.5 | 0.000749 | 0.7823 | +1 | 9 |
| - | - | 1.525E+05 | 958.5 | - | - | 0 | - |
| - | - | 4.391E+04 | 959.5 | - | - | 0 | - |
| - | - | 6200 | 960.5 | - | - | 0 | - |
| - | - | 8.614E+04 | 970.5 | - | - | 0 | - |
| - | - | 4.254E+04 | 971.5 | - | - | 0 | - |
| - | - | 1.25E+04 | 972.5 | - | - | 0 | - |
| - | - | 2093 | 973.5 | - | - | 0 | - |
| - | - | 4879 | 974.5 | - | - | 0 | - |
| - | - | 1877 | 975.5 | - | - | 0 | - |
| - | - | 3621 | 984.5 | - | - | 0 | - |
| - | - | 2574 | 985.5 | - | - | 0 | - |
| - | - | 1.476E+04 | 986.6 | - | - | 0 | - |
| - | - | 9351 | 987.6 | - | - | 0 | - |
| - | - | 2729 | 988.6 | - | - | 0 | - |
| - | - | 8157 | 995.5 | - | - | 0 | - |
| - | - | 4759 | 996.5 | - | - | 0 | - |
| - | - | 1634 | 997.5 | - | - | 0 | - |
| - | - | 2254 | 1010 | - | - | 0 | - |
| - | - | 5221 | 1012 | - | - | 0 | - |
| - | - | 1.009E+04 | 1013 | - | - | 0 | - |
| - | - | 3.458E+05 | 1014 | - | - | 0 | - |
| - | - | 1.966E+05 | 1015 | - | - | 0 | - |
| - | - | 6.454E+04 | 1016 | - | - | 0 | - |
| - | - | 1.014E+04 | 1017 | - | - | 0 | - |
| - | - | 1114 | 1027 | - | - | 0 | - |
| - | - | 5708 | 1028 | - | - | 0 | - |
| - | - | 1.535E+04 | 1029 | - | - | 0 | - |
| - | - | 2.717E+05 | 1030 | - | - | 0 | - |
| - | - | 1.059E+06 | 1031 | - | - | 0 | - |
| - | - | 5.796E+05 | 1032 | - | - | 0 | - |
| - | - | 1.769E+05 | 1033 | - | - | 0 | - |
| - | - | 2.221E+04 | 1034 | - | - | 0 | - |
| - | - | 2004 | 1046 | - | - | 0 | - |
| - | - | 1318 | 1063 | - | - | 0 | - |
| - | - | 1932 | 1064 | - | - | 0 | - |
| - | - | 2809 | 1079 | - | - | 0 | - |
| - | - | 1751 | 1080 | - | - | 0 | - |
| - | - | 1099 | 1081 | - | - | 0 | - |
| - | - | 5366 | 1096 | - | - | 0 | - |
| - | - | 3315 | 1097 | - | - | 0 | - |
| - | - | 1317 | 1141 | - | - | 0 | - |
| - | - | 1.098E+04 | 1143 | - | - | 0 | - |
| - | - | 7499 | 1144 | - | - | 0 | - |
| - | - | 2361 | 1145 | - | - | 0 | - |
| - | - | 1243 | 1211 | - | - | 0 | - |
| - | - | 1302 | 1269 | - | - | 0 | - |
| - | - | 8796 | 1271 | - | - | 0 | - |
| - | - | 5713 | 1272 | - | - | 0 | - |
| - | - | 3007 | 1273 | - | - | 0 | - |
| - | - | 981.8 | 1283 | - | - | 0 | - |
| - | - | 1108 | 1287 | - | - | 0 | - |
| - | - | 957.4 | 1288 | - | - | 0 | - |
| - | - | 1772 | 1298 | - | - | 0 | - |
| - | - | 1506 | 1299 | - | - | 0 | - |
| - | - | 1786 | 1304 | - | - | 0 | - |
| - | - | 1336 | 1306 | - | - | 0 | - |
| - | - | 1218 | 1337 | - | - | 0 | - |
| - | - | 1041 | 1353 | - | - | 0 | - |
| - | - | 2496 | 1354 | - | - | 0 | - |
| - | - | 1563 | 1355 | - | - | 0 | - |
| - | - | 1387 | 1382 | - | - | 0 | - |
| - | - | 1014 | 1395 | - | - | 0 | - |
| - | - | 1259 | 1396 | - | - | 0 | - |
| - | - | 3200 | 1397 | - | - | 0 | - |
| - | - | 3410 | 1398 | - | - | 0 | - |
| - | - | 2772 | 1399 | - | - | 0 | - |
| - | - | 1943 | 1412 | - | - | 0 | - |
| - | - | 1641 | 1413 | - | - | 0 | - |
| - | - | 2031 | 1426 | - | - | 0 | - |
| - | - | 3083 | 1428 | - | - | 0 | - |
| - | - | 2147 | 1429 | - | - | 0 | - |
| - | - | 1282 | 1430 | - | - | 0 | - |
| - | - | 1135 | 1441 | - | - | 0 | - |
| - | - | 3511 | 1484 | - | - | 0 | - |
| - | - | 5438 | 1485 | - | - | 0 | - |
| - | - | 5928 | 1486 | - | - | 0 | - |
| - | - | 1335 | 1487 | - | - | 0 | - |
| - | - | 1533 | 1498 | - | - | 0 | - |
| - | - | 2325 | 1499 | - | - | 0 | - |
| - | - | 2320 | 1500 | - | - | 0 | - |
| - | - | 1233 | 1509 | - | - | 0 | - |
| - | - | 1616 | 1510 | - | - | 0 | - |
| - | - | 1506 | 1511 | - | - | 0 | - |
| - | - | 1163 | 1516 | - | - | 0 | - |
| - | - | 5905 | 1524 | - | - | 0 | - |
| - | - | 8564 | 1525 | - | - | 0 | - |
| - | - | 1.268E+04 | 1526 | - | - | 0 | - |
| - | - | 2.719E+04 | 1527 | - | - | 0 | - |
| - | - | 2.019E+04 | 1528 | - | - | 0 | - |
| - | - | 8426 | 1529 | - | - | 0 | - |
| - | - | 4394 | 1530 | - | - | 0 | - |
| - | - | 1467 | 1531 | - | - | 0 | - |
| - | - | 5878 | 1541 | - | - | 0 | - |
| - | - | 2.102E+04 | 1542 | - | - | 0 | - |
| - | - | 1.524E+04 | 1543 | - | - | 0 | - |
| - | - | 3.264E+04 | 1544 | - | - | 0 | - |
| - | - | 2.464E+04 | 1545 | - | - | 0 | - |
| - | - | 1.015E+04 | 1546 | - | - | 0 | - |
| - | - | 4870 | 1547 | - | - | 0 | - |
| - | - | 2183 | 1548 | - | - | 0 | - |

m/z Charge Intensity FragmentType MassShift Position
120.08126831054688 0 1878.0388
120.9655990600586 0 425.44806
122.35568237304688 0 593.24615
123.05572509765625 0 536.8993
125.10768127441406 0 3607.2312
129.10269165039062 0 903.5159
131.0706329345703 0 1422.9626
133.08079528808594 0 1042.0598
133.08631896972656 0 17936.559
139.46348571777344 0 619.2663
140.08226013183594 0 11463.876
141.32200622558594 0 600.13916
142.12303161621094 0 12748.876
143.12640380859375 0 1168.8262
148.9554901123047 0 777.7441
152.1072998046875 0 6052.0664
153.11099243164062 0 752.19135
155.09315490722656 0 4969.1514
157.10853576660156 0 889.31537
163.0717010498047 0 2025.1605
169.13397216796875 0 9680.147
170.13735961914062 0 583.4367
171.07705688476562 0 839.6973
172.69766235351562 0 533.48755
173.8200225830078 0 624.2128
175.09715270996094 0 1817.1018
177.11256408691406 0 13938.915
178.11599731445312 0 1319.558
183.11407470703125 0 676.3299
187.10818481445312 0 197263.77 y 8
187.14456176757812 0 85947.04
188.11158752441406 0 16423.445
188.14019775390625 0 589.79144
188.14791870117188 0 8225.288
189.08712768554688 0 1128.656
194.12933349609375 0 1310.383
195.13699340820312 0 3745.6533
197.12893676757812 0 11301.664
198.1328887939453 0 898.6052
201.12374877929688 0 3424.2695
213.16006469726562 0 6915.5933
215.1394805908203 0 88942.375 c Ammonia loss 1
216.14285278320312 0 9196.749
221.1039581298828 0 1148.1158
221.13876342773438 0 4903.105
223.1446075439453 0 19670.676
224.103271484375 0 1106.7341
234.1480255126953 0 738.61005
239.11427307128906 0 2017.1865
239.1504669189453 0 3753.7617
241.1551513671875 0 37979.06
245.12889099121094 0 961.5006
251.1507110595703 0 1394.0155
252.1584930419922 0 8173.507
256.1772766113281 0 1058.6401
260.1514587402344 0 718.4527
265.1650085449219 0 2326.352
273.2039794921875 0 2047.8258
276.1559753417969 0 16625.174
283.17462158203125 0 919.9734
284.16094970703125 0 11594.526
285.1648254394531 0 1493.1831
294.1555480957031 0 748.3931
295.1622314453125 0 1271.2957
296.1722106933594 0 16212.052
297.1751403808594 0 2408.9622
299.2193298339844 0 995.6875
302.1717834472656 0 1009.3232 c Ammonia loss 2
304.1758117675781 0 670.42346
308.16339111328125 0 934.37317
309.1915588378906 0 2277.4097
310.15338134765625 0 731.49036
310.2132568359375 0 1100.8508
322.1636657714844 0 1037.404
323.16998291015625 0 1007.17017
325.16571044921875 0 1136.3052
326.1361389160156 0 1043.4625
326.24444580078125 0 1317.7916
327.2024230957031 0 1800.1968
334.1767883300781 0 9826.347 y 7
335.1798095703125 0 1948.6263
336.2289733886719 0 39823.402
339.1910705566406 0 15581.557
339.6833801269531 0 912.91156
340.1947326660156 0 3107.434
341.1819152832031 0 1046.0116
343.1773376464844 0 1381.1984
348.6873779296875 0 766.9212 c Ammonia loss 6
349.18865966796875 0 827.25854
352.1993103027344 0 862.3172
353.15972900390625 0 1360.0017
353.195068359375 0 1324.8733
353.2179260253906 0 923.264
354.1632385253906 0 834.6533
354.2393493652344 0 22077.746
359.1936950683594 0 1728.9348 c Ammonia loss 3
363.6824951171875 0 2851.1582
366.18194580078125 0 938.8289
371.1731872558594 0 2975.4622
371.2275390625 0 2246.2043
375.2008972167969 0 867.0978
380.1816711425781 0 2365.6418
381.1883544921875 0 10288.232
382.19390869140625 0 3261.835
383.204345703125 0 12364.716
384.20770263671875 0 2367.0889
386.1828918457031 0 3161.406
387.18597412109375 0 1139.4207
388.2096252441406 0 1587.4901
390.2153625488281 0 956.95856
391.7416687011719 0 2660.0417
392.2423095703125 0 878.79114
395.2045593261719 0 1792.7809
396.1977844238281 0 812.8732
397.21435546875 0 1592.245
398.2154541015625 0 2400.874
399.2010498046875 0 4325.8438 y Water loss 2
399.7034606933594 0 2261.6338
400.2059326171875 0 841.38873
401.2626953125 0 10927.483
402.2661437988281 0 2605.601
403.21923828125 0 862.5806
404.2121276855469 0 4203.357
405.2010192871094 0 1431.692
406.1838073730469 0 1751.3843
408.2032470703125 0 2657.512 y 2
408.22589111328125 0 4077.0024
408.7278747558594 0 2059.7222
413.2168273925781 0 3486.9495
413.2770690917969 0 1591.4855
413.7178039550781 0 2275.3057
414.20849609375 0 2527.841
414.7085266113281 0 2266.0283
415.2545166015625 0 3176.8643
422.22271728515625 0 7557.2505 c Ammonia loss 7
422.7239990234375 0 1912.2317
423.2611083984375 0 4236.596
427.2598876953125 0 1300.6261
427.7626953125 0 757.136
430.2673645019531 0 2488.3977
435.22637939453125 0 2069.8184 y 6
437.216796875 0 885.4105
438.2194519042969 0 1156.7103
438.73779296875 0 827.3302
440.2395935058594 0 5036.9434
440.71978759765625 0 3368.8079
441.2199401855469 0 1530.9202
441.2716979980469 0 24509.656
445.196533203125 0 2507.3818
446.2019348144531 0 860.0087
447.21038818359375 0 1028.0873
448.73406982421875 0 891.24255
449.2371520996094 0 1078.8243
449.72607421875 0 1422.0686 y Water loss 1
450.2271423339844 0 880.4762
451.2256164550781 0 1373.0255
452.7381591796875 0 1851.4524
456.75225830078125 0 1912.3049
457.2536926269531 0 1300.4331
458.2625427246094 0 2229.6816 c Ammonia loss 4
458.7310485839844 0 6767.23 y 1
459.2347717285156 0 1915.2332
459.2808532714844 0 48083.258
459.7328796386719 0 1670.8438
460.28436279296875 0 10407.782
461.7433776855469 0 2934.7373
462.24322509765625 0 1878.1833
464.2259826660156 0 3502.7446
465.2294006347656 0 1135.5441
470.2374267578125 0 1112.4989
470.74945068359375 0 4573.812 c Ammonia loss 8
471.2490234375 0 3885.729
471.75048828125 0 872.55493
473.2591247558594 0 856.9804
475.7403869628906 0 782.86884
481.22943115234375 0 2189.7908
482.2364501953125 0 21077.764
483.2392578125 0 5332.232
483.80224609375 0 4770.22
484.252685546875 0 3406.8777
484.302978515625 0 2383.76
485.25323486328125 0 1354.4884
488.2581481933594 0 987.16394
488.7497253417969 0 1326.5021
497.261962890625 0 13730.779
497.7630920410156 0 8631.152
498.29254150390625 0 13720.586
498.7657775878906 0 904.3393
499.2667541503906 0 14266.275
500.26947021484375 0 4895.143
501.2763671875 0 1002.77203
506.26727294921875 0 18839.576
506.7689514160156 0 9405.553
507.2311706542969 0 3798.436
507.27069091796875 0 3221.5195
508.2394714355469 0 811.09576
509.25018310546875 0 1364.6616
514.2846069335938 0 3317.8716
515.27294921875 0 19386.74
515.3173828125 0 1360.9249
515.7739868164062 0 13494.155
516.2080078125 0 1485.3973
516.3038940429688 0 38689.777
521.7838745117188 0 1485.4724
524.2626953125 0 2680.1917
525.2734375 0 1472.0177
526.2542114257812 0 3783.0137
527.2612915039062 0 6017.637
528.262939453125 0 1606.481
532.8055419921875 0 914.4176
538.254638671875 0 4360.591 z Water loss 5
539.2610473632812 0 3652.3716
539.8106079101562 0 4887.9775
540.31201171875 0 2783.8337
540.8142700195312 0 1662.4576
542.2722778320312 0 7602.9985
543.2767333984375 0 2280.6804
548.3236694335938 0 9866.925
548.8251953125 0 5645.014
549.3165893554688 0 1326.181
553.3222045898438 0 5896.2617
554.2680053710938 0 1103.7344 y Water loss 5
554.3207397460938 0 884.05133
555.2777099609375 0 1893.3647
556.2650756835938 0 28322.586 z 5
557.2722778320312 0 150809.88
558.2754516601562 0 48283.45
559.2786865234375 0 7458.179
567.3258666992188 0 2764.0435
568.2835693359375 0 6910.1553
568.3338623046875 0 18369.56
569.2863159179688 0 1665.1693
569.3406372070312 0 9438.348
570.2675170898438 0 853.17426
570.3458251953125 0 1785.9691
571.276123046875 0 8098.6226
572.2832641601562 0 24869.414 y 5
573.2860107421875 0 9086.126
574.2897338867188 0 1318.0836
577.3101806640625 0 1272.8632
583.2896728515625 0 4192.1045
584.2899780273438 0 1692.6754
585.2806396484375 0 2288.5625
586.2984619140625 0 5705.162
587.3016357421875 0 2765.1958
593.280029296875 0 1167.8671
593.3280639648438 0 1350.2397
594.3312377929688 0 2822.1597 c Water loss 5
595.3215942382812 0 11573.085 c Ammonia loss 5
596.3226928710938 0 4879.0347
597.3359985351562 0 4648.5376
598.3357543945312 0 879.1392
600.2747802734375 0 933.32697
601.3095703125 0 4597.7793
602.3168334960938 0 1719.577
605.3359375 0 900.8388
606.337890625 0 871.2143
610.33056640625 0 2083.8467
611.29248046875 0 18762.188
611.33984375 0 75266.56
612.3466796875 0 237158.28 c 5
613.349853515625 0 71503.58
614.2943725585938 0 1769.1514
614.3519897460938 0 12704.421
615.3553466796875 0 927.9435
619.3507690429688 0 1004.60144
624.34228515625 0 880.5845
626.3741455078125 0 12176.686
627.379638671875 0 2340.8806
629.3051147460938 0 51343.375
630.3076171875 0 18028.86
631.3103637695312 0 4780.7944
640.3104858398438 0 5178.607 w 4
640.843017578125 0 1059.4583
641.3115844726562 0 1057.2523
643.8519287109375 0 730.66364
649.8607177734375 0 1246.6859
650.3611450195312 0 936.7711
651.3695068359375 0 6168.307
651.8609008789062 0 1871.3782
652.3719482421875 0 2382.7869
654.3245849609375 0 1225.117
655.333251953125 0 37120.562 z 4
656.3397827148438 0 58544.293
657.3431396484375 0 19843.496
658.3449096679688 0 3783.838
660.3516235351562 0 1518.8983
668.3799438476562 0 1106.8942
669.324462890625 0 1367.242
669.3809814453125 0 7076.216
670.3417358398438 0 3136.9897
670.3894653320312 0 2513.673
671.35107421875 0 10080.872 y 4
672.3555908203125 0 3428.718
676.3854370117188 0 1335.0015
677.3861694335938 0 1300.0464
678.3583374023438 0 6530.9004
679.3585815429688 0 2756.7148
680.3626098632812 0 792.3838
683.3577880859375 0 1263.6145
684.3529052734375 0 1382.1547
686.3309326171875 0 868.5992
688.3641967773438 0 1042.8722
690.8802490234375 0 1994.1835
694.3403930664062 0 4934.905 z Water loss 3
695.33203125 0 1135.8071
696.3688354492188 0 12551.158 c Ammonia loss 6
697.3744506835938 0 4942.21
697.8753051757812 0 860.4441
698.3819580078125 0 11339.641
698.8869018554688 0 23104.566
699.3883666992188 0 11450.331
699.89208984375 0 5499.6035
700.3916015625 0 1211.6831
705.892822265625 0 1597.5702
710.361083984375 0 1219.657 y Water loss 3
711.37646484375 0 2865.9624
712.3511352539062 0 17952.643 z 3
712.385498046875 0 19891.906
713.393310546875 0 162007.23 c 6
714.3959350585938 0 58808.242
715.3994140625 0 12487.938
716.3992919921875 0 1301.1714
726.3569946289062 0 2102.9453
727.3652954101562 0 4553.6655
728.3731689453125 0 28520.385 y 3
729.3765258789062 0 9940.934
730.3535766601562 0 19311.35
731.3561401367188 0 5838.2344
732.3580932617188 0 1396.8058
740.3391723632812 0 1617.8381
742.40673828125 0 3963.0637
742.9111328125 0 3512.0188
743.41162109375 0 2558.6313
744.3671875 0 1154.565
749.4098510742188 0 1695.621
750.4020385742188 0 2339.6665
750.9053344726562 0 1626.9293
751.4017333984375 0 1226.1422
751.9051513671875 0 1082.6199
754.9011840820312 0 1107.2194
756.4044799804688 0 1697.4183
762.4000244140625 0 1329.9885
762.910400390625 0 3928.3474
763.404296875 0 11721.205
763.90625 0 7590.609
764.40478515625 0 4306.2524
764.9043579101562 0 1677.4658
765.4088134765625 0 993.0644
770.41650390625 0 2642.246
770.9187622070312 0 8568.504
771.4135131835938 0 11142.523
771.9177856445312 0 13281.647
772.416748046875 0 8116.5176
772.9171752929688 0 2103.1538
773.4151000976562 0 1954.4033
774.424560546875 0 992.4569
781.3788452148438 0 9958.762 z Water loss 2
782.3825073242188 0 5306.951 w 2
782.4689331054688 0 1260.5498
783.38232421875 0 1581.5012
783.479248046875 0 874.6369
789.4057006835938 0 2083.985
790.4064331054688 0 1106.8705
797.392822265625 0 11227.329 y Water loss 2
798.3870239257812 0 9026.689
799.3875732421875 0 134330.94 z 2
800.3910522460938 0 62536.777
801.3941650390625 0 16865.258
802.389892578125 0 1772.0409
807.4157104492188 0 13232.152
808.4185180664062 0 5627.42
809.4151000976562 0 2094.9072
813.3930053710938 0 1045.31
814.3890991210938 0 1304.7897
815.4061279296875 0 146902.6 y 2
816.4093627929688 0 64657.37
817.4118041992188 0 16851.004
818.4146118164062 0 1936.1351
825.4263305664062 0 44410.85
826.4290771484375 0 22083.879
827.430419921875 0 5546.1714
839.4434204101562 0 2017.2397
841.42333984375 0 1956.8606
843.4365234375 0 89845.55 c Ammonia loss 7
844.4397583007812 0 38810.85
845.4414672851562 0 11132.507
846.4445190429688 0 998.83295
880.428955078125 0 3462.4277
881.4329223632812 0 2140.6633
882.4246215820312 0 7131.2954 z Water loss 1
883.4279174804688 0 4531.811 w 1
885.4210815429688 0 1238.4421 w 1
887.4865112304688 0 909.20917
898.4252319335938 0 13883.486 y Water loss 1
899.4276733398438 0 21080.441
900.4332885742188 0 64999.13 z 1
901.4367065429688 0 29868.453
902.44091796875 0 9408.276
903.4471435546875 0 1020.8705
908.4301147460938 0 1712.2814
909.4286499023438 0 977.26276
913.5020141601562 0 2855.3105
916.452880859375 0 53394.47 y 1
917.4566650390625 0 31295.506
918.460205078125 0 10926.016
919.4666748046875 0 2045.8141
926.4373779296875 0 6970.3726
927.4397583007812 0 3156.3318
951.3693237304688 0 4415.3804
952.3741455078125 0 2187.5408
953.3778686523438 0 954.4504
955.4983520507812 0 2228.2336
956.4993286132812 0 1062.7218
957.5159912109375 0 262933.84 c 8
958.5187377929688 0 152493.62
959.5213623046875 0 43911.82
960.5238647460938 0 6199.9927
970.4640502929688 0 86138.42
971.4663696289062 0 42539.36
972.4678344726562 0 12498.496
973.4779663085938 0 2093.2456
974.479248046875 0 4879.044
975.4797973632812 0 1877.3665
984.5400390625 0 3621.4805
985.5328979492188 0 2574.027
986.5534057617188 0 14755.985
987.55712890625 0 9351.479
988.560546875 0 2728.8574
995.5084228515625 0 8156.759
996.5120849609375 0 4759.013
997.5184326171875 0 1633.6244
1010.4627075195312 0 2253.948
1011.5037841796875 0 5220.9365
1012.5250854492188 0 10090.559
1013.518310546875 0 345802.34
1014.5215454101562 0 196641.69
1015.52392578125 0 64537.438
1016.5274047851562 0 10141.332
1026.5042724609375 0 1114.4327
1027.513671875 0 5707.7495
1028.52734375 0 15348.857
1029.537109375 0 271687.16
1030.5445556640625 0 1059261.5
1031.54736328125 0 579596.6
1032.55029296875 0 176878.58
1033.5528564453125 0 22207.65
1045.5081787109375 0 2003.648
1062.528076171875 0 1318.061
1063.5968017578125 0 1932.423
1078.61181640625 0 2809.2786
1079.6212158203125 0 1751.1383
1080.6195068359375 0 1099.1053
1095.6370849609375 0 5366.135
1096.6434326171875 0 3315.0493
1140.5755615234375 0 1317.3541
1142.5712890625 0 10983.06
1143.5740966796875 0 7498.616
1144.5743408203125 0 2360.6836
1210.6844482421875 0 1242.9857
1268.6439208984375 0 1301.6213
1270.6282958984375 0 8795.664
1271.6324462890625 0 5713.147
1272.639404296875 0 3007.3523
1282.69482421875 0 981.7781
1286.6514892578125 0 1108.2454
1287.6517333984375 0 957.3907
1297.709716796875 0 1771.9259
1298.709716796875 0 1505.5836
1303.73095703125 0 1786.0709
1305.732421875 0 1335.6747
1336.7174072265625 0 1217.6329
1352.7607421875 0 1041.4894
1353.7642822265625 0 2495.6294
1354.7662353515625 0 1563.0454
1381.7705078125 0 1387.3296
1394.76220703125 0 1014.42926
1395.7652587890625 0 1258.754
1396.782470703125 0 3200.1553
1397.787353515625 0 3410.2065
1398.7845458984375 0 2772.34
1411.7894287109375 0 1943.1317
1412.7861328125 0 1641.0215
1425.7359619140625 0 2031.0267
1427.7327880859375 0 3082.6133
1428.7418212890625 0 2146.975
1429.76171875 0 1281.6235
1440.81396484375 0 1135.0126
1483.781005859375 0 3510.8691
1484.794921875 0 5438.2476
1485.8009033203125 0 5928.1914
1486.80712890625 0 1334.7637
1497.8284912109375 0 1532.9202
1498.8143310546875 0 2325.1746
1499.8106689453125 0 2319.867
1508.8157958984375 0 1233.3212
1509.7744140625 0 1615.9294
1510.78955078125 0 1506.2485
1515.8216552734375 0 1163.3317
1523.8170166015625 0 5905.2607
1524.8128662109375 0 8563.957
1525.8157958984375 0 12681.121
1526.8114013671875 0 27191.977
1527.810302734375 0 20188.795
1528.8138427734375 0 8426.09
1529.81494140625 0 4393.826
1530.8150634765625 0 1467.1727
1540.827880859375 0 5877.7197
1541.83154296875 0 21023.184
1542.830078125 0 15243.866
1543.8310546875 0 32642.977
1544.834228515625 0 24644.668
1545.8363037109375 0 10152.389
1546.833740234375 0 4869.987
1547.824951171875 0 2182.6108

Spectrum Details

|  |  |
| --- | --- |
| Matched peaks? Matched peaks The total absolute number of peaks matched. Additionally in brackets the total fraction of peaks matched and the total number of peaks is shown. | 43 (8.35% of 515) |
| FDR? FDR The false discovery rate estimated for this peptide. It is calculated by matching all theoretical fragments with a non-integer shift with the raw peaks for this spectrum. This is done with 40 different shifts. The resulting percentage is the average number of annotated peaks over the number of annotated peaks with the correct spectrum. | 0.06% |
| Satellite FDR? Satellite FDR See the FDR for details on its calculation. This satellite ion specific FDR only contains the satellite ions (d/w) for I/L/J positions. | - |
| PSM Score? PSM Score The PSM Score as given by Hecklib to this annotated spectrum. It is shown with three significant figures. | 492 |

#### Spectrum 5425? Spectrum 5425 The raw spectrum of this peptide as annotated by Hecklib. The fragments are coloured according to ion type (see legend). Any peaks with a star '\*' as text can be hovered over to see the full details, first the ion type second the mass shift type. By hovering over the amino acids in the peptide or ions in the legend the corresponding peaks are highlighted. By toggling the 'Unassigned' label you can turn the background (unassigned) peaks on or off in the plot. By updating the slider in the Ion legend you can update the spectrum to only show the top X% of the peaks with labels. The top X% means any peak that is within X% of the highest intensity. By dragging in the spectrum you can zoom in to a specific part of the spectrum and use 'Zoom Out' to get back to the original zoom level. The annotation of the spectrum is based on the given sequence in the peptides file and is done with different software so inconsistencies are likely. The peaks are annotated based on the given sequence, with 20 ppm tolerance.

Copy Data

##### Spectrum 5425 (TSV)

###### Preview

```
Loading example...
```

*Click on the button to copy the data to your clipboard.*

Mz MinMz MaxIntensity Max

WidthHeightPeptide font sizePeptide stroke widthSpectrum font sizeSpectrum stroke widthCompact peptide

Ion legend

wxyz

abcd

OtherUnassignedIon^Charge^_Position_Show for top:%

JTSGVHTFPA

07.70e+41.54e+52.31e+53.08e+5

Zoom Out

a^+1^_2_a^+1^_2_y^+1^_2_a^+1^_2_b^+1^_2_b^+1^_2_a^+1^_3_b^+1^_3_y^+1^_3_b^+2^_7_b^+2^_7_b^+1^_4_b^+2^_7_b^+1^_4_y^+2^_8_b^+2^_8_b^+2^_8_y^+1^_4_b^+1^_5_y^+2^_9_b^+1^_5_y^+2^_9_b^+2^_9_b^+2^_9_\*\*y^+1^_5_y^+1^_5_b^+1^_6_b^+1^_6_y^+1^_6_y^+1^_6_b^+1^_7_b^+1^_7_y^+1^_7_y^+1^_7_y^+1^_8_y^+1^_8_b^+1^_8_b^+1^_8_y^+1^_9_y^+1^_9_

0665133019952660

Fragment Matches Table

Show background peaks

| Position | Ion type | Intensity | mz Theoretical | mz Error (Th) | mz Error (ppm) | Charge | Series Number |
| --- | --- | --- | --- | --- | --- | --- | --- |
| - | - | 1.537E+04 | 120.1 | - | - | 0 | - |
| - | - | 1599 | 121.1 | - | - | 0 | - |
| - | - | 1349 | 122.1 | - | - | 0 | - |
| - | - | 368.7 | 124.1 | - | - | 0 | - |
| - | - | 373.5 | 125.1 | - | - | 0 | - |
| - | - | 2.067E+04 | 125.1 | - | - | 0 | - |
| - | - | 494.3 | 126.1 | - | - | 0 | - |
| - | - | 719.2 | 126.1 | - | - | 0 | - |
| - | - | 1584 | 126.1 | - | - | 0 | - |
| - | - | 336.9 | 128 | - | - | 0 | - |
| - | - | 482.8 | 128.1 | - | - | 0 | - |
| - | - | 457.9 | 129.1 | - | - | 0 | - |
| - | - | 5634 | 129.1 | - | - | 0 | - |
| - | - | 543.7 | 130.1 | - | - | 0 | - |
| - | - | 1239 | 130.1 | - | - | 0 | - |
| - | - | 447.3 | 131.1 | - | - | 0 | - |
| - | - | 1148 | 132 | - | - | 0 | - |
| - | - | 487.8 | 132.1 | - | - | 0 | - |
| - | - | 1569 | 133.1 | - | - | 0 | - |
| - | - | 3009 | 133.1 | - | - | 0 | - |
| - | - | 4465 | 136.1 | - | - | 0 | - |
| - | - | 630.8 | 137.1 | - | - | 0 | - |
| - | - | 1824 | 138.1 | - | - | 0 | - |
| - | - | 575.7 | 139.1 | - | - | 0 | - |
| - | - | 825.2 | 140.1 | - | - | 0 | - |
| - | - | 2173 | 141.1 | - | - | 0 | - |
| - | - | 6.893E+04 | 142.1 | - | - | 0 | - |
| - | - | 948.2 | 143.1 | - | - | 0 | - |
| - | - | 5604 | 143.1 | - | - | 0 | - |
| - | - | 590.5 | 144.1 | - | - | 0 | - |
| - | - | 3536 | 145.1 | - | - | 0 | - |
| - | - | 675.2 | 146.1 | - | - | 0 | - |
| - | - | 742.6 | 147.1 | - | - | 0 | - |
| - | - | 1888 | 150.1 | - | - | 0 | - |
| - | - | 470.1 | 151.4 | - | - | 0 | - |
| - | - | 1016 | 152.1 | - | - | 0 | - |
| - | - | 752.8 | 152.1 | - | - | 0 | - |
| - | - | 3.928E+04 | 152.1 | - | - | 0 | - |
| - | - | 794.3 | 153.1 | - | - | 0 | - |
| - | - | 3132 | 153.1 | - | - | 0 | - |
| - | - | 633.3 | 154.1 | - | - | 0 | - |
| - | - | 509.1 | 154.1 | - | - | 0 | - |
| - | - | 812 | 155.1 | - | - | 0 | - |
| - | - | 5152 | 155.1 | - | - | 0 | - |
| - | - | 3715 | 156.1 | - | - | 0 | - |
| - | - | 3663 | 157.1 | - | - | 0 | - |
| - | - | 788.9 | 157.1 | - | - | 0 | - |
| - | - | 551.5 | 157.1 | - | - | 0 | - |
| - | - | 2308 | 158.1 | - | - | 0 | - |
| - | - | 472.8 | 158.1 | - | - | 0 | - |
| - | - | 3936 | 161.1 | - | - | 0 | - |
| - | - | 1.147E+04 | 163.1 | - | - | 0 | - |
| - | - | 1445 | 164.1 | - | - | 0 | - |
| - | - | 1241 | 165.1 | - | - | 0 | - |
| - | - | 573.5 | 165.1 | - | - | 0 | - |
| - | - | 994.4 | 166.1 | - | - | 0 | - |
| - | - | 2442 | 166.1 | - | - | 0 | - |
| - | - | 5850 | 166.1 | - | - | 0 | - |
| - | - | 1236 | 167.1 | - | - | 0 | - |
| - | - | 747 | 169.1 | - | - | 0 | - |
| 2 | a | 2.366E+04 | 169.1 | 0.000341 | 2.016 | +1 | 2 |
| 2 | a | 662.3 | 170.1 | 0.0008835 | 5.194 | +1 | 2 |
| - | - | 2096 | 170.1 | - | - | 0 | - |
| - | - | 7056 | 171.1 | - | - | 0 | - |
| - | - | 1429 | 171.1 | - | - | 0 | - |
| - | - | 457.3 | 172.1 | - | - | 0 | - |
| - | - | 1460 | 174.1 | - | - | 0 | - |
| - | - | 7999 | 175.1 | - | - | 0 | - |
| - | - | 778.3 | 175.1 | - | - | 0 | - |
| - | - | 1661 | 176.1 | - | - | 0 | - |
| - | - | 1041 | 176.1 | - | - | 0 | - |
| - | - | 560.7 | 176.1 | - | - | 0 | - |
| - | - | 1763 | 177.1 | - | - | 0 | - |
| - | - | 4035 | 178.1 | - | - | 0 | - |
| - | - | 2025 | 180.1 | - | - | 0 | - |
| - | - | 801.8 | 181.1 | - | - | 0 | - |
| - | - | 880.6 | 183.1 | - | - | 0 | - |
| - | - | 892.9 | 183.1 | - | - | 0 | - |
| - | - | 1280 | 185.1 | - | - | 0 | - |
| - | - | 556.7 | 185.1 | - | - | 0 | - |
| - | - | 608.9 | 185.1 | - | - | 0 | - |
| - | - | 1747 | 185.2 | - | - | 0 | - |
| 9 | y | 3.048E+05 | 187.1 | 0.0004203 | 2.246 | +1 | 2 |
| 2 | a | 1.405E+05 | 187.1 | 0.0003812 | 2.037 | +1 | 2 |
| - | - | 2.587E+04 | 188.1 | - | - | 0 | - |
| - | - | 1.114E+04 | 188.1 | - | - | 0 | - |
| - | - | 3912 | 189.1 | - | - | 0 | - |
| - | - | 1925 | 189.1 | - | - | 0 | - |
| - | - | 580.7 | 191.1 | - | - | 0 | - |
| - | - | 1.288E+04 | 193.1 | - | - | 0 | - |
| - | - | 4692 | 194.1 | - | - | 0 | - |
| - | - | 1099 | 194.1 | - | - | 0 | - |
| - | - | 1.156E+04 | 195.1 | - | - | 0 | - |
| - | - | 975.5 | 195.1 | - | - | 0 | - |
| - | - | 1503 | 195.1 | - | - | 0 | - |
| - | - | 972.1 | 195.1 | - | - | 0 | - |
| 2 | b | 2.619E+04 | 197.1 | 0.0003452 | 1.751 | +1 | 2 |
| - | - | 676.5 | 198.1 | - | - | 0 | - |
| - | - | 2317 | 198.1 | - | - | 0 | - |
| - | - | 871.5 | 199.1 | - | - | 0 | - |
| - | - | 1125 | 199.1 | - | - | 0 | - |
| - | - | 649.3 | 201.1 | - | - | 0 | - |
| - | - | 808.5 | 201.1 | - | - | 0 | - |
| - | - | 882.8 | 205.1 | - | - | 0 | - |
| - | - | 935.1 | 207.1 | - | - | 0 | - |
| - | - | 549.9 | 207.1 | - | - | 0 | - |
| - | - | 683.5 | 209.1 | - | - | 0 | - |
| - | - | 3469 | 209.1 | - | - | 0 | - |
| - | - | 1252 | 210.1 | - | - | 0 | - |
| - | - | 3767 | 211.1 | - | - | 0 | - |
| - | - | 614.4 | 211.1 | - | - | 0 | - |
| - | - | 1215 | 212.1 | - | - | 0 | - |
| - | - | 2022 | 213.2 | - | - | 0 | - |
| 2 | b | 9.107E+04 | 215.1 | 0.0003396 | 1.579 | +1 | 2 |
| - | - | 8907 | 216.1 | - | - | 0 | - |
| - | - | 2701 | 217.1 | - | - | 0 | - |
| - | - | 735.4 | 217.1 | - | - | 0 | - |
| - | - | 1208 | 219.1 | - | - | 0 | - |
| - | - | 1221 | 219.1 | - | - | 0 | - |
| - | - | 1387 | 220.1 | - | - | 0 | - |
| - | - | 3.554E+04 | 221.1 | - | - | 0 | - |
| - | - | 2621 | 221.1 | - | - | 0 | - |
| - | - | 2748 | 222.1 | - | - | 0 | - |
| - | - | 1901 | 223.1 | - | - | 0 | - |
| - | - | 1509 | 223.1 | - | - | 0 | - |
| - | - | 1587 | 226.1 | - | - | 0 | - |
| - | - | 851.3 | 226.2 | - | - | 0 | - |
| - | - | 620.7 | 227.1 | - | - | 0 | - |
| - | - | 7365 | 228.1 | - | - | 0 | - |
| - | - | 1255 | 231.1 | - | - | 0 | - |
| - | - | 1339 | 231.1 | - | - | 0 | - |
| - | - | 2508 | 233.2 | - | - | 0 | - |
| - | - | 909.6 | 235.1 | - | - | 0 | - |
| - | - | 3546 | 237.1 | - | - | 0 | - |
| - | - | 8498 | 237.1 | - | - | 0 | - |
| - | - | 4264 | 238.1 | - | - | 0 | - |
| - | - | 3.882E+04 | 239.1 | - | - | 0 | - |
| - | - | 1213 | 239.1 | - | - | 0 | - |
| - | - | 4076 | 239.2 | - | - | 0 | - |
| - | - | 3442 | 240.1 | - | - | 0 | - |
| - | - | 658.7 | 240.1 | - | - | 0 | - |
| - | - | 759.8 | 241.1 | - | - | 0 | - |
| - | - | 1734 | 241.2 | - | - | 0 | - |
| - | - | 3780 | 244.1 | - | - | 0 | - |
| - | - | 2969 | 245.1 | - | - | 0 | - |
| - | - | 5071 | 246.1 | - | - | 0 | - |
| - | - | 662 | 247.1 | - | - | 0 | - |
| - | - | 696.1 | 248.2 | - | - | 0 | - |
| - | - | 1973 | 249.1 | - | - | 0 | - |
| - | - | 1553 | 249.1 | - | - | 0 | - |
| - | - | 727.7 | 250.1 | - | - | 0 | - |
| - | - | 1360 | 251.2 | - | - | 0 | - |
| - | - | 595.8 | 254.1 | - | - | 0 | - |
| - | - | 800.8 | 255.1 | - | - | 0 | - |
| - | - | 6831 | 256.1 | - | - | 0 | - |
| - | - | 1036 | 256.2 | - | - | 0 | - |
| 3 | a | 701.1 | 257.1 | 0.00197 | 7.661 | +1 | 3 |
| - | - | 1044 | 259.1 | - | - | 0 | - |
| - | - | 791.2 | 261.2 | - | - | 0 | - |
| - | - | 988.3 | 265.1 | - | - | 0 | - |
| - | - | 852.2 | 266.1 | - | - | 0 | - |
| - | - | 719.6 | 266.2 | - | - | 0 | - |
| - | - | 1634 | 267.1 | - | - | 0 | - |
| - | - | 1022 | 269.2 | - | - | 0 | - |
| - | - | 648.4 | 275.1 | - | - | 0 | - |
| - | - | 7464 | 276.1 | - | - | 0 | - |
| - | - | 833 | 277.1 | - | - | 0 | - |
| - | - | 785.7 | 277.1 | - | - | 0 | - |
| - | - | 880.2 | 281.2 | - | - | 0 | - |
| - | - | 605.1 | 282.1 | - | - | 0 | - |
| - | - | 737 | 282.1 | - | - | 0 | - |
| - | - | 764.9 | 283.1 | - | - | 0 | - |
| - | - | 2441 | 284.1 | - | - | 0 | - |
| 3 | b | 1.313E+04 | 284.2 | 0.000406 | 1.429 | +1 | 3 |
| - | - | 1175 | 285.2 | - | - | 0 | - |
| - | - | 804.3 | 289.2 | - | - | 0 | - |
| - | - | 1.154E+04 | 294.2 | - | - | 0 | - |
| - | - | 935.3 | 294.2 | - | - | 0 | - |
| - | - | 2243 | 295.2 | - | - | 0 | - |
| - | - | 830.5 | 296.1 | - | - | 0 | - |
| - | - | 841.1 | 302.2 | - | - | 0 | - |
| - | - | 1340 | 303.1 | - | - | 0 | - |
| - | - | 680.7 | 309.2 | - | - | 0 | - |
| - | - | 1088 | 311.2 | - | - | 0 | - |
| - | - | 539.9 | 312.2 | - | - | 0 | - |
| - | - | 1833 | 314.2 | - | - | 0 | - |
| - | - | 735.1 | 315.7 | - | - | 0 | - |
| - | - | 5479 | 317.2 | - | - | 0 | - |
| - | - | 802.5 | 317.7 | - | - | 0 | - |
| - | - | 775.2 | 318.2 | - | - | 0 | - |
| - | - | 4633 | 320.2 | - | - | 0 | - |
| - | - | 912.8 | 321.7 | - | - | 0 | - |
| - | - | 1260 | 322.2 | - | - | 0 | - |
| - | - | 855.2 | 322.2 | - | - | 0 | - |
| - | - | 1727 | 323.2 | - | - | 0 | - |
| - | - | 1.419E+04 | 324.1 | - | - | 0 | - |
| - | - | 646.7 | 324.2 | - | - | 0 | - |
| - | - | 2871 | 325.1 | - | - | 0 | - |
| - | - | 2834 | 327.2 | - | - | 0 | - |
| - | - | 881 | 329.1 | - | - | 0 | - |
| - | - | 2182 | 330.7 | - | - | 0 | - |
| - | - | 1009 | 333.7 | - | - | 0 | - |
| 8 | y | 1.154E+04 | 334.2 | 0.0005336 | 1.597 | +1 | 3 |
| - | - | 4446 | 335.2 | - | - | 0 | - |
| - | - | 1596 | 336.2 | - | - | 0 | - |
| - | - | 783 | 337.2 | - | - | 0 | - |
| - | - | 3756 | 338.2 | - | - | 0 | - |
| 7 | b | 2538 | 339.7 | 7.618E-05 | 0.2243 | +2 | 7 |
| 7 | b | 1.292E+04 | 340.2 | 0.003338 | 9.814 | +2 | 7 |
| 4 | b | 5879 | 341.2 | 0.0004278 | 1.254 | +1 | 4 |
| - | - | 7913 | 342.2 | - | - | 0 | - |
| - | - | 909.8 | 342.2 | - | - | 0 | - |
| - | - | 1259 | 343.2 | - | - | 0 | - |
| - | - | 3887 | 345.2 | - | - | 0 | - |
| - | - | 841.8 | 346.2 | - | - | 0 | - |
| - | - | 799.2 | 347.7 | - | - | 0 | - |
| 7 | b | 1906 | 348.7 | 0.0002621 | 0.7516 | +2 | 7 |
| - | - | 830.3 | 349.2 | - | - | 0 | - |
| - | - | 1.045E+04 | 350.2 | - | - | 0 | - |
| - | - | 818.2 | 350.2 | - | - | 0 | - |
| - | - | 1651 | 351.2 | - | - | 0 | - |
| - | - | 675.3 | 352.2 | - | - | 0 | - |
| - | - | 1.634E+04 | 353.2 | - | - | 0 | - |
| - | - | 947.6 | 353.2 | - | - | 0 | - |
| - | - | 1876 | 354.2 | - | - | 0 | - |
| - | - | 1311 | 354.7 | - | - | 0 | - |
| - | - | 856.9 | 356.7 | - | - | 0 | - |
| - | - | 2202 | 358.2 | - | - | 0 | - |
| 4 | b | 2913 | 359.2 | 0.001074 | 2.991 | +1 | 4 |
| - | - | 890.2 | 360.2 | - | - | 0 | - |
| - | - | 5764 | 363.2 | - | - | 0 | - |
| - | - | 877.4 | 363.2 | - | - | 0 | - |
| - | - | 1645 | 363.7 | - | - | 0 | - |
| - | - | 1085 | 364.2 | - | - | 0 | - |
| - | - | 831.7 | 365.2 | - | - | 0 | - |
| - | - | 1262 | 365.2 | - | - | 0 | - |
| - | - | 665.6 | 365.7 | - | - | 0 | - |
| - | - | 906.1 | 367.2 | - | - | 0 | - |
| - | - | 7179 | 368.2 | - | - | 0 | - |
| - | - | 1638 | 369.2 | - | - | 0 | - |
| - | - | 852.5 | 369.2 | - | - | 0 | - |
| - | - | 973.7 | 371.2 | - | - | 0 | - |
| - | - | 1026 | 375.2 | - | - | 0 | - |
| - | - | 994 | 375.2 | - | - | 0 | - |
| - | - | 7835 | 377.2 | - | - | 0 | - |
| - | - | 1476 | 378.2 | - | - | 0 | - |
| - | - | 3.109E+04 | 381.2 | - | - | 0 | - |
| - | - | 5277 | 382.2 | - | - | 0 | - |
| - | - | 651.3 | 382.7 | - | - | 0 | - |
| - | - | 1023 | 383.2 | - | - | 0 | - |
| - | - | 1239 | 383.2 | - | - | 0 | - |
| - | - | 2690 | 383.2 | - | - | 0 | - |
| - | - | 900.4 | 384.2 | - | - | 0 | - |
| - | - | 1.664E+04 | 386.2 | - | - | 0 | - |
| - | - | 1151 | 386.2 | - | - | 0 | - |
| - | - | 2613 | 387.2 | - | - | 0 | - |
| - | - | 1618 | 388.2 | - | - | 0 | - |
| - | - | 804.7 | 390.2 | - | - | 0 | - |
| - | - | 1176 | 390.7 | - | - | 0 | - |
| - | - | 1141 | 391.2 | - | - | 0 | - |
| - | - | 2061 | 392.2 | - | - | 0 | - |
| - | - | 639.2 | 393.2 | - | - | 0 | - |
| - | - | 7941 | 395.2 | - | - | 0 | - |
| - | - | 1138 | 396.2 | - | - | 0 | - |
| - | - | 2470 | 396.2 | - | - | 0 | - |
| - | - | 769.2 | 396.2 | - | - | 0 | - |
| - | - | 1997 | 397.2 | - | - | 0 | - |
| 3 | y | 3112 | 399.2 | 0.0005972 | 1.496 | +2 | 8 |
| - | - | 859.9 | 399.2 | - | - | 0 | - |
| - | - | 711.3 | 399.7 | - | - | 0 | - |
| - | - | 1801 | 399.7 | - | - | 0 | - |
| - | - | 876.4 | 400.2 | - | - | 0 | - |
| - | - | 646.6 | 403.7 | - | - | 0 | - |
| - | - | 3043 | 404.2 | - | - | 0 | - |
| - | - | 1008 | 404.7 | - | - | 0 | - |
| - | - | 1129 | 405.2 | - | - | 0 | - |
| - | - | 5707 | 408.2 | - | - | 0 | - |
| - | - | 2583 | 408.7 | - | - | 0 | - |
| - | - | 1206 | 409.2 | - | - | 0 | - |
| - | - | 2017 | 410.2 | - | - | 0 | - |
| - | - | 1202 | 412.3 | - | - | 0 | - |
| 8 | b | 5352 | 413.2 | 0.0003238 | 0.7836 | +2 | 8 |
| - | - | 1472 | 413.7 | - | - | 0 | - |
| - | - | 988 | 414.2 | - | - | 0 | - |
| - | - | 1849 | 414.2 | - | - | 0 | - |
| - | - | 961.7 | 414.7 | - | - | 0 | - |
| - | - | 3652 | 418.2 | - | - | 0 | - |
| - | - | 874.4 | 418.7 | - | - | 0 | - |
| - | - | 947.8 | 419.2 | - | - | 0 | - |
| - | - | 755.8 | 419.2 | - | - | 0 | - |
| - | - | 1633 | 420.2 | - | - | 0 | - |
| 8 | b | 3470 | 422.2 | 0.0005957 | 1.411 | +2 | 8 |
| - | - | 1247 | 422.7 | - | - | 0 | - |
| - | - | 769.9 | 427.6 | - | - | 0 | - |
| - | - | 754.5 | 427.9 | - | - | 0 | - |
| - | - | 3845 | 430.3 | - | - | 0 | - |
| - | - | 1073 | 431.3 | - | - | 0 | - |
| 7 | y | 1199 | 435.2 | 0.003148 | 7.233 | +1 | 4 |
| - | - | 6634 | 436.2 | - | - | 0 | - |
| - | - | 5484 | 437.2 | - | - | 0 | - |
| - | - | 8122 | 438.2 | - | - | 0 | - |
| - | - | 862.5 | 438.2 | - | - | 0 | - |
| - | - | 1838 | 439.2 | - | - | 0 | - |
| - | - | 882.2 | 439.2 | - | - | 0 | - |
| - | - | 765.9 | 439.7 | - | - | 0 | - |
| 5 | b | 3198 | 440.3 | 0.0002687 | 0.6104 | +1 | 5 |
| - | - | 3088 | 440.7 | - | - | 0 | - |
| - | - | 1249 | 441.2 | - | - | 0 | - |
| - | - | 862.5 | 441.3 | - | - | 0 | - |
| - | - | 7245 | 446.2 | - | - | 0 | - |
| - | - | 1760 | 447.2 | - | - | 0 | - |
| - | - | 1599 | 447.7 | - | - | 0 | - |
| - | - | 877.2 | 448.7 | - | - | 0 | - |
| 2 | y | 1669 | 449.7 | 0.000527 | 1.172 | +2 | 9 |
| - | - | 2188 | 452.7 | - | - | 0 | - |
| - | - | 749.8 | 453.2 | - | - | 0 | - |
| - | - | 1.507E+04 | 454.2 | - | - | 0 | - |
| - | - | 3446 | 455.2 | - | - | 0 | - |
| - | - | 925.5 | 455.3 | - | - | 0 | - |
| - | - | 739 | 456.2 | - | - | 0 | - |
| - | - | 899.6 | 456.3 | - | - | 0 | - |
| - | - | 1491 | 456.7 | - | - | 0 | - |
| - | - | 1610 | 457.3 | - | - | 0 | - |
| 5 | b | 2803 | 458.3 | 6.137E-05 | 0.1339 | +1 | 5 |
| 2 | y | 3348 | 458.7 | 0.00028 | 0.6104 | +2 | 9 |
| - | - | 2051 | 459.2 | - | - | 0 | - |
| - | - | 723.9 | 459.3 | - | - | 0 | - |
| - | - | 1284 | 459.7 | - | - | 0 | - |
| - | - | 709.8 | 461.2 | - | - | 0 | - |
| - | - | 2709 | 461.3 | - | - | 0 | - |
| 9 | b | 2415 | 461.7 | 0.0003396 | 0.7354 | +2 | 9 |
| - | - | 2182 | 462.2 | - | - | 0 | - |
| - | - | 3.866E+04 | 464.2 | - | - | 0 | - |
| - | - | 1.062E+04 | 465.2 | - | - | 0 | - |
| - | - | 1231 | 466.2 | - | - | 0 | - |
| - | - | 1562 | 467.2 | - | - | 0 | - |
| - | - | 858.3 | 468.2 | - | - | 0 | - |
| 9 | b | 3940 | 470.7 | 0.0006114 | 1.299 | +2 | 9 |
| - | - | 2292 | 471.3 | - | - | 0 | - |
| - | - | 659.4 | 471.7 | - | - | 0 | - |
| - | - | 3136 | 473.3 | - | - | 0 | - |
| - | - | 1363 | 474.2 | - | - | 0 | - |
| - | - | 685.4 | 474.8 | - | - | 0 | - |
| - | - | 654 | 475.2 | - | - | 0 | - |
| - | - | 1879 | 475.3 | - | - | 0 | - |
| - | - | 1203 | 475.8 | - | - | 0 | - |
| - | - | 5.655E+04 | 482.2 | - | - | 0 | - |
| - | - | 1.536E+04 | 483.2 | - | - | 0 | - |
| - | - | 3026 | 484.2 | - | - | 0 | - |
| - | - | 2526 | 485.3 | - | - | 0 | - |
| - | - | 628.6 | 486.2 | - | - | 0 | - |
| - | - | 1135 | 488.3 | - | - | 0 | - |
| - | - | 806.2 | 488.7 | - | - | 0 | - |
| - | - | 1471 | 489.6 | - | - | 0 | - |
| - | - | 653.4 | 489.9 | - | - | 0 | - |
| - | - | 1044 | 490.3 | - | - | 0 | - |
| - | - | 2981 | 492.2 | - | - | 0 | - |
| - | - | 638.6 | 493.3 | - | - | 0 | - |
| - | - | 2086 | 496.3 | - | - | 0 | - |
| - | - | 1.045E+04 | 497.3 | - | - | 0 | - |
| - | - | 7373 | 497.8 | - | - | 0 | - |
| - | - | 1764 | 498.3 | - | - | 0 | - |
| - | - | 1508 | 499.3 | - | - | 0 | - |
| - | - | 967.4 | 503.2 | - | - | 0 | - |
| 0 | Precursor | 1.07E+04 | 506.3 | 3.164E-05 | 0.0625 | +2 | -1 |
| - | - | 6868 | 506.8 | - | - | 0 | - |
| - | - | 2099 | 507.3 | - | - | 0 | - |
| - | - | 903.9 | 508.6 | - | - | 0 | - |
| - | - | 927.7 | 511.3 | - | - | 0 | - |
| - | - | 1037 | 512.2 | - | - | 0 | - |
| - | - | 2668 | 514.3 | - | - | 0 | - |
| 0 | Precursor | 1.007E+04 | 515.3 | 0.0005759 | 1.118 | +2 | -1 |
| - | - | 5892 | 515.8 | - | - | 0 | - |
| - | - | 1948 | 516.3 | - | - | 0 | - |
| - | - | 1617 | 521.2 | - | - | 0 | - |
| - | - | 1111 | 521.8 | - | - | 0 | - |
| - | - | 8156 | 524.3 | - | - | 0 | - |
| - | - | 2874 | 525.3 | - | - | 0 | - |
| - | - | 7378 | 525.8 | - | - | 0 | - |
| - | - | 4016 | 526.3 | - | - | 0 | - |
| - | - | 2091 | 526.8 | - | - | 0 | - |
| - | - | 845.7 | 528.3 | - | - | 0 | - |
| - | - | 2485 | 529.3 | - | - | 0 | - |
| - | - | 1129 | 531.3 | - | - | 0 | - |
| - | - | 2167 | 537.3 | - | - | 0 | - |
| - | - | 1176 | 538.3 | - | - | 0 | - |
| - | - | 3252 | 539.3 | - | - | 0 | - |
| - | - | 2.777E+04 | 539.8 | - | - | 0 | - |
| - | - | 902.2 | 540.3 | - | - | 0 | - |
| - | - | 1.555E+04 | 540.3 | - | - | 0 | - |
| - | - | 4801 | 540.8 | - | - | 0 | - |
| - | - | 714.4 | 541.8 | - | - | 0 | - |
| - | - | 9272 | 542.3 | - | - | 0 | - |
| - | - | 2905 | 543.3 | - | - | 0 | - |
| - | - | 782.6 | 544.3 | - | - | 0 | - |
| - | - | 5810 | 547.3 | - | - | 0 | - |
| - | - | 1895 | 548.3 | - | - | 0 | - |
| - | - | 667.6 | 548.3 | - | - | 0 | - |
| - | - | 1129 | 548.8 | - | - | 0 | - |
| - | - | 1889 | 549.3 | - | - | 0 | - |
| - | - | 669.8 | 550.3 | - | - | 0 | - |
| - | - | 739.6 | 553.3 | - | - | 0 | - |
| 6 | y | 7631 | 554.3 | 0.0007298 | 1.317 | +1 | 5 |
| - | - | 4118 | 555.3 | - | - | 0 | - |
| - | - | 1061 | 556.3 | - | - | 0 | - |
| - | - | 3607 | 557.3 | - | - | 0 | - |
| - | - | 1155 | 558.3 | - | - | 0 | - |
| - | - | 1038 | 559.3 | - | - | 0 | - |
| - | - | 625.6 | 563.6 | - | - | 0 | - |
| - | - | 1.517E+04 | 565.3 | - | - | 0 | - |
| - | - | 6089 | 566.3 | - | - | 0 | - |
| - | - | 2539 | 567.3 | - | - | 0 | - |
| - | - | 5496 | 567.3 | - | - | 0 | - |
| - | - | 2025 | 568.3 | - | - | 0 | - |
| - | - | 1248 | 571.3 | - | - | 0 | - |
| 6 | y | 1.578E+04 | 572.3 | 8.189E-06 | 0.01431 | +1 | 5 |
| - | - | 5606 | 573.3 | - | - | 0 | - |
| - | - | 1985 | 574.3 | - | - | 0 | - |
| - | - | 1673 | 574.8 | - | - | 0 | - |
| - | - | 2780 | 575.3 | - | - | 0 | - |
| 6 | b | 2521 | 577.3 | 0.0006784 | 1.175 | +1 | 6 |
| - | - | 1144 | 578.3 | - | - | 0 | - |
| - | - | 2.072E+04 | 583.3 | - | - | 0 | - |
| - | - | 6635 | 584.3 | - | - | 0 | - |
| - | - | 8410 | 585.3 | - | - | 0 | - |
| - | - | 3038 | 586.3 | - | - | 0 | - |
| - | - | 719.5 | 589.3 | - | - | 0 | - |
| - | - | 7035 | 593.3 | - | - | 0 | - |
| - | - | 2487 | 594.3 | - | - | 0 | - |
| 6 | b | 1.172E+04 | 595.3 | 0.0001705 | 0.2864 | +1 | 6 |
| - | - | 2816 | 596.3 | - | - | 0 | - |
| - | - | 1.747E+04 | 601.3 | - | - | 0 | - |
| - | - | 5049 | 602.3 | - | - | 0 | - |
| - | - | 676.1 | 603.3 | - | - | 0 | - |
| - | - | 5.883E+04 | 611.3 | - | - | 0 | - |
| - | - | 2.105E+04 | 612.3 | - | - | 0 | - |
| - | - | 3808 | 613.3 | - | - | 0 | - |
| - | - | 810.8 | 618.9 | - | - | 0 | - |
| - | - | 944.5 | 624.3 | - | - | 0 | - |
| - | - | 712.4 | 626.3 | - | - | 0 | - |
| - | - | 1592 | 626.8 | - | - | 0 | - |
| - | - | 1865 | 627.4 | - | - | 0 | - |
| - | - | 1391 | 627.9 | - | - | 0 | - |
| - | - | 7.086E+04 | 629.3 | - | - | 0 | - |
| - | - | 2.456E+04 | 630.3 | - | - | 0 | - |
| - | - | 1279 | 630.4 | - | - | 0 | - |
| - | - | 4580 | 631.3 | - | - | 0 | - |
| - | - | 784.5 | 632.3 | - | - | 0 | - |
| - | - | 825.6 | 633.3 | - | - | 0 | - |
| - | - | 1222 | 634.3 | - | - | 0 | - |
| - | - | 850.7 | 640.3 | - | - | 0 | - |
| - | - | 1.474E+04 | 640.8 | - | - | 0 | - |
| - | - | 1.057E+04 | 641.3 | - | - | 0 | - |
| - | - | 2820 | 641.8 | - | - | 0 | - |
| - | - | 2318 | 642.3 | - | - | 0 | - |
| - | - | 948.3 | 642.8 | - | - | 0 | - |
| - | - | 1265 | 643.3 | - | - | 0 | - |
| - | - | 1538 | 650.3 | - | - | 0 | - |
| - | - | 2087 | 650.4 | - | - | 0 | - |
| - | - | 1785 | 651.4 | - | - | 0 | - |
| - | - | 3519 | 652.3 | - | - | 0 | - |
| 5 | y | 1838 | 653.3 | 0.002323 | 3.556 | +1 | 6 |
| - | - | 1025 | 654.3 | - | - | 0 | - |
| - | - | 1843 | 658.3 | - | - | 0 | - |
| - | - | 4677 | 660.3 | - | - | 0 | - |
| - | - | 1761 | 661.3 | - | - | 0 | - |
| - | - | 1353 | 666.3 | - | - | 0 | - |
| - | - | 1321 | 668.3 | - | - | 0 | - |
| - | - | 3498 | 668.4 | - | - | 0 | - |
| - | - | 1210 | 669.4 | - | - | 0 | - |
| - | - | 729.7 | 670.4 | - | - | 0 | - |
| 5 | y | 3332 | 671.4 | 0.0001848 | 0.2753 | +1 | 6 |
| - | - | 788.9 | 672.4 | - | - | 0 | - |
| - | - | 3180 | 676.3 | - | - | 0 | - |
| - | - | 9026 | 676.4 | - | - | 0 | - |
| - | - | 7882 | 676.9 | - | - | 0 | - |
| - | - | 1183 | 677.3 | - | - | 0 | - |
| - | - | 2749 | 677.4 | - | - | 0 | - |
| - | - | 697.7 | 677.9 | - | - | 0 | - |
| 7 | b | 1.709E+04 | 678.4 | 7.805E-05 | 0.1151 | +1 | 7 |
| - | - | 6943 | 679.4 | - | - | 0 | - |
| - | - | 1553 | 680.4 | - | - | 0 | - |
| - | - | 824.7 | 682.3 | - | - | 0 | - |
| - | - | 723.4 | 683.3 | - | - | 0 | - |
| - | - | 5000 | 684.3 | - | - | 0 | - |
| - | - | 2188 | 685.3 | - | - | 0 | - |
| - | - | 4242 | 686.3 | - | - | 0 | - |
| - | - | 1417 | 687.3 | - | - | 0 | - |
| - | - | 880.7 | 688.4 | - | - | 0 | - |
| - | - | 7904 | 690.4 | - | - | 0 | - |
| - | - | 7353 | 690.9 | - | - | 0 | - |
| - | - | 3148 | 691.4 | - | - | 0 | - |
| - | - | 671.5 | 692.4 | - | - | 0 | - |
| - | - | 1.004E+04 | 694.3 | - | - | 0 | - |
| - | - | 2666 | 695.3 | - | - | 0 | - |
| 7 | b | 1.755E+04 | 696.4 | 0.0002668 | 0.3831 | +1 | 7 |
| - | - | 7358 | 697.4 | - | - | 0 | - |
| - | - | 1845 | 698.4 | - | - | 0 | - |
| - | - | 6453 | 702.4 | - | - | 0 | - |
| - | - | 1989 | 703.4 | - | - | 0 | - |
| - | - | 1383 | 708.3 | - | - | 0 | - |
| 4 | y | 1771 | 710.4 | 0.002783 | 3.918 | +1 | 7 |
| - | - | 1095 | 711.4 | - | - | 0 | - |
| - | - | 2.255E+04 | 712.3 | - | - | 0 | - |
| - | - | 8232 | 713.3 | - | - | 0 | - |
| - | - | 1368 | 714.3 | - | - | 0 | - |
| - | - | 3440 | 722.3 | - | - | 0 | - |
| - | - | 1709 | 723.3 | - | - | 0 | - |
| - | - | 716.9 | 725.4 | - | - | 0 | - |
| - | - | 5390 | 726.4 | - | - | 0 | - |
| - | - | 1803 | 727.4 | - | - | 0 | - |
| 4 | y | 1.114E+04 | 728.4 | 0.001141 | 1.566 | +1 | 7 |
| - | - | 4665 | 729.4 | - | - | 0 | - |
| - | - | 3.042E+04 | 730.4 | - | - | 0 | - |
| - | - | 1.065E+04 | 731.4 | - | - | 0 | - |
| - | - | 2892 | 732.4 | - | - | 0 | - |
| - | - | 3660 | 740.3 | - | - | 0 | - |
| - | - | 1456 | 741.3 | - | - | 0 | - |
| - | - | 1475 | 742.4 | - | - | 0 | - |
| - | - | 1480 | 743.4 | - | - | 0 | - |
| - | - | 2408 | 744.4 | - | - | 0 | - |
| - | - | 1130 | 745.4 | - | - | 0 | - |
| - | - | 944.6 | 746.4 | - | - | 0 | - |
| - | - | 822.5 | 753.4 | - | - | 0 | - |
| - | - | 770 | 754.4 | - | - | 0 | - |
| - | - | 1950 | 763.4 | - | - | 0 | - |
| - | - | 1869 | 764.4 | - | - | 0 | - |
| - | - | 940.4 | 765.4 | - | - | 0 | - |
| - | - | 3286 | 771.4 | - | - | 0 | - |
| - | - | 690 | 772.4 | - | - | 0 | - |
| - | - | 1772 | 779.4 | - | - | 0 | - |
| - | - | 812.4 | 780.4 | - | - | 0 | - |
| - | - | 3033 | 781.4 | - | - | 0 | - |
| - | - | 1261 | 782.4 | - | - | 0 | - |
| - | - | 2852 | 789.4 | - | - | 0 | - |
| - | - | 2228 | 790.4 | - | - | 0 | - |
| 3 | y | 1.109E+04 | 797.4 | 0.004983 | 6.25 | +1 | 8 |
| - | - | 6556 | 798.4 | - | - | 0 | - |
| - | - | 1.077E+04 | 799.4 | - | - | 0 | - |
| - | - | 3868 | 800.4 | - | - | 0 | - |
| - | - | 1143 | 801.4 | - | - | 0 | - |
| - | - | 1.556E+04 | 807.4 | - | - | 0 | - |
| - | - | 7031 | 808.4 | - | - | 0 | - |
| - | - | 1646 | 809.4 | - | - | 0 | - |
| - | - | 1017 | 813.4 | - | - | 0 | - |
| 3 | y | 1.123E+05 | 815.4 | 0.0005223 | 0.6405 | +1 | 8 |
| - | - | 5.378E+04 | 816.4 | - | - | 0 | - |
| - | - | 1.254E+04 | 817.4 | - | - | 0 | - |
| - | - | 798.6 | 818.4 | - | - | 0 | - |
| - | - | 1427 | 819.5 | - | - | 0 | - |
| - | - | 1168 | 820.5 | - | - | 0 | - |
| 8 | b | 5.111E+04 | 825.4 | 0.000926 | 1.122 | +1 | 8 |
| - | - | 2.303E+04 | 826.4 | - | - | 0 | - |
| - | - | 7230 | 827.4 | - | - | 0 | - |
| - | - | 867.9 | 828.4 | - | - | 0 | - |
| - | - | 1063 | 839.4 | - | - | 0 | - |
| - | - | 979.1 | 841.4 | - | - | 0 | - |
| - | - | 756.8 | 842.4 | - | - | 0 | - |
| 8 | b | 7.155E+04 | 843.4 | 0.001115 | 1.322 | +1 | 8 |
| - | - | 3.041E+04 | 844.4 | - | - | 0 | - |
| - | - | 9085 | 845.4 | - | - | 0 | - |
| - | - | 1046 | 846.4 | - | - | 0 | - |
| - | - | 717.9 | 854.4 | - | - | 0 | - |
| - | - | 4346 | 880.4 | - | - | 0 | - |
| - | - | 2418 | 881.4 | - | - | 0 | - |
| - | - | 934.3 | 882.4 | - | - | 0 | - |
| 2 | y | 9566 | 898.4 | 0.0003366 | 0.3747 | +1 | 9 |
| - | - | 5230 | 899.4 | - | - | 0 | - |
| - | - | 1015 | 900.4 | - | - | 0 | - |
| - | - | 982.6 | 908.4 | - | - | 0 | - |
| - | - | 845.5 | 914.4 | - | - | 0 | - |
| 2 | y | 4.686E+04 | 916.5 | 0.001197 | 1.306 | +1 | 9 |
| - | - | 2.4E+04 | 917.5 | - | - | 0 | - |
| - | - | 7694 | 918.5 | - | - | 0 | - |
| - | - | 991.1 | 919.5 | - | - | 0 | - |
| - | - | 5186 | 926.4 | - | - | 0 | - |
| - | - | 2886 | 927.4 | - | - | 0 | - |
| - | - | 917.2 | 958.5 | - | - | 0 | - |
| - | - | 640.2 | 1078 | - | - | 0 | - |
| - | - | 648.8 | 1146 | - | - | 0 | - |
| - | - | 626.9 | 1255 | - | - | 0 | - |
| - | - | 633.4 | 1853 | - | - | 0 | - |
| - | - | 706.4 | 2633 | - | - | 0 | - |

m/z Charge Intensity FragmentType MassShift Position
120.08110809326172 0 15367.677
121.08465576171875 0 1598.6136
122.07170104980469 0 1349.1136
124.08722686767578 0 368.7025
125.06344604492188 0 373.54263
125.10765838623047 0 20671.783
126.05547332763672 0 494.26706
126.1026840209961 0 719.23535
126.111083984375 0 1584.3873
128.02313232421875 0 336.94275
128.10731506347656 0 482.8273
129.0659637451172 0 457.85895
129.10255432128906 0 5634.449
130.05026245117188 0 543.71497
130.06532287597656 0 1239.0015
131.07046508789062 0 447.27454
132.04458618164062 0 1148.284
132.14535522460938 0 487.77243
133.06103515625 0 1568.5133
133.08621215820312 0 3008.7488
136.07606506347656 0 4465.1616
137.0795135498047 0 630.8296
138.06658935546875 0 1824.3801
139.08682250976562 0 575.74445
140.1437530517578 0 825.1765
141.1025390625 0 2172.6426
142.12301635742188 0 68931.945
143.12025451660156 0 948.1764
143.1262969970703 0 5604.355
144.10214233398438 0 590.4983
145.06112670898438 0 3535.8174
146.06011962890625 0 675.2307
147.0652313232422 0 742.5875
150.06666564941406 0 1888.3582
151.3741455078125 0 470.0785
152.07070922851562 0 1015.7593
152.08245849609375 0 752.8443
152.10733032226562 0 39276.754
153.10325622558594 0 794.319
153.11068725585938 0 3131.7317
154.08694458007812 0 633.27966
154.09739685058594 0 509.08575
155.08193969726562 0 811.95184
155.09303283691406 0 5151.665
156.0769805908203 0 3714.551
157.07640075683594 0 3662.908
157.09759521484375 0 788.9356
157.1332244873047 0 551.45917
158.060302734375 0 2308.096
158.08035278320312 0 472.8135
161.09243774414062 0 3935.8472
163.07164001464844 0 11467.217
164.11856079101562 0 1444.8225
165.10284423828125 0 1240.7428
165.1133270263672 0 573.4617
166.0538330078125 0 994.3679
166.0614776611328 0 2442.3013
166.09780883789062 0 5849.5146
167.09320068359375 0 1236.026
169.0974578857422 0 746.9737
169.13388061523438 0 23657.203 a Water loss 1
170.11843872070312 0 662.28357 a Ammonia loss 1
170.13723754882812 0 2095.7444
171.07681274414062 0 7055.793
171.11325073242188 0 1428.8035
172.10862731933594 0 457.3225
174.0550994873047 0 1459.6544
175.0869903564453 0 7998.6475
175.09579467773438 0 778.3287
176.08229064941406 0 1660.9503
176.09033203125 0 1040.9979
176.11878967285156 0 560.73206
177.11253356933594 0 1763.1589
178.13421630859375 0 4034.693
180.07704162597656 0 2024.6224
181.0612030029297 0 801.80994
183.05575561523438 0 880.6169
183.1494140625 0 892.87524
185.09242248535156 0 1280.1278
185.10450744628906 0 556.6715
185.12852478027344 0 608.8948
185.1652374267578 0 1747.3024
187.10813903808594 0 304804.25 y 8
187.1444854736328 0 140505.2 a 1
188.11148071289062 0 25871.768
188.14772033691406 0 11144.404
189.0873260498047 0 3911.86
189.11328125 0 1924.5394
191.09255981445312 0 580.65405
193.1086883544922 0 12884.672
194.0927276611328 0 4692.453
194.11233520507812 0 1099.3184
195.08798217773438 0 11562.531
195.09706115722656 0 975.4613
195.11273193359375 0 1503.0522
195.1234588623047 0 972.0523
197.12879943847656 0 26187.209 b Water loss 1
198.12330627441406 0 676.543
198.13238525390625 0 2317.3909
199.0709991455078 0 871.50867
199.1082763671875 0 1124.9768
201.0872039794922 0 649.25256
201.12315368652344 0 808.49243
205.1085662841797 0 882.7672
207.08795166015625 0 935.13806
207.12319946289062 0 549.89197
209.09225463867188 0 683.51624
209.1399383544922 0 3468.776
210.08749389648438 0 1251.9346
211.11936950683594 0 3767.2322
211.14459228515625 0 614.3846
212.1150665283203 0 1214.6792
213.16009521484375 0 2021.5844
215.1393585205078 0 91067.36 b 1
216.14279174804688 0 8907.255
217.13397216796875 0 2700.636
217.14459228515625 0 735.4008
219.0883331298828 0 1208.4365
219.12374877929688 0 1220.5225
220.10845947265625 0 1386.7451
221.10360717773438 0 35540.14
221.12901306152344 0 2620.9448
222.1070098876953 0 2747.6543
223.08291625976562 0 1901.1931
223.1441192626953 0 1509.4668
226.1190643310547 0 1586.8044
226.15541076660156 0 851.3422
227.1030731201172 0 620.66785
228.09832763671875 0 7364.9204
231.1132354736328 0 1255.2898
231.12469482421875 0 1338.6946
233.16497802734375 0 2507.8582
235.11923217773438 0 909.57654
237.09869384765625 0 3545.7502
237.1348876953125 0 8497.9375
238.13023376464844 0 4263.7744
239.11424255371094 0 38817.984
239.1379852294922 0 1213.4565
239.1508026123047 0 4076.305
240.11749267578125 0 3441.5737
240.13168334960938 0 658.69366
241.12013244628906 0 759.8192
241.15478515625 0 1734.0205
244.12957763671875 0 3779.5417
245.1290740966797 0 2969.4846
246.1089630126953 0 5070.854
247.1089324951172 0 662.00354
248.1509246826172 0 696.0691
249.0983123779297 0 1972.5708
249.1234588623047 0 1553.0962
250.1021728515625 0 727.70703
251.1506805419922 0 1359.881
254.11293029785156 0 595.8066
255.1456756591797 0 800.8031
256.1437683105469 0 6830.615
256.1776123046875 0 1036.0747
257.1476135253906 0 701.08307 a Ammonia loss 2
259.11993408203125 0 1043.7484
261.158935546875 0 791.222
265.13037109375 0 988.2501
266.1490173339844 0 852.2234
266.1634216308594 0 719.62396
267.109375 0 1633.5375
269.18609619140625 0 1022.4641
275.149658203125 0 648.41846
276.1457824707031 0 7463.7505
277.130615234375 0 832.99634
277.1487121582031 0 785.6786
281.16064453125 0 880.21423
282.1199035644531 0 605.1377
282.1445617675781 0 737.0188
283.1431884765625 0 764.87994
284.1403503417969 0 2440.5518
284.160888671875 0 13130.65 b Water loss 2
285.16326904296875 0 1174.9789
289.1923828125 0 804.26074
294.1563720703125 0 11542.507
294.1734619140625 0 935.27783
295.1596984863281 0 2242.8035
296.149658203125 0 830.46356
302.1618957519531 0 841.093
303.0578918457031 0 1339.6746
309.1559753417969 0 680.7457
311.1826171875 0 1087.6238
312.16796875 0 539.85406
314.16180419921875 0 1832.9666
315.6824645996094 0 735.09796
317.1826477050781 0 5479.0923
317.6703796386719 0 802.51776
318.1869201660156 0 775.1823
320.17169189453125 0 4632.7266
321.67218017578125 0 912.7924
322.1502380371094 0 1259.5325
322.16937255859375 0 855.2353
323.17169189453125 0 1726.6035
324.14593505859375 0 14193.345
324.1797180175781 0 646.73785
325.1490173339844 0 2870.8848
327.1671142578125 0 2834.454
329.1481018066406 0 880.9896
330.6773681640625 0 2181.9375
333.672607421875 0 1008.51495
334.1766662597656 0 11538.531 y 7
335.1817932128906 0 4445.701
336.1668701171875 0 1595.9043
337.1986389160156 0 783.00574
338.1824645996094 0 3756.391
339.6820373535156 0 2537.733 b Water loss 6
340.1774597167969 0 12915.161 b Ammonia loss 6
341.1815185546875 0 5879.111 b Water loss 3
342.1565856933594 0 7912.9585
342.18060302734375 0 909.7531
343.1604309082031 0 1259.0728
345.1766052246094 0 3887.4155
346.177978515625 0 841.79596
347.6703796386719 0 799.2019
348.6871337890625 0 1906.032 b 6
349.1957092285156 0 830.3235
350.16119384765625 0 10453.231
350.1829833984375 0 818.2484
351.1637878417969 0 1650.8999
352.2009582519531 0 675.3482
353.19342041015625 0 16340.692
353.2180480957031 0 947.5714
354.1958923339844 0 1875.706
354.676025390625 0 1311.4196
356.67425537109375 0 856.91736
358.1873474121094 0 2201.8044
359.1914367675781 0 2913.4072 b 3
360.1932067871094 0 890.21747
363.1777038574219 0 5764.4014
363.20233154296875 0 877.3894
363.68182373046875 0 1645.4414
364.1810302734375 0 1084.5913
365.1577453613281 0 831.7072
365.19219970703125 0 1261.8898
365.68048095703125 0 665.63513
367.2102966308594 0 906.0681
368.1718444824219 0 7179.1846
369.1752624511719 0 1638.4667
369.2313537597656 0 852.5367
371.1718444824219 0 973.65753
375.16778564453125 0 1025.647
375.1990966796875 0 994.0454
377.1936950683594 0 7835.1104
378.19708251953125 0 1475.7336
381.1883544921875 0 31085.492
382.1916198730469 0 5276.9565
382.6990051269531 0 651.3042
383.1685485839844 0 1022.87
383.19512939453125 0 1238.8733
383.2276306152344 0 2689.9905
384.2340087890625 0 900.3924
386.1825866699219 0 16635.975
386.2102355957031 0 1151.426
387.18511962890625 0 2612.691
388.2090759277344 0 1618.2266
390.2140808105469 0 804.7352
390.7142333984375 0 1175.5015
391.2048034667969 0 1140.6128
392.2035217285156 0 2060.5315
393.1802673339844 0 639.1898
395.2041015625 0 7940.8633
396.16619873046875 0 1138.2135
396.2030029296875 0 2470.1777
396.2337341308594 0 769.2491
397.2120056152344 0 1997.3594
399.2000732421875 0 3111.9849 y Water loss 2
399.2247619628906 0 859.9305
399.69647216796875 0 711.32153
399.721435546875 0 1800.8917
400.20843505859375 0 876.3503
403.6756896972656 0 646.6118
404.2113037109375 0 3042.947
404.7121276855469 0 1007.65466
405.2005920410156 0 1128.995
408.22479248046875 0 5707.3574
408.7256774902344 0 2582.506
409.2251281738281 0 1205.7982
410.2154846191406 0 2016.8914
412.2576599121094 0 1202.092
413.2166442871094 0 5351.8477 b Water loss 7
413.7180480957031 0 1472.1523
414.1783447265625 0 987.9537
414.2068786621094 0 1849.4148
414.7093811035156 0 961.65265
418.2216491699219 0 3652.2336
418.74566650390625 0 874.44104
419.20147705078125 0 947.8135
419.2497863769531 0 755.8186
420.2008361816406 0 1632.6561
422.2221984863281 0 3470.034 b 7
422.72210693359375 0 1246.8293
427.5708312988281 0 769.85516
427.8994140625 0 754.4561
430.26641845703125 0 3844.564
431.2701721191406 0 1072.5077
435.2269592285156 0 1198.9526 y 6
436.2306823730469 0 6633.8164
437.2148132324219 0 5483.5996
438.2106628417969 0 8122.0757
438.2434997558594 0 862.5029
439.21331787109375 0 1838.4253
439.24517822265625 0 882.18463
439.7310485839844 0 765.86676
440.2500915527344 0 3197.8162 b Water loss 4
440.7200622558594 0 3087.781
441.2211608886719 0 1249.2229
441.2535095214844 0 862.466
446.2116394042969 0 7245.3477
447.2138671875 0 1760.1621
447.74444580078125 0 1599.2222
448.73443603515625 0 877.2428
449.72503662109375 0 1668.9614 y Water loss 1
452.7380065917969 0 2187.763
453.2408142089844 0 749.8296
454.2411804199219 0 15066.772
455.2444152832031 0 3446.2385
455.27630615234375 0 925.475
456.2454833984375 0 738.9844
456.27764892578125 0 899.59894
456.7497253417969 0 1491.3135
457.25238037109375 0 1609.7324
458.260986328125 0 2802.9192 b 4
458.7300720214844 0 3347.5544 y 1
459.2306823730469 0 2051.4692
459.2651062011719 0 723.93866
459.7341613769531 0 1284.1129
461.2374267578125 0 709.78894
461.2913513183594 0 2709.4963
461.7430419921875 0 2414.6208 b Water loss 8
462.2447509765625 0 2182.3523
464.2255859375 0 38658.777
465.2286071777344 0 10618.915
466.2323303222656 0 1231.3513
467.23919677734375 0 1561.5895
468.2425231933594 0 858.3378
470.74859619140625 0 3939.8662 b 8
471.2503662109375 0 2292.0273
471.7497253417969 0 659.3922
473.2592468261719 0 3136.365
474.2364501953125 0 1363.063
474.7956237792969 0 685.3981
475.2497863769531 0 653.95624
475.2901916503906 0 1878.564
475.790283203125 0 1202.5461
482.2361755371094 0 56547.12
483.23834228515625 0 15364.748
484.24176025390625 0 3026.332
485.25054931640625 0 2526.145
486.2113342285156 0 628.5569
488.2596740722656 0 1135.1298
488.74908447265625 0 806.162
489.6011657714844 0 1470.5896
489.9341735839844 0 653.4063
490.2779846191406 0 1043.8508
492.2201232910156 0 2981.4695
493.25592041015625 0 638.6139
496.2658996582031 0 2085.9666
497.2615966796875 0 10445.248
497.7629699707031 0 7373.286
498.26226806640625 0 1763.6552
499.2645263671875 0 1508.172
503.2362060546875 0 967.4274
506.2665100097656 0 10699.947 Precursor Water loss
506.7680358886719 0 6867.897
507.2702941894531 0 2099.1965
508.6065979003906 0 903.8744
511.262451171875 0 927.7427
512.222900390625 0 1037.0905
514.2777099609375 0 2667.5547
515.2723999023438 0 10073.211 Precursor
515.7733154296875 0 5892.3735
516.2758178710938 0 1947.5062
521.2466430664062 0 1616.5099
521.7852783203125 0 1110.721
524.26171875 0 8155.877
525.2647094726562 0 2874.2148
525.8120727539062 0 7377.5947
526.3128051757812 0 4015.7522
526.814208984375 0 2090.795
528.2577514648438 0 845.65594
529.2535400390625 0 2484.8381
531.3005981445312 0 1129.3276
537.2779541015625 0 2167.3982
538.2584838867188 0 1176.4175
539.2586059570312 0 3251.9949
539.8095703125 0 27768.94
540.2655029296875 0 902.1596
540.3108520507812 0 15548.051
540.8123779296875 0 4801.415
541.8135986328125 0 714.35126
542.27197265625 0 9272.174
543.2740478515625 0 2905.3513
544.2789916992188 0 782.63245
547.2630004882812 0 5810.3047
548.26416015625 0 1894.7582
548.31982421875 0 667.6271
548.8231811523438 0 1129.3524
549.3178100585938 0 1888.642
550.3029174804688 0 669.75854
553.2783813476562 0 739.59106
554.2728881835938 0 7630.9517 y Water loss 5
555.2833862304688 0 4118.036
556.2900390625 0 1060.7334
557.2832641601562 0 3606.5242
558.2847290039062 0 1155.1553
559.2969970703125 0 1038.2428
563.61572265625 0 625.62225
565.2739868164062 0 15173.93
566.275390625 0 6089.4224
567.2708129882812 0 2539.0952
567.325439453125 0 5495.6357
568.326171875 0 2024.6316
571.2875366210938 0 1247.5632
572.28271484375 0 15780.236 y 5
573.2857055664062 0 5605.7173
574.3200073242188 0 1984.7844
574.8209838867188 0 1673.3301
575.2589721679688 0 2780.0715
577.30859375 0 2521.4553 b Water loss 5
578.31494140625 0 1144.1167
583.29150390625 0 20717.777
584.2904052734375 0 6634.6484
585.2802734375 0 8410.397
586.281982421875 0 3038.3015
589.2991943359375 0 719.4809
593.2794799804688 0 7034.7227
594.2824096679688 0 2487.481
595.3200073242188 0 11720.3125 b 5
596.3218383789062 0 2816.2627
601.30908203125 0 17468.486
602.3128051757812 0 5048.686
603.3174438476562 0 676.1477
611.2936401367188 0 58831.52
612.2965087890625 0 21054.283
613.2975463867188 0 3808.186
618.8502197265625 0 810.7731
624.3473510742188 0 944.53326
626.3179931640625 0 712.3968
626.8494873046875 0 1591.9324
627.3519897460938 0 1864.5409
627.8513793945312 0 1391.2477
629.30419921875 0 70857.875
630.3070678710938 0 24555.549
630.3571166992188 0 1279.2478
631.3082275390625 0 4580.3364
632.3458251953125 0 784.4778
633.3384399414062 0 825.5563
634.3281860351562 0 1221.6467
640.3223876953125 0 850.67395
640.8464965820312 0 14741.217
641.3475952148438 0 10572.1455
641.8490600585938 0 2819.8584
642.3464965820312 0 2317.8203
642.8460693359375 0 948.28613
643.3427124023438 0 1265.4359
650.3019409179688 0 1538.4933
650.363037109375 0 2086.521
651.3506469726562 0 1785.3373
652.34228515625 0 3519.4836
653.3428955078125 0 1838.3497 y Water loss 4
654.3411865234375 0 1025.1362
658.3289184570312 0 1843.0317
660.3453979492188 0 4676.5103
661.3493041992188 0 1761.1787
666.333740234375 0 1352.7848
668.3125610351562 0 1321.1152
668.3720092773438 0 3498.3804
669.3768310546875 0 1210.409
670.3674926757812 0 729.7206
671.3509521484375 0 3331.7908 y 4
672.3540649414062 0 788.8517
676.3203735351562 0 3180.2524
676.3826904296875 0 9026.235
676.8841552734375 0 7882.4517
677.3222045898438 0 1182.9712
677.385986328125 0 2749.0005
677.8868408203125 0 697.67334
678.3568725585938 0 17087.59 b Water loss 6
679.3599853515625 0 6943.0884
680.362548828125 0 1552.9309
682.3303833007812 0 824.69
683.3258666992188 0 723.35223
684.3463134765625 0 5000.487
685.3468017578125 0 2188.1167
686.3252563476562 0 4241.5283
687.3343505859375 0 1417.127
688.3602905273438 0 880.6508
690.3805541992188 0 7904.2524
690.8817138671875 0 7352.942
691.3836669921875 0 3147.5474
692.38134765625 0 671.4988
694.3304443359375 0 10043.916
695.3324584960938 0 2666.3628
696.3672485351562 0 17550.082 b 6
697.3692016601562 0 7358.028
698.3681640625 0 1844.7622
702.3562622070312 0 6453.224
703.3607177734375 0 1988.6227
708.3436279296875 0 1383.3026
710.3592529296875 0 1771.3016 y Water loss 3
711.3622436523438 0 1094.9081
712.340576171875 0 22548.23
713.3438110351562 0 8232.395
714.3458251953125 0 1367.6195
722.3247680664062 0 3439.8022
723.3267211914062 0 1709.2374
725.3792724609375 0 716.93524
726.3573608398438 0 5390.4307
727.3612060546875 0 1802.6785
728.3714599609375 0 11142.839 y 3
729.3738403320312 0 4665.067
730.3515625 0 30423.494
731.354248046875 0 10650.34
732.358154296875 0 2892.473
740.3360595703125 0 3660.3591
741.3414306640625 0 1456.3562
742.3875122070312 0 1474.6611
743.391357421875 0 1480.1143
744.3665771484375 0 2408.3298
745.375244140625 0 1129.5585
746.3680419921875 0 944.6276
753.4003295898438 0 822.4636
754.4033813476562 0 769.98804
763.3884887695312 0 1949.9407
764.3784790039062 0 1869.1848
765.3715209960938 0 940.36365
771.409912109375 0 3285.762
772.4144287109375 0 689.9572
779.4174194335938 0 1771.758
780.412109375 0 812.4036
781.4002075195312 0 3033.243
782.4041748046875 0 1261.4094
789.40185546875 0 2851.987
790.4028930664062 0 2227.8247
797.3990478515625 0 11093.1 y Water loss 2
798.4056396484375 0 6555.7217
799.4091186523438 0 10765.804
800.4129028320312 0 3868.0818
801.416015625 0 1142.9136
807.4142456054688 0 15555.549
808.4172973632812 0 7031.341
809.4086303710938 0 1646.2058
813.3921508789062 0 1016.949
815.4051513671875 0 112264.21 y 2
816.4078979492188 0 53777.82
817.4100341796875 0 12538.421
818.4071655273438 0 798.63293
819.4566650390625 0 1426.7654
820.4553833007812 0 1167.9751
825.4244384765625 0 51111.992 b Water loss 7
826.4276123046875 0 23031.994
827.425537109375 0 7229.643
828.4212036132812 0 867.87445
839.4444580078125 0 1063.2767
841.4173583984375 0 979.0774
842.4217529296875 0 756.7937
843.434814453125 0 71545.195 b 7
844.4383544921875 0 30407.857
845.4398193359375 0 9085.467
846.4402465820312 0 1046.3958
854.4203491210938 0 717.9004
880.4298706054688 0 4345.811
881.4313354492188 0 2418.02
882.42041015625 0 934.26025
898.44140625 0 9566.337 y Water loss 1
899.4425659179688 0 5230.266
900.4437866210938 0 1014.6745
908.4251098632812 0 982.5514
914.4352416992188 0 845.49396
916.4511108398438 0 46862.387 y 1
917.4536743164062 0 24004.357
918.4566040039062 0 7693.783
919.462646484375 0 991.0792
926.4363403320312 0 5185.9077
927.4360961914062 0 2886.0835
958.5003662109375 0 917.2271
1077.6500244140625 0 640.2207
1145.5283203125 0 648.81616
1254.6739501953125 0 626.94293
1852.9644775390625 0 633.38165
2633.291015625 0 706.35583

Spectrum Details

|  |  |
| --- | --- |
| Matched peaks? Matched peaks The total absolute number of peaks matched. Additionally in brackets the total fraction of peaks matched and the total number of peaks is shown. | 42 (7.20% of 583) |
| FDR? FDR The false discovery rate estimated for this peptide. It is calculated by matching all theoretical fragments with a non-integer shift with the raw peaks for this spectrum. This is done with 40 different shifts. The resulting percentage is the average number of annotated peaks over the number of annotated peaks with the correct spectrum. | 0.00% |
| Satellite FDR? Satellite FDR See the FDR for details on its calculation. This satellite ion specific FDR only contains the satellite ions (d/w) for I/L/J positions. | - |
| PSM Score? PSM Score The PSM Score as given by Hecklib to this annotated spectrum. It is shown with three significant figures. | 476 |

#### Reverse Lookup? Reverse Lookup All places where this read could be placed.

| Group | Segment | Template | Template Part | Read Part | Score | Unique |
| --- | --- | --- | --- | --- | --- | --- |
| Homo sapiens Heavy Chain | IGHC | IGHG1 | [45..55] | [0..10] | 80 | False |
| Homo sapiens Heavy Chain | IGHC | IGHG3 | [45..55] | [0..10] | 80 | False |
| Homo sapiens Heavy Chain | IGHC | IGHG2 | [45..55] | [0..10] | 80 | False |
| Homo sapiens Heavy Chain | IGHC | IGHG4 | [45..55] | [0..10] | 80 | False |

| Recombined | Template Part | Read Part | Score | Unique |
| --- | --- | --- | --- | --- |
| REC-0-1 | [167..177] | [0..10] | 80 | True |

#### Meta Information from Multiple reads

##### Number of combined reads

3

##### Intensity

0.9254

##### TotalArea

1.389E+09

##### Changes to the peptide sequence

JTSGVHTFPA

L→JNo support for either Leucine or Isoleucine based on side chain ions (Position: 1)

#### Positional Score

Copy Data

##### Positional Score (TSV)

###### Preview

```
Loading example...
```

*Click on the button to copy the data to your clipboard.*

100123456789

Label Value
"0" 0.333
"1" 0.333
"2" 0.33
"3" 0.317
"4" 0.32
"5" 0.277
"6" 0.317
"7" 0.33
"8" 0.297
"9" 0.31

#### Meta Information from PEAKS

##### Scan Identifier

F2:5220

##### Original sequence

L

T

S

G

V

H

T

F

P

A

##### Posttranslational Modifications

##### Source File

D:\separate\_stitch\_analyses\xle-disambiguation\raw\20210323\_F1\_UM1\_Peng0013\_SA\_F59\_ingel\_3ug\_TL.raw

##### Fraction

2

##### Scan Feature

F2:4691

##### De Novo Score

98

##### ConfidenceScore

98

### m/z

515.2727

##### Mass

1028.5291

##### Charge

2

##### Retention Time

28

##### Predicted Retention Time

-

##### Area

6.947E+08

##### Parts Per Million

1.8

##### Fragmentation mode

ETHCD

##### Originating file

01 D:\separate\_stitch\_analyses\xle-disambiguation\20210325\_F59\_3ug\_DENOVO\_12.csv

#### Meta Information from PEAKS

##### Scan Identifier

F2:5282

##### Original sequence

L

T

S

G

V

H

T

F

P

A

##### Posttranslational Modifications

##### Source File

D:\separate\_stitch\_analyses\xle-disambiguation\raw\20210323\_F1\_UM1\_Peng0013\_SA\_F59\_ingel\_3ug\_TL.raw

##### Fraction

2

##### Scan Feature

F2:4691

##### De Novo Score

98

##### ConfidenceScore

98

### m/z

515.2727

##### Mass

1028.5291

##### Charge

2

##### Retention Time

28

##### Predicted Retention Time

-

##### Area

6.947E+08

##### Parts Per Million

1.8

##### Fragmentation mode

ETHCD

##### Originating file

01 D:\separate\_stitch\_analyses\xle-disambiguation\20210325\_F59\_3ug\_DENOVO\_12.csv

#### Meta Information from PEAKS

##### Scan Identifier

F2:5425

##### Original sequence

L

T

S

G

V

H

T

F

P

A

##### Posttranslational Modifications

##### Source File

D:\separate\_stitch\_analyses\xle-disambiguation\raw\20210323\_F1\_UM1\_Peng0013\_SA\_F59\_ingel\_3ug\_TL.raw

##### Fraction

2

##### Scan Feature

-

##### De Novo Score

95

##### ConfidenceScore

95

### m/z

515.2729

##### Mass

1028.5291

##### Charge

2

##### Retention Time

29.45

##### Predicted Retention Time

-

##### Area

0

##### Parts Per Million

2.1

##### Fragmentation mode

HCD

##### Originating file

01 D:\separate\_stitch\_analyses\xle-disambiguation\20210325\_F59\_3ug\_DENOVO\_12.csv
