## Supplementary material for "A handle on mass coincidence errors in *de novo* sequencing of antibodies by bottom-up proteomics": Combined_074.html

Details Combined\_074 | Stitch OverviewUndefined

### Read Combined\_074

#### Sequence (length=10)

JVKDYFPEPV

#### Spectrum 8316? Spectrum 8316 The raw spectrum of this peptide as annotated by Hecklib. The fragments are coloured according to ion type (see legend). Any peaks with a star '\*' as text can be hovered over to see the full details, first the ion type second the mass shift type. By hovering over the amino acids in the peptide or ions in the legend the corresponding peaks are highlighted. By toggling the 'Unassigned' label you can turn the background (unassigned) peaks on or off in the plot. By updating the slider in the Ion legend you can update the spectrum to only show the top X% of the peaks with labels. The top X% means any peak that is within X% of the highest intensity. By dragging in the spectrum you can zoom in to a specific part of the spectrum and use 'Zoom Out' to get back to the original zoom level. The annotation of the spectrum is based on the given sequence in the peptides file and is done with different software so inconsistencies are likely. The peaks are annotated based on the given sequence, with 20 ppm tolerance.

Copy Data

##### Spectrum 8316 (TSV)

###### Preview

```
Loading example...
```

*Click on the button to copy the data to your clipboard.*

Mz MinMz MaxIntensity Max

WidthHeightPeptide font sizePeptide stroke widthSpectrum font sizeSpectrum stroke widthCompact peptide

Ion legend

wxyz

abcd

OtherUnassignedIon^Charge^_Position_Show for top:%

JVKDYFPEPV

04.78e+59.57e+51.43e+61.91e+6

Zoom Out

y^+1^_2_c^+2^_5_y^+1^_3_c^+1^_3_y^+1^_3_c^+1^_3_c^+2^_6_y^+1^_4_c^+2^_7_y^+1^_4_c^+1^_4_c^+1^_4_c^+2^_8_c^+2^_9_y^+2^_9_z^+1^_5_y^+1^_5_y^+1^_5_c^+1^_5_c^+1^_5_y^+1^_6_z^+1^_6_y^+1^_6_c^+1^_6_c^+1^_7_y^+1^_7_c^+1^_7_w^+1^_8_y^+1^_8_z^+1^_8_c^+1^_8_y^+1^_8_y^+1^_9_y^+1^_9_z^+1^_9_y^+1^_9_c^+1^_9_

03136279401253

Fragment Matches Table

Show background peaks

| Position | Ion type | Intensity | mz Theoretical | mz Error (Th) | mz Error (ppm) | Charge | Series Number |
| --- | --- | --- | --- | --- | --- | --- | --- |
| - | - | 6287 | 120.1 | - | - | 0 | - |
| - | - | 395.9 | 121.1 | - | - | 0 | - |
| - | - | 3.716E+04 | 129.1 | - | - | 0 | - |
| - | - | 1550 | 130.1 | - | - | 0 | - |
| - | - | 1288 | 136.1 | - | - | 0 | - |
| - | - | 387.7 | 138.4 | - | - | 0 | - |
| - | - | 425.5 | 148.2 | - | - | 0 | - |
| - | - | 693.5 | 148.9 | - | - | 0 | - |
| - | - | 477.6 | 158.9 | - | - | 0 | - |
| - | - | 1.304E+04 | 169.1 | - | - | 0 | - |
| - | - | 1044 | 170.1 | - | - | 0 | - |
| - | - | 503.3 | 171.9 | - | - | 0 | - |
| - | - | 439.5 | 172.5 | - | - | 0 | - |
| - | - | 2942 | 173.4 | - | - | 0 | - |
| - | - | 776.1 | 181.1 | - | - | 0 | - |
| - | - | 600.8 | 181.1 | - | - | 0 | - |
| - | - | 501.8 | 182.1 | - | - | 0 | - |
| - | - | 717.7 | 183.2 | - | - | 0 | - |
| - | - | 1.457E+05 | 185.2 | - | - | 0 | - |
| - | - | 1.564E+04 | 186.2 | - | - | 0 | - |
| - | - | 736.8 | 187.2 | - | - | 0 | - |
| - | - | 622.5 | 191.1 | - | - | 0 | - |
| - | - | 5105 | 197.1 | - | - | 0 | - |
| - | - | 679.7 | 198.1 | - | - | 0 | - |
| - | - | 7740 | 199.1 | - | - | 0 | - |
| - | - | 608.1 | 200.1 | - | - | 0 | - |
| - | - | 713.6 | 201.1 | - | - | 0 | - |
| - | - | 1820 | 203.1 | - | - | 0 | - |
| - | - | 6444 | 209.1 | - | - | 0 | - |
| - | - | 748.9 | 210.1 | - | - | 0 | - |
| - | - | 3112 | 213.1 | - | - | 0 | - |
| - | - | 8.568E+04 | 213.2 | - | - | 0 | - |
| - | - | 1.194E+04 | 214.2 | - | - | 0 | - |
| 9 | y | 1.894E+06 | 215.1 | 0.00095 | 4.416 | +1 | 2 |
| - | - | 2.062E+05 | 216.1 | - | - | 0 | - |
| - | - | 1.312E+04 | 217.1 | - | - | 0 | - |
| - | - | 5.919E+04 | 227.1 | - | - | 0 | - |
| - | - | 5316 | 228.1 | - | - | 0 | - |
| - | - | 3616 | 228.2 | - | - | 0 | - |
| - | - | 1826 | 231.1 | - | - | 0 | - |
| - | - | 9145 | 244.1 | - | - | 0 | - |
| - | - | 2352 | 245.1 | - | - | 0 | - |
| - | - | 1545 | 251.1 | - | - | 0 | - |
| - | - | 904.5 | 255.1 | - | - | 0 | - |
| - | - | 849.7 | 261.2 | - | - | 0 | - |
| - | - | 1426 | 278.2 | - | - | 0 | - |
| - | - | 5219 | 279.1 | - | - | 0 | - |
| - | - | 699.2 | 280.1 | - | - | 0 | - |
| - | - | 1253 | 280.2 | - | - | 0 | - |
| - | - | 926.5 | 283.1 | - | - | 0 | - |
| - | - | 3998 | 301.2 | - | - | 0 | - |
| 5 | c | 1698 | 310.2 | 0.001724 | 5.557 | +2 | 5 |
| - | - | 842.9 | 310.2 | - | - | 0 | - |
| - | - | 2800 | 311.1 | - | - | 0 | - |
| - | - | 702.3 | 312.1 | - | - | 0 | - |
| - | - | 1044 | 315.3 | - | - | 0 | - |
| - | - | 1160 | 320.2 | - | - | 0 | - |
| - | - | 3.751E+04 | 324.2 | - | - | 0 | - |
| - | - | 6981 | 325.2 | - | - | 0 | - |
| 8 | y | 1.932E+04 | 326.2 | 0.001255 | 3.847 | +1 | 3 |
| - | - | 4184 | 327.2 | - | - | 0 | - |
| - | - | 2186 | 340.2 | - | - | 0 | - |
| - | - | 1214 | 341.2 | - | - | 0 | - |
| 3 | c | 1.034E+04 | 341.3 | 0.001569 | 4.598 | +1 | 3 |
| - | - | 1511 | 342.3 | - | - | 0 | - |
| - | - | 1767 | 343.2 | - | - | 0 | - |
| 8 | y | 2.242E+04 | 344.2 | 0.00128 | 3.719 | +1 | 3 |
| - | - | 1205 | 344.2 | - | - | 0 | - |
| - | - | 1141 | 344.7 | - | - | 0 | - |
| - | - | 2989 | 345.2 | - | - | 0 | - |
| - | - | 888 | 353.2 | - | - | 0 | - |
| - | - | 2313 | 355.2 | - | - | 0 | - |
| - | - | 1610 | 356.3 | - | - | 0 | - |
| - | - | 1.039E+05 | 357.3 | - | - | 0 | - |
| 3 | c | 2.534E+04 | 358.3 | 0.002153 | 6.008 | +1 | 3 |
| - | - | 1208 | 358.7 | - | - | 0 | - |
| - | - | 2215 | 359.3 | - | - | 0 | - |
| - | - | 785.5 | 362.2 | - | - | 0 | - |
| - | - | 5146 | 369.7 | - | - | 0 | - |
| - | - | 2028 | 370.2 | - | - | 0 | - |
| - | - | 2253 | 371.2 | - | - | 0 | - |
| - | - | 1396 | 373.2 | - | - | 0 | - |
| - | - | 1.399E+04 | 374.2 | - | - | 0 | - |
| - | - | 3001 | 375.2 | - | - | 0 | - |
| - | - | 690.8 | 377.2 | - | - | 0 | - |
| - | - | 1341 | 379.2 | - | - | 0 | - |
| - | - | 928.4 | 381.1 | - | - | 0 | - |
| - | - | 674.1 | 383.2 | - | - | 0 | - |
| 6 | c | 4046 | 383.7 | 0.0009946 | 2.592 | +2 | 6 |
| - | - | 2474 | 384.2 | - | - | 0 | - |
| - | - | 1394 | 389.2 | - | - | 0 | - |
| - | - | 626.2 | 390.7 | - | - | 0 | - |
| - | - | 3001 | 391.2 | - | - | 0 | - |
| - | - | 3273 | 392.2 | - | - | 0 | - |
| - | - | 1328 | 395.2 | - | - | 0 | - |
| - | - | 739.3 | 398.2 | - | - | 0 | - |
| - | - | 1589 | 403.2 | - | - | 0 | - |
| - | - | 1016 | 405.2 | - | - | 0 | - |
| - | - | 620.8 | 406.2 | - | - | 0 | - |
| - | - | 1.33E+04 | 407.2 | - | - | 0 | - |
| - | - | 708.5 | 407.2 | - | - | 0 | - |
| - | - | 2912 | 408.2 | - | - | 0 | - |
| - | - | 794.6 | 411.3 | - | - | 0 | - |
| - | - | 1.91E+04 | 418.2 | - | - | 0 | - |
| - | - | 1.266E+04 | 418.7 | - | - | 0 | - |
| - | - | 3481 | 419.2 | - | - | 0 | - |
| 7 | y | 4.618E+04 | 423.2 | 0.001805 | 4.266 | +1 | 4 |
| - | - | 1.331E+04 | 424.2 | - | - | 0 | - |
| - | - | 1747 | 425.2 | - | - | 0 | - |
| - | - | 2.134E+04 | 426.2 | - | - | 0 | - |
| - | - | 5379 | 427.2 | - | - | 0 | - |
| - | - | 1200 | 429.3 | - | - | 0 | - |
| - | - | 668.1 | 430.3 | - | - | 0 | - |
| 7 | c | 2.645E+04 | 432.2 | 0.001743 | 4.032 | +2 | 7 |
| - | - | 1.375E+04 | 432.7 | - | - | 0 | - |
| - | - | 3280 | 433.2 | - | - | 0 | - |
| - | - | 5163 | 438.3 | - | - | 0 | - |
| - | - | 1934 | 438.3 | - | - | 0 | - |
| - | - | 984.8 | 439.2 | - | - | 0 | - |
| - | - | 2116 | 439.3 | - | - | 0 | - |
| - | - | 908.8 | 440.2 | - | - | 0 | - |
| 7 | y | 2.038E+05 | 441.2 | 0.001922 | 4.355 | +1 | 4 |
| - | - | 5.095E+04 | 442.2 | - | - | 0 | - |
| - | - | 7564 | 443.2 | - | - | 0 | - |
| - | - | 848.3 | 443.3 | - | - | 0 | - |
| - | - | 896.1 | 447.2 | - | - | 0 | - |
| - | - | 2485 | 452.2 | - | - | 0 | - |
| - | - | 1309 | 454.3 | - | - | 0 | - |
| 4 | c | 3.361E+04 | 456.3 | 0.001848 | 4.05 | +1 | 4 |
| - | - | 1662 | 457.2 | - | - | 0 | - |
| - | - | 9085 | 457.3 | - | - | 0 | - |
| - | - | 1435 | 458.3 | - | - | 0 | - |
| - | - | 693.8 | 464.3 | - | - | 0 | - |
| - | - | 1402 | 467.3 | - | - | 0 | - |
| - | - | 1.037E+04 | 470.2 | - | - | 0 | - |
| - | - | 3793 | 471.2 | - | - | 0 | - |
| - | - | 2.002E+04 | 472.3 | - | - | 0 | - |
| 4 | c | 4.059E+04 | 473.3 | 0.001483 | 3.133 | +1 | 4 |
| - | - | 983.3 | 473.8 | - | - | 0 | - |
| - | - | 1133 | 474.3 | - | - | 0 | - |
| - | - | 9442 | 474.3 | - | - | 0 | - |
| - | - | 1079 | 475.3 | - | - | 0 | - |
| - | - | 7248 | 482.3 | - | - | 0 | - |
| - | - | 1.455E+04 | 482.8 | - | - | 0 | - |
| - | - | 1.176E+04 | 483.3 | - | - | 0 | - |
| - | - | 2194 | 483.8 | - | - | 0 | - |
| - | - | 3043 | 487.8 | - | - | 0 | - |
| - | - | 2129 | 488.3 | - | - | 0 | - |
| - | - | 3237 | 488.3 | - | - | 0 | - |
| - | - | 1104 | 488.8 | - | - | 0 | - |
| - | - | 766.2 | 489.2 | - | - | 0 | - |
| - | - | 986.7 | 489.3 | - | - | 0 | - |
| - | - | 2140 | 491.3 | - | - | 0 | - |
| - | - | 620.5 | 493.9 | - | - | 0 | - |
| - | - | 637.2 | 495.3 | - | - | 0 | - |
| - | - | 735.5 | 495.7 | - | - | 0 | - |
| 8 | c | 2.501E+04 | 496.8 | 0.002236 | 4.501 | +2 | 8 |
| - | - | 1.65E+04 | 497.3 | - | - | 0 | - |
| - | - | 6427 | 497.8 | - | - | 0 | - |
| - | - | 657.4 | 502.3 | - | - | 0 | - |
| - | - | 1677 | 506.3 | - | - | 0 | - |
| - | - | 1071 | 507.3 | - | - | 0 | - |
| - | - | 3255 | 509.2 | - | - | 0 | - |
| - | - | 2108 | 510.2 | - | - | 0 | - |
| - | - | 6337 | 511.3 | - | - | 0 | - |
| - | - | 2698 | 512.3 | - | - | 0 | - |
| - | - | 1119 | 521.3 | - | - | 0 | - |
| - | - | 2344 | 522.3 | - | - | 0 | - |
| - | - | 838.5 | 522.8 | - | - | 0 | - |
| - | - | 3194 | 523.2 | - | - | 0 | - |
| - | - | 2271 | 523.3 | - | - | 0 | - |
| - | - | 813.5 | 524.2 | - | - | 0 | - |
| - | - | 997.6 | 524.3 | - | - | 0 | - |
| - | - | 2181 | 526.3 | - | - | 0 | - |
| - | - | 999.1 | 527.3 | - | - | 0 | - |
| - | - | 3514 | 531.3 | - | - | 0 | - |
| - | - | 1887 | 531.8 | - | - | 0 | - |
| - | - | 793.1 | 532.3 | - | - | 0 | - |
| - | - | 8285 | 534.3 | - | - | 0 | - |
| - | - | 2742 | 535.3 | - | - | 0 | - |
| - | - | 5486 | 536.3 | - | - | 0 | - |
| - | - | 856.9 | 536.8 | - | - | 0 | - |
| - | - | 3558 | 537.2 | - | - | 0 | - |
| - | - | 1937 | 538.2 | - | - | 0 | - |
| - | - | 7678 | 539.3 | - | - | 0 | - |
| - | - | 2085 | 540.3 | - | - | 0 | - |
| - | - | 1177 | 540.3 | - | - | 0 | - |
| 9 | c | 2.879E+04 | 545.3 | 0.001855 | 3.402 | +2 | 9 |
| - | - | 1.699E+04 | 545.8 | - | - | 0 | - |
| - | - | 6706 | 546.3 | - | - | 0 | - |
| - | - | 973.4 | 546.8 | - | - | 0 | - |
| 2 | y | 797.6 | 547.3 | 0.001346 | 2.459 | +2 | 9 |
| - | - | 1275 | 548.3 | - | - | 0 | - |
| - | - | 2563 | 549.3 | - | - | 0 | - |
| - | - | 890.4 | 550.3 | - | - | 0 | - |
| - | - | 761.2 | 551.3 | - | - | 0 | - |
| - | - | 2688 | 553.3 | - | - | 0 | - |
| 6 | z | 9.409E+04 | 554.3 | 0.01032 | 18.61 | +1 | 5 |
| - | - | 3.076E+04 | 555.3 | - | - | 0 | - |
| - | - | 5319 | 556.3 | - | - | 0 | - |
| - | - | 943 | 558.3 | - | - | 0 | - |
| - | - | 1069 | 567.3 | - | - | 0 | - |
| - | - | 1.646E+04 | 567.4 | - | - | 0 | - |
| - | - | 5493 | 568.4 | - | - | 0 | - |
| 6 | y | 1849 | 570.3 | 0.005199 | 9.117 | +1 | 5 |
| - | - | 618.2 | 571.3 | - | - | 0 | - |
| - | - | 824 | 574.3 | - | - | 0 | - |
| - | - | 3199 | 580.8 | - | - | 0 | - |
| - | - | 2372 | 581.3 | - | - | 0 | - |
| - | - | 1372 | 581.8 | - | - | 0 | - |
| - | - | 1125 | 582.3 | - | - | 0 | - |
| - | - | 1973 | 585.3 | - | - | 0 | - |
| - | - | 2795 | 587.3 | - | - | 0 | - |
| 6 | y | 1.066E+05 | 588.3 | 0.002386 | 4.056 | +1 | 5 |
| - | - | 3.561E+04 | 589.3 | - | - | 0 | - |
| - | - | 6199 | 590.3 | - | - | 0 | - |
| - | - | 1.337E+04 | 591.4 | - | - | 0 | - |
| - | - | 9565 | 592.4 | - | - | 0 | - |
| - | - | 654.1 | 593.3 | - | - | 0 | - |
| - | - | 3222 | 593.4 | - | - | 0 | - |
| - | - | 5787 | 594.8 | - | - | 0 | - |
| - | - | 4745 | 595.3 | - | - | 0 | - |
| - | - | 2312 | 595.8 | - | - | 0 | - |
| - | - | 2466 | 597.3 | - | - | 0 | - |
| - | - | 1081 | 598.3 | - | - | 0 | - |
| - | - | 1842 | 599.3 | - | - | 0 | - |
| - | - | 3227 | 601.3 | - | - | 0 | - |
| - | - | 839.9 | 602.3 | - | - | 0 | - |
| - | - | 3836 | 603.4 | - | - | 0 | - |
| - | - | 2.842E+04 | 603.8 | - | - | 0 | - |
| - | - | 2.52E+04 | 604.3 | - | - | 0 | - |
| - | - | 8860 | 604.8 | - | - | 0 | - |
| - | - | 1191 | 605.3 | - | - | 0 | - |
| - | - | 1091 | 610.3 | - | - | 0 | - |
| - | - | 2284 | 615.3 | - | - | 0 | - |
| - | - | 874.5 | 617.3 | - | - | 0 | - |
| 5 | c | 8.748E+04 | 619.3 | 0.002606 | 4.208 | +1 | 5 |
| - | - | 3.462E+04 | 620.4 | - | - | 0 | - |
| - | - | 6619 | 621.4 | - | - | 0 | - |
| - | - | 930.7 | 624.3 | - | - | 0 | - |
| - | - | 3921 | 633.3 | - | - | 0 | - |
| - | - | 6.689E+04 | 635.4 | - | - | 0 | - |
| 5 | c | 8.9E+04 | 636.4 | 0.001265 | 1.987 | +1 | 5 |
| - | - | 2.585E+04 | 637.4 | - | - | 0 | - |
| - | - | 4986 | 638.3 | - | - | 0 | - |
| - | - | 3830 | 638.4 | - | - | 0 | - |
| - | - | 1783 | 639.3 | - | - | 0 | - |
| - | - | 598 | 644.2 | - | - | 0 | - |
| - | - | 1.084E+04 | 651.3 | - | - | 0 | - |
| - | - | 1154 | 651.4 | - | - | 0 | - |
| - | - | 1.675E+04 | 652.3 | - | - | 0 | - |
| - | - | 3821 | 652.3 | - | - | 0 | - |
| - | - | 7544 | 653.3 | - | - | 0 | - |
| - | - | 1.116E+04 | 653.3 | - | - | 0 | - |
| - | - | 4354 | 654.3 | - | - | 0 | - |
| - | - | 1000 | 655.3 | - | - | 0 | - |
| - | - | 4043 | 663.3 | - | - | 0 | - |
| - | - | 4217 | 664.4 | - | - | 0 | - |
| - | - | 1541 | 665.4 | - | - | 0 | - |
| - | - | 1874 | 667.3 | - | - | 0 | - |
| - | - | 856 | 668.3 | - | - | 0 | - |
| - | - | 1399 | 669.4 | - | - | 0 | - |
| - | - | 1.502E+04 | 681.3 | - | - | 0 | - |
| - | - | 3.868E+04 | 682.4 | - | - | 0 | - |
| - | - | 718 | 683.3 | - | - | 0 | - |
| - | - | 1.626E+04 | 683.4 | - | - | 0 | - |
| - | - | 2817 | 684.4 | - | - | 0 | - |
| - | - | 1543 | 695.3 | - | - | 0 | - |
| - | - | 2132 | 703.4 | - | - | 0 | - |
| - | - | 923 | 704.4 | - | - | 0 | - |
| - | - | 1245 | 705.3 | - | - | 0 | - |
| - | - | 1386 | 706.3 | - | - | 0 | - |
| - | - | 1090 | 714.4 | - | - | 0 | - |
| - | - | 2544 | 714.4 | - | - | 0 | - |
| - | - | 1541 | 715.4 | - | - | 0 | - |
| - | - | 4876 | 716.4 | - | - | 0 | - |
| - | - | 1916 | 717.4 | - | - | 0 | - |
| - | - | 1214 | 720.4 | - | - | 0 | - |
| - | - | 2516 | 721.4 | - | - | 0 | - |
| - | - | 799.8 | 725.4 | - | - | 0 | - |
| 5 | y | 982 | 733.4 | 0.0137 | 18.68 | +1 | 6 |
| 5 | z | 1.051E+04 | 735.3 | 0.00252 | 3.427 | +1 | 6 |
| - | - | 7000 | 736.4 | - | - | 0 | - |
| - | - | 2852 | 737.4 | - | - | 0 | - |
| - | - | 2.746E+04 | 738.4 | - | - | 0 | - |
| - | - | 6.616E+04 | 739.4 | - | - | 0 | - |
| - | - | 6.26E+04 | 740.4 | - | - | 0 | - |
| - | - | 2.067E+04 | 741.4 | - | - | 0 | - |
| - | - | 3677 | 742.4 | - | - | 0 | - |
| - | - | 1253 | 744.3 | - | - | 0 | - |
| - | - | 1.168E+04 | 748.4 | - | - | 0 | - |
| - | - | 5699 | 749.4 | - | - | 0 | - |
| - | - | 2817 | 750.4 | - | - | 0 | - |
| 5 | y | 1.14E+04 | 751.4 | 0.003572 | 4.754 | +1 | 6 |
| - | - | 1.27E+04 | 752.4 | - | - | 0 | - |
| - | - | 5793 | 753.4 | - | - | 0 | - |
| - | - | 733.2 | 754.4 | - | - | 0 | - |
| - | - | 877.6 | 760.4 | - | - | 0 | - |
| - | - | 9645 | 762.3 | - | - | 0 | - |
| - | - | 4472 | 763.4 | - | - | 0 | - |
| - | - | 1406 | 764.4 | - | - | 0 | - |
| - | - | 1876 | 765.4 | - | - | 0 | - |
| - | - | 1579 | 765.4 | - | - | 0 | - |
| 6 | c | 3.563E+05 | 766.4 | 0.003284 | 4.285 | +1 | 6 |
| - | - | 1.71E+05 | 767.4 | - | - | 0 | - |
| - | - | 4.173E+04 | 768.4 | - | - | 0 | - |
| - | - | 4861 | 769.4 | - | - | 0 | - |
| - | - | 2445 | 778.4 | - | - | 0 | - |
| - | - | 1640 | 779.4 | - | - | 0 | - |
| - | - | 1.978E+05 | 780.4 | - | - | 0 | - |
| - | - | 8.767E+04 | 781.4 | - | - | 0 | - |
| - | - | 2.248E+04 | 782.4 | - | - | 0 | - |
| - | - | 2546 | 783.4 | - | - | 0 | - |
| - | - | 845.6 | 784.4 | - | - | 0 | - |
| - | - | 1201 | 790.3 | - | - | 0 | - |
| - | - | 905.7 | 800.4 | - | - | 0 | - |
| - | - | 3981 | 811.4 | - | - | 0 | - |
| - | - | 2233 | 812.4 | - | - | 0 | - |
| - | - | 2164 | 817.4 | - | - | 0 | - |
| - | - | 843.3 | 822.4 | - | - | 0 | - |
| - | - | 4453 | 825.2 | - | - | 0 | - |
| - | - | 3934 | 827.4 | - | - | 0 | - |
| - | - | 2877 | 828.4 | - | - | 0 | - |
| - | - | 4774 | 829.4 | - | - | 0 | - |
| - | - | 1832 | 830.4 | - | - | 0 | - |
| - | - | 3705 | 835.5 | - | - | 0 | - |
| - | - | 7026 | 836.5 | - | - | 0 | - |
| - | - | 3265 | 837.5 | - | - | 0 | - |
| - | - | 776.3 | 838.5 | - | - | 0 | - |
| - | - | 1.053E+04 | 845.4 | - | - | 0 | - |
| - | - | 5447 | 846.4 | - | - | 0 | - |
| - | - | 2306 | 847.5 | - | - | 0 | - |
| - | - | 8217 | 850.4 | - | - | 0 | - |
| - | - | 1.158E+04 | 851.4 | - | - | 0 | - |
| - | - | 5903 | 852.4 | - | - | 0 | - |
| - | - | 2618 | 853.4 | - | - | 0 | - |
| - | - | 925.6 | 860.4 | - | - | 0 | - |
| - | - | 1176 | 861.4 | - | - | 0 | - |
| - | - | 3149 | 862.5 | - | - | 0 | - |
| 7 | c | 3.13E+04 | 863.5 | 0.002888 | 3.345 | +1 | 7 |
| - | - | 1.532E+04 | 864.5 | - | - | 0 | - |
| - | - | 1584 | 865.4 | - | - | 0 | - |
| - | - | 5049 | 865.5 | - | - | 0 | - |
| 4 | y | 6449 | 866.4 | 0.003118 | 3.599 | +1 | 7 |
| - | - | 3199 | 867.4 | - | - | 0 | - |
| - | - | 1101 | 868.4 | - | - | 0 | - |
| - | - | 4316 | 872.4 | - | - | 0 | - |
| - | - | 2953 | 873.4 | - | - | 0 | - |
| - | - | 1170 | 875.4 | - | - | 0 | - |
| - | - | 1.039E+04 | 877.4 | - | - | 0 | - |
| - | - | 4988 | 878.4 | - | - | 0 | - |
| - | - | 5.231E+04 | 879.5 | - | - | 0 | - |
| 7 | c | 2.16E+05 | 880.5 | 0.00289 | 3.282 | +1 | 7 |
| - | - | 1.068E+05 | 881.5 | - | - | 0 | - |
| - | - | 3.03E+04 | 882.5 | - | - | 0 | - |
| - | - | 2908 | 883.5 | - | - | 0 | - |
| - | - | 5646 | 889.4 | - | - | 0 | - |
| - | - | 3211 | 890.4 | - | - | 0 | - |
| - | - | 955.5 | 891.5 | - | - | 0 | - |
| - | - | 2747 | 893.4 | - | - | 0 | - |
| - | - | 932.4 | 894.5 | - | - | 0 | - |
| - | - | 4907 | 895.5 | - | - | 0 | - |
| - | - | 2078 | 896.5 | - | - | 0 | - |
| - | - | 1043 | 897.5 | - | - | 0 | - |
| - | - | 1074 | 898.2 | - | - | 0 | - |
| - | - | 1005 | 904.4 | - | - | 0 | - |
| - | - | 1025 | 905.4 | - | - | 0 | - |
| - | - | 1219 | 905.9 | - | - | 0 | - |
| - | - | 2.479E+04 | 906.5 | - | - | 0 | - |
| - | - | 2535 | 906.5 | - | - | 0 | - |
| - | - | 1366 | 906.9 | - | - | 0 | - |
| - | - | 2.17E+04 | 907.4 | - | - | 0 | - |
| - | - | 1.099E+04 | 908.4 | - | - | 0 | - |
| - | - | 978.1 | 908.5 | - | - | 0 | - |
| - | - | 2672 | 909.4 | - | - | 0 | - |
| - | - | 3258 | 919.4 | - | - | 0 | - |
| 3 | w | 1894 | 920.4 | 0.000733 | 0.7964 | +1 | 8 |
| - | - | 1632 | 921.4 | - | - | 0 | - |
| - | - | 3590 | 933.5 | - | - | 0 | - |
| - | - | 7.538E+04 | 934.5 | - | - | 0 | - |
| - | - | 4.518E+04 | 935.5 | - | - | 0 | - |
| - | - | 1.329E+04 | 936.5 | - | - | 0 | - |
| - | - | 2246 | 937.5 | - | - | 0 | - |
| - | - | 3702 | 946.5 | - | - | 0 | - |
| - | - | 3871 | 947.5 | - | - | 0 | - |
| - | - | 2466 | 948.5 | - | - | 0 | - |
| - | - | 2626 | 956.5 | - | - | 0 | - |
| - | - | 1927 | 957.5 | - | - | 0 | - |
| - | - | 1092 | 959.5 | - | - | 0 | - |
| - | - | 5.783E+04 | 964.5 | - | - | 0 | - |
| - | - | 4.169E+04 | 965.5 | - | - | 0 | - |
| - | - | 1.759E+04 | 966.5 | - | - | 0 | - |
| - | - | 3420 | 967.5 | - | - | 0 | - |
| - | - | 1340 | 971.5 | - | - | 0 | - |
| - | - | 3.703E+04 | 974.5 | - | - | 0 | - |
| - | - | 2.201E+04 | 975.5 | - | - | 0 | - |
| 3 | y | 1.88E+04 | 976.5 | 0.01082 | 11.08 | +1 | 8 |
| - | - | 1.004E+04 | 977.5 | - | - | 0 | - |
| 3 | z | 1.744E+05 | 978.5 | 0.003966 | 4.054 | +1 | 8 |
| - | - | 1.024E+05 | 979.5 | - | - | 0 | - |
| - | - | 3.44E+04 | 980.5 | - | - | 0 | - |
| - | - | 5141 | 981.5 | - | - | 0 | - |
| - | - | 1.503E+04 | 990.6 | - | - | 0 | - |
| - | - | 1.134E+04 | 991.6 | - | - | 0 | - |
| 8 | c | 7.632E+05 | 992.5 | 0.00418 | 4.211 | +1 | 8 |
| - | - | 4.658E+05 | 993.5 | - | - | 0 | - |
| 3 | y | 2.696E+05 | 994.5 | 0.01594 | 16.03 | +1 | 8 |
| - | - | 1.188E+05 | 995.5 | - | - | 0 | - |
| - | - | 3.512E+04 | 996.5 | - | - | 0 | - |
| - | - | 3913 | 997.5 | - | - | 0 | - |
| - | - | 1397 | 1002 | - | - | 0 | - |
| - | - | 1051 | 1021 | - | - | 0 | - |
| - | - | 2742 | 1034 | - | - | 0 | - |
| - | - | 1148 | 1035 | - | - | 0 | - |
| - | - | 1157 | 1037 | - | - | 0 | - |
| - | - | 960 | 1047 | - | - | 0 | - |
| - | - | 1192 | 1048 | - | - | 0 | - |
| - | - | 8700 | 1063 | - | - | 0 | - |
| - | - | 5956 | 1064 | - | - | 0 | - |
| - | - | 2431 | 1065 | - | - | 0 | - |
| - | - | 1205 | 1075 | - | - | 0 | - |
| 2 | y | 1710 | 1076 | 0.000269 | 0.2501 | +1 | 9 |
| 2 | y | 3138 | 1077 | 0.02065 | 19.18 | +1 | 9 |
| 2 | z | 4.713E+04 | 1078 | 0.004156 | 3.857 | +1 | 9 |
| - | - | 2.931E+04 | 1079 | - | - | 0 | - |
| - | - | 9930 | 1080 | - | - | 0 | - |
| - | - | 2456 | 1081 | - | - | 0 | - |
| - | - | 958.7 | 1091 | - | - | 0 | - |
| - | - | 1731 | 1092 | - | - | 0 | - |
| 2 | y | 2.006E+04 | 1094 | 0.004719 | 4.315 | +1 | 9 |
| - | - | 1.115E+04 | 1095 | - | - | 0 | - |
| - | - | 4266 | 1096 | - | - | 0 | - |
| - | - | 1702 | 1100 | - | - | 0 | - |
| - | - | 1582 | 1101 | - | - | 0 | - |
| - | - | 1326 | 1102 | - | - | 0 | - |
| - | - | 1208 | 1106 | - | - | 0 | - |
| 9 | c | 2.395E+05 | 1107 | 0.004212 | 3.807 | +1 | 9 |
| - | - | 1.655E+05 | 1108 | - | - | 0 | - |
| - | - | 6.182E+04 | 1109 | - | - | 0 | - |
| - | - | 9681 | 1110 | - | - | 0 | - |
| - | - | 1274 | 1117 | - | - | 0 | - |
| - | - | 1451 | 1118 | - | - | 0 | - |
| - | - | 1.101E+04 | 1119 | - | - | 0 | - |
| - | - | 6713 | 1120 | - | - | 0 | - |
| - | - | 3298 | 1121 | - | - | 0 | - |
| - | - | 1204 | 1124 | - | - | 0 | - |
| - | - | 1366 | 1134 | - | - | 0 | - |
| - | - | 2.611E+04 | 1136 | - | - | 0 | - |
| - | - | 1.754E+04 | 1137 | - | - | 0 | - |
| - | - | 7005 | 1138 | - | - | 0 | - |
| - | - | 1077 | 1139 | - | - | 0 | - |
| - | - | 4080 | 1146 | - | - | 0 | - |
| - | - | 7267 | 1147 | - | - | 0 | - |
| - | - | 4.83E+04 | 1148 | - | - | 0 | - |
| - | - | 3.52E+04 | 1149 | - | - | 0 | - |
| - | - | 1.348E+04 | 1150 | - | - | 0 | - |
| - | - | 2961 | 1151 | - | - | 0 | - |
| - | - | 3.129E+04 | 1152 | - | - | 0 | - |
| - | - | 2.156E+04 | 1153 | - | - | 0 | - |
| - | - | 8780 | 1154 | - | - | 0 | - |
| - | - | 1317 | 1155 | - | - | 0 | - |
| - | - | 4.895E+04 | 1162 | - | - | 0 | - |
| - | - | 4.53E+04 | 1163 | - | - | 0 | - |
| - | - | 7.389E+04 | 1164 | - | - | 0 | - |
| - | - | 4.472E+04 | 1165 | - | - | 0 | - |
| - | - | 1.475E+04 | 1166 | - | - | 0 | - |
| - | - | 2471 | 1167 | - | - | 0 | - |
| - | - | 2237 | 1172 | - | - | 0 | - |
| - | - | 1620 | 1173 | - | - | 0 | - |
| - | - | 3.457E+04 | 1180 | - | - | 0 | - |
| - | - | 2.553E+04 | 1181 | - | - | 0 | - |
| - | - | 1.031E+04 | 1182 | - | - | 0 | - |
| - | - | 1541 | 1183 | - | - | 0 | - |
| - | - | 1657 | 1189 | - | - | 0 | - |
| - | - | 1.253E+04 | 1190 | - | - | 0 | - |
| - | - | 2.611E+05 | 1191 | - | - | 0 | - |
| - | - | 1.971E+05 | 1192 | - | - | 0 | - |
| - | - | 7.941E+04 | 1193 | - | - | 0 | - |
| - | - | 1.481E+04 | 1194 | - | - | 0 | - |
| - | - | 3224 | 1205 | - | - | 0 | - |
| - | - | 1.121E+04 | 1206 | - | - | 0 | - |
| - | - | 3.257E+05 | 1207 | - | - | 0 | - |
| - | - | 1.309E+06 | 1208 | - | - | 0 | - |
| - | - | 9.142E+05 | 1209 | - | - | 0 | - |
| - | - | 3.434E+05 | 1210 | - | - | 0 | - |
| - | - | 5.905E+04 | 1211 | - | - | 0 | - |
| - | - | 2257 | 1223 | - | - | 0 | - |
| - | - | 1011 | 1224 | - | - | 0 | - |
| - | - | 3394 | 1240 | - | - | 0 | - |
| - | - | 2389 | 1241 | - | - | 0 | - |

m/z Charge Intensity FragmentType MassShift Position
120.08134460449219 0 6286.6055
121.08432006835938 0 395.8706
129.1028289794922 0 37162.54
130.10614013671875 0 1550.3977
136.07632446289062 0 1287.9532
138.4123077392578 0 387.70462
148.1997528076172 0 425.5316
148.94764709472656 0 693.4892
158.93199157714844 0 477.57053
169.13421630859375 0 13044.899
170.1377410888672 0 1043.5449
171.9104766845703 0 503.33124
172.5270538330078 0 439.48102
173.43959045410156 0 2941.665
181.0973663330078 0 776.13385
181.1346435546875 0 600.80505
182.0828399658203 0 501.78696
183.15016174316406 0 717.7488
185.1656036376953 0 145699.33
186.16897583007812 0 15636.487
187.1715087890625 0 736.8495
191.10227966308594 0 622.487
197.12913513183594 0 5105.073
198.13235473632812 0 679.6867
199.10842895507812 0 7740.464
200.1114501953125 0 608.10406
201.12429809570312 0 713.63824
203.10362243652344 0 1820.2169
209.09280395507812 0 6443.929
210.095458984375 0 748.93964
213.12416076660156 0 3111.58
213.1607208251953 0 85683.484
214.16429138183594 0 11940.14
215.1399688720703 0 1894352.5 y 8
216.14317321777344 0 206182.77
217.14527893066406 0 13119.781
227.10353088378906 0 59189.254
228.10690307617188 0 5316.148
228.1715545654297 0 3616.3523
231.14987182617188 0 1826.4829
244.13009643554688 0 9144.723
245.13011169433594 0 2352.22
251.10302734375 0 1545.4949
255.0983123779297 0 904.53925
261.1585693359375 0 849.7216
278.15142822265625 0 1426.1882
279.0986022949219 0 5219.1255
280.10302734375 0 699.2326
280.1658935546875 0 1253.0381
283.1449279785156 0 926.5304
301.2121887207031 0 3998.4995
310.1778564453125 0 1698.0054 c Ammonia loss 4
310.2130432128906 0 842.8745
311.1402893066406 0 2800.078
312.1435241699219 0 702.30023
315.2759704589844 0 1044.1879
320.19793701171875 0 1159.5745
324.1567687988281 0 37513.715
325.1602783203125 0 6981.278
326.17230224609375 0 19322.506 y Water loss 7
327.1753845214844 0 4184.466
340.1875 0 2186.456
341.1839599609375 0 1213.9622
341.25628662109375 0 10340.894 c Ammonia loss 2
342.25921630859375 0 1511.1682
343.1981506347656 0 1766.8503
344.1828918457031 0 22421.357 y 7
344.2059631347656 0 1204.7546
344.7060852050781 0 1141.0831
345.1859436035156 0 2989.1738
353.21978759765625 0 887.96716
355.1993408203125 0 2313.159
356.267822265625 0 1609.556
357.27471923828125 0 103939.19
358.27911376953125 0 25340.59 c 2
358.70330810546875 0 1208.0402
359.28375244140625 0 2215.4226
362.1733093261719 0 785.54297
369.71441650390625 0 5145.688
370.21514892578125 0 2027.7803
371.1937255859375 0 2252.6448
373.188720703125 0 1395.6984
374.17236328125 0 13994.255
375.1748046875 0 3000.581
377.2221984863281 0 690.7576
379.1995544433594 0 1341.17
381.14715576171875 0 928.3994
383.19317626953125 0 674.12683
383.7113342285156 0 4045.8262 c Ammonia loss 5
384.2132263183594 0 2474.0093
389.1835632324219 0 1393.896
390.6809387207031 0 626.24457
391.1986999511719 0 3001.1094
392.1830749511719 0 3273.4832
395.2293701171875 0 1327.9122
398.17425537109375 0 739.2781
403.23577880859375 0 1588.522
405.21539306640625 0 1015.9089
406.21453857421875 0 620.83295
407.19415283203125 0 13303.614
407.2245788574219 0 708.45013
408.1962585449219 0 2912.4932
411.2607421875 0 794.5636
418.2411193847656 0 19099.924
418.74249267578125 0 12657.49
419.2442932128906 0 3481.488
423.2256164550781 0 46181.65 y Water loss 6
424.2283020019531 0 13308.234
425.2303161621094 0 1746.8
426.1677551269531 0 21341.115
427.17120361328125 0 5379.2896
429.2947998046875 0 1199.7516
430.3051452636719 0 668.1315
432.23846435546875 0 26449.77 c Ammonia loss 6
432.7402648925781 0 13746.2705
433.2411804199219 0 3280.3523
438.2729797363281 0 5163.1367
438.310302734375 0 1934.3969
439.208740234375 0 984.8078
439.25714111328125 0 2115.9272
440.22467041015625 0 908.793
441.2362976074219 0 203766.33 y 6
442.239501953125 0 50954.242
443.2417907714844 0 7563.5215
443.29913330078125 0 848.2693
447.2276916503906 0 896.0509
452.2156982421875 0 2484.7102
454.2669372558594 0 1309.1641
456.28350830078125 0 33606.85 c Ammonia loss 3
457.2483825683594 0 1661.5967
457.2867431640625 0 9085.445
458.2867431640625 0 1434.6968
464.25567626953125 0 693.80054
467.2655944824219 0 1401.5024
470.2266540527344 0 10365.533
471.2279357910156 0 3793.4348
472.3023986816406 0 20022.281
473.3096923828125 0 40588.766 c 3
473.7537536621094 0 983.25574
474.25628662109375 0 1133.1395
474.31298828125 0 9442.4375
475.3131408691406 0 1079.0312
482.2628173828125 0 7248.026
482.7626953125 0 14549.513
483.2642822265625 0 11760.647
483.764404296875 0 2194.2568
487.7545471191406 0 3042.831
488.2557067871094 0 2129.4175
488.3240966796875 0 3236.6736
488.7541809082031 0 1104.0162
489.2445373535156 0 766.2243
489.3288269042969 0 986.67773
491.2516174316406 0 2139.7483
493.851806640625 0 620.49426
495.2972717285156 0 637.1818
495.74932861328125 0 735.5335
496.76025390625 0 25006.047 c Ammonia loss 7
497.2617492675781 0 16502.613
497.762451171875 0 6426.6113
502.2711486816406 0 657.4372
506.2624816894531 0 1676.9978
507.2654113769531 0 1070.7612
509.2411193847656 0 3255.2195
510.2456359863281 0 2107.6392
511.2564697265625 0 6336.811
512.2610473632812 0 2697.824
521.282470703125 0 1118.8893
522.2845458984375 0 2343.5544
522.78515625 0 838.54767
523.2210083007812 0 3193.6023
523.293212890625 0 2270.7502
524.224609375 0 813.49133
524.2962646484375 0 997.579
526.2677612304688 0 2181.472
527.2763061523438 0 999.12555
531.2883911132812 0 3513.7705
531.7875366210938 0 1886.7231
532.2881469726562 0 793.1192
534.258056640625 0 8284.728
535.2611083984375 0 2741.8152
536.2530517578125 0 5486.2236
536.7830810546875 0 856.852
537.2399291992188 0 3558.0154
538.2405395507812 0 1936.9667
539.2518920898438 0 7678.45
540.2548828125 0 2084.9795
540.3063354492188 0 1176.7052
545.2862548828125 0 28786.357 c Ammonia loss 8
545.7882080078125 0 16992.174
546.2898559570312 0 6705.6035
546.7875366210938 0 973.41437
547.283203125 0 797.593 y 1
548.2733154296875 0 1275.2073
549.34228515625 0 2562.6282
550.3447875976562 0 890.441
551.2868041992188 0 761.174
553.3365478515625 0 2688.0637
554.26318359375 0 94087.46 z Water loss 5
555.2659912109375 0 30755.387
556.26953125 0 5318.714
558.3308715820312 0 942.9879
567.3059692382812 0 1069.4879
567.3524780273438 0 16460.127
568.3549194335938 0 5493.198
570.2974243164062 0 1848.849 y Water loss 5
571.28369140625 0 618.1613
574.3380737304688 0 824.04584
580.82421875 0 3198.8188
581.323974609375 0 2371.66
581.8278198242188 0 1371.7423
582.2551879882812 0 1124.7675
585.3414306640625 0 1972.7024
587.29638671875 0 2795.3162
588.30517578125 0 106608.63 y 5
589.3082885742188 0 35607
590.3104248046875 0 6199.1265
591.3522338867188 0 13374.38
592.3579711914062 0 9564.516
593.30615234375 0 654.1449
593.3623657226562 0 3221.9001
594.8209838867188 0 5786.7485
595.3214721679688 0 4744.784
595.8230590820312 0 2312.4065
597.290771484375 0 2465.9795
598.2955322265625 0 1081.0682
599.2849731445312 0 1841.5088
601.3369750976562 0 3226.7734
602.3391723632812 0 839.92316
603.3536376953125 0 3835.7432
603.8264770507812 0 28417.03
604.32763671875 0 25203.898
604.8290405273438 0 8860.413
605.3305053710938 0 1190.6315
610.3229370117188 0 1091.4204
615.2811889648438 0 2284.0544
617.2893676757812 0 874.48175
619.3475952148438 0 87481.5 c Ammonia loss 4
620.3505859375 0 34617.156
621.3526611328125 0 6618.869
624.2706909179688 0 930.67035
633.2906494140625 0 3921.1067
635.3663940429688 0 66886.31
636.372802734375 0 88996.44 c 4
637.37646484375 0 25847.51
638.32080078125 0 4986.463
638.3795776367188 0 3830.332
639.3242797851562 0 1782.5186
644.1674194335938 0 598.0355
651.3165283203125 0 10838.224
651.3812255859375 0 1154.2194
652.2636108398438 0 16751.955
652.32177734375 0 3820.68
653.2664794921875 0 7544.184
653.3311157226562 0 11160.578
654.335205078125 0 4353.96
655.3414306640625 0 1000.0153
663.3169555664062 0 4043.1426
664.3697509765625 0 4217.243
665.3717041015625 0 1540.8175
667.3466796875 0 1873.6445
668.3431396484375 0 856.012
669.4014282226562 0 1398.6249
681.3267822265625 0 15020.6045
682.380126953125 0 38684.953
683.320556640625 0 717.96204
683.3831787109375 0 16256.55
684.3843383789062 0 2817.136
695.3440551757812 0 1542.8353
703.3724975585938 0 2132.153
704.375244140625 0 923.0125
705.26611328125 0 1245.3495
706.2725830078125 0 1386.4124
714.3582763671875 0 1090.0728
714.421630859375 0 2543.8853
715.425048828125 0 1540.8281
716.40185546875 0 4875.522
717.402099609375 0 1915.7444
720.4105834960938 0 1213.6401
721.3984985351562 0 2515.7756
725.3941040039062 0 799.82196
733.3418579101562 0 982.00635 y Water loss 4
735.3499145507812 0 10506.19 z 4
736.3556518554688 0 7000.457
737.359619140625 0 2852.3464
738.4214477539062 0 27455.225
739.4281616210938 0 66160.56
740.4345703125 0 62604.508
741.438232421875 0 20673.24
742.4398193359375 0 3677.266
744.343017578125 0 1253.2443
748.4053955078125 0 11675.865
749.4032592773438 0 5698.762
750.3675537109375 0 2816.835
751.3696899414062 0 11404.541 y 4
752.3670654296875 0 12703.966
753.3690795898438 0 5792.821
754.3568115234375 0 733.1951
760.3526000976562 0 877.5804
762.3491821289062 0 9645.408
763.3528442382812 0 4472.143
764.4075927734375 0 1406.3463
765.3524169921875 0 1876.0096
765.4097290039062 0 1579.3876
766.4166870117188 0 356273.7 c Ammonia loss 5
767.419677734375 0 170973.34
768.4224853515625 0 41732.574
769.4247436523438 0 4861.4126
778.3687133789062 0 2445.3684
779.37548828125 0 1639.884
780.359130859375 0 197750.17
781.3624267578125 0 87674.2
782.365234375 0 22479.516
783.3665771484375 0 2546.1096
784.4270629882812 0 845.59033
790.3379516601562 0 1200.7709
800.4461669921875 0 905.69916
811.43994140625 0 3981.456
812.4429931640625 0 2232.8013
817.4470825195312 0 2163.523
822.44189453125 0 843.33655
825.2326049804688 0 4452.5054
827.4305419921875 0 3934.4749
828.4380493164062 0 2877.4663
829.4471435546875 0 4773.74
830.4478149414062 0 1831.7925
835.4769897460938 0 3704.666
836.4824829101562 0 7025.525
837.484130859375 0 3264.5713
838.4864501953125 0 776.3169
845.4449462890625 0 10532.524
846.447998046875 0 5447.142
847.452392578125 0 2306.0632
850.4375610351562 0 8217.126
851.38525390625 0 11581.474
852.3890380859375 0 5902.5386
853.3886108398438 0 2618.2947
860.42822265625 0 925.60144
861.4108276367188 0 1175.8885
862.4611206054688 0 3148.6992
863.4690551757812 0 31300.002 c Ammonia loss 6
864.471435546875 0 15319.617
865.3924560546875 0 1583.9794
865.4758911132812 0 5049.4863
866.3961791992188 0 6448.9463 y 3
867.401123046875 0 3199.4473
868.3970947265625 0 1100.9652
872.429931640625 0 4315.904
873.431640625 0 2952.7253
875.4225463867188 0 1170.285
877.4138793945312 0 10392.932
878.4200439453125 0 4987.998
879.4885864257812 0 52307.69
880.49560546875 0 215957.4 c 6
881.4986572265625 0 106799.72
882.5014038085938 0 30296.11
883.501220703125 0 2908.4229
889.410888671875 0 5646.455
890.4136962890625 0 3210.8777
891.4842529296875 0 955.52936
893.4446411132812 0 2747.1235
894.4524536132812 0 932.4452
895.4577026367188 0 4907.1733
896.4637451171875 0 2077.9163
897.4662475585938 0 1043.1483
898.1597900390625 0 1073.76
904.4401245117188 0 1004.9492
905.3859252929688 0 1025.062
905.8952026367188 0 1218.6685
906.4515380859375 0 24791.557
906.5353393554688 0 2534.6196
906.8984375 0 1365.7068
907.435302734375 0 21695.727
908.4321899414062 0 10994.117
908.529296875 0 978.1466
909.4320068359375 0 2672.3945
919.3983154296875 0 3258.1768
920.4028930664062 0 1893.6768 w 2
921.4335327148438 0 1632.4797
933.4761962890625 0 3589.8235
934.4827270507812 0 75379.43
935.4852905273438 0 45180.59
936.4882202148438 0 13285.945
937.4910888671875 0 2245.9854
946.5059204101562 0 3701.7505
947.5028686523438 0 3871.0337
948.5029907226562 0 2465.7207
956.489990234375 0 2625.803
957.4874267578125 0 1926.9985
959.4581298828125 0 1091.5481
964.517578125 0 57834.11
965.5221557617188 0 41688.418
966.5252685546875 0 17590.133
967.5315551757812 0 3420.3186
971.5010375976562 0 1339.593
974.5018310546875 0 37026.883
975.5043334960938 0 22008.703
976.48828125 0 18796.31 y Water loss 2
977.4829711914062 0 10037.026
978.4732666015625 0 174350.11 z 2
979.4764404296875 0 102374.305
980.4794921875 0 34402.152
981.4832153320312 0 5140.709
990.5571899414062 0 15028.76
991.5599365234375 0 11341.681
992.512939453125 0 763169.7 c Ammonia loss 7
993.5157470703125 0 465792
994.5039672851562 0 269590.06 y 2
995.4984741210938 0 118843.91
996.4974365234375 0 35116.617
997.5021362304688 0 3913.1238
1001.5780029296875 0 1396.9062
1021.476806640625 0 1050.6476
1033.54833984375 0 2742.2485
1034.5660400390625 0 1147.9626
1036.565185546875 0 1156.5417
1046.5428466796875 0 960.0177
1047.5546875 0 1192.1592
1062.5753173828125 0 8699.619
1063.5810546875 0 5955.6274
1064.5826416015625 0 2430.8367
1074.6343994140625 0 1204.9552
1075.546142578125 0 1709.7031 y Water loss 1
1076.550537109375 0 3138.2302 y Ammonia loss 1
1077.5418701171875 0 47130.777 z 1
1078.544677734375 0 29305.342
1079.5482177734375 0 9930.285
1080.5570068359375 0 2455.8638
1090.632080078125 0 958.67645
1091.5718994140625 0 1731.1869
1093.5611572265625 0 20055.453 y 1
1094.5640869140625 0 11146.461
1095.5654296875 0 4266.412
1099.5941162109375 0 1702.4479
1100.603759765625 0 1581.5093
1101.615478515625 0 1325.6992
1105.568359375 0 1208.3004
1106.59228515625 0 239501.5 c 8
1107.5947265625 0 165452.53
1108.597412109375 0 61818.613
1109.598388671875 0 9680.968
1116.602294921875 0 1274.411
1117.65283203125 0 1450.5016
1118.6064453125 0 11007.093
1119.6075439453125 0 6713.3975
1120.614501953125 0 3297.9985
1123.5479736328125 0 1203.5251
1133.626953125 0 1365.782
1135.6309814453125 0 26113.65
1136.631591796875 0 17540.402
1137.6336669921875 0 7004.5874
1138.6414794921875 0 1077.0029
1145.6446533203125 0 4079.8003
1146.640625 0 7267.2593
1147.612548828125 0 48298.74
1148.5966796875 0 35197.7
1149.619384765625 0 13483.858
1150.6024169921875 0 2960.753
1151.589111328125 0 31286.648
1152.5927734375 0 21556.1
1153.5955810546875 0 8780.322
1154.5867919921875 0 1316.8606
1161.6470947265625 0 48947.33
1162.6507568359375 0 45299.96
1163.6597900390625 0 73890.445
1164.663818359375 0 44717.855
1165.6661376953125 0 14749.14
1166.657470703125 0 2471.391
1171.630126953125 0 2237.4658
1172.6385498046875 0 1619.8713
1179.656982421875 0 34565.664
1180.65966796875 0 25533.393
1181.6627197265625 0 10307.514
1182.6602783203125 0 1540.5459
1188.6048583984375 0 1656.94
1189.634521484375 0 12526.883
1190.6268310546875 0 261141.5
1191.6298828125 0 197092.86
1192.63232421875 0 79408.39
1193.634765625 0 14812.265
1204.629150390625 0 3224.4949
1205.634033203125 0 11210.02
1206.6451416015625 0 325735.53
1207.65234375 0 1308700.4
1208.6551513671875 0 914234.8
1209.658203125 0 343373.6
1210.6605224609375 0 59046.11
1222.61328125 0 2256.7722
1223.59814453125 0 1010.57544
1239.6405029296875 0 3393.9746
1240.6505126953125 0 2388.506

Spectrum Details

|  |  |
| --- | --- |
| Matched peaks? Matched peaks The total absolute number of peaks matched. Additionally in brackets the total fraction of peaks matched and the total number of peaks is shown. | 37 (7.55% of 490) |
| FDR? FDR The false discovery rate estimated for this peptide. It is calculated by matching all theoretical fragments with a non-integer shift with the raw peaks for this spectrum. This is done with 40 different shifts. The resulting percentage is the average number of annotated peaks over the number of annotated peaks with the correct spectrum. | 3.60% |
| Satellite FDR? Satellite FDR See the FDR for details on its calculation. This satellite ion specific FDR only contains the satellite ions (d/w) for I/L/J positions. | - |
| PSM Score? PSM Score The PSM Score as given by Hecklib to this annotated spectrum. It is shown with three significant figures. | 440 |

#### Spectrum 8595? Spectrum 8595 The raw spectrum of this peptide as annotated by Hecklib. The fragments are coloured according to ion type (see legend). Any peaks with a star '\*' as text can be hovered over to see the full details, first the ion type second the mass shift type. By hovering over the amino acids in the peptide or ions in the legend the corresponding peaks are highlighted. By toggling the 'Unassigned' label you can turn the background (unassigned) peaks on or off in the plot. By updating the slider in the Ion legend you can update the spectrum to only show the top X% of the peaks with labels. The top X% means any peak that is within X% of the highest intensity. By dragging in the spectrum you can zoom in to a specific part of the spectrum and use 'Zoom Out' to get back to the original zoom level. The annotation of the spectrum is based on the given sequence in the peptides file and is done with different software so inconsistencies are likely. The peaks are annotated based on the given sequence, with 20 ppm tolerance.

Copy Data

##### Spectrum 8595 (TSV)

###### Preview

```
Loading example...
```

*Click on the button to copy the data to your clipboard.*

Mz MinMz MaxIntensity Max

WidthHeightPeptide font sizePeptide stroke widthSpectrum font sizeSpectrum stroke widthCompact peptide

Ion legend

wxyz

abcd

OtherUnassignedIon^Charge^_Position_Show for top:%

JVKDYFPEPV

06.40e+41.28e+51.92e+52.56e+5

Zoom Out

y^+1^_2_y^+1^_3_c^+1^_3_y^+1^_4_c^+2^_7_y^+1^_4_c^+1^_4_c^+1^_4_c^+2^_9_y^+1^_5_c^+1^_5_c^+1^_5_z^+1^_6_y^+1^_6_c^+1^_6_c^+1^_7_y^+1^_7_c^+1^_7_z^+1^_8_c^+1^_8_y^+1^_8_z^+1^_9_c^+1^_9_

0825165124763302

Fragment Matches Table

Show background peaks

| Position | Ion type | Intensity | mz Theoretical | mz Error (Th) | mz Error (ppm) | Charge | Series Number |
| --- | --- | --- | --- | --- | --- | --- | --- |
| - | - | 573.3 | 120.1 | - | - | 0 | - |
| - | - | 1946 | 129.1 | - | - | 0 | - |
| - | - | 471.9 | 133.8 | - | - | 0 | - |
| - | - | 496.4 | 148.9 | - | - | 0 | - |
| - | - | 493.1 | 148.9 | - | - | 0 | - |
| - | - | 685.1 | 148.9 | - | - | 0 | - |
| - | - | 710.5 | 148.9 | - | - | 0 | - |
| - | - | 1185 | 148.9 | - | - | 0 | - |
| - | - | 1266 | 148.9 | - | - | 0 | - |
| - | - | 1637 | 148.9 | - | - | 0 | - |
| - | - | 3331 | 148.9 | - | - | 0 | - |
| - | - | 4413 | 149 | - | - | 0 | - |
| - | - | 2497 | 149 | - | - | 0 | - |
| - | - | 1176 | 149 | - | - | 0 | - |
| - | - | 1184 | 149 | - | - | 0 | - |
| - | - | 854.8 | 149 | - | - | 0 | - |
| - | - | 693 | 149 | - | - | 0 | - |
| - | - | 606.2 | 149 | - | - | 0 | - |
| - | - | 658.8 | 149 | - | - | 0 | - |
| - | - | 424 | 149 | - | - | 0 | - |
| - | - | 560.6 | 149 | - | - | 0 | - |
| - | - | 422.7 | 149 | - | - | 0 | - |
| - | - | 445.5 | 149.1 | - | - | 0 | - |
| - | - | 452.4 | 151.2 | - | - | 0 | - |
| - | - | 474.5 | 156.4 | - | - | 0 | - |
| - | - | 456.3 | 168.9 | - | - | 0 | - |
| - | - | 596.5 | 169.1 | - | - | 0 | - |
| - | - | 465.6 | 173.1 | - | - | 0 | - |
| - | - | 6215 | 185.2 | - | - | 0 | - |
| - | - | 555.2 | 187.1 | - | - | 0 | - |
| - | - | 3448 | 213.2 | - | - | 0 | - |
| 9 | y | 8.06E+04 | 215.1 | 0.0002023 | 0.9402 | +1 | 2 |
| - | - | 9222 | 216.1 | - | - | 0 | - |
| - | - | 2273 | 227.1 | - | - | 0 | - |
| - | - | 710.8 | 301.2 | - | - | 0 | - |
| - | - | 507.4 | 311.1 | - | - | 0 | - |
| - | - | 1697 | 324.2 | - | - | 0 | - |
| 8 | y | 978.9 | 326.2 | 0.0006751 | 2.07 | +1 | 3 |
| - | - | 1.75E+04 | 357.3 | - | - | 0 | - |
| 3 | c | 4193 | 358.3 | 0.00319 | 8.904 | +1 | 3 |
| - | - | 1068 | 374.2 | - | - | 0 | - |
| - | - | 894.1 | 407.2 | - | - | 0 | - |
| - | - | 647.7 | 418.7 | - | - | 0 | - |
| 7 | y | 1574 | 423.2 | 8.68E-05 | 0.2051 | +1 | 4 |
| - | - | 1175 | 426.2 | - | - | 0 | - |
| 7 | c | 1143 | 432.2 | 0.0005831 | 1.349 | +2 | 7 |
| 7 | y | 8784 | 441.2 | 0.000579 | 1.312 | +1 | 4 |
| - | - | 2005 | 442.2 | - | - | 0 | - |
| 4 | c | 1734 | 456.3 | 0.0005051 | 1.107 | +1 | 4 |
| - | - | 655 | 470.2 | - | - | 0 | - |
| - | - | 3921 | 472.3 | - | - | 0 | - |
| 4 | c | 6194 | 473.3 | 1.804E-05 | 0.03811 | +1 | 4 |
| - | - | 1645 | 474.3 | - | - | 0 | - |
| - | - | 584.3 | 483.3 | - | - | 0 | - |
| - | - | 1070 | 497.3 | - | - | 0 | - |
| - | - | 973 | 511.3 | - | - | 0 | - |
| - | - | 817.1 | 543.3 | - | - | 0 | - |
| 9 | c | 1301 | 545.3 | 0.001441 | 2.643 | +2 | 9 |
| - | - | 3867 | 554.3 | - | - | 0 | - |
| - | - | 1129 | 555.3 | - | - | 0 | - |
| - | - | 792.7 | 587.3 | - | - | 0 | - |
| 6 | y | 8255 | 588.3 | 0.0003718 | 0.6321 | +1 | 5 |
| - | - | 2345 | 589.3 | - | - | 0 | - |
| - | - | 647.7 | 590.3 | - | - | 0 | - |
| - | - | 1162 | 591.3 | - | - | 0 | - |
| - | - | 897.3 | 603.8 | - | - | 0 | - |
| - | - | 1155 | 604.3 | - | - | 0 | - |
| 5 | c | 3061 | 619.3 | 0.0009583 | 1.547 | +1 | 5 |
| - | - | 1363 | 620.3 | - | - | 0 | - |
| - | - | 578.8 | 631.8 | - | - | 0 | - |
| - | - | 1.066E+04 | 635.4 | - | - | 0 | - |
| 5 | c | 1.662E+04 | 636.4 | 0.0008105 | 1.274 | +1 | 5 |
| - | - | 5070 | 637.4 | - | - | 0 | - |
| - | - | 825.4 | 638.4 | - | - | 0 | - |
| - | - | 969.1 | 681.3 | - | - | 0 | - |
| - | - | 987.3 | 682.4 | - | - | 0 | - |
| - | - | 687.5 | 683.4 | - | - | 0 | - |
| - | - | 2359 | 706.4 | - | - | 0 | - |
| - | - | 1081 | 707.4 | - | - | 0 | - |
| - | - | 897.3 | 717.4 | - | - | 0 | - |
| - | - | 589.5 | 718.4 | - | - | 0 | - |
| 5 | z | 1005 | 735.3 | 0.0006282 | 0.8543 | +1 | 6 |
| - | - | 1199 | 736.4 | - | - | 0 | - |
| - | - | 1468 | 738.4 | - | - | 0 | - |
| - | - | 1.142E+04 | 739.4 | - | - | 0 | - |
| - | - | 1.089E+04 | 740.4 | - | - | 0 | - |
| - | - | 4291 | 741.4 | - | - | 0 | - |
| - | - | 675.2 | 750.4 | - | - | 0 | - |
| 5 | y | 1572 | 751.4 | 0.0006398 | 0.8515 | +1 | 6 |
| - | - | 1061 | 752.4 | - | - | 0 | - |
| - | - | 784.1 | 762.3 | - | - | 0 | - |
| 6 | c | 1.553E+04 | 766.4 | 0.0002559 | 0.3339 | +1 | 6 |
| - | - | 7045 | 767.4 | - | - | 0 | - |
| - | - | 1403 | 768.4 | - | - | 0 | - |
| - | - | 8294 | 780.4 | - | - | 0 | - |
| - | - | 4372 | 781.4 | - | - | 0 | - |
| - | - | 1410 | 782.4 | - | - | 0 | - |
| - | - | 1101 | 787.5 | - | - | 0 | - |
| - | - | 614.2 | 807.4 | - | - | 0 | - |
| - | - | 983.8 | 836.5 | - | - | 0 | - |
| - | - | 948.5 | 837.5 | - | - | 0 | - |
| - | - | 1843 | 851.4 | - | - | 0 | - |
| - | - | 1352 | 852.4 | - | - | 0 | - |
| 7 | c | 1306 | 863.5 | 0.002483 | 2.875 | +1 | 7 |
| 4 | y | 964 | 866.4 | 0.004705 | 5.43 | +1 | 7 |
| - | - | 1393 | 872.4 | - | - | 0 | - |
| - | - | 945.1 | 879.4 | - | - | 0 | - |
| - | - | 1.136E+04 | 879.5 | - | - | 0 | - |
| 7 | c | 3.674E+04 | 880.5 | 0.0007114 | 0.808 | +1 | 7 |
| - | - | 1.915E+04 | 881.5 | - | - | 0 | - |
| - | - | 5049 | 882.5 | - | - | 0 | - |
| - | - | 4860 | 906.4 | - | - | 0 | - |
| - | - | 3106 | 907.4 | - | - | 0 | - |
| - | - | 1147 | 919.4 | - | - | 0 | - |
| - | - | 977.6 | 933.5 | - | - | 0 | - |
| - | - | 1.455E+04 | 934.5 | - | - | 0 | - |
| - | - | 8980 | 935.5 | - | - | 0 | - |
| - | - | 1931 | 936.5 | - | - | 0 | - |
| - | - | 2941 | 964.5 | - | - | 0 | - |
| - | - | 3564 | 965.5 | - | - | 0 | - |
| - | - | 1551 | 966.5 | - | - | 0 | - |
| - | - | 1811 | 974.5 | - | - | 0 | - |
| - | - | 845.1 | 975.5 | - | - | 0 | - |
| 3 | z | 3.088E+04 | 978.5 | 6.006E-05 | 0.06138 | +1 | 8 |
| - | - | 1.752E+04 | 979.5 | - | - | 0 | - |
| - | - | 5601 | 980.5 | - | - | 0 | - |
| - | - | 891.5 | 981.5 | - | - | 0 | - |
| - | - | 3121 | 990.6 | - | - | 0 | - |
| - | - | 3091 | 991.6 | - | - | 0 | - |
| 8 | c | 3.199E+04 | 992.5 | 3.178E-05 | 0.03202 | +1 | 8 |
| - | - | 1.84E+04 | 993.5 | - | - | 0 | - |
| 3 | y | 1.149E+04 | 994.5 | 0.009595 | 9.648 | +1 | 8 |
| - | - | 5264 | 995.5 | - | - | 0 | - |
| - | - | 1792 | 996.5 | - | - | 0 | - |
| - | - | 1541 | 1029 | - | - | 0 | - |
| - | - | 875.4 | 1054 | - | - | 0 | - |
| - | - | 1948 | 1063 | - | - | 0 | - |
| - | - | 1163 | 1064 | - | - | 0 | - |
| 2 | z | 8508 | 1078 | 5.517E-06 | 0.00512 | +1 | 9 |
| - | - | 5485 | 1079 | - | - | 0 | - |
| - | - | 1752 | 1080 | - | - | 0 | - |
| - | - | 744.9 | 1090 | - | - | 0 | - |
| 9 | c | 4.447E+04 | 1107 | 0.0007925 | 0.7162 | +1 | 9 |
| - | - | 3.088E+04 | 1108 | - | - | 0 | - |
| - | - | 1.136E+04 | 1109 | - | - | 0 | - |
| - | - | 1929 | 1110 | - | - | 0 | - |
| - | - | 1496 | 1116 | - | - | 0 | - |
| - | - | 2178 | 1119 | - | - | 0 | - |
| - | - | 769.1 | 1121 | - | - | 0 | - |
| - | - | 4404 | 1136 | - | - | 0 | - |
| - | - | 3988 | 1137 | - | - | 0 | - |
| - | - | 1568 | 1138 | - | - | 0 | - |
| - | - | 729.7 | 1146 | - | - | 0 | - |
| - | - | 1027 | 1147 | - | - | 0 | - |
| - | - | 8833 | 1148 | - | - | 0 | - |
| - | - | 7094 | 1149 | - | - | 0 | - |
| - | - | 3431 | 1150 | - | - | 0 | - |
| - | - | 5847 | 1152 | - | - | 0 | - |
| - | - | 4561 | 1153 | - | - | 0 | - |
| - | - | 1717 | 1154 | - | - | 0 | - |
| - | - | 8910 | 1162 | - | - | 0 | - |
| - | - | 9124 | 1163 | - | - | 0 | - |
| - | - | 1.446E+04 | 1164 | - | - | 0 | - |
| - | - | 8313 | 1165 | - | - | 0 | - |
| - | - | 2370 | 1166 | - | - | 0 | - |
| - | - | 6965 | 1180 | - | - | 0 | - |
| - | - | 5123 | 1181 | - | - | 0 | - |
| - | - | 1111 | 1182 | - | - | 0 | - |
| - | - | 2319 | 1190 | - | - | 0 | - |
| - | - | 4.912E+04 | 1191 | - | - | 0 | - |
| - | - | 3.376E+04 | 1192 | - | - | 0 | - |
| - | - | 1.223E+04 | 1193 | - | - | 0 | - |
| - | - | 2202 | 1194 | - | - | 0 | - |
| - | - | 1750 | 1205 | - | - | 0 | - |
| - | - | 2320 | 1206 | - | - | 0 | - |
| - | - | 6.27E+04 | 1207 | - | - | 0 | - |
| - | - | 2.535E+05 | 1208 | - | - | 0 | - |
| - | - | 1.688E+05 | 1209 | - | - | 0 | - |
| - | - | 6.315E+04 | 1210 | - | - | 0 | - |
| - | - | 1.007E+04 | 1211 | - | - | 0 | - |
| - | - | 739.3 | 1766 | - | - | 0 | - |
| - | - | 701 | 1812 | - | - | 0 | - |
| - | - | 883.3 | 3034 | - | - | 0 | - |
| - | - | 763 | 3269 | - | - | 0 | - |

m/z Charge Intensity FragmentType MassShift Position
120.08065032958984 0 573.2552
129.10247802734375 0 1945.8154
133.78009033203125 0 471.89395
148.89056396484375 0 496.42886
148.89768981933594 0 493.1404
148.9053192138672 0 685.0977
148.9127197265625 0 710.473
148.91964721679688 0 1185.3622
148.92681884765625 0 1266.0385
148.93411254882812 0 1636.7231
148.94186401367188 0 3330.591
148.95852661132812 0 4412.8057
148.96630859375 0 2496.9487
148.9736785888672 0 1175.9534
148.98110961914062 0 1184.095
148.988037109375 0 854.79895
148.9954376220703 0 693.00104
149.00291442871094 0 606.2254
149.0098114013672 0 658.8436
149.0246124267578 0 423.97617
149.031494140625 0 560.5817
149.04632568359375 0 422.6609
149.11837768554688 0 445.51254
151.19471740722656 0 452.41568
156.4000701904297 0 474.52838
168.89187622070312 0 456.27295
169.13429260253906 0 596.5127
173.09207153320312 0 465.63803
185.1651611328125 0 6214.6973
187.10791015625 0 555.15283
213.1600799560547 0 3447.8157
215.13922119140625 0 80602.88 y 8
216.14263916015625 0 9221.846
227.10289001464844 0 2273.323
301.211181640625 0 710.8065
311.1374816894531 0 507.44846
324.1554870605469 0 1696.7034
326.1717224121094 0 978.8623 y Water loss 7
357.2735290527344 0 17500.666
358.278076171875 0 4192.6763 c 2
374.1709289550781 0 1067.8689
407.1917724609375 0 894.12335
418.7416687011719 0 647.65045
423.2237243652344 0 1573.7119 y Water loss 6
426.1668701171875 0 1174.9197
432.2373046875 0 1142.5393 c Ammonia loss 6
441.2349548339844 0 8783.581 y 6
442.23895263671875 0 2005.4421
456.28216552734375 0 1734.4414 c Ammonia loss 3
470.222900390625 0 654.9974
472.301513671875 0 3920.5908
473.3082275390625 0 6194.225 c 3
474.3124694824219 0 1645.3112
483.2604675292969 0 584.27844
497.26031494140625 0 1069.5612
511.25244140625 0 973.0377
543.313232421875 0 817.07385
545.282958984375 0 1301.3702 c Ammonia loss 8
554.2607421875 0 3867.0378
555.262939453125 0 1129.286
587.2937622070312 0 792.68896
588.3031616210938 0 8254.723 y 5
589.30517578125 0 2345.1497
590.3034057617188 0 647.72205
591.34814453125 0 1161.8365
603.8234252929688 0 897.32574
604.3260498046875 0 1155.1879
619.345947265625 0 3060.9785 c Ammonia loss 4
620.3494262695312 0 1363.4869
631.7743530273438 0 578.82086
635.364013671875 0 10656.397
636.3707275390625 0 16623.19 c 4
637.3735961914062 0 5069.567
638.3795166015625 0 825.3869
681.3262939453125 0 969.1046
682.378173828125 0 987.27594
683.3767700195312 0 687.50055
706.3768920898438 0 2358.7856
707.3779296875 0 1080.6315
717.38134765625 0 897.33057
718.3772583007812 0 589.52826
735.3480224609375 0 1005.19476 z 4
736.3538208007812 0 1198.5125
738.4205932617188 0 1467.958
739.4263916015625 0 11415.074
740.431884765625 0 10886.215
741.4341430664062 0 4290.669
750.3547973632812 0 675.18567
751.365478515625 0 1571.6716 y 4
752.36083984375 0 1060.5305
762.336669921875 0 784.1078
766.4131469726562 0 15525.679 c Ammonia loss 5
767.4163208007812 0 7044.6187
768.4212646484375 0 1402.665
780.356201171875 0 8294.408
781.3599853515625 0 4371.573
782.36181640625 0 1409.6072
787.470458984375 0 1100.9247
807.39208984375 0 614.1885
836.4785766601562 0 983.759
837.48291015625 0 948.4715
851.3832397460938 0 1843.3585
852.3807983398438 0 1351.9998
863.4636840820312 0 1306.1412 c Ammonia loss 6
866.3977661132812 0 963.9501 y 3
872.420166015625 0 1393.4482
879.398193359375 0 945.11115
879.4857788085938 0 11364.961
880.4920043945312 0 36737.234 c 6
881.4951171875 0 19148.506
882.4981079101562 0 5049.231
906.4486083984375 0 4859.857
907.4462280273438 0 3106.2078
919.3898315429688 0 1147.0896
933.477294921875 0 977.63776
934.4789428710938 0 14545.575
935.481201171875 0 8979.815
936.485107421875 0 1930.7621
964.5133666992188 0 2941.0557
965.520751953125 0 3564.3135
966.5234375 0 1550.6971
974.494873046875 0 1811.2927
975.4948120117188 0 845.0949
978.4693603515625 0 30876.979 z 2
979.4724731445312 0 17518.932
980.474853515625 0 5600.7847
981.4736938476562 0 891.4755
990.552734375 0 3121.058
991.5545043945312 0 3091.4988
992.5087280273438 0 31991.027 c Ammonia loss 7
993.5115356445312 0 18400.627
994.4976196289062 0 11488.53 y 2
995.4959106445312 0 5264.1606
996.493896484375 0 1791.8885
1028.6163330078125 0 1540.9119
1053.56103515625 0 875.4028
1062.5692138671875 0 1948.4198
1063.5750732421875 0 1162.7908
1077.5377197265625 0 8508.203 z 1
1078.5386962890625 0 5484.7188
1079.5430908203125 0 1752.2152
1089.62353515625 0 744.90717
1106.5872802734375 0 44472.8 c 8
1107.59130859375 0 30879.506
1108.59375 0 11357.083
1109.5928955078125 0 1928.9453
1115.6434326171875 0 1495.8226
1118.6053466796875 0 2178.1826
1120.581298828125 0 769.09985
1135.625 0 4403.6934
1136.6234130859375 0 3987.9324
1137.62109375 0 1567.8601
1145.64453125 0 729.71576
1146.6356201171875 0 1026.5391
1147.599853515625 0 8832.685
1148.5921630859375 0 7093.9326
1149.5814208984375 0 3430.9836
1151.584716796875 0 5846.56
1152.5858154296875 0 4561.1714
1153.5904541015625 0 1716.7432
1161.6417236328125 0 8910.46
1162.6461181640625 0 9124.165
1163.6556396484375 0 14462.011
1164.660400390625 0 8313.076
1165.664794921875 0 2370.1157
1179.6527099609375 0 6965.361
1180.6534423828125 0 5122.761
1181.6495361328125 0 1111.4656
1189.6275634765625 0 2318.8115
1190.62109375 0 49120.09
1191.624267578125 0 33755.043
1192.6297607421875 0 12229.2
1193.6318359375 0 2201.835
1204.658935546875 0 1750.3875
1205.631591796875 0 2319.915
1206.6392822265625 0 62698.97
1207.6470947265625 0 253503.42
1208.650146484375 0 168838.27
1209.6529541015625 0 63151.598
1210.6571044921875 0 10068.999
1765.8890380859375 0 739.33325
1811.8299560546875 0 700.9808
3033.823486328125 0 883.3351
3268.9814453125 0 763.0087

Spectrum Details

|  |  |
| --- | --- |
| Matched peaks? Matched peaks The total absolute number of peaks matched. Additionally in brackets the total fraction of peaks matched and the total number of peaks is shown. | 23 (12.50% of 184) |
| FDR? FDR The false discovery rate estimated for this peptide. It is calculated by matching all theoretical fragments with a non-integer shift with the raw peaks for this spectrum. This is done with 40 different shifts. The resulting percentage is the average number of annotated peaks over the number of annotated peaks with the correct spectrum. | 2.28% |
| Satellite FDR? Satellite FDR See the FDR for details on its calculation. This satellite ion specific FDR only contains the satellite ions (d/w) for I/L/J positions. | - |
| PSM Score? PSM Score The PSM Score as given by Hecklib to this annotated spectrum. It is shown with three significant figures. | 268 |

#### Spectrum 8799? Spectrum 8799 The raw spectrum of this peptide as annotated by Hecklib. The fragments are coloured according to ion type (see legend). Any peaks with a star '\*' as text can be hovered over to see the full details, first the ion type second the mass shift type. By hovering over the amino acids in the peptide or ions in the legend the corresponding peaks are highlighted. By toggling the 'Unassigned' label you can turn the background (unassigned) peaks on or off in the plot. By updating the slider in the Ion legend you can update the spectrum to only show the top X% of the peaks with labels. The top X% means any peak that is within X% of the highest intensity. By dragging in the spectrum you can zoom in to a specific part of the spectrum and use 'Zoom Out' to get back to the original zoom level. The annotation of the spectrum is based on the given sequence in the peptides file and is done with different software so inconsistencies are likely. The peaks are annotated based on the given sequence, with 20 ppm tolerance.

Copy Data

##### Spectrum 8799 (TSV)

###### Preview

```
Loading example...
```

*Click on the button to copy the data to your clipboard.*

Mz MinMz MaxIntensity Max

WidthHeightPeptide font sizePeptide stroke widthSpectrum font sizeSpectrum stroke widthCompact peptide

Ion legend

wxyz

abcd

OtherUnassignedIon^Charge^_Position_Show for top:%

JVKDYFPEPV

03.15e+46.29e+49.44e+41.26e+5

Zoom Out

y^+1^_2_y^+1^_3_c^+1^_3_y^+1^_4_y^+1^_4_c^+1^_4_c^+1^_4_c^+2^_8_y^+1^_5_c^+1^_5_c^+1^_5_z^+1^_6_y^+1^_6_c^+1^_6_c^+1^_7_z^+1^_8_c^+1^_8_y^+1^_8_z^+1^_9_c^+1^_9_

0578115517332311

Fragment Matches Table

Show background peaks

| Position | Ion type | Intensity | mz Theoretical | mz Error (Th) | mz Error (ppm) | Charge | Series Number |
| --- | --- | --- | --- | --- | --- | --- | --- |
| - | - | 391.5 | 124.1 | - | - | 0 | - |
| - | - | 900.9 | 129.1 | - | - | 0 | - |
| - | - | 435 | 162.3 | - | - | 0 | - |
| - | - | 587 | 173.1 | - | - | 0 | - |
| - | - | 1502 | 173.5 | - | - | 0 | - |
| - | - | 3187 | 185.2 | - | - | 0 | - |
| - | - | 1625 | 213.2 | - | - | 0 | - |
| 9 | y | 4.002E+04 | 215.1 | 0.0002023 | 0.9402 | +1 | 2 |
| - | - | 4611 | 216.1 | - | - | 0 | - |
| - | - | 1050 | 227.1 | - | - | 0 | - |
| - | - | 517.1 | 228.1 | - | - | 0 | - |
| - | - | 546.1 | 259.6 | - | - | 0 | - |
| - | - | 635.7 | 275.3 | - | - | 0 | - |
| - | - | 515 | 298.5 | - | - | 0 | - |
| - | - | 620.1 | 324.2 | - | - | 0 | - |
| 8 | y | 602.2 | 326.2 | 0.000332 | 1.018 | +1 | 3 |
| - | - | 8705 | 357.3 | - | - | 0 | - |
| 3 | c | 2980 | 358.3 | 0.002641 | 7.371 | +1 | 3 |
| - | - | 553 | 369.3 | - | - | 0 | - |
| 7 | y | 1347 | 423.2 | 0.001134 | 2.679 | +1 | 4 |
| - | - | 745.7 | 426.2 | - | - | 0 | - |
| 7 | y | 3855 | 441.2 | 0.0007316 | 1.658 | +1 | 4 |
| - | - | 1033 | 442.2 | - | - | 0 | - |
| 4 | c | 806.6 | 456.3 | 0.0009934 | 2.177 | +1 | 4 |
| - | - | 1632 | 472.3 | - | - | 0 | - |
| 4 | c | 3691 | 473.3 | 0.0006284 | 1.328 | +1 | 4 |
| - | - | 800.3 | 474.3 | - | - | 0 | - |
| 8 | c | 829.9 | 496.8 | 0.001503 | 3.026 | +2 | 8 |
| - | - | 2278 | 554.3 | - | - | 0 | - |
| - | - | 617.6 | 554.6 | - | - | 0 | - |
| - | - | 693.5 | 555.3 | - | - | 0 | - |
| - | - | 563.1 | 587.3 | - | - | 0 | - |
| 6 | y | 4880 | 588.3 | 0.000677 | 1.151 | +1 | 5 |
| - | - | 964.9 | 589.3 | - | - | 0 | - |
| - | - | 661.1 | 604.3 | - | - | 0 | - |
| 5 | c | 1959 | 619.3 | 0.0008117 | 1.311 | +1 | 5 |
| - | - | 5704 | 635.4 | - | - | 0 | - |
| 5 | c | 6812 | 636.4 | 0.001299 | 2.041 | +1 | 5 |
| - | - | 1808 | 637.4 | - | - | 0 | - |
| 5 | z | 988.1 | 735.3 | 0.005084 | 6.913 | +1 | 6 |
| - | - | 841.9 | 738.4 | - | - | 0 | - |
| - | - | 5136 | 739.4 | - | - | 0 | - |
| - | - | 5736 | 740.4 | - | - | 0 | - |
| - | - | 2359 | 741.4 | - | - | 0 | - |
| 5 | y | 1055 | 751.4 | 0.002532 | 3.37 | +1 | 6 |
| 6 | c | 7831 | 766.4 | 0.0001948 | 0.2542 | +1 | 6 |
| - | - | 2800 | 767.4 | - | - | 0 | - |
| - | - | 801.5 | 770.9 | - | - | 0 | - |
| - | - | 4396 | 780.4 | - | - | 0 | - |
| - | - | 2050 | 781.4 | - | - | 0 | - |
| - | - | 1556 | 851.4 | - | - | 0 | - |
| - | - | 739.1 | 852.4 | - | - | 0 | - |
| - | - | 721.6 | 872.4 | - | - | 0 | - |
| - | - | 5443 | 879.5 | - | - | 0 | - |
| 7 | c | 2.113E+04 | 880.5 | 0.0007725 | 0.8773 | +1 | 7 |
| - | - | 9346 | 881.5 | - | - | 0 | - |
| - | - | 2459 | 882.5 | - | - | 0 | - |
| - | - | 2196 | 906.5 | - | - | 0 | - |
| - | - | 1020 | 907.4 | - | - | 0 | - |
| - | - | 7088 | 934.5 | - | - | 0 | - |
| - | - | 3555 | 935.5 | - | - | 0 | - |
| - | - | 986.1 | 936.5 | - | - | 0 | - |
| - | - | 1602 | 964.5 | - | - | 0 | - |
| - | - | 1416 | 965.5 | - | - | 0 | - |
| - | - | 814.1 | 966.5 | - | - | 0 | - |
| - | - | 861.5 | 974.5 | - | - | 0 | - |
| 3 | z | 1.471E+04 | 978.5 | 0.0002451 | 0.2505 | +1 | 8 |
| - | - | 9075 | 979.5 | - | - | 0 | - |
| - | - | 2293 | 980.5 | - | - | 0 | - |
| - | - | 1640 | 990.6 | - | - | 0 | - |
| - | - | 929.2 | 991.6 | - | - | 0 | - |
| 8 | c | 1.633E+04 | 992.5 | 9.281E-05 | 0.09351 | +1 | 8 |
| - | - | 8605 | 993.5 | - | - | 0 | - |
| 3 | y | 5144 | 994.5 | 0.01063 | 10.69 | +1 | 8 |
| - | - | 2040 | 995.5 | - | - | 0 | - |
| 2 | z | 4563 | 1078 | 0.0007269 | 0.6746 | +1 | 9 |
| - | - | 3485 | 1079 | - | - | 0 | - |
| - | - | 1415 | 1080 | - | - | 0 | - |
| - | - | 839.6 | 1090 | - | - | 0 | - |
| 9 | c | 2.314E+04 | 1107 | 0.001159 | 1.047 | +1 | 9 |
| - | - | 1.434E+04 | 1108 | - | - | 0 | - |
| - | - | 4196 | 1109 | - | - | 0 | - |
| - | - | 969.9 | 1110 | - | - | 0 | - |
| - | - | 1158 | 1119 | - | - | 0 | - |
| - | - | 2330 | 1136 | - | - | 0 | - |
| - | - | 1685 | 1137 | - | - | 0 | - |
| - | - | 4115 | 1148 | - | - | 0 | - |
| - | - | 3397 | 1149 | - | - | 0 | - |
| - | - | 1873 | 1150 | - | - | 0 | - |
| - | - | 3238 | 1152 | - | - | 0 | - |
| - | - | 1974 | 1153 | - | - | 0 | - |
| - | - | 5125 | 1162 | - | - | 0 | - |
| - | - | 4554 | 1163 | - | - | 0 | - |
| - | - | 7399 | 1164 | - | - | 0 | - |
| - | - | 4698 | 1165 | - | - | 0 | - |
| - | - | 1530 | 1166 | - | - | 0 | - |
| - | - | 3140 | 1180 | - | - | 0 | - |
| - | - | 2485 | 1181 | - | - | 0 | - |
| - | - | 1043 | 1182 | - | - | 0 | - |
| - | - | 1828 | 1190 | - | - | 0 | - |
| - | - | 2.559E+04 | 1191 | - | - | 0 | - |
| - | - | 1.832E+04 | 1192 | - | - | 0 | - |
| - | - | 7151 | 1193 | - | - | 0 | - |
| - | - | 1923 | 1194 | - | - | 0 | - |
| - | - | 1574 | 1206 | - | - | 0 | - |
| - | - | 3.205E+04 | 1207 | - | - | 0 | - |
| - | - | 1.246E+05 | 1208 | - | - | 0 | - |
| - | - | 8.372E+04 | 1209 | - | - | 0 | - |
| - | - | 3.143E+04 | 1210 | - | - | 0 | - |
| - | - | 5241 | 1211 | - | - | 0 | - |
| - | - | 849 | 1813 | - | - | 0 | - |
| - | - | 770.5 | 2042 | - | - | 0 | - |
| - | - | 845.5 | 2288 | - | - | 0 | - |

m/z Charge Intensity FragmentType MassShift Position
124.06621551513672 0 391.48618
129.10231018066406 0 900.90247
162.27328491210938 0 434.9809
173.09228515625 0 587.028
173.45164489746094 0 1501.7083
185.1650848388672 0 3187.2974
213.15965270996094 0 1624.7411
215.13922119140625 0 40017.95 y 8
216.14242553710938 0 4610.7666
227.1026611328125 0 1049.8281
228.106201171875 0 517.0995
259.57244873046875 0 546.13245
275.3149719238281 0 635.6981
298.5101013183594 0 514.9709
324.15728759765625 0 620.141
326.17071533203125 0 602.17975 y Water loss 7
357.2736511230469 0 8704.989
358.27862548828125 0 2979.6787 c 2
369.3236999511719 0 552.9749
423.2249450683594 0 1346.5281 y Water loss 6
426.16851806640625 0 745.736
441.235107421875 0 3854.9712 y 6
442.2382507324219 0 1032.6007
456.28265380859375 0 806.6386 c Ammonia loss 3
472.2998046875 0 1632.44
473.308837890625 0 3691.424 c 3
474.3110046386719 0 800.28235
496.759521484375 0 829.91315 c Ammonia loss 7
554.26123046875 0 2277.5928
554.5965576171875 0 617.59607
555.2628173828125 0 693.53925
587.2874145507812 0 563.1136
588.303466796875 0 4880.3804 y 5
589.3084716796875 0 964.8712
604.306640625 0 661.0798
619.3441772460938 0 1959.1472 c Ammonia loss 4
635.3644409179688 0 5704.2485
636.3702392578125 0 6812.048 c 4
637.3748168945312 0 1808.0203
735.3524780273438 0 988.1466 z 4
738.4157104492188 0 841.9068
739.4260864257812 0 5136.0845
740.4315185546875 0 5736.1914
741.4375610351562 0 2358.6223
751.3635864257812 0 1054.5883 y 4
766.4132080078125 0 7831.4937 c Ammonia loss 5
767.4166259765625 0 2800.0986
770.9293212890625 0 801.5095
780.3564453125 0 4396.0337
781.3597412109375 0 2050.1155
851.3812255859375 0 1555.8793
852.3873291015625 0 739.0591
872.4234008789062 0 721.56854
879.4843139648438 0 5442.6646
880.491943359375 0 21128.078 c 6
881.4963989257812 0 9345.526
882.4996948242188 0 2459.3916
906.4505615234375 0 2196.2805
907.4384155273438 0 1020.2681
934.47900390625 0 7087.5493
935.4808959960938 0 3555.3462
936.479736328125 0 986.1298
964.5109252929688 0 1602.4268
965.5195922851562 0 1416.454
966.5137939453125 0 814.1095
974.4994506835938 0 861.4563
978.4690551757812 0 14712.715 z 2
979.4727783203125 0 9075.081
980.4742431640625 0 2293.1335
990.555419921875 0 1639.9855
991.56103515625 0 929.1687
992.5086669921875 0 16326.011 c Ammonia loss 7
993.5117797851562 0 8605.475
994.4986572265625 0 5143.911 y 2
995.4964599609375 0 2040.1276
1077.5369873046875 0 4562.788 z 1
1078.5423583984375 0 3484.799
1079.5474853515625 0 1414.8531
1089.616943359375 0 839.58936
1106.5869140625 0 23142.205 c 8
1107.5911865234375 0 14343.637
1108.5936279296875 0 4196.286
1109.5853271484375 0 969.9457
1118.6063232421875 0 1157.9146
1135.6258544921875 0 2329.5203
1136.633056640625 0 1684.6349
1147.6114501953125 0 4114.75
1148.6214599609375 0 3396.8699
1149.592041015625 0 1872.5709
1151.5848388671875 0 3238.0188
1152.590087890625 0 1973.8822
1161.64013671875 0 5124.701
1162.6446533203125 0 4553.7866
1163.6561279296875 0 7399.079
1164.6566162109375 0 4697.9697
1165.6644287109375 0 1529.6018
1179.650634765625 0 3139.704
1180.65869140625 0 2485.3452
1181.661376953125 0 1043.4309
1189.63427734375 0 1828.4128
1190.6209716796875 0 25592.074
1191.623779296875 0 18315.172
1192.628173828125 0 7151.094
1193.6343994140625 0 1922.5984
1205.625732421875 0 1574.4279
1206.638916015625 0 32047.715
1207.6468505859375 0 124554.64
1208.6497802734375 0 83724.76
1209.653076171875 0 31432.941
1210.6566162109375 0 5241.212
1812.8687744140625 0 849.0236
2042.4332275390625 0 770.4525
2288.02734375 0 845.46954

Spectrum Details

|  |  |
| --- | --- |
| Matched peaks? Matched peaks The total absolute number of peaks matched. Additionally in brackets the total fraction of peaks matched and the total number of peaks is shown. | 20 (17.70% of 113) |
| FDR? FDR The false discovery rate estimated for this peptide. It is calculated by matching all theoretical fragments with a non-integer shift with the raw peaks for this spectrum. This is done with 40 different shifts. The resulting percentage is the average number of annotated peaks over the number of annotated peaks with the correct spectrum. | 1.79% |
| Satellite FDR? Satellite FDR See the FDR for details on its calculation. This satellite ion specific FDR only contains the satellite ions (d/w) for I/L/J positions. | - |
| PSM Score? PSM Score The PSM Score as given by Hecklib to this annotated spectrum. It is shown with three significant figures. | 234 |

#### Spectrum 8526? Spectrum 8526 The raw spectrum of this peptide as annotated by Hecklib. The fragments are coloured according to ion type (see legend). Any peaks with a star '\*' as text can be hovered over to see the full details, first the ion type second the mass shift type. By hovering over the amino acids in the peptide or ions in the legend the corresponding peaks are highlighted. By toggling the 'Unassigned' label you can turn the background (unassigned) peaks on or off in the plot. By updating the slider in the Ion legend you can update the spectrum to only show the top X% of the peaks with labels. The top X% means any peak that is within X% of the highest intensity. By dragging in the spectrum you can zoom in to a specific part of the spectrum and use 'Zoom Out' to get back to the original zoom level. The annotation of the spectrum is based on the given sequence in the peptides file and is done with different software so inconsistencies are likely. The peaks are annotated based on the given sequence, with 20 ppm tolerance.

Copy Data

##### Spectrum 8526 (TSV)

###### Preview

```
Loading example...
```

*Click on the button to copy the data to your clipboard.*

Mz MinMz MaxIntensity Max

WidthHeightPeptide font sizePeptide stroke widthSpectrum font sizeSpectrum stroke widthCompact peptide

Ion legend

wxyz

abcd

OtherUnassignedIon^Charge^_Position_Show for top:%

JVKDYFPEPV

08.26e+41.65e+52.48e+53.30e+5

Zoom Out

y^+1^_2_y^+1^_3_y^+1^_3_c^+1^_3_z^+2^_6_c^+2^_6_y^+1^_4_c^+2^_7_y^+1^_4_c^+1^_4_c^+1^_4_c^+2^_8_c^+2^_9_y^+1^_5_c^+1^_5_c^+1^_5_z^+1^_6_y^+1^_6_c^+1^_6_c^+1^_7_y^+1^_7_c^+1^_7_y^+1^_8_y^+1^_8_z^+1^_8_c^+1^_8_y^+1^_8_z^+1^_9_y^+1^_9_c^+1^_9_

0766153322993065

Fragment Matches Table

Show background peaks

| Position | Ion type | Intensity | mz Theoretical | mz Error (Th) | mz Error (ppm) | Charge | Series Number |
| --- | --- | --- | --- | --- | --- | --- | --- |
| - | - | 1424 | 120.1 | - | - | 0 | - |
| - | - | 366.7 | 122.5 | - | - | 0 | - |
| - | - | 1983 | 129.1 | - | - | 0 | - |
| - | - | 962.3 | 133.1 | - | - | 0 | - |
| - | - | 563.9 | 136.1 | - | - | 0 | - |
| - | - | 385.8 | 143.7 | - | - | 0 | - |
| - | - | 424.6 | 146.5 | - | - | 0 | - |
| - | - | 405.1 | 158 | - | - | 0 | - |
| - | - | 618.7 | 173.4 | - | - | 0 | - |
| - | - | 646.6 | 177.1 | - | - | 0 | - |
| - | - | 6580 | 185.2 | - | - | 0 | - |
| - | - | 791.8 | 186.2 | - | - | 0 | - |
| - | - | 659.1 | 199.1 | - | - | 0 | - |
| - | - | 580.5 | 207.1 | - | - | 0 | - |
| - | - | 611.4 | 209.1 | - | - | 0 | - |
| - | - | 4988 | 213.2 | - | - | 0 | - |
| 9 | y | 1.019E+05 | 215.1 | 0.0003244 | 1.508 | +1 | 2 |
| - | - | 565.9 | 215.9 | - | - | 0 | - |
| - | - | 9850 | 216.1 | - | - | 0 | - |
| - | - | 653.8 | 217.1 | - | - | 0 | - |
| - | - | 1038 | 219.1 | - | - | 0 | - |
| - | - | 3267 | 227.1 | - | - | 0 | - |
| - | - | 5815 | 229.2 | - | - | 0 | - |
| - | - | 1404 | 243.1 | - | - | 0 | - |
| - | - | 603.4 | 287.1 | - | - | 0 | - |
| - | - | 951.2 | 301.2 | - | - | 0 | - |
| - | - | 811.4 | 304.2 | - | - | 0 | - |
| - | - | 2205 | 324.2 | - | - | 0 | - |
| - | - | 521.2 | 325.2 | - | - | 0 | - |
| 8 | y | 946.8 | 326.2 | 0.000492 | 1.508 | +1 | 3 |
| - | - | 1384 | 343.2 | - | - | 0 | - |
| 8 | y | 1068 | 344.2 | 0.001345 | 3.907 | +1 | 3 |
| - | - | 2.551E+04 | 357.3 | - | - | 0 | - |
| 3 | c | 5240 | 358.3 | 0.003312 | 9.245 | +1 | 3 |
| 5 | z | 605.1 | 359.2 | 0.002202 | 6.132 | +2 | 6 |
| - | - | 849.4 | 360.2 | - | - | 0 | - |
| - | - | 1295 | 361.2 | - | - | 0 | - |
| - | - | 953.3 | 374.2 | - | - | 0 | - |
| 6 | c | 709.1 | 392.2 | 0.003857 | 9.834 | +2 | 6 |
| - | - | 877.5 | 407.2 | - | - | 0 | - |
| - | - | 1544 | 418.2 | - | - | 0 | - |
| 7 | y | 2001 | 423.2 | 0.001378 | 3.256 | +1 | 4 |
| - | - | 1406 | 426.2 | - | - | 0 | - |
| 7 | c | 905.9 | 432.2 | 0.000546 | 1.263 | +2 | 7 |
| - | - | 978.4 | 432.7 | - | - | 0 | - |
| 7 | y | 1.02E+04 | 441.2 | 0.0008231 | 1.865 | +1 | 4 |
| - | - | 3106 | 442.2 | - | - | 0 | - |
| 4 | c | 2136 | 456.3 | 0.0009019 | 1.977 | +1 | 4 |
| - | - | 625.8 | 467.3 | - | - | 0 | - |
| - | - | 712.5 | 468.2 | - | - | 0 | - |
| - | - | 595.9 | 470.2 | - | - | 0 | - |
| - | - | 3446 | 472.3 | - | - | 0 | - |
| 4 | c | 8162 | 473.3 | 0.0003232 | 0.6829 | +1 | 4 |
| - | - | 2007 | 474.3 | - | - | 0 | - |
| - | - | 930.7 | 483.3 | - | - | 0 | - |
| - | - | 593.8 | 485.3 | - | - | 0 | - |
| - | - | 670.2 | 486.2 | - | - | 0 | - |
| - | - | 2325 | 491.3 | - | - | 0 | - |
| - | - | 1114 | 492.3 | - | - | 0 | - |
| - | - | 662.5 | 495.7 | - | - | 0 | - |
| 8 | c | 1296 | 496.8 | 0.0005878 | 1.183 | +2 | 8 |
| - | - | 697.5 | 497.3 | - | - | 0 | - |
| - | - | 1215 | 511.3 | - | - | 0 | - |
| - | - | 659.2 | 536.2 | - | - | 0 | - |
| - | - | 1064 | 543.3 | - | - | 0 | - |
| - | - | 716 | 544.3 | - | - | 0 | - |
| 9 | c | 1072 | 545.3 | 0.001868 | 3.426 | +2 | 9 |
| - | - | 1404 | 545.8 | - | - | 0 | - |
| - | - | 708.2 | 552.2 | - | - | 0 | - |
| - | - | 5306 | 554.3 | - | - | 0 | - |
| - | - | 1417 | 555.3 | - | - | 0 | - |
| - | - | 584.5 | 563.9 | - | - | 0 | - |
| - | - | 859.6 | 567.3 | - | - | 0 | - |
| - | - | 1189 | 585.3 | - | - | 0 | - |
| 6 | y | 1.138E+04 | 588.3 | 0.0003718 | 0.6321 | +1 | 5 |
| - | - | 4611 | 589.3 | - | - | 0 | - |
| - | - | 846.3 | 590.3 | - | - | 0 | - |
| - | - | 1506 | 592.4 | - | - | 0 | - |
| - | - | 2322 | 603.3 | - | - | 0 | - |
| - | - | 963.6 | 603.8 | - | - | 0 | - |
| - | - | 1869 | 604.3 | - | - | 0 | - |
| - | - | 578.3 | 613 | - | - | 0 | - |
| 5 | c | 4892 | 619.3 | 0.001141 | 1.843 | +1 | 5 |
| - | - | 1727 | 620.3 | - | - | 0 | - |
| - | - | 1.437E+04 | 635.4 | - | - | 0 | - |
| 5 | c | 1.811E+04 | 636.4 | 0.0009326 | 1.465 | +1 | 5 |
| - | - | 5467 | 637.4 | - | - | 0 | - |
| - | - | 1038 | 638.4 | - | - | 0 | - |
| - | - | 565.2 | 645.3 | - | - | 0 | - |
| - | - | 1475 | 645.3 | - | - | 0 | - |
| - | - | 835.1 | 646.4 | - | - | 0 | - |
| - | - | 611.4 | 651.3 | - | - | 0 | - |
| - | - | 829 | 652.3 | - | - | 0 | - |
| - | - | 606.9 | 654.3 | - | - | 0 | - |
| - | - | 851.1 | 681.3 | - | - | 0 | - |
| - | - | 1804 | 682.4 | - | - | 0 | - |
| - | - | 981.5 | 683.4 | - | - | 0 | - |
| - | - | 603.1 | 723.8 | - | - | 0 | - |
| - | - | 1271 | 732.4 | - | - | 0 | - |
| - | - | 1133 | 733.4 | - | - | 0 | - |
| 5 | z | 2088 | 735.3 | 0.0009944 | 1.352 | +1 | 6 |
| - | - | 1635 | 736.4 | - | - | 0 | - |
| - | - | 1806 | 738.4 | - | - | 0 | - |
| - | - | 1.446E+04 | 739.4 | - | - | 0 | - |
| - | - | 1.541E+04 | 740.4 | - | - | 0 | - |
| - | - | 4826 | 741.4 | - | - | 0 | - |
| - | - | 1215 | 742.4 | - | - | 0 | - |
| - | - | 908.4 | 747.4 | - | - | 0 | - |
| - | - | 868.5 | 750.4 | - | - | 0 | - |
| 5 | y | 2268 | 751.4 | 0.0005199 | 0.6919 | +1 | 6 |
| - | - | 1520 | 752.4 | - | - | 0 | - |
| 6 | c | 2.082E+04 | 766.4 | 0.0001714 | 0.2236 | +1 | 6 |
| - | - | 1.024E+04 | 767.4 | - | - | 0 | - |
| - | - | 1998 | 768.4 | - | - | 0 | - |
| - | - | 9264 | 780.4 | - | - | 0 | - |
| - | - | 4741 | 781.4 | - | - | 0 | - |
| - | - | 1078 | 782.4 | - | - | 0 | - |
| - | - | 762.5 | 815.4 | - | - | 0 | - |
| - | - | 1142 | 832.5 | - | - | 0 | - |
| - | - | 1596 | 836.5 | - | - | 0 | - |
| - | - | 609.6 | 837.5 | - | - | 0 | - |
| - | - | 1615 | 842.5 | - | - | 0 | - |
| - | - | 707.2 | 843.5 | - | - | 0 | - |
| - | - | 1422 | 844.5 | - | - | 0 | - |
| - | - | 950.4 | 845.5 | - | - | 0 | - |
| - | - | 752.7 | 846.4 | - | - | 0 | - |
| - | - | 4142 | 851.4 | - | - | 0 | - |
| - | - | 1312 | 852.4 | - | - | 0 | - |
| 7 | c | 1797 | 863.5 | 0.0001023 | 0.1184 | +1 | 7 |
| - | - | 1470 | 864.5 | - | - | 0 | - |
| 4 | y | 1547 | 866.4 | 0.0002391 | 0.2759 | +1 | 7 |
| - | - | 1109 | 872.4 | - | - | 0 | - |
| - | - | 705.6 | 873.4 | - | - | 0 | - |
| - | - | 1.348E+04 | 879.5 | - | - | 0 | - |
| 7 | c | 4.819E+04 | 880.5 | 0.0004062 | 0.4614 | +1 | 7 |
| - | - | 2.471E+04 | 881.5 | - | - | 0 | - |
| - | - | 5585 | 882.5 | - | - | 0 | - |
| - | - | 795.5 | 889.4 | - | - | 0 | - |
| - | - | 6366 | 906.4 | - | - | 0 | - |
| - | - | 3147 | 907.4 | - | - | 0 | - |
| - | - | 720 | 908.4 | - | - | 0 | - |
| - | - | 1.72E+04 | 934.5 | - | - | 0 | - |
| - | - | 1.009E+04 | 935.5 | - | - | 0 | - |
| - | - | 3288 | 936.5 | - | - | 0 | - |
| - | - | 1155 | 945.5 | - | - | 0 | - |
| - | - | 848.4 | 946.5 | - | - | 0 | - |
| - | - | 3561 | 964.5 | - | - | 0 | - |
| - | - | 4675 | 965.5 | - | - | 0 | - |
| - | - | 1742 | 966.5 | - | - | 0 | - |
| - | - | 1851 | 974.5 | - | - | 0 | - |
| - | - | 1287 | 975.5 | - | - | 0 | - |
| 3 | y | 2004 | 976.5 | 0.003192 | 3.269 | +1 | 8 |
| 3 | y | 1629 | 977.5 | 0.01088 | 11.13 | +1 | 8 |
| 3 | z | 3.895E+04 | 978.5 | 0.001037 | 1.059 | +1 | 8 |
| - | - | 2.25E+04 | 979.5 | - | - | 0 | - |
| - | - | 7910 | 980.5 | - | - | 0 | - |
| - | - | 1580 | 981.5 | - | - | 0 | - |
| - | - | 4092 | 990.6 | - | - | 0 | - |
| - | - | 2835 | 991.6 | - | - | 0 | - |
| 8 | c | 4.3E+04 | 992.5 | 0.0003955 | 0.3985 | +1 | 8 |
| - | - | 2.671E+04 | 993.5 | - | - | 0 | - |
| 3 | y | 1.478E+04 | 994.5 | 0.01149 | 11.55 | +1 | 8 |
| - | - | 6503 | 995.5 | - | - | 0 | - |
| - | - | 1860 | 996.5 | - | - | 0 | - |
| - | - | 1844 | 1056 | - | - | 0 | - |
| - | - | 1987 | 1063 | - | - | 0 | - |
| - | - | 1533 | 1064 | - | - | 0 | - |
| - | - | 770.8 | 1071 | - | - | 0 | - |
| 2 | z | 1.144E+04 | 1078 | 0.0003717 | 0.345 | +1 | 9 |
| - | - | 7125 | 1079 | - | - | 0 | - |
| - | - | 2498 | 1080 | - | - | 0 | - |
| - | - | 763.8 | 1090 | - | - | 0 | - |
| - | - | 1007 | 1091 | - | - | 0 | - |
| - | - | 1017 | 1092 | - | - | 0 | - |
| - | - | 1837 | 1093 | - | - | 0 | - |
| 2 | y | 2293 | 1094 | 0.005696 | 5.208 | +1 | 9 |
| - | - | 1163 | 1095 | - | - | 0 | - |
| - | - | 1081 | 1105 | - | - | 0 | - |
| 9 | c | 5.598E+04 | 1107 | 6.006E-05 | 0.05428 | +1 | 9 |
| - | - | 3.823E+04 | 1108 | - | - | 0 | - |
| - | - | 1.297E+04 | 1109 | - | - | 0 | - |
| - | - | 1441 | 1110 | - | - | 0 | - |
| - | - | 2204 | 1119 | - | - | 0 | - |
| - | - | 1656 | 1120 | - | - | 0 | - |
| - | - | 6377 | 1136 | - | - | 0 | - |
| - | - | 4366 | 1137 | - | - | 0 | - |
| - | - | 1432 | 1138 | - | - | 0 | - |
| - | - | 1104 | 1146 | - | - | 0 | - |
| - | - | 2387 | 1147 | - | - | 0 | - |
| - | - | 1.125E+04 | 1148 | - | - | 0 | - |
| - | - | 9262 | 1149 | - | - | 0 | - |
| - | - | 4077 | 1150 | - | - | 0 | - |
| - | - | 8924 | 1152 | - | - | 0 | - |
| - | - | 5473 | 1153 | - | - | 0 | - |
| - | - | 1910 | 1154 | - | - | 0 | - |
| - | - | 738 | 1161 | - | - | 0 | - |
| - | - | 1.106E+04 | 1162 | - | - | 0 | - |
| - | - | 1.042E+04 | 1163 | - | - | 0 | - |
| - | - | 1.645E+04 | 1164 | - | - | 0 | - |
| - | - | 1.015E+04 | 1165 | - | - | 0 | - |
| - | - | 3885 | 1166 | - | - | 0 | - |
| - | - | 799.6 | 1167 | - | - | 0 | - |
| - | - | 967.3 | 1172 | - | - | 0 | - |
| - | - | 7063 | 1180 | - | - | 0 | - |
| - | - | 5015 | 1181 | - | - | 0 | - |
| - | - | 1955 | 1182 | - | - | 0 | - |
| - | - | 971.8 | 1183 | - | - | 0 | - |
| - | - | 3183 | 1189 | - | - | 0 | - |
| - | - | 6416 | 1190 | - | - | 0 | - |
| - | - | 7.014E+04 | 1191 | - | - | 0 | - |
| - | - | 4.961E+04 | 1192 | - | - | 0 | - |
| - | - | 1.835E+04 | 1193 | - | - | 0 | - |
| - | - | 4342 | 1194 | - | - | 0 | - |
| - | - | 2810 | 1205 | - | - | 0 | - |
| - | - | 9212 | 1206 | - | - | 0 | - |
| - | - | 8.621E+04 | 1207 | - | - | 0 | - |
| - | - | 3.272E+05 | 1208 | - | - | 0 | - |
| - | - | 2.112E+05 | 1209 | - | - | 0 | - |
| - | - | 7.963E+04 | 1210 | - | - | 0 | - |
| - | - | 1.301E+04 | 1211 | - | - | 0 | - |
| - | - | 768.6 | 1992 | - | - | 0 | - |
| - | - | 783.3 | 3034 | - | - | 0 | - |
| - | - | 720.1 | 3035 | - | - | 0 | - |

m/z Charge Intensity FragmentType MassShift Position
120.08111572265625 0 1423.8317
122.52171325683594 0 366.6997
129.10247802734375 0 1982.6208
133.08621215820312 0 962.34955
136.07594299316406 0 563.923
143.7473602294922 0 385.83868
146.51641845703125 0 424.56332
158.04730224609375 0 405.1353
173.4381866455078 0 618.7355
177.11233520507812 0 646.6058
185.16510009765625 0 6579.9756
186.1682586669922 0 791.80273
199.1077117919922 0 659.13153
207.1107940673828 0 580.5089
209.09197998046875 0 611.4395
213.16018676757812 0 4988.271
215.13934326171875 0 101921.32 y 8
215.90476989746094 0 565.8801
216.14259338378906 0 9849.524
217.14480590820312 0 653.7615
219.1343231201172 0 1038.1699
227.10304260253906 0 3266.766
229.1550750732422 0 5814.618
243.13426208496094 0 1404.1515
287.13848876953125 0 603.356
301.21124267578125 0 951.2421
304.1652526855469 0 811.404
324.1561584472656 0 2205.1965
325.1622009277344 0 521.1615
326.1715393066406 0 946.7506 y Water loss 7
343.177001953125 0 1383.8474
344.1802673339844 0 1068.399 y 7
357.27374267578125 0 25507.373
358.2779541015625 0 5239.8105 c 2
359.17425537109375 0 605.055 z Water loss 4
360.17498779296875 0 849.3888
361.18768310546875 0 1295.4886
374.17047119140625 0 953.31433
392.2197570800781 0 709.0559 c 5
407.1946716308594 0 877.4868
418.2388610839844 0 1544.0032
423.2251892089844 0 2000.6152 y Water loss 6
426.1678466796875 0 1406.0272
432.2361755371094 0 905.9051 c Ammonia loss 6
432.7377014160156 0 978.4225
441.2351989746094 0 10197.153 y 6
442.2389831542969 0 3105.6265
456.2825622558594 0 2135.9329 c Ammonia loss 3
467.264404296875 0 625.76154
468.2242431640625 0 712.53864
470.2231140136719 0 595.91
472.3014221191406 0 3445.8794
473.30853271484375 0 8162.4023 c 3
474.3113098144531 0 2006.6045
483.2627258300781 0 930.67896
485.30767822265625 0 593.78284
486.2327880859375 0 670.1586
491.2705993652344 0 2325.2217
492.27838134765625 0 1113.6907
495.6523132324219 0 662.5235
496.75860595703125 0 1295.9587 c Ammonia loss 7
497.2636413574219 0 697.5169
511.2554626464844 0 1215.4879
536.248779296875 0 659.1627
543.3154296875 0 1063.5286
544.318603515625 0 716.03375
545.2825317382812 0 1072.2905 c Ammonia loss 8
545.786376953125 0 1403.8701
552.23779296875 0 708.22095
554.2611083984375 0 5306.2275
555.2660522460938 0 1417.2628
563.9158325195312 0 584.529
567.3497924804688 0 859.60095
585.3026733398438 0 1189.0188
588.3031616210938 0 11378.152 y 5
589.3065185546875 0 4610.963
590.308349609375 0 846.3328
592.3590087890625 0 1505.5111
603.3125 0 2321.8499
603.8214721679688 0 963.64404
604.3197021484375 0 1868.5234
613.0093994140625 0 578.3052
619.3461303710938 0 4892.2515 c Ammonia loss 4
620.3470458984375 0 1727.1329
635.3646850585938 0 14368.835
636.37060546875 0 18113.367 c 4
637.3740844726562 0 5467.3667
638.3822021484375 0 1037.5837
645.2852783203125 0 565.2328
645.3447265625 0 1474.982
646.3519897460938 0 835.05396
651.3194580078125 0 611.4161
652.2595825195312 0 829.0101
654.335205078125 0 606.87994
681.3226928710938 0 851.0722
682.3800659179688 0 1804.3497
683.3831176757812 0 981.49835
723.8109130859375 0 603.14185
732.374755859375 0 1270.604
733.3809814453125 0 1133.1205
735.348388671875 0 2088.3413 z 4
736.3523559570312 0 1634.6987
738.4186401367188 0 1805.7006
739.4263305664062 0 14462.088
740.4324951171875 0 15413.225
741.4356079101562 0 4825.8867
742.4388427734375 0 1214.5294
747.3684692382812 0 908.3749
750.3662719726562 0 868.50464
751.3666381835938 0 2268.2842 y 4
752.3692016601562 0 1519.7725
766.41357421875 0 20816.143 c Ammonia loss 5
767.4171142578125 0 10240.023
768.4201049804688 0 1998.2542
780.3568725585938 0 9264.414
781.3591918945312 0 4740.86
782.3621215820312 0 1077.6406
815.42626953125 0 762.45685
832.4533081054688 0 1142.0775
836.48291015625 0 1596.3433
837.4833374023438 0 609.6435
842.4541625976562 0 1615.1885
843.4515380859375 0 707.16895
844.4708251953125 0 1421.6482
845.4572143554688 0 950.4101
846.4375 0 752.6577
851.3826293945312 0 4141.856
852.3861694335938 0 1311.5
863.466064453125 0 1796.5707 c Ammonia loss 6
864.46923828125 0 1470.4427
866.392822265625 0 1547.4088 y 3
872.4246215820312 0 1109.3229
873.4353637695312 0 705.61304
879.485107421875 0 13480.983
880.4923095703125 0 48188.77 c 6
881.4955444335938 0 24709.361
882.4981689453125 0 5585.07
889.4047241210938 0 795.453
906.4480590820312 0 6366.2603
907.4470825195312 0 3147.1675
908.4305419921875 0 719.9781
934.479248046875 0 17197.68
935.4822998046875 0 10094.689
936.4862060546875 0 3288.265
945.517578125 0 1155.1488
946.5089111328125 0 848.40753
964.5133666992188 0 3560.6257
965.5184326171875 0 4675.421
966.5231323242188 0 1742.2471
974.4964599609375 0 1851.3843
975.5001831054688 0 1287.0875
976.4806518554688 0 2003.8705 y Water loss 2
977.4723510742188 0 1629.3384 y Ammonia loss 2
978.4703369140625 0 38947.324 z 2
979.47314453125 0 22496.373
980.4766845703125 0 7909.8066
981.4832763671875 0 1580.3407
990.555908203125 0 4092.395
991.555908203125 0 2835.4966
992.5091552734375 0 43000.06 c Ammonia loss 7
993.5123901367188 0 26706.9
994.49951171875 0 14781.814 y 2
995.4945068359375 0 6502.8706
996.4937133789062 0 1859.9603
1055.5111083984375 0 1844.2754
1062.56884765625 0 1987.4127
1063.5762939453125 0 1533.2441
1071.498779296875 0 770.8295
1077.5380859375 0 11443.205 z 1
1078.5416259765625 0 7124.8975
1079.5435791015625 0 2498.395
1089.6226806640625 0 763.83124
1090.5587158203125 0 1006.5993
1091.55419921875 0 1016.8117
1092.5628662109375 0 1837.2294
1093.5621337890625 0 2293.0706 y 1
1094.56591796875 0 1162.9492
1105.4276123046875 0 1081.284
1106.5880126953125 0 55975.094 c 8
1107.59130859375 0 38228.203
1108.59375 0 12972.263
1109.5899658203125 0 1440.6761
1118.59521484375 0 2203.8276
1119.593017578125 0 1656.2192
1135.6258544921875 0 6377.176
1136.628173828125 0 4366.4624
1137.6357421875 0 1432.1752
1145.643798828125 0 1103.726
1146.6329345703125 0 2386.627
1147.5941162109375 0 11250.401
1148.59033203125 0 9262.445
1149.5986328125 0 4077.3936
1151.586181640625 0 8923.893
1152.5899658203125 0 5473.2734
1153.5897216796875 0 1909.6727
1160.6292724609375 0 738.0304
1161.6429443359375 0 11060.283
1162.646728515625 0 10415.021
1163.655029296875 0 16448.121
1164.6600341796875 0 10154.598
1165.6630859375 0 3884.6404
1166.6585693359375 0 799.5564
1171.6280517578125 0 967.2599
1179.65380859375 0 7062.9995
1180.655517578125 0 5014.514
1181.65966796875 0 1955.32
1182.646728515625 0 971.8078
1188.60400390625 0 3182.544
1189.6234130859375 0 6415.813
1190.6219482421875 0 70135.93
1191.624267578125 0 49607.45
1192.62744140625 0 18346.941
1193.62890625 0 4341.507
1204.622802734375 0 2810.0688
1205.6280517578125 0 9212.039
1206.639404296875 0 86211.13
1207.647705078125 0 327222.12
1208.650634765625 0 211226.11
1209.653076171875 0 79630.695
1210.6544189453125 0 13011.481
1991.8818359375 0 768.5663
3034.243408203125 0 783.3251
3034.908447265625 0 720.121

Spectrum Details

|  |  |
| --- | --- |
| Matched peaks? Matched peaks The total absolute number of peaks matched. Additionally in brackets the total fraction of peaks matched and the total number of peaks is shown. | 30 (13.45% of 223) |
| FDR? FDR The false discovery rate estimated for this peptide. It is calculated by matching all theoretical fragments with a non-integer shift with the raw peaks for this spectrum. This is done with 40 different shifts. The resulting percentage is the average number of annotated peaks over the number of annotated peaks with the correct spectrum. | 2.30% |
| Satellite FDR? Satellite FDR See the FDR for details on its calculation. This satellite ion specific FDR only contains the satellite ions (d/w) for I/L/J positions. | - |
| PSM Score? PSM Score The PSM Score as given by Hecklib to this annotated spectrum. It is shown with three significant figures. | 341 |

#### Spectrum 8462? Spectrum 8462 The raw spectrum of this peptide as annotated by Hecklib. The fragments are coloured according to ion type (see legend). Any peaks with a star '\*' as text can be hovered over to see the full details, first the ion type second the mass shift type. By hovering over the amino acids in the peptide or ions in the legend the corresponding peaks are highlighted. By toggling the 'Unassigned' label you can turn the background (unassigned) peaks on or off in the plot. By updating the slider in the Ion legend you can update the spectrum to only show the top X% of the peaks with labels. The top X% means any peak that is within X% of the highest intensity. By dragging in the spectrum you can zoom in to a specific part of the spectrum and use 'Zoom Out' to get back to the original zoom level. The annotation of the spectrum is based on the given sequence in the peptides file and is done with different software so inconsistencies are likely. The peaks are annotated based on the given sequence, with 20 ppm tolerance.

Copy Data

##### Spectrum 8462 (TSV)

###### Preview

```
Loading example...
```

*Click on the button to copy the data to your clipboard.*

Mz MinMz MaxIntensity Max

WidthHeightPeptide font sizePeptide stroke widthSpectrum font sizeSpectrum stroke widthCompact peptide

Ion legend

wxyz

abcd

OtherUnassignedIon^Charge^_Position_Show for top:%

JVKDYFPEPV

02.26e+54.51e+56.77e+59.02e+5

Zoom Out

a^+1^_2_b^+1^_2_y^+1^_2_b^+2^_5_b^+1^_3_y^+1^_3_b^+1^_3_y^+1^_3_b^+2^_6_b^+2^_7_b^+2^_7_b^+1^_4_y^+1^_4_b^+1^_4_b^+2^_8_b^+2^_8_b^+2^_9_b^+2^_9_y^+1^_5_y^+1^_5_\*b^+1^_5_\*b^+1^_5_b^+1^_6_b^+1^_6_y^+1^_6_b^+1^_6_b^+1^_7_b^+1^_7_b^+1^_7_b^+1^_8_b^+1^_8_y^+1^_8_b^+1^_8_y^+1^_8_y^+1^_9_

043286512971730

Fragment Matches Table

Show background peaks

| Position | Ion type | Intensity | mz Theoretical | mz Error (Th) | mz Error (ppm) | Charge | Series Number |
| --- | --- | --- | --- | --- | --- | --- | --- |
| - | - | 3.656E+04 | 120.1 | - | - | 0 | - |
| - | - | 394.4 | 121.1 | - | - | 0 | - |
| - | - | 2939 | 121.1 | - | - | 0 | - |
| - | - | 381.6 | 127 | - | - | 0 | - |
| - | - | 1080 | 127.1 | - | - | 0 | - |
| - | - | 797.3 | 129.1 | - | - | 0 | - |
| - | - | 6.941E+04 | 129.1 | - | - | 0 | - |
| - | - | 488.3 | 130.1 | - | - | 0 | - |
| - | - | 1010 | 130.1 | - | - | 0 | - |
| - | - | 4569 | 130.1 | - | - | 0 | - |
| - | - | 505.7 | 131 | - | - | 0 | - |
| - | - | 553.9 | 132.1 | - | - | 0 | - |
| - | - | 785.3 | 132.1 | - | - | 0 | - |
| - | - | 410.2 | 132.8 | - | - | 0 | - |
| - | - | 761 | 133.1 | - | - | 0 | - |
| - | - | 595.7 | 133.1 | - | - | 0 | - |
| - | - | 434 | 134.1 | - | - | 0 | - |
| - | - | 1.59E+04 | 136.1 | - | - | 0 | - |
| - | - | 767.5 | 137.1 | - | - | 0 | - |
| - | - | 2079 | 139.1 | - | - | 0 | - |
| - | - | 561.4 | 140.1 | - | - | 0 | - |
| - | - | 974.1 | 141.1 | - | - | 0 | - |
| - | - | 522.4 | 143.1 | - | - | 0 | - |
| - | - | 675.7 | 147 | - | - | 0 | - |
| - | - | 828.8 | 149 | - | - | 0 | - |
| - | - | 573.9 | 153.1 | - | - | 0 | - |
| - | - | 803.6 | 155.1 | - | - | 0 | - |
| - | - | 1027 | 155.1 | - | - | 0 | - |
| - | - | 534.3 | 155.5 | - | - | 0 | - |
| - | - | 808.1 | 156.1 | - | - | 0 | - |
| - | - | 1026 | 157.1 | - | - | 0 | - |
| - | - | 649.5 | 159.1 | - | - | 0 | - |
| - | - | 669.9 | 159.1 | - | - | 0 | - |
| - | - | 1245 | 165.1 | - | - | 0 | - |
| - | - | 466.2 | 166.3 | - | - | 0 | - |
| - | - | 1732 | 167.1 | - | - | 0 | - |
| - | - | 2054 | 167.1 | - | - | 0 | - |
| - | - | 1839 | 168.1 | - | - | 0 | - |
| - | - | 1122 | 169.1 | - | - | 0 | - |
| - | - | 527.2 | 169.1 | - | - | 0 | - |
| - | - | 1.879E+04 | 169.1 | - | - | 0 | - |
| - | - | 2309 | 170.1 | - | - | 0 | - |
| - | - | 447.1 | 170.9 | - | - | 0 | - |
| - | - | 4928 | 172.1 | - | - | 0 | - |
| - | - | 833.2 | 172.1 | - | - | 0 | - |
| - | - | 1117 | 173.1 | - | - | 0 | - |
| - | - | 1154 | 173.1 | - | - | 0 | - |
| - | - | 950.3 | 173.4 | - | - | 0 | - |
| - | - | 2212 | 174.1 | - | - | 0 | - |
| - | - | 787.5 | 175.1 | - | - | 0 | - |
| - | - | 495.2 | 177.1 | - | - | 0 | - |
| - | - | 506.4 | 180.7 | - | - | 0 | - |
| - | - | 7299 | 181.1 | - | - | 0 | - |
| - | - | 4978 | 181.1 | - | - | 0 | - |
| - | - | 556.9 | 182.1 | - | - | 0 | - |
| - | - | 600.9 | 182.1 | - | - | 0 | - |
| - | - | 922.6 | 182.1 | - | - | 0 | - |
| - | - | 661.5 | 182.1 | - | - | 0 | - |
| - | - | 1769 | 183.1 | - | - | 0 | - |
| - | - | 2718 | 183.1 | - | - | 0 | - |
| - | - | 734.1 | 184.1 | - | - | 0 | - |
| - | - | 1707 | 185.1 | - | - | 0 | - |
| - | - | 1723 | 185.1 | - | - | 0 | - |
| 2 | a | 7.739E+04 | 185.2 | 0.000413 | 2.23 | +1 | 2 |
| - | - | 2999 | 186.1 | - | - | 0 | - |
| - | - | 7233 | 186.2 | - | - | 0 | - |
| - | - | 1296 | 187.1 | - | - | 0 | - |
| - | - | 748.1 | 187.1 | - | - | 0 | - |
| - | - | 1506 | 188.1 | - | - | 0 | - |
| - | - | 3278 | 188.1 | - | - | 0 | - |
| - | - | 899.9 | 190.1 | - | - | 0 | - |
| - | - | 918.1 | 191.1 | - | - | 0 | - |
| - | - | 6611 | 195.1 | - | - | 0 | - |
| - | - | 931.1 | 197.1 | - | - | 0 | - |
| - | - | 3404 | 197.1 | - | - | 0 | - |
| - | - | 1168 | 197.2 | - | - | 0 | - |
| - | - | 842.8 | 198.1 | - | - | 0 | - |
| - | - | 2.518E+04 | 199.1 | - | - | 0 | - |
| - | - | 1778 | 200.1 | - | - | 0 | - |
| - | - | 2105 | 201.1 | - | - | 0 | - |
| - | - | 752.8 | 202.1 | - | - | 0 | - |
| - | - | 2052 | 203.1 | - | - | 0 | - |
| - | - | 1852 | 203.2 | - | - | 0 | - |
| - | - | 2138 | 204.1 | - | - | 0 | - |
| - | - | 1.634E+04 | 209.1 | - | - | 0 | - |
| - | - | 582.2 | 209.1 | - | - | 0 | - |
| - | - | 1083 | 210.1 | - | - | 0 | - |
| - | - | 1418 | 210.2 | - | - | 0 | - |
| - | - | 1769 | 211.1 | - | - | 0 | - |
| - | - | 2976 | 212.1 | - | - | 0 | - |
| - | - | 766.7 | 213.1 | - | - | 0 | - |
| - | - | 3714 | 213.1 | - | - | 0 | - |
| 2 | b | 2.876E+04 | 213.2 | 0.0004629 | 2.172 | +1 | 2 |
| - | - | 2831 | 214.2 | - | - | 0 | - |
| 9 | y | 8.931E+05 | 215.1 | 0.0005227 | 2.43 | +1 | 2 |
| - | - | 9.095E+04 | 216.1 | - | - | 0 | - |
| - | - | 4086 | 217.1 | - | - | 0 | - |
| - | - | 6752 | 217.1 | - | - | 0 | - |
| - | - | 808.8 | 223.1 | - | - | 0 | - |
| - | - | 1481 | 225.1 | - | - | 0 | - |
| - | - | 2.03E+04 | 226.1 | - | - | 0 | - |
| - | - | 965.7 | 226.2 | - | - | 0 | - |
| - | - | 6.64E+04 | 227.1 | - | - | 0 | - |
| - | - | 1748 | 227.1 | - | - | 0 | - |
| - | - | 6884 | 228.1 | - | - | 0 | - |
| - | - | 6708 | 228.2 | - | - | 0 | - |
| - | - | 849.2 | 229.1 | - | - | 0 | - |
| - | - | 4586 | 229.2 | - | - | 0 | - |
| - | - | 1641 | 230.2 | - | - | 0 | - |
| - | - | 707.9 | 231.1 | - | - | 0 | - |
| - | - | 1.538E+04 | 231.1 | - | - | 0 | - |
| - | - | 2435 | 232.2 | - | - | 0 | - |
| - | - | 960.4 | 233.1 | - | - | 0 | - |
| - | - | 2061 | 233.2 | - | - | 0 | - |
| - | - | 1534 | 238.1 | - | - | 0 | - |
| - | - | 2837 | 238.2 | - | - | 0 | - |
| - | - | 760.5 | 239.2 | - | - | 0 | - |
| - | - | 789.4 | 240.1 | - | - | 0 | - |
| - | - | 651.3 | 243.1 | - | - | 0 | - |
| - | - | 3.35E+04 | 244.1 | - | - | 0 | - |
| - | - | 1018 | 245.1 | - | - | 0 | - |
| - | - | 8423 | 245.1 | - | - | 0 | - |
| - | - | 1238 | 246.1 | - | - | 0 | - |
| - | - | 583.2 | 246.9 | - | - | 0 | - |
| - | - | 2879 | 247.1 | - | - | 0 | - |
| - | - | 1500 | 249.2 | - | - | 0 | - |
| - | - | 2.329E+04 | 251.1 | - | - | 0 | - |
| - | - | 2770 | 252.1 | - | - | 0 | - |
| - | - | 3768 | 254.1 | - | - | 0 | - |
| - | - | 1167 | 254.2 | - | - | 0 | - |
| - | - | 2014 | 256.2 | - | - | 0 | - |
| - | - | 826 | 257.1 | - | - | 0 | - |
| - | - | 694 | 259.1 | - | - | 0 | - |
| - | - | 817.3 | 261.1 | - | - | 0 | - |
| - | - | 600.2 | 261.1 | - | - | 0 | - |
| - | - | 3480 | 261.2 | - | - | 0 | - |
| - | - | 1055 | 262.1 | - | - | 0 | - |
| - | - | 677.6 | 262.2 | - | - | 0 | - |
| - | - | 707.6 | 265.2 | - | - | 0 | - |
| - | - | 628.3 | 266.2 | - | - | 0 | - |
| - | - | 954 | 269.2 | - | - | 0 | - |
| - | - | 727.7 | 270.2 | - | - | 0 | - |
| - | - | 5833 | 272.1 | - | - | 0 | - |
| - | - | 929 | 273.1 | - | - | 0 | - |
| - | - | 880 | 276.2 | - | - | 0 | - |
| - | - | 1430 | 277.2 | - | - | 0 | - |
| - | - | 1030 | 278.2 | - | - | 0 | - |
| - | - | 2.305E+04 | 279.1 | - | - | 0 | - |
| - | - | 3479 | 280.1 | - | - | 0 | - |
| - | - | 1249 | 280.2 | - | - | 0 | - |
| - | - | 1.088E+04 | 283.1 | - | - | 0 | - |
| - | - | 699.1 | 283.2 | - | - | 0 | - |
| - | - | 1450 | 284.1 | - | - | 0 | - |
| - | - | 637.3 | 286.1 | - | - | 0 | - |
| - | - | 2901 | 287.2 | - | - | 0 | - |
| - | - | 589 | 292.1 | - | - | 0 | - |
| - | - | 900.6 | 292.2 | - | - | 0 | - |
| - | - | 1438 | 294.2 | - | - | 0 | - |
| - | - | 987.5 | 296.1 | - | - | 0 | - |
| - | - | 1965 | 301.2 | - | - | 0 | - |
| - | - | 3335 | 303.2 | - | - | 0 | - |
| - | - | 1513 | 304.2 | - | - | 0 | - |
| - | - | 674.7 | 306.1 | - | - | 0 | - |
| - | - | 773.4 | 306.2 | - | - | 0 | - |
| - | - | 1860 | 308.2 | - | - | 0 | - |
| - | - | 1116 | 309.2 | - | - | 0 | - |
| - | - | 1006 | 310.1 | - | - | 0 | - |
| 5 | b | 2658 | 310.2 | 0.001022 | 3.294 | +2 | 5 |
| - | - | 805 | 310.2 | - | - | 0 | - |
| - | - | 7589 | 311.1 | - | - | 0 | - |
| - | - | 1539 | 312.1 | - | - | 0 | - |
| - | - | 921 | 312.2 | - | - | 0 | - |
| - | - | 1883 | 313.2 | - | - | 0 | - |
| - | - | 2205 | 315.2 | - | - | 0 | - |
| - | - | 652.5 | 320.2 | - | - | 0 | - |
| - | - | 640.9 | 321.1 | - | - | 0 | - |
| - | - | 663.2 | 322.2 | - | - | 0 | - |
| - | - | 1346 | 323.2 | - | - | 0 | - |
| - | - | 2353 | 323.2 | - | - | 0 | - |
| - | - | 1.488E+04 | 324.2 | - | - | 0 | - |
| 3 | b | 681.1 | 324.2 | 0.001812 | 5.589 | +1 | 3 |
| - | - | 2598 | 325.2 | - | - | 0 | - |
| - | - | 2941 | 325.2 | - | - | 0 | - |
| - | - | 1048 | 325.2 | - | - | 0 | - |
| 8 | y | 8820 | 326.2 | 0.0006141 | 1.883 | +1 | 3 |
| - | - | 692.6 | 326.2 | - | - | 0 | - |
| - | - | 1785 | 327.2 | - | - | 0 | - |
| - | - | 1037 | 328.1 | - | - | 0 | - |
| - | - | 1087 | 328.2 | - | - | 0 | - |
| - | - | 2263 | 329.2 | - | - | 0 | - |
| - | - | 552.6 | 330.2 | - | - | 0 | - |
| - | - | 1455 | 331.2 | - | - | 0 | - |
| - | - | 1526 | 335.2 | - | - | 0 | - |
| - | - | 612.1 | 336.2 | - | - | 0 | - |
| - | - | 2793 | 337.2 | - | - | 0 | - |
| - | - | 1149 | 337.2 | - | - | 0 | - |
| - | - | 802.2 | 338.2 | - | - | 0 | - |
| - | - | 1284 | 339.2 | - | - | 0 | - |
| - | - | 5052 | 340.2 | - | - | 0 | - |
| - | - | 5023 | 341.2 | - | - | 0 | - |
| 3 | b | 5919 | 341.3 | 0.0008063 | 2.363 | +1 | 3 |
| - | - | 1213 | 342.1 | - | - | 0 | - |
| - | - | 995.7 | 342.2 | - | - | 0 | - |
| - | - | 963.9 | 342.3 | - | - | 0 | - |
| - | - | 8285 | 343.2 | - | - | 0 | - |
| - | - | 1288 | 344.2 | - | - | 0 | - |
| 8 | y | 5508 | 344.2 | 0.000639 | 1.856 | +1 | 3 |
| - | - | 998.5 | 344.2 | - | - | 0 | - |
| - | - | 1740 | 344.7 | - | - | 0 | - |
| - | - | 1223 | 345.2 | - | - | 0 | - |
| - | - | 4876 | 346.2 | - | - | 0 | - |
| - | - | 4200 | 347.2 | - | - | 0 | - |
| - | - | 1412 | 347.7 | - | - | 0 | - |
| - | - | 1305 | 349.2 | - | - | 0 | - |
| - | - | 520.1 | 353.2 | - | - | 0 | - |
| - | - | 5365 | 353.2 | - | - | 0 | - |
| - | - | 2456 | 353.2 | - | - | 0 | - |
| - | - | 1147 | 354.2 | - | - | 0 | - |
| - | - | 751.2 | 354.2 | - | - | 0 | - |
| - | - | 4565 | 355.2 | - | - | 0 | - |
| - | - | 2785 | 355.2 | - | - | 0 | - |
| - | - | 1644 | 356.2 | - | - | 0 | - |
| - | - | 875.5 | 356.2 | - | - | 0 | - |
| - | - | 694.1 | 358.2 | - | - | 0 | - |
| - | - | 671.3 | 359.1 | - | - | 0 | - |
| - | - | 755.9 | 360.2 | - | - | 0 | - |
| - | - | 667.9 | 361.7 | - | - | 0 | - |
| - | - | 1888 | 362.2 | - | - | 0 | - |
| - | - | 677.4 | 363.2 | - | - | 0 | - |
| - | - | 875.8 | 363.2 | - | - | 0 | - |
| - | - | 718.8 | 365.2 | - | - | 0 | - |
| - | - | 931.9 | 367.2 | - | - | 0 | - |
| - | - | 1102 | 369.2 | - | - | 0 | - |
| - | - | 6160 | 369.7 | - | - | 0 | - |
| - | - | 727.4 | 370.2 | - | - | 0 | - |
| - | - | 3565 | 370.2 | - | - | 0 | - |
| - | - | 1214 | 370.7 | - | - | 0 | - |
| - | - | 7229 | 371.2 | - | - | 0 | - |
| - | - | 2057 | 372.2 | - | - | 0 | - |
| - | - | 1252 | 372.2 | - | - | 0 | - |
| - | - | 633.3 | 372.2 | - | - | 0 | - |
| - | - | 2426 | 373.2 | - | - | 0 | - |
| - | - | 1.233E+04 | 374.2 | - | - | 0 | - |
| - | - | 1659 | 375.2 | - | - | 0 | - |
| - | - | 680.9 | 375.2 | - | - | 0 | - |
| - | - | 645.5 | 376.2 | - | - | 0 | - |
| - | - | 6917 | 379.2 | - | - | 0 | - |
| - | - | 819.7 | 380.2 | - | - | 0 | - |
| - | - | 2810 | 381.1 | - | - | 0 | - |
| - | - | 785.3 | 382.1 | - | - | 0 | - |
| - | - | 998.8 | 383.2 | - | - | 0 | - |
| - | - | 2866 | 383.2 | - | - | 0 | - |
| 6 | b | 1133 | 383.7 | 0.0007504 | 1.956 | +2 | 6 |
| - | - | 7852 | 389.2 | - | - | 0 | - |
| - | - | 1671 | 390.2 | - | - | 0 | - |
| - | - | 983.4 | 390.2 | - | - | 0 | - |
| - | - | 1210 | 391.2 | - | - | 0 | - |
| - | - | 1.46E+04 | 391.2 | - | - | 0 | - |
| - | - | 2884 | 392.2 | - | - | 0 | - |
| - | - | 1675 | 392.2 | - | - | 0 | - |
| - | - | 2952 | 403.2 | - | - | 0 | - |
| - | - | 1151 | 404.2 | - | - | 0 | - |
| - | - | 736.6 | 406.2 | - | - | 0 | - |
| - | - | 1.533E+04 | 407.2 | - | - | 0 | - |
| - | - | 866.5 | 408.2 | - | - | 0 | - |
| - | - | 3700 | 408.2 | - | - | 0 | - |
| - | - | 962.5 | 408.2 | - | - | 0 | - |
| - | - | 1208 | 409.2 | - | - | 0 | - |
| - | - | 1864 | 411.3 | - | - | 0 | - |
| - | - | 785.3 | 415.2 | - | - | 0 | - |
| - | - | 1017 | 417.2 | - | - | 0 | - |
| - | - | 9575 | 418.2 | - | - | 0 | - |
| - | - | 8398 | 418.7 | - | - | 0 | - |
| - | - | 1203 | 419.2 | - | - | 0 | - |
| - | - | 1213 | 419.2 | - | - | 0 | - |
| - | - | 711.7 | 420.3 | - | - | 0 | - |
| - | - | 704.5 | 421.2 | - | - | 0 | - |
| - | - | 774.6 | 422.2 | - | - | 0 | - |
| 7 | b | 1.688E+04 | 423.2 | 0.006738 | 15.92 | +2 | 7 |
| - | - | 8114 | 424.2 | - | - | 0 | - |
| - | - | 1604 | 425.2 | - | - | 0 | - |
| - | - | 2.125E+04 | 426.2 | - | - | 0 | - |
| - | - | 5160 | 427.2 | - | - | 0 | - |
| - | - | 729.5 | 428.2 | - | - | 0 | - |
| 7 | b | 5764 | 432.2 | 0.0005831 | 1.349 | +2 | 7 |
| - | - | 3471 | 432.7 | - | - | 0 | - |
| - | - | 1109 | 433.2 | - | - | 0 | - |
| - | - | 1466 | 434.2 | - | - | 0 | - |
| - | - | 614.6 | 434.3 | - | - | 0 | - |
| - | - | 3759 | 435.2 | - | - | 0 | - |
| - | - | 652.1 | 436.2 | - | - | 0 | - |
| - | - | 1631 | 436.2 | - | - | 0 | - |
| - | - | 786.6 | 436.3 | - | - | 0 | - |
| 4 | b | 8376 | 438.3 | 0.001457 | 3.324 | +1 | 4 |
| - | - | 1554 | 438.3 | - | - | 0 | - |
| - | - | 1729 | 439.3 | - | - | 0 | - |
| - | - | 1348 | 440.3 | - | - | 0 | - |
| 7 | y | 5.389E+04 | 441.2 | 0.0008536 | 1.935 | +1 | 4 |
| - | - | 1.077E+04 | 442.2 | - | - | 0 | - |
| - | - | 2132 | 443.2 | - | - | 0 | - |
| - | - | 1575 | 445.2 | - | - | 0 | - |
| - | - | 5170 | 452.2 | - | - | 0 | - |
| - | - | 862.2 | 452.3 | - | - | 0 | - |
| - | - | 792.7 | 453.2 | - | - | 0 | - |
| - | - | 910.5 | 454.2 | - | - | 0 | - |
| - | - | 1843 | 454.3 | - | - | 0 | - |
| 4 | b | 2.775E+04 | 456.3 | 0.0009324 | 2.043 | +1 | 4 |
| - | - | 1686 | 457.2 | - | - | 0 | - |
| - | - | 7814 | 457.3 | - | - | 0 | - |
| - | - | 1003 | 458.3 | - | - | 0 | - |
| - | - | 823.9 | 460.2 | - | - | 0 | - |
| - | - | 888.8 | 464.2 | - | - | 0 | - |
| - | - | 1531 | 464.3 | - | - | 0 | - |
| - | - | 966.1 | 465.3 | - | - | 0 | - |
| - | - | 1440 | 468.2 | - | - | 0 | - |
| - | - | 1043 | 469.2 | - | - | 0 | - |
| - | - | 1.434E+04 | 470.2 | - | - | 0 | - |
| - | - | 4620 | 471.2 | - | - | 0 | - |
| - | - | 1542 | 473.2 | - | - | 0 | - |
| - | - | 687.6 | 473.2 | - | - | 0 | - |
| - | - | 1009 | 473.3 | - | - | 0 | - |
| - | - | 788.4 | 474.2 | - | - | 0 | - |
| - | - | 1520 | 476.3 | - | - | 0 | - |
| - | - | 1483 | 478.3 | - | - | 0 | - |
| - | - | 780.1 | 480.2 | - | - | 0 | - |
| - | - | 7931 | 482.3 | - | - | 0 | - |
| - | - | 4905 | 482.8 | - | - | 0 | - |
| - | - | 877.5 | 483.2 | - | - | 0 | - |
| - | - | 4949 | 483.3 | - | - | 0 | - |
| - | - | 983.4 | 483.8 | - | - | 0 | - |
| - | - | 1199 | 486.2 | - | - | 0 | - |
| - | - | 867.9 | 487.3 | - | - | 0 | - |
| 8 | b | 1382 | 488.2 | 0.007271 | 14.89 | +2 | 8 |
| - | - | 2867 | 488.3 | - | - | 0 | - |
| - | - | 1551 | 490.3 | - | - | 0 | - |
| - | - | 764.6 | 491.2 | - | - | 0 | - |
| - | - | 726.6 | 492.2 | - | - | 0 | - |
| - | - | 684.4 | 495.2 | - | - | 0 | - |
| 8 | b | 5790 | 496.8 | 0.0007099 | 1.429 | +2 | 8 |
| - | - | 2636 | 497.3 | - | - | 0 | - |
| - | - | 1414 | 497.8 | - | - | 0 | - |
| - | - | 1380 | 498.2 | - | - | 0 | - |
| - | - | 935.3 | 500.2 | - | - | 0 | - |
| - | - | 1242 | 500.3 | - | - | 0 | - |
| - | - | 853 | 502.3 | - | - | 0 | - |
| - | - | 1837 | 504.2 | - | - | 0 | - |
| - | - | 1035 | 504.3 | - | - | 0 | - |
| - | - | 804 | 505.2 | - | - | 0 | - |
| - | - | 3186 | 506.3 | - | - | 0 | - |
| - | - | 856.9 | 507.3 | - | - | 0 | - |
| - | - | 3085 | 508.3 | - | - | 0 | - |
| - | - | 4153 | 509.2 | - | - | 0 | - |
| - | - | 1916 | 510.2 | - | - | 0 | - |
| - | - | 1219 | 516.2 | - | - | 0 | - |
| - | - | 1767 | 518.2 | - | - | 0 | - |
| - | - | 1643 | 518.3 | - | - | 0 | - |
| - | - | 764.4 | 519.2 | - | - | 0 | - |
| - | - | 937 | 520.3 | - | - | 0 | - |
| - | - | 1714 | 521.3 | - | - | 0 | - |
| - | - | 627.9 | 522.3 | - | - | 0 | - |
| - | - | 2204 | 523.2 | - | - | 0 | - |
| - | - | 825.4 | 523.3 | - | - | 0 | - |
| - | - | 704 | 524.2 | - | - | 0 | - |
| - | - | 752.4 | 525.2 | - | - | 0 | - |
| - | - | 1.25E+04 | 526.3 | - | - | 0 | - |
| - | - | 2564 | 527.3 | - | - | 0 | - |
| - | - | 749.6 | 531.3 | - | - | 0 | - |
| - | - | 855.8 | 531.3 | - | - | 0 | - |
| - | - | 644.3 | 531.8 | - | - | 0 | - |
| - | - | 6833 | 534.3 | - | - | 0 | - |
| - | - | 1353 | 535.3 | - | - | 0 | - |
| - | - | 7678 | 536.3 | - | - | 0 | - |
| 9 | b | 664.2 | 536.8 | 0.006523 | 12.15 | +2 | 9 |
| - | - | 2626 | 537.2 | - | - | 0 | - |
| - | - | 4357 | 539.3 | - | - | 0 | - |
| - | - | 1239 | 540.3 | - | - | 0 | - |
| - | - | 1493 | 540.3 | - | - | 0 | - |
| - | - | 1167 | 541.3 | - | - | 0 | - |
| 9 | b | 5049 | 545.3 | 0.00039 | 0.7152 | +2 | 9 |
| - | - | 2988 | 545.8 | - | - | 0 | - |
| - | - | 935.8 | 548.3 | - | - | 0 | - |
| - | - | 2214 | 549.3 | - | - | 0 | - |
| - | - | 1180 | 551.3 | - | - | 0 | - |
| - | - | 674.8 | 552.3 | - | - | 0 | - |
| - | - | 2361 | 553.3 | - | - | 0 | - |
| - | - | 5.457E+04 | 554.3 | - | - | 0 | - |
| - | - | 1.64E+04 | 555.3 | - | - | 0 | - |
| - | - | 2695 | 556.3 | - | - | 0 | - |
| - | - | 4616 | 558.3 | - | - | 0 | - |
| - | - | 1152 | 559.3 | - | - | 0 | - |
| - | - | 1015 | 564.2 | - | - | 0 | - |
| - | - | 1021 | 567.3 | - | - | 0 | - |
| - | - | 8157 | 567.4 | - | - | 0 | - |
| - | - | 2880 | 568.4 | - | - | 0 | - |
| - | - | 2073 | 569.3 | - | - | 0 | - |
| - | - | 812 | 569.4 | - | - | 0 | - |
| 6 | y | 965.1 | 570.3 | 0.001049 | 1.839 | +1 | 5 |
| - | - | 616.7 | 571.3 | - | - | 0 | - |
| - | - | 692.7 | 572.3 | - | - | 0 | - |
| - | - | 1436 | 574.3 | - | - | 0 | - |
| - | - | 878.4 | 575.3 | - | - | 0 | - |
| - | - | 1017 | 580.3 | - | - | 0 | - |
| - | - | 883.9 | 580.8 | - | - | 0 | - |
| - | - | 3673 | 582.3 | - | - | 0 | - |
| - | - | 1737 | 583.3 | - | - | 0 | - |
| - | - | 698.1 | 584.3 | - | - | 0 | - |
| - | - | 1300 | 585.3 | - | - | 0 | - |
| 6 | y | 1.755E+04 | 588.3 | 0.000677 | 1.151 | +1 | 5 |
| - | - | 6735 | 589.3 | - | - | 0 | - |
| - | - | 665.9 | 590.3 | - | - | 0 | - |
| - | - | 9857 | 591.4 | - | - | 0 | - |
| - | - | 3110 | 592.4 | - | - | 0 | - |
| 0 | Precursor | 694.4 | 595.3 | 0.01012 | 17.01 | +2 | -1 |
| - | - | 3623 | 597.3 | - | - | 0 | - |
| - | - | 961.8 | 598.3 | - | - | 0 | - |
| - | - | 1790 | 599.3 | - | - | 0 | - |
| 5 | b | 2310 | 601.3 | 0.00133 | 2.212 | +1 | 5 |
| - | - | 1186 | 602.3 | - | - | 0 | - |
| - | - | 4148 | 603.4 | - | - | 0 | - |
| 0 | Precursor | 3258 | 603.8 | 0.0008787 | 1.455 | +2 | -1 |
| - | - | 2272 | 604.3 | - | - | 0 | - |
| - | - | 1240 | 605.3 | - | - | 0 | - |
| - | - | 1053 | 608.3 | - | - | 0 | - |
| - | - | 1823 | 615.3 | - | - | 0 | - |
| - | - | 1598 | 617.3 | - | - | 0 | - |
| 5 | b | 3.134E+04 | 619.3 | 0.0008973 | 1.449 | +1 | 5 |
| - | - | 1.179E+04 | 620.3 | - | - | 0 | - |
| - | - | 1798 | 621.4 | - | - | 0 | - |
| - | - | 3335 | 623.3 | - | - | 0 | - |
| - | - | 944 | 624.3 | - | - | 0 | - |
| - | - | 1747 | 624.3 | - | - | 0 | - |
| - | - | 2772 | 625.3 | - | - | 0 | - |
| - | - | 1242 | 629.3 | - | - | 0 | - |
| - | - | 783.4 | 631.3 | - | - | 0 | - |
| - | - | 4634 | 633.3 | - | - | 0 | - |
| - | - | 1883 | 634.3 | - | - | 0 | - |
| - | - | 1061 | 635.3 | - | - | 0 | - |
| - | - | 936.9 | 636.3 | - | - | 0 | - |
| - | - | 1167 | 636.4 | - | - | 0 | - |
| - | - | 1018 | 638.3 | - | - | 0 | - |
| - | - | 772.7 | 639.4 | - | - | 0 | - |
| - | - | 980.9 | 646.4 | - | - | 0 | - |
| - | - | 7211 | 651.3 | - | - | 0 | - |
| - | - | 1146 | 651.4 | - | - | 0 | - |
| - | - | 6389 | 652.3 | - | - | 0 | - |
| - | - | 2284 | 652.3 | - | - | 0 | - |
| - | - | 1832 | 653.3 | - | - | 0 | - |
| - | - | 7263 | 653.3 | - | - | 0 | - |
| - | - | 3065 | 654.3 | - | - | 0 | - |
| - | - | 2439 | 663.3 | - | - | 0 | - |
| - | - | 814.7 | 664.3 | - | - | 0 | - |
| - | - | 2149 | 664.4 | - | - | 0 | - |
| - | - | 806.1 | 665.4 | - | - | 0 | - |
| - | - | 2712 | 667.3 | - | - | 0 | - |
| - | - | 1270 | 668.4 | - | - | 0 | - |
| - | - | 903.1 | 669.4 | - | - | 0 | - |
| - | - | 8213 | 681.3 | - | - | 0 | - |
| - | - | 2032 | 682.3 | - | - | 0 | - |
| - | - | 1.919E+04 | 682.4 | - | - | 0 | - |
| - | - | 1003 | 683.3 | - | - | 0 | - |
| - | - | 6418 | 683.4 | - | - | 0 | - |
| - | - | 1102 | 684.4 | - | - | 0 | - |
| - | - | 878.7 | 695.3 | - | - | 0 | - |
| - | - | 708.2 | 696.3 | - | - | 0 | - |
| - | - | 731.2 | 700.4 | - | - | 0 | - |
| - | - | 666.5 | 702.4 | - | - | 0 | - |
| - | - | 2288 | 703.4 | - | - | 0 | - |
| - | - | 1350 | 714.4 | - | - | 0 | - |
| - | - | 4962 | 716.4 | - | - | 0 | - |
| - | - | 1717 | 717.4 | - | - | 0 | - |
| - | - | 1824 | 720.4 | - | - | 0 | - |
| - | - | 3353 | 721.4 | - | - | 0 | - |
| - | - | 2340 | 722.4 | - | - | 0 | - |
| - | - | 1585 | 734.4 | - | - | 0 | - |
| - | - | 1119 | 735.3 | - | - | 0 | - |
| - | - | 816.2 | 737.3 | - | - | 0 | - |
| - | - | 1.402E+04 | 738.4 | - | - | 0 | - |
| - | - | 5762 | 739.4 | - | - | 0 | - |
| - | - | 1432 | 740.4 | - | - | 0 | - |
| - | - | 906.8 | 742.3 | - | - | 0 | - |
| - | - | 1159 | 744.3 | - | - | 0 | - |
| 6 | b | 6621 | 748.4 | 0.0004944 | 0.6606 | +1 | 6 |
| 6 | b | 3516 | 749.4 | 0.01476 | 19.69 | +1 | 6 |
| - | - | 2141 | 750.4 | - | - | 0 | - |
| 5 | y | 1649 | 751.4 | 0.008149 | 10.85 | +1 | 6 |
| - | - | 1.088E+04 | 752.4 | - | - | 0 | - |
| - | - | 4697 | 753.4 | - | - | 0 | - |
| - | - | 1779 | 754.4 | - | - | 0 | - |
| - | - | 1437 | 760.4 | - | - | 0 | - |
| - | - | 754.5 | 761.4 | - | - | 0 | - |
| - | - | 6678 | 762.3 | - | - | 0 | - |
| - | - | 3418 | 763.3 | - | - | 0 | - |
| - | - | 1267 | 764.4 | - | - | 0 | - |
| - | - | 1044 | 765.3 | - | - | 0 | - |
| 6 | b | 1.12E+05 | 766.4 | 0.0007817 | 1.02 | +1 | 6 |
| - | - | 5.266E+04 | 767.4 | - | - | 0 | - |
| - | - | 1141 | 767.5 | - | - | 0 | - |
| - | - | 1.311E+04 | 768.4 | - | - | 0 | - |
| - | - | 1330 | 769.4 | - | - | 0 | - |
| - | - | 812 | 778.4 | - | - | 0 | - |
| - | - | 6.884E+04 | 780.4 | - | - | 0 | - |
| - | - | 3.146E+04 | 781.4 | - | - | 0 | - |
| - | - | 7597 | 782.4 | - | - | 0 | - |
| - | - | 883.4 | 783.4 | - | - | 0 | - |
| - | - | 938.1 | 783.4 | - | - | 0 | - |
| - | - | 1702 | 784.4 | - | - | 0 | - |
| - | - | 954.8 | 785.4 | - | - | 0 | - |
| - | - | 727.9 | 799.4 | - | - | 0 | - |
| - | - | 1253 | 811.4 | - | - | 0 | - |
| - | - | 3167 | 817.4 | - | - | 0 | - |
| - | - | 1686 | 818.4 | - | - | 0 | - |
| - | - | 1265 | 822.4 | - | - | 0 | - |
| - | - | 1257 | 827.4 | - | - | 0 | - |
| - | - | 1038 | 828.4 | - | - | 0 | - |
| - | - | 3714 | 829.4 | - | - | 0 | - |
| - | - | 1231 | 830.4 | - | - | 0 | - |
| - | - | 2069 | 835.5 | - | - | 0 | - |
| - | - | 1159 | 836.5 | - | - | 0 | - |
| 7 | b | 5429 | 845.5 | 0.01426 | 16.86 | +1 | 7 |
| 7 | b | 3530 | 846.4 | 0.004535 | 5.358 | +1 | 7 |
| - | - | 1625 | 847.4 | - | - | 0 | - |
| - | - | 4007 | 850.4 | - | - | 0 | - |
| - | - | 2917 | 851.4 | - | - | 0 | - |
| - | - | 1757 | 852.4 | - | - | 0 | - |
| - | - | 1474 | 861.4 | - | - | 0 | - |
| 7 | b | 9917 | 863.5 | 0.0005081 | 0.5884 | +1 | 7 |
| - | - | 4607 | 864.5 | - | - | 0 | - |
| - | - | 982.6 | 865.5 | - | - | 0 | - |
| - | - | 3599 | 877.4 | - | - | 0 | - |
| - | - | 1542 | 878.4 | - | - | 0 | - |
| - | - | 7019 | 879.4 | - | - | 0 | - |
| - | - | 3870 | 880.4 | - | - | 0 | - |
| - | - | 1451 | 881.4 | - | - | 0 | - |
| - | - | 2802 | 889.4 | - | - | 0 | - |
| - | - | 1011 | 890.4 | - | - | 0 | - |
| - | - | 1739 | 893.4 | - | - | 0 | - |
| - | - | 656.6 | 894.4 | - | - | 0 | - |
| - | - | 1209 | 895.5 | - | - | 0 | - |
| - | - | 7055 | 907.4 | - | - | 0 | - |
| - | - | 4104 | 908.4 | - | - | 0 | - |
| - | - | 929.9 | 909.4 | - | - | 0 | - |
| - | - | 1101 | 928.5 | - | - | 0 | - |
| - | - | 2207 | 946.5 | - | - | 0 | - |
| - | - | 2608 | 947.5 | - | - | 0 | - |
| - | - | 1616 | 948.5 | - | - | 0 | - |
| - | - | 1898 | 956.5 | - | - | 0 | - |
| - | - | 1745 | 957.5 | - | - | 0 | - |
| - | - | 859 | 958.5 | - | - | 0 | - |
| - | - | 2.284E+04 | 964.5 | - | - | 0 | - |
| - | - | 1.233E+04 | 965.5 | - | - | 0 | - |
| - | - | 3574 | 966.5 | - | - | 0 | - |
| 8 | b | 1.172E+04 | 974.5 | 3.486E-05 | 0.03577 | +1 | 8 |
| 8 | b | 6523 | 975.5 | 0.01608 | 16.48 | +1 | 8 |
| 3 | y | 7195 | 976.5 | 0.005206 | 5.332 | +1 | 8 |
| - | - | 625.3 | 977.4 | - | - | 0 | - |
| - | - | 3015 | 977.5 | - | - | 0 | - |
| - | - | 1348 | 978.5 | - | - | 0 | - |
| - | - | 866.6 | 990.5 | - | - | 0 | - |
| 8 | b | 2.014E+05 | 992.5 | 0.0001513 | 0.1525 | +1 | 8 |
| - | - | 1.229E+05 | 993.5 | - | - | 0 | - |
| 3 | y | 7.362E+04 | 994.5 | 0.01082 | 10.88 | +1 | 8 |
| - | - | 3.35E+04 | 995.5 | - | - | 0 | - |
| - | - | 9417 | 996.5 | - | - | 0 | - |
| - | - | 1727 | 997.5 | - | - | 0 | - |
| 2 | y | 4750 | 1094 | 0.0005301 | 0.4847 | +1 | 9 |
| - | - | 3480 | 1095 | - | - | 0 | - |
| - | - | 1160 | 1096 | - | - | 0 | - |
| - | - | 744.3 | 1712 | - | - | 0 | - |

m/z Charge Intensity FragmentType MassShift Position
120.0811538696289 0 36563.62
121.06529998779297 0 394.3606
121.0844955444336 0 2939.3406
127.04984283447266 0 381.5974
127.08709716796875 0 1079.8042
129.06639099121094 0 797.32336
129.10264587402344 0 69409.08
130.05007934570312 0 488.33987
130.06570434570312 0 1010.1256
130.10598754882812 0 4568.8613
131.045166015625 0 505.73276
132.08111572265625 0 553.93225
132.10223388671875 0 785.3329
132.7752685546875 0 410.20193
133.06106567382812 0 761.0011
133.0865020751953 0 595.6678
134.09698486328125 0 433.99265
136.07608032226562 0 15904.34
137.0795135498047 0 767.4844
139.0869598388672 0 2079.4023
140.0902862548828 0 561.40894
141.10260009765625 0 974.1441
143.08203125 0 522.357
147.0444793701172 0 675.67737
148.9532928466797 0 828.7505
153.06646728515625 0 573.9167
155.0821075439453 0 803.60974
155.11822509765625 0 1027.2715
155.50643920898438 0 534.2746
156.1136932373047 0 808.1099
157.0974884033203 0 1025.7631
159.09213256835938 0 649.485
159.1131134033203 0 669.87146
165.1028594970703 0 1245.397
166.32249450683594 0 466.2391
167.08184814453125 0 1732.0403
167.11831665039062 0 2054.0833
168.10235595703125 0 1839.2708
169.09759521484375 0 1121.5122
169.10597229003906 0 527.22906
169.13392639160156 0 18790.402
170.1372528076172 0 2308.9077
170.91209411621094 0 447.11874
172.11233520507812 0 4927.75
172.14488220214844 0 833.2235
173.0921630859375 0 1117.3772
173.1288604736328 0 1153.7372
173.4384307861328 0 950.30365
174.1280517578125 0 2212.2498
175.08651733398438 0 787.52057
177.10275268554688 0 495.24652
180.72451782226562 0 506.36874
181.09759521484375 0 7298.9595
181.1339874267578 0 4978.3994
182.08187866210938 0 556.8696
182.1007537841797 0 600.87195
182.129150390625 0 922.6462
182.13719177246094 0 661.45074
183.1132049560547 0 1768.6357
183.14959716796875 0 2718.0437
184.10873413085938 0 734.123
185.09222412109375 0 1706.6198
185.12887573242188 0 1723.4637
185.16525268554688 0 77390.586 a 1
186.1281280517578 0 2999.0513
186.16860961914062 0 7233.3066
187.10809326171875 0 1296.0653
187.1441192626953 0 748.1263
188.1073455810547 0 1505.8806
188.14378356933594 0 3278.0708
190.12303161621094 0 899.91766
191.08224487304688 0 918.1375
195.11314392089844 0 6610.922
197.09266662597656 0 931.13824
197.12884521484375 0 3403.8486
197.16567993164062 0 1167.7059
198.1240997314453 0 842.7746
199.1081085205078 0 25184.254
200.1109161376953 0 1777.6123
201.12359619140625 0 2104.6047
202.1225128173828 0 752.8024
203.1032257080078 0 2051.5188
203.15499877929688 0 1852.3218
204.1385955810547 0 2137.567
209.09243774414062 0 16336.655
209.12855529785156 0 582.1775
210.0966796875 0 1082.7865
210.1603240966797 0 1417.6909
211.1443634033203 0 1769.3043
212.13963317871094 0 2975.7322
213.08755493164062 0 766.73175
213.1238555908203 0 3714.124
213.16021728515625 0 28760.258 b 1
214.16354370117188 0 2831.4497
215.13954162597656 0 893076.25 y 8
216.14279174804688 0 90947.086
217.13365173339844 0 4085.596
217.14466857910156 0 6751.945
223.10800170898438 0 808.75085
225.0874786376953 0 1480.6135
226.11911010742188 0 20303.258
226.15597534179688 0 965.70917
227.10308837890625 0 66396.74
227.1232452392578 0 1748.1697
228.1064453125 0 6883.5596
228.17120361328125 0 6708.0903
229.1072540283203 0 849.19244
229.15509033203125 0 4585.517
230.18692016601562 0 1641.2799
231.0987548828125 0 707.9253
231.14964294433594 0 15383.575
232.1531219482422 0 2434.6091
233.0924530029297 0 960.4402
233.16546630859375 0 2061.3962
238.123046875 0 1533.553
238.15562438964844 0 2837.1494
239.1589813232422 0 760.46924
240.13356018066406 0 789.4263
243.13453674316406 0 651.2976
244.12965393066406 0 33497.8
245.1144256591797 0 1018.1718
245.12974548339844 0 8423.493
246.1334686279297 0 1237.9175
246.88633728027344 0 583.1981
247.14471435546875 0 2878.8442
249.16000366210938 0 1500.3483
251.1031036376953 0 23291.516
252.10638427734375 0 2769.5974
254.11375427246094 0 3767.5024
254.15048217773438 0 1167.1996
256.16607666015625 0 2014.1166
257.12890625 0 826.0354
259.1429748535156 0 694
261.0878601074219 0 817.33374
261.1226806640625 0 600.17377
261.1589660644531 0 3479.7524
262.07098388671875 0 1054.6405
262.15936279296875 0 677.5504
265.1562805175781 0 707.57227
266.1502380371094 0 628.25165
269.1855163574219 0 953.956
270.1805725097656 0 727.6685
272.1244201660156 0 5833.384
273.12786865234375 0 929.0393
276.17144775390625 0 879.99963
277.15509033203125 0 1430.464
278.15069580078125 0 1029.6373
279.0980224609375 0 23050.752
280.1015930175781 0 3478.5745
280.1656799316406 0 1249.0656
283.1445617675781 0 10877.6875
283.17315673828125 0 699.1228
284.1478576660156 0 1450.4258
286.1385192871094 0 637.33344
287.2121276855469 0 2900.505
292.12890625 0 588.96466
292.1667785644531 0 900.6013
294.1817321777344 0 1437.5653
296.12469482421875 0 987.5114
301.15496826171875 0 1965.3052
303.20660400390625 0 3335.199
304.1661376953125 0 1513.3145
306.1468200683594 0 674.7142
306.2183837890625 0 773.3762
308.1607971191406 0 1860.1588
309.1568603515625 0 1115.8693
310.1403503417969 0 1006.0259
310.1771545410156 0 2657.528 b 4
310.2120056152344 0 804.9571
311.1397705078125 0 7589.395
312.1431884765625 0 1538.8325
312.19189453125 0 921.0063
313.192138671875 0 1883.2301
315.2079162597656 0 2204.706
320.1612548828125 0 652.5035
321.12408447265625 0 640.9263
322.1756591796875 0 663.21063
323.1719055175781 0 1346.0668
323.2445068359375 0 2352.7837
324.1559753417969 0 14875.404
324.22998046875 0 681.1281 b Ammonia loss 2
325.16009521484375 0 2598.1719
325.187744140625 0 2941.2021
325.2237854003906 0 1048.467
326.1716613769531 0 8820.357 y Water loss 7
326.1922912597656 0 692.6497
327.1740417480469 0 1785.1969
328.14813232421875 0 1036.5519
328.1681213378906 0 1086.52
329.1504821777344 0 2263.0312
330.18359375 0 552.64575
331.2022399902344 0 1454.7852
335.1745910644531 0 1525.9688
336.17657470703125 0 612.12683
337.1514892578125 0 2793.103
337.18817138671875 0 1148.808
338.1520690917969 0 802.1878
339.2035827636719 0 1284.1388
340.1875 0 5052.5
341.1834716796875 0 5022.522
341.2555236816406 0 5918.503 b 2
342.1304016113281 0 1213.0905
342.1855773925781 0 995.7452
342.2591247558594 0 963.9299
343.1982116699219 0 8285.141
344.1614074707031 0 1288.2506
344.1822509765625 0 5507.618 y 7
344.20233154296875 0 998.53296
344.7054138183594 0 1740.4268
345.1875 0 1223.3938
346.1767883300781 0 4875.802
347.2027587890625 0 4199.9214
347.7043151855469 0 1411.7834
349.1906433105469 0 1305.0713
353.1539001464844 0 520.11145
353.1824035644531 0 5364.999
353.21917724609375 0 2455.9343
354.185546875 0 1146.5808
354.2213134765625 0 751.18256
355.1617736816406 0 4564.6196
355.1978454589844 0 2784.9973
356.1631164550781 0 1643.9845
356.1998291015625 0 875.53033
358.21148681640625 0 694.11414
359.12249755859375 0 671.32416
360.2271728515625 0 755.85205
361.6967468261719 0 667.87994
362.1719055175781 0 1887.9246
363.1768493652344 0 677.39087
363.2029724121094 0 875.7892
365.1827087402344 0 718.79236
367.1986389160156 0 931.94727
369.1766662597656 0 1102.3844
369.71337890625 0 6159.5957
370.1793212890625 0 727.4463
370.21466064453125 0 3565.4568
370.7156677246094 0 1213.9468
371.1929626464844 0 7229.2314
372.1562194824219 0 2056.957
372.19549560546875 0 1252.3143
372.2237548828125 0 633.292
373.1873779296875 0 2426.154
374.1716613769531 0 12325.417
375.17437744140625 0 1658.645
375.2408142089844 0 680.8872
376.15350341796875 0 645.4824
379.19805908203125 0 6916.775
380.1980285644531 0 819.672
381.14459228515625 0 2810.4714
382.14874267578125 0 785.2838
383.158203125 0 998.80676
383.193115234375 0 2866.276
383.7110900878906 0 1133.0785 b 5
389.1825866699219 0 7851.6523
390.16510009765625 0 1670.9268
390.1890869140625 0 983.3766
391.16748046875 0 1210.3251
391.1980895996094 0 14597.085
392.1811828613281 0 2883.9736
392.2031555175781 0 1675.3715
403.2337341308594 0 2952.0469
404.238037109375 0 1150.8885
406.2109680175781 0 736.5975
407.1932373046875 0 15325.468
408.15606689453125 0 866.49524
408.1957702636719 0 3699.816
408.2236633300781 0 962.5461
409.2049255371094 0 1207.8518
411.2607421875 0 1864.1753
415.2274169921875 0 785.2794
417.17889404296875 0 1017.28723
418.24017333984375 0 9574.75
418.7419128417969 0 8397.833
419.1935119628906 0 1202.7506
419.2416687011719 0 1212.9513
420.29736328125 0 711.7391
421.2452392578125 0 704.5327
422.20672607421875 0 774.6218
423.2247009277344 0 16877.574 b Water loss 6
424.2239074707031 0 8114.498
425.226318359375 0 1603.7483
426.1667785644531 0 21253.615
427.1700744628906 0 5159.6357
428.1739807128906 0 729.48505
432.2373046875 0 5764.3716 b 6
432.7388000488281 0 3470.8975
433.24114990234375 0 1108.6887
434.2048645019531 0 1465.8143
434.2767028808594 0 614.5636
435.1881408691406 0 3758.6226
436.15167236328125 0 652.1474
436.188720703125 0 1630.8087
436.25628662109375 0 786.6183
438.2725524902344 0 8375.862 b Water loss 3
438.3091735839844 0 1553.6849
439.27691650390625 0 1729.2485
440.2503967285156 0 1347.7345
441.2352294921875 0 53892.72 y 6
442.23834228515625 0 10765.068
443.2380676269531 0 2131.95
445.244873046875 0 1575.1737
452.2151184082031 0 5169.962
452.28704833984375 0 862.1748
453.2176208496094 0 792.7177
454.231201171875 0 910.4697
454.2684020996094 0 1842.577
456.2825927734375 0 27749.8 b 3
457.2461242675781 0 1686.372
457.2852478027344 0 7813.839
458.2836608886719 0 1002.5497
460.24530029296875 0 823.93353
464.215576171875 0 888.7651
464.2503967285156 0 1531.3597
465.2541198730469 0 966.1345
468.24530029296875 0 1439.5569
469.2458801269531 0 1043.2449
470.2253112792969 0 14340.095
471.2278747558594 0 4619.712
473.20263671875 0 1542.4663
473.2391357421875 0 687.6008
473.3070068359375 0 1009.01227
474.24090576171875 0 788.36163
476.25140380859375 0 1520.4407
478.26690673828125 0 1482.9146
480.2111511230469 0 780.0808
482.2618408203125 0 7930.8066
482.76171875 0 4905.4277
483.22686767578125 0 877.4945
483.2642822265625 0 4949.28
483.7640380859375 0 983.3794
486.23455810546875 0 1199.199
487.2543640136719 0 867.93805
488.25201416015625 0 1382.019 b Ammonia loss 7
488.3242492675781 0 2866.9666
490.2677001953125 0 1551.1101
491.21124267578125 0 764.5875
492.21795654296875 0 726.6362
495.2273864746094 0 684.42883
496.75872802734375 0 5789.67 b 7
497.259765625 0 2635.7212
497.7601013183594 0 1414.4622
498.2214660644531 0 1379.8525
500.21490478515625 0 935.3212
500.2530517578125 0 1241.6349
502.26708984375 0 853.0287
504.2468566894531 0 1837.2871
504.31817626953125 0 1034.5696
505.1939392089844 0 803.97894
506.2616882324219 0 3185.9902
507.2656555175781 0 856.90753
508.2557373046875 0 3085.2388
509.2407531738281 0 4152.5835
510.24298095703125 0 1915.8154
516.2469482421875 0 1218.7867
518.226318359375 0 1766.9514
518.2625122070312 0 1642.7444
519.2293090820312 0 764.3778
520.2787475585938 0 937.03265
521.2752075195312 0 1714.3612
522.2821044921875 0 627.9166
523.2195434570312 0 2204.103
523.2921142578125 0 825.3598
524.2227783203125 0 704.00885
525.2362670898438 0 752.3787
526.2669677734375 0 12497.028
527.2703857421875 0 2563.998
531.2846069335938 0 749.6073
531.3301391601562 0 855.7991
531.7884521484375 0 644.3266
534.2567138671875 0 6833.072
535.2597045898438 0 1352.6646
536.2503662109375 0 7678.4233
536.7776489257812 0 664.24396 b Ammonia loss 8
537.2432861328125 0 2625.997
539.2505493164062 0 4357.0137
540.254150390625 0 1238.9387
540.3038330078125 0 1493.3073
541.3033447265625 0 1167.4255
545.2847900390625 0 5049.0156 b 8
545.7862548828125 0 2987.6416
548.2720336914062 0 935.7861
549.3392944335938 0 2214.045
551.2820434570312 0 1179.8059
552.28857421875 0 674.8172
553.3363037109375 0 2361.39
554.261474609375 0 54573.785
555.2645263671875 0 16403.615
556.2671508789062 0 2694.7349
558.3291625976562 0 4615.8213
559.3320922851562 0 1151.5822
564.244873046875 0 1015.28394
567.3131103515625 0 1021.49506
567.3510131835938 0 8157.2036
568.3539428710938 0 2880.1367
569.2953491210938 0 2073.0579
569.35498046875 0 812.0282
570.2932739257812 0 965.0839 y Water loss 5
571.2866821289062 0 616.7146
572.2689819335938 0 692.71484
574.328857421875 0 1436.1642
575.322509765625 0 878.43604
580.3229370117188 0 1016.618
580.82080078125 0 883.9278
582.2562866210938 0 3672.5244
583.259033203125 0 1737.1123
584.3091430664062 0 698.0715
585.336181640625 0 1300.2466
588.303466796875 0 17545.38 y 5
589.3070678710938 0 6735.2354
590.3099975585938 0 665.9217
591.350830078125 0 9856.773
592.3544921875 0 3109.575
595.3207397460938 0 694.3927 Precursor Ammonia loss
597.2882690429688 0 3623.2227
598.2909545898438 0 961.8343
599.2816772460938 0 1789.9137
601.3357543945312 0 2310.4788 b Water loss 4
602.334228515625 0 1186.0731
603.3500366210938 0 4148.019
603.8247680664062 0 3257.8567 Precursor
604.32470703125 0 2272.426
605.2952270507812 0 1239.6066
608.3110961914062 0 1053.1708
615.2775268554688 0 1823.1245
617.2922973632812 0 1598.2681
619.3458862304688 0 31343.455 b 4
620.3484497070312 0 11793.596
621.3502807617188 0 1798.4408
623.3209228515625 0 3334.7712
624.26806640625 0 944.02893
624.3234252929688 0 1747.1212
625.3349609375 0 2771.9143
629.3298950195312 0 1242.326
631.3056030273438 0 783.4137
633.2882690429688 0 4633.5137
634.2938842773438 0 1883.0532
635.318115234375 0 1061.3162
636.3170776367188 0 936.9246
636.3744506835938 0 1167.4547
638.2760620117188 0 1018.32544
639.3519897460938 0 772.7368
646.3564453125 0 980.89557
651.3143310546875 0 7210.5347
651.379638671875 0 1146.1248
652.26171875 0 6389.189
652.3192138671875 0 2283.719
653.2649536132812 0 1831.8405
653.329345703125 0 7263.496
654.3323974609375 0 3064.836
663.3141479492188 0 2438.7256
664.3093872070312 0 814.65027
664.36767578125 0 2149.3953
665.3681640625 0 806.06146
667.347900390625 0 2712.157
668.3519287109375 0 1269.5706
669.4014892578125 0 903.1041
681.3242797851562 0 8212.879
682.3251953125 0 2032.1312
682.3779907226562 0 19189.84
683.317138671875 0 1003.4275
683.3807983398438 0 6417.6294
684.3804321289062 0 1101.958
695.3397216796875 0 878.723
696.3333129882812 0 708.2147
700.4002685546875 0 731.2295
702.37890625 0 666.48895
703.3693237304688 0 2287.9277
714.4180297851562 0 1349.6049
716.3995361328125 0 4961.845
717.400390625 0 1717.421
720.405029296875 0 1824.2631
721.3949584960938 0 3353.3252
722.3947143554688 0 2340.3142
734.354248046875 0 1585.4202
735.3446655273438 0 1119.1956
737.3490600585938 0 816.1588
738.4191284179688 0 14016.989
739.4219360351562 0 5762.4165
740.4237670898438 0 1432.1016
742.34228515625 0 906.8164
744.3385009765625 0 1158.6083
748.40234375 0 6620.866 b Water loss 5
749.401611328125 0 3515.9258 b Ammonia loss 5
750.3880004882812 0 2141.0105
751.374267578125 0 1649.2151 y 4
752.3621215820312 0 10883.852
753.3652954101562 0 4697.018
754.366943359375 0 1778.5127
760.3568115234375 0 1436.648
761.3507690429688 0 754.51495
762.3468627929688 0 6677.5894
763.3490600585938 0 3417.7478
764.4055786132812 0 1267.2059
765.347900390625 0 1043.7668
766.4141845703125 0 111959.73 b 5
767.4169921875 0 52664.113
767.5005493164062 0 1140.7577
768.4198608398438 0 13113.959
769.4263305664062 0 1329.8102
778.3765258789062 0 811.98987
780.35693359375 0 68836.805
781.3599243164062 0 31462.188
782.3628540039062 0 7596.5244
783.359375 0 883.352
783.4395751953125 0 938.0937
784.425048828125 0 1701.853
785.4314575195312 0 954.8068
799.4359741210938 0 727.8677
811.4326782226562 0 1252.7208
817.4470825195312 0 3167.4668
818.4478759765625 0 1685.5668
822.438232421875 0 1264.9664
827.429443359375 0 1257.2921
828.4371948242188 0 1038.264
829.4449462890625 0 3714.1084
830.4469604492188 0 1230.8777
835.470458984375 0 2068.8208
836.4723510742188 0 1158.7773
845.4413452148438 0 5428.742 b Water loss 6
846.4441528320312 0 3530.438 b Ammonia loss 6
847.449951171875 0 1624.6256
850.4346923828125 0 4006.6028
851.4354248046875 0 2917.4478
852.4360961914062 0 1757.1283
861.4130859375 0 1474.0459
863.4666748046875 0 9916.73 b 6
864.4697265625 0 4607.2324
865.4689331054688 0 982.631
877.41162109375 0 3599.4175
878.4129028320312 0 1541.8323
879.4251098632812 0 7019.1865
880.4281005859375 0 3870.4036
881.4353637695312 0 1451.1538
889.4119262695312 0 2801.7163
890.4110107421875 0 1010.8433
893.4420166015625 0 1739.3512
894.4371948242188 0 656.5829
895.4547729492188 0 1208.7767
907.4190673828125 0 7055.201
908.4214477539062 0 4103.535
909.4220581054688 0 929.8514
928.4928588867188 0 1101.29
946.5056762695312 0 2206.5845
947.4935302734375 0 2607.7234
948.5001831054688 0 1615.749
956.4876708984375 0 1898.4598
957.483154296875 0 1744.9017
958.486083984375 0 858.9912
964.5134887695312 0 22840.035
965.5172119140625 0 12327.661
966.5193481445312 0 3574.1448
974.4982299804688 0 11720.239 b Water loss 7
975.498291015625 0 6522.76 b Ammonia loss 7
976.482666015625 0 7195.023 y Water loss 2
977.3609008789062 0 625.32544
977.4820556640625 0 3015.246
978.482177734375 0 1347.7225
990.4904174804688 0 866.5712
992.5089111328125 0 201369.8 b 7
993.5120239257812 0 122896.945
994.4988403320312 0 73616.18 y 2
995.49462890625 0 33496.945
996.4942016601562 0 9416.556
997.4951782226562 0 1726.8801
1093.555908203125 0 4749.9927 y 1
1094.5589599609375 0 3479.664
1095.562255859375 0 1159.66
1712.379638671875 0 744.2931

Spectrum Details

|  |  |
| --- | --- |
| Matched peaks? Matched peaks The total absolute number of peaks matched. Additionally in brackets the total fraction of peaks matched and the total number of peaks is shown. | 37 (6.51% of 568) |
| FDR? FDR The false discovery rate estimated for this peptide. It is calculated by matching all theoretical fragments with a non-integer shift with the raw peaks for this spectrum. This is done with 40 different shifts. The resulting percentage is the average number of annotated peaks over the number of annotated peaks with the correct spectrum. | 1.03% |
| Satellite FDR? Satellite FDR See the FDR for details on its calculation. This satellite ion specific FDR only contains the satellite ions (d/w) for I/L/J positions. | - |
| PSM Score? PSM Score The PSM Score as given by Hecklib to this annotated spectrum. It is shown with three significant figures. | 440 |

#### Spectrum 8375? Spectrum 8375 The raw spectrum of this peptide as annotated by Hecklib. The fragments are coloured according to ion type (see legend). Any peaks with a star '\*' as text can be hovered over to see the full details, first the ion type second the mass shift type. By hovering over the amino acids in the peptide or ions in the legend the corresponding peaks are highlighted. By toggling the 'Unassigned' label you can turn the background (unassigned) peaks on or off in the plot. By updating the slider in the Ion legend you can update the spectrum to only show the top X% of the peaks with labels. The top X% means any peak that is within X% of the highest intensity. By dragging in the spectrum you can zoom in to a specific part of the spectrum and use 'Zoom Out' to get back to the original zoom level. The annotation of the spectrum is based on the given sequence in the peptides file and is done with different software so inconsistencies are likely. The peaks are annotated based on the given sequence, with 20 ppm tolerance.

Copy Data

##### Spectrum 8375 (TSV)

###### Preview

```
Loading example...
```

*Click on the button to copy the data to your clipboard.*

Mz MinMz MaxIntensity Max

WidthHeightPeptide font sizePeptide stroke widthSpectrum font sizeSpectrum stroke widthCompact peptide

Ion legend

wxyz

abcd

OtherUnassignedIon^Charge^_Position_Show for top:%

JVKDYFPEPV

01.28e+62.57e+63.85e+65.14e+6

Zoom Out

a^+1^_2_b^+1^_2_y^+1^_2_b^+2^_5_y^+1^_3_b^+1^_3_y^+1^_3_b^+2^_6_b^+2^_7_b^+2^_7_b^+1^_4_b^+1^_4_y^+1^_4_b^+1^_4_b^+2^_8_b^+2^_8_b^+2^_9_y^+1^_5_\*\*b^+1^_5_\*b^+1^_5_b^+1^_6_b^+1^_6_y^+1^_6_b^+1^_6_b^+1^_7_b^+1^_7_b^+1^_7_b^+1^_8_b^+1^_8_y^+1^_8_b^+1^_8_y^+1^_8_b^+1^_9_y^+1^_9_

0856171125673423

Fragment Matches Table

Show background peaks

| Position | Ion type | Intensity | mz Theoretical | mz Error (Th) | mz Error (ppm) | Charge | Series Number |
| --- | --- | --- | --- | --- | --- | --- | --- |
| - | - | 1.97E+05 | 120.1 | - | - | 0 | - |
| - | - | 1.674E+04 | 121.1 | - | - | 0 | - |
| - | - | 2495 | 128.1 | - | - | 0 | - |
| - | - | 3.533E+05 | 129.1 | - | - | 0 | - |
| - | - | 3460 | 130.1 | - | - | 0 | - |
| - | - | 2.132E+04 | 130.1 | - | - | 0 | - |
| - | - | 3950 | 134.1 | - | - | 0 | - |
| - | - | 2320 | 134.9 | - | - | 0 | - |
| - | - | 7.855E+04 | 136.1 | - | - | 0 | - |
| - | - | 4342 | 137.1 | - | - | 0 | - |
| - | - | 1.075E+04 | 139.1 | - | - | 0 | - |
| - | - | 3781 | 147 | - | - | 0 | - |
| - | - | 4979 | 155.1 | - | - | 0 | - |
| - | - | 5151 | 156.1 | - | - | 0 | - |
| - | - | 8155 | 165.1 | - | - | 0 | - |
| - | - | 8284 | 167.1 | - | - | 0 | - |
| - | - | 1.123E+04 | 167.1 | - | - | 0 | - |
| - | - | 1.052E+04 | 168.1 | - | - | 0 | - |
| - | - | 1.171E+05 | 169.1 | - | - | 0 | - |
| - | - | 1.015E+04 | 170.1 | - | - | 0 | - |
| - | - | 2.871E+04 | 172.1 | - | - | 0 | - |
| - | - | 4231 | 172.1 | - | - | 0 | - |
| - | - | 1.06E+04 | 173.4 | - | - | 0 | - |
| - | - | 7632 | 174.1 | - | - | 0 | - |
| - | - | 2916 | 175.1 | - | - | 0 | - |
| - | - | 5044 | 177.1 | - | - | 0 | - |
| - | - | 3.764E+04 | 181.1 | - | - | 0 | - |
| - | - | 2.291E+04 | 181.1 | - | - | 0 | - |
| - | - | 2874 | 182.1 | - | - | 0 | - |
| - | - | 6687 | 183.1 | - | - | 0 | - |
| - | - | 1.759E+04 | 183.1 | - | - | 0 | - |
| - | - | 5179 | 184.1 | - | - | 0 | - |
| - | - | 1.035E+04 | 185.1 | - | - | 0 | - |
| 2 | a | 4.533E+05 | 185.2 | 0.0003519 | 1.901 | +1 | 2 |
| - | - | 1.789E+04 | 186.1 | - | - | 0 | - |
| - | - | 4.7E+04 | 186.2 | - | - | 0 | - |
| - | - | 4618 | 187.1 | - | - | 0 | - |
| - | - | 8513 | 188.1 | - | - | 0 | - |
| - | - | 1.711E+04 | 188.1 | - | - | 0 | - |
| - | - | 3970 | 190.1 | - | - | 0 | - |
| - | - | 9169 | 191.1 | - | - | 0 | - |
| - | - | 3.041E+04 | 195.1 | - | - | 0 | - |
| - | - | 3169 | 196.1 | - | - | 0 | - |
| - | - | 1.607E+04 | 197.1 | - | - | 0 | - |
| - | - | 3641 | 197.2 | - | - | 0 | - |
| - | - | 7208 | 198.1 | - | - | 0 | - |
| - | - | 1.432E+05 | 199.1 | - | - | 0 | - |
| - | - | 9821 | 200.1 | - | - | 0 | - |
| - | - | 4053 | 201.1 | - | - | 0 | - |
| - | - | 5250 | 202.1 | - | - | 0 | - |
| - | - | 6078 | 203.2 | - | - | 0 | - |
| - | - | 1E+04 | 204.1 | - | - | 0 | - |
| - | - | 9.107E+04 | 209.1 | - | - | 0 | - |
| - | - | 6961 | 210.1 | - | - | 0 | - |
| - | - | 5877 | 210.2 | - | - | 0 | - |
| - | - | 2549 | 210.8 | - | - | 0 | - |
| - | - | 3430 | 211.1 | - | - | 0 | - |
| - | - | 6913 | 211.1 | - | - | 0 | - |
| - | - | 9814 | 212.1 | - | - | 0 | - |
| - | - | 4565 | 213.1 | - | - | 0 | - |
| - | - | 2.245E+04 | 213.1 | - | - | 0 | - |
| 2 | b | 1.689E+05 | 213.2 | 0.0003866 | 1.814 | +1 | 2 |
| - | - | 1.979E+04 | 214.2 | - | - | 0 | - |
| 9 | y | 5.088E+06 | 215.1 | 0.0004312 | 2.004 | +1 | 2 |
| - | - | 5.061E+05 | 216.1 | - | - | 0 | - |
| - | - | 2.019E+04 | 217.1 | - | - | 0 | - |
| - | - | 3.705E+04 | 217.1 | - | - | 0 | - |
| - | - | 4504 | 218.1 | - | - | 0 | - |
| - | - | 6920 | 225.1 | - | - | 0 | - |
| - | - | 1.21E+05 | 226.1 | - | - | 0 | - |
| - | - | 3.548E+05 | 227.1 | - | - | 0 | - |
| - | - | 1.186E+04 | 227.1 | - | - | 0 | - |
| - | - | 4.14E+04 | 228.1 | - | - | 0 | - |
| - | - | 3.66E+04 | 228.2 | - | - | 0 | - |
| - | - | 3599 | 229.1 | - | - | 0 | - |
| - | - | 3172 | 229.2 | - | - | 0 | - |
| - | - | 4344 | 229.2 | - | - | 0 | - |
| - | - | 4108 | 230.2 | - | - | 0 | - |
| - | - | 8.459E+04 | 231.1 | - | - | 0 | - |
| - | - | 1.103E+04 | 232.2 | - | - | 0 | - |
| - | - | 5625 | 233.1 | - | - | 0 | - |
| - | - | 8084 | 233.2 | - | - | 0 | - |
| - | - | 5267 | 237.1 | - | - | 0 | - |
| - | - | 7995 | 238.1 | - | - | 0 | - |
| - | - | 1.411E+04 | 238.2 | - | - | 0 | - |
| - | - | 4544 | 243.1 | - | - | 0 | - |
| - | - | 1.989E+05 | 244.1 | - | - | 0 | - |
| - | - | 4258 | 245.1 | - | - | 0 | - |
| - | - | 6415 | 245.1 | - | - | 0 | - |
| - | - | 4.728E+04 | 245.1 | - | - | 0 | - |
| - | - | 4823 | 246.1 | - | - | 0 | - |
| - | - | 2.395E+04 | 247.1 | - | - | 0 | - |
| - | - | 5810 | 249.2 | - | - | 0 | - |
| - | - | 1.31E+05 | 251.1 | - | - | 0 | - |
| - | - | 1.471E+04 | 252.1 | - | - | 0 | - |
| - | - | 2.15E+04 | 254.1 | - | - | 0 | - |
| - | - | 7730 | 254.2 | - | - | 0 | - |
| - | - | 1.023E+04 | 256.2 | - | - | 0 | - |
| - | - | 3308 | 257.1 | - | - | 0 | - |
| - | - | 3561 | 261.1 | - | - | 0 | - |
| - | - | 1.748E+04 | 261.2 | - | - | 0 | - |
| - | - | 3283 | 262.1 | - | - | 0 | - |
| - | - | 3393 | 268.5 | - | - | 0 | - |
| - | - | 4075 | 269.2 | - | - | 0 | - |
| - | - | 3.411E+04 | 272.1 | - | - | 0 | - |
| - | - | 4888 | 277.2 | - | - | 0 | - |
| - | - | 5722 | 278.2 | - | - | 0 | - |
| - | - | 1.376E+05 | 279.1 | - | - | 0 | - |
| - | - | 1.453E+04 | 280.1 | - | - | 0 | - |
| - | - | 5025 | 280.2 | - | - | 0 | - |
| - | - | 5.702E+04 | 283.1 | - | - | 0 | - |
| - | - | 9787 | 284.1 | - | - | 0 | - |
| - | - | 1.888E+04 | 287.2 | - | - | 0 | - |
| - | - | 4600 | 288.2 | - | - | 0 | - |
| - | - | 4424 | 294.2 | - | - | 0 | - |
| - | - | 4365 | 295.1 | - | - | 0 | - |
| - | - | 3677 | 296.2 | - | - | 0 | - |
| - | - | 3199 | 297.1 | - | - | 0 | - |
| - | - | 1.175E+04 | 301.2 | - | - | 0 | - |
| - | - | 1.772E+04 | 303.2 | - | - | 0 | - |
| - | - | 9191 | 304.2 | - | - | 0 | - |
| - | - | 4942 | 308.2 | - | - | 0 | - |
| - | - | 4037 | 309.2 | - | - | 0 | - |
| - | - | 5474 | 310.1 | - | - | 0 | - |
| 5 | b | 1.401E+04 | 310.2 | 0.0003199 | 1.031 | +2 | 5 |
| - | - | 4.253E+04 | 311.1 | - | - | 0 | - |
| - | - | 6382 | 312.1 | - | - | 0 | - |
| - | - | 7249 | 312.2 | - | - | 0 | - |
| - | - | 6990 | 313.2 | - | - | 0 | - |
| - | - | 7684 | 315.2 | - | - | 0 | - |
| - | - | 6234 | 323.2 | - | - | 0 | - |
| - | - | 1.591E+04 | 323.2 | - | - | 0 | - |
| - | - | 7.934E+04 | 324.2 | - | - | 0 | - |
| - | - | 3309 | 324.2 | - | - | 0 | - |
| - | - | 1.25E+04 | 325.2 | - | - | 0 | - |
| - | - | 1.501E+04 | 325.2 | - | - | 0 | - |
| 8 | y | 4.894E+04 | 326.2 | 0.0006446 | 1.976 | +1 | 3 |
| - | - | 1.281E+04 | 327.2 | - | - | 0 | - |
| - | - | 7947 | 328.1 | - | - | 0 | - |
| - | - | 9026 | 328.2 | - | - | 0 | - |
| - | - | 1.343E+04 | 329.2 | - | - | 0 | - |
| - | - | 5521 | 330.2 | - | - | 0 | - |
| - | - | 9249 | 331.2 | - | - | 0 | - |
| - | - | 9903 | 335.2 | - | - | 0 | - |
| - | - | 2.277E+04 | 337.2 | - | - | 0 | - |
| - | - | 6612 | 337.2 | - | - | 0 | - |
| - | - | 3244 | 339.1 | - | - | 0 | - |
| - | - | 6972 | 339.2 | - | - | 0 | - |
| - | - | 3.426E+04 | 340.2 | - | - | 0 | - |
| - | - | 4128 | 340.3 | - | - | 0 | - |
| - | - | 2.859E+04 | 341.2 | - | - | 0 | - |
| 3 | b | 2.838E+04 | 341.3 | 0.000318 | 0.932 | +1 | 3 |
| - | - | 5560 | 342.1 | - | - | 0 | - |
| - | - | 3376 | 342.2 | - | - | 0 | - |
| - | - | 7809 | 342.3 | - | - | 0 | - |
| - | - | 3.71E+04 | 343.2 | - | - | 0 | - |
| - | - | 8369 | 344.2 | - | - | 0 | - |
| 8 | y | 2.714E+04 | 344.2 | 0.0006085 | 1.768 | +1 | 3 |
| - | - | 5939 | 344.2 | - | - | 0 | - |
| - | - | 9111 | 344.7 | - | - | 0 | - |
| - | - | 8427 | 345.2 | - | - | 0 | - |
| - | - | 6015 | 345.2 | - | - | 0 | - |
| - | - | 3.181E+04 | 346.2 | - | - | 0 | - |
| - | - | 4572 | 347.2 | - | - | 0 | - |
| - | - | 1.953E+04 | 347.2 | - | - | 0 | - |
| - | - | 8552 | 347.7 | - | - | 0 | - |
| - | - | 8337 | 349.2 | - | - | 0 | - |
| - | - | 2.576E+04 | 353.2 | - | - | 0 | - |
| - | - | 1.565E+04 | 353.2 | - | - | 0 | - |
| - | - | 5022 | 354.2 | - | - | 0 | - |
| - | - | 3959 | 354.2 | - | - | 0 | - |
| - | - | 2.221E+04 | 355.2 | - | - | 0 | - |
| - | - | 1.662E+04 | 355.2 | - | - | 0 | - |
| - | - | 7667 | 356.2 | - | - | 0 | - |
| - | - | 4417 | 356.2 | - | - | 0 | - |
| - | - | 5379 | 357.2 | - | - | 0 | - |
| - | - | 6384 | 358.2 | - | - | 0 | - |
| - | - | 5048 | 358.7 | - | - | 0 | - |
| - | - | 5041 | 361.2 | - | - | 0 | - |
| - | - | 9046 | 362.2 | - | - | 0 | - |
| - | - | 4117 | 364.2 | - | - | 0 | - |
| - | - | 9648 | 367.2 | - | - | 0 | - |
| - | - | 6145 | 369.2 | - | - | 0 | - |
| - | - | 3512 | 369.2 | - | - | 0 | - |
| - | - | 4.216E+04 | 369.7 | - | - | 0 | - |
| - | - | 1.909E+04 | 370.2 | - | - | 0 | - |
| - | - | 3.737E+04 | 371.2 | - | - | 0 | - |
| - | - | 7494 | 372.2 | - | - | 0 | - |
| - | - | 6617 | 372.2 | - | - | 0 | - |
| - | - | 5328 | 372.2 | - | - | 0 | - |
| - | - | 8666 | 373.2 | - | - | 0 | - |
| - | - | 6.35E+04 | 374.2 | - | - | 0 | - |
| - | - | 1.482E+04 | 375.2 | - | - | 0 | - |
| - | - | 4658 | 376.2 | - | - | 0 | - |
| - | - | 3.221E+04 | 379.2 | - | - | 0 | - |
| - | - | 5698 | 380.2 | - | - | 0 | - |
| - | - | 1.223E+04 | 381.1 | - | - | 0 | - |
| - | - | 4230 | 382.1 | - | - | 0 | - |
| - | - | 6485 | 383.2 | - | - | 0 | - |
| - | - | 1.271E+04 | 383.2 | - | - | 0 | - |
| 6 | b | 6468 | 383.7 | 0.0005923 | 1.544 | +2 | 6 |
| - | - | 4462 | 384.3 | - | - | 0 | - |
| - | - | 4.632E+04 | 389.2 | - | - | 0 | - |
| - | - | 8582 | 390.2 | - | - | 0 | - |
| - | - | 6132 | 391.2 | - | - | 0 | - |
| - | - | 9.045E+04 | 391.2 | - | - | 0 | - |
| - | - | 1.118E+04 | 392.2 | - | - | 0 | - |
| - | - | 1.379E+04 | 392.2 | - | - | 0 | - |
| - | - | 4874 | 398.2 | - | - | 0 | - |
| - | - | 4008 | 401.2 | - | - | 0 | - |
| - | - | 1.843E+04 | 403.2 | - | - | 0 | - |
| - | - | 7834 | 406.2 | - | - | 0 | - |
| - | - | 8.762E+04 | 407.2 | - | - | 0 | - |
| - | - | 8259 | 408.2 | - | - | 0 | - |
| - | - | 2.247E+04 | 408.2 | - | - | 0 | - |
| - | - | 4890 | 411.3 | - | - | 0 | - |
| - | - | 6597 | 417.2 | - | - | 0 | - |
| - | - | 6.469E+04 | 418.2 | - | - | 0 | - |
| - | - | 2.797E+04 | 418.7 | - | - | 0 | - |
| - | - | 8846 | 419.2 | - | - | 0 | - |
| - | - | 1.06E+04 | 419.2 | - | - | 0 | - |
| - | - | 3636 | 420.3 | - | - | 0 | - |
| 7 | b | 1.114E+05 | 423.2 | 0.006799 | 16.07 | +2 | 7 |
| - | - | 5.002E+04 | 424.2 | - | - | 0 | - |
| - | - | 9687 | 425.2 | - | - | 0 | - |
| - | - | 1.192E+05 | 426.2 | - | - | 0 | - |
| - | - | 2.36E+04 | 427.2 | - | - | 0 | - |
| 7 | b | 3.066E+04 | 432.2 | 0.0004 | 0.9254 | +2 | 7 |
| - | - | 1.405E+04 | 432.7 | - | - | 0 | - |
| - | - | 3781 | 433.2 | - | - | 0 | - |
| - | - | 5874 | 434.2 | - | - | 0 | - |
| - | - | 3808 | 434.3 | - | - | 0 | - |
| - | - | 2.286E+04 | 435.2 | - | - | 0 | - |
| - | - | 4342 | 436.2 | - | - | 0 | - |
| - | - | 1.199E+04 | 436.2 | - | - | 0 | - |
| - | - | 4979 | 437.2 | - | - | 0 | - |
| 4 | b | 4.417E+04 | 438.3 | 0.0007549 | 1.722 | +1 | 4 |
| - | - | 1.176E+04 | 438.3 | - | - | 0 | - |
| 4 | b | 1.001E+04 | 439.3 | 0.000687 | 1.564 | +1 | 4 |
| - | - | 6212 | 440.3 | - | - | 0 | - |
| 7 | y | 3.017E+05 | 441.2 | 0.000701 | 1.589 | +1 | 4 |
| - | - | 7.003E+04 | 442.2 | - | - | 0 | - |
| - | - | 9381 | 443.2 | - | - | 0 | - |
| - | - | 5257 | 443.3 | - | - | 0 | - |
| - | - | 4364 | 445.2 | - | - | 0 | - |
| - | - | 3.313E+04 | 452.2 | - | - | 0 | - |
| - | - | 5158 | 453.2 | - | - | 0 | - |
| - | - | 6658 | 454.2 | - | - | 0 | - |
| - | - | 6798 | 454.3 | - | - | 0 | - |
| 4 | b | 1.717E+05 | 456.3 | 0.0007493 | 1.642 | +1 | 4 |
| - | - | 1.483E+04 | 457.2 | - | - | 0 | - |
| - | - | 4.542E+04 | 457.3 | - | - | 0 | - |
| - | - | 5352 | 458.3 | - | - | 0 | - |
| - | - | 3886 | 461.2 | - | - | 0 | - |
| - | - | 8652 | 464.2 | - | - | 0 | - |
| - | - | 1.051E+04 | 468.2 | - | - | 0 | - |
| - | - | 7.883E+04 | 470.2 | - | - | 0 | - |
| - | - | 6357 | 470.3 | - | - | 0 | - |
| - | - | 2.087E+04 | 471.2 | - | - | 0 | - |
| - | - | 5799 | 472.2 | - | - | 0 | - |
| - | - | 7270 | 473.2 | - | - | 0 | - |
| - | - | 4240 | 473.3 | - | - | 0 | - |
| - | - | 6724 | 476.3 | - | - | 0 | - |
| - | - | 1.026E+04 | 478.3 | - | - | 0 | - |
| - | - | 5823 | 482.2 | - | - | 0 | - |
| - | - | 5.088E+04 | 482.3 | - | - | 0 | - |
| - | - | 2.73E+04 | 482.8 | - | - | 0 | - |
| - | - | 2.265E+04 | 483.3 | - | - | 0 | - |
| - | - | 5001 | 487.3 | - | - | 0 | - |
| 8 | b | 1.079E+04 | 488.2 | 0.007454 | 15.27 | +2 | 8 |
| - | - | 1.459E+04 | 488.3 | - | - | 0 | - |
| - | - | 6866 | 489.3 | - | - | 0 | - |
| - | - | 5265 | 492.2 | - | - | 0 | - |
| - | - | 4524 | 493.3 | - | - | 0 | - |
| - | - | 6975 | 495.2 | - | - | 0 | - |
| - | - | 4415 | 495.3 | - | - | 0 | - |
| 8 | b | 3.664E+04 | 496.8 | 0.0001606 | 0.3233 | +2 | 8 |
| - | - | 1.586E+04 | 497.3 | - | - | 0 | - |
| - | - | 8506 | 497.8 | - | - | 0 | - |
| - | - | 6909 | 498.2 | - | - | 0 | - |
| - | - | 4710 | 500.2 | - | - | 0 | - |
| - | - | 5187 | 500.3 | - | - | 0 | - |
| - | - | 5876 | 502.3 | - | - | 0 | - |
| - | - | 1.112E+04 | 504.2 | - | - | 0 | - |
| - | - | 6188 | 504.3 | - | - | 0 | - |
| - | - | 1.4E+04 | 506.3 | - | - | 0 | - |
| - | - | 1.918E+04 | 508.3 | - | - | 0 | - |
| - | - | 2.913E+04 | 509.2 | - | - | 0 | - |
| - | - | 8944 | 510.2 | - | - | 0 | - |
| - | - | 4009 | 511.2 | - | - | 0 | - |
| - | - | 6586 | 516.2 | - | - | 0 | - |
| - | - | 1.017E+04 | 518.2 | - | - | 0 | - |
| - | - | 8340 | 518.3 | - | - | 0 | - |
| - | - | 4264 | 519.2 | - | - | 0 | - |
| - | - | 8323 | 520.3 | - | - | 0 | - |
| - | - | 6994 | 521.3 | - | - | 0 | - |
| - | - | 1.269E+04 | 523.2 | - | - | 0 | - |
| - | - | 4207 | 523.3 | - | - | 0 | - |
| - | - | 4076 | 524.3 | - | - | 0 | - |
| - | - | 6.603E+04 | 526.3 | - | - | 0 | - |
| - | - | 2.621E+04 | 527.3 | - | - | 0 | - |
| - | - | 3936 | 528.3 | - | - | 0 | - |
| - | - | 5803 | 531.3 | - | - | 0 | - |
| - | - | 3.916E+04 | 534.3 | - | - | 0 | - |
| - | - | 1.136E+04 | 535.3 | - | - | 0 | - |
| - | - | 4.098E+04 | 536.3 | - | - | 0 | - |
| - | - | 1.724E+04 | 537.2 | - | - | 0 | - |
| - | - | 6667 | 538.2 | - | - | 0 | - |
| - | - | 2.733E+04 | 539.3 | - | - | 0 | - |
| - | - | 8090 | 540.3 | - | - | 0 | - |
| - | - | 3866 | 541.3 | - | - | 0 | - |
| 9 | b | 2.951E+04 | 545.3 | 0.0002679 | 0.4913 | +2 | 9 |
| - | - | 8962 | 545.8 | - | - | 0 | - |
| - | - | 6808 | 546.3 | - | - | 0 | - |
| - | - | 6747 | 548.3 | - | - | 0 | - |
| - | - | 6872 | 549.3 | - | - | 0 | - |
| - | - | 6738 | 550.3 | - | - | 0 | - |
| - | - | 1.198E+04 | 551.3 | - | - | 0 | - |
| - | - | 1.921E+04 | 553.3 | - | - | 0 | - |
| - | - | 3.108E+05 | 554.3 | - | - | 0 | - |
| - | - | 9.764E+04 | 555.3 | - | - | 0 | - |
| - | - | 1.487E+04 | 556.3 | - | - | 0 | - |
| - | - | 2.222E+04 | 558.3 | - | - | 0 | - |
| - | - | 8937 | 564.2 | - | - | 0 | - |
| - | - | 6259 | 567.3 | - | - | 0 | - |
| - | - | 3.994E+04 | 567.4 | - | - | 0 | - |
| - | - | 1.863E+04 | 568.4 | - | - | 0 | - |
| - | - | 4632 | 569.3 | - | - | 0 | - |
| - | - | 7857 | 571.3 | - | - | 0 | - |
| - | - | 5007 | 572.3 | - | - | 0 | - |
| - | - | 4677 | 573.3 | - | - | 0 | - |
| - | - | 6749 | 574.3 | - | - | 0 | - |
| - | - | 7590 | 579.3 | - | - | 0 | - |
| - | - | 8923 | 580.8 | - | - | 0 | - |
| - | - | 5266 | 581.3 | - | - | 0 | - |
| - | - | 2.375E+04 | 582.3 | - | - | 0 | - |
| - | - | 4891 | 583.3 | - | - | 0 | - |
| - | - | 4535 | 585.3 | - | - | 0 | - |
| - | - | 4012 | 586.3 | - | - | 0 | - |
| 6 | y | 1.102E+05 | 588.3 | 0.0002498 | 0.4246 | +1 | 5 |
| - | - | 3.862E+04 | 589.3 | - | - | 0 | - |
| - | - | 8896 | 590.3 | - | - | 0 | - |
| - | - | 5.75E+04 | 591.4 | - | - | 0 | - |
| - | - | 2.375E+04 | 592.4 | - | - | 0 | - |
| 0 | Precursor | 5128 | 594.8 | 0.001034 | 1.739 | +2 | -1 |
| 0 | Precursor | 6843 | 595.3 | 0.008233 | 13.83 | +2 | -1 |
| - | - | 1.785E+04 | 597.3 | - | - | 0 | - |
| - | - | 8163 | 598.3 | - | - | 0 | - |
| - | - | 9562 | 599.3 | - | - | 0 | - |
| 5 | b | 1.401E+04 | 601.3 | 0.0008061 | 1.34 | +1 | 5 |
| - | - | 8622 | 602.3 | - | - | 0 | - |
| - | - | 2.436E+04 | 603.4 | - | - | 0 | - |
| 0 | Precursor | 1.708E+04 | 603.8 | 0.0006346 | 1.051 | +2 | -1 |
| - | - | 1.208E+04 | 604.3 | - | - | 0 | - |
| - | - | 5652 | 604.8 | - | - | 0 | - |
| - | - | 9336 | 605.3 | - | - | 0 | - |
| - | - | 4296 | 606.4 | - | - | 0 | - |
| - | - | 3802 | 608.3 | - | - | 0 | - |
| - | - | 1.311E+04 | 615.3 | - | - | 0 | - |
| - | - | 9496 | 617.3 | - | - | 0 | - |
| - | - | 4111 | 618.3 | - | - | 0 | - |
| - | - | 4136 | 618.4 | - | - | 0 | - |
| 5 | b | 1.838E+05 | 619.3 | 0.0005921 | 0.956 | +1 | 5 |
| - | - | 6.642E+04 | 620.3 | - | - | 0 | - |
| - | - | 9497 | 621.4 | - | - | 0 | - |
| - | - | 1.494E+04 | 623.3 | - | - | 0 | - |
| - | - | 5119 | 624.3 | - | - | 0 | - |
| - | - | 1.726E+04 | 625.3 | - | - | 0 | - |
| - | - | 8509 | 626.3 | - | - | 0 | - |
| - | - | 6179 | 629.3 | - | - | 0 | - |
| - | - | 7550 | 631.3 | - | - | 0 | - |
| - | - | 3.034E+04 | 633.3 | - | - | 0 | - |
| - | - | 7010 | 634.3 | - | - | 0 | - |
| - | - | 8112 | 635.3 | - | - | 0 | - |
| - | - | 5687 | 636.3 | - | - | 0 | - |
| - | - | 6274 | 636.4 | - | - | 0 | - |
| - | - | 4947 | 637.3 | - | - | 0 | - |
| - | - | 7041 | 638.3 | - | - | 0 | - |
| - | - | 6492 | 638.3 | - | - | 0 | - |
| - | - | 4511 | 649.3 | - | - | 0 | - |
| - | - | 3790 | 650.3 | - | - | 0 | - |
| - | - | 3.879E+04 | 651.3 | - | - | 0 | - |
| - | - | 1.019E+04 | 651.4 | - | - | 0 | - |
| - | - | 3.4E+04 | 652.3 | - | - | 0 | - |
| - | - | 1.205E+04 | 652.3 | - | - | 0 | - |
| - | - | 1.544E+04 | 653.3 | - | - | 0 | - |
| - | - | 4.286E+04 | 653.3 | - | - | 0 | - |
| - | - | 1.501E+04 | 654.3 | - | - | 0 | - |
| - | - | 1.183E+04 | 663.3 | - | - | 0 | - |
| - | - | 1.902E+04 | 664.4 | - | - | 0 | - |
| - | - | 5253 | 665.4 | - | - | 0 | - |
| - | - | 1.248E+04 | 667.3 | - | - | 0 | - |
| - | - | 5639 | 668.4 | - | - | 0 | - |
| - | - | 6931 | 669.4 | - | - | 0 | - |
| - | - | 4.493E+04 | 681.3 | - | - | 0 | - |
| - | - | 1.13E+04 | 682.3 | - | - | 0 | - |
| - | - | 9.59E+04 | 682.4 | - | - | 0 | - |
| - | - | 4.112E+04 | 683.4 | - | - | 0 | - |
| - | - | 7241 | 684.4 | - | - | 0 | - |
| - | - | 6126 | 688.4 | - | - | 0 | - |
| - | - | 1.146E+04 | 703.4 | - | - | 0 | - |
| - | - | 4839 | 704.4 | - | - | 0 | - |
| - | - | 5432 | 714.4 | - | - | 0 | - |
| - | - | 2.793E+04 | 716.4 | - | - | 0 | - |
| - | - | 1.124E+04 | 717.4 | - | - | 0 | - |
| - | - | 7083 | 720.4 | - | - | 0 | - |
| - | - | 1.941E+04 | 721.4 | - | - | 0 | - |
| - | - | 1.192E+04 | 722.4 | - | - | 0 | - |
| - | - | 3834 | 732.4 | - | - | 0 | - |
| - | - | 5774 | 734.4 | - | - | 0 | - |
| - | - | 5684 | 735.3 | - | - | 0 | - |
| - | - | 4405 | 737.4 | - | - | 0 | - |
| - | - | 7.831E+04 | 738.4 | - | - | 0 | - |
| - | - | 4.199E+04 | 739.4 | - | - | 0 | - |
| - | - | 7206 | 740.4 | - | - | 0 | - |
| - | - | 4924 | 742.3 | - | - | 0 | - |
| - | - | 4340 | 744.3 | - | - | 0 | - |
| 6 | b | 4.069E+04 | 748.4 | 0.0004334 | 0.5791 | +1 | 6 |
| 6 | b | 1.793E+04 | 749.4 | 0.01464 | 19.53 | +1 | 6 |
| - | - | 1.081E+04 | 750.4 | - | - | 0 | - |
| - | - | 3865 | 751.3 | - | - | 0 | - |
| 5 | y | 8598 | 751.4 | 0.007783 | 10.36 | +1 | 6 |
| - | - | 6.466E+04 | 752.4 | - | - | 0 | - |
| - | - | 2.38E+04 | 753.4 | - | - | 0 | - |
| - | - | 4660 | 754.4 | - | - | 0 | - |
| - | - | 4566 | 760.4 | - | - | 0 | - |
| - | - | 2.683E+04 | 762.3 | - | - | 0 | - |
| - | - | 1.088E+04 | 763.3 | - | - | 0 | - |
| 6 | b | 6.352E+05 | 766.4 | 0.0002324 | 0.3032 | +1 | 6 |
| - | - | 2.895E+05 | 767.4 | - | - | 0 | - |
| - | - | 6.79E+04 | 768.4 | - | - | 0 | - |
| - | - | 9121 | 769.4 | - | - | 0 | - |
| - | - | 5301 | 778.4 | - | - | 0 | - |
| - | - | 3.895E+05 | 780.4 | - | - | 0 | - |
| - | - | 1.769E+05 | 781.4 | - | - | 0 | - |
| - | - | 6426 | 781.4 | - | - | 0 | - |
| - | - | 4.538E+04 | 782.4 | - | - | 0 | - |
| - | - | 4948 | 783.4 | - | - | 0 | - |
| - | - | 9372 | 784.4 | - | - | 0 | - |
| - | - | 6287 | 785.4 | - | - | 0 | - |
| - | - | 6262 | 808.3 | - | - | 0 | - |
| - | - | 8610 | 811.4 | - | - | 0 | - |
| - | - | 1.22E+04 | 817.4 | - | - | 0 | - |
| - | - | 8133 | 818.4 | - | - | 0 | - |
| - | - | 4511 | 822.4 | - | - | 0 | - |
| - | - | 4788 | 823.4 | - | - | 0 | - |
| - | - | 1.211E+04 | 827.4 | - | - | 0 | - |
| - | - | 7537 | 828.4 | - | - | 0 | - |
| - | - | 1.582E+04 | 829.4 | - | - | 0 | - |
| - | - | 9099 | 830.4 | - | - | 0 | - |
| - | - | 3906 | 832.4 | - | - | 0 | - |
| - | - | 7554 | 835.5 | - | - | 0 | - |
| - | - | 4335 | 836.5 | - | - | 0 | - |
| 7 | b | 2.913E+04 | 845.5 | 0.01407 | 16.65 | +1 | 7 |
| 7 | b | 1.89E+04 | 846.4 | 0.004535 | 5.358 | +1 | 7 |
| - | - | 4692 | 847.5 | - | - | 0 | - |
| - | - | 2.467E+04 | 850.4 | - | - | 0 | - |
| - | - | 1.559E+04 | 851.4 | - | - | 0 | - |
| - | - | 6669 | 852.4 | - | - | 0 | - |
| - | - | 4152 | 861.4 | - | - | 0 | - |
| 7 | b | 4.971E+04 | 863.5 | 8.085E-05 | 0.09364 | +1 | 7 |
| - | - | 1.977E+04 | 864.5 | - | - | 0 | - |
| - | - | 9187 | 865.5 | - | - | 0 | - |
| - | - | 2.059E+04 | 877.4 | - | - | 0 | - |
| - | - | 5934 | 878.4 | - | - | 0 | - |
| - | - | 4.172E+04 | 879.4 | - | - | 0 | - |
| - | - | 2.676E+04 | 880.4 | - | - | 0 | - |
| - | - | 6448 | 881.4 | - | - | 0 | - |
| - | - | 1.085E+04 | 889.4 | - | - | 0 | - |
| - | - | 9903 | 893.4 | - | - | 0 | - |
| - | - | 6355 | 894.4 | - | - | 0 | - |
| - | - | 1.048E+04 | 895.5 | - | - | 0 | - |
| - | - | 4038 | 896.5 | - | - | 0 | - |
| - | - | 3.242E+04 | 907.4 | - | - | 0 | - |
| - | - | 1.963E+04 | 908.4 | - | - | 0 | - |
| - | - | 1.016E+04 | 946.5 | - | - | 0 | - |
| - | - | 1.531E+04 | 947.5 | - | - | 0 | - |
| - | - | 5907 | 949.5 | - | - | 0 | - |
| - | - | 6383 | 956.5 | - | - | 0 | - |
| - | - | 5512 | 957.5 | - | - | 0 | - |
| - | - | 1.362E+05 | 964.5 | - | - | 0 | - |
| - | - | 5615 | 965.4 | - | - | 0 | - |
| - | - | 7.519E+04 | 965.5 | - | - | 0 | - |
| - | - | 2.661E+04 | 966.5 | - | - | 0 | - |
| - | - | 4679 | 974.4 | - | - | 0 | - |
| 8 | b | 7.47E+04 | 974.5 | 0.001247 | 1.28 | +1 | 8 |
| 8 | b | 3.824E+04 | 975.5 | 0.01755 | 17.99 | +1 | 8 |
| 3 | y | 3.547E+04 | 976.5 | 0.004779 | 4.894 | +1 | 8 |
| - | - | 2.129E+04 | 977.5 | - | - | 0 | - |
| - | - | 4814 | 978.5 | - | - | 0 | - |
| - | - | 5384 | 991.5 | - | - | 0 | - |
| 8 | b | 1.131E+06 | 992.5 | 0.000337 | 0.3395 | +1 | 8 |
| - | - | 6.717E+05 | 993.5 | - | - | 0 | - |
| 3 | y | 4.232E+05 | 994.5 | 0.01069 | 10.75 | +1 | 8 |
| - | - | 1.877E+05 | 995.5 | - | - | 0 | - |
| - | - | 5.379E+04 | 996.5 | - | - | 0 | - |
| - | - | 1.009E+04 | 997.5 | - | - | 0 | - |
| 9 | b | 3988 | 1090 | 0.001709 | 1.568 | +1 | 9 |
| 2 | y | 2.942E+04 | 1094 | 0.000408 | 0.3731 | +1 | 9 |
| - | - | 1.889E+04 | 1095 | - | - | 0 | - |
| - | - | 4311 | 1108 | - | - | 0 | - |
| - | - | 3661 | 1361 | - | - | 0 | - |
| - | - | 3956 | 2410 | - | - | 0 | - |
| - | - | 4997 | 3226 | - | - | 0 | - |
| - | - | 4139 | 3389 | - | - | 0 | - |

m/z Charge Intensity FragmentType MassShift Position
120.08111572265625 0 196970.47
121.08450317382812 0 16743.502
128.07107543945312 0 2494.9011
129.10255432128906 0 353279.5
130.10072326660156 0 3459.669
130.10594177246094 0 21319.496
134.09657287597656 0 3949.6743
134.91490173339844 0 2319.9602
136.07598876953125 0 78545.85
137.079345703125 0 4342.135
139.08685302734375 0 10745.767
147.04446411132812 0 3780.864
155.11839294433594 0 4978.5645
156.07733154296875 0 5151.441
165.10235595703125 0 8154.6084
167.08180236816406 0 8284.168
167.11822509765625 0 11234.2
168.10211181640625 0 10515.851
169.1338653564453 0 117077.27
170.1371612548828 0 10149.555
172.11245727539062 0 28712.852
172.1449432373047 0 4230.932
173.43911743164062 0 10597.633
174.12832641601562 0 7632.388
175.10821533203125 0 2915.9448
177.10240173339844 0 5043.717
181.09750366210938 0 37639.492
181.13372802734375 0 22906.436
182.13723754882812 0 2874.4124
183.1127166748047 0 6686.9336
183.1495361328125 0 17585.09
184.1084442138672 0 5179.058
185.1289825439453 0 10345.808
185.16519165039062 0 453333.9 a 1
186.12783813476562 0 17893.113
186.1685333251953 0 46999.977
187.13134765625 0 4617.8267
188.107421875 0 8513.254
188.14364624023438 0 17111.842
190.1234130859375 0 3970.0903
191.0821075439453 0 9168.79
195.1130828857422 0 30411.197
196.11630249023438 0 3169.34
197.1287841796875 0 16069.722
197.1648712158203 0 3641.4932
198.12396240234375 0 7208.452
199.10801696777344 0 143225.66
200.11099243164062 0 9821.419
201.12403869628906 0 4052.8096
202.12278747558594 0 5249.556
203.15512084960938 0 6077.5083
204.13856506347656 0 10002.202
209.09239196777344 0 91066.266
210.09564208984375 0 6960.6055
210.16041564941406 0 5877.1543
210.79127502441406 0 2549.1921
211.09829711914062 0 3430.412
211.1442108154297 0 6912.711
212.13983154296875 0 9813.827
213.08741760253906 0 4564.596
213.1236114501953 0 22451.402
213.16014099121094 0 168880.56 b 1
214.16348266601562 0 19788.287
215.1394500732422 0 5088432 y 8
216.1426544189453 0 506058.06
217.1334991455078 0 20191.72
217.14479064941406 0 37052.27
218.1470947265625 0 4504.299
225.08761596679688 0 6919.6167
226.11892700195312 0 121019.47
227.10299682617188 0 354777.66
227.12327575683594 0 11855.704
228.10633850097656 0 41401.062
228.17100524902344 0 36604.35
229.10671997070312 0 3598.8857
229.15481567382812 0 3171.612
229.17416381835938 0 4344.1934
230.1859893798828 0 4107.6147
231.14952087402344 0 84586.67
232.15292358398438 0 11034.266
233.09188842773438 0 5625.497
233.16551208496094 0 8084.2056
237.08665466308594 0 5266.518
238.12283325195312 0 7995.3613
238.1555633544922 0 14106.833
243.09872436523438 0 4543.7876
244.1295623779297 0 198923.1
245.091552734375 0 4258.0405
245.11387634277344 0 6415.029
245.1298370361328 0 47283.664
246.1326904296875 0 4823.0312
247.14451599121094 0 23950.697
249.1601104736328 0 5810.2417
251.10301208496094 0 131026.555
252.10630798339844 0 14709.1
254.11398315429688 0 21496.777
254.15089416503906 0 7730.0977
256.1658020019531 0 10231.662
257.1295471191406 0 3308.4531
261.086669921875 0 3560.696
261.1579284667969 0 17476.309
262.0716552734375 0 3283.0745
268.5059814453125 0 3392.5435
269.1865539550781 0 4075.1726
272.1243591308594 0 34105.61
277.1537170410156 0 4888.2817
278.1500244140625 0 5721.9087
279.0979309082031 0 137620.34
280.10076904296875 0 14531.794
280.1661682128906 0 5025.476
283.14434814453125 0 57019.746
284.1479187011719 0 9786.688
287.21209716796875 0 18882.041
288.215087890625 0 4600.304
294.1819152832031 0 4424.288
295.1432800292969 0 4365.2334
296.1797790527344 0 3676.9075
297.10821533203125 0 3198.5317
301.1551513671875 0 11751.665
303.2070007324219 0 17721.367
304.1667785644531 0 9191.273
308.1611328125 0 4942.2456
309.191650390625 0 4037.4756
310.1405334472656 0 5473.916
310.17645263671875 0 14005.901 b 4
311.1394958496094 0 42525.324
312.1423034667969 0 6382.132
312.1923828125 0 7249.133
313.1915588378906 0 6990.39
315.2073059082031 0 7684.214
323.16973876953125 0 6234.449
323.2445373535156 0 15914.198
324.1559753417969 0 79342.15
324.24786376953125 0 3308.7932
325.1593017578125 0 12503.856
325.1878356933594 0 15014.014
326.17169189453125 0 48944.293 y Water loss 7
327.1744384765625 0 12814.552
328.14971923828125 0 7946.798
328.16717529296875 0 9026.18
329.1502685546875 0 13430.051
330.217529296875 0 5521.391
331.201904296875 0 9249.073
335.1733703613281 0 9902.733
337.1513671875 0 22768.895
337.187255859375 0 6612.4263
339.1361999511719 0 3243.6682
339.2035217285156 0 6971.5537
340.18719482421875 0 34261.656
340.30206298828125 0 4128.2153
341.1829528808594 0 28591.354
341.2550354003906 0 28380.92 b 2
342.13031005859375 0 5560.413
342.1853942871094 0 3375.9973
342.2589416503906 0 7809.4536
343.19805908203125 0 37104.004
344.160888671875 0 8369.292
344.1822204589844 0 27142.734 y 7
344.2026672363281 0 5939.4893
344.7049865722656 0 9111.462
345.1865539550781 0 8426.934
345.2090759277344 0 6015.3604
346.1765441894531 0 31814.598
347.1800842285156 0 4571.7085
347.2028503417969 0 19525.639
347.7047119140625 0 8551.731
349.1914367675781 0 8337.217
353.1825866699219 0 25760.975
353.2189025878906 0 15645.224
354.1856384277344 0 5022.148
354.22119140625 0 3959.4934
355.1611328125 0 22206.43
355.1983642578125 0 16619.996
356.1628112792969 0 7667.2324
356.1986999511719 0 4416.946
357.21258544921875 0 5378.907
358.21075439453125 0 6384.1123
358.7035827636719 0 5047.8115
361.1860046386719 0 5041.4136
362.17108154296875 0 9046.265
364.1860656738281 0 4116.854
367.1983947753906 0 9647.9795
369.17657470703125 0 6145.139
369.2472839355469 0 3511.6953
369.7130432128906 0 42163.613
370.2149353027344 0 19088.146
371.1929016113281 0 37372.566
372.1561584472656 0 7494.4155
372.197265625 0 6617.273
372.2254638671875 0 5328.4087
373.1866455078125 0 8666.483
374.1712341308594 0 63499.76
375.1748046875 0 14818.777
376.15093994140625 0 4658.259
379.1982116699219 0 32212.404
380.20257568359375 0 5697.781
381.14471435546875 0 12226.464
382.14697265625 0 4229.575
383.1560363769531 0 6485.0723
383.19256591796875 0 12712.351
383.7097473144531 0 6468.095 b 5
384.296875 0 4461.8657
389.18243408203125 0 46319.266
390.1651611328125 0 8582.274
391.1687927246094 0 6132.4336
391.19793701171875 0 90445.9
392.1797180175781 0 11178.375
392.2015380859375 0 13792.854
398.1712646484375 0 4874.061
401.18359375 0 4008.122
403.2347717285156 0 18434.965
406.2088928222656 0 7833.9194
407.1928405761719 0 87615.32
408.1569519042969 0 8258.578
408.19549560546875 0 22472.271
411.2591857910156 0 4889.7812
417.1761169433594 0 6597.2207
418.2398681640625 0 64692.758
418.7410888671875 0 27969.953
419.1925964355469 0 8845.688
419.2431335449219 0 10598.317
420.3016662597656 0 3636.4565
423.2246398925781 0 111393.99 b Water loss 6
424.22381591796875 0 50022.96
425.2247314453125 0 9686.8
426.16650390625 0 119190.12
427.169189453125 0 23599.639
432.23712158203125 0 30664.395 b 6
432.73876953125 0 14051.089
433.2381591796875 0 3781.3914
434.2044372558594 0 5873.837
434.2766418457031 0 3808.2205
435.18804931640625 0 22859.998
436.1549987792969 0 4342.2217
436.1896667480469 0 11988.008
437.2391052246094 0 4979.3984
438.2718505859375 0 44166.02 b Water loss 3
438.3079528808594 0 11764.531
439.25579833984375 0 10014.117 b Ammonia loss 3
440.25177001953125 0 6212.491
441.2350769042969 0 301744.66 y 6
442.2379150390625 0 70034.125
443.23858642578125 0 9381.298
443.3015441894531 0 5256.701
445.2458190917969 0 4363.864
452.214599609375 0 33132.027
453.2191467285156 0 5157.586
454.230712890625 0 6658.2666
454.26617431640625 0 6797.588
456.28240966796875 0 171708.23 b 3
457.2451477050781 0 14833.201
457.28509521484375 0 45422.535
458.28936767578125 0 5351.738
461.2395935058594 0 3885.6084
464.2497863769531 0 8651.561
468.2452697753906 0 10507.628
470.2253723144531 0 78826.58
470.30224609375 0 6356.6064
471.22723388671875 0 20871.436
472.2204284667969 0 5798.8403
473.20361328125 0 7269.722
473.3094787597656 0 4240.012
476.2519836425781 0 6723.5806
478.26763916015625 0 10256.805
482.22564697265625 0 5822.7583
482.26171875 0 50875.84
482.76080322265625 0 27299.016
483.26336669921875 0 22653.305
487.25592041015625 0 5001.3013
488.252197265625 0 10791.117 b Ammonia loss 7
488.3236083984375 0 14589.742
489.3280944824219 0 6866.3657
492.2141418457031 0 5264.9033
493.28155517578125 0 4524.1885
495.2230529785156 0 6975.178
495.2942810058594 0 4414.5312
496.7581787109375 0 36640.027 b 7
497.2599792480469 0 15857.861
497.7603759765625 0 8506.058
498.220703125 0 6909.218
500.21343994140625 0 4709.706
500.25091552734375 0 5187.414
502.2674560546875 0 5876.4985
504.2452392578125 0 11117.711
504.3187255859375 0 6187.793
506.2626037597656 0 13995.545
508.2553405761719 0 19182.104
509.2410583496094 0 29130.43
510.2426452636719 0 8943.742
511.247314453125 0 4008.9639
516.2459716796875 0 6586.4746
518.2249755859375 0 10167.443
518.26220703125 0 8340.244
519.2272338867188 0 4264.413
520.27587890625 0 8323.211
521.2753295898438 0 6993.862
523.2197265625 0 12691.22
523.289306640625 0 4207.2153
524.2908325195312 0 4075.9521
526.266357421875 0 66027
527.2701416015625 0 26207.723
528.2695922851562 0 3936.1594
531.2862548828125 0 5803.075
534.256103515625 0 39155.812
535.2587280273438 0 11358.376
536.2503051757812 0 40975.71
537.2374267578125 0 17238.684
538.2391357421875 0 6666.9927
539.2509765625 0 27333.04
540.30419921875 0 8089.8696
541.3079223632812 0 3865.627
545.28466796875 0 29507.908 b 8
545.7858276367188 0 8961.782
546.2886962890625 0 6808.0513
548.2702026367188 0 6747.2607
549.3399047851562 0 6872.257
550.3439331054688 0 6737.5894
551.281982421875 0 11982.615
553.3355102539062 0 19214.97
554.2611694335938 0 310844.3
555.2640991210938 0 97641.8
556.266845703125 0 14869.059
558.328857421875 0 22216.268
564.24365234375 0 8936.574
567.3099365234375 0 6258.6714
567.350830078125 0 39942.555
568.353515625 0 18630.916
569.2915649414062 0 4632.4775
571.293212890625 0 7856.85
572.2723388671875 0 5006.726
573.3380737304688 0 4677.37
574.326171875 0 6748.955
579.2780151367188 0 7590.331
580.8219604492188 0 8922.907
581.32080078125 0 5266.415
582.256103515625 0 23746.611
583.2608032226562 0 4890.588
585.333984375 0 4534.734
586.322509765625 0 4011.8574
588.3030395507812 0 110167.766 y 5
589.3057250976562 0 38620.203
590.3006591796875 0 8896.472
591.3505249023438 0 57498.957
592.353759765625 0 23749.436
594.8196411132812 0 5128.4517 Precursor Water loss
595.31884765625 0 6843.1475 Precursor Ammonia loss
597.287353515625 0 17845.377
598.2919921875 0 8163.163
599.282470703125 0 9561.841
601.3336181640625 0 14011.248 b Water loss 4
602.33447265625 0 8622.26
603.3507080078125 0 24363.98
603.8245239257812 0 17075.42 Precursor
604.3239135742188 0 12078.173
604.8245239257812 0 5652.137
605.2954711914062 0 9335.605
606.3565063476562 0 4295.819
608.3114013671875 0 3801.82
615.2780151367188 0 13109.999
617.2920532226562 0 9496.23
618.2836303710938 0 4110.8145
618.3648681640625 0 4136.4966
619.3455810546875 0 183762.25 b 4
620.3485107421875 0 66415.42
621.3504028320312 0 9497.024
623.3191528320312 0 14940.625
624.267333984375 0 5118.5825
625.3350219726562 0 17261.857
626.3364868164062 0 8509.172
629.3289794921875 0 6178.796
631.3099365234375 0 7549.884
633.28955078125 0 30339.594
634.2897338867188 0 7010.0986
635.3175659179688 0 8112.2207
636.3134765625 0 5686.8726
636.37451171875 0 6273.622
637.301025390625 0 4947.497
638.2759399414062 0 7041.0527
638.325927734375 0 6492.288
649.3338623046875 0 4511.4155
650.3175659179688 0 3790.0496
651.314453125 0 38793.457
651.3873901367188 0 10194.922
652.2613525390625 0 33997.082
652.3177490234375 0 12051.497
653.2646484375 0 15438.6
653.3291625976562 0 42856.754
654.3341674804688 0 15014.405
663.3153686523438 0 11830.01
664.3677368164062 0 19021.74
665.370361328125 0 5252.5874
667.3457641601562 0 12480.902
668.3505859375 0 5639.2324
669.3970947265625 0 6931.409
681.3244018554688 0 44927.797
682.3253784179688 0 11300.07
682.3779296875 0 95900.76
683.3798828125 0 41121.07
684.3812866210938 0 7241.4204
688.4039916992188 0 6125.7485
703.3693237304688 0 11461.705
704.369873046875 0 4839.175
714.4144287109375 0 5432.4077
716.3986206054688 0 27929.953
717.399658203125 0 11242.9
720.4049072265625 0 7083.2715
721.395751953125 0 19413.553
722.3919677734375 0 11916.259
732.3587036132812 0 3833.843
734.3538818359375 0 5774.2075
735.3363647460938 0 5683.678
737.3572998046875 0 4405.188
738.418701171875 0 78312.26
739.4216918945312 0 41989.41
740.425048828125 0 7205.735
742.3447875976562 0 4924.123
744.343017578125 0 4339.8022
748.4024047851562 0 40687.465 b Water loss 5
749.4014892578125 0 17926.344 b Ammonia loss 5
750.3855590820312 0 10812.35
751.2722778320312 0 3864.6714
751.3739013671875 0 8597.814 y 4
752.3621215820312 0 64661.254
753.36474609375 0 23802.115
754.36669921875 0 4660.3145
760.3622436523438 0 4566.0317
762.3460693359375 0 26832.84
763.3483276367188 0 10877.028
766.4136352539062 0 635170.6 b 5
767.41650390625 0 289453.7
768.4193115234375 0 67895.98
769.4232177734375 0 9121.096
778.3694458007812 0 5300.951
780.3565063476562 0 389509.8
781.3597412109375 0 176887.88
781.4414672851562 0 6426.3623
782.362060546875 0 45377.64
783.361572265625 0 4948.392
784.423828125 0 9371.509
785.4271850585938 0 6287.395
808.34765625 0 6262.0864
811.4334716796875 0 8610.485
817.4474487304688 0 12201.211
818.44384765625 0 8132.6475
822.4410400390625 0 4510.565
823.447509765625 0 4788.325
827.427001953125 0 12111.082
828.43212890625 0 7537.1685
829.4454345703125 0 15815.029
830.4439697265625 0 9099.115
832.4112548828125 0 3905.7249
835.4707641601562 0 7553.9893
836.4813842773438 0 4335.2134
845.4415283203125 0 29126.406 b Water loss 6
846.4441528320312 0 18897.615 b Ammonia loss 6
847.4505004882812 0 4691.6504
850.4356689453125 0 24672.453
851.4349365234375 0 15585.069
852.4318237304688 0 6669.0386
861.4086303710938 0 4152.1304
863.4662475585938 0 49706.863 b 6
864.46923828125 0 19770.64
865.4669799804688 0 9187.384
877.41259765625 0 20589.13
878.4091796875 0 5934.336
879.423583984375 0 41716.234
880.42822265625 0 26762.535
881.4259033203125 0 6447.6533
889.40771484375 0 10850.632
893.4402465820312 0 9903.152
894.4424438476562 0 6354.5493
895.4559326171875 0 10481.023
896.4608764648438 0 4037.7021
907.4195556640625 0 32416.451
908.4213256835938 0 19633.488
946.5015258789062 0 10161.061
947.4957885742188 0 15306.759
949.5017700195312 0 5906.7104
956.4921264648438 0 6383.1323
957.4812622070312 0 5512.168
964.5133056640625 0 136238.67
965.4118041992188 0 5614.591
965.5171508789062 0 75193.98
966.5182495117188 0 26612.498
974.3833618164062 0 4678.724
974.4969482421875 0 74704.195 b Water loss 7
975.499755859375 0 38242.816 b Ammonia loss 7
976.4822387695312 0 35465.62 y Water loss 2
977.4811401367188 0 21286.346
978.4794311523438 0 4814.0347
991.497802734375 0 5384.4575
992.5084228515625 0 1130724 b 7
993.5114135742188 0 671650.56
994.4987182617188 0 423165.94 y 2
995.4942626953125 0 187748.97
996.4934692382812 0 53794.32
997.4949340820312 0 10085.139
1089.563232421875 0 3988.3333 b 8
1093.5560302734375 0 29416.914 y 1
1094.5599365234375 0 18892.113
1107.564208984375 0 4311.394
1361.0574951171875 0 3660.9363
2409.72314453125 0 3956.2458
3226.327392578125 0 4997.1626
3389.08642578125 0 4139.289

Spectrum Details

|  |  |
| --- | --- |
| Matched peaks? Matched peaks The total absolute number of peaks matched. Additionally in brackets the total fraction of peaks matched and the total number of peaks is shown. | 37 (7.33% of 505) |
| FDR? FDR The false discovery rate estimated for this peptide. It is calculated by matching all theoretical fragments with a non-integer shift with the raw peaks for this spectrum. This is done with 40 different shifts. The resulting percentage is the average number of annotated peaks over the number of annotated peaks with the correct spectrum. | 0.90% |
| Satellite FDR? Satellite FDR See the FDR for details on its calculation. This satellite ion specific FDR only contains the satellite ions (d/w) for I/L/J positions. | - |
| PSM Score? PSM Score The PSM Score as given by Hecklib to this annotated spectrum. It is shown with three significant figures. | 440 |

#### Spectrum 9005? Spectrum 9005 The raw spectrum of this peptide as annotated by Hecklib. The fragments are coloured according to ion type (see legend). Any peaks with a star '\*' as text can be hovered over to see the full details, first the ion type second the mass shift type. By hovering over the amino acids in the peptide or ions in the legend the corresponding peaks are highlighted. By toggling the 'Unassigned' label you can turn the background (unassigned) peaks on or off in the plot. By updating the slider in the Ion legend you can update the spectrum to only show the top X% of the peaks with labels. The top X% means any peak that is within X% of the highest intensity. By dragging in the spectrum you can zoom in to a specific part of the spectrum and use 'Zoom Out' to get back to the original zoom level. The annotation of the spectrum is based on the given sequence in the peptides file and is done with different software so inconsistencies are likely. The peaks are annotated based on the given sequence, with 20 ppm tolerance.

Copy Data

##### Spectrum 9005 (TSV)

###### Preview

```
Loading example...
```

*Click on the button to copy the data to your clipboard.*

Mz MinMz MaxIntensity Max

WidthHeightPeptide font sizePeptide stroke widthSpectrum font sizeSpectrum stroke widthCompact peptide

Ion legend

wxyz

abcd

OtherUnassignedIon^Charge^_Position_Show for top:%

JVKDYFPEPV

02.16e+44.31e+46.47e+48.62e+4

Zoom Out

y^+1^_2_c^+1^_3_y^+1^_4_c^+1^_4_c^+1^_4_y^+1^_5_c^+1^_5_c^+1^_5_z^+1^_6_c^+1^_6_c^+1^_7_c^+1^_7_z^+1^_8_c^+1^_8_y^+1^_8_z^+1^_9_c^+1^_9_

0591118217732364

Fragment Matches Table

Show background peaks

| Position | Ion type | Intensity | mz Theoretical | mz Error (Th) | mz Error (ppm) | Charge | Series Number |
| --- | --- | --- | --- | --- | --- | --- | --- |
| - | - | 386.8 | 120.9 | - | - | 0 | - |
| - | - | 417.3 | 121.3 | - | - | 0 | - |
| - | - | 481.4 | 123.6 | - | - | 0 | - |
| - | - | 671.6 | 129.1 | - | - | 0 | - |
| - | - | 438 | 131.4 | - | - | 0 | - |
| - | - | 452.1 | 133.1 | - | - | 0 | - |
| - | - | 457.3 | 143.3 | - | - | 0 | - |
| - | - | 499.4 | 147.8 | - | - | 0 | - |
| - | - | 404.1 | 147.9 | - | - | 0 | - |
| - | - | 390.3 | 152 | - | - | 0 | - |
| - | - | 387.8 | 162.4 | - | - | 0 | - |
| - | - | 685.3 | 173.1 | - | - | 0 | - |
| - | - | 1049 | 173.5 | - | - | 0 | - |
| - | - | 451.6 | 181.9 | - | - | 0 | - |
| - | - | 1385 | 185.2 | - | - | 0 | - |
| - | - | 450.6 | 186.1 | - | - | 0 | - |
| - | - | 444.5 | 192.3 | - | - | 0 | - |
| - | - | 443.5 | 193.7 | - | - | 0 | - |
| - | - | 750.6 | 213.2 | - | - | 0 | - |
| 9 | y | 2.836E+04 | 215.1 | 0.000126 | 0.5856 | +1 | 2 |
| - | - | 3178 | 216.1 | - | - | 0 | - |
| - | - | 598.7 | 223.9 | - | - | 0 | - |
| - | - | 626.3 | 227.1 | - | - | 0 | - |
| - | - | 483.8 | 264.8 | - | - | 0 | - |
| - | - | 559.4 | 281.5 | - | - | 0 | - |
| - | - | 6350 | 357.3 | - | - | 0 | - |
| 3 | c | 1930 | 358.3 | 0.003709 | 10.35 | +1 | 3 |
| - | - | 669.5 | 383.1 | - | - | 0 | - |
| - | - | 661.8 | 391.9 | - | - | 0 | - |
| - | - | 733.3 | 397.2 | - | - | 0 | - |
| - | - | 590.8 | 401.2 | - | - | 0 | - |
| - | - | 646.6 | 415.5 | - | - | 0 | - |
| 7 | y | 2367 | 441.2 | 0.0004569 | 1.036 | +1 | 4 |
| - | - | 638.1 | 442.2 | - | - | 0 | - |
| 4 | c | 919.8 | 456.3 | 0.00215 | 4.712 | +1 | 4 |
| 4 | c | 2106 | 473.3 | 0.0002927 | 0.6184 | +1 | 4 |
| - | - | 701.7 | 482.3 | - | - | 0 | - |
| - | - | 725.8 | 487.2 | - | - | 0 | - |
| - | - | 571.2 | 519.3 | - | - | 0 | - |
| - | - | 1292 | 554.3 | - | - | 0 | - |
| - | - | 590.9 | 567.8 | - | - | 0 | - |
| - | - | 540.2 | 569.3 | - | - | 0 | - |
| - | - | 744.7 | 580.3 | - | - | 0 | - |
| 6 | y | 3380 | 588.3 | 0.0004826 | 0.8204 | +1 | 5 |
| - | - | 1098 | 589.3 | - | - | 0 | - |
| - | - | 636.5 | 592.4 | - | - | 0 | - |
| - | - | 658.2 | 604.3 | - | - | 0 | - |
| 5 | c | 1313 | 619.3 | 0.003314 | 5.351 | +1 | 5 |
| - | - | 3923 | 635.4 | - | - | 0 | - |
| 5 | c | 4933 | 636.4 | 0.0008715 | 1.37 | +1 | 5 |
| - | - | 1740 | 637.4 | - | - | 0 | - |
| 5 | z | 846.2 | 735.3 | 0.0006893 | 0.9373 | +1 | 6 |
| - | - | 675.2 | 736.4 | - | - | 0 | - |
| - | - | 3097 | 739.4 | - | - | 0 | - |
| - | - | 3288 | 740.4 | - | - | 0 | - |
| - | - | 1917 | 741.4 | - | - | 0 | - |
| 6 | c | 5658 | 766.4 | 1.174E-05 | 0.01532 | +1 | 6 |
| - | - | 2757 | 767.4 | - | - | 0 | - |
| - | - | 1819 | 780.4 | - | - | 0 | - |
| - | - | 1869 | 781.4 | - | - | 0 | - |
| - | - | 672.5 | 831.9 | - | - | 0 | - |
| - | - | 1090 | 851.4 | - | - | 0 | - |
| 7 | c | 726.2 | 863.5 | 0.003072 | 3.557 | +1 | 7 |
| - | - | 3715 | 879.5 | - | - | 0 | - |
| - | - | 835.6 | 880.4 | - | - | 0 | - |
| 7 | c | 1.323E+04 | 880.5 | 0.0001011 | 0.1148 | +1 | 7 |
| - | - | 6108 | 881.5 | - | - | 0 | - |
| - | - | 1783 | 882.5 | - | - | 0 | - |
| - | - | 699.7 | 906.4 | - | - | 0 | - |
| - | - | 911.9 | 907.4 | - | - | 0 | - |
| - | - | 627.7 | 909.2 | - | - | 0 | - |
| - | - | 4447 | 934.5 | - | - | 0 | - |
| - | - | 2401 | 935.5 | - | - | 0 | - |
| - | - | 737.6 | 936.5 | - | - | 0 | - |
| - | - | 809.2 | 964.5 | - | - | 0 | - |
| - | - | 1212 | 965.5 | - | - | 0 | - |
| 3 | z | 9035 | 978.5 | 0.0002451 | 0.2505 | +1 | 8 |
| - | - | 6145 | 979.5 | - | - | 0 | - |
| - | - | 1647 | 980.5 | - | - | 0 | - |
| - | - | 1421 | 990.6 | - | - | 0 | - |
| - | - | 722 | 991.6 | - | - | 0 | - |
| 8 | c | 1.217E+04 | 992.5 | 3.178E-05 | 0.03202 | +1 | 8 |
| - | - | 8104 | 993.5 | - | - | 0 | - |
| 3 | y | 3755 | 994.5 | 0.01198 | 12.04 | +1 | 8 |
| - | - | 2248 | 995.5 | - | - | 0 | - |
| 2 | z | 2597 | 1078 | 0.0001166 | 0.1082 | +1 | 9 |
| - | - | 2034 | 1079 | - | - | 0 | - |
| 9 | c | 1.47E+04 | 1107 | 0.001281 | 1.157 | +1 | 9 |
| - | - | 8604 | 1108 | - | - | 0 | - |
| - | - | 2444 | 1109 | - | - | 0 | - |
| - | - | 904.5 | 1119 | - | - | 0 | - |
| - | - | 1022 | 1136 | - | - | 0 | - |
| - | - | 640.7 | 1137 | - | - | 0 | - |
| - | - | 3468 | 1148 | - | - | 0 | - |
| - | - | 2313 | 1149 | - | - | 0 | - |
| - | - | 896.3 | 1150 | - | - | 0 | - |
| - | - | 2252 | 1152 | - | - | 0 | - |
| - | - | 1924 | 1153 | - | - | 0 | - |
| - | - | 2826 | 1162 | - | - | 0 | - |
| - | - | 2684 | 1163 | - | - | 0 | - |
| - | - | 5019 | 1164 | - | - | 0 | - |
| - | - | 2543 | 1165 | - | - | 0 | - |
| - | - | 821.8 | 1166 | - | - | 0 | - |
| - | - | 2082 | 1180 | - | - | 0 | - |
| - | - | 660.3 | 1181 | - | - | 0 | - |
| - | - | 1452 | 1189 | - | - | 0 | - |
| - | - | 970.5 | 1190 | - | - | 0 | - |
| - | - | 1.739E+04 | 1191 | - | - | 0 | - |
| - | - | 1.319E+04 | 1192 | - | - | 0 | - |
| - | - | 4497 | 1193 | - | - | 0 | - |
| - | - | 794.4 | 1205 | - | - | 0 | - |
| - | - | 1994 | 1206 | - | - | 0 | - |
| - | - | 2.309E+04 | 1207 | - | - | 0 | - |
| - | - | 8.539E+04 | 1208 | - | - | 0 | - |
| - | - | 5.653E+04 | 1209 | - | - | 0 | - |
| - | - | 1.992E+04 | 1210 | - | - | 0 | - |
| - | - | 3981 | 1211 | - | - | 0 | - |
| - | - | 620.1 | 1241 | - | - | 0 | - |
| - | - | 716.4 | 2104 | - | - | 0 | - |
| - | - | 711.5 | 2341 | - | - | 0 | - |

m/z Charge Intensity FragmentType MassShift Position
120.88493347167969 0 386.84543
121.30936431884766 0 417.34116
123.56439208984375 0 481.4475
129.10250854492188 0 671.5733
131.37515258789062 0 438.03558
133.08578491210938 0 452.0523
143.2843780517578 0 457.31802
147.8327178955078 0 499.44806
147.8662567138672 0 404.0923
152.03709411621094 0 390.3178
162.42189025878906 0 387.84775
173.09149169921875 0 685.3157
173.45263671875 0 1049.1084
181.9421844482422 0 451.62137
185.1650390625 0 1385.0134
186.0881805419922 0 450.61212
192.28990173339844 0 444.49442
193.7004852294922 0 443.4674
213.15994262695312 0 750.63336
215.13914489746094 0 28360.682 y 8
216.14259338378906 0 3177.7766
223.9035186767578 0 598.70483
227.102783203125 0 626.2558
264.7906799316406 0 483.84262
281.4610595703125 0 559.43536
357.2734680175781 0 6350.367
358.2775573730469 0 1929.5623 c 2
383.12384033203125 0 669.4703
391.9351501464844 0 661.758
397.2458190917969 0 733.29016
401.239501953125 0 590.83307
415.51226806640625 0 646.62823
441.2348327636719 0 2366.796 y 6
442.23638916015625 0 638.1018
456.2795104980469 0 919.78186 c Ammonia loss 3
473.3085021972656 0 2106.2563 c 3
482.2624206542969 0 701.6788
487.223388671875 0 725.7652
519.29833984375 0 571.17505
554.2606201171875 0 1291.9454
567.8067016601562 0 590.895
569.282958984375 0 540.17834
580.2693481445312 0 744.7177
588.3023071289062 0 3380.24 y 5
589.3048706054688 0 1098.0773
592.357421875 0 636.49304
604.328857421875 0 658.1857
619.3416748046875 0 1312.8679 c Ammonia loss 4
635.3643798828125 0 3923.0388
636.3706665039062 0 4933.4834 c 4
637.373046875 0 1740.0398
735.3480834960938 0 846.2085 z 4
736.3543701171875 0 675.1956
739.4285888671875 0 3097.0737
740.4312744140625 0 3287.8665
741.4347534179688 0 1916.7643
766.4133911132812 0 5658.0215 c Ammonia loss 5
767.41650390625 0 2756.8625
780.3543701171875 0 1818.78
781.3590087890625 0 1869.3344
831.8983764648438 0 672.4581
851.3807983398438 0 1089.749
863.46923828125 0 726.1527 c Ammonia loss 6
879.4847412109375 0 3715.3171
880.3961791992188 0 835.59924
880.4926147460938 0 13232.211 c 6
881.4956665039062 0 6107.804
882.496826171875 0 1783.4915
906.4424438476562 0 699.7393
907.44091796875 0 911.9233
909.1639404296875 0 627.7069
934.4791870117188 0 4447.2617
935.4819946289062 0 2401.0095
936.4830932617188 0 737.5588
964.5165405273438 0 809.2047
965.5167236328125 0 1212.231
978.4690551757812 0 9034.534 z 2
979.471923828125 0 6145.187
980.4735107421875 0 1647.0608
990.5518188476562 0 1421.0679
991.559814453125 0 721.9864
992.5087280273438 0 12174.1875 c Ammonia loss 7
993.5112915039062 0 8103.7505
994.5 0 3755.2607 y 2
995.4972534179688 0 2248.2957
1077.53759765625 0 2596.5059 z 1
1078.5377197265625 0 2033.8997
1106.5867919921875 0 14704.71 c 8
1107.5908203125 0 8603.985
1108.5931396484375 0 2444.1692
1118.5970458984375 0 904.50885
1135.626708984375 0 1021.5096
1136.6202392578125 0 640.7043
1147.5804443359375 0 3467.8694
1148.6201171875 0 2312.7305
1149.620361328125 0 896.3078
1151.5889892578125 0 2251.8767
1152.5946044921875 0 1923.9255
1161.6395263671875 0 2826.2415
1162.6497802734375 0 2683.6323
1163.65673828125 0 5019.462
1164.6590576171875 0 2542.6792
1165.66552734375 0 821.83435
1179.6610107421875 0 2082.273
1180.6556396484375 0 660.2622
1188.601806640625 0 1452.1262
1189.6192626953125 0 970.52356
1190.6201171875 0 17386.834
1191.6236572265625 0 13188.582
1192.625 0 4496.7144
1204.6043701171875 0 794.3699
1205.6114501953125 0 1993.9504
1206.638671875 0 23093.98
1207.6463623046875 0 85393.49
1208.649658203125 0 56528.79
1209.652099609375 0 19916.184
1210.6536865234375 0 3980.6653
1240.62109375 0 620.0669
2104.290283203125 0 716.37915
2340.87255859375 0 711.52515

Spectrum Details

|  |  |
| --- | --- |
| Matched peaks? Matched peaks The total absolute number of peaks matched. Additionally in brackets the total fraction of peaks matched and the total number of peaks is shown. | 17 (14.17% of 120) |
| FDR? FDR The false discovery rate estimated for this peptide. It is calculated by matching all theoretical fragments with a non-integer shift with the raw peaks for this spectrum. This is done with 40 different shifts. The resulting percentage is the average number of annotated peaks over the number of annotated peaks with the correct spectrum. | 1.96% |
| Satellite FDR? Satellite FDR See the FDR for details on its calculation. This satellite ion specific FDR only contains the satellite ions (d/w) for I/L/J positions. | - |
| PSM Score? PSM Score The PSM Score as given by Hecklib to this annotated spectrum. It is shown with three significant figures. | 200 |

#### Reverse Lookup? Reverse Lookup All places where this read could be placed.

| Group | Segment | Template | Template Part | Read Part | Score | Unique |
| --- | --- | --- | --- | --- | --- | --- |
| Homo sapiens Heavy Chain | IGHC | IGHG1 | [27..37] | [0..10] | 80 | False |
| Homo sapiens Heavy Chain | IGHC | IGHG3 | [27..37] | [0..10] | 80 | False |
| Homo sapiens Heavy Chain | IGHC | IGHG2 | [27..37] | [0..10] | 80 | False |
| Homo sapiens Heavy Chain | IGHC | IGHG4 | [27..37] | [0..10] | 80 | False |

| Recombined | Template Part | Read Part | Score | Unique |
| --- | --- | --- | --- | --- |
| REC-0-1 | [149..159] | [0..10] | 80 | True |

#### Meta Information from Multiple reads

##### Number of combined reads

7

##### Intensity

0.9022

##### TotalArea

1.149E+09

##### Changes to the peptide sequence

JVKDYFPEPV

L→JNo support for either Leucine or Isoleucine based on side chain ions (Position: 1)

#### Positional Score

Copy Data

##### Positional Score (TSV)

###### Preview

```
Loading example...
```

*Click on the button to copy the data to your clipboard.*

100123456789

Label Value
"0" 0.703
"1" 0.693
"2" 0.659
"3" 0.684
"4" 0.707
"5" 0.683
"6" 0.689
"7" 0.707
"8" 0.691
"9" 0.697

#### Meta Information from PEAKS

##### Scan Identifier

F1:8316

##### Original sequence

L

V

K

D

Y

F

P

E

P

V

##### Posttranslational Modifications

##### Source File

D:\separate\_stitch\_analyses\xle-disambiguation\raw\20210323\_F1\_UM1\_Peng0013\_SA\_F59\_ingel\_3ug\_ELA.raw

##### Fraction

1

##### Scan Feature

F1:8542

##### De Novo Score

99

##### ConfidenceScore

99

### m/z

603.8257

##### Mass

1205.6331

##### Charge

2

##### Retention Time

45.61

##### Predicted Retention Time

-

##### Area

2.872E+08

##### Parts Per Million

3.1

##### Fragmentation mode

ETHCD

##### Originating file

01 D:\separate\_stitch\_analyses\xle-disambiguation\20210325\_F59\_3ug\_DENOVO\_12.csv

#### Meta Information from PEAKS

##### Scan Identifier

F1:8595

##### Original sequence

L

V

K

D

Y

F

P

E

P

V

##### Posttranslational Modifications

##### Source File

D:\separate\_stitch\_analyses\xle-disambiguation\raw\20210323\_F1\_UM1\_Peng0013\_SA\_F59\_ingel\_3ug\_ELA.raw

##### Fraction

1

##### Scan Feature

-

##### De Novo Score

98

##### ConfidenceScore

98

### m/z

603.8249

##### Mass

1205.6331

##### Charge

2

##### Retention Time

47.18

##### Predicted Retention Time

-

##### Area

0

##### Parts Per Million

1.8

##### Fragmentation mode

ETHCD

##### Originating file

01 D:\separate\_stitch\_analyses\xle-disambiguation\20210325\_F59\_3ug\_DENOVO\_12.csv

#### Meta Information from PEAKS

##### Scan Identifier

F1:8799

##### Original sequence

L

V

K

D

Y

F

P

E

P

V

##### Posttranslational Modifications

##### Source File

D:\separate\_stitch\_analyses\xle-disambiguation\raw\20210323\_F1\_UM1\_Peng0013\_SA\_F59\_ingel\_3ug\_ELA.raw

##### Fraction

1

##### Scan Feature

-

##### De Novo Score

97

##### ConfidenceScore

97

### m/z

603.8249

##### Mass

1205.6331

##### Charge

2

##### Retention Time

48.37

##### Predicted Retention Time

-

##### Area

0

##### Parts Per Million

1.8

##### Fragmentation mode

ETHCD

##### Originating file

01 D:\separate\_stitch\_analyses\xle-disambiguation\20210325\_F59\_3ug\_DENOVO\_12.csv

#### Meta Information from PEAKS

##### Scan Identifier

F1:8526

##### Original sequence

L

V

K

D

Y

F

P

E

P

V

##### Posttranslational Modifications

##### Source File

D:\separate\_stitch\_analyses\xle-disambiguation\raw\20210323\_F1\_UM1\_Peng0013\_SA\_F59\_ingel\_3ug\_ELA.raw

##### Fraction

1

##### Scan Feature

F1:8542

##### De Novo Score

97

##### ConfidenceScore

97

### m/z

603.8257

##### Mass

1205.6331

##### Charge

2

##### Retention Time

45.61

##### Predicted Retention Time

-

##### Area

2.872E+08

##### Parts Per Million

3.1

##### Fragmentation mode

ETHCD

##### Originating file

01 D:\separate\_stitch\_analyses\xle-disambiguation\20210325\_F59\_3ug\_DENOVO\_12.csv

#### Meta Information from PEAKS

##### Scan Identifier

F1:8462

##### Original sequence

L

V

K

D

Y

F

P

E

P

V

##### Posttranslational Modifications

##### Source File

D:\separate\_stitch\_analyses\xle-disambiguation\raw\20210323\_F1\_UM1\_Peng0013\_SA\_F59\_ingel\_3ug\_ELA.raw

##### Fraction

1

##### Scan Feature

F1:8542

##### De Novo Score

96

##### ConfidenceScore

96

### m/z

603.8257

##### Mass

1205.6331

##### Charge

2

##### Retention Time

45.61

##### Predicted Retention Time

-

##### Area

2.872E+08

##### Parts Per Million

3.1

##### Fragmentation mode

HCD

##### Originating file

01 D:\separate\_stitch\_analyses\xle-disambiguation\20210325\_F59\_3ug\_DENOVO\_12.csv

#### Meta Information from PEAKS

##### Scan Identifier

F1:8375

##### Original sequence

L

V

K

D

Y

F

P

E

P

V

##### Posttranslational Modifications

##### Source File

D:\separate\_stitch\_analyses\xle-disambiguation\raw\20210323\_F1\_UM1\_Peng0013\_SA\_F59\_ingel\_3ug\_ELA.raw

##### Fraction

1

##### Scan Feature

F1:8542

##### De Novo Score

96

##### ConfidenceScore

96

### m/z

603.8257

##### Mass

1205.6331

##### Charge

2

##### Retention Time

45.61

##### Predicted Retention Time

-

##### Area

2.872E+08

##### Parts Per Million

3.1

##### Fragmentation mode

HCD

##### Originating file

01 D:\separate\_stitch\_analyses\xle-disambiguation\20210325\_F59\_3ug\_DENOVO\_12.csv

#### Meta Information from PEAKS

##### Scan Identifier

F1:9005

##### Original sequence

L

V

K

D

Y

F

P

E

P

V

##### Posttranslational Modifications

##### Source File

D:\separate\_stitch\_analyses\xle-disambiguation\raw\20210323\_F1\_UM1\_Peng0013\_SA\_F59\_ingel\_3ug\_ELA.raw

##### Fraction

1

##### Scan Feature

-

##### De Novo Score

96

##### ConfidenceScore

96

### m/z

603.8259

##### Mass

1205.6331

##### Charge

2

##### Retention Time

49.58

##### Predicted Retention Time

-

##### Area

0

##### Parts Per Million

3.5

##### Fragmentation mode

ETHCD

##### Originating file

01 D:\separate\_stitch\_analyses\xle-disambiguation\20210325\_F59\_3ug\_DENOVO\_12.csv
