## Supplementary material for "A handle on mass coincidence errors in *de novo* sequencing of antibodies by bottom-up proteomics": Combined_075.html

Details Combined\_075 | Stitch OverviewUndefined

### Read Combined\_075

#### Sequence (length=9)

VLQSSGJYS

#### Spectrum 5480? Spectrum 5480 The raw spectrum of this peptide as annotated by Hecklib. The fragments are coloured according to ion type (see legend). Any peaks with a star '\*' as text can be hovered over to see the full details, first the ion type second the mass shift type. By hovering over the amino acids in the peptide or ions in the legend the corresponding peaks are highlighted. By toggling the 'Unassigned' label you can turn the background (unassigned) peaks on or off in the plot. By updating the slider in the Ion legend you can update the spectrum to only show the top X% of the peaks with labels. The top X% means any peak that is within X% of the highest intensity. By dragging in the spectrum you can zoom in to a specific part of the spectrum and use 'Zoom Out' to get back to the original zoom level. The annotation of the spectrum is based on the given sequence in the peptides file and is done with different software so inconsistencies are likely. The peaks are annotated based on the given sequence, with 20 ppm tolerance.

Copy Data

##### Spectrum 5480 (TSV)

###### Preview

```
Loading example...
```

*Click on the button to copy the data to your clipboard.*

Mz MinMz MaxIntensity Max

WidthHeightPeptide font sizePeptide stroke widthSpectrum font sizeSpectrum stroke widthCompact peptide

Ion legend

wxyz

abcd

OtherUnassignedIon^Charge^_Position_Show for top:%

VLQSSGJYS

09.91e+31.98e+42.97e+43.96e+4

Zoom Out

y^+1^_2_y^+1^_2_c^+1^_3_c^+1^_3_c^+1^_4_c^+1^_4_y^+1^_5_c^+1^_5_y^+1^_5_c^+1^_5_c^+1^_6_c^+1^_6_c^+1^_6_z^+1^_6_y^+1^_6_c^+1^_7_c^+1^_7_z^+1^_7_y^+1^_7_z^+1^_7_y^+1^_7_w^+1^_8_z^+1^_8_z^+1^_8_c^+1^_8_y^+1^_8_c^+1^_8_

0816163324493266

Fragment Matches Table

Show background peaks

| Position | Ion type | Intensity | mz Theoretical | mz Error (Th) | mz Error (ppm) | Charge | Series Number |
| --- | --- | --- | --- | --- | --- | --- | --- |
| - | - | 377.1 | 125.7 | - | - | 0 | - |
| - | - | 547.6 | 128.1 | - | - | 0 | - |
| - | - | 997.5 | 131.1 | - | - | 0 | - |
| - | - | 7146 | 133.1 | - | - | 0 | - |
| - | - | 520.8 | 134.1 | - | - | 0 | - |
| - | - | 1.638E+04 | 136.1 | - | - | 0 | - |
| - | - | 420.1 | 136.7 | - | - | 0 | - |
| - | - | 983.6 | 137.1 | - | - | 0 | - |
| - | - | 863.8 | 146.1 | - | - | 0 | - |
| - | - | 488.5 | 149 | - | - | 0 | - |
| - | - | 2591 | 149 | - | - | 0 | - |
| - | - | 701.2 | 167.1 | - | - | 0 | - |
| - | - | 1390 | 175.1 | - | - | 0 | - |
| - | - | 421.9 | 176.8 | - | - | 0 | - |
| - | - | 4598 | 177.1 | - | - | 0 | - |
| - | - | 655.7 | 178.1 | - | - | 0 | - |
| - | - | 5392 | 185.2 | - | - | 0 | - |
| - | - | 524.5 | 190.4 | - | - | 0 | - |
| - | - | 7553 | 213.2 | - | - | 0 | - |
| - | - | 706.7 | 216.1 | - | - | 0 | - |
| - | - | 730.3 | 219.1 | - | - | 0 | - |
| - | - | 4590 | 221.1 | - | - | 0 | - |
| - | - | 2342 | 221.1 | - | - | 0 | - |
| - | - | 818.1 | 223.1 | - | - | 0 | - |
| - | - | 533.3 | 223.8 | - | - | 0 | - |
| - | - | 1506 | 225 | - | - | 0 | - |
| - | - | 1330 | 233.1 | - | - | 0 | - |
| - | - | 612 | 235.2 | - | - | 0 | - |
| - | - | 6054 | 239.1 | - | - | 0 | - |
| - | - | 1086 | 240.1 | - | - | 0 | - |
| - | - | 1894 | 242.1 | - | - | 0 | - |
| 8 | y | 737.4 | 251.1 | 0.0005313 | 2.116 | +1 | 2 |
| - | - | 1046 | 265.2 | - | - | 0 | - |
| - | - | 522.9 | 265.8 | - | - | 0 | - |
| 8 | y | 6243 | 269.1 | 8.39E-06 | 0.03118 | +1 | 2 |
| - | - | 798.5 | 270.1 | - | - | 0 | - |
| - | - | 850.6 | 281.1 | - | - | 0 | - |
| - | - | 1524 | 283.1 | - | - | 0 | - |
| - | - | 7102 | 295.1 | - | - | 0 | - |
| - | - | 2193 | 296.1 | - | - | 0 | - |
| - | - | 658.3 | 297.1 | - | - | 0 | - |
| - | - | 1747 | 299.1 | - | - | 0 | - |
| - | - | 688.8 | 301.1 | - | - | 0 | - |
| - | - | 700.1 | 311.1 | - | - | 0 | - |
| - | - | 896.2 | 313.1 | - | - | 0 | - |
| - | - | 621 | 314.1 | - | - | 0 | - |
| - | - | 1045 | 314.2 | - | - | 0 | - |
| - | - | 1809 | 315.2 | - | - | 0 | - |
| - | - | 637.8 | 317.2 | - | - | 0 | - |
| - | - | 660.1 | 332.2 | - | - | 0 | - |
| - | - | 786.1 | 340.2 | - | - | 0 | - |
| - | - | 762.5 | 341 | - | - | 0 | - |
| 3 | c | 4882 | 341.2 | 8.245E-05 | 0.2416 | +1 | 3 |
| - | - | 800.9 | 342.2 | - | - | 0 | - |
| - | - | 3176 | 345.2 | - | - | 0 | - |
| - | - | 858.9 | 347.2 | - | - | 0 | - |
| - | - | 2899 | 355.1 | - | - | 0 | - |
| - | - | 4500 | 357.2 | - | - | 0 | - |
| 3 | c | 2.927E+04 | 358.2 | 0.00013 | 0.3628 | +1 | 3 |
| - | - | 1768 | 359 | - | - | 0 | - |
| - | - | 4704 | 359.2 | - | - | 0 | - |
| - | - | 567.5 | 360.2 | - | - | 0 | - |
| - | - | 1308 | 361.2 | - | - | 0 | - |
| - | - | 1439 | 365.2 | - | - | 0 | - |
| - | - | 8787 | 369.1 | - | - | 0 | - |
| - | - | 3753 | 370.1 | - | - | 0 | - |
| - | - | 1693 | 371.1 | - | - | 0 | - |
| - | - | 820.7 | 371.2 | - | - | 0 | - |
| - | - | 1198 | 377.2 | - | - | 0 | - |
| - | - | 1288 | 384.3 | - | - | 0 | - |
| - | - | 1193 | 389.2 | - | - | 0 | - |
| - | - | 1473 | 395.2 | - | - | 0 | - |
| - | - | 596.5 | 396.2 | - | - | 0 | - |
| - | - | 1337 | 398.1 | - | - | 0 | - |
| - | - | 4986 | 402.3 | - | - | 0 | - |
| - | - | 708.7 | 403.3 | - | - | 0 | - |
| - | - | 1176 | 415.3 | - | - | 0 | - |
| - | - | 595.2 | 416.3 | - | - | 0 | - |
| - | - | 1408 | 426.3 | - | - | 0 | - |
| - | - | 615.9 | 428.2 | - | - | 0 | - |
| 4 | c | 1849 | 428.3 | 0.0006163 | 1.439 | +1 | 4 |
| - | - | 2570 | 429.1 | - | - | 0 | - |
| - | - | 726.8 | 429.3 | - | - | 0 | - |
| - | - | 1153 | 430.3 | - | - | 0 | - |
| - | - | 1562 | 441.2 | - | - | 0 | - |
| - | - | 9428 | 444.3 | - | - | 0 | - |
| 4 | c | 3.156E+04 | 445.3 | 9.868E-05 | 0.2216 | +1 | 4 |
| - | - | 6436 | 446.3 | - | - | 0 | - |
| - | - | 1120 | 447.3 | - | - | 0 | - |
| - | - | 1346 | 456.2 | - | - | 0 | - |
| - | - | 2312 | 458.2 | - | - | 0 | - |
| - | - | 686.5 | 458.3 | - | - | 0 | - |
| - | - | 767.6 | 459.2 | - | - | 0 | - |
| - | - | 1.758E+04 | 459.3 | - | - | 0 | - |
| - | - | 4380 | 460.3 | - | - | 0 | - |
| - | - | 871.4 | 461.3 | - | - | 0 | - |
| - | - | 729.6 | 464.8 | - | - | 0 | - |
| - | - | 728.3 | 473.2 | - | - | 0 | - |
| - | - | 1391 | 475.3 | - | - | 0 | - |
| - | - | 1076 | 476.2 | - | - | 0 | - |
| - | - | 843.6 | 476.3 | - | - | 0 | - |
| - | - | 2281 | 476.3 | - | - | 0 | - |
| - | - | 9099 | 477.2 | - | - | 0 | - |
| - | - | 825.7 | 477.3 | - | - | 0 | - |
| - | - | 2428 | 478.2 | - | - | 0 | - |
| - | - | 897.8 | 478.2 | - | - | 0 | - |
| - | - | 1366 | 497.3 | - | - | 0 | - |
| 5 | y | 1675 | 508.2 | 7.721E-05 | 0.1519 | +1 | 5 |
| 5 | c | 3311 | 515.3 | 8.202E-05 | 0.1592 | +1 | 5 |
| - | - | 3871 | 525.2 | - | - | 0 | - |
| 5 | y | 2107 | 526.3 | 0.001731 | 3.289 | +1 | 5 |
| - | - | 1853 | 531.3 | - | - | 0 | - |
| 5 | c | 1494 | 532.3 | 0.001992 | 3.742 | +1 | 5 |
| - | - | 653.8 | 536.3 | - | - | 0 | - |
| - | - | 939.1 | 545.3 | - | - | 0 | - |
| - | - | 1116 | 546.3 | - | - | 0 | - |
| - | - | 1443 | 554.3 | - | - | 0 | - |
| - | - | 937.1 | 555.3 | - | - | 0 | - |
| - | - | 868.6 | 556.3 | - | - | 0 | - |
| - | - | 967.8 | 557.3 | - | - | 0 | - |
| - | - | 1112 | 570.3 | - | - | 0 | - |
| 6 | c | 2855 | 571.3 | 0.01051 | 18.4 | +1 | 6 |
| 6 | c | 4570 | 572.3 | 0.00169 | 2.952 | +1 | 6 |
| - | - | 1207 | 573.3 | - | - | 0 | - |
| - | - | 763.3 | 586.3 | - | - | 0 | - |
| - | - | 2953 | 588.3 | - | - | 0 | - |
| 6 | c | 1.655E+04 | 589.3 | 0.0006532 | 1.108 | +1 | 6 |
| - | - | 4682 | 590.3 | - | - | 0 | - |
| - | - | 763.1 | 594.3 | - | - | 0 | - |
| 4 | z | 1308 | 597.3 | 0.0001425 | 0.2386 | +1 | 6 |
| - | - | 1100 | 598.3 | - | - | 0 | - |
| - | - | 2604 | 612.3 | - | - | 0 | - |
| 4 | y | 9904 | 613.3 | 0.0001898 | 0.3095 | +1 | 6 |
| - | - | 2248 | 614.3 | - | - | 0 | - |
| - | - | 667.3 | 620.3 | - | - | 0 | - |
| - | - | 830.7 | 638.3 | - | - | 0 | - |
| - | - | 1027 | 640.4 | - | - | 0 | - |
| - | - | 2437 | 657.4 | - | - | 0 | - |
| - | - | 1430 | 658.4 | - | - | 0 | - |
| - | - | 932.4 | 659.4 | - | - | 0 | - |
| - | - | 1808 | 667.4 | - | - | 0 | - |
| - | - | 1442 | 668.4 | - | - | 0 | - |
| - | - | 3228 | 669.4 | - | - | 0 | - |
| - | - | 2930 | 670.4 | - | - | 0 | - |
| - | - | 664.7 | 671.4 | - | - | 0 | - |
| - | - | 572.6 | 684.4 | - | - | 0 | - |
| 7 | c | 1.296E+04 | 685.4 | 0.0002211 | 0.3226 | +1 | 7 |
| - | - | 4187 | 686.4 | - | - | 0 | - |
| - | - | 2082 | 687.4 | - | - | 0 | - |
| - | - | 965.2 | 690.3 | - | - | 0 | - |
| 7 | c | 2.165E+04 | 702.4 | 0.0002684 | 0.3821 | +1 | 7 |
| - | - | 8074 | 703.4 | - | - | 0 | - |
| - | - | 1567 | 704.4 | - | - | 0 | - |
| - | - | 1075 | 706.3 | - | - | 0 | - |
| 3 | z | 3131 | 707.3 | 0.0001207 | 0.1706 | +1 | 7 |
| - | - | 735 | 707.4 | - | - | 0 | - |
| - | - | 842.5 | 708.3 | - | - | 0 | - |
| - | - | 606 | 720.4 | - | - | 0 | - |
| 3 | y | 803.8 | 723.3 | 0.0009615 | 1.329 | +1 | 7 |
| 3 | z | 5039 | 725.3 | 6.525E-05 | 0.08997 | +1 | 7 |
| - | - | 2398 | 726.3 | - | - | 0 | - |
| - | - | 4173 | 733.4 | - | - | 0 | - |
| - | - | 1926 | 734.4 | - | - | 0 | - |
| - | - | 1090 | 735.4 | - | - | 0 | - |
| 3 | y | 2892 | 741.3 | 0.0001126 | 0.1518 | +1 | 7 |
| - | - | 1060 | 742.3 | - | - | 0 | - |
| - | - | 2476 | 765.4 | - | - | 0 | - |
| - | - | 1203 | 766.4 | - | - | 0 | - |
| - | - | 1022 | 776.4 | - | - | 0 | - |
| - | - | 634.8 | 778.4 | - | - | 0 | - |
| 2 | w | 4325 | 795.4 | 6.494E-05 | 0.08164 | +1 | 8 |
| - | - | 1309 | 796.4 | - | - | 0 | - |
| - | - | 815.3 | 797.4 | - | - | 0 | - |
| - | - | 1153 | 803.4 | - | - | 0 | - |
| 2 | z | 1457 | 820.4 | 0.0003834 | 0.4673 | +1 | 8 |
| - | - | 6956 | 821.5 | - | - | 0 | - |
| - | - | 3099 | 822.5 | - | - | 0 | - |
| - | - | 710.7 | 823.5 | - | - | 0 | - |
| - | - | 6397 | 832.4 | - | - | 0 | - |
| - | - | 2637 | 833.4 | - | - | 0 | - |
| - | - | 875.4 | 834.4 | - | - | 0 | - |
| 2 | z | 9873 | 838.4 | 0.0002824 | 0.3368 | +1 | 8 |
| - | - | 4195 | 839.4 | - | - | 0 | - |
| - | - | 1491 | 840.4 | - | - | 0 | - |
| 8 | c | 3908 | 848.5 | 0.0009886 | 1.165 | +1 | 8 |
| - | - | 2009 | 849.5 | - | - | 0 | - |
| - | - | 692.1 | 850.5 | - | - | 0 | - |
| - | - | 1701 | 853.4 | - | - | 0 | - |
| 2 | y | 734.3 | 854.4 | 0.003122 | 3.654 | +1 | 8 |
| 8 | c | 3.925E+04 | 865.5 | 5.021E-05 | 0.05802 | +1 | 8 |
| - | - | 1.605E+04 | 866.5 | - | - | 0 | - |
| - | - | 6491 | 867.5 | - | - | 0 | - |
| - | - | 745.3 | 868.5 | - | - | 0 | - |
| - | - | 6320 | 881.4 | - | - | 0 | - |
| - | - | 2296 | 882.4 | - | - | 0 | - |
| - | - | 919.7 | 883.4 | - | - | 0 | - |
| - | - | 1077 | 908.5 | - | - | 0 | - |
| - | - | 5142 | 909.5 | - | - | 0 | - |
| - | - | 1646 | 910.5 | - | - | 0 | - |
| - | - | 1007 | 911.5 | - | - | 0 | - |
| - | - | 1158 | 919.5 | - | - | 0 | - |
| - | - | 955.8 | 926.5 | - | - | 0 | - |
| - | - | 801.3 | 927.5 | - | - | 0 | - |
| - | - | 7851 | 937.5 | - | - | 0 | - |
| - | - | 3636 | 938.5 | - | - | 0 | - |
| - | - | 1524 | 951.5 | - | - | 0 | - |
| - | - | 839 | 952.5 | - | - | 0 | - |
| - | - | 1.334E+04 | 953.5 | - | - | 0 | - |
| - | - | 2.187E+04 | 954.5 | - | - | 0 | - |
| - | - | 1.006E+04 | 955.5 | - | - | 0 | - |
| - | - | 2891 | 956.5 | - | - | 0 | - |
| - | - | 703.2 | 1288 | - | - | 0 | - |
| - | - | 771.2 | 1397 | - | - | 0 | - |
| - | - | 666.3 | 1398 | - | - | 0 | - |
| - | - | 782.5 | 1399 | - | - | 0 | - |
| - | - | 744.1 | 1432 | - | - | 0 | - |
| - | - | 797.8 | 2739 | - | - | 0 | - |
| - | - | 748.2 | 3072 | - | - | 0 | - |
| - | - | 652.8 | 3234 | - | - | 0 | - |

m/z Charge Intensity FragmentType MassShift Position
125.65748596191406 0 377.12683
128.08206176757812 0 547.5737
131.0706024169922 0 997.5173
133.0860595703125 0 7145.6445
134.0899200439453 0 520.8133
136.0758056640625 0 16378.448
136.70286560058594 0 420.1282
137.07913208007812 0 983.555
146.093017578125 0 863.8151
148.95509338378906 0 488.4911
149.044921875 0 2590.6667
167.0552520751953 0 701.2389
175.09642028808594 0 1390.4403
176.82127380371094 0 421.86456
177.11221313476562 0 4598.2705
178.1160888671875 0 655.7424
185.16482543945312 0 5391.749
190.44277954101562 0 524.52924
213.1597900390625 0 7553.1724
216.0982208251953 0 706.72876
219.12269592285156 0 730.34814
221.08433532714844 0 4590.3022
221.1385040283203 0 2341.7473
223.0633087158203 0 818.1014
223.8359832763672 0 533.2519
225.0430450439453 0 1506.4945
233.1245880126953 0 1329.7499
235.19923400878906 0 612.0361
239.09503173828125 0 6054.3306
240.0963134765625 0 1086.3003
242.14990234375 0 1894.4495
251.10316467285156 0 737.3602 y Water loss 7
265.16461181640625 0 1045.931
265.7780456542969 0 522.86414
269.1131896972656 0 6243.0156 y 7
270.11663818359375 0 798.5047
281.0511169433594 0 850.64136
283.1107177734375 0 1523.656
295.1033630371094 0 7102.145
296.1038818359375 0 2193.2998
297.100341796875 0 658.301
299.062255859375 0 1746.9111
301.0594787597656 0 688.7566
311.1064758300781 0 700.0659
313.11407470703125 0 896.1569
314.11279296875 0 621.04315
314.2311706542969 0 1044.8585
315.2389831542969 0 1809.305
317.18170166015625 0 637.822
332.2322692871094 0 660.1011
340.21185302734375 0 786.1062
341.0206604003906 0 762.52466
341.2184143066406 0 4882.15 c Ammonia loss 2
342.2216796875 0 800.90485
345.1768493652344 0 3176.0835
347.19287109375 0 858.89264
355.0697326660156 0 2898.8406
357.2372131347656 0 4499.698
358.2447509765625 0 29267.195 c 2
359.0282287597656 0 1768.4817
359.2474670410156 0 4704.203
360.2467346191406 0 567.5084
361.16650390625 0 1308.3085
365.20001220703125 0 1438.537
369.121826171875 0 8786.865
370.1231689453125 0 3753.1746
371.119873046875 0 1692.5604
371.2265625 0 820.6764
377.1778564453125 0 1198.2137
384.26031494140625 0 1288.4586
389.1612548828125 0 1193.435
395.2276306152344 0 1472.976
396.232421875 0 596.4983
398.1232604980469 0 1337.1022
402.27117919921875 0 4985.893
403.2721252441406 0 708.71765
415.2552490234375 0 1176.3086
416.2569580078125 0 595.21
426.2586975097656 0 1407.5963
428.2181396484375 0 615.9268
428.2509765625 0 1848.7087 c Ammonia loss 3
429.0896911621094 0 2569.8186
429.25360107421875 0 726.7915
430.2658996582031 0 1153.2566
441.2336730957031 0 1562.2574
444.2695617675781 0 9428.125
445.2770080566406 0 31558.232 c 3
446.2799987792969 0 6435.7026
447.2827453613281 0 1120.2699
456.20941162109375 0 1345.5453
458.22509765625 0 2311.7415
458.25933837890625 0 686.4899
459.2264099121094 0 767.62744
459.2803649902344 0 17584.21
460.2842712402344 0 4380.364
461.2829895019531 0 871.3625
464.7688903808594 0 729.6298
473.2347717285156 0 728.3488
475.2500305175781 0 1391.088
476.1856384277344 0 1076.2112
476.26904296875 0 843.60516
476.3066101074219 0 2280.9575
477.1751403808594 0 9099.018
477.3121032714844 0 825.74774
478.1778564453125 0 2427.5063
478.2111511230469 0 897.7573
497.27197265625 0 1366.3433
508.2401123046875 0 1675.472 y Water loss 4
515.282470703125 0 3310.7502 c Ammonia loss 4
525.2429809570312 0 3870.584
526.2490234375 0 2106.7017 y 4
531.30224609375 0 1853.409
532.3069458007812 0 1493.5394 c 4
536.280029296875 0 653.80084
545.3181762695312 0 939.0876
546.3265380859375 0 1116.4476
554.2959594726562 0 1443.2445
555.283935546875 0 937.093
556.2808837890625 0 868.64496
557.3300170898438 0 967.7819
570.30078125 0 1111.849
571.309326171875 0 2854.8203 c Water loss 5
572.3055419921875 0 4570.2485 c Ammonia loss 5
573.3063354492188 0 1206.6011
586.3197021484375 0 763.2566
588.3336181640625 0 2953.4436
589.3310546875 0 16548.797 c 5
590.3336181640625 0 4681.5435
594.2628173828125 0 763.07837
597.263916015625 0 1307.8933 z 3
598.2706909179688 0 1099.8588
612.2763061523438 0 2603.8938
613.2825927734375 0 9904.072 y 3
614.2861328125 0 2247.9907
620.282470703125 0 667.34937
638.3158569335938 0 830.71515
640.3696899414062 0 1027.3066
657.3931274414062 0 2436.836
658.3973999023438 0 1430.3013
659.4091186523438 0 932.42596
667.376220703125 0 1808.0527
668.3663330078125 0 1441.8376
669.3704833984375 0 3227.8604
670.375732421875 0 2929.909
671.37939453125 0 664.6605
684.3899536132812 0 572.6326
685.3876953125 0 12959.01 c Ammonia loss 6
686.3887329101562 0 4186.6724
687.4010009765625 0 2081.808
690.3082885742188 0 965.23346
702.4147338867188 0 21652.861 c 6
703.417236328125 0 8073.883
704.4174194335938 0 1567.3312
706.3089599609375 0 1075.031
707.3119506835938 0 3131.4768 z Water loss 2
707.3750610351562 0 735.00134
708.3168334960938 0 842.54315
720.3975830078125 0 605.9523
723.329833984375 0 803.7676 y Water loss 2
725.3225708007812 0 5038.665 z 2
726.3253173828125 0 2397.7812
733.36474609375 0 4173.171
734.3694458007812 0 1925.6364
735.3724365234375 0 1090.3707
741.3412475585938 0 2891.9744 y 2
742.344482421875 0 1059.6625
765.401123046875 0 2476.3542
766.407470703125 0 1202.7911
776.3668823242188 0 1021.79407
778.40380859375 0 634.84174
795.3519897460938 0 4324.7217 w 1
796.3579711914062 0 1309.4258
797.3549194335938 0 815.3404
803.4486083984375 0 1153.1115
820.395751953125 0 1457.4576 z Water loss 1
821.464599609375 0 6955.68
822.467529296875 0 3098.6953
823.4732055664062 0 710.65784
832.4328002929688 0 6396.887
833.437255859375 0 2636.89
834.4375610351562 0 875.4322
838.406982421875 0 9872.512 z 1
839.4105224609375 0 4194.7573
840.4190673828125 0 1490.5488
848.4502563476562 0 3907.966 c Ammonia loss 7
849.4501953125 0 2008.6403
850.4619750976562 0 692.086
853.4187622070312 0 1700.7223
854.4223022460938 0 734.3476 y 1
865.4778442382812 0 39248.977 c 7
866.48046875 0 16048.174
867.48291015625 0 6491.3716
868.489990234375 0 745.2579
881.4149169921875 0 6320.2227
882.419677734375 0 2296.3008
883.4171142578125 0 919.6682
908.4970703125 0 1076.7306
909.4810180664062 0 5141.6094
910.4808959960938 0 1646.1073
911.4830932617188 0 1007.24915
919.46044921875 0 1157.9114
926.5103149414062 0 955.8326
927.515869140625 0 801.2502
937.4751586914062 0 7850.897
938.47802734375 0 3636.244
951.4578247070312 0 1523.952
952.4671630859375 0 839.0424
953.4935913085938 0 13337.045
954.4990234375 0 21872.61
955.5006103515625 0 10059.372
956.4974975585938 0 2890.745
1287.6943359375 0 703.15
1396.7017822265625 0 771.17194
1397.7454833984375 0 666.2927
1398.72314453125 0 782.4718
1431.71484375 0 744.1364
2739.22900390625 0 797.7995
3072.343017578125 0 748.1517
3233.53369140625 0 652.8047

Spectrum Details

|  |  |
| --- | --- |
| Matched peaks? Matched peaks The total absolute number of peaks matched. Additionally in brackets the total fraction of peaks matched and the total number of peaks is shown. | 27 (12.33% of 219) |
| FDR? FDR The false discovery rate estimated for this peptide. It is calculated by matching all theoretical fragments with a non-integer shift with the raw peaks for this spectrum. This is done with 40 different shifts. The resulting percentage is the average number of annotated peaks over the number of annotated peaks with the correct spectrum. | 1.68% |
| Satellite FDR? Satellite FDR See the FDR for details on its calculation. This satellite ion specific FDR only contains the satellite ions (d/w) for I/L/J positions. | 7.14% |
| PSM Score? PSM Score The PSM Score as given by Hecklib to this annotated spectrum. It is shown with three significant figures. | 368 |

#### Spectrum 5352? Spectrum 5352 The raw spectrum of this peptide as annotated by Hecklib. The fragments are coloured according to ion type (see legend). Any peaks with a star '\*' as text can be hovered over to see the full details, first the ion type second the mass shift type. By hovering over the amino acids in the peptide or ions in the legend the corresponding peaks are highlighted. By toggling the 'Unassigned' label you can turn the background (unassigned) peaks on or off in the plot. By updating the slider in the Ion legend you can update the spectrum to only show the top X% of the peaks with labels. The top X% means any peak that is within X% of the highest intensity. By dragging in the spectrum you can zoom in to a specific part of the spectrum and use 'Zoom Out' to get back to the original zoom level. The annotation of the spectrum is based on the given sequence in the peptides file and is done with different software so inconsistencies are likely. The peaks are annotated based on the given sequence, with 20 ppm tolerance.

Copy Data

##### Spectrum 5352 (TSV)

###### Preview

```
Loading example...
```

*Click on the button to copy the data to your clipboard.*

Mz MinMz MaxIntensity Max

WidthHeightPeptide font sizePeptide stroke widthSpectrum font sizeSpectrum stroke widthCompact peptide

Ion legend

wxyz

abcd

OtherUnassignedIon^Charge^_Position_Show for top:%

VLQSSGJYS

02.00e+44.01e+46.01e+48.01e+4

Zoom Out

y^+1^_2_y^+1^_2_c^+1^_3_c^+1^_3_y^+2^_7_y^+1^_4_c^+1^_4_c^+1^_4_c^+1^_4_y^+1^_5_c^+1^_5_y^+1^_5_c^+1^_5_c^+1^_6_c^+1^_6_c^+1^_6_y^+1^_6_z^+1^_6_y^+1^_6_c^+1^_7_c^+1^_7_z^+1^_7_y^+1^_7_z^+1^_7_y^+1^_7_w^+1^_8_z^+1^_8_z^+1^_8_c^+1^_8_y^+1^_8_c^+1^_8_

0656131119672623

Fragment Matches Table

Show background peaks

| Position | Ion type | Intensity | mz Theoretical | mz Error (Th) | mz Error (ppm) | Charge | Series Number |
| --- | --- | --- | --- | --- | --- | --- | --- |
| - | - | 457.4 | 120.1 | - | - | 0 | - |
| - | - | 376.4 | 121.2 | - | - | 0 | - |
| - | - | 437.1 | 124 | - | - | 0 | - |
| - | - | 368.2 | 125.6 | - | - | 0 | - |
| - | - | 390.6 | 127.5 | - | - | 0 | - |
| - | - | 911.6 | 128.1 | - | - | 0 | - |
| - | - | 579.8 | 129.1 | - | - | 0 | - |
| - | - | 1636 | 131.1 | - | - | 0 | - |
| - | - | 1522 | 131.1 | - | - | 0 | - |
| - | - | 2.888E+04 | 133.1 | - | - | 0 | - |
| - | - | 2627 | 134.1 | - | - | 0 | - |
| - | - | 2.935E+04 | 136.1 | - | - | 0 | - |
| - | - | 1965 | 137.1 | - | - | 0 | - |
| - | - | 1646 | 146.1 | - | - | 0 | - |
| - | - | 1155 | 149 | - | - | 0 | - |
| - | - | 1372 | 149 | - | - | 0 | - |
| - | - | 448.5 | 151.6 | - | - | 0 | - |
| - | - | 424.7 | 157 | - | - | 0 | - |
| - | - | 497.7 | 165.1 | - | - | 0 | - |
| - | - | 479.9 | 166.1 | - | - | 0 | - |
| - | - | 738.2 | 167.1 | - | - | 0 | - |
| - | - | 1206 | 173.4 | - | - | 0 | - |
| - | - | 1703 | 175.1 | - | - | 0 | - |
| - | - | 2.116E+04 | 177.1 | - | - | 0 | - |
| - | - | 2402 | 178.1 | - | - | 0 | - |
| - | - | 6349 | 185.2 | - | - | 0 | - |
| - | - | 546.3 | 186.2 | - | - | 0 | - |
| - | - | 432.9 | 187.1 | - | - | 0 | - |
| - | - | 893 | 212.1 | - | - | 0 | - |
| - | - | 599.8 | 212.2 | - | - | 0 | - |
| - | - | 4906 | 213.2 | - | - | 0 | - |
| - | - | 834.5 | 214.2 | - | - | 0 | - |
| - | - | 487.5 | 215.1 | - | - | 0 | - |
| - | - | 1149 | 216.1 | - | - | 0 | - |
| - | - | 1248 | 219.1 | - | - | 0 | - |
| - | - | 3757 | 221.1 | - | - | 0 | - |
| - | - | 5773 | 221.1 | - | - | 0 | - |
| - | - | 949.7 | 222.1 | - | - | 0 | - |
| - | - | 871.1 | 223.1 | - | - | 0 | - |
| - | - | 1123 | 225 | - | - | 0 | - |
| - | - | 2639 | 225.1 | - | - | 0 | - |
| - | - | 2231 | 233.1 | - | - | 0 | - |
| - | - | 1547 | 237.1 | - | - | 0 | - |
| - | - | 6252 | 239.1 | - | - | 0 | - |
| - | - | 1048 | 239.1 | - | - | 0 | - |
| - | - | 1070 | 240.1 | - | - | 0 | - |
| - | - | 2.264E+04 | 242.2 | - | - | 0 | - |
| - | - | 531 | 247.1 | - | - | 0 | - |
| 8 | y | 1277 | 251.1 | 0.0002871 | 1.143 | +1 | 2 |
| - | - | 563.5 | 256.3 | - | - | 0 | - |
| - | - | 3262 | 265.2 | - | - | 0 | - |
| - | - | 689.2 | 266.2 | - | - | 0 | - |
| 8 | y | 1.085E+04 | 269.1 | 2.213E-05 | 0.08222 | +1 | 2 |
| - | - | 627.1 | 270.1 | - | - | 0 | - |
| - | - | 903.3 | 283.1 | - | - | 0 | - |
| - | - | 1396 | 283.2 | - | - | 0 | - |
| - | - | 542.7 | 285 | - | - | 0 | - |
| - | - | 6069 | 295.1 | - | - | 0 | - |
| - | - | 2038 | 296.1 | - | - | 0 | - |
| - | - | 723.8 | 297.1 | - | - | 0 | - |
| - | - | 488.3 | 297.9 | - | - | 0 | - |
| - | - | 2535 | 299.1 | - | - | 0 | - |
| - | - | 629 | 302.6 | - | - | 0 | - |
| - | - | 2784 | 309.2 | - | - | 0 | - |
| - | - | 654.9 | 310.2 | - | - | 0 | - |
| - | - | 781.3 | 311.1 | - | - | 0 | - |
| - | - | 1175 | 313.1 | - | - | 0 | - |
| - | - | 7050 | 313.2 | - | - | 0 | - |
| - | - | 1856 | 314.2 | - | - | 0 | - |
| - | - | 3715 | 315.2 | - | - | 0 | - |
| - | - | 1020 | 316.2 | - | - | 0 | - |
| - | - | 969.4 | 317.2 | - | - | 0 | - |
| - | - | 784.7 | 327.2 | - | - | 0 | - |
| - | - | 1754 | 327.2 | - | - | 0 | - |
| - | - | 681.7 | 328.2 | - | - | 0 | - |
| - | - | 1561 | 340.2 | - | - | 0 | - |
| 3 | c | 8393 | 341.2 | 0.0004792 | 1.404 | +1 | 3 |
| - | - | 1128 | 342.2 | - | - | 0 | - |
| - | - | 3493 | 345.2 | - | - | 0 | - |
| - | - | 2486 | 347.2 | - | - | 0 | - |
| - | - | 686.4 | 353.2 | - | - | 0 | - |
| - | - | 1425 | 353.2 | - | - | 0 | - |
| - | - | 1601 | 355.1 | - | - | 0 | - |
| - | - | 6068 | 357.2 | - | - | 0 | - |
| 3 | c | 4.782E+04 | 358.2 | 6.895E-05 | 0.1925 | +1 | 3 |
| - | - | 1705 | 359 | - | - | 0 | - |
| - | - | 7837 | 359.2 | - | - | 0 | - |
| - | - | 1005 | 360.2 | - | - | 0 | - |
| 3 | y | 899.1 | 362.2 | 0.001586 | 4.379 | +2 | 7 |
| - | - | 1539 | 365.2 | - | - | 0 | - |
| - | - | 8210 | 369.1 | - | - | 0 | - |
| - | - | 3228 | 370.1 | - | - | 0 | - |
| - | - | 945.4 | 371.1 | - | - | 0 | - |
| - | - | 1471 | 371.1 | - | - | 0 | - |
| - | - | 4045 | 371.2 | - | - | 0 | - |
| - | - | 586.2 | 372.2 | - | - | 0 | - |
| - | - | 2294 | 377.2 | - | - | 0 | - |
| - | - | 2126 | 384.3 | - | - | 0 | - |
| - | - | 581 | 397.2 | - | - | 0 | - |
| - | - | 6232 | 402.3 | - | - | 0 | - |
| - | - | 1374 | 403.3 | - | - | 0 | - |
| - | - | 630.5 | 414.2 | - | - | 0 | - |
| - | - | 5512 | 415.3 | - | - | 0 | - |
| - | - | 1624 | 416.3 | - | - | 0 | - |
| 6 | y | 1030 | 421.2 | 0.0008857 | 2.103 | +1 | 4 |
| - | - | 878 | 426.2 | - | - | 0 | - |
| - | - | 2342 | 426.3 | - | - | 0 | - |
| 4 | c | 700.3 | 427.3 | 0.0007502 | 1.756 | +1 | 4 |
| - | - | 857.8 | 428.2 | - | - | 0 | - |
| 4 | c | 3476 | 428.3 | 0.0001585 | 0.3702 | +1 | 4 |
| - | - | 1194 | 429.1 | - | - | 0 | - |
| - | - | 1816 | 430.3 | - | - | 0 | - |
| - | - | 794.7 | 441.3 | - | - | 0 | - |
| - | - | 1.633E+04 | 444.3 | - | - | 0 | - |
| 4 | c | 4.734E+04 | 445.3 | 3.765E-05 | 0.08455 | +1 | 4 |
| - | - | 9848 | 446.3 | - | - | 0 | - |
| - | - | 867.2 | 447.3 | - | - | 0 | - |
| - | - | 730 | 455.2 | - | - | 0 | - |
| - | - | 1916 | 456.2 | - | - | 0 | - |
| - | - | 997.5 | 457.2 | - | - | 0 | - |
| - | - | 3126 | 458.2 | - | - | 0 | - |
| - | - | 604 | 458.3 | - | - | 0 | - |
| - | - | 7.932E+04 | 459.3 | - | - | 0 | - |
| - | - | 2.313E+04 | 460.3 | - | - | 0 | - |
| - | - | 886.5 | 461.2 | - | - | 0 | - |
| - | - | 3900 | 461.3 | - | - | 0 | - |
| - | - | 1302 | 471.3 | - | - | 0 | - |
| - | - | 825.4 | 473.2 | - | - | 0 | - |
| - | - | 3237 | 475.3 | - | - | 0 | - |
| - | - | 596.6 | 476.3 | - | - | 0 | - |
| - | - | 798.2 | 476.3 | - | - | 0 | - |
| - | - | 1.156E+04 | 476.3 | - | - | 0 | - |
| - | - | 3587 | 477.3 | - | - | 0 | - |
| - | - | 753.7 | 477.8 | - | - | 0 | - |
| - | - | 3559 | 478.3 | - | - | 0 | - |
| - | - | 807.2 | 478.3 | - | - | 0 | - |
| - | - | 629 | 480.2 | - | - | 0 | - |
| - | - | 2436 | 497.3 | - | - | 0 | - |
| - | - | 1024 | 507.2 | - | - | 0 | - |
| 5 | y | 2314 | 508.2 | 0.001054 | 2.073 | +1 | 5 |
| - | - | 827.1 | 509.2 | - | - | 0 | - |
| - | - | 1232 | 513.3 | - | - | 0 | - |
| 5 | c | 4215 | 515.3 | 0.0002041 | 0.3961 | +1 | 5 |
| - | - | 1296 | 516.3 | - | - | 0 | - |
| - | - | 6970 | 525.2 | - | - | 0 | - |
| 5 | y | 2610 | 526.3 | 0.002097 | 3.985 | +1 | 5 |
| - | - | 1437 | 527.2 | - | - | 0 | - |
| - | - | 1989 | 531.3 | - | - | 0 | - |
| 5 | c | 2677 | 532.3 | 0.0005271 | 0.9903 | +1 | 5 |
| - | - | 914.8 | 533.3 | - | - | 0 | - |
| - | - | 1439 | 545.3 | - | - | 0 | - |
| - | - | 1364 | 546.3 | - | - | 0 | - |
| - | - | 2596 | 554.3 | - | - | 0 | - |
| - | - | 1840 | 555.3 | - | - | 0 | - |
| - | - | 1845 | 557.3 | - | - | 0 | - |
| - | - | 656.1 | 558.3 | - | - | 0 | - |
| - | - | 2399 | 570.3 | - | - | 0 | - |
| 6 | c | 4018 | 571.3 | 0.01039 | 18.18 | +1 | 6 |
| 6 | c | 7473 | 572.3 | 0.0009572 | 1.672 | +1 | 6 |
| - | - | 2746 | 573.3 | - | - | 0 | - |
| - | - | 727.8 | 586.3 | - | - | 0 | - |
| - | - | 3954 | 588.3 | - | - | 0 | - |
| 6 | c | 2.222E+04 | 589.3 | 0.000287 | 0.4869 | +1 | 6 |
| - | - | 6758 | 590.3 | - | - | 0 | - |
| - | - | 1457 | 591.3 | - | - | 0 | - |
| - | - | 1558 | 594.3 | - | - | 0 | - |
| 4 | y | 913.9 | 595.3 | 0.001159 | 1.946 | +1 | 6 |
| 4 | z | 1182 | 597.3 | 0.001302 | 2.18 | +1 | 6 |
| - | - | 1041 | 598.3 | - | - | 0 | - |
| - | - | 3740 | 612.3 | - | - | 0 | - |
| 4 | y | 1.593E+04 | 613.3 | 0.0003729 | 0.6081 | +1 | 6 |
| - | - | 5561 | 614.3 | - | - | 0 | - |
| - | - | 618.5 | 615.3 | - | - | 0 | - |
| - | - | 607.8 | 615.4 | - | - | 0 | - |
| - | - | 854.2 | 619.3 | - | - | 0 | - |
| - | - | 1735 | 620.3 | - | - | 0 | - |
| - | - | 564.9 | 636.3 | - | - | 0 | - |
| - | - | 1661 | 638.3 | - | - | 0 | - |
| - | - | 769.7 | 639.3 | - | - | 0 | - |
| - | - | 732.3 | 640.4 | - | - | 0 | - |
| - | - | 1436 | 649.4 | - | - | 0 | - |
| - | - | 791.1 | 650.4 | - | - | 0 | - |
| - | - | 1078 | 652.4 | - | - | 0 | - |
| - | - | 588.3 | 656.3 | - | - | 0 | - |
| - | - | 3064 | 657.4 | - | - | 0 | - |
| - | - | 1945 | 658.4 | - | - | 0 | - |
| - | - | 1758 | 659.4 | - | - | 0 | - |
| - | - | 2697 | 667.4 | - | - | 0 | - |
| - | - | 2904 | 668.4 | - | - | 0 | - |
| - | - | 5571 | 669.4 | - | - | 0 | - |
| - | - | 6048 | 670.4 | - | - | 0 | - |
| - | - | 1435 | 671.4 | - | - | 0 | - |
| - | - | 1574 | 684.4 | - | - | 0 | - |
| 7 | c | 1.926E+04 | 685.4 | 0.00016 | 0.2335 | +1 | 7 |
| - | - | 8123 | 686.4 | - | - | 0 | - |
| - | - | 4084 | 687.4 | - | - | 0 | - |
| - | - | 1401 | 688.4 | - | - | 0 | - |
| - | - | 1729 | 690.3 | - | - | 0 | - |
| - | - | 752.3 | 691.3 | - | - | 0 | - |
| - | - | 674.9 | 697.4 | - | - | 0 | - |
| 7 | c | 3.418E+04 | 702.4 | 3.678E-05 | 0.05236 | +1 | 7 |
| - | - | 1.195E+04 | 703.4 | - | - | 0 | - |
| - | - | 2731 | 704.4 | - | - | 0 | - |
| - | - | 662.2 | 705.3 | - | - | 0 | - |
| - | - | 1813 | 706.3 | - | - | 0 | - |
| 3 | z | 4861 | 707.3 | 6.242E-05 | 0.08825 | +1 | 7 |
| - | - | 989.3 | 708.3 | - | - | 0 | - |
| - | - | 687.7 | 710 | - | - | 0 | - |
| - | - | 640.7 | 713.9 | - | - | 0 | - |
| 3 | y | 1166 | 723.3 | 0.003677 | 5.084 | +1 | 7 |
| 3 | z | 9776 | 725.3 | 0.001103 | 1.52 | +1 | 7 |
| - | - | 4519 | 726.3 | - | - | 0 | - |
| - | - | 864.2 | 727.3 | - | - | 0 | - |
| - | - | 6166 | 733.4 | - | - | 0 | - |
| - | - | 3568 | 734.4 | - | - | 0 | - |
| 3 | y | 4397 | 741.3 | 0.0004177 | 0.5635 | +1 | 7 |
| - | - | 1704 | 742.3 | - | - | 0 | - |
| - | - | 5320 | 765.4 | - | - | 0 | - |
| - | - | 1879 | 766.4 | - | - | 0 | - |
| - | - | 1607 | 776.4 | - | - | 0 | - |
| - | - | 1747 | 790.4 | - | - | 0 | - |
| 2 | w | 7404 | 795.4 | 0.0002402 | 0.3021 | +1 | 8 |
| - | - | 2934 | 796.4 | - | - | 0 | - |
| - | - | 1985 | 803.4 | - | - | 0 | - |
| - | - | 914.1 | 804.4 | - | - | 0 | - |
| 2 | z | 2055 | 820.4 | 0.00197 | 2.402 | +1 | 8 |
| - | - | 1.014E+04 | 821.5 | - | - | 0 | - |
| - | - | 722.2 | 822.4 | - | - | 0 | - |
| - | - | 4913 | 822.5 | - | - | 0 | - |
| - | - | 919.5 | 823.5 | - | - | 0 | - |
| - | - | 899.8 | 825.5 | - | - | 0 | - |
| - | - | 739.6 | 830.4 | - | - | 0 | - |
| - | - | 837.6 | 831.4 | - | - | 0 | - |
| - | - | 9274 | 832.4 | - | - | 0 | - |
| - | - | 4741 | 833.4 | - | - | 0 | - |
| - | - | 1672 | 834.4 | - | - | 0 | - |
| 2 | z | 1.435E+04 | 838.4 | 9.928E-05 | 0.1184 | +1 | 8 |
| - | - | 6074 | 839.4 | - | - | 0 | - |
| - | - | 2299 | 840.4 | - | - | 0 | - |
| 8 | c | 7242 | 848.5 | 0.0006834 | 0.8055 | +1 | 8 |
| - | - | 3893 | 849.5 | - | - | 0 | - |
| - | - | 1533 | 850.5 | - | - | 0 | - |
| - | - | 3069 | 853.4 | - | - | 0 | - |
| 2 | y | 2345 | 854.4 | 0.005136 | 6.011 | +1 | 8 |
| - | - | 680.3 | 855.4 | - | - | 0 | - |
| 8 | c | 5.81E+04 | 865.5 | 0.0007432 | 0.8588 | +1 | 8 |
| - | - | 2.552E+04 | 866.5 | - | - | 0 | - |
| - | - | 8534 | 867.5 | - | - | 0 | - |
| - | - | 1518 | 868.5 | - | - | 0 | - |
| - | - | 1.025E+04 | 881.4 | - | - | 0 | - |
| - | - | 4264 | 882.4 | - | - | 0 | - |
| - | - | 924.1 | 883.4 | - | - | 0 | - |
| - | - | 1818 | 891.5 | - | - | 0 | - |
| - | - | 706.7 | 892.4 | - | - | 0 | - |
| - | - | 940.7 | 898.4 | - | - | 0 | - |
| - | - | 652.2 | 906.3 | - | - | 0 | - |
| - | - | 801 | 908.5 | - | - | 0 | - |
| - | - | 6570 | 909.5 | - | - | 0 | - |
| - | - | 3175 | 910.5 | - | - | 0 | - |
| - | - | 1209 | 911.5 | - | - | 0 | - |
| - | - | 1391 | 919.5 | - | - | 0 | - |
| - | - | 970.8 | 920.5 | - | - | 0 | - |
| - | - | 1486 | 926.5 | - | - | 0 | - |
| - | - | 1037 | 927.5 | - | - | 0 | - |
| - | - | 1030 | 935.5 | - | - | 0 | - |
| - | - | 799.2 | 936.5 | - | - | 0 | - |
| - | - | 1.286E+04 | 937.5 | - | - | 0 | - |
| - | - | 5322 | 938.5 | - | - | 0 | - |
| - | - | 1646 | 939.5 | - | - | 0 | - |
| - | - | 789.6 | 951.5 | - | - | 0 | - |
| - | - | 1153 | 952.5 | - | - | 0 | - |
| - | - | 2.21E+04 | 953.5 | - | - | 0 | - |
| - | - | 3.731E+04 | 954.5 | - | - | 0 | - |
| - | - | 1.462E+04 | 955.5 | - | - | 0 | - |
| - | - | 3768 | 956.5 | - | - | 0 | - |
| - | - | 683 | 1305 | - | - | 0 | - |
| - | - | 901.7 | 1319 | - | - | 0 | - |
| - | - | 756.7 | 1320 | - | - | 0 | - |
| - | - | 689.2 | 1355 | - | - | 0 | - |
| - | - | 836.6 | 1370 | - | - | 0 | - |
| - | - | 688.8 | 1394 | - | - | 0 | - |
| - | - | 767.3 | 1398 | - | - | 0 | - |
| - | - | 674.6 | 1411 | - | - | 0 | - |
| - | - | 790.4 | 1413 | - | - | 0 | - |
| - | - | 938.9 | 1416 | - | - | 0 | - |
| - | - | 866.2 | 1417 | - | - | 0 | - |
| - | - | 829.1 | 1429 | - | - | 0 | - |
| - | - | 725.9 | 1430 | - | - | 0 | - |
| - | - | 1240 | 1431 | - | - | 0 | - |
| - | - | 1082 | 1432 | - | - | 0 | - |
| - | - | 676.4 | 1537 | - | - | 0 | - |
| - | - | 693.7 | 1711 | - | - | 0 | - |
| - | - | 833.1 | 2597 | - | - | 0 | - |

m/z Charge Intensity FragmentType MassShift Position
120.08099365234375 0 457.39148
121.19173431396484 0 376.424
124.01115417480469 0 437.13513
125.60460662841797 0 368.23557
127.4601821899414 0 390.62762
128.08209228515625 0 911.55927
129.06593322753906 0 579.76556
131.0706329345703 0 1635.8339
131.11813354492188 0 1521.589
133.086181640625 0 28880.957
134.0895538330078 0 2626.978
136.075927734375 0 29345.713
137.0792694091797 0 1964.5671
146.092529296875 0 1645.8683
148.95449829101562 0 1155.0969
149.04519653320312 0 1371.7769
151.6209716796875 0 448.45258
157.03550720214844 0 424.69687
165.1026611328125 0 497.6821
166.08702087402344 0 479.85147
167.05593872070312 0 738.185
173.43898010253906 0 1205.7782
175.09657287597656 0 1703.0511
177.11231994628906 0 21161.445
178.1158447265625 0 2402.18
185.1650848388672 0 6349.0522
186.16883850097656 0 546.32745
187.14364624023438 0 432.85123
212.13963317871094 0 892.97797
212.1522216796875 0 599.7928
213.15988159179688 0 4905.7656
214.15444946289062 0 834.52185
215.11354064941406 0 487.53064
216.09799194335938 0 1148.8188
219.12269592285156 0 1247.7976
221.08457946777344 0 3757.418
221.13845825195312 0 5773.031
222.1421661376953 0 949.68304
223.06365966796875 0 871.10693
225.04315185546875 0 1123.3412
225.12359619140625 0 2639.0938
233.12484741210938 0 2231.1208
237.1232147216797 0 1547.4124
239.09507751464844 0 6251.912
239.14901733398438 0 1048.4513
240.09591674804688 0 1070.4738
242.15016174316406 0 22639.104
247.10679626464844 0 530.95264
251.10292053222656 0 1277.2345 y Water loss 7
256.3260192871094 0 563.4796
265.1647033691406 0 3262.17
266.1689453125 0 689.24554
269.11322021484375 0 10847.714 y 7
270.11663818359375 0 627.14514
283.1109313964844 0 903.27966
283.17529296875 0 1396.0164
285.0113830566406 0 542.74207
295.10345458984375 0 6069.3706
296.10479736328125 0 2037.5642
297.0992431640625 0 723.78864
297.94793701171875 0 488.26334
299.06182861328125 0 2534.7966
302.58868408203125 0 629.02783
309.19122314453125 0 2784.067
310.19390869140625 0 654.90063
311.1057434082031 0 781.25726
313.1125793457031 0 1174.685
313.1875 0 7050.0547
314.2319641113281 0 1855.8881
315.2394714355469 0 3715.0256
316.24188232421875 0 1019.5741
317.1830139160156 0 969.37683
327.16644287109375 0 784.7085
327.2021484375 0 1754.4429
328.1516418457031 0 681.7382
340.21026611328125 0 1560.9446
341.21881103515625 0 8392.868 c Ammonia loss 2
342.222412109375 0 1127.6678
345.1769714355469 0 3493.3613
347.192626953125 0 2485.5623
353.1659851074219 0 686.38544
353.21697998046875 0 1424.5945
355.0699462890625 0 1600.8871
357.2369384765625 0 6068.04
358.24481201171875 0 47816.684 c 2
359.0287170410156 0 1705.3335
359.24761962890625 0 7836.799
360.2495422363281 0 1004.88745
362.1674499511719 0 899.1258 y Water loss 2
365.1828308105469 0 1538.543
369.1219482421875 0 8209.743
370.1226501464844 0 3227.8015
371.09869384765625 0 945.36914
371.1210632324219 0 1471.4774
371.22784423828125 0 4044.5437
372.2325134277344 0 586.1875
377.17767333984375 0 2294.401
384.2606201171875 0 2125.9854
397.2418518066406 0 581.03033
402.271240234375 0 6231.95
403.27398681640625 0 1373.869
414.20037841796875 0 630.5313
415.25421142578125 0 5512.0757
416.2567443847656 0 1624.4877
421.207275390625 0 1030.3231 y Water loss 5
426.23089599609375 0 877.9501
426.25982666015625 0 2342.45
427.2655944824219 0 700.27295 c Water loss 3
428.2173156738281 0 857.7876
428.2505187988281 0 3475.7476 c Ammonia loss 3
429.08953857421875 0 1193.5216
430.2656555175781 0 1816.4337
441.2690734863281 0 794.68365
444.26959228515625 0 16330.383
445.2769470214844 0 47338.453 c 3
446.2798767089844 0 9848.458
447.2823791503906 0 867.16266
455.2244873046875 0 729.9899
456.20880126953125 0 1916.419
457.212158203125 0 997.54254
458.2239074707031 0 3125.6938
458.2564392089844 0 604.0412
459.2804260253906 0 79319.836
460.2838439941406 0 23128.982
461.2393493652344 0 886.5244
461.28570556640625 0 3899.7512
471.2920837402344 0 1301.877
473.2364196777344 0 825.4132
475.2511901855469 0 3236.7786
476.2511291503906 0 596.5976
476.2665710449219 0 798.1863
476.3067932128906 0 11555.517
477.3107604980469 0 3587.448
477.8225402832031 0 753.741
478.266357421875 0 3558.8047
478.3053894042969 0 807.2114
480.240478515625 0 629.02167
497.27276611328125 0 2435.5938
507.23260498046875 0 1023.93243
508.2391357421875 0 2314.1328 y Water loss 4
509.24542236328125 0 827.07697
513.293212890625 0 1231.6918
515.2825927734375 0 4215.215 c Ammonia loss 4
516.2880249023438 0 1295.7139
525.2431030273438 0 6970.0903
526.2486572265625 0 2610.4094 y 4
527.2479248046875 0 1436.8431
531.3024291992188 0 1989.424
532.3084106445312 0 2676.9912 c 4
533.3103637695312 0 914.79114
545.31689453125 0 1438.6302
546.325439453125 0 1364.3699
554.2944946289062 0 2596.0652
555.2800903320312 0 1840.2008
557.3286743164062 0 1845.1879
558.3324584960938 0 656.091
570.2999267578125 0 2399.174
571.3094482421875 0 4018.4368 c Water loss 5
572.3048095703125 0 7472.5376 c Ammonia loss 5
573.3074340820312 0 2745.8848
586.3162841796875 0 727.7556
588.3350830078125 0 3954.322
589.3306884765625 0 22215.64 c 5
590.3341064453125 0 6757.8975
591.334716796875 0 1456.5409
594.2655029296875 0 1558.0159
595.2733764648438 0 913.918 y Water loss 3
597.2627563476562 0 1181.7871 z 3
598.273193359375 0 1041.1237
612.2759399414062 0 3739.832
613.2824096679688 0 15932.119 y 3
614.2861938476562 0 5560.704
615.283447265625 0 618.46826
615.3505859375 0 607.8073
619.2730102539062 0 854.20746
620.279296875 0 1735.0789
636.2960815429688 0 564.9117
638.3148193359375 0 1660.9259
639.3226318359375 0 769.66534
640.3692016601562 0 732.3465
649.365966796875 0 1436.311
650.3660278320312 0 791.1456
652.36181640625 0 1077.6703
656.3358154296875 0 588.29285
657.3925170898438 0 3064.3833
658.3983154296875 0 1944.6405
659.4061279296875 0 1758.0077
667.3768920898438 0 2696.7063
668.3657836914062 0 2903.8074
669.3695678710938 0 5570.638
670.3758544921875 0 6048.4043
671.3720092773438 0 1435.2042
684.3851928710938 0 1574.3907
685.3877563476562 0 19255.117 c Ammonia loss 6
686.3905639648438 0 8122.7886
687.400146484375 0 4084.081
688.4031372070312 0 1401.0864
690.307373046875 0 1728.813
691.3109741210938 0 752.26764
697.441162109375 0 674.9118
702.4144287109375 0 34180.242 c 6
703.4170532226562 0 11950.004
704.4193115234375 0 2730.6477
705.3204956054688 0 662.2017
706.3052978515625 0 1813.3019
707.3121337890625 0 4860.77 z Water loss 2
708.3136596679688 0 989.27325
710.0383911132812 0 687.65485
713.8781127929688 0 640.6532
723.33447265625 0 1165.5853 y Water loss 2
725.321533203125 0 9776.278 z 2
726.3262939453125 0 4519.483
727.32421875 0 864.2485
733.3635864257812 0 6165.5425
734.36669921875 0 3568.0107
741.3409423828125 0 4396.8076 y 2
742.3441162109375 0 1703.8035
765.4016723632812 0 5320.1206
766.4054565429688 0 1879.3323
776.3695068359375 0 1607.1984
790.3948364257812 0 1747.0897
795.3516845703125 0 7403.5 w 1
796.3547973632812 0 2933.5017
803.4497680664062 0 1985.0522
804.4407958984375 0 914.11993
820.3941650390625 0 2055.3591 z Water loss 1
821.4641723632812 0 10135.458
822.3950805664062 0 722.1944
822.4674072265625 0 4912.8296
823.4599609375 0 919.4543
825.5365600585938 0 899.77985
830.4442138671875 0 739.59436
831.432861328125 0 837.6461
832.4323120117188 0 9273.689
833.4361572265625 0 4741.009
834.4400634765625 0 1671.983
838.4067993164062 0 14354.611 z 1
839.40966796875 0 6073.892
840.4105224609375 0 2298.537
848.4505615234375 0 7241.994 c Ammonia loss 7
849.454345703125 0 3893.0552
850.4585571289062 0 1533.314
853.4161376953125 0 3068.7783
854.4202880859375 0 2345.45 y 1
855.42724609375 0 680.26666
865.47705078125 0 58101.23 c 7
866.4796142578125 0 25517.771
867.4835205078125 0 8533.599
868.4864501953125 0 1518.3782
881.4142456054688 0 10245.481
882.41796875 0 4264.4077
883.4105224609375 0 924.1055
891.4735107421875 0 1817.5361
892.4495239257812 0 706.6913
898.43505859375 0 940.7497
906.3356323242188 0 652.1995
908.503662109375 0 801.0112
909.4805908203125 0 6570.3384
910.481689453125 0 3174.819
911.487548828125 0 1209.015
919.4635620117188 0 1391.196
920.4710083007812 0 970.8318
926.508544921875 0 1486.4896
927.5028686523438 0 1037.1046
935.4508056640625 0 1030.021
936.4803466796875 0 799.17676
937.4746704101562 0 12855.011
938.4773559570312 0 5321.9634
939.4801025390625 0 1646.4
951.453857421875 0 789.64777
952.47802734375 0 1152.7949
953.4924926757812 0 22099.883
954.4995727539062 0 37312.8
955.5032348632812 0 14617.298
956.5055541992188 0 3768.1492
1304.7047119140625 0 682.96924
1318.68994140625 0 901.6974
1319.7142333984375 0 756.653
1354.7027587890625 0 689.2313
1369.7093505859375 0 836.5766
1393.71240234375 0 688.83374
1397.7069091796875 0 767.30743
1410.774169921875 0 674.60626
1412.6976318359375 0 790.3652
1415.6988525390625 0 938.88983
1416.679931640625 0 866.235
1428.7435302734375 0 829.07465
1429.7269287109375 0 725.94794
1430.6954345703125 0 1240.214
1431.7164306640625 0 1081.7704
1537.2491455078125 0 676.362
1711.209716796875 0 693.65857
2596.827880859375 0 833.134

Spectrum Details

|  |  |
| --- | --- |
| Matched peaks? Matched peaks The total absolute number of peaks matched. Additionally in brackets the total fraction of peaks matched and the total number of peaks is shown. | 31 (10.58% of 293) |
| FDR? FDR The false discovery rate estimated for this peptide. It is calculated by matching all theoretical fragments with a non-integer shift with the raw peaks for this spectrum. This is done with 40 different shifts. The resulting percentage is the average number of annotated peaks over the number of annotated peaks with the correct spectrum. | 1.77% |
| Satellite FDR? Satellite FDR See the FDR for details on its calculation. This satellite ion specific FDR only contains the satellite ions (d/w) for I/L/J positions. | 9.52% |
| PSM Score? PSM Score The PSM Score as given by Hecklib to this annotated spectrum. It is shown with three significant figures. | 429 |

#### Reverse Lookup? Reverse Lookup All places where this read could be placed.

| Group | Segment | Template | Template Part | Read Part | Score | Unique |
| --- | --- | --- | --- | --- | --- | --- |
| Homo sapiens Heavy Chain | IGHC | IGHG1 | [55..64] | [0..9] | 72 | False |
| Homo sapiens Heavy Chain | IGHC | IGHG3 | [55..64] | [0..9] | 72 | False |
| Homo sapiens Heavy Chain | IGHC | IGHG2 | [55..64] | [0..9] | 72 | False |
| Homo sapiens Heavy Chain | IGHC | IGHG4 | [55..64] | [0..9] | 72 | False |

| Recombined | Template Part | Read Part | Score | Unique |
| --- | --- | --- | --- | --- |
| REC-0-1 | [177..186] | [0..9] | 72 | True |

#### Meta Information from Multiple reads

##### Number of combined reads

2

##### Intensity

1

##### TotalArea

0

##### Changes to the peptide sequence

VLQSSGJYS

L→JNo support for either Leucine or Isoleucine based on side chain ions (Position: 7)

#### Positional Score

Copy Data

##### Positional Score (TSV)

###### Preview

```
Loading example...
```

*Click on the button to copy the data to your clipboard.*

00012345678

Label Value
"0" 0
"1" 0
"2" 0
"3" 0
"4" 0
"5" 0
"6" 0
"7" 0
"8" 0

#### Meta Information from PEAKS

##### Scan Identifier

F2:5480

##### Original sequence

V

L

Q

S

S

G

L

Y

S

##### Posttranslational Modifications

##### Source File

D:\separate\_stitch\_analyses\xle-disambiguation\raw\20210323\_F1\_UM1\_Peng0013\_SA\_F59\_ingel\_3ug\_TL.raw

##### Fraction

2

##### Scan Feature

-

##### De Novo Score

98

##### ConfidenceScore

98

### m/z

477.2519

##### Mass

952.4865

##### Charge

2

##### Retention Time

29.76

##### Predicted Retention Time

-

##### Area

0

##### Parts Per Million

2.8

##### Fragmentation mode

ETHCD

##### Originating file

01 D:\separate\_stitch\_analyses\xle-disambiguation\20210325\_F59\_3ug\_DENOVO\_12.csv

#### Meta Information from PEAKS

##### Scan Identifier

F2:5352

##### Original sequence

V

L

Q

S

S

G

L

Y

S

##### Posttranslational Modifications

##### Source File

D:\separate\_stitch\_analyses\xle-disambiguation\raw\20210323\_F1\_UM1\_Peng0013\_SA\_F59\_ingel\_3ug\_TL.raw

##### Fraction

2

##### Scan Feature

-

##### De Novo Score

98

##### ConfidenceScore

98

### m/z

477.2516

##### Mass

952.4865

##### Charge

2

##### Retention Time

29.02

##### Predicted Retention Time

-

##### Area

0

##### Parts Per Million

2.2

##### Fragmentation mode

ETHCD

##### Originating file

01 D:\separate\_stitch\_analyses\xle-disambiguation\20210325\_F59\_3ug\_DENOVO\_12.csv
