## Supplementary material for "A handle on mass coincidence errors in *de novo* sequencing of antibodies by bottom-up proteomics": Combined_076.html

Details Combined\_076 | Stitch OverviewUndefined

### Read Combined\_076

#### Sequence (length=9)

VFGGGTKJT

#### Spectrum 4540? Spectrum 4540 The raw spectrum of this peptide as annotated by Hecklib. The fragments are coloured according to ion type (see legend). Any peaks with a star '\*' as text can be hovered over to see the full details, first the ion type second the mass shift type. By hovering over the amino acids in the peptide or ions in the legend the corresponding peaks are highlighted. By toggling the 'Unassigned' label you can turn the background (unassigned) peaks on or off in the plot. By updating the slider in the Ion legend you can update the spectrum to only show the top X% of the peaks with labels. The top X% means any peak that is within X% of the highest intensity. By dragging in the spectrum you can zoom in to a specific part of the spectrum and use 'Zoom Out' to get back to the original zoom level. The annotation of the spectrum is based on the given sequence in the peptides file and is done with different software so inconsistencies are likely. The peaks are annotated based on the given sequence, with 20 ppm tolerance.

Copy Data

##### Spectrum 4540 (TSV)

###### Preview

```
Loading example...
```

*Click on the button to copy the data to your clipboard.*

Mz MinMz MaxIntensity Max

WidthHeightPeptide font sizePeptide stroke widthSpectrum font sizeSpectrum stroke widthCompact peptide

Ion legend

wxyz

abcd

OtherUnassignedIon^Charge^_Position_Show for top:%

VFGGGTKJT

05.38e+61.08e+71.61e+72.15e+7

Zoom Out

b^+2^_4_y^+1^_2_a^+1^_2_y^+2^_4_y^+1^_2_b^+1^_2_y^+2^_5_y^+2^_5_y^+2^_6_y^+2^_6_b^+1^_3_y^+2^_7_b^+2^_7_b^+2^_7_y^+1^_3_y^+1^_3_b^+1^_4_y^+1^_3_b^+2^_8_b^+2^_8_y^+2^_8_y^+2^_8_b^+1^_5_\*\*y^+1^_4_y^+1^_4_b^+1^_6_y^+1^_5_b^+1^_6_y^+1^_5_y^+1^_6_y^+1^_6_y^+1^_7_b^+1^_7_y^+1^_7_b^+1^_7_b^+1^_8_b^+1^_8_y^+1^_8_y^+1^_8_

0768153623043072

Fragment Matches Table

Show background peaks

| Position | Ion type | Intensity | mz Theoretical | mz Error (Th) | mz Error (ppm) | Charge | Series Number |
| --- | --- | --- | --- | --- | --- | --- | --- |
| - | - | 2.129E+07 | 120.1 | - | - | 0 | - |
| - | - | 3.568E+04 | 121.1 | - | - | 0 | - |
| - | - | 6.027E+04 | 121.1 | - | - | 0 | - |
| - | - | 1.674E+06 | 121.1 | - | - | 0 | - |
| - | - | 5545 | 122.1 | - | - | 0 | - |
| - | - | 5.372E+04 | 122.1 | - | - | 0 | - |
| - | - | 5086 | 124.1 | - | - | 0 | - |
| - | - | 2.105E+04 | 125.1 | - | - | 0 | - |
| - | - | 1.181E+05 | 127.1 | - | - | 0 | - |
| - | - | 1.492E+04 | 127.1 | - | - | 0 | - |
| - | - | 6401 | 128.1 | - | - | 0 | - |
| - | - | 5819 | 128.1 | - | - | 0 | - |
| - | - | 2.288E+04 | 128.1 | - | - | 0 | - |
| - | - | 1.127E+07 | 129.1 | - | - | 0 | - |
| - | - | 1.989E+04 | 130.1 | - | - | 0 | - |
| - | - | 1.146E+04 | 130.1 | - | - | 0 | - |
| - | - | 6.738E+04 | 130.1 | - | - | 0 | - |
| - | - | 6.759E+05 | 130.1 | - | - | 0 | - |
| - | - | 2.013E+04 | 131 | - | - | 0 | - |
| - | - | 5.152E+05 | 131.1 | - | - | 0 | - |
| - | - | 1.461E+04 | 131.1 | - | - | 0 | - |
| - | - | 1.361E+04 | 132.1 | - | - | 0 | - |
| - | - | 1.088E+05 | 132.1 | - | - | 0 | - |
| - | - | 1.347E+04 | 132.1 | - | - | 0 | - |
| - | - | 8837 | 133.1 | - | - | 0 | - |
| - | - | 4259 | 137.1 | - | - | 0 | - |
| - | - | 5.317E+04 | 139.1 | - | - | 0 | - |
| - | - | 1.133E+04 | 139.1 | - | - | 0 | - |
| - | - | 5338 | 140.1 | - | - | 0 | - |
| - | - | 6447 | 140.1 | - | - | 0 | - |
| - | - | 3.187E+04 | 140.1 | - | - | 0 | - |
| - | - | 6891 | 140.1 | - | - | 0 | - |
| - | - | 2.202E+05 | 141.1 | - | - | 0 | - |
| - | - | 5.016E+04 | 141.1 | - | - | 0 | - |
| - | - | 1.439E+04 | 142.1 | - | - | 0 | - |
| - | - | 3.432E+04 | 142.1 | - | - | 0 | - |
| - | - | 5792 | 143 | - | - | 0 | - |
| - | - | 3.598E+04 | 143.1 | - | - | 0 | - |
| - | - | 5822 | 143.1 | - | - | 0 | - |
| - | - | 4673 | 143.1 | - | - | 0 | - |
| - | - | 5931 | 146.1 | - | - | 0 | - |
| - | - | 3.679E+04 | 146.1 | - | - | 0 | - |
| - | - | 5641 | 146.5 | - | - | 0 | - |
| - | - | 7691 | 147.1 | - | - | 0 | - |
| - | - | 4.05E+04 | 148.1 | - | - | 0 | - |
| - | - | 7725 | 148.9 | - | - | 0 | - |
| - | - | 8102 | 148.9 | - | - | 0 | - |
| - | - | 1.934E+04 | 148.9 | - | - | 0 | - |
| - | - | 3.719E+04 | 149 | - | - | 0 | - |
| - | - | 2.1E+04 | 149 | - | - | 0 | - |
| - | - | 9665 | 149 | - | - | 0 | - |
| - | - | 7865 | 149 | - | - | 0 | - |
| - | - | 6450 | 149 | - | - | 0 | - |
| - | - | 5186 | 149 | - | - | 0 | - |
| - | - | 5344 | 149.1 | - | - | 0 | - |
| - | - | 4964 | 149.1 | - | - | 0 | - |
| - | - | 3.687E+04 | 151.1 | - | - | 0 | - |
| - | - | 6807 | 151.1 | - | - | 0 | - |
| - | - | 5654 | 152.1 | - | - | 0 | - |
| - | - | 1.159E+05 | 152.1 | - | - | 0 | - |
| - | - | 7.089E+04 | 153.1 | - | - | 0 | - |
| - | - | 3.335E+04 | 153.1 | - | - | 0 | - |
| - | - | 7603 | 153.1 | - | - | 0 | - |
| - | - | 3.994E+04 | 154.1 | - | - | 0 | - |
| - | - | 7407 | 154.1 | - | - | 0 | - |
| - | - | 8.947E+04 | 155.1 | - | - | 0 | - |
| - | - | 3.237E+05 | 155.1 | - | - | 0 | - |
| - | - | 3846 | 156.1 | - | - | 0 | - |
| - | - | 4509 | 156.1 | - | - | 0 | - |
| - | - | 4418 | 156.1 | - | - | 0 | - |
| - | - | 2.887E+04 | 156.1 | - | - | 0 | - |
| - | - | 9381 | 157.1 | - | - | 0 | - |
| - | - | 1.093E+05 | 158.1 | - | - | 0 | - |
| - | - | 3.893E+05 | 159.1 | - | - | 0 | - |
| - | - | 2.335E+04 | 160.1 | - | - | 0 | - |
| - | - | 1.075E+04 | 165.1 | - | - | 0 | - |
| - | - | 7.783E+04 | 167.1 | - | - | 0 | - |
| - | - | 1.47E+04 | 168.1 | - | - | 0 | - |
| - | - | 8748 | 168.1 | - | - | 0 | - |
| - | - | 6882 | 168.2 | - | - | 0 | - |
| - | - | 4332 | 169.1 | - | - | 0 | - |
| - | - | 2.819E+04 | 169.1 | - | - | 0 | - |
| - | - | 1.549E+04 | 169.1 | - | - | 0 | - |
| - | - | 9869 | 169.2 | - | - | 0 | - |
| - | - | 2.052E+05 | 170.1 | - | - | 0 | - |
| - | - | 1.078E+05 | 171.1 | - | - | 0 | - |
| - | - | 1.579E+04 | 171.1 | - | - | 0 | - |
| - | - | 1.111E+04 | 171.1 | - | - | 0 | - |
| - | - | 2.346E+04 | 171.1 | - | - | 0 | - |
| - | - | 4657 | 171.2 | - | - | 0 | - |
| - | - | 5.238E+05 | 172.1 | - | - | 0 | - |
| - | - | 1.207E+04 | 172.1 | - | - | 0 | - |
| - | - | 3.098E+04 | 173.1 | - | - | 0 | - |
| - | - | 4182 | 173.1 | - | - | 0 | - |
| - | - | 4313 | 174.1 | - | - | 0 | - |
| - | - | 1.967E+04 | 174.1 | - | - | 0 | - |
| - | - | 4063 | 175.1 | - | - | 0 | - |
| - | - | 2.01E+05 | 176.1 | - | - | 0 | - |
| - | - | 7.01E+05 | 177.1 | - | - | 0 | - |
| - | - | 2.681E+04 | 177.1 | - | - | 0 | - |
| - | - | 5456 | 178.1 | - | - | 0 | - |
| - | - | 5.705E+04 | 178.1 | - | - | 0 | - |
| - | - | 4260 | 179.1 | - | - | 0 | - |
| - | - | 1.643E+04 | 179.2 | - | - | 0 | - |
| - | - | 1.71E+04 | 180.1 | - | - | 0 | - |
| - | - | 1.14E+04 | 180.1 | - | - | 0 | - |
| - | - | 1.869E+04 | 181.1 | - | - | 0 | - |
| 4 | b | 3597 | 181.1 | 0.001418 | 7.828 | +2 | 4 |
| - | - | 5398 | 181.1 | - | - | 0 | - |
| - | - | 1.895E+04 | 181.2 | - | - | 0 | - |
| - | - | 5045 | 182.1 | - | - | 0 | - |
| - | - | 1.638E+04 | 182.1 | - | - | 0 | - |
| - | - | 7640 | 183.1 | - | - | 0 | - |
| - | - | 8.861E+04 | 183.1 | - | - | 0 | - |
| - | - | 3.019E+04 | 184.1 | - | - | 0 | - |
| - | - | 7369 | 184.1 | - | - | 0 | - |
| - | - | 1.658E+04 | 184.1 | - | - | 0 | - |
| - | - | 3.848E+04 | 185.1 | - | - | 0 | - |
| - | - | 1.663E+05 | 185.1 | - | - | 0 | - |
| - | - | 1.445E+05 | 186.1 | - | - | 0 | - |
| - | - | 1.027E+04 | 186.1 | - | - | 0 | - |
| - | - | 5034 | 187.1 | - | - | 0 | - |
| - | - | 4268 | 187.1 | - | - | 0 | - |
| - | - | 1.248E+04 | 187.1 | - | - | 0 | - |
| - | - | 1.351E+04 | 187.1 | - | - | 0 | - |
| - | - | 5770 | 188.1 | - | - | 0 | - |
| - | - | 9.145E+04 | 188.1 | - | - | 0 | - |
| - | - | 1.29E+04 | 188.1 | - | - | 0 | - |
| - | - | 2.781E+04 | 189.1 | - | - | 0 | - |
| - | - | 5677 | 190.1 | - | - | 0 | - |
| - | - | 2.249E+04 | 192.1 | - | - | 0 | - |
| - | - | 3474 | 194.1 | - | - | 0 | - |
| - | - | 6.007E+04 | 194.1 | - | - | 0 | - |
| - | - | 7524 | 195.1 | - | - | 0 | - |
| - | - | 4.911E+04 | 195.2 | - | - | 0 | - |
| - | - | 5.219E+04 | 196.1 | - | - | 0 | - |
| - | - | 1.128E+04 | 196.1 | - | - | 0 | - |
| - | - | 3887 | 196.2 | - | - | 0 | - |
| - | - | 8355 | 196.2 | - | - | 0 | - |
| - | - | 1.313E+04 | 197.1 | - | - | 0 | - |
| - | - | 7239 | 197.1 | - | - | 0 | - |
| - | - | 1.35E+04 | 197.1 | - | - | 0 | - |
| - | - | 5.038E+05 | 197.2 | - | - | 0 | - |
| - | - | 3.039E+05 | 198.1 | - | - | 0 | - |
| - | - | 8988 | 198.1 | - | - | 0 | - |
| - | - | 4.741E+04 | 198.2 | - | - | 0 | - |
| - | - | 4595 | 198.2 | - | - | 0 | - |
| - | - | 2.358E+04 | 199.1 | - | - | 0 | - |
| - | - | 5083 | 199.1 | - | - | 0 | - |
| - | - | 4925 | 199.2 | - | - | 0 | - |
| - | - | 8882 | 200.1 | - | - | 0 | - |
| - | - | 1.568E+04 | 200.1 | - | - | 0 | - |
| - | - | 4.76E+04 | 201.1 | - | - | 0 | - |
| - | - | 1.156E+04 | 202.1 | - | - | 0 | - |
| - | - | 4.949E+04 | 203.1 | - | - | 0 | - |
| - | - | 1.292E+04 | 203.2 | - | - | 0 | - |
| - | - | 4311 | 204.1 | - | - | 0 | - |
| - | - | 1.244E+06 | 205.1 | - | - | 0 | - |
| - | - | 1.517E+05 | 206.1 | - | - | 0 | - |
| - | - | 1.265E+04 | 206.1 | - | - | 0 | - |
| - | - | 6766 | 206.1 | - | - | 0 | - |
| - | - | 7559 | 207.1 | - | - | 0 | - |
| - | - | 2.326E+05 | 207.2 | - | - | 0 | - |
| - | - | 4125 | 208.1 | - | - | 0 | - |
| - | - | 2.085E+05 | 208.1 | - | - | 0 | - |
| - | - | 2.36E+04 | 208.2 | - | - | 0 | - |
| - | - | 1.614E+05 | 209.1 | - | - | 0 | - |
| - | - | 1.539E+04 | 209.1 | - | - | 0 | - |
| - | - | 8873 | 209.2 | - | - | 0 | - |
| - | - | 9.494E+04 | 210.1 | - | - | 0 | - |
| - | - | 6637 | 210.1 | - | - | 0 | - |
| - | - | 1.406E+04 | 210.1 | - | - | 0 | - |
| - | - | 4512 | 210.1 | - | - | 0 | - |
| - | - | 8642 | 211.1 | - | - | 0 | - |
| - | - | 4534 | 211.1 | - | - | 0 | - |
| - | - | 8505 | 211.1 | - | - | 0 | - |
| - | - | 7295 | 212.1 | - | - | 0 | - |
| - | - | 1.047E+06 | 212.1 | - | - | 0 | - |
| - | - | 3.766E+04 | 213.1 | - | - | 0 | - |
| - | - | 1.048E+05 | 213.1 | - | - | 0 | - |
| - | - | 1.694E+04 | 214.1 | - | - | 0 | - |
| - | - | 8.722E+04 | 214.2 | - | - | 0 | - |
| - | - | 2.001E+04 | 215.1 | - | - | 0 | - |
| - | - | 1.788E+05 | 215.1 | - | - | 0 | - |
| 8 | y | 1.837E+05 | 215.1 | 0.001194 | 5.55 | +1 | 2 |
| - | - | 7505 | 215.2 | - | - | 0 | - |
| - | - | 2.211E+05 | 216.1 | - | - | 0 | - |
| - | - | 1.232E+04 | 216.1 | - | - | 0 | - |
| - | - | 2.056E+04 | 216.1 | - | - | 0 | - |
| - | - | 1.024E+05 | 217.1 | - | - | 0 | - |
| - | - | 1.052E+04 | 218.1 | - | - | 0 | - |
| 2 | a | 1.873E+07 | 219.1 | 0.00114 | 5.202 | +1 | 2 |
| - | - | 2.447E+06 | 220.2 | - | - | 0 | - |
| - | - | 3.224E+04 | 221.1 | - | - | 0 | - |
| - | - | 1.44E+05 | 221.2 | - | - | 0 | - |
| - | - | 1.985E+04 | 222.1 | - | - | 0 | - |
| - | - | 5468 | 222.2 | - | - | 0 | - |
| - | - | 1.557E+04 | 223.1 | - | - | 0 | - |
| - | - | 2.133E+04 | 224.1 | - | - | 0 | - |
| - | - | 3.594E+05 | 224.2 | - | - | 0 | - |
| - | - | 8144 | 225.1 | - | - | 0 | - |
| - | - | 3809 | 225.1 | - | - | 0 | - |
| - | - | 1.377E+05 | 225.1 | - | - | 0 | - |
| - | - | 9.924E+04 | 225.2 | - | - | 0 | - |
| - | - | 4.47E+04 | 225.2 | - | - | 0 | - |
| - | - | 2.919E+04 | 226.1 | - | - | 0 | - |
| - | - | 1.849E+04 | 226.1 | - | - | 0 | - |
| - | - | 1.269E+04 | 226.1 | - | - | 0 | - |
| - | - | 1.538E+04 | 226.2 | - | - | 0 | - |
| - | - | 7.995E+05 | 227.1 | - | - | 0 | - |
| - | - | 4.263E+05 | 228.1 | - | - | 0 | - |
| - | - | 6.695E+04 | 228.1 | - | - | 0 | - |
| - | - | 3.664E+04 | 228.1 | - | - | 0 | - |
| - | - | 2.356E+04 | 228.1 | - | - | 0 | - |
| - | - | 9.822E+04 | 229.1 | - | - | 0 | - |
| - | - | 1.175E+04 | 229.1 | - | - | 0 | - |
| - | - | 2.159E+06 | 230.2 | - | - | 0 | - |
| - | - | 4.122E+04 | 231.1 | - | - | 0 | - |
| - | - | 2.26E+05 | 231.2 | - | - | 0 | - |
| 6 | y | 8172 | 231.6 | 0.001662 | 7.175 | +2 | 4 |
| - | - | 2.345E+04 | 232.2 | - | - | 0 | - |
| - | - | 8853 | 233.1 | - | - | 0 | - |
| - | - | 3.976E+04 | 233.1 | - | - | 0 | - |
| 8 | y | 3.821E+05 | 233.1 | 0.001265 | 5.425 | +1 | 2 |
| - | - | 1.279E+04 | 233.2 | - | - | 0 | - |
| - | - | 6.605E+04 | 234.1 | - | - | 0 | - |
| - | - | 4.039E+04 | 234.2 | - | - | 0 | - |
| - | - | 6738 | 235.1 | - | - | 0 | - |
| - | - | 5498 | 235.2 | - | - | 0 | - |
| - | - | 4882 | 236.2 | - | - | 0 | - |
| - | - | 1.342E+05 | 237.1 | - | - | 0 | - |
| - | - | 4687 | 237.1 | - | - | 0 | - |
| - | - | 5798 | 237.1 | - | - | 0 | - |
| - | - | 1.815E+04 | 238.1 | - | - | 0 | - |
| - | - | 1.343E+04 | 238.1 | - | - | 0 | - |
| - | - | 9870 | 238.1 | - | - | 0 | - |
| - | - | 1.43E+05 | 240.1 | - | - | 0 | - |
| - | - | 2.02E+04 | 240.2 | - | - | 0 | - |
| - | - | 1.875E+04 | 241.1 | - | - | 0 | - |
| - | - | 7.129E+04 | 241.2 | - | - | 0 | - |
| - | - | 4.473E+05 | 242.2 | - | - | 0 | - |
| - | - | 9.732E+04 | 243.1 | - | - | 0 | - |
| - | - | 5.523E+04 | 243.1 | - | - | 0 | - |
| - | - | 3.829E+04 | 243.2 | - | - | 0 | - |
| - | - | 4.957E+04 | 243.2 | - | - | 0 | - |
| - | - | 6173 | 243.7 | - | - | 0 | - |
| - | - | 4.511E+04 | 244.1 | - | - | 0 | - |
| - | - | 4.588E+05 | 245.1 | - | - | 0 | - |
| - | - | 2.106E+04 | 245.2 | - | - | 0 | - |
| - | - | 3E+04 | 246.1 | - | - | 0 | - |
| 2 | b | 4.65E+06 | 247.1 | 0.001373 | 5.556 | +1 | 2 |
| - | - | 6.717E+05 | 248.1 | - | - | 0 | - |
| - | - | 1.201E+04 | 249.1 | - | - | 0 | - |
| - | - | 4.356E+04 | 249.2 | - | - | 0 | - |
| - | - | 1.206E+04 | 250.2 | - | - | 0 | - |
| - | - | 9761 | 250.2 | - | - | 0 | - |
| 5 | y | 9.74E+04 | 251.2 | 0.003131 | 12.47 | +2 | 5 |
| - | - | 1.737E+04 | 252.1 | - | - | 0 | - |
| - | - | 1.317E+04 | 252.2 | - | - | 0 | - |
| - | - | 4.398E+04 | 252.2 | - | - | 0 | - |
| - | - | 8875 | 253.1 | - | - | 0 | - |
| - | - | 6.463E+04 | 254.1 | - | - | 0 | - |
| - | - | 1.057E+04 | 254.2 | - | - | 0 | - |
| - | - | 1.171E+06 | 255.1 | - | - | 0 | - |
| - | - | 1.26E+05 | 256.1 | - | - | 0 | - |
| - | - | 4658 | 256.2 | - | - | 0 | - |
| - | - | 1.174E+04 | 257.1 | - | - | 0 | - |
| - | - | 2.768E+04 | 257.2 | - | - | 0 | - |
| - | - | 6737 | 258.1 | - | - | 0 | - |
| - | - | 5046 | 258.2 | - | - | 0 | - |
| - | - | 5.017E+04 | 259.1 | - | - | 0 | - |
| - | - | 4497 | 260.1 | - | - | 0 | - |
| 5 | y | 1.364E+04 | 260.2 | 0.001932 | 7.426 | +2 | 5 |
| - | - | 7.836E+04 | 260.2 | - | - | 0 | - |
| - | - | 8.176E+04 | 261.1 | - | - | 0 | - |
| - | - | 6761 | 261.2 | - | - | 0 | - |
| - | - | 8526 | 261.2 | - | - | 0 | - |
| - | - | 3.455E+05 | 262.1 | - | - | 0 | - |
| - | - | 7636 | 262.2 | - | - | 0 | - |
| - | - | 4.21E+04 | 263.1 | - | - | 0 | - |
| - | - | 1.335E+04 | 263.2 | - | - | 0 | - |
| - | - | 2.983E+04 | 264.2 | - | - | 0 | - |
| - | - | 5654 | 265.1 | - | - | 0 | - |
| - | - | 1.967E+04 | 265.1 | - | - | 0 | - |
| - | - | 1.823E+04 | 266.2 | - | - | 0 | - |
| - | - | 1.276E+04 | 268.2 | - | - | 0 | - |
| - | - | 1.37E+06 | 269.2 | - | - | 0 | - |
| - | - | 1.628E+05 | 270.2 | - | - | 0 | - |
| - | - | 1.592E+05 | 270.2 | - | - | 0 | - |
| - | - | 2.63E+04 | 271.1 | - | - | 0 | - |
| - | - | 3.319E+04 | 271.1 | - | - | 0 | - |
| - | - | 1.37E+04 | 271.2 | - | - | 0 | - |
| - | - | 1.842E+04 | 271.2 | - | - | 0 | - |
| - | - | 8013 | 271.2 | - | - | 0 | - |
| - | - | 1.508E+04 | 272.1 | - | - | 0 | - |
| - | - | 1.949E+05 | 272.1 | - | - | 0 | - |
| - | - | 2.002E+06 | 273.1 | - | - | 0 | - |
| - | - | 2.689E+05 | 274.1 | - | - | 0 | - |
| - | - | 8988 | 274.6 | - | - | 0 | - |
| - | - | 3.872E+04 | 275.1 | - | - | 0 | - |
| - | - | 1.726E+04 | 275.2 | - | - | 0 | - |
| - | - | 8010 | 279.1 | - | - | 0 | - |
| - | - | 5552 | 279.2 | - | - | 0 | - |
| 4 | y | 7.65E+04 | 279.7 | 0.001609 | 5.755 | +2 | 6 |
| - | - | 2.08E+04 | 280.2 | - | - | 0 | - |
| - | - | 1.881E+04 | 280.2 | - | - | 0 | - |
| - | - | 5353 | 280.7 | - | - | 0 | - |
| - | - | 1.699E+04 | 281.2 | - | - | 0 | - |
| - | - | 4140 | 281.2 | - | - | 0 | - |
| - | - | 4.624E+04 | 281.2 | - | - | 0 | - |
| - | - | 9925 | 281.2 | - | - | 0 | - |
| - | - | 1.202E+05 | 282.2 | - | - | 0 | - |
| - | - | 1.017E+04 | 282.2 | - | - | 0 | - |
| - | - | 1.421E+04 | 283.1 | - | - | 0 | - |
| - | - | 7360 | 283.1 | - | - | 0 | - |
| - | - | 1.028E+04 | 283.2 | - | - | 0 | - |
| - | - | 8007 | 283.2 | - | - | 0 | - |
| - | - | 5675 | 284.1 | - | - | 0 | - |
| - | - | 2.375E+04 | 284.2 | - | - | 0 | - |
| - | - | 5.592E+04 | 285.2 | - | - | 0 | - |
| - | - | 1.238E+04 | 286.2 | - | - | 0 | - |
| - | - | 3.915E+05 | 287.2 | - | - | 0 | - |
| - | - | 1.109E+04 | 287.2 | - | - | 0 | - |
| - | - | 2.331E+04 | 288.1 | - | - | 0 | - |
| - | - | 4.886E+04 | 288.2 | - | - | 0 | - |
| 4 | y | 2.613E+04 | 288.7 | 0.001362 | 4.72 | +2 | 6 |
| - | - | 9233 | 289.2 | - | - | 0 | - |
| - | - | 1.644E+04 | 290.1 | - | - | 0 | - |
| - | - | 7.596E+04 | 290.1 | - | - | 0 | - |
| - | - | 1.958E+04 | 290.2 | - | - | 0 | - |
| - | - | 1.365E+05 | 291.1 | - | - | 0 | - |
| - | - | 1.763E+04 | 292.1 | - | - | 0 | - |
| - | - | 1.179E+04 | 293.2 | - | - | 0 | - |
| - | - | 5.314E+04 | 295.2 | - | - | 0 | - |
| - | - | 4383 | 296.2 | - | - | 0 | - |
| - | - | 3.018E+04 | 297.2 | - | - | 0 | - |
| - | - | 1.928E+04 | 297.2 | - | - | 0 | - |
| - | - | 9315 | 298.2 | - | - | 0 | - |
| - | - | 1.187E+04 | 299.2 | - | - | 0 | - |
| - | - | 2.467E+04 | 299.2 | - | - | 0 | - |
| - | - | 2.996E+04 | 299.2 | - | - | 0 | - |
| - | - | 5.558E+04 | 300.1 | - | - | 0 | - |
| - | - | 2.073E+04 | 300.2 | - | - | 0 | - |
| - | - | 2.004E+04 | 301.1 | - | - | 0 | - |
| - | - | 2.683E+04 | 303.2 | - | - | 0 | - |
| - | - | 2.865E+04 | 304.1 | - | - | 0 | - |
| 3 | b | 6.214E+05 | 304.2 | 0.00179 | 5.886 | +1 | 3 |
| - | - | 1.091E+05 | 305.2 | - | - | 0 | - |
| - | - | 9827 | 306.2 | - | - | 0 | - |
| - | - | 4.351E+04 | 307.2 | - | - | 0 | - |
| 3 | y | 8.027E+04 | 308.2 | 0.0002419 | 0.7849 | +2 | 7 |
| - | - | 7.752E+04 | 308.2 | - | - | 0 | - |
| - | - | 9834 | 308.2 | - | - | 0 | - |
| - | - | 2.149E+04 | 308.7 | - | - | 0 | - |
| - | - | 1.636E+04 | 309.2 | - | - | 0 | - |
| - | - | 6629 | 309.2 | - | - | 0 | - |
| - | - | 1.235E+04 | 309.2 | - | - | 0 | - |
| - | - | 2.155E+04 | 310.2 | - | - | 0 | - |
| - | - | 5302 | 311.1 | - | - | 0 | - |
| - | - | 1.524E+04 | 311.2 | - | - | 0 | - |
| - | - | 1.35E+04 | 311.2 | - | - | 0 | - |
| - | - | 1.48E+04 | 313.2 | - | - | 0 | - |
| - | - | 4.779E+04 | 313.2 | - | - | 0 | - |
| - | - | 1.66E+04 | 314.2 | - | - | 0 | - |
| 7 | b | 1.739E+04 | 315.2 | 0.001186 | 3.761 | +2 | 7 |
| - | - | 7824 | 315.2 | - | - | 0 | - |
| - | - | 3.051E+04 | 315.2 | - | - | 0 | - |
| - | - | 1.072E+04 | 315.7 | - | - | 0 | - |
| - | - | 2.347E+04 | 316.2 | - | - | 0 | - |
| - | - | 6969 | 316.2 | - | - | 0 | - |
| - | - | 7369 | 317.2 | - | - | 0 | - |
| - | - | 7.968E+04 | 317.2 | - | - | 0 | - |
| - | - | 8319 | 317.2 | - | - | 0 | - |
| - | - | 2.84E+04 | 317.7 | - | - | 0 | - |
| - | - | 4.986E+04 | 318.1 | - | - | 0 | - |
| - | - | 1.814E+05 | 319.1 | - | - | 0 | - |
| - | - | 2.98E+04 | 320.1 | - | - | 0 | - |
| - | - | 3.998E+04 | 321.2 | - | - | 0 | - |
| - | - | 2.904E+04 | 321.2 | - | - | 0 | - |
| - | - | 7564 | 322.2 | - | - | 0 | - |
| - | - | 7.644E+04 | 322.2 | - | - | 0 | - |
| - | - | 2.525E+04 | 322.7 | - | - | 0 | - |
| - | - | 9.189E+04 | 323.2 | - | - | 0 | - |
| 7 | b | 6.107E+04 | 324.2 | 0.0009691 | 2.989 | +2 | 7 |
| - | - | 1.163E+04 | 324.7 | - | - | 0 | - |
| - | - | 5998 | 325.1 | - | - | 0 | - |
| - | - | 2.179E+05 | 325.2 | - | - | 0 | - |
| - | - | 9.758E+05 | 326.2 | - | - | 0 | - |
| - | - | 3.96E+04 | 326.2 | - | - | 0 | - |
| - | - | 1.204E+05 | 327.2 | - | - | 0 | - |
| - | - | 9010 | 327.2 | - | - | 0 | - |
| - | - | 5754 | 327.7 | - | - | 0 | - |
| - | - | 1.71E+04 | 328.1 | - | - | 0 | - |
| - | - | 1.789E+04 | 328.2 | - | - | 0 | - |
| - | - | 1.725E+04 | 328.2 | - | - | 0 | - |
| - | - | 8152 | 328.2 | - | - | 0 | - |
| - | - | 2.805E+04 | 329.1 | - | - | 0 | - |
| - | - | 2.122E+04 | 329.2 | - | - | 0 | - |
| - | - | 9279 | 329.2 | - | - | 0 | - |
| - | - | 2.539E+04 | 330.2 | - | - | 0 | - |
| - | - | 2.104E+04 | 330.2 | - | - | 0 | - |
| - | - | 1.18E+04 | 331.1 | - | - | 0 | - |
| - | - | 5.503E+04 | 331.2 | - | - | 0 | - |
| - | - | 2.666E+04 | 331.7 | - | - | 0 | - |
| - | - | 4.907E+04 | 333.2 | - | - | 0 | - |
| - | - | 9108 | 334.2 | - | - | 0 | - |
| - | - | 1.935E+04 | 336.2 | - | - | 0 | - |
| - | - | 1.458E+04 | 336.2 | - | - | 0 | - |
| - | - | 6372 | 336.2 | - | - | 0 | - |
| - | - | 6941 | 337.2 | - | - | 0 | - |
| - | - | 1.22E+04 | 337.2 | - | - | 0 | - |
| - | - | 3.477E+04 | 338.2 | - | - | 0 | - |
| - | - | 3.024E+04 | 338.2 | - | - | 0 | - |
| - | - | 2.512E+05 | 339.2 | - | - | 0 | - |
| - | - | 2.037E+04 | 340.2 | - | - | 0 | - |
| - | - | 5.731E+04 | 340.2 | - | - | 0 | - |
| - | - | 1.363E+05 | 341.2 | - | - | 0 | - |
| - | - | 6213 | 341.2 | - | - | 0 | - |
| - | - | 2.593E+04 | 342.2 | - | - | 0 | - |
| 7 | y | 3.876E+05 | 343.2 | 0.002072 | 6.035 | +1 | 3 |
| - | - | 1.701E+05 | 344.2 | - | - | 0 | - |
| 7 | y | 3.485E+04 | 344.2 | 0.00118 | 3.427 | +1 | 3 |
| - | - | 5.997E+04 | 344.2 | - | - | 0 | - |
| - | - | 1.563E+04 | 345.2 | - | - | 0 | - |
| - | - | 3.337E+04 | 345.2 | - | - | 0 | - |
| - | - | 8646 | 345.2 | - | - | 0 | - |
| - | - | 5022 | 345.2 | - | - | 0 | - |
| - | - | 1.29E+04 | 347.1 | - | - | 0 | - |
| - | - | 2.01E+04 | 347.2 | - | - | 0 | - |
| - | - | 7333 | 347.2 | - | - | 0 | - |
| - | - | 2.283E+05 | 348.2 | - | - | 0 | - |
| - | - | 4.018E+04 | 349.2 | - | - | 0 | - |
| - | - | 6520 | 350.2 | - | - | 0 | - |
| - | - | 8059 | 350.2 | - | - | 0 | - |
| - | - | 7052 | 351.2 | - | - | 0 | - |
| - | - | 1.614E+04 | 353.2 | - | - | 0 | - |
| - | - | 5.552E+04 | 354.2 | - | - | 0 | - |
| - | - | 6094 | 354.2 | - | - | 0 | - |
| - | - | 1.076E+04 | 354.3 | - | - | 0 | - |
| - | - | 6500 | 355.2 | - | - | 0 | - |
| - | - | 1.22E+04 | 355.2 | - | - | 0 | - |
| - | - | 1.545E+04 | 356.2 | - | - | 0 | - |
| - | - | 1.217E+04 | 356.2 | - | - | 0 | - |
| - | - | 4.186E+04 | 356.2 | - | - | 0 | - |
| - | - | 1.356E+05 | 357.2 | - | - | 0 | - |
| - | - | 5.015E+04 | 357.2 | - | - | 0 | - |
| - | - | 6689 | 357.2 | - | - | 0 | - |
| - | - | 4558 | 357.7 | - | - | 0 | - |
| - | - | 2.973E+04 | 358.2 | - | - | 0 | - |
| - | - | 9.213E+04 | 358.2 | - | - | 0 | - |
| - | - | 1.753E+04 | 359.2 | - | - | 0 | - |
| - | - | 8922 | 360.2 | - | - | 0 | - |
| 4 | b | 6.539E+04 | 361.2 | 0.003428 | 9.492 | +1 | 4 |
| 7 | y | 1.402E+06 | 361.2 | 0.002066 | 5.719 | +1 | 3 |
| - | - | 1.119E+04 | 362.2 | - | - | 0 | - |
| - | - | 2.544E+05 | 362.2 | - | - | 0 | - |
| - | - | 5010 | 362.7 | - | - | 0 | - |
| - | - | 2.966E+04 | 363.3 | - | - | 0 | - |
| - | - | 4.674E+04 | 364.2 | - | - | 0 | - |
| - | - | 3.805E+05 | 365.2 | - | - | 0 | - |
| - | - | 9.101E+04 | 366.2 | - | - | 0 | - |
| - | - | 4.958E+04 | 366.2 | - | - | 0 | - |
| - | - | 1.308E+05 | 366.7 | - | - | 0 | - |
| - | - | 1.525E+04 | 367.2 | - | - | 0 | - |
| - | - | 6677 | 367.2 | - | - | 0 | - |
| - | - | 5.672E+04 | 367.2 | - | - | 0 | - |
| - | - | 8122 | 367.7 | - | - | 0 | - |
| - | - | 8590 | 367.7 | - | - | 0 | - |
| - | - | 2.138E+05 | 368.2 | - | - | 0 | - |
| - | - | 4.363E+04 | 369.2 | - | - | 0 | - |
| - | - | 2.14E+04 | 371.2 | - | - | 0 | - |
| 8 | b | 4.492E+04 | 371.7 | 0.001817 | 4.888 | +2 | 8 |
| - | - | 1.988E+04 | 372.2 | - | - | 0 | - |
| - | - | 1.178E+04 | 372.2 | - | - | 0 | - |
| - | - | 4.689E+04 | 372.3 | - | - | 0 | - |
| - | - | 1.943E+04 | 372.7 | - | - | 0 | - |
| - | - | 1.847E+04 | 373.2 | - | - | 0 | - |
| - | - | 1.167E+04 | 373.3 | - | - | 0 | - |
| - | - | 1.538E+05 | 374.2 | - | - | 0 | - |
| - | - | 8.009E+04 | 375.2 | - | - | 0 | - |
| - | - | 8250 | 375.2 | - | - | 0 | - |
| - | - | 3.409E+04 | 376.2 | - | - | 0 | - |
| - | - | 8961 | 376.2 | - | - | 0 | - |
| - | - | 7467 | 376.3 | - | - | 0 | - |
| - | - | 1.386E+04 | 376.7 | - | - | 0 | - |
| - | - | 1.824E+04 | 376.7 | - | - | 0 | - |
| - | - | 1.005E+04 | 377.2 | - | - | 0 | - |
| - | - | 6840 | 378.2 | - | - | 0 | - |
| 8 | b | 3.025E+04 | 380.7 | 0.001692 | 4.445 | +2 | 8 |
| - | - | 1.127E+04 | 381.2 | - | - | 0 | - |
| 2 | y | 2.758E+05 | 381.7 | 0.002416 | 6.33 | +2 | 8 |
| - | - | 1.077E+05 | 382.2 | - | - | 0 | - |
| - | - | 2.639E+05 | 382.2 | - | - | 0 | - |
| - | - | 3.224E+04 | 382.7 | - | - | 0 | - |
| - | - | 1.899E+06 | 383.2 | - | - | 0 | - |
| - | - | 5.5E+04 | 383.3 | - | - | 0 | - |
| - | - | 2.015E+04 | 384.2 | - | - | 0 | - |
| - | - | 3.276E+05 | 384.2 | - | - | 0 | - |
| - | - | 2.836E+04 | 385.2 | - | - | 0 | - |
| - | - | 3.981E+04 | 385.2 | - | - | 0 | - |
| - | - | 9573 | 385.3 | - | - | 0 | - |
| - | - | 5936 | 386.2 | - | - | 0 | - |
| - | - | 1.252E+05 | 386.2 | - | - | 0 | - |
| - | - | 5074 | 386.7 | - | - | 0 | - |
| - | - | 4.939E+04 | 387.2 | - | - | 0 | - |
| - | - | 4844 | 388.2 | - | - | 0 | - |
| - | - | 5.758E+04 | 390.2 | - | - | 0 | - |
| 2 | y | 2.012E+06 | 390.7 | 0.002261 | 5.787 | +2 | 8 |
| - | - | 8.152E+05 | 391.2 | - | - | 0 | - |
| - | - | 1.972E+05 | 391.7 | - | - | 0 | - |
| - | - | 1.93E+05 | 392.2 | - | - | 0 | - |
| - | - | 1.913E+04 | 392.2 | - | - | 0 | - |
| - | - | 4.451E+04 | 393.2 | - | - | 0 | - |
| - | - | 6897 | 393.3 | - | - | 0 | - |
| - | - | 6928 | 394.2 | - | - | 0 | - |
| - | - | 3.517E+04 | 394.2 | - | - | 0 | - |
| - | - | 5.09E+04 | 395.2 | - | - | 0 | - |
| - | - | 1.26E+05 | 395.7 | - | - | 0 | - |
| - | - | 4.12E+04 | 396.2 | - | - | 0 | - |
| - | - | 7650 | 396.2 | - | - | 0 | - |
| - | - | 1.09E+04 | 396.7 | - | - | 0 | - |
| - | - | 4.248E+05 | 400.3 | - | - | 0 | - |
| - | - | 1.944E+06 | 401.2 | - | - | 0 | - |
| - | - | 7.557E+04 | 401.3 | - | - | 0 | - |
| - | - | 5.803E+05 | 402.2 | - | - | 0 | - |
| - | - | 3.562E+05 | 402.2 | - | - | 0 | - |
| - | - | 8379 | 402.3 | - | - | 0 | - |
| - | - | 1.222E+05 | 403.2 | - | - | 0 | - |
| - | - | 4.579E+04 | 403.2 | - | - | 0 | - |
| - | - | 1.468E+04 | 404.2 | - | - | 0 | - |
| - | - | 1.693E+04 | 404.2 | - | - | 0 | - |
| - | - | 6503 | 407.2 | - | - | 0 | - |
| - | - | 6709 | 409.3 | - | - | 0 | - |
| - | - | 6093 | 410.2 | - | - | 0 | - |
| - | - | 2.127E+04 | 411.2 | - | - | 0 | - |
| - | - | 2.897E+04 | 411.3 | - | - | 0 | - |
| - | - | 1.009E+04 | 412.2 | - | - | 0 | - |
| - | - | 6843 | 412.3 | - | - | 0 | - |
| - | - | 5.696E+04 | 413.3 | - | - | 0 | - |
| - | - | 1.141E+04 | 414.3 | - | - | 0 | - |
| - | - | 4933 | 417.2 | - | - | 0 | - |
| - | - | 4885 | 417.3 | - | - | 0 | - |
| 5 | b | 3.294E+04 | 418.2 | 0.002656 | 6.35 | +1 | 5 |
| - | - | 9043 | 418.2 | - | - | 0 | - |
| - | - | 9129 | 418.3 | - | - | 0 | - |
| - | - | 3.494E+04 | 419.2 | - | - | 0 | - |
| - | - | 3.346E+05 | 420.2 | - | - | 0 | - |
| - | - | 6.812E+04 | 421.2 | - | - | 0 | - |
| - | - | 3590 | 421.2 | - | - | 0 | - |
| - | - | 7.067E+04 | 421.3 | - | - | 0 | - |
| - | - | 9325 | 422.2 | - | - | 0 | - |
| - | - | 1.225E+04 | 422.2 | - | - | 0 | - |
| - | - | 6292 | 422.3 | - | - | 0 | - |
| - | - | 2.961E+04 | 423.2 | - | - | 0 | - |
| - | - | 6218 | 424.2 | - | - | 0 | - |
| - | - | 1.602E+04 | 424.3 | - | - | 0 | - |
| - | - | 1.057E+04 | 426.3 | - | - | 0 | - |
| - | - | 7615 | 429.1 | - | - | 0 | - |
| - | - | 1.564E+05 | 429.3 | - | - | 0 | - |
| - | - | 1.582E+04 | 430.2 | - | - | 0 | - |
| - | - | 3.704E+04 | 430.3 | - | - | 0 | - |
| 0 | Precursor | 1.325E+05 | 431.2 | 0.002694 | 6.248 | +2 | -1 |
| - | - | 5.151E+04 | 431.7 | - | - | 0 | - |
| - | - | 1.686E+04 | 432.2 | - | - | 0 | - |
| - | - | 1.47E+04 | 433.3 | - | - | 0 | - |
| - | - | 2.267E+04 | 434.3 | - | - | 0 | - |
| - | - | 3.202E+04 | 437.2 | - | - | 0 | - |
| - | - | 6951 | 438.2 | - | - | 0 | - |
| - | - | 5.07E+05 | 439.3 | - | - | 0 | - |
| 0 | Precursor | 2.384E+05 | 440.3 | 0.003088 | 7.015 | +2 | -1 |
| - | - | 1.359E+05 | 440.8 | - | - | 0 | - |
| - | - | 6.558E+04 | 441.3 | - | - | 0 | - |
| - | - | 4912 | 441.3 | - | - | 0 | - |
| - | - | 1.347E+04 | 442.2 | - | - | 0 | - |
| - | - | 1.722E+04 | 442.3 | - | - | 0 | - |
| - | - | 7493 | 443.2 | - | - | 0 | - |
| 6 | y | 5.984E+04 | 444.3 | 0.002885 | 6.495 | +1 | 4 |
| - | - | 1.591E+04 | 445.3 | - | - | 0 | - |
| - | - | 6418 | 446.3 | - | - | 0 | - |
| - | - | 7604 | 447.2 | - | - | 0 | - |
| - | - | 1.312E+04 | 447.2 | - | - | 0 | - |
| - | - | 4610 | 448.5 | - | - | 0 | - |
| - | - | 1.672E+04 | 448.6 | - | - | 0 | - |
| - | - | 2.526E+04 | 448.6 | - | - | 0 | - |
| - | - | 3.799E+04 | 448.7 | - | - | 0 | - |
| - | - | 4821 | 448.7 | - | - | 0 | - |
| - | - | 6843 | 449.3 | - | - | 0 | - |
| - | - | 8138 | 449.6 | - | - | 0 | - |
| - | - | 2.916E+04 | 450.3 | - | - | 0 | - |
| - | - | 8.299E+04 | 451.3 | - | - | 0 | - |
| - | - | 1.685E+05 | 452.3 | - | - | 0 | - |
| - | - | 3.324E+04 | 453.3 | - | - | 0 | - |
| - | - | 5574 | 454.3 | - | - | 0 | - |
| - | - | 5686 | 455.2 | - | - | 0 | - |
| - | - | 3.261E+04 | 456.2 | - | - | 0 | - |
| - | - | 8.034E+05 | 457.3 | - | - | 0 | - |
| - | - | 1.856E+05 | 458.3 | - | - | 0 | - |
| - | - | 2.497E+04 | 459.3 | - | - | 0 | - |
| - | - | 1.967E+04 | 460.3 | - | - | 0 | - |
| - | - | 5.893E+04 | 461.3 | - | - | 0 | - |
| - | - | 1.495E+04 | 462.3 | - | - | 0 | - |
| 6 | y | 3.235E+05 | 462.3 | 0.002819 | 6.098 | +1 | 4 |
| - | - | 7.727E+04 | 463.3 | - | - | 0 | - |
| - | - | 1.436E+04 | 464.3 | - | - | 0 | - |
| - | - | 5856 | 466.2 | - | - | 0 | - |
| - | - | 1.74E+04 | 467.3 | - | - | 0 | - |
| - | - | 6394 | 468.3 | - | - | 0 | - |
| - | - | 1.008E+05 | 468.3 | - | - | 0 | - |
| - | - | 6112 | 469.2 | - | - | 0 | - |
| - | - | 6.02E+04 | 469.3 | - | - | 0 | - |
| - | - | 1.897E+04 | 470.2 | - | - | 0 | - |
| - | - | 1.281E+05 | 470.3 | - | - | 0 | - |
| - | - | 8038 | 470.3 | - | - | 0 | - |
| - | - | 5785 | 471.2 | - | - | 0 | - |
| - | - | 3.24E+04 | 471.3 | - | - | 0 | - |
| - | - | 6168 | 472.3 | - | - | 0 | - |
| - | - | 4.033E+04 | 473.3 | - | - | 0 | - |
| - | - | 2.135E+04 | 474.2 | - | - | 0 | - |
| - | - | 9035 | 475.2 | - | - | 0 | - |
| - | - | 1.276E+04 | 475.3 | - | - | 0 | - |
| - | - | 3.204E+05 | 478.3 | - | - | 0 | - |
| - | - | 6.727E+04 | 479.3 | - | - | 0 | - |
| - | - | 2.046E+04 | 480.3 | - | - | 0 | - |
| - | - | 1.425E+04 | 481.1 | - | - | 0 | - |
| - | - | 8669 | 481.2 | - | - | 0 | - |
| - | - | 9157 | 483.2 | - | - | 0 | - |
| - | - | 2.291E+04 | 483.3 | - | - | 0 | - |
| - | - | 6447 | 484.3 | - | - | 0 | - |
| - | - | 9317 | 485.3 | - | - | 0 | - |
| - | - | 1.263E+06 | 486.3 | - | - | 0 | - |
| - | - | 2.931E+05 | 487.3 | - | - | 0 | - |
| - | - | 1.273E+04 | 488.3 | - | - | 0 | - |
| - | - | 3.825E+04 | 488.3 | - | - | 0 | - |
| - | - | 5689 | 489.3 | - | - | 0 | - |
| - | - | 4.701E+04 | 491.3 | - | - | 0 | - |
| - | - | 1.247E+04 | 492.3 | - | - | 0 | - |
| - | - | 2.044E+04 | 495.2 | - | - | 0 | - |
| - | - | 2.541E+06 | 496.3 | - | - | 0 | - |
| - | - | 6.219E+05 | 497.3 | - | - | 0 | - |
| - | - | 9.226E+04 | 498.3 | - | - | 0 | - |
| 6 | b | 1.576E+05 | 501.2 | 0.002987 | 5.959 | +1 | 6 |
| 5 | y | 8.235E+04 | 501.3 | 0.00312 | 6.223 | +1 | 5 |
| - | - | 5.012E+04 | 502.3 | - | - | 0 | - |
| - | - | 1.804E+04 | 502.3 | - | - | 0 | - |
| - | - | 1.067E+04 | 503.3 | - | - | 0 | - |
| - | - | 6913 | 503.3 | - | - | 0 | - |
| - | - | 1.212E+04 | 504.3 | - | - | 0 | - |
| - | - | 4914 | 505.3 | - | - | 0 | - |
| - | - | 5311 | 506.3 | - | - | 0 | - |
| - | - | 8.31E+04 | 512.3 | - | - | 0 | - |
| - | - | 2.274E+04 | 513.3 | - | - | 0 | - |
| - | - | 4731 | 513.3 | - | - | 0 | - |
| - | - | 6.09E+06 | 514.3 | - | - | 0 | - |
| - | - | 1.544E+06 | 515.3 | - | - | 0 | - |
| - | - | 7829 | 515.5 | - | - | 0 | - |
| - | - | 2.442E+05 | 516.3 | - | - | 0 | - |
| - | - | 1.414E+04 | 517.3 | - | - | 0 | - |
| - | - | 8563 | 518.3 | - | - | 0 | - |
| 6 | b | 7.502E+04 | 519.3 | 0.00408 | 7.857 | +1 | 6 |
| 5 | y | 4.714E+05 | 519.3 | 0.002962 | 5.703 | +1 | 5 |
| - | - | 2.176E+04 | 520.3 | - | - | 0 | - |
| - | - | 1.157E+05 | 520.3 | - | - | 0 | - |
| - | - | 6047 | 521.3 | - | - | 0 | - |
| - | - | 2.112E+04 | 521.3 | - | - | 0 | - |
| - | - | 1.8E+04 | 524.3 | - | - | 0 | - |
| - | - | 2.901E+05 | 530.3 | - | - | 0 | - |
| - | - | 9.838E+04 | 531.3 | - | - | 0 | - |
| - | - | 1.013E+05 | 532.3 | - | - | 0 | - |
| - | - | 2.976E+04 | 533.3 | - | - | 0 | - |
| - | - | 8997 | 540.3 | - | - | 0 | - |
| - | - | 2.634E+04 | 540.3 | - | - | 0 | - |
| - | - | 5.345E+05 | 548.3 | - | - | 0 | - |
| - | - | 1.624E+05 | 549.3 | - | - | 0 | - |
| - | - | 2.688E+04 | 550.3 | - | - | 0 | - |
| - | - | 7650 | 552.3 | - | - | 0 | - |
| - | - | 7740 | 553.3 | - | - | 0 | - |
| - | - | 7522 | 553.4 | - | - | 0 | - |
| - | - | 1.625E+04 | 554.3 | - | - | 0 | - |
| - | - | 2.578E+04 | 558.3 | - | - | 0 | - |
| 4 | y | 1.285E+05 | 558.3 | 0.003232 | 5.789 | +1 | 6 |
| - | - | 4.511E+04 | 559.3 | - | - | 0 | - |
| - | - | 9394 | 560.3 | - | - | 0 | - |
| - | - | 5666 | 562.3 | - | - | 0 | - |
| - | - | 5905 | 569.3 | - | - | 0 | - |
| - | - | 1.515E+04 | 570.3 | - | - | 0 | - |
| - | - | 5.439E+04 | 571.3 | - | - | 0 | - |
| - | - | 1.497E+05 | 571.4 | - | - | 0 | - |
| - | - | 1.573E+04 | 572.3 | - | - | 0 | - |
| - | - | 4.113E+04 | 572.4 | - | - | 0 | - |
| - | - | 8220 | 573.4 | - | - | 0 | - |
| 4 | y | 1.554E+06 | 576.3 | 0.003348 | 5.81 | +1 | 6 |
| - | - | 3.932E+05 | 577.3 | - | - | 0 | - |
| - | - | 8.044E+04 | 578.3 | - | - | 0 | - |
| - | - | 2.347E+04 | 579.3 | - | - | 0 | - |
| - | - | 1.276E+04 | 580.3 | - | - | 0 | - |
| - | - | 7619 | 581.3 | - | - | 0 | - |
| - | - | 6293 | 585.3 | - | - | 0 | - |
| - | - | 3.381E+04 | 587.4 | - | - | 0 | - |
| - | - | 1.353E+04 | 588.4 | - | - | 0 | - |
| - | - | 2.776E+04 | 589.3 | - | - | 0 | - |
| - | - | 6228 | 590.3 | - | - | 0 | - |
| - | - | 1.972E+05 | 597.3 | - | - | 0 | - |
| - | - | 8.645E+04 | 598.3 | - | - | 0 | - |
| - | - | 5.375E+04 | 599.3 | - | - | 0 | - |
| - | - | 1.421E+04 | 600.3 | - | - | 0 | - |
| - | - | 1.681E+04 | 611.3 | - | - | 0 | - |
| - | - | 5430 | 612.3 | - | - | 0 | - |
| - | - | 1.82E+04 | 613.4 | - | - | 0 | - |
| 3 | y | 8.973E+05 | 615.3 | 0.003558 | 5.782 | +1 | 7 |
| - | - | 3.19E+05 | 616.4 | - | - | 0 | - |
| - | - | 8.11E+04 | 617.4 | - | - | 0 | - |
| - | - | 1.198E+04 | 618.3 | - | - | 0 | - |
| - | - | 5.801E+04 | 625.3 | - | - | 0 | - |
| - | - | 2.643E+04 | 626.3 | - | - | 0 | - |
| 7 | b | 1.124E+05 | 629.3 | 0.003544 | 5.631 | +1 | 7 |
| - | - | 3.271E+04 | 630.3 | - | - | 0 | - |
| - | - | 6218 | 631.4 | - | - | 0 | - |
| 3 | y | 1.396E+07 | 633.4 | 0.003735 | 5.898 | +1 | 7 |
| - | - | 4.242E+06 | 634.4 | - | - | 0 | - |
| - | - | 8.581E+05 | 635.4 | - | - | 0 | - |
| - | - | 4.541E+04 | 636.4 | - | - | 0 | - |
| - | - | 6.357E+05 | 643.4 | - | - | 0 | - |
| - | - | 2.178E+05 | 644.4 | - | - | 0 | - |
| - | - | 4.341E+04 | 645.4 | - | - | 0 | - |
| 7 | b | 3.184E+05 | 647.4 | 0.003538 | 5.466 | +1 | 7 |
| - | - | 9.581E+04 | 648.4 | - | - | 0 | - |
| - | - | 1.973E+04 | 649.4 | - | - | 0 | - |
| - | - | 1.265E+04 | 653.3 | - | - | 0 | - |
| - | - | 1.196E+06 | 661.4 | - | - | 0 | - |
| - | - | 4.263E+05 | 662.4 | - | - | 0 | - |
| - | - | 7.84E+04 | 663.4 | - | - | 0 | - |
| - | - | 5315 | 664.4 | - | - | 0 | - |
| - | - | 1.789E+04 | 671.4 | - | - | 0 | - |
| - | - | 8865 | 672.4 | - | - | 0 | - |
| - | - | 3.102E+04 | 679.4 | - | - | 0 | - |
| - | - | 1.321E+04 | 680.4 | - | - | 0 | - |
| - | - | 5042 | 689.4 | - | - | 0 | - |
| - | - | 4895 | 698.4 | - | - | 0 | - |
| - | - | 5617 | 714.4 | - | - | 0 | - |
| - | - | 4522 | 715.4 | - | - | 0 | - |
| - | - | 8400 | 716.4 | - | - | 0 | - |
| - | - | 3.723E+04 | 718.4 | - | - | 0 | - |
| - | - | 1.502E+04 | 719.4 | - | - | 0 | - |
| - | - | 7579 | 724.4 | - | - | 0 | - |
| - | - | 7789 | 725.4 | - | - | 0 | - |
| - | - | 5.797E+04 | 732.4 | - | - | 0 | - |
| - | - | 2.123E+04 | 733.4 | - | - | 0 | - |
| - | - | 1.156E+04 | 736.4 | - | - | 0 | - |
| 8 | b | 1.562E+05 | 742.4 | 0.003953 | 5.324 | +1 | 8 |
| - | - | 5.769E+04 | 743.4 | - | - | 0 | - |
| - | - | 4.793E+04 | 744.4 | - | - | 0 | - |
| - | - | 1.755E+04 | 745.4 | - | - | 0 | - |
| - | - | 6843 | 746.4 | - | - | 0 | - |
| 8 | b | 4.619E+05 | 760.4 | 0.003825 | 5.03 | +1 | 8 |
| - | - | 1.97E+05 | 761.4 | - | - | 0 | - |
| 2 | y | 1.774E+05 | 762.4 | 0.007959 | 10.44 | +1 | 8 |
| - | - | 8.662E+04 | 763.4 | - | - | 0 | - |
| - | - | 2.031E+04 | 764.4 | - | - | 0 | - |
| - | - | 7261 | 772.4 | - | - | 0 | - |
| - | - | 8806 | 778.4 | - | - | 0 | - |
| - | - | 6070 | 779.5 | - | - | 0 | - |
| 2 | y | 3.325E+06 | 780.4 | 0.003742 | 4.795 | +1 | 8 |
| - | - | 1.328E+06 | 781.4 | - | - | 0 | - |
| - | - | 3.487E+05 | 782.4 | - | - | 0 | - |
| - | - | 6185 | 782.5 | - | - | 0 | - |
| - | - | 1.893E+04 | 783.4 | - | - | 0 | - |
| - | - | 5.418E+04 | 790.4 | - | - | 0 | - |
| - | - | 2.526E+04 | 791.4 | - | - | 0 | - |
| - | - | 7330 | 792.4 | - | - | 0 | - |
| - | - | 1.089E+04 | 1293 | - | - | 0 | - |
| - | - | 4693 | 1308 | - | - | 0 | - |
| - | - | 4730 | 1308 | - | - | 0 | - |
| - | - | 4428 | 1557 | - | - | 0 | - |
| - | - | 1.763E+04 | 1779 | - | - | 0 | - |
| - | - | 2.684E+04 | 1780 | - | - | 0 | - |
| - | - | 5012 | 1818 | - | - | 0 | - |
| - | - | 6372 | 1818 | - | - | 0 | - |
| - | - | 6336 | 2389 | - | - | 0 | - |
| - | - | 1.331E+04 | 2389 | - | - | 0 | - |
| - | - | 5011 | 3042 | - | - | 0 | - |

m/z Charge Intensity FragmentType MassShift Position
120.08136749267578 0 21287488
121.0694351196289 0 35679.58
121.07929229736328 0 60267.984
121.08456420898438 0 1674138.5
122.08174133300781 0 5545.387
122.08795928955078 0 53724.2
124.08724975585938 0 5085.539
125.07179260253906 0 21050.46
127.0510025024414 0 118056.1
127.08760833740234 0 14918.546
128.0540008544922 0 6400.902
128.0720672607422 0 5819.266
128.10775756835938 0 22880.662
129.10299682617188 0 11265332
130.0503387451172 0 19893.318
130.06578063964844 0 11458.841
130.1002655029297 0 67382.08
130.10623168945312 0 675949.6
131.04971313476562 0 20132.229
131.08216857910156 0 515161.5
131.1075897216797 0 14606.774
132.0773468017578 0 13606.745
132.08143615722656 0 108756.766
132.08599853515625 0 13465.143
133.08505249023438 0 8837.145
137.10855102539062 0 4259.199
139.0873260498047 0 53174.895
139.123779296875 0 11328.291
140.08328247070312 0 5337.8535
140.0907745361328 0 6447.4
140.1078643798828 0 31869.814
140.14425659179688 0 6890.806
141.06666564941406 0 220247.9
141.10311889648438 0 50158.68
142.07008361816406 0 14392.195
142.12356567382812 0 34323.84
143.04608154296875 0 5792.287
143.08230590820312 0 35979.117
143.11907958984375 0 5822.3276
143.1267547607422 0 4672.677
146.0611572265625 0 5930.82
146.129638671875 0 36793.684
146.49412536621094 0 5641.1
147.1136932373047 0 7690.7373
148.07652282714844 0 40495.42
148.9243621826172 0 7725.4194
148.93167114257812 0 8101.5034
148.93910217285156 0 19339.574
148.95547485351562 0 37187.945
148.96331787109375 0 21001.152
148.9706573486328 0 9665.099
148.97801208496094 0 7864.8667
148.98545837402344 0 6449.9844
148.99319458007812 0 5185.5166
149.07974243164062 0 5344.0835
149.10777282714844 0 4964.2573
151.08740234375 0 36867.7
151.12400817871094 0 6807.1597
152.08338928222656 0 5653.803
152.1442108154297 0 115851.34
153.06671142578125 0 70885.43
153.10308837890625 0 33346.633
153.1479034423828 0 7603.2505
154.0619354248047 0 39944.23
154.06918334960938 0 7406.87
155.08236694335938 0 89472.49
155.1187286376953 0 323672.16
156.0784912109375 0 3846.0576
156.08546447753906 0 4509.4116
156.1158447265625 0 4417.6294
156.1220703125 0 28874.672
157.09783935546875 0 9380.557
158.09329223632812 0 109345.52
159.0772705078125 0 389254.9
160.0804901123047 0 23352.838
165.1033172607422 0 10747.685
167.1187744140625 0 77830.67
168.11431884765625 0 14698.647
168.1217803955078 0 8748.318
168.1503448486328 0 6882.3726
169.06210327148438 0 4331.876
169.09800720214844 0 28185.252
169.13446044921875 0 15494.591
169.1710662841797 0 9869.417
170.09335327148438 0 205185.19
171.07737731933594 0 107818.055
171.0887908935547 0 15793.766
171.09666442871094 0 11114.88
171.11373901367188 0 23460.7
171.15074157714844 0 4656.8403
172.0726318359375 0 523755.3
172.109130859375 0 12065.073
173.07594299316406 0 30982.256
173.09291076660156 0 4182.0015
174.07688903808594 0 4312.8477
174.12864685058594 0 19669.984
175.08810424804688 0 4062.555
176.10781860351562 0 201042.22
177.10317993164062 0 700964.25
177.11085510253906 0 26807.266
178.09927368164062 0 5456.269
178.10659790039062 0 57047.387
179.118896484375 0 4260.244
179.1550750732422 0 16425.307
180.0778045654297 0 17100.484
180.11404418945312 0 11395.865
181.06182861328125 0 18687.295
181.09857177734375 0 3596.5847 b 3
181.1095428466797 0 5397.7695
181.1710968017578 0 18953.717
182.0570526123047 0 5044.8154
182.09347534179688 0 16379.083
183.08905029296875 0 7640.389
183.11378479003906 0 88610.42
184.07276916503906 0 30189.682
184.1172332763672 0 7368.7
184.14556884765625 0 16580.682
185.0930938720703 0 38483.34
185.12945556640625 0 166272.7
186.12472534179688 0 144450.2
186.13291931152344 0 10265.562
187.0869140625 0 5033.6284
187.1080780029297 0 4267.9355
187.1243438720703 0 12478.644
187.14517211914062 0 13509.825
188.0723419189453 0 5769.953
188.1040496826172 0 91447.31
188.14437866210938 0 12898.238
189.102783203125 0 27805.799
190.08326721191406 0 5676.6343
192.07794189453125 0 22490.807
194.0944061279297 0 3474.2886
194.12979125976562 0 60068.566
195.13377380371094 0 7523.955
195.15028381347656 0 49109.496
196.10897827148438 0 52194.914
196.11758422851562 0 11282.802
196.1546630859375 0 3887.0867
196.18136596679688 0 8354.899
197.0929718017578 0 13131.184
197.1037139892578 0 7239.415
197.12948608398438 0 13501.878
197.16586303710938 0 503847.2
198.0883331298828 0 303859.56
198.1239776611328 0 8987.786
198.16920471191406 0 47409.78
198.1967315673828 0 4595.3867
199.091552734375 0 23582.004
199.14547729492188 0 5082.788
199.17149353027344 0 4924.6836
200.0675811767578 0 8881.705
200.1038818359375 0 15684.942
201.09922790527344 0 47595.066
202.12403869628906 0 11562.193
203.1189422607422 0 49487.293
203.1552276611328 0 12915.333
204.12203979492188 0 4310.7466
205.09820556640625 0 1243591.5
206.1016387939453 0 151731.6
206.12985229492188 0 12652.959
206.13954162597656 0 6766.243
207.10397338867188 0 7558.7793
207.15025329589844 0 232586.11
208.0726318359375 0 4124.8447
208.10916137695312 0 208506.25
208.15380859375 0 23603.441
209.10443115234375 0 161446.11
209.12925720214844 0 15386.021
209.16600036621094 0 8872.532
210.08843994140625 0 94936.04
210.09906005859375 0 6636.7593
210.1077117919922 0 14058.59
210.12440490722656 0 4512.351
211.08274841308594 0 8642.413
211.09327697753906 0 4534.018
211.12060546875 0 8505.408
212.06784057617188 0 7294.9463
212.14048767089844 0 1046986
213.1246337890625 0 37655.93
213.1438751220703 0 104780.65
214.11965942382812 0 16938.566
214.19256591796875 0 87215.14
215.08290100097656 0 20011.516
215.1154327392578 0 178753.56
215.1402130126953 0 183671.58 y Water loss 7
215.1964874267578 0 7505.1875
216.09912109375 0 221128.14
216.1176300048828 0 12322.678
216.14393615722656 0 20559.258
217.09872436523438 0 102354.12
218.10223388671875 0 10519.122
219.15032958984375 0 18731308 a 1
220.15347290039062 0 2447324.5
221.12933349609375 0 32239.059
221.1564178466797 0 143980.02
222.1248779296875 0 19851.223
222.1613311767578 0 5468.416
223.14578247070312 0 15573.509
224.14059448242188 0 21329.861
224.17686462402344 0 359399.38
225.1001739501953 0 8144.3486
225.1248779296875 0 3809.049
225.1357879638672 0 137716.86
225.16094970703125 0 99236.53
225.18038940429688 0 44700.7
226.08331298828125 0 29188.33
226.11978149414062 0 18485.205
226.1393280029297 0 12693.846
226.16419982910156 0 15383.313
227.1150665283203 0 799472.6
228.09906005859375 0 426306.56
228.11050415039062 0 66946.11
228.119384765625 0 36641.44
228.13534545898438 0 23556.799
229.09434509277344 0 98220.12
229.13113403320312 0 11745.409
230.15113830566406 0 2158980
231.11412048339844 0 41221.438
231.1536865234375 0 225966.1
231.6514129638672 0 8172.258 y 5
232.15501403808594 0 23445.414
233.0931396484375 0 8853.488
233.1293487548828 0 39764.094
233.15084838867188 0 382057.12 y 7
233.16525268554688 0 12792.32
234.12489318847656 0 66048.18
234.15432739257812 0 40389.98
235.12826538085938 0 6737.506
235.15626525878906 0 5497.9688
236.1768798828125 0 4882.057
237.09950256347656 0 134220
237.12257385253906 0 4686.6035
237.13748168945312 0 5798.091
238.08352661132812 0 18148.121
238.1030731201172 0 13430.148
238.1306915283203 0 9870.449
240.1356658935547 0 143013.61
240.17184448242188 0 20204.719
241.13917541503906 0 18746.12
241.1560821533203 0 71287.72
242.1876678466797 0 447322.72
243.11415100097656 0 97324
243.14683532714844 0 55228.1
243.17184448242188 0 38294.75
243.19091796875 0 49573.76
243.65516662597656 0 6172.7744
244.1096649169922 0 45114.164
245.12583923339844 0 458843.34
245.16566467285156 0 21055.535
246.12875366210938 0 29995.625
247.14547729492188 0 4650401 b 1
248.14878845214844 0 671731.1
249.1251220703125 0 12013.076
249.1514434814453 0 43561.746
250.15599060058594 0 12063.424
250.1936492919922 0 9760.749
251.15206909179688 0 97404.1 y Water loss 4
252.1354217529297 0 17369.863
252.15484619140625 0 13165.112
252.17202758789062 0 43980.66
253.131591796875 0 8875.472
254.12612915039062 0 64632.285
254.15138244628906 0 10569.116
255.11019897460938 0 1171218.6
256.11309814453125 0 126018.766
256.1656188964844 0 4658.0054
257.115966796875 0 11744.727
257.1624755859375 0 27676.662
258.1466979980469 0 6736.855
258.162841796875 0 5046.2886
259.1455993652344 0 50171.742
260.1472473144531 0 4497.3486
260.16241455078125 0 13641.964 y 4
260.1982116699219 0 78359.24
261.1248474121094 0 81756.555
261.1592712402344 0 6760.7686
261.2016906738281 0 8525.79
262.12005615234375 0 345505.2
262.1931457519531 0 7636.462
263.12347412109375 0 42102.695
263.1764831542969 0 13347.447
264.1722106933594 0 29831.7
265.0968017578125 0 5654.319
265.1307067871094 0 19674.83
266.1507568359375 0 18229.67
268.1667785644531 0 12756.327
269.16229248046875 0 1370089.8
270.1654357910156 0 162846.2
270.18389892578125 0 159197.03
271.1093444824219 0 26298.426
271.14141845703125 0 33190.598
271.1680603027344 0 13703.676
271.18585205078125 0 18416.666
271.2139587402344 0 8012.5195
272.1043701171875 0 15078.792
272.1369934082031 0 194899
273.120849609375 0 2001900
274.122802734375 0 268906.66
274.646484375 0 8988.263
275.1248779296875 0 38724.477
275.17645263671875 0 17255.691
279.1482238769531 0 8009.617
279.21923828125 0 5551.5894
279.66754150390625 0 76501.45 y Water loss 3
280.16900634765625 0 20800.867
280.2035217285156 0 18808.504
280.6678771972656 0 5352.727
281.16229248046875 0 16993.873
281.1826171875 0 4139.514
281.1988830566406 0 46239.46
281.234375 0 9925.182
282.15771484375 0 120189.64
282.1842956542969 0 10170.621
283.1059875488281 0 14209.186
283.1419982910156 0 7359.8086
283.16064453125 0 10278.341
283.17779541015625 0 8007.2393
284.1029357910156 0 5675.231
284.1624450683594 0 23749.242
285.1573791503906 0 55924.934
286.15863037109375 0 12379.604
287.1730041503906 0 391528.75
287.21246337890625 0 11089.624
288.1357727050781 0 23306.84
288.1758728027344 0 48859.75
288.6725769042969 0 26134.922 y 3
289.17620849609375 0 9233.345
290.1156311035156 0 16437.014
290.1478271484375 0 75959.03
290.1885681152344 0 19580.469
291.1470642089844 0 136451
292.14996337890625 0 17634.65
293.1987609863281 0 11787.393
295.1782531738281 0 53141.094
296.1787414550781 0 4382.7954
297.157470703125 0 30180.562
297.23052978515625 0 19278.104
298.2140808105469 0 9315.102
299.1739807128906 0 11874.93
299.2094421386719 0 24665.205
299.24591064453125 0 29957.016
300.1358642578125 0 55581.14
300.1683349609375 0 20730.902
301.1319885253906 0 20040.166
303.18304443359375 0 26832.049
304.14404296875 0 28651.62
304.1673583984375 0 621427.25 b 2
305.17034912109375 0 109083.06
306.1741638183594 0 9826.902
307.2144775390625 0 43511.965
308.1764221191406 0 80268.56 y Water loss 2
308.1988220214844 0 77518.75
308.2189636230469 0 9834.259
308.6794738769531 0 21492.277
309.1571044921875 0 16363.812
309.1773376464844 0 6629.116
309.2010192871094 0 12353.381
310.15289306640625 0 21547.848
311.1368103027344 0 5301.7666
311.1732177734375 0 15243.321
311.20928955078125 0 13495.503
313.1525573730469 0 14804.317
313.188720703125 0 47787.42
314.1859130859375 0 16600.72
315.17510986328125 0 17389.064 b Water loss 6
315.2080078125 0 7823.513
315.2411804199219 0 30507.514
315.6773986816406 0 10721.336
316.167236328125 0 23466.875
316.2438659667969 0 6969.0405
317.1635437011719 0 7368.7544
317.1898498535156 0 79683.836
317.22137451171875 0 8319.493
317.6922302246094 0 28399.094
318.1467590332031 0 49856.71
319.1421813964844 0 181355.7
320.1449279785156 0 29803.852
321.1688537597656 0 39980.605
321.1939697265625 0 29044.105
322.15283203125 0 7563.799
322.18365478515625 0 76444.414
322.6844787597656 0 25246.38
323.17352294921875 0 91891.125
324.18017578125 0 61073.258 b 6
324.6827392578125 0 11631.051
325.12322998046875 0 5997.714
325.225341796875 0 217888.12
326.1842346191406 0 975839.5
326.2296142578125 0 39604.832
327.1868896484375 0 120363.92
327.21002197265625 0 9010.066
327.6754455566406 0 5754.089
328.131103515625 0 17102.814
328.1638488769531 0 17891.055
328.1893310546875 0 17252.62
328.23748779296875 0 8152.441
329.1264343261719 0 28047.164
329.1625671386719 0 21215.076
329.1846923828125 0 9279.303
330.158203125 0 25390.459
330.1832275390625 0 21035.914
331.1424865722656 0 11802.3545
331.1886901855469 0 55033.81
331.6905517578125 0 26658.195
333.1939697265625 0 49071.266
334.1975402832031 0 9108.125
336.1680908203125 0 19346.23
336.18328857421875 0 14577.262
336.2412109375 0 6371.7686
337.1527404785156 0 6941.2695
337.2255554199219 0 12202.188
338.1839904785156 0 34766.496
338.2209167480469 0 30243.27
339.17962646484375 0 251208.12
340.1816711425781 0 20369.088
340.199951171875 0 57307.5
341.18389892578125 0 136267.92
341.20184326171875 0 6213.3535
342.18414306640625 0 25930.615
343.2360534667969 0 387641.25 y Water loss 6
344.19482421875 0 170095.27
344.21917724609375 0 34848.867 y Ammonia loss 6
344.2394104003906 0 59970.008
345.15771484375 0 15633.682
345.1964416503906 0 33369.496
345.220947265625 0 8646.268
345.2449035644531 0 5021.67
347.1371765136719 0 12901.052
347.183837890625 0 20102.016
347.20843505859375 0 7333.157
348.1687927246094 0 228332.81
349.171630859375 0 40183.96
350.182373046875 0 6520.336
350.2201232910156 0 8059.182
351.1678466796875 0 7052.1846
353.2212219238281 0 16136.821
354.17950439453125 0 55523.1
354.22235107421875 0 6094.476
354.25164794921875 0 10757.979
355.1826477050781 0 6499.7983
355.2369079589844 0 12199.701
356.1737365722656 0 15445.637
356.1956481933594 0 12174.753
356.231201171875 0 41856.61
357.1579895019531 0 135586.22
357.190185546875 0 50153.19
357.2325744628906 0 6688.714
357.7214660644531 0 4558.1006
358.1587219238281 0 29731.846
358.2108459472656 0 92125.86
359.2127380371094 0 17533.717
360.1945495605469 0 8921.9795
361.1904602050781 0 65392.12 b 3
361.2466125488281 0 1402039.1 y 6
362.19342041015625 0 11190.535
362.2496337890625 0 254432.36
362.7142639160156 0 5010.0015
363.25201416015625 0 29662.291
364.2364196777344 0 46735.824
365.1951904296875 0 380546.47
366.17901611328125 0 91009.88
366.1992492675781 0 49575.586
366.7259521484375 0 130824.29
367.1799011230469 0 15250.606
367.2035827636719 0 6676.7524
367.2274169921875 0 56718.395
367.7129211425781 0 8121.543
367.7311706542969 0 8589.568
368.1949768066406 0 213814.5
369.19805908203125 0 43631.293
371.2314453125 0 21395.84
371.7177734375 0 44921.53 b Water loss 7
372.18939208984375 0 19881.94
372.21844482421875 0 11778.833
372.2628173828125 0 46891.383
372.708984375 0 19433.605
373.1904296875 0 18468.236
373.2665710449219 0 11667.578
374.18463134765625 0 153753.97
375.16961669921875 0 80091.32
375.19091796875 0 8250.329
376.1663513183594 0 34085.414
376.2006530761719 0 8961.155
376.2847900390625 0 7466.97
376.7152099609375 0 13857.154
376.7463684082031 0 18237.508
377.2327880859375 0 10050.038
378.21490478515625 0 6840.103
380.7229309082031 0 30248.174 b 7
381.2246398925781 0 11273.33
381.7132873535156 0 275824.22 y Water loss 1
382.21453857421875 0 107743.22
382.2470703125 0 263930.06
382.7149353027344 0 32238.371
383.2060241699219 0 1898640.5
383.2513427734375 0 55002.723
384.1679992675781 0 20148.166
384.2088623046875 0 327639
385.1521301269531 0 28362.135
385.21337890625 0 39813.176
385.25897216796875 0 9572.898
386.1575927734375 0 5935.631
386.2056579589844 0 125229.77
386.7065124511719 0 5074.2104
387.20611572265625 0 49388.324
388.2054138183594 0 4843.7695
390.2157897949219 0 57583.848
390.7184143066406 0 2011834.6 y 1
391.2199401855469 0 815223.56
391.7213439941406 0 197216.27
392.1950988769531 0 192971.94
392.2231750488281 0 19133.352
393.19610595703125 0 44511.973
393.2607421875 0 6896.6353
394.1995544433594 0 6927.5337
394.24725341796875 0 35173.65
395.2430114746094 0 50900.926
395.71063232421875 0 126009.36
396.2122802734375 0 41203.2
396.2441711425781 0 7649.6953
396.71490478515625 0 10902.957
400.2579040527344 0 424830.22
401.2166748046875 0 1944056.2
401.26226806640625 0 75571.016
402.1796569824219 0 580308.56
402.2194519042969 0 356185.56
402.26470947265625 0 8379.338
403.1827087402344 0 122192.47
403.221435546875 0 45789.25
404.1851501464844 0 14680.601
404.2312927246094 0 16926.238
407.2403869628906 0 6503.4277
409.25152587890625 0 6708.582
410.2417297363281 0 6093.335
411.2012939453125 0 21266.264
411.2735900878906 0 28974.889
412.1651306152344 0 10091.568
412.2553405761719 0 6842.934
413.25341796875 0 56963.203
414.2574768066406 0 11411.497
417.22198486328125 0 4933.4688
417.2503356933594 0 4885.3296
418.2111511230469 0 32942.55 b 4
418.2424621582031 0 9043.065
418.27093505859375 0 9128.671
419.2073059082031 0 34944.35
420.19036865234375 0 334563.9
421.1932373046875 0 68123.32
421.22650146484375 0 3589.6406
421.25860595703125 0 70667.875
422.1943054199219 0 9324.643
422.2423095703125 0 12250.16
422.26849365234375 0 6292.1587
423.2385559082031 0 29612.459
424.239501953125 0 6217.536
424.26995849609375 0 16023.989
426.27276611328125 0 10574.149
429.1164855957031 0 7615.3784
429.2846984863281 0 156383.28
430.1745300292969 0 15819.147
430.28729248046875 0 37040.445
431.2477722167969 0 132497.53 Precursor Water loss
431.748779296875 0 51506.465
432.2494812011719 0 16861.145
433.2579040527344 0 14699.492
434.2540283203125 0 22666.994
437.2172546386719 0 32016.441
438.2170715332031 0 6950.7314
439.26910400390625 0 507044.78
440.2534484863281 0 238430.22 Precursor
440.75445556640625 0 135884.72
441.25140380859375 0 65584.67
441.2835388183594 0 4911.8687
442.2494812011719 0 13466.64
442.2802429199219 0 17222.2
443.2435302734375 0 7493.381
444.2845458984375 0 59836.094 y Water loss 5
445.2877197265625 0 15914.906
446.27984619140625 0 6417.682
447.20269775390625 0 7604.027
447.2383117675781 0 13116.575
448.5225830078125 0 4610.1323
448.591552734375 0 16721.45
448.613037109375 0 25256.285
448.6522216796875 0 37989.082
448.68878173828125 0 4821.159
449.2538146972656 0 6842.9355
449.6268005371094 0 8137.5166
450.2848815917969 0 29157.168
451.26904296875 0 82985.125
452.2649230957031 0 168501.28
453.2674865722656 0 33235.008
454.2682800292969 0 5573.8555
455.24212646484375 0 5686.207
456.22674560546875 0 32614.129
457.2796630859375 0 803375.06
458.2825927734375 0 185608.62
459.2839660644531 0 24970.506
460.2678527832031 0 19673.326
461.25335693359375 0 58933.41
462.25860595703125 0 14947.922
462.2950439453125 0 323479.94 y 5
463.2980651855469 0 77274.36
464.300537109375 0 14357.397
466.2416076660156 0 5856.4937
467.26458740234375 0 17400.346
468.2616271972656 0 6394.18
468.29595947265625 0 100770.88
469.2421875 0 6111.9985
469.2808532714844 0 60203.4
470.24237060546875 0 18972.904
470.2760009765625 0 128054.086
470.31085205078125 0 8037.966
471.2451171875 0 5784.556
471.2782287597656 0 32402.799
472.27655029296875 0 6167.716
473.2539367675781 0 40333.1
474.2402648925781 0 21345.21
475.23480224609375 0 9034.6455
475.2920837402344 0 12761.646
478.2801513671875 0 320442.94
479.2651672363281 0 67272.984
480.26763916015625 0 20456.396
481.1451721191406 0 14253.991
481.1877136230469 0 8668.561
483.2362365722656 0 9156.618
483.2955017089844 0 22908.803
484.2540588378906 0 6446.947
485.2796630859375 0 9316.949
486.30657958984375 0 1262741.9
487.3094787597656 0 293066.5
488.2531433105469 0 12730.017
488.3124694824219 0 38249.25
489.2558898925781 0 5689.0586
491.26422119140625 0 47007.3
492.2675476074219 0 12472.731
495.2393493652344 0 20443.488
496.2909240722656 0 2540946.8
497.2934875488281 0 621931
498.2957458496094 0 92255.09
501.24859619140625 0 157556.84 b Water loss 5
501.3062438964844 0 82354.47 y Water loss 4
502.25506591796875 0 50121.844
502.3080749511719 0 18036.408
503.2585144042969 0 10669.456
503.30145263671875 0 6912.9116
504.2621765136719 0 12119.379
505.2773132324219 0 4913.935
506.2775573730469 0 5310.83
512.264892578125 0 83095.305
513.2660522460938 0 22741.14
513.30810546875 0 4730.917
514.3013916015625 0 6089883
515.3043212890625 0 1543546.9
515.5014038085938 0 7828.7217
516.306640625 0 244199.3
517.3090209960938 0 14135.155
518.273681640625 0 8562.691
519.26025390625 0 75020.41 b 5
519.316650390625 0 471375.06 y 4
520.2626953125 0 21761.596
520.3200073242188 0 115709.28
521.2706298828125 0 6046.6567
521.322509765625 0 21115.352
524.2852172851562 0 18001.156
530.2752075195312 0 290149.53
531.2781982421875 0 98383.695
532.3121948242188 0 101270.47
533.314697265625 0 29760.57
540.2625122070312 0 8996.66
540.3170776367188 0 26339.629
548.2857666015625 0 534533.25
549.2891845703125 0 162396.58
550.2910766601562 0 26882.953
552.3163452148438 0 7650.0923
553.310302734375 0 7739.995
553.35205078125 0 7522.3726
554.3317260742188 0 16246.046
558.271240234375 0 25776.055
558.3278198242188 0 128516.42 y Water loss 3
559.3300170898438 0 45105.01
560.330078125 0 9393.889
562.3026123046875 0 5665.6953
569.3457641601562 0 5905.0474
570.3263549804688 0 15146.331
571.3209228515625 0 54386.598
571.3599243164062 0 149664
572.322998046875 0 15726.223
572.363037109375 0 41131.41
573.3673706054688 0 8220.288
576.3385009765625 0 1554311.5 y 3
577.34130859375 0 393235.8
578.34375 0 80439.41
579.3297729492188 0 23473.162
580.3187255859375 0 12758.525
581.3237915039062 0 7619.2295
585.3160400390625 0 6293.335
587.3539428710938 0 33809.105
588.3563232421875 0 13526.866
589.3348388671875 0 27755.393
590.3360595703125 0 6228.327
597.3389892578125 0 197248.19
598.3338012695312 0 86450.77
599.3327026367188 0 53747.047
600.3343505859375 0 14207.926
611.3328247070312 0 16813.129
612.3362426757812 0 5430.214
613.370361328125 0 18204.943
615.349609375 0 897269.1 y Water loss 2
616.3513793945312 0 319004.44
617.3502807617188 0 81097.766
618.3486328125 0 11982.685
625.3487548828125 0 58009.9
626.3463134765625 0 26425.273
629.3441162109375 0 112381.94 b Water loss 6
630.346923828125 0 32709.152
631.3510131835938 0 6218.4717
633.3603515625 0 13955515 y 2
634.363037109375 0 4242355
635.365478515625 0 858091.06
636.3678588867188 0 45408.66
643.359375 0 635748.94
644.3617553710938 0 217798.4
645.3645629882812 0 43406.703
647.3546752929688 0 318439.94 b 6
648.357666015625 0 95807.62
649.3580322265625 0 19726.2
653.34423828125 0 12645.584
661.3704223632812 0 1195821.1
662.3731689453125 0 426346.34
663.3759155273438 0 78397.41
664.3790893554688 0 5315.1006
671.3539428710938 0 17892.428
672.358154296875 0 8865.051
679.3807983398438 0 31024.568
680.386474609375 0 13212.064
689.3726806640625 0 5041.5083
698.4083862304688 0 4895.4907
714.4317626953125 0 5617.3804
715.4178466796875 0 4522.17
716.4138793945312 0 8399.587
718.4264526367188 0 37228.543
719.4301147460938 0 15019.012
724.4191284179688 0 7578.535
725.4140625 0 7788.889
732.4439086914062 0 57971.617
733.4466552734375 0 21229.861
736.4021606445312 0 11555.984
742.4285888671875 0 156180.27 b Water loss 7
743.4310302734375 0 57688.52
744.4105834960938 0 47934.277
745.4043579101562 0 17554.312
746.3941650390625 0 6842.9746
760.4390258789062 0 461887.66 b 7
761.4418334960938 0 197048.3
762.4224243164062 0 177427.19 y Water loss 1
763.4205932617188 0 86622.91
764.421142578125 0 20305.096
772.4059448242188 0 7261.3667
778.4464111328125 0 8806.022
779.4572143554688 0 6069.897
780.4287719726562 0 3324503.5 y 1
781.4317626953125 0 1328282.2
782.4341430664062 0 348701.97
782.5230712890625 0 6184.542
783.4359130859375 0 18926.389
790.4130249023438 0 54180.016
791.416259765625 0 25258.396
792.4163818359375 0 7329.7305
1292.581787109375 0 10887.114
1308.268798828125 0 4692.635
1308.423583984375 0 4730.204
1557.480712890625 0 4427.5044
1779.3353271484375 0 17625.242
1779.651123046875 0 26837.65
1817.5548095703125 0 5012.384
1817.79345703125 0 6371.686
2388.74853515625 0 6336.4673
2389.22314453125 0 13314.978
3042.030517578125 0 5011.2417

Spectrum Details

|  |  |
| --- | --- |
| Matched peaks? Matched peaks The total absolute number of peaks matched. Additionally in brackets the total fraction of peaks matched and the total number of peaks is shown. | 41 (5.26% of 780) |
| FDR? FDR The false discovery rate estimated for this peptide. It is calculated by matching all theoretical fragments with a non-integer shift with the raw peaks for this spectrum. This is done with 40 different shifts. The resulting percentage is the average number of annotated peaks over the number of annotated peaks with the correct spectrum. | 0.52% |
| Satellite FDR? Satellite FDR See the FDR for details on its calculation. This satellite ion specific FDR only contains the satellite ions (d/w) for I/L/J positions. | - |
| PSM Score? PSM Score The PSM Score as given by Hecklib to this annotated spectrum. It is shown with three significant figures. | 421 |

#### Spectrum 4777? Spectrum 4777 The raw spectrum of this peptide as annotated by Hecklib. The fragments are coloured according to ion type (see legend). Any peaks with a star '\*' as text can be hovered over to see the full details, first the ion type second the mass shift type. By hovering over the amino acids in the peptide or ions in the legend the corresponding peaks are highlighted. By toggling the 'Unassigned' label you can turn the background (unassigned) peaks on or off in the plot. By updating the slider in the Ion legend you can update the spectrum to only show the top X% of the peaks with labels. The top X% means any peak that is within X% of the highest intensity. By dragging in the spectrum you can zoom in to a specific part of the spectrum and use 'Zoom Out' to get back to the original zoom level. The annotation of the spectrum is based on the given sequence in the peptides file and is done with different software so inconsistencies are likely. The peaks are annotated based on the given sequence, with 20 ppm tolerance.

Copy Data

##### Spectrum 4777 (TSV)

###### Preview

```
Loading example...
```

*Click on the button to copy the data to your clipboard.*

Mz MinMz MaxIntensity Max

WidthHeightPeptide font sizePeptide stroke widthSpectrum font sizeSpectrum stroke widthCompact peptide

Ion legend

wxyz

abcd

OtherUnassignedIon^Charge^_Position_Show for top:%

VFGGGTKJT

05.43e+41.09e+51.63e+52.17e+5

Zoom Out

y^+1^_1_b^+2^_4_y^+1^_2_a^+1^_2_y^+1^_2_b^+1^_2_y^+2^_5_b^+1^_3_y^+1^_3_y^+1^_3_y^+2^_8_y^+2^_8_b^+1^_5_\*\*y^+1^_4_b^+1^_6_b^+1^_6_y^+1^_5_y^+1^_6_y^+1^_6_y^+1^_7_b^+1^_7_y^+1^_7_b^+1^_7_b^+1^_8_b^+1^_8_y^+1^_8_y^+1^_8_

0769153823073076

Fragment Matches Table

Show background peaks

| Position | Ion type | Intensity | mz Theoretical | mz Error (Th) | mz Error (ppm) | Charge | Series Number |
| --- | --- | --- | --- | --- | --- | --- | --- |
| 9 | y | 1.125E+04 | 120.1 | 0.0003297 | 2.746 | +1 | 1 |
| - | - | 2.149E+05 | 120.1 | - | - | 0 | - |
| - | - | 1.794E+04 | 121.1 | - | - | 0 | - |
| - | - | 525.9 | 122.1 | - | - | 0 | - |
| - | - | 421.8 | 123.1 | - | - | 0 | - |
| - | - | 480.9 | 124.1 | - | - | 0 | - |
| - | - | 898.6 | 125.1 | - | - | 0 | - |
| - | - | 2669 | 127.1 | - | - | 0 | - |
| - | - | 942.6 | 127.1 | - | - | 0 | - |
| - | - | 633.6 | 128.1 | - | - | 0 | - |
| - | - | 624.6 | 128.1 | - | - | 0 | - |
| - | - | 366.5 | 129.1 | - | - | 0 | - |
| - | - | 1.249E+05 | 129.1 | - | - | 0 | - |
| - | - | 2528 | 130.1 | - | - | 0 | - |
| - | - | 1103 | 130.1 | - | - | 0 | - |
| - | - | 8095 | 130.1 | - | - | 0 | - |
| - | - | 454.4 | 131 | - | - | 0 | - |
| - | - | 5651 | 131.1 | - | - | 0 | - |
| - | - | 927.6 | 132.1 | - | - | 0 | - |
| - | - | 922.9 | 132.1 | - | - | 0 | - |
| - | - | 427.9 | 133.1 | - | - | 0 | - |
| - | - | 4360 | 136.1 | - | - | 0 | - |
| - | - | 440.8 | 138.2 | - | - | 0 | - |
| - | - | 952.9 | 139.1 | - | - | 0 | - |
| - | - | 729.6 | 140.1 | - | - | 0 | - |
| - | - | 2113 | 141.1 | - | - | 0 | - |
| - | - | 1081 | 141.1 | - | - | 0 | - |
| - | - | 1861 | 143.1 | - | - | 0 | - |
| - | - | 422.9 | 143.9 | - | - | 0 | - |
| - | - | 732.5 | 145.1 | - | - | 0 | - |
| - | - | 2717 | 146.1 | - | - | 0 | - |
| - | - | 706 | 148 | - | - | 0 | - |
| - | - | 683.2 | 148.1 | - | - | 0 | - |
| - | - | 1073 | 149 | - | - | 0 | - |
| - | - | 622.9 | 151.1 | - | - | 0 | - |
| - | - | 822.4 | 152.1 | - | - | 0 | - |
| - | - | 921.8 | 152.1 | - | - | 0 | - |
| - | - | 979.7 | 153.1 | - | - | 0 | - |
| - | - | 1889 | 153.1 | - | - | 0 | - |
| - | - | 586.6 | 153.1 | - | - | 0 | - |
| - | - | 1069 | 154.1 | - | - | 0 | - |
| - | - | 1461 | 155.1 | - | - | 0 | - |
| - | - | 948.1 | 155.1 | - | - | 0 | - |
| - | - | 3787 | 155.1 | - | - | 0 | - |
| - | - | 1325 | 157.1 | - | - | 0 | - |
| - | - | 1223 | 158.1 | - | - | 0 | - |
| - | - | 3007 | 159.1 | - | - | 0 | - |
| - | - | 2386 | 159.1 | - | - | 0 | - |
| - | - | 6871 | 159.1 | - | - | 0 | - |
| - | - | 447.6 | 159.4 | - | - | 0 | - |
| - | - | 478.7 | 160.1 | - | - | 0 | - |
| - | - | 894.3 | 162.1 | - | - | 0 | - |
| - | - | 1101 | 167.1 | - | - | 0 | - |
| - | - | 480.5 | 168.1 | - | - | 0 | - |
| - | - | 914.9 | 168.1 | - | - | 0 | - |
| - | - | 700.7 | 169.1 | - | - | 0 | - |
| - | - | 534 | 169.1 | - | - | 0 | - |
| - | - | 1698 | 170.1 | - | - | 0 | - |
| - | - | 1972 | 171.1 | - | - | 0 | - |
| - | - | 933.1 | 171.1 | - | - | 0 | - |
| - | - | 5661 | 172.1 | - | - | 0 | - |
| - | - | 849.6 | 173.1 | - | - | 0 | - |
| - | - | 6350 | 173.1 | - | - | 0 | - |
| - | - | 1373 | 173.1 | - | - | 0 | - |
| - | - | 3816 | 173.4 | - | - | 0 | - |
| - | - | 1393 | 174.1 | - | - | 0 | - |
| - | - | 489.5 | 174.1 | - | - | 0 | - |
| - | - | 1970 | 175.1 | - | - | 0 | - |
| - | - | 2179 | 176.1 | - | - | 0 | - |
| - | - | 6205 | 177.1 | - | - | 0 | - |
| - | - | 774.5 | 177.1 | - | - | 0 | - |
| - | - | 680.3 | 178.1 | - | - | 0 | - |
| 4 | b | 1034 | 181.1 | 0.0001665 | 0.9192 | +2 | 4 |
| - | - | 503.2 | 183.1 | - | - | 0 | - |
| - | - | 2558 | 183.1 | - | - | 0 | - |
| - | - | 1938 | 185.1 | - | - | 0 | - |
| - | - | 2151 | 185.1 | - | - | 0 | - |
| - | - | 2088 | 186.1 | - | - | 0 | - |
| - | - | 4255 | 187.1 | - | - | 0 | - |
| - | - | 1221 | 188.1 | - | - | 0 | - |
| - | - | 540.7 | 189.1 | - | - | 0 | - |
| - | - | 562.7 | 189.1 | - | - | 0 | - |
| - | - | 673.9 | 191.1 | - | - | 0 | - |
| - | - | 646.3 | 191.2 | - | - | 0 | - |
| - | - | 1151 | 195.1 | - | - | 0 | - |
| - | - | 1148 | 196.1 | - | - | 0 | - |
| - | - | 827.8 | 197.1 | - | - | 0 | - |
| - | - | 4548 | 197.2 | - | - | 0 | - |
| - | - | 3562 | 198.1 | - | - | 0 | - |
| - | - | 538.4 | 198.2 | - | - | 0 | - |
| - | - | 627 | 199.1 | - | - | 0 | - |
| - | - | 1176 | 201.1 | - | - | 0 | - |
| - | - | 769.7 | 201.1 | - | - | 0 | - |
| - | - | 3767 | 203.1 | - | - | 0 | - |
| - | - | 1.203E+04 | 205.1 | - | - | 0 | - |
| - | - | 498.5 | 205.4 | - | - | 0 | - |
| - | - | 1246 | 206.1 | - | - | 0 | - |
| - | - | 2307 | 207.1 | - | - | 0 | - |
| - | - | 1933 | 208.1 | - | - | 0 | - |
| - | - | 486.8 | 208.2 | - | - | 0 | - |
| - | - | 1629 | 209.1 | - | - | 0 | - |
| - | - | 1420 | 210.1 | - | - | 0 | - |
| - | - | 818 | 211.1 | - | - | 0 | - |
| - | - | 778.1 | 212.1 | - | - | 0 | - |
| - | - | 9601 | 212.1 | - | - | 0 | - |
| - | - | 588.6 | 213.1 | - | - | 0 | - |
| - | - | 551.4 | 213.1 | - | - | 0 | - |
| - | - | 700.4 | 213.1 | - | - | 0 | - |
| - | - | 3597 | 213.2 | - | - | 0 | - |
| - | - | 1015 | 214.2 | - | - | 0 | - |
| - | - | 822.5 | 215.1 | - | - | 0 | - |
| 8 | y | 2271 | 215.1 | 0.0004006 | 1.862 | +1 | 2 |
| - | - | 1878 | 216.1 | - | - | 0 | - |
| - | - | 1050 | 217.1 | - | - | 0 | - |
| - | - | 541.5 | 218.2 | - | - | 0 | - |
| 2 | a | 1.65E+05 | 219.1 | 0.0004533 | 2.068 | +1 | 2 |
| - | - | 562.8 | 220.1 | - | - | 0 | - |
| - | - | 2.029E+04 | 220.2 | - | - | 0 | - |
| - | - | 1163 | 221.2 | - | - | 0 | - |
| - | - | 590.2 | 223.2 | - | - | 0 | - |
| - | - | 624.3 | 224.1 | - | - | 0 | - |
| - | - | 3647 | 224.2 | - | - | 0 | - |
| - | - | 1671 | 225.1 | - | - | 0 | - |
| - | - | 1062 | 225.2 | - | - | 0 | - |
| - | - | 727.9 | 226.1 | - | - | 0 | - |
| - | - | 2102 | 226.2 | - | - | 0 | - |
| - | - | 1141 | 227.1 | - | - | 0 | - |
| - | - | 7927 | 227.1 | - | - | 0 | - |
| - | - | 3603 | 228.1 | - | - | 0 | - |
| - | - | 662.2 | 229.1 | - | - | 0 | - |
| - | - | 2.004E+04 | 230.2 | - | - | 0 | - |
| - | - | 2041 | 231.2 | - | - | 0 | - |
| 8 | y | 2931 | 233.1 | 0.0005019 | 2.153 | +1 | 2 |
| - | - | 666.1 | 233.2 | - | - | 0 | - |
| - | - | 1209 | 234.1 | - | - | 0 | - |
| - | - | 842.7 | 237.1 | - | - | 0 | - |
| - | - | 586.8 | 238.1 | - | - | 0 | - |
| - | - | 751.5 | 239.2 | - | - | 0 | - |
| - | - | 2344 | 240.1 | - | - | 0 | - |
| - | - | 2434 | 241.2 | - | - | 0 | - |
| - | - | 680.3 | 242.1 | - | - | 0 | - |
| - | - | 4134 | 242.2 | - | - | 0 | - |
| - | - | 590.6 | 243.1 | - | - | 0 | - |
| - | - | 4660 | 245.1 | - | - | 0 | - |
| 2 | b | 4.113E+04 | 247.1 | 0.0004575 | 1.851 | +1 | 2 |
| - | - | 5829 | 248.1 | - | - | 0 | - |
| 5 | y | 1860 | 251.2 | 0.003299 | 13.14 | +2 | 5 |
| - | - | 685.2 | 253.1 | - | - | 0 | - |
| - | - | 761.5 | 254.1 | - | - | 0 | - |
| - | - | 9573 | 255.1 | - | - | 0 | - |
| - | - | 1610 | 256.1 | - | - | 0 | - |
| - | - | 4041 | 262.1 | - | - | 0 | - |
| - | - | 563.9 | 266.1 | - | - | 0 | - |
| - | - | 1350 | 268.1 | - | - | 0 | - |
| - | - | 565.8 | 268.2 | - | - | 0 | - |
| - | - | 1.206E+04 | 269.2 | - | - | 0 | - |
| - | - | 1175 | 270.2 | - | - | 0 | - |
| - | - | 1160 | 270.2 | - | - | 0 | - |
| - | - | 1666 | 272.1 | - | - | 0 | - |
| - | - | 1.812E+04 | 273.1 | - | - | 0 | - |
| - | - | 775.3 | 273.1 | - | - | 0 | - |
| - | - | 2217 | 274.1 | - | - | 0 | - |
| - | - | 773.1 | 280.1 | - | - | 0 | - |
| - | - | 648.8 | 280.2 | - | - | 0 | - |
| - | - | 681 | 282.2 | - | - | 0 | - |
| - | - | 4106 | 287.2 | - | - | 0 | - |
| - | - | 1243 | 288.2 | - | - | 0 | - |
| - | - | 776.8 | 290.1 | - | - | 0 | - |
| - | - | 1185 | 291.1 | - | - | 0 | - |
| - | - | 597.5 | 295.2 | - | - | 0 | - |
| - | - | 1199 | 301.1 | - | - | 0 | - |
| 3 | b | 5281 | 304.2 | 0.0002035 | 0.669 | +1 | 3 |
| - | - | 1107 | 305.2 | - | - | 0 | - |
| - | - | 716.9 | 311.7 | - | - | 0 | - |
| - | - | 655.8 | 318.1 | - | - | 0 | - |
| - | - | 1996 | 319.1 | - | - | 0 | - |
| - | - | 4041 | 319.7 | - | - | 0 | - |
| - | - | 1575 | 320.2 | - | - | 0 | - |
| - | - | 964.6 | 323.2 | - | - | 0 | - |
| - | - | 1669 | 325.2 | - | - | 0 | - |
| - | - | 1478 | 325.7 | - | - | 0 | - |
| - | - | 7461 | 326.2 | - | - | 0 | - |
| - | - | 975.3 | 326.7 | - | - | 0 | - |
| - | - | 720.1 | 333.1 | - | - | 0 | - |
| - | - | 664.5 | 333.2 | - | - | 0 | - |
| - | - | 1014 | 334.7 | - | - | 0 | - |
| - | - | 604.5 | 335.2 | - | - | 0 | - |
| - | - | 1878 | 339.2 | - | - | 0 | - |
| - | - | 943.4 | 340.2 | - | - | 0 | - |
| - | - | 1055 | 341.2 | - | - | 0 | - |
| 7 | y | 2791 | 343.2 | 0.0006372 | 1.857 | +1 | 3 |
| - | - | 2267 | 344.2 | - | - | 0 | - |
| - | - | 876.7 | 346.1 | - | - | 0 | - |
| - | - | 569.9 | 347.1 | - | - | 0 | - |
| - | - | 1863 | 348.2 | - | - | 0 | - |
| - | - | 1416 | 357.2 | - | - | 0 | - |
| - | - | 949.1 | 358.2 | - | - | 0 | - |
| - | - | 875 | 361.2 | - | - | 0 | - |
| 7 | y | 1.186E+04 | 361.2 | 0.0003264 | 0.9036 | +1 | 3 |
| - | - | 2055 | 362.2 | - | - | 0 | - |
| - | - | 3195 | 365.2 | - | - | 0 | - |
| - | - | 1064 | 366.2 | - | - | 0 | - |
| - | - | 751.2 | 366.7 | - | - | 0 | - |
| - | - | 2146 | 368.2 | - | - | 0 | - |
| - | - | 605 | 369.2 | - | - | 0 | - |
| - | - | 1346 | 374.2 | - | - | 0 | - |
| 2 | y | 2086 | 381.7 | 0.0003412 | 0.8938 | +2 | 8 |
| - | - | 1130 | 382.2 | - | - | 0 | - |
| - | - | 2041 | 382.2 | - | - | 0 | - |
| - | - | 804.4 | 383.2 | - | - | 0 | - |
| - | - | 1.708E+04 | 383.2 | - | - | 0 | - |
| - | - | 3869 | 384.2 | - | - | 0 | - |
| - | - | 692.4 | 385.2 | - | - | 0 | - |
| - | - | 673.4 | 390.2 | - | - | 0 | - |
| 2 | y | 1.43E+04 | 390.7 | 0.0003994 | 1.022 | +2 | 8 |
| - | - | 5734 | 391.2 | - | - | 0 | - |
| - | - | 961.5 | 391.7 | - | - | 0 | - |
| - | - | 1468 | 392.2 | - | - | 0 | - |
| - | - | 929.5 | 395.7 | - | - | 0 | - |
| - | - | 678.2 | 396.2 | - | - | 0 | - |
| - | - | 2596 | 400.3 | - | - | 0 | - |
| - | - | 1.786E+04 | 401.2 | - | - | 0 | - |
| - | - | 784.5 | 401.3 | - | - | 0 | - |
| - | - | 5917 | 402.2 | - | - | 0 | - |
| - | - | 3375 | 402.2 | - | - | 0 | - |
| - | - | 1200 | 403.2 | - | - | 0 | - |
| - | - | 818.5 | 413.3 | - | - | 0 | - |
| 5 | b | 718.3 | 418.2 | 0.001527 | 3.65 | +1 | 5 |
| - | - | 845.4 | 419.2 | - | - | 0 | - |
| - | - | 2359 | 420.2 | - | - | 0 | - |
| - | - | 744.8 | 420.2 | - | - | 0 | - |
| - | - | 856.5 | 421.2 | - | - | 0 | - |
| - | - | 993 | 421.8 | - | - | 0 | - |
| - | - | 610.2 | 422.2 | - | - | 0 | - |
| - | - | 1025 | 429.3 | - | - | 0 | - |
| 0 | Precursor | 1209 | 431.2 | 0.0009243 | 2.143 | +2 | -1 |
| - | - | 1695 | 438.2 | - | - | 0 | - |
| - | - | 869.2 | 439.2 | - | - | 0 | - |
| - | - | 4006 | 439.3 | - | - | 0 | - |
| - | - | 8463 | 439.8 | - | - | 0 | - |
| 0 | Precursor | 1717 | 440.3 | 0.001349 | 3.064 | +2 | -1 |
| - | - | 1112 | 440.8 | - | - | 0 | - |
| - | - | 2817 | 440.8 | - | - | 0 | - |
| - | - | 1013 | 441.3 | - | - | 0 | - |
| - | - | 837 | 451.3 | - | - | 0 | - |
| - | - | 1591 | 452.3 | - | - | 0 | - |
| - | - | 7896 | 457.3 | - | - | 0 | - |
| - | - | 1386 | 458.3 | - | - | 0 | - |
| - | - | 664.1 | 461.3 | - | - | 0 | - |
| 6 | y | 3184 | 462.3 | 0.000408 | 0.8825 | +1 | 4 |
| - | - | 712.7 | 463.3 | - | - | 0 | - |
| - | - | 1194 | 466.2 | - | - | 0 | - |
| - | - | 852.5 | 468.3 | - | - | 0 | - |
| - | - | 1263 | 470.3 | - | - | 0 | - |
| - | - | 945.7 | 473.3 | - | - | 0 | - |
| - | - | 3571 | 478.3 | - | - | 0 | - |
| - | - | 716.7 | 479.3 | - | - | 0 | - |
| - | - | 1.161E+04 | 486.3 | - | - | 0 | - |
| - | - | 2096 | 487.3 | - | - | 0 | - |
| - | - | 2.335E+04 | 496.3 | - | - | 0 | - |
| - | - | 6073 | 497.3 | - | - | 0 | - |
| - | - | 727.2 | 498.3 | - | - | 0 | - |
| 6 | b | 1633 | 501.2 | 0.0002784 | 0.5555 | +1 | 6 |
| - | - | 755.5 | 512.3 | - | - | 0 | - |
| - | - | 5.716E+04 | 514.3 | - | - | 0 | - |
| - | - | 480.9 | 515.3 | - | - | 0 | - |
| - | - | 1.447E+04 | 515.3 | - | - | 0 | - |
| - | - | 2091 | 516.3 | - | - | 0 | - |
| 6 | b | 1036 | 519.3 | 0.001761 | 3.391 | +1 | 6 |
| 5 | y | 5483 | 519.3 | 0.0003371 | 0.6491 | +1 | 5 |
| - | - | 946.3 | 520.3 | - | - | 0 | - |
| - | - | 2615 | 530.3 | - | - | 0 | - |
| - | - | 1151 | 532.3 | - | - | 0 | - |
| - | - | 5501 | 548.3 | - | - | 0 | - |
| - | - | 790 | 548.3 | - | - | 0 | - |
| - | - | 1519 | 549.3 | - | - | 0 | - |
| 4 | y | 1694 | 558.3 | 0.0008516 | 1.525 | +1 | 6 |
| - | - | 935.2 | 571.3 | - | - | 0 | - |
| - | - | 1170 | 571.4 | - | - | 0 | - |
| 4 | y | 1.448E+04 | 576.3 | 0.0004798 | 0.8325 | +1 | 6 |
| - | - | 4241 | 577.3 | - | - | 0 | - |
| - | - | 1020 | 587.4 | - | - | 0 | - |
| - | - | 1950 | 597.3 | - | - | 0 | - |
| - | - | 720.8 | 598.3 | - | - | 0 | - |
| - | - | 785.4 | 598.3 | - | - | 0 | - |
| 3 | y | 8793 | 615.3 | 0.0005671 | 0.9216 | +1 | 7 |
| - | - | 2759 | 616.3 | - | - | 0 | - |
| - | - | 969.5 | 625.3 | - | - | 0 | - |
| 7 | b | 895.8 | 629.3 | 6.494E-05 | 0.1032 | +1 | 7 |
| - | - | 721.8 | 631.3 | - | - | 0 | - |
| 3 | y | 1.411E+05 | 633.4 | 0.0006835 | 1.079 | +1 | 7 |
| - | - | 4.362E+04 | 634.4 | - | - | 0 | - |
| - | - | 8979 | 635.4 | - | - | 0 | - |
| - | - | 6632 | 643.4 | - | - | 0 | - |
| - | - | 2508 | 644.4 | - | - | 0 | - |
| 7 | b | 3099 | 647.4 | 0.0003069 | 0.474 | +1 | 7 |
| - | - | 1040 | 648.4 | - | - | 0 | - |
| - | - | 1.326E+04 | 661.4 | - | - | 0 | - |
| - | - | 3528 | 662.4 | - | - | 0 | - |
| - | - | 1244 | 663.4 | - | - | 0 | - |
| - | - | 633.4 | 721.3 | - | - | 0 | - |
| 8 | b | 1436 | 742.4 | 0.0003809 | 0.513 | +1 | 8 |
| 8 | b | 4642 | 760.4 | 0.0002034 | 0.2674 | +1 | 8 |
| - | - | 2269 | 761.4 | - | - | 0 | - |
| 2 | y | 2282 | 762.4 | 0.0005736 | 0.7523 | +1 | 8 |
| 2 | y | 3.667E+04 | 780.4 | 1.865E-05 | 0.0239 | +1 | 8 |
| - | - | 1.606E+04 | 781.4 | - | - | 0 | - |
| - | - | 4902 | 782.4 | - | - | 0 | - |
| - | - | 868.8 | 790.4 | - | - | 0 | - |
| - | - | 576.1 | 1510 | - | - | 0 | - |
| - | - | 689.2 | 1983 | - | - | 0 | - |
| - | - | 649.9 | 2374 | - | - | 0 | - |
| - | - | 655.9 | 2418 | - | - | 0 | - |
| - | - | 929.4 | 3045 | - | - | 0 | - |

m/z Charge Intensity FragmentType MassShift Position
120.06584930419922 0 11248.502 y 8
120.08120727539062 0 214869.05
121.08448791503906 0 17936.9
122.07188415527344 0 525.8578
123.11692810058594 0 421.8061
124.08731079101562 0 480.9118
125.0714111328125 0 898.6464
127.05047607421875 0 2668.9336
127.0870132446289 0 942.6315
128.09478759765625 0 633.58185
128.1074676513672 0 624.55286
129.05496215820312 0 366.51236
129.1026611328125 0 124887.28
130.0655059814453 0 2527.549
130.10031127929688 0 1103.4556
130.10598754882812 0 8094.8286
131.04942321777344 0 454.4078
131.08184814453125 0 5651.3105
132.08114624023438 0 927.55884
132.1023712158203 0 922.89484
133.0614013671875 0 427.8742
136.07601928710938 0 4360.2905
138.1524200439453 0 440.8498
139.08697509765625 0 952.868
140.08241271972656 0 729.61865
141.0662078857422 0 2112.853
141.1025848388672 0 1081.148
143.08177185058594 0 1861.148
143.88719177246094 0 422.88608
145.0609893798828 0 732.53156
146.0604248046875 0 2717.135
148.0396728515625 0 706.00256
148.07591247558594 0 683.17523
148.9536590576172 0 1072.7306
151.08676147460938 0 622.91534
152.07098388671875 0 822.4336
152.14376831054688 0 921.7622
153.0665283203125 0 979.7448
153.07728576660156 0 1888.5812
153.10263061523438 0 586.6084
154.06185913085938 0 1068.9363
155.0821533203125 0 1460.9597
155.09298706054688 0 948.0609
155.1182403564453 0 3787.346
157.09744262695312 0 1325.246
158.09262084960938 0 1223.3202
159.07679748535156 0 3007.1287
159.09193420410156 0 2385.5151
159.1131134033203 0 6870.582
159.37197875976562 0 447.57275
160.0960693359375 0 478.65073
162.0552978515625 0 894.28613
167.1182861328125 0 1101.1055
168.0775909423828 0 480.45648
168.1138458251953 0 914.9443
169.09779357910156 0 700.67413
169.133056640625 0 533.9532
170.09251403808594 0 1698.0779
171.07688903808594 0 1971.61
171.11312866210938 0 933.0883
172.07208251953125 0 5660.888
173.07118225097656 0 849.63696
173.09237670898438 0 6350.1313
173.12901306152344 0 1373.0405
173.44032287597656 0 3816.3284
174.05528259277344 0 1393.4253
174.134765625 0 489.45407
175.08694458007812 0 1969.7426
176.1072998046875 0 2179.2285
177.10264587402344 0 6205.07
177.1105194091797 0 774.51184
178.1059112548828 0 680.25287
181.09732055664062 0 1033.7612 b 3
183.07687377929688 0 503.1671
183.11314392089844 0 2557.6335
185.0923614501953 0 1938.0552
185.1288604736328 0 2150.6719
186.12403869628906 0 2088.0562
187.1081085205078 0 4255.4556
188.10342407226562 0 1220.7231
189.0662078857422 0 540.70905
189.10171508789062 0 562.69885
191.0818328857422 0 673.94073
191.15455627441406 0 646.2791
195.14907836914062 0 1150.9916
196.10816955566406 0 1147.9113
197.0925750732422 0 827.83185
197.16522216796875 0 4548.0396
198.08770751953125 0 3561.9314
198.16807556152344 0 538.3624
199.10809326171875 0 626.95087
201.0877685546875 0 1176.2906
201.12355041503906 0 769.7121
203.10304260253906 0 3766.947
205.09754943847656 0 12029.283
205.44078063964844 0 498.5485
206.10089111328125 0 1245.9387
207.14955139160156 0 2306.9602
208.10838317871094 0 1933.4424
208.1531219482422 0 486.78244
209.10365295410156 0 1629.3657
210.0879364013672 0 1420.2925
211.10906982421875 0 818.01746
212.1028289794922 0 778.1196
212.13975524902344 0 9600.729
213.0869140625 0 588.6075
213.1249542236328 0 551.42676
213.142822265625 0 700.4277
213.16036987304688 0 3597.1792
214.19174194335938 0 1015.3542
215.1148223876953 0 822.52515
215.13941955566406 0 2271.389 y Water loss 7
216.0982208251953 0 1877.7169
217.0977783203125 0 1050.2198
218.15028381347656 0 541.5031
219.14964294433594 0 165013.6 a 1
220.05978393554688 0 562.7506
220.15292358398438 0 20292.645
221.15626525878906 0 1162.9843
223.15570068359375 0 590.24335
224.14016723632812 0 624.3127
224.1761474609375 0 3646.8125
225.13455200195312 0 1671.3372
225.16049194335938 0 1062.0386
226.08328247070312 0 727.85486
226.15533447265625 0 2101.8188
227.1022186279297 0 1140.6116
227.1143341064453 0 7927.0576
228.0982666015625 0 3602.87
229.09300231933594 0 662.18396
230.15036010742188 0 20035.883
231.1525421142578 0 2041.3699
233.15008544921875 0 2930.905 y 7
233.162841796875 0 666.08386
234.1244354248047 0 1209.039
237.0994110107422 0 842.68024
238.1182098388672 0 586.8037
239.15090942382812 0 751.5079
240.13455200195312 0 2344.39
241.1549072265625 0 2433.8152
242.11398315429688 0 680.25665
242.18667602539062 0 4134.202
243.11282348632812 0 590.6421
245.1251678466797 0 4660.048
247.14456176757812 0 41130.023 b 1
248.1479034423828 0 5828.6943
251.1519012451172 0 1859.6188 y Water loss 4
253.13157653808594 0 685.15717
254.12498474121094 0 761.4933
255.10922241210938 0 9572.828
256.1114196777344 0 1609.791
262.118896484375 0 4040.5188
266.12451171875 0 563.94586
268.1441345214844 0 1350.4412
268.1656494140625 0 565.8141
269.1611328125 0 12062.412
270.1649475097656 0 1174.5387
270.181884765625 0 1160.4797
272.1360778808594 0 1665.6083
273.1195983886719 0 18123.623
273.1336975097656 0 775.3249
274.12164306640625 0 2216.6052
280.1288146972656 0 773.10236
280.1648864746094 0 648.8323
282.1808166503906 0 681.01135
287.1718444824219 0 4105.9395
288.1749572753906 0 1242.6985
290.14678955078125 0 776.84717
291.1451721191406 0 1185.0513
295.1781921386719 0 597.5289
301.1298522949219 0 1198.9408
304.165771484375 0 5281.375 b 2
305.16961669921875 0 1107.4872
311.66162109375 0 716.9041
318.1441650390625 0 655.7565
319.14056396484375 0 1995.8414
319.6588439941406 0 4040.8208
320.16009521484375 0 1574.6576
323.17236328125 0 964.6104
325.2235107421875 0 1669.048
325.65838623046875 0 1477.5364
326.1828918457031 0 7460.6387
326.66363525390625 0 975.2549
333.125244140625 0 720.094
333.19183349609375 0 664.5097
334.663818359375 0 1014.1394
335.167236328125 0 604.4876
339.17913818359375 0 1877.8986
340.1999206542969 0 943.3762
341.1827392578125 0 1055.2771
343.234619140625 0 2791.144 y Water loss 6
344.1930236816406 0 2266.6553
346.1147155761719 0 876.7076
347.1439208984375 0 569.9417
348.1669006347656 0 1862.9495
357.15509033203125 0 1416.217
358.2081298828125 0 949.0729
361.2189025878906 0 874.9637
361.244873046875 0 11858.723 y 6
362.2480163574219 0 2054.6882
365.1933898925781 0 3194.9783
366.1755676269531 0 1063.7668
366.7243347167969 0 751.2317
368.193115234375 0 2146.493
369.15655517578125 0 605.03595
374.18212890625 0 1346.1692
381.7112121582031 0 2086.447 y Water loss 1
382.211181640625 0 1129.6935
382.2457580566406 0 2040.9464
383.1756591796875 0 804.4392
383.2041320800781 0 17082.621
384.20745849609375 0 3868.5151
385.20916748046875 0 692.3757
390.21380615234375 0 673.41595
390.716552734375 0 14304.035 y 1
391.2181396484375 0 5734.4175
391.72039794921875 0 961.45416
392.1938171386719 0 1468.3771
395.7093811035156 0 929.4782
396.21087646484375 0 678.22437
400.2554016113281 0 2596.0415
401.21490478515625 0 17864.459
401.2593688964844 0 784.5469
402.17779541015625 0 5917.346
402.2173156738281 0 3374.608
403.18084716796875 0 1200.3154
413.2536926269531 0 818.4587
418.21002197265625 0 718.29553 b 4
419.2022399902344 0 845.3946
420.1878967285156 0 2359.372
420.21942138671875 0 744.8084
421.18951416015625 0 856.52686
421.8326721191406 0 992.97235
422.198974609375 0 610.21313
429.2838439941406 0 1025.0411
431.2460021972656 0 1209.0836 Precursor Water loss
438.2290344238281 0 1695.4171
439.2354736328125 0 869.21173
439.2668762207031 0 4006.3547
439.8439636230469 0 8462.968
440.251708984375 0 1716.6001 Precursor
440.75433349609375 0 1112.0831
440.8449401855469 0 2817.0134
441.2553405761719 0 1013.14655
451.2688903808594 0 836.9743
452.26312255859375 0 1590.7551
457.2776794433594 0 7896.1353
458.2803039550781 0 1385.615
461.2517395019531 0 664.0964
462.2926330566406 0 3183.7996 y 5
463.296875 0 712.723
466.2251892089844 0 1193.7258
468.2879943847656 0 852.5478
470.27276611328125 0 1262.8735
473.2510070800781 0 945.69916
478.2782287597656 0 3570.6304
479.27886962890625 0 716.73456
486.303955078125 0 11614.334
487.30645751953125 0 2096.344
496.2883605957031 0 23346.293
497.29107666015625 0 6072.9277
498.29290771484375 0 727.15784
501.2453308105469 0 1633.0775 b Water loss 5
512.2650146484375 0 755.5144
514.2987670898438 0 57155.56
515.2625732421875 0 480.9179
515.3015747070312 0 14472.486
516.3043212890625 0 2091.102
519.2579345703125 0 1035.9994 b 5
519.3140258789062 0 5483.3994 y 4
520.3178100585938 0 946.29144
530.2728881835938 0 2614.5374
532.310302734375 0 1150.8226
548.2825317382812 0 5501.135
548.3218994140625 0 789.9943
549.28662109375 0 1518.9374
558.325439453125 0 1693.7925 y Water loss 3
571.31494140625 0 935.22906
571.3594970703125 0 1169.9896
576.3356323242188 0 14477.744 y 3
577.3385009765625 0 4241.3984
587.3509521484375 0 1019.80206
597.3341064453125 0 1950.3574
598.2713012695312 0 720.80347
598.3323364257812 0 785.39886
615.3466186523438 0 8792.794 y Water loss 2
616.3478393554688 0 2759.051
625.346435546875 0 969.4693
629.3406372070312 0 895.7595 b Water loss 6
631.3414306640625 0 721.8445
633.3572998046875 0 141113.75 y 2
634.360107421875 0 43624.32
635.3623657226562 0 8978.62
643.3553466796875 0 6632.406
644.3594360351562 0 2507.5166
647.350830078125 0 3098.5403 b 6
648.3558349609375 0 1039.7804
661.3670654296875 0 13258.488
662.37060546875 0 3527.5234
663.3720092773438 0 1243.9382
721.33984375 0 633.36224
742.4242553710938 0 1436.1521 b Water loss 7
760.4349975585938 0 4642.0103 b 7
761.4386596679688 0 2268.9116
762.4150390625 0 2282.3042 y Water loss 1
780.425048828125 0 36673.55 y 1
781.4281616210938 0 16063.934
782.431396484375 0 4902.1836
790.4031372070312 0 868.7575
1509.9884033203125 0 576.10046
1983.390625 0 689.1595
2373.86328125 0 649.8642
2417.75048828125 0 655.93274
3045.419921875 0 929.4024

Spectrum Details

|  |  |
| --- | --- |
| Matched peaks? Matched peaks The total absolute number of peaks matched. Additionally in brackets the total fraction of peaks matched and the total number of peaks is shown. | 29 (9.24% of 314) |
| FDR? FDR The false discovery rate estimated for this peptide. It is calculated by matching all theoretical fragments with a non-integer shift with the raw peaks for this spectrum. This is done with 40 different shifts. The resulting percentage is the average number of annotated peaks over the number of annotated peaks with the correct spectrum. | 0.33% |
| Satellite FDR? Satellite FDR See the FDR for details on its calculation. This satellite ion specific FDR only contains the satellite ions (d/w) for I/L/J positions. | - |
| PSM Score? PSM Score The PSM Score as given by Hecklib to this annotated spectrum. It is shown with three significant figures. | 355 |

#### Spectrum 4605? Spectrum 4605 The raw spectrum of this peptide as annotated by Hecklib. The fragments are coloured according to ion type (see legend). Any peaks with a star '\*' as text can be hovered over to see the full details, first the ion type second the mass shift type. By hovering over the amino acids in the peptide or ions in the legend the corresponding peaks are highlighted. By toggling the 'Unassigned' label you can turn the background (unassigned) peaks on or off in the plot. By updating the slider in the Ion legend you can update the spectrum to only show the top X% of the peaks with labels. The top X% means any peak that is within X% of the highest intensity. By dragging in the spectrum you can zoom in to a specific part of the spectrum and use 'Zoom Out' to get back to the original zoom level. The annotation of the spectrum is based on the given sequence in the peptides file and is done with different software so inconsistencies are likely. The peaks are annotated based on the given sequence, with 20 ppm tolerance.

Copy Data

##### Spectrum 4605 (TSV)

###### Preview

```
Loading example...
```

*Click on the button to copy the data to your clipboard.*

Mz MinMz MaxIntensity Max

WidthHeightPeptide font sizePeptide stroke widthSpectrum font sizeSpectrum stroke widthCompact peptide

Ion legend

wxyz

abcd

OtherUnassignedIon^Charge^_Position_Show for top:%

VFGGGTKJT

01.03e+52.06e+53.09e+54.12e+5

Zoom Out

y^+1^_1_y^+1^_2_y^+1^_2_y^+2^_7_y^+2^_7_c^+2^_7_z^+1^_3_y^+1^_3_z^+1^_3_y^+1^_3_c^+1^_4_c^+2^_8_y^+2^_8_z^+2^_8_y^+2^_8_w^+1^_4_w^+1^_4_c^+1^_5_y^+1^_4_z^+1^_4_y^+1^_4_z^+1^_5_y^+1^_5_z^+1^_5_c^+1^_6_y^+1^_5_c^+1^_6_z^+1^_6_y^+1^_6_y^+1^_6_z^+1^_6_y^+1^_6_z^+1^_7_y^+1^_7_z^+1^_7_y^+1^_7_c^+1^_7_c^+1^_7_z^+1^_8_c^+1^_8_z^+1^_8_c^+1^_8_y^+1^_8_

0770154123113081

Fragment Matches Table

Show background peaks

| Position | Ion type | Intensity | mz Theoretical | mz Error (Th) | mz Error (ppm) | Charge | Series Number |
| --- | --- | --- | --- | --- | --- | --- | --- |
| 9 | y | 1.388E+04 | 120.1 | 0.0001847 | 1.539 | +1 | 1 |
| - | - | 6.749E+04 | 120.1 | - | - | 0 | - |
| - | - | 6165 | 121.1 | - | - | 0 | - |
| - | - | 2.239E+04 | 129.1 | - | - | 0 | - |
| - | - | 1499 | 130.1 | - | - | 0 | - |
| - | - | 441.6 | 138.4 | - | - | 0 | - |
| - | - | 798.1 | 144.1 | - | - | 0 | - |
| - | - | 961.5 | 153.1 | - | - | 0 | - |
| - | - | 408.9 | 158.1 | - | - | 0 | - |
| - | - | 541.7 | 165.2 | - | - | 0 | - |
| - | - | 501.9 | 168.6 | - | - | 0 | - |
| - | - | 451.4 | 170.5 | - | - | 0 | - |
| - | - | 566 | 172.1 | - | - | 0 | - |
| - | - | 1079 | 173.1 | - | - | 0 | - |
| - | - | 574.7 | 176.6 | - | - | 0 | - |
| - | - | 2153 | 177.1 | - | - | 0 | - |
| - | - | 442.7 | 177.7 | - | - | 0 | - |
| - | - | 519.8 | 186.1 | - | - | 0 | - |
| - | - | 2856 | 187.1 | - | - | 0 | - |
| - | - | 1176 | 187.1 | - | - | 0 | - |
| - | - | 813.3 | 188.1 | - | - | 0 | - |
| - | - | 744 | 191.2 | - | - | 0 | - |
| - | - | 1116 | 197.2 | - | - | 0 | - |
| - | - | 649.4 | 199.2 | - | - | 0 | - |
| - | - | 8014 | 205.1 | - | - | 0 | - |
| - | - | 572.8 | 206.1 | - | - | 0 | - |
| - | - | 1003 | 210.1 | - | - | 0 | - |
| - | - | 1116 | 212.1 | - | - | 0 | - |
| 8 | y | 4875 | 215.1 | 0.0001412 | 0.6565 | +1 | 2 |
| - | - | 2.189E+05 | 219.1 | - | - | 0 | - |
| - | - | 2.96E+04 | 220.2 | - | - | 0 | - |
| - | - | 815.9 | 221.1 | - | - | 0 | - |
| - | - | 1615 | 221.2 | - | - | 0 | - |
| - | - | 531.2 | 227.2 | - | - | 0 | - |
| - | - | 607.4 | 228.1 | - | - | 0 | - |
| - | - | 701.5 | 230.1 | - | - | 0 | - |
| - | - | 5276 | 230.2 | - | - | 0 | - |
| 8 | y | 5971 | 233.1 | 0.0002577 | 1.105 | +1 | 2 |
| - | - | 687.8 | 234.2 | - | - | 0 | - |
| - | - | 903.9 | 242.2 | - | - | 0 | - |
| - | - | 542.1 | 244.2 | - | - | 0 | - |
| - | - | 1047 | 245.1 | - | - | 0 | - |
| - | - | 570.5 | 245.2 | - | - | 0 | - |
| - | - | 9.01E+04 | 247.1 | - | - | 0 | - |
| - | - | 1.289E+04 | 248.1 | - | - | 0 | - |
| - | - | 1790 | 255.1 | - | - | 0 | - |
| - | - | 1957 | 262.1 | - | - | 0 | - |
| - | - | 657.2 | 263.1 | - | - | 0 | - |
| - | - | 1166 | 269.2 | - | - | 0 | - |
| - | - | 562.3 | 270.4 | - | - | 0 | - |
| - | - | 827.8 | 272.1 | - | - | 0 | - |
| - | - | 3425 | 272.2 | - | - | 0 | - |
| - | - | 9631 | 273.1 | - | - | 0 | - |
| - | - | 675.8 | 273.2 | - | - | 0 | - |
| - | - | 1226 | 274.1 | - | - | 0 | - |
| - | - | 1388 | 287.2 | - | - | 0 | - |
| - | - | 1239 | 289.2 | - | - | 0 | - |
| - | - | 2895 | 290.1 | - | - | 0 | - |
| - | - | 597.3 | 299.2 | - | - | 0 | - |
| - | - | 828 | 300.2 | - | - | 0 | - |
| - | - | 4167 | 301.2 | - | - | 0 | - |
| - | - | 907.3 | 302.2 | - | - | 0 | - |
| - | - | 3481 | 302.2 | - | - | 0 | - |
| - | - | 617.3 | 303.2 | - | - | 0 | - |
| - | - | 1.097E+04 | 304.2 | - | - | 0 | - |
| - | - | 1791 | 305.2 | - | - | 0 | - |
| 3 | y | 732.9 | 308.2 | 0.002322 | 7.533 | +2 | 7 |
| - | - | 908.1 | 310.2 | - | - | 0 | - |
| - | - | 1927 | 311.2 | - | - | 0 | - |
| 3 | y | 785.8 | 317.2 | 0.004058 | 12.79 | +2 | 7 |
| - | - | 727 | 319.1 | - | - | 0 | - |
| - | - | 598.9 | 322.2 | - | - | 0 | - |
| 7 | c | 691.5 | 324.2 | 0.0009996 | 3.083 | +2 | 7 |
| - | - | 2096 | 326.2 | - | - | 0 | - |
| - | - | 580 | 326.3 | - | - | 0 | - |
| - | - | 662.7 | 327.2 | - | - | 0 | - |
| 7 | z | 565.6 | 327.2 | 0.0002005 | 0.6127 | +1 | 3 |
| - | - | 2689 | 328.2 | - | - | 0 | - |
| - | - | 732.1 | 328.2 | - | - | 0 | - |
| - | - | 2674 | 329.2 | - | - | 0 | - |
| - | - | 688.4 | 331.2 | - | - | 0 | - |
| - | - | 599.5 | 333.2 | - | - | 0 | - |
| - | - | 694.1 | 341.2 | - | - | 0 | - |
| - | - | 1006 | 342.2 | - | - | 0 | - |
| 7 | y | 1669 | 343.2 | 0.0008814 | 2.568 | +1 | 3 |
| - | - | 2618 | 344.2 | - | - | 0 | - |
| 7 | z | 3.628E+04 | 345.2 | 0.0001601 | 0.4638 | +1 | 3 |
| - | - | 1230 | 345.2 | - | - | 0 | - |
| - | - | 9.1E+04 | 346.2 | - | - | 0 | - |
| - | - | 1.405E+04 | 347.2 | - | - | 0 | - |
| - | - | 1457 | 348.2 | - | - | 0 | - |
| - | - | 2860 | 358.2 | - | - | 0 | - |
| - | - | 2141 | 358.3 | - | - | 0 | - |
| - | - | 8007 | 359.2 | - | - | 0 | - |
| - | - | 636.3 | 359.3 | - | - | 0 | - |
| - | - | 552.2 | 360.2 | - | - | 0 | - |
| - | - | 700.5 | 360.2 | - | - | 0 | - |
| - | - | 1173 | 360.2 | - | - | 0 | - |
| - | - | 2379 | 361.2 | - | - | 0 | - |
| 7 | y | 1.912E+04 | 361.2 | 3.978E-05 | 0.1101 | +1 | 3 |
| - | - | 3088 | 362.2 | - | - | 0 | - |
| - | - | 843.9 | 365.2 | - | - | 0 | - |
| - | - | 1792 | 366.7 | - | - | 0 | - |
| - | - | 684.2 | 367.2 | - | - | 0 | - |
| - | - | 5719 | 367.2 | - | - | 0 | - |
| - | - | 792.2 | 367.7 | - | - | 0 | - |
| - | - | 1661 | 368.2 | - | - | 0 | - |
| - | - | 809.2 | 368.2 | - | - | 0 | - |
| - | - | 1199 | 371.7 | - | - | 0 | - |
| - | - | 1036 | 372.2 | - | - | 0 | - |
| - | - | 622.6 | 377.2 | - | - | 0 | - |
| 4 | c | 1332 | 378.2 | 0.0008733 | 2.309 | +1 | 4 |
| 8 | c | 1126 | 380.7 | 0.0008102 | 2.128 | +2 | 8 |
| 2 | y | 3151 | 381.7 | 9.705E-05 | 0.2542 | +2 | 8 |
| - | - | 1736 | 382.2 | - | - | 0 | - |
| - | - | 752.5 | 382.2 | - | - | 0 | - |
| 2 | z | 962.2 | 382.7 | 0.00613 | 16.02 | +2 | 8 |
| - | - | 8714 | 383.2 | - | - | 0 | - |
| - | - | 1075 | 384.2 | - | - | 0 | - |
| - | - | 3240 | 385.2 | - | - | 0 | - |
| - | - | 7810 | 385.2 | - | - | 0 | - |
| - | - | 2971 | 386.2 | - | - | 0 | - |
| - | - | 1016 | 386.2 | - | - | 0 | - |
| - | - | 1850 | 389.7 | - | - | 0 | - |
| - | - | 2294 | 390.2 | - | - | 0 | - |
| 2 | y | 4.185E+04 | 390.7 | 2.693E-06 | 0.006891 | +2 | 8 |
| - | - | 1.611E+04 | 391.2 | - | - | 0 | - |
| - | - | 3981 | 391.7 | - | - | 0 | - |
| - | - | 2289 | 395.7 | - | - | 0 | - |
| - | - | 1007 | 396.2 | - | - | 0 | - |
| - | - | 808.7 | 398.7 | - | - | 0 | - |
| - | - | 2169 | 400.3 | - | - | 0 | - |
| - | - | 2.098E+04 | 401.2 | - | - | 0 | - |
| - | - | 997.2 | 401.3 | - | - | 0 | - |
| - | - | 1881 | 402.2 | - | - | 0 | - |
| - | - | 3355 | 402.2 | - | - | 0 | - |
| - | - | 1.553E+04 | 402.3 | - | - | 0 | - |
| - | - | 8386 | 403.2 | - | - | 0 | - |
| - | - | 2423 | 403.3 | - | - | 0 | - |
| - | - | 1550 | 404.2 | - | - | 0 | - |
| - | - | 1648 | 414.3 | - | - | 0 | - |
| - | - | 5202 | 415.3 | - | - | 0 | - |
| - | - | 1122 | 416.2 | - | - | 0 | - |
| - | - | 1154 | 416.3 | - | - | 0 | - |
| - | - | 2262 | 418.2 | - | - | 0 | - |
| - | - | 1725 | 420.2 | - | - | 0 | - |
| - | - | 1417 | 421.7 | - | - | 0 | - |
| - | - | 1533 | 421.8 | - | - | 0 | - |
| - | - | 1143 | 422.2 | - | - | 0 | - |
| - | - | 621.3 | 422.7 | - | - | 0 | - |
| - | - | 1588 | 424.3 | - | - | 0 | - |
| - | - | 841.2 | 425.2 | - | - | 0 | - |
| - | - | 823.7 | 429.2 | - | - | 0 | - |
| 6 | w | 1425 | 429.3 | 0.003714 | 8.651 | +1 | 4 |
| 6 | w | 3431 | 431.3 | 0.003078 | 7.137 | +1 | 4 |
| - | - | 895.3 | 431.7 | - | - | 0 | - |
| - | - | 933.1 | 433.2 | - | - | 0 | - |
| - | - | 4023 | 434.2 | - | - | 0 | - |
| 5 | c | 1964 | 435.2 | 0.001249 | 2.871 | +1 | 5 |
| - | - | 1369 | 439.2 | - | - | 0 | - |
| - | - | 2157 | 439.3 | - | - | 0 | - |
| - | - | 1741 | 439.8 | - | - | 0 | - |
| - | - | 7508 | 440.3 | - | - | 0 | - |
| - | - | 3506 | 440.8 | - | - | 0 | - |
| - | - | 3005 | 441.3 | - | - | 0 | - |
| - | - | 846.3 | 441.3 | - | - | 0 | - |
| - | - | 1.743E+04 | 442.3 | - | - | 0 | - |
| - | - | 3281 | 443.3 | - | - | 0 | - |
| - | - | 1.261E+04 | 444.3 | - | - | 0 | - |
| 6 | y | 2.193E+04 | 445.3 | 1.005E-05 | 0.02257 | +1 | 4 |
| 6 | z | 1.131E+04 | 446.3 | 0.001742 | 3.903 | +1 | 4 |
| - | - | 1.691E+04 | 447.2 | - | - | 0 | - |
| - | - | 1.798E+05 | 447.3 | - | - | 0 | - |
| - | - | 5416 | 448.2 | - | - | 0 | - |
| - | - | 3.987E+04 | 448.3 | - | - | 0 | - |
| - | - | 852.4 | 449.2 | - | - | 0 | - |
| - | - | 842.7 | 449.3 | - | - | 0 | - |
| - | - | 4702 | 449.3 | - | - | 0 | - |
| - | - | 1302 | 455.3 | - | - | 0 | - |
| - | - | 6109 | 457.3 | - | - | 0 | - |
| - | - | 967.9 | 458.3 | - | - | 0 | - |
| - | - | 1118 | 459.3 | - | - | 0 | - |
| - | - | 4373 | 459.3 | - | - | 0 | - |
| - | - | 969.7 | 460.2 | - | - | 0 | - |
| - | - | 712.8 | 460.3 | - | - | 0 | - |
| - | - | 604.5 | 461.2 | - | - | 0 | - |
| - | - | 2430 | 461.3 | - | - | 0 | - |
| 6 | y | 6000 | 462.3 | 0.0005991 | 1.296 | +1 | 4 |
| - | - | 1252 | 463.3 | - | - | 0 | - |
| - | - | 3866 | 470.3 | - | - | 0 | - |
| - | - | 676.4 | 471.3 | - | - | 0 | - |
| - | - | 753.9 | 471.3 | - | - | 0 | - |
| - | - | 8808 | 472.3 | - | - | 0 | - |
| - | - | 2538 | 473.3 | - | - | 0 | - |
| - | - | 1003 | 478.3 | - | - | 0 | - |
| - | - | 652.5 | 484.2 | - | - | 0 | - |
| 5 | z | 1252 | 485.3 | 0.001014 | 2.089 | +1 | 5 |
| - | - | 5883 | 486.2 | - | - | 0 | - |
| - | - | 6875 | 486.3 | - | - | 0 | - |
| - | - | 1328 | 487.2 | - | - | 0 | - |
| - | - | 998 | 487.3 | - | - | 0 | - |
| - | - | 623.3 | 488.3 | - | - | 0 | - |
| - | - | 2286 | 491.3 | - | - | 0 | - |
| - | - | 729.6 | 492.3 | - | - | 0 | - |
| - | - | 1206 | 493.3 | - | - | 0 | - |
| - | - | 1.474E+04 | 496.3 | - | - | 0 | - |
| - | - | 3339 | 497.3 | - | - | 0 | - |
| - | - | 1752 | 498.3 | - | - | 0 | - |
| - | - | 2860 | 498.3 | - | - | 0 | - |
| - | - | 1042 | 498.8 | - | - | 0 | - |
| - | - | 1.479E+04 | 499.3 | - | - | 0 | - |
| - | - | 3361 | 500.3 | - | - | 0 | - |
| - | - | 7102 | 501.2 | - | - | 0 | - |
| - | - | 2188 | 502.2 | - | - | 0 | - |
| 5 | y | 899.1 | 502.3 | 0.0003969 | 0.7902 | +1 | 5 |
| 5 | z | 1.247E+04 | 503.3 | 0.0001039 | 0.2064 | +1 | 5 |
| - | - | 1.066E+05 | 504.3 | - | - | 0 | - |
| - | - | 2.442E+05 | 504.3 | - | - | 0 | - |
| - | - | 2.53E+04 | 505.3 | - | - | 0 | - |
| - | - | 5.884E+04 | 505.3 | - | - | 0 | - |
| - | - | 4240 | 506.3 | - | - | 0 | - |
| - | - | 9657 | 506.3 | - | - | 0 | - |
| - | - | 3375 | 512.3 | - | - | 0 | - |
| - | - | 972.9 | 513.3 | - | - | 0 | - |
| - | - | 6.719E+04 | 514.3 | - | - | 0 | - |
| - | - | 1.579E+04 | 515.3 | - | - | 0 | - |
| - | - | 1.521E+04 | 516.3 | - | - | 0 | - |
| - | - | 9562 | 517.3 | - | - | 0 | - |
| - | - | 3096 | 517.3 | - | - | 0 | - |
| 6 | c | 5319 | 518.3 | 0.003115 | 6.011 | +1 | 6 |
| - | - | 1.592E+04 | 518.3 | - | - | 0 | - |
| - | - | 5425 | 519.3 | - | - | 0 | - |
| 5 | y | 2.97E+04 | 519.3 | 0.0007005 | 1.349 | +1 | 5 |
| - | - | 1516 | 520.3 | - | - | 0 | - |
| - | - | 6897 | 520.3 | - | - | 0 | - |
| - | - | 1399 | 521.3 | - | - | 0 | - |
| - | - | 759.2 | 525.2 | - | - | 0 | - |
| - | - | 897.3 | 527.3 | - | - | 0 | - |
| - | - | 1071 | 530.3 | - | - | 0 | - |
| - | - | 925.4 | 532.3 | - | - | 0 | - |
| - | - | 3.238E+04 | 535.3 | - | - | 0 | - |
| 6 | c | 1.226E+05 | 536.3 | 0.0002523 | 0.4705 | +1 | 6 |
| - | - | 3.028E+04 | 537.3 | - | - | 0 | - |
| - | - | 5446 | 538.3 | - | - | 0 | - |
| 4 | z | 3143 | 542.3 | 7.765E-05 | 0.1432 | +1 | 6 |
| - | - | 3623 | 543.3 | - | - | 0 | - |
| - | - | 1984 | 543.3 | - | - | 0 | - |
| - | - | 1132 | 544.3 | - | - | 0 | - |
| - | - | 827.5 | 547.3 | - | - | 0 | - |
| - | - | 5780 | 548.3 | - | - | 0 | - |
| - | - | 731.9 | 549.3 | - | - | 0 | - |
| - | - | 7915 | 555.3 | - | - | 0 | - |
| - | - | 750 | 556.3 | - | - | 0 | - |
| - | - | 2424 | 556.3 | - | - | 0 | - |
| 4 | y | 2049 | 558.3 | 0.002322 | 4.159 | +1 | 6 |
| 4 | y | 992.5 | 559.3 | 0.008169 | 14.61 | +1 | 6 |
| 4 | z | 2.666E+04 | 560.3 | 9.986E-05 | 0.1782 | +1 | 6 |
| - | - | 7270 | 561.3 | - | - | 0 | - |
| - | - | 1.197E+05 | 561.3 | - | - | 0 | - |
| - | - | 1705 | 562.3 | - | - | 0 | - |
| - | - | 3.1E+04 | 562.3 | - | - | 0 | - |
| - | - | 5807 | 563.3 | - | - | 0 | - |
| - | - | 1449 | 571.4 | - | - | 0 | - |
| - | - | 1.054E+04 | 574.3 | - | - | 0 | - |
| - | - | 2539 | 575.3 | - | - | 0 | - |
| - | - | 1.297E+04 | 575.3 | - | - | 0 | - |
| 4 | y | 7.708E+04 | 576.3 | 0.0003747 | 0.6501 | +1 | 6 |
| - | - | 1.994E+04 | 577.3 | - | - | 0 | - |
| - | - | 4440 | 578.3 | - | - | 0 | - |
| - | - | 2450 | 587.4 | - | - | 0 | - |
| - | - | 852.5 | 588.4 | - | - | 0 | - |
| - | - | 715.9 | 589.3 | - | - | 0 | - |
| - | - | 757.7 | 591.3 | - | - | 0 | - |
| - | - | 1948 | 597.3 | - | - | 0 | - |
| - | - | 923.2 | 598.3 | - | - | 0 | - |
| 3 | z | 4817 | 599.3 | 0.0003011 | 0.5025 | +1 | 7 |
| - | - | 2171 | 600.3 | - | - | 0 | - |
| 3 | y | 1.198E+04 | 615.3 | 0.0001043 | 0.1695 | +1 | 7 |
| - | - | 3931 | 616.3 | - | - | 0 | - |
| 3 | z | 4.278E+04 | 617.3 | 1.563E-06 | 0.002532 | +1 | 7 |
| - | - | 2.284E+04 | 618.3 | - | - | 0 | - |
| - | - | 7178 | 619.4 | - | - | 0 | - |
| - | - | 4919 | 620.4 | - | - | 0 | - |
| - | - | 1676 | 621.4 | - | - | 0 | - |
| - | - | 1181 | 629.3 | - | - | 0 | - |
| - | - | 1064 | 630.3 | - | - | 0 | - |
| - | - | 3123 | 631.3 | - | - | 0 | - |
| - | - | 2903 | 632.3 | - | - | 0 | - |
| 3 | y | 3.973E+05 | 633.4 | 0.0001953 | 0.3083 | +1 | 7 |
| - | - | 1.198E+05 | 634.4 | - | - | 0 | - |
| - | - | 2.43E+04 | 635.4 | - | - | 0 | - |
| - | - | 1252 | 636.4 | - | - | 0 | - |
| - | - | 1091 | 639.8 | - | - | 0 | - |
| - | - | 2825 | 643.4 | - | - | 0 | - |
| - | - | 1222 | 644.4 | - | - | 0 | - |
| - | - | 2417 | 646.3 | - | - | 0 | - |
| 7 | c | 8815 | 647.4 | 0.0007341 | 1.134 | +1 | 7 |
| - | - | 2838 | 648.4 | - | - | 0 | - |
| - | - | 886 | 649.4 | - | - | 0 | - |
| - | - | 699.8 | 649.8 | - | - | 0 | - |
| - | - | 820.4 | 650.4 | - | - | 0 | - |
| - | - | 1533 | 659.8 | - | - | 0 | - |
| - | - | 6091 | 660.3 | - | - | 0 | - |
| - | - | 1.477E+04 | 661.4 | - | - | 0 | - |
| - | - | 5451 | 661.8 | - | - | 0 | - |
| - | - | 2469 | 662.3 | - | - | 0 | - |
| - | - | 4866 | 662.4 | - | - | 0 | - |
| - | - | 9805 | 663.4 | - | - | 0 | - |
| 7 | c | 2.644E+05 | 664.4 | 6.053E-05 | 0.09111 | +1 | 7 |
| - | - | 9.131E+04 | 665.4 | - | - | 0 | - |
| - | - | 1.923E+04 | 666.4 | - | - | 0 | - |
| - | - | 1503 | 667.4 | - | - | 0 | - |
| - | - | 653.8 | 672.9 | - | - | 0 | - |
| - | - | 3456 | 690.4 | - | - | 0 | - |
| - | - | 1977 | 691.4 | - | - | 0 | - |
| - | - | 1223 | 693.3 | - | - | 0 | - |
| - | - | 885.8 | 701.4 | - | - | 0 | - |
| - | - | 4263 | 702.4 | - | - | 0 | - |
| - | - | 1199 | 703.4 | - | - | 0 | - |
| - | - | 1962 | 708.3 | - | - | 0 | - |
| - | - | 705.1 | 709.3 | - | - | 0 | - |
| - | - | 3373 | 718.4 | - | - | 0 | - |
| - | - | 882.6 | 719.4 | - | - | 0 | - |
| - | - | 865.2 | 721.3 | - | - | 0 | - |
| - | - | 764.2 | 722.3 | - | - | 0 | - |
| - | - | 1004 | 732.4 | - | - | 0 | - |
| - | - | 2.102E+04 | 733.4 | - | - | 0 | - |
| - | - | 7891 | 734.4 | - | - | 0 | - |
| - | - | 1768 | 735.5 | - | - | 0 | - |
| - | - | 1849 | 742.4 | - | - | 0 | - |
| - | - | 1115 | 743.4 | - | - | 0 | - |
| - | - | 645.5 | 744.4 | - | - | 0 | - |
| - | - | 1397 | 745.4 | - | - | 0 | - |
| 2 | z | 4867 | 746.4 | 0.0006819 | 0.9136 | +1 | 8 |
| - | - | 1475 | 747.4 | - | - | 0 | - |
| - | - | 651 | 750.3 | - | - | 0 | - |
| - | - | 1484 | 750.3 | - | - | 0 | - |
| 8 | c | 1.569E+04 | 760.4 | 0.001119 | 1.471 | +1 | 8 |
| - | - | 5431 | 761.4 | - | - | 0 | - |
| - | - | 8585 | 762.4 | - | - | 0 | - |
| - | - | 3284 | 763.5 | - | - | 0 | - |
| 2 | z | 5.698E+04 | 764.4 | 0.0004834 | 0.6323 | +1 | 8 |
| - | - | 2.167E+04 | 765.4 | - | - | 0 | - |
| - | - | 6019 | 766.4 | - | - | 0 | - |
| - | - | 2496 | 775.4 | - | - | 0 | - |
| - | - | 1062 | 776.5 | - | - | 0 | - |
| 8 | c | 4.082E+05 | 777.5 | 1.907E-05 | 0.02453 | +1 | 8 |
| - | - | 1.588E+05 | 778.5 | - | - | 0 | - |
| - | - | 4.071E+04 | 779.5 | - | - | 0 | - |
| 2 | y | 6.823E+04 | 780.4 | 0.0004086 | 0.5236 | +1 | 8 |
| - | - | 2.663E+04 | 781.4 | - | - | 0 | - |
| - | - | 6600 | 782.4 | - | - | 0 | - |
| - | - | 821.2 | 789.4 | - | - | 0 | - |
| - | - | 2069 | 790.4 | - | - | 0 | - |
| - | - | 1776 | 792.4 | - | - | 0 | - |
| - | - | 1367 | 801.5 | - | - | 0 | - |
| - | - | 1966 | 802.5 | - | - | 0 | - |
| - | - | 1912 | 807.4 | - | - | 0 | - |
| - | - | 4583 | 808.3 | - | - | 0 | - |
| - | - | 1083 | 809.3 | - | - | 0 | - |
| - | - | 1076 | 816.5 | - | - | 0 | - |
| - | - | 707.6 | 817.5 | - | - | 0 | - |
| - | - | 857.5 | 818.5 | - | - | 0 | - |
| - | - | 1091 | 822.4 | - | - | 0 | - |
| - | - | 3244 | 823.4 | - | - | 0 | - |
| - | - | 2679 | 824.4 | - | - | 0 | - |
| - | - | 1190 | 825.4 | - | - | 0 | - |
| - | - | 4872 | 834.5 | - | - | 0 | - |
| - | - | 1920 | 835.5 | - | - | 0 | - |
| - | - | 2140 | 842.3 | - | - | 0 | - |
| - | - | 3559 | 843.3 | - | - | 0 | - |
| - | - | 2665 | 844.3 | - | - | 0 | - |
| - | - | 878.2 | 845.3 | - | - | 0 | - |
| - | - | 4135 | 845.5 | - | - | 0 | - |
| - | - | 938.3 | 846.3 | - | - | 0 | - |
| - | - | 839.7 | 846.5 | - | - | 0 | - |
| - | - | 963.5 | 847.3 | - | - | 0 | - |
| - | - | 1165 | 848.3 | - | - | 0 | - |
| - | - | 5312 | 848.5 | - | - | 0 | - |
| - | - | 1212 | 849.5 | - | - | 0 | - |
| - | - | 1483 | 852.5 | - | - | 0 | - |
| - | - | 900.7 | 860.3 | - | - | 0 | - |
| - | - | 1151 | 861.3 | - | - | 0 | - |
| - | - | 3204 | 861.5 | - | - | 0 | - |
| - | - | 3069 | 862.3 | - | - | 0 | - |
| - | - | 6209 | 862.5 | - | - | 0 | - |
| - | - | 3.23E+05 | 863.5 | - | - | 0 | - |
| - | - | 1.391E+05 | 864.5 | - | - | 0 | - |
| - | - | 4.022E+04 | 865.5 | - | - | 0 | - |
| - | - | 3034 | 866.5 | - | - | 0 | - |
| - | - | 914 | 869.4 | - | - | 0 | - |
| - | - | 1216 | 871.4 | - | - | 0 | - |
| - | - | 1123 | 871.9 | - | - | 0 | - |
| - | - | 1232 | 872.4 | - | - | 0 | - |
| - | - | 2059 | 872.9 | - | - | 0 | - |
| - | - | 792.8 | 873.5 | - | - | 0 | - |
| - | - | 914.9 | 877.3 | - | - | 0 | - |
| - | - | 1492 | 877.5 | - | - | 0 | - |
| - | - | 1445 | 878.3 | - | - | 0 | - |
| - | - | 4695 | 878.5 | - | - | 0 | - |
| - | - | 2.553E+05 | 879.5 | - | - | 0 | - |
| - | - | 2.184E+05 | 880.5 | - | - | 0 | - |
| - | - | 7.962E+04 | 881.5 | - | - | 0 | - |
| - | - | 1013 | 882.3 | - | - | 0 | - |
| - | - | 1.653E+04 | 882.5 | - | - | 0 | - |
| - | - | 1410 | 883.3 | - | - | 0 | - |
| - | - | 1005 | 883.5 | - | - | 0 | - |
| - | - | 896.6 | 1076 | - | - | 0 | - |
| - | - | 832.4 | 1282 | - | - | 0 | - |
| - | - | 1690 | 1289 | - | - | 0 | - |
| - | - | 844.9 | 1290 | - | - | 0 | - |
| - | - | 1186 | 1299 | - | - | 0 | - |
| - | - | 2426 | 1300 | - | - | 0 | - |
| - | - | 1463 | 1301 | - | - | 0 | - |
| - | - | 960.7 | 1302 | - | - | 0 | - |
| - | - | 2106 | 1303 | - | - | 0 | - |
| - | - | 1785 | 1304 | - | - | 0 | - |
| - | - | 827.6 | 1307 | - | - | 0 | - |
| - | - | 900.2 | 1317 | - | - | 0 | - |
| - | - | 1194 | 1319 | - | - | 0 | - |
| - | - | 806.4 | 1320 | - | - | 0 | - |
| - | - | 1867 | 1321 | - | - | 0 | - |
| - | - | 784.3 | 1322 | - | - | 0 | - |
| - | - | 1976 | 1323 | - | - | 0 | - |
| - | - | 4219 | 1324 | - | - | 0 | - |
| - | - | 2544 | 1325 | - | - | 0 | - |
| - | - | 910.1 | 1745 | - | - | 0 | - |
| - | - | 629.8 | 2613 | - | - | 0 | - |
| - | - | 619.9 | 3051 | - | - | 0 | - |

m/z Charge Intensity FragmentType MassShift Position
120.06570434570312 0 13884.312 y 8
120.08100128173828 0 67487.21
121.0843276977539 0 6165.322
129.10243225097656 0 22386.178
130.10586547851562 0 1499.4053
138.4063262939453 0 441.5602
144.12619018554688 0 798.0849
153.0771942138672 0 961.4987
158.1381378173828 0 408.88727
165.18092346191406 0 541.7362
168.6046905517578 0 501.92838
170.49765014648438 0 451.41956
172.07200622558594 0 566.03937
173.09182739257812 0 1079.333
176.61012268066406 0 574.6829
177.10256958007812 0 2153.0142
177.66595458984375 0 442.74734
186.1232452392578 0 519.82086
187.1078338623047 0 2856.3477
187.14456176757812 0 1176.4617
188.13941955566406 0 813.3111
191.15460205078125 0 743.9626
197.16517639160156 0 1116.2173
199.16952514648438 0 649.3572
205.09719848632812 0 8014.0586
206.10092163085938 0 572.8032
210.1493682861328 0 1003.0224
212.1392059326172 0 1116.3497
215.13916015625 0 4875.196 y Water loss 7
219.14939880371094 0 218850.16
220.15267944335938 0 29604.057
221.09185791015625 0 815.918
221.15626525878906 0 1615.2507
227.16273498535156 0 531.18207
228.09768676757812 0 607.39636
230.11700439453125 0 701.54663
230.15005493164062 0 5275.7544
233.14984130859375 0 5971.4497 y 7
234.15426635742188 0 687.7832
242.18597412109375 0 903.86066
244.20252990722656 0 542.08563
245.12445068359375 0 1047.1448
245.17245483398438 0 570.48975
247.14427185058594 0 90103.375
248.14755249023438 0 12886.1875
255.10865783691406 0 1790.0818
262.11846923828125 0 1957.0581
263.1226806640625 0 657.172
269.1615905761719 0 1165.8446
270.3630065917969 0 562.2595
272.1356506347656 0 827.82007
272.1607666015625 0 3425.1572
273.1193542480469 0 9630.865
273.16485595703125 0 675.80725
274.12255859375 0 1225.58
287.17169189453125 0 1388.1466
289.1630554199219 0 1239.1538
290.1461181640625 0 2895.1287
299.22149658203125 0 597.2971
300.2290954589844 0 828.01105
301.187255859375 0 4167.2725
302.1727600097656 0 907.28046
302.19439697265625 0 3480.8074
303.19781494140625 0 617.289
304.1657409667969 0 10972.468
305.1691589355469 0 1790.7855
308.1789855957031 0 732.9324 y Water loss 2
310.212890625 0 908.1115
311.1717834472656 0 1926.6276
317.1860046386719 0 785.8027 y 2
319.1402893066406 0 726.99426
322.1824951171875 0 598.8881
324.1802062988281 0 691.5431 c Ammonia loss 6
326.1827087402344 0 2096.0713
326.3380432128906 0 580.0142
327.18365478515625 0 662.7393
327.2150573730469 0 565.56604 z Water loss 6
328.1745300292969 0 2688.9968
328.2222900390625 0 732.1259
329.181640625 0 2673.6013
331.1867980957031 0 688.3875
333.19305419921875 0 599.45593
341.2055358886719 0 694.0735
342.2051696777344 0 1005.7901
343.23486328125 0 1669.0353 y Water loss 6
344.1929626464844 0 2618.4434
345.2259826660156 0 36281.027 z 6
345.2467956542969 0 1230.1875
346.2335205078125 0 90997.49
347.2367858886719 0 14047.338
348.2387390136719 0 1456.926
358.2087097167969 0 2860.1204
358.2576599121094 0 2141.0974
359.2161865234375 0 8006.5967
359.25994873046875 0 636.2515
360.1697692871094 0 552.20825
360.2160339355469 0 700.4966
360.23846435546875 0 1172.5005
361.1869812011719 0 2379.3
361.2445068359375 0 19118.834 y 6
362.2480163574219 0 3087.6057
365.1935729980469 0 843.8823
366.7235412597656 0 1792.2759
367.1636657714844 0 684.2358
367.23333740234375 0 5718.567
367.7132873535156 0 792.1816
368.1929016113281 0 1661.0676
368.2365417480469 0 809.15137
371.7140808105469 0 1199.4857
372.2232971191406 0 1036.346
377.2059020996094 0 622.6299
378.21270751953125 0 1332.495 c 3
380.7204284667969 0 1125.9048 c Ammonia loss 7
381.7109680175781 0 3151.1145 y Water loss 1
382.2121887207031 0 1736.4058
382.24041748046875 0 752.468
382.7129211425781 0 962.2133 z 1
383.2032775878906 0 8714.176
384.2074890136719 0 1075.3324
385.19561767578125 0 3239.7515
385.2444763183594 0 7810.147
386.20208740234375 0 2970.8503
386.24755859375 0 1015.82385
389.66949462890625 0 1849.5608
390.2136535644531 0 2294.4785
390.7161560058594 0 41850.78 y 1
391.2177734375 0 16114.469
391.7189636230469 0 3980.67
395.7080993652344 0 2288.9512
396.211181640625 0 1007.3594
398.65789794921875 0 808.72534
400.255859375 0 2169.4695
401.2143859863281 0 20980.832
401.2771911621094 0 997.2311
402.17718505859375 0 1881.178
402.2178649902344 0 3354.916
402.27117919921875 0 15527.625
403.22979736328125 0 8386.115
403.27484130859375 0 2423.1292
404.2322692871094 0 1550.1171
414.2708740234375 0 1648.1093
415.2787170410156 0 5202.341
416.239013671875 0 1121.9639
416.28228759765625 0 1154.4991
418.2088317871094 0 2261.8848
420.1893310546875 0 1724.7281
421.6613464355469 0 1417.3169
421.83245849609375 0 1533.2523
422.1643981933594 0 1142.8518
422.6646728515625 0 621.2912
424.2541809082031 0 1587.7765
425.1615905761719 0 841.1685
429.2190246582031 0 823.74854
429.27447509765625 0 1425.4952 w 5
431.2469482421875 0 3430.6726 w 5
431.7480773925781 0 895.2574
433.1798400878906 0 933.09644
434.2273254394531 0 4022.5781
435.2337951660156 0 1964.4429 c 4
439.2021789550781 0 1368.6179
439.2671203613281 0 2156.839
439.8442687988281 0 1740.6426
440.25042724609375 0 7508.1323
440.7524719238281 0 3505.7258
441.256103515625 0 3004.9219
441.2936706542969 0 846.3411
442.2571105957031 0 17429.354
443.2597351074219 0 3280.8457
444.2586669921875 0 12613.086
445.26568603515625 0 21926.16 y Ammonia loss 5
446.2717590332031 0 11306.996 z 5
447.2328796386719 0 16909.283
447.2816162109375 0 179846.8
448.2386779785156 0 5416.384
448.2845458984375 0 39865.51
449.24029541015625 0 852.368
449.2509460449219 0 842.72864
449.2870788574219 0 4701.7397
455.262939453125 0 1301.8495
457.2769470214844 0 6108.9453
458.2819519042969 0 967.8765
459.2583312988281 0 1117.604
459.2939453125 0 4372.7363
460.2476501464844 0 969.65826
460.2920227050781 0 712.81464
461.2478942871094 0 604.4881
461.2851257324219 0 2430.2083
462.2916259765625 0 5999.763 y 5
463.2951354980469 0 1252.1742
470.2620544433594 0 3866.313
471.2615051269531 0 676.4178
471.29534912109375 0 753.8741
472.3004150390625 0 8807.596
473.30438232421875 0 2538.2651
478.27593994140625 0 1003.14355
484.24285888671875 0 652.486
485.28338623046875 0 1251.9154 z Water loss 4
486.2437744140625 0 5882.7554
486.3006591796875 0 6874.5625
487.24700927734375 0 1327.5931
487.3045654296875 0 998.04126
488.3081359863281 0 623.2909
491.26080322265625 0 2285.627
492.2653503417969 0 729.64197
493.2777404785156 0 1205.7166
496.28814697265625 0 14735.211
497.291259765625 0 3338.973
498.2795104980469 0 1751.5414
498.3160705566406 0 2859.8718
498.79180908203125 0 1042.3289
499.27618408203125 0 14792.611
500.27972412109375 0 3361.0964
501.24542236328125 0 7101.882
502.2478332519531 0 2188.289
502.28753662109375 0 899.08856 y Ammonia loss 4
503.29486083984375 0 12472.873 z 4
504.2544250488281 0 106567.49
504.3028564453125 0 244232.61
505.25775146484375 0 25303.713
505.3059387207031 0 58843.25
506.2602844238281 0 4239.541
506.3082275390625 0 9657.071
512.2832641601562 0 3374.7144
513.2861938476562 0 972.905
514.29833984375 0 67186.44
515.3012084960938 0 15790.554
516.3126831054688 0 15205.556
517.2618408203125 0 9561.727
517.316162109375 0 3095.793
518.26904296875 0 5319.185 c Water loss 5
518.3065795898438 0 15924.89
519.2595825195312 0 5424.721
519.31298828125 0 29697.074 y 4
520.2618408203125 0 1515.6593
520.316650390625 0 6897.298
521.318359375 0 1398.5751
525.2269897460938 0 759.16956
527.3059692382812 0 897.29486
530.273681640625 0 1071.2074
532.3088989257812 0 925.4384
535.2747802734375 0 32379.957
536.282470703125 0 122634.39 c 5
537.2853393554688 0 30278.506
538.28857421875 0 5445.6875
542.3057861328125 0 3143.3572 z Water loss 3
543.264892578125 0 3622.5068
543.3137817382812 0 1984.1671
544.2679443359375 0 1131.8807
547.3076782226562 0 827.48254
548.2828369140625 0 5779.5737
549.2887573242188 0 731.93066
555.3375854492188 0 7914.98
556.27392578125 0 750.01337
556.3385009765625 0 2423.664
558.322265625 0 2049.4553 y Water loss 3
559.3167724609375 0 992.4516 y Ammonia loss 3
560.3165283203125 0 26663.914 z 3
561.2763671875 0 7270.008
561.323974609375 0 119719.14
562.2807006835938 0 1704.6141
562.3267822265625 0 31001.875
563.3294677734375 0 5807.039
571.3545532226562 0 1448.6122
574.2828369140625 0 10536.393
575.2838134765625 0 2539.2795
575.3275146484375 0 12970.871
576.3347778320312 0 77077.914 y 3
577.33740234375 0 19940.86
578.3400268554688 0 4439.884
587.3504028320312 0 2450.3289
588.3526000976562 0 852.52545
589.339111328125 0 715.8848
591.3363647460938 0 757.66693
597.3336791992188 0 1948.3224
598.3335571289062 0 923.16077
599.3270263671875 0 4817.197 z Water loss 2
600.3291625976562 0 2170.9583
615.345947265625 0 11982.311 y Water loss 2
616.3489379882812 0 3930.8967
617.337890625 0 42779.324 z 2
618.343017578125 0 22844.312
619.3544921875 0 7177.586
620.3648681640625 0 4919.2603
621.36962890625 0 1676.4445
629.3423461914062 0 1180.604
630.345458984375 0 1064.1688
631.3427124023438 0 3122.567
632.34716796875 0 2902.9912
633.3568115234375 0 397339.03 y 2
634.3594970703125 0 119803.22
635.3612670898438 0 24299.96
636.3576049804688 0 1252.3346
639.8079833984375 0 1091.3217
643.3558959960938 0 2824.5078
644.3572998046875 0 1221.9874
646.3452758789062 0 2417.2654
647.3504028320312 0 8815.217 c Ammonia loss 6
648.353515625 0 2837.5852
649.3518676757812 0 885.9663
649.8414306640625 0 699.80316
650.3506469726562 0 820.3528
659.8394165039062 0 1532.543
660.3436279296875 0 6090.9976
661.3655395507812 0 14769.405
661.82373046875 0 5451.125
662.3201904296875 0 2469.2798
662.3687133789062 0 4866.2393
663.3810424804688 0 9805.406
664.3777465820312 0 264364 c 6
665.380615234375 0 91312.39
666.3828125 0 19227.967
667.3834838867188 0 1502.5267
672.93994140625 0 653.8112
690.3935546875 0 3456.1243
691.3973999023438 0 1976.781
693.3292236328125 0 1223.1503
701.4287109375 0 885.7567
702.4067993164062 0 4262.6943
703.4140625 0 1198.8787
708.3428955078125 0 1962.3977
709.3466796875 0 705.14966
718.4349975585938 0 3373.3286
719.4364013671875 0 882.6393
721.3494873046875 0 865.1692
722.3438110351562 0 764.20044
732.434814453125 0 1003.6422
733.4476318359375 0 21017.701
734.4496459960938 0 7891.457
735.4507446289062 0 1767.7311
742.427001953125 0 1849.0581
743.4322509765625 0 1115.1185
744.405517578125 0 645.5049
745.4120483398438 0 1397.0458
746.3964233398438 0 4867.075 z Water loss 1
747.4005737304688 0 1475.1016
750.2526245117188 0 650.9824
750.336181640625 0 1484.3258
760.43408203125 0 15687.181 c Ammonia loss 7
761.4362182617188 0 5431.161
762.4461669921875 0 8585.01
763.4519653320312 0 3283.632
764.4058227539062 0 56982.707 z 1
765.4086303710938 0 21674.758
766.410888671875 0 6018.6353
775.4456176757812 0 2496.0706
776.4526977539062 0 1061.5802
777.4617309570312 0 408215.66 c 7
778.4642944335938 0 158829.14
779.4668579101562 0 40709.797
780.4246215820312 0 68227.5 y 1
781.4273681640625 0 26628.389
782.43017578125 0 6600.35
789.4061889648438 0 821.1753
790.4089965820312 0 2068.8513
792.4006958007812 0 1775.5009
801.4730834960938 0 1367.0118
802.461669921875 0 1965.6233
807.41357421875 0 1911.8417
808.341552734375 0 4583.3496
809.3404541015625 0 1082.5223
816.4934692382812 0 1076.3458
817.4716186523438 0 707.6294
818.4761962890625 0 857.5428
822.3512573242188 0 1090.6049
823.3650512695312 0 3244.2996
824.43701171875 0 2679.0696
825.4412841796875 0 1190.3839
834.4934692382812 0 4872.483
835.4957275390625 0 1920.1072
842.3157958984375 0 2140.0774
843.3211059570312 0 3558.8281
844.3229370117188 0 2665.4766
845.3176879882812 0 878.20593
845.4633178710938 0 4135.3975
846.3143310546875 0 938.3337
846.4656982421875 0 839.71643
847.3215942382812 0 963.5488
848.3279418945312 0 1165.2095
848.4524536132812 0 5311.6074
849.4627685546875 0 1211.7319
852.50830078125 0 1482.7047
860.2930297851562 0 900.6838
861.3055419921875 0 1151.297
861.4569091796875 0 3203.8938
862.3107299804688 0 3068.7205
862.4816284179688 0 6208.526
863.4743041992188 0 323029.4
864.4772338867188 0 139084.97
865.4796752929688 0 40217.71
866.4800415039062 0 3033.7878
869.44580078125 0 914.0299
871.4413452148438 0 1215.9517
871.9346313476562 0 1122.9362
872.4343872070312 0 1232.2253
872.945068359375 0 2058.6248
873.452880859375 0 792.77563
877.3082275390625 0 914.86053
877.475341796875 0 1491.9384
878.3005981445312 0 1444.5853
878.4824829101562 0 4694.9614
879.49267578125 0 255280.83
880.4981689453125 0 218429.77
881.5010986328125 0 79619.805
882.3034057617188 0 1012.7572
882.505126953125 0 16534.145
883.2989501953125 0 1409.572
883.509033203125 0 1004.5073
1075.5826416015625 0 896.59546
1281.6571044921875 0 832.3688
1288.6019287109375 0 1690.2112
1289.6126708984375 0 844.862
1298.6978759765625 0 1186.118
1299.68701171875 0 2425.7324
1300.6771240234375 0 1463.0308
1301.66748046875 0 960.73096
1302.65576171875 0 2105.6853
1303.6656494140625 0 1785.3728
1306.641357421875 0 827.58514
1316.7047119140625 0 900.2471
1318.694580078125 0 1194.126
1319.6990966796875 0 806.35693
1320.6822509765625 0 1867.2936
1321.658203125 0 784.29004
1322.6407470703125 0 1975.6696
1323.6441650390625 0 4218.7505
1324.654541015625 0 2543.6406
1744.8973388671875 0 910.06287
2612.806884765625 0 629.8007
3050.944091796875 0 619.93506

Spectrum Details

|  |  |
| --- | --- |
| Matched peaks? Matched peaks The total absolute number of peaks matched. Additionally in brackets the total fraction of peaks matched and the total number of peaks is shown. | 43 (10.02% of 429) |
| FDR? FDR The false discovery rate estimated for this peptide. It is calculated by matching all theoretical fragments with a non-integer shift with the raw peaks for this spectrum. This is done with 40 different shifts. The resulting percentage is the average number of annotated peaks over the number of annotated peaks with the correct spectrum. | 0.50% |
| Satellite FDR? Satellite FDR See the FDR for details on its calculation. This satellite ion specific FDR only contains the satellite ions (d/w) for I/L/J positions. | - |
| PSM Score? PSM Score The PSM Score as given by Hecklib to this annotated spectrum. It is shown with three significant figures. | 528 |

#### Spectrum 4698? Spectrum 4698 The raw spectrum of this peptide as annotated by Hecklib. The fragments are coloured according to ion type (see legend). Any peaks with a star '\*' as text can be hovered over to see the full details, first the ion type second the mass shift type. By hovering over the amino acids in the peptide or ions in the legend the corresponding peaks are highlighted. By toggling the 'Unassigned' label you can turn the background (unassigned) peaks on or off in the plot. By updating the slider in the Ion legend you can update the spectrum to only show the top X% of the peaks with labels. The top X% means any peak that is within X% of the highest intensity. By dragging in the spectrum you can zoom in to a specific part of the spectrum and use 'Zoom Out' to get back to the original zoom level. The annotation of the spectrum is based on the given sequence in the peptides file and is done with different software so inconsistencies are likely. The peaks are annotated based on the given sequence, with 20 ppm tolerance.

Copy Data

##### Spectrum 4698 (TSV)

###### Preview

```
Loading example...
```

*Click on the button to copy the data to your clipboard.*

Mz MinMz MaxIntensity Max

WidthHeightPeptide font sizePeptide stroke widthSpectrum font sizeSpectrum stroke widthCompact peptide

Ion legend

wxyz

abcd

OtherUnassignedIon^Charge^_Position_Show for top:%

VFGGGTKJT

03.85e+47.71e+41.16e+51.54e+5

Zoom Out

y^+1^_1_y^+1^_2_y^+1^_2_y^+2^_7_z^+1^_3_y^+1^_3_c^+1^_4_y^+2^_8_y^+2^_8_w^+1^_4_c^+1^_5_y^+1^_4_z^+1^_4_y^+1^_4_z^+1^_5_c^+1^_6_y^+1^_5_c^+1^_6_z^+1^_6_z^+1^_6_y^+1^_6_z^+1^_7_y^+1^_7_z^+1^_7_y^+1^_7_c^+1^_7_c^+1^_7_z^+1^_8_c^+1^_8_z^+1^_8_c^+1^_8_y^+1^_8_

0505100915142019

Fragment Matches Table

Show background peaks

| Position | Ion type | Intensity | mz Theoretical | mz Error (Th) | mz Error (ppm) | Charge | Series Number |
| --- | --- | --- | --- | --- | --- | --- | --- |
| 9 | y | 3691 | 120.1 | 0.0002381 | 1.983 | +1 | 1 |
| - | - | 1.913E+04 | 120.1 | - | - | 0 | - |
| - | - | 1240 | 121.1 | - | - | 0 | - |
| - | - | 352.3 | 127.1 | - | - | 0 | - |
| - | - | 7992 | 129.1 | - | - | 0 | - |
| - | - | 775 | 130.1 | - | - | 0 | - |
| - | - | 401.4 | 137.7 | - | - | 0 | - |
| - | - | 415.2 | 142.8 | - | - | 0 | - |
| - | - | 447.4 | 148.9 | - | - | 0 | - |
| - | - | 470.1 | 148.9 | - | - | 0 | - |
| - | - | 822.2 | 148.9 | - | - | 0 | - |
| - | - | 1036 | 148.9 | - | - | 0 | - |
| - | - | 1003 | 148.9 | - | - | 0 | - |
| - | - | 1508 | 148.9 | - | - | 0 | - |
| - | - | 3160 | 148.9 | - | - | 0 | - |
| - | - | 4820 | 149 | - | - | 0 | - |
| - | - | 3187 | 149 | - | - | 0 | - |
| - | - | 1430 | 149 | - | - | 0 | - |
| - | - | 1035 | 149 | - | - | 0 | - |
| - | - | 925.6 | 149 | - | - | 0 | - |
| - | - | 819.5 | 149 | - | - | 0 | - |
| - | - | 544.6 | 149 | - | - | 0 | - |
| - | - | 470.5 | 149 | - | - | 0 | - |
| - | - | 722.9 | 149 | - | - | 0 | - |
| - | - | 853.6 | 153.1 | - | - | 0 | - |
| - | - | 518.7 | 155.1 | - | - | 0 | - |
| - | - | 417.4 | 161.2 | - | - | 0 | - |
| - | - | 914.6 | 173.1 | - | - | 0 | - |
| - | - | 628.2 | 177.1 | - | - | 0 | - |
| - | - | 541.6 | 182.9 | - | - | 0 | - |
| - | - | 708.3 | 187.1 | - | - | 0 | - |
| - | - | 615.2 | 188.1 | - | - | 0 | - |
| - | - | 2931 | 205.1 | - | - | 0 | - |
| 8 | y | 1086 | 215.1 | 0.0002786 | 1.295 | +1 | 2 |
| - | - | 514 | 215.7 | - | - | 0 | - |
| - | - | 6.913E+04 | 219.1 | - | - | 0 | - |
| - | - | 8943 | 220.2 | - | - | 0 | - |
| - | - | 2241 | 230.2 | - | - | 0 | - |
| 8 | y | 2050 | 233.1 | 0.0003645 | 1.563 | +1 | 2 |
| - | - | 2.653E+04 | 247.1 | - | - | 0 | - |
| - | - | 3935 | 248.1 | - | - | 0 | - |
| - | - | 820.7 | 249.2 | - | - | 0 | - |
| - | - | 620.6 | 255.1 | - | - | 0 | - |
| - | - | 990 | 272.2 | - | - | 0 | - |
| - | - | 3423 | 273.1 | - | - | 0 | - |
| - | - | 532.1 | 280.1 | - | - | 0 | - |
| - | - | 1130 | 290.1 | - | - | 0 | - |
| - | - | 1766 | 301.2 | - | - | 0 | - |
| - | - | 975.8 | 302.2 | - | - | 0 | - |
| - | - | 3494 | 304.2 | - | - | 0 | - |
| - | - | 567.8 | 305.2 | - | - | 0 | - |
| - | - | 1315 | 311.2 | - | - | 0 | - |
| 3 | y | 901.4 | 317.2 | 0.0006708 | 2.115 | +2 | 7 |
| - | - | 1049 | 326.2 | - | - | 0 | - |
| - | - | 685.2 | 328.2 | - | - | 0 | - |
| - | - | 874.3 | 328.2 | - | - | 0 | - |
| - | - | 1354 | 329.2 | - | - | 0 | - |
| - | - | 771.4 | 340.3 | - | - | 0 | - |
| - | - | 621.5 | 341.1 | - | - | 0 | - |
| 7 | z | 1.507E+04 | 345.2 | 0.0003737 | 1.083 | +1 | 3 |
| - | - | 541 | 345.2 | - | - | 0 | - |
| - | - | 3.658E+04 | 346.2 | - | - | 0 | - |
| - | - | 5721 | 347.2 | - | - | 0 | - |
| - | - | 991.3 | 358.2 | - | - | 0 | - |
| - | - | 652.3 | 358.3 | - | - | 0 | - |
| - | - | 3347 | 359.2 | - | - | 0 | - |
| 7 | y | 6226 | 361.2 | 0.0001313 | 0.3635 | +1 | 3 |
| - | - | 1284 | 362.2 | - | - | 0 | - |
| - | - | 1958 | 367.2 | - | - | 0 | - |
| - | - | 625.4 | 368.2 | - | - | 0 | - |
| 4 | c | 605.9 | 378.2 | 0.001171 | 3.097 | +1 | 4 |
| 2 | y | 1386 | 381.7 | 0.0005548 | 1.453 | +2 | 8 |
| - | - | 649.6 | 382.2 | - | - | 0 | - |
| - | - | 2962 | 383.2 | - | - | 0 | - |
| - | - | 1041 | 385.2 | - | - | 0 | - |
| - | - | 3142 | 385.2 | - | - | 0 | - |
| - | - | 713.5 | 386.2 | - | - | 0 | - |
| - | - | 617 | 386.2 | - | - | 0 | - |
| 2 | y | 1.013E+04 | 390.7 | 0.0003384 | 0.8661 | +2 | 8 |
| - | - | 4469 | 391.2 | - | - | 0 | - |
| - | - | 6507 | 401.2 | - | - | 0 | - |
| - | - | 1075 | 402.2 | - | - | 0 | - |
| - | - | 6619 | 402.3 | - | - | 0 | - |
| - | - | 2823 | 403.2 | - | - | 0 | - |
| - | - | 1417 | 403.3 | - | - | 0 | - |
| - | - | 1055 | 415.3 | - | - | 0 | - |
| - | - | 736.3 | 418.2 | - | - | 0 | - |
| - | - | 1300 | 421.8 | - | - | 0 | - |
| 6 | w | 1055 | 431.3 | 0.003566 | 8.269 | +1 | 4 |
| - | - | 1678 | 434.2 | - | - | 0 | - |
| 5 | c | 1101 | 435.2 | 0.001951 | 4.483 | +1 | 5 |
| - | - | 670.2 | 439.3 | - | - | 0 | - |
| - | - | 3009 | 439.8 | - | - | 0 | - |
| - | - | 2293 | 440.3 | - | - | 0 | - |
| - | - | 695.9 | 440.8 | - | - | 0 | - |
| - | - | 1218 | 440.8 | - | - | 0 | - |
| - | - | 706 | 441.2 | - | - | 0 | - |
| - | - | 1159 | 441.3 | - | - | 0 | - |
| - | - | 7274 | 442.3 | - | - | 0 | - |
| - | - | 1098 | 443.3 | - | - | 0 | - |
| - | - | 4678 | 444.3 | - | - | 0 | - |
| 6 | y | 8592 | 445.3 | 4.057E-05 | 0.09111 | +1 | 4 |
| 6 | z | 4846 | 446.3 | 0.001528 | 3.425 | +1 | 4 |
| - | - | 6131 | 447.2 | - | - | 0 | - |
| - | - | 7.269E+04 | 447.3 | - | - | 0 | - |
| - | - | 2211 | 448.2 | - | - | 0 | - |
| - | - | 1.465E+04 | 448.3 | - | - | 0 | - |
| - | - | 2569 | 449.3 | - | - | 0 | - |
| - | - | 787.7 | 455.3 | - | - | 0 | - |
| - | - | 2131 | 457.3 | - | - | 0 | - |
| - | - | 1781 | 459.3 | - | - | 0 | - |
| - | - | 563.5 | 460.3 | - | - | 0 | - |
| 6 | y | 2265 | 462.3 | 0.0005991 | 1.296 | +1 | 4 |
| - | - | 2007 | 470.3 | - | - | 0 | - |
| - | - | 3306 | 472.3 | - | - | 0 | - |
| - | - | 920 | 473.3 | - | - | 0 | - |
| - | - | 2203 | 486.2 | - | - | 0 | - |
| - | - | 2400 | 486.3 | - | - | 0 | - |
| - | - | 726.9 | 487.2 | - | - | 0 | - |
| - | - | 712.9 | 487.3 | - | - | 0 | - |
| - | - | 4694 | 496.3 | - | - | 0 | - |
| - | - | 1162 | 498.3 | - | - | 0 | - |
| - | - | 5509 | 499.3 | - | - | 0 | - |
| - | - | 721 | 500.3 | - | - | 0 | - |
| - | - | 1674 | 501.2 | - | - | 0 | - |
| - | - | 945.6 | 502.2 | - | - | 0 | - |
| 5 | z | 5352 | 503.3 | 0.0004759 | 0.9456 | +1 | 5 |
| - | - | 3.885E+04 | 504.3 | - | - | 0 | - |
| - | - | 9.552E+04 | 504.3 | - | - | 0 | - |
| - | - | 9796 | 505.3 | - | - | 0 | - |
| - | - | 2.339E+04 | 505.3 | - | - | 0 | - |
| - | - | 1506 | 506.3 | - | - | 0 | - |
| - | - | 3677 | 506.3 | - | - | 0 | - |
| - | - | 1480 | 512.3 | - | - | 0 | - |
| - | - | 832.6 | 513.3 | - | - | 0 | - |
| - | - | 1.902E+04 | 514.3 | - | - | 0 | - |
| - | - | 4949 | 515.3 | - | - | 0 | - |
| - | - | 4654 | 516.3 | - | - | 0 | - |
| - | - | 3548 | 517.3 | - | - | 0 | - |
| - | - | 1052 | 517.3 | - | - | 0 | - |
| 6 | c | 2006 | 518.3 | 0.003482 | 6.718 | +1 | 6 |
| - | - | 5796 | 518.3 | - | - | 0 | - |
| - | - | 1604 | 519.3 | - | - | 0 | - |
| 5 | y | 8839 | 519.3 | 0.0008226 | 1.584 | +1 | 5 |
| - | - | 2941 | 520.3 | - | - | 0 | - |
| - | - | 1.086E+04 | 535.3 | - | - | 0 | - |
| 6 | c | 4.764E+04 | 536.3 | 5.285E-05 | 0.09854 | +1 | 6 |
| - | - | 1.211E+04 | 537.3 | - | - | 0 | - |
| - | - | 2033 | 538.3 | - | - | 0 | - |
| 4 | z | 1180 | 542.3 | 0.0005049 | 0.931 | +1 | 6 |
| - | - | 1648 | 543.3 | - | - | 0 | - |
| - | - | 1416 | 548.3 | - | - | 0 | - |
| - | - | 2741 | 555.3 | - | - | 0 | - |
| - | - | 960.2 | 556.3 | - | - | 0 | - |
| - | - | 661.8 | 559.3 | - | - | 0 | - |
| - | - | 955.8 | 559.3 | - | - | 0 | - |
| 4 | z | 1.197E+04 | 560.3 | 0.000344 | 0.6139 | +1 | 6 |
| - | - | 3159 | 561.3 | - | - | 0 | - |
| - | - | 4.496E+04 | 561.3 | - | - | 0 | - |
| - | - | 505.9 | 562.3 | - | - | 0 | - |
| - | - | 1.216E+04 | 562.3 | - | - | 0 | - |
| - | - | 2529 | 563.3 | - | - | 0 | - |
| - | - | 738.4 | 571.4 | - | - | 0 | - |
| - | - | 3924 | 574.3 | - | - | 0 | - |
| - | - | 1025 | 575.3 | - | - | 0 | - |
| - | - | 3747 | 575.3 | - | - | 0 | - |
| 4 | y | 2.615E+04 | 576.3 | 8.491E-06 | 0.01473 | +1 | 6 |
| - | - | 8236 | 577.3 | - | - | 0 | - |
| - | - | 1411 | 578.3 | - | - | 0 | - |
| 3 | z | 1903 | 599.3 | 0.0008505 | 1.419 | +1 | 7 |
| 3 | y | 4254 | 615.3 | 0.0002264 | 0.3679 | +1 | 7 |
| - | - | 1666 | 616.3 | - | - | 0 | - |
| 3 | z | 1.359E+04 | 617.3 | 0.0001205 | 0.1952 | +1 | 7 |
| - | - | 8575 | 618.3 | - | - | 0 | - |
| - | - | 2639 | 619.4 | - | - | 0 | - |
| - | - | 2235 | 620.4 | - | - | 0 | - |
| - | - | 1211 | 632.3 | - | - | 0 | - |
| 3 | y | 1.255E+05 | 633.4 | 0.0003173 | 0.501 | +1 | 7 |
| - | - | 3.84E+04 | 634.4 | - | - | 0 | - |
| - | - | 7850 | 635.4 | - | - | 0 | - |
| - | - | 588.5 | 646.3 | - | - | 0 | - |
| 7 | c | 3238 | 647.4 | 6.273E-05 | 0.0969 | +1 | 7 |
| - | - | 849.8 | 648.4 | - | - | 0 | - |
| - | - | 829.9 | 659.3 | - | - | 0 | - |
| - | - | 954.3 | 659.8 | - | - | 0 | - |
| - | - | 2999 | 660.3 | - | - | 0 | - |
| - | - | 4243 | 661.4 | - | - | 0 | - |
| - | - | 2508 | 662.4 | - | - | 0 | - |
| - | - | 3461 | 663.4 | - | - | 0 | - |
| 7 | c | 1.04E+05 | 664.4 | 0.0003047 | 0.4586 | +1 | 7 |
| - | - | 3.487E+04 | 665.4 | - | - | 0 | - |
| - | - | 7896 | 666.4 | - | - | 0 | - |
| - | - | 765 | 690.3 | - | - | 0 | - |
| - | - | 1172 | 690.4 | - | - | 0 | - |
| - | - | 623.4 | 701.4 | - | - | 0 | - |
| - | - | 919.1 | 702.4 | - | - | 0 | - |
| - | - | 1024 | 718.4 | - | - | 0 | - |
| - | - | 860.3 | 721.3 | - | - | 0 | - |
| - | - | 696 | 732.4 | - | - | 0 | - |
| - | - | 9065 | 733.4 | - | - | 0 | - |
| - | - | 2280 | 734.5 | - | - | 0 | - |
| - | - | 659 | 742.4 | - | - | 0 | - |
| 2 | z | 2428 | 746.4 | 0.0006819 | 0.9136 | +1 | 8 |
| 8 | c | 4849 | 760.4 | 0.001607 | 2.114 | +1 | 8 |
| - | - | 1797 | 761.4 | - | - | 0 | - |
| - | - | 3122 | 762.4 | - | - | 0 | - |
| - | - | 1574 | 763.5 | - | - | 0 | - |
| 2 | z | 1.991E+04 | 764.4 | 0.0004223 | 0.5525 | +1 | 8 |
| - | - | 7088 | 765.4 | - | - | 0 | - |
| - | - | 2329 | 766.4 | - | - | 0 | - |
| - | - | 1086 | 775.4 | - | - | 0 | - |
| 8 | c | 1.527E+05 | 777.5 | 0.000103 | 0.1325 | +1 | 8 |
| - | - | 6.168E+04 | 778.5 | - | - | 0 | - |
| - | - | 1.51E+04 | 779.5 | - | - | 0 | - |
| 2 | y | 1.926E+04 | 780.4 | 0.0002628 | 0.3367 | +1 | 8 |
| - | - | 6580 | 781.4 | - | - | 0 | - |
| - | - | 2264 | 782.4 | - | - | 0 | - |
| - | - | 1197 | 802.5 | - | - | 0 | - |
| - | - | 827.8 | 817.5 | - | - | 0 | - |
| - | - | 700.5 | 823.4 | - | - | 0 | - |
| - | - | 1533 | 824.4 | - | - | 0 | - |
| - | - | 2148 | 834.5 | - | - | 0 | - |
| - | - | 657.2 | 835.4 | - | - | 0 | - |
| - | - | 1082 | 845.4 | - | - | 0 | - |
| - | - | 1012 | 845.5 | - | - | 0 | - |
| - | - | 1857 | 848.4 | - | - | 0 | - |
| - | - | 1205 | 849.5 | - | - | 0 | - |
| - | - | 754.9 | 853.5 | - | - | 0 | - |
| - | - | 696.6 | 854.4 | - | - | 0 | - |
| - | - | 1159 | 861.5 | - | - | 0 | - |
| - | - | 2565 | 862.3 | - | - | 0 | - |
| - | - | 2083 | 862.5 | - | - | 0 | - |
| - | - | 1.192E+05 | 863.5 | - | - | 0 | - |
| - | - | 5.342E+04 | 864.5 | - | - | 0 | - |
| - | - | 789.6 | 865.4 | - | - | 0 | - |
| - | - | 1.463E+04 | 865.5 | - | - | 0 | - |
| - | - | 1390 | 866.5 | - | - | 0 | - |
| - | - | 2654 | 878.5 | - | - | 0 | - |
| - | - | 732.8 | 879 | - | - | 0 | - |
| - | - | 9.698E+04 | 879.5 | - | - | 0 | - |
| - | - | 8.063E+04 | 880.5 | - | - | 0 | - |
| - | - | 1232 | 881.4 | - | - | 0 | - |
| - | - | 2.753E+04 | 881.5 | - | - | 0 | - |
| - | - | 992.7 | 882.4 | - | - | 0 | - |
| - | - | 6240 | 882.5 | - | - | 0 | - |
| - | - | 821.5 | 883.4 | - | - | 0 | - |
| - | - | 1075 | 962.5 | - | - | 0 | - |
| - | - | 729.1 | 1229 | - | - | 0 | - |
| - | - | 1341 | 1303 | - | - | 0 | - |
| - | - | 2389 | 1304 | - | - | 0 | - |
| - | - | 882.8 | 1305 | - | - | 0 | - |
| - | - | 796.9 | 1319 | - | - | 0 | - |
| - | - | 2225 | 1320 | - | - | 0 | - |
| - | - | 1142 | 1321 | - | - | 0 | - |
| - | - | 849.2 | 1322 | - | - | 0 | - |
| - | - | 701.5 | 1323 | - | - | 0 | - |
| - | - | 846.8 | 1999 | - | - | 0 | - |

m/z Charge Intensity FragmentType MassShift Position
120.06575775146484 0 3691.097 y 8
120.08104705810547 0 19126.94
121.0844497680664 0 1239.5117
127.08662414550781 0 352.2893
129.10252380371094 0 7991.7144
130.10597229003906 0 775.0325
137.71043395996094 0 401.4151
142.77149963378906 0 415.20816
148.88992309570312 0 447.3941
148.904052734375 0 470.0596
148.9114227294922 0 822.18726
148.91859436035156 0 1035.9349
148.92568969726562 0 1003.4796
148.93264770507812 0 1508.0035
148.94052124023438 0 3160.1707
148.95704650878906 0 4820.1094
148.96484375 0 3187.409
148.97238159179688 0 1430.2678
148.9792938232422 0 1034.9691
148.98681640625 0 925.6141
148.9936981201172 0 819.5044
149.0006561279297 0 544.6187
149.00856018066406 0 470.52936
149.02354431152344 0 722.8957
153.0777130126953 0 853.61053
155.09320068359375 0 518.65125
161.2008056640625 0 417.41766
173.0920867919922 0 914.56775
177.10252380371094 0 628.22
182.90550231933594 0 541.5522
187.10797119140625 0 708.2593
188.14041137695312 0 615.1581
205.09764099121094 0 2930.5215
215.13929748535156 0 1085.658 y Water loss 7
215.7398681640625 0 514.0387
219.1494598388672 0 69127.35
220.15277099609375 0 8942.792
230.15013122558594 0 2240.8188
233.1499481201172 0 2050.0645 y 7
247.14434814453125 0 26529.744
248.14781188964844 0 3935.37
249.15089416503906 0 820.7001
255.108154296875 0 620.5929
272.1597900390625 0 989.96106
273.11932373046875 0 3423.2122
280.0957336425781 0 532.0538
290.14630126953125 0 1130.3721
301.1876220703125 0 1765.7158
302.19439697265625 0 975.80884
304.1665954589844 0 3494.0906
305.1693420410156 0 567.7772
311.17095947265625 0 1315.1267
317.1826171875 0 901.37335 y 2
326.18304443359375 0 1049.4507
328.1744384765625 0 685.18097
328.2227783203125 0 874.2602
329.1819152832031 0 1353.5978
340.25933837890625 0 771.3725
341.1001892089844 0 621.5174
345.2261962890625 0 15071.617 z 6
345.24798583984375 0 540.9821
346.23370361328125 0 36579.234
347.23675537109375 0 5721.01
358.208251953125 0 991.3334
358.2572937011719 0 652.32965
359.2163391113281 0 3347.4067
361.2444152832031 0 6226.017 y 6
362.2476806640625 0 1283.6495
367.2335205078125 0 1958.4808
368.2381591796875 0 625.35126
378.2147521972656 0 605.9466 c 3
381.71142578125 0 1386.4684 y Water loss 1
382.2127380371094 0 649.59204
383.2036437988281 0 2961.78
385.1958923339844 0 1041.4266
385.2449035644531 0 3142.061
386.2033386230469 0 713.49994
386.2450256347656 0 617.0418
390.71649169921875 0 10134.993 y 1
391.2179260253906 0 4468.7334
401.21453857421875 0 6506.9736
402.21923828125 0 1074.8046
402.2712707519531 0 6619.497
403.2301940917969 0 2823.2385
403.2743835449219 0 1416.7433
415.2795104980469 0 1055.144
418.2080078125 0 736.3399
421.833984375 0 1300.1064
431.2464599609375 0 1055.3499 w 5
434.2279357910156 0 1677.7344
435.23309326171875 0 1101.166 c 4
439.2647705078125 0 670.24
439.8442077636719 0 3008.9685
440.25128173828125 0 2293.1516
440.7542419433594 0 695.8563
440.8460388183594 0 1217.7219
441.2197265625 0 705.9711
441.2554931640625 0 1159.1963
442.2569274902344 0 7273.5435
443.25201416015625 0 1097.6423
444.2584228515625 0 4678.4346
445.2657165527344 0 8591.86 y Ammonia loss 5
446.27197265625 0 4845.9863 z 5
447.2336120605469 0 6131.457
447.2818603515625 0 72690.336
448.2390441894531 0 2211.2905
448.2850036621094 0 14646.159
449.28741455078125 0 2569.4075
455.26055908203125 0 787.7038
457.2767639160156 0 2131.2788
459.2937316894531 0 1781.3835
460.2955322265625 0 563.52496
462.2916259765625 0 2264.5503 y 5
470.2610778808594 0 2007.0674
472.3005676269531 0 3305.9446
473.30279541015625 0 920.0132
486.2439270019531 0 2202.607
486.3018798828125 0 2400.4368
487.249267578125 0 726.9104
487.3043518066406 0 712.9476
496.2882995605469 0 4694.4097
498.31781005859375 0 1162.4769
499.2763366699219 0 5508.575
500.28045654296875 0 721.02405
501.2460632324219 0 1674.2977
502.2486572265625 0 945.5584
503.2954406738281 0 5351.709 z 4
504.2545471191406 0 38848.92
504.3030090332031 0 95523.98
505.2580871582031 0 9796.374
505.30633544921875 0 23386.672
506.2611083984375 0 1505.6389
506.3079833984375 0 3677.2415
512.2826538085938 0 1480.4292
513.287841796875 0 832.588
514.2985229492188 0 19020.48
515.3016357421875 0 4949.0474
516.3128051757812 0 4654.1494
517.2616577148438 0 3548.2815
517.3126831054688 0 1051.9635
518.2686767578125 0 2005.904 c Water loss 5
518.3064575195312 0 5795.995
519.2589721679688 0 1604.4052
519.3128662109375 0 8839.22 y 4
520.3155517578125 0 2941.2249
535.2750244140625 0 10855.145
536.2827758789062 0 47637.26 c 5
537.2860107421875 0 12108.646
538.2882080078125 0 2032.9438
542.3053588867188 0 1180.2919 z Water loss 3
543.2656860351562 0 1647.5096
548.2801513671875 0 1415.9469
555.3372192382812 0 2740.6702
556.3416748046875 0 960.167
559.2675170898438 0 661.75977
559.3211669921875 0 955.8366
560.3167724609375 0 11971.856 z 3
561.2760009765625 0 3158.995
561.3243408203125 0 44955.125
562.278564453125 0 505.8629
562.3272094726562 0 12162.856
563.3300170898438 0 2528.57
571.3578491210938 0 738.3538
574.2833251953125 0 3923.8647
575.2865600585938 0 1024.996
575.3280029296875 0 3747.2893
576.3351440429688 0 26145.885 y 3
577.3383178710938 0 8235.629
578.3412475585938 0 1411.178
599.3264770507812 0 1903.053 z Water loss 2
615.3458251953125 0 4254.0454 y Water loss 2
616.3470458984375 0 1666.4779
617.3380126953125 0 13585.629 z 2
618.3436279296875 0 8574.829
619.353515625 0 2638.6697
620.3622436523438 0 2235.2856
632.34619140625 0 1211.274
633.35693359375 0 125466.57 y 2
634.35986328125 0 38395.43
635.3616333007812 0 7849.98
646.3406372070312 0 588.52313
647.35107421875 0 3238.1113 c Ammonia loss 6
648.3552856445312 0 849.76276
659.3466186523438 0 829.9185
659.8457641601562 0 954.31146
660.3455200195312 0 2999.2908
661.3649291992188 0 4242.978
662.3656616210938 0 2507.5222
663.3812866210938 0 3460.607
664.3779907226562 0 104014.55 c 6
665.3807983398438 0 34869.51
666.3829956054688 0 7895.5513
690.3322143554688 0 764.9563
690.3914184570312 0 1171.8528
701.4288940429688 0 623.3593
702.40380859375 0 919.055
718.4365234375 0 1024.4827
721.3486938476562 0 860.27155
732.4412231445312 0 696.0373
733.4480590820312 0 9064.588
734.45166015625 0 2280.1584
742.4276733398438 0 659.0074
746.3964233398438 0 2427.8467 z Water loss 1
760.43359375 0 4849.094 c Ammonia loss 7
761.4385986328125 0 1796.5208
762.4495239257812 0 3121.9705
763.4528198242188 0 1573.7847
764.4058837890625 0 19910.967 z 1
765.4090576171875 0 7087.889
766.4124145507812 0 2328.868
775.445068359375 0 1086.282
777.4618530273438 0 152671.38 c 7
778.464599609375 0 61682.465
779.4671630859375 0 15104.771
780.42529296875 0 19260.523 y 1
781.4284057617188 0 6579.9214
782.431640625 0 2264.387
802.4611206054688 0 1197.1755
817.4794921875 0 827.8248
823.3696899414062 0 700.5401
824.4395751953125 0 1533.3088
834.4955444335938 0 2148.3677
835.3831787109375 0 657.15643
845.3671875 0 1081.5139
845.4629516601562 0 1012.43146
848.4479370117188 0 1856.9663
849.4547119140625 0 1205.0011
853.519287109375 0 754.87836
854.4285888671875 0 696.6349
861.45947265625 0 1158.7249
862.3079223632812 0 2565.0698
862.48681640625 0 2083.0767
863.4745483398438 0 119232.664
864.4775390625 0 53417.805
865.3941650390625 0 789.6392
865.4804077148438 0 14633.591
866.478515625 0 1389.9801
878.4804077148438 0 2653.788
878.9839477539062 0 732.7796
879.4931030273438 0 96984.82
880.4983520507812 0 80629.01
881.4180908203125 0 1232.3097
881.501953125 0 27526.62
882.4107055664062 0 992.73193
882.5053100585938 0 6240.05
883.383056640625 0 821.4744
962.5004272460938 0 1075.3596
1229.2325439453125 0 729.0745
1302.6719970703125 0 1341.2947
1303.6722412109375 0 2388.8384
1304.67822265625 0 882.7652
1318.6917724609375 0 796.88904
1319.6910400390625 0 2224.5088
1320.7037353515625 0 1141.7279
1321.6866455078125 0 849.1749
1322.688720703125 0 701.5492
1998.9569091796875 0 846.75073

Spectrum Details

|  |  |
| --- | --- |
| Matched peaks? Matched peaks The total absolute number of peaks matched. Additionally in brackets the total fraction of peaks matched and the total number of peaks is shown. | 32 (12.45% of 257) |
| FDR? FDR The false discovery rate estimated for this peptide. It is calculated by matching all theoretical fragments with a non-integer shift with the raw peaks for this spectrum. This is done with 40 different shifts. The resulting percentage is the average number of annotated peaks over the number of annotated peaks with the correct spectrum. | 0.22% |
| Satellite FDR? Satellite FDR See the FDR for details on its calculation. This satellite ion specific FDR only contains the satellite ions (d/w) for I/L/J positions. | ∞ |
| PSM Score? PSM Score The PSM Score as given by Hecklib to this annotated spectrum. It is shown with three significant figures. | 402 |

#### Spectrum 4927? Spectrum 4927 The raw spectrum of this peptide as annotated by Hecklib. The fragments are coloured according to ion type (see legend). Any peaks with a star '\*' as text can be hovered over to see the full details, first the ion type second the mass shift type. By hovering over the amino acids in the peptide or ions in the legend the corresponding peaks are highlighted. By toggling the 'Unassigned' label you can turn the background (unassigned) peaks on or off in the plot. By updating the slider in the Ion legend you can update the spectrum to only show the top X% of the peaks with labels. The top X% means any peak that is within X% of the highest intensity. By dragging in the spectrum you can zoom in to a specific part of the spectrum and use 'Zoom Out' to get back to the original zoom level. The annotation of the spectrum is based on the given sequence in the peptides file and is done with different software so inconsistencies are likely. The peaks are annotated based on the given sequence, with 20 ppm tolerance.

Copy Data

##### Spectrum 4927 (TSV)

###### Preview

```
Loading example...
```

*Click on the button to copy the data to your clipboard.*

Mz MinMz MaxIntensity Max

WidthHeightPeptide font sizePeptide stroke widthSpectrum font sizeSpectrum stroke widthCompact peptide

Ion legend

wxyz

abcd

OtherUnassignedIon^Charge^_Position_Show for top:%

VFGGGTKJT

03.90e+47.79e+41.17e+51.56e+5

Zoom Out

y^+1^_1_y^+1^_2_a^+1^_2_y^+1^_2_b^+1^_2_y^+2^_5_b^+1^_3_y^+2^_7_y^+1^_3_y^+1^_3_y^+2^_8_y^+2^_8_\*\*y^+1^_4_b^+1^_6_y^+1^_5_y^+1^_6_y^+1^_6_y^+1^_7_b^+1^_7_y^+1^_7_b^+1^_7_b^+1^_8_b^+1^_8_y^+1^_8_y^+1^_8_

0769153923083077

Fragment Matches Table

Show background peaks

| Position | Ion type | Intensity | mz Theoretical | mz Error (Th) | mz Error (ppm) | Charge | Series Number |
| --- | --- | --- | --- | --- | --- | --- | --- |
| 9 | y | 9061 | 120.1 | 0.0003297 | 2.746 | +1 | 1 |
| - | - | 1.543E+05 | 120.1 | - | - | 0 | - |
| - | - | 407.6 | 121.1 | - | - | 0 | - |
| - | - | 771.2 | 121.1 | - | - | 0 | - |
| - | - | 1.401E+04 | 121.1 | - | - | 0 | - |
| - | - | 635.9 | 122.1 | - | - | 0 | - |
| - | - | 375.2 | 123.1 | - | - | 0 | - |
| - | - | 391.3 | 124.1 | - | - | 0 | - |
| - | - | 678.6 | 125.1 | - | - | 0 | - |
| - | - | 551.4 | 126.1 | - | - | 0 | - |
| - | - | 590.6 | 126.1 | - | - | 0 | - |
| - | - | 1694 | 127.1 | - | - | 0 | - |
| - | - | 1030 | 127.1 | - | - | 0 | - |
| - | - | 514.5 | 128.1 | - | - | 0 | - |
| - | - | 749.8 | 128.1 | - | - | 0 | - |
| - | - | 832.6 | 129.1 | - | - | 0 | - |
| - | - | 9.114E+04 | 129.1 | - | - | 0 | - |
| - | - | 598.8 | 130.1 | - | - | 0 | - |
| - | - | 1573 | 130.1 | - | - | 0 | - |
| - | - | 827.5 | 130.1 | - | - | 0 | - |
| - | - | 6206 | 130.1 | - | - | 0 | - |
| - | - | 408.2 | 130.2 | - | - | 0 | - |
| - | - | 615.7 | 131 | - | - | 0 | - |
| - | - | 405.1 | 131.1 | - | - | 0 | - |
| - | - | 4195 | 131.1 | - | - | 0 | - |
| - | - | 455.7 | 131.2 | - | - | 0 | - |
| - | - | 1529 | 132.1 | - | - | 0 | - |
| - | - | 1462 | 132.1 | - | - | 0 | - |
| - | - | 758 | 133.1 | - | - | 0 | - |
| - | - | 625.9 | 134 | - | - | 0 | - |
| - | - | 450.9 | 136 | - | - | 0 | - |
| - | - | 4043 | 136.1 | - | - | 0 | - |
| - | - | 688.2 | 139.1 | - | - | 0 | - |
| - | - | 755.1 | 140.1 | - | - | 0 | - |
| - | - | 507.7 | 140.1 | - | - | 0 | - |
| - | - | 1840 | 141.1 | - | - | 0 | - |
| - | - | 1114 | 141.1 | - | - | 0 | - |
| - | - | 1339 | 146.1 | - | - | 0 | - |
| - | - | 419.1 | 146.3 | - | - | 0 | - |
| - | - | 568.1 | 148.9 | - | - | 0 | - |
| - | - | 538.6 | 148.9 | - | - | 0 | - |
| - | - | 759.2 | 148.9 | - | - | 0 | - |
| - | - | 1146 | 148.9 | - | - | 0 | - |
| - | - | 1617 | 148.9 | - | - | 0 | - |
| - | - | 3223 | 148.9 | - | - | 0 | - |
| - | - | 4861 | 149 | - | - | 0 | - |
| - | - | 2998 | 149 | - | - | 0 | - |
| - | - | 1452 | 149 | - | - | 0 | - |
| - | - | 954.1 | 149 | - | - | 0 | - |
| - | - | 759.5 | 149 | - | - | 0 | - |
| - | - | 532.5 | 149 | - | - | 0 | - |
| - | - | 910.5 | 149 | - | - | 0 | - |
| - | - | 967.4 | 149 | - | - | 0 | - |
| - | - | 453.7 | 149.1 | - | - | 0 | - |
| - | - | 650.7 | 151.1 | - | - | 0 | - |
| - | - | 1027 | 152.1 | - | - | 0 | - |
| - | - | 557 | 152.1 | - | - | 0 | - |
| - | - | 928.3 | 153.1 | - | - | 0 | - |
| - | - | 1647 | 153.1 | - | - | 0 | - |
| - | - | 612.7 | 153.1 | - | - | 0 | - |
| - | - | 459.6 | 154 | - | - | 0 | - |
| - | - | 916 | 154.1 | - | - | 0 | - |
| - | - | 1019 | 154.1 | - | - | 0 | - |
| - | - | 1154 | 155.1 | - | - | 0 | - |
| - | - | 1121 | 155.1 | - | - | 0 | - |
| - | - | 2699 | 155.1 | - | - | 0 | - |
| - | - | 473.9 | 156.1 | - | - | 0 | - |
| - | - | 1069 | 158.1 | - | - | 0 | - |
| - | - | 2905 | 159.1 | - | - | 0 | - |
| - | - | 972 | 159.1 | - | - | 0 | - |
| - | - | 549.6 | 162.1 | - | - | 0 | - |
| - | - | 1064 | 166.1 | - | - | 0 | - |
| - | - | 573.1 | 166.1 | - | - | 0 | - |
| - | - | 527.8 | 166.1 | - | - | 0 | - |
| - | - | 569.4 | 167.1 | - | - | 0 | - |
| - | - | 1236 | 170.1 | - | - | 0 | - |
| - | - | 1971 | 171.1 | - | - | 0 | - |
| - | - | 742.8 | 171.1 | - | - | 0 | - |
| - | - | 4583 | 172.1 | - | - | 0 | - |
| - | - | 858.5 | 173.1 | - | - | 0 | - |
| - | - | 3124 | 173.1 | - | - | 0 | - |
| - | - | 1788 | 173.1 | - | - | 0 | - |
| - | - | 1210 | 174.1 | - | - | 0 | - |
| - | - | 672.7 | 175.1 | - | - | 0 | - |
| - | - | 735.4 | 176.1 | - | - | 0 | - |
| - | - | 4085 | 177.1 | - | - | 0 | - |
| - | - | 642.7 | 178 | - | - | 0 | - |
| - | - | 493.8 | 180.1 | - | - | 0 | - |
| - | - | 554.5 | 182.1 | - | - | 0 | - |
| - | - | 1214 | 183.1 | - | - | 0 | - |
| - | - | 687.8 | 185.1 | - | - | 0 | - |
| - | - | 940.1 | 185.1 | - | - | 0 | - |
| - | - | 1010 | 185.1 | - | - | 0 | - |
| - | - | 1580 | 186.1 | - | - | 0 | - |
| - | - | 1635 | 187.1 | - | - | 0 | - |
| - | - | 666.3 | 191.2 | - | - | 0 | - |
| - | - | 599.4 | 194.1 | - | - | 0 | - |
| - | - | 1224 | 195.1 | - | - | 0 | - |
| - | - | 576.8 | 195.2 | - | - | 0 | - |
| - | - | 527.1 | 197.1 | - | - | 0 | - |
| - | - | 3537 | 197.2 | - | - | 0 | - |
| - | - | 1791 | 198.1 | - | - | 0 | - |
| - | - | 710.3 | 200.1 | - | - | 0 | - |
| - | - | 1143 | 201.1 | - | - | 0 | - |
| - | - | 715.4 | 201.1 | - | - | 0 | - |
| - | - | 852.5 | 202.1 | - | - | 0 | - |
| - | - | 554.8 | 202.1 | - | - | 0 | - |
| - | - | 652.7 | 204.1 | - | - | 0 | - |
| - | - | 7025 | 205.1 | - | - | 0 | - |
| - | - | 1432 | 206.1 | - | - | 0 | - |
| - | - | 1405 | 207.1 | - | - | 0 | - |
| - | - | 1326 | 208.1 | - | - | 0 | - |
| - | - | 1478 | 209.1 | - | - | 0 | - |
| - | - | 800.6 | 210.1 | - | - | 0 | - |
| - | - | 6631 | 212.1 | - | - | 0 | - |
| - | - | 1158 | 213.1 | - | - | 0 | - |
| - | - | 526.1 | 214.1 | - | - | 0 | - |
| - | - | 742.5 | 214.2 | - | - | 0 | - |
| - | - | 910.8 | 215.1 | - | - | 0 | - |
| 8 | y | 1401 | 215.1 | 0.0002633 | 1.224 | +1 | 2 |
| - | - | 1057 | 216.1 | - | - | 0 | - |
| - | - | 623.7 | 217 | - | - | 0 | - |
| - | - | 661.3 | 217.1 | - | - | 0 | - |
| 2 | a | 1.137E+05 | 219.1 | 0.0004075 | 1.859 | +1 | 2 |
| - | - | 1.523E+04 | 220.2 | - | - | 0 | - |
| - | - | 877.5 | 221.2 | - | - | 0 | - |
| - | - | 674 | 222.1 | - | - | 0 | - |
| - | - | 603.8 | 223.2 | - | - | 0 | - |
| - | - | 2222 | 224.2 | - | - | 0 | - |
| - | - | 764.4 | 225.1 | - | - | 0 | - |
| - | - | 792.2 | 225.2 | - | - | 0 | - |
| - | - | 748.2 | 226.1 | - | - | 0 | - |
| - | - | 2066 | 226.2 | - | - | 0 | - |
| - | - | 4782 | 227.1 | - | - | 0 | - |
| - | - | 1629 | 228.1 | - | - | 0 | - |
| - | - | 569.8 | 229.1 | - | - | 0 | - |
| - | - | 874 | 229.2 | - | - | 0 | - |
| - | - | 1.152E+04 | 230.2 | - | - | 0 | - |
| - | - | 1663 | 231.2 | - | - | 0 | - |
| - | - | 547.8 | 232.2 | - | - | 0 | - |
| 8 | y | 1758 | 233.1 | 0.0004866 | 2.087 | +1 | 2 |
| - | - | 1195 | 233.2 | - | - | 0 | - |
| - | - | 813.4 | 234.1 | - | - | 0 | - |
| - | - | 527.7 | 237.1 | - | - | 0 | - |
| - | - | 1210 | 240.1 | - | - | 0 | - |
| - | - | 638.7 | 240.1 | - | - | 0 | - |
| - | - | 2118 | 242.2 | - | - | 0 | - |
| - | - | 775.8 | 243.1 | - | - | 0 | - |
| - | - | 586.1 | 243.2 | - | - | 0 | - |
| - | - | 3809 | 245.1 | - | - | 0 | - |
| 2 | b | 3.227E+04 | 247.1 | 0.0004422 | 1.789 | +1 | 2 |
| - | - | 4474 | 248.1 | - | - | 0 | - |
| 5 | y | 1689 | 251.2 | 0.004474 | 17.81 | +2 | 5 |
| - | - | 755 | 252.1 | - | - | 0 | - |
| - | - | 1286 | 253.2 | - | - | 0 | - |
| - | - | 8862 | 255.1 | - | - | 0 | - |
| - | - | 913.6 | 261.1 | - | - | 0 | - |
| - | - | 566.6 | 261.2 | - | - | 0 | - |
| - | - | 2654 | 262.1 | - | - | 0 | - |
| - | - | 6797 | 269.2 | - | - | 0 | - |
| - | - | 806.2 | 270.2 | - | - | 0 | - |
| - | - | 707.6 | 271.1 | - | - | 0 | - |
| - | - | 1422 | 272.1 | - | - | 0 | - |
| - | - | 1.306E+04 | 273.1 | - | - | 0 | - |
| - | - | 549.6 | 273.1 | - | - | 0 | - |
| - | - | 1327 | 274.1 | - | - | 0 | - |
| - | - | 1893 | 287.2 | - | - | 0 | - |
| - | - | 928.9 | 290.1 | - | - | 0 | - |
| - | - | 611.3 | 290.2 | - | - | 0 | - |
| - | - | 1076 | 291.1 | - | - | 0 | - |
| - | - | 735.6 | 301.1 | - | - | 0 | - |
| 3 | b | 3636 | 304.2 | 0.0004171 | 1.371 | +1 | 3 |
| 3 | y | 637.8 | 317.2 | 0.005523 | 17.41 | +2 | 7 |
| - | - | 1233 | 319.1 | - | - | 0 | - |
| - | - | 1065 | 325.2 | - | - | 0 | - |
| - | - | 5798 | 326.2 | - | - | 0 | - |
| - | - | 1747 | 339.2 | - | - | 0 | - |
| - | - | 1099 | 341.2 | - | - | 0 | - |
| 7 | y | 1908 | 343.2 | 0.0004846 | 1.412 | +1 | 3 |
| - | - | 1312 | 344.2 | - | - | 0 | - |
| - | - | 745.2 | 348.2 | - | - | 0 | - |
| - | - | 885.6 | 357.2 | - | - | 0 | - |
| - | - | 636.4 | 358.2 | - | - | 0 | - |
| 7 | y | 8323 | 361.2 | 5.178E-05 | 0.1433 | +1 | 3 |
| - | - | 1363 | 362.2 | - | - | 0 | - |
| - | - | 3138 | 365.2 | - | - | 0 | - |
| - | - | 702.8 | 366.2 | - | - | 0 | - |
| - | - | 572.1 | 366.7 | - | - | 0 | - |
| - | - | 1416 | 368.2 | - | - | 0 | - |
| - | - | 784.5 | 375.2 | - | - | 0 | - |
| 2 | y | 1121 | 381.7 | 0.0006659 | 1.744 | +2 | 8 |
| - | - | 681 | 382.2 | - | - | 0 | - |
| - | - | 2084 | 382.2 | - | - | 0 | - |
| - | - | 1.288E+04 | 383.2 | - | - | 0 | - |
| - | - | 1984 | 384.2 | - | - | 0 | - |
| - | - | 701.7 | 387.2 | - | - | 0 | - |
| 2 | y | 1.305E+04 | 390.7 | 0.0003079 | 0.788 | +2 | 8 |
| - | - | 4423 | 391.2 | - | - | 0 | - |
| - | - | 1035 | 391.7 | - | - | 0 | - |
| - | - | 819.7 | 392.2 | - | - | 0 | - |
| - | - | 2335 | 400.3 | - | - | 0 | - |
| - | - | 1.254E+04 | 401.2 | - | - | 0 | - |
| - | - | 3940 | 402.2 | - | - | 0 | - |
| - | - | 1337 | 402.2 | - | - | 0 | - |
| - | - | 800.5 | 403.2 | - | - | 0 | - |
| - | - | 702.9 | 419.2 | - | - | 0 | - |
| - | - | 2128 | 420.2 | - | - | 0 | - |
| - | - | 1107 | 421.8 | - | - | 0 | - |
| - | - | 645.1 | 429.3 | - | - | 0 | - |
| - | - | 730.1 | 430.3 | - | - | 0 | - |
| 0 | Precursor | 873.8 | 431.2 | 0.001107 | 2.568 | +2 | -1 |
| - | - | 672.7 | 439.2 | - | - | 0 | - |
| - | - | 2638 | 439.3 | - | - | 0 | - |
| - | - | 8897 | 439.8 | - | - | 0 | - |
| 0 | Precursor | 1752 | 440.3 | 0.0008909 | 2.024 | +2 | -1 |
| - | - | 947.4 | 440.8 | - | - | 0 | - |
| - | - | 3462 | 440.8 | - | - | 0 | - |
| - | - | 1213 | 441.3 | - | - | 0 | - |
| - | - | 586.6 | 450.3 | - | - | 0 | - |
| - | - | 1050 | 452.3 | - | - | 0 | - |
| - | - | 733.2 | 456.2 | - | - | 0 | - |
| - | - | 5203 | 457.3 | - | - | 0 | - |
| - | - | 1219 | 458.3 | - | - | 0 | - |
| 6 | y | 2056 | 462.3 | 0.0007131 | 1.543 | +1 | 4 |
| - | - | 666.9 | 463.3 | - | - | 0 | - |
| - | - | 1324 | 470.3 | - | - | 0 | - |
| - | - | 1925 | 478.3 | - | - | 0 | - |
| - | - | 629 | 480.3 | - | - | 0 | - |
| - | - | 7599 | 486.3 | - | - | 0 | - |
| - | - | 1966 | 487.3 | - | - | 0 | - |
| - | - | 1731 | 490.3 | - | - | 0 | - |
| - | - | 811.3 | 491.3 | - | - | 0 | - |
| - | - | 1.69E+04 | 496.3 | - | - | 0 | - |
| - | - | 3715 | 497.3 | - | - | 0 | - |
| 6 | b | 1042 | 501.2 | 0.001835 | 3.661 | +1 | 6 |
| - | - | 587.9 | 506.2 | - | - | 0 | - |
| - | - | 4.259E+04 | 514.3 | - | - | 0 | - |
| - | - | 1.073E+04 | 515.3 | - | - | 0 | - |
| - | - | 1616 | 516.3 | - | - | 0 | - |
| 5 | y | 3242 | 519.3 | 2.914E-05 | 0.05612 | +1 | 5 |
| - | - | 833 | 520.3 | - | - | 0 | - |
| - | - | 984.3 | 522.3 | - | - | 0 | - |
| - | - | 1765 | 530.3 | - | - | 0 | - |
| - | - | 918.3 | 532.3 | - | - | 0 | - |
| - | - | 3381 | 548.3 | - | - | 0 | - |
| - | - | 1098 | 549.3 | - | - | 0 | - |
| 4 | y | 718.6 | 558.3 | 0.000247 | 0.4424 | +1 | 6 |
| - | - | 688 | 571.3 | - | - | 0 | - |
| - | - | 828.9 | 571.4 | - | - | 0 | - |
| 4 | y | 1.01E+04 | 576.3 | 5.254E-05 | 0.09117 | +1 | 6 |
| - | - | 3052 | 577.3 | - | - | 0 | - |
| - | - | 633.3 | 587.3 | - | - | 0 | - |
| - | - | 1425 | 597.3 | - | - | 0 | - |
| 3 | y | 7535 | 615.3 | 0.0005671 | 0.9216 | +1 | 7 |
| - | - | 1926 | 616.4 | - | - | 0 | - |
| - | - | 761 | 617.4 | - | - | 0 | - |
| 7 | b | 1141 | 629.3 | 0.001156 | 1.836 | +1 | 7 |
| - | - | 660.6 | 630.3 | - | - | 0 | - |
| - | - | 647.4 | 631.3 | - | - | 0 | - |
| 3 | y | 1.012E+05 | 633.4 | 0.0006225 | 0.9829 | +1 | 7 |
| - | - | 3.079E+04 | 634.4 | - | - | 0 | - |
| - | - | 6495 | 635.4 | - | - | 0 | - |
| - | - | 677.5 | 636.4 | - | - | 0 | - |
| - | - | 4239 | 643.4 | - | - | 0 | - |
| - | - | 1619 | 644.4 | - | - | 0 | - |
| 7 | b | 2837 | 647.4 | 5.934E-05 | 0.09167 | +1 | 7 |
| - | - | 1023 | 648.4 | - | - | 0 | - |
| - | - | 865.6 | 659.3 | - | - | 0 | - |
| - | - | 8553 | 661.4 | - | - | 0 | - |
| - | - | 3306 | 662.4 | - | - | 0 | - |
| - | - | 796.8 | 663.4 | - | - | 0 | - |
| 8 | b | 976.3 | 742.4 | 0.004197 | 5.653 | +1 | 8 |
| 8 | b | 3821 | 760.4 | 0.0002644 | 0.3477 | +1 | 8 |
| - | - | 1168 | 761.4 | - | - | 0 | - |
| 2 | y | 1392 | 762.4 | 0.0008177 | 1.073 | +1 | 8 |
| 2 | y | 2.545E+04 | 780.4 | 0.0001407 | 0.1803 | +1 | 8 |
| - | - | 1.135E+04 | 781.4 | - | - | 0 | - |
| - | - | 3269 | 782.4 | - | - | 0 | - |
| - | - | 836.5 | 788.4 | - | - | 0 | - |
| - | - | 959.6 | 790.4 | - | - | 0 | - |
| - | - | 613.1 | 1061 | - | - | 0 | - |
| - | - | 619.6 | 1219 | - | - | 0 | - |
| - | - | 603.6 | 1594 | - | - | 0 | - |
| - | - | 657.9 | 1861 | - | - | 0 | - |
| - | - | 723.7 | 2133 | - | - | 0 | - |
| - | - | 646.9 | 2414 | - | - | 0 | - |
| - | - | 839 | 3047 | - | - | 0 | - |

m/z Charge Intensity FragmentType MassShift Position
120.06584930419922 0 9060.634 y 8
120.08118438720703 0 154344.8
121.06926727294922 0 407.63217
121.07923889160156 0 771.2429
121.0844955444336 0 14009.39
122.0876235961914 0 635.8926
123.11699676513672 0 375.1556
124.08714294433594 0 391.26065
125.07152557373047 0 678.5868
126.05545806884766 0 551.37103
126.10287475585938 0 590.6308
127.05057525634766 0 1694.2313
127.0871353149414 0 1029.907
128.08197021484375 0 514.5008
128.10726928710938 0 749.81116
129.06607055664062 0 832.6488
129.10263061523438 0 91143.35
130.0504608154297 0 598.8455
130.06549072265625 0 1572.6471
130.10025024414062 0 827.4603
130.10595703125 0 6205.8765
130.1694793701172 0 408.20657
131.04515075683594 0 615.6539
131.07029724121094 0 405.0817
131.08180236816406 0 4194.9556
131.17303466796875 0 455.74814
132.0812530517578 0 1528.9019
132.10218811035156 0 1462.2122
133.06114196777344 0 758.04095
134.0271453857422 0 625.8794
136.01893615722656 0 450.92422
136.0760498046875 0 4042.7576
139.08734130859375 0 688.2208
140.08213806152344 0 755.0729
140.1070556640625 0 507.65204
141.06629943847656 0 1839.7812
141.10279846191406 0 1113.6532
146.06057739257812 0 1338.7202
146.32325744628906 0 419.1416
148.89688110351562 0 568.0842
148.91168212890625 0 538.55286
148.91888427734375 0 759.15344
148.9261474609375 0 1145.9587
148.933349609375 0 1616.6412
148.94097900390625 0 3222.649
148.9575958251953 0 4860.924
148.96539306640625 0 2997.7227
148.97280883789062 0 1452.2572
148.97982788085938 0 954.11237
148.98765563964844 0 759.4585
148.99484252929688 0 532.46625
149.001953125 0 910.48987
149.02369689941406 0 967.44916
149.0531463623047 0 453.691
151.08663940429688 0 650.66736
152.0709228515625 0 1027.4037
152.14349365234375 0 557.02966
153.06613159179688 0 928.30444
153.07733154296875 0 1646.7466
153.1023406982422 0 612.67236
154.0233612060547 0 459.59772
154.0613555908203 0 916.0374
154.09793090820312 0 1018.51746
155.08157348632812 0 1153.786
155.0930938720703 0 1120.9459
155.11817932128906 0 2698.7544
156.07749938964844 0 473.89453
158.09275817871094 0 1069.1337
159.07684326171875 0 2905.4673
159.0918731689453 0 971.9857
162.05551147460938 0 549.61676
166.0535888671875 0 1063.8868
166.0611572265625 0 573.1428
166.09779357910156 0 527.81885
167.11795043945312 0 569.4145
170.09278869628906 0 1235.8977
171.07664489746094 0 1971.1675
171.11300659179688 0 742.7611
172.072021484375 0 4582.5547
173.07139587402344 0 858.5028
173.09251403808594 0 3123.7808
173.1289825439453 0 1788.4078
174.05538940429688 0 1210.1696
175.08648681640625 0 672.7454
176.10780334472656 0 735.3689
177.1025848388672 0 4084.8152
178.0499267578125 0 642.6939
180.07762145996094 0 493.84918
182.0928497314453 0 554.51324
183.1129150390625 0 1214.1384
185.0557861328125 0 687.83905
185.09278869628906 0 940.1419
185.12857055664062 0 1010.4861
186.124267578125 0 1580.492
187.10829162597656 0 1635.3323
191.1542205810547 0 666.33594
194.12901306152344 0 599.4331
195.1129150390625 0 1224.323
195.15008544921875 0 576.77106
197.12887573242188 0 527.0599
197.16517639160156 0 3537.3792
198.0880584716797 0 1791.1427
200.10382080078125 0 710.30566
201.0874786376953 0 1142.5287
201.1232452392578 0 715.38074
202.08258056640625 0 852.53925
202.1079559326172 0 554.76807
204.13438415527344 0 652.6779
205.09750366210938 0 7025.0757
206.10092163085938 0 1432.3418
207.14964294433594 0 1405.0804
208.1084442138672 0 1326.2168
209.10336303710938 0 1478.1986
210.08767700195312 0 800.5622
212.13966369628906 0 6631.4316
213.12344360351562 0 1157.8309
214.11898803710938 0 526.13684
214.1911163330078 0 742.4848
215.1145477294922 0 910.8478
215.1392822265625 0 1400.5455 y Water loss 7
216.0981903076172 0 1056.9215
217.03611755371094 0 623.6929
217.0970001220703 0 661.3064
219.14959716796875 0 113740.836 a 1
220.15289306640625 0 15232.093
221.1566619873047 0 877.5316
222.1244354248047 0 674.00616
223.15516662597656 0 603.7669
224.17611694335938 0 2221.9482
225.13470458984375 0 764.44086
225.1599578857422 0 792.19617
226.11883544921875 0 748.2374
226.15516662597656 0 2065.6877
227.11422729492188 0 4782.364
228.0981903076172 0 1629.2971
229.093017578125 0 569.78925
229.1553192138672 0 874.03406
230.15040588378906 0 11518.195
231.15325927734375 0 1662.9191
232.18321228027344 0 547.80646
233.1500701904297 0 1758.3942 y 7
233.16470336914062 0 1195.3633
234.12359619140625 0 813.3618
237.0991668701172 0 527.6783
240.13467407226562 0 1210.4935
240.1467742919922 0 638.72955
242.18670654296875 0 2118.0295
243.14593505859375 0 775.8304
243.1900634765625 0 586.1269
245.12477111816406 0 3809.3262
247.14454650878906 0 32274.773 b 1
248.1481475830078 0 4473.6416
251.15072631835938 0 1689.2871 y Water loss 4
252.13491821289062 0 754.9622
253.1664581298828 0 1285.9456
255.10922241210938 0 8862.073
261.12396240234375 0 913.64307
261.1610412597656 0 566.60406
262.1185302734375 0 2654.3284
269.16119384765625 0 6797.441
270.17987060546875 0 806.23425
271.1073303222656 0 707.6435
272.13525390625 0 1421.7223
273.11962890625 0 13057.246
273.1344299316406 0 549.5989
274.12030029296875 0 1326.6053
287.1717834472656 0 1892.5521
290.1464538574219 0 928.91833
290.1864929199219 0 611.29865
291.1470031738281 0 1075.5236
301.12994384765625 0 735.5975
304.1659851074219 0 3635.912 b 2
317.1874694824219 0 637.7652 y 2
319.1397705078125 0 1233.0594
325.2236633300781 0 1064.988
326.18292236328125 0 5797.71
339.17791748046875 0 1746.7903
341.18310546875 0 1099.2349
343.2344665527344 0 1908.2234 y Water loss 6
344.19268798828125 0 1312.2538
348.16583251953125 0 745.19275
357.156494140625 0 885.6038
358.2080383300781 0 636.38025
361.2445983886719 0 8323.306 y 6
362.2471008300781 0 1363.4485
365.19268798828125 0 3138.0334
366.1753845214844 0 702.8244
366.7264099121094 0 572.06036
368.193359375 0 1415.9308
375.16162109375 0 784.4926
381.710205078125 0 1120.9904 y Water loss 1
382.21453857421875 0 680.9973
382.2456970214844 0 2084.4634
383.2041015625 0 12877.23
384.2076416015625 0 1983.6226
387.20220947265625 0 701.6555
390.7164611816406 0 13053.006 y 1
391.21820068359375 0 4423.1797
391.7193298339844 0 1035.1273
392.1931457519531 0 819.70984
400.2565002441406 0 2334.608
401.2149658203125 0 12535.337
402.177978515625 0 3939.8098
402.2178649902344 0 1337.3875
403.18035888671875 0 800.51434
419.20574951171875 0 702.89734
420.18829345703125 0 2127.6982
421.83258056640625 0 1107.0099
429.28082275390625 0 645.1465
430.28857421875 0 730.07794
431.2461853027344 0 873.8214 Precursor Water loss
439.1726379394531 0 672.683
439.26544189453125 0 2638.433
439.8441467285156 0 8896.647
440.2512512207031 0 1752.2764 Precursor
440.7548522949219 0 947.36316
440.8445129394531 0 3461.8357
441.25653076171875 0 1212.6204
450.28338623046875 0 586.6482
452.2628173828125 0 1050.1818
456.22332763671875 0 733.21075
457.27764892578125 0 5203.0615
458.2798156738281 0 1218.8575
462.2929382324219 0 2056.2498 y 5
463.29461669921875 0 666.8539
470.2738037109375 0 1323.6821
478.2781677246094 0 1925.1323
480.2607421875 0 628.9898
486.3041687011719 0 7598.648
487.30645751953125 0 1966.3641
490.2752380371094 0 1731.0529
491.2763366699219 0 811.34064
496.2881774902344 0 16896.605
497.2910461425781 0 3714.6987
501.2437744140625 0 1041.9484 b Water loss 5
506.2325439453125 0 587.8589
514.2987670898438 0 42590.99
515.3016967773438 0 10730.286
516.3043823242188 0 1615.6989
519.3136596679688 0 3241.8047 y 4
520.3148193359375 0 833.0248
522.287109375 0 984.2505
530.2742919921875 0 1764.7787
532.3082885742188 0 918.2505
548.282958984375 0 3381.3228
549.2857666015625 0 1098.4817
558.3243408203125 0 718.6417 y Water loss 3
571.3145751953125 0 688.0208
571.3597412109375 0 828.9134
576.335205078125 0 10101.076 y 3
577.3379516601562 0 3051.525
587.348876953125 0 633.33954
597.3368530273438 0 1424.7488
615.3466186523438 0 7534.924 y Water loss 2
616.3500366210938 0 1925.9395
617.355224609375 0 761.0071
629.3394165039062 0 1141.4431 b Water loss 6
630.3408203125 0 660.6248
631.3255004882812 0 647.3652
633.3572387695312 0 101163.766 y 2
634.3603515625 0 30788.783
635.361572265625 0 6494.6006
636.3591918945312 0 677.4631
643.3568115234375 0 4239.0293
644.3587036132812 0 1619.2761
647.3511962890625 0 2836.9734 b 6
648.3573608398438 0 1022.53644
659.31689453125 0 865.60016
661.3663940429688 0 8553.36
662.369873046875 0 3305.5903
663.3740234375 0 796.789
742.4288330078125 0 976.2735 b Water loss 7
760.4349365234375 0 3820.881 b 7
761.4420166015625 0 1167.6034
762.415283203125 0 1391.6713 y Water loss 1
780.4251708984375 0 25453.266 y 1
781.42822265625 0 11353.093
782.4318237304688 0 3269.18
788.3626708984375 0 836.472
790.4090576171875 0 959.60077
1060.8607177734375 0 613.0872
1219.4566650390625 0 619.5688
1594.147216796875 0 603.58746
1861.033203125 0 657.94653
2132.991455078125 0 723.661
2414.223876953125 0 646.9244
3046.755615234375 0 838.99664

Spectrum Details

|  |  |
| --- | --- |
| Matched peaks? Matched peaks The total absolute number of peaks matched. Additionally in brackets the total fraction of peaks matched and the total number of peaks is shown. | 27 (9.41% of 287) |
| FDR? FDR The false discovery rate estimated for this peptide. It is calculated by matching all theoretical fragments with a non-integer shift with the raw peaks for this spectrum. This is done with 40 different shifts. The resulting percentage is the average number of annotated peaks over the number of annotated peaks with the correct spectrum. | 0.44% |
| Satellite FDR? Satellite FDR See the FDR for details on its calculation. This satellite ion specific FDR only contains the satellite ions (d/w) for I/L/J positions. | - |
| PSM Score? PSM Score The PSM Score as given by Hecklib to this annotated spectrum. It is shown with three significant figures. | 313 |

#### Spectrum 4999? Spectrum 4999 The raw spectrum of this peptide as annotated by Hecklib. The fragments are coloured according to ion type (see legend). Any peaks with a star '\*' as text can be hovered over to see the full details, first the ion type second the mass shift type. By hovering over the amino acids in the peptide or ions in the legend the corresponding peaks are highlighted. By toggling the 'Unassigned' label you can turn the background (unassigned) peaks on or off in the plot. By updating the slider in the Ion legend you can update the spectrum to only show the top X% of the peaks with labels. The top X% means any peak that is within X% of the highest intensity. By dragging in the spectrum you can zoom in to a specific part of the spectrum and use 'Zoom Out' to get back to the original zoom level. The annotation of the spectrum is based on the given sequence in the peptides file and is done with different software so inconsistencies are likely. The peaks are annotated based on the given sequence, with 20 ppm tolerance.

Copy Data

##### Spectrum 4999 (TSV)

###### Preview

```
Loading example...
```

*Click on the button to copy the data to your clipboard.*

Mz MinMz MaxIntensity Max

WidthHeightPeptide font sizePeptide stroke widthSpectrum font sizeSpectrum stroke widthCompact peptide

Ion legend

wxyz

abcd

OtherUnassignedIon^Charge^_Position_Show for top:%

VFGGGTKJT

09.52e+31.90e+42.86e+43.81e+4

Zoom Out

y^+1^_1_z^+1^_3_y^+1^_3_y^+2^_8_y^+1^_4_z^+1^_4_z^+1^_5_c^+1^_6_y^+1^_5_c^+1^_6_z^+1^_6_y^+1^_6_z^+1^_7_y^+1^_7_z^+1^_7_y^+1^_7_c^+1^_7_z^+1^_8_c^+1^_8_z^+1^_8_c^+1^_8_y^+1^_8_

0812162524373250

Fragment Matches Table

Show background peaks

| Position | Ion type | Intensity | mz Theoretical | mz Error (Th) | mz Error (ppm) | Charge | Series Number |
| --- | --- | --- | --- | --- | --- | --- | --- |
| 9 | y | 1207 | 120.1 | 0.0002839 | 2.365 | +1 | 1 |
| - | - | 4993 | 120.1 | - | - | 0 | - |
| - | - | 380.8 | 124 | - | - | 0 | - |
| - | - | 386.9 | 126.8 | - | - | 0 | - |
| - | - | 2414 | 129.1 | - | - | 0 | - |
| - | - | 995.9 | 133.1 | - | - | 0 | - |
| - | - | 407.1 | 136.1 | - | - | 0 | - |
| - | - | 379.4 | 151.6 | - | - | 0 | - |
| - | - | 1073 | 153.1 | - | - | 0 | - |
| - | - | 791.7 | 155.1 | - | - | 0 | - |
| - | - | 2360 | 173.4 | - | - | 0 | - |
| - | - | 545.9 | 205.1 | - | - | 0 | - |
| - | - | 598.9 | 212.1 | - | - | 0 | - |
| - | - | 1383 | 213.1 | - | - | 0 | - |
| - | - | 1.613E+04 | 219.1 | - | - | 0 | - |
| - | - | 2246 | 220.2 | - | - | 0 | - |
| - | - | 671.1 | 229.2 | - | - | 0 | - |
| - | - | 673.6 | 231.2 | - | - | 0 | - |
| - | - | 6550 | 247.1 | - | - | 0 | - |
| - | - | 756.1 | 248.1 | - | - | 0 | - |
| - | - | 809.5 | 273.1 | - | - | 0 | - |
| - | - | 568.1 | 294.1 | - | - | 0 | - |
| - | - | 678.9 | 322.1 | - | - | 0 | - |
| - | - | 1089 | 327.2 | - | - | 0 | - |
| 7 | z | 4474 | 345.2 | 0.0001601 | 0.4638 | +1 | 3 |
| - | - | 958.4 | 346.2 | - | - | 0 | - |
| - | - | 9747 | 346.2 | - | - | 0 | - |
| - | - | 1200 | 347.2 | - | - | 0 | - |
| - | - | 1060 | 359.2 | - | - | 0 | - |
| 7 | y | 1819 | 361.2 | 3.978E-05 | 0.1101 | +1 | 3 |
| - | - | 1137 | 385.2 | - | - | 0 | - |
| - | - | 546.1 | 386.2 | - | - | 0 | - |
| 2 | y | 2577 | 390.7 | 0.0001194 | 0.3055 | +2 | 8 |
| - | - | 1198 | 391.2 | - | - | 0 | - |
| - | - | 1642 | 401.2 | - | - | 0 | - |
| - | - | 1802 | 402.3 | - | - | 0 | - |
| - | - | 789.4 | 403.8 | - | - | 0 | - |
| - | - | 1004 | 421.8 | - | - | 0 | - |
| - | - | 586.8 | 422.8 | - | - | 0 | - |
| - | - | 829.1 | 434.2 | - | - | 0 | - |
| - | - | 3527 | 439.8 | - | - | 0 | - |
| - | - | 564.9 | 440.3 | - | - | 0 | - |
| - | - | 1147 | 440.8 | - | - | 0 | - |
| - | - | 620.7 | 441.3 | - | - | 0 | - |
| - | - | 885.4 | 441.3 | - | - | 0 | - |
| - | - | 1894 | 442.3 | - | - | 0 | - |
| - | - | 1881 | 444.3 | - | - | 0 | - |
| 6 | y | 2406 | 445.3 | 0.0002237 | 0.5023 | +1 | 4 |
| 6 | z | 923.9 | 446.3 | 0.00162 | 3.63 | +1 | 4 |
| - | - | 1633 | 447.2 | - | - | 0 | - |
| - | - | 1.86E+04 | 447.3 | - | - | 0 | - |
| - | - | 4072 | 448.3 | - | - | 0 | - |
| - | - | 574.3 | 457.2 | - | - | 0 | - |
| - | - | 1217 | 472.3 | - | - | 0 | - |
| - | - | 624.9 | 486.2 | - | - | 0 | - |
| - | - | 1715 | 496.3 | - | - | 0 | - |
| - | - | 1642 | 499.3 | - | - | 0 | - |
| - | - | 730.6 | 501.2 | - | - | 0 | - |
| 5 | z | 1107 | 503.3 | 1.235E-05 | 0.02453 | +1 | 5 |
| - | - | 9390 | 504.3 | - | - | 0 | - |
| - | - | 2.671E+04 | 504.3 | - | - | 0 | - |
| - | - | 3311 | 505.3 | - | - | 0 | - |
| - | - | 6304 | 505.3 | - | - | 0 | - |
| - | - | 692.9 | 506.3 | - | - | 0 | - |
| - | - | 5139 | 514.3 | - | - | 0 | - |
| - | - | 1409 | 515.3 | - | - | 0 | - |
| - | - | 1298 | 516.3 | - | - | 0 | - |
| - | - | 921.7 | 517.3 | - | - | 0 | - |
| 6 | c | 706.4 | 518.3 | 0.006839 | 13.19 | +1 | 6 |
| - | - | 1326 | 518.3 | - | - | 0 | - |
| 5 | y | 3034 | 519.3 | 0.0008226 | 1.584 | +1 | 5 |
| - | - | 672.6 | 520.3 | - | - | 0 | - |
| - | - | 3356 | 535.3 | - | - | 0 | - |
| 6 | c | 1.228E+04 | 536.3 | 0.0001913 | 0.3567 | +1 | 6 |
| - | - | 3150 | 537.3 | - | - | 0 | - |
| - | - | 646.7 | 538.3 | - | - | 0 | - |
| 4 | z | 2755 | 560.3 | 0.0003274 | 0.5843 | +1 | 6 |
| - | - | 769.7 | 561.3 | - | - | 0 | - |
| - | - | 1.252E+04 | 561.3 | - | - | 0 | - |
| - | - | 2525 | 562.3 | - | - | 0 | - |
| - | - | 944.1 | 574.3 | - | - | 0 | - |
| 4 | y | 7011 | 576.3 | 0.0003137 | 0.5442 | +1 | 6 |
| - | - | 1594 | 577.3 | - | - | 0 | - |
| 3 | z | 961.5 | 599.3 | 0.001827 | 3.048 | +1 | 7 |
| 3 | y | 1280 | 615.3 | 0.001055 | 1.715 | +1 | 7 |
| 3 | z | 3051 | 617.3 | 0.0005509 | 0.8923 | +1 | 7 |
| - | - | 1765 | 618.3 | - | - | 0 | - |
| - | - | 1023 | 619.4 | - | - | 0 | - |
| - | - | 793.2 | 632.3 | - | - | 0 | - |
| 3 | y | 3.102E+04 | 633.4 | 7.32E-05 | 0.1156 | +1 | 7 |
| - | - | 1.01E+04 | 634.4 | - | - | 0 | - |
| - | - | 2074 | 635.4 | - | - | 0 | - |
| - | - | 899.3 | 659.9 | - | - | 0 | - |
| - | - | 737 | 660.3 | - | - | 0 | - |
| - | - | 1546 | 661.4 | - | - | 0 | - |
| - | - | 1623 | 663.4 | - | - | 0 | - |
| 7 | c | 2.793E+04 | 664.4 | 0.0001826 | 0.2748 | +1 | 7 |
| - | - | 9414 | 665.4 | - | - | 0 | - |
| - | - | 2419 | 666.4 | - | - | 0 | - |
| - | - | 1611 | 716.2 | - | - | 0 | - |
| - | - | 848.9 | 717.2 | - | - | 0 | - |
| - | - | 1890 | 733.4 | - | - | 0 | - |
| - | - | 905.6 | 734.5 | - | - | 0 | - |
| 2 | z | 1229 | 746.4 | 0.004933 | 6.61 | +1 | 8 |
| 8 | c | 850 | 760.4 | 0.002523 | 3.317 | +1 | 8 |
| - | - | 1014 | 762.5 | - | - | 0 | - |
| 2 | z | 4983 | 764.4 | 0.0002392 | 0.3129 | +1 | 8 |
| - | - | 1695 | 765.4 | - | - | 0 | - |
| 8 | c | 3.771E+04 | 777.5 | 1.907E-05 | 0.02453 | +1 | 8 |
| - | - | 1.738E+04 | 778.5 | - | - | 0 | - |
| - | - | 4046 | 779.5 | - | - | 0 | - |
| 2 | y | 4427 | 780.4 | 4.238E-05 | 0.05431 | +1 | 8 |
| - | - | 2001 | 781.4 | - | - | 0 | - |
| - | - | 1138 | 789.4 | - | - | 0 | - |
| - | - | 687.8 | 805.4 | - | - | 0 | - |
| - | - | 670.3 | 820.4 | - | - | 0 | - |
| - | - | 595.8 | 836.3 | - | - | 0 | - |
| - | - | 667.5 | 840.5 | - | - | 0 | - |
| - | - | 3.152E+04 | 863.5 | - | - | 0 | - |
| - | - | 1007 | 864.4 | - | - | 0 | - |
| - | - | 1.398E+04 | 864.5 | - | - | 0 | - |
| - | - | 4415 | 865.5 | - | - | 0 | - |
| - | - | 883.2 | 877.4 | - | - | 0 | - |
| - | - | 3743 | 878.4 | - | - | 0 | - |
| - | - | 2.36E+04 | 879.5 | - | - | 0 | - |
| - | - | 2.004E+04 | 880.5 | - | - | 0 | - |
| - | - | 891.6 | 881.4 | - | - | 0 | - |
| - | - | 7516 | 881.5 | - | - | 0 | - |
| - | - | 627.3 | 882.4 | - | - | 0 | - |
| - | - | 1690 | 882.5 | - | - | 0 | - |
| - | - | 708.7 | 956.6 | - | - | 0 | - |
| - | - | 758.4 | 1189 | - | - | 0 | - |
| - | - | 752.9 | 1288 | - | - | 0 | - |
| - | - | 1121 | 1302 | - | - | 0 | - |
| - | - | 1228 | 1303 | - | - | 0 | - |
| - | - | 1494 | 1304 | - | - | 0 | - |
| - | - | 739.9 | 1305 | - | - | 0 | - |
| - | - | 998.9 | 1318 | - | - | 0 | - |
| - | - | 1145 | 1319 | - | - | 0 | - |
| - | - | 2095 | 1320 | - | - | 0 | - |
| - | - | 1941 | 1321 | - | - | 0 | - |
| - | - | 605.6 | 1402 | - | - | 0 | - |
| - | - | 611.5 | 1422 | - | - | 0 | - |
| - | - | 660.5 | 2219 | - | - | 0 | - |
| - | - | 715.6 | 3218 | - | - | 0 | - |

m/z Charge Intensity FragmentType MassShift Position
120.06580352783203 0 1207.4208 y 8
120.08097076416016 0 4993.076
124.02474212646484 0 380.7984
126.82208251953125 0 386.934
129.10235595703125 0 2413.5723
133.08624267578125 0 995.90015
136.0751953125 0 407.05356
151.62965393066406 0 379.3969
153.07713317871094 0 1072.7052
155.0924835205078 0 791.72565
173.4400634765625 0 2359.9407
205.09796142578125 0 545.92615
212.13870239257812 0 598.9068
213.12367248535156 0 1383.02
219.1493377685547 0 16130.764
220.1526336669922 0 2246.1235
229.1547393798828 0 671.0727
231.16993713378906 0 673.57385
247.1441650390625 0 6550.2314
248.14785766601562 0 756.14136
273.1189270019531 0 809.50385
294.0758972167969 0 568.0616
322.10723876953125 0 678.857
327.2017517089844 0 1088.5764
345.2259826660156 0 4474.4287 z 6
346.2113037109375 0 958.44147
346.23358154296875 0 9746.546
347.2366943359375 0 1200.1677
359.21588134765625 0 1060.0941
361.2445068359375 0 1819.3204 y 6
385.24395751953125 0 1137.4286
386.2006530761719 0 546.13153
390.7160339355469 0 2577.4863 y 1
391.2182312011719 0 1198.1989
401.21527099609375 0 1641.9087
402.2711486816406 0 1802.2765
403.8240051269531 0 789.414
421.8330383300781 0 1003.8047
422.8336486816406 0 586.7602
434.2278747558594 0 829.0641
439.8440856933594 0 3527.0146
440.2513427734375 0 564.88824
440.8450012207031 0 1146.515
441.2601318359375 0 620.73083
441.2979431152344 0 885.36536
442.2579650878906 0 1894.4706
444.2572021484375 0 1881.0734
445.2658996582031 0 2406.1985 y Ammonia loss 5
446.2718811035156 0 923.8802 z 5
447.23388671875 0 1633.122
447.2818908691406 0 18598.4
448.2850341796875 0 4071.603
457.1685485839844 0 574.3111
472.3008117675781 0 1217.307
486.2437438964844 0 624.88293
496.28759765625 0 1714.8643
499.2750244140625 0 1642.1791
501.24627685546875 0 730.63336
503.2949523925781 0 1106.7676 z 4
504.2545166015625 0 9389.74
504.30291748046875 0 26708.467
505.2578430175781 0 3311.2898
505.3058776855469 0 6303.7617
506.3087158203125 0 692.9238
514.2984619140625 0 5139.407
515.3023681640625 0 1409.067
516.3109130859375 0 1298.0275
517.262451171875 0 921.7339
518.2653198242188 0 706.4243 c Water loss 5
518.3053588867188 0 1325.5948
519.3128662109375 0 3033.5103 y 4
520.3145751953125 0 672.56055
535.2745971679688 0 3355.5264
536.2825317382812 0 12277.156 c 5
537.2853393554688 0 3150.0806
538.2841186523438 0 646.71906
560.3161010742188 0 2755.0037 z 3
561.2766723632812 0 769.7273
561.324462890625 0 12523.286
562.326171875 0 2524.5383
574.2839965820312 0 944.09863
576.3348388671875 0 7011.302 y 3
577.339599609375 0 1594.3527
599.3255004882812 0 961.5476 z Water loss 2
615.3471069335938 0 1279.5165 y Water loss 2
617.3373413085938 0 3050.5054 z 2
618.3447265625 0 1764.891
619.3543090820312 0 1023.4969
632.3477783203125 0 793.206
633.356689453125 0 31018.06 y 2
634.3599243164062 0 10101.14
635.3595581054688 0 2073.9355
659.8511962890625 0 899.25543
660.3449096679688 0 736.96356
661.3649291992188 0 1545.5851
663.3800659179688 0 1622.8292
664.3778686523438 0 27926.467 c 6
665.3807983398438 0 9413.596
666.384521484375 0 2419.128
716.2301025390625 0 1611.1436
717.2254638671875 0 848.91797
733.4489135742188 0 1889.6715
734.4547119140625 0 905.56024
746.3908081054688 0 1228.8785 z Water loss 1
760.4326782226562 0 850.0011 c Ammonia loss 7
762.4503173828125 0 1013.80634
764.4060668945312 0 4983.4556 z 1
765.4107055664062 0 1695.4958
777.4617309570312 0 37706.56 c 7
778.46435546875 0 17380.164
779.466796875 0 4046.4292
780.4249877929688 0 4426.9546 y 1
781.4265747070312 0 2000.8058
789.4012451171875 0 1137.655
805.3851928710938 0 687.8167
820.4036865234375 0 670.3273
836.3292846679688 0 595.8222
840.5151977539062 0 667.478
863.474365234375 0 31516.482
864.3897094726562 0 1006.8569
864.4778442382812 0 13983.182
865.4810180664062 0 4414.5215
877.421875 0 883.24774
878.4302368164062 0 3742.721
879.4932250976562 0 23599.826
880.4982299804688 0 20039.453
881.4180297851562 0 891.6114
881.5010986328125 0 7516.0884
882.3875122070312 0 627.2704
882.5062866210938 0 1689.5286
956.5667724609375 0 708.6867
1188.591796875 0 758.4032
1287.68408203125 0 752.8595
1301.6898193359375 0 1121.1638
1302.6763916015625 0 1227.8217
1303.6795654296875 0 1493.9542
1304.693603515625 0 739.86334
1317.677490234375 0 998.8818
1318.69189453125 0 1144.9327
1319.695556640625 0 2095.2139
1320.7081298828125 0 1940.9501
1402.356689453125 0 605.6067
1422.07275390625 0 611.4676
2218.84423828125 0 660.5192
3217.511962890625 0 715.55023

Spectrum Details

|  |  |
| --- | --- |
| Matched peaks? Matched peaks The total absolute number of peaks matched. Additionally in brackets the total fraction of peaks matched and the total number of peaks is shown. | 22 (15.17% of 145) |
| FDR? FDR The false discovery rate estimated for this peptide. It is calculated by matching all theoretical fragments with a non-integer shift with the raw peaks for this spectrum. This is done with 40 different shifts. The resulting percentage is the average number of annotated peaks over the number of annotated peaks with the correct spectrum. | 0.54% |
| Satellite FDR? Satellite FDR See the FDR for details on its calculation. This satellite ion specific FDR only contains the satellite ions (d/w) for I/L/J positions. | - |
| PSM Score? PSM Score The PSM Score as given by Hecklib to this annotated spectrum. It is shown with three significant figures. | 270 |

#### Reverse Lookup? Reverse Lookup All places where this read could be placed.

| Group | Segment | Template | Template Part | Read Part | Score | Unique |
| --- | --- | --- | --- | --- | --- | --- |
| Homo sapiens Light Chain | IGLJ | IGLJ2 | [1..10] | [0..9] | 72 | True |

| Recombined | Template Part | Read Part | Score | Unique |
| --- | --- | --- | --- | --- |
| REC-0-1\_002 | [100..109] | [0..9] | 72 | True |

#### Meta Information from Multiple reads

##### Number of combined reads

6

##### Intensity

0.9568

##### TotalArea

1.068E+09

##### Changes to the peptide sequence

VFGGGTKJT

L→JNo support for either Leucine or Isoleucine based on side chain ions (Position: 8)

#### Positional Score

Copy Data

##### Positional Score (TSV)

###### Preview

```
Loading example...
```

*Click on the button to copy the data to your clipboard.*

10012345678

Label Value
"0" 0.667
"1" 0.667
"2" 0.667
"3" 0.638
"4" 0.585
"5" 0.622
"6" 0.662
"7" 0.665
"8" 0.665

#### Meta Information from PEAKS

##### Scan Identifier

F1:4540

##### Original sequence

V

F

G

G

G

T

K

L

T

##### Posttranslational Modifications

##### Source File

D:\separate\_stitch\_analyses\xle-disambiguation\raw\20210323\_F1\_UM1\_Peng0013\_SA\_F59\_ingel\_3ug\_ELA.raw

##### Fraction

1

##### Scan Feature

F1:1900

##### De Novo Score

99

##### ConfidenceScore

99

### m/z

440.2511

##### Mass

878.4861

##### Charge

2

##### Retention Time

24.06

##### Predicted Retention Time

-

##### Area

5.338E+08

##### Parts Per Million

1.8

##### Fragmentation mode

HCD

##### Originating file

01 D:\separate\_stitch\_analyses\xle-disambiguation\20210325\_F59\_3ug\_DENOVO\_12.csv

#### Meta Information from PEAKS

##### Scan Identifier

F1:4777

##### Original sequence

V

F

G

G

G

T

K

L

T

##### Posttranslational Modifications

##### Source File

D:\separate\_stitch\_analyses\xle-disambiguation\raw\20210323\_F1\_UM1\_Peng0013\_SA\_F59\_ingel\_3ug\_ELA.raw

##### Fraction

1

##### Scan Feature

-

##### De Novo Score

98

##### ConfidenceScore

98

### m/z

440.2511

##### Mass

878.4861

##### Charge

2

##### Retention Time

25.43

##### Predicted Retention Time

-

##### Area

0

##### Parts Per Million

1.7

##### Fragmentation mode

HCD

##### Originating file

01 D:\separate\_stitch\_analyses\xle-disambiguation\20210325\_F59\_3ug\_DENOVO\_12.csv

#### Meta Information from PEAKS

##### Scan Identifier

F1:4605

##### Original sequence

V

F

G

G

G

T

K

L

T

##### Posttranslational Modifications

##### Source File

D:\separate\_stitch\_analyses\xle-disambiguation\raw\20210323\_F1\_UM1\_Peng0013\_SA\_F59\_ingel\_3ug\_ELA.raw

##### Fraction

1

##### Scan Feature

F1:1900

##### De Novo Score

97

##### ConfidenceScore

97

### m/z

440.2511

##### Mass

878.4861

##### Charge

2

##### Retention Time

24.06

##### Predicted Retention Time

-

##### Area

5.338E+08

##### Parts Per Million

1.8

##### Fragmentation mode

ETHCD

##### Originating file

01 D:\separate\_stitch\_analyses\xle-disambiguation\20210325\_F59\_3ug\_DENOVO\_12.csv

#### Meta Information from PEAKS

##### Scan Identifier

F1:4698

##### Original sequence

V

F

G

G

G

T

K

L

T

##### Posttranslational Modifications

##### Source File

D:\separate\_stitch\_analyses\xle-disambiguation\raw\20210323\_F1\_UM1\_Peng0013\_SA\_F59\_ingel\_3ug\_ELA.raw

##### Fraction

1

##### Scan Feature

-

##### De Novo Score

97

##### ConfidenceScore

97

### m/z

440.2513

##### Mass

878.4861

##### Charge

2

##### Retention Time

24.97

##### Predicted Retention Time

-

##### Area

0

##### Parts Per Million

2.1

##### Fragmentation mode

ETHCD

##### Originating file

01 D:\separate\_stitch\_analyses\xle-disambiguation\20210325\_F59\_3ug\_DENOVO\_12.csv

#### Meta Information from PEAKS

##### Scan Identifier

F1:4927

##### Original sequence

V

F

G

G

G

T

K

L

T

##### Posttranslational Modifications

##### Source File

D:\separate\_stitch\_analyses\xle-disambiguation\raw\20210323\_F1\_UM1\_Peng0013\_SA\_F59\_ingel\_3ug\_ELA.raw

##### Fraction

1

##### Scan Feature

-

##### De Novo Score

97

##### ConfidenceScore

97

### m/z

440.2514

##### Mass

878.4861

##### Charge

2

##### Retention Time

26.26

##### Predicted Retention Time

-

##### Area

0

##### Parts Per Million

2.4

##### Fragmentation mode

HCD

##### Originating file

01 D:\separate\_stitch\_analyses\xle-disambiguation\20210325\_F59\_3ug\_DENOVO\_12.csv

#### Meta Information from PEAKS

##### Scan Identifier

F1:4999

##### Original sequence

V

F

G

G

G

T

K

L

T

##### Posttranslational Modifications

##### Source File

D:\separate\_stitch\_analyses\xle-disambiguation\raw\20210323\_F1\_UM1\_Peng0013\_SA\_F59\_ingel\_3ug\_ELA.raw

##### Fraction

1

##### Scan Feature

-

##### De Novo Score

95

##### ConfidenceScore

95

### m/z

440.2512

##### Mass

878.4861

##### Charge

2

##### Retention Time

26.67

##### Predicted Retention Time

-

##### Area

0

##### Parts Per Million

1.9

##### Fragmentation mode

ETHCD

##### Originating file

01 D:\separate\_stitch\_analyses\xle-disambiguation\20210325\_F59\_3ug\_DENOVO\_12.csv
